## Supplementary material for "Reversible m^6^Am methylation of snRNA by FTO controls morphine reward and tolerance without altering analgesia": Table S1

### Table S1. m6A_methylation

| chr_A_pos | NAc_GFP_ | NAc_GFP_ | NAc_GFP_ | NAc_GFP_ | NAc_GFP_ | NAc_GFP_ | NAc_GFP_ | NAc_GFP_ | NAc_GFP_ | NAc_GFP_ | NAc_Cre_1 | NAc_Cre_1 | NAc_Cre_2 | NAc_Cre_2 | NAc_Cre_3 | NAc_Cre_3 | NAc_Cre_4 | NAc_Cre_4 | NAc_Cre_n | NAc_Cre_merged_m6A_percent |
|---|---|---|---|---|---|---|---|---|---|---|---|---|---|---|---|---|---|---|---|---|
| 1_3270485 | 3 | 0 | 4 | 0.25 | 3 | 0 | 7 | 0.2857 | 17 | 0.1765 | 3 | 0 | 3 | 0 | 5 | 0 | 4 | 0.25 | 15 | 0.0667 |
| 1_3270753 | 5 | 0.2 | 6 | 0.6667 | 5 | 0.4 | 6 | 0.3333 | 22 | 0.4091 | 3 | 0.6667 | 4 | 0.25 | 5 | 0 | 4 | 0.5 | 16 | 0.3125 |
| 1_3274138 | NA | NA | 1 | 0 | NA | NA | 1 | 1 | 2 | 0.5 | 1 | 0 | NA | NA | 2 | 0.5 | 2 | 0 | 5 | 0.2 |
| 1_3274156 | 1 | 0 | 1 | 0 | NA | NA | 1 | 1 | 3 | 0.3333 | 1 | 1 | 1 | 0 | 3 | 0.3333 | NA | NA | 5 | 0.4 |
| 1_3275110 | 1 | 0 | 1 | 1 | NA | NA | 1 | 1 | 3 | 0.6667 | 1 | 0 | 1 | 1 | 3 | 0.6667 | NA | NA | 5 | 0.6 |
| 1_3275165 | 1 | 0 | 1 | 0 | NA | NA | 1 | 1 | 3 | 0.3333 | NA | NA | 1 | 1 | 1 | 0 | NA | NA | 2 | 0.5 |
| 1_3275322 | 1 | 0 | 1 | 1 | NA | NA | NA | NA | 2 | 0.5 | 1 | 0 | 1 | 1 | 2 | 0 | NA | NA | 4 | 0.25 |
| 1_3275479 | NA | NA | 1 | 1 | NA | NA | 1 | 1 | 2 | 1 | NA | NA | 1 | 1 | 2 | 1 | NA | NA | 3 | 1 |
| 1_3275537 | NA | NA | 1 | 0 | NA | NA | 1 | 1 | 2 | 0.5 | 1 | 0 | 1 | 0 | 2 | 0.5 | NA | NA | 4 | 0.25 |
| 1_3275663 | NA | NA | 1 | 0 | NA | NA | 1 | 1 | 2 | 0.5 | 1 | 1 | 1 | 1 | 2 | 1 | NA | NA | 4 | 1 |
| 1_3277255 | NA | NA | 1 | 1 | NA | NA | 1 | 1 | 2 | 1 | 1 | 1 | NA | NA | NA | NA | NA | NA | 1 | 1 |
| 1_3277355 | 1 | 0 | 1 | 1 | NA | NA | 1 | 1 | 3 | 0.6667 | 1 | 1 | NA | NA | NA | NA | NA | NA | 1 | 1 |
| 1_3277458 | 1 | 0 | 1 | 1 | NA | NA | 1 | 0 | 3 | 0.3333 | 1 | 1 | NA | NA | NA | NA | NA | NA | 1 | 1 |
| 1_3277530 | NA | NA | 1 | 1 | NA | NA | 1 | 0 | 2 | 0.5 | 1 | 1 | NA | NA | NA | NA | NA | NA | 1 | 1 |
| 1_3284083 | 1 | 1 | 1 | 1 | NA | NA | 1 | 1 | 3 | 1 | 1 | 1 | NA | NA | NA | NA | NA | NA | 1 | 1 |
| 1_3284151 | NA | NA | 1 | 1 | NA | NA | 1 | 0 | 2 | 0.5 | 1 | 1 | NA | NA | NA | NA | NA | NA | 1 | 1 |
| 1_4561697 | 2 | 0.5 | 2 | 0 | 2 | 0 | 1 | 1 | 7 | 0.2857 | NA | NA | 4 | 0.75 | NA | NA | 2 | 0 | 6 | 0.5 |
| 1_4561751 | 2 | 0.5 | 3 | 0.6667 | 2 | 0.5 | 1 | 1 | 8 | 0.625 | NA | NA | 4 | 0.5 | NA | NA | 1 | 0 | 5 | 0.4 |
| 1_4561797 | NA | NA | NA | NA | 2 | 0 | 1 | 1 | 3 | 0.3333 | NA | NA | NA | NA | NA | NA | 1 | 1 | 1 | 1 |
| 1_4562041 | 2 | 1 | 1 | 1 | 2 | 1 | 1 | 1 | 6 | 1 | NA | NA | 4 | 1 | NA | NA | 2 | 1 | 6 | 1 |
| 1_4562092 | 2 | 1 | 3 | 1 | 1 | 0 | 1 | 1 | 7 | 0.8571 | NA | NA | 3 | 1 | NA | NA | 2 | 1 | 5 | 1 |
| 1_4562108 | 1 | 0 | 2 | 0.5 | 1 | 0 | 1 | 1 | 5 | 0.4 | NA | NA | 4 | 0.5 | NA | NA | 2 | 0.5 | 6 | 0.5 |
| 1_4562426 | 1 | 1 | 1 | 1 | 2 | 1 | 1 | 1 | 5 | 1 | NA | NA | 4 | 1 | NA | NA | NA | NA | 4 | 1 |
| 1_4562468 | 1 | 0 | 2 | 1 | 2 | 0 | 1 | 1 | 6 | 0.5 | NA | NA | 4 | 0.75 | NA | NA | 1 | 0 | 5 | 0.6 |
| 1_4562501 | 2 | 0.5 | 3 | 0.3333 | 2 | 0.5 | 1 | 1 | 8 | 0.5 | NA | NA | 4 | 0.25 | NA | NA | 1 | 0 | 5 | 0.2 |
| 1_4562531 | 2 | 1 | 3 | 0.3333 | NA | NA | 1 | 0 | 6 | 0.5 | NA | NA | 3 | 0.3333 | NA | NA | NA | NA | 3 | 0.3333 |
| 1_4562575 | 2 | 1 | 2 | 1 | 1 | 1 | 1 | 1 | 6 | 1 | NA | NA | 4 | 1 | NA | NA | 1 | 1 | 5 | 1 |
| 1_4562585 | 2 | 1 | 2 | 1 | 2 | 1 | 1 | 1 | 7 | 1 | NA | NA | 3 | 1 | NA | NA | 1 | 1 | 4 | 1 |
| 1_4846666 | 6 | 0.5 | 11 | 0.3636 | 9 | 0.5556 | 2 | 0 | 28 | 0.4286 | 8 | 0.25 | 8 | 0.375 | 8 | 0.375 | 9 | 0.2222 | 33 | 0.303 |
| 1_4846686 | 6 | 0 | 11 | 0.6364 | 5 | 0.4 | 4 | 0.75 | 26 | 0.4615 | 8 | 0.5 | 6 | 0.8333 | 6 | 0.1667 | 9 | 0.3333 | 29 | 0.4483 |
| 1_4846825 | 2 | 1 | 4 | 0.25 | 6 | 0.6667 | 2 | 0.5 | 14 | 0.5714 | 6 | 0.5 | 6 | 1 | 8 | 0.625 | 5 | 0.6 | 25 | 0.68 |
| 1_4915308 | 6 | 0 | 3 | 0.3333 | NA | NA | 3 | 0 | 12 | 0.0833 | 4 | 0 | 2 | 0 | 5 | 0 | 6 | 0.1667 | 17 | 0.0588 |
| 1_4915320 | 6 | 0.1667 | 3 | 0 | NA | NA | 4 | 0.25 | 13 | 0.1538 | 4 | 0.25 | NA | NA | 4 | 0 | 5 | 0 | 13 | 0.0769 |
| 1_4967192 | 10 | 0 | 13 | 0.2308 | 14 | 0.2857 | 16 | 0.1875 | 53 | 0.1887 | 10 | 0.3 | 16 | 0.125 | 18 | 0.1111 | 20 | 0.25 | 64 | 0.1875 |
| 1_4967415 | 11 | 0.2727 | 16 | 0.3125 | 14 | 0.1429 | 16 | 0.125 | 57 | 0.2105 | 10 | 0.2 | 16 | 0.125 | 15 | 0.1333 | 23 | 0.2609 | 64 | 0.1875 |
| 1_4967480 | 10 | 0 | 15 | 0.0667 | 15 | 0.2 | 16 | 0 | 56 | 0.0714 | 11 | 0 | 15 | 0.0667 | 20 | 0.15 | 22 | 0.0455 | 68 | 0.0735 |
| 1_4976025 | NA | NA | NA | NA | NA | NA | 1 | 1 | 1 | 1 | NA | NA | NA | NA | 1 | 1 | 1 | 1 | 2 | 1 |
| 1_4980589 | 2 | 0.5 | NA | NA | 1 | 0 | 1 | 1 | 4 | 0.5 | 3 | 1 | 4 | 0.75 | 6 | 1 | 3 | 1 | 16 | 0.9375 |
| 1_4980647 | 3 | 0.6667 | NA | NA | 1 | 0 | NA | NA | 4 | 0.5 | 3 | 1 | 2 | 1 | 4 | 0.75 | 3 | 0.6667 | 12 | 0.8333 |
| 1_4980755 | 2 | 0.5 | NA | NA | 1 | 0 | 1 | 0 | 4 | 0.25 | 2 | 0 | 2 | 0 | 5 | 0.2 | 3 | 0.6667 | 12 | 0.25 |
| 1_4980776 | 3 | 1 | NA | NA | 1 | 0 | 1 | 1 | 5 | 0.8 | 4 | 0.5 | 4 | 0.75 | 7 | 0.4286 | 2 | 0 | 17 | 0.4706 |
| 1_5669019 | 3 | 0 | 1 | 0 | 1 | 1 | 3 | 0.3333 | 8 | 0.25 | 1 | 0 | 2 | 0 | 1 | 1 | 2 | 0 | 6 | 0.1667 |
| 1_5669058 | 2 | 1 | 1 | 1 | 1 | 1 | 3 | 1 | 7 | 1 | 1 | 1 | 3 | 0.6667 | 1 | 1 | 2 | 1 | 7 | 0.8571 |
| 1_5672661 | 3 | 1 | 1 | 0 | 1 | 1 | 3 | 0.3333 | 8 | 0.625 | 1 | 1 | 3 | 0.6667 | 1 | 1 | NA | NA | 5 | 0.8 |
| 1_5672726 | 3 | 1 | NA | NA | NA | NA | 3 | 1 | 6 | 1 | 1 | 1 | 3 | 1 | 1 | 1 | 1 | 1 | 6 | 1 |
| 1_5672871 | 3 | 0.3333 | 1 | 1 | 2 | 0.5 | 5 | 0 | 11 | 0.2727 | 1 | 0 | 2 | 1 | NA | NA | 2 | 0 | 5 | 0.4 |
| 1_5672892 | 3 | 1 | 1 | 1 | 1 | 1 | 5 | 1 | 10 | 1 | 1 | 0 | 3 | 1 | 1 | 1 | 2 | 1 | 7 | 0.8571 |
| 1_5673119 | 2 | 0 | 1 | 0 | 2 | 1 | 4 | 0 | 9 | 0.2222 | 1 | 1 | 3 | 0.6667 | 1 | 1 | 1 | 1 | 6 | 0.8333 |
| 1_5673125 | 3 | 0.6667 | NA | NA | 2 | 0 | 5 | 0.4 | 10 | 0.4 | 1 | 1 | 3 | 0 | 1 | 1 | 2 | 0 | 7 | 0.2857 |
| 1_5673200 | 3 | 0.3333 | 1 | 0 | 1 | 0 | 5 | 0 | 10 | 0.1 | 1 | 0 | 3 | 0.3333 | 1 | 0 | 2 | 0 | 7 | 0.1429 |
| 1_5673244 | 1 | 0 | 1 | 1 | 1 | 1 | 5 | 1 | 8 | 0.875 | 1 | 1 | 2 | 1 | NA | NA | 2 | 1 | 5 | 1 |
| 1_5673288 | 2 | 0.5 | 1 | 1 | 2 | 1 | 5 | 0.8 | 10 | 0.8 | 1 | 1 | 3 | 0.3333 | 1 | 0 | 2 | 0.5 | 7 | 0.4286 |
| 1_5673410 | 2 | 0.5 | 1 | 1 | 2 | 1 | 3 | 0.6667 | 8 | 0.75 | 1 | 0 | 3 | 0.6667 | 1 | 1 | 2 | 1 | 7 | 0.7143 |
| 1_5673419 | 3 | 1 | 1 | 1 | 2 | 1 | 3 | 1 | 9 | 1 | 1 | 1 | 2 | 1 | 1 | 1 | 2 | 1 | 6 | 1 |
| 1_5673552 | 3 | 1 | 1 | 1 | 2 | 1 | 4 | 1 | 10 | 1 | 1 | 1 | 3 | 0.6667 | 1 | 1 | 2 | 1 | 7 | 0.8571 |
| 1_5673594 | 3 | 1 | 1 | 1 | 2 | 1 | 5 | 1 | 11 | 1 | 1 | 1 | 3 | 0.6667 | 1 | 1 | 2 | 1 | 7 | 0.8571 |
| 1_5673604 | 2 | 0.5 | 1 | 1 | 2 | 0 | 5 | 0.6 | 10 | 0.5 | 1 | 1 | 3 | 0 | 1 | 0 | 2 | 0.5 | 7 | 0.2857 |
| 1_5673656 | 1 | 0 | 2 | 0.5 | 2 | 0 | 4 | 0 | 9 | 0.1111 | 1 | 1 | 3 | 0 | 1 | 0 | 2 | 0 | 7 | 0.1429 |
| 1_5673728 | 1 | 1 | 2 | 1 | NA | NA | 5 | 1 | 8 | 1 | 1 | 1 | 3 | 1 | 1 | 1 | 2 | 1 | 7 | 1 |
| 1_5675855 | 5 | 0 | 3 | 0.3333 | 2 | 0 | 7 | 0.2857 | 17 | 0.1765 | NA | NA | 5 | 0.4 | 2 | 0.5 | 2 | 0 | 9 | 0.3333 |
| 1_5675866 | 3 | 0.3333 | 2 | 0.5 | 2 | 1 | 6 | 0.5 | 13 | 0.5385 | 1 | 0 | 6 | 0.5 | 1 | 1 | 2 | 0.5 | 10 | 0.5 |
| 1_6308592 | 1 | 1 | 1 | 0 | NA | NA | 1 | 0 | 3 | 0.3333 | 1 | 0 | NA | NA | 1 | 0 | 3 | 0.3333 | 5 | 0.2 |
| 1_6310320 | 1 | 1 | 1 | 0 | NA | NA | NA | NA | 2 | 0.5 | 1 | 1 | NA | NA | 1 | 1 | 3 | 0.3333 | 5 | 0.6 |
| 1_6310355 | 1 | 0 | 1 | 0 | NA | NA | 1 | 1 | 3 | 0.3333 | 1 | 0 | NA | NA | 2 | 0.5 | 3 | 0 | 6 | 0.1667 |

| | | | | | | | | | | | | | | | | | | | | |
|---|---|---|---|---|---|---|---|---|---|---|---|---|---|---|---|---|---|---|---|---|
| 1_6310434 | 1 | 1 | 1 | 0 | NA | NA | 1 | 1 | 3 | 0.6667 | 1 | 1 | NA | NA | 2 | 0 | 2 | 0.5 | 5 | 0.4 |
| 1_6310493 | 1 | 0 | 1 | 1 | NA | NA | 1 | 0 | 3 | 0.3333 | 1 | 0 | 1 | 0 | 2 | 1 | 2 | 1 | 6 | 0.6667 |
| 1_6310529 | 1 | 1 | 1 | 1 | NA | NA | 1 | 0 | 3 | 0.6667 | 1 | 1 | 1 | 0 | 2 | 0.5 | 3 | 0 | 7 | 0.2857 |
| 1_6318738 | 3 | 0.3333 | 1 | 1 | NA | NA | 2 | 0.5 | 6 | 0.5 | 2 | 0 | 1 | 0 | 2 | 1 | 3 | 0.6667 | 8 | 0.5 |
| 1_6318771 | 3 | 0.3333 | 1 | 0 | NA | NA | 2 | 0.5 | 6 | 0.3333 | 2 | 0.5 | NA | NA | 3 | 0 | 3 | 0 | 8 | 0.125 |
| 1_6318999 | 3 | 0.6667 | 1 | 0 | 1 | 0 | 2 | 0.5 | 7 | 0.4286 | 2 | 0 | 1 | 0 | 3 | 0.3333 | 4 | 0.25 | 10 | 0.2 |
| 1_6319049 | 1 | 1 | 1 | 0 | 1 | 0 | 2 | 0.5 | 5 | 0.4 | 2 | 0 | 1 | 1 | 1 | 0 | 2 | 0.5 | 6 | 0.3333 |
| 1_6319066 | 3 | 0.3333 | 1 | 0 | 1 | 0 | 2 | 0 | 7 | 0.1429 | 2 | 1 | 1 | 1 | 3 | 0.6667 | 4 | 0.75 | 10 | 0.8 |
| 1_6319074 | 4 | 0.5 | 1 | 0 | 1 | 0 | 2 | 0.5 | 8 | 0.375 | 2 | 0 | NA | NA | 3 | 0 | 3 | 0.6667 | 8 | 0.25 |
| 1_6319177 | 4 | 0.25 | 1 | 0 | 1 | 0 | 2 | 0 | 8 | 0.125 | 2 | 0 | 1 | 0 | 4 | 0.5 | 4 | 0.5 | 11 | 0.3636 |
| 1_6319230 | 4 | 0 | 1 | 0 | 1 | 0 | 2 | 1 | 8 | 0.25 | 2 | 0.5 | NA | NA | 4 | 0 | 4 | 1 | 10 | 0.5 |
| 1_6319308 | 4 | 0 | 1 | 1 | 1 | 0 | 2 | 0 | 8 | 0.125 | 2 | 0 | 1 | 0 | 4 | 0.25 | 4 | 0.25 | 11 | 0.1818 |
| 1_6319429 | 3 | 0.6667 | 1 | 0 | 1 | 0 | 2 | 0 | 7 | 0.2857 | 2 | 0 | 1 | 1 | 4 | 0 | 3 | 0.6667 | 10 | 0.3 |
| 1_6319725 | 4 | 0.25 | 2 | 0.5 | 2 | 0 | 2 | 0 | 10 | 0.2 | 2 | 0 | 1 | 0 | 4 | 0.25 | 5 | 0 | 12 | 0.0833 |
| 1_6319759 | 3 | 0 | 2 | 0 | 3 | 0.6667 | 2 | 0 | 10 | 0.2 | 1 | 1 | NA | NA | 5 | 0 | 5 | 0.2 | 11 | 0.1818 |
| 1_6319859 | 4 | 0 | 2 | 0.5 | 3 | 0.3333 | 2 | 0 | 11 | 0.1818 | 2 | 0 | 1 | 0 | 5 | 0.2 | 4 | 0 | 12 | 0.0833 |
| 1_6319874 | 3 | 0 | 2 | 0.5 | 3 | 0 | 2 | 0 | 10 | 0.1 | 1 | 0 | 1 | 0 | 5 | 0.4 | 5 | 0.4 | 12 | 0.3333 |
| 1_6319958 | 3 | 0 | 3 | 0.3333 | 3 | 0.3333 | 1 | 1 | 10 | 0.3 | 2 | 0 | 1 | 0 | 5 | 0 | 5 | 0.2 | 13 | 0.0769 |
| 1_6320083 | 3 | 0.6667 | 3 | 0 | 2 | 0 | 2 | 0.5 | 10 | 0.3 | 2 | 0.5 | NA | NA | 4 | 0 | 5 | 0.2 | 11 | 0.1818 |
| 1_6320121 | 4 | 0.5 | 3 | 0.3333 | 3 | 0 | 1 | 0 | 11 | 0.2727 | 2 | 1 | 1 | 0 | 4 | 0.25 | 4 | 0.5 | 11 | 0.4545 |
| 1_6320224 | 5 | 0.4 | 3 | 0.6667 | 3 | 0.6667 | 2 | 0.5 | 13 | 0.5385 | 2 | 1 | 1 | 1 | 5 | 0.4 | 5 | 0.6 | 13 | 0.6154 |
| 1_6320254 | 4 | 0 | 4 | 0 | 2 | 0.5 | 2 | 0.5 | 12 | 0.1667 | 1 | 1 | 1 | 0 | 3 | 0.3333 | 5 | 0.4 | 10 | 0.4 |
| 1_6320300 | 5 | 0.4 | 4 | 0.25 | 3 | 0 | 2 | 0 | 14 | 0.2143 | 2 | 0 | 1 | 1 | 5 | 0.2 | 5 | 0.4 | 13 | 0.3077 |
| 1_6320332 | 4 | 0.5 | 4 | 0.75 | 3 | 0.3333 | 2 | 0.5 | 13 | 0.5385 | 2 | 0 | 1 | 1 | 5 | 0.4 | 5 | 0.2 | 13 | 0.3077 |
| 1_6344658 | 9 | 0.7778 | 4 | 0.75 | 3 | 0.6667 | 3 | 0 | 19 | 0.6316 | 4 | 0.5 | NA | NA | 4 | 0.25 | 6 | 0.6667 | 14 | 0.5 |
| 1_6344769 | 8 | 0 | 6 | 0.1667 | 2 | 0 | 3 | 0 | 19 | 0.0526 | 4 | 0 | 1 | 0 | 4 | 0.25 | 4 | 0.25 | 13 | 0.1538 |
| 1_6345205 | 9 | 0.4444 | 5 | 0 | 2 | 0 | 3 | 0 | 19 | 0.2105 | 4 | 0 | 1 | 0 | 5 | 0.6 | 4 | 0.5 | 14 | 0.3571 |
| 1_6345233 | 10 | 0.6 | 6 | 0.8333 | 2 | 0.5 | 3 | 0.3333 | 21 | 0.619 | 4 | 0.5 | 1 | 0 | 6 | 0.6667 | 2 | 0.5 | 13 | 0.5385 |
| 1_6345259 | 7 | 0.4286 | 2 | 0 | NA | NA | 3 | 0.3333 | 12 | 0.3333 | 4 | 0 | 1 | 0 | 5 | 0.2 | 4 | 0 | 14 | 0.0714 |
| 1_6873048 | NA | NA | NA | NA | NA | NA | 2 | 1 | 2 | 1 | NA | NA | NA | NA | 1 | 1 | NA | NA | 1 | 1 |
| 1_6873243 | NA | NA | NA | NA | NA | NA | 2 | 0.5 | 2 | 0.5 | NA | NA | 1 | 1 | 1 | 0 | NA | NA | 2 | 0.5 |
| 1_6928757 | NA | NA | NA | NA | NA | NA | 2 | 0.5 | 2 | 0.5 | NA | NA | 1 | 1 | 1 | 1 | NA | NA | 2 | 1 |
| 1_6928865 | NA | NA | NA | NA | NA | NA | 2 | 0.5 | 2 | 0.5 | NA | NA | 1 | 1 | 1 | 0 | NA | NA | 2 | 0.5 |
| 1_7190570 | 4 | 0.25 | 7 | 0.1429 | 5 | 0 | 10 | 0.1 | 26 | 0.1154 | 2 | 0.5 | 7 | 0.1429 | 4 | 0.25 | 3 | 0.3333 | 16 | 0.25 |
| 1_7190608 | 4 | 0.5 | 5 | 0 | 4 | 0.25 | 8 | 0.375 | 21 | 0.2857 | 3 | 0 | 7 | 0.5714 | 3 | 0.3333 | 3 | 0.3333 | 16 | 0.375 |
| 1_7190675 | 4 | 0 | 7 | 0.1429 | 4 | 0 | 10 | 0.2 | 25 | 0.12 | 2 | 0 | 6 | 0.1667 | 4 | 0.25 | 3 | 0 | 15 | 0.1333 |
| 1_7190690 | 3 | 0.6667 | 7 | 0.5714 | 5 | 0.4 | 10 | 0.7 | 25 | 0.6 | 2 | 0 | 7 | 0.5714 | 4 | 0.5 | 3 | 0.3333 | 16 | 0.4375 |
| 1_7230845 | NA | NA | 1 | 1 | 1 | 1 | 1 | 0 | 3 | 0.6667 | NA | NA | 1 | 1 | 1 | 0 | 1 | 0 | 3 | 0.3333 |
| 1_7239802 | 5 | 0.2 | 10 | 0.1 | 11 | 0.2727 | 12 | 0.1667 | 38 | 0.1842 | 2 | 0.5 | 11 | 0.4545 | 7 | 0.1429 | 7 | 0.4286 | 27 | 0.3704 |
| 1_7239878 | 4 | 0 | 6 | 0 | 10 | 0.2 | 7 | 0 | 27 | 0.0741 | 3 | 0 | 10 | 0.2 | 4 | 0 | 8 | 0.125 | 25 | 0.12 |
| 1_7239934 | 5 | 0 | 10 | 0.1 | 12 | 0 | 10 | 0.1 | 37 | 0.0541 | 3 | 0.3333 | 9 | 0.3333 | 6 | 0 | 8 | 0.125 | 26 | 0.1923 |
| 1_7240086 | 6 | 0.1667 | 12 | 0.0833 | 11 | 0.1818 | 11 | 0 | 40 | 0.1 | 3 | 0 | 12 | 0.0833 | 9 | 0.4444 | 9 | 0.4444 | 33 | 0.2727 |
| 1_7240099 | 6 | 0.3333 | 12 | 0.1667 | 10 | 0.1 | 12 | 0.25 | 40 | 0.2 | 3 | 0 | 11 | 0.1818 | 8 | 0.25 | 7 | 0.1429 | 29 | 0.1724 |
| 1_7240106 | 6 | 0.3333 | 10 | 0.3 | 8 | 0.375 | 9 | 0.3333 | 33 | 0.3333 | 3 | 0 | 10 | 0.4 | 8 | 0.375 | 7 | 0.2857 | 28 | 0.3214 |
| 1_7240136 | 5 | 0 | 7 | 0.1429 | 11 | 0.3636 | 9 | 0 | 32 | 0.1562 | 3 | 0.3333 | 11 | 0.2727 | 7 | 0.1429 | 8 | 0.125 | 29 | 0.2069 |
| 1_7240171 | 5 | 0.4 | 11 | 0.6364 | 11 | 0.6364 | 11 | 0.4545 | 38 | 0.5526 | 4 | 0.5 | 12 | 0.8333 | 9 | 0.7778 | 9 | 0.6667 | 34 | 0.7353 |
| 1_7240348 | 6 | 0 | 10 | 0.1 | 9 | 0.1111 | 11 | 0.4545 | 36 | 0.1944 | 4 | 0 | 11 | 0.2727 | 8 | 0 | 9 | 0 | 32 | 0.0938 |
| 1_7241767 | 10 | 0 | 17 | 0 | 13 | 0.1538 | 14 | 0.1429 | 54 | 0.0741 | 9 | 0.1111 | 12 | 0.0833 | 11 | 0.0909 | 10 | 0.1 | 42 | 0.0952 |
| 1_7242802 | 12 | 0.8333 | 18 | 0.7778 | 22 | 0.8182 | 20 | 0.75 | 72 | 0.7917 | 10 | 0.8 | 19 | 0.9474 | 16 | 0.625 | 13 | 0.6923 | 58 | 0.7759 |
| 1_7243099 | 2 | 0.5 | 2 | 0 | 2 | 0 | 3 | 0 | 9 | 0.1111 | 2 | 0 | 4 | 0.5 | 5 | 0 | 2 | 0 | 13 | 0.1538 |
| 1_7243108 | 2 | 0 | 2 | 0 | 4 | 0.25 | 3 | 0.3333 | 11 | 0.1818 | 2 | 0.5 | 4 | 0.25 | 5 | 0.2 | 2 | 0 | 13 | 0.2308 |
| 1_7243426 | 3 | 0.3333 | 1 | 0 | 3 | 0 | 3 | 0 | 10 | 0.1 | 2 | 0 | 4 | 0.25 | 6 | 0.1667 | 2 | 0 | 14 | 0.1429 |
| 1_7243694 | 3 | 0 | 2 | 0.5 | 4 | 0.25 | 3 | 0.3333 | 12 | 0.25 | 1 | 0 | 4 | 0.25 | 6 | 0.1667 | 2 | 0.5 | 13 | 0.2308 |
| 1_8430445 | 2 | 0.5 | 1 | 0 | 2 | 0 | 5 | 0 | 10 | 0.1 | 3 | 0 | 2 | 0.5 | NA | NA | NA | NA | 5 | 0.2 |
| 1_8432599 | 1 | 1 | NA | NA | 1 | 0 | 4 | 0.25 | 6 | 0.3333 | 2 | 0.5 | 2 | 0.5 | 1 | 0 | NA | NA | 5 | 0.4 |
| 1_8433560 | 2 | 0 | NA | NA | 1 | 1 | 4 | 0 | 7 | 0.1429 | 1 | 0 | 2 | 1 | 2 | 0.5 | NA | NA | 5 | 0.6 |
| 1_8433604 | 2 | 0.5 | NA | NA | 1 | 1 | 3 | 1 | 6 | 0.8333 | 1 | 0 | 2 | 1 | 2 | 0.5 | NA | NA | 5 | 0.6 |
| 1_8433637 | 2 | 0.5 | 1 | 0 | 1 | 0 | 3 | 0 | 7 | 0.1429 | 1 | 0 | 2 | 0.5 | 2 | 0 | NA | NA | 5 | 0.2 |
| 1_8433781 | 1 | 0 | 1 | 1 | 1 | 1 | 3 | 1 | 6 | 0.8333 | NA | NA | 2 | 1 | 2 | 0.5 | NA | NA | 4 | 0.75 |
| 1_9616381 | 1 | 0 | 4 | 0.25 | NA | NA | 1 | 0 | 6 | 0.1667 | 3 | 0 | 4 | 0.25 | 3 | 0 | 1 | 0 | 11 | 0.0909 |
| 1_9616527 | 1 | 1 | 4 | 0.25 | NA | NA | 3 | 0 | 8 | 0.25 | 3 | 0 | 4 | 0.5 | 3 | 0 | 3 | 0.3333 | 13 | 0.2308 |
| 1_9616548 | 1 | 1 | 5 | 0 | NA | NA | 3 | 0 | 9 | 0.1111 | 4 | 0 | 5 | 0.2 | 3 | 0 | 3 | 0 | 15 | 0.0667 |
| 1_9616702 | 1 | 1 | 6 | 1 | NA | NA | 3 | 1 | 10 | 1 | 4 | 1 | 5 | 1 | 3 | 1 | 3 | 0.6667 | 15 | 0.9333 |
| 1_9616862 | NA | NA | 6 | 0.3333 | NA | NA | 2 | 0 | 8 | 0.25 | 4 | 0.25 | 6 | 0.1667 | 3 | 0.3333 | 3 | 0 | 16 | 0.1875 |
| 1_9616882 | 1 | 1 | 5 | 1 | NA | NA | 3 | 1 | 9 | 1 | 3 | 1 | 4 | 1 | 2 | 1 | 3 | 1 | 12 | 1 |

| | | | | | | | | | | | | | | | | | | | | |
|---|---|---|---|---|---|---|---|---|---|---|---|---|---|---|---|---|---|---|---|---|
| 1_9616932 | NA | NA | 4 | 1 | NA | NA | 3 | 1 | 7 | 1 | 4 | 1 | 6 | 1 | 3 | 1 | 3 | 1 | 16 | 1 |
| 1_9616939 | 1 | 1 | 5 | 0 | NA | NA | 2 | 0.5 | 8 | 0.25 | 3 | 0 | 3 | 0.3333 | 2 | 0 | 3 | 0.3333 | 11 | 0.1818 |
| 1_9616995 | 1 | 0 | 5 | 0.6 | NA | NA | 3 | 1 | 9 | 0.6667 | 5 | 0.6 | 5 | 0.4 | 3 | 1 | 4 | 0.5 | 17 | 0.5882 |
| 1_9617001 | 1 | 0 | 5 | 0.4 | NA | NA | 2 | 0.5 | 8 | 0.375 | 4 | 0.5 | 5 | 0.4 | 2 | 0.5 | 3 | 0 | 14 | 0.3571 |
| 1_9617083 | 1 | 1 | 5 | 0.4 | NA | NA | 2 | 0.5 | 8 | 0.5 | 3 | 0.3333 | 5 | 0.2 | 3 | 0.3333 | 4 | 0 | 15 | 0.2 |
| 1_9646490 | 7 | 0.1429 | 2 | 0.5 | 4 | 0.25 | 3 | 0.3333 | 16 | 0.25 | 1 | 0 | 1 | 0 | 8 | 0.125 | 6 | 0.1667 | 16 | 0.125 |
| 1_9671664 | 5 | 0.4 | 17 | 0.2353 | 11 | 0.2727 | 6 | 0.1667 | 39 | 0.2564 | 11 | 0.1818 | 7 | 0 | 4 | 0 | 16 | 0.3125 | 38 | 0.1842 |
| 1_9695059 | 8 | 0.125 | 21 | 0.2381 | 15 | 0.2667 | 5 | 0 | 49 | 0.2041 | 13 | 0 | 9 | 0 | 6 | 0.1667 | 24 | 0.0833 | 52 | 0.0577 |
| 1_9695184 | 5 | 0.6 | 25 | 0.68 | 12 | 0.75 | 6 | 0.8333 | 48 | 0.7083 | 13 | 0.6154 | 10 | 0.9 | 5 | 0.6 | 23 | 0.913 | 51 | 0.8039 |
| 1_9695190 | 8 | 0.875 | 22 | 0.9091 | 16 | 0.9375 | 5 | 1 | 51 | 0.9216 | 13 | 0.9231 | 8 | 0.625 | 6 | 1 | 20 | 0.85 | 47 | 0.8511 |
| 1_9695305 | 7 | 0.4286 | 19 | 0.4737 | 14 | 0.5714 | 5 | 1 | 45 | 0.5556 | 10 | 0.5 | 8 | 0.625 | 6 | 0.8333 | 17 | 0.4706 | 41 | 0.561 |
| 1_9695391 | 8 | 0 | 24 | 0.25 | 12 | 0 | 7 | 0.1429 | 51 | 0.1373 | 12 | 0.25 | 7 | 0.2857 | 5 | 0.2 | 20 | 0.25 | 44 | 0.25 |
| 1_9695919 | 12 | 0 | 26 | 0.1154 | 12 | 0.0833 | 7 | 0 | 57 | 0.0702 | 17 | 0.0588 | 9 | 0.2222 | 8 | 0 | 22 | 0.0909 | 56 | 0.0893 |
| 1_9696172 | 10 | 0 | 27 | 0.0741 | 11 | 0.1818 | 5 | 0 | 53 | 0.0755 | 13 | 0.0769 | 8 | 0.125 | 6 | 0.1667 | 18 | 0.1111 | 45 | 0.1111 |
| 1_9696651 | 20 | 0.4 | 34 | 0.4706 | 18 | 0.2222 | 7 | 0.2857 | 79 | 0.3797 | 21 | 0.1429 | 14 | 0.2143 | 10 | 0.3 | 33 | 0.2424 | 78 | 0.2179 |
| 1_9696864 | 23 | 0.2174 | 25 | 0.4 | 15 | 0.2667 | 5 | 0.4 | 68 | 0.3088 | 19 | 0.2632 | 12 | 0 | 8 | 0.25 | 28 | 0.3214 | 67 | 0.2388 |
| 1_9697053 | 22 | 0.0455 | 37 | 0.0811 | 18 | 0.0556 | 5 | 0 | 82 | 0.061 | 23 | 0.087 | 9 | 0 | 8 | 0 | 31 | 0.0645 | 71 | 0.0563 |
| 1_9793476 | 4 | 0.25 | 2 | 0 | 6 | 0.1667 | 10 | 0.1 | 22 | 0.1364 | 4 | 0.5 | 5 | 0.2 | 9 | 0.2222 | 10 | 0 | 28 | 0.1786 |
| 1_9794886 | NA | NA | 2 | 1 | 4 | 0.25 | 7 | 0.2857 | 13 | 0.3846 | 2 | 0 | 1 | 0 | 5 | 0.4 | 5 | 0.6 | 13 | 0.3846 |
| 1_9794899 | NA | NA | 1 | 0 | 4 | 0.25 | 9 | 0.4444 | 14 | 0.3571 | 3 | 0.3333 | 3 | 0.3333 | 7 | 0 | 8 | 0.375 | 21 | 0.2381 |
| 1_9794913 | NA | NA | 2 | 1 | 3 | 1 | 8 | 0.875 | 13 | 0.9231 | 2 | 1 | 3 | 1 | 7 | 0.8571 | 7 | 1 | 19 | 0.9474 |
| 1_9795011 | NA | NA | 1 | 1 | 3 | 1 | 6 | 0.8333 | 10 | 0.9 | 3 | 1 | 3 | 1 | 7 | 0.8571 | 6 | 1 | 19 | 0.9474 |
| 1_9795201 | NA | NA | 2 | 0 | 4 | 0.75 | 8 | 0.5 | 14 | 0.5 | 1 | 1 | 3 | 1 | 7 | 0.7143 | 6 | 0.3333 | 17 | 0.6471 |
| 1_9795273 | NA | NA | 2 | 0.5 | 4 | 0.75 | 7 | 0.7143 | 13 | 0.6923 | 1 | 1 | 3 | 0.6667 | 7 | 0.8571 | 5 | 0.8 | 16 | 0.8125 |
| 1_9795358 | NA | NA | 1 | 0 | 4 | 0.25 | 5 | 0 | 10 | 0.1 | 2 | 0 | 3 | 0 | 7 | 0 | 6 | 0.1667 | 18 | 0.0556 |
| 1_9795472 | NA | NA | NA | NA | 2 | 0.5 | 9 | 0.6667 | 11 | 0.6364 | 1 | 1 | 3 | 0.3333 | 6 | 1 | 8 | 0.625 | 18 | 0.7222 |
| 1_9795499 | NA | NA | 2 | 1 | 1 | 1 | 4 | 0.5 | 7 | 0.7143 | 1 | 1 | 2 | 1 | 4 | 0.75 | 4 | 0.5 | 11 | 0.7273 |
| 1_9816152 | NA | NA | 2 | 1 | 2 | 0.5 | 5 | 0.8 | 9 | 0.7778 | 1 | 0 | 2 | 0 | 5 | 1 | 4 | 1 | 12 | 0.75 |
| 1_9816242 | NA | NA | 2 | 0 | 2 | 0.5 | 5 | 0.8 | 9 | 0.5556 | 1 | 1 | 1 | 1 | 6 | 0.8333 | 5 | 0.8 | 13 | 0.8462 |
| 1_9816284 | NA | NA | 2 | 0 | 1 | 1 | 4 | 0.25 | 7 | 0.2857 | 1 | 0 | 2 | 0.5 | 6 | 0.3333 | 5 | 0 | 14 | 0.2143 |
| 1_9816297 | NA | NA | 2 | 1 | 1 | 0 | 2 | 1 | 5 | 0.8 | 1 | 1 | 2 | 0.5 | 5 | 0.8 | 4 | 0.5 | 12 | 0.6667 |
| 1_9816342 | NA | NA | 2 | 0 | 2 | 0 | 5 | 0.4 | 9 | 0.2222 | 1 | 0 | 2 | 0 | 6 | 0.1667 | 6 | 0.1667 | 15 | 0.1333 |
| 1_9816422 | NA | NA | 1 | 0 | 2 | 0.5 | 5 | 0.6 | 8 | 0.5 | 1 | 0 | 2 | 0.5 | 4 | 1 | 4 | 0.75 | 11 | 0.7273 |
| 1_9816501 | NA | NA | 2 | 0 | 2 | 0 | 5 | 0.2 | 9 | 0.1111 | 1 | 1 | 2 | 0 | 5 | 0.2 | 6 | 0 | 14 | 0.1429 |
| 1_9816799 | NA | NA | 2 | 0.5 | 1 | 1 | 4 | 0 | 7 | 0.2857 | 1 | 1 | 1 | 1 | 5 | 0 | 5 | 0.2 | 12 | 0.25 |
| 1_9816915 | NA | NA | 2 | 1 | NA | NA | 4 | 0 | 6 | 0.3333 | NA | NA | 1 | 0 | 5 | 0.2 | 5 | 0 | 11 | 0.0909 |
| 1_9816954 | NA | NA | 2 | 1 | NA | NA | 4 | 0.75 | 6 | 0.8333 | NA | NA | NA | NA | 4 | 0.75 | 5 | 0.6 | 9 | 0.6667 |
| 1_9816997 | NA | NA | 2 | 1 | 1 | 1 | 5 | 1 | 8 | 1 | NA | NA | 1 | 1 | 3 | 1 | 4 | 1 | 8 | 1 |
| 1_9817058 | NA | NA | NA | NA | 1 | 1 | 1 | 1 | 2 | 1 | NA | NA | 1 | 1 | 4 | 1 | 2 | 1 | 7 | 1 |
| 1_9817117 | NA | NA | 2 | 1 | NA | NA | 5 | 1 | 7 | 1 | NA | NA | 1 | 1 | 5 | 0.8 | 4 | 0.5 | 10 | 0.7 |
| 1_9817237 | NA | NA | 2 | 0.5 | NA | NA | 4 | 1 | 6 | 0.8333 | NA | NA | 1 | 1 | 5 | 0.6 | 4 | 0.5 | 10 | 0.6 |
| 1_9817582 | NA | NA | 2 | 1 | 1 | 1 | 4 | 1 | 7 | 1 | NA | NA | NA | NA | 2 | 1 | 5 | 1 | 7 | 1 |
| 1_9817609 | NA | NA | 1 | 1 | NA | NA | 4 | 0.25 | 5 | 0.4 | NA | NA | NA | NA | 4 | 0.5 | 3 | 0.3333 | 7 | 0.4286 |
| 1_9817734 | NA | NA | 1 | 1 | NA | NA | 4 | 1 | 5 | 1 | NA | NA | NA | NA | 4 | 1 | 2 | 1 | 6 | 1 |
| 1_9817917 | NA | NA | 1 | 0 | NA | NA | 2 | 1 | 3 | 0.6667 | NA | NA | NA | NA | 4 | 0.75 | 3 | 1 | 7 | 0.8571 |
| 1_9817966 | NA | NA | NA | NA | NA | NA | 4 | 0.5 | 4 | 0.5 | NA | NA | NA | NA | 2 | 1 | 3 | 1 | 5 | 1 |
| 1_9818040 | NA | NA | NA | NA | NA | NA | 3 | 0.3333 | 3 | 0.3333 | NA | NA | NA | NA | 2 | 0.5 | 3 | 0.3333 | 5 | 0.4 |
| 1_9935435 | 3 | 0 | 2 | 0 | 3 | 0.3333 | 1 | 0 | 9 | 0.1111 | 1 | 0 | 3 | 0 | NA | NA | 2 | 0.5 | 6 | 0.1667 |
| 1_9969024 | 4 | 0.25 | 8 | 0.125 | 5 | 0 | 5 | 0.2 | 22 | 0.1364 | 1 | 0 | 5 | 0 | 1 | 0 | 3 | 0.3333 | 10 | 0.1 |
| 1_9969102 | 5 | 0.2 | 8 | 0.375 | 5 | 0.6 | 6 | 0.5 | 24 | 0.4167 | 1 | 0 | 6 | 0.1667 | 1 | 1 | 3 | 0 | 11 | 0.1818 |
| 1_9969116 | 3 | 1 | 8 | 0.75 | 5 | 0.6 | 6 | 0.6667 | 22 | 0.7273 | 1 | 1 | 4 | 0.25 | 1 | 1 | 3 | 1 | 9 | 0.6667 |
| 1_9969124 | 4 | 0.25 | 7 | 0 | 4 | 0.25 | 4 | 0.25 | 19 | 0.1579 | 1 | 0 | 6 | 0.1667 | 1 | 0 | 4 | 0.25 | 12 | 0.1667 |
| 1_9969142 | 4 | 0 | 6 | 0.1667 | 5 | 0.2 | 5 | 0 | 20 | 0.1 | 1 | 0 | 5 | 0.4 | 1 | 1 | 3 | 0.6667 | 10 | 0.5 |
| 1_9969302 | 5 | 0.2 | 6 | 0.3333 | 5 | 0 | 6 | 0.1667 | 22 | 0.1818 | 1 | 0 | 6 | 0.3333 | 1 | 0 | 4 | 0.25 | 12 | 0.25 |
| 1_9969800 | 5 | 0.2 | 7 | 0 | 7 | 0.4286 | 6 | 0.5 | 25 | 0.28 | 1 | 1 | 8 | 0.375 | 1 | 1 | 6 | 0.6667 | 16 | 0.5625 |
| 1_1009525 | 20 | 0.05 | 15 | 0.4667 | 16 | 0.0625 | 9 | 0 | 60 | 0.15 | 5 | 0.2 | 8 | 0.125 | 9 | 0.2222 | 12 | 0.1667 | 34 | 0.1765 |
| 1_1010790 | 17 | 0.1176 | 12 | 0.0833 | 9 | 0 | 4 | 0 | 42 | 0.0714 | 3 | 0 | 6 | 0 | 8 | 0.125 | 9 | 0.1111 | 26 | 0.0769 |
| 1_1010828 | NA | NA | 1 | 0 | 1 | 1 | NA | NA | 2 | 0.5 | NA | NA | NA | NA | NA | NA | 1 | 1 | 1 | 1 |
| 1_1013674 | NA | NA | NA | NA | NA | NA | 1 | 1 | 1 | 1 | NA | NA | NA | NA | NA | NA | 1 | 1 | 1 | 1 |
| 1_1020801 | 10 | 0.2 | 7 | 0.2857 | 11 | 0.3636 | 8 | 0 | 36 | 0.2222 | 6 | 0.3333 | 11 | 0.0909 | 6 | 0.3333 | 5 | 0 | 28 | 0.1786 |
| 1_1020819 | 10 | 0.7 | 6 | 0.5 | 15 | 0.6667 | 8 | 0.5 | 39 | 0.6154 | 5 | 0.2 | 14 | 0.2857 | 6 | 0.5 | 9 | 0.5556 | 34 | 0.3824 |
| 1_1020822 | 9 | 0 | 7 | 0.1429 | 14 | 0.0714 | 9 | 0 | 39 | 0.0513 | 4 | 0 | 16 | 0.0625 | 6 | 0.1667 | 8 | 0.125 | 34 | 0.0882 |
| 1_1020839 | 9 | 0.2222 | 7 | 0 | 14 | 0.0714 | 8 | 0 | 38 | 0.0789 | 4 | 0 | 14 | 0.1429 | 4 | 0 | 7 | 0 | 29 | 0.069 |
| 1_1020872 | 8 | 0.75 | 4 | 0.25 | 11 | 0.6364 | 7 | 0.5714 | 30 | 0.6 | 5 | 0.4 | 9 | 0.4444 | 3 | 0.6667 | 8 | 0.5 | 25 | 0.48 |
| 1_1020907 | 7 | 0.7143 | 4 | 0.25 | 11 | 0.3636 | 6 | 0.5 | 28 | 0.4643 | 4 | 0.5 | 11 | 0.2727 | 2 | 0 | 8 | 0.375 | 25 | 0.32 |

| | | | | | | | | | | | | | | | | | | | | |
|---|---|---|---|---|---|---|---|---|---|---|---|---|---|---|---|---|---|---|---|---|
| 1_1024313 | 4 | 0 | 1 | 0 | 6 | 0 | 3 | 0.3333 | 14 | 0.0714 | 2 | 0.5 | 4 | 0 | 1 | 0 | 3 | 0 | 10 | 0.1 |
| 1_1136810 | NA | NA | 3 | 0.3333 | 2 | 0.5 | 1 | 1 | 6 | 0.5 | NA | NA | 1 | 0 | 2 | 1 | 2 | 0 | 5 | 0.4 |
| 1_1166785 | 4 | 0.5 | 2 | 0 | NA | NA | 7 | 0.4286 | 13 | 0.3846 | 3 | 0 | 3 | 0.3333 | 1 | 0 | 8 | 0.25 | 15 | 0.2 |
| 1_1166806 | 5 | 0.2 | 2 | 0 | 1 | 0 | 6 | 0.1667 | 14 | 0.1429 | 3 | 0 | 3 | 0 | 1 | 0 | 8 | 0.25 | 15 | 0.1333 |
| 1_1166811 | 5 | 0 | 2 | 0 | 1 | 0 | 5 | 0.2 | 13 | 0.0769 | 2 | 0.5 | 3 | 0 | 1 | 0 | 8 | 0 | 14 | 0.0714 |
| 1_1166820 | 5 | 0.4 | 2 | 0 | 1 | 0 | 5 | 0.4 | 13 | 0.3077 | 4 | 0.25 | 3 | 0 | 1 | 1 | 8 | 0.25 | 16 | 0.25 |
| 1_1167120 | 6 | 0.1667 | 2 | 0 | 4 | 0 | 3 | 0.3333 | 15 | 0.1333 | 2 | 0.5 | 2 | 0 | 5 | 0 | 4 | 0 | 13 | 0.0769 |
| 1_1167124 | 6 | 0.1667 | 2 | 0 | 4 | 0 | 2 | 0.5 | 14 | 0.1429 | 2 | 0.5 | 2 | 0.5 | 5 | 0.2 | 3 | 0 | 12 | 0.25 |
| 1_1167136 | 6 | 0.1667 | 3 | 0 | 4 | 0 | 3 | 0 | 16 | 0.0625 | 2 | 0.5 | 2 | 0 | 5 | 0 | 3 | 0 | 12 | 0.0833 |
| 1_1167145 | 5 | 0.4 | 2 | 0 | 4 | 0.25 | 3 | 0 | 14 | 0.2143 | 1 | 1 | 1 | 0 | 5 | 0.2 | 4 | 0 | 11 | 0.1818 |
| 1_1292917 | 1 | 1 | 2 | 1 | NA | NA | 6 | 1 | 9 | 1 | 2 | 1 | 4 | 0.75 | 3 | 1 | 2 | 0.5 | 11 | 0.8182 |
| 1_1292917 | 1 | 1 | 2 | 1 | NA | NA | 5 | 1 | 8 | 1 | 1 | 1 | 4 | 1 | 2 | 0.5 | 2 | 0.5 | 9 | 0.7778 |
| 1_1292921 | NA | NA | 1 | 0 | NA | NA | 3 | 0.3333 | 4 | 0.25 | NA | NA | 2 | 0.5 | 1 | 0 | 1 | 0 | 4 | 0.25 |
| 1_1292925 | 1 | 1 | 2 | 1 | NA | NA | 4 | 1 | 7 | 1 | 2 | 1 | 6 | 1 | 3 | 1 | 2 | 1 | 13 | 1 |
| 1_1292927 | 1 | 0 | 1 | 1 | NA | NA | 4 | 0.5 | 6 | 0.5 | 2 | 0.5 | 4 | 0.25 | 3 | 0.3333 | 1 | 0 | 10 | 0.3 |
| 1_1292940 | 1 | 0 | 3 | 0 | NA | NA | 6 | 0.1667 | 10 | 0.1 | 2 | 0.5 | 3 | 0 | 3 | 0 | 2 | 0 | 10 | 0.1 |
| 1_1292943 | 1 | 0 | 3 | 0.3333 | NA | NA | 5 | 0.2 | 9 | 0.2222 | 2 | 0.5 | 7 | 0 | 3 | 0 | 3 | 0.3333 | 15 | 0.1333 |
| 1_1292945 | 1 | 0 | 3 | 0.6667 | NA | NA | 7 | 0.4286 | 11 | 0.4545 | 2 | 0.5 | 7 | 0.5714 | 4 | 0.5 | 3 | 0.6667 | 16 | 0.5625 |
| 1_1292948 | 2 | 0.5 | 2 | 1 | NA | NA | 7 | 0.4286 | 11 | 0.5455 | 1 | 1 | 6 | 0.3333 | 4 | 0.75 | 1 | 0 | 12 | 0.5 |
| 1_1292949 | 1 | 0 | 3 | 1 | NA | NA | 5 | 1 | 9 | 0.8889 | 2 | 1 | 6 | 0.8333 | 3 | 0 | 2 | 1 | 13 | 0.6923 |
| 1_1292964 | 1 | 0 | NA | NA | NA | NA | 7 | 0.1429 | 8 | 0.125 | NA | NA | 5 | 0.2 | 3 | 0 | 3 | 0 | 11 | 0.0909 |
| 1_1292969 | 2 | 0.5 | 2 | 0 | 1 | 0 | 4 | 0 | 9 | 0.1111 | 2 | 0 | 5 | 0.2 | 3 | 0 | 3 | 0.3333 | 13 | 0.1538 |
| 1_1292969 | 2 | 0.5 | 2 | 0.5 | 1 | 1 | 5 | 0.8 | 10 | 0.7 | NA | NA | 3 | 0.6667 | 4 | 0.25 | 2 | 0 | 9 | 0.3333 |
| 1_1292990 | 3 | 0.3333 | 3 | 0 | NA | NA | 5 | 0 | 11 | 0.0909 | 2 | 0 | 6 | 0.1667 | 2 | 0 | 2 | 0 | 12 | 0.0833 |
| 1_1292992 | 2 | 1 | 3 | 0 | 1 | 1 | 6 | 0.1667 | 12 | 0.3333 | 2 | 0 | 8 | 0.5 | 4 | 0.25 | 5 | 0 | 19 | 0.2632 |
| 1_1292998 | 3 | 0.6667 | 3 | 0.6667 | 1 | 1 | 7 | 0.2857 | 14 | 0.5 | 3 | 0 | 7 | 0.2857 | 4 | 0.5 | 4 | 0 | 18 | 0.2222 |
| 1_1293002 | 3 | 1 | 3 | 0.6667 | NA | NA | 7 | 0.7143 | 13 | 0.7692 | 3 | 0.3333 | 6 | 1 | 4 | 1 | 5 | 1 | 18 | 0.8889 |
| 1_1293003 | 3 | 0 | 3 | 0 | 1 | 1 | 7 | 0.2857 | 14 | 0.2143 | 2 | 0 | 5 | 0.6 | 3 | 0 | 3 | 0 | 13 | 0.2308 |
| 1_1293013 | 3 | 0.3333 | 3 | 0.3333 | 1 | 1 | 7 | 0 | 14 | 0.2143 | 2 | 0 | 7 | 0.2857 | 2 | 0 | 5 | 0.4 | 16 | 0.25 |
| 1_1293043 | 4 | 0.25 | 5 | 0.4 | 2 | 0 | 7 | 0.2857 | 18 | 0.2778 | 3 | 0 | 7 | 0.1429 | 4 | 0 | 5 | 0 | 19 | 0.0526 |
| 1_1293047 | 4 | 0 | 5 | 0 | 2 | 0 | 7 | 0.4286 | 18 | 0.1667 | 2 | 0 | 6 | 0.3333 | 3 | 0 | 5 | 0.2 | 16 | 0.1875 |
| 1_1293094 | 4 | 1 | NA | NA | 1 | 0 | 2 | 1 | 7 | 0.8571 | 1 | 1 | 2 | 1 | 1 | 1 | 1 | 0 | 5 | 0.8 |
| 1_1293686 | NA | NA | 1 | 0 | 1 | 0 | 1 | 1 | 3 | 0.3333 | 1 | 0 | 3 | 0 | 3 | 0.3333 | 1 | 0 | 8 | 0.125 |
| 1_1294147 | NA | NA | 1 | 1 | NA | NA | NA | NA | 1 | 1 | 1 | 1 | 1 | 1 | 2 | 0.5 | 1 | 1 | 5 | 0.8 |
| 1_1294191 | NA | NA | 1 | 1 | NA | NA | NA | NA | 1 | 1 | NA | NA | 1 | 1 | 2 | 1 | 1 | 1 | 4 | 1 |
| 1_1294196 | NA | NA | 1 | 1 | NA | NA | NA | NA | 1 | 1 | 1 | 1 | 1 | 1 | 2 | 0.5 | 1 | 0 | 5 | 0.6 |
| 1_1294198 | NA | NA | 1 | 1 | NA | NA | NA | NA | 1 | 1 | 1 | 1 | 1 | 0 | 2 | 1 | 1 | 0 | 5 | 0.6 |
| 1_1294212 | NA | NA | 1 | 1 | NA | NA | NA | NA | 1 | 1 | 1 | 0 | NA | NA | 2 | 0 | 1 | 1 | 4 | 0.25 |
| 1_1294216 | NA | NA | 1 | 1 | NA | NA | NA | NA | 1 | 1 | 1 | 1 | 1 | 1 | 1 | 1 | 1 | 1 | 4 | 1 |
| 1_1294225 | NA | NA | 1 | 1 | NA | NA | NA | NA | 1 | 1 | 1 | 0 | 1 | 1 | 2 | 0.5 | 1 | 1 | 5 | 0.6 |
| 1_1321065 | 6 | 0 | 10 | 0.1 | 13 | 0 | 9 | 0.2222 | 38 | 0.0789 | 3 | 0 | 5 | 0 | 7 | 0 | 6 | 0.3333 | 21 | 0.0952 |
| 1_1321196 | 3 | 0.3333 | 7 | 0 | 7 | 0.1429 | 11 | 0.0909 | 28 | 0.1071 | 2 | 0 | 4 | 0 | 9 | 0.1111 | 4 | 0.5 | 19 | 0.1579 |
| 1_1321253 | 2 | 1 | 6 | 0.5 | 5 | 0.2 | 10 | 0.5 | 23 | 0.4783 | 2 | 0.5 | 3 | 0.6667 | 8 | 0.375 | 2 | 0.5 | 15 | 0.4667 |
| 1_1321260 | 2 | 0 | 5 | 0.4 | 7 | 0 | 11 | 0.1818 | 25 | 0.16 | 2 | 0 | 3 | 0 | 9 | 0.1111 | 3 | 0.3333 | 17 | 0.1176 |
| 1_1321270 | 2 | 1 | 4 | 0 | 6 | 0.3333 | 9 | 0.4444 | 21 | 0.381 | 2 | 0.5 | 3 | 0.6667 | 8 | 0.5 | 3 | 1 | 16 | 0.625 |
| 1_1321285 | 1 | 0 | 3 | 0 | 6 | 0.1667 | 5 | 0.4 | 15 | 0.2 | 1 | 0 | 2 | 0 | 7 | 0 | 2 | 0.5 | 12 | 0.0833 |
| 1_1321287 | 1 | 1 | 3 | 0.6667 | 6 | 0.5 | 6 | 0.5 | 16 | 0.5625 | 2 | 0.5 | 4 | 0.5 | 8 | 0.375 | 1 | 0 | 15 | 0.4 |
| 1_1321293 | 1 | 0 | 2 | 0 | 6 | 0.3333 | 9 | 0.3333 | 18 | 0.2778 | 1 | 1 | 1 | 0 | 7 | 0 | 2 | 0 | 11 | 0.0909 |
| 1_1321297 | 2 | 0 | 3 | 0 | 6 | 0 | 8 | 0.375 | 19 | 0.1579 | 2 | 0.5 | 3 | 0 | 8 | 0 | 3 | 0 | 16 | 0.0625 |
| 1_1323523 | 2 | 0.5 | NA | NA | 1 | 1 | 5 | 0 | 8 | 0.25 | 1 | 0 | 1 | 0 | 3 | 0.3333 | 2 | 0 | 7 | 0.1429 |
| 1_1324438 | 2 | 0 | NA | NA | 1 | 1 | 3 | 0 | 6 | 0.1667 | 1 | 0 | 1 | 0 | 2 | 0 | 2 | 0.5 | 6 | 0.1667 |
| 1_1324451 | 2 | 0 | NA | NA | 1 | 1 | 3 | 0.3333 | 6 | 0.3333 | 1 | 0 | 1 | 0 | 1 | 1 | 2 | 0.5 | 5 | 0.4 |
| 1_1324459 | 2 | 1 | NA | NA | 1 | 0 | 3 | 1 | 6 | 0.8333 | 1 | 1 | NA | NA | 2 | 0.5 | 2 | 0.5 | 5 | 0.6 |
| 1_1324461 | 2 | 0.5 | NA | NA | 1 | 0 | 3 | 0.6667 | 6 | 0.5 | 1 | 1 | NA | NA | 1 | 0 | 2 | 1 | 4 | 0.75 |
| 1_1324462 | 1 | 1 | NA | NA | 1 | 1 | 3 | 0.3333 | 5 | 0.6 | 1 | 1 | NA | NA | 2 | 1 | 2 | 0.5 | 5 | 0.8 |
| 1_1324469 | 1 | 1 | NA | NA | NA | NA | 3 | 0.6667 | 4 | 0.75 | 1 | 1 | 1 | 1 | 1 | 1 | 2 | 1 | 5 | 1 |
| 1_1324470 | 2 | 0.5 | NA | NA | NA | NA | 3 | 1 | 5 | 0.8 | 1 | 0 | 1 | 1 | 1 | 0 | 2 | 1 | 5 | 0.6 |
| 1_1324481 | 2 | 0.5 | NA | NA | NA | NA | 2 | 0.5 | 4 | 0.5 | NA | NA | 1 | 0 | 2 | 0 | 2 | 1 | 5 | 0.4 |
| 1_1324493 | 2 | 1 | NA | NA | NA | NA | 3 | 1 | 5 | 1 | NA | NA | NA | NA | 2 | 1 | 1 | 1 | 3 | 1 |
| 1_1324494 | 2 | 1 | NA | NA | NA | NA | 3 | 0.6667 | 5 | 0.8 | NA | NA | 1 | 1 | 2 | 1 | NA | NA | 3 | 1 |
| 1_1324495 | 2 | 0 | NA | NA | NA | NA | 3 | 0.6667 | 5 | 0.4 | NA | NA | NA | NA | 1 | 0 | 1 | 1 | 2 | 0.5 |
| 1_1324503 | 2 | 0 | NA | NA | NA | NA | 3 | 0.6667 | 5 | 0.4 | NA | NA | 1 | 1 | 2 | 0.5 | 1 | 0 | 4 | 0.5 |
| 1_1324545 | 1 | 1 | NA | NA | NA | NA | 2 | 0.5 | 3 | 0.6667 | NA | NA | 1 | 1 | 2 | 0 | NA | NA | 3 | 0.3333 |
| 1_1363506 | 3 | 0.3333 | 3 | 0.3333 | 4 | 0 | 5 | 0.2 | 15 | 0.2 | 3 | 0 | 6 | 0 | 1 | 0 | 6 | 0.1667 | 16 | 0.0625 |
| 1_1368700 | 1 | 0 | 2 | 0.5 | NA | NA | NA | NA | 3 | 0.3333 | 2 | 0.5 | NA | NA | 3 | 0.3333 | NA | NA | 5 | 0.4 |

| | | | | | | | | | | | | | | | | | | | | |
|---|---|---|---|---|---|---|---|---|---|---|---|---|---|---|---|---|---|---|---|---|
| 1_1368724 | 1 | 0 | 2 | 0.5 | NA | NA | NA | NA | 3 | 0.3333 | 2 | 0 | NA | NA | 3 | 0.3333 | NA | NA | 5 | 0.2 |
| 1_1369531 | 1 | 0 | 2 | 0.5 | 2 | 0 | NA | NA | 5 | 0.2 | 1 | 0 | NA | NA | 4 | 0 | 2 | 0.5 | 7 | 0.1429 |
| 1_1369656 | 1 | 0 | 1 | 0 | 2 | 0.5 | NA | NA | 4 | 0.25 | 2 | 0 | NA | NA | 4 | 0.5 | 2 | 0 | 8 | 0.25 |
| 1_1423984 | 2 | 1 | 2 | 0 | NA | NA | 1 | 0 | 5 | 0.4 | NA | NA | 3 | 0.6667 | 1 | 0 | 1 | 1 | 5 | 0.6 |
| 1_1424056 | 2 | 0.5 | 2 | 1 | NA | NA | 1 | 1 | 5 | 0.8 | NA | NA | 3 | 0.6667 | NA | NA | 1 | 1 | 4 | 0.75 |
| 1_1424059 | 1 | 1 | 2 | 1 | NA | NA | 1 | 0 | 4 | 0.75 | NA | NA | 3 | 0.3333 | NA | NA | 1 | 0 | 4 | 0.25 |
| 1_1424065 | 2 | 1 | 1 | 1 | NA | NA | 1 | 1 | 4 | 1 | NA | NA | 2 | 1 | NA | NA | NA | NA | 2 | 1 |
| 1_1424076 | 2 | 0.5 | NA | NA | NA | NA | 1 | 0 | 3 | 0.3333 | NA | NA | 2 | 1 | NA | NA | 1 | 1 | 3 | 1 |
| 1_1424078 | 2 | 0.5 | 2 | 0 | NA | NA | 1 | 0 | 5 | 0.2 | NA | NA | 2 | 0.5 | NA | NA | 1 | 1 | 3 | 0.6667 |
| 1_1424083 | 2 | 0.5 | 2 | 0 | NA | NA | 1 | 0 | 5 | 0.2 | NA | NA | 3 | 0.3333 | NA | NA | 1 | 1 | 4 | 0.5 |
| 1_1424084 | 1 | 0 | 2 | 0.5 | NA | NA | 1 | 1 | 4 | 0.5 | NA | NA | 3 | 0.3333 | NA | NA | 1 | 1 | 4 | 0.5 |
| 1_1424097 | 2 | 1 | 2 | 1 | NA | NA | 1 | 1 | 5 | 1 | NA | NA | 3 | 0.6667 | NA | NA | 1 | 0 | 4 | 0.5 |
| 1_1424105 | 2 | 1 | 2 | 1 | NA | NA | 1 | 1 | 5 | 1 | NA | NA | 2 | 1 | NA | NA | 1 | 1 | 3 | 1 |
| 1_1482398 | NA | NA | NA | NA | NA | NA | 2 | 0.5 | 2 | 0.5 | NA | NA | 1 | 1 | NA | NA | NA | NA | 1 | 1 |
| 1_1482401 | NA | NA | NA | NA | NA | NA | 2 | 0.5 | 2 | 0.5 | NA | NA | 1 | 1 | NA | NA | NA | NA | 1 | 1 |
| 1_1482439 | NA | NA | NA | NA | NA | NA | 2 | 1 | 2 | 1 | NA | NA | 1 | 1 | NA | NA | NA | NA | 1 | 1 |
| 1_1482442 | NA | NA | NA | NA | NA | NA | 1 | 1 | 1 | 1 | NA | NA | 1 | 1 | NA | NA | NA | NA | 1 | 1 |
| 1_1482447 | NA | NA | NA | NA | NA | NA | 2 | 0.5 | 2 | 0.5 | NA | NA | 1 | 1 | NA | NA | NA | NA | 1 | 1 |
| 1_1578174 | NA | NA | NA | NA | 1 | 1 | NA | NA | 1 | 1 | NA | NA | 1 | 1 | NA | NA | NA | NA | 1 | 1 |
| 1_1578188 | NA | NA | NA | NA | 1 | 1 | NA | NA | 1 | 1 | NA | NA | 1 | 1 | NA | NA | NA | NA | 1 | 1 |
| 1_1578214 | NA | NA | NA | NA | 1 | 1 | NA | NA | 1 | 1 | NA | NA | 1 | 1 | NA | NA | NA | NA | 1 | 1 |
| 1_1578228 | NA | NA | NA | NA | 1 | 1 | NA | NA | 1 | 1 | NA | NA | 1 | 1 | NA | NA | NA | NA | 1 | 1 |
| 1_1591328 | 2 | 0 | 3 | 0.3333 | 3 | 1 | 2 | 1 | 10 | 0.6 | NA | NA | 1 | 1 | NA | NA | NA | NA | 1 | 1 |
| 1_1591330 | 2 | 0 | 3 | 0 | 3 | 0.3333 | 2 | 0 | 10 | 0.1 | NA | NA | 1 | 1 | NA | NA | NA | NA | 1 | 1 |
| 1_1617620 | NA | NA | NA | NA | 1 | 1 | NA | NA | 1 | 1 | NA | NA | 1 | 1 | NA | NA | NA | NA | 1 | 1 |
| 1_1617624 | NA | NA | NA | NA | 1 | 1 | NA | NA | 1 | 1 | NA | NA | 1 | 1 | NA | NA | NA | NA | 1 | 1 |
| 1_1629917 | 2 | 0 | 8 | 0.5 | 4 | 0.25 | 1 | 1 | 15 | 0.4 | NA | NA | NA | NA | 5 | 1 | NA | NA | 5 | 1 |
| 1_1629918 | 3 | 0 | 7 | 0.1429 | 5 | 0.2 | NA | NA | 15 | 0.1333 | 1 | 0 | 2 | 0.5 | 4 | 0.5 | NA | NA | 7 | 0.4286 |
| 1_1629937 | 4 | 0.5 | 7 | 0.5714 | 4 | 0.25 | NA | NA | 15 | 0.4667 | NA | NA | 2 | 0 | 5 | 0.8 | 1 | 0 | 8 | 0.5 |
| 1_1629979 | 3 | 0 | 7 | 0 | 3 | 0.3333 | 1 | 0 | 14 | 0.0714 | NA | NA | 2 | 0 | 5 | 0.6 | 1 | 0 | 8 | 0.375 |
| 1_1629991 | 3 | 0.3333 | 6 | 0.5 | 4 | 0.75 | 1 | 0 | 14 | 0.5 | NA | NA | 2 | 0 | 4 | 0.5 | 1 | 0 | 7 | 0.2857 |
| 1_1630002 | 3 | 0 | 7 | 0.4286 | 3 | 0.3333 | 1 | 0 | 14 | 0.2857 | NA | NA | 2 | 0.5 | 4 | 0.5 | NA | NA | 6 | 0.5 |
| 1_1641461 | 13 | 0.0769 | 20 | 0.05 | 16 | 0.0625 | 10 | 0.1 | 59 | 0.0678 | 9 | 0.1111 | 10 | 0.3 | 15 | 0.1333 | 8 | 0.125 | 42 | 0.1667 |
| 1_1641493 | 11 | 0.8182 | 20 | 0.7 | 16 | 0.6875 | 11 | 0.8182 | 58 | 0.7414 | 5 | 0.8 | 9 | 1 | 15 | 0.8 | 8 | 0.875 | 37 | 0.8649 |
| 1_1641495 | 11 | 0.3636 | 19 | 0.4211 | 16 | 0.1875 | 11 | 0.1818 | 57 | 0.2982 | 5 | 0 | 9 | 0.3333 | 16 | 0.375 | 9 | 0.6667 | 39 | 0.3846 |
| 1_1641513 | 9 | 0.2222 | 22 | 0.3182 | 15 | 0.4667 | 11 | 0.3636 | 57 | 0.3509 | 6 | 0.5 | 11 | 0.7273 | 16 | 0.5 | 7 | 0.8571 | 40 | 0.625 |
| 1_1641588 | 7 | 0 | 17 | 0.0588 | 6 | 0.1667 | 10 | 0.1 | 40 | 0.075 | 4 | 0 | 7 | 0.4286 | 12 | 0.0833 | 6 | 0 | 29 | 0.1379 |
| 1_1641589 | NA | NA | 3 | 0.3333 | NA | NA | NA | NA | 3 | 0.3333 | 1 | 0 | 1 | 1 | 1 | 1 | 1 | 0 | 4 | 0.5 |
| 1_1641604 | 6 | 0.1667 | 14 | 0.1429 | 7 | 0 | 7 | 0.2857 | 34 | 0.1471 | 4 | 0.25 | 4 | 0 | 12 | 0.0833 | 4 | 0.25 | 24 | 0.125 |
| 1_1641605 | 10 | 0.3 | 21 | 0.2381 | 9 | 0.4444 | 11 | 0 | 51 | 0.2353 | 3 | 0 | 8 | 0.125 | 17 | 0.1176 | 8 | 0 | 36 | 0.0833 |
| 1_1641612 | 9 | 0.2222 | 15 | 0.0667 | 10 | 0.2 | 12 | 0.3333 | 46 | 0.1957 | 3 | 0 | 6 | 0 | 15 | 0.2 | 6 | 0.1667 | 30 | 0.1333 |
| 1_1664092 | 19 | 0.1053 | 14 | 0.0714 | 8 | 0.25 | 12 | 0.25 | 53 | 0.1509 | 12 | 0.1667 | 13 | 0.1538 | 16 | 0.0625 | 12 | 0 | 53 | 0.0943 |
| 1_1664117 | 17 | 0.1176 | 11 | 0 | 7 | 0.1429 | 12 | 0.0833 | 47 | 0.0851 | 8 | 0.125 | 10 | 0.2 | 14 | 0 | 10 | 0.1 | 42 | 0.0952 |
| 1_1664142 | 15 | 0.2 | 12 | 0.4167 | 8 | 0.375 | 9 | 0.4444 | 44 | 0.3409 | 9 | 0.3333 | 10 | 0.3 | 13 | 0.1538 | 9 | 0.6667 | 41 | 0.3415 |
| 1_1670954 | 2 | 1 | 1 | 0 | 3 | 0.3333 | 1 | 0 | 7 | 0.4286 | NA | NA | 1 | 0 | 1 | 0 | 1 | 1 | 3 | 0.3333 |
| 1_1671332 | 18 | 0.2778 | 18 | 0.3333 | 19 | 0.3158 | 24 | 0.5 | 79 | 0.3671 | 14 | 0.4286 | 10 | 0.5 | 21 | 0.2857 | 16 | 0.4375 | 61 | 0.3934 |
| 1_1671349 | 19 | 0.1053 | 20 | 0.05 | 29 | 0.1034 | 31 | 0.129 | 99 | 0.101 | 20 | 0.15 | 21 | 0.1429 | 28 | 0.0714 | 20 | 0.15 | 89 | 0.1236 |
| 1_1671354 | 21 | 0 | 23 | 0.2174 | 33 | 0.2424 | 31 | 0.1935 | 108 | 0.1759 | 20 | 0.3 | 27 | 0.1481 | 27 | 0.1481 | 18 | 0.0556 | 92 | 0.163 |
| 1_1674739 | 13 | 0.2308 | 11 | 0.4545 | 7 | 0.4286 | 7 | 0.2857 | 38 | 0.3421 | 5 | 0.4 | 7 | 0.4286 | 10 | 0.1 | 5 | 0.6 | 27 | 0.3333 |
| 1_1674747 | 12 | 0.1667 | 10 | 0 | 7 | 0 | 8 | 0.125 | 37 | 0.0811 | 5 | 0 | 8 | 0.125 | 11 | 0 | 5 | 0.2 | 29 | 0.069 |
| 1_1674756 | 13 | 0.6154 | 11 | 0.9091 | 8 | 0.875 | 6 | 1 | 38 | 0.8158 | 5 | 0.8 | 6 | 0.8333 | 9 | 0.6667 | 4 | 0.25 | 24 | 0.6667 |
| 1_1674762 | 11 | 0.5455 | 11 | 0.4545 | 7 | 0.4286 | 7 | 0.5714 | 36 | 0.5 | 7 | 0.2857 | 9 | 0.4444 | 9 | 0.3333 | 5 | 0.4 | 30 | 0.3667 |
| 1_1674765 | 11 | 0.2727 | 10 | 0.2 | 9 | 0.1111 | 8 | 0.125 | 38 | 0.1842 | 5 | 0.2 | 10 | 0.3 | 10 | 0.1 | 6 | 0.1667 | 31 | 0.1935 |
| 1_1674777 | 13 | 0 | 12 | 0.0833 | 9 | 0.2222 | 9 | 0.1111 | 43 | 0.093 | 7 | 0.1429 | 10 | 0 | 10 | 0.2 | 7 | 0 | 34 | 0.0882 |
| 1_1706682 | 2 | 0.5 | 2 | 0 | 4 | 0 | NA | NA | 8 | 0.125 | 3 | 0.3333 | 3 | 0.6667 | 2 | 0 | 1 | 0 | 9 | 0.3333 |
| 1_1706709 | 2 | 0 | 2 | 0.5 | 4 | 0.25 | NA | NA | 8 | 0.25 | 3 | 0.6667 | 2 | 0.5 | 2 | 0 | NA | NA | 7 | 0.4286 |
| 1_1706718 | 1 | 0 | NA | NA | 1 | 1 | NA | NA | 2 | 0.5 | 2 | 0.5 | 3 | 0.6667 | 1 | 1 | NA | NA | 6 | 0.6667 |
| 1_1707418 | 2 | 0 | 2 | 0 | 3 | 0.3333 | NA | NA | 7 | 0.1429 | 2 | 0.5 | 3 | 0 | 1 | 0 | NA | NA | 6 | 0.1667 |
| 1_1707420 | 2 | 0.5 | 2 | 0 | 2 | 0 | NA | NA | 6 | 0.1667 | 2 | 1 | 3 | 0 | 1 | 0 | NA | NA | 6 | 0.3333 |
| 1_1707435 | 2 | 1 | 1 | 0 | 2 | 0 | NA | NA | 5 | 0.4 | 2 | 0 | 2 | 0.5 | 1 | 0 | NA | NA | 5 | 0.2 |
| 1_1707449 | 2 | 0 | 1 | 0 | 2 | 0.5 | NA | NA | 5 | 0.2 | 2 | 0.5 | 2 | 0 | 1 | 0 | NA | NA | 5 | 0.2 |
| 1_1707453 | 2 | 1 | NA | NA | 3 | 0.6667 | NA | NA | 5 | 0.8 | 1 | 1 | 3 | 0.6667 | 1 | 1 | NA | NA | 5 | 0.8 |
| 1_1716160 | NA | NA | NA | NA | 1 | 1 | NA | NA | 1 | 1 | 2 | 1 | 2 | 1 | NA | NA | NA | NA | 4 | 1 |
| 1_1719341 | 1 | 1 | NA | NA | NA | NA | NA | NA | 1 | 1 | NA | NA | 2 | 0.5 | NA | NA | NA | NA | 2 | 0.5 |
| 1_1719345 | 1 | 1 | NA | NA | NA | NA | NA | NA | 1 | 1 | NA | NA | 1 | 1 | NA | NA | NA | NA | 1 | 1 |

| | | | | | | | | | | | | | | | | | | | | |
|---|---|---|---|---|---|---|---|---|---|---|---|---|---|---|---|---|---|---|---|---|
| 1_1723146 | 5 | 0.2 | 3 | 0 | 3 | 0 | 8 | 0 | 19 | 0.0526 | 4 | 0 | 9 | 0 | 2 | 0.5 | 3 | 0.3333 | 18 | 0.1111 |
| 1_1723148 | 5 | 0 | 2 | 0 | 4 | 0.25 | 7 | 0 | 18 | 0.0556 | 4 | 0.25 | 8 | 0.25 | 1 | 0 | 3 | 0.3333 | 16 | 0.25 |
| 1_1723153 | 5 | 0.6 | 3 | 0.6667 | 4 | 0.5 | 8 | 0.5 | 20 | 0.55 | 6 | 0.6667 | 8 | 0.375 | 1 | 0 | 3 | 0.6667 | 18 | 0.5 |
| 1_1723397 | 10 | 0.1 | 8 | 0.125 | 12 | 0.0833 | 10 | 0 | 40 | 0.075 | 9 | 0 | 17 | 0.1176 | 7 | 0 | 6 | 0 | 39 | 0.0513 |
| 1_1723409 | 9 | 0.1111 | 8 | 0.125 | 10 | 0 | 10 | 0 | 37 | 0.0541 | 7 | 0.1429 | 17 | 0.0588 | 7 | 0 | 5 | 0 | 36 | 0.0556 |
| 1_1806487 | 1 | 0 | NA | NA | 1 | 0 | 2 | 1 | 4 | 0.5 | NA | NA | NA | NA | 1 | 1 | NA | NA | 1 | 1 |
| 1_1806492 | 1 | 1 | NA | NA | 1 | 1 | 2 | 1 | 4 | 1 | NA | NA | NA | NA | 1 | 1 | NA | NA | 1 | 1 |
| 1_1806506 | 1 | 1 | NA | NA | 1 | 1 | 1 | 1 | 3 | 1 | NA | NA | NA | NA | 1 | 1 | NA | NA | 1 | 1 |
| 1_1806509 | 1 | 1 | NA | NA | 1 | 1 | 1 | 1 | 3 | 1 | NA | NA | NA | NA | 1 | 1 | NA | NA | 1 | 1 |
| 1_1806527 | 1 | 1 | NA | NA | 1 | 1 | 2 | 1 | 4 | 1 | NA | NA | NA | NA | 1 | 1 | NA | NA | 1 | 1 |
| 1_1806539 | 1 | 1 | NA | NA | NA | NA | 2 | 1 | 3 | 1 | NA | NA | NA | NA | 1 | 1 | NA | NA | 1 | 1 |
| 1_1806543 | 1 | 1 | NA | NA | 1 | 1 | 1 | 1 | 3 | 1 | NA | NA | NA | NA | 1 | 1 | NA | NA | 1 | 1 |
| 1_1806561 | 1 | 1 | NA | NA | NA | NA | NA | NA | 1 | 1 | NA | NA | NA | NA | 1 | 1 | NA | NA | 1 | 1 |
| 1_1806568 | 1 | 1 | NA | NA | 1 | 1 | 1 | 1 | 3 | 1 | NA | NA | NA | NA | 1 | 1 | NA | NA | 1 | 1 |
| 1_1806581 | 1 | 1 | NA | NA | 1 | 0 | 1 | 0 | 3 | 0.3333 | NA | NA | NA | NA | 1 | 1 | NA | NA | 1 | 1 |
| 1_2100493 | 6 | 1 | 11 | 0.6364 | 8 | 0.625 | 5 | 0.8 | 30 | 0.7333 | 10 | 0.8 | 1 | 1 | 3 | 1 | 4 | 0.75 | 18 | 0.8333 |
| 1_2100508 | 6 | 0.8333 | 9 | 0.7778 | 8 | 0.875 | 6 | 0.6667 | 29 | 0.7931 | 10 | 0.9 | 1 | 1 | 3 | 0.3333 | 5 | 1 | 19 | 0.8421 |
| 1_2100528 | 6 | 0.8333 | 10 | 1 | 9 | 1 | 5 | 1 | 30 | 0.9667 | 11 | 1 | 1 | 1 | 2 | 1 | 4 | 1 | 18 | 1 |
| 1_2100551 | 9 | 0.2222 | 11 | 0.0909 | 9 | 0.2222 | 7 | 0 | 36 | 0.1389 | 15 | 0.0667 | 1 | 0 | 4 | 0 | 4 | 0.25 | 24 | 0.0833 |
| 1_2100567 | 9 | 0.2222 | 12 | 0.25 | 10 | 0.4 | 7 | 0.4286 | 38 | 0.3158 | 16 | 0.25 | 2 | 0.5 | 3 | 0 | 6 | 0.1667 | 27 | 0.2222 |
| 1_2100569 | 11 | 0.8182 | 14 | 0.7857 | 11 | 0.7273 | 9 | 0.8889 | 45 | 0.8 | 16 | 0.5625 | 1 | 0 | 5 | 0.8 | 6 | 0.6667 | 28 | 0.6071 |
| 1_2100575 | 9 | 0.5556 | 13 | 0.9231 | 10 | 0.7 | 7 | 0.7143 | 39 | 0.7436 | 11 | 0.7273 | 2 | 1 | 2 | 0.5 | 4 | 1 | 19 | 0.7895 |
| 1_2100588 | 8 | 0.625 | 14 | 0.8571 | 9 | 1 | 6 | 0.6667 | 37 | 0.8108 | 14 | 0.8571 | 2 | 0.5 | 5 | 0.8 | 6 | 0.8333 | 27 | 0.8148 |
| 1_2100589 | 6 | 0.6667 | 9 | 1 | 9 | 0.6667 | 5 | 0.8 | 29 | 0.7931 | 11 | 0.7273 | NA | NA | NA | NA | 5 | 0.8 | 16 | 0.75 |
| 1_2100595 | 12 | 0.5833 | 16 | 0.4375 | 13 | 0.3846 | 9 | 0.3333 | 50 | 0.44 | 17 | 0.4118 | 2 | 0.5 | 8 | 0.375 | 8 | 0.375 | 35 | 0.4 |
| 1_2100596 | 12 | 0 | 14 | 0.0714 | 13 | 0.0769 | 9 | 0.2222 | 48 | 0.0833 | 17 | 0.1176 | 2 | 0 | 6 | 0 | 8 | 0 | 33 | 0.0606 |
| 1_2100598 | 12 | 0.4167 | 10 | 0.6 | 13 | 0.3846 | 8 | 0.625 | 43 | 0.4884 | 14 | 0.5 | 2 | 0.5 | 6 | 0.6667 | 8 | 0.25 | 30 | 0.4667 |
| 1_2100605 | 12 | 0.0833 | 16 | 0.1875 | 12 | 0.1667 | 9 | 0.1111 | 49 | 0.1429 | 16 | 0.125 | 2 | 0 | 5 | 0 | 9 | 0.3333 | 32 | 0.1562 |
| 1_2100626 | 14 | 0.1429 | 13 | 0.1538 | 17 | 0.0588 | 8 | 0 | 52 | 0.0962 | 14 | 0.3571 | 2 | 0 | 7 | 0.1429 | 10 | 0 | 33 | 0.1818 |
| 1_2100645 | 14 | 0.0714 | 16 | 0.125 | 19 | 0 | 8 | 0 | 57 | 0.0526 | 17 | 0.1765 | 3 | 0.3333 | 6 | 0.1667 | 10 | 0 | 36 | 0.1389 |
| 1_2100646 | 14 | 0.1429 | 13 | 0 | 17 | 0.1765 | 10 | 0.2 | 54 | 0.1296 | 16 | 0.0625 | 3 | 0 | 6 | 0 | 10 | 0.1 | 35 | 0.0571 |
| 1_2100647 | 11 | 1 | 16 | 0.9375 | 18 | 1 | 10 | 1 | 55 | 0.9818 | 16 | 1 | 2 | 1 | 7 | 1 | 10 | 1 | 35 | 1 |
| 1_2100654 | 15 | 0.1333 | 10 | 0.2 | 17 | 0.2941 | 9 | 0.1111 | 51 | 0.1961 | 15 | 0.2667 | 3 | 0.3333 | 5 | 0.2 | 10 | 0.3 | 33 | 0.2727 |
| 1_2100655 | 10 | 1 | 16 | 0.9375 | 16 | 0.875 | 8 | 0.875 | 50 | 0.92 | 15 | 1 | 2 | 1 | 6 | 1 | 8 | 1 | 31 | 1 |
| 1_2100662 | 14 | 0.2143 | 15 | 0.2 | 18 | 0.0556 | 9 | 0.2222 | 56 | 0.1607 | 17 | 0.2353 | 4 | 0.25 | 7 | 0.4286 | 9 | 0.1111 | 37 | 0.2432 |
| 1_2100663 | 13 | 0 | 14 | 0.0714 | 19 | 0.1579 | 9 | 0.1111 | 55 | 0.0909 | 15 | 0.1333 | 4 | 0 | 6 | 0 | 11 | 0 | 36 | 0.0556 |
| 1_2100663 | 13 | 1 | 13 | 1 | 18 | 1 | 9 | 1 | 53 | 1 | 18 | 0.9444 | 4 | 0.75 | 6 | 1 | 10 | 1 | 38 | 0.9474 |
| 1_2100680 | 9 | 0.8889 | 13 | 0.8462 | 12 | 0.8333 | 3 | 1 | 37 | 0.8649 | 10 | 0.7 | NA | NA | 5 | 1 | 6 | 0.8333 | 21 | 0.8095 |
| 1_2100687 | 15 | 0.2667 | 16 | 0.125 | 19 | 0.1579 | 10 | 0.2 | 60 | 0.1833 | 17 | 0.1765 | 4 | 0.25 | 8 | 0.25 | 11 | 0 | 40 | 0.15 |
| 1_2100692 | 15 | 1 | 16 | 0.9375 | 18 | 1 | 10 | 1 | 59 | 0.9831 | 18 | 0.9444 | 4 | 1 | 8 | 1 | 11 | 1 | 41 | 0.9756 |
| 1_2100694 | 11 | 0.9091 | 14 | 0.7143 | 17 | 0.8235 | 5 | 0.6 | 47 | 0.7872 | 16 | 0.8125 | 2 | 1 | 6 | 1 | 8 | 0.875 | 32 | 0.875 |
| 1_2100695 | 10 | 0 | 10 | 0.2 | 15 | 0.0667 | 10 | 0.2 | 45 | 0.1111 | 13 | 0.0769 | 5 | 0.6 | 9 | 0.1111 | 9 | 0 | 36 | 0.1389 |
| 1_2100702 | 15 | 0.9333 | 17 | 0.8824 | 19 | 0.8947 | 9 | 1 | 60 | 0.9167 | 18 | 0.9444 | 4 | 1 | 9 | 1 | 10 | 0.9 | 41 | 0.9512 |
| 1_2100706 | 10 | 1 | 16 | 0.6875 | 15 | 0.6 | 8 | 0.75 | 49 | 0.7347 | 14 | 0.8571 | 2 | 1 | 9 | 0.7778 | 8 | 0.875 | 33 | 0.8485 |
| 1_2100712 | 14 | 0.6429 | 17 | 0.4706 | 16 | 0.5625 | 10 | 0.3 | 57 | 0.5088 | 19 | 0.6316 | 5 | 0.4 | 8 | 0.5 | 11 | 0.1818 | 43 | 0.4651 |
| 1_2100718 | 15 | 0.0667 | 17 | 0.1765 | 18 | 0.0556 | 8 | 0 | 58 | 0.0862 | 17 | 0 | 5 | 0 | 6 | 0 | 10 | 0.2 | 38 | 0.0526 |
| 1_2100737 | 13 | 0.9231 | 15 | 1 | 15 | 0.8667 | 8 | 1 | 51 | 0.9412 | 17 | 1 | 4 | 1 | 6 | 1 | 11 | 0.9091 | 38 | 0.9737 |
| 1_2100738 | 12 | 0.8333 | 12 | 1 | 13 | 0.6154 | 8 | 0.5 | 45 | 0.7556 | 17 | 0.8824 | 5 | 0.4 | 7 | 0.5714 | 5 | 0.6 | 34 | 0.7059 |
| 1_2100751 | 16 | 0.8125 | 16 | 0.75 | 18 | 0.6667 | 8 | 0.75 | 58 | 0.7414 | 18 | 0.8889 | 3 | 0.6667 | 8 | 0.5 | 12 | 0.75 | 41 | 0.7561 |
| 1_2100753 | 15 | 0.2 | 19 | 0.1053 | 17 | 0.0588 | 9 | 0.1111 | 60 | 0.1167 | 20 | 0.15 | 5 | 0.2 | 8 | 0.25 | 13 | 0.0769 | 46 | 0.1522 |
| 1_2100757 | 15 | 0.6 | 17 | 0.5294 | 17 | 0.6471 | 8 | 0.375 | 57 | 0.5614 | 17 | 0.7059 | 5 | 0.8 | 10 | 0.7 | 10 | 0.8 | 42 | 0.7381 |
| 1_2100761 | 16 | 0.25 | 18 | 0.0556 | 19 | 0.2105 | 11 | 0.0909 | 64 | 0.1562 | 18 | 0.0556 | 6 | 0.5 | 9 | 0.1111 | 11 | 0.1818 | 44 | 0.1591 |
| 1_2100766 | 11 | 0.8182 | 17 | 0.6471 | 14 | 0.5 | 7 | 0.1429 | 49 | 0.5714 | 14 | 0.5714 | 4 | 0.5 | 12 | 0.6667 | 6 | 0.5 | 36 | 0.5833 |
| 1_2100770 | 14 | 0.0714 | 20 | 0.05 | 14 | 0.1429 | 7 | 0 | 55 | 0.0727 | 15 | 0.0667 | 4 | 0.25 | 14 | 0 | 9 | 0.3333 | 42 | 0.119 |
| 1_2100791 | 13 | 0.0769 | 17 | 0.2353 | 15 | 0.1333 | 7 | 0 | 52 | 0.1346 | 12 | 0.1667 | 4 | 0.25 | 12 | 0 | 11 | 0.0909 | 39 | 0.1026 |
| 1_2100792 | 14 | 0.9286 | 16 | 0.9375 | 13 | 1 | 5 | 1 | 48 | 0.9583 | 11 | 1 | 3 | 1 | 15 | 1 | 10 | 0.9 | 39 | 0.9744 |
| 1_2100793 | 14 | 0.6429 | 19 | 0.5263 | 14 | 0.5 | 5 | 0.6 | 52 | 0.5577 | 14 | 0.6429 | 4 | 0.75 | 15 | 0.6 | 11 | 0.5455 | 44 | 0.6136 |
| 1_2100803 | 13 | 1 | 17 | 0.7647 | 14 | 0.9286 | 7 | 0.7143 | 51 | 0.8627 | 14 | 0.7857 | 2 | 1 | 13 | 0.9231 | 12 | 0.9167 | 41 | 0.878 |
| 1_2100821 | 7 | 0.5714 | 16 | 0.5625 | 13 | 0.6154 | 8 | 0.375 | 44 | 0.5455 | 12 | 0.5833 | 3 | 0.6667 | 12 | 0.4167 | 9 | 0.5556 | 36 | 0.5278 |
| 1_2100828 | 14 | 0.2143 | 18 | 0.2222 | 17 | 0.2353 | 9 | 0.3333 | 58 | 0.2414 | 15 | 0.2667 | 4 | 0 | 13 | 0.2308 | 11 | 0.1818 | 43 | 0.2093 |
| 1_2100833 | 14 | 0.6429 | 14 | 0.3571 | 18 | 0.6111 | 8 | 0.5 | 54 | 0.537 | 13 | 0.6923 | 5 | 0.4 | 13 | 0.6923 | 11 | 0.6364 | 42 | 0.6429 |
| 1_2100845 | 11 | 0.0909 | 14 | 0 | 14 | 0.0714 | 8 | 0.125 | 47 | 0.0638 | 15 | 0.0667 | 4 | 0 | 16 | 0 | 9 | 0.2222 | 44 | 0.0682 |
| 1_2100845 | 13 | 1 | 15 | 1 | 16 | 1 | 8 | 1 | 52 | 1 | 12 | 1 | 5 | 1 | 14 | 0.8571 | 11 | 1 | 42 | 0.9524 |
| 1_2100846 | 16 | 0.5 | 17 | 0.3529 | 19 | 0.6316 | 10 | 0.5 | 62 | 0.5 | 15 | 0.6 | 5 | 0.8 | 15 | 0.5333 | 11 | 0.7273 | 46 | 0.6304 |
| 1_2100850 | 16 | 0.875 | 19 | 0.9474 | 19 | 0.8947 | 10 | 0.7 | 64 | 0.875 | 13 | 0.7692 | 5 | 0.8 | 15 | 0.8667 | 12 | 0.75 | 45 | 0.8 |

| | | | | | | | | | | | | | | | | | | | | |
|---|---|---|---|---|---|---|---|---|---|---|---|---|---|---|---|---|---|---|---|---|
| 1_21009052 | 10 | 0.3 | 13 | 0.5385 | 17 | 0.5882 | 7 | 0.7143 | 47 | 0.5319 | 11 | 0.3636 | 4 | 0.5 | 10 | 0.5 | 10 | 0.5 | 35 | 0.4571 |
| 1_21009154 | 11 | 0 | 18 | 0 | 20 | 0.05 | 8 | 0.25 | 57 | 0.0526 | 10 | 0 | 6 | 0.1667 | 12 | 0.0833 | 10 | 0.1 | 38 | 0.0789 |
| 1_21009346 | 8 | 0.25 | 12 | 0.1667 | 9 | 0.1111 | 6 | 0.1667 | 35 | 0.1714 | 9 | 0.3333 | 2 | 0 | 7 | 0.1429 | 7 | 0.1429 | 25 | 0.2 |
| 1_21299665 | NA | NA | 1 | 1 | NA | NA | 2 | 0 | 3 | 0.3333 | NA | NA | 1 | 1 | 3 | 0.6667 | 1 | 0 | 5 | 0.6 |
| 1_21299740 | NA | NA | 2 | 0.5 | 1 | 1 | 2 | 1 | 5 | 0.8 | NA | NA | NA | NA | 3 | 0.6667 | 1 | 1 | 4 | 0.75 |
| 1_21299781 | NA | NA | 2 | 1 | 1 | 1 | 1 | 0 | 4 | 0.75 | NA | NA | 1 | 1 | 3 | 0.3333 | 1 | 1 | 5 | 0.6 |
| 1_21299809 | NA | NA | 1 | 0 | 1 | 0 | 1 | 1 | 3 | 0.3333 | NA | NA | 1 | 0 | 1 | 1 | 1 | 0 | 3 | 0.3333 |
| 1_21299833 | 2 | 0 | 2 | 1 | 6 | 0.1667 | 7 | 0.2857 | 17 | 0.2941 | 3 | 0.3333 | 3 | 0 | 5 | 0.2 | 4 | 0 | 15 | 0.1333 |
| 1_21299862 | 2 | 0 | 3 | 0.3333 | 6 | 0.1667 | 5 | 0 | 16 | 0.125 | 3 | 0 | 3 | 0.6667 | 4 | 0.5 | 4 | 0.25 | 14 | 0.3571 |
| 1_21300133 | 3 | 0 | 3 | 0 | 7 | 0.1429 | 6 | 0.1667 | 19 | 0.1053 | 3 | 0 | 3 | 0 | 6 | 0.1667 | 5 | 0.4 | 17 | 0.1765 |
| 1_21469655 | 6 | 0.3333 | 12 | 0.3333 | 4 | 0.25 | 7 | 0.4286 | 29 | 0.3448 | 2 | 0.5 | 10 | 0.4 | 7 | 0.4286 | 6 | 0.3333 | 25 | 0.4 |
| 1_21469733 | 7 | 0.7143 | 10 | 1 | 4 | 1 | 6 | 0.3333 | 27 | 0.7778 | 2 | 1 | 8 | 0.75 | 6 | 0.1667 | 6 | 0.6667 | 22 | 0.5909 |
| 1_21469769 | 7 | 0 | 11 | 0.3636 | 4 | 0.25 | 5 | 0 | 27 | 0.1852 | 1 | 0 | 7 | 0 | 7 | 0.1429 | 6 | 0.5 | 21 | 0.1905 |
| 1_21469850 | 7 | 0 | 8 | 0.125 | 2 | 0.5 | 6 | 0 | 23 | 0.087 | 1 | 0 | 4 | 0 | 6 | 0.1667 | 3 | 0.3333 | 14 | 0.1429 |
| 1_21469928 | 7 | 1 | 12 | 1 | 4 | 1 | 7 | 1 | 30 | 1 | 2 | 1 | 8 | 1 | 6 | 1 | 5 | 1 | 21 | 1 |
| 1_21470003 | 6 | 1 | 10 | 1 | 3 | 1 | 6 | 1 | 25 | 1 | 2 | 1 | 7 | 1 | 6 | 0.8333 | 5 | 1 | 20 | 0.95 |
| 1_21470143 | 6 | 1 | 10 | 1 | 1 | 1 | 6 | 1 | 23 | 1 | 2 | 1 | 7 | 1 | 4 | 1 | 4 | 1 | 17 | 1 |
| 1_21470153 | 4 | 0.25 | 6 | 0 | 1 | 0 | 5 | 0.6 | 16 | 0.25 | 1 | 0 | 4 | 0.5 | 3 | 0.3333 | 5 | 0.4 | 13 | 0.3846 |
| 1_21470345 | 5 | 0.2 | 6 | 0.3333 | 3 | 0 | 2 | 0.5 | 16 | 0.25 | 1 | 0 | 4 | 0.5 | 5 | 0.2 | 4 | 0.25 | 14 | 0.2857 |
| 1_21470811 | 4 | 0.25 | 10 | 0.1 | 1 | 0 | 6 | 0 | 21 | 0.0952 | 1 | 0 | 6 | 0 | 6 | 0.1667 | 5 | 0 | 18 | 0.0556 |
| 1_21471480 | 5 | 0.4 | 6 | 0.3333 | 1 | 0 | 4 | 0.25 | 16 | 0.3125 | NA | NA | 4 | 0.5 | 4 | 0.5 | 1 | 1 | 9 | 0.5556 |
| 1_21471812 | 3 | 0 | 8 | 0.75 | 1 | 0 | 4 | 0.5 | 16 | 0.5 | 1 | 0 | 3 | 0.3333 | 5 | 0.4 | 3 | 0.6667 | 12 | 0.4167 |
| 1_21471884 | 4 | 0 | 7 | 0.2857 | 1 | 1 | 4 | 0 | 16 | 0.1875 | 1 | 0 | 3 | 0 | 4 | 0.25 | 3 | 0 | 11 | 0.0909 |
| 1_21471966 | 7 | 0 | 8 | 0.125 | 3 | 0 | 6 | 0.1667 | 24 | 0.0833 | 4 | 0 | 6 | 0 | 5 | 0.2 | 4 | 0 | 19 | 0.0526 |
| 1_21472014 | 9 | 0.1111 | 9 | 0.2222 | 3 | 0 | 6 | 0.1667 | 27 | 0.1481 | 4 | 0 | 8 | 0.375 | 4 | 0.5 | 6 | 0.3333 | 22 | 0.3182 |
| 1_21472391 | 6 | 0.3333 | 8 | 0.375 | 4 | 0.5 | 3 | 1 | 21 | 0.4762 | 1 | 1 | 4 | 0.25 | 5 | 0.6 | 3 | 0.3333 | 13 | 0.4615 |
| 1_21472396 | 7 | 0 | 10 | 0.2 | 4 | 0 | 4 | 0 | 25 | 0.08 | 2 | 0.5 | 8 | 0.25 | 6 | 0.1667 | 4 | 0 | 20 | 0.2 |
| 1_21472430 | 10 | 0.3 | 8 | 0 | 4 | 0 | 4 | 0.25 | 26 | 0.1538 | 2 | 0 | 7 | 0.2857 | 6 | 0.1667 | 4 | 0.25 | 19 | 0.2105 |
| 1_21472448 | 4 | 0.5 | 7 | 1 | 3 | 0.6667 | 2 | 1 | 16 | 0.8125 | 2 | 1 | 5 | 0.4 | 3 | 0 | 2 | 0.5 | 12 | 0.4167 |
| 1_21472473 | 8 | 0.875 | 9 | 1 | 4 | 1 | 4 | 1 | 25 | 0.96 | 2 | 1 | 7 | 0.8571 | 5 | 1 | 3 | 1 | 17 | 0.9412 |
| 1_21472484 | 8 | 0.75 | 6 | 0.6667 | 4 | 0.75 | 4 | 0.75 | 22 | 0.7273 | 2 | 1 | 6 | 0.8333 | 4 | 1 | 4 | 0.75 | 16 | 0.875 |
| 1_21472539 | 8 | 0 | 6 | 0.1667 | 3 | 0 | 3 | 0 | 20 | 0.05 | 1 | 0 | 7 | 0.1429 | 5 | 0 | 3 | 0.3333 | 16 | 0.125 |
| 1_21472545 | 9 | 0.7778 | 9 | 0.8889 | 4 | 0.75 | 4 | 0.75 | 26 | 0.8077 | 2 | 1 | 7 | 0.8571 | 5 | 0.6 | 4 | 0.75 | 18 | 0.7778 |
| 1_21472658 | 5 | 0.6 | 7 | 0.5714 | 1 | 0 | 3 | 0.3333 | 16 | 0.5 | 1 | 1 | 3 | 1 | 4 | 0.75 | 3 | 0 | 11 | 0.6364 |
| 1_21472663 | 5 | 1 | 7 | 1 | 4 | 0.5 | 2 | 1 | 18 | 0.8889 | 2 | 1 | 7 | 0.8571 | 2 | 1 | 2 | 1 | 13 | 0.9231 |
| 1_21472746 | 9 | 0.6667 | 9 | 0.5556 | 4 | 0.5 | 3 | 0.3333 | 25 | 0.56 | 2 | 1 | 7 | 0.5714 | 6 | 0.6667 | 4 | 0.5 | 19 | 0.6316 |
| 1_21472784 | 8 | 0.75 | 8 | 0.875 | 3 | 1 | 3 | 0.6667 | 22 | 0.8182 | 1 | 1 | 7 | 0.8571 | 5 | 1 | 3 | 1 | 16 | 0.9375 |
| 1_21472894 | 7 | 0.1429 | 7 | 0 | 3 | 0 | 2 | 0 | 19 | 0.0526 | NA | NA | 4 | 0.25 | 5 | 0 | 4 | 0 | 13 | 0.0769 |
| 1_21472994 | 5 | 0.2 | 6 | 0.3333 | 3 | 0 | 2 | 0.5 | 16 | 0.25 | NA | NA | 6 | 0.1667 | 4 | 0 | 3 | 0 | 13 | 0.0769 |
| 1_21473085 | 5 | 1 | 5 | 1 | 2 | 1 | 2 | 1 | 14 | 1 | NA | NA | 7 | 0.8571 | 2 | 1 | 4 | 1 | 13 | 0.9231 |
| 1_21473192 | 8 | 0.375 | 5 | 0.2 | 4 | 0.5 | 2 | 0.5 | 19 | 0.3684 | NA | NA | 7 | 0.1429 | 6 | 0.1667 | 4 | 0.5 | 17 | 0.2353 |
| 1_21473219 | 8 | 0.125 | 4 | 0 | 3 | 0 | 2 | 0 | 17 | 0.0588 | NA | NA | 7 | 0.1429 | 5 | 0.2 | 4 | 0 | 16 | 0.125 |
| 1_21473226 | 8 | 0.125 | 5 | 0.2 | 3 | 0 | 2 | 0 | 18 | 0.1111 | NA | NA | 7 | 0.1429 | 5 | 0 | 4 | 0 | 16 | 0.0625 |
| 1_21473230 | 4 | 0 | 4 | 0.25 | 4 | 0.25 | 1 | 0 | 13 | 0.1538 | NA | NA | 7 | 0 | 4 | 0.5 | 3 | 0 | 14 | 0.1429 |
| 1_22356657 | 9 | 0.1111 | 17 | 0.1176 | 9 | 0.1111 | 7 | 0 | 42 | 0.0952 | 11 | 0.1818 | 6 | 0 | 5 | 0 | 3 | 0 | 25 | 0.08 |
| 1_22356907 | 12 | 0.1667 | 18 | 0.0556 | 12 | 0 | 9 | 0.2222 | 51 | 0.098 | 9 | 0.2222 | 8 | 0.5 | 6 | 0.1667 | 5 | 0.2 | 28 | 0.2857 |
| 1_22357727 | 8 | 0.125 | 18 | 0.0556 | 11 | 0.0909 | 8 | 0.125 | 45 | 0.0889 | 10 | 0 | 6 | 0 | 5 | 0.2 | 5 | 0.4 | 26 | 0.1154 |
| 1_22358557 | 6 | 0.1667 | 12 | 0 | 5 | 0.2 | 4 | 0.25 | 27 | 0.1111 | 6 | 0.3333 | 4 | 0.25 | 4 | 0 | 3 | 0.3333 | 17 | 0.2353 |
| 1_22358595 | 7 | 0 | 15 | 0.0667 | 8 | 0 | 5 | 0.2 | 35 | 0.0571 | 6 | 0.1667 | 5 | 0.2 | 4 | 0 | 4 | 0.25 | 19 | 0.1579 |
| 1_22358790 | 5 | 0.6 | 12 | 0.5 | 5 | 0.8 | 4 | 0 | 26 | 0.5 | 6 | 0 | 5 | 0.6 | 4 | 0.5 | 3 | 0.3333 | 18 | 0.3333 |
| 1_22497467 | NA | NA | 1 | 1 | NA | NA | NA | NA | 1 | 1 | NA | NA | 1 | 1 | NA | NA | NA | NA | 1 | 1 |
| 1_22573055 | 3 | 0.3333 | 2 | 0 | 4 | 0.25 | 1 | 0 | 10 | 0.2 | 3 | 0.3333 | NA | NA | NA | NA | 1 | 0 | 4 | 0.25 |
| 1_22573077 | 3 | 0 | 2 | 0 | 4 | 0.25 | 1 | 0 | 10 | 0.1 | 3 | 0.3333 | NA | NA | NA | NA | 1 | 1 | 4 | 0.5 |
| 1_22573157 | 2 | 0 | 2 | 1 | 3 | 0 | 1 | 1 | 8 | 0.375 | 2 | 0 | NA | NA | NA | NA | 1 | 1 | 3 | 0.3333 |
| 1_22844975 | NA | NA | 1 | 0 | NA | NA | 2 | 0.5 | 3 | 0.3333 | 1 | 1 | NA | NA | NA | NA | 1 | 0 | 2 | 0.5 |
| 1_23408674 | 3 | 1 | 10 | 1 | 6 | 1 | 10 | 1 | 29 | 1 | 3 | 1 | 9 | 1 | 7 | 1 | 8 | 1 | 27 | 1 |
| 1_23408717 | 1 | 1 | 10 | 1 | 6 | 0.8333 | 10 | 1 | 27 | 0.963 | 1 | 1 | 8 | 1 | 6 | 0.8333 | 6 | 1 | 21 | 0.9524 |
| 1_23408781 | NA | NA | NA | NA | NA | NA | 1 | 1 | 1 | 1 | NA | NA | 1 | 1 | NA | NA | NA | NA | 1 | 1 |
| 1_23408788 | NA | NA | NA | NA | NA | NA | 1 | 1 | 1 | 1 | NA | NA | NA | NA | 1 | 1 | 1 | 1 | 2 | 1 |
| 1_23408789 | NA | NA | 2 | 1 | NA | NA | NA | NA | 2 | 1 | NA | NA | NA | NA | 1 | 1 | 1 | 1 | 2 | 1 |
| 1_23408790 | NA | NA | NA | NA | 2 | 1 | NA | NA | 2 | 1 | NA | NA | 1 | 1 | NA | NA | NA | NA | 1 | 1 |
| 1_23408791 | NA | NA | 1 | 1 | NA | NA | 1 | 1 | 2 | 1 | 1 | 1 | 1 | 1 | NA | NA | NA | NA | 2 | 1 |
| 1_23408792 | NA | NA | NA | NA | 1 | 1 | 1 | 1 | 2 | 1 | NA | NA | NA | NA | NA | NA | 1 | 1 | 1 | 1 |
| 1_23408795 | NA | NA | NA | NA | NA | NA | 1 | 1 | 1 | 1 | NA | NA | NA | NA | 1 | 1 | NA | NA | 1 | 1 |
| 1_23408844 | 2 | 0 | 9 | 0 | 5 | 0.2 | 9 | 0.1111 | 25 | 0.08 | 2 | 0 | 8 | 0.25 | 5 | 0.2 | 4 | 0 | 19 | 0.1579 |

| | | | | | | | | | | | | | | | | | | | | |
|---|---|---|---|---|---|---|---|---|---|---|---|---|---|---|---|---|---|---|---|---|
| 1_2340891 | 2 | 1 | 9 | 1 | 6 | 1 | 8 | 1 | 25 | 1 | 1 | 1 | 9 | 1 | 5 | 1 | 6 | 1 | 21 | 1 |
| 1_2340903 | 1 | 1 | 8 | 0.875 | 5 | 0.8 | 7 | 0.7143 | 21 | 0.8095 | 2 | 1 | 9 | 0.6667 | 5 | 0.8 | 6 | 0.6667 | 22 | 0.7273 |
| 1_2340908 | 1 | 0 | 8 | 0.25 | 4 | 0 | 7 | 0.4286 | 20 | 0.25 | 3 | 0 | 5 | 0.2 | 6 | 0.1667 | 4 | 0.25 | 18 | 0.1667 |
| 1_2340917 | 1 | 1 | 9 | 1 | 5 | 1 | 8 | 1 | 23 | 1 | 3 | 1 | 8 | 1 | 7 | 1 | 6 | 1 | 24 | 1 |
| 1_2340926 | NA | NA | 5 | 0.8 | 4 | 1 | 3 | 1 | 12 | 0.9167 | 2 | 1 | 5 | 1 | 7 | 0.8571 | 4 | 1 | 18 | 0.9444 |
| 1_2340931 | 1 | 1 | 7 | 0.5714 | 5 | 0.4 | 8 | 0.25 | 21 | 0.4286 | 2 | 1 | 7 | 0.2857 | 5 | 0.4 | 5 | 0.2 | 19 | 0.3684 |
| 1_2340935 | NA | NA | 7 | 0.7143 | 4 | 0.5 | 5 | 1 | 16 | 0.75 | 2 | 0.5 | 6 | 0.5 | 6 | 1 | 5 | 1 | 19 | 0.7895 |
| 1_2388443 | 2 | 0.5 | NA | NA | NA | NA | NA | NA | 2 | 0.5 | 1 | 0 | 1 | 0 | 2 | 1 | NA | NA | 4 | 0.5 |
| 1_2388464 | 1 | 1 | NA | NA | NA | NA | NA | NA | 1 | 1 | 2 | 0.5 | 1 | 1 | 2 | 1 | NA | NA | 5 | 0.8 |
| 1_2388741 | 9 | 0.1111 | 18 | 0.1111 | 8 | 0.125 | 8 | 0 | 43 | 0.093 | 8 | 0.125 | 12 | 0.0833 | 9 | 0.1111 | 11 | 0 | 40 | 0.075 |
| 1_2388747 | 8 | 0.375 | 18 | 0.5 | 10 | 0.2 | 7 | 0.1429 | 43 | 0.3488 | 8 | 0.5 | 11 | 0.4545 | 10 | 0.4 | 14 | 0.1429 | 43 | 0.3488 |
| 1_2388753 | 8 | 0.375 | 19 | 0.5263 | 8 | 0.5 | 7 | 0.2857 | 42 | 0.4524 | 7 | 0.2857 | 10 | 0.5 | 11 | 0.4545 | 10 | 0.5 | 38 | 0.4474 |
| 1_2403529 | 7 | 0 | 6 | 0.1667 | 5 | 0 | 6 | 0.5 | 24 | 0.1667 | 9 | 0.5556 | 10 | 0.3 | 5 | 0 | 3 | 0.3333 | 27 | 0.3333 |
| 1_2405031 | 1 | 1 | 1 | 1 | 1 | 0 | NA | NA | 3 | 0.6667 | NA | NA | NA | NA | NA | NA | 5 | 0.2 | 5 | 0.2 |
| 1_2405135 | 2 | 1 | 4 | 0.75 | 2 | 0.5 | 2 | 0.5 | 10 | 0.7 | 2 | 1 | NA | NA | 2 | 0.5 | 4 | 0.25 | 8 | 0.5 |
| 1_2406746 | 1 | 1 | 3 | 0.3333 | 1 | 1 | 2 | 0.5 | 7 | 0.5714 | 1 | 0 | NA | NA | 2 | 1 | 1 | 0 | 4 | 0.5 |
| 1_2406749 | 1 | 0 | 2 | 0 | 1 | 1 | 2 | 0.5 | 6 | 0.3333 | 1 | 0 | NA | NA | 3 | 0 | 2 | 0.5 | 6 | 0.1667 |
| 1_2406750 | 1 | 1 | 1 | 0 | NA | NA | 2 | 0 | 4 | 0.25 | 1 | 0 | NA | NA | 2 | 0 | 1 | 1 | 4 | 0.25 |
| 1_2406788 | 1 | 0 | 3 | 0.3333 | NA | NA | 2 | 0 | 6 | 0.1667 | 1 | 0 | NA | NA | 3 | 0 | 2 | 0.5 | 6 | 0.1667 |
| 1_2406811 | 1 | 0 | 3 | 0.3333 | 1 | 0 | 2 | 0.5 | 7 | 0.2857 | 1 | 0 | NA | NA | 3 | 0 | 2 | 0.5 | 6 | 0.1667 |
| 1_2406815 | 1 | 1 | 3 | 1 | 1 | 1 | 2 | 0.5 | 7 | 0.8571 | 1 | 0 | NA | NA | 3 | 1 | 2 | 0.5 | 6 | 0.6667 |
| 1_2406816 | 1 | 1 | 1 | 1 | 1 | 1 | 2 | 0.5 | 5 | 0.8 | 1 | 1 | NA | NA | 3 | 1 | 2 | 0.5 | 6 | 0.8333 |
| 1_2406826 | 1 | 0 | 2 | 0.5 | 1 | 0 | 2 | 0.5 | 6 | 0.3333 | 1 | 1 | NA | NA | 2 | 0.5 | 2 | 1 | 5 | 0.8 |
| 1_2406832 | NA | NA | 3 | 1 | NA | NA | 1 | 1 | 4 | 1 | 1 | 1 | NA | NA | 1 | 1 | 2 | 0.5 | 4 | 0.75 |
| 1_2406849 | NA | NA | 2 | 0.5 | 1 | 0 | 2 | 0.5 | 5 | 0.4 | 1 | 0 | NA | NA | 2 | 0.5 | 2 | 1 | 5 | 0.6 |
| 1_2406876 | NA | NA | 2 | 0.5 | NA | NA | 2 | 0.5 | 4 | 0.5 | 1 | 0 | NA | NA | 2 | 1 | 2 | 0.5 | 5 | 0.6 |
| 1_2406883 | NA | NA | 1 | 1 | NA | NA | 2 | 0 | 3 | 0.3333 | NA | NA | NA | NA | 1 | 0 | 2 | 0.5 | 3 | 0.3333 |
| 1_2406906 | NA | NA | 1 | 1 | NA | NA | NA | NA | 1 | 1 | NA | NA | NA | NA | 1 | 0 | 1 | 1 | 2 | 0.5 |
| 1_2480178 | 8 | 0.125 | 16 | 0.125 | 16 | 0.0625 | 16 | 0.1875 | 56 | 0.125 | 7 | 0 | 8 | 0.375 | 11 | 0.0909 | 10 | 0.4 | 36 | 0.2222 |
| 1_2480190 | 7 | 0.5714 | 10 | 0.8 | 13 | 0.8462 | 12 | 0.75 | 42 | 0.7619 | 5 | 0.6 | 6 | 1 | 10 | 0.6 | 9 | 0.5556 | 30 | 0.6667 |
| 1_2480234 | 5 | 0.6 | 9 | 0.6667 | 7 | 0.1429 | 12 | 0.4167 | 33 | 0.4545 | 6 | 0 | 7 | 0.1429 | 4 | 0.25 | 7 | 0.5714 | 24 | 0.25 |
| 1_2480249 | 5 | 0.2 | 12 | 0.1667 | 9 | 0.2222 | 14 | 0.2857 | 40 | 0.225 | 7 | 0 | 9 | 0.1111 | 5 | 0.4 | 6 | 0.3333 | 27 | 0.1852 |
| 1_2480258 | 5 | 0.2 | 12 | 0.5 | 11 | 0.1818 | 13 | 0.2308 | 41 | 0.2927 | 6 | 0 | 9 | 0.3333 | 5 | 0.4 | 7 | 0.1429 | 27 | 0.2222 |
| 1_2480260 | 5 | 0 | 11 | 0.0909 | 10 | 0 | 14 | 0.0714 | 40 | 0.05 | 6 | 0.1667 | 8 | 0 | 5 | 0.2 | 6 | 0.1667 | 25 | 0.12 |
| 1_2480265 | 3 | 0.6667 | 8 | 0.75 | 9 | 0.5556 | 8 | 0.625 | 28 | 0.6429 | 6 | 0.5 | 7 | 0.5714 | 3 | 0.6667 | 5 | 1 | 21 | 0.6667 |
| 1_2480276 | 6 | 0.3333 | 4 | 0.25 | 7 | 0 | 7 | 0.2857 | 24 | 0.2083 | 5 | 0.2 | 6 | 0.6667 | 3 | 0.6667 | 4 | 0.5 | 18 | 0.5 |
| 1_2480409 | 6 | 0 | 10 | 0 | 9 | 0.2222 | 11 | 0.0909 | 36 | 0.0833 | 5 | 0 | 8 | 0.125 | 6 | 0.1667 | 7 | 0.1429 | 26 | 0.1154 |
| 1_2480449 | 7 | 0.2857 | 11 | 0.6364 | 8 | 0.25 | 11 | 0.3636 | 37 | 0.4054 | 5 | 0 | 10 | 0.2 | 7 | 0.2857 | 5 | 0 | 27 | 0.1481 |
| 1_2480461 | 7 | 0.4286 | 10 | 0.2 | 7 | 0.1429 | 9 | 0.1111 | 33 | 0.2121 | 7 | 0.1429 | 12 | 0.25 | 7 | 0.1429 | 6 | 0.1667 | 32 | 0.1875 |
| 1_2510714 | 10 | 0.4 | 18 | 0.2222 | 19 | 0.2632 | 14 | 0.2143 | 61 | 0.2623 | 8 | 0.25 | 13 | 0.2308 | 11 | 0.2727 | 9 | 0.3333 | 41 | 0.2683 |
| 1_2510716 | 11 | 0.1818 | 16 | 0.4375 | 16 | 0.4375 | 13 | 0.0769 | 56 | 0.3036 | 9 | 0.3333 | 13 | 0.3846 | 11 | 0.5455 | 7 | 0.2857 | 40 | 0.4 |
| 1_2510716 | 9 | 0.8889 | 15 | 0.6 | 15 | 0.8 | 10 | 0.9 | 49 | 0.7755 | 8 | 0.875 | 11 | 0.6364 | 9 | 0.6667 | 8 | 0.875 | 36 | 0.75 |
| 1_2510719 | 10 | 0.5 | 18 | 0.1111 | 21 | 0.1905 | 10 | 0.4 | 59 | 0.2542 | 8 | 0.25 | 14 | 0.2143 | 11 | 0.7273 | 8 | 0.5 | 41 | 0.4146 |
| 1_2510720 | 11 | 0.0909 | 17 | 0.2941 | 13 | 0.1538 | 13 | 0.0769 | 54 | 0.1667 | 7 | 0.1429 | 13 | 0 | 11 | 0.1818 | 5 | 0 | 36 | 0.0833 |
| 1_2510722 | 11 | 0.4545 | 19 | 0.5263 | 21 | 0.381 | 13 | 0.3846 | 64 | 0.4375 | 9 | 0.5556 | 14 | 0.5714 | 10 | 0.3 | 8 | 0.875 | 41 | 0.561 |
| 1_2510723 | 10 | 0.3 | 18 | 0.7222 | 14 | 0.5714 | 11 | 0.8182 | 53 | 0.6226 | 6 | 0.1667 | 12 | 0.5 | 10 | 0.6 | 7 | 0.7143 | 35 | 0.5143 |
| 1_2510740 | 7 | 0.1429 | 16 | 0.3125 | 18 | 0.1667 | 10 | 0.3 | 51 | 0.2353 | 7 | 0.1429 | 14 | 0.2143 | 10 | 0.3 | 8 | 0.25 | 39 | 0.2308 |
| 1_2513289 | 7 | 0.8571 | 13 | 0.6154 | 17 | 0.8235 | 9 | 0.6667 | 46 | 0.7391 | 6 | 0.8333 | 11 | 0.5455 | 5 | 0.6 | 6 | 1 | 28 | 0.7143 |
| 1_2513297 | 9 | 0.2222 | 11 | 0 | 13 | 0.1538 | 8 | 0.125 | 41 | 0.122 | 5 | 0 | 8 | 0.25 | 6 | 0.1667 | 4 | 0.25 | 23 | 0.1739 |
| 1_2513300 | 5 | 1 | 10 | 0.9 | 14 | 1 | 9 | 0.7778 | 38 | 0.9211 | 7 | 1 | 11 | 0.8182 | 6 | 1 | 5 | 0.8 | 29 | 0.8966 |
| 1_2513301 | 7 | 0.7143 | 11 | 0.8182 | 13 | 0.9231 | 9 | 0.8889 | 40 | 0.85 | 7 | 1 | 12 | 0.9167 | 5 | 1 | 6 | 0.8333 | 30 | 0.9333 |
| 1_2513316 | 7 | 0.8571 | 12 | 1 | 11 | 0.7273 | 9 | 0.6667 | 39 | 0.8205 | 7 | 0.7143 | 10 | 0.7 | 4 | 1 | 5 | 0.4 | 26 | 0.6923 |
| 1_2513320 | 5 | 1 | 11 | 0.7273 | 14 | 0.9286 | 8 | 0.875 | 38 | 0.8684 | 7 | 1 | 10 | 1 | 5 | 1 | 5 | 0.8 | 27 | 0.963 |
| 1_2513322 | 6 | 0 | 12 | 0 | 13 | 0.1538 | 9 | 0.1111 | 40 | 0.075 | 6 | 0 | 9 | 0.2222 | 5 | 0 | 5 | 0 | 25 | 0.08 |
| 1_2513324 | 7 | 1 | 12 | 0.9167 | 14 | 0.9286 | 9 | 1 | 42 | 0.9524 | 6 | 0.8333 | 8 | 1 | 5 | 0.8 | 6 | 0.8333 | 25 | 0.88 |
| 1_2513325 | 4 | 0.5 | 11 | 0.3636 | 7 | 0.7143 | 4 | 0.75 | 26 | 0.5385 | 4 | 0.25 | 7 | 0.2857 | 4 | 0 | 3 | 1 | 18 | 0.3333 |
| 1_2513326 | 5 | 0 | 12 | 0.3333 | 13 | 0.4615 | 7 | 0.5714 | 37 | 0.3784 | 6 | 0.8333 | 9 | 0.4444 | 5 | 0.2 | 5 | 0.2 | 25 | 0.44 |
| 1_2578291 | NA | NA | NA | NA | 2 | 1 | NA | NA | 2 | 1 | NA | NA | 1 | 0 | 1 | 1 | NA | NA | 2 | 0.5 |
| 1_3084161 | 2 | 1 | 2 | 1 | 4 | 0.75 | NA | NA | 8 | 0.875 | NA | NA | 4 | 0.75 | 2 | 1 | 5 | 0.8 | 11 | 0.8182 |
| 1_3084179 | 1 | 0 | 1 | 1 | 4 | 0.75 | 1 | 1 | 7 | 0.7143 | NA | NA | 5 | 0.6 | 2 | 1 | 5 | 0.4 | 12 | 0.5833 |
| 1_3084241 | 5 | 0.2 | 10 | 0.2 | 7 | 0.2857 | 7 | 0.2857 | 29 | 0.2414 | 6 | 0.6667 | 7 | 0.5714 | 8 | 0.375 | 7 | 0.7143 | 28 | 0.5714 |
| 1_3084277 | 6 | 0.3333 | 8 | 0.5 | 3 | 0.6667 | 5 | 0.6 | 22 | 0.5 | 4 | 0 | 3 | 0 | 3 | 0.3333 | 4 | 0.25 | 14 | 0.1429 |
| 1_3084282 | 7 | 1 | 10 | 1 | 6 | 1 | 8 | 1 | 31 | 1 | 5 | 1 | 6 | 1 | 7 | 1 | 8 | 1 | 26 | 1 |
| 1_3084287 | 5 | 0 | 11 | 0.2727 | 5 | 0.4 | 6 | 0 | 27 | 0.1852 | 4 | 0.25 | 6 | 0.5 | 7 | 0.2857 | 6 | 0.3333 | 23 | 0.3478 |
| 1_3084291 | 6 | 0.3333 | 11 | 0.1818 | 7 | 0.2857 | 7 | 0.2857 | 31 | 0.2581 | 3 | 0 | 5 | 0 | 7 | 0.4286 | 8 | 0.375 | 23 | 0.2609 |

| | | | | | | | | | | | | | | | | | | | | |
|---|---|---|---|---|---|---|---|---|---|---|---|---|---|---|---|---|---|---|---|---|
| 1_30842917 | 6 | 0 | 12 | 0.0833 | 6 | 0.1667 | 7 | 0.1429 | 31 | 0.0968 | 5 | 0.2 | 6 | 0.1667 | 7 | 0.1429 | 8 | 0 | 26 | 0.1154 |
| 1_30842938 | 4 | 0.5 | 10 | 0.7 | 5 | 0.6 | 5 | 0.4 | 24 | 0.5833 | 3 | 0.3333 | 4 | 1 | 6 | 0.5 | 6 | 0.3333 | 19 | 0.5263 |
| 1_30842968 | 6 | 0 | 11 | 0.0909 | 7 | 0 | 6 | 0.3333 | 30 | 0.1 | 5 | 0.4 | 3 | 0.3333 | 6 | 0 | 6 | 0 | 20 | 0.15 |
| 1_30843010 | 2 | 0 | 7 | 0.2857 | 4 | 0.5 | 6 | 0.5 | 19 | 0.3684 | 2 | 0 | 3 | 0.6667 | 4 | 0.75 | 4 | 0.75 | 13 | 0.6154 |
| 1_30843133 | 4 | 0.5 | 10 | 0.2 | 7 | 0.2857 | 6 | 0.6667 | 27 | 0.3704 | 4 | 0 | 5 | 0.2 | 6 | 0.5 | 5 | 0.2 | 20 | 0.25 |
| 1_30843139 | 4 | 0.25 | 11 | 0.1818 | 6 | 0.5 | 6 | 0.1667 | 27 | 0.2593 | 4 | 0.5 | 4 | 0.5 | 7 | 0.4286 | 5 | 0.8 | 20 | 0.55 |
| 1_30843250 | 4 | 0 | 10 | 0.3 | 5 | 0.4 | 7 | 0.1429 | 26 | 0.2308 | 2 | 0 | 4 | 0.5 | 6 | 0.3333 | 6 | 0.3333 | 18 | 0.3333 |
| 1_30843292 | 5 | 0.4 | 10 | 0 | 7 | 0 | 7 | 0 | 29 | 0.069 | 3 | 0 | 4 | 0 | 6 | 0 | 6 | 0.1667 | 19 | 0.0526 |
| 1_30843310 | 4 | 0.5 | 9 | 0.2222 | 6 | 0.6667 | 6 | 0.3333 | 25 | 0.4 | 2 | 1 | 3 | 1 | 6 | 0.6667 | 6 | 0.5 | 17 | 0.7059 |
| 1_30843330 | 5 | 0 | 10 | 0.1 | 6 | 0.1667 | 7 | 0.2857 | 28 | 0.1429 | 2 | 0 | 4 | 0 | 6 | 0 | 6 | 0.1667 | 18 | 0.0556 |
| 1_30843541 | 5 | 0.2 | 5 | 0.2 | 2 | 0 | 4 | 0.25 | 16 | 0.1875 | 1 | 0 | 3 | 0 | 3 | 0.3333 | 2 | 0 | 9 | 0.1111 |
| 1_30843549 | 3 | 0 | 8 | 0.25 | 5 | 0.2 | 4 | 0.25 | 20 | 0.2 | 1 | 0 | 2 | 0.5 | 5 | 0.8 | 4 | 0.25 | 12 | 0.5 |
| 1_30843625 | 3 | 1 | 5 | 1 | 4 | 0.75 | 4 | 0.75 | 16 | 0.875 | NA | NA | 3 | 1 | 3 | 0.6667 | 3 | 1 | 9 | 0.8889 |
| 1_30844051 | 3 | 0 | 6 | 0.5 | 4 | 0.25 | 3 | 1 | 16 | 0.4375 | NA | NA | 3 | 0.3333 | 4 | 0 | 3 | 0.6667 | 10 | 0.3 |
| 1_30844186 | 1 | 1 | 2 | 0.5 | 2 | 1 | 2 | 1 | 7 | 0.8571 | NA | NA | 3 | 1 | 2 | 1 | 3 | 1 | 8 | 1 |
| 1_30844196 | 1 | 0 | 3 | 0.6667 | 1 | 1 | 1 | 1 | 6 | 0.6667 | NA | NA | 3 | 0.6667 | 4 | 1 | 3 | 1 | 10 | 0.9 |
| 1_30844239 | 1 | 1 | 3 | 1 | 1 | 1 | 1 | 1 | 6 | 1 | NA | NA | 2 | 1 | 4 | 1 | 2 | 0.5 | 8 | 0.875 |
| 1_30844250 | NA | NA | 3 | 0.3333 | 1 | 0 | 2 | 0.5 | 6 | 0.3333 | NA | NA | 2 | 0 | 4 | 0.25 | 3 | 0.6667 | 9 | 0.3333 |
| 1_30844400 | 1 | 0 | 3 | 0.6667 | 2 | 0.5 | 2 | 1 | 8 | 0.625 | NA | NA | 3 | 1 | 4 | 0.5 | 3 | 1 | 10 | 0.8 |
| 1_30844557 | 1 | 0 | 2 | 0.5 | 2 | 0.5 | 2 | 0 | 7 | 0.2857 | NA | NA | 3 | 0.3333 | 3 | 0.3333 | 2 | 0 | 8 | 0.25 |
| 1_30844672 | 1 | 1 | 1 | 1 | 1 | 1 | 2 | 1 | 5 | 1 | NA | NA | 3 | 1 | 3 | 1 | 3 | 1 | 9 | 1 |
| 1_30844834 | 1 | 1 | 1 | 1 | 2 | 1 | 1 | 0 | 5 | 0.8 | NA | NA | 3 | 1 | 4 | 1 | 3 | 1 | 10 | 1 |
| 1_30981013 | 16 | 0.1875 | 10 | 0.1 | 14 | 0.1429 | 15 | 0.2 | 55 | 0.1636 | 9 | 0.1111 | 9 | 0.2222 | 16 | 0.1875 | 14 | 0.1429 | 48 | 0.1667 |
| 1_30981267 | 13 | 0 | 11 | 0 | 15 | 0.1333 | 15 | 0.0667 | 54 | 0.0556 | 8 | 0.125 | 9 | 0.3333 | 20 | 0.05 | 17 | 0.0588 | 54 | 0.1111 |
| 1_30981293 | 12 | 0.0833 | 11 | 0.1818 | 15 | 0.0667 | 15 | 0 | 53 | 0.0755 | 9 | 0.1111 | 7 | 0 | 22 | 0.1818 | 16 | 0.25 | 54 | 0.1667 |
| 1_30981873 | 8 | 0.125 | 8 | 0.125 | 12 | 0.0833 | 11 | 0.0909 | 39 | 0.1026 | 10 | 0.2 | 9 | 0 | 18 | 0 | 12 | 0.25 | 49 | 0.102 |
| 1_30982089 | 9 | 0.8889 | 7 | 0.8571 | 12 | 0.9167 | 12 | 0.9167 | 40 | 0.9 | 8 | 1 | 8 | 1 | 17 | 0.8824 | 12 | 1 | 45 | 0.9556 |
| 1_30982238 | 9 | 0.8889 | 10 | 0.3 | 12 | 0.9167 | 12 | 0.9167 | 43 | 0.7674 | 9 | 0.8889 | 8 | 0.625 | 19 | 0.9474 | 12 | 0.8333 | 48 | 0.8542 |
| 1_30982250 | 8 | 0.375 | 8 | 0.375 | 11 | 0.2727 | 12 | 0.4167 | 39 | 0.359 | 9 | 0 | 6 | 0.3333 | 18 | 0.2778 | 14 | 0.1429 | 47 | 0.1915 |
| 1_30982285 | 7 | 0.4286 | 8 | 0.375 | 11 | 0.7273 | 13 | 0.6154 | 39 | 0.5641 | 8 | 0.5 | 8 | 0.5 | 15 | 0.4 | 10 | 0.6 | 41 | 0.4878 |
| 1_30985354 | 9 | 0.1111 | 9 | 0.1111 | 8 | 0.125 | 10 | 0.3 | 36 | 0.1667 | 8 | 0.5 | 7 | 0.1429 | 15 | 0.2 | 12 | 0.1667 | 42 | 0.2381 |
| 1_30985456 | 7 | 0 | 8 | 0.125 | 9 | 0.4444 | 10 | 0.2 | 34 | 0.2059 | 8 | 0.125 | 6 | 0.6667 | 14 | 0.5 | 13 | 0.2308 | 41 | 0.3659 |
| 1_30985567 | 9 | 0.4444 | 6 | 0.3333 | 9 | 0.3333 | 10 | 0 | 34 | 0.2647 | 7 | 0.2857 | 7 | 0.1429 | 12 | 0 | 12 | 0.4167 | 38 | 0.2105 |
| 1_30985624 | 7 | 0.7143 | 6 | 0.1667 | 8 | 0.75 | 9 | 0.7778 | 30 | 0.6333 | 6 | 0.6667 | 4 | 0.5 | 11 | 0.7273 | 11 | 1 | 32 | 0.7812 |
| 1_32696653 | NA | NA | NA | NA | 2 | 0.5 | 3 | 0 | 5 | 0.2 | NA | NA | 2 | 0 | NA | NA | 5 | 0.2 | 7 | 0.1429 |
| 1_32696802 | NA | NA | NA | NA | 2 | 0 | 2 | 0.5 | 4 | 0.25 | NA | NA | 1 | 0 | NA | NA | 5 | 0.4 | 6 | 0.3333 |
| 1_32696827 | NA | NA | NA | NA | 2 | 0 | 2 | 0.5 | 4 | 0.25 | NA | NA | 2 | 0 | NA | NA | 4 | 0.25 | 6 | 0.1667 |
| 1_32696842 | NA | NA | NA | NA | 2 | 0 | 3 | 0.3333 | 5 | 0.2 | NA | NA | NA | NA | NA | NA | 4 | 0.5 | 4 | 0.5 |
| 1_33493195 | NA | NA | 1 | 1 | NA | NA | 2 | 0.5 | 3 | 0.6667 | NA | NA | 1 | 0 | 1 | 0 | 3 | 0.3333 | 5 | 0.2 |
| 1_33778429 | 2 | 0.5 | 2 | 0 | 1 | 0 | 4 | 0 | 9 | 0.1111 | 1 | 0 | 1 | 0 | 1 | 0 | 4 | 0.5 | 7 | 0.2857 |
| 1_33778474 | 2 | 1 | 1 | 1 | 1 | 1 | 5 | 0.8 | 9 | 0.8889 | NA | NA | 2 | 0 | 1 | 0 | 3 | 0.6667 | 6 | 0.3333 |
| 1_33778489 | 2 | 0.5 | 1 | 1 | 1 | 0 | 5 | 0.8 | 9 | 0.6667 | 1 | 1 | 2 | 0.5 | 1 | 0 | 4 | 0.5 | 8 | 0.5 |
| 1_33778578 | 1 | 0 | 1 | 1 | 1 | 1 | 5 | 0.2 | 8 | 0.375 | 1 | 0 | 2 | 0 | 1 | 1 | 5 | 0.8 | 9 | 0.5556 |
| 1_33778652 | 2 | 1 | 1 | 1 | 1 | 1 | 4 | 1 | 8 | 1 | 1 | 0 | 2 | 1 | 1 | 1 | 5 | 0.8 | 9 | 0.7778 |
| 1_33778702 | 1 | 1 | 2 | 1 | 2 | 1 | 5 | 1 | 10 | 1 | 1 | 1 | 2 | 1 | 1 | 1 | 4 | 1 | 8 | 1 |
| 1_33778753 | 2 | 0 | 2 | 0 | 1 | 0 | 5 | 0.2 | 10 | 0.1 | 1 | 0 | 2 | 0 | 1 | 0 | 5 | 0.2 | 9 | 0.1111 |
| 1_33778767 | 2 | 0.5 | 1 | 1 | 1 | 1 | 4 | 0.5 | 8 | 0.625 | NA | NA | 2 | 0.5 | 1 | 0 | 5 | 0.2 | 8 | 0.25 |
| 1_33778801 | 2 | 1 | 2 | 1 | 1 | 1 | 5 | 1 | 10 | 1 | 1 | 1 | 1 | 1 | 1 | 1 | 4 | 1 | 7 | 1 |
| 1_33778878 | NA | NA | 2 | 0.5 | NA | NA | 4 | 0.75 | 6 | 0.6667 | 1 | 1 | 2 | 0.5 | 1 | 1 | 4 | 0.75 | 8 | 0.75 |
| 1_33778899 | 2 | 0.5 | 2 | 1 | 1 | 1 | 4 | 0.75 | 9 | 0.7778 | 1 | 1 | 2 | 1 | 1 | 1 | 4 | 1 | 8 | 1 |
| 1_33778909 | 2 | 1 | 2 | 0 | NA | NA | 4 | 0.5 | 8 | 0.5 | 1 | 0 | 2 | 0.5 | 1 | 0 | 4 | 0.25 | 8 | 0.25 |
| 1_33778964 | 2 | 1 | 2 | 1 | 1 | 1 | 5 | 1 | 10 | 1 | NA | NA | 2 | 1 | 1 | 1 | 5 | 1 | 8 | 1 |
| 1_33779098 | 2 | 1 | 1 | 0 | 1 | 0 | 4 | 0.25 | 8 | 0.375 | 1 | 0 | 2 | 0 | 1 | 1 | 5 | 0.2 | 9 | 0.2222 |
| 1_33779116 | 1 | 1 | 1 | 0 | NA | NA | 2 | 0.5 | 4 | 0.5 | NA | NA | 1 | 0 | 1 | 0 | 3 | 0.3333 | 5 | 0.2 |
| 1_33779122 | 2 | 0.5 | 2 | 0.5 | 1 | 0 | 5 | 0.4 | 10 | 0.4 | 1 | 0 | 2 | 1 | 1 | 1 | 5 | 0.2 | 9 | 0.4444 |
| 1_33779245 | 1 | 0 | 2 | 0 | 1 | 1 | 4 | 0 | 8 | 0.125 | 1 | 0 | 1 | 0 | NA | NA | 5 | 0.2 | 7 | 0.1429 |
| 1_33779403 | 1 | 1 | 1 | 0 | 1 | 1 | 6 | 0 | 9 | 0.2222 | 1 | 1 | 2 | 0 | 1 | 1 | 5 | 0.4 | 9 | 0.4444 |
| 1_33779421 | 1 | 0 | 2 | 0.5 | NA | NA | 7 | 0.2857 | 10 | 0.3 | 1 | 0 | 2 | 0 | 1 | 0 | 6 | 0.1667 | 10 | 0.1 |
| 1_33779564 | 2 | 1 | 2 | 0.5 | 1 | 1 | 7 | 0.1429 | 12 | 0.4167 | 1 | 1 | 2 | 0.5 | 1 | 0 | 6 | 0.5 | 10 | 0.5 |
| 1_33780625 | NA | NA | 3 | 0.3333 | NA | NA | 4 | 0.25 | 7 | 0.2857 | 1 | 0 | 1 | 1 | 1 | 1 | 6 | 0.6667 | 9 | 0.6667 |
| 1_33785471 | 2 | 1 | 3 | 1 | NA | NA | NA | NA | 5 | 1 | 2 | 1 | NA | NA | 2 | 1 | 1 | 1 | 5 | 1 |
| 1_33785476 | 2 | 0 | 1 | 1 | NA | NA | NA | NA | 3 | 0.3333 | 2 | 1 | NA | NA | 2 | 0.5 | NA | NA | 4 | 0.75 |
| 1_33785494 | 1 | 0 | 2 | 1 | NA | NA | NA | NA | 3 | 0.6667 | 1 | 0 | NA | NA | 2 | 0.5 | 1 | 1 | 4 | 0.5 |
| 1_33785710 | 2 | 1 | 1 | 0 | NA | NA | NA | NA | 3 | 0.6667 | 2 | 1 | NA | NA | 1 | 0 | 1 | 0 | 4 | 0.5 |
| 1_33785782 | 2 | 0.5 | 3 | 0 | NA | NA | NA | NA | 5 | 0.2 | 1 | 0 | NA | NA | 1 | 0 | 1 | 1 | 3 | 0.3333 |

| | | | | | | | | | | | | | | | | | | | | |
|---|---|---|---|---|---|---|---|---|---|---|---|---|---|---|---|---|---|---|---|---|
| 1_33785795 | 1 | 0 | 3 | 0.6667 | NA | NA | NA | NA | 4 | 0.5 | 2 | 0.5 | NA | NA | 1 | 0 | 1 | 1 | 4 | 0.5 |
| 1_33815809 | NA | NA | NA | NA | NA | NA | 1 | 1 | 1 | 1 | 2 | 1 | 2 | 1 | 2 | 0 | 1 | 1 | 7 | 0.7143 |
| 1_33816117 | NA | NA | NA | NA | NA | NA | 1 | 1 | 1 | 1 | 3 | 0 | 2 | 0.5 | 1 | 1 | 1 | 1 | 7 | 0.4286 |
| 1_33816612 | NA | NA | NA | NA | NA | NA | 1 | 1 | 1 | 1 | 2 | 1 | 1 | 0 | 1 | 1 | NA | NA | 4 | 0.75 |
| 1_33816797 | NA | NA | NA | NA | NA | NA | 1 | 1 | 1 | 1 | 2 | 0 | 2 | 0.5 | 1 | 0 | 1 | 0 | 6 | 0.1667 |
| 1_33832192 | NA | NA | NA | NA | NA | NA | 2 | 0.5 | 2 | 0.5 | 1 | 0 | 3 | 0 | 2 | 0.5 | 2 | 0 | 8 | 0.125 |
| 1_33839949 | NA | NA | 4 | 0.75 | 1 | 1 | NA | NA | 5 | 0.8 | NA | NA | NA | NA | NA | NA | 1 | 1 | 1 | 1 |
| 1_33840197 | NA | NA | 2 | 1 | 1 | 1 | 1 | 1 | 4 | 1 | NA | NA | NA | NA | NA | NA | 1 | 1 | 1 | 1 |
| 1_33840566 | NA | NA | 2 | 1 | 1 | 1 | 1 | 1 | 4 | 1 | NA | NA | NA | NA | NA | NA | 1 | 1 | 1 | 1 |
| 1_33840624 | NA | NA | 2 | 1 | 1 | 1 | 1 | 1 | 4 | 1 | NA | NA | NA | NA | NA | NA | 1 | 1 | 1 | 1 |
| 1_33840671 | NA | NA | 3 | 1 | 1 | 1 | 1 | 0 | 5 | 0.8 | NA | NA | NA | NA | NA | NA | 1 | 1 | 1 | 1 |
| 1_33840881 | NA | NA | 3 | 1 | 1 | 1 | 1 | 0 | 5 | 0.8 | NA | NA | NA | NA | NA | NA | 1 | 1 | 1 | 1 |
| 1_33840888 | NA | NA | 3 | 0.3333 | 1 | 0 | 1 | 0 | 5 | 0.2 | NA | NA | NA | NA | NA | NA | 1 | 1 | 1 | 1 |
| 1_33841026 | NA | NA | 2 | 1 | 1 | 1 | 1 | 0 | 4 | 0.75 | NA | NA | NA | NA | NA | NA | 1 | 1 | 1 | 1 |
| 1_33841052 | NA | NA | 2 | 1 | 1 | 1 | 1 | 0 | 4 | 0.75 | NA | NA | NA | NA | NA | NA | 1 | 1 | 1 | 1 |
| 1_33841193 | NA | NA | 3 | 0.6667 | 1 | 0 | 1 | 1 | 5 | 0.6 | NA | NA | NA | NA | NA | NA | 1 | 1 | 1 | 1 |
| 1_33841232 | NA | NA | 2 | 1 | 1 | 0 | NA | NA | 3 | 0.6667 | NA | NA | NA | NA | NA | NA | 1 | 1 | 1 | 1 |
| 1_33841556 | NA | NA | 3 | 1 | 1 | 1 | 1 | 1 | 5 | 1 | NA | NA | NA | NA | NA | NA | 1 | 1 | 1 | 1 |
| 1_33841565 | NA | NA | 3 | 1 | 1 | 1 | 1 | 1 | 5 | 1 | NA | NA | NA | NA | NA | NA | 1 | 1 | 1 | 1 |
| 1_33892112 | 7 | 0.2857 | 8 | 0 | 7 | 0 | 12 | 0.25 | 34 | 0.1471 | 6 | 0.1667 | 9 | 0.4444 | 9 | 0 | 9 | 0 | 33 | 0.1515 |
| 1_33892334 | 11 | 0.8182 | 8 | 0.75 | 10 | 0.9 | 12 | 0.9167 | 41 | 0.8537 | 10 | 0.9 | 12 | 0.8333 | 7 | 1 | 8 | 0.875 | 37 | 0.8919 |
| 1_33892374 | 9 | 0.6667 | 7 | 0.5714 | 9 | 0.1111 | 12 | 0.3333 | 37 | 0.4054 | 8 | 0.25 | 8 | 0.5 | 9 | 0.2222 | 8 | 0.125 | 33 | 0.2727 |
| 1_34267024 | 1 | 1 | NA | NA | NA | NA | 1 | 0 | 2 | 0.5 | 2 | 1 | 2 | 0 | 1 | 1 | NA | NA | 5 | 0.6 |
| 1_34267814 | 1 | 0 | 1 | 1 | NA | NA | 1 | 1 | 3 | 0.6667 | 1 | 1 | 1 | 0 | NA | NA | NA | NA | 2 | 0.5 |
| 1_34268113 | 1 | 0 | 1 | 1 | 1 | 1 | 1 | 1 | 4 | 0.75 | 3 | 0.6667 | 2 | 0.5 | 1 | 0 | NA | NA | 6 | 0.5 |
| 1_34299319 | 1 | 0 | 2 | 0 | 2 | 0.5 | 2 | 0 | 7 | 0.1429 | 3 | 0 | 3 | 0.3333 | 5 | 0 | 3 | 0 | 14 | 0.0714 |
| 1_34346507 | 8 | 0.125 | 9 | 0.3333 | 9 | 0.1111 | 14 | 0.0714 | 40 | 0.15 | 13 | 0.2308 | 12 | 0.0833 | 11 | 0.1818 | 11 | 0.2727 | 47 | 0.1915 |
| 1_34346557 | 7 | 0.2857 | 11 | 0.1818 | 10 | 0.2 | 14 | 0.1429 | 42 | 0.1905 | 11 | 0.1818 | 11 | 0.2727 | 12 | 0 | 13 | 0.1538 | 47 | 0.1489 |
| 1_34346565 | 7 | 0 | 12 | 0.3333 | 11 | 0.0909 | 15 | 0.3333 | 45 | 0.2222 | 12 | 0.1667 | 13 | 0.2308 | 12 | 0 | 13 | 0.2308 | 50 | 0.16 |
| 1_34346891 | 10 | 0.3 | 17 | 0.2353 | 13 | 0.4615 | 17 | 0.3529 | 57 | 0.3333 | 10 | 0.2 | 10 | 0.2 | 13 | 0.6923 | 15 | 0.3333 | 48 | 0.375 |
| 1_34346902 | 7 | 0.7143 | 11 | 0.7273 | 9 | 0.8889 | 11 | 0.9091 | 38 | 0.8158 | 5 | 0.6 | 8 | 0.625 | 9 | 0.6667 | 11 | 0.7273 | 33 | 0.6667 |
| 1_34347375 | 10 | 0.2 | 11 | 0.1818 | 14 | 0.1429 | 20 | 0.3 | 55 | 0.2182 | 13 | 0.0769 | 11 | 0.2727 | 12 | 0.1667 | 13 | 0.2308 | 49 | 0.1837 |
| 1_34347397 | 10 | 0.5 | 18 | 0.7778 | 16 | 0.375 | 19 | 0.3158 | 63 | 0.4921 | 11 | 0.7273 | 11 | 0.4545 | 15 | 0.4667 | 16 | 0.625 | 53 | 0.566 |
| 1_34347542 | 9 | 0.4444 | 13 | 0.0769 | 7 | 0.2857 | 9 | 0.2222 | 38 | 0.2368 | 6 | 0 | 7 | 0 | 11 | 0.2727 | 6 | 0.1667 | 30 | 0.1333 |
| 1_34349072 | NA | NA | NA | NA | NA | NA | 1 | 1 | 1 | 1 | NA | NA | 1 | 1 | NA | NA | NA | NA | 1 | 1 |
| 1_34483601 | 3 | 0.3333 | 3 | 1 | 2 | 0 | 2 | 0.5 | 10 | 0.5 | 1 | 0 | 5 | 0.6 | 10 | 0.7 | 5 | 0.4 | 21 | 0.5714 |
| 1_34483719 | 3 | 0.6667 | 4 | 0.25 | 1 | 0 | 2 | 0 | 10 | 0.3 | 1 | 0 | 3 | 0 | 9 | 0.2222 | 4 | 0 | 17 | 0.1176 |
| 1_34483756 | 4 | 0 | 5 | 0.4 | 1 | 0 | 2 | 0 | 12 | 0.1667 | 1 | 1 | 5 | 0.2 | 9 | 0.4444 | 4 | 0.25 | 19 | 0.3684 |
| 1_34483764 | 3 | 0 | 5 | 0 | 2 | 0.5 | 2 | 0 | 12 | 0.0833 | 1 | 0 | 5 | 0 | 8 | 0 | 5 | 0.2 | 19 | 0.0526 |
| 1_34483771 | 4 | 0.5 | 5 | 0.8 | 1 | 0 | 2 | 1 | 12 | 0.6667 | 1 | 1 | 4 | 0.75 | 9 | 0.6667 | 5 | 0.6 | 19 | 0.6842 |
| 1_34483854 | 3 | 0 | 4 | 0 | 1 | 1 | 1 | 0 | 9 | 0.1111 | 1 | 0 | 3 | 0 | 7 | 0 | 5 | 0.2 | 16 | 0.0625 |
| 1_34626030 | 1 | 1 | 1 | 1 | NA | NA | NA | NA | 2 | 1 | NA | NA | 1 | 1 | NA | NA | NA | NA | 1 | 1 |
| 1_34626268 | 1 | 1 | 1 | 0 | NA | NA | NA | NA | 2 | 0.5 | NA | NA | 2 | 0 | 1 | 0 | 1 | 1 | 4 | 0.25 |
| 1_34626370 | 1 | 1 | 1 | 1 | NA | NA | NA | NA | 2 | 1 | NA | NA | 2 | 0.5 | 1 | 0 | 1 | 0 | 4 | 0.25 |
| 1_34626577 | 1 | 1 | 1 | 1 | NA | NA | NA | NA | 2 | 1 | NA | NA | 2 | 1 | 1 | 1 | NA | NA | 3 | 1 |
| 1_34626670 | NA | NA | 1 | 1 | NA | NA | NA | NA | 1 | 1 | NA | NA | 2 | 0.5 | 1 | 1 | 1 | 1 | 4 | 0.75 |
| 1_34626982 | 1 | 0 | 1 | 1 | NA | NA | 1 | 1 | 3 | 0.6667 | NA | NA | 3 | 0.6667 | 1 | 1 | NA | NA | 4 | 0.75 |
| 1_34627010 | 1 | 1 | 1 | 1 | NA | NA | 1 | 1 | 3 | 1 | NA | NA | 2 | 1 | 1 | 1 | NA | NA | 3 | 1 |
| 1_34627060 | 1 | 1 | NA | NA | NA | NA | 1 | 1 | 2 | 1 | 1 | 1 | 2 | 0.5 | NA | NA | 1 | 1 | 4 | 0.75 |
| 1_34627238 | NA | NA | 1 | 1 | NA | NA | NA | NA | 1 | 1 | 1 | 0 | 3 | 1 | 1 | 1 | 1 | 0 | 6 | 0.6667 |
| 1_34627677 | 1 | 0 | 2 | 0.5 | NA | NA | 1 | 0 | 4 | 0.25 | 1 | 0 | 3 | 1 | NA | NA | 1 | 0 | 5 | 0.6 |
| 1_34627948 | 1 | 0 | 2 | 0.5 | NA | NA | 1 | 0 | 4 | 0.25 | 2 | 0.5 | 3 | 0.3333 | 1 | 0 | 1 | 0 | 7 | 0.2857 |
| 1_34627975 | 1 | 1 | 2 | 0 | NA | NA | 1 | 1 | 4 | 0.5 | 2 | 0 | 3 | 0.6667 | NA | NA | 1 | 0 | 6 | 0.3333 |
| 1_34628172 | NA | NA | 2 | 1 | NA | NA | 1 | 1 | 3 | 1 | 2 | 0.5 | 2 | 1 | 1 | 1 | 1 | 1 | 6 | 0.8333 |
| 1_34628244 | 1 | 1 | 3 | 1 | 1 | 1 | 1 | 1 | 6 | 1 | 2 | 1 | 3 | 1 | 1 | 1 | 1 | 1 | 7 | 1 |
| 1_34628271 | 1 | 1 | 3 | 1 | 1 | 1 | 1 | 1 | 6 | 1 | 1 | 1 | 2 | 1 | 1 | 1 | 1 | 1 | 5 | 1 |
| 1_34628365 | 1 | 1 | 3 | 0.6667 | 1 | 0 | 1 | 1 | 6 | 0.6667 | 2 | 0.5 | 3 | 0.6667 | 2 | 0 | 2 | 1 | 9 | 0.5556 |
| 1_34628502 | 2 | 1 | 4 | 1 | 2 | 1 | NA | NA | 8 | 1 | 2 | 1 | 3 | 1 | 1 | 1 | NA | NA | 6 | 1 |
| 1_34628537 | 2 | 0 | 3 | 0.3333 | 2 | 1 | 1 | 1 | 8 | 0.5 | 1 | 1 | 3 | 0 | 1 | 0 | 1 | 1 | 6 | 0.3333 |
| 1_34628675 | 2 | 1 | 3 | 1 | 2 | 0 | 1 | 1 | 8 | 0.75 | 3 | 1 | 4 | 0.5 | 1 | 0 | 1 | 1 | 9 | 0.6667 |
| 1_34628768 | 1 | 0 | 4 | 0.75 | 1 | 1 | NA | NA | 6 | 0.6667 | 3 | 1 | 4 | 0.75 | 1 | 1 | 1 | 0 | 9 | 0.7778 |
| 1_34629104 | 2 | 0 | 4 | 0.75 | 1 | 0 | 1 | 1 | 8 | 0.5 | 2 | 1 | 4 | 0.25 | 2 | 0.5 | 3 | 0.3333 | 11 | 0.4545 |
| 1_34629166 | 1 | 1 | 5 | 0.8 | 2 | 1 | 1 | 1 | 9 | 0.8889 | 4 | 1 | 2 | 0.5 | 2 | 1 | 2 | 0.5 | 10 | 0.8 |
| 1_34629244 | 2 | 1 | 6 | 1 | 4 | 1 | 1 | 1 | 13 | 1 | 4 | 1 | 5 | 1 | 2 | 1 | 2 | 1 | 13 | 1 |
| 1_34629264 | 2 | 0 | 5 | 0.2 | 2 | 0 | NA | NA | 9 | 0.1111 | 4 | 0 | 5 | 0 | 1 | 0 | 2 | 0.5 | 12 | 0.0833 |

| | | | | | | | | | | | | | | | | | | | | |
|---|---|---|---|---|---|---|---|---|---|---|---|---|---|---|---|---|---|---|---|---|
| 1_3462927 | 2 | 0.5 | 6 | 1 | 4 | 0.5 | 1 | 1 | 13 | 0.7692 | 4 | 0.75 | 4 | 0.25 | 2 | 1 | 2 | 0 | 12 | 0.5 |
| 1_3462942 | 2 | 0 | 6 | 0.1667 | 3 | 0 | 2 | 0 | 13 | 0.0769 | 4 | 0.25 | 4 | 0 | 2 | 0 | 2 | 0 | 12 | 0.0833 |
| 1_3462943 | 2 | 0 | 5 | 0.6 | 2 | 0.5 | 2 | 0.5 | 11 | 0.4545 | 4 | 0.25 | 4 | 0.5 | 2 | 0 | 2 | 0.5 | 12 | 0.3333 |
| 1_3462951 | 2 | 0 | 6 | 0.6667 | 4 | 0.25 | 2 | 0.5 | 14 | 0.4286 | 4 | 0.5 | 5 | 0.6 | 3 | 0.3333 | 2 | 0.5 | 14 | 0.5 |
| 1_3462952 | 2 | 0 | 4 | 0.5 | 3 | 0.3333 | 1 | 1 | 10 | 0.4 | 3 | 0 | 4 | 0.75 | 2 | 0.5 | NA | NA | 9 | 0.4444 |
| 1_3462957 | 2 | 0 | 5 | 0.2 | 4 | 0 | 2 | 0 | 13 | 0.0769 | 4 | 0 | 5 | 0.2 | 4 | 0 | 3 | 0 | 16 | 0.0625 |
| 1_3462960 | 2 | 0 | 6 | 0 | 3 | 0 | 2 | 0.5 | 13 | 0.0769 | 4 | 0 | 5 | 0 | 4 | 0.25 | 2 | 0 | 15 | 0.0667 |
| 1_3462963 | 2 | 0 | 6 | 0 | 4 | 0.25 | 2 | 0.5 | 14 | 0.1429 | 3 | 0.3333 | 4 | 0 | 4 | 0 | 3 | 0 | 14 | 0.0714 |
| 1_3462964 | 1 | 0 | 6 | 0.3333 | 3 | 0.3333 | 2 | 0.5 | 12 | 0.3333 | 2 | 0.5 | 4 | 0.25 | 2 | 0 | 2 | 0 | 10 | 0.2 |
| 1_3462972 | 2 | 0.5 | 6 | 1 | 3 | 0.6667 | 2 | 1 | 13 | 0.8462 | 4 | 0.75 | 5 | 0.8 | 3 | 1 | 2 | 0.5 | 14 | 0.7857 |
| 1_3462981 | 1 | 0 | 6 | 0.3333 | 2 | 0.5 | 2 | 0.5 | 11 | 0.3636 | 3 | 0.3333 | 6 | 0.5 | 2 | 0.5 | 2 | 0.5 | 13 | 0.4615 |
| 1_3463006 | NA | NA | 1 | 1 | NA | NA | 1 | 0 | 2 | 0.5 | 1 | 1 | NA | NA | NA | NA | NA | NA | 1 | 1 |
| 1_3463007 | NA | NA | 1 | 0 | NA | NA | 1 | 1 | 2 | 0.5 | 1 | 1 | NA | NA | NA | NA | NA | NA | 1 | 1 |
| 1_3484684 | 14 | 0.0714 | 18 | 0.0556 | 17 | 0.0588 | 15 | 0.0667 | 64 | 0.0625 | 13 | 0 | 13 | 0.1538 | 16 | 0.0625 | 13 | 0 | 55 | 0.0545 |
| 1_3484713 | NA | NA | 1 | 1 | 1 | 1 | NA | NA | 2 | 1 | NA | NA | 1 | 0 | NA | NA | 1 | 1 | 2 | 0.5 |
| 1_3484771 | NA | NA | 2 | 0.5 | 1 | 0 | 1 | 1 | 4 | 0.5 | NA | NA | 2 | 0 | NA | NA | 1 | 1 | 3 | 0.3333 |
| 1_3484806 | NA | NA | 3 | 0.6667 | 1 | 1 | 1 | 1 | 5 | 0.8 | NA | NA | 2 | 0.5 | NA | NA | 1 | 1 | 3 | 0.6667 |
| 1_3484900 | NA | NA | NA | NA | 1 | 1 | 1 | 0 | 2 | 0.5 | NA | NA | 2 | 0 | 1 | 1 | NA | NA | 3 | 0.3333 |
| 1_3484924 | 1 | 0 | NA | NA | 1 | 1 | NA | NA | 2 | 0.5 | NA | NA | 3 | 0.3333 | NA | NA | NA | NA | 3 | 0.3333 |
| 1_3485094 | 20 | 0.7 | 23 | 0.5652 | 29 | 0.6897 | 20 | 0.7 | 92 | 0.663 | 18 | 0.6111 | 24 | 0.5833 | 21 | 0.8095 | 14 | 0.7857 | 77 | 0.6883 |
| 1_3485100 | 15 | 1 | 18 | 0.8889 | 25 | 0.76 | 17 | 0.8824 | 75 | 0.8667 | 15 | 0.8 | 14 | 1 | 14 | 0.7143 | 14 | 1 | 57 | 0.8772 |
| 1_3485101 | 8 | 0.25 | 12 | 0.1667 | 11 | 0.1818 | 10 | 0.1 | 41 | 0.1707 | 14 | 0.2143 | 10 | 0.2 | 10 | 0.2 | 8 | 0.125 | 42 | 0.1905 |
| 1_3485109 | 18 | 0.1667 | 24 | 0.1667 | 25 | 0.16 | 23 | 0.3043 | 90 | 0.2 | 22 | 0.0909 | 22 | 0.2273 | 16 | 0.25 | 17 | 0.2353 | 77 | 0.1948 |
| 1_3485110 | 20 | 0.5 | 22 | 0.4091 | 26 | 0.4231 | 20 | 0.4 | 88 | 0.4318 | 20 | 0.3 | 23 | 0.4783 | 25 | 0.56 | 15 | 0.7333 | 83 | 0.506 |
| 1_3485119 | 19 | 1 | 23 | 0.9565 | 27 | 1 | 26 | 0.9615 | 95 | 0.9789 | 21 | 0.9048 | 31 | 0.9677 | 22 | 0.9545 | 18 | 0.8889 | 92 | 0.9348 |
| 1_3485165 | 23 | 0.5652 | 28 | 0.4643 | 23 | 0.5652 | 21 | 0.5714 | 95 | 0.5368 | 23 | 0.3913 | 28 | 0.5357 | 24 | 0.625 | 20 | 0.45 | 95 | 0.5053 |
| 1_3485166 | 16 | 0.75 | 18 | 0.7222 | 25 | 0.68 | 14 | 0.5714 | 73 | 0.6849 | 19 | 0.8947 | 21 | 0.619 | 14 | 0.7143 | 16 | 0.625 | 70 | 0.7143 |
| 1_3485173 | 24 | 0.5 | 30 | 0.4333 | 31 | 0.3548 | 29 | 0.4138 | 114 | 0.4211 | 27 | 0.2963 | 31 | 0.3226 | 24 | 0.5 | 22 | 0.4091 | 104 | 0.375 |
| 1_3485417 | 19 | 0.0526 | 18 | 0.1667 | 12 | 0.0833 | 15 | 0.1333 | 64 | 0.1094 | 8 | 0.125 | 13 | 0 | 19 | 0.1579 | 14 | 0 | 54 | 0.0741 |
| 1_3485565 | 9 | 0.2222 | 7 | 0.4286 | 6 | 0.1667 | 7 | 0 | 29 | 0.2069 | 3 | 0 | 8 | 0.25 | 4 | 0.25 | 7 | 0.1429 | 22 | 0.1818 |
| 1_3485591 | 8 | 0.25 | 6 | 0 | 3 | 0 | 7 | 0 | 24 | 0.0833 | 3 | 0.3333 | 7 | 0.1429 | 5 | 0.2 | 6 | 0 | 21 | 0.1429 |
| 1_3485598 | 8 | 0.125 | 8 | 0.375 | 7 | 0.2857 | 7 | 0.4286 | 30 | 0.3 | 3 | 0 | 10 | 0.2 | 6 | 0.1667 | 8 | 0.375 | 27 | 0.2222 |
| 1_3485601 | 9 | 0.2222 | 5 | 0.2 | 5 | 0 | 10 | 0 | 29 | 0.1034 | 3 | 0 | 8 | 0.125 | 6 | 0 | 9 | 0.3333 | 26 | 0.1538 |
| 1_3485602 | 8 | 0.625 | 9 | 0.2222 | 7 | 0.2857 | 8 | 0.125 | 32 | 0.3125 | 2 | 0 | 8 | 0.5 | 6 | 0.3333 | 7 | 0 | 23 | 0.2609 |
| 1_3485612 | 8 | 0.5 | 9 | 0.1111 | 6 | 0.1667 | 9 | 0 | 32 | 0.1875 | 2 | 0.5 | 9 | 0.3333 | 5 | 0.4 | 8 | 0.25 | 24 | 0.3333 |
| 1_3485616 | 7 | 0.1429 | 8 | 0.125 | 8 | 0.25 | 8 | 0.125 | 31 | 0.1613 | 3 | 0.3333 | 8 | 0.375 | 5 | 0 | 6 | 0.1667 | 22 | 0.2273 |
| 1_3485629 | 7 | 0.7143 | 10 | 0.8 | 7 | 0.5714 | 9 | 0.4444 | 33 | 0.6364 | 3 | 0.3333 | 4 | 0.5 | 2 | 0.5 | 9 | 0.6667 | 18 | 0.5556 |
| 1_3485632 | 7 | 0.7143 | 10 | 0.8 | 5 | 0.2 | 8 | 0.625 | 30 | 0.6333 | 3 | 0.3333 | 5 | 0.4 | 5 | 0.6 | 8 | 0.5 | 21 | 0.4762 |
| 1_3485637 | 7 | 0.2857 | 7 | 0.4286 | 7 | 0.1429 | 8 | 0.125 | 29 | 0.2414 | 3 | 0 | 5 | 0.2 | 4 | 0 | 7 | 0.2857 | 19 | 0.1579 |
| 1_3485641 | 6 | 0.8333 | 9 | 0.2222 | 9 | 0 | 7 | 0.2857 | 31 | 0.2903 | 3 | 0.3333 | 5 | 0.6 | 5 | 0.6 | 8 | 0.375 | 21 | 0.4762 |
| 1_3610819 | 1 | 1 | NA | NA | 4 | 0.5 | 3 | 0 | 8 | 0.375 | 2 | 0.5 | 4 | 0 | 3 | 0.3333 | 1 | 0 | 10 | 0.2 |
| 1_3614275 | 2 | 0.5 | 2 | 1 | 3 | 0.6667 | 3 | 1 | 10 | 0.8 | 2 | 0.5 | 5 | 0.6 | 3 | 1 | 2 | 1 | 12 | 0.75 |
| 1_3614306 | 4 | 0.75 | 3 | 1 | 6 | 0.6667 | 4 | 0.75 | 17 | 0.7647 | 2 | 1 | 6 | 0.6667 | 6 | 1 | 3 | 0.6667 | 17 | 0.8235 |
| 1_3614309 | 3 | 0.6667 | 3 | 1 | 6 | 0.6667 | 3 | 1 | 15 | 0.8 | 2 | 1 | 6 | 0.8333 | 7 | 0.7143 | 4 | 0.75 | 19 | 0.7895 |
| 1_3614311 | 3 | 0 | 3 | 0 | 5 | 0 | 2 | 1 | 13 | 0.1538 | 1 | 0 | 6 | 0.1667 | 5 | 0.2 | 4 | 0.25 | 16 | 0.1875 |
| 1_3614312 | 3 | 1 | 3 | 1 | 6 | 1 | 5 | 0.6 | 17 | 0.8824 | 2 | 1 | 6 | 1 | 6 | 1 | 4 | 0.75 | 18 | 0.9444 |
| 1_3614315 | 1 | 1 | 4 | 0.5 | 6 | 0.3333 | 4 | 0.5 | 15 | 0.4667 | NA | NA | 4 | 0.25 | 5 | 0.6 | 1 | 1 | 10 | 0.5 |
| 1_3614323 | 3 | 0 | 4 | 0 | 5 | 0.2 | 5 | 0 | 17 | 0.0588 | 2 | 0.5 | 7 | 0.1429 | 5 | 0.6 | 4 | 0 | 18 | 0.2778 |
| 1_3614327 | 3 | 1 | 4 | 1 | 4 | 1 | 5 | 1 | 16 | 1 | 2 | 1 | 5 | 1 | 5 | 1 | 4 | 0.75 | 16 | 0.9375 |
| 1_3614342 | 1 | 1 | 5 | 0.8 | 6 | 0.3333 | 3 | 1 | 15 | 0.6667 | 2 | 0 | 4 | 0.75 | 7 | 0.8571 | 4 | 0.5 | 17 | 0.6471 |
| 1_3614347 | 3 | 0.3333 | 5 | 0.2 | 3 | 0.3333 | 5 | 0.2 | 16 | 0.25 | 2 | 0 | 7 | 0 | 6 | 0 | 5 | 0.2 | 20 | 0.05 |
| 1_3614350 | 2 | 1 | 5 | 1 | 6 | 1 | 4 | 1 | 17 | 1 | 1 | 1 | 4 | 0.5 | 4 | 1 | 6 | 1 | 15 | 0.8667 |
| 1_3614353 | 1 | 0 | 5 | 0.2 | 5 | 0.2 | 5 | 0.2 | 16 | 0.1875 | 2 | 0 | 5 | 0 | 6 | 0.1667 | 5 | 0 | 18 | 0.0556 |
| 1_3614378 | 3 | 0 | 4 | 0 | 5 | 0.2 | 8 | 0.25 | 20 | 0.15 | 2 | 0.5 | 7 | 0.1429 | 7 | 0 | 7 | 0.1429 | 23 | 0.1304 |
| 1_3614388 | 3 | 0.6667 | 5 | 0.6 | 4 | 0.5 | 6 | 0.5 | 18 | 0.5556 | 2 | 0 | 4 | 0.75 | 7 | 0.5714 | 5 | 0.8 | 18 | 0.6111 |
| 1_3614409 | 6 | 0.1667 | 6 | 0.3333 | 7 | 0 | 9 | 0.1111 | 28 | 0.1429 | 4 | 0.25 | 5 | 0 | 7 | 0.2857 | 9 | 0.1111 | 25 | 0.16 |
| 1_3614459 | 7 | 0 | 6 | 0 | 9 | 0 | 11 | 0.2727 | 33 | 0.0909 | 5 | 0 | 7 | 0.1429 | 7 | 0.2857 | 10 | 0 | 29 | 0.1034 |
| 1_3614517 | 6 | 0.6667 | 12 | 0.5833 | 10 | 1 | 11 | 0.6364 | 39 | 0.7179 | 7 | 0.4286 | 11 | 0.5455 | 10 | 0.6 | 12 | 0.75 | 40 | 0.6 |
| 1_3614522 | 8 | 0.375 | 12 | 0.4167 | 11 | 0.3636 | 9 | 0.2222 | 40 | 0.35 | 5 | 0.2 | 9 | 0.2222 | 11 | 0.3636 | 11 | 0.1818 | 36 | 0.25 |
| 1_3614523 | 8 | 0.125 | 13 | 0.3077 | 10 | 0.1 | 8 | 0.25 | 39 | 0.2051 | 6 | 0.3333 | 11 | 0 | 9 | 0.3333 | 8 | 0.25 | 34 | 0.2059 |
| 1_3614529 | 7 | 0.2857 | 12 | 0.25 | 10 | 0.3 | 10 | 0 | 39 | 0.2051 | 6 | 0.1667 | 9 | 0 | 9 | 0.1111 | 11 | 0.1818 | 35 | 0.1143 |
| 1_3614530 | 6 | 0.5 | 15 | 0.7333 | 12 | 0.9167 | 10 | 0.9 | 43 | 0.7907 | 6 | 0.8333 | 11 | 0.4545 | 12 | 0.5833 | 11 | 0.6364 | 40 | 0.6 |
| 1_3614532 | 10 | 0.6 | 12 | 0.4167 | 11 | 0.3636 | 10 | 0.6 | 43 | 0.4884 | 7 | 0.5714 | 12 | 0.3333 | 10 | 0.4 | 9 | 0.2222 | 38 | 0.3684 |
| 1_3614533 | 7 | 0.8571 | 11 | 0.7273 | 10 | 0.8 | 9 | 0.6667 | 37 | 0.7568 | 6 | 0.8333 | 6 | 0.8333 | 10 | 0.8 | 9 | 0.7778 | 31 | 0.8065 |
| 1_3617931 | 1 | 0 | 1 | 1 | NA | NA | NA | NA | 2 | 0.5 | 2 | 0 | 2 | 0 | 1 | 1 | NA | NA | 5 | 0.2 |

| | | | | | | | | | | | | | | | | | | | | |
|---|---|---|---|---|---|---|---|---|---|---|---|---|---|---|---|---|---|---|---|---|
| 1_3617943 | 1 | 0 | 1 | 1 | NA | NA | NA | NA | 2 | 0.5 | 1 | 1 | 1 | 1 | 1 | 1 | NA | NA | 3 | 1 |
| 1_3617979 | 1 | 1 | 1 | 1 | NA | NA | NA | NA | 2 | 1 | 1 | 1 | 2 | 0.5 | 1 | 0 | NA | NA | 4 | 0.5 |
| 1_3618234 | 1 | 0 | 1 | 1 | NA | NA | NA | NA | 2 | 0.5 | 1 | 0 | 1 | 0 | 1 | 0 | 2 | 0.5 | 5 | 0.2 |
| 1_3618260 | 5 | 0.2 | 6 | 0.3333 | 1 | 0 | 5 | 0 | 17 | 0.1765 | 4 | 0.25 | 4 | 0.25 | 2 | 0 | 4 | 0.25 | 14 | 0.2143 |
| 1_3618308 | 3 | 1 | 5 | 0.8 | 1 | 1 | 5 | 1 | 14 | 0.9286 | 2 | 1 | 3 | 1 | 2 | 1 | 7 | 0.7143 | 14 | 0.8571 |
| 1_3618317 | 1 | 0 | 3 | 1 | 1 | 1 | 6 | 0.6667 | 11 | 0.7273 | 1 | 1 | NA | NA | 2 | 0.5 | 3 | 0.6667 | 6 | 0.6667 |
| 1_3618326 | 3 | 0 | 4 | 0.25 | 1 | 0 | 6 | 0.1667 | 14 | 0.1429 | 2 | 0 | 3 | 0 | 2 | 0.5 | 6 | 0.1667 | 13 | 0.1538 |
| 1_3618326 | 3 | 0.6667 | 5 | 1 | 1 | 1 | 5 | 1 | 14 | 0.9286 | 2 | 0.5 | 3 | 1 | 2 | 0.5 | 7 | 0.7143 | 14 | 0.7143 |
| 1_3635828 | NA | NA | 1 | 0 | 2 | 0.5 | NA | NA | 3 | 0.3333 | 1 | 0 | NA | NA | NA | NA | 2 | 1 | 3 | 0.6667 |
| 1_3635837 | NA | NA | 1 | 1 | 2 | 1 | NA | NA | 3 | 1 | 1 | 1 | NA | NA | NA | NA | 1 | 1 | 2 | 1 |
| 1_3635841 | NA | NA | 1 | 1 | 1 | 1 | NA | NA | 2 | 1 | 1 | 1 | NA | NA | NA | NA | 2 | 1 | 3 | 1 |
| 1_3635857 | NA | NA | 1 | 1 | 2 | 1 | NA | NA | 3 | 1 | 1 | 1 | NA | NA | NA | NA | 2 | 1 | 3 | 1 |
| 1_3635861 | NA | NA | 1 | 1 | 1 | 1 | NA | NA | 2 | 1 | 1 | 1 | NA | NA | NA | NA | 2 | 0 | 3 | 0.3333 |
| 1_3635887 | NA | NA | NA | NA | 2 | 0.5 | NA | NA | 2 | 0.5 | 1 | 1 | NA | NA | 1 | 1 | 1 | 1 | 3 | 1 |
| 1_3635891 | NA | NA | NA | NA | 1 | 1 | NA | NA | 1 | 1 | 1 | 1 | NA | NA | NA | NA | 1 | 1 | 2 | 1 |
| 1_3635919 | NA | NA | NA | NA | 2 | 1 | NA | NA | 2 | 1 | 1 | 1 | NA | NA | 1 | 1 | 1 | 1 | 3 | 1 |
| 1_3635927 | NA | NA | 1 | 0 | 1 | 1 | NA | NA | 2 | 0.5 | 1 | 1 | 1 | 0 | NA | NA | 1 | 1 | 3 | 0.6667 |
| 1_3635947 | NA | NA | 1 | 0 | 3 | 1 | NA | NA | 4 | 0.75 | NA | NA | 1 | 0 | 1 | 1 | 2 | 0 | 4 | 0.25 |
| 1_3635957 | NA | NA | 1 | 0 | 2 | 0.5 | NA | NA | 3 | 0.3333 | 1 | 1 | 1 | 0 | NA | NA | 2 | 1 | 4 | 0.75 |
| 1_3635958 | NA | NA | 1 | 0 | 1 | 1 | NA | NA | 2 | 0.5 | NA | NA | 1 | 1 | NA | NA | 1 | 1 | 2 | 1 |
| 1_3637506 | 3 | 0.6667 | 4 | 0.25 | 5 | 0.2 | 10 | 0.1 | 22 | 0.2273 | 6 | 0 | 7 | 0.1429 | 5 | 0.4 | 5 | 0.2 | 23 | 0.1739 |
| 1_3637508 | 4 | 0.25 | 5 | 0.6 | 5 | 0.4 | 10 | 0.5 | 24 | 0.4583 | 6 | 0.3333 | 4 | 0.25 | 5 | 0.8 | 3 | 0.6667 | 18 | 0.5 |
| 1_3637515 | 4 | 0.5 | 5 | 0.2 | 5 | 0.6 | 10 | 0.3 | 24 | 0.375 | 8 | 0.125 | 7 | 0.4286 | 3 | 0 | 4 | 0.25 | 22 | 0.2273 |
| 1_3637517 | 2 | 1 | 3 | 1 | 4 | 0.75 | 9 | 0.8889 | 18 | 0.8889 | 4 | 1 | 6 | 0.6667 | 4 | 0.5 | 4 | 0.75 | 18 | 0.7222 |
| 1_3637521 | 3 | 0 | 1 | 1 | 3 | 0.3333 | 10 | 0.3 | 17 | 0.2941 | 8 | 0.125 | 4 | 0.25 | 3 | 0.3333 | 2 | 0 | 17 | 0.1765 |
| 1_3637522 | 1 | 0 | 3 | 0.6667 | 4 | 0.5 | 9 | 0.5556 | 17 | 0.5294 | 7 | 1 | 7 | 0.5714 | 2 | 0 | 6 | 0.6667 | 22 | 0.6818 |
| 1_3637525 | 4 | 0.25 | 2 | 0 | 5 | 0 | 10 | 0.2 | 21 | 0.1429 | 6 | 0.1667 | 6 | 0.1667 | 2 | 0.5 | 3 | 0 | 17 | 0.1765 |
| 1_3637544 | 4 | 0 | 6 | 0.1667 | 5 | 0 | 9 | 0.1111 | 24 | 0.0833 | 7 | 0.2857 | 6 | 0.1667 | 5 | 0 | 3 | 0 | 21 | 0.1429 |
| 1_3637569 | 3 | 0 | 5 | 0.4 | 3 | 0 | 10 | 0 | 21 | 0.0952 | 5 | 0 | 7 | 0.1429 | 4 | 0 | 3 | 0 | 19 | 0.0526 |
| 1_3645803 | 1 | 1 | 4 | 1 | NA | NA | 1 | 1 | 6 | 1 | 2 | 1 | 3 | 1 | 1 | 0 | 1 | 1 | 7 | 0.8571 |
| 1_3645871 | 1 | 0 | 4 | 0.25 | NA | NA | 1 | 0 | 6 | 0.1667 | 1 | 1 | 2 | 0 | 1 | 1 | 1 | 0 | 5 | 0.4 |
| 1_3645898 | 1 | 0 | 4 | 0.5 | NA | NA | NA | NA | 5 | 0.4 | 1 | 1 | 1 | 0 | 1 | 1 | 1 | 1 | 4 | 0.75 |
| 1_3645945 | 1 | 0 | 3 | 0 | NA | NA | 1 | 1 | 5 | 0.2 | 1 | 0 | NA | NA | 1 | 1 | NA | NA | 2 | 0.5 |
| 1_3645956 | 1 | 0 | 4 | 0.25 | NA | NA | 1 | 1 | 6 | 0.3333 | 1 | 0 | NA | NA | 1 | 1 | 1 | 1 | 3 | 0.6667 |
| 1_3645991 | NA | NA | 3 | 0.3333 | NA | NA | 1 | 1 | 4 | 0.5 | NA | NA | NA | NA | 1 | 1 | 1 | 1 | 2 | 1 |
| 1_3646227 | 5 | 0.4 | 1 | 0 | 4 | 0.25 | 4 | 0 | 14 | 0.2143 | 3 | 0.6667 | 3 | 0.3333 | 5 | 0 | 5 | 0 | 16 | 0.1875 |
| 1_3646252 | 3 | 0.3333 | 1 | 0 | 5 | 0.6 | 6 | 0 | 15 | 0.2667 | 3 | 0.3333 | 4 | 0 | 4 | 0.25 | 6 | 0.1667 | 17 | 0.1765 |
| 1_3646256 | 5 | 1 | 1 | 1 | 6 | 1 | 4 | 0.75 | 16 | 0.9375 | 4 | 0.75 | 2 | 1 | 5 | 1 | 5 | 0.8 | 16 | 0.875 |
| 1_3646257 | 2 | 1 | 1 | 1 | 7 | 0.8571 | 7 | 0.8571 | 17 | 0.8824 | 3 | 1 | 1 | 1 | 5 | 0.6 | 5 | 1 | 14 | 0.8571 |
| 1_3646266 | 5 | 0.8 | 2 | 1 | 5 | 0.6 | 7 | 0.2857 | 19 | 0.5789 | 4 | 0.5 | 3 | 0.3333 | 6 | 0.6667 | 4 | 0.25 | 17 | 0.4706 |
| 1_3646272 | 5 | 0.2 | 1 | 0 | 4 | 0.25 | 6 | 0 | 16 | 0.125 | 4 | 0.25 | 3 | 0 | 6 | 0.1667 | 4 | 0 | 17 | 0.1176 |
| 1_3646274 | 3 | 0 | 2 | 0 | 3 | 0 | 6 | 0.1667 | 14 | 0.0714 | 2 | 0.5 | 2 | 0 | 5 | 0 | 2 | 0.5 | 11 | 0.1818 |
| 1_3646284 | 5 | 0.6 | 1 | 1 | 4 | 0.5 | 7 | 0.5714 | 17 | 0.5882 | 4 | 0.5 | 3 | 0.6667 | 5 | 0.6 | 4 | 0.75 | 16 | 0.625 |
| 1_3646290 | 5 | 0.6 | 2 | 1 | 2 | 1 | 7 | 0.8571 | 16 | 0.8125 | 2 | 0.5 | 1 | 0 | 3 | 1 | 5 | 0.4 | 11 | 0.5455 |
| 1_3646300 | 6 | 0.1667 | 2 | 0 | 3 | 0.3333 | 7 | 0.1429 | 18 | 0.1667 | 3 | 0.3333 | 3 | 0 | 6 | 0.6667 | 3 | 0.3333 | 15 | 0.4 |
| 1_3646308 | 4 | 0 | 2 | 0.5 | 3 | 0 | 5 | 0.2 | 14 | 0.1429 | 4 | 0.5 | 1 | 0 | 6 | 0.1667 | 3 | 0.6667 | 14 | 0.3571 |
| 1_3646313 | 5 | 1 | 2 | 1 | 4 | 1 | 5 | 0.8 | 16 | 0.9375 | 4 | 0.75 | 1 | 1 | 6 | 1 | 4 | 1 | 15 | 0.9333 |
| 1_3646314 | 4 | 1 | 1 | 1 | 2 | 1 | 5 | 0.6 | 12 | 0.8333 | 4 | 1 | 2 | 1 | 5 | 0.8 | 3 | 0.6667 | 14 | 0.8571 |
| 1_3646320 | 6 | 0.8333 | 1 | 1 | 4 | 1 | 6 | 0.5 | 17 | 0.7647 | 3 | 0.3333 | 1 | 1 | 6 | 0.6667 | 4 | 0.75 | 14 | 0.6429 |
| 1_3646329 | 5 | 0.2 | 1 | 0 | 2 | 0 | 4 | 0 | 12 | 0.0833 | 3 | 0.3333 | NA | NA | 6 | 0 | 3 | 0 | 12 | 0.0833 |
| 1_3646335 | 5 | 0.4 | 1 | 1 | 4 | 0.75 | 6 | 0.5 | 16 | 0.5625 | 3 | 0.3333 | 1 | 1 | 4 | 0.5 | 4 | 0.75 | 12 | 0.5833 |
| 1_3651173 | 1 | 1 | NA | NA | NA | NA | NA | NA | 1 | 1 | NA | NA | NA | NA | 1 | 1 | NA | NA | 1 | 1 |
| 1_3651182 | 1 | 1 | NA | NA | NA | NA | NA | NA | 1 | 1 | NA | NA | NA | NA | 1 | 1 | NA | NA | 1 | 1 |
| 1_3651187 | 1 | 1 | NA | NA | NA | NA | NA | NA | 1 | 1 | NA | NA | NA | NA | 1 | 1 | NA | NA | 1 | 1 |
| 1_3651194 | 1 | 1 | NA | NA | NA | NA | NA | NA | 1 | 1 | NA | NA | NA | NA | 1 | 1 | NA | NA | 1 | 1 |
| 1_3651204 | 1 | 1 | NA | NA | NA | NA | 1 | 1 | 2 | 1 | NA | NA | NA | NA | 1 | 1 | NA | NA | 1 | 1 |
| 1_3654558 | 2 | 0.5 | 1 | 1 | NA | NA | NA | NA | 3 | 0.6667 | NA | NA | NA | NA | 1 | 1 | NA | NA | 1 | 1 |
| 1_3654602 | 2 | 0.5 | 1 | 0 | 1 | 0 | 1 | 0 | 5 | 0.2 | NA | NA | NA | NA | 2 | 0.5 | NA | NA | 2 | 0.5 |
| 1_3654606 | 2 | 0.5 | 1 | 1 | 1 | 0 | NA | NA | 4 | 0.5 | NA | NA | 1 | 1 | 1 | 0 | NA | NA | 2 | 0.5 |
| 1_3654624 | 1 | 0 | 1 | 1 | 1 | 0 | 1 | 0 | 4 | 0.25 | NA | NA | NA | NA | 1 | 1 | NA | NA | 1 | 1 |
| 1_3654669 | 2 | 0 | 1 | 1 | NA | NA | 2 | 0 | 5 | 0.2 | 1 | 1 | 1 | 0 | NA | NA | NA | NA | 2 | 0.5 |
| 1_3654780 | 2 | 0.5 | 1 | 1 | 3 | 0.3333 | 3 | 0 | 9 | 0.3333 | 2 | 1 | 1 | 0 | 1 | 0 | NA | NA | 4 | 0.5 |
| 1_3655109 | NA | NA | 1 | 0 | NA | NA | 1 | 1 | 2 | 0.5 | 1 | 1 | 2 | 0.5 | NA | NA | NA | NA | 3 | 0.6667 |
| 1_3655122 | 1 | 1 | 1 | 1 | NA | NA | 2 | 1 | 4 | 1 | 1 | 1 | 3 | 0.3333 | NA | NA | NA | NA | 4 | 0.5 |
| 1_3655197 | 1 | 1 | 1 | 1 | NA | NA | 3 | 0.3333 | 5 | 0.6 | 1 | 0 | 2 | 0.5 | NA | NA | NA | NA | 3 | 0.3333 |

| | | | | | | | | | | | | | | | | | | | | |
|---|---|---|---|---|---|---|---|---|---|---|---|---|---|---|---|---|---|---|---|---|
| 1_3655199 | 1 | 1 | 1 | 1 | NA | NA | 3 | 1 | 5 | 1 | 2 | 1 | 3 | 0.6667 | NA | NA | NA | NA | 5 | 0.8 |
| 1_3655209 | 1 | 1 | 1 | 1 | NA | NA | 2 | 0 | 4 | 0.5 | 2 | 0.5 | 3 | 0.3333 | NA | NA | NA | NA | 5 | 0.4 |
| 1_3655210 | 1 | 1 | NA | NA | NA | NA | 3 | 0.6667 | 4 | 0.75 | 2 | 0.5 | 3 | 1 | NA | NA | NA | NA | 5 | 0.8 |
| 1_3655213 | 1 | 0 | 1 | 1 | NA | NA | 2 | 0.5 | 4 | 0.5 | 2 | 0.5 | 2 | 1 | NA | NA | NA | NA | 4 | 0.75 |
| 1_3656434 | 1 | 0 | 2 | 1 | 2 | 0.5 | 2 | 0 | 7 | 0.4286 | 1 | 1 | 4 | 0.5 | 4 | 0.75 | NA | NA | 9 | 0.6667 |
| 1_3656454 | NA | NA | 3 | 0.3333 | 3 | 0 | 4 | 0.5 | 10 | 0.3 | 1 | 1 | 4 | 0.25 | 4 | 0.25 | NA | NA | 9 | 0.3333 |
| 1_3656521 | 1 | 1 | 3 | 0.3333 | 3 | 0 | 5 | 0 | 12 | 0.1667 | 2 | 0 | 6 | 0 | 8 | 0.125 | 1 | 0 | 17 | 0.0588 |
| 1_3656525 | 1 | 0 | 3 | 0 | 3 | 0 | 5 | 0.4 | 12 | 0.1667 | 2 | 0 | 7 | 0.1429 | 9 | 0 | 1 | 0 | 19 | 0.0526 |
| 1_3656598 | 1 | 0 | 2 | 0.5 | 4 | 0.5 | 5 | 0.4 | 12 | 0.4167 | 2 | 0.5 | 6 | 0.1667 | 6 | 0.6667 | 2 | 0 | 16 | 0.375 |
| 1_3656603 | 2 | 0 | 4 | 0.75 | 5 | 0.4 | 5 | 0.6 | 16 | 0.5 | 1 | 1 | 8 | 0.25 | 9 | 0.4444 | 2 | 0.5 | 20 | 0.4 |
| 1_3656604 | NA | NA | 4 | 0 | 4 | 0 | 6 | 0.5 | 14 | 0.2143 | 1 | 0 | 7 | 0.1429 | 10 | 0.6 | 2 | 0 | 20 | 0.35 |
| 1_3656611 | 1 | 0 | 3 | 0.3333 | 5 | 0.2 | 4 | 0.5 | 13 | 0.3077 | 2 | 1 | 6 | 1 | 9 | 0.6667 | 1 | 0 | 18 | 0.7778 |
| 1_3656628 | 1 | 0 | 2 | 1 | 3 | 0 | 5 | 0.4 | 11 | 0.3636 | 2 | 0 | 6 | 0.1667 | 9 | 0.2222 | 2 | 0 | 19 | 0.1579 |
| 1_3657829 | 2 | 0.5 | 1 | 0 | 2 | 0 | 1 | 0 | 6 | 0.1667 | 1 | 0 | 1 | 1 | NA | NA | NA | NA | 2 | 0.5 |
| 1_3657842 | 3 | 0.3333 | 1 | 0 | 2 | 0 | 1 | 0 | 7 | 0.1429 | 1 | 0 | 1 | 0 | NA | NA | 1 | 1 | 3 | 0.3333 |
| 1_3658847 | 1 | 1 | 1 | 1 | NA | NA | NA | NA | 2 | 1 | NA | NA | NA | NA | 1 | 1 | NA | NA | 1 | 1 |
| 1_3658850 | 1 | 1 | 1 | 1 | NA | NA | NA | NA | 2 | 1 | NA | NA | NA | NA | 1 | 1 | NA | NA | 1 | 1 |
| 1_3658877 | 1 | 1 | 1 | 1 | NA | NA | NA | NA | 2 | 1 | NA | NA | NA | NA | 1 | 1 | NA | NA | 1 | 1 |
| 1_3658893 | 1 | 1 | 1 | 1 | NA | NA | NA | NA | 2 | 1 | NA | NA | NA | NA | 1 | 1 | NA | NA | 1 | 1 |
| 1_3658901 | 1 | 1 | 1 | 0 | NA | NA | NA | NA | 2 | 0.5 | NA | NA | NA | NA | 1 | 1 | NA | NA | 1 | 1 |
| 1_3659869 | NA | NA | 1 | 1 | NA | NA | NA | NA | 1 | 1 | NA | NA | NA | NA | 1 | 1 | NA | NA | 1 | 1 |
| 1_3673763 | 10 | 0 | 20 | 0.15 | 10 | 0 | 16 | 0.375 | 56 | 0.1607 | 14 | 0.2857 | 10 | 0 | 12 | 0.1667 | 10 | 0.1 | 46 | 0.1522 |
| 1_3673766 | 12 | 0.25 | 20 | 0.05 | 10 | 0.1 | 12 | 0.0833 | 54 | 0.1111 | 12 | 0.0833 | 8 | 0 | 8 | 0.125 | 9 | 0.1111 | 37 | 0.0811 |
| 1_3673828 | 14 | 0.1429 | 19 | 0.3158 | 15 | 0.2 | 17 | 0.2353 | 65 | 0.2308 | 22 | 0.1818 | 15 | 0.0667 | 17 | 0.0588 | 13 | 0.0769 | 67 | 0.1045 |
| 1_3673863 | 21 | 0.1429 | 27 | 0.0741 | 17 | 0 | 20 | 0.1 | 85 | 0.0824 | 23 | 0 | 16 | 0.125 | 21 | 0.0476 | 11 | 0.0909 | 71 | 0.0563 |
| 1_3673864 | 26 | 0.0769 | 29 | 0.1379 | 27 | 0.1481 | 25 | 0.24 | 107 | 0.1495 | 31 | 0.129 | 23 | 0.0435 | 24 | 0.0417 | 16 | 0.0625 | 94 | 0.0745 |
| 1_3673881 | 27 | 0.1481 | 32 | 0.0938 | 27 | 0.0741 | 28 | 0 | 114 | 0.0789 | 31 | 0.129 | 24 | 0.0417 | 24 | 0.0417 | 17 | 0.1176 | 96 | 0.0833 |
| 1_3673891 | 26 | 0.1923 | 25 | 0.24 | 22 | 0.1818 | 29 | 0.2759 | 102 | 0.2255 | 25 | 0.16 | 21 | 0.2381 | 23 | 0.2174 | 13 | 0.2308 | 82 | 0.2073 |
| 1_3673914 | 17 | 0.5294 | 20 | 0.15 | 17 | 0.3529 | 18 | 0.3333 | 72 | 0.3333 | 24 | 0.4167 | 16 | 0.375 | 13 | 0.3077 | 7 | 0 | 60 | 0.3333 |
| 1_3676325 | NA | NA | 1 | 1 | NA | NA | 1 | 0 | 2 | 0.5 | 1 | 1 | NA | NA | NA | NA | NA | NA | 1 | 1 |
| 1_3676346 | NA | NA | 1 | 1 | NA | NA | NA | NA | 1 | 1 | 1 | 1 | NA | NA | NA | NA | NA | NA | 1 | 1 |
| 1_3676350 | NA | NA | NA | NA | NA | NA | 1 | 1 | 1 | 1 | 1 | 1 | NA | NA | NA | NA | NA | NA | 1 | 1 |
| 1_3676366 | NA | NA | 1 | 0 | NA | NA | 1 | 1 | 2 | 0.5 | 1 | 1 | NA | NA | NA | NA | NA | NA | 1 | 1 |
| 1_3683156 | 4 | 0.5 | 12 | 0.5833 | 5 | 0.2 | 8 | 0.75 | 29 | 0.5517 | 3 | 0.6667 | 8 | 0.375 | 4 | 0.5 | 8 | 0.625 | 23 | 0.5217 |
| 1_3683180 | 4 | 0.75 | 11 | 0.9091 | 5 | 0.6 | 7 | 0.8571 | 27 | 0.8148 | 2 | 0.5 | 6 | 1 | 4 | 1 | 6 | 0.8333 | 18 | 0.8889 |
| 1_3683181 | 5 | 0.2 | 10 | 0.1 | 5 | 0.2 | 8 | 0.25 | 28 | 0.1786 | 2 | 0.5 | 7 | 0.1429 | 3 | 0 | 6 | 0.1667 | 18 | 0.1667 |
| 1_3683209 | 4 | 0 | 11 | 0.1818 | 5 | 0.4 | 6 | 0.3333 | 26 | 0.2308 | 2 | 0 | 8 | 0.25 | 3 | 0 | 6 | 0 | 19 | 0.1053 |
| 1_3684704 | 2 | 0.5 | 5 | 0 | 1 | 0 | 3 | 0 | 11 | 0.0909 | 2 | 0.5 | 3 | 0 | 3 | 0 | 3 | 0 | 11 | 0.0909 |
| 1_3684707 | 2 | 0 | 6 | 0 | 1 | 0 | 5 | 0.2 | 14 | 0.0714 | 2 | 0 | 4 | 0.25 | 2 | 0.5 | 4 | 0.25 | 12 | 0.25 |
| 1_3684731 | 2 | 0 | 7 | 0.5714 | 1 | 0 | 5 | 0.2 | 15 | 0.3333 | 1 | 0 | 4 | 0.25 | 1 | 1 | 3 | 0 | 9 | 0.2222 |
| 1_3684737 | 1 | 0 | 6 | 0.1667 | 1 | 0 | 5 | 0.6 | 13 | 0.3077 | 1 | 0 | 4 | 0.5 | 2 | 1 | 3 | 0.6667 | 10 | 0.6 |
| 1_3684739 | 2 | 0 | 6 | 0.1667 | 1 | 0 | 5 | 0 | 14 | 0.0714 | 1 | 0 | 2 | 0.5 | 1 | 0 | 3 | 0 | 7 | 0.1429 |
| 1_3684742 | 2 | 0.5 | 6 | 0.1667 | 1 | 0 | 3 | 0 | 12 | 0.1667 | 1 | 1 | 4 | 0 | 1 | 0 | 3 | 0.3333 | 9 | 0.2222 |
| 1_3744256 | 2 | 0 | 3 | 0.3333 | 2 | 0 | 1 | 0 | 8 | 0.125 | 1 | 0 | NA | NA | 3 | 0 | 4 | 0.25 | 8 | 0.125 |
| 1_3744258 | NA | NA | 4 | 0.5 | 2 | 0.5 | 1 | 0 | 7 | 0.4286 | 1 | 0 | NA | NA | 2 | 0.5 | 3 | 0.3333 | 6 | 0.3333 |
| 1_3744278 | 2 | 1 | 3 | 0.6667 | 2 | 1 | 1 | 0 | 8 | 0.75 | NA | NA | NA | NA | 2 | 0.5 | 3 | 0.6667 | 5 | 0.6 |
| 1_3744289 | 1 | 0 | 3 | 0.6667 | 2 | 0.5 | 1 | 1 | 7 | 0.5714 | 1 | 1 | NA | NA | 3 | 1 | 3 | 0.6667 | 7 | 0.8571 |
| 1_3744295 | 2 | 0.5 | 4 | 0.25 | 2 | 0 | 2 | 0 | 10 | 0.2 | 1 | 0 | NA | NA | 3 | 0.3333 | 3 | 0.6667 | 7 | 0.4286 |
| 1_3744318 | 2 | 0 | 4 | 0.25 | 2 | 0 | 2 | 0.5 | 10 | 0.2 | 1 | 0 | 1 | 0 | 3 | 0.3333 | 1 | 0 | 6 | 0.1667 |
| 1_3744731 | 4 | 0.75 | 12 | 0.75 | 9 | 0.7778 | 5 | 0.8 | 30 | 0.7667 | 4 | 0.75 | 6 | 0.8333 | 5 | 0.4 | 7 | 0.2857 | 22 | 0.5455 |
| 1_3744734 | 3 | 0 | 9 | 0 | 6 | 0.3333 | 7 | 0.1429 | 25 | 0.12 | 2 | 0 | 4 | 0.25 | 5 | 0.2 | 7 | 0.1429 | 18 | 0.1667 |
| 1_3744772 | 5 | 0.2 | 14 | 0.5 | 11 | 0.2727 | 7 | 0.4286 | 37 | 0.3784 | 4 | 0.25 | 7 | 0.4286 | 5 | 0.6 | 9 | 0.2222 | 25 | 0.36 |
| 1_3744854 | 9 | 0.5556 | 17 | 0.2941 | 11 | 0.0909 | 6 | 0.3333 | 43 | 0.3023 | 6 | 0.3333 | 6 | 0.3333 | 6 | 0.1667 | 9 | 0.2222 | 27 | 0.2593 |
| 1_3744873 | 12 | 0.0833 | 22 | 0.1364 | 13 | 0 | 9 | 0.1111 | 56 | 0.0893 | 10 | 0 | 9 | 0.2222 | 4 | 0 | 11 | 0.0909 | 34 | 0.0882 |
| 1_3744925 | 16 | 0.25 | 21 | 0.1429 | 13 | 0 | 11 | 0.3636 | 61 | 0.1803 | 15 | 0.2 | 12 | 0 | 6 | 0 | 16 | 0.3125 | 49 | 0.1633 |
| 1_3744929 | 14 | 0.2857 | 21 | 0.1429 | 16 | 0 | 12 | 0 | 63 | 0.1111 | 13 | 0.1538 | 11 | 0.0909 | 8 | 0 | 14 | 0.2143 | 46 | 0.1304 |
| 1_3744930 | 17 | 0.4118 | 23 | 0.5217 | 16 | 0.1875 | 12 | 0.5 | 68 | 0.4118 | 16 | 0.3125 | 11 | 0.3636 | 8 | 0.25 | 16 | 0.25 | 51 | 0.2941 |
| 1_3744935 | 15 | 0.4 | 24 | 0.5417 | 13 | 0.3846 | 12 | 0.4167 | 64 | 0.4531 | 15 | 0.5333 | 12 | 0.5 | 7 | 0.2857 | 17 | 0.4118 | 51 | 0.451 |
| 1_3744961 | 15 | 1 | 21 | 0.8571 | 16 | 0.875 | 10 | 0.9 | 62 | 0.9032 | 15 | 0.9333 | 12 | 0.9167 | 9 | 0.8889 | 20 | 0.85 | 56 | 0.8929 |
| 1_3744969 | 15 | 0.4 | 19 | 0.1579 | 15 | 0.1333 | 10 | 0.4 | 59 | 0.2542 | 16 | 0.1875 | 9 | 0.4444 | 10 | 0.3 | 19 | 0.3684 | 54 | 0.3148 |
| 1_3745843 | 10 | 0.5 | 3 | 0.6667 | 5 | 0.2 | 8 | 0.5 | 26 | 0.4615 | 8 | 0.5 | 9 | 0.2222 | 9 | 0.1111 | 6 | 0.6667 | 32 | 0.3438 |
| 1_3745884 | 9 | 0.3333 | 3 | 0.3333 | 7 | 0.2857 | 10 | 0.5 | 29 | 0.3793 | 7 | 0.4286 | 9 | 0.4444 | 10 | 0.2 | 8 | 0.25 | 34 | 0.3235 |
| 1_3745904 | 9 | 0.6667 | 3 | 0.3333 | 7 | 0.7143 | 7 | 0.7143 | 26 | 0.6538 | 8 | 0.5 | 9 | 0.6667 | 10 | 0.5 | 7 | 0.8571 | 34 | 0.6176 |
| 1_3745939 | 7 | 0 | 3 | 0.6667 | 7 | 0.2857 | 8 | 0 | 25 | 0.16 | 8 | 0 | 8 | 0.25 | 7 | 0.5714 | 7 | 0 | 30 | 0.2 |
| 1_3745946 | 5 | 1 | 4 | 1 | 9 | 1 | 10 | 1 | 28 | 1 | 8 | 1 | 8 | 0.875 | 8 | 1 | 7 | 1 | 31 | 0.9677 |

| | | | | | | | | | | | | | | | | | | | | |
|---|---|---|---|---|---|---|---|---|---|---|---|---|---|---|---|---|---|---|---|---|
| 1_3745954( | 7 | 0.1429 | 4 | 0 | 9 | 0 | 9 | 0.1111 | 29 | 0.069 | 7 | 0.1429 | 8 | 0.125 | 8 | 0.125 | 8 | 0.125 | 31 | 0.129 |
| 1_3747051 | 8 | 1 | 7 | 0.5714 | 5 | 1 | 6 | 0.8333 | 26 | 0.8462 | 4 | 0.75 | 12 | 0.6667 | 4 | 1 | 5 | 0.8 | 25 | 0.76 |
| 1_3747057 | 9 | 0.1111 | 8 | 0 | 5 | 0 | 7 | 0.1429 | 29 | 0.069 | 4 | 0 | 10 | 0.1 | 5 | 0.2 | 6 | 0.1667 | 25 | 0.12 |
| 1_3747785 | 11 | 0 | 9 | 0.2222 | 7 | 0.5714 | 10 | 0.6 | 37 | 0.3243 | 6 | 0.6667 | 12 | 0.25 | 8 | 0.375 | 10 | 0.3 | 36 | 0.3611 |
| 1_3747791 | 12 | 0.1667 | 9 | 0.3333 | 7 | 0.4286 | 10 | 0.3 | 38 | 0.2895 | 7 | 0.5714 | 11 | 0.3636 | 6 | 0.5 | 10 | 0.2 | 34 | 0.3824 |
| 1_3747915 | 3 | 0.3333 | 2 | 0 | 2 | 0 | 1 | 0 | 8 | 0.125 | 1 | 0 | 7 | 0 | 3 | 0.3333 | 3 | 0 | 14 | 0.0714 |
| 1_3747955 | 1 | 0 | 2 | 0 | 2 | 0.5 | 1 | 0 | 6 | 0.1667 | 1 | 1 | 3 | 0 | 3 | 0.3333 | 2 | 0 | 9 | 0.2222 |
| 1_3747977 | 2 | 0.5 | 1 | 0 | 1 | 1 | 1 | 0 | 5 | 0.4 | 1 | 1 | 3 | 0.6667 | 2 | 1 | 2 | 0.5 | 8 | 0.75 |
| 1_3748360 | 1 | 1 | 1 | 1 | NA | NA | NA | NA | 2 | 1 | NA | NA | NA | NA | NA | NA | 1 | 1 | 1 | 1 |
| 1_3765084 | 47 | 0.8298 | 33 | 0.8788 | 37 | 0.7027 | 30 | 0.8 | 147 | 0.8027 | 23 | 0.7826 | 38 | 0.7632 | 45 | 0.8222 | 39 | 0.8974 | 145 | 0.8207 |
| 1_3765119 | 48 | 0.1042 | 34 | 0.0588 | 38 | 0.0263 | 33 | 0 | 153 | 0.0523 | 22 | 0.0455 | 43 | 0.0233 | 45 | 0.0889 | 42 | 0.0476 | 152 | 0.0526 |
| 1_3765124 | 43 | 0.186 | 32 | 0.2812 | 37 | 0.2703 | 29 | 0.1724 | 141 | 0.227 | 22 | 0.0455 | 39 | 0.2051 | 45 | 0.4 | 34 | 0.2059 | 140 | 0.2429 |
| 1_3765125 | 40 | 0.95 | 21 | 0.9524 | 35 | 0.9143 | 28 | 0.8929 | 124 | 0.9274 | 18 | 1 | 33 | 0.8788 | 42 | 0.9524 | 36 | 0.9167 | 129 | 0.9302 |
| 1_3765128 | 35 | 0.0857 | 25 | 0.16 | 29 | 0.1724 | 24 | 0.0833 | 113 | 0.1239 | 18 | 0 | 32 | 0.0938 | 35 | 0.0571 | 31 | 0.0323 | 116 | 0.0517 |
| 1_3765130 | 36 | 0 | 27 | 0.1111 | 27 | 0.1111 | 27 | 0.1481 | 117 | 0.0855 | 18 | 0.0556 | 37 | 0.0541 | 34 | 0.1176 | 36 | 0.0556 | 125 | 0.072 |
| 1_3766290 | 35 | 0.6857 | 27 | 0.7037 | 33 | 0.8485 | 17 | 0.6471 | 112 | 0.7321 | 15 | 0.5333 | 30 | 0.6667 | 35 | 0.7143 | 36 | 0.7778 | 116 | 0.6983 |
| 1_3766302 | 26 | 0.1154 | 22 | 0.1364 | 28 | 0 | 15 | 0.0667 | 91 | 0.0769 | 13 | 0.1538 | 24 | 0.0417 | 28 | 0.0357 | 25 | 0.16 | 90 | 0.0889 |
| 1_3766309 | 19 | 0 | 19 | 0.1053 | 24 | 0.1667 | 13 | 0.2308 | 75 | 0.12 | 4 | 0.5 | 21 | 0.0476 | 26 | 0.1923 | 24 | 0.125 | 75 | 0.1467 |
| 1_3766318 | 27 | 0.5926 | 23 | 0.8261 | 29 | 0.8276 | 17 | 0.5294 | 96 | 0.7083 | 11 | 0.7273 | 25 | 0.72 | 28 | 0.6786 | 29 | 0.6207 | 93 | 0.6774 |
| 1_3766324 | 28 | 0.0714 | 22 | 0.1818 | 32 | 0.0625 | 14 | 0.1429 | 96 | 0.1042 | 12 | 0.0833 | 26 | 0.1923 | 27 | 0.0741 | 29 | 0.1034 | 94 | 0.117 |
| 1_3766330 | 27 | 0.1111 | 23 | 0.1304 | 26 | 0.0769 | 16 | 0.125 | 92 | 0.1087 | 12 | 0.0833 | 24 | 0.0833 | 28 | 0.1786 | 28 | 0.1429 | 92 | 0.1304 |
| 1_3766333 | 23 | 0.4348 | 17 | 0.2941 | 19 | 0.4211 | 8 | 0.75 | 67 | 0.4328 | 5 | 0.8 | 16 | 0.375 | 23 | 0.3913 | 23 | 0.4783 | 67 | 0.4478 |
| 1_3766334( | NA | NA | 2 | 0.5 | 1 | 1 | NA | NA | 3 | 0.6667 | 1 | 0 | 2 | 0.5 | 1 | 0 | 2 | 0 | 6 | 0.1667 |
| 1_3766336 | 26 | 0.2308 | 21 | 0.2857 | 29 | 0.3448 | 15 | 0.2667 | 91 | 0.2857 | 8 | 0.5 | 26 | 0.2692 | 30 | 0.2 | 29 | 0.2759 | 93 | 0.2688 |
| 1_3766336 | 27 | 0.1852 | 22 | 0.0909 | 26 | 0.0769 | 15 | 0.3333 | 90 | 0.1556 | 11 | 0.0909 | 27 | 0.1111 | 31 | 0.0645 | 30 | 0.3333 | 99 | 0.1616 |
| 1_3766386 | 19 | 0.1053 | 17 | 0 | 23 | 0.0435 | 11 | 0.0909 | 70 | 0.0571 | 8 | 0 | 20 | 0 | 23 | 0.0435 | 18 | 0.1667 | 69 | 0.058 |
| 1_3766390 | 23 | 0.2609 | 22 | 0.0909 | 24 | 0.2917 | 11 | 0.1818 | 80 | 0.2125 | 8 | 0 | 20 | 0.3 | 21 | 0.1905 | 19 | 0.2105 | 68 | 0.2059 |
| 1_3766394 | 24 | 0.4167 | 21 | 0.5714 | 25 | 0.36 | 10 | 0.4 | 80 | 0.4375 | 8 | 0.25 | 21 | 0.3333 | 20 | 0.35 | 21 | 0.4286 | 70 | 0.3571 |
| 1_3766406 | 20 | 0.65 | 17 | 0.6471 | 20 | 0.8 | 10 | 0.7 | 67 | 0.7015 | 9 | 0.7778 | 18 | 0.6667 | 20 | 0.75 | 18 | 0.7778 | 65 | 0.7385 |
| 1_3766413 | 19 | 0.1579 | 19 | 0.2632 | 23 | 0.087 | 10 | 0 | 71 | 0.1408 | 8 | 0 | 17 | 0.1765 | 21 | 0.0952 | 15 | 0.0667 | 61 | 0.0984 |
| 1_3766419 | 21 | 0.0476 | 11 | 0.1818 | 23 | 0.1739 | 11 | 0 | 66 | 0.1061 | 7 | 0.1429 | 17 | 0.1765 | 16 | 0 | 16 | 0.0625 | 56 | 0.0893 |
| 1_3766425 | 20 | 0.1 | 15 | 0.1333 | 22 | 0.1364 | 8 | 0.25 | 65 | 0.1385 | 6 | 0 | 17 | 0 | 21 | 0.1429 | 12 | 0.1667 | 56 | 0.0893 |
| 1_3766432 | 14 | 0.1429 | 12 | 0.0833 | 18 | 0.1111 | 8 | 0 | 52 | 0.0962 | 7 | 0.1429 | 18 | 0.1111 | 14 | 0 | 12 | 0.1667 | 51 | 0.098 |
| 1_3766438 | 19 | 0.1053 | 11 | 0.0909 | 17 | 0.1765 | 7 | 0.1429 | 54 | 0.1296 | 8 | 0.125 | 17 | 0.0588 | 18 | 0.1667 | 8 | 0.125 | 51 | 0.1176 |
| 1_3766444 | 17 | 0.3529 | 10 | 0.3 | 13 | 0.3846 | 6 | 0.3333 | 46 | 0.3478 | 8 | 0.375 | 15 | 0.3333 | 16 | 0.0625 | 9 | 0.4444 | 48 | 0.2708 |
| 1_3766444 | 3 | 0.3333 | 1 | 0 | 4 | 0 | 1 | 0 | 9 | 0.1111 | 1 | 1 | NA | NA | NA | NA | 1 | 0 | 2 | 0.5 |
| 1_3766449 | 21 | 0.2381 | 11 | 0.2727 | 18 | 0.2222 | 6 | 0.3333 | 56 | 0.25 | 9 | 0 | 19 | 0.1579 | 21 | 0.2381 | 12 | 0 | 61 | 0.1311 |
| 1_3766450 | 21 | 0.0476 | 12 | 0.0833 | 18 | 0.0556 | 5 | 0.2 | 56 | 0.0714 | 9 | 0.1111 | 19 | 0.2105 | 19 | 0.1579 | 12 | 0.25 | 59 | 0.1864 |
| 1_3766459 | 19 | 0.5263 | 8 | 0.625 | 16 | 0.75 | 5 | 0.4 | 48 | 0.6042 | 7 | 0.5714 | 16 | 0.4375 | 12 | 0.5 | 10 | 0.6 | 45 | 0.5111 |
| 1_3766464 | 14 | 0.9286 | 7 | 1 | 14 | 0.7857 | 6 | 0.6667 | 41 | 0.8537 | 7 | 1 | 18 | 0.8333 | 12 | 0.8333 | 9 | 0.7778 | 46 | 0.8478 |
| 1_3766489 | 21 | 0.0952 | 9 | 0.1111 | 17 | 0.0588 | 6 | 0 | 53 | 0.0755 | 7 | 0.2857 | 16 | 0.0625 | 12 | 0.1667 | 11 | 0 | 46 | 0.1087 |
| 1_3766500 | 21 | 0.5714 | 10 | 0.7 | 17 | 0.5294 | 6 | 0.5 | 54 | 0.5741 | 7 | 0.8571 | 16 | 0.3125 | 12 | 0.25 | 12 | 0.5833 | 47 | 0.4468 |
| 1_3766778 | 1 | 0 | 3 | 0.3333 | NA | NA | NA | NA | 4 | 0.25 | 1 | 1 | NA | NA | 1 | 0 | NA | NA | 2 | 0.5 |
| 1_3771557 | 2 | 1 | 3 | 0.6667 | 3 | 1 | NA | NA | 8 | 0.875 | 1 | 1 | 2 | 0.5 | 5 | 0.8 | 3 | 1 | 11 | 0.8182 |
| 1_3771564 | 2 | 0.5 | 3 | 0.3333 | 2 | 0.5 | NA | NA | 7 | 0.4286 | 1 | 1 | 1 | 1 | 4 | 0.25 | 3 | 0.3333 | 9 | 0.4444 |
| 1_3771574 | 2 | 0 | 3 | 0.3333 | 3 | 0.3333 | NA | NA | 8 | 0.25 | 1 | 0 | 2 | 0.5 | 4 | 0.25 | 2 | 0.5 | 9 | 0.3333 |
| 1_3787331 | 1 | 0 | 1 | 0 | NA | NA | 1 | 1 | 3 | 0.3333 | NA | NA | NA | NA | 1 | 1 | 3 | 0.6667 | 4 | 0.75 |
| 1_3787338 | 1 | 0 | 1 | 1 | NA | NA | 1 | 0 | 3 | 0.3333 | NA | NA | NA | NA | 2 | 0 | 3 | 0.3333 | 5 | 0.2 |
| 1_3791474 | 1 | 1 | NA | NA | NA | NA | NA | NA | 1 | 1 | NA | NA | 1 | 1 | NA | NA | NA | NA | 1 | 1 |
| 1_3791477 | 1 | 1 | NA | NA | NA | NA | NA | NA | 1 | 1 | NA | NA | 1 | 1 | NA | NA | NA | NA | 1 | 1 |
| 1_3791499 | 1 | 1 | NA | NA | NA | NA | NA | NA | 1 | 1 | NA | NA | 1 | 1 | NA | NA | 1 | 1 | 2 | 1 |
| 1_3791511 | 1 | 1 | NA | NA | NA | NA | NA | NA | 1 | 1 | NA | NA | 1 | 1 | 1 | 1 | 1 | 1 | 3 | 1 |
| 1_3791522 | 1 | 1 | NA | NA | NA | NA | NA | NA | 1 | 1 | NA | NA | 1 | 1 | 1 | 0 | 1 | 0 | 3 | 0.3333 |
| 1_3791828 | NA | NA | 1 | 1 | NA | NA | 4 | 0.5 | 5 | 0.6 | 2 | 1 | NA | NA | NA | NA | 1 | 0 | 3 | 0.6667 |
| 1_3791838 | NA | NA | 2 | 0 | NA | NA | 3 | 0.3333 | 5 | 0.2 | 2 | 0.5 | NA | NA | NA | NA | 1 | 0 | 3 | 0.3333 |
| 1_3793724 | 20 | 0.3 | 15 | 0.1333 | 14 | 0.3571 | 21 | 0.5238 | 70 | 0.3429 | 17 | 0.2353 | 20 | 0.3 | 15 | 0.2 | 19 | 0.2632 | 71 | 0.2535 |
| 1_3802490 | 2 | 0.5 | NA | NA | NA | NA | NA | NA | 2 | 0.5 | 1 | 0 | NA | NA | 1 | 1 | 4 | 0 | 6 | 0.1667 |
| 1_3802497 | 2 | 0.5 | NA | NA | NA | NA | NA | NA | 2 | 0.5 | 1 | 1 | NA | NA | 1 | 1 | 4 | 0.25 | 6 | 0.5 |
| 1_3802632 | 2 | 0.5 | 2 | 0 | NA | NA | NA | NA | 4 | 0.25 | NA | NA | 1 | 0 | 4 | 0.25 | 4 | 0.25 | 9 | 0.2222 |
| 1_3802674 | 23 | 0.2174 | 9 | 0.2222 | 8 | 0.125 | 9 | 0 | 49 | 0.1633 | 8 | 0.125 | 10 | 0.5 | 11 | 0.2727 | 12 | 0.25 | 41 | 0.2927 |
| 1_3802681 | 20 | 0.15 | 9 | 0 | 8 | 0 | 10 | 0.2 | 47 | 0.1064 | 6 | 0.1667 | 6 | 0.1667 | 8 | 0.625 | 10 | 0.1 | 30 | 0.2667 |
| 1_3803385 | 3 | 0 | NA | NA | 1 | 0 | 1 | 1 | 5 | 0.2 | NA | NA | NA | NA | 1 | 1 | 2 | 0.5 | 3 | 0.6667 |
| 1_3803464 | 3 | 0.6667 | NA | NA | NA | NA | 1 | 0 | 4 | 0.5 | NA | NA | NA | NA | 1 | 1 | 2 | 0.5 | 3 | 0.6667 |
| 1_3803481 | 17 | 0.8235 | 7 | 0.5714 | 3 | 0.6667 | 9 | 0.7778 | 36 | 0.75 | 4 | 0.5 | 6 | 0.8333 | 8 | 0.5 | 10 | 0.8 | 28 | 0.6786 |
| 1_3805807 | NA | NA | 2 | 0.5 | NA | NA | NA | NA | 2 | 0.5 | NA | NA | NA | NA | 1 | 1 | NA | NA | 1 | 1 |

| | | | | | | | | | | | | | | | | | | | | |
|---|---|---|---|---|---|---|---|---|---|---|---|---|---|---|---|---|---|---|---|---|
| 1_38090718 | 9 | 0.2222 | 8 | 0 | 9 | 0.1111 | 3 | 0 | 29 | 0.1034 | 3 | 0 | 2 | 0 | 5 | 0.2 | 8 | 0.25 | 18 | 0.1667 |
| 1_38090748 | 9 | 0 | 8 | 0.125 | 6 | 0.1667 | 3 | 0 | 26 | 0.0769 | 2 | 0 | 1 | 0 | 4 | 0 | 9 | 0.1111 | 16 | 0.0625 |
| 1_38090809 | 9 | 0.1111 | 9 | 0.4444 | 8 | 0.125 | 3 | 0.3333 | 29 | 0.2414 | 3 | 0 | 2 | 0 | 3 | 0 | 8 | 0.25 | 16 | 0.125 |
| 1_38090878 | 9 | 0.8889 | 9 | 0.8889 | 8 | 0.875 | 3 | 1 | 29 | 0.8966 | 3 | 1 | 1 | 1 | 5 | 0.8 | 9 | 0.8889 | 18 | 0.8889 |
| 1_38090994 | 10 | 0.1 | 9 | 0 | 11 | 0.2727 | 3 | 0.3333 | 33 | 0.1515 | 3 | 0 | 2 | 0 | 4 | 0.5 | 10 | 0.3 | 19 | 0.2632 |
| 1_38091134 | 3 | 1 | NA | NA | 2 | 1 | 1 | 0 | 6 | 0.8333 | NA | NA | NA | NA | 1 | 1 | 6 | 0.5 | 7 | 0.5714 |
| 1_38091748 | 2 | 1 | 1 | 0 | 1 | 0 | 1 | 0 | 5 | 0.4 | NA | NA | NA | NA | 1 | 0 | 7 | 0.5714 | 8 | 0.5 |
| 1_38091775 | NA | NA | 1 | 0 | NA | NA | 1 | 1 | 2 | 0.5 | NA | NA | NA | NA | NA | NA | 6 | 0.3333 | 6 | 0.3333 |
| 1_38092441 | 1 | 0 | 2 | 0.5 | 1 | 0 | NA | NA | 4 | 0.25 | NA | NA | NA | NA | NA | NA | 5 | 0.2 | 5 | 0.2 |
| 1_38092529 | 1 | 1 | 3 | 0.3333 | 1 | 0 | NA | NA | 5 | 0.4 | NA | NA | 1 | 0 | 1 | 0 | 5 | 0.4 | 7 | 0.2857 |
| 1_38093043 | 4 | 0 | 4 | 0.25 | 4 | 0 | 1 | 0 | 13 | 0.0769 | 3 | 0.3333 | 1 | 0 | 2 | 0 | 5 | 0 | 11 | 0.0909 |
| 1_38127134 | 1 | 0 | 2 | 0 | 1 | 1 | NA | NA | 4 | 0.25 | 2 | 0 | NA | NA | 1 | 1 | NA | NA | 3 | 0.3333 |
| 1_38127160 | 1 | 1 | 2 | 0.5 | 1 | 0 | NA | NA | 4 | 0.5 | 2 | 1 | NA | NA | 1 | 1 | NA | NA | 3 | 1 |
| 1_38127194 | NA | NA | 2 | 0.5 | 1 | 0 | NA | NA | 3 | 0.3333 | 2 | 0 | NA | NA | 1 | 1 | NA | NA | 3 | 0.3333 |
| 1_38127231 | 1 | 1 | 2 | 1 | 1 | 0 | NA | NA | 4 | 0.75 | 2 | 0.5 | NA | NA | 1 | 0 | NA | NA | 3 | 0.3333 |
| 1_38127274 | NA | NA | 1 | 1 | 1 | 0 | NA | NA | 2 | 0.5 | 2 | 1 | NA | NA | 1 | 1 | NA | NA | 3 | 1 |
| 1_38127345 | 1 | 0 | 2 | 0.5 | 1 | 0 | NA | NA | 4 | 0.25 | 2 | 0.5 | NA | NA | 1 | 1 | NA | NA | 3 | 0.6667 |
| 1_38127376 | 1 | 1 | 2 | 1 | 1 | 1 | NA | NA | 4 | 1 | 2 | 0.5 | NA | NA | 1 | 1 | NA | NA | 3 | 0.6667 |
| 1_38127450 | NA | NA | 2 | 1 | NA | NA | NA | NA | 2 | 1 | 2 | 0.5 | NA | NA | 1 | 1 | NA | NA | 3 | 0.6667 |
| 1_38127547 | 1 | 1 | 2 | 1 | NA | NA | NA | NA | 3 | 1 | 2 | 1 | NA | NA | NA | NA | NA | NA | 2 | 1 |
| 1_38127578 | 1 | 1 | 2 | 1 | NA | NA | NA | NA | 3 | 1 | 1 | 1 | NA | NA | NA | NA | NA | NA | 1 | 1 |
| 1_38127704 | 1 | 1 | 2 | 0.5 | NA | NA | NA | NA | 3 | 0.6667 | 2 | 1 | NA | NA | NA | NA | NA | NA | 2 | 1 |
| 1_38218471 | 4 | 0.75 | 3 | 0.6667 | 3 | 1 | 4 | 0.75 | 14 | 0.7857 | 2 | 1 | 2 | 0.5 | 3 | 0.6667 | 3 | 0.3333 | 10 | 0.6 |
| 1_38218509 | 5 | 0.6 | 4 | 0.75 | 2 | 0.5 | 4 | 0.75 | 15 | 0.6667 | 2 | 0 | 2 | 1 | 3 | 0.3333 | 2 | 1 | 9 | 0.5556 |
| 1_38218527 | 4 | 0.25 | 4 | 0 | 2 | 0 | 4 | 0.25 | 14 | 0.1429 | 2 | 0 | 2 | 1 | 3 | 0.3333 | 3 | 0.3333 | 10 | 0.4 |
| 1_38248663 | 3 | 0.3333 | 1 | 1 | 1 | 1 | 3 | 0.6667 | 8 | 0.625 | NA | NA | 2 | 0.5 | 1 | 1 | 2 | 0.5 | 5 | 0.6 |
| 1_38248713 | 3 | 0.6667 | 1 | 1 | 1 | 1 | 3 | 0.6667 | 8 | 0.75 | NA | NA | 2 | 1 | 1 | 1 | 2 | 0 | 5 | 0.6 |
| 1_38248824 | 3 | 0.6667 | 1 | 0 | 1 | 0 | 3 | 0.6667 | 8 | 0.5 | NA | NA | 2 | 1 | 1 | 0 | 2 | 1 | 5 | 0.8 |
| 1_38248844 | 3 | 1 | 1 | 1 | 1 | 0 | 2 | 0.5 | 7 | 0.7143 | NA | NA | 2 | 0.5 | 1 | 1 | 2 | 0.5 | 5 | 0.6 |
| 1_38249070 | 2 | 0.5 | 1 | 0 | 1 | 1 | 3 | 0.6667 | 7 | 0.5714 | NA | NA | 2 | 1 | 1 | 1 | 2 | 0 | 5 | 0.6 |
| 1_38249200 | 2 | 0 | 1 | 0 | NA | NA | 3 | 0.3333 | 6 | 0.1667 | NA | NA | 2 | 0 | 1 | 1 | 2 | 0 | 5 | 0.2 |
| 1_38574230 | 2 | 1 | NA | NA | NA | NA | 2 | 0 | 4 | 0.5 | NA | NA | 1 | 0 | 1 | 0 | 1 | 1 | 3 | 0.3333 |
| 1_38832912 | 22 | 0.0909 | 15 | 0.1333 | 9 | 0 | 32 | 0.0625 | 78 | 0.0769 | 14 | 0.0714 | 22 | 0.0455 | 17 | 0 | 14 | 0.1429 | 67 | 0.0597 |
| 1_38832960 | 16 | 0.25 | 16 | 0 | 10 | 0 | 30 | 0.1 | 72 | 0.0972 | 13 | 0.0769 | 20 | 0.1 | 13 | 0 | 11 | 0.0909 | 57 | 0.0702 |
| 1_38833799 | 13 | 0.1538 | 9 | 0.1111 | 6 | 0 | 17 | 0.1765 | 45 | 0.1333 | 8 | 0.125 | 17 | 0.1176 | 11 | 0.1818 | 11 | 0.0909 | 47 | 0.1277 |
| 1_38833953 | 14 | 0.2857 | 11 | 0.1818 | 8 | 0.25 | 20 | 0.15 | 53 | 0.2075 | 9 | 0.2222 | 17 | 0.2353 | 14 | 0.2143 | 10 | 0 | 50 | 0.18 |
| 1_38835983 | 10 | 0.5 | 8 | 0.25 | 4 | 0.75 | 12 | 0.4167 | 34 | 0.4412 | 9 | 0.3333 | 8 | 0.625 | 6 | 0.5 | 5 | 0.6 | 28 | 0.5 |
| 1_38836189 | 6 | 0 | 8 | 0.125 | 5 | 0 | 9 | 0.1111 | 28 | 0.0714 | 7 | 0.1429 | 5 | 0 | 7 | 0.1429 | 7 | 0 | 26 | 0.0769 |
| 1_38836638 | 3 | 0.6667 | 8 | 1 | 4 | 1 | 11 | 0.9091 | 26 | 0.9231 | 5 | 1 | 5 | 1 | 9 | 0.8889 | 6 | 1 | 25 | 0.96 |
| 1_38836659 | 4 | 0.5 | 9 | 0 | 5 | 0 | 10 | 0.1 | 28 | 0.1071 | 5 | 0 | 6 | 0.3333 | 9 | 0.2222 | 7 | 0 | 27 | 0.1481 |
| 1_38836684 | 3 | 1 | 8 | 1 | 4 | 0.75 | 11 | 0.9091 | 26 | 0.9231 | 5 | 1 | 6 | 1 | 9 | 1 | 7 | 0.8571 | 27 | 0.963 |
| 1_38836689 | 3 | 0 | 5 | 0.2 | 2 | 0 | 7 | 0.1429 | 17 | 0.1176 | 4 | 0 | 5 | 0.2 | 8 | 0 | 4 | 0.25 | 21 | 0.0952 |
| 1_38836755 | 4 | 0.5 | 7 | 0.5714 | 4 | 0.5 | 12 | 0.75 | 27 | 0.6296 | 3 | 0.3333 | 7 | 0.5714 | 9 | 0.5556 | 5 | 0.6 | 24 | 0.5417 |
| 1_38836809 | 4 | 0 | 6 | 0.1667 | 4 | 0 | 12 | 0.1667 | 26 | 0.1154 | 6 | 0.1667 | 6 | 0 | 8 | 0.125 | 6 | 0 | 26 | 0.0769 |
| 1_38836854 | 4 | 0 | 6 | 0.3333 | 3 | 0.3333 | 12 | 0.25 | 25 | 0.24 | 5 | 0.4 | 7 | 0.2857 | 8 | 0.125 | 6 | 0.5 | 26 | 0.3077 |
| 1_38836938 | 3 | 1 | 7 | 1 | 4 | 0.75 | 12 | 1 | 26 | 0.9615 | 4 | 1 | 5 | 1 | 9 | 0.8889 | 4 | 1 | 22 | 0.9545 |
| 1_38836944 | 4 | 0 | 7 | 0 | 3 | 0.3333 | 10 | 0.1 | 24 | 0.0833 | 6 | 0 | 7 | 0.2857 | 9 | 0 | 6 | 0.1667 | 28 | 0.1071 |
| 1_38836993 | 4 | 0.75 | 7 | 1 | 3 | 0.6667 | 9 | 1 | 23 | 0.913 | 6 | 1 | 6 | 1 | 8 | 1 | 4 | 1 | 24 | 1 |
| 1_38837057 | 4 | 1 | 5 | 0.6 | 2 | 0.5 | 11 | 0.7273 | 22 | 0.7273 | 3 | 0.6667 | 7 | 0.7143 | 6 | 0.6667 | 6 | 0.8333 | 22 | 0.7273 |
| 1_38903145 | 14 | 0.5714 | 10 | 0.7 | 4 | 0.5 | 11 | 0.6364 | 39 | 0.6154 | 4 | 0.5 | 5 | 0.4 | 15 | 0.3333 | 10 | 0.8 | 34 | 0.5 |
| 1_38903153 | 11 | 0.9091 | 13 | 0.9231 | 6 | 0.8333 | 12 | 0.75 | 42 | 0.8571 | 4 | 0.75 | 4 | 0.75 | 13 | 0.7692 | 11 | 0.9091 | 32 | 0.8125 |
| 1_38903183 | 12 | 0.4167 | 12 | 0.0833 | 4 | 0 | 7 | 0.1429 | 35 | 0.2 | 5 | 0.2 | 5 | 0.2 | 13 | 0.3077 | 10 | 0.4 | 33 | 0.303 |
| 1_38903301 | 14 | 0.7143 | 13 | 0.4615 | 6 | 0.5 | 8 | 1 | 41 | 0.6585 | 3 | 1 | 4 | 1 | 16 | 0.5625 | 11 | 0.5455 | 34 | 0.6471 |
| 1_38903575 | 16 | 0.0625 | 13 | 0.0769 | 5 | 0.2 | 10 | 0 | 44 | 0.0682 | 1 | 1 | 5 | 0.2 | 15 | 0 | 13 | 0 | 34 | 0.0588 |
| 1_38903652 | 15 | 0.8 | 9 | 0.8889 | 3 | 1 | 8 | 1 | 35 | 0.8857 | 1 | 1 | 2 | 1 | 8 | 0.875 | 12 | 0.9167 | 23 | 0.913 |
| 1_38903704 | 15 | 0.4667 | 11 | 0.2727 | 6 | 0.3333 | 9 | 0.4444 | 41 | 0.3902 | NA | NA | 4 | 0.25 | 15 | 0.3333 | 10 | 0.2 | 29 | 0.2759 |
| 1_38903728 | 15 | 0.3333 | 13 | 0.8462 | 5 | 0.4 | 11 | 0.4545 | 44 | 0.5227 | 1 | 1 | 5 | 1 | 15 | 0.6667 | 12 | 0.6667 | 33 | 0.7273 |
| 1_38903891 | 12 | 0.1667 | 13 | 0.2308 | 5 | 0.2 | 10 | 0.2 | 40 | 0.2 | 1 | 0 | 4 | 0.5 | 13 | 0.0769 | 11 | 0.0909 | 29 | 0.1379 |
| 1_38903932 | 15 | 0.4667 | 11 | 0.5455 | 5 | 0.4 | 10 | 0.3 | 41 | 0.439 | 1 | 0 | 4 | 0.25 | 14 | 0.2143 | 12 | 0.5 | 31 | 0.3226 |
| 1_38903939 | 14 | 0.7857 | 8 | 0.875 | 6 | 1 | 7 | 0.8571 | 35 | 0.8571 | 1 | 1 | 3 | 0.6667 | 10 | 0.9 | 10 | 1 | 24 | 0.9167 |
| 1_38904016 | 16 | 0.375 | 12 | 0.3333 | 5 | 0.6 | 9 | 0.1111 | 42 | 0.3333 | 1 | 1 | 2 | 0 | 11 | 0.5455 | 11 | 0.0909 | 25 | 0.32 |
| 1_38904040 | 16 | 0.25 | 12 | 0.1667 | 6 | 0.3333 | 11 | 0.0909 | 45 | 0.2 | 1 | 1 | 3 | 0 | 11 | 0.2727 | 12 | 0.3333 | 27 | 0.2963 |
| 1_38904067 | 9 | 0.3333 | 9 | 0 | 3 | 0 | 6 | 0.1667 | 27 | 0.1481 | 1 | 0 | NA | NA | 11 | 0.0909 | 8 | 0.125 | 20 | 0.1 |
| 1_38904091 | 13 | 0.3077 | 12 | 0.5833 | 5 | 0.4 | 10 | 0.4 | 40 | 0.425 | 1 | 0 | 3 | 0 | 12 | 0.5833 | 9 | 0.4444 | 25 | 0.44 |
| 1_38904198 | 16 | 0.375 | 13 | 0.3077 | 6 | 0.5 | 11 | 0.2727 | 46 | 0.3478 | 1 | 0 | 3 | 0.6667 | 13 | 0.3846 | 11 | 0.4545 | 28 | 0.4286 |

| | | | | | | | | | | | | | | | | | | | | |
|---|---|---|---|---|---|---|---|---|---|---|---|---|---|---|---|---|---|---|---|---|
| 1_3890438 | 15 | 0.7333 | 12 | 0.5833 | 6 | 0.5 | 10 | 0.8 | 43 | 0.6744 | 1 | 1 | 2 | 0.5 | 13 | 0.8462 | 10 | 0.5 | 26 | 0.6923 |
| 1_3890462 | 13 | 0.7692 | 9 | 0.8889 | 3 | 0.6667 | 9 | 1 | 34 | 0.8529 | 1 | 1 | 3 | 1 | 10 | 1 | 9 | 0.7778 | 23 | 0.913 |
| 1_3890468 | 7 | 0.5714 | 8 | 0.125 | 6 | 0.6667 | 11 | 0.1818 | 32 | 0.3438 | 1 | 0 | 3 | 0 | 11 | 0.4545 | 8 | 0.25 | 23 | 0.3043 |
| 1_3890475 | 13 | 0 | 10 | 0.2 | 6 | 0 | 11 | 0.1818 | 40 | 0.1 | 1 | 0 | 3 | 0 | 10 | 0.2 | 12 | 0 | 26 | 0.0769 |
| 1_3890479 | 11 | 0.0909 | 8 | 0 | 5 | 0 | 10 | 0.1 | 34 | 0.0588 | 1 | 0 | 2 | 0 | 10 | 0.1 | 9 | 0.1111 | 22 | 0.0909 |
| 1_3890480 | 12 | 0.25 | 10 | 0.2 | 5 | 0 | 8 | 0.25 | 35 | 0.2 | NA | NA | 2 | 0 | 12 | 0.25 | 8 | 0.375 | 22 | 0.2727 |
| 1_3890484 | 14 | 0.7143 | 10 | 0.5 | 6 | 0.6667 | 8 | 0.75 | 38 | 0.6579 | 1 | 1 | 3 | 1 | 10 | 0.7 | 10 | 0.7 | 24 | 0.75 |
| 1_3890487 | 12 | 0.25 | 11 | 0.3636 | 5 | 0.4 | 9 | 0.1111 | 37 | 0.2703 | 1 | 1 | 2 | 0.5 | 10 | 0.3 | 10 | 0.6 | 23 | 0.4783 |
| 1_3890495 | 12 | 0.1667 | 9 | 0.4444 | 5 | 0.4 | 10 | 0.3 | 36 | 0.3056 | 1 | 0 | 2 | 0.5 | 9 | 0.4444 | 11 | 0.0909 | 23 | 0.2609 |
| 1_3890496 | 5 | 0.8 | 6 | 0.3333 | 3 | 0.3333 | 4 | 0.75 | 18 | 0.5556 | 1 | 0 | NA | NA | 4 | 0.25 | 5 | 0.2 | 10 | 0.2 |
| 1_3890506 | 14 | 1 | 11 | 0.9091 | 5 | 1 | 9 | 0.8889 | 39 | 0.9487 | 1 | 1 | 1 | 1 | 9 | 0.8889 | 10 | 0.9 | 21 | 0.9048 |
| 1_3903440 | 3 | 0.3333 | 5 | 0.2 | 5 | 0.2 | 6 | 0 | 19 | 0.1579 | 7 | 0.1429 | 5 | 0.4 | 10 | 0 | 9 | 0.2222 | 31 | 0.1613 |
| 1_3903534 | NA | NA | 6 | 0 | 2 | 0.5 | 5 | 0.4 | 13 | 0.2308 | 7 | 0.2857 | 3 | 0.6667 | 7 | 0.1429 | 9 | 0.2222 | 26 | 0.2692 |
| 1_3903538 | 3 | 0 | 5 | 0 | 5 | 0.2 | 7 | 0.2857 | 20 | 0.15 | 6 | 0 | 4 | 0 | 10 | 0.2 | 8 | 0.25 | 28 | 0.1429 |
| 1_3903551 | 3 | 0 | 5 | 0.2 | 5 | 0.4 | 7 | 0.4286 | 20 | 0.3 | 8 | 0.125 | 6 | 0.1667 | 11 | 0.0909 | 10 | 0 | 35 | 0.0857 |
| 1_3903553 | 2 | 0.5 | 6 | 0.1667 | 5 | 0 | 6 | 0.5 | 19 | 0.2632 | 7 | 0 | 6 | 0.3333 | 9 | 0 | 10 | 0.1 | 32 | 0.0938 |
| 1_3903567 | 2 | 0 | 7 | 0 | 5 | 0.2 | 9 | 0.2222 | 23 | 0.1304 | 8 | 0.375 | 6 | 0 | 10 | 0 | 11 | 0.0909 | 35 | 0.1143 |
| 1_3940122 | 8 | 0.25 | 6 | 0.3333 | 6 | 0.3333 | 3 | 0 | 23 | 0.2609 | 2 | 1 | 6 | 0.3333 | 10 | 0.6 | 4 | 0.25 | 22 | 0.5 |
| 1_3940128 | 11 | 0.0909 | 8 | 0.125 | 6 | 0 | 5 | 0 | 30 | 0.0667 | 2 | 0 | 7 | 0.1429 | 10 | 0.2 | 4 | 0.25 | 23 | 0.1739 |
| 1_3940151 | 9 | 0.6667 | 6 | 0.1667 | 3 | 0.6667 | 1 | 1 | 19 | 0.5263 | 2 | 0.5 | 7 | 0.2857 | 9 | 0.6667 | 2 | 0.5 | 20 | 0.5 |
| 1_3940176 | 9 | 0.2222 | 8 | 0.375 | 6 | 0 | 6 | 0.1667 | 29 | 0.2069 | 2 | 1 | 6 | 0.5 | 11 | 0.2727 | 6 | 0 | 25 | 0.32 |
| 1_3940186 | 11 | 0.3636 | 8 | 0.375 | 8 | 0.125 | 6 | 0.1667 | 33 | 0.2727 | 2 | 0.5 | 8 | 0.375 | 12 | 0.25 | 5 | 0.2 | 27 | 0.2963 |
| 1_3940191 | 10 | 0.1 | 6 | 0.3333 | 5 | 0 | 6 | 0.1667 | 27 | 0.1481 | 2 | 0.5 | 9 | 0.3333 | 13 | 0.1538 | 6 | 0.3333 | 30 | 0.2667 |
| 1_3941117 | 4 | 0.5 | 3 | 0.3333 | 3 | 0.6667 | 8 | 0.25 | 18 | 0.3889 | 5 | 0.4 | 1 | 0 | 8 | 0.375 | 5 | 0.6 | 19 | 0.4211 |
| 1_3941126 | 6 | 0.5 | 3 | 0.3333 | 3 | 0.3333 | 7 | 0.2857 | 19 | 0.3684 | 2 | 0.5 | 4 | 0.25 | 7 | 0.4286 | 4 | 0.25 | 17 | 0.3529 |
| 1_3941130 | 5 | 0.6 | 3 | 0.3333 | 3 | 1 | 4 | 0.25 | 15 | 0.5333 | 4 | 0.5 | 5 | 0.8 | 5 | 0.2 | 4 | 0.75 | 18 | 0.5556 |
| 1_3941145 | 5 | 0.6 | 4 | 0.75 | 2 | 0.5 | 5 | 1 | 16 | 0.75 | 4 | 0.25 | 4 | 0.25 | 6 | 0.3333 | 5 | 0.6 | 19 | 0.3684 |
| 1_3941159 | 5 | 0 | 3 | 0 | 2 | 0 | 5 | 0.2 | 15 | 0.0667 | 2 | 0 | 4 | 0 | 4 | 0.25 | 5 | 0.4 | 15 | 0.2 |
| 1_3941160 | 5 | 1 | 4 | 1 | 2 | 1 | 8 | 0.875 | 19 | 0.9474 | 4 | 1 | 5 | 1 | 7 | 0.7143 | 6 | 0.8333 | 22 | 0.8636 |
| 1_3941162 | 4 | 0.5 | 3 | 0 | 2 | 0.5 | 5 | 0.2 | 14 | 0.2857 | 3 | 1 | 2 | 0 | 6 | 0.6667 | 4 | 0.75 | 15 | 0.6667 |
| 1_3941167 | 4 | 1 | 4 | 0.75 | 2 | 1 | 7 | 0.7143 | 17 | 0.8235 | 2 | 0 | 4 | 1 | 5 | 0.8 | 5 | 0.8 | 16 | 0.75 |
| 1_3942040 | 3 | 0.3333 | 2 | 0 | NA | NA | 6 | 0.1667 | 11 | 0.1818 | 3 | 0.3333 | 5 | 0 | 4 | 0.25 | 5 | 0 | 17 | 0.1176 |
| 1_3957525 | 3 | 0.3333 | 3 | 0 | 2 | 0 | NA | NA | 8 | 0.125 | 2 | 0.5 | 2 | 0.5 | 3 | 0 | NA | NA | 7 | 0.2857 |
| 1_3958422 | 6 | 0.5 | 4 | 1 | 5 | 0.2 | 3 | 0.3333 | 18 | 0.5 | 2 | 1 | 5 | 0.8 | 5 | 0.6 | 1 | 0 | 13 | 0.6923 |
| 1_3958433 | 6 | 0 | 4 | 0.5 | 5 | 0.2 | 3 | 0.3333 | 18 | 0.2222 | 1 | 1 | 6 | 0 | 6 | 0 | 2 | 0 | 15 | 0.0667 |
| 1_3958436 | 6 | 0 | 2 | 0.5 | 3 | 0 | 3 | 0 | 14 | 0.0714 | 2 | 0 | 4 | 0.25 | 5 | 0 | 2 | 0 | 13 | 0.0769 |
| 1_3958444 | 5 | 0 | 2 | 0.5 | 3 | 0 | 2 | 0 | 12 | 0.0833 | 2 | 0 | 5 | 0 | 4 | 0 | 2 | 0.5 | 13 | 0.0769 |
| 1_3958549 | 2 | 0.5 | NA | NA | 3 | 0 | 1 | 1 | 6 | 0.3333 | 1 | 0 | 1 | 0 | 1 | 1 | 2 | 0 | 5 | 0.2 |
| 1_3958562 | 1 | 0 | 1 | 1 | 2 | 1 | 1 | 0 | 5 | 0.6 | 2 | 1 | 1 | 1 | 3 | 1 | 2 | 0 | 8 | 0.75 |
| 1_3958584 | 1 | 0 | 2 | 1 | 3 | 0.6667 | 1 | 0 | 7 | 0.5714 | 2 | 1 | 1 | 0 | 4 | 1 | 2 | 0 | 9 | 0.6667 |
| 1_3959093 | 3 | 0 | 4 | 0 | 3 | 0 | 5 | 0.2 | 15 | 0.0667 | 3 | 0 | 11 | 0.0909 | 7 | 0 | 6 | 0.1667 | 27 | 0.0741 |
| 1_3959138 | 4 | 0.25 | 4 | 0.25 | 3 | 0.3333 | 6 | 0.5 | 17 | 0.3529 | 3 | 0 | 10 | 0.5 | 6 | 0.3333 | 5 | 0.6 | 24 | 0.4167 |
| 1_3959145 | 1 | 1 | 4 | 0.5 | 2 | 1 | 6 | 0.8333 | 13 | 0.7692 | 3 | 0.6667 | 9 | 0.8889 | 6 | 0.8333 | 5 | 1 | 23 | 0.8696 |
| 1_3959150 | 4 | 0.25 | 2 | 0.5 | 3 | 0 | 5 | 0.6 | 14 | 0.3571 | 3 | 0.6667 | 10 | 0.5 | 5 | 0.2 | 6 | 0.1667 | 24 | 0.375 |
| 1_3965770 | 5 | 0.6 | 6 | 0.6667 | 3 | 0.6667 | 4 | 1 | 18 | 0.7222 | NA | NA | 7 | 0.5714 | NA | NA | 6 | 1 | 13 | 0.7692 |
| 1_3965782 | 5 | 0.8 | 8 | 0.5 | 6 | 0.6667 | 5 | 0.6 | 24 | 0.625 | 2 | 1 | 8 | 0.625 | 2 | 1 | 6 | 1 | 18 | 0.8333 |
| 1_3965789 | 5 | 1 | 7 | 1 | 5 | 0.6 | 6 | 1 | 23 | 0.913 | 2 | 1 | 8 | 1 | 2 | 0.5 | 5 | 0.8 | 17 | 0.8824 |
| 1_3965791 | 5 | 0.8 | 7 | 1 | 6 | 0.8333 | 6 | 0.6667 | 24 | 0.8333 | 2 | 0.5 | 8 | 0.875 | 2 | 1 | 5 | 1 | 17 | 0.8824 |
| 1_3965795 | 5 | 1 | 6 | 1 | 4 | 1 | 6 | 1 | 21 | 1 | 2 | 1 | 7 | 0.8571 | 2 | 1 | 5 | 1 | 16 | 0.9375 |
| 1_3965802 | 1 | 1 | 7 | 0.8571 | 6 | 0.6667 | 5 | 1 | 19 | 0.8421 | 2 | 1 | 7 | 0.7143 | 2 | 1 | 5 | 1 | 16 | 0.875 |
| 1_3965814 | 3 | 1 | 5 | 1 | 4 | 1 | 4 | 1 | 16 | 1 | 1 | 1 | 6 | 1 | 2 | 1 | 4 | 1 | 13 | 1 |
| 1_3965819 | 4 | 0.25 | 5 | 0.2 | 6 | 0 | 5 | 0.2 | 20 | 0.15 | 2 | 0.5 | 6 | 0 | 2 | 0 | 4 | 0.25 | 14 | 0.1429 |
| 1_3965822 | 4 | 1 | 7 | 1 | 6 | 1 | 5 | 1 | 22 | 1 | 2 | 1 | 7 | 1 | 2 | 1 | 3 | 1 | 14 | 1 |
| 1_3965845 | 3 | 0.3333 | 6 | 0.1667 | 6 | 0.1667 | 4 | 0 | 19 | 0.1579 | 2 | 0 | 5 | 0.2 | 1 | 1 | 3 | 0 | 11 | 0.1818 |
| 1_3965846 | 3 | 0.3333 | 5 | 0.2 | 6 | 0.1667 | 5 | 0 | 19 | 0.1579 | 2 | 0 | 5 | 0.2 | 2 | 0 | 2 | 0 | 11 | 0.0909 |
| 1_3965881 | 3 | 0 | 6 | 0.1667 | 6 | 0.1667 | 4 | 0 | 19 | 0.1053 | 2 | 0 | 4 | 0.25 | 2 | 0 | 1 | 0 | 9 | 0.1111 |
| 1_3965887 | 3 | 0.3333 | 4 | 0.5 | 6 | 0.5 | 5 | 0.2 | 18 | 0.3889 | 2 | 0.5 | 5 | 0.2 | 2 | 0.5 | 2 | 0 | 11 | 0.2727 |
| 1_3965903 | 2 | 0.5 | 5 | 0.6 | 4 | 0.5 | 5 | 0 | 16 | 0.375 | 2 | 0.5 | 4 | 0.25 | 2 | 0 | 2 | 0.5 | 10 | 0.3 |
| 1_3966117 | 1 | 1 | 3 | 0.3333 | NA | NA | 3 | 0.3333 | 7 | 0.4286 | 1 | 0 | 2 | 0.5 | 2 | 0 | 1 | 1 | 6 | 0.3333 |
| 1_3966212 | NA | NA | 2 | 0.5 | NA | NA | 1 | 0 | 3 | 0.3333 | NA | NA | 1 | 1 | 1 | 1 | 1 | 0 | 3 | 0.6667 |
| 1_3966219 | 1 | 0 | 2 | 0.5 | NA | NA | 2 | 0.5 | 5 | 0.4 | 1 | 1 | 3 | 0 | 1 | 1 | 1 | 1 | 6 | 0.5 |
| 1_3966220 | NA | NA | 2 | 0.5 | NA | NA | 2 | 0.5 | 4 | 0.5 | 1 | 0 | 3 | 0.3333 | 1 | 0 | 2 | 0 | 7 | 0.1429 |
| 1_3969440 | NA | NA | 1 | 1 | NA | NA | NA | NA | 1 | 1 | 1 | 1 | NA | NA | NA | NA | NA | NA | 1 | 1 |
| 1_3969448 | NA | NA | 1 | 1 | NA | NA | NA | NA | 1 | 1 | 2 | 1 | NA | NA | NA | NA | NA | NA | 2 | 1 |
| 1_3969463 | NA | NA | 1 | 1 | NA | NA | NA | NA | 1 | 1 | 2 | 1 | NA | NA | NA | NA | NA | NA | 2 | 1 |

| | | | | | | | | | | | | | | | | | | | | |
|---|---|---|---|---|---|---|---|---|---|---|---|---|---|---|---|---|---|---|---|---|
| 1_40063852 | 2 | 0 | 1 | 0 | 1 | 1 | 1 | 1 | 5 | 0.4 | 1 | 0 | 2 | 0.5 | 2 | 0.5 | 5 | 0.4 | 10 | 0.4 |
| 1_40063978 | 3 | 0.3333 | 1 | 0 | NA | NA | 1 | 0 | 5 | 0.2 | 1 | 0 | 2 | 0 | 2 | 0 | 5 | 0.2 | 10 | 0.1 |
| 1_40064124 | 2 | 1 | NA | NA | 1 | 0 | 1 | 1 | 4 | 0.75 | NA | NA | NA | NA | 3 | 1 | 3 | 0.6667 | 6 | 0.8333 |
| 1_40064159 | 1 | 1 | NA | NA | 1 | 1 | NA | NA | 2 | 1 | 1 | 1 | NA | NA | NA | NA | 3 | 1 | 4 | 1 |
| 1_40065207 | 8 | 0 | 13 | 0.0769 | 8 | 0 | 7 | 0.1429 | 36 | 0.0556 | 10 | 0.2 | 4 | 0 | 8 | 0 | 6 | 0 | 28 | 0.0714 |
| 1_40088599 | NA | NA | NA | NA | NA | NA | 1 | 1 | 1 | 1 | NA | NA | NA | NA | 1 | 1 | NA | NA | 1 | 1 |
| 1_40089025 | NA | NA | NA | NA | NA | NA | 1 | 1 | 1 | 1 | NA | NA | NA | NA | 1 | 1 | NA | NA | 1 | 1 |
| 1_40352507 | NA | NA | 1 | 1 | 1 | 1 | 1 | 1 | 3 | 1 | 1 | 1 | NA | NA | NA | NA | NA | NA | 1 | 1 |
| 1_40352579 | NA | NA | 1 | 1 | 1 | 1 | 1 | 1 | 3 | 1 | 1 | 1 | NA | NA | NA | NA | NA | NA | 1 | 1 |
| 1_40352705 | NA | NA | 1 | 1 | 1 | 1 | 1 | 1 | 3 | 1 | 1 | 1 | NA | NA | NA | NA | NA | NA | 1 | 1 |
| 1_40352719 | NA | NA | 1 | 1 | 1 | 0 | 1 | 0 | 3 | 0.3333 | 1 | 1 | NA | NA | NA | NA | NA | NA | 1 | 1 |
| 1_40352883 | NA | NA | NA | NA | NA | NA | 1 | 1 | 1 | 1 | 1 | 1 | NA | NA | NA | NA | NA | NA | 1 | 1 |
| 1_40353855 | NA | NA | 1 | 0 | 1 | 0 | 1 | 1 | 3 | 0.3333 | 1 | 1 | 1 | 0 | NA | NA | NA | NA | 2 | 0.5 |
| 1_40353876 | NA | NA | 1 | 1 | 1 | 1 | 1 | 1 | 3 | 1 | 1 | 1 | 1 | 1 | NA | NA | NA | NA | 2 | 1 |
| 1_40354410 | NA | NA | 1 | 0 | 1 | 1 | 1 | 0 | 3 | 0.3333 | NA | NA | 1 | 1 | NA | NA | NA | NA | 1 | 1 |
| 1_40354913 | NA | NA | 1 | 0 | 1 | 1 | 1 | 0 | 3 | 0.3333 | 1 | 1 | 1 | 0 | NA | NA | NA | NA | 2 | 0.5 |
| 1_40807029 | 2 | 1 | NA | NA | NA | NA | NA | NA | 2 | 1 | 1 | 0 | NA | NA | 3 | 0.6667 | NA | NA | 4 | 0.5 |
| 1_40807206 | 2 | 1 | NA | NA | NA | NA | NA | NA | 2 | 1 | 1 | 1 | NA | NA | 3 | 0.6667 | NA | NA | 4 | 0.75 |
| 1_40807339 | 2 | 1 | NA | NA | NA | NA | NA | NA | 2 | 1 | 1 | 1 | NA | NA | 2 | 1 | NA | NA | 3 | 1 |
| 1_40807348 | 2 | 1 | NA | NA | NA | NA | NA | NA | 2 | 1 | NA | NA | NA | NA | 2 | 1 | NA | NA | 2 | 1 |
| 1_40807411 | 1 | 1 | NA | NA | NA | NA | NA | NA | 1 | 1 | 1 | 1 | NA | NA | 3 | 0.6667 | NA | NA | 4 | 0.75 |
| 1_40807425 | 2 | 0.5 | NA | NA | NA | NA | NA | NA | 2 | 0.5 | 1 | 0 | NA | NA | 3 | 0.6667 | NA | NA | 4 | 0.5 |
| 1_40811654 | 1 | 1 | NA | NA | NA | NA | NA | NA | 1 | 1 | NA | NA | 1 | 1 | 1 | 1 | 1 | 1 | 3 | 1 |
| 1_40811726 | 1 | 1 | NA | NA | NA | NA | NA | NA | 1 | 1 | NA | NA | NA | NA | 1 | 1 | 1 | 1 | 2 | 1 |
| 1_40811737 | 1 | 1 | NA | NA | NA | NA | NA | NA | 1 | 1 | NA | NA | 1 | 1 | 1 | 0 | 1 | 1 | 3 | 0.6667 |
| 1_40811806 | 1 | 1 | NA | NA | NA | NA | NA | NA | 1 | 1 | NA | NA | 1 | 1 | 1 | 0 | 1 | 0 | 3 | 0.3333 |
| 1_40811817 | 1 | 1 | NA | NA | NA | NA | NA | NA | 1 | 1 | NA | NA | 1 | 1 | 1 | 0 | 1 | 1 | 3 | 0.6667 |
| 1_42687458 | 8 | 0.375 | 17 | 0.1176 | 12 | 0.25 | 10 | 0.2 | 47 | 0.2128 | 10 | 0.2 | 16 | 0 | 15 | 0.2 | 13 | 0.0769 | 54 | 0.1111 |
| 1_42687550 | 5 | 0.2 | 12 | 0.0833 | 10 | 0.1 | 8 | 0.25 | 35 | 0.1429 | 8 | 0.375 | 14 | 0.1429 | 15 | 0.0667 | 8 | 0.25 | 45 | 0.1778 |
| 1_42735607 | 1 | 1 | NA | NA | 1 | 1 | NA | NA | 2 | 1 | 1 | 1 | 1 | 1 | NA | NA | NA | NA | 2 | 1 |
| 1_42735676 | 1 | 1 | NA | NA | 1 | 0 | NA | NA | 2 | 0.5 | 1 | 0 | 1 | 1 | NA | NA | NA | NA | 2 | 0.5 |
| 1_42735782 | 1 | 1 | NA | NA | 1 | 0 | NA | NA | 2 | 0.5 | NA | NA | 1 | 1 | NA | NA | NA | NA | 1 | 1 |
| 1_42736848 | 3 | 0.3333 | 1 | 1 | 4 | 0.75 | 3 | 0.6667 | 11 | 0.6364 | 2 | 0.5 | 3 | 0.6667 | NA | NA | NA | NA | 5 | 0.6 |
| 1_42737090 | 4 | 0 | 3 | 0.3333 | 6 | 0.1667 | 4 | 0 | 17 | 0.1176 | 2 | 0.5 | 3 | 0.3333 | NA | NA | 1 | 1 | 6 | 0.5 |
| 1_42737172 | 5 | 0.2 | 2 | 0 | 6 | 0.5 | 5 | 0.2 | 18 | 0.2778 | 3 | 0 | 3 | 0.3333 | NA | NA | 3 | 0 | 9 | 0.1111 |
| 1_42737240 | 6 | 0.1667 | 3 | 0 | 10 | 0.1 | 4 | 0 | 23 | 0.087 | 4 | 0 | 6 | 0.1667 | 1 | 0 | 3 | 0.3333 | 14 | 0.1429 |
| 1_42737411 | 4 | 0 | 3 | 0.3333 | 9 | 0.2222 | 4 | 0.25 | 20 | 0.2 | 3 | 0.3333 | 7 | 0.8571 | 1 | 1 | 4 | 0.25 | 15 | 0.6 |
| 1_42737459 | 5 | 0 | 2 | 0 | 9 | 0 | 5 | 0.4 | 21 | 0.0952 | 2 | 0.5 | 7 | 0.1429 | NA | NA | 3 | 0 | 12 | 0.1667 |
| 1_42738034 | 5 | 0.6 | 5 | 1 | 8 | 0.75 | 5 | 0 | 23 | 0.6087 | 5 | 0.8 | 7 | 0.5714 | 1 | 1 | 3 | 0.6667 | 16 | 0.6875 |
| 1_42738066 | 7 | 0.1429 | 6 | 0.3333 | 12 | 0.0833 | 4 | 0.25 | 29 | 0.1724 | 7 | 0.2857 | 5 | 0.2 | NA | NA | 6 | 0 | 18 | 0.1667 |
| 1_42738085 | 5 | 0.6 | 9 | 0.6667 | 12 | 0.75 | 6 | 0.8333 | 32 | 0.7188 | 5 | 0.6 | 8 | 0.75 | NA | NA | 6 | 0.8333 | 19 | 0.7368 |
| 1_42738113 | 5 | 0.8 | 7 | 0.8571 | 13 | 0.5385 | 7 | 0.4286 | 32 | 0.625 | 8 | 0.75 | 9 | 0.8889 | 1 | 1 | 6 | 0.5 | 24 | 0.75 |
| 1_42738147 | 6 | 1 | 6 | 1 | 13 | 1 | 7 | 1 | 32 | 1 | 8 | 1 | 7 | 0.8571 | 2 | 1 | 6 | 1 | 23 | 0.9565 |
| 1_42738204 | 4 | 0.75 | 6 | 0.8333 | 11 | 0.6364 | 7 | 0.4286 | 28 | 0.6429 | 8 | 0.625 | 7 | 0.8571 | 3 | 0.3333 | 6 | 0.8333 | 24 | 0.7083 |
| 1_42738361 | 6 | 0 | 7 | 0.2857 | 13 | 0.1538 | 9 | 0 | 35 | 0.1143 | 7 | 0.2857 | 9 | 0.1111 | 3 | 0.3333 | 5 | 0.4 | 24 | 0.25 |
| 1_42738420 | 5 | 0 | 7 | 0.2857 | 11 | 0.1818 | 8 | 0.375 | 31 | 0.2258 | 8 | 0.375 | 7 | 0.4286 | 3 | 0.3333 | 6 | 0.3333 | 24 | 0.375 |
| 1_42738478 | 6 | 0.3333 | 5 | 0.2 | 13 | 0.3077 | 9 | 0.1111 | 33 | 0.2424 | 5 | 0 | 9 | 0.1111 | 3 | 0.3333 | 4 | 0.25 | 21 | 0.1429 |
| 1_42738491 | 7 | 1 | 7 | 1 | 12 | 1 | 8 | 1 | 34 | 1 | 8 | 1 | 8 | 1 | 2 | 1 | 5 | 0.8 | 23 | 0.9565 |
| 1_42738528 | 6 | 0 | 6 | 0.1667 | 13 | 0.0769 | 8 | 0 | 33 | 0.0606 | 7 | 0.2857 | 6 | 0.1667 | 1 | 0 | 5 | 0 | 19 | 0.1579 |
| 1_42738565 | 7 | 0 | 7 | 0.1429 | 7 | 0.1429 | 6 | 0 | 27 | 0.0741 | 5 | 0.2 | 4 | 0 | NA | NA | 5 | 0 | 14 | 0.0714 |
| 1_42738608 | 8 | 0.875 | 9 | 0.7778 | 13 | 0.6923 | 7 | 0.8571 | 37 | 0.7838 | 3 | 0.6667 | 9 | 0.8889 | 3 | 1 | 4 | 0.75 | 19 | 0.8421 |
| 1_42738648 | 8 | 0 | 8 | 0 | 12 | 0.1667 | 10 | 0 | 38 | 0.0526 | 8 | 0.125 | 10 | 0 | 3 | 0.3333 | 7 | 0 | 28 | 0.0714 |
| 1_42738768 | 8 | 0.125 | 6 | 0.1667 | 13 | 0 | 8 | 0.125 | 35 | 0.0857 | 5 | 0.2 | 6 | 0 | 3 | 0 | 6 | 0.1667 | 20 | 0.1 |
| 1_42738815 | 7 | 0.4286 | 7 | 0.4286 | 12 | 0.4167 | 12 | 0.25 | 38 | 0.3684 | 8 | 0.25 | 9 | 0.2222 | 4 | 0.5 | 5 | 0.2 | 26 | 0.2692 |
| 1_42738832 | 8 | 0.375 | 9 | 0.6667 | 12 | 0.6667 | 12 | 0.5 | 41 | 0.561 | 7 | 0.8571 | 9 | 0.7778 | 3 | 0.6667 | 7 | 0.5714 | 26 | 0.7308 |
| 1_42738896 | 7 | 1 | 3 | 1 | 12 | 0.9167 | 9 | 1 | 31 | 0.9677 | 7 | 0.8571 | 9 | 1 | 2 | 1 | 7 | 1 | 25 | 0.96 |
| 1_42738924 | 10 | 0.2 | 10 | 0 | 16 | 0.25 | 12 | 0.3333 | 48 | 0.2083 | 9 | 0.4444 | 10 | 0.3 | 4 | 0 | 8 | 0.25 | 31 | 0.2903 |
| 1_42738950 | 6 | 0.3333 | 7 | 0.1429 | 13 | 0.2308 | 9 | 0.1111 | 35 | 0.2 | 9 | 0.4444 | 9 | 0.4444 | 5 | 0 | 6 | 0 | 29 | 0.2759 |
| 1_42739057 | 10 | 0.9 | 5 | 1 | 13 | 1 | 10 | 1 | 38 | 0.9737 | 8 | 1 | 6 | 1 | 4 | 1 | 6 | 1 | 24 | 1 |
| 1_42739136 | 7 | 0.5714 | 10 | 0.7 | 17 | 0.2941 | 11 | 0.2727 | 45 | 0.4222 | 11 | 0.4545 | 9 | 0.7778 | 6 | 0.1667 | 9 | 0.3333 | 35 | 0.4571 |
| 1_42739170 | 9 | 0.8889 | 5 | 0.8 | 10 | 0.8 | 9 | 0.8889 | 33 | 0.8485 | 6 | 0.8333 | 8 | 1 | 3 | 1 | 6 | 1 | 23 | 0.9565 |
| 1_42739261 | 12 | 0.75 | 9 | 1 | 16 | 0.9375 | 12 | 0.75 | 49 | 0.8571 | 11 | 0.9091 | 8 | 1 | 6 | 1 | 10 | 1 | 35 | 0.9714 |
| 1_42783108 | NA | NA | NA | NA | 1 | 1 | NA | NA | 1 | 1 | NA | NA | NA | NA | NA | NA | 1 | 1 | 1 | 1 |
| 1_42783328 | NA | NA | NA | NA | 1 | 1 | NA | NA | 1 | 1 | NA | NA | NA | NA | NA | NA | 1 | 1 | 1 | 1 |
| 1_42783519 | NA | NA | NA | NA | 1 | 1 | NA | NA | 1 | 1 | NA | NA | 1 | 1 | NA | NA | 1 | 1 | 2 | 1 |

| | | | | | | | | | | | | | | | | | | | | |
|---|---|---|---|---|---|---|---|---|---|---|---|---|---|---|---|---|---|---|---|---|
| 1_42992193 | NA | NA | 1 | 1 | 1 | 0 | NA | NA | 2 | 0.5 | NA | NA | 1 | 1 | NA | NA | NA | NA | 1 | 1 |
| 1_42993010 | NA | NA | 2 | 1 | 1 | 1 | NA | NA | 3 | 1 | 1 | 0 | 1 | 1 | 1 | 0 | NA | NA | 3 | 0.3333 |
| 1_42993018 | NA | NA | 2 | 0 | 1 | 1 | NA | NA | 3 | 0.3333 | 1 | 1 | 1 | 1 | 1 | 0 | NA | NA | 3 | 0.6667 |
| 1_42993769 | NA | NA | 2 | 0.5 | 1 | 0 | NA | NA | 3 | 0.3333 | 1 | 0 | 1 | 1 | NA | NA | NA | NA | 2 | 0.5 |
| 1_42993824 | NA | NA | 3 | 0.6667 | 1 | 1 | NA | NA | 4 | 0.75 | 1 | 1 | 1 | 1 | NA | NA | NA | NA | 2 | 1 |
| 1_43074050 | 3 | 0 | 1 | 0 | 1 | 0 | 2 | 0.5 | 7 | 0.1429 | 1 | 1 | NA | NA | NA | NA | 1 | 0 | 2 | 0.5 |
| 1_43074264 | 2 | 0 | 1 | 0 | 1 | 0 | 2 | 0.5 | 6 | 0.1667 | NA | NA | NA | NA | NA | NA | 1 | 1 | 1 | 1 |
| 1_43086845 | 5 | 0 | 2 | 0.5 | 1 | 0 | 2 | 0 | 10 | 0.1 | 3 | 0 | 4 | 0 | 1 | 0 | 6 | 0.1667 | 14 | 0.0714 |
| 1_43087067 | 5 | 0 | 2 | 0.5 | 1 | 0 | 2 | 0 | 10 | 0.1 | NA | NA | 3 | 0 | 1 | 0 | 5 | 0.2 | 9 | 0.1111 |
| 1_43087097 | 5 | 0 | 2 | 0 | 1 | 0 | 2 | 0.5 | 10 | 0.1 | 2 | 0 | 3 | 0.3333 | 2 | 0 | 5 | 0.4 | 12 | 0.25 |
| 1_43087282 | 5 | 0.2 | 1 | 0 | 1 | 0 | 1 | 0 | 8 | 0.125 | 2 | 0 | 3 | 0.3333 | 2 | 0 | 5 | 0.2 | 12 | 0.1667 |
| 1_43087620 | 4 | 1 | 1 | 1 | 1 | 1 | 2 | 1 | 8 | 1 | 1 | 1 | 2 | 0 | 2 | 0.5 | 5 | 0.6 | 10 | 0.5 |
| 1_43087993 | 3 | 0.6667 | 1 | 1 | 1 | 0 | 2 | 0.5 | 7 | 0.5714 | 1 | 0 | 2 | 0 | 2 | 0 | 5 | 0.2 | 10 | 0.1 |
| 1_43088412 | 3 | 1 | 1 | 1 | 1 | 0 | NA | NA | 5 | 0.8 | NA | NA | 3 | 0.3333 | 2 | 0.5 | 3 | 0.3333 | 8 | 0.375 |
| 1_43088529 | 2 | 0.5 | 1 | 0 | 1 | 0 | 1 | 0 | 5 | 0.2 | NA | NA | 1 | 0 | 2 | 0 | 5 | 0.2 | 8 | 0.125 |
| 1_43088625 | 2 | 1 | NA | NA | 1 | 0 | 1 | 1 | 4 | 0.75 | NA | NA | 3 | 1 | 1 | 1 | 5 | 0.2 | 9 | 0.5556 |
| 1_43088691 | 3 | 1 | NA | NA | 1 | 1 | 1 | 1 | 5 | 1 | NA | NA | 2 | 0 | 1 | 1 | 3 | 0.6667 | 6 | 0.5 |
| 1_43088824 | 3 | 0.6667 | NA | NA | 1 | 0 | 1 | 1 | 5 | 0.6 | NA | NA | 1 | 1 | 1 | 1 | 4 | 1 | 6 | 1 |
| 1_43090797 | 3 | 1 | NA | NA | NA | NA | 1 | 1 | 4 | 1 | NA | NA | NA | NA | 1 | 1 | 2 | 1 | 3 | 1 |
| 1_43090908 | 2 | 1 | NA | NA | 1 | 1 | 1 | 1 | 4 | 1 | NA | NA | NA | NA | 2 | 1 | 4 | 1 | 6 | 1 |
| 1_43091031 | 2 | 0 | NA | NA | 1 | 0 | 1 | 1 | 4 | 0.25 | NA | NA | NA | NA | 1 | 0 | 2 | 0.5 | 3 | 0.3333 |
| 1_43114500 | 2 | 0 | NA | NA | 1 | 1 | 1 | 0 | 4 | 0.25 | NA | NA | 1 | 0 | 1 | 1 | 3 | 0 | 5 | 0.2 |
| 1_43114632 | 2 | 1 | NA | NA | 1 | 0 | 1 | 1 | 4 | 0.75 | NA | NA | 1 | 1 | 1 | 1 | 3 | 0.6667 | 5 | 0.8 |
| 1_43114660 | 1 | 1 | NA | NA | 1 | 1 | 1 | 1 | 3 | 1 | NA | NA | 1 | 1 | 1 | 1 | 2 | 1 | 4 | 1 |
| 1_43114782 | 2 | 0.5 | NA | NA | 1 | 1 | 1 | 0 | 4 | 0.5 | NA | NA | 1 | 1 | 1 | 0 | 2 | 0.5 | 4 | 0.5 |
| 1_43114921 | 2 | 1 | NA | NA | NA | NA | 1 | 1 | 3 | 1 | NA | NA | 1 | 1 | 1 | 1 | 2 | 1 | 4 | 1 |
| 1_43131996 | 1 | 1 | NA | NA | NA | NA | NA | NA | 1 | 1 | NA | NA | 1 | 1 | NA | NA | 1 | 0 | 2 | 0.5 |
| 1_43150373 | 1 | 0 | NA | NA | 3 | 0.3333 | 4 | 0 | 8 | 0.125 | NA | NA | 2 | 1 | 5 | 0.2 | NA | NA | 7 | 0.4286 |
| 1_43150400 | 1 | 0 | NA | NA | 3 | 0.3333 | 4 | 0.25 | 8 | 0.25 | NA | NA | 2 | 1 | 5 | 0.8 | NA | NA | 7 | 0.8571 |
| 1_43150418 | 1 | 0 | NA | NA | 3 | 0.3333 | 4 | 0.25 | 8 | 0.25 | NA | NA | 1 | 0 | 5 | 0.2 | NA | NA | 6 | 0.1667 |
| 1_43152492 | 1 | 1 | NA | NA | 3 | 0 | 4 | 0 | 8 | 0.125 | NA | NA | 2 | 1 | 4 | 1 | NA | NA | 6 | 1 |
| 1_43154813 | 2 | 0 | 2 | 0 | 2 | 0 | 9 | 0.1111 | 15 | 0.0667 | 2 | 0 | 3 | 0 | 7 | 0.2857 | 2 | 0 | 14 | 0.1429 |
| 1_43154825 | 3 | 0 | 2 | 0 | 2 | 0.5 | 10 | 0.1 | 17 | 0.1176 | 2 | 0 | 4 | 0 | 8 | 0.125 | 2 | 0 | 16 | 0.0625 |
| 1_43154843 | 4 | 0 | 2 | 0.5 | 3 | 0 | 9 | 0 | 18 | 0.0556 | 2 | 0 | 4 | 0.25 | 9 | 0.2222 | 2 | 0.5 | 17 | 0.2353 |
| 1_43162358 | 11 | 0 | 8 | 0.25 | 7 | 0.1429 | 1 | 0 | 27 | 0.1111 | 9 | 0.1111 | 7 | 0.1429 | 1 | 0 | 6 | 0 | 23 | 0.087 |
| 1_43162511 | 8 | 0.25 | 9 | 0.2222 | 6 | 0.3333 | 1 | 1 | 24 | 0.2917 | 6 | 0 | 6 | 0 | 1 | 0 | 7 | 0.1429 | 20 | 0.05 |
| 1_43162693 | 8 | 0.25 | 10 | 0.1 | 4 | 0.5 | 1 | 0 | 23 | 0.2174 | 8 | 0 | 6 | 0 | 1 | 0 | 7 | 0.2857 | 22 | 0.0909 |
| 1_43593455 | 1 | 1 | 2 | 0.5 | 2 | 0 | 2 | 0 | 7 | 0.2857 | 2 | 0.5 | 2 | 1 | 3 | 0.3333 | 2 | 0.5 | 9 | 0.5556 |
| 1_43593486 | 1 | 1 | 2 | 0.5 | 2 | 0 | 2 | 0 | 7 | 0.2857 | 2 | 0 | 2 | 0 | 3 | 0.3333 | 1 | 0 | 8 | 0.125 |
| 1_43593543 | 1 | 1 | 1 | 1 | 1 | 1 | 1 | 1 | 4 | 1 | 2 | 1 | 1 | 1 | 1 | 1 | NA | NA | 4 | 1 |
| 1_43593729 | 1 | 1 | 2 | 1 | 2 | 0.5 | 1 | 1 | 6 | 0.8333 | 1 | 1 | 3 | 1 | 4 | 0.75 | 2 | 1 | 10 | 0.9 |
| 1_43608375 | 2 | 0 | 4 | 0.25 | 3 | 0 | 2 | 0.5 | 11 | 0.1818 | 2 | 0.5 | 3 | 0.3333 | 4 | 0 | 1 | 0 | 10 | 0.2 |
| 1_43608819 | 3 | 0.3333 | 3 | 1 | 5 | 1 | 2 | 0.5 | 13 | 0.7692 | 3 | 0.6667 | 2 | 1 | 4 | 0.25 | 2 | 1 | 11 | 0.6364 |
| 1_43608831 | 3 | 1 | 4 | 0.5 | 5 | 0.8 | 2 | 1 | 14 | 0.7857 | 3 | 0.3333 | 2 | 0.5 | 4 | 0.5 | 2 | 1 | 11 | 0.5455 |
| 1_43608879 | 3 | 0 | 4 | 0 | 5 | 0.2 | 1 | 0 | 13 | 0.0769 | 3 | 0 | 2 | 0 | 3 | 0 | 2 | 0.5 | 10 | 0.1 |
| 1_43609071 | 3 | 0 | 4 | 0.25 | 5 | 0 | 3 | 0 | 15 | 0.0667 | 4 | 0.25 | 4 | 0.5 | 5 | 0 | 1 | 0 | 14 | 0.2143 |
| 1_43609550 | 2 | 0 | 4 | 0 | 3 | 0 | 4 | 0.25 | 13 | 0.0769 | 4 | 0 | 3 | 0 | 6 | 0 | 2 | 0.5 | 15 | 0.0667 |
| 1_43788917 | 1 | 0 | 3 | 0 | 5 | 0.2 | 1 | 0 | 10 | 0.1 | 2 | 0 | 6 | 0 | 3 | 0 | 4 | 0.25 | 15 | 0.0667 |
| 1_43789250 | 3 | 0 | 4 | 0.5 | 4 | 0 | 1 | 0 | 12 | 0.1667 | 5 | 0 | 6 | 0 | 3 | 0.3333 | 4 | 0.5 | 18 | 0.1667 |
| 1_43789308 | 3 | 0 | 3 | 0.3333 | 3 | 0 | NA | NA | 9 | 0.1111 | 4 | 0 | 6 | 0.1667 | 3 | 0 | 4 | 0 | 17 | 0.0588 |
| 1_43789342 | 3 | 0.6667 | 3 | 1 | 3 | 1 | 1 | 0 | 10 | 0.8 | 3 | 0.6667 | 5 | 0.8 | 2 | 0.5 | 4 | 0.75 | 14 | 0.7143 |
| 1_43789367 | 3 | 0.6667 | 2 | 0.5 | 4 | 0.25 | 1 | 1 | 10 | 0.5 | 5 | 0.8 | 5 | 0.2 | 3 | 0.3333 | 3 | 1 | 16 | 0.5625 |
| 1_43928740 | 2 | 0 | 2 | 0.5 | 7 | 0 | 3 | 0 | 14 | 0.0714 | 3 | 0.3333 | 6 | 0.1667 | 2 | 0 | 3 | 0 | 14 | 0.1429 |
| 1_44126060 | 2 | 1 | 1 | 0 | 1 | 0 | 1 | 1 | 5 | 0.6 | NA | NA | 2 | 0 | 1 | 1 | NA | NA | 3 | 0.3333 |
| 1_44157751 | 4 | 0.5 | 1 | 0 | NA | NA | NA | NA | 5 | 0.4 | NA | NA | 1 | 1 | NA | NA | 1 | 0 | 2 | 0.5 |
| 1_44165468 | 2 | 0 | 2 | 0 | 1 | 0 | 2 | 0.5 | 7 | 0.1429 | NA | NA | NA | NA | NA | NA | 3 | 0.6667 | 3 | 0.6667 |
| 1_44165534 | 2 | 0.5 | 2 | 0.5 | 2 | 0 | 2 | 0 | 8 | 0.25 | NA | NA | 1 | 1 | NA | NA | 4 | 0 | 5 | 0.2 |
| 1_44165593 | 1 | 0 | 3 | 0 | 2 | 0.5 | 2 | 0 | 8 | 0.125 | NA | NA | NA | NA | NA | NA | 3 | 0.6667 | 3 | 0.6667 |
| 1_44165729 | 2 | 1 | 3 | 0.6667 | 2 | 1 | 2 | 0.5 | 9 | 0.7778 | NA | NA | 1 | 1 | NA | NA | 5 | 0.8 | 6 | 0.8333 |
| 1_44165804 | 2 | 0 | 2 | 0.5 | 2 | 0 | 2 | 0 | 8 | 0.125 | NA | NA | 1 | 0 | NA | NA | 5 | 0.2 | 6 | 0.1667 |
| 1_44182047 | 2 | 1 | 3 | 0.3333 | 2 | 0.5 | 2 | 0.5 | 9 | 0.5556 | 1 | 1 | 2 | 0.5 | 1 | 0 | 7 | 0.5714 | 11 | 0.5455 |
| 1_44182076 | 2 | 1 | 2 | 1 | 1 | 1 | 2 | 1 | 7 | 1 | 1 | 1 | 3 | 1 | 1 | 1 | 7 | 0.5714 | 12 | 0.75 |
| 1_44182154 | 2 | 1 | 3 | 0.6667 | 2 | 0.5 | 2 | 1 | 9 | 0.7778 | 2 | 0.5 | 3 | 1 | 1 | 1 | 7 | 0.7143 | 13 | 0.7692 |
| 1_44182217 | 3 | 0 | 3 | 0.3333 | 2 | 0 | 2 | 0 | 10 | 0.1 | 2 | 0.5 | 3 | 0.3333 | 1 | 1 | 7 | 0.2857 | 13 | 0.3846 |
| 1_44182259 | 3 | 0.6667 | 2 | 0 | 2 | 0 | 2 | 0 | 9 | 0.2222 | 2 | 0 | 3 | 0 | 1 | 0 | 7 | 0.1429 | 13 | 0.0769 |

| | | | | | | | | | | | | | | | | | | | | |
|---|---|---|---|---|---|---|---|---|---|---|---|---|---|---|---|---|---|---|---|---|
| 1_44182298 | 3 | 0.6667 | 2 | 1 | 2 | 0.5 | 2 | 0.5 | 9 | 0.6667 | 2 | 0.5 | 3 | 0.6667 | NA | NA | 7 | 0.4286 | 12 | 0.5 |
| 1_44182354 | 3 | 0.6667 | 3 | 1 | 2 | 1 | 2 | 1 | 10 | 0.9 | 2 | 0.5 | 3 | 1 | 1 | 1 | 7 | 1 | 13 | 0.9231 |
| 1_44183174 | 3 | 0 | 3 | 0.3333 | 2 | 0 | 2 | 0 | 10 | 0.1 | 2 | 0 | 5 | 0 | 4 | 0 | 7 | 0.1429 | 18 | 0.0556 |
| 1_44183460 | 3 | 0.3333 | 4 | 0 | 2 | 0 | 2 | 0.5 | 11 | 0.1818 | 2 | 0.5 | 5 | 0.2 | 4 | 0 | 8 | 0.25 | 19 | 0.2105 |
| 1_44183588 | 2 | 0.5 | 4 | 0.75 | 2 | 0.5 | 2 | 0 | 10 | 0.5 | 2 | 0.5 | 4 | 0.75 | 4 | 0.25 | 7 | 0.5714 | 17 | 0.5294 |
| 1_44206430 | NA | NA | NA | NA | 3 | 0.6667 | 1 | 0 | 4 | 0.5 | NA | NA | NA | NA | 2 | 0.5 | 1 | 0 | 3 | 0.3333 |
| 1_44206474 | NA | NA | 1 | 0 | 3 | 0.6667 | 1 | 1 | 5 | 0.6 | NA | NA | NA | NA | 2 | 0.5 | 1 | 0 | 3 | 0.3333 |
| 1_44206482 | NA | NA | NA | NA | 3 | 0.6667 | 1 | 1 | 4 | 0.75 | NA | NA | NA | NA | 2 | 0.5 | 1 | 0 | 3 | 0.3333 |
| 1_44206634 | NA | NA | 1 | 1 | 3 | 0 | 1 | 0 | 5 | 0.2 | NA | NA | NA | NA | 1 | 0 | 1 | 1 | 2 | 0.5 |
| 1_44206759 | 1 | 1 | 1 | 1 | 1 | 1 | 1 | 1 | 4 | 1 | NA | NA | NA | NA | 2 | 0.5 | 1 | 1 | 3 | 0.6667 |
| 1_44212679 | 1 | 1 | 1 | 0 | 3 | 0 | 2 | 0 | 7 | 0.1429 | NA | NA | NA | NA | 2 | 0 | 1 | 1 | 3 | 0.3333 |
| 1_44219838 | 2 | 0 | NA | NA | 3 | 1 | 2 | 1 | 7 | 0.7143 | NA | NA | 2 | 0.5 | 3 | 0.6667 | 5 | 0.2 | 10 | 0.4 |
| 1_44219901 | 2 | 0.5 | 1 | 0 | 4 | 0.5 | 2 | 0.5 | 9 | 0.4444 | 1 | 0 | 1 | 1 | 3 | 0.6667 | 5 | 0.4 | 10 | 0.5 |
| 1_44220088 | 2 | 0 | 1 | 0 | 4 | 0.25 | 2 | 0 | 9 | 0.1111 | 1 | 0 | 2 | 0 | 3 | 0 | 4 | 0.25 | 10 | 0.1 |
| 1_44220159 | 2 | 0 | NA | NA | 3 | 0.6667 | 1 | 0 | 6 | 0.3333 | 1 | 1 | 2 | 1 | 2 | 1 | 4 | 0.25 | 9 | 0.6667 |
| 1_44220171 | 2 | 0.5 | 1 | 1 | 4 | 0.75 | 2 | 0.5 | 9 | 0.6667 | 2 | 1 | 2 | 0.5 | 1 | 1 | 2 | 0 | 7 | 0.5714 |
| 1_44220200 | 2 | 0 | 1 | 0 | 4 | 0.5 | 1 | 0 | 8 | 0.25 | 1 | 0 | 2 | 0.5 | 3 | 1 | 5 | 0.4 | 11 | 0.5455 |
| 1_44220206 | 2 | 0 | 1 | 0 | 4 | 0.25 | 2 | 0 | 9 | 0.1111 | 2 | 0.5 | 2 | 0 | 3 | 0 | 4 | 0 | 11 | 0.0909 |
| 1_45415184 | NA | NA | NA | NA | NA | NA | 1 | 1 | 1 | 1 | 1 | 1 | NA | NA | NA | NA | NA | NA | 1 | 1 |
| 1_45862578 | 4 | 0.25 | 5 | 0 | 3 | 0 | 7 | 0.2857 | 19 | 0.1579 | 3 | 0 | 3 | 0 | 6 | 0.1667 | 5 | 0.2 | 17 | 0.1176 |
| 1_45947705 | 1 | 0 | 1 | 1 | 1 | 1 | 1 | 0 | 4 | 0.5 | 1 | 1 | 1 | 1 | 1 | 1 | NA | NA | 3 | 1 |
| 1_45947753 | 1 | 0 | 1 | 1 | NA | NA | 1 | 0 | 3 | 0.3333 | 1 | 1 | 1 | 0 | 1 | 0 | NA | NA | 3 | 0.3333 |
| 1_45947865 | NA | NA | 1 | 1 | 1 | 0 | 1 | 0 | 3 | 0.3333 | 1 | 1 | 1 | 0 | 1 | 0 | NA | NA | 3 | 0.3333 |
| 1_45948680 | NA | NA | NA | NA | 1 | 1 | 1 | 0 | 2 | 0.5 | 1 | 1 | NA | NA | 1 | 1 | NA | NA | 2 | 1 |
| 1_45948783 | NA | NA | 1 | 1 | 1 | 1 | NA | NA | 2 | 1 | 1 | 0 | NA | NA | 1 | 1 | NA | NA | 2 | 0.5 |
| 1_45950320 | NA | NA | NA | NA | 1 | 0 | 1 | 1 | 2 | 0.5 | 1 | 1 | NA | NA | 1 | 0 | NA | NA | 2 | 0.5 |
| 1_45950337 | NA | NA | NA | NA | 1 | 1 | 1 | 1 | 2 | 1 | 1 | 1 | NA | NA | 1 | 1 | NA | NA | 2 | 1 |
| 1_45950382 | NA | NA | NA | NA | 1 | 1 | 1 | 1 | 2 | 1 | 1 | 1 | NA | NA | NA | NA | NA | NA | 1 | 1 |
| 1_45950477 | NA | NA | NA | NA | 1 | 1 | 1 | 0 | 2 | 0.5 | 1 | 1 | NA | NA | 1 | 1 | NA | NA | 2 | 1 |
| 1_45950626 | NA | NA | NA | NA | 1 | 1 | 1 | 0 | 2 | 0.5 | 1 | 1 | NA | NA | 1 | 0 | NA | NA | 2 | 0.5 |
| 1_46848763 | 6 | 0.6667 | 10 | 0.8 | 15 | 0.5333 | 9 | 0.5556 | 40 | 0.625 | 8 | 0.875 | 14 | 0.5714 | 15 | 0.6667 | 7 | 0.7143 | 44 | 0.6818 |
| 1_46848903 | 5 | 0.2 | 13 | 0.2308 | 15 | 0.0667 | 11 | 0 | 44 | 0.1136 | 7 | 0.1429 | 13 | 0 | 17 | 0.1176 | 7 | 0.1429 | 44 | 0.0909 |
| 1_46848974 | 6 | 0.8333 | 9 | 0.8889 | 15 | 0.9333 | 12 | 0.9167 | 42 | 0.9048 | 8 | 0.625 | 12 | 0.9167 | 15 | 0.9333 | 6 | 1 | 41 | 0.878 |
| 1_46849188 | 4 | 1 | 13 | 0.9231 | 10 | 0.9 | 10 | 0.4 | 37 | 0.7838 | 9 | 0.6667 | 13 | 0.9231 | 12 | 0.6667 | 5 | 0.6 | 39 | 0.7436 |
| 1_46858352 | 4 | 0.5 | 11 | 0.0909 | 12 | 0.0833 | 8 | 0.125 | 35 | 0.1429 | 7 | 0 | 11 | 0.1818 | 16 | 0.3125 | 3 | 0.3333 | 37 | 0.2162 |
| 1_46858454 | 3 | 0.6667 | 9 | 0.7778 | 10 | 0.8 | 2 | 1 | 24 | 0.7917 | 6 | 0.8333 | 6 | 1 | 14 | 0.8571 | 3 | 0.6667 | 29 | 0.8621 |
| 1_46858473 | 4 | 0.25 | 10 | 0.3 | 9 | 0.5556 | 6 | 0.6667 | 29 | 0.4483 | 6 | 0.8333 | 7 | 0.2857 | 10 | 0.3 | 2 | 0.5 | 25 | 0.44 |
| 1_46865334 | 3 | 0.6667 | 11 | 0.0909 | 9 | 0.2222 | 9 | 0.1111 | 32 | 0.1875 | 7 | 0 | 10 | 0 | 14 | 0.1429 | 3 | 0 | 34 | 0.0588 |
| 1_46871798 | 3 | 0 | 8 | 0.25 | 9 | 0.2222 | 5 | 0.6 | 25 | 0.28 | 5 | 0.2 | 8 | 0.25 | 14 | 0.2857 | 3 | 0 | 30 | 0.2333 |
| 1_46871810 | 3 | 0 | 10 | 0.2 | 12 | 0.0833 | 9 | 0.2222 | 34 | 0.1471 | 5 | 0 | 9 | 0.1111 | 16 | 0.125 | 3 | 0 | 33 | 0.0909 |
| 1_46874389 | 3 | 0 | 10 | 0.1 | 11 | 0.0909 | 6 | 0.1667 | 30 | 0.1 | 3 | 0 | 10 | 0.2 | 13 | 0 | 2 | 0 | 28 | 0.0714 |
| 1_46874413 | 1 | 1 | 10 | 0.7 | 10 | 0.6 | 7 | 0.5714 | 28 | 0.6429 | 3 | 0.6667 | 7 | 0.8571 | 9 | 0.6667 | 1 | 0 | 20 | 0.7 |
| 1_46874422 | 3 | 0.6667 | 10 | 0.4 | 11 | 0.0909 | 8 | 0.375 | 32 | 0.3125 | 3 | 0.3333 | 9 | 0.4444 | 12 | 0.5 | 2 | 0.5 | 26 | 0.4615 |
| 1_46874497 | 3 | 0.3333 | 9 | 0.1111 | 9 | 0.1111 | 8 | 0.125 | 29 | 0.1379 | 4 | 0 | 7 | 0.1429 | 11 | 0.5455 | 3 | 0.3333 | 25 | 0.32 |
| 1_46874524 | 2 | 0.5 | 6 | 0.1667 | 6 | 0.5 | 5 | 0.6 | 19 | 0.4211 | 3 | 0.6667 | 7 | 0.5714 | 8 | 0.375 | 2 | 0.5 | 20 | 0.5 |
| 1_46874652 | 3 | 0.6667 | 10 | 0.1 | 9 | 0.1111 | 7 | 0.1429 | 29 | 0.1724 | 4 | 1 | 9 | 0.4444 | 13 | 0.3846 | 3 | 0.3333 | 29 | 0.4828 |
| 1_46874705 | 3 | 0 | 10 | 0 | 9 | 0.1111 | 7 | 0.2857 | 29 | 0.1034 | 4 | 0.25 | 7 | 0 | 10 | 0.3 | 2 | 0 | 23 | 0.1739 |
| 1_46874963 | 2 | 0 | 8 | 0.5 | 5 | 0.2 | 6 | 0.8333 | 21 | 0.4762 | 2 | 0.5 | 7 | 0.4286 | 9 | 0.3333 | 3 | 0.3333 | 21 | 0.381 |
| 1_46874993 | 2 | 1 | 9 | 0.6667 | 9 | 0.5556 | 6 | 0.6667 | 26 | 0.6538 | 3 | 1 | 7 | 0.5714 | 11 | 0.6364 | 3 | 1 | 24 | 0.7083 |
| 1_46875038 | 3 | 0 | 10 | 0.1 | 9 | 0.2222 | 6 | 0.1667 | 28 | 0.1429 | 3 | 0.3333 | 7 | 0.1429 | 12 | 0.3333 | 3 | 0 | 25 | 0.24 |
| 1_46875049 | 3 | 0.3333 | 9 | 0.7778 | 8 | 0.75 | 4 | 0.5 | 24 | 0.6667 | 3 | 0.6667 | 7 | 0.4286 | 11 | 0.6364 | 3 | 0.6667 | 24 | 0.5833 |
| 1_46875103 | 1 | 1 | 6 | 0.8333 | 3 | 0.6667 | 4 | 0.5 | 14 | 0.7143 | NA | NA | 6 | 1 | 8 | 1 | 2 | 1 | 16 | 1 |
| 1_46875104 | NA | NA | 1 | 1 | 1 | 1 | 2 | 0.5 | 4 | 0.75 | NA | NA | 1 | 1 | 1 | 1 | NA | NA | 2 | 1 |
| 1_46875255 | 2 | 0 | 9 | 0.4444 | 9 | 0.1111 | 4 | 0.25 | 24 | 0.25 | 1 | 0 | 7 | 0.2857 | 11 | 0.1818 | 2 | 0.5 | 21 | 0.2381 |
| 1_50966887 | 2 | 0 | 3 | 0.3333 | 6 | 0.3333 | 5 | 0.2 | 16 | 0.25 | 2 | 0 | 10 | 0.3 | 8 | 0.375 | 7 | 0.1429 | 27 | 0.2593 |
| 1_50967146 | 2 | 0.5 | 4 | 0.5 | 5 | 0.4 | 3 | 0.3333 | 14 | 0.4286 | 2 | 0.5 | 4 | 1 | 6 | 0.5 | 7 | 0.5714 | 19 | 0.6316 |
| 1_51224682 | 6 | 0.6667 | 11 | 0.6364 | 9 | 0.3333 | 9 | 0.6667 | 35 | 0.5714 | 3 | 1 | 10 | 0.6 | 8 | 0.625 | 10 | 0.4 | 31 | 0.5806 |
| 1_51224704 | 9 | 0.1111 | 12 | 0.1667 | 10 | 0.3 | 9 | 0.1111 | 40 | 0.175 | 3 | 0 | 10 | 0.1 | 11 | 0 | 10 | 0.4 | 34 | 0.1471 |
| 1_51340289 | 1 | 1 | 1 | 0 | NA | NA | NA | NA | 2 | 0.5 | 1 | 1 | 1 | 1 | 3 | 1 | NA | NA | 5 | 1 |
| 1_51340527 | 2 | 0 | 1 | 1 | NA | NA | NA | NA | 3 | 0.3333 | 1 | 0 | 1 | 0 | 3 | 0.3333 | NA | NA | 5 | 0.2 |
| 1_51340666 | 2 | 1 | 1 | 1 | NA | NA | 1 | 1 | 4 | 1 | 1 | 1 | NA | NA | 3 | 1 | NA | NA | 4 | 1 |
| 1_51340688 | 2 | 1 | 1 | 0 | NA | NA | 2 | 0.5 | 5 | 0.6 | 1 | 0 | 1 | 1 | 3 | 0 | NA | NA | 5 | 0.2 |
| 1_51340723 | 2 | 1 | NA | NA | NA | NA | 1 | 1 | 3 | 1 | NA | NA | 1 | 1 | 4 | 0.75 | NA | NA | 5 | 0.8 |
| 1_51340851 | 2 | 0.5 | 1 | 1 | NA | NA | 1 | 0 | 4 | 0.5 | 1 | 1 | NA | NA | 3 | 0.3333 | NA | NA | 4 | 0.5 |
| 1_51341069 | 2 | 0.5 | 1 | 0 | NA | NA | 2 | 0 | 5 | 0.2 | 1 | 0 | 1 | 1 | 4 | 0 | NA | NA | 6 | 0.1667 |

| | | | | | | | | | | | | | | | | | | | | |
|---|---|---|---|---|---|---|---|---|---|---|---|---|---|---|---|---|---|---|---|---|
| 1_5134115 | 1 | 1 | 1 | 1 | NA | NA | 2 | 0.5 | 4 | 0.75 | NA | NA | 1 | 0 | 3 | 1 | NA | NA | 4 | 0.75 |
| 1_5134185 | 2 | 0 | 2 | 0 | 1 | 0 | 3 | 0.3333 | 8 | 0.125 | 1 | 0 | NA | NA | 4 | 0.25 | 2 | 0 | 7 | 0.1429 |
| 1_5134187 | NA | NA | 2 | 1 | NA | NA | 2 | 1 | 4 | 1 | 1 | 1 | NA | NA | 2 | 0.5 | 2 | 1 | 5 | 0.8 |
| 1_5134195 | 1 | 0 | 2 | 0 | 1 | 1 | 3 | 0.3333 | 7 | 0.2857 | 1 | 1 | NA | NA | 2 | 0.5 | 2 | 1 | 5 | 0.8 |
| 1_5151035 | NA | NA | 2 | 0.5 | 1 | 0 | 2 | 0.5 | 5 | 0.4 | NA | NA | NA | NA | 1 | 1 | 1 | 0 | 2 | 0.5 |
| 1_5179030 | 3 | 0.3333 | 2 | 0 | 5 | 0.4 | 2 | 0 | 12 | 0.25 | 1 | 1 | 5 | 0.6 | 6 | 0.5 | 2 | 0 | 14 | 0.5 |
| 1_5219862 | 3 | 1 | 1 | 1 | NA | NA | 2 | 0.5 | 6 | 0.8333 | 2 | 1 | NA | NA | NA | NA | 3 | 1 | 5 | 1 |
| 1_5219872 | 3 | 0.3333 | 1 | 0 | NA | NA | 2 | 0.5 | 6 | 0.3333 | 3 | 0.6667 | NA | NA | 1 | 1 | 3 | 0.6667 | 7 | 0.7143 |
| 1_5219881 | 3 | 0.6667 | NA | NA | NA | NA | 2 | 0.5 | 5 | 0.6 | 3 | 0.3333 | NA | NA | NA | NA | 2 | 1 | 5 | 0.6 |
| 1_5219905 | 2 | 0.5 | 2 | 0 | NA | NA | NA | NA | 4 | 0.25 | 2 | 0 | NA | NA | 2 | 0.5 | 3 | 0.6667 | 7 | 0.4286 |
| 1_5220279 | 18 | 0.0556 | 36 | 0.0278 | 23 | 0.0435 | 16 | 0.125 | 93 | 0.0538 | 20 | 0.1 | 15 | 0.0667 | 20 | 0.05 | 24 | 0.1667 | 79 | 0.1013 |
| 1_5220306 | 16 | 0.0625 | 30 | 0.0333 | 19 | 0.1053 | 14 | 0.1429 | 79 | 0.0759 | 20 | 0.1 | 13 | 0.0769 | 18 | 0.0556 | 25 | 0.12 | 76 | 0.0921 |
| 1_5220375 | 8 | 0.125 | 22 | 0.1364 | 14 | 0.1429 | 7 | 0 | 51 | 0.1176 | 13 | 0.0769 | 9 | 0.1111 | 10 | 0.1 | 19 | 0.1053 | 51 | 0.098 |
| 1_5220490 | 6 | 0 | 17 | 0 | 8 | 0.125 | 5 | 0.2 | 36 | 0.0556 | 9 | 0 | 4 | 0 | 8 | 0.125 | 18 | 0.0556 | 39 | 0.0513 |
| 1_5220504 | 5 | 0.2 | 17 | 0.2353 | 7 | 0.1429 | 4 | 0.25 | 33 | 0.2121 | 6 | 0.3333 | 4 | 0.25 | 8 | 0.25 | 16 | 0.1875 | 34 | 0.2353 |
| 1_5220526 | 4 | 0 | 13 | 0.1538 | 9 | 0.1111 | 4 | 0 | 30 | 0.1 | 6 | 0.1667 | 3 | 0.3333 | 7 | 0.1429 | 13 | 0.0769 | 29 | 0.1379 |
| 1_5220527 | 3 | 0 | 12 | 0.0833 | 7 | 0.1429 | 4 | 0 | 26 | 0.0769 | 5 | 0 | 3 | 0.3333 | 6 | 0 | 10 | 0.1 | 24 | 0.0833 |
| 1_5220530 | 3 | 0 | 12 | 0.1667 | 9 | 0.3333 | 3 | 1 | 27 | 0.2963 | 5 | 0.4 | 2 | 0.5 | 6 | 0.5 | 13 | 0.3077 | 26 | 0.3846 |
| 1_5220534 | 4 | 0.5 | 10 | 0.4 | 3 | 0.6667 | 3 | 0.3333 | 20 | 0.45 | 3 | 0.3333 | 3 | 0.3333 | 7 | 0.4286 | 11 | 0.0909 | 24 | 0.25 |
| 1_5220543 | 4 | 0 | 14 | 0.2143 | 8 | 0.25 | 4 | 0 | 30 | 0.1667 | 4 | 0 | 5 | 0.2 | 7 | 0.4286 | 14 | 0.1429 | 30 | 0.2 |
| 1_5222519 | 4 | 0.25 | 2 | 0 | 7 | 0 | 3 | 0 | 16 | 0.0625 | 3 | 0 | 2 | 0 | 3 | 0 | 9 | 0.2222 | 17 | 0.1176 |
| 1_5222522 | 4 | 0 | 1 | 0 | 4 | 0 | 3 | 0.6667 | 12 | 0.1667 | 3 | 0 | 3 | 0.3333 | 4 | 0.25 | 5 | 0.4 | 15 | 0.2667 |
| 1_5222691 | 2 | 1 | 1 | 0 | 6 | 0.1667 | 2 | 0.5 | 11 | 0.3636 | 3 | 0 | 4 | 0 | 3 | 0.3333 | 7 | 0.1429 | 17 | 0.1176 |
| 1_5222693 | 2 | 0.5 | 1 | 0 | 5 | 0.4 | 2 | 0.5 | 10 | 0.4 | 3 | 0.3333 | 4 | 0.25 | 1 | 1 | 6 | 0.5 | 14 | 0.4286 |
| 1_5222694 | 2 | 0 | 1 | 0 | 3 | 0.3333 | 1 | 0 | 7 | 0.1429 | 3 | 0 | 4 | 0 | 2 | 0 | 7 | 0.1429 | 16 | 0.0625 |
| 1_5227225 | 2 | 0.5 | 5 | 0.6 | 1 | 0 | 1 | 0 | 9 | 0.4444 | 2 | 1 | 2 | 1 | 1 | 0 | 4 | 1 | 9 | 0.8889 |
| 1_5249822 | 2 | 0 | 5 | 0.6 | 1 | 1 | 1 | 1 | 9 | 0.5556 | 4 | 0.5 | 3 | 0.3333 | NA | NA | 2 | 0.5 | 9 | 0.4444 |
| 1_5249827 | 2 | 0.5 | 5 | 0.4 | 1 | 0 | 1 | 0 | 9 | 0.3333 | 4 | 0.25 | 3 | 0 | NA | NA | 2 | 0 | 9 | 0.1111 |
| 1_5249844 | 2 | 0.5 | 4 | 0.25 | 1 | 0 | 1 | 0 | 8 | 0.25 | 4 | 0.5 | 4 | 0 | 1 | 0 | 1 | 1 | 10 | 0.3 |
| 1_5252925 | 2 | 0.5 | 5 | 1 | 1 | 0 | NA | NA | 8 | 0.75 | 2 | 0.5 | 3 | 0.6667 | NA | NA | NA | NA | 5 | 0.6 |
| 1_5252941 | 2 | 0 | 4 | 0.75 | 1 | 0 | NA | NA | 7 | 0.4286 | 3 | 0.6667 | 3 | 0.6667 | NA | NA | NA | NA | 6 | 0.6667 |
| 1_5252942 | 2 | 0 | 4 | 0.5 | 1 | 0 | NA | NA | 7 | 0.2857 | 3 | 0.3333 | 3 | 0.3333 | NA | NA | NA | NA | 6 | 0.3333 |
| 1_5252963 | 2 | 1 | 4 | 0.5 | 1 | 1 | NA | NA | 7 | 0.7143 | 3 | 1 | 1 | 1 | NA | NA | NA | NA | 4 | 1 |
| 1_5252967 | 2 | 1 | 3 | 1 | 1 | 1 | NA | NA | 6 | 1 | 3 | 1 | 2 | 0 | NA | NA | NA | NA | 5 | 0.6 |
| 1_5252987 | 2 | 0 | 4 | 0.5 | 1 | 0 | NA | NA | 7 | 0.2857 | 3 | 0 | 1 | 1 | NA | NA | NA | NA | 4 | 0.25 |
| 1_5268906 | NA | NA | NA | NA | NA | NA | 2 | 0.5 | 2 | 0.5 | 1 | 1 | NA | NA | NA | NA | NA | NA | 1 | 1 |
| 1_5268910 | NA | NA | NA | NA | NA | NA | 2 | 1 | 2 | 1 | 1 | 1 | NA | NA | NA | NA | NA | NA | 1 | 1 |
| 1_5268922 | NA | NA | NA | NA | NA | NA | 2 | 0.5 | 2 | 0.5 | 1 | 1 | NA | NA | NA | NA | NA | NA | 1 | 1 |
| 1_5268923 | NA | NA | NA | NA | NA | NA | 1 | 1 | 1 | 1 | 1 | 1 | NA | NA | NA | NA | NA | NA | 1 | 1 |
| 1_5269553 | 9 | 0.1111 | 19 | 0 | 16 | 0.125 | 7 | 0 | 51 | 0.0588 | 8 | 0 | 8 | 0 | 7 | 0.2857 | 5 | 0 | 28 | 0.0714 |
| 1_5269582 | 8 | 0 | 16 | 0.25 | 15 | 0 | 6 | 0.3333 | 45 | 0.1333 | 8 | 0.25 | 6 | 0.5 | 7 | 0.1429 | 6 | 0.3333 | 27 | 0.2963 |
| 1_5269593 | 6 | 0.1667 | 20 | 0.1 | 16 | 0 | 6 | 0.1667 | 48 | 0.0833 | 6 | 0.1667 | 5 | 0.2 | 6 | 0 | 6 | 0.1667 | 23 | 0.1304 |
| 1_5269600 | 7 | 0.1429 | 16 | 0.1875 | 13 | 0.0769 | 5 | 0.6 | 41 | 0.1951 | 5 | 0 | 6 | 0 | 2 | 0.5 | 3 | 0.6667 | 16 | 0.1875 |
| 1_5269617 | 6 | 0 | 17 | 0.3529 | 15 | 0.4 | 4 | 0.5 | 42 | 0.3333 | 7 | 0.4286 | 6 | 0.5 | 5 | 0.6 | 5 | 0.4 | 23 | 0.4783 |
| 1_5269629 | 6 | 0 | 16 | 0 | 16 | 0.125 | 5 | 0.4 | 43 | 0.093 | 6 | 0.3333 | 2 | 0 | 4 | 0.25 | 5 | 0 | 17 | 0.1765 |
| 1_5269635 | 5 | 0.2 | 13 | 0.0769 | 15 | 0.0667 | 3 | 0 | 36 | 0.0833 | 5 | 0 | 4 | 0 | 4 | 0.25 | 3 | 0.3333 | 16 | 0.125 |
| 1_5269641 | 6 | 0.1667 | 17 | 0.2941 | 16 | 0 | 4 | 0.25 | 43 | 0.1628 | 6 | 0.1667 | 4 | 0 | 5 | 0 | 5 | 0.2 | 20 | 0.1 |
| 1_5269673 | 5 | 0 | 11 | 0.0909 | 14 | 0.0714 | 5 | 0 | 35 | 0.0571 | 5 | 0.2 | 3 | 0.3333 | 4 | 0 | 4 | 0.25 | 16 | 0.1875 |
| 1_5269698 | 3 | 0 | 10 | 0.3 | 12 | 0.1667 | 5 | 0.2 | 30 | 0.2 | 5 | 0.2 | 1 | 0 | 4 | 0.25 | 4 | 0 | 14 | 0.1429 |
| 1_5269709 | 3 | 0.3333 | 11 | 0.5455 | 12 | 0.5 | 5 | 0.4 | 31 | 0.4839 | 4 | 0.25 | 2 | 1 | 4 | 0.75 | 4 | 0.5 | 14 | 0.5714 |
| 1_5269713 | 2 | 0 | 9 | 0.2222 | 10 | 0.2 | 3 | 0 | 24 | 0.1667 | 2 | 0 | 3 | 0.6667 | 4 | 0 | 4 | 0.25 | 13 | 0.2308 |
| 1_5269723 | 3 | 0 | 8 | 0.25 | 13 | 0.1538 | 5 | 0 | 29 | 0.1379 | 4 | 0 | 3 | 0.3333 | 3 | 0.3333 | 4 | 0 | 14 | 0.1429 |
| 1_5269754 | 2 | 0.5 | 9 | 0.4444 | 11 | 0.3636 | 4 | 0.75 | 26 | 0.4615 | 2 | 1 | 1 | 1 | 3 | 0 | 3 | 0.6667 | 9 | 0.5556 |
| 1_5274742 | 3 | 0.6667 | 5 | 0.8 | 6 | 0.8333 | 4 | 0.25 | 18 | 0.6667 | 3 | 1 | NA | NA | 1 | 1 | 1 | 0 | 5 | 0.8 |
| 1_5274754 | 3 | 1 | 6 | 0.8333 | 4 | 0.75 | 3 | 0.6667 | 16 | 0.8125 | 3 | 1 | 2 | 1 | 1 | 1 | 3 | 1 | 9 | 1 |
| 1_5274758 | 3 | 0 | 6 | 0.1667 | 5 | 0.8 | 3 | 0.6667 | 17 | 0.4118 | 3 | 0.6667 | 2 | 1 | 1 | 0 | 3 | 0 | 9 | 0.4444 |
| 1_5274761 | 3 | 0.3333 | 6 | 0.5 | 6 | 0.3333 | 4 | 0.5 | 19 | 0.4211 | 3 | 0 | 1 | 1 | 2 | 0 | 3 | 0.6667 | 9 | 0.3333 |
| 1_5274775 | 2 | 0 | 5 | 0.6 | 4 | 0.5 | 4 | 0 | 15 | 0.3333 | 2 | 1 | 1 | 0 | 2 | 0.5 | 3 | 0.6667 | 8 | 0.625 |
| 1_5274811 | 2 | 1 | 5 | 0.8 | 5 | 1 | 3 | 1 | 15 | 0.9333 | 3 | 1 | 1 | 1 | 2 | 1 | 2 | 1 | 8 | 1 |
| 1_5274816 | 2 | 0 | 6 | 0 | 5 | 0.2 | 4 | 0 | 17 | 0.0588 | 3 | 0.3333 | 1 | 0 | 2 | 0 | 2 | 0 | 8 | 0.125 |
| 1_5274831 | 2 | 0.5 | 4 | 0.5 | 2 | 0.5 | 1 | 1 | 9 | 0.5556 | 3 | 1 | NA | NA | 2 | 0 | 2 | 1 | 7 | 0.7143 |
| 1_5274833 | 2 | 1 | 5 | 0.2 | 3 | 0.3333 | 3 | 1 | 13 | 0.5385 | 3 | 1 | NA | NA | 1 | 0 | 2 | 0 | 6 | 0.5 |
| 1_5274854 | 2 | 1 | 6 | 1 | 4 | 1 | 4 | 1 | 16 | 1 | 3 | 1 | 1 | 1 | 1 | 1 | 2 | 1 | 7 | 1 |
| 1_5274857 | 1 | 0 | 6 | 0.1667 | 3 | 0 | 3 | 0 | 13 | 0.0769 | 2 | 0 | NA | NA | 2 | 0 | 2 | 0.5 | 6 | 0.1667 |
| 1_5274868 | 2 | 0.5 | 6 | 0 | 3 | 0 | 3 | 0 | 14 | 0.0714 | 3 | 0 | 1 | 1 | 2 | 0.5 | 2 | 0 | 8 | 0.25 |

| | | | | | | | | | | | | | | | | | | | | |
|---|---|---|---|---|---|---|---|---|---|---|---|---|---|---|---|---|---|---|---|---|
| 1_5274869 | 3 | 1 | 6 | 0.8333 | 4 | 0.75 | 3 | 1 | 16 | 0.875 | 3 | 1 | 1 | 1 | 2 | 1 | 2 | 1 | 8 | 1 |
| 1_5274872 | 2 | 0.5 | 6 | 0.5 | 3 | 0 | 1 | 0 | 12 | 0.3333 | 3 | 0 | NA | NA | 2 | 1 | 2 | 0.5 | 7 | 0.4286 |
| 1_5282530 | 2 | 0.5 | 1 | 0 | 2 | 0.5 | 3 | 0.3333 | 8 | 0.375 | NA | NA | 1 | 0 | 1 | 1 | 1 | 0 | 3 | 0.3333 |
| 1_5282872 | 4 | 0.25 | 2 | 0 | NA | NA | 4 | 0.5 | 10 | 0.3 | 2 | 0.5 | 2 | 0 | 2 | 0.5 | 2 | 0 | 8 | 0.25 |
| 1_5282880 | 5 | 0.8 | 2 | 0.5 | 1 | 1 | 4 | 0.75 | 12 | 0.75 | 2 | 1 | 2 | 0.5 | 3 | 0.6667 | 1 | 1 | 8 | 0.75 |
| 1_5282882 | 4 | 1 | 2 | 1 | 1 | 1 | 1 | 1 | 8 | 1 | 1 | 1 | 2 | 1 | 3 | 1 | 2 | 1 | 8 | 1 |
| 1_5282883 | 4 | 0.5 | 2 | 0 | 1 | 1 | 5 | 0 | 12 | 0.25 | 2 | 0.5 | 3 | 0.3333 | 3 | 0 | 2 | 0 | 10 | 0.2 |
| 1_5282888 | 4 | 0.25 | 2 | 0 | 1 | 1 | 5 | 0 | 12 | 0.1667 | 2 | 0 | 3 | 0.3333 | 3 | 0 | 2 | 0 | 10 | 0.1 |
| 1_5282893 | 4 | 1 | 3 | 1 | 1 | 1 | 3 | 1 | 11 | 1 | 1 | 1 | 2 | 0.5 | 2 | 1 | 2 | 1 | 7 | 0.8571 |
| 1_5282899 | 5 | 0.2 | 3 | 0 | 1 | 0 | 3 | 0.3333 | 12 | 0.1667 | 2 | 0 | 3 | 0 | 3 | 0.3333 | 1 | 0 | 9 | 0.1111 |
| 1_5282902 | 5 | 0.2 | 2 | 0.5 | 1 | 0 | 3 | 0.3333 | 11 | 0.2727 | 2 | 1 | 3 | 0.3333 | 3 | 0.3333 | 1 | 0 | 9 | 0.4444 |
| 1_5282908 | 4 | 0.75 | 2 | 1 | 1 | 1 | 4 | 1 | 11 | 0.9091 | 2 | 1 | 2 | 1 | 2 | 1 | 2 | 1 | 8 | 1 |
| 1_5282909 | 4 | 0.5 | 3 | 0.6667 | 1 | 1 | 3 | 1 | 11 | 0.7273 | 1 | 1 | 1 | 0 | 2 | 0 | 2 | 0.5 | 6 | 0.3333 |
| 1_5282911 | 4 | 0.5 | 3 | 0 | 1 | 0 | 4 | 0.25 | 12 | 0.25 | 2 | 0 | 2 | 0 | 3 | 1 | 1 | 1 | 8 | 0.5 |
| 1_5282921 | 4 | 0 | 3 | 0.3333 | 1 | 0 | 3 | 0.6667 | 11 | 0.2727 | 1 | 1 | 2 | 1 | 2 | 0.5 | 2 | 0.5 | 7 | 0.7143 |
| 1_5282926 | 5 | 0.6 | 3 | 0.6667 | 1 | 1 | 4 | 0.5 | 13 | 0.6154 | 2 | 0 | 2 | 1 | 3 | 0.3333 | 2 | 0.5 | 9 | 0.4444 |
| 1_5282928 | 5 | 0 | 3 | 0.3333 | 1 | 1 | 4 | 0.25 | 13 | 0.2308 | 2 | 0.5 | 2 | 0 | 3 | 0.3333 | 2 | 1 | 9 | 0.4444 |
| 1_5295948 | NA | NA | 2 | 0 | 1 | 0 | 2 | 0.5 | 5 | 0.2 | NA | NA | 1 | 1 | 4 | 0 | 3 | 0 | 8 | 0.125 |
| 1_5295965 | NA | NA | 2 | 1 | 1 | 1 | 1 | 1 | 4 | 1 | NA | NA | 1 | 1 | 4 | 0.75 | 3 | 0.6667 | 8 | 0.75 |
| 1_5295981 | NA | NA | 1 | 0 | 1 | 1 | 3 | 0.3333 | 5 | 0.4 | NA | NA | 1 | 0 | 4 | 0.5 | 2 | 0.5 | 7 | 0.4286 |
| 1_5296208 | NA | NA | 3 | 0 | 6 | 0.3333 | NA | NA | 9 | 0.2222 | 4 | 0 | 3 | 0.3333 | 1 | 0 | NA | NA | 8 | 0.125 |
| 1_5296242 | NA | NA | 2 | 0.5 | 4 | 0 | NA | NA | 6 | 0.1667 | 2 | 0 | 2 | 0 | 1 | 1 | NA | NA | 5 | 0.2 |
| 1_5296457 | 2 | 1 | 2 | 0 | 4 | 0.25 | NA | NA | 8 | 0.375 | 1 | 0 | 1 | 1 | NA | NA | NA | NA | 2 | 0.5 |
| 1_5296467 | 2 | 1 | 2 | 1 | 3 | 1 | NA | NA | 7 | 1 | 1 | 1 | 1 | 1 | NA | NA | NA | NA | 2 | 1 |
| 1_5296486 | 2 | 0.5 | 2 | 1 | 4 | 0.75 | NA | NA | 8 | 0.75 | 1 | 1 | NA | NA | NA | NA | NA | NA | 1 | 1 |
| 1_5322850 | 1 | 0 | 1 | 1 | 1 | 0 | NA | NA | 3 | 0.3333 | 1 | 0 | NA | NA | NA | NA | 1 | 1 | 2 | 0.5 |
| 1_5324595 | 1 | 1 | NA | NA | NA | NA | NA | NA | 1 | 1 | 1 | 1 | NA | NA | NA | NA | 1 | 1 | 2 | 1 |
| 1_5324622 | 1 | 1 | NA | NA | NA | NA | NA | NA | 1 | 1 | 1 | 1 | NA | NA | NA | NA | 1 | 1 | 2 | 1 |
| 1_5324626 | 1 | 1 | NA | NA | NA | NA | NA | NA | 1 | 1 | 1 | 1 | NA | NA | NA | NA | 1 | 0 | 2 | 0.5 |
| 1_5334814 | 3 | 0.3333 | NA | NA | 2 | 0 | 1 | 1 | 6 | 0.3333 | 2 | 0 | 4 | 0 | 2 | 0.5 | NA | NA | 8 | 0.125 |
| 1_5334818 | 3 | 0.3333 | NA | NA | 2 | 0.5 | 2 | 0 | 7 | 0.2857 | 1 | 0 | 4 | 0.25 | 3 | 0.3333 | 1 | 0 | 9 | 0.2222 |
| 1_5335719 | 2 | 1 | NA | NA | 3 | 0.6667 | 3 | 1 | 8 | 0.875 | NA | NA | 4 | 0.75 | NA | NA | NA | NA | 4 | 0.75 |
| 1_5335729 | 2 | 0 | 1 | 0 | 4 | 0 | 2 | 0.5 | 9 | 0.1111 | NA | NA | 4 | 0.25 | NA | NA | NA | NA | 4 | 0.25 |
| 1_5335925 | 2 | 0.5 | 1 | 1 | 2 | 0.5 | 3 | 1 | 8 | 0.75 | 1 | 1 | 3 | 0.6667 | NA | NA | NA | NA | 4 | 0.75 |
| 1_5338394 | 6 | 0 | 5 | 0.2 | 5 | 0 | 3 | 0 | 19 | 0.0526 | 3 | 0.6667 | 4 | 0 | 1 | 0 | 1 | 0 | 9 | 0.2222 |
| 1_5338401 | 6 | 0.3333 | 4 | 0 | 4 | 0.5 | 3 | 0 | 17 | 0.2353 | 4 | 0 | 4 | 0.25 | 1 | 1 | 3 | 0 | 12 | 0.1667 |
| 1_5338628 | 5 | 0.8 | 3 | 0 | 2 | 0.5 | 4 | 0.75 | 14 | 0.5714 | 3 | 0.6667 | 4 | 0.75 | 1 | 0 | 2 | 0.5 | 10 | 0.6 |
| 1_5338633 | 4 | 0.5 | 2 | 0 | 2 | 0 | 4 | 0 | 12 | 0.1667 | 3 | 0.3333 | 3 | 0 | NA | NA | 3 | 0 | 9 | 0.1111 |
| 1_5338636 | 5 | 0.4 | 2 | 0.5 | 2 | 0.5 | 4 | 0.5 | 13 | 0.4615 | 3 | 0.3333 | 3 | 0 | 1 | 0 | 3 | 0.3333 | 10 | 0.2 |
| 1_5338640 | 4 | 0 | 3 | 0.3333 | 2 | 0 | 4 | 0.25 | 13 | 0.1538 | 3 | 0.6667 | 4 | 0 | 1 | 0 | 1 | 0 | 9 | 0.2222 |
| 1_5338668 | 2 | 0 | 3 | 0 | 2 | 0 | 4 | 0.5 | 11 | 0.1818 | 2 | 0.5 | 2 | 0.5 | NA | NA | 3 | 0 | 7 | 0.2857 |
| 1_5338682 | 3 | 1 | 3 | 1 | 2 | 1 | 4 | 0.5 | 12 | 0.8333 | 3 | 1 | 4 | 1 | NA | NA | 2 | 0.5 | 9 | 0.8889 |
| 1_5338698 | 3 | 0.3333 | 3 | 0 | 2 | 0.5 | 4 | 0 | 12 | 0.1667 | 3 | 0 | 4 | 0.25 | NA | NA | 3 | 0 | 10 | 0.1 |
| 1_5338721 | 3 | 0.6667 | 3 | 0 | 2 | 0.5 | 3 | 0.3333 | 11 | 0.3636 | 3 | 0.6667 | 4 | 0.25 | NA | NA | 2 | 0.5 | 9 | 0.4444 |
| 1_5338744 | 2 | 0 | 1 | 0 | 2 | 0.5 | 4 | 0.5 | 9 | 0.3333 | 3 | 0.3333 | 2 | 0.5 | NA | NA | 2 | 0 | 7 | 0.2857 |
| 1_5379479 | 1 | 0 | 2 | 0 | NA | NA | 1 | 1 | 4 | 0.25 | NA | NA | 1 | 1 | NA | NA | NA | NA | 1 | 1 |
| 1_5379652 | 3 | 0 | 1 | 0 | 1 | 0 | 2 | 0.5 | 7 | 0.1429 | 1 | 0 | 2 | 0.5 | NA | NA | NA | NA | 3 | 0.3333 |
| 1_5379662 | 2 | 1 | 1 | 1 | 2 | 1 | 2 | 0.5 | 7 | 0.8571 | 1 | 1 | 2 | 1 | 1 | 1 | NA | NA | 4 | 1 |
| 1_5379667 | 3 | 1 | 2 | 1 | 1 | 1 | 2 | 1 | 8 | 1 | 1 | 1 | 1 | 1 | 1 | 1 | NA | NA | 3 | 1 |
| 1_5385085 | 3 | 0 | 1 | 0 | 3 | 0.3333 | 4 | 0 | 11 | 0.0909 | 5 | 0.4 | 6 | 0.1667 | 1 | 0 | 1 | 0 | 13 | 0.2308 |
| 1_5385129 | 2 | 0 | 1 | 0 | 1 | 0 | 3 | 0.3333 | 7 | 0.1429 | 4 | 0.25 | 5 | 0 | NA | NA | NA | NA | 9 | 0.1111 |
| 1_5385184 | 1 | 0 | 1 | 0 | 2 | 0 | 3 | 0.3333 | 7 | 0.1429 | 4 | 0.25 | 4 | 0.25 | NA | NA | 1 | 0 | 9 | 0.2222 |
| 1_5385206 | 1 | 1 | 1 | 1 | NA | NA | 3 | 1 | 5 | 1 | 4 | 0.75 | 4 | 0.75 | NA | NA | 1 | 1 | 9 | 0.7778 |
| 1_5385207 | 1 | 1 | 1 | 1 | 1 | 1 | 2 | 0.5 | 5 | 0.8 | 3 | 0.3333 | 4 | 0.75 | NA | NA | 1 | 1 | 8 | 0.625 |
| 1_5385208 | 1 | 1 | 1 | 0 | 1 | 0 | 3 | 0.3333 | 6 | 0.3333 | 3 | 0.3333 | 4 | 0.25 | NA | NA | 1 | 0 | 8 | 0.25 |
| 1_5385222 | 1 | 1 | 1 | 1 | 1 | 1 | 3 | 1 | 6 | 1 | 3 | 1 | 4 | 1 | NA | NA | 1 | 1 | 8 | 1 |
| 1_5385227 | 1 | 1 | 1 | 1 | 1 | 1 | 2 | 1 | 5 | 1 | 3 | 1 | 4 | 1 | NA | NA | NA | NA | 7 | 1 |
| 1_5385231 | 1 | 1 | 1 | 1 | 2 | 0.5 | NA | NA | 4 | 0.75 | 2 | 1 | 3 | 0.6667 | NA | NA | 1 | 1 | 6 | 0.8333 |
| 1_5385239 | 1 | 1 | NA | NA | 1 | 0 | 2 | 1 | 4 | 0.75 | 4 | 0.75 | 4 | 1 | NA | NA | 1 | 1 | 9 | 0.8889 |
| 1_5385242 | 1 | 1 | 1 | 0 | 1 | 0 | 2 | 0.5 | 5 | 0.4 | 1 | 1 | 2 | 0.5 | NA | NA | NA | NA | 3 | 0.6667 |
| 1_5396546 | NA | NA | 1 | 1 | NA | NA | NA | NA | 1 | 1 | 1 | 1 | NA | NA | NA | NA | NA | NA | 1 | 1 |
| 1_5396549 | NA | NA | 1 | 1 | NA | NA | NA | NA | 1 | 1 | 1 | 1 | NA | NA | NA | NA | NA | NA | 1 | 1 |
| 1_5396566 | NA | NA | 1 | 1 | NA | NA | NA | NA | 1 | 1 | 1 | 1 | NA | NA | NA | NA | NA | NA | 1 | 1 |
| 1_5396570 | NA | NA | 1 | 1 | NA | NA | NA | NA | 1 | 1 | 1 | 1 | NA | NA | NA | NA | NA | NA | 1 | 1 |
| 1_5443540 | 6 | 0 | 2 | 0.5 | 2 | 0 | 2 | 0 | 12 | 0.0833 | 5 | 0 | 6 | 0 | 4 | 0.25 | 2 | 0 | 17 | 0.0588 |

| | | | | | | | | | | | | | | | | | | | | |
|---|---|---|---|---|---|---|---|---|---|---|---|---|---|---|---|---|---|---|---|---|
| 1_54435787 | 3 | 0.6667 | 1 | 1 | 1 | 1 | 2 | 1 | 7 | 0.8571 | 3 | 0.3333 | 5 | 0.8 | 3 | 1 | 2 | 0.5 | 13 | 0.6923 |
| 1_54436133 | 3 | 0.6667 | NA | NA | NA | NA | 1 | 1 | 4 | 0.75 | 3 | 0 | 3 | 1 | NA | NA | 2 | 0 | 8 | 0.375 |
| 1_54436732 | 2 | 0.5 | 1 | 0 | NA | NA | 2 | 0 | 5 | 0.2 | 2 | 0 | 2 | 0.5 | 2 | 0 | 2 | 0.5 | 8 | 0.25 |
| 1_54436899 | 2 | 0 | 3 | 0.3333 | 2 | 0 | 3 | 0.3333 | 10 | 0.2 | 5 | 0 | 6 | 0 | 3 | 0.3333 | 4 | 0.25 | 18 | 0.1111 |
| 1_54436991 | 2 | 0 | 2 | 0 | 2 | 0 | 3 | 0.6667 | 9 | 0.2222 | 3 | 0.3333 | 6 | 0.3333 | 2 | 0.5 | 4 | 0.25 | 15 | 0.3333 |
| 1_54437010 | 2 | 0 | 3 | 0.6667 | 2 | 0 | 2 | 0.5 | 9 | 0.3333 | 5 | 0 | 7 | 0.4286 | 3 | 0.3333 | 3 | 0.3333 | 18 | 0.2778 |
| 1_54519625 | NA | NA | 1 | 1 | NA | NA | 2 | 1 | 3 | 1 | NA | NA | NA | NA | 1 | 1 | 1 | 0 | 2 | 0.5 |
| 1_54519863 | NA | NA | 1 | 0 | NA | NA | 2 | 1 | 3 | 0.6667 | NA | NA | NA | NA | 1 | 1 | 1 | 1 | 2 | 1 |
| 1_54684655 | 1 | 0 | 1 | 1 | NA | NA | 1 | 0 | 3 | 0.3333 | 1 | 1 | NA | NA | NA | NA | NA | NA | 1 | 1 |
| 1_55024793 | 4 | 0.25 | 7 | 0 | 3 | 0.3333 | 4 | 0 | 18 | 0.1111 | 2 | 0 | 2 | 0.5 | 3 | 0 | 5 | 0 | 12 | 0.0833 |
| 1_55025288 | 3 | 0.3333 | 3 | 0 | 1 | 1 | 3 | 0 | 10 | 0.2 | 2 | 0 | 3 | 0.3333 | 1 | 0 | 3 | 0 | 9 | 0.1111 |
| 1_55026310 | 15 | 0.0667 | 27 | 0.1852 | 18 | 0.2778 | 25 | 0.28 | 85 | 0.2118 | 18 | 0.2222 | 18 | 0 | 18 | 0.1111 | 22 | 0.1818 | 76 | 0.1316 |
| 1_55026511 | 13 | 0.3077 | 28 | 0.3571 | 20 | 0.5 | 27 | 0.3704 | 88 | 0.3864 | 17 | 0.4706 | 15 | 0.4 | 19 | 0.3158 | 21 | 0.381 | 72 | 0.3889 |
| 1_55027216 | 10 | 0.1 | 24 | 0.0833 | 16 | 0.125 | 20 | 0.1 | 70 | 0.1 | 9 | 0.2222 | 13 | 0.1538 | 15 | 0.0667 | 17 | 0.1176 | 54 | 0.1296 |
| 1_55054447 | 1 | 0 | 3 | 0.3333 | 3 | 0.3333 | 4 | 0 | 11 | 0.1818 | 3 | 0.3333 | 2 | 0 | NA | NA | 4 | 0 | 9 | 0.1111 |
| 1_55105790 | 1 | 1 | NA | NA | NA | NA | NA | NA | 1 | 1 | NA | NA | NA | NA | 1 | 1 | NA | NA | 1 | 1 |
| 1_55110899 | 4 | 0 | 5 | 0.4 | 8 | 0 | 8 | 0.25 | 25 | 0.16 | 2 | 0.5 | 3 | 0 | 1 | 0 | 10 | 0.1 | 16 | 0.125 |
| 1_55110915 | 4 | 0.25 | 8 | 0.25 | 8 | 0.375 | 8 | 0.375 | 28 | 0.3214 | 2 | 0 | 3 | 0.6667 | 1 | 0 | 10 | 0.4 | 16 | 0.375 |
| 1_55111107 | 4 | 0 | 5 | 0.2 | 6 | 0.5 | 4 | 0.75 | 19 | 0.3684 | NA | NA | 3 | 0.6667 | 1 | 0 | 7 | 0.2857 | 11 | 0.3636 |
| 1_55111113 | 5 | 1 | 7 | 1 | 9 | 0.8889 | 8 | 1 | 29 | 0.9655 | 2 | 1 | 4 | 0.75 | 1 | 1 | 11 | 0.9091 | 18 | 0.8889 |
| 1_55111152 | 5 | 0.8 | 6 | 0.1667 | 9 | 0.6667 | 7 | 0.4286 | 27 | 0.5185 | 1 | 0 | 4 | 0 | NA | NA | 10 | 0.3 | 15 | 0.2 |
| 1_55117124 | 30 | 0.2 | 22 | 0.0455 | 24 | 0.2083 | 17 | 0.1765 | 93 | 0.1613 | 11 | 0.0909 | 24 | 0.1667 | 19 | 0.0526 | 18 | 0.2222 | 72 | 0.1389 |
| 1_55117224 | 30 | 0.3 | 23 | 0.0435 | 19 | 0.3684 | 14 | 0.2143 | 86 | 0.2326 | 11 | 0.1818 | 24 | 0.2917 | 16 | 0.25 | 17 | 0.2941 | 68 | 0.2647 |
| 1_55117414 | 12 | 0.5 | 15 | 0.2667 | 11 | 0.1818 | 11 | 0.3636 | 49 | 0.3265 | 6 | 0.3333 | 13 | 0.5385 | 10 | 0.9 | 10 | 0.4 | 39 | 0.5641 |
| 1_55117505 | 28 | 0.0357 | 23 | 0.1739 | 19 | 0.0526 | 16 | 0.5 | 86 | 0.1628 | 9 | 0.1111 | 22 | 0.1818 | 22 | 0.2273 | 15 | 0.0667 | 68 | 0.1618 |
| 1_55192218 | 15 | 0 | 8 | 0.25 | 13 | 0 | 4 | 0 | 40 | 0.05 | 4 | 0.25 | 7 | 0.1429 | 8 | 0.25 | 13 | 0 | 32 | 0.125 |
| 1_55193793 | 18 | 0.1111 | 6 | 0.1667 | 8 | 0.125 | 5 | 0 | 37 | 0.1081 | 7 | 0.1429 | 6 | 0.1667 | 5 | 0 | 12 | 0 | 30 | 0.0667 |
| 1_55209460 | 3 | 0.6667 | 2 | 0 | 1 | 0 | 4 | 0 | 10 | 0.2 | 4 | 0 | 1 | 1 | 3 | 0.6667 | 2 | 0.5 | 10 | 0.4 |
| 1_55209503 | 3 | 0 | 2 | 0.5 | 2 | 0 | 4 | 0 | 11 | 0.0909 | 4 | 0 | 1 | 0 | 2 | 0 | 1 | 1 | 8 | 0.125 |
| 1_55209658 | 3 | 0.3333 | 2 | 0 | 2 | 0 | 4 | 0 | 11 | 0.0909 | 3 | 0.6667 | 1 | 0 | 3 | 0 | 2 | 0 | 9 | 0.2222 |
| 1_55209998 | 3 | 0.3333 | 2 | 0 | 2 | 0 | 4 | 0 | 11 | 0.0909 | 3 | 0.3333 | 1 | 0 | 3 | 0 | 1 | 0 | 8 | 0.125 |
| 1_55210685 | 2 | 0 | 2 | 0 | 2 | 0.5 | 4 | 0 | 10 | 0.1 | 1 | 0 | 1 | 0 | 3 | 0.3333 | NA | NA | 5 | 0.2 |
| 1_55210922 | 4 | 0 | 4 | 0 | 5 | 0.2 | 3 | 0.3333 | 16 | 0.125 | 4 | 0 | 2 | 0 | 5 | 0.2 | 1 | 0 | 12 | 0.0833 |
| 1_55211014 | 2 | 0.5 | 4 | 1 | 5 | 0.8 | 5 | 1 | 16 | 0.875 | 3 | 1 | 1 | 1 | 4 | 0.75 | 1 | 1 | 9 | 0.8889 |
| 1_55211026 | 3 | 0 | 3 | 0.3333 | 5 | 0 | 5 | 0.2 | 16 | 0.125 | 4 | 0.25 | 1 | 0 | 4 | 0.25 | 2 | 0 | 11 | 0.1818 |
| 1_55245601 | 1 | 0 | 4 | 0.25 | 3 | 0 | 4 | 0 | 12 | 0.0833 | 4 | 0.25 | 2 | 0.5 | 3 | 0 | 2 | 0 | 11 | 0.1818 |
| 1_55265407 | 1 | 0 | 3 | 0.3333 | 4 | 0.25 | 2 | 0.5 | 10 | 0.3 | 3 | 0.3333 | 1 | 1 | 2 | 1 | 2 | 0 | 8 | 0.5 |
| 1_55265426 | 1 | 0 | 2 | 0 | 3 | 0.6667 | 2 | 0.5 | 8 | 0.375 | 2 | 0 | 2 | 0 | 1 | 1 | 1 | 0 | 6 | 0.1667 |
| 1_55274655 | 2 | 0.5 | NA | NA | 2 | 0.5 | NA | NA | 4 | 0.5 | 1 | 0 | 1 | 1 | 1 | 0 | 1 | 0 | 4 | 0.25 |
| 1_55277455 | 2 | 1 | 1 | 1 | 1 | 1 | NA | NA | 4 | 1 | NA | NA | 1 | 1 | 1 | 1 | NA | NA | 2 | 1 |
| 1_55277525 | 2 | 1 | 1 | 0 | NA | NA | NA | NA | 3 | 0.6667 | NA | NA | 1 | 0 | 1 | 1 | NA | NA | 2 | 0.5 |
| 1_55277564 | 2 | 1 | 1 | 1 | NA | NA | NA | NA | 3 | 1 | NA | NA | 1 | 1 | 1 | 1 | NA | NA | 2 | 1 |
| 1_55277714 | 1 | 0 | 1 | 1 | 1 | 0 | NA | NA | 3 | 0.3333 | NA | NA | NA | NA | 1 | 1 | NA | NA | 1 | 1 |
| 1_55277774 | 1 | 1 | NA | NA | NA | NA | NA | NA | 1 | 1 | NA | NA | 1 | 1 | NA | NA | NA | NA | 1 | 1 |
| 1_55277864 | 2 | 1 | 1 | 1 | 1 | 1 | NA | NA | 4 | 1 | NA | NA | 1 | 1 | 1 | 1 | NA | NA | 2 | 1 |
| 1_55277941 | 3 | 1 | 1 | 1 | 1 | 1 | 1 | 1 | 6 | 1 | NA | NA | 1 | 1 | 1 | 1 | NA | NA | 2 | 1 |
| 1_55277991 | 3 | 0.6667 | 1 | 0 | 1 | 1 | 1 | 0 | 6 | 0.5 | NA | NA | 1 | 1 | 1 | 1 | NA | NA | 2 | 1 |
| 1_55278054 | 3 | 1 | 1 | 1 | NA | NA | 1 | 1 | 5 | 1 | NA | NA | 1 | 1 | NA | NA | NA | NA | 1 | 1 |
| 1_55278056 | NA | NA | NA | NA | 1 | 1 | NA | NA | 1 | 1 | NA | NA | NA | NA | 1 | 1 | NA | NA | 1 | 1 |
| 1_55278087 | 2 | 1 | 1 | 0 | 1 | 1 | 1 | 1 | 5 | 0.8 | NA | NA | 1 | 1 | 1 | 0 | NA | NA | 2 | 0.5 |
| 1_55278114 | 3 | 1 | 1 | 0 | 1 | 0 | NA | NA | 5 | 0.6 | NA | NA | 1 | 0 | 1 | 1 | NA | NA | 2 | 0.5 |
| 1_55278119 | 3 | 1 | 1 | 0 | NA | NA | 1 | 1 | 5 | 0.8 | NA | NA | NA | NA | 1 | 1 | NA | NA | 1 | 1 |
| 1_55278130 | 3 | 1 | 1 | 1 | 1 | 1 | 1 | 1 | 6 | 1 | NA | NA | 1 | 1 | 1 | 1 | NA | NA | 2 | 1 |
| 1_55278154 | 3 | 0.6667 | 1 | 1 | 1 | 0 | 1 | 0 | 6 | 0.5 | NA | NA | 1 | 0 | 1 | 1 | NA | NA | 2 | 0.5 |
| 1_55278162 | 3 | 0.6667 | 1 | 0 | 1 | 1 | 1 | 0 | 6 | 0.5 | NA | NA | 1 | 0 | 1 | 1 | NA | NA | 2 | 0.5 |
| 1_55278177 | 2 | 1 | 1 | 1 | 1 | 1 | 1 | 1 | 5 | 1 | NA | NA | 1 | 1 | 1 | 0 | NA | NA | 2 | 0.5 |
| 1_55735139 | 2 | 1 | 1 | 0 | 2 | 0.5 | NA | NA | 5 | 0.6 | NA | NA | NA | NA | NA | NA | 1 | 1 | 1 | 1 |
| 1_55735223 | 2 | 0.5 | 2 | 1 | 2 | 0.5 | NA | NA | 6 | 0.6667 | NA | NA | NA | NA | NA | NA | 2 | 0.5 | 2 | 0.5 |
| 1_55735268 | 2 | 1 | 2 | 1 | 2 | 0.5 | NA | NA | 6 | 0.8333 | NA | NA | NA | NA | NA | NA | 1 | 1 | 1 | 1 |
| 1_55735590 | 2 | 1 | 2 | 0.5 | 2 | 1 | NA | NA | 6 | 0.8333 | NA | NA | NA | NA | NA | NA | 2 | 1 | 2 | 1 |
| 1_55735823 | 2 | 0 | 3 | 0.3333 | 2 | 0.5 | NA | NA | 7 | 0.2857 | NA | NA | NA | NA | NA | NA | 2 | 0.5 | 2 | 0.5 |
| 1_55736205 | 2 | 0 | 4 | 0.5 | 2 | 0.5 | NA | NA | 8 | 0.375 | NA | NA | NA | NA | 1 | 1 | 2 | 0.5 | 3 | 0.6667 |
| 1_55736423 | 2 | 0.5 | 4 | 0.5 | 2 | 0.5 | NA | NA | 8 | 0.5 | NA | NA | NA | NA | 1 | 0 | 2 | 0.5 | 3 | 0.3333 |
| 1_55736466 | 2 | 1 | 4 | 1 | 2 | 1 | NA | NA | 8 | 1 | NA | NA | NA | NA | NA | NA | 2 | 1 | 2 | 1 |
| 1_55736552 | 1 | 1 | 4 | 0.5 | 2 | 0 | NA | NA | 7 | 0.4286 | NA | NA | NA | NA | 1 | 0 | 2 | 0.5 | 3 | 0.3333 |

| | | | | | | | | | | | | | | | | | | | | |
|---|---|---|---|---|---|---|---|---|---|---|---|---|---|---|---|---|---|---|---|---|
| 1_5573663 | 2 | 1 | 3 | 1 | 2 | 0.5 | NA | NA | 7 | 0.8571 | NA | NA | NA | NA | 1 | 0 | 2 | 1 | 3 | 0.6667 |
| 1_5573664 | 2 | 0.5 | 3 | 1 | 2 | 0.5 | NA | NA | 7 | 0.7143 | NA | NA | NA | NA | 1 | 1 | 1 | 1 | 2 | 1 |
| 1_5573685 | 2 | 0 | 4 | 0.25 | 2 | 0.5 | NA | NA | 8 | 0.25 | NA | NA | NA | NA | 1 | 0 | 3 | 0.6667 | 4 | 0.5 |
| 1_5573695 | 1 | 1 | 2 | 1 | 1 | 1 | NA | NA | 4 | 1 | NA | NA | NA | NA | 1 | 0 | 2 | 0.5 | 3 | 0.3333 |
| 1_5573696 | 2 | 0.5 | 3 | 0.6667 | 2 | 1 | NA | NA | 7 | 0.7143 | NA | NA | NA | NA | 1 | 1 | 3 | 0.6667 | 4 | 0.75 |
| 1_5573700 | 2 | 0 | 3 | 0.3333 | 2 | 0 | NA | NA | 7 | 0.1429 | NA | NA | NA | NA | 1 | 0 | 3 | 0.3333 | 4 | 0.25 |
| 1_5573718 | 2 | 0 | 3 | 0.3333 | 2 | 0 | NA | NA | 7 | 0.1429 | NA | NA | NA | NA | 1 | 1 | 3 | 0 | 4 | 0.25 |
| 1_5579052 | 1 | 1 | 3 | 1 | 4 | 1 | NA | NA | 8 | 1 | 1 | 1 | NA | NA | 1 | 0 | 3 | 1 | 5 | 0.8 |
| 1_5579161 | 2 | 0 | 3 | 0 | 5 | 0 | 1 | 1 | 11 | 0.0909 | 1 | 0 | NA | NA | 5 | 0.2 | 4 | 0 | 10 | 0.1 |
| 1_5683390 | NA | NA | 2 | 0.5 | 1 | 0 | NA | NA | 3 | 0.3333 | NA | NA | NA | NA | NA | NA | 1 | 1 | 1 | 1 |
| 1_5742196 | 3 | 0.3333 | 1 | 0 | 2 | 0 | 5 | 0 | 11 | 0.0909 | 2 | 0 | 1 | 0 | 4 | 0.25 | 3 | 0 | 10 | 0.1 |
| 1_5742205 | 3 | 0 | NA | NA | 4 | 0.25 | 5 | 0 | 12 | 0.0833 | 3 | 0 | 2 | 0 | 4 | 0 | 3 | 0.3333 | 12 | 0.0833 |
| 1_5742209 | 2 | 0.5 | 1 | 0 | 3 | 0.3333 | 5 | 0.4 | 11 | 0.3636 | 2 | 0 | 1 | 0 | 3 | 0.6667 | 1 | 0 | 7 | 0.2857 |
| 1_5742210 | 3 | 1 | 1 | 1 | 3 | 1 | 5 | 1 | 12 | 1 | 3 | 1 | NA | NA | 4 | 1 | 3 | 1 | 10 | 1 |
| 1_5742216 | 2 | 0.5 | 1 | 1 | 3 | 0.6667 | 4 | 0.75 | 10 | 0.7 | 3 | 0.3333 | 2 | 0.5 | 3 | 0.3333 | 3 | 1 | 11 | 0.5455 |
| 1_5742218 | 2 | 0 | 1 | 0 | 2 | 0 | 4 | 0.25 | 9 | 0.1111 | 3 | 0.3333 | 2 | 0 | 4 | 0 | 2 | 0 | 11 | 0.0909 |
| 1_5742222 | 3 | 0.3333 | 1 | 0 | 3 | 1 | 5 | 0.2 | 12 | 0.4167 | 2 | 0 | 1 | 1 | 3 | 0.6667 | 2 | 1 | 8 | 0.625 |
| 1_5742227 | 3 | 0.6667 | 1 | 1 | 4 | 0.5 | 4 | 0.75 | 12 | 0.6667 | 3 | 1 | 3 | 1 | 4 | 0.75 | 3 | 0.3333 | 13 | 0.7692 |
| 1_5742241 | 2 | 0.5 | 1 | 0 | 4 | 0.25 | 4 | 0.5 | 11 | 0.3636 | 2 | 0 | 3 | 1 | 4 | 0 | 2 | 0 | 11 | 0.2727 |
| 1_5742241 | 3 | 0.3333 | 1 | 0 | 3 | 0 | 5 | 0.6 | 12 | 0.3333 | 2 | 0.5 | 3 | 0.3333 | 3 | 0 | 2 | 0 | 10 | 0.2 |
| 1_5742244 | 2 | 0 | NA | NA | 3 | 1 | 3 | 1 | 8 | 0.75 | 2 | 1 | 2 | 1 | 3 | 0.6667 | 2 | 0.5 | 9 | 0.7778 |
| 1_5742265 | 3 | 0.3333 | 1 | 0 | 4 | 0 | 4 | 0 | 12 | 0.0833 | 3 | 0 | 3 | 0 | 3 | 0.3333 | 4 | 0.25 | 13 | 0.1538 |
| 1_5742272 | 3 | 0.6667 | 1 | 0 | 3 | 0.6667 | 5 | 0.8 | 12 | 0.6667 | 3 | 0.6667 | 3 | 0.6667 | 4 | 0.25 | 4 | 0.5 | 14 | 0.5 |
| 1_5742302 | 3 | 0 | 1 | 0 | 3 | 0 | 5 | 0.2 | 12 | 0.0833 | 3 | 0 | 3 | 0 | 4 | 0.25 | 4 | 0 | 14 | 0.0714 |
| 1_5742425 | 2 | 0 | 2 | 0 | 2 | 0.5 | 5 | 0.2 | 11 | 0.1818 | 2 | 0.5 | 5 | 0.2 | 5 | 0.4 | 3 | 0 | 15 | 0.2667 |
| 1_5742429 | 3 | 0 | 2 | 1 | 2 | 0 | 5 | 0.4 | 12 | 0.3333 | 3 | 0.6667 | 4 | 0 | 5 | 0.8 | 4 | 0.25 | 16 | 0.4375 |
| 1_5742432 | 3 | 0 | 1 | 0 | 3 | 0 | 5 | 0.2 | 12 | 0.0833 | 2 | 0 | 3 | 0 | 5 | 0.2 | 4 | 0 | 14 | 0.0714 |
| 1_5742747 | 3 | 0.3333 | 2 | 0 | 2 | 0.5 | 2 | 0 | 9 | 0.2222 | NA | NA | 1 | 0 | 5 | 0 | 2 | 0.5 | 8 | 0.125 |
| 1_5742766 | 3 | 0 | 1 | 0 | 2 | 1 | 2 | 0 | 8 | 0.25 | NA | NA | 2 | 0 | 5 | 0.2 | 3 | 0.3333 | 10 | 0.2 |
| 1_5742771 | 2 | 0.5 | 1 | 0 | 2 | 0.5 | 2 | 0.5 | 7 | 0.4286 | NA | NA | 1 | 1 | 5 | 0.6 | 2 | 0 | 8 | 0.5 |
| 1_5744600 | 4 | 0.25 | 2 | 0.5 | 2 | 0.5 | 2 | 0 | 10 | 0.3 | 2 | 0.5 | NA | NA | 1 | 0 | 2 | 0 | 5 | 0.2 |
| 1_5744630 | 5 | 0.8 | 2 | 1 | 3 | 1 | 2 | 0.5 | 12 | 0.8333 | 3 | 1 | 1 | 1 | 1 | 1 | 1 | 1 | 6 | 1 |
| 1_5745492 | 7 | 0.5714 | 1 | 0 | 2 | 0.5 | 3 | 0 | 13 | 0.3846 | 3 | 0.6667 | NA | NA | 1 | 0 | 2 | 0 | 6 | 0.3333 |
| 1_5745647 | 2 | 0.5 | 1 | 0 | 1 | 0 | 1 | 0 | 5 | 0.2 | 1 | 0 | 2 | 0.5 | NA | NA | 2 | 0 | 5 | 0.2 |
| 1_5798234 | 3 | 0.3333 | 3 | 0 | 1 | 0 | 3 | 0 | 10 | 0.1 | 1 | 0 | 5 | 0.2 | NA | NA | NA | NA | 6 | 0.1667 |
| 1_5798598 | 3 | 0.3333 | 2 | 0 | 1 | 0 | 3 | 0.3333 | 9 | 0.2222 | 2 | 0.5 | 6 | 0 | NA | NA | 1 | 0 | 9 | 0.1111 |
| 1_5799534 | NA | NA | 2 | 0.5 | 2 | 1 | 2 | 1 | 6 | 0.8333 | 1 | 0 | NA | NA | 1 | 1 | 2 | 0.5 | 4 | 0.5 |
| 1_5799550 | NA | NA | 2 | 0.5 | 2 | 0 | 2 | 0.5 | 6 | 0.3333 | 2 | 0.5 | 1 | 1 | 1 | 0 | 2 | 0.5 | 6 | 0.5 |
| 1_5799562 | NA | NA | 2 | 1 | 2 | 1 | 2 | 1 | 6 | 1 | 2 | 1 | 1 | 0 | 1 | 1 | 3 | 0.6667 | 7 | 0.7143 |
| 1_5799587 | NA | NA | 2 | 0.5 | 2 | 1 | 1 | 1 | 5 | 0.8 | 2 | 1 | 1 | 1 | 1 | 1 | 2 | 1 | 6 | 1 |
| 1_5799594 | NA | NA | NA | NA | 1 | 1 | 1 | 0 | 2 | 0.5 | 1 | 1 | 1 | 0 | 1 | 1 | NA | NA | 3 | 0.6667 |
| 1_5844500 | 14 | 0.2143 | 14 | 0.1429 | 16 | 0.125 | 8 | 0.25 | 52 | 0.1731 | 15 | 0.1333 | 18 | 0.0556 | 17 | 0.2941 | 15 | 0.3333 | 65 | 0.2 |
| 1_5844542 | 11 | 0.0909 | 12 | 0 | 17 | 0.1176 | 6 | 0 | 46 | 0.0652 | 17 | 0.0588 | 16 | 0.1875 | 16 | 0.0625 | 12 | 0 | 61 | 0.082 |
| 1_5845132 | 9 | 0 | 14 | 0.2143 | 12 | 0 | 18 | 0.1111 | 53 | 0.0943 | 8 | 0 | 19 | 0.0526 | 23 | 0.1739 | 17 | 0.1176 | 67 | 0.1045 |
| 1_5845952 | NA | NA | 9 | 0.3333 | 4 | 0 | 6 | 0.1667 | 19 | 0.2105 | 8 | 0.125 | 9 | 0.3333 | 12 | 0.25 | 7 | 0 | 36 | 0.1944 |
| 1_5847035 | 4 | 0.5 | 7 | 0.4286 | 1 | 1 | 3 | 0.3333 | 15 | 0.4667 | 7 | 0.4286 | 3 | 1 | 5 | 0.4 | NA | NA | 15 | 0.5333 |
| 1_5847039 | 4 | 0.25 | 6 | 0.3333 | 1 | 1 | 2 | 0 | 13 | 0.3077 | 6 | 0.5 | 4 | 0.5 | 4 | 0.5 | NA | NA | 14 | 0.5 |
| 1_5848417 | NA | NA | NA | NA | NA | NA | 1 | 1 | 1 | 1 | 1 | 1 | NA | NA | NA | NA | NA | NA | 1 | 1 |
| 1_5848692 | 2 | 1 | 1 | 1 | 1 | 1 | 3 | 1 | 7 | 1 | 1 | 1 | 2 | 1 | 2 | 1 | 4 | 1 | 9 | 1 |
| 1_5848696 | 2 | 1 | 1 | 1 | 1 | 1 | 4 | 1 | 8 | 1 | 1 | 1 | 3 | 1 | 2 | 0.5 | 4 | 0.75 | 10 | 0.8 |
| 1_5850030 | 2 | 0.5 | 1 | 1 | NA | NA | 1 | 1 | 4 | 0.75 | NA | NA | NA | NA | 1 | 1 | NA | NA | 1 | 1 |
| 1_5850058 | 3 | 1 | 2 | 1 | 1 | 1 | 3 | 1 | 9 | 1 | 1 | 1 | 4 | 1 | 2 | 1 | 5 | 0.8 | 12 | 0.9167 |
| 1_5850064 | 3 | 0.3333 | 2 | 0 | 1 | 1 | 3 | 0.3333 | 9 | 0.3333 | 1 | 0 | 4 | 0.5 | 1 | 0 | 5 | 0.2 | 11 | 0.2727 |
| 1_5850075 | 1 | 1 | NA | NA | 1 | 1 | 3 | 1 | 5 | 1 | NA | NA | 2 | 0.5 | 1 | 1 | 1 | 1 | 4 | 0.75 |
| 1_5850082 | 1 | 1 | NA | NA | 1 | 0 | 3 | 1 | 5 | 0.8 | NA | NA | 2 | 0.5 | 1 | 0 | 1 | 0 | 4 | 0.25 |
| 1_5850092 | 1 | 1 | NA | NA | 1 | 0 | 3 | 0.3333 | 5 | 0.4 | NA | NA | 2 | 1 | 1 | 1 | 1 | 0 | 4 | 0.75 |
| 1_5850108 | 1 | 0 | NA | NA | 1 | 1 | 3 | 0 | 5 | 0.2 | NA | NA | 2 | 0.5 | 1 | 0 | 1 | 0 | 4 | 0.25 |
| 1_5850109 | 1 | 1 | NA | NA | 1 | 1 | 3 | 1 | 5 | 1 | NA | NA | 2 | 0.5 | 1 | 0 | 1 | 1 | 4 | 0.5 |
| 1_5850110 | 1 | 1 | NA | NA | 1 | 1 | 3 | 0.6667 | 5 | 0.8 | NA | NA | 3 | 0.6667 | 1 | 1 | 1 | 1 | 5 | 0.8 |
| 1_5850119 | 1 | 0 | NA | NA | 1 | 1 | 2 | 0.5 | 4 | 0.5 | NA | NA | 3 | 0.3333 | 1 | 0 | 1 | 0 | 5 | 0.2 |
| 1_5850269 | NA | NA | NA | NA | 1 | 1 | 1 | 0 | 2 | 0.5 | NA | NA | 1 | 1 | NA | NA | NA | NA | 1 | 1 |
| 1_5850285 | 6 | 0.1667 | 1 | 1 | 1 | 0 | 5 | 0.2 | 13 | 0.2308 | 2 | 0 | 1 | 0 | 6 | 0 | 7 | 0.5714 | 16 | 0.25 |
| 1_5850288 | 6 | 0.8333 | 1 | 1 | 1 | 1 | 4 | 0.75 | 12 | 0.8333 | 2 | 0.5 | 1 | 1 | 7 | 0.5714 | 7 | 0.8571 | 17 | 0.7059 |
| 1_5850294 | 5 | 0 | 1 | 0 | 1 | 1 | 5 | 0.4 | 12 | 0.25 | 1 | 0 | 1 | 0 | 5 | 0.8 | 7 | 1 | 14 | 0.7857 |
| 1_5850308 | 4 | 0.75 | 1 | 1 | 1 | 1 | 4 | 0.5 | 10 | 0.7 | 2 | 1 | 1 | 1 | 6 | 0.5 | 6 | 0.8333 | 15 | 0.7333 |

| | | | | | | | | | | | | | | | | | | | | |
|---|---|---|---|---|---|---|---|---|---|---|---|---|---|---|---|---|---|---|---|---|
| 1_58503123 | 4 | 0 | 1 | 0 | NA | NA | 4 | 0.25 | 9 | 0.1111 | 2 | 0.5 | NA | NA | 7 | 0.1429 | 6 | 0.1667 | 15 | 0.2 |
| 1_58503216 | 4 | 1 | 1 | 1 | NA | NA | 5 | 0.8 | 10 | 0.9 | 2 | 1 | 1 | 1 | 4 | 1 | 4 | 1 | 11 | 1 |
| 1_58503344 | 2 | 0.5 | 1 | 1 | 1 | 1 | 5 | 0.4 | 9 | 0.5556 | 2 | 0.5 | 1 | 0 | 5 | 1 | 4 | 0.5 | 12 | 0.6667 |
| 1_58562349 | 15 | 0.0667 | 11 | 0.0909 | 16 | 0.0625 | 12 | 0.0833 | 54 | 0.0741 | 10 | 0 | 9 | 0 | 13 | 0.0769 | 7 | 0.1429 | 39 | 0.0513 |
| 1_58562504 | 15 | 0.0667 | 12 | 0.0833 | 15 | 0.2 | 12 | 0.1667 | 54 | 0.1296 | 5 | 0 | 9 | 0.2222 | 11 | 0.1818 | 9 | 0.1111 | 34 | 0.1471 |
| 1_58562873 | 15 | 0 | 10 | 0.2 | 15 | 0.1333 | 11 | 0 | 51 | 0.0784 | 3 | 0.3333 | 10 | 0 | 11 | 0 | 9 | 0.2222 | 33 | 0.0909 |
| 1_58563258 | 14 | 0 | 9 | 0.3333 | 15 | 0.0667 | 12 | 0.1667 | 50 | 0.12 | 3 | 0 | 8 | 0.125 | 12 | 0 | 8 | 0.25 | 31 | 0.0968 |
| 1_58568038 | 2 | 0 | 3 | 0.3333 | 6 | 0 | 5 | 0.4 | 16 | 0.1875 | 1 | 0 | 1 | 0 | 3 | 0.3333 | 2 | 0.5 | 7 | 0.2857 |
| 1_58568076 | 1 | 0 | 3 | 0.3333 | 5 | 0.2 | 4 | 0.5 | 13 | 0.3077 | 1 | 1 | 2 | 0 | 3 | 0 | 2 | 0 | 8 | 0.125 |
| 1_58568084 | 2 | 0 | 2 | 0 | 4 | 0.25 | 5 | 0.2 | 13 | 0.1538 | 1 | 0 | 2 | 0 | 3 | 0.6667 | 2 | 0.5 | 8 | 0.375 |
| 1_58568353 | 2 | 0 | 3 | 0 | 4 | 0 | 4 | 0.25 | 13 | 0.0769 | 1 | 0 | 1 | 0 | 4 | 0.25 | 1 | 0 | 7 | 0.1429 |
| 1_58568418 | 2 | 0 | 3 | 0.3333 | 1 | 0 | 4 | 0.75 | 10 | 0.4 | 1 | 1 | NA | NA | 1 | 0 | 1 | 0 | 3 | 0.3333 |
| 1_58568516 | 2 | 0.5 | 2 | 0 | 3 | 0 | 4 | 0.5 | 11 | 0.2727 | 1 | 1 | 1 | 0 | 1 | 0 | NA | NA | 3 | 0.3333 |
| 1_58569037 | 2 | 1 | 2 | 0.5 | 2 | 1 | 3 | 0.6667 | 9 | 0.7778 | NA | NA | NA | NA | 1 | 1 | NA | NA | 1 | 1 |
| 1_58569068 | 1 | 1 | 3 | 1 | 2 | 1 | 2 | 1 | 8 | 1 | NA | NA | 1 | 1 | 1 | 1 | NA | NA | 2 | 1 |
| 1_58569253 | 2 | 0 | 3 | 0.6667 | 2 | 0 | 4 | 0 | 11 | 0.1818 | 1 | 1 | 1 | 0 | 1 | 0 | NA | NA | 3 | 0.3333 |
| 1_58569320 | 2 | 0 | 3 | 1 | 2 | 0 | 3 | 0.3333 | 10 | 0.4 | 1 | 1 | NA | NA | 1 | 0 | NA | NA | 2 | 0.5 |
| 1_58765695 | NA | NA | 1 | 1 | NA | NA | NA | NA | 1 | 1 | NA | NA | 1 | 1 | NA | NA | NA | NA | 1 | 1 |
| 1_58940288 | 8 | 0.375 | 20 | 0.6 | 12 | 0.5833 | 12 | 0.3333 | 52 | 0.5 | 12 | 0.25 | 19 | 0.5789 | 8 | 0.25 | 15 | 0.4 | 54 | 0.4074 |
| 1_58940345 | 6 | 0.8333 | 19 | 0.5789 | 8 | 0.625 | 12 | 0.6667 | 45 | 0.6444 | 11 | 0.3636 | 16 | 0.6875 | 6 | 0.8333 | 12 | 0.5833 | 45 | 0.6 |
| 1_58940742 | 5 | 0.2 | 13 | 0 | 8 | 0.125 | 10 | 0.2 | 36 | 0.1111 | 7 | 0.1429 | 17 | 0.1176 | 5 | 0 | 14 | 0 | 43 | 0.0698 |
| 1_58940760 | 5 | 0.2 | 17 | 0.4118 | 10 | 0.2 | 13 | 0.1538 | 45 | 0.2667 | 7 | 0.4286 | 18 | 0.3889 | 6 | 0.6667 | 13 | 0.0769 | 44 | 0.3409 |
| 1_58941963 | 3 | 1 | 6 | 1 | 7 | 0.8571 | 9 | 0.5556 | 25 | 0.8 | 4 | 0.75 | 10 | 0.8 | 3 | 0.6667 | 11 | 0.8182 | 28 | 0.7857 |
| 1_58942096 | 4 | 1 | 10 | 0.9 | 8 | 0.875 | 6 | 0.6667 | 28 | 0.8571 | 4 | 0.75 | 12 | 0.8333 | 2 | 1 | 10 | 1 | 28 | 0.8929 |
| 1_58942112 | 1 | 1 | 10 | 0.8 | 7 | 0.8571 | 5 | 0.8 | 23 | 0.8261 | 3 | 0.6667 | 9 | 0.7778 | 3 | 1 | 9 | 1 | 24 | 0.875 |
| 1_58942312 | 3 | 0.3333 | 8 | 0.125 | 7 | 0 | 4 | 0 | 22 | 0.0909 | 3 | 0 | 8 | 0.125 | 3 | 0.3333 | 10 | 0 | 24 | 0.0833 |
| 1_58942338 | 2 | 0.5 | 9 | 0.2222 | 8 | 0 | 4 | 0 | 23 | 0.1304 | 4 | 0.5 | 10 | 0.1 | 3 | 0.6667 | 9 | 0.2222 | 26 | 0.2692 |
| 1_58942351 | 2 | 0.5 | 9 | 0.6667 | 7 | 0.2857 | 4 | 1 | 22 | 0.5909 | 4 | 0.75 | 10 | 0.6 | 3 | 1 | 11 | 0.9091 | 28 | 0.7857 |
| 1_58942377 | 3 | 0.3333 | 9 | 0.2222 | 6 | 0.1667 | 4 | 0.5 | 22 | 0.2727 | 4 | 0 | 12 | 0 | 3 | 0.3333 | 10 | 0.6 | 29 | 0.2414 |
| 1_58942446 | 2 | 0 | 8 | 0 | 9 | 0.2222 | 5 | 0 | 24 | 0.0833 | 4 | 0.25 | 10 | 0.2 | 3 | 0 | 10 | 0 | 27 | 0.1111 |
| 1_58942468 | 2 | 0.5 | 8 | 0.5 | 8 | 0.5 | 3 | 1 | 21 | 0.5714 | 4 | 0.5 | 9 | 0.3333 | 3 | 0.6667 | 10 | 0.4 | 26 | 0.4231 |
| 1_58942557 | 3 | 1 | 9 | 1 | 9 | 1 | 5 | 1 | 26 | 1 | 4 | 1 | 12 | 0.9167 | 2 | 1 | 10 | 0.9 | 28 | 0.9286 |
| 1_58942650 | 2 | 0.5 | 4 | 0.75 | 4 | 0 | 1 | 1 | 11 | 0.4545 | 3 | 0.6667 | 7 | 0.4286 | 1 | 0 | 3 | 0.6667 | 14 | 0.5 |
| 1_58942759 | 2 | 1 | 8 | 1 | 7 | 1 | 4 | 1 | 21 | 1 | 3 | 1 | 10 | 1 | 3 | 1 | 7 | 1 | 23 | 1 |
| 1_58942910 | 2 | 0.5 | 7 | 1 | 7 | 0.5714 | 5 | 1 | 21 | 0.8095 | 2 | 0.5 | 11 | 1 | 2 | 1 | 10 | 0.8 | 25 | 0.88 |
| 1_58942924 | 2 | 0.5 | 8 | 0.5 | 7 | 0.5714 | 4 | 1 | 21 | 0.619 | 2 | 0.5 | 8 | 0.75 | 3 | 0.6667 | 9 | 0.6667 | 22 | 0.6818 |
| 1_58942951 | 1 | 0 | 8 | 0.125 | 8 | 0.125 | 4 | 0 | 21 | 0.0952 | 3 | 0 | 11 | 0.1818 | 3 | 0 | 10 | 0 | 27 | 0.0741 |
| 1_58943222 | 2 | 1 | 7 | 0.8571 | 7 | 0.5714 | 5 | 0.6 | 21 | 0.7143 | 3 | 1 | 6 | 0.8333 | 2 | 0.5 | 7 | 0.2857 | 18 | 0.6111 |
| 1_58943257 | 2 | 1 | 7 | 0.2857 | 7 | 0.4286 | 3 | 0.6667 | 19 | 0.4737 | 3 | 0.3333 | 6 | 1 | 2 | 0.5 | 6 | 1 | 17 | 0.8235 |
| 1_58943279 | 2 | 0 | 8 | 0 | 6 | 0 | 3 | 0.3333 | 19 | 0.0526 | 2 | 0 | 6 | 0.1667 | 1 | 0 | 7 | 0.1429 | 16 | 0.125 |
| 1_58947639 | NA | NA | NA | NA | 1 | 1 | NA | NA | 1 | 1 | 2 | 0.5 | NA | NA | NA | NA | NA | NA | 2 | 0.5 |
| 1_58947819 | 1 | 0 | 5 | 0.2 | 5 | 0 | 5 | 0.4 | 16 | 0.1875 | 2 | 0.5 | 7 | 0.4286 | 1 | 1 | 7 | 0.2857 | 17 | 0.4118 |
| 1_59033576 | 4 | 0.5 | 6 | 0.3333 | 3 | 0 | 3 | 0 | 16 | 0.25 | 2 | 0.5 | 4 | 0.5 | 1 | 0 | 4 | 0.75 | 11 | 0.5455 |
| 1_59033675 | 4 | 0 | 7 | 0.4286 | 3 | 0.3333 | 4 | 0 | 18 | 0.2222 | 2 | 0 | 3 | 0 | NA | NA | 4 | 0.25 | 9 | 0.1111 |
| 1_59033877 | 5 | 0.4 | 7 | 0.5714 | 3 | 0.3333 | 4 | 0 | 19 | 0.3684 | 2 | 0.5 | 4 | 0.75 | 1 | 1 | 4 | 0.5 | 11 | 0.6364 |
| 1_59034154 | 5 | 0.2 | 6 | 0.1667 | 3 | 0 | 5 | 0 | 19 | 0.1053 | 1 | 0 | 5 | 0.4 | 2 | 0 | 4 | 0.25 | 12 | 0.25 |
| 1_59202047 | NA | NA | 1 | 1 | NA | NA | NA | NA | 1 | 1 | NA | NA | NA | NA | 1 | 1 | NA | NA | 1 | 1 |
| 1_59202101 | NA | NA | 5 | 0.6 | 3 | 0 | 1 | 0 | 9 | 0.3333 | 3 | 0 | 3 | 0.6667 | NA | NA | 1 | 0 | 7 | 0.2857 |
| 1_59202231 | 1 | 0 | 7 | 0.8571 | 4 | 0.5 | 1 | 0 | 13 | 0.6154 | 5 | 0.6 | 3 | 0.6667 | 1 | 1 | 4 | 0.75 | 13 | 0.6923 |
| 1_59202245 | 1 | 0 | 5 | 0.2 | 4 | 0.5 | 1 | 1 | 11 | 0.3636 | 2 | 0.5 | 4 | 0.5 | 2 | 0.5 | 4 | 0.75 | 12 | 0.5833 |
| 1_59202399 | 1 | 0 | 5 | 0 | 2 | 0.5 | 1 | 0 | 9 | 0.1111 | 4 | 0 | 4 | 0.25 | 2 | 0 | 4 | 0.25 | 14 | 0.1429 |
| 1_59202458 | NA | NA | 1 | 0 | 2 | 1 | NA | NA | 3 | 0.6667 | 1 | 1 | 4 | 1 | 1 | 1 | 3 | 1 | 9 | 1 |
| 1_59202542 | 1 | 0 | 3 | 0.6667 | 3 | 0 | 1 | 0 | 8 | 0.25 | 3 | 0.3333 | 4 | 0.5 | 1 | 1 | 4 | 0.25 | 12 | 0.4167 |
| 1_59202662 | 1 | 0 | 5 | 0.2 | 3 | 0 | NA | NA | 9 | 0.1111 | 4 | 0 | 3 | 0.3333 | 1 | 0 | 3 | 0.3333 | 11 | 0.1818 |
| 1_59202672 | 1 | 0 | 4 | 0.5 | 3 | 0.3333 | 1 | 0 | 9 | 0.3333 | 5 | 0 | 3 | 0 | 1 | 0 | 4 | 0.25 | 13 | 0.0769 |
| 1_59202894 | NA | NA | 6 | 0.8333 | 2 | 0.5 | NA | NA | 8 | 0.75 | 2 | 1 | 2 | 0.5 | NA | NA | 2 | 0 | 6 | 0.5 |
| 1_59203140 | NA | NA | 4 | 0.75 | 2 | 0 | NA | NA | 6 | 0.5 | 1 | 1 | 2 | 1 | 1 | 1 | 4 | 0.25 | 8 | 0.625 |
| 1_59203227 | NA | NA | 4 | 1 | 1 | 1 | NA | NA | 5 | 1 | 2 | 1 | 2 | 0.5 | NA | NA | 4 | 0.75 | 8 | 0.75 |
| 1_59203248 | NA | NA | 3 | 0.3333 | 1 | 0 | NA | NA | 4 | 0.25 | 2 | 0.5 | 2 | 0.5 | 1 | 1 | 4 | 0 | 9 | 0.3333 |
| 1_59522945 | 1 | 1 | NA | NA | NA | NA | NA | NA | 1 | 1 | 1 | 1 | NA | NA | NA | NA | NA | NA | 1 | 1 |
| 1_59523905 | NA | NA | NA | NA | NA | NA | 1 | 1 | 1 | 1 | 2 | 1 | NA | NA | NA | NA | NA | NA | 2 | 1 |
| 1_59524063 | 1 | 0 | NA | NA | NA | NA | 1 | 1 | 2 | 0.5 | 2 | 1 | NA | NA | NA | NA | NA | NA | 2 | 1 |
| 1_59524076 | 1 | 0 | NA | NA | NA | NA | 1 | 1 | 2 | 0.5 | 2 | 0.5 | NA | NA | NA | NA | NA | NA | 2 | 0.5 |
| 1_59524155 | 1 | 0 | NA | NA | NA | NA | 1 | 1 | 2 | 0.5 | 1 | 1 | NA | NA | NA | NA | NA | NA | 1 | 1 |
| 1_59525951 | 2 | 0 | 1 | 0 | 1 | 0 | 3 | 0.3333 | 7 | 0.1429 | 2 | 0.5 | NA | NA | 2 | 0 | 1 | 0 | 5 | 0.2 |

| | | | | | | | | | | | | | | | | | | | | |
|---|---|---|---|---|---|---|---|---|---|---|---|---|---|---|---|---|---|---|---|---|
| 1_5966936 | 1 | 0 | 2 | 0.5 | NA | NA | NA | NA | 3 | 0.3333 | 1 | 1 | NA | NA | 1 | 1 | 2 | 0 | 4 | 0.5 |
| 1_5967528 | 1 | 1 | 1 | 0 | 1 | 0 | 2 | 0.5 | 5 | 0.4 | 1 | 0 | NA | NA | 1 | 0 | 2 | 0.5 | 4 | 0.25 |
| 1_5967531 | 2 | 0 | 2 | 0 | 2 | 0.5 | 2 | 0 | 8 | 0.125 | 1 | 1 | NA | NA | 2 | 0.5 | 2 | 0 | 5 | 0.4 |
| 1_5967549 | 1 | 0 | 2 | 1 | 2 | 0 | 2 | 0 | 7 | 0.2857 | 1 | 0 | NA | NA | 2 | 0.5 | 2 | 0 | 5 | 0.2 |
| 1_5967869 | 23 | 0.1304 | 18 | 0.1667 | 15 | 0.0667 | 23 | 0.1739 | 79 | 0.1392 | 13 | 0.1538 | 23 | 0.1304 | 22 | 0.1364 | 27 | 0 | 85 | 0.0941 |
| 1_5967889 | 24 | 0.0417 | 22 | 0.2273 | 20 | 0 | 28 | 0.0714 | 94 | 0.0851 | 15 | 0 | 25 | 0 | 27 | 0.0741 | 26 | 0.1538 | 93 | 0.0645 |
| 1_5967894 | 17 | 0.9412 | 14 | 0.8571 | 13 | 0.6923 | 22 | 0.5909 | 66 | 0.7576 | 12 | 0.75 | 16 | 0.6875 | 21 | 0.7619 | 19 | 0.6842 | 68 | 0.7206 |
| 1_5967895 | 19 | 0.8947 | 15 | 0.6667 | 13 | 0.6923 | 21 | 0.6667 | 68 | 0.7353 | 12 | 0.75 | 16 | 0.9375 | 22 | 0.6364 | 20 | 0.7 | 70 | 0.7429 |
| 1_5967899 | 24 | 0.9583 | 19 | 1 | 16 | 0.875 | 27 | 0.963 | 86 | 0.9535 | 14 | 0.9286 | 20 | 0.85 | 26 | 0.8846 | 25 | 0.88 | 85 | 0.8824 |
| 1_5967909 | 27 | 0.6296 | 22 | 0.6364 | 17 | 0.7059 | 29 | 0.6207 | 95 | 0.6421 | 16 | 0.625 | 24 | 0.625 | 26 | 0.8077 | 29 | 0.6552 | 95 | 0.6842 |
| 1_5980354 | NA | NA | 2 | 0.5 | NA | NA | NA | NA | 2 | 0.5 | NA | NA | NA | NA | 1 | 1 | 1 | 0 | 2 | 0.5 |
| 1_5990656 | 3 | 0 | 6 | 0 | 1 | 0 | 3 | 0.3333 | 13 | 0.0769 | 3 | 0 | 2 | 0 | 4 | 0.25 | 1 | 0 | 10 | 0.1 |
| 1_5990690 | 3 | 0 | 4 | 0 | 1 | 0 | 3 | 0.3333 | 11 | 0.0909 | 4 | 0 | 2 | 0.5 | 5 | 0.2 | NA | NA | 11 | 0.1818 |
| 1_5990693 | 3 | 0.3333 | 6 | 0.1667 | 1 | 0 | 2 | 0 | 12 | 0.1667 | 3 | 0.3333 | 2 | 0 | 5 | 0 | NA | NA | 10 | 0.1 |
| 1_5990706 | 1 | 1 | 5 | 0.8 | NA | NA | 2 | 0.5 | 8 | 0.75 | 4 | 0.25 | 2 | 1 | 4 | 0.5 | 1 | 0 | 11 | 0.4545 |
| 1_5990707 | 3 | 1 | 4 | 0.75 | 1 | 0 | 3 | 0.6667 | 11 | 0.7273 | 4 | 0.5 | 1 | 1 | 4 | 0.75 | 1 | 1 | 10 | 0.7 |
| 1_5990715 | 3 | 0 | 6 | 0 | 1 | 1 | 3 | 0 | 13 | 0.0769 | 4 | 0 | 2 | 0 | 6 | 0.1667 | 1 | 0 | 13 | 0.0769 |
| 1_5990717 | 3 | 1 | 5 | 0.6 | 1 | 1 | 3 | 0.3333 | 12 | 0.6667 | 4 | 0.75 | 2 | 0.5 | 6 | 0.1667 | 1 | 1 | 13 | 0.4615 |
| 1_5990721 | 3 | 0 | 5 | 0.2 | 1 | 0 | 3 | 0.3333 | 12 | 0.1667 | 4 | 0 | 2 | 0 | 5 | 0.2 | 1 | 0 | 12 | 0.0833 |
| 1_5990724 | 3 | 0 | 6 | 0.3333 | 1 | 0 | 3 | 0.3333 | 13 | 0.2308 | 4 | 0.25 | 2 | 0.5 | 6 | 0.3333 | NA | NA | 12 | 0.3333 |
| 1_5990726 | 3 | 0 | 6 | 0.3333 | NA | NA | 3 | 0.3333 | 12 | 0.25 | 4 | 0.25 | 2 | 0.5 | 6 | 0 | 1 | 1 | 13 | 0.2308 |
| 1_5990731 | 2 | 0 | 5 | 0.6 | 1 | 0 | 1 | 0 | 9 | 0.3333 | 4 | 0 | 1 | 0 | 6 | 0.1667 | 1 | 0 | 12 | 0.0833 |
| 1_5990732 | 1 | 0 | 1 | 0 | 1 | 0 | 3 | 0.3333 | 6 | 0.1667 | 3 | 0.6667 | 2 | 0 | 4 | 0.5 | 1 | 0 | 10 | 0.4 |
| 1_5990751 | 3 | 0.6667 | 4 | 0.5 | 1 | 0 | 2 | 1 | 10 | 0.6 | 5 | 1 | 3 | 0.6667 | 4 | 0.75 | 2 | 0.5 | 14 | 0.7857 |
| 1_5990764 | 2 | 0 | 6 | 0 | 1 | 0 | 3 | 0.3333 | 12 | 0.0833 | 4 | 0.25 | 3 | 0 | 4 | 0 | 1 | 0 | 12 | 0.0833 |
| 1_5990951 | 3 | 0.6667 | 2 | 1 | 2 | 0.5 | 3 | 0.6667 | 10 | 0.7 | 5 | 0.6 | 1 | 0 | 6 | 0.8333 | 3 | 1 | 15 | 0.7333 |
| 1_5990970 | 4 | 0 | 6 | 0.1667 | 2 | 0 | 3 | 0.3333 | 15 | 0.1333 | 5 | 0.2 | 3 | 0 | 4 | 0.25 | 2 | 0 | 14 | 0.1429 |
| 1_5990971 | 4 | 0.5 | 4 | 0.5 | 1 | 0 | 2 | 0.5 | 11 | 0.4545 | 3 | 0.6667 | 3 | 0 | 6 | 0.1667 | 1 | 0 | 13 | 0.2308 |
| 1_5990976 | 4 | 0 | 6 | 0.3333 | 2 | 0.5 | 1 | 0 | 13 | 0.2308 | 4 | 0.5 | 3 | 0.6667 | 6 | 0.3333 | 2 | 1 | 15 | 0.5333 |
| 1_5991030 | 4 | 0 | 7 | 0.1429 | 3 | 0 | 5 | 0.2 | 19 | 0.1053 | 5 | 0.4 | 3 | 0.3333 | 6 | 0.1667 | 3 | 0 | 17 | 0.2353 |
| 1_5991571 | 21 | 0.2381 | 15 | 0.0667 | 8 | 0.25 | 24 | 0 | 68 | 0.1176 | 9 | 0 | 15 | 0 | 17 | 0.0588 | 14 | 0.2143 | 55 | 0.0727 |
| 1_5991628 | 26 | 0.0769 | 14 | 0.2857 | 11 | 0.1818 | 23 | 0.087 | 74 | 0.1351 | 10 | 0.3 | 16 | 0.1875 | 19 | 0.4211 | 18 | 0.1667 | 63 | 0.2698 |
| 1_5991667 | 22 | 0.1364 | 15 | 0.0667 | 11 | 0 | 21 | 0.0952 | 69 | 0.087 | 11 | 0 | 18 | 0 | 23 | 0.087 | 18 | 0.1111 | 70 | 0.0571 |
| 1_6002081 | 3 | 0 | 5 | 0 | 2 | 0 | 3 | 0.3333 | 13 | 0.0769 | 2 | 0 | 4 | 0 | 1 | 0 | 3 | 0.3333 | 10 | 0.1 |
| 1_6002092 | 3 | 0 | 6 | 0.3333 | 1 | 1 | 3 | 0.3333 | 13 | 0.3077 | 2 | 0 | 5 | 0.2 | 1 | 1 | 2 | 0.5 | 10 | 0.3 |
| 1_6002093 | 2 | 1 | 5 | 1 | 2 | 1 | 3 | 1 | 12 | 1 | 1 | 1 | 4 | 1 | 1 | 1 | 2 | 1 | 8 | 1 |
| 1_6002100 | 3 | 0 | 6 | 0.1667 | 2 | 0.5 | 2 | 0 | 13 | 0.1538 | 1 | 0 | 5 | 0.2 | 1 | 0 | 3 | 0 | 10 | 0.1 |
| 1_6002102 | 3 | 0.3333 | 4 | 1 | 2 | 0.5 | 3 | 0.3333 | 12 | 0.5833 | 1 | 1 | 5 | 0.6 | 1 | 1 | 4 | 0.75 | 11 | 0.7273 |
| 1_6002114 | 3 | 1 | 4 | 0.25 | 1 | 0 | 2 | 1 | 10 | 0.6 | 1 | 1 | 5 | 0.6 | 1 | 1 | 2 | 0.5 | 9 | 0.6667 |
| 1_6002117 | 2 | 1 | 6 | 1 | 2 | 1 | 3 | 1 | 13 | 1 | 1 | 1 | 5 | 1 | 1 | 1 | 5 | 1 | 12 | 1 |
| 1_6002121 | 5 | 0.2 | 6 | 0 | 1 | 0 | 3 | 0.3333 | 15 | 0.1333 | 2 | 0.5 | 5 | 0 | 1 | 0 | 2 | 0 | 10 | 0.1 |
| 1_6002140 | 1 | 0 | 4 | 1 | 2 | 1 | 4 | 0.5 | 11 | 0.7273 | 1 | 1 | 1 | 1 | 2 | 1 | 4 | 0.5 | 8 | 0.75 |
| 1_6002142 | 1 | 1 | 4 | 0 | 2 | 0 | 4 | 0 | 11 | 0.0909 | 1 | 0 | 3 | 0 | 1 | 0 | 3 | 0.3333 | 8 | 0.125 |
| 1_6002145 | 2 | 0.5 | 3 | 1 | 2 | 1 | 3 | 1 | 10 | 0.9 | NA | NA | 3 | 0.6667 | 2 | 1 | 2 | 1 | 7 | 0.8571 |
| 1_6002178 | 3 | 0 | 6 | 0 | 3 | 0.3333 | 3 | 0 | 15 | 0.0667 | 3 | 0.3333 | 3 | 0.3333 | 2 | 0 | 5 | 0.2 | 13 | 0.2308 |
| 1_6002181 | 4 | 0 | 7 | 0.1429 | 2 | 0.5 | 4 | 0 | 17 | 0.1176 | 2 | 0 | 3 | 0 | 3 | 0 | 6 | 0.1667 | 14 | 0.0714 |
| 1_6002195 | 4 | 0.25 | 7 | 0.7143 | 2 | 0.5 | 4 | 0 | 17 | 0.4118 | 2 | 0 | 3 | 0.3333 | 3 | 0.6667 | 6 | 0.5 | 14 | 0.4286 |
| 1_6002195 | 3 | 0 | 7 | 0.1429 | 1 | 1 | 4 | 0.25 | 15 | 0.2 | 2 | 0.5 | 3 | 0 | 2 | 0.5 | 5 | 0 | 12 | 0.1667 |
| 1_6002404 | 9 | 0 | 9 | 0.1111 | 9 | 0.2222 | 9 | 0.1111 | 36 | 0.1111 | 2 | 0 | 7 | 0 | 5 | 0.4 | 7 | 0 | 21 | 0.0952 |
| 1_6002442 | 9 | 0.2222 | 10 | 0.1 | 9 | 0.3333 | 11 | 0.0909 | 39 | 0.1795 | 3 | 0.3333 | 11 | 0.1818 | 8 | 0.25 | 10 | 0.1 | 32 | 0.1875 |
| 1_6002895 | NA | NA | 2 | 0.5 | 3 | 0.3333 | 1 | 0 | 6 | 0.3333 | 1 | 0 | 2 | 0.5 | 2 | 0 | 2 | 0 | 7 | 0.1429 |
| 1_6002898 | NA | NA | 2 | 0 | 4 | 0.25 | 1 | 0 | 7 | 0.1429 | 1 | 0 | 2 | 0 | 2 | 0 | 5 | 0.4 | 10 | 0.2 |
| 1_6002941 | NA | NA | 2 | 1 | 1 | 0 | 2 | 1 | 5 | 0.8 | NA | NA | 1 | 0 | 2 | 1 | 2 | 0 | 5 | 0.4 |
| 1_6002950 | NA | NA | 2 | 0 | 3 | 0.3333 | 1 | 1 | 6 | 0.3333 | 1 | 1 | 1 | 0 | 2 | 0 | 2 | 0.5 | 6 | 0.3333 |
| 1_6002957 | NA | NA | 2 | 0 | 3 | 0 | 2 | 0.5 | 7 | 0.1429 | 1 | 0 | 1 | 0 | 1 | 1 | 3 | 0 | 6 | 0.1667 |
| 1_6002960 | NA | NA | 2 | 0.5 | 3 | 0.3333 | 2 | 1 | 7 | 0.5714 | 1 | 0 | 2 | 0.5 | 2 | 1 | NA | NA | 5 | 0.6 |
| 1_6002964 | NA | NA | 2 | 0 | 3 | 0.3333 | 2 | 0 | 7 | 0.1429 | 1 | 1 | 2 | 0 | 2 | 0 | 3 | 0 | 8 | 0.125 |
| 1_6002970 | NA | NA | 2 | 0.5 | 2 | 0.5 | 2 | 1 | 6 | 0.6667 | 1 | 1 | 2 | 0.5 | NA | NA | 3 | 0.6667 | 6 | 0.6667 |
| 1_6002983 | 3 | 0 | 4 | 0 | 3 | 0.3333 | 1 | 0 | 11 | 0.0909 | 1 | 0 | 2 | 0.5 | 1 | 0 | 2 | 0.5 | 6 | 0.3333 |
| 1_6003009 | 3 | 1 | 3 | 1 | 3 | 1 | 1 | 1 | 10 | 1 | 2 | 1 | 1 | 1 | 1 | 1 | NA | NA | 4 | 1 |
| 1_6003012 | 3 | 1 | 2 | 1 | 3 | 1 | 1 | 1 | 9 | 1 | 1 | 0 | 2 | 0.5 | 2 | 1 | 2 | 1 | 7 | 0.7143 |
| 1_6003026 | 3 | 0.6667 | 2 | 0.5 | 4 | 0.5 | 1 | 0 | 10 | 0.5 | 1 | 0 | 2 | 0.5 | 2 | 0.5 | 1 | 1 | 6 | 0.5 |
| 1_6003031 | 2 | 1 | 2 | 1 | 4 | 0.5 | 2 | 1 | 10 | 0.8 | 2 | 0 | 2 | 0.5 | 2 | 1 | 2 | 1 | 8 | 0.625 |
| 1_6003034 | 3 | 0 | 1 | 1 | 2 | 0.5 | 1 | 0 | 7 | 0.2857 | 1 | 0 | 2 | 0 | 2 | 0.5 | 2 | 0.5 | 7 | 0.2857 |
| 1_6011722 | 3 | 0.3333 | 2 | 0 | NA | NA | 3 | 0.3333 | 8 | 0.25 | 1 | 1 | 1 | 0 | 1 | 0 | 1 | 0 | 4 | 0.25 |

| | | | | | | | | | | | | | | | | | | | | |
|---|---|---|---|---|---|---|---|---|---|---|---|---|---|---|---|---|---|---|---|---|
| 1_60117302 | 3 | 0.6667 | 2 | 0.5 | NA | NA | 3 | 0.3333 | 8 | 0.5 | 2 | 0.5 | 2 | 0 | 1 | 0 | 2 | 0 | 7 | 0.1429 |
| 1_60187154 | 2 | 1 | NA | NA | NA | NA | 1 | 1 | 3 | 1 | NA | NA | NA | NA | 1 | 1 | 1 | 1 | 2 | 1 |
| 1_60187301 | 2 | 0.5 | NA | NA | NA | NA | 1 | 1 | 3 | 0.6667 | NA | NA | NA | NA | 1 | 0 | 1 | 1 | 2 | 0.5 |
| 1_60189821 | 2 | 1 | NA | NA | 1 | 1 | 1 | 1 | 4 | 1 | NA | NA | NA | NA | 1 | 1 | 1 | 1 | 2 | 1 |
| 1_60190505 | 3 | 0 | 1 | 0 | 2 | 0.5 | 1 | 0 | 7 | 0.1429 | 1 | 0 | NA | NA | 1 | 0 | 2 | 0.5 | 4 | 0.25 |
| 1_60190986 | 3 | 0 | 1 | 0 | 2 | 0.5 | 1 | 0 | 7 | 0.1429 | 1 | 0 | NA | NA | 2 | 0 | 2 | 0.5 | 5 | 0.2 |
| 1_60191916 | 2 | 0 | 2 | 0.5 | 2 | 0 | 1 | 1 | 7 | 0.2857 | 1 | 0 | NA | NA | 3 | 0.6667 | 2 | 0 | 6 | 0.3333 |
| 1_60426394 | 5 | 0.2 | 6 | 0.3333 | 2 | 0.5 | 1 | 0 | 14 | 0.2857 | NA | NA | 1 | 1 | 2 | 0 | 2 | 0 | 5 | 0.2 |
| 1_60516040 | 7 | 1 | 10 | 0.8 | 12 | 1 | 14 | 1 | 43 | 0.9535 | 8 | 1 | 9 | 0.8889 | 10 | 0.8 | 11 | 1 | 38 | 0.9211 |
| 1_60516226 | 8 | 0.625 | 6 | 1 | 8 | 1 | 10 | 1 | 32 | 0.9062 | 11 | 0.6364 | 10 | 1 | 8 | 1 | 12 | 0.9167 | 41 | 0.878 |
| 1_60516298 | 8 | 0.25 | 10 | 0.2 | 14 | 0.2143 | 13 | 0.2308 | 45 | 0.2222 | 12 | 0.0833 | 10 | 0.2 | 11 | 0.1818 | 14 | 0.2143 | 47 | 0.1702 |
| 1_60516333 | 8 | 0.625 | 10 | 0.9 | 13 | 0.8462 | 13 | 0.8462 | 44 | 0.8182 | 11 | 0.5455 | 10 | 1 | 13 | 0.8462 | 12 | 1 | 46 | 0.8478 |
| 1_60516355 | 10 | 0.1 | 9 | 0.1111 | 13 | 0 | 14 | 0.1429 | 46 | 0.087 | 12 | 0 | 10 | 0.4 | 12 | 0.0833 | 13 | 0.0769 | 47 | 0.1277 |
| 1_60516361 | 7 | 0.8571 | 4 | 0.75 | 3 | 1 | 4 | 0.75 | 18 | 0.8333 | 8 | 0.75 | 3 | 0.6667 | 6 | 1 | 6 | 0.8333 | 23 | 0.8261 |
| 1_60516365 | 9 | 0.2222 | 11 | 0.3636 | 11 | 0.7273 | 12 | 0.3333 | 43 | 0.4186 | 12 | 0.25 | 8 | 0.625 | 11 | 0.4545 | 8 | 0.125 | 39 | 0.359 |
| 1_60516440 | 8 | 0.5 | 11 | 0.5455 | 14 | 0.6429 | 13 | 0.3846 | 46 | 0.5217 | 11 | 0.2727 | 9 | 0.5556 | 11 | 0.5455 | 12 | 0.8333 | 43 | 0.5581 |
| 1_60516491 | 10 | 0.4 | 12 | 0.6667 | 15 | 0.5333 | 12 | 0.75 | 49 | 0.5918 | 13 | 0.4615 | 9 | 0.6667 | 12 | 0.5833 | 13 | 0.7692 | 47 | 0.617 |
| 1_60516991 | 11 | 0.0909 | 13 | 0.1538 | 16 | 0.125 | 9 | 0.1111 | 49 | 0.1224 | 8 | 0.125 | 10 | 0 | 10 | 0.1 | 11 | 0.0909 | 39 | 0.0769 |
| 1_60517116 | 13 | 0.1538 | 14 | 0.1429 | 18 | 0.0556 | 15 | 0.0667 | 60 | 0.1 | 15 | 0.1333 | 12 | 0 | 16 | 0 | 17 | 0.1176 | 60 | 0.0667 |
| 1_60517788 | 18 | 0.1111 | 14 | 0.0714 | 21 | 0.0952 | 17 | 0 | 70 | 0.0714 | 15 | 0.2 | 11 | 0.2727 | 18 | 0.0556 | 17 | 0.0588 | 61 | 0.1311 |
| 1_60517906 | 19 | 0.1579 | 20 | 0.2 | 19 | 0 | 22 | 0.0455 | 80 | 0.1 | 15 | 0.1333 | 16 | 0 | 25 | 0.16 | 27 | 0.2222 | 83 | 0.1446 |
| 1_60517959 | 19 | 0.1053 | 20 | 0.2 | 23 | 0.087 | 23 | 0.0435 | 85 | 0.1059 | 17 | 0 | 14 | 0.1429 | 24 | 0.125 | 27 | 0.1852 | 82 | 0.122 |
| 1_60517970 | 21 | 0.1905 | 18 | 0.1667 | 21 | 0.3333 | 26 | 0.1154 | 86 | 0.1977 | 17 | 0.1765 | 15 | 0.2 | 23 | 0.2174 | 25 | 0.24 | 80 | 0.2125 |
| 1_60518171 | 24 | 0.5 | 16 | 0.25 | 19 | 0.4737 | 17 | 0.1765 | 76 | 0.3684 | 13 | 0.3077 | 13 | 0.2308 | 21 | 0.2857 | 19 | 0.2105 | 66 | 0.2576 |
| 1_60518251 | 24 | 0.0833 | 18 | 0.0556 | 19 | 0.0526 | 23 | 0.0435 | 84 | 0.0595 | 14 | 0.0714 | 15 | 0 | 22 | 0.1364 | 24 | 0.2083 | 75 | 0.12 |
| 1_60518684 | 29 | 0.2414 | 21 | 0.0476 | 27 | 0.1111 | 29 | 0.2759 | 106 | 0.1792 | 16 | 0.125 | 19 | 0.1579 | 25 | 0.28 | 25 | 0.24 | 85 | 0.2118 |
| 1_60518689 | 16 | 0.0625 | 14 | 0.0714 | 17 | 0 | 20 | 0.2 | 67 | 0.0896 | 12 | 0.1667 | 13 | 0.0769 | 12 | 0 | 21 | 0.0952 | 58 | 0.0862 |
| 1_60518943 | 22 | 0.1818 | 18 | 0.1667 | 18 | 0.2222 | 23 | 0.0435 | 81 | 0.1481 | 17 | 0.1176 | 14 | 0.0714 | 21 | 0.2381 | 27 | 0.1111 | 79 | 0.1392 |
| 1_60519038 | 20 | 0.2 | 17 | 0.0588 | 19 | 0.0526 | 25 | 0.08 | 81 | 0.0988 | 17 | 0.0588 | 14 | 0 | 21 | 0.1429 | 27 | 0.037 | 79 | 0.0633 |
| 1_60520841 | 1 | 1 | 3 | 0.3333 | 1 | 1 | 1 | 0 | 6 | 0.5 | 1 | 0 | 1 | 0 | 2 | 0 | 6 | 0.1667 | 10 | 0.1 |
| 1_60522981 | 2 | 0.5 | 2 | 0.5 | 1 | 1 | 6 | 0.3333 | 11 | 0.4545 | NA | NA | 3 | 0.6667 | 2 | 0.5 | 3 | 1 | 8 | 0.75 |
| 1_60523231 | 1 | 0 | 3 | 0.3333 | 1 | 0 | 7 | 0 | 12 | 0.0833 | NA | NA | 2 | 0 | 3 | 0.3333 | 3 | 0 | 8 | 0.125 |
| 1_60523264 | 3 | 0.3333 | 4 | 0 | 1 | 0 | 8 | 0.125 | 16 | 0.125 | NA | NA | 4 | 0.25 | 3 | 0 | 3 | 0.3333 | 10 | 0.2 |
| 1_60524499 | 1 | 0 | 4 | 0.25 | NA | NA | 4 | 0.25 | 9 | 0.2222 | NA | NA | 3 | 0.3333 | 2 | 0 | 1 | 0 | 6 | 0.1667 |
| 1_60527452 | 1 | 1 | NA | NA | NA | NA | 1 | 0 | 2 | 0.5 | NA | NA | 3 | 1 | 1 | 1 | 2 | 1 | 6 | 1 |
| 1_60527460 | 1 | 1 | NA | NA | NA | NA | 2 | 1 | 3 | 1 | NA | NA | 3 | 1 | 1 | 1 | 2 | 1 | 6 | 1 |
| 1_60527490 | NA | NA | NA | NA | NA | NA | 1 | 1 | 1 | 1 | NA | NA | 3 | 1 | 1 | 1 | 2 | 0 | 6 | 0.6667 |
| 1_60527588 | NA | NA | NA | NA | NA | NA | 1 | 1 | 1 | 1 | NA | NA | NA | NA | 1 | 1 | 1 | 1 | 2 | 1 |
| 1_60527701 | NA | NA | NA | NA | NA | NA | 2 | 1 | 2 | 1 | NA | NA | 2 | 1 | 1 | 1 | 1 | 1 | 4 | 1 |
| 1_60527795 | NA | NA | NA | NA | NA | NA | 2 | 0.5 | 2 | 0.5 | NA | NA | 1 | 0 | 1 | 1 | 2 | 0 | 4 | 0.25 |
| 1_60527829 | NA | NA | NA | NA | NA | NA | 1 | 1 | 1 | 1 | NA | NA | 1 | 0 | 1 | 1 | 1 | 0 | 3 | 0.3333 |
| 1_60528000 | NA | NA | NA | NA | NA | NA | 1 | 1 | 1 | 1 | NA | NA | 2 | 1 | 1 | 1 | 1 | 1 | 4 | 1 |
| 1_60528609 | NA | NA | NA | NA | NA | NA | 1 | 1 | 1 | 1 | NA | NA | 2 | 1 | NA | NA | NA | NA | 2 | 1 |
| 1_60744407 | NA | NA | 3 | 0.3333 | 4 | 0 | 1 | 0 | 8 | 0.125 | 3 | 0.6667 | 4 | 0.25 | 1 | 0 | 3 | 0 | 11 | 0.2727 |
| 1_60744689 | 2 | 0 | 7 | 0 | 7 | 0.1429 | 3 | 0 | 19 | 0.0526 | 6 | 0.1667 | 7 | 0 | 5 | 0.2 | 3 | 0 | 21 | 0.0952 |
| 1_61763155 | NA | NA | NA | NA | NA | NA | 1 | 1 | 1 | 1 | NA | NA | NA | NA | NA | NA | 2 | 1 | 2 | 1 |
| 1_62743076 | NA | NA | 1 | 1 | NA | NA | NA | NA | 1 | 1 | NA | NA | 1 | 0 | 2 | 0.5 | NA | NA | 3 | 0.3333 |
| 1_62743308 | NA | NA | 1 | 1 | NA | NA | NA | NA | 1 | 1 | NA | NA | 2 | 1 | 2 | 1 | NA | NA | 4 | 1 |
| 1_62835194 | 2 | 1 | 1 | 1 | NA | NA | 2 | 1 | 5 | 1 | NA | NA | 3 | 1 | 1 | 1 | NA | NA | 4 | 1 |
| 1_62835990 | 2 | 0.5 | 1 | 1 | NA | NA | 2 | 0.5 | 5 | 0.6 | 1 | 0 | 4 | 0.25 | 2 | 0 | NA | NA | 7 | 0.1429 |
| 1_62854894 | NA | NA | 2 | 1 | 1 | 0 | 1 | 1 | 4 | 0.75 | NA | NA | NA | NA | 1 | 1 | NA | NA | 1 | 1 |
| 1_62854944 | NA | NA | 2 | 1 | 1 | 1 | 1 | 1 | 4 | 1 | NA | NA | NA | NA | 1 | 1 | NA | NA | 1 | 1 |
| 1_62855012 | NA | NA | 2 | 0.5 | 1 | 0 | 1 | 1 | 4 | 0.5 | NA | NA | NA | NA | 2 | 1 | NA | NA | 2 | 1 |
| 1_62855062 | NA | NA | 2 | 0.5 | 1 | 1 | 1 | 1 | 4 | 0.75 | NA | NA | NA | NA | 2 | 1 | NA | NA | 2 | 1 |
| 1_62856329 | NA | NA | 2 | 0.5 | 2 | 1 | 4 | 1 | 8 | 0.875 | NA | NA | NA | NA | 2 | 0.5 | NA | NA | 2 | 0.5 |
| 1_62857731 | 1 | 0 | 4 | 0 | 3 | 0 | 5 | 0.4 | 13 | 0.1538 | 3 | 0 | 2 | 0 | 3 | 0.3333 | NA | NA | 8 | 0.125 |
| 1_63087992 | 2 | 0 | 2 | 0.5 | 2 | 0 | 4 | 0.25 | 10 | 0.2 | 2 | 0.5 | 3 | 0 | 2 | 0 | 2 | 0 | 9 | 0.1111 |
| 1_63088480 | 2 | 0.5 | 1 | 0 | 1 | 0 | 1 | 0 | 5 | 0.2 | 1 | 1 | 2 | 0 | 2 | 0 | 1 | 0 | 6 | 0.1667 |
| 1_63089549 | 1 | 0 | 1 | 1 | NA | NA | 3 | 1 | 5 | 0.8 | NA | NA | 2 | 0.5 | 2 | 0 | NA | NA | 4 | 0.25 |
| 1_63153632 | NA | NA | 1 | 1 | NA | NA | NA | NA | 1 | 1 | NA | NA | NA | NA | 1 | 1 | NA | NA | 1 | 1 |
| 1_63153866 | 1 | 0 | 1 | 1 | NA | NA | NA | NA | 2 | 0.5 | NA | NA | 1 | 0 | 1 | 1 | NA | NA | 2 | 0.5 |
| 1_63182948 | 35 | 0.1714 | 25 | 0.2 | 22 | 0.2273 | 22 | 0.2727 | 104 | 0.2115 | 21 | 0.2381 | 11 | 0.2727 | 20 | 0.2 | 23 | 0.1304 | 75 | 0.2 |
| 1_63202912 | 29 | 0.0345 | 17 | 0.1176 | 9 | 0.1111 | 12 | 0 | 67 | 0.0597 | 14 | 0 | 9 | 0.1111 | 13 | 0.0769 | 18 | 0.0556 | 54 | 0.0556 |
| 1_63216206 | NA | NA | 3 | 0.6667 | 2 | 0 | 5 | 0 | 10 | 0.2 | 1 | 1 | 4 | 0.25 | 1 | 0 | NA | NA | 6 | 0.3333 |
| 1_63218728 | 46 | 0.087 | 48 | 0.0625 | 38 | 0.0526 | 45 | 0.0889 | 177 | 0.0734 | 32 | 0 | 58 | 0.1034 | 47 | 0.0638 | 37 | 0.1351 | 174 | 0.0805 |

| | | | | | | | | | | | | | | | | | | | | |
|---|---|---|---|---|---|---|---|---|---|---|---|---|---|---|---|---|---|---|---|---|
| 1_6334603 | 1 | 1 | NA | NA | NA | NA | 1 | 1 | 2 | 1 | NA | NA | NA | NA | 1 | 1 | NA | NA | 1 | 1 |
| 1_6334656 | 1 | 1 | NA | NA | NA | NA | 1 | 1 | 2 | 1 | NA | NA | NA | NA | 2 | 1 | NA | NA | 2 | 1 |
| 1_6334696 | 1 | 1 | NA | NA | NA | NA | 1 | 1 | 2 | 1 | 1 | 1 | NA | NA | 2 | 1 | NA | NA | 3 | 1 |
| 1_6334731 | 1 | 1 | NA | NA | NA | NA | 1 | 1 | 2 | 1 | 1 | 0 | NA | NA | 2 | 1 | 1 | 1 | 4 | 0.75 |
| 1_6334771 | 1 | 1 | NA | NA | NA | NA | 1 | 1 | 2 | 1 | 2 | 1 | NA | NA | 2 | 1 | 1 | 1 | 5 | 1 |
| 1_6334781 | 1 | 0 | NA | NA | NA | NA | 1 | 1 | 2 | 0.5 | 2 | 0.5 | NA | NA | 2 | 0 | 1 | 0 | 5 | 0.2 |
| 1_6334788 | 1 | 0 | NA | NA | NA | NA | 1 | 1 | 2 | 0.5 | 2 | 0.5 | NA | NA | 2 | 0.5 | 1 | 0 | 5 | 0.4 |
| 1_6334795 | 1 | 1 | NA | NA | NA | NA | 1 | 1 | 2 | 1 | 1 | 1 | NA | NA | 2 | 1 | 1 | 1 | 4 | 1 |
| 1_6334823 | 1 | 1 | NA | NA | NA | NA | 1 | 1 | 2 | 1 | 2 | 1 | NA | NA | 1 | 1 | 1 | 1 | 4 | 1 |
| 1_6334891 | 1 | 0 | 1 | 1 | NA | NA | 1 | 0 | 3 | 0.3333 | 2 | 0.5 | NA | NA | 2 | 0 | 1 | 1 | 5 | 0.4 |
| 1_6334899 | 2 | 0.5 | 1 | 1 | NA | NA | 1 | 0 | 4 | 0.5 | 2 | 1 | NA | NA | 3 | 0 | 1 | 1 | 6 | 0.5 |
| 1_6334916 | 1 | 1 | 2 | 0.5 | NA | NA | 1 | 1 | 4 | 0.75 | 2 | 1 | NA | NA | 3 | 1 | NA | NA | 5 | 1 |
| 1_6334996 | 1 | 0 | 3 | 0.3333 | NA | NA | 1 | 0 | 5 | 0.2 | 1 | 0 | NA | NA | 2 | 0.5 | NA | NA | 3 | 0.3333 |
| 1_6335205 | 1 | 0 | 5 | 0.2 | NA | NA | 2 | 0 | 8 | 0.125 | 1 | 0 | 2 | 0.5 | 7 | 0.1429 | 1 | 0 | 11 | 0.1818 |
| 1_6335213 | 1 | 0 | 5 | 0.4 | NA | NA | 2 | 0 | 8 | 0.25 | 2 | 0 | 3 | 1 | 6 | 0.5 | 1 | 0 | 12 | 0.5 |
| 1_6335350 | 1 | 1 | 7 | 0.2857 | NA | NA | 4 | 0.5 | 12 | 0.4167 | 4 | 0.5 | 4 | 0.5 | 7 | 0.5714 | 2 | 0.5 | 17 | 0.5294 |
| 1_6348704 | 2 | 0.5 | 4 | 0 | 1 | 0 | 4 | 0.25 | 11 | 0.1818 | 2 | 0 | 3 | 0 | 5 | 0.2 | 3 | 0.6667 | 13 | 0.2308 |
| 1_6363189 | 9 | 0.6667 | 11 | 0.9091 | 3 | 1 | 5 | 0.6 | 28 | 0.7857 | 2 | 0.5 | 9 | 0.8889 | 6 | 0.8333 | 10 | 0.8 | 27 | 0.8148 |
| 1_6363191 | 13 | 0.1538 | 11 | 0.1818 | 2 | 0 | 7 | 0 | 33 | 0.1212 | 3 | 0 | 12 | 0.0833 | 7 | 0.2857 | 12 | 0.0833 | 34 | 0.1176 |
| 1_6363192 | 14 | 0.4286 | 13 | 0.3846 | 3 | 0 | 7 | 0.2857 | 37 | 0.3514 | 4 | 0 | 10 | 0.2 | 6 | 0.1667 | 11 | 0 | 31 | 0.0968 |
| 1_6363199 | 10 | 0.1 | 12 | 0.0833 | 2 | 0.5 | 8 | 0.25 | 32 | 0.1562 | 3 | 0 | 8 | 0.25 | 8 | 0.125 | 12 | 0.1667 | 31 | 0.1613 |
| 1_6363204 | 14 | 0.3571 | 12 | 0.25 | 2 | 0.5 | 9 | 0.2222 | 37 | 0.2973 | 4 | 0 | 10 | 0.2 | 8 | 0.125 | 13 | 0.0769 | 35 | 0.1143 |
| 1_6363213 | 8 | 0.125 | 6 | 0.5 | 2 | 0 | 5 | 0.4 | 21 | 0.2857 | 2 | 0 | 6 | 0.1667 | 3 | 0 | 5 | 0 | 16 | 0.0625 |
| 1_6363215 | 9 | 0 | 9 | 0.2222 | 2 | 0 | 6 | 0 | 26 | 0.0769 | 3 | 0 | 6 | 0.1667 | 4 | 0.5 | 6 | 0 | 19 | 0.1579 |
| 1_6363222 | 11 | 0.6364 | 9 | 0.7778 | 2 | 0.5 | 5 | 0.8 | 27 | 0.7037 | 3 | 1 | 4 | 0.75 | 4 | 1 | 7 | 0.7143 | 18 | 0.8333 |
| 1_6363227 | 10 | 0.8 | 9 | 0.8889 | NA | NA | 4 | 1 | 23 | 0.8696 | 2 | 0.5 | 5 | 0.4 | 5 | 0.4 | 8 | 0.75 | 20 | 0.55 |
| 1_6363249 | 11 | 0.9091 | 10 | 1 | 1 | 1 | 5 | 1 | 27 | 0.963 | 6 | 0.8333 | 6 | 0.6667 | 6 | 0.8333 | 6 | 1 | 24 | 0.8333 |
| 1_6363253 | 10 | 0 | 6 | 0 | 1 | 0 | 4 | 0.5 | 21 | 0.0952 | 7 | 0.2857 | 4 | 0.5 | 5 | 0.2 | 6 | 0.1667 | 22 | 0.2727 |
| 1_6363256 | 11 | 0.0909 | 12 | 0.25 | 2 | 0 | 4 | 0.5 | 29 | 0.2069 | 7 | 0.2857 | 5 | 0.2 | 6 | 0.3333 | 9 | 0.2222 | 27 | 0.2593 |
| 1_6363294 | 14 | 0 | 12 | 0.4167 | 5 | 0 | 6 | 0 | 37 | 0.1351 | 5 | 0 | 6 | 0.3333 | 6 | 0 | 7 | 0 | 24 | 0.0833 |
| 1_6363343 | 16 | 0 | 12 | 0.0833 | 7 | 0 | 7 | 0.2857 | 42 | 0.0714 | 6 | 0 | 6 | 0.1667 | 6 | 0 | 10 | 0.2 | 28 | 0.1071 |
| 1_6363378 | 17 | 0.1765 | 14 | 0.3571 | 7 | 0.1429 | 8 | 0 | 46 | 0.1957 | 6 | 0 | 7 | 0 | 7 | 0.1429 | 10 | 0.3 | 30 | 0.1333 |
| 1_6363379 | 17 | 0.1176 | 12 | 0.0833 | 4 | 0 | 6 | 0 | 39 | 0.0769 | 3 | 0 | 8 | 0.375 | 7 | 0.1429 | 12 | 0.5833 | 30 | 0.3667 |
| 1_6363502 | 23 | 0.1304 | 22 | 0.1818 | 19 | 0.1053 | 9 | 0.3333 | 73 | 0.1644 | 8 | 0.25 | 11 | 0.0909 | 9 | 0.3333 | 17 | 0.2353 | 45 | 0.2222 |
| 1_6377067 | 1 | 0 | 3 | 0.3333 | 2 | 0 | NA | NA | 6 | 0.1667 | NA | NA | NA | NA | 2 | 0.5 | NA | NA | 2 | 0.5 |
| 1_6377093 | 1 | 0 | 4 | 1 | 2 | 1 | NA | NA | 7 | 0.8571 | NA | NA | NA | NA | 2 | 0.5 | NA | NA | 2 | 0.5 |
| 1_6377097 | 1 | 1 | 4 | 0.5 | 2 | 0.5 | NA | NA | 7 | 0.5714 | NA | NA | NA | NA | 2 | 0.5 | NA | NA | 2 | 0.5 |
| 1_6377100 | 1 | 1 | 4 | 0.75 | 2 | 0.5 | NA | NA | 7 | 0.7143 | NA | NA | NA | NA | 2 | 1 | NA | NA | 2 | 1 |
| 1_6377111 | 1 | 1 | 4 | 0.75 | 2 | 1 | NA | NA | 7 | 0.8571 | NA | NA | NA | NA | 2 | 1 | NA | NA | 2 | 1 |
| 1_6377121 | 1 | 0 | 4 | 0 | 1 | 1 | NA | NA | 6 | 0.1667 | NA | NA | NA | NA | 2 | 0.5 | NA | NA | 2 | 0.5 |
| 1_6407020 | 1 | 0 | 3 | 0 | 3 | 0.3333 | NA | NA | 7 | 0.1429 | 2 | 0 | 3 | 0 | NA | NA | 2 | 0.5 | 7 | 0.1429 |
| 1_6407231 | NA | NA | 2 | 0.5 | 1 | 0 | NA | NA | 3 | 0.3333 | NA | NA | 2 | 0 | NA | NA | 1 | 1 | 3 | 0.3333 |
| 1_6407247 | NA | NA | 2 | 0.5 | 1 | 0 | NA | NA | 3 | 0.3333 | NA | NA | 1 | 0 | 1 | 1 | 1 | 0 | 3 | 0.3333 |
| 1_6407271 | NA | NA | 1 | 1 | NA | NA | NA | NA | 1 | 1 | NA | NA | NA | NA | 1 | 1 | 1 | 0 | 2 | 0.5 |
| 1_6407379 | NA | NA | 1 | 1 | NA | NA | NA | NA | 1 | 1 | NA | NA | NA | NA | 1 | 1 | NA | NA | 1 | 1 |
| 1_6407420 | 1 | 1 | NA | NA | NA | NA | NA | NA | 1 | 1 | NA | NA | NA | NA | 1 | 1 | NA | NA | 1 | 1 |
| 1_6463644 | NA | NA | 3 | 0.3333 | 2 | 0 | 1 | 1 | 6 | 0.3333 | NA | NA | 1 | 1 | NA | NA | 1 | 0 | 2 | 0.5 |
| 1_6463648 | NA | NA | 2 | 0.5 | 4 | 0.5 | 1 | 1 | 7 | 0.5714 | 1 | 1 | 1 | 0 | 1 | 0 | 1 | 1 | 4 | 0.5 |
| 1_6463651 | 1 | 1 | 3 | 1 | 4 | 0.75 | 1 | 1 | 9 | 0.8889 | 1 | 1 | 2 | 1 | 1 | 1 | 1 | 1 | 5 | 1 |
| 1_6463657 | NA | NA | 2 | 1 | 2 | 1 | 1 | 1 | 5 | 1 | 1 | 1 | NA | NA | 1 | 1 | 1 | 1 | 3 | 1 |
| 1_6463660 | 1 | 1 | 3 | 1 | 4 | 0.75 | 1 | 1 | 9 | 0.8889 | 1 | 1 | 2 | 0.5 | 1 | 1 | 1 | 0 | 5 | 0.6 |
| 1_6463660 | 1 | 0 | 3 | 0.6667 | 3 | 0.3333 | 1 | 0 | 8 | 0.375 | 1 | 0 | 2 | 0.5 | 1 | 0 | 1 | 0 | 5 | 0.2 |
| 1_6463662 | 1 | 1 | 1 | 0 | 2 | 0.5 | 1 | 0 | 5 | 0.4 | 1 | 1 | 1 | 0 | 1 | 0 | NA | NA | 3 | 0.3333 |
| 1_6463664 | 1 | 1 | 2 | 0.5 | 4 | 0.5 | 1 | 1 | 8 | 0.625 | 1 | 1 | 2 | 1 | 1 | 0 | NA | NA | 4 | 0.75 |
| 1_6463671 | NA | NA | 2 | 0.5 | 3 | 0.3333 | 1 | 1 | 6 | 0.5 | 1 | 0 | 1 | 1 | NA | NA | 1 | 0 | 3 | 0.3333 |
| 1_6463677 | 1 | 0 | 3 | 0.3333 | 4 | 0.25 | 1 | 0 | 9 | 0.2222 | 1 | 1 | 1 | 0 | 1 | 0 | 1 | 0 | 4 | 0.25 |
| 1_6463694 | 1 | 1 | 2 | 0 | 4 | 1 | 1 | 0 | 8 | 0.625 | 1 | 0 | 2 | 0 | 1 | 1 | NA | NA | 4 | 0.25 |
| 1_6463815 | 1 | 0 | 4 | 0 | 5 | 0.4 | 2 | 0.5 | 12 | 0.25 | 1 | 0 | 1 | 0 | 2 | 0.5 | 1 | 0 | 5 | 0.2 |
| 1_6463819 | NA | NA | 3 | 0.3333 | 5 | 0.8 | 1 | 1 | 9 | 0.6667 | 1 | 0 | 1 | 1 | 1 | 0 | NA | NA | 3 | 0.3333 |
| 1_6463836 | 2 | 0 | 5 | 0 | 5 | 0.2 | 2 | 0 | 14 | 0.0714 | 2 | 0 | NA | NA | 2 | 0 | 2 | 0.5 | 6 | 0.1667 |
| 1_6463964 | 3 | 0 | 5 | 0.2 | 5 | 0 | 2 | 0.5 | 15 | 0.1333 | 2 | 0 | 1 | 0 | 4 | 0.25 | 3 | 0 | 10 | 0.1 |
| 1_6464670 | 1 | 1 | 3 | 0.3333 | 2 | 0.5 | 1 | 1 | 7 | 0.5714 | NA | NA | 7 | 0.4286 | 1 | 1 | 2 | 1 | 10 | 0.6 |
| 1_6464690 | 1 | 1 | 2 | 0 | 3 | 1 | 1 | 0 | 7 | 0.5714 | NA | NA | 6 | 0.8333 | NA | NA | 2 | 1 | 8 | 0.875 |
| 1_6464701 | NA | NA | 2 | 0.5 | 3 | 0.3333 | 1 | 0 | 6 | 0.3333 | NA | NA | 6 | 0.1667 | NA | NA | 1 | 0 | 7 | 0.1429 |
| 1_6464707 | NA | NA | 2 | 1 | 3 | 1 | 1 | 1 | 6 | 1 | NA | NA | 7 | 1 | NA | NA | 2 | 1 | 9 | 1 |

| | | | | | | | | | | | | | | | | | | | | |
|---|---|---|---|---|---|---|---|---|---|---|---|---|---|---|---|---|---|---|---|---|
| 1_6464709 | NA | NA | 1 | 0 | 3 | 0.3333 | 1 | 1 | 5 | 0.4 | NA | NA | 8 | 0.25 | NA | NA | 2 | 0.5 | 10 | 0.3 |
| 1_6464717 | NA | NA | 2 | 0 | 2 | 0.5 | 1 | 1 | 5 | 0.4 | NA | NA | 7 | 0.2857 | NA | NA | 2 | 0.5 | 9 | 0.3333 |
| 1_6464718 | NA | NA | 2 | 0 | 1 | 1 | 1 | 0 | 4 | 0.25 | NA | NA | 8 | 0.125 | NA | NA | 2 | 0.5 | 10 | 0.2 |
| 1_6471917 | 1 | 1 | NA | NA | NA | NA | NA | NA | 1 | 1 | NA | NA | NA | NA | 1 | 1 | NA | NA | 1 | 1 |
| 1_6476271 | NA | NA | 1 | 0 | NA | NA | 3 | 0.6667 | 4 | 0.5 | 1 | 1 | 5 | 0.6 | NA | NA | 1 | 1 | 7 | 0.7143 |
| 1_6476283 | NA | NA | 1 | 1 | 1 | 1 | 2 | 0.5 | 4 | 0.75 | 1 | 1 | 4 | 1 | NA | NA | 1 | 1 | 6 | 1 |
| 1_6476284 | NA | NA | 1 | 0 | 1 | 1 | 3 | 0.3333 | 5 | 0.4 | 1 | 1 | 5 | 0 | NA | NA | 1 | 0 | 7 | 0.1429 |
| 1_6476289 | NA | NA | 1 | 0 | NA | NA | 3 | 0.3333 | 4 | 0.25 | 1 | 0 | 5 | 0.2 | NA | NA | 1 | 0 | 7 | 0.1429 |
| 1_6476292 | NA | NA | NA | NA | 1 | 1 | 3 | 0.6667 | 4 | 0.75 | 1 | 1 | 5 | 1 | NA | NA | 1 | 1 | 7 | 1 |
| 1_6476293 | NA | NA | NA | NA | 1 | 1 | 3 | 0.3333 | 4 | 0.5 | 1 | 0 | 5 | 0.4 | NA | NA | 1 | 0 | 7 | 0.2857 |
| 1_6476293 | NA | NA | 1 | 1 | 1 | 0 | 2 | 0.5 | 4 | 0.5 | 1 | 0 | 4 | 0.25 | NA | NA | 1 | 1 | 6 | 0.3333 |
| 1_6476296 | NA | NA | 1 | 0 | 1 | 1 | 2 | 1 | 4 | 0.75 | 1 | 1 | 5 | 0.8 | NA | NA | 1 | 0 | 7 | 0.7143 |
| 1_6476461 | 1 | 0 | 1 | 1 | 3 | 0.6667 | 1 | 1 | 6 | 0.6667 | 2 | 0.5 | 4 | 0 | 3 | 0.3333 | NA | NA | 9 | 0.2222 |
| 1_6477011 | 1 | 1 | NA | NA | NA | NA | 1 | 0 | 2 | 0.5 | 1 | 0 | 1 | 0 | 1 | 1 | 1 | 1 | 4 | 0.5 |
| 1_6482842 | 4 | 0 | 2 | 0 | 4 | 0 | 6 | 0.1667 | 16 | 0.0625 | 3 | 0 | 5 | 0.2 | 2 | 0 | 6 | 0 | 16 | 0.0625 |
| 1_6482869 | 5 | 0.2 | 2 | 0 | 4 | 0 | 5 | 0 | 16 | 0.0625 | 3 | 0 | 7 | 0 | 1 | 1 | 5 | 0 | 16 | 0.0625 |
| 1_6483309 | 3 | 0 | 1 | 0 | NA | NA | 3 | 0.6667 | 7 | 0.2857 | 1 | 0 | NA | NA | 2 | 1 | NA | NA | 3 | 0.6667 |
| 1_6483333 | 3 | 0.3333 | 1 | 0 | NA | NA | 3 | 0 | 7 | 0.1429 | 1 | 0 | NA | NA | 2 | 1 | NA | NA | 3 | 0.6667 |
| 1_6483344 | 2 | 1 | 1 | 1 | NA | NA | 3 | 0.6667 | 6 | 0.8333 | 1 | 1 | NA | NA | 1 | 1 | NA | NA | 2 | 1 |
| 1_6483347 | 2 | 1 | 1 | 0 | NA | NA | 2 | 0.5 | 5 | 0.6 | 1 | 0 | NA | NA | 1 | 1 | NA | NA | 2 | 0.5 |
| 1_6483358 | 3 | 1 | 1 | 0 | NA | NA | 3 | 0.6667 | 7 | 0.7143 | 1 | 0 | NA | NA | 2 | 0.5 | NA | NA | 3 | 0.3333 |
| 1_6483364 | 2 | 0.5 | 1 | 1 | NA | NA | 2 | 0.5 | 5 | 0.6 | 1 | 1 | NA | NA | 2 | 0 | NA | NA | 3 | 0.3333 |
| 1_6483377 | 2 | 1 | 1 | 0 | NA | NA | 3 | 0.6667 | 6 | 0.6667 | 1 | 1 | NA | NA | 2 | 1 | NA | NA | 3 | 1 |
| 1_6483380 | 2 | 0 | 1 | 0 | NA | NA | 2 | 0.5 | 5 | 0.2 | 1 | 0 | NA | NA | 2 | 1 | NA | NA | 3 | 0.6667 |
| 1_6483382 | 1 | 1 | 1 | 1 | NA | NA | 3 | 0.3333 | 5 | 0.6 | 1 | 1 | NA | NA | 1 | 0 | NA | NA | 2 | 0.5 |
| 1_6483400 | 2 | 1 | 1 | 0 | NA | NA | 3 | 0.6667 | 6 | 0.6667 | NA | NA | NA | NA | 2 | 1 | NA | NA | 2 | 1 |
| 1_6496099 | 1 | 1 | 1 | 1 | NA | NA | 1 | 0 | 3 | 0.6667 | 1 | 0 | NA | NA | 1 | 1 | NA | NA | 2 | 0.5 |
| 1_6496115 | 2 | 1 | 1 | 1 | NA | NA | 2 | 0.5 | 5 | 0.8 | 1 | 1 | NA | NA | 1 | 1 | NA | NA | 2 | 1 |
| 1_6496125 | NA | NA | 1 | 1 | NA | NA | 2 | 0.5 | 3 | 0.6667 | 1 | 0 | NA | NA | 1 | 1 | NA | NA | 2 | 0.5 |
| 1_6496135 | 1 | 1 | 1 | 1 | NA | NA | 1 | 1 | 3 | 1 | 1 | 1 | NA | NA | 1 | 1 | NA | NA | 2 | 1 |
| 1_6497699 | 2 | 0 | 1 | 1 | NA | NA | 1 | 0 | 4 | 0.25 | 1 | 1 | NA | NA | NA | NA | NA | NA | 1 | 1 |
| 1_6497706 | 2 | 0.5 | 1 | 0 | NA | NA | 1 | 1 | 4 | 0.5 | 1 | 1 | NA | NA | 1 | 1 | NA | NA | 2 | 1 |
| 1_6497724 | 2 | 0.5 | 1 | 1 | NA | NA | 1 | 1 | 4 | 0.75 | 1 | 0 | NA | NA | 1 | 1 | NA | NA | 2 | 0.5 |
| 1_6497733 | 2 | 1 | 1 | 1 | NA | NA | 1 | 1 | 4 | 1 | 1 | 0 | NA | NA | 1 | 1 | NA | NA | 2 | 0.5 |
| 1_6519867 | 5 | 0 | 10 | 0 | 5 | 0.4 | 5 | 0.4 | 25 | 0.16 | 7 | 0 | 8 | 0.125 | 7 | 0.1429 | 4 | 0 | 26 | 0.0769 |
| 1_6528556 | NA | NA | NA | NA | 2 | 0.5 | NA | NA | 2 | 0.5 | 2 | 1 | 1 | 1 | NA | NA | NA | NA | 3 | 1 |
| 1_6528575 | NA | NA | NA | NA | 2 | 0.5 | 1 | 1 | 3 | 0.6667 | 2 | 0.5 | 1 | 1 | NA | NA | NA | NA | 3 | 0.6667 |
| 1_6528580 | NA | NA | NA | NA | 2 | 0.5 | 1 | 0 | 3 | 0.3333 | 2 | 1 | 1 | 1 | NA | NA | NA | NA | 3 | 1 |
| 1_6528587 | NA | NA | 1 | 1 | 2 | 0.5 | 1 | 1 | 4 | 0.75 | 2 | 0.5 | 1 | 0 | NA | NA | NA | NA | 3 | 0.3333 |
| 1_6528595 | NA | NA | 1 | 0 | 3 | 0.3333 | NA | NA | 4 | 0.25 | 1 | 0 | 1 | 1 | NA | NA | NA | NA | 2 | 0.5 |
| 1_6528609 | NA | NA | 1 | 1 | 3 | 1 | 1 | 1 | 5 | 1 | 2 | 0.5 | NA | NA | NA | NA | NA | NA | 2 | 0.5 |
| 1_6528619 | NA | NA | 1 | 0 | 3 | 0.6667 | NA | NA | 4 | 0.5 | 1 | 1 | 1 | 0 | NA | NA | NA | NA | 2 | 0.5 |
| 1_6531284 | NA | NA | 2 | 0.5 | 4 | 0.5 | 2 | 1 | 8 | 0.625 | 2 | 1 | 1 | 0 | 1 | 0 | 2 | 0.5 | 6 | 0.5 |
| 1_6531294 | NA | NA | 2 | 1 | 4 | 0.25 | 3 | 0.6667 | 9 | 0.5556 | 2 | 1 | 1 | 1 | NA | NA | 1 | 1 | 4 | 1 |
| 1_6531297 | NA | NA | 2 | 0 | 3 | 0 | 3 | 0.3333 | 8 | 0.125 | 1 | 0 | 1 | 0 | 1 | 0 | 2 | 0.5 | 5 | 0.2 |
| 1_6531302 | NA | NA | 2 | 0.5 | 4 | 0.5 | 2 | 0.5 | 8 | 0.5 | 1 | 0 | 1 | 0 | 1 | 1 | 2 | 0.5 | 5 | 0.4 |
| 1_6531306 | NA | NA | 2 | 0.5 | 4 | 0.25 | 3 | 0 | 9 | 0.2222 | 1 | 0 | 1 | 1 | NA | NA | 2 | 0 | 4 | 0.25 |
| 1_6531408 | NA | NA | 3 | 0.3333 | 4 | 0.25 | 3 | 0 | 10 | 0.2 | 2 | 0 | 2 | 0 | 2 | 0 | 4 | 0.25 | 10 | 0.1 |
| 1_6542782 | 1 | 1 | NA | NA | NA | NA | NA | NA | 1 | 1 | 1 | 1 | 1 | 1 | NA | NA | NA | NA | 2 | 1 |
| 1_6641976 | NA | NA | 4 | 0 | 5 | 0.2 | 3 | 0.3333 | 12 | 0.1667 | 2 | 1 | 5 | 0.4 | 2 | 0.5 | 3 | 0 | 12 | 0.4167 |
| 1_6645179 | 1 | 0 | 6 | 0.3333 | 5 | 0.2 | 3 | 0.6667 | 15 | 0.3333 | 3 | 0.3333 | 5 | 0.2 | 3 | 0.3333 | 7 | 0.2857 | 18 | 0.2778 |
| 1_6645189 | 1 | 1 | 7 | 1 | 4 | 1 | 3 | 1 | 15 | 1 | 3 | 1 | 5 | 1 | 3 | 1 | 6 | 1 | 17 | 1 |
| 1_6645203 | 1 | 0 | 7 | 0.1429 | 2 | 0 | 5 | 0.2 | 15 | 0.1333 | 3 | 0.3333 | 4 | 0 | 4 | 0.5 | 7 | 0.1429 | 18 | 0.2222 |
| 1_6645211 | 1 | 0 | 8 | 0.25 | 4 | 0 | 3 | 0 | 16 | 0.125 | 3 | 0 | 4 | 0.25 | 5 | 0 | 8 | 0 | 20 | 0.05 |
| 1_6645271 | 3 | 0 | 10 | 0.7 | 7 | 0.4286 | 8 | 0.625 | 28 | 0.5357 | 4 | 0.5 | 5 | 0.2 | 8 | 0.375 | 10 | 0.7 | 27 | 0.4815 |
| 1_6645289 | 3 | 0 | 9 | 0.1111 | 7 | 0.1429 | 8 | 0 | 27 | 0.0741 | 4 | 0.5 | 5 | 0 | 9 | 0 | 10 | 0 | 28 | 0.0714 |
| 1_6645312 | 3 | 0.6667 | 11 | 0.4545 | 7 | 0.2857 | 9 | 0.2222 | 30 | 0.3667 | 4 | 0.5 | 6 | 0.5 | 9 | 0.5556 | 11 | 0.1818 | 30 | 0.4 |
| 1_6645344 | 4 | 0.25 | 12 | 0.25 | 7 | 0 | 8 | 0.5 | 31 | 0.2581 | 4 | 0.25 | 7 | 0.1429 | 9 | 0.2222 | 13 | 0.0769 | 33 | 0.1515 |
| 1_6645395 | 3 | 0.3333 | 13 | 0.0769 | 9 | 0 | 8 | 0.125 | 33 | 0.0909 | 7 | 0.1429 | 7 | 0 | 13 | 0.1538 | 15 | 0.2 | 42 | 0.1429 |
| 1_6645412 | 3 | 0.3333 | 11 | 0.5455 | 8 | 0.625 | 7 | 0.4286 | 29 | 0.5172 | 5 | 1 | 8 | 0.375 | 12 | 0.5 | 13 | 0.5385 | 38 | 0.5526 |
| 1_6645428 | 5 | 0.6 | 15 | 0.4 | 12 | 0.3333 | 10 | 0.5 | 42 | 0.4286 | 8 | 0.375 | 7 | 0.4286 | 14 | 0.2857 | 19 | 0.3158 | 48 | 0.3333 |
| 1_6645429 | 4 | 0.75 | 10 | 0.7 | 10 | 0.6 | 9 | 0.8889 | 33 | 0.7273 | 7 | 0.8571 | 8 | 0.75 | 13 | 0.6923 | 18 | 0.7778 | 46 | 0.7609 |
| 1_6645460 | 7 | 0 | 17 | 0.1176 | 14 | 0 | 11 | 0.0909 | 49 | 0.0612 | 8 | 0.125 | 11 | 0 | 17 | 0.1176 | 23 | 0.0435 | 59 | 0.0678 |
| 1_6645474 | 8 | 0 | 17 | 0 | 18 | 0.1667 | 15 | 0.2 | 58 | 0.1034 | 11 | 0.0909 | 12 | 0.0833 | 20 | 0.05 | 22 | 0.2273 | 65 | 0.1231 |
| 1_6645487 | 8 | 0 | 13 | 0.2308 | 16 | 0.1875 | 14 | 0.0714 | 51 | 0.1373 | 10 | 0.1 | 9 | 0.4444 | 19 | 0.1053 | 24 | 0.125 | 62 | 0.1613 |

| | | | | | | | | | | | | | | | | | | | | |
|---|---|---|---|---|---|---|---|---|---|---|---|---|---|---|---|---|---|---|---|---|
| 1_6647836 | 21 | 0.4762 | 46 | 0.3478 | 45 | 0.4222 | 36 | 0.3333 | 148 | 0.3851 | 30 | 0.2333 | 36 | 0.3056 | 42 | 0.381 | 49 | 0.4082 | 157 | 0.3439 |
| 1_6647839 | 19 | 0.7895 | 46 | 0.587 | 43 | 0.6512 | 32 | 0.4688 | 140 | 0.6071 | 29 | 0.7586 | 36 | 0.5556 | 39 | 0.6923 | 48 | 0.5833 | 152 | 0.6382 |
| 1_6647978 | 35 | 0.0571 | 69 | 0.0435 | 59 | 0.0847 | 45 | 0.1111 | 208 | 0.0721 | 40 | 0.025 | 40 | 0.025 | 57 | 0.1228 | 60 | 0.1167 | 197 | 0.0812 |
| 1_6670841 | 3 | 0.3333 | 3 | 0.3333 | 2 | 0 | 1 | 0 | 9 | 0.2222 | 2 | 0 | NA | NA | 2 | 0 | 2 | 1 | 6 | 0.3333 |
| 1_6673463 | 4 | 0 | 12 | 0.3333 | 7 | 0.4286 | 4 | 0.5 | 27 | 0.3333 | 3 | 0.6667 | 4 | 0.25 | 5 | 0.2 | 8 | 0.375 | 20 | 0.35 |
| 1_6673472 | 4 | 1 | 11 | 0.8182 | 7 | 0.8571 | 5 | 0.8 | 27 | 0.8519 | 3 | 1 | 2 | 1 | 7 | 1 | 8 | 0.875 | 20 | 0.95 |
| 1_6673479 | 4 | 0.75 | 10 | 0.6 | 6 | 0.6667 | 5 | 0.6 | 25 | 0.64 | 3 | 0.6667 | 4 | 1 | 9 | 0.7778 | 8 | 0.625 | 24 | 0.75 |
| 1_6673495 | 5 | 0.4 | 9 | 0.5556 | 8 | 0.25 | 6 | 0.5 | 28 | 0.4286 | 3 | 0 | 3 | 0.6667 | 9 | 0.3333 | 11 | 0.6364 | 26 | 0.4615 |
| 1_6673503 | 5 | 0.2 | 12 | 0.1667 | 8 | 0 | 6 | 0 | 31 | 0.0968 | 3 | 0.3333 | 4 | 0 | 9 | 0.1111 | 12 | 0.25 | 28 | 0.1786 |
| 1_6673531 | 5 | 0 | 12 | 0.25 | 8 | 0.125 | 7 | 0.1429 | 32 | 0.1562 | 2 | 0 | 4 | 0 | 8 | 0.375 | 13 | 0.4615 | 27 | 0.3333 |
| 1_6673587 | 6 | 0.1667 | 12 | 0.0833 | 12 | 0 | 10 | 0 | 40 | 0.05 | 3 | 0.3333 | 3 | 0.3333 | 13 | 0 | 14 | 0.0714 | 33 | 0.0909 |
| 1_6673643 | 6 | 0 | 13 | 0.0769 | 12 | 0.1667 | 10 | 0 | 41 | 0.0732 | 4 | 0.25 | 5 | 0.2 | 9 | 0.1111 | 16 | 0.0625 | 34 | 0.1176 |
| 1_6673658 | 5 | 0.2 | 13 | 0.2308 | 13 | 0.0769 | 12 | 0.4167 | 43 | 0.2326 | 2 | 0 | 5 | 0 | 10 | 0.3 | 16 | 0.25 | 33 | 0.2121 |
| 1_6673684 | 6 | 0.3333 | 15 | 0.2 | 13 | 0.3077 | 14 | 0.4286 | 48 | 0.3125 | 5 | 0.2 | 6 | 0.8333 | 13 | 0.2308 | 19 | 0.3684 | 43 | 0.3721 |
| 1_6673688 | 6 | 0.3333 | 15 | 0.5333 | 11 | 0.3636 | 15 | 0.4 | 47 | 0.4255 | 4 | 0.25 | 7 | 0.8571 | 16 | 0.5 | 17 | 0.5882 | 44 | 0.5682 |
| 1_6673697 | 6 | 0.1667 | 17 | 0.1765 | 15 | 0.1333 | 16 | 0.125 | 54 | 0.1481 | 5 | 0.2 | 7 | 0.1429 | 16 | 0 | 16 | 0.125 | 44 | 0.0909 |
| 1_6673762 | 9 | 0.2222 | 16 | 0.125 | 14 | 0.0714 | 18 | 0.1111 | 57 | 0.1228 | 7 | 0 | 7 | 0.1429 | 14 | 0.0714 | 22 | 0.1364 | 50 | 0.1 |
| 1_6674033 | 1 | 0 | 2 | 0.5 | 1 | 0 | 1 | 1 | 5 | 0.4 | 1 | 1 | NA | NA | 2 | 1 | NA | NA | 3 | 1 |
| 1_6674045 | 1 | 0 | 2 | 0.5 | 1 | 0 | 1 | 0 | 5 | 0.2 | 1 | 0 | NA | NA | 2 | 0.5 | NA | NA | 3 | 0.3333 |
| 1_6674047 | 1 | 1 | 2 | 1 | 1 | 1 | 1 | 0 | 5 | 0.8 | NA | NA | NA | NA | 2 | 1 | NA | NA | 2 | 1 |
| 1_6674060 | NA | NA | 2 | 0.5 | 1 | 0 | NA | NA | 3 | 0.3333 | 1 | 1 | NA | NA | 1 | 1 | NA | NA | 2 | 1 |
| 1_6674066 | NA | NA | 1 | 1 | 1 | 0 | NA | NA | 2 | 0.5 | 1 | 0 | NA | NA | 1 | 1 | NA | NA | 2 | 0.5 |
| 1_6675709 | 2 | 1 | 6 | 0.5 | 2 | 1 | 3 | 0.6667 | 13 | 0.6923 | 1 | 1 | 3 | 0.3333 | 4 | 0.75 | 1 | 1 | 9 | 0.6667 |
| 1_6676008 | 1 | 0 | 1 | 0 | NA | NA | 3 | 0.3333 | 5 | 0.2 | 1 | 0 | 1 | 1 | 1 | 0 | 2 | 0 | 5 | 0.2 |
| 1_6684027 | NA | NA | NA | NA | 1 | 1 | NA | NA | 1 | 1 | NA | NA | NA | NA | NA | NA | 2 | 0.5 | 2 | 0.5 |
| 1_6684046 | 1 | 0 | NA | NA | 1 | 1 | NA | NA | 2 | 0.5 | NA | NA | NA | NA | NA | NA | 2 | 1 | 2 | 1 |
| 1_6684069 | 1 | 0 | NA | NA | 1 | 1 | NA | NA | 2 | 0.5 | NA | NA | NA | NA | NA | NA | 2 | 0.5 | 2 | 0.5 |
| 1_6684088 | 1 | 1 | NA | NA | 1 | 0 | NA | NA | 2 | 0.5 | NA | NA | NA | NA | NA | NA | 2 | 0.5 | 2 | 0.5 |
| 1_6704034 | 7 | 0.1429 | 3 | 0 | 5 | 0 | 6 | 0.1667 | 21 | 0.0952 | 4 | 0 | 3 | 0 | 3 | 0.3333 | 2 | 0 | 12 | 0.0833 |
| 1_6704123 | 5 | 0.2 | 3 | 0 | 4 | 0.25 | 6 | 0 | 18 | 0.1111 | 3 | 0.3333 | 4 | 0.25 | 4 | 0.25 | 2 | 0 | 13 | 0.2308 |
| 1_6704128 | 3 | 0.6667 | 3 | 0.6667 | 4 | 0.25 | 5 | 0.2 | 15 | 0.4 | 2 | 0 | 5 | 0.2 | 3 | 0.3333 | 2 | 0.5 | 12 | 0.25 |
| 1_6704151 | 5 | 0.4 | 3 | 0.6667 | 4 | 0.25 | 6 | 0 | 18 | 0.2778 | 2 | 0.5 | 4 | 0 | 5 | 0 | 2 | 0 | 13 | 0.0769 |
| 1_6704154 | 5 | 0 | 5 | 0 | 4 | 0 | 6 | 0.3333 | 20 | 0.1 | 1 | 0 | 3 | 0.6667 | 4 | 0 | NA | NA | 8 | 0.25 |
| 1_6704155 | 4 | 0 | 5 | 0 | 4 | 0.25 | 7 | 0 | 20 | 0.05 | 2 | 0.5 | 5 | 0 | 5 | 0.4 | 2 | 0 | 14 | 0.2143 |
| 1_6704159 | 5 | 0.4 | 4 | 0 | 4 | 0 | 6 | 0.3333 | 19 | 0.2105 | 2 | 0 | 5 | 0 | 4 | 0.25 | 2 | 0 | 13 | 0.0769 |
| 1_6704254 | 30 | 0.0333 | 24 | 0.0833 | 26 | 0.0385 | 37 | 0.0541 | 117 | 0.0513 | 16 | 0.0625 | 19 | 0.1579 | 24 | 0.125 | 19 | 0.1053 | 78 | 0.1154 |
| 1_6704264 | 30 | 0.1667 | 24 | 0.0417 | 20 | 0.15 | 33 | 0.1515 | 107 | 0.1308 | 16 | 0.1875 | 20 | 0.05 | 25 | 0 | 15 | 0.0667 | 76 | 0.0658 |
| 1_6807152 | 5 | 0.2 | 6 | 0 | 10 | 0.3 | 7 | 0.2857 | 28 | 0.2143 | 6 | 0 | 4 | 0.25 | 3 | 0 | 6 | 0.1667 | 19 | 0.1053 |
| 1_6807458 | NA | NA | 3 | 0.3333 | 2 | 0.5 | 2 | 0 | 7 | 0.2857 | 1 | 1 | 1 | 1 | 1 | 0 | 4 | 0.5 | 7 | 0.5714 |
| 1_6807773 | NA | NA | 1 | 1 | 1 | 0 | 2 | 0 | 4 | 0.25 | NA | NA | NA | NA | 1 | 1 | NA | NA | 1 | 1 |
| 1_6807793 | NA | NA | 2 | 1 | 1 | 1 | 2 | 0.5 | 5 | 0.8 | NA | NA | NA | NA | 1 | 1 | NA | NA | 1 | 1 |
| 1_6807932 | NA | NA | 2 | 0.5 | NA | NA | NA | NA | 2 | 0.5 | NA | NA | NA | NA | 1 | 1 | NA | NA | 1 | 1 |
| 1_6957077 | 1 | 1 | 2 | 0 | NA | NA | 2 | 0.5 | 5 | 0.4 | NA | NA | NA | NA | NA | NA | 1 | 1 | 1 | 1 |
| 1_7076805 | 1 | 1 | 1 | 0 | NA | NA | NA | NA | 2 | 0.5 | 1 | 0 | NA | NA | NA | NA | 1 | 1 | 2 | 0.5 |
| 1_7076807 | 1 | 1 | 1 | 0 | NA | NA | NA | NA | 2 | 0.5 | NA | NA | NA | NA | NA | NA | 1 | 1 | 1 | 1 |
| 1_7076818 | 1 | 1 | 1 | 1 | NA | NA | NA | NA | 2 | 1 | 1 | 1 | NA | NA | NA | NA | 1 | 1 | 2 | 1 |
| 1_7161833 | 3 | 0 | 4 | 0.25 | 3 | 0.3333 | 5 | 0.2 | 15 | 0.2 | 2 | 0.5 | 3 | 0.3333 | 6 | 0.1667 | 1 | 0 | 12 | 0.25 |
| 1_7162491 | 7 | 0 | 8 | 0.125 | 4 | 0.25 | 4 | 0.5 | 23 | 0.1739 | 5 | 0.2 | 3 | 0.6667 | 2 | 0 | 5 | 0 | 15 | 0.2 |
| 1_7162495 | 7 | 0 | 5 | 0.2 | 4 | 0 | 4 | 0 | 20 | 0.05 | 5 | 0.2 | 3 | 0 | 2 | 0 | 5 | 0 | 15 | 0.0667 |
| 1_7162518 | 6 | 0.1667 | 3 | 0 | 3 | 0 | 3 | 0 | 15 | 0.0667 | 5 | 0.2 | 1 | 0 | 1 | 0 | 4 | 0.25 | 11 | 0.1818 |
| 1_7163670 | 3 | 0.3333 | 3 | 0 | 2 | 0 | 2 | 0.5 | 10 | 0.2 | 4 | 0 | 2 | 0 | NA | NA | 4 | 0.5 | 10 | 0.2 |
| 1_7219922 | 1 | 1 | NA | NA | NA | NA | 2 | 0.5 | 3 | 0.6667 | NA | NA | NA | NA | 1 | 1 | NA | NA | 1 | 1 |
| 1_7219972 | 1 | 1 | NA | NA | NA | NA | 2 | 1 | 3 | 1 | NA | NA | NA | NA | 1 | 1 | NA | NA | 1 | 1 |
| 1_7219989 | NA | NA | NA | NA | NA | NA | 2 | 1 | 2 | 1 | NA | NA | NA | NA | 1 | 1 | NA | NA | 1 | 1 |
| 1_7220007 | 1 | 1 | NA | NA | NA | NA | NA | NA | 1 | 1 | NA | NA | NA | NA | 1 | 1 | NA | NA | 1 | 1 |
| 1_7234006 | 1 | 1 | 1 | 1 | NA | NA | 1 | 1 | 3 | 1 | 3 | 0.6667 | NA | NA | NA | NA | 1 | 1 | 4 | 0.75 |
| 1_7234016 | 1 | 1 | 1 | 1 | NA | NA | 1 | 1 | 3 | 1 | 3 | 1 | NA | NA | NA | NA | 1 | 0 | 4 | 0.75 |
| 1_7234031 | 1 | 1 | 1 | 0 | NA | NA | 1 | 0 | 3 | 0.3333 | 3 | 0.3333 | NA | NA | NA | NA | 1 | 0 | 4 | 0.25 |
| 1_7234036 | 1 | 1 | 1 | 1 | NA | NA | 1 | 1 | 3 | 1 | 3 | 1 | NA | NA | NA | NA | 1 | 1 | 4 | 1 |
| 1_7234038 | 1 | 1 | NA | NA | NA | NA | 1 | 1 | 2 | 1 | 3 | 1 | NA | NA | NA | NA | 1 | 1 | 4 | 1 |
| 1_7234043 | 1 | 1 | NA | NA | NA | NA | 1 | 0 | 2 | 0.5 | 2 | 1 | NA | NA | NA | NA | 1 | 1 | 3 | 1 |
| 1_7234047 | 1 | 0 | 1 | 1 | NA | NA | 1 | 1 | 3 | 0.6667 | 3 | 1 | NA | NA | NA | NA | 1 | 1 | 4 | 1 |
| 1_7234056 | 1 | 0 | 1 | 1 | NA | NA | 1 | 1 | 3 | 0.6667 | 3 | 1 | NA | NA | NA | NA | NA | NA | 3 | 1 |
| 1_7234067 | 1 | 0 | 1 | 1 | NA | NA | NA | NA | 2 | 0.5 | 1 | 1 | NA | NA | NA | NA | 1 | 0 | 2 | 0.5 |
| 1_7234069 | 1 | 1 | 1 | 1 | NA | NA | NA | NA | 2 | 1 | 3 | 1 | NA | NA | NA | NA | NA | NA | 3 | 1 |

| | | | | | | | | | | | | | | | | | | | | |
|---|---|---|---|---|---|---|---|---|---|---|---|---|---|---|---|---|---|---|---|---|
| 1_7234088 | 1 | 0 | 1 | 0 | NA | NA | 1 | 1 | 3 | 0.3333 | 3 | 1 | NA | NA | NA | NA | 1 | 0 | 4 | 0.75 |
| 1_7234208 | 1 | 0 | 3 | 0.6667 | NA | NA | 2 | 0.5 | 6 | 0.5 | 5 | 0.4 | 1 | 1 | 1 | 1 | 1 | 0 | 8 | 0.5 |
| 1_7234683 | 1 | 0 | NA | NA | 2 | 0.5 | 1 | 1 | 4 | 0.5 | NA | NA | NA | NA | NA | NA | 1 | 1 | 1 | 1 |
| 1_7246638 | 4 | 0.5 | 3 | 0.3333 | 4 | 1 | 2 | 0.5 | 13 | 0.6154 | 2 | 0.5 | 2 | 0.5 | 3 | 0 | 3 | 0.6667 | 10 | 0.4 |
| 1_7246641 | 5 | 0.4 | 2 | 0 | 4 | 0.5 | 2 | 0.5 | 13 | 0.3846 | 3 | 0.3333 | 2 | 0 | 3 | 0 | 2 | 0.5 | 10 | 0.2 |
| 1_7246802 | 3 | 0.3333 | NA | NA | 2 | 0.5 | NA | NA | 5 | 0.4 | 2 | 0.5 | 1 | 0 | 1 | 1 | 3 | 0.3333 | 7 | 0.4286 |
| 1_7246806 | 3 | 0.3333 | NA | NA | 2 | 0.5 | NA | NA | 5 | 0.4 | 2 | 0.5 | NA | NA | NA | NA | 3 | 1 | 5 | 0.8 |
| 1_7246808 | 2 | 1 | NA | NA | 2 | 0.5 | NA | NA | 4 | 0.75 | 2 | 0.5 | NA | NA | 1 | 0 | 2 | 0.5 | 5 | 0.4 |
| 1_7257383 | 1 | 1 | NA | NA | 3 | 1 | NA | NA | 4 | 1 | 1 | 1 | NA | NA | 1 | 1 | 2 | 1 | 4 | 1 |
| 1_7257386 | 2 | 1 | NA | NA | 3 | 1 | NA | NA | 5 | 1 | 2 | 0.5 | NA | NA | 1 | 1 | 3 | 1 | 6 | 0.8333 |
| 1_7257387 | 2 | 0 | NA | NA | 2 | 0.5 | NA | NA | 4 | 0.25 | 1 | 1 | NA | NA | NA | NA | 3 | 0.6667 | 4 | 0.75 |
| 1_7257423 | 1 | 1 | NA | NA | 2 | 1 | NA | NA | 3 | 1 | 1 | 1 | NA | NA | 1 | 1 | 2 | 1 | 4 | 1 |
| 1_7257433 | 1 | 0 | NA | NA | 2 | 1 | NA | NA | 3 | 0.6667 | 1 | 0 | NA | NA | NA | NA | 2 | 1 | 3 | 0.6667 |
| 1_7257437 | 1 | 0 | NA | NA | 2 | 0.5 | NA | NA | 3 | 0.3333 | 1 | 0 | NA | NA | 1 | 0 | 2 | 0.5 | 4 | 0.25 |
| 1_7257447 | 1 | 0 | NA | NA | 1 | 1 | NA | NA | 2 | 0.5 | 1 | 1 | NA | NA | 1 | 1 | 2 | 0.5 | 4 | 0.75 |
| 1_7257466 | 1 | 1 | NA | NA | 2 | 0.5 | NA | NA | 3 | 0.6667 | 1 | 0 | NA | NA | 1 | 0 | 2 | 0.5 | 4 | 0.25 |
| 1_7257512 | 1 | 1 | NA | NA | 2 | 0.5 | NA | NA | 3 | 0.6667 | 1 | 1 | NA | NA | 1 | 0 | 1 | 0 | 3 | 0.3333 |
| 1_7262510 | 5 | 0.4 | 3 | 1 | 3 | 0.6667 | 4 | 0.5 | 15 | 0.6 | 1 | 1 | 1 | 0 | 1 | 1 | 4 | 0.25 | 7 | 0.4286 |
| 1_7262527 | 4 | 0.25 | 3 | 0.6667 | 3 | 0 | 2 | 0 | 12 | 0.25 | NA | NA | 1 | 0 | 2 | 0 | 4 | 0.25 | 7 | 0.1429 |
| 1_7262530 | 3 | 0 | 2 | 0 | 2 | 0.5 | 2 | 0 | 9 | 0.1111 | 1 | 0 | 1 | 1 | 2 | 0 | 5 | 0.6 | 9 | 0.4444 |
| 1_7262537 | 4 | 0.25 | 2 | 0.5 | 2 | 0.5 | 3 | 1 | 11 | 0.5455 | 1 | 1 | 1 | 1 | 1 | 1 | 3 | 1 | 6 | 1 |
| 1_7262540 | 5 | 0.2 | 3 | 0.6667 | 4 | 0.5 | 2 | 0 | 14 | 0.3571 | 1 | 1 | 1 | 1 | 2 | 0.5 | 4 | 0.25 | 8 | 0.5 |
| 1_7262548 | 4 | 0.5 | 4 | 0.5 | 4 | 0.5 | 4 | 0.5 | 16 | 0.5 | 1 | 1 | NA | NA | 2 | 0 | 5 | 0.4 | 8 | 0.375 |
| 1_7267219 | 8 | 0.875 | 6 | 0.6667 | 9 | 0.8889 | 10 | 0.6 | 33 | 0.7576 | 2 | 0.5 | 7 | 1 | 6 | 0.8333 | 11 | 0.7273 | 26 | 0.8077 |
| 1_7267265 | 4 | 0.5 | 3 | 0 | 2 | 0.5 | 1 | 0 | 10 | 0.3 | 2 | 0.5 | 4 | 0 | 2 | 0.5 | 2 | 0 | 10 | 0.2 |
| 1_7267425 | 6 | 0.1667 | 3 | 0 | 4 | 0.25 | 2 | 0.5 | 15 | 0.2 | 3 | 0 | 6 | 0.1667 | 6 | 0 | 1 | 0 | 16 | 0.0625 |
| 1_7267466 | 7 | 0.5714 | 3 | 0.3333 | 4 | 0.5 | 3 | 1 | 17 | 0.5882 | 4 | 0.75 | 7 | 0.4286 | 7 | 0.1429 | 3 | 0 | 21 | 0.3333 |
| 1_7267468 | 8 | 0.375 | 2 | 0 | 4 | 0 | 3 | 0.3333 | 17 | 0.2353 | 4 | 0 | 6 | 0.1667 | 7 | 0.4286 | 3 | 0 | 20 | 0.2 |
| 1_7267471 | 7 | 0.7143 | 3 | 0.3333 | 4 | 0.25 | 3 | 0.3333 | 17 | 0.4706 | 4 | 0.75 | 6 | 0.1667 | 7 | 0.5714 | 2 | 0 | 19 | 0.4211 |
| 1_7267472 | 6 | 0.6667 | 2 | 0.5 | 4 | 0.5 | 3 | 1 | 15 | 0.6667 | 3 | 1 | 7 | 0.5714 | 6 | 0.5 | 3 | 0.3333 | 19 | 0.5789 |
| 1_7267481 | 1 | 1 | NA | NA | 1 | 0 | 1 | 1 | 3 | 0.6667 | 1 | 1 | 1 | 0 | 3 | 0.6667 | NA | NA | 5 | 0.6 |
| 1_7267486 | 2 | 1 | NA | NA | NA | NA | 1 | 1 | 3 | 1 | 1 | 1 | NA | NA | 2 | 1 | NA | NA | 3 | 1 |
| 1_7289128 | 16 | 0.125 | 8 | 0.125 | 13 | 0 | 3 | 0 | 40 | 0.075 | 6 | 0.1667 | 5 | 0 | 7 | 0 | 7 | 0.2857 | 25 | 0.12 |
| 1_7289775 | 4 | 0.25 | 7 | 0.1429 | 13 | 0 | 9 | 0.2222 | 33 | 0.1212 | 19 | 0.2105 | 3 | 0 | 4 | 0.5 | 5 | 0.4 | 31 | 0.2581 |
| 1_7289777 | 3 | 0 | 8 | 0.375 | 12 | 0.3333 | 11 | 0.2727 | 34 | 0.2941 | 20 | 0.3 | 3 | 0.3333 | 4 | 0.25 | 5 | 0.2 | 32 | 0.2812 |
| 1_7289800 | 4 | 0.25 | 8 | 0 | 14 | 0.0714 | 11 | 0.1818 | 37 | 0.1081 | 19 | 0.2105 | 3 | 0.3333 | 3 | 0.3333 | 5 | 0 | 30 | 0.2 |
| 1_7289805 | 4 | 0.75 | 8 | 0.375 | 13 | 0.6923 | 10 | 0.6 | 35 | 0.6 | 20 | 0.65 | 3 | 0.6667 | 3 | 1 | 5 | 0.6 | 31 | 0.6774 |
| 1_7289817 | 3 | 0 | 7 | 0 | 13 | 0.0769 | 10 | 0.1 | 33 | 0.0606 | 17 | 0.1176 | 3 | 0 | 3 | 0.3333 | 5 | 0.2 | 28 | 0.1429 |
| 1_7289829 | 3 | 1 | 4 | 0.5 | 13 | 0.4615 | 7 | 0.4286 | 27 | 0.5185 | 17 | 0.5294 | 3 | 0 | 3 | 0.6667 | 4 | 0.25 | 27 | 0.4444 |
| 1_7289832 | 3 | 0.6667 | 5 | 0.2 | 12 | 0.5833 | 10 | 0.6 | 30 | 0.5333 | 20 | 0.4 | 3 | 0 | 3 | 0.6667 | 6 | 0.1667 | 32 | 0.3438 |
| 1_7289838 | 2 | 0.5 | 7 | 0 | 12 | 0 | 7 | 0.1429 | 28 | 0.0714 | 20 | 0.1 | 2 | 0.5 | 2 | 0 | 5 | 0 | 29 | 0.1034 |
| 1_7289839 | 3 | 0 | 8 | 0.125 | 14 | 0.0714 | 7 | 0.1429 | 32 | 0.0938 | 19 | 0.1053 | 3 | 0 | 3 | 0 | 5 | 0 | 30 | 0.0667 |
| 1_7289869 | 2 | 1 | 6 | 0.6667 | 10 | 0.5 | 8 | 0.375 | 26 | 0.5385 | 14 | 0.3571 | 1 | 0 | 1 | 0 | 4 | 0.75 | 20 | 0.4 |
| 1_7289869 | 2 | 0.5 | 8 | 0.125 | 12 | 0 | 8 | 0.125 | 30 | 0.1 | 19 | 0.1579 | 2 | 0 | 2 | 0.5 | 5 | 0 | 28 | 0.1429 |
| 1_7289874 | 2 | 0.5 | 7 | 0 | 13 | 0.1538 | 7 | 0 | 29 | 0.1034 | 21 | 0.1429 | 2 | 0 | 2 | 0 | 5 | 0.2 | 30 | 0.1333 |
| 1_7289875 | 2 | 0.5 | 7 | 0 | 14 | 0.1429 | 8 | 0.125 | 31 | 0.129 | 19 | 0.1579 | 2 | 0.5 | 2 | 1 | 5 | 0 | 28 | 0.2143 |
| 1_7289886 | 1 | 0 | 7 | 0 | 14 | 0.1429 | 6 | 0 | 28 | 0.0714 | 18 | 0.2778 | 2 | 0 | 2 | 0 | 4 | 0 | 26 | 0.1923 |
| 1_7289901 | 2 | 1 | 6 | 0.1667 | 13 | 0.3077 | 7 | 0.1429 | 28 | 0.2857 | 15 | 0.4 | 1 | 0 | 2 | 0 | 4 | 0.25 | 22 | 0.3182 |
| 1_7289902 | 2 | 1 | 5 | 0.6 | 11 | 0.7273 | 6 | 0.6667 | 24 | 0.7083 | 15 | 0.7333 | 2 | 0.5 | 2 | 1 | 4 | 0.75 | 23 | 0.7391 |
| 1_7289922 | 2 | 1 | 5 | 0.8 | 11 | 1 | 8 | 1 | 26 | 0.9615 | 17 | 1 | 2 | 1 | 1 | 1 | 4 | 1 | 24 | 1 |
| 1_7289929 | 2 | 0.5 | 5 | 0.8 | 12 | 0.75 | 8 | 0.5 | 27 | 0.6667 | 17 | 0.8235 | 1 | 1 | 1 | 1 | 4 | 0.25 | 23 | 0.7391 |
| 1_7289930 | 1 | 0 | 4 | 0 | 9 | 0.3333 | 6 | 0.1667 | 20 | 0.2 | 15 | 0.2667 | NA | NA | 2 | 0 | 4 | 0.5 | 21 | 0.2857 |
| 1_7289934 | 1 | 0 | 5 | 0.8 | 10 | 0.7 | 8 | 0.875 | 24 | 0.75 | 13 | 0.6154 | 1 | 1 | 2 | 0 | 4 | 0.25 | 20 | 0.5 |
| 1_7289937 | 2 | 1 | 5 | 0.2 | 11 | 0.1818 | 8 | 0.375 | 26 | 0.3077 | 15 | 0.5333 | 2 | 0.5 | NA | NA | 4 | 0.25 | 21 | 0.4762 |
| 1_7289946 | 2 | 0 | 4 | 0 | 13 | 0.0769 | 8 | 0.125 | 27 | 0.0741 | 13 | 0.3846 | 2 | 0 | 2 | 0 | 3 | 0 | 20 | 0.25 |
| 1_7289949 | 1 | 1 | 5 | 1 | 12 | 1 | 5 | 1 | 23 | 1 | 14 | 1 | 2 | 1 | 2 | 1 | 2 | 1 | 20 | 1 |
| 1_7289949 | 2 | 0 | 5 | 0.2 | 13 | 0.0769 | 8 | 0.125 | 28 | 0.1071 | 16 | 0.25 | 2 | 0.5 | 2 | 0.5 | 3 | 0.3333 | 23 | 0.3043 |
| 1_7289953 | 2 | 1 | 6 | 1 | 11 | 0.8182 | 6 | 1 | 25 | 0.92 | 16 | 0.875 | 2 | 1 | 1 | 1 | 4 | 1 | 23 | 0.913 |
| 1_7289965 | 2 | 0 | 6 | 0.1667 | 12 | 0.25 | 5 | 0 | 25 | 0.16 | 13 | 0.5385 | 1 | 0 | 2 | 0 | 3 | 0 | 19 | 0.3684 |
| 1_7289975 | 2 | 0.5 | 5 | 0.8 | 13 | 0.0769 | 5 | 0.4 | 25 | 0.32 | 14 | 0.4286 | 2 | 0 | 2 | 0 | 4 | 0.5 | 22 | 0.3636 |
| 1_7289986 | 1 | 0 | 4 | 0 | 13 | 0.0769 | 5 | 0.2 | 23 | 0.087 | 13 | 0.0769 | 2 | 0 | 2 | 0 | 3 | 0 | 20 | 0.05 |
| 1_7289986 | 2 | 0 | 5 | 0 | 14 | 0.1429 | 6 | 0 | 27 | 0.0741 | 14 | 0.1429 | 2 | 0 | 2 | 0 | 3 | 0 | 21 | 0.0952 |
| 1_7289993 | 2 | 0 | 3 | 0 | 9 | 0.1111 | 6 | 0 | 20 | 0.05 | 12 | 0 | 2 | 0 | 1 | 0 | 3 | 0.3333 | 18 | 0.0556 |
| 1_7290028 | 1 | 0 | 1 | 0 | 6 | 0.3333 | 3 | 0 | 11 | 0.1818 | 11 | 0.0909 | 1 | 0 | 1 | 0 | 3 | 0 | 16 | 0.0625 |
| 1_7290079 | 1 | 0 | 2 | 0 | 7 | 0.1429 | 5 | 0 | 15 | 0.0667 | 11 | 0.0909 | NA | NA | 1 | 0 | 2 | 0 | 14 | 0.0714 |

| | | | | | | | | | | | | | | | | | | | | |
|---|---|---|---|---|---|---|---|---|---|---|---|---|---|---|---|---|---|---|---|---|
| 1_72901183 | 1 | 0 | 2 | 0 | 8 | 0.625 | 4 | 0 | 15 | 0.3333 | 9 | 0.1111 | NA | NA | 1 | 0 | 2 | 0 | 12 | 0.0833 |
| 1_72901314 | 1 | 0 | 2 | 0 | 6 | 0.1667 | 5 | 0.4 | 14 | 0.2143 | 8 | 0.25 | NA | NA | 1 | 0 | 2 | 0.5 | 11 | 0.2727 |
| 1_72901398 | 1 | 0 | 2 | 0.5 | 8 | 0.125 | 4 | 0.5 | 15 | 0.2667 | 9 | 0.4444 | NA | NA | 1 | 0 | 2 | 0 | 12 | 0.3333 |
| 1_72901576 | 1 | 1 | 2 | 1 | 7 | 0.1429 | 2 | 0.5 | 12 | 0.4167 | 9 | 0.8889 | NA | NA | 1 | 0 | 2 | 0 | 12 | 0.6667 |
| 1_72913110 | NA | NA | NA | NA | 4 | 0 | 4 | 0.25 | 8 | 0.125 | 1 | 0 | NA | NA | NA | NA | 1 | 1 | 2 | 0.5 |
| 1_72913470 | 1 | 0 | 1 | 0 | 3 | 0 | 4 | 0.25 | 9 | 0.1111 | 3 | 0.3333 | NA | NA | NA | NA | 1 | 0 | 4 | 0.25 |
| 1_73952667 | NA | NA | 5 | 0.2 | NA | NA | 1 | 0 | 6 | 0.1667 | 2 | 1 | 3 | 0.3333 | 1 | 0 | 2 | 0 | 8 | 0.375 |
| 1_73952889 | NA | NA | 3 | 0.3333 | NA | NA | 2 | 0 | 5 | 0.2 | 1 | 0 | 2 | 0.5 | 1 | 0 | 2 | 1 | 6 | 0.5 |
| 1_73953168 | NA | NA | 1 | 1 | 1 | 0 | 3 | 0 | 5 | 0.2 | 2 | 0.5 | 4 | 0 | NA | NA | 1 | 0 | 7 | 0.1429 |
| 1_73953221 | NA | NA | 2 | 0 | 1 | 1 | 3 | 0 | 6 | 0.1667 | 2 | 1 | 3 | 0.3333 | 1 | 0 | 2 | 0 | 8 | 0.375 |
| 1_73953395 | NA | NA | 2 | 1 | NA | NA | 2 | 0 | 4 | 0.5 | 1 | 1 | 3 | 0.6667 | 1 | 1 | 1 | 1 | 6 | 0.8333 |
| 1_73953591 | NA | NA | 2 | 1 | NA | NA | 3 | 0.6667 | 5 | 0.8 | 1 | 1 | 3 | 1 | 1 | 1 | 2 | 1 | 7 | 1 |
| 1_73953628 | NA | NA | 1 | 1 | NA | NA | 2 | 0 | 3 | 0.3333 | 1 | 0 | 3 | 0.3333 | 1 | 0 | 2 | 0 | 7 | 0.1429 |
| 1_73963888 | NA | NA | NA | NA | NA | NA | 1 | 1 | 1 | 1 | NA | NA | 1 | 1 | NA | NA | 2 | 0.5 | 3 | 0.6667 |
| 1_74306997 | 29 | 0.069 | 30 | 0.1 | 26 | 0.1923 | 28 | 0.0714 | 113 | 0.1062 | 26 | 0.2308 | 29 | 0.2414 | 24 | 0.2917 | 24 | 0.1667 | 103 | 0.233 |
| 1_74307127 | 41 | 0.6341 | 47 | 0.7872 | 40 | 0.8 | 47 | 0.7234 | 175 | 0.7371 | 41 | 0.878 | 43 | 0.814 | 44 | 0.7045 | 36 | 0.8333 | 164 | 0.8049 |
| 1_74319286 | 17 | 0.6471 | 21 | 0.4762 | 19 | 0.3684 | 19 | 0.7368 | 76 | 0.5526 | 14 | 0.8571 | 27 | 0.5556 | 15 | 0.4 | 26 | 0.4615 | 82 | 0.5488 |
| 1_74326471 | 12 | 0.0833 | 6 | 0.1667 | 4 | 0 | 6 | 0.1667 | 28 | 0.1071 | 6 | 0.3333 | 9 | 0.2222 | 4 | 0 | 1 | 0 | 20 | 0.2 |
| 1_74326478 | 13 | 0.1538 | 5 | 0.6 | 3 | 1 | 7 | 0.1429 | 28 | 0.3214 | 6 | 0.1667 | 6 | 0.5 | 5 | 0.6 | 1 | 0 | 18 | 0.3889 |
| 1_74327823 | 4 | 0.5 | 6 | 0.3333 | 8 | 0.25 | 5 | 0 | 23 | 0.2609 | 9 | 0.2222 | 3 | 0.3333 | 3 | 0 | 3 | 0.6667 | 18 | 0.2778 |
| 1_74328071 | 4 | 0 | 7 | 0.4286 | 7 | 0.1429 | 5 | 0.2 | 23 | 0.2174 | 9 | 0.1111 | 4 | 0 | 4 | 0.25 | 3 | 0.3333 | 20 | 0.15 |
| 1_74328497 | 2 | 0.5 | 3 | 0.3333 | 5 | 0.2 | 3 | 0 | 13 | 0.2308 | 8 | 0.25 | NA | NA | 2 | 0 | 1 | 1 | 11 | 0.2727 |
| 1_74328535 | 2 | 1 | 4 | 0.75 | 6 | 1 | 2 | 1 | 14 | 0.9286 | 6 | 0.1667 | 1 | 0 | 2 | 1 | 3 | 0.6667 | 12 | 0.4167 |
| 1_74391071 | 15 | 0.4667 | 11 | 0.2727 | 12 | 0.3333 | 11 | 0.7273 | 49 | 0.449 | 9 | 0.4444 | 16 | 0.375 | 16 | 0.5 | 9 | 0.6667 | 50 | 0.48 |
| 1_74391263 | 16 | 0.1875 | 11 | 0 | 12 | 0.25 | 14 | 0.1429 | 53 | 0.1509 | 11 | 0.1818 | 19 | 0.0526 | 16 | 0.125 | 11 | 0 | 57 | 0.0877 |
| 1_74391354 | 15 | 0.2667 | 11 | 0.4545 | 10 | 0.1 | 17 | 0 | 53 | 0.1887 | 9 | 0.2222 | 16 | 0.125 | 14 | 0.0714 | 11 | 0.1818 | 50 | 0.14 |
| 1_74391398 | 8 | 0.375 | 10 | 0.6 | 9 | 0.6667 | 14 | 0.3571 | 41 | 0.4878 | 11 | 0.4545 | 12 | 0.1667 | 11 | 0.3636 | 8 | 0.75 | 42 | 0.4048 |
| 1_74391603 | 16 | 0.1875 | 12 | 0.1667 | 13 | 0.1538 | 17 | 0.1176 | 58 | 0.1552 | 11 | 0.2727 | 20 | 0.1 | 19 | 0.1053 | 13 | 0.2308 | 63 | 0.1587 |
| 1_74391619 | 15 | 0.2667 | 13 | 0.0769 | 13 | 0.0769 | 17 | 0.0588 | 58 | 0.1207 | 11 | 0.2727 | 19 | 0.1053 | 19 | 0.0526 | 13 | 0.0769 | 62 | 0.1129 |
| 1_74391651 | 14 | 0.2857 | 13 | 0.1538 | 13 | 0.0769 | 15 | 0.2667 | 55 | 0.2 | 9 | 0 | 17 | 0.1765 | 20 | 0.2 | 10 | 0.3 | 56 | 0.1786 |
| 1_74391689 | 13 | 0.4615 | 14 | 0.5714 | 12 | 0.5 | 13 | 0.4615 | 52 | 0.5 | 9 | 0.6667 | 17 | 0.4706 | 16 | 0.625 | 12 | 0.5 | 54 | 0.5556 |
| 1_74391895 | 12 | 0.3333 | 16 | 0.375 | 14 | 0.3571 | 19 | 0.2105 | 61 | 0.3115 | 13 | 0.3846 | 19 | 0.2105 | 19 | 0.2105 | 14 | 0.2143 | 65 | 0.2462 |
| 1_74392448 | 16 | 0.0625 | 16 | 0.125 | 14 | 0 | 19 | 0.1579 | 65 | 0.0923 | 12 | 0.3333 | 21 | 0 | 18 | 0.2222 | 18 | 0.2778 | 69 | 0.1884 |
| 1_74392471 | 15 | 0.3333 | 14 | 0.2143 | 13 | 0.1538 | 16 | 0.25 | 58 | 0.2414 | 12 | 0.25 | 18 | 0.1667 | 19 | 0.2105 | 16 | 0.25 | 65 | 0.2154 |
| 1_74392757 | 20 | 0.1 | 18 | 0.2778 | 13 | 0.2308 | 22 | 0.0909 | 73 | 0.1644 | 18 | 0.1667 | 29 | 0.1379 | 24 | 0.2083 | 17 | 0 | 88 | 0.1364 |
| 1_74424967 | 1 | 1 | NA | NA | NA | NA | NA | NA | 1 | 1 | NA | NA | NA | NA | 1 | 1 | NA | NA | 1 | 1 |
| 1_74434802 | 2 | 0.5 | 4 | 0.25 | 5 | 0.8 | 3 | 0.3333 | 14 | 0.5 | 6 | 0.3333 | 4 | 0.5 | 4 | 0.5 | 3 | 0.6667 | 17 | 0.4706 |
| 1_74434896 | 3 | 0.3333 | 4 | 0.5 | 6 | 0.5 | 3 | 0.6667 | 16 | 0.5 | 6 | 0.3333 | 4 | 0 | 6 | 0.5 | 3 | 0 | 19 | 0.2632 |
| 1_74435064 | 3 | 0.6667 | 5 | 0 | 6 | 0 | 4 | 0 | 18 | 0.1111 | 8 | 0.125 | 4 | 0.25 | 4 | 0 | 3 | 0 | 19 | 0.1053 |
| 1_74477064 | NA | NA | 3 | 0.6667 | 1 | 1 | 2 | 0.5 | 6 | 0.6667 | NA | NA | NA | NA | 1 | 1 | 3 | 0.6667 | 4 | 0.75 |
| 1_74477115 | 1 | 0 | 3 | 0 | 1 | 0 | 3 | 0.3333 | 8 | 0.125 | NA | NA | NA | NA | 1 | 0 | 3 | 0.3333 | 4 | 0.25 |
| 1_74477646 | NA | NA | 2 | 0.5 | 1 | 0 | 1 | 1 | 4 | 0.5 | NA | NA | NA | NA | 1 | 0 | 1 | 1 | 2 | 0.5 |
| 1_74478466 | NA | NA | 1 | 1 | 1 | 1 | 1 | 0 | 3 | 0.6667 | NA | NA | NA | NA | 1 | 1 | 1 | 1 | 2 | 1 |
| 1_74568118 | 2 | 0.5 | 3 | 1 | 1 | 1 | 3 | 1 | 9 | 0.8889 | 1 | 1 | 3 | 1 | 2 | 1 | 4 | 0.5 | 10 | 0.8 |
| 1_74568166 | 2 | 0.5 | 3 | 0 | 1 | 0 | 3 | 0.3333 | 9 | 0.2222 | 2 | 0 | 4 | 0 | 2 | 0 | 6 | 0.1667 | 14 | 0.0714 |
| 1_74568377 | 2 | 0 | 3 | 0 | 1 | 0 | 3 | 0.3333 | 9 | 0.1111 | 2 | 0 | 2 | 0 | 2 | 0 | 5 | 0.2 | 11 | 0.0909 |
| 1_74568426 | 2 | 0 | 2 | 0 | 1 | 0 | 7 | 0.2857 | 12 | 0.1667 | 2 | 0 | 3 | 0.3333 | 2 | 0 | 5 | 0 | 12 | 0.0833 |
| 1_74568520 | 2 | 0 | 3 | 0 | 1 | 0 | 5 | 0.6 | 11 | 0.2727 | 3 | 0 | 4 | 0.5 | 3 | 0 | 6 | 0.1667 | 16 | 0.1875 |
| 1_74568758 | 1 | 0 | 1 | 0 | NA | NA | 6 | 0.1667 | 8 | 0.125 | 2 | 0 | 2 | 0 | 2 | 0 | 4 | 0.25 | 10 | 0.1 |
| 1_74569081 | 1 | 0 | 2 | 0 | NA | NA | 6 | 0.8333 | 9 | 0.5556 | 4 | 0.25 | 1 | 0 | 2 | 0 | 3 | 0.3333 | 10 | 0.2 |
| 1_74569382 | 1 | 0 | 2 | 0 | NA | NA | 9 | 0.1111 | 12 | 0.0833 | 6 | 0 | 3 | 0.3333 | 2 | 0 | 3 | 0 | 14 | 0.0714 |
| 1_74569549 | 2 | 0 | 2 | 0.5 | 1 | 1 | 9 | 0.1111 | 14 | 0.2143 | 6 | 0 | 3 | 0.3333 | 2 | 0 | 4 | 0.25 | 15 | 0.1333 |
| 1_74569754 | 2 | 0.5 | 2 | 0 | 2 | 0.5 | 10 | 0.7 | 16 | 0.5625 | 6 | 0.5 | 3 | 0.3333 | 2 | 0 | 4 | 0.75 | 15 | 0.4667 |
| 1_74569842 | 2 | 0 | 2 | 0 | 2 | 0 | 10 | 0.3 | 16 | 0.1875 | 6 | 0.1667 | 3 | 0.3333 | 1 | 0 | 4 | 0 | 14 | 0.1429 |
| 1_74588506 | 2 | 0 | 2 | 0 | 1 | 1 | 1 | 0 | 6 | 0.1667 | 2 | 0 | NA | NA | 1 | 0 | 3 | 0.6667 | 6 | 0.3333 |
| 1_74599003 | NA | NA | 1 | 1 | NA | NA | NA | NA | 1 | 1 | 1 | 1 | NA | NA | 1 | 1 | NA | NA | 2 | 1 |
| 1_74605056 | 4 | 0.5 | 2 | 0.5 | 3 | 0.6667 | 3 | 0 | 12 | 0.4167 | 4 | 0 | NA | NA | 2 | 0 | 4 | 0.25 | 10 | 0.1 |
| 1_74605094 | 3 | 0 | 2 | 0.5 | 3 | 0 | 2 | 0.5 | 10 | 0.2 | 3 | 0 | NA | NA | 2 | 0.5 | 4 | 0 | 9 | 0.1111 |
| 1_74605812 | 4 | 0 | 4 | 0.25 | NA | NA | 4 | 0.25 | 12 | 0.1667 | 2 | 0 | 4 | 0 | 1 | 0 | 4 | 0.25 | 11 | 0.0909 |
| 1_74605836 | 2 | 1 | 4 | 0.75 | NA | NA | 3 | 1 | 9 | 0.8889 | 1 | 0 | 4 | 1 | 2 | 1 | 3 | 0.6667 | 10 | 0.8 |
| 1_74605848 | 4 | 1 | 4 | 1 | NA | NA | 5 | 1 | 13 | 1 | 2 | 1 | 4 | 1 | 2 | 1 | 5 | 0.8 | 13 | 0.9231 |
| 1_74605892 | 4 | 1 | 3 | 0.6667 | NA | NA | 5 | 0.6 | 12 | 0.75 | 2 | 0.5 | 4 | 1 | 2 | 1 | 5 | 0.4 | 13 | 0.6923 |
| 1_74605960 | 3 | 0.6667 | 3 | 1 | NA | NA | 6 | 1 | 12 | 0.9167 | 2 | 0.5 | 4 | 1 | 2 | 1 | 5 | 1 | 13 | 0.9231 |
| 1_74605979 | 3 | 1 | 3 | 0.3333 | NA | NA | 5 | 0.4 | 11 | 0.5455 | 1 | 1 | 4 | 0.75 | 1 | 1 | 4 | 0.75 | 10 | 0.8 |
| 1_74606049 | 2 | 1 | 1 | 1 | NA | NA | 5 | 1 | 8 | 1 | 2 | 0.5 | 1 | 0 | 2 | 1 | 4 | 1 | 9 | 0.7778 |

| | | | | | | | | | | | | | | | | | | | | |
|---|---|---|---|---|---|---|---|---|---|---|---|---|---|---|---|---|---|---|---|---|
| 1_7460628( | 4 | 0.5 | 1 | 1 | NA | NA | 6 | 1 | 11 | 0.8182 | 2 | 1 | 3 | 1 | 2 | 1 | 4 | 1 | 11 | 1 |
| 1_7460895! | 1 | 1 | 1 | 1 | NA | NA | 3 | 0.3333 | 5 | 0.6 | 2 | 1 | 3 | 0.6667 | 1 | 0 | 2 | 0.5 | 8 | 0.625 |
| 1_7460899 | 1 | 0 | 1 | 1 | NA | NA | 5 | 0.8 | 7 | 0.7143 | 2 | 1 | 3 | 1 | 2 | 1 | 2 | 1 | 9 | 1 |
| 1_7460903 | 1 | 1 | 2 | 1 | NA | NA | 3 | 1 | 6 | 1 | 2 | 0.5 | 2 | 0.5 | 1 | 1 | 2 | 1 | 7 | 0.7143 |
| 1_7460911 | 2 | 0 | 2 | 0.5 | NA | NA | 5 | 0 | 9 | 0.1111 | 1 | 1 | 3 | 0 | 2 | 0 | 1 | 0 | 7 | 0.1429 |
| 1_7460927! | 2 | 0 | 2 | 0 | NA | NA | 4 | 0.75 | 8 | 0.375 | 2 | 1 | 3 | 0 | 2 | 0.5 | 1 | 0 | 8 | 0.375 |
| 1_7460958 | 2 | 0.5 | 1 | 0 | NA | NA | 2 | 0 | 5 | 0.2 | 2 | 0.5 | 1 | 0 | 1 | 0 | NA | NA | 4 | 0.25 |
| 1_7460960 | 2 | 0 | 1 | 0 | NA | NA | 2 | 1 | 5 | 0.4 | 2 | 0.5 | 2 | 0 | 1 | 0 | NA | NA | 5 | 0.2 |
| 1_7460969 | 2 | 0.5 | 2 | 1 | NA | NA | 1 | 1 | 5 | 0.8 | 2 | 0.5 | 2 | 1 | 1 | 1 | 1 | 1 | 6 | 0.8333 |
| 1_7460979( | NA | NA | 2 | 0.5 | NA | NA | 2 | 0.5 | 4 | 0.5 | 2 | 0.5 | 1 | 1 | 1 | 0 | 1 | 1 | 5 | 0.6 |
| 1_7460995( | 2 | 0 | 2 | 1 | NA | NA | 1 | 1 | 5 | 0.6 | 2 | 1 | 2 | 1 | 1 | 1 | 1 | 1 | 6 | 1 |
| 1_7460995! | 2 | 0.5 | 2 | 0 | NA | NA | 1 | 0 | 5 | 0.2 | 2 | 0.5 | 2 | 0 | 1 | 1 | 1 | 1 | 6 | 0.5 |
| 1_7461008 | 2 | 1 | NA | NA | NA | NA | 1 | 1 | 3 | 1 | 2 | 1 | 2 | 0.5 | NA | NA | 1 | 1 | 5 | 0.8 |
| 1_7461017 | 2 | 1 | 1 | 1 | NA | NA | 1 | 1 | 4 | 1 | 2 | 0.5 | 2 | 0.5 | 1 | 1 | NA | NA | 5 | 0.6 |
| 1_7461026 | 2 | 0.5 | 1 | 0 | NA | NA | 1 | 1 | 4 | 0.5 | 2 | 0 | 2 | 0.5 | 1 | 0 | NA | NA | 5 | 0.2 |
| 1_7461081 | 1 | 1 | 1 | 1 | NA | NA | 1 | 1 | 3 | 1 | 2 | 1 | 1 | 0 | 1 | 1 | NA | NA | 4 | 0.75 |
| 1_7461131( | 1 | 0 | 2 | 0.5 | NA | NA | 1 | 0 | 4 | 0.25 | 2 | 1 | NA | NA | NA | NA | NA | NA | 2 | 1 |
| 1_7461150 | NA | NA | 2 | 0.5 | NA | NA | 1 | 1 | 3 | 0.6667 | 2 | 1 | NA | NA | 1 | 1 | NA | NA | 3 | 1 |
| 1_7461516 | 1 | 1 | 2 | 1 | NA | NA | 1 | 1 | 4 | 1 | 1 | 1 | NA | NA | 1 | 1 | NA | NA | 2 | 1 |
| 1_7462476 | 1 | 1 | 2 | 0.5 | NA | NA | NA | NA | 3 | 0.6667 | 1 | 1 | NA | NA | 1 | 1 | NA | NA | 2 | 1 |
| 1_7462873 | NA | NA | 5 | 0.8 | 1 | 0 | 5 | 0.8 | 11 | 0.7273 | NA | NA | 2 | 0.5 | 1 | 0 | 3 | 0.6667 | 6 | 0.5 |
| 1_7462875 | NA | NA | 5 | 0.8 | 2 | 0.5 | 6 | 0.3333 | 13 | 0.5385 | NA | NA | 2 | 1 | 1 | 1 | 3 | 0.6667 | 6 | 0.8333 |
| 1_7462888 | NA | NA | 5 | 0 | 2 | 0.5 | 5 | 0 | 12 | 0.0833 | 1 | 0 | 2 | 0.5 | 1 | 0 | 3 | 0 | 7 | 0.1429 |
| 1_7462918 | 1 | 0 | 6 | 0.1667 | 1 | 0 | 7 | 0.1429 | 15 | 0.1333 | 1 | 0 | 2 | 0.5 | 2 | 0.5 | 3 | 0 | 8 | 0.25 |
| 1_7462945 | 1 | 0 | 5 | 0.2 | 2 | 0 | 7 | 0.1429 | 15 | 0.1333 | 1 | 0 | 2 | 0 | 2 | 0.5 | 3 | 0 | 8 | 0.125 |
| 1_7463017 | 2 | 0 | 4 | 0 | 1 | 0 | 7 | 0.1429 | 14 | 0.0714 | 1 | 0 | 1 | 0 | 3 | 0.3333 | 3 | 0 | 8 | 0.125 |
| 1_7463110 | 4 | 0.75 | 6 | 0.3333 | 2 | 0 | 6 | 0.5 | 18 | 0.4444 | 1 | 0 | 2 | 0 | 3 | 0 | 5 | 0.4 | 11 | 0.1818 |
| 1_7463126 | 3 | 0 | 5 | 0.2 | 3 | 0 | 4 | 0 | 15 | 0.0667 | NA | NA | 2 | 0 | 2 | 0 | 3 | 0.3333 | 7 | 0.1429 |
| 1_7463140( | NA | NA | 1 | 1 | 2 | 0 | 1 | 0 | 4 | 0.25 | NA | NA | 1 | 0 | 1 | 0 | 2 | 0.5 | 4 | 0.25 |
| 1_7463311 | 5 | 0.8 | 5 | 0.8 | 3 | 1 | 5 | 0.8 | 18 | 0.8333 | 2 | 0.5 | 7 | 0.4286 | 4 | 0.75 | 7 | 0.5714 | 20 | 0.55 |
| 1_7463312 | 6 | 0.3333 | 4 | 0 | 3 | 0.3333 | 5 | 0.8 | 18 | 0.3889 | 3 | 0.6667 | 7 | 0.1429 | 5 | 0.8 | 6 | 0.6667 | 21 | 0.5238 |
| 1_7463321 | 6 | 0 | 5 | 0.6 | 4 | 0.25 | 5 | 0.6 | 20 | 0.35 | 2 | 0.5 | 5 | 0 | 2 | 1 | 7 | 0 | 16 | 0.1875 |
| 1_7463321! | 7 | 0.4286 | 6 | 0 | 4 | 0.75 | 5 | 0.2 | 22 | 0.3182 | 2 | 0.5 | 6 | 0.5 | 5 | 0.2 | 8 | 0.375 | 21 | 0.381 |
| 1_7463325( | 2 | 1 | 3 | 0.6667 | 3 | 1 | 4 | 0.75 | 12 | 0.8333 | 2 | 1 | 5 | 0.6 | 3 | 0.6667 | 3 | 1 | 13 | 0.7692 |
| 1_7463340 | 8 | 0.125 | 4 | 0 | 2 | 0 | 5 | 0.2 | 19 | 0.1053 | 2 | 0.5 | 5 | 0 | 5 | 0.2 | 5 | 0 | 17 | 0.1176 |
| 1_7464048 | 4 | 0 | 3 | 0.3333 | 1 | 0 | 4 | 0.25 | 12 | 0.1667 | NA | NA | 2 | 0 | 4 | 0 | 3 | 0.3333 | 9 | 0.1111 |
| 1_7473684 | 1 | 0 | 2 | 0.5 | NA | NA | NA | NA | 3 | 0.3333 | NA | NA | NA | NA | 2 | 0.5 | 1 | 1 | 3 | 0.6667 |
| 1_7473765! | NA | NA | 4 | 0.25 | 1 | 0 | NA | NA | 5 | 0.2 | NA | NA | NA | NA | 2 | 0 | 1 | 1 | 3 | 0.3333 |
| 1_7473770 | NA | NA | 4 | 0.25 | NA | NA | NA | NA | 4 | 0.25 | NA | NA | NA | NA | 1 | 1 | 1 | 0 | 2 | 0.5 |
| 1_7473874 | NA | NA | 6 | 0.3333 | 1 | 0 | NA | NA | 7 | 0.2857 | NA | NA | NA | NA | 2 | 1 | 1 | 0 | 3 | 0.6667 |
| 1_7473988( | NA | NA | 4 | 0 | 4 | 0.5 | NA | NA | 8 | 0.25 | NA | NA | NA | NA | 1 | 1 | 1 | 0 | 2 | 0.5 |
| 1_7477692 | 1 | 1 | NA | NA | NA | NA | 1 | 0 | 2 | 0.5 | NA | NA | NA | NA | NA | NA | 1 | 1 | 1 | 1 |
| 1_7481137 | NA | NA | 1 | 1 | NA | NA | NA | NA | 1 | 1 | 1 | 1 | NA | NA | NA | NA | NA | NA | 1 | 1 |
| 1_7482367 | 1 | 0 | 1 | 1 | NA | NA | NA | NA | 2 | 0.5 | 1 | 1 | NA | NA | NA | NA | NA | NA | 1 | 1 |
| 1_7482381 | 1 | 1 | 1 | 1 | NA | NA | NA | NA | 2 | 1 | 1 | 1 | NA | NA | NA | NA | NA | NA | 1 | 1 |
| 1_7484258 | NA | NA | 1 | 1 | 1 | 1 | 1 | 1 | 3 | 1 | NA | NA | 2 | 1 | NA | NA | NA | NA | 2 | 1 |
| 1_7484294! | NA | NA | 1 | 1 | 2 | 1 | NA | NA | 3 | 1 | NA | NA | 2 | 0.5 | NA | NA | NA | NA | 2 | 0.5 |
| 1_7484318 | NA | NA | 1 | 1 | 2 | 1 | 1 | 0 | 4 | 0.75 | NA | NA | 2 | 1 | NA | NA | NA | NA | 2 | 1 |
| 1_7484330( | NA | NA | 1 | 0 | 3 | 0.6667 | 1 | 1 | 5 | 0.6 | NA | NA | 1 | 1 | NA | NA | NA | NA | 1 | 1 |
| 1_7484331 | NA | NA | 1 | 1 | 3 | 1 | 1 | 1 | 5 | 1 | NA | NA | 2 | 1 | NA | NA | NA | NA | 2 | 1 |
| 1_7489456 | 20 | 0.05 | 23 | 0.0435 | 37 | 0.0811 | 19 | 0 | 99 | 0.0505 | 25 | 0.08 | 25 | 0.04 | 24 | 0.0833 | 25 | 0 | 99 | 0.0505 |
| 1_7489490 | 37 | 0.1622 | 39 | 0.1282 | 43 | 0.1395 | 25 | 0.04 | 144 | 0.125 | 39 | 0.0513 | 31 | 0.0968 | 39 | 0.1026 | 32 | 0.1562 | 141 | 0.0993 |
| 1_7489502 | 33 | 0.0606 | 35 | 0.1143 | 44 | 0.1136 | 30 | 0.0667 | 142 | 0.0915 | 43 | 0.093 | 32 | 0.125 | 42 | 0.1429 | 40 | 0.025 | 157 | 0.0955 |
| 1_7489531 | 40 | 0.275 | 40 | 0.3 | 55 | 0.2909 | 33 | 0.3636 | 168 | 0.3036 | 47 | 0.2553 | 36 | 0.4722 | 43 | 0.2558 | 45 | 0.4 | 171 | 0.3392 |
| 1_7489534 | 32 | 1 | 32 | 0.9688 | 40 | 0.975 | 23 | 1 | 127 | 0.9843 | 33 | 1 | 30 | 1 | 29 | 0.9655 | 32 | 1 | 124 | 0.9919 |
| 1_7489535 | 29 | 0.9655 | 24 | 1 | 39 | 0.9231 | 23 | 1 | 115 | 0.9652 | 38 | 0.9737 | 32 | 0.9062 | 30 | 0.9667 | 34 | 0.9706 | 134 | 0.9552 |
| 1_7489535! | 29 | 0.6552 | 31 | 0.7742 | 39 | 0.8205 | 24 | 0.8333 | 123 | 0.7724 | 38 | 0.8421 | 29 | 0.7931 | 39 | 0.8462 | 35 | 0.9429 | 141 | 0.8582 |
| 1_7489543 | 38 | 0.1316 | 44 | 0.1591 | 51 | 0.0784 | 30 | 0.2 | 163 | 0.135 | 40 | 0.1 | 40 | 0.125 | 43 | 0.186 | 43 | 0.1163 | 166 | 0.1325 |
| 1_7489605 | 51 | 0.098 | 59 | 0.2034 | 69 | 0.087 | 41 | 0.122 | 220 | 0.1273 | 55 | 0.1273 | 48 | 0.125 | 57 | 0.1053 | 53 | 0.0943 | 213 | 0.1127 |
| 1_7489605 | 38 | 0.0526 | 39 | 0.0513 | 49 | 0.0408 | 34 | 0.0588 | 160 | 0.05 | 40 | 0.075 | 35 | 0.0857 | 40 | 0.025 | 38 | 0.1053 | 153 | 0.0719 |
| 1_7489628 | 46 | 0.1304 | 59 | 0.0678 | 63 | 0.0476 | 44 | 0.0909 | 212 | 0.0802 | 59 | 0.0678 | 45 | 0.1111 | 54 | 0.0185 | 57 | 0.0351 | 215 | 0.0558 |
| 1_7489647 | 55 | 0.4182 | 68 | 0.3529 | 73 | 0.4247 | 40 | 0.4 | 236 | 0.3983 | 60 | 0.4333 | 49 | 0.5918 | 59 | 0.5932 | 52 | 0.4808 | 220 | 0.5227 |
| 1_7489656 | 32 | 0.1562 | 43 | 0.186 | 48 | 0.1458 | 24 | 0.0417 | 147 | 0.1429 | 36 | 0.1389 | 40 | 0.2 | 36 | 0.2222 | 37 | 0.2703 | 149 | 0.2081 |
| 1_7496418 | NA | NA | NA | NA | NA | NA | 1 | 1 | 1 | 1 | NA | NA | NA | NA | NA | NA | 1 | 1 | 1 | 1 |
| 1_7500663! | NA | NA | 1 | 0 | 1 | 1 | NA | NA | 2 | 0.5 | NA | NA | 2 | 1 | 1 | 0 | NA | NA | 3 | 0.6667 |

| | | | | | | | | | | | | | | | | | | | | |
|---|---|---|---|---|---|---|---|---|---|---|---|---|---|---|---|---|---|---|---|---|
| 1_75006729 | NA | NA | 1 | 1 | 1 | 1 | 1 | 0 | 3 | 0.6667 | NA | NA | 2 | 1 | 1 | 1 | NA | NA | 3 | 1 |
| 1_75112827 | 4 | 0 | 4 | 0.25 | 3 | 0 | 2 | 0 | 13 | 0.0769 | 4 | 0 | 1 | 0 | 5 | 0.2 | 2 | 0 | 12 | 0.0833 |
| 1_75113241 | 3 | 0 | 3 | 0.3333 | 1 | 0 | 1 | 0 | 8 | 0.125 | 4 | 0.5 | 2 | 0 | 7 | 0.1429 | 1 | 0 | 14 | 0.2143 |
| 1_75113362 | 3 | 0.3333 | 5 | 0 | 3 | 0 | 3 | 0.3333 | 14 | 0.1429 | 4 | 0 | 2 | 0 | 6 | 0 | 2 | 0.5 | 14 | 0.0714 |
| 1_75123209 | 7 | 0 | 3 | 0 | 8 | 0.125 | 15 | 0.3333 | 33 | 0.1818 | 7 | 0.2857 | 9 | 0.3333 | 11 | 0.4545 | 8 | 0.125 | 35 | 0.3143 |
| 1_75123476 | 20 | 0.85 | 14 | 0.5 | 21 | 0.7143 | 12 | 0.6667 | 67 | 0.7015 | 11 | 0.5455 | 15 | 0.7333 | 21 | 0.6667 | 12 | 0.5833 | 59 | 0.6441 |
| 1_75123530 | 15 | 0.8667 | 12 | 0.8333 | 20 | 0.95 | 11 | 0.8182 | 58 | 0.8793 | 10 | 1 | 11 | 1 | 17 | 0.8235 | 11 | 0.7273 | 49 | 0.8776 |
| 1_75123588 | 23 | 0.3043 | 16 | 0.25 | 25 | 0.12 | 14 | 0.2143 | 78 | 0.2179 | 15 | 0.1333 | 18 | 0.1111 | 24 | 0.125 | 16 | 0.0625 | 73 | 0.1096 |
| 1_75123867 | 27 | 0.1852 | 16 | 0.3125 | 27 | 0.1481 | 17 | 0.1765 | 87 | 0.1954 | 17 | 0.1176 | 18 | 0.1667 | 26 | 0.1154 | 21 | 0.0476 | 82 | 0.1098 |
| 1_75124058 | 29 | 0.1034 | 21 | 0.1429 | 29 | 0.1034 | 18 | 0.1111 | 97 | 0.1134 | 18 | 0 | 19 | 0.0526 | 31 | 0.129 | 22 | 0.0455 | 90 | 0.0667 |
| 1_75124417 | 28 | 0 | 19 | 0.1053 | 32 | 0.125 | 17 | 0.1765 | 96 | 0.0938 | 19 | 0 | 23 | 0.2174 | 30 | 0.3333 | 24 | 0.0833 | 96 | 0.1771 |
| 1_75147959 | 2 | 0 | 2 | 0.5 | 4 | 0 | 2 | 0 | 10 | 0.1 | 2 | 0 | 6 | 0.1667 | 2 | 0 | 2 | 0 | 12 | 0.0833 |
| 1_75148073 | 1 | 1 | 3 | 0 | 3 | 1 | 1 | 0 | 8 | 0.5 | 1 | 1 | 5 | 0.6 | 4 | 0 | 4 | 0 | 14 | 0.2857 |
| 1_75148254 | 2 | 0 | 3 | 0 | 4 | 0.25 | 1 | 0 | 10 | 0.1 | 2 | 0.5 | 5 | 0.6 | 3 | 0 | 3 | 0 | 13 | 0.3077 |
| 1_75148490 | 1 | 0 | 2 | 0 | NA | NA | 1 | 1 | 4 | 0.25 | 3 | 0 | 2 | 0 | 1 | 1 | 1 | 0 | 7 | 0.1429 |
| 1_75157515 | 15 | 0 | 16 | 0.0625 | 11 | 0.0909 | 13 | 0.0769 | 55 | 0.0545 | 14 | 0.0714 | 17 | 0.0588 | 24 | 0 | 11 | 0.1818 | 66 | 0.0606 |
| 1_75157587 | 17 | 0.1176 | 18 | 0.2222 | 13 | 0.2308 | 14 | 0.2143 | 62 | 0.1935 | 16 | 0.0625 | 15 | 0 | 22 | 0.0909 | 11 | 0.1818 | 64 | 0.0781 |
| 1_75157709 | 15 | 0.6 | 16 | 0.5625 | 10 | 0.3 | 11 | 0.4545 | 52 | 0.5 | 10 | 0.2 | 12 | 0.4167 | 21 | 0.6667 | 9 | 0.6667 | 52 | 0.5192 |
| 1_75158185 | 15 | 0.0667 | 12 | 0.0833 | 10 | 0 | 12 | 0.1667 | 49 | 0.0816 | 11 | 0 | 17 | 0.1765 | 18 | 0.1111 | 10 | 0.1 | 56 | 0.1071 |
| 1_75158319 | 13 | 0.2308 | 12 | 0.25 | 8 | 0.125 | 12 | 0.1667 | 45 | 0.2 | 10 | 0.2 | 16 | 0.3125 | 18 | 0.1111 | 11 | 0.4545 | 55 | 0.2545 |
| 1_75158349 | 12 | 0.4167 | 11 | 0.4545 | 9 | 0.3333 | 13 | 0.2308 | 45 | 0.3556 | 10 | 0.1 | 11 | 0.3636 | 12 | 0.6667 | 8 | 0.25 | 41 | 0.3659 |
| 1_75158385 | 14 | 0.6429 | 12 | 0.75 | 11 | 0.5455 | 12 | 0.8333 | 49 | 0.6939 | 12 | 0.5 | 13 | 0.6923 | 16 | 0.6875 | 11 | 0.6364 | 52 | 0.6346 |
| 1_75158490 | 12 | 0.5833 | 10 | 0.3 | 9 | 0.6667 | 13 | 0.6154 | 44 | 0.5455 | 11 | 0.4545 | 16 | 0.5625 | 16 | 0.625 | 9 | 0.4444 | 52 | 0.5385 |
| 1_75162844 | 9 | 0.1111 | 9 | 0.2222 | 4 | 0.25 | 8 | 0.375 | 30 | 0.2333 | 9 | 0 | 9 | 0.2222 | 8 | 0.25 | 5 | 0.4 | 31 | 0.1935 |
| 1_75162979 | 4 | 0 | 8 | 0.25 | 4 | 0 | 5 | 0.2 | 21 | 0.1429 | 8 | 0.375 | 9 | 0.1111 | 5 | 0.2 | 4 | 0 | 26 | 0.1923 |
| 1_75163325 | 6 | 0.3333 | 6 | 0 | 3 | 0 | 7 | 0.2857 | 22 | 0.1818 | 10 | 0.1 | 11 | 0.1818 | 6 | 0.1667 | 3 | 0.6667 | 30 | 0.2 |
| 1_75163449 | 5 | 0 | 6 | 0.1667 | 3 | 0 | 5 | 0 | 19 | 0.0526 | 8 | 0.125 | 10 | 0 | 6 | 0.1667 | 3 | 0 | 27 | 0.0741 |
| 1_75163479 | 5 | 0.2 | 7 | 0.2857 | 2 | 0 | 5 | 0 | 19 | 0.1579 | 7 | 0 | 10 | 0.2 | 5 | 0 | 1 | 0 | 23 | 0.087 |
| 1_75168985 | 1 | 0 | 2 | 0.5 | NA | NA | NA | NA | 3 | 0.3333 | NA | NA | NA | NA | NA | NA | 3 | 0.6667 | 3 | 0.6667 |
| 1_75173699 | NA | NA | NA | NA | NA | NA | 1 | 1 | 1 | 1 | 1 | 1 | NA | NA | NA | NA | NA | NA | 1 | 1 |
| 1_75174296 | 1 | 1 | 1 | 1 | 1 | 1 | 1 | 1 | 4 | 1 | 2 | 0.5 | NA | NA | NA | NA | NA | NA | 2 | 0.5 |
| 1_75174564 | 1 | 0 | 2 | 0 | 2 | 1 | 2 | 0 | 7 | 0.2857 | NA | NA | 1 | 0 | 2 | 0 | 4 | 0.5 | 7 | 0.2857 |
| 1_75174976 | 1 | 1 | 1 | 0 | 1 | 1 | 1 | 0 | 4 | 0.5 | 2 | 0.5 | NA | NA | NA | NA | NA | NA | 2 | 0.5 |
| 1_75175302 | 1 | 1 | NA | NA | 1 | 1 | 1 | 1 | 3 | 1 | 1 | 1 | NA | NA | NA | NA | NA | NA | 1 | 1 |
| 1_75175680 | NA | NA | 1 | 0 | 1 | 1 | NA | NA | 2 | 0.5 | 2 | 0.5 | NA | NA | NA | NA | NA | NA | 2 | 0.5 |
| 1_75175730 | NA | NA | 1 | 0 | 1 | 1 | NA | NA | 2 | 0.5 | 1 | 1 | NA | NA | NA | NA | NA | NA | 1 | 1 |
| 1_75175871 | 1 | 0 | 2 | 0.5 | 2 | 0 | 3 | 0 | 8 | 0.125 | NA | NA | 1 | 1 | 2 | 0 | 4 | 0 | 7 | 0.1429 |
| 1_75175993 | NA | NA | 1 | 1 | 1 | 0 | NA | NA | 2 | 0.5 | 2 | 0.5 | NA | NA | NA | NA | NA | NA | 2 | 0.5 |
| 1_75176050 | NA | NA | 1 | 1 | NA | NA | NA | NA | 1 | 1 | 2 | 0.5 | NA | NA | NA | NA | 1 | 1 | 3 | 0.6667 |
| 1_75186287 | NA | NA | 1 | 1 | 1 | 1 | NA | NA | 2 | 1 | 2 | 0.5 | NA | NA | NA | NA | NA | NA | 2 | 0.5 |
| 1_75188611 | 5 | 0.2 | 6 | 0.1667 | 10 | 0.2 | 4 | 0.75 | 25 | 0.28 | 3 | 0 | 2 | 0 | 2 | 0 | 6 | 0.5 | 13 | 0.2308 |
| 1_75189586 | 4 | 0.25 | 8 | 0 | 12 | 0 | 6 | 0.3333 | 30 | 0.1 | 3 | 0 | 6 | 0.1667 | 2 | 0 | 6 | 0.1667 | 17 | 0.1176 |
| 1_75189861 | 5 | 0.2 | 7 | 0 | 11 | 0.1818 | 7 | 0.2857 | 30 | 0.1667 | 4 | 0.25 | 6 | 0.3333 | 4 | 0 | 7 | 0 | 21 | 0.1429 |
| 1_75190609 | 4 | 0.5 | 7 | 0.7143 | 13 | 0.5385 | 7 | 0.7143 | 31 | 0.6129 | 4 | 0.5 | 6 | 0.8333 | 4 | 0.25 | 5 | 0.4 | 19 | 0.5263 |
| 1_75190655 | 5 | 0.4 | 5 | 0.2 | 9 | 0.4444 | 7 | 0.4286 | 26 | 0.3846 | 4 | 0.5 | 4 | 0.25 | 4 | 0.25 | 8 | 0.5 | 20 | 0.4 |
| 1_75190751 | 6 | 0.1667 | 8 | 0.25 | 14 | 0.2857 | 7 | 0.5714 | 35 | 0.3143 | 5 | 0.4 | 5 | 0 | 6 | 0.3333 | 8 | 0.375 | 24 | 0.2917 |
| 1_75190924 | 1 | 0 | NA | NA | NA | NA | 2 | 1 | 3 | 0.6667 | NA | NA | 2 | 0.5 | 1 | 1 | NA | NA | 3 | 0.6667 |
| 1_75191004 | 2 | 0.5 | NA | NA | NA | NA | NA | NA | 2 | 0.5 | NA | NA | 1 | 1 | NA | NA | 2 | 0 | 3 | 0.3333 |
| 1_75191126 | 2 | 1 | NA | NA | NA | NA | NA | NA | 2 | 1 | NA | NA | NA | NA | NA | NA | 1 | 1 | 1 | 1 |
| 1_75191187 | 2 | 1 | NA | NA | NA | NA | NA | NA | 2 | 1 | NA | NA | 1 | 1 | NA | NA | 1 | 1 | 2 | 1 |
| 1_75191750 | 23 | 0.6957 | 23 | 0.8696 | 25 | 0.84 | 17 | 0.7647 | 88 | 0.7955 | 29 | 0.7931 | 13 | 0.7692 | 9 | 0.8889 | 15 | 0.8 | 66 | 0.803 |
| 1_75191816 | 34 | 0.6471 | 48 | 0.5625 | 35 | 0.2857 | 32 | 0.7188 | 149 | 0.5503 | 44 | 0.4773 | 20 | 0.55 | 22 | 0.4545 | 20 | 0.4 | 106 | 0.4717 |
| 1_75191883 | 31 | 0.9677 | 48 | 1 | 42 | 1 | 27 | 0.9259 | 148 | 0.9797 | 47 | 0.9787 | 23 | 1 | 23 | 1 | 25 | 1 | 118 | 0.9915 |
| 1_75191965 | 34 | 0.3824 | 48 | 0.4167 | 39 | 0.5128 | 29 | 0.4483 | 150 | 0.44 | 44 | 0.5 | 23 | 0.3043 | 18 | 0.2222 | 20 | 0.45 | 105 | 0.4 |
| 1_75191977 | 38 | 0.2895 | 50 | 0.54 | 42 | 0.3095 | 34 | 0.2647 | 164 | 0.3659 | 45 | 0.2444 | 28 | 0.2857 | 20 | 0.4 | 23 | 0.3478 | 116 | 0.3017 |
| 1_75192003 | 34 | 0.9706 | 48 | 0.9792 | 42 | 0.9762 | 35 | 0.9143 | 159 | 0.9623 | 45 | 0.9556 | 29 | 0.9655 | 23 | 0.913 | 28 | 0.9286 | 125 | 0.944 |
| 1_75192025 | 34 | 0.7647 | 44 | 0.8864 | 42 | 0.8333 | 34 | 0.8824 | 154 | 0.8442 | 41 | 0.7805 | 27 | 0.6667 | 16 | 0.75 | 26 | 0.8462 | 110 | 0.7636 |
| 1_75192074 | 30 | 0.3333 | 33 | 0.2727 | 28 | 0.2143 | 25 | 0.12 | 116 | 0.2414 | 40 | 0.275 | 24 | 0.2917 | 17 | 0.3529 | 21 | 0.381 | 102 | 0.3137 |
| 1_75192110 | 32 | 0.0312 | 42 | 0.0952 | 33 | 0.1515 | 29 | 0.0345 | 136 | 0.0809 | 40 | 0.1 | 28 | 0.0714 | 19 | 0 | 25 | 0.24 | 112 | 0.1071 |
| 1_75192177 | 24 | 0.2917 | 35 | 0.1714 | 28 | 0.1429 | 26 | 0 | 113 | 0.1504 | 32 | 0.0625 | 23 | 0.0435 | 11 | 0.0909 | 17 | 0.2941 | 83 | 0.1084 |
| 1_75192212 | 22 | 0.0455 | 34 | 0.0882 | 28 | 0.0714 | 22 | 0.0455 | 106 | 0.066 | 30 | 0.1 | 20 | 0.1 | 17 | 0.1176 | 30 | 0.1333 | 97 | 0.1134 |
| 1_75192426 | 47 | 0.6596 | 68 | 0.4412 | 48 | 0.4583 | 40 | 0.65 | 203 | 0.5369 | 63 | 0.4762 | 44 | 0.6136 | 38 | 0.5526 | 44 | 0.5227 | 189 | 0.5344 |
| 1_75192438 | 46 | 0.3043 | 73 | 0.3014 | 54 | 0.4259 | 45 | 0.3111 | 218 | 0.3349 | 67 | 0.2239 | 46 | 0.4348 | 37 | 0.2703 | 51 | 0.3333 | 201 | 0.3085 |
| 1_75192579 | 43 | 1 | 60 | 1 | 46 | 1 | 41 | 1 | 190 | 1 | 63 | 1 | 45 | 0.9778 | 33 | 1 | 33 | 1 | 174 | 0.9943 |
| 1_75192696 | 47 | 0.0851 | 72 | 0.0972 | 52 | 0.1538 | 47 | 0.1064 | 218 | 0.1101 | 68 | 0.0882 | 54 | 0.1111 | 34 | 0.0588 | 44 | 0.0455 | 200 | 0.08 |

| | | | | | | | | | | | | | | | | | | | | |
|---|---|---|---|---|---|---|---|---|---|---|---|---|---|---|---|---|---|---|---|---|
| 1_7519285( | 41 | 0.0976 | 60 | 0.0333 | 45 | 0.0222 | 45 | 0.0667 | 191 | 0.0524 | 64 | 0.0781 | 48 | 0.0417 | 35 | 0.0857 | 37 | 0.0541 | 184 | 0.0652 |
| 1_7519286' | 43 | 0.8605 | 64 | 0.8281 | 46 | 0.913 | 45 | 0.8 | 198 | 0.8485 | 62 | 0.7903 | 46 | 0.7826 | 34 | 0.8824 | 39 | 0.8462 | 181 | 0.8177 |
| 1_7519299' | 40 | 0.075 | 59 | 0.1525 | 46 | 0.1522 | 37 | 0.1622 | 182 | 0.1374 | 51 | 0.2157 | 50 | 0.24 | 32 | 0.0938 | 36 | 0.1667 | 169 | 0.1893 |
| 1_7519301 | 32 | 0.9688 | 57 | 0.9298 | 44 | 0.8636 | 37 | 0.973 | 170 | 0.9294 | 45 | 0.9111 | 41 | 0.9512 | 30 | 0.9333 | 38 | 0.9474 | 154 | 0.9351 |
| 1_7519302' | 26 | 0.1923 | 45 | 0.2222 | 33 | 0.2121 | 23 | 0.3043 | 127 | 0.2283 | 43 | 0.1628 | 32 | 0.2812 | 24 | 0.25 | 25 | 0.24 | 124 | 0.2258 |
| 1_7519313 | 36 | 0.3889 | 60 | 0.2833 | 40 | 0.275 | 39 | 0.1538 | 175 | 0.2743 | 48 | 0.2917 | 39 | 0.2821 | 32 | 0.3125 | 34 | 0.2353 | 153 | 0.281 |
| 1_7519322( | 34 | 0.0882 | 60 | 0.0667 | 44 | 0.0682 | 40 | 0.1 | 178 | 0.0787 | 51 | 0.1176 | 42 | 0.0952 | 28 | 0.1429 | 33 | 0.0606 | 154 | 0.1039 |
| 1_7519333( | 1 | 1 | NA | NA | NA | NA | NA | NA | 1 | 1 | NA | NA | 1 | 1 | NA | NA | NA | NA | 1 | 1 |
| 1_7519342' | 1 | 1 | NA | NA | NA | NA | NA | NA | 1 | 1 | NA | NA | 1 | 1 | NA | NA | NA | NA | 1 | 1 |
| 1_7522073 | 2 | 0 | 1 | 0 | 6 | 0.5 | 2 | 0.5 | 11 | 0.3636 | 1 | 0 | NA | NA | 1 | 1 | NA | NA | 2 | 0.5 |
| 1_7522081 | 2 | 0 | 2 | 0 | 7 | 0.1429 | 3 | 0.3333 | 14 | 0.1429 | 1 | 1 | 1 | 0 | 2 | 0.5 | NA | NA | 4 | 0.5 |
| 1_7522207 | 10 | 0.2 | 14 | 0.0714 | 10 | 0.1 | 9 | 0.3333 | 43 | 0.1628 | 10 | 0 | 4 | 0 | 8 | 0.125 | 7 | 0.1429 | 29 | 0.069 |
| 1_7522209 | 5 | 0.2 | 13 | 0.1538 | 10 | 0.2 | 8 | 0.375 | 36 | 0.2222 | 7 | 0.2857 | 5 | 0.2 | 4 | 0 | 7 | 0.2857 | 23 | 0.2174 |
| 1_7523466 | 31 | 0.4516 | 21 | 0.381 | 24 | 0.5 | 27 | 0.5926 | 103 | 0.4854 | 26 | 0.4231 | 29 | 0.3103 | 12 | 0.5 | 22 | 0.5 | 89 | 0.4157 |
| 1_7523480 | 21 | 0.6667 | 11 | 0.6364 | 15 | 0.5333 | 20 | 0.6 | 67 | 0.6119 | 16 | 0.5 | 20 | 0.55 | 9 | 0.5556 | 12 | 0.5 | 57 | 0.5263 |
| 1_7523481 | 27 | 0.7037 | 19 | 0.7895 | 22 | 0.5909 | 23 | 0.7826 | 91 | 0.7143 | 17 | 0.7647 | 22 | 0.5455 | 8 | 0.625 | 11 | 0.8182 | 58 | 0.6724 |
| 1_7536459 | 3 | 1 | 2 | 0.5 | 2 | 0.5 | 1 | 1 | 8 | 0.75 | 3 | 1 | 2 | 0.5 | 1 | 1 | 6 | 0.5 | 12 | 0.6667 |
| 1_7536466 | 4 | 0.25 | 3 | 0 | 2 | 0.5 | 2 | 0.5 | 11 | 0.2727 | 3 | 0.3333 | 3 | 0 | 1 | 0 | 5 | 0 | 12 | 0.0833 |
| 1_7536483 | 3 | 0.3333 | 3 | 0 | 3 | 0.3333 | 1 | 0 | 10 | 0.2 | 3 | 0.3333 | 3 | 0 | 2 | 0.5 | 7 | 0.2857 | 15 | 0.2667 |
| 1_7536510 | 4 | 0.75 | 2 | 0.5 | 3 | 0.6667 | 1 | 1 | 10 | 0.7 | 3 | 0.3333 | 2 | 0.5 | 1 | 0 | 6 | 0.1667 | 12 | 0.25 |
| 1_7536537 | 4 | 1 | 1 | 1 | 2 | 0.5 | 1 | 1 | 8 | 0.875 | 4 | 0.75 | 4 | 1 | 1 | 1 | 7 | 0.8571 | 16 | 0.875 |
| 1_7536550 | 3 | 0.3333 | 3 | 0 | 5 | 0.2 | 2 | 0.5 | 13 | 0.2308 | 4 | 0 | 3 | 0.3333 | 2 | 0.5 | 7 | 0.4286 | 16 | 0.3125 |
| 1_7536558 | 4 | 1 | 4 | 1 | 5 | 0.8 | 2 | 1 | 15 | 0.9333 | 5 | 1 | 4 | 1 | 2 | 1 | 8 | 1 | 19 | 1 |
| 1_7536565 | 2 | 0 | 4 | 0.25 | 5 | 0.2 | 2 | 0 | 13 | 0.1538 | 5 | 0.2 | 4 | 0.25 | 2 | 0 | 8 | 0.375 | 19 | 0.2632 |
| 1_7536568 | 4 | 0 | 4 | 0.25 | 4 | 0 | 2 | 0 | 14 | 0.0714 | 3 | 0.6667 | 3 | 0.6667 | 1 | 0 | 6 | 0.5 | 13 | 0.5385 |
| 1_7536569 | 4 | 0 | 2 | 0.5 | 5 | 0.2 | 2 | 0 | 13 | 0.1538 | 4 | 0 | 4 | 0.25 | 1 | 0 | 6 | 0.3333 | 15 | 0.2 |
| 1_7538055 | 7 | 0.2857 | 8 | 0.375 | 6 | 0.6667 | 7 | 0.5714 | 28 | 0.4643 | 9 | 0.2222 | 10 | 0.5 | 4 | 0 | 9 | 0.4444 | 32 | 0.3438 |
| 1_7538067 | 6 | 0.3333 | 8 | 0.625 | 6 | 0.3333 | 5 | 0.2 | 25 | 0.4 | 7 | 0.5714 | 9 | 0.3333 | 4 | 0.75 | 10 | 0.6 | 30 | 0.5333 |
| 1_7539848 | 1 | 1 | 1 | 1 | NA | NA | NA | NA | 2 | 1 | NA | NA | NA | NA | 1 | 1 | NA | NA | 1 | 1 |
| 1_7539850 | 1 | 1 | 1 | 1 | NA | NA | NA | NA | 2 | 1 | NA | NA | 1 | 1 | NA | NA | 1 | 1 | 2 | 1 |
| 1_7539917 | 1 | 1 | 1 | 1 | NA | NA | NA | NA | 2 | 1 | NA | NA | 1 | 1 | 1 | 1 | 1 | 1 | 3 | 1 |
| 1_7539993 | 1 | 1 | 1 | 0 | NA | NA | NA | NA | 2 | 0.5 | NA | NA | 1 | 0 | 1 | 0 | 2 | 0.5 | 4 | 0.25 |
| 1_7539999 | 1 | 1 | 1 | 0 | NA | NA | NA | NA | 2 | 0.5 | NA | NA | 1 | 1 | 1 | 0 | 2 | 0 | 4 | 0.25 |
| 1_7540618 | 3 | 0 | 4 | 0 | NA | NA | 2 | 0.5 | 9 | 0.1111 | 1 | 0 | 3 | 0 | 2 | 1 | 6 | 0.1667 | 12 | 0.25 |
| 1_7540815 | 8 | 0.75 | 5 | 0.8 | 1 | 1 | 3 | 1 | 17 | 0.8235 | 3 | 1 | 6 | 1 | 2 | 1 | 6 | 0.8333 | 17 | 0.9412 |
| 1_7540818 | 8 | 0.125 | 4 | 0 | 1 | 0 | 1 | 0 | 14 | 0.0714 | 2 | 0.5 | 5 | 0 | 2 | 0 | 6 | 0.3333 | 15 | 0.2 |
| 1_7540826 | 7 | 0.8571 | 5 | 0.6 | 2 | 0.5 | 3 | 0.6667 | 17 | 0.7059 | 3 | 1 | 6 | 0.6667 | 2 | 1 | 8 | 0.75 | 19 | 0.7895 |
| 1_7540827 | 8 | 0 | 5 | 0.2 | 2 | 0 | 3 | 0 | 18 | 0.0556 | 3 | 0 | 5 | 0.4 | 2 | 0 | 5 | 0.2 | 15 | 0.2 |
| 1_7540849 | 11 | 0.0909 | 6 | 0.3333 | 3 | 0 | 3 | 0.3333 | 23 | 0.1739 | 5 | 0.2 | 7 | 0 | 2 | 0 | 7 | 0.2857 | 21 | 0.1429 |
| 1_7541281 | 2 | 0 | 3 | 0.3333 | NA | NA | 2 | 0 | 7 | 0.1429 | 1 | 0 | 1 | 0 | 1 | 0 | 1 | 1 | 4 | 0.25 |
| 1_7541958 | 4 | 0 | 11 | 0.0909 | 1 | 0 | 5 | 0.2 | 21 | 0.0952 | 4 | 0.25 | 2 | 0 | 2 | 0 | 4 | 0.25 | 12 | 0.1667 |
| 1_7541967( | 3 | 0.6667 | 10 | 0.6 | 1 | 0 | 9 | 0.5556 | 23 | 0.5652 | 5 | 0.6 | 3 | 0.6667 | 6 | 0.6667 | 4 | 0.25 | 18 | 0.5556 |
| 1_7541972 | 3 | 0.3333 | 9 | 0.2222 | 2 | 0 | 8 | 0.125 | 22 | 0.1818 | 5 | 0 | 3 | 0 | 6 | 0.3333 | 6 | 0.1667 | 20 | 0.15 |
| 1_7541981 | 5 | 0 | 11 | 0.0909 | 2 | 0 | 8 | 0.125 | 26 | 0.0769 | 5 | 0 | 3 | 0.3333 | 5 | 0 | 5 | 0.4 | 18 | 0.1667 |
| 1_7542766 | 1 | 1 | 1 | 0 | NA | NA | 1 | 0 | 3 | 0.3333 | 1 | 1 | NA | NA | 3 | 0.6667 | 5 | 1 | 9 | 0.8889 |
| 1_7542778 | 1 | 0 | 1 | 0 | NA | NA | 1 | 1 | 3 | 0.3333 | 1 | 0 | NA | NA | 3 | 0 | 5 | 0.4 | 9 | 0.2222 |
| 1_7545035( | 2 | 0 | NA | NA | 1 | 0 | 1 | 1 | 4 | 0.25 | 3 | 0.3333 | NA | NA | 6 | 0.1667 | 3 | 0.3333 | 12 | 0.25 |
| 1_7545123 | 10 | 0.6 | 10 | 0.4 | 13 | 0.4615 | 4 | 0.25 | 37 | 0.4595 | 15 | 0.4 | 14 | 0.3571 | 14 | 0.6429 | 14 | 0.2143 | 57 | 0.4035 |
| 1_7545146 | 10 | 0.3 | 9 | 0.5556 | 14 | 0.4286 | 3 | 0.6667 | 36 | 0.4444 | 12 | 0.3333 | 13 | 0.3077 | 16 | 0.5 | 12 | 0.4167 | 53 | 0.3962 |
| 1_7545147 | 12 | 1 | 11 | 0.6364 | 13 | 0.6154 | 3 | 0.6667 | 39 | 0.7436 | 13 | 0.8462 | 13 | 0.9231 | 15 | 0.8 | 12 | 0.8333 | 53 | 0.8491 |
| 1_7545148 | 8 | 0.875 | 8 | 0.875 | 12 | 1 | 2 | 1 | 30 | 0.9333 | 9 | 1 | 11 | 1 | 15 | 1 | 13 | 0.9231 | 48 | 0.9792 |
| 1_7545155 | 13 | 0.0769 | 10 | 0.2 | 12 | 0.1667 | 3 | 0 | 38 | 0.1316 | 13 | 0.3077 | 10 | 0.2 | 11 | 0.2727 | 13 | 0.1538 | 47 | 0.234 |
| 1_7545158 | 12 | 0.25 | 9 | 0.3333 | 11 | 0.7273 | 3 | 0 | 35 | 0.4 | 13 | 0.3077 | 11 | 0.4545 | 16 | 0.4375 | 13 | 0.5385 | 53 | 0.434 |
| 1_7545162 | 10 | 0.2 | 9 | 0.1111 | 13 | 0.1538 | 3 | 0.3333 | 35 | 0.1714 | 10 | 0.4 | 11 | 0.1818 | 13 | 0.1538 | 11 | 0.2727 | 45 | 0.2444 |
| 1_7545163 | 11 | 0 | 9 | 0.1111 | 12 | 0.0833 | 3 | 0 | 35 | 0.0571 | 11 | 0.1818 | 10 | 0.2 | 16 | 0.0625 | 10 | 0 | 47 | 0.1064 |
| 1_7545171 | 13 | 0.0769 | 11 | 0.1818 | 14 | 0.3571 | 3 | 0.6667 | 41 | 0.2439 | 12 | 0.4167 | 14 | 0.2143 | 16 | 0.3125 | 14 | 0.5 | 56 | 0.3571 |
| 1_7545206 | 9 | 0.2222 | 9 | 0.1111 | 12 | 0.1667 | 1 | 0 | 31 | 0.1613 | 9 | 0.1111 | 12 | 0.0833 | 15 | 0.2 | 12 | 0.5 | 48 | 0.2292 |
| 1_7545211 | 10 | 0.7 | 5 | 0.8 | 10 | 0.8 | 1 | 1 | 26 | 0.7692 | 6 | 0.8333 | 11 | 0.7273 | 13 | 0.9231 | 9 | 1 | 39 | 0.8718 |
| 1_7545233 | 8 | 0.25 | 10 | 0.4 | 10 | 0.4 | 2 | 0.5 | 30 | 0.3667 | 6 | 0.5 | 13 | 0.3846 | 11 | 0.4545 | 8 | 0.375 | 38 | 0.4211 |
| 1_7545238( | 8 | 0.375 | 8 | 0.125 | 8 | 0.5 | 2 | 1 | 26 | 0.3846 | 6 | 0.6667 | 9 | 0.4444 | 14 | 0.5 | 4 | 0.75 | 33 | 0.5455 |
| 1_7545252 | 10 | 0 | 10 | 0.1 | 12 | 0.1667 | 1 | 0 | 33 | 0.0909 | 5 | 0 | 10 | 0.1 | 13 | 0.1538 | 8 | 0 | 36 | 0.0833 |
| 1_7545254 | 10 | 0.1 | 6 | 0 | 9 | 0.4444 | 2 | 0 | 27 | 0.1852 | 6 | 0.1667 | 11 | 0.0909 | 14 | 0.0714 | 8 | 0.125 | 39 | 0.1026 |
| 1_7545265 | 9 | 0.2222 | 7 | 0.2857 | 6 | 0.3333 | 2 | 0.5 | 24 | 0.2917 | 5 | 0.2 | 13 | 0.3077 | 10 | 0.1 | 8 | 0.375 | 36 | 0.25 |
| 1_7545274 | 5 | 0.4 | 6 | 0.3333 | 4 | 0 | 1 | 0 | 16 | 0.25 | 4 | 0.25 | 9 | 0.1111 | 10 | 0.3 | 4 | 0.25 | 27 | 0.2222 |
| 1_7545275 | 8 | 0.125 | 8 | 0.375 | 8 | 0.25 | 1 | 0 | 25 | 0.24 | 6 | 0.1667 | 11 | 0.4545 | 15 | 0.6 | 9 | 0.6667 | 41 | 0.5122 |

| | | | | | | | | | | | | | | | | | | | | |
|---|---|---|---|---|---|---|---|---|---|---|---|---|---|---|---|---|---|---|---|---|
| 1_7545408 | 5 | 0.8 | 5 | 0.4 | 8 | 0.25 | 1 | 0 | 19 | 0.4211 | 4 | 0.25 | 10 | 0.5 | 12 | 0.3333 | 7 | 0.4286 | 33 | 0.3939 |
| 1_7545414 | 4 | 0.25 | 3 | 0 | 3 | 0 | NA | NA | 10 | 0.1 | 4 | 0 | 3 | 0 | 5 | 0.4 | 6 | 0.1667 | 18 | 0.1667 |
| 1_7545415 | 7 | 0.8571 | 6 | 0.8333 | 7 | 0.7143 | 1 | 1 | 21 | 0.8095 | 4 | 0.75 | 9 | 0.8889 | 10 | 0.9 | 6 | 0.8333 | 29 | 0.8621 |
| 1_7545419 | 4 | 0.5 | 6 | 0.5 | 6 | 0.1667 | 1 | 0 | 17 | 0.3529 | 2 | 0.5 | 7 | 0.1429 | 8 | 0.5 | 8 | 0.5 | 25 | 0.4 |
| 1_7545420 | 8 | 0.125 | 6 | 0 | 7 | 0.2857 | 2 | 0 | 23 | 0.1304 | 4 | 0 | 10 | 0.1 | 12 | 0.0833 | 7 | 0.2857 | 33 | 0.1212 |
| 1_7545425 | 8 | 0.25 | 5 | 0.2 | 7 | 0.4286 | NA | NA | 20 | 0.3 | 2 | 0 | 10 | 0.4 | 11 | 0.3636 | 8 | 0.375 | 31 | 0.3548 |
| 1_7545435 | 3 | 0.3333 | 4 | 0.75 | 4 | 0.75 | 1 | 1 | 12 | 0.6667 | 2 | 0.5 | 9 | 1 | 8 | 0.75 | 6 | 1 | 25 | 0.88 |
| 1_7545561 | 4 | 0.25 | 5 | 0.2 | 4 | 0 | 1 | 0 | 14 | 0.1429 | 2 | 0 | 5 | 0.2 | 5 | 0 | 5 | 0 | 17 | 0.0588 |
| 1_7545949 | 5 | 0 | 5 | 0 | 3 | 0.3333 | 6 | 0.3333 | 19 | 0.1579 | 4 | 0.25 | 3 | 0.3333 | 9 | 0.5556 | 4 | 0.25 | 20 | 0.4 |
| 1_7545962 | 5 | 0.6 | 9 | 0.3333 | 5 | 0.2 | 4 | 0.5 | 23 | 0.3913 | 6 | 0.1667 | 3 | 0.6667 | 7 | 0 | 4 | 0.25 | 20 | 0.2 |
| 1_7546156 | 7 | 0.8571 | 8 | 0.5 | 7 | 0.8571 | 9 | 0.6667 | 31 | 0.7097 | 9 | 0.8889 | 6 | 0.6667 | 7 | 0.8571 | 7 | 0.5714 | 29 | 0.7586 |
| 1_7546164 | 6 | 0.6667 | 8 | 0.625 | 5 | 0.6 | 4 | 1 | 23 | 0.6957 | 9 | 0.7778 | 4 | 0.5 | 9 | 0.7778 | 5 | 0.8 | 27 | 0.7407 |
| 1_7546183 | 7 | 0.1429 | 9 | 0.1111 | 7 | 0 | 5 | 0.4 | 28 | 0.1429 | 7 | 0.4286 | 4 | 0 | 5 | 0 | 4 | 0.25 | 20 | 0.2 |
| 1_7546723 | NA | NA | 2 | 1 | NA | NA | 2 | 0 | 4 | 0.5 | NA | NA | 1 | 1 | 1 | 1 | 1 | 1 | 3 | 1 |
| 1_7546757 | NA | NA | 1 | 1 | NA | NA | 2 | 1 | 3 | 1 | NA | NA | 1 | 1 | 1 | 1 | NA | NA | 2 | 1 |
| 1_7546761 | NA | NA | 2 | 0 | NA | NA | 1 | 1 | 3 | 0.3333 | NA | NA | 1 | 0 | NA | NA | 1 | 1 | 2 | 0.5 |
| 1_7547325 | NA | NA | 2 | 1 | NA | NA | NA | NA | 2 | 1 | 1 | 0 | NA | NA | 1 | 1 | NA | NA | 2 | 0.5 |
| 1_7547331 | NA | NA | 2 | 1 | NA | NA | NA | NA | 2 | 1 | NA | NA | NA | NA | 1 | 1 | NA | NA | 1 | 1 |
| 1_7548193 | NA | NA | 3 | 0.6667 | NA | NA | 1 | 1 | 4 | 0.75 | 1 | 0 | 1 | 0 | 1 | 1 | NA | NA | 3 | 0.3333 |
| 1_7548248 | 1 | 1 | 2 | 0 | 1 | 0 | 1 | 0 | 5 | 0.2 | 1 | 1 | 1 | 0 | NA | NA | NA | NA | 2 | 0.5 |
| 1_7548276 | 1 | 1 | 2 | 0.5 | NA | NA | 1 | 0 | 4 | 0.5 | 1 | 1 | 1 | 1 | NA | NA | NA | NA | 2 | 1 |
| 1_7548294 | 1 | 1 | 2 | 0 | 1 | 0 | 1 | 0 | 5 | 0.2 | 2 | 0.5 | 1 | 1 | NA | NA | NA | NA | 3 | 0.6667 |
| 1_7548295 | 1 | 1 | 2 | 1 | 1 | 1 | 1 | 1 | 5 | 1 | 2 | 1 | 1 | 1 | NA | NA | NA | NA | 3 | 1 |
| 1_7548298 | NA | NA | 2 | 0.5 | 1 | 1 | 1 | 0 | 4 | 0.5 | 1 | 1 | 1 | 0 | 1 | 0 | NA | NA | 3 | 0.3333 |
| 1_7548309 | 1 | 1 | 2 | 0 | NA | NA | 1 | 1 | 4 | 0.5 | 1 | 1 | NA | NA | 1 | 1 | NA | NA | 2 | 1 |
| 1_7548323 | 1 | 1 | 2 | 0.5 | 1 | 1 | 1 | 1 | 5 | 0.8 | 2 | 1 | 1 | 1 | 1 | 1 | NA | NA | 4 | 1 |
| 1_7548330 | 1 | 0 | 2 | 0.5 | NA | NA | 1 | 0 | 4 | 0.25 | 2 | 1 | 1 | 1 | 1 | 0 | NA | NA | 4 | 0.75 |
| 1_7548374 | NA | NA | 1 | 0 | 2 | 0.5 | 1 | 1 | 4 | 0.5 | 2 | 0.5 | 1 | 0 | NA | NA | NA | NA | 3 | 0.3333 |
| 1_7548388 | 1 | 1 | 1 | 0 | 2 | 0.5 | 1 | 1 | 5 | 0.6 | 2 | 0.5 | 2 | 1 | 3 | 1 | NA | NA | 7 | 0.8571 |
| 1_7548394 | 1 | 1 | 3 | 1 | 2 | 0 | 1 | 0 | 7 | 0.5714 | 2 | 0 | 1 | 0 | 3 | 0.6667 | NA | NA | 6 | 0.3333 |
| 1_7548576 | NA | NA | 3 | 0 | 6 | 0.1667 | 4 | 0.75 | 13 | 0.3077 | 2 | 0.5 | 4 | 0.5 | 4 | 0.25 | 1 | 0 | 11 | 0.3636 |
| 1_7548608 | 2 | 0 | 4 | 0.25 | 6 | 0.1667 | 4 | 0 | 16 | 0.125 | 2 | 0 | 6 | 0.1667 | 6 | 0 | 1 | 0 | 15 | 0.0667 |
| 1_7548621 | 1 | 1 | 3 | 0.3333 | 3 | 0 | 3 | 0.6667 | 10 | 0.4 | 2 | 0.5 | 4 | 0.25 | 4 | 0.75 | 2 | 0.5 | 12 | 0.5 |
| 1_7548627 | 1 | 0 | 2 | 0 | 5 | 0.2 | 3 | 0 | 11 | 0.0909 | 2 | 0 | 4 | 0 | 6 | 0.1667 | 1 | 0 | 13 | 0.0769 |
| 1_7548647 | 3 | 0 | 4 | 0.25 | 7 | 0 | 4 | 0 | 18 | 0.0556 | 2 | 0 | 7 | 0 | 6 | 0.3333 | 2 | 0 | 17 | 0.1176 |
| 1_7548653 | 3 | 0.6667 | 4 | 0.5 | 7 | 0.2857 | 5 | 0 | 19 | 0.3158 | 2 | 0 | 6 | 0.3333 | 6 | 0.5 | 2 | 0.5 | 16 | 0.375 |
| 1_7548688 | 2 | 0 | 4 | 0.25 | 6 | 0.1667 | 4 | 0 | 16 | 0.125 | 2 | 0.5 | 8 | 0.125 | 5 | 0.2 | 3 | 0 | 18 | 0.1667 |
| 1_7548690 | 3 | 1 | 4 | 0 | 7 | 0 | 5 | 0.2 | 19 | 0.2105 | 1 | 0 | 7 | 0 | 6 | 0.1667 | 2 | 0 | 16 | 0.0625 |
| 1_7551302 | 1 | 0 | 1 | 0 | NA | NA | 4 | 0.25 | 6 | 0.1667 | 3 | 1 | 1 | 0 | 6 | 0.5 | 2 | 0.5 | 12 | 0.5833 |
| 1_7551340 | 1 | 0 | 2 | 0.5 | 2 | 0 | 5 | 0 | 10 | 0.1 | 4 | 0 | 1 | 0 | 8 | 0.125 | 2 | 0 | 15 | 0.0667 |
| 1_7552403 | 4 | 0 | 2 | 0 | 3 | 0.3333 | 3 | 0 | 12 | 0.0833 | NA | NA | NA | NA | 3 | 0 | 3 | 0.3333 | 6 | 0.1667 |
| 1_7553355 | 10 | 0 | 14 | 0.0714 | 16 | 0.0625 | 12 | 0.0833 | 52 | 0.0577 | 7 | 0 | 12 | 0 | 17 | 0.0588 | 10 | 0.2 | 46 | 0.0652 |
| 1_7553567 | 13 | 0 | 16 | 0.125 | 17 | 0.1176 | 17 | 0.1765 | 63 | 0.1111 | 10 | 0.2 | 18 | 0.0556 | 17 | 0.2353 | 13 | 0.3077 | 58 | 0.1897 |
| 1_7553776 | NA | NA | NA | NA | NA | NA | 1 | 1 | 1 | 1 | 1 | 1 | NA | NA | NA | NA | 1 | 0 | 2 | 0.5 |
| 1_7734413 | 8 | 0.75 | 12 | 0.75 | 14 | 0.5714 | 12 | 0.5 | 46 | 0.6304 | 12 | 0.5 | 10 | 0.5 | 13 | 0.5385 | 12 | 0.6667 | 47 | 0.5532 |
| 1_7734526 | 7 | 0.1429 | 6 | 0.3333 | 6 | 0.1667 | 12 | 0.3333 | 31 | 0.2581 | 8 | 0.375 | 8 | 0.125 | 9 | 0.2222 | 11 | 0.3636 | 36 | 0.2778 |
| 1_7734531 | 6 | 0.6667 | 6 | 0.8333 | 9 | 0.7778 | 12 | 0.75 | 33 | 0.7576 | 8 | 0.875 | 5 | 0.8 | 11 | 0.8182 | 10 | 0.8 | 34 | 0.8235 |
| 1_7734540 | 6 | 0 | 6 | 0 | 8 | 0.125 | 12 | 0.1667 | 32 | 0.0938 | 7 | 0.2857 | 8 | 0.125 | 9 | 0 | 11 | 0.4545 | 35 | 0.2286 |
| 1_7734545 | 6 | 0.8333 | 5 | 0.8 | 4 | 0.75 | 7 | 0.8571 | 22 | 0.8182 | 7 | 1 | 7 | 0.8571 | 9 | 0.8889 | 7 | 1 | 30 | 0.9333 |
| 1_7734584 | 8 | 1 | 6 | 1 | 7 | 0.7143 | 9 | 0.7778 | 30 | 0.8667 | 8 | 0.875 | 6 | 1 | 10 | 0.8 | 8 | 0.875 | 32 | 0.875 |
| 1_7734588 | 7 | 0.1429 | 5 | 0.2 | 6 | 0 | 3 | 0 | 21 | 0.0952 | 7 | 0.1429 | 5 | 0.2 | 10 | 0.1 | 8 | 0 | 30 | 0.1 |
| 1_7734594 | 7 | 0.2857 | 6 | 0.6667 | 7 | 0.2857 | 8 | 0.625 | 28 | 0.4643 | 8 | 0.625 | 7 | 0.5714 | 9 | 0.6667 | 9 | 0.5556 | 33 | 0.6061 |
| 1_7734601 | 8 | 0.5 | 4 | 0.25 | 6 | 0.5 | 6 | 0.3333 | 24 | 0.4167 | 7 | 0.4286 | 5 | 0.6 | 9 | 0.3333 | 9 | 0.5556 | 30 | 0.4667 |
| 1_7734617 | 8 | 0.375 | 6 | 0.1667 | 5 | 0.2 | 8 | 0.25 | 27 | 0.2593 | 7 | 0.4286 | 6 | 0 | 8 | 0 | 8 | 0.125 | 29 | 0.1379 |
| 1_7734965 | 8 | 0.125 | 4 | 0.25 | 5 | 0 | 6 | 0 | 23 | 0.087 | 8 | 0.125 | 6 | 0 | 7 | 0.5714 | 8 | 0 | 29 | 0.1724 |
| 1_7734971 | 6 | 1 | 3 | 0.6667 | 4 | 0.5 | 5 | 0.8 | 18 | 0.7778 | 4 | 0.5 | 6 | 0.5 | 9 | 0.8889 | 5 | 0.4 | 24 | 0.625 |
| 1_7734972 | 8 | 0 | 2 | 0.5 | 3 | 0 | 4 | 0 | 17 | 0.0588 | 5 | 0.2 | 6 | 0 | 5 | 0.2 | 7 | 0.1429 | 23 | 0.1304 |
| 1_7734973 | 8 | 0.125 | 4 | 0 | 4 | 0.25 | 6 | 0 | 22 | 0.0909 | 8 | 0 | 6 | 0 | 9 | 0.1111 | 7 | 0.1429 | 30 | 0.0667 |
| 1_7734974 | 5 | 0.4 | 4 | 0.75 | 2 | 0.5 | 4 | 0.75 | 15 | 0.6 | 4 | 1 | 3 | 0.3333 | 7 | 0.4286 | 6 | 0.5 | 20 | 0.55 |
| 1_7734979 | 6 | 0.6667 | 2 | 1 | 4 | 0.25 | 5 | 0.6 | 17 | 0.5882 | 6 | 0.5 | 2 | 1 | 7 | 0.4286 | 3 | 1 | 18 | 0.6111 |
| 1_7734981 | 6 | 0.1667 | 4 | 0.75 | 3 | 0.6667 | 5 | 0.4 | 18 | 0.4444 | 8 | 0.5 | 4 | 0.5 | 7 | 0.5714 | 7 | 0.4286 | 26 | 0.5 |
| 1_7734990 | 6 | 1 | 4 | 1 | 4 | 0.75 | 4 | 0.75 | 18 | 0.8889 | 8 | 1 | 4 | 1 | 9 | 1 | 6 | 0.8333 | 27 | 0.963 |
| 1_7735007 | 3 | 0 | 3 | 0.3333 | 3 | 0.6667 | 2 | 0 | 11 | 0.2727 | 8 | 0.625 | 5 | 0.4 | 7 | 0.5714 | 6 | 0.6667 | 26 | 0.5769 |
| 1_7735012 | 3 | 0.6667 | 3 | 0 | 3 | 0.6667 | 4 | 0.75 | 13 | 0.5385 | 8 | 0.625 | 5 | 0.6 | 7 | 0.5714 | 7 | 0.7143 | 27 | 0.6296 |
| 1_7735023 | 3 | 0 | 4 | 0 | 3 | 0 | 3 | 0.3333 | 13 | 0.0769 | 7 | 0 | 2 | 0 | 3 | 0.3333 | 3 | 0 | 15 | 0.0667 |

| | | | | | | | | | | | | | | | | | | | | |
|---|---|---|---|---|---|---|---|---|---|---|---|---|---|---|---|---|---|---|---|---|
| 1_7735025 | 3 | 0 | 3 | 0 | 3 | 0.3333 | 4 | 0 | 13 | 0.0769 | 8 | 0.125 | 4 | 0.25 | 7 | 0 | 6 | 0.1667 | 25 | 0.12 |
| 1_7735030 | 4 | 0.75 | 3 | 0.6667 | 3 | 1 | 4 | 0.5 | 14 | 0.7143 | 6 | 0.6667 | 4 | 0.25 | 5 | 0.6 | 4 | 1 | 19 | 0.6316 |
| 1_7735034 | 2 | 0 | 3 | 0 | 2 | 0 | 4 | 0.5 | 11 | 0.1818 | 3 | 0 | 3 | 0.3333 | 7 | 0.4286 | 4 | 0.25 | 17 | 0.2941 |
| 1_7735035 | 4 | 1 | 2 | 1 | 3 | 1 | 4 | 0.5 | 13 | 0.8462 | 7 | 0.8571 | 4 | 0.75 | 7 | 0.8571 | 6 | 0.8333 | 24 | 0.8333 |
| 1_7740333 | 2 | 0 | 1 | 0 | 2 | 0.5 | 1 | 1 | 6 | 0.3333 | 3 | 0.6667 | NA | NA | 2 | 0.5 | 1 | 1 | 6 | 0.6667 |
| 1_7748353 | NA | NA | NA | NA | 2 | 0.5 | 1 | 1 | 3 | 0.6667 | 1 | 1 | NA | NA | 1 | 1 | NA | NA | 2 | 1 |
| 1_7748365 | 1 | 0 | 1 | 1 | 2 | 1 | NA | NA | 4 | 0.75 | 1 | 1 | NA | NA | 1 | 1 | NA | NA | 2 | 1 |
| 1_7748368 | NA | NA | NA | NA | 2 | 1 | NA | NA | 2 | 1 | 1 | 1 | NA | NA | 1 | 1 | NA | NA | 2 | 1 |
| 1_7748374 | 1 | 0 | 1 | 0 | 2 | 0.5 | NA | NA | 4 | 0.25 | 1 | 0 | NA | NA | 1 | 1 | NA | NA | 2 | 0.5 |
| 1_7819198 | 1 | 1 | NA | NA | NA | NA | NA | NA | 1 | 1 | 2 | 0.5 | NA | NA | NA | NA | NA | NA | 2 | 0.5 |
| 1_7819249 | NA | NA | 1 | 1 | NA | NA | NA | NA | 1 | 1 | 1 | 1 | NA | NA | NA | NA | NA | NA | 1 | 1 |
| 1_7819282 | 1 | 1 | 1 | 1 | NA | NA | NA | NA | 2 | 1 | 1 | 1 | NA | NA | NA | NA | NA | NA | 1 | 1 |
| 1_7839372 | NA | NA | 1 | 1 | NA | NA | NA | NA | 1 | 1 | NA | NA | NA | NA | 1 | 1 | 1 | 0 | 2 | 0.5 |
| 1_7839422 | NA | NA | 1 | 1 | NA | NA | NA | NA | 1 | 1 | NA | NA | NA | NA | 1 | 1 | NA | NA | 1 | 1 |
| 1_7839428 | NA | NA | 1 | 1 | NA | NA | NA | NA | 1 | 1 | NA | NA | NA | NA | 1 | 1 | 1 | 1 | 2 | 1 |
| 1_7839452 | NA | NA | 1 | 1 | NA | NA | NA | NA | 1 | 1 | NA | NA | NA | NA | 1 | 1 | 1 | 1 | 2 | 1 |
| 1_7839453 | NA | NA | 1 | 1 | NA | NA | NA | NA | 1 | 1 | NA | NA | NA | NA | 1 | 1 | 1 | 1 | 2 | 1 |
| 1_7839458 | NA | NA | 1 | 1 | NA | NA | 1 | 1 | 2 | 1 | NA | NA | NA | NA | 1 | 1 | NA | NA | 1 | 1 |
| 1_7839465 | NA | NA | 1 | 1 | NA | NA | NA | NA | 1 | 1 | NA | NA | NA | NA | 1 | 1 | NA | NA | 1 | 1 |
| 1_7839492 | NA | NA | NA | NA | NA | NA | 1 | 1 | 1 | 1 | NA | NA | NA | NA | 1 | 1 | 1 | 1 | 2 | 1 |
| 1_7847578 | NA | NA | NA | NA | 1 | 1 | NA | NA | 1 | 1 | NA | NA | 1 | 1 | NA | NA | NA | NA | 1 | 1 |
| 1_7847616 | NA | NA | NA | NA | 1 | 1 | NA | NA | 1 | 1 | NA | NA | 1 | 1 | NA | NA | NA | NA | 1 | 1 |
| 1_7847621 | NA | NA | NA | NA | 1 | 1 | NA | NA | 1 | 1 | NA | NA | 1 | 1 | NA | NA | NA | NA | 1 | 1 |
| 1_7864399 | 1 | 1 | NA | NA | NA | NA | NA | NA | 1 | 1 | NA | NA | NA | NA | NA | NA | 1 | 1 | 1 | 1 |
| 1_7864400 | 1 | 1 | NA | NA | NA | NA | NA | NA | 1 | 1 | NA | NA | NA | NA | NA | NA | 1 | 1 | 1 | 1 |
| 1_7864421 | 1 | 1 | NA | NA | NA | NA | NA | NA | 1 | 1 | NA | NA | NA | NA | NA | NA | 1 | 1 | 1 | 1 |
| 1_7865954 | 20 | 0.45 | 23 | 0.4783 | 23 | 0.4348 | 13 | 0.3846 | 79 | 0.443 | 12 | 0.25 | 7 | 0.5714 | 16 | 0.5 | 20 | 0.65 | 55 | 0.5091 |
| 1_7865965 | 21 | 0.2857 | 27 | 0.3333 | 26 | 0.3077 | 16 | 0.1875 | 90 | 0.2889 | 13 | 0.0769 | 8 | 0.125 | 18 | 0.2778 | 23 | 0.4348 | 62 | 0.2742 |
| 1_7868403 | 31 | 0.5161 | 35 | 0.5143 | 31 | 0.3871 | 20 | 0.7 | 117 | 0.5128 | 14 | 0.5714 | 9 | 0.3333 | 22 | 0.5455 | 31 | 0.4516 | 76 | 0.4868 |
| 1_7868410 | 29 | 0.069 | 41 | 0.0976 | 35 | 0.0857 | 22 | 0 | 127 | 0.0709 | 16 | 0.125 | 14 | 0.1429 | 28 | 0.1786 | 32 | 0.0938 | 90 | 0.1333 |
| 1_7868426 | 23 | 0.1304 | 36 | 0.0278 | 31 | 0.0968 | 19 | 0.1053 | 109 | 0.0826 | 19 | 0.1579 | 12 | 0.3333 | 24 | 0.0833 | 34 | 0.0882 | 89 | 0.1348 |
| 1_7868427 | 29 | 0.1724 | 38 | 0.0789 | 36 | 0.1111 | 21 | 0 | 124 | 0.0968 | 22 | 0.1818 | 14 | 0.0714 | 28 | 0.1071 | 35 | 0.0286 | 99 | 0.0909 |
| 1_7868435 | 28 | 0.6429 | 41 | 0.7805 | 33 | 0.8182 | 19 | 0.6842 | 121 | 0.7438 | 18 | 0.7778 | 13 | 0.7692 | 29 | 0.7241 | 33 | 0.697 | 93 | 0.7312 |
| 1_7868437 | 30 | 0.6667 | 43 | 0.3953 | 33 | 0.4242 | 18 | 0.6111 | 124 | 0.5 | 21 | 0.5238 | 15 | 0.4 | 27 | 0.4444 | 34 | 0.5882 | 97 | 0.5052 |
| 1_7879597 | 1 | 1 | NA | NA | NA | NA | NA | NA | 1 | 1 | NA | NA | NA | NA | NA | NA | 1 | 1 | 1 | 1 |
| 1_7941297 | 41 | 0.4146 | 62 | 0.2742 | 44 | 0.2727 | 46 | 0.1739 | 193 | 0.2798 | 36 | 0.2778 | 47 | 0.2553 | 41 | 0.3171 | 44 | 0.25 | 168 | 0.2738 |
| 1_7941298 | 53 | 0.2264 | 75 | 0.2533 | 60 | 0.2333 | 60 | 0.3167 | 248 | 0.2581 | 44 | 0.2045 | 63 | 0.2381 | 55 | 0.2727 | 57 | 0.3158 | 219 | 0.2603 |
| 1_7941303 | 35 | 0.8857 | 56 | 0.8214 | 46 | 0.7174 | 48 | 0.8333 | 185 | 0.8108 | 30 | 0.8 | 41 | 0.8293 | 42 | 0.7857 | 42 | 0.9286 | 155 | 0.8387 |
| 1_7941341 | 44 | 0.9773 | 59 | 0.9661 | 50 | 0.98 | 51 | 1 | 204 | 0.9804 | 40 | 0.95 | 52 | 0.9615 | 46 | 0.9565 | 50 | 0.96 | 188 | 0.9574 |
| 1_7941371 | 22 | 0.2727 | 17 | 0.2353 | 23 | 0.1304 | 18 | 0.2222 | 80 | 0.2125 | 17 | 0.2353 | 25 | 0.24 | 20 | 0.35 | 24 | 0.375 | 86 | 0.3023 |
| 1_7941391 | 41 | 0.0244 | 47 | 0.0851 | 41 | 0.0732 | 47 | 0.0851 | 176 | 0.0682 | 36 | 0.1944 | 45 | 0.0667 | 39 | 0.0513 | 40 | 0.15 | 160 | 0.1125 |
| 1_7941394 | 43 | 0.8837 | 48 | 0.6042 | 42 | 0.8333 | 45 | 0.7333 | 178 | 0.7584 | 36 | 0.8889 | 48 | 0.75 | 39 | 0.7436 | 40 | 0.9 | 163 | 0.816 |
| 1_7941415 | 33 | 0.2121 | 35 | 0.1429 | 30 | 0.2 | 35 | 0.0857 | 133 | 0.1579 | 22 | 0.2727 | 35 | 0.3143 | 31 | 0.2903 | 34 | 0.2353 | 122 | 0.2787 |
| 1_7941425 | 35 | 0.3714 | 46 | 0.3261 | 34 | 0.2941 | 41 | 0.3902 | 156 | 0.3462 | 30 | 0.2667 | 45 | 0.3333 | 37 | 0.4324 | 38 | 0.4474 | 150 | 0.3733 |
| 1_7941427 | 34 | 0.0588 | 43 | 0.2093 | 29 | 0.1034 | 40 | 0.1 | 146 | 0.1233 | 32 | 0.1875 | 43 | 0.2326 | 36 | 0.1389 | 35 | 0.1714 | 146 | 0.1849 |
| 1_7941429 | 36 | 0.0833 | 45 | 0.1333 | 34 | 0.0294 | 41 | 0.0244 | 156 | 0.0705 | 29 | 0.1034 | 45 | 0.0222 | 36 | 0.0556 | 33 | 0.1212 | 143 | 0.0699 |
| 1_7975565 | 5 | 0.2 | 5 | 0.4 | 2 | 0 | 3 | 0.3333 | 15 | 0.2667 | 2 | 0.5 | 3 | 0.3333 | 5 | 0.2 | 4 | 0 | 14 | 0.2143 |
| 1_7975566 | 5 | 0 | 5 | 0.6 | 2 | 0 | 3 | 0 | 15 | 0.2 | 2 | 0.5 | 5 | 0 | 4 | 0.5 | 3 | 0.3333 | 14 | 0.2857 |
| 1_7975575 | 6 | 1 | 4 | 0.75 | 1 | 1 | 3 | 0.6667 | 14 | 0.8571 | 2 | 1 | 6 | 0.3333 | 5 | 0.8 | 3 | 0.6667 | 16 | 0.625 |
| 1_7975576 | 6 | 0.8333 | 4 | 0.75 | 1 | 1 | 3 | 1 | 14 | 0.8571 | 2 | 1 | 6 | 0.8333 | 3 | 1 | 4 | 1 | 15 | 0.9333 |
| 1_7975591 | 5 | 0.2 | 5 | 0.2 | 2 | 0.5 | 3 | 0.3333 | 15 | 0.2667 | 2 | 0 | 5 | 0 | 5 | 0.2 | 4 | 0 | 16 | 0.0625 |
| 1_7975597 | 7 | 0.1429 | 5 | 0 | 2 | 0.5 | 3 | 0 | 17 | 0.1176 | 2 | 0 | 6 | 0.3333 | 6 | 0.3333 | 3 | 0 | 17 | 0.2353 |
| 1_7975727 | 6 | 0.1667 | 3 | 0 | NA | NA | 3 | 0 | 12 | 0.0833 | 1 | 0 | 3 | 0 | 5 | 0.4 | 3 | 0 | 12 | 0.1667 |
| 1_7975879 | 7 | 0 | 5 | 0 | 2 | 0 | 6 | 0.1667 | 20 | 0.05 | 3 | 0.3333 | 6 | 0 | 7 | 0.1429 | 3 | 0 | 19 | 0.1053 |
| 1_7977144 | 1 | 0 | NA | NA | 2 | 1 | 1 | 1 | 4 | 0.75 | NA | NA | 2 | 0.5 | NA | NA | 2 | 0.5 | 4 | 0.5 |
| 1_7977159 | 1 | 0 | NA | NA | 2 | 0 | 1 | 1 | 4 | 0.25 | NA | NA | 2 | 0.5 | NA | NA | 2 | 0 | 4 | 0.25 |
| 1_7979461 | 26 | 0.0385 | 29 | 0.0345 | 32 | 0.0938 | 25 | 0.04 | 112 | 0.0536 | 14 | 0.0714 | 25 | 0.04 | 31 | 0.0968 | 29 | 0.069 | 99 | 0.0707 |
| 1_7979921 | 27 | 0.4074 | 29 | 0.1724 | 31 | 0.2258 | 27 | 0.0741 | 114 | 0.2193 | 15 | 0.2 | 24 | 0.2917 | 31 | 0.3226 | 30 | 0.1333 | 100 | 0.24 |
| 1_8024427 | 2 | 0 | 8 | 0.375 | 9 | 0.4444 | 10 | 0.5 | 29 | 0.4138 | 8 | 0.125 | 11 | 0.1818 | 9 | 0.1111 | 11 | 0.4545 | 39 | 0.2308 |
| 1_8024428 | 4 | 0 | 7 | 0.2857 | 11 | 0.1818 | 11 | 0.3636 | 33 | 0.2424 | 7 | 0.1429 | 11 | 0 | 10 | 0 | 13 | 0.6923 | 41 | 0.2439 |
| 1_8024475 | 5 | 0.2 | 18 | 0.1111 | 19 | 0.1053 | 16 | 0.25 | 58 | 0.1552 | 12 | 0.0833 | 17 | 0.2941 | 15 | 0.0667 | 22 | 0.3182 | 66 | 0.2121 |
| 1_8024482 | 5 | 1 | 18 | 1 | 18 | 1 | 18 | 0.9444 | 59 | 0.9831 | 9 | 1 | 18 | 1 | 15 | 1 | 16 | 0.9375 | 58 | 0.9828 |
| 1_8024484 | 3 | 1 | 17 | 0.8824 | 18 | 1 | 18 | 0.9444 | 56 | 0.9464 | 8 | 1 | 16 | 1 | 14 | 1 | 19 | 0.9474 | 57 | 0.9825 |
| 1_8024491 | 3 | 0 | 15 | 0 | 13 | 0.2308 | 13 | 0.0769 | 44 | 0.0909 | 6 | 0 | 12 | 0.1667 | 14 | 0.2143 | 21 | 0.0952 | 53 | 0.1321 |
| 1_8024493 | 6 | 1 | 18 | 0.9444 | 17 | 0.8824 | 17 | 1 | 58 | 0.9483 | 10 | 0.9 | 15 | 1 | 16 | 0.75 | 22 | 0.9091 | 63 | 0.8889 |

| | | | | | | | | | | | | | | | | | | | | |
|---|---|---|---|---|---|---|---|---|---|---|---|---|---|---|---|---|---|---|---|---|
| 1_80479387 | 2 | 0.5 | 4 | 0.75 | 1 | 1 | 4 | 0.75 | 11 | 0.7273 | 3 | 0 | 4 | 0.25 | 5 | 0.8 | 4 | 0.75 | 16 | 0.5 |
| 1_81218932 | 1 | 1 | NA | NA | 1 | 0 | 1 | 1 | 3 | 0.6667 | NA | NA | 1 | 1 | NA | NA | 1 | 1 | 2 | 1 |
| 1_81218989 | 1 | 1 | NA | NA | 1 | 1 | 2 | 1 | 4 | 1 | NA | NA | 1 | 1 | NA | NA | 1 | 1 | 2 | 1 |
| 1_81315049 | 1 | 1 | 1 | 0 | 1 | 1 | 2 | 0 | 5 | 0.4 | NA | NA | 1 | 1 | NA | NA | NA | NA | 1 | 1 |
| 1_81318232 | 3 | 0.3333 | 2 | 0.5 | 3 | 0.3333 | 9 | 0.1111 | 17 | 0.2353 | NA | NA | 3 | 0.3333 | 1 | 0 | 3 | 0.3333 | 7 | 0.2857 |
| 1_81318843 | 2 | 0 | 2 | 0.5 | 3 | 0 | 5 | 0.2 | 12 | 0.1667 | NA | NA | 6 | 0.1667 | 2 | 0 | 2 | 0.5 | 10 | 0.2 |
| 1_82729897 | 34 | 0.2941 | 33 | 0.3333 | 33 | 0.2727 | 24 | 0.2083 | 124 | 0.2823 | 27 | 0.2222 | 26 | 0.2308 | 33 | 0.2121 | 33 | 0.2424 | 119 | 0.2269 |
| 1_82872790 | 10 | 0.6 | 9 | 0.3333 | 4 | 0 | 3 | 0 | 26 | 0.3462 | 4 | 0.75 | 6 | 0.3333 | 7 | 0.2857 | 3 | 0.6667 | 20 | 0.45 |
| 1_82873478 | 7 | 0.2857 | 7 | 0 | 5 | 0 | 1 | 1 | 20 | 0.15 | 1 | 0 | 5 | 0.2 | 4 | 0 | 4 | 0.25 | 14 | 0.1429 |
| 1_82873488 | 8 | 0 | 8 | 0.25 | 5 | 0.2 | 1 | 0 | 22 | 0.1364 | 2 | 0 | 5 | 0 | 6 | 0.1667 | 4 | 0 | 17 | 0.0588 |
| 1_82873547 | 8 | 0.375 | 8 | 0.25 | 4 | 0 | 2 | 0 | 22 | 0.2273 | 2 | 0 | 4 | 0.25 | 6 | 0 | 5 | 0.2 | 17 | 0.1176 |
| 1_82873795 | 9 | 0.2222 | 7 | 0.1429 | 5 | 0.2 | 3 | 0 | 24 | 0.1667 | 3 | 0.3333 | 5 | 0.2 | 5 | 0 | 5 | 0.2 | 18 | 0.1667 |
| 1_82873838 | 8 | 0.375 | 5 | 0.2 | 4 | 0 | 3 | 0.6667 | 20 | 0.3 | 3 | 0 | 6 | 0.1667 | 6 | 0.1667 | 5 | 0.4 | 20 | 0.2 |
| 1_82873986 | 8 | 0 | 5 | 0 | 2 | 0 | 3 | 0.3333 | 18 | 0.0556 | 2 | 0 | 6 | 0.1667 | 5 | 0.2 | 5 | 0.2 | 18 | 0.1667 |
| 1_83233831 | 6 | 0.3333 | 19 | 0.5263 | 12 | 0.4167 | 7 | 0.5714 | 44 | 0.4773 | 9 | 0.4444 | 2 | 0.5 | 7 | 0.1429 | 5 | 0.4 | 23 | 0.3478 |
| 1_83234242 | 4 | 0.5 | 12 | 0 | 8 | 0.125 | 6 | 0 | 30 | 0.1 | 8 | 0.125 | 1 | 0 | 5 | 0 | 6 | 0 | 20 | 0.05 |
| 1_83234773 | 6 | 0.1667 | 13 | 0.1538 | 8 | 0.375 | 7 | 0.2857 | 34 | 0.2353 | 7 | 0.4286 | 1 | 0 | 5 | 0.4 | 4 | 0 | 17 | 0.2941 |
| 1_83234857 | 6 | 0 | 10 | 0.1 | 7 | 0 | 7 | 0.1429 | 30 | 0.0667 | 6 | 0 | 1 | 1 | 5 | 0 | 5 | 0 | 17 | 0.0588 |
| 1_83234870 | 5 | 0 | 14 | 0.0714 | 8 | 0.375 | 8 | 0.125 | 35 | 0.1429 | 8 | 0.125 | 1 | 0 | 5 | 0.4 | 5 | 0.4 | 19 | 0.2632 |
| 1_83234886 | 6 | 0.1667 | 14 | 0.0714 | 9 | 0.3333 | 7 | 0.1429 | 36 | 0.1667 | 9 | 0.1111 | 1 | 0 | 5 | 0.4 | 6 | 0.1667 | 21 | 0.1905 |
| 1_83243729 | 1 | 1 | 3 | 0 | 1 | 0 | 1 | 0 | 6 | 0.1667 | 2 | 0.5 | 1 | 0 | 1 | 0 | 1 | 0 | 5 | 0.2 |
| 1_83253455 | 5 | 0.6 | 9 | 0.4444 | 6 | 0.5 | 3 | 1 | 23 | 0.5652 | 7 | 0.5714 | 1 | 0 | 3 | 0.6667 | 4 | 0.75 | 15 | 0.6 |
| 1_83253490 | 6 | 0.6667 | 11 | 0.4545 | 5 | 0.8 | 4 | 0.75 | 26 | 0.6154 | 6 | 0.6667 | 1 | 1 | 4 | 0.5 | 4 | 0.75 | 15 | 0.6667 |
| 1_83253719 | 6 | 0.3333 | 9 | 0.8889 | 7 | 0.4286 | 4 | 0.25 | 26 | 0.5385 | 3 | 0.6667 | 1 | 1 | 4 | 0.5 | 4 | 0.5 | 12 | 0.5833 |
| 1_83253831 | 4 | 0.25 | 8 | 0.25 | 5 | 0.4 | 4 | 0.5 | 21 | 0.3333 | 4 | 0 | 1 | 0 | 3 | 0.3333 | 4 | 0.25 | 12 | 0.1667 |
| 1_83253885 | 6 | 0 | 8 | 0.375 | 5 | 0 | 3 | 0.3333 | 22 | 0.1818 | 5 | 0 | 1 | 0 | 3 | 0 | 4 | 0.25 | 13 | 0.0769 |
| 1_83253901 | 6 | 1 | 7 | 0.8571 | 5 | 0.8 | 3 | 1 | 21 | 0.9048 | 2 | 1 | NA | NA | 3 | 1 | 3 | 0.6667 | 8 | 0.875 |
| 1_83253996 | 5 | 0.4 | 7 | 0 | 5 | 0 | 1 | 0 | 18 | 0.1111 | 5 | 0.2 | 1 | 0 | 1 | 1 | 2 | 0 | 9 | 0.2222 |
| 1_83254088 | 5 | 0.4 | 6 | 0.3333 | 4 | 0.25 | 3 | 0.6667 | 18 | 0.3889 | 4 | 0.75 | 1 | 1 | 2 | 0.5 | 2 | 0 | 9 | 0.5556 |
| 1_83254135 | 4 | 0 | 8 | 0.125 | 5 | 0.2 | 1 | 0 | 18 | 0.1111 | 3 | 0.6667 | 1 | 0 | 2 | 0 | NA | NA | 6 | 0.3333 |
| 1_83254191 | 5 | 0 | 8 | 0.125 | 5 | 0.2 | 3 | 0 | 21 | 0.0952 | 5 | 0.2 | 1 | 0 | 3 | 0.3333 | 1 | 0 | 10 | 0.2 |
| 1_83254524 | 3 | 1 | 4 | 0.25 | 5 | 0.6 | 1 | 1 | 13 | 0.6154 | 4 | 0.5 | 1 | 1 | 2 | 1 | 2 | 0.5 | 9 | 0.6667 |
| 1_83254594 | 4 | 0.75 | 5 | 0.2 | 4 | 0.25 | 1 | 1 | 14 | 0.4286 | 4 | 0.5 | 1 | 1 | 2 | 0 | 1 | 0 | 8 | 0.375 |
| 1_83254987 | 3 | 0.6667 | 3 | 0.3333 | 4 | 0.25 | 2 | 0 | 12 | 0.3333 | 4 | 0.5 | 1 | 0 | 2 | 0 | 1 | 0 | 8 | 0.25 |
| 1_83255393 | 1 | 1 | 1 | 1 | 1 | 0 | 2 | 0 | 5 | 0.4 | 2 | 0 | NA | NA | 2 | 0 | 1 | 1 | 5 | 0.2 |
| 1_83255647 | 2 | 0 | 4 | 0.5 | 3 | 0.3333 | 2 | 0 | 11 | 0.2727 | 2 | 0.5 | NA | NA | 2 | 1 | 1 | 0 | 5 | 0.6 |
| 1_83255750 | 2 | 0 | 2 | 0 | 2 | 0.5 | 1 | 1 | 7 | 0.2857 | 3 | 0.3333 | 1 | 1 | NA | NA | NA | NA | 4 | 0.5 |
| 1_83255921 | 2 | 0.5 | 2 | 0.5 | 2 | 0.5 | NA | NA | 6 | 0.5 | 2 | 0.5 | 1 | 0 | NA | NA | 1 | 0 | 4 | 0.25 |
| 1_83256142 | 2 | 0 | 4 | 0.25 | 3 | 0 | 1 | 0 | 10 | 0.1 | 3 | 0.3333 | 1 | 0 | NA | NA | 1 | 0 | 5 | 0.2 |
| 1_84014078 | 3 | 0 | 3 | 0.3333 | 2 | 0 | 4 | 0 | 12 | 0.0833 | 1 | 0 | 1 | 0 | 1 | 0 | 3 | 0.3333 | 6 | 0.1667 |
| 1_84014205 | 7 | 0.5714 | 9 | 0.3333 | 7 | 0.2857 | 6 | 0 | 29 | 0.3103 | 1 | 0 | 6 | 0.1667 | 1 | 0 | 9 | 0.2222 | 17 | 0.1765 |
| 1_84014499 | 7 | 0.2857 | 9 | 0.2222 | 6 | 0.1667 | 7 | 0.4286 | 29 | 0.2759 | 1 | 0 | 6 | 0.5 | 2 | 0 | 9 | 0.4444 | 18 | 0.3889 |
| 1_84014608 | 6 | 0.3333 | 8 | 0.5 | 6 | 0.5 | 5 | 0.2 | 25 | 0.4 | 1 | 0 | 6 | 0 | NA | NA | 8 | 0.125 | 15 | 0.0667 |
| 1_84014712 | 6 | 0.1667 | 8 | 0 | 6 | 0.1667 | 6 | 0.1667 | 26 | 0.1154 | 1 | 0 | 5 | 0.2 | 1 | 0 | 9 | 0.2222 | 16 | 0.1875 |
| 1_84014732 | 5 | 1 | 7 | 0.7143 | 4 | 1 | 4 | 1 | 20 | 0.9 | 1 | 1 | 5 | 0.8 | NA | NA | 8 | 0.875 | 14 | 0.8571 |
| 1_84014782 | 5 | 0.6 | 7 | 0.1429 | 5 | 0.6 | 3 | 0.6667 | 20 | 0.45 | 1 | 1 | 5 | 0.2 | NA | NA | 6 | 0.3333 | 12 | 0.3333 |
| 1_84014802 | 6 | 0.5 | 8 | 0.625 | 6 | 0.1667 | 5 | 0.2 | 25 | 0.4 | 1 | 0 | 6 | 0.5 | 1 | 0 | 8 | 0.5 | 16 | 0.4375 |
| 1_84014831 | 4 | 0 | 8 | 0.375 | 6 | 0.3333 | 5 | 0.2 | 23 | 0.2609 | 1 | 0 | 5 | 0 | 1 | 0 | 6 | 0.3333 | 13 | 0.1538 |
| 1_84014912 | 3 | 0.3333 | 8 | 0.125 | 6 | 0 | 5 | 0 | 22 | 0.0909 | 1 | 0 | 4 | 0 | 1 | 0 | 8 | 0.125 | 14 | 0.0714 |
| 1_84014979 | 3 | 1 | 8 | 0.75 | 6 | 1 | 3 | 1 | 20 | 0.9 | 1 | 1 | 5 | 1 | 1 | 1 | 8 | 0.875 | 15 | 0.9333 |
| 1_84015002 | 3 | 0 | 7 | 0.1429 | 6 | 0 | 5 | 0.2 | 21 | 0.0952 | 1 | 0 | 4 | 0 | 1 | 0 | 8 | 0.25 | 14 | 0.1429 |
| 1_84015271 | 3 | 1 | 6 | 0.8333 | 5 | 0.6 | 4 | 0.75 | 18 | 0.7778 | 1 | 1 | 5 | 0.8 | 1 | 1 | 3 | 1 | 10 | 0.9 |
| 1_84015275 | 3 | 0.3333 | 8 | 0.625 | 5 | 0.8 | 4 | 0.75 | 20 | 0.65 | NA | NA | 5 | 0.4 | NA | NA | 2 | 0.5 | 7 | 0.4286 |
| 1_84015288 | 4 | 0.5 | 8 | 0.875 | 5 | 0.8 | 5 | 0.8 | 22 | 0.7727 | 1 | 1 | 4 | 0.75 | 1 | 0 | 6 | 0.6667 | 12 | 0.6667 |
| 1_84015405 | 4 | 1 | 8 | 1 | 5 | 1 | 5 | 1 | 22 | 1 | 1 | 1 | 3 | 1 | 1 | 1 | 7 | 1 | 12 | 1 |
| 1_84015516 | 3 | 1 | 8 | 0.75 | 4 | 0.75 | 4 | 0.5 | 19 | 0.7368 | 1 | 0 | 2 | 0 | 1 | 1 | 4 | 0.75 | 8 | 0.5 |
| 1_84015577 | 4 | 1 | 6 | 1 | 5 | 0.8 | 4 | 1 | 19 | 0.9474 | 1 | 1 | 2 | 1 | 1 | 1 | 6 | 1 | 10 | 1 |
| 1_84015659 | 4 | 1 | 6 | 1 | 5 | 1 | 5 | 1 | 20 | 1 | 1 | 1 | 2 | 1 | NA | NA | 5 | 1 | 8 | 1 |
| 1_84015747 | 3 | 0.3333 | 6 | 0.8333 | 4 | 1 | 5 | 0.4 | 18 | 0.6667 | 1 | 1 | 2 | 0.5 | 1 | 1 | 5 | 0.8 | 9 | 0.7778 |
| 1_84015841 | 4 | 0.25 | 6 | 0.1667 | 3 | 0.3333 | 5 | 0.2 | 18 | 0.2222 | 1 | 0 | 2 | 0.5 | 1 | 0 | 4 | 0.25 | 8 | 0.25 |
| 1_84015918 | 3 | 1 | 6 | 1 | 4 | 1 | 5 | 1 | 18 | 1 | 1 | 1 | 2 | 1 | NA | NA | 5 | 1 | 8 | 1 |
| 1_84348194 | 23 | 0.1739 | 41 | 0.3171 | 20 | 0.15 | 31 | 0.3226 | 115 | 0.2609 | 25 | 0.28 | 20 | 0.45 | 20 | 0.3 | 24 | 0.2083 | 89 | 0.3034 |
| 1_84699093 | 5 | 0.6 | 5 | 0.6 | 4 | 0.75 | 3 | 0.6667 | 17 | 0.6471 | 4 | 0.75 | 5 | 0.6 | 5 | 0.4 | 4 | 1 | 18 | 0.6667 |
| 1_84699141 | 5 | 0 | 5 | 0.2 | 4 | 0 | 3 | 0.3333 | 17 | 0.1176 | 5 | 0.2 | 4 | 0 | 7 | 0.2857 | 5 | 0.2 | 21 | 0.1905 |
| 1_84699161 | 5 | 0 | 5 | 0.2 | 4 | 0.25 | 3 | 0 | 17 | 0.1176 | 5 | 0 | 5 | 0.4 | 7 | 0 | 5 | 0 | 22 | 0.0909 |

| | | | | | | | | | | | | | | | | | | | | |
|---|---|---|---|---|---|---|---|---|---|---|---|---|---|---|---|---|---|---|---|---|
| 1_8469918 | 5 | 0.4 | 5 | 0.2 | 4 | 0 | 3 | 0 | 17 | 0.1765 | 4 | 0 | 4 | 0.25 | 6 | 0 | 4 | 0 | 18 | 0.0556 |
| 1_8469920 | 4 | 0.5 | 5 | 0 | 4 | 0.5 | 2 | 0 | 15 | 0.2667 | 1 | 1 | 3 | 0.3333 | 5 | 0.2 | 3 | 0.3333 | 12 | 0.3333 |
| 1_8470091 | 4 | 0.25 | 7 | 0 | 9 | 0.1111 | 4 | 0 | 24 | 0.0833 | 3 | 0 | 6 | 0.3333 | 7 | 0 | 2 | 0 | 18 | 0.1111 |
| 1_8470098 | 3 | 0 | 5 | 0.2 | 6 | 0.1667 | 3 | 0 | 17 | 0.1176 | 2 | 0 | 4 | 0.25 | 7 | 0 | 2 | 0 | 15 | 0.0667 |
| 1_8470219 | 12 | 0.75 | 13 | 0.7692 | 11 | 0.7273 | 4 | 1 | 40 | 0.775 | 6 | 0.6667 | 12 | 0.9167 | 13 | 0.6923 | 9 | 0.5556 | 40 | 0.725 |
| 1_8470238 | 13 | 0.0769 | 13 | 0 | 12 | 0.1667 | 5 | 0 | 43 | 0.0698 | 5 | 0 | 11 | 0.1818 | 12 | 0.25 | 9 | 0.3333 | 37 | 0.2162 |
| 1_8470239 | 11 | 0.3636 | 12 | 0.1667 | 12 | 0.25 | 5 | 0.2 | 40 | 0.25 | 4 | 0.5 | 11 | 0.0909 | 10 | 0.3 | 7 | 0.1429 | 32 | 0.2188 |
| 1_8470241 | 10 | 0.6 | 12 | 0.5833 | 9 | 0.8889 | 5 | 0.8 | 36 | 0.6944 | 4 | 1 | 11 | 0.4545 | 10 | 0.7 | 5 | 0.2 | 30 | 0.5667 |
| 1_8470243 | 12 | 0.3333 | 12 | 0.5 | 9 | 0.6667 | 3 | 1 | 36 | 0.5278 | 5 | 0.8 | 10 | 0.6 | 10 | 0.7 | 9 | 0.5556 | 34 | 0.6471 |
| 1_8470244 | 11 | 0.0909 | 12 | 0.1667 | 13 | 0.3077 | 5 | 0.2 | 41 | 0.1951 | 5 | 0.6 | 12 | 0.4167 | 12 | 0.1667 | 9 | 0.1111 | 38 | 0.2895 |
| 1_8477128 | 1 | 1 | 2 | 0 | 1 | 0 | NA | NA | 4 | 0.25 | 1 | 0 | 3 | 0.6667 | 3 | 0 | 1 | 0 | 8 | 0.25 |
| 1_8477131 | 1 | 1 | 1 | 1 | NA | NA | NA | NA | 2 | 1 | NA | NA | 3 | 0.3333 | 1 | 1 | 1 | 0 | 5 | 0.4 |
| 1_8477139 | 1 | 0 | 2 | 0.5 | NA | NA | NA | NA | 3 | 0.3333 | NA | NA | 2 | 0 | 4 | 0 | 1 | 1 | 7 | 0.1429 |
| 1_8477169 | 1 | 0 | 2 | 0 | 1 | 1 | NA | NA | 4 | 0.25 | 1 | 1 | 3 | 0.3333 | 3 | 1 | 1 | 0 | 8 | 0.625 |
| 1_8477180 | 1 | 1 | 2 | 1 | 1 | 1 | NA | NA | 4 | 1 | NA | NA | 3 | 1 | 3 | 0.6667 | 1 | 1 | 7 | 0.8571 |
| 1_8477190 | 1 | 0 | 1 | 1 | 1 | 1 | NA | NA | 3 | 0.6667 | NA | NA | 2 | 0.5 | 3 | 0.3333 | 1 | 0 | 6 | 0.3333 |
| 1_8477194 | 1 | 0 | 2 | 0.5 | 1 | 0 | NA | NA | 4 | 0.25 | NA | NA | 3 | 1 | 3 | 0.6667 | 1 | 1 | 7 | 0.8571 |
| 1_8488150 | 1 | 1 | 3 | 0.6667 | 1 | 0 | 1 | 1 | 6 | 0.6667 | NA | NA | NA | NA | NA | NA | 3 | 0.6667 | 3 | 0.6667 |
| 1_8488156 | NA | NA | 3 | 1 | NA | NA | 1 | 1 | 4 | 1 | NA | NA | NA | NA | NA | NA | 3 | 1 | 3 | 1 |
| 1_8577717 | 5 | 0.8 | 9 | 0.8889 | 10 | 0.8 | 5 | 0.6 | 29 | 0.7931 | 6 | 0.1667 | 9 | 0.8889 | 8 | 0.625 | 19 | 0.7895 | 42 | 0.6905 |
| 1_8577717 | 9 | 0.8889 | 14 | 0.9286 | 16 | 0.8125 | 14 | 1 | 53 | 0.9057 | 10 | 1 | 15 | 0.9333 | 15 | 0.8667 | 36 | 0.8611 | 76 | 0.8947 |
| 1_8577720 | 10 | 0.4 | 14 | 0.2143 | 16 | 0.375 | 14 | 0.6429 | 54 | 0.4074 | 10 | 0.3 | 15 | 0.7333 | 14 | 0.4286 | 32 | 0.5312 | 71 | 0.5211 |
| 1_8577738 | 12 | 0.9167 | 14 | 0.9286 | 14 | 1 | 14 | 1 | 54 | 0.963 | 10 | 1 | 15 | 1 | 15 | 1 | 31 | 0.9355 | 71 | 0.9718 |
| 1_8577751 | 15 | 0.2667 | 16 | 0.375 | 17 | 0.2941 | 15 | 0.2667 | 63 | 0.3016 | 10 | 0.2 | 15 | 0.2667 | 15 | 0.1333 | 33 | 0.3939 | 73 | 0.2877 |
| 1_8577752 | 15 | 0.1333 | 16 | 0.0625 | 15 | 0.0667 | 15 | 0.0667 | 61 | 0.082 | 9 | 0 | 16 | 0.1875 | 16 | 0.0625 | 33 | 0.1212 | 74 | 0.1081 |
| 1_8577793 | 20 | 0.05 | 13 | 0.1538 | 16 | 0 | 12 | 0.3333 | 61 | 0.1148 | 8 | 0 | 17 | 0.0588 | 10 | 0.2 | 31 | 0.129 | 66 | 0.1061 |
| 1_8577870 | 21 | 0.1905 | 16 | 0.0625 | 15 | 0.0667 | 13 | 0.0769 | 65 | 0.1077 | 11 | 0.0909 | 15 | 0.1333 | 20 | 0.15 | 34 | 0.2059 | 80 | 0.1625 |
| 1_8583564 | 118 | 0.1186 | 82 | 0.1341 | 79 | 0.1646 | 119 | 0.1092 | 398 | 0.1281 | 77 | 0.1039 | 135 | 0.1037 | 125 | 0.12 | 118 | 0.1186 | 455 | 0.1121 |
| 1_8583575 | 209 | 0.1962 | 161 | 0.2484 | 141 | 0.2199 | 210 | 0.2143 | 721 | 0.2178 | 144 | 0.2778 | 242 | 0.2231 | 209 | 0.2584 | 198 | 0.2879 | 793 | 0.2585 |
| 1_8583581 | 158 | 0.0506 | 130 | 0.0923 | 112 | 0.1071 | 167 | 0.0838 | 567 | 0.0811 | 119 | 0.1092 | 177 | 0.1186 | 164 | 0.1341 | 144 | 0.1389 | 604 | 0.1258 |
| 1_8583616 | NA | NA | NA | NA | 1 | 1 | NA | NA | 1 | 1 | NA | NA | 1 | 1 | NA | NA | NA | NA | 1 | 1 |
| 1_8585920 | NA | NA | NA | NA | 1 | 1 | NA | NA | 1 | 1 | 1 | 1 | NA | NA | NA | NA | NA | NA | 1 | 1 |
| 1_8586932 | 1 | 1 | NA | NA | NA | NA | NA | NA | 1 | 1 | NA | NA | NA | NA | NA | NA | 2 | 1 | 2 | 1 |
| 1_8586933 | 1 | 1 | NA | NA | NA | NA | NA | NA | 1 | 1 | NA | NA | NA | NA | NA | NA | 2 | 1 | 2 | 1 |
| 1_8586950 | 1 | 1 | NA | NA | NA | NA | NA | NA | 1 | 1 | NA | NA | NA | NA | NA | NA | 2 | 1 | 2 | 1 |
| 1_8587065 | 1 | 1 | NA | NA | NA | NA | NA | NA | 1 | 1 | 1 | 0 | NA | NA | NA | NA | 2 | 0.5 | 3 | 0.3333 |
| 1_8598082 | NA | NA | 1 | 1 | 2 | 1 | NA | NA | 3 | 1 | NA | NA | 1 | 1 | 1 | 0 | NA | NA | 2 | 0.5 |
| 1_8598085 | 1 | 1 | 1 | 1 | 2 | 1 | NA | NA | 4 | 1 | NA | NA | NA | NA | 1 | 1 | NA | NA | 1 | 1 |
| 1_8598090 | 1 | 0 | NA | NA | 2 | 0.5 | NA | NA | 3 | 0.3333 | NA | NA | NA | NA | 1 | 1 | NA | NA | 1 | 1 |
| 1_8598091 | NA | NA | 1 | 1 | 1 | 1 | NA | NA | 2 | 1 | NA | NA | 1 | 1 | 1 | 1 | NA | NA | 2 | 1 |
| 1_8598098 | 1 | 1 | 1 | 1 | 2 | 0.5 | NA | NA | 4 | 0.75 | NA | NA | 1 | 1 | 2 | 0.5 | NA | NA | 3 | 0.6667 |
| 1_8598124 | NA | NA | 1 | 1 | 3 | 0 | NA | NA | 4 | 0.25 | 1 | 0 | 1 | 0 | 2 | 0.5 | NA | NA | 4 | 0.25 |
| 1_8598139 | NA | NA | 1 | 1 | 4 | 0.25 | NA | NA | 5 | 0.4 | 1 | 0 | 1 | 0 | 2 | 1 | NA | NA | 4 | 0.5 |
| 1_8598145 | NA | NA | NA | NA | 4 | 0.25 | NA | NA | 4 | 0.25 | 1 | 0 | 1 | 0 | 2 | 0.5 | NA | NA | 4 | 0.25 |
| 1_8598161 | NA | NA | NA | NA | 4 | 0.25 | NA | NA | 4 | 0.25 | 1 | 0 | NA | NA | 2 | 1 | NA | NA | 3 | 0.6667 |
| 1_8598281 | NA | NA | 1 | 1 | 3 | 0 | NA | NA | 4 | 0.25 | 1 | 1 | NA | NA | 1 | 1 | NA | NA | 2 | 1 |
| 1_8627271 | 3 | 0 | 3 | 0.3333 | 1 | 1 | 3 | 0 | 10 | 0.2 | NA | NA | 2 | 0 | 1 | 0 | 3 | 0.3333 | 6 | 0.1667 |
| 1_8627473 | NA | NA | NA | NA | 1 | 1 | 2 | 0.5 | 3 | 0.6667 | NA | NA | 2 | 0 | 4 | 0.75 | 2 | 0.5 | 8 | 0.5 |
| 1_8627521 | NA | NA | 3 | 0.6667 | 2 | 0 | 3 | 0 | 8 | 0.25 | NA | NA | 2 | 0 | 4 | 0.5 | 1 | 0 | 7 | 0.2857 |
| 1_8627789 | 2 | 0.5 | 4 | 0.25 | 5 | 0.2 | 5 | 0 | 16 | 0.1875 | 2 | 0 | 3 | 0.3333 | 1 | 0 | NA | NA | 6 | 0.1667 |
| 1_8627791 | 1 | 0 | 2 | 0 | 5 | 0.2 | 3 | 0 | 11 | 0.0909 | NA | NA | 3 | 0.3333 | 3 | 0 | 1 | 0 | 7 | 0.1429 |
| 1_8627821 | 7 | 0.4286 | 6 | 0.5 | 9 | 0.5556 | 7 | 0.7143 | 29 | 0.5517 | 2 | 1 | 5 | 0.6 | 4 | 0.25 | 3 | 0.3333 | 14 | 0.5 |
| 1_8627822 | 18 | 0.0556 | 24 | 0.2083 | 23 | 0.2174 | 35 | 0.1429 | 100 | 0.16 | 12 | 0 | 31 | 0.129 | 31 | 0.0968 | 29 | 0.1724 | 103 | 0.1165 |
| 1_8627823 | 19 | 0.2632 | 20 | 0.1 | 22 | 0.2727 | 37 | 0.1081 | 98 | 0.1735 | 11 | 0 | 25 | 0.2 | 30 | 0.1 | 24 | 0.125 | 90 | 0.1222 |
| 1_8627824 | 24 | 0.8333 | 27 | 0.8889 | 30 | 0.8333 | 45 | 0.8667 | 126 | 0.8571 | 13 | 0.8462 | 31 | 0.8387 | 40 | 0.975 | 30 | 0.9333 | 114 | 0.9123 |
| 1_8627829 | 21 | 1 | 24 | 0.9583 | 25 | 0.96 | 33 | 0.9091 | 103 | 0.9515 | 10 | 1 | 28 | 1 | 32 | 0.9688 | 27 | 1 | 97 | 0.9897 |
| 1_8627866 | 13 | 0.3077 | 16 | 0.5 | 22 | 0.2273 | 31 | 0.2903 | 82 | 0.3171 | 8 | 0.375 | 21 | 0.1429 | 21 | 0.2381 | 20 | 0.45 | 70 | 0.2857 |
| 1_8628729 | NA | NA | NA | NA | 1 | 1 | NA | NA | 1 | 1 | NA | NA | 2 | 1 | NA | NA | NA | NA | 2 | 1 |
| 1_8628731 | NA | NA | NA | NA | 1 | 0 | 1 | 1 | 2 | 0.5 | NA | NA | 2 | 0.5 | NA | NA | NA | NA | 2 | 0.5 |
| 1_8645794 | 39 | 0.0256 | 79 | 0.038 | 72 | 0.0972 | 51 | 0.1176 | 241 | 0.0705 | 44 | 0.1136 | 52 | 0.0385 | 50 | 0.02 | 42 | 0.0714 | 188 | 0.0585 |
| 1_8645810 | 45 | 0.0444 | 93 | 0.0753 | 77 | 0.1429 | 53 | 0.0943 | 268 | 0.0933 | 54 | 0.0926 | 58 | 0.1207 | 59 | 0.0847 | 49 | 0.1429 | 220 | 0.1091 |
| 1_8647079 | 9 | 0.1111 | 8 | 0.25 | 7 | 0 | 11 | 0.1818 | 35 | 0.1429 | 4 | 0 | 15 | 0.0667 | 13 | 0.2308 | 8 | 0.375 | 40 | 0.175 |
| 1_8647103 | 11 | 0.6364 | 8 | 0.375 | 4 | 0.5 | 11 | 0.5455 | 34 | 0.5294 | 3 | 0.6667 | 11 | 0.4545 | 11 | 0.5455 | 7 | 0.5714 | 32 | 0.5312 |
| 1_8647113 | 9 | 0 | 8 | 0.5 | 5 | 0 | 9 | 0 | 31 | 0.129 | 3 | 0 | 12 | 0.0833 | 14 | 0.2143 | 6 | 0.3333 | 35 | 0.1714 |
| 1_8647117 | 10 | 0.4 | 7 | 0.5714 | 5 | 0.2 | 7 | 0.4286 | 29 | 0.4138 | 4 | 0.75 | 14 | 0.4286 | 15 | 0.4 | 5 | 0.2 | 38 | 0.4211 |

| | | | | | | | | | | | | | | | | | | | | |
|---|---|---|---|---|---|---|---|---|---|---|---|---|---|---|---|---|---|---|---|---|
| 1_8653313 | NA | NA | 5 | 0 | 5 | 0.2 | 3 | 0.3333 | 13 | 0.1538 | 1 | 0 | 1 | 0 | 6 | 0.1667 | 2 | 0.5 | 10 | 0.2 |
| 1_8653379 | 1 | 0 | 4 | 0 | 5 | 0.2 | 4 | 0 | 14 | 0.0714 | 1 | 0 | 1 | 0 | 6 | 0 | 3 | 0.3333 | 11 | 0.0909 |
| 1_8653388 | 1 | 0 | 4 | 0.75 | 5 | 0.8 | 4 | 0.75 | 14 | 0.7143 | 1 | 0 | 1 | 0 | 6 | 0.8333 | 3 | 0.6667 | 11 | 0.6364 |
| 1_8653389 | 1 | 0 | 3 | 0.6667 | 4 | 0 | 3 | 0 | 11 | 0.1818 | 1 | 0 | 1 | 0 | 6 | 0.1667 | 2 | 0 | 10 | 0.1 |
| 1_8653390 | 1 | 0 | 5 | 0 | 5 | 0.2 | 4 | 0 | 15 | 0.0667 | 1 | 0 | 1 | 0 | 6 | 0 | 3 | 0.3333 | 11 | 0.0909 |
| 1_8659424 | 3 | 0.3333 | 2 | 0.5 | 1 | 0 | NA | NA | 6 | 0.3333 | 2 | 0 | NA | NA | 1 | 0 | 3 | 0.3333 | 6 | 0.1667 |
| 1_8697746 | 4 | 0.5 | 2 | 0.5 | 2 | 0.5 | 2 | 0.5 | 10 | 0.5 | 5 | 0.4 | 3 | 0.3333 | 4 | 0.5 | 1 | 1 | 13 | 0.4615 |
| 1_8697752 | 4 | 0.75 | 2 | 1 | 2 | 0 | 2 | 0 | 10 | 0.5 | 6 | 0 | 2 | 0 | 6 | 0.1667 | 1 | 0 | 15 | 0.0667 |
| 1_8707550 | 2 | 0 | 1 | 0 | 2 | 0 | 9 | 0.2222 | 14 | 0.1429 | 1 | 1 | 9 | 0.5556 | 4 | 0.5 | 7 | 0.4286 | 21 | 0.5238 |
| 1_8707574 | 2 | 1 | NA | NA | 3 | 0.3333 | 12 | 0.4167 | 17 | 0.4706 | 1 | 0 | 6 | 0.1667 | 2 | 1 | 10 | 0.1 | 19 | 0.2105 |
| 1_8708208 | 3 | 0 | NA | NA | 3 | 0 | 11 | 0.0909 | 17 | 0.0588 | NA | NA | 4 | 0.5 | 1 | 0 | 6 | 0.1667 | 11 | 0.2727 |
| 1_8708233 | 2 | 1 | NA | NA | 2 | 0 | 6 | 0.5 | 10 | 0.5 | NA | NA | 3 | 0.6667 | 3 | 1 | 6 | 0.8333 | 12 | 0.8333 |
| 1_8708262 | 2 | 0.5 | NA | NA | 2 | 0 | 9 | 0.2222 | 13 | 0.2308 | NA | NA | 2 | 0.5 | 2 | 0.5 | 3 | 0 | 7 | 0.2857 |
| 1_8716575 | 2 | 0.5 | 2 | 1 | NA | NA | 2 | 1 | 6 | 0.8333 | 1 | 1 | 1 | 1 | NA | NA | 2 | 1 | 4 | 1 |
| 1_8716581 | 1 | 1 | 2 | 1 | NA | NA | 2 | 0.5 | 5 | 0.8 | 1 | 1 | NA | NA | NA | NA | 2 | 0.5 | 3 | 0.6667 |
| 1_8716592 | 1 | 1 | 2 | 1 | 1 | 0 | 2 | 1 | 6 | 0.8333 | 1 | 1 | 1 | 0 | 1 | 1 | 3 | 1 | 6 | 0.8333 |
| 1_8716617 | 2 | 1 | NA | NA | 1 | 0 | 2 | 1 | 5 | 0.8 | 1 | 0 | 1 | 1 | 1 | 1 | 2 | 1 | 5 | 0.8 |
| 1_8716634 | 2 | 0.5 | 1 | 0 | 1 | 0 | 1 | 0 | 5 | 0.2 | 1 | 1 | 2 | 0 | 1 | 1 | 2 | 0 | 6 | 0.3333 |
| 1_8716637 | 2 | 0.5 | 2 | 1 | 1 | 0 | 2 | 1 | 7 | 0.7143 | 1 | 1 | 1 | 0 | 2 | 0.5 | 2 | 0.5 | 6 | 0.5 |
| 1_8716655 | 2 | 1 | 2 | 1 | 1 | 1 | 2 | 1 | 7 | 1 | NA | NA | 2 | 1 | 2 | 1 | 1 | 1 | 5 | 1 |
| 1_8716664 | 1 | 0 | 2 | 0.5 | 1 | 0 | 2 | 0 | 6 | 0.1667 | 1 | 0 | 2 | 0.5 | 2 | 0 | 1 | 0 | 6 | 0.1667 |
| 1_8716673 | 2 | 0.5 | 1 | 1 | 1 | 1 | 1 | 1 | 5 | 0.8 | NA | NA | 1 | 0 | 1 | 1 | 2 | 1 | 4 | 0.75 |
| 1_8716714 | 2 | 0.5 | 3 | 0 | 1 | 0 | 2 | 0 | 8 | 0.125 | 1 | 0 | 2 | 0 | 2 | 1 | 1 | 0 | 6 | 0.3333 |
| 1_8716715 | 2 | 0.5 | 4 | 1 | 1 | 1 | 3 | 0.6667 | 10 | 0.8 | 1 | 0 | 2 | 0.5 | 1 | 1 | 2 | 1 | 6 | 0.6667 |
| 1_8719232 | 6 | 0.3333 | 2 | 0 | 1 | 0 | 3 | 0 | 12 | 0.1667 | NA | NA | 3 | 0.3333 | 1 | 1 | 2 | 0 | 6 | 0.3333 |
| 1_8719233 | 4 | 0.5 | 2 | 0 | 1 | 0 | 3 | 0.3333 | 10 | 0.3 | NA | NA | 2 | 0.5 | NA | NA | 2 | 0.5 | 4 | 0.5 |
| 1_8721730 | 2 | 0 | 3 | 0.3333 | 1 | 0 | 3 | 0 | 9 | 0.1111 | 1 | 0 | 4 | 0.25 | 1 | 0 | 1 | 1 | 7 | 0.2857 |
| 1_8723784 | 6 | 0 | 2 | 0.5 | 2 | 1 | 3 | 0.3333 | 13 | 0.3077 | 1 | 0 | 5 | 0.2 | 2 | 0 | 2 | 0 | 10 | 0.1 |
| 1_8723803 | 6 | 0.3333 | 3 | 0.3333 | 2 | 0.5 | 3 | 0.3333 | 14 | 0.3571 | 1 | 0 | 4 | 0.25 | 2 | 0 | 2 | 0 | 9 | 0.1111 |
| 1_8723822 | 5 | 0 | 3 | 0.3333 | 4 | 0.25 | 4 | 0 | 16 | 0.125 | 1 | 0 | 5 | 0.2 | 2 | 0 | 2 | 0 | 10 | 0.1 |
| 1_8723836 | 4 | 0 | 3 | 0.6667 | 4 | 0.25 | 4 | 0.5 | 15 | 0.3333 | 1 | 0 | 5 | 0 | 2 | 0 | 2 | 0.5 | 10 | 0.1 |
| 1_8737692 | 3 | 0.3333 | 4 | 0.25 | 1 | 1 | 3 | 0.3333 | 11 | 0.3636 | 2 | 0.5 | 3 | 0.3333 | 2 | 0.5 | NA | NA | 7 | 0.4286 |
| 1_8737697 | 3 | 0.3333 | 5 | 0.2 | 1 | 0 | 4 | 0 | 13 | 0.1538 | 2 | 0 | 3 | 0.6667 | 4 | 0 | NA | NA | 9 | 0.2222 |
| 1_8737705 | 3 | 0.6667 | 3 | 0.6667 | 1 | 0 | 2 | 1 | 9 | 0.6667 | 2 | 1 | 2 | 0.5 | 2 | 0.5 | 1 | 1 | 7 | 0.7143 |
| 1_8737707 | 3 | 0.3333 | 5 | 0.2 | 1 | 1 | 4 | 0.25 | 13 | 0.3077 | 1 | 1 | 3 | 0 | 3 | 0 | 1 | 1 | 8 | 0.25 |
| 1_8737732 | 4 | 0.25 | 5 | 0.2 | 1 | 0 | 4 | 0.5 | 14 | 0.2857 | 2 | 0 | 3 | 0.3333 | 4 | 0.5 | 2 | 0.5 | 11 | 0.3636 |
| 1_8737735 | 4 | 0.5 | 3 | 0.6667 | 1 | 0 | 3 | 0.6667 | 11 | 0.5455 | 2 | 0.5 | 2 | 0.5 | 3 | 0.6667 | 2 | 0.5 | 9 | 0.5556 |
| 1_8737757 | 3 | 0 | 6 | 0.1667 | 1 | 0 | 3 | 0.3333 | 13 | 0.1538 | 2 | 0 | 2 | 0.5 | 4 | 0.25 | 2 | 0 | 10 | 0.2 |
| 1_8737760 | 4 | 0.5 | 7 | 0.4286 | 1 | 1 | 4 | 1 | 16 | 0.625 | 2 | 0.5 | 3 | 0.3333 | 4 | 0.5 | 2 | 0 | 11 | 0.3636 |
| 1_8737763 | 4 | 0.25 | 7 | 0.2857 | 1 | 0 | 5 | 0.2 | 17 | 0.2353 | 1 | 1 | 3 | 0 | 4 | 0.25 | 2 | 0 | 10 | 0.2 |
| 1_8737767 | 4 | 0 | 6 | 0.6667 | 2 | 0.5 | 5 | 0.2 | 17 | 0.3529 | 2 | 0.5 | 2 | 0 | 4 | 0.5 | 2 | 0.5 | 10 | 0.4 |
| 1_8737777 | 5 | 0.4 | 7 | 0.4286 | 2 | 1 | 3 | 0 | 17 | 0.4118 | 2 | 0.5 | 3 | 0 | 3 | 0.6667 | 2 | 0 | 10 | 0.3 |
| 1_8737790 | 4 | 0 | 5 | 0.2 | 2 | 0 | 5 | 0.2 | 16 | 0.125 | 2 | 0.5 | 3 | 0.6667 | 3 | 0.3333 | 2 | 0 | 10 | 0.4 |
| 1_8737805 | 5 | 0.6 | 7 | 0.5714 | 2 | 0.5 | 5 | 0.4 | 19 | 0.5263 | 3 | 0.6667 | 1 | 1 | 3 | 0.6667 | 3 | 0.3333 | 10 | 0.6 |
| 1_8737806 | 5 | 0 | 7 | 0 | 3 | 0 | 5 | 0.2 | 20 | 0.05 | 3 | 0.3333 | 2 | 1 | 4 | 0.25 | 3 | 0 | 12 | 0.3333 |
| 1_8737818 | 5 | 0 | 7 | 0.1429 | 3 | 0 | 5 | 0 | 20 | 0.05 | 3 | 0 | 3 | 0 | 5 | 0.2 | 3 | 0 | 14 | 0.0714 |
| 1_8737832 | 6 | 0.5 | 8 | 0.5 | 2 | 1 | 4 | 0.75 | 20 | 0.6 | 3 | 0.6667 | 3 | 1 | 5 | 0.4 | 4 | 0.25 | 15 | 0.5333 |
| 1_8740491 | 57 | 0.1404 | 38 | 0.1579 | 46 | 0.2174 | 58 | 0.0862 | 199 | 0.1457 | 35 | 0.1143 | 62 | 0.2097 | 75 | 0.2267 | 45 | 0.2 | 217 | 0.1982 |
| 1_8740498 | 60 | 0.1 | 38 | 0.0526 | 44 | 0.1591 | 62 | 0.0323 | 204 | 0.0833 | 39 | 0.1026 | 66 | 0.1818 | 77 | 0.0779 | 51 | 0.1765 | 233 | 0.133 |
| 1_8752248 | NA | NA | NA | NA | 2 | 1 | NA | NA | 2 | 1 | 1 | 0 | 1 | 1 | NA | NA | NA | NA | 2 | 0.5 |
| 1_8752251 | NA | NA | NA | NA | 2 | 0.5 | NA | NA | 2 | 0.5 | 1 | 0 | 1 | 1 | NA | NA | NA | NA | 2 | 0.5 |
| 1_8752434 | NA | NA | 1 | 1 | 2 | 0 | 1 | 0 | 4 | 0.25 | 2 | 0.5 | 1 | 1 | NA | NA | NA | NA | 3 | 0.6667 |
| 1_8752446 | NA | NA | 2 | 0.5 | 2 | 0.5 | 1 | 0 | 5 | 0.4 | 2 | 0.5 | 1 | 0 | NA | NA | NA | NA | 3 | 0.3333 |
| 1_8752448 | NA | NA | 2 | 0.5 | 2 | 0.5 | 1 | 1 | 5 | 0.6 | 2 | 0.5 | 1 | 1 | NA | NA | NA | NA | 3 | 0.6667 |
| 1_8752449 | NA | NA | 1 | 0 | 2 | 1 | 1 | 1 | 4 | 0.75 | 1 | 0 | 1 | 1 | NA | NA | NA | NA | 2 | 0.5 |
| 1_8752460 | NA | NA | 1 | 0 | 1 | 1 | 1 | 0 | 3 | 0.3333 | 1 | 0 | 1 | 1 | NA | NA | NA | NA | 2 | 0.5 |
| 1_8752470 | NA | NA | 1 | 0 | 2 | 0.5 | 1 | 0 | 4 | 0.25 | 1 | 0 | 1 | 1 | NA | NA | NA | NA | 2 | 0.5 |
| 1_8752482 | NA | NA | 2 | 0 | 2 | 0.5 | 1 | 0 | 5 | 0.2 | 2 | 0 | 1 | 1 | NA | NA | NA | NA | 3 | 0.3333 |
| 1_8752495 | NA | NA | 1 | 0 | 2 | 0.5 | 1 | 0 | 4 | 0.25 | 2 | 0.5 | NA | NA | NA | NA | NA | NA | 2 | 0.5 |
| 1_8752506 | NA | NA | 2 | 0 | 1 | 1 | 1 | 0 | 4 | 0.25 | 2 | 0 | 1 | 1 | NA | NA | NA | NA | 3 | 0.3333 |
| 1_8764565 | 1 | 0 | NA | NA | 3 | 0.6667 | 1 | 0 | 5 | 0.4 | NA | NA | 2 | 0.5 | 1 | 0 | NA | NA | 3 | 0.3333 |
| 1_8764574 | 1 | 0 | NA | NA | 4 | 0.25 | 2 | 0 | 7 | 0.1429 | NA | NA | 2 | 0.5 | 1 | 0 | NA | NA | 3 | 0.3333 |
| 1_8764713 | 1 | 0 | NA | NA | 4 | 0.5 | 3 | 0.3333 | 8 | 0.375 | NA | NA | 1 | 1 | 1 | 0 | NA | NA | 2 | 0.5 |
| 1_8764731 | 2 | 1 | NA | NA | 3 | 0.6667 | 2 | 0.5 | 7 | 0.7143 | 1 | 1 | NA | NA | NA | NA | NA | NA | 1 | 1 |
| 1_8764736 | 1 | 1 | NA | NA | 4 | 1 | 3 | 1 | 8 | 1 | 1 | 1 | 1 | 1 | 1 | 0 | NA | NA | 3 | 0.6667 |

| | | | | | | | | | | | | | | | | | | | | |
|---|---|---|---|---|---|---|---|---|---|---|---|---|---|---|---|---|---|---|---|---|
| 1_8764750 | 2 | 0.5 | NA | NA | 4 | 0 | 3 | 0 | 9 | 0.1111 | 1 | 0 | 2 | 0.5 | NA | NA | NA | NA | 3 | 0.3333 |
| 1_8764761 | 2 | 0.5 | NA | NA | 4 | 0 | 4 | 0.25 | 10 | 0.2 | 1 | 1 | 2 | 0 | NA | NA | NA | NA | 3 | 0.3333 |
| 1_8764783 | 1 | 1 | NA | NA | 4 | 1 | 3 | 0.3333 | 8 | 0.75 | NA | NA | 1 | 1 | NA | NA | NA | NA | 1 | 1 |
| 1_8771913 | 4 | 0.5 | 11 | 0.7273 | 2 | 0.5 | 5 | 1 | 22 | 0.7273 | 3 | 0.6667 | 6 | 0.8333 | 4 | 0.5 | 7 | 0.7143 | 20 | 0.7 |
| 1_8771914 | 5 | 0.2 | 16 | 0.1875 | 3 | 0 | 6 | 0 | 30 | 0.1333 | 4 | 0 | 7 | 0.1429 | 5 | 0 | 8 | 0.125 | 24 | 0.0833 |
| 1_8771919 | 6 | 0.1667 | 13 | 0.1538 | 2 | 0 | 5 | 0 | 26 | 0.1154 | 3 | 0 | 7 | 0.1429 | 5 | 0 | 9 | 0.2222 | 24 | 0.125 |
| 1_8771931 | 6 | 0.3333 | 12 | 0.25 | 1 | 0 | 6 | 0 | 25 | 0.2 | 4 | 0.5 | 8 | 0.25 | 3 | 0 | 7 | 0.4286 | 22 | 0.3182 |
| 1_8771932 | 5 | 0.2 | 15 | 0.2667 | 2 | 0 | 6 | 0.5 | 28 | 0.2857 | 6 | 0.3333 | 8 | 0.125 | 6 | 0.3333 | 8 | 0 | 28 | 0.1786 |
| 1_8771962 | 6 | 0.3333 | 16 | 0.25 | 3 | 0 | 7 | 0.1429 | 32 | 0.2188 | 6 | 0.3333 | 6 | 0.5 | 6 | 0.1667 | 10 | 0.6 | 28 | 0.4286 |
| 1_8785379 | 3 | 0 | 3 | 0.3333 | 1 | 0 | 2 | 0 | 9 | 0.1111 | 1 | 0 | NA | NA | 2 | 0.5 | 2 | 0.5 | 5 | 0.4 |
| 1_8787023 | 9 | 0.3333 | 8 | 0.375 | 6 | 0.1667 | 6 | 0.6667 | 29 | 0.3793 | 6 | 0 | 7 | 0.1429 | 8 | 0.25 | 6 | 0.1667 | 27 | 0.1481 |
| 1_8787025 | 9 | 0.4444 | 8 | 0.125 | 5 | 0.6 | 7 | 0.4286 | 29 | 0.3793 | 5 | 0.6 | 7 | 0.4286 | 8 | 0.625 | 6 | 0.3333 | 26 | 0.5 |
| 1_8787030 | 8 | 0 | 9 | 0.4444 | 6 | 0.6667 | 7 | 0.2857 | 30 | 0.3333 | 6 | 0.1667 | 7 | 0.1429 | 8 | 0.375 | 5 | 0.4 | 26 | 0.2692 |
| 1_8787041 | 9 | 0.5556 | 8 | 0.125 | 7 | 0.4286 | 7 | 0.4286 | 31 | 0.3871 | 7 | 0.7143 | 7 | 0.7143 | 9 | 0.4444 | 6 | 0.6667 | 29 | 0.6207 |
| 1_8787051 | 8 | 0 | 5 | 0 | 6 | 0.1667 | 6 | 0.1667 | 25 | 0.08 | 8 | 0.125 | 3 | 0.3333 | 8 | 0.125 | 7 | 0.1429 | 26 | 0.1538 |
| 1_8787086 | 12 | 0.3333 | 9 | 0.3333 | 7 | 0.2857 | 7 | 0.5714 | 35 | 0.3714 | 9 | 0.5556 | 6 | 0.1667 | 12 | 0.3333 | 9 | 0.4444 | 36 | 0.3889 |
| 1_8787091 | 9 | 0.2222 | 7 | 0 | 7 | 0.1429 | 4 | 0 | 27 | 0.1111 | 9 | 0 | 5 | 0.4 | 7 | 0 | 5 | 0.2 | 26 | 0.1154 |
| 1_8787113 | 15 | 0.4 | 9 | 0.2222 | 2 | 0.5 | 6 | 0 | 32 | 0.2812 | 9 | 0.6667 | 6 | 0.5 | 8 | 0.625 | 7 | 0.1429 | 30 | 0.5 |
| 1_8787128 | 13 | 0.3077 | 8 | 0.125 | 6 | 0.1667 | 7 | 0.1429 | 34 | 0.2059 | 13 | 0.3077 | 8 | 0.25 | 11 | 0.3636 | 9 | 0.1111 | 41 | 0.2683 |
| 1_8787129 | 11 | 0.1818 | 9 | 0.1111 | 4 | 0.25 | 5 | 0.2 | 29 | 0.1724 | 9 | 0 | 8 | 0.125 | 10 | 0 | 11 | 0.0909 | 38 | 0.0526 |
| 1_8787191 | 15 | 0 | 11 | 0.0909 | 4 | 0 | 9 | 0.1111 | 39 | 0.0513 | 11 | 0 | 7 | 0.1429 | 8 | 0.125 | 13 | 0 | 39 | 0.0513 |
| 1_8787383 | NA | NA | 2 | 1 | 1 | 1 | 1 | 0 | 4 | 0.75 | NA | NA | 3 | 1 | 2 | 1 | NA | NA | 5 | 1 |
| 1_8787403 | NA | NA | 2 | 0 | 1 | 1 | 1 | 1 | 4 | 0.5 | NA | NA | 2 | 1 | 2 | 0.5 | NA | NA | 4 | 0.75 |
| 1_8787410 | NA | NA | 2 | 0 | 2 | 0.5 | 1 | 1 | 5 | 0.4 | NA | NA | 3 | 0.3333 | 2 | 0 | NA | NA | 5 | 0.2 |
| 1_8787423 | NA | NA | NA | NA | 2 | 0.5 | 1 | 1 | 3 | 0.6667 | NA | NA | 3 | 0.3333 | 2 | 1 | NA | NA | 5 | 0.6 |
| 1_8813258 | 2 | 0.5 | 2 | 0 | NA | NA | NA | NA | 4 | 0.25 | NA | NA | 1 | 0 | 1 | 1 | NA | NA | 2 | 0.5 |
| 1_8813261 | 2 | 1 | 1 | 1 | 1 | 1 | NA | NA | 4 | 1 | NA | NA | 1 | 0 | 1 | 1 | NA | NA | 2 | 0.5 |
| 1_8813264 | 2 | 1 | 2 | 1 | 1 | 1 | NA | NA | 5 | 1 | NA | NA | 1 | 1 | 1 | 1 | NA | NA | 2 | 1 |
| 1_8813265 | 2 | 1 | 1 | 1 | 1 | 1 | NA | NA | 4 | 1 | NA | NA | 1 | 1 | NA | NA | NA | NA | 1 | 1 |
| 1_8813266 | 2 | 1 | 1 | 1 | 1 | 1 | NA | NA | 4 | 1 | NA | NA | 1 | 1 | 1 | 1 | NA | NA | 2 | 1 |
| 1_8818671 | NA | NA | 1 | 1 | NA | NA | 2 | 0 | 3 | 0.3333 | 1 | 1 | 1 | 1 | NA | NA | 1 | 1 | 3 | 1 |
| 1_8819162 | 3 | 0 | 3 | 0 | 2 | 0 | 3 | 0.3333 | 11 | 0.0909 | 3 | 0 | 4 | 0.25 | 4 | 0 | 2 | 0 | 13 | 0.0769 |
| 1_8819186 | NA | NA | NA | NA | NA | NA | 1 | 1 | 1 | 1 | NA | NA | 1 | 1 | NA | NA | NA | NA | 1 | 1 |
| 1_8819201 | 2 | 0 | 3 | 0 | 2 | 0 | 2 | 1 | 9 | 0.2222 | 3 | 0.3333 | 2 | 0.5 | 4 | 0 | 1 | 1 | 10 | 0.3 |
| 1_8819202 | 2 | 0.5 | 2 | 0 | 2 | 0.5 | 2 | 0.5 | 8 | 0.375 | 2 | 0 | 3 | 0.3333 | 4 | 0 | 2 | 0 | 11 | 0.0909 |
| 1_8819223 | 2 | 1 | 2 | 0 | 2 | 0.5 | 2 | 0.5 | 8 | 0.5 | 3 | 0.3333 | 4 | 0.5 | 2 | 0.5 | 1 | 0 | 10 | 0.4 |
| 1_8819233 | 2 | 0 | NA | NA | 1 | 1 | 3 | 0.3333 | 6 | 0.3333 | 1 | 0 | 5 | 0.2 | 1 | 0 | 2 | 0.5 | 9 | 0.2222 |
| 1_8819281 | 2 | 0.5 | 2 | 1 | 2 | 1 | 2 | 1 | 8 | 0.875 | 3 | 1 | 5 | 1 | 4 | 1 | 2 | 1 | 14 | 1 |
| 1_8819286 | 2 | 1 | 2 | 1 | 1 | 0 | 2 | 1 | 7 | 0.8571 | 3 | 1 | 4 | 1 | 4 | 1 | 2 | 1 | 13 | 1 |
| 1_8819300 | 2 | 1 | 2 | 1 | 1 | 1 | 1 | 1 | 6 | 1 | 3 | 1 | 4 | 1 | 4 | 0.75 | 2 | 1 | 13 | 0.9231 |
| 1_8819302 | 1 | 0 | 2 | 1 | 2 | 0.5 | 2 | 1 | 7 | 0.7143 | 2 | 1 | 5 | 1 | 4 | 0.75 | 2 | 1 | 13 | 0.9231 |
| 1_8819311 | 1 | 0 | 1 | 1 | 2 | 0.5 | 2 | 1 | 6 | 0.6667 | 3 | 1 | 5 | 1 | 3 | 0.6667 | 2 | 1 | 13 | 0.9231 |
| 1_8819323 | 1 | 0 | 1 | 1 | NA | NA | 2 | 0.5 | 4 | 0.5 | 2 | 1 | 4 | 1 | 2 | 1 | 1 | 1 | 9 | 1 |
| 1_8819361 | NA | NA | 1 | 0 | 2 | 0.5 | NA | NA | 3 | 0.3333 | 1 | 0 | 4 | 0.25 | 4 | 0.25 | NA | NA | 9 | 0.2222 |
| 1_8819379 | 1 | 1 | 1 | 0 | 2 | 1 | NA | NA | 4 | 0.75 | 2 | 1 | 4 | 1 | 2 | 0.5 | 1 | 0 | 9 | 0.7778 |
| 1_8819380 | 1 | 1 | 1 | 0 | 1 | 1 | 1 | 1 | 4 | 0.75 | 2 | 0.5 | 5 | 0.8 | 3 | 0.6667 | 1 | 0 | 11 | 0.6364 |
| 1_8819389 | 1 | 0 | NA | NA | 2 | 1 | 1 | 1 | 4 | 0.75 | 2 | 0.5 | 4 | 0.5 | 3 | 0.3333 | 1 | 1 | 10 | 0.5 |
| 1_8819393 | 1 | 0 | 1 | 1 | 2 | 0.5 | 1 | 1 | 5 | 0.6 | 2 | 0.5 | 3 | 1 | 4 | 0.5 | 1 | 1 | 10 | 0.7 |
| 1_8862757 | 2 | 0.5 | 1 | 1 | 3 | 1 | 2 | 0.5 | 8 | 0.75 | NA | NA | 4 | 0.5 | 4 | 1 | 1 | 1 | 9 | 0.7778 |
| 1_8862792 | 1 | 0 | 3 | 0 | 3 | 0.3333 | 2 | 0 | 9 | 0.1111 | 1 | 0 | 4 | 0 | 5 | 0 | 2 | 0.5 | 12 | 0.0833 |
| 1_8862792 | 1 | 0 | 3 | 0.3333 | 1 | 0 | 3 | 0 | 8 | 0.125 | 1 | 0 | 3 | 0 | 3 | 0.3333 | 2 | 1 | 9 | 0.3333 |
| 1_8862802 | 2 | 1 | NA | NA | 2 | 0 | 2 | 1 | 6 | 0.6667 | 1 | 1 | 2 | 0.5 | 4 | 0.5 | 1 | 1 | 8 | 0.625 |
| 1_8862808 | 1 | 1 | 1 | 1 | 1 | 0 | 3 | 1 | 6 | 0.8333 | 1 | 1 | 4 | 0.75 | 4 | 1 | 2 | 1 | 11 | 0.9091 |
| 1_8862813 | 1 | 0 | 1 | 1 | 2 | 0 | 3 | 0.6667 | 7 | 0.4286 | NA | NA | 4 | 0 | 4 | 0.25 | 2 | 0.5 | 10 | 0.2 |
| 1_8862814 | 1 | 0 | 1 | 1 | 2 | 0 | 3 | 0.3333 | 7 | 0.2857 | 1 | 1 | 3 | 1 | 3 | 0.3333 | NA | NA | 7 | 0.7143 |
| 1_8862823 | 1 | 0 | 2 | 0.5 | 3 | 0 | 1 | 0 | 7 | 0.1429 | 1 | 1 | 2 | 0.5 | 5 | 0.2 | 2 | 1 | 10 | 0.5 |
| 1_8862833 | 1 | 1 | 2 | 1 | 1 | 1 | 1 | 0 | 5 | 0.8 | 1 | 1 | 2 | 1 | 1 | 1 | NA | NA | 4 | 1 |
| 1_8862844 | 1 | 1 | 2 | 0.5 | 2 | 0 | 1 | 0 | 6 | 0.3333 | 1 | 0 | 1 | 0 | 1 | 1 | NA | NA | 3 | 0.3333 |
| 1_8862857 | 1 | 1 | 2 | 1 | 2 | 1 | 1 | 1 | 6 | 1 | 1 | 1 | 1 | 1 | 2 | 1 | NA | NA | 4 | 1 |
| 1_8862864 | 1 | 1 | 2 | 1 | 1 | 1 | 1 | 1 | 5 | 1 | 1 | 1 | 1 | 0 | 2 | 0.5 | NA | NA | 4 | 0.5 |
| 1_8862865 | 1 | 1 | 1 | 1 | 1 | 1 | 1 | 1 | 4 | 1 | NA | NA | NA | NA | 1 | 1 | NA | NA | 1 | 1 |
| 1_8862871 | 1 | 0 | 2 | 0.5 | 1 | 0 | 1 | 0 | 5 | 0.2 | 1 | 0 | NA | NA | 2 | 0.5 | NA | NA | 3 | 0.3333 |
| 1_8862886 | 1 | 1 | 1 | 1 | 1 | 0 | 1 | 0 | 4 | 0.5 | 1 | 1 | NA | NA | 2 | 1 | NA | NA | 3 | 1 |
| 1_8907134 | NA | NA | 1 | 0 | 1 | 1 | NA | NA | 2 | 0.5 | 1 | 1 | NA | NA | 2 | 0.5 | 1 | 0 | 4 | 0.5 |
| 1_8907159 | NA | NA | 1 | 0 | 1 | 1 | NA | NA | 2 | 0.5 | 2 | 0.5 | NA | NA | 2 | 0 | 1 | 0 | 5 | 0.2 |

| | | | | | | | | | | | | | | | | | | | | |
|---|---|---|---|---|---|---|---|---|---|---|---|---|---|---|---|---|---|---|---|---|
| 1_89071825 | NA | NA | 1 | 1 | 1 | 1 | NA | NA | 2 | 1 | 2 | 0.5 | NA | NA | 2 | 0.5 | 1 | 1 | 5 | 0.6 |
| 1_89072244 | 1 | 0 | 2 | 0.5 | 1 | 0 | 1 | 0 | 5 | 0.2 | 1 | 1 | NA | NA | 3 | 0.6667 | 1 | 0 | 5 | 0.6 |
| 1_89072406 | 1 | 1 | 1 | 1 | 1 | 1 | NA | NA | 3 | 1 | 2 | 0.5 | NA | NA | 3 | 0.6667 | 1 | 1 | 6 | 0.6667 |
| 1_89072478 | 1 | 0 | 2 | 0.5 | 2 | 0 | 1 | 0 | 6 | 0.1667 | 2 | 0.5 | NA | NA | 2 | 0 | 1 | 0 | 5 | 0.2 |
| 1_89072618 | 1 | 0 | 2 | 0 | 2 | 0.5 | 1 | 0 | 6 | 0.1667 | 2 | 0.5 | 2 | 0.5 | 3 | 0 | 1 | 0 | 8 | 0.25 |
| 1_89072627 | 1 | 0 | 2 | 0.5 | 2 | 0 | 1 | 0 | 6 | 0.1667 | 2 | 0 | 2 | 0.5 | 3 | 0 | 1 | 0 | 8 | 0.125 |
| 1_89072638 | 1 | 1 | 2 | 0.5 | 1 | 1 | 1 | 1 | 5 | 0.8 | 1 | 1 | 2 | 1 | 2 | 0.5 | 1 | 1 | 6 | 0.8333 |
| 1_89072678 | 1 | 0 | 2 | 0 | 2 | 0 | 1 | 1 | 6 | 0.1667 | 2 | 0 | 1 | 0 | 1 | 1 | 1 | 0 | 5 | 0.2 |
| 1_89073140 | 1 | 0 | 2 | 0.5 | 2 | 0 | 1 | 1 | 6 | 0.3333 | 1 | 0 | 2 | 0 | 2 | 0.5 | 1 | 0 | 6 | 0.1667 |
| 1_89073235 | 2 | 1 | 2 | 0.5 | 2 | 0.5 | 1 | 1 | 7 | 0.7143 | 2 | 1 | 1 | 0 | 3 | 0.3333 | 1 | 1 | 7 | 0.5714 |
| 1_89073245 | 1 | 1 | 2 | 0.5 | 2 | 0 | 1 | 0 | 6 | 0.3333 | 1 | 0 | 2 | 0 | 2 | 0.5 | 1 | 0 | 6 | 0.1667 |
| 1_89073428 | 2 | 1 | 2 | 1 | 2 | 1 | 1 | 1 | 7 | 1 | 2 | 0.5 | 1 | 1 | 2 | 1 | 1 | 0 | 6 | 0.6667 |
| 1_89073444 | 2 | 0 | 2 | 0.5 | 2 | 0 | 1 | 0 | 7 | 0.1429 | 1 | 0 | 1 | 1 | 1 | 0 | 1 | 0 | 4 | 0.25 |
| 1_89073525 | 2 | 1 | 2 | 1 | 2 | 0 | 1 | 1 | 7 | 0.7143 | 2 | 1 | 1 | 1 | 2 | 0.5 | 1 | 0 | 6 | 0.6667 |
| 1_89073573 | 2 | 0.5 | 2 | 0.5 | 2 | 0.5 | 1 | 1 | 7 | 0.5714 | 3 | 0.6667 | 2 | 1 | 2 | 0.5 | 1 | 0 | 8 | 0.625 |
| 1_89080993 | 3 | 1 | 3 | 1 | 2 | 1 | 1 | 1 | 9 | 1 | 3 | 1 | 1 | 1 | 4 | 1 | 1 | 1 | 9 | 1 |
| 1_89081009 | 3 | 0 | 2 | 0 | 2 | 0.5 | 1 | 1 | 8 | 0.25 | 3 | 0 | 2 | 0.5 | 2 | 0.5 | 1 | 0 | 8 | 0.25 |
| 1_89081018 | 2 | 0.5 | 3 | 0.6667 | 1 | 0 | NA | NA | 6 | 0.5 | 2 | 1 | 2 | 0 | 2 | 0.5 | 1 | 1 | 7 | 0.5714 |
| 1_89081051 | 4 | 0 | 2 | 0.5 | 1 | 0 | NA | NA | 7 | 0.1429 | 3 | 0.3333 | 3 | 0.3333 | 3 | 0.6667 | 1 | 1 | 10 | 0.5 |
| 1_89081060 | 3 | 0 | 3 | 1 | 2 | 1 | NA | NA | 8 | 0.625 | 3 | 0.6667 | 3 | 0.3333 | 2 | 0 | 1 | 0 | 9 | 0.3333 |
| 1_89081109 | 4 | 0.75 | 3 | 1 | 2 | 1 | 1 | 1 | 10 | 0.9 | 3 | 1 | 3 | 1 | 3 | 1 | 1 | 1 | 10 | 1 |
| 1_89081130 | 4 | 1 | 2 | 1 | 2 | 1 | 1 | 1 | 9 | 1 | 3 | 1 | 3 | 0.6667 | 3 | 0.6667 | 1 | 1 | 10 | 0.8 |
| 1_89081152 | 3 | 1 | 2 | 1 | 2 | 1 | 1 | 1 | 8 | 1 | 3 | 0.6667 | 3 | 0.3333 | 3 | 1 | 1 | 1 | 10 | 0.7 |
| 1_89081173 | 4 | 0.5 | 3 | 1 | 2 | 1 | 1 | 1 | 10 | 0.8 | 3 | 1 | 3 | 1 | 2 | 1 | NA | NA | 8 | 1 |
| 1_89081277 | 3 | 0.6667 | 2 | 1 | 1 | 1 | 1 | 1 | 7 | 0.8571 | 3 | 0.6667 | 2 | 1 | 4 | 0.75 | 1 | 1 | 10 | 0.8 |
| 1_89081284 | 2 | 1 | 4 | 1 | 2 | 1 | 1 | 1 | 9 | 1 | 2 | 1 | 3 | 1 | 4 | 1 | 1 | 1 | 10 | 1 |
| 1_89081291 | 4 | 0.25 | 3 | 0.3333 | 2 | 0.5 | 1 | 0 | 10 | 0.3 | 2 | 0 | 2 | 0.5 | 3 | 0.3333 | 1 | 0 | 8 | 0.25 |
| 1_89081462 | 1 | 1 | 3 | 1 | 2 | 1 | 1 | 1 | 7 | 1 | 2 | 1 | 4 | 0.75 | 1 | 1 | NA | NA | 7 | 0.8571 |
| 1_89508719 | NA | NA | NA | NA | NA | NA | 3 | 1 | 3 | 1 | NA | NA | NA | NA | 1 | 1 | NA | NA | 1 | 1 |
| 1_89815352 | 2 | 0 | 2 | 0.5 | 7 | 0.2857 | 7 | 0.2857 | 18 | 0.2778 | 3 | 0.3333 | 4 | 0.75 | 2 | 0.5 | 4 | 0.5 | 13 | 0.5385 |
| 1_89816555 | 2 | 0 | 2 | 0 | 7 | 0 | 6 | 0.1667 | 17 | 0.0588 | 3 | 0 | 4 | 0 | 2 | 0 | 3 | 0.3333 | 12 | 0.0833 |
| 1_89816582 | NA | NA | 1 | 0 | 5 | 0.4 | 5 | 0.2 | 11 | 0.2727 | 2 | 0 | 4 | 0.25 | 1 | 1 | 4 | 0.25 | 11 | 0.2727 |
| 1_89816760 | 2 | 0.5 | 2 | 0.5 | 7 | 0.1429 | 6 | 0.1667 | 17 | 0.2353 | 3 | 0 | 4 | 0.25 | 2 | 0 | 4 | 0 | 13 | 0.0769 |
| 1_89816766 | 3 | 1 | 2 | 1 | 6 | 1 | 5 | 0.6 | 16 | 0.875 | 2 | 1 | 4 | 1 | 2 | 0.5 | 3 | 0.6667 | 11 | 0.8182 |
| 1_89816810 | 1 | 1 | 1 | 1 | 6 | 0.8333 | 5 | 0.6 | 13 | 0.7692 | 2 | 1 | 2 | 1 | 1 | 1 | 2 | 1 | 7 | 1 |
| 1_89816845 | 3 | 0.3333 | 2 | 0 | 8 | 0.125 | 7 | 0.1429 | 20 | 0.15 | 2 | 0.5 | 2 | 0 | 2 | 0 | 4 | 0.25 | 10 | 0.2 |
| 1_89816888 | 2 | 0.5 | 3 | 0.3333 | 8 | 0.25 | 7 | 0.1429 | 20 | 0.25 | 3 | 0.3333 | 4 | 0.25 | 1 | 0 | 4 | 0 | 12 | 0.1667 |
| 1_89817558 | 1 | 0 | 3 | 1 | 8 | 0.625 | 4 | 0.5 | 16 | 0.625 | 2 | 1 | 5 | 0.8 | 3 | 1 | 4 | 0.75 | 14 | 0.8571 |
| 1_89817719 | NA | NA | 3 | 0.3333 | 5 | 0 | 3 | 0 | 11 | 0.0909 | 3 | 0 | 3 | 0 | NA | NA | 4 | 0.25 | 10 | 0.1 |
| 1_89817834 | NA | NA | 4 | 0.25 | 4 | 0 | 2 | 0 | 10 | 0.1 | 3 | 0.3333 | 4 | 0 | NA | NA | 3 | 0 | 10 | 0.1 |
| 1_89818108 | NA | NA | 5 | 0.2 | 5 | 0.4 | 6 | 0 | 16 | 0.1875 | 4 | 0 | 4 | 0 | NA | NA | 4 | 0.25 | 12 | 0.0833 |
| 1_89818183 | NA | NA | 5 | 0.6 | 3 | 0.6667 | 4 | 0.5 | 12 | 0.5833 | 2 | 0.5 | 4 | 0.5 | NA | NA | 3 | 0.3333 | 9 | 0.4444 |
| 1_89818205 | NA | NA | 5 | 0 | 5 | 0.2 | 6 | 0 | 16 | 0.0625 | 3 | 0 | 4 | 0.25 | NA | NA | 4 | 0.25 | 11 | 0.1818 |
| 1_89818211 | NA | NA | 5 | 0.2 | 4 | 0 | 6 | 0 | 15 | 0.0667 | 3 | 0.3333 | 4 | 0 | NA | NA | 4 | 0 | 11 | 0.0909 |
| 1_89818224 | NA | NA | 4 | 0.25 | 4 | 0.5 | 6 | 0.6667 | 14 | 0.5 | 3 | 0 | 4 | 0.5 | NA | NA | 3 | 0 | 10 | 0.2 |
| 1_89818287 | NA | NA | 2 | 0.5 | 5 | 0.6 | 5 | 0.2 | 12 | 0.4167 | 4 | 0.5 | 5 | 0.2 | NA | NA | 3 | 0.3333 | 12 | 0.3333 |
| 1_89818362 | 1 | 0 | 5 | 0.2 | 5 | 0 | 6 | 0.1667 | 17 | 0.1176 | 3 | 0.3333 | 6 | 0 | NA | NA | 4 | 0 | 13 | 0.0769 |
| 1_89818386 | 1 | 0 | 5 | 0.6 | 5 | 1 | 5 | 1 | 16 | 0.8125 | 3 | 1 | 5 | 1 | NA | NA | 3 | 1 | 11 | 1 |
| 1_89818421 | NA | NA | 4 | 0 | 4 | 0.5 | 4 | 0.25 | 12 | 0.25 | 2 | 0.5 | 5 | 0.2 | NA | NA | 2 | 0.5 | 9 | 0.3333 |
| 1_89818446 | 1 | 0 | 6 | 0 | 5 | 0 | 5 | 0.2 | 17 | 0.0588 | 5 | 0.2 | 6 | 0.1667 | NA | NA | 4 | 0 | 15 | 0.1333 |
| 1_89818542 | NA | NA | 4 | 0.25 | 4 | 0.25 | 4 | 0 | 12 | 0.1667 | 5 | 0.4 | 6 | 0.1667 | 1 | 1 | 4 | 0.5 | 16 | 0.375 |
| 1_89818599 | 1 | 0 | 7 | 0.4286 | 3 | 0.3333 | 5 | 0.6 | 16 | 0.4375 | 4 | 0.5 | 4 | 0.25 | 1 | 1 | 3 | 0.6667 | 12 | 0.5 |
| 1_89818764 | 1 | 0 | 7 | 0.2857 | 6 | 0.1667 | 6 | 0.1667 | 20 | 0.2 | 4 | 0.75 | 6 | 0.5 | 3 | 0.3333 | 4 | 0.25 | 17 | 0.4706 |
| 1_89819093 | 2 | 0 | 6 | 0.1667 | 6 | 0 | 6 | 0 | 20 | 0.05 | 3 | 0 | 5 | 0.2 | 3 | 0 | 3 | 0 | 14 | 0.0714 |
| 1_89819193 | 2 | 0 | 7 | 0.1429 | 4 | 0 | 7 | 0.1429 | 20 | 0.1 | 5 | 0.2 | 6 | 0 | 3 | 0.3333 | 3 | 0 | 17 | 0.1176 |
| 1_89821946 | 7 | 0.2857 | 8 | 0.25 | 5 | 0 | 9 | 0.2222 | 29 | 0.2069 | 9 | 0.1111 | 7 | 0 | 7 | 0 | 8 | 0.125 | 31 | 0.0645 |
| 1_89822428 | 7 | 0 | 11 | 0.1818 | 10 | 0 | 11 | 0 | 39 | 0.0513 | 11 | 0.0909 | 8 | 0 | 10 | 0 | 8 | 0.125 | 37 | 0.0541 |
| 1_89822806 | 10 | 0.3 | 11 | 0 | 11 | 0.0909 | 10 | 0 | 42 | 0.0952 | 11 | 0.0909 | 7 | 0 | 10 | 0.2 | 9 | 0.1111 | 37 | 0.1081 |
| 1_90141562 | 1 | 0 | 2 | 0 | 3 | 0.3333 | NA | NA | 6 | 0.1667 | NA | NA | 4 | 0 | 1 | 0 | 1 | 1 | 6 | 0.1667 |
| 1_90142366 | 2 | 1 | 2 | 1 | 5 | 1 | NA | NA | 9 | 1 | 1 | 1 | 4 | 1 | 2 | 1 | 1 | 1 | 8 | 1 |
| 1_90142596 | 2 | 1 | 1 | 1 | 5 | 0.8 | NA | NA | 8 | 0.875 | 1 | 1 | 5 | 0.8 | 2 | 0 | 1 | 1 | 9 | 0.6667 |
| 1_90142621 | 1 | 0 | 2 | 0.5 | 4 | 0.25 | NA | NA | 7 | 0.2857 | 1 | 1 | 3 | 0.6667 | 2 | 0.5 | 1 | 0 | 7 | 0.5714 |
| 1_90142658 | 2 | 0 | 2 | 0.5 | 4 | 1 | NA | NA | 8 | 0.625 | 1 | 1 | 5 | 0.8 | 1 | 0 | 1 | 1 | 8 | 0.75 |
| 1_90142939 | 2 | 0.5 | 2 | 0.5 | 6 | 0.8333 | NA | NA | 10 | 0.7 | NA | NA | 5 | 0.8 | 2 | 1 | 1 | 0 | 8 | 0.75 |
| 1_90142953 | 2 | 1 | 1 | 1 | 6 | 1 | NA | NA | 9 | 1 | 1 | 1 | 5 | 1 | 2 | 1 | 1 | 1 | 9 | 1 |

| | | | | | | | | | | | | | | | | | | | | |
|---|---|---|---|---|---|---|---|---|---|---|---|---|---|---|---|---|---|---|---|---|
| 1_90143002 | 2 | 0.5 | 2 | 0 | 6 | 0.3333 | NA | NA | 10 | 0.3 | 1 | 0 | 4 | 0.5 | 1 | 0 | 1 | 0 | 7 | 0.2857 |
| 1_90143025 | 1 | 0 | 2 | 0 | 5 | 0.2 | NA | NA | 8 | 0.125 | 1 | 0 | 5 | 0.2 | 2 | 0.5 | 1 | 0 | 9 | 0.2222 |
| 1_90143146 | 2 | 0.5 | 2 | 0.5 | 5 | 0.2 | NA | NA | 9 | 0.3333 | NA | NA | 5 | 0.2 | 2 | 0.5 | 1 | 1 | 8 | 0.375 |
| 1_90265918 | 1 | 1 | NA | NA | 1 | 1 | NA | NA | 2 | 1 | 2 | 0.5 | 1 | 0 | 1 | 0 | NA | NA | 4 | 0.25 |
| 1_90265925 | 1 | 0 | NA | NA | 1 | 1 | NA | NA | 2 | 0.5 | 2 | 0 | 1 | 1 | 1 | 0 | NA | NA | 4 | 0.25 |
| 1_90266092 | 1 | 1 | NA | NA | 2 | 1 | NA | NA | 3 | 1 | 2 | 1 | 1 | 1 | 1 | 1 | NA | NA | 4 | 1 |
| 1_90378991 | NA | NA | NA | NA | 1 | 1 | NA | NA | 1 | 1 | 1 | 0 | NA | NA | 1 | 1 | NA | NA | 2 | 0.5 |
| 1_90379079 | 1 | 1 | 1 | 1 | 1 | 1 | NA | NA | 3 | 1 | 1 | 1 | NA | NA | 1 | 1 | NA | NA | 2 | 1 |
| 1_90379179 | 1 | 1 | NA | NA | 1 | 1 | NA | NA | 2 | 1 | 2 | 1 | NA | NA | 1 | 1 | NA | NA | 3 | 1 |
| 1_90380108 | 2 | 0 | 1 | 0 | 1 | 1 | NA | NA | 4 | 0.25 | 2 | 1 | NA | NA | 1 | 1 | NA | NA | 3 | 1 |
| 1_90381108 | NA | NA | 2 | 0.5 | NA | NA | NA | NA | 2 | 0.5 | 2 | 0.5 | NA | NA | NA | NA | NA | NA | 2 | 0.5 |
| 1_90382446 | 1 | 1 | 2 | 0 | 1 | 0 | NA | NA | 4 | 0.25 | 2 | 0.5 | NA | NA | NA | NA | NA | NA | 2 | 0.5 |
| 1_90433061 | NA | NA | NA | NA | 1 | 1 | NA | NA | 1 | 1 | NA | NA | 1 | 1 | NA | NA | NA | NA | 1 | 1 |
| 1_90433336 | NA | NA | NA | NA | 1 | 1 | NA | NA | 1 | 1 | 1 | 1 | 1 | 1 | NA | NA | NA | NA | 2 | 1 |
| 1_90433427 | NA | NA | NA | NA | 1 | 1 | NA | NA | 1 | 1 | 1 | 1 | NA | NA | NA | NA | NA | NA | 1 | 1 |
| 1_90433609 | NA | NA | 1 | 1 | 1 | 1 | NA | NA | 2 | 1 | NA | NA | 1 | 1 | NA | NA | NA | NA | 1 | 1 |
| 1_90540028 | 15 | 0.1333 | 19 | 0.2632 | 18 | 0.1111 | 17 | 0.0588 | 69 | 0.1449 | 11 | 0.2727 | 13 | 0.1538 | 23 | 0.1739 | 10 | 0.5 | 57 | 0.2456 |
| 1_90540054 | 12 | 0.25 | 17 | 0.2353 | 16 | 0.0625 | 16 | 0.125 | 61 | 0.1639 | 10 | 0.1 | 10 | 0.1 | 17 | 0.1176 | 7 | 0 | 44 | 0.0909 |
| 1_90540224 | 15 | 0.9333 | 18 | 0.8333 | 20 | 1 | 17 | 0.8235 | 70 | 0.9 | 11 | 0.8182 | 10 | 0.9 | 16 | 0.8125 | 10 | 0.8 | 47 | 0.8298 |
| 1_90540229 | 10 | 0.9 | 16 | 0.8125 | 13 | 0.9231 | 15 | 0.9333 | 54 | 0.8889 | 11 | 0.9091 | 11 | 0.8182 | 21 | 0.6667 | 6 | 1 | 49 | 0.7959 |
| 1_90540341 | 18 | 1 | 16 | 1 | 19 | 0.8947 | 17 | 1 | 70 | 0.9714 | 12 | 0.9167 | 14 | 0.8571 | 19 | 0.9474 | 9 | 1 | 54 | 0.9259 |
| 1_90540457 | 13 | 0.7692 | 14 | 0.6429 | 14 | 0.6429 | 15 | 0.8667 | 56 | 0.7321 | 9 | 0.8889 | 13 | 0.6923 | 14 | 0.7857 | 7 | 0.8571 | 43 | 0.7907 |
| 1_90540524 | 16 | 0.1875 | 17 | 0 | 22 | 0.0455 | 14 | 0 | 69 | 0.058 | 10 | 0 | 12 | 0.1667 | 20 | 0.05 | 10 | 0.1 | 52 | 0.0769 |
| 1_90540528 | 12 | 0.6667 | 17 | 0.4706 | 17 | 0.2941 | 17 | 0.5294 | 63 | 0.4762 | 11 | 0.3636 | 13 | 0.4615 | 16 | 0.5625 | 9 | 0.5556 | 49 | 0.4898 |
| 1_90540544 | 12 | 0.5 | 15 | 0.6667 | 18 | 0.3333 | 17 | 0.5294 | 62 | 0.5 | 12 | 0.5833 | 15 | 0.6667 | 19 | 0.7368 | 8 | 0.5 | 54 | 0.6481 |
| 1_90540659 | 18 | 0.8333 | 20 | 0.85 | 22 | 0.9545 | 20 | 0.85 | 80 | 0.875 | 13 | 1 | 16 | 0.9375 | 21 | 0.9048 | 8 | 1 | 58 | 0.9483 |
| 1_91005607 | NA | NA | NA | NA | NA | NA | 1 | 1 | 1 | 1 | NA | NA | 2 | 0 | NA | NA | 1 | 1 | 3 | 0.3333 |
| 1_91055950 | 7 | 0.7143 | 4 | 0.75 | 4 | 0.75 | 8 | 0.625 | 23 | 0.6957 | 3 | 0.3333 | 11 | 0.7273 | 7 | 0.8571 | 7 | 0.8571 | 28 | 0.75 |
| 1_91056167 | 7 | 0.5714 | 4 | 1 | 5 | 0.8 | 5 | 0.8 | 21 | 0.7619 | 5 | 0.6 | 10 | 0.8 | 6 | 1 | 4 | 1 | 25 | 0.84 |
| 1_91056208 | 8 | 0.5 | 4 | 0.25 | 5 | 0.2 | 7 | 0.2857 | 24 | 0.3333 | 3 | 0.6667 | 12 | 0.25 | 8 | 0.625 | 6 | 0.3333 | 29 | 0.4138 |
| 1_91056231 | 9 | 0.6667 | 4 | 1 | 5 | 0.2 | 9 | 0.5556 | 27 | 0.5926 | 5 | 0.6 | 12 | 0.75 | 9 | 0.4444 | 5 | 0.4 | 31 | 0.5806 |
| 1_91056320 | 8 | 0.375 | 4 | 0 | 5 | 0.4 | 9 | 0.3333 | 26 | 0.3077 | 5 | 0.8 | 9 | 0.3333 | 8 | 0.25 | 6 | 0.3333 | 28 | 0.3929 |
| 1_91056447 | 10 | 0.3 | 4 | 0.25 | 5 | 0.2 | 6 | 0.5 | 25 | 0.32 | 6 | 0.3333 | 12 | 0.4167 | 11 | 0.3636 | 7 | 0.4286 | 36 | 0.3889 |
| 1_91056483 | 9 | 0.5556 | 4 | 0.75 | 4 | 0.75 | 9 | 0.5556 | 26 | 0.6154 | 6 | 0.8333 | 12 | 0.6667 | 10 | 0.5 | 9 | 0.5556 | 37 | 0.6216 |
| 1_91151186 | 34 | 0.6176 | 18 | 0.5556 | 38 | 0.6316 | 22 | 0.5455 | 112 | 0.5982 | 19 | 0.7368 | 29 | 0.4483 | 32 | 0.5938 | 27 | 0.3704 | 107 | 0.5234 |
| 1_91151342 | 29 | 0.1034 | 17 | 0 | 36 | 0.0833 | 22 | 0 | 104 | 0.0577 | 19 | 0.1053 | 29 | 0.1034 | 35 | 0.1143 | 27 | 0.1111 | 110 | 0.1091 |
| 1_91151675 | 2 | 1 | NA | NA | NA | NA | NA | NA | 2 | 1 | 1 | 0 | NA | NA | NA | NA | 4 | 0.5 | 5 | 0.4 |
| 1_91151681 | 2 | 0.5 | NA | NA | 1 | 0 | NA | NA | 3 | 0.3333 | 1 | 0 | NA | NA | NA | NA | 4 | 0.25 | 5 | 0.2 |
| 1_91151728 | 2 | 0 | NA | NA | 2 | 0.5 | NA | NA | 4 | 0.25 | 1 | 0 | NA | NA | NA | NA | 4 | 0.25 | 5 | 0.2 |
| 1_91151903 | 2 | 0 | NA | NA | 2 | 0.5 | NA | NA | 4 | 0.25 | 1 | 0 | NA | NA | NA | NA | 4 | 0.25 | 5 | 0.2 |
| 1_91152195 | 1 | 1 | NA | NA | 2 | 0 | NA | NA | 3 | 0.3333 | 1 | 0 | NA | NA | NA | NA | 3 | 0.3333 | 4 | 0.25 |
| 1_91152249 | 2 | 1 | NA | NA | 2 | 1 | NA | NA | 4 | 1 | 1 | 1 | NA | NA | NA | NA | 4 | 1 | 5 | 1 |
| 1_91152373 | 2 | 0.5 | NA | NA | 1 | 1 | NA | NA | 3 | 0.6667 | 1 | 1 | NA | NA | NA | NA | 4 | 0.5 | 5 | 0.6 |
| 1_91152692 | 1 | 0 | NA | NA | 2 | 0.5 | 1 | 1 | 4 | 0.5 | NA | NA | NA | NA | NA | NA | 4 | 0.5 | 4 | 0.5 |
| 1_91152784 | 1 | 1 | NA | NA | 2 | 0.5 | 1 | 1 | 4 | 0.75 | NA | NA | NA | NA | NA | NA | 3 | 0.6667 | 3 | 0.6667 |
| 1_91213104 | 7 | 1 | 7 | 0.8571 | 9 | 1 | 7 | 0.7143 | 30 | 0.9 | 6 | 1 | 8 | 0.875 | 8 | 0.75 | 8 | 0.875 | 30 | 0.8667 |
| 1_91213372 | 5 | 0.6 | 3 | 0.6667 | 5 | 1 | 5 | 0.6 | 18 | 0.7222 | 5 | 0.8 | 6 | 0.6667 | 5 | 0.8 | 5 | 0.8 | 21 | 0.7619 |
| 1_91213443 | 7 | 0.1429 | 8 | 0 | 9 | 0.2222 | 6 | 0.1667 | 30 | 0.1333 | 5 | 0 | 7 | 0.2857 | 9 | 0 | 9 | 0 | 30 | 0.0667 |
| 1_91213457 | 7 | 0.2857 | 8 | 0.375 | 7 | 0.7143 | 6 | 0.6667 | 28 | 0.5 | 6 | 0.8333 | 6 | 0.6667 | 8 | 0.5 | 8 | 0.375 | 28 | 0.5714 |
| 1_91213490 | 7 | 0.8571 | 7 | 0.5714 | 8 | 1 | 4 | 0.5 | 26 | 0.7692 | 5 | 0.8 | 6 | 0.6667 | 6 | 0.8333 | 7 | 0.5714 | 24 | 0.7083 |
| 1_91213717 | 2 | 0 | 3 | 0 | 3 | 0.3333 | 2 | 0 | 10 | 0.1 | 2 | 0 | 3 | 0.3333 | 3 | 0 | 2 | 0 | 10 | 0.1 |
| 1_91214000 | 2 | 0 | 4 | 0.25 | 3 | 0 | 2 | 0 | 11 | 0.0909 | 2 | 0 | 3 | 0 | 3 | 0 | 2 | 0.5 | 10 | 0.1 |
| 1_91214066 | 2 | 0 | 5 | 0.2 | 3 | 0 | 2 | 0 | 12 | 0.0833 | 2 | 0 | 3 | 0.3333 | 4 | 0 | 2 | 0 | 11 | 0.0909 |
| 1_91247983 | NA | NA | 3 | 1 | 5 | 0.8 | NA | NA | 8 | 0.875 | 1 | 0 | NA | NA | 2 | 0.5 | 3 | 0.3333 | 6 | 0.3333 |
| 1_91248152 | 3 | 0 | 2 | 0 | 5 | 0.2 | NA | NA | 10 | 0.1 | 1 | 0 | 2 | 0.5 | 2 | 0 | 3 | 0 | 8 | 0.125 |
| 1_91248170 | 3 | 0.3333 | 2 | 0 | 5 | 0.4 | NA | NA | 10 | 0.3 | 1 | 0 | 3 | 0.3333 | 1 | 0 | 3 | 0.6667 | 8 | 0.375 |
| 1_91248778 | 2 | 0 | 2 | 0.5 | 4 | 0.25 | NA | NA | 8 | 0.25 | 1 | 0 | 1 | 0 | 1 | 0 | 4 | 0.25 | 7 | 0.1429 |
| 1_91303670 | 8 | 0.125 | 6 | 0 | 6 | 0.1667 | 10 | 0.1 | 30 | 0.1 | 6 | 0.1667 | 5 | 0 | 11 | 0.2727 | 7 | 0 | 29 | 0.1379 |
| 1_91339763 | 2 | 0 | 4 | 0.25 | 3 | 0.3333 | NA | NA | 9 | 0.2222 | NA | NA | NA | NA | 1 | 1 | 1 | 0 | 2 | 0.5 |
| 1_91343839 | 5 | 0.4 | 5 | 0.2 | 2 | 0 | 2 | 0 | 14 | 0.2143 | NA | NA | 5 | 0.2 | 5 | 0 | 5 | 0 | 15 | 0.0667 |
| 1_91343908 | 6 | 0 | 7 | 0 | 2 | 0 | 2 | 0.5 | 17 | 0.0588 | 3 | 0 | 3 | 0.3333 | 5 | 0 | 6 | 0 | 17 | 0.0588 |
| 1_91343924 | 6 | 0 | 6 | 0.1667 | 2 | 0 | 3 | 0 | 17 | 0.0588 | 3 | 0 | 6 | 0.3333 | 4 | 0 | 6 | 0 | 19 | 0.1053 |
| 1_91344116 | 4 | 0 | 4 | 0.25 | 2 | 0.5 | NA | NA | 10 | 0.2 | 3 | 0 | 5 | 0 | 5 | 0.2 | 6 | 0.1667 | 19 | 0.1053 |
| 1_91344278 | 4 | 1 | 5 | 0.6 | 1 | 1 | 1 | 1 | 11 | 0.8182 | 3 | 1 | 4 | 1 | 5 | 1 | 6 | 0.8333 | 18 | 0.9444 |
| 1_91344357 | 5 | 0.4 | 3 | 0.6667 | 1 | 1 | 2 | 1 | 11 | 0.6364 | 3 | 0.6667 | 4 | 0.5 | 4 | 0.75 | 7 | 0.5714 | 18 | 0.6111 |

| | | | | | | | | | | | | | | | | | | | | |
|---|---|---|---|---|---|---|---|---|---|---|---|---|---|---|---|---|---|---|---|---|
| 1_91344422 | 6 | 1 | 4 | 1 | 2 | 1 | NA | NA | 12 | 1 | 3 | 1 | 5 | 1 | 4 | 1 | 7 | 1 | 19 | 1 |
| 1_91344514 | 5 | 0 | 5 | 0.4 | 1 | 0 | NA | NA | 11 | 0.1818 | 3 | 0.6667 | 5 | 0.4 | 4 | 0.25 | 7 | 0.1429 | 19 | 0.3158 |
| 1_91344735 | 5 | 0.2 | 5 | 0.2 | 2 | 1 | 1 | 0 | 13 | 0.3077 | 4 | 0.75 | 5 | 0.6 | 4 | 0.5 | 7 | 0.2857 | 20 | 0.5 |
| 1_91344745 | 3 | 1 | 3 | 1 | 1 | 1 | NA | NA | 7 | 1 | 3 | 1 | 5 | 1 | 4 | 1 | 6 | 1 | 18 | 1 |
| 1_91344774 | 5 | 0.4 | 5 | 0.2 | 2 | 0 | 1 | 1 | 13 | 0.3077 | 3 | 0.3333 | 4 | 0.25 | 4 | 0 | 7 | 0.4286 | 18 | 0.2778 |
| 1_91344924 | 4 | 0.25 | 5 | 0.4 | 2 | 0.5 | NA | NA | 11 | 0.3636 | 3 | 0.3333 | 6 | 0.5 | 4 | 0.75 | 7 | 0.7143 | 20 | 0.6 |
| 1_91344968 | 3 | 1 | 5 | 1 | 2 | 1 | 1 | 1 | 11 | 1 | 2 | 1 | 6 | 1 | 4 | 0.75 | 6 | 1 | 18 | 0.9444 |
| 1_91345155 | 4 | 1 | 5 | 1 | 2 | 1 | 2 | 1 | 13 | 1 | 2 | 1 | 6 | 0.8333 | 3 | 0.6667 | 6 | 1 | 17 | 0.8824 |
| 1_91345227 | 2 | 0.5 | 4 | 0 | NA | NA | 1 | 0 | 7 | 0.1429 | 1 | 0 | 4 | 0 | 3 | 0 | 6 | 0.1667 | 14 | 0.0714 |
| 1_91345344 | 4 | 0.75 | 5 | 0.6 | 2 | 0.5 | 1 | 1 | 12 | 0.6667 | 2 | 0 | 6 | 0.6667 | 3 | 0.6667 | 7 | 0.2857 | 18 | 0.4444 |
| 1_91345351 | 4 | 0.5 | 4 | 0.25 | 1 | 1 | 1 | 1 | 10 | 0.5 | 1 | 0 | 3 | 0.3333 | 3 | 0.3333 | 6 | 0.3333 | 13 | 0.3077 |
| 1_91345436 | 4 | 0.5 | 4 | 0.5 | 2 | 0.5 | 1 | 1 | 11 | 0.5455 | 2 | 0.5 | 5 | 0.4 | 3 | 0.6667 | 5 | 1 | 15 | 0.6667 |
| 1_91345465 | 4 | 0 | 4 | 0 | 2 | 0 | 1 | 1 | 11 | 0.0909 | 2 | 0 | 3 | 0 | 3 | 0.3333 | 5 | 0 | 13 | 0.0769 |
| 1_91345515 | 3 | 1 | 5 | 1 | 2 | 1 | 1 | 0 | 11 | 0.9091 | 2 | 1 | 5 | 1 | 3 | 1 | 7 | 0.8571 | 17 | 0.9412 |
| 1_91345668 | 4 | 0.5 | 4 | 0 | 2 | 0.5 | 1 | 0 | 11 | 0.2727 | 2 | 0.5 | 6 | 0.1667 | 4 | 0.25 | 4 | 0.25 | 16 | 0.25 |
| 1_91351461 | 2 | 0 | 2 | 0 | 1 | 1 | 1 | 0 | 6 | 0.1667 | 1 | 0 | 4 | 0 | 4 | 0.25 | 4 | 0.25 | 13 | 0.1538 |
| 1_91351666 | 2 | 0.5 | 2 | 0 | 1 | 0 | 1 | 0 | 6 | 0.1667 | 1 | 1 | 4 | 0 | 4 | 0.25 | 4 | 0.25 | 13 | 0.2308 |
| 1_91352121 | 2 | 1 | 2 | 0 | 2 | 0 | 1 | 0 | 7 | 0.2857 | 1 | 1 | 2 | 1 | 4 | 0.5 | 3 | 0.6667 | 10 | 0.7 |
| 1_91455899 | NA | NA | NA | NA | 3 | 0.3333 | 1 | 0 | 4 | 0.25 | 1 | 1 | 1 | 0 | NA | NA | 2 | 0.5 | 4 | 0.5 |
| 1_91456640 | 1 | 0 | 1 | 1 | 3 | 0.6667 | 1 | 1 | 6 | 0.6667 | 1 | 1 | 1 | 1 | NA | NA | 2 | 0 | 4 | 0.5 |
| 1_91482663 | 1 | 1 | NA | NA | NA | NA | NA | NA | 1 | 1 | NA | NA | NA | NA | 1 | 1 | NA | NA | 1 | 1 |
| 1_91482747 | 1 | 1 | NA | NA | NA | NA | NA | NA | 1 | 1 | NA | NA | NA | NA | 1 | 1 | NA | NA | 1 | 1 |
| 1_91856710 | 4 | 0 | 4 | 0.5 | 4 | 0.25 | 7 | 0 | 19 | 0.1579 | 6 | 0.3333 | 3 | 0 | 5 | 0.2 | 5 | 0 | 19 | 0.1579 |
| 1_91857413 | 3 | 0 | 4 | 0.75 | 5 | 0.2 | 7 | 0.2857 | 19 | 0.3158 | 6 | 0.5 | 3 | 0 | 5 | 0.2 | 5 | 0.4 | 19 | 0.3158 |
| 1_91857441 | 3 | 0 | 5 | 0 | 3 | 0 | 6 | 0.1667 | 17 | 0.0588 | 6 | 0.1667 | 3 | 0 | 5 | 0 | 5 | 0 | 19 | 0.0526 |
| 1_91858688 | 1 | 0 | 3 | 0.6667 | NA | NA | 6 | 0.3333 | 10 | 0.4 | 5 | 0.2 | 3 | 0.3333 | 4 | 0.25 | 3 | 0.3333 | 15 | 0.2667 |
| 1_91860269 | 1 | 1 | 2 | 0.5 | 1 | 0 | 4 | 0 | 8 | 0.25 | 2 | 0.5 | 2 | 0 | 4 | 0 | 3 | 0.3333 | 11 | 0.1818 |
| 1_91860295 | 1 | 0 | 3 | 0.3333 | 1 | 0 | 3 | 0 | 8 | 0.125 | 3 | 0.3333 | 2 | 0 | 2 | 0 | 2 | 0 | 9 | 0.1111 |
| 1_91860305 | 1 | 0 | 1 | 0 | 1 | 1 | 2 | 0 | 5 | 0.2 | 1 | 0 | 2 | 0.5 | 3 | 0.6667 | 2 | 0 | 8 | 0.375 |
| 1_91860324 | 1 | 1 | 3 | 1 | 1 | 0 | 4 | 1 | 9 | 0.8889 | 4 | 0.75 | 2 | 1 | 4 | 0.75 | 3 | 1 | 13 | 0.8462 |
| 1_91860444 | 1 | 0 | 1 | 0 | NA | NA | 4 | 0.25 | 6 | 0.1667 | 3 | 0.3333 | 2 | 0 | 3 | 0 | 2 | 0 | 10 | 0.1 |
| 1_91860449 | NA | NA | 1 | 1 | NA | NA | 3 | 0.6667 | 4 | 0.75 | 2 | 1 | 1 | 1 | 4 | 0.75 | 1 | 1 | 8 | 0.875 |
| 1_91860503 | 1 | 1 | 2 | 1 | 1 | 0 | 3 | 0.3333 | 7 | 0.5714 | 4 | 0.75 | 1 | 1 | 3 | 0.3333 | 2 | 1 | 10 | 0.7 |
| 1_91860544 | 1 | 1 | 2 | 0.5 | 1 | 1 | 3 | 0.3333 | 7 | 0.5714 | 2 | 0.5 | 1 | 0 | 4 | 0.75 | 3 | 1 | 10 | 0.7 |
| 1_91860576 | 1 | 0 | 1 | 1 | NA | NA | 3 | 0 | 5 | 0.2 | 3 | 0.3333 | 1 | 1 | 1 | 0 | 1 | 0 | 6 | 0.3333 |
| 1_91860676 | NA | NA | 2 | 1 | 1 | 1 | 3 | 0.6667 | 6 | 0.8333 | 3 | 0.6667 | 1 | 0 | 3 | 0.6667 | 2 | 0.5 | 9 | 0.5556 |
| 1_91860687 | 1 | 0 | 2 | 1 | 1 | 0 | 3 | 0.6667 | 7 | 0.5714 | 3 | 0.3333 | 1 | 0 | 3 | 1 | 2 | 0.5 | 9 | 0.5556 |
| 1_91860691 | 1 | 1 | 1 | 0 | 1 | 0 | 2 | 0 | 5 | 0.2 | 3 | 0.6667 | 1 | 0 | 3 | 0 | 3 | 0 | 10 | 0.2 |
| 1_91860730 | NA | NA | 2 | 0 | NA | NA | 2 | 0.5 | 4 | 0.25 | 3 | 0.3333 | 1 | 0 | 3 | 0 | 3 | 0 | 10 | 0.1 |
| 1_91860857 | 1 | 0 | 1 | 1 | NA | NA | 3 | 0.3333 | 5 | 0.4 | 3 | 0.3333 | 1 | 0 | 2 | 0 | 3 | 0.6667 | 9 | 0.3333 |
| 1_91860961 | NA | NA | 2 | 0.5 | NA | NA | 2 | 0.5 | 4 | 0.5 | 3 | 0.3333 | 1 | 0 | 2 | 1 | 2 | 0.5 | 8 | 0.5 |
| 1_92367250 | 26 | 0 | 39 | 0.0513 | 28 | 0.1429 | 35 | 0.1429 | 128 | 0.0859 | 38 | 0.1053 | 27 | 0.1852 | 31 | 0.129 | 23 | 0.3478 | 119 | 0.1765 |
| 1_92367358 | 34 | 0.1471 | 40 | 0.225 | 32 | 0.1875 | 35 | 0.2286 | 141 | 0.1986 | 41 | 0.1463 | 24 | 0.125 | 32 | 0.125 | 29 | 0.2069 | 126 | 0.1508 |
| 1_92367498 | 25 | 0.08 | 34 | 0.1471 | 21 | 0.0952 | 24 | 0.125 | 104 | 0.1154 | 29 | 0.1379 | 17 | 0.1765 | 23 | 0.1739 | 24 | 0.2083 | 93 | 0.172 |
| 1_92564893 | 25 | 0.04 | 36 | 0.1389 | 20 | 0.15 | 36 | 0.1389 | 117 | 0.1197 | 32 | 0.0625 | 34 | 0.2059 | 29 | 0.1034 | 20 | 0.1 | 115 | 0.1217 |
| 1_92564964 | 34 | 0.3529 | 37 | 0.2973 | 27 | 0.3704 | 34 | 0.3529 | 132 | 0.3409 | 36 | 0.25 | 35 | 0.4286 | 30 | 0.4 | 25 | 0.36 | 126 | 0.3571 |
| 1_92759666 | 4 | 0 | 3 | 0 | 4 | 0.25 | 2 | 0 | 13 | 0.0769 | NA | NA | NA | NA | 4 | 0 | 3 | 0.3333 | 7 | 0.1429 |
| 1_92782649 | 5 | 0 | 4 | 0 | 5 | 0.2 | 3 | 0 | 17 | 0.0588 | NA | NA | 1 | 1 | 9 | 0 | 5 | 0 | 15 | 0.0667 |
| 1_92782811 | 6 | 0 | 5 | 0 | 4 | 0.25 | 2 | 0 | 17 | 0.0588 | NA | NA | 1 | 0 | 8 | 0.125 | 5 | 0 | 14 | 0.0714 |
| 1_92782820 | 2 | 0 | 3 | 0.3333 | 5 | 0.2 | 1 | 1 | 11 | 0.2727 | NA | NA | NA | NA | 5 | 1 | 2 | 0.5 | 7 | 0.8571 |
| 1_92786306 | 10 | 0.8 | 10 | 1 | 6 | 1 | 7 | 0.8571 | 33 | 0.9091 | 1 | 1 | 8 | 1 | 11 | 0.9091 | 8 | 1 | 28 | 0.9643 |
| 1_92786332 | 8 | 0.125 | 6 | 0.1667 | 7 | 0.1429 | 6 | 0.1667 | 27 | 0.1481 | 1 | 0 | 8 | 0.25 | 10 | 0.1 | 5 | 0.2 | 24 | 0.1667 |
| 1_92786519 | 10 | 0.3 | 5 | 0.6 | 7 | 0.2857 | 6 | 0.1667 | 28 | 0.3214 | 1 | 0 | 6 | 0.3333 | 10 | 0.4 | 3 | 0 | 20 | 0.3 |
| 1_92786674 | 9 | 0.1111 | 9 | 0.3333 | 5 | 0.2 | 7 | 0 | 30 | 0.1667 | 2 | 1 | 9 | 0.2222 | 9 | 0.2222 | 9 | 0.4444 | 29 | 0.3448 |
| 1_92786861 | 15 | 0 | 12 | 0.25 | 9 | 0.1111 | 9 | 0.2222 | 45 | 0.1333 | 1 | 0 | 11 | 0.1818 | 13 | 0.2308 | 9 | 0.2222 | 34 | 0.2059 |
| 1_92787034 | 9 | 0.4444 | 9 | 0.4444 | 10 | 0.7 | 9 | 0.4444 | 37 | 0.5135 | 1 | 0 | 10 | 0.4 | 11 | 0.1818 | 7 | 0.4286 | 29 | 0.3103 |
| 1_92787787 | 17 | 0.0588 | 14 | 0.0714 | 16 | 0 | 15 | 0.1333 | 62 | 0.0645 | 6 | 0 | 15 | 0.0667 | 17 | 0.1176 | 13 | 0.0769 | 51 | 0.0784 |
| 1_92787820 | 7 | 0 | 13 | 0.3077 | 11 | 0.0909 | 10 | 0.1 | 41 | 0.1463 | 5 | 0 | 10 | 0.2 | 10 | 0.1 | 7 | 0.2857 | 32 | 0.1562 |
| 1_92835547 | 4 | 0.5 | 1 | 1 | 1 | 1 | 4 | 0.75 | 10 | 0.7 | 5 | 0.2 | NA | NA | 6 | 1 | 2 | 1 | 13 | 0.6923 |
| 1_92835726 | 3 | 0.6667 | 3 | 0 | 2 | 0.5 | 5 | 0.2 | 13 | 0.3077 | 6 | 0.1667 | NA | NA | 6 | 0 | 2 | 0 | 14 | 0.0714 |
| 1_92835748 | 3 | 0.3333 | 3 | 0.6667 | 2 | 0 | 4 | 0.25 | 12 | 0.3333 | 5 | 0.2 | NA | NA | 6 | 0.1667 | 2 | 0 | 13 | 0.1538 |
| 1_92839096 | 3 | 0.3333 | 1 | 0 | 2 | 0 | 1 | 0 | 7 | 0.1429 | NA | NA | NA | NA | 3 | 0 | 2 | 0.5 | 5 | 0.2 |
| 1_92847468 | 5 | 0 | 2 | 0 | 3 | 0 | 4 | 0.25 | 14 | 0.0714 | 2 | 0 | 1 | 1 | 4 | 0 | 1 | 0 | 8 | 0.125 |
| 1_92847495 | 6 | 0.3333 | 2 | 0 | 2 | 0 | 4 | 0.75 | 14 | 0.3571 | 1 | 0 | 1 | 1 | 5 | 0.6 | 2 | 0 | 9 | 0.4444 |
| 1_92847501 | 4 | 0.25 | 3 | 0 | 2 | 0.5 | 3 | 0 | 12 | 0.1667 | 2 | 0.5 | 1 | 1 | 4 | 0.25 | 2 | 0 | 9 | 0.3333 |

| | | | | | | | | | | | | | | | | | | | | |
|---|---|---|---|---|---|---|---|---|---|---|---|---|---|---|---|---|---|---|---|---|
| 1_92847589 | 6 | 0.6667 | 2 | 0 | 3 | 0.6667 | 3 | 0 | 14 | 0.4286 | 1 | 0 | 1 | 0 | 3 | 0.3333 | 2 | 0 | 7 | 0.1429 |
| 1_92847647 | 5 | 0.6 | 1 | 1 | 2 | 0.5 | 4 | 0.75 | 12 | 0.6667 | 2 | 1 | 1 | 1 | 3 | 1 | 2 | 1 | 8 | 1 |
| 1_92847683 | 6 | 0.8333 | 2 | 1 | 3 | 1 | 3 | 1 | 14 | 0.9286 | 2 | 1 | 1 | 1 | 5 | 1 | 2 | 0.5 | 10 | 0.9 |
| 1_92847731 | 7 | 0.7143 | 4 | 0.25 | 3 | 1 | 5 | 0.8 | 19 | 0.6842 | 2 | 0.5 | 1 | 1 | 4 | 0.75 | 2 | 0.5 | 9 | 0.6667 |
| 1_92847754 | 5 | 0 | 3 | 0.3333 | 2 | 0.5 | 1 | 0 | 11 | 0.1818 | 2 | 1 | 1 | 1 | 4 | 0.25 | 2 | 0.5 | 9 | 0.5556 |
| 1_92847838 | 6 | 0.1667 | 3 | 0 | 2 | 0 | 3 | 0.6667 | 14 | 0.2143 | 2 | 0.5 | 1 | 0 | 5 | 0.4 | 2 | 0.5 | 10 | 0.4 |
| 1_92847888 | 4 | 0 | 1 | 0 | 1 | 0 | 2 | 0.5 | 8 | 0.125 | 1 | 0 | NA | NA | 3 | 1 | 1 | 1 | 5 | 0.8 |
| 1_92848032 | 7 | 0.2857 | 3 | 1 | 3 | 0.6667 | 4 | 0.25 | 17 | 0.4706 | 2 | 0.5 | 1 | 1 | 5 | 0.6 | NA | NA | 8 | 0.625 |
| 1_92848039 | 7 | 0.1429 | 4 | 0 | 4 | 0 | 5 | 0 | 20 | 0.05 | 1 | 0 | 1 | 0 | 6 | 0.3333 | 1 | 0 | 9 | 0.2222 |
| 1_92848053 | 5 | 0.2 | 4 | 0.75 | 2 | 0.5 | 5 | 0.6 | 16 | 0.5 | 2 | 1 | 1 | 1 | 2 | 0.5 | 1 | 1 | 6 | 0.8333 |
| 1_92848097 | 6 | 0.1667 | 4 | 0.75 | 3 | 1 | 5 | 0.4 | 18 | 0.5 | 2 | 0.5 | 1 | 1 | 5 | 0.6 | 2 | 0 | 10 | 0.5 |
| 1_92848248 | 7 | 0.1429 | 3 | 0.3333 | 3 | 0 | 4 | 0.25 | 17 | 0.1765 | 1 | 0 | 1 | 0 | 5 | 0 | 1 | 1 | 8 | 0.125 |
| 1_92848260 | 5 | 0 | 4 | 0.25 | 2 | 0.5 | 4 | 0.25 | 15 | 0.2 | 2 | 0.5 | 1 | 0 | 4 | 0 | 2 | 0 | 9 | 0.1111 |
| 1_92848269 | 6 | 0.6667 | 3 | 0.3333 | 2 | 0.5 | 4 | 0.25 | 15 | 0.4667 | 2 | 1 | 1 | 1 | 5 | 0.6 | 2 | 0.5 | 10 | 0.7 |
| 1_92868181 | 1 | 0 | 3 | 0.3333 | 1 | 0 | 4 | 0 | 9 | 0.1111 | 1 | 0 | 1 | 1 | 5 | 0.2 | 5 | 0.2 | 12 | 0.25 |
| 1_92868581 | 1 | 0 | 2 | 0.5 | 1 | 0 | 3 | 0.3333 | 7 | 0.2857 | 1 | 0 | 1 | 0 | 1 | 0 | 3 | 0.3333 | 6 | 0.1667 |
| 1_92875379 | 1 | 1 | 2 | 0.5 | NA | NA | 4 | 0.75 | 7 | 0.7143 | 2 | 0.5 | 1 | 0 | 4 | 0.75 | 3 | 1 | 10 | 0.7 |
| 1_92875585 | 1 | 0 | 2 | 0 | 2 | 0 | 4 | 0.25 | 9 | 0.1111 | 4 | 0.25 | 1 | 1 | 5 | 0.2 | 3 | 0.3333 | 13 | 0.3077 |
| 1_92943450 | 88 | 0.2045 | 79 | 0.1899 | 70 | 0.1429 | 59 | 0.2034 | 296 | 0.1858 | 72 | 0.3194 | 53 | 0.2642 | 77 | 0.3636 | 44 | 0.2045 | 246 | 0.3008 |
| 1_92943509 | 83 | 0.1687 | 78 | 0.1538 | 73 | 0.137 | 68 | 0.2794 | 302 | 0.1821 | 69 | 0.2319 | 48 | 0.25 | 71 | 0.2394 | 47 | 0.1702 | 235 | 0.2255 |
| 1_92943543 | 80 | 0.0375 | 79 | 0.1139 | 67 | 0.0299 | 64 | 0.0156 | 290 | 0.0517 | 71 | 0.0704 | 51 | 0.0392 | 68 | 0.1471 | 47 | 0.0638 | 237 | 0.0844 |
| 1_92945139 | 31 | 0.129 | 31 | 0.129 | 34 | 0.0294 | 38 | 0.0526 | 134 | 0.0821 | 34 | 0.0588 | 24 | 0.0833 | 35 | 0.0857 | 26 | 0.0385 | 119 | 0.0672 |
| 1_92945166 | 33 | 0.0303 | 39 | 0.1795 | 36 | 0.1111 | 40 | 0.15 | 148 | 0.1216 | 33 | 0.1212 | 25 | 0.04 | 35 | 0.0286 | 25 | 0.12 | 118 | 0.0763 |
| 1_92945813 | 28 | 0.1071 | 24 | 0.0417 | 23 | 0.087 | 24 | 0.2083 | 99 | 0.1111 | 23 | 0.1304 | 16 | 0.25 | 35 | 0.2286 | 18 | 0.2778 | 92 | 0.2174 |
| 1_92946162 | 24 | 0.1667 | 23 | 0.0435 | 18 | 0.1111 | 25 | 0.16 | 90 | 0.1222 | 17 | 0.1176 | 19 | 0.0526 | 28 | 0.1071 | 12 | 0.3333 | 76 | 0.1316 |
| 1_93225271 | NA | NA | NA | NA | NA | NA | 1 | 1 | 1 | 1 | NA | NA | NA | NA | NA | NA | 1 | 1 | 1 | 1 |
| 1_93225625 | NA | NA | NA | NA | NA | NA | 1 | 1 | 1 | 1 | NA | NA | NA | NA | NA | NA | 1 | 1 | 1 | 1 |
| 1_93226036 | NA | NA | NA | NA | NA | NA | 1 | 1 | 1 | 1 | NA | NA | NA | NA | NA | NA | 1 | 1 | 1 | 1 |
| 1_93228640 | 2 | 0.5 | 1 | 1 | NA | NA | 1 | 0 | 4 | 0.5 | 2 | 0.5 | 1 | 0 | NA | NA | 1 | 1 | 4 | 0.5 |
| 1_93228962 | 1 | 1 | 3 | 1 | 4 | 1 | 1 | 0 | 9 | 0.8889 | 2 | 1 | 4 | 0.5 | 2 | 0.5 | NA | NA | 8 | 0.625 |
| 1_93229131 | 1 | 0 | 3 | 0 | 4 | 0.75 | 1 | 1 | 9 | 0.4444 | 2 | 0.5 | 3 | 0 | 2 | 0.5 | NA | NA | 7 | 0.2857 |
| 1_93229269 | 1 | 1 | 2 | 0.5 | 2 | 0.5 | 1 | 1 | 6 | 0.6667 | NA | NA | 1 | 1 | 1 | 1 | NA | NA | 2 | 1 |
| 1_93229310 | 1 | 1 | 3 | 0.6667 | 1 | 0 | 1 | 1 | 6 | 0.6667 | NA | NA | 3 | 0.3333 | 2 | 0.5 | NA | NA | 5 | 0.4 |
| 1_93229348 | NA | NA | 2 | 0 | 2 | 0 | 1 | 1 | 5 | 0.2 | NA | NA | 3 | 0.3333 | 2 | 0.5 | NA | NA | 5 | 0.4 |
| 1_93229431 | NA | NA | 3 | 0.6667 | 2 | 0.5 | 1 | 0 | 6 | 0.5 | NA | NA | 2 | 0.5 | 2 | 0 | NA | NA | 4 | 0.25 |
| 1_93229497 | 1 | 0 | 2 | 0.5 | 2 | 0.5 | 1 | 0 | 6 | 0.3333 | NA | NA | 3 | 1 | 2 | 1 | NA | NA | 5 | 1 |
| 1_93232391 | 1 | 1 | 2 | 0.5 | 1 | 0 | 1 | 0 | 5 | 0.4 | NA | NA | 3 | 0.3333 | 2 | 0 | NA | NA | 5 | 0.2 |
| 1_93232460 | 1 | 1 | 1 | 1 | 1 | 1 | 1 | 1 | 4 | 1 | NA | NA | 3 | 0.6667 | 1 | 0 | NA | NA | 4 | 0.5 |
| 1_93232548 | 1 | 1 | NA | NA | 2 | 0.5 | NA | NA | 3 | 0.6667 | NA | NA | 2 | 1 | NA | NA | NA | NA | 2 | 1 |
| 1_93292828 | 15 | 0.6 | 10 | 0.8 | 19 | 0.7368 | 13 | 0.7692 | 57 | 0.7193 | 10 | 0.9 | 14 | 0.6429 | 10 | 0.9 | 12 | 0.75 | 46 | 0.7826 |
| 1_93293110 | 6 | 0.3333 | 5 | 0.2 | 9 | 0.2222 | 4 | 0.75 | 24 | 0.3333 | 4 | 0.5 | 5 | 0.2 | 3 | 0.6667 | 6 | 0.5 | 18 | 0.4444 |
| 1_93293137 | 6 | 0 | 6 | 0.1667 | 8 | 0.5 | 3 | 0.3333 | 23 | 0.2609 | 4 | 0.5 | 4 | 0 | 3 | 1 | 4 | 0.5 | 15 | 0.4667 |
| 1_93293168 | 6 | 0 | 6 | 0.1667 | 9 | 0.1111 | 3 | 0.3333 | 24 | 0.125 | 4 | 0 | 5 | 0.2 | 3 | 0 | 7 | 0.1429 | 19 | 0.1053 |
| 1_93293430 | 3 | 1 | 5 | 0.6 | 5 | 1 | 4 | 1 | 17 | 0.8824 | 4 | 0.75 | 3 | 1 | NA | NA | 3 | 1 | 10 | 0.9 |
| 1_93293590 | 3 | 0.3333 | 2 | 0.5 | 6 | 0.3333 | 3 | 0.3333 | 14 | 0.3571 | 4 | 0 | 4 | 0.5 | 1 | 0 | 4 | 0.5 | 13 | 0.3077 |
| 1_93293594 | 3 | 0.3333 | 4 | 0.25 | 6 | 0.3333 | 3 | 0.3333 | 16 | 0.3125 | 2 | 0.5 | 3 | 0 | 1 | 1 | 1 | 1 | 7 | 0.4286 |
| 1_93293662 | NA | NA | 5 | 0.2 | 1 | 1 | 1 | 1 | 7 | 0.4286 | 2 | 1 | 1 | 0 | 2 | 0 | 1 | 0 | 6 | 0.3333 |
| 1_93293813 | 4 | 1 | 5 | 0.6 | 8 | 0.75 | 3 | 0.6667 | 20 | 0.75 | 3 | 1 | 4 | 0.5 | 2 | 0.5 | 2 | 0.5 | 11 | 0.6364 |
| 1_93293846 | 4 | 1 | 7 | 0.8571 | 9 | 0.8889 | 3 | 0.6667 | 23 | 0.8696 | 3 | 1 | 6 | 0.3333 | 3 | 0.6667 | 4 | 0.75 | 16 | 0.625 |
| 1_93293941 | 5 | 1 | 6 | 0.6667 | 6 | 0.8333 | 4 | 0.75 | 21 | 0.8095 | 4 | 1 | 6 | 0.6667 | 2 | 1 | 6 | 0.8333 | 18 | 0.8333 |
| 1_93294000 | 5 | 0.8 | 7 | 0.1429 | 6 | 0.1667 | 3 | 0.6667 | 21 | 0.381 | 3 | 0 | 4 | 0 | 2 | 0.5 | 4 | 0.25 | 13 | 0.1538 |
| 1_93294031 | 4 | 0.25 | 7 | 0.2857 | 7 | 0.4286 | 4 | 0.25 | 22 | 0.3182 | 3 | 0.6667 | 7 | 0.1429 | 3 | 0.3333 | 5 | 0.6 | 18 | 0.3889 |
| 1_93294108 | 6 | 1 | 7 | 0.8571 | 9 | 0.8889 | 4 | 1 | 26 | 0.9231 | 2 | 1 | 7 | 1 | 4 | 1 | 6 | 1 | 19 | 1 |
| 1_93294488 | 4 | 0.25 | 6 | 0 | 8 | 0 | 5 | 0.2 | 23 | 0.087 | 4 | 0 | 7 | 0.1429 | 2 | 0.5 | 5 | 0 | 18 | 0.1111 |
| 1_93294517 | 5 | 0.2 | 6 | 0.3333 | 8 | 0.25 | 3 | 0 | 22 | 0.2273 | 3 | 0 | 6 | 0 | 2 | 0 | 5 | 0.2 | 16 | 0.0625 |
| 1_93294686 | 4 | 0 | 5 | 0 | 8 | 0.25 | 3 | 0 | 20 | 0.1 | 3 | 0.3333 | 7 | 0 | 2 | 0 | 5 | 0.2 | 17 | 0.1176 |
| 1_93294755 | 5 | 0.2 | 6 | 0.1667 | 6 | 0 | 5 | 0.2 | 22 | 0.1364 | 4 | 0 | 7 | 0.4286 | 2 | 0 | 6 | 0.1667 | 19 | 0.2105 |
| 1_93294946 | 5 | 0.2 | 7 | 0.4286 | 8 | 0.125 | 4 | 0.25 | 24 | 0.25 | 4 | 0.5 | 7 | 0 | 3 | 0 | 6 | 0.5 | 20 | 0.25 |
| 1_93300657 | 5 | 0.6 | 3 | 0.6667 | 4 | 0.75 | 5 | 1 | 17 | 0.7647 | NA | NA | 5 | 1 | 1 | 1 | 2 | 1 | 8 | 1 |
| 1_93300680 | 4 | 0.25 | 5 | 0.2 | 3 | 0.3333 | 6 | 0.1667 | 18 | 0.2222 | NA | NA | 4 | 0.5 | 1 | 0 | 1 | 0 | 6 | 0.3333 |
| 1_93300795 | 5 | 0 | 5 | 0 | 4 | 0.25 | 5 | 0.2 | 19 | 0.1053 | 1 | 0 | 5 | 0.2 | 3 | 0 | 2 | 0 | 11 | 0.0909 |
| 1_93300831 | 5 | 0.6 | 5 | 0.8 | 5 | 0.6 | 6 | 1 | 21 | 0.7619 | 1 | 0 | 5 | 0.8 | 3 | 1 | 2 | 1 | 11 | 0.8182 |
| 1_93300930 | 5 | 0.4 | 5 | 1 | 4 | 0.75 | 5 | 1 | 19 | 0.7895 | 1 | 1 | 5 | 0.8 | 3 | 1 | 3 | 1 | 12 | 0.9167 |
| 1_93300986 | 5 | 0 | 4 | 0.25 | 4 | 0.25 | 3 | 0 | 16 | 0.125 | 2 | 0 | 5 | 0.2 | 3 | 0.3333 | 3 | 0 | 13 | 0.1538 |
| 1_93301126 | 5 | 0 | 6 | 0.3333 | 5 | 0 | 6 | 0.3333 | 22 | 0.1818 | 1 | 1 | 6 | 0.3333 | 3 | 0 | 4 | 0.75 | 14 | 0.4286 |

| | | | | | | | | | | | | | | | | | | | | |
|---|---|---|---|---|---|---|---|---|---|---|---|---|---|---|---|---|---|---|---|---|
| 1_9333390 | 6 | 0.1667 | 7 | 0 | 11 | 0.1818 | 10 | 0.1 | 34 | 0.1176 | 4 | 0 | 12 | 0.3333 | 8 | 0.125 | 8 | 0 | 32 | 0.1562 |
| 1_9333550 | 9 | 0.1111 | 9 | 0.1111 | 7 | 0 | 9 | 0.2222 | 34 | 0.1176 | 6 | 0.1667 | 9 | 0.1111 | 8 | 0 | 10 | 0 | 33 | 0.0606 |
| 1_9333567 | 7 | 0 | 13 | 0.2308 | 8 | 0 | 8 | 0.125 | 36 | 0.1111 | 9 | 0.2222 | 8 | 0.25 | 4 | 0.25 | 5 | 0.2 | 26 | 0.2308 |
| 1_9333581 | 9 | 0.2222 | 13 | 0.2308 | 9 | 0.2222 | 10 | 0.1 | 41 | 0.1951 | 7 | 0 | 9 | 0.1111 | 8 | 0.25 | 8 | 0.125 | 32 | 0.125 |
| 1_9333588 | 9 | 0 | 14 | 0.0714 | 7 | 0.1429 | 8 | 0.125 | 38 | 0.0789 | 7 | 0.2857 | 7 | 0 | 7 | 0 | 8 | 0.5 | 29 | 0.2069 |
| 1_9334127 | 3 | 0 | 7 | 0.1429 | 5 | 0.2 | 3 | 0 | 18 | 0.1111 | 4 | 0.25 | 2 | 0 | 4 | 0 | 4 | 0 | 14 | 0.0714 |
| 1_9335882 | NA | NA | 5 | 0 | 3 | 0.3333 | NA | NA | 8 | 0.125 | 2 | 0 | 1 | 0 | 3 | 0 | 1 | 1 | 7 | 0.1429 |
| 1_9336493 | NA | NA | 5 | 0.2 | 1 | 0 | NA | NA | 6 | 0.1667 | 1 | 0 | 2 | 0.5 | 3 | 0 | 2 | 0.5 | 8 | 0.25 |
| 1_9352085 | NA | NA | 2 | 0.5 | NA | NA | 1 | 1 | 3 | 0.6667 | NA | NA | 3 | 0.3333 | 1 | 0 | NA | NA | 4 | 0.25 |
| 1_9354910 | 4 | 0.75 | NA | NA | 3 | 0.6667 | 3 | 0.6667 | 10 | 0.7 | 1 | 1 | NA | NA | NA | NA | NA | NA | 1 | 1 |
| 1_9354985 | 32 | 0.2188 | 33 | 0.3333 | 26 | 0.2308 | 31 | 0.4194 | 122 | 0.3033 | 22 | 0.1818 | 24 | 0.375 | 22 | 0.2273 | 25 | 0.08 | 93 | 0.2151 |
| 1_9354988 | 31 | 0.0968 | 38 | 0 | 27 | 0.0741 | 30 | 0.0667 | 126 | 0.0556 | 22 | 0.0909 | 28 | 0.0714 | 24 | 0 | 26 | 0.0769 | 100 | 0.06 |
| 1_9354992 | 28 | 0.1071 | 38 | 0.1579 | 27 | 0.1111 | 25 | 0.32 | 118 | 0.1695 | 24 | 0.1667 | 26 | 0.2308 | 24 | 0.125 | 25 | 0.16 | 99 | 0.1717 |
| 1_9354997 | 33 | 0 | 39 | 0.1026 | 31 | 0.0323 | 31 | 0.0968 | 134 | 0.0597 | 26 | 0.0769 | 29 | 0.069 | 31 | 0.129 | 29 | 0.1724 | 115 | 0.113 |
| 1_9355009 | 30 | 0.4 | 36 | 0.5833 | 26 | 0.6154 | 32 | 0.5938 | 124 | 0.5484 | 27 | 0.5926 | 29 | 0.5172 | 26 | 0.7308 | 27 | 0.5926 | 109 | 0.6055 |
| 1_9355013 | 34 | 0.5882 | 37 | 0.5946 | 28 | 0.8214 | 32 | 0.6875 | 131 | 0.6641 | 27 | 0.7037 | 26 | 0.6538 | 29 | 0.7241 | 27 | 0.5556 | 109 | 0.6606 |
| 1_9355538 | 20 | 0.1 | 19 | 0.0526 | 16 | 0.0625 | 19 | 0.0526 | 74 | 0.0676 | 14 | 0.1429 | 16 | 0 | 13 | 0.1538 | 16 | 0.125 | 59 | 0.1017 |
| 1_9361339 | 3 | 0 | 9 | 0 | 4 | 0.25 | 1 | 0 | 17 | 0.0588 | 4 | 0.25 | 1 | 0 | 5 | 0.2 | 2 | 0 | 12 | 0.1667 |
| 1_9362183 | 7 | 0 | 8 | 0.125 | 5 | 0 | 2 | 0.5 | 22 | 0.0909 | 2 | 0 | 1 | 0 | 3 | 0.3333 | 1 | 0 | 7 | 0.1429 |
| 1_9362326 | 13 | 0 | 18 | 0.1111 | 15 | 0.1333 | 6 | 0 | 52 | 0.0769 | 11 | 0.0909 | 6 | 0 | 7 | 0 | 10 | 0.1 | 34 | 0.0588 |
| 1_9367803 | 1 | 1 | NA | NA | 2 | 0.5 | 2 | 0 | 5 | 0.4 | 3 | 0.3333 | 2 | 0 | 3 | 0.6667 | NA | NA | 8 | 0.375 |
| 1_9367849 | 1 | 0 | NA | NA | 1 | 1 | 2 | 0 | 4 | 0.25 | 1 | 0 | 1 | 0 | 2 | 0.5 | NA | NA | 4 | 0.25 |
| 1_9367858 | 1 | 1 | NA | NA | 1 | 0 | 2 | 0 | 4 | 0.25 | 2 | 0 | 1 | 0 | 2 | 0.5 | NA | NA | 5 | 0.2 |
| 1_9367864 | 1 | 0 | NA | NA | 1 | 1 | 1 | 1 | 3 | 0.6667 | 3 | 0 | 1 | 0 | 2 | 0.5 | NA | NA | 6 | 0.1667 |
| 1_9371554 | 8 | 0 | 4 | 0 | 3 | 0 | 9 | 0.3333 | 24 | 0.125 | 1 | 0 | 9 | 0.1111 | 3 | 0 | 6 | 0 | 19 | 0.0526 |
| 1_9371564 | 8 | 0.5 | 4 | 0.75 | 3 | 0.3333 | 10 | 0.4 | 25 | 0.48 | 1 | 1 | 11 | 0.6364 | 3 | 0.6667 | 7 | 0.2857 | 22 | 0.5455 |
| 1_9371565 | 8 | 1 | 5 | 1 | 1 | 0 | 9 | 1 | 23 | 0.9565 | 1 | 1 | 13 | 0.9231 | 3 | 0.6667 | 8 | 1 | 25 | 0.92 |
| 1_9371569 | 9 | 0.4444 | 3 | 0.6667 | 2 | 0.5 | 10 | 0.5 | 24 | 0.5 | 1 | 1 | 12 | 0.6667 | 3 | 0.3333 | 7 | 0.8571 | 23 | 0.6957 |
| 1_9371590 | 6 | 0.3333 | 2 | 0 | 3 | 0.6667 | 4 | 0.5 | 15 | 0.4 | NA | NA | 6 | 0.5 | NA | NA | 4 | 0.25 | 10 | 0.4 |
| 1_9371617 | 5 | 0 | 3 | 0.3333 | 5 | 0 | 4 | 0 | 17 | 0.0588 | NA | NA | 7 | 0.2857 | NA | NA | 4 | 0 | 11 | 0.1818 |
| 1_9372051 | 1 | 0 | 5 | 0.4 | 1 | 1 | 6 | 0.8333 | 13 | 0.6154 | 3 | 0.3333 | 6 | 0.8333 | 5 | 0.4 | 2 | 0.5 | 16 | 0.5625 |
| 1_9372766 | NA | NA | 2 | 0.5 | 1 | 0 | NA | NA | 3 | 0.3333 | NA | NA | 1 | 0 | 2 | 0.5 | NA | NA | 3 | 0.3333 |
| 1_9374712 | NA | NA | 1 | 0 | 1 | 1 | 1 | 0 | 3 | 0.3333 | 1 | 1 | NA | NA | NA | NA | NA | NA | 1 | 1 |
| 1_9374822 | 1 | 0 | 1 | 1 | 1 | 0 | 2 | 0 | 5 | 0.2 | 1 | 1 | NA | NA | 1 | 0 | NA | NA | 2 | 0.5 |
| 1_9377849 | NA | NA | 1 | 0 | NA | NA | 2 | 0.5 | 3 | 0.3333 | 1 | 0 | NA | NA | 3 | 0.3333 | NA | NA | 4 | 0.25 |
| 1_9377852 | NA | NA | 1 | 1 | NA | NA | 2 | 1 | 3 | 1 | 1 | 1 | NA | NA | 2 | 0.5 | NA | NA | 3 | 0.6667 |
| 1_9377858 | NA | NA | 1 | 0 | NA | NA | 2 | 0.5 | 3 | 0.3333 | 1 | 0 | 1 | 1 | 3 | 0.3333 | NA | NA | 5 | 0.4 |
| 1_9377864 | NA | NA | 2 | 0.5 | NA | NA | 2 | 0 | 4 | 0.25 | 1 | 0 | 1 | 1 | 2 | 0.5 | NA | NA | 4 | 0.5 |
| 1_9377865 | NA | NA | 1 | 1 | NA | NA | 2 | 0.5 | 3 | 0.6667 | 1 | 1 | 1 | 1 | 3 | 1 | NA | NA | 5 | 1 |
| 1_9377869 | NA | NA | 2 | 1 | NA | NA | 2 | 0.5 | 4 | 0.75 | 1 | 0 | 1 | 1 | 2 | 0.5 | NA | NA | 4 | 0.5 |
| 1_9377917 | NA | NA | 2 | 0 | NA | NA | 2 | 0.5 | 4 | 0.25 | NA | NA | NA | NA | 3 | 0.3333 | 1 | 0 | 4 | 0.25 |
| 1_9377942 | NA | NA | 2 | 1 | NA | NA | 2 | 1 | 4 | 1 | 2 | 0.5 | 1 | 1 | 2 | 0.5 | 1 | 1 | 6 | 0.6667 |
| 1_9377949 | NA | NA | 2 | 0 | NA | NA | 2 | 0.5 | 4 | 0.25 | 2 | 0 | 1 | 0 | 3 | 0.3333 | 2 | 0 | 8 | 0.125 |
| 1_9377950 | NA | NA | NA | NA | NA | NA | 2 | 1 | 2 | 1 | 2 | 0 | 1 | 0 | 2 | 0.5 | 1 | 1 | 6 | 0.3333 |
| 1_9377964 | NA | NA | 2 | 0.5 | NA | NA | 1 | 1 | 3 | 0.6667 | 2 | 0.5 | 1 | 0 | 3 | 0.3333 | 2 | 1 | 8 | 0.5 |
| 1_9377978 | NA | NA | 2 | 0.5 | NA | NA | 2 | 0.5 | 4 | 0.5 | 2 | 0.5 | 1 | 0 | 3 | 0 | 2 | 0 | 8 | 0.125 |
| 1_9377985 | NA | NA | 2 | 0.5 | NA | NA | 2 | 0.5 | 4 | 0.5 | 1 | 1 | NA | NA | 2 | 1 | 2 | 0.5 | 5 | 0.8 |
| 1_9395239 | NA | NA | 1 | 1 | 2 | 0.5 | NA | NA | 3 | 0.6667 | NA | NA | 1 | 1 | NA | NA | NA | NA | 1 | 1 |
| 1_9395290 | 1 | 1 | 1 | 0 | 2 | 0 | NA | NA | 4 | 0.25 | NA | NA | 1 | 1 | NA | NA | NA | NA | 1 | 1 |
| 1_9395301 | 1 | 0 | 2 | 0 | 2 | 0.5 | NA | NA | 5 | 0.2 | NA | NA | 1 | 1 | NA | NA | NA | NA | 1 | 1 |
| 1_9395332 | 1 | 0 | 1 | 0 | 1 | 1 | NA | NA | 3 | 0.3333 | NA | NA | 1 | 1 | NA | NA | NA | NA | 1 | 1 |
| 1_9526184 | 18 | 0.2222 | 14 | 0 | 10 | 0.1 | 16 | 0 | 58 | 0.0862 | 6 | 0 | 12 | 0.0833 | 17 | 0.1176 | 22 | 0.1364 | 57 | 0.1053 |
| 1_9551546 | 1 | 1 | 1 | 0 | 2 | 0 | 2 | 0 | 6 | 0.1667 | 2 | 1 | 1 | 1 | 3 | 0 | 2 | 0.5 | 8 | 0.5 |
| 1_9551892 | 1 | 1 | 1 | 1 | NA | NA | NA | NA | 2 | 1 | NA | NA | NA | NA | 2 | 1 | 2 | 0.5 | 4 | 0.75 |
| 1_9551902 | NA | NA | 1 | 1 | NA | NA | NA | NA | 1 | 1 | NA | NA | NA | NA | 1 | 1 | 2 | 1 | 3 | 1 |
| 1_9551921 | 1 | 1 | 1 | 1 | NA | NA | NA | NA | 2 | 1 | NA | NA | NA | NA | 2 | 0.5 | 1 | 1 | 3 | 0.6667 |
| 1_9551929 | 1 | 1 | NA | NA | NA | NA | NA | NA | 1 | 1 | NA | NA | NA | NA | 2 | 1 | 1 | 1 | 3 | 1 |
| 1_9551938 | NA | NA | 1 | 1 | NA | NA | NA | NA | 1 | 1 | NA | NA | NA | NA | 2 | 1 | 2 | 1 | 4 | 1 |
| 1_9551947 | NA | NA | 1 | 1 | NA | NA | NA | NA | 1 | 1 | NA | NA | NA | NA | 2 | 0 | 2 | 0.5 | 4 | 0.25 |
| 1_9551953 | NA | NA | 1 | 1 | NA | NA | NA | NA | 1 | 1 | NA | NA | NA | NA | 2 | 1 | 2 | 1 | 4 | 1 |
| 1_9555527 | 1 | 1 | NA | NA | NA | NA | NA | NA | 1 | 1 | 1 | 0 | NA | NA | 2 | 0 | 2 | 1 | 5 | 0.4 |
| 1_9555538 | 1 | 1 | NA | NA | NA | NA | NA | NA | 1 | 1 | 1 | 0 | NA | NA | 1 | 1 | 2 | 0.5 | 4 | 0.5 |
| 1_9559500 | 1 | 1 | NA | NA | NA | NA | NA | NA | 1 | 1 | NA | NA | NA | NA | 3 | 0.6667 | 2 | 0.5 | 5 | 0.6 |
| 1_9674664 | NA | NA | NA | NA | 1 | 1 | NA | NA | 1 | 1 | 1 | 1 | 1 | 1 | NA | NA | NA | NA | 2 | 1 |
| 1_9674679 | NA | NA | NA | NA | 1 | 1 | NA | NA | 1 | 1 | NA | NA | 1 | 1 | NA | NA | NA | NA | 1 | 1 |

| | | | | | | | | | | | | | | | | | | | | |
|---|---|---|---|---|---|---|---|---|---|---|---|---|---|---|---|---|---|---|---|---|
| 1_9680845 | NA | NA | 1 | 1 | NA | NA | NA | NA | 1 | 1 | NA | NA | NA | NA | 1 | 1 | NA | NA | 1 | 1 |
| 1_9680842 | NA | NA | 1 | 1 | NA | NA | NA | NA | 1 | 1 | NA | NA | NA | NA | 1 | 1 | NA | NA | 1 | 1 |
| 1_9757196 | 6 | 0.1667 | 12 | 0.0833 | 2 | 0 | 3 | 0 | 23 | 0.087 | 1 | 1 | 7 | 0 | 3 | 0 | 8 | 0 | 19 | 0.0526 |
| 1_9757297 | 6 | 0.1667 | 5 | 0 | NA | NA | 3 | 0.3333 | 14 | 0.1429 | 3 | 0 | 5 | 0.2 | 3 | 0 | 8 | 0.125 | 19 | 0.1053 |
| 1_9757320 | 5 | 0.4 | 3 | 0.3333 | NA | NA | 2 | 0 | 10 | 0.3 | 3 | 0.3333 | 3 | 0.6667 | 1 | 0 | 8 | 0.125 | 15 | 0.2667 |
| 1_9757345 | 4 | 0 | 4 | 0.5 | 1 | 0 | 2 | 0 | 11 | 0.1818 | 2 | 1 | 5 | 0.2 | 1 | 0 | 5 | 0.2 | 13 | 0.3077 |
| 1_9757346 | 3 | 0 | 3 | 0.6667 | NA | NA | 2 | 0.5 | 8 | 0.375 | 3 | 0.6667 | 5 | 0.6 | 2 | 1 | 6 | 0.1667 | 16 | 0.5 |
| 1_9757353 | 3 | 0.3333 | 4 | 0.5 | NA | NA | 1 | 0 | 8 | 0.375 | 3 | 0.3333 | 4 | 0 | 1 | 1 | 4 | 0 | 12 | 0.1667 |
| 1_9757355 | 4 | 1 | 4 | 1 | NA | NA | 2 | 1 | 10 | 1 | 3 | 1 | 4 | 1 | 1 | 1 | 5 | 1 | 13 | 1 |
| 1_9757363 | 2 | 1 | 3 | 1 | 1 | 1 | 2 | 1 | 8 | 1 | 1 | 1 | 3 | 1 | 1 | 1 | 2 | 1 | 7 | 1 |
| 1_9757366 | 4 | 0 | 5 | 0.2 | NA | NA | 2 | 0 | 11 | 0.0909 | 3 | 0.3333 | 5 | 0.2 | 2 | 0 | 6 | 0 | 16 | 0.125 |
| 1_9757373 | 4 | 0 | 5 | 0.2 | 1 | 0 | 1 | 0 | 11 | 0.0909 | 2 | 0 | 5 | 0.4 | 2 | 0.5 | 6 | 0.3333 | 15 | 0.3333 |
| 1_9757376 | 4 | 1 | 5 | 1 | 1 | 1 | 2 | 1 | 12 | 1 | 2 | 1 | 5 | 1 | 2 | 1 | 5 | 1 | 14 | 1 |
| 1_9757377 | 5 | 0.8 | 4 | 1 | NA | NA | 2 | 0 | 11 | 0.7273 | 2 | 0.5 | 5 | 0.8 | 2 | 0.5 | 5 | 0.8 | 14 | 0.7143 |
| 1_9757385 | 4 | 0.75 | 5 | 0.2 | NA | NA | 1 | 0 | 10 | 0.4 | 2 | 0.5 | 4 | 0.75 | 2 | 0 | 6 | 0.3333 | 14 | 0.4286 |
| 1_9757389 | 4 | 1 | 4 | 1 | 1 | 1 | 2 | 1 | 11 | 1 | 2 | 1 | 5 | 1 | 1 | 1 | 4 | 1 | 12 | 1 |
| 1_9757392 | 5 | 1 | 5 | 1 | NA | NA | 1 | 1 | 11 | 1 | 2 | 1 | 5 | 1 | 2 | 1 | 6 | 0.6667 | 15 | 0.8667 |
| 1_9758863 | NA | NA | 1 | 1 | NA | NA | 1 | 1 | 2 | 1 | NA | NA | NA | NA | 1 | 1 | NA | NA | 1 | 1 |
| 1_9758871 | 1 | 1 | 1 | 1 | NA | NA | 1 | 1 | 3 | 1 | NA | NA | NA | NA | 2 | 1 | NA | NA | 2 | 1 |
| 1_9763414 | 1 | 0 | 3 | 0 | 4 | 0 | 5 | 0.2 | 13 | 0.0769 | 2 | 0 | 2 | 0 | 3 | 0.6667 | 3 | 0 | 10 | 0.2 |
| 1_9763437 | 1 | 0 | 3 | 0.3333 | 2 | 0.5 | 4 | 0 | 10 | 0.2 | 2 | 0 | 3 | 0 | 3 | 0.3333 | 4 | 0.25 | 12 | 0.1667 |
| 1_9763459 | 1 | 0 | 3 | 0.3333 | 4 | 0 | 5 | 0 | 13 | 0.0769 | 2 | 0.5 | 4 | 0 | 3 | 0 | 5 | 0 | 14 | 0.0714 |
| 1_9763500 | 1 | 0 | 3 | 0 | 4 | 0 | 5 | 0.2 | 13 | 0.0769 | 2 | 0.5 | 4 | 0 | 3 | 0 | 4 | 0 | 13 | 0.0769 |
| 1_9763512 | 1 | 0 | 3 | 0.3333 | 4 | 0 | 3 | 0 | 11 | 0.0909 | 2 | 0 | 4 | 0.25 | 3 | 0.3333 | 3 | 0 | 12 | 0.1667 |
| 1_9763513 | 1 | 0 | 3 | 0 | 4 | 0.25 | 3 | 0 | 11 | 0.0909 | 2 | 0 | 3 | 0 | 3 | 0 | 4 | 0.25 | 12 | 0.0833 |
| 1_9763536 | 1 | 0 | 3 | 0 | 3 | 0 | 3 | 0.3333 | 10 | 0.1 | 2 | 0.5 | 4 | 0.25 | 3 | 0 | 3 | 0 | 12 | 0.1667 |
| 1_9763540 | 1 | 1 | 3 | 0.6667 | 4 | 1 | 4 | 0.75 | 12 | 0.8333 | 2 | 0.5 | 4 | 0.75 | 2 | 0.5 | 4 | 0.75 | 12 | 0.6667 |
| 1_9768929 | 1 | 0 | NA | NA | 2 | 1 | NA | NA | 3 | 0.6667 | 1 | 0 | 1 | 1 | NA | NA | NA | NA | 2 | 0.5 |
| 1_9771372 | 1 | 0 | 1 | 0 | 1 | 1 | 3 | 0 | 6 | 0.1667 | 1 | 0 | 1 | 0 | NA | NA | 1 | 1 | 3 | 0.3333 |
| 1_9772015 | 2 | 0.5 | 1 | 1 | 1 | 1 | 3 | 1 | 7 | 0.8571 | 1 | 1 | 2 | 0.5 | NA | NA | 2 | 1 | 5 | 0.8 |
| 1_9772017 | NA | NA | 1 | 1 | 1 | 1 | 2 | 1 | 4 | 1 | 1 | 1 | 1 | 1 | NA | NA | 2 | 0.5 | 4 | 0.75 |
| 1_9772024 | 2 | 0.5 | 1 | 0 | 1 | 0 | 4 | 1 | 8 | 0.625 | 1 | 1 | 2 | 0.5 | NA | NA | 2 | 0 | 5 | 0.4 |
| 1_9774941 | 16 | 0.0625 | 31 | 0.1935 | 6 | 0.1667 | 15 | 0.1333 | 68 | 0.1471 | 18 | 0.2778 | 12 | 0.25 | 11 | 0.5455 | 18 | 0.1667 | 59 | 0.2881 |
| 1_9775368 | 10 | 0.1 | 21 | 0.1429 | 5 | 0 | 12 | 0.0833 | 48 | 0.1042 | 16 | 0.125 | 8 | 0.25 | 9 | 0 | 15 | 0.0667 | 48 | 0.1042 |
| 1_9779210 | 5 | 0.2 | 13 | 0.0769 | 4 | 0.5 | 9 | 0 | 31 | 0.129 | 8 | 0.25 | 5 | 0.2 | 4 | 0 | 11 | 0 | 28 | 0.1071 |
| 1_1004178 | 3 | 0.3333 | 3 | 0 | 2 | 0.5 | 1 | 0 | 9 | 0.2222 | NA | NA | NA | NA | 1 | 0 | 3 | 0.6667 | 4 | 0.5 |
| 1_1048618 | NA | NA | 1 | 1 | NA | NA | NA | NA | 1 | 1 | NA | NA | NA | NA | 1 | 0 | 1 | 1 | 2 | 0.5 |
| 1_1048618 | NA | NA | 1 | 1 | NA | NA | NA | NA | 1 | 1 | NA | NA | NA | NA | 1 | 1 | NA | NA | 1 | 1 |
| 1_1049216 | NA | NA | NA | NA | NA | NA | 1 | 1 | 1 | 1 | NA | NA | NA | NA | 1 | 0 | 1 | 1 | 2 | 0.5 |
| 1_1049217 | NA | NA | NA | NA | NA | NA | 1 | 1 | 1 | 1 | NA | NA | NA | NA | 1 | 1 | 1 | 0 | 2 | 0.5 |
| 1_1049219 | NA | NA | 1 | 1 | NA | NA | 1 | 1 | 2 | 1 | NA | NA | NA | NA | 1 | 1 | 1 | 1 | 2 | 1 |
| 1_1049220 | NA | NA | 1 | 1 | NA | NA | 1 | 1 | 2 | 1 | NA | NA | NA | NA | 1 | 1 | 1 | 1 | 2 | 1 |
| 1_1049220 | NA | NA | 1 | 1 | NA | NA | 1 | 1 | 2 | 1 | NA | NA | NA | NA | NA | NA | 1 | 1 | 1 | 1 |
| 1_1049220 | NA | NA | 1 | 1 | NA | NA | NA | NA | 1 | 1 | NA | NA | NA | NA | 1 | 1 | NA | NA | 1 | 1 |
| 1_1052056 | 2 | 1 | NA | NA | NA | NA | 1 | 0 | 3 | 0.6667 | NA | NA | 1 | 1 | 1 | 1 | NA | NA | 2 | 1 |
| 1_1052117 | 1 | 1 | NA | NA | NA | NA | NA | NA | 1 | 1 | NA | NA | NA | NA | NA | NA | 1 | 1 | 1 | 1 |
| 1_1052118 | 1 | 1 | NA | NA | NA | NA | NA | NA | 1 | 1 | NA | NA | NA | NA | NA | NA | 1 | 1 | 1 | 1 |
| 1_1052122 | 1 | 1 | NA | NA | NA | NA | NA | NA | 1 | 1 | NA | NA | NA | NA | NA | NA | 1 | 1 | 1 | 1 |
| 1_1054495 | 2 | 0.5 | 1 | 1 | 1 | 1 | 2 | 1 | 6 | 0.8333 | NA | NA | NA | NA | 1 | 1 | 1 | 1 | 2 | 1 |
| 1_1054496 | 2 | 0 | 1 | 0 | 1 | 1 | 2 | 1 | 6 | 0.5 | NA | NA | NA | NA | 1 | 0 | 1 | 1 | 2 | 0.5 |
| 1_1054497 | 2 | 0 | NA | NA | 1 | 1 | 1 | 0 | 4 | 0.25 | NA | NA | NA | NA | 1 | 0 | 1 | 1 | 2 | 0.5 |
| 1_1055917 | 2 | 0 | 5 | 0.2 | 1 | 0 | NA | NA | 8 | 0.125 | NA | NA | 2 | 0.5 | 1 | 0 | 1 | 0 | 4 | 0.25 |
| 1_1055919 | 2 | 1 | 3 | 0 | NA | NA | NA | NA | 5 | 0.4 | NA | NA | 3 | 0.3333 | NA | NA | 2 | 0.5 | 5 | 0.4 |
| 1_1056681 | 2 | 0 | 5 | 0 | 2 | 0 | 4 | 0.25 | 13 | 0.0769 | NA | NA | 6 | 0.1667 | 1 | 0 | 7 | 0 | 14 | 0.0714 |
| 1_1056812 | 2 | 0 | 5 | 0 | 2 | 0.5 | 4 | 0 | 13 | 0.0769 | NA | NA | 5 | 0 | 1 | 0 | 5 | 0.2 | 11 | 0.0909 |
| 1_1056813 | 3 | 0 | 7 | 0.1429 | 3 | 0.3333 | 3 | 0 | 16 | 0.125 | 1 | 1 | 8 | 0.375 | 4 | 0 | 9 | 0.2222 | 22 | 0.2727 |
| 1_1056813 | 4 | 0 | 8 | 0.25 | 3 | 0.3333 | 5 | 0 | 20 | 0.15 | 1 | 0 | 9 | 0.1111 | 4 | 0 | 9 | 0.1111 | 23 | 0.087 |
| 1_1056813 | 4 | 0.5 | 8 | 0.125 | 4 | 0.25 | 5 | 0 | 21 | 0.1905 | 1 | 0 | 10 | 0.1 | 3 | 0.6667 | 7 | 0.2857 | 21 | 0.2381 |
| 1_1057551 | NA | NA | 1 | 1 | NA | NA | NA | NA | 1 | 1 | 1 | 1 | NA | NA | NA | NA | NA | NA | 1 | 1 |
| 1_1057552 | NA | NA | 1 | 1 | NA | NA | NA | NA | 1 | 1 | 1 | 1 | NA | NA | NA | NA | NA | NA | 1 | 1 |
| 1_1057724 | NA | NA | 1 | 1 | NA | NA | NA | NA | 1 | 1 | 1 | 1 | NA | NA | NA | NA | NA | NA | 1 | 1 |
| 1_1057725 | NA | NA | 1 | 1 | NA | NA | NA | NA | 1 | 1 | 1 | 1 | NA | NA | NA | NA | NA | NA | 1 | 1 |
| 1_1059581 | 1 | 1 | 1 | 1 | 1 | 1 | 2 | 1 | 5 | 1 | 4 | 1 | NA | NA | NA | NA | 1 | 1 | 5 | 1 |
| 1_1059584 | NA | NA | 2 | 1 | 2 | 1 | 2 | 0.5 | 6 | 0.8333 | 3 | 1 | 1 | 0 | 1 | 1 | 1 | 1 | 6 | 0.8333 |
| 1_1059585 | NA | NA | 2 | 0.5 | 2 | 0.5 | 2 | 0 | 6 | 0.3333 | 4 | 0.75 | 1 | 0 | 1 | 1 | 1 | 0 | 7 | 0.5714 |

| | | | | | | | | | | | | | | | | | | | | |
|---|---|---|---|---|---|---|---|---|---|---|---|---|---|---|---|---|---|---|---|---|
| 1_10595861 | NA | NA | 2 | 1 | 2 | 1 | 1 | 1 | 5 | 1 | 4 | 1 | 1 | 0 | 1 | 1 | 1 | 1 | 7 | 0.8571 |
| 1_10595872 | 1 | 1 | 2 | 1 | 2 | 1 | 2 | 1 | 7 | 1 | 4 | 1 | 2 | 1 | 1 | 1 | 1 | 1 | 8 | 1 |
| 1_10595874 | 1 | 1 | 2 | 1 | 2 | 1 | 2 | 1 | 7 | 1 | 4 | 1 | 2 | 1 | 1 | 1 | NA | NA | 7 | 1 |
| 1_10595891 | 1 | 0 | 2 | 1 | 2 | 1 | 2 | 1 | 7 | 0.8571 | 1 | 1 | 2 | 0.5 | 1 | 1 | 1 | 1 | 5 | 0.8 |
| 1_10595892 | 1 | 1 | 2 | 0.5 | 2 | 1 | 2 | 1 | 7 | 0.8571 | 2 | 1 | 2 | 1 | 1 | 1 | 1 | 1 | 6 | 1 |
| 1_10595898 | 1 | 1 | NA | NA | NA | NA | 1 | 0 | 2 | 0.5 | 3 | 1 | 1 | 1 | NA | NA | 1 | 1 | 5 | 1 |
| 1_10595988 | 2 | 0 | 2 | 0.5 | 2 | 0 | 2 | 0.5 | 8 | 0.25 | 4 | 0 | 2 | 0 | 1 | 1 | 1 | 0 | 8 | 0.125 |
| 1_10596011 | 1 | 0 | 2 | 0 | 2 | 0.5 | 2 | 0 | 7 | 0.1429 | 3 | 0.3333 | 2 | 0 | 1 | 0 | 1 | 0 | 7 | 0.1429 |
| 1_10596013 | NA | NA | 2 | 0.5 | 2 | 0.5 | 2 | 0 | 6 | 0.3333 | 3 | 0 | 2 | 1 | NA | NA | NA | NA | 5 | 0.4 |
| 1_10596055 | NA | NA | 2 | 0.5 | 2 | 0.5 | 2 | 0 | 6 | 0.3333 | 4 | 0.5 | 2 | 0 | 1 | 0 | 1 | 1 | 8 | 0.375 |
| 1_10596061 | NA | NA | 1 | 1 | 2 | 1 | 2 | 0 | 5 | 0.6 | 3 | 0.6667 | 1 | 0 | 1 | 1 | 1 | 1 | 6 | 0.6667 |
| 1_10596062 | NA | NA | 2 | 0 | 2 | 0 | 2 | 0.5 | 6 | 0.1667 | 4 | 0.25 | 2 | 0 | 1 | 1 | 1 | 0 | 8 | 0.25 |
| 1_10596091 | 1 | 1 | 2 | 0.5 | 1 | 0 | 2 | 0 | 6 | 0.3333 | 4 | 0.25 | 2 | 0 | 1 | 1 | 1 | 0 | 8 | 0.25 |
| 1_10596120 | 1 | 0 | 2 | 0.5 | 2 | 0 | 2 | 0 | 7 | 0.1429 | 3 | 0 | 2 | 0.5 | NA | NA | 1 | 1 | 6 | 0.3333 |
| 1_10609778 | NA | NA | 1 | 1 | NA | NA | NA | NA | 1 | 1 | NA | NA | NA | NA | 1 | 1 | NA | NA | 1 | 1 |
| 1_10610017 | 3 | 0.3333 | NA | NA | 1 | 1 | 1 | 0 | 5 | 0.4 | 2 | 0 | NA | NA | 1 | 1 | 2 | 0 | 5 | 0.2 |
| 1_10610023 | 4 | 0.5 | NA | NA | 1 | 1 | 4 | 0.5 | 9 | 0.5556 | 4 | 0.75 | 1 | 1 | 1 | 1 | 2 | 1 | 8 | 0.875 |
| 1_10610032 | 3 | 0 | NA | NA | NA | NA | 4 | 0.25 | 7 | 0.1429 | 3 | 0.3333 | 1 | 0 | 1 | 0 | 2 | 0.5 | 7 | 0.2857 |
| 1_10610060 | 4 | 1 | NA | NA | 1 | 1 | 3 | 1 | 8 | 1 | 2 | 0.5 | 1 | 0 | 2 | 1 | 1 | 1 | 6 | 0.6667 |
| 1_10610072 | 5 | 0 | NA | NA | 2 | 0.5 | 5 | 0 | 12 | 0.0833 | 4 | 0.25 | 2 | 0.5 | 2 | 0 | 1 | 0 | 9 | 0.2222 |
| 1_10610099 | 4 | 0.25 | 1 | 0 | 3 | 0 | 7 | 0.1429 | 15 | 0.1333 | 5 | 0.4 | 1 | 0 | 2 | 0 | 3 | 0.3333 | 11 | 0.2727 |
| 1_10610108 | 5 | 0.2 | NA | NA | 2 | 0 | 8 | 0.25 | 15 | 0.2 | 5 | 0 | 2 | 0 | 3 | 0.3333 | 3 | 0 | 13 | 0.0769 |
| 1_10610114 | 4 | 0.25 | 1 | 0 | 2 | 0 | 7 | 0.2857 | 14 | 0.2143 | 3 | 0 | 1 | 0 | 3 | 0.6667 | 3 | 0.6667 | 10 | 0.4 |
| 1_10631996 | 3 | 0 | 3 | 0 | 4 | 0.25 | 10 | 0.1 | 20 | 0.1 | 4 | 0.25 | 6 | 0 | 8 | 0.125 | 4 | 0.25 | 22 | 0.1364 |
| 1_10632000 | 4 | 0.25 | 5 | 0.2 | 7 | 0.4286 | 17 | 0.3529 | 33 | 0.3333 | 6 | 0.3333 | 8 | 0.375 | 9 | 0.1111 | 5 | 0.6 | 28 | 0.3214 |
| 1_10632015 | 4 | 0 | 5 | 0.2 | 9 | 0.1111 | 14 | 0.1429 | 32 | 0.125 | 6 | 0.1667 | 9 | 0.1111 | 9 | 0 | 6 | 0.1667 | 30 | 0.1 |
| 1_10632024 | 5 | 0.8 | 6 | 0.8333 | 10 | 0.6 | 14 | 0.6429 | 35 | 0.6857 | 4 | 0.5 | 9 | 0.4444 | 11 | 0.6364 | 4 | 0.25 | 28 | 0.5 |
| 1_10632031 | 6 | 1 | 5 | 1 | 8 | 1 | 14 | 1 | 33 | 1 | 6 | 1 | 8 | 1 | 9 | 1 | 6 | 1 | 29 | 1 |
| 1_10632051 | 6 | 0.1667 | 5 | 0.2 | 8 | 0.125 | 17 | 0.1765 | 36 | 0.1667 | 4 | 0 | 8 | 0.25 | 8 | 0 | 7 | 0.1429 | 27 | 0.1111 |
| 1_10632052 | 5 | 0 | 6 | 0.1667 | 10 | 0.2 | 14 | 0.0714 | 35 | 0.1143 | 5 | 0.2 | 7 | 0.1429 | 11 | 0 | 5 | 0 | 28 | 0.0714 |
| 1_10632057 | 5 | 0 | 6 | 0.3333 | 9 | 0.2222 | 16 | 0.0625 | 36 | 0.1389 | 6 | 0 | 11 | 0.0909 | 12 | 0.1667 | 7 | 0 | 36 | 0.0833 |
| 1_10632061 | 4 | 0.5 | 4 | 0.75 | 5 | 0.2 | 12 | 0.6667 | 25 | 0.56 | 4 | 0.75 | 8 | 0.5 | 13 | 0.4615 | 6 | 0.8333 | 31 | 0.5806 |
| 1_10632070 | 6 | 0.1667 | 6 | 0 | 8 | 0.125 | 15 | 0.4 | 35 | 0.2286 | 5 | 0 | 11 | 0.2727 | 12 | 0 | 7 | 0.4286 | 35 | 0.1714 |
| 1_10632072 | 6 | 0.8333 | 6 | 0.5 | 9 | 0.5556 | 13 | 0.5385 | 34 | 0.5882 | 5 | 0.4 | 9 | 0.7778 | 11 | 0.3636 | 7 | 0.4286 | 32 | 0.5 |
| 1_10632104 | 4 | 1 | 6 | 0.8333 | 10 | 1 | 17 | 0.8824 | 37 | 0.9189 | 6 | 0.8333 | 13 | 0.8462 | 12 | 0.8333 | 7 | 1 | 38 | 0.8684 |
| 1_10632109 | 2 | 0 | 4 | 0.25 | 7 | 0.1429 | 14 | 0 | 27 | 0.0741 | 4 | 0.5 | 9 | 0.3333 | 9 | 0.1111 | 3 | 0 | 25 | 0.24 |
| 1_10632110 | 3 | 0.3333 | 1 | 1 | 3 | 0.3333 | 5 | 0.6 | 12 | 0.5 | 1 | 1 | 7 | 0 | 2 | 0.5 | 3 | 0 | 13 | 0.1538 |
| 1_10632111 | 2 | 1 | 5 | 1 | 9 | 1 | 8 | 0.875 | 24 | 0.9583 | 4 | 0.75 | 8 | 1 | 6 | 1 | 4 | 1 | 22 | 0.9545 |
| 1_10632117 | 7 | 0.2857 | 7 | 0.1429 | 11 | 0.1818 | 18 | 0.2778 | 43 | 0.2326 | 5 | 0 | 13 | 0.1538 | 12 | 0.1667 | 6 | 0.1667 | 36 | 0.1389 |
| 1_10632159 | 6 | 0 | 8 | 0.125 | 9 | 0 | 17 | 0.0588 | 40 | 0.05 | 8 | 0.125 | 10 | 0.1 | 14 | 0.0714 | 7 | 0.1429 | 39 | 0.1026 |
| 1_10632167 | 8 | 0.125 | 9 | 0.1111 | 11 | 0.0909 | 17 | 0.1765 | 45 | 0.1333 | 8 | 0 | 14 | 0.2143 | 13 | 0.0769 | 7 | 0.1429 | 42 | 0.119 |
| 1_10632182 | 7 | 0.4286 | 10 | 0.4 | 10 | 0.4 | 16 | 0.4375 | 43 | 0.4186 | 9 | 0.3333 | 15 | 0.3333 | 14 | 0.4286 | 8 | 0.125 | 46 | 0.3261 |
| 1_10632193 | 6 | 0.3333 | 8 | 0 | 11 | 0 | 13 | 0 | 38 | 0.0526 | 6 | 0.1667 | 13 | 0 | 12 | 0.1667 | 8 | 0.375 | 39 | 0.1538 |
| 1_10646670 | 6 | 0.3333 | NA | NA | 3 | 0.3333 | NA | NA | 9 | 0.3333 | 2 | 0 | 3 | 0 | 5 | 0.4 | 2 | 0 | 12 | 0.1667 |
| 1_10646711 | 6 | 0 | 1 | 0 | 2 | 0.5 | NA | NA | 9 | 0.1111 | 1 | 0 | 1 | 0 | 4 | 0 | 3 | 0.3333 | 9 | 0.1111 |
| 1_10646899 | 1 | 0 | 1 | 0 | 2 | 0.5 | NA | NA | 4 | 0.25 | NA | NA | 2 | 0 | 1 | 1 | 1 | 0 | 4 | 0.25 |
| 1_10646981 | 1 | 1 | NA | NA | 2 | 1 | NA | NA | 3 | 1 | NA | NA | 1 | 1 | 1 | 1 | NA | NA | 2 | 1 |
| 1_10646983 | 1 | 0 | NA | NA | 2 | 1 | NA | NA | 3 | 0.6667 | NA | NA | 1 | 1 | 1 | 0 | NA | NA | 2 | 0.5 |
| 1_10646989 | 2 | 1 | NA | NA | 1 | 1 | NA | NA | 3 | 1 | NA | NA | NA | NA | 1 | 1 | NA | NA | 1 | 1 |
| 1_10646992 | 1 | 1 | NA | NA | 2 | 1 | NA | NA | 3 | 1 | NA | NA | NA | NA | 1 | 1 | NA | NA | 1 | 1 |
| 1_10647022 | 1 | 1 | NA | NA | 2 | 0.5 | NA | NA | 3 | 0.6667 | NA | NA | NA | NA | 1 | 1 | NA | NA | 1 | 1 |
| 1_10647033 | 2 | 1 | NA | NA | 1 | 1 | NA | NA | 3 | 1 | NA | NA | 1 | 1 | NA | NA | NA | NA | 1 | 1 |
| 1_10647034 | 2 | 1 | NA | NA | 1 | 0 | NA | NA | 3 | 0.6667 | NA | NA | 1 | 1 | NA | NA | NA | NA | 1 | 1 |
| 1_10647036 | NA | NA | NA | NA | 2 | 1 | NA | NA | 2 | 1 | NA | NA | 1 | 1 | NA | NA | NA | NA | 1 | 1 |
| 1_10647043 | 2 | 0.5 | NA | NA | 2 | 1 | NA | NA | 4 | 0.75 | NA | NA | 1 | 1 | NA | NA | NA | NA | 1 | 1 |
| 1_10664021 | 1 | 0 | NA | NA | 1 | 1 | NA | NA | 2 | 0.5 | NA | NA | 1 | 1 | NA | NA | NA | NA | 1 | 1 |
| 1_10664065 | NA | NA | NA | NA | 2 | 1 | NA | NA | 2 | 1 | NA | NA | 1 | 1 | NA | NA | NA | NA | 1 | 1 |
| 1_10664088 | NA | NA | NA | NA | 1 | 1 | NA | NA | 1 | 1 | 1 | 1 | 1 | 1 | NA | NA | NA | NA | 2 | 1 |
| 1_10664132 | NA | NA | NA | NA | 1 | 1 | NA | NA | 1 | 1 | 1 | 1 | 1 | 0 | NA | NA | NA | NA | 2 | 0.5 |
| 1_10664134 | NA | NA | NA | NA | 1 | 1 | NA | NA | 1 | 1 | 1 | 0 | 1 | 1 | NA | NA | NA | NA | 2 | 0.5 |
| 1_10664144 | NA | NA | NA | NA | 1 | 1 | NA | NA | 1 | 1 | 1 | 1 | 1 | 1 | NA | NA | NA | NA | 2 | 1 |
| 1_10664916 | 2 | 0 | 2 | 0 | 2 | 0 | 4 | 0.25 | 10 | 0.1 | 2 | 0 | 2 | 0 | 1 | 0 | 3 | 0.3333 | 8 | 0.125 |
| 1_10665206 | 3 | 0 | 1 | 0 | 3 | 0 | 6 | 0.1667 | 13 | 0.0769 | 3 | 0.3333 | 2 | 0.5 | 2 | 0 | 3 | 0 | 10 | 0.2 |
| 1_10665242 | 2 | 0.5 | 1 | 0 | 1 | 0 | 5 | 0.4 | 9 | 0.3333 | 1 | 0 | 3 | 0.3333 | 2 | 0.5 | 4 | 0.25 | 10 | 0.3 |
| 1_10665247 | 2 | 0.5 | 2 | 0.5 | 2 | 0 | 5 | 0.8 | 11 | 0.5455 | 1 | 0 | 3 | 0.6667 | 2 | 0.5 | 4 | 0.75 | 10 | 0.6 |

| | | | | | | | | | | | | | | | | | | | | |
|---|---|---|---|---|---|---|---|---|---|---|---|---|---|---|---|---|---|---|---|---|
| 1_1066525 | 2 | 0 | 4 | 0 | 3 | 0.6667 | 5 | 0.2 | 14 | 0.2143 | 1 | 0 | 3 | 0.3333 | 2 | 0 | 4 | 0.25 | 10 | 0.2 |
| 1_1066525 | 2 | 1 | 4 | 0.25 | 3 | 0.3333 | 4 | 0.5 | 13 | 0.4615 | 1 | 0 | 2 | 0.5 | 2 | 1 | 3 | 0.6667 | 8 | 0.625 |
| 1_1066525 | 2 | 0 | 3 | 0.3333 | 3 | 0.3333 | 5 | 0 | 13 | 0.1538 | NA | NA | 2 | 0 | 3 | 0 | 3 | 0.6667 | 8 | 0.25 |
| 1_1066526 | 2 | 1 | 5 | 0.8 | 3 | 0.6667 | 5 | 1 | 15 | 0.8667 | 1 | 1 | 3 | 1 | 3 | 0.6667 | 4 | 1 | 11 | 0.9091 |
| 1_1066526 | 2 | 0 | 4 | 0.25 | 2 | 0.5 | 5 | 0.4 | 13 | 0.3077 | 1 | 0 | 3 | 0 | 2 | 0 | 4 | 0.25 | 10 | 0.1 |
| 1_1066935 | 1 | 1 | NA | NA | 1 | 1 | 1 | 0 | 3 | 0.6667 | NA | NA | 1 | 1 | NA | NA | NA | NA | 1 | 1 |
| 1_1066940 | 1 | 1 | NA | NA | 1 | 1 | NA | NA | 2 | 1 | NA | NA | 1 | 1 | NA | NA | NA | NA | 1 | 1 |
| 1_1066991 | 6 | 0.1667 | 7 | 0.1429 | NA | NA | 7 | 0.7143 | 20 | 0.35 | 4 | 0.5 | 4 | 0 | 1 | 0 | 3 | 0.3333 | 12 | 0.25 |
| 1_1066992 | 6 | 0 | 5 | 0.2 | 2 | 0 | 7 | 0 | 20 | 0.05 | 3 | 0 | 3 | 0 | 3 | 0 | 4 | 0.25 | 13 | 0.0769 |
| 1_1066994 | 6 | 0.5 | 4 | 0.5 | 3 | 0.6667 | 6 | 0.5 | 19 | 0.5263 | 2 | 1 | 4 | 0.75 | 4 | 0.5 | 3 | 1 | 13 | 0.7692 |
| 1_1066995 | 5 | 0 | 5 | 0.2 | 2 | 0 | 7 | 0 | 19 | 0.0526 | 2 | 0 | 4 | 0.25 | 4 | 0.25 | 4 | 0 | 14 | 0.1429 |
| 1_1066999 | 5 | 0.2 | 5 | 0.6 | 1 | 0 | 5 | 0.4 | 16 | 0.375 | 2 | 1 | 3 | 0 | 4 | 0.25 | 3 | 0.6667 | 12 | 0.4167 |
| 1_1066999 | 4 | 0.25 | 5 | 0.2 | 1 | 0 | 4 | 0 | 14 | 0.1429 | 1 | 0 | 3 | 0 | 3 | 0 | 3 | 0.3333 | 10 | 0.1 |
| 1_1067000 | 4 | 0.5 | 4 | 0 | 1 | 0 | 4 | 0 | 13 | 0.1538 | 1 | 0 | 3 | 0 | 2 | 0 | 4 | 0.25 | 10 | 0.1 |
| 1_1067001 | 5 | 0.2 | 3 | 0 | NA | NA | 5 | 0 | 13 | 0.0769 | 2 | 0 | 3 | 0 | 3 | 0 | 4 | 0.25 | 12 | 0.0833 |
| 1_1067001 | 5 | 0.2 | 5 | 0 | 1 | 0 | 5 | 0 | 16 | 0.0625 | 2 | 0 | 4 | 0.25 | 3 | 0 | 4 | 0.25 | 13 | 0.1538 |
| 1_1067001 | 5 | 0.2 | 3 | 0.3333 | 1 | 0 | 5 | 0.2 | 14 | 0.2143 | 1 | 0 | 4 | 0 | 3 | 0 | 4 | 0.75 | 12 | 0.25 |
| 1_1067002 | 4 | 0 | 5 | 0 | 1 | 0 | 5 | 0.6 | 15 | 0.2 | 2 | 0.5 | 4 | 0 | 3 | 0.3333 | 4 | 0.5 | 13 | 0.3077 |
| 1_1099113 | 1 | 1 | NA | NA | NA | NA | NA | NA | 1 | 1 | NA | NA | 1 | 0 | NA | NA | 2 | 1 | 3 | 0.6667 |
| 1_1099114 | 1 | 1 | NA | NA | NA | NA | NA | NA | 1 | 1 | NA | NA | 1 | 0 | NA | NA | 2 | 0.5 | 3 | 0.3333 |
| 1_1099115 | 1 | 1 | NA | NA | NA | NA | NA | NA | 1 | 1 | NA | NA | 1 | 0 | NA | NA | 3 | 0.6667 | 4 | 0.5 |
| 1_1099218 | 1 | 1 | NA | NA | NA | NA | NA | NA | 1 | 1 | NA | NA | 1 | 1 | NA | NA | 3 | 1 | 4 | 1 |
| 1_1099218 | 1 | 1 | NA | NA | NA | NA | NA | NA | 1 | 1 | NA | NA | 1 | 1 | 1 | 1 | 3 | 1 | 5 | 1 |
| 1_1099220 | 1 | 1 | NA | NA | NA | NA | NA | NA | 1 | 1 | NA | NA | NA | NA | 1 | 1 | 3 | 0.3333 | 4 | 0.5 |
| 1_1099259 | 1 | 0 | 1 | 0 | 1 | 1 | 2 | 0 | 5 | 0.2 | NA | NA | 1 | 0 | 1 | 0 | 3 | 0.3333 | 5 | 0.2 |
| 1_1099260 | 1 | 0 | 1 | 1 | NA | NA | 2 | 1 | 4 | 0.75 | NA | NA | 1 | 1 | NA | NA | 4 | 0.75 | 5 | 0.8 |
| 1_1099261 | 1 | 1 | 1 | 0 | 1 | 0 | 2 | 0 | 5 | 0.2 | NA | NA | 1 | 0 | 1 | 0 | 4 | 0.25 | 6 | 0.1667 |
| 1_1099262 | 1 | 0 | 1 | 1 | 1 | 0 | 2 | 1 | 5 | 0.6 | NA | NA | 1 | 1 | 1 | 0 | 4 | 0.5 | 6 | 0.5 |
| 1_1108166 | 1 | 0 | NA | NA | 3 | 0.6667 | 6 | 0 | 10 | 0.2 | 1 | 1 | NA | NA | 1 | 1 | 3 | 0.3333 | 5 | 0.6 |
| 1_1108166 | 1 | 0 | NA | NA | 3 | 0.3333 | 4 | 0 | 8 | 0.125 | 1 | 0 | 1 | 1 | 1 | 0 | 2 | 0.5 | 5 | 0.4 |
| 1_1108171 | 1 | 0 | NA | NA | 3 | 0.3333 | 3 | 0.3333 | 7 | 0.2857 | 1 | 0 | 1 | 1 | 1 | 0 | 2 | 0 | 5 | 0.2 |
| 1_1108174 | 1 | 1 | NA | NA | 1 | 1 | 4 | 0.75 | 6 | 0.8333 | NA | NA | 1 | 1 | 1 | 1 | 2 | 1 | 4 | 1 |
| 1_1108175 | 1 | 1 | NA | NA | 2 | 1 | 4 | 0.5 | 7 | 0.7143 | NA | NA | 1 | 0 | 1 | 1 | 2 | 0 | 4 | 0.25 |
| 1_1108824 | NA | NA | NA | NA | 1 | 1 | 1 | 0 | 2 | 0.5 | NA | NA | 1 | 1 | 1 | 0 | 1 | 0 | 3 | 0.3333 |
| 1_1117865 | NA | NA | 3 | 1 | NA | NA | 5 | 0.4 | 8 | 0.625 | NA | NA | 2 | 1 | 3 | 0.3333 | 1 | 1 | 6 | 0.6667 |
| 1_1117865 | NA | NA | 3 | 0 | NA | NA | 5 | 0.4 | 8 | 0.25 | NA | NA | 3 | 0 | 3 | 0.3333 | 1 | 0 | 7 | 0.1429 |
| 1_1117866 | NA | NA | 3 | 1 | NA | NA | 5 | 1 | 8 | 1 | NA | NA | 3 | 0.3333 | 3 | 0.3333 | 1 | 1 | 7 | 0.4286 |
| 1_1117868 | NA | NA | 3 | 0.3333 | NA | NA | 4 | 0.75 | 7 | 0.5714 | NA | NA | 3 | 0 | NA | NA | 1 | 1 | 4 | 0.25 |
| 1_1117869 | NA | NA | 2 | 1 | NA | NA | 4 | 1 | 6 | 1 | NA | NA | 3 | 1 | 2 | 1 | 1 | 1 | 6 | 1 |
| 1_1117869 | NA | NA | 3 | 0.6667 | NA | NA | 4 | 1 | 7 | 0.8571 | NA | NA | 3 | 1 | 2 | 1 | 1 | 1 | 6 | 1 |
| 1_1117869 | NA | NA | 3 | 0.6667 | NA | NA | 3 | 0.3333 | 6 | 0.5 | NA | NA | 2 | 0 | 2 | 1 | 1 | 1 | 5 | 0.6 |
| 1_1117874 | NA | NA | 3 | 0 | NA | NA | 4 | 0.5 | 7 | 0.2857 | NA | NA | 3 | 0.3333 | 2 | 0 | 1 | 0 | 6 | 0.1667 |
| 1_1117875 | NA | NA | 3 | 1 | NA | NA | 2 | 0.5 | 5 | 0.8 | NA | NA | 2 | 1 | 1 | 0 | 1 | 1 | 4 | 0.75 |
| 1_1117876 | NA | NA | 3 | 0.3333 | NA | NA | 4 | 1 | 7 | 0.7143 | NA | NA | 3 | 1 | 2 | 1 | 1 | 0 | 6 | 0.8333 |
| 1_1117876 | NA | NA | 3 | 1 | NA | NA | 3 | 0.3333 | 6 | 0.6667 | NA | NA | 3 | 1 | 2 | 1 | 1 | 1 | 6 | 1 |
| 1_1117877 | NA | NA | 2 | 0 | NA | NA | 4 | 0.25 | 6 | 0.1667 | NA | NA | 3 | 0 | 2 | 0 | 1 | 1 | 6 | 0.1667 |
| 1_1117878 | NA | NA | 3 | 1 | NA | NA | 2 | 0.5 | 5 | 0.8 | NA | NA | 2 | 1 | 1 | 1 | 1 | 1 | 4 | 1 |
| 1_1117878 | NA | NA | 2 | 0.5 | NA | NA | 4 | 0.25 | 6 | 0.3333 | NA | NA | 3 | 0.6667 | 2 | 0 | 1 | 1 | 6 | 0.5 |
| 1_1117880 | NA | NA | 2 | 1 | NA | NA | 4 | 1 | 6 | 1 | NA | NA | 3 | 1 | 2 | 0.5 | 1 | 1 | 6 | 0.8333 |
| 1_1117880 | NA | NA | 3 | 0.3333 | NA | NA | 4 | 0.75 | 7 | 0.5714 | NA | NA | 3 | 0.6667 | 1 | 0 | 1 | 1 | 5 | 0.6 |
| 1_1117883 | NA | NA | 1 | 1 | NA | NA | 4 | 0.5 | 5 | 0.6 | NA | NA | 2 | 0.5 | 2 | 0 | 1 | 1 | 5 | 0.4 |
| 1_1117883 | NA | NA | 3 | 0 | NA | NA | 3 | 0.3333 | 6 | 0.1667 | NA | NA | 3 | 0.3333 | 2 | 0 | 1 | 0 | 6 | 0.1667 |
| 1_1117884 | NA | NA | 3 | 0.3333 | NA | NA | 2 | 0.5 | 5 | 0.4 | NA | NA | 3 | 0.3333 | 2 | 0.5 | 1 | 0 | 6 | 0.3333 |
| 1_1117885 | NA | NA | 3 | 0 | NA | NA | 2 | 0.5 | 5 | 0.2 | NA | NA | 2 | 1 | 2 | 0 | 1 | 1 | 5 | 0.6 |
| 1_1117888 | NA | NA | 3 | 0.3333 | NA | NA | 3 | 0.3333 | 6 | 0.3333 | NA | NA | 2 | 0.5 | 1 | 0 | NA | NA | 3 | 0.3333 |
| 1_1117889 | NA | NA | 1 | 1 | NA | NA | 2 | 1 | 3 | 1 | NA | NA | 2 | 0.5 | 1 | 1 | NA | NA | 3 | 0.6667 |
| 1_1117890 | NA | NA | 2 | 1 | NA | NA | 3 | 0 | 5 | 0.4 | NA | NA | 2 | 1 | 1 | 1 | NA | NA | 3 | 1 |
| 1_1117891 | NA | NA | 2 | 0.5 | NA | NA | 2 | 0.5 | 4 | 0.5 | NA | NA | 2 | 0.5 | NA | NA | NA | NA | 2 | 0.5 |
| 1_1117892 | NA | NA | 2 | 1 | NA | NA | 2 | 1 | 4 | 1 | NA | NA | 2 | 1 | NA | NA | NA | NA | 2 | 1 |
| 1_1117897 | NA | NA | 1 | 1 | NA | NA | 2 | 0 | 3 | 0.3333 | NA | NA | 2 | 0.5 | NA | NA | NA | NA | 2 | 0.5 |
| 1_1117900 | NA | NA | NA | NA | NA | NA | 2 | 0.5 | 2 | 0.5 | NA | NA | 1 | 1 | NA | NA | NA | NA | 1 | 1 |
| 1_1117900 | NA | NA | 1 | 1 | NA | NA | 1 | 1 | 2 | 1 | NA | NA | 1 | 1 | NA | NA | NA | NA | 1 | 1 |
| 1_1117910 | 1 | 0 | 1 | 1 | NA | NA | 2 | 0.5 | 4 | 0.5 | NA | NA | 1 | 1 | NA | NA | NA | NA | 1 | 1 |
| 1_1117911 | 1 | 0 | 1 | 1 | NA | NA | 2 | 1 | 4 | 0.75 | NA | NA | 2 | 1 | NA | NA | NA | NA | 2 | 1 |
| 1_1117911 | 1 | 1 | NA | NA | NA | NA | 2 | 1 | 3 | 1 | NA | NA | 2 | 0.5 | NA | NA | NA | NA | 2 | 0.5 |

| | | | | | | | | | | | | | | | | | | | | |
|---|---|---|---|---|---|---|---|---|---|---|---|---|---|---|---|---|---|---|---|---|
| 1_1123063( | NA | NA | NA | NA | 1 | 1 | NA | NA | 1 | 1 | NA | NA | 1 | 1 | NA | NA | NA | NA | 1 | 1 |
| 1_1123069 | NA | NA | NA | NA | 1 | 1 | NA | NA | 1 | 1 | NA | NA | 2 | 1 | NA | NA | NA | NA | 2 | 1 |
| 1_1123072 | NA | NA | NA | NA | 1 | 1 | NA | NA | 1 | 1 | NA | NA | 2 | 1 | NA | NA | NA | NA | 2 | 1 |
| 1_1123077( | NA | NA | NA | NA | 1 | 1 | NA | NA | 1 | 1 | NA | NA | 1 | 1 | NA | NA | NA | NA | 1 | 1 |
| 1_1123080 | NA | NA | NA | NA | 1 | 1 | NA | NA | 1 | 1 | NA | NA | 1 | 1 | NA | NA | NA | NA | 1 | 1 |
| 1_1123236 | 1 | 1 | NA | NA | 2 | 1 | 2 | 1 | 5 | 1 | NA | NA | 3 | 1 | 2 | 1 | 1 | 1 | 6 | 1 |
| 1_1123236 | 1 | 1 | NA | NA | 2 | 1 | NA | NA | 3 | 1 | NA | NA | 3 | 1 | 2 | 1 | 1 | 1 | 6 | 1 |
| 1_1123237 | 1 | 1 | NA | NA | 2 | 1 | 2 | 1 | 5 | 1 | NA | NA | 3 | 1 | 2 | 1 | 1 | 1 | 6 | 1 |
| 1_1123238 | 1 | 0 | NA | NA | NA | NA | 2 | 1 | 3 | 0.6667 | NA | NA | 2 | 0.5 | 2 | 1 | NA | NA | 4 | 0.75 |
| 1_1123240 | 1 | 1 | NA | NA | 2 | 0.5 | 1 | 0 | 4 | 0.5 | NA | NA | 2 | 1 | 2 | 1 | 1 | 0 | 5 | 0.8 |
| 1_1123240 | 1 | 0 | NA | NA | 2 | 0.5 | 1 | 0 | 4 | 0.25 | NA | NA | 2 | 1 | 2 | 1 | NA | NA | 4 | 1 |
| 1_1123252 | 1 | 0 | NA | NA | 1 | 1 | NA | NA | 2 | 0.5 | NA | NA | NA | NA | NA | NA | 1 | 1 | 1 | 1 |
| 1_1123320 | 1 | 0 | NA | NA | 2 | 0.5 | NA | NA | 3 | 0.3333 | NA | NA | 1 | 1 | NA | NA | NA | NA | 1 | 1 |
| 1_1123324 | 1 | 1 | NA | NA | 1 | 1 | NA | NA | 2 | 1 | NA | NA | 1 | 1 | NA | NA | NA | NA | 1 | 1 |
| 1_1155930 | NA | NA | NA | NA | 1 | 0 | 2 | 0.5 | 3 | 0.3333 | NA | NA | NA | NA | NA | NA | 1 | 1 | 1 | 1 |
| 1_1155937( | 1 | 0 | NA | NA | 1 | 0 | 2 | 0.5 | 4 | 0.25 | NA | NA | NA | NA | 1 | 1 | 1 | 0 | 2 | 0.5 |
| 1_1155969 | NA | NA | NA | NA | 1 | 1 | 2 | 1 | 3 | 1 | NA | NA | NA | NA | 1 | 1 | NA | NA | 1 | 1 |
| 1_1155970 | NA | NA | NA | NA | 1 | 1 | 2 | 1 | 3 | 1 | NA | NA | NA | NA | 1 | 1 | NA | NA | 1 | 1 |
| 1_1155971 | NA | NA | NA | NA | 1 | 1 | 2 | 0.5 | 3 | 0.6667 | NA | NA | NA | NA | 1 | 1 | NA | NA | 1 | 1 |
| 1_1155971( | NA | NA | NA | NA | 1 | 1 | 2 | 1 | 3 | 1 | NA | NA | NA | NA | 1 | 1 | NA | NA | 1 | 1 |
| 1_1165083 | NA | NA | 2 | 1 | 2 | 1 | 2 | 1 | 6 | 1 | 1 | 1 | 1 | 1 | 2 | 1 | NA | NA | 4 | 1 |
| 1_1165083 | NA | NA | 2 | 0.5 | 2 | 1 | 2 | 0.5 | 6 | 0.6667 | 1 | 1 | 1 | 1 | 2 | 1 | NA | NA | 4 | 1 |
| 1_1165084 | NA | NA | 2 | 0.5 | 3 | 0 | 2 | 0 | 7 | 0.1429 | 1 | 0 | 2 | 1 | 2 | 0.5 | NA | NA | 5 | 0.6 |
| 1_1182229( | NA | NA | NA | NA | 2 | 1 | 2 | 0.5 | 4 | 0.75 | NA | NA | NA | NA | 2 | 1 | NA | NA | 2 | 1 |
| 1_1182264 | 8 | 0 | 8 | 0.125 | 6 | 0.1667 | 7 | 0 | 29 | 0.069 | 4 | 0.25 | 8 | 0.25 | 6 | 0 | 3 | 0 | 21 | 0.1429 |
| 1_1182282 | 4 | 0 | 6 | 0 | 3 | 0 | 6 | 0.1667 | 19 | 0.0526 | 5 | 0 | 6 | 0 | 3 | 0 | 2 | 0.5 | 16 | 0.0625 |
| 1_1182607 | 2 | 1 | 2 | 1 | 7 | 0.8571 | 2 | 0.5 | 13 | 0.8462 | 2 | 1 | 3 | 0.6667 | 1 | 1 | 3 | 0.6667 | 9 | 0.7778 |
| 1_1182607( | 1 | 0 | 2 | 0.5 | 9 | 0.6667 | 2 | 1 | 14 | 0.6429 | 2 | 1 | 3 | 0.3333 | 1 | 1 | 2 | 1 | 8 | 0.75 |
| 1_1182608 | 2 | 0 | 2 | 0.5 | 9 | 0.3333 | 2 | 0 | 15 | 0.2667 | 2 | 1 | 4 | 0 | 1 | 0 | 4 | 0.25 | 11 | 0.2727 |
| 1_1182612( | 1 | 1 | 2 | 0.5 | 9 | 0.5556 | 2 | 0 | 14 | 0.5 | 2 | 0.5 | 3 | 0 | NA | NA | 3 | 0.3333 | 8 | 0.25 |
| 1_1183161 | NA | NA | NA | NA | NA | NA | 1 | 1 | 1 | 1 | 1 | 1 | NA | NA | NA | NA | NA | NA | 1 | 1 |
| 1_1183162 | NA | NA | NA | NA | NA | NA | 1 | 1 | 1 | 1 | 1 | 1 | NA | NA | NA | NA | NA | NA | 1 | 1 |
| 1_1183162 | NA | NA | NA | NA | NA | NA | 1 | 1 | 1 | 1 | 1 | 1 | NA | NA | NA | NA | NA | NA | 1 | 1 |
| 1_1183475 | 1 | 0 | 2 | 0.5 | NA | NA | 1 | 1 | 4 | 0.5 | 1 | 0 | NA | NA | 1 | 1 | NA | NA | 2 | 0.5 |
| 1_1183475 | 1 | 0 | 2 | 0 | NA | NA | 1 | 1 | 4 | 0.25 | 1 | 1 | NA | NA | 1 | 0 | NA | NA | 2 | 0.5 |
| 1_1183869 | 1 | 0 | 4 | 0.25 | 3 | 0 | 3 | 0.3333 | 11 | 0.1818 | NA | NA | 1 | 0 | 3 | 0.3333 | 2 | 0 | 6 | 0.1667 |
| 1_1185348 | 1 | 1 | 3 | 0.6667 | 2 | 1 | 5 | 1 | 11 | 0.9091 | 5 | 1 | 1 | 1 | 4 | 1 | 1 | 1 | 11 | 1 |
| 1_1185350 | 2 | 1 | 3 | 0.3333 | 1 | 0 | 5 | 0.6 | 11 | 0.5455 | 4 | 0.75 | 2 | 0.5 | 7 | 0.4286 | 2 | 0 | 15 | 0.4667 |
| 1_1185360 | 3 | 0 | 6 | 0.1667 | 3 | 0.3333 | 7 | 0 | 19 | 0.1053 | 5 | 0.6 | 4 | 0.5 | 9 | 0.1111 | 3 | 0.3333 | 21 | 0.3333 |
| 1_1185364 | 3 | 0 | 7 | 0.2857 | 5 | 0 | 6 | 0.1667 | 21 | 0.1429 | 5 | 0.2 | 7 | 0.1429 | 8 | 0.25 | 3 | 0 | 23 | 0.1739 |
| 1_1185365 | 3 | 0.6667 | 7 | 0.7143 | 6 | 1 | 7 | 1 | 23 | 0.8696 | 5 | 1 | 6 | 1 | 9 | 0.8889 | 3 | 1 | 23 | 0.9565 |
| 1_1185365 | 3 | 0.6667 | 8 | 0.875 | 6 | 0.8333 | 6 | 1 | 23 | 0.8696 | 6 | 1 | 7 | 1 | 9 | 1 | 3 | 1 | 25 | 1 |
| 1_1185366( | 1 | 1 | 7 | 0.8571 | 5 | 0.8 | 7 | 0.8571 | 20 | 0.85 | 6 | 0.8333 | 6 | 0.6667 | 8 | 1 | 3 | 1 | 23 | 0.8696 |
| 1_1185366 | 2 | 0 | 3 | 0 | 4 | 0.75 | 6 | 0.1667 | 15 | 0.2667 | 4 | 0 | 6 | 0.5 | 7 | 0 | 3 | 0.3333 | 20 | 0.2 |
| 1_1185366 | 3 | 0.3333 | 7 | 0.1429 | 6 | 0 | 6 | 0 | 22 | 0.0909 | 4 | 0.25 | 6 | 0 | 9 | 0.1111 | 3 | 0 | 22 | 0.0909 |
| 1_1185367 | 4 | 0 | 5 | 0.2 | 5 | 0 | 7 | 0.1429 | 21 | 0.0952 | 4 | 0.25 | 7 | 0 | 6 | 0 | 2 | 0 | 19 | 0.0526 |
| 1_1185367 | 4 | 0.25 | 9 | 0.1111 | 6 | 0.3333 | 9 | 0.1111 | 28 | 0.1786 | 5 | 0.4 | 7 | 0.4286 | 9 | 0 | 4 | 0.5 | 25 | 0.28 |
| 1_1185368 | 4 | 0.25 | 8 | 0.125 | 6 | 0.1667 | 7 | 0.1429 | 25 | 0.16 | 7 | 0.2857 | 5 | 0.6 | 8 | 0.25 | 3 | 0.3333 | 23 | 0.3478 |
| 1_1185369 | 4 | 0.75 | 10 | 0.7 | 6 | 0.8333 | 9 | 0.7778 | 29 | 0.7586 | 8 | 0.75 | 5 | 0.8 | 9 | 0.7778 | 5 | 0.8 | 27 | 0.7778 |
| 1_1185369 | 5 | 0.2 | 9 | 0 | 6 | 0.3333 | 7 | 0.1429 | 27 | 0.1481 | 8 | 0.25 | 6 | 0 | 9 | 0.1111 | 6 | 0.3333 | 29 | 0.1724 |
| 1_1185369 | 5 | 0.4 | 9 | 0.2222 | 5 | 0.2 | 9 | 0.6667 | 28 | 0.3929 | 6 | 0.5 | 6 | 0.1667 | 10 | 0.4 | 5 | 0.4 | 27 | 0.3704 |
| 1_1193446( | NA | NA | 1 | 1 | 1 | 1 | NA | NA | 2 | 1 | NA | NA | NA | NA | NA | NA | 1 | 1 | 1 | 1 |
| 1_1193448( | NA | NA | 1 | 1 | 1 | 1 | NA | NA | 2 | 1 | NA | NA | NA | NA | NA | NA | 1 | 1 | 1 | 1 |
| 1_1193451 | NA | NA | 1 | 1 | NA | NA | NA | NA | 1 | 1 | NA | NA | NA | NA | NA | NA | 1 | 1 | 1 | 1 |
| 1_1193452( | NA | NA | 1 | 1 | NA | NA | NA | NA | 1 | 1 | NA | NA | NA | NA | NA | NA | 1 | 1 | 1 | 1 |
| 1_1193452 | NA | NA | 1 | 1 | NA | NA | NA | NA | 1 | 1 | NA | NA | NA | NA | NA | NA | 1 | 1 | 1 | 1 |
| 1_1193454 | NA | NA | 1 | 1 | NA | NA | NA | NA | 1 | 1 | NA | NA | NA | NA | NA | NA | 1 | 1 | 1 | 1 |
| 1_1193991( | 2 | 0 | 3 | 0.3333 | 3 | 0 | 2 | 0 | 10 | 0.1 | 2 | 1 | 1 | 0 | NA | NA | 2 | 0 | 5 | 0.4 |
| 1_1193993 | 2 | 1 | 2 | 1 | 3 | 0.6667 | 1 | 0 | 8 | 0.75 | 1 | 1 | 1 | 1 | NA | NA | 2 | 1 | 4 | 1 |
| 1_1193994 | 2 | 0 | 2 | 0 | 3 | 0 | 2 | 0.5 | 9 | 0.1111 | 2 | 0.5 | 1 | 0 | NA | NA | 1 | 0 | 4 | 0.25 |
| 1_1194542( | 1 | 0 | 2 | 1 | 1 | 0 | 1 | 1 | 5 | 0.6 | 2 | 0.5 | 2 | 0.5 | 2 | 0 | 1 | 1 | 7 | 0.4286 |
| 1_1194546( | 2 | 1 | 4 | 0.75 | NA | NA | 1 | 1 | 7 | 0.8571 | 2 | 0.5 | 3 | 1 | 2 | 0.5 | 2 | 1 | 9 | 0.7778 |
| 1_1194547 | 1 | 0 | 1 | 1 | 1 | 1 | 1 | 0 | 4 | 0.5 | 2 | 0.5 | 1 | 0 | 2 | 0 | NA | NA | 5 | 0.2 |
| 1_1194548( | 2 | 0 | 3 | 0.3333 | NA | NA | 1 | 0 | 6 | 0.1667 | 2 | 0 | 3 | 0 | 2 | 0.5 | 2 | 0.5 | 9 | 0.2222 |
| 1_1194548 | 2 | 0 | 4 | 0.25 | 1 | 1 | 1 | 0 | 8 | 0.25 | 2 | 0.5 | 3 | 0 | 1 | 0 | 2 | 0 | 8 | 0.125 |

| | | | | | | | | | | | | | | | | | | | | |
|---|---|---|---|---|---|---|---|---|---|---|---|---|---|---|---|---|---|---|---|---|
| 1_11945494 | 2 | 1 | 4 | 1 | 1 | 1 | 2 | 1 | 9 | 1 | 2 | 1 | 2 | 1 | 2 | 1 | 3 | 1 | 9 | 1 |
| 1_11945500 | 2 | 0.5 | 4 | 1 | 1 | 1 | 2 | 1 | 9 | 0.8889 | 1 | 1 | 2 | 1 | 2 | 1 | 3 | 1 | 8 | 1 |
| 1_11945527 | 2 | 0.5 | 3 | 0.3333 | 1 | 0 | 2 | 0.5 | 8 | 0.375 | 3 | 0.3333 | 2 | 1 | 2 | 0.5 | 3 | 0.6667 | 10 | 0.6 |
| 1_11945528 | 2 | 1 | 3 | 0.6667 | 1 | 1 | 2 | 1 | 8 | 0.875 | 2 | 1 | 3 | 1 | 3 | 0.6667 | 2 | 1 | 10 | 0.9 |
| 1_11945532 | 1 | 1 | 4 | 1 | 1 | 1 | 2 | 1 | 8 | 1 | 1 | 1 | 3 | 0.6667 | 3 | 0.3333 | 3 | 1 | 10 | 0.7 |
| 1_11945542 | 3 | 0 | 5 | 0.4 | 1 | 0 | 2 | 0.5 | 11 | 0.2727 | 2 | 0.5 | 2 | 0 | 3 | 0 | 3 | 0.3333 | 10 | 0.2 |
| 1_11945546 | 3 | 0 | 4 | 0.5 | 1 | 0 | 3 | 0.3333 | 11 | 0.2727 | 3 | 0.6667 | 3 | 0.3333 | 3 | 0 | 3 | 0.3333 | 12 | 0.3333 |
| 1_11945547 | 3 | 1 | 5 | 1 | NA | NA | 3 | 1 | 11 | 1 | 2 | 1 | 2 | 1 | 2 | 1 | 3 | 1 | 9 | 1 |
| 1_11945550 | 3 | 0.6667 | 4 | 0.75 | NA | NA | 1 | 0 | 8 | 0.625 | 1 | 1 | 2 | 0 | 3 | 0.6667 | 2 | 0.5 | 8 | 0.5 |
| 1_11945558 | 2 | 0.5 | 4 | 0.5 | 1 | 1 | 2 | 0.5 | 9 | 0.5556 | 3 | 0.3333 | 3 | 0.6667 | 3 | 0 | 3 | 0.3333 | 12 | 0.3333 |
| 1_11945581 | 1 | 0 | 6 | 0 | 1 | 1 | 3 | 0 | 11 | 0.0909 | 3 | 0.3333 | 3 | 0 | 3 | 0 | 3 | 0 | 12 | 0.0833 |
| 1_11958037 | 5 | 0 | 15 | 0.0667 | 8 | 0.125 | 11 | 0 | 39 | 0.0513 | 6 | 0 | 6 | 0 | 9 | 0 | 13 | 0.1538 | 34 | 0.0588 |
| 1_11958053 | 3 | 0.3333 | 12 | 0.25 | 5 | 0.6 | 8 | 0.375 | 28 | 0.3571 | 5 | 0.6 | 3 | 0.3333 | 7 | 0.4286 | 12 | 0.1667 | 27 | 0.3333 |
| 1_11958214 | 2 | 0 | 6 | 0 | 3 | 0.3333 | 5 | 0 | 16 | 0.0625 | 3 | 0 | 2 | 0 | 4 | 0 | 9 | 0.1111 | 18 | 0.0556 |
| 1_11958270 | 2 | 0 | 6 | 0 | 3 | 0.3333 | 5 | 0 | 16 | 0.0625 | 3 | 0 | 1 | 0 | 4 | 0 | 11 | 0.0909 | 19 | 0.0526 |
| 1_11958398 | 2 | 0.5 | 6 | 0 | 4 | 0 | 4 | 0 | 16 | 0.0625 | 4 | 0 | 1 | 0 | 4 | 0 | 10 | 0.1 | 19 | 0.0526 |
| 1_11958400 | 2 | 0.5 | 6 | 0.1667 | 4 | 0.25 | 4 | 0 | 16 | 0.1875 | 2 | 0 | NA | NA | 4 | 0.25 | 8 | 0.25 | 14 | 0.2143 |
| 1_11958412 | 2 | 0.5 | 5 | 0 | 3 | 0.3333 | 3 | 0 | 13 | 0.1538 | 2 | 0 | 1 | 0 | 2 | 0.5 | 9 | 0.2222 | 14 | 0.2143 |
| 1_11958665 | 2 | 0 | 4 | 0.75 | 1 | 1 | NA | NA | 7 | 0.5714 | 3 | 0.6667 | 2 | 1 | 3 | 0.6667 | 3 | 1 | 11 | 0.8182 |
| 1_11958668 | 2 | 0 | 4 | 0.25 | 1 | 0 | NA | NA | 7 | 0.1429 | 3 | 0.3333 | 2 | 1 | 3 | 0.3333 | 5 | 0.4 | 13 | 0.4615 |
| 1_11958697 | 1 | 0 | 3 | 0.3333 | 1 | 0 | NA | NA | 5 | 0.2 | 2 | 0 | 2 | 0 | 3 | 0.3333 | 3 | 0 | 10 | 0.1 |
| 1_11958697 | 2 | 1 | 3 | 0.6667 | 1 | 0 | NA | NA | 6 | 0.6667 | 3 | 0.6667 | 2 | 1 | 3 | 0.6667 | 2 | 0.5 | 10 | 0.7 |
| 1_11958704 | 1 | 1 | 4 | 0.75 | 1 | 0 | NA | NA | 6 | 0.6667 | 3 | 0.6667 | 2 | 0.5 | 3 | 0.6667 | 2 | 0.5 | 10 | 0.6 |
| 1_11958705 | 2 | 0 | 4 | 0.5 | 1 | 1 | NA | NA | 7 | 0.4286 | 2 | 1 | 2 | 1 | 3 | 0.6667 | 5 | 0.6 | 12 | 0.75 |
| 1_11958734 | NA | NA | 4 | 1 | 1 | 1 | NA | NA | 5 | 1 | 2 | 1 | 1 | 1 | 2 | 1 | 1 | 1 | 6 | 1 |
| 1_11958734 | 1 | 1 | NA | NA | NA | NA | NA | NA | 1 | 1 | NA | NA | NA | NA | NA | NA | 1 | 1 | 1 | 1 |
| 1_11958737 | 1 | 1 | 3 | 0.3333 | 1 | 1 | NA | NA | 5 | 0.6 | 4 | 0 | 1 | 0 | 3 | 0.3333 | 3 | 0.3333 | 11 | 0.1818 |
| 1_11958744 | 1 | 1 | 3 | 1 | 1 | 0 | NA | NA | 5 | 0.8 | 4 | 0.5 | NA | NA | 3 | 1 | 3 | 0.6667 | 10 | 0.7 |
| 1_11958746 | 1 | 1 | 3 | 0.6667 | NA | NA | NA | NA | 4 | 0.75 | 4 | 1 | 1 | 1 | 3 | 0.6667 | 3 | 1 | 11 | 0.9091 |
| 1_11958753 | NA | NA | 3 | 1 | 1 | 1 | NA | NA | 4 | 1 | 3 | 0.6667 | 1 | 0 | 3 | 1 | 4 | 1 | 11 | 0.8182 |
| 1_11958754 | 1 | 0 | 4 | 0.25 | 1 | 0 | NA | NA | 6 | 0.1667 | 3 | 0.3333 | 1 | 0 | 2 | 0 | 3 | 0.6667 | 9 | 0.3333 |
| 1_11983705 | 1 | 0 | 2 | 0.5 | 3 | 0 | NA | NA | 6 | 0.1667 | NA | NA | 2 | 0.5 | 1 | 0 | NA | NA | 3 | 0.3333 |
| 1_11983741 | NA | NA | 2 | 1 | 3 | 0.6667 | NA | NA | 5 | 0.8 | NA | NA | 2 | 0.5 | 1 | 1 | NA | NA | 3 | 0.6667 |
| 1_11983751 | 2 | 0.5 | 2 | 1 | 1 | 0 | 2 | 0.5 | 7 | 0.5714 | 1 | 1 | 2 | 0.5 | 3 | 1 | 1 | 0 | 7 | 0.7143 |
| 1_11983758 | 1 | 0 | 2 | 1 | 2 | 1 | 2 | 1 | 7 | 0.8571 | 1 | 1 | 2 | 0.5 | 2 | 0.5 | 2 | 1 | 7 | 0.7143 |
| 1_11983768 | NA | NA | 2 | 1 | 1 | 1 | 2 | 1 | 5 | 1 | 1 | 1 | 2 | 0.5 | 3 | 0.6667 | 2 | 1 | 8 | 0.75 |
| 1_11983779 | 1 | 1 | 1 | 1 | NA | NA | 1 | 1 | 3 | 1 | NA | NA | 2 | 0.5 | 2 | 1 | NA | NA | 4 | 0.75 |
| 1_11983801 | NA | NA | 2 | 1 | 1 | 1 | 2 | 1 | 5 | 1 | NA | NA | 2 | 0.5 | 3 | 0.6667 | 2 | 1 | 7 | 0.7143 |
| 1_11983807 | 1 | 1 | 2 | 1 | 2 | 0.5 | 2 | 1 | 7 | 0.8571 | 1 | 0 | 2 | 0.5 | 3 | 0.6667 | 2 | 0.5 | 8 | 0.5 |
| 1_11983817 | 1 | 0 | 2 | 0 | 2 | 0 | 1 | 1 | 6 | 0.1667 | 1 | 1 | 2 | 0 | 3 | 0 | 2 | 0 | 8 | 0.125 |
| 1_11983832 | NA | NA | 1 | 1 | 1 | 1 | 2 | 0.5 | 4 | 0.75 | 1 | 0 | 1 | 0 | 1 | 1 | 1 | 1 | 4 | 0.5 |
| 1_11983846 | 1 | 1 | 2 | 1 | 2 | 1 | 1 | 1 | 6 | 1 | NA | NA | 2 | 0.5 | 2 | 1 | 1 | 1 | 5 | 0.8 |
| 1_11983863 | NA | NA | 1 | 1 | 2 | 1 | 1 | 1 | 4 | 1 | NA | NA | 1 | 0 | 2 | 0.5 | 1 | 1 | 4 | 0.5 |
| 1_11983882 | 1 | 1 | 2 | 0.5 | 2 | 0 | 2 | 0 | 7 | 0.2857 | NA | NA | 2 | 0 | 3 | 0.6667 | 1 | 0 | 6 | 0.3333 |
| 1_11999541 | 2 | 1 | 1 | 0 | 1 | 1 | NA | NA | 4 | 0.75 | NA | NA | 1 | 1 | NA | NA | NA | NA | 1 | 1 |
| 1_11999543 | 2 | 1 | NA | NA | 1 | 1 | NA | NA | 3 | 1 | NA | NA | 1 | 1 | NA | NA | NA | NA | 1 | 1 |
| 1_11999545 | 2 | 1 | 1 | 1 | 1 | 1 | NA | NA | 4 | 1 | NA | NA | 1 | 1 | NA | NA | NA | NA | 1 | 1 |
| 1_11999584 | 1 | 1 | 1 | 1 | 1 | 1 | NA | NA | 3 | 1 | NA | NA | 1 | 1 | NA | NA | NA | NA | 1 | 1 |
| 1_12015487 | NA | NA | NA | NA | 1 | 1 | NA | NA | 1 | 1 | NA | NA | NA | NA | 1 | 1 | 1 | 0 | 2 | 0.5 |
| 1_12027062 | 1 | 1 | NA | NA | NA | NA | 1 | 1 | 2 | 1 | 1 | 1 | NA | NA | NA | NA | NA | NA | 1 | 1 |
| 1_12123276 | 8 | 0.125 | 6 | 0.1667 | 4 | 0 | 9 | 0.2222 | 27 | 0.1481 | 8 | 0.375 | 5 | 0.2 | 9 | 0.1111 | 4 | 0.25 | 26 | 0.2308 |
| 1_12123278 | 8 | 0.625 | 7 | 0.8571 | 5 | 0.6 | 12 | 0.6667 | 32 | 0.6875 | 7 | 0.5714 | 4 | 0.25 | 9 | 0.6667 | 5 | 0.4 | 25 | 0.52 |
| 1_12123496 | 6 | 0.1667 | 2 | 0 | 6 | 0 | 9 | 0.1111 | 23 | 0.087 | 8 | 0.5 | 5 | 0 | 5 | 0.2 | 2 | 0 | 20 | 0.25 |
| 1_12124743 | 7 | 0.1429 | 4 | 0.5 | 6 | 0 | 6 | 0.1667 | 23 | 0.1739 | 9 | 0.3333 | 5 | 0.6 | 6 | 0.1667 | 3 | 0.3333 | 23 | 0.3478 |
| 1_12124758 | 7 | 0.8571 | 3 | 0.6667 | 5 | 0.6 | 5 | 1 | 20 | 0.8 | 7 | 0.5714 | 3 | 0.6667 | 5 | 0.8 | 3 | 1 | 18 | 0.7222 |
| 1_12142906 | 1 | 1 | 1 | 0 | NA | NA | NA | NA | 2 | 0.5 | NA | NA | NA | NA | 2 | 1 | 1 | 1 | 3 | 1 |
| 1_12142915 | 1 | 1 | 1 | 1 | NA | NA | NA | NA | 2 | 1 | NA | NA | NA | NA | 2 | 1 | 1 | 1 | 3 | 1 |
| 1_12143363 | 3 | 0.3333 | NA | NA | 2 | 0 | 2 | 0 | 7 | 0.1429 | NA | NA | 1 | 0 | 4 | 0.25 | 2 | 0 | 7 | 0.1429 |
| 1_12147778 | NA | NA | NA | NA | 1 | 1 | NA | NA | 1 | 1 | NA | NA | 1 | 1 | NA | NA | NA | NA | 1 | 1 |
| 1_12148165 | 7 | 0.7143 | 4 | 0.25 | 2 | 0 | 2 | 0 | 15 | 0.4 | 3 | 0 | 4 | 0.25 | 3 | 0 | 4 | 0.5 | 14 | 0.2143 |
| 1_12148169 | 7 | 0.1429 | 4 | 0.25 | 2 | 0 | 2 | 0 | 15 | 0.1333 | 3 | 0 | 5 | 0 | 3 | 0.3333 | 4 | 0.25 | 15 | 0.1333 |
| 1_12148170 | 6 | 1 | 3 | 0.6667 | 2 | 1 | 2 | 0.5 | 13 | 0.8462 | 3 | 0.6667 | 4 | 1 | 3 | 0.6667 | 4 | 0.75 | 14 | 0.7857 |
| 1_12149379 | 2 | 0 | 1 | 0 | 1 | 0 | 2 | 0.5 | 6 | 0.1667 | 2 | 0 | 3 | 0.6667 | 2 | 0 | 3 | 0 | 10 | 0.2 |
| 1_12326031 | 2 | 0.5 | 5 | 0.2 | 12 | 0.25 | 7 | 0.1429 | 26 | 0.2308 | 7 | 0.1429 | 10 | 0 | 4 | 0.5 | 3 | 0.6667 | 24 | 0.2083 |
| 1_12326183 | 5 | 0 | 12 | 0 | 21 | 0.0476 | 14 | 0.1429 | 52 | 0.0577 | 6 | 0 | 11 | 0.0909 | 10 | 0 | 10 | 0.1 | 37 | 0.0541 |

| | | | | | | | | | | | | | | | | | | | | |
|---|---|---|---|---|---|---|---|---|---|---|---|---|---|---|---|---|---|---|---|---|
| 1_12326183 | 4 | 0 | 8 | 0.375 | 14 | 0.1429 | 15 | 0.4 | 41 | 0.2683 | 6 | 0.1667 | 9 | 0.1111 | 4 | 0 | 6 | 0.5 | 25 | 0.2 |
| 1_12326184 | 5 | 0.2 | 12 | 0.1667 | 20 | 0 | 14 | 0.0714 | 51 | 0.0784 | 5 | 0 | 11 | 0 | 10 | 0.3 | 11 | 0.0909 | 37 | 0.1081 |
| 1_12326191 | 3 | 1 | 10 | 1 | 19 | 0.8947 | 13 | 1 | 45 | 0.9556 | 6 | 1 | 10 | 0.9 | 7 | 0.8571 | 11 | 0.9091 | 34 | 0.9118 |
| 1_12326193 | 5 | 0.8 | 10 | 0.4 | 18 | 0.2778 | 16 | 0.5625 | 49 | 0.449 | 5 | 0.4 | 9 | 0.3333 | 7 | 0.5714 | 10 | 0.4 | 31 | 0.4194 |
| 1_12326197 | 5 | 0.8 | 10 | 1 | 16 | 0.875 | 14 | 0.9286 | 45 | 0.9111 | 6 | 1 | 10 | 0.9 | 8 | 0.75 | 8 | 0.75 | 32 | 0.8438 |
| 1_12397345 | 1 | 0 | 3 | 0.3333 | 3 | 0 | 2 | 0 | 9 | 0.1111 | 1 | 1 | 2 | 0 | NA | NA | 1 | 1 | 4 | 0.5 |
| 1_12397360 | 2 | 0 | 3 | 0.6667 | 7 | 0 | 5 | 0.2 | 17 | 0.1765 | 1 | 0 | 2 | 1 | 2 | 0 | NA | NA | 5 | 0.4 |
| 1_12532109 | 34 | 0.2059 | 36 | 0.2222 | 21 | 0.1905 | 34 | 0.2353 | 125 | 0.216 | 23 | 0.5217 | 34 | 0.1765 | 35 | 0.2 | 25 | 0.08 | 117 | 0.2308 |
| 1_12532110 | 31 | 0.0968 | 37 | 0.1081 | 22 | 0.0455 | 29 | 0.2069 | 119 | 0.1176 | 23 | 0.0435 | 32 | 0.125 | 32 | 0.125 | 21 | 0.1429 | 108 | 0.1111 |
| 1_12532127 | 30 | 0.2333 | 37 | 0.3784 | 20 | 0.3 | 30 | 0.3 | 117 | 0.3077 | 20 | 0.3 | 37 | 0.2703 | 33 | 0.3939 | 24 | 0.4583 | 114 | 0.3509 |
| 1_12532133 | 26 | 0.0385 | 34 | 0.1176 | 20 | 0.1 | 25 | 0.16 | 105 | 0.1048 | 19 | 0.2105 | 33 | 0.0606 | 28 | 0.2143 | 20 | 0 | 100 | 0.12 |
| 1_12552283 | 1 | 1 | 4 | 0 | 2 | 0.5 | 1 | 0 | 8 | 0.25 | 3 | 0.3333 | 1 | 0 | 1 | 1 | 3 | 0 | 8 | 0.25 |
| 1_12552289 | NA | NA | 4 | 0.25 | 2 | 0 | 2 | 0 | 8 | 0.125 | 2 | 0 | 1 | 0 | 1 | 1 | NA | NA | 4 | 0.25 |
| 1_12552290 | 1 | 1 | 4 | 0.5 | 2 | 0.5 | 2 | 0 | 9 | 0.4444 | 3 | 0.6667 | 1 | 0 | 1 | 0 | 1 | 0 | 6 | 0.3333 |
| 1_12579589 | 1 | 1 | 1 | 0 | NA | NA | 1 | 0 | 3 | 0.3333 | 1 | 0 | NA | NA | 1 | 0 | 2 | 0.5 | 4 | 0.25 |
| 1_12579647 | NA | NA | NA | NA | 1 | 1 | NA | NA | 1 | 1 | 1 | 1 | 1 | 1 | 1 | 0 | 1 | 1 | 4 | 0.75 |
| 1_12579662 | NA | NA | 1 | 0 | 1 | 1 | NA | NA | 2 | 0.5 | 1 | 0 | 1 | 1 | NA | NA | 1 | 0 | 3 | 0.3333 |
| 1_12579700 | NA | NA | 1 | 0 | 1 | 1 | NA | NA | 2 | 0.5 | 1 | 1 | NA | NA | 1 | 0 | 1 | 0 | 3 | 0.3333 |
| 1_12579990 | 4 | 1 | 8 | 1 | 3 | 0.6667 | 3 | 1 | 18 | 0.9444 | 7 | 1 | 10 | 0.8 | 3 | 1 | 6 | 1 | 26 | 0.9231 |
| 1_12580005 | 11 | 0.2727 | 8 | 0.375 | 4 | 0.25 | 6 | 0.1667 | 29 | 0.2759 | 7 | 0.4286 | 12 | 0.25 | 11 | 0.4545 | 11 | 0.1818 | 41 | 0.3171 |
| 1_12580006 | 10 | 0.1 | 8 | 0.25 | 4 | 0 | 5 | 0 | 27 | 0.1111 | 8 | 0 | 15 | 0.0667 | 10 | 0 | 11 | 0.1818 | 44 | 0.0682 |
| 1_12580009 | 12 | 0.0833 | 7 | 0 | 3 | 0 | 6 | 0.3333 | 28 | 0.1071 | 6 | 0.1667 | 15 | 0.3333 | 13 | 0.1538 | 12 | 0.25 | 46 | 0.2391 |
| 1_12580028 | 11 | 0.5455 | 7 | 0.7143 | 4 | 1 | 5 | 0.6 | 27 | 0.6667 | 7 | 0.7143 | 13 | 0.7692 | 13 | 0.9231 | 10 | 0.6 | 43 | 0.7674 |
| 1_12580030 | 13 | 0.5385 | 8 | 1 | 4 | 0.5 | 5 | 0.8 | 30 | 0.7 | 7 | 0.8571 | 15 | 0.6 | 12 | 0.6667 | 10 | 0.9 | 44 | 0.7273 |
| 1_12580054 | 10 | 1 | 6 | 1 | 4 | 1 | 4 | 1 | 24 | 1 | 6 | 1 | 12 | 1 | 12 | 1 | 10 | 1 | 40 | 1 |
| 1_12580054 | 14 | 0.2143 | 7 | 0.1429 | 5 | 0.2 | 6 | 0.1667 | 32 | 0.1875 | 5 | 0 | 14 | 0.0714 | 13 | 0.1538 | 10 | 0.3 | 42 | 0.1429 |
| 1_12580060 | 10 | 0.1 | 5 | 0.8 | 2 | 0 | 3 | 0 | 20 | 0.25 | 6 | 0.5 | 8 | 0.25 | 11 | 0.4545 | 7 | 0.5714 | 32 | 0.4375 |
| 1_12580063 | 11 | 1 | 7 | 0.8571 | 3 | 0.3333 | 4 | 1 | 25 | 0.88 | 5 | 1 | 12 | 0.8333 | 12 | 0.9167 | 8 | 0.875 | 37 | 0.8919 |
| 1_12580067 | 10 | 0.2 | 6 | 0.5 | NA | NA | 3 | 0 | 19 | 0.2632 | 2 | 1 | 9 | 0.4444 | 7 | 0.1429 | 6 | 0 | 24 | 0.2917 |
| 1_12580069 | 13 | 0.9231 | 7 | 1 | 2 | 1 | 4 | 1 | 26 | 0.9615 | 6 | 1 | 11 | 0.9091 | 9 | 1 | 10 | 0.7 | 36 | 0.8889 |
| 1_12580090 | 12 | 0.3333 | 5 | 0 | 1 | 0 | 4 | 0.5 | 22 | 0.2727 | 5 | 0 | 13 | 0.5385 | 13 | 0.6154 | 8 | 0.375 | 39 | 0.4615 |
| 1_12580121 | 13 | 0.3077 | 6 | 0.5 | NA | NA | 4 | 0 | 23 | 0.3043 | 5 | 0.4 | 10 | 0.2 | 12 | 0.1667 | 11 | 0.1818 | 38 | 0.2105 |
| 1_12723445 | 1 | 0 | 1 | 1 | NA | NA | 2 | 0.5 | 4 | 0.5 | NA | NA | 1 | 1 | 1 | 0 | NA | NA | 2 | 0.5 |
| 1_12741045 | 1 | 1 | 3 | 1 | NA | NA | 3 | 1 | 7 | 1 | NA | NA | 1 | 1 | 1 | 1 | NA | NA | 2 | 1 |
| 1_12741048 | 1 | 1 | 4 | 0.25 | NA | NA | 2 | 0.5 | 7 | 0.4286 | NA | NA | 1 | 0 | 2 | 0.5 | NA | NA | 3 | 0.3333 |
| 1_12741054 | NA | NA | 4 | 1 | NA | NA | 3 | 0.6667 | 7 | 0.8571 | NA | NA | 1 | 1 | 1 | 1 | NA | NA | 2 | 1 |
| 1_12741079 | 3 | 0.3333 | 4 | 0.25 | 1 | 0 | 3 | 0.3333 | 11 | 0.2727 | NA | NA | 2 | 0.5 | 3 | 0.3333 | NA | NA | 5 | 0.4 |
| 1_12741084 | 3 | 1 | 4 | 0.75 | 1 | 1 | 3 | 1 | 11 | 0.9091 | NA | NA | 2 | 1 | 2 | 1 | NA | NA | 4 | 1 |
| 1_12741134 | 4 | 0 | 4 | 0.25 | 2 | 0 | 4 | 0 | 14 | 0.0714 | 1 | 0 | 4 | 0 | 4 | 0 | 2 | 0.5 | 11 | 0.0909 |
| 1_12741225 | 5 | 0 | 5 | 0 | 3 | 0.3333 | 4 | 0.25 | 17 | 0.1176 | 2 | 0.5 | 8 | 0 | 4 | 0.25 | 3 | 0 | 17 | 0.1176 |
| 1_12741569 | 2 | 0 | 3 | 0 | 2 | 0.5 | NA | NA | 7 | 0.1429 | 1 | 0 | 3 | 0.3333 | 2 | 0 | 2 | 0 | 8 | 0.125 |
| 1_12741599 | 4 | 0.25 | 5 | 0.2 | 1 | 0 | NA | NA | 10 | 0.2 | 3 | 0.6667 | 5 | 0.4 | 2 | 0 | 1 | 1 | 11 | 0.4545 |
| 1_12741906 | 2 | 0 | 4 | 0.25 | 5 | 0.4 | 2 | 0.5 | 13 | 0.3077 | 1 | 0 | NA | NA | 1 | 1 | 1 | 0 | 3 | 0.3333 |
| 1_12741916 | 2 | 0.5 | 3 | 0 | 4 | 0 | 2 | 0.5 | 11 | 0.1818 | 1 | 0 | NA | NA | 1 | 0 | 1 | 1 | 3 | 0.3333 |
| 1_12772992 | 3 | 0.3333 | 5 | 0.4 | 1 | 0 | 5 | 0.2 | 14 | 0.2857 | 1 | 0 | 2 | 0 | 3 | 1 | 3 | 0.3333 | 9 | 0.4444 |
| 1_12772994 | 3 | 0.3333 | 2 | 0 | 1 | 0 | 6 | 0 | 12 | 0.0833 | 1 | 0 | 2 | 1 | 3 | 0 | 3 | 0 | 9 | 0.2222 |
| 1_12773019 | 3 | 0.3333 | 4 | 0.5 | 2 | 1 | 4 | 0.75 | 13 | 0.6154 | 1 | 0 | 2 | 0.5 | 2 | 0.5 | 2 | 1 | 7 | 0.5714 |
| 1_12773020 | 3 | 0.3333 | 5 | 0.2 | 2 | 0 | 7 | 0 | 17 | 0.1176 | 1 | 0 | 3 | 0.3333 | 3 | 0 | 3 | 0 | 10 | 0.1 |
| 1_12773026 | 2 | 1 | 5 | 0.8 | 3 | 1 | 7 | 0.7143 | 17 | 0.8235 | 1 | 1 | 3 | 1 | 4 | 0.75 | 3 | 1 | 11 | 0.9091 |
| 1_12773030 | 1 | 1 | 4 | 0.5 | 3 | 0.3333 | 5 | 0.6 | 13 | 0.5385 | NA | NA | 2 | 1 | 4 | 0.5 | NA | NA | 6 | 0.6667 |
| 1_12773046 | 3 | 1 | 3 | 0.6667 | 3 | 0.3333 | 6 | 0.6667 | 15 | 0.6667 | 1 | 0 | 4 | 0.25 | 3 | 0.3333 | 2 | 0.5 | 10 | 0.3 |
| 1_12773050 | 2 | 0 | 4 | 0.25 | 3 | 0.6667 | 3 | 0 | 12 | 0.25 | NA | NA | 3 | 0 | 3 | 0.3333 | NA | NA | 6 | 0.1667 |
| 1_12773059 | 2 | 0.5 | 5 | 0 | 2 | 0.5 | 4 | 0.5 | 13 | 0.3077 | 1 | 0 | 4 | 0.75 | 4 | 0 | 2 | 1 | 11 | 0.4545 |
| 1_12773072 | 3 | 0 | 2 | 0 | 4 | 0.25 | 5 | 0.2 | 14 | 0.1429 | 1 | 0 | 4 | 0.25 | 4 | 0 | 2 | 0 | 11 | 0.0909 |
| 1_12773114 | 3 | 1 | 4 | 0.5 | 5 | 1 | 7 | 0.7143 | 19 | 0.7895 | 1 | 1 | 4 | 1 | 4 | 1 | 2 | 0.5 | 11 | 0.9091 |
| 1_12773116 | 1 | 0 | 4 | 0.25 | 5 | 0.4 | 6 | 0 | 16 | 0.1875 | 1 | 0 | 3 | 0.3333 | 4 | 0 | 2 | 0 | 10 | 0.1 |
| 1_12773116 | 3 | 1 | 4 | 0.75 | 4 | 1 | 5 | 0.8 | 16 | 0.875 | 1 | 0 | 3 | 1 | 3 | 0.6667 | 2 | 0 | 9 | 0.5556 |
| 1_12773123 | 3 | 1 | 6 | 1 | 5 | 0.8 | 2 | 1 | 16 | 0.9375 | 1 | 1 | 4 | 1 | 5 | 1 | 1 | 1 | 11 | 1 |
| 1_12773124 | 3 | 1 | 6 | 0.6667 | 5 | 1 | 6 | 0.8333 | 20 | 0.85 | 1 | 1 | 4 | 1 | 5 | 0.8 | 3 | 1 | 13 | 0.9231 |
| 1_12773126 | 3 | 1 | 6 | 0.8333 | 5 | 1 | 7 | 1 | 21 | 0.9524 | 1 | 1 | 3 | 1 | 5 | 1 | 2 | 1 | 11 | 1 |
| 1_12773129 | 3 | 0.6667 | 6 | 0.3333 | 3 | 0 | 6 | 0 | 18 | 0.2222 | 1 | 0 | 3 | 0 | 4 | 0.5 | 3 | 0.6667 | 11 | 0.3636 |
| 1_12773147 | 2 | 1 | 6 | 0.8333 | 5 | 1 | 6 | 0.8333 | 19 | 0.8947 | 1 | 1 | 3 | 1 | 4 | 1 | 1 | 1 | 9 | 1 |
| 1_12773150 | 3 | 0.6667 | 5 | 0.2 | 6 | 0.8333 | 5 | 0.2 | 19 | 0.4737 | 1 | 0 | 4 | 0.5 | 5 | 0.2 | 2 | 0.5 | 12 | 0.3333 |
| 1_12773192 | 3 | 0 | 6 | 0.3333 | 6 | 0 | 7 | 0 | 22 | 0.0909 | 1 | 0 | 4 | 0 | 6 | 0 | 4 | 0.25 | 15 | 0.0667 |
| 1_12773266 | 4 | 0.25 | 4 | 0 | 4 | 0.25 | 5 | 0.4 | 17 | 0.2353 | NA | NA | 4 | 0.5 | 7 | 0 | 4 | 0.25 | 15 | 0.2 |

| | | | | | | | | | | | | | | | | | | | | |
|---|---|---|---|---|---|---|---|---|---|---|---|---|---|---|---|---|---|---|---|---|
| 1_1277326( | 4 | 0.25 | 5 | 0 | 5 | 0 | 4 | 0 | 18 | 0.0556 | NA | NA | 4 | 0 | 7 | 0 | 4 | 0.25 | 15 | 0.0667 |
| 1_1277334( | 4 | 0.5 | 5 | 0.6 | 6 | 0.6667 | 8 | 0.375 | 23 | 0.5217 | NA | NA | 4 | 0.5 | 7 | 0.8571 | 4 | 0.5 | 15 | 0.6667 |
| 1_1277335: | 3 | 0.3333 | 3 | 0.3333 | 4 | 0.75 | 6 | 0.5 | 16 | 0.5 | 1 | 1 | 3 | 1 | 6 | 0.5 | 4 | 0.5 | 14 | 0.6429 |
| 1_1277335 | 4 | 0 | 3 | 0.3333 | 3 | 0 | 6 | 0 | 16 | 0.0625 | 1 | 0 | 3 | 0.3333 | 6 | 0.1667 | 4 | 0.25 | 14 | 0.2143 |
| 1_1277335 | 4 | 0.25 | 3 | 0 | 6 | 0.1667 | 5 | 0 | 18 | 0.1111 | 1 | 0 | 3 | 0.6667 | 6 | 0 | 4 | 0 | 14 | 0.1429 |
| 1_1277338 | 3 | 1 | 1 | 1 | 3 | 0.6667 | 6 | 0.8333 | 13 | 0.8462 | 1 | 1 | 1 | 1 | 5 | 0.8 | 4 | 1 | 11 | 0.9091 |
| 1_1277349 | 4 | 0 | 1 | 1 | 3 | 0 | 6 | 0 | 14 | 0.0714 | NA | NA | 3 | 0 | 5 | 0.2 | 3 | 0 | 11 | 0.0909 |
| 1_1277354 | 5 | 0.2 | 1 | 1 | 1 | 0 | 6 | 0 | 13 | 0.1538 | NA | NA | 5 | 0.2 | 5 | 0 | 4 | 0 | 14 | 0.0714 |
| 1_1277373 | 1 | 1 | NA | NA | NA | NA | NA | NA | 1 | 1 | NA | NA | NA | NA | NA | NA | 1 | 1 | 1 | 1 |
| 1_1277375 | 1 | 1 | NA | NA | NA | NA | NA | NA | 1 | 1 | NA | NA | NA | NA | NA | NA | 1 | 1 | 1 | 1 |
| 1_1277377 | 1 | 1 | NA | NA | NA | NA | NA | NA | 1 | 1 | NA | NA | NA | NA | NA | NA | 1 | 1 | 1 | 1 |
| 1_1277429 | NA | NA | NA | NA | NA | NA | 2 | 0.5 | 2 | 0.5 | 1 | 1 | 2 | 0 | 1 | 1 | 3 | 0.6667 | 7 | 0.5714 |
| 1_1277452( | NA | NA | NA | NA | NA | NA | 2 | 1 | 2 | 1 | NA | NA | 1 | 1 | 1 | 1 | 1 | 1 | 3 | 1 |
| 1_1277452: | NA | NA | NA | NA | NA | NA | 2 | 0.5 | 2 | 0.5 | 1 | 1 | 1 | 1 | 1 | 1 | 1 | 0 | 4 | 0.75 |
| 1_1277452 | NA | NA | NA | NA | NA | NA | 2 | 1 | 2 | 1 | 1 | 1 | 1 | 1 | NA | NA | 2 | 0.5 | 4 | 0.75 |
| 1_1277498 | NA | NA | NA | NA | NA | NA | 2 | 1 | 2 | 1 | 1 | 1 | 1 | 1 | NA | NA | 2 | 1 | 4 | 1 |
| 1_1277503 | NA | NA | NA | NA | NA | NA | 1 | 1 | 1 | 1 | NA | NA | 1 | 1 | NA | NA | NA | NA | 1 | 1 |
| 1_1277504 | NA | NA | NA | NA | NA | NA | 1 | 1 | 1 | 1 | NA | NA | 1 | 1 | NA | NA | 1 | 1 | 2 | 1 |
| 1_1278477 | 1 | 1 | NA | NA | NA | NA | NA | NA | 1 | 1 | 1 | 1 | NA | NA | NA | NA | NA | NA | 1 | 1 |
| 1_1278481 | 1 | 1 | NA | NA | NA | NA | NA | NA | 1 | 1 | 1 | 1 | NA | NA | NA | NA | NA | NA | 1 | 1 |
| 1_1278703( | 5 | 0.6 | 5 | 0.8 | 9 | 0.6667 | 4 | 0.75 | 23 | 0.6957 | 2 | 0.5 | 11 | 0.6364 | 9 | 0.7778 | 7 | 0.7143 | 29 | 0.6897 |
| 1_1278703( | 6 | 0.3333 | 8 | 0.125 | 7 | 0.2857 | 6 | 0 | 27 | 0.1852 | 1 | 0 | 10 | 0.5 | 10 | 0.5 | 9 | 0.2222 | 30 | 0.4 |
| 1_1278704 | 4 | 0.5 | 9 | 0.7778 | 8 | 0.375 | 5 | 1 | 26 | 0.6538 | 5 | 1 | 10 | 0.9 | 8 | 0.875 | 6 | 0.3333 | 29 | 0.7931 |
| 1_1278705 | 6 | 0 | 7 | 0.1429 | 10 | 0 | 7 | 0.1429 | 30 | 0.0667 | 5 | 0 | 10 | 0 | 9 | 0.1111 | 7 | 0.2857 | 31 | 0.0968 |
| 1_1278706 | 5 | 0.4 | 10 | 0.5 | 8 | 0.125 | 5 | 0.2 | 28 | 0.3214 | 5 | 0.6 | 11 | 0.4545 | 11 | 0.3636 | 7 | 0.1429 | 34 | 0.3824 |
| 1_1278707 | 7 | 0.1429 | 10 | 0 | 9 | 0 | 7 | 0.1429 | 33 | 0.0606 | 6 | 0 | 10 | 0.1 | 13 | 0 | 10 | 0.1 | 39 | 0.0513 |
| 1_1278714 | 5 | 0.6 | 10 | 0.4 | 9 | 0.6667 | 9 | 0.2222 | 33 | 0.4545 | 7 | 0.5714 | 14 | 0.6429 | 12 | 0.5 | 11 | 0.5455 | 44 | 0.5682 |
| 1_1278823 | 1 | 0 | 1 | 1 | 1 | 1 | NA | NA | 3 | 0.6667 | NA | NA | NA | NA | NA | NA | 2 | 1 | 2 | 1 |
| 1_1278825 | 1 | 0 | 1 | 0 | 2 | 1 | NA | NA | 4 | 0.5 | NA | NA | NA | NA | NA | NA | 2 | 0.5 | 2 | 0.5 |
| 1_1278876 | 1 | 1 | 1 | 1 | 1 | 1 | NA | NA | 3 | 1 | NA | NA | NA | NA | NA | NA | 2 | 1 | 2 | 1 |
| 1_1279042 | NA | NA | 1 | 1 | NA | NA | NA | NA | 1 | 1 | NA | NA | NA | NA | NA | NA | 1 | 1 | 1 | 1 |
| 1_1281642 | 10 | 0.2 | 13 | 0.6154 | 8 | 0.125 | 18 | 0.3333 | 49 | 0.3469 | 15 | 0.4 | 14 | 0.3571 | 11 | 0.5455 | 9 | 0.5556 | 49 | 0.449 |
| 1_1281643 | 14 | 0 | 19 | 0.1053 | 10 | 0 | 14 | 0.0714 | 57 | 0.0526 | 14 | 0.0714 | 17 | 0.0588 | 16 | 0 | 13 | 0.1538 | 60 | 0.0667 |
| 1_1281644 | 10 | 0.1 | 15 | 0 | 10 | 0.1 | 12 | 0.25 | 47 | 0.1064 | 10 | 0.1 | 11 | 0.0909 | 12 | 0.1667 | 7 | 0 | 40 | 0.1 |
| 1_1281645 | 13 | 0.6923 | 19 | 0.2105 | 9 | 0 | 17 | 0.4118 | 58 | 0.3448 | 16 | 0.1875 | 15 | 0.2667 | 13 | 0.2308 | 13 | 0.3077 | 57 | 0.2456 |
| 1_1281645 | 14 | 0.7857 | 24 | 0.7083 | 12 | 0.5833 | 17 | 0.7059 | 67 | 0.7015 | 17 | 0.7059 | 20 | 0.65 | 17 | 0.8235 | 15 | 0.8667 | 69 | 0.7536 |
| 1_1281646 | 14 | 0.2143 | 24 | 0.1667 | 11 | 0 | 19 | 0.1579 | 68 | 0.1471 | 17 | 0.3529 | 18 | 0.0556 | 19 | 0.1579 | 14 | 0.0714 | 68 | 0.1618 |
| 1_1281648 | 18 | 0.1111 | 26 | 0.1154 | 13 | 0.1538 | 22 | 0.0909 | 79 | 0.1139 | 19 | 0 | 21 | 0.1905 | 20 | 0 | 14 | 0 | 74 | 0.0541 |
| 1_1281653 | 13 | 0.1538 | 22 | 0.1364 | 6 | 0.1667 | 19 | 0.0526 | 60 | 0.1167 | 13 | 0.1538 | 18 | 0 | 17 | 0.1176 | 11 | 0.0909 | 59 | 0.0847 |
| 1_1282025 | 3 | 0 | 4 | 0 | 5 | 0.2 | 1 | 0 | 13 | 0.0769 | 3 | 0.3333 | 5 | 0.2 | 3 | 0 | 4 | 0 | 15 | 0.1333 |
| 1_1282047 | 6 | 0.1667 | 7 | 0.1429 | 9 | 0.1111 | 4 | 0 | 26 | 0.1154 | 3 | 0.3333 | 8 | 0.25 | 10 | 0.1 | 5 | 0 | 26 | 0.1538 |
| 1_1282047 | 6 | 0 | 6 | 0.1667 | 8 | 0 | 4 | 0.25 | 24 | 0.0833 | 3 | 0 | 9 | 0.2222 | 10 | 0 | 6 | 0.1667 | 28 | 0.1071 |
| 1_1282048 | 7 | 0.1429 | 5 | 0.2 | 8 | 0.25 | 4 | 0.25 | 24 | 0.2083 | 4 | 0.25 | 9 | 0.4444 | 10 | 0.2 | 5 | 0.4 | 28 | 0.3214 |
| 1_1282056 | 2 | 0 | 3 | 0.3333 | 2 | 0 | 2 | 0 | 9 | 0.1111 | 2 | 0.5 | 3 | 0.3333 | 2 | 0 | NA | NA | 7 | 0.2857 |
| 1_1282059 | 3 | 0.3333 | 3 | 0 | 1 | 0 | 2 | 0 | 9 | 0.1111 | 2 | 0 | 2 | 0.5 | 2 | 0 | NA | NA | 6 | 0.1667 |
| 1_1282595 | 1 | 0 | 1 | 0 | 2 | 1 | NA | NA | 4 | 0.5 | 1 | 1 | 1 | 0 | NA | NA | NA | NA | 2 | 0.5 |
| 1_1282596 | 1 | 0 | 1 | 0 | 2 | 0.5 | NA | NA | 4 | 0.25 | 1 | 0 | 1 | 0 | NA | NA | 1 | 1 | 3 | 0.3333 |
| 1_1301459 | NA | NA | NA | NA | 1 | 1 | NA | NA | 1 | 1 | NA | NA | NA | NA | 3 | 0.3333 | NA | NA | 3 | 0.3333 |
| 1_1301460 | NA | NA | NA | NA | 1 | 1 | NA | NA | 1 | 1 | NA | NA | NA | NA | 3 | 0.3333 | NA | NA | 3 | 0.3333 |
| 1_1301460 | NA | NA | NA | NA | 1 | 1 | NA | NA | 1 | 1 | NA | NA | NA | NA | 3 | 1 | NA | NA | 3 | 1 |
| 1_1301462 | NA | NA | NA | NA | 1 | 1 | NA | NA | 1 | 1 | NA | NA | NA | NA | 3 | 0.6667 | NA | NA | 3 | 0.6667 |
| 1_1301462 | NA | NA | NA | NA | 1 | 1 | NA | NA | 1 | 1 | NA | NA | NA | NA | 2 | 0.5 | NA | NA | 2 | 0.5 |
| 1_1306174 | 1 | 0 | 1 | 0 | 1 | 1 | 4 | 0.25 | 7 | 0.2857 | NA | NA | 1 | 0 | 7 | 0.2857 | 7 | 0.2857 | 15 | 0.2667 |
| 1_1306181( | 1 | 0 | NA | NA | 1 | 0 | 2 | 0.5 | 4 | 0.25 | NA | NA | NA | NA | 7 | 0.8571 | 4 | 0.75 | 11 | 0.8182 |
| 1_1306181 | 1 | 1 | NA | NA | 1 | 0 | 2 | 0 | 4 | 0.25 | NA | NA | 1 | 0 | 8 | 0 | 4 | 0.25 | 13 | 0.0769 |
| 1_1306184 | 1 | 0 | NA | NA | 1 | 0 | 3 | 0.6667 | 5 | 0.4 | NA | NA | 1 | 1 | 3 | 0.3333 | 3 | 0 | 7 | 0.2857 |
| 1_1306185 | 1 | 1 | NA | NA | 1 | 1 | 3 | 1 | 5 | 1 | NA | NA | 1 | 1 | 5 | 0.8 | 4 | 0.5 | 10 | 0.7 |
| 1_1306185 | 1 | 0 | NA | NA | 1 | 0 | 3 | 1 | 5 | 0.6 | NA | NA | 1 | 0 | 5 | 1 | 4 | 0.25 | 10 | 0.6 |
| 1_1306185 | NA | NA | NA | NA | 1 | 0 | 3 | 0.6667 | 4 | 0.5 | NA | NA | 1 | 1 | 4 | 0.25 | 4 | 0 | 9 | 0.2222 |
| 1_1306187 | 1 | 1 | NA | NA | 1 | 1 | 2 | 0.5 | 4 | 0.75 | NA | NA | 1 | 1 | 4 | 0.5 | 3 | 1 | 8 | 0.75 |
| 1_1306187 | 1 | 1 | NA | NA | NA | NA | 1 | 1 | 2 | 1 | NA | NA | 1 | 0 | 5 | 0.6 | 4 | 0.5 | 10 | 0.5 |
| 1_1306216 | 1 | 0 | NA | NA | 1 | 1 | 1 | 0 | 3 | 0.3333 | NA | NA | NA | NA | 3 | 0.3333 | 1 | 0 | 4 | 0.25 |
| 1_1306254 | 1 | 1 | NA | NA | NA | NA | 1 | 0 | 2 | 0.5 | NA | NA | NA | NA | 1 | 0 | 1 | 1 | 2 | 0.5 |
| 1_1306467 | NA | NA | 1 | 1 | NA | NA | NA | NA | 1 | 1 | NA | NA | NA | NA | 1 | 0 | 2 | 0.5 | 3 | 0.3333 |
| 1_1306470 | NA | NA | 1 | 1 | NA | NA | NA | NA | 1 | 1 | NA | NA | 1 | 1 | 2 | 0.5 | 3 | 1 | 6 | 0.8333 |

| | | | | | | | | | | | | | | | | | | | | |
|---|---|---|---|---|---|---|---|---|---|---|---|---|---|---|---|---|---|---|---|---|
| 1_13064709 | NA | NA | 1 | 0 | NA | NA | 1 | 1 | 2 | 0.5 | NA | NA | 1 | 1 | 2 | 1 | 3 | 0 | 6 | 0.5 |
| 1_13064711 | 1 | 1 | 1 | 1 | NA | NA | 1 | 1 | 3 | 1 | NA | NA | 1 | 1 | 2 | 1 | 3 | 0.3333 | 6 | 0.6667 |
| 1_13064724 | 1 | 1 | 1 | 0 | NA | NA | 1 | 0 | 3 | 0.3333 | NA | NA | 1 | 1 | 2 | 0 | 1 | 0 | 4 | 0.25 |
| 1_13064728 | NA | NA | 1 | 1 | NA | NA | 1 | 0 | 2 | 0.5 | NA | NA | 1 | 0 | 2 | 1 | 2 | 0 | 5 | 0.4 |
| 1_13064728 | 1 | 1 | 1 | 1 | NA | NA | NA | NA | 2 | 1 | NA | NA | 1 | 1 | 1 | 1 | 3 | 0.3333 | 5 | 0.6 |
| 1_13064742 | NA | NA | 1 | 1 | NA | NA | 1 | 1 | 2 | 1 | NA | NA | 1 | 1 | 2 | 0.5 | 3 | 0.6667 | 6 | 0.6667 |
| 1_13064751 | 1 | 1 | 1 | 1 | NA | NA | 1 | 1 | 3 | 1 | NA | NA | 1 | 1 | 2 | 0.5 | 2 | 1 | 5 | 0.8 |
| 1_13064765 | 1 | 0 | 1 | 1 | NA | NA | 2 | 0.5 | 4 | 0.5 | NA | NA | 1 | 1 | 2 | 0 | 3 | 0 | 6 | 0.1667 |
| 1_13064783 | 1 | 1 | 1 | 0 | NA | NA | 2 | 0.5 | 4 | 0.5 | NA | NA | NA | NA | 2 | 1 | 3 | 0 | 5 | 0.4 |
| 1_13067039 | 1 | 1 | NA | NA | NA | NA | 1 | 1 | 2 | 1 | NA | NA | 2 | 1 | NA | NA | NA | NA | 2 | 1 |
| 1_13067104 | 1 | 1 | NA | NA | NA | NA | NA | NA | 1 | 1 | NA | NA | 2 | 1 | NA | NA | NA | NA | 2 | 1 |
| 1_13067122 | 1 | 1 | NA | NA | NA | NA | 1 | 1 | 2 | 1 | NA | NA | 3 | 1 | NA | NA | NA | NA | 3 | 1 |
| 1_13067134 | 1 | 1 | NA | NA | NA | NA | 1 | 1 | 2 | 1 | NA | NA | 2 | 0.5 | NA | NA | NA | NA | 2 | 0.5 |
| 1_13067145 | 1 | 0 | NA | NA | NA | NA | 1 | 1 | 2 | 0.5 | NA | NA | 2 | 0.5 | 1 | 0 | NA | NA | 3 | 0.3333 |
| 1_13067148 | 1 | 1 | NA | NA | NA | NA | 1 | 1 | 2 | 1 | NA | NA | 3 | 0.6667 | NA | NA | NA | NA | 3 | 0.6667 |
| 1_13067164 | 1 | 1 | NA | NA | NA | NA | 1 | 1 | 2 | 1 | NA | NA | 2 | 1 | NA | NA | NA | NA | 2 | 1 |
| 1_13067179 | 1 | 0 | NA | NA | NA | NA | 1 | 1 | 2 | 0.5 | NA | NA | 3 | 0.6667 | NA | NA | NA | NA | 3 | 0.6667 |
| 1_13067198 | 1 | 1 | NA | NA | NA | NA | 1 | 1 | 2 | 1 | NA | NA | 3 | 0.6667 | NA | NA | NA | NA | 3 | 0.6667 |
| 1_13067208 | 1 | 1 | NA | NA | NA | NA | 1 | 1 | 2 | 1 | NA | NA | 2 | 0.5 | NA | NA | NA | NA | 2 | 0.5 |
| 1_13067226 | 1 | 1 | NA | NA | NA | NA | NA | NA | 1 | 1 | NA | NA | 2 | 1 | NA | NA | NA | NA | 2 | 1 |
| 1_13098150 | 4 | 0 | 6 | 0 | 2 | 0 | 5 | 0.2 | 17 | 0.0588 | 3 | 0 | 3 | 0 | 2 | 0.5 | 3 | 0 | 11 | 0.0909 |
| 1_13098174 | 3 | 0.3333 | 3 | 0.3333 | 2 | 0 | 4 | 0 | 12 | 0.1667 | 3 | 0.3333 | 3 | 0 | 1 | 1 | 2 | 0 | 9 | 0.2222 |
| 1_13098177 | 3 | 0.3333 | 3 | 0.3333 | 2 | 0 | 4 | 0 | 12 | 0.1667 | 3 | 0.3333 | 2 | 0.5 | 2 | 0 | 2 | 0.5 | 9 | 0.3333 |
| 1_13098221 | 4 | 0 | 2 | 0 | 2 | 0.5 | 5 | 0.2 | 13 | 0.1538 | 3 | 0.3333 | 3 | 0 | 2 | 0 | 2 | 0 | 10 | 0.1 |
| 1_13098275 | 4 | 0.75 | 2 | 0.5 | 2 | 0.5 | 5 | 0.6 | 13 | 0.6154 | 3 | 0.6667 | 2 | 1 | 1 | 1 | 2 | 0.5 | 8 | 0.75 |
| 1_13098276 | 4 | 0.25 | 3 | 0 | 2 | 0 | 3 | 0 | 12 | 0.0833 | 3 | 0.3333 | 2 | 0 | 2 | 0 | 1 | 0 | 8 | 0.125 |
| 1_13098280 | 4 | 0 | 4 | 0.5 | 2 | 0.5 | 4 | 0 | 14 | 0.2143 | 3 | 0 | 1 | 0 | 2 | 0 | 2 | 0.5 | 8 | 0.125 |
| 1_13105619 | NA | NA | 1 | 0 | 1 | 1 | 1 | 1 | 3 | 0.6667 | 2 | 0.5 | 2 | 0.5 | NA | NA | 1 | 1 | 5 | 0.6 |
| 1_13105627 | NA | NA | 1 | 0 | 1 | 1 | 1 | 0 | 3 | 0.3333 | 2 | 0 | 2 | 0.5 | NA | NA | 1 | 0 | 5 | 0.2 |
| 1_13105629 | NA | NA | 2 | 0 | 2 | 0.5 | 1 | 0 | 5 | 0.2 | 2 | 0.5 | 1 | 0 | NA | NA | 1 | 1 | 4 | 0.5 |
| 1_13105639 | NA | NA | 2 | 1 | 2 | 0.5 | 1 | 1 | 5 | 0.8 | 3 | 1 | 2 | 1 | NA | NA | 1 | 1 | 6 | 1 |
| 1_13105640 | NA | NA | 2 | 0.5 | 2 | 1 | NA | NA | 4 | 0.75 | 1 | 1 | 2 | 0 | NA | NA | 1 | 1 | 4 | 0.5 |
| 1_13105652 | NA | NA | 2 | 1 | 1 | 1 | NA | NA | 3 | 1 | 2 | 1 | 2 | 1 | NA | NA | 1 | 1 | 5 | 1 |
| 1_13105657 | NA | NA | NA | NA | 1 | 1 | NA | NA | 1 | 1 | 1 | 1 | 1 | 1 | NA | NA | 1 | 1 | 3 | 1 |
| 1_13105670 | NA | NA | 2 | 0 | 1 | 1 | NA | NA | 3 | 0.3333 | 3 | 0.3333 | 1 | 0 | NA | NA | 1 | 0 | 5 | 0.2 |
| 1_13105673 | NA | NA | 2 | 1 | 1 | 1 | 1 | 1 | 4 | 1 | 3 | 1 | 1 | 1 | NA | NA | 1 | 1 | 5 | 1 |
| 1_13105688 | NA | NA | 2 | 0.5 | 1 | 0 | 1 | 0 | 4 | 0.25 | 3 | 0 | 1 | 1 | NA | NA | 1 | 0 | 5 | 0.2 |
| 1_13105696 | NA | NA | 2 | 0.5 | 1 | 1 | 1 | 1 | 4 | 0.75 | 2 | 0.5 | 1 | 1 | NA | NA | 1 | 1 | 4 | 0.75 |
| 1_13105697 | NA | NA | 2 | 1 | 1 | 1 | NA | NA | 3 | 1 | 3 | 1 | NA | NA | NA | NA | 1 | 1 | 4 | 1 |
| 1_13105729 | NA | NA | 2 | 0.5 | 1 | 0 | NA | NA | 3 | 0.3333 | 3 | 0.6667 | 1 | 0 | NA | NA | 1 | 0 | 5 | 0.4 |
| 1_13105730 | NA | NA | 1 | 1 | 1 | 1 | NA | NA | 2 | 1 | 3 | 1 | 1 | 1 | NA | NA | 1 | 1 | 5 | 1 |
| 1_13105737 | NA | NA | 2 | 0.5 | 1 | 1 | 1 | 0 | 4 | 0.5 | 2 | 0 | NA | NA | NA | NA | 1 | 1 | 3 | 0.3333 |
| 1_13105751 | NA | NA | 1 | 1 | 1 | 1 | NA | NA | 2 | 1 | 1 | 0 | NA | NA | NA | NA | 1 | 1 | 2 | 0.5 |
| 1_13110496 | NA | NA | 1 | 1 | 2 | 0 | NA | NA | 3 | 0.3333 | 3 | 1 | 3 | 0.6667 | 2 | 0 | 3 | 0.3333 | 11 | 0.5455 |
| 1_13110513 | NA | NA | 1 | 1 | 2 | 0 | NA | NA | 3 | 0.3333 | 3 | 0.3333 | 3 | 0.3333 | 2 | 0 | 2 | 0 | 10 | 0.2 |
| 1_13110576 | NA | NA | 1 | 1 | 2 | 1 | NA | NA | 3 | 1 | 1 | 1 | 1 | 1 | 2 | 1 | 2 | 1 | 6 | 1 |
| 1_13110604 | NA | NA | 1 | 0 | 2 | 1 | NA | NA | 3 | 0.6667 | 1 | 1 | 1 | 0 | 1 | 1 | 2 | 0.5 | 5 | 0.6 |
| 1_13110737 | NA | NA | NA | NA | 1 | 0 | 1 | 1 | 2 | 0.5 | NA | NA | NA | NA | NA | NA | 1 | 1 | 1 | 1 |
| 1_13110744 | NA | NA | NA | NA | NA | NA | 1 | 1 | 1 | 1 | NA | NA | NA | NA | NA | NA | 1 | 1 | 1 | 1 |
| 1_13110745 | NA | NA | NA | NA | 1 | 1 | NA | NA | 1 | 1 | NA | NA | NA | NA | NA | NA | 1 | 1 | 1 | 1 |
| 1_13110752 | NA | NA | NA | NA | 1 | 0 | 1 | 1 | 2 | 0.5 | NA | NA | NA | NA | NA | NA | 1 | 1 | 1 | 1 |
| 1_13110768 | NA | NA | NA | NA | 1 | 1 | NA | NA | 1 | 1 | NA | NA | NA | NA | NA | NA | 1 | 1 | 1 | 1 |
| 1_13121430 | 2 | 0 | 7 | 0 | 3 | 0.3333 | 4 | 0.25 | 16 | 0.125 | 5 | 0.2 | 2 | 0 | 9 | 0.2222 | 4 | 0 | 20 | 0.15 |
| 1_13121442 | 2 | 0 | 5 | 0.2 | 4 | 0.5 | 4 | 0 | 15 | 0.2 | 3 | 0 | 1 | 0 | 8 | 0.25 | 4 | 0.5 | 16 | 0.25 |
| 1_13121444 | 2 | 0 | 6 | 0.1667 | 3 | 0 | 3 | 0 | 14 | 0.0714 | 4 | 0.25 | 2 | 0 | 8 | 0.375 | 2 | 0 | 16 | 0.25 |
| 1_13121445 | 2 | 1 | 7 | 1 | 3 | 1 | 3 | 1 | 15 | 1 | 4 | 0.75 | 1 | 1 | 10 | 0.9 | 3 | 1 | 18 | 0.8889 |
| 1_13121448 | 1 | 0 | 5 | 0.2 | 2 | 0 | 3 | 0 | 11 | 0.0909 | 4 | 0 | 1 | 0 | 5 | 0 | 3 | 0.3333 | 13 | 0.0769 |
| 1_13121457 | 2 | 0 | 7 | 0.5714 | 3 | 0.6667 | 4 | 0.25 | 16 | 0.4375 | 5 | 0.6 | 2 | 0 | 10 | 0.6 | 2 | 1 | 19 | 0.5789 |
| 1_13121466 | 2 | 1 | 3 | 1 | 3 | 0.6667 | 4 | 1 | 12 | 0.9167 | 5 | 1 | 1 | 0 | 7 | 1 | 3 | 1 | 16 | 0.9375 |
| 1_13121497 | 2 | 0 | 5 | 0.2 | NA | NA | 2 | 1 | 9 | 0.3333 | 3 | 0.6667 | 1 | 0 | 5 | 0.2 | 3 | 0.3333 | 12 | 0.3333 |
| 1_13121498 | 1 | 0 | 6 | 0.5 | 2 | 0 | 3 | 1 | 12 | 0.5 | 2 | 0.5 | 2 | 0 | 7 | 0.5714 | 4 | 0.5 | 15 | 0.4667 |
| 1_13121628 | 1 | 0 | 4 | 0.25 | 4 | 0 | 3 | 0 | 12 | 0.0833 | 3 | 0.6667 | 2 | 0 | 7 | 0.1429 | 2 | 0 | 14 | 0.2143 |
| 1_13121643 | 1 | 0 | 3 | 0 | 5 | 0.2 | 3 | 0.3333 | 12 | 0.1667 | 3 | 0 | 2 | 0 | 6 | 0.1667 | 2 | 0 | 13 | 0.0769 |
| 1_13121645 | 1 | 0 | 4 | 0 | 5 | 0 | 2 | 0.5 | 12 | 0.0833 | 3 | 0 | 2 | 0.5 | 6 | 0 | 1 | 0 | 12 | 0.0833 |
| 1_13121675 | 1 | 0 | 4 | 0.5 | 3 | 0 | 2 | 0.5 | 10 | 0.3 | 1 | 0 | 1 | 0 | 4 | 0.25 | NA | NA | 6 | 0.1667 |

| | | | | | | | | | | | | | | | | | | | | |
|---|---|---|---|---|---|---|---|---|---|---|---|---|---|---|---|---|---|---|---|---|
| 1_13121702 | 1 | 1 | 4 | 0.5 | 2 | 0.5 | 4 | 0.75 | 11 | 0.6364 | 2 | 1 | 2 | 1 | 6 | 0.6667 | 2 | 1 | 12 | 0.8333 |
| 1_13121706 | 1 | 0 | 4 | 0 | 3 | 0.6667 | 4 | 0.25 | 12 | 0.25 | 2 | 0 | 2 | 0.5 | 6 | 0 | 3 | 0 | 13 | 0.0769 |
| 1_13121726 | 1 | 1 | 2 | 0.5 | 3 | 0.6667 | 3 | 1 | 9 | 0.7778 | NA | NA | 2 | 1 | 5 | 0.8 | 2 | 0.5 | 9 | 0.7778 |
| 1_13121730 | 1 | 1 | 2 | 1 | 3 | 0.6667 | 4 | 0.75 | 10 | 0.8 | 1 | 1 | 2 | 0.5 | 5 | 1 | 3 | 0.6667 | 11 | 0.8182 |
| 1_13122050 | 1 | 0 | 2 | 0.5 | 1 | 0 | 4 | 0.5 | 8 | 0.375 | 1 | 0 | 2 | 1 | 5 | 0.2 | 1 | 0 | 9 | 0.3333 |
| 1_13122321 | 3 | 0.3333 | 1 | 0 | 1 | 0 | 2 | 0 | 7 | 0.1429 | 2 | 0 | 2 | 0.5 | 3 | 0.3333 | 2 | 0 | 9 | 0.2222 |
| 1_13145308 | 1 | 1 | NA | NA | NA | NA | 1 | 0 | 2 | 0.5 | NA | NA | 2 | 0.5 | 1 | 1 | 1 | 0 | 4 | 0.5 |
| 1_13145622 | 1 | 1 | NA | NA | 1 | 0 | NA | NA | 2 | 0.5 | 1 | 1 | NA | NA | NA | NA | NA | NA | 1 | 1 |
| 1_13145632 | 1 | 1 | NA | NA | 1 | 1 | NA | NA | 2 | 1 | 1 | 1 | NA | NA | NA | NA | NA | NA | 1 | 1 |
| 1_13163972 | 1 | 1 | NA | NA | NA | NA | NA | NA | 1 | 1 | NA | NA | 1 | 1 | 2 | 1 | NA | NA | 3 | 1 |
| 1_13164027 | 1 | 1 | NA | NA | NA | NA | NA | NA | 1 | 1 | NA | NA | 1 | 1 | 1 | 1 | NA | NA | 2 | 1 |
| 1_13164040 | 1 | 1 | NA | NA | NA | NA | NA | NA | 1 | 1 | NA | NA | 1 | 1 | 1 | 1 | NA | NA | 2 | 1 |
| 1_13164069 | 1 | 1 | NA | NA | NA | NA | NA | NA | 1 | 1 | NA | NA | 1 | 1 | 1 | 0 | NA | NA | 2 | 0.5 |
| 1_13164097 | 1 | 1 | NA | NA | NA | NA | NA | NA | 1 | 1 | NA | NA | 1 | 1 | 2 | 1 | NA | NA | 3 | 1 |
| 1_13164103 | 1 | 1 | NA | NA | NA | NA | NA | NA | 1 | 1 | NA | NA | 1 | 1 | 1 | 0 | NA | NA | 2 | 0.5 |
| 1_13175777 | 2 | 0.5 | 3 | 0 | 1 | 0 | 1 | 0 | 7 | 0.1429 | 1 | 0 | 1 | 0 | 3 | 0 | 4 | 0.25 | 9 | 0.1111 |
| 1_13175796 | 2 | 0.5 | 4 | 0.75 | 1 | 1 | 1 | 1 | 8 | 0.75 | 1 | 1 | 1 | 1 | 2 | 0.5 | 3 | 0.6667 | 7 | 0.7143 |
| 1_13175801 | 3 | 0.6667 | 4 | 0.25 | 2 | 0 | 1 | 1 | 10 | 0.4 | 2 | 0 | NA | NA | 2 | 0.5 | 4 | 0.5 | 8 | 0.375 |
| 1_13175814 | 3 | 1 | 5 | 1 | 2 | 1 | 1 | 1 | 11 | 1 | 2 | 1 | 1 | 1 | 3 | 0.6667 | 2 | 1 | 8 | 0.875 |
| 1_13175828 | 4 | 0.5 | 5 | 0.2 | NA | NA | NA | NA | 9 | 0.3333 | 2 | 0 | 1 | 0 | 3 | 0.3333 | 4 | 0 | 10 | 0.1 |
| 1_13175838 | 4 | 1 | 3 | 0.6667 | 2 | 1 | 1 | 1 | 10 | 0.9 | 2 | 0.5 | 1 | 0 | 4 | 0.75 | 4 | 1 | 11 | 0.7273 |
| 1_13175846 | 4 | 0.5 | 5 | 0.4 | 1 | 0 | 1 | 0 | 11 | 0.3636 | 2 | 0 | 1 | 1 | 5 | 0.2 | 4 | 0.75 | 12 | 0.4167 |
| 1_13175865 | 3 | 1 | 2 | 1 | 2 | 0.5 | 1 | 1 | 8 | 0.875 | 1 | 0 | NA | NA | 4 | 1 | 4 | 0.75 | 9 | 0.7778 |
| 1_13175869 | 2 | 0.5 | 5 | 0.2 | 1 | 0 | 1 | 1 | 9 | 0.3333 | 2 | 0 | 1 | 1 | 5 | 0.8 | 3 | 1 | 11 | 0.7273 |
| 1_13177440 | 5 | 0.2 | 6 | 0.3333 | 6 | 0.3333 | 3 | 0 | 20 | 0.25 | 1 | 1 | 5 | 0.4 | 7 | 0.4286 | 7 | 0.4286 | 20 | 0.45 |
| 1_13177446 | 6 | 0.1667 | 7 | 0.2857 | 6 | 0.3333 | 2 | 0 | 21 | 0.2381 | 1 | 0 | 4 | 0 | 8 | 0.125 | 7 | 0.2857 | 20 | 0.15 |
| 1_13177469 | 6 | 0 | 5 | 0 | 6 | 0 | 3 | 0.3333 | 20 | 0.05 | 2 | 0 | 4 | 0.25 | 8 | 0.125 | 7 | 0.1429 | 21 | 0.0952 |
| 1_13177476 | 7 | 0.2857 | 6 | 0.1667 | 6 | 0.3333 | 2 | 0 | 21 | 0.2381 | 2 | 0 | 4 | 0.25 | 7 | 0 | 5 | 0 | 18 | 0.0556 |
| 1_13177569 | 11 | 0.0909 | 10 | 0.1 | 9 | 0.2222 | 5 | 0 | 35 | 0.1143 | 5 | 0 | 7 | 0.1429 | 10 | 0.3 | 7 | 0 | 29 | 0.1379 |
| 1_13177603 | 12 | 0.0833 | 10 | 0.2 | 10 | 0.3 | 6 | 0.1667 | 38 | 0.1842 | 5 | 0 | 8 | 0.125 | 10 | 0.1 | 7 | 0.5714 | 30 | 0.2 |
| 1_13177622 | 9 | 0.2222 | 7 | 0.1429 | 10 | 0.3 | 4 | 0 | 30 | 0.2 | 6 | 0.1667 | 8 | 0 | 11 | 0.1818 | 5 | 0.6 | 30 | 0.2 |
| 1_13177649 | 16 | 0.125 | 11 | 0.0909 | 11 | 0.3636 | 4 | 0 | 42 | 0.1667 | 7 | 0.1429 | 7 | 0 | 13 | 0.1538 | 7 | 0.1429 | 34 | 0.1176 |
| 1_13180030 | NA | NA | 1 | 1 | 3 | 0 | NA | NA | 4 | 0.25 | 1 | 0 | 1 | 1 | NA | NA | 1 | 0 | 3 | 0.3333 |
| 1_13180045 | NA | NA | 1 | 0 | 3 | 1 | NA | NA | 4 | 0.75 | 1 | 1 | NA | NA | 1 | 0 | 2 | 0.5 | 4 | 0.5 |
| 1_13180047 | NA | NA | 1 | 0 | 3 | 0.6667 | NA | NA | 4 | 0.5 | 1 | 0 | NA | NA | 1 | 1 | 2 | 0 | 4 | 0.25 |
| 1_13185899 | 1 | 1 | 2 | 0.5 | 4 | 0 | 6 | 0.6667 | 13 | 0.4615 | NA | NA | 9 | 0.2222 | 5 | 0.4 | 8 | 0.5 | 22 | 0.3636 |
| 1_13185901 | 1 | 0 | 2 | 0 | 5 | 0 | 6 | 0.1667 | 14 | 0.0714 | NA | NA | 9 | 0.2222 | 5 | 0.2 | 9 | 0.1111 | 23 | 0.1739 |
| 1_13185925 | 1 | 0 | 3 | 0 | 3 | 0 | 7 | 0.1429 | 14 | 0.0714 | NA | NA | 10 | 0.4 | 6 | 0.1667 | 8 | 0.125 | 24 | 0.25 |
| 1_13185942 | 1 | 0 | 3 | 0 | 6 | 0.3333 | 8 | 0.125 | 18 | 0.1667 | NA | NA | 11 | 0.2727 | 6 | 0.1667 | 6 | 0.1667 | 23 | 0.2174 |
| 1_13186033 | 1 | 0 | 5 | 0.2 | 8 | 0 | 3 | 0.3333 | 17 | 0.1176 | NA | NA | 5 | 0.4 | 6 | 0.1667 | 5 | 0 | 16 | 0.1875 |
| 1_13186382 | 12 | 0.0833 | 15 | 0 | 16 | 0.25 | 17 | 0 | 60 | 0.0833 | 9 | 0.1111 | 19 | 0.1053 | 10 | 0.2 | 15 | 0.1333 | 53 | 0.1321 |
| 1_13194577 | 1 | 1 | NA | NA | NA | NA | NA | NA | 1 | 1 | NA | NA | 1 | 1 | 1 | 1 | NA | NA | 2 | 1 |
| 1_13194600 | 1 | 1 | NA | NA | NA | NA | NA | NA | 1 | 1 | NA | NA | 1 | 1 | 1 | 0 | NA | NA | 2 | 0.5 |
| 1_13195089 | NA | NA | 4 | 0.25 | 4 | 0 | NA | NA | 8 | 0.125 | 2 | 0.5 | NA | NA | NA | NA | 1 | 0 | 3 | 0.3333 |
| 1_13195140 | 1 | 1 | NA | NA | NA | NA | NA | NA | 1 | 1 | NA | NA | 1 | 1 | 1 | 1 | NA | NA | 2 | 1 |
| 1_13195151 | 1 | 1 | NA | NA | NA | NA | NA | NA | 1 | 1 | NA | NA | NA | NA | 1 | 1 | NA | NA | 1 | 1 |
| 1_13196006 | 4 | 0 | 3 | 0 | 2 | 0 | 4 | 0.25 | 13 | 0.0769 | 1 | 0 | 4 | 0.5 | 2 | 0 | 2 | 1 | 9 | 0.4444 |
| 1_13196019 | 4 | 0 | 3 | 0 | 2 | 0.5 | 6 | 0 | 15 | 0.0667 | 2 | 0 | 3 | 0 | 2 | 1 | 3 | 0 | 10 | 0.2 |
| 1_13196496 | 6 | 0.1667 | 6 | 0 | 2 | 0 | 1 | 0 | 15 | 0.0667 | 2 | 0.5 | 3 | 0 | 1 | 0 | 4 | 0.25 | 10 | 0.2 |
| 1_13204145 | NA | NA | 10 | 0.1 | 8 | 0.125 | 1 | 0 | 19 | 0.1053 | 6 | 0.1667 | 5 | 0 | 3 | 0 | 1 | 1 | 15 | 0.1333 |
| 1_13204154 | NA | NA | 9 | 0.2222 | 10 | 0.3 | 1 | 0 | 20 | 0.25 | 6 | 0.1667 | 5 | 0 | 1 | 1 | 1 | 0 | 13 | 0.1538 |
| 1_13204193 | NA | NA | 12 | 0.3333 | 11 | 0.3636 | 1 | 0 | 24 | 0.3333 | 7 | 0.5714 | 4 | 0.5 | 3 | 0.6667 | 2 | 1 | 16 | 0.625 |
| 1_13204220 | NA | NA | 12 | 0.25 | 12 | 0.1667 | 1 | 0 | 25 | 0.2 | 7 | 0.2857 | 4 | 0 | 3 | 0.3333 | 2 | 0 | 16 | 0.1875 |
| 1_13204224 | NA | NA | 11 | 0.2727 | 11 | 0.1818 | 2 | 0 | 24 | 0.2083 | 7 | 0.1429 | 4 | 0.5 | 2 | 0 | 2 | 0 | 15 | 0.2 |
| 1_13204230 | NA | NA | 12 | 0.3333 | 11 | 0.4545 | 2 | 0 | 25 | 0.36 | 6 | 0 | 4 | 0 | 2 | 0.5 | 2 | 0.5 | 14 | 0.1429 |
| 1_13204257 | NA | NA | 8 | 0 | 5 | 0.4 | 2 | 0 | 15 | 0.1333 | 2 | 0 | 4 | 0.5 | 1 | 0 | NA | NA | 7 | 0.2857 |
| 1_13204262 | NA | NA | 11 | 0.3636 | 6 | 0.8333 | 2 | 0.5 | 19 | 0.5263 | 5 | 0.4 | 4 | 1 | 3 | 0.3333 | 2 | 0 | 14 | 0.5 |
| 1_13213535 | NA | NA | 1 | 1 | NA | NA | NA | NA | 1 | 1 | NA | NA | 1 | 1 | NA | NA | NA | NA | 1 | 1 |
| 1_13213541 | NA | NA | 1 | 1 | NA | NA | NA | NA | 1 | 1 | NA | NA | 1 | 1 | NA | NA | NA | NA | 1 | 1 |
| 1_13213552 | NA | NA | 1 | 1 | NA | NA | NA | NA | 1 | 1 | NA | NA | 1 | 0 | NA | NA | 1 | 1 | 2 | 0.5 |
| 1_13213554 | NA | NA | 1 | 1 | NA | NA | NA | NA | 1 | 1 | NA | NA | 1 | 0 | NA | NA | 1 | 1 | 2 | 0.5 |
| 1_13222696 | 1 | 1 | 3 | 0.3333 | 1 | 1 | 4 | 0.25 | 9 | 0.4444 | 4 | 0.25 | NA | NA | 1 | 0 | 1 | 1 | 6 | 0.3333 |
| 1_13222703 | 1 | 1 | 4 | 1 | 2 | 1 | 4 | 1 | 11 | 1 | 4 | 1 | NA | NA | 1 | 1 | 1 | 1 | 6 | 1 |
| 1_13222715 | 1 | 1 | 4 | 0.25 | 2 | 1 | 4 | 0.25 | 11 | 0.4545 | 3 | 0.6667 | 1 | 0 | 1 | 1 | 2 | 0.5 | 7 | 0.5714 |
| 1_13222719 | 1 | 0 | 3 | 0.3333 | 2 | 0 | 5 | 0.2 | 11 | 0.1818 | 3 | 0 | 2 | 0 | 1 | 0 | 1 | 1 | 7 | 0.1429 |

| | | | | | | | | | | | | | | | | | | | | |
|---|---|---|---|---|---|---|---|---|---|---|---|---|---|---|---|---|---|---|---|---|
| 1_13223245 | 1 | 0 | NA | NA | 2 | 0.5 | 5 | 0.6 | 8 | 0.5 | 3 | 0.3333 | NA | NA | 2 | 0.5 | 3 | 0.6667 | 8 | 0.5 |
| 1_13223248 | 1 | 1 | 5 | 1 | 2 | 1 | 7 | 1 | 15 | 1 | 5 | 1 | 2 | 1 | 2 | 1 | 3 | 1 | 12 | 1 |
| 1_13223252 | 1 | 0 | 3 | 0.3333 | 2 | 0 | 2 | 0 | 8 | 0.125 | 5 | 0 | 1 | 0 | 1 | 0 | 3 | 0.3333 | 10 | 0.1 |
| 1_13223260 | NA | NA | 3 | 1 | 2 | 1 | 7 | 0.8571 | 12 | 0.9167 | 4 | 1 | 2 | 1 | 2 | 0.5 | 3 | 1 | 11 | 0.9091 |
| 1_13223264 | 1 | 0 | 5 | 0.2 | 1 | 0 | 7 | 0.4286 | 14 | 0.2857 | 4 | 0.25 | 2 | 0 | 2 | 0 | 2 | 0 | 10 | 0.1 |
| 1_13223274 | 1 | 1 | 5 | 0.8 | 2 | 1 | 6 | 1 | 14 | 0.9286 | 5 | 1 | 3 | 1 | 2 | 1 | 1 | 1 | 11 | 1 |
| 1_13223308 | NA | NA | 4 | 0.75 | 2 | 0.5 | 6 | 0.6667 | 12 | 0.6667 | 3 | 0.3333 | 3 | 0.3333 | 2 | 0 | 1 | 0 | 9 | 0.2222 |
| 1_13223322 | 1 | 1 | 4 | 0.25 | 1 | 1 | 5 | 0.2 | 11 | 0.3636 | 3 | 0.6667 | 3 | 0.3333 | 2 | 0.5 | 1 | 0 | 9 | 0.4444 |
| 1_13228409 | 30 | 0.1333 | 40 | 0.075 | 44 | 0.1818 | 34 | 0.1176 | 148 | 0.1284 | 34 | 0.0294 | 24 | 0.125 | 22 | 0.1818 | 29 | 0.069 | 109 | 0.0917 |
| 1_13228442 | 27 | 0.2963 | 41 | 0.1951 | 37 | 0.1622 | 31 | 0.2258 | 136 | 0.2132 | 31 | 0.129 | 23 | 0.2609 | 22 | 0.1364 | 27 | 0.2222 | 103 | 0.1845 |
| 1_13228466 | 23 | 0.087 | 33 | 0.0303 | 33 | 0.1212 | 24 | 0.0833 | 113 | 0.0796 | 27 | 0.0741 | 22 | 0.0455 | 20 | 0.1 | 23 | 0.1739 | 92 | 0.0978 |
| 1_13228468 | 24 | 0.25 | 34 | 0.1176 | 34 | 0.1765 | 25 | 0.36 | 117 | 0.2137 | 24 | 0.0833 | 23 | 0.1304 | 19 | 0.2105 | 23 | 0.1304 | 89 | 0.1348 |
| 1_13228496 | 20 | 0.8 | 25 | 0.84 | 28 | 0.9643 | 23 | 1 | 96 | 0.9062 | 22 | 0.8182 | 19 | 0.8947 | 14 | 0.9286 | 22 | 0.9091 | 77 | 0.8831 |
| 1_13228503 | 23 | 0.6087 | 31 | 0.7419 | 24 | 0.6667 | 24 | 0.6667 | 102 | 0.6765 | 20 | 0.65 | 17 | 0.8235 | 17 | 0.6471 | 23 | 0.6957 | 77 | 0.7013 |
| 1_13228513 | 20 | 0.1 | 22 | 0.1818 | 23 | 0.1739 | 21 | 0.1905 | 86 | 0.1628 | 17 | 0.1765 | 15 | 0.1333 | 15 | 0.0667 | 16 | 0.25 | 63 | 0.1587 |
| 1_13228514 | 25 | 0.16 | 29 | 0.069 | 29 | 0.1034 | 25 | 0.16 | 108 | 0.1204 | 20 | 0.2 | 21 | 0 | 17 | 0 | 22 | 0.0455 | 80 | 0.0625 |
| 1_13228551 | 21 | 0.0476 | 23 | 0.087 | 22 | 0.0455 | 20 | 0.05 | 86 | 0.0581 | 20 | 0 | 16 | 0.0625 | 14 | 0.1429 | 19 | 0.1053 | 69 | 0.0725 |
| 1_13228805 | 17 | 0.3529 | 21 | 0.1429 | 17 | 0.1765 | 20 | 0.15 | 75 | 0.2 | 16 | 0.25 | 13 | 0.0769 | 14 | 0.2857 | 16 | 0.125 | 59 | 0.1864 |
| 1_13228807 | 19 | 0.1053 | 24 | 0.0833 | 21 | 0.1429 | 19 | 0.1579 | 83 | 0.1205 | 20 | 0.05 | 13 | 0.0769 | 15 | 0.2667 | 18 | 0.1667 | 66 | 0.1364 |
| 1_13228809 | 18 | 0.1667 | 21 | 0.0476 | 18 | 0.0556 | 19 | 0.2632 | 76 | 0.1316 | 16 | 0.25 | 15 | 0.1333 | 15 | 0.0667 | 16 | 0.25 | 62 | 0.1774 |
| 1_13228812 | 18 | 0.3889 | 21 | 0.6667 | 20 | 0.45 | 19 | 0.4211 | 78 | 0.4872 | 16 | 0.5625 | 11 | 0.3636 | 13 | 0.3846 | 17 | 0.8824 | 57 | 0.5789 |
| 1_13228814 | 20 | 0.8 | 21 | 0.7619 | 19 | 0.7895 | 18 | 0.7778 | 78 | 0.7821 | 15 | 0.8667 | 14 | 0.7857 | 12 | 0.75 | 18 | 0.8889 | 59 | 0.8305 |
| 1_13228833 | 18 | 0.1667 | 19 | 0.3684 | 20 | 0.1 | 17 | 0.3529 | 74 | 0.2432 | 16 | 0.1875 | 13 | 0 | 13 | 0.3846 | 15 | 0.2667 | 57 | 0.2105 |
| 1_13228835 | 19 | 0.1053 | 16 | 0.125 | 21 | 0.2381 | 14 | 0.0714 | 70 | 0.1429 | 11 | 0.1818 | 10 | 0 | 13 | 0.1538 | 15 | 0.1333 | 49 | 0.1224 |
| 1_13228863 | 13 | 0.6154 | 12 | 0.5 | 11 | 0.2727 | 10 | 0.4 | 46 | 0.4565 | 10 | 0.5 | 7 | 0.2857 | 10 | 0.3 | 6 | 0.8333 | 33 | 0.4545 |
| 1_13228872 | 16 | 0.125 | 12 | 0.0833 | 15 | 0.1333 | 14 | 0 | 57 | 0.0877 | 15 | 0.1333 | 10 | 0.4 | 11 | 0.1818 | 11 | 0.1818 | 47 | 0.2128 |
| 1_13228888 | 17 | 0 | 14 | 0.3571 | 14 | 0.2857 | 12 | 0.1667 | 57 | 0.193 | 12 | 0.1667 | 8 | 0.25 | 10 | 0.2 | 9 | 0.4444 | 39 | 0.2564 |
| 1_13228889 | 16 | 0 | 14 | 0.1429 | 16 | 0 | 14 | 0.1429 | 60 | 0.0667 | 12 | 0 | 8 | 0.5 | 9 | 0.1111 | 10 | 0.1 | 39 | 0.1538 |
| 1_13230853 | 15 | 0.4667 | 10 | 0.4 | 10 | 0.4 | 10 | 0.2 | 45 | 0.3778 | 8 | 0.375 | 7 | 0.7143 | 6 | 0.3333 | 8 | 0.5 | 29 | 0.4828 |
| 1_13230877 | 16 | 0.5 | 9 | 0.4444 | 10 | 0.9 | 11 | 0.6364 | 46 | 0.6087 | 10 | 0.7 | 5 | 0.6 | 6 | 0.5 | 7 | 0.5714 | 28 | 0.6071 |
| 1_13231895 | 10 | 0.8 | 9 | 0.4444 | 7 | 0.1429 | 11 | 0.8182 | 37 | 0.5946 | 6 | 0.3333 | 4 | 0.5 | 4 | 0 | 5 | 0.4 | 19 | 0.3158 |
| 1_13231905 | 7 | 0.1429 | 8 | 0.625 | 9 | 0.6667 | 12 | 0.5833 | 36 | 0.5278 | 4 | 0.5 | 5 | 0.6 | 3 | 0.6667 | 5 | 0.6 | 17 | 0.5882 |
| 1_13231912 | 7 | 0.2857 | 10 | 0.4 | 9 | 0.5556 | 13 | 0.3846 | 39 | 0.4103 | 5 | 0.2 | 5 | 0.4 | 3 | 0.3333 | 5 | 0.8 | 18 | 0.4444 |
| 1_13231917 | 6 | 0 | 10 | 0 | 8 | 0 | 12 | 0.1667 | 36 | 0.0556 | 5 | 0 | 3 | 0 | 2 | 0 | 5 | 0.4 | 15 | 0.1333 |
| 1_13231918 | 8 | 0.125 | 7 | 0.1429 | 7 | 0.1429 | 11 | 0.0909 | 33 | 0.1212 | 1 | 0 | 4 | 0 | 2 | 0.5 | 4 | 0 | 11 | 0.0909 |
| 1_13236204 | NA | NA | 1 | 1 | NA | NA | NA | NA | 1 | 1 | NA | NA | 3 | 0.6667 | NA | NA | NA | NA | 3 | 0.6667 |
| 1_13237730 | 1 | 1 | 2 | 0 | NA | NA | NA | NA | 3 | 0.3333 | 2 | 1 | 1 | 0 | NA | NA | NA | NA | 3 | 0.6667 |
| 1_13237734 | 1 | 1 | 2 | 1 | NA | NA | NA | NA | 3 | 1 | NA | NA | 3 | 1 | NA | NA | NA | NA | 3 | 1 |
| 1_13237736 | 1 | 1 | 2 | 0.5 | NA | NA | NA | NA | 3 | 0.6667 | 1 | 1 | 3 | 1 | NA | NA | NA | NA | 4 | 1 |
| 1_13239119 | 4 | 0.5 | 6 | 0.3333 | 3 | 0 | 3 | 0.3333 | 16 | 0.3125 | 3 | 0.6667 | 6 | 0.1667 | 1 | 0 | NA | NA | 10 | 0.3 |
| 1_13239123 | 4 | 1 | 3 | 1 | 2 | 0.5 | 3 | 1 | 12 | 0.9167 | 2 | 0.5 | 4 | 1 | 1 | 1 | NA | NA | 7 | 0.8571 |
| 1_13239135 | 3 | 0.6667 | 7 | 0.5714 | 3 | 1 | 2 | 1 | 15 | 0.7333 | 2 | 1 | 3 | 1 | 1 | 1 | NA | NA | 6 | 1 |
| 1_13239146 | 4 | 0.75 | 7 | 0.7143 | 2 | 1 | 3 | 1 | 16 | 0.8125 | 4 | 1 | 5 | 0.8 | 1 | 0 | NA | NA | 10 | 0.8 |
| 1_13239151 | 4 | 0 | 7 | 0.2857 | 2 | 0 | 2 | 0 | 15 | 0.1333 | 3 | 0.3333 | 5 | 0.2 | 1 | 0 | NA | NA | 9 | 0.2222 |
| 1_13239218 | 3 | 0 | 7 | 0.2857 | 5 | 0.2 | 4 | 0.5 | 19 | 0.2632 | 4 | 0.25 | 7 | 0.2857 | 1 | 0 | NA | NA | 12 | 0.25 |
| 1_13239227 | 1 | 0 | 6 | 0.3333 | 4 | 0 | 4 | 0 | 15 | 0.1333 | 4 | 0 | 6 | 0.1667 | 1 | 0 | NA | NA | 11 | 0.0909 |
| 1_13239338 | 3 | 0 | 7 | 0.1429 | 5 | 0 | 4 | 0.25 | 19 | 0.1053 | 5 | 0 | 15 | 0.0667 | 1 | 1 | NA | NA | 21 | 0.0952 |
| 1_13239424 | 9 | 0.2222 | 6 | 0.3333 | 5 | 0.2 | 4 | 0.25 | 24 | 0.25 | 5 | 0 | 7 | 0.2857 | 3 | 0 | 1 | 0 | 16 | 0.125 |
| 1_13243158 | 13 | 1 | 4 | 1 | 2 | 1 | 5 | 1 | 24 | 1 | 2 | 1 | 4 | 1 | 3 | 1 | 3 | 0.6667 | 12 | 0.9167 |
| 1_13243184 | 11 | 0.2727 | 5 | 0.4 | 1 | 0 | 7 | 0.2857 | 24 | 0.2917 | 4 | 0.25 | 2 | 0.5 | 5 | 0 | 3 | 0 | 14 | 0.1429 |
| 1_13243192 | 14 | 0.8571 | 6 | 0.6667 | 3 | 0.6667 | 9 | 0.5556 | 32 | 0.7188 | 5 | 0.4 | 5 | 0.8 | 6 | 0.3333 | 4 | 0.25 | 20 | 0.45 |
| 1_13243196 | 14 | 0.2857 | 6 | 0.5 | 3 | 0.3333 | 8 | 0.125 | 31 | 0.2903 | 5 | 0.4 | 5 | 0.6 | 5 | 0 | 4 | 0 | 19 | 0.2632 |
| 1_13243207 | 14 | 0.2857 | 7 | 0.1429 | 3 | 0 | 9 | 0.1111 | 33 | 0.1818 | 5 | 0 | 5 | 0 | 6 | 0.1667 | 4 | 0.25 | 20 | 0.1 |
| 1_13243236 | 11 | 0.2727 | 6 | 0.1667 | 3 | 0.6667 | 11 | 0.3636 | 31 | 0.3226 | 5 | 0.6 | 5 | 0.6 | 6 | 0.8333 | 3 | 0.3333 | 19 | 0.6316 |
| 1_13243504 | NA | NA | NA | NA | 1 | 1 | NA | NA | 1 | 1 | NA | NA | 2 | 1 | 1 | 0 | NA | NA | 3 | 0.6667 |
| 1_13243515 | NA | NA | NA | NA | 1 | 1 | NA | NA | 1 | 1 | 1 | 1 | 2 | 1 | 2 | 0.5 | NA | NA | 5 | 0.8 |
| 1_13243520 | NA | NA | NA | NA | 1 | 1 | NA | NA | 1 | 1 | 1 | 0 | 2 | 0.5 | 2 | 0 | NA | NA | 5 | 0.2 |
| 1_13243549 | NA | NA | NA | NA | 1 | 1 | NA | NA | 1 | 1 | 1 | 0 | 1 | 0 | 2 | 0.5 | NA | NA | 4 | 0.25 |
| 1_13243568 | NA | NA | NA | NA | 1 | 1 | NA | NA | 1 | 1 | 1 | 0 | 2 | 1 | 2 | 0.5 | NA | NA | 5 | 0.6 |
| 1_13243571 | NA | NA | NA | NA | 1 | 1 | NA | NA | 1 | 1 | 1 | 1 | NA | NA | 2 | 1 | NA | NA | 3 | 1 |
| 1_13243797 | NA | NA | 1 | 1 | NA | NA | NA | NA | 1 | 1 | NA | NA | NA | NA | 1 | 1 | NA | NA | 1 | 1 |
| 1_13243957 | 9 | 0.4444 | 17 | 0.1765 | 11 | 0.1818 | 6 | 0.1667 | 43 | 0.2326 | 14 | 0.2143 | 12 | 0.1667 | 5 | 0 | 9 | 0.2222 | 40 | 0.175 |
| 1_13243965 | 7 | 0.1429 | 14 | 0.1429 | 11 | 0.1818 | 4 | 0.25 | 36 | 0.1667 | 9 | 0 | 12 | 0.1667 | 5 | 0 | 9 | 0.2222 | 35 | 0.1143 |
| 1_13244063 | 2 | 0.5 | 7 | 0.2857 | 5 | 0.4 | 3 | 0.3333 | 17 | 0.3529 | 2 | 0.5 | 5 | 0.2 | 2 | 0.5 | 3 | 0.3333 | 12 | 0.3333 |
| 1_13244073 | 3 | 0.3333 | 10 | 0.2 | 10 | 0 | 2 | 0 | 25 | 0.12 | 6 | 0.1667 | 7 | 0 | 6 | 0.5 | 7 | 0.1429 | 26 | 0.1923 |

| | | | | | | | | | | | | | | | | | | | | |
|---|---|---|---|---|---|---|---|---|---|---|---|---|---|---|---|---|---|---|---|---|
| 1_13244129 | 2 | 1 | 9 | 0.6667 | 5 | 0.4 | 1 | 1 | 17 | 0.6471 | 2 | 0.5 | 5 | 0.4 | 4 | 0.75 | 7 | 0.4286 | 18 | 0.5 |
| 1_13244157 | 1 | 0 | 8 | 0.375 | 7 | 0.2857 | 2 | 0 | 18 | 0.2778 | 3 | 0 | 4 | 0.25 | 3 | 0.6667 | 7 | 0.1429 | 17 | 0.2353 |
| 1_13244159 | 3 | 0.3333 | 9 | 0.2222 | 7 | 0.1429 | 4 | 0 | 23 | 0.1739 | 2 | 0 | 5 | 0 | 5 | 0 | 6 | 0.1667 | 18 | 0.0556 |
| 1_13244167 | 1 | 1 | 7 | 0.7143 | 8 | 0.375 | NA | NA | 16 | 0.5625 | 2 | 0.5 | 5 | 0.8 | 5 | 0.6 | 4 | 0.25 | 16 | 0.5625 |
| 1_13244191 | 2 | 1 | 9 | 0.8889 | 6 | 0.6667 | 2 | 1 | 19 | 0.8421 | 2 | 1 | 3 | 1 | 3 | 1 | 5 | 1 | 13 | 1 |
| 1_13244197 | 1 | 1 | 8 | 0.25 | 5 | 0.2 | NA | NA | 14 | 0.2857 | 2 | 0 | 2 | 0.5 | 3 | 0.3333 | 5 | 0.4 | 12 | 0.3333 |
| 1_13244211 | 1 | 1 | 8 | 0.5 | 7 | 0.7143 | 2 | 0.5 | 18 | 0.6111 | 3 | 0 | 3 | 0.6667 | 3 | 1 | 5 | 0.4 | 14 | 0.5 |
| 1_13244224 | 1 | 1 | 7 | 1 | 6 | 1 | 2 | 1 | 16 | 1 | 3 | 1 | 3 | 1 | 3 | 1 | 6 | 1 | 15 | 1 |
| 1_13244225 | NA | NA | 5 | 0 | 7 | 0.1429 | 1 | 0 | 13 | 0.0769 | 3 | 0.3333 | 2 | 0 | 3 | 0.3333 | 6 | 0.1667 | 14 | 0.2143 |
| 1_13244241 | 2 | 0.5 | 7 | 0.7143 | 7 | 0.8571 | 1 | 1 | 17 | 0.7647 | 3 | 0 | 2 | 1 | 4 | 1 | 6 | 0.5 | 15 | 0.6 |
| 1_13244248 | 2 | 1 | 5 | 0.8 | 5 | 1 | 1 | 1 | 13 | 0.9231 | 2 | 1 | 2 | 1 | 3 | 1 | 4 | 1 | 11 | 1 |
| 1_13244253 | 2 | 0.5 | 6 | 1 | 6 | 0.6667 | 1 | 0 | 15 | 0.7333 | 3 | 1 | 2 | 1 | 1 | 0 | 5 | 0.8 | 11 | 0.8182 |
| 1_13244265 | 1 | 1 | 6 | 0.3333 | 5 | 0.4 | NA | NA | 12 | 0.4167 | 3 | 0.6667 | 2 | 0.5 | 2 | 0.5 | 2 | 0 | 9 | 0.4444 |
| 1_13244272 | 1 | 0 | 4 | 0.25 | 6 | 0.1667 | NA | NA | 11 | 0.1818 | 1 | 0 | NA | NA | 2 | 0 | 3 | 0.3333 | 6 | 0.1667 |
| 1_13244281 | 1 | 0 | 5 | 0.6 | 5 | 0.2 | 2 | 0.5 | 13 | 0.3846 | 3 | 0.6667 | 2 | 0 | 2 | 0.5 | 4 | 0.25 | 11 | 0.3636 |
| 1_13244286 | 1 | 1 | 6 | 0.6667 | 3 | 0.6667 | 1 | 1 | 11 | 0.7273 | 3 | 0 | 2 | 1 | 2 | 1 | 4 | 0.25 | 11 | 0.4545 |
| 1_13244296 | 1 | 0 | 5 | 0.2 | 5 | 0 | 2 | 0 | 13 | 0.0769 | 3 | 0.3333 | 2 | 0.5 | 2 | 0 | 4 | 0 | 11 | 0.1818 |
| 1_13244298 | 1 | 0 | 6 | 0.3333 | 4 | 0.25 | 2 | 0 | 13 | 0.2308 | 3 | 0.6667 | 2 | 0 | 3 | 0.3333 | 3 | 0 | 11 | 0.2727 |
| 1_13244299 | 1 | 1 | 6 | 0.8333 | 4 | 0.5 | 2 | 0.5 | 13 | 0.6923 | 3 | 0.6667 | 1 | 1 | 2 | 1 | 2 | 0.5 | 8 | 0.75 |
| 1_13244304 | 2 | 0.5 | 6 | 0.6667 | 5 | 0.4 | 2 | 0.5 | 15 | 0.5333 | 3 | 1 | 2 | 1 | 3 | 1 | 4 | 0.75 | 12 | 0.9167 |
| 1_13244315 | 1 | 0 | 6 | 0 | 5 | 0.2 | 2 | 0 | 14 | 0.0714 | 3 | 0 | 2 | 0 | 3 | 0.3333 | 4 | 0 | 12 | 0.0833 |
| 1_13244322 | 1 | 0 | 5 | 0.8 | 3 | 0.3333 | 1 | 0 | 10 | 0.5 | 3 | 0 | 1 | 1 | 3 | 0.3333 | 1 | 0 | 8 | 0.25 |
| 1_13244324 | 2 | 0.5 | 6 | 0.3333 | 4 | 0 | 2 | 0.5 | 14 | 0.2857 | 3 | 0 | 2 | 0 | 2 | 0 | 1 | 1 | 8 | 0.125 |
| 1_13249255 | 9 | 0 | 5 | 0 | 8 | 0.125 | 13 | 0.1538 | 35 | 0.0857 | 8 | 0.125 | 3 | 0 | 8 | 0.125 | 10 | 0 | 29 | 0.069 |
| 1_13249256 | 7 | 0.7143 | 4 | 0.5 | 8 | 0.625 | 10 | 0.7 | 29 | 0.6552 | 6 | 0.6667 | 5 | 0.8 | 7 | 0.5714 | 9 | 0.7778 | 27 | 0.7037 |
| 1_13249257 | 8 | 0.375 | 3 | 0.6667 | 8 | 0.375 | 15 | 0.4 | 34 | 0.4118 | 8 | 0.75 | 5 | 0.6 | 7 | 0.2857 | 9 | 0.4444 | 29 | 0.5172 |
| 1_13249259 | 8 | 0.5 | 5 | 0.8 | 6 | 0.3333 | 14 | 0.3571 | 33 | 0.4545 | 8 | 0.125 | 6 | 0.5 | 8 | 0.375 | 10 | 0.4 | 32 | 0.3438 |
| 1_13249311 | 7 | 0.1429 | 5 | 0.4 | 6 | 0.1667 | 13 | 0.1538 | 31 | 0.1935 | 8 | 0 | 4 | 0.25 | 6 | 0.1667 | 10 | 0.1 | 28 | 0.1071 |
| 1_13249331 | 2 | 0.5 | 1 | 0 | 4 | 0.25 | 3 | 0.3333 | 10 | 0.3 | 4 | 0.5 | 2 | 0 | 2 | 0 | 5 | 0.4 | 13 | 0.3077 |
| 1_13249642 | 2 | 0.5 | 1 | 0 | 4 | 0.25 | 4 | 0.5 | 11 | 0.3636 | 1 | 1 | 2 | 1 | 1 | 1 | 1 | 1 | 5 | 1 |
| 1_13249669 | 2 | 0 | 2 | 0.5 | 3 | 0.3333 | 4 | 0.25 | 11 | 0.2727 | 1 | 0 | 2 | 0 | 1 | 1 | 2 | 0.5 | 6 | 0.3333 |
| 1_13249678 | 2 | 0.5 | 1 | 0 | 3 | 0 | 4 | 0.25 | 10 | 0.2 | 1 | 1 | 2 | 0 | 1 | 0 | 2 | 0.5 | 6 | 0.3333 |
| 1_13249689 | 2 | 1 | 1 | 0 | 4 | 0.5 | 4 | 0.75 | 11 | 0.6364 | NA | NA | 2 | 0 | 1 | 1 | 2 | 1 | 5 | 0.6 |
| 1_13249703 | 4 | 0.5 | 5 | 0.6 | 8 | 0.625 | 6 | 0.8333 | 23 | 0.6522 | 6 | 0.3333 | 14 | 0.6429 | 5 | 0.6 | 5 | 0.8 | 30 | 0.6 |
| 1_13249704 | 5 | 0.6 | 5 | 0 | 8 | 0.125 | 6 | 0.1667 | 24 | 0.2083 | 6 | 0 | 14 | 0.1429 | 5 | 0 | 6 | 0.1667 | 31 | 0.0968 |
| 1_13249722 | 5 | 1 | 4 | 0.5 | 8 | 0.75 | 6 | 0.8333 | 23 | 0.7826 | 7 | 0.7143 | 13 | 0.7692 | 5 | 0.8 | 7 | 0.7143 | 32 | 0.75 |
| 1_13249732 | 5 | 1 | 6 | 1 | 8 | 0.875 | 6 | 1 | 25 | 0.96 | 7 | 0.7143 | 15 | 1 | 4 | 1 | 7 | 1 | 33 | 0.9394 |
| 1_13249744 | 5 | 0.8 | 4 | 1 | 8 | 0.875 | 6 | 0.8333 | 23 | 0.8696 | 7 | 0.8571 | 15 | 0.8 | 5 | 0.8 | 7 | 0.8571 | 34 | 0.8235 |
| 1_13249748 | 4 | 0.5 | 3 | 0 | 6 | 0.6667 | 6 | 0.6667 | 19 | 0.5263 | 7 | 0.5714 | 9 | 0.5556 | 3 | 0.3333 | 5 | 0.8 | 24 | 0.5833 |
| 1_13249806 | 2 | 1 | 5 | 0.6 | 6 | 0.8333 | 1 | 0 | 14 | 0.7143 | 4 | 0.25 | 10 | 0.2 | 4 | 0 | 5 | 0.4 | 23 | 0.2174 |
| 1_13249823 | 2 | 1 | 7 | 0.7143 | 7 | 0.7143 | 3 | 1 | 19 | 0.7895 | 5 | 0.8 | 10 | 0.7 | 2 | 0.5 | 6 | 0.6667 | 23 | 0.6957 |
| 1_13249830 | 2 | 1 | 5 | 0.2 | 6 | 0.6667 | 3 | 0.6667 | 16 | 0.5625 | 6 | 0.6667 | 12 | 0.6667 | 2 | 0.5 | 5 | 0.4 | 25 | 0.6 |
| 1_13252976 | 1 | 0 | 1 | 0 | 1 | 0 | 2 | 0.5 | 5 | 0.2 | 5 | 0 | 7 | 0.1429 | 3 | 0 | 3 | 0 | 18 | 0.0556 |
| 1_13280824 | 4 | 0.25 | 4 | 0 | 2 | 0 | 2 | 0.5 | 12 | 0.1667 | 7 | 0.2857 | 6 | 0.1667 | 7 | 0 | 3 | 0 | 23 | 0.1304 |
| 1_13286479 | 7 | 0.4286 | 5 | 0 | 5 | 0 | 5 | 0 | 22 | 0.1364 | 8 | 0 | 5 | 0 | 11 | 0.1818 | 10 | 0.1 | 34 | 0.0882 |
| 1_13286483 | 7 | 0.5714 | 5 | 0.6 | 5 | 0.6 | 5 | 0.2 | 22 | 0.5 | 9 | 0 | 8 | 0.25 | 11 | 0.0909 | 11 | 0.2727 | 39 | 0.1538 |
| 1_13286486 | 6 | 0.5 | 7 | 0.4286 | 3 | 0.3333 | 6 | 0.1667 | 22 | 0.3636 | 9 | 0.3333 | 7 | 0.1429 | 9 | 0.2222 | 6 | 0.1667 | 31 | 0.2258 |
| 1_13286491 | 6 | 0.5 | 8 | 0.5 | 5 | 0.4 | 5 | 0.2 | 24 | 0.4167 | 10 | 0.4 | 8 | 0 | 12 | 0.3333 | 12 | 0.0833 | 42 | 0.2143 |
| 1_13286519 | 10 | 0.3 | 8 | 0.5 | 5 | 0.4 | 7 | 0.7143 | 30 | 0.4667 | 8 | 0.125 | 9 | 0 | 11 | 0.5455 | 11 | 0.4545 | 39 | 0.3077 |
| 1_13286523 | 8 | 0.125 | 7 | 0.1429 | 7 | 0.1429 | 7 | 0 | 29 | 0.1034 | 9 | 0 | 7 | 0 | 11 | 0.0909 | 12 | 0.25 | 39 | 0.1026 |
| 1_13286526 | 11 | 0.0909 | 8 | 0.125 | 7 | 0.1429 | 7 | 0 | 33 | 0.0909 | 11 | 0.0909 | 8 | 0.125 | 12 | 0.0833 | 13 | 0.0769 | 44 | 0.0909 |
| 1_13286544 | 15 | 0.6 | 8 | 0.5 | 8 | 0.375 | 7 | 0.5714 | 38 | 0.5263 | 11 | 0.6364 | 7 | 0.5714 | 11 | 0.5455 | 13 | 0.4615 | 42 | 0.5476 |
| 1_13286555 | 10 | 0.6 | 7 | 0.8571 | 6 | 0.6667 | 5 | 1 | 28 | 0.75 | 7 | 0.7143 | 4 | 1 | 10 | 0.7 | 7 | 0.5714 | 28 | 0.7143 |
| 1_13286555 | 9 | 0.7778 | 5 | 0.8 | 3 | 0.3333 | 2 | 1 | 19 | 0.7368 | 7 | 0.8571 | 1 | 1 | 5 | 0.8 | 5 | 0.8 | 18 | 0.8333 |
| 1_13286564 | 15 | 0.6667 | 8 | 0.875 | 7 | 0.4286 | 5 | 0.8 | 35 | 0.6857 | 11 | 0.5455 | 8 | 0.875 | 13 | 0.6154 | 14 | 0.5714 | 46 | 0.6304 |
| 1_13286576 | 13 | 0.1538 | 9 | 0.1111 | 9 | 0 | 7 | 0 | 38 | 0.0789 | 12 | 0 | 10 | 0.1 | 11 | 0.2727 | 14 | 0.2143 | 47 | 0.1489 |
| 1_13286577 | 17 | 0.2353 | 9 | 0.2222 | 9 | 0.2222 | 8 | 0.25 | 43 | 0.2326 | 12 | 0.25 | 12 | 0 | 13 | 0.3077 | 17 | 0.1176 | 54 | 0.1667 |
| 1_13286580 | 18 | 0.1667 | 8 | 0 | 9 | 0 | 8 | 0.25 | 43 | 0.1163 | 12 | 0.25 | 12 | 0.0833 | 12 | 0 | 15 | 0.2 | 51 | 0.1373 |
| 1_13286580 | 16 | 0.6875 | 9 | 0.8889 | 9 | 0.7778 | 6 | 0.8333 | 40 | 0.775 | 12 | 1 | 12 | 0.6667 | 14 | 0.8571 | 15 | 0.8 | 53 | 0.8302 |
| 1_13286582 | 18 | 0.2222 | 10 | 0.5 | 8 | 0.25 | 8 | 0.25 | 44 | 0.2955 | 11 | 0.1818 | 11 | 0.5455 | 13 | 0.3846 | 18 | 0.3889 | 53 | 0.3774 |
| 1_13286588 | 16 | 0.3125 | 8 | 0.125 | 7 | 0.4286 | 8 | 0.25 | 39 | 0.2821 | 12 | 0.25 | 10 | 0.1 | 14 | 0.2143 | 16 | 0.0625 | 52 | 0.1538 |
| 1_13286595 | 19 | 0.3158 | 10 | 0.5 | 9 | 0.5556 | 8 | 0.125 | 46 | 0.3696 | 11 | 0.1818 | 12 | 0.3333 | 14 | 0.1429 | 18 | 0.1667 | 55 | 0.2 |
| 1_13286599 | 21 | 0.1905 | 10 | 0.1 | 9 | 0.1111 | 9 | 0 | 49 | 0.1224 | 11 | 0 | 13 | 0.0769 | 14 | 0.1429 | 17 | 0 | 55 | 0.0545 |
| 1_13286605 | 21 | 0.6667 | 8 | 0.5 | 7 | 0.7143 | 7 | 0.5714 | 43 | 0.6279 | 11 | 0.7273 | 10 | 0.8 | 13 | 0.9231 | 18 | 0.6667 | 52 | 0.7692 |
| 1_13286629 | 22 | 0.0909 | 12 | 0.0833 | 9 | 0.3333 | 6 | 0 | 49 | 0.1224 | 11 | 0.3636 | 12 | 0.25 | 15 | 0.3333 | 13 | 0.0769 | 51 | 0.2549 |

| | | | | | | | | | | | | | | | | | | | | |
|---|---|---|---|---|---|---|---|---|---|---|---|---|---|---|---|---|---|---|---|---|
| 1_13286643 | 11 | 0.9091 | 7 | 0.8571 | 5 | 1 | 8 | 1 | 31 | 0.9355 | 12 | 0.9167 | 7 | 0.8571 | 11 | 0.9091 | 9 | 0.7778 | 39 | 0.8718 |
| 1_13286645 | 22 | 0.0455 | 13 | 0 | 9 | 0.1111 | 9 | 0.2222 | 53 | 0.0755 | 11 | 0.1818 | 13 | 0.2308 | 15 | 0.0667 | 18 | 0.0556 | 57 | 0.1228 |
| 1_13286650 | 22 | 0.4545 | 10 | 0.5 | 9 | 0.5556 | 7 | 0.4286 | 48 | 0.4792 | 13 | 0.4615 | 13 | 0.3846 | 15 | 0.3333 | 17 | 0.5294 | 58 | 0.431 |
| 1_13286651 | 22 | 0.9091 | 12 | 0.75 | 8 | 0.75 | 8 | 0.75 | 50 | 0.82 | 14 | 0.6429 | 13 | 0.9231 | 13 | 0.6923 | 15 | 0.8667 | 55 | 0.7818 |
| 1_13286652 | 23 | 1 | 13 | 0.9231 | 9 | 1 | 7 | 1 | 52 | 0.9808 | 15 | 1 | 11 | 1 | 15 | 1 | 16 | 1 | 57 | 1 |
| 1_13286663 | 24 | 0.5417 | 12 | 0.4167 | 11 | 0.3636 | 7 | 0.8571 | 54 | 0.5185 | 15 | 0.2 | 12 | 0.5833 | 16 | 0.25 | 16 | 0.625 | 59 | 0.4068 |
| 1_13286677 | 20 | 1 | 13 | 1 | 11 | 1 | 10 | 1 | 54 | 1 | 17 | 1 | 13 | 1 | 19 | 1 | 14 | 1 | 63 | 1 |
| 1_13286681 | 21 | 0.5238 | 15 | 0.5333 | 13 | 0.6923 | 10 | 0.4 | 59 | 0.5424 | 13 | 0.4615 | 13 | 0.3846 | 18 | 0.5556 | 16 | 0.5 | 60 | 0.4833 |
| 1_13286710 | 22 | 0.0455 | 14 | 0.1429 | 15 | 0.0667 | 9 | 0.1111 | 60 | 0.0833 | 20 | 0.15 | 15 | 0.0667 | 20 | 0 | 18 | 0.0556 | 73 | 0.0685 |
| 1_13286711 | 23 | 0.7391 | 15 | 0.9333 | 16 | 0.875 | 11 | 0.9091 | 65 | 0.8462 | 18 | 0.6667 | 17 | 0.4706 | 19 | 0.7368 | 18 | 0.6667 | 72 | 0.6389 |
| 1_13286732 | 22 | 1 | 17 | 1 | 15 | 0.9333 | 15 | 1 | 69 | 0.9855 | 22 | 1 | 18 | 1 | 21 | 0.9524 | 18 | 0.9444 | 79 | 0.9747 |
| 1_13286761 | 20 | 0.45 | 15 | 0.4667 | 18 | 0.5556 | 15 | 0.3333 | 68 | 0.4559 | 16 | 0.5625 | 18 | 0.5556 | 20 | 0.3 | 20 | 0.3 | 74 | 0.4189 |
| 1_13291327 | 1 | 0 | 2 | 0.5 | NA | NA | 1 | 0 | 4 | 0.25 | 1 | 0 | 3 | 0.3333 | 1 | 0 | NA | NA | 5 | 0.2 |
| 1_13291413 | 5 | 0 | 4 | 0 | 7 | 0 | 6 | 0.3333 | 22 | 0.0909 | 2 | 0 | 8 | 0.25 | 6 | 0 | 4 | 0 | 20 | 0.1 |
| 1_13291497 | 5 | 0 | 3 | 0.3333 | 5 | 0 | 7 | 0 | 20 | 0.05 | 4 | 0 | 5 | 0 | 6 | 0.1667 | 4 | 0 | 19 | 0.0526 |
| 1_13291560 | 6 | 0 | NA | NA | 5 | 0 | 6 | 0.3333 | 17 | 0.1176 | 4 | 0 | 6 | 0.3333 | 7 | 0.1429 | 4 | 0.25 | 21 | 0.1905 |
| 1_13291607 | 5 | 0 | NA | NA | 4 | 0 | 5 | 0.4 | 14 | 0.1429 | 3 | 0 | 4 | 0 | 4 | 0 | 3 | 0.3333 | 14 | 0.0714 |
| 1_13291653 | 3 | 0 | NA | NA | 3 | 0 | 6 | 0.3333 | 12 | 0.1667 | 3 | 0.3333 | 4 | 0.25 | 2 | 0.5 | 3 | 0 | 12 | 0.25 |
| 1_13291691 | 3 | 0.3333 | NA | NA | 1 | 0 | 3 | 0 | 7 | 0.1429 | 1 | 0 | 3 | 0 | 3 | 0 | 2 | 0.5 | 9 | 0.1111 |
| 1_13291946 | 1 | 1 | NA | NA | NA | NA | 2 | 0 | 3 | 0.3333 | 1 | 1 | 2 | 0.5 | 1 | 0 | NA | NA | 4 | 0.5 |
| 1_13291952 | 1 | 1 | NA | NA | NA | NA | 1 | 0 | 2 | 0.5 | 1 | 1 | 1 | 0 | 1 | 1 | NA | NA | 3 | 0.6667 |
| 1_13291977 | 1 | 0 | NA | NA | NA | NA | 2 | 0.5 | 3 | 0.3333 | NA | NA | 1 | 1 | 1 | 0 | NA | NA | 2 | 0.5 |
| 1_13291980 | 1 | 1 | NA | NA | NA | NA | 2 | 1 | 3 | 1 | 1 | 1 | 1 | 1 | NA | NA | NA | NA | 2 | 1 |
| 1_13291985 | 1 | 0 | NA | NA | NA | NA | 1 | 1 | 2 | 0.5 | 1 | 0 | 1 | 1 | 1 | 1 | NA | NA | 3 | 0.6667 |
| 1_13303386 | 1 | 0 | NA | NA | 2 | 0.5 | NA | NA | 3 | 0.3333 | 1 | 1 | 1 | 0 | 1 | 0 | 1 | 1 | 4 | 0.5 |
| 1_13303436 | 6 | 0.5 | 1 | 0 | 3 | 0.6667 | 2 | 0.5 | 12 | 0.5 | 2 | 0 | 2 | 0.5 | 2 | 0 | 2 | 1 | 8 | 0.375 |
| 1_13303444 | 4 | 0.5 | NA | NA | 1 | 1 | NA | NA | 5 | 0.6 | 2 | 0.5 | 1 | 1 | 2 | 0 | 2 | 1 | 7 | 0.5714 |
| 1_13303457 | 5 | 0.8 | 1 | 1 | 3 | 1 | 1 | 1 | 10 | 0.9 | 2 | 0.5 | 2 | 1 | 2 | 1 | 2 | 1 | 8 | 0.875 |
| 1_13303474 | 5 | 0.4 | 1 | 1 | 2 | 0.5 | 2 | 0 | 10 | 0.4 | 2 | 0.5 | 1 | 0 | 2 | 0 | 2 | 1 | 7 | 0.4286 |
| 1_13303539 | 6 | 0.3333 | 2 | 0.5 | 5 | 0.2 | 2 | 0.5 | 15 | 0.3333 | 1 | 0 | 2 | 0.5 | 3 | 0.3333 | 1 | 1 | 7 | 0.4286 |
| 1_13305925 | 3 | 0.3333 | 1 | 0 | 1 | 0 | 1 | 0 | 6 | 0.1667 | NA | NA | NA | NA | 1 | 1 | NA | NA | 1 | 1 |
| 1_13305936 | 3 | 0.3333 | 1 | 0 | NA | NA | 1 | 0 | 5 | 0.2 | NA | NA | NA | NA | 1 | 1 | NA | NA | 1 | 1 |
| 1_13305965 | 4 | 0.5 | 1 | 0 | 1 | 0 | 1 | 0 | 7 | 0.2857 | NA | NA | 1 | 1 | 1 | 1 | 1 | 0 | 3 | 0.6667 |
| 1_13305990 | 5 | 0.8 | 1 | 1 | 1 | 0 | NA | NA | 7 | 0.7143 | NA | NA | 2 | 0.5 | 1 | 1 | 1 | 0 | 4 | 0.5 |
| 1_13305994 | 5 | 0.2 | 1 | 0 | 1 | 0 | 1 | 0 | 8 | 0.125 | NA | NA | 2 | 0.5 | 1 | 0 | 1 | 0 | 4 | 0.25 |
| 1_13305996 | 5 | 0.6 | 1 | 0 | 1 | 0 | 1 | 0 | 8 | 0.375 | NA | NA | 2 | 0 | 1 | 1 | 1 | 1 | 4 | 0.5 |
| 1_13306006 | 5 | 1 | 1 | 0 | 1 | 0 | 1 | 0 | 8 | 0.625 | NA | NA | 2 | 0.5 | 1 | 1 | NA | NA | 3 | 0.6667 |
| 1_13306006 | 4 | 0.25 | 1 | 0 | NA | NA | 1 | 0 | 6 | 0.1667 | NA | NA | 2 | 0.5 | NA | NA | 1 | 0 | 3 | 0.3333 |
| 1_13306015 | 5 | 0.8 | NA | NA | 1 | 1 | 1 | 1 | 7 | 0.8571 | NA | NA | 2 | 0.5 | NA | NA | 1 | 1 | 3 | 0.6667 |
| 1_13306046 | 5 | 0.6 | 1 | 1 | 1 | 1 | 1 | 0 | 8 | 0.625 | NA | NA | 2 | 1 | 1 | 1 | 1 | 1 | 4 | 1 |
| 1_13306070 | 5 | 0.4 | 1 | 1 | 2 | 0.5 | 1 | 0 | 9 | 0.4444 | NA | NA | 2 | 0.5 | 1 | 1 | 2 | 0 | 5 | 0.4 |
| 1_13306074 | 6 | 0.6667 | 1 | 0 | 2 | 1 | 1 | 0 | 10 | 0.6 | NA | NA | 2 | 1 | NA | NA | 1 | 0 | 3 | 0.6667 |
| 1_13306084 | 6 | 0.6667 | 1 | 1 | 2 | 0.5 | 2 | 1 | 11 | 0.7273 | NA | NA | 2 | 0.5 | 1 | 0 | 3 | 0.3333 | 6 | 0.3333 |
| 1_13306087 | 5 | 0.8 | 1 | 1 | 1 | 1 | NA | NA | 7 | 0.8571 | NA | NA | 2 | 0.5 | 1 | 1 | 1 | 0 | 4 | 0.5 |
| 1_13306090 | 5 | 0.2 | 1 | 0 | 1 | 0 | 2 | 0 | 9 | 0.1111 | NA | NA | 2 | 0 | 1 | 0 | 2 | 0.5 | 5 | 0.2 |
| 1_13306095 | 6 | 0.3333 | 1 | 0 | 2 | 1 | 2 | 0.5 | 11 | 0.4545 | NA | NA | 2 | 1 | 1 | 0 | 3 | 0 | 6 | 0.3333 |
| 1_13306097 | 5 | 0.8 | 1 | 0 | 2 | 0.5 | 1 | 1 | 9 | 0.6667 | NA | NA | 2 | 0.5 | 1 | 0 | 3 | 0.6667 | 6 | 0.5 |
| 1_13306101 | 6 | 0.1667 | 1 | 1 | 2 | 0.5 | 3 | 0.6667 | 12 | 0.4167 | NA | NA | 2 | 0 | 1 | 1 | 3 | 0 | 6 | 0.1667 |
| 1_13306110 | 3 | 1 | NA | NA | 2 | 1 | 2 | 1 | 7 | 1 | NA | NA | 2 | 0.5 | 1 | 1 | 3 | 1 | 6 | 0.8333 |
| 1_13306112 | 5 | 0.6 | 1 | 0 | 2 | 0 | 4 | 0.5 | 12 | 0.4167 | NA | NA | 2 | 0.5 | 1 | 0 | 3 | 0 | 6 | 0.1667 |
| 1_13306119 | 6 | 1 | 1 | 1 | 2 | 1 | 4 | 1 | 13 | 1 | NA | NA | 2 | 1 | 1 | 1 | 3 | 1 | 6 | 1 |
| 1_13306446 | 5 | 0.2 | 1 | 0 | 2 | 1 | 4 | 0.25 | 12 | 0.3333 | NA | NA | 3 | 0.6667 | 2 | 0.5 | 2 | 0.5 | 7 | 0.5714 |
| 1_13306457 | 4 | 1 | 1 | 1 | 2 | 1 | 3 | 1 | 10 | 1 | NA | NA | 2 | 1 | 3 | 1 | 4 | 0.75 | 9 | 0.8889 |
| 1_13306459 | 6 | 1 | 1 | 1 | 2 | 1 | 6 | 1 | 15 | 1 | NA | NA | 4 | 1 | 3 | 1 | 4 | 1 | 11 | 1 |
| 1_13306468 | 6 | 0.5 | 2 | 0 | 2 | 0 | 4 | 0.5 | 14 | 0.3571 | NA | NA | 3 | 0.6667 | 3 | 0 | 2 | 0 | 8 | 0.25 |
| 1_13306470 | 6 | 0.8333 | 1 | 0 | 1 | 0 | 6 | 0.5 | 14 | 0.5714 | NA | NA | 3 | 0.6667 | 1 | 1 | 3 | 1 | 7 | 0.8571 |
| 1_13306519 | 5 | 0 | 2 | 0 | 2 | 0 | 6 | 0.1667 | 15 | 0.0667 | 1 | 0 | 5 | 0.4 | 3 | 0 | 4 | 0 | 13 | 0.1538 |
| 1_13306596 | 9 | 0 | 4 | 0 | 7 | 0.1429 | 13 | 0.3077 | 33 | 0.1515 | 1 | 0 | 7 | 0 | 4 | 0 | 7 | 0.2857 | 19 | 0.1053 |
| 1_13306696 | 10 | 0.1 | 4 | 0.25 | 7 | 0 | 13 | 0.1538 | 34 | 0.1176 | 3 | 0 | 8 | 0 | 3 | 0.6667 | 4 | 0 | 18 | 0.1111 |
| 1_13306714 | 11 | 0 | 4 | 0 | 8 | 0 | 14 | 0.2143 | 37 | 0.0811 | 4 | 0 | 10 | 0 | 3 | 0.6667 | 10 | 0 | 27 | 0.0741 |
| 1_13320014 | NA | NA | NA | NA | NA | NA | 1 | 1 | 1 | 1 | NA | NA | 1 | 0 | 1 | 1 | 1 | 0 | 3 | 0.3333 |
| 1_13322785 | NA | NA | NA | NA | 1 | 0 | 2 | 0.5 | 3 | 0.3333 | 1 | 0 | 2 | 0 | 1 | 0 | 2 | 0.5 | 6 | 0.1667 |
| 1_13322786 | NA | NA | NA | NA | 1 | 1 | 1 | 0 | 2 | 0.5 | 1 | 0 | 2 | 0.5 | 2 | 0.5 | 1 | 1 | 6 | 0.5 |
| 1_13322834 | 1 | 0 | 2 | 0.5 | 1 | 1 | 2 | 1 | 6 | 0.6667 | 1 | 0 | 2 | 0.5 | 3 | 0.6667 | 2 | 0 | 8 | 0.375 |
| 1_13322870 | 1 | 0 | 2 | 0 | 1 | 1 | 2 | 0 | 6 | 0.1667 | 1 | 0 | 4 | 0.25 | 3 | 0 | 3 | 0.3333 | 11 | 0.1818 |

| | | | | | | | | | | | | | | | | | | | | |
|---|---|---|---|---|---|---|---|---|---|---|---|---|---|---|---|---|---|---|---|---|
| 1_13322883 | 1 | 1 | 2 | 1 | 1 | 1 | 2 | 1 | 6 | 1 | 1 | 1 | 4 | 1 | 3 | 1 | 3 | 1 | 11 | 1 |
| 1_13322891 | 3 | 1 | 2 | 1 | 1 | 1 | 2 | 1 | 8 | 1 | 1 | 1 | 4 | 1 | 4 | 1 | 3 | 1 | 12 | 1 |
| 1_13322894 | 3 | 0.6667 | 2 | 0 | 1 | 0 | 2 | 0 | 8 | 0.25 | 1 | 0 | 4 | 0.25 | 3 | 0 | 2 | 0.5 | 10 | 0.2 |
| 1_13322902 | 3 | 1 | 1 | 0 | 1 | 1 | 2 | 1 | 7 | 0.8571 | 1 | 1 | 1 | 0 | 3 | 1 | 3 | 1 | 8 | 0.875 |
| 1_13322914 | 3 | 0.6667 | 2 | 0.5 | 1 | 1 | 1 | 0 | 7 | 0.5714 | 1 | 1 | 2 | 1 | 4 | 0.5 | 4 | 0.75 | 11 | 0.7273 |
| 1_13322973 | 3 | 0 | 2 | 0.5 | 2 | 0.5 | 2 | 0 | 9 | 0.2222 | 1 | 0 | 5 | 0 | 3 | 0.3333 | 4 | 0.25 | 13 | 0.1538 |
| 1_13322979 | 2 | 0 | 2 | 0.5 | 2 | 0 | 2 | 0 | 8 | 0.125 | 1 | 0 | 5 | 0.2 | 4 | 0 | 4 | 0 | 14 | 0.0714 |
| 1_13331128 | 1 | 0 | NA | NA | 1 | 1 | 1 | 1 | 3 | 0.6667 | NA | NA | 1 | 1 | 1 | 1 | NA | NA | 2 | 1 |
| 1_13331144 | NA | NA | NA | NA | NA | NA | 1 | 1 | 1 | 1 | NA | NA | 1 | 1 | NA | NA | NA | NA | 1 | 1 |
| 1_13354788 | 20 | 0.4 | 13 | 0.4615 | 21 | 0.7143 | 10 | 0.3 | 64 | 0.5 | 6 | 0.3333 | 11 | 0.6364 | 13 | 0.5385 | 11 | 0.4545 | 41 | 0.5122 |
| 1_13354792 | 19 | 0.4737 | 15 | 0.4 | 20 | 0.5 | 9 | 0.5556 | 63 | 0.4762 | 9 | 0.6667 | 10 | 0.5 | 12 | 0.5 | 11 | 0.4545 | 42 | 0.5238 |
| 1_13354797 | 16 | 0.75 | 12 | 0.9167 | 18 | 0.7778 | 8 | 0.625 | 54 | 0.7778 | 6 | 0.8333 | 8 | 0.625 | 10 | 0.5 | 7 | 0.5714 | 31 | 0.6129 |
| 1_13354805 | 18 | 0.7778 | 15 | 0.8667 | 16 | 0.75 | 5 | 0.8 | 54 | 0.7963 | 8 | 0.875 | 8 | 0.75 | 9 | 0.6667 | 10 | 0.9 | 35 | 0.8 |
| 1_13354808 | 14 | 0.2857 | 12 | 0.25 | 10 | 0.2 | 9 | 0.2222 | 45 | 0.2444 | 7 | 0 | 8 | 0.25 | 9 | 0.5556 | 5 | 0.2 | 29 | 0.2759 |
| 1_13354814 | 19 | 0.7368 | 13 | 0.9231 | 16 | 0.875 | 8 | 0.75 | 56 | 0.8214 | 9 | 1 | 10 | 0.5 | 11 | 0.7273 | 11 | 0.8182 | 41 | 0.7561 |
| 1_13354817 | 17 | 0.1176 | 15 | 0.1333 | 19 | 0.1053 | 8 | 0.125 | 59 | 0.1186 | 8 | 0.125 | 10 | 0.5 | 10 | 0 | 11 | 0.1818 | 39 | 0.2051 |
| 1_13354836 | 18 | 0.8333 | 13 | 0.4615 | 17 | 0.7647 | 8 | 0.75 | 56 | 0.7143 | 7 | 0.5714 | 6 | 0.3333 | 9 | 0.8889 | 7 | 0.2857 | 29 | 0.5517 |
| 1_13354840 | 10 | 0.1 | 5 | 0.2 | 11 | 0.0909 | 1 | 0 | 27 | 0.1111 | 4 | 0 | 4 | 0 | 6 | 0.1667 | 2 | 0 | 16 | 0.0625 |
| 1_13354848 | 18 | 0.0556 | 16 | 0 | 18 | 0.1667 | 7 | 0.2857 | 59 | 0.1017 | 9 | 0.2222 | 10 | 0.1 | 11 | 0 | 9 | 0 | 39 | 0.0769 |
| 1_13354872 | 17 | 0.0588 | 9 | 0 | 16 | 0.1875 | 4 | 0 | 46 | 0.087 | 5 | 0 | 10 | 0.1 | 10 | 0 | 7 | 0.1429 | 32 | 0.0625 |
| 1_13354873 | 14 | 0.2143 | 10 | 0 | 16 | 0.1875 | 7 | 0 | 47 | 0.1277 | 7 | 0.2857 | 10 | 0 | 10 | 0.2 | 10 | 0.2 | 37 | 0.1622 |
| 1_13354882 | 18 | 0.2222 | 14 | 0 | 17 | 0.1176 | 8 | 0.125 | 57 | 0.1228 | 8 | 0 | 9 | 0.1111 | 9 | 0 | 8 | 0.25 | 34 | 0.0882 |
| 1_13354885 | 16 | 0.0625 | 13 | 0.0769 | 16 | 0.5 | 8 | 0.375 | 53 | 0.2453 | 6 | 0.1667 | 10 | 0.1 | 8 | 0.25 | 8 | 0.25 | 32 | 0.1875 |
| 1_13354887 | 18 | 0 | 12 | 0 | 18 | 0.1111 | 8 | 0.125 | 56 | 0.0536 | 8 | 0.125 | 9 | 0 | 10 | 0.1 | 8 | 0 | 35 | 0.0571 |
| 1_13354897 | 14 | 0.6429 | 10 | 0.6 | 18 | 0.8333 | 7 | 0.5714 | 49 | 0.6939 | 6 | 0.5 | 10 | 0.5 | 7 | 0.5714 | 9 | 0.6667 | 32 | 0.5625 |
| 1_13354900 | 14 | 0.7143 | 12 | 0.6667 | 13 | 0.9231 | 6 | 0.5 | 45 | 0.7333 | 8 | 0.625 | 9 | 0.6667 | 8 | 0.75 | 6 | 0.5 | 31 | 0.6452 |
| 1_13354907 | 14 | 0.5714 | 10 | 0.9 | 12 | 0.8333 | 6 | 0.6667 | 42 | 0.7381 | 6 | 0.5 | 8 | 0.75 | 10 | 0.9 | 7 | 0.7143 | 31 | 0.7419 |
| 1_13354911 | 14 | 0.9286 | 11 | 0.9091 | 19 | 0.8947 | 6 | 1 | 50 | 0.92 | 8 | 1 | 10 | 0.8 | 8 | 0.875 | 7 | 1 | 33 | 0.9091 |
| 1_13354921 | 17 | 0.7059 | 9 | 0.8889 | 12 | 1 | 6 | 0.8333 | 44 | 0.8409 | 6 | 0.8333 | 8 | 0.75 | 9 | 0.7778 | 5 | 0.6 | 28 | 0.75 |
| 1_13354922 | 7 | 0.5714 | 4 | 1 | 7 | 0.8571 | 4 | 1 | 22 | 0.8182 | 3 | 1 | 4 | 0.75 | 6 | 0.8333 | 2 | 1 | 15 | 0.8667 |
| 1_13354926 | 17 | 1 | 11 | 1 | 16 | 1 | 8 | 1 | 52 | 1 | 6 | 1 | 10 | 1 | 9 | 1 | 7 | 1 | 32 | 1 |
| 1_13354928 | 16 | 0.0625 | 12 | 0.0833 | 16 | 0.125 | 7 | 0.2857 | 51 | 0.1176 | 7 | 0.2857 | 9 | 0 | 9 | 0.2222 | 7 | 0.1429 | 32 | 0.1562 |
| 1_13354938 | 15 | 0.5333 | 10 | 0.1 | 15 | 0.4 | 7 | 0.4286 | 47 | 0.383 | 6 | 0.3333 | 10 | 0.3 | 9 | 0.5556 | 8 | 0.5 | 33 | 0.4242 |
| 1_13355468 | 8 | 0.375 | 6 | 0.1667 | 9 | 0.1111 | 5 | 0.2 | 28 | 0.2143 | 5 | 0.2 | 5 | 0.2 | 6 | 0.1667 | 5 | 0 | 21 | 0.1429 |
| 1_13355472 | 8 | 0.375 | 7 | 0.4286 | 9 | 0 | 7 | 0.2857 | 31 | 0.2581 | 6 | 0.1667 | 4 | 0.25 | 7 | 0.1429 | 5 | 0.2 | 22 | 0.1818 |
| 1_13355478 | 9 | 0.2222 | 7 | 0 | 10 | 0 | 7 | 0.2857 | 33 | 0.1212 | 5 | 0.2 | 4 | 0 | 6 | 0 | 5 | 0 | 20 | 0.05 |
| 1_13355486 | 9 | 0.1111 | 6 | 0.1667 | 10 | 0.3 | 7 | 0.2857 | 32 | 0.2188 | 5 | 0.2 | 4 | 0 | 4 | 0.5 | 4 | 0.25 | 17 | 0.2353 |
| 1_13358192 | NA | NA | 1 | 0 | 4 | 0.75 | 2 | 1 | 7 | 0.7143 | 1 | 0 | 1 | 0 | 1 | 1 | 2 | 0.5 | 5 | 0.4 |
| 1_13363074 | 9 | 0.2222 | 21 | 0.1429 | 14 | 0.1429 | 12 | 0.1667 | 56 | 0.1607 | 11 | 0.1818 | 5 | 0 | 9 | 0.4444 | 14 | 0.1429 | 39 | 0.2051 |
| 1_13363102 | 7 | 0 | 23 | 0.2609 | 14 | 0.2143 | 12 | 0.1667 | 56 | 0.1964 | 13 | 0.1538 | 6 | 0.1667 | 12 | 0.25 | 13 | 0.0769 | 44 | 0.1591 |
| 1_13363306 | 3 | 0 | 9 | 0.1111 | 7 | 0.1429 | 5 | 0.4 | 24 | 0.1667 | 6 | 0 | 1 | 1 | 8 | 0.25 | 6 | 0.6667 | 21 | 0.3333 |
| 1_13363386 | 3 | 0 | 11 | 0.2727 | 7 | 0.1429 | 3 | 0 | 24 | 0.1667 | 4 | 0.25 | 1 | 1 | 7 | 0.1429 | 4 | 0 | 16 | 0.1875 |
| 1_13363387 | 1 | 0 | 11 | 0.0909 | 7 | 0.2857 | 3 | 0.3333 | 22 | 0.1818 | 4 | 0.25 | 1 | 0 | 7 | 0.1429 | 4 | 0.25 | 16 | 0.1875 |
| 1_13363412 | 2 | 0.5 | 10 | 0.5 | 6 | 0.1667 | 3 | 0 | 21 | 0.3333 | 5 | 0.2 | 1 | 0 | 6 | 0.1667 | 2 | 0.5 | 14 | 0.2143 |
| 1_13363426 | 2 | 0 | 10 | 0.1 | 5 | 0 | 3 | 0 | 20 | 0.05 | 4 | 0 | 1 | 0 | 7 | 0.2857 | 2 | 0 | 14 | 0.1429 |
| 1_13363428 | NA | NA | 11 | 0 | 4 | 0 | 4 | 0.25 | 19 | 0.0526 | 4 | 0.25 | 1 | 0 | 6 | 0 | 2 | 0.5 | 13 | 0.1538 |
| 1_13363435 | 2 | 1 | 11 | 1 | 5 | 1 | 4 | 1 | 22 | 1 | 5 | 1 | 1 | 1 | 7 | 0.8571 | 2 | 1 | 15 | 0.9333 |
| 1_13363442 | 2 | 1 | 9 | 1 | 3 | 1 | 3 | 1 | 17 | 1 | 3 | 0.6667 | 1 | 1 | 3 | 1 | 1 | 1 | 8 | 0.875 |
| 1_13363446 | 2 | 0 | 9 | 0 | 2 | 0.5 | 4 | 0 | 17 | 0.0588 | 2 | 0 | 1 | 0 | 7 | 0.1429 | NA | NA | 10 | 0.1 |
| 1_13363449 | 2 | 1 | 6 | 1 | 3 | 1 | 3 | 1 | 14 | 1 | 2 | 1 | 1 | 1 | 7 | 1 | NA | NA | 10 | 1 |
| 1_13363465 | 1 | 0 | 11 | 0.0909 | 3 | 0 | 3 | 0 | 18 | 0.0556 | 4 | 0 | 1 | 0 | 5 | 0.2 | 1 | 0 | 11 | 0.0909 |
| 1_13363472 | 1 | 0 | 8 | 0.25 | 2 | 0 | 4 | 0 | 15 | 0.1333 | 4 | 0.25 | 1 | 1 | 6 | 0 | NA | NA | 11 | 0.1818 |
| 1_13365757 | 1 | 0 | 6 | 0.3333 | NA | NA | 2 | 0 | 9 | 0.2222 | 3 | 0.3333 | 1 | 0 | 2 | 0.5 | 1 | 0 | 7 | 0.2857 |
| 1_13368084 | NA | NA | 2 | 1 | NA | NA | 1 | 0 | 3 | 0.6667 | 2 | 0.5 | 1 | 0 | 1 | 0 | NA | NA | 4 | 0.25 |
| 1_13368093 | NA | NA | 2 | 0.5 | NA | NA | 1 | 1 | 3 | 0.6667 | 2 | 1 | 1 | 1 | 1 | 1 | NA | NA | 4 | 1 |
| 1_13368108 | NA | NA | 2 | 1 | NA | NA | NA | NA | 2 | 1 | 2 | 1 | 1 | 0 | 1 | 1 | NA | NA | 4 | 0.75 |
| 1_13368122 | NA | NA | 1 | 1 | NA | NA | NA | NA | 1 | 1 | 2 | 1 | 1 | 1 | 1 | 1 | NA | NA | 4 | 1 |
| 1_13368135 | NA | NA | NA | NA | NA | NA | 1 | 1 | 1 | 1 | 2 | 1 | 1 | 0 | 1 | 1 | NA | NA | 4 | 0.75 |
| 1_13384032 | 6 | 0.8333 | 3 | 1 | 4 | 0.5 | 1 | 1 | 14 | 0.7857 | NA | NA | NA | NA | 1 | 1 | NA | NA | 1 | 1 |
| 1_13384038 | 5 | 0.4 | 3 | 0.6667 | 4 | 0.25 | NA | NA | 12 | 0.4167 | NA | NA | NA | NA | 2 | 0.5 | NA | NA | 2 | 0.5 |
| 1_13384232 | 4 | 0.5 | 3 | 0.3333 | 3 | 0 | 1 | 1 | 11 | 0.3636 | NA | NA | NA | NA | 1 | 1 | NA | NA | 1 | 1 |
| 1_13384233 | 4 | 0.25 | 2 | 0.5 | 3 | 0 | NA | NA | 9 | 0.2222 | NA | NA | NA | NA | 1 | 1 | NA | NA | 1 | 1 |
| 1_13384256 | 4 | 0 | 3 | 0 | 3 | 0.3333 | 1 | 0 | 11 | 0.0909 | NA | NA | NA | NA | 1 | 1 | NA | NA | 1 | 1 |
| 1_13384264 | 2 | 0 | 2 | 0 | 3 | 0.3333 | 1 | 0 | 8 | 0.125 | NA | NA | NA | NA | 1 | 1 | NA | NA | 1 | 1 |
| 1_13384275 | 4 | 0.25 | 2 | 0 | 3 | 0.3333 | NA | NA | 9 | 0.2222 | NA | NA | NA | NA | 1 | 1 | NA | NA | 1 | 1 |

| | | | | | | | | | | | | | | | | | | | | |
|---|---|---|---|---|---|---|---|---|---|---|---|---|---|---|---|---|---|---|---|---|
| 1_13396802 | 3 | 0.3333 | NA | NA | 1 | 0 | 1 | 0 | 5 | 0.2 | NA | NA | NA | NA | NA | NA | 2 | 0.5 | 2 | 0.5 |
| 1_13396844 | 5 | 0.4 | 1 | 1 | 2 | 1 | 2 | 0.5 | 10 | 0.6 | NA | NA | NA | NA | NA | NA | 2 | 1 | 2 | 1 |
| 1_13396851 | 6 | 0.6667 | 1 | 0 | 1 | 1 | 2 | 0 | 10 | 0.5 | NA | NA | NA | NA | NA | NA | 2 | 0.5 | 2 | 0.5 |
| 1_13396864 | 7 | 0.1429 | 1 | 0 | 2 | 0 | 1 | 0 | 11 | 0.0909 | 1 | 1 | NA | NA | NA | NA | 2 | 0 | 3 | 0.3333 |
| 1_13396885 | 5 | 0.2 | 2 | 0.5 | 3 | 1 | 1 | 1 | 11 | 0.5455 | 1 | 1 | NA | NA | 1 | 1 | 1 | 1 | 3 | 1 |
| 1_13397448 | 7 | 0.1429 | 2 | 0.5 | 6 | 0.5 | 1 | 0 | 16 | 0.3125 | 1 | 0 | 1 | 1 | 1 | 0 | 3 | 0 | 6 | 0.1667 |
| 1_13400311 | 3 | 0 | 3 | 0 | 3 | 0.3333 | 4 | 0 | 13 | 0.0769 | 2 | 0.5 | 1 | 1 | 4 | 0 | 1 | 0 | 8 | 0.25 |
| 1_13400325 | 3 | 0 | 4 | 0 | 2 | 0.5 | 2 | 0.5 | 11 | 0.1818 | 3 | 0 | 1 | 0 | 4 | 0.25 | 1 | 0 | 9 | 0.1111 |
| 1_13400441 | 2 | 1 | NA | NA | 3 | 0.3333 | 2 | 1 | 7 | 0.7143 | 2 | 0.5 | NA | NA | 4 | 0.5 | 2 | 1 | 8 | 0.625 |
| 1_13400486 | 1 | 1 | NA | NA | 1 | 1 | 1 | 1 | 3 | 1 | 2 | 1 | NA | NA | 3 | 1 | 2 | 1 | 7 | 1 |
| 1_13400502 | 1 | 1 | NA | NA | 1 | 1 | NA | NA | 2 | 1 | 2 | 1 | NA | NA | 3 | 1 | 1 | 1 | 6 | 1 |
| 1_1340051[illegible] | NA | NA | NA | NA | 2 | 1 | 1 | 1 | 3 | 1 | NA | NA | NA | NA | 3 | 0.6667 | 2 | 1 | 5 | 0.8 |
| 1_13400516 | 1 | 0 | NA | NA | 2 | 0 | 2 | 0.5 | 5 | 0.2 | 2 | 0.5 | NA | NA | 3 | 0.3333 | 2 | 0.5 | 7 | 0.4286 |
| 1_13400532 | 1 | 1 | NA | NA | 2 | 1 | 2 | 1 | 5 | 1 | 1 | 1 | NA | NA | 3 | 0.3333 | 2 | 1 | 6 | 0.6667 |
| 1_13400536 | 1 | 0 | NA | NA | 2 | 0 | 2 | 0.5 | 5 | 0.2 | 2 | 0.5 | NA | NA | 3 | 0 | 2 | 0 | 7 | 0.1429 |
| 1_13412706 | 12 | 0.0833 | 14 | 0.2857 | 4 | 0.75 | 9 | 0.3333 | 39 | 0.2821 | 8 | 0 | 6 | 0.1667 | 12 | 0.25 | 12 | 0.25 | 38 | 0.1842 |
| 1_13412724 | 11 | 0.1818 | 13 | 0 | 4 | 0 | 13 | 0.0769 | 41 | 0.0732 | 10 | 0.1 | 6 | 0 | 12 | 0.0833 | 12 | 0.25 | 40 | 0.125 |
| 1_13412737 | 10 | 0.1 | 12 | 0.0833 | 4 | 0.25 | 11 | 0.0909 | 37 | 0.1081 | 6 | 0.1667 | 5 | 0 | 9 | 0.3333 | 11 | 0.4545 | 31 | 0.2903 |
| 1_13412852 | 9 | 0.1111 | 9 | 0.2222 | 2 | 0.5 | 10 | 0.3 | 30 | 0.2333 | 6 | 0.1667 | 2 | 0 | 9 | 0.2222 | 7 | 0.2857 | 24 | 0.2083 |
| 1_13412872 | 9 | 0 | 9 | 0.1111 | 2 | 0.5 | 8 | 0 | 28 | 0.0714 | 7 | 0.1429 | 3 | 0.3333 | 8 | 0.125 | 7 | 0 | 25 | 0.12 |
| 1_13412937 | 10 | 0.6 | 9 | 0 | 3 | 0.3333 | 12 | 0.5 | 34 | 0.3824 | 9 | 0.7778 | 3 | 0.3333 | 6 | 0.3333 | 5 | 0 | 23 | 0.4348 |
| 1_13412943 | 8 | 0.375 | 10 | 0.3 | 3 | 0.3333 | 10 | 0.7 | 31 | 0.4516 | 8 | 0.625 | 4 | 0.25 | 4 | 0.5 | 4 | 0 | 20 | 0.4 |
| 1_13412949 | 3 | 1 | 3 | 1 | 2 | 1 | 6 | 1 | 14 | 1 | 3 | 1 | 2 | 0.5 | 2 | 1 | 3 | 1 | 10 | 0.9 |
| 1_13412960 | 7 | 0.7143 | 8 | 0.75 | 3 | 0.3333 | 11 | 0.8182 | 29 | 0.7241 | 7 | 0.7143 | 3 | 0.6667 | 6 | 0.5 | 4 | 0.5 | 20 | 0.6 |
| 1_13412969 | 8 | 0.125 | 9 | 0 | 2 | 0.5 | 12 | 0.0833 | 31 | 0.0968 | 9 | 0 | 2 | 0 | 6 | 0.3333 | 6 | 0 | 23 | 0.087 |
| 1_13412970 | 9 | 0.3333 | 8 | 0.375 | 3 | 0 | 12 | 0.1667 | 32 | 0.25 | 7 | 0.1429 | 4 | 0 | 6 | 0.5 | 6 | 0.3333 | 23 | 0.2609 |
| 1_13412986 | 9 | 0.6667 | 6 | 0.8333 | 2 | 1 | 9 | 0.7778 | 26 | 0.7692 | 5 | 0.8 | 2 | 0.5 | 3 | 1 | 5 | 1 | 15 | 0.8667 |
| 1_13412987 | 7 | 1 | 5 | 1 | 2 | 1 | 11 | 0.9091 | 25 | 0.96 | 5 | 1 | 2 | 1 | 4 | 1 | 5 | 1 | 16 | 1 |
| 1_13412994 | 6 | 0.8333 | 6 | 0.8333 | 2 | 1 | 8 | 1 | 22 | 0.9091 | 5 | 0.4 | 1 | 0 | 4 | 0.5 | 4 | 0.75 | 14 | 0.5 |
| 1_13413004 | 6 | 0.6667 | 7 | 1 | 2 | 1 | 8 | 0.875 | 23 | 0.8696 | 4 | 0.75 | NA | NA | 4 | 1 | 4 | 1 | 12 | 0.9167 |
| 1_13413010 | 8 | 0.125 | 7 | 0.1429 | 2 | 0 | 10 | 0.1 | 27 | 0.1111 | 4 | 0.25 | 1 | 0 | 4 | 0.25 | 5 | 0.2 | 14 | 0.2143 |
| 1_13413030 | 7 | 0.4286 | 6 | 0 | 2 | 0.5 | 8 | 0.375 | 23 | 0.3043 | 4 | 0 | 1 | 0 | 4 | 0.75 | 4 | 0.75 | 13 | 0.4615 |
| 1_13413049 | 5 | 0.6 | 6 | 0.8333 | 2 | 1 | 10 | 1 | 23 | 0.8696 | 3 | 1 | 1 | 1 | 4 | 1 | 4 | 0.75 | 12 | 0.9167 |
| 1_13413053 | 6 | 0.3333 | 2 | 0.5 | 2 | 0.5 | 4 | 0.5 | 14 | 0.4286 | 2 | 0.5 | 1 | 0 | 4 | 0.5 | 1 | 0 | 8 | 0.375 |
| 1_13413056 | 8 | 1 | 6 | 1 | 2 | 1 | 8 | 1 | 24 | 1 | 3 | 1 | NA | NA | 4 | 0.75 | 3 | 1 | 10 | 0.9 |
| 1_13413069 | 6 | 0.8333 | 4 | 0.75 | 2 | 0.5 | 10 | 0.6 | 22 | 0.6818 | 3 | 1 | NA | NA | 3 | 1 | 2 | 0.5 | 8 | 0.875 |
| 1_13413081 | 5 | 0.2 | 6 | 0.1667 | 2 | 0.5 | 8 | 0.5 | 21 | 0.3333 | 3 | 0.6667 | NA | NA | 3 | 0 | 2 | 0.5 | 8 | 0.375 |
| 1_13413090 | 4 | 0.75 | 4 | 1 | 2 | 1 | 5 | 0.8 | 15 | 0.8667 | 3 | 1 | NA | NA | 2 | 1 | NA | NA | 5 | 1 |
| 1_13416249 | 1 | 0 | 3 | 0 | 1 | 0 | 2 | 0.5 | 7 | 0.1429 | 1 | 1 | NA | NA | NA | NA | 1 | 0 | 2 | 0.5 |
| 1_13416309 | 2 | 1 | NA | NA | NA | NA | NA | NA | 2 | 1 | 1 | 1 | NA | NA | 1 | 0 | NA | NA | 2 | 0.5 |
| 1_13422558 | 1 | 0 | 4 | 0.25 | 1 | 0 | 2 | 0 | 8 | 0.125 | 2 | 0 | 2 | 0 | 3 | 0 | 2 | 0.5 | 9 | 0.1111 |
| 1_13422591 | 2 | 0 | 4 | 0 | 1 | 0 | 2 | 0.5 | 9 | 0.1111 | 2 | 0 | 3 | 0.3333 | 1 | 0 | 2 | 0 | 8 | 0.125 |
| 1_13422608 | NA | NA | 3 | 0 | NA | NA | 3 | 0.3333 | 6 | 0.1667 | 3 | 0.6667 | 2 | 0 | 2 | 0.5 | 1 | 1 | 8 | 0.5 |
| 1_13422610 | 2 | 1 | 4 | 0.75 | NA | NA | 2 | 0.5 | 8 | 0.75 | 3 | 0.6667 | 2 | 0.5 | 2 | 0.5 | 2 | 0.5 | 9 | 0.5556 |
| 1_13422611 | 3 | 1 | 6 | 1 | 3 | 0.6667 | 6 | 0.8333 | 18 | 0.8889 | 3 | 1 | 5 | 1 | 4 | 1 | 2 | 1 | 14 | 1 |
| 1_13422615 | 7 | 0.8571 | 5 | 0.8 | 4 | 0.25 | 6 | 0.8333 | 22 | 0.7273 | 3 | 0.3333 | 5 | 0.6 | 4 | 0.75 | 1 | 1 | 13 | 0.6154 |
| 1_13422619 | 4 | 0.75 | 2 | 0.5 | 2 | 0 | 3 | 0.3333 | 11 | 0.4545 | 1 | 0 | 4 | 0.5 | 1 | 0 | 1 | 1 | 7 | 0.4286 |
| 1_13422623 | 8 | 0.375 | 5 | 0 | 5 | 0 | 6 | 0 | 24 | 0.125 | 4 | 0 | 3 | 0.3333 | 4 | 0.5 | 2 | 0.5 | 13 | 0.3077 |
| 1_13422652 | 7 | 0.2857 | 3 | 0.6667 | 3 | 0 | 4 | 0.25 | 17 | 0.2941 | 3 | 0.6667 | 5 | 0.2 | 5 | 0.4 | 2 | 0.5 | 15 | 0.4 |
| 1_13422654 | 8 | 0.25 | 5 | 0 | 5 | 0.2 | 5 | 0.6 | 23 | 0.2609 | 4 | 0.25 | 3 | 0 | 4 | 0.25 | 2 | 0 | 13 | 0.1538 |
| 1_13425515 | 3 | 0.3333 | 2 | 0 | 1 | 0 | NA | NA | 6 | 0.1667 | 1 | 0 | 1 | 1 | 2 | 0 | 1 | 0 | 5 | 0.2 |
| 1_13426045 | 2 | 0 | 2 | 1 | NA | NA | 1 | 1 | 5 | 0.6 | NA | NA | 1 | 1 | 1 | 1 | 1 | 1 | 3 | 1 |
| 1_13427537 | 5 | 0 | 9 | 0 | 6 | 0.1667 | 6 | 0.1667 | 26 | 0.0769 | 7 | 0.1429 | 8 | 0.125 | 7 | 0.2857 | 2 | 0 | 24 | 0.1667 |
| 1_13427556 | 3 | 0 | 11 | 0.1818 | 6 | 0.1667 | 7 | 0.1429 | 27 | 0.1481 | 5 | 0.2 | 6 | 0.1667 | 6 | 0 | 2 | 0.5 | 19 | 0.1579 |
| 1_13428334 | 1 | 1 | 7 | 0.5714 | 3 | 0.3333 | 5 | 0.2 | 16 | 0.4375 | 2 | 0 | 3 | 0 | 3 | 0 | 2 | 0.5 | 10 | 0.1 |
| 1_13433916 | 3 | 0 | 5 | 0 | NA | NA | 3 | 0.6667 | 11 | 0.1818 | 3 | 0 | 3 | 0 | 7 | 0.1429 | 1 | 0 | 14 | 0.0714 |
| 1_13435942 | 16 | 0.375 | 15 | 0 | 14 | 0.2857 | 14 | 0 | 59 | 0.1695 | 12 | 0.0833 | 13 | 0.0769 | 20 | 0.1 | 16 | 0.0625 | 61 | 0.082 |
| 1_13435952 | 15 | 0 | 15 | 0 | 14 | 0.2143 | 16 | 0.0625 | 60 | 0.0667 | 11 | 0.1818 | 12 | 0.1667 | 19 | 0.1579 | 16 | 0 | 58 | 0.1207 |
| 1_13435969 | 10 | 0.7 | 8 | 0.625 | 7 | 0.5714 | 9 | 0.6667 | 34 | 0.6471 | 7 | 0.7143 | 8 | 0.75 | 12 | 0.5 | 8 | 0.625 | 35 | 0.6286 |
| 1_13435984 | 19 | 0.0526 | 17 | 0 | 17 | 0.0588 | 15 | 0.2 | 68 | 0.0735 | 11 | 0.3636 | 13 | 0.0769 | 21 | 0.0952 | 15 | 0.1333 | 60 | 0.15 |
| 1_13436004 | 18 | 0.0556 | 20 | 0 | 17 | 0.1765 | 18 | 0.0556 | 73 | 0.0685 | 10 | 0.3 | 14 | 0.0714 | 23 | 0.087 | 16 | 0.25 | 63 | 0.1587 |
| 1_13436081 | 1 | 0 | 1 | 1 | 2 | 0 | 2 | 0 | 6 | 0.1667 | 1 | 0 | 1 | 1 | NA | NA | NA | NA | 2 | 0.5 |
| 1_13436100 | NA | NA | 1 | 0 | NA | NA | 2 | 0.5 | 3 | 0.3333 | 1 | 1 | 1 | 1 | 2 | 0 | NA | NA | 4 | 0.5 |
| 1_13441719 | 2 | 1 | 1 | 1 | 3 | 0.6667 | 3 | 0.3333 | 9 | 0.6667 | 1 | 1 | NA | NA | 2 | 1 | 3 | 1 | 6 | 1 |
| 1_13441722 | 3 | 0.3333 | 1 | 0 | 2 | 0 | 4 | 0.25 | 10 | 0.2 | 2 | 0 | 3 | 0 | 2 | 0 | 4 | 0.5 | 11 | 0.1818 |

| | | | | | | | | | | | | | | | | | | | | |
|---|---|---|---|---|---|---|---|---|---|---|---|---|---|---|---|---|---|---|---|---|
| 1_13441724 | 2 | 0 | 1 | 0 | 3 | 0.3333 | 4 | 0.5 | 10 | 0.3 | 2 | 1 | 3 | 0.3333 | 1 | 1 | 1 | 1 | 7 | 0.7143 |
| 1_13441789 | 5 | 0.8 | 1 | 1 | 8 | 0.5 | 6 | 0.1667 | 20 | 0.5 | 2 | 0.5 | 5 | 0.2 | 2 | 0.5 | 3 | 0.6667 | 12 | 0.4167 |
| 1_13441795 | 5 | 0 | 2 | 0.5 | 7 | 0 | 6 | 0 | 20 | 0.05 | 2 | 0 | 4 | 0 | 2 | 0.5 | 4 | 0 | 12 | 0.0833 |
| 1_13441836 | 4 | 0.5 | 2 | 0.5 | 7 | 0.1429 | 8 | 0.125 | 21 | 0.2381 | 3 | 0.3333 | 6 | 0.3333 | 3 | 0 | 5 | 0.8 | 17 | 0.4118 |
| 1_13441859 | 1 | 0 | 1 | 0 | 5 | 0.4 | 7 | 0.2857 | 14 | 0.2857 | 3 | 0 | 6 | 0.1667 | 1 | 0 | 4 | 0.25 | 14 | 0.1429 |
| 1_13443397 | 1 | 1 | 7 | 1 | 3 | 0.6667 | NA | NA | 11 | 0.9091 | 3 | 0.3333 | 3 | 1 | 5 | 1 | 3 | 0.6667 | 14 | 0.7857 |
| 1_13443398 | 1 | 1 | 6 | 0.5 | 2 | 1 | NA | NA | 9 | 0.6667 | 3 | 0.6667 | 2 | 1 | 6 | 1 | 3 | 0.6667 | 14 | 0.8571 |
| 1_13443405 | NA | NA | 3 | 1 | NA | NA | NA | NA | 3 | 1 | NA | NA | NA | NA | 1 | 1 | 1 | 1 | 2 | 1 |
| 1_13443409 | 2 | 0 | 5 | 0.6 | 3 | 0.3333 | NA | NA | 10 | 0.4 | 3 | 0.3333 | 3 | 0.6667 | 4 | 0.5 | 3 | 0.3333 | 13 | 0.4615 |
| 1_13443411 | 2 | 0 | 7 | 0.2857 | 3 | 0.3333 | NA | NA | 12 | 0.25 | 5 | 0.2 | 3 | 0 | 6 | 0 | 2 | 0 | 16 | 0.0625 |
| 1_13443412 | 1 | 1 | 7 | 0.8571 | 3 | 0.6667 | NA | NA | 11 | 0.8182 | 4 | 0.75 | 3 | 1 | 5 | 1 | 2 | 1 | 14 | 0.9286 |
| 1_13443415 | 2 | 0.5 | 7 | 0.4286 | 3 | 0 | NA | NA | 12 | 0.3333 | 4 | 0.5 | 3 | 0 | 6 | 0.5 | 3 | 0.3333 | 16 | 0.375 |
| 1_13443419 | 2 | 0.5 | 6 | 0 | 2 | 0 | NA | NA | 10 | 0.1 | 5 | 0 | 1 | 0 | 6 | 0.1667 | 3 | 0 | 15 | 0.0667 |
| 1_13443429 | 2 | 0.5 | 6 | 0 | 2 | 0 | NA | NA | 10 | 0.1 | 3 | 0.3333 | 3 | 0 | 5 | 0.2 | 3 | 0.6667 | 14 | 0.2857 |
| 1_13443496 | 2 | 0.5 | 7 | 0.1429 | 1 | 0 | NA | NA | 10 | 0.2 | 5 | 0.4 | 5 | 0.6 | 7 | 0.5714 | 3 | 0.6667 | 20 | 0.55 |
| 1_13443503 | 3 | 0 | 7 | 0.1429 | 2 | 0.5 | NA | NA | 12 | 0.1667 | 6 | 0 | 4 | 0 | 7 | 0.1429 | 2 | 1 | 19 | 0.1579 |
| 1_13443519 | 4 | 0.75 | 7 | 0.7143 | 3 | 0.6667 | NA | NA | 14 | 0.7143 | 6 | 1 | 5 | 0.8 | 8 | 1 | 3 | 1 | 22 | 0.9545 |
| 1_13443541 | 6 | 0 | 9 | 0.2222 | 4 | 0 | NA | NA | 19 | 0.1053 | 5 | 0.2 | 5 | 0.2 | 8 | 0.25 | 2 | 0 | 20 | 0.2 |
| 1_13443546 | 6 | 0.1667 | 10 | 0.1 | 4 | 0 | NA | NA | 20 | 0.1 | 7 | 0.1429 | 5 | 0.4 | 8 | 0.125 | 3 | 0 | 23 | 0.1739 |
| 1_13454562 | 2 | 0 | 10 | 0.2 | 1 | 0 | 1 | 0 | 14 | 0.1429 | 3 | 0 | 3 | 0.3333 | 4 | 0.25 | 2 | 0 | 12 | 0.1667 |
| 1_13454566 | 2 | 0.5 | 3 | 0.3333 | NA | NA | 2 | 1 | 7 | 0.5714 | 3 | 0.3333 | 2 | 0.5 | 4 | 0.25 | 1 | 0 | 10 | 0.3 |
| 1_13455260 | 1 | 1 | 7 | 1 | 2 | 1 | 5 | 1 | 15 | 1 | 2 | 1 | 1 | 1 | 4 | 1 | 4 | 0.75 | 11 | 0.9091 |
| 1_13455280 | 2 | 0 | 10 | 0 | 3 | 0.3333 | 5 | 0 | 20 | 0.05 | 3 | 0.6667 | 3 | 0.3333 | 4 | 0 | 3 | 0 | 13 | 0.2308 |
| 1_13455280 | 2 | 1 | 8 | 0.375 | 3 | 0.3333 | 3 | 0.3333 | 16 | 0.4375 | 3 | 0 | 1 | 1 | 4 | 0.25 | 3 | 0 | 11 | 0.1818 |
| 1_13455288 | 2 | 0.5 | 7 | 0.7143 | 2 | 1 | 2 | 1 | 13 | 0.7692 | 3 | 0.6667 | 2 | 0.5 | 4 | 0.5 | 5 | 0.4 | 14 | 0.5 |
| 1_13455832 | 3 | 1 | 13 | 0.8462 | 3 | 0.6667 | 5 | 0.4 | 24 | 0.75 | 4 | 0.75 | 4 | 0.5 | 6 | 0.8333 | 7 | 0.7143 | 21 | 0.7143 |
| 1_13455833 | 1 | 0 | 8 | 1 | 3 | 1 | 3 | 0.3333 | 15 | 0.8 | 3 | 1 | 4 | 1 | 4 | 0.75 | 5 | 1 | 16 | 0.9375 |
| 1_13455836 | 3 | 0 | 11 | 0.1818 | 4 | 0 | 5 | 0.2 | 23 | 0.1304 | 4 | 0.25 | 4 | 0.25 | 5 | 0.2 | 7 | 0 | 20 | 0.15 |
| 1_13455838 | 4 | 0 | 10 | 0.2 | 4 | 0 | 5 | 0 | 23 | 0.087 | 3 | 0 | 4 | 0.25 | 7 | 0.2857 | 6 | 0.1667 | 20 | 0.2 |
| 1_13455843 | 4 | 0.25 | 8 | 0.375 | 3 | 0 | 4 | 0.5 | 19 | 0.3158 | 3 | 0.6667 | 4 | 0.25 | 6 | 0.6667 | 6 | 0.3333 | 19 | 0.4737 |
| 1_13455853 | 2 | 0 | 10 | 0.1 | 1 | 0 | 2 | 0 | 15 | 0.0667 | 3 | 0 | 3 | 0 | 4 | 0.25 | 3 | 0 | 13 | 0.0769 |
| 1_13455854 | 2 | 0.5 | 8 | 0.125 | NA | NA | 4 | 0 | 14 | 0.1429 | 2 | 0 | 2 | 0 | 5 | 0.2 | 3 | 0 | 12 | 0.0833 |
| 1_13455868 | 3 | 1 | 8 | 0.625 | 1 | 0 | 3 | 0.6667 | 15 | 0.6667 | 4 | 1 | 3 | 1 | 5 | 0.6 | NA | NA | 12 | 0.8333 |
| 1_13455883 | 3 | 1 | 10 | 1 | 1 | 1 | 2 | 1 | 16 | 1 | 4 | 1 | 3 | 1 | 5 | 1 | 3 | 1 | 15 | 1 |
| 1_13455889 | 3 | 0.3333 | 8 | 0.125 | NA | NA | 2 | 0.5 | 13 | 0.2308 | 3 | 0.3333 | 3 | 0.3333 | 4 | 0.25 | 3 | 0.6667 | 13 | 0.3846 |
| 1_13455897 | 4 | 1 | 10 | 0.9 | 3 | 0.3333 | 6 | 0.6667 | 23 | 0.7826 | 4 | 0.75 | 5 | 0.8 | 6 | 0.8333 | 5 | 0.8 | 20 | 0.8 |
| 1_13455912 | 2 | 0 | 9 | 0.2222 | 4 | 0 | 4 | 0 | 19 | 0.1053 | 4 | 0.25 | 5 | 0 | 5 | 0.2 | 5 | 0 | 19 | 0.1053 |
| 1_13455914 | 2 | 1 | 13 | 1 | 4 | 1 | 7 | 1 | 26 | 1 | 6 | 1 | 5 | 1 | 7 | 1 | 6 | 1 | 24 | 1 |
| 1_13455919 | 5 | 1 | 10 | 0.7 | 5 | 0.6 | 6 | 0.5 | 26 | 0.6923 | 5 | 0.6 | 3 | 1 | 7 | 1 | 8 | 0.375 | 23 | 0.6957 |
| 1_13455925 | 5 | 1 | 11 | 1 | 3 | 0.6667 | 5 | 0.8 | 24 | 0.9167 | 6 | 1 | 5 | 1 | 7 | 0.8571 | 5 | 0.8 | 23 | 0.913 |
| 1_13455927 | 4 | 0 | 7 | 0.1429 | 4 | 0 | 4 | 0.25 | 19 | 0.1053 | 6 | 0 | 4 | 0.25 | 5 | 0.4 | 7 | 0 | 22 | 0.1364 |
| 1_13455929 | 5 | 0 | 12 | 0.1667 | 5 | 0 | 5 | 0.2 | 27 | 0.1111 | 6 | 0.3333 | 4 | 0.5 | 8 | 0.125 | 8 | 0.125 | 26 | 0.2308 |
| 1_13455935 | 3 | 0 | 10 | 0.6 | 3 | 0.3333 | 6 | 0.5 | 22 | 0.4545 | 3 | 1 | 5 | 0.6 | 6 | 0.5 | 5 | 0.6 | 19 | 0.6316 |
| 1_13455977 | 4 | 0.5 | 5 | 0.6 | 5 | 0.4 | 7 | 0.2857 | 21 | 0.4286 | 4 | 0.5 | 5 | 0.4 | 7 | 0.1429 | 9 | 0.5556 | 25 | 0.4 |
| 1_13456031 | NA | NA | 3 | 0 | 2 | 0 | 7 | 0.2857 | 12 | 0.1667 | 1 | 0 | 2 | 0.5 | 5 | 0.2 | 3 | 0 | 11 | 0.1818 |
| 1_13467548 | 1 | 1 | 1 | 1 | 4 | 1 | 1 | 1 | 7 | 1 | NA | NA | NA | NA | 3 | 0.6667 | NA | NA | 3 | 0.6667 |
| 1_13467559 | 1 | 0 | 1 | 0 | 4 | 0.75 | NA | NA | 6 | 0.5 | NA | NA | NA | NA | 3 | 0.6667 | NA | NA | 3 | 0.6667 |
| 1_13467573 | 1 | 0 | 1 | 1 | 2 | 0 | 1 | 0 | 5 | 0.2 | NA | NA | NA | NA | 2 | 1 | NA | NA | 2 | 1 |
| 1_13467577 | 1 | 1 | 1 | 0 | 2 | 0 | NA | NA | 4 | 0.25 | NA | NA | NA | NA | 3 | 0.6667 | NA | NA | 3 | 0.6667 |
| 1_13467580 | 1 | 1 | 1 | 1 | 4 | 0.75 | 1 | 0 | 7 | 0.7143 | NA | NA | NA | NA | 2 | 1 | NA | NA | 2 | 1 |
| 1_13467595 | 1 | 1 | 1 | 1 | 5 | 1 | 2 | 1 | 9 | 1 | NA | NA | NA | NA | 3 | 0.6667 | NA | NA | 3 | 0.6667 |
| 1_13467609 | 1 | 1 | 1 | 1 | 3 | 1 | 1 | 1 | 6 | 1 | NA | NA | NA | NA | 2 | 1 | NA | NA | 2 | 1 |
| 1_13467609 | 1 | 1 | NA | NA | 4 | 0.5 | 1 | 0 | 6 | 0.5 | NA | NA | NA | NA | 3 | 1 | NA | NA | 3 | 1 |
| 1_13467617 | 1 | 1 | 1 | 1 | 4 | 1 | 2 | 1 | 8 | 1 | NA | NA | NA | NA | 3 | 1 | NA | NA | 3 | 1 |
| 1_13467620 | 1 | 0 | NA | NA | 4 | 0.25 | 2 | 0.5 | 7 | 0.2857 | NA | NA | NA | NA | 1 | 1 | NA | NA | 1 | 1 |
| 1_13467622 | 1 | 1 | 1 | 1 | 4 | 1 | 2 | 1 | 8 | 1 | NA | NA | NA | NA | 2 | 1 | NA | NA | 2 | 1 |
| 1_13467645 | 1 | 1 | 1 | 1 | 3 | 1 | 2 | 1 | 7 | 1 | NA | NA | NA | NA | 3 | 1 | NA | NA | 3 | 1 |
| 1_13467653 | NA | NA | 1 | 1 | 5 | 0.6 | 2 | 0.5 | 8 | 0.625 | NA | NA | NA | NA | 3 | 1 | NA | NA | 3 | 1 |
| 1_13467654 | 1 | 1 | 1 | 0 | 5 | 0.4 | 2 | 0 | 9 | 0.3333 | NA | NA | NA | NA | 2 | 0.5 | NA | NA | 2 | 0.5 |
| 1_13467659 | 1 | 0 | 1 | 0 | 4 | 0.75 | 2 | 0 | 8 | 0.375 | NA | NA | NA | NA | 3 | 1 | 1 | 0 | 4 | 0.75 |
| 1_13467667 | NA | NA | 1 | 1 | 5 | 0.8 | 2 | 0.5 | 8 | 0.75 | NA | NA | NA | NA | 2 | 1 | 1 | 1 | 3 | 1 |
| 1_13467707 | NA | NA | 1 | 0 | NA | NA | 1 | 1 | 2 | 0.5 | NA | NA | NA | NA | 2 | 0.5 | NA | NA | 2 | 0.5 |
| 1_13467859 | 2 | 0.5 | 1 | 1 | 1 | 0 | 1 | 0 | 5 | 0.4 | 1 | 0 | NA | NA | 3 | 0 | 1 | 1 | 5 | 0.2 |
| 1_13467913 | 4 | 0.5 | NA | NA | 3 | 0.3333 | 2 | 0.5 | 9 | 0.4444 | 2 | 0.5 | 1 | 0 | 3 | 0 | NA | NA | 6 | 0.1667 |
| 1_13467915 | 5 | 0.2 | 2 | 1 | 3 | 0 | 3 | 0 | 13 | 0.2308 | 2 | 0 | 1 | 0 | 4 | 0.25 | 1 | 0 | 8 | 0.125 |

| | | | | | | | | | | | | | | | | | | | | |
|---|---|---|---|---|---|---|---|---|---|---|---|---|---|---|---|---|---|---|---|---|
| 1_1346794 | 4 | 0.5 | 2 | 0.5 | 1 | 1 | 4 | 0 | 11 | 0.3636 | 2 | 0 | 2 | 0 | 3 | 0.3333 | NA | NA | 7 | 0.1429 |
| 1_1346805 | 5 | 0.2 | 3 | 0 | 4 | 0 | 5 | 0.2 | 17 | 0.1176 | 4 | 0.25 | 4 | 0 | 4 | 0 | 1 | 0 | 13 | 0.0769 |
| 1_1346806 | 4 | 0.25 | 3 | 0 | 5 | 0.2 | 5 | 0.2 | 17 | 0.1765 | 4 | 0 | 4 | 0.25 | 3 | 0.3333 | 1 | 0 | 12 | 0.1667 |
| 1_1346825 | 8 | 0.875 | 11 | 1 | 3 | 0.6667 | 12 | 0.8333 | 34 | 0.8824 | 5 | 1 | 7 | 0.8571 | 6 | 0.6667 | 9 | 0.8889 | 27 | 0.8519 |
| 1_1346829 | 8 | 0.5 | 9 | 0.5556 | 4 | 0.75 | 11 | 0.4545 | 32 | 0.5312 | 5 | 1 | 8 | 0.75 | 5 | 0.6 | 10 | 1 | 28 | 0.8571 |
| 1_1346830 | 8 | 0.75 | 8 | 0.5 | 4 | 0.5 | 7 | 0.5714 | 27 | 0.5926 | 3 | 1 | 8 | 0.5 | 5 | 0.6 | 9 | 0.4444 | 25 | 0.56 |
| 1_1346830 | 8 | 0.25 | 8 | 0.125 | 4 | 0 | 9 | 0.2222 | 29 | 0.1724 | 5 | 0 | 8 | 0.125 | 5 | 0 | 9 | 0.4444 | 27 | 0.1852 |
| 1_1346831 | 8 | 0.875 | 6 | 0.8333 | 4 | 1 | 8 | 0.625 | 26 | 0.8077 | 5 | 1 | 8 | 0.75 | 5 | 0.4 | 9 | 0.6667 | 27 | 0.7037 |
| 1_1346832 | 6 | 0.6667 | 5 | 0.4 | 2 | 0 | 6 | 0.5 | 19 | 0.4737 | 5 | 0.4 | 6 | 0.3333 | 6 | 0.8333 | 5 | 0.8 | 22 | 0.5909 |
| 1_1346833 | 6 | 0.3333 | 9 | 0.3333 | 2 | 0.5 | 7 | 0.1429 | 24 | 0.2917 | 4 | 0.75 | 7 | 0.4286 | 5 | 0 | 8 | 0.375 | 24 | 0.375 |
| 1_1346833 | 5 | 0 | 4 | 0 | 2 | 0 | 1 | 1 | 12 | 0.0833 | 3 | 0.3333 | 3 | 0 | 4 | 0.25 | 3 | 0 | 13 | 0.1538 |
| 1_1346835 | 5 | 0.6 | 8 | 1 | 1 | 0 | 7 | 0.5714 | 21 | 0.7143 | 4 | 0.5 | 5 | 1 | 4 | 1 | 7 | 0.8571 | 20 | 0.85 |
| 1_1346840 | 4 | 0 | 7 | 0.4286 | 1 | 0 | 7 | 0 | 19 | 0.1579 | 2 | 0 | 5 | 0.2 | 4 | 0 | 5 | 0 | 16 | 0.0625 |
| 1_1346847 | 4 | 0.25 | 4 | 0 | 1 | 0 | 5 | 0 | 14 | 0.0714 | 1 | 0 | 4 | 0 | 4 | 0.25 | 2 | 0 | 11 | 0.0909 |
| 1_1346853 | 3 | 0 | 4 | 0 | 1 | 0 | 4 | 0.25 | 12 | 0.0833 | 3 | 0 | 4 | 0.25 | 3 | 0 | 3 | 0 | 13 | 0.0769 |
| 1_1346862 | 2 | 0.5 | 1 | 0 | NA | NA | 2 | 1 | 5 | 0.6 | 3 | 0.3333 | 2 | 0.5 | 3 | 0.3333 | 1 | 0 | 9 | 0.3333 |
| 1_1346868 | 1 | 0 | 2 | 0.5 | NA | NA | 1 | 1 | 4 | 0.5 | 3 | 0.6667 | 2 | 0.5 | 3 | 0.6667 | 1 | 1 | 9 | 0.6667 |
| 1_1346869 | 2 | 0.5 | 2 | 0.5 | 1 | 0 | 1 | 0 | 6 | 0.3333 | 3 | 0.3333 | 1 | 0 | 3 | 0 | 1 | 0 | 8 | 0.125 |
| 1_1346869 | 2 | 0.5 | 1 | 0 | 1 | 0 | 1 | 1 | 5 | 0.4 | 2 | 0.5 | 2 | 0.5 | 2 | 0.5 | 1 | 0 | 7 | 0.4286 |
| 1_1346870 | 2 | 0.5 | 3 | 0.3333 | 1 | 0 | 1 | 0 | 7 | 0.2857 | 2 | 0 | 2 | 0.5 | 3 | 0.3333 | 1 | 0 | 8 | 0.25 |
| 1_1346875 | 1 | 1 | 3 | 0.6667 | 1 | 0 | 1 | 0 | 6 | 0.5 | 1 | 1 | 2 | 1 | 2 | 0.5 | 1 | 1 | 6 | 0.8333 |
| 1_1346876 | 2 | 0.5 | 2 | 0 | 1 | 0 | 1 | 0 | 6 | 0.1667 | 1 | 0 | 1 | 0 | 2 | 0.5 | 1 | 0 | 5 | 0.2 |
| 1_1346876 | 1 | 1 | 3 | 0.3333 | 1 | 1 | 1 | 1 | 6 | 0.6667 | 1 | 1 | 1 | 0 | 2 | 1 | 1 | 0 | 5 | 0.6 |
| 1_1346877 | 2 | 0.5 | 2 | 0.5 | 1 | 0 | 1 | 1 | 6 | 0.5 | 1 | 1 | 2 | 1 | 1 | 1 | NA | NA | 4 | 1 |
| 1_1346880 | 2 | 0 | 3 | 0.3333 | 1 | 1 | 1 | 0 | 7 | 0.2857 | 1 | 1 | 2 | 0.5 | 2 | 1 | NA | NA | 5 | 0.8 |
| 1_1350376 | 2 | 1 | 1 | 1 | 1 | 0 | 1 | 0 | 5 | 0.6 | NA | NA | 1 | 0 | 1 | 1 | NA | NA | 2 | 0.5 |
| 1_1350715 | NA | NA | NA | NA | 1 | 1 | NA | NA | 1 | 1 | NA | NA | NA | NA | NA | NA | 1 | 1 | 1 | 1 |
| 1_1350729 | 1 | 1 | 1 | 1 | 1 | 1 | NA | NA | 3 | 1 | NA | NA | NA | NA | NA | NA | 1 | 1 | 1 | 1 |
| 1_1350747 | 14 | 0.2857 | 16 | 0.0625 | 16 | 0.3125 | 15 | 0.2667 | 61 | 0.2295 | 13 | 0.3846 | 15 | 0.2 | 20 | 0.2 | 23 | 0.2609 | 71 | 0.2535 |
| 1_1350881 | 22 | 0.2727 | 25 | 0.32 | 29 | 0.1724 | 22 | 0.3636 | 98 | 0.2755 | 38 | 0.3947 | 20 | 0.4 | 20 | 0.4 | 24 | 0.375 | 102 | 0.3922 |
| 1_1350881 | 25 | 0.32 | 30 | 0.3333 | 35 | 0.2857 | 27 | 0.1852 | 117 | 0.2821 | 39 | 0.2564 | 20 | 0.25 | 20 | 0.3 | 19 | 0.2632 | 98 | 0.2653 |
| 1_1350882 | 28 | 0.1429 | 32 | 0.125 | 38 | 0.1316 | 30 | 0.0667 | 128 | 0.1172 | 46 | 0.1304 | 30 | 0.1333 | 26 | 0.1154 | 29 | 0.2069 | 131 | 0.145 |
| 1_1350884 | 29 | 0.1724 | 27 | 0.0741 | 29 | 0.069 | 26 | 0.1154 | 111 | 0.1081 | 42 | 0.2143 | 26 | 0.1923 | 24 | 0.1667 | 25 | 0.04 | 117 | 0.1624 |
| 1_1350884 | 32 | 1 | 30 | 0.9333 | 36 | 0.9722 | 27 | 1 | 125 | 0.976 | 43 | 0.9535 | 28 | 0.9286 | 24 | 1 | 30 | 1 | 125 | 0.968 |
| 1_1350887 | 25 | 0.2 | 30 | 0.2 | 34 | 0.2059 | 27 | 0.1481 | 116 | 0.1897 | 36 | 0.1944 | 24 | 0.2083 | 22 | 0.1818 | 25 | 0.16 | 107 | 0.1869 |
| 1_1350887 | 27 | 0.2593 | 32 | 0.1562 | 38 | 0.2105 | 28 | 0.25 | 125 | 0.216 | 38 | 0.2368 | 26 | 0.2692 | 24 | 0.1667 | 30 | 0.1667 | 118 | 0.2119 |
| 1_1350888 | 28 | 0.3929 | 32 | 0.5 | 38 | 0.3421 | 23 | 0.3913 | 121 | 0.405 | 34 | 0.2353 | 26 | 0.3462 | 22 | 0.4091 | 27 | 0.4444 | 109 | 0.3486 |
| 1_1350889 | 29 | 0.4483 | 30 | 0.3 | 34 | 0.3235 | 25 | 0.4 | 118 | 0.3644 | 38 | 0.3684 | 24 | 0.4583 | 23 | 0.4783 | 27 | 0.2593 | 112 | 0.3839 |
| 1_1350891 | 30 | 0.0667 | 25 | 0.08 | 30 | 0 | 24 | 0.0833 | 109 | 0.055 | 31 | 0.0323 | 21 | 0.0476 | 23 | 0.0435 | 22 | 0.0909 | 97 | 0.0515 |
| 1_1350892 | 28 | 0.9286 | 29 | 0.931 | 28 | 1 | 24 | 1 | 109 | 0.9633 | 34 | 0.9412 | 25 | 1 | 19 | 1 | 25 | 1 | 103 | 0.9806 |
| 1_1350892 | 25 | 0.84 | 23 | 0.8261 | 28 | 0.9643 | 24 | 0.875 | 100 | 0.88 | 32 | 0.7812 | 18 | 0.9444 | 15 | 0.8667 | 20 | 0.8 | 85 | 0.8353 |
| 1_1350892 | 25 | 0.4 | 26 | 0.3846 | 24 | 0.5 | 20 | 0.3 | 95 | 0.4 | 33 | 0.3939 | 20 | 0.3 | 16 | 0.25 | 21 | 0.381 | 90 | 0.3444 |
| 1_1350947 | 18 | 1 | 20 | 0.9 | 24 | 0.8333 | 24 | 0.875 | 86 | 0.8953 | 29 | 0.931 | 23 | 0.7826 | 22 | 0.8636 | 18 | 0.8889 | 92 | 0.8696 |
| 1_1350949 | 17 | 0.2941 | 18 | 0.2222 | 18 | 0.4444 | 19 | 0.3158 | 72 | 0.3194 | 26 | 0.4615 | 19 | 0.3684 | 15 | 0.4 | 13 | 0.4615 | 73 | 0.4247 |
| 1_1350950 | 18 | 0.6111 | 21 | 0.2381 | 24 | 0.375 | 23 | 0.3043 | 86 | 0.3721 | 29 | 0.4483 | 22 | 0.2273 | 19 | 0.1579 | 21 | 0.3333 | 91 | 0.3077 |
| 1_1350951 | 12 | 0.25 | 20 | 0.45 | 19 | 0.6316 | 21 | 0.7143 | 72 | 0.5417 | 26 | 0.5769 | 18 | 0.4444 | 15 | 0.3333 | 15 | 0.4 | 74 | 0.4595 |
| 1_1351906 | 14 | 0.1429 | 10 | 0.3 | 9 | 0.3333 | 8 | 0.25 | 41 | 0.2439 | 5 | 0.2 | 8 | 0.25 | 11 | 0.3636 | 11 | 0.3636 | 35 | 0.3143 |
| 1_1351907 | 15 | 0.0667 | 10 | 0.2 | 11 | 0.2727 | 8 | 0.125 | 44 | 0.1591 | 6 | 0 | 6 | 0.1667 | 9 | 0.1111 | 13 | 0.3077 | 34 | 0.1765 |
| 1_1351930 | 10 | 0.1 | 9 | 0 | 7 | 0.1429 | 8 | 0 | 34 | 0.0588 | 6 | 0 | 7 | 0.2857 | 8 | 0 | 11 | 0 | 32 | 0.0625 |
| 1_1352113 | 8 | 0.25 | 8 | 0 | 4 | 0 | 8 | 0.25 | 28 | 0.1429 | 3 | 0 | 7 | 0 | 5 | 0.2 | 8 | 0.5 | 23 | 0.2174 |
| 1_1352116 | 8 | 0.125 | 9 | 0.2222 | 3 | 0.3333 | 6 | 0 | 26 | 0.1538 | 2 | 0.5 | 5 | 0.2 | 4 | 0 | 8 | 0 | 19 | 0.1053 |
| 1_1352218 | 1 | 1 | NA | NA | 1 | 1 | 1 | 1 | 3 | 1 | NA | NA | 2 | 0.5 | 1 | 1 | NA | NA | 3 | 0.6667 |
| 1_1352226 | 1 | 0 | NA | NA | 1 | 0 | 1 | 1 | 3 | 0.3333 | NA | NA | 1 | 0 | 2 | 1 | NA | NA | 3 | 0.6667 |
| 1_1352228 | 1 | 1 | NA | NA | 1 | 1 | 1 | 1 | 3 | 1 | NA | NA | 1 | 1 | 2 | 1 | NA | NA | 3 | 1 |
| 1_1352229 | 4 | 0.5 | 1 | 0 | 2 | 0 | 1 | 1 | 8 | 0.375 | 1 | 1 | NA | NA | 2 | 0.5 | 1 | 0 | 4 | 0.5 |
| 1_1352231 | 4 | 0 | NA | NA | 3 | 0 | 1 | 1 | 8 | 0.125 | NA | NA | NA | NA | 3 | 0.3333 | 1 | 1 | 4 | 0.5 |
| 1_1352236 | 2 | 0 | 1 | 0 | 3 | 0.3333 | NA | NA | 6 | 0.1667 | NA | NA | NA | NA | 3 | 0.3333 | 1 | 0 | 4 | 0.25 |
| 1_1352292 | 6 | 0.1667 | 12 | 0.0833 | 9 | 0.1111 | 9 | 0.2222 | 36 | 0.1389 | 10 | 0.3 | 15 | 0.2 | 12 | 0.3333 | 12 | 0.1667 | 49 | 0.2449 |
| 1_1352293 | 8 | 0.125 | 13 | 0.1538 | 13 | 0.2308 | 9 | 0.1111 | 43 | 0.1628 | 8 | 0.125 | 11 | 0.1818 | 15 | 0 | 12 | 0.25 | 46 | 0.1304 |
| 1_1352295 | 10 | 0.6 | 13 | 0.6923 | 19 | 0.7895 | 11 | 0.4545 | 53 | 0.6604 | 16 | 0.625 | 20 | 0.9 | 18 | 0.7778 | 16 | 0.8125 | 70 | 0.7857 |
| 1_1352295 | 10 | 0.2 | 16 | 0.25 | 22 | 0.4091 | 12 | 0.5833 | 60 | 0.3667 | 16 | 0.375 | 21 | 0.619 | 19 | 0.5789 | 14 | 0.5714 | 70 | 0.5429 |
| 1_1352295 | 4 | 0.25 | 8 | 0 | 12 | 0.25 | 8 | 0.25 | 32 | 0.1875 | 11 | 0.0909 | 16 | 0.0625 | 16 | 0.25 | 10 | 0.2 | 53 | 0.1509 |
| 1_1352296 | 9 | 0.7778 | 14 | 0.2143 | 22 | 0.4091 | 13 | 0.5385 | 58 | 0.4483 | 14 | 0.7143 | 21 | 0.381 | 17 | 0.7059 | 16 | 0.4375 | 68 | 0.5441 |
| 1_1352987 | 2 | 0 | 2 | 0.5 | 1 | 0 | NA | NA | 5 | 0.2 | 1 | 0 | 1 | 1 | 1 | 0 | NA | NA | 3 | 0.3333 |
| 1_1352989 | 8 | 0.625 | 14 | 0.2857 | 8 | 0.5 | 4 | 0.25 | 34 | 0.4118 | 8 | 0.375 | 5 | 0.2 | 5 | 0.8 | 6 | 0.5 | 24 | 0.4583 |

| | | | | | | | | | | | | | | | | | | | | |
|---|---|---|---|---|---|---|---|---|---|---|---|---|---|---|---|---|---|---|---|---|
| 1_13529893 | 10 | 0.2 | 21 | 0.1905 | 11 | 0.1818 | 6 | 0.5 | 48 | 0.2292 | 13 | 0.1538 | 11 | 0 | 8 | 0.25 | 10 | 0.1 | 42 | 0.119 |
| 1_13529904 | 7 | 1 | 18 | 0.8889 | 9 | 0.7778 | 6 | 0.8333 | 40 | 0.875 | 10 | 0.9 | 6 | 1 | 5 | 0.8 | 6 | 1 | 27 | 0.9259 |
| 1_13529914 | 11 | 0.6364 | 17 | 0.1176 | 10 | 0.7 | 6 | 0.1667 | 44 | 0.3864 | 12 | 0.6667 | 10 | 0.5 | 7 | 0.7143 | 8 | 0.625 | 37 | 0.6216 |
| 1_13531019 | 5 | 0.2 | 8 | 0 | 5 | 0.2 | 7 | 0.1429 | 25 | 0.12 | 4 | 0 | 3 | 0 | 8 | 0.125 | 3 | 0 | 18 | 0.0556 |
| 1_13531074 | 3 | 0 | 5 | 0 | 3 | 0.3333 | 7 | 0 | 18 | 0.0556 | 3 | 0 | 3 | 0 | 7 | 0.2857 | 1 | 0 | 14 | 0.1429 |
| 1_13531090 | 3 | 0.3333 | 6 | 0.1667 | 3 | 0 | 6 | 0.5 | 18 | 0.2778 | 3 | 0 | 3 | 0.3333 | 6 | 0.3333 | 1 | 0 | 13 | 0.2308 |
| 1_13531099 | 2 | 0 | 5 | 0.4 | 2 | 0.5 | 7 | 0.4286 | 16 | 0.375 | 2 | 0.5 | 3 | 0 | 7 | 0.5714 | 1 | 0 | 13 | 0.3846 |
| 1_13531103 | 3 | 0.6667 | 4 | 1 | 3 | 0.6667 | 7 | 0.5714 | 17 | 0.7059 | 3 | 0.3333 | 3 | 1 | 6 | 0.8333 | 1 | 0 | 13 | 0.6923 |
| 1_13531110 | 2 | 0.5 | 4 | 0.5 | 3 | 0 | 5 | 0.6 | 14 | 0.4286 | 3 | 0.6667 | 3 | 0.3333 | 6 | 0.3333 | NA | NA | 12 | 0.4167 |
| 1_13531172 | 4 | 0 | 4 | 0 | 3 | 0 | 6 | 0.1667 | 17 | 0.0588 | 3 | 0 | 3 | 0 | 5 | 0.2 | 1 | 0 | 12 | 0.0833 |
| 1_13531174 | 4 | 0.5 | 4 | 0.25 | 3 | 0.3333 | 7 | 0.2857 | 18 | 0.3333 | 3 | 0.3333 | 2 | 0.5 | 5 | 0 | 1 | 0 | 11 | 0.1818 |
| 1_13531207 | 3 | 0 | 4 | 0.25 | 1 | 0 | 5 | 0 | 13 | 0.0769 | 2 | 0 | 3 | 0 | 3 | 0.3333 | 1 | 0 | 9 | 0.1111 |
| 1_13531271 | 5 | 0.2 | 7 | 0 | 3 | 0.6667 | 11 | 0.0909 | 26 | 0.1538 | 3 | 0.3333 | 7 | 0 | 6 | 0.1667 | 13 | 0.0769 | 29 | 0.1034 |
| 1_13531282 | 3 | 0 | 5 | 0.2 | 2 | 0 | 9 | 0.1111 | 19 | 0.1053 | 2 | 0 | 6 | 0.1667 | 6 | 0.1667 | 12 | 0.1667 | 26 | 0.1538 |
| 1_13531288 | 5 | 1 | 6 | 0.8333 | 1 | 0 | 10 | 0.7 | 22 | 0.7727 | 1 | 1 | 6 | 0.6667 | 6 | 0.8333 | 9 | 0.8889 | 22 | 0.8182 |
| 1_13531292 | 4 | 0.5 | 6 | 0.6667 | 3 | 0.6667 | 6 | 1 | 19 | 0.7368 | 3 | 1 | 6 | 0.8333 | 5 | 1 | 9 | 0.5556 | 23 | 0.7826 |
| 1_13531297 | 5 | 0.4 | 5 | 0.2 | 2 | 0.5 | 7 | 0.2857 | 19 | 0.3158 | 3 | 0.3333 | 6 | 0.1667 | 6 | 0 | 8 | 0.375 | 23 | 0.2174 |
| 1_13531298 | 6 | 0.6667 | 7 | 0.8571 | 2 | 1 | 9 | 0.7778 | 24 | 0.7917 | 3 | 1 | 8 | 1 | 7 | 0.7143 | 12 | 0.8333 | 30 | 0.8667 |
| 1_13531305 | 2 | 0.5 | 4 | 1 | 1 | 1 | 3 | 0.3333 | 10 | 0.7 | 1 | 1 | 5 | 0.6 | 3 | 0.3333 | 9 | 0.2222 | 18 | 0.3889 |
| 1_13531311 | 3 | 0 | 8 | 0.25 | 2 | 0 | 9 | 0.4444 | 22 | 0.2727 | 3 | 0.3333 | 6 | 0.5 | 6 | 0.3333 | 11 | 0.4545 | 26 | 0.4231 |
| 1_13531724 | 2 | 0 | 5 | 0.2 | 3 | 0 | 8 | 0.375 | 18 | 0.2222 | 1 | 0 | 2 | 0 | 2 | 0 | 10 | 0.1 | 15 | 0.0667 |
| 1_13536258 | 4 | 0 | 6 | 0 | 6 | 0.1667 | NA | NA | 16 | 0.0625 | 3 | 0 | 4 | 0.25 | 4 | 0 | 2 | 0.5 | 13 | 0.1538 |
| 1_13536394 | 3 | 0 | 3 | 0 | 5 | 0.2 | NA | NA | 11 | 0.0909 | 3 | 0 | 5 | 0.2 | 3 | 0 | 1 | 0 | 12 | 0.0833 |
| 1_13536560 | 2 | 0.5 | 3 | 0 | 5 | 0.4 | NA | NA | 10 | 0.3 | 2 | 1 | 2 | 0 | 3 | 0 | NA | NA | 7 | 0.2857 |
| 1_13536568 | 2 | 0.5 | 2 | 0 | 5 | 0.2 | NA | NA | 9 | 0.2222 | 2 | 0 | 2 | 0 | 3 | 0.3333 | NA | NA | 7 | 0.1429 |
| 1_13536576 | 2 | 0 | 3 | 0 | 5 | 0.2 | NA | NA | 10 | 0.1 | 3 | 0.6667 | 2 | 0.5 | 3 | 0 | 1 | 0 | 9 | 0.3333 |
| 1_13536607 | 1 | 0 | 3 | 0.3333 | 3 | 0.3333 | NA | NA | 7 | 0.2857 | 5 | 0.6 | 3 | 0 | 3 | 0 | 3 | 0.3333 | 14 | 0.2857 |
| 1_13536635 | 1 | 0 | 6 | 0 | 4 | 0 | 1 | 1 | 12 | 0.0833 | 6 | 0.5 | 4 | 0.25 | 6 | 0 | 3 | 0.3333 | 19 | 0.2632 |
| 1_13536637 | 1 | 1 | 6 | 0.6667 | 4 | 1 | 1 | 1 | 12 | 0.8333 | 7 | 1 | 4 | 0.75 | 7 | 0.7143 | 4 | 0.75 | 22 | 0.8182 |
| 1_13536638 | 2 | 0.5 | 5 | 0.2 | 4 | 0.25 | 1 | 0 | 12 | 0.25 | 7 | 0.4286 | 4 | 0.75 | 6 | 0.3333 | 2 | 0 | 19 | 0.4211 |
| 1_13536648 | 1 | 1 | 4 | 0 | 5 | 0.2 | 1 | 0 | 11 | 0.1818 | 7 | 0.1429 | 4 | 1 | 7 | 0.4286 | 4 | 0 | 22 | 0.3636 |
| 1_13536663 | 1 | 0 | 5 | 0.6 | 5 | 0.8 | 1 | 1 | 12 | 0.6667 | 8 | 0.375 | 4 | 1 | 7 | 0.5714 | 4 | 0.25 | 23 | 0.5217 |
| 1_13536673 | 1 | 0 | 5 | 0.4 | 5 | 0.2 | 1 | 0 | 12 | 0.25 | 7 | 0.5714 | 2 | 0.5 | 6 | 0.1667 | 3 | 0.3333 | 18 | 0.3889 |
| 1_13536678 | 1 | 1 | 5 | 0.4 | 5 | 0.2 | 1 | 1 | 12 | 0.4167 | 7 | 0.2857 | 3 | 0.3333 | 5 | 0.4 | 3 | 0.3333 | 18 | 0.3333 |
| 1_13536702 | 1 | 0 | 5 | 1 | 4 | 0.5 | NA | NA | 10 | 0.7 | 8 | 0.625 | 3 | 1 | 7 | 0.7143 | 3 | 0.6667 | 21 | 0.7143 |
| 1_13536705 | 1 | 1 | 4 | 0.25 | 4 | 0 | NA | NA | 9 | 0.2222 | 7 | 0.1429 | 4 | 0 | 7 | 0.1429 | 4 | 0 | 22 | 0.0909 |
| 1_13536707 | 2 | 1 | 3 | 0.6667 | 4 | 0.5 | NA | NA | 9 | 0.6667 | 6 | 0.3333 | 4 | 1 | 6 | 0.3333 | 3 | 0.3333 | 19 | 0.4737 |
| 1_13536709 | 1 | 1 | 3 | 0 | 4 | 0.5 | NA | NA | 8 | 0.375 | 7 | 0.1429 | 4 | 0.5 | 7 | 0 | 4 | 0.25 | 22 | 0.1818 |
| 1_13536719 | 1 | 0 | 4 | 0.25 | 4 | 0 | NA | NA | 9 | 0.1111 | 7 | 0.2857 | 3 | 0 | 6 | 0 | 3 | 0.3333 | 19 | 0.1579 |
| 1_13536728 | NA | NA | 4 | 0.5 | 5 | 0.2 | NA | NA | 9 | 0.3333 | 7 | 0.7143 | 4 | 1 | 7 | 0.7143 | 3 | 0.6667 | 21 | 0.7619 |
| 1_13536768 | 1 | 1 | 3 | 0.6667 | 4 | 0.25 | NA | NA | 8 | 0.5 | 5 | 0.6 | 4 | 0.75 | 6 | 0.6667 | 3 | 0.6667 | 18 | 0.6667 |
| 1_13536780 | 2 | 0.5 | 3 | 0 | 3 | 0 | NA | NA | 8 | 0.125 | 6 | 0.1667 | 3 | 0.3333 | 7 | 0.5714 | 3 | 1 | 19 | 0.4737 |
| 1_13536787 | 2 | 0.5 | 3 | 0.3333 | 4 | 0 | NA | NA | 9 | 0.2222 | 5 | 0.2 | 4 | 0.25 | 7 | 0.4286 | 3 | 0.6667 | 19 | 0.3684 |
| 1_13536801 | 1 | 0 | 3 | 0 | 4 | 0.25 | NA | NA | 8 | 0.125 | 6 | 0.1667 | 4 | 0 | 7 | 0 | 2 | 0 | 19 | 0.0526 |
| 1_13536813 | 1 | 0 | 3 | 0.3333 | 3 | 0.3333 | NA | NA | 7 | 0.2857 | 5 | 0.4 | 2 | 0.5 | 4 | 0.25 | 2 | 0.5 | 13 | 0.3846 |
| 1_13536823 | 1 | 0 | 2 | 0.5 | 4 | 0.5 | NA | NA | 7 | 0.4286 | 6 | 0.6667 | 3 | 0.6667 | 3 | 0.6667 | 3 | 1 | 15 | 0.7333 |
| 1_13536833 | 1 | 0 | 2 | 0.5 | 3 | 0.3333 | NA | NA | 6 | 0.3333 | 6 | 0.5 | 4 | 0.5 | 6 | 0.1667 | 3 | 0.6667 | 19 | 0.4211 |
| 1_13536836 | 1 | 1 | 2 | 0.5 | 4 | 0 | NA | NA | 7 | 0.2857 | 4 | 0 | 4 | 0 | 6 | 0.1667 | 3 | 0 | 17 | 0.0588 |
| 1_13536872 | 1 | 1 | 1 | 0 | 4 | 0.75 | 2 | 0.5 | 8 | 0.625 | 4 | 0.75 | 2 | 0.5 | 1 | 0 | 2 | 0 | 9 | 0.4444 |
| 1_13536886 | 1 | 0 | 1 | 0 | 4 | 0 | 3 | 0.3333 | 9 | 0.1111 | 4 | 0 | 5 | 0.2 | 2 | 0 | 2 | 0 | 13 | 0.0769 |
| 1_13536897 | 1 | 1 | 1 | 1 | 4 | 0.75 | 3 | 1 | 9 | 0.8889 | 4 | 0.75 | 5 | 0.8 | 2 | 1 | 1 | 1 | 12 | 0.8333 |
| 1_13536901 | NA | NA | NA | NA | NA | NA | 1 | 1 | 1 | 1 | 4 | 0.25 | 1 | 0 | 1 | 0 | NA | NA | 6 | 0.1667 |
| 1_13536906 | 1 | 0 | NA | NA | 3 | 0.3333 | 3 | 0 | 7 | 0.1429 | 5 | 0 | 5 | 0 | 2 | 0.5 | 2 | 0 | 14 | 0.0714 |
| 1_13536907 | 1 | 0 | 1 | 1 | 3 | 0 | 2 | 0 | 7 | 0.1429 | 4 | 0 | 5 | 0.2 | 2 | 0 | 2 | 0 | 13 | 0.0769 |
| 1_13536908 | 1 | 0 | 1 | 1 | 3 | 0 | 2 | 0 | 7 | 0.1429 | 4 | 0 | 4 | 0 | 1 | 0 | 2 | 0.5 | 11 | 0.0909 |
| 1_13536912 | NA | NA | 1 | 1 | 3 | 0.6667 | 1 | 1 | 5 | 0.8 | 3 | 1 | NA | NA | 1 | 0 | NA | NA | 4 | 0.75 |
| 1_13536921 | 1 | 0 | 1 | 0 | 3 | 0.3333 | 3 | 0.3333 | 8 | 0.25 | 5 | 0.2 | 3 | 0 | 2 | 0 | 2 | 0 | 12 | 0.0833 |
| 1_13536925 | 1 | 1 | 1 | 1 | 3 | 1 | 3 | 0.6667 | 8 | 0.875 | 5 | 0.8 | 2 | 1 | 1 | 1 | 1 | 1 | 9 | 0.8889 |
| 1_13536926 | 1 | 0 | NA | NA | 3 | 0.3333 | 3 | 0.3333 | 7 | 0.2857 | 5 | 0 | 2 | 0.5 | 2 | 0 | 1 | 0 | 10 | 0.1 |
| 1_13536931 | 1 | 0 | 1 | 0 | 3 | 0.3333 | 2 | 0.5 | 7 | 0.2857 | 5 | 0.2 | 2 | 0 | 2 | 0.5 | 2 | 0 | 11 | 0.1818 |
| 1_13536945 | 2 | 0.5 | 1 | 0 | 3 | 0.3333 | 3 | 0.6667 | 9 | 0.4444 | 4 | 0.25 | 3 | 0.6667 | 2 | 0.5 | 2 | 0 | 11 | 0.3636 |
| 1_13539832 | NA | NA | NA | NA | 1 | 1 | 2 | 0.5 | 3 | 0.6667 | 3 | 0.6667 | 1 | 1 | NA | NA | NA | NA | 4 | 0.75 |
| 1_13539843 | NA | NA | NA | NA | 1 | 1 | 2 | 0 | 3 | 0.3333 | 3 | 0.6667 | 1 | 1 | NA | NA | NA | NA | 4 | 0.75 |
| 1_13552747 | 1 | 1 | NA | NA | NA | NA | NA | NA | 1 | 1 | NA | NA | NA | NA | NA | NA | 1 | 1 | 1 | 1 |
| 1_13567898 | 24 | 0.375 | 27 | 0.2963 | 30 | 0.5 | 22 | 0.5909 | 103 | 0.4369 | 20 | 0.65 | 13 | 0.6154 | 20 | 0.3 | 21 | 0.4286 | 74 | 0.4865 |

| | | | | | | | | | | | | | | | | | | | | |
|---|---|---|---|---|---|---|---|---|---|---|---|---|---|---|---|---|---|---|---|---|
| 1_1356945 | 6 | 0.5 | 5 | 0 | 7 | 0 | 8 | 0.375 | 26 | 0.2308 | 3 | 0 | 2 | 0 | 6 | 0.1667 | 4 | 0.25 | 15 | 0.1333 |
| 1_1356945 | 5 | 0.8 | 4 | 0.75 | 6 | 0.8333 | 5 | 0.6 | 20 | 0.75 | 2 | 0.5 | 1 | 1 | 5 | 0.6 | 4 | 1 | 12 | 0.75 |
| 1_1356963 | 6 | 1 | 5 | 1 | 6 | 0.3333 | 9 | 0.7778 | 26 | 0.7692 | 4 | 1 | 2 | 0 | 6 | 1 | 6 | 0.6667 | 18 | 0.7778 |
| 1_1356963 | 4 | 1 | 4 | 1 | 6 | 1 | 7 | 1 | 21 | 1 | 3 | 1 | 2 | 0.5 | 4 | 1 | 5 | 1 | 14 | 0.9286 |
| 1_1356964 | 7 | 0.5714 | 5 | 0.6 | 6 | 0.1667 | 7 | 0.2857 | 25 | 0.4 | 3 | 0 | 2 | 1 | 4 | 0.5 | 6 | 0 | 15 | 0.2667 |
| 1_1356964 | 7 | 0.2857 | 4 | 0.5 | 6 | 0 | 6 | 0.3333 | 23 | 0.2609 | 3 | 0 | 2 | 0.5 | 5 | 0.2 | 6 | 0.1667 | 16 | 0.1875 |
| 1_1356964 | 5 | 1 | 5 | 0.8 | 6 | 1 | 9 | 0.7778 | 25 | 0.88 | 2 | 1 | 2 | 0.5 | 6 | 1 | 5 | 0.8 | 15 | 0.8667 |
| 1_1356965 | 7 | 0.1429 | 4 | 0 | 5 | 0.6 | 9 | 0.3333 | 25 | 0.28 | 4 | 0.25 | 2 | 0.5 | 6 | 0 | 5 | 0.2 | 17 | 0.1765 |
| 1_1356965 | 7 | 0.5714 | 5 | 0.6 | 6 | 0.6667 | 8 | 0.625 | 26 | 0.6154 | 3 | 0.6667 | 2 | 1 | 6 | 0.8333 | 7 | 0.7143 | 18 | 0.7778 |
| 1_1356966 | 7 | 0.4286 | 5 | 0.6 | 5 | 0.4 | 9 | 0.4444 | 26 | 0.4615 | 2 | 0 | 2 | 0.5 | 6 | 1 | 6 | 0.6667 | 16 | 0.6875 |
| 1_1356966 | 5 | 1 | 3 | 1 | 5 | 1 | 8 | 1 | 21 | 1 | 3 | 1 | NA | NA | 3 | 1 | 4 | 1 | 10 | 1 |
| 1_1359621 | 6 | 0.5 | 9 | 0.2222 | 7 | 0.1429 | 11 | 0.0909 | 33 | 0.2121 | 6 | 0 | 10 | 0 | 9 | 0.3333 | 7 | 0.7143 | 32 | 0.25 |
| 1_1359624 | 5 | 0.6 | 10 | 0.9 | 9 | 0.6667 | 11 | 0.8182 | 35 | 0.7714 | 7 | 0.4286 | 10 | 0.6 | 11 | 0.1818 | 9 | 0.7778 | 37 | 0.4865 |
| 1_1359626 | 7 | 0.1429 | 11 | 0.2727 | 7 | 0 | 12 | 0.1667 | 37 | 0.1622 | 7 | 0.1429 | 9 | 0.2222 | 9 | 0.4444 | 8 | 0.125 | 33 | 0.2424 |
| 1_1361018 | 1 | 0 | 3 | 0 | 1 | 1 | 1 | 0 | 6 | 0.1667 | 2 | 0.5 | 2 | 0 | 2 | 0.5 | 2 | 0 | 8 | 0.25 |
| 1_1361020 | 1 | 1 | 4 | 0 | NA | NA | NA | NA | 5 | 0.2 | 2 | 0 | 2 | 0.5 | 3 | 0 | 2 | 0.5 | 9 | 0.2222 |
| 1_1361030 | 1 | 0 | 3 | 0.6667 | 1 | 0 | 1 | 1 | 6 | 0.5 | 2 | 0.5 | 2 | 0.5 | 2 | 0.5 | 3 | 0.3333 | 9 | 0.4444 |
| 1_1361031 | 1 | 0 | 3 | 0.3333 | 1 | 0 | 1 | 1 | 6 | 0.3333 | 1 | 1 | 2 | 0 | 2 | 1 | 3 | 0.6667 | 8 | 0.625 |
| 1_1361033 | 1 | 0 | 2 | 0 | 1 | 0 | 3 | 0.3333 | 7 | 0.1429 | 2 | 0 | 3 | 0 | 2 | 0.5 | 4 | 0 | 11 | 0.0909 |
| 1_1361034 | 1 | 1 | 4 | 0.25 | 1 | 0 | 2 | 1 | 8 | 0.5 | 2 | 1 | 3 | 0.6667 | 3 | 1 | 3 | 1 | 11 | 0.9091 |
| 1_1361035 | NA | NA | 3 | 0.3333 | 1 | 0 | 2 | 0.5 | 6 | 0.3333 | 1 | 0 | 2 | 0.5 | 3 | 0.6667 | 2 | 0.5 | 8 | 0.5 |
| 1_1361036 | 1 | 1 | 4 | 0.75 | 1 | 0 | 3 | 0.6667 | 9 | 0.6667 | 2 | 1 | 4 | 1 | 3 | 0.6667 | 5 | 0.8 | 14 | 0.8571 |
| 1_1361037 | 1 | 1 | 4 | 1 | 1 | 1 | 2 | 1 | 8 | 1 | 1 | 1 | 3 | 0 | 3 | 1 | 3 | 1 | 10 | 0.7 |
| 1_1361039 | 2 | 1 | 2 | 0.5 | 1 | 1 | 3 | 0.6667 | 8 | 0.75 | 2 | 0 | 2 | 0 | 3 | 0.6667 | 4 | 0.5 | 11 | 0.3636 |
| 1_1361041 | 2 | 0 | 4 | 0.25 | 1 | 0 | 3 | 0 | 10 | 0.1 | 2 | 0.5 | 5 | 0.6 | 4 | 0.5 | 4 | 0 | 15 | 0.4 |
| 1_1361043 | 2 | 0 | 3 | 0.3333 | 1 | 0 | 3 | 0 | 9 | 0.1111 | 2 | 0 | 4 | 0 | 3 | 0.3333 | 4 | 0 | 13 | 0.0769 |
| 1_1361043 | 2 | 0 | 5 | 0.2 | NA | NA | 2 | 1 | 9 | 0.3333 | 2 | 1 | 5 | 0.6 | 2 | 1 | 4 | 0.5 | 13 | 0.6923 |
| 1_1361046 | 2 | 0 | 5 | 0.2 | 2 | 0.5 | 3 | 0 | 12 | 0.1667 | 2 | 0 | 5 | 0 | 6 | 0 | 5 | 0.2 | 18 | 0.0556 |
| 1_1361046 | 2 | 1 | 4 | 1 | 1 | 1 | 4 | 0.75 | 11 | 0.9091 | 2 | 1 | 3 | 1 | 6 | 0.8333 | 6 | 1 | 17 | 0.9412 |
| 1_1361047 | 2 | 0 | 5 | 0.4 | 2 | 0.5 | 3 | 0.3333 | 12 | 0.3333 | 2 | 0 | 5 | 0.4 | 6 | 0.1667 | 6 | 0 | 19 | 0.1579 |
| 1_1361047 | 2 | 1 | 3 | 1 | 2 | 1 | 4 | 0.75 | 11 | 0.9091 | 1 | 1 | 2 | 1 | 5 | 0.8 | 5 | 0.8 | 13 | 0.8462 |
| 1_1361047 | 1 | 0 | NA | NA | 1 | 0 | 2 | 1 | 4 | 0.5 | 2 | 1 | 2 | 0.5 | 3 | 0.6667 | 3 | 1 | 10 | 0.8 |
| 1_1361049 | 2 | 0 | 6 | 0.5 | 2 | 0 | 4 | 0.5 | 14 | 0.3571 | 1 | 0 | 5 | 0 | 7 | 0.5714 | 7 | 0.2857 | 20 | 0.3 |
| 1_1361050 | 3 | 0.3333 | 6 | 0.3333 | 2 | 0 | 4 | 0 | 15 | 0.2 | 2 | 0 | 4 | 0 | 6 | 0 | 6 | 0.1667 | 18 | 0.0556 |
| 1_1361874 | NA | NA | 1 | 1 | 1 | 1 | NA | NA | 2 | 1 | 1 | 0 | 3 | 1 | 2 | 1 | NA | NA | 6 | 0.8333 |
| 1_1361875 | NA | NA | NA | NA | 1 | 1 | 1 | 0 | 2 | 0.5 | 1 | 1 | 1 | 1 | 1 | 1 | NA | NA | 3 | 1 |
| 1_1361876 | NA | NA | NA | NA | 1 | 1 | NA | NA | 1 | 1 | 1 | 0 | 3 | 0.6667 | 2 | 1 | NA | NA | 6 | 0.6667 |
| 1_1361961 | 8 | 0.75 | 11 | 0.7273 | 12 | 0.75 | 11 | 0.7273 | 42 | 0.7381 | 5 | 0.8 | 13 | 0.6154 | 7 | 0.7143 | 13 | 0.6923 | 38 | 0.6842 |
| 1_1361962 | 9 | 0.3333 | 13 | 0.6154 | 13 | 0.7692 | 18 | 0.4444 | 53 | 0.5472 | 5 | 0.8 | 10 | 0.5 | 7 | 0.5714 | 12 | 0.5833 | 34 | 0.5882 |
| 1_1361966 | 8 | 0.125 | 7 | 0.2857 | 8 | 0.125 | 12 | 0.0833 | 35 | 0.1429 | 4 | 0.25 | 7 | 0 | 6 | 0.1667 | 9 | 0.1111 | 26 | 0.1154 |
| 1_1361966 | 10 | 0.1 | 11 | 0.0909 | 13 | 0.0769 | 18 | 0.1111 | 52 | 0.0962 | 7 | 0.4286 | 12 | 0.0833 | 8 | 0.25 | 12 | 0.3333 | 39 | 0.2564 |
| 1_1361982 | 7 | 0.1429 | 7 | 0.4286 | 10 | 0 | 11 | 0 | 35 | 0.1143 | 6 | 0.1667 | 4 | 0 | 10 | 0.2 | 11 | 0 | 31 | 0.0968 |
| 1_1361982 | 8 | 0.5 | 6 | 0.5 | 9 | 0.3333 | 12 | 0.4167 | 35 | 0.4286 | 5 | 0.2 | 5 | 0.4 | 9 | 0.5556 | 13 | 0.7692 | 32 | 0.5625 |
| 1_1361983 | 7 | 0 | 9 | 0.4444 | 11 | 0.0909 | 13 | 0.1538 | 40 | 0.175 | 6 | 0.1667 | 5 | 0 | 11 | 0.0909 | 15 | 0.1333 | 37 | 0.1081 |
| 1_1361984 | 4 | 0.75 | 7 | 0.7143 | 9 | 1 | 7 | 0.7143 | 27 | 0.8148 | 5 | 1 | 4 | 0.5 | 8 | 0.625 | 11 | 0.9091 | 28 | 0.7857 |
| 1_1361984 | 6 | 0.1667 | 8 | 0.625 | 6 | 0 | 8 | 0.625 | 28 | 0.3929 | 4 | 0 | 4 | 0 | 9 | 0.3333 | 10 | 0.4 | 27 | 0.2593 |
| 1_1361985 | 7 | 0 | 10 | 0.2 | 9 | 0.2222 | 12 | 0.0833 | 38 | 0.1316 | 5 | 0 | 6 | 0.3333 | 10 | 0.1 | 12 | 0.1667 | 33 | 0.1515 |
| 1_1361986 | 2 | 0.5 | 6 | 0.8333 | 4 | 0.75 | 3 | 0.6667 | 15 | 0.7333 | 2 | 1 | 2 | 1 | 5 | 1 | 4 | 1 | 13 | 1 |
| 1_1361986 | 2 | 0 | NA | NA | 1 | 1 | NA | NA | 3 | 0.3333 | 1 | 0 | NA | NA | 3 | 0.6667 | 1 | 0 | 5 | 0.4 |
| 1_1361986 | 1 | 0 | NA | NA | 1 | 1 | 1 | 0 | 3 | 0.3333 | NA | NA | 2 | 0.5 | NA | NA | 2 | 0.5 | 4 | 0.5 |
| 1_1361986 | 6 | 0.3333 | 9 | 0.3333 | 12 | 0.1667 | 13 | 0.4615 | 40 | 0.325 | 6 | 0.1667 | 7 | 0.4286 | 10 | 0.1 | 14 | 0.2857 | 37 | 0.2432 |
| 1_1361986 | 6 | 0.1667 | 6 | 0.3333 | 7 | 0.4286 | 11 | 0.2727 | 30 | 0.3 | 5 | 0.2 | 5 | 0 | 8 | 0.25 | 12 | 0.3333 | 30 | 0.2333 |
| 1_1361987 | 6 | 1 | 9 | 0.6667 | 8 | 0.875 | 10 | 0.9 | 33 | 0.8485 | 6 | 0.6667 | 5 | 0.4 | 9 | 0.6667 | 12 | 0.8333 | 32 | 0.6875 |
| 1_1361987 | 5 | 0.6 | 7 | 0.7143 | 5 | 0.4 | 7 | 0.4286 | 24 | 0.5417 | 4 | 0.25 | 3 | 0.3333 | 8 | 0.625 | 7 | 0.1429 | 22 | 0.3636 |
| 1_1361987 | 3 | 0 | 5 | 0.2 | 6 | 0.3333 | 4 | 0.25 | 18 | 0.2222 | 4 | 0.5 | 4 | 0.25 | 7 | 0.2857 | 7 | 0.2857 | 22 | 0.3182 |
| 1_1361987 | 7 | 0 | 9 | 0.2222 | 7 | 0.1429 | 10 | 0.2 | 33 | 0.1515 | 5 | 0.2 | 6 | 0 | 9 | 0.1111 | 12 | 0.0833 | 32 | 0.0938 |
| 1_1362082 | 7 | 0.1429 | 7 | 0.1429 | 5 | 0.4 | 8 | 0.25 | 27 | 0.2222 | 4 | 0.25 | 4 | 1 | 7 | 0.5714 | 10 | 0.5 | 25 | 0.56 |
| 1_1362082 | 7 | 1 | 7 | 0.8571 | 5 | 0.8 | 8 | 1 | 27 | 0.9259 | 5 | 0.8 | 3 | 1 | 7 | 1 | 11 | 1 | 26 | 0.9615 |
| 1_1362083 | 6 | 0.6667 | 8 | 1 | 4 | 0.5 | 6 | 0.8333 | 24 | 0.7917 | 5 | 0.8 | 2 | 1 | 7 | 1 | 10 | 0.7 | 24 | 0.8333 |
| 1_1362083 | 7 | 1 | 7 | 0.8571 | 4 | 0.75 | 7 | 1 | 25 | 0.92 | 5 | 0.6 | 3 | 0.6667 | 5 | 1 | 10 | 1 | 23 | 0.8696 |
| 1_1362084 | 5 | 0 | 6 | 0.1667 | 4 | 0 | 8 | 0.25 | 23 | 0.1304 | 5 | 0.6 | 3 | 0 | 6 | 0.3333 | 9 | 0.1111 | 23 | 0.2609 |
| 1_1362085 | 5 | 0.4 | 6 | 0.1667 | 3 | 0 | 6 | 0.3333 | 20 | 0.25 | 3 | 0.3333 | 3 | 0.3333 | 5 | 0.2 | 8 | 0.25 | 19 | 0.2632 |
| 1_1362085 | 2 | 0 | 6 | 0.1667 | 3 | 0 | 5 | 0.4 | 16 | 0.1875 | 3 | 0 | 2 | 0 | 4 | 0 | 9 | 0.1111 | 18 | 0.0556 |
| 1_1362085 | 5 | 1 | 6 | 1 | 5 | 1 | 6 | 1 | 22 | 1 | 5 | 1 | 4 | 0.75 | 5 | 0.6 | 8 | 1 | 22 | 0.8636 |
| 1_1362088 | 2 | 0.5 | 6 | 0.5 | 5 | 0.4 | 7 | 0.2857 | 20 | 0.4 | 5 | 0.2 | 4 | 1 | 3 | 0.3333 | 8 | 0.75 | 20 | 0.6 |

| | | | | | | | | | | | | | | | | | | | | |
|---|---|---|---|---|---|---|---|---|---|---|---|---|---|---|---|---|---|---|---|---|
| 1_13620905 | 2 | 0 | 6 | 0 | 4 | 0 | 7 | 0.1429 | 19 | 0.0526 | 5 | 0 | 2 | 0.5 | 2 | 0 | 6 | 0 | 15 | 0.0667 |
| 1_13620934 | 2 | 0.5 | 4 | 0.25 | 5 | 0 | 8 | 0.25 | 19 | 0.2105 | 5 | 0.2 | 2 | 0 | 3 | 0 | 4 | 0.25 | 14 | 0.1429 |
| 1_13620955 | 1 | 0 | 2 | 0.5 | 4 | 0.25 | 7 | 0.5714 | 14 | 0.4286 | 2 | 0.5 | 2 | 1 | 2 | 0.5 | 3 | 0.3333 | 9 | 0.5556 |
| 1_13620969 | 1 | 0 | 4 | 0 | 4 | 0.25 | 4 | 0.25 | 13 | 0.1538 | 4 | 0 | 2 | 0 | 2 | 0 | 2 | 0.5 | 10 | 0.1 |
| 1_13620997 | 1 | 0 | 2 | 0 | 1 | 0 | 5 | 0.6 | 9 | 0.3333 | 3 | 0 | 2 | 0.5 | 2 | 0.5 | 2 | 0 | 9 | 0.2222 |
| 1_13620997 | NA | NA | 2 | 1 | 2 | 1 | 4 | 1 | 8 | 1 | 2 | 1 | 2 | 1 | 2 | 1 | 3 | 1 | 9 | 1 |
| 1_13655331 | 1 | 1 | NA | NA | NA | NA | 1 | 1 | 2 | 1 | 1 | 1 | 2 | 0 | 2 | 0 | 2 | 1 | 7 | 0.4286 |
| 1_13655362 | 1 | 1 | 1 | 1 | 1 | 1 | 2 | 1 | 5 | 1 | 1 | 1 | 5 | 1 | 2 | 1 | 3 | 1 | 11 | 1 |
| 1_13655393 | 1 | 1 | 1 | 0 | 2 | 0.5 | 2 | 0.5 | 6 | 0.5 | 1 | 1 | 6 | 0.5 | 2 | 0 | 3 | 0.3333 | 12 | 0.4167 |
| 1_13655401 | 1 | 0 | 1 | 1 | 2 | 0.5 | 2 | 0 | 6 | 0.3333 | 1 | 0 | 5 | 0.4 | 2 | 0 | 3 | 0.3333 | 11 | 0.2727 |
| 1_13655402 | 1 | 1 | 1 | 1 | 1 | 0 | 1 | 0 | 4 | 0.5 | 1 | 0 | 6 | 0.3333 | 3 | 0.3333 | 3 | 0.6667 | 13 | 0.3846 |
| 1_13655405 | 1 | 1 | 1 | 1 | 2 | 0 | 2 | 0 | 6 | 0.3333 | 1 | 1 | 5 | 0.2 | 3 | 0 | 2 | 0 | 11 | 0.1818 |
| 1_13655408 | 1 | 1 | 1 | 1 | 1 | 1 | 2 | 1 | 5 | 1 | 1 | 0 | 4 | 1 | 2 | 1 | 3 | 1 | 10 | 0.9 |
| 1_13655427 | 1 | 1 | 1 | 1 | 2 | 1 | 2 | 1 | 6 | 1 | 1 | 0 | 5 | 0.8 | 3 | 0.6667 | 3 | 1 | 12 | 0.75 |
| 1_13655431 | 1 | 0 | 1 | 0 | 3 | 0.3333 | 1 | 0 | 6 | 0.1667 | 1 | 1 | 4 | 0 | 3 | 0 | 3 | 0 | 11 | 0.0909 |
| 1_13655438 | NA | NA | NA | NA | 3 | 1 | 2 | 1 | 5 | 1 | 1 | 1 | 4 | 0.75 | 2 | 1 | 3 | 1 | 10 | 0.9 |
| 1_13655468 | NA | NA | 1 | 0 | 1 | 0 | 2 | 0.5 | 4 | 0.25 | NA | NA | 4 | 0.5 | 3 | 0.6667 | 2 | 0.5 | 9 | 0.5556 |
| 1_13655474 | 1 | 1 | 1 | 0 | 2 | 1 | 2 | 1 | 6 | 0.8333 | 1 | 1 | 4 | 0.5 | 3 | 1 | 3 | 0.6667 | 11 | 0.7273 |
| 1_13655476 | 1 | 1 | 1 | 1 | 3 | 0.3333 | 2 | 1 | 7 | 0.7143 | NA | NA | 2 | 0.5 | 3 | 1 | 2 | 0.5 | 7 | 0.7143 |
| 1_13655487 | NA | NA | 1 | 0 | 3 | 0.3333 | 3 | 0 | 7 | 0.1429 | 1 | 0 | 4 | 0 | 3 | 0 | 3 | 0.3333 | 11 | 0.0909 |
| 1_13655493 | NA | NA | 1 | 0 | 2 | 0.5 | 1 | 1 | 4 | 0.5 | 1 | 0 | 2 | 0.5 | 2 | 0.5 | 1 | 0 | 6 | 0.3333 |
| 1_13655507 | NA | NA | 1 | 1 | 3 | 1 | 3 | 0.6667 | 7 | 0.8571 | 1 | 1 | 6 | 1 | 4 | 1 | 2 | 1 | 13 | 1 |
| 1_13655513 | NA | NA | 1 | 0 | 2 | 0.5 | 1 | 1 | 4 | 0.5 | 1 | 0 | 2 | 0 | 4 | 0.5 | 2 | 0.5 | 9 | 0.3333 |
| 1_13655533 | NA | NA | NA | NA | 3 | 0.6667 | 3 | 0 | 6 | 0.3333 | 1 | 1 | 4 | 0.5 | 5 | 0.6 | 3 | 1 | 13 | 0.6923 |
| 1_13655534 | NA | NA | 1 | 1 | 3 | 0.6667 | 3 | 1 | 7 | 0.8571 | 1 | 1 | 6 | 0.8333 | 5 | 0.4 | 3 | 0.3333 | 15 | 0.6 |
| 1_13655537 | 1 | 0 | 1 | 1 | 2 | 0.5 | 3 | 0 | 7 | 0.2857 | 1 | 0 | 4 | 0.75 | 5 | 0.2 | 3 | 1 | 13 | 0.5385 |
| 1_13655547 | 2 | 0.5 | 3 | 0.6667 | 2 | 1 | 3 | 1 | 10 | 0.8 | 1 | 1 | 6 | 0.3333 | 4 | 0.75 | 3 | 0 | 14 | 0.4286 |
| 1_13655550 | 1 | 1 | 3 | 0.6667 | 3 | 0.6667 | 2 | 1 | 9 | 0.7778 | NA | NA | 2 | 1 | 4 | 0.75 | 2 | 1 | 8 | 0.875 |
| 1_13655555 | 2 | 0 | 2 | 0 | 3 | 0.3333 | 1 | 1 | 8 | 0.25 | 1 | 0 | 4 | 0.5 | 5 | 0.2 | 2 | 0.5 | 12 | 0.3333 |
| 1_13655557 | 1 | 1 | 3 | 1 | 2 | 0.5 | 2 | 0.5 | 8 | 0.75 | NA | NA | 4 | 0.75 | 3 | 0.6667 | 2 | 0.5 | 9 | 0.6667 |
| 1_13655559 | 1 | 1 | 3 | 1 | 3 | 1 | 2 | 1 | 9 | 1 | 1 | 1 | 5 | 0.6 | 5 | 0.6 | 3 | 0.6667 | 14 | 0.6429 |
| 1_13655560 | 1 | 1 | 3 | 1 | 3 | 0.6667 | 3 | 0.6667 | 10 | 0.8 | 1 | 1 | 4 | 1 | 5 | 1 | 3 | 1 | 13 | 1 |
| 1_13655567 | 2 | 0.5 | 3 | 0 | 3 | 0 | 3 | 0 | 11 | 0.0909 | 1 | 1 | 6 | 0 | 5 | 0 | 3 | 0.3333 | 15 | 0.1333 |
| 1_13655592 | 2 | 0 | 5 | 0 | 5 | 0.2 | 2 | 0 | 14 | 0.0714 | 2 | 0 | 5 | 0.2 | 4 | 0 | 2 | 0 | 13 | 0.0769 |
| 1_13655606 | 1 | 1 | 5 | 0 | 5 | 0 | 3 | 0 | 14 | 0.0714 | 2 | 0.5 | 5 | 0.2 | 5 | 0 | 3 | 0 | 15 | 0.1333 |
| 1_13655700 | 3 | 0.3333 | 7 | 0 | 8 | 0 | 5 | 0.2 | 23 | 0.087 | 1 | 1 | 5 | 0.4 | 6 | 0 | 4 | 0.25 | 16 | 0.25 |
| 1_13655725 | 3 | 0.6667 | 10 | 0.6 | 10 | 0.7 | 4 | 0.5 | 27 | 0.6296 | 3 | 1 | 7 | 0.2857 | 5 | 0.2 | 4 | 0.5 | 19 | 0.4211 |
| 1_13655761 | 3 | 0 | 11 | 0 | 10 | 0.2 | 6 | 0.1667 | 30 | 0.1 | 4 | 0 | 9 | 0 | 8 | 0.25 | 3 | 0 | 24 | 0.0833 |
| 1_13841211 | 11 | 0 | 9 | 0 | 1 | 1 | 6 | 0.1667 | 27 | 0.0741 | 4 | 0 | 1 | 1 | 4 | 0.25 | 8 | 0.125 | 17 | 0.1765 |
| 1_13841268 | 10 | 0.1 | 7 | 0 | 2 | 0.5 | 8 | 0 | 27 | 0.0741 | 3 | 0 | 2 | 0 | 4 | 0 | 10 | 0.4 | 19 | 0.2105 |
| 1_13841294 | 10 | 0.2 | 6 | 0.1667 | 2 | 0 | 8 | 0 | 26 | 0.1154 | 3 | 0.3333 | 2 | 0 | 4 | 0.25 | 8 | 0.125 | 17 | 0.1765 |
| 1_13841404 | 8 | 0 | 6 | 0.1667 | 2 | 0.5 | 8 | 0.125 | 24 | 0.125 | 3 | 0.3333 | 1 | 0 | 3 | 0.6667 | 8 | 0.25 | 15 | 0.3333 |
| 1_13841422 | 7 | 0 | 6 | 0 | 2 | 0 | 7 | 0.2857 | 22 | 0.0909 | 3 | 0.3333 | 1 | 1 | 2 | 0 | 8 | 0.125 | 14 | 0.2143 |
| 1_13841432 | 6 | 0 | 5 | 0.2 | 2 | 0 | 6 | 0.3333 | 19 | 0.1579 | 2 | 0.5 | 1 | 0 | 2 | 0 | 7 | 0.4286 | 12 | 0.3333 |
| 1_13841433 | 6 | 0 | 6 | 0.1667 | 1 | 0 | 7 | 0 | 20 | 0.05 | 3 | 0 | 1 | 1 | 1 | 0 | 7 | 0.4286 | 12 | 0.3333 |
| 1_13841438 | 6 | 0.1667 | 6 | 0.5 | 2 | 0.5 | 8 | 0.125 | 22 | 0.2727 | 3 | 0.3333 | 1 | 0 | 1 | 0 | 6 | 0.1667 | 11 | 0.1818 |
| 1_13841439 | 6 | 0 | 5 | 0.2 | 2 | 0 | 5 | 0 | 18 | 0.0556 | 3 | 0.3333 | NA | NA | 2 | 0 | 5 | 0 | 10 | 0.1 |
| 1_13841447 | 3 | 0.3333 | 5 | 1 | 1 | 1 | 5 | 0.8 | 14 | 0.7857 | 2 | 1 | 1 | 1 | 2 | 1 | 6 | 0.8333 | 11 | 0.9091 |
| 1_13909774 | NA | NA | NA | NA | NA | NA | 2 | 1 | 2 | 1 | NA | NA | NA | NA | NA | NA | 1 | 1 | 1 | 1 |
| 1_13909791 | NA | NA | NA | NA | NA | NA | 2 | 0.5 | 2 | 0.5 | NA | NA | NA | NA | NA | NA | 1 | 1 | 1 | 1 |
| 1_13909794 | NA | NA | NA | NA | NA | NA | 2 | 1 | 2 | 1 | NA | NA | NA | NA | NA | NA | 1 | 1 | 1 | 1 |
| 1_13909799 | NA | NA | NA | NA | NA | NA | 2 | 0.5 | 2 | 0.5 | NA | NA | NA | NA | NA | NA | 1 | 1 | 1 | 1 |
| 1_13909839 | NA | NA | NA | NA | NA | NA | 2 | 1 | 2 | 1 | NA | NA | NA | NA | NA | NA | 1 | 1 | 1 | 1 |
| 1_13909844 | NA | NA | NA | NA | NA | NA | 2 | 1 | 2 | 1 | NA | NA | NA | NA | NA | NA | 1 | 1 | 1 | 1 |
| 1_13909856 | NA | NA | NA | NA | NA | NA | 2 | 1 | 2 | 1 | NA | NA | NA | NA | NA | NA | 1 | 1 | 1 | 1 |
| 1_13909868 | NA | NA | NA | NA | NA | NA | 2 | 1 | 2 | 1 | NA | NA | NA | NA | NA | NA | 1 | 1 | 1 | 1 |
| 1_13909995 | NA | NA | NA | NA | NA | NA | 2 | 0.5 | 2 | 0.5 | NA | NA | 1 | 1 | NA | NA | 1 | 0 | 2 | 0.5 |
| 1_13910335 | NA | NA | NA | NA | NA | NA | 3 | 0.3333 | 3 | 0.3333 | 1 | 1 | 2 | 0.5 | 1 | 1 | 1 | 1 | 5 | 0.8 |
| 1_13910364 | NA | NA | NA | NA | NA | NA | 3 | 0.6667 | 3 | 0.6667 | 3 | 0.3333 | 1 | 0 | 1 | 1 | 1 | 1 | 6 | 0.5 |
| 1_13937721 | 1 | 1 | 1 | 0 | 2 | 0.5 | 1 | 1 | 5 | 0.6 | NA | NA | NA | NA | 1 | 0 | 1 | 1 | 2 | 0.5 |
| 1_13937927 | 2 | 0.5 | 1 | 0 | 3 | 0 | 1 | 1 | 7 | 0.2857 | 1 | 1 | NA | NA | 2 | 0 | 2 | 1 | 5 | 0.6 |
| 1_13937944 | 2 | 0.5 | 2 | 0 | 3 | 0 | 1 | 0 | 8 | 0.125 | NA | NA | NA | NA | 2 | 0 | 3 | 1 | 5 | 0.6 |
| 1_13937949 | 2 | 0.5 | 1 | 1 | 3 | 0.3333 | 1 | 1 | 7 | 0.5714 | 1 | 0 | NA | NA | 2 | 1 | 1 | 1 | 4 | 0.75 |
| 1_13937961 | 2 | 0.5 | 2 | 0.5 | 2 | 0.5 | 1 | 1 | 7 | 0.5714 | NA | NA | NA | NA | 3 | 0.6667 | 1 | 0 | 4 | 0.5 |
| 1_13937964 | 1 | 1 | 1 | 1 | 1 | 0 | 1 | 1 | 4 | 0.75 | 1 | 1 | NA | NA | 3 | 0.6667 | 2 | 0.5 | 6 | 0.6667 |

| | | | | | | | | | | | | | | | | | | | | |
|---|---|---|---|---|---|---|---|---|---|---|---|---|---|---|---|---|---|---|---|---|
| 1_1393803 | NA | NA | 3 | 0.3333 | 5 | 0 | NA | NA | 8 | 0.125 | 1 | 0 | NA | NA | 3 | 0 | 4 | 0.5 | 8 | 0.25 |
| 1_1393805 | NA | NA | 3 | 0 | 4 | 0.5 | NA | NA | 7 | 0.2857 | 1 | 0 | NA | NA | 3 | 1 | 2 | 1 | 6 | 0.8333 |
| 1_1404509 | 1 | 1 | NA | NA | 1 | 1 | NA | NA | 2 | 1 | 3 | 0.6667 | 2 | 1 | NA | NA | NA | NA | 5 | 0.8 |
| 1_1405374 | NA | NA | 1 | 1 | 1 | 0 | NA | NA | 2 | 0.5 | 3 | 0.6667 | 2 | 0.5 | 1 | 1 | 1 | 1 | 7 | 0.7143 |
| 1_1405412 | NA | NA | NA | NA | 2 | 0.5 | 1 | 0 | 3 | 0.3333 | 1 | 1 | NA | NA | 2 | 0 | 1 | 0 | 4 | 0.25 |
| 1_1405413 | NA | NA | NA | NA | 2 | 1 | 1 | 1 | 3 | 1 | 1 | 1 | NA | NA | 1 | 0 | NA | NA | 2 | 0.5 |
| 1_1405413 | NA | NA | NA | NA | 2 | 1 | 1 | 1 | 3 | 1 | 1 | 1 | NA | NA | NA | NA | 1 | 0 | 2 | 0.5 |
| 1_1405414 | NA | NA | NA | NA | 2 | 1 | 1 | 1 | 3 | 1 | 1 | 1 | NA | NA | 2 | 1 | 1 | 1 | 4 | 1 |
| 1_1434833 | NA | NA | 1 | 0 | 2 | 0.5 | 1 | 0 | 4 | 0.25 | NA | NA | 2 | 0.5 | 1 | 0 | 4 | 0.5 | 7 | 0.4286 |
| 1_1434838 | 1 | 0 | 3 | 0 | 6 | 0.1667 | 2 | 0 | 12 | 0.0833 | 2 | 0 | 2 | 0 | 2 | 0 | 3 | 0.3333 | 9 | 0.1111 |
| 1_1434840 | 1 | 1 | 3 | 0 | 5 | 0.4 | 3 | 0 | 12 | 0.25 | 2 | 0.5 | 1 | 0 | NA | NA | 1 | 0 | 4 | 0.25 |
| 1_1434841 | 1 | 1 | 3 | 0 | 5 | 0 | 3 | 0 | 12 | 0.0833 | 2 | 0.5 | 1 | 0 | 1 | 0 | 3 | 0.3333 | 7 | 0.2857 |
| 1_1434841 | 1 | 0 | 2 | 0 | 5 | 0.4 | 3 | 0.6667 | 11 | 0.3636 | 2 | 0 | 1 | 0 | NA | NA | 3 | 0.3333 | 6 | 0.1667 |
| 1_1434841 | NA | NA | 2 | 0.5 | 5 | 0.4 | 3 | 0.3333 | 10 | 0.4 | 1 | 0 | 1 | 1 | 1 | 0 | 2 | 0.5 | 5 | 0.4 |
| 1_1435166 | 1 | 0 | 5 | 0 | 2 | 0.5 | NA | NA | 8 | 0.125 | NA | NA | NA | NA | 1 | 0 | 1 | 1 | 2 | 0.5 |
| 1_1435168 | 1 | 0 | 5 | 0 | 2 | 0.5 | 1 | 0 | 9 | 0.1111 | NA | NA | NA | NA | 1 | 0 | 3 | 0.3333 | 4 | 0.25 |
| 1_1435169 | 1 | 1 | 5 | 1 | 2 | 0.5 | 1 | 0 | 9 | 0.7778 | NA | NA | NA | NA | 1 | 1 | 3 | 1 | 4 | 1 |
| 1_1435169 | 1 | 0 | 6 | 0.6667 | 2 | 1 | 2 | 1 | 11 | 0.7273 | NA | NA | NA | NA | 1 | 0 | 2 | 1 | 3 | 0.6667 |
| 1_1435219 | 1 | 1 | 4 | 1 | 1 | 1 | 1 | 1 | 7 | 1 | NA | NA | NA | NA | 1 | 1 | 2 | 1 | 3 | 1 |
| 1_1435220 | 1 | 1 | 3 | 0.6667 | NA | NA | 1 | 0 | 5 | 0.6 | NA | NA | NA | NA | NA | NA | 2 | 0.5 | 2 | 0.5 |
| 1_1435220 | 1 | 0 | 7 | 0.2857 | 2 | 0 | 2 | 0.5 | 12 | 0.25 | NA | NA | NA | NA | 1 | 0 | 3 | 0.3333 | 4 | 0.25 |
| 1_1435221 | 1 | 1 | 6 | 0.3333 | 2 | 0 | 1 | 0 | 10 | 0.3 | NA | NA | NA | NA | 1 | 0 | 3 | 0.6667 | 4 | 0.5 |
| 1_1435221 | NA | NA | 6 | 0.1667 | 2 | 0.5 | 2 | 0 | 10 | 0.2 | NA | NA | NA | NA | 1 | 0 | 3 | 0.6667 | 4 | 0.5 |
| 1_1435222 | 1 | 1 | 7 | 0.5714 | 2 | 0.5 | 1 | 1 | 11 | 0.6364 | NA | NA | NA | NA | 1 | 0 | 2 | 0.5 | 3 | 0.3333 |
| 1_1435222 | 1 | 1 | 7 | 0.8571 | 2 | 1 | NA | NA | 10 | 0.9 | NA | NA | NA | NA | 1 | 1 | 2 | 1 | 3 | 1 |
| 1_1435222 | NA | NA | 7 | 1 | 3 | 1 | 1 | 1 | 11 | 1 | NA | NA | NA | NA | 1 | 1 | 3 | 1 | 4 | 1 |
| 1_1435223 | 1 | 0 | 7 | 0.5714 | 4 | 0.5 | 1 | 0 | 13 | 0.4615 | NA | NA | NA | NA | 1 | 0 | 3 | 0.3333 | 4 | 0.25 |
| 1_1435223 | NA | NA | 6 | 0.6667 | 4 | 0.5 | 2 | 1 | 12 | 0.6667 | NA | NA | NA | NA | NA | NA | 3 | 0.6667 | 3 | 0.6667 |
| 1_1435223 | 1 | 1 | 6 | 0.6667 | 4 | 1 | 2 | 0.5 | 13 | 0.7692 | NA | NA | NA | NA | 1 | 0 | 3 | 0.6667 | 4 | 0.5 |
| 1_1435225 | 1 | 1 | 8 | 0.125 | 4 | 0.25 | 2 | 0 | 15 | 0.2 | NA | NA | NA | NA | NA | NA | 3 | 0.3333 | 3 | 0.3333 |
| 1_1435225 | 1 | 0 | 8 | 0.25 | 4 | 0.25 | 2 | 0 | 15 | 0.2 | NA | NA | NA | NA | NA | NA | 3 | 0.3333 | 3 | 0.3333 |
| 1_1435225 | 1 | 1 | 7 | 1 | 3 | 1 | 2 | 1 | 13 | 1 | NA | NA | NA | NA | 1 | 1 | 3 | 1 | 4 | 1 |
| 1_1435226 | 1 | 1 | 6 | 1 | 3 | 1 | 1 | 1 | 11 | 1 | NA | NA | NA | NA | 1 | 1 | 3 | 1 | 4 | 1 |
| 1_1435226 | 1 | 0 | 10 | 0.1 | 4 | 0.25 | 2 | 1 | 17 | 0.2353 | NA | NA | NA | NA | 1 | 0 | 3 | 0.3333 | 4 | 0.25 |
| 1_1435226 | 1 | 0 | 7 | 0.1429 | 4 | 0.75 | 2 | 1 | 14 | 0.4286 | NA | NA | NA | NA | 1 | 0 | 3 | 1 | 4 | 0.75 |
| 1_1435227 | 1 | 1 | 5 | 1 | 4 | 1 | NA | NA | 10 | 1 | NA | NA | NA | NA | 1 | 1 | 1 | 1 | 2 | 1 |
| 1_1435228 | NA | NA | 9 | 1 | 2 | 1 | 2 | 1 | 13 | 1 | NA | NA | NA | NA | 1 | 1 | 3 | 1 | 4 | 1 |
| 1_1435230 | 2 | 1 | 8 | 1 | 1 | 1 | 1 | 0 | 12 | 0.9167 | 2 | 0.5 | NA | NA | 1 | 1 | 2 | 0.5 | 5 | 0.6 |
| 1_1435230 | 3 | 0.3333 | 9 | 0.2222 | 3 | 0.3333 | 2 | 0.5 | 17 | 0.2941 | NA | NA | NA | NA | 1 | 0 | 3 | 0.6667 | 4 | 0.5 |
| 1_1435231 | 3 | 0 | 10 | 0.1 | 3 | 0.3333 | 2 | 0 | 18 | 0.1111 | 1 | 0 | NA | NA | 1 | 1 | 3 | 0 | 5 | 0.2 |
| 1_1435235 | 3 | 0 | 10 | 0.2 | 5 | 0 | 1 | 0 | 19 | 0.1053 | 3 | 0.3333 | NA | NA | 1 | 0 | 3 | 0 | 7 | 0.1429 |
| 1_1436214 | NA | NA | 1 | 1 | NA | NA | 2 | 0.5 | 3 | 0.6667 | 1 | 0 | 1 | 1 | NA | NA | 1 | 1 | 3 | 0.6667 |
| 1_1436222 | 14 | 0.0714 | 12 | 0.1667 | 12 | 0 | 7 | 0.1429 | 45 | 0.0889 | 9 | 0.1111 | 9 | 0.3333 | 7 | 0.2857 | 7 | 0.1429 | 32 | 0.2188 |
| 1_1436223 | 14 | 0.2143 | 10 | 0.6 | 13 | 0.2308 | 9 | 0.2222 | 46 | 0.3043 | 11 | 0.1818 | 10 | 0.1 | 6 | 0.1667 | 5 | 0.6 | 32 | 0.2188 |
| 1_1436224 | 11 | 0.1818 | 12 | 0.25 | 9 | 0.1111 | 6 | 0.3333 | 38 | 0.2105 | 10 | 0.1 | 6 | 0.1667 | 6 | 0.1667 | 5 | 0.4 | 27 | 0.1852 |
| 1_1436225 | 12 | 0 | 10 | 0.1 | 10 | 0.1 | 6 | 0 | 38 | 0.0526 | 12 | 0.1667 | 10 | 0.1 | 6 | 0.1667 | 6 | 0.1667 | 34 | 0.1471 |
| 1_1436227 | 3 | 0.3333 | 2 | 0.5 | 4 | 0 | 4 | 0.25 | 13 | 0.2308 | 2 | 0 | 4 | 0.5 | 2 | 0 | 4 | 0.5 | 12 | 0.3333 |
| 1_1436229 | 3 | 0 | 2 | 0.5 | 3 | 0 | 2 | 0 | 10 | 0.1 | 1 | 0 | 3 | 0 | 1 | 0 | 3 | 0.3333 | 8 | 0.125 |
| 1_1436237 | 4 | 0.25 | 2 | 0.5 | 4 | 0.25 | 3 | 0 | 13 | 0.2308 | 3 | 0.3333 | 5 | 0.4 | 1 | 1 | 5 | 0.2 | 14 | 0.3571 |
| 1_1436238 | 3 | 0.3333 | 1 | 0 | 3 | 0.6667 | 4 | 0.25 | 11 | 0.3636 | 3 | 0 | 4 | 0.5 | 2 | 0.5 | 5 | 0 | 14 | 0.2143 |
| 1_1436267 | 9 | 0.7778 | 7 | 0.5714 | 7 | 0.4286 | 8 | 0.25 | 31 | 0.5161 | 1 | 0 | 2 | 0 | 6 | 0.6667 | 4 | 0.5 | 13 | 0.4615 |
| 1_1436273 | 7 | 0 | 7 | 0 | 6 | 0.1667 | 7 | 0.1429 | 27 | 0.0741 | 2 | 0 | 1 | 0 | 6 | 0.1667 | 3 | 0 | 12 | 0.0833 |
| 1_1436274 | 8 | 0 | 7 | 0 | 6 | 0 | 8 | 0.375 | 29 | 0.1034 | 2 | 0 | 1 | 0 | 6 | 0.1667 | 4 | 0 | 13 | 0.0769 |
| 1_1436278 | 7 | 0.4286 | 5 | 0.2 | 5 | 0.2 | 6 | 0.1667 | 23 | 0.2609 | 2 | 0 | 1 | 0 | 6 | 0 | 3 | 0.3333 | 12 | 0.0833 |
| 1_1436330 | 1 | 0 | NA | NA | 2 | 0 | 1 | 1 | 4 | 0.25 | NA | NA | NA | NA | 1 | 1 | NA | NA | 1 | 1 |
| 1_1436332 | 2 | 0 | NA | NA | 1 | 1 | 2 | 0.5 | 5 | 0.4 | NA | NA | NA | NA | 1 | 1 | NA | NA | 1 | 1 |
| 1_1436533 | NA | NA | 1 | 0 | 2 | 1 | 1 | 0 | 4 | 0.5 | 1 | 1 | 1 | 1 | 1 | 1 | 1 | 1 | 4 | 1 |
| 1_1436533 | NA | NA | 1 | 1 | 1 | 1 | 1 | 1 | 3 | 1 | 1 | 1 | 1 | 1 | 1 | 0 | NA | NA | 3 | 0.6667 |
| 1_1436534 | NA | NA | 1 | 1 | 1 | 1 | 1 | 1 | 3 | 1 | NA | NA | 1 | 1 | NA | NA | 1 | 1 | 2 | 1 |
| 1_1436534 | NA | NA | 1 | 1 | 2 | 1 | NA | NA | 3 | 1 | NA | NA | 1 | 1 | 1 | 1 | NA | NA | 2 | 1 |
| 1_1436536 | NA | NA | 1 | 1 | 3 | 0.6667 | 1 | 1 | 5 | 0.8 | 1 | 0 | 1 | 1 | 1 | 1 | 1 | 1 | 4 | 0.75 |
| 1_1436537 | NA | NA | 1 | 1 | 1 | 1 | NA | NA | 2 | 1 | 1 | 1 | 1 | 1 | 1 | 1 | NA | NA | 3 | 1 |
| 1_1436538 | NA | NA | 1 | 0 | 3 | 0.3333 | 1 | 0 | 5 | 0.2 | 1 | 0 | 1 | 0 | 1 | 1 | 1 | 0 | 4 | 0.25 |
| 1_1436538 | NA | NA | 1 | 0 | 3 | 1 | 1 | 1 | 5 | 0.8 | 1 | 1 | 1 | 1 | 1 | 0 | 1 | 1 | 4 | 0.75 |
| 1_1436539 | NA | NA | 1 | 1 | 3 | 1 | 1 | 1 | 5 | 1 | 1 | 1 | 1 | 1 | 1 | 1 | 1 | 1 | 4 | 1 |

| | | | | | | | | | | | | | | | | | | | | |
|---|---|---|---|---|---|---|---|---|---|---|---|---|---|---|---|---|---|---|---|---|
| 1_14365419 | NA | NA | NA | NA | 3 | 1 | 1 | 0 | 4 | 0.75 | 1 | 0 | 1 | 0 | 1 | 1 | NA | NA | 3 | 0.3333 |
| 1_14365693 | NA | NA | NA | NA | 3 | 0.3333 | 1 | 0 | 4 | 0.25 | 1 | 0 | 1 | 0 | NA | NA | 1 | 1 | 3 | 0.3333 |
| 1_14368251 | 2 | 0.5 | 8 | 0.5 | 9 | 0.6667 | 7 | 0 | 26 | 0.4231 | 4 | 0.75 | 10 | 0.4 | 8 | 0.375 | 5 | 0.2 | 27 | 0.4074 |
| 1_14368253 | 3 | 0.6667 | 12 | 0.25 | 10 | 0.5 | 8 | 0.375 | 33 | 0.3939 | 5 | 0.8 | 12 | 0.3333 | 9 | 0.3333 | 8 | 0.125 | 34 | 0.3529 |
| 1_14368254 | 3 | 0.3333 | 11 | 0.1818 | 11 | 0 | 5 | 0 | 30 | 0.1 | 4 | 0 | 10 | 0.1 | 7 | 0 | 6 | 0.1667 | 27 | 0.0741 |
| 1_14368255 | 2 | 0 | 11 | 0.1818 | 11 | 0.2727 | 6 | 0.1667 | 30 | 0.2 | 4 | 0.25 | 12 | 0.25 | 6 | 0.1667 | 7 | 0.1429 | 29 | 0.2069 |
| 1_14387653 | 11 | 0.0909 | 4 | 0.25 | 8 | 0.375 | 12 | 0.0833 | 35 | 0.1714 | 4 | 0 | 12 | 0.5 | 10 | 0.2 | 5 | 0.6 | 31 | 0.3548 |
| 1_14387665 | 8 | 0.25 | 5 | 0 | 8 | 0.125 | 10 | 0.4 | 31 | 0.2258 | 5 | 0.4 | 8 | 0.5 | 10 | 0.3 | 5 | 0.2 | 28 | 0.3571 |
| 1_14387728 | 12 | 0.3333 | 16 | 0.25 | 13 | 0.3077 | 16 | 0.375 | 57 | 0.3158 | 8 | 0.875 | 14 | 0.5 | 16 | 0.5625 | 7 | 0.4286 | 45 | 0.5778 |
| 1_14387733 | 3 | 0.3333 | 1 | 1 | 2 | 1 | 3 | 0.6667 | 9 | 0.6667 | NA | NA | 2 | 0.5 | 6 | 0.8333 | 1 | 0 | 9 | 0.6667 |
| 1_14387733 | 2 | 0.5 | 6 | 0.5 | 2 | 1 | 5 | 0.6 | 15 | 0.6 | NA | NA | NA | NA | 3 | 1 | 1 | 0 | 4 | 0.75 |
| 1_14387733 | NA | NA | 3 | 0.3333 | 1 | 0 | 2 | 1 | 6 | 0.5 | 1 | 0 | 4 | 0.75 | 1 | 1 | 1 | 1 | 7 | 0.7143 |
| 1_14387733 | 1 | 1 | 1 | 0 | 1 | 1 | 1 | 1 | 4 | 0.75 | NA | NA | NA | NA | 1 | 1 | NA | NA | 1 | 1 |
| 1_14387739 | 12 | 1 | 13 | 1 | 13 | 1 | 19 | 1 | 57 | 1 | 7 | 0.8571 | 17 | 1 | 20 | 1 | 9 | 0.8889 | 53 | 0.9623 |
| 1_14387746 | 12 | 0.8333 | 16 | 0.75 | 13 | 0.8462 | 19 | 0.7895 | 60 | 0.8 | 7 | 0.4286 | 16 | 0.9375 | 19 | 0.6842 | 9 | 0.5556 | 51 | 0.7059 |
| 1_14387760 | 13 | 0.1538 | 11 | 0.1818 | 11 | 0.1818 | 17 | 0.1176 | 52 | 0.1538 | 6 | 0.1667 | 16 | 0.3125 | 19 | 0.2105 | 8 | 0.375 | 49 | 0.2653 |
| 1_14387763 | 1 | 1 | NA | NA | NA | NA | NA | NA | 1 | 1 | NA | NA | 1 | 1 | NA | NA | NA | NA | 1 | 1 |
| 1_14387791 | 9 | 0.2222 | 12 | 0 | 8 | 0.25 | 13 | 0.1538 | 42 | 0.1429 | 4 | 0.25 | 13 | 0.0769 | 11 | 0.0909 | 6 | 0.3333 | 34 | 0.1471 |
| 1_14639061 | 1 | 1 | NA | NA | NA | NA | 2 | 0 | 3 | 0.3333 | 2 | 0.5 | NA | NA | NA | NA | 1 | 0 | 3 | 0.3333 |
| 1_14662759 | 2 | 0.5 | NA | NA | 1 | 0 | 3 | 0.6667 | 6 | 0.5 | 2 | 0.5 | NA | NA | NA | NA | 1 | 0 | 3 | 0.3333 |
| 1_14670737 | 1 | 0 | NA | NA | 1 | 1 | 3 | 0.3333 | 5 | 0.4 | 2 | 0.5 | NA | NA | NA | NA | 1 | 0 | 3 | 0.3333 |
| 1_14677697 | 2 | 1 | 1 | 1 | 1 | 1 | 2 | 1 | 6 | 1 | 2 | 1 | NA | NA | NA | NA | 1 | 0 | 3 | 0.6667 |
| 1_14677734 | 2 | 1 | 1 | 1 | 1 | 1 | 4 | 1 | 8 | 1 | 2 | 1 | NA | NA | NA | NA | 1 | 1 | 3 | 1 |
| 1_14677738 | 2 | 1 | 1 | 1 | 1 | 1 | 4 | 1 | 8 | 1 | NA | NA | NA | NA | NA | NA | 1 | 1 | 1 | 1 |
| 1_14677739 | 2 | 1 | NA | NA | 1 | 1 | 3 | 0.3333 | 6 | 0.6667 | 2 | 1 | NA | NA | NA | NA | 1 | 0 | 3 | 0.6667 |
| 1_14677742 | 2 | 0.5 | 1 | 0 | 1 | 1 | 2 | 0.5 | 6 | 0.5 | 2 | 0 | NA | NA | NA | NA | 1 | 1 | 3 | 0.3333 |
| 1_14677759 | 3 | 0.6667 | 2 | 1 | 1 | 0 | 5 | 0.8 | 11 | 0.7273 | 2 | 1 | NA | NA | NA | NA | 1 | 1 | 3 | 1 |
| 1_14677761 | 3 | 1 | 2 | 1 | 1 | 1 | 5 | 0.8 | 11 | 0.9091 | 2 | 1 | NA | NA | NA | NA | 1 | 1 | 3 | 1 |
| 1_14677764 | 3 | 1 | 2 | 1 | 1 | 1 | 5 | 1 | 11 | 1 | 2 | 1 | NA | NA | NA | NA | 1 | 1 | 3 | 1 |
| 1_14677767 | 3 | 1 | 2 | 1 | 1 | 1 | 5 | 0.8 | 11 | 0.9091 | 2 | 0.5 | NA | NA | NA | NA | 1 | 1 | 3 | 0.6667 |
| 1_14677769 | 2 | 1 | 2 | 1 | 1 | 1 | 4 | 0.75 | 9 | 0.8889 | 2 | 0.5 | NA | NA | NA | NA | NA | NA | 2 | 0.5 |
| 1_14677774 | 3 | 1 | 2 | 1 | 1 | 1 | 3 | 1 | 9 | 1 | 1 | 1 | NA | NA | NA | NA | 1 | 1 | 2 | 1 |
| 1_14677800 | 3 | 1 | 2 | 1 | 1 | 1 | 5 | 1 | 11 | 1 | 3 | 0.6667 | NA | NA | NA | NA | NA | NA | 3 | 0.6667 |
| 1_14970572 | 1 | 1 | NA | NA | 2 | 0.5 | NA | NA | 3 | 0.6667 | NA | NA | 2 | 0.5 | NA | NA | NA | NA | 2 | 0.5 |
| 1_14998118 | 2 | 1 | NA | NA | NA | NA | 1 | 1 | 3 | 1 | 1 | 1 | 1 | 1 | NA | NA | NA | NA | 2 | 1 |
| 1_14998127 | 2 | 1 | NA | NA | NA | NA | 1 | 1 | 3 | 1 | 1 | 1 | NA | NA | NA | NA | NA | NA | 1 | 1 |
| 1_14998149 | 2 | 0.5 | NA | NA | NA | NA | 1 | 0 | 3 | 0.3333 | 1 | 1 | 1 | 0 | NA | NA | NA | NA | 2 | 0.5 |
| 1_15023725 | 1 | 1 | 8 | 0.125 | 5 | 0.2 | 9 | 0.5556 | 23 | 0.3478 | 3 | 0.3333 | 2 | 0.5 | 7 | 0.4286 | 7 | 0.2857 | 19 | 0.3684 |
| 1_15023807 | 1 | 1 | 8 | 0.5 | 6 | 0.1667 | 8 | 0.375 | 23 | 0.3913 | 1 | 0 | NA | NA | 5 | 0.2 | 7 | 0.2857 | 13 | 0.2308 |
| 1_15023816 | 2 | 0.5 | 7 | 0.4286 | 4 | 0.5 | 9 | 0.4444 | 22 | 0.4545 | 1 | 1 | NA | NA | 5 | 0.4 | 7 | 0.7143 | 13 | 0.6154 |
| 1_15023845 | 3 | 0.3333 | 7 | 0.1429 | 5 | 0.4 | 6 | 0 | 21 | 0.1905 | 1 | 0 | 1 | 1 | 4 | 0.5 | 6 | 0 | 12 | 0.25 |
| 1_15030515 | 5 | 0 | 5 | 0.2 | NA | NA | 4 | 0 | 14 | 0.0714 | 1 | 0 | 3 | 0 | 8 | 0.125 | 3 | 0.3333 | 15 | 0.1333 |
| 1_15095043 | 1 | 1 | 1 | 1 | 1 | 1 | NA | NA | 3 | 1 | NA | NA | 1 | 1 | NA | NA | NA | NA | 1 | 1 |
| 1_15095048 | 1 | 1 | NA | NA | NA | NA | NA | NA | 1 | 1 | 1 | 0 | 1 | 1 | NA | NA | NA | NA | 2 | 0.5 |
| 1_15121987 | NA | NA | NA | NA | 1 | 1 | NA | NA | 1 | 1 | 2 | 0.5 | 1 | 1 | NA | NA | NA | NA | 3 | 0.6667 |
| 1_15121995 | 1 | 1 | NA | NA | 1 | 0 | NA | NA | 2 | 0.5 | 1 | 1 | 2 | 1 | NA | NA | NA | NA | 3 | 1 |
| 1_15122517 | 9 | 0.1111 | 9 | 0.4444 | 9 | 0.3333 | 13 | 0.1538 | 40 | 0.25 | 7 | 0 | 5 | 0.2 | 8 | 0 | 12 | 0.1667 | 32 | 0.0938 |
| 1_15123113 | 6 | 0.6667 | 3 | 0.3333 | 2 | 0.5 | 2 | 1 | 13 | 0.6154 | 5 | 0.2 | 2 | 0.5 | 5 | 0.6 | 5 | 0.4 | 17 | 0.4118 |
| 1_15123119 | 5 | 0.8 | 3 | 0.6667 | 3 | 0.6667 | 1 | 0 | 12 | 0.6667 | 5 | 0.6 | 1 | 1 | 3 | 1 | 4 | 0.75 | 13 | 0.7692 |
| 1_15123164 | 5 | 0.8 | 2 | 0.5 | 3 | 0.3333 | 2 | 0.5 | 12 | 0.5833 | 5 | 0.2 | 2 | 0.5 | 4 | 0.75 | 4 | 0.25 | 15 | 0.4 |
| 1_15130474 | 1 | 0 | NA | NA | 2 | 0 | 3 | 0.6667 | 6 | 0.3333 | 1 | 1 | 3 | 0.3333 | 1 | 0 | 2 | 0.5 | 7 | 0.4286 |
| 1_15133368 | 4 | 0.25 | 3 | 0.3333 | 5 | 0.6 | 6 | 0.3333 | 18 | 0.3889 | 4 | 1 | 5 | 0.4 | 2 | 0 | 4 | 0 | 15 | 0.4 |
| 1_15134543 | 6 | 0 | 4 | 0 | 5 | 0.4 | 4 | 0.25 | 19 | 0.1579 | 5 | 0.2 | 6 | 0 | 6 | 0.1667 | 2 | 0.5 | 19 | 0.1579 |
| 1_15134688 | 5 | 0.2 | 2 | 0 | 4 | 0.5 | 4 | 0.25 | 15 | 0.2667 | 2 | 0 | 4 | 0.25 | 6 | 0 | 2 | 0 | 14 | 0.0714 |
| 1_15134690 | 4 | 0 | 3 | 0.3333 | 2 | 0.5 | 2 | 0 | 11 | 0.1818 | 2 | 0.5 | 4 | 0.75 | 4 | 0 | 2 | 0 | 12 | 0.3333 |
| 1_15134698 | 3 | 0 | 3 | 0.3333 | 3 | 0.6667 | 4 | 0.5 | 13 | 0.3846 | 1 | 1 | 2 | 0 | 6 | 0.5 | 2 | 0.5 | 11 | 0.4545 |
| 1_15134703 | 5 | 0.2 | 1 | 1 | 2 | 1 | 3 | 0.6667 | 11 | 0.5455 | 2 | 0 | 3 | 0.3333 | 4 | 0 | 2 | 0.5 | 11 | 0.1818 |
| 1_15134704 | 5 | 0.4 | 3 | 0.6667 | 2 | 0 | 4 | 0.25 | 14 | 0.3571 | 2 | 0.5 | 3 | 0.6667 | 6 | 0.6667 | 1 | 1 | 12 | 0.6667 |
| 1_15134710 | 5 | 0.2 | 3 | 0.6667 | 3 | 0.6667 | 3 | 0 | 14 | 0.3571 | 2 | 0 | 3 | 0.3333 | 6 | 0.1667 | 1 | 0 | 12 | 0.1667 |
| 1_15134897 | 4 | 0.25 | 3 | 0.3333 | 3 | 0 | NA | NA | 10 | 0.2 | 2 | 0.5 | 4 | 0.25 | 5 | 0 | NA | NA | 11 | 0.1818 |
| 1_15169421 | 2 | 1 | NA | NA | NA | NA | 1 | 1 | 3 | 1 | 2 | 0.5 | NA | NA | NA | NA | NA | NA | 2 | 0.5 |
| 1_15169431 | 2 | 1 | NA | NA | NA | NA | 1 | 1 | 3 | 1 | 2 | 0.5 | NA | NA | NA | NA | NA | NA | 2 | 0.5 |
| 1_15169433 | 1 | 0 | NA | NA | NA | NA | 1 | 1 | 2 | 0.5 | 2 | 0.5 | NA | NA | NA | NA | NA | NA | 2 | 0.5 |
| 1_15169434 | 3 | 0.6667 | NA | NA | NA | NA | 1 | 0 | 4 | 0.5 | 1 | 1 | NA | NA | NA | NA | NA | NA | 1 | 1 |
| 1_15169439 | 2 | 1 | NA | NA | NA | NA | 1 | 1 | 3 | 1 | 2 | 1 | NA | NA | NA | NA | NA | NA | 2 | 1 |

| | | | | | | | | | | | | | | | | | | | | |
|---|---|---|---|---|---|---|---|---|---|---|---|---|---|---|---|---|---|---|---|---|
| 1_1516944 | 3 | 0.6667 | NA | NA | NA | NA | 1 | 1 | 4 | 0.75 | 3 | 0.3333 | NA | NA | NA | NA | NA | NA | 3 | 0.3333 |
| 1_1516944 | 2 | 1 | NA | NA | NA | NA | 1 | 1 | 3 | 1 | 3 | 0.6667 | NA | NA | NA | NA | NA | NA | 3 | 0.6667 |
| 1_1516944 | 3 | 1 | NA | NA | NA | NA | 1 | 1 | 4 | 1 | 3 | 1 | NA | NA | NA | NA | NA | NA | 3 | 1 |
| 1_1516945 | 2 | 0.5 | NA | NA | NA | NA | 1 | 0 | 3 | 0.3333 | 2 | 0.5 | NA | NA | NA | NA | NA | NA | 2 | 0.5 |
| 1_1516946 | 3 | 0 | NA | NA | NA | NA | 1 | 1 | 4 | 0.25 | 2 | 0.5 | NA | NA | NA | NA | NA | NA | 2 | 0.5 |
| 1_1516946 | 3 | 0.6667 | NA | NA | NA | NA | 1 | 0 | 4 | 0.5 | 2 | 0.5 | NA | NA | NA | NA | NA | NA | 2 | 0.5 |
| 1_1516947 | 1 | 1 | NA | NA | NA | NA | NA | NA | 1 | 1 | 3 | 0.3333 | NA | NA | NA | NA | NA | NA | 3 | 0.3333 |
| 1_1516947 | 2 | 0.5 | NA | NA | NA | NA | NA | NA | 2 | 0.5 | 3 | 1 | NA | NA | NA | NA | NA | NA | 3 | 1 |
| 1_1516948 | 2 | 1 | NA | NA | NA | NA | NA | NA | 2 | 1 | 1 | 1 | NA | NA | NA | NA | NA | NA | 1 | 1 |
| 1_1516955 | 3 | 0.3333 | NA | NA | NA | NA | 2 | 0 | 5 | 0.2 | 3 | 0 | 1 | 1 | NA | NA | NA | NA | 4 | 0.25 |
| 1_1516975 | 3 | 0 | 6 | 0 | 3 | 0.3333 | 4 | 0.25 | 16 | 0.125 | 6 | 0.1667 | 7 | 0.2857 | 3 | 0 | 2 | 0 | 18 | 0.1667 |
| 1_1517604 | 3 | 1 | 3 | 0.3333 | 6 | 0.5 | 1 | 0 | 13 | 0.5385 | 2 | 0.5 | 3 | 0.6667 | NA | NA | NA | NA | 5 | 0.6 |
| 1_1517610 | 2 | 0 | NA | NA | 5 | 0.2 | 1 | 0 | 8 | 0.125 | 2 | 0.5 | 1 | 0 | NA | NA | NA | NA | 3 | 0.3333 |
| 1_1517669 | 1 | 0 | 2 | 0 | 2 | 0.5 | 1 | 0 | 6 | 0.1667 | NA | NA | 1 | 0 | 2 | 0.5 | NA | NA | 3 | 0.3333 |
| 1_1517670 | 1 | 0 | 2 | 0.5 | 2 | 1 | 2 | 1 | 7 | 0.7143 | NA | NA | 2 | 1 | 2 | 0.5 | NA | NA | 4 | 0.75 |
| 1_1517671 | 1 | 1 | 2 | 1 | 1 | 1 | 2 | 1 | 6 | 1 | NA | NA | 2 | 1 | 2 | 0.5 | NA | NA | 4 | 0.75 |
| 1_1517671 | 1 | 0 | 1 | 1 | NA | NA | 2 | 0.5 | 4 | 0.5 | NA | NA | 2 | 1 | 2 | 0.5 | NA | NA | 4 | 0.75 |
| 1_1517673 | NA | NA | 2 | 1 | NA | NA | 2 | 1 | 4 | 1 | NA | NA | 2 | 1 | 1 | 0 | NA | NA | 3 | 0.6667 |
| 1_1517675 | NA | NA | 1 | 1 | NA | NA | 2 | 0 | 3 | 0.3333 | NA | NA | 2 | 1 | 1 | 0 | NA | NA | 3 | 0.6667 |
| 1_1517684 | 5 | 0 | 7 | 0.2857 | 2 | 1 | 6 | 0.1667 | 20 | 0.25 | 3 | 0.3333 | 1 | 1 | 3 | 0.6667 | 3 | 0 | 10 | 0.4 |
| 1_1517686 | 5 | 0 | 3 | 0.3333 | 1 | 0 | 5 | 0.6 | 14 | 0.2857 | 2 | 0.5 | 1 | 0 | 2 | 0 | 2 | 0 | 7 | 0.1429 |
| 1_1517687 | 5 | 0 | 6 | 0.5 | 1 | 0 | 6 | 0.1667 | 18 | 0.2222 | 3 | 0.3333 | 1 | 0 | 2 | 0.5 | 1 | 1 | 7 | 0.4286 |
| 1_1517689 | 3 | 0.3333 | 5 | 0.8 | NA | NA | 2 | 0.5 | 10 | 0.6 | 1 | 1 | 2 | 1 | 2 | 0.5 | 2 | 0 | 7 | 0.5714 |
| 1_1517689 | 4 | 0 | 7 | 0.2857 | 1 | 0 | 5 | 0.2 | 17 | 0.1765 | 2 | 0.5 | 2 | 0 | 2 | 0.5 | 2 | 0.5 | 8 | 0.375 |
| 1_1517690 | 4 | 0.5 | 6 | 0.5 | NA | NA | 5 | 0.4 | 15 | 0.4667 | 3 | 0.6667 | 3 | 0.6667 | 2 | 0.5 | 1 | 1 | 9 | 0.6667 |
| 1_1522466 | 7 | 0.1429 | 8 | 0.125 | 3 | 0.3333 | NA | NA | 18 | 0.1667 | 3 | 0 | 5 | 0 | 6 | 0.1667 | 4 | 0 | 18 | 0.0556 |
| 1_1522466 | 6 | 0.3333 | 8 | 0.125 | 4 | 0.25 | NA | NA | 18 | 0.2222 | 4 | 0.25 | 5 | 0.2 | 6 | 0.3333 | 4 | 0.25 | 19 | 0.2632 |
| 1_1522468 | 5 | 0 | 4 | 0.5 | 3 | 0 | NA | NA | 12 | 0.1667 | 4 | 0.25 | 4 | 0 | 4 | 0 | 4 | 0 | 16 | 0.0625 |
| 1_1523925 | 9 | 0.1111 | 3 | 0 | NA | NA | 3 | 0.3333 | 15 | 0.1333 | 7 | 0 | 8 | 0.125 | 3 | 0.3333 | 4 | 0 | 22 | 0.0909 |
| 1_1523930 | 9 | 0.3333 | 4 | 0.5 | NA | NA | 6 | 0.5 | 19 | 0.4211 | 8 | 0.375 | 10 | 0.3 | 3 | 0.3333 | 5 | 0 | 26 | 0.2692 |
| 1_1523930 | 10 | 0 | 4 | 0.25 | NA | NA | 7 | 0.1429 | 21 | 0.0952 | 12 | 0.0833 | 10 | 0.1 | 4 | 0 | 6 | 0 | 32 | 0.0625 |
| 1_1523931 | 9 | 0.1111 | 4 | 0.25 | NA | NA | 4 | 0.5 | 17 | 0.2353 | 7 | 0.4286 | 8 | 0.25 | 1 | 0 | 8 | 0.25 | 24 | 0.2917 |
| 1_1523932 | 9 | 0.4444 | 4 | 0.25 | NA | NA | 7 | 0.1429 | 20 | 0.3 | 8 | 0.125 | 9 | 0 | 2 | 0 | 8 | 0.25 | 27 | 0.1111 |
| 1_1523936 | 8 | 0.125 | 10 | 0.1 | 5 | 0 | 7 | 0.1429 | 30 | 0.1 | 15 | 0 | 10 | 0.1 | 12 | 0.0833 | 10 | 0.1 | 47 | 0.0638 |
| 1_1523936 | 13 | 0.5385 | 11 | 0.6364 | 7 | 0.8571 | 6 | 0.8333 | 37 | 0.6757 | 15 | 0.3333 | 13 | 0.5385 | 13 | 0.8462 | 11 | 0.7273 | 52 | 0.5962 |
| 1_1523937 | 11 | 0.3636 | 10 | 0.3 | 7 | 0.8571 | 6 | 0.3333 | 34 | 0.4412 | 15 | 0.2667 | 11 | 0.2727 | 12 | 0.3333 | 11 | 0.3636 | 49 | 0.3061 |
| 1_1523938 | 10 | 0.4 | 8 | 0.5 | 4 | 0.25 | 7 | 0.4286 | 29 | 0.4138 | 7 | 0.5714 | 11 | 0.7273 | 8 | 0.5 | 8 | 0.75 | 34 | 0.6471 |
| 1_1523938 | 9 | 1 | 10 | 1 | 6 | 1 | 5 | 1 | 30 | 1 | 11 | 0.9091 | 9 | 1 | 10 | 1 | 12 | 1 | 42 | 0.9762 |
| 1_1523939 | 8 | 0.125 | 9 | 0.1111 | 6 | 0.1667 | 5 | 0 | 28 | 0.1071 | 7 | 0.2857 | 8 | 0 | 6 | 0.1667 | 9 | 0.1111 | 30 | 0.1333 |
| 1_1523941 | 11 | 0.1818 | 10 | 0.2 | 2 | 0 | 7 | 0.4286 | 30 | 0.2333 | 9 | 0 | 11 | 0.0909 | 8 | 0.125 | 11 | 0 | 39 | 0.0513 |
| 1_1523942 | 12 | 0.25 | 7 | 0.2857 | 5 | 0.2 | 7 | 0.1429 | 31 | 0.2258 | 7 | 0 | 9 | 0.1111 | 9 | 0 | 10 | 0.1 | 35 | 0.0571 |
| 1_1523943 | 13 | 0 | 12 | 0.1667 | 6 | 0.1667 | 8 | 0.125 | 39 | 0.1026 | 11 | 0 | 12 | 0.1667 | 8 | 0.125 | 11 | 0.0909 | 42 | 0.0952 |
| 1_1523943 | 13 | 0.0769 | 11 | 0.0909 | 5 | 0 | 8 | 0.375 | 37 | 0.1351 | 11 | 0.0909 | 10 | 0.3 | 8 | 0.125 | 10 | 0.2 | 39 | 0.1795 |
| 1_1523944 | 12 | 1 | 11 | 1 | 5 | 1 | 7 | 1 | 35 | 1 | 9 | 1 | 11 | 1 | 7 | 1 | 9 | 0.7778 | 36 | 0.9444 |
| 1_1523944 | 12 | 0.5 | 9 | 0.4444 | 5 | 0.2 | 6 | 0.5 | 32 | 0.4375 | 10 | 0.3 | 12 | 0.5833 | 5 | 0.8 | 11 | 0.6364 | 38 | 0.5526 |
| 1_1523945 | 13 | 0.0769 | 9 | 0.3333 | 6 | 0.1667 | 8 | 0.25 | 36 | 0.1944 | 9 | 0.1111 | 12 | 0.3333 | 6 | 0.3333 | 9 | 0.3333 | 36 | 0.2778 |
| 1_1523947 | 14 | 0.7857 | 11 | 1 | 5 | 0.8 | 7 | 0.7143 | 37 | 0.8378 | 10 | 0.9 | 12 | 0.9167 | 6 | 1 | 9 | 0.7778 | 37 | 0.8919 |
| 1_1523948 | 13 | 0.6923 | 8 | 0.625 | 3 | 0.3333 | 7 | 0.7143 | 31 | 0.6452 | 8 | 0.5 | 12 | 0.75 | 6 | 1 | 10 | 0.6 | 36 | 0.6944 |
| 1_1523948 | 15 | 0.2 | 11 | 0.2727 | 5 | 0.2 | 9 | 0.1111 | 40 | 0.2 | 10 | 0.1 | 13 | 0.0769 | 6 | 0.1667 | 10 | 0.1 | 39 | 0.1026 |
| 1_1523948 | 15 | 0.0667 | 8 | 0.25 | 5 | 0 | 7 | 0.1429 | 35 | 0.1143 | 9 | 0 | 11 | 0.1818 | 3 | 0 | 8 | 0 | 31 | 0.0645 |
| 1_1523948 | 10 | 0.7 | 8 | 0.625 | 5 | 0.2 | 6 | 0.6667 | 29 | 0.5862 | 5 | 0.8 | 9 | 0.7778 | 2 | 0 | 7 | 0.2857 | 23 | 0.5652 |
| 1_1523948 | 13 | 1 | 10 | 0.9 | 4 | 1 | 6 | 1 | 33 | 0.9697 | 10 | 0.9 | 10 | 0.8 | 5 | 0.6 | 9 | 1 | 34 | 0.8529 |
| 1_1523949 | 14 | 0.2143 | 10 | 0.2 | 2 | 0 | 6 | 0 | 32 | 0.1562 | 10 | 0.1 | 11 | 0.0909 | 6 | 0 | 10 | 0.3 | 37 | 0.1351 |
| 1_1524622 | 4 | 0 | 4 | 0 | 1 | 0 | 4 | 0.25 | 13 | 0.0769 | 4 | 0.25 | 8 | 0 | 4 | 0 | 2 | 0 | 18 | 0.0556 |
| 1_1526417 | 2 | 0.5 | 1 | 1 | NA | NA | 2 | 0 | 5 | 0.4 | 1 | 1 | 3 | 0.6667 | NA | NA | NA | NA | 4 | 0.75 |
| 1_1526426 | 13 | 0.0769 | 17 | 0.1765 | 7 | 0.1429 | 14 | 0.2143 | 51 | 0.1569 | 15 | 0.2 | 16 | 0.25 | 9 | 0.2222 | 16 | 0.3125 | 56 | 0.25 |
| 1_1526501 | 17 | 0.1765 | 20 | 0.1 | 10 | 0.3 | 15 | 0.0667 | 62 | 0.1452 | 20 | 0.3 | 21 | 0.2381 | 11 | 0 | 20 | 0.15 | 72 | 0.1944 |
| 1_1526504 | 17 | 0.0588 | 20 | 0.15 | 12 | 0 | 12 | 0.1667 | 61 | 0.0984 | 21 | 0.1429 | 21 | 0.1429 | 12 | 0.1667 | 22 | 0.1818 | 76 | 0.1579 |
| 1_1526510 | 20 | 0.15 | 29 | 0.1034 | 16 | 0.125 | 16 | 0.1875 | 81 | 0.1358 | 22 | 0 | 22 | 0.1364 | 14 | 0.1429 | 25 | 0.12 | 83 | 0.0964 |
| 1_1527129 | 8 | 0.125 | 5 | 0.2 | 6 | 0.3333 | 4 | 0 | 23 | 0.1739 | 3 | 0 | 6 | 0.1667 | 7 | 0.2857 | 6 | 0.1667 | 22 | 0.1818 |
| 1_1527134 | 8 | 0.125 | 13 | 0.1538 | 8 | 0 | 7 | 0.1429 | 36 | 0.1111 | 5 | 0 | 10 | 0.2 | 12 | 0 | 9 | 0 | 36 | 0.0556 |
| 1_1527137 | 9 | 0.8889 | 15 | 0.2 | 6 | 0.5 | 6 | 0.6667 | 36 | 0.5 | 8 | 0.625 | 7 | 0.5714 | 13 | 0.5385 | 8 | 0.5 | 36 | 0.5556 |
| 1_1527137 | 10 | 0.7 | 13 | 0.8462 | 5 | 0.8 | 7 | 1 | 35 | 0.8286 | 7 | 0.7143 | 5 | 0.8 | 12 | 0.9167 | 9 | 0.7778 | 33 | 0.8182 |
| 1_1527137 | 8 | 0.875 | 13 | 0.6923 | 4 | 1 | 6 | 0.6667 | 31 | 0.7742 | 7 | 0.8571 | 8 | 0.875 | 9 | 0.8889 | 8 | 0.75 | 32 | 0.8438 |
| 1_1527138 | 10 | 0.8 | 14 | 0.7857 | 5 | 0.8 | 5 | 1 | 34 | 0.8235 | 8 | 0.875 | 8 | 0.75 | 9 | 0.7778 | 9 | 0.7778 | 34 | 0.7941 |

| | | | | | | | | | | | | | | | | | | | | |
|---|---|---|---|---|---|---|---|---|---|---|---|---|---|---|---|---|---|---|---|---|
| 1_1527146 | 7 | 0 | 13 | 0 | 5 | 0.2 | 5 | 0.2 | 30 | 0.0667 | 5 | 0 | 5 | 0.2 | 9 | 0 | 5 | 0.4 | 24 | 0.125 |
| 1_1527148 | 4 | 0.25 | 11 | 0.5455 | 3 | 1 | 2 | 0.5 | 20 | 0.55 | 4 | 0.25 | 4 | 1 | 5 | 0.4 | 3 | 0.3333 | 16 | 0.5 |
| 1_1527149 | 8 | 0.75 | 9 | 0.5556 | 1 | 0 | 5 | 0.2 | 23 | 0.5217 | 4 | 0 | 3 | 0.6667 | 4 | 1 | 5 | 0.8 | 16 | 0.625 |
| 1_1527149 | 6 | 0.1667 | 8 | 0 | 1 | 0 | 3 | 0 | 18 | 0.0556 | 4 | 0 | 3 | 0 | 5 | 0 | 6 | 0.1667 | 18 | 0.0556 |
| 1_1527200 | 4 | 0.5 | 5 | 0.6 | 1 | 0 | 3 | 0 | 13 | 0.3846 | 3 | 1 | 3 | 0.3333 | 3 | 0.3333 | 2 | 0.5 | 11 | 0.5455 |
| 1_1527211 | 3 | 0 | 4 | 0.5 | 1 | 0 | 2 | 0.5 | 10 | 0.3 | 2 | 0.5 | 3 | 0 | 3 | 0 | 1 | 0 | 9 | 0.1111 |
| 1_1527244 | NA | NA | 2 | 0 | 1 | 0 | 3 | 0.6667 | 6 | 0.3333 | 1 | 1 | NA | NA | NA | NA | NA | NA | 1 | 1 |
| 1_1529915 | NA | NA | 2 | 0.5 | NA | NA | 1 | 0 | 3 | 0.3333 | 4 | 0.75 | 4 | 0.75 | 1 | 0 | 2 | 0.5 | 11 | 0.6364 |
| 1_1529916 | 1 | 0 | 2 | 0 | 1 | 0 | 2 | 0.5 | 6 | 0.1667 | 3 | 0 | 3 | 0.6667 | 1 | 0 | 4 | 0.25 | 11 | 0.2727 |
| 1_1529916 | 1 | 1 | 2 | 0 | 1 | 0 | 2 | 0 | 6 | 0.1667 | 4 | 0.5 | 5 | 0 | 2 | 0 | 3 | 0 | 14 | 0.1429 |
| 1_1529928 | 2 | 0.5 | 8 | 0 | 7 | 0 | 3 | 0 | 20 | 0.05 | 7 | 0.2857 | 7 | 0 | 5 | 0.2 | 5 | 0 | 24 | 0.125 |
| 1_1529944 | 7 | 0 | 9 | 0 | 13 | 0.1538 | 10 | 0.1 | 39 | 0.0769 | 6 | 0.1667 | 10 | 0.1 | 11 | 0 | 7 | 0.1429 | 34 | 0.0882 |
| 1_1529986 | 2 | 0.5 | 1 | 0 | NA | NA | 3 | 0.3333 | 6 | 0.3333 | NA | NA | 2 | 0.5 | 2 | 1 | NA | NA | 4 | 0.75 |
| 1_1529989 | 2 | 0.5 | 1 | 0 | 1 | 0 | 2 | 0.5 | 6 | 0.3333 | NA | NA | 2 | 0.5 | 1 | 1 | NA | NA | 3 | 0.6667 |
| 1_1529995 | 2 | 0.5 | 1 | 0 | 1 | 0 | 3 | 0 | 7 | 0.1429 | NA | NA | 2 | 0.5 | NA | NA | NA | NA | 2 | 0.5 |
| 1_1529996 | 2 | 1 | NA | NA | 1 | 0 | 3 | 0.6667 | 6 | 0.6667 | NA | NA | 1 | 1 | 1 | 0 | NA | NA | 2 | 0.5 |
| 1_1529997 | 2 | 1 | NA | NA | 1 | 1 | 3 | 0.6667 | 6 | 0.8333 | NA | NA | 1 | 1 | 1 | 0 | NA | NA | 2 | 0.5 |
| 1_1530961 | 4 | 0 | 5 | 0.2 | 2 | 0 | 1 | 0 | 12 | 0.0833 | 1 | 0 | NA | NA | 4 | 0.25 | 2 | 0 | 7 | 0.1429 |
| 1_1530963 | 5 | 0.8 | 5 | 0.2 | 1 | 0 | NA | NA | 11 | 0.4545 | 1 | 0 | NA | NA | 3 | 0.3333 | 2 | 0.5 | 6 | 0.3333 |
| 1_1530964 | 5 | 0.2 | 4 | 0.25 | 2 | 0 | 1 | 1 | 12 | 0.25 | NA | NA | NA | NA | 3 | 0.3333 | 2 | 0 | 5 | 0.2 |
| 1_1530968 | 4 | 0.5 | 5 | 0.2 | 1 | 0 | 1 | 1 | 11 | 0.3636 | 1 | 1 | NA | NA | 3 | 0.3333 | 2 | 0 | 6 | 0.3333 |
| 1_1530969 | 4 | 0.25 | 2 | 0.5 | 1 | 0 | NA | NA | 7 | 0.2857 | 1 | 1 | NA | NA | 2 | 0.5 | 2 | 0.5 | 5 | 0.6 |
| 1_1530969 | 4 | 0.75 | 5 | 0.6 | 1 | 0 | 1 | 1 | 11 | 0.6364 | 1 | 1 | NA | NA | 3 | 0.6667 | 1 | 0 | 5 | 0.6 |
| 1_1530970 | 2 | 0 | 3 | 0.3333 | 2 | 0 | 1 | 0 | 8 | 0.125 | 1 | 1 | NA | NA | 1 | 0 | 1 | 0 | 3 | 0.3333 |
| 1_1530971 | 3 | 0.3333 | 5 | 0 | 2 | 0 | 1 | 0 | 11 | 0.0909 | 1 | 1 | NA | NA | 3 | 0.6667 | 3 | 0.3333 | 7 | 0.5714 |
| 1_1530973 | 3 | 1 | 5 | 0.6 | 2 | 0 | 1 | 1 | 11 | 0.6364 | 1 | 1 | NA | NA | 3 | 0.6667 | 3 | 0.6667 | 7 | 0.7143 |
| 1_1530973 | 4 | 0.75 | 5 | 1 | 2 | 0.5 | 1 | 1 | 12 | 0.8333 | 1 | 0 | NA | NA | 3 | 1 | 3 | 1 | 7 | 0.8571 |
| 1_1535685 | 8 | 0.75 | 2 | 1 | 5 | 0.6 | 4 | 0 | 19 | 0.5789 | 4 | 0.75 | 4 | 0.25 | 2 | 1 | 4 | 0.5 | 14 | 0.5714 |
| 1_1535686 | 8 | 0.75 | 2 | 0.5 | 5 | 0.6 | 4 | 1 | 19 | 0.7368 | 5 | 1 | 4 | 0.25 | 2 | 1 | 3 | 0.6667 | 14 | 0.7143 |
| 1_1535688 | 8 | 0.75 | 1 | 1 | 5 | 1 | 5 | 0.6 | 19 | 0.7895 | 4 | 1 | 4 | 0.25 | 2 | 0.5 | 7 | 0.5714 | 17 | 0.5882 |
| 1_1535694 | 8 | 0.375 | 3 | 0.3333 | 4 | 0 | 4 | 0.75 | 19 | 0.3684 | 3 | 0.3333 | 4 | 0 | 3 | 0 | 8 | 0.375 | 18 | 0.2222 |
| 1_1535696 | 10 | 0 | 3 | 0 | 5 | 0.2 | 7 | 0.4286 | 25 | 0.16 | 4 | 0 | 4 | 0 | 4 | 0.25 | 8 | 0.125 | 20 | 0.1 |
| 1_1535697 | 10 | 0.5 | 3 | 0.3333 | 4 | 0.75 | 7 | 0.7143 | 24 | 0.5833 | 4 | 0.75 | 3 | 0.3333 | 4 | 0.25 | 9 | 0.6667 | 20 | 0.55 |
| 1_1535697 | 9 | 0.3333 | 3 | 1 | 4 | 0.75 | 7 | 0.5714 | 23 | 0.5652 | 4 | 0.5 | 3 | 0.3333 | 4 | 0.5 | 7 | 0.5714 | 18 | 0.5 |
| 1_1535697 | 8 | 0.75 | 3 | 1 | 5 | 1 | 7 | 1 | 23 | 0.913 | 3 | 1 | 4 | 0.5 | 3 | 1 | 8 | 1 | 18 | 0.8889 |
| 1_1535698 | 9 | 0.1111 | 3 | 0 | 5 | 0 | 8 | 0.125 | 25 | 0.08 | 4 | 0.5 | 4 | 0 | 6 | 0 | 9 | 0.1111 | 23 | 0.1304 |
| 1_1535699 | 7 | 0.1429 | 2 | 0 | 6 | 0 | 6 | 0.1667 | 21 | 0.0952 | 5 | 0.2 | 3 | 0 | 7 | 0 | 8 | 0.125 | 23 | 0.087 |
| 1_1535699 | 8 | 0.375 | 2 | 0 | 6 | 0.5 | 6 | 0.5 | 22 | 0.4091 | 4 | 0.25 | 3 | 0.3333 | 6 | 0.1667 | 7 | 0.5714 | 20 | 0.35 |
| 1_1535701 | 8 | 0.25 | 3 | 0.3333 | 4 | 0.5 | 6 | 0.3333 | 21 | 0.3333 | 4 | 0.25 | 5 | 0.2 | 7 | 0.2857 | 9 | 0.3333 | 25 | 0.28 |
| 1_1535704 | 9 | 0.2222 | 3 | 0 | 6 | 0.1667 | 8 | 0.375 | 26 | 0.2308 | 5 | 0.6 | 6 | 0.1667 | 7 | 0.2857 | 10 | 0.4 | 28 | 0.3571 |
| 1_1535704 | 9 | 0.6667 | 3 | 0.6667 | 5 | 0.4 | 10 | 0.4 | 27 | 0.5185 | 5 | 0.6 | 5 | 0.4 | 7 | 0.1429 | 10 | 0.2 | 27 | 0.2963 |
| 1_1535713 | 6 | 0.5 | 3 | 0 | 9 | 0.5556 | 14 | 0.2857 | 32 | 0.375 | 5 | 0.2 | 9 | 0.5556 | 8 | 0.375 | 10 | 0.4 | 32 | 0.4062 |
| 1_1535714 | 8 | 0.125 | 3 | 0.3333 | 11 | 0.0909 | 16 | 0.125 | 38 | 0.1316 | 5 | 0 | 8 | 0.125 | 8 | 0 | 10 | 0.2 | 31 | 0.0968 |
| 1_1535714 | 6 | 0.1667 | 2 | 0.5 | 10 | 0 | 16 | 0.1875 | 34 | 0.1471 | 6 | 0.3333 | 8 | 0.25 | 8 | 0.375 | 10 | 0.2 | 32 | 0.2812 |
| 1_1535714 | 7 | 0.2857 | 3 | 0.3333 | 9 | 0.1111 | 16 | 0 | 35 | 0.1143 | 5 | 0 | 6 | 0 | 9 | 0 | 11 | 0.1818 | 31 | 0.0645 |
| 1_1535715 | 8 | 0.875 | 4 | 0.75 | 8 | 0.875 | 16 | 0.6875 | 36 | 0.7778 | 5 | 0.6 | 8 | 1 | 7 | 0.8571 | 11 | 0.7273 | 31 | 0.8065 |
| 1_1535717 | 8 | 0.5 | 3 | 0.6667 | 8 | 0.375 | 11 | 0.8182 | 30 | 0.6 | 5 | 0.8 | 8 | 0.625 | 10 | 0.7 | 10 | 0.3 | 33 | 0.5758 |
| 1_1535718 | 8 | 0.125 | 5 | 0 | 11 | 0.0909 | 16 | 0 | 40 | 0.05 | 6 | 0.1667 | 11 | 0.0909 | 11 | 0 | 9 | 0.2222 | 37 | 0.1081 |
| 1_1535719 | 9 | 0.2222 | 3 | 0 | 9 | 0 | 12 | 0.1667 | 33 | 0.1212 | 4 | 0 | 9 | 0.1111 | 10 | 0.2 | 11 | 0.0909 | 34 | 0.1176 |
| 1_1535719 | 11 | 0.9091 | 3 | 0.6667 | 11 | 0.4545 | 13 | 0.7692 | 38 | 0.7105 | 4 | 0.5 | 9 | 0.4444 | 9 | 0.4444 | 12 | 0.4167 | 34 | 0.4412 |
| 1_1535721 | 7 | 0.2857 | 6 | 0 | 11 | 0 | 16 | 0.1875 | 40 | 0.125 | 6 | 0.1667 | 11 | 0 | 10 | 0.1 | 10 | 0 | 37 | 0.0541 |
| 1_1535722 | 9 | 0.5556 | 6 | 0.6667 | 12 | 0.3333 | 15 | 0.5333 | 42 | 0.5 | 6 | 0.6667 | 11 | 0.6364 | 11 | 0.4545 | 13 | 0.3846 | 41 | 0.5122 |
| 1_1535722 | 11 | 0.8182 | 7 | 0.2857 | 13 | 0.3846 | 17 | 0.4118 | 48 | 0.4792 | 6 | 0.3333 | 12 | 0.25 | 11 | 0.3636 | 14 | 0.2857 | 43 | 0.3023 |
| 1_1535726 | 4 | 0.25 | 5 | 0 | 10 | 0.1 | 14 | 0 | 33 | 0.0606 | 5 | 0.2 | 13 | 0 | 10 | 0 | 9 | 0.1111 | 37 | 0.0541 |
| 1_1535728 | 12 | 0.0833 | 9 | 0.2222 | 12 | 0.0833 | 18 | 0.2222 | 51 | 0.1569 | 6 | 0.5 | 16 | 0.0625 | 14 | 0.1429 | 15 | 0.0667 | 51 | 0.1373 |
| 1_1535730 | 10 | 0.5 | 9 | 0.2222 | 11 | 0.4545 | 18 | 0.3889 | 48 | 0.3958 | 5 | 0.6 | 15 | 0.4 | 13 | 0.3077 | 13 | 0.4615 | 46 | 0.413 |
| 1_1535731 | 13 | 0.3077 | 9 | 0.3333 | 12 | 0.25 | 19 | 0.3684 | 53 | 0.3208 | 6 | 0.3333 | 17 | 0.2353 | 16 | 0.25 | 16 | 0.25 | 55 | 0.2545 |
| 1_1535731 | 14 | 0.0714 | 10 | 0.2 | 11 | 0.3636 | 18 | 0.0556 | 53 | 0.1509 | 8 | 0.625 | 14 | 0.2143 | 15 | 0.2 | 15 | 0.4 | 52 | 0.3269 |
| 1_1535731 | 11 | 0.0909 | 5 | 0 | 8 | 0.25 | 14 | 0 | 38 | 0.0789 | 6 | 0.1667 | 9 | 0.1111 | 9 | 0 | 8 | 0 | 32 | 0.0625 |
| 1_1536194 | 3 | 0.6667 | 2 | 0.5 | 1 | 1 | 1 | 1 | 7 | 0.7143 | 1 | 0 | 1 | 1 | 1 | 1 | NA | NA | 3 | 0.6667 |
| 1_1536197 | 3 | 0.3333 | 2 | 0 | 1 | 0 | 1 | 0 | 7 | 0.1429 | 1 | 1 | 1 | 0 | 1 | 0 | NA | NA | 3 | 0.3333 |
| 1_1536197 | 3 | 1 | 2 | 0 | 1 | 1 | 1 | 1 | 7 | 0.7143 | 1 | 1 | 1 | 0 | 1 | 0 | NA | NA | 3 | 0.3333 |
| 1_1536198 | 3 | 0.6667 | 1 | 1 | 1 | 1 | 1 | 0 | 6 | 0.6667 | 1 | 1 | 1 | 1 | 1 | 1 | NA | NA | 3 | 1 |
| 1_1536198 | 2 | 0.5 | 1 | 0 | 1 | 1 | 1 | 1 | 5 | 0.6 | 1 | 0 | 1 | 1 | 1 | 0 | NA | NA | 3 | 0.3333 |
| 1_1536200 | 3 | 0.6667 | 1 | 1 | 1 | 1 | 1 | 1 | 6 | 0.8333 | 1 | 1 | 1 | 1 | 1 | 1 | NA | NA | 3 | 1 |

| | | | | | | | | | | | | | | | | | | | | |
|---|---|---|---|---|---|---|---|---|---|---|---|---|---|---|---|---|---|---|---|---|
| 1_1536207 | 3 | 0.3333 | 2 | 0.5 | 1 | 0 | 1 | 0 | 7 | 0.2857 | 1 | 0 | 1 | 1 | NA | NA | NA | NA | 2 | 0.5 |
| 1_1536208 | 3 | 0 | 1 | 0 | 2 | 0.5 | 1 | 0 | 7 | 0.1429 | 1 | 0 | 1 | 0 | 1 | 1 | NA | NA | 3 | 0.3333 |
| 1_1536307 | NA | NA | NA | NA | NA | NA | 1 | 1 | 1 | 1 | 1 | 0 | NA | NA | 1 | 1 | NA | NA | 2 | 0.5 |
| 1_1536308 | NA | NA | NA | NA | NA | NA | 1 | 1 | 1 | 1 | 1 | 0 | NA | NA | 1 | 1 | NA | NA | 2 | 0.5 |
| 1_1536388 | 1 | 1 | NA | NA | NA | NA | NA | NA | 1 | 1 | NA | NA | 1 | 1 | NA | NA | NA | NA | 1 | 1 |
| 1_1537837 | 96 | 0.6458 | 107 | 0.5234 | 107 | 0.5607 | 80 | 0.575 | 390 | 0.5744 | 117 | 0.5983 | 89 | 0.618 | 83 | 0.6386 | 81 | 0.6296 | 370 | 0.6189 |
| 1_1537839 | 85 | 0.3412 | 90 | 0.3778 | 97 | 0.2784 | 62 | 0.2742 | 334 | 0.3204 | 101 | 0.4059 | 76 | 0.3684 | 64 | 0.25 | 59 | 0.4576 | 300 | 0.3733 |
| 1_1537839 | 108 | 0.963 | 111 | 0.8919 | 121 | 0.9421 | 78 | 0.8462 | 418 | 0.9163 | 128 | 0.8906 | 103 | 0.8835 | 80 | 0.95 | 89 | 0.8989 | 400 | 0.9025 |
| 1_1537839 | 111 | 0.1532 | 105 | 0.1143 | 115 | 0.0609 | 85 | 0.0824 | 416 | 0.1034 | 125 | 0.144 | 102 | 0.098 | 78 | 0.1026 | 89 | 0 | 394 | 0.0914 |
| 1_1537842 | 106 | 0.1132 | 108 | 0.1759 | 108 | 0.213 | 85 | 0.1529 | 407 | 0.1646 | 126 | 0.1746 | 97 | 0.1546 | 84 | 0.2024 | 85 | 0.1647 | 392 | 0.1735 |
| 1_1537851 | 279 | 0.0609 | 270 | 0.0852 | 257 | 0.0506 | 201 | 0.0647 | 1007 | 0.0655 | 249 | 0.0723 | 232 | 0.0776 | 210 | 0.0571 | 229 | 0.0568 | 920 | 0.0663 |
| 1_1537853 | 265 | 0.0981 | 277 | 0.0975 | 261 | 0.092 | 214 | 0.0888 | 1017 | 0.0944 | 255 | 0.0863 | 246 | 0.0894 | 216 | 0.0787 | 235 | 0.0809 | 952 | 0.084 |
| 1_1542668 | 18 | 0.0556 | 35 | 0.0857 | 20 | 0.05 | 27 | 0.037 | 100 | 0.06 | 13 | 0.1538 | 20 | 0 | 15 | 0.2 | 21 | 0.0476 | 69 | 0.087 |
| 1_1542673 | 14 | 0.2143 | 26 | 0.1154 | 20 | 0.15 | 19 | 0.1053 | 79 | 0.1392 | 9 | 0.1111 | 12 | 0.0833 | 14 | 0.2143 | 14 | 0.0714 | 49 | 0.1224 |
| 1_1542673 | 2 | 0 | 5 | 0.4 | NA | NA | 2 | 0 | 9 | 0.2222 | 1 | 0 | NA | NA | NA | NA | 3 | 0.3333 | 4 | 0.25 |
| 1_1542680 | 15 | 0.1333 | 24 | 0.125 | 17 | 0 | 16 | 0.0625 | 72 | 0.0833 | 8 | 0 | 11 | 0 | 14 | 0.1429 | 14 | 0.0714 | 47 | 0.0638 |
| 1_1542683 | 10 | 0 | 22 | 0.0909 | 11 | 0 | 17 | 0.0588 | 60 | 0.05 | 8 | 0 | 7 | 0 | 13 | 0.0769 | 15 | 0.1333 | 43 | 0.0698 |
| 1_1542731 | 3 | 0.3333 | 5 | 0.2 | 1 | 0 | NA | NA | 9 | 0.2222 | 2 | 0 | NA | NA | 6 | 0.1667 | 1 | 0 | 9 | 0.1111 |
| 1_1542736 | 2 | 1 | 5 | 0.8 | NA | NA | 1 | 1 | 8 | 0.875 | 2 | 1 | 1 | 1 | 3 | 1 | NA | NA | 6 | 1 |
| 1_1542737 | 2 | 0 | 4 | 0.5 | NA | NA | NA | NA | 6 | 0.3333 | 1 | 0 | NA | NA | 4 | 0.5 | NA | NA | 5 | 0.4 |
| 1_1542737 | 1 | 1 | 4 | 0.25 | NA | NA | 1 | 0 | 6 | 0.3333 | 1 | 0 | 1 | 0 | 3 | 0.3333 | NA | NA | 5 | 0.2 |
| 1_1542738 | 1 | 1 | 5 | 1 | NA | NA | 1 | 0 | 7 | 0.8571 | 1 | 1 | NA | NA | 3 | 1 | NA | NA | 4 | 1 |
| 1_1542739 | 2 | 0.5 | 5 | 0.6 | NA | NA | 1 | 1 | 8 | 0.625 | 1 | 0 | 1 | 0 | 2 | 0.5 | NA | NA | 4 | 0.25 |
| 1_1542739 | 2 | 0.5 | 3 | 0.6667 | NA | NA | 1 | 0 | 6 | 0.5 | 1 | 1 | NA | NA | 2 | 0 | NA | NA | 3 | 0.3333 |
| 1_1542739 | 2 | 0.5 | 4 | 0.5 | NA | NA | 1 | 0 | 7 | 0.4286 | 1 | 1 | NA | NA | 3 | 0.6667 | NA | NA | 4 | 0.75 |
| 1_1542741 | 2 | 0.5 | 3 | 0 | NA | NA | NA | NA | 5 | 0.2 | 1 | 0 | NA | NA | 2 | 0.5 | NA | NA | 3 | 0.3333 |
| 1_1542743 | 2 | 0 | 2 | 1 | NA | NA | NA | NA | 4 | 0.5 | 1 | 1 | 1 | 0 | 2 | 1 | NA | NA | 4 | 0.75 |
| 1_1542743 | 2 | 1 | 3 | 0.6667 | NA | NA | NA | NA | 5 | 0.8 | 1 | 1 | NA | NA | 2 | 1 | NA | NA | 3 | 1 |
| 1_1542743 | 2 | 0.5 | 3 | 0.6667 | NA | NA | NA | NA | 5 | 0.6 | 1 | 0 | 1 | 0 | 2 | 1 | NA | NA | 4 | 0.5 |
| 1_1542834 | 1 | 0 | 2 | 0.5 | NA | NA | NA | NA | 3 | 0.3333 | NA | NA | 1 | 0 | 2 | 0.5 | NA | NA | 3 | 0.3333 |
| 1_1549734 | 1 | 1 | 7 | 0.4286 | 8 | 0.5 | 4 | 0.75 | 20 | 0.55 | 4 | 0.75 | 8 | 0.625 | 4 | 0.75 | 6 | 0.3333 | 22 | 0.5909 |
| 1_1549734 | NA | NA | 5 | 0.6 | 8 | 0.75 | 4 | 1 | 17 | 0.7647 | 6 | 0.5 | 5 | 0.4 | 4 | 0.5 | 4 | 0.5 | 19 | 0.4737 |
| 1_1549735 | NA | NA | 6 | 0 | 8 | 0.125 | 4 | 0.25 | 18 | 0.1111 | 6 | 0.1667 | 7 | 0.2857 | 4 | 0.25 | 4 | 0.5 | 21 | 0.2857 |
| 1_1549736 | 1 | 1 | 6 | 1 | 5 | 1 | 2 | 1 | 14 | 1 | 6 | 1 | 4 | 1 | 4 | 0.75 | 4 | 1 | 18 | 0.9444 |
| 1_1549736 | NA | NA | 4 | 0.75 | 8 | 0.625 | 2 | 1 | 14 | 0.7143 | 7 | 0.7143 | 4 | 0.25 | 3 | 0.6667 | 5 | 0.6 | 19 | 0.5789 |
| 1_1549737 | NA | NA | 5 | 1 | 8 | 1 | 3 | 1 | 16 | 1 | 5 | 1 | 5 | 1 | 4 | 1 | 6 | 1 | 20 | 1 |
| 1_1549737 | NA | NA | 5 | 0.2 | 8 | 0.125 | 3 | 0.3333 | 16 | 0.1875 | 6 | 0.1667 | 5 | 0.2 | 4 | 0.25 | 5 | 0 | 20 | 0.15 |
| 1_1549738 | NA | NA | 6 | 0.8333 | 8 | 0.875 | 2 | 1 | 16 | 0.875 | 4 | 0.75 | 5 | 0.6 | 3 | 0.6667 | 4 | 1 | 16 | 0.75 |
| 1_1549738 | NA | NA | 6 | 1 | 8 | 0.75 | 3 | 1 | 17 | 0.8824 | 5 | 1 | 4 | 0.5 | 4 | 0.75 | 4 | 1 | 17 | 0.8235 |
| 1_1549739 | NA | NA | 5 | 1 | 7 | 1 | 3 | 1 | 15 | 1 | 6 | 1 | 5 | 1 | 4 | 1 | 4 | 1 | 19 | 1 |
| 1_1549739 | NA | NA | 6 | 0.6667 | 8 | 0.75 | 3 | 1 | 17 | 0.7647 | 4 | 0.25 | 4 | 0.5 | 4 | 0.5 | 3 | 1 | 15 | 0.5333 |
| 1_1549741 | NA | NA | 5 | 1 | 5 | 1 | 3 | 1 | 13 | 1 | 2 | 1 | 4 | 1 | 3 | 1 | 4 | 1 | 13 | 1 |
| 1_1549741 | NA | NA | 5 | 0.6 | 3 | 0 | 2 | 0 | 10 | 0.3 | 4 | 0.25 | 3 | 0 | 3 | 0.3333 | 4 | 0 | 14 | 0.1429 |
| 1_1549742 | NA | NA | 6 | 0.3333 | 6 | 0.6667 | 3 | 0.3333 | 15 | 0.4667 | 5 | 0.2 | 4 | 0 | 3 | 0.6667 | 5 | 0 | 17 | 0.1765 |
| 1_1549742 | NA | NA | 6 | 0.1667 | 6 | 0.3333 | 3 | 0.6667 | 15 | 0.3333 | 6 | 1 | 5 | 0.4 | 4 | 0.5 | 4 | 0.5 | 19 | 0.6316 |
| 1_1549743 | NA | NA | 3 | 0.6667 | 4 | 0.5 | 3 | 0.3333 | 10 | 0.5 | 6 | 0.5 | 5 | 0.4 | 4 | 0.5 | 4 | 0.5 | 19 | 0.4737 |
| 1_1549747 | NA | NA | 5 | 0.6 | 2 | 1 | 3 | 1 | 10 | 0.8 | 4 | 0.75 | 5 | 0.6 | 4 | 0.25 | 2 | 0.5 | 15 | 0.5333 |
| 1_1549749 | NA | NA | 5 | 0.2 | 3 | 0 | NA | NA | 8 | 0.125 | 2 | 0 | 4 | 0.25 | 3 | 0 | NA | NA | 9 | 0.1111 |
| 1_1549750 | NA | NA | 4 | 0 | 2 | 0 | 2 | 0.5 | 8 | 0.125 | 4 | 0 | 2 | 0 | 4 | 0.25 | 2 | 0 | 12 | 0.0833 |
| 1_1550048 | NA | NA | 2 | 1 | NA | NA | NA | NA | 2 | 1 | NA | NA | NA | NA | 1 | 1 | NA | NA | 1 | 1 |
| 1_1550778 | 14 | 0.2143 | 12 | 0.0833 | 6 | 0.1667 | 7 | 0.4286 | 39 | 0.2051 | 11 | 0.0909 | 6 | 0 | 5 | 0.2 | 9 | 0.1111 | 31 | 0.0968 |
| 1_1550779 | 14 | 0.5 | 7 | 0.1429 | 7 | 0.2857 | 7 | 0.5714 | 35 | 0.4 | 12 | 0.3333 | 6 | 0.3333 | 5 | 0.6 | 9 | 0.3333 | 32 | 0.375 |
| 1_1550780 | 11 | 0.7273 | 9 | 0.6667 | 8 | 0.625 | 5 | 0.4 | 33 | 0.6364 | 13 | 0.6154 | 7 | 0.4286 | 4 | 0.75 | 9 | 0.6667 | 33 | 0.6061 |
| 1_1550780 | 14 | 0.7857 | 9 | 0.8889 | 7 | 0.4286 | 7 | 0.8571 | 37 | 0.7568 | 12 | 0.8333 | 6 | 0.5 | 6 | 0.5 | 10 | 0.9 | 34 | 0.7353 |
| 1_1550780 | 13 | 0.9231 | 10 | 0.9 | 7 | 0.8571 | 7 | 0.8571 | 37 | 0.8919 | 12 | 0.9167 | 6 | 1 | 3 | 1 | 9 | 0.8889 | 30 | 0.9333 |
| 1_1550780 | 9 | 0 | 5 | 0 | 6 | 0.1667 | 4 | 0.25 | 24 | 0.0833 | 12 | 0.0833 | 4 | 0 | 4 | 0.25 | 4 | 0 | 24 | 0.0833 |
| 1_1550781 | 12 | 0.5833 | 9 | 0.2222 | 6 | 0.1667 | 6 | 0.3333 | 33 | 0.3636 | 13 | 0.6154 | 6 | 0.3333 | 6 | 0.5 | 8 | 0.125 | 33 | 0.4242 |
| 1_1550784 | 12 | 0.1667 | 14 | 0.0714 | 8 | 0.125 | 8 | 0.125 | 42 | 0.119 | 12 | 0.0833 | 7 | 0.1429 | 7 | 0 | 10 | 0 | 36 | 0.0556 |
| 1_1550784 | 9 | 0.6667 | 12 | 1 | 7 | 0.7143 | 5 | 0.8 | 33 | 0.8182 | 10 | 0.9 | 6 | 0.8333 | 5 | 0.8 | 6 | 0.6667 | 27 | 0.8148 |
| 1_1550786 | 9 | 0.3333 | 10 | 0 | 6 | 0.1667 | 4 | 0 | 29 | 0.1379 | 12 | 0.1667 | 6 | 0 | 5 | 0.6 | 6 | 0.1667 | 29 | 0.2069 |
| 1_1550787 | 12 | 0.0833 | 11 | 0.0909 | 8 | 0 | 6 | 0 | 37 | 0.0541 | 13 | 0.3077 | 6 | 0 | 6 | 0 | 8 | 0 | 33 | 0.1212 |
| 1_1550798 | 1 | 1 | 1 | 0 | 1 | 0 | 1 | 0 | 4 | 0.25 | 2 | 0 | NA | NA | 1 | 1 | NA | NA | 3 | 0.3333 |
| 1_1550801 | NA | NA | 1 | 1 | 1 | 1 | 1 | 1 | 3 | 1 | 2 | 0.5 | NA | NA | 1 | 0 | NA | NA | 3 | 0.3333 |
| 1_1550805 | 1 | 1 | 1 | 1 | 1 | 0 | 1 | 0 | 4 | 0.5 | 2 | 0.5 | NA | NA | 1 | 1 | NA | NA | 3 | 0.6667 |
| 1_1550807 | 1 | 1 | 1 | 1 | NA | NA | 1 | 0 | 3 | 0.6667 | 1 | 1 | NA | NA | 1 | 1 | NA | NA | 2 | 1 |

| | | | | | | | | | | | | | | | | | | | | |
|---|---|---|---|---|---|---|---|---|---|---|---|---|---|---|---|---|---|---|---|---|
| 1_15515165 | 9 | 0.5556 | 9 | 0.8889 | 15 | 0.6667 | 12 | 0.6667 | 45 | 0.6889 | 5 | 0.8 | 10 | 0.6 | 16 | 0.8125 | 18 | 0.7778 | 49 | 0.7551 |
| 1_15515211 | 12 | 0.5833 | 6 | 0.5 | 14 | 0.5714 | 13 | 0.4615 | 45 | 0.5333 | 4 | 0.5 | 10 | 0.5 | 14 | 0.4286 | 14 | 0.7143 | 42 | 0.5476 |
| 1_15515224 | 4 | 0.75 | 5 | 0 | 8 | 0.375 | 12 | 0.5833 | 29 | 0.4483 | 3 | 0 | 4 | 0.25 | 9 | 0.5556 | 9 | 0.4444 | 25 | 0.4 |
| 1_15515231 | 9 | 0.4444 | 6 | 0.1667 | 11 | 0.3636 | 15 | 0.4667 | 41 | 0.3902 | 3 | 0.3333 | 10 | 0.5 | 13 | 0.7692 | 14 | 0.6429 | 40 | 0.625 |
| 1_15515238 | 10 | 0.5 | 3 | 0 | 13 | 0.4615 | 9 | 0.2222 | 35 | 0.3714 | 2 | 0.5 | 8 | 0 | 11 | 0.2727 | 14 | 0.4286 | 35 | 0.2857 |
| 1_15515260 | 9 | 0.1111 | 6 | 0.1667 | 11 | 0.1818 | 9 | 0.1111 | 35 | 0.1429 | 3 | 0 | 10 | 0.1 | 11 | 0.1818 | 15 | 0.2 | 39 | 0.1538 |
| 1_15515261 | 10 | 0.5 | 6 | 0.5 | 12 | 0.8333 | 7 | 0.5714 | 35 | 0.6286 | 3 | 0.6667 | 7 | 0.4286 | 9 | 0.7778 | 14 | 0.8571 | 33 | 0.7273 |
| 1_15515418 | 3 | 0 | 3 | 0.3333 | 9 | 0.1111 | 4 | 0.5 | 19 | 0.2105 | NA | NA | 9 | 0 | 6 | 0.1667 | 7 | 0.2857 | 22 | 0.1364 |
| 1_15515566 | 7 | 0.2857 | 7 | 0.2857 | 13 | 0.1538 | 7 | 0.5714 | 34 | 0.2941 | 4 | 0.75 | 10 | 0.6 | 10 | 0.2 | 10 | 0.2 | 34 | 0.3824 |
| 1_15515609 | 6 | 0.1667 | 4 | 0.25 | 11 | 0.1818 | 3 | 0 | 24 | 0.1667 | 3 | 0 | 8 | 0.375 | 10 | 0.1 | 6 | 0.3333 | 27 | 0.2222 |
| 1_15515621 | 1 | 1 | 5 | 0.8 | 6 | 1 | 3 | 1 | 15 | 0.9333 | 4 | 1 | 5 | 1 | 9 | 0.6667 | 5 | 0.8 | 23 | 0.8261 |
| 1_15515629 | 3 | 0 | 5 | 0.6 | 7 | 0.5714 | 6 | 0.5 | 21 | 0.4762 | 4 | 0.75 | 4 | 0.5 | 10 | 0.6 | 4 | 0.25 | 22 | 0.5455 |
| 1_15515636 | 3 | 0 | 5 | 0.4 | 7 | 0 | 5 | 0.4 | 20 | 0.2 | 4 | 0.25 | 2 | 0 | 9 | 0.3333 | 2 | 0 | 17 | 0.2353 |
| 1_15515637 | 3 | 0.6667 | 5 | 1 | 7 | 1 | 4 | 0.75 | 19 | 0.8947 | 3 | 1 | 3 | 1 | 9 | 1 | 3 | 0.6667 | 18 | 0.9444 |
| 1_15515650 | 3 | 0 | 5 | 0.2 | 6 | 0 | 4 | 0 | 18 | 0.0556 | 4 | 0 | 4 | 0 | 9 | 0.2222 | 3 | 0 | 20 | 0.1 |
| 1_15515650 | 3 | 0.6667 | 4 | 0.25 | 6 | 0.8333 | 5 | 0.4 | 18 | 0.5556 | 3 | 0.3333 | 3 | 0 | 10 | 0.3 | 1 | 0 | 17 | 0.2353 |
| 1_15515661 | 3 | 1 | 5 | 0.8 | 6 | 1 | 4 | 1 | 18 | 0.9444 | 4 | 0.75 | 3 | 1 | 10 | 0.9 | 3 | 1 | 20 | 0.9 |
| 1_15515664 | 3 | 1 | 3 | 0.3333 | 5 | 0.6 | 4 | 0.5 | 15 | 0.6 | 4 | 0.75 | 1 | 1 | 7 | 0.2857 | 2 | 0.5 | 14 | 0.5 |
| 1_15543443 | 17 | 0 | 15 | 0.0667 | 10 | 0.1 | 11 | 0.0909 | 53 | 0.0566 | 9 | 0 | 10 | 0.1 | 5 | 0.2 | 8 | 0 | 32 | 0.0625 |
| 1_15546326 | 1 | 0 | NA | NA | 1 | 1 | NA | NA | 2 | 0.5 | 1 | 0 | 1 | 0 | NA | NA | 1 | 1 | 3 | 0.3333 |
| 1_15546754 | 1 | 1 | NA | NA | NA | NA | 1 | 0 | 2 | 0.5 | 1 | 1 | 1 | 0 | NA | NA | NA | NA | 2 | 0.5 |
| 1_15546784 | 1 | 1 | NA | NA | NA | NA | 1 | 1 | 2 | 1 | 1 | 0 | 1 | 1 | NA | NA | NA | NA | 2 | 0.5 |
| 1_15546805 | 1 | 0 | NA | NA | NA | NA | 1 | 1 | 2 | 0.5 | 1 | 1 | 1 | 1 | NA | NA | NA | NA | 2 | 1 |
| 1_15546827 | NA | NA | NA | NA | NA | NA | 1 | 1 | 1 | 1 | 1 | 0 | 1 | 1 | NA | NA | NA | NA | 2 | 0.5 |
| 1_15556288 | 9 | 0.4444 | 9 | 0.2222 | 8 | 0.25 | 7 | 0.4286 | 33 | 0.3333 | 4 | 0.5 | 2 | 0.5 | 4 | 0.5 | 4 | 0.25 | 14 | 0.4286 |
| 1_15556290 | 9 | 0.4444 | 11 | 0.1818 | 8 | 0.25 | 7 | 0 | 35 | 0.2286 | 4 | 0.25 | 3 | 0 | 4 | 0.25 | 4 | 0.25 | 15 | 0.2 |
| 1_15556297 | 6 | 0.1667 | 9 | 0.2222 | 7 | 0.1429 | 6 | 0.3333 | 28 | 0.2143 | 4 | 0 | 3 | 0 | 4 | 0 | 4 | 0.25 | 15 | 0.0667 |
| 1_15565443 | NA | NA | 2 | 1 | 2 | 0.5 | 1 | 0 | 5 | 0.6 | NA | NA | 1 | 1 | 4 | 0.25 | NA | NA | 5 | 0.4 |
| 1_15565560 | NA | NA | 2 | 0.5 | NA | NA | NA | NA | 2 | 0.5 | 1 | 0 | NA | NA | 3 | 0.3333 | NA | NA | 4 | 0.25 |
| 1_15572102 | 3 | 1 | 2 | 1 | 4 | 0.75 | 4 | 1 | 13 | 0.9231 | 2 | 1 | 4 | 0.75 | 4 | 1 | 2 | 1 | 12 | 0.9167 |
| 1_15572130 | 4 | 0.25 | 2 | 0.5 | 5 | 0.4 | 4 | 0.25 | 15 | 0.3333 | 4 | 1 | 3 | 0.6667 | 4 | 0 | 4 | 0 | 15 | 0.4 |
| 1_15572221 | 3 | 0 | 2 | 0 | 3 | 0 | 3 | 0.3333 | 11 | 0.0909 | 2 | 0 | 2 | 0 | 3 | 0 | 4 | 0.25 | 11 | 0.0909 |
| 1_15572257 | 3 | 0.6667 | 1 | 1 | 2 | 1 | NA | NA | 6 | 0.8333 | 2 | 1 | 1 | 1 | 2 | 1 | 3 | 1 | 8 | 1 |
| 1_15572261 | 3 | 1 | 2 | 1 | 4 | 0.5 | 3 | 1 | 12 | 0.8333 | 2 | 1 | 2 | 1 | 3 | 1 | 3 | 0.3333 | 10 | 0.8 |
| 1_15572278 | 2 | 0.5 | 2 | 1 | 3 | 0.3333 | 2 | 0.5 | 9 | 0.5556 | 2 | 1 | 1 | 1 | 3 | 0.6667 | 3 | 0.3333 | 9 | 0.6667 |
| 1_15572280 | 3 | 0 | 1 | 1 | 3 | 0 | 1 | 0 | 8 | 0.125 | 2 | 0 | 1 | 0 | 2 | 0.5 | 4 | 0 | 9 | 0.1111 |
| 1_15572281 | 3 | 1 | 2 | 1 | 4 | 0.75 | 3 | 1 | 12 | 0.9167 | 2 | 1 | 1 | 1 | 3 | 1 | 4 | 1 | 10 | 1 |
| 1_15572285 | 3 | 0.3333 | 2 | 1 | 4 | 0.25 | 3 | 0 | 12 | 0.3333 | 2 | 0 | 1 | 1 | 2 | 0 | 2 | 0 | 7 | 0.1429 |
| 1_15572296 | 1 | 1 | 2 | 1 | 3 | 0.6667 | 1 | 1 | 7 | 0.8571 | 2 | 0.5 | 1 | 1 | 3 | 1 | 3 | 1 | 9 | 0.8889 |
| 1_15572302 | 3 | 0.3333 | 2 | 0.5 | 3 | 0 | 3 | 0 | 11 | 0.1818 | 2 | 0 | 1 | 1 | 3 | 0 | 2 | 0 | 8 | 0.125 |
| 1_15572303 | 3 | 0 | 2 | 0.5 | 4 | 0 | 3 | 0 | 12 | 0.0833 | 1 | 0 | 1 | 0 | 2 | 0.5 | 2 | 0 | 6 | 0.1667 |
| 1_15572309 | 2 | 0.5 | 2 | 1 | 4 | 1 | 1 | 1 | 9 | 0.8889 | 1 | 1 | 1 | 1 | 3 | 1 | 3 | 1 | 8 | 1 |
| 1_15572310 | 3 | 1 | 2 | 1 | 4 | 1 | 2 | 1 | 11 | 1 | 2 | 1 | 1 | 1 | 1 | 1 | 3 | 1 | 7 | 1 |
| 1_15572311 | 2 | 0 | 2 | 1 | 3 | 0.3333 | 3 | 0.6667 | 10 | 0.5 | 2 | 0.5 | NA | NA | 3 | 0.6667 | 3 | 0.6667 | 8 | 0.625 |
| 1_15572315 | 3 | 0.3333 | 2 | 0.5 | 4 | 0 | 1 | 0 | 10 | 0.2 | 1 | 0 | 1 | 0 | 3 | 0.3333 | 3 | 0.6667 | 8 | 0.375 |
| 1_15572338 | 3 | 0 | 2 | 0.5 | 1 | 1 | 2 | 0 | 8 | 0.25 | 2 | 0 | 1 | 1 | 3 | 0.3333 | 3 | 0 | 9 | 0.2222 |
| 1_15572345 | 3 | 0 | 2 | 0.5 | 3 | 0 | 2 | 0 | 10 | 0.1 | 2 | 0.5 | 1 | 1 | 3 | 0.3333 | 2 | 0 | 8 | 0.375 |
| 1_15572347 | 3 | 0 | 2 | 0 | 4 | 0.25 | 2 | 0 | 11 | 0.0909 | 2 | 1 | 1 | 1 | 2 | 1 | 3 | 0 | 8 | 0.625 |
| 1_15572351 | 3 | 0 | 1 | 0 | 4 | 0.5 | 2 | 0 | 10 | 0.2 | 1 | 0 | 1 | 1 | 2 | 0.5 | 3 | 0.3333 | 7 | 0.4286 |
| 1_15572361 | 3 | 1 | 2 | 0.5 | 4 | 0.5 | 1 | 0 | 10 | 0.6 | 2 | 1 | NA | NA | 2 | 1 | 2 | 1 | 6 | 1 |
| 1_15572368 | 3 | 0 | 2 | 0.5 | 2 | 0.5 | 1 | 1 | 8 | 0.375 | 2 | 0.5 | NA | NA | 2 | 1 | 2 | 0.5 | 6 | 0.6667 |
| 1_15572385 | 2 | 0.5 | 2 | 0 | 3 | 0.6667 | 1 | 0 | 8 | 0.375 | 2 | 1 | NA | NA | 2 | 1 | 2 | 1 | 6 | 1 |
| 1_15572386 | 3 | 0.6667 | 2 | 1 | 4 | 1 | NA | NA | 9 | 0.8889 | 2 | 1 | NA | NA | 2 | 1 | 3 | 1 | 7 | 1 |
| 1_15572391 | 2 | 0.5 | 2 | 0.5 | 4 | 0 | 1 | 0 | 9 | 0.2222 | 2 | 0.5 | NA | NA | 2 | 0 | 2 | 0 | 6 | 0.1667 |
| 1_15572396 | 2 | 0 | 2 | 0 | 3 | 0 | 1 | 1 | 8 | 0.125 | 2 | 0.5 | NA | NA | 2 | 0.5 | 3 | 0 | 7 | 0.2857 |
| 1_15572400 | NA | NA | 1 | 1 | 2 | 0.5 | NA | NA | 3 | 0.6667 | 1 | 1 | NA | NA | 2 | 1 | 2 | 0.5 | 5 | 0.8 |
| 1_15572400 | 2 | 0 | 2 | 0 | 4 | 0.25 | 1 | 0 | 9 | 0.1111 | 2 | 0.5 | NA | NA | 2 | 0 | 2 | 0 | 6 | 0.1667 |
| 1_15572406 | 2 | 1 | 2 | 0.5 | 4 | 0.5 | 2 | 0.5 | 10 | 0.6 | 2 | 1 | NA | NA | 2 | 0 | 3 | 0 | 7 | 0.2857 |
| 1_15572412 | 1 | 1 | 1 | 1 | 2 | 1 | 1 | 1 | 5 | 1 | 1 | 1 | NA | NA | 2 | 1 | 2 | 1 | 5 | 1 |
| 1_15572416 | 2 | 1 | 2 | 1 | 3 | 1 | 2 | 1 | 9 | 1 | 2 | 1 | NA | NA | 2 | 1 | 3 | 1 | 7 | 1 |
| 1_15572418 | 2 | 1 | 2 | 1 | 3 | 1 | 2 | 0.5 | 9 | 0.8889 | 2 | 1 | NA | NA | 2 | 1 | 3 | 1 | 7 | 1 |
| 1_15572420 | 1 | 0 | 2 | 0.5 | 3 | 1 | 1 | 0 | 7 | 0.5714 | 2 | 1 | NA | NA | 2 | 0.5 | 2 | 1 | 6 | 0.8333 |
| 1_15572428 | 2 | 1 | 2 | 0.5 | 3 | 0.3333 | 2 | 0.5 | 9 | 0.5556 | 2 | 0.5 | NA | NA | 2 | 0 | 3 | 0.6667 | 7 | 0.4286 |
| 1_15572438 | 2 | 0 | 2 | 0.5 | 3 | 0 | 1 | 0 | 8 | 0.125 | 2 | 0 | NA | NA | 2 | 0.5 | 2 | 0 | 6 | 0.1667 |
| 1_15573778 | 1 | 1 | 1 | 0 | 1 | 0 | 1 | 0 | 4 | 0.25 | 1 | 1 | NA | NA | 1 | 1 | 1 | 0 | 3 | 0.6667 |

| | | | | | | | | | | | | | | | | | | | | |
|---|---|---|---|---|---|---|---|---|---|---|---|---|---|---|---|---|---|---|---|---|
| 1_1557380 | NA | NA | 1 | 0 | 1 | 0 | 1 | 1 | 3 | 0.3333 | 1 | 0 | NA | NA | 1 | 1 | 1 | 0 | 3 | 0.3333 |
| 1_1557381 | NA | NA | 1 | 0 | 1 | 1 | 1 | 0 | 3 | 0.3333 | NA | NA | NA | NA | 1 | 0 | 1 | 1 | 2 | 0.5 |
| 1_1557383 | NA | NA | NA | NA | NA | NA | 1 | 1 | 1 | 1 | 1 | 0 | NA | NA | 1 | 1 | 1 | 1 | 3 | 0.6667 |
| 1_1557384 | NA | NA | NA | NA | NA | NA | 1 | 1 | 1 | 1 | 1 | 0 | NA | NA | 1 | 0 | 1 | 1 | 3 | 0.3333 |
| 1_1557386 | NA | NA | NA | NA | NA | NA | 1 | 1 | 1 | 1 | 1 | 0 | NA | NA | 1 | 1 | 1 | 1 | 3 | 0.6667 |
| 1_1558806 | 3 | 0 | 6 | 0.1667 | 5 | 0 | NA | NA | 14 | 0.0714 | 6 | 0 | 1 | 0 | 7 | 0 | 4 | 0.25 | 18 | 0.0556 |
| 1_1558822 | 2 | 0.5 | 4 | 0 | 2 | 0 | NA | NA | 8 | 0.125 | 2 | 0 | 1 | 0 | 3 | 0 | 3 | 0.3333 | 9 | 0.1111 |
| 1_1558825 | 4 | 0.5 | 1 | 0 | 1 | 0 | 2 | 0 | 8 | 0.25 | 2 | 0 | 2 | 0.5 | 4 | 0.5 | 1 | 1 | 9 | 0.4444 |
| 1_1558826 | 6 | 0.8333 | 3 | 0.6667 | 1 | 1 | 1 | 1 | 11 | 0.8182 | 3 | 1 | 2 | 0 | 3 | 1 | 3 | 1 | 11 | 0.8182 |
| 1_1558826 | 6 | 0.1667 | 2 | 0 | 1 | 0 | 2 | 0 | 11 | 0.0909 | 3 | 0 | 2 | 0.5 | 4 | 0.5 | 3 | 0.3333 | 12 | 0.3333 |
| 1_1558826 | 3 | 0.6667 | 3 | 1 | NA | NA | 1 | 0 | 7 | 0.7143 | 3 | 1 | 1 | 1 | 3 | 0.6667 | 1 | 1 | 8 | 0.875 |
| 1_1558827 | 4 | 1 | 2 | 1 | 1 | 1 | 1 | 1 | 8 | 1 | 3 | 1 | 2 | 1 | 4 | 0.75 | 3 | 1 | 12 | 0.9167 |
| 1_1558828 | 3 | 0.3333 | 3 | 0 | 1 | 1 | 1 | 0 | 8 | 0.25 | 3 | 1 | 1 | 1 | 4 | 0.25 | 2 | 0.5 | 10 | 0.6 |
| 1_1558830 | 4 | 0.75 | 3 | 0.3333 | 1 | 1 | 2 | 0 | 10 | 0.5 | 3 | 0 | 2 | 0.5 | 4 | 0.5 | 2 | 1 | 11 | 0.4545 |
| 1_1558830 | 5 | 0.2 | 3 | 0 | 1 | 0 | 2 | 0.5 | 11 | 0.1818 | 3 | 0 | 2 | 0 | 3 | 0.3333 | 3 | 0 | 11 | 0.0909 |
| 1_1558830 | 5 | 1 | 3 | 1 | 1 | 1 | 2 | 1 | 11 | 1 | 2 | 1 | 2 | 1 | 4 | 0.5 | 2 | 1 | 10 | 0.8 |
| 1_1558831 | 3 | 0 | 2 | 0 | 1 | 0 | 2 | 0.5 | 8 | 0.125 | 2 | 0.5 | 2 | 0 | 3 | 0 | 3 | 0 | 10 | 0.1 |
| 1_1558833 | 4 | 1 | 3 | 1 | NA | NA | 2 | 1 | 9 | 1 | 3 | 1 | 2 | 1 | 3 | 1 | 3 | 1 | 11 | 1 |
| 1_1558833 | 5 | 0.8 | 2 | 0.5 | NA | NA | 2 | 0.5 | 9 | 0.6667 | 3 | 0.6667 | 2 | 0.5 | 3 | 0.6667 | 2 | 0 | 10 | 0.5 |
| 1_1559115 | 4 | 0.25 | 2 | 0 | NA | NA | 1 | 1 | 7 | 0.2857 | 1 | 1 | 2 | 0 | 3 | 0.3333 | 2 | 0 | 8 | 0.25 |
| 1_1559270 | 5 | 0 | 6 | 0.3333 | 11 | 0 | 3 | 0.3333 | 25 | 0.12 | 7 | 0.1429 | 5 | 0 | 3 | 0 | 2 | 0 | 17 | 0.0588 |
| 1_1559271 | 4 | 0 | 2 | 0.5 | 9 | 0.2222 | 3 | 0 | 18 | 0.1667 | 7 | 0.2857 | 4 | 0.75 | 1 | 0 | 2 | 0.5 | 14 | 0.4286 |
| 1_1559271 | 4 | 0.25 | 6 | 0 | 12 | 0.5 | 4 | 0.25 | 26 | 0.3077 | 6 | 0.3333 | 5 | 0.4 | 4 | 0.5 | 3 | 0.6667 | 18 | 0.4444 |
| 1_1559272 | 4 | 0.75 | 4 | 0.75 | 6 | 1 | 4 | 1 | 18 | 0.8889 | 7 | 0.7143 | 5 | 1 | 2 | 0 | 1 | 1 | 15 | 0.7333 |
| 1_1559273 | 5 | 0.8 | 5 | 1 | 10 | 0.9 | 4 | 1 | 24 | 0.9167 | 8 | 1 | 5 | 1 | 2 | 1 | 3 | 1 | 18 | 1 |
| 1_1559273 | 4 | 1 | 6 | 1 | 11 | 1 | 4 | 1 | 25 | 1 | 5 | 1 | 6 | 1 | 3 | 1 | 3 | 1 | 17 | 1 |
| 1_1559273 | 3 | 1 | 5 | 0.6 | 10 | 0.5 | 1 | 1 | 19 | 0.6316 | 8 | 0.5 | 4 | 0.75 | 3 | 0.6667 | 2 | 1 | 17 | 0.6471 |
| 1_1559273 | 4 | 1 | 5 | 1 | 11 | 1 | 4 | 1 | 24 | 1 | 8 | 0.875 | 5 | 1 | 3 | 0.6667 | 1 | 1 | 17 | 0.8824 |
| 1_1559274 | 4 | 1 | 7 | 0.8571 | 13 | 1 | 4 | 1 | 28 | 0.9643 | 9 | 1 | 5 | 1 | 3 | 1 | 3 | 1 | 20 | 1 |
| 1_1559274 | 5 | 0 | 4 | 0.25 | 8 | 0.125 | 5 | 0.2 | 22 | 0.1364 | 8 | 0.125 | 6 | 0.1667 | 2 | 0 | 2 | 0 | 18 | 0.1111 |
| 1_1559275 | 4 | 0.75 | 7 | 0.7143 | 12 | 0.75 | 5 | 0.2 | 28 | 0.6429 | 9 | 0.6667 | 7 | 0.4286 | 3 | 0.6667 | 2 | 0.5 | 21 | 0.5714 |
| 1_1559275 | 3 | 0 | 6 | 0.3333 | 13 | 0.1538 | 4 | 0 | 26 | 0.1538 | 7 | 0.1429 | 7 | 0.1429 | 3 | 0 | 2 | 0 | 19 | 0.1053 |
| 1_1559275 | 6 | 0.8333 | 6 | 1 | 12 | 0.9167 | 4 | 1 | 28 | 0.9286 | 8 | 1 | 6 | 0.8333 | 3 | 1 | 2 | 1 | 19 | 0.9474 |
| 1_1559275 | 5 | 0 | 6 | 0 | 12 | 0.0833 | 5 | 0.2 | 28 | 0.0714 | 8 | 0.125 | 7 | 0.2857 | 2 | 0 | 1 | 0 | 18 | 0.1667 |
| 1_1559277 | 5 | 0 | 8 | 0.25 | 13 | 0.1538 | 5 | 0 | 31 | 0.129 | 8 | 0.375 | 5 | 0 | 3 | 0.3333 | 3 | 0 | 19 | 0.2105 |
| 1_1559287 | 4 | 0.25 | 1 | 0 | 6 | 0 | 2 | 0.5 | 13 | 0.1538 | 5 | 0.2 | 2 | 0 | 2 | 0 | 1 | 0 | 10 | 0.1 |
| 1_1559375 | 1 | 0 | NA | NA | NA | NA | 2 | 0.5 | 3 | 0.3333 | 1 | 0 | NA | NA | 1 | 0 | 1 | 1 | 3 | 0.3333 |
| 1_1559407 | 1 | 0 | NA | NA | NA | NA | 1 | 1 | 2 | 0.5 | 2 | 0 | NA | NA | 1 | 1 | NA | NA | 3 | 0.3333 |
| 1_1559408 | 1 | 1 | 1 | 1 | NA | NA | 2 | 1 | 4 | 1 | 2 | 1 | NA | NA | 1 | 1 | 1 | 0 | 4 | 0.75 |
| 1_1559408 | 1 | 1 | 1 | 1 | NA | NA | 2 | 0.5 | 4 | 0.75 | 1 | 1 | NA | NA | 1 | 0 | 1 | 0 | 3 | 0.3333 |
| 1_1559411 | 1 | 1 | 1 | 1 | 1 | 1 | 2 | 1 | 5 | 1 | 1 | 1 | NA | NA | 1 | 1 | 1 | 1 | 3 | 1 |
| 1_1559411 | 1 | 0 | NA | NA | 1 | 0 | 2 | 1 | 4 | 0.5 | 2 | 1 | NA | NA | NA | NA | 1 | 1 | 3 | 1 |
| 1_1559412 | 1 | 1 | 1 | 0 | 1 | 1 | 2 | 1 | 5 | 0.8 | 2 | 1 | NA | NA | NA | NA | 1 | 0 | 3 | 0.6667 |
| 1_1559413 | 1 | 0 | 1 | 0 | 1 | 1 | 2 | 0.5 | 5 | 0.4 | 2 | 1 | NA | NA | 1 | 1 | NA | NA | 3 | 1 |
| 1_1559413 | NA | NA | 1 | 1 | 1 | 1 | 1 | 1 | 3 | 1 | 2 | 1 | NA | NA | 1 | 1 | 1 | 1 | 4 | 1 |
| 1_1559415 | 1 | 1 | 1 | 1 | 1 | 1 | 2 | 1 | 5 | 1 | 2 | 1 | NA | NA | 1 | 1 | 1 | 1 | 4 | 1 |
| 1_1559415 | NA | NA | NA | NA | NA | NA | 1 | 1 | 1 | 1 | 2 | 0.5 | NA | NA | 1 | 0 | 1 | 1 | 4 | 0.5 |
| 1_1559415 | 1 | 0 | 1 | 1 | 1 | 1 | 2 | 1 | 5 | 0.8 | 2 | 0.5 | NA | NA | 1 | 1 | 1 | 1 | 4 | 0.75 |
| 1_1559419 | 1 | 1 | 1 | 0 | 1 | 1 | 2 | 0 | 5 | 0.4 | 2 | 1 | NA | NA | NA | NA | 1 | 0 | 3 | 0.6667 |
| 1_1559527 | NA | NA | 1 | 0 | 2 | 0.5 | NA | NA | 3 | 0.3333 | NA | NA | 1 | 0 | 2 | 0.5 | NA | NA | 3 | 0.3333 |
| 1_1559527 | NA | NA | 1 | 1 | 1 | 0 | NA | NA | 2 | 0.5 | NA | NA | 1 | 0 | 2 | 0.5 | NA | NA | 3 | 0.3333 |
| 1_1559535 | NA | NA | 1 | 0 | 1 | 1 | NA | NA | 2 | 0.5 | NA | NA | 1 | 0 | 2 | 0.5 | NA | NA | 3 | 0.3333 |
| 1_1559548 | NA | NA | NA | NA | 1 | 1 | NA | NA | 1 | 1 | NA | NA | 1 | 1 | NA | NA | NA | NA | 1 | 1 |
| 1_1560755 | NA | NA | NA | NA | NA | NA | 1 | 1 | 1 | 1 | NA | NA | NA | NA | NA | NA | 3 | 1 | 3 | 1 |
| 1_1560762 | NA | NA | NA | NA | NA | NA | 1 | 1 | 1 | 1 | NA | NA | NA | NA | NA | NA | 2 | 1 | 2 | 1 |
| 1_1560764 | NA | NA | NA | NA | NA | NA | 1 | 1 | 1 | 1 | NA | NA | NA | NA | NA | NA | 2 | 0.5 | 2 | 0.5 |
| 1_1560772 | NA | NA | NA | NA | NA | NA | 1 | 1 | 1 | 1 | NA | NA | NA | NA | NA | NA | 1 | 1 | 1 | 1 |
| 1_1560776 | NA | NA | NA | NA | NA | NA | 1 | 1 | 1 | 1 | NA | NA | NA | NA | NA | NA | 1 | 1 | 1 | 1 |
| 1_1562523 | 2 | 0.5 | 1 | 1 | 5 | 0.8 | 3 | 1 | 11 | 0.8182 | 2 | 1 | NA | NA | NA | NA | 1 | 0 | 3 | 0.6667 |
| 1_1562526 | NA | NA | 1 | 0 | 4 | 0.25 | 4 | 0.5 | 9 | 0.3333 | 2 | 0.5 | NA | NA | NA | NA | NA | NA | 2 | 0.5 |
| 1_1562528 | NA | NA | 1 | 1 | 3 | 0.6667 | 3 | 0.6667 | 7 | 0.7143 | 3 | 0.6667 | NA | NA | NA | NA | 1 | 0 | 4 | 0.5 |
| 1_1562545 | NA | NA | NA | NA | 2 | 0.5 | 1 | 0 | 3 | 0.3333 | 3 | 0.3333 | NA | NA | NA | NA | NA | NA | 3 | 0.3333 |
| 1_1562578 | NA | NA | 1 | 0 | 3 | 0 | 4 | 0.25 | 8 | 0.125 | NA | NA | 1 | 0 | 1 | 1 | 1 | 0 | 3 | 0.3333 |
| 1_1562583 | 1 | 1 | NA | NA | 4 | 0.5 | 2 | 0.5 | 7 | 0.5714 | NA | NA | 1 | 1 | 1 | 0 | 2 | 1 | 4 | 0.75 |
| 1_1562584 | NA | NA | 1 | 0 | 3 | 0.3333 | 2 | 0.5 | 6 | 0.3333 | NA | NA | 1 | 0 | 1 | 1 | 2 | 0 | 4 | 0.25 |

| | | | | | | | | | | | | | | | | | | | | |
|---|---|---|---|---|---|---|---|---|---|---|---|---|---|---|---|---|---|---|---|---|
| 1_15625864 | NA | NA | NA | NA | 4 | 0.25 | 2 | 1 | 6 | 0.5 | NA | NA | 1 | 0 | 1 | 1 | 1 | 1 | 3 | 0.6667 |
| 1_15646893 | 1 | 1 | NA | NA | NA | NA | NA | NA | 1 | 1 | 1 | 1 | NA | NA | NA | NA | 1 | 0 | 2 | 0.5 |
| 1_15646918 | 1 | 1 | NA | NA | NA | NA | NA | NA | 1 | 1 | 2 | 1 | 1 | 1 | NA | NA | NA | NA | 3 | 1 |
| 1_15646936 | 1 | 1 | NA | NA | NA | NA | NA | NA | 1 | 1 | 1 | 1 | 1 | 1 | NA | NA | 1 | 1 | 3 | 1 |
| 1_15646977 | 1 | 1 | NA | NA | NA | NA | NA | NA | 1 | 1 | 2 | 0 | 1 | 1 | NA | NA | 1 | 0 | 4 | 0.25 |
| 1_15647012 | 1 | 1 | NA | NA | NA | NA | NA | NA | 1 | 1 | 2 | 1 | 1 | 0 | NA | NA | 1 | 1 | 4 | 0.75 |
| 1_15647022 | 1 | 1 | NA | NA | NA | NA | NA | NA | 1 | 1 | 2 | 0.5 | 1 | 1 | NA | NA | 1 | 0 | 4 | 0.5 |
| 1_15647034 | 1 | 1 | NA | NA | NA | NA | 1 | 1 | 2 | 1 | 2 | 1 | 2 | 1 | NA | NA | 1 | 1 | 5 | 1 |
| 1_15647037 | 1 | 1 | NA | NA | NA | NA | 2 | 1 | 3 | 1 | 2 | 1 | 2 | 1 | 1 | 1 | 1 | 0 | 6 | 0.8333 |
| 1_15647258 | 1 | 0 | NA | NA | 2 | 0.5 | 4 | 0 | 7 | 0.1429 | 3 | 0.3333 | NA | NA | 2 | 0 | 1 | 0 | 6 | 0.1667 |
| 1_15648144 | NA | NA | 2 | 0.5 | NA | NA | NA | NA | 2 | 0.5 | NA | NA | 1 | 0 | NA | NA | 2 | 0.5 | 3 | 0.3333 |
| 1_15648158 | NA | NA | 2 | 0.5 | NA | NA | NA | NA | 2 | 0.5 | NA | NA | NA | NA | NA | NA | 2 | 0.5 | 2 | 0.5 |
| 1_15648264 | 2 | 0 | 3 | 0 | 1 | 1 | NA | NA | 6 | 0.1667 | 1 | 0 | 1 | 1 | NA | NA | 2 | 0 | 4 | 0.25 |
| 1_15648280 | 2 | 0 | 3 | 0.3333 | 1 | 0 | NA | NA | 6 | 0.1667 | 1 | 0 | 1 | 0 | NA | NA | 1 | 1 | 3 | 0.3333 |
| 1_15648280 | 1 | 1 | 3 | 0.3333 | 1 | 1 | NA | NA | 5 | 0.6 | 1 | 1 | 1 | 1 | NA | NA | 1 | 0 | 3 | 0.6667 |
| 1_15648292 | 2 | 1 | 3 | 1 | 1 | 1 | NA | NA | 6 | 1 | 1 | 1 | 1 | 1 | NA | NA | 1 | 1 | 3 | 1 |
| 1_15648303 | 1 | 1 | 2 | 0 | 1 | 1 | NA | NA | 4 | 0.5 | 1 | 0 | 1 | 0 | NA | NA | 2 | 0.5 | 4 | 0.25 |
| 1_15648308 | 1 | 1 | 2 | 0.5 | 1 | 1 | NA | NA | 4 | 0.75 | 1 | 0 | 1 | 1 | NA | NA | 2 | 0.5 | 4 | 0.5 |
| 1_15648309 | 1 | 1 | 2 | 0 | NA | NA | NA | NA | 3 | 0.3333 | 1 | 0 | 1 | 1 | NA | NA | 2 | 0 | 4 | 0.25 |
| 1_15648322 | 2 | 1 | 1 | 0 | 1 | 1 | NA | NA | 4 | 0.75 | 1 | 1 | 1 | 0 | NA | NA | 1 | 1 | 3 | 0.6667 |
| 1_15648325 | 2 | 0.5 | 3 | 0.6667 | NA | NA | NA | NA | 5 | 0.6 | 1 | 1 | 1 | 0 | NA | NA | 1 | 1 | 3 | 0.6667 |
| 1_15650631 | 5 | 0.4 | 8 | 0.375 | 8 | 0 | 13 | 0.4615 | 34 | 0.3235 | 8 | 0 | 4 | 0.25 | 7 | 0.2857 | 8 | 0 | 27 | 0.1111 |
| 1_15650647 | 8 | 0.75 | 8 | 0.625 | 9 | 0.5556 | 14 | 0.5 | 39 | 0.5897 | 9 | 0.4444 | 7 | 0.4286 | 6 | 0.6667 | 9 | 0.6667 | 31 | 0.5484 |
| 1_15650741 | 5 | 0 | 5 | 0.2 | 5 | 0.4 | 12 | 0.5833 | 27 | 0.3704 | 6 | 0.1667 | 6 | 0.3333 | 2 | 0.5 | 5 | 0.2 | 19 | 0.2632 |
| 1_15650825 | 1 | 0 | 6 | 0.3333 | 6 | 0.1667 | 9 | 0.2222 | 22 | 0.2273 | 7 | 0.2857 | 4 | 0.25 | 2 | 0 | 8 | 0 | 21 | 0.1429 |
| 1_15650848 | 2 | 1 | 7 | 0.1429 | 6 | 0.8333 | 9 | 0.3333 | 24 | 0.4583 | 6 | 0 | 3 | 0 | 2 | 0.5 | 4 | 0.25 | 15 | 0.1333 |
| 1_15650859 | 2 | 0 | 6 | 0.1667 | 6 | 0.1667 | 9 | 0.2222 | 23 | 0.1739 | 6 | 0 | 2 | 0.5 | 2 | 0 | 6 | 0.1667 | 16 | 0.125 |
| 1_15650873 | 2 | 0 | 5 | 0.6 | 3 | 0.3333 | 6 | 0.5 | 16 | 0.4375 | 7 | 0.1429 | 3 | 0.3333 | 2 | 0 | 5 | 0.4 | 17 | 0.2353 |
| 1_15650881 | 2 | 0.5 | 3 | 0.3333 | 6 | 0.8333 | 6 | 1 | 17 | 0.7647 | 5 | 0.4 | 3 | 0.6667 | NA | NA | 4 | 0.75 | 12 | 0.5833 |
| 1_15650893 | 1 | 0 | 5 | 0 | 6 | 0.3333 | 5 | 0.4 | 17 | 0.2353 | 7 | 0.2857 | 3 | 0.3333 | 2 | 0 | 3 | 0 | 15 | 0.2 |
| 1_15650895 | 1 | 1 | 4 | 1 | 4 | 1 | 7 | 1 | 16 | 1 | 6 | 1 | 3 | 1 | 1 | 1 | 4 | 1 | 14 | 1 |
| 1_15650908 | 1 | 0 | 4 | 0.25 | 6 | 0.3333 | 7 | 0 | 18 | 0.1667 | 6 | 0.1667 | 3 | 0 | 1 | 0 | 5 | 0.4 | 15 | 0.2 |
| 1_15650909 | 1 | 0 | 4 | 0.5 | 5 | 1 | 6 | 0.8333 | 16 | 0.75 | 6 | 0.6667 | 2 | 1 | 1 | 1 | 5 | 1 | 14 | 0.8571 |
| 1_15650918 | 1 | 1 | 5 | 0.6 | 5 | 0.6 | 8 | 0.5 | 19 | 0.5789 | 5 | 0.4 | 3 | 1 | 1 | 1 | 5 | 1 | 14 | 0.7857 |
| 1_15653329 | 1 | 0 | 2 | 0.5 | 4 | 0.25 | 6 | 0.3333 | 13 | 0.3077 | 1 | 1 | 2 | 0 | 1 | 1 | 4 | 0.25 | 8 | 0.375 |
| 1_15653344 | 2 | 0.5 | NA | NA | 4 | 0.75 | 5 | 1 | 11 | 0.8182 | 2 | 1 | 2 | 0.5 | NA | NA | 3 | 0.6667 | 7 | 0.7143 |
| 1_15653347 | 2 | 0 | 1 | 0 | 2 | 0.5 | 3 | 0 | 8 | 0.125 | 1 | 0 | 2 | 0.5 | 1 | 0 | 4 | 0 | 8 | 0.125 |
| 1_15653350 | 2 | 0.5 | 1 | 0 | 4 | 0.25 | 5 | 0 | 12 | 0.1667 | 2 | 0 | 2 | 0 | 1 | 0 | 5 | 0.2 | 10 | 0.1 |
| 1_15653353 | 1 | 0 | 1 | 1 | 3 | 0.3333 | 6 | 0.5 | 11 | 0.4545 | 3 | 0.3333 | 2 | 0.5 | 1 | 1 | 4 | 0.75 | 10 | 0.6 |
| 1_15663203 | 3 | 0.3333 | 4 | 0 | 2 | 0 | 6 | 0 | 15 | 0.0667 | 5 | 0.2 | 3 | 0 | 5 | 0.6 | 7 | 0.1429 | 20 | 0.25 |
| 1_15663204 | 3 | 0 | 5 | 0.2 | 2 | 0 | 6 | 0.3333 | 16 | 0.1875 | 5 | 0.2 | 3 | 0 | 6 | 0 | 7 | 0.1429 | 21 | 0.0952 |
| 1_15663348 | 2 | 0.5 | 3 | 0 | 1 | 0 | 3 | 1 | 9 | 0.4444 | 1 | 1 | 2 | 0.5 | 4 | 0.5 | 3 | 0.3333 | 10 | 0.5 |
| 1_15663369 | 2 | 0.5 | 2 | 0 | 1 | 0 | 2 | 0.5 | 7 | 0.2857 | 2 | 0.5 | 3 | 0 | 5 | 0 | 2 | 0 | 12 | 0.0833 |
| 1_15663379 | 1 | 1 | 1 | 0 | NA | NA | 1 | 1 | 3 | 0.6667 | 2 | 1 | 3 | 0 | 4 | 0.25 | 2 | 0 | 11 | 0.2727 |
| 1_15663385 | 1 | 0 | NA | NA | NA | NA | 1 | 1 | 2 | 0.5 | 2 | 0.5 | 2 | 0 | 3 | 0.3333 | NA | NA | 7 | 0.2857 |
| 1_15663651 | 1 | 1 | 1 | 1 | NA | NA | NA | NA | 2 | 1 | 1 | 0 | 1 | 1 | NA | NA | 2 | 0 | 4 | 0.25 |
| 1_15663653 | 1 | 0 | 1 | 1 | 1 | 1 | NA | NA | 3 | 0.6667 | 1 | 1 | 1 | 1 | 3 | 0.6667 | 2 | 1 | 7 | 0.8571 |
| 1_15696342 | 1 | 0 | 12 | 0.25 | 5 | 0.2 | 5 | 0 | 23 | 0.1739 | 3 | 0.3333 | 7 | 0.1429 | 4 | 0 | 7 | 0.2857 | 21 | 0.1905 |
| 1_15696344 | 3 | 0 | 14 | 0.0714 | 6 | 0.1667 | 6 | 0.1667 | 29 | 0.1034 | 4 | 0 | 7 | 0.1429 | 4 | 0 | 9 | 0.3333 | 24 | 0.1667 |
| 1_15696460 | 1 | 0 | 9 | 0.1111 | 3 | 0 | 6 | 0 | 19 | 0.0526 | 2 | 0 | 4 | 0 | 3 | 0 | 8 | 0.125 | 17 | 0.0588 |
| 1_15696756 | NA | NA | 3 | 1 | 2 | 0 | NA | NA | 5 | 0.6 | NA | NA | 3 | 1 | 1 | 0 | 2 | 1 | 6 | 0.8333 |
| 1_15696769 | NA | NA | 3 | 0.3333 | 2 | 1 | 1 | 0 | 6 | 0.5 | NA | NA | 3 | 0.6667 | 1 | 0 | 2 | 1 | 6 | 0.6667 |
| 1_15696811 | NA | NA | 2 | 1 | 1 | 1 | NA | NA | 3 | 1 | NA | NA | 2 | 1 | 1 | 0 | 1 | 0 | 4 | 0.5 |
| 1_15696826 | NA | NA | 1 | 1 | 1 | 1 | NA | NA | 2 | 1 | NA | NA | 1 | 1 | 1 | 1 | 1 | 1 | 3 | 1 |
| 1_15698468 | NA | NA | 1 | 1 | 1 | 1 | NA | NA | 2 | 1 | NA | NA | NA | NA | NA | NA | 1 | 1 | 1 | 1 |
| 1_15698483 | NA | NA | 1 | 1 | NA | NA | NA | NA | 1 | 1 | NA | NA | 2 | 0 | 1 | 0 | 1 | 1 | 4 | 0.25 |
| 1_15700354 | NA | NA | 1 | 1 | NA | NA | NA | NA | 1 | 1 | NA | NA | 2 | 1 | 1 | 1 | NA | NA | 3 | 1 |
| 1_15700370 | NA | NA | 1 | 1 | NA | NA | NA | NA | 1 | 1 | NA | NA | 1 | 1 | NA | NA | NA | NA | 1 | 1 |
| 1_15724759 | 7 | 0.4286 | 1 | 0 | 2 | 1 | 1 | 1 | 11 | 0.5455 | 2 | 0 | 3 | 0.3333 | 1 | 1 | 4 | 0.5 | 10 | 0.4 |
| 1_15724769 | 7 | 0.2857 | 1 | 1 | 6 | 0.5 | 1 | 1 | 15 | 0.4667 | 4 | 0.5 | 3 | 0.3333 | 1 | 0 | 4 | 0.5 | 12 | 0.4167 |
| 1_15807286 | 6 | 0.1667 | 2 | 0 | 4 | 0 | 2 | 0.5 | 14 | 0.1429 | 9 | 0.1111 | 6 | 0.1667 | 2 | 0 | 3 | 0 | 20 | 0.1 |
| 1_15807305 | 10 | 0.1 | 13 | 0.2308 | 13 | 0 | 9 | 0.1111 | 45 | 0.1111 | 18 | 0.2778 | 12 | 0.1667 | 7 | 0.2857 | 9 | 0.2222 | 46 | 0.2391 |
| 1_15807338 | 9 | 0.6667 | 9 | 0.5556 | 10 | 0.6 | 8 | 0.875 | 36 | 0.6667 | 12 | 0.5833 | 8 | 0.5 | 6 | 1 | 6 | 0.8333 | 32 | 0.6875 |
| 1_15807343 | 6 | 0 | 9 | 0.4444 | 11 | 0.2727 | 6 | 0.3333 | 32 | 0.2812 | 15 | 0.2 | 10 | 0.4 | 7 | 0.5714 | 6 | 0.5 | 38 | 0.3684 |
| 1_15807343 | 9 | 0.8889 | 10 | 0.8 | 11 | 0.9091 | 6 | 0.6667 | 36 | 0.8333 | 16 | 1 | 8 | 1 | 8 | 0.75 | 6 | 1 | 38 | 0.9474 |

| | | | | | | | | | | | | | | | | | | | | |
|---|---|---|---|---|---|---|---|---|---|---|---|---|---|---|---|---|---|---|---|---|
| 1_15807361 | 9 | 0.5556 | 12 | 0.5 | 9 | 0.6667 | 7 | 0.4286 | 37 | 0.5405 | 14 | 0.4286 | 10 | 0.6 | 9 | 0.6667 | 7 | 0.7143 | 40 | 0.575 |
| 1_15807362 | 7 | 0.8571 | 7 | 0.7143 | 6 | 0.8333 | 7 | 0.5714 | 27 | 0.7407 | 13 | 1 | 9 | 1 | 6 | 1 | 4 | 0.75 | 32 | 0.9688 |
| 1_15807366 | 8 | 0.75 | 11 | 0.7273 | 10 | 0.8 | 7 | 0.5714 | 36 | 0.7222 | 15 | 0.6 | 11 | 0.6364 | 8 | 0.75 | 7 | 0.8571 | 41 | 0.6829 |
| 1_15807378 | 8 | 0.5 | 11 | 0.5455 | 9 | 0.4444 | 7 | 0.7143 | 35 | 0.5429 | 15 | 0.5333 | 10 | 0.4 | 7 | 0.4286 | 6 | 0.6667 | 38 | 0.5 |
| 1_15807394 | 8 | 0.125 | 12 | 0.0833 | 8 | 0 | 7 | 0.1429 | 35 | 0.0857 | 15 | 0.1333 | 10 | 0.2 | 7 | 0.4286 | 6 | 0.1667 | 38 | 0.2105 |
| 1_15807397 | 8 | 1 | 13 | 0.9231 | 9 | 1 | 7 | 1 | 37 | 0.973 | 15 | 0.9333 | 10 | 1 | 7 | 1 | 6 | 1 | 38 | 0.9737 |
| 1_15807400 | 6 | 1 | 10 | 0.9 | 9 | 0.8889 | 6 | 1 | 31 | 0.9355 | 11 | 0.8182 | 8 | 0.75 | 5 | 0.6 | 6 | 1 | 30 | 0.8 |
| 1_15807411 | 6 | 0.5 | 12 | 0.0833 | 8 | 0.375 | 4 | 0.25 | 30 | 0.2667 | 11 | 0.2727 | 9 | 0 | 4 | 0.25 | 6 | 0.3333 | 30 | 0.2 |
| 1_15807414 | 5 | 0.2 | 13 | 0.0769 | 7 | 0.1429 | 5 | 0 | 30 | 0.1 | 14 | 0.0714 | 5 | 0.2 | 5 | 0 | 7 | 0 | 31 | 0.0645 |
| 1_15807419 | 6 | 0.1667 | 11 | 0.0909 | 7 | 0.2857 | 3 | 0 | 27 | 0.1481 | 12 | 0.1667 | 7 | 0 | 3 | 0 | 6 | 0.1667 | 28 | 0.1071 |
| 1_15807421 | 8 | 0 | 12 | 0.3333 | 8 | 0 | 6 | 0.3333 | 34 | 0.1765 | 15 | 0.4 | 8 | 0.25 | 7 | 0.1429 | 7 | 0.2857 | 37 | 0.2973 |
| 1_15807422 | 4 | 0 | 12 | 0.25 | 6 | 0 | 5 | 0 | 27 | 0.1111 | 13 | 0.0769 | 7 | 0.1429 | 7 | 0 | 5 | 0 | 32 | 0.0625 |
| 1_15807423 | 6 | 1 | 12 | 1 | 8 | 1 | 5 | 0.8 | 31 | 0.9677 | 13 | 1 | 9 | 1 | 7 | 1 | 5 | 1 | 34 | 1 |
| 1_15807427 | 7 | 1 | 12 | 0.9167 | 7 | 1 | 6 | 0.8333 | 32 | 0.9375 | 13 | 0.8462 | 9 | 1 | 7 | 1 | 7 | 1 | 36 | 0.9444 |
| 1_15807428 | 8 | 0.125 | 12 | 0.1667 | 7 | 0.4286 | 5 | 0 | 32 | 0.1875 | 14 | 0.2143 | 8 | 0.125 | 7 | 0.1429 | 7 | 0.4286 | 36 | 0.2222 |
| 1_15807438 | 6 | 0.3333 | 11 | 0.1818 | 4 | 0 | 2 | 1 | 23 | 0.2609 | 13 | 0.2308 | 5 | 0.8 | 5 | 0.6 | 3 | 0 | 26 | 0.3846 |
| 1_15807458 | 6 | 0 | 9 | 0.3333 | 6 | 0 | 6 | 0 | 27 | 0.1111 | 13 | 0.2308 | 9 | 0 | 6 | 0 | 4 | 0.75 | 32 | 0.1875 |
| 1_15807461 | 5 | 0.2 | 6 | 0.3333 | 4 | 0 | 5 | 0 | 20 | 0.15 | 7 | 0.4286 | 7 | 0.2857 | 5 | 0 | 4 | 0.25 | 23 | 0.2609 |
| 1_15807465 | 5 | 1 | 8 | 0.75 | 5 | 0.8 | 5 | 1 | 23 | 0.8696 | 12 | 0.75 | 8 | 0.75 | 5 | 1 | 5 | 0.8 | 30 | 0.8 |
| 1_15807470 | 5 | 0.2 | 8 | 0.125 | 4 | 0 | 6 | 0.1667 | 23 | 0.1304 | 10 | 0.1 | 8 | 0.125 | 5 | 0.2 | 4 | 0.25 | 27 | 0.1481 |
| 1_15807700 | 4 | 0.5 | 6 | 0.3333 | 6 | 0.3333 | 5 | 0.2 | 21 | 0.3333 | 11 | 0.2727 | 7 | 0.1429 | 5 | 0.6 | 4 | 0.25 | 27 | 0.2963 |
| 1_15818321 | 1 | 1 | 4 | 0.5 | 2 | 0.5 | 3 | 0.3333 | 10 | 0.5 | 4 | 0 | 4 | 0.25 | 4 | 0.5 | 3 | 0.3333 | 15 | 0.2667 |
| 1_15818329 | 1 | 1 | 3 | 0.3333 | 2 | 0.5 | 2 | 0.5 | 8 | 0.5 | 2 | 0.5 | 4 | 0.75 | 4 | 0.5 | 2 | 0.5 | 12 | 0.5833 |
| 1_15818349 | 1 | 1 | 2 | 0.5 | 1 | 1 | 3 | 0.6667 | 7 | 0.7143 | 3 | 0 | 4 | 1 | 3 | 1 | 2 | 1 | 12 | 0.75 |
| 1_15818350 | 1 | 0 | NA | NA | 2 | 0.5 | 3 | 0.3333 | 6 | 0.3333 | 3 | 0.6667 | 4 | 0 | 1 | 0 | 3 | 0 | 11 | 0.1818 |
| 1_15818369 | 1 | 0 | 2 | 0 | 1 | 0 | 3 | 0.3333 | 7 | 0.1429 | 2 | 0 | 3 | 0 | 2 | 0 | 2 | 0.5 | 9 | 0.1111 |
| 1_15832993 | 2 | 0 | 4 | 0.25 | 2 | 0.5 | 1 | 1 | 9 | 0.3333 | 2 | 0 | 2 | 0.5 | 1 | 1 | 5 | 0.2 | 10 | 0.3 |
| 1_15851613 | 10 | 0.6 | 8 | 0.75 | 6 | 0.5 | 7 | 0.5714 | 31 | 0.6129 | 5 | 0.4 | 9 | 0.7778 | 8 | 1 | 7 | 0.7143 | 29 | 0.7586 |
| 1_15851622 | 8 | 0.875 | 10 | 0.8 | 5 | 0.6 | 6 | 0.8333 | 29 | 0.7931 | 8 | 1 | 7 | 1 | 9 | 0.8889 | 4 | 1 | 28 | 0.9643 |
| 1_15851622 | 9 | 0.8889 | 5 | 0.6 | 4 | 1 | 5 | 0.8 | 23 | 0.8261 | 7 | 0.7143 | 4 | 1 | 3 | 1 | 7 | 1 | 21 | 0.9048 |
| 1_15851624 | 11 | 0.0909 | 14 | 0.2143 | 6 | 0.1667 | 8 | 0.25 | 39 | 0.1795 | 10 | 0 | 16 | 0.1875 | 11 | 0.2727 | 8 | 0 | 45 | 0.1333 |
| 1_15851629 | 11 | 0.2727 | 12 | 0.1667 | 7 | 0.1429 | 7 | 0.1429 | 37 | 0.1892 | 10 | 0.1 | 12 | 0.0833 | 11 | 0.1818 | 8 | 0.125 | 41 | 0.122 |
| 1_15851630 | 11 | 0.0909 | 13 | 0.0769 | 7 | 0 | 7 | 0.1429 | 38 | 0.0789 | 10 | 0.3 | 13 | 0.2308 | 12 | 0.4167 | 7 | 0.4286 | 42 | 0.3333 |
| 1_15851631 | 10 | 0.2 | 11 | 0.2727 | 7 | 0.1429 | 8 | 0.125 | 36 | 0.1944 | 10 | 0.1 | 13 | 0.3077 | 12 | 0.1667 | 4 | 0 | 39 | 0.1795 |
| 1_15851633 | 10 | 0.3 | 11 | 0.1818 | 7 | 0.2857 | 7 | 0.2857 | 35 | 0.2571 | 8 | 0.25 | 13 | 0.4615 | 12 | 0.25 | 7 | 0.1429 | 40 | 0.3 |
| 1_15851634 | 8 | 0.125 | 7 | 0.2857 | 6 | 0.5 | 5 | 0 | 26 | 0.2308 | 9 | 0.1111 | 8 | 0.375 | 11 | 0.0909 | 6 | 0.3333 | 34 | 0.2059 |
| 1_15851640 | 9 | 0.3333 | 10 | 0.5 | 6 | 0.5 | 8 | 0.375 | 33 | 0.4242 | 7 | 0.5714 | 11 | 0.8182 | 13 | 1 | 7 | 0.7143 | 38 | 0.8158 |
| 1_15851641 | 6 | 0.1667 | 10 | 0 | 6 | 0 | 5 | 0.2 | 27 | 0.0741 | 7 | 0.2857 | 12 | 0.3333 | 7 | 0 | 6 | 0 | 32 | 0.1875 |
| 1_15851675 | 6 | 0.3333 | 12 | 0.1667 | 8 | 0.5 | 7 | 0.7143 | 33 | 0.3939 | 8 | 0.125 | 10 | 0.3 | 13 | 0.1538 | 8 | 0.375 | 39 | 0.2308 |
| 1_15851690 | 11 | 0.1818 | 13 | 0.1538 | 11 | 0.3636 | 9 | 0.1111 | 44 | 0.2045 | 9 | 0.4444 | 17 | 0.4118 | 14 | 0.3571 | 10 | 0.3 | 50 | 0.38 |
| 1_15851695 | 11 | 0.7273 | 13 | 0.9231 | 10 | 0.8 | 10 | 0.7 | 44 | 0.7955 | 10 | 0.8 | 20 | 0.75 | 16 | 0.875 | 11 | 0.8182 | 57 | 0.807 |
| 1_15851696 | 8 | 0.25 | 9 | 0.2222 | 7 | 0.2857 | 4 | 0.25 | 28 | 0.25 | 6 | 0.3333 | 16 | 0.375 | 10 | 0.6 | 7 | 0.5714 | 39 | 0.4615 |
| 1_15851700 | 12 | 0.3333 | 13 | 0.3846 | 9 | 0.4444 | 10 | 0.4 | 44 | 0.3864 | 11 | 0.1818 | 19 | 0.4737 | 15 | 0.6 | 11 | 0.7273 | 56 | 0.5 |
| 1_15851725 | 11 | 0.3636 | 18 | 0.3333 | 13 | 0.3846 | 10 | 0.2 | 52 | 0.3269 | 12 | 0.25 | 22 | 0.4545 | 18 | 0.4444 | 11 | 0.4545 | 63 | 0.4127 |
| 1_15851752 | 12 | 0.4167 | 17 | 0.4706 | 13 | 0.5385 | 12 | 0.4167 | 54 | 0.463 | 12 | 0.5 | 23 | 0.6522 | 19 | 0.5263 | 13 | 0.8462 | 67 | 0.6269 |
| 1_15851757 | 9 | 0 | 18 | 0.0556 | 11 | 0.1818 | 11 | 0 | 49 | 0.0612 | 11 | 0.2727 | 24 | 0 | 18 | 0.1111 | 11 | 0.0909 | 64 | 0.0938 |
| 1_15851761 | 13 | 0.0769 | 19 | 0.1579 | 11 | 0.2727 | 13 | 0.0769 | 56 | 0.1429 | 11 | 0.0909 | 25 | 0.2 | 20 | 0.2 | 13 | 0.1538 | 69 | 0.1739 |
| 1_15851762 | 13 | 0 | 20 | 0.25 | 12 | 0 | 13 | 0.0769 | 58 | 0.1034 | 12 | 0.0833 | 25 | 0.04 | 19 | 0.1579 | 13 | 0 | 69 | 0.0725 |
| 1_15851837 | 9 | 0.1111 | 13 | 0.2308 | 11 | 0.4545 | 9 | 0.2222 | 42 | 0.2619 | 11 | 0.0909 | 21 | 0.4762 | 17 | 0.2353 | 8 | 0.25 | 57 | 0.2982 |
| 1_15851838 | 12 | 0.0833 | 21 | 0.1905 | 11 | 0.0909 | 14 | 0.1429 | 58 | 0.1379 | 12 | 0 | 27 | 0.037 | 18 | 0.1667 | 12 | 0.0833 | 69 | 0.0725 |
| 1_15851847 | 18 | 0.1111 | 25 | 0.16 | 15 | 0.0667 | 14 | 0 | 72 | 0.0972 | 13 | 0.0769 | 27 | 0.1111 | 23 | 0.087 | 15 | 0.1333 | 78 | 0.1026 |
| 1_15851851 | 16 | 0.1875 | 23 | 0.0435 | 16 | 0 | 11 | 0 | 66 | 0.0606 | 9 | 0.2222 | 22 | 0 | 22 | 0.0455 | 13 | 0.0769 | 66 | 0.0606 |
| 1_15851887 | 19 | 0.5789 | 23 | 0.5217 | 18 | 0.4444 | 14 | 0.3571 | 74 | 0.4865 | 14 | 0.3571 | 20 | 0.45 | 29 | 0.5517 | 15 | 0.7333 | 78 | 0.5256 |
| 1_15851894 | 13 | 0.3846 | 18 | 0.3889 | 16 | 0.5 | 11 | 0.2727 | 58 | 0.3966 | 15 | 0.3333 | 20 | 0.35 | 22 | 0.3636 | 14 | 0.5714 | 71 | 0.3944 |
| 1_15915620 | 9 | 0.1111 | 13 | 0.3077 | 7 | 0 | 7 | 0.2857 | 36 | 0.1944 | 7 | 0.1429 | 6 | 0.3333 | 4 | 0.25 | 12 | 0.3333 | 29 | 0.2759 |
| 1_15917255 | 10 | 0.6 | 12 | 0.5833 | 7 | 0.7143 | 7 | 0.7143 | 36 | 0.6389 | 4 | 0.25 | 7 | 0.8571 | 7 | 0.5714 | 12 | 0.6667 | 30 | 0.6333 |
| 1_15917260 | 7 | 0.7143 | 7 | 0.8571 | 7 | 0.8571 | 6 | 0.8333 | 27 | 0.8148 | 5 | 0.6 | 3 | 0.6667 | 6 | 0.6667 | 12 | 0.9167 | 26 | 0.7692 |
| 1_15917323 | 1 | 1 | NA | NA | NA | NA | 2 | 0 | 3 | 0.3333 | NA | NA | 2 | 1 | 1 | 0 | 1 | 0 | 4 | 0.5 |
| 1_15967788 | NA | NA | NA | NA | NA | NA | 1 | 1 | 1 | 1 | NA | NA | NA | NA | NA | NA | 1 | 1 | 1 | 1 |
| 1_15967980 | NA | NA | NA | NA | NA | NA | 1 | 1 | 1 | 1 | NA | NA | NA | NA | NA | NA | 1 | 1 | 1 | 1 |
| 1_15967983 | NA | NA | NA | NA | NA | NA | 1 | 1 | 1 | 1 | NA | NA | NA | NA | 1 | 1 | 1 | 1 | 2 | 1 |
| 1_15967989 | NA | NA | NA | NA | NA | NA | 2 | 0.5 | 2 | 0.5 | NA | NA | NA | NA | 1 | 1 | 1 | 0 | 2 | 0.5 |
| 1_15974684 | NA | NA | NA | NA | 1 | 1 | NA | NA | 1 | 1 | NA | NA | NA | NA | 1 | 1 | NA | NA | 1 | 1 |
| 1_15975199 | 1 | 0 | 1 | 0 | 3 | 0.6667 | 3 | 0.3333 | 8 | 0.375 | 1 | 0 | NA | NA | NA | NA | 2 | 0.5 | 3 | 0.3333 |
| 1_15975250 | 2 | 0 | 1 | 0 | 2 | 0 | 2 | 0.5 | 7 | 0.1429 | 1 | 1 | 1 | 0 | 3 | 0 | 2 | 0.5 | 7 | 0.2857 |

| | | | | | | | | | | | | | | | | | | | | |
|---|---|---|---|---|---|---|---|---|---|---|---|---|---|---|---|---|---|---|---|---|
| 1_1597528 | 1 | 0 | 1 | 0 | 2 | 0.5 | 1 | 1 | 5 | 0.4 | 3 | 0.3333 | 1 | 0 | 2 | 0.5 | 2 | 0.5 | 8 | 0.375 |
| 1_1597529 | 2 | 0 | NA | NA | 3 | 0 | 2 | 0.5 | 7 | 0.1429 | 2 | 0.5 | 1 | 0 | NA | NA | 2 | 0 | 5 | 0.2 |
| 1_1597542 | 4 | 0.25 | 3 | 0.3333 | 2 | 0.5 | 3 | 0.6667 | 12 | 0.4167 | 3 | 0 | 2 | 1 | 1 | 0 | 2 | 0 | 8 | 0.25 |
| 1_1597542 | 3 | 0.3333 | 3 | 0 | 1 | 1 | 3 | 0.3333 | 10 | 0.3 | 2 | 0.5 | 2 | 0 | 1 | 0 | 2 | 0.5 | 7 | 0.2857 |
| 1_1597544 | 4 | 1 | 4 | 1 | 4 | 1 | 3 | 1 | 15 | 1 | 3 | 0.6667 | 1 | 1 | 1 | 1 | 2 | 1 | 7 | 0.8571 |
| 1_1597544 | 4 | 0.5 | 3 | 0.3333 | 3 | 1 | 2 | 0.5 | 12 | 0.5833 | 3 | 0.3333 | 2 | 1 | 1 | 1 | 2 | 1 | 8 | 0.75 |
| 1_1597546 | 3 | 0.6667 | 4 | 0.75 | 3 | 0.6667 | 1 | 1 | 11 | 0.7273 | 2 | 1 | 2 | 0.5 | 1 | 1 | 2 | 0.5 | 7 | 0.7143 |
| 1_1597546 | 3 | 0.6667 | 2 | 0.5 | 2 | 0 | 2 | 0.5 | 9 | 0.4444 | 3 | 0.6667 | 2 | 0.5 | 1 | 0 | 1 | 1 | 7 | 0.5714 |
| 1_1597546 | 4 | 0.25 | 3 | 0 | 2 | 0 | 2 | 0 | 11 | 0.0909 | 3 | 0.6667 | 2 | 0 | 1 | 0 | 1 | 0 | 7 | 0.2857 |
| 1_1597547 | 4 | 1 | 4 | 0.25 | 4 | 0.75 | 3 | 1 | 15 | 0.7333 | 3 | 0.3333 | 2 | 0.5 | 1 | 1 | 2 | 0.5 | 8 | 0.5 |
| 1_1597548 | 3 | 0.3333 | 3 | 0.3333 | 3 | 0.3333 | 4 | 0.5 | 13 | 0.3846 | 2 | 1 | 2 | 0 | 1 | 0 | 1 | 0 | 6 | 0.3333 |
| 1_1597550 | 4 | 0.25 | 6 | 0 | 2 | 0 | 4 | 0.25 | 16 | 0.125 | 4 | 0.25 | 2 | 0 | 1 | 0 | 2 | 0 | 9 | 0.1111 |
| 1_1597550 | 4 | 0.25 | 5 | 0.2 | 4 | 0.5 | 5 | 0.6 | 18 | 0.3889 | 4 | 0 | 2 | 0 | 1 | 0 | 3 | 0.6667 | 10 | 0.2 |
| 1_1597554 | 3 | 0.3333 | 7 | 0.1429 | 3 | 0 | 4 | 0 | 17 | 0.1176 | 4 | 0.25 | 1 | 1 | 1 | 0 | 2 | 0 | 8 | 0.25 |
| 1_1597557 | 3 | 0 | 5 | 0.6 | 4 | 0.25 | 6 | 0.5 | 18 | 0.3889 | 3 | 0.3333 | 1 | 0 | 3 | 0.3333 | 3 | 0.6667 | 10 | 0.4 |
| 1_1597559 | 4 | 0.25 | 5 | 0.2 | 3 | 0.3333 | 5 | 0 | 17 | 0.1765 | 2 | 0 | 2 | 0 | 2 | 0 | 3 | 0.3333 | 9 | 0.1111 |
| 1_1597560 | 5 | 0.2 | 6 | 0.3333 | 4 | 0.25 | 6 | 0.3333 | 21 | 0.2857 | 5 | 0.2 | 2 | 0 | 3 | 0 | 2 | 0 | 12 | 0.0833 |
| 1_1597564 | 5 | 0 | 6 | 0 | 5 | 0.4 | 6 | 0.1667 | 22 | 0.1364 | 4 | 0 | 1 | 0 | 3 | 0 | 2 | 0.5 | 10 | 0.1 |
| 1_1597566 | 6 | 0.1667 | 7 | 0.1429 | 4 | 0 | 6 | 0 | 23 | 0.087 | 6 | 0.1667 | 3 | 0 | 3 | 0 | 3 | 0 | 15 | 0.0667 |
| 1_1597567 | 5 | 0.2 | 7 | 0.8571 | 3 | 0.3333 | 3 | 0.6667 | 18 | 0.5556 | 6 | 0.5 | 3 | 0.3333 | 3 | 0.6667 | 4 | 0.75 | 16 | 0.5625 |
| 1_1597568 | 7 | 0.2857 | 8 | 0.125 | 5 | 0.2 | 6 | 0.1667 | 26 | 0.1923 | 5 | 0.2 | 3 | 0 | 3 | 0 | 3 | 0 | 14 | 0.0714 |
| 1_1597572 | 9 | 0.2222 | 9 | 0.1111 | 6 | 0.1667 | 5 | 0 | 29 | 0.1379 | 6 | 0.1667 | 2 | 0 | 4 | 0.25 | 4 | 0 | 16 | 0.125 |
| 1_1597576 | 9 | 0.1111 | 14 | 0 | 6 | 0.1667 | 5 | 0.2 | 34 | 0.0882 | 5 | 0 | 4 | 0 | 4 | 0.25 | 4 | 0 | 17 | 0.0588 |
| 1_1597581 | 14 | 0.2143 | 14 | 0.0714 | 6 | 0 | 7 | 0.2857 | 41 | 0.1463 | 7 | 0.4286 | 5 | 0.2 | 7 | 0 | 4 | 0.25 | 23 | 0.2174 |
| 1_1597581 | 14 | 0.1429 | 15 | 0.1333 | 7 | 0 | 6 | 0.3333 | 42 | 0.1429 | 6 | 0.1667 | 5 | 0 | 7 | 0 | 3 | 0.3333 | 21 | 0.0952 |
| 1_1597589 | 17 | 0.1176 | 17 | 0.2353 | 8 | 0.25 | 9 | 0.3333 | 51 | 0.2157 | 8 | 0.125 | 7 | 0.1429 | 8 | 0.125 | 5 | 0.2 | 28 | 0.1429 |
| 1_1597591 | 16 | 0.125 | 18 | 0.2778 | 10 | 0.3 | 12 | 0.25 | 56 | 0.2321 | 11 | 0.1818 | 10 | 0.2 | 10 | 0.1 | 7 | 0.2857 | 38 | 0.1842 |
| 1_1599052 | NA | NA | 1 | 1 | 1 | 1 | NA | NA | 2 | 1 | NA | NA | 1 | 1 | NA | NA | NA | NA | 1 | 1 |
| 1_1599054 | NA | NA | 1 | 0 | 1 | 1 | NA | NA | 2 | 0.5 | NA | NA | 1 | 1 | NA | NA | NA | NA | 1 | 1 |
| 1_1599056 | NA | NA | 1 | 0 | 1 | 1 | NA | NA | 2 | 0.5 | NA | NA | 1 | 1 | NA | NA | NA | NA | 1 | 1 |
| 1_1599058 | NA | NA | 1 | 0 | 1 | 1 | NA | NA | 2 | 0.5 | NA | NA | 1 | 1 | NA | NA | NA | NA | 1 | 1 |
| 1_1599059 | NA | NA | 1 | 1 | 1 | 1 | NA | NA | 2 | 1 | NA | NA | 1 | 1 | NA | NA | NA | NA | 1 | 1 |
| 1_1599059 | NA | NA | 1 | 1 | 1 | 1 | NA | NA | 2 | 1 | NA | NA | 1 | 1 | NA | NA | NA | NA | 1 | 1 |
| 1_1600272 | 13 | 0 | 14 | 0.1429 | 11 | 0.2727 | 17 | 0.0588 | 55 | 0.1091 | 17 | 0 | 12 | 0.1667 | 19 | 0 | 9 | 0.2222 | 57 | 0.0702 |
| 1_1600310 | 19 | 0.3158 | 17 | 0.1176 | 24 | 0.0833 | 18 | 0.3889 | 78 | 0.2179 | 11 | 0.0909 | 27 | 0.0741 | 16 | 0.0625 | 11 | 0.3636 | 65 | 0.1231 |
| 1_1600311 | 20 | 0.5 | 15 | 0.5333 | 15 | 0.6 | 18 | 0.7222 | 68 | 0.5882 | 10 | 0.4 | 20 | 0.6 | 15 | 0.4 | 12 | 0.4167 | 57 | 0.4737 |
| 1_1600473 | 4 | 0.25 | 16 | 0.0625 | 6 | 0.1667 | 8 | 0.125 | 34 | 0.1176 | 6 | 0.3333 | 7 | 0 | 2 | 0.5 | 10 | 0 | 25 | 0.12 |
| 1_1600511 | 2 | 0 | 5 | 0 | 1 | 0 | 4 | 0.25 | 12 | 0.0833 | 3 | 0.3333 | 2 | 0 | 3 | 0 | 4 | 0 | 12 | 0.0833 |
| 1_1600516 | 2 | 0.5 | 8 | 0.125 | 1 | 0 | 3 | 0 | 14 | 0.1429 | 3 | 0.3333 | 2 | 0.5 | 1 | 0 | 3 | 0.3333 | 9 | 0.3333 |
| 1_1600516 | 2 | 0.5 | 8 | 0 | 1 | 0 | 4 | 0 | 15 | 0.0667 | 3 | 0 | 1 | 0 | 1 | 0 | 4 | 0.25 | 9 | 0.1111 |
| 1_1600516 | 2 | 1 | 8 | 0.875 | 2 | 0.5 | 2 | 0.5 | 14 | 0.7857 | 3 | 1 | 2 | 1 | 2 | 0.5 | 4 | 0.75 | 11 | 0.8182 |
| 1_1600518 | 2 | 1 | 8 | 0.875 | 1 | 1 | 2 | 0.5 | 13 | 0.8462 | NA | NA | 1 | 0 | 1 | 1 | 4 | 1 | 6 | 0.8333 |
| 1_1600518 | 2 | 1 | 7 | 0.8571 | 1 | 1 | 2 | 0.5 | 12 | 0.8333 | 2 | 1 | 1 | 0 | 1 | 1 | 3 | 0.6667 | 7 | 0.7143 |
| 1_1600541 | 2 | 1 | 8 | 0 | 1 | 1 | 2 | 0.5 | 13 | 0.3077 | 2 | 1 | 1 | 0 | 1 | 1 | 4 | 0.5 | 8 | 0.625 |
| 1_1600541 | 1 | 0 | 8 | 0.125 | 1 | 0 | 2 | 0 | 12 | 0.0833 | 2 | 0.5 | 1 | 0 | 2 | 0 | 4 | 0 | 9 | 0.1111 |
| 1_1600846 | NA | NA | 1 | 1 | NA | NA | 1 | 1 | 2 | 1 | NA | NA | 3 | 1 | 1 | 1 | NA | NA | 4 | 1 |
| 1_1600846 | NA | NA | 1 | 1 | NA | NA | 1 | 1 | 2 | 1 | NA | NA | 2 | 1 | 1 | 1 | NA | NA | 3 | 1 |
| 1_1604027 | 11 | 0.3636 | 3 | 0 | 4 | 0 | 9 | 0.2222 | 27 | 0.2222 | 3 | 0.3333 | 8 | 0.125 | 7 | 0.1429 | 6 | 0.5 | 24 | 0.25 |
| 1_1604030 | 12 | 0.5 | 2 | 0 | 5 | 0.2 | 5 | 0.6 | 24 | 0.4167 | 3 | 0.3333 | 7 | 0.2857 | 5 | 0.6 | 5 | 0.6 | 20 | 0.45 |
| 1_1604042 | 10 | 0.8 | 2 | 0 | 3 | 0.3333 | 4 | 0.5 | 19 | 0.5789 | 2 | 0.5 | 4 | 0.25 | 4 | 0.25 | 5 | 0.6 | 15 | 0.4 |
| 1_1604043 | 10 | 0.3 | 2 | 0 | 3 | 0.3333 | 4 | 0.75 | 19 | 0.3684 | 2 | 0.5 | 3 | 0 | 4 | 0.25 | 4 | 0.25 | 13 | 0.2308 |
| 1_1604043 | 9 | 0.7778 | 2 | 0 | 2 | 1 | 5 | 1 | 18 | 0.7778 | 1 | 1 | 3 | 0.3333 | 3 | 1 | 3 | 1 | 10 | 0.8 |
| 1_1604044 | 6 | 1 | 2 | 0 | 4 | 1 | 5 | 1 | 17 | 0.8824 | 2 | 1 | 3 | 0.6667 | 4 | 1 | 4 | 1 | 13 | 0.9231 |
| 1_1604046 | 7 | 0.1429 | 1 | 0 | 2 | 0 | 2 | 0 | 12 | 0.0833 | 1 | 0 | 3 | 0 | 2 | 0 | 3 | 0.3333 | 9 | 0.1111 |
| 1_1604047 | 7 | 0.5714 | 2 | 0.5 | 3 | 0.6667 | 5 | 0.6 | 17 | 0.5882 | 1 | 1 | 3 | 1 | 2 | 0.5 | 4 | 0.25 | 10 | 0.6 |
| 1_1604049 | 7 | 1 | 2 | 0.5 | 2 | 1 | 4 | 1 | 15 | 0.9333 | 1 | 1 | 2 | 1 | 2 | 1 | 4 | 0.75 | 9 | 0.8889 |
| 1_1604050 | 5 | 0.4 | 2 | 1 | 2 | 0.5 | 4 | 0.75 | 13 | 0.6154 | 1 | 1 | 3 | 0.6667 | 2 | 0 | 2 | 1 | 8 | 0.625 |
| 1_1604051 | 5 | 1 | 1 | 1 | 1 | 1 | 4 | 1 | 11 | 1 | 1 | 0 | 3 | 0.6667 | 2 | 0 | 3 | 0.6667 | 9 | 0.4444 |
| 1_1604051 | 4 | 1 | 1 | 1 | 2 | 1 | 3 | 1 | 10 | 1 | NA | NA | 2 | 1 | 2 | 1 | 2 | 1 | 6 | 1 |
| 1_1604052 | 5 | 0.2 | 2 | 0 | 2 | 0 | 3 | 0 | 12 | 0.0833 | 1 | 0 | 3 | 0.6667 | 2 | 0.5 | 4 | 0 | 10 | 0.3 |
| 1_1604053 | 5 | 1 | 2 | 1 | 2 | 1 | 3 | 1 | 12 | 1 | 1 | 1 | 3 | 1 | 2 | 1 | 4 | 0.75 | 10 | 0.9 |
| 1_1604053 | 4 | 0.25 | 2 | 0 | 2 | 0 | 3 | 0.3333 | 11 | 0.1818 | 1 | 0 | 3 | 0 | 1 | 0 | 3 | 0.3333 | 8 | 0.125 |
| 1_1604054 | 5 | 0.6 | 2 | 0.5 | 2 | 0.5 | 4 | 0.75 | 13 | 0.6154 | 1 | 1 | 3 | 0.6667 | 2 | 0.5 | 4 | 1 | 10 | 0.8 |
| 1_1604054 | 1 | 1 | 1 | 1 | 1 | 1 | 1 | 1 | 4 | 1 | 1 | 1 | 1 | 0 | NA | NA | 3 | 0.6667 | 5 | 0.6 |
| 1_1604066 | NA | NA | NA | NA | NA | NA | 2 | 0.5 | 2 | 0.5 | NA | NA | 1 | 0 | NA | NA | 1 | 1 | 2 | 0.5 |

| | | | | | | | | | | | | | | | | | | | | |
|---|---|---|---|---|---|---|---|---|---|---|---|---|---|---|---|---|---|---|---|---|
| 1_1604067 | NA | NA | NA | NA | NA | NA | 1 | 1 | 1 | 1 | NA | NA | 3 | 1 | NA | NA | 1 | 1 | 4 | 1 |
| 1_1604068 | 3 | 1 | 2 | 1 | 1 | 1 | 2 | 1 | 8 | 1 | 1 | 1 | 3 | 1 | NA | NA | 3 | 1 | 7 | 1 |
| 1_1604068 | 4 | 0.5 | 2 | 0.5 | 1 | 0 | 2 | 1 | 9 | 0.5556 | NA | NA | 3 | 0.6667 | NA | NA | 3 | 0.6667 | 6 | 0.6667 |
| 1_1604070 | 2 | 0 | 1 | 0 | NA | NA | 3 | 0.3333 | 6 | 0.1667 | 1 | 0 | 2 | 0.5 | NA | NA | 3 | 0 | 6 | 0.1667 |
| 1_1604070 | 4 | 0.75 | NA | NA | 1 | 1 | 3 | 1 | 8 | 0.875 | 1 | 1 | 2 | 1 | NA | NA | 3 | 1 | 6 | 1 |
| 1_1604070 | 4 | 0 | 1 | 1 | 1 | 1 | 3 | 0.3333 | 9 | 0.3333 | 1 | 0 | 3 | 0 | NA | NA | 3 | 0.6667 | 7 | 0.2857 |
| 1_1604070 | 4 | 0.75 | NA | NA | 1 | 1 | 2 | 0 | 7 | 0.5714 | 1 | 0 | 2 | 0.5 | NA | NA | 1 | 1 | 4 | 0.5 |
| 1_1604071 | 4 | 0.25 | 1 | 0 | 1 | 1 | 1 | 1 | 7 | 0.4286 | NA | NA | 2 | 0.5 | NA | NA | 3 | 0.3333 | 5 | 0.4 |
| 1_1608320 | 1 | 0 | NA | NA | 1 | 1 | 1 | 0 | 3 | 0.3333 | NA | NA | 2 | 1 | NA | NA | 1 | 0 | 3 | 0.6667 |
| 1_1608322 | 1 | 0 | NA | NA | 1 | 1 | 1 | 0 | 3 | 0.3333 | NA | NA | 3 | 0 | 1 | 1 | 1 | 0 | 5 | 0.2 |
| 1_1608326 | NA | NA | NA | NA | 1 | 1 | 1 | 0 | 2 | 0.5 | NA | NA | 3 | 0 | 1 | 1 | 1 | 1 | 5 | 0.4 |
| 1_1608474 | 2 | 1 | NA | NA | 1 | 1 | NA | NA | 3 | 1 | NA | NA | NA | NA | 1 | 1 | NA | NA | 1 | 1 |
| 1_1608596 | 2 | 0.5 | NA | NA | NA | NA | NA | NA | 2 | 0.5 | NA | NA | NA | NA | 1 | 1 | NA | NA | 1 | 1 |
| 1_1608596 | 2 | 0.5 | NA | NA | NA | NA | NA | NA | 2 | 0.5 | NA | NA | NA | NA | 1 | 1 | NA | NA | 1 | 1 |
| 1_1608691 | 1 | 1 | 2 | 1 | 1 | 1 | 4 | 0.5 | 8 | 0.75 | 3 | 1 | 1 | 1 | 3 | 0.6667 | 2 | 1 | 9 | 0.8889 |
| 1_1608692 | 2 | 0.5 | 2 | 0 | 1 | 0 | 4 | 0.25 | 9 | 0.2222 | 4 | 0.25 | 1 | 1 | 3 | 0.3333 | 2 | 1 | 10 | 0.5 |
| 1_1608692 | 2 | 0 | 3 | 0 | 1 | 0 | 3 | 0.3333 | 9 | 0.1111 | 4 | 0.25 | 1 | 0 | 3 | 0 | 2 | 0.5 | 10 | 0.2 |
| 1_1608694 | 2 | 0.5 | 3 | 0 | 1 | 0 | 4 | 0 | 10 | 0.1 | 4 | 0.25 | 1 | 0 | 2 | 0 | 1 | 0 | 8 | 0.125 |
| 1_1608976 | 1 | 1 | 2 | 0.5 | NA | NA | NA | NA | 3 | 0.6667 | 1 | 1 | 2 | 1 | 2 | 0.5 | NA | NA | 5 | 0.8 |
| 1_1608977 | 1 | 0 | 2 | 1 | NA | NA | NA | NA | 3 | 0.6667 | 1 | 0 | 1 | 1 | 2 | 0 | NA | NA | 4 | 0.25 |
| 1_1608979 | 1 | 1 | 1 | 1 | NA | NA | NA | NA | 2 | 1 | 1 | 1 | 2 | 1 | 1 | 1 | NA | NA | 4 | 1 |
| 1_1609162 | 3 | 0.3333 | 1 | 0 | 3 | 0.3333 | 4 | 0.25 | 11 | 0.2727 | 2 | 0 | NA | NA | 1 | 0 | 5 | 0.2 | 8 | 0.125 |
| 1_1609164 | 1 | 1 | 1 | 0 | 1 | 1 | 3 | 0.6667 | 6 | 0.6667 | 2 | 1 | NA | NA | NA | NA | 3 | 1 | 5 | 1 |
| 1_1609165 | 1 | 1 | 1 | 0 | 2 | 1 | 4 | 1 | 8 | 0.875 | 2 | 1 | NA | NA | 1 | 1 | 2 | 1 | 5 | 1 |
| 1_1609165 | 2 | 0 | 1 | 0 | 3 | 0 | 4 | 0.75 | 10 | 0.3 | 2 | 0.5 | NA | NA | 1 | 0 | 4 | 0.5 | 7 | 0.4286 |
| 1_1609166 | 1 | 0 | 1 | 0 | 1 | 0 | 4 | 0.75 | 7 | 0.4286 | 2 | 0.5 | NA | NA | NA | NA | 3 | 0.3333 | 5 | 0.4 |
| 1_1609168 | 2 | 0 | 1 | 0 | 2 | 0 | 4 | 0.25 | 9 | 0.1111 | 2 | 0 | NA | NA | NA | NA | 3 | 0.3333 | 5 | 0.2 |
| 1_1609179 | 1 | 0 | NA | NA | 2 | 0.5 | 1 | 0 | 4 | 0.25 | 1 | 0 | NA | NA | NA | NA | 2 | 0.5 | 3 | 0.3333 |
| 1_1609180 | 2 | 0 | 1 | 1 | 2 | 0.5 | 3 | 0 | 8 | 0.25 | 1 | 0 | NA | NA | NA | NA | 3 | 0.3333 | 4 | 0.25 |
| 1_1609369 | 1 | 1 | NA | NA | NA | NA | NA | NA | 1 | 1 | 1 | 1 | NA | NA | NA | NA | 1 | 0 | 2 | 0.5 |
| 1_1609370 | 1 | 1 | NA | NA | NA | NA | 1 | 1 | 2 | 1 | 1 | 1 | NA | NA | NA | NA | 2 | 1 | 3 | 1 |
| 1_1609787 | 1 | 0 | 7 | 0.4286 | 4 | 0.25 | 5 | 0.4 | 17 | 0.3529 | 2 | 0.5 | 2 | 0 | NA | NA | 5 | 0.2 | 9 | 0.2222 |
| 1_1609787 | 1 | 0 | 7 | 0.5714 | 4 | 0.25 | 6 | 0.5 | 18 | 0.4444 | 2 | 0.5 | 2 | 0 | 1 | 0 | 7 | 0.5714 | 12 | 0.4167 |
| 1_1609954 | 3 | 0 | 9 | 0.1111 | 6 | 0 | 8 | 0.125 | 26 | 0.0769 | 5 | 0 | 3 | 0 | 2 | 0.5 | 8 | 0.25 | 18 | 0.1667 |
| 1_1609954 | 3 | 1 | 7 | 1 | 4 | 1 | 3 | 1 | 17 | 1 | 2 | 1 | 2 | 1 | 2 | 1 | 9 | 0.8889 | 15 | 0.9333 |
| 1_1609955 | 3 | 0.3333 | 9 | 0.6667 | 6 | 0.6667 | 8 | 0.875 | 26 | 0.6923 | 5 | 0.4 | 3 | 1 | 3 | 0 | 9 | 0.4444 | 20 | 0.45 |
| 1_1609970 | 2 | 0 | 10 | 0.1 | 6 | 0 | 8 | 0.125 | 26 | 0.0769 | 3 | 0.3333 | 5 | 0.2 | 5 | 0.2 | 9 | 0 | 22 | 0.1364 |
| 1_1609977 | 7 | 0 | 16 | 0.0625 | 6 | 0.1667 | 10 | 0 | 39 | 0.0513 | 7 | 0.2857 | 8 | 0.125 | 5 | 0.2 | 12 | 0.25 | 32 | 0.2188 |
| 1_1609980 | 7 | 0 | 17 | 0.1765 | 7 | 0.1429 | 14 | 0 | 45 | 0.0889 | 6 | 0 | 13 | 0 | 6 | 0.3333 | 11 | 0 | 36 | 0.0556 |
| 1_1609986 | NA | NA | NA | NA | NA | NA | 1 | 1 | 1 | 1 | 1 | 1 | 1 | 1 | 2 | 1 | NA | NA | 4 | 1 |
| 1_1610688 | 54 | 0.037 | 34 | 0.1176 | 49 | 0.0612 | 38 | 0.1053 | 175 | 0.0743 | 40 | 0.125 | 37 | 0.0811 | 33 | 0.0909 | 45 | 0.0444 | 155 | 0.0839 |
| 1_1610689 | 67 | 0.3582 | 40 | 0.675 | 52 | 0.4423 | 42 | 0.4286 | 201 | 0.4577 | 54 | 0.4074 | 41 | 0.5854 | 38 | 0.6842 | 55 | 0.5273 | 188 | 0.5372 |
| 1_1610690 | 64 | 0.2969 | 40 | 0.275 | 49 | 0.3265 | 42 | 0.3571 | 195 | 0.3128 | 53 | 0.2075 | 42 | 0.5 | 41 | 0.3902 | 51 | 0.3137 | 187 | 0.3422 |
| 1_1616454 | 3 | 0.3333 | 3 | 0 | NA | NA | NA | NA | 6 | 0.1667 | 3 | 0.3333 | 4 | 0 | 4 | 0.25 | 7 | 0 | 18 | 0.1111 |
| 1_1616459 | 1 | 0 | NA | NA | NA | NA | 1 | 1 | 2 | 0.5 | 2 | 0.5 | 2 | 1 | 2 | 0.5 | 4 | 0.5 | 10 | 0.6 |
| 1_1616459 | 1 | 1 | NA | NA | NA | NA | 1 | 1 | 2 | 1 | 2 | 1 | 2 | 1 | 2 | 1 | 4 | 1 | 10 | 1 |
| 1_1616460 | 1 | 1 | NA | NA | NA | NA | 1 | 1 | 2 | 1 | 1 | 1 | 1 | 1 | 2 | 1 | 3 | 1 | 7 | 1 |
| 1_1616461 | 1 | 1 | NA | NA | NA | NA | NA | NA | 1 | 1 | NA | NA | NA | NA | NA | NA | 2 | 1 | 2 | 1 |
| 1_1616461 | 1 | 1 | NA | NA | NA | NA | 1 | 1 | 2 | 1 | NA | NA | 1 | 1 | 3 | 1 | 3 | 0.6667 | 7 | 0.8571 |
| 1_1616461 | 1 | 1 | NA | NA | NA | NA | NA | NA | 1 | 1 | 1 | 1 | 1 | 1 | 2 | 1 | 3 | 1 | 7 | 1 |
| 1_1616462 | 1 | 0 | NA | NA | NA | NA | 1 | 1 | 2 | 0.5 | 1 | 1 | 2 | 0.5 | 3 | 0.3333 | 4 | 0.5 | 10 | 0.5 |
| 1_1617981 | 2 | 0.5 | 2 | 0.5 | 1 | 0 | 2 | 0.5 | 7 | 0.4286 | 2 | 0.5 | NA | NA | 1 | 0 | NA | NA | 3 | 0.3333 |
| 1_1617982 | 3 | 0.6667 | 3 | 0.6667 | 1 | 1 | 3 | 0.6667 | 10 | 0.7 | 2 | 0.5 | NA | NA | 1 | 1 | 1 | 1 | 4 | 0.75 |
| 1_1617982 | 3 | 1 | 2 | 1 | 1 | 1 | 3 | 1 | 9 | 1 | 1 | 1 | NA | NA | NA | NA | NA | NA | 1 | 1 |
| 1_1617984 | 3 | 0.3333 | 3 | 0.6667 | 1 | 1 | 3 | 0.3333 | 10 | 0.5 | 1 | 0 | NA | NA | 1 | 1 | 1 | 1 | 3 | 0.6667 |
| 1_1617984 | 2 | 0.5 | 3 | 0.6667 | 1 | 0 | 4 | 0.75 | 10 | 0.6 | 2 | 0 | NA | NA | 1 | 1 | 1 | 1 | 4 | 0.5 |
| 1_1617985 | 4 | 0.25 | 3 | 0 | NA | NA | 2 | 0.5 | 9 | 0.2222 | 2 | 0 | NA | NA | 1 | 1 | 1 | 0 | 4 | 0.25 |
| 1_1617986 | 3 | 0.6667 | 4 | 0.75 | NA | NA | 3 | 0.6667 | 10 | 0.7 | 2 | 0 | NA | NA | 2 | 1 | 1 | 1 | 5 | 0.6 |
| 1_1617988 | 3 | 0.3333 | 3 | 0.6667 | 1 | 1 | 4 | 0.75 | 11 | 0.6364 | 2 | 0.5 | NA | NA | 2 | 1 | 1 | 1 | 5 | 0.8 |
| 1_1617989 | 4 | 0.75 | 4 | 0.75 | 1 | 1 | 5 | 0.6 | 14 | 0.7143 | 2 | 0.5 | NA | NA | 2 | 1 | 1 | 0 | 5 | 0.6 |
| 1_1617989 | 4 | 0 | 4 | 0 | 1 | 0 | 5 | 0.4 | 14 | 0.1429 | 2 | 0.5 | NA | NA | 2 | 0 | 1 | 0 | 5 | 0.2 |
| 1_1617989 | 4 | 1 | 2 | 1 | 1 | 1 | 4 | 1 | 11 | 1 | 2 | 0.5 | NA | NA | 2 | 1 | 1 | 1 | 5 | 0.8 |
| 1_1617990 | 2 | 0 | 2 | 0.5 | 1 | 0 | 4 | 0 | 9 | 0.1111 | 1 | 0 | NA | NA | 2 | 0.5 | 1 | 1 | 4 | 0.5 |
| 1_1617990 | 4 | 0.25 | 5 | 0.8 | 1 | 0 | 5 | 0.8 | 15 | 0.6 | 2 | 0.5 | NA | NA | 2 | 0 | 1 | 0 | 5 | 0.2 |
| 1_1618169 | 12 | 0.5 | 6 | 0.5 | 7 | 0.2857 | 14 | 0.2857 | 39 | 0.3846 | 5 | 0.2 | 10 | 0.4 | 8 | 0.5 | 2 | 0 | 25 | 0.36 |

| | | | | | | | | | | | | | | | | | | | | |
|---|---|---|---|---|---|---|---|---|---|---|---|---|---|---|---|---|---|---|---|---|
| 1_16181716 | 10 | 0.1 | 4 | 0.25 | 6 | 0.1667 | 13 | 0 | 33 | 0.0909 | 5 | 0 | 12 | 0 | 7 | 0.2857 | 2 | 0 | 26 | 0.0769 |
| 1_16181738 | 10 | 0.4 | 4 | 0.5 | 6 | 0.1667 | 10 | 0 | 30 | 0.2333 | 5 | 0.2 | 11 | 0.2727 | 8 | 0.25 | 1 | 0 | 25 | 0.24 |
| 1_16181869 | 6 | 0 | 2 | 0.5 | 4 | 0 | 8 | 0 | 20 | 0.05 | 3 | 0 | 5 | 0 | 8 | 0.125 | 1 | 0 | 17 | 0.0588 |
| 1_16181872 | 4 | 0.25 | 1 | 0 | 3 | 0 | 6 | 0 | 14 | 0.0714 | 1 | 0 | 1 | 0 | 6 | 0.5 | 1 | 0 | 9 | 0.3333 |
| 1_16181908 | 4 | 0.25 | 2 | 1 | 5 | 0.2 | 8 | 0.125 | 19 | 0.2632 | 2 | 1 | 5 | 0.6 | 6 | 1 | 1 | 1 | 14 | 0.8571 |
| 1_16181949 | 2 | 1 | 1 | 1 | 5 | 0.8 | 6 | 0.8333 | 14 | 0.8571 | 3 | 1 | 5 | 1 | 5 | 1 | 1 | 1 | 14 | 1 |
| 1_16181959 | 3 | 0.3333 | 1 | 0 | 3 | 0 | 5 | 0.4 | 12 | 0.25 | 3 | 0.3333 | 4 | 0 | 6 | 0.8333 | 1 | 0 | 14 | 0.4286 |
| 1_16183381 | 1 | 1 | 4 | 0.25 | NA | NA | NA | NA | 5 | 0.4 | NA | NA | 1 | 0 | NA | NA | 1 | 1 | 2 | 0.5 |
| 1_16183385 | 1 | 0 | 4 | 0.75 | NA | NA | NA | NA | 5 | 0.6 | NA | NA | 1 | 1 | NA | NA | NA | NA | 1 | 1 |
| 1_16183850 | 2 | 0 | 4 | 0.25 | 3 | 0 | 6 | 0.1667 | 15 | 0.1333 | 2 | 1 | 5 | 0 | 5 | 0.2 | 1 | 0 | 13 | 0.2308 |
| 1_16236116 | NA | NA | NA | NA | NA | NA | 1 | 1 | 1 | 1 | NA | NA | 2 | 0.5 | 2 | 0.5 | NA | NA | 4 | 0.5 |
| 1_16236146 | 1 | 1 | 1 | 1 | NA | NA | NA | NA | 2 | 1 | 1 | 0 | 2 | 0 | 2 | 0.5 | NA | NA | 5 | 0.2 |
| 1_16236147 | NA | NA | 1 | 1 | NA | NA | 2 | 0 | 3 | 0.3333 | NA | NA | 2 | 0 | 3 | 0.3333 | 1 | 0 | 6 | 0.1667 |
| 1_16236176 | 1 | 1 | 1 | 1 | NA | NA | 2 | 1 | 4 | 1 | 1 | 0 | 2 | 0.5 | 3 | 1 | 1 | 1 | 7 | 0.7143 |
| 1_16236180 | 1 | 0 | 1 | 1 | NA | NA | 1 | 1 | 3 | 0.6667 | 1 | 1 | 2 | 0 | 3 | 1 | 1 | 0 | 7 | 0.5714 |
| 1_16236180 | 1 | 1 | 1 | 0 | NA | NA | 1 | 1 | 3 | 0.6667 | 1 | 0 | 1 | 1 | 2 | 0 | NA | NA | 4 | 0.25 |
| 1_16236192 | 1 | 0 | 1 | 1 | NA | NA | 2 | 0 | 4 | 0.25 | NA | NA | 2 | 0 | 3 | 0 | 1 | 1 | 6 | 0.1667 |
| 1_16236207 | 1 | 1 | 1 | 1 | NA | NA | 2 | 1 | 4 | 1 | 1 | 1 | 2 | 0 | 2 | 0 | NA | NA | 5 | 0.2 |
| 1_16237343 | NA | NA | 1 | 1 | NA | NA | NA | NA | 1 | 1 | NA | NA | 1 | 1 | NA | NA | 1 | 1 | 2 | 1 |
| 1_16237383 | 1 | 0 | 1 | 1 | NA | NA | NA | NA | 2 | 0.5 | 1 | 0 | 1 | 0 | 1 | 0 | 1 | 1 | 4 | 0.25 |
| 1_16237389 | 1 | 0 | 1 | 1 | NA | NA | NA | NA | 2 | 0.5 | 1 | 0 | 1 | 0 | 1 | 0 | 1 | 1 | 4 | 0.25 |
| 1_16237393 | 1 | 0 | 1 | 1 | NA | NA | NA | NA | 2 | 0.5 | 1 | 1 | 1 | 0 | 1 | 0 | 1 | 0 | 4 | 0.25 |
| 1_16242218 | 5 | 0 | 5 | 0 | 9 | 0.1111 | 5 | 0.2 | 24 | 0.0833 | 3 | 0 | 8 | 0.25 | 9 | 0 | 5 | 0 | 25 | 0.08 |
| 1_16246689 | 4 | 0.25 | 2 | 0 | 5 | 0.6 | 4 | 0.25 | 15 | 0.3333 | 1 | 0 | 3 | 0.3333 | 1 | 0 | 2 | 0.5 | 7 | 0.2857 |
| 1_16246702 | 5 | 0 | 2 | 0.5 | 5 | 0 | 5 | 0 | 17 | 0.0588 | 2 | 0 | 3 | 0 | 1 | 0 | 2 | 0.5 | 8 | 0.125 |
| 1_16246709 | 4 | 0 | 2 | 0 | 3 | 0.3333 | 5 | 0.2 | 14 | 0.1429 | 2 | 0.5 | 3 | 0.3333 | 2 | 0 | 3 | 0.6667 | 10 | 0.4 |
| 1_16246717 | 5 | 0.2 | 1 | 1 | 5 | 0 | 5 | 0.4 | 16 | 0.25 | 2 | 0.5 | 2 | 0 | 1 | 0 | 3 | 0 | 8 | 0.125 |
| 1_16246718 | 6 | 0.1667 | 2 | 0 | 5 | 0 | 5 | 0 | 18 | 0.0556 | 2 | 0 | 3 | 0 | 2 | 0 | 3 | 0.3333 | 10 | 0.1 |
| 1_16247603 | 6 | 0 | 3 | 0 | 6 | 0.1667 | 4 | 0 | 19 | 0.0526 | 2 | 1 | 4 | 0 | 3 | 0.3333 | 5 | 0 | 14 | 0.2143 |
| 1_16247628 | 7 | 1 | 3 | 1 | 5 | 0.8 | 5 | 1 | 20 | 0.95 | 2 | 1 | 4 | 1 | 3 | 1 | 6 | 0.8333 | 15 | 0.9333 |
| 1_16247635 | 6 | 1 | 3 | 1 | 5 | 0.8 | 5 | 1 | 19 | 0.9474 | 2 | 0.5 | 3 | 1 | 2 | 1 | 5 | 1 | 12 | 0.9167 |
| 1_16247638 | 5 | 0.4 | 2 | 0.5 | 3 | 1 | 3 | 0.6667 | 13 | 0.6154 | 1 | 1 | 5 | 0.6 | 2 | 0 | 4 | 0.5 | 12 | 0.5 |
| 1_16247639 | 7 | 0.4286 | 2 | 0 | 5 | 0.2 | 5 | 0.8 | 19 | 0.4211 | 1 | 0 | 5 | 0.2 | 2 | 0.5 | 6 | 0.8333 | 14 | 0.5 |
| 1_16247649 | 6 | 0.6667 | 4 | 0.25 | 5 | 0.8 | 6 | 1 | 21 | 0.7143 | 3 | 1 | 5 | 0.6 | 2 | 0.5 | 6 | 0.3333 | 16 | 0.5625 |
| 1_16247651 | 5 | 0.2 | 4 | 0.25 | 4 | 0.25 | 6 | 0.3333 | 19 | 0.2632 | 3 | 0 | 5 | 0 | 3 | 0.6667 | 7 | 0.2857 | 18 | 0.2222 |
| 1_16247663 | 8 | 0.25 | 4 | 0.25 | 6 | 0.1667 | 6 | 0.3333 | 24 | 0.25 | 3 | 0.3333 | 4 | 0 | 3 | 0 | 6 | 0.1667 | 16 | 0.125 |
| 1_16247668 | 6 | 0 | 4 | 0.25 | 5 | 0.4 | 4 | 0.5 | 19 | 0.2632 | 2 | 0 | 4 | 0.5 | 3 | 0 | 5 | 0 | 14 | 0.1429 |
| 1_16247672 | 7 | 0.7143 | 4 | 0 | 6 | 0.3333 | 7 | 0.1429 | 24 | 0.3333 | 3 | 0 | 5 | 0.2 | 3 | 0.6667 | 7 | 0.7143 | 18 | 0.4444 |
| 1_16247692 | 6 | 0.1667 | 3 | 0.3333 | 5 | 0.8 | 7 | 0.4286 | 21 | 0.4286 | 4 | 0.25 | 5 | 0.4 | 3 | 0.6667 | 6 | 0.1667 | 18 | 0.3333 |
| 1_16247695 | 6 | 0.6667 | 3 | 0.6667 | 6 | 0.8333 | 9 | 0.7778 | 24 | 0.75 | 4 | 0.75 | 5 | 1 | 3 | 0.6667 | 6 | 1 | 18 | 0.8889 |
| 1_16249857 | 1 | 0 | 1 | 1 | NA | NA | NA | NA | 2 | 0.5 | 1 | 1 | 1 | 0 | NA | NA | NA | NA | 2 | 0.5 |
| 1_16249967 | 12 | 0 | 17 | 0.0588 | 13 | 0.0769 | 13 | 0.1538 | 55 | 0.0727 | 13 | 0.0769 | 12 | 0.1667 | 12 | 0.0833 | 8 | 0 | 45 | 0.0889 |
| 1_16249977 | 14 | 0.3571 | 15 | 0.1333 | 13 | 0.1538 | 11 | 0.4545 | 53 | 0.2642 | 13 | 0.3077 | 11 | 0.4545 | 12 | 0.25 | 9 | 0.2222 | 45 | 0.3111 |
| 1_16250011 | 12 | 0.4167 | 12 | 0.1667 | 11 | 0 | 7 | 0 | 42 | 0.1667 | 10 | 0 | 10 | 0.1 | 12 | 0 | 8 | 0.375 | 40 | 0.1 |
| 1_16250065 | 8 | 0.25 | 10 | 0.3 | 8 | 0.125 | 7 | 0.4286 | 33 | 0.2727 | 8 | 0.375 | 5 | 0.4 | 7 | 0.1429 | 5 | 0.2 | 25 | 0.28 |
| 1_16250086 | 8 | 0 | 9 | 0.2222 | 10 | 0 | 7 | 0 | 34 | 0.0588 | 8 | 0.125 | 7 | 0 | 9 | 0 | 7 | 0.1429 | 31 | 0.0645 |
| 1_16250101 | 6 | 0.5 | 7 | 0.1429 | 6 | 0.1667 | 7 | 0.7143 | 26 | 0.3846 | 5 | 0.4 | 6 | 0.3333 | 5 | 0.2 | 5 | 0.6 | 21 | 0.381 |
| 1_16250102 | 7 | 0.4286 | 7 | 0.4286 | 8 | 0.375 | 6 | 0.8333 | 28 | 0.5 | 4 | 1 | 5 | 0.6 | 6 | 0.5 | 5 | 0.6 | 20 | 0.65 |
| 1_16250113 | 8 | 0.375 | 9 | 0.3333 | 7 | 0.7143 | 8 | 0.75 | 32 | 0.5312 | 7 | 0.5714 | 4 | 1 | 6 | 0.5 | 6 | 0.6667 | 23 | 0.6522 |
| 1_16252511 | 5 | 0.2 | 2 | 0 | 3 | 0 | 3 | 0 | 13 | 0.0769 | 3 | 0 | 4 | 0 | 3 | 0 | 1 | 1 | 11 | 0.0909 |
| 1_16252575 | 3 | 0.3333 | NA | NA | 2 | 0 | 2 | 0.5 | 7 | 0.2857 | 1 | 0 | 3 | 0 | 2 | 0.5 | 1 | 1 | 7 | 0.2857 |
| 1_16253227 | 2 | 0 | NA | NA | 2 | 0.5 | 2 | 0.5 | 6 | 0.3333 | NA | NA | 1 | 1 | 1 | 1 | 1 | 0 | 3 | 0.6667 |
| 1_16253258 | 2 | 0 | NA | NA | 2 | 0.5 | 2 | 0 | 6 | 0.1667 | NA | NA | 1 | 0 | 1 | 1 | 1 | 0 | 3 | 0.3333 |
| 1_16253289 | 2 | 0 | NA | NA | 2 | 0.5 | 2 | 0 | 6 | 0.1667 | NA | NA | NA | NA | 1 | 1 | 1 | 0 | 2 | 0.5 |
| 1_16265760 | 2 | 0 | 1 | 1 | NA | NA | NA | NA | 3 | 0.3333 | NA | NA | 1 | 1 | NA | NA | 2 | 0.5 | 3 | 0.6667 |
| 1_16270450 | 1 | 0 | 1 | 1 | NA | NA | 1 | 1 | 3 | 0.6667 | 1 | 1 | NA | NA | NA | NA | NA | NA | 1 | 1 |
| 1_16307345 | NA | NA | 1 | 1 | 1 | 0 | NA | NA | 2 | 0.5 | 1 | 1 | NA | NA | 1 | 0 | 1 | 0 | 3 | 0.3333 |
| 1_16307569 | NA | NA | 1 | 1 | NA | NA | NA | NA | 1 | 1 | 1 | 1 | NA | NA | NA | NA | 1 | 0 | 2 | 0.5 |
| 1_16307588 | NA | NA | 1 | 1 | NA | NA | NA | NA | 1 | 1 | 1 | 1 | NA | NA | NA | NA | 1 | 1 | 2 | 1 |
| 1_16321384 | NA | NA | 2 | 0.5 | 1 | 1 | NA | NA | 3 | 0.6667 | NA | NA | 1 | 0 | 2 | 1 | 2 | 1 | 5 | 0.8 |
| 1_16321387 | NA | NA | 2 | 0.5 | 1 | 0 | NA | NA | 3 | 0.3333 | NA | NA | 1 | 0 | 2 | 0.5 | 1 | 0 | 4 | 0.25 |
| 1_16321391 | NA | NA | 2 | 0.5 | 1 | 0 | NA | NA | 3 | 0.3333 | NA | NA | NA | NA | 3 | 0.6667 | 1 | 1 | 4 | 0.75 |
| 1_16321395 | NA | NA | 1 | 1 | 1 | 0 | NA | NA | 2 | 0.5 | NA | NA | 1 | 0 | 2 | 1 | 2 | 1 | 5 | 0.8 |
| 1_16321396 | NA | NA | 2 | 0 | 1 | 1 | NA | NA | 3 | 0.3333 | NA | NA | 1 | 1 | 3 | 0.6667 | 1 | 0 | 5 | 0.6 |
| 1_16352728 | NA | NA | NA | NA | 1 | 1 | NA | NA | 1 | 1 | 1 | 1 | NA | NA | NA | NA | NA | NA | 1 | 1 |

| | | | | | | | | | | | | | | | | | | | | |
|---|---|---|---|---|---|---|---|---|---|---|---|---|---|---|---|---|---|---|---|---|
| 1_1637434 | 18 | 0.1667 | 26 | 0.3846 | 19 | 0.1579 | 23 | 0.2609 | 86 | 0.2558 | 16 | 0.1875 | 19 | 0.3158 | 17 | 0.2353 | 13 | 0.2308 | 65 | 0.2462 |
| 1_1637441 | 30 | 0.2 | 37 | 0.3514 | 24 | 0.375 | 31 | 0.3226 | 122 | 0.3115 | 20 | 0.3 | 23 | 0.3913 | 26 | 0.3846 | 14 | 0.3571 | 83 | 0.3614 |
| 1_1637795 | 1 | 1 | NA | NA | 1 | 0 | 2 | 0.5 | 4 | 0.5 | 2 | 1 | 1 | 1 | NA | NA | 1 | 1 | 4 | 1 |
| 1_1637797 | 2 | 0.5 | NA | NA | 2 | 0.5 | 3 | 1 | 7 | 0.7143 | 2 | 1 | 1 | 0 | NA | NA | 2 | 0.5 | 5 | 0.6 |
| 1_1637798 | 2 | 0 | NA | NA | 1 | 1 | 3 | 0.3333 | 6 | 0.3333 | 2 | 0.5 | 1 | 0 | NA | NA | 2 | 0.5 | 5 | 0.4 |
| 1_1637799 | 2 | 0 | NA | NA | 2 | 0 | 1 | 1 | 5 | 0.2 | 2 | 0 | 1 | 0 | NA | NA | 2 | 0.5 | 5 | 0.2 |
| 1_1637799 | 3 | 1 | NA | NA | 2 | 1 | 3 | 0.3333 | 8 | 0.75 | 2 | 1 | NA | NA | NA | NA | 2 | 1 | 4 | 1 |
| 1_1637800 | 3 | 1 | NA | NA | 1 | 1 | 3 | 1 | 7 | 1 | 2 | 1 | 1 | 1 | NA | NA | 2 | 1 | 5 | 1 |
| 1_1637801 | 3 | 0.6667 | NA | NA | 1 | 1 | 3 | 1 | 7 | 0.8571 | 2 | 0 | 1 | 1 | NA | NA | 3 | 0.3333 | 6 | 0.3333 |
| 1_1637811 | 2 | 0.5 | NA | NA | 1 | 0 | 3 | 0.3333 | 6 | 0.3333 | 2 | 0 | 1 | 0 | NA | NA | 3 | 0.3333 | 6 | 0.1667 |
| 1_1637811 | 2 | 0.5 | NA | NA | 2 | 0.5 | 3 | 0 | 7 | 0.2857 | 2 | 0 | 1 | 1 | NA | NA | 3 | 0.3333 | 6 | 0.3333 |
| 1_1637812 | 3 | 1 | NA | NA | 1 | 1 | 3 | 0.6667 | 7 | 0.8571 | 2 | 0.5 | 1 | 1 | NA | NA | 2 | 1 | 5 | 0.8 |
| 1_1637812 | 2 | 0.5 | NA | NA | 2 | 0.5 | 2 | 0.5 | 6 | 0.5 | 1 | 1 | 1 | 1 | NA | NA | 1 | 0 | 3 | 0.6667 |
| 1_1638237 | NA | NA | 1 | 1 | 1 | 1 | NA | NA | 2 | 1 | 1 | 0 | NA | NA | 1 | 1 | NA | NA | 2 | 0.5 |
| 1_1638239 | NA | NA | 1 | 1 | 1 | 1 | NA | NA | 2 | 1 | 1 | 0 | NA | NA | 1 | 1 | NA | NA | 2 | 0.5 |
| 1_1638241 | NA | NA | 1 | 0 | 1 | 1 | NA | NA | 2 | 0.5 | 1 | 0 | NA | NA | 1 | 1 | NA | NA | 2 | 0.5 |
| 1_1638242 | NA | NA | 1 | 1 | 1 | 1 | NA | NA | 2 | 1 | 1 | 1 | NA | NA | 1 | 1 | NA | NA | 2 | 1 |
| 1_1638244 | NA | NA | 1 | 1 | 1 | 1 | NA | NA | 2 | 1 | 1 | 1 | NA | NA | 1 | 1 | NA | NA | 2 | 1 |
| 1_1638246 | NA | NA | 1 | 1 | 1 | 0 | NA | NA | 2 | 0.5 | 1 | 1 | NA | NA | 1 | 1 | NA | NA | 2 | 1 |
| 1_1640844 | 1 | 1 | 2 | 1 | NA | NA | 2 | 1 | 5 | 1 | 1 | 1 | 2 | 1 | 1 | 1 | 1 | 1 | 5 | 1 |
| 1_1640844 | 1 | 1 | 1 | 1 | NA | NA | 1 | 1 | 3 | 1 | 1 | 1 | 1 | 1 | 1 | 1 | 1 | 1 | 4 | 1 |
| 1_1640845 | 1 | 1 | 2 | 1 | NA | NA | 1 | 1 | 4 | 1 | 1 | 1 | 1 | 1 | 1 | 1 | 1 | 1 | 4 | 1 |
| 1_1640848 | 1 | 1 | 2 | 0.5 | NA | NA | 2 | 1 | 5 | 0.8 | 1 | 0 | 2 | 0.5 | 2 | 0.5 | 1 | 0 | 6 | 0.3333 |
| 1_1640897 | 1 | 0 | 2 | 0 | NA | NA | 2 | 0.5 | 5 | 0.2 | 1 | 0 | 3 | 0 | 1 | 1 | 1 | 0 | 6 | 0.1667 |
| 1_1640911 | 1 | 0 | 2 | 0 | NA | NA | 2 | 0.5 | 5 | 0.2 | NA | NA | 3 | 0.3333 | 1 | 0 | 1 | 1 | 5 | 0.4 |
| 1_1640913 | 1 | 0 | 3 | 0.3333 | NA | NA | 2 | 1 | 6 | 0.5 | 1 | 1 | 2 | 1 | 2 | 1 | 2 | 0.5 | 7 | 0.8571 |
| 1_1640913 | 1 | 1 | 3 | 0 | NA | NA | 2 | 1 | 6 | 0.5 | 1 | 0 | 3 | 0.3333 | 2 | 1 | 2 | 0.5 | 8 | 0.5 |
| 1_1640918 | 2 | 0 | 4 | 0.25 | NA | NA | 2 | 1 | 8 | 0.375 | 1 | 0 | 3 | 0.3333 | 2 | 0.5 | 3 | 0.6667 | 9 | 0.4444 |
| 1_1640921 | 2 | 0 | 5 | 0 | NA | NA | 2 | 0.5 | 9 | 0.1111 | 1 | 0 | 3 | 0 | 2 | 0 | 3 | 0.6667 | 9 | 0.2222 |
| 1_1640927 | 2 | 0 | 5 | 0 | 1 | 0 | 3 | 0.6667 | 11 | 0.1818 | 1 | 1 | 3 | 0.6667 | 3 | 0 | 3 | 0.6667 | 10 | 0.5 |
| 1_1640928 | 2 | 0 | 3 | 0 | NA | NA | 3 | 0.3333 | 8 | 0.125 | 2 | 0 | 3 | 0.3333 | 2 | 0 | 2 | 0 | 9 | 0.1111 |
| 1_1641076 | 3 | 0 | 3 | 0 | 2 | 0 | 4 | 0.25 | 12 | 0.0833 | 3 | 0.6667 | 1 | 0 | 2 | 0 | NA | NA | 6 | 0.3333 |
| 1_1641077 | 5 | 0.2 | 2 | 0.5 | 2 | 0 | 5 | 0 | 14 | 0.1429 | 3 | 0.3333 | 1 | 0 | 3 | 0 | NA | NA | 7 | 0.1429 |
| 1_1641077 | 5 | 0.2 | 3 | 0 | 2 | 0 | 4 | 0.25 | 14 | 0.1429 | 3 | 0 | 1 | 1 | 2 | 0 | NA | NA | 6 | 0.1667 |
| 1_1641077 | 5 | 0.2 | 3 | 0.3333 | 2 | 0 | 5 | 0 | 15 | 0.1333 | 3 | 0 | 1 | 0 | 3 | 0.3333 | NA | NA | 7 | 0.1429 |
| 1_1641078 | 5 | 0.4 | 3 | 0.6667 | 2 | 1 | 5 | 0.6 | 15 | 0.6 | 3 | 0.6667 | 1 | 1 | 2 | 0.5 | NA | NA | 6 | 0.6667 |
| 1_1641079 | 5 | 0.6 | 3 | 0.3333 | 1 | 0 | 5 | 0 | 14 | 0.2857 | 2 | 0 | 1 | 0 | 3 | 0.3333 | NA | NA | 6 | 0.1667 |
| 1_1641080 | 3 | 0.3333 | 3 | 0.6667 | 2 | 0.5 | 4 | 0.5 | 12 | 0.5 | 2 | 0 | 1 | 0 | 3 | 0.3333 | NA | NA | 6 | 0.1667 |
| 1_1641233 | 3 | 0.3333 | 1 | 1 | 6 | 0.1667 | 2 | 1 | 12 | 0.4167 | 3 | 0.6667 | 2 | 0 | 3 | 1 | 3 | 0.3333 | 11 | 0.5455 |
| 1_1641299 | 3 | 0 | 1 | 0 | 5 | 0.2 | 2 | 0 | 11 | 0.0909 | 3 | 0.3333 | 1 | 0 | 2 | 0.5 | 4 | 0 | 10 | 0.2 |
| 1_1641300 | 4 | 0 | 2 | 0.5 | 5 | 0 | 3 | 0 | 14 | 0.0714 | 3 | 0.3333 | 1 | 0 | 2 | 0 | 5 | 0.4 | 11 | 0.2727 |
| 1_1642647 | 4 | 0.25 | 2 | 0 | 3 | 0 | 2 | 0 | 11 | 0.0909 | 7 | 0.1429 | 3 | 0 | 1 | 0 | 2 | 0 | 13 | 0.0769 |
| 1_1642654 | 3 | 0 | 8 | 0.125 | 6 | 0 | 3 | 0 | 20 | 0.05 | 7 | 0.1429 | NA | NA | 2 | 0 | 2 | 0 | 11 | 0.0909 |
| 1_1642657 | 281 | 0.0783 | 332 | 0.1145 | 243 | 0.0782 | 245 | 0.1061 | 1101 | 0.0954 | 223 | 0.0807 | 223 | 0.1031 | 221 | 0.0905 | 235 | 0.0468 | 902 | 0.0798 |
| 1_1642657 | 254 | 0.0827 | 303 | 0.132 | 235 | 0.1787 | 227 | 0.1189 | 1019 | 0.1276 | 211 | 0.1422 | 219 | 0.1279 | 204 | 0.1373 | 221 | 0.1222 | 855 | 0.1322 |
| 1_1642659 | 219 | 0.1735 | 241 | 0.2033 | 178 | 0.191 | 184 | 0.1304 | 822 | 0.1764 | 168 | 0.1548 | 163 | 0.2025 | 179 | 0.1564 | 186 | 0.1667 | 696 | 0.1695 |
| 1_1642659 | 190 | 0.1842 | 202 | 0.2228 | 162 | 0.2222 | 159 | 0.2516 | 713 | 0.2188 | 143 | 0.2238 | 157 | 0.2229 | 146 | 0.2466 | 165 | 0.2485 | 611 | 0.2357 |
| 1_1642865 | NA | NA | NA | NA | 1 | 1 | NA | NA | 1 | 1 | NA | NA | 2 | 1 | NA | NA | 1 | 1 | 3 | 1 |
| 1_1642866 | NA | NA | NA | NA | 1 | 1 | NA | NA | 1 | 1 | NA | NA | 1 | 1 | NA | NA | 1 | 0 | 2 | 0.5 |
| 1_1642868 | NA | NA | NA | NA | 1 | 1 | NA | NA | 1 | 1 | NA | NA | 2 | 0.5 | NA | NA | 1 | 0 | 3 | 0.3333 |
| 1_1642897 | NA | NA | NA | NA | 1 | 1 | NA | NA | 1 | 1 | NA | NA | NA | NA | NA | NA | 2 | 1 | 2 | 1 |
| 1_1642913 | NA | NA | 1 | 0 | 2 | 0.5 | 1 | 1 | 4 | 0.5 | NA | NA | 2 | 0 | 1 | 1 | 3 | 0.3333 | 6 | 0.3333 |
| 1_1650023 | 2 | 0 | 4 | 0 | 3 | 0 | 4 | 0.25 | 13 | 0.0769 | 2 | 0 | 2 | 0 | 5 | 0 | 5 | 0.2 | 14 | 0.0714 |
| 1_1650054 | 1 | 0 | 3 | 0 | 2 | 0 | 3 | 0.3333 | 9 | 0.1111 | 2 | 0 | NA | NA | 3 | 0.3333 | 3 | 0 | 8 | 0.125 |
| 1_1650059 | NA | NA | 3 | 0 | 1 | 0 | 2 | 1 | 6 | 0.3333 | 2 | 0 | NA | NA | 3 | 0.3333 | 3 | 0.3333 | 8 | 0.25 |
| 1_1650061 | 1 | 0 | 2 | 0.5 | 1 | 1 | 1 | 0 | 5 | 0.4 | 2 | 0 | NA | NA | 3 | 0.3333 | 3 | 0 | 8 | 0.125 |
| 1_1650062 | 1 | 1 | 4 | 0.25 | 2 | 1 | 4 | 0.25 | 11 | 0.4545 | 1 | 1 | 2 | 0 | 3 | 0.3333 | 2 | 0.5 | 8 | 0.375 |
| 1_1650065 | 1 | 0 | 3 | 0.3333 | 2 | 0 | 3 | 0.3333 | 9 | 0.2222 | 2 | 0.5 | 2 | 0 | 3 | 0.3333 | 2 | 0 | 9 | 0.2222 |
| 1_1650066 | 1 | 1 | 3 | 1 | 2 | 1 | 3 | 1 | 9 | 1 | 2 | 0.5 | 1 | 1 | 3 | 1 | NA | NA | 6 | 0.8333 |
| 1_1650067 | 1 | 0 | 2 | 1 | 1 | 1 | 2 | 0.5 | 6 | 0.6667 | 2 | 0 | 2 | 0.5 | 2 | 0 | 2 | 0 | 8 | 0.125 |
| 1_1650426 | 15 | 0.0667 | 6 | 0.3333 | 10 | 0 | 17 | 0.1176 | 48 | 0.1042 | 2 | 0 | 12 | 0.0833 | 8 | 0.125 | 13 | 0 | 35 | 0.0571 |
| 1_1650429 | 14 | 0.1429 | 8 | 0.25 | 6 | 0.3333 | 15 | 0.2 | 43 | 0.2093 | 3 | 0.3333 | 13 | 0.1538 | 8 | 0.125 | 12 | 0.0833 | 36 | 0.1389 |
| 1_1650429 | 15 | 0.1333 | 8 | 0 | 5 | 0 | 15 | 0.2667 | 43 | 0.1395 | 2 | 0 | 12 | 0.25 | 8 | 0.125 | 12 | 0.25 | 34 | 0.2059 |
| 1_1650430 | 15 | 0.2667 | 7 | 0.2857 | 7 | 0.2857 | 16 | 0.25 | 45 | 0.2667 | 2 | 0 | 12 | 0.1667 | 8 | 0.125 | 11 | 0.3636 | 33 | 0.2121 |
| 1_1650430 | 14 | 0.5 | 8 | 0.25 | 7 | 0.2857 | 14 | 0.2143 | 43 | 0.3256 | 2 | 0 | 9 | 0.1111 | 8 | 0.125 | 12 | 0.1667 | 31 | 0.129 |

| | | | | | | | | | | | | | | | | | | | | |
|---|---|---|---|---|---|---|---|---|---|---|---|---|---|---|---|---|---|---|---|---|
| 1_16504327 | 15 | 0.0667 | 6 | 0 | 7 | 0.4286 | 12 | 0.3333 | 40 | 0.2 | 3 | 0.3333 | 11 | 0.2727 | 7 | 0.2857 | 13 | 0.3077 | 34 | 0.2941 |
| 1_16504329 | 13 | 0.4615 | 8 | 0.25 | 7 | 0.4286 | 10 | 0.2 | 38 | 0.3421 | 1 | 0 | 10 | 0.4 | 7 | 0.2857 | 11 | 0.4545 | 29 | 0.3793 |
| 1_16504333 | 10 | 0.9 | 7 | 0.7143 | 6 | 0.5 | 10 | 0.9 | 33 | 0.7879 | 1 | 1 | 10 | 0.7 | 6 | 0.6667 | 11 | 0.8182 | 28 | 0.75 |
| 1_16517853 | 7 | 0.1429 | 9 | 0.1111 | 7 | 0.2857 | 5 | 0.2 | 28 | 0.1786 | 7 | 0.2857 | 13 | 0.1538 | 9 | 0.1111 | 8 | 0.125 | 37 | 0.1622 |
| 1_16517859 | 4 | 0.5 | 7 | 0.2857 | 5 | 0.6 | 4 | 0.5 | 20 | 0.45 | 5 | 0.4 | 13 | 0.2308 | 6 | 0.1667 | 7 | 0.2857 | 31 | 0.2581 |
| 1_16517867 | 8 | 0.25 | 12 | 0.25 | 6 | 0.5 | 5 | 0 | 31 | 0.2581 | 6 | 0 | 12 | 0.0833 | 7 | 0.4286 | 6 | 0.1667 | 31 | 0.1613 |
| 1_16517872 | 10 | 0.4 | 12 | 0.1667 | 6 | 0.1667 | 5 | 0.2 | 33 | 0.2424 | 6 | 0.3333 | 13 | 0.1538 | 10 | 0.4 | 8 | 0.125 | 37 | 0.2432 |
| 1_16517873 | 10 | 0 | 12 | 0 | 8 | 0.25 | 4 | 0 | 34 | 0.0588 | 7 | 0 | 14 | 0.0714 | 10 | 0 | 8 | 0.125 | 39 | 0.0513 |
| 1_16541995 | NA | NA | 5 | 0.2 | 3 | 0.3333 | 2 | 0 | 10 | 0.2 | 3 | 0.3333 | NA | NA | 3 | 0 | NA | NA | 6 | 0.1667 |
| 1_16542013 | NA | NA | 5 | 0.2 | 3 | 0.6667 | 1 | 1 | 9 | 0.4444 | 3 | 0 | NA | NA | 3 | 0.3333 | NA | NA | 6 | 0.1667 |
| 1_16542016 | NA | NA | 4 | 0.75 | 3 | 0.6667 | 2 | 0.5 | 9 | 0.6667 | 2 | 0.5 | NA | NA | 3 | 1 | NA | NA | 5 | 0.8 |
| 1_16542040 | NA | NA | 3 | 0 | 1 | 0 | 1 | 1 | 5 | 0.2 | 2 | 0.5 | NA | NA | 2 | 0.5 | NA | NA | 4 | 0.5 |
| 1_16542064 | NA | NA | 5 | 0 | 2 | 0.5 | 2 | 0 | 9 | 0.1111 | 3 | 0 | NA | NA | 2 | 0.5 | NA | NA | 5 | 0.2 |
| 1_16542083 | NA | NA | 5 | 0.2 | 2 | 0 | 1 | 0 | 8 | 0.125 | 3 | 0.3333 | NA | NA | 2 | 0.5 | NA | NA | 5 | 0.4 |
| 1_16542106 | NA | NA | 5 | 0 | 2 | 0.5 | 1 | 1 | 8 | 0.25 | 3 | 0.3333 | NA | NA | 2 | 0.5 | NA | NA | 5 | 0.4 |
| 1_16542107 | NA | NA | 5 | 0.8 | 1 | 1 | 2 | 1 | 8 | 0.875 | 3 | 0.6667 | NA | NA | 2 | 0.5 | NA | NA | 5 | 0.6 |
| 1_16547716 | 1 | 0 | 1 | 1 | NA | NA | 1 | 0 | 3 | 0.3333 | 1 | 1 | NA | NA | NA | NA | NA | NA | 1 | 1 |
| 1_16547719 | 2 | 0 | 1 | 0 | NA | NA | 1 | 1 | 4 | 0.25 | 1 | 1 | NA | NA | NA | NA | NA | NA | 1 | 1 |
| 1_16547739 | 2 | 0.5 | 1 | 1 | NA | NA | 1 | 1 | 4 | 0.75 | NA | NA | NA | NA | 1 | 1 | NA | NA | 1 | 1 |
| 1_16547754 | 1 | 1 | 1 | 1 | NA | NA | 1 | 1 | 3 | 1 | 1 | 1 | NA | NA | 1 | 1 | NA | NA | 2 | 1 |
| 1_16548354 | 2 | 1 | NA | NA | NA | NA | NA | NA | 2 | 1 | NA | NA | NA | NA | 1 | 1 | NA | NA | 1 | 1 |
| 1_16559832 | 2 | 0 | 2 | 0 | 4 | 0 | 3 | 0.3333 | 11 | 0.0909 | 1 | 0 | 1 | 1 | 4 | 0 | 4 | 0.25 | 10 | 0.2 |
| 1_16560143 | 2 | 0 | 2 | 0.5 | 7 | 0 | 2 | 0 | 13 | 0.0769 | 4 | 0.25 | 2 | 0 | 5 | 0.2 | 3 | 0 | 14 | 0.1429 |
| 1_16560209 | 2 | 0 | 1 | 0 | 2 | 0 | 4 | 0.25 | 9 | 0.1111 | 4 | 0.5 | 3 | 0 | 4 | 0.25 | 3 | 0.3333 | 14 | 0.2857 |
| 1_16560273 | 4 | 0.25 | 1 | 0 | 6 | 0 | 4 | 0 | 15 | 0.0667 | 6 | 0.1667 | 3 | 0.3333 | 7 | 0.2857 | 5 | 0.2 | 21 | 0.2381 |
| 1_16560280 | 3 | 0 | 1 | 0 | 6 | 0.3333 | 4 | 0.25 | 14 | 0.2143 | 7 | 0 | 3 | 0 | 5 | 0.2 | 5 | 0.4 | 20 | 0.15 |
| 1_16569438 | 1 | 0 | 1 | 1 | 2 | 0 | NA | NA | 4 | 0.25 | 2 | 0.5 | 1 | 0 | NA | NA | NA | NA | 3 | 0.3333 |
| 1_16613645 | NA | NA | 2 | 0.5 | NA | NA | NA | NA | 2 | 0.5 | NA | NA | 1 | 1 | 1 | 0 | 1 | 0 | 3 | 0.3333 |
| 1_16613837 | NA | NA | 2 | 1 | NA | NA | NA | NA | 2 | 1 | NA | NA | 2 | 0.5 | 2 | 0 | 1 | 0 | 5 | 0.2 |
| 1_16613866 | NA | NA | 3 | 0.6667 | 1 | 1 | NA | NA | 4 | 0.75 | 1 | 0 | 2 | 1 | 1 | 1 | 1 | 0 | 5 | 0.6 |
| 1_16613882 | NA | NA | 3 | 1 | 1 | 1 | NA | NA | 4 | 1 | 1 | 1 | 2 | 1 | 2 | 1 | 1 | 1 | 6 | 1 |
| 1_16613895 | NA | NA | 2 | 0.5 | 1 | 1 | NA | NA | 3 | 0.6667 | 1 | 1 | 2 | 0.5 | 1 | 0 | NA | NA | 4 | 0.5 |
| 1_16613897 | NA | NA | 2 | 0.5 | 1 | 1 | NA | NA | 3 | 0.6667 | 1 | 1 | 2 | 0 | 2 | 1 | 1 | 0 | 6 | 0.5 |
| 1_16613904 | NA | NA | 3 | 1 | 1 | 0 | NA | NA | 4 | 0.75 | 1 | 1 | 1 | 1 | 1 | 1 | 1 | 1 | 4 | 1 |
| 1_16613904 | NA | NA | 3 | 0.3333 | 1 | 1 | NA | NA | 4 | 0.5 | 1 | 0 | 2 | 0.5 | NA | NA | 1 | 0 | 4 | 0.25 |
| 1_16613907 | NA | NA | 3 | 0.3333 | NA | NA | NA | NA | 3 | 0.3333 | 2 | 0 | 1 | 1 | 2 | 0.5 | 1 | 1 | 6 | 0.5 |
| 1_16613927 | NA | NA | 3 | 1 | 1 | 0 | NA | NA | 4 | 0.75 | 1 | 1 | 2 | 0.5 | 2 | 0.5 | 1 | 1 | 6 | 0.6667 |
| 1_16613949 | NA | NA | 2 | 1 | 1 | 1 | 1 | 1 | 4 | 1 | 2 | 1 | 2 | 1 | 1 | 0 | 1 | 1 | 6 | 0.8333 |
| 1_16613955 | NA | NA | 3 | 1 | 1 | 1 | 1 | 0 | 5 | 0.8 | NA | NA | 1 | 1 | 1 | 1 | 1 | 1 | 3 | 1 |
| 1_16613956 | NA | NA | 3 | 0.3333 | 1 | 0 | 1 | 0 | 5 | 0.2 | 2 | 0.5 | 2 | 0 | 3 | 0.3333 | 1 | 0 | 8 | 0.25 |
| 1_16613958 | NA | NA | 2 | 1 | 2 | 1 | 1 | 0 | 5 | 0.8 | 1 | 1 | NA | NA | 2 | 1 | 1 | 1 | 4 | 1 |
| 1_16613973 | NA | NA | 3 | 0.3333 | 1 | 1 | 1 | 0 | 5 | 0.4 | 2 | 1 | 2 | 0.5 | 3 | 0.6667 | 1 | 1 | 8 | 0.75 |
| 1_16613976 | NA | NA | 3 | 1 | 1 | 1 | NA | NA | 4 | 1 | 1 | 1 | 1 | 1 | 2 | 1 | 1 | 1 | 5 | 1 |
| 1_16613981 | NA | NA | 3 | 0.3333 | 2 | 1 | 1 | 1 | 6 | 0.6667 | 2 | 1 | 2 | 1 | 2 | 0.5 | 1 | 0 | 7 | 0.7143 |
| 1_16613993 | NA | NA | 2 | 0.5 | 2 | 1 | 1 | 0 | 5 | 0.6 | 1 | 0 | 3 | 1 | 2 | 1 | 1 | 1 | 7 | 0.8571 |
| 1_16613997 | NA | NA | 3 | 1 | 2 | 1 | 1 | 1 | 6 | 1 | 2 | 1 | 2 | 1 | 1 | 1 | 1 | 1 | 6 | 1 |
| 1_16613998 | NA | NA | 1 | 1 | NA | NA | 1 | 1 | 2 | 1 | 2 | 0.5 | 1 | 1 | 2 | 1 | 1 | 1 | 6 | 0.8333 |
| 1_16614009 | NA | NA | 4 | 0.25 | 2 | 0.5 | 1 | 0 | 7 | 0.2857 | 2 | 0.5 | 2 | 1 | 3 | 0.6667 | NA | NA | 7 | 0.7143 |
| 1_16614024 | NA | NA | 5 | 0.4 | 2 | 0.5 | 1 | 0 | 8 | 0.375 | 2 | 0 | 3 | 1 | 1 | 0 | NA | NA | 6 | 0.5 |
| 1_16614202 | 2 | 0 | 8 | 0.125 | 2 | 0 | 1 | 0 | 13 | 0.0769 | 3 | 0 | 3 | 0.3333 | 2 | 0 | 3 | 0.3333 | 11 | 0.1818 |
| 1_16614261 | 2 | 0.5 | 7 | 0.1429 | 1 | 0 | 2 | 0 | 12 | 0.1667 | 4 | 0.25 | 2 | 0 | 2 | 0.5 | 3 | 0.3333 | 11 | 0.2727 |
| 1_16622027 | 6 | 0.8333 | 9 | 0.6667 | NA | NA | 9 | 0.8889 | 24 | 0.7917 | 6 | 0.6667 | 8 | 0.625 | 10 | 0.8 | 5 | 0.4 | 29 | 0.6552 |
| 1_16622031 | 6 | 0 | 9 | 0.4444 | NA | NA | 9 | 0.2222 | 24 | 0.25 | 6 | 0 | 7 | 0.2857 | 8 | 0.5 | 5 | 0.4 | 26 | 0.3077 |
| 1_16622036 | 7 | 0 | 10 | 0 | NA | NA | 9 | 0.2222 | 26 | 0.0769 | 6 | 0.1667 | 8 | 0 | 10 | 0.2 | 5 | 0 | 29 | 0.1034 |
| 1_16622125 | NA | NA | 2 | 0.5 | NA | NA | NA | NA | 2 | 0.5 | 1 | 1 | NA | NA | 3 | 0 | NA | NA | 4 | 0.25 |
| 1_16622249 | 1 | 0 | 4 | 0.25 | 2 | 0 | 4 | 0.25 | 11 | 0.1818 | 1 | 0 | 4 | 0.25 | 6 | 0 | 2 | 0.5 | 13 | 0.1538 |
| 1_16622263 | 1 | 0 | 5 | 0.2 | 2 | 0 | 4 | 0.25 | 12 | 0.1667 | 2 | 0 | 5 | 0.2 | 6 | 0.1667 | 2 | 0 | 15 | 0.1333 |
| 1_16622305 | 1 | 0 | 6 | 0 | 2 | 0 | 5 | 0.4 | 14 | 0.1429 | 3 | 0.3333 | 6 | 0.3333 | 4 | 0 | 5 | 0.2 | 18 | 0.2222 |
| 1_16622375 | 1 | 0 | 6 | 0 | 2 | 0.5 | 4 | 0 | 13 | 0.0769 | 2 | 0 | 5 | 0 | 3 | 0 | 4 | 0.25 | 14 | 0.0714 |
| 1_16622418 | 1 | 0 | 4 | 0.25 | 2 | 0.5 | 4 | 0 | 11 | 0.1818 | 1 | 0 | 6 | 0 | 3 | 0 | 3 | 0.6667 | 13 | 0.1538 |
| 1_16622427 | 1 | 1 | 4 | 0 | 2 | 0.5 | 2 | 0.5 | 9 | 0.3333 | 1 | 0 | 6 | 0.5 | 2 | 1 | 4 | 0.5 | 13 | 0.5385 |
| 1_16622558 | 1 | 0 | 3 | 0.3333 | 1 | 0 | 3 | 0.3333 | 8 | 0.25 | 1 | 0 | 1 | 0 | NA | NA | 4 | 0.25 | 6 | 0.1667 |
| 1_16622592 | 1 | 0 | 2 | 0 | 1 | 0 | 3 | 0.3333 | 7 | 0.1429 | 1 | 0 | 3 | 0.3333 | NA | NA | 5 | 0.2 | 9 | 0.2222 |
| 1_16622625 | 1 | 1 | 2 | 0.5 | 1 | 0 | 3 | 1 | 7 | 0.7143 | 2 | 1 | 3 | 0.6667 | NA | NA | 3 | 0 | 8 | 0.5 |
| 1_16622626 | 1 | 1 | 3 | 1 | 1 | 1 | 3 | 1 | 8 | 1 | 2 | 1 | 3 | 1 | NA | NA | 4 | 1 | 9 | 1 |

| | | | | | | | | | | | | | | | | | | | | |
|---|---|---|---|---|---|---|---|---|---|---|---|---|---|---|---|---|---|---|---|---|
| 1_16622627 | 1 | 1 | 4 | 0.75 | 1 | 0 | 3 | 0.6667 | 9 | 0.6667 | 2 | 1 | 2 | 1 | NA | NA | 4 | 0.5 | 8 | 0.75 |
| 1_16622631 | 1 | 0 | 3 | 0.3333 | 1 | 0 | 2 | 0.5 | 7 | 0.2857 | 1 | 1 | NA | NA | NA | NA | 3 | 0.6667 | 4 | 0.75 |
| 1_16622642 | 1 | 0 | 3 | 0.6667 | 1 | 0 | NA | NA | 5 | 0.4 | 2 | 1 | 3 | 0.6667 | NA | NA | 2 | 1 | 7 | 0.8571 |
| 1_16622650 | 1 | 0 | 4 | 0.5 | 1 | 0 | 2 | 0 | 8 | 0.25 | 1 | 1 | 3 | 0.3333 | NA | NA | 5 | 0 | 9 | 0.2222 |
| 1_16622662 | NA | NA | 3 | 1 | NA | NA | 2 | 1 | 5 | 1 | 1 | 1 | 1 | 1 | NA | NA | 4 | 1 | 6 | 1 |
| 1_16622662 | 1 | 1 | 1 | 0 | NA | NA | 2 | 0 | 4 | 0.25 | NA | NA | 3 | 0.6667 | NA | NA | 3 | 0.6667 | 6 | 0.6667 |
| 1_16622665 | 1 | 1 | 1 | 1 | 1 | 0 | 3 | 0.6667 | 6 | 0.6667 | 1 | 1 | 3 | 0.3333 | NA | NA | 2 | 0 | 6 | 0.3333 |
| 1_16622672 | NA | NA | 3 | 0.3333 | NA | NA | 3 | 0 | 6 | 0.1667 | 1 | 0 | 3 | 0.3333 | NA | NA | 4 | 0.25 | 8 | 0.25 |
| 1_16622675 | 1 | 0 | 2 | 0.5 | 1 | 0 | 3 | 0 | 7 | 0.1429 | 1 | 0 | 2 | 0.5 | NA | NA | 3 | 0.6667 | 6 | 0.5 |
| 1_16622676 | 1 | 1 | 2 | 1 | 1 | 1 | 2 | 1 | 6 | 1 | 1 | 1 | 3 | 1 | NA | NA | 3 | 1 | 7 | 1 |
| 1_16622696 | 1 | 1 | 2 | 1 | NA | NA | 2 | 1 | 5 | 1 | NA | NA | 2 | 1 | NA | NA | 1 | 1 | 3 | 1 |
| 1_16622720 | NA | NA | 2 | 1 | NA | NA | 1 | 0 | 3 | 0.6667 | NA | NA | 1 | 1 | NA | NA | 2 | 1 | 3 | 1 |
| 1_16622726 | 1 | 1 | 2 | 0.5 | NA | NA | 3 | 0.3333 | 6 | 0.5 | NA | NA | 2 | 0.5 | NA | NA | 2 | 0 | 4 | 0.25 |
| 1_16622739 | 1 | 1 | 1 | 0 | NA | NA | 1 | 1 | 3 | 0.6667 | NA | NA | 2 | 0.5 | NA | NA | 2 | 0.5 | 4 | 0.5 |
| 1_16622760 | NA | NA | NA | NA | NA | NA | 1 | 1 | 1 | 1 | NA | NA | 1 | 1 | NA | NA | 2 | 1 | 3 | 1 |
| 1_16623665 | 1 | 0 | NA | NA | NA | NA | 3 | 0.6667 | 4 | 0.5 | NA | NA | 1 | 1 | NA | NA | 1 | 1 | 2 | 1 |
| 1_16682667 | 1 | 0 | 3 | 0.3333 | NA | NA | 1 | 0 | 5 | 0.2 | NA | NA | 3 | 0 | 1 | 0 | 4 | 0.25 | 8 | 0.125 |
| 1_16682668 | 1 | 1 | 3 | 0 | NA | NA | 2 | 0.5 | 6 | 0.3333 | NA | NA | 2 | 0 | 1 | 0 | 4 | 0.25 | 7 | 0.1429 |
| 1_16690615 | 3 | 0.3333 | 1 | 0 | 1 | 1 | 3 | 0 | 8 | 0.25 | NA | NA | 2 | 0 | 2 | 0 | 1 | 1 | 5 | 0.2 |
| 1_16690620 | 3 | 1 | 1 | 1 | 1 | 0 | 3 | 1 | 8 | 0.875 | NA | NA | 3 | 1 | 2 | 1 | 1 | 1 | 6 | 1 |
| 1_16690630 | 3 | 0 | 1 | 0 | 1 | 0 | 2 | 0.5 | 7 | 0.1429 | NA | NA | 3 | 0.3333 | 2 | 0.5 | 1 | 0 | 6 | 0.3333 |
| 1_16690633 | 3 | 0.3333 | 1 | 1 | 1 | 0 | 2 | 1 | 7 | 0.5714 | NA | NA | 3 | 0.6667 | 2 | 1 | 1 | 1 | 6 | 0.8333 |
| 1_16690642 | 2 | 1 | 2 | 1 | 1 | 1 | 3 | 1 | 8 | 1 | NA | NA | 2 | 0.5 | 1 | 1 | 1 | 1 | 4 | 0.75 |
| 1_16690644 | 2 | 0.5 | 2 | 1 | NA | NA | 2 | 0.5 | 6 | 0.6667 | NA | NA | 3 | 0.6667 | 2 | 0.5 | 1 | 1 | 6 | 0.6667 |
| 1_16690648 | 3 | 1 | 2 | 1 | 1 | 1 | 2 | 1 | 8 | 1 | NA | NA | 3 | 1 | 2 | 0.5 | 1 | 1 | 6 | 0.8333 |
| 1_16690659 | 2 | 0.5 | 2 | 0 | 1 | 0 | 3 | 0.6667 | 8 | 0.375 | NA | NA | 2 | 0 | 3 | 0.6667 | 1 | 0 | 6 | 0.3333 |
| 1_16690663 | 3 | 1 | 2 | 1 | NA | NA | 4 | 1 | 9 | 1 | NA | NA | 1 | 1 | 1 | 0 | 1 | 1 | 3 | 0.6667 |
| 1_16690665 | NA | NA | 1 | 0 | NA | NA | 4 | 0.25 | 5 | 0.2 | NA | NA | 2 | 0 | 3 | 0.3333 | 1 | 0 | 6 | 0.1667 |
| 1_16690667 | 3 | 0.3333 | 2 | 0 | 1 | 0 | 4 | 0.25 | 10 | 0.2 | NA | NA | 3 | 0.3333 | 3 | 0 | 1 | 0 | 7 | 0.1429 |
| 1_16690670 | 3 | 0.6667 | 2 | 0.5 | 1 | 0 | 4 | 0.5 | 10 | 0.5 | NA | NA | 4 | 0.25 | 2 | 0 | 1 | 0 | 7 | 0.1429 |
| 1_16690672 | 3 | 0 | 2 | 0.5 | 1 | 0 | 4 | 0 | 10 | 0.1 | NA | NA | 4 | 0.25 | 2 | 0 | 1 | 0 | 7 | 0.1429 |
| 1_16690685 | 3 | 1 | 1 | 1 | NA | NA | 4 | 1 | 8 | 1 | NA | NA | 4 | 1 | 2 | 0.5 | 1 | 0 | 7 | 0.7143 |
| 1_16690691 | 1 | 1 | NA | NA | NA | NA | 4 | 0.75 | 5 | 0.8 | NA | NA | 4 | 0.75 | 1 | 1 | NA | NA | 5 | 0.8 |
| 1_16690696 | 2 | 0 | 1 | 0 | NA | NA | 4 | 0.25 | 7 | 0.1429 | NA | NA | 4 | 0 | 2 | 0.5 | 1 | 0 | 7 | 0.1429 |
| 1_16690706 | 1 | 1 | NA | NA | NA | NA | 1 | 1 | 2 | 1 | NA | NA | NA | NA | 1 | 1 | NA | NA | 1 | 1 |
| 1_16690721 | 1 | 0 | NA | NA | NA | NA | 1 | 1 | 2 | 0.5 | NA | NA | NA | NA | 1 | 1 | NA | NA | 1 | 1 |
| 1_16690724 | 1 | 0 | NA | NA | NA | NA | 1 | 1 | 2 | 0.5 | NA | NA | 1 | 1 | 1 | 1 | NA | NA | 2 | 1 |
| 1_16691565 | 1 | 1 | 2 | 0 | NA | NA | 2 | 1 | 5 | 0.6 | NA | NA | 5 | 1 | 2 | 1 | 1 | 0 | 8 | 0.875 |
| 1_16691566 | 2 | 0 | 2 | 0.5 | NA | NA | 2 | 0 | 6 | 0.1667 | NA | NA | 5 | 0 | 2 | 0.5 | 1 | 0 | 8 | 0.125 |
| 1_16691579 | 2 | 1 | 3 | 0.6667 | NA | NA | 1 | 1 | 6 | 0.8333 | NA | NA | 3 | 1 | 2 | 0.5 | 1 | 1 | 6 | 0.8333 |
| 1_16691591 | 2 | 0.5 | 3 | 0.6667 | NA | NA | 2 | 1 | 7 | 0.7143 | NA | NA | 5 | 1 | 1 | 1 | 1 | 1 | 7 | 1 |
| 1_16691609 | 2 | 1 | 2 | 1 | NA | NA | 1 | 1 | 5 | 1 | NA | NA | 3 | 1 | 2 | 1 | 1 | 1 | 6 | 1 |
| 1_16691609 | 2 | 0 | 3 | 0.3333 | NA | NA | 1 | 0 | 6 | 0.1667 | NA | NA | 3 | 0 | 2 | 0.5 | 1 | 0 | 6 | 0.1667 |
| 1_16691757 | 2 | 0 | 3 | 0 | NA | NA | 3 | 0.3333 | 8 | 0.125 | NA | NA | 6 | 0.3333 | 3 | 0.6667 | 1 | 1 | 10 | 0.5 |
| 1_16691829 | 2 | 0 | 3 | 0.6667 | NA | NA | 3 | 0.3333 | 8 | 0.375 | NA | NA | 3 | 0.3333 | 5 | 0.2 | 1 | 0 | 9 | 0.2222 |
| 1_16691845 | 2 | 1 | 4 | 0.75 | NA | NA | 3 | 1 | 9 | 0.8889 | NA | NA | 5 | 1 | 2 | 1 | 1 | 1 | 8 | 1 |
| 1_16705370 | 1 | 1 | 3 | 0 | NA | NA | 3 | 0 | 7 | 0.1429 | 1 | 0 | 1 | 1 | 2 | 0.5 | NA | NA | 4 | 0.5 |
| 1_16705373 | NA | NA | 4 | 0.25 | NA | NA | 3 | 0.6667 | 7 | 0.4286 | 1 | 0 | NA | NA | 2 | 0.5 | NA | NA | 3 | 0.3333 |
| 1_16705376 | 1 | 0 | 4 | 0 | NA | NA | 3 | 1 | 8 | 0.375 | 1 | 0 | 1 | 1 | 2 | 1 | NA | NA | 4 | 0.75 |
| 1_16705378 | 1 | 0 | 2 | 0 | NA | NA | 3 | 0.3333 | 6 | 0.1667 | NA | NA | 1 | 0 | 2 | 0.5 | NA | NA | 3 | 0.3333 |
| 1_16705390 | 2 | 0.5 | 4 | 0.25 | NA | NA | 2 | 0 | 8 | 0.25 | 1 | 0 | NA | NA | 2 | 0.5 | NA | NA | 3 | 0.3333 |
| 1_16705395 | 2 | 0.5 | 3 | 0 | NA | NA | 2 | 0.5 | 7 | 0.2857 | 1 | 0 | 1 | 1 | 1 | 0 | NA | NA | 3 | 0.3333 |
| 1_16719974 | 1 | 1 | 4 | 0.5 | NA | NA | 3 | 0.3333 | 8 | 0.5 | NA | NA | NA | NA | 1 | 0 | 2 | 0.5 | 3 | 0.3333 |
| 1_16742609 | 4 | 1 | 3 | 0.3333 | 13 | 0.6923 | 5 | 0.8 | 25 | 0.72 | 3 | 1 | 10 | 0.7 | 8 | 0.625 | 6 | 0.8333 | 27 | 0.7407 |
| 1_16742630 | 4 | 0.5 | 4 | 0.25 | 12 | 0 | 6 | 0.1667 | 26 | 0.1538 | 2 | 0.5 | 9 | 0.4444 | 10 | 0.4 | 5 | 0.2 | 26 | 0.3846 |
| 1_16746699 | 10 | 0.2 | 8 | 0 | 17 | 0.1176 | 9 | 0.1111 | 44 | 0.1136 | 5 | 0 | 13 | 0 | 16 | 0.1875 | 9 | 0 | 43 | 0.0698 |
| 1_16798606 | 2 | 0 | 9 | 0.2222 | 8 | 0.125 | 6 | 0.5 | 25 | 0.24 | 6 | 0.3333 | 3 | 0.3333 | 5 | 0.4 | 5 | 0.2 | 19 | 0.3158 |
| 1_16798612 | 1 | 1 | 9 | 1 | 5 | 0.8 | 5 | 0.8 | 20 | 0.9 | 6 | 0.8333 | 2 | 1 | 4 | 0.75 | 5 | 0.6 | 17 | 0.7647 |
| 1_16825942 | 1 | 1 | 2 | 0.5 | NA | NA | 2 | 0 | 5 | 0.4 | 1 | 1 | NA | NA | 1 | 1 | NA | NA | 2 | 1 |
| 1_16952025 | 3 | 0.3333 | 1 | 1 | 4 | 0 | 9 | 0.1111 | 17 | 0.1765 | 2 | 0 | 3 | 0 | 7 | 0 | 5 | 0.2 | 17 | 0.0588 |
| 1_16952048 | 2 | 0 | NA | NA | 2 | 0 | 7 | 0.1429 | 11 | 0.0909 | 1 | 0 | 4 | 0.25 | 5 | 0.2 | 5 | 0 | 15 | 0.1333 |
| 1_16952274 | 8 | 0.25 | 5 | 0.6 | 7 | 0.2857 | 12 | 0.25 | 32 | 0.3125 | 6 | 0.1667 | 5 | 0.2 | 9 | 0.3333 | 9 | 0.2222 | 29 | 0.2414 |
| 1_16956890 | 1 | 1 | 2 | 0 | 1 | 0 | NA | NA | 4 | 0.25 | NA | NA | 3 | 0 | 3 | 0.3333 | 1 | 0 | 7 | 0.1429 |
| 1_16956997 | 25 | 0.08 | 37 | 0.1081 | 40 | 0.15 | 27 | 0.1481 | 129 | 0.124 | 14 | 0.2143 | 37 | 0.1892 | 33 | 0.1212 | 27 | 0.1852 | 111 | 0.1712 |
| 1_16957022 | 29 | 0.1034 | 36 | 0.1944 | 42 | 0.1667 | 35 | 0.1143 | 142 | 0.1479 | 15 | 0.1333 | 38 | 0.1316 | 32 | 0.1875 | 27 | 0.2222 | 112 | 0.1696 |

| | | | | | | | | | | | | | | | | | | | | |
|---|---|---|---|---|---|---|---|---|---|---|---|---|---|---|---|---|---|---|---|---|
| 1_1695704( | 32 | 0.125 | 33 | 0.0303 | 37 | 0.1081 | 33 | 0.0606 | 135 | 0.0815 | 16 | 0.0625 | 35 | 0.0286 | 32 | 0.0312 | 28 | 0.1071 | 111 | 0.0541 |
| 1_1695708( | 29 | 0.5517 | 28 | 0.5714 | 34 | 0.5294 | 32 | 0.6562 | 123 | 0.5772 | 13 | 0.6923 | 34 | 0.4412 | 26 | 0.5 | 25 | 0.6 | 98 | 0.5306 |
| 1_1695711! | 25 | 0.08 | 28 | 0 | 36 | 0.0833 | 31 | 0.0645 | 120 | 0.0583 | 12 | 0.0833 | 37 | 0.0811 | 30 | 0 | 20 | 0.05 | 99 | 0.0505 |
| 1_1695712' | 23 | 0.913 | 27 | 0.963 | 37 | 0.8649 | 30 | 0.9 | 117 | 0.906 | 10 | 0.9 | 32 | 0.9062 | 29 | 0.931 | 24 | 0.9583 | 95 | 0.9263 |
| 1_1695712( | 19 | 0.4737 | 29 | 0.3103 | 34 | 0.5 | 27 | 0.4074 | 109 | 0.422 | 11 | 0.5455 | 31 | 0.4516 | 27 | 0.4444 | 24 | 0.3333 | 93 | 0.4301 |
| 1_1695714; | 16 | 0.375 | 24 | 0.375 | 24 | 0.2083 | 21 | 0.381 | 85 | 0.3294 | 8 | 0.375 | 25 | 0.28 | 20 | 0.2 | 20 | 0.2 | 73 | 0.2466 |
| 1_1697615( | NA | NA | 1 | 1 | 3 | 1 | NA | NA | 4 | 1 | 2 | 1 | 1 | 1 | 2 | 1 | 3 | 1 | 8 | 1 |
| 1_1697617; | 1 | 1 | 2 | 0.5 | 3 | 1 | NA | NA | 6 | 0.8333 | 2 | 1 | 1 | 1 | 2 | 1 | 4 | 1 | 9 | 1 |
| 1_1697619; | 1 | 1 | 3 | 0.6667 | 3 | 1 | NA | NA | 7 | 0.8571 | 2 | 1 | 1 | 1 | 2 | 0.5 | 4 | 1 | 9 | 0.8889 |
| 1_1697620; | 1 | 0 | 1 | 0 | 3 | 0.6667 | NA | NA | 5 | 0.4 | 3 | 0.6667 | NA | NA | 1 | 1 | 2 | 0.5 | 6 | 0.6667 |
| 1_1697638; | NA | NA | NA | NA | 1 | 1 | NA | NA | 1 | 1 | NA | NA | NA | NA | NA | NA | 2 | 0.5 | 2 | 0.5 |
| 1_1697639! | NA | NA | NA | NA | 1 | 1 | NA | NA | 1 | 1 | NA | NA | NA | NA | NA | NA | 1 | 1 | 1 | 1 |
| 1_1697641' | NA | NA | NA | NA | 1 | 1 | NA | NA | 1 | 1 | NA | NA | NA | NA | NA | NA | 2 | 1 | 2 | 1 |
| 1_1697802; | 1 | 0 | 3 | 0.3333 | 2 | 0.5 | NA | NA | 6 | 0.3333 | 2 | 0 | 1 | 1 | NA | NA | 1 | 0 | 4 | 0.25 |
| 1_1697802( | 1 | 0 | 3 | 0.3333 | 2 | 0 | NA | NA | 6 | 0.1667 | 2 | 0.5 | 1 | 0 | NA | NA | 1 | 0 | 4 | 0.25 |
| 1_1697803! | 1 | 1 | 3 | 0.6667 | 1 | 0 | NA | NA | 5 | 0.6 | 2 | 0.5 | 1 | 1 | NA | NA | 1 | 0 | 4 | 0.5 |
| 1_1697804( | 1 | 1 | 3 | 1 | 2 | 0.5 | NA | NA | 6 | 0.8333 | 2 | 1 | 1 | 1 | NA | NA | 1 | 1 | 4 | 1 |
| 1_1698012; | 1 | 0 | NA | NA | NA | NA | 2 | 0.5 | 3 | 0.3333 | 1 | 1 | NA | NA | NA | NA | NA | NA | 1 | 1 |
| 1_1699695! | 1 | 0 | 3 | 0.3333 | 4 | 0.5 | 3 | 0 | 11 | 0.2727 | 3 | 0 | 4 | 0.25 | 3 | 0.6667 | 2 | 0 | 12 | 0.25 |
| 1_1699697! | 1 | 0 | 6 | 0.5 | 7 | 0 | 6 | 0.1667 | 20 | 0.2 | 5 | 0.4 | 6 | 0.1667 | 6 | 0.1667 | 3 | 0.3333 | 20 | 0.25 |
| 1_1699698! | 1 | 0 | 6 | 0 | 6 | 0.1667 | 7 | 0.2857 | 20 | 0.15 | 5 | 0 | 4 | 0.25 | 6 | 0.1667 | 4 | 0.5 | 19 | 0.2105 |
| 1_1699699' | 1 | 1 | 6 | 0 | 6 | 0.6667 | 7 | 0.5714 | 20 | 0.45 | 5 | 0.8 | 6 | 0.3333 | 5 | 0.2 | 4 | 0.5 | 20 | 0.45 |
| 1_1699699; | 1 | 0 | 7 | 0.1429 | 4 | 0.25 | 5 | 0.2 | 17 | 0.1765 | 5 | 0.6 | 6 | 0.3333 | 7 | 0.2857 | 4 | 0.25 | 22 | 0.3636 |
| 1_1699699! | 1 | 0 | 5 | 0 | 6 | 0.1667 | 7 | 0.1429 | 19 | 0.1053 | 4 | 0.25 | 5 | 0.2 | 7 | 0 | 4 | 0.25 | 20 | 0.15 |
| 1_1699699! | NA | NA | 4 | 0.25 | 1 | 0 | 3 | 0 | 8 | 0.125 | 2 | 0 | 4 | 0.25 | 7 | 0 | 3 | 0.3333 | 16 | 0.125 |
| 1_1699943! | NA | NA | 6 | 0.1667 | 3 | 0.3333 | 4 | 0.75 | 13 | 0.3846 | 4 | 0.25 | 2 | 1 | 4 | 0.25 | 1 | 0 | 11 | 0.3636 |
| 1_1699943; | NA | NA | 4 | 0.25 | 3 | 0.6667 | 3 | 0.6667 | 10 | 0.5 | 4 | 0.75 | 2 | 0 | 3 | 0.3333 | 2 | 0 | 11 | 0.3636 |
| 1_1700211! | 27 | 0.2963 | 30 | 0.1 | 33 | 0.1818 | 31 | 0.129 | 121 | 0.1736 | 6 | 0.1667 | 24 | 0.125 | 28 | 0.0714 | 21 | 0.1429 | 79 | 0.1139 |
| 1_1700261' | 3 | 0.3333 | 11 | 0 | 9 | 0.4444 | 8 | 0.375 | 31 | 0.2581 | 2 | 0 | 8 | 0 | 10 | 0.1 | 8 | 0.25 | 28 | 0.1071 |
| 1_1700272! | 2 | 0 | 6 | 0.3333 | 4 | 0.25 | 4 | 0 | 16 | 0.1875 | 1 | 1 | 4 | 0.25 | 8 | 0.125 | 6 | 0.3333 | 19 | 0.2632 |
| 1_1700274; | 3 | 1 | 6 | 1 | 6 | 0.8333 | 6 | 1 | 21 | 0.9524 | NA | NA | 5 | 0.8 | 7 | 0.8571 | 6 | 0.8333 | 18 | 0.8333 |
| 1_1701264( | NA | NA | NA | NA | 1 | 1 | 1 | 1 | 2 | 1 | 2 | 0.5 | NA | NA | 1 | 0 | NA | NA | 3 | 0.3333 |
| 1_1701266' | NA | NA | NA | NA | 1 | 1 | 1 | 1 | 2 | 1 | 2 | 0.5 | NA | NA | 1 | 0 | NA | NA | 3 | 0.3333 |
| 1_1701281' | NA | NA | NA | NA | 1 | 1 | NA | NA | 1 | 1 | 2 | 0.5 | NA | NA | 1 | 1 | NA | NA | 3 | 0.6667 |
| 1_1701282( | NA | NA | NA | NA | 1 | 1 | 1 | 1 | 2 | 1 | 2 | 0.5 | NA | NA | 1 | 0 | NA | NA | 3 | 0.3333 |
| 1_1701282; | NA | NA | NA | NA | 1 | 0 | 1 | 1 | 2 | 0.5 | 2 | 0 | NA | NA | 1 | 1 | NA | NA | 3 | 0.3333 |
| 1_1701287( | NA | NA | NA | NA | 1 | 1 | NA | NA | 1 | 1 | 1 | 1 | NA | NA | 1 | 1 | NA | NA | 2 | 1 |
| 1_1701289; | NA | NA | NA | NA | 1 | 1 | 1 | 1 | 2 | 1 | 1 | 1 | NA | NA | 1 | 1 | NA | NA | 2 | 1 |
| 1_1701290· | NA | NA | NA | NA | 1 | 1 | 1 | 1 | 2 | 1 | 1 | 1 | NA | NA | 1 | 1 | NA | NA | 2 | 1 |
| 1_1701302; | NA | NA | NA | NA | 1 | 1 | NA | NA | 1 | 1 | 1 | 1 | NA | NA | 1 | 1 | NA | NA | 2 | 1 |
| 1_1701424' | 2 | 0 | 6 | 0.3333 | 6 | 0 | 3 | 0 | 17 | 0.1176 | 5 | 0 | 3 | 0.3333 | 1 | 0 | 2 | 0 | 11 | 0.0909 |
| 1_1701431· | 9 | 0.1111 | 9 | 0 | 8 | 0 | 9 | 0.1111 | 35 | 0.0571 | 11 | 0 | 10 | 0.2 | 8 | 0 | 10 | 0 | 39 | 0.0513 |
| 1_1701432' | 10 | 0.2 | 8 | 0.125 | 10 | 0.1 | 9 | 0.1111 | 37 | 0.1351 | 9 | 0.2222 | 11 | 0 | 7 | 0 | 9 | 0.1111 | 36 | 0.0833 |
| 1_1701432! | 10 | 0.3 | 8 | 0.375 | 9 | 0.3333 | 11 | 0.1818 | 38 | 0.2895 | 7 | 0.4286 | 10 | 0 | 9 | 0.1111 | 10 | 0.2 | 36 | 0.1667 |
| 1_1701433; | 10 | 0.9 | 7 | 0.4286 | 10 | 0.4 | 9 | 0.6667 | 36 | 0.6111 | 9 | 0.6667 | 9 | 1 | 10 | 0.6 | 10 | 0.7 | 38 | 0.7368 |
| 1_1701433· | 9 | 1 | 9 | 1 | 9 | 1 | 11 | 0.9091 | 38 | 0.9737 | 9 | 1 | 7 | 0.7143 | 10 | 1 | 6 | 0.8333 | 32 | 0.9062 |
| 1_1701433; | 8 | 0.125 | 8 | 0.125 | 9 | 0.2222 | 7 | 0.1429 | 32 | 0.1562 | 10 | 0.2 | 10 | 0.3 | 7 | 0.1429 | 10 | 0 | 37 | 0.1622 |
| 1_1701434! | 7 | 0.8571 | 4 | 1 | 5 | 1 | 8 | 0.875 | 24 | 0.9167 | 8 | 1 | 6 | 1 | 5 | 0.8 | 9 | 1 | 28 | 0.9643 |
| 1_1701434( | 7 | 0.4286 | 6 | 0.6667 | 5 | 0.6 | 10 | 0.5 | 28 | 0.5357 | 7 | 0.5714 | 8 | 0.625 | 10 | 0.7 | 5 | 0.6 | 30 | 0.6333 |
| 1_1701436; | 8 | 0.375 | 7 | 0.5714 | 7 | 0.5714 | 9 | 0.5556 | 31 | 0.5161 | 7 | 0.7143 | 8 | 0.75 | 9 | 0.6667 | 6 | 0.6667 | 30 | 0.7 |
| 1_1701437! | 8 | 0.375 | 7 | 1 | 7 | 0.2857 | 9 | 0.4444 | 31 | 0.5161 | 7 | 0.2857 | 8 | 0.625 | 10 | 0.3 | 6 | 0.6667 | 31 | 0.4516 |
| 1_1701439· | 7 | 0 | 4 | 0.5 | 6 | 0.3333 | 5 | 0.4 | 22 | 0.2727 | 5 | 0.4 | 7 | 0.2857 | 10 | 0.3 | 4 | 0.25 | 26 | 0.3077 |
| 1_1701455' | 1 | 0 | 3 | 0.3333 | 2 | 0 | 2 | 0 | 8 | 0.125 | 4 | 0 | 4 | 0 | 3 | 0.3333 | NA | NA | 11 | 0.0909 |
| 1_1701456; | 4 | 0 | 4 | 0 | 4 | 0.25 | 5 | 0.2 | 17 | 0.1176 | 4 | 0 | 5 | 0.4 | 6 | 0 | 2 | 0 | 17 | 0.1176 |
| 1_1701460( | 3 | 0.3333 | 3 | 0 | 2 | 0 | 6 | 0.5 | 14 | 0.2857 | 3 | 0.3333 | 4 | 0 | 3 | 0 | 2 | 0 | 12 | 0.0833 |
| 1_1701461; | 1 | 0 | 3 | 0 | 2 | 0 | 5 | 0.2 | 11 | 0.0909 | 3 | 0 | 4 | 0.25 | 3 | 0.3333 | 2 | 0 | 12 | 0.1667 |
| 1_1701462! | 2 | 0 | 3 | 0.3333 | 5 | 0.6 | 6 | 0.8333 | 16 | 0.5625 | 3 | 0.6667 | 3 | 0.6667 | 3 | 1 | 2 | 0 | 11 | 0.6364 |
| 1_1701463; | 2 | 0 | 4 | 0.75 | 4 | 0.75 | 5 | 0.8 | 15 | 0.6667 | 4 | 0.5 | 2 | 0 | 3 | 0.6667 | 2 | 0 | 11 | 0.3636 |
| 1_1701463; | 2 | 0.5 | 4 | 0.75 | 5 | 0.4 | 6 | 0.5 | 17 | 0.5294 | 4 | 0.75 | 3 | 0.6667 | 3 | 1 | 2 | 0 | 12 | 0.6667 |
| 1_1704965; | NA | NA | 1 | 0 | 1 | 0 | 1 | 1 | 3 | 0.3333 | NA | NA | NA | NA | NA | NA | 1 | 1 | 1 | 1 |
| 1_1704966( | NA | NA | 1 | 1 | 1 | 1 | 1 | 1 | 3 | 1 | NA | NA | NA | NA | NA | NA | 1 | 1 | 1 | 1 |
| 1_1704967; | NA | NA | 1 | 0 | 1 | 1 | 1 | 1 | 3 | 0.6667 | NA | NA | NA | NA | NA | NA | 1 | 1 | 1 | 1 |
| 1_1704967! | NA | NA | 1 | 1 | 1 | 1 | 1 | 0 | 3 | 0.6667 | NA | NA | NA | NA | NA | NA | 1 | 1 | 1 | 1 |
| 1_1705095( | NA | NA | 1 | 1 | 1 | 1 | 1 | 1 | 3 | 1 | NA | NA | NA | NA | NA | NA | 1 | 1 | 1 | 1 |
| 1_1705096( | NA | NA | NA | NA | 1 | 1 | 1 | 0 | 2 | 0.5 | NA | NA | NA | NA | NA | NA | 1 | 1 | 1 | 1 |

| | | | | | | | | | | | | | | | | | | | | |
|---|---|---|---|---|---|---|---|---|---|---|---|---|---|---|---|---|---|---|---|---|
| 1_17050970 | NA | NA | 1 | 1 | 1 | 1 | 1 | 1 | 3 | 1 | NA | NA | NA | NA | NA | NA | 1 | 1 | 1 | 1 |
| 1_17050981 | 1 | 1 | NA | NA | NA | NA | 1 | 1 | 2 | 1 | NA | NA | NA | NA | NA | NA | 1 | 1 | 1 | 1 |
| 1_17051013 | 1 | 0 | 1 | 1 | 1 | 0 | NA | NA | 3 | 0.3333 | NA | NA | NA | NA | NA | NA | 1 | 1 | 1 | 1 |
| 1_17053305 | 4 | 0.25 | 1 | 0 | 4 | 0 | 10 | 0 | 19 | 0.0526 | 4 | 0 | 4 | 0 | 6 | 0.1667 | 6 | 0 | 20 | 0.05 |
| 1_17053327 | 3 | 0 | 1 | 0 | 1 | 1 | 4 | 0 | 9 | 0.1111 | 2 | 0 | 3 | 0 | 4 | 0 | 3 | 0.3333 | 12 | 0.0833 |
| 1_17053542 | 4 | 0 | 3 | 0 | 4 | 0 | 8 | 0.125 | 19 | 0.0526 | 1 | 0 | 2 | 0 | 3 | 0 | 5 | 0.2 | 11 | 0.0909 |
| 1_17053608 | 2 | 0 | 3 | 0 | 5 | 0 | 8 | 0.5 | 18 | 0.2222 | 1 | 1 | 1 | 0 | 2 | 0 | 3 | 0.3333 | 7 | 0.2857 |
| 1_17053733 | 3 | 0 | 2 | 0 | 2 | 0 | 4 | 0.25 | 11 | 0.0909 | 1 | 0 | 1 | 0 | 1 | 0 | 4 | 0.25 | 7 | 0.1429 |
| 1_17053737 | 3 | 0.3333 | 3 | 0.3333 | 2 | 1 | 4 | 0.5 | 12 | 0.5 | 1 | 1 | 3 | 0.3333 | 2 | 0.5 | 3 | 0.3333 | 9 | 0.4444 |
| 1_17053742 | 2 | 0 | 3 | 0.6667 | 2 | 0.5 | 4 | 0.75 | 11 | 0.5455 | 1 | 1 | 3 | 0.6667 | 2 | 1 | 3 | 0.6667 | 9 | 0.7778 |
| 1_17053743 | 1 | 0 | 2 | 0.5 | 1 | 1 | 2 | 0 | 6 | 0.3333 | 1 | 1 | 3 | 0.3333 | 1 | 1 | 2 | 0.5 | 7 | 0.5714 |
| 1_17053761 | 3 | 0 | 3 | 0 | NA | NA | 4 | 0.25 | 10 | 0.1 | 1 | 1 | 2 | 0 | NA | NA | 3 | 0.6667 | 6 | 0.5 |
| 1_17070187 | 1 | 0 | 1 | 1 | 3 | 0.6667 | 1 | 1 | 6 | 0.6667 | NA | NA | 1 | 0 | 5 | 0.8 | 3 | 0.3333 | 9 | 0.5556 |
| 1_17070188 | 1 | 0 | 1 | 1 | 4 | 0.5 | 1 | 0 | 7 | 0.4286 | 1 | 0 | 2 | 0.5 | 5 | 0.4 | 3 | 0 | 11 | 0.2727 |
| 1_17070197 | 1 | 0 | 1 | 0 | 4 | 0.25 | 1 | 0 | 7 | 0.1429 | 2 | 0 | 3 | 0.3333 | 5 | 0.4 | 4 | 0.25 | 14 | 0.2857 |
| 1_17070197 | 1 | 0 | 1 | 0 | 2 | 0.5 | 1 | 0 | 5 | 0.2 | 2 | 0.5 | 3 | 0.3333 | 3 | 0 | 4 | 0.25 | 12 | 0.25 |
| 1_17070203 | 1 | 0 | 1 | 0 | 4 | 0.75 | 1 | 0 | 7 | 0.4286 | 2 | 0 | 3 | 0 | 5 | 0.2 | 3 | 0 | 13 | 0.0769 |
| 1_17070723 | 1 | 0 | 1 | 0 | 3 | 0.6667 | 1 | 0 | 6 | 0.3333 | 2 | 0.5 | 3 | 0 | 4 | 0.5 | 3 | 0 | 12 | 0.25 |
| 1_17078843 | 1 | 1 | 1 | 1 | NA | NA | NA | NA | 2 | 1 | 1 | 1 | 1 | 1 | NA | NA | NA | NA | 2 | 1 |
| 1_17087884 | 2 | 0.5 | 7 | 0.5714 | 2 | 0.5 | 2 | 1 | 13 | 0.6154 | 4 | 0.5 | 3 | 0.3333 | NA | NA | 3 | 0 | 10 | 0.3 |
| 1_17087897 | NA | NA | 5 | 0.2 | 2 | 0 | 1 | 0 | 8 | 0.125 | 4 | 0 | 2 | 0.5 | NA | NA | 2 | 0 | 8 | 0.125 |
| 1_17087899 | 1 | 1 | 6 | 0.6667 | 2 | 1 | 1 | 1 | 10 | 0.8 | 5 | 0.4 | 3 | 0.6667 | 1 | 0 | 3 | 0.3333 | 12 | 0.4167 |
| 1_17087934 | 1 | 0 | 7 | 0.4286 | 2 | 0 | 2 | 0 | 12 | 0.25 | 4 | 0.5 | 2 | 0 | 1 | 0 | 2 | 0 | 9 | 0.2222 |
| 1_17088157 | 1 | 0 | 7 | 0.1429 | 2 | 0 | 2 | 0 | 12 | 0.0833 | 4 | 0 | 2 | 0.5 | 1 | 0 | 2 | 0 | 9 | 0.1111 |
| 1_17100189 | 38 | 0.0526 | 62 | 0.0645 | 81 | 0.037 | 34 | 0.0882 | 215 | 0.0558 | 30 | 0.0333 | 83 | 0.0482 | 91 | 0.0989 | 59 | 0.0678 | 263 | 0.0684 |
| 1_17104381 | NA | NA | 1 | 1 | NA | NA | NA | NA | 1 | 1 | NA | NA | NA | NA | NA | NA | 1 | 1 | 1 | 1 |
| 1_17104405 | NA | NA | 1 | 1 | NA | NA | 1 | 1 | 2 | 1 | NA | NA | NA | NA | NA | NA | 1 | 1 | 1 | 1 |
| 1_17104528 | NA | NA | 1 | 1 | NA | NA | NA | NA | 1 | 1 | 1 | 1 | NA | NA | 1 | 1 | NA | NA | 2 | 1 |
| 1_17104549 | 4 | 0 | 4 | 0.25 | 2 | 0 | 5 | 0 | 15 | 0.0667 | 4 | 0 | 4 | 0.25 | 3 | 0 | 1 | 0 | 12 | 0.0833 |
| 1_17104553 | 4 | 0.25 | 4 | 0 | 2 | 1 | 4 | 0.25 | 14 | 0.2857 | 4 | 0.5 | 4 | 0.5 | 3 | 0.6667 | 1 | 1 | 12 | 0.5833 |
| 1_17104563 | 3 | 0 | 4 | 0.5 | 1 | 0 | 4 | 0 | 12 | 0.1667 | 4 | 0.5 | 4 | 0.25 | 3 | 0.3333 | 1 | 0 | 12 | 0.3333 |
| 1_17104565 | 4 | 1 | 4 | 0.75 | 2 | 1 | 3 | 1 | 13 | 0.9231 | 3 | 1 | 4 | 0.5 | 2 | 1 | 1 | 1 | 10 | 0.8 |
| 1_17104570 | 4 | 1 | 4 | 0.75 | 1 | 0 | 3 | 0.3333 | 12 | 0.6667 | 4 | 0.5 | 3 | 0.3333 | 3 | 0.6667 | 1 | 1 | 11 | 0.5455 |
| 1_17104575 | 4 | 0 | 3 | 0 | NA | NA | 4 | 0.25 | 11 | 0.0909 | 3 | 0.6667 | 3 | 0 | 2 | 0 | 1 | 0 | 9 | 0.2222 |
| 1_17104595 | 4 | 0.25 | 4 | 0 | 2 | 0 | 4 | 0 | 14 | 0.0714 | 3 | 0 | 3 | 0.3333 | 2 | 0 | 1 | 0 | 9 | 0.1111 |
| 1_17104601 | 4 | 0.25 | 4 | 0 | 2 | 0.5 | 3 | 0 | 13 | 0.1538 | 4 | 0.25 | 3 | 0.3333 | 3 | 0.3333 | 1 | 0 | 11 | 0.2727 |
| 1_17104608 | 3 | 0 | 2 | 0.5 | NA | NA | 1 | 0 | 6 | 0.1667 | 2 | 0.5 | 2 | 0 | 3 | 0 | NA | NA | 7 | 0.1429 |
| 1_17104641 | 4 | 0.75 | 3 | 0.6667 | 2 | 0.5 | 2 | 0.5 | 11 | 0.6364 | 6 | 0.8333 | 2 | 1 | 4 | 1 | NA | NA | 12 | 0.9167 |
| 1_17104641 | 4 | 0.75 | 3 | 0.3333 | 2 | 1 | 4 | 0.75 | 13 | 0.6923 | 5 | 0.8 | 3 | 1 | 3 | 1 | NA | NA | 11 | 0.9091 |
| 1_17104656 | 4 | 0.75 | 2 | 1 | 1 | 1 | 4 | 0.75 | 11 | 0.8182 | 4 | 1 | 3 | 1 | 3 | 1 | NA | NA | 10 | 1 |
| 1_17104681 | 14 | 0.3571 | 16 | 0.5625 | 10 | 0.6 | 9 | 0 | 49 | 0.4082 | 16 | 0.4375 | 18 | 0.4444 | 13 | 0.6154 | 10 | 0.4 | 57 | 0.4737 |
| 1_17104692 | 9 | 0.3333 | 9 | 0.5556 | 9 | 0.2222 | 8 | 0.5 | 35 | 0.4 | 10 | 0.2 | 14 | 0.3571 | 17 | 0.3529 | 10 | 0.7 | 51 | 0.3922 |
| 1_17104702 | 13 | 0.7692 | 15 | 0.8 | 11 | 0.8182 | 10 | 0.8 | 49 | 0.7959 | 10 | 0.8 | 15 | 0.7333 | 19 | 0.6842 | 9 | 0.5556 | 53 | 0.6981 |
| 1_17105727 | 5 | 0 | 1 | 0 | 6 | 0.3333 | 4 | 0.25 | 16 | 0.1875 | 6 | 0 | 4 | 0 | 4 | 0 | 3 | 0.6667 | 17 | 0.1176 |
| 1_17108668 | 1 | 1 | NA | NA | 1 | 0 | NA | NA | 2 | 0.5 | 1 | 0 | 1 | 0 | NA | NA | 1 | 1 | 3 | 0.3333 |
| 1_17108679 | 1 | 1 | NA | NA | 1 | 1 | NA | NA | 2 | 1 | 1 | 1 | 1 | 1 | NA | NA | 1 | 1 | 3 | 1 |
| 1_17108729 | 1 | 0 | NA | NA | 2 | 0.5 | NA | NA | 3 | 0.3333 | 1 | 0 | 2 | 0.5 | NA | NA | 1 | 0 | 4 | 0.25 |
| 1_17108905 | 1 | 1 | NA | NA | NA | NA | NA | NA | 1 | 1 | 1 | 1 | NA | NA | NA | NA | 1 | 1 | 2 | 1 |
| 1_17110397 | 4 | 0.75 | 4 | 0.5 | 2 | 0.5 | 4 | 0.5 | 14 | 0.5714 | 7 | 0.5714 | 3 | 0.3333 | 7 | 0.7143 | 2 | 0.5 | 19 | 0.5789 |
| 1_17110407 | 8 | 0.5 | 7 | 0 | 4 | 0 | 4 | 0 | 23 | 0.1739 | 9 | 0.1111 | 7 | 0.2857 | 9 | 0 | 4 | 0.5 | 29 | 0.1724 |
| 1_17110415 | 7 | 0.1429 | 9 | 0.1111 | 3 | 0 | 4 | 0.5 | 23 | 0.1739 | 9 | 0.6667 | 7 | 0.1429 | 9 | 0.1111 | 3 | 0.6667 | 28 | 0.3571 |
| 1_17110425 | 6 | 0.6667 | 4 | 1 | 3 | 0.6667 | 4 | 0.25 | 17 | 0.6471 | 9 | 0.5556 | 5 | 0.8 | 7 | 0.7143 | 4 | 0.75 | 25 | 0.68 |
| 1_17110858 | NA | NA | 1 | 1 | 2 | 0 | 2 | 0 | 5 | 0.2 | NA | NA | NA | NA | NA | NA | 1 | 1 | 1 | 1 |
| 1_17111400 | 2 | 0.5 | 2 | 0.5 | 2 | 0.5 | 3 | 0.6667 | 9 | 0.5556 | 4 | 0.75 | 3 | 0.3333 | 5 | 0.2 | 5 | 0.6 | 17 | 0.4706 |
| 1_17111417 | 2 | 1 | 1 | 0 | 1 | 0 | 4 | 0.75 | 8 | 0.625 | 4 | 0.75 | 3 | 0.6667 | 4 | 0.5 | 4 | 1 | 15 | 0.7333 |
| 1_17111420 | 1 | 0 | 3 | 0 | 1 | 0 | 4 | 0.25 | 9 | 0.1111 | 2 | 0.5 | 3 | 0.6667 | 2 | 0.5 | 4 | 0 | 11 | 0.3636 |
| 1_17111432 | 1 | 1 | 2 | 0 | 1 | 1 | 3 | 0.6667 | 7 | 0.5714 | 2 | 0 | 3 | 0 | 3 | 0.3333 | 3 | 0 | 11 | 0.0909 |
| 1_17111448 | 1 | 1 | 2 | 0 | 2 | 0 | 5 | 0 | 10 | 0.1 | 4 | 0 | 3 | 0 | 3 | 0.6667 | 4 | 0.25 | 14 | 0.2143 |
| 1_17111629 | 22 | 0.0455 | 17 | 0.2941 | 15 | 0.0667 | 19 | 0.1053 | 73 | 0.1233 | 14 | 0.1429 | 20 | 0.15 | 14 | 0 | 15 | 0.0667 | 63 | 0.0952 |
| 1_17111634 | 21 | 0.0952 | 19 | 0.1579 | 13 | 0.0769 | 19 | 0.1053 | 72 | 0.1111 | 11 | 0.1818 | 22 | 0.0455 | 15 | 0 | 15 | 0.1333 | 63 | 0.0794 |
| 1_17111643 | 21 | 0.0952 | 16 | 0.1875 | 16 | 0 | 18 | 0.0556 | 71 | 0.0845 | 10 | 0 | 20 | 0.1 | 14 | 0.0714 | 15 | 0.2667 | 59 | 0.1186 |
| 1_17111644 | 18 | 0 | 15 | 0 | 15 | 0.0667 | 18 | 0.1667 | 66 | 0.0606 | 12 | 0.0833 | 18 | 0.1111 | 16 | 0.125 | 11 | 0.1818 | 57 | 0.1228 |
| 1_17111646 | 16 | 0.0625 | 14 | 0.1429 | 16 | 0.0625 | 17 | 0 | 63 | 0.0635 | 11 | 0.0909 | 17 | 0.1176 | 14 | 0.0714 | 12 | 0.1667 | 54 | 0.1111 |
| 1_17116606 | 2 | 0.5 | 3 | 0.6667 | 5 | 0.2 | 3 | 1 | 13 | 0.5385 | 2 | 0.5 | 2 | 0.5 | 2 | 0.5 | 2 | 1 | 8 | 0.625 |
| 1_17116847 | 2 | 0 | 5 | 0 | 4 | 0.75 | 2 | 0.5 | 13 | 0.3077 | 2 | 0.5 | 3 | 0 | 1 | 0 | 2 | 0 | 8 | 0.125 |

| | | | | | | | | | | | | | | | | | | | | |
|---|---|---|---|---|---|---|---|---|---|---|---|---|---|---|---|---|---|---|---|---|
| 1_1711685 | NA | NA | 1 | 1 | NA | NA | 2 | 1 | 3 | 1 | 1 | 1 | NA | NA | NA | NA | NA | NA | 1 | 1 |
| 1_1711686 | 2 | 0 | 5 | 0.2 | 4 | 0.25 | 4 | 0 | 15 | 0.1333 | 2 | 0 | 3 | 0 | 2 | 0 | 2 | 0.5 | 9 | 0.1111 |
| 1_1711686 | 3 | 0.6667 | 5 | 0.8 | 3 | 0.6667 | 3 | 1 | 14 | 0.7857 | 2 | 1 | 1 | 1 | 1 | 1 | 2 | 0.5 | 6 | 0.8333 |
| 1_1711687 | 3 | 1 | 5 | 0.8 | 5 | 0.8 | 4 | 1 | 17 | 0.8824 | 2 | 1 | 3 | 0.6667 | 2 | 1 | 2 | 1 | 9 | 0.8889 |
| 1_1711687 | 3 | 0.3333 | 5 | 0 | 4 | 0 | 4 | 0 | 16 | 0.0625 | 2 | 0 | 2 | 0.5 | 2 | 0 | 2 | 0.5 | 8 | 0.25 |
| 1_1711687 | 3 | 0.3333 | 5 | 0.2 | 5 | 0.8 | 3 | 1 | 16 | 0.5625 | 2 | 0.5 | 2 | 0.5 | 1 | 1 | 2 | 1 | 7 | 0.7143 |
| 1_1711687 | 2 | 1 | 5 | 1 | 5 | 1 | 3 | 1 | 15 | 1 | 2 | 1 | 2 | 1 | 2 | 1 | 2 | 1 | 8 | 1 |
| 1_1711690 | 3 | 0.3333 | 6 | 0.1667 | 4 | 0 | 5 | 0.4 | 18 | 0.2222 | 2 | 0 | 2 | 0 | 2 | 0.5 | 1 | 0 | 7 | 0.1429 |
| 1_1711691 | 1 | 0 | 4 | 0.25 | 1 | 0 | 4 | 0.25 | 10 | 0.2 | 2 | 0.5 | 1 | 0 | 1 | 0 | 3 | 0.6667 | 7 | 0.4286 |
| 1_1711698 | 4 | 0 | 4 | 0 | 5 | 0.2 | 5 | 0.2 | 18 | 0.1111 | 3 | 0.3333 | 3 | 0 | 2 | 0.5 | NA | NA | 8 | 0.25 |
| 1_1711699 | 6 | 0.6667 | 5 | 0.4 | 7 | 0.7143 | 8 | 0.75 | 26 | 0.6538 | 4 | 0.5 | 3 | 0.3333 | 3 | 1 | 3 | 0.6667 | 13 | 0.6154 |
| 1_1711699 | 8 | 0.625 | 5 | 0.4 | 5 | 0.8 | 7 | 0.5714 | 25 | 0.6 | 4 | 1 | 4 | 0.5 | 2 | 1 | 2 | 1 | 12 | 0.8333 |
| 1_1711700 | 9 | 0.1111 | 5 | 0.2 | 6 | 0 | 9 | 0.1111 | 29 | 0.1034 | 4 | 0 | 4 | 0 | 2 | 0.5 | 3 | 0 | 13 | 0.0769 |
| 1_1711700 | 9 | 0 | 5 | 0 | 6 | 0.1667 | 8 | 0.125 | 28 | 0.0714 | 4 | 0.25 | 4 | 0.25 | 2 | 0.5 | 3 | 0 | 13 | 0.2308 |
| 1_1711729 | 1 | 0 | NA | NA | 1 | 0 | 2 | 1 | 4 | 0.5 | NA | NA | NA | NA | 1 | 1 | 1 | 0 | 2 | 0.5 |
| 1_1711854 | 25 | 0.08 | 29 | 0.069 | 23 | 0.0435 | 12 | 0.0833 | 89 | 0.0674 | 18 | 0.1111 | 24 | 0.0417 | 14 | 0.0714 | 21 | 0.1905 | 77 | 0.1039 |
| 1_1712144 | 1 | 1 | NA | NA | NA | NA | NA | NA | 1 | 1 | NA | NA | NA | NA | 1 | 1 | 1 | 1 | 2 | 1 |
| 1_1712149 | 1 | 1 | NA | NA | NA | NA | NA | NA | 1 | 1 | NA | NA | NA | NA | 1 | 0 | 1 | 1 | 2 | 0.5 |
| 1_1712456 | 13 | 0.3077 | 6 | 0.1667 | 5 | 0.4 | 7 | 0.5714 | 31 | 0.3548 | 8 | 0.125 | 5 | 0.2 | 1 | 0 | 7 | 0.7143 | 21 | 0.3333 |
| 1_1712456 | 14 | 0.3571 | 4 | 0.25 | 5 | 0.4 | 5 | 0.8 | 28 | 0.4286 | 9 | 0.4444 | 6 | 0.3333 | 4 | 0.5 | 5 | 0.6 | 24 | 0.4583 |
| 1_1712457 | 16 | 0.8125 | 8 | 0.5 | 7 | 0.8571 | 6 | 0.8333 | 37 | 0.7568 | 9 | 0.6667 | 7 | 0.8571 | 5 | 0.8 | 7 | 0.4286 | 28 | 0.6786 |
| 1_1712457 | 16 | 0.9375 | 7 | 1 | 7 | 0.8571 | 5 | 0.6 | 35 | 0.8857 | 9 | 0.8889 | 8 | 0.75 | 6 | 0.8333 | 7 | 1 | 30 | 0.8667 |
| 1_1712458 | 13 | 0.9231 | 4 | 1 | 5 | 0.8 | 7 | 0.4286 | 29 | 0.7931 | 4 | 1 | 7 | 1 | 5 | 0.8 | 8 | 0.625 | 24 | 0.8333 |
| 1_1712459 | 14 | 0.8571 | 6 | 1 | 7 | 0.7143 | 8 | 0.5 | 35 | 0.7714 | 9 | 0.8889 | 8 | 0.375 | 7 | 0.5714 | 10 | 0.8 | 34 | 0.6765 |
| 1_1712459 | 12 | 0.5833 | 8 | 0.5 | 7 | 0.7143 | 7 | 0.2857 | 34 | 0.5294 | 7 | 0.5714 | 8 | 0.5 | 7 | 0.7143 | 10 | 0.4 | 32 | 0.5312 |
| 1_1712460 | 18 | 0.5556 | 8 | 0.25 | 7 | 0.1429 | 7 | 0.4286 | 40 | 0.4 | 8 | 0.5 | 8 | 0.25 | 9 | 0.3333 | 12 | 0.5833 | 37 | 0.4324 |
| 1_1712462 | 17 | 0.1176 | 8 | 0.375 | 9 | 0.2222 | 8 | 0.375 | 42 | 0.2381 | 11 | 0.1818 | 8 | 0.125 | 9 | 0 | 8 | 0.25 | 36 | 0.1389 |
| 1_1712908 | NA | NA | 3 | 0.6667 | 1 | 0 | 3 | 0 | 7 | 0.2857 | 3 | 0.6667 | 2 | 0.5 | 1 | 0 | 1 | 0 | 7 | 0.4286 |
| 1_1712910 | NA | NA | 3 | 1 | 2 | 1 | 3 | 1 | 8 | 1 | 2 | 1 | 2 | 1 | 1 | 1 | 1 | 1 | 6 | 1 |
| 1_1712912 | NA | NA | 3 | 1 | 2 | 1 | 3 | 1 | 8 | 1 | 3 | 0.6667 | 2 | 1 | NA | NA | 1 | 1 | 6 | 0.8333 |
| 1_1712912 | NA | NA | 3 | 0.6667 | 1 | 0 | 1 | 1 | 5 | 0.6 | 2 | 0 | 1 | 1 | 1 | 1 | 1 | 0 | 5 | 0.4 |
| 1_1712913 | NA | NA | 3 | 0.3333 | 1 | 0 | 3 | 0.3333 | 7 | 0.2857 | 3 | 0.3333 | 2 | 0.5 | NA | NA | 1 | 0 | 6 | 0.3333 |
| 1_1712913 | NA | NA | 3 | 0.6667 | 2 | 0 | 3 | 0 | 8 | 0.25 | 3 | 0.3333 | 2 | 0 | 1 | 1 | 1 | 0 | 7 | 0.2857 |
| 1_1713314 | 2 | 1 | 1 | 0 | NA | NA | 2 | 1 | 5 | 0.8 | NA | NA | 1 | 1 | NA | NA | NA | NA | 1 | 1 |
| 1_1713319 | 3 | 1 | 2 | 0.5 | 1 | 1 | 1 | 1 | 7 | 0.8571 | 1 | 1 | 1 | 1 | NA | NA | NA | NA | 2 | 1 |
| 1_1713322 | 1 | 0 | 2 | 0.5 | 2 | 0.5 | 1 | 1 | 6 | 0.5 | NA | NA | 1 | 1 | 1 | 0 | NA | NA | 2 | 0.5 |
| 1_1718321 | NA | NA | 1 | 1 | 1 | 1 | 1 | 0 | 3 | 0.6667 | NA | NA | NA | NA | 1 | 1 | 1 | 0 | 2 | 0.5 |
| 1_1718323 | NA | NA | NA | NA | 1 | 0 | 1 | 1 | 2 | 0.5 | NA | NA | NA | NA | 1 | 1 | 2 | 1 | 3 | 1 |
| 1_1718940 | 14 | 0.2857 | 5 | 0.2 | 9 | 0.1111 | 8 | 0.25 | 36 | 0.2222 | 11 | 0.2727 | 7 | 0.5714 | 7 | 0.5714 | 4 | 0 | 29 | 0.3793 |
| 1_1718940 | 12 | 0.4167 | 3 | 0.3333 | 10 | 0.5 | 6 | 0.3333 | 31 | 0.4194 | 9 | 0.7778 | 7 | 0.4286 | 6 | 0.3333 | 4 | 0 | 26 | 0.4615 |
| 1_1718941 | 8 | 0.75 | 4 | 0.5 | 4 | 0.5 | 5 | 0.8 | 21 | 0.6667 | 10 | 0.9 | 4 | 0.5 | 3 | 0.3333 | 3 | 0.3333 | 20 | 0.65 |
| 1_1718941 | 9 | 1 | 5 | 0.8 | 8 | 1 | 5 | 1 | 27 | 0.963 | 10 | 0.9 | 5 | 1 | 6 | 0.8333 | 3 | 1 | 24 | 0.9167 |
| 1_1718942 | 11 | 0.5455 | 3 | 0.6667 | 6 | 0.3333 | 7 | 0.4286 | 27 | 0.4815 | 5 | 0.8 | 4 | 0.5 | 4 | 0.25 | 4 | 0.75 | 17 | 0.5882 |
| 1_1718942 | 10 | 0.6 | 5 | 0.2 | 9 | 0.2222 | 3 | 0 | 27 | 0.3333 | 10 | 0.1 | 6 | 0.1667 | 5 | 0 | 3 | 0.3333 | 24 | 0.125 |
| 1_1719619 | 5 | 0.6 | 7 | 0 | 2 | 0.5 | 6 | 0.5 | 20 | 0.35 | 8 | 0.375 | 5 | 0.2 | 9 | 0.3333 | 6 | 0 | 28 | 0.25 |
| 1_1719619 | 7 | 0.4286 | 7 | 0.7143 | 3 | 0.3333 | 5 | 0.6 | 22 | 0.5455 | 9 | 0.5556 | 8 | 0.625 | 13 | 0.5385 | 4 | 0.5 | 34 | 0.5588 |
| 1_1719624 | 9 | 0.3333 | 11 | 0.1818 | 2 | 0 | 10 | 0.1 | 32 | 0.1875 | 12 | 0.0833 | 6 | 0.1667 | 16 | 0.25 | 7 | 0 | 41 | 0.1463 |
| 1_1719636 | 2 | 0.5 | 2 | 0 | 2 | 0 | 1 | 0 | 7 | 0.1429 | 2 | 0.5 | 3 | 0 | NA | NA | NA | NA | 5 | 0.2 |
| 1_1719636 | 2 | 0 | 2 | 1 | 1 | 1 | 1 | 1 | 6 | 0.6667 | 3 | 0.3333 | 3 | 0.6667 | 1 | 0 | NA | NA | 7 | 0.4286 |
| 1_1719637 | 2 | 0 | 5 | 0.2 | 1 | 0 | 1 | 0 | 9 | 0.1111 | 4 | 0.5 | 3 | 0 | 1 | 0 | NA | NA | 8 | 0.25 |
| 1_1719637 | 2 | 0 | 4 | 0.75 | 2 | 0 | 1 | 1 | 9 | 0.4444 | 3 | 0.6667 | 3 | 0 | 1 | 0 | NA | NA | 7 | 0.2857 |
| 1_1719638 | 2 | 0 | 4 | 0.5 | 2 | 0.5 | 1 | 0 | 9 | 0.3333 | 4 | 0.5 | 2 | 0 | 1 | 0 | NA | NA | 7 | 0.2857 |
| 1_1719663 | NA | NA | NA | NA | 1 | 1 | NA | NA | 1 | 1 | 1 | 0 | 2 | 0.5 | NA | NA | NA | NA | 3 | 0.3333 |
| 1_1719716 | 2 | 1 | NA | NA | NA | NA | NA | NA | 2 | 1 | 2 | 0.5 | NA | NA | NA | NA | 1 | 1 | 3 | 0.6667 |
| 1_1719717 | 3 | 1 | NA | NA | NA | NA | NA | NA | 3 | 1 | 2 | 0.5 | 1 | 0 | NA | NA | NA | NA | 3 | 0.3333 |
| 1_1719718 | 3 | 0.6667 | NA | NA | NA | NA | NA | NA | 3 | 0.6667 | 2 | 1 | 1 | 0 | NA | NA | NA | NA | 3 | 0.6667 |
| 1_1720000 | 4 | 1 | 3 | 1 | 1 | 1 | 4 | 0.75 | 12 | 0.9167 | 3 | 0.6667 | 2 | 1 | 1 | 1 | 2 | 0.5 | 8 | 0.75 |
| 1_1720000 | 4 | 0.25 | 3 | 0 | 1 | 0 | 4 | 0.5 | 12 | 0.25 | 3 | 0 | 3 | 0 | 1 | 1 | 2 | 0 | 9 | 0.1111 |
| 1_1720001 | 3 | 0 | 2 | 0 | 2 | 0.5 | 3 | 0 | 10 | 0.1 | 2 | 0 | 3 | 0.6667 | 1 | 0 | 2 | 0 | 8 | 0.25 |
| 1_1720002 | 4 | 0.25 | 3 | 0 | 2 | 0.5 | 3 | 0 | 12 | 0.1667 | 3 | 0 | 3 | 0.6667 | 1 | 0 | 3 | 0.3333 | 10 | 0.3 |
| 1_1720003 | 4 | 0.25 | 4 | 0 | 2 | 0 | 4 | 0 | 14 | 0.0714 | 3 | 0.3333 | 2 | 0 | 1 | 1 | 1 | 1 | 7 | 0.4286 |
| 1_1720005 | 4 | 0.25 | 2 | 0.5 | 3 | 0.3333 | 2 | 0.5 | 11 | 0.3636 | 2 | 0.5 | 3 | 0.3333 | 2 | 0.5 | 2 | 0 | 9 | 0.3333 |
| 1_1720223 | 7 | 1 | 8 | 0.75 | 6 | 0.8333 | 7 | 0.7143 | 28 | 0.8214 | 3 | 1 | 7 | 1 | 5 | 0.8 | 8 | 0.875 | 23 | 0.913 |
| 1_1720223 | 6 | 0.1667 | 10 | 0 | 5 | 0 | 9 | 0.1111 | 30 | 0.0667 | 3 | 0 | 5 | 0 | 5 | 0.6 | 8 | 0 | 21 | 0.1429 |
| 1_1720224 | 7 | 0.8571 | 10 | 0.6 | 8 | 0.875 | 9 | 0.6667 | 34 | 0.7353 | 5 | 0.6 | 6 | 0.3333 | 5 | 0.6 | 10 | 0.8 | 26 | 0.6154 |

| | | | | | | | | | | | | | | | | | | | | |
|---|---|---|---|---|---|---|---|---|---|---|---|---|---|---|---|---|---|---|---|---|
| 1_17202355 | 8 | 0 | 13 | 0 | 9 | 0 | 7 | 0.2857 | 37 | 0.0541 | 9 | 0.1111 | 6 | 0.1667 | 7 | 0 | 7 | 0 | 29 | 0.069 |
| 1_17202455 | 42 | 0.0476 | 26 | 0.0769 | 32 | 0.0938 | 43 | 0.1395 | 143 | 0.0909 | 33 | 0.0909 | 39 | 0.1795 | 37 | 0 | 27 | 0 | 136 | 0.0735 |
| 1_17202538 | 43 | 0.1163 | 39 | 0.1026 | 37 | 0.0811 | 47 | 0.0426 | 166 | 0.0843 | 52 | 0.0962 | 52 | 0.0577 | 50 | 0.08 | 31 | 0.129 | 185 | 0.0865 |
| 1_17209998 | 6 | 0.1667 | 9 | 0 | 9 | 0.1111 | 6 | 0.1667 | 30 | 0.1 | 5 | 0 | 9 | 0.1111 | 4 | 0.5 | 6 | 0 | 24 | 0.125 |
| 1_17210036 | 81 | 0.0741 | 86 | 0.0581 | 58 | 0.1034 | 73 | 0.0685 | 298 | 0.0738 | 62 | 0 | 62 | 0.0323 | 53 | 0.0755 | 52 | 0.1346 | 229 | 0.0568 |
| 1_17210044 | 78 | 0.141 | 84 | 0.131 | 49 | 0.102 | 67 | 0.1045 | 278 | 0.1223 | 49 | 0.0612 | 51 | 0.0784 | 49 | 0.102 | 48 | 0.2292 | 197 | 0.1168 |
| 1_17211235 | 18 | 0.1111 | 20 | 0.2 | 25 | 0.12 | 22 | 0.0909 | 85 | 0.1294 | 17 | 0.1765 | 25 | 0.04 | 19 | 0 | 10 | 0.2 | 71 | 0.0845 |
| 1_17214381 | 14 | 0.0714 | 14 | 0.1429 | 12 | 0 | 18 | 0.0556 | 58 | 0.069 | 5 | 0.2 | 8 | 0 | 12 | 0.0833 | 12 | 0 | 37 | 0.0541 |
| 1_17214401 | 16 | 0.5 | 16 | 0.375 | 14 | 0.5 | 24 | 0.4583 | 70 | 0.4571 | 10 | 0.5 | 14 | 0.5714 | 16 | 0.4375 | 17 | 0.7059 | 57 | 0.5614 |
| 1_17214514 | 15 | 0.5333 | 12 | 0.3333 | 13 | 0.4615 | 21 | 0.3333 | 61 | 0.4098 | 11 | 0.1818 | 11 | 0.2727 | 17 | 0.3529 | 19 | 0.3684 | 58 | 0.3103 |
| 1_17214522 | 16 | 0.25 | 13 | 0.3846 | 15 | 0.5333 | 22 | 0.3636 | 66 | 0.3788 | 8 | 0.25 | 13 | 0.3846 | 15 | 0.2667 | 19 | 0.5263 | 55 | 0.3818 |
| 1_17214566 | 19 | 0.3684 | 16 | 0.125 | 14 | 0.2143 | 21 | 0.1905 | 70 | 0.2286 | 8 | 0.125 | 15 | 0.4667 | 17 | 0.3529 | 18 | 0.3333 | 58 | 0.3448 |
| 1_17214576 | 20 | 0.15 | 11 | 0.0909 | 14 | 0.1429 | 22 | 0.1364 | 67 | 0.1343 | 11 | 0 | 5 | 0.2 | 12 | 0.0833 | 12 | 0.0833 | 40 | 0.075 |
| 1_17214578 | 25 | 0.16 | 20 | 0.15 | 18 | 0.1111 | 26 | 0.0385 | 89 | 0.1124 | 11 | 0.1818 | 14 | 0.1429 | 18 | 0.1667 | 17 | 0.1176 | 60 | 0.15 |
| 1_17214582 | 27 | 0.1481 | 23 | 0.087 | 18 | 0 | 28 | 0.1429 | 96 | 0.1042 | 11 | 0.0909 | 16 | 0 | 22 | 0.1364 | 18 | 0.2778 | 67 | 0.1343 |
| 1_17214646 | 27 | 0.0741 | 23 | 0.087 | 18 | 0.0556 | 35 | 0.0286 | 103 | 0.0583 | 12 | 0.0833 | 15 | 0.0667 | 25 | 0.08 | 20 | 0.1 | 72 | 0.0833 |
| 1_17214716 | 30 | 0.1 | 33 | 0.1212 | 24 | 0.1667 | 40 | 0.125 | 127 | 0.126 | 18 | 0.2778 | 24 | 0.2917 | 27 | 0.1481 | 25 | 0.16 | 94 | 0.2128 |
| 1_17214828 | 4 | 0 | 9 | 0.1111 | 5 | 0 | 2 | 0.5 | 20 | 0.1 | 4 | 0.25 | 3 | 0 | 1 | 0 | 8 | 0.125 | 16 | 0.125 |
| 1_17214839 | 4 | 0.25 | 6 | 0.1667 | 6 | 0.5 | 3 | 0 | 19 | 0.2632 | 5 | 0.2 | 3 | 0.3333 | 1 | 1 | 8 | 0 | 17 | 0.1765 |
| 1_17214851 | 4 | 0 | 5 | 0.2 | 7 | 0.1429 | 4 | 0 | 20 | 0.1 | 4 | 0.5 | 3 | 0.3333 | 1 | 0 | 9 | 0.2222 | 17 | 0.2941 |
| 1_17214865 | 4 | 0.5 | 7 | 0 | 7 | 0.2857 | 5 | 0.2 | 23 | 0.2174 | 4 | 0 | 4 | 0 | 1 | 0 | 10 | 0.2 | 19 | 0.1053 |
| 1_17214866 | 4 | 0.75 | 6 | 0.5 | 6 | 0.8333 | 5 | 0.6 | 21 | 0.6667 | 4 | 0.25 | 4 | 0.75 | 1 | 1 | 9 | 0.4444 | 18 | 0.5 |
| 1_17214872 | 4 | 0.25 | 6 | 0.3333 | 4 | 0.25 | 4 | 0.5 | 18 | 0.3333 | 4 | 0.25 | 2 | 0 | 1 | 1 | 10 | 0.3 | 17 | 0.2941 |
| 1_17214880 | 2 | 0 | 6 | 0.1667 | 2 | 0 | 5 | 0.2 | 15 | 0.1333 | 3 | 0 | 3 | 0 | NA | NA | 8 | 0.25 | 14 | 0.1429 |
| 1_17214887 | 2 | 0.5 | 5 | 0 | 4 | 0 | 5 | 0.4 | 16 | 0.1875 | 2 | 0.5 | 4 | 0 | 1 | 0 | 9 | 0 | 16 | 0.0625 |
| 1_17214888 | 4 | 0 | 6 | 0.1667 | 4 | 0 | 5 | 0.4 | 19 | 0.1579 | 1 | 0 | 3 | 0.3333 | 1 | 1 | 7 | 0.4286 | 12 | 0.4167 |
| 1_17214889 | 4 | 0.75 | 6 | 0.5 | 3 | 0.6667 | 3 | 0.3333 | 16 | 0.5625 | 2 | 0.5 | 4 | 0.5 | 1 | 0 | 7 | 0.5714 | 14 | 0.5 |
| 1_17214906 | 4 | 0.25 | 4 | 0 | 3 | 0 | 4 | 0.25 | 15 | 0.1333 | 3 | 0.3333 | 3 | 0.3333 | 1 | 1 | 9 | 0 | 16 | 0.1875 |
| 1_17214913 | 2 | 0 | 5 | 0.2 | 4 | 0 | 4 | 0 | 15 | 0.0667 | 3 | 0 | 3 | 0 | 1 | 0 | 8 | 0.375 | 15 | 0.2 |
| 1_17214918 | 2 | 0 | 4 | 0 | 4 | 0 | 5 | 0.4 | 15 | 0.1333 | 1 | 1 | 3 | 0.3333 | 1 | 1 | 8 | 0.125 | 13 | 0.3077 |
| 1_17214946 | 3 | 0.3333 | 4 | 0 | 1 | 1 | 5 | 0 | 13 | 0.1538 | 2 | 0 | 3 | 0.6667 | 1 | 0 | 7 | 0 | 13 | 0.1538 |
| 1_17214967 | 2 | 0.5 | 3 | 0 | 1 | 0 | 4 | 0.25 | 10 | 0.2 | 2 | 0 | 2 | 0.5 | NA | NA | 5 | 0 | 9 | 0.1111 |
| 1_17214997 | 1 | 0 | 3 | 0.3333 | 1 | 1 | 4 | 0.25 | 9 | 0.3333 | 2 | 0 | 2 | 0.5 | NA | NA | 2 | 0 | 6 | 0.1667 |
| 1_17215017 | 1 | 1 | 1 | 1 | 1 | 1 | 2 | 0 | 5 | 0.6 | 1 | 1 | 1 | 1 | 1 | 1 | 3 | 0 | 6 | 0.5 |
| 1_17215038 | NA | NA | 2 | 1 | 1 | 1 | 2 | 1 | 5 | 1 | 1 | 1 | 1 | 1 | 1 | 1 | 1 | 0 | 4 | 0.75 |
| 1_17215043 | NA | NA | 1 | 1 | 1 | 1 | 2 | 1 | 4 | 1 | 1 | 1 | 1 | 0 | NA | NA | 2 | 1 | 4 | 0.75 |
| 1_17215065 | NA | NA | 1 | 0 | 1 | 1 | 1 | 1 | 3 | 0.6667 | 1 | 1 | NA | NA | 1 | 0 | NA | NA | 2 | 0.5 |
| 1_17219677 | NA | NA | 7 | 1 | 8 | 0.625 | 4 | 1 | 19 | 0.8421 | 4 | 1 | 5 | 0.8 | 1 | 1 | 1 | 1 | 11 | 0.9091 |
| 1_17219678 | 1 | 1 | 7 | 0.7143 | 5 | 0.6 | 3 | 1 | 16 | 0.75 | 4 | 0.5 | 5 | 1 | 1 | 1 | 2 | 1 | 12 | 0.8333 |
| 1_17219719 | 1 | 1 | 9 | 0.7778 | 9 | 0.4444 | 4 | 0 | 23 | 0.5217 | 4 | 0.25 | 5 | 0.6 | 3 | 0.6667 | 2 | 0.5 | 14 | 0.5 |
| 1_17219720 | 1 | 0 | 10 | 0.1 | 9 | 0.1111 | 3 | 0 | 23 | 0.087 | 4 | 0 | 5 | 0 | 2 | 0.5 | 2 | 0 | 13 | 0.0769 |
| 1_17219722 | 1 | 0 | 5 | 0 | 6 | 0.1667 | 3 | 0.3333 | 15 | 0.1333 | 2 | 0.5 | 4 | 0 | 3 | 0 | 2 | 0 | 11 | 0.0909 |
| 1_17219724 | NA | NA | 9 | 0.8889 | 7 | 0.8571 | 3 | 1 | 19 | 0.8947 | 3 | 0.6667 | 3 | 0.6667 | 1 | 1 | 1 | 1 | 8 | 0.75 |
| 1_17219731 | 2 | 1 | 9 | 0.8889 | 9 | 1 | 4 | 1 | 24 | 0.9583 | 3 | 0.6667 | 4 | 0.75 | 2 | 1 | 2 | 1 | 11 | 0.8182 |
| 1_17219734 | 2 | 1 | 9 | 0.8889 | 8 | 0.625 | 4 | 0.75 | 23 | 0.7826 | 5 | 1 | 5 | 1 | 1 | 1 | 3 | 0.3333 | 14 | 0.8571 |
| 1_17219771 | 2 | 0.5 | 11 | 0.1818 | 9 | 0 | 4 | 0 | 26 | 0.1154 | 6 | 0.1667 | 5 | 0.2 | 6 | 0.1667 | 4 | 0.5 | 21 | 0.2381 |
| 1_17219784 | NA | NA | 6 | 1 | 5 | 1 | 1 | 1 | 12 | 1 | 4 | 1 | 2 | 1 | 3 | 0 | 4 | 0.75 | 13 | 0.6923 |
| 1_17219793 | 2 | 1 | 10 | 0.7 | 7 | 0.5714 | 5 | 0.4 | 24 | 0.625 | 5 | 0.8 | 5 | 0.4 | 4 | 0.25 | 3 | 0.3333 | 17 | 0.4706 |
| 1_17219794 | 4 | 0.5 | 9 | 0.7778 | 9 | 1 | 5 | 0.8 | 27 | 0.8148 | 4 | 1 | 5 | 1 | 5 | 0.6 | 4 | 1 | 18 | 0.8889 |
| 1_17219795 | 4 | 0.75 | 8 | 0.625 | 6 | 0.5 | 4 | 0.5 | 22 | 0.5909 | 5 | 0.2 | 4 | 0.5 | 4 | 1 | 3 | 0.3333 | 16 | 0.5 |
| 1_17219799 | 4 | 0.5 | 11 | 0.3636 | 8 | 0.625 | 5 | 0 | 28 | 0.3929 | 6 | 0.3333 | 4 | 0.25 | 6 | 0.1667 | 5 | 0 | 21 | 0.1905 |
| 1_17219804 | 1 | 0 | 8 | 0.375 | 7 | 0.1429 | 3 | 0.3333 | 19 | 0.2632 | 4 | 0.5 | 5 | 0.6 | 4 | 0.5 | 3 | 0.3333 | 16 | 0.5 |
| 1_17219819 | 5 | 1 | 12 | 1 | 9 | 1 | 5 | 1 | 31 | 1 | 5 | 1 | 4 | 1 | 7 | 1 | 4 | 1 | 20 | 1 |
| 1_17219828 | 4 | 0.75 | 7 | 0.8571 | 6 | 0.8333 | 3 | 1 | 20 | 0.85 | 3 | 0.6667 | 2 | 1 | 4 | 1 | 4 | 0.75 | 13 | 0.8462 |
| 1_17219829 | 4 | 0 | 11 | 0.0909 | 8 | 0.125 | 5 | 0.2 | 28 | 0.1071 | 6 | 0.1667 | 4 | 0 | 4 | 0.25 | 4 | 0.25 | 18 | 0.1667 |
| 1_17219900 | 4 | 0.25 | 9 | 0.1111 | 7 | 0.2857 | 3 | 0 | 23 | 0.1739 | 6 | 0 | 5 | 0.4 | 5 | 0.4 | 2 | 0 | 18 | 0.2222 |
| 1_17219909 | 8 | 0.375 | 13 | 0.5385 | 12 | 0.0833 | 6 | 0.1667 | 39 | 0.3077 | 9 | 0.1111 | 7 | 0.1429 | 7 | 0.4286 | 6 | 0.1667 | 29 | 0.2069 |
| 1_17219920 | 7 | 0.5714 | 12 | 0.5 | 11 | 0.4545 | 6 | 0.5 | 36 | 0.5 | 8 | 0.125 | 7 | 0.1429 | 8 | 0.375 | 6 | 0 | 29 | 0.1724 |
| 1_17219937 | 8 | 0.875 | 13 | 0.7692 | 14 | 0.7143 | 6 | 0.6667 | 41 | 0.7561 | 9 | 0.7778 | 6 | 0.8333 | 6 | 0.8333 | 5 | 1 | 26 | 0.8462 |
| 1_17219944 | 7 | 0.1429 | 11 | 0.1818 | 11 | 0.0909 | 7 | 0.2857 | 36 | 0.1667 | 9 | 0.2222 | 6 | 0 | 7 | 0 | 7 | 0 | 29 | 0.069 |
| 1_17219946 | 8 | 0.375 | 13 | 0.1538 | 13 | 0.0769 | 6 | 0.3333 | 40 | 0.2 | 8 | 0.25 | 7 | 0.2857 | 6 | 0 | 6 | 0.1667 | 27 | 0.1852 |
| 1_17219951 | 7 | 0.8571 | 11 | 1 | 11 | 1 | 6 | 0.8333 | 35 | 0.9429 | 7 | 0.8571 | 6 | 1 | 7 | 0.8571 | 6 | 0.8333 | 26 | 0.8846 |
| 1_17219952 | 7 | 0.8571 | 12 | 1 | 8 | 0.875 | 4 | 0.75 | 31 | 0.9032 | 7 | 0.7143 | 3 | 1 | 7 | 0.7143 | 4 | 0.75 | 21 | 0.7619 |
| 1_17219971 | 8 | 0.125 | 13 | 0.1538 | 13 | 0.1538 | 7 | 0.4286 | 41 | 0.1951 | 7 | 0 | 7 | 0.1429 | 8 | 0.5 | 8 | 0.375 | 30 | 0.2667 |
| 1_17219993 | 7 | 0.2857 | 14 | 0.1429 | 13 | 0.2308 | 7 | 0.4286 | 41 | 0.2439 | 7 | 0.1429 | 7 | 0.2857 | 5 | 0.4 | 6 | 0.1667 | 25 | 0.24 |

| | | | | | | | | | | | | | | | | | | | | |
|---|---|---|---|---|---|---|---|---|---|---|---|---|---|---|---|---|---|---|---|---|
| 1_1722004 | 12 | 0.5 | 17 | 0.2941 | 15 | 0.5333 | 8 | 0.375 | 52 | 0.4231 | 10 | 0.5 | 9 | 0.4444 | 8 | 0.375 | 13 | 0.4615 | 40 | 0.45 |
| 1_1722015 | 18 | 0.3333 | 17 | 0 | 20 | 0.2 | 9 | 0 | 64 | 0.1562 | 14 | 0 | 12 | 0 | 12 | 0.25 | 18 | 0.1111 | 56 | 0.0893 |
| 1_1722043 | NA | NA | 1 | 1 | NA | NA | 1 | 1 | 2 | 1 | 1 | 1 | NA | NA | NA | NA | 1 | 0 | 2 | 0.5 |
| 1_1722043 | NA | NA | 1 | 1 | NA | NA | 1 | 1 | 2 | 1 | NA | NA | NA | NA | 1 | 1 | 2 | 0.5 | 3 | 0.6667 |
| 1_1722044 | NA | NA | 1 | 0 | NA | NA | 1 | 1 | 2 | 0.5 | 1 | 0 | NA | NA | 1 | 1 | 2 | 0 | 4 | 0.25 |
| 1_1722047 | NA | NA | 1 | 1 | NA | NA | 1 | 0 | 2 | 0.5 | 1 | 0 | NA | NA | 1 | 1 | 2 | 0.5 | 4 | 0.5 |
| 1_1722051 | NA | NA | 1 | 0 | NA | NA | 1 | 1 | 2 | 0.5 | 1 | 1 | NA | NA | 1 | 1 | 2 | 0.5 | 4 | 0.75 |
| 1_1722052 | NA | NA | NA | NA | NA | NA | 2 | 0.5 | 2 | 0.5 | 1 | 0 | 1 | 1 | 1 | 1 | 2 | 0 | 5 | 0.4 |
| 1_1722052 | NA | NA | 1 | 1 | NA | NA | 2 | 1 | 3 | 1 | 1 | 1 | 1 | 1 | 1 | 1 | 2 | 0 | 5 | 0.6 |
| 1_1722053 | NA | NA | 1 | 1 | NA | NA | 2 | 1 | 3 | 1 | 1 | 0 | NA | NA | 1 | 0 | 1 | 1 | 3 | 0.3333 |
| 1_1722055 | NA | NA | 1 | 1 | NA | NA | 2 | 0.5 | 3 | 0.6667 | 1 | 0 | NA | NA | NA | NA | 2 | 1 | 3 | 0.6667 |
| 1_1722055 | NA | NA | 1 | 1 | NA | NA | 2 | 1 | 3 | 1 | 1 | 1 | 1 | 1 | 1 | 1 | 2 | 1 | 5 | 1 |
| 1_1722056 | NA | NA | 1 | 1 | NA | NA | 1 | 1 | 2 | 1 | NA | NA | 1 | 1 | NA | NA | 1 | 1 | 2 | 1 |
| 1_1722056 | NA | NA | NA | NA | NA | NA | 2 | 1 | 2 | 1 | 1 | 1 | 1 | 1 | 1 | 1 | 2 | 1 | 5 | 1 |
| 1_1722063 | NA | NA | NA | NA | NA | NA | 2 | 0.5 | 2 | 0.5 | 1 | 0 | 1 | 0 | 1 | 0 | 2 | 0.5 | 5 | 0.2 |
| 1_1723038 | 2 | 1 | NA | NA | NA | NA | 1 | 1 | 3 | 1 | NA | NA | NA | NA | 1 | 1 | NA | NA | 1 | 1 |
| 1_1723038 | 2 | 0.5 | NA | NA | NA | NA | 1 | 1 | 3 | 0.6667 | NA | NA | NA | NA | 1 | 1 | NA | NA | 1 | 1 |
| 1_1723058 | 2 | 1 | NA | NA | NA | NA | 1 | 0 | 3 | 0.6667 | NA | NA | NA | NA | 1 | 1 | NA | NA | 1 | 1 |
| 1_1723059 | 2 | 0 | 1 | 1 | NA | NA | 1 | 0 | 4 | 0.25 | NA | NA | NA | NA | 1 | 1 | NA | NA | 1 | 1 |
| 1_1723247 | 1 | 1 | NA | NA | NA | NA | 1 | 1 | 2 | 1 | NA | NA | 1 | 0 | 1 | 1 | NA | NA | 2 | 0.5 |
| 1_1723247 | 1 | 1 | NA | NA | NA | NA | 1 | 0 | 2 | 0.5 | NA | NA | 1 | 0 | 1 | 1 | NA | NA | 2 | 0.5 |
| 1_1723249 | 1 | 1 | NA | NA | NA | NA | 1 | 0 | 2 | 0.5 | NA | NA | 1 | 1 | 1 | 1 | NA | NA | 2 | 1 |
| 1_1723250 | 1 | 1 | NA | NA | NA | NA | 1 | 1 | 2 | 1 | NA | NA | 1 | 1 | 1 | 1 | NA | NA | 2 | 1 |
| 1_1723261 | 1 | 0 | NA | NA | NA | NA | 1 | 1 | 2 | 0.5 | NA | NA | 2 | 0 | 1 | 1 | NA | NA | 3 | 0.3333 |
| 1_1723337 | 2 | 0.5 | 2 | 0 | 2 | 0 | 3 | 0 | 9 | 0.1111 | NA | NA | 2 | 0 | 2 | 0 | 2 | 0.5 | 6 | 0.1667 |
| 1_1723512 | NA | NA | 1 | 1 | NA | NA | NA | NA | 1 | 1 | NA | NA | 1 | 1 | NA | NA | NA | NA | 1 | 1 |
| 1_1723512 | NA | NA | 1 | 1 | NA | NA | NA | NA | 1 | 1 | NA | NA | 1 | 1 | NA | NA | NA | NA | 1 | 1 |
| 1_1723513 | NA | NA | 1 | 1 | NA | NA | NA | NA | 1 | 1 | NA | NA | 1 | 1 | NA | NA | NA | NA | 1 | 1 |
| 1_1723515 | NA | NA | 1 | 1 | NA | NA | NA | NA | 1 | 1 | NA | NA | 1 | 1 | NA | NA | NA | NA | 1 | 1 |
| 1_1723516 | NA | NA | 1 | 1 | NA | NA | NA | NA | 1 | 1 | NA | NA | 1 | 1 | NA | NA | NA | NA | 1 | 1 |
| 1_1724587 | NA | NA | NA | NA | NA | NA | 2 | 0.5 | 2 | 0.5 | 1 | 1 | NA | NA | NA | NA | NA | NA | 1 | 1 |
| 1_1724590 | NA | NA | NA | NA | NA | NA | 2 | 1 | 2 | 1 | 1 | 1 | 1 | 1 | 1 | 0 | NA | NA | 3 | 0.6667 |
| 1_1724591 | NA | NA | NA | NA | NA | NA | 2 | 0.5 | 2 | 0.5 | 2 | 1 | 1 | 0 | 1 | 1 | NA | NA | 4 | 0.75 |
| 1_1731599 | 9 | 0.2222 | 12 | 0 | 12 | 0.1667 | 3 | 0.3333 | 36 | 0.1389 | 8 | 0.125 | 12 | 0.3333 | 9 | 0.5556 | 7 | 0.1429 | 36 | 0.3056 |
| 1_1731600 | 8 | 0.75 | 9 | 0.6667 | 12 | 0.5833 | 2 | 0.5 | 31 | 0.6452 | 4 | 0.75 | 11 | 0.8182 | 8 | 0.75 | 7 | 0.7143 | 30 | 0.7667 |
| 1_1731600 | 8 | 0.75 | 13 | 0.6154 | 9 | 0.6667 | 3 | 0.6667 | 33 | 0.6667 | 7 | 0.7143 | 11 | 0.7273 | 7 | 0.7143 | 8 | 0.875 | 33 | 0.7576 |
| 1_1731600 | 9 | 0.6667 | 12 | 0.4167 | 9 | 0.2222 | 3 | 0.6667 | 33 | 0.4545 | 6 | 0.6667 | 10 | 0.3 | 8 | 0.5 | 8 | 0.25 | 32 | 0.4062 |
| 1_1731602 | 6 | 0 | 9 | 0.1111 | 10 | 0.2 | 2 | 0 | 27 | 0.1111 | 7 | 0.1429 | 7 | 0 | 8 | 0.125 | 7 | 0.1429 | 29 | 0.1034 |
| 1_1731604 | 7 | 0.4286 | 10 | 0.5 | 8 | 0.375 | 1 | 0 | 26 | 0.4231 | 6 | 0.6667 | 6 | 0.1667 | 7 | 0.8571 | 4 | 0.75 | 23 | 0.6087 |
| 1_1731604 | 7 | 0 | 9 | 0.1111 | 11 | 0.0909 | 2 | 0 | 29 | 0.069 | 6 | 0.1667 | 10 | 0.1 | 9 | 0 | 7 | 0 | 32 | 0.0625 |
| 1_1731611 | 6 | 0 | 11 | 0.2727 | 9 | 0 | 2 | 0 | 28 | 0.1071 | 4 | 0 | 10 | 0.2 | 6 | 0.5 | 6 | 0 | 26 | 0.1923 |
| 1_1731623 | 3 | 0 | 8 | 0 | 9 | 0.4444 | 1 | 0 | 21 | 0.1905 | 6 | 0.1667 | 8 | 0.375 | 4 | 0 | 4 | 0 | 22 | 0.1818 |
| 1_1731637 | 20 | 0.25 | 26 | 0.5 | 8 | 0.625 | 19 | 0.4211 | 73 | 0.4247 | 15 | 0.6667 | 23 | 0.6087 | 17 | 0.4706 | 14 | 0.1429 | 69 | 0.4928 |
| 1_1731637 | 24 | 0.0833 | 31 | 0.1613 | 15 | 0.0667 | 21 | 0.1429 | 91 | 0.1209 | 18 | 0.3333 | 33 | 0.1515 | 23 | 0.087 | 17 | 0 | 91 | 0.1429 |
| 1_1731645 | 13 | 0.9231 | 15 | 0.8667 | 4 | 1 | 13 | 0.6923 | 45 | 0.8444 | 11 | 0.9091 | 19 | 0.8947 | 12 | 0.75 | 6 | 0.6667 | 48 | 0.8333 |
| 1_1731645 | 15 | 0.8 | 21 | 0.7143 | 5 | 0.8 | 13 | 0.6154 | 54 | 0.7222 | 13 | 0.6923 | 25 | 0.76 | 14 | 0.6429 | 6 | 1 | 58 | 0.7414 |
| 1_1732927 | NA | NA | 2 | 1 | NA | NA | NA | NA | 2 | 1 | 1 | 1 | NA | NA | NA | NA | NA | NA | 1 | 1 |
| 1_1732927 | NA | NA | 2 | 0.5 | NA | NA | NA | NA | 2 | 0.5 | 1 | 1 | NA | NA | NA | NA | NA | NA | 1 | 1 |
| 1_1732930 | NA | NA | 2 | 0.5 | NA | NA | NA | NA | 2 | 0.5 | 1 | 1 | 1 | 0 | NA | NA | NA | NA | 2 | 0.5 |
| 1_1732930 | 1 | 1 | 2 | 0.5 | NA | NA | NA | NA | 3 | 0.6667 | 1 | 1 | 1 | 1 | NA | NA | NA | NA | 2 | 1 |
| 1_1743298 | NA | NA | 1 | 0 | 1 | 1 | NA | NA | 2 | 0.5 | 1 | 0 | 1 | 1 | 1 | 1 | NA | NA | 3 | 0.6667 |
| 1_1743302 | 1 | 1 | 1 | 1 | 1 | 1 | NA | NA | 3 | 1 | 1 | 0 | 1 | 1 | 3 | 0.6667 | NA | NA | 5 | 0.6 |
| 1_1743303 | 1 | 1 | 1 | 1 | 1 | 1 | NA | NA | 3 | 1 | 1 | 1 | 1 | 0 | 3 | 0.3333 | NA | NA | 5 | 0.4 |
| 1_1743304 | NA | NA | 1 | 1 | NA | NA | NA | NA | 1 | 1 | NA | NA | NA | NA | 3 | 0.6667 | 1 | 1 | 4 | 0.75 |
| 1_1743305 | 1 | 0 | 1 | 1 | 1 | 0 | NA | NA | 3 | 0.3333 | 1 | 0 | 1 | 0 | 2 | 0.5 | 1 | 0 | 5 | 0.2 |
| 1_1743309 | 2 | 0.5 | NA | NA | NA | NA | NA | NA | 2 | 0.5 | 1 | 1 | 1 | 0 | 3 | 0.6667 | 1 | 0 | 6 | 0.5 |
| 1_1744360 | 2 | 0 | 1 | 1 | 1 | 0 | NA | NA | 4 | 0.25 | 2 | 0.5 | 2 | 1 | 2 | 0.5 | 2 | 1 | 8 | 0.75 |
| 1_1744360 | 2 | 0.5 | 1 | 1 | 1 | 1 | NA | NA | 4 | 0.75 | 2 | 0.5 | 2 | 0.5 | 2 | 0.5 | 2 | 1 | 8 | 0.625 |
| 1_1744361 | 3 | 0.6667 | 1 | 1 | 1 | 1 | NA | NA | 5 | 0.8 | 1 | 0 | 2 | 0.5 | 3 | 0.6667 | 2 | 1 | 8 | 0.625 |
| 1_1744361 | 3 | 0.3333 | 1 | 1 | 1 | 1 | NA | NA | 5 | 0.6 | 2 | 0 | 2 | 1 | 3 | 0.3333 | 1 | 1 | 8 | 0.5 |
| 1_1744363 | 3 | 1 | 1 | 0 | 1 | 0 | NA | NA | 5 | 0.6 | 2 | 0.5 | 2 | 0.5 | 3 | 1 | 3 | 0.6667 | 10 | 0.7 |
| 1_1744363 | 2 | 0 | 1 | 1 | 1 | 0 | NA | NA | 4 | 0.25 | 2 | 0 | 2 | 0 | 3 | 0 | 3 | 0.3333 | 10 | 0.1 |
| 1_1744364 | 3 | 0.3333 | 1 | 1 | 2 | 0.5 | NA | NA | 6 | 0.5 | 3 | 1 | 1 | 0 | 2 | 0.5 | 2 | 1 | 8 | 0.75 |
| 1_1744365 | 2 | 1 | 1 | 0 | 1 | 0 | NA | NA | 4 | 0.5 | 2 | 1 | 1 | 1 | 2 | 1 | 3 | 0.6667 | 8 | 0.875 |
| 1_1744366 | 3 | 0.6667 | 1 | 0 | 2 | 0 | NA | NA | 6 | 0.3333 | 2 | 0 | 2 | 0 | 4 | 0.25 | 3 | 0.3333 | 11 | 0.1818 |

| | | | | | | | | | | | | | | | | | | | | |
|---|---|---|---|---|---|---|---|---|---|---|---|---|---|---|---|---|---|---|---|---|
| 1_17466160 | 2 | 1 | 6 | 0.5 | 3 | 0.6667 | 2 | 1 | 13 | 0.6923 | 3 | 0.6667 | 11 | 0.5455 | 4 | 0.5 | 3 | 0.6667 | 21 | 0.5714 |
| 1_17466169 | 1 | 0 | 7 | 0.4286 | 3 | 0.6667 | NA | NA | 11 | 0.4545 | 2 | 1 | 8 | 0.125 | 3 | 0.6667 | 4 | 0 | 17 | 0.2941 |
| 1_17466181 | 1 | 1 | 6 | 0 | 1 | 0 | 1 | 0 | 9 | 0.1111 | 2 | 0 | 9 | 0.1111 | 3 | 0 | 4 | 0.25 | 18 | 0.1111 |
| 1_17466251 | 2 | 0 | 5 | 0.2 | 3 | 0 | 1 | 0 | 11 | 0.0909 | 2 | 0.5 | 6 | 0.1667 | 3 | 0 | 3 | 0.3333 | 14 | 0.2143 |
| 1_17466258 | 1 | 0 | 3 | 0.6667 | 3 | 0.6667 | 1 | 0 | 8 | 0.5 | 2 | 0.5 | 4 | 0 | 2 | 0.5 | 3 | 0.3333 | 11 | 0.2727 |
| 1_17466279 | 2 | 0.5 | 4 | 0 | 3 | 0 | 1 | 0 | 10 | 0.1 | 2 | 0 | 5 | 0.2 | 3 | 0 | 3 | 0 | 13 | 0.0769 |
| 1_17466291 | 1 | 1 | 3 | 0.3333 | 3 | 1 | 1 | 0 | 8 | 0.625 | 2 | 1 | 6 | 0.6667 | 3 | 0 | 3 | 0.6667 | 14 | 0.5714 |
| 1_17466324 | 1 | 0 | 4 | 0 | 3 | 0.3333 | 1 | 0 | 9 | 0.1111 | 2 | 0 | 5 | 0.2 | 2 | 0 | 1 | 0 | 10 | 0.1 |
| 1_17466328 | NA | NA | 4 | 0 | 2 | 0.5 | 1 | 0 | 7 | 0.1429 | 2 | 0 | 5 | 0.2 | 2 | 0 | 2 | 0 | 11 | 0.0909 |
| 1_17466350 | 1 | 0 | 3 | 0.3333 | 2 | 1 | 1 | 0 | 7 | 0.4286 | 1 | 0 | 5 | 1 | 1 | 0 | 2 | 0 | 9 | 0.5556 |
| 1_17466364 | 1 | 1 | 1 | 1 | 2 | 0.5 | 1 | 1 | 5 | 0.8 | 2 | 0.5 | 5 | 0.4 | 2 | 1 | 2 | 1 | 11 | 0.6364 |
| 1_17466369 | 1 | 0 | 3 | 0.6667 | 3 | 0.6667 | 1 | 1 | 8 | 0.625 | 1 | 1 | 5 | 0.6 | 1 | 1 | 2 | 0.5 | 9 | 0.6667 |
| 1_17466371 | 1 | 0 | 3 | 0.3333 | 3 | 0 | 1 | 0 | 8 | 0.125 | 2 | 1 | 4 | 0.75 | 1 | 1 | 2 | 0.5 | 9 | 0.7778 |
| 1_17466373 | 1 | 1 | 2 | 1 | 3 | 1 | 1 | 1 | 7 | 1 | 2 | 1 | 5 | 1 | 1 | 1 | 2 | 1 | 10 | 1 |
| 1_17466374 | 1 | 1 | 2 | 1 | 3 | 1 | 1 | 1 | 7 | 1 | 2 | 1 | 3 | 1 | 1 | 1 | 2 | 1 | 8 | 1 |
| 1_17466377 | NA | NA | 1 | 1 | 3 | 1 | NA | NA | 4 | 1 | NA | NA | 5 | 0.8 | NA | NA | 2 | 1 | 7 | 0.8571 |
| 1_17466389 | 1 | 1 | 1 | 1 | 2 | 1 | 1 | 1 | 5 | 1 | 2 | 1 | 4 | 1 | 1 | 1 | 2 | 1 | 9 | 1 |
| 1_17466391 | 1 | 0 | 2 | 0.5 | 3 | 0.3333 | 1 | 1 | 7 | 0.4286 | 2 | 1 | 5 | 1 | 1 | 1 | 2 | 0.5 | 10 | 0.9 |
| 1_17466393 | NA | NA | 2 | 1 | 3 | 1 | 1 | 0 | 6 | 0.8333 | 2 | 1 | 4 | 0.5 | 1 | 1 | 2 | 1 | 9 | 0.7778 |
| 1_17466407 | 1 | 1 | 2 | 0.5 | 2 | 0 | 1 | 1 | 6 | 0.5 | 1 | 0 | 5 | 0.2 | 1 | 1 | 2 | 0.5 | 9 | 0.3333 |
| 1_17466408 | 1 | 1 | 2 | 0 | 2 | 0 | NA | NA | 5 | 0.2 | 2 | 0 | 4 | 0.25 | NA | NA | 2 | 0 | 8 | 0.125 |
| 1_17466418 | NA | NA | 1 | 0 | 2 | 0.5 | 1 | 1 | 4 | 0.5 | 1 | 1 | 5 | 0.2 | 1 | 1 | 1 | 0 | 8 | 0.375 |
| 1_17466435 | 1 | 1 | 2 | 0.5 | 1 | 1 | 1 | 1 | 5 | 0.8 | 2 | 0.5 | 5 | 0.6 | 1 | 1 | NA | NA | 8 | 0.625 |
| 1_17466436 | 1 | 1 | 2 | 0 | 3 | 0.3333 | 1 | 1 | 7 | 0.4286 | 2 | 0.5 | 4 | 0.5 | 1 | 0 | 2 | 0.5 | 9 | 0.4444 |
| 1_17466450 | 1 | 0 | 2 | 0.5 | 2 | 0 | 1 | 0 | 6 | 0.1667 | 2 | 0 | 4 | 0.5 | 1 | 1 | 1 | 1 | 8 | 0.5 |
| 1_17466463 | NA | NA | 1 | 1 | 3 | 0.3333 | 1 | 1 | 5 | 0.6 | 2 | 0 | 4 | 0.5 | 1 | 1 | 1 | 1 | 8 | 0.5 |
| 1_17488680 | 13 | 0.5385 | 5 | 0.6 | 10 | 0.4 | 10 | 0.3 | 38 | 0.4474 | 7 | 0.2857 | 8 | 0.375 | 8 | 0.375 | 3 | 0.6667 | 26 | 0.3846 |
| 1_17488688 | 11 | 0 | 4 | 0.5 | 11 | 0.1818 | 11 | 0 | 37 | 0.1081 | 4 | 0 | 5 | 0.2 | 10 | 0 | 1 | 0 | 20 | 0.05 |
| 1_17488695 | 12 | 0.1667 | 6 | 0.6667 | 10 | 0.1 | 11 | 0.1818 | 39 | 0.2308 | 5 | 0.4 | 7 | 0.1429 | 8 | 0.25 | 2 | 0.5 | 22 | 0.2727 |
| 1_17488703 | 12 | 0.0833 | 4 | 0.25 | 7 | 0 | 10 | 0.2 | 33 | 0.1212 | 5 | 0.2 | 5 | 0.2 | 9 | 0 | 2 | 0 | 21 | 0.0952 |
| 1_17488719 | 11 | 0.8182 | 6 | 0.8333 | 8 | 0.5 | 11 | 0.9091 | 36 | 0.7778 | 4 | 1 | 5 | 0.4 | 10 | 0.9 | 2 | 1 | 21 | 0.8095 |
| 1_17542808 | 5 | 0 | 4 | 0.5 | 3 | 0 | 2 | 0 | 14 | 0.1429 | NA | NA | 4 | 0.25 | 1 | 0 | 4 | 0.5 | 9 | 0.3333 |
| 1_17549010 | NA | NA | 1 | 1 | 2 | 0 | 1 | 1 | 4 | 0.5 | 1 | 0 | 2 | 0.5 | 1 | 1 | NA | NA | 4 | 0.5 |
| 1_17549019 | NA | NA | NA | NA | 1 | 1 | 1 | 1 | 2 | 1 | 1 | 1 | 2 | 1 | 1 | 1 | NA | NA | 4 | 1 |
| 1_17549037 | NA | NA | 1 | 0 | 1 | 1 | NA | NA | 2 | 0.5 | 1 | 0 | 2 | 0 | 1 | 1 | NA | NA | 4 | 0.25 |
| 1_17579021 | 1 | 0 | 2 | 1 | 2 | 0.5 | 1 | 0 | 6 | 0.5 | NA | NA | 1 | 1 | NA | NA | 1 | 0 | 2 | 0.5 |
| 1_17579135 | 1 | 1 | 3 | 1 | 1 | 1 | 1 | 1 | 6 | 1 | NA | NA | NA | NA | NA | NA | 2 | 0.5 | 2 | 0.5 |
| 1_17579138 | 1 | 1 | 3 | 0.3333 | 3 | 0 | 1 | 1 | 8 | 0.375 | NA | NA | NA | NA | NA | NA | 2 | 0.5 | 2 | 0.5 |
| 1_17579152 | 1 | 1 | 3 | 0.6667 | 3 | 1 | 2 | 1 | 9 | 0.8889 | NA | NA | NA | NA | NA | NA | 2 | 1 | 2 | 1 |
| 1_17610619 | NA | NA | NA | NA | NA | NA | 1 | 1 | 1 | 1 | NA | NA | NA | NA | NA | NA | 1 | 1 | 1 | 1 |
| 1_17656250 | 3 | 0 | 5 | 0.2 | 2 | 1 | 6 | 0.1667 | 16 | 0.25 | 2 | 1 | 3 | 0.3333 | 4 | 0.25 | 7 | 0.2857 | 16 | 0.375 |
| 1_17656267 | 3 | 0 | 3 | 0.3333 | 2 | 0 | 6 | 0 | 14 | 0.0714 | 3 | 0.3333 | 2 | 0 | 4 | 0 | 7 | 0.1429 | 16 | 0.125 |
| 1_17656270 | 2 | 0 | 4 | 0 | 2 | 0 | 5 | 0.2 | 13 | 0.0769 | 3 | 0.6667 | 2 | 0.5 | 4 | 0.25 | 6 | 0 | 15 | 0.2667 |
| 1_17656284 | 1 | 0 | 4 | 0.75 | 2 | 0.5 | 5 | 0.8 | 12 | 0.6667 | 2 | 1 | 2 | 1 | 4 | 0.75 | 7 | 0.8571 | 15 | 0.8667 |
| 1_17656290 | 2 | 0 | 4 | 0.25 | 2 | 0 | 6 | 0 | 14 | 0.0714 | 3 | 0 | 2 | 0 | 4 | 0.25 | 5 | 0 | 14 | 0.0714 |
| 1_17656322 | 3 | 1 | 5 | 1 | 2 | 1 | 6 | 0.5 | 16 | 0.8125 | 2 | 1 | 1 | 1 | 4 | 0.75 | 6 | 0.8333 | 13 | 0.8462 |
| 1_17656351 | 3 | 0.6667 | 4 | 0.25 | 2 | 1 | 5 | 0.4 | 14 | 0.5 | 2 | 1 | 1 | 0 | 4 | 0.5 | 5 | 0.8 | 12 | 0.6667 |
| 1_17658280 | 2 | 0 | 1 | 0 | 1 | 0 | 4 | 0.25 | 8 | 0.125 | NA | NA | 1 | 0 | 3 | 0.3333 | 4 | 0.25 | 8 | 0.25 |
| 1_17658282 | NA | NA | 3 | 0.6667 | 1 | 0 | 4 | 0.75 | 8 | 0.625 | NA | NA | 1 | 0 | 2 | 0.5 | 2 | 1 | 5 | 0.6 |
| 1_17658291 | NA | NA | 3 | 1 | NA | NA | 4 | 1 | 7 | 1 | 2 | 1 | 1 | 0 | 3 | 0.6667 | 3 | 0.6667 | 9 | 0.6667 |
| 1_17658293 | NA | NA | 4 | 0.25 | 1 | 0 | 4 | 0.5 | 9 | 0.3333 | 1 | 0 | 1 | 0 | 3 | 0 | 4 | 0.5 | 9 | 0.2222 |
| 1_17658300 | NA | NA | 2 | 1 | NA | NA | 3 | 0.6667 | 5 | 0.8 | 1 | 0 | 1 | 1 | 1 | 1 | 3 | 0.6667 | 6 | 0.6667 |
| 1_17658311 | NA | NA | 1 | 1 | 1 | 0 | 1 | 0 | 3 | 0.3333 | 1 | 0 | 1 | 0 | 3 | 0.3333 | 3 | 0.3333 | 8 | 0.25 |
| 1_17658317 | NA | NA | 3 | 0.3333 | 1 | 0 | 3 | 0.3333 | 7 | 0.2857 | 1 | 0 | 1 | 0 | 3 | 0.3333 | 4 | 0.5 | 9 | 0.3333 |
| 1_17658324 | NA | NA | 3 | 0.3333 | 1 | 0 | 4 | 0.25 | 8 | 0.25 | 1 | 0 | 1 | 0 | 2 | 0.5 | 4 | 0.25 | 8 | 0.25 |
| 1_17658334 | NA | NA | 3 | 0.3333 | 1 | 0 | 4 | 0.5 | 8 | 0.375 | 1 | 0 | 1 | 0 | 2 | 0 | 4 | 0.25 | 8 | 0.125 |
| 1_17658362 | NA | NA | 2 | 1 | 1 | 0 | 4 | 0.5 | 7 | 0.5714 | 1 | 1 | NA | NA | 2 | 0.5 | 4 | 0.5 | 7 | 0.5714 |
| 1_17658373 | NA | NA | 2 | 0.5 | 1 | 0 | 4 | 0.75 | 7 | 0.5714 | 1 | 1 | NA | NA | 2 | 1 | 4 | 0.5 | 7 | 0.7143 |
| 1_17658376 | NA | NA | 2 | 0 | 1 | 0 | 4 | 0.25 | 7 | 0.1429 | 1 | 0 | NA | NA | 2 | 0.5 | 4 | 0.5 | 7 | 0.4286 |
| 1_17658385 | NA | NA | 2 | 0 | 1 | 0 | 4 | 0.25 | 7 | 0.1429 | 1 | 0 | NA | NA | 2 | 0 | 3 | 0.3333 | 6 | 0.1667 |
| 1_17658402 | NA | NA | 2 | 0 | 1 | 0 | 3 | 1 | 6 | 0.5 | 1 | 0 | NA | NA | 1 | 0 | 4 | 0.75 | 6 | 0.5 |
| 1_17658412 | NA | NA | 2 | 0.5 | 1 | 1 | 4 | 0 | 7 | 0.2857 | 1 | 1 | NA | NA | 1 | 1 | 4 | 1 | 6 | 1 |
| 1_17658420 | NA | NA | 2 | 0 | 1 | 0 | 3 | 0.6667 | 6 | 0.3333 | 1 | 1 | NA | NA | 1 | 0 | 4 | 0.25 | 6 | 0.3333 |
| 1_17659699 | NA | NA | NA | NA | NA | NA | 2 | 0.5 | 2 | 0.5 | NA | NA | NA | NA | NA | NA | 2 | 0.5 | 2 | 0.5 |
| 1_17660215 | NA | NA | 2 | 0 | 1 | 0 | 2 | 0.5 | 5 | 0.2 | 1 | 0 | NA | NA | 1 | 0 | 4 | 0.25 | 6 | 0.1667 |

| | | | | | | | | | | | | | | | | | | | | |
|---|---|---|---|---|---|---|---|---|---|---|---|---|---|---|---|---|---|---|---|---|
| 1_1766022 | NA | NA | 2 | 0 | 1 | 1 | 3 | 0 | 6 | 0.1667 | 1 | 0 | NA | NA | 1 | 0 | 3 | 0.3333 | 5 | 0.2 |
| 1_1766023 | NA | NA | 2 | 0.5 | 1 | 0 | 3 | 0.3333 | 6 | 0.3333 | 1 | 0 | NA | NA | 1 | 0 | 2 | 0.5 | 4 | 0.25 |
| 1_1768478 | 1 | 1 | 3 | 0.6667 | 3 | 1 | 1 | 0 | 8 | 0.75 | NA | NA | NA | NA | NA | NA | 1 | 1 | 1 | 1 |
| 1_1768478 | 9 | 0 | 6 | 0 | 11 | 0.1818 | 11 | 0 | 37 | 0.0541 | 9 | 0.1111 | 8 | 0 | 4 | 0.25 | 11 | 0.2727 | 32 | 0.1562 |
| 1_1768478 | 9 | 0 | 3 | 0 | 10 | 0.1 | 10 | 0.2 | 32 | 0.0938 | 5 | 0 | 7 | 0.2857 | 4 | 0 | 6 | 0 | 22 | 0.0909 |
| 1_1768479 | 9 | 0 | 5 | 0.2 | 11 | 0.3636 | 10 | 0.2 | 35 | 0.2 | 9 | 0.1111 | 7 | 0.4286 | 5 | 0.2 | 11 | 0 | 32 | 0.1562 |
| 1_1768479 | 8 | 0.125 | 6 | 0.1667 | 10 | 0.1 | 12 | 0.0833 | 36 | 0.1111 | 9 | 0 | 8 | 0.125 | 5 | 0 | 9 | 0.1111 | 31 | 0.0645 |
| 1_1768490 | 4 | 0 | 3 | 0 | 3 | 0 | 5 | 0.2 | 15 | 0.0667 | 4 | 0 | 5 | 0.2 | 1 | 0 | 5 | 0 | 15 | 0.0667 |
| 1_1768491 | 5 | 0 | 3 | 0 | 3 | 0 | 7 | 0.1429 | 18 | 0.0556 | 3 | 0 | 6 | 0 | 1 | 0 | 9 | 0.1111 | 19 | 0.0526 |
| 1_1768511 | 7 | 0.2857 | 2 | 0 | 7 | 0.1429 | 6 | 0.3333 | 22 | 0.2273 | 10 | 0.3 | 8 | 0.375 | 3 | 0.6667 | 4 | 0.5 | 25 | 0.4 |
| 1_1768516 | 5 | 0.2 | 3 | 0 | 9 | 0.2222 | 5 | 0.2 | 22 | 0.1818 | 9 | 0.1111 | 7 | 0.2857 | 3 | 0 | 6 | 0.3333 | 25 | 0.2 |
| 1_1768517 | 6 | 0.1667 | 5 | 0 | 8 | 0 | 5 | 0.2 | 24 | 0.0833 | 7 | 0.2857 | 7 | 0.2857 | 3 | 0 | 5 | 0.2 | 22 | 0.2273 |
| 1_1768522 | 5 | 0.6 | 5 | 1 | 10 | 0.4 | 2 | 1 | 22 | 0.6364 | 6 | 0.8333 | 5 | 1 | 2 | 1 | 3 | 0.6667 | 16 | 0.875 |
| 1_1768523 | 4 | 0.25 | 4 | 0.25 | 8 | 0.125 | 4 | 0.5 | 20 | 0.25 | 7 | 0.1429 | 5 | 0.2 | 4 | 0.25 | 2 | 0.5 | 18 | 0.2222 |
| 1_1768527 | 3 | 0.3333 | 3 | 0 | 8 | 0.375 | 3 | 0 | 17 | 0.2353 | 3 | 0.3333 | 5 | 0 | 3 | 0.6667 | 1 | 1 | 12 | 0.3333 |
| 1_1768528 | 3 | 0.6667 | 2 | 0.5 | 7 | 0.5714 | 2 | 1 | 14 | 0.6429 | 7 | 0.2857 | 3 | 0.6667 | 3 | 0 | 1 | 0 | 14 | 0.2857 |
| 1_1768528 | 2 | 1 | 3 | 0.6667 | 7 | 0.5714 | 3 | 0.3333 | 15 | 0.6 | 5 | 0.6 | 2 | 0.5 | 3 | 0 | 1 | 0 | 11 | 0.3636 |
| 1_1768528 | 3 | 0.3333 | 3 | 0.6667 | 8 | 0.125 | 2 | 0.5 | 16 | 0.3125 | 7 | 0.5714 | 4 | 0.5 | 3 | 0.3333 | 1 | 1 | 15 | 0.5333 |
| 1_1772747 | 3 | 1 | 4 | 0.5 | 2 | 1 | 5 | 0.6 | 14 | 0.7143 | 2 | 0.5 | 4 | 0.75 | 5 | 0.6 | 5 | 0.4 | 16 | 0.5625 |
| 1_1772748 | 4 | 0 | 12 | 0.0833 | 5 | 0.2 | 6 | 0.1667 | 27 | 0.1111 | 2 | 0.5 | 8 | 0 | 9 | 0.1111 | 7 | 0.2857 | 26 | 0.1538 |
| 1_1772748 | 4 | 0 | 10 | 0.4 | 4 | 0 | 7 | 0.1429 | 25 | 0.2 | 2 | 0 | 6 | 0.1667 | 6 | 0 | 6 | 0.1667 | 20 | 0.1 |
| 1_1772748 | 3 | 1 | 13 | 0.8462 | 5 | 1 | 7 | 0.8571 | 28 | 0.8929 | 3 | 1 | 7 | 1 | 10 | 0.6 | 7 | 0.5714 | 27 | 0.7407 |
| 1_1772749 | 3 | 1 | 10 | 1 | 3 | 1 | 6 | 1 | 22 | 1 | 3 | 0.3333 | 5 | 0.8 | 7 | 0.8571 | 6 | 0.8333 | 21 | 0.7619 |
| 1_1772749 | 5 | 0 | 11 | 0 | 5 | 0.2 | 7 | 0.1429 | 28 | 0.0714 | 4 | 0.25 | 5 | 0.2 | 7 | 0 | 6 | 0 | 22 | 0.0909 |
| 1_1772751 | 4 | 0.25 | 10 | 0.3 | 6 | 0.3333 | 5 | 0.2 | 25 | 0.28 | 1 | 0 | 8 | 0.25 | 5 | 0 | 8 | 0 | 22 | 0.0909 |
| 1_1772751 | 4 | 1 | 10 | 1 | 5 | 0.4 | 6 | 0.8333 | 25 | 0.84 | 4 | 1 | 8 | 1 | 9 | 0.8889 | 8 | 0.875 | 29 | 0.931 |
| 1_1772752 | 5 | 0.2 | 12 | 0.25 | 6 | 0 | 6 | 0 | 29 | 0.1379 | 5 | 0.4 | 9 | 0.1111 | 10 | 0 | 6 | 0.5 | 30 | 0.2 |
| 1_1772753 | 4 | 0.75 | 6 | 0.8333 | 4 | 0.5 | 2 | 1 | 16 | 0.75 | 4 | 0.5 | 3 | 1 | 3 | 1 | 4 | 1 | 14 | 0.8571 |
| 1_1772753 | 4 | 0 | 7 | 0.2857 | 4 | 0.5 | 4 | 0.25 | 19 | 0.2632 | 5 | 0.2 | 4 | 0 | 6 | 0.1667 | 4 | 0 | 19 | 0.1053 |
| 1_1772754 | 3 | 0.6667 | 5 | 0.4 | 2 | 0.5 | 3 | 0 | 13 | 0.3846 | 3 | 0.3333 | 3 | 0.3333 | 5 | 0.4 | 3 | 0.3333 | 14 | 0.3571 |
| 1_1772755 | 7 | 0.8571 | 11 | 0.7273 | 5 | 1 | 6 | 0.6667 | 29 | 0.7931 | 5 | 0.6 | 8 | 0.625 | 9 | 0.8889 | 5 | 1 | 27 | 0.7778 |
| 1_1772755 | 6 | 0.5 | 10 | 0.4 | 3 | 0 | 6 | 0.1667 | 25 | 0.32 | 5 | 0.2 | 5 | 0.2 | 9 | 0.2222 | 5 | 0.2 | 24 | 0.2083 |
| 1_1772756 | 10 | 1 | 13 | 1 | 5 | 1 | 6 | 1 | 34 | 1 | 6 | 0.8333 | 6 | 1 | 9 | 1 | 8 | 0.875 | 29 | 0.931 |
| 1_1772756 | 8 | 0.5 | 13 | 0.5385 | 5 | 0.4 | 7 | 0.7143 | 33 | 0.5455 | 5 | 0.2 | 8 | 0.625 | 7 | 0.7143 | 8 | 0.75 | 28 | 0.6071 |
| 1_1772756 | 10 | 0.3 | 13 | 0.2308 | 6 | 0.1667 | 8 | 0.125 | 37 | 0.2162 | 6 | 0 | 9 | 0.2222 | 10 | 0.2 | 8 | 0.25 | 33 | 0.1818 |
| 1_1772757 | 5 | 0.6 | 11 | 0.3636 | 5 | 0.4 | 7 | 0.5714 | 28 | 0.4643 | 5 | 0.4 | 7 | 0.2857 | 8 | 0.625 | 6 | 0.5 | 26 | 0.4615 |
| 1_1772759 | 10 | 0.9 | 15 | 0.9333 | 6 | 0.8333 | 10 | 0.9 | 41 | 0.9024 | 5 | 0.8 | 10 | 1 | 11 | 0.7273 | 8 | 1 | 34 | 0.8824 |
| 1_1772759 | 7 | 1 | 10 | 1 | 3 | 1 | 4 | 1 | 24 | 1 | 3 | 1 | 4 | 1 | 9 | 1 | 6 | 1 | 22 | 1 |
| 1_1772761 | 13 | 0.2308 | 16 | 0.1875 | 3 | 0 | 10 | 0.2 | 42 | 0.1905 | 7 | 0.2857 | 9 | 0.3333 | 9 | 0.4444 | 8 | 0.625 | 33 | 0.4242 |
| 1_1772761 | 10 | 0.7 | 14 | 0.4286 | 6 | 0.6667 | 7 | 0.1429 | 37 | 0.4865 | 6 | 0.8333 | 9 | 0.8889 | 9 | 0.5556 | 7 | 0.1429 | 31 | 0.6129 |
| 1_1772761 | 11 | 1 | 18 | 0.9444 | 6 | 1 | 9 | 1 | 44 | 0.9773 | 6 | 1 | 7 | 1 | 10 | 1 | 10 | 1 | 33 | 1 |
| 1_1772762 | 12 | 0.5 | 16 | 0.25 | 5 | 0.4 | 10 | 0.5 | 43 | 0.3953 | 7 | 0.4286 | 9 | 0.3333 | 10 | 0.5 | 11 | 0.4545 | 37 | 0.4324 |
| 1_1772762 | 9 | 0.7778 | 13 | 0.9231 | 6 | 0.8333 | 8 | 1 | 36 | 0.8889 | 3 | 1 | 9 | 0.7778 | 10 | 0.9 | 6 | 0.8333 | 28 | 0.8571 |
| 1_1772763 | 8 | 0.5 | 17 | 0.4706 | 5 | 0.8 | 8 | 0.375 | 38 | 0.5 | 6 | 0.1667 | 10 | 0.3 | 9 | 0.2222 | 9 | 0.3333 | 34 | 0.2647 |
| 1_1772767 | 13 | 0.0769 | 22 | 0.0909 | 8 | 0 | 8 | 0.125 | 51 | 0.0784 | 7 | 0 | 9 | 0 | 8 | 0.125 | 11 | 0.0909 | 35 | 0.0571 |
| 1_1772768 | 11 | 0.4545 | 16 | 0.4375 | 7 | 0.8571 | 10 | 0.4 | 44 | 0.5 | 8 | 0.625 | 8 | 0.375 | 9 | 0.6667 | 7 | 0.7143 | 32 | 0.5938 |
| 1_1772768 | 12 | 0.3333 | 21 | 0.1905 | 6 | 0.1667 | 11 | 0.2727 | 50 | 0.24 | 9 | 0.3333 | 8 | 0.125 | 11 | 0.3636 | 12 | 0.1667 | 40 | 0.25 |
| 1_1772775 | 17 | 0.6471 | 21 | 0.9524 | 8 | 0.75 | 9 | 0.8889 | 55 | 0.8182 | 9 | 0.7778 | 7 | 0.8571 | 15 | 0.7333 | 17 | 1 | 48 | 0.8542 |
| 1_1772775 | 18 | 0.6111 | 26 | 0.5769 | 9 | 0.4444 | 12 | 0.6667 | 65 | 0.5846 | 8 | 0.875 | 11 | 0.5455 | 14 | 0.6429 | 14 | 0.7857 | 47 | 0.7021 |
| 1_1772776 | 17 | 0.1765 | 27 | 0.2222 | 8 | 0.125 | 12 | 0.25 | 64 | 0.2031 | 10 | 0.2 | 13 | 0.1538 | 13 | 0 | 16 | 0.25 | 52 | 0.1538 |
| 1_1772778 | 18 | 0.5556 | 30 | 0.6667 | 12 | 0.6667 | 12 | 0.6667 | 72 | 0.6389 | 10 | 0.6 | 15 | 0.5333 | 15 | 0.4667 | 17 | 0.5882 | 57 | 0.5439 |
| 1_1772780 | 22 | 0.5455 | 36 | 0.6667 | 16 | 0.8125 | 12 | 0.75 | 86 | 0.6744 | 11 | 0.4545 | 15 | 0.7333 | 16 | 0.625 | 18 | 0.5556 | 60 | 0.6 |
| 1_1775912 | 13 | 0.2308 | 5 | 0 | 11 | 0 | 10 | 0 | 39 | 0.0769 | 16 | 0.0625 | 15 | 0.0667 | 12 | 0.0833 | 8 | 0 | 51 | 0.0588 |
| 1_1775915 | 11 | 0 | 4 | 0.25 | 8 | 0.125 | 9 | 0 | 32 | 0.0625 | 16 | 0.125 | 13 | 0.0769 | 10 | 0.1 | 8 | 0.125 | 47 | 0.1064 |
| 1_1775915 | 11 | 0.7273 | 3 | 1 | 7 | 0.8571 | 9 | 0.6667 | 30 | 0.7667 | 12 | 0.6667 | 14 | 0.7857 | 10 | 0.8 | 5 | 0.6 | 41 | 0.7317 |
| 1_1775917 | 9 | 0.8889 | 4 | 0.5 | 8 | 0.375 | 10 | 0.4 | 31 | 0.5484 | 16 | 0.3125 | 14 | 0.6429 | 10 | 0.3 | 6 | 0.1667 | 46 | 0.3913 |
| 1_1775917 | 11 | 0.6364 | 4 | 1 | 9 | 0.5556 | 9 | 0.6667 | 33 | 0.6667 | 16 | 0.8125 | 15 | 0.6 | 10 | 0.3 | 6 | 0.6667 | 47 | 0.617 |
| 1_1778119 | NA | NA | 1 | 1 | NA | NA | NA | NA | 1 | 1 | NA | NA | NA | NA | NA | NA | 1 | 1 | 1 | 1 |
| 1_1778120 | NA | NA | 1 | 1 | NA | NA | NA | NA | 1 | 1 | NA | NA | NA | NA | NA | NA | 1 | 1 | 1 | 1 |
| 1_1778152 | NA | NA | 1 | 0 | NA | NA | 1 | 1 | 2 | 0.5 | NA | NA | NA | NA | NA | NA | 1 | 1 | 1 | 1 |
| 1_1778156 | NA | NA | 1 | 1 | NA | NA | 1 | 1 | 2 | 1 | NA | NA | NA | NA | 1 | 1 | 1 | 1 | 2 | 1 |
| 1_1778156 | NA | NA | 1 | 0 | NA | NA | 1 | 1 | 2 | 0.5 | NA | NA | NA | NA | NA | NA | 1 | 1 | 1 | 1 |
| 1_1778157 | NA | NA | 1 | 1 | NA | NA | NA | NA | 1 | 1 | NA | NA | NA | NA | NA | NA | 1 | 1 | 1 | 1 |
| 1_1778157 | NA | NA | 1 | 0 | NA | NA | 1 | 1 | 2 | 0.5 | NA | NA | NA | NA | 1 | 0 | 1 | 1 | 2 | 0.5 |
| 1_1778160 | NA | NA | 1 | 0 | NA | NA | 1 | 1 | 2 | 0.5 | NA | NA | NA | NA | 1 | 1 | 1 | 1 | 2 | 1 |

| | | | | | | | | | | | | | | | | | | | | |
|---|---|---|---|---|---|---|---|---|---|---|---|---|---|---|---|---|---|---|---|---|
| 1_17792622 | NA | NA | 1 | 0 | 1 | 1 | NA | NA | 2 | 0.5 | NA | NA | 2 | 0.5 | NA | NA | 2 | 0.5 | 4 | 0.5 |
| 1_17793141 | NA | NA | 1 | 1 | 1 | 0 | NA | NA | 2 | 0.5 | NA | NA | 2 | 0.5 | NA | NA | 2 | 0 | 4 | 0.25 |
| 1_17793147 | 1 | 1 | 1 | 1 | 1 | 1 | NA | NA | 3 | 1 | NA | NA | 1 | 0 | NA | NA | 1 | 1 | 2 | 0.5 |
| 1_17793896 | 1 | 1 | NA | NA | 1 | 0 | NA | NA | 2 | 0.5 | NA | NA | 2 | 0 | NA | NA | 2 | 0.5 | 4 | 0.25 |
| 1_17798909 | 1 | 1 | 1 | 1 | NA | NA | NA | NA | 2 | 1 | 1 | 1 | NA | NA | NA | NA | 1 | 1 | 2 | 1 |
| 1_17798911 | 1 | 1 | 1 | 1 | NA | NA | NA | NA | 2 | 1 | 1 | 1 | NA | NA | NA | NA | 1 | 1 | 2 | 1 |
| 1_17798914 | 1 | 0 | 1 | 1 | NA | NA | NA | NA | 2 | 0.5 | NA | NA | NA | NA | NA | NA | 1 | 1 | 1 | 1 |
| 1_17798929 | 1 | 1 | 1 | 1 | NA | NA | NA | NA | 2 | 1 | 1 | 1 | NA | NA | NA | NA | 1 | 1 | 2 | 1 |
| 1_17798959 | 1 | 1 | 1 | 0 | NA | NA | NA | NA | 2 | 0.5 | 1 | 1 | NA | NA | NA | NA | 1 | 1 | 2 | 1 |
| 1_17798972 | 1 | 1 | 1 | 1 | NA | NA | NA | NA | 2 | 1 | 1 | 1 | NA | NA | NA | NA | NA | NA | 1 | 1 |
| 1_17798986 | 1 | 0 | 1 | 1 | NA | NA | NA | NA | 2 | 0.5 | 1 | 1 | NA | NA | NA | NA | NA | NA | 1 | 1 |
| 1_17798991 | 1 | 1 | 1 | 0 | NA | NA | NA | NA | 2 | 0.5 | 1 | 1 | NA | NA | NA | NA | 1 | 1 | 2 | 1 |
| 1_17798993 | 1 | 1 | 1 | 1 | NA | NA | NA | NA | 2 | 1 | 1 | 1 | NA | NA | NA | NA | 1 | 1 | 2 | 1 |
| 1_17798997 | 1 | 1 | 1 | 0 | NA | NA | NA | NA | 2 | 0.5 | 1 | 1 | NA | NA | NA | NA | 1 | 1 | 2 | 1 |
| 1_17799001 | NA | NA | 1 | 1 | NA | NA | NA | NA | 1 | 1 | NA | NA | NA | NA | NA | NA | 1 | 1 | 1 | 1 |
| 1_17807701 | 2 | 0 | 3 | 0.3333 | NA | NA | 6 | 0.3333 | 11 | 0.2727 | 2 | 0 | 2 | 1 | NA | NA | 1 | 0 | 5 | 0.4 |
| 1_17807705 | 2 | 1 | NA | NA | NA | NA | 6 | 0.5 | 8 | 0.625 | 2 | 1 | 1 | 1 | NA | NA | 2 | 0 | 5 | 0.6 |
| 1_17807737 | 2 | 1 | 5 | 0.4 | NA | NA | 6 | 1 | 13 | 0.7692 | 2 | 1 | 3 | 1 | NA | NA | 3 | 0 | 8 | 0.625 |
| 1_17808010 | 7 | 0.2857 | 4 | 0.5 | 1 | 0 | 10 | 0.3 | 22 | 0.3182 | 4 | 0.5 | 7 | 0.2857 | 5 | 0.4 | 1 | 1 | 17 | 0.4118 |
| 1_17814878 | 2 | 0 | 6 | 0.3333 | 5 | 0 | 7 | 0 | 20 | 0.1 | NA | NA | 3 | 0 | 1 | 0 | 1 | 1 | 5 | 0.2 |
| 1_17814883 | 2 | 0 | 5 | 0.2 | 4 | 0 | 7 | 0.1429 | 18 | 0.1111 | NA | NA | 3 | 0.3333 | 2 | 0 | NA | NA | 5 | 0.2 |
| 1_17814892 | 2 | 0 | 6 | 0.1667 | 5 | 0 | 6 | 0 | 19 | 0.0526 | NA | NA | 3 | 0.3333 | 2 | 0 | 1 | 0 | 6 | 0.1667 |
| 1_17814904 | 2 | 0.5 | 6 | 0.3333 | 5 | 0 | 7 | 0.4286 | 20 | 0.3 | NA | NA | 3 | 0.3333 | 2 | 0.5 | 1 | 1 | 6 | 0.5 |
| 1_17814995 | 2 | 0 | 5 | 0.2 | 5 | 0.2 | 7 | 0.1429 | 19 | 0.1579 | NA | NA | 1 | 0 | 1 | 1 | 1 | 1 | 3 | 0.6667 |
| 1_17815179 | 1 | 0 | 2 | 0 | 4 | 0 | 3 | 0.3333 | 10 | 0.1 | NA | NA | 2 | 0 | 1 | 0 | 1 | 1 | 4 | 0.25 |
| 1_17815297 | 1 | 0 | 4 | 0.25 | 5 | 0.2 | 2 | 0.5 | 12 | 0.25 | 2 | 0 | 3 | 0 | 3 | 0 | 4 | 0.25 | 12 | 0.0833 |
| 1_17815305 | 2 | 0 | 4 | 0.25 | 5 | 0 | 2 | 0 | 13 | 0.0769 | 2 | 0 | 3 | 0 | 2 | 0 | 4 | 0.25 | 11 | 0.0909 |
| 1_17815690 | 26 | 0.7692 | 35 | 0.8571 | 23 | 0.8261 | 47 | 0.7234 | 131 | 0.7863 | 16 | 0.75 | 28 | 0.8214 | 37 | 0.6216 | 30 | 0.6 | 111 | 0.6847 |
| 1_17816829 | NA | NA | 2 | 1 | NA | NA | 1 | 1 | 3 | 1 | NA | NA | NA | NA | NA | NA | 1 | 1 | 1 | 1 |
| 1_17876607 | 1 | 1 | 1 | 1 | NA | NA | NA | NA | 2 | 1 | NA | NA | 1 | 1 | NA | NA | NA | NA | 1 | 1 |
| 1_17934790 | 2 | 0.5 | 1 | 1 | 2 | 0.5 | NA | NA | 5 | 0.6 | 3 | 0.3333 | NA | NA | 1 | 1 | 1 | 0 | 5 | 0.4 |
| 1_17935619 | 1 | 0 | 1 | 1 | 2 | 0.5 | 1 | 0 | 5 | 0.4 | NA | NA | 2 | 1 | 1 | 0 | NA | NA | 3 | 0.6667 |
| 1_17935621 | 5 | 0.6 | 5 | 0.2 | 3 | 0.6667 | 3 | 0.3333 | 16 | 0.4375 | NA | NA | 6 | 0.1667 | 5 | 0.4 | 3 | 0.6667 | 14 | 0.3571 |
| 1_17935645 | 6 | 0.3333 | 2 | 0 | 3 | 0 | 4 | 0 | 15 | 0.1333 | NA | NA | 6 | 0 | 4 | 0.25 | 3 | 0 | 13 | 0.0769 |
| 1_17935663 | 4 | 0 | 4 | 0 | 3 | 0.3333 | 2 | 0.5 | 13 | 0.1538 | NA | NA | 6 | 0.1667 | 5 | 0 | 3 | 0 | 14 | 0.0714 |
| 1_17935667 | 5 | 1 | 3 | 1 | 3 | 1 | 4 | 1 | 15 | 1 | NA | NA | 4 | 0.75 | 5 | 1 | 3 | 1 | 12 | 0.9167 |
| 1_17935675 | 5 | 0.2 | 3 | 0.3333 | 4 | 0 | 4 | 0 | 16 | 0.125 | NA | NA | 4 | 0.25 | 5 | 0 | 3 | 0 | 12 | 0.0833 |
| 1_17935682 | 3 | 1 | 3 | 0.3333 | 3 | 0 | 3 | 1 | 12 | 0.5833 | NA | NA | 4 | 0 | 2 | 1 | 3 | 0.3333 | 9 | 0.3333 |
| 1_17935682 | 1 | 1 | NA | NA | NA | NA | NA | NA | 1 | 1 | NA | NA | 2 | 0 | 2 | 1 | NA | NA | 4 | 0.5 |
| 1_17937359 | 3 | 0 | 1 | 1 | 3 | 0 | 3 | 0.3333 | 10 | 0.2 | NA | NA | 4 | 0.25 | 4 | 0 | 2 | 0.5 | 10 | 0.2 |
| 1_17937370 | 3 | 0.3333 | 1 | 1 | 2 | 0.5 | 3 | 0.3333 | 9 | 0.4444 | NA | NA | 3 | 0.3333 | 4 | 0.5 | 2 | 0.5 | 9 | 0.4444 |
| 1_17940715 | 1 | 1 | NA | NA | NA | NA | NA | NA | 1 | 1 | 1 | 0 | 1 | 1 | NA | NA | NA | NA | 2 | 0.5 |
| 1_17943752 | 1 | 0 | 1 | 1 | 1 | 0 | NA | NA | 3 | 0.3333 | NA | NA | 2 | 0.5 | 1 | 0 | NA | NA | 3 | 0.3333 |
| 1_17943763 | 2 | 0.5 | 1 | 0 | 2 | 0 | 1 | 0 | 6 | 0.1667 | 1 | 0 | 2 | 0.5 | NA | NA | NA | NA | 3 | 0.3333 |
| 1_17943765 | 2 | 0.5 | 1 | 1 | 3 | 0.3333 | 1 | 1 | 7 | 0.5714 | 2 | 1 | 3 | 1 | 2 | 1 | NA | NA | 7 | 1 |
| 1_17943776 | 2 | 1 | 1 | 1 | 1 | 1 | 1 | 0 | 5 | 0.8 | 2 | 1 | 2 | 1 | 2 | 0.5 | NA | NA | 6 | 0.8333 |
| 1_17943781 | 1 | 1 | 1 | 1 | 2 | 0 | 1 | 1 | 5 | 0.6 | 2 | 0 | 3 | 0.3333 | 2 | 0.5 | NA | NA | 7 | 0.2857 |
| 1_17943790 | 2 | 1 | NA | NA | 2 | 1 | 1 | 1 | 5 | 1 | 1 | 1 | 3 | 1 | 2 | 0.5 | NA | NA | 6 | 0.8333 |
| 1_17943799 | 2 | 0 | 1 | 0 | 3 | 0.3333 | 1 | 1 | 7 | 0.2857 | 2 | 0 | 3 | 0.3333 | 2 | 0.5 | 1 | 1 | 8 | 0.375 |
| 1_17945263 | 2 | 0.5 | 1 | 1 | 2 | 0 | 1 | 0 | 6 | 0.3333 | 2 | 0 | 3 | 0.6667 | 3 | 0.3333 | 1 | 0 | 9 | 0.3333 |
| 1_17945268 | 2 | 1 | 1 | 1 | 3 | 1 | NA | NA | 6 | 1 | 2 | 0.5 | 3 | 0.6667 | 3 | 1 | 2 | 1 | 10 | 0.8 |
| 1_17945270 | 1 | 1 | 1 | 1 | 3 | 1 | NA | NA | 5 | 1 | 2 | 1 | 3 | 1 | 3 | 0.6667 | 1 | 1 | 9 | 0.8889 |
| 1_17945274 | 2 | 0.5 | 1 | 1 | 4 | 0.75 | NA | NA | 7 | 0.7143 | 2 | 0 | 4 | 0 | 3 | 0.6667 | 1 | 0 | 10 | 0.2 |
| 1_17945282 | 2 | 0.5 | NA | NA | 3 | 0 | 1 | 0 | 6 | 0.1667 | 2 | 0.5 | 4 | 0.5 | 3 | 0.3333 | 1 | 0 | 10 | 0.4 |
| 1_17945288 | 3 | 0.3333 | 1 | 1 | 5 | 0.6 | 1 | 0 | 10 | 0.5 | 2 | 0.5 | 5 | 0.4 | 3 | 0.3333 | 2 | 1 | 12 | 0.5 |
| 1_17945295 | 2 | 1 | 1 | 1 | 2 | 1 | 1 | 1 | 6 | 1 | 2 | 1 | 3 | 0.6667 | 3 | 1 | 2 | 0.5 | 10 | 0.8 |
| 1_17945315 | 3 | 1 | 1 | 1 | 3 | 0.6667 | 1 | 0 | 8 | 0.75 | 1 | 1 | 4 | 0.75 | 3 | 0.3333 | 2 | 0.5 | 10 | 0.6 |
| 1_17945338 | 4 | 0.25 | NA | NA | 4 | 0.5 | 1 | 0 | 9 | 0.3333 | 1 | 1 | 4 | 0.5 | 2 | 0.5 | 2 | 0 | 9 | 0.4444 |
| 1_17945341 | 3 | 0.3333 | 1 | 0 | 1 | 1 | 2 | 0 | 7 | 0.2857 | NA | NA | 6 | 0.5 | 3 | 0.6667 | 2 | 0 | 11 | 0.4545 |
| 1_17945343 | 4 | 0.25 | 1 | 0 | 3 | 0.3333 | 2 | 0 | 10 | 0.2 | 2 | 1 | 5 | 0.2 | 2 | 0.5 | 2 | 0 | 11 | 0.3636 |
| 1_17945385 | 8 | 0 | 1 | 0 | 6 | 0.1667 | 2 | 0 | 17 | 0.0588 | 2 | 0.5 | 7 | 0 | 3 | 0 | 2 | 0 | 14 | 0.0714 |
| 1_17945396 | 7 | 0.1429 | 1 | 0 | 6 | 0 | NA | NA | 14 | 0.0714 | 2 | 0.5 | 4 | 0.25 | 4 | 0 | 2 | 0 | 12 | 0.1667 |
| 1_17945418 | 8 | 0 | 1 | 1 | 7 | 0 | 2 | 0 | 18 | 0.0556 | 2 | 0 | 8 | 0.125 | 4 | 0 | 3 | 0 | 17 | 0.0588 |
| 1_17945423 | 9 | 0.1111 | 1 | 0 | 5 | 0 | 2 | 0 | 17 | 0.0588 | 3 | 0 | 8 | 0 | 3 | 0 | 2 | 0.5 | 16 | 0.0625 |
| 1_17945436 | 9 | 0 | 1 | 0 | 6 | 0.1667 | 2 | 0 | 18 | 0.0556 | 3 | 0 | 8 | 0.125 | 4 | 0 | 2 | 0 | 17 | 0.0588 |

| | | | | | | | | | | | | | | | | | | | | |
|---|---|---|---|---|---|---|---|---|---|---|---|---|---|---|---|---|---|---|---|---|
| 1_1794544 | 6 | 0.1667 | 1 | 0 | 7 | 0 | 2 | 0 | 16 | 0.0625 | 3 | 0.3333 | 8 | 0.125 | 1 | 0 | 2 | 0 | 14 | 0.1429 |
| 1_1794546 | 9 | 0 | NA | NA | 7 | 0.2857 | 2 | 0 | 18 | 0.1111 | 3 | 0 | 5 | 0.2 | 3 | 0 | 2 | 0 | 13 | 0.0769 |
| 1_1794548 | 6 | 0.1667 | 1 | 0 | 5 | 0.2 | 2 | 0 | 14 | 0.1429 | 3 | 0.6667 | 5 | 0.4 | 4 | 0 | 2 | 0 | 14 | 0.2857 |
| 1_1794549 | 9 | 0.1111 | 1 | 0 | 8 | 0 | 2 | 0 | 20 | 0.05 | 4 | 0 | 7 | 0.1429 | 4 | 0 | 3 | 0 | 18 | 0.0556 |
| 1_1794549 | 9 | 0.3333 | 1 | 0 | 6 | 0.3333 | 2 | 0.5 | 18 | 0.3333 | 4 | 0.75 | 6 | 0.1667 | 2 | 0 | 2 | 0 | 14 | 0.2857 |
| 1_1795143 | 16 | 0.0625 | 17 | 0.1176 | 9 | 0.1111 | 9 | 0.1111 | 51 | 0.098 | 13 | 0 | 13 | 0 | 9 | 0.2222 | 15 | 0.1333 | 50 | 0.08 |
| 1_1795143 | 13 | 0.3846 | 18 | 0.5 | 10 | 0.6 | 9 | 0.2222 | 50 | 0.44 | 12 | 0.3333 | 13 | 0.3846 | 9 | 0.5556 | 16 | 0.5 | 50 | 0.44 |
| 1_1795144 | 10 | 0 | 13 | 0.1538 | 11 | 0.0909 | 5 | 0 | 39 | 0.0769 | 13 | 0.0769 | 9 | 0.1111 | 7 | 0 | 10 | 0.1 | 39 | 0.0769 |
| 1_1795146 | 16 | 0.125 | 15 | 0.2 | 13 | 0.0769 | 6 | 0 | 50 | 0.12 | 9 | 0 | 11 | 0 | 11 | 0 | 18 | 0.2778 | 49 | 0.102 |
| 1_1795147 | 16 | 0.1875 | 14 | 0.1429 | 11 | 0.4545 | 4 | 0 | 45 | 0.2222 | 10 | 0.3 | 12 | 0.25 | 11 | 0.2727 | 13 | 0.1538 | 46 | 0.2391 |
| 1_1795727 | 2 | 0 | 2 | 0.5 | 3 | 0 | 5 | 0.4 | 12 | 0.25 | 5 | 0 | 2 | 0 | 4 | 0.25 | 4 | 0.25 | 15 | 0.1333 |
| 1_1795728 | 2 | 0 | 4 | 0 | 5 | 0 | 3 | 0.3333 | 14 | 0.0714 | 3 | 0.3333 | 2 | 0 | 4 | 0 | 5 | 0 | 14 | 0.0714 |
| 1_1795730 | 2 | 0 | 5 | 0 | 5 | 0.2 | 3 | 0.3333 | 15 | 0.1333 | 4 | 0.25 | 1 | 0 | 3 | 0 | 5 | 0 | 13 | 0.0769 |
| 1_1795730 | 3 | 0.3333 | 5 | 0.4 | 5 | 0.2 | 2 | 0.5 | 15 | 0.3333 | 4 | 0.25 | NA | NA | 5 | 0.2 | 5 | 0.2 | 14 | 0.2143 |
| 1_1795731 | 3 | 0 | 5 | 0 | 5 | 0.2 | 3 | 0.3333 | 16 | 0.125 | 4 | 0 | 1 | 0 | 4 | 0.25 | 6 | 0.1667 | 15 | 0.1333 |
| 1_1795732 | 2 | 0 | 5 | 0.8 | 4 | 0.25 | 4 | 0.5 | 15 | 0.4667 | 3 | 0 | 2 | 0.5 | 5 | 0.2 | 6 | 0.3333 | 16 | 0.25 |
| 1_1795732 | 3 | 1 | 5 | 1 | 4 | 1 | 4 | 1 | 16 | 1 | 2 | 1 | 1 | 1 | 5 | 0.4 | 5 | 1 | 13 | 0.7692 |
| 1_1795735 | 3 | 0 | 5 | 0.6 | 5 | 0 | 4 | 0 | 17 | 0.1765 | 3 | 0 | 2 | 0 | 5 | 0.8 | 6 | 0.1667 | 16 | 0.3125 |
| 1_1795735 | 3 | 0 | 4 | 0.5 | 4 | 0 | 4 | 0 | 15 | 0.1333 | 3 | 0 | 1 | 0 | 5 | 0.2 | 6 | 0.1667 | 15 | 0.1333 |
| 1_1795739 | 1 | 0 | 3 | 0.6667 | 1 | 1 | 1 | 1 | 6 | 0.6667 | NA | NA | 2 | 0 | 2 | 0.5 | 4 | 0.25 | 8 | 0.25 |
| 1_1795739 | 3 | 0 | 4 | 0.25 | 3 | 0.3333 | 4 | 0 | 14 | 0.1429 | 1 | 0 | 2 | 0 | 3 | 0.3333 | 5 | 0.2 | 11 | 0.1818 |
| 1_1795799 | 2 | 0.5 | 2 | 0.5 | 3 | 1 | 3 | 0.6667 | 10 | 0.7 | NA | NA | 2 | 1 | 3 | 0.6667 | 4 | 0.5 | 9 | 0.6667 |
| 1_1795799 | 3 | 0 | 2 | 0.5 | 3 | 0.3333 | 4 | 0.25 | 12 | 0.25 | NA | NA | 1 | 0 | 3 | 0.6667 | 6 | 0.1667 | 10 | 0.3 |
| 1_1795799 | 3 | 0.3333 | 2 | 1 | 1 | 1 | 4 | 0.75 | 10 | 0.7 | NA | NA | 1 | 1 | 2 | 0.5 | 3 | 0.6667 | 6 | 0.6667 |
| 1_1795800 | 3 | 0.3333 | 2 | 0 | 3 | 0.3333 | 3 | 0.6667 | 11 | 0.3636 | NA | NA | 2 | 0 | 2 | 0.5 | 6 | 0.5 | 10 | 0.4 |
| 1_1795803 | 3 | 0.3333 | 2 | 1 | 3 | 1 | 3 | 1 | 11 | 0.8182 | NA | NA | 2 | 1 | 1 | 0 | 4 | 0.25 | 7 | 0.4286 |
| 1_1795805 | 3 | 0.6667 | 1 | 1 | 3 | 0.6667 | 2 | 1 | 9 | 0.7778 | NA | NA | 2 | 1 | 3 | 0.6667 | 4 | 0.75 | 9 | 0.7778 |
| 1_1795807 | 3 | 0.3333 | 1 | 0 | 3 | 0 | 3 | 0 | 10 | 0.1 | NA | NA | 2 | 0 | 3 | 0.3333 | 4 | 0 | 9 | 0.1111 |
| 1_1795809 | 2 | 0 | 1 | 0 | 3 | 0.3333 | 3 | 0.3333 | 9 | 0.2222 | NA | NA | 2 | 0 | 1 | 0 | 4 | 0.25 | 7 | 0.1429 |
| 1_1795810 | 2 | 1 | 1 | 1 | 3 | 1 | 3 | 1 | 9 | 1 | NA | NA | 2 | 0.5 | 2 | 1 | 4 | 1 | 8 | 0.875 |
| 1_1795812 | 1 | 1 | 1 | 1 | 3 | 0.6667 | 2 | 0.5 | 7 | 0.7143 | NA | NA | 2 | 1 | 2 | 0.5 | 4 | 1 | 8 | 0.875 |
| 1_1795812 | 1 | 0 | 1 | 0 | 3 | 0.3333 | 1 | 0 | 6 | 0.1667 | NA | NA | 2 | 0.5 | 2 | 0 | 4 | 0.25 | 8 | 0.25 |
| 1_1795814 | 1 | 0 | 1 | 1 | 3 | 0.3333 | 2 | 0 | 7 | 0.2857 | NA | NA | 1 | 0 | 2 | 0.5 | 3 | 0 | 6 | 0.1667 |
| 1_1795814 | 1 | 1 | 1 | 1 | 3 | 0.6667 | 2 | 1 | 7 | 0.8571 | NA | NA | 2 | 1 | 2 | 1 | 3 | 0.6667 | 7 | 0.8571 |
| 1_1795815 | 1 | 1 | 1 | 0 | 3 | 0.6667 | 1 | 0 | 6 | 0.5 | NA | NA | 2 | 0.5 | 2 | 1 | 4 | 0.25 | 8 | 0.5 |
| 1_1797876 | NA | NA | 2 | 0.5 | NA | NA | NA | NA | 2 | 0.5 | NA | NA | NA | NA | 1 | 0 | 1 | 1 | 2 | 0.5 |
| 1_1797881 | NA | NA | 2 | 1 | NA | NA | NA | NA | 2 | 1 | NA | NA | NA | NA | 2 | 0.5 | 2 | 0.5 | 4 | 0.5 |
| 1_1797882 | NA | NA | 2 | 1 | NA | NA | NA | NA | 2 | 1 | 1 | 0 | NA | NA | 3 | 0.6667 | 1 | 1 | 5 | 0.6 |
| 1_1799721 | 2 | 0 | 6 | 0.1667 | 7 | 0.4286 | 7 | 0.2857 | 22 | 0.2727 | 2 | 0.5 | 8 | 0.5 | 6 | 0.3333 | 7 | 0.1429 | 23 | 0.3478 |
| 1_1799721 | 2 | 0.5 | 7 | 0.5714 | 8 | 0.25 | 6 | 0.1667 | 23 | 0.3478 | 4 | 0.25 | 8 | 0.5 | 7 | 0.4286 | 6 | 0.1667 | 25 | 0.36 |
| 1_1799722 | 2 | 0.5 | 7 | 0.7143 | 8 | 0.5 | 7 | 0.5714 | 24 | 0.5833 | 3 | 0.3333 | 7 | 0.8571 | 5 | 0.8 | 6 | 0.5 | 21 | 0.6667 |
| 1_1799896 | 3 | 0.6667 | 7 | 0.2857 | 10 | 0.2 | 8 | 0.125 | 28 | 0.25 | 4 | 0.25 | 8 | 0.125 | 8 | 0.25 | 5 | 0 | 25 | 0.16 |
| 1_1799897 | 3 | 0.6667 | 9 | 0.1111 | 10 | 0.1 | 6 | 0 | 28 | 0.1429 | 3 | 0.3333 | 11 | 0.1818 | 8 | 0.25 | 5 | 0 | 27 | 0.1852 |
| 1_1799897 | 6 | 0.5 | 12 | 0.1667 | 7 | 0.2857 | 8 | 0 | 33 | 0.2121 | 5 | 0.2 | 9 | 0.1111 | 9 | 0.2222 | 9 | 0 | 32 | 0.125 |
| 1_1799897 | 5 | 0.6 | 13 | 0.6154 | 12 | 0.3333 | 11 | 0.5455 | 41 | 0.5122 | 6 | 0.3333 | 11 | 0.4545 | 9 | 0.6667 | 9 | 0.3333 | 35 | 0.4571 |
| 1_1799898 | 7 | 0.2857 | 15 | 0.2667 | 12 | 0.0833 | 12 | 0.0833 | 46 | 0.1739 | 6 | 0.1667 | 10 | 0 | 10 | 0.2 | 8 | 0 | 34 | 0.0882 |
| 1_1799907 | 7 | 0.1429 | 10 | 0.4 | 6 | 0.6667 | 8 | 0.25 | 31 | 0.3548 | 4 | 0.5 | 5 | 0.2 | 8 | 0.75 | 4 | 0.5 | 21 | 0.5238 |
| 1_1799908 | 8 | 0 | 10 | 0.2 | 6 | 0 | 11 | 0.3636 | 35 | 0.1714 | 5 | 0 | 3 | 0.3333 | 6 | 0.1667 | 7 | 0 | 21 | 0.0952 |
| 1_1799941 | 1 | 1 | 3 | 0.3333 | 2 | 0 | 3 | 0 | 9 | 0.2222 | 2 | 0 | 4 | 0.25 | 6 | 0 | 3 | 0 | 15 | 0.0667 |
| 1_1799943 | 3 | 0 | 3 | 0 | 2 | 0 | 5 | 0.2 | 13 | 0.0769 | 4 | 0 | 4 | 0.25 | 5 | 0 | 3 | 0 | 16 | 0.0625 |
| 1_1799943 | 5 | 0 | 4 | 0.25 | 2 | 0 | 5 | 0 | 16 | 0.0625 | 5 | 0 | 6 | 0 | 6 | 0.1667 | 3 | 0 | 20 | 0.05 |
| 1_1799943 | 4 | 0.75 | 3 | 0.3333 | 2 | 0.5 | 5 | 0.6 | 14 | 0.5714 | 5 | 0.8 | 4 | 0.5 | 7 | 0.4286 | 2 | 0.5 | 18 | 0.5556 |
| 1_1799943 | 4 | 0.5 | 3 | 0.3333 | 1 | 0 | 5 | 0 | 13 | 0.2308 | 4 | 0.25 | 5 | 0 | 6 | 0.1667 | 1 | 0 | 16 | 0.125 |
| 1_1799945 | 5 | 0 | 3 | 0.3333 | 2 | 0 | 4 | 0 | 14 | 0.0714 | 4 | 0.25 | 2 | 0 | 4 | 0 | 3 | 0 | 13 | 0.0769 |
| 1_1799945 | 3 | 0 | 3 | 0 | 2 | 0.5 | 4 | 0 | 12 | 0.0833 | 4 | 0 | 4 | 0 | 6 | 0.1667 | 2 | 0 | 16 | 0.0625 |
| 1_1800065 | 1 | 1 | 2 | 1 | NA | NA | 2 | 0.5 | 5 | 0.8 | 1 | 0 | 2 | 1 | 2 | 1 | 1 | 1 | 6 | 0.8333 |
| 1_1800066 | 1 | 0 | 2 | 0 | NA | NA | 2 | 0.5 | 5 | 0.2 | 1 | 0 | 4 | 0.25 | 2 | 0 | 1 | 0 | 8 | 0.125 |
| 1_1800068 | 1 | 0 | 1 | 0 | NA | NA | 2 | 0.5 | 4 | 0.25 | 2 | 0 | 5 | 0 | 2 | 0.5 | 1 | 0 | 10 | 0.1 |
| 1_1800546 | 3 | 0.6667 | 3 | 0 | 2 | 0.5 | 1 | 1 | 9 | 0.4444 | 3 | 0 | 2 | 0.5 | 2 | 0 | 4 | 0.5 | 11 | 0.2727 |
| 1_1800549 | 3 | 0.3333 | 2 | 0.5 | 3 | 0 | 1 | 0 | 9 | 0.2222 | 3 | 0.3333 | 2 | 0.5 | 2 | 0.5 | 4 | 0.25 | 11 | 0.3636 |
| 1_1800551 | 2 | 1 | 2 | 1 | 3 | 1 | NA | NA | 7 | 1 | 3 | 1 | 2 | 1 | 2 | 1 | NA | NA | 7 | 1 |
| 1_1800552 | 3 | 0.6667 | 2 | 0 | 3 | 0.6667 | NA | NA | 8 | 0.5 | 2 | 1 | 2 | 0.5 | 1 | 1 | 3 | 0.3333 | 8 | 0.625 |
| 1_1801547 | NA | NA | 1 | 0 | 1 | 1 | 2 | 1 | 4 | 0.75 | 2 | 1 | NA | NA | 1 | 1 | NA | NA | 3 | 1 |
| 1_1801547 | NA | NA | 1 | 1 | 1 | 1 | 2 | 1 | 4 | 1 | 1 | 1 | NA | NA | 1 | 1 | NA | NA | 2 | 1 |
| 1_1801548 | NA | NA | NA | NA | 1 | 1 | 2 | 0.5 | 3 | 0.6667 | 2 | 0 | NA | NA | 1 | 1 | NA | NA | 3 | 0.3333 |

| | | | | | | | | | | | | | | | | | | | | |
|---|---|---|---|---|---|---|---|---|---|---|---|---|---|---|---|---|---|---|---|---|
| 1_18015518 | NA | NA | 1 | 1 | 1 | 0 | 2 | 1 | 4 | 0.75 | 1 | 0 | NA | NA | 1 | 1 | NA | NA | 2 | 0.5 |
| 1_18016144 | NA | NA | 1 | 1 | NA | NA | NA | NA | 1 | 1 | 1 | 1 | NA | NA | NA | NA | NA | NA | 1 | 1 |
| 1_18025018 | NA | NA | 1 | 1 | NA | NA | 1 | 1 | 2 | 1 | 4 | 0 | NA | NA | 1 | 0 | 1 | 1 | 6 | 0.1667 |
| 1_18025072 | NA | NA | 1 | 1 | NA | NA | 1 | 0 | 2 | 0.5 | 4 | 0.5 | NA | NA | 1 | 1 | 1 | 0 | 6 | 0.5 |
| 1_18026040 | 10 | 0.3 | 15 | 0.4 | 8 | 0.375 | 5 | 0.6 | 38 | 0.3947 | 6 | 0.3333 | 9 | 0.4444 | 4 | 0.5 | 9 | 0.3333 | 28 | 0.3929 |
| 1_18026081 | 11 | 0.0909 | 11 | 0.2727 | 10 | 0 | 4 | 0 | 36 | 0.1111 | 8 | 0.25 | 11 | 0.1818 | 3 | 0 | 6 | 0.5 | 28 | 0.25 |
| 1_18026163 | 8 | 0.125 | 7 | 0 | 6 | 0.1667 | 3 | 0 | 24 | 0.0833 | 8 | 0.25 | 4 | 0 | 2 | 0 | 5 | 0.2 | 19 | 0.1579 |
| 1_18026202 | 9 | 0 | 9 | 0.2222 | 11 | 0.2727 | 5 | 0.4 | 34 | 0.2059 | 8 | 0.25 | 3 | 0 | 3 | 0 | 4 | 0.25 | 18 | 0.1667 |
| 1_18026387 | 4 | 0 | 3 | 0 | 8 | 0.125 | 2 | 0 | 17 | 0.0588 | 5 | 0 | 3 | 0.3333 | 2 | 0.5 | 2 | 0 | 12 | 0.1667 |
| 1_18026408 | 4 | 1 | 3 | 0.3333 | 6 | 0.8333 | 1 | 0 | 14 | 0.7143 | 2 | 0.5 | 3 | 1 | 1 | 1 | 3 | 0.3333 | 9 | 0.6667 |
| 1_18026432 | 7 | 0.1429 | 7 | 0 | 7 | 0.4286 | 2 | 0 | 23 | 0.1739 | 6 | 0.1667 | 5 | 0.2 | 5 | 0.2 | 3 | 0 | 19 | 0.1579 |
| 1_18026454 | 7 | 0.1429 | 7 | 0.4286 | 6 | 0.3333 | 2 | 0 | 22 | 0.2727 | 4 | 1 | 6 | 1 | 4 | 0.5 | 4 | 0.5 | 18 | 0.7778 |
| 1_18026461 | 3 | 1 | 4 | 0.75 | 3 | 0.3333 | NA | NA | 10 | 0.7 | 2 | 1 | 6 | 0.6667 | 3 | 0.3333 | 2 | 0.5 | 13 | 0.6154 |
| 1_18026462 | 5 | 0 | 6 | 0.1667 | 7 | 0 | 2 | 0 | 20 | 0.05 | 4 | 0 | 4 | 0.25 | 5 | 0 | 3 | 0.3333 | 16 | 0.125 |
| 1_18026462 | 5 | 0.4 | 7 | 0.5714 | 5 | 0.8 | 2 | 0.5 | 19 | 0.5789 | 5 | 0.6 | 6 | 0.8333 | 3 | 0.6667 | 4 | 0.25 | 18 | 0.6111 |
| 1_18026468 | 7 | 0.1429 | 8 | 0.25 | 8 | 0.375 | 2 | 0 | 25 | 0.24 | 4 | 0.5 | 6 | 0.3333 | 5 | 0.8 | 6 | 0.1667 | 21 | 0.4286 |
| 1_18026478 | 9 | 1 | 9 | 1 | 11 | 0.9091 | 2 | 0.5 | 31 | 0.9355 | 6 | 1 | 10 | 1 | 7 | 1 | 8 | 1 | 31 | 1 |
| 1_18026487 | 9 | 0.8889 | 7 | 0.7143 | 8 | 1 | 2 | 1 | 26 | 0.8846 | 6 | 1 | 10 | 0.9 | 7 | 1 | 8 | 1 | 31 | 0.9677 |
| 1_18026492 | 8 | 0.875 | 7 | 1 | 9 | 0.7778 | 2 | 0.5 | 26 | 0.8462 | 7 | 0.8571 | 10 | 0.9 | 7 | 0.8571 | 8 | 0.875 | 32 | 0.875 |
| 1_18026492 | 7 | 0.4286 | 7 | 0.1429 | 10 | 0.4 | 2 | 0 | 26 | 0.3077 | 7 | 0.4286 | 10 | 0.3 | 7 | 0.1429 | 7 | 0.7143 | 31 | 0.3871 |
| 1_18026498 | 8 | 1 | 7 | 1 | 12 | 1 | 2 | 1 | 29 | 1 | 6 | 1 | 12 | 1 | 7 | 1 | 8 | 1 | 33 | 1 |
| 1_18026551 | 5 | 0.2 | 6 | 0.5 | 7 | 0 | 2 | 0 | 20 | 0.2 | 2 | 0 | 10 | 0.3 | 7 | 0 | 5 | 0.4 | 24 | 0.2083 |
| 1_18026566 | 4 | 0 | 5 | 0 | 9 | 0.1111 | 1 | 0 | 19 | 0.0526 | 1 | 0 | 6 | 0.1667 | 7 | 0 | 5 | 0 | 19 | 0.0526 |
| 1_18026582 | 3 | 0.3333 | 5 | 0.4 | 5 | 0.2 | 1 | 0 | 14 | 0.2857 | 2 | 0 | 9 | 0.4444 | 6 | 0.5 | 4 | 0.25 | 21 | 0.381 |
| 1_18055380 | 2 | 0.5 | NA | NA | NA | NA | NA | NA | 2 | 0.5 | NA | NA | 2 | 1 | 3 | 0.6667 | 1 | 1 | 6 | 0.8333 |
| 1_18057441 | 2 | 0.5 | NA | NA | 2 | 0 | 4 | 0.5 | 8 | 0.375 | 1 | 0 | 5 | 0.4 | 5 | 0.2 | 3 | 0.3333 | 14 | 0.2857 |
| 1_18057453 | 2 | 0.5 | 1 | 0 | NA | NA | 4 | 0.25 | 7 | 0.2857 | 1 | 1 | 5 | 0.4 | 5 | 0.8 | 2 | 0.5 | 13 | 0.6154 |
| 1_18057454 | 1 | 1 | 1 | 1 | 1 | 1 | 4 | 1 | 7 | 1 | 1 | 1 | 6 | 1 | 7 | 0.8571 | 2 | 1 | 16 | 0.9375 |
| 1_18057457 | 2 | 0.5 | 1 | 1 | 2 | 1 | 4 | 0.5 | 9 | 0.6667 | 1 | 1 | 5 | 0.8 | 6 | 1 | 3 | 0.6667 | 15 | 0.8667 |
| 1_18057457 | 2 | 0.5 | 1 | 1 | 2 | 0 | 2 | 0 | 7 | 0.2857 | 1 | 0 | 6 | 0 | 7 | 0.1429 | 3 | 0 | 17 | 0.0588 |
| 1_18057980 | 2 | 0 | 1 | 1 | 2 | 1 | 4 | 0.75 | 9 | 0.6667 | 1 | 0 | 4 | 0.25 | 6 | 0.5 | 3 | 1 | 14 | 0.5 |
| 1_18057999 | 2 | 0 | NA | NA | 1 | 1 | 5 | 0.2 | 8 | 0.25 | 1 | 0 | 5 | 0.4 | 7 | 0.1429 | 3 | 0 | 16 | 0.1875 |
| 1_18058005 | 3 | 0.6667 | 1 | 1 | 2 | 1 | 4 | 1 | 10 | 0.9 | 1 | 1 | 6 | 1 | 7 | 1 | 4 | 1 | 18 | 1 |
| 1_18058008 | 3 | 0.3333 | 1 | 1 | 2 | 1 | 5 | 0.8 | 11 | 0.7273 | 1 | 0 | 5 | 0.8 | 7 | 0.7143 | 4 | 1 | 17 | 0.7647 |
| 1_18058012 | 3 | 1 | 1 | 0 | 1 | 0 | 5 | 0 | 10 | 0.3 | 2 | 0 | 5 | 0.2 | 6 | 0 | 4 | 0.25 | 17 | 0.1176 |
| 1_18058036 | 3 | 0.3333 | 1 | 0 | 1 | 0 | 5 | 0 | 10 | 0.1 | 2 | 0 | 4 | 0.25 | 6 | 0 | 5 | 0 | 17 | 0.0588 |
| 1_18058057 | 2 | 0 | 2 | 0.5 | 2 | 0 | 6 | 0 | 12 | 0.0833 | 2 | 0 | 6 | 0.3333 | 7 | 0 | 5 | 0.4 | 20 | 0.2 |
| 1_18058066 | 2 | 0.5 | 2 | 0.5 | 1 | 0 | 6 | 0 | 11 | 0.1818 | 2 | 0 | 6 | 0.1667 | 5 | 0 | 5 | 0.4 | 18 | 0.1667 |
| 1_18058146 | 2 | 0 | 4 | 0.25 | 4 | 0 | 8 | 0 | 18 | 0.0556 | 3 | 0.3333 | 7 | 0.1429 | 6 | 0 | 4 | 0 | 20 | 0.1 |
| 1_18058210 | NA | NA | 1 | 1 | NA | NA | NA | NA | 1 | 1 | 1 | 0 | 1 | 1 | NA | NA | NA | NA | 2 | 0.5 |
| 1_18061932 | NA | NA | NA | NA | NA | NA | 1 | 1 | 1 | 1 | NA | NA | 2 | 1 | NA | NA | NA | NA | 2 | 1 |
| 1_18061969 | NA | NA | NA | NA | NA | NA | 1 | 1 | 1 | 1 | NA | NA | 2 | 0.5 | NA | NA | NA | NA | 2 | 0.5 |
| 1_18061986 | NA | NA | NA | NA | NA | NA | 1 | 1 | 1 | 1 | NA | NA | 2 | 0.5 | NA | NA | NA | NA | 2 | 0.5 |
| 1_18063033 | 63 | 0.1905 | 55 | 0.0727 | 39 | 0.1795 | 65 | 0.0923 | 222 | 0.1306 | 57 | 0.1579 | 48 | 0.1458 | 55 | 0.1455 | 68 | 0.1618 | 228 | 0.1535 |
| 1_18069018 | 2 | 0.5 | 2 | 1 | 1 | 1 | 1 | 1 | 6 | 0.8333 | NA | NA | 1 | 1 | 1 | 1 | NA | NA | 2 | 1 |
| 1_18069372 | 2 | 0.5 | 2 | 1 | 1 | 1 | NA | NA | 5 | 0.8 | 1 | 1 | 2 | 1 | 3 | 1 | 1 | 1 | 7 | 1 |
| 1_18069373 | 2 | 1 | 1 | 1 | NA | NA | 1 | 0 | 4 | 0.75 | 2 | 1 | 2 | 1 | 3 | 1 | 1 | 1 | 8 | 1 |
| 1_18069375 | 2 | 0.5 | 1 | 0 | 1 | 0 | 1 | 0 | 5 | 0.2 | 2 | 0 | 2 | 0 | 4 | 0.25 | 1 | 0 | 9 | 0.1111 |
| 1_18069380 | 2 | 0.5 | 2 | 1 | 1 | 1 | 1 | 1 | 6 | 0.8333 | 2 | 1 | 2 | 0.5 | 4 | 1 | 1 | 1 | 9 | 0.8889 |
| 1_18069381 | 2 | 0 | 2 | 0.5 | 1 | 0 | 1 | 0 | 6 | 0.1667 | 1 | 1 | 2 | 1 | 3 | 0 | 1 | 0 | 7 | 0.4286 |
| 1_18069389 | 2 | 1 | 2 | 1 | 1 | 1 | 1 | 1 | 6 | 1 | 2 | 1 | 2 | 0.5 | 3 | 0.6667 | 1 | 0 | 8 | 0.625 |
| 1_18069399 | 2 | 0.5 | 2 | 1 | 1 | 0 | 1 | 0 | 6 | 0.5 | 2 | 0 | 2 | 0 | 3 | 0 | 1 | 1 | 8 | 0.125 |
| 1_18069408 | 2 | 1 | 2 | 0.5 | 1 | 1 | 1 | 0 | 6 | 0.6667 | 2 | 0.5 | 2 | 0.5 | 3 | 0 | 1 | 0 | 8 | 0.25 |
| 1_18073437 | 5 | 0.2 | 2 | 0 | 3 | 0 | 2 | 0.5 | 12 | 0.1667 | 2 | 0.5 | 3 | 0 | 1 | 0 | 4 | 0 | 10 | 0.1 |
| 1_18073521 | 4 | 1 | 3 | 0.6667 | 2 | 0.5 | 2 | 0.5 | 11 | 0.7273 | 3 | 0.6667 | 3 | 0.6667 | 2 | 1 | 7 | 0.5714 | 15 | 0.6667 |
| 1_18080183 | NA | NA | 2 | 0 | 6 | 0.5 | NA | NA | 8 | 0.375 | 3 | 0.6667 | 4 | 0.5 | 2 | 0.5 | NA | NA | 9 | 0.5556 |
| 1_18080229 | 2 | 0 | 4 | 0 | 4 | 0.5 | NA | NA | 10 | 0.2 | 4 | 0 | 3 | 0.3333 | 2 | 0.5 | 1 | 0 | 10 | 0.2 |
| 1_18091494 | 1 | 1 | 2 | 1 | 1 | 1 | 3 | 1 | 7 | 1 | 1 | 1 | NA | NA | NA | NA | 2 | 0.5 | 3 | 0.6667 |
| 1_18091726 | NA | NA | 1 | 1 | 2 | 0 | 1 | 0 | 4 | 0.25 | 1 | 1 | NA | NA | 1 | 0 | NA | NA | 2 | 0.5 |
| 1_18091800 | 2 | 0 | 1 | 1 | 4 | 0.25 | 1 | 0 | 8 | 0.25 | 2 | 0.5 | NA | NA | 1 | 0 | 1 | 0 | 4 | 0.25 |
| 1_18091856 | 2 | 1 | 1 | 0 | 2 | 1 | 1 | 1 | 6 | 0.8333 | 1 | 1 | NA | NA | 1 | 1 | NA | NA | 2 | 1 |
| 1_18091856 | 2 | 1 | 1 | 1 | 1 | 0 | 1 | 1 | 5 | 0.8 | 2 | 0.5 | NA | NA | 1 | 0 | NA | NA | 3 | 0.3333 |
| 1_18091872 | 3 | 0.3333 | 1 | 0 | 2 | 0 | 1 | 0 | 7 | 0.1429 | 2 | 0.5 | NA | NA | NA | NA | NA | NA | 2 | 0.5 |
| 1_18091879 | 2 | 1 | 1 | 1 | 1 | 0 | 1 | 0 | 5 | 0.6 | 2 | 1 | NA | NA | NA | NA | NA | NA | 2 | 1 |
| 1_18091883 | 2 | 1 | 1 | 1 | 1 | 1 | 1 | 1 | 5 | 1 | 2 | 0.5 | NA | NA | 1 | 1 | NA | NA | 3 | 0.6667 |

| | | | | | | | | | | | | | | | | | | | | |
|---|---|---|---|---|---|---|---|---|---|---|---|---|---|---|---|---|---|---|---|---|
| 1_1809189( | 2 | 0 | 1 | 1 | 2 | 0 | 1 | 0 | 6 | 0.1667 | 2 | 0.5 | NA | NA | 1 | 0 | NA | NA | 3 | 0.3333 |
| 1_1809191! | NA | NA | 1 | 1 | 1 | 0 | 1 | 0 | 3 | 0.3333 | 2 | 1 | NA | NA | 1 | 1 | NA | NA | 3 | 1 |
| 1_1809192: | 2 | 0 | 1 | 1 | 2 | 0.5 | 1 | 0 | 6 | 0.3333 | 1 | 1 | NA | NA | 1 | 0 | NA | NA | 2 | 0.5 |
| 1_1809193( | 1 | 1 | NA | NA | 2 | 0 | NA | NA | 3 | 0.3333 | 2 | 0.5 | NA | NA | 1 | 1 | NA | NA | 3 | 0.6667 |
| 1_1809194 | 1 | 1 | NA | NA | 1 | 0 | NA | NA | 2 | 0.5 | 2 | 0 | NA | NA | 1 | 1 | NA | NA | 3 | 0.3333 |
| 1_1809197: | 1 | 1 | NA | NA | 1 | 1 | NA | NA | 2 | 1 | 2 | 0.5 | NA | NA | NA | NA | NA | NA | 2 | 0.5 |
| 1_1809417! | 3 | 0 | 1 | 1 | 2 | 0 | 1 | 0 | 7 | 0.1429 | 1 | 0 | NA | NA | 1 | 1 | NA | NA | 2 | 0.5 |
| 1_1809957! | 4 | 0.25 | 4 | 0.25 | 4 | 0 | 2 | 0 | 14 | 0.1429 | 6 | 0 | NA | NA | 6 | 0.3333 | 2 | 0 | 14 | 0.1429 |
| 1_1810011( | 4 | 0.25 | 9 | 0.5556 | 6 | 0.5 | 6 | 0.3333 | 25 | 0.44 | 5 | 0.4 | 8 | 0.25 | 12 | 0.1667 | 4 | 0.25 | 29 | 0.2414 |
| 1_1810021 | 2 | 0 | 3 | 0.3333 | 5 | 0 | 5 | 0 | 15 | 0.0667 | 4 | 0 | 7 | 0 | 4 | 0.25 | 2 | 0 | 17 | 0.0588 |
| 1_1810028( | 5 | 0 | 9 | 0 | 8 | 0.25 | 5 | 0 | 27 | 0.0741 | 5 | 0 | 7 | 0.1429 | 10 | 0.1 | 2 | 0 | 24 | 0.0833 |
| 1_1810048( | 2 | 0 | 9 | 0 | 10 | 0.4 | 5 | 0 | 26 | 0.1538 | 6 | 0 | 7 | 0 | 7 | 0 | 6 | 0.3333 | 26 | 0.0769 |
| 1_1810049: | 1 | 0 | 9 | 0.2222 | 10 | 0.3 | 5 | 0 | 25 | 0.2 | 3 | 0.3333 | 7 | 0 | 7 | 0.4286 | 4 | 0 | 21 | 0.1905 |
| 1_1810049! | 3 | 0 | 10 | 0 | 12 | 0 | 7 | 0.2857 | 32 | 0.0625 | 6 | 0.3333 | 6 | 0 | 9 | 0 | 5 | 0 | 26 | 0.0769 |
| 1_1810050! | 2 | 1 | 6 | 1 | 7 | 1 | 6 | 0.8333 | 21 | 0.9524 | 5 | 0.8 | 4 | 1 | 7 | 1 | 5 | 1 | 21 | 0.9524 |
| 1_1810051 | 2 | 1 | 8 | 1 | 10 | 1 | 7 | 1 | 27 | 1 | 5 | 0.8 | 6 | 0.6667 | 8 | 0.875 | 6 | 1 | 25 | 0.84 |
| 1_1810051! | 2 | 1 | 8 | 0.75 | 10 | 0.6 | 7 | 0.4286 | 27 | 0.6296 | 6 | 0.6667 | 7 | 0.7143 | 9 | 0.6667 | 5 | 0.6 | 27 | 0.6667 |
| 1_1810052 | 1 | 0 | 5 | 0.4 | 9 | 0.3333 | 6 | 0.1667 | 21 | 0.2857 | 4 | 0.5 | 5 | 0 | 9 | 0.3333 | 5 | 0.2 | 23 | 0.2609 |
| 1_1810052 | 2 | 1 | 5 | 0.8 | 4 | 1 | 6 | 0.6667 | 17 | 0.8235 | 5 | 1 | 5 | 0.6 | 9 | 0.7778 | 5 | 0.8 | 24 | 0.7917 |
| 1_1810052 | 2 | 0 | 8 | 0.375 | 9 | 0.1111 | 7 | 0 | 26 | 0.1538 | 6 | 0.1667 | 7 | 0.2857 | 8 | 0 | 5 | 0.4 | 26 | 0.1923 |
| 1_1810052 | 2 | 1 | 7 | 0.7143 | 9 | 1 | 6 | 0.8333 | 24 | 0.875 | 6 | 0.8333 | 7 | 0.8571 | 7 | 1 | 3 | 1 | 23 | 0.913 |
| 1_1810052! | 2 | 1 | 7 | 0 | 9 | 0.1111 | 5 | 0 | 23 | 0.1304 | 6 | 0.1667 | 6 | 0.3333 | 6 | 0 | 6 | 0.1667 | 24 | 0.1667 |
| 1_1810390( | 1 | 0 | 2 | 0 | 4 | 0.5 | 5 | 0 | 12 | 0.1667 | 2 | 0.5 | 4 | 0 | 4 | 0.25 | 1 | 0 | 11 | 0.1818 |
| 1_1810395! | NA | NA | NA | NA | 1 | 1 | NA | NA | 1 | 1 | 1 | 1 | NA | NA | NA | NA | 1 | 0 | 2 | 0.5 |
| 1_1811804! | 3 | 0 | 6 | 0.3333 | 2 | 0.5 | 1 | 0 | 12 | 0.25 | 1 | 0 | 1 | 0 | 2 | 0.5 | 1 | 0 | 5 | 0.2 |
| 1_1811806! | 3 | 0.3333 | 6 | 0.6667 | 3 | 0.6667 | 1 | 0 | 13 | 0.5385 | 3 | 0.3333 | 1 | 1 | 1 | 1 | 4 | 0.25 | 9 | 0.4444 |
| 1_1811807! | 2 | 1 | 7 | 1 | 3 | 1 | 1 | 0 | 13 | 0.9231 | 3 | 1 | 1 | 1 | 3 | 0.3333 | 5 | 1 | 12 | 0.8333 |
| 1_1811809: | 2 | 0.5 | 5 | 0 | 4 | 0.5 | 1 | 0 | 12 | 0.25 | 2 | 0.5 | 1 | 1 | 1 | 1 | 5 | 0.4 | 9 | 0.5556 |
| 1_1812869 | 5 | 0 | 11 | 0.1818 | 6 | 0 | 2 | 0.5 | 24 | 0.125 | 7 | 0 | 1 | 1 | 3 | 0.3333 | 6 | 0.1667 | 17 | 0.1765 |
| 1_1812870! | 5 | 0.8 | 10 | 0.5 | 5 | 0.2 | 2 | 0.5 | 22 | 0.5 | 8 | 0.25 | 2 | 0.5 | 4 | 0 | 5 | 0.2 | 19 | 0.2105 |
| 1_1812871 | 5 | 1 | 7 | 1 | 8 | 1 | 1 | 1 | 21 | 1 | 6 | 0.8333 | 1 | 1 | 3 | 1 | 4 | 1 | 14 | 0.9286 |
| 1_1812871! | 4 | 0 | 10 | 0 | 8 | 0.375 | 1 | 0 | 23 | 0.1304 | 6 | 0.1667 | 2 | 0.5 | 2 | 0.5 | 5 | 0.4 | 15 | 0.3333 |
| 1_1812871! | 5 | 1 | 10 | 1 | 7 | 1 | 2 | 1 | 24 | 1 | 7 | 1 | 2 | 1 | 3 | 1 | 6 | 1 | 18 | 1 |
| 1_1812872: | 5 | 0.2 | 12 | 0 | 7 | 0.1429 | 2 | 0 | 26 | 0.0769 | 9 | 0 | 2 | 0.5 | 4 | 0 | 5 | 0.2 | 20 | 0.1 |
| 1_1812872 | 5 | 0.8 | 11 | 0.9091 | 8 | 0.875 | 2 | 1 | 26 | 0.8846 | 9 | 0.8889 | 3 | 1 | 5 | 0.8 | 6 | 1 | 23 | 0.913 |
| 1_1812872( | 5 | 1 | 10 | 0.8 | 8 | 0.875 | 1 | 1 | 24 | 0.875 | 9 | 1 | 3 | 1 | 4 | 1 | 4 | 1 | 20 | 1 |
| 1_1812873: | 5 | 1 | 11 | 0.9091 | 9 | 0.8889 | 2 | 1 | 27 | 0.9259 | 9 | 1 | 3 | 1 | 3 | 1 | 8 | 1 | 23 | 1 |
| 1_1812874: | 4 | 1 | 10 | 1 | 7 | 0.8571 | 1 | 1 | 22 | 0.9545 | 5 | 1 | 2 | 0.5 | 2 | 1 | 6 | 0.8333 | 15 | 0.8667 |
| 1_1812876 | 2 | 1 | 7 | 0.7143 | 5 | 1 | 1 | 1 | 15 | 0.8667 | 6 | 0.6667 | 3 | 1 | 2 | 1 | 6 | 1 | 17 | 0.8824 |
| 1_1812877! | 2 | 0.5 | 12 | 0.0833 | 5 | 0.2 | 2 | 0.5 | 21 | 0.1905 | 5 | 0.4 | 3 | 0 | 4 | 0.25 | 7 | 0 | 19 | 0.1579 |
| 1_1812877! | 1 | 0 | 12 | 0.25 | 6 | 0.5 | NA | NA | 19 | 0.3158 | 7 | 0.2857 | 4 | 0 | 4 | 0.25 | 8 | 0.25 | 23 | 0.2174 |
| 1_1812878 | 3 | 0 | 16 | 0.125 | 6 | 0.1667 | 2 | 0 | 27 | 0.1111 | 9 | 0.2222 | 4 | 0 | 4 | 0 | 8 | 0.125 | 25 | 0.12 |
| 1_1816443( | NA | NA | NA | NA | 4 | 0 | 3 | 0.3333 | 7 | 0.1429 | NA | NA | 2 | 0 | NA | NA | 2 | 1 | 4 | 0.5 |
| 1_1817241: | 9 | 0.1111 | 5 | 0.4 | 6 | 0 | 7 | 0.1429 | 27 | 0.1481 | 3 | 0 | 11 | 0.0909 | 6 | 0 | 5 | 0.2 | 25 | 0.08 |
| 1_1817241: | 8 | 0.75 | 4 | 0.5 | 6 | 1 | 6 | 0.8333 | 24 | 0.7917 | 3 | 1 | 11 | 0.9091 | 4 | 0.25 | 5 | 1 | 23 | 0.8261 |
| 1_1817242( | 9 | 0.5556 | 6 | 0 | 6 | 0.5 | 7 | 0.2857 | 28 | 0.3571 | 3 | 0.6667 | 11 | 0.1818 | 6 | 0.3333 | 6 | 0 | 26 | 0.2308 |
| 1_1817243! | 7 | 0.1429 | 6 | 0.1667 | 8 | 0.25 | 6 | 0 | 27 | 0.1481 | 2 | 0 | 11 | 0.1818 | 5 | 0.2 | 5 | 0.2 | 23 | 0.1739 |
| 1_1817245: | 8 | 0.375 | 4 | 0 | 5 | 0.4 | 6 | 0.6667 | 23 | 0.3913 | 3 | 0.3333 | 9 | 0.2222 | 6 | 0.5 | 5 | 0.2 | 23 | 0.3043 |
| 1_1817247 | 7 | 0.1429 | 5 | 0 | 5 | 0.6 | 6 | 0.1667 | 23 | 0.2174 | 3 | 0 | 7 | 0.5714 | 6 | 0 | 4 | 0.5 | 20 | 0.3 |
| 1_1817247 | 6 | 0 | 3 | 0.3333 | 5 | 0.2 | 6 | 0 | 20 | 0.1 | 3 | 0.3333 | 8 | 0.25 | 6 | 0 | 4 | 0.25 | 21 | 0.1905 |
| 1_1817247! | 7 | 0 | 5 | 0.6 | 4 | 0 | 3 | 0 | 19 | 0.1579 | 3 | 0 | 3 | 0.3333 | 2 | 0 | NA | NA | 8 | 0.125 |
| 1_1817257( | 4 | 0 | 3 | 0 | 3 | 0 | 4 | 0.25 | 14 | 0.0714 | 3 | 0 | 1 | 0 | 4 | 0.25 | 3 | 0 | 11 | 0.0909 |
| 1_1817306 | 7 | 0 | 2 | 0 | 2 | 0 | 8 | 0.125 | 19 | 0.0526 | 3 | 0 | 5 | 0.2 | 6 | 0.1667 | 4 | 0 | 18 | 0.1111 |
| 1_1817306! | 8 | 0.625 | 2 | 0.5 | 1 | 1 | 8 | 0.75 | 19 | 0.6842 | 1 | 1 | 6 | 0.8333 | 7 | 0.5714 | 4 | 0.75 | 18 | 0.7222 |
| 1_1817307! | 8 | 0.125 | 2 | 0 | 3 | 0.6667 | 9 | 0.1111 | 22 | 0.1818 | 3 | 0 | 4 | 0.25 | 8 | 0.375 | 4 | 0 | 19 | 0.2105 |
| 1_1817309( | 5 | 0.6 | 2 | 1 | 3 | 1 | 7 | 0.7143 | 17 | 0.7647 | 3 | 1 | 4 | 0.5 | 7 | 0.8571 | 4 | 1 | 18 | 0.8333 |
| 1_1817309: | 6 | 0 | 3 | 0.3333 | 3 | 0 | 10 | 0.2 | 22 | 0.1364 | 1 | 0 | 2 | 0 | 7 | 0.1429 | 5 | 0 | 15 | 0.0667 |
| 1_1817309! | 4 | 0.5 | 1 | 1 | 1 | 1 | 5 | 0.8 | 11 | 0.7273 | 2 | 1 | 4 | 1 | 4 | 0.5 | 5 | 0.8 | 15 | 0.8 |
| 1_1817309! | 5 | 0.4 | 2 | 0.5 | 2 | 1 | 9 | 0.2222 | 18 | 0.3889 | 3 | 0.6667 | 3 | 1 | 6 | 0.5 | 3 | 0.3333 | 15 | 0.6 |
| 1_1817310: | 3 | 0.6667 | 2 | 0.5 | 3 | 0.6667 | 7 | 0.4286 | 15 | 0.5333 | 3 | 1 | 3 | 0.3333 | 7 | 0.5714 | 4 | 0.75 | 17 | 0.6471 |
| 1_1817310( | 7 | 0.1429 | 2 | 0 | 3 | 0.3333 | 9 | 0 | 21 | 0.0952 | 3 | 0 | 4 | 0 | 8 | 0.125 | 5 | 0 | 20 | 0.05 |
| 1_1817311 | 5 | 0.2 | 2 | 0 | 2 | 0.5 | 8 | 0 | 17 | 0.1176 | 4 | 0 | 4 | 0 | 8 | 0.375 | 7 | 0 | 23 | 0.1304 |
| 1_1817322( | 9 | 0 | 1 | 0 | 7 | 0.2857 | 10 | 0.1 | 27 | 0.1111 | 5 | 0.2 | 6 | 0 | 9 | 0.2222 | 8 | 0 | 28 | 0.1071 |
| 1_1817326 | 4 | 0 | 3 | 0.3333 | 3 | 0.3333 | 8 | 0.125 | 18 | 0.1667 | 4 | 0.5 | 8 | 0.25 | 8 | 0 | 2 | 0 | 22 | 0.1818 |
| 1_1817329( | 4 | 0.25 | 2 | 0.5 | 5 | 0.2 | 7 | 0.2857 | 18 | 0.2778 | 5 | 0 | 5 | 0.2 | 6 | 0.3333 | 4 | 0 | 20 | 0.15 |

| | | | | | | | | | | | | | | | | | | | | |
|---|---|---|---|---|---|---|---|---|---|---|---|---|---|---|---|---|---|---|---|---|
| 1_18173322 | 4 | 0 | 4 | 0.25 | 7 | 0 | 10 | 0.4 | 25 | 0.2 | 3 | 0.3333 | 6 | 0 | 5 | 0 | 6 | 0.5 | 20 | 0.2 |
| 1_18173327 | 7 | 1 | 2 | 1 | 6 | 1 | 8 | 1 | 23 | 1 | 4 | 1 | 6 | 1 | 6 | 0.6667 | 5 | 1 | 21 | 0.9048 |
| 1_18173328 | 6 | 0.1667 | 2 | 0 | 6 | 0.3333 | 10 | 0 | 24 | 0.125 | 6 | 0 | 8 | 0.125 | 5 | 0.2 | 4 | 0 | 23 | 0.087 |
| 1_18173334 | 6 | 0.3333 | 2 | 0.5 | 5 | 0.2 | 9 | 0.3333 | 22 | 0.3182 | 4 | 0 | 7 | 0.2857 | 4 | 0 | 4 | 0.25 | 19 | 0.1579 |
| 1_18173340 | 7 | 1 | 3 | 0.6667 | 7 | 0.8571 | 9 | 0.6667 | 26 | 0.8077 | 6 | 0.8333 | 8 | 0.75 | 5 | 1 | 4 | 0.75 | 23 | 0.8261 |
| 1_18174928 | 5 | 0.2 | 2 | 0 | 5 | 0 | 8 | 0 | 20 | 0.05 | 3 | 0 | 7 | 0 | 3 | 0 | 5 | 0.2 | 18 | 0.0556 |
| 1_18174956 | 4 | 0.25 | NA | NA | 3 | 0.6667 | 6 | 0.5 | 13 | 0.4615 | NA | NA | 4 | 0.5 | NA | NA | 1 | 0 | 5 | 0.4 |
| 1_18174978 | 3 | 0.3333 | NA | NA | 1 | 1 | 4 | 0.5 | 8 | 0.5 | NA | NA | NA | NA | NA | NA | 1 | 1 | 1 | 1 |
| 1_18174979 | NA | NA | NA | NA | NA | NA | 1 | 1 | 1 | 1 | NA | NA | 1 | 1 | NA | NA | NA | NA | 1 | 1 |
| 1_18174992 | 3 | 1 | 1 | 1 | 3 | 1 | 5 | 0.8 | 12 | 0.9167 | 1 | 1 | 3 | 1 | NA | NA | 1 | 1 | 5 | 1 |
| 1_18185700 | NA | NA | NA | NA | NA | NA | 1 | 1 | 1 | 1 | NA | NA | NA | NA | NA | NA | 1 | 1 | 1 | 1 |
| 1_18195907 | 17 | 0.4706 | 21 | 0.3333 | 21 | 0.4286 | 26 | 0.3462 | 85 | 0.3882 | 15 | 0.4667 | 22 | 0.4545 | 20 | 0.55 | 16 | 0.4375 | 73 | 0.4795 |
| 1_18195912 | 19 | 0.2105 | 26 | 0.1538 | 21 | 0.1905 | 29 | 0 | 95 | 0.1263 | 14 | 0.2143 | 24 | 0.0833 | 19 | 0.1053 | 16 | 0.125 | 73 | 0.1233 |
| 1_18195922 | 20 | 0.25 | 26 | 0.3077 | 20 | 0.1 | 28 | 0.25 | 94 | 0.234 | 16 | 0.3125 | 25 | 0.24 | 18 | 0.1667 | 15 | 0.2 | 74 | 0.2297 |
| 1_18195950 | 11 | 0 | 14 | 0.2857 | 8 | 0.125 | 17 | 0.1176 | 50 | 0.14 | 8 | 0 | 9 | 0.3333 | 7 | 0.1429 | 7 | 0.1429 | 31 | 0.1613 |
| 1_18195959 | 9 | 0.1111 | 14 | 0.2143 | 5 | 0 | 15 | 0 | 43 | 0.093 | 9 | 0 | 5 | 0.2 | 7 | 0 | 6 | 0.1667 | 27 | 0.0741 |
| 1_18195968 | 10 | 0.1 | 15 | 0.2667 | 7 | 0.4286 | 15 | 0.2667 | 47 | 0.2553 | 8 | 0.125 | 7 | 0.2857 | 6 | 0.5 | 6 | 0.6667 | 27 | 0.3704 |
| 1_18195971 | 9 | 0 | 13 | 0.4615 | 7 | 0.4286 | 16 | 0.3125 | 45 | 0.3111 | 10 | 0.8 | 10 | 0.3 | 7 | 0.4286 | 5 | 0.2 | 32 | 0.4688 |
| 1_18210397 | 13 | 0.3846 | 21 | 0.5238 | 14 | 0.3571 | 11 | 0.4545 | 59 | 0.4407 | 12 | 0.5833 | 12 | 0.25 | 13 | 0.4615 | 12 | 0.5833 | 49 | 0.4694 |
| 1_18210397 | 11 | 0 | 21 | 0.1429 | 13 | 0.1538 | 13 | 0.0769 | 58 | 0.1034 | 10 | 0 | 13 | 0.2308 | 15 | 0.0667 | 12 | 0.0833 | 50 | 0.1 |
| 1_18210438 | 10 | 0.2 | 19 | 0.3158 | 15 | 0.2667 | 14 | 0.1429 | 58 | 0.2414 | 11 | 0.1818 | 14 | 0.2857 | 16 | 0.375 | 15 | 0.2667 | 56 | 0.2857 |
| 1_18210661 | 9 | 0.3333 | 21 | 0.1905 | 13 | 0.4615 | 12 | 0.25 | 55 | 0.2909 | 11 | 0.1818 | 12 | 0.3333 | 15 | 0.1333 | 14 | 0.0714 | 52 | 0.1731 |
| 1_18210664 | 9 | 0.8889 | 19 | 0.7368 | 13 | 1 | 15 | 0.8 | 56 | 0.8393 | 11 | 0.7273 | 14 | 0.8571 | 15 | 0.8667 | 14 | 0.9286 | 54 | 0.8519 |
| 1_18210667 | 8 | 0.5 | 10 | 0.5 | 9 | 0.6667 | 9 | 0.7778 | 36 | 0.6111 | 8 | 0.5 | 9 | 0.5556 | 9 | 0.7778 | 13 | 0.6923 | 39 | 0.641 |
| 1_18210677 | 8 | 0.75 | 18 | 0.6667 | 10 | 0.7 | 13 | 0.4615 | 49 | 0.6327 | 10 | 0.5 | 9 | 0.7778 | 14 | 0.9286 | 12 | 0.5833 | 45 | 0.7111 |
| 1_18210705 | 8 | 0 | 14 | 0.3571 | 11 | 0.1818 | 12 | 0.0833 | 45 | 0.1778 | 11 | 0.2727 | 10 | 0.3 | 12 | 0.0833 | 12 | 0.25 | 45 | 0.2222 |
| 1_18210711 | 5 | 1 | 13 | 0.7692 | 9 | 1 | 12 | 0.8333 | 39 | 0.8718 | 11 | 0.8182 | 9 | 1 | 9 | 1 | 13 | 0.7692 | 42 | 0.881 |
| 1_18210717 | 4 | 0.75 | 13 | 0.9231 | 8 | 1 | 9 | 1 | 34 | 0.9412 | 7 | 0.8571 | 4 | 1 | 8 | 1 | 9 | 1 | 28 | 0.9643 |
| 1_18214162 | 1 | 1 | 2 | 1 | 3 | 0 | 1 | 0 | 7 | 0.4286 | 2 | 0.5 | NA | NA | 6 | 0.6667 | 3 | 0.6667 | 11 | 0.6364 |
| 1_18214203 | NA | NA | 4 | 0.25 | 2 | 0 | 2 | 0 | 8 | 0.125 | 1 | 0 | 1 | 0 | 7 | 0 | 5 | 0.4 | 14 | 0.1429 |
| 1_18214311 | 1 | 0 | 5 | 0.2 | 4 | 0.75 | 2 | 0 | 12 | 0.3333 | 1 | 0 | 2 | 0.5 | 7 | 0.1429 | 5 | 0.4 | 15 | 0.2667 |
| 1_18214350 | 1 | 0 | 3 | 0 | 6 | 0 | 3 | 0.3333 | 13 | 0.0769 | NA | NA | 2 | 0 | 7 | 0 | 7 | 0.1429 | 16 | 0.0625 |
| 1_18214361 | 1 | 0 | 3 | 0.3333 | 6 | 0.5 | 3 | 0 | 13 | 0.3077 | NA | NA | 2 | 0 | 6 | 0.6667 | 7 | 0.7143 | 15 | 0.6 |
| 1_18214392 | NA | NA | 2 | 0.5 | 6 | 0.8333 | 3 | 0.6667 | 11 | 0.7273 | NA | NA | 1 | 0 | 6 | 0.8333 | 6 | 1 | 13 | 0.8462 |
| 1_18214405 | 1 | 1 | 1 | 1 | 5 | 1 | 3 | 1 | 10 | 1 | NA | NA | 2 | 1 | 4 | 1 | 6 | 1 | 12 | 1 |
| 1_18214412 | 1 | 1 | 3 | 1 | 7 | 0.5714 | 3 | 0.3333 | 14 | 0.6429 | NA | NA | 5 | 0.2 | 6 | 0.5 | 7 | 0.7143 | 18 | 0.5 |
| 1_18214427 | NA | NA | 3 | 0.6667 | 6 | 0.1667 | 2 | 0 | 11 | 0.2727 | NA | NA | 5 | 0.4 | 6 | 0.6667 | 7 | 0 | 18 | 0.3333 |
| 1_18214434 | NA | NA | 2 | 1 | 6 | 0.5 | 3 | 0.6667 | 11 | 0.6364 | NA | NA | 4 | 0 | 6 | 0.1667 | 6 | 0.6667 | 16 | 0.3125 |
| 1_18214449 | NA | NA | 3 | 1 | 5 | 1 | 3 | 1 | 11 | 1 | NA | NA | 5 | 1 | 4 | 1 | 6 | 1 | 15 | 1 |
| 1_18214456 | NA | NA | 3 | 0.6667 | 4 | 1 | 3 | 1 | 10 | 0.9 | NA | NA | 4 | 0.25 | 5 | 0.8 | 6 | 0.6667 | 15 | 0.6 |
| 1_18214462 | NA | NA | 3 | 0.3333 | 5 | 0.2 | 2 | 0 | 10 | 0.2 | NA | NA | 3 | 0 | 3 | 0 | 5 | 0.2 | 11 | 0.0909 |
| 1_18214465 | NA | NA | 3 | 1 | 4 | 0.25 | 2 | 0.5 | 9 | 0.5556 | NA | NA | 5 | 0.4 | 5 | 0.6 | 6 | 0.8333 | 16 | 0.625 |
| 1_18214473 | NA | NA | 2 | 0.5 | 5 | 0.6 | 1 | 0 | 8 | 0.5 | NA | NA | 4 | 0.25 | 5 | 0.4 | 6 | 0.3333 | 15 | 0.3333 |
| 1_18227391 | 5 | 0.4 | 2 | 0.5 | 1 | 0 | 2 | 1 | 10 | 0.5 | NA | NA | 1 | 0 | 2 | 0.5 | 1 | 0 | 4 | 0.25 |
| 1_18227628 | 8 | 1 | 2 | 0.5 | 2 | 0.5 | 2 | 1 | 14 | 0.8571 | 1 | 1 | 1 | 1 | 2 | 0 | 2 | 1 | 6 | 0.6667 |
| 1_18227630 | 8 | 0.375 | 2 | 0 | 1 | 0 | 2 | 0 | 13 | 0.2308 | 1 | 0 | 1 | 0 | 3 | 0.3333 | 2 | 0 | 7 | 0.1429 |
| 1_18227637 | 4 | 1 | 2 | 1 | 1 | 1 | 2 | 1 | 9 | 1 | 1 | 1 | 1 | 1 | 1 | 1 | 2 | 1 | 5 | 1 |
| 1_18227645 | 5 | 0 | 2 | 0 | 3 | 0 | 2 | 0.5 | 12 | 0.0833 | 1 | 0 | 1 | 0 | 3 | 0 | 1 | 1 | 6 | 0.1667 |
| 1_18227649 | 5 | 0.6 | 2 | 0.5 | 1 | 0 | 1 | 1 | 9 | 0.5556 | 1 | 1 | NA | NA | 2 | 0 | NA | NA | 3 | 0.3333 |
| 1_18227655 | 8 | 0 | 2 | 0.5 | 2 | 0 | 1 | 0 | 13 | 0.0769 | 1 | 0 | 1 | 0 | 3 | 0 | 2 | 0.5 | 7 | 0.1429 |
| 1_18228937 | 8 | 0.75 | 4 | 0.5 | 3 | 0.3333 | 2 | 0.5 | 17 | 0.5882 | 2 | 0.5 | 4 | 0.25 | 4 | 0.25 | 3 | 0.3333 | 13 | 0.3077 |
| 1_18228943 | 8 | 0.375 | 4 | 0.75 | 3 | 0 | 1 | 1 | 16 | 0.4375 | 2 | 0 | 4 | 0.25 | 4 | 0 | 3 | 0.3333 | 13 | 0.1538 |
| 1_18228954 | 9 | 0.4444 | 4 | 0.25 | 3 | 0.6667 | 2 | 1 | 18 | 0.5 | 1 | 1 | 3 | 0.6667 | 4 | 0 | 2 | 0.5 | 10 | 0.4 |
| 1_18228955 | 9 | 1 | 3 | 1 | 3 | 0.6667 | 2 | 1 | 17 | 0.9412 | 2 | 1 | 4 | 1 | 4 | 1 | 3 | 1 | 13 | 1 |
| 1_18228963 | 8 | 0.625 | 4 | 1 | 3 | 1 | 2 | 1 | 17 | 0.8235 | 1 | 1 | 1 | 1 | 4 | 0.5 | 3 | 0.6667 | 9 | 0.6667 |
| 1_18228973 | 8 | 0.5 | 5 | 0.4 | 2 | 0 | 1 | 0 | 16 | 0.375 | 2 | 1 | 3 | 0.3333 | 5 | 0.4 | 3 | 0.6667 | 13 | 0.5385 |
| 1_18228975 | 9 | 0.1111 | 4 | 0 | 3 | 0 | 2 | 0 | 18 | 0.0556 | 2 | 0.5 | 4 | 0 | 5 | 0 | 4 | 0 | 15 | 0.0667 |
| 1_18229497 | 13 | 0.4615 | 7 | 0.2857 | 7 | 0.1429 | 9 | 0.2222 | 36 | 0.3056 | 8 | 0.125 | 8 | 0.125 | 11 | 0.4545 | 6 | 0.5 | 33 | 0.303 |
| 1_18229532 | 12 | 0.25 | 9 | 0.1111 | 7 | 0.2857 | 10 | 0 | 38 | 0.1579 | 6 | 0.1667 | 7 | 0.4286 | 12 | 0.25 | 6 | 0 | 31 | 0.2258 |
| 1_18229534 | 11 | 0.3636 | 9 | 0.1111 | 5 | 0.6 | 9 | 0.3333 | 34 | 0.3235 | 5 | 0.4 | 8 | 0.125 | 11 | 0.1818 | 6 | 0 | 30 | 0.1667 |
| 1_18229563 | 11 | 0.7273 | 7 | 0.4286 | 5 | 0.6 | 8 | 0.875 | 31 | 0.6774 | 6 | 0.5 | 5 | 0.8 | 10 | 0.5 | 5 | 0.4 | 26 | 0.5385 |
| 1_18229569 | 12 | 0.1667 | 7 | 0 | 6 | 0 | 7 | 0.1429 | 32 | 0.0938 | 5 | 0.4 | 4 | 0 | 8 | 0 | 5 | 0 | 22 | 0.0909 |
| 1_18229570 | 10 | 0.1 | 6 | 0 | 6 | 0 | 7 | 0.1429 | 29 | 0.069 | 4 | 0.25 | 7 | 0.1429 | 10 | 0.2 | 5 | 0.2 | 26 | 0.1923 |
| 1_18229570 | 9 | 0.1111 | 7 | 0.4286 | 6 | 0.3333 | 9 | 0.3333 | 31 | 0.2903 | 6 | 0 | 4 | 0 | 10 | 0.4 | 5 | 0.2 | 25 | 0.2 |
| 1_18231750 | NA | NA | 2 | 0.5 | 1 | 0 | NA | NA | 3 | 0.3333 | NA | NA | 1 | 0 | 2 | 0.5 | NA | NA | 3 | 0.3333 |

| | | | | | | | | | | | | | | | | | | | | |
|---|---|---|---|---|---|---|---|---|---|---|---|---|---|---|---|---|---|---|---|---|
| 1_18231762 | NA | NA | 2 | 1 | 1 | 1 | NA | NA | 3 | 1 | 2 | 0.5 | 1 | 1 | 1 | 1 | NA | NA | 4 | 0.75 |
| 1_18231770 | NA | NA | 2 | 1 | 1 | 1 | 1 | 1 | 4 | 1 | 2 | 1 | 1 | 0 | 2 | 1 | NA | NA | 5 | 0.8 |
| 1_18231784 | NA | NA | 2 | 0.5 | 1 | 0 | 1 | 0 | 4 | 0.25 | 2 | 1 | 1 | 0 | 3 | 0.3333 | NA | NA | 6 | 0.5 |
| 1_18231801 | NA | NA | 2 | 0 | 1 | 1 | 1 | 1 | 4 | 0.5 | 2 | 1 | 1 | 1 | 3 | 1 | NA | NA | 6 | 1 |
| 1_18231813 | NA | NA | 2 | 0.5 | 1 | 0 | 1 | 0 | 4 | 0.25 | 2 | 0.5 | NA | NA | 3 | 0.6667 | NA | NA | 5 | 0.6 |
| 1_18231818 | NA | NA | 1 | 0 | 1 | 0 | 1 | 1 | 3 | 0.3333 | 2 | 0.5 | NA | NA | 3 | 0 | NA | NA | 5 | 0.2 |
| 1_18231833 | NA | NA | 1 | 0 | 1 | 1 | 1 | 1 | 3 | 0.6667 | 3 | 1 | NA | NA | 2 | 1 | NA | NA | 5 | 1 |
| 1_18231839 | NA | NA | 2 | 0.5 | 1 | 1 | 2 | 1 | 5 | 0.8 | 3 | 1 | NA | NA | 3 | 1 | NA | NA | 6 | 1 |
| 1_18231842 | NA | NA | 2 | 0 | 1 | 0 | 2 | 0.5 | 5 | 0.2 | 3 | 0 | NA | NA | 5 | 0.2 | NA | NA | 8 | 0.125 |
| 1_18231942 | 1 | 0 | 5 | 0.4 | 2 | 0.5 | 2 | 0 | 10 | 0.3 | 2 | 0 | 1 | 1 | 3 | 0 | NA | NA | 6 | 0.1667 |
| 1_18231966 | NA | NA | 3 | 0.3333 | 3 | 0 | 2 | 0 | 8 | 0.125 | 3 | 0 | 1 | 0 | 3 | 0.6667 | NA | NA | 7 | 0.2857 |
| 1_18231976 | 1 | 0 | 3 | 0.6667 | 2 | 0 | 2 | 0.5 | 8 | 0.375 | 3 | 0.3333 | 1 | 0 | 3 | 0 | NA | NA | 7 | 0.1429 |
| 1_18231977 | 2 | 0.5 | 5 | 0.4 | 3 | 0.3333 | 2 | 1 | 12 | 0.5 | 3 | 0.6667 | NA | NA | 3 | 0.3333 | NA | NA | 6 | 0.5 |
| 1_18231979 | 2 | 1 | 4 | 1 | 3 | 1 | 2 | 1 | 11 | 1 | 3 | 1 | 1 | 1 | 4 | 1 | NA | NA | 8 | 1 |
| 1_18232019 | 2 | 0.5 | 3 | 0.3333 | 3 | 0.6667 | NA | NA | 8 | 0.5 | 2 | 0.5 | 1 | 1 | 2 | 1 | NA | NA | 5 | 0.8 |
| 1_18232022 | 2 | 1 | 2 | 1 | 2 | 1 | NA | NA | 6 | 1 | 1 | 1 | 1 | 1 | 1 | 1 | NA | NA | 3 | 1 |
| 1_18232044 | 2 | 0 | 3 | 0.3333 | 3 | 0 | NA | NA | 8 | 0.125 | 1 | 0 | 1 | 1 | 2 | 1 | NA | NA | 4 | 0.75 |
| 1_18232061 | 2 | 1 | 3 | 1 | 3 | 1 | NA | NA | 8 | 1 | 2 | 0.5 | 1 | 1 | 2 | 1 | NA | NA | 5 | 0.8 |
| 1_18232068 | 1 | 0 | 3 | 0.3333 | 3 | 0.6667 | NA | NA | 7 | 0.4286 | 1 | 0 | 1 | 1 | 2 | 0 | NA | NA | 4 | 0.25 |
| 1_18232068 | 2 | 0.5 | 3 | 0.6667 | 3 | 0.6667 | NA | NA | 8 | 0.625 | 1 | 1 | 1 | 1 | 1 | 1 | NA | NA | 3 | 1 |
| 1_18232076 | 1 | 1 | NA | NA | 1 | 1 | NA | NA | 2 | 1 | NA | NA | 1 | 1 | 1 | 1 | NA | NA | 2 | 1 |
| 1_18232077 | 2 | 0.5 | 2 | 1 | 1 | 0 | NA | NA | 5 | 0.6 | 1 | 0 | 1 | 1 | 1 | 1 | NA | NA | 3 | 0.6667 |
| 1_18232100 | 2 | 0 | 2 | 0.5 | 3 | 0 | NA | NA | 7 | 0.1429 | 2 | 1 | 1 | 0 | 2 | 0.5 | 1 | 0 | 6 | 0.5 |
| 1_18232125 | 1 | 0 | 3 | 0.6667 | 3 | 0.3333 | NA | NA | 7 | 0.4286 | 1 | 0 | NA | NA | 1 | 1 | NA | NA | 2 | 0.5 |
| 1_18232125 | 2 | 1 | 3 | 1 | 3 | 1 | NA | NA | 8 | 1 | 2 | 1 | 1 | 1 | 1 | 1 | 1 | 0 | 5 | 0.8 |
| 1_18232128 | 2 | 0 | 2 | 0.5 | 1 | 0 | NA | NA | 5 | 0.2 | 2 | 0 | 1 | 1 | 2 | 0 | 1 | 0 | 6 | 0.1667 |
| 1_18232131 | 2 | 0 | 3 | 0.3333 | 2 | 0 | NA | NA | 7 | 0.1429 | 2 | 0.5 | NA | NA | NA | NA | 1 | 0 | 3 | 0.3333 |
| 1_18232134 | 2 | 1 | 1 | 1 | 1 | 1 | NA | NA | 4 | 1 | 1 | 1 | 1 | 1 | 1 | 1 | 1 | 1 | 4 | 1 |
| 1_18232151 | NA | NA | 1 | 1 | 1 | 0 | NA | NA | 2 | 0.5 | NA | NA | 1 | 1 | NA | NA | NA | NA | 1 | 1 |
| 1_18232170 | NA | NA | 1 | 1 | 1 | 0 | NA | NA | 2 | 0.5 | NA | NA | NA | NA | 1 | 1 | NA | NA | 1 | 1 |
| 1_18232268 | NA | NA | 1 | 1 | 1 | 1 | 1 | 0 | 3 | 0.6667 | NA | NA | NA | NA | 1 | 1 | NA | NA | 1 | 1 |
| 1_18232287 | NA | NA | 1 | 1 | NA | NA | 1 | 0 | 2 | 0.5 | NA | NA | NA | NA | 1 | 1 | NA | NA | 1 | 1 |
| 1_18259268 | 9 | 0.1111 | 8 | 0.25 | 3 | 0 | 9 | 0 | 29 | 0.1034 | 4 | 0 | 7 | 0.1429 | 6 | 0.1667 | 5 | 0.2 | 22 | 0.1364 |
| 1_18271498 | 12 | 0.1667 | 10 | 0 | 2 | 0 | 12 | 0 | 36 | 0.0556 | 5 | 0 | 11 | 0.1818 | 9 | 0.1111 | 7 | 0 | 32 | 0.0938 |
| 1_18271964 | 12 | 0.3333 | 7 | 0.4286 | 2 | 0 | 11 | 0.1818 | 32 | 0.2812 | 6 | 0.1667 | 9 | 0.3333 | 7 | 0.2857 | 7 | 0.1429 | 29 | 0.2414 |
| 1_18272286 | 15 | 0.4 | 12 | 0.4167 | 2 | 0 | 17 | 0.2353 | 46 | 0.3261 | 11 | 0.5455 | 13 | 0.4615 | 12 | 0.5 | 10 | 0.5 | 46 | 0.5 |
| 1_18272287 | 18 | 0.2778 | 13 | 0.3846 | 4 | 0 | 17 | 0.2941 | 52 | 0.2885 | 11 | 0.5455 | 16 | 0.1875 | 14 | 0.4286 | 11 | 0.2727 | 52 | 0.3462 |
| 1_18272290 | 12 | 0.5833 | 14 | 0.7857 | 3 | 0.6667 | 18 | 0.6667 | 47 | 0.6809 | 10 | 0.7 | 14 | 0.6429 | 10 | 0.6 | 10 | 0.7 | 44 | 0.6591 |
| 1_18272293 | 18 | 0.9444 | 13 | 0.9231 | 3 | 0.6667 | 19 | 0.7895 | 53 | 0.8679 | 10 | 1 | 16 | 0.9375 | 14 | 0.8571 | 11 | 0.7273 | 51 | 0.8824 |
| 1_18272298 | 14 | 0.7143 | 13 | 0.8462 | 4 | 0.5 | 12 | 0.8333 | 43 | 0.7674 | 11 | 0.5455 | 12 | 0.5833 | 14 | 0.7857 | 9 | 0.8889 | 46 | 0.6957 |
| 1_18272302 | 17 | 0.2353 | 14 | 0.5714 | 3 | 0.3333 | 12 | 0 | 46 | 0.2826 | 12 | 0.1667 | 14 | 0 | 15 | 0.4 | 10 | 0.2 | 51 | 0.1961 |
| 1_18272308 | 18 | 0.2778 | 14 | 0.3571 | 3 | 0 | 18 | 0.3333 | 53 | 0.3019 | 11 | 0.4545 | 16 | 0.4375 | 17 | 0.5294 | 12 | 0.5833 | 56 | 0.5 |
| 1_18272318 | 18 | 1 | 12 | 0.8333 | 4 | 0.75 | 16 | 1 | 50 | 0.94 | 11 | 0.7273 | 14 | 0.7857 | 18 | 0.8889 | 10 | 0.8 | 53 | 0.8113 |
| 1_18272319 | 19 | 0.6842 | 12 | 0.8333 | 4 | 0.75 | 19 | 0.6842 | 54 | 0.7222 | 12 | 0.8333 | 14 | 0.7143 | 20 | 0.7 | 10 | 0.5 | 56 | 0.6964 |
| 1_18272331 | 21 | 0.8095 | 13 | 0.9231 | 4 | 0.75 | 20 | 0.7 | 58 | 0.7931 | 12 | 0.75 | 16 | 0.5625 | 16 | 0.75 | 12 | 0.5833 | 56 | 0.6607 |
| 1_18272332 | 19 | 0.0526 | 14 | 0.0714 | 4 | 0 | 21 | 0.1429 | 58 | 0.0862 | 11 | 0.0909 | 17 | 0.1765 | 16 | 0.125 | 12 | 0.1667 | 56 | 0.1429 |
| 1_18272334 | 14 | 0.1429 | 8 | 0.25 | 3 | 0 | 11 | 0.3636 | 36 | 0.2222 | 10 | 0.2 | 11 | 0 | 14 | 0.1429 | 8 | 0.375 | 43 | 0.1628 |
| 1_18272349 | 20 | 0.3 | 15 | 0.2667 | 5 | 0.2 | 20 | 0.4 | 60 | 0.3167 | 11 | 0.1818 | 19 | 0.1579 | 20 | 0.55 | 14 | 0.2857 | 64 | 0.3125 |
| 1_18272351 | 20 | 0.25 | 15 | 0.1333 | 5 | 0.2 | 23 | 0.0435 | 63 | 0.1429 | 12 | 0.0833 | 17 | 0.0588 | 21 | 0.0476 | 15 | 0.1333 | 65 | 0.0769 |
| 1_18272405 | 21 | 0.0952 | 16 | 0 | 5 | 0 | 20 | 0.1 | 62 | 0.0645 | 11 | 0.1818 | 20 | 0 | 24 | 0.0417 | 9 | 0.1111 | 64 | 0.0625 |
| 1_18286794 | NA | NA | 1 | 1 | 2 | 1 | 2 | 1 | 5 | 1 | NA | NA | 1 | 1 | NA | NA | NA | NA | 1 | 1 |
| 1_18305803 | 5 | 0.2 | 4 | 0.25 | 6 | 0.1667 | 10 | 0.4 | 25 | 0.28 | 8 | 0.375 | 10 | 0.4 | 6 | 0.3333 | 4 | 0.25 | 28 | 0.3571 |
| 1_18305828 | 9 | 0.1111 | 4 | 0.25 | 8 | 0.125 | 11 | 0.2727 | 32 | 0.1875 | 6 | 0.5 | 12 | 0.25 | 6 | 0.1667 | 7 | 0.1429 | 31 | 0.2581 |
| 1_18305929 | 6 | 0.3333 | 5 | 0.4 | 6 | 0.6667 | 6 | 0.6667 | 23 | 0.5217 | 8 | 0.5 | 12 | 0.5833 | 4 | 0.75 | 5 | 0.4 | 29 | 0.5517 |
| 1_18305965 | 7 | 0.1429 | 5 | 0 | 5 | 0.2 | 7 | 0 | 24 | 0.0833 | 6 | 0.1667 | 9 | 0.1111 | 3 | 0.6667 | 4 | 0 | 22 | 0.1818 |
| 1_18305966 | 7 | 0.1429 | 5 | 0 | 6 | 0 | 7 | 0.2857 | 25 | 0.12 | 6 | 0.1667 | 10 | 0.3 | 3 | 0 | 5 | 0.2 | 24 | 0.2083 |
| 1_18305971 | 5 | 0 | 5 | 0.2 | 5 | 0.2 | 7 | 0.1429 | 22 | 0.1364 | 6 | 0.1667 | 8 | 0.25 | 3 | 0 | 3 | 0.3333 | 20 | 0.2 |
| 1_18305987 | 6 | 0.3333 | 4 | 0.5 | 5 | 0.8 | 6 | 0 | 21 | 0.381 | 6 | 0.1667 | 7 | 0.5714 | 3 | 0.3333 | 4 | 1 | 20 | 0.5 |
| 1_18305993 | 5 | 0.8 | 4 | 0.25 | 6 | 0.5 | 7 | 0.5714 | 22 | 0.5455 | 6 | 0.5 | 10 | 1 | 3 | 0.6667 | 3 | 0.3333 | 22 | 0.7273 |
| 1_18310346 | 4 | 0.75 | 6 | 0.8333 | 3 | 0.6667 | 3 | 1 | 16 | 0.8125 | 2 | 1 | 5 | 1 | 4 | 0.75 | 3 | 0.3333 | 14 | 0.7857 |
| 1_18310360 | 7 | 0.2857 | 6 | 0.1667 | 3 | 0.6667 | 4 | 0.25 | 20 | 0.3 | 2 | 0.5 | 5 | 0 | 7 | 0.1429 | 2 | 0.5 | 16 | 0.1875 |
| 1_18310363 | 6 | 0.1667 | 7 | 0 | 3 | 0 | 3 | 0 | 19 | 0.0526 | 2 | 0 | 6 | 0.1667 | 6 | 0.3333 | 3 | 0 | 17 | 0.1765 |
| 1_18310375 | 6 | 0.5 | 7 | 0.5714 | 4 | 0.75 | 6 | 0.6667 | 23 | 0.6087 | 2 | 0.5 | 6 | 0.3333 | 7 | 0.5714 | 3 | 0 | 18 | 0.3889 |
| 1_18310377 | 8 | 0.125 | 6 | 0.1667 | 4 | 0.25 | 4 | 0 | 22 | 0.1364 | 2 | 0 | 6 | 0 | 6 | 0.3333 | 2 | 0 | 16 | 0.125 |
| 1_18310378 | 9 | 0.1111 | 8 | 0.125 | 4 | 0 | 5 | 0 | 26 | 0.0769 | 2 | 0 | 6 | 0.5 | 6 | 0.1667 | 2 | 0 | 16 | 0.25 |

| | | | | | | | | | | | | | | | | | | | | |
|---|---|---|---|---|---|---|---|---|---|---|---|---|---|---|---|---|---|---|---|---|
| 1_1831038 | 8 | 0.625 | 9 | 0.3333 | 3 | 0 | 6 | 0.3333 | 26 | 0.3846 | 2 | 0.5 | 6 | 0.3333 | 6 | 0.3333 | 2 | 0.5 | 16 | 0.375 |
| 1_1831039 | 3 | 0.6667 | 6 | 0.6667 | 2 | 1 | 4 | 0.5 | 15 | 0.6667 | 1 | 1 | 5 | 0.6 | 6 | 0.8333 | 2 | 1 | 14 | 0.7857 |
| 1_1831039 | 9 | 1 | 9 | 0.7778 | 4 | 1 | 3 | 1 | 25 | 0.92 | 1 | 1 | 4 | 1 | 7 | 1 | 2 | 0.5 | 14 | 0.9286 |
| 1_1831040 | 7 | 0.4286 | 8 | 0.25 | 5 | 0.2 | 6 | 0.5 | 26 | 0.3462 | 2 | 0.5 | 6 | 0.3333 | 6 | 0.1667 | 3 | 0.3333 | 17 | 0.2941 |
| 1_1831043 | 9 | 0.5556 | 9 | 0.3333 | 5 | 0.2 | 7 | 0.4286 | 30 | 0.4 | 2 | 0.5 | 5 | 0.2 | 7 | 0.1429 | 2 | 0 | 16 | 0.1875 |
| 1_1831044 | 6 | 0.3333 | 8 | 0.375 | 4 | 0.5 | 7 | 0 | 25 | 0.28 | 2 | 1 | 6 | 0.1667 | 3 | 0 | 3 | 0 | 14 | 0.2143 |
| 1_1831055 | 2 | 0 | NA | NA | 1 | 0 | 2 | 0.5 | 5 | 0.2 | 1 | 0 | 1 | 0 | 1 | 1 | 1 | 0 | 4 | 0.25 |
| 1_1831059 | 3 | 0 | 1 | 1 | 3 | 0 | 2 | 0 | 9 | 0.1111 | 1 | 0 | 2 | 0 | 1 | 1 | 1 | 0 | 5 | 0.2 |
| 1_1831077 | 10 | 0.2 | 12 | 0.5833 | 11 | 0.1818 | 10 | 0.4 | 43 | 0.3488 | 7 | 0.5714 | 8 | 0.125 | 12 | 0.3333 | 7 | 0.1429 | 34 | 0.2941 |
| 1_1831081 | 9 | 1 | 10 | 1 | 11 | 0.9091 | 10 | 0.9 | 40 | 0.95 | 7 | 0.7143 | 9 | 0.8889 | 11 | 0.7273 | 6 | 0.8333 | 33 | 0.7879 |
| 1_1831081 | 8 | 0.5 | 10 | 0.5 | 11 | 0.2727 | 10 | 0.2 | 39 | 0.359 | 6 | 0.3333 | 7 | 0.1429 | 13 | 0.1538 | 6 | 0.1667 | 32 | 0.1875 |
| 1_1831082 | 9 | 0.5556 | 9 | 0.1111 | 9 | 0 | 8 | 0 | 35 | 0.1714 | 6 | 0.1667 | 6 | 0.1667 | 12 | 0.3333 | 6 | 0.3333 | 30 | 0.2667 |
| 1_1831082 | 9 | 0.6667 | 9 | 0.7778 | 9 | 0.6667 | 9 | 0.6667 | 36 | 0.6944 | 5 | 0.6 | 7 | 1 | 12 | 0.6667 | 6 | 0.8333 | 30 | 0.7667 |
| 1_1831084 | 10 | 0.1 | 11 | 0.0909 | 11 | 0.4545 | 10 | 0.3 | 42 | 0.2381 | 5 | 0.6 | 6 | 0.3333 | 11 | 0.1818 | 5 | 0.6 | 27 | 0.3704 |
| 1_1831376 | 3 | 0 | 2 | 0 | 1 | 0 | 2 | 0.5 | 8 | 0.125 | 1 | 0 | 2 | 0.5 | 2 | 0.5 | 3 | 0 | 8 | 0.25 |
| 1_1831377 | 2 | 0.5 | 2 | 1 | NA | NA | 1 | 1 | 5 | 0.8 | 2 | 1 | 1 | 1 | 2 | 0.5 | 3 | 0.6667 | 8 | 0.75 |
| 1_1831380 | 2 | 0.5 | 1 | 1 | 2 | 1 | 2 | 0.5 | 7 | 0.7143 | 2 | 0.5 | 2 | 0 | 2 | 0 | 1 | 1 | 7 | 0.2857 |
| 1_1831383 | 1 | 0 | 1 | 0 | 2 | 0.5 | 2 | 0 | 6 | 0.1667 | 1 | 0 | 2 | 1 | 2 | 0 | 1 | 0 | 6 | 0.3333 |
| 1_1831384 | 2 | 0.5 | 1 | 0 | 2 | 0 | 2 | 0.5 | 7 | 0.2857 | 1 | 1 | NA | NA | 2 | 1 | 2 | 0.5 | 5 | 0.8 |
| 1_1831388 | 2 | 0 | 1 | 1 | 1 | 1 | 1 | 1 | 5 | 0.6 | 1 | 1 | 2 | 0 | 2 | 0.5 | 2 | 0 | 7 | 0.2857 |
| 1_1831389 | 2 | 0 | 1 | 1 | NA | NA | NA | NA | 3 | 0.3333 | 1 | 0 | 2 | 0.5 | 1 | 0 | 1 | 1 | 5 | 0.4 |
| 1_1831390 | 1 | 1 | NA | NA | 1 | 0 | NA | NA | 2 | 0.5 | 1 | 0 | 1 | 1 | 1 | 0 | 1 | 0 | 4 | 0.25 |
| 1_1831392 | 1 | 0 | 1 | 1 | 1 | 0 | NA | NA | 3 | 0.3333 | 1 | 1 | 1 | 1 | NA | NA | 1 | 1 | 3 | 1 |
| 1_1831393 | NA | NA | 1 | 0 | 1 | 1 | NA | NA | 2 | 0.5 | 1 | 1 | 1 | 0 | NA | NA | 1 | 0 | 3 | 0.3333 |
| 1_1831393 | 1 | 1 | 1 | 0 | 1 | 0 | NA | NA | 3 | 0.3333 | 1 | 1 | 1 | 1 | NA | NA | 1 | 1 | 3 | 1 |
| 1_1831399 | 1 | 1 | 1 | 1 | 1 | 0 | NA | NA | 3 | 0.6667 | 1 | 1 | NA | NA | NA | NA | NA | NA | 1 | 1 |
| 1_1831401 | 1 | 1 | NA | NA | 1 | 1 | NA | NA | 2 | 1 | 1 | 1 | NA | NA | NA | NA | NA | NA | 1 | 1 |
| 1_1831904 | 1 | 0 | 2 | 1 | NA | NA | 3 | 0.6667 | 6 | 0.6667 | NA | NA | 4 | 1 | 1 | 1 | NA | NA | 5 | 1 |
| 1_1831951 | NA | NA | NA | NA | NA | NA | 1 | 1 | 1 | 1 | NA | NA | NA | NA | 1 | 1 | NA | NA | 1 | 1 |
| 1_1831952 | NA | NA | NA | NA | NA | NA | 1 | 1 | 1 | 1 | NA | NA | NA | NA | 1 | 1 | NA | NA | 1 | 1 |
| 1_1831952 | NA | NA | NA | NA | NA | NA | 1 | 1 | 1 | 1 | NA | NA | NA | NA | 1 | 1 | NA | NA | 1 | 1 |
| 1_1831955 | NA | NA | NA | NA | 1 | 0 | 1 | 1 | 2 | 0.5 | NA | NA | NA | NA | 1 | 1 | NA | NA | 1 | 1 |
| 1_1834088 | 1 | 1 | NA | NA | NA | NA | NA | NA | 1 | 1 | NA | NA | NA | NA | 1 | 1 | NA | NA | 1 | 1 |
| 1_1834094 | 1 | 1 | NA | NA | NA | NA | NA | NA | 1 | 1 | NA | NA | NA | NA | 1 | 1 | NA | NA | 1 | 1 |
| 1_1834094 | 1 | 1 | NA | NA | NA | NA | NA | NA | 1 | 1 | NA | NA | NA | NA | 1 | 1 | NA | NA | 1 | 1 |
| 1_1834095 | 1 | 1 | NA | NA | NA | NA | NA | NA | 1 | 1 | NA | NA | NA | NA | 1 | 1 | NA | NA | 1 | 1 |
| 1_1837696 | NA | NA | 1 | 1 | 1 | 0 | NA | NA | 2 | 0.5 | NA | NA | NA | NA | NA | NA | 1 | 1 | 1 | 1 |
| 1_1837697 | NA | NA | 1 | 1 | 1 | 1 | NA | NA | 2 | 1 | 1 | 1 | NA | NA | NA | NA | 1 | 1 | 2 | 1 |
| 1_1838067 | NA | NA | 1 | 1 | 1 | 0 | NA | NA | 2 | 0.5 | 1 | 1 | NA | NA | NA | NA | NA | NA | 1 | 1 |
| 1_1838069 | NA | NA | 1 | 1 | 1 | 1 | NA | NA | 2 | 1 | 1 | 1 | NA | NA | NA | NA | 1 | 1 | 2 | 1 |
| 1_1838069 | NA | NA | 1 | 0 | 1 | 1 | NA | NA | 2 | 0.5 | 1 | 0 | NA | NA | NA | NA | 1 | 1 | 2 | 0.5 |
| 1_1838071 | NA | NA | 1 | 1 | NA | NA | NA | NA | 1 | 1 | 1 | 1 | NA | NA | NA | NA | 1 | 1 | 2 | 1 |
| 1_1838072 | NA | NA | 1 | 1 | NA | NA | NA | NA | 1 | 1 | NA | NA | NA | NA | NA | NA | 1 | 1 | 1 | 1 |
| 1_1838072 | NA | NA | 1 | 1 | NA | NA | NA | NA | 1 | 1 | NA | NA | NA | NA | NA | NA | 1 | 1 | 1 | 1 |
| 1_1838073 | NA | NA | 1 | 1 | NA | NA | NA | NA | 1 | 1 | 1 | 1 | NA | NA | NA | NA | 1 | 1 | 2 | 1 |
| 1_1844594 | NA | NA | NA | NA | NA | NA | 1 | 1 | 1 | 1 | 1 | 1 | NA | NA | NA | NA | NA | NA | 1 | 1 |
| 1_1844599 | NA | NA | NA | NA | NA | NA | 1 | 1 | 1 | 1 | 1 | 1 | NA | NA | NA | NA | NA | NA | 1 | 1 |
| 1_1846043 | 1 | 1 | 2 | 1 | NA | NA | NA | NA | 3 | 1 | 2 | 0 | NA | NA | NA | NA | 2 | 0.5 | 4 | 0.25 |
| 1_1846044 | 1 | 0 | 2 | 0.5 | NA | NA | NA | NA | 3 | 0.3333 | 3 | 0 | NA | NA | NA | NA | 2 | 1 | 5 | 0.4 |
| 1_1846044 | 1 | 0 | 1 | 1 | NA | NA | NA | NA | 2 | 0.5 | NA | NA | NA | NA | NA | NA | 1 | 1 | 1 | 1 |
| 1_1846052 | 1 | 0 | 1 | 1 | NA | NA | NA | NA | 2 | 0.5 | 2 | 0.5 | NA | NA | NA | NA | 1 | 0 | 3 | 0.3333 |
| 1_1846291 | 13 | 0.2308 | 9 | 0.2222 | 9 | 0.2222 | 9 | 0.2222 | 40 | 0.225 | 8 | 0.375 | 16 | 0.125 | 15 | 0.2667 | 8 | 0.375 | 47 | 0.2553 |
| 1_1846293 | 11 | 0.0909 | 10 | 0.1 | 6 | 0 | 8 | 0 | 35 | 0.0571 | 9 | 0 | 13 | 0.1538 | 9 | 0.1111 | 4 | 0 | 35 | 0.0857 |
| 1_1846294 | 9 | 0.3333 | 10 | 0.2 | 8 | 0.375 | 8 | 0.25 | 35 | 0.2857 | 7 | 0.2857 | 16 | 0.25 | 13 | 0.3077 | 7 | 0.2857 | 43 | 0.2791 |
| 1_1846294 | 10 | 0.1 | 9 | 0 | 7 | 0.1429 | 5 | 0.2 | 31 | 0.0968 | 7 | 0.2857 | 12 | 0.0833 | 13 | 0 | 7 | 0.1429 | 39 | 0.1026 |
| 1_1846295 | 10 | 0.2 | 9 | 0.2222 | 7 | 0.5714 | 5 | 0.4 | 31 | 0.3226 | 6 | 0.8333 | 12 | 0.25 | 10 | 0.5 | 6 | 0.1667 | 34 | 0.4118 |
| 1_1846296 | 8 | 0.25 | 7 | 0.4286 | 4 | 0.5 | 5 | 0.6 | 24 | 0.4167 | 6 | 0 | 8 | 0.25 | 10 | 0.2 | 6 | 0.3333 | 30 | 0.2 |
| 1_1846296 | 10 | 1 | 9 | 1 | 8 | 1 | 8 | 1 | 35 | 1 | 8 | 1 | 12 | 1 | 13 | 1 | 6 | 1 | 39 | 1 |
| 1_1846297 | 7 | 0 | 9 | 0 | 5 | 0.2 | 5 | 0.2 | 26 | 0.0769 | 7 | 0.1429 | 13 | 0.0769 | 12 | 0.0833 | 4 | 0 | 36 | 0.0833 |
| 1_1846299 | 6 | 0.8333 | 8 | 0.875 | 4 | 1 | 6 | 1 | 24 | 0.9167 | 7 | 1 | 12 | 1 | 11 | 1 | 6 | 1 | 36 | 1 |
| 1_1846300 | 6 | 0.1667 | 9 | 0.1111 | 7 | 0 | 6 | 0.1667 | 28 | 0.1071 | 7 | 0.1429 | 11 | 0 | 11 | 0.2727 | 6 | 0 | 35 | 0.1143 |
| 1_1846301 | 7 | 1 | 9 | 0.8889 | 7 | 0.7143 | 7 | 1 | 30 | 0.9 | 6 | 1 | 13 | 1 | 12 | 1 | 6 | 1 | 37 | 1 |
| 1_1846302 | 6 | 0 | 8 | 0.25 | 6 | 0.1667 | 6 | 0.1667 | 26 | 0.1538 | 4 | 0.25 | 10 | 0.1 | 9 | 0 | 7 | 0 | 30 | 0.0667 |
| 1_1846307 | 4 | 1 | 12 | 1 | 6 | 1 | 11 | 1 | 33 | 1 | 11 | 1 | 12 | 0.9167 | 12 | 1 | 6 | 1 | 41 | 0.9756 |
| 1_1846307 | 8 | 1 | 13 | 1 | 8 | 1 | 11 | 1 | 40 | 1 | 11 | 1 | 14 | 0.9286 | 16 | 1 | 7 | 1 | 48 | 0.9792 |

| | | | | | | | | | | | | | | | | | | | | |
|---|---|---|---|---|---|---|---|---|---|---|---|---|---|---|---|---|---|---|---|---|
| 1_1846308 | 7 | 0.4286 | 10 | 0.3 | 7 | 0.5714 | 5 | 0.8 | 29 | 0.4828 | 12 | 0.4167 | 11 | 0.0909 | 16 | 0.25 | 7 | 0.1429 | 46 | 0.2391 |
| 1_1846309 | 4 | 0.75 | 9 | 0.7778 | 5 | 1 | 9 | 0.7778 | 27 | 0.8148 | 11 | 1 | 9 | 1 | 13 | 0.8462 | 6 | 0.5 | 39 | 0.8718 |
| 1_1846309 | 7 | 0.2857 | 12 | 0.3333 | 7 | 0.2857 | 11 | 0.1818 | 37 | 0.2703 | 12 | 0.4167 | 13 | 0.1538 | 18 | 0.3333 | 7 | 0.2857 | 50 | 0.3 |
| 1_1849833 | NA | NA | 5 | 0 | 1 | 0 | 2 | 0.5 | 8 | 0.125 | 3 | 0 | 3 | 0.3333 | 3 | 0 | 1 | 0 | 10 | 0.1 |
| 1_1849959 | NA | NA | 8 | 0.125 | 2 | 0 | 4 | 0 | 14 | 0.0714 | 3 | 0 | 5 | 0.2 | 3 | 0 | 3 | 0 | 14 | 0.0714 |
| 1_1850163 | NA | NA | 4 | 0.25 | 2 | 0 | NA | NA | 6 | 0.1667 | 1 | 0 | 7 | 0.1429 | NA | NA | 3 | 0 | 11 | 0.0909 |
| 1_1850187 | NA | NA | 4 | 0.5 | 2 | 0.5 | NA | NA | 6 | 0.5 | NA | NA | 3 | 0.6667 | 1 | 0 | NA | NA | 4 | 0.5 |
| 1_1850189 | 4 | 0 | 1 | 1 | 11 | 0 | 3 | 0 | 19 | 0.0526 | 2 | 0 | 3 | 0.3333 | 2 | 0 | 3 | 0 | 10 | 0.1 |
| 1_1850192 | 5 | 0 | 3 | 0.3333 | 11 | 0.2727 | 4 | 0 | 23 | 0.1739 | 2 | 0 | 1 | 0 | 2 | 0.5 | 2 | 0 | 7 | 0.1429 |
| 1_1850192 | 5 | 0 | 1 | 0 | 10 | 0 | 4 | 0.25 | 20 | 0.05 | 2 | 0.5 | 1 | 0 | 1 | 0 | 3 | 0.3333 | 7 | 0.2857 |
| 1_1850851 | NA | NA | NA | NA | NA | NA | 1 | 1 | 1 | 1 | 1 | 1 | NA | NA | NA | NA | NA | NA | 1 | 1 |
| 1_1850852 | NA | NA | NA | NA | NA | NA | 1 | 1 | 1 | 1 | 1 | 1 | NA | NA | NA | NA | NA | NA | 1 | 1 |
| 1_1850857 | NA | NA | NA | NA | 1 | 0 | 1 | 1 | 2 | 0.5 | 1 | 1 | NA | NA | NA | NA | NA | NA | 1 | 1 |
| 1_1850887 | 3 | 0.3333 | 8 | 0.625 | 5 | 0 | 4 | 0.5 | 20 | 0.4 | 7 | 0.2857 | 6 | 0.3333 | 6 | 0.3333 | 2 | 0 | 21 | 0.2857 |
| 1_1850888 | 6 | 0.3333 | 7 | 0 | 5 | 0 | 4 | 0 | 22 | 0.0909 | 7 | 0 | 7 | 0.2857 | 7 | 0.1429 | 2 | 0 | 23 | 0.1304 |
| 1_1850888 | 6 | 0.1667 | 6 | 0.3333 | 5 | 0.2 | 3 | 1 | 20 | 0.35 | 4 | 1 | 6 | 0.1667 | 7 | 0.4286 | 2 | 0 | 19 | 0.4211 |
| 1_1850889 | 6 | 0.1667 | 8 | 0.125 | 6 | 0 | 5 | 0 | 25 | 0.08 | 7 | 0.2857 | 7 | 0 | 9 | 0.3333 | 4 | 0 | 27 | 0.1852 |
| 1_1851336 | 3 | 0.3333 | 5 | 0 | 4 | 0 | 6 | 0 | 18 | 0.0556 | 3 | 0 | 4 | 0 | 4 | 0.25 | 2 | 0 | 13 | 0.0769 |
| 1_1851337 | 3 | 0.6667 | 7 | 0.5714 | 4 | 0.5 | 5 | 0 | 19 | 0.4211 | 4 | 0.75 | 3 | 0.6667 | 6 | 0.5 | 2 | 1 | 15 | 0.6667 |
| 1_1851392 | 3 | 0 | 8 | 0.125 | 5 | 0 | 6 | 0.1667 | 22 | 0.0909 | 3 | 0.3333 | 4 | 0.25 | 7 | 0.2857 | 2 | 0 | 16 | 0.25 |
| 1_1851398 | 3 | 0.3333 | 8 | 0.125 | 5 | 0.2 | 7 | 0.4286 | 23 | 0.2609 | 5 | 0.2 | 7 | 0.1429 | 7 | 0.4286 | 3 | 0.3333 | 22 | 0.2727 |
| 1_1851603 | 7 | 0.1429 | 9 | 0.2222 | 7 | 0.5714 | 10 | 0.6 | 33 | 0.3939 | 6 | 0.6667 | 11 | 0.2727 | 13 | 0.3846 | 5 | 0.6 | 35 | 0.4286 |
| 1_1851605 | 6 | 0 | 6 | 0 | 4 | 0.25 | 10 | 0.4 | 26 | 0.1923 | 6 | 0.3333 | 12 | 0.1667 | 12 | 0.4167 | 4 | 0.5 | 34 | 0.3235 |
| 1_1851964 | 1 | 1 | NA | NA | 1 | 1 | 2 | 1 | 4 | 1 | 1 | 1 | NA | NA | 1 | 1 | NA | NA | 2 | 1 |
| 1_1851966 | 1 | 1 | NA | NA | 1 | 1 | 2 | 1 | 4 | 1 | 1 | 1 | NA | NA | 1 | 1 | NA | NA | 2 | 1 |
| 1_1851966 | 1 | 0 | NA | NA | 1 | 0 | 2 | 1 | 4 | 0.5 | 1 | 0 | NA | NA | 1 | 1 | NA | NA | 2 | 0.5 |
| 1_1851967 | 2 | 1 | NA | NA | 2 | 1 | 2 | 0.5 | 6 | 0.8333 | 1 | 1 | NA | NA | 1 | 0 | NA | NA | 2 | 0.5 |
| 1_1851967 | 2 | 0 | NA | NA | 2 | 0 | 3 | 0.6667 | 7 | 0.2857 | 1 | 1 | NA | NA | 1 | 0 | NA | NA | 2 | 0.5 |
| 1_1851967 | 2 | 0 | NA | NA | 2 | 1 | 3 | 0 | 7 | 0.2857 | 1 | 0 | NA | NA | 2 | 0.5 | NA | NA | 3 | 0.3333 |
| 1_1851969 | 1 | 0 | 1 | 1 | 2 | 0 | 2 | 0.5 | 6 | 0.3333 | 1 | 1 | NA | NA | 2 | 0 | NA | NA | 3 | 0.3333 |
| 1_1851969 | 2 | 0.5 | 1 | 1 | 2 | 1 | 3 | 1 | 8 | 0.875 | 1 | 1 | NA | NA | 2 | 1 | NA | NA | 3 | 1 |
| 1_1851969 | 2 | 1 | 1 | 0 | 1 | 1 | 1 | 1 | 5 | 0.8 | NA | NA | NA | NA | 2 | 1 | NA | NA | 2 | 1 |
| 1_1851970 | 2 | 0.5 | NA | NA | 2 | 1 | 3 | 1 | 7 | 0.8571 | 1 | 1 | NA | NA | 2 | 1 | NA | NA | 3 | 1 |
| 1_1851970 | 2 | 0.5 | 1 | 1 | 2 | 0.5 | 2 | 0.5 | 7 | 0.5714 | 1 | 0 | NA | NA | 1 | 1 | NA | NA | 2 | 0.5 |
| 1_1851971 | 2 | 0.5 | 1 | 1 | 1 | 1 | 3 | 0.3333 | 7 | 0.5714 | 1 | 0 | NA | NA | 2 | 0.5 | NA | NA | 3 | 0.3333 |
| 1_1851972 | 2 | 0.5 | 1 | 0 | 2 | 0 | 2 | 0 | 7 | 0.1429 | 1 | 1 | NA | NA | 1 | 0 | 1 | 0 | 3 | 0.3333 |
| 1_1863567 | 1 | 0 | 2 | 1 | 1 | 0 | 1 | 1 | 5 | 0.6 | NA | NA | 1 | 0 | 1 | 1 | NA | NA | 2 | 0.5 |
| 1_1863569 | 2 | 1 | 1 | 1 | 1 | 1 | 1 | 1 | 5 | 1 | NA | NA | 1 | 1 | 2 | 1 | NA | NA | 3 | 1 |
| 1_1863570 | 1 | 1 | 1 | 1 | NA | NA | 1 | 1 | 3 | 1 | NA | NA | NA | NA | 2 | 1 | NA | NA | 2 | 1 |
| 1_1863571 | 2 | 1 | 1 | 1 | NA | NA | 1 | 1 | 4 | 1 | NA | NA | 1 | 1 | 1 | 1 | NA | NA | 2 | 1 |
| 1_1863573 | 1 | 0 | 1 | 1 | NA | NA | 1 | 0 | 3 | 0.3333 | NA | NA | 1 | 0 | 2 | 0.5 | NA | NA | 3 | 0.3333 |
| 1_1863573 | 2 | 0.5 | 1 | 1 | NA | NA | 1 | 1 | 4 | 0.75 | NA | NA | 1 | 0 | 2 | 0.5 | NA | NA | 3 | 0.3333 |
| 1_1863574 | 2 | 0 | 1 | 1 | NA | NA | 1 | 0 | 4 | 0.25 | NA | NA | NA | NA | 2 | 0.5 | NA | NA | 2 | 0.5 |
| 1_1863574 | 1 | 1 | 1 | 0 | NA | NA | NA | NA | 2 | 0.5 | NA | NA | 1 | 1 | 1 | 0 | NA | NA | 2 | 0.5 |
| 1_1863575 | 1 | 1 | NA | NA | NA | NA | NA | NA | 1 | 1 | NA | NA | NA | NA | 2 | 1 | NA | NA | 2 | 1 |
| 1_1864367 | 1 | 1 | NA | NA | NA | NA | 1 | 0 | 2 | 0.5 | NA | NA | NA | NA | 1 | 1 | NA | NA | 1 | 1 |
| 1_1864533 | 1 | 0 | 1 | 0 | 3 | 0.3333 | NA | NA | 5 | 0.2 | 1 | 0 | 1 | 0 | NA | NA | 2 | 0.5 | 4 | 0.25 |
| 1_1864534 | 1 | 1 | 2 | 0.5 | 2 | 0.5 | NA | NA | 5 | 0.6 | 1 | 0 | 1 | 1 | 1 | 0 | 2 | 0.5 | 5 | 0.4 |
| 1_1864535 | 1 | 1 | 1 | 1 | 3 | 1 | NA | NA | 5 | 1 | 1 | 1 | 1 | 1 | 1 | 0 | 1 | 1 | 4 | 0.75 |
| 1_1864536 | NA | NA | 1 | 1 | 2 | 0 | NA | NA | 3 | 0.3333 | 1 | 1 | 1 | 1 | 1 | 0 | 2 | 0 | 5 | 0.4 |
| 1_1864718 | 1 | 1 | NA | NA | 4 | 0.75 | NA | NA | 5 | 0.8 | 1 | 1 | 1 | 1 | 1 | 1 | 1 | 1 | 4 | 1 |
| 1_1864719 | 1 | 0 | NA | NA | 4 | 0.25 | NA | NA | 5 | 0.2 | NA | NA | 1 | 1 | 1 | 0 | 2 | 0 | 4 | 0.25 |
| 1_1869697 | NA | NA | NA | NA | 2 | 0.5 | NA | NA | 2 | 0.5 | NA | NA | 1 | 0 | 1 | 1 | NA | NA | 2 | 0.5 |
| 1_1870806 | 2 | 0.5 | 1 | 0 | 2 | 0 | 1 | 0 | 6 | 0.1667 | NA | NA | 1 | 1 | 2 | 0.5 | NA | NA | 3 | 0.6667 |
| 1_1870807 | 2 | 1 | 1 | 1 | 2 | 1 | 3 | 1 | 8 | 1 | NA | NA | 1 | 1 | 2 | 1 | NA | NA | 3 | 1 |
| 1_1870807 | 2 | 0 | 1 | 0 | 2 | 0.5 | 3 | 0 | 8 | 0.125 | NA | NA | NA | NA | 2 | 0.5 | NA | NA | 2 | 0.5 |
| 1_1870809 | 2 | 1 | 1 | 1 | 2 | 0.5 | 3 | 0.6667 | 8 | 0.75 | NA | NA | NA | NA | 2 | 0.5 | NA | NA | 2 | 0.5 |
| 1_1870810 | 2 | 0 | 1 | 1 | 2 | 0.5 | 4 | 0.25 | 9 | 0.3333 | NA | NA | 1 | 0 | 1 | 1 | NA | NA | 2 | 0.5 |
| 1_1870812 | 2 | 0.5 | NA | NA | 2 | 0.5 | 4 | 0.25 | 8 | 0.375 | NA | NA | 1 | 0 | 2 | 0.5 | NA | NA | 3 | 0.3333 |
| 1_1870812 | 2 | 0.5 | 1 | 0 | 2 | 0.5 | 3 | 0.3333 | 8 | 0.375 | NA | NA | 1 | 0 | 2 | 1 | NA | NA | 3 | 0.6667 |
| 1_1870814 | 2 | 0.5 | 2 | 0.5 | 2 | 0.5 | 4 | 0 | 10 | 0.3 | NA | NA | 1 | 1 | 2 | 0.5 | NA | NA | 3 | 0.6667 |
| 1_1870816 | 2 | 0 | 2 | 0.5 | 1 | 0 | 4 | 0.25 | 9 | 0.2222 | NA | NA | 1 | 0 | 2 | 0.5 | NA | NA | 3 | 0.3333 |
| 1_1870819 | 3 | 0 | 2 | 0 | 2 | 0 | 4 | 0.75 | 11 | 0.2727 | NA | NA | 1 | 0 | 2 | 0.5 | NA | NA | 3 | 0.3333 |
| 1_1870820 | 3 | 1 | 2 | 0 | 2 | 1 | 4 | 0.5 | 11 | 0.6364 | NA | NA | 1 | 0 | 1 | 1 | NA | NA | 2 | 0.5 |
| 1_1870822 | 3 | 0.6667 | 2 | 0 | 3 | 0.6667 | 4 | 0.5 | 12 | 0.5 | NA | NA | 1 | 1 | 2 | 0 | NA | NA | 3 | 0.3333 |

| | | | | | | | | | | | | | | | | | | | | |
|---|---|---|---|---|---|---|---|---|---|---|---|---|---|---|---|---|---|---|---|---|
| 1_1870822 | 3 | 1 | 2 | 0.5 | 3 | 1 | 2 | 1 | 10 | 0.9 | NA | NA | 1 | 1 | 1 | 1 | NA | NA | 2 | 1 |
| 1_1870823 | 1 | 1 | 2 | 1 | 4 | 1 | 4 | 1 | 11 | 1 | NA | NA | 1 | 1 | 2 | 1 | NA | NA | 3 | 1 |
| 1_1870825 | NA | NA | 2 | 0.5 | 2 | 0.5 | 3 | 0.3333 | 7 | 0.4286 | NA | NA | 1 | 1 | 1 | 0 | NA | NA | 2 | 0.5 |
| 1_1870825 | 3 | 0 | 2 | 0 | 3 | 0.3333 | 3 | 0 | 11 | 0.0909 | NA | NA | 1 | 1 | 2 | 1 | NA | NA | 3 | 1 |
| 1_1870825 | 3 | 0.3333 | 2 | 0 | 4 | 1 | 3 | 0.3333 | 12 | 0.5 | NA | NA | 1 | 1 | 2 | 0.5 | NA | NA | 3 | 0.6667 |
| 1_1870834 | 2 | 0 | 2 | 0 | 5 | 0.4 | 3 | 0.6667 | 12 | 0.3333 | 1 | 0 | 1 | 1 | 2 | 0 | NA | NA | 4 | 0.25 |
| 1_1877758 | NA | NA | 1 | 1 | NA | NA | 1 | 1 | 2 | 1 | 1 | 1 | NA | NA | 1 | 1 | NA | NA | 2 | 1 |
| 1_1879433 | NA | NA | 1 | 1 | 1 | 1 | NA | NA | 2 | 1 | 1 | 1 | NA | NA | NA | NA | NA | NA | 1 | 1 |
| 1_1879433 | NA | NA | 2 | 0 | 1 | 1 | NA | NA | 3 | 0.3333 | 1 | 1 | NA | NA | 1 | 1 | NA | NA | 2 | 1 |
| 1_1879434 | NA | NA | 2 | 1 | 1 | 1 | 1 | 1 | 4 | 1 | 2 | 1 | NA | NA | 1 | 1 | NA | NA | 3 | 1 |
| 1_1879438 | NA | NA | 2 | 0.5 | NA | NA | 1 | 0 | 3 | 0.3333 | 1 | 1 | NA | NA | 1 | 0 | NA | NA | 2 | 0.5 |
| 1_1879439 | NA | NA | 2 | 1 | 1 | 1 | 1 | 1 | 4 | 1 | 2 | 1 | NA | NA | 1 | 1 | NA | NA | 3 | 1 |
| 1_1879439 | NA | NA | 2 | 1 | 1 | 1 | 1 | 1 | 4 | 1 | 3 | 0.6667 | NA | NA | 1 | 0 | NA | NA | 4 | 0.5 |
| 1_1879443 | NA | NA | 3 | 0.6667 | 2 | 1 | 1 | 0 | 6 | 0.6667 | 3 | 0.6667 | NA | NA | 1 | 0 | NA | NA | 4 | 0.5 |
| 1_1879446 | NA | NA | NA | NA | 1 | 0 | 1 | 1 | 2 | 0.5 | 3 | 0.6667 | NA | NA | 1 | 0 | NA | NA | 4 | 0.5 |
| 1_1879447 | NA | NA | 2 | 0.5 | 2 | 1 | 2 | 1 | 6 | 0.8333 | 3 | 0.3333 | NA | NA | NA | NA | NA | NA | 3 | 0.3333 |
| 1_1879448 | NA | NA | 2 | 0.5 | 3 | 0.3333 | 2 | 0.5 | 7 | 0.4286 | 4 | 0.5 | NA | NA | 1 | 0 | NA | NA | 5 | 0.4 |
| 1_1879450 | 1 | 0 | 2 | 0.5 | 3 | 1 | 1 | 1 | 7 | 0.7143 | 4 | 0.25 | NA | NA | 1 | 0 | NA | NA | 5 | 0.2 |
| 1_1879451 | 1 | 1 | 3 | 0.3333 | 3 | 0.3333 | 2 | 0.5 | 9 | 0.4444 | 3 | 0.6667 | 1 | 1 | 1 | 0 | NA | NA | 5 | 0.6 |
| 1_1879456 | 1 | 0 | 3 | 0.6667 | 3 | 0 | 2 | 0 | 9 | 0.2222 | 4 | 0 | 1 | 0 | 1 | 1 | NA | NA | 6 | 0.1667 |
| 1_1879457 | NA | NA | 3 | 0.3333 | 3 | 0.3333 | 2 | 1 | 8 | 0.5 | 2 | 0 | 1 | 1 | NA | NA | NA | NA | 3 | 0.3333 |
| 1_1879465 | 2 | 0 | 4 | 0.5 | 4 | 0 | 2 | 0.5 | 12 | 0.25 | 4 | 0 | 2 | 0 | 1 | 1 | 1 | 0 | 8 | 0.125 |
| 1_1879466 | 2 | 0.5 | 5 | 0 | 5 | 0.4 | 2 | 0.5 | 14 | 0.2857 | 4 | 0.25 | 2 | 0 | 1 | 0 | 2 | 0.5 | 9 | 0.2222 |
| 1_1879467 | 2 | 0.5 | 6 | 0.5 | 4 | 0.75 | 2 | 0.5 | 14 | 0.5714 | 5 | 0.2 | 2 | 0.5 | 1 | 0 | 2 | 0.5 | 10 | 0.3 |
| 1_1879467 | 2 | 0 | 5 | 0.2 | 5 | 0.2 | 2 | 0 | 14 | 0.0714 | 4 | 0.25 | 2 | 0 | 1 | 0 | 2 | 0 | 9 | 0.1111 |
| 1_1887035 | 6 | 0.3333 | 12 | 0.4167 | 4 | 0.25 | 12 | 0.1667 | 34 | 0.2941 | 4 | 0.25 | 6 | 0.6667 | 6 | 0.3333 | 7 | 0.1429 | 23 | 0.3478 |
| 1_1887036 | 6 | 0.8333 | 14 | 0.5 | 5 | 0.2 | 14 | 0.4286 | 39 | 0.4872 | 5 | 0 | 6 | 0.6667 | 7 | 0.2857 | 8 | 0.5 | 26 | 0.3846 |
| 1_1887036 | 7 | 0.4286 | 12 | 0.75 | 3 | 0.3333 | 15 | 0.2 | 37 | 0.4324 | 5 | 0.6 | 5 | 0 | 7 | 0.2857 | 9 | 0.2222 | 26 | 0.2692 |
| 1_1887039 | 5 | 0.4 | 11 | 0.0909 | 4 | 0.25 | 10 | 0.1 | 30 | 0.1667 | 4 | 0.25 | 5 | 0.2 | 6 | 0.1667 | 6 | 0.1667 | 21 | 0.1905 |
| 1_1887041 | 7 | 0.1429 | 13 | 0.2308 | 5 | 0 | 14 | 0.2857 | 39 | 0.2051 | 4 | 0 | 6 | 0 | 9 | 0.2222 | 7 | 0.1429 | 26 | 0.1154 |
| 1_1887043 | 5 | 0.2 | 9 | 0.8889 | 2 | 0.5 | 10 | 0.3 | 26 | 0.5 | 3 | 0.6667 | 5 | 0.6 | 6 | 0.6667 | 7 | 0.2857 | 21 | 0.5238 |
| 1_1887045 | 3 | 0 | 3 | 0.3333 | 3 | 0 | 6 | 0 | 15 | 0.0667 | 3 | 0 | 3 | 0.3333 | 6 | 0 | 5 | 0.2 | 17 | 0.1176 |
| 1_1887045 | 6 | 0.3333 | 9 | 0.3333 | 4 | 0.5 | 10 | 0.2 | 29 | 0.3103 | 4 | 0.25 | 5 | 0.2 | 7 | 0.4286 | 9 | 0.3333 | 25 | 0.32 |
| 1_1887046 | 6 | 0.8333 | 7 | 0.1429 | 4 | 1 | 13 | 0.3077 | 30 | 0.4667 | 4 | 0.5 | 5 | 0.6 | 8 | 0.625 | 8 | 0.625 | 25 | 0.6 |
| 1_1887047 | 5 | 0 | 9 | 0.2222 | 5 | 0.2 | 14 | 0.0714 | 33 | 0.1212 | 4 | 0 | 5 | 0 | 7 | 0.1429 | 8 | 0.25 | 24 | 0.125 |
| 1_1887047 | 4 | 0.5 | 8 | 0.875 | 5 | 0.2 | 14 | 0.5 | 31 | 0.5484 | 3 | 0.3333 | 2 | 0.5 | 8 | 0.25 | 7 | 0.5714 | 20 | 0.4 |
| 1_1887047 | 5 | 0.8 | 8 | 0.375 | 5 | 0.8 | 14 | 0.5 | 32 | 0.5625 | 4 | 0.75 | 3 | 0.6667 | 9 | 0.5556 | 8 | 0.75 | 24 | 0.6667 |
| 1_1887047 | 5 | 0.2 | 8 | 0.75 | 5 | 0.8 | 14 | 0.8571 | 32 | 0.7188 | 3 | 1 | 3 | 1 | 7 | 0.8571 | 9 | 0.8889 | 22 | 0.9091 |
| 1_1889407 | 7 | 0.5714 | 10 | 0.8 | 10 | 0.7 | 11 | 0.6364 | 38 | 0.6842 | 9 | 0.5556 | 13 | 0.7692 | 10 | 0.6 | 5 | 0.4 | 37 | 0.6216 |
| 1_1889408 | 6 | 0 | 7 | 0 | 11 | 0.3636 | 11 | 0 | 35 | 0.1143 | 9 | 0 | 10 | 0.2 | 9 | 0.1111 | 7 | 0.1429 | 35 | 0.1143 |
| 1_1889409 | 6 | 0.1667 | 6 | 0.1667 | 10 | 0.4 | 10 | 0.2 | 32 | 0.25 | 9 | 0.2222 | 11 | 0.0909 | 9 | 0.3333 | 7 | 0.2857 | 36 | 0.2222 |
| 1_1889410 | 5 | 0.6 | 8 | 0.75 | 11 | 0.5455 | 11 | 0.6364 | 35 | 0.6286 | 7 | 0.7143 | 9 | 0.7778 | 10 | 0.6 | 9 | 0.4444 | 35 | 0.6286 |
| 1_1889410 | 4 | 0.25 | 6 | 0.6667 | 10 | 0.3 | 9 | 0.4444 | 29 | 0.4138 | 8 | 0.125 | 8 | 0.25 | 5 | 0 | 7 | 0.1429 | 28 | 0.1429 |
| 1_1889414 | 6 | 0 | 5 | 0.4 | 7 | 0.2857 | 8 | 0.5 | 26 | 0.3077 | 10 | 0.2 | 10 | 0.4 | 7 | 0.5714 | 7 | 0.5714 | 34 | 0.4118 |
| 1_1889417 | 5 | 0.4 | 6 | 0.5 | 7 | 0.5714 | 9 | 0.3333 | 27 | 0.4444 | 9 | 0.3333 | 6 | 0.5 | 8 | 0.5 | 8 | 0.5 | 31 | 0.4516 |
| 1_1889417 | 4 | 0.75 | 7 | 0.5714 | 7 | 0.7143 | 6 | 0.5 | 24 | 0.625 | 5 | 0.8 | 2 | 1 | 4 | 1 | 8 | 0.25 | 19 | 0.6316 |
| 1_1889419 | 5 | 1 | 6 | 1 | 6 | 1 | 8 | 1 | 25 | 1 | 9 | 1 | 8 | 1 | 6 | 1 | 9 | 1 | 32 | 1 |
| 1_1889419 | 4 | 0 | 5 | 0.8 | 5 | 0.6 | 6 | 0.3333 | 20 | 0.45 | 7 | 0.4286 | 4 | 0.75 | 6 | 0.5 | 8 | 0.625 | 25 | 0.56 |
| 1_1889420 | 4 | 0.25 | 5 | 1 | 6 | 0.6667 | 8 | 0.625 | 23 | 0.6522 | 8 | 0.625 | 9 | 0.8889 | 7 | 0.5714 | 9 | 0.8889 | 33 | 0.7576 |
| 1_1889421 | 3 | 0 | 4 | 0 | 3 | 0 | 7 | 0.1429 | 17 | 0.0588 | 8 | 0 | 5 | 0.2 | 5 | 0.2 | 7 | 0 | 25 | 0.08 |
| 1_1889421 | 6 | 0.1667 | 6 | 0.5 | 5 | 0.2 | 7 | 0.1429 | 24 | 0.25 | 8 | 0.125 | 9 | 0.1111 | 7 | 0 | 9 | 0.2222 | 33 | 0.1212 |
| 1_1889421 | 5 | 0.2 | 5 | 0.2 | 4 | 0 | 5 | 0 | 19 | 0.1053 | 5 | 0 | 5 | 0 | 2 | 0 | 6 | 0.1667 | 18 | 0.0556 |
| 1_1889421 | 1 | 1 | 3 | 1 | 4 | 1 | 4 | 0.75 | 12 | 0.9167 | 2 | 1 | 3 | 0.6667 | 2 | 1 | 5 | 0.8 | 12 | 0.8333 |
| 1_1889422 | 5 | 0.2 | 3 | 0 | 6 | 0.1667 | 5 | 0 | 19 | 0.1053 | 5 | 0 | 4 | 0 | 6 | 0.1667 | 6 | 0.1667 | 21 | 0.0952 |
| 1_1890721 | 4 | 0.5 | 4 | 0 | 3 | 0 | 6 | 0 | 17 | 0.1176 | 6 | 0.1667 | 6 | 0 | 3 | 0 | 8 | 0.25 | 23 | 0.1304 |
| 1_1890757 | 1 | 0 | 1 | 0 | 1 | 0 | 5 | 0.2 | 8 | 0.125 | 3 | 0 | 3 | 0.3333 | 2 | 0 | 5 | 0 | 13 | 0.0769 |
| 1_1890759 | 1 | 0 | NA | NA | NA | NA | 3 | 0.3333 | 4 | 0.25 | NA | NA | 3 | 0 | NA | NA | 3 | 0.3333 | 6 | 0.1667 |
| 1_1890765 | 1 | 1 | 2 | 0.5 | 1 | 1 | NA | NA | 4 | 0.75 | NA | NA | 1 | 1 | 1 | 1 | NA | NA | 2 | 1 |
| 1_1890767 | 1 | 1 | 2 | 1 | 1 | 1 | NA | NA | 4 | 1 | NA | NA | 1 | 1 | 2 | 0.5 | 1 | 0 | 4 | 0.5 |
| 1_1890767 | 1 | 0 | 1 | 1 | 1 | 1 | NA | NA | 3 | 0.6667 | NA | NA | 1 | 1 | 2 | 0 | 1 | 0 | 4 | 0.25 |
| 1_1890768 | 1 | 0 | 2 | 0.5 | 1 | 1 | NA | NA | 4 | 0.5 | NA | NA | 1 | 1 | 2 | 1 | 1 | 0 | 4 | 0.75 |
| 1_1890769 | 1 | 1 | 4 | 0.75 | 1 | 1 | NA | NA | 6 | 0.8333 | NA | NA | 1 | 1 | 1 | 0 | 1 | 0 | 3 | 0.3333 |
| 1_1893746 | NA | NA | 1 | 1 | NA | NA | NA | NA | 1 | 1 | NA | NA | NA | NA | 1 | 1 | NA | NA | 1 | 1 |
| 1_1893747 | NA | NA | 1 | 1 | NA | NA | NA | NA | 1 | 1 | NA | NA | NA | NA | 1 | 1 | NA | NA | 1 | 1 |
| 1_1896129 | 7 | 0.7143 | 10 | 0.7 | 10 | 0.8 | 8 | 0.625 | 35 | 0.7143 | 11 | 0.4545 | 2 | 0.5 | 4 | 0.75 | 4 | 0.5 | 21 | 0.5238 |

| | | | | | | | | | | | | | | | | | | | | |
|---|---|---|---|---|---|---|---|---|---|---|---|---|---|---|---|---|---|---|---|---|
| 1_19050529 | 3 | 0.3333 | NA | NA | NA | NA | 4 | 0 | 7 | 0.1429 | NA | NA | 1 | 0 | 2 | 0.5 | 1 | 0 | 4 | 0.25 |
| 1_19050531 | 4 | 0.75 | 2 | 1 | 2 | 1 | 7 | 1 | 15 | 0.9333 | 4 | 0.75 | 6 | 1 | 4 | 1 | 4 | 1 | 18 | 0.9444 |
| 1_19050533 | 4 | 0 | 3 | 0.6667 | 2 | 0 | 5 | 0.4 | 14 | 0.2857 | 2 | 0.5 | 3 | 0 | 4 | 0.5 | 5 | 0 | 14 | 0.2143 |
| 1_19050533 | 1 | 0 | NA | NA | NA | NA | 2 | 0.5 | 3 | 0.3333 | NA | NA | 3 | 0.6667 | 1 | 0 | NA | NA | 4 | 0.5 |
| 1_19050537 | 4 | 0 | 3 | 0.3333 | 2 | 0 | 5 | 0 | 14 | 0.0714 | 3 | 0.3333 | 6 | 0 | 6 | 0 | 5 | 0 | 20 | 0.05 |
| 1_19050538 | 5 | 0.8 | 3 | 0 | 2 | 1 | 7 | 0.7143 | 17 | 0.6471 | 4 | 0.5 | 6 | 0.5 | 6 | 0.6667 | 3 | 1 | 19 | 0.6316 |
| 1_19050540 | 2 | 0 | 2 | 0 | 2 | 0 | 5 | 0.4 | 11 | 0.1818 | 3 | 0.3333 | 4 | 0.75 | 4 | 0.25 | 2 | 0 | 13 | 0.3846 |
| 1_19050542 | 4 | 0 | 3 | 0 | 1 | 0 | 7 | 0.1429 | 15 | 0.0667 | 4 | 0 | 6 | 0.1667 | 6 | 0 | 4 | 0 | 20 | 0.05 |
| 1_19050545 | 5 | 1 | 3 | 1 | 2 | 1 | 6 | 1 | 16 | 1 | 2 | 1 | 7 | 1 | 5 | 1 | 4 | 1 | 18 | 1 |
| 1_19050549 | 4 | 0.5 | 3 | 0.6667 | 2 | 1 | 7 | 0.2857 | 16 | 0.5 | 4 | 0.75 | 7 | 0.5714 | 6 | 0.3333 | 4 | 0.75 | 21 | 0.5714 |
| 1_19050551 | 5 | 0.4 | 2 | 0.5 | 2 | 1 | 7 | 0.5714 | 16 | 0.5625 | 4 | 0.25 | 6 | 0.1667 | 6 | 0.3333 | 4 | 0.25 | 20 | 0.25 |
| 1_19050568 | 5 | 1 | 1 | 0 | 2 | 1 | 8 | 0.75 | 16 | 0.8125 | 4 | 1 | 6 | 1 | 6 | 0.8333 | 4 | 1 | 20 | 0.95 |
| 1_19050569 | 4 | 0.75 | 3 | 1 | 2 | 0.5 | 7 | 0.1429 | 16 | 0.5 | 4 | 0.75 | 7 | 0.5714 | 6 | 1 | 4 | 0.75 | 21 | 0.7619 |
| 1_19052833 | NA | NA | 3 | 1 | NA | NA | NA | NA | 3 | 1 | NA | NA | 1 | 1 | NA | NA | NA | NA | 1 | 1 |
| 1_19052853 | NA | NA | 2 | 0.5 | NA | NA | NA | NA | 2 | 0.5 | NA | NA | 1 | 1 | NA | NA | NA | NA | 1 | 1 |
| 1_19053057 | NA | NA | 2 | 0.5 | NA | NA | 1 | 0 | 3 | 0.3333 | NA | NA | 1 | 1 | NA | NA | NA | NA | 1 | 1 |
| 1_19053070 | NA | NA | 2 | 0.5 | NA | NA | 1 | 1 | 3 | 0.6667 | NA | NA | 1 | 1 | NA | NA | NA | NA | 1 | 1 |
| 1_19053093 | 4 | 0 | 3 | 0 | NA | NA | 8 | 0.125 | 15 | 0.0667 | 2 | 0.5 | 4 | 0 | 3 | 0 | 2 | 0.5 | 11 | 0.1818 |
| 1_19053120 | NA | NA | 1 | 1 | NA | NA | 1 | 1 | 2 | 1 | NA | NA | 1 | 1 | NA | NA | NA | NA | 1 | 1 |
| 1_19053136 | 4 | 0.25 | 3 | 0.3333 | NA | NA | 8 | 0.375 | 15 | 0.3333 | 2 | 0 | 4 | 0.25 | 3 | 0.6667 | 2 | 0 | 11 | 0.2727 |
| 1_19053143 | 3 | 0.3333 | 3 | 0.3333 | NA | NA | 8 | 0.25 | 14 | 0.2857 | 2 | 0.5 | 3 | 0.3333 | 2 | 0 | 2 | 0 | 9 | 0.2222 |
| 1_19053176 | 4 | 1 | 2 | 1 | NA | NA | 8 | 0.875 | 14 | 0.9286 | 2 | 1 | 4 | 0.75 | 2 | 1 | 2 | 1 | 10 | 0.9 |
| 1_19053180 | 3 | 0 | 2 | 0.5 | NA | NA | 7 | 0.5714 | 12 | 0.4167 | NA | NA | 2 | 0 | 1 | 1 | 1 | 0 | 4 | 0.25 |
| 1_19053189 | 4 | 1 | 2 | 1 | NA | NA | 7 | 0.8571 | 13 | 0.9231 | 1 | 1 | 4 | 0.75 | 2 | 1 | 2 | 1 | 9 | 0.8889 |
| 1_19053192 | 3 | 0.3333 | 2 | 0 | NA | NA | 6 | 0.1667 | 11 | 0.1818 | 1 | 0 | 3 | 0 | 2 | 0.5 | 2 | 0 | 8 | 0.125 |
| 1_19053204 | 4 | 0.75 | 1 | 1 | NA | NA | 7 | 0.5714 | 12 | 0.6667 | 1 | 0 | 3 | 0.6667 | 2 | 0.5 | 2 | 0.5 | 8 | 0.5 |
| 1_19053217 | 4 | 1 | 1 | 1 | NA | NA | 6 | 1 | 11 | 1 | 2 | 1 | 3 | 0.6667 | 2 | 1 | 2 | 1 | 9 | 0.8889 |
| 1_19053220 | 3 | 0.6667 | 1 | 0 | NA | NA | 4 | 0.25 | 8 | 0.375 | 2 | 0 | 2 | 0 | 1 | 0 | 2 | 1 | 7 | 0.2857 |
| 1_19053238 | 3 | 0 | 1 | 0 | NA | NA | 5 | 0.2 | 9 | 0.1111 | 2 | 0 | 3 | 0 | 2 | 0 | 2 | 0.5 | 9 | 0.1111 |
| 1_19053243 | 2 | 0.5 | 1 | 1 | NA | NA | 5 | 0.8 | 8 | 0.75 | 2 | 1 | 3 | 0.6667 | 2 | 1 | 2 | 1 | 9 | 0.8889 |
| 1_19053271 | 1 | 0 | 1 | 0 | NA | NA | 2 | 0.5 | 4 | 0.25 | 2 | 0.5 | NA | NA | 1 | 0 | 2 | 0.5 | 5 | 0.4 |
| 1_19066518 | 4 | 0.75 | 1 | 0 | 3 | 0.3333 | NA | NA | 8 | 0.5 | 1 | 0 | 2 | 1 | 6 | 0 | 1 | 0 | 10 | 0.2 |
| 1_19066523 | 4 | 0.75 | NA | NA | 2 | 1 | 1 | 0 | 7 | 0.7143 | 1 | 1 | 2 | 0.5 | 5 | 0.8 | 1 | 1 | 9 | 0.7778 |
| 1_19067320 | 4 | 0.25 | 1 | 0 | 7 | 0 | 2 | 0 | 14 | 0.0714 | NA | NA | 5 | 0 | 6 | 0.1667 | 1 | 0 | 12 | 0.0833 |
| 1_19067320 | 4 | 0.5 | 3 | 0 | 7 | 0.1429 | 3 | 0.3333 | 17 | 0.2353 | NA | NA | 5 | 0.6 | 7 | 0.1429 | 2 | 1 | 14 | 0.4286 |
| 1_19067331 | 4 | 0 | 3 | 0.3333 | 7 | 0.4286 | 3 | 0 | 17 | 0.2353 | 1 | 0 | 7 | 0 | 6 | 0.3333 | 1 | 0 | 15 | 0.1333 |
| 1_19067341 | 3 | 0.6667 | 3 | 0.3333 | 7 | 0.2857 | 3 | 1 | 16 | 0.5 | NA | NA | 4 | 0 | 5 | 0.6 | 1 | 0 | 10 | 0.3 |
| 1_19067718 | 5 | 0.2 | 3 | 0 | 6 | 0 | 3 | 0 | 17 | 0.0588 | 2 | 0 | 8 | 0.125 | 8 | 0.375 | 3 | 0 | 21 | 0.1905 |
| 1_19067721 | 5 | 0.6 | 3 | 1 | 6 | 0.1667 | 3 | 0.3333 | 17 | 0.4706 | 1 | 1 | 5 | 0.6 | 7 | 0.5714 | 3 | 1 | 16 | 0.6875 |
| 1_19067728 | 6 | 0 | 3 | 0.3333 | 6 | 0.1667 | 3 | 0 | 18 | 0.1111 | 2 | 0 | 6 | 0 | 7 | 0.1429 | 3 | 0 | 18 | 0.0556 |
| 1_19067730 | 6 | 0.5 | 2 | 0 | 7 | 0.8571 | 2 | 1 | 17 | 0.6471 | 2 | 1 | 7 | 0.7143 | 6 | 0.6667 | 3 | 1 | 18 | 0.7778 |
| 1_19067748 | 5 | 0 | 3 | 0.3333 | 6 | 0.5 | 2 | 0.5 | 16 | 0.3125 | 2 | 0 | 8 | 0.25 | 8 | 0.375 | 3 | 0.6667 | 21 | 0.3333 |
| 1_19067751 | 6 | 0.1667 | 2 | 0 | 6 | 0 | 3 | 0 | 17 | 0.0588 | 2 | 0 | 7 | 0.1429 | 7 | 0 | 3 | 0 | 19 | 0.0526 |
| 1_19073938 | NA | NA | NA | NA | 1 | 1 | 1 | 0 | 2 | 0.5 | NA | NA | 1 | 1 | NA | NA | 1 | 0 | 2 | 0.5 |
| 1_19075780 | NA | NA | 1 | 1 | 1 | 1 | NA | NA | 2 | 1 | NA | NA | NA | NA | 1 | 1 | NA | NA | 1 | 1 |
| 1_19075790 | NA | NA | 1 | 1 | NA | NA | NA | NA | 1 | 1 | NA | NA | NA | NA | 1 | 1 | NA | NA | 1 | 1 |
| 1_19077803 | 1 | 0 | 2 | 1 | NA | NA | NA | NA | 3 | 0.6667 | NA | NA | 1 | 1 | NA | NA | 1 | 1 | 2 | 1 |
| 1_19077806 | 3 | 0 | 4 | 0 | 2 | 0.5 | 2 | 0 | 11 | 0.0909 | NA | NA | 2 | 0 | NA | NA | 2 | 0.5 | 4 | 0.25 |
| 1_19077817 | 2 | 0.5 | 4 | 0.5 | 2 | 1 | 1 | 1 | 9 | 0.6667 | NA | NA | 2 | 1 | NA | NA | 2 | 0 | 4 | 0.5 |
| 1_19108211 | 2 | 0.5 | 1 | 1 | 1 | 0 | 2 | 0 | 6 | 0.3333 | 1 | 0 | NA | NA | 2 | 1 | 1 | 0 | 4 | 0.5 |
| 1_19108212 | 2 | 0 | 1 | 0 | 1 | 1 | 2 | 0.5 | 6 | 0.3333 | 1 | 1 | NA | NA | 2 | 0 | 1 | 0 | 4 | 0.25 |
| 1_19108222 | 2 | 1 | 1 | 1 | 1 | 1 | 2 | 0.5 | 6 | 0.8333 | 1 | 1 | 1 | 1 | 2 | 0.5 | 1 | 1 | 5 | 0.8 |
| 1_19108232 | 1 | 1 | 1 | 1 | 2 | 1 | 2 | 1 | 6 | 1 | 1 | 1 | 1 | 1 | 2 | 1 | 1 | 1 | 5 | 1 |
| 1_19108470 | 7 | 0 | 18 | 0.2778 | 11 | 0 | 9 | 0 | 45 | 0.1111 | 5 | 0.2 | 15 | 0.2 | 15 | 0.1333 | 13 | 0.2308 | 48 | 0.1875 |
| 1_19108503 | 7 | 0.7143 | 18 | 0.8889 | 9 | 0.7778 | 9 | 0.7778 | 43 | 0.814 | 6 | 0.8333 | 15 | 0.9333 | 15 | 1 | 12 | 0.8333 | 48 | 0.9167 |
| 1_19110478 | 3 | 0.3333 | 10 | 0.5 | 7 | 0.2857 | 6 | 0.1667 | 26 | 0.3462 | 2 | 0.5 | 10 | 0.4 | 11 | 0.3636 | 8 | 0.375 | 31 | 0.3871 |
| 1_19112895 | 1 | 0 | 4 | 0.25 | 2 | 0 | 5 | 0.2 | 12 | 0.1667 | 2 | 0.5 | 8 | 0.375 | 5 | 0 | 3 | 0.3333 | 18 | 0.2778 |
| 1_19127108 | NA | NA | 1 | 1 | NA | NA | NA | NA | 1 | 1 | 1 | 1 | NA | NA | NA | NA | NA | NA | 1 | 1 |
| 1_19127133 | NA | NA | 1 | 1 | NA | NA | NA | NA | 1 | 1 | 2 | 1 | NA | NA | NA | NA | 1 | 1 | 3 | 1 |
| 1_19127292 | NA | NA | 1 | 1 | NA | NA | NA | NA | 1 | 1 | 2 | 0.5 | NA | NA | NA | NA | 1 | 1 | 3 | 0.6667 |
| 1_19135335 | 2 | 0 | 3 | 0.3333 | 3 | 0.3333 | 3 | 0.6667 | 11 | 0.3636 | 3 | 0.6667 | 3 | 0.6667 | 4 | 1 | 3 | 0 | 13 | 0.6154 |
| 1_19135343 | 2 | 0.5 | 3 | 0.6667 | 3 | 0.6667 | 3 | 0.6667 | 11 | 0.6364 | 3 | 1 | 3 | 0.3333 | 3 | 0.6667 | 2 | 0.5 | 11 | 0.6364 |
| 1_19135344 | 2 | 0 | 3 | 0 | 3 | 0.3333 | 3 | 0.3333 | 11 | 0.1818 | 3 | 0.3333 | 3 | 0 | 5 | 0.2 | 3 | 0 | 14 | 0.1429 |
| 1_19135347 | 2 | 0 | 3 | 0 | 3 | 0.3333 | 3 | 0 | 11 | 0.0909 | 3 | 0 | 4 | 0 | 5 | 0 | 4 | 0.25 | 16 | 0.0625 |
| 1_19135361 | 3 | 0.3333 | 3 | 0.3333 | 3 | 0.3333 | 3 | 0 | 12 | 0.25 | 2 | 0 | 4 | 0 | 5 | 0.4 | 4 | 0 | 15 | 0.1333 |

| | | | | | | | | | | | | | | | | | | | | |
|---|---|---|---|---|---|---|---|---|---|---|---|---|---|---|---|---|---|---|---|---|
| 1_1913543 | 3 | 0.3333 | 3 | 0 | 4 | 0.5 | 6 | 0.1667 | 16 | 0.25 | 3 | 0.3333 | 3 | 0.3333 | 4 | 0.75 | 4 | 0.25 | 14 | 0.4286 |
| 1_1913545 | 4 | 0.25 | 3 | 0 | 4 | 0 | 6 | 0 | 17 | 0.0588 | 3 | 0 | 4 | 0 | 3 | 0.3333 | 4 | 0 | 14 | 0.0714 |
| 1_1913545 | 4 | 0.5 | 3 | 0.3333 | 4 | 0 | 4 | 0.5 | 15 | 0.3333 | 2 | 0 | 3 | 0.3333 | 3 | 0.3333 | 4 | 0.5 | 12 | 0.3333 |
| 1_1913546 | 4 | 0.25 | 4 | 0.5 | 3 | 0 | 6 | 0.8333 | 17 | 0.4706 | 4 | 0 | 3 | 0.3333 | 4 | 0.5 | 5 | 0.6 | 16 | 0.375 |
| 1_1913548 | 1 | 1 | 1 | 0 | NA | NA | NA | NA | 2 | 0.5 | 1 | 1 | NA | NA | NA | NA | NA | NA | 1 | 1 |
| 1_1913555 | 1 | 1 | 1 | 0 | NA | NA | NA | NA | 2 | 0.5 | 2 | 0.5 | NA | NA | NA | NA | NA | NA | 2 | 0.5 |
| 1_1914516 | 2 | 0.5 | 3 | 0 | 3 | 0.3333 | 4 | 0.25 | 12 | 0.25 | 1 | 1 | NA | NA | 4 | 0 | 2 | 0.5 | 7 | 0.2857 |
| 1_1914516 | 2 | 1 | 4 | 1 | 4 | 0.5 | 6 | 0.6667 | 16 | 0.75 | 3 | 0.6667 | 1 | 1 | 5 | 0.6 | 4 | 0.5 | 13 | 0.6154 |
| 1_1915106 | 2 | 0.5 | 3 | 0.3333 | 7 | 0.4286 | 8 | 0.625 | 20 | 0.5 | 2 | 0.5 | 2 | 0.5 | 8 | 0.25 | 8 | 0.25 | 20 | 0.3 |
| 1_1915106 | 2 | 0.5 | 3 | 1 | 6 | 0.8333 | 4 | 1 | 15 | 0.8667 | 1 | 1 | 2 | 1 | 6 | 1 | 8 | 1 | 17 | 1 |
| 1_1915106 | 2 | 0.5 | 3 | 0.3333 | 8 | 0.375 | 7 | 0.4286 | 20 | 0.4 | 3 | 0.6667 | 2 | 0 | 7 | 0.2857 | 10 | 0.3 | 22 | 0.3182 |
| 1_1915107 | NA | NA | 3 | 0.3333 | 9 | 0.1111 | 6 | 0.1667 | 18 | 0.1667 | 3 | 0.3333 | NA | NA | 6 | 0 | 8 | 0.625 | 17 | 0.3529 |
| 1_1915108 | 1 | 1 | 3 | 1 | 5 | 0.8 | 5 | 0.8 | 14 | 0.8571 | 1 | 1 | 1 | 1 | 4 | 1 | 9 | 0.8889 | 15 | 0.9333 |
| 1_1915109 | NA | NA | 5 | 0.4 | 8 | 0.125 | 8 | 0.375 | 21 | 0.2857 | 4 | 0 | 1 | 0 | 7 | 0.2857 | 8 | 0.125 | 20 | 0.15 |
| 1_1915109 | NA | NA | 4 | 0 | 8 | 0.125 | 7 | 0.1429 | 19 | 0.1053 | 4 | 0 | 1 | 0 | 6 | 0.3333 | 9 | 0.2222 | 20 | 0.2 |
| 1_1915110 | 1 | 0 | 4 | 0.5 | 8 | 0.25 | 7 | 0.1429 | 20 | 0.25 | 4 | 0.25 | 1 | 1 | 5 | 0 | 9 | 0 | 19 | 0.1053 |
| 1_1915124 | 4 | 0.25 | 7 | 0.1429 | 9 | 0 | 11 | 0 | 31 | 0.0645 | 8 | 0 | 2 | 0 | 10 | 0 | 11 | 0.1818 | 31 | 0.0645 |
| 1_1915135 | 5 | 0.2 | 15 | 0.2 | 15 | 0.1333 | 13 | 0.1538 | 48 | 0.1667 | 8 | 0.125 | 5 | 0.2 | 9 | 0.4444 | 14 | 0.0714 | 36 | 0.1944 |
| 1_1915136 | 5 | 0.4 | 15 | 0.6667 | 16 | 0.6875 | 16 | 0.75 | 52 | 0.6731 | 7 | 0.7143 | 4 | 1 | 12 | 0.6667 | 13 | 0.7692 | 36 | 0.75 |
| 1_1915137 | 4 | 0 | 15 | 0.0667 | 16 | 0.0625 | 16 | 0.1875 | 51 | 0.098 | 9 | 0 | 5 | 0.2 | 13 | 0.2308 | 14 | 0.1429 | 41 | 0.1463 |
| 1_1915138 | 5 | 0.2 | 16 | 0.125 | 17 | 0.4706 | 18 | 0.5556 | 56 | 0.375 | 9 | 0.3333 | 5 | 0.2 | 16 | 0.5625 | 16 | 0.5625 | 46 | 0.4783 |
| 1_1915142 | 7 | 0 | 15 | 0.2 | 17 | 0.2353 | 16 | 0.0625 | 55 | 0.1455 | 9 | 0.1111 | 7 | 0.1429 | 16 | 0.1875 | 17 | 0.1765 | 49 | 0.1633 |
| 1_1915142 | 7 | 0 | 16 | 0.0625 | 17 | 0.0588 | 20 | 0.1 | 60 | 0.0667 | 10 | 0.1 | 8 | 0.25 | 17 | 0.2353 | 17 | 0.0588 | 52 | 0.1538 |
| 1_1915157 | 15 | 0.2 | 22 | 0.0909 | 20 | 0.2 | 21 | 0.1429 | 78 | 0.1538 | 13 | 0.2308 | 11 | 0.2727 | 18 | 0.1111 | 19 | 0.1579 | 61 | 0.1803 |
| 1_1916411 | 2 | 0.5 | NA | NA | NA | NA | 3 | 0.6667 | 5 | 0.6 | NA | NA | 2 | 0 | NA | NA | 2 | 1 | 4 | 0.5 |
| 1_1916412 | 3 | 0.3333 | NA | NA | NA | NA | 3 | 0.6667 | 6 | 0.5 | NA | NA | 2 | 1 | NA | NA | 2 | 0.5 | 4 | 0.75 |
| 1_1916413 | 3 | 0.6667 | NA | NA | NA | NA | 3 | 1 | 6 | 0.8333 | NA | NA | 3 | 1 | NA | NA | 1 | 1 | 4 | 1 |
| 1_1916415 | 3 | 0.6667 | NA | NA | NA | NA | 3 | 0.3333 | 6 | 0.5 | NA | NA | 3 | 1 | NA | NA | 2 | 1 | 5 | 1 |
| 1_1916416 | 2 | 1 | NA | NA | NA | NA | 3 | 1 | 5 | 1 | NA | NA | 2 | 0.5 | NA | NA | 3 | 1 | 5 | 0.8 |
| 1_1916416 | 3 | 0.6667 | NA | NA | NA | NA | 3 | 0.6667 | 6 | 0.6667 | NA | NA | 3 | 0.6667 | NA | NA | 2 | 0 | 5 | 0.4 |
| 1_1916417 | 3 | 0.6667 | NA | NA | NA | NA | 3 | 1 | 6 | 0.8333 | NA | NA | 3 | 1 | NA | NA | 2 | 1 | 5 | 1 |
| 1_1916417 | 5 | 1 | NA | NA | NA | NA | 3 | 1 | 8 | 1 | NA | NA | 3 | 0.6667 | NA | NA | 3 | 1 | 6 | 0.8333 |
| 1_1916418 | 4 | 1 | NA | NA | NA | NA | 3 | 1 | 7 | 1 | NA | NA | 3 | 1 | NA | NA | 2 | 1 | 5 | 1 |
| 1_1916419 | 5 | 0.2 | NA | NA | NA | NA | 2 | 0 | 7 | 0.1429 | 1 | 0 | 4 | 0.5 | NA | NA | 3 | 0 | 8 | 0.25 |
| 1_1916419 | 4 | 1 | NA | NA | NA | NA | 2 | 1 | 6 | 1 | 1 | 1 | 4 | 1 | NA | NA | 3 | 1 | 8 | 1 |
| 1_1916420 | 5 | 0.2 | 1 | 0 | NA | NA | 1 | 0 | 7 | 0.1429 | 1 | 0 | 4 | 0.25 | NA | NA | 3 | 0.3333 | 8 | 0.25 |
| 1_1916420 | 4 | 1 | NA | NA | NA | NA | 2 | 1 | 6 | 1 | 1 | 1 | 3 | 0.6667 | NA | NA | 3 | 1 | 7 | 0.8571 |
| 1_1916442 | 5 | 0.4 | 6 | 0.5 | 2 | 1 | 4 | 0.5 | 17 | 0.5294 | 4 | 0 | 4 | 0.5 | 3 | 1 | 2 | 1 | 13 | 0.5385 |
| 1_1916443 | 5 | 0 | 6 | 0.1667 | 2 | 0 | 5 | 0 | 18 | 0.0556 | 3 | 0 | 4 | 0.25 | 3 | 0 | 2 | 0 | 12 | 0.0833 |
| 1_1916447 | 6 | 0.8333 | 7 | 0.5714 | 2 | 1 | 4 | 0.5 | 19 | 0.6842 | 3 | 0.6667 | 5 | 0.2 | 4 | 0.75 | 3 | 0.6667 | 15 | 0.5333 |
| 1_1917581 | 5 | 0 | 7 | 0.1429 | 4 | 0.25 | 3 | 0.3333 | 19 | 0.1579 | 13 | 0 | 5 | 0 | 1 | 0 | 3 | 0.6667 | 22 | 0.0909 |
| 1_1917843 | 2 | 0.5 | 2 | 0.5 | 2 | 0 | 4 | 0.75 | 10 | 0.5 | 1 | 0 | 3 | 0 | 1 | 1 | 1 | 0 | 6 | 0.1667 |
| 1_1917849 | 1 | 1 | 1 | 1 | 1 | 1 | 3 | 1 | 6 | 1 | 1 | 1 | 2 | 1 | 1 | 1 | NA | NA | 4 | 1 |
| 1_1917850 | NA | NA | 1 | 1 | 1 | 1 | 2 | 1 | 4 | 1 | NA | NA | 1 | 0 | 1 | 1 | NA | NA | 2 | 0.5 |
| 1_1917851 | 1 | 1 | 1 | 1 | 1 | 0 | 2 | 1 | 5 | 0.8 | 1 | 0 | 2 | 1 | 1 | 0 | NA | NA | 4 | 0.5 |
| 1_1920198 | 2 | 0.5 | 1 | 1 | NA | NA | NA | NA | 3 | 0.6667 | NA | NA | 3 | 0.3333 | 2 | 0.5 | 1 | 1 | 6 | 0.5 |
| 1_1920199 | 2 | 0.5 | 1 | 1 | NA | NA | NA | NA | 3 | 0.6667 | NA | NA | 3 | 0.6667 | 2 | 1 | 2 | 0.5 | 7 | 0.7143 |
| 1_1920200 | 2 | 1 | 1 | 1 | NA | NA | NA | NA | 3 | 1 | NA | NA | 3 | 0.6667 | 3 | 0.6667 | 2 | 0.5 | 8 | 0.625 |
| 1_1921876 | 4 | 0.5 | 2 | 0 | NA | NA | NA | NA | 6 | 0.3333 | NA | NA | 5 | 0.2 | 3 | 0 | 2 | 0 | 10 | 0.1 |
| 1_1921877 | 4 | 0 | 2 | 0.5 | NA | NA | NA | NA | 6 | 0.1667 | NA | NA | 5 | 0 | 3 | 0 | 2 | 0.5 | 10 | 0.1 |
| 1_1921877 | 4 | 1 | 2 | 1 | NA | NA | NA | NA | 6 | 1 | NA | NA | 5 | 1 | 3 | 1 | 2 | 1 | 10 | 1 |
| 1_1921881 | 4 | 0.5 | 2 | 0.5 | NA | NA | NA | NA | 6 | 0.5 | NA | NA | 7 | 0.8571 | 4 | 0.75 | 3 | 0.6667 | 14 | 0.7857 |
| 1_1921884 | 4 | 0.25 | 3 | 0 | NA | NA | NA | NA | 7 | 0.1429 | NA | NA | 6 | 0 | 4 | 0.25 | 3 | 0 | 13 | 0.0769 |
| 1_1921884 | 5 | 1 | 3 | 1 | NA | NA | NA | NA | 8 | 1 | NA | NA | 7 | 1 | 4 | 1 | 3 | 1 | 14 | 1 |
| 1_1921884 | 5 | 0.2 | 2 | 0.5 | NA | NA | NA | NA | 7 | 0.2857 | NA | NA | 5 | 0.4 | 4 | 0.5 | 3 | 0.3333 | 12 | 0.4167 |
| 1_1921885 | 5 | 0.2 | 3 | 0 | NA | NA | NA | NA | 8 | 0.125 | NA | NA | 6 | 0.1667 | 2 | 0 | 3 | 0.3333 | 11 | 0.1818 |
| 1_1921886 | 4 | 0 | 3 | 0.3333 | 1 | 0 | NA | NA | 8 | 0.125 | NA | NA | 6 | 0 | 3 | 0.6667 | 2 | 0.5 | 11 | 0.2727 |
| 1_1921887 | 5 | 1 | 3 | 1 | 1 | 1 | 1 | 1 | 10 | 1 | NA | NA | 6 | 1 | 4 | 1 | 2 | 1 | 12 | 1 |
| 1_1921888 | 5 | 0 | 3 | 0.3333 | 1 | 1 | 1 | 0 | 10 | 0.2 | NA | NA | 7 | 0.1429 | 4 | 0.25 | 2 | 0 | 13 | 0.1538 |
| 1_1921888 | 5 | 1 | 3 | 0.6667 | 1 | 1 | 1 | 1 | 10 | 0.9 | NA | NA | 7 | 1 | 4 | 0.75 | 3 | 1 | 14 | 0.9286 |
| 1_1921889 | 4 | 0.5 | 3 | 0.6667 | 2 | 0 | NA | NA | 9 | 0.4444 | NA | NA | 5 | 0.4 | 3 | 0.3333 | 3 | 0.3333 | 11 | 0.3636 |
| 1_1921889 | 4 | 1 | 3 | 1 | 1 | 1 | 2 | 1 | 10 | 1 | NA | NA | 6 | 1 | 3 | 1 | 3 | 1 | 12 | 1 |
| 1_1921889 | 4 | 1 | 3 | 1 | 2 | 0.5 | 2 | 1 | 11 | 0.9091 | NA | NA | 6 | 0.8333 | 4 | 1 | 3 | 1 | 13 | 0.9231 |
| 1_1921890 | 3 | 1 | 2 | 1 | 1 | 1 | 1 | 1 | 7 | 1 | NA | NA | 5 | 1 | 4 | 0.75 | 2 | 1 | 11 | 0.9091 |
| 1_1921891 | 2 | 1 | 3 | 1 | 1 | 1 | 2 | 1 | 8 | 1 | NA | NA | 4 | 0.75 | 3 | 1 | 2 | 0.5 | 9 | 0.7778 |

| | | | | | | | | | | | | | | | | | | | | |
|---|---|---|---|---|---|---|---|---|---|---|---|---|---|---|---|---|---|---|---|---|
| 1_19218917 | 1 | 1 | 3 | 0.6667 | 2 | 0.5 | 1 | 0 | 7 | 0.5714 | NA | NA | 6 | 0 | 3 | 0 | 2 | 0.5 | 11 | 0.0909 |
| 1_19218957 | 2 | 0 | 3 | 0.3333 | 2 | 0 | 3 | 0.3333 | 10 | 0.2 | 1 | 0 | 6 | 0.1667 | 4 | 0.25 | 2 | 0 | 13 | 0.1538 |
| 1_19218991 | 3 | 0 | 4 | 0.25 | 4 | 0 | 4 | 0 | 15 | 0.0667 | 1 | 0 | 8 | 0.125 | 6 | 0.3333 | 4 | 0 | 19 | 0.1579 |
| 1_19219008 | 2 | 0 | 4 | 0 | 4 | 0.25 | 3 | 0 | 13 | 0.0769 | 1 | 0 | 6 | 0.1667 | 6 | 0.3333 | 2 | 0 | 15 | 0.2 |
| 1_19219047 | 4 | 0.25 | 5 | 0.2 | 4 | 0 | 5 | 0 | 18 | 0.1111 | 2 | 0 | 8 | 0.25 | 7 | 0.2857 | 4 | 0 | 21 | 0.1905 |
| 1_19219068 | 4 | 0 | 5 | 0 | 5 | 0.2 | 5 | 0 | 19 | 0.0526 | 3 | 0 | 9 | 0 | 7 | 0.2857 | 4 | 0 | 23 | 0.087 |
| 1_19219114 | 3 | 0 | 4 | 0 | 5 | 0 | 5 | 0.2 | 17 | 0.0588 | 2 | 0 | 6 | 0.1667 | 8 | 0.125 | 2 | 0 | 18 | 0.1111 |
| 1_19219177 | 9 | 0.4444 | 6 | 0.5 | 6 | 0.6667 | 6 | 0.3333 | 27 | 0.4815 | 2 | 0 | 13 | 0.4615 | 9 | 0.5556 | 5 | 0.4 | 29 | 0.4483 |
| 1_19219183 | 11 | 0.1818 | 6 | 0.3333 | 6 | 0.1667 | 6 | 0.3333 | 29 | 0.2414 | 3 | 0.3333 | 12 | 0.25 | 9 | 0.2222 | 4 | 0 | 28 | 0.2143 |
| 1_19219191 | 7 | 0.8571 | 4 | 1 | 5 | 1 | 5 | 0.8 | 21 | 0.9048 | 2 | 1 | 10 | 0.7 | 8 | 0.625 | 5 | 0.6 | 25 | 0.68 |
| 1_19219200 | 11 | 0.2727 | 6 | 0.1667 | 7 | 0.4286 | 6 | 0.1667 | 30 | 0.2667 | 2 | 0.5 | 15 | 0.3333 | 8 | 0.375 | 7 | 0 | 32 | 0.2812 |
| 1_19219206 | 10 | 0.6 | 7 | 0.2857 | 6 | 0.5 | 5 | 0 | 28 | 0.3929 | 4 | 0.25 | 14 | 0.2857 | 6 | 0.6667 | 8 | 0.25 | 32 | 0.3438 |
| 1_19219209 | 12 | 0.3333 | 7 | 0.7143 | 7 | 0.4286 | 5 | 0.4 | 31 | 0.4516 | 4 | 0.25 | 17 | 0.2941 | 9 | 0.5556 | 7 | 0.2857 | 37 | 0.3514 |
| 1_19219211 | 13 | 0.7692 | 7 | 1 | 6 | 1 | 6 | 0.8333 | 32 | 0.875 | 3 | 1 | 14 | 0.8571 | 9 | 0.8889 | 6 | 0.8333 | 32 | 0.875 |
| 1_19219216 | 12 | 0.1667 | 6 | 0.3333 | 8 | 0 | 5 | 0 | 31 | 0.129 | 3 | 1 | 14 | 0.2143 | 6 | 0.1667 | 7 | 0.2857 | 30 | 0.3 |
| 1_19219234 | 12 | 0.1667 | 9 | 0 | 9 | 0 | 6 | 0 | 36 | 0.0556 | 5 | 0.2 | 16 | 0.0625 | 8 | 0 | 7 | 0 | 36 | 0.0556 |
| 1_19219557 | NA | NA | 1 | 1 | NA | NA | NA | NA | 1 | 1 | NA | NA | NA | NA | 2 | 0.5 | NA | NA | 2 | 0.5 |
| 1_19219602 | NA | NA | 1 | 1 | NA | NA | NA | NA | 1 | 1 | NA | NA | NA | NA | 1 | 1 | NA | NA | 1 | 1 |
| 1_19252721 | 7 | 0.7143 | 5 | 0.4 | 3 | 0.6667 | 4 | 1 | 19 | 0.6842 | NA | NA | 6 | 0.8333 | 4 | 0.5 | NA | NA | 10 | 0.7 |
| 1_19252741 | 7 | 0 | 5 | 0 | 4 | 0.25 | 4 | 0.25 | 20 | 0.1 | 1 | 0 | 5 | 0.2 | 2 | 0 | 1 | 0 | 9 | 0.1111 |
| 1_19252751 | 6 | 0.1667 | 5 | 0.2 | 4 | 0 | 4 | 0.25 | 19 | 0.1579 | 1 | 0 | 6 | 0.5 | 2 | 0.5 | 2 | 0 | 11 | 0.3636 |
| 1_19252822 | 5 | 0.4 | 5 | 0.4 | 3 | 1 | 4 | 0.75 | 17 | 0.5882 | 1 | 1 | 2 | 0 | 5 | 0.2 | NA | NA | 8 | 0.25 |
| 1_19252834 | 5 | 0.2 | 5 | 0.2 | 3 | 0 | 5 | 0.2 | 18 | 0.1667 | 1 | 0 | 2 | 0 | 4 | 0.25 | NA | NA | 7 | 0.1429 |
| 1_19252871 | 4 | 0 | 5 | 0 | 4 | 0.5 | 4 | 0 | 17 | 0.1176 | 1 | 1 | 3 | 0.3333 | 3 | 0 | 2 | 0 | 9 | 0.2222 |
| 1_19252877 | 4 | 0 | 2 | 0 | 4 | 0.25 | 4 | 0.25 | 14 | 0.1429 | 1 | 0 | 2 | 0.5 | 2 | 0 | 2 | 0 | 7 | 0.1429 |
| 1_19252879 | 4 | 1 | 4 | 1 | 4 | 1 | 4 | 1 | 16 | 1 | NA | NA | 2 | 1 | 3 | 1 | 2 | 1 | 7 | 1 |
| 1_19252884 | 4 | 1 | 4 | 0.75 | 4 | 0.75 | 2 | 1 | 14 | 0.8571 | 1 | 1 | 2 | 1 | 4 | 0.75 | 2 | 0.5 | 9 | 0.7778 |
| 1_19252901 | 3 | 1 | 2 | 1 | 3 | 1 | 4 | 1 | 12 | 1 | NA | NA | 2 | 1 | 4 | 1 | 2 | 1 | 8 | 1 |
| 1_19252930 | 2 | 1 | 4 | 1 | 3 | 1 | 3 | 1 | 12 | 1 | NA | NA | NA | NA | 3 | 1 | 2 | 1 | 5 | 1 |
| 1_19253869 | NA | NA | NA | NA | NA | NA | 1 | 1 | 1 | 1 | NA | NA | 1 | 1 | NA | NA | NA | NA | 1 | 1 |
| 1_19253869 | NA | NA | NA | NA | NA | NA | 1 | 1 | 1 | 1 | NA | NA | 1 | 1 | NA | NA | NA | NA | 1 | 1 |
| 1_19253892 | NA | NA | NA | NA | 1 | 1 | 1 | 1 | 2 | 1 | NA | NA | 1 | 1 | NA | NA | NA | NA | 1 | 1 |
| 1_19253904 | NA | NA | NA | NA | 1 | 1 | 1 | 0 | 2 | 0.5 | NA | NA | 1 | 1 | NA | NA | NA | NA | 1 | 1 |
| 1_19257391 | 4 | 0.5 | 2 | 0.5 | 1 | 0 | 1 | 0 | 8 | 0.375 | 2 | 0.5 | 1 | 1 | 2 | 0.5 | NA | NA | 5 | 0.6 |
| 1_19257703 | NA | NA | 2 | 0 | 2 | 0.5 | NA | NA | 4 | 0.25 | 2 | 0 | 1 | 1 | NA | NA | NA | NA | 3 | 0.3333 |
| 1_19257753 | NA | NA | 1 | 0 | 2 | 1 | NA | NA | 3 | 0.6667 | 2 | 0.5 | 1 | 1 | NA | NA | NA | NA | 3 | 0.6667 |
| 1_19257812 | NA | NA | 3 | 0 | 2 | 0.5 | 1 | 0 | 6 | 0.1667 | 2 | 0.5 | 1 | 0 | NA | NA | NA | NA | 3 | 0.3333 |
| 1_19257817 | NA | NA | 3 | 0.3333 | 2 | 0 | 1 | 1 | 6 | 0.3333 | 2 | 0.5 | 1 | 0 | NA | NA | NA | NA | 3 | 0.3333 |
| 1_19257823 | NA | NA | 3 | 0 | 2 | 0.5 | NA | NA | 5 | 0.2 | 2 | 1 | 1 | 1 | NA | NA | NA | NA | 3 | 1 |
| 1_19257938 | NA | NA | 2 | 1 | 2 | 0.5 | 1 | 1 | 5 | 0.8 | 1 | 1 | NA | NA | NA | NA | NA | NA | 1 | 1 |
| 1_19257944 | NA | NA | 3 | 0.6667 | 2 | 0.5 | 1 | 0 | 6 | 0.5 | 1 | 1 | NA | NA | NA | NA | NA | NA | 1 | 1 |
| 1_19257965 | NA | NA | 2 | 0.5 | 1 | 1 | 1 | 1 | 4 | 0.75 | 1 | 0 | 1 | 1 | NA | NA | NA | NA | 2 | 0.5 |
| 1_19257967 | NA | NA | 3 | 1 | 2 | 1 | 1 | 1 | 6 | 1 | 1 | 1 | 1 | 1 | NA | NA | NA | NA | 2 | 1 |
| 1_19257978 | NA | NA | 2 | 1 | 1 | 1 | 1 | 1 | 4 | 1 | 1 | 1 | NA | NA | NA | NA | NA | NA | 1 | 1 |
| 1_19278704 | 1 | 0 | 1 | 1 | 3 | 0.6667 | 2 | 1 | 7 | 0.7143 | NA | NA | 1 | 1 | 2 | 1 | 1 | 1 | 4 | 1 |
| 1_19278851 | NA | NA | NA | NA | 1 | 0 | 2 | 0.5 | 3 | 0.3333 | NA | NA | 1 | 0 | 1 | 1 | 1 | 0 | 3 | 0.3333 |
| 1_19278882 | NA | NA | NA | NA | 1 | 1 | 2 | 0.5 | 3 | 0.6667 | NA | NA | 1 | 0 | 1 | 1 | 1 | 1 | 3 | 0.6667 |
| 1_19278966 | 1 | 0 | 1 | 1 | 2 | 0.5 | 4 | 1 | 8 | 0.75 | NA | NA | 2 | 0.5 | 2 | 0.5 | NA | NA | 4 | 0.5 |
| 1_19278981 | 1 | 0 | 1 | 0 | 3 | 0.6667 | 6 | 0.8333 | 11 | 0.6364 | NA | NA | 2 | 1 | 2 | 0.5 | 2 | 0 | 6 | 0.5 |
| 1_19278982 | 1 | 1 | 1 | 1 | 2 | 1 | 6 | 0.8333 | 10 | 0.9 | NA | NA | 2 | 1 | 1 | 1 | 2 | 1 | 5 | 1 |
| 1_19278983 | 1 | 1 | 1 | 1 | 2 | 1 | 6 | 1 | 10 | 1 | NA | NA | 2 | 1 | 2 | 1 | NA | NA | 4 | 1 |
| 1_19279005 | NA | NA | 1 | 1 | 1 | 1 | 5 | 0.8 | 7 | 0.8571 | NA | NA | 1 | 1 | 1 | 1 | NA | NA | 2 | 1 |
| 1_19280267 | NA | NA | 1 | 1 | 1 | 1 | 4 | 0 | 6 | 0.3333 | NA | NA | 1 | 0 | 1 | 1 | 1 | 1 | 3 | 0.6667 |
| 1_19280278 | NA | NA | 1 | 0 | 1 | 0 | 3 | 0.3333 | 5 | 0.2 | NA | NA | 1 | 1 | NA | NA | 1 | 1 | 2 | 1 |
| 1_19285660 | 1 | 1 | NA | NA | NA | NA | NA | NA | 1 | 1 | NA | NA | 1 | 1 | NA | NA | NA | NA | 1 | 1 |
| 1_19285669 | 1 | 1 | NA | NA | NA | NA | NA | NA | 1 | 1 | NA | NA | 1 | 1 | NA | NA | NA | NA | 1 | 1 |
| 1_19285687 | 1 | 1 | NA | NA | NA | NA | NA | NA | 1 | 1 | NA | NA | 1 | 1 | NA | NA | NA | NA | 1 | 1 |
| 1_19285703 | 1 | 1 | NA | NA | NA | NA | NA | NA | 1 | 1 | NA | NA | 1 | 1 | NA | NA | NA | NA | 1 | 1 |
| 1_19285712 | 1 | 1 | NA | NA | NA | NA | NA | NA | 1 | 1 | NA | NA | 1 | 1 | NA | NA | NA | NA | 1 | 1 |
| 1_19285732 | 2 | 1 | NA | NA | NA | NA | NA | NA | 2 | 1 | NA | NA | 1 | 1 | NA | NA | NA | NA | 1 | 1 |
| 1_19285735 | 2 | 0.5 | NA | NA | NA | NA | NA | NA | 2 | 0.5 | NA | NA | 1 | 1 | NA | NA | NA | NA | 1 | 1 |
| 1_19290412 | 2 | 0 | NA | NA | 3 | 0.3333 | 2 | 0 | 7 | 0.1429 | 4 | 0 | 2 | 0 | 1 | 1 | NA | NA | 7 | 0.1429 |
| 1_19290433 | 3 | 0.6667 | NA | NA | 3 | 0.6667 | 2 | 1 | 8 | 0.75 | 3 | 0.6667 | 1 | 1 | 1 | 1 | 1 | 1 | 6 | 0.8333 |
| 1_19295464 | 1 | 1 | NA | NA | 4 | 1 | 2 | 1 | 7 | 1 | 1 | 1 | 1 | 1 | NA | NA | 1 | 1 | 3 | 1 |
| 1_19295468 | 1 | 1 | 1 | 1 | 4 | 0 | 2 | 0 | 8 | 0.25 | NA | NA | 1 | 1 | NA | NA | NA | NA | 1 | 1 |

| | | | | | | | | | | | | | | | | | | | | |
|---|---|---|---|---|---|---|---|---|---|---|---|---|---|---|---|---|---|---|---|---|
| 1_1929547 | 1 | 1 | 1 | 1 | 3 | 0.3333 | 2 | 1 | 7 | 0.7143 | 1 | 0 | 1 | 1 | NA | NA | 1 | 1 | 3 | 0.6667 |
| 1_1930260 | 1 | 0 | NA | NA | 1 | 1 | 1 | 1 | 3 | 0.6667 | 1 | 1 | 3 | 1 | 1 | 0 | 1 | 1 | 6 | 0.8333 |
| 1_1930288 | 5 | 0.2 | 2 | 0.5 | 8 | 0.125 | 7 | 0.1429 | 22 | 0.1818 | 1 | 0 | 9 | 0.3333 | 9 | 0.5556 | 5 | 0 | 24 | 0.3333 |
| 1_1930288 | 5 | 0.6 | 3 | 0.6667 | 9 | 0.4444 | 6 | 0.6667 | 23 | 0.5652 | 1 | 1 | 11 | 0.7273 | 8 | 0.75 | 3 | 0.6667 | 23 | 0.7391 |
| 1_1930288 | 5 | 0.2 | 3 | 0 | 9 | 0.2222 | 6 | 0.1667 | 23 | 0.1739 | 1 | 0 | 9 | 0.1111 | 7 | 0.1429 | 2 | 0 | 19 | 0.1053 |
| 1_1930290 | 3 | 0 | 2 | 0 | 6 | 0.3333 | 7 | 0 | 18 | 0.1111 | NA | NA | 10 | 0.3 | 7 | 0.1429 | 3 | 0.3333 | 20 | 0.25 |
| 1_1930291 | 3 | 0 | 3 | 0.3333 | 5 | 0.2 | 4 | 0 | 15 | 0.1333 | 1 | 0 | 6 | 0.1667 | 9 | 0.1111 | 3 | 0 | 19 | 0.1053 |
| 1_1943025 | NA | NA | NA | NA | NA | NA | 1 | 1 | 1 | 1 | NA | NA | 2 | 0.5 | 1 | 1 | NA | NA | 3 | 0.6667 |
| 1_1943026 | NA | NA | NA | NA | NA | NA | 1 | 1 | 1 | 1 | NA | NA | 1 | 1 | 1 | 1 | NA | NA | 2 | 1 |
| 1_1943026 | NA | NA | NA | NA | NA | NA | 1 | 1 | 1 | 1 | NA | NA | 2 | 0 | 1 | 1 | NA | NA | 3 | 0.3333 |
| 1_1943262 | NA | NA | NA | NA | NA | NA | 1 | 1 | 1 | 1 | NA | NA | 1 | 1 | NA | NA | NA | NA | 1 | 1 |
| 1_1943262 | NA | NA | NA | NA | NA | NA | 1 | 1 | 1 | 1 | NA | NA | 2 | 0.5 | 1 | 1 | NA | NA | 3 | 0.6667 |
| 1_1943267 | NA | NA | NA | NA | NA | NA | 1 | 1 | 1 | 1 | NA | NA | 2 | 1 | 1 | 0 | NA | NA | 3 | 0.6667 |
| 1_1943268 | 1 | 1 | NA | NA | NA | NA | 1 | 0 | 2 | 0.5 | NA | NA | 3 | 0.3333 | 1 | 0 | NA | NA | 4 | 0.25 |
| 1_1943271 | 1 | 0 | NA | NA | NA | NA | 1 | 1 | 2 | 0.5 | 1 | 0 | 2 | 0.5 | 1 | 0 | NA | NA | 4 | 0.25 |
| 1_1943318 | 1 | 0 | NA | NA | NA | NA | 1 | 1 | 2 | 0.5 | 1 | 0 | 3 | 0.6667 | 1 | 0 | NA | NA | 5 | 0.4 |
| 1_1944924 | 2 | 0 | 1 | 0 | 7 | 0.2857 | 3 | 0.3333 | 13 | 0.2308 | 2 | 0.5 | 5 | 0.2 | 4 | 0 | 4 | 0.25 | 15 | 0.2 |
| 1_1944944 | 2 | 1 | 1 | 1 | 5 | 1 | 3 | 1 | 11 | 1 | 2 | 1 | 4 | 1 | 4 | 0.75 | 3 | 1 | 13 | 0.9231 |
| 1_1944945 | 2 | 0 | 3 | 0 | 7 | 0.1429 | 3 | 0 | 15 | 0.0667 | 2 | 0 | 6 | 0.1667 | 3 | 0 | 4 | 0 | 15 | 0.0667 |
| 1_1944946 | 2 | 0.5 | 3 | 1 | 6 | 1 | 3 | 1 | 14 | 0.9286 | 2 | 1 | 5 | 1 | 2 | 1 | 4 | 0.5 | 13 | 0.8462 |
| 1_1944946 | 2 | 0.5 | 3 | 1 | 7 | 0.4286 | 5 | 0.6 | 17 | 0.5882 | 2 | 0.5 | 6 | 0.6667 | 4 | 1 | 4 | 1 | 16 | 0.8125 |
| 1_1944947 | 4 | 1 | 3 | 0.6667 | 7 | 0.7143 | 4 | 1 | 18 | 0.8333 | 1 | 1 | 5 | 1 | 5 | 0.4 | 3 | 1 | 14 | 0.7857 |
| 1_1944948 | 4 | 0.75 | 2 | 1 | 7 | 0.7143 | 5 | 1 | 18 | 0.8333 | 2 | 1 | 3 | 1 | 5 | 0.4 | 4 | 0.75 | 14 | 0.7143 |
| 1_1944948 | 4 | 0 | 1 | 0 | 5 | 0 | 5 | 0.2 | 15 | 0.0667 | 1 | 0 | 4 | 0.25 | 3 | 0 | 4 | 0.25 | 12 | 0.1667 |
| 1_1944950 | 3 | 0.6667 | 1 | 1 | 1 | 0 | 1 | 1 | 6 | 0.6667 | 1 | 1 | 3 | 0.6667 | 1 | 1 | 2 | 0.5 | 7 | 0.7143 |
| 1_1944951 | 3 | 0 | 2 | 0 | 2 | 0 | 2 | 0.5 | 9 | 0.1111 | NA | NA | 3 | 0.3333 | NA | NA | 2 | 0.5 | 5 | 0.4 |
| 1_1944952 | 3 | 0.6667 | 2 | 0.5 | 2 | 0.5 | 3 | 0.6667 | 10 | 0.6 | 1 | 1 | 4 | 1 | 2 | 0.5 | 4 | 0.5 | 11 | 0.7273 |
| 1_1944952 | 3 | 0.6667 | 2 | 0 | 2 | 0 | 3 | 0.6667 | 10 | 0.4 | 1 | 1 | 3 | 0 | 1 | 1 | 3 | 0.3333 | 8 | 0.375 |
| 1_1944953 | 3 | 0.6667 | 2 | 1 | NA | NA | 3 | 0.6667 | 8 | 0.75 | NA | NA | 4 | 0.5 | 2 | 1 | 3 | 0.3333 | 9 | 0.5556 |
| 1_1944954 | 1 | 1 | 2 | 1 | 2 | 1 | 4 | 1 | 9 | 1 | 2 | 1 | 4 | 1 | 2 | 1 | 4 | 1 | 12 | 1 |
| 1_1944954 | 3 | 1 | 2 | 1 | 2 | 1 | 4 | 1 | 11 | 1 | 2 | 1 | 4 | 1 | 2 | 1 | 2 | 1 | 10 | 1 |
| 1_1944955 | 3 | 0.3333 | 2 | 0.5 | 2 | 0 | 4 | 0.25 | 11 | 0.2727 | 2 | 0.5 | 4 | 0.25 | 2 | 0.5 | 3 | 0.3333 | 11 | 0.3636 |
| 1_1944960 | 5 | 0.4 | 3 | 0.6667 | 2 | 0.5 | 6 | 0.5 | 16 | 0.5 | 2 | 0.5 | 3 | 0.3333 | 3 | 0.6667 | 6 | 0.5 | 14 | 0.5 |
| 1_1944961 | 6 | 0.5 | 4 | 0.25 | 2 | 0 | 8 | 0.25 | 20 | 0.3 | 5 | 0.4 | 4 | 0.5 | 3 | 0.3333 | 5 | 0.2 | 17 | 0.3529 |
| 1_1944961 | 4 | 0.75 | 3 | 1 | 1 | 1 | 5 | 1 | 13 | 0.9231 | 4 | 0.75 | 4 | 1 | 3 | 0.6667 | 5 | 0.6 | 16 | 0.75 |
| 1_1944963 | 6 | 0.3333 | 4 | 0 | 4 | 0 | 8 | 0 | 22 | 0.0909 | 5 | 0 | 4 | 0 | 3 | 0.3333 | 6 | 0.3333 | 18 | 0.1667 |
| 1_1944964 | 7 | 0.2857 | 4 | 0 | 4 | 0 | 8 | 0 | 23 | 0.087 | 5 | 0.4 | 5 | 0.2 | 2 | 0.5 | 6 | 0.3333 | 18 | 0.3333 |
| 1_1944965 | 5 | 0.6 | 2 | 0.5 | 3 | 0.3333 | 6 | 0.3333 | 16 | 0.4375 | 5 | 0 | 2 | 1 | 3 | 0.3333 | 6 | 0.3333 | 16 | 0.3125 |
| 1_1944965 | 5 | 0.4 | 3 | 0.6667 | 3 | 0.3333 | 5 | 0.6 | 16 | 0.5 | 4 | 0.5 | 3 | 0.6667 | 3 | 1 | 4 | 0.5 | 14 | 0.6429 |
| 1_1944966 | 7 | 0.5714 | 4 | 0.25 | 4 | 0.25 | 7 | 0.1429 | 22 | 0.3182 | 5 | 0.4 | 4 | 0.75 | 3 | 1 | 5 | 0.6 | 17 | 0.6471 |
| 1_1944967 | 3 | 1 | 2 | 1 | 3 | 0.6667 | 5 | 0.8 | 13 | 0.8462 | 4 | 0.75 | 4 | 1 | 2 | 1 | 4 | 0.75 | 14 | 0.8571 |
| 1_1944974 | 7 | 0 | 4 | 0 | 3 | 0 | 8 | 0.25 | 22 | 0.0909 | 6 | 0.3333 | 7 | 0 | 7 | 0.1429 | 6 | 0.1667 | 26 | 0.1538 |
| 1_1944974 | 7 | 0.2857 | 8 | 0.375 | 5 | 0.4 | 10 | 0.2 | 30 | 0.3 | 7 | 0.1429 | 8 | 0.25 | 7 | 0.2857 | 7 | 0.1429 | 29 | 0.2069 |
| 1_1944988 | 10 | 0.2 | 8 | 0.25 | 9 | 0.5556 | 12 | 0.25 | 39 | 0.3077 | 8 | 0.5 | 11 | 0.4545 | 10 | 0.3 | 8 | 0.25 | 37 | 0.3784 |
| 1_1946424 | 2 | 1 | 1 | 1 | 3 | 0.6667 | 2 | 1 | 8 | 0.875 | 3 | 0.6667 | 2 | 1 | 1 | 1 | NA | NA | 6 | 0.8333 |
| 1_1946425 | 4 | 0.25 | 1 | 0 | 4 | 0.25 | 3 | 1 | 12 | 0.4167 | 2 | 1 | 2 | 0.5 | 1 | 0 | NA | NA | 5 | 0.6 |
| 1_1946434 | 2 | 0 | 1 | 0 | 4 | 0.25 | 3 | 0.3333 | 10 | 0.2 | 3 | 0.3333 | 2 | 0 | 1 | 0 | NA | NA | 6 | 0.1667 |
| 1_1947185 | 1 | 0 | 1 | 0 | 3 | 0.3333 | 4 | 0.25 | 9 | 0.2222 | 1 | 0 | 1 | 0 | 2 | 1 | 3 | 0 | 7 | 0.2857 |
| 1_1947185 | 1 | 0 | 2 | 0.5 | 3 | 0.3333 | 4 | 0.25 | 10 | 0.3 | 1 | 0 | NA | NA | 3 | 0.3333 | 3 | 0 | 7 | 0.1429 |
| 1_1947186 | 1 | 1 | 2 | 0.5 | 3 | 1 | 3 | 0.6667 | 9 | 0.7778 | 1 | 1 | NA | NA | 3 | 1 | 2 | 0 | 6 | 0.6667 |
| 1_1947189 | 1 | 0 | 1 | 0 | 4 | 0.25 | 4 | 0.25 | 10 | 0.2 | 1 | 0 | 2 | 0 | 4 | 0.5 | 3 | 0 | 10 | 0.2 |
| 1_1947194 | 2 | 0 | 2 | 0 | 6 | 0 | 10 | 0.1 | 20 | 0.05 | 5 | 0 | 5 | 0.2 | 6 | 0.1667 | 3 | 0 | 19 | 0.1053 |
| 1_1947194 | 2 | 0 | 2 | 0 | 5 | 0.2 | 9 | 0.5556 | 18 | 0.3333 | 3 | 0.3333 | 6 | 0 | 7 | 0.1429 | 2 | 0.5 | 18 | 0.1667 |
| 1_1947195 | 2 | 0 | 2 | 0 | 5 | 0 | 11 | 0.1818 | 20 | 0.1 | 5 | 0.4 | 6 | 0.1667 | 7 | 0.2857 | 3 | 0 | 21 | 0.2381 |
| 1_1947197 | 2 | 1 | 2 | 0.5 | 7 | 0.5714 | 11 | 0.5455 | 22 | 0.5909 | 5 | 0.6 | 6 | 1 | 7 | 0.8571 | 3 | 0 | 21 | 0.7143 |
| 1_1947198 | 2 | 0 | 2 | 1 | 7 | 0.8571 | 10 | 0.5 | 21 | 0.619 | 4 | 0.5 | 6 | 0.5 | 7 | 0.5714 | 1 | 0 | 18 | 0.5 |
| 1_1947219 | NA | NA | NA | NA | 2 | 1 | NA | NA | 2 | 1 | NA | NA | NA | NA | 1 | 1 | NA | NA | 1 | 1 |
| 1_1947792 | 2 | 0.5 | 3 | 0.6667 | 1 | 1 | 1 | 1 | 7 | 0.7143 | NA | NA | 2 | 1 | 1 | 1 | 2 | 0.5 | 5 | 0.8 |
| 1_1947792 | 1 | 1 | 2 | 1 | 1 | 1 | 1 | 0 | 5 | 0.8 | NA | NA | 2 | 1 | 1 | 1 | 2 | 1 | 5 | 1 |
| 1_1947792 | NA | NA | 2 | 1 | 1 | 1 | 1 | 1 | 4 | 1 | NA | NA | 1 | 1 | 1 | 1 | 3 | 1 | 5 | 1 |
| 1_1947792 | 2 | 0 | 3 | 1 | 1 | 1 | 1 | 1 | 7 | 0.7143 | NA | NA | 2 | 1 | 1 | 1 | 3 | 0.3333 | 6 | 0.6667 |
| 1_1947794 | 2 | 1 | 2 | 1 | 1 | 1 | 1 | 1 | 6 | 1 | NA | NA | 1 | 1 | 1 | 1 | 3 | 1 | 5 | 1 |
| 1_1947795 | 2 | 1 | 3 | 0 | 1 | 0 | 1 | 0 | 7 | 0.2857 | NA | NA | 2 | 0 | 1 | 0 | 3 | 0.3333 | 6 | 0.1667 |
| 1_1947795 | 2 | 0 | 2 | 0 | 1 | 0 | 1 | 1 | 6 | 0.1667 | NA | NA | 2 | 0 | 1 | 0 | 3 | 0.3333 | 6 | 0.1667 |
| 1_1947798 | NA | NA | 2 | 1 | NA | NA | 1 | 0 | 3 | 0.6667 | NA | NA | 2 | 1 | NA | NA | 3 | 1 | 5 | 1 |

| | | | | | | | | | | | | | | | | | | | | |
|---|---|---|---|---|---|---|---|---|---|---|---|---|---|---|---|---|---|---|---|---|
| 1_19477992 | 1 | 0 | 3 | 0 | 1 | 1 | 1 | 0 | 6 | 0.1667 | NA | NA | 2 | 1 | NA | NA | 3 | 0 | 5 | 0.4 |
| 1_19478001 | 1 | 0 | 3 | 0.6667 | 1 | 1 | 1 | 0 | 6 | 0.5 | NA | NA | 2 | 0 | NA | NA | 3 | 0.3333 | 5 | 0.2 |
| 1_19478004 | 1 | 1 | 3 | 0 | 1 | 1 | 1 | 0 | 6 | 0.3333 | NA | NA | 2 | 0.5 | NA | NA | 3 | 0 | 5 | 0.2 |
| 1_19478038 | 1 | 0 | 3 | 0.3333 | 1 | 0 | 1 | 0 | 6 | 0.1667 | NA | NA | 2 | 0 | NA | NA | 2 | 0.5 | 4 | 0.25 |
| 1_19478074 | 1 | 1 | 3 | 1 | 1 | 1 | NA | NA | 5 | 1 | NA | NA | 2 | 1 | NA | NA | 3 | 1 | 5 | 1 |
| 1_19478082 | 1 | 1 | 3 | 1 | 1 | 1 | 1 | 1 | 6 | 1 | NA | NA | 2 | 1 | NA | NA | 3 | 0.6667 | 5 | 0.8 |
| 1_19478085 | 1 | 1 | 3 | 0.6667 | 1 | 1 | 1 | 1 | 6 | 0.8333 | NA | NA | 2 | 1 | NA | NA | 3 | 1 | 5 | 1 |
| 1_19478089 | 1 | 1 | 3 | 1 | 1 | 1 | 1 | 1 | 6 | 1 | NA | NA | 2 | 1 | NA | NA | 2 | 1 | 4 | 1 |
| 1_19478107 | 2 | 0.5 | 3 | 0.6667 | 1 | 1 | 4 | 1 | 10 | 0.8 | NA | NA | 3 | 0.3333 | 1 | 1 | 3 | 1 | 7 | 0.7143 |
| 1_19478110 | 3 | 0 | 3 | 0.3333 | 2 | 1 | 4 | 0.25 | 12 | 0.3333 | NA | NA | 2 | 0.5 | 1 | 1 | 4 | 0 | 7 | 0.2857 |
| 1_19479997 | 3 | 0.6667 | 3 | 0.6667 | 1 | 0 | 5 | 0.2 | 12 | 0.4167 | 1 | 1 | 4 | 0 | 1 | 0 | 2 | 0.5 | 8 | 0.25 |
| 1_19480012 | 5 | 0 | 3 | 0 | 2 | 0 | 6 | 0.1667 | 16 | 0.0625 | 2 | 0.5 | 4 | 0 | 1 | 0 | 4 | 0 | 11 | 0.0909 |
| 1_19481360 | NA | NA | 2 | 0.5 | NA | NA | NA | NA | 2 | 0.5 | NA | NA | 1 | 1 | 1 | 0 | 1 | 1 | 3 | 0.6667 |
| 10_3913498 | 1 | 1 | NA | NA | NA | NA | NA | NA | 1 | 1 | 1 | 1 | NA | NA | 1 | 1 | NA | NA | 2 | 1 |
| 10_3913551 | 1 | 1 | NA | NA | NA | NA | NA | NA | 1 | 1 | 2 | 1 | NA | NA | 1 | 1 | NA | NA | 3 | 1 |
| 10_3913716 | 1 | 1 | NA | NA | NA | NA | NA | NA | 1 | 1 | 2 | 1 | NA | NA | NA | NA | NA | NA | 2 | 1 |
| 10_3913779 | 1 | 1 | NA | NA | NA | NA | NA | NA | 1 | 1 | 1 | 1 | NA | NA | 1 | 0 | NA | NA | 2 | 0.5 |
| 10_3913893 | 1 | 1 | NA | NA | NA | NA | NA | NA | 1 | 1 | 2 | 1 | NA | NA | 1 | 1 | NA | NA | 3 | 1 |
| 10_3914010 | 1 | 1 | NA | NA | NA | NA | NA | NA | 1 | 1 | 2 | 0.5 | NA | NA | NA | NA | NA | NA | 2 | 0.5 |
| 10_3914340 | 1 | 1 | NA | NA | NA | NA | NA | NA | 1 | 1 | 2 | 1 | NA | NA | 1 | 0 | NA | NA | 3 | 0.6667 |
| 10_3914350 | 1 | 1 | NA | NA | NA | NA | NA | NA | 1 | 1 | 2 | 1 | NA | NA | 1 | 0 | NA | NA | 3 | 0.6667 |
| 10_3914462 | 1 | 1 | NA | NA | NA | NA | NA | NA | 1 | 1 | 1 | 1 | NA | NA | 1 | 0 | NA | NA | 2 | 0.5 |
| 10_3915696 | 2 | 0.5 | NA | NA | NA | NA | NA | NA | 2 | 0.5 | 2 | 0.5 | NA | NA | 1 | 1 | NA | NA | 3 | 0.6667 |
| 10_3916580 | 2 | 0.5 | NA | NA | NA | NA | NA | NA | 2 | 0.5 | 3 | 0.3333 | NA | NA | 1 | 0 | NA | NA | 4 | 0.25 |
| 10_3920726 | 1 | 1 | NA | NA | NA | NA | NA | NA | 1 | 1 | 1 | 1 | NA | NA | NA | NA | NA | NA | 1 | 1 |
| 10_3923127 | 1 | 1 | NA | NA | NA | NA | NA | NA | 1 | 1 | 1 | 1 | NA | NA | NA | NA | NA | NA | 1 | 1 |
| 10_4105912 | 1 | 0 | 1 | 1 | 2 | 0.5 | 1 | 1 | 5 | 0.6 | NA | NA | 1 | 1 | NA | NA | 2 | 0.5 | 3 | 0.6667 |
| 10_4105935 | 2 | 1 | 1 | 0 | 2 | 0.5 | 1 | 0 | 6 | 0.5 | NA | NA | 1 | 0 | NA | NA | 2 | 0.5 | 3 | 0.3333 |
| 10_4106082 | 2 | 0.5 | 1 | 0 | 2 | 0 | 1 | 0 | 6 | 0.1667 | NA | NA | 1 | 0 | 1 | 0 | 3 | 0.6667 | 5 | 0.4 |
| 10_4304950 | 1 | 1 | NA | NA | NA | NA | NA | NA | 1 | 1 | NA | NA | 1 | 1 | NA | NA | NA | NA | 1 | 1 |
| 10_4305475 | 1 | 0 | 1 | 1 | NA | NA | NA | NA | 2 | 0.5 | NA | NA | 1 | 1 | NA | NA | NA | NA | 1 | 1 |
| 10_4306629 | 2 | 1 | 2 | 0.5 | NA | NA | NA | NA | 4 | 0.75 | NA | NA | 1 | 1 | NA | NA | NA | NA | 1 | 1 |
| 10_4308672 | 2 | 0.5 | 3 | 0.3333 | NA | NA | 1 | 0 | 6 | 0.3333 | NA | NA | 1 | 1 | NA | NA | NA | NA | 1 | 1 |
| 10_4309206 | 7 | 0.8571 | 4 | 0.5 | NA | NA | 2 | 0.5 | 13 | 0.6923 | NA | NA | 3 | 0.3333 | NA | NA | NA | NA | 3 | 0.3333 |
| 10_4317225 | NA | NA | 3 | 0.6667 | 3 | 0 | 3 | 0.3333 | 9 | 0.3333 | 1 | 0 | 1 | 0 | 1 | 0 | 1 | 1 | 4 | 0.25 |
| 10_4317945 | NA | NA | 2 | 0 | 1 | 1 | 3 | 0 | 6 | 0.1667 | NA | NA | NA | NA | 1 | 0 | 3 | 0.3333 | 4 | 0.25 |
| 10_4318325 | NA | NA | 1 | 0 | 1 | 0 | 1 | 1 | 3 | 0.3333 | NA | NA | 1 | 0 | 1 | 0 | 3 | 0.3333 | 5 | 0.2 |
| 10_4318422 | NA | NA | 1 | 1 | NA | NA | 3 | 1 | 4 | 1 | NA | NA | 1 | 1 | NA | NA | 2 | 1 | 3 | 1 |
| 10_4318458 | NA | NA | 1 | 1 | 1 | 1 | 3 | 1 | 5 | 1 | NA | NA | 1 | 1 | 1 | 1 | 2 | 1 | 4 | 1 |
| 10_4318484 | NA | NA | 1 | 0 | 1 | 1 | 1 | 1 | 3 | 0.6667 | NA | NA | 1 | 1 | 1 | 1 | 3 | 1 | 5 | 1 |
| 10_4318497 | NA | NA | 1 | 0 | 1 | 1 | 3 | 0 | 5 | 0.2 | NA | NA | 1 | 0 | 1 | 0 | 3 | 0.6667 | 5 | 0.4 |
| 10_4318545 | NA | NA | 1 | 1 | 1 | 1 | 3 | 1 | 5 | 1 | NA | NA | 1 | 1 | 1 | 1 | 1 | 1 | 3 | 1 |
| 10_4318556 | NA | NA | 1 | 1 | 1 | 1 | 2 | 1 | 4 | 1 | NA | NA | 1 | 1 | 1 | 1 | 2 | 1 | 4 | 1 |
| 10_4318712 | NA | NA | 1 | 1 | 1 | 1 | 2 | 1 | 4 | 1 | NA | NA | 1 | 1 | 1 | 1 | 2 | 1 | 4 | 1 |
| 10_4319410 | NA | NA | 1 | 1 | 1 | 1 | 1 | 1 | 3 | 1 | NA | NA | 1 | 1 | NA | NA | 1 | 1 | 2 | 1 |
| 10_4319457 | NA | NA | NA | NA | 1 | 1 | 1 | 0 | 2 | 0.5 | NA | NA | 1 | 0 | 1 | 0 | 1 | 1 | 3 | 0.3333 |
| 10_4319505 | NA | NA | NA | NA | 1 | 1 | 1 | 1 | 2 | 1 | NA | NA | 1 | 1 | 1 | 1 | 1 | 0 | 3 | 0.6667 |
| 10_4319563 | NA | NA | NA | NA | 1 | 0 | 1 | 1 | 2 | 0.5 | NA | NA | 1 | 1 | 1 | 0 | 1 | 0 | 3 | 0.3333 |
| 10_4324722 | NA | NA | NA | NA | 1 | 1 | NA | NA | 1 | 1 | NA | NA | 1 | 0 | 1 | 1 | 1 | 0 | 3 | 0.3333 |
| 10_4345973 | 1 | 1 | 2 | 0.5 | 1 | 1 | NA | NA | 4 | 0.75 | NA | NA | 1 | 1 | 1 | 1 | 1 | 1 | 3 | 1 |
| 10_4353176 | 4 | 0.25 | 3 | 0 | NA | NA | 5 | 0 | 12 | 0.0833 | NA | NA | 6 | 0 | 5 | 0.2 | 2 | 0 | 13 | 0.0769 |
| 10_4353201 | 4 | 0.75 | 4 | 0.5 | 2 | 1 | 5 | 0.2 | 15 | 0.5333 | 1 | 1 | 5 | 0.4 | 4 | 0.25 | 2 | 0 | 12 | 0.3333 |
| 10_4377368 | 3 | 1 | 4 | 1 | NA | NA | 4 | 0.75 | 11 | 0.9091 | 1 | 1 | 3 | 1 | 1 | 1 | 2 | 0.5 | 7 | 0.8571 |
| 10_4377396 | 2 | 0.5 | 4 | 0.75 | NA | NA | 4 | 0.5 | 10 | 0.6 | 1 | 1 | 4 | 0.75 | NA | NA | 2 | 0 | 7 | 0.5714 |
| 10_4377438 | 2 | 1 | 4 | 1 | NA | NA | 5 | 0.4 | 11 | 0.7273 | 1 | 1 | 3 | 0.6667 | 2 | 1 | 2 | 0.5 | 8 | 0.75 |
| 10_4377542 | 2 | 0 | 4 | 0 | NA | NA | 5 | 0.2 | 11 | 0.0909 | 1 | 0 | 4 | 0.25 | 1 | 0 | 1 | 0 | 7 | 0.1429 |
| 10_4377585 | 2 | 0 | 4 | 0 | NA | NA | 4 | 0.25 | 10 | 0.1 | 1 | 0 | 3 | 0 | 1 | 1 | 2 | 0 | 7 | 0.1429 |
| 10_4377606 | 2 | 1 | 4 | 0.5 | NA | NA | 5 | 0.8 | 11 | 0.7273 | 1 | 0 | 4 | 0.75 | 1 | 1 | 2 | 0.5 | 8 | 0.625 |
| 10_4389548 | 3 | 0 | 2 | 0.5 | NA | NA | NA | NA | 5 | 0.2 | 2 | 0.5 | 3 | 0.6667 | 4 | 0 | 1 | 0 | 10 | 0.3 |
| 10_4403587 | 3 | 0 | 2 | 0.5 | NA | NA | 1 | 0 | 6 | 0.1667 | 1 | 0 | 3 | 0 | 5 | 0 | 4 | 0.5 | 13 | 0.1538 |
| 10_4403684 | 3 | 0.3333 | 2 | 0 | NA | NA | 1 | 0 | 6 | 0.1667 | 1 | 1 | 4 | 0.75 | 5 | 0.6 | 4 | 0.25 | 14 | 0.5714 |
| 10_4403710 | 3 | 0.6667 | 2 | 0.5 | NA | NA | 1 | 0 | 6 | 0.5 | 1 | 1 | 4 | 1 | 5 | 0.8 | 4 | 1 | 14 | 0.9286 |
| 10_4403919 | 3 | 1 | 1 | 0 | 1 | 1 | 1 | 0 | 6 | 0.6667 | 1 | 1 | 4 | 1 | 5 | 0.6 | 5 | 1 | 15 | 0.8667 |
| 10_4403975 | 2 | 0.5 | 2 | 0 | 1 | 0 | 1 | 0 | 6 | 0.1667 | 1 | 0 | 4 | 0.5 | 4 | 0 | 4 | 0.75 | 13 | 0.3846 |
| 10_4404114 | 2 | 1 | 2 | 1 | 1 | 1 | 1 | 1 | 6 | 1 | 2 | 0.5 | 4 | 1 | 5 | 0.8 | 5 | 0.8 | 16 | 0.8125 |

| | | | | | | | | | | | | | | | | | | | | |
|---|---|---|---|---|---|---|---|---|---|---|---|---|---|---|---|---|---|---|---|---|
| 10_440426( | 3 | 0 | 1 | 0 | 1 | 1 | 1 | 0 | 6 | 0.1667 | 1 | 0 | 4 | 0.25 | 4 | 0.25 | 5 | 0.2 | 14 | 0.2143 |
| 10_440481( | 3 | 0 | 3 | 0.3333 | 2 | 0 | 1 | 0 | 9 | 0.1111 | 1 | 0 | 5 | 0 | 5 | 0 | 5 | 0.2 | 16 | 0.0625 |
| 10_440482' | 3 | 0 | 3 | 0.3333 | 1 | 0 | 1 | 1 | 8 | 0.25 | 1 | 0 | 5 | 0.6 | 5 | 0 | 6 | 0 | 17 | 0.1765 |
| 10_440483( | 3 | 0.3333 | 3 | 0.3333 | 1 | 1 | 1 | 0 | 8 | 0.375 | 2 | 0 | 5 | 0.6 | 5 | 0.4 | 4 | 0.25 | 16 | 0.375 |
| 10_440496' | 3 | 0.3333 | 3 | 0.6667 | 2 | 0.5 | 1 | 1 | 9 | 0.5556 | 1 | 0 | 6 | 0.8333 | 4 | 0.75 | 6 | 0.3333 | 17 | 0.5882 |
| 10_440498' | 4 | 0 | 3 | 0.3333 | 2 | 0 | 1 | 0 | 10 | 0.1 | 2 | 0 | 6 | 0.3333 | 6 | 0 | 6 | 0.1667 | 20 | 0.15 |
| 10_497049 | 21 | 0.0952 | 23 | 0.0435 | 20 | 0.05 | 24 | 0.0417 | 88 | 0.0568 | 14 | 0.0714 | 17 | 0.1765 | 27 | 0.1481 | 22 | 0.0455 | 80 | 0.1125 |
| 10_497067' | 18 | 0.1111 | 22 | 0.1364 | 19 | 0.0526 | 18 | 0.3889 | 77 | 0.1688 | 12 | 0.0833 | 12 | 0.1667 | 23 | 0.2609 | 17 | 0.2941 | 64 | 0.2188 |
| 10_497076' | 16 | 0.125 | 19 | 0.0526 | 13 | 0.3077 | 19 | 0.1579 | 67 | 0.1493 | 7 | 0 | 15 | 0.1333 | 19 | 0.1053 | 15 | 0.1333 | 56 | 0.1071 |
| 10_497079' | 14 | 0.0714 | 17 | 0.1176 | 15 | 0.0667 | 20 | 0.05 | 66 | 0.0758 | 8 | 0.25 | 16 | 0.1875 | 22 | 0.0455 | 18 | 0.0556 | 64 | 0.1094 |
| 10_497118 | 6 | 0.8333 | 8 | 0.875 | 4 | 1 | 4 | 0.75 | 22 | 0.8636 | 3 | 0.6667 | 6 | 1 | 7 | 0.4286 | 4 | 0.75 | 20 | 0.7 |
| 10_497126 | 6 | 0.6667 | 8 | 0.875 | 4 | 1 | 3 | 0.6667 | 21 | 0.8095 | 3 | 1 | 4 | 0.75 | 5 | 0.8 | 4 | 1 | 16 | 0.875 |
| 10_497130 | 5 | 1 | 5 | 1 | 4 | 1 | 2 | 1 | 16 | 1 | 2 | 1 | 3 | 1 | 6 | 1 | 3 | 1 | 14 | 1 |
| 10_497141 | 5 | 0 | 6 | 0 | 3 | 0.6667 | 2 | 0 | 16 | 0.125 | 1 | 0 | 4 | 0.75 | 7 | 0 | 3 | 0.6667 | 15 | 0.3333 |
| 10_497145 | 5 | 0.8 | 6 | 0.8333 | 4 | 1 | 3 | 1 | 18 | 0.8889 | 2 | 1 | 3 | 0.6667 | 5 | 0.8 | 4 | 1 | 14 | 0.8571 |
| 10_497148' | 5 | 1 | 6 | 0.8333 | 4 | 0.75 | 3 | 1 | 18 | 0.8889 | 3 | 1 | 3 | 1 | 6 | 0.8333 | 4 | 1 | 16 | 0.9375 |
| 10_497166 | 3 | 0.3333 | 4 | 1 | 3 | 0 | 2 | 0.5 | 12 | 0.5 | 2 | 1 | 1 | 0 | 4 | 0.5 | 3 | 0 | 10 | 0.4 |
| 10_497170 | 2 | 0 | 4 | 0.25 | 3 | 0 | 1 | 0 | 10 | 0.1 | 3 | 0.3333 | 1 | 0 | 6 | 0 | 3 | 0 | 13 | 0.0769 |
| 10_497171 | 2 | 0 | 4 | 0.5 | 3 | 1 | 2 | 0.5 | 11 | 0.5455 | 1 | 1 | 2 | 0.5 | 5 | 0 | 3 | 0.3333 | 11 | 0.2727 |
| 10_497174' | 2 | 0 | 4 | 0.25 | 3 | 0.3333 | 2 | 0 | 11 | 0.1818 | 2 | 0.5 | 1 | 0 | 4 | 0.25 | 2 | 1 | 9 | 0.4444 |
| 10_497208 | 2 | 0 | 4 | 0 | 4 | 0 | 2 | 0.5 | 12 | 0.0833 | 2 | 0 | 2 | 0 | 4 | 0.25 | 1 | 0 | 9 | 0.1111 |
| 10_497212' | NA | NA | 4 | 0.25 | 4 | 0.5 | 2 | 1 | 10 | 0.5 | 2 | 0.5 | 2 | 1 | 3 | 1 | 1 | 0 | 8 | 0.75 |
| 10_528709 | 17 | 0.1765 | 29 | 0.3448 | 14 | 0.5 | 14 | 0.2143 | 74 | 0.3108 | 12 | 0.4167 | 20 | 0.3 | 19 | 0.2105 | 17 | 0.3529 | 68 | 0.3088 |
| 10_528714 | 16 | 0.1875 | 28 | 0.25 | 15 | 0.2667 | 8 | 0.125 | 67 | 0.2239 | 10 | 0.1 | 16 | 0.4375 | 17 | 0.2941 | 14 | 0.3571 | 57 | 0.3158 |
| 10_528724 | 14 | 0.7143 | 29 | 0.5517 | 10 | 0.5 | 8 | 0.75 | 61 | 0.6066 | 8 | 0.75 | 15 | 0.6667 | 15 | 0.7333 | 13 | 0.6923 | 51 | 0.7059 |
| 10_528727 | 15 | 0.1333 | 31 | 0 | 14 | 0.1429 | 14 | 0 | 74 | 0.0541 | 11 | 0.0909 | 21 | 0.0476 | 17 | 0.1176 | 19 | 0.1053 | 68 | 0.0882 |
| 10_528770 | 10 | 0 | 26 | 0.1923 | 9 | 0.1111 | 9 | 0 | 54 | 0.1111 | 9 | 0.2222 | 13 | 0 | 12 | 0.1667 | 12 | 0.0833 | 46 | 0.1087 |
| 10_528852 | 7 | 0.5714 | 23 | 0.2174 | 7 | 0.5714 | 11 | 0.5455 | 48 | 0.3958 | 5 | 0.4 | 10 | 0.6 | 15 | 0.4 | 15 | 0.4 | 45 | 0.4444 |
| 10_528893' | 6 | 0.3333 | 11 | 0 | 4 | 0 | 3 | 0 | 24 | 0.0833 | 6 | 0 | 7 | 0.1429 | 12 | 0.0833 | 10 | 0 | 35 | 0.0571 |
| 10_528894 | 7 | 0.5714 | 23 | 0.6957 | 8 | 0.375 | 7 | 0.7143 | 45 | 0.6222 | 7 | 0.7143 | 10 | 0.3 | 14 | 0.4286 | 13 | 0.6154 | 44 | 0.5 |
| 10_528903 | 7 | 0.1429 | 18 | 0.3333 | 6 | 0.3333 | 6 | 0.5 | 37 | 0.3243 | 4 | 0.25 | 9 | 0.1111 | 16 | 0.1875 | 10 | 0.3 | 39 | 0.2051 |
| 10_528905 | 5 | 1 | 20 | 0.95 | 4 | 0.75 | 7 | 1 | 36 | 0.9444 | 3 | 1 | 5 | 1 | 14 | 0.8571 | 11 | 0.9091 | 33 | 0.9091 |
| 10_574938 | NA | NA | 1 | 1 | NA | NA | NA | NA | 1 | 1 | NA | NA | 1 | 1 | NA | NA | 1 | 1 | 2 | 1 |
| 10_575199 | NA | NA | 1 | 1 | NA | NA | NA | NA | 1 | 1 | NA | NA | NA | NA | NA | NA | 1 | 1 | 1 | 1 |
| 10_575200 | NA | NA | 1 | 1 | NA | NA | NA | NA | 1 | 1 | NA | NA | NA | NA | NA | NA | 1 | 1 | 1 | 1 |
| 10_575225 | NA | NA | 1 | 1 | NA | NA | NA | NA | 1 | 1 | NA | NA | NA | NA | NA | NA | 1 | 1 | 1 | 1 |
| 10_575233 | NA | NA | 1 | 1 | NA | NA | NA | NA | 1 | 1 | NA | NA | NA | NA | NA | NA | 1 | 1 | 1 | 1 |
| 10_576309' | 1 | 1 | 1 | 1 | NA | NA | 1 | 1 | 3 | 1 | 1 | 1 | 1 | 1 | 2 | 1 | 1 | 1 | 5 | 1 |
| 10_576321 | 1 | 1 | 1 | 1 | NA | NA | 1 | 0 | 3 | 0.6667 | 1 | 0 | 1 | 1 | 4 | 0.5 | 2 | 0.5 | 8 | 0.5 |
| 10_576325 | 1 | 1 | 2 | 1 | NA | NA | 1 | 1 | 4 | 1 | 1 | 1 | 1 | 1 | 4 | 1 | 2 | 0.5 | 8 | 0.875 |
| 10_576329' | 1 | 1 | 2 | 0.5 | NA | NA | 1 | 0 | 4 | 0.5 | 1 | 0 | 1 | 1 | 3 | 0 | 2 | 0 | 7 | 0.1429 |
| 10_577597 | 7 | 0.1429 | 10 | 0.1 | 6 | 0.1667 | 7 | 0.1429 | 30 | 0.1333 | 5 | 0.2 | 3 | 0 | 11 | 0.1818 | 13 | 0 | 32 | 0.0938 |
| 10_577839 | 1 | 0 | 4 | 0 | 4 | 0.25 | 4 | 0 | 13 | 0.0769 | 2 | 0 | 2 | 0 | 5 | 0.2 | 3 | 0 | 12 | 0.0833 |
| 10_577863 | 1 | 1 | 3 | 0.6667 | 4 | 0.25 | 4 | 0 | 12 | 0.3333 | 2 | 0 | 2 | 0 | 6 | 0.1667 | 3 | 0 | 13 | 0.0769 |
| 10_577871' | 1 | 0 | 3 | 0.3333 | 4 | 0 | 4 | 0.25 | 12 | 0.1667 | 2 | 0.5 | 2 | 0 | 5 | 0 | 3 | 0 | 12 | 0.0833 |
| 10_577922 | 1 | 0 | 3 | 0 | 4 | 0 | 4 | 0.25 | 12 | 0.0833 | 2 | 0 | 2 | 0 | 4 | 0.25 | 3 | 0.3333 | 11 | 0.1818 |
| 10_578044' | NA | NA | 3 | 0.3333 | 3 | 0.3333 | 2 | 0.5 | 8 | 0.375 | NA | NA | 1 | 0 | 1 | 0 | 3 | 0.6667 | 5 | 0.4 |
| 10_578215' | 1 | 0 | 1 | 1 | 3 | 0 | 3 | 0 | 8 | 0.125 | NA | NA | 1 | 0 | 2 | 0 | 4 | 0.5 | 7 | 0.2857 |
| 10_578255 | 1 | 0 | 2 | 1 | 3 | 0.6667 | 1 | 1 | 7 | 0.7143 | NA | NA | 1 | 0 | 2 | 1 | 4 | 0.75 | 7 | 0.7143 |
| 10_578261 | 1 | 0 | 2 | 0.5 | 2 | 0 | 1 | 0 | 6 | 0.1667 | NA | NA | 2 | 0.5 | 2 | 0 | 4 | 0.25 | 8 | 0.25 |
| 10_578284' | NA | NA | 1 | 1 | 2 | 0.5 | 1 | 0 | 4 | 0.5 | NA | NA | 1 | 0 | 2 | 0.5 | 4 | 0.75 | 7 | 0.5714 |
| 10_578301' | NA | NA | 1 | 1 | 2 | 1 | 1 | 1 | 4 | 1 | NA | NA | 1 | 0 | 2 | 0.5 | 5 | 0.8 | 8 | 0.625 |
| 10_680270 | NA | NA | 1 | 1 | NA | NA | NA | NA | 1 | 1 | NA | NA | 2 | 0.5 | 1 | 1 | NA | NA | 3 | 0.6667 |
| 10_683691 | 2 | 0 | 7 | 0 | 3 | 0 | 4 | 0.25 | 16 | 0.0625 | 3 | 0.3333 | 2 | 0 | 1 | 0 | 1 | 0 | 7 | 0.1429 |
| 10_683955 | 1 | 0 | 2 | 0.5 | 2 | 0 | 1 | 1 | 6 | 0.3333 | 2 | 0.5 | NA | NA | NA | NA | NA | NA | 2 | 0.5 |
| 10_683996 | NA | NA | 1 | 1 | 2 | 0 | 1 | 1 | 4 | 0.5 | 2 | 0.5 | NA | NA | 1 | 0 | NA | NA | 3 | 0.3333 |
| 10_683999 | NA | NA | NA | NA | 2 | 1 | NA | NA | 2 | 1 | 1 | 1 | NA | NA | 1 | 1 | NA | NA | 2 | 1 |
| 10_684006 | NA | NA | 1 | 1 | 2 | 0 | NA | NA | 3 | 0.3333 | 1 | 1 | NA | NA | 1 | 1 | NA | NA | 2 | 1 |
| 10_684013 | NA | NA | 2 | 1 | 2 | 1 | 1 | 1 | 5 | 1 | 1 | 1 | NA | NA | 1 | 0 | NA | NA | 2 | 0.5 |
| 10_684036 | NA | NA | 2 | 0 | 2 | 0 | 1 | 1 | 5 | 0.2 | 2 | 0.5 | NA | NA | 1 | 0 | NA | NA | 3 | 0.3333 |
| 10_684054 | NA | NA | 2 | 0.5 | 2 | 0.5 | 1 | 1 | 5 | 0.6 | 2 | 1 | NA | NA | NA | NA | NA | NA | 2 | 1 |
| 10_684056 | NA | NA | 2 | 0.5 | 2 | 0.5 | 1 | 1 | 5 | 0.6 | 2 | 0.5 | NA | NA | NA | NA | NA | NA | 2 | 0.5 |
| 10_686980 | NA | NA | 1 | 1 | 2 | 0 | 1 | 1 | 4 | 0.5 | 2 | 1 | NA | NA | 1 | 1 | NA | NA | 3 | 1 |
| 10_706963 | 1 | 1 | 1 | 0 | NA | NA | 1 | 0 | 3 | 0.3333 | 2 | 0 | NA | NA | 1 | 1 | NA | NA | 3 | 0.3333 |
| 10_706992 | 1 | 1 | 1 | 1 | NA | NA | 1 | 1 | 3 | 1 | 2 | 0.5 | NA | NA | 1 | 1 | NA | NA | 3 | 0.6667 |

| | | | | | | | | | | | | | | | | | | | | |
|---|---|---|---|---|---|---|---|---|---|---|---|---|---|---|---|---|---|---|---|---|
| 10_706993 | 1 | 1 | NA | NA | NA | NA | 1 | 1 | 2 | 1 | 1 | 1 | NA | NA | 1 | 1 | NA | NA | 2 | 1 |
| 10_707001 | 1 | 1 | 1 | 1 | NA | NA | 1 | 0 | 3 | 0.6667 | 1 | 0 | NA | NA | 1 | 1 | NA | NA | 2 | 0.5 |
| 10_707006 | 1 | 1 | 1 | 1 | NA | NA | 1 | 1 | 3 | 1 | 1 | 1 | NA | NA | 1 | 1 | NA | NA | 2 | 1 |
| 10_707021 | 1 | 1 | 1 | 1 | NA | NA | 1 | 1 | 3 | 1 | 1 | 0 | NA | NA | 1 | 1 | NA | NA | 2 | 0.5 |
| 10_707022 | 1 | 1 | 1 | 1 | NA | NA | 1 | 1 | 3 | 1 | 1 | 1 | NA | NA | 1 | 1 | NA | NA | 2 | 1 |
| 10_707029 | NA | NA | 1 | 1 | NA | NA | 1 | 1 | 2 | 1 | 1 | 1 | NA | NA | 1 | 0 | NA | NA | 2 | 0.5 |
| 10_707046 | 1 | 1 | NA | NA | NA | NA | 1 | 0 | 2 | 0.5 | 1 | 0 | NA | NA | 1 | 1 | NA | NA | 2 | 0.5 |
| 10_746583 | 7 | 0.1429 | 5 | 0 | 5 | 0.2 | 8 | 0 | 25 | 0.08 | 3 | 0 | 6 | 0.1667 | 10 | 0.2 | 6 | 0.5 | 25 | 0.24 |
| 10_746595 | 6 | 0.8333 | 3 | 0.6667 | 5 | 0.4 | 9 | 0.7778 | 23 | 0.6957 | 4 | 0.5 | 5 | 0.6 | 10 | 0.6 | 7 | 0.5714 | 26 | 0.5769 |
| 10_749952 | 13 | 0.6923 | 7 | 0.5714 | 6 | 0.3333 | 7 | 0.7143 | 33 | 0.6061 | 12 | 0.5833 | 14 | 0.6429 | 15 | 0.4667 | 14 | 0.6429 | 55 | 0.5818 |
| 10_749963 | 11 | 0.2727 | 6 | 0.5 | 7 | 0.2857 | 9 | 0.4444 | 33 | 0.3636 | 13 | 0.4615 | 14 | 0.2857 | 16 | 0.1875 | 12 | 0.3333 | 55 | 0.3091 |
| 10_749964 | 15 | 0.1333 | 9 | 0 | 9 | 0.1111 | 11 | 0.2727 | 44 | 0.1364 | 14 | 0.2143 | 15 | 0.1333 | 16 | 0 | 17 | 0.1176 | 62 | 0.1129 |
| 10_749966 | 13 | 0.0769 | 7 | 0.1429 | 9 | 0.2222 | 8 | 0.25 | 37 | 0.1622 | 12 | 0.0833 | 14 | 0.0714 | 18 | 0.1667 | 13 | 0.3077 | 57 | 0.1579 |
| 10_749973 | 9 | 0.4444 | 9 | 0.4444 | 6 | 0.1667 | 9 | 0.4444 | 33 | 0.3939 | 11 | 0.5455 | 8 | 0.375 | 11 | 0.4545 | 14 | 0.5 | 44 | 0.4773 |
| 10_750016 | 15 | 0.2 | 13 | 0.1538 | 11 | 0.0909 | 11 | 0.1818 | 50 | 0.16 | 15 | 0.1333 | 21 | 0.0476 | 19 | 0 | 15 | 0.0667 | 70 | 0.0571 |
| 10_750021 | 17 | 0.4706 | 13 | 0.8462 | 9 | 0.5556 | 10 | 0.7 | 49 | 0.6327 | 13 | 0.6923 | 18 | 0.6111 | 19 | 0.6316 | 15 | 0.6667 | 65 | 0.6462 |
| 10_750663 | 14 | 0.9286 | 24 | 0.9167 | 21 | 0.9524 | 18 | 0.9444 | 77 | 0.9351 | 10 | 1 | 11 | 1 | 22 | 0.9091 | 12 | 0.9167 | 55 | 0.9455 |
| 10_751368 | 5 | 0.2 | 6 | 0.1667 | 1 | 0 | 4 | 0 | 16 | 0.125 | 1 | 0 | 3 | 0.3333 | 7 | 0.4286 | 3 | 0 | 14 | 0.2857 |
| 10_753900 | 10 | 0 | 13 | 0.1538 | 11 | 0.1818 | 9 | 0.1111 | 43 | 0.1163 | 4 | 0 | 6 | 0 | 16 | 0.125 | 7 | 0 | 33 | 0.0606 |
| 10_753910 | 2 | 1 | 1 | 1 | 1 | 1 | 3 | 0.6667 | 7 | 0.8571 | 1 | 1 | NA | NA | 6 | 0.6667 | 4 | 0.25 | 11 | 0.5455 |
| 10_753914 | 1 | 1 | 2 | 0.5 | 1 | 0 | 2 | 1 | 6 | 0.6667 | NA | NA | NA | NA | 6 | 0.6667 | 4 | 0.75 | 10 | 0.7 |
| 10_753922 | 2 | 1 | 2 | 0.5 | 1 | 1 | 2 | 0.5 | 7 | 0.7143 | NA | NA | NA | NA | 6 | 0.1667 | 4 | 0.25 | 10 | 0.2 |
| 10_755446 | 2 | 0.5 | NA | NA | 1 | 1 | 2 | 0 | 5 | 0.4 | 1 | 0 | NA | NA | 2 | 0.5 | 1 | 0 | 4 | 0.25 |
| 10_755606 | 1 | 1 | NA | NA | NA | NA | NA | NA | 1 | 1 | NA | NA | NA | NA | NA | NA | 1 | 1 | 1 | 1 |
| 10_755644 | 1 | 1 | NA | NA | NA | NA | NA | NA | 1 | 1 | NA | NA | NA | NA | NA | NA | 1 | 1 | 1 | 1 |
| 10_755658 | 1 | 1 | NA | NA | 1 | 1 | NA | NA | 2 | 1 | NA | NA | 1 | 1 | NA | NA | 1 | 1 | 2 | 1 |
| 10_757757 | NA | NA | 1 | 1 | 1 | 1 | NA | NA | 2 | 1 | NA | NA | NA | NA | 1 | 1 | 2 | 1 | 3 | 1 |
| 10_757771 | NA | NA | NA | NA | 1 | 1 | NA | NA | 1 | 1 | NA | NA | NA | NA | 1 | 1 | 2 | 0.5 | 3 | 0.6667 |
| 10_757797 | NA | NA | 1 | 0 | 1 | 1 | NA | NA | 2 | 0.5 | NA | NA | NA | NA | 2 | 0.5 | 2 | 1 | 4 | 0.75 |
| 10_757802 | NA | NA | NA | NA | 1 | 1 | NA | NA | 1 | 1 | NA | NA | NA | NA | 2 | 1 | NA | NA | 2 | 1 |
| 10_757815 | NA | NA | 1 | 1 | 1 | 1 | NA | NA | 2 | 1 | NA | NA | NA | NA | 2 | 1 | 2 | 1 | 4 | 1 |
| 10_757840 | NA | NA | 1 | 0 | 1 | 1 | 1 | 1 | 3 | 0.6667 | 1 | 1 | 1 | 0 | 1 | 0 | 2 | 0.5 | 5 | 0.4 |
| 10_757845 | NA | NA | 1 | 1 | 1 | 0 | 1 | 1 | 3 | 0.6667 | 1 | 1 | NA | NA | 2 | 1 | 2 | 1 | 5 | 1 |
| 10_757862 | NA | NA | NA | NA | 1 | 1 | NA | NA | 1 | 1 | 1 | 1 | 1 | 0 | 2 | 0 | 2 | 0.5 | 6 | 0.3333 |
| 10_757866 | NA | NA | 1 | 1 | 1 | 1 | NA | NA | 2 | 1 | 1 | 0 | 2 | 1 | 2 | 0.5 | 1 | 0 | 6 | 0.5 |
| 10_758131 | NA | NA | 1 | 1 | 1 | 1 | NA | NA | 2 | 1 | NA | NA | 2 | 1 | 2 | 1 | 2 | 1 | 6 | 1 |
| 10_758134 | NA | NA | 1 | 1 | 1 | 1 | 1 | 1 | 3 | 1 | NA | NA | 2 | 1 | 2 | 1 | 1 | 1 | 5 | 1 |
| 10_758141 | NA | NA | 1 | 1 | 1 | 1 | 1 | 0 | 3 | 0.6667 | NA | NA | 2 | 1 | 2 | 1 | 2 | 1 | 6 | 1 |
| 10_758156 | NA | NA | 1 | 1 | 1 | 1 | 1 | 1 | 3 | 1 | 1 | 1 | 2 | 0.5 | 2 | 1 | 2 | 1 | 7 | 0.8571 |
| 10_758164 | NA | NA | 1 | 1 | 1 | 1 | NA | NA | 2 | 1 | 1 | 1 | 3 | 0.6667 | 2 | 0.5 | 2 | 0.5 | 8 | 0.625 |
| 10_758841 | 1 | 1 | 1 | 0 | 1 | 0 | NA | NA | 3 | 0.3333 | 2 | 1 | 2 | 0 | 2 | 0.5 | 1 | 0 | 7 | 0.4286 |
| 10_758844 | 1 | 0 | 2 | 0.5 | 1 | 0 | 1 | 0 | 5 | 0.2 | 2 | 0.5 | 2 | 0 | 2 | 0.5 | 2 | 0.5 | 8 | 0.375 |
| 10_758855 | 1 | 1 | 1 | 1 | 1 | 0 | 1 | 0 | 4 | 0.5 | 2 | 1 | 3 | 1 | 1 | 1 | 2 | 1 | 8 | 1 |
| 10_758863 | 1 | 0 | 1 | 1 | 1 | 1 | NA | NA | 3 | 0.6667 | 2 | 0.5 | 2 | 1 | 2 | 1 | 2 | 0.5 | 8 | 0.75 |
| 10_758899 | 1 | 0 | 1 | 1 | NA | NA | 1 | 0 | 3 | 0.3333 | 2 | 0 | 3 | 0.6667 | 1 | 1 | 1 | 0 | 7 | 0.4286 |
| 10_759060 | 3 | 0.3333 | 1 | 0 | 1 | 0 | 5 | 0.2 | 10 | 0.2 | 3 | 0 | 1 | 0 | 6 | 0.1667 | 5 | 0.4 | 15 | 0.2 |
| 10_759110 | 2 | 0 | 3 | 0.3333 | 1 | 0 | 5 | 0.2 | 11 | 0.1818 | 3 | 0 | 3 | 0.3333 | 7 | 0.1429 | 4 | 0 | 17 | 0.1176 |
| 10_759115 | 2 | 0.5 | 4 | 0.5 | 1 | 0 | 5 | 0.4 | 12 | 0.4167 | 3 | 0 | 3 | 0 | 5 | 0.4 | 6 | 0.3333 | 17 | 0.2353 |
| 10_759123 | 3 | 0 | 4 | 0.5 | NA | NA | 5 | 0.4 | 12 | 0.3333 | 3 | 0 | 2 | 0 | 6 | 0.3333 | 6 | 0.1667 | 17 | 0.1765 |
| 10_759142 | 4 | 0.25 | 3 | 0 | 1 | 0 | 5 | 0.2 | 13 | 0.1538 | 3 | 0 | 3 | 0.3333 | 5 | 0.2 | 5 | 0.6 | 16 | 0.3125 |
| 10_764389 | 5 | 0 | 5 | 0 | 2 | 0.5 | 3 | 0 | 15 | 0.0667 | 2 | 0.5 | 7 | 0.1429 | 2 | 0 | 2 | 0 | 13 | 0.1538 |
| 10_764396 | 6 | 0.3333 | 6 | 0.3333 | 4 | 0.25 | 3 | 0.6667 | 19 | 0.3684 | 1 | 0 | 8 | 0.5 | 2 | 0 | 3 | 0 | 14 | 0.2857 |
| 10_764404 | 6 | 0 | 5 | 0.2 | 4 | 0.5 | 2 | 0 | 17 | 0.1765 | 1 | 0 | 8 | 0.125 | 2 | 0 | 3 | 0 | 14 | 0.0714 |
| 10_764420 | 6 | 0 | 5 | 0 | 4 | 0 | 3 | 0.3333 | 18 | 0.0556 | 2 | 0 | 8 | 0.125 | 2 | 0.5 | 3 | 0.3333 | 15 | 0.2 |
| 10_764422 | 4 | 1 | 5 | 1 | 3 | 1 | 2 | 1 | 14 | 1 | 2 | 0.5 | 5 | 0.8 | 2 | 1 | 3 | 1 | 12 | 0.8333 |
| 10_766848 | 1 | 1 | NA | NA | NA | NA | NA | NA | 1 | 1 | NA | NA | NA | NA | 1 | 1 | NA | NA | 1 | 1 |
| 10_769691 | 4 | 0 | 6 | 0.1667 | NA | NA | 8 | 0.25 | 18 | 0.1667 | 2 | 0 | 4 | 0.25 | 2 | 0.5 | 3 | 0.6667 | 11 | 0.3636 |
| 10_769698 | 2 | 0.5 | 7 | 1 | NA | NA | 6 | 1 | 15 | 0.9333 | NA | NA | 4 | 1 | 2 | 1 | 3 | 1 | 9 | 1 |
| 10_769705 | 4 | 0.25 | 6 | 0 | NA | NA | 8 | 0.375 | 18 | 0.2222 | 1 | 1 | 4 | 0 | 3 | 0.6667 | 3 | 0.6667 | 11 | 0.4545 |
| 10_769751 | 5 | 0 | 10 | 0.3 | NA | NA | 8 | 0.5 | 23 | 0.3043 | 2 | 0 | 4 | 0.25 | 3 | 0.6667 | 4 | 0.25 | 13 | 0.3077 |
| 10_778199 | 3 | 0.3333 | 6 | 0.3333 | 4 | 0 | 5 | 0 | 18 | 0.1667 | 5 | 0 | 3 | 0 | 2 | 0 | 8 | 0.125 | 18 | 0.0556 |
| 10_778200 | 6 | 0.1667 | 7 | 0.4286 | 5 | 0.4 | 5 | 0.4 | 23 | 0.3478 | 6 | 0.1667 | 3 | 0 | 2 | 0 | 9 | 0.3333 | 20 | 0.2 |
| 10_778204 | 6 | 0 | 6 | 0.1667 | 5 | 0.4 | 5 | 0 | 22 | 0.1364 | 5 | 0.2 | 3 | 0 | 2 | 0 | 9 | 0.1111 | 19 | 0.1053 |
| 10_778208 | 5 | 1 | 8 | 0.75 | 4 | 0.75 | 5 | 0.2 | 22 | 0.6818 | 5 | 0.4 | 3 | 0 | 2 | 1 | 9 | 0.7778 | 19 | 0.5789 |
| 10_778218 | 5 | 0.6 | 7 | 0.5714 | 4 | 0.25 | 5 | 0 | 21 | 0.381 | 4 | 0.75 | 3 | 0 | 2 | 0 | 6 | 0.5 | 15 | 0.4 |

| | | | | | | | | | | | | | | | | | | | | |
|---|---|---|---|---|---|---|---|---|---|---|---|---|---|---|---|---|---|---|---|---|
| 10_778229 | 5 | 0.2 | 5 | 0 | 4 | 0 | 4 | 0 | 18 | 0.0556 | 6 | 0.1667 | 3 | 0 | 2 | 0 | 8 | 0.25 | 19 | 0.1579 |
| 10_778244 | 4 | 0.25 | 5 | 0.4 | 2 | 0 | 5 | 0.2 | 16 | 0.25 | 6 | 0 | 2 | 0 | 2 | 0.5 | 8 | 0.25 | 18 | 0.1667 |
| 10_778255 | 5 | 0 | 6 | 0.3333 | 4 | 0 | 5 | 0 | 20 | 0.1 | 6 | 0.1667 | 3 | 0 | 2 | 0 | 6 | 0 | 17 | 0.0588 |
| 10_778279 | 3 | 0.6667 | 4 | 0.5 | 2 | 0.5 | 3 | 0.3333 | 12 | 0.5 | 5 | 0.6 | 2 | 0 | 1 | 0 | 7 | 0.4286 | 15 | 0.4 |
| 10_778280 | 2 | 0.5 | 3 | 0.6667 | 1 | 0 | 4 | 0.5 | 10 | 0.5 | 4 | 0.25 | 2 | 0 | 1 | 1 | 7 | 0.7143 | 14 | 0.5 |
| 10_778281 | 4 | 0.25 | 2 | 0 | 3 | 0 | 3 | 0 | 12 | 0.0833 | 6 | 0.1667 | 2 | 0 | 2 | 0 | 6 | 0 | 16 | 0.0625 |
| 10_778288 | 4 | 0.5 | 5 | 0 | 3 | 0 | 3 | 0.3333 | 15 | 0.2 | 5 | 0.4 | 2 | 0 | 2 | 0.5 | 8 | 0.125 | 17 | 0.2353 |
| 10_778292 | 4 | 0.75 | 4 | 1 | 3 | 1 | 4 | 1 | 15 | 0.9333 | 3 | 1 | 2 | 0.5 | 2 | 1 | 6 | 0.6667 | 13 | 0.7692 |
| 10_778299 | 4 | 0.25 | 5 | 0.4 | 3 | 0 | 4 | 0.25 | 16 | 0.25 | 4 | 0.25 | 2 | 0 | 2 | 0 | 6 | 0 | 14 | 0.0714 |
| 10_778308 | 3 | 0.3333 | 3 | 0.6667 | 3 | 0.3333 | 4 | 0.25 | 13 | 0.3846 | 5 | 0.2 | 1 | 0 | 1 | 0 | 6 | 0.5 | 13 | 0.3077 |
| 10_778322 | 3 | 1 | 3 | 0.6667 | 3 | 1 | 4 | 0.5 | 13 | 0.7692 | 3 | 0.6667 | 2 | 0.5 | 2 | 0.5 | 6 | 0.8333 | 13 | 0.6923 |
| 10_779491 | 4 | 0 | 5 | 0 | 2 | 0.5 | 4 | 0 | 15 | 0.0667 | 3 | 0 | 2 | 0 | 2 | 0 | 4 | 0.25 | 11 | 0.0909 |
| 10_779497 | 3 | 0.3333 | 3 | 0.3333 | 2 | 0 | 4 | 0.75 | 12 | 0.4167 | 3 | 0.3333 | 1 | 1 | 2 | 0 | 4 | 0.25 | 10 | 0.3 |
| 10_779502 | 2 | 1 | 1 | 1 | 2 | 1 | 1 | 1 | 6 | 1 | 4 | 0.75 | 1 | 1 | 1 | 1 | NA | NA | 6 | 0.8333 |
| 10_779514 | 3 | 0 | 3 | 0 | 2 | 0 | 4 | 0.25 | 12 | 0.0833 | 4 | 0.25 | NA | NA | 2 | 0 | 2 | 0 | 8 | 0.125 |
| 10_779524 | 1 | 0 | 4 | 0.5 | 2 | 0.5 | 3 | 0.3333 | 10 | 0.4 | 3 | 0 | 1 | 0 | 2 | 0 | 3 | 0.3333 | 9 | 0.1111 |
| 10_779552 | 2 | 0 | 5 | 0.2 | 2 | 0 | 4 | 0 | 13 | 0.0769 | 2 | 0.5 | 1 | 0 | 2 | 0.5 | 4 | 0.25 | 9 | 0.3333 |
| 10_779571 | 1 | 0 | 4 | 0.5 | 2 | 0.5 | 2 | 1 | 9 | 0.5556 | 1 | 1 | NA | NA | 1 | 1 | 3 | 0.3333 | 5 | 0.6 |
| 10_779571 | 1 | 0 | 5 | 0.4 | 2 | 0 | 2 | 0 | 10 | 0.2 | 2 | 0 | 1 | 0 | 1 | 0 | 3 | 0.3333 | 7 | 0.1429 |
| 10_779592 | NA | NA | 4 | 1 | 2 | 1 | 1 | 1 | 7 | 1 | NA | NA | NA | NA | NA | NA | 3 | 1 | 3 | 1 |
| 10_779597 | NA | NA | 4 | 0.5 | 2 | 0 | 2 | 0 | 8 | 0.25 | 2 | 0.5 | 1 | 0 | 1 | 1 | 2 | 1 | 6 | 0.6667 |
| 10_779599 | NA | NA | 5 | 0.4 | 2 | 0.5 | 2 | 1 | 9 | 0.5556 | 2 | 1 | 1 | 0 | 1 | 0 | 2 | 0.5 | 6 | 0.5 |
| 10_779612 | NA | NA | 2 | 0.5 | 2 | 1 | 1 | 1 | 5 | 0.8 | 1 | 1 | 1 | 1 | NA | NA | 1 | 1 | 3 | 1 |
| 10_779617 | NA | NA | 5 | 0.6 | 2 | 1 | 2 | 1 | 9 | 0.7778 | NA | NA | NA | NA | 1 | 1 | 1 | 1 | 2 | 1 |
| 10_779623 | NA | NA | 5 | 0.8 | 2 | 0.5 | 2 | 1 | 9 | 0.7778 | 2 | 0 | 1 | 0 | 1 | 0 | 1 | 1 | 5 | 0.2 |
| 10_808123 | 3 | 0.3333 | 4 | 0.5 | 3 | 0.3333 | 2 | 1 | 12 | 0.5 | 3 | 0.3333 | 4 | 0.25 | 3 | 0 | 5 | 0.6 | 15 | 0.3333 |
| 10_808134 | 3 | 0.3333 | 4 | 0.25 | 3 | 0.3333 | 2 | 0.5 | 12 | 0.3333 | 1 | 0 | 3 | 0.3333 | 4 | 0.25 | 2 | 0.5 | 10 | 0.3 |
| 10_808184 | 2 | 0 | 3 | 0.3333 | 1 | 0 | 1 | 0 | 7 | 0.1429 | 1 | 0 | 3 | 0 | 3 | 0 | 5 | 0.4 | 12 | 0.1667 |
| 10_808237 | 2 | 0 | 3 | 0.3333 | NA | NA | 2 | 0.5 | 7 | 0.2857 | 1 | 0 | 4 | 0 | 3 | 0.3333 | 4 | 0 | 12 | 0.0833 |
| 10_808304 | 1 | 1 | 3 | 0 | 1 | 1 | 1 | 0 | 6 | 0.3333 | NA | NA | 3 | 0.3333 | 2 | 1 | 2 | 0.5 | 7 | 0.5714 |
| 10_808307 | 2 | 1 | 3 | 0.6667 | 1 | 1 | 1 | 1 | 7 | 0.8571 | 1 | 1 | 4 | 1 | 2 | 1 | 2 | 1 | 9 | 1 |
| 10_808328 | 2 | 1 | 1 | 1 | 1 | 1 | 2 | 1 | 6 | 1 | 1 | 1 | 3 | 1 | 2 | 1 | 1 | 1 | 7 | 1 |
| 10_808331 | 2 | 0.5 | 2 | 0 | 1 | 0 | 1 | 0 | 6 | 0.1667 | 1 | 0 | 3 | 0 | 1 | 0 | 2 | 0.5 | 7 | 0.1429 |
| 10_859830 | 5 | 0.2 | 13 | 0.0769 | 3 | 0 | 5 | 0 | 26 | 0.0769 | 7 | 0.1429 | 1 | 0 | 6 | 0 | 3 | 0 | 17 | 0.0588 |
| 10_859838 | 5 | 0.4 | 14 | 0.4286 | 4 | 0.25 | 3 | 0 | 26 | 0.3462 | 7 | 0.1429 | 2 | 0.5 | 6 | 0.3333 | 4 | 0.5 | 19 | 0.3158 |
| 10_859941 | 4 | 0.25 | 11 | 0 | NA | NA | 3 | 0 | 18 | 0.0556 | 5 | 0.2 | 2 | 0 | 4 | 0 | 2 | 0 | 13 | 0.0769 |
| 10_860069 | 4 | 0.25 | 6 | 0.1667 | 3 | 0 | 4 | 0.5 | 17 | 0.2353 | 3 | 0.3333 | 1 | 0 | 5 | 0 | 3 | 0 | 12 | 0.0833 |
| 10_860073 | 4 | 1 | 6 | 0.6667 | 4 | 0.5 | 4 | 0.75 | 18 | 0.7222 | 3 | 0.6667 | 1 | 1 | 4 | 0.75 | 2 | 1 | 10 | 0.8 |
| 10_860079 | 4 | 0.5 | 6 | 0.3333 | 2 | 0 | 5 | 0 | 17 | 0.2353 | 2 | 0 | 1 | 0 | 4 | 0.25 | 1 | 1 | 8 | 0.25 |
| 10_860082 | 3 | 0.3333 | 6 | 0.1667 | 2 | 0 | 5 | 0.2 | 16 | 0.1875 | 2 | 0 | 1 | 0 | 5 | 0.2 | 2 | 0 | 10 | 0.1 |
| 10_860084 | 2 | 1 | 5 | 1 | 3 | 0.6667 | 1 | 1 | 11 | 0.9091 | 3 | 0.6667 | 1 | 1 | 4 | 0.75 | 2 | 0.5 | 10 | 0.7 |
| 10_860098 | 3 | 0 | 5 | 0 | 3 | 0.3333 | 4 | 0.25 | 15 | 0.1333 | 3 | 0 | NA | NA | 5 | 0.2 | 3 | 0.3333 | 11 | 0.1818 |
| 10_860099 | 4 | 0.25 | 4 | 0.25 | 2 | 0 | 5 | 0.2 | 15 | 0.2 | 3 | 0 | NA | NA | 4 | 0.75 | 2 | 0 | 9 | 0.3333 |
| 10_860103 | 3 | 1 | 5 | 0 | 1 | 0 | 5 | 0.4 | 14 | 0.3571 | 2 | 0 | 1 | 0 | 4 | 0.75 | 3 | 0 | 10 | 0.3 |
| 10_860113 | 3 | 0.6667 | 5 | 0.8 | 1 | 1 | 4 | 1 | 13 | 0.8462 | 3 | 1 | 1 | 1 | 5 | 0.8 | 2 | 0.5 | 11 | 0.8182 |
| 10_860114 | 2 | 0.5 | 3 | 0.6667 | 2 | 1 | 1 | 1 | 8 | 0.75 | 1 | 1 | 1 | 1 | 4 | 1 | 3 | 1 | 9 | 1 |
| 10_860128 | 4 | 0.5 | 3 | 0.3333 | 2 | 0 | 4 | 0.5 | 13 | 0.3846 | 2 | 0 | 1 | 0 | 4 | 0.25 | 3 | 0 | 10 | 0.1 |
| 10_860129 | 4 | 1 | 4 | 0.5 | 2 | 1 | 5 | 0.2 | 15 | 0.6 | 2 | 1 | 1 | 1 | 5 | 0.6 | 3 | 0.3333 | 11 | 0.6364 |
| 10_860131 | 2 | 0 | 3 | 0 | 3 | 0.3333 | 4 | 0.5 | 12 | 0.25 | 2 | 0 | 1 | 0 | 5 | 0.6 | 2 | 0 | 10 | 0.3 |
| 10_860132 | 4 | 0.5 | 4 | 0.75 | 2 | 0.5 | 4 | 0 | 14 | 0.4286 | 1 | 1 | 1 | 0 | 5 | 0.6 | 3 | 0.3333 | 10 | 0.5 |
| 10_860521 | 4 | 0 | 3 | 0.3333 | 2 | 0 | 4 | 0 | 13 | 0.0769 | 1 | 0 | 1 | 0 | 4 | 0.25 | 3 | 0 | 9 | 0.1111 |
| 10_860527 | 3 | 0.3333 | 2 | 0.5 | 2 | 0 | 4 | 0 | 11 | 0.1818 | 1 | 0 | 1 | 0 | 3 | 0 | 3 | 0.3333 | 8 | 0.125 |
| 10_860530 | 3 | 0.6667 | 3 | 1 | 1 | 0 | 2 | 0.5 | 9 | 0.6667 | 1 | 1 | 1 | 1 | 4 | 0.75 | 3 | 0.3333 | 9 | 0.6667 |
| 10_860549 | 3 | 0 | 2 | 0 | 1 | 0 | 3 | 0.3333 | 9 | 0.1111 | 1 | 1 | 1 | 0 | 4 | 0.75 | 3 | 0.6667 | 9 | 0.6667 |
| 10_860555 | 3 | 1 | 2 | 1 | 1 | 1 | 3 | 1 | 9 | 1 | 1 | 1 | 1 | 1 | 3 | 1 | 3 | 1 | 8 | 1 |
| 10_860568 | 2 | 0.5 | 3 | 0 | 3 | 0 | 2 | 0 | 10 | 0.1 | 1 | 0 | 1 | 0 | 3 | 0.3333 | 1 | 0 | 6 | 0.1667 |
| 10_860600 | 3 | 1 | 2 | 0 | 2 | 0.5 | 2 | 0.5 | 9 | 0.5556 | 1 | 1 | NA | NA | 3 | 1 | 2 | 1 | 6 | 1 |
| 10_860603 | 3 | 0.6667 | 2 | 0.5 | 2 | 1 | 3 | 0.6667 | 10 | 0.7 | 1 | 0 | 1 | 1 | 4 | 0.75 | 2 | 0.5 | 8 | 0.625 |
| 10_862700 | 1 | 0 | 1 | 0 | 1 | 1 | 2 | 0.5 | 5 | 0.4 | NA | NA | 1 | 0 | 1 | 1 | 1 | 0 | 3 | 0.3333 |
| 10_949877 | 1 | 1 | NA | NA | 2 | 0 | NA | NA | 3 | 0.3333 | 1 | 1 | 1 | 0 | 1 | 0 | 1 | 0 | 4 | 0.25 |
| 10_949891 | 2 | 0 | NA | NA | 2 | 0.5 | NA | NA | 4 | 0.25 | 1 | 1 | 1 | 1 | 1 | 0 | 1 | 1 | 4 | 0.75 |
| 10_963165 | 6 | 0.3333 | 6 | 0 | 3 | 0.3333 | 5 | 0 | 20 | 0.15 | 2 | 0 | 3 | 0 | 6 | 0 | 7 | 0.1429 | 18 | 0.0556 |
| 10_963612 | 5 | 0.6 | 7 | 0.7143 | 3 | 0.6667 | 6 | 1 | 21 | 0.7619 | NA | NA | 1 | 1 | 6 | 0.6667 | 6 | 0.8333 | 13 | 0.7692 |
| 10_963617 | 6 | 1 | 4 | 0.5 | 3 | 1 | 6 | 0.8333 | 19 | 0.8421 | NA | NA | 2 | 1 | 3 | 0.6667 | 8 | 0.875 | 13 | 0.8462 |
| 10_963622 | 4 | 1 | 6 | 1 | 3 | 1 | 8 | 1 | 21 | 1 | NA | NA | 1 | 1 | 4 | 1 | 10 | 0.9 | 15 | 0.9333 |

| | | | | | | | | | | | | | | | | | | | | |
|---|---|---|---|---|---|---|---|---|---|---|---|---|---|---|---|---|---|---|---|---|
| 10_9636318 | 6 | 0.8333 | 5 | 0.8 | 3 | 1 | 5 | 1 | 19 | 0.8947 | NA | NA | 1 | 1 | 5 | 0.8 | 8 | 0.875 | 14 | 0.8571 |
| 10_9636336 | 6 | 0.8333 | 6 | 0.3333 | 2 | 0.5 | 7 | 0.8571 | 21 | 0.6667 | NA | NA | 2 | 1 | 5 | 0.6 | 12 | 0.5833 | 19 | 0.6316 |
| 10_9636364 | 6 | 1 | 7 | 1 | 2 | 1 | 7 | 0.8571 | 22 | 0.9545 | NA | NA | 3 | 1 | 7 | 0.8571 | 11 | 0.8182 | 21 | 0.8571 |
| 10_9636395 | 6 | 0.3333 | 7 | 0 | 1 | 0 | 6 | 0.5 | 20 | 0.25 | NA | NA | 3 | 0.3333 | 6 | 0 | 10 | 0.1 | 19 | 0.1053 |
| 10_9636422 | 6 | 0.8333 | 7 | 1 | 2 | 1 | 5 | 1 | 20 | 0.95 | NA | NA | 2 | 1 | 5 | 1 | 9 | 1 | 16 | 1 |
| 10_9636484 | 5 | 0.2 | 7 | 0.2857 | 1 | 0 | 6 | 0.3333 | 19 | 0.2632 | 1 | 0 | 2 | 0.5 | 4 | 0.25 | 8 | 0.25 | 15 | 0.2667 |
| 10_9636493 | 4 | 0.25 | 6 | 0.1667 | 1 | 0 | 3 | 0 | 14 | 0.1429 | NA | NA | 1 | 1 | 6 | 0 | 8 | 0.125 | 15 | 0.1333 |
| 10_9636545 | 6 | 0.5 | 7 | 0.5714 | 2 | 1 | 6 | 0.6667 | 21 | 0.619 | NA | NA | 3 | 0.3333 | 5 | 0.8 | 12 | 0.4167 | 20 | 0.5 |
| 10_9636648 | 5 | 0.6 | 6 | 0.3333 | 2 | 0.5 | 6 | 0.3333 | 19 | 0.4211 | NA | NA | 1 | 0 | 6 | 0.5 | 9 | 0.2222 | 16 | 0.3125 |
| 10_9776494 | 3 | 1 | 1 | 1 | NA | NA | 2 | 0.5 | 6 | 0.8333 | NA | NA | 1 | 1 | NA | NA | 3 | 1 | 4 | 1 |
| 10_9776736 | 2 | 1 | NA | NA | NA | NA | 1 | 1 | 3 | 1 | NA | NA | 1 | 1 | NA | NA | 2 | 0.5 | 3 | 0.6667 |
| 10_1042195 | 1 | 1 | NA | NA | NA | NA | NA | NA | 1 | 1 | NA | NA | NA | NA | 1 | 1 | NA | NA | 1 | 1 |
| 10_1056222 | 7 | 0.2857 | 4 | 0 | 1 | 0 | 3 | 0 | 15 | 0.1333 | NA | NA | 4 | 0 | 2 | 0 | 3 | 0.3333 | 9 | 0.1111 |
| 10_1056352 | 4 | 0.25 | 4 | 0 | 1 | 0 | 2 | 0.5 | 11 | 0.1818 | 1 | 0 | 3 | 0 | 2 | 0.5 | 2 | 0.5 | 8 | 0.25 |
| 10_1056359 | 3 | 1 | 3 | 1 | 1 | 0 | 2 | 1 | 9 | 0.8889 | 1 | 0 | 2 | 0.5 | 1 | 1 | NA | NA | 4 | 0.5 |
| 10_1056367 | 3 | 1 | 3 | 0.6667 | 1 | 0 | 1 | 1 | 8 | 0.75 | 2 | 1 | NA | NA | NA | NA | NA | NA | 2 | 1 |
| 10_1056378 | 2 | 1 | 2 | 1 | 1 | 0 | 1 | 1 | 6 | 0.8333 | 1 | 1 | 3 | 1 | 2 | 1 | 1 | 1 | 7 | 1 |
| 10_1056408 | 3 | 0.6667 | 3 | 0.6667 | 1 | 1 | NA | NA | 7 | 0.7143 | NA | NA | 2 | 1 | 2 | 0.5 | 2 | 1 | 6 | 0.8333 |
| 10_1056409 | 4 | 1 | 3 | 1 | 1 | 1 | 1 | 1 | 9 | 1 | 1 | 1 | 2 | 1 | 2 | 1 | NA | NA | 5 | 1 |
| 10_1056413 | 3 | 0 | 4 | 0.75 | 1 | 0 | 1 | 0 | 9 | 0.3333 | 1 | 0 | 2 | 0.5 | 2 | 1 | 2 | 0 | 7 | 0.4286 |
| 10_1056415 | 4 | 1 | 4 | 0.75 | 1 | 1 | 1 | 0 | 10 | 0.8 | 1 | 1 | 2 | 1 | 2 | 1 | 2 | 1 | 7 | 1 |
| 10_1056416 | 3 | 0.6667 | 4 | 0.75 | 1 | 1 | 1 | 1 | 9 | 0.7778 | 1 | 1 | 2 | 0.5 | 2 | 0 | 2 | 1 | 7 | 0.5714 |
| 10_1056435 | 2 | 0.5 | 2 | 0.5 | 1 | 0 | 1 | 0 | 6 | 0.3333 | 1 | 0 | NA | NA | 2 | 0.5 | 2 | 0.5 | 5 | 0.4 |
| 10_1056441 | 2 | 0.5 | 2 | 0 | 1 | 0 | 1 | 0 | 6 | 0.1667 | 1 | 0 | 2 | 0 | 2 | 0.5 | 1 | 1 | 6 | 0.3333 |
| 10_1056443 | 2 | 1 | 2 | 1 | 1 | 0 | 1 | 1 | 6 | 0.8333 | 1 | 1 | 1 | 1 | 2 | 0.5 | 1 | 1 | 5 | 0.8 |
| 10_1056443 | 2 | 1 | 1 | 1 | 1 | 1 | 1 | 1 | 5 | 1 | 1 | 1 | 2 | 1 | 1 | 1 | 2 | 1 | 6 | 1 |
| 10_1056461 | 3 | 0 | 2 | 0.5 | 1 | 0 | 1 | 0 | 7 | 0.1429 | NA | NA | 2 | 0.5 | 2 | 0 | 2 | 0.5 | 6 | 0.3333 |
| 10_1056473 | 2 | 1 | 2 | 1 | NA | NA | NA | NA | 4 | 1 | NA | NA | 1 | 1 | 1 | 1 | 2 | 1 | 4 | 1 |
| 10_1056512 | 1 | 1 | 2 | 0 | 1 | 1 | 1 | 0 | 5 | 0.4 | NA | NA | 1 | 0 | 1 | 0 | 1 | 1 | 3 | 0.3333 |
| 10_1056514 | NA | NA | 2 | 0.5 | 1 | 0 | 1 | 0 | 4 | 0.25 | NA | NA | NA | NA | 1 | 1 | 1 | 1 | 2 | 1 |
| 10_1056542 | 1 | 0 | 2 | 1 | 1 | 0 | 1 | 0 | 5 | 0.4 | NA | NA | 1 | 0 | 1 | 1 | NA | NA | 2 | 0.5 |
| 10_1056558 | 1 | 0 | 2 | 0 | 1 | 1 | NA | NA | 4 | 0.25 | NA | NA | 1 | 1 | 1 | 1 | NA | NA | 2 | 1 |
| 10_1059534 | NA | NA | 1 | 1 | 1 | 1 | NA | NA | 2 | 1 | NA | NA | 1 | 1 | NA | NA | NA | NA | 1 | 1 |
| 10_1059577 | NA | NA | 1 | 1 | 1 | 1 | NA | NA | 2 | 1 | NA | NA | NA | NA | 1 | 1 | NA | NA | 1 | 1 |
| 10_1102777 | NA | NA | NA | NA | 2 | 0.5 | NA | NA | 2 | 0.5 | NA | NA | NA | NA | NA | NA | 1 | 1 | 1 | 1 |
| 10_1104039 | NA | NA | 1 | 1 | 2 | 1 | NA | NA | 3 | 1 | NA | NA | 1 | 1 | NA | NA | NA | NA | 1 | 1 |
| 10_1109025 | 2 | 0.5 | 1 | 0 | 9 | 0 | 6 | 0 | 18 | 0.0556 | NA | NA | 5 | 0 | 4 | 0 | 6 | 0.1667 | 15 | 0.0667 |
| 10_1109187 | NA | NA | NA | NA | 1 | 0 | 2 | 0.5 | 3 | 0.3333 | NA | NA | 1 | 0 | 2 | 0 | 5 | 0.2 | 8 | 0.125 |
| 10_1116531 | NA | NA | 1 | 1 | NA | NA | NA | NA | 1 | 1 | NA | NA | NA | NA | 1 | 1 | NA | NA | 1 | 1 |
| 10_1116610 | NA | NA | 1 | 1 | NA | NA | NA | NA | 1 | 1 | NA | NA | NA | NA | 1 | 1 | NA | NA | 1 | 1 |
| 10_1116637 | NA | NA | 1 | 1 | NA | NA | NA | NA | 1 | 1 | NA | NA | NA | NA | 1 | 1 | NA | NA | 1 | 1 |
| 10_1116665 | NA | NA | 1 | 1 | NA | NA | NA | NA | 1 | 1 | NA | NA | NA | NA | 1 | 1 | NA | NA | 1 | 1 |
| 10_1116678 | NA | NA | 1 | 1 | NA | NA | NA | NA | 1 | 1 | NA | NA | NA | NA | 1 | 1 | NA | NA | 1 | 1 |
| 10_1117099 | NA | NA | 1 | 1 | NA | NA | NA | NA | 1 | 1 | NA | NA | NA | NA | 1 | 1 | NA | NA | 1 | 1 |
| 10_1117104 | NA | NA | 1 | 1 | NA | NA | NA | NA | 1 | 1 | NA | NA | NA | NA | 1 | 1 | NA | NA | 1 | 1 |
| 10_1117110 | NA | NA | 1 | 1 | NA | NA | NA | NA | 1 | 1 | NA | NA | NA | NA | 1 | 1 | NA | NA | 1 | 1 |
| 10_1132445 | NA | NA | 3 | 0 | 2 | 0.5 | 1 | 0 | 6 | 0.1667 | 1 | 0 | NA | NA | 1 | 0 | 1 | 1 | 3 | 0.3333 |
| 10_1133294 | NA | NA | 3 | 0 | 1 | 1 | 1 | 0 | 5 | 0.2 | 1 | 1 | NA | NA | NA | NA | NA | NA | 1 | 1 |
| 10_1133311 | NA | NA | 3 | 0.6667 | 1 | 1 | 1 | 0 | 5 | 0.6 | 1 | 0 | NA | NA | 1 | 1 | 1 | 1 | 3 | 0.6667 |
| 10_1133320 | NA | NA | 3 | 1 | 2 | 0.5 | 1 | 1 | 6 | 0.8333 | 1 | 1 | NA | NA | 1 | 1 | 1 | 1 | 3 | 1 |
| 10_1133324 | NA | NA | 2 | 1 | 2 | 1 | NA | NA | 4 | 1 | 1 | 1 | NA | NA | NA | NA | NA | NA | 1 | 1 |
| 10_1148533 | NA | NA | NA | NA | 1 | 1 | NA | NA | 1 | 1 | NA | NA | 1 | 1 | NA | NA | NA | NA | 1 | 1 |
| 10_1225833 | 5 | 0.2 | 5 | 0.2 | 4 | 0 | 7 | 0 | 21 | 0.0952 | 2 | 0 | 5 | 0 | 9 | 0 | 4 | 0.5 | 20 | 0.1 |
| 10_1225925 | 4 | 0 | 4 | 0 | 4 | 0 | 8 | 0.125 | 20 | 0.05 | 3 | 0 | 6 | 0.1667 | 7 | 0 | 4 | 0 | 20 | 0.05 |
| 10_1225977 | 3 | 0 | 2 | 0 | 3 | 0.3333 | 5 | 0 | 13 | 0.0769 | 2 | 0.5 | 4 | 0 | 2 | 0 | 4 | 0 | 12 | 0.0833 |
| 10_1228173 | 4 | 0.25 | 2 | 0 | 3 | 0.3333 | 2 | 0 | 11 | 0.1818 | 3 | 0.3333 | 4 | 0 | 4 | 0.25 | 2 | 0.5 | 13 | 0.2308 |
| 10_1249036 | NA | NA | NA | NA | 1 | 1 | NA | NA | 1 | 1 | NA | NA | NA | NA | 2 | 0 | 3 | 0.3333 | 5 | 0.2 |
| 10_1249051 | NA | NA | NA | NA | 1 | 1 | NA | NA | 1 | 1 | NA | NA | NA | NA | 2 | 0.5 | 2 | 0 | 4 | 0.25 |
| 10_1288460 | 9 | 0.2222 | 7 | 0.5714 | 17 | 0.1765 | 13 | 0.1538 | 46 | 0.2391 | 11 | 0.0909 | 13 | 0.2308 | 17 | 0.1765 | 16 | 0.1875 | 57 | 0.1754 |
| 10_1288462 | 9 | 0.4444 | 11 | 0.2727 | 19 | 0.2632 | 13 | 0.3077 | 52 | 0.3077 | 11 | 0.0909 | 11 | 0.4545 | 17 | 0.2353 | 14 | 0.5 | 53 | 0.3208 |
| 10_1288475 | 10 | 0.9 | 12 | 1 | 18 | 0.8333 | 15 | 0.9333 | 55 | 0.9091 | 13 | 0.8462 | 13 | 0.6923 | 20 | 1 | 15 | 0.9333 | 61 | 0.8852 |
| 10_1288478 | 8 | 0.875 | 5 | 0.8 | 12 | 0.9167 | 10 | 0.9 | 35 | 0.8857 | 9 | 0.8889 | 7 | 0.7143 | 17 | 0.8824 | 12 | 1 | 45 | 0.8889 |
| 10_1288482 | 10 | 0.1 | 11 | 0.0909 | 18 | 0.0556 | 13 | 0.1538 | 52 | 0.0962 | 14 | 0 | 10 | 0.1 | 20 | 0.05 | 15 | 0.0667 | 59 | 0.0508 |
| 10_1300328 | NA | NA | 1 | 1 | NA | NA | 1 | 1 | 2 | 1 | 1 | 1 | NA | NA | NA | NA | 1 | 1 | 2 | 1 |
| 10_1300506 | 1 | 0 | 2 | 0.5 | NA | NA | 2 | 1 | 5 | 0.6 | 2 | 0 | NA | NA | 1 | 1 | 2 | 0 | 5 | 0.2 |

| | | | | | | | | | | | | | | | | | | | | |
|---|---|---|---|---|---|---|---|---|---|---|---|---|---|---|---|---|---|---|---|---|
| 10_1300537 | 1 | 0 | 2 | 0 | NA | NA | 3 | 0.3333 | 6 | 0.1667 | 1 | 0 | NA | NA | 1 | 0 | 2 | 0.5 | 4 | 0.25 |
| 10_1300539 | 1 | 0 | 2 | 1 | NA | NA | 2 | 1 | 5 | 0.8 | 2 | 1 | NA | NA | 1 | 1 | 2 | 1 | 5 | 1 |
| 10_1300557 | 1 | 0 | 1 | 0 | NA | NA | 3 | 0.3333 | 5 | 0.2 | 2 | 0.5 | NA | NA | 1 | 0 | 2 | 0 | 5 | 0.2 |
| 10_1300567 | 2 | 1 | 2 | 0.5 | NA | NA | 3 | 1 | 7 | 0.8571 | 2 | 0.5 | NA | NA | 1 | 1 | 2 | 0.5 | 5 | 0.6 |
| 10_1300599 | 2 | 0.5 | 2 | 0 | NA | NA | 3 | 0 | 7 | 0.1429 | 2 | 0 | 1 | 1 | 3 | 0.3333 | 2 | 0 | 8 | 0.25 |
| 10_1300611 | 2 | 0.5 | 1 | 0 | NA | NA | 3 | 1 | 6 | 0.6667 | 2 | 0.5 | 1 | 1 | 3 | 0.6667 | 2 | 0.5 | 8 | 0.625 |
| 10_1300617 | 2 | 1 | 1 | 1 | NA | NA | 4 | 0.5 | 7 | 0.7143 | 2 | 0.5 | 1 | 0 | 3 | 1 | 2 | 0 | 8 | 0.5 |
| 10_1300626 | 2 | 0 | 1 | 0 | NA | NA | 4 | 0.25 | 7 | 0.1429 | 2 | 0 | 2 | 0.5 | 3 | 0.3333 | 2 | 0 | 9 | 0.2222 |
| 10_1300632 | 2 | 0.5 | 1 | 0 | NA | NA | 4 | 0.5 | 7 | 0.4286 | 2 | 0 | 2 | 0.5 | 3 | 0 | 2 | 0 | 9 | 0.1111 |
| 10_1300634 | 2 | 0.5 | 1 | 1 | NA | NA | 4 | 0.5 | 7 | 0.5714 | 2 | 1 | 2 | 1 | 3 | 0.3333 | 2 | 0.5 | 9 | 0.6667 |
| 10_1300640 | 3 | 0.3333 | 1 | 0 | NA | NA | 4 | 0.5 | 8 | 0.375 | 2 | 0 | 1 | 1 | 3 | 0.6667 | 2 | 0.5 | 8 | 0.5 |
| 10_1300651 | 3 | 0.3333 | 2 | 0 | NA | NA | 4 | 0 | 9 | 0.1111 | 2 | 0 | 1 | 1 | 3 | 0.3333 | 1 | 0 | 7 | 0.2857 |
| 10_1300669 | 2 | 0.5 | 2 | 0 | NA | NA | 4 | 0 | 8 | 0.125 | 2 | 0 | 2 | 0 | 2 | 0 | 2 | 0.5 | 8 | 0.125 |
| 10_1300675 | 3 | 0.6667 | 2 | 0 | NA | NA | 4 | 0.25 | 9 | 0.3333 | 2 | 0 | 2 | 0 | 3 | 0.3333 | 2 | 0 | 9 | 0.1111 |
| 10_1300686 | 2 | 0 | 1 | 1 | NA | NA | 3 | 0.6667 | 6 | 0.5 | 1 | 1 | 2 | 0 | 3 | 0.6667 | 1 | 0 | 7 | 0.4286 |
| 10_1300695 | 3 | 0 | 2 | 0.5 | NA | NA | 4 | 0 | 9 | 0.1111 | 2 | 0 | 3 | 0 | 2 | 0.5 | 2 | 0 | 9 | 0.1111 |
| 10_1300697 | 3 | 0 | 2 | 0.5 | NA | NA | 4 | 0.5 | 9 | 0.3333 | 1 | 1 | 2 | 0 | 3 | 0.3333 | 2 | 0.5 | 8 | 0.375 |
| 10_1300703 | 3 | 0.3333 | 2 | 0 | NA | NA | 4 | 0.75 | 9 | 0.4444 | 2 | 0.5 | 2 | 0 | 3 | 0.3333 | 2 | 0 | 9 | 0.2222 |
| 10_1300709 | 3 | 0 | 1 | 0 | NA | NA | 4 | 0.5 | 8 | 0.25 | 1 | 1 | 3 | 0 | 3 | 0 | 2 | 0 | 9 | 0.1111 |
| 10_1300715 | 3 | 0.3333 | 2 | 0 | NA | NA | 4 | 0.5 | 9 | 0.3333 | 2 | 0.5 | 3 | 0 | 4 | 0 | 2 | 0.5 | 11 | 0.1818 |
| 10_1300721 | 3 | 0.3333 | 1 | 0 | NA | NA | 4 | 0 | 8 | 0.125 | 2 | 1 | 3 | 0 | 4 | 0 | 2 | 0 | 11 | 0.1818 |
| 10_1300724 | 3 | 0 | 2 | 0 | NA | NA | 4 | 0.25 | 9 | 0.1111 | 2 | 0 | 3 | 0.3333 | 4 | 0.25 | 2 | 0 | 11 | 0.1818 |
| 10_1300726 | 3 | 0 | 2 | 0.5 | NA | NA | 4 | 0.5 | 9 | 0.3333 | 2 | 0 | 3 | 0 | 4 | 0.25 | 3 | 0 | 12 | 0.0833 |
| 10_1305462 | NA | NA | NA | NA | 3 | 0.3333 | 1 | 0 | 4 | 0.25 | 1 | 1 | 3 | 0.3333 | 4 | 0.5 | 3 | 0 | 11 | 0.3636 |
| 10_1308946 | NA | NA | 2 | 1 | NA | NA | NA | NA | 2 | 1 | NA | NA | 1 | 1 | 2 | 0 | 1 | 0 | 4 | 0.25 |
| 10_1308951 | NA | NA | 3 | 0.6667 | NA | NA | 1 | 1 | 4 | 0.75 | NA | NA | 1 | 0 | 2 | 0.5 | 1 | 0 | 4 | 0.25 |
| 10_1308958 | NA | NA | 4 | 0.5 | 1 | 1 | NA | NA | 5 | 0.6 | NA | NA | 1 | 1 | 2 | 1 | NA | NA | 3 | 1 |
| 10_1308964 | NA | NA | 3 | 0.6667 | 1 | 1 | 1 | 0 | 5 | 0.6 | NA | NA | 1 | 1 | 2 | 1 | 1 | 0 | 4 | 0.75 |
| 10_1312923 | NA | NA | 3 | 1 | NA | NA | 1 | 0 | 4 | 0.75 | NA | NA | 1 | 0 | 2 | 0.5 | NA | NA | 3 | 0.3333 |
| 10_1312936 | NA | NA | 3 | 1 | NA | NA | NA | NA | 3 | 1 | NA | NA | 1 | 0 | 1 | 1 | NA | NA | 2 | 0.5 |
| 10_1338154 | 6 | 0 | 5 | 0.4 | 5 | 0.4 | 2 | 0 | 18 | 0.2222 | 1 | 0 | NA | NA | 3 | 0.3333 | 3 | 0 | 7 | 0.1429 |
| 10_1338158 | 7 | 0.5714 | 5 | 0.4 | 4 | 0.25 | 2 | 0.5 | 18 | 0.4444 | 1 | 0 | 2 | 0 | 4 | 0.25 | 3 | 0.3333 | 10 | 0.2 |
| 10_1339056 | 7 | 0.8571 | 5 | 0.6 | 6 | 0.6667 | 4 | 0.5 | 22 | 0.6818 | 1 | 1 | 3 | 0.6667 | 4 | 0.75 | 3 | 0.6667 | 11 | 0.7273 |
| 10_1339098 | 7 | 1 | 4 | 0.75 | 6 | 1 | 8 | 0.875 | 25 | 0.92 | 1 | 1 | 3 | 1 | 4 | 0.5 | 3 | 0.6667 | 11 | 0.7273 |
| 10_1339100 | 7 | 0.1429 | 6 | 0.1667 | 6 | 0 | 8 | 0.625 | 27 | 0.2593 | 1 | 0 | 3 | 0 | 3 | 0.3333 | 3 | 0.3333 | 10 | 0.2 |
| 10_1339111 | 6 | 1 | 2 | 0.5 | 6 | 0.6667 | 7 | 0.4286 | 21 | 0.6667 | 2 | 1 | 3 | 0.6667 | 2 | 0.5 | 3 | 0.3333 | 10 | 0.6 |
| 10_1339113 | 7 | 0.8571 | 6 | 1 | 7 | 0.8571 | 9 | 1 | 29 | 0.931 | 2 | 1 | 2 | 1 | 3 | 1 | 3 | 1 | 10 | 1 |
| 10_1339122 | 7 | 0.4286 | 3 | 1 | 7 | 0 | 6 | 0.1667 | 23 | 0.3043 | 3 | 0.3333 | 3 | 0 | 3 | 0.3333 | 4 | 0 | 13 | 0.1538 |
| 10_1339129 | 5 | 0.4 | 4 | 0.25 | 1 | 0 | 2 | 0 | 12 | 0.25 | 3 | 0 | 1 | 0 | 2 | 0.5 | 2 | 0 | 8 | 0.125 |
| 10_1340022 | 2 | 1 | 1 | 0 | 7 | 0.8571 | 7 | 0.4286 | 17 | 0.6471 | 2 | 0.5 | 4 | 1 | 4 | 0.75 | 4 | 0.25 | 14 | 0.6429 |
| 10_1340026 | 2 | 1 | 1 | 1 | 8 | 1 | 7 | 0.8571 | 18 | 0.9444 | 2 | 1 | 5 | 1 | 4 | 1 | 5 | 0.8 | 16 | 0.9375 |
| 10_1340060 | 1 | 1 | 2 | 0 | 7 | 0.4286 | 6 | 0.3333 | 16 | 0.375 | 3 | 0.6667 | 4 | 0.25 | 3 | 0.3333 | 4 | 0.5 | 14 | 0.4286 |
| 10_1340068 | 1 | 0 | 2 | 0 | 7 | 0 | 8 | 0.125 | 18 | 0.0556 | 3 | 0 | 4 | 0 | 4 | 0 | 5 | 0.2 | 16 | 0.0625 |
| 10_1343881 | 1 | 0 | 2 | 0.5 | NA | NA | 1 | 0 | 4 | 0.25 | NA | NA | 2 | 0.5 | 1 | 0 | 2 | 0 | 5 | 0.2 |
| 10_1343922 | 1 | 0 | 2 | 0.5 | NA | NA | 2 | 0 | 5 | 0.2 | NA | NA | 2 | 0 | 1 | 0 | 2 | 0.5 | 5 | 0.2 |
| 10_1343932 | 1 | 1 | 2 | 0 | NA | NA | 2 | 0 | 5 | 0.2 | NA | NA | 2 | 0 | 1 | 0 | 2 | 0.5 | 5 | 0.2 |
| 10_1352322 | 6 | 0.6667 | 8 | 1 | 5 | 0.8 | 4 | 0.75 | 23 | 0.8261 | 5 | 0.8 | 5 | 0.4 | 12 | 0.6667 | 2 | 0.5 | 24 | 0.625 |
| 10_1352328 | 10 | 0.2 | 8 | 0.125 | 8 | 0.375 | 5 | 0.2 | 31 | 0.2258 | 5 | 0.4 | 10 | 0.1 | 12 | 0.0833 | 4 | 0 | 31 | 0.129 |
| 10_1400319 | NA | NA | 1 | 1 | NA | NA | 1 | 1 | 2 | 1 | NA | NA | NA | NA | 1 | 1 | NA | NA | 1 | 1 |
| 10_1400340 | 1 | 1 | 1 | 1 | NA | NA | 1 | 1 | 3 | 1 | NA | NA | NA | NA | 1 | 1 | NA | NA | 1 | 1 |
| 10_1400341 | NA | NA | 1 | 1 | NA | NA | 1 | 0 | 2 | 0.5 | NA | NA | NA | NA | 2 | 0.5 | NA | NA | 2 | 0.5 |
| 10_1400349 | 1 | 0 | 1 | 1 | NA | NA | 1 | 0 | 3 | 0.3333 | NA | NA | NA | NA | 2 | 0.5 | NA | NA | 2 | 0.5 |
| 10_1400381 | 1 | 0 | 1 | 1 | 2 | 0.5 | 1 | 1 | 5 | 0.6 | NA | NA | NA | NA | 2 | 0.5 | NA | NA | 2 | 0.5 |
| 10_1400431 | 1 | 1 | 1 | 1 | 2 | 1 | 1 | 1 | 5 | 1 | NA | NA | NA | NA | 2 | 1 | NA | NA | 2 | 1 |
| 10_1400459 | 1 | 1 | 2 | 1 | 2 | 1 | 1 | 0 | 6 | 0.8333 | 1 | 0 | NA | NA | 3 | 1 | NA | NA | 4 | 0.75 |
| 10_1400501 | 1 | 1 | 1 | 1 | 4 | 0.75 | 1 | 1 | 7 | 0.8571 | 1 | 0 | NA | NA | 4 | 1 | 1 | 1 | 6 | 0.8333 |
| 10_1400541 | 1 | 0 | 2 | 0.5 | 4 | 0 | 1 | 1 | 8 | 0.25 | 1 | 0 | 1 | 0 | 3 | 0.3333 | 1 | 0 | 6 | 0.1667 |
| 10_1400586 | 1 | 0 | 2 | 0 | 4 | 0.25 | 1 | 0 | 8 | 0.125 | 1 | 1 | 2 | 0 | 3 | 0 | 1 | 0 | 7 | 0.1429 |
| 10_1400615 | 1 | 0 | 3 | 0.3333 | 4 | 0 | NA | NA | 8 | 0.125 | 2 | 0.5 | 3 | 0 | 3 | 0 | 2 | 0.5 | 10 | 0.2 |
| 10_1400640 | 1 | 1 | 2 | 0 | 4 | 0.75 | 1 | 0 | 8 | 0.5 | 1 | 1 | 1 | 0 | 2 | 0.5 | 2 | 1 | 6 | 0.6667 |
| 10_1400685 | NA | NA | 5 | 1 | 2 | 1 | 1 | 1 | 8 | 1 | 2 | 0.5 | 1 | 0 | 4 | 1 | 2 | 1 | 9 | 0.7778 |
| 10_1400715 | 1 | 0 | 4 | 0.25 | 5 | 0 | 1 | 0 | 11 | 0.0909 | 1 | 0 | 2 | 0 | 4 | 0.25 | 4 | 0 | 11 | 0.0909 |
| 10_1400744 | 1 | 0 | 6 | 0.5 | 4 | 0.75 | 2 | 1 | 13 | 0.6154 | NA | NA | 1 | 0 | 6 | 0.6667 | 4 | 0.5 | 11 | 0.5455 |
| 10_1400749 | 1 | 0 | 5 | 0.4 | 5 | 0.2 | 2 | 0.5 | 13 | 0.3077 | 1 | 1 | 3 | 1 | 5 | 0.6 | 4 | 0.5 | 13 | 0.6923 |
| 10_1400760 | 1 | 1 | 5 | 0 | 5 | 0.2 | 2 | 0 | 13 | 0.1538 | 1 | 0 | 4 | 0.25 | 4 | 0.25 | 5 | 0.2 | 14 | 0.2143 |

| | | | | | | | | | | | | | | | | | | | | |
|---|---|---|---|---|---|---|---|---|---|---|---|---|---|---|---|---|---|---|---|---|
| 10_1400778 | 1 | 0 | 6 | 0.3333 | 5 | 0.4 | 2 | 0 | 14 | 0.2857 | 2 | 0 | 4 | 0.25 | 5 | 0.4 | 5 | 0.4 | 16 | 0.3125 |
| 10_1400781 | 1 | 0 | 6 | 0.1667 | 5 | 0.2 | 2 | 0 | 14 | 0.1429 | 3 | 0 | 4 | 0.25 | 6 | 0 | 4 | 0 | 17 | 0.0588 |
| 10_1400806 | 1 | 1 | 7 | 0.4286 | 5 | 0.2 | 3 | 0.3333 | 16 | 0.375 | 2 | 0.5 | 3 | 0.3333 | 8 | 0.5 | 4 | 0.5 | 17 | 0.4706 |
| 10_1400828 | 2 | 0 | 7 | 0.2857 | 8 | 0.25 | 5 | 0.2 | 22 | 0.2273 | 3 | 0.3333 | 4 | 0.5 | 8 | 0.125 | 5 | 0.4 | 20 | 0.3 |
| 10_1400832 | 2 | 0.5 | 5 | 0.2 | 8 | 0.125 | 4 | 0 | 19 | 0.1579 | 2 | 0.5 | 5 | 0 | 8 | 0 | 3 | 0 | 18 | 0.0556 |
| 10_1401907 | 2 | 1 | 5 | 0.8 | 6 | 0.6667 | 7 | 1 | 20 | 0.85 | 3 | 1 | 3 | 0.6667 | 8 | 0.875 | 8 | 0.875 | 22 | 0.8636 |
| 10_1401914 | 2 | 1 | 10 | 1 | 8 | 0.75 | 7 | 0.8571 | 27 | 0.8889 | 3 | 1 | 5 | 0.8 | 10 | 0.9 | 10 | 0.9 | 28 | 0.8929 |
| 10_1401920 | 2 | 1 | 10 | 0.3 | 8 | 0.375 | 7 | 0.2857 | 27 | 0.3704 | 4 | 0.5 | 4 | 0.75 | 12 | 0.5 | 10 | 0.5 | 30 | 0.5333 |
| 10_1401921 | 2 | 0 | 8 | 0.5 | 7 | 0.1429 | 7 | 0 | 24 | 0.2083 | 4 | 0.5 | 4 | 0.25 | 11 | 0.3636 | 10 | 0.1 | 29 | 0.2759 |
| 10_1401929 | 1 | 1 | 9 | 1 | 8 | 0.75 | 6 | 0.3333 | 24 | 0.75 | 3 | 0.3333 | 4 | 0.75 | 8 | 0.75 | 10 | 0.5 | 25 | 0.6 |
| 10_1401954 | 1 | 0 | 9 | 1 | 7 | 1 | 8 | 1 | 25 | 0.96 | 4 | 1 | 5 | 1 | 7 | 1 | 10 | 1 | 26 | 1 |
| 10_1401965 | 2 | 0 | 8 | 0.25 | 7 | 0.4286 | 9 | 0.3333 | 26 | 0.3077 | 4 | 0.75 | 3 | 0.6667 | 11 | 0.4545 | 10 | 0.7 | 28 | 0.6071 |
| 10_1402472 | 2 | 0.5 | 9 | 0.2222 | 8 | 0.375 | 7 | 0.1429 | 26 | 0.2692 | 4 | 0 | 4 | 0 | 12 | 0.5 | 9 | 0 | 29 | 0.2069 |
| 10_1402476 | 2 | 1 | 11 | 0.7273 | 8 | 0.375 | 9 | 0.7778 | 30 | 0.6667 | 4 | 0.25 | 5 | 0.4 | 13 | 0.4615 | 11 | 0.4545 | 33 | 0.4242 |
| 10_1402499 | 2 | 0.5 | 8 | 0 | 6 | 0.1667 | 7 | 0.1429 | 23 | 0.1304 | 4 | 0 | 5 | 0 | 11 | 0.1818 | 9 | 0.1111 | 29 | 0.1034 |
| 10_1402506 | 3 | 0.3333 | 10 | 0 | 6 | 0.1667 | 8 | 0.125 | 27 | 0.1111 | 4 | 0.25 | 5 | 0.4 | 12 | 0.0833 | 10 | 0.1 | 31 | 0.1613 |
| 10_1402507 | 3 | 0 | 12 | 0.3333 | 9 | 0.1111 | 9 | 0.1111 | 33 | 0.1818 | 5 | 0 | 5 | 0 | 14 | 0.1429 | 12 | 0.25 | 36 | 0.1389 |
| 10_1402510 | 3 | 0 | 13 | 0.1538 | 8 | 0.125 | 7 | 0.4286 | 31 | 0.1935 | 5 | 0 | 3 | 0 | 13 | 0.2308 | 11 | 0.4545 | 32 | 0.25 |
| 10_1402511 | 2 | 0 | 12 | 0.1667 | 8 | 0 | 8 | 0 | 30 | 0.0667 | 5 | 0 | 6 | 0.3333 | 13 | 0.0769 | 11 | 0 | 35 | 0.0857 |
| 10_1402512 | 1 | 1 | 12 | 0.9167 | 8 | 1 | 8 | 0.75 | 29 | 0.8966 | 4 | 1 | 5 | 0.8 | 12 | 1 | 10 | 1 | 31 | 0.9677 |
| 10_1402521 | 3 | 0.3333 | 10 | 0.2 | 9 | 0 | 7 | 0.4286 | 29 | 0.2069 | 4 | 0 | 5 | 0.2 | 13 | 0.3846 | 10 | 0.2 | 32 | 0.25 |
| 10_1402524 | 2 | 1 | 11 | 0.9091 | 5 | 0.8 | 7 | 1 | 25 | 0.92 | 4 | 1 | 4 | 1 | 8 | 1 | 7 | 1 | 23 | 1 |
| 10_1402525 | NA | NA | NA | NA | 1 | 1 | 1 | 1 | 2 | 1 | NA | NA | NA | NA | 1 | 1 | 1 | 1 | 2 | 1 |
| 10_1402530 | 3 | 0.6667 | 12 | 0.5 | 8 | 0.125 | 9 | 0.6667 | 32 | 0.4688 | 5 | 0.6 | 6 | 0.6667 | 13 | 0.7692 | 10 | 0.8 | 34 | 0.7353 |
| 10_1402532 | 3 | 1 | 11 | 1 | 9 | 1 | 9 | 0.8889 | 32 | 0.9688 | 5 | 1 | 5 | 1 | 15 | 0.8 | 11 | 1 | 36 | 0.9167 |
| 10_1402534 | 3 | 1 | 12 | 1 | 7 | 1 | 9 | 1 | 31 | 1 | 3 | 1 | 6 | 1 | 13 | 0.8462 | 10 | 1 | 32 | 0.9375 |
| 10_1402540 | 3 | 1 | 12 | 1 | 9 | 1 | 9 | 1 | 33 | 1 | 5 | 1 | 6 | 1 | 15 | 1 | 11 | 1 | 37 | 1 |
| 10_1402545 | 3 | 0.3333 | 13 | 0.1538 | 10 | 0 | 9 | 0 | 35 | 0.0857 | 6 | 0.3333 | 6 | 0.1667 | 15 | 0.1333 | 11 | 0.0909 | 38 | 0.1579 |
| 10_1402549 | 2 | 1 | 11 | 1 | 10 | 1 | 8 | 1 | 31 | 1 | 6 | 1 | 5 | 1 | 12 | 1 | 11 | 1 | 34 | 1 |
| 10_1402584 | 5 | 0.6 | 13 | 0.3846 | 6 | 0.8333 | 9 | 0.7778 | 33 | 0.6061 | 5 | 0.8 | 5 | 0.8 | 15 | 0.3333 | 7 | 0.5714 | 32 | 0.5312 |
| 10_1402592 | 6 | 0 | 13 | 0.0769 | 10 | 0.1 | 9 | 0.2222 | 38 | 0.1053 | 6 | 0.1667 | 7 | 0.1429 | 15 | 0 | 10 | 0 | 38 | 0.0526 |
| 10_1402643 | 6 | 0.5 | 15 | 0.1333 | 13 | 0.2308 | 7 | 0.2857 | 41 | 0.2439 | 5 | 0.2 | 5 | 0.4 | 15 | 0.3333 | 11 | 0.5455 | 36 | 0.3889 |
| 10_1402650 | 7 | 1 | 13 | 1 | 9 | 1 | 6 | 1 | 35 | 1 | 4 | 1 | 7 | 0.8571 | 14 | 1 | 11 | 1 | 36 | 0.9722 |
| 10_1402661 | 8 | 0.5 | 14 | 0.4286 | 14 | 0.5 | 9 | 0.5556 | 45 | 0.4889 | 5 | 0.4 | 6 | 0.6667 | 18 | 0.3333 | 13 | 0.3077 | 42 | 0.381 |
| 10_1402662 | 7 | 0.7143 | 15 | 0.6667 | 9 | 0.8889 | 4 | 1 | 35 | 0.7714 | 3 | 0.3333 | 7 | 0.8571 | 17 | 0.7647 | 11 | 0.7273 | 38 | 0.7368 |
| 10_1402665 | 7 | 0.8571 | 11 | 0.9091 | 12 | 0.8333 | 9 | 1 | 39 | 0.8974 | 4 | 1 | 6 | 1 | 19 | 1 | 15 | 0.9333 | 44 | 0.9773 |
| 10_1428073 | NA | NA | NA | NA | 2 | 1 | 1 | 1 | 3 | 1 | NA | NA | 2 | 1 | NA | NA | NA | NA | 2 | 1 |
| 10_1428074 | NA | NA | NA | NA | 2 | 1 | 1 | 1 | 3 | 1 | NA | NA | 2 | 1 | NA | NA | NA | NA | 2 | 1 |
| 10_1428077 | NA | NA | NA | NA | 2 | 0.5 | 1 | 1 | 3 | 0.6667 | 1 | 1 | 2 | 0 | NA | NA | NA | NA | 3 | 0.3333 |
| 10_1428087 | NA | NA | NA | NA | 1 | 1 | 1 | 1 | 2 | 1 | NA | NA | 1 | 1 | NA | NA | NA | NA | 1 | 1 |
| 10_1428105 | NA | NA | NA | NA | 1 | 1 | NA | NA | 1 | 1 | 1 | 1 | NA | NA | NA | NA | NA | NA | 1 | 1 |
| 10_1434307 | NA | NA | NA | NA | NA | NA | 1 | 1 | 1 | 1 | 1 | 1 | 1 | 1 | NA | NA | NA | NA | 2 | 1 |
| 10_1434313 | NA | NA | NA | NA | NA | NA | 1 | 1 | 1 | 1 | 1 | 1 | NA | NA | NA | NA | NA | NA | 1 | 1 |
| 10_1434321 | NA | NA | NA | NA | NA | NA | 1 | 1 | 1 | 1 | 1 | 0 | 1 | 1 | NA | NA | NA | NA | 2 | 0.5 |
| 10_1434322 | NA | NA | NA | NA | NA | NA | 1 | 1 | 1 | 1 | NA | NA | 1 | 1 | NA | NA | NA | NA | 1 | 1 |
| 10_1453114 | 7 | 0 | 8 | 0.125 | 8 | 0 | 11 | 0.0909 | 34 | 0.0588 | 6 | 0 | 8 | 0 | 7 | 0 | 11 | 0.1818 | 32 | 0.0625 |
| 10_1453132 | 10 | 0.1 | 9 | 0.3333 | 8 | 0.125 | 11 | 0.2727 | 38 | 0.2105 | 6 | 0.3333 | 7 | 0.2857 | 7 | 0 | 12 | 0.3333 | 32 | 0.25 |
| 10_1453136 | 10 | 0 | 8 | 0.125 | 9 | 0 | 12 | 0.0833 | 39 | 0.0513 | 6 | 0.1667 | 8 | 0 | 5 | 0.2 | 9 | 0 | 28 | 0.0714 |
| 10_1453140 | 6 | 0.1667 | 9 | 0.1111 | 6 | 0 | 10 | 0.2 | 31 | 0.129 | 5 | 0.4 | 7 | 0.1429 | 6 | 0.1667 | 9 | 0.3333 | 27 | 0.2593 |
| 10_1455035 | NA | NA | 2 | 0.5 | NA | NA | NA | NA | 2 | 0.5 | NA | NA | 1 | 1 | NA | NA | NA | NA | 1 | 1 |
| 10_1463572 | 1 | 1 | 3 | 0.3333 | 2 | 1 | NA | NA | 6 | 0.6667 | 3 | 0 | NA | NA | NA | NA | 2 | 0.5 | 5 | 0.2 |
| 10_1463588 | 1 | 0 | 4 | 0 | 3 | 0.3333 | NA | NA | 8 | 0.125 | 4 | 0.25 | NA | NA | NA | NA | 3 | 0 | 7 | 0.1429 |
| 10_1463594 | 1 | 1 | 4 | 0 | 4 | 0.25 | NA | NA | 9 | 0.2222 | 5 | 0.2 | NA | NA | NA | NA | 3 | 0.3333 | 8 | 0.25 |
| 10_1463595 | 1 | 0 | 4 | 0 | 4 | 0.25 | NA | NA | 9 | 0.1111 | 5 | 0.2 | NA | NA | NA | NA | 3 | 0 | 8 | 0.125 |
| 10_1464260 | 1 | 0 | 5 | 0 | 4 | 0.25 | NA | NA | 10 | 0.1 | 4 | 0 | NA | NA | 1 | 0 | 3 | 0.3333 | 8 | 0.125 |
| 10_1464271 | NA | NA | 6 | 1 | 5 | 0.8 | 1 | 1 | 12 | 0.9167 | 7 | 0.8571 | NA | NA | 1 | 0 | 7 | 0.7143 | 15 | 0.7333 |
| 10_1464621 | 1 | 1 | 8 | 0.75 | 5 | 0.4 | 1 | 1 | 15 | 0.6667 | 7 | 0.7143 | NA | NA | 1 | 0 | 9 | 0.7778 | 17 | 0.7059 |
| 10_1464624 | 1 | 0 | 10 | 0.1 | 5 | 0 | 2 | 0 | 18 | 0.0556 | 7 | 0.2857 | NA | NA | 1 | 0 | 9 | 0.2222 | 17 | 0.2353 |
| 10_1464627 | 1 | 1 | 10 | 0.9 | 5 | 0.8 | 2 | 1 | 18 | 0.8889 | 6 | 0.8333 | NA | NA | 1 | 1 | 7 | 1 | 14 | 0.9286 |
| 10_1464637 | 1 | 1 | 11 | 0.4545 | 5 | 0.6 | 2 | 0 | 19 | 0.4737 | 7 | 0.1429 | NA | NA | NA | NA | 9 | 0.4444 | 16 | 0.3125 |
| 10_1464643 | 2 | 0 | 11 | 0.2727 | 4 | 0.75 | 2 | 0.5 | 19 | 0.3684 | 8 | 0.125 | NA | NA | 1 | 1 | 8 | 0.5 | 17 | 0.3529 |
| 10_1464664 | 2 | 0 | 11 | 0.0909 | 5 | 0 | 2 | 0 | 20 | 0.05 | 8 | 0 | NA | NA | 1 | 0 | 9 | 0.1111 | 18 | 0.0556 |
| 10_1759992 | 15 | 0.9333 | 15 | 0.9333 | 11 | 1 | 11 | 0.9091 | 52 | 0.9423 | 5 | 0.8 | 13 | 0.9231 | 13 | 0.9231 | 9 | 1 | 40 | 0.925 |
| 10_1760013 | 17 | 0.5882 | 16 | 0.4375 | 12 | 0.3333 | 12 | 0.3333 | 57 | 0.4386 | 7 | 0.2857 | 17 | 0.4706 | 14 | 0.3571 | 12 | 0.25 | 50 | 0.36 |
| 10_1760034 | 15 | 1 | 17 | 0.9412 | 10 | 1 | 10 | 1 | 52 | 0.9808 | 7 | 1 | 14 | 1 | 10 | 0.9 | 8 | 1 | 39 | 0.9744 |

| | | | | | | | | | | | | | | | | | | | | |
|---|---|---|---|---|---|---|---|---|---|---|---|---|---|---|---|---|---|---|---|---|
| 10_176004( | 18 | 0.5 | 13 | 0.6154 | 12 | 0.5833 | 11 | 0.7273 | 54 | 0.5926 | 7 | 0.7143 | 10 | 0.8 | 12 | 0.6667 | 8 | 0.5 | 37 | 0.6757 |
| 10_176004: | 12 | 1 | 12 | 0.9167 | 11 | 0.9091 | 7 | 1 | 42 | 0.9524 | 4 | 1 | 10 | 0.8 | 10 | 0.8 | 4 | 1 | 28 | 0.8571 |
| 10_176004( | 17 | 1 | 18 | 0.9444 | 12 | 1 | 10 | 1 | 57 | 0.9825 | 8 | 1 | 16 | 1 | 14 | 1 | 11 | 1 | 49 | 1 |
| 10_176005( | 19 | 0.1579 | 18 | 0.0556 | 12 | 0.0833 | 13 | 0.1538 | 62 | 0.1129 | 8 | 0.125 | 15 | 0.3333 | 14 | 0.1429 | 11 | 0.0909 | 48 | 0.1875 |
| 10_176006( | 19 | 0.2105 | 13 | 0.4615 | 12 | 0 | 7 | 0.2857 | 51 | 0.2353 | 8 | 0.375 | 12 | 0.25 | 11 | 0.1818 | 7 | 0.4286 | 38 | 0.2895 |
| 10_176006! | 21 | 0.1429 | 20 | 0.05 | 15 | 0.0667 | 13 | 0 | 69 | 0.0725 | 9 | 0.1111 | 16 | 0.0625 | 14 | 0 | 11 | 0.1818 | 50 | 0.08 |
| 10_176006( | 22 | 0.2273 | 17 | 0.1176 | 15 | 0.0667 | 13 | 0 | 67 | 0.1194 | 9 | 0.2222 | 15 | 0.0667 | 12 | 0.1667 | 11 | 0.0909 | 47 | 0.1277 |
| 10_176006 | 18 | 0.3333 | 16 | 0.4375 | 13 | 0.3077 | 12 | 0.5 | 59 | 0.3898 | 9 | 0 | 14 | 0.2857 | 14 | 0.2143 | 8 | 0.5 | 45 | 0.2444 |
| 10_176009 | 24 | 0.25 | 19 | 0.3158 | 15 | 0.2667 | 12 | 0.1667 | 70 | 0.2571 | 7 | 0.1429 | 15 | 0 | 14 | 0.2143 | 10 | 0.1 | 46 | 0.1087 |
| 10_176010 | 27 | 0.5556 | 19 | 0.5789 | 17 | 0.6471 | 13 | 0.6154 | 76 | 0.5921 | 9 | 0.6667 | 20 | 0.5 | 13 | 0.4615 | 11 | 0.5455 | 53 | 0.5283 |
| 10_176011( | 21 | 0.1429 | 16 | 0.0625 | 15 | 0.1333 | 14 | 0.1429 | 66 | 0.1212 | 6 | 0.1667 | 17 | 0.1176 | 15 | 0 | 11 | 0.0909 | 49 | 0.0816 |
| 10_177192! | NA | NA | NA | NA | 1 | 1 | NA | NA | 1 | 1 | 1 | 1 | NA | NA | 1 | 0 | NA | NA | 2 | 0.5 |
| 10_177750( | 5 | 0.4 | 3 | 0.6667 | 2 | 0 | 2 | 0 | 12 | 0.3333 | 1 | 0 | 1 | 0 | 2 | 0 | 7 | 0.1429 | 11 | 0.0909 |
| 10_177750 | 5 | 0.2 | 3 | 0 | 2 | 0 | 2 | 0 | 12 | 0.0833 | 1 | 0 | 1 | 0 | 2 | 0.5 | 6 | 0 | 10 | 0.1 |
| 10_177780 | 1 | 1 | NA | NA | NA | NA | 1 | 0 | 2 | 0.5 | 1 | 0 | NA | NA | 1 | 1 | 1 | 0 | 3 | 0.3333 |
| 10_177907 | 1 | 0 | NA | NA | NA | NA | 1 | 1 | 2 | 0.5 | 1 | 0 | 1 | 1 | 1 | 1 | 1 | 0 | 4 | 0.5 |
| 10_177908( | 1 | 1 | NA | NA | NA | NA | 1 | 0 | 2 | 0.5 | 1 | 1 | 1 | 1 | NA | NA | 1 | 1 | 3 | 1 |
| 10_177908: | NA | NA | NA | NA | NA | NA | 1 | 1 | 1 | 1 | NA | NA | NA | NA | 1 | 1 | NA | NA | 1 | 1 |
| 10_177908( | 1 | 1 | NA | NA | NA | NA | 1 | 0 | 2 | 0.5 | 1 | 0 | 1 | 1 | NA | NA | NA | NA | 2 | 0.5 |
| 10_177909! | NA | NA | NA | NA | NA | NA | 1 | 1 | 1 | 1 | 1 | 1 | NA | NA | 1 | 0 | 1 | 1 | 3 | 0.6667 |
| 10_177910 | 1 | 1 | NA | NA | NA | NA | 1 | 1 | 2 | 1 | 1 | 1 | 1 | 1 | NA | NA | 1 | 1 | 3 | 1 |
| 10_177912! | NA | NA | NA | NA | NA | NA | 1 | 1 | 1 | 1 | 1 | 0 | NA | NA | 1 | 1 | NA | NA | 2 | 0.5 |
| 10_177913 | 1 | 1 | NA | NA | NA | NA | 1 | 0 | 2 | 0.5 | 1 | 1 | 1 | 1 | NA | NA | NA | NA | 2 | 1 |
| 10_177914 | 1 | 1 | NA | NA | NA | NA | 1 | 1 | 2 | 1 | 1 | 1 | 1 | 1 | NA | NA | NA | NA | 2 | 1 |
| 10_177915! | 1 | 1 | NA | NA | NA | NA | 1 | 0 | 2 | 0.5 | 1 | 0 | 1 | 1 | 1 | 0 | NA | NA | 3 | 0.3333 |
| 10_178873: | 1 | 0 | 4 | 0 | 2 | 0.5 | NA | NA | 7 | 0.1429 | 1 | 0 | 3 | 0.3333 | 3 | 0 | 5 | 0 | 12 | 0.0833 |
| 10_178874 | 1 | 1 | 3 | 0 | 2 | 0 | NA | NA | 6 | 0.1667 | 1 | 0 | 3 | 0.3333 | 4 | 0.5 | 3 | 0 | 11 | 0.2727 |
| 10_180007 | 3 | 0 | 2 | 0 | 5 | 0.2 | 7 | 0.2857 | 17 | 0.1765 | 3 | 0 | 4 | 0 | 4 | 0 | 2 | 1 | 13 | 0.1538 |
| 10_180007( | 4 | 0.25 | 2 | 0 | 6 | 0.1667 | 8 | 0 | 20 | 0.1 | 4 | 0 | 5 | 0 | 6 | 0.1667 | 4 | 0 | 19 | 0.0526 |
| 10_180008 | 2 | 0.5 | 2 | 0.5 | 3 | 0.6667 | 6 | 1 | 13 | 0.7692 | 3 | 0.6667 | 5 | 0.6 | 1 | 1 | 2 | 1 | 11 | 0.7273 |
| 10_180896: | NA | NA | 4 | 0.25 | 1 | 0 | 6 | 0.1667 | 11 | 0.1818 | 2 | 0 | 4 | 0 | 8 | 0.125 | 6 | 0 | 20 | 0.05 |
| 10_180899: | NA | NA | 6 | 0 | 2 | 0.5 | 6 | 0 | 14 | 0.0714 | 3 | 0 | 5 | 0 | 7 | 0.2857 | 4 | 0 | 19 | 0.1053 |
| 10_184072( | NA | NA | 4 | 0.75 | 1 | 0 | NA | NA | 5 | 0.6 | 1 | 1 | NA | NA | NA | NA | 1 | 0 | 2 | 0.5 |
| 10_184075! | 1 | 1 | 1 | 0 | 1 | 1 | NA | NA | 3 | 0.6667 | 1 | 1 | NA | NA | NA | NA | 1 | 1 | 2 | 1 |
| 10_184076 | NA | NA | 3 | 1 | NA | NA | NA | NA | 3 | 1 | 1 | 1 | NA | NA | NA | NA | 1 | 1 | 2 | 1 |
| 10_184077( | 2 | 0 | 4 | 0.25 | 1 | 0 | NA | NA | 7 | 0.1429 | 1 | 0 | NA | NA | NA | NA | 1 | 1 | 2 | 0.5 |
| 10_184078( | 2 | 0.5 | 4 | 0.5 | 1 | 1 | NA | NA | 7 | 0.5714 | 1 | 1 | NA | NA | NA | NA | 1 | 1 | 2 | 1 |
| 10_184081! | 2 | 0 | 3 | 0.3333 | 2 | 0.5 | NA | NA | 7 | 0.2857 | 1 | 1 | NA | NA | NA | NA | 1 | 0 | 2 | 0.5 |
| 10_184092 | 3 | 0.3333 | 6 | 0.3333 | 4 | 0.5 | NA | NA | 13 | 0.3846 | 1 | 1 | NA | NA | NA | NA | 1 | 0 | 2 | 0.5 |
| 10_184581 | 8 | 0 | 7 | 0 | 6 | 0.3333 | 7 | 0.2857 | 28 | 0.1429 | 4 | 0.25 | 6 | 0.3333 | 10 | 0.1 | 11 | 0.0909 | 31 | 0.1613 |
| 10_184584! | 6 | 0 | 7 | 0 | 4 | 0.5 | 7 | 0.1429 | 24 | 0.125 | 4 | 0 | 6 | 0.3333 | 9 | 0 | 11 | 0 | 30 | 0.0667 |
| 10_184594! | 5 | 0 | 5 | 0.2 | 3 | 0 | 6 | 0.1667 | 19 | 0.1053 | 3 | 0.3333 | 4 | 0.5 | 8 | 0 | 8 | 0.125 | 23 | 0.1739 |
| 10_184597( | 4 | 0.25 | 4 | 0 | 3 | 0.6667 | 4 | 0.5 | 15 | 0.3333 | 2 | 1 | 3 | 0.3333 | 6 | 0.3333 | 7 | 0.4286 | 18 | 0.4444 |
| 10_184601 | 2 | 0 | 5 | 1 | 3 | 0.3333 | 3 | 0.6667 | 13 | 0.6154 | 1 | 1 | 3 | 0.3333 | 7 | 0.8571 | 6 | 0.6667 | 17 | 0.7059 |
| 10_184635( | 2 | 0 | 4 | 0 | NA | NA | 1 | 1 | 7 | 0.1429 | 1 | 0 | 1 | 0 | 2 | 0 | 6 | 0.1667 | 10 | 0.1 |
| 10_184644( | 1 | 0 | 2 | 0.5 | 1 | 0 | NA | NA | 4 | 0.25 | 1 | 0 | 1 | 0 | 2 | 0 | 4 | 0.25 | 8 | 0.125 |
| 10_184647 | 2 | 0.5 | 1 | 0 | 1 | 0 | NA | NA | 4 | 0.25 | 1 | 1 | 1 | 0 | NA | NA | 4 | 0 | 6 | 0.1667 |
| 10_184651: | 1 | 1 | 1 | 1 | NA | NA | NA | NA | 2 | 1 | NA | NA | 1 | 0 | NA | NA | 2 | 1 | 3 | 0.6667 |
| 10_184652: | NA | NA | 1 | 1 | NA | NA | NA | NA | 1 | 1 | NA | NA | 1 | 0 | 1 | 0 | 2 | 0.5 | 4 | 0.25 |
| 10_184653: | NA | NA | 1 | 1 | NA | NA | NA | NA | 1 | 1 | NA | NA | 1 | 1 | 1 | 1 | 2 | 1 | 4 | 1 |
| 10_184654( | NA | NA | 1 | 1 | NA | NA | NA | NA | 1 | 1 | NA | NA | 1 | 0 | 1 | 0 | 3 | 0.3333 | 5 | 0.2 |
| 10_187318( | 3 | 0.3333 | 1 | 0 | 1 | 1 | 1 | 0 | 6 | 0.3333 | 1 | 1 | 3 | 0 | 4 | 0.5 | NA | NA | 8 | 0.375 |
| 10_187323 | 3 | 0.3333 | 1 | 0 | 1 | 0 | 2 | 0 | 7 | 0.1429 | 1 | 0 | 3 | 0.3333 | 3 | 0 | NA | NA | 7 | 0.1429 |
| 10_188770( | NA | NA | 1 | 1 | 1 | 0 | NA | NA | 2 | 0.5 | 1 | 1 | NA | NA | 1 | 1 | NA | NA | 2 | 1 |
| 10_188775 | NA | NA | 1 | 1 | 1 | 1 | NA | NA | 2 | 1 | 1 | 1 | NA | NA | 1 | 0 | NA | NA | 2 | 0.5 |
| 10_188776: | NA | NA | 1 | 1 | 1 | 1 | NA | NA | 2 | 1 | 1 | 0 | NA | NA | 1 | 1 | NA | NA | 2 | 0.5 |
| 10_188777: | NA | NA | 1 | 1 | NA | NA | NA | NA | 1 | 1 | 1 | 1 | NA | NA | 1 | 1 | NA | NA | 2 | 1 |
| 10_188781( | NA | NA | 1 | 1 | 1 | 1 | NA | NA | 2 | 1 | 1 | 1 | NA | NA | 1 | 1 | NA | NA | 2 | 1 |
| 10_188782! | NA | NA | 1 | 1 | NA | NA | NA | NA | 1 | 1 | 1 | 1 | NA | NA | 1 | 1 | NA | NA | 2 | 1 |
| 10_188783 | NA | NA | 1 | 1 | 1 | 1 | NA | NA | 2 | 1 | 1 | 1 | NA | NA | 1 | 1 | NA | NA | 2 | 1 |
| 10_188786( | NA | NA | 1 | 1 | NA | NA | NA | NA | 1 | 1 | 1 | 1 | NA | NA | 1 | 0 | NA | NA | 2 | 0.5 |
| 10_194850 | 2 | 0 | 2 | 0 | 5 | 0.4 | 2 | 1 | 11 | 0.3636 | 2 | 0 | 3 | 0 | 3 | 0.3333 | 4 | 0.25 | 12 | 0.1667 |
| 10_194852 | 3 | 1 | 3 | 1 | 5 | 1 | 2 | 1 | 13 | 1 | 3 | 1 | 3 | 1 | 2 | 1 | 5 | 1 | 13 | 1 |
| 10_194853: | 2 | 0.5 | 3 | 0 | 4 | 0 | 2 | 0 | 11 | 0.0909 | 3 | 0.6667 | 3 | 0 | 3 | 0 | 5 | 0 | 14 | 0.1429 |
| 10_194854( | 2 | 1 | 3 | 0.6667 | 5 | 0.4 | 2 | 1 | 12 | 0.6667 | 3 | 1 | 3 | 1 | 3 | 0.3333 | 4 | 0.75 | 13 | 0.7692 |

| | | | | | | | | | | | | | | | | | | | | |
|---|---|---|---|---|---|---|---|---|---|---|---|---|---|---|---|---|---|---|---|---|
| 10_194854 | 2 | 0 | 2 | 0 | 5 | 0 | 2 | 0.5 | 11 | 0.0909 | 3 | 0.6667 | 3 | 0.6667 | 3 | 0 | 5 | 0.2 | 14 | 0.3571 |
| 10_194855 | 3 | 0.6667 | 3 | 0 | 5 | 0.2 | 1 | 1 | 12 | 0.3333 | 2 | 0 | 3 | 0.6667 | 2 | 0.5 | 5 | 0.4 | 12 | 0.4167 |
| 10_194856 | 4 | 0.5 | 2 | 0 | 5 | 0.4 | 2 | 0 | 13 | 0.3077 | 3 | 0.3333 | 2 | 0.5 | 3 | 0.6667 | 5 | 0.2 | 13 | 0.3846 |
| 10_194858 | 2 | 0.5 | 1 | 0 | 7 | 0.2857 | 2 | 0 | 12 | 0.25 | 4 | 0.25 | 3 | 0 | 3 | 0.6667 | 3 | 0.3333 | 13 | 0.3077 |
| 10_196347 | NA | NA | 1 | 1 | NA | NA | 1 | 0 | 2 | 0.5 | NA | NA | 2 | 0.5 | NA | NA | NA | NA | 2 | 0.5 |
| 10_196349 | NA | NA | 1 | 0 | 1 | 1 | 1 | 1 | 3 | 0.6667 | NA | NA | 2 | 0.5 | NA | NA | NA | NA | 2 | 0.5 |
| 10_196350 | NA | NA | 1 | 0 | 1 | 1 | 1 | 0 | 3 | 0.3333 | NA | NA | 1 | 1 | NA | NA | NA | NA | 1 | 1 |
| 10_196351 | NA | NA | 1 | 1 | 1 | 1 | 1 | 0 | 3 | 0.6667 | NA | NA | 2 | 0.5 | NA | NA | NA | NA | 2 | 0.5 |
| 10_196351 | NA | NA | 1 | 1 | 1 | 1 | 1 | 1 | 3 | 1 | NA | NA | 2 | 1 | NA | NA | NA | NA | 2 | 1 |
| 10_196352 | NA | NA | 1 | 1 | 1 | 1 | 1 | 1 | 3 | 1 | NA | NA | 2 | 1 | NA | NA | NA | NA | 2 | 1 |
| 10_196352 | NA | NA | 1 | 1 | 1 | 1 | 1 | 1 | 3 | 1 | NA | NA | 1 | 1 | NA | NA | NA | NA | 1 | 1 |
| 10_196353 | NA | NA | 1 | 1 | 1 | 0 | 1 | 0 | 3 | 0.3333 | NA | NA | 2 | 0.5 | NA | NA | NA | NA | 2 | 0.5 |
| 10_196354 | NA | NA | 1 | 1 | 1 | 0 | NA | NA | 2 | 0.5 | NA | NA | 2 | 0.5 | NA | NA | NA | NA | 2 | 0.5 |
| 10_196354 | NA | NA | 1 | 1 | 1 | 1 | NA | NA | 2 | 1 | NA | NA | 2 | 1 | NA | NA | NA | NA | 2 | 1 |
| 10_196355 | NA | NA | 1 | 1 | 1 | 1 | 1 | 1 | 3 | 1 | NA | NA | 2 | 1 | NA | NA | NA | NA | 2 | 1 |
| 10_196355 | NA | NA | 1 | 1 | 1 | 0 | 1 | 1 | 3 | 0.6667 | NA | NA | 1 | 1 | NA | NA | NA | NA | 1 | 1 |
| 10_197238 | 11 | 0.0909 | 4 | 0 | 8 | 0.25 | 7 | 0.2857 | 30 | 0.1667 | 1 | 0 | 12 | 0 | 8 | 0.125 | 3 | 0.3333 | 24 | 0.0833 |
| 10_197243 | 10 | 1 | 3 | 1 | 5 | 1 | 7 | 0.8571 | 25 | 0.96 | 2 | 0.5 | 10 | 0.9 | 6 | 0.8333 | 3 | 1 | 21 | 0.8571 |
| 10_197243 | 10 | 0.1 | 3 | 0 | 7 | 0.1429 | 8 | 0.375 | 28 | 0.1786 | 2 | 0 | 10 | 0.2 | 8 | 0.25 | 4 | 0.25 | 24 | 0.2083 |
| 10_197244 | 9 | 0.7778 | 4 | 0.75 | 7 | 0.8571 | 7 | 0.8571 | 27 | 0.8148 | 2 | 0.5 | 6 | 0.5 | 7 | 0.7143 | 4 | 1 | 19 | 0.6842 |
| 10_197244 | 9 | 0 | 4 | 0 | 6 | 0.1667 | 8 | 0.25 | 27 | 0.1111 | 1 | 0 | 9 | 0 | 6 | 0.1667 | 4 | 0.25 | 20 | 0.1 |
| 10_197245 | 6 | 0 | 2 | 1 | 5 | 0.2 | 4 | 0.25 | 17 | 0.2353 | 1 | 0 | 6 | 0.3333 | 3 | 0.3333 | 3 | 0.3333 | 13 | 0.3077 |
| 10_197246 | 6 | 0.1667 | 3 | 0 | 7 | 0.4286 | 9 | 0.2222 | 25 | 0.24 | 1 | 1 | 2 | 0 | 3 | 0 | 3 | 0.3333 | 9 | 0.2222 |
| 10_197248 | 5 | 0.8 | 3 | 0 | 5 | 0.4 | 7 | 0.1429 | 20 | 0.35 | 1 | 0 | 2 | 0.5 | 4 | 0.5 | 3 | 0.6667 | 10 | 0.5 |
| 10_197249 | 2 | 0.5 | 2 | 0 | 5 | 0.2 | 4 | 0.25 | 13 | 0.2308 | 1 | 1 | 2 | 0 | 3 | 0 | 1 | 1 | 7 | 0.2857 |
| 10_197251 | 5 | 1 | 2 | 1 | 5 | 0.8 | 7 | 1 | 19 | 0.9474 | 1 | 0 | 3 | 1 | 4 | 1 | 1 | 0 | 9 | 0.7778 |
| 10_197251 | 5 | 0.6 | 2 | 1 | 4 | 0.5 | 7 | 0.7143 | 18 | 0.6667 | 1 | 1 | 3 | 0.3333 | 4 | 0.5 | 3 | 0.3333 | 11 | 0.4545 |
| 10_197252 | 3 | 0.3333 | 2 | 0 | 5 | 0.2 | 5 | 0.4 | 15 | 0.2667 | 1 | 0 | 2 | 0 | 2 | 0.5 | 1 | 0 | 6 | 0.1667 |
| 10_197253 | 4 | 0 | 2 | 0 | 5 | 0.2 | 8 | 0.375 | 19 | 0.2105 | 1 | 1 | 2 | 0.5 | 3 | 0.6667 | 3 | 0.3333 | 9 | 0.5556 |
| 10_197261 | 1 | 1 | NA | NA | NA | NA | NA | NA | 1 | 1 | 1 | 1 | NA | NA | 1 | 1 | NA | NA | 2 | 1 |
| 10_199940 | 3 | 0 | 1 | 0 | NA | NA | 3 | 0.3333 | 7 | 0.1429 | 2 | 0 | 6 | 0 | 3 | 0.3333 | 4 | 0 | 15 | 0.0667 |
| 10_200173 | 3 | 0.3333 | 2 | 0 | 2 | 0 | 5 | 0 | 12 | 0.0833 | 2 | 0 | 6 | 0.1667 | 5 | 0 | 5 | 0 | 18 | 0.0556 |
| 10_200173 | 3 | 0 | 1 | 0 | 2 | 0 | 7 | 0.1429 | 13 | 0.0769 | 2 | 0.5 | 7 | 0 | 5 | 0 | 6 | 0 | 20 | 0.05 |
| 10_200173 | 3 | 1 | 2 | 1 | 1 | 1 | 7 | 1 | 13 | 1 | 2 | 0.5 | 6 | 1 | 4 | 1 | 5 | 0.6 | 17 | 0.8235 |
| 10_200174 | 4 | 0 | 2 | 0.5 | 2 | 0 | 8 | 0.25 | 16 | 0.1875 | 1 | 0 | 6 | 0.1667 | 3 | 0 | 7 | 0.1429 | 17 | 0.1176 |
| 10_200174 | 4 | 0.25 | 1 | 0 | 2 | 0 | 7 | 0 | 14 | 0.0714 | 2 | 0 | 7 | 0.1429 | 5 | 0 | 7 | 0.1429 | 21 | 0.0952 |
| 10_200175 | 4 | 1 | 2 | 1 | 2 | 1 | 7 | 0.8571 | 15 | 0.9333 | 1 | 1 | 7 | 0.8571 | 5 | 1 | 5 | 0.8 | 18 | 0.8889 |
| 10_200175 | 4 | 0.25 | 2 | 0.5 | 2 | 0 | 7 | 0 | 15 | 0.1333 | 2 | 0 | 7 | 0.1429 | 5 | 0 | 7 | 0.4286 | 21 | 0.1905 |
| 10_200176 | 4 | 0.5 | 4 | 0.25 | 2 | 0.5 | 8 | 0.5 | 18 | 0.4444 | 2 | 0 | 7 | 0.4286 | 4 | 0.5 | 7 | 0.5714 | 20 | 0.45 |
| 10_200178 | 5 | 0.2 | 4 | 0.75 | 2 | 0 | 8 | 0.375 | 19 | 0.3684 | 2 | 0.5 | 7 | 0.1429 | 5 | 0 | 7 | 0.4286 | 21 | 0.2381 |
| 10_200178 | 5 | 0.2 | 3 | 0 | 2 | 0 | 7 | 0.2857 | 17 | 0.1765 | 2 | 0 | 7 | 0.2857 | 3 | 0 | 5 | 0.2 | 17 | 0.1765 |
| 10_201558 | 4 | 0 | 7 | 0.1429 | 8 | 0.25 | 10 | 0.1 | 29 | 0.1379 | 3 | 0.3333 | 4 | 0 | 9 | 0 | 4 | 0.25 | 20 | 0.1 |
| 10_201558 | 4 | 1 | 8 | 0.5 | 9 | 0.2222 | 12 | 0.5833 | 33 | 0.5152 | 4 | 0.25 | 4 | 0 | 7 | 0.4286 | 5 | 0.2 | 20 | 0.25 |
| 10_201560 | 2 | 1 | 8 | 0.75 | 9 | 0.7778 | 8 | 0.875 | 27 | 0.8148 | 3 | 0.6667 | 4 | 1 | 7 | 0.4286 | 6 | 0.6667 | 20 | 0.65 |
| 10_201561 | 4 | 0 | 6 | 0 | 10 | 0.1 | 9 | 0.1111 | 29 | 0.069 | 4 | 0 | 3 | 0 | 5 | 0.2 | 4 | 0 | 16 | 0.0625 |
| 10_201562 | 6 | 0.3333 | 8 | 0 | 10 | 0.1 | 13 | 0.2308 | 37 | 0.1622 | 3 | 0 | 4 | 0 | 8 | 0.125 | 4 | 0.25 | 19 | 0.1053 |
| 10_201563 | 6 | 0 | 9 | 0.1111 | 9 | 0.1111 | 11 | 0 | 35 | 0.0571 | 4 | 0.5 | 4 | 0 | 6 | 0.1667 | 5 | 0.2 | 19 | 0.2105 |
| 10_201564 | 5 | 0.8 | 9 | 0.2222 | 10 | 0.4 | 11 | 0.2727 | 35 | 0.3714 | 5 | 0.4 | 3 | 0 | 7 | 0.2857 | 5 | 0.6 | 20 | 0.35 |
| 10_201564 | 6 | 0 | 8 | 0.125 | 10 | 0.1 | 12 | 0 | 36 | 0.0556 | 4 | 0.25 | 3 | 0.3333 | 8 | 0 | 6 | 0.1667 | 21 | 0.1429 |
| 10_201564 | 5 | 0.2 | 8 | 0 | 12 | 0.25 | 13 | 0.3077 | 38 | 0.2105 | 5 | 0.2 | 4 | 0 | 7 | 0.5714 | 7 | 0.2857 | 23 | 0.3043 |
| 10_201567 | 5 | 0.2 | 10 | 0.1 | 11 | 0.4545 | 11 | 0.5455 | 37 | 0.3514 | 7 | 0.2857 | 5 | 0.6 | 8 | 0.25 | 6 | 0.1667 | 26 | 0.3077 |
| 10_201568 | 7 | 0 | 8 | 0 | 12 | 0.3333 | 11 | 0.0909 | 38 | 0.1316 | 6 | 0.5 | 4 | 0.5 | 7 | 0.1429 | 6 | 0 | 23 | 0.2609 |
| 10_201568 | 7 | 0.4286 | 10 | 0.3 | 13 | 0.5385 | 13 | 0.3077 | 43 | 0.3953 | 6 | 0.5 | 7 | 0.5714 | 8 | 0.125 | 9 | 0.3333 | 30 | 0.3667 |
| 10_201569 | 7 | 0 | 10 | 0 | 11 | 0.1818 | 12 | 0 | 40 | 0.05 | 7 | 0.1429 | 7 | 0.1429 | 8 | 0 | 9 | 0.1111 | 31 | 0.0968 |
| 10_201991 | 3 | 0.3333 | 10 | 0.1 | NA | NA | 7 | 0.4286 | 20 | 0.25 | 2 | 0 | 1 | 0 | 6 | 0.3333 | 2 | 0.5 | 11 | 0.2727 |
| 10_202011 | 6 | 0 | 13 | 0 | 1 | 0 | 10 | 0.2 | 30 | 0.0667 | 5 | 0.2 | 3 | 0 | 10 | 0 | 2 | 0 | 20 | 0.05 |
| 10_202012 | 6 | 0.3333 | 13 | 0.4615 | 2 | 0.5 | 6 | 0.3333 | 27 | 0.4074 | 5 | 0.2 | 2 | 0.5 | 8 | 0.5 | 2 | 0 | 17 | 0.3529 |
| 10_202012 | 6 | 0.6667 | 12 | 1 | 2 | 1 | 8 | 0.875 | 28 | 0.8929 | 5 | 0.4 | 3 | 1 | 9 | 0.7778 | 2 | 1 | 19 | 0.7368 |
| 10_202155 | 12 | 0.5833 | 16 | 0.625 | 8 | 0.375 | 13 | 0.6154 | 49 | 0.5714 | 9 | 0.4444 | 7 | 0.4286 | 14 | 0.3571 | 6 | 0.6667 | 36 | 0.4444 |
| 10_202164 | 9 | 0 | 7 | 0.2857 | 7 | 0 | 12 | 0.25 | 35 | 0.1429 | 6 | 0.1667 | 5 | 0.2 | 10 | 0.2 | 9 | 0.2222 | 30 | 0.2 |
| 10_202181 | 13 | 0.1538 | 15 | 0.1333 | 13 | 0.0769 | 20 | 0.2 | 61 | 0.1475 | 5 | 0 | 10 | 0.2 | 17 | 0.2353 | 14 | 0.2857 | 46 | 0.2174 |
| 10_202201 | NA | NA | 1 | 1 | NA | NA | NA | NA | 1 | 1 | 3 | 1 | 2 | 1 | 1 | 1 | 3 | 1 | 9 | 1 |
| 10_202201 | NA | NA | 1 | 1 | 1 | 1 | NA | NA | 2 | 1 | 3 | 1 | 2 | 1 | 2 | 1 | 3 | 1 | 10 | 1 |
| 10_202728 | 11 | 0 | 11 | 0.0909 | 13 | 0.0769 | 17 | 0.0588 | 52 | 0.0577 | 2 | 0 | 22 | 0.1364 | 20 | 0.05 | 8 | 0 | 52 | 0.0769 |
| 10_202729 | 16 | 0 | 10 | 0 | 15 | 0.1333 | 19 | 0.1053 | 60 | 0.0667 | 2 | 0 | 20 | 0.2 | 21 | 0.1429 | 9 | 0.1111 | 52 | 0.1538 |

| | | | | | | | | | | | | | | | | | | | | |
|---|---|---|---|---|---|---|---|---|---|---|---|---|---|---|---|---|---|---|---|---|
| 10_2027404 | 2 | 0.5 | 1 | 1 | 3 | 0.3333 | 3 | 0.3333 | 9 | 0.4444 | 1 | 0 | 6 | 0.3333 | 2 | 0.5 | 2 | 0.5 | 11 | 0.3636 |
| 10_2027408 | 5 | 1 | 1 | 1 | 3 | 0.6667 | 3 | 1 | 12 | 0.9167 | 1 | 1 | 6 | 0.8333 | 2 | 1 | 2 | 1 | 11 | 0.9091 |
| 10_2027419 | 4 | 0.5 | 1 | 1 | 2 | 0.5 | 2 | 0.5 | 9 | 0.5556 | 1 | 1 | 6 | 0.1667 | 2 | 0.5 | 2 | 0.5 | 11 | 0.3636 |
| 10_2027430 | 5 | 0 | 1 | 0 | 2 | 0.5 | 2 | 0 | 10 | 0.1 | 1 | 0 | 6 | 0 | 2 | 0.5 | 1 | 0 | 10 | 0.1 |
| 10_2027466 | 4 | 0.25 | 1 | 1 | 1 | 0 | 2 | 0 | 8 | 0.25 | 1 | 0 | 6 | 0.5 | 2 | 1 | NA | NA | 9 | 0.5556 |
| 10_2027518 | 3 | 0 | 1 | 0 | 1 | 1 | 2 | 0 | 7 | 0.1429 | NA | NA | 5 | 0.2 | 1 | 1 | 1 | 0 | 7 | 0.2857 |
| 10_2027545 | 3 | 0.6667 | 3 | 0 | 2 | 0.5 | 2 | 0 | 10 | 0.3 | 1 | 0 | 4 | 0.25 | 2 | 0 | 3 | 0.3333 | 10 | 0.2 |
| 10_2027558 | 2 | 1 | 3 | 0.6667 | 2 | 1 | 2 | 1 | 9 | 0.8889 | NA | NA | 3 | 1 | 1 | 1 | 2 | 1 | 6 | 1 |
| 10_2027564 | 2 | 1 | 1 | 1 | 2 | 1 | 2 | 1 | 7 | 1 | 1 | 1 | 4 | 1 | 1 | 1 | 2 | 1 | 8 | 1 |
| 10_2027564 | 3 | 1 | 2 | 0.5 | 2 | 1 | 2 | 0.5 | 9 | 0.7778 | 1 | 1 | 4 | 0.25 | 2 | 0.5 | 2 | 0 | 9 | 0.3333 |
| 10_2027566 | 3 | 0.6667 | 2 | 0.5 | 2 | 0.5 | 2 | 0.5 | 9 | 0.5556 | 1 | 0 | 5 | 0.6 | 2 | 1 | 2 | 0.5 | 10 | 0.6 |
| 10_2027568 | 3 | 0.3333 | 2 | 0 | 2 | 1 | 2 | 0.5 | 9 | 0.4444 | 1 | 0 | 5 | 0.4 | 2 | 0.5 | 2 | 0 | 10 | 0.3 |
| 10_2027577 | 1 | 1 | 1 | 0 | 1 | 0 | 1 | 1 | 4 | 0.5 | NA | NA | 2 | 0.5 | 2 | 1 | NA | NA | 4 | 0.75 |
| 10_2027578 | 2 | 1 | 1 | 1 | 2 | 1 | 2 | 1 | 7 | 1 | 1 | 1 | 5 | 1 | 2 | 1 | 1 | 1 | 9 | 1 |
| 10_2027582 | 2 | 1 | 1 | 1 | 1 | 1 | 2 | 1 | 6 | 1 | 2 | 1 | 5 | 0.8 | 2 | 1 | 1 | 1 | 10 | 0.9 |
| 10_2027586 | 1 | 1 | 1 | 0 | 2 | 0.5 | 2 | 1 | 6 | 0.6667 | 2 | 0.5 | 5 | 0.8 | 1 | 1 | 1 | 0 | 9 | 0.6667 |
| 10_2027619 | 2 | 0 | 1 | 0 | 2 | 0.5 | 1 | 0 | 6 | 0.1667 | 2 | 0.5 | 4 | 0 | 1 | 0 | 1 | 0 | 8 | 0.125 |
| 10_2042375 | NA | NA | NA | NA | 1 | 1 | NA | NA | 1 | 1 | 1 | 1 | NA | NA | NA | NA | NA | NA | 1 | 1 |
| 10_2083560 | 6 | 0.6667 | 8 | 0.125 | 5 | 0.4 | 15 | 0.3333 | 34 | 0.3529 | 3 | 0.3333 | 10 | 0.2 | 7 | 0.2857 | 7 | 0.4286 | 27 | 0.2963 |
| 10_2083576 | 6 | 0.1667 | 8 | 0.125 | 5 | 0 | 16 | 0.0625 | 35 | 0.0857 | 3 | 0.3333 | 10 | 0 | 7 | 0.4286 | 7 | 0 | 27 | 0.1481 |
| 10_2083970 | 7 | 0 | 10 | 0 | 6 | 0.3333 | 18 | 0.0556 | 41 | 0.0732 | 3 | 0 | 10 | 0.1 | 12 | 0.0833 | 8 | 0 | 33 | 0.0606 |
| 10_2095526 | 22 | 0.0909 | 13 | 0.0769 | 12 | 0.0833 | 36 | 0.1111 | 83 | 0.0964 | 11 | 0.0909 | 19 | 0.1053 | 27 | 0.0741 | 25 | 0.12 | 82 | 0.0976 |
| 10_2095558 | 27 | 0.0741 | 15 | 0.0667 | 15 | 0.1333 | 37 | 0.0541 | 94 | 0.0745 | 14 | 0.0714 | 21 | 0.1429 | 28 | 0.0714 | 21 | 0.0476 | 84 | 0.0833 |
| 10_2118529 | NA | NA | 1 | 1 | NA | NA | NA | NA | 1 | 1 | NA | NA | 1 | 1 | NA | NA | NA | NA | 1 | 1 |
| 10_2118626 | NA | NA | 1 | 1 | NA | NA | NA | NA | 1 | 1 | NA | NA | 1 | 1 | NA | NA | NA | NA | 1 | 1 |
| 10_2118627 | NA | NA | 1 | 1 | NA | NA | NA | NA | 1 | 1 | NA | NA | 1 | 1 | NA | NA | NA | NA | 1 | 1 |
| 10_2118631 | NA | NA | 1 | 1 | NA | NA | NA | NA | 1 | 1 | NA | NA | 1 | 1 | NA | NA | NA | NA | 1 | 1 |
| 10_2121763 | 2 | 0.5 | 1 | 0 | 1 | 0 | 6 | 0.3333 | 10 | 0.3 | 1 | 0 | 4 | 0.5 | 2 | 0 | 4 | 0.25 | 11 | 0.2727 |
| 10_2124430 | 6 | 0.5 | 5 | 0.6 | 6 | 0.5 | 8 | 0.5 | 25 | 0.52 | 2 | 0.5 | 5 | 0.4 | 5 | 0.2 | 7 | 0.5714 | 19 | 0.4211 |
| 10_2124439 | 4 | 0 | 7 | 0.4286 | 6 | 0.1667 | 9 | 0.1111 | 26 | 0.1923 | 3 | 0.3333 | 5 | 0 | 4 | 0.25 | 7 | 0.1429 | 19 | 0.1579 |
| 10_2124444 | 6 | 0.3333 | 6 | 0.5 | 6 | 0.3333 | 8 | 0.5 | 26 | 0.4231 | 4 | 0.5 | 5 | 0.2 | 5 | 0.6 | 8 | 0.375 | 22 | 0.4091 |
| 10_2124446 | 5 | 0.2 | 6 | 0.1667 | 6 | 0.1667 | 9 | 0.1111 | 26 | 0.1538 | 3 | 0 | 5 | 0.2 | 5 | 0.2 | 7 | 0.1429 | 20 | 0.15 |
| 10_2124448 | 5 | 0.8 | 7 | 0.5714 | 6 | 0.6667 | 7 | 0.5714 | 25 | 0.64 | 3 | 0.6667 | 4 | 0.5 | 4 | 1 | 6 | 0.6667 | 17 | 0.7059 |
| 10_2124456 | 4 | 0 | 4 | 0.25 | 6 | 0 | 9 | 0.1111 | 23 | 0.087 | 3 | 0 | 5 | 0.2 | 4 | 0 | 5 | 0.2 | 17 | 0.1176 |
| 10_2187475 | 40 | 0.875 | 33 | 1 | 37 | 0.8919 | 40 | 0.925 | 150 | 0.92 | 33 | 0.8788 | 21 | 0.9524 | 25 | 0.88 | 33 | 0.9091 | 112 | 0.9018 |
| 10_2187480 | 40 | 0.125 | 37 | 0.1081 | 39 | 0.1026 | 39 | 0.1026 | 155 | 0.1097 | 32 | 0.125 | 22 | 0.1364 | 24 | 0 | 31 | 0.0968 | 109 | 0.0917 |
| 10_2187481 | 28 | 0.8929 | 32 | 0.875 | 31 | 0.871 | 31 | 0.9355 | 122 | 0.8934 | 21 | 0.7619 | 16 | 0.875 | 20 | 0.95 | 29 | 0.8621 | 86 | 0.8605 |
| 10_2187501 | 43 | 0.093 | 39 | 0.1282 | 42 | 0.0952 | 41 | 0.0976 | 165 | 0.103 | 35 | 0.0571 | 24 | 0.0417 | 25 | 0.08 | 36 | 0.0556 | 120 | 0.0583 |
| 10_2187528 | 43 | 0.2558 | 40 | 0.2 | 40 | 0.175 | 41 | 0.1463 | 164 | 0.1951 | 36 | 0.1389 | 24 | 0.1667 | 24 | 0.1667 | 38 | 0.2368 | 122 | 0.1803 |
| 10_2187530 | 36 | 0.4722 | 28 | 0.6786 | 29 | 0.5172 | 32 | 0.5938 | 125 | 0.56 | 26 | 0.5385 | 17 | 0.4706 | 20 | 0.6 | 28 | 0.6071 | 91 | 0.5604 |
| 10_2187534 | 48 | 0.1667 | 42 | 0.1905 | 46 | 0.1087 | 41 | 0.0732 | 177 | 0.1356 | 36 | 0.0833 | 28 | 0.1071 | 25 | 0.2 | 42 | 0.1667 | 131 | 0.1374 |
| 10_2187547 | 47 | 0.2766 | 43 | 0.2326 | 45 | 0.2444 | 43 | 0.186 | 178 | 0.236 | 33 | 0.2424 | 24 | 0.25 | 28 | 0.2857 | 40 | 0.3 | 125 | 0.272 |
| 10_2191581 | NA | NA | NA | NA | 1 | 1 | NA | NA | 1 | 1 | NA | NA | NA | NA | 1 | 1 | NA | NA | 1 | 1 |
| 10_2191587 | NA | NA | NA | NA | 1 | 1 | NA | NA | 1 | 1 | NA | NA | NA | NA | 1 | 1 | NA | NA | 1 | 1 |
| 10_2191614 | NA | NA | NA | NA | 1 | 1 | NA | NA | 1 | 1 | NA | NA | NA | NA | 1 | 1 | NA | NA | 1 | 1 |
| 10_2191623 | NA | NA | NA | NA | 1 | 1 | NA | NA | 1 | 1 | NA | NA | NA | NA | 1 | 1 | NA | NA | 1 | 1 |
| 10_2193085 | NA | NA | 3 | 0.3333 | NA | NA | NA | NA | 3 | 0.3333 | 1 | 0 | 1 | 1 | NA | NA | 2 | 1 | 4 | 0.75 |
| 10_2257819 | 1 | 0 | 1 | 1 | NA | NA | 1 | 1 | 3 | 0.6667 | 1 | 1 | NA | NA | NA | NA | NA | NA | 1 | 1 |
| 10_2257849 | 1 | 0 | 1 | 1 | NA | NA | 1 | 0 | 3 | 0.3333 | 1 | 1 | NA | NA | NA | NA | NA | NA | 1 | 1 |
| 10_2258152 | 6 | 0.1667 | 9 | 0.1111 | 6 | 0 | 11 | 0.0909 | 32 | 0.0938 | 2 | 0 | 10 | 0.2 | 7 | 0.2857 | 7 | 0.1429 | 26 | 0.1923 |
| 10_2258160 | 4 | 0.5 | 9 | 0.8889 | 5 | 1 | 12 | 0.75 | 30 | 0.8 | 3 | 0 | 11 | 0.6364 | 6 | 0.6667 | 7 | 0.2857 | 27 | 0.4815 |
| 10_2258163 | 3 | 0 | 4 | 0.5 | 3 | 0.3333 | 5 | 0.8 | 15 | 0.4667 | 3 | 0 | 5 | 0.6 | 3 | 0.6667 | 3 | 0.3333 | 14 | 0.4286 |
| 10_2258165 | 7 | 0.8571 | 10 | 1 | 5 | 0.6 | 12 | 0.8333 | 34 | 0.8529 | 3 | 1 | 10 | 0.9 | 6 | 1 | 6 | 0.8333 | 25 | 0.92 |
| 10_2258170 | 7 | 0 | 8 | 0.125 | 6 | 0.1667 | 12 | 0 | 33 | 0.0606 | 2 | 0 | 8 | 0.25 | 6 | 0 | 6 | 0 | 22 | 0.0909 |
| 10_2258791 | 4 | 0.25 | 5 | 0 | 3 | 0 | 8 | 0.125 | 20 | 0.1 | 1 | 0 | 7 | 0.1429 | 3 | 0 | 4 | 0 | 15 | 0.0667 |
| 10_2370260 | 1 | 0 | 1 | 1 | NA | NA | NA | NA | 2 | 0.5 | 1 | 1 | 2 | 0.5 | 1 | 0 | 1 | 1 | 5 | 0.6 |
| 10_2370261 | 1 | 0 | 2 | 0.5 | NA | NA | NA | NA | 3 | 0.3333 | 1 | 1 | 3 | 0.3333 | 2 | 0 | NA | NA | 6 | 0.3333 |
| 10_2370279 | 1 | 0 | 2 | 0.5 | NA | NA | NA | NA | 3 | 0.3333 | 1 | 1 | 3 | 0.3333 | 2 | 0.5 | 1 | 0 | 7 | 0.4286 |
| 10_2406426 | 9 | 0.1111 | 15 | 0.1333 | 13 | 0.1538 | 17 | 0.1765 | 54 | 0.1481 | 6 | 0.3333 | 12 | 0.4167 | 10 | 0 | 11 | 0.3636 | 39 | 0.2821 |
| 10_2417755 | NA | NA | NA | NA | NA | NA | 2 | 0.5 | 2 | 0.5 | 1 | 1 | 1 | 1 | NA | NA | NA | NA | 2 | 1 |
| 10_2417776 | NA | NA | NA | NA | 1 | 0 | 2 | 0.5 | 3 | 0.3333 | 1 | 1 | 2 | 0.5 | NA | NA | NA | NA | 3 | 0.6667 |
| 10_2447345 | 2 | 0 | 7 | 0.4286 | 4 | 0.5 | 5 | 0.6 | 18 | 0.4444 | 3 | 0.3333 | 3 | 0.6667 | 3 | 0.3333 | 3 | 0 | 12 | 0.3333 |
| 10_2447352 | 4 | 0.5 | 9 | 0.6667 | 4 | 0.5 | 4 | 0.75 | 21 | 0.619 | 3 | 0.6667 | 4 | 0.75 | 3 | 0 | 2 | 0.5 | 12 | 0.5 |
| 10_2447353 | 3 | 0.6667 | 6 | 0.6667 | 4 | 0.5 | 5 | 0.6 | 18 | 0.6111 | 3 | 1 | 4 | 1 | 3 | 0.3333 | 2 | 0 | 12 | 0.6667 |
| 10_2447354 | 5 | 0.2 | 9 | 0 | 5 | 0.4 | 6 | 0.1667 | 25 | 0.16 | 3 | 0.6667 | 4 | 0 | 3 | 0.3333 | 3 | 0 | 13 | 0.2308 |

| | | | | | | | | | | | | | | | | | | | | |
|---|---|---|---|---|---|---|---|---|---|---|---|---|---|---|---|---|---|---|---|---|
| 10_2447355 | 4 | 0.25 | 4 | 0.75 | 5 | 0.8 | 5 | 0.2 | 18 | 0.5 | 3 | 0.3333 | 4 | 0.25 | 2 | 0.5 | 2 | 0.5 | 11 | 0.3636 |
| 10_2478871 | 1 | 0 | 5 | 0 | 5 | 0 | 6 | 0.1667 | 17 | 0.0588 | 6 | 0 | 6 | 0 | 2 | 0 | 5 | 0.2 | 19 | 0.0526 |
| 10_2478874 | 1 | 0 | 5 | 0 | 5 | 0 | 6 | 0.1667 | 17 | 0.0588 | 5 | 0.2 | 6 | 0 | 2 | 0.5 | 5 | 0.2 | 18 | 0.1667 |
| 10_2478876 | 2 | 1 | 5 | 0.8 | 5 | 0.4 | 5 | 0.6 | 17 | 0.6471 | 5 | 0.8 | 6 | 0.5 | 2 | 0.5 | 3 | 0.6667 | 16 | 0.625 |
| 10_2504605 | 1 | 1 | 5 | 0.4 | 6 | 1 | 2 | 1 | 14 | 0.7857 | 1 | 0 | 4 | 1 | 2 | 1 | 2 | 0.5 | 9 | 0.7778 |
| 10_2504619 | 1 | 0 | 4 | 0 | 6 | 0.3333 | 4 | 0 | 15 | 0.1333 | 2 | 0 | 4 | 0 | 2 | 0.5 | 2 | 0 | 10 | 0.1 |
| 10_2504620 | 1 | 1 | 2 | 1 | 6 | 1 | 4 | 1 | 13 | 1 | 2 | 0.5 | 1 | 1 | 1 | 1 | 1 | 1 | 5 | 0.8 |
| 10_2504660 | NA | NA | 3 | 0.3333 | 6 | 0.5 | 3 | 0 | 12 | 0.3333 | 2 | 0.5 | 3 | 0 | 1 | 1 | 1 | 0 | 7 | 0.2857 |
| 10_2504662 | NA | NA | 2 | 1 | 5 | 0 | 4 | 0.5 | 11 | 0.3636 | 2 | 0.5 | 4 | 0.25 | 1 | 0 | 1 | 1 | 8 | 0.375 |
| 10_2504665 | NA | NA | 2 | 0.5 | 5 | 0.6 | 4 | 0.5 | 11 | 0.5455 | 3 | 0.3333 | 3 | 0.6667 | 1 | 1 | NA | NA | 7 | 0.5714 |
| 10_2504666 | NA | NA | 3 | 1 | 6 | 1 | 4 | 1 | 13 | 1 | 3 | 1 | 4 | 1 | 1 | 1 | 1 | 1 | 9 | 1 |
| 10_2504669 | NA | NA | 3 | 0 | 5 | 0.6 | 3 | 0.3333 | 11 | 0.3636 | 2 | 0.5 | 4 | 0.5 | 1 | 0 | 1 | 0 | 8 | 0.375 |
| 10_2504670 | NA | NA | 4 | 0.25 | 6 | 0.1667 | 4 | 0.5 | 14 | 0.2857 | 3 | 0.6667 | 4 | 0.25 | 1 | 1 | 1 | 0 | 9 | 0.4444 |
| 10_2504679 | NA | NA | 4 | 0.5 | 4 | 0 | 4 | 0.25 | 12 | 0.25 | 3 | 0.3333 | 4 | 0.25 | 1 | 0 | 1 | 1 | 9 | 0.3333 |
| 10_2504690 | NA | NA | 4 | 0 | 6 | 0.5 | 4 | 0.25 | 14 | 0.2857 | 2 | 0.5 | 3 | 0 | 1 | 1 | NA | NA | 6 | 0.3333 |
| 10_2504690 | NA | NA | 4 | 0.75 | 5 | 0.8 | 4 | 0.75 | 13 | 0.7692 | 3 | 0.6667 | 4 | 0.75 | 1 | 1 | NA | NA | 8 | 0.75 |
| 10_2537787 | 1 | 0 | 2 | 0.5 | 1 | 0 | 3 | 1 | 7 | 0.5714 | 3 | 0.3333 | 3 | 0 | 2 | 0 | 3 | 0 | 11 | 0.0909 |
| 10_2537812 | NA | NA | 1 | 0 | 4 | 0.25 | NA | NA | 5 | 0.2 | 5 | 0.2 | 1 | 1 | 2 | 0 | 1 | 1 | 9 | 0.3333 |
| 10_2539885 | 3 | 0 | 8 | 0.125 | 10 | 0 | 7 | 0.5714 | 28 | 0.1786 | 5 | 0 | 7 | 0.1429 | 9 | 0 | 8 | 0.125 | 29 | 0.069 |
| 10_2586852 | NA | NA | NA | NA | 2 | 0.5 | NA | NA | 2 | 0.5 | NA | NA | 1 | 1 | NA | NA | NA | NA | 1 | 1 |
| 10_2586881 | NA | NA | NA | NA | 2 | 1 | NA | NA | 2 | 1 | NA | NA | 1 | 1 | NA | NA | NA | NA | 1 | 1 |
| 10_2586884 | NA | NA | NA | NA | 2 | 1 | NA | NA | 2 | 1 | NA | NA | 1 | 1 | NA | NA | NA | NA | 1 | 1 |
| 10_2586889 | NA | NA | NA | NA | 2 | 0.5 | NA | NA | 2 | 0.5 | NA | NA | 1 | 1 | NA | NA | NA | NA | 1 | 1 |
| 10_2586897 | NA | NA | NA | NA | 2 | 1 | NA | NA | 2 | 1 | NA | NA | 1 | 1 | NA | NA | NA | NA | 1 | 1 |
| 10_2586915 | NA | NA | NA | NA | 2 | 1 | NA | NA | 2 | 1 | NA | NA | 1 | 1 | NA | NA | NA | NA | 1 | 1 |
| 10_2586957 | NA | NA | NA | NA | 2 | 1 | NA | NA | 2 | 1 | NA | NA | 1 | 1 | NA | NA | NA | NA | 1 | 1 |
| 10_2615061 | 3 | 1 | 3 | 0.3333 | 2 | 0 | NA | NA | 8 | 0.5 | 3 | 1 | NA | NA | 5 | 1 | 1 | 1 | 9 | 1 |
| 10_2615064 | 1 | 0 | 3 | 0.3333 | 2 | 0 | NA | NA | 6 | 0.1667 | 1 | 1 | 1 | 0 | 6 | 0.1667 | 1 | 0 | 9 | 0.2222 |
| 10_2615068 | 4 | 0.75 | 4 | 0.25 | 2 | 0.5 | NA | NA | 10 | 0.5 | 2 | 1 | 1 | 1 | 7 | 0.8571 | 2 | 0.5 | 12 | 0.8333 |
| 10_2615079 | 3 | 0 | 4 | 0.25 | 2 | 0 | NA | NA | 9 | 0.1111 | 3 | 0.3333 | 2 | 0 | 6 | 0.1667 | 2 | 0 | 13 | 0.1538 |
| 10_2615098 | 3 | 0.3333 | 3 | 0.3333 | NA | NA | NA | NA | 6 | 0.3333 | 1 | 0 | 1 | 1 | 3 | 0.6667 | 2 | 0.5 | 7 | 0.5714 |
| 10_2615110 | 3 | 1 | 3 | 0.6667 | 2 | 0.5 | NA | NA | 8 | 0.75 | 3 | 0.3333 | 1 | 1 | 7 | 0.7143 | 2 | 1 | 13 | 0.6923 |
| 10_2615126 | 3 | 0 | 3 | 0.6667 | 2 | 0 | NA | NA | 8 | 0.25 | 3 | 0.3333 | 1 | 0 | 5 | 0.4 | 2 | 0.5 | 11 | 0.3636 |
| 10_2615131 | 2 | 1 | 3 | 0.3333 | 2 | 0 | NA | NA | 7 | 0.4286 | 2 | 0.5 | 1 | 0 | 4 | 0.25 | 2 | 0.5 | 9 | 0.3333 |
| 10_2615145 | 3 | 0.6667 | 3 | 0.6667 | 2 | 0.5 | NA | NA | 8 | 0.625 | 3 | 0.6667 | NA | NA | 5 | 1 | 1 | 1 | 9 | 0.8889 |
| 10_2615156 | 2 | 1 | 1 | 0 | 2 | 0.5 | NA | NA | 5 | 0.6 | 3 | 1 | NA | NA | 6 | 0.8333 | 2 | 1 | 11 | 0.9091 |
| 10_2615170 | 2 | 1 | 3 | 0 | 2 | 0 | NA | NA | 7 | 0.2857 | 2 | 1 | NA | NA | 1 | 1 | 2 | 0 | 5 | 0.6 |
| 10_2615177 | 3 | 0.3333 | 2 | 0 | 2 | 0 | NA | NA | 7 | 0.1429 | 2 | 0.5 | NA | NA | 3 | 0.6667 | 1 | 1 | 6 | 0.6667 |
| 10_2615195 | 2 | 1 | 2 | 1 | 2 | 1 | NA | NA | 6 | 1 | 2 | 1 | NA | NA | 5 | 0.6 | 2 | 1 | 9 | 0.7778 |
| 10_2615202 | 2 | 1 | 2 | 0.5 | 2 | 0.5 | NA | NA | 6 | 0.6667 | 1 | 1 | NA | NA | 3 | 1 | NA | NA | 4 | 1 |
| 10_2615212 | 1 | 1 | 2 | 0.5 | NA | NA | NA | NA | 3 | 0.6667 | 2 | 1 | NA | NA | 3 | 1 | 2 | 1 | 7 | 1 |
| 10_2615615 | 1 | 1 | 2 | 0 | 1 | 0 | NA | NA | 4 | 0.25 | 1 | 0 | NA | NA | 4 | 0.75 | 2 | 0 | 7 | 0.4286 |
| 10_2662343 | 1 | 1 | 1 | 1 | NA | NA | NA | NA | 2 | 1 | 1 | 1 | NA | NA | NA | NA | NA | NA | 1 | 1 |
| 10_2662361 | 1 | 1 | 1 | 0 | NA | NA | NA | NA | 2 | 0.5 | 1 | 1 | NA | NA | NA | NA | NA | NA | 1 | 1 |
| 10_2662366 | 1 | 1 | 1 | 0 | NA | NA | NA | NA | 2 | 0.5 | 1 | 1 | NA | NA | NA | NA | NA | NA | 1 | 1 |
| 10_2679303 | NA | NA | NA | NA | NA | NA | 3 | 0.6667 | 3 | 0.6667 | 1 | 1 | 4 | 0.25 | 2 | 0 | NA | NA | 7 | 0.2857 |
| 10_2679386 | NA | NA | NA | NA | NA | NA | 4 | 0.5 | 4 | 0.5 | 1 | 0 | 4 | 0.25 | 1 | 0 | NA | NA | 6 | 0.1667 |
| 10_2679386 | NA | NA | NA | NA | NA | NA | 4 | 0.25 | 4 | 0.25 | 1 | 1 | 3 | 0 | 2 | 0 | NA | NA | 6 | 0.1667 |
| 10_2685753 | 4 | 0.25 | 2 | 0.5 | NA | NA | 4 | 0 | 10 | 0.2 | 1 | 0 | 2 | 0.5 | NA | NA | NA | NA | 3 | 0.3333 |
| 10_2847202 | 1 | 0 | 4 | 0.25 | 2 | 0 | 3 | 0.3333 | 10 | 0.2 | 4 | 0 | NA | NA | 2 | 0.5 | 3 | 0 | 9 | 0.1111 |
| 10_2901956 | NA | NA | 2 | 1 | 1 | 1 | NA | NA | 3 | 1 | NA | NA | NA | NA | 1 | 1 | NA | NA | 1 | 1 |
| 10_2907235 | 2 | 1 | 4 | 1 | 1 | 0 | 5 | 0.6 | 12 | 0.75 | 1 | 0 | NA | NA | 4 | 0.5 | 3 | 0 | 8 | 0.25 |
| 10_2907237 | 2 | 0 | 5 | 0.4 | NA | NA | 3 | 0.3333 | 10 | 0.3 | 1 | 1 | NA | NA | 4 | 0.5 | 3 | 0.3333 | 8 | 0.5 |
| 10_2907240 | 2 | 0 | 3 | 0 | 1 | 0 | 6 | 0.3333 | 12 | 0.1667 | 1 | 0 | 1 | 0 | 3 | 0.3333 | 3 | 0.6667 | 8 | 0.375 |
| 10_2907252 | 1 | 0 | 5 | 0 | NA | NA | 7 | 0.4286 | 13 | 0.2308 | 1 | 0 | NA | NA | 4 | 0.25 | 3 | 0.3333 | 8 | 0.25 |
| 10_2907325 | 2 | 0.5 | 6 | 0 | 2 | 0 | 7 | 0.1429 | 17 | 0.1176 | 3 | 0.3333 | 2 | 0.5 | 5 | 0.4 | 4 | 0.25 | 14 | 0.3571 |
| 10_2907326 | 3 | 0.6667 | 5 | 0.8 | 2 | 0 | 6 | 0.5 | 16 | 0.5625 | 1 | 0 | 2 | 0.5 | 5 | 0.8 | 3 | 0.3333 | 11 | 0.5455 |
| 10_2907518 | 2 | 0.5 | 6 | 0.3333 | 5 | 0 | 3 | 0.3333 | 16 | 0.25 | 2 | 0 | 3 | 0 | 2 | 0 | 5 | 0.2 | 12 | 0.0833 |
| 10_2907519 | 3 | 0.3333 | 8 | 0.75 | 4 | 0.5 | 7 | 0.5714 | 22 | 0.5909 | 3 | 0.6667 | 3 | 0.6667 | 4 | 0.5 | 6 | 0.5 | 16 | 0.5625 |
| 10_2907521 | 2 | 0 | 8 | 0 | 6 | 0.3333 | 7 | 0.2857 | 23 | 0.1739 | 3 | 0.6667 | 3 | 0.3333 | 5 | 0 | 7 | 0.2857 | 18 | 0.2778 |
| 10_2907537 | 1 | 1 | 5 | 0.4 | 3 | 0.3333 | 4 | 0.25 | 13 | 0.3846 | 2 | 0.5 | 2 | 1 | 2 | 0 | 5 | 0.8 | 11 | 0.6364 |
| 10_2907539 | NA | NA | 5 | 0.8 | 3 | 0.6667 | 4 | 1 | 12 | 0.8333 | 2 | 1 | 2 | 1 | 2 | 0 | 5 | 0.6 | 11 | 0.6364 |
| 10_2907608 | NA | NA | NA | NA | 1 | 0 | 1 | 1 | 2 | 0.5 | NA | NA | NA | NA | NA | NA | 1 | 1 | 1 | 1 |
| 10_2908754 | NA | NA | 2 | 0.5 | 1 | 0 | NA | NA | 3 | 0.3333 | NA | NA | 2 | 0.5 | 1 | 1 | 1 | 0 | 4 | 0.5 |
| 10_2908771 | NA | NA | 3 | 0.3333 | 1 | 0 | 1 | 0 | 5 | 0.2 | 1 | 0 | 3 | 0 | 1 | 0 | 1 | 1 | 6 | 0.1667 |

| | | | | | | | | | | | | | | | | | | | | |
|---|---|---|---|---|---|---|---|---|---|---|---|---|---|---|---|---|---|---|---|---|
| 10_290877 | NA | NA | 3 | 0.6667 | 1 | 0 | 1 | 0 | 5 | 0.4 | NA | NA | 3 | 0.6667 | 1 | 1 | 1 | 1 | 5 | 0.8 |
| 10_290878 | NA | NA | 3 | 0.3333 | 1 | 0 | 2 | 0 | 6 | 0.1667 | 1 | 0 | 3 | 0.3333 | NA | NA | NA | NA | 4 | 0.25 |
| 10_290880 | NA | NA | 3 | 0.3333 | NA | NA | NA | NA | 3 | 0.3333 | NA | NA | 2 | 0.5 | 1 | 1 | 1 | 1 | 4 | 0.75 |
| 10_290880 | NA | NA | 2 | 1 | 1 | 1 | 1 | 0 | 4 | 0.75 | NA | NA | 1 | 1 | 1 | 1 | 1 | 1 | 3 | 1 |
| 10_290881 | NA | NA | 2 | 0 | 1 | 1 | 1 | 0 | 4 | 0.25 | NA | NA | 2 | 0.5 | 1 | 1 | 1 | 1 | 4 | 0.75 |
| 10_290940 | 1 | 0 | 3 | 0.3333 | 1 | 0 | 2 | 0.5 | 7 | 0.2857 | NA | NA | 2 | 1 | 1 | 0 | 1 | 0 | 4 | 0.5 |
| 10_290941 | 3 | 0 | 5 | 0 | 2 | 0 | 8 | 0.125 | 18 | 0.0556 | 3 | 0.3333 | 3 | 0 | 3 | 0 | 3 | 0 | 12 | 0.0833 |
| 10_291002 | 3 | 0 | 7 | 0 | 4 | 0.25 | 9 | 0.1111 | 23 | 0.087 | 4 | 0.25 | 3 | 0 | 6 | 0.1667 | 4 | 0.25 | 17 | 0.1765 |
| 10_291003 | 2 | 1 | 9 | 0.8889 | 5 | 0.8 | 8 | 1 | 24 | 0.9167 | 4 | 0.75 | 2 | 1 | 7 | 1 | 5 | 1 | 18 | 0.9444 |
| 10_291005 | 3 | 0.3333 | 7 | 0.7143 | 4 | 0.5 | 9 | 0.4444 | 23 | 0.5217 | 1 | 1 | 2 | 0.5 | 5 | 0.8 | 3 | 0.6667 | 11 | 0.7273 |
| 10_291005 | 2 | 0 | 8 | 0.5 | 5 | 0.4 | 9 | 0.2222 | 24 | 0.3333 | 4 | 0.25 | 2 | 0.5 | 5 | 0.8 | 2 | 1 | 13 | 0.6154 |
| 10_291006 | 2 | 0 | 10 | 0.3 | 4 | 0.25 | 6 | 0 | 22 | 0.1818 | 4 | 0.25 | 2 | 0 | 6 | 0.8333 | 5 | 0 | 17 | 0.3529 |
| 10_291006 | 3 | 0 | 7 | 0.5714 | 4 | 0.5 | 7 | 0.2857 | 21 | 0.381 | 4 | 0.5 | 1 | 0 | 7 | 0.7143 | 4 | 0.25 | 16 | 0.5 |
| 10_291007 | 3 | 1 | 9 | 0.8889 | 3 | 1 | 8 | 1 | 23 | 0.9565 | 1 | 0 | 2 | 1 | 4 | 0.75 | 3 | 1 | 10 | 0.8 |
| 10_291007 | 3 | 0 | 9 | 0.2222 | 5 | 0.2 | 10 | 0.3 | 27 | 0.2222 | 4 | 0 | 3 | 0 | 7 | 0.4286 | 5 | 0 | 19 | 0.1579 |
| 10_291008 | 2 | 0.5 | 6 | 0.5 | 5 | 0.6 | 9 | 0.7778 | 22 | 0.6364 | 3 | 0.6667 | 2 | 1 | 4 | 0.25 | 5 | 1 | 14 | 0.7143 |
| 10_291009 | 2 | 1 | 8 | 0.875 | 4 | 1 | 5 | 0.8 | 19 | 0.8947 | 2 | 1 | 3 | 0.6667 | 4 | 0.5 | 4 | 1 | 13 | 0.7692 |
| 10_291011 | 3 | 0.3333 | 8 | 0.375 | 4 | 0.5 | 11 | 0.3636 | 26 | 0.3846 | 5 | 0.4 | 3 | 0 | 6 | 0.1667 | 6 | 0.3333 | 20 | 0.25 |
| 10_291011 | 3 | 0 | 9 | 0 | 3 | 0 | 8 | 0.5 | 23 | 0.1739 | 5 | 0.6 | 3 | 0 | 7 | 0.5714 | 6 | 0 | 21 | 0.3333 |
| 10_291033 | 4 | 1 | 7 | 0.7143 | 5 | 1 | 12 | 0.8333 | 28 | 0.8571 | 7 | 0.8571 | 3 | 1 | 9 | 1 | 6 | 1 | 25 | 0.96 |
| 10_291034 | 3 | 0.3333 | 8 | 0.75 | 3 | 0.6667 | 9 | 0.7778 | 23 | 0.6957 | 5 | 0.6 | 4 | 1 | 7 | 0.7143 | 4 | 0.75 | 20 | 0.75 |
| 10_291034 | 3 | 0.6667 | 10 | 0.6 | 4 | 0.25 | 12 | 0.75 | 29 | 0.6207 | 8 | 0.625 | 5 | 0.6 | 7 | 0.5714 | 8 | 0.5 | 28 | 0.5714 |
| 10_291034 | 3 | 0 | 7 | 0.1429 | 5 | 0.2 | 10 | 0.1 | 25 | 0.12 | 7 | 0 | 1 | 0 | 6 | 0 | 9 | 0.2222 | 23 | 0.087 |
| 10_291036 | 6 | 0.5 | 9 | 0.7778 | 7 | 0.5714 | 11 | 0.7273 | 33 | 0.6667 | 7 | 0.7143 | 5 | 0.8 | 8 | 1 | 8 | 0.75 | 28 | 0.8214 |
| 10_291039 | 2 | 0 | 10 | 0.1 | 7 | 0.1429 | 8 | 0 | 27 | 0.0741 | 3 | 0 | 2 | 0 | 4 | 0.25 | 5 | 0 | 14 | 0.0714 |
| 10_291040 | 3 | 0 | 8 | 0 | 6 | 0.1667 | 3 | 0 | 20 | 0.05 | 1 | 0 | 3 | 0 | 3 | 0.3333 | 3 | 0 | 10 | 0.1 |
| 10_291043 | 3 | 0 | 7 | 0.1429 | 6 | 0 | 3 | 0 | 19 | 0.0526 | 3 | 0 | 4 | 0 | 5 | 0 | 5 | 0.2 | 17 | 0.0588 |
| 10_291056 | NA | NA | 3 | 0.3333 | 2 | 0 | 2 | 0 | 7 | 0.1429 | 4 | 0.25 | 1 | 0 | 4 | 0 | 3 | 0 | 12 | 0.0833 |
| 10_292203 | 1 | 0 | 1 | 0 | 1 | 1 | NA | NA | 3 | 0.3333 | 1 | 0 | NA | NA | 5 | 0.6 | NA | NA | 6 | 0.5 |
| 10_292203 | 1 | 1 | 1 | 0 | 2 | 0 | NA | NA | 4 | 0.25 | 1 | 0 | NA | NA | 3 | 0.3333 | NA | NA | 4 | 0.25 |
| 10_292204 | 2 | 0.5 | 3 | 0.6667 | 2 | 1 | 2 | 1 | 9 | 0.7778 | 1 | 1 | 4 | 0.25 | 4 | 1 | 1 | 0 | 10 | 0.6 |
| 10_292206 | 3 | 0.6667 | 3 | 0.6667 | 2 | 1 | 2 | 0.5 | 10 | 0.7 | 1 | 0 | 3 | 1 | 4 | 0.5 | 1 | 0 | 9 | 0.5556 |
| 10_292208 | 2 | 0 | 2 | 0 | 2 | 0 | 2 | 0.5 | 8 | 0.125 | 1 | 0 | 5 | 0.4 | 3 | 0.6667 | 1 | 0 | 10 | 0.4 |
| 10_292228 | 5 | 0 | 9 | 0.2222 | 5 | 0 | 7 | 0.1429 | 26 | 0.1154 | 4 | 0.25 | 7 | 0.1429 | 8 | 0.125 | 5 | 0.2 | 24 | 0.1667 |
| 10_292228 | 4 | 0 | 9 | 0 | 5 | 0.2 | 9 | 0.1111 | 27 | 0.0741 | 6 | 0.3333 | 5 | 0.2 | 6 | 0.3333 | 6 | 0.1667 | 23 | 0.2609 |
| 10_292228 | 6 | 0.6667 | 10 | 1 | 5 | 0.6 | 5 | 1 | 26 | 0.8462 | 6 | 0.6667 | 6 | 1 | 8 | 0.875 | 5 | 0.8 | 25 | 0.84 |
| 10_292229 | 5 | 0.4 | 9 | 0.6667 | 4 | 0.5 | 4 | 0.5 | 22 | 0.5455 | 5 | 0.6 | 6 | 0.5 | 5 | 0.2 | 5 | 0.4 | 21 | 0.4286 |
| 10_292229 | 4 | 1 | 8 | 1 | 5 | 0.8 | 6 | 1 | 23 | 0.9565 | 4 | 1 | 6 | 1 | 8 | 1 | 6 | 1 | 24 | 1 |
| 10_292231 | 6 | 0.8333 | 9 | 1 | 5 | 0.8 | 7 | 1 | 27 | 0.9259 | 4 | 1 | 6 | 1 | 8 | 0.875 | 4 | 0.75 | 22 | 0.9091 |
| 10_292232 | 5 | 0.2 | 9 | 0.3333 | 5 | 0.8 | 4 | 0.5 | 23 | 0.4348 | 4 | 1 | 5 | 1 | 8 | 0.625 | 5 | 0.6 | 22 | 0.7727 |
| 10_292232 | 4 | 0.5 | 7 | 0.8571 | 5 | 1 | 5 | 0.6 | 21 | 0.7619 | 5 | 0.8 | 5 | 1 | 7 | 1 | 5 | 0.6 | 22 | 0.8636 |
| 10_292233 | 4 | 0.75 | 8 | 1 | 4 | 1 | 7 | 1 | 23 | 0.9565 | 4 | 1 | 5 | 1 | 7 | 1 | 6 | 1 | 22 | 1 |
| 10_292233 | 4 | 0.75 | 9 | 0.8889 | 4 | 1 | 6 | 0.8333 | 23 | 0.8696 | 4 | 0.5 | 5 | 1 | 7 | 1 | 6 | 1 | 22 | 0.9091 |
| 10_292234 | 4 | 0.5 | 8 | 0 | 5 | 0.6 | 7 | 0.1429 | 24 | 0.25 | 3 | 0.3333 | 3 | 0.3333 | 3 | 0 | 5 | 0.2 | 14 | 0.2143 |
| 10_292234 | 5 | 0 | 9 | 0.4444 | 4 | 0.25 | 7 | 0.4286 | 25 | 0.32 | 4 | 0.25 | 5 | 0 | 7 | 0.2857 | 6 | 0.3333 | 22 | 0.2273 |
| 10_292236 | 4 | 0.25 | 7 | 0.4286 | 4 | 0.75 | 4 | 0.25 | 19 | 0.4211 | 3 | 0.6667 | 5 | 0.8 | 5 | 0.6 | 5 | 0.8 | 18 | 0.7222 |
| 10_292237 | 5 | 0 | 9 | 0.2222 | 4 | 0.25 | 7 | 0 | 25 | 0.12 | 4 | 0.25 | 5 | 0.4 | 7 | 0 | 5 | 0 | 21 | 0.1429 |
| 10_292237 | 5 | 0.2 | 7 | 0.1429 | 4 | 0 | 6 | 0 | 22 | 0.0909 | 4 | 0 | 4 | 0.25 | 5 | 0.2 | 6 | 0.3333 | 19 | 0.2105 |
| 10_293285 | NA | NA | 1 | 1 | NA | NA | NA | NA | 1 | 1 | 1 | 1 | NA | NA | NA | NA | NA | NA | 1 | 1 |
| 10_293300 | NA | NA | 3 | 1 | NA | NA | NA | NA | 3 | 1 | 1 | 1 | 2 | 1 | 2 | 1 | 3 | 1 | 8 | 1 |
| 10_293301 | NA | NA | 3 | 0.6667 | NA | NA | 1 | 1 | 4 | 0.75 | 1 | 0 | 1 | 1 | 2 | 0.5 | 3 | 0.6667 | 7 | 0.5714 |
| 10_304111 | 2 | 0.5 | 2 | 0 | NA | NA | 2 | 0 | 6 | 0.1667 | 1 | 1 | 1 | 0 | NA | NA | 3 | 0.3333 | 5 | 0.4 |
| 10_304792 | 4 | 0 | 5 | 0.2 | 4 | 0.25 | 7 | 0 | 20 | 0.1 | 2 | 0 | 6 | 0.1667 | 8 | 0 | 4 | 0 | 20 | 0.05 |
| 10_304795 | 5 | 0 | 4 | 0.25 | 4 | 0.5 | 7 | 0.1429 | 20 | 0.2 | 2 | 0 | 4 | 0.5 | 6 | 0.1667 | 4 | 0.25 | 16 | 0.25 |
| 10_304799 | 5 | 0.2 | 3 | 0 | 2 | 0 | 6 | 0 | 16 | 0.0625 | NA | NA | 3 | 0 | 7 | 0 | 4 | 0.25 | 14 | 0.0714 |
| 10_304843 | 2 | 0 | 2 | 0 | NA | NA | 4 | 0.25 | 8 | 0.125 | 1 | 0 | 1 | 1 | 2 | 0 | 1 | 1 | 5 | 0.4 |
| 10_305218 | 6 | 0.1667 | 10 | 0.1 | 7 | 0.1429 | 5 | 0 | 28 | 0.1071 | 2 | 0 | 9 | 0.2222 | 6 | 0.1667 | 1 | 0 | 18 | 0.1667 |
| 10_305218 | 6 | 0 | 10 | 0.3 | 7 | 0.2857 | 4 | 0.25 | 27 | 0.2222 | 3 | 0 | 7 | 0 | 7 | 0.2857 | 3 | 0 | 20 | 0.1 |
| 10_305232 | 4 | 1 | 7 | 1 | 3 | 1 | 2 | 1 | 16 | 1 | 1 | 1 | 6 | 1 | 4 | 1 | 4 | 1 | 15 | 1 |
| 10_305234 | 4 | 1 | 4 | 1 | 3 | 1 | NA | NA | 11 | 1 | NA | NA | 4 | 1 | 6 | 0.8333 | 3 | 1 | 13 | 0.9231 |
| 10_305235 | 4 | 0.25 | 8 | 0.25 | 3 | 0 | 1 | 0 | 16 | 0.1875 | NA | NA | 6 | 0.1667 | 6 | 0 | 4 | 0 | 16 | 0.0625 |
| 10_305235 | 3 | 1 | 6 | 0.8333 | 3 | 1 | 1 | 1 | 13 | 0.9231 | NA | NA | 6 | 0.8333 | 4 | 0.75 | 3 | 1 | 13 | 0.8462 |
| 10_305235 | 4 | 0.75 | 5 | 0.2 | 3 | 0.6667 | 2 | 0 | 14 | 0.4286 | NA | NA | 6 | 0.1667 | 6 | 0.6667 | 4 | 0.5 | 16 | 0.4375 |
| 10_305235 | 4 | 0.5 | 7 | 0.1429 | 3 | 0 | 1 | 0 | 15 | 0.2 | NA | NA | 5 | 0.2 | 5 | 0.2 | 4 | 0 | 14 | 0.1429 |
| 10_305668 | 4 | 0.25 | 5 | 0.2 | 2 | 0 | 2 | 0.5 | 13 | 0.2308 | NA | NA | 4 | 0 | 2 | 0.5 | 1 | 1 | 7 | 0.2857 |

| | | | | | | | | | | | | | | | | | | | | |
|---|---|---|---|---|---|---|---|---|---|---|---|---|---|---|---|---|---|---|---|---|
| 10_3056692 | 3 | 0.3333 | 5 | 0 | 2 | 0 | 1 | 0 | 11 | 0.0909 | NA | NA | 3 | 0.6667 | 2 | 0 | 1 | 0 | 6 | 0.3333 |
| 10_3056696 | 4 | 0.5 | 5 | 0.2 | 1 | 0 | NA | NA | 10 | 0.3 | NA | NA | 3 | 0 | 1 | 1 | 1 | 0 | 5 | 0.2 |
| 10_3056701 | 3 | 0.3333 | 6 | 0.3333 | 1 | 0 | 2 | 0 | 12 | 0.25 | NA | NA | 3 | 0.3333 | 2 | 0 | 1 | 0 | 6 | 0.1667 |
| 10_3056712 | 4 | 0.25 | 6 | 0.3333 | 1 | 1 | NA | NA | 11 | 0.3636 | NA | NA | 2 | 0 | 1 | 0 | 2 | 0.5 | 5 | 0.2 |
| 10_3056725 | 2 | 0.5 | 5 | 1 | NA | NA | NA | NA | 7 | 0.8571 | NA | NA | 2 | 1 | 1 | 1 | 1 | 1 | 4 | 1 |
| 10_3056727 | 4 | 0.25 | 5 | 0.4 | 1 | 0 | NA | NA | 10 | 0.3 | NA | NA | 4 | 0.25 | 1 | 0 | 2 | 0.5 | 7 | 0.2857 |
| 10_3056741 | 4 | 0.5 | 3 | 0.6667 | 1 | 1 | NA | NA | 8 | 0.625 | NA | NA | 4 | 0.5 | 1 | 1 | 1 | 0 | 6 | 0.5 |
| 10_3056743 | 2 | 1 | 4 | 0.75 | 1 | 1 | NA | NA | 7 | 0.8571 | NA | NA | 3 | 1 | 1 | 1 | 1 | 0 | 5 | 0.8 |
| 10_3056745 | 3 | 0.3333 | 5 | 0.8 | 1 | 0 | NA | NA | 9 | 0.5556 | NA | NA | 3 | 0.3333 | NA | NA | 1 | 0 | 4 | 0.25 |
| 10_3056765 | 3 | 0.3333 | 5 | 0.2 | 1 | 0 | NA | NA | 9 | 0.2222 | NA | NA | 4 | 0.25 | 1 | 0 | 1 | 1 | 6 | 0.3333 |
| 10_3070849 | NA | NA | 4 | 0.5 | 2 | 1 | 2 | 0.5 | 8 | 0.625 | 3 | 0.6667 | NA | NA | 2 | 1 | 1 | 1 | 6 | 0.8333 |
| 10_3070890 | NA | NA | 4 | 1 | 2 | 0.5 | 1 | 1 | 7 | 0.8571 | 3 | 1 | 1 | 1 | 2 | 0.5 | 1 | 1 | 7 | 0.8571 |
| 10_3070902 | NA | NA | 4 | 0.25 | 2 | 0 | 1 | 0 | 7 | 0.1429 | 2 | 1 | 1 | 0 | 2 | 0 | 1 | 0 | 6 | 0.3333 |
| 10_3070910 | NA | NA | 4 | 0.25 | 2 | 0 | 2 | 0 | 8 | 0.125 | 2 | 0 | 1 | 1 | 2 | 0.5 | 1 | 0 | 6 | 0.3333 |
| 10_3070915 | NA | NA | 4 | 0.5 | 2 | 0 | 1 | 1 | 7 | 0.4286 | 3 | 0.3333 | 1 | 1 | 2 | 0 | 1 | 1 | 7 | 0.4286 |
| 10_3070916 | NA | NA | 4 | 0.75 | 2 | 0.5 | 2 | 1 | 8 | 0.75 | 3 | 0.6667 | NA | NA | 2 | 0 | 1 | 1 | 6 | 0.5 |
| 10_3070919 | NA | NA | 4 | 0.25 | 2 | 0 | 2 | 0 | 8 | 0.125 | 3 | 0.6667 | 1 | 0 | 2 | 0 | 1 | 0 | 7 | 0.2857 |
| 10_3070935 | NA | NA | 2 | 1 | 2 | 0 | 1 | 1 | 5 | 0.6 | 1 | 0 | 1 | 1 | 1 | 1 | NA | NA | 3 | 0.6667 |
| 10_3070941 | NA | NA | 3 | 0.3333 | 2 | 0.5 | 1 | 0 | 6 | 0.3333 | 2 | 1 | 1 | 0 | 1 | 0 | NA | NA | 4 | 0.5 |
| 10_3070943 | NA | NA | 4 | 0.5 | 2 | 0 | 1 | 0 | 7 | 0.2857 | 2 | 0 | 1 | 0 | 2 | 0.5 | 1 | 1 | 6 | 0.3333 |
| 10_3070953 | NA | NA | 3 | 0 | 2 | 0.5 | 2 | 0 | 7 | 0.1429 | 3 | 0.3333 | 1 | 0 | 2 | 0 | 1 | 0 | 7 | 0.1429 |
| 10_3070959 | NA | NA | 1 | 1 | 1 | 0 | 2 | 0 | 4 | 0.25 | 2 | 0 | NA | NA | 1 | 1 | 1 | 0 | 4 | 0.25 |
| 10_3070970 | NA | NA | 2 | 0.5 | 1 | 1 | 1 | 1 | 4 | 0.75 | NA | NA | NA | NA | 2 | 1 | NA | NA | 2 | 1 |
| 10_3070975 | NA | NA | 1 | 0 | 2 | 0.5 | 2 | 0.5 | 5 | 0.4 | 1 | 1 | 1 | 1 | 2 | 0.5 | 1 | 1 | 5 | 0.8 |
| 10_3070978 | NA | NA | 3 | 1 | 2 | 1 | 2 | 0 | 7 | 0.7143 | 2 | 1 | 1 | 0 | 2 | 0.5 | 1 | 1 | 6 | 0.6667 |
| 10_3071014 | NA | NA | 1 | 1 | 2 | 1 | 2 | 1 | 5 | 1 | NA | NA | NA | NA | 2 | 1 | 1 | 1 | 3 | 1 |
| 10_3120410 | 7 | 0 | 2 | 0 | 3 | 0 | 4 | 0.5 | 16 | 0.125 | 2 | 0 | 4 | 0.5 | 2 | 0 | 1 | 0 | 9 | 0.2222 |
| 10_3120859 | 3 | 0 | 5 | 0.2 | 5 | 0 | 4 | 0 | 17 | 0.0588 | 2 | 0 | 6 | 0.3333 | 3 | 0.3333 | 5 | 0 | 16 | 0.1875 |
| 10_3128954 | NA | NA | 1 | 1 | 2 | 1 | 1 | 1 | 4 | 1 | NA | NA | NA | NA | 1 | 1 | 2 | 1 | 3 | 1 |
| 10_3137073 | 2 | 0.5 | 1 | 0 | NA | NA | 1 | 0 | 4 | 0.25 | 2 | 1 | NA | NA | 2 | 0 | 1 | 1 | 5 | 0.6 |
| 10_3137361 | 1 | 0 | 1 | 1 | NA | NA | 1 | 0 | 3 | 0.3333 | 2 | 0.5 | NA | NA | NA | NA | 1 | 0 | 3 | 0.3333 |
| 10_3137383 | NA | NA | 1 | 1 | NA | NA | 1 | 0 | 2 | 0.5 | 2 | 0.5 | NA | NA | 1 | 0 | 1 | 0 | 4 | 0.25 |
| 10_3137836 | 1 | 0 | 1 | 1 | NA | NA | 1 | 0 | 3 | 0.3333 | NA | NA | NA | NA | 1 | 0 | 1 | 1 | 2 | 0.5 |
| 10_3137840 | NA | NA | 1 | 1 | NA | NA | NA | NA | 1 | 1 | 1 | 0 | NA | NA | 2 | 0 | 1 | 1 | 4 | 0.25 |
| 10_3137855 | 1 | 1 | NA | NA | NA | NA | 1 | 0 | 2 | 0.5 | 2 | 0.5 | NA | NA | 4 | 0.25 | 3 | 0.6667 | 9 | 0.4444 |
| 10_3137867 | 1 | 1 | NA | NA | NA | NA | 1 | 1 | 2 | 1 | 1 | 0 | NA | NA | 3 | 0.3333 | 2 | 0.5 | 6 | 0.3333 |
| 10_3141079 | 1 | 1 | 1 | 0 | NA | NA | 1 | 1 | 3 | 0.6667 | NA | NA | NA | NA | 3 | 0.3333 | 2 | 0 | 5 | 0.2 |
| 10_3148490 | 1 | 0 | NA | NA | NA | NA | 1 | 1 | 2 | 0.5 | NA | NA | NA | NA | 2 | 0.5 | 1 | 0 | 3 | 0.3333 |
| 10_3148551 | NA | NA | NA | NA | NA | NA | 1 | 1 | 1 | 1 | NA | NA | NA | NA | 1 | 1 | NA | NA | 1 | 1 |
| 10_3156679 | 3 | 0.3333 | 1 | 0 | 2 | 0 | 3 | 0.3333 | 9 | 0.2222 | 4 | 0 | 4 | 0.5 | 1 | 1 | NA | NA | 9 | 0.3333 |
| 10_3156692 | 2 | 0 | 1 | 1 | 1 | 0 | 3 | 0 | 7 | 0.1429 | 3 | 0.3333 | 4 | 0 | 2 | 0.5 | NA | NA | 9 | 0.2222 |
| 10_3156700 | 3 | 0.6667 | 1 | 0 | 2 | 0.5 | 2 | 1 | 8 | 0.625 | 3 | 0.3333 | 3 | 0.6667 | 1 | 0 | NA | NA | 7 | 0.4286 |
| 10_3156702 | 2 | 1 | 1 | 1 | 2 | 1 | 3 | 1 | 8 | 1 | 4 | 0.75 | 4 | 1 | 1 | 1 | NA | NA | 9 | 0.8889 |
| 10_3156720 | 1 | 0 | 1 | 0 | 2 | 0.5 | 2 | 0.5 | 6 | 0.3333 | 4 | 0 | 2 | 0.5 | 1 | 0 | NA | NA | 7 | 0.1429 |
| 10_3156764 | 1 | 1 | 1 | 0 | 1 | 0 | 3 | 0.3333 | 6 | 0.3333 | 2 | 0.5 | 3 | 0.6667 | 1 | 0 | NA | NA | 6 | 0.5 |
| 10_3276665 | 1 | 1 | NA | NA | 1 | 0 | NA | NA | 2 | 0.5 | NA | NA | 1 | 1 | NA | NA | NA | NA | 1 | 1 |
| 10_3276669 | 1 | 1 | NA | NA | 1 | 0 | NA | NA | 2 | 0.5 | NA | NA | 1 | 1 | NA | NA | NA | NA | 1 | 1 |
| 10_3276670 | 1 | 1 | NA | NA | 1 | 1 | NA | NA | 2 | 1 | NA | NA | 1 | 1 | NA | NA | NA | NA | 1 | 1 |
| 10_3276674 | 1 | 1 | NA | NA | NA | NA | NA | NA | 1 | 1 | NA | NA | 1 | 1 | NA | NA | NA | NA | 1 | 1 |
| 10_3276682 | 1 | 1 | NA | NA | 1 | 1 | NA | NA | 2 | 1 | NA | NA | 1 | 1 | NA | NA | NA | NA | 1 | 1 |
| 10_3276695 | 1 | 1 | NA | NA | 1 | 1 | NA | NA | 2 | 1 | NA | NA | 1 | 1 | NA | NA | NA | NA | 1 | 1 |
| 10_3276718 | 1 | 1 | NA | NA | 1 | 1 | NA | NA | 2 | 1 | NA | NA | 1 | 1 | NA | NA | NA | NA | 1 | 1 |
| 10_3276727 | 2 | 1 | NA | NA | 1 | 1 | NA | NA | 3 | 1 | NA | NA | 1 | 1 | NA | NA | NA | NA | 1 | 1 |
| 10_3276733 | 2 | 1 | NA | NA | 1 | 0 | NA | NA | 3 | 0.6667 | 1 | 1 | 1 | 1 | NA | NA | NA | NA | 2 | 1 |
| 10_3276737 | 2 | 1 | NA | NA | 1 | 1 | NA | NA | 3 | 1 | 1 | 1 | 1 | 1 | NA | NA | NA | NA | 2 | 1 |
| 10_3276743 | 1 | 1 | NA | NA | NA | NA | NA | NA | 1 | 1 | 1 | 1 | NA | NA | NA | NA | NA | NA | 1 | 1 |
| 10_3276762 | 1 | 1 | NA | NA | NA | NA | NA | NA | 1 | 1 | 1 | 0 | 1 | 1 | NA | NA | NA | NA | 2 | 0.5 |
| 10_3276773 | 2 | 0.5 | NA | NA | 1 | 0 | NA | NA | 3 | 0.3333 | 1 | 1 | 1 | 1 | NA | NA | NA | NA | 2 | 1 |
| 10_3338872 | 3 | 0 | 5 | 0 | 3 | 0 | 7 | 0.1429 | 18 | 0.0556 | 1 | 0 | 1 | 1 | 4 | 0 | 5 | 0 | 11 | 0.0909 |
| 10_3338892 | 4 | 0.25 | 10 | 0.3 | 7 | 0 | 9 | 0.1111 | 30 | 0.1667 | 4 | 0.25 | 2 | 1 | 5 | 0.6 | 8 | 0.625 | 19 | 0.5789 |
| 10_3338894 | 4 | 1 | 9 | 1 | 7 | 1 | 8 | 1 | 28 | 1 | 4 | 1 | 2 | 0.5 | 4 | 0.75 | 8 | 1 | 18 | 0.8889 |
| 10_3338910 | 4 | 0 | 8 | 0.125 | 7 | 0 | 8 | 0.125 | 27 | 0.0741 | 3 | 0 | 2 | 0 | 3 | 0.3333 | 6 | 0 | 14 | 0.0714 |
| 10_3338916 | 4 | 0.25 | 6 | 0.5 | 7 | 0.2857 | 7 | 0.8571 | 24 | 0.5 | 2 | 0.5 | 1 | 0 | 4 | 0 | 6 | 0.1667 | 13 | 0.1538 |
| 10_3338919 | 4 | 0.5 | 9 | 0.3333 | 6 | 0.1667 | 8 | 0.375 | 27 | 0.3333 | 3 | 0.3333 | 2 | 0 | 4 | 0.25 | 7 | 0.1429 | 16 | 0.1875 |
| 10_3338925 | 3 | 1 | 6 | 0.8333 | 5 | 0.8 | 8 | 0.625 | 22 | 0.7727 | 3 | 1 | 2 | 1 | 4 | 1 | 7 | 0.8571 | 16 | 0.9375 |

| | | | | | | | | | | | | | | | | | | | | |
|---|---|---|---|---|---|---|---|---|---|---|---|---|---|---|---|---|---|---|---|---|
| 10_334718 | 5 | 0 | 4 | 0 | 6 | 0.6667 | 7 | 0.1429 | 22 | 0.2273 | 1 | 0 | 2 | 0 | 3 | 0 | 6 | 0.1667 | 12 | 0.0833 |
| 10_334985 | 3 | 0.6667 | 5 | 0.4 | 6 | 0.1667 | 6 | 0.1667 | 20 | 0.3 | 2 | 0 | 1 | 0 | 2 | 0.5 | 6 | 0.3333 | 11 | 0.2727 |
| 10_334986 | 3 | 0 | 5 | 0.2 | 6 | 0.1667 | 6 | 0 | 20 | 0.1 | 2 | 0 | 1 | 0 | 1 | 0 | 6 | 0.1667 | 10 | 0.1 |
| 10_334988 | 2 | 0.5 | 4 | 1 | 6 | 1 | 3 | 0.6667 | 15 | 0.8667 | 1 | 0 | 1 | 1 | 1 | 1 | 5 | 1 | 8 | 0.875 |
| 10_334988 | 2 | 1 | 4 | 0.5 | 6 | 1 | 4 | 0.5 | 16 | 0.75 | 2 | 1 | 1 | 0 | 1 | 1 | 6 | 1 | 10 | 0.9 |
| 10_334988 | 3 | 0.3333 | 5 | 0 | 6 | 0 | 5 | 0 | 19 | 0.0526 | 2 | 0 | 1 | 0 | 1 | 0 | 5 | 0.4 | 9 | 0.2222 |
| 10_334988 | 3 | 0 | 4 | 0 | 6 | 0.1667 | 4 | 0 | 17 | 0.0588 | 2 | 0 | 1 | 1 | NA | NA | 4 | 0.25 | 7 | 0.2857 |
| 10_334988 | 3 | 0.3333 | 4 | 0.5 | 5 | 0.2 | 5 | 0 | 17 | 0.2353 | 2 | 0 | 1 | 1 | 1 | 0 | 4 | 0 | 8 | 0.125 |
| 10_334988 | 2 | 1 | 5 | 0.4 | 6 | 0.8333 | 4 | 0.5 | 17 | 0.6471 | 2 | 0.5 | 1 | 1 | 1 | 1 | 5 | 0.4 | 9 | 0.5556 |
| 10_334989 | 3 | 0.3333 | 4 | 0.5 | 6 | 0.6667 | 5 | 0.2 | 18 | 0.4444 | 2 | 0.5 | 1 | 0 | 1 | 0 | 5 | 0.4 | 9 | 0.3333 |
| 10_334989 | 3 | 1 | 4 | 1 | 5 | 1 | 4 | 1 | 16 | 1 | 2 | 1 | 1 | 0 | 1 | 1 | 5 | 0.6 | 9 | 0.6667 |
| 10_334989 | 1 | 1 | 5 | 0.6 | 4 | 0.5 | 5 | 0.6 | 15 | 0.6 | 2 | 0.5 | 1 | 0 | 1 | 1 | 5 | 0.8 | 9 | 0.6667 |
| 10_337952 | 2 | 0.5 | 1 | 0 | 2 | 0 | 1 | 0 | 6 | 0.1667 | 3 | 0 | 1 | 1 | 3 | 0 | NA | NA | 7 | 0.1429 |
| 10_338253 | NA | NA | NA | NA | NA | NA | 1 | 1 | 1 | 1 | 1 | 1 | NA | NA | NA | NA | NA | NA | 1 | 1 |
| 10_338640 | 1 | 0 | 1 | 0 | 2 | 0.5 | 2 | 0.5 | 6 | 0.3333 | 1 | 0 | 1 | 1 | 1 | 1 | 1 | 0 | 4 | 0.5 |
| 10_338725 | 10 | 0.2 | 8 | 0.25 | 14 | 0.1429 | 7 | 0.2857 | 39 | 0.2051 | 6 | 0 | 3 | 0 | 11 | 0.0909 | 10 | 0.2 | 30 | 0.1 |
| 10_338727 | 14 | 0.3571 | 8 | 0.625 | 15 | 0.4667 | 5 | 0.6 | 42 | 0.4762 | 6 | 0.5 | 3 | 1 | 10 | 0.4 | 10 | 0.7 | 29 | 0.5862 |
| 10_338727 | 14 | 0.0714 | 11 | 0 | 15 | 0.0667 | 7 | 0.2857 | 47 | 0.0851 | 6 | 0 | 3 | 0.3333 | 11 | 0.0909 | 10 | 0 | 30 | 0.0667 |
| 10_339674 | NA | NA | NA | NA | NA | NA | 1 | 1 | 1 | 1 | NA | NA | 2 | 1 | 1 | 0 | NA | NA | 3 | 0.6667 |
| 10_339675 | NA | NA | NA | NA | NA | NA | 1 | 1 | 1 | 1 | NA | NA | 2 | 1 | 2 | 1 | NA | NA | 4 | 1 |
| 10_339679 | NA | NA | NA | NA | NA | NA | 1 | 1 | 1 | 1 | NA | NA | 2 | 1 | 2 | 1 | NA | NA | 4 | 1 |
| 10_339679 | NA | NA | NA | NA | NA | NA | 1 | 1 | 1 | 1 | NA | NA | 2 | 1 | 2 | 1 | NA | NA | 4 | 1 |
| 10_339679 | NA | NA | NA | NA | NA | NA | 1 | 1 | 1 | 1 | NA | NA | 2 | 1 | 2 | 0.5 | NA | NA | 4 | 0.75 |
| 10_339723 | NA | NA | NA | NA | NA | NA | 1 | 1 | 1 | 1 | NA | NA | NA | NA | 1 | 1 | NA | NA | 1 | 1 |
| 10_340021 | 1 | 1 | NA | NA | 1 | 1 | NA | NA | 2 | 1 | NA | NA | NA | NA | NA | NA | 1 | 1 | 1 | 1 |
| 10_340023 | 1 | 1 | NA | NA | 1 | 1 | NA | NA | 2 | 1 | NA | NA | NA | NA | NA | NA | 1 | 1 | 1 | 1 |
| 10_340276 | NA | NA | 1 | 1 | 1 | 0 | 1 | 1 | 3 | 0.6667 | NA | NA | 1 | 1 | 1 | 1 | 2 | 0.5 | 4 | 0.75 |
| 10_340277 | NA | NA | 1 | 0 | 1 | 0 | 1 | 1 | 3 | 0.3333 | NA | NA | 1 | 1 | 1 | 0 | 1 | 0 | 3 | 0.3333 |
| 10_340278 | NA | NA | 1 | 1 | 1 | 1 | NA | NA | 2 | 1 | NA | NA | 1 | 1 | 1 | 1 | 2 | 1 | 4 | 1 |
| 10_340283 | NA | NA | 1 | 1 | NA | NA | NA | NA | 1 | 1 | NA | NA | 1 | 1 | 1 | 1 | 1 | 1 | 3 | 1 |
| 10_340283 | NA | NA | 1 | 1 | NA | NA | NA | NA | 1 | 1 | NA | NA | NA | NA | 1 | 1 | 2 | 1 | 3 | 1 |
| 10_340284 | NA | NA | 1 | 1 | NA | NA | NA | NA | 1 | 1 | NA | NA | NA | NA | 1 | 1 | 1 | 0 | 2 | 0.5 |
| 10_340284 | NA | NA | 1 | 1 | NA | NA | NA | NA | 1 | 1 | NA | NA | NA | NA | NA | NA | 1 | 1 | 1 | 1 |
| 10_340285 | NA | NA | 1 | 1 | NA | NA | NA | NA | 1 | 1 | NA | NA | 1 | 1 | 1 | 0 | 1 | 1 | 3 | 0.6667 |
| 10_340287 | NA | NA | 1 | 1 | NA | NA | NA | NA | 1 | 1 | NA | NA | 1 | 1 | 1 | 1 | 1 | 1 | 3 | 1 |
| 10_340289 | NA | NA | 1 | 1 | NA | NA | NA | NA | 1 | 1 | NA | NA | NA | NA | 1 | 1 | 1 | 1 | 2 | 1 |
| 10_340293 | NA | NA | 1 | 1 | NA | NA | NA | NA | 1 | 1 | NA | NA | 1 | 1 | NA | NA | 1 | 0 | 2 | 0.5 |
| 10_340295 | NA | NA | 1 | 1 | NA | NA | NA | NA | 1 | 1 | NA | NA | 1 | 1 | NA | NA | NA | NA | 1 | 1 |
| 10_341588 | 16 | 0.5625 | 17 | 0.5294 | 20 | 0.55 | 16 | 0.5 | 69 | 0.5362 | 6 | 0.5 | 19 | 0.4737 | 23 | 0.3913 | 16 | 0.5625 | 64 | 0.4688 |
| 10_341588 | 16 | 1 | 18 | 1 | 22 | 1 | 18 | 0.9444 | 74 | 0.9865 | 9 | 0.8889 | 26 | 0.9615 | 23 | 0.913 | 17 | 1 | 75 | 0.9467 |
| 10_341589 | 23 | 0 | 20 | 0.1 | 26 | 0.0385 | 17 | 0.2941 | 86 | 0.093 | 13 | 0.1538 | 27 | 0.037 | 24 | 0 | 20 | 0.1 | 84 | 0.0595 |
| 10_341589 | 26 | 0.0769 | 21 | 0.0952 | 25 | 0.16 | 20 | 0.1 | 92 | 0.1087 | 12 | 0.0833 | 27 | 0.0741 | 27 | 0.037 | 19 | 0.1053 | 85 | 0.0706 |
| 10_341590 | 27 | 1 | 21 | 0.9524 | 26 | 0.9231 | 21 | 0.9524 | 95 | 0.9579 | 13 | 1 | 27 | 0.9259 | 31 | 0.9355 | 18 | 1 | 89 | 0.9551 |
| 10_341590 | 24 | 0.375 | 20 | 0.55 | 26 | 0.6538 | 21 | 0.4286 | 91 | 0.5055 | 16 | 0.5 | 25 | 0.48 | 33 | 0.6667 | 20 | 0.65 | 94 | 0.5851 |
| 10_341592 | 22 | 0.2727 | 18 | 0.1111 | 25 | 0.24 | 19 | 0.2632 | 84 | 0.2262 | 16 | 0.1875 | 24 | 0.25 | 29 | 0.3103 | 18 | 0.2222 | 87 | 0.2529 |
| 10_341592 | 25 | 0.12 | 21 | 0.2381 | 28 | 0.0357 | 21 | 0 | 95 | 0.0947 | 18 | 0.1667 | 24 | 0.1667 | 31 | 0.0968 | 21 | 0.1429 | 94 | 0.1383 |
| 10_341594 | 24 | 1 | 20 | 0.95 | 25 | 1 | 15 | 1 | 84 | 0.9881 | 19 | 1 | 18 | 0.9444 | 30 | 1 | 19 | 1 | 86 | 0.9884 |
| 10_341595 | 28 | 0.8214 | 22 | 0.7273 | 23 | 0.913 | 21 | 0.9048 | 94 | 0.8404 | 22 | 0.8636 | 25 | 0.8 | 29 | 0.6897 | 19 | 0.8421 | 95 | 0.7895 |
| 10_341595 | 23 | 0.9565 | 21 | 1 | 28 | 1 | 22 | 0.9091 | 94 | 0.9681 | 19 | 1 | 23 | 0.9565 | 28 | 1 | 22 | 1 | 92 | 0.9891 |
| 10_341595 | 24 | 1 | 24 | 1 | 28 | 1 | 22 | 0.9545 | 98 | 0.9898 | 19 | 0.8947 | 23 | 1 | 32 | 0.9375 | 21 | 1 | 95 | 0.9579 |
| 10_341596 | 30 | 0.9333 | 24 | 0.9583 | 25 | 1 | 21 | 1 | 100 | 0.97 | 24 | 0.9583 | 26 | 1 | 29 | 0.931 | 23 | 0.9565 | 102 | 0.9608 |
| 10_341596 | 33 | 0.9394 | 25 | 0.96 | 26 | 1 | 23 | 1 | 107 | 0.972 | 24 | 0.875 | 27 | 0.9259 | 32 | 0.9375 | 24 | 1 | 107 | 0.9346 |
| 10_341597 | 29 | 0.2414 | 21 | 0.0952 | 27 | 0.1852 | 18 | 0.1667 | 95 | 0.1789 | 22 | 0.1364 | 24 | 0.0833 | 33 | 0.1212 | 26 | 0.1154 | 105 | 0.1143 |
| 10_341597 | 31 | 0.8387 | 23 | 0.8261 | 26 | 0.7308 | 20 | 0.85 | 100 | 0.81 | 21 | 0.5238 | 22 | 0.6818 | 29 | 0.7586 | 26 | 0.5769 | 98 | 0.6429 |
| 10_341598 | 24 | 0.9167 | 24 | 0.9583 | 24 | 0.9167 | 20 | 1 | 92 | 0.9457 | 21 | 1 | 24 | 1 | 29 | 1 | 26 | 1 | 100 | 1 |
| 10_341605 | 36 | 0.2222 | 36 | 0.2778 | 32 | 0.2188 | 24 | 0.2917 | 128 | 0.25 | 21 | 0.3333 | 32 | 0.2812 | 32 | 0.2188 | 30 | 0.3 | 115 | 0.2783 |
| 10_341606 | 46 | 0.3261 | 40 | 0.35 | 39 | 0.3846 | 31 | 0.1935 | 156 | 0.3205 | 28 | 0.2143 | 41 | 0.4146 | 44 | 0.25 | 32 | 0.25 | 145 | 0.2897 |
| 10_341735 | 13 | 0 | 10 | 0.1 | 6 | 0.1667 | 9 | 0 | 38 | 0.0526 | 1 | 0 | 5 | 0 | 9 | 0.1111 | 5 | 0 | 20 | 0.05 |
| 10_341740 | 13 | 0.0769 | 11 | 0.0909 | 6 | 0 | 9 | 0 | 39 | 0.0513 | 3 | 0 | 10 | 0.2 | 10 | 0 | 8 | 0 | 31 | 0.0645 |
| 10_341740 | 17 | 0.8824 | 12 | 0.9167 | 7 | 0.8571 | 10 | 0.9 | 46 | 0.8913 | 5 | 1 | 9 | 0.7778 | 9 | 1 | 8 | 1 | 31 | 0.9355 |
| 10_341743 | 20 | 0.3 | 14 | 0 | 9 | 0.1111 | 11 | 0.0909 | 54 | 0.1481 | 6 | 0.1667 | 16 | 0.125 | 16 | 0.1875 | 9 | 0.1111 | 47 | 0.1489 |
| 10_341745 | 20 | 0.05 | 15 | 0.0667 | 10 | 0.1 | 17 | 0.0588 | 62 | 0.0645 | 10 | 0.1 | 18 | 0.0556 | 15 | 0.1333 | 11 | 0.1818 | 54 | 0.1111 |
| 10_341749 | 25 | 0.84 | 14 | 0.9286 | 12 | 0.8333 | 17 | 0.7059 | 68 | 0.8235 | 8 | 0.625 | 18 | 0.3889 | 14 | 0.7143 | 13 | 0.8462 | 53 | 0.6226 |
| 10_341750 | 29 | 0.3448 | 18 | 0.2778 | 16 | 0.1875 | 23 | 0.3043 | 86 | 0.2907 | 11 | 0.2727 | 24 | 0.4167 | 23 | 0.1304 | 16 | 0.25 | 74 | 0.2703 |
| 10_341750 | 32 | 0.125 | 19 | 0.1053 | 17 | 0 | 25 | 0.12 | 93 | 0.0968 | 14 | 0.0714 | 25 | 0.12 | 23 | 0.0435 | 22 | 0.1364 | 84 | 0.0952 |

| | | | | | | | | | | | | | | | | | | | | |
|---|---|---|---|---|---|---|---|---|---|---|---|---|---|---|---|---|---|---|---|---|
| 10_341751( | 26 | 0.6923 | 16 | 0.6875 | 12 | 0.8333 | 18 | 0.7222 | 72 | 0.7222 | 12 | 0.75 | 22 | 0.6364 | 19 | 0.8421 | 22 | 0.5455 | 75 | 0.68 |
| 10_341751! | 32 | 0.9062 | 21 | 0.9048 | 15 | 0.9333 | 23 | 1 | 91 | 0.9341 | 13 | 1 | 21 | 0.9048 | 23 | 1 | 21 | 0.9048 | 78 | 0.9487 |
| 10_341752; | 37 | 0.973 | 23 | 1 | 15 | 1 | 25 | 0.96 | 100 | 0.98 | 13 | 1 | 25 | 0.96 | 27 | 1 | 24 | 0.875 | 89 | 0.9551 |
| 10_341754ł | 31 | 0.0968 | 18 | 0.0556 | 15 | 0 | 18 | 0.0556 | 82 | 0.061 | 11 | 0 | 24 | 0.125 | 21 | 0.0476 | 22 | 0.1364 | 78 | 0.0897 |
| 10_341754! | 32 | 0.5938 | 19 | 0.5263 | 15 | 0.3333 | 18 | 0.1667 | 84 | 0.4405 | 14 | 0.3571 | 26 | 0.4231 | 22 | 0.4091 | 19 | 0.4737 | 81 | 0.4198 |
| 10_341761‹ | 43 | 0.4651 | 26 | 0.3846 | 22 | 0.3636 | 27 | 0.2593 | 118 | 0.3814 | 18 | 0.2778 | 30 | 0.2667 | 28 | 0.4643 | 28 | 0.2857 | 104 | 0.3269 |
| 10_341763( | 47 | 0.0851 | 34 | 0.0588 | 25 | 0.12 | 33 | 0 | 139 | 0.0647 | 19 | 0.2105 | 34 | 0.0588 | 30 | 0.1333 | 33 | 0.1212 | 116 | 0.1207 |
| 10_341764‹ | 46 | 0.1304 | 33 | 0.0303 | 24 | 0 | 29 | 0.069 | 132 | 0.0682 | 21 | 0.0952 | 34 | 0.1176 | 32 | 0.0312 | 32 | 0.125 | 119 | 0.0924 |
| 10_341765; | 44 | 0.1591 | 34 | 0.0882 | 21 | 0.1905 | 27 | 0.037 | 126 | 0.119 | 22 | 0 | 34 | 0.1471 | 35 | 0.0857 | 34 | 0.2353 | 125 | 0.128 |
| 10_341769' | 65 | 0.1692 | 56 | 0.1964 | 35 | 0.2286 | 54 | 0.2407 | 210 | 0.2048 | 33 | 0.1515 | 45 | 0.2667 | 54 | 0.2407 | 53 | 0.2453 | 185 | 0.2324 |
| 10_341769; | 54 | 0.2222 | 43 | 0.1163 | 33 | 0.0909 | 43 | 0.186 | 173 | 0.1618 | 34 | 0.2059 | 41 | 0.1951 | 50 | 0.18 | 44 | 0.1818 | 169 | 0.1893 |
| 10_341770; | 68 | 0.2941 | 58 | 0.2586 | 38 | 0.3421 | 53 | 0.2264 | 217 | 0.2765 | 32 | 0.2812 | 49 | 0.2041 | 54 | 0.2593 | 50 | 0.36 | 185 | 0.2757 |
| 10_341770; | 60 | 0.3 | 54 | 0.2778 | 37 | 0.2432 | 47 | 0.3191 | 198 | 0.2879 | 26 | 0.3846 | 42 | 0.3333 | 58 | 0.2414 | 49 | 0.4286 | 175 | 0.3371 |
| 10_341770; | 66 | 0.0455 | 60 | 0.0833 | 39 | 0.0513 | 55 | 0.0909 | 220 | 0.0682 | 34 | 0 | 49 | 0.1224 | 60 | 0.0833 | 54 | 0.0556 | 197 | 0.0711 |
| 10_341771! | 62 | 0.5161 | 63 | 0.4444 | 41 | 0.4146 | 56 | 0.3929 | 222 | 0.4459 | 31 | 0.4516 | 52 | 0.4808 | 58 | 0.2759 | 51 | 0.4118 | 192 | 0.3958 |
| 10_341771! | 63 | 0.3175 | 53 | 0.2642 | 34 | 0.2353 | 54 | 0.2407 | 204 | 0.2696 | 27 | 0.2593 | 46 | 0.3261 | 55 | 0.2545 | 48 | 0.3333 | 176 | 0.2955 |
| 10_355850' | NA | NA | 2 | 1 | NA | NA | NA | NA | 2 | 1 | NA | NA | 2 | 0.5 | 2 | 0 | NA | NA | 4 | 0.25 |
| 10_355851! | NA | NA | 2 | 0.5 | 1 | 0 | 1 | 1 | 4 | 0.5 | NA | NA | 2 | 0.5 | 2 | 1 | NA | NA | 4 | 0.75 |
| 10_355852; | NA | NA | 2 | 0.5 | NA | NA | 1 | 1 | 3 | 0.6667 | NA | NA | 1 | 1 | 1 | 0 | NA | NA | 2 | 0.5 |
| 10_355852! | NA | NA | 2 | 0.5 | NA | NA | 1 | 0 | 3 | 0.3333 | NA | NA | 1 | 1 | 1 | 1 | NA | NA | 2 | 1 |
| 10_355863( | NA | NA | NA | NA | NA | NA | 1 | 1 | 1 | 1 | NA | NA | NA | NA | 1 | 1 | NA | NA | 1 | 1 |
| 10_367088‹ | 2 | 0.5 | NA | NA | 2 | 0.5 | NA | NA | 4 | 0.5 | NA | NA | 1 | 1 | NA | NA | NA | NA | 1 | 1 |
| 10_367091( | 2 | 1 | NA | NA | 2 | 1 | NA | NA | 4 | 1 | NA | NA | 1 | 1 | NA | NA | NA | NA | 1 | 1 |
| 10_367091' | 2 | 1 | NA | NA | 2 | 1 | NA | NA | 4 | 1 | NA | NA | 1 | 1 | NA | NA | NA | NA | 1 | 1 |
| 10_367094! | 2 | 1 | NA | NA | 2 | 1 | NA | NA | 4 | 1 | NA | NA | 1 | 1 | NA | NA | NA | NA | 1 | 1 |
| 10_367094! | 2 | 0.5 | NA | NA | 1 | 0 | NA | NA | 3 | 0.3333 | NA | NA | 1 | 1 | NA | NA | NA | NA | 1 | 1 |
| 10_367097( | 2 | 0.5 | NA | NA | 2 | 0 | NA | NA | 4 | 0.25 | NA | NA | 1 | 1 | NA | NA | NA | NA | 1 | 1 |
| 10_367097' | 2 | 0.5 | NA | NA | 2 | 0 | NA | NA | 4 | 0.25 | NA | NA | 2 | 0.5 | NA | NA | NA | NA | 2 | 0.5 |
| 10_368776( | 15 | 0.4667 | 16 | 0.4375 | 15 | 0.6667 | 14 | 0.5 | 60 | 0.5167 | 12 | 0.5 | 4 | 0.5 | 15 | 0.4 | 19 | 0.7895 | 50 | 0.58 |
| 10_370093( | 7 | 0.1429 | 2 | 0 | 2 | 0 | 5 | 0 | 16 | 0.0625 | 3 | 0 | 1 | 1 | 2 | 0 | 1 | 1 | 7 | 0.2857 |
| 10_370097( | 14 | 0 | 12 | 0.0833 | 12 | 0.0833 | 19 | 0.1579 | 57 | 0.0877 | 10 | 0 | 17 | 0.0588 | 13 | 0.0769 | 9 | 0.1111 | 49 | 0.0612 |
| 10_370097ł | 5 | 0.2 | 1 | 1 | 3 | 0 | 7 | 0.2857 | 16 | 0.25 | 3 | 0 | 6 | 0.8333 | 4 | 0.25 | 3 | 0.3333 | 16 | 0.4375 |
| 10_370103( | 11 | 0.1818 | 11 | 0.0909 | 8 | 0 | 17 | 0.3529 | 47 | 0.1915 | 5 | 0.6 | 16 | 0.1875 | 13 | 0.1538 | 6 | 0 | 40 | 0.2 |
| 10_370103( | 11 | 0 | 10 | 0.1 | 8 | 0 | 17 | 0.2353 | 46 | 0.1087 | 5 | 0.2 | 14 | 0.0714 | 11 | 0 | 9 | 0 | 39 | 0.0513 |
| 10_370105; | 9 | 0.1111 | 12 | 0.0833 | 8 | 0 | 13 | 0.1538 | 42 | 0.0952 | 5 | 0.2 | 12 | 0 | 9 | 0.2222 | 7 | 0.1429 | 33 | 0.1212 |
| 10_370105! | 10 | 0.1 | 12 | 0.0833 | 9 | 0 | 14 | 0.1429 | 45 | 0.0889 | 5 | 0 | 14 | 0.0714 | 13 | 0.0769 | 7 | 0 | 39 | 0.0513 |
| 10_370110! | 35 | 0.0286 | 47 | 0.1064 | 31 | 0.0968 | 43 | 0.0233 | 156 | 0.0641 | 26 | 0.0769 | 35 | 0.0857 | 29 | 0.2759 | 32 | 0.0625 | 122 | 0.123 |
| 10_370111‹ | 34 | 0.8235 | 45 | 0.7778 | 31 | 0.5806 | 41 | 0.7073 | 151 | 0.7285 | 26 | 0.7308 | 33 | 0.7879 | 24 | 0.75 | 30 | 0.7667 | 113 | 0.7611 |
| 10_370111( | 22 | 0.0909 | 34 | 0.0882 | 22 | 0 | 36 | 0.0833 | 114 | 0.0702 | 23 | 0.0435 | 30 | 0.1333 | 20 | 0.2 | 27 | 0.1481 | 100 | 0.13 |
| 10_370112‹ | 29 | 0.0345 | 39 | 0.0513 | 28 | 0.1071 | 37 | 0.027 | 133 | 0.0526 | 23 | 0.087 | 28 | 0.1786 | 25 | 0.08 | 23 | 0 | 99 | 0.0909 |
| 10_370120; | 25 | 0.48 | 21 | 0.4286 | 15 | 0.4667 | 26 | 0.4231 | 87 | 0.4483 | 16 | 0.25 | 17 | 0.5294 | 13 | 0.4615 | 16 | 0.5 | 62 | 0.4355 |
| 10_370120' | 17 | 0.9412 | 14 | 1 | 13 | 0.9231 | 23 | 0.9565 | 67 | 0.9552 | 11 | 1 | 12 | 1 | 9 | 1 | 11 | 0.9091 | 43 | 0.9767 |
| 10_394098ł | 14 | 0.0714 | 16 | 0.0625 | 10 | 0.4 | 13 | 0.1538 | 53 | 0.1509 | 7 | 0.2857 | 10 | 0.3 | 10 | 0.1 | 8 | 0.25 | 35 | 0.2286 |
| 10_394398; | 15 | 0.0667 | 19 | 0.1579 | 9 | 0 | 12 | 0.0833 | 55 | 0.0909 | 6 | 0 | 11 | 0 | 10 | 0.1 | 8 | 0.25 | 35 | 0.0857 |
| 10_394399' | 15 | 0.6 | 17 | 0.8824 | 10 | 0.6 | 15 | 0.7333 | 57 | 0.7193 | 8 | 0.875 | 13 | 0.7692 | 10 | 0.4 | 9 | 0.7778 | 40 | 0.7 |
| 10_394400ł | 19 | 0.5263 | 20 | 0.55 | 12 | 0.5 | 14 | 0.5714 | 65 | 0.5385 | 9 | 0.6667 | 13 | 0.7692 | 12 | 0.25 | 8 | 0.375 | 42 | 0.5238 |
| 10_394400! | 20 | 0.25 | 19 | 0.3158 | 12 | 0.3333 | 16 | 0.0625 | 67 | 0.2388 | 10 | 0.1 | 15 | 0.0667 | 11 | 0.0909 | 9 | 0 | 45 | 0.0667 |
| 10_394401; | 21 | 0.2381 | 18 | 0.4444 | 11 | 0.2727 | 15 | 0.2667 | 65 | 0.3077 | 10 | 0.2 | 14 | 0.1429 | 11 | 0.0909 | 10 | 0.7 | 45 | 0.2667 |
| 10_394401ł | 20 | 0.25 | 19 | 0.1579 | 12 | 0.25 | 16 | 0.1875 | 67 | 0.209 | 11 | 0 | 14 | 0.1429 | 14 | 0 | 11 | 0.2727 | 50 | 0.1 |
| 10_394405! | 7 | 0 | 3 | 0.3333 | 2 | 0 | 5 | 0.2 | 17 | 0.1176 | 3 | 0 | 3 | 0 | 3 | 0 | 2 | 0.5 | 11 | 0.0909 |
| 10_394406( | 6 | 0.3333 | 3 | 0.3333 | 2 | 1 | 5 | 0.2 | 16 | 0.375 | 3 | 0.3333 | 2 | 0.5 | 2 | 0.5 | 2 | 0.5 | 9 | 0.4444 |
| 10_394407( | 6 | 0.5 | 2 | 0.5 | 2 | 0 | 5 | 0 | 15 | 0.2667 | 3 | 0 | 3 | 0.6667 | 3 | 0 | 3 | 0.3333 | 12 | 0.25 |
| 10_394410ł | 7 | 0.1429 | 4 | 0 | 2 | 0 | 6 | 0 | 19 | 0.0526 | 2 | 0 | 3 | 0.3333 | 3 | 0 | 1 | 0 | 9 | 0.1111 |
| 10_394975‹ | 2 | 0 | NA | NA | 1 | 1 | 1 | 0 | 4 | 0.25 | NA | NA | 1 | 1 | NA | NA | 1 | 0 | 2 | 0.5 |
| 10_394980! | 3 | 0.6667 | 3 | 1 | 1 | 1 | 2 | 1 | 9 | 0.8889 | NA | NA | NA | NA | 1 | 1 | 1 | 1 | 2 | 1 |
| 10_394981; | 3 | 1 | 2 | 0.5 | NA | NA | 1 | 1 | 6 | 0.8333 | NA | NA | 1 | 1 | 1 | 1 | 1 | 0 | 3 | 0.6667 |
| 10_394982( | 2 | 0 | 3 | 0 | NA | NA | 2 | 0.5 | 7 | 0.1429 | NA | NA | 2 | 0.5 | 1 | 1 | 1 | 0 | 4 | 0.5 |
| 10_394982( | 4 | 0 | 3 | 0.3333 | 1 | 0 | 2 | 0.5 | 10 | 0.2 | NA | NA | 2 | 0 | 1 | 1 | 1 | 1 | 4 | 0.5 |
| 10_394982! | 4 | 1 | 3 | 1 | 1 | 1 | 1 | 1 | 9 | 1 | NA | NA | 2 | 1 | 1 | 1 | 1 | 1 | 4 | 1 |
| 10_394983‹ | 4 | 0.5 | 3 | 0.6667 | 1 | 1 | 2 | 0.5 | 10 | 0.6 | NA | NA | 1 | 1 | 1 | 1 | NA | NA | 2 | 1 |
| 10_394984! | 3 | 0.6667 | 3 | 0.3333 | 1 | 1 | 2 | 0 | 9 | 0.4444 | NA | NA | 2 | 0.5 | 1 | 0 | 1 | 1 | 4 | 0.5 |
| 10_394985( | 3 | 0 | 3 | 0.3333 | 1 | 0 | 2 | 0.5 | 9 | 0.2222 | NA | NA | 2 | 0 | NA | NA | 1 | 1 | 3 | 0.3333 |
| 10_394985ł | 4 | 0.25 | 1 | 0 | NA | NA | 2 | 1 | 7 | 0.4286 | NA | NA | 2 | 0 | 1 | 1 | 1 | 0 | 4 | 0.25 |
| 10_395021' | NA | NA | NA | NA | NA | NA | 1 | 1 | 1 | 1 | NA | NA | 1 | 1 | NA | NA | 1 | 1 | 2 | 1 |
| 10_395024( | NA | NA | NA | NA | NA | NA | 1 | 1 | 1 | 1 | NA | NA | NA | NA | NA | NA | 1 | 1 | 1 | 1 |

| | | | | | | | | | | | | | | | | | | | | |
|---|---|---|---|---|---|---|---|---|---|---|---|---|---|---|---|---|---|---|---|---|
| 10_395025 | NA | NA | NA | NA | NA | NA | 1 | 1 | 1 | 1 | NA | NA | 1 | 1 | NA | NA | 1 | 1 | 2 | 1 |
| 10_395137 | 1 | 1 | NA | NA | NA | NA | NA | NA | 1 | 1 | 1 | 0 | 1 | 1 | NA | NA | NA | NA | 2 | 0.5 |
| 10_395137 | 1 | 1 | NA | NA | NA | NA | NA | NA | 1 | 1 | NA | NA | 1 | 1 | NA | NA | NA | NA | 1 | 1 |
| 10_396996 | 1 | 1 | 1 | 1 | NA | NA | 1 | 1 | 3 | 1 | NA | NA | NA | NA | NA | NA | 1 | 1 | 1 | 1 |
| 10_397000 | 1 | 1 | 1 | 0 | NA | NA | 1 | 1 | 3 | 0.6667 | NA | NA | NA | NA | NA | NA | 1 | 1 | 1 | 1 |
| 10_397006 | NA | NA | 1 | 0 | NA | NA | 1 | 1 | 2 | 0.5 | NA | NA | NA | NA | NA | NA | 2 | 0.5 | 2 | 0.5 |
| 10_397008 | 1 | 1 | 1 | 0 | NA | NA | 1 | 0 | 3 | 0.3333 | NA | NA | NA | NA | NA | NA | 2 | 1 | 2 | 1 |
| 10_397011 | 1 | 0 | 2 | 1 | NA | NA | 1 | 1 | 4 | 0.75 | NA | NA | NA | NA | NA | NA | 1 | 1 | 1 | 1 |
| 10_397042 | 1 | 1 | 2 | 0.5 | NA | NA | 3 | 0.6667 | 6 | 0.6667 | NA | NA | NA | NA | NA | NA | 2 | 0.5 | 2 | 0.5 |
| 10_397043 | 2 | 1 | 4 | 0.75 | NA | NA | 3 | 0.3333 | 9 | 0.6667 | NA | NA | 1 | 1 | NA | NA | 1 | 1 | 2 | 1 |
| 10_397044 | 3 | 1 | 4 | 1 | NA | NA | 3 | 1 | 10 | 1 | NA | NA | 1 | 1 | NA | NA | 2 | 1 | 3 | 1 |
| 10_397044 | 2 | 0.5 | 4 | 0.5 | NA | NA | 4 | 0.75 | 10 | 0.6 | NA | NA | 1 | 1 | NA | NA | 1 | 0 | 2 | 0.5 |
| 10_397406 | 2 | 0.5 | 5 | 0 | 1 | 0 | 5 | 0.2 | 13 | 0.1538 | NA | NA | 4 | 0.25 | NA | NA | 3 | 0 | 7 | 0.1429 |
| 10_397502 | 3 | 0.6667 | 7 | 0.5714 | 3 | 0.3333 | 7 | 0.5714 | 20 | 0.55 | NA | NA | 3 | 0.6667 | 2 | 1 | 3 | 0.6667 | 8 | 0.75 |
| 10_397503 | 3 | 0 | 7 | 0.2857 | 2 | 0 | 8 | 0.125 | 20 | 0.15 | NA | NA | 4 | 0 | 3 | 0.3333 | 4 | 0 | 11 | 0.0909 |
| 10_397504 | 2 | 0 | 8 | 0.75 | 3 | 0.3333 | 6 | 0.6667 | 19 | 0.5789 | NA | NA | 3 | 0.3333 | 3 | 0.3333 | 3 | 1 | 9 | 0.5556 |
| 10_397505 | 1 | 0 | 8 | 0.125 | 2 | 0 | 9 | 0.1111 | 20 | 0.1 | NA | NA | 3 | 0 | 3 | 0 | 4 | 0.25 | 10 | 0.1 |
| 10_397673 | 2 | 1 | 1 | 0 | NA | NA | NA | NA | 3 | 0.6667 | NA | NA | NA | NA | NA | NA | 2 | 1 | 2 | 1 |
| 10_397673 | 1 | 1 | 1 | 1 | NA | NA | NA | NA | 2 | 1 | 1 | 1 | NA | NA | NA | NA | 1 | 1 | 2 | 1 |
| 10_397675 | 2 | 0.5 | 1 | 1 | NA | NA | NA | NA | 3 | 0.6667 | 1 | 1 | NA | NA | NA | NA | 1 | 1 | 2 | 1 |
| 10_397676 | 2 | 1 | 1 | 1 | NA | NA | NA | NA | 3 | 1 | NA | NA | NA | NA | NA | NA | 2 | 1 | 2 | 1 |
| 10_397676 | 1 | 1 | 1 | 1 | NA | NA | NA | NA | 2 | 1 | NA | NA | NA | NA | NA | NA | 1 | 1 | 1 | 1 |
| 10_397676 | 2 | 1 | 1 | 0 | NA | NA | NA | NA | 3 | 0.6667 | 1 | 0 | NA | NA | NA | NA | 2 | 0.5 | 3 | 0.3333 |
| 10_397677 | 2 | 1 | 1 | 0 | NA | NA | NA | NA | 3 | 0.6667 | 1 | 0 | NA | NA | NA | NA | 2 | 1 | 3 | 0.6667 |
| 10_397677 | 2 | 1 | 1 | 1 | NA | NA | NA | NA | 3 | 1 | 1 | 1 | NA | NA | NA | NA | 1 | 1 | 2 | 1 |
| 10_397678 | 2 | 0.5 | 1 | 1 | NA | NA | NA | NA | 3 | 0.6667 | 1 | 1 | NA | NA | NA | NA | 2 | 1 | 3 | 1 |
| 10_397681 | 2 | 1 | NA | NA | NA | NA | NA | NA | 2 | 1 | NA | NA | NA | NA | NA | NA | 2 | 1 | 2 | 1 |
| 10_397684 | 2 | 0.5 | NA | NA | NA | NA | NA | NA | 2 | 0.5 | 1 | 1 | NA | NA | NA | NA | NA | NA | 1 | 1 |
| 10_397686 | 2 | 0.5 | NA | NA | NA | NA | NA | NA | 2 | 0.5 | 1 | 1 | NA | NA | NA | NA | 1 | 0 | 2 | 0.5 |
| 10_397686 | 2 | 1 | NA | NA | NA | NA | NA | NA | 2 | 1 | 1 | 1 | NA | NA | NA | NA | 1 | 1 | 2 | 1 |
| 10_397687 | 2 | 1 | NA | NA | NA | NA | NA | NA | 2 | 1 | 1 | 1 | NA | NA | NA | NA | 1 | 1 | 2 | 1 |
| 10_399126 | NA | NA | 1 | 1 | NA | NA | NA | NA | 1 | 1 | NA | NA | NA | NA | NA | NA | 1 | 1 | 1 | 1 |
| 10_399129 | NA | NA | 1 | 1 | NA | NA | NA | NA | 1 | 1 | NA | NA | NA | NA | NA | NA | 1 | 1 | 1 | 1 |
| 10_399130 | NA | NA | 1 | 1 | NA | NA | NA | NA | 1 | 1 | NA | NA | NA | NA | NA | NA | 1 | 1 | 1 | 1 |
| 10_400997 | 2 | 0 | 4 | 0.5 | 3 | 0.3333 | 2 | 0.5 | 11 | 0.3636 | 1 | 0 | 2 | 0 | NA | NA | 2 | 1 | 5 | 0.4 |
| 10_401239 | NA | NA | 2 | 0.5 | 1 | 1 | 1 | 0 | 4 | 0.5 | 3 | 0 | 3 | 0.3333 | 1 | 0 | 1 | 0 | 8 | 0.125 |
| 10_401249 | NA | NA | 2 | 0 | NA | NA | 1 | 1 | 3 | 0.3333 | 4 | 0.25 | 1 | 0 | 1 | 0 | 2 | 0.5 | 8 | 0.25 |
| 10_401249 | NA | NA | 2 | 0 | NA | NA | 1 | 1 | 3 | 0.3333 | 4 | 0.25 | 1 | 0 | 1 | 1 | 2 | 0 | 8 | 0.25 |
| 10_401252 | NA | NA | 2 | 0.5 | NA | NA | 1 | 0 | 3 | 0.3333 | 4 | 0.25 | NA | NA | 1 | 0 | 1 | 0 | 6 | 0.1667 |
| 10_401253 | NA | NA | 2 | 0 | NA | NA | 1 | 1 | 3 | 0.3333 | 4 | 0.25 | NA | NA | 1 | 1 | NA | NA | 5 | 0.4 |
| 10_401253 | NA | NA | 2 | 0 | NA | NA | 1 | 1 | 3 | 0.3333 | 4 | 0.25 | NA | NA | NA | NA | 1 | 0 | 5 | 0.2 |
| 10_401254 | NA | NA | 2 | 0 | NA | NA | 1 | 1 | 3 | 0.3333 | 4 | 0 | NA | NA | 1 | 1 | 1 | 1 | 6 | 0.3333 |
| 10_401255 | 1 | 0 | 6 | 0.3333 | 1 | 0 | 2 | 0.5 | 10 | 0.3 | 4 | 1 | 3 | 0.6667 | 2 | 0.5 | 3 | 0.3333 | 12 | 0.6667 |
| 10_401255 | 2 | 0.5 | 7 | 0.5714 | 1 | 0 | 4 | 0 | 14 | 0.3571 | 3 | 0.6667 | 3 | 0 | 1 | 0 | 4 | 0.25 | 11 | 0.2727 |
| 10_401257 | NA | NA | 5 | 0.2 | 1 | 0 | 4 | 0 | 10 | 0.1 | 2 | 0 | 3 | 0 | 3 | 0.3333 | 3 | 0.3333 | 11 | 0.1818 |
| 10_401635 | 11 | 0 | 22 | 0.1818 | 11 | 0.0909 | 14 | 0.0714 | 58 | 0.1034 | 15 | 0.2667 | 21 | 0.0952 | 17 | 0.0588 | 15 | 0.0667 | 68 | 0.1176 |
| 10_401638 | 15 | 0.4 | 14 | 0.9286 | 13 | 0.6923 | 14 | 0.8571 | 56 | 0.7143 | 17 | 0.7647 | 14 | 0.6429 | 16 | 0.625 | 15 | 0.8 | 62 | 0.7097 |
| 10_401639 | 16 | 0.5 | 21 | 0.9048 | 15 | 0.6 | 16 | 0.6875 | 68 | 0.6912 | 14 | 0.7143 | 20 | 0.65 | 18 | 0.8333 | 18 | 0.8333 | 70 | 0.7571 |
| 10_401640 | 17 | 0.6471 | 21 | 0.7619 | 14 | 0.5714 | 16 | 0.875 | 68 | 0.7206 | 17 | 0.4118 | 19 | 0.7895 | 21 | 0.7619 | 17 | 0.7647 | 74 | 0.6892 |
| 10_401640 | 16 | 0.4375 | 21 | 0.5238 | 15 | 0.4667 | 18 | 0.4444 | 70 | 0.4714 | 16 | 0.625 | 19 | 0.6842 | 16 | 0.5 | 17 | 0.5294 | 68 | 0.5882 |
| 10_401641 | 14 | 0.3571 | 19 | 0.3684 | 14 | 0.1429 | 16 | 0.375 | 63 | 0.3175 | 19 | 0.0526 | 20 | 0.25 | 17 | 0.1765 | 12 | 0.25 | 68 | 0.1765 |
| 10_401644 | 15 | 0.1333 | 20 | 0.15 | 12 | 0 | 13 | 0.2308 | 60 | 0.1333 | 18 | 0.1111 | 15 | 0.2 | 14 | 0.1429 | 15 | 0.3333 | 62 | 0.1935 |
| 10_401645 | 16 | 0.1875 | 21 | 0.1429 | 14 | 0.2143 | 15 | 0.1333 | 66 | 0.1667 | 19 | 0 | 18 | 0.1667 | 13 | 0.0769 | 12 | 0.1667 | 62 | 0.0968 |
| 10_401649 | 16 | 0.4375 | 23 | 0.2174 | 16 | 0.3125 | 18 | 0.2778 | 73 | 0.3014 | 16 | 0.25 | 19 | 0.2105 | 23 | 0.3478 | 15 | 0.4667 | 73 | 0.3151 |
| 10_401650 | 16 | 0.0625 | 21 | 0.1429 | 17 | 0.2353 | 19 | 0.0526 | 73 | 0.1233 | 18 | 0.2222 | 18 | 0.2222 | 23 | 0.1304 | 15 | 0.1333 | 74 | 0.1757 |
| 10_401650 | 15 | 0.1333 | 21 | 0.2381 | 18 | 0.1667 | 16 | 0.0625 | 70 | 0.1571 | 18 | 0.1667 | 17 | 0.2353 | 21 | 0.1905 | 13 | 0.2308 | 69 | 0.2029 |
| 10_401662 | 11 | 0 | 15 | 0.0667 | 16 | 0.1875 | 14 | 0.1429 | 56 | 0.1071 | 11 | 0.0909 | 10 | 0.1 | 16 | 0.0625 | 11 | 0.0909 | 48 | 0.0833 |
| 10_401779 | 7 | 0.7143 | 13 | 0.3846 | 10 | 0.3 | 12 | 0 | 42 | 0.3095 | 10 | 0.2 | 10 | 0.5 | 16 | 0.25 | 13 | 0.3077 | 49 | 0.3061 |
| 10_401780 | 11 | 0.0909 | 13 | 0.2308 | 12 | 0.1667 | 15 | 0.0667 | 51 | 0.1373 | 12 | 0.25 | 12 | 0.25 | 15 | 0.1333 | 13 | 0.1538 | 52 | 0.1923 |
| 10_401781 | 8 | 0.25 | 9 | 0.3333 | 10 | 0.1 | 13 | 0.1538 | 40 | 0.2 | 9 | 0.4444 | 11 | 0.5455 | 15 | 0.2 | 10 | 0.4 | 45 | 0.3778 |
| 10_401781 | 10 | 0.1 | 14 | 0.5 | 12 | 0.25 | 15 | 0.3333 | 51 | 0.3137 | 11 | 0.3636 | 12 | 0 | 18 | 0.5 | 11 | 0.2727 | 52 | 0.3077 |
| 10_402255 | 1 | 1 | NA | NA | 3 | 0.3333 | NA | NA | 4 | 0.5 | 1 | 0 | 1 | 0 | 1 | 1 | NA | NA | 3 | 0.3333 |
| 10_403558 | 1 | 1 | 1 | 1 | 8 | 0.625 | 3 | 0.6667 | 13 | 0.6923 | 4 | 0.75 | 1 | 1 | 2 | 0.5 | NA | NA | 7 | 0.7143 |
| 10_403558 | 1 | 0 | 2 | 0 | 9 | 0.1111 | 2 | 0 | 14 | 0.0714 | 6 | 0.1667 | 4 | 0 | 3 | 0.3333 | 2 | 0 | 15 | 0.1333 |
| 10_403558 | 1 | 0 | 1 | 1 | 7 | 1 | 1 | 1 | 10 | 0.9 | 5 | 1 | 2 | 1 | 4 | 0.75 | 2 | 0.5 | 13 | 0.8462 |

| | | | | | | | | | | | | | | | | | | | | |
|---|---|---|---|---|---|---|---|---|---|---|---|---|---|---|---|---|---|---|---|---|
| 10_4035589 | 1 | 0 | 2 | 0 | 9 | 0.1111 | 3 | 0 | 15 | 0.0667 | 6 | 0.1667 | 4 | 0 | 5 | 0 | 1 | 0 | 16 | 0.0625 |
| 10_4035593 | NA | NA | 1 | 1 | 8 | 1 | 3 | 1 | 12 | 1 | 5 | 1 | 3 | 1 | 5 | 1 | 2 | 1 | 15 | 1 |
| 10_4035595 | 1 | 0 | 1 | 0 | 7 | 0.2857 | 2 | 0 | 11 | 0.1818 | 3 | 0.3333 | 3 | 0.6667 | 4 | 0.75 | 1 | 1 | 11 | 0.6364 |
| 10_4035601 | 1 | 1 | 2 | 1 | 10 | 0.8 | 3 | 0.3333 | 16 | 0.75 | 6 | 0.3333 | 4 | 1 | 4 | 1 | 2 | 1 | 16 | 0.75 |
| 10_4035618 | 2 | 1 | 1 | 1 | 6 | 1 | 1 | 1 | 10 | 1 | 6 | 0.8333 | 4 | 1 | 5 | 1 | 1 | 1 | 16 | 0.9375 |
| 10_4035624 | 3 | 0.3333 | 2 | 0 | 9 | 0.2222 | 4 | 0.5 | 18 | 0.2778 | 6 | 0.5 | 4 | 0.75 | 4 | 0.75 | 3 | 0.3333 | 17 | 0.5882 |
| 10_4035624 | 2 | 0 | 2 | 0 | 8 | 0 | 4 | 0.25 | 16 | 0.0625 | 4 | 0 | 3 | 0.3333 | 4 | 0 | 3 | 0.3333 | 14 | 0.1429 |
| 10_4035628 | 2 | 1 | 2 | 0.5 | 11 | 0.8182 | 3 | 1 | 18 | 0.8333 | 5 | 0.8 | 4 | 1 | 5 | 1 | 3 | 1 | 17 | 0.9412 |
| 10_4035634 | 1 | 0 | 1 | 0 | 7 | 0.1429 | NA | NA | 9 | 0.1111 | 2 | 0.5 | 2 | 0 | 2 | 0 | 2 | 0 | 8 | 0.125 |
| 10_4035635 | 3 | 0.3333 | 2 | 0.5 | 8 | 0.375 | 3 | 0 | 16 | 0.3125 | 5 | 0 | 3 | 0.3333 | 3 | 0 | 3 | 0 | 14 | 0.0714 |
| 10_4035637 | 2 | 0 | 1 | 0 | 8 | 0.125 | 2 | 0 | 13 | 0.0769 | 4 | 0 | 3 | 0 | 4 | 0 | 2 | 1 | 13 | 0.1538 |
| 10_4035641 | 1 | 1 | NA | NA | 12 | 0.75 | 4 | 1 | 17 | 0.8235 | 4 | 1 | 3 | 1 | 3 | 1 | 3 | 1 | 13 | 1 |
| 10_4035658 | 2 | 0 | 2 | 0.5 | 10 | 0.3 | 5 | 0.6 | 19 | 0.3684 | 6 | 0 | 4 | 0.75 | 4 | 0.75 | 3 | 0.3333 | 17 | 0.4118 |
| 10_4035662 | 3 | 0.3333 | 2 | 0.5 | 13 | 0.3077 | 3 | 0 | 21 | 0.2857 | 4 | 0.5 | 3 | 0 | 4 | 0.25 | 2 | 0.5 | 13 | 0.3077 |
| 10_4035676 | 3 | 0.6667 | 2 | 0 | 10 | 0.7 | 4 | 0.75 | 19 | 0.6316 | 6 | 0.5 | 3 | 0.6667 | 4 | 0.75 | 2 | 0.5 | 15 | 0.6 |
| 10_4035685 | 3 | 0.3333 | 2 | 0.5 | 11 | 0.2727 | 5 | 0.4 | 21 | 0.3333 | 5 | 0.4 | 4 | 0.5 | 5 | 0.6 | 4 | 0 | 18 | 0.3889 |
| 10_4035888 | 7 | 0.1429 | 4 | 0 | 10 | 0.2 | 7 | 0.1429 | 28 | 0.1429 | 5 | 0.2 | 5 | 0.4 | 7 | 0.4286 | 7 | 0 | 24 | 0.25 |
| 10_4035969 | 10 | 0 | 9 | 0.1111 | 10 | 0 | 8 | 0.125 | 37 | 0.0541 | 6 | 0 | 4 | 0 | 10 | 0.1 | 8 | 0.25 | 28 | 0.1071 |
| 10_4035974 | 8 | 0.375 | 8 | 0.25 | 15 | 0.3333 | 8 | 0.375 | 39 | 0.3333 | 7 | 0 | 3 | 0.3333 | 9 | 0.4444 | 7 | 0.2857 | 26 | 0.2692 |
| 10_4052568 | NA | NA | 2 | 0.5 | 1 | 0 | 1 | 0 | 4 | 0.25 | NA | NA | NA | NA | 1 | 1 | NA | NA | 1 | 1 |
| 10_4081243 | 43 | 0.3953 | 33 | 0.3939 | 18 | 0.2778 | 34 | 0.3235 | 128 | 0.3594 | 29 | 0.2759 | 36 | 0.3056 | 38 | 0.4474 | 31 | 0.3548 | 134 | 0.3507 |
| 10_4081378 | 48 | 0.2917 | 33 | 0.5455 | 19 | 0.1579 | 34 | 0.2941 | 134 | 0.3358 | 34 | 0.3824 | 42 | 0.4524 | 40 | 0.3 | 36 | 0.5 | 152 | 0.4079 |
| 10_4081381 | 48 | 0.3333 | 35 | 0.2286 | 22 | 0.4091 | 39 | 0.4103 | 144 | 0.3403 | 35 | 0.2857 | 40 | 0.35 | 40 | 0.525 | 35 | 0.3714 | 150 | 0.3867 |
| 10_4081384 | 42 | 0.7619 | 25 | 0.48 | 17 | 0.7059 | 31 | 0.5161 | 115 | 0.6261 | 32 | 0.5 | 36 | 0.5278 | 40 | 0.6 | 31 | 0.4516 | 139 | 0.5252 |
| 10_4081386 | 39 | 0.1282 | 31 | 0.129 | 17 | 0.1176 | 33 | 0.1515 | 120 | 0.1333 | 33 | 0.1212 | 40 | 0.175 | 37 | 0.1892 | 33 | 0.1212 | 143 | 0.1538 |
| 10_4081421 | 58 | 0.2759 | 41 | 0.2439 | 29 | 0.2759 | 47 | 0.1702 | 175 | 0.24 | 42 | 0.2143 | 48 | 0.2083 | 44 | 0.2045 | 43 | 0.2791 | 177 | 0.226 |
| 10_4094616 | 11 | 0 | 5 | 0 | 7 | 0.1429 | 21 | 0.0952 | 44 | 0.0682 | 7 | 0.2857 | 18 | 0.2222 | 15 | 0.0667 | 11 | 0 | 51 | 0.1373 |
| 10_4094624 | 13 | 0.0769 | 5 | 0 | 10 | 0.2 | 21 | 0.2857 | 49 | 0.1837 | 7 | 0 | 20 | 0.05 | 17 | 0.1765 | 12 | 0.0833 | 56 | 0.0893 |
| 10_4094633 | 10 | 0.1 | 5 | 0.6 | 7 | 0.1429 | 18 | 0.1667 | 40 | 0.2 | 5 | 0.2 | 16 | 0.125 | 10 | 0.3 | 5 | 0.2 | 36 | 0.1944 |
| 10_4094634 | 10 | 1 | 5 | 0.2 | 9 | 0.8889 | 17 | 0.9412 | 41 | 0.8537 | 6 | 0.8333 | 14 | 0.9286 | 16 | 0.8125 | 9 | 1 | 45 | 0.8889 |
| 10_4094636 | 12 | 1 | 5 | 0.8 | 8 | 1 | 19 | 1 | 44 | 0.9773 | 7 | 1 | 17 | 1 | 15 | 0.9333 | 8 | 1 | 47 | 0.9787 |
| 10_4094638 | 11 | 0.4545 | 5 | 0.2 | 9 | 0 | 19 | 0.3158 | 44 | 0.2727 | 7 | 0.5714 | 20 | 0.3 | 16 | 0.5 | 11 | 0.4545 | 54 | 0.4259 |
| 10_4094640 | 10 | 1 | 4 | 0.75 | 3 | 1 | 15 | 0.9333 | 32 | 0.9375 | 5 | 1 | 15 | 1 | 10 | 1 | 7 | 1 | 37 | 1 |
| 10_4094661 | 11 | 0 | 4 | 0 | 8 | 0 | 17 | 0.1176 | 40 | 0.05 | 6 | 0.3333 | 16 | 0.0625 | 15 | 0.1333 | 10 | 0 | 47 | 0.1064 |
| 10_4094666 | 13 | 0.4615 | 4 | 0 | 6 | 0.1667 | 19 | 0.4211 | 42 | 0.3571 | 7 | 0.4286 | 19 | 0.3684 | 13 | 0.3077 | 10 | 0.3 | 49 | 0.3469 |
| 10_4094674 | 9 | 1 | 4 | 0.75 | 7 | 0.8571 | 12 | 0.9167 | 32 | 0.9062 | 7 | 1 | 17 | 1 | 10 | 1 | 9 | 0.8889 | 43 | 0.9767 |
| 10_4094695 | 5 | 1 | 4 | 1 | 7 | 1 | 14 | 1 | 30 | 1 | 6 | 1 | 15 | 1 | 9 | 0.8889 | 7 | 1 | 37 | 0.973 |
| 10_4094699 | 8 | 1 | 4 | 1 | 7 | 1 | 17 | 0.9412 | 36 | 0.9722 | 6 | 1 | 14 | 1 | 10 | 1 | 8 | 1 | 38 | 1 |
| 10_4094703 | 8 | 0.125 | 3 | 0.6667 | 8 | 0.375 | 15 | 0.3333 | 34 | 0.3235 | 6 | 0.5 | 15 | 0.2667 | 9 | 0.2222 | 7 | 0.2857 | 37 | 0.2973 |
| 10_4094708 | 7 | 0 | 4 | 0.25 | 7 | 0 | 16 | 0.0625 | 34 | 0.0588 | 6 | 0 | 14 | 0.2143 | 8 | 0 | 8 | 0.125 | 36 | 0.1111 |
| 10_4094716 | 6 | 0.8333 | 3 | 1 | 5 | 1 | 15 | 0.9333 | 29 | 0.931 | 6 | 1 | 14 | 0.9286 | 8 | 1 | 7 | 0.8571 | 35 | 0.9429 |
| 10_4094732 | 6 | 1 | 3 | 1 | 5 | 1 | 13 | 1 | 27 | 1 | 3 | 1 | 13 | 1 | 5 | 1 | 6 | 1 | 27 | 1 |
| 10_4094738 | 6 | 0.6667 | 3 | 0.6667 | 8 | 0.625 | 13 | 0.5385 | 30 | 0.6 | 6 | 0.5 | 13 | 0.4615 | 5 | 0.4 | 8 | 0.5 | 32 | 0.4688 |
| 10_4094743 | 8 | 0.25 | 4 | 0.25 | 7 | 0.1429 | 15 | 0.2 | 34 | 0.2059 | 5 | 0.4 | 12 | 0.3333 | 7 | 0.1429 | 7 | 0 | 31 | 0.2258 |
| 10_4106444 | 3 | 0.6667 | 3 | 0 | 5 | 0.8 | 3 | 1 | 14 | 0.6429 | 7 | 0.2857 | 6 | 0.5 | 6 | 0.3333 | 4 | 1 | 23 | 0.4783 |
| 10_4106450 | 1 | 1 | 4 | 0 | 3 | 0 | 2 | 0 | 10 | 0.1 | 6 | 0 | 5 | 0.2 | 7 | 0.2857 | 4 | 0 | 22 | 0.1364 |
| 10_4106456 | 2 | 0.5 | 4 | 0.5 | 5 | 0 | 3 | 0.6667 | 14 | 0.3571 | 6 | 0.5 | 5 | 0.6 | 7 | 0.7143 | 4 | 0.5 | 22 | 0.5909 |
| 10_4132753 | NA | NA | 1 | 1 | 1 | 1 | NA | NA | 2 | 1 | NA | NA | NA | NA | 3 | 1 | 1 | 1 | 4 | 1 |
| 10_4132764 | NA | NA | 2 | 0.5 | NA | NA | NA | NA | 2 | 0.5 | NA | NA | NA | NA | 3 | 1 | 1 | 0 | 4 | 0.75 |
| 10_4132768 | NA | NA | 2 | 0 | 1 | 1 | NA | NA | 3 | 0.3333 | 1 | 0 | NA | NA | 3 | 0.3333 | 2 | 0.5 | 6 | 0.3333 |
| 10_4132770 | NA | NA | 2 | 0.5 | 1 | 0 | 1 | 0 | 4 | 0.25 | 1 | 0 | NA | NA | 4 | 0.25 | 2 | 0 | 7 | 0.1429 |
| 10_4132782 | NA | NA | 1 | 1 | NA | NA | NA | NA | 1 | 1 | NA | NA | NA | NA | 3 | 0.3333 | 1 | 1 | 4 | 0.5 |
| 10_4132785 | NA | NA | 3 | 1 | 1 | 1 | NA | NA | 4 | 1 | 1 | 1 | NA | NA | 4 | 0.5 | 2 | 0.5 | 7 | 0.5714 |
| 10_4132800 | NA | NA | 3 | 0.6667 | 1 | 1 | NA | NA | 4 | 0.75 | 1 | 1 | NA | NA | 4 | 0.75 | 1 | 1 | 6 | 0.8333 |
| 10_4134050 | 1 | 1 | 3 | 0.6667 | NA | NA | NA | NA | 4 | 0.75 | 1 | 1 | 1 | 1 | 3 | 1 | 1 | 0 | 6 | 0.8333 |
| 10_4134059 | 1 | 0 | 2 | 1 | NA | NA | NA | NA | 3 | 0.6667 | 1 | 1 | 1 | 0 | 2 | 1 | 1 | 1 | 5 | 0.8 |
| 10_4134079 | 2 | 1 | 4 | 1 | NA | NA | 1 | 1 | 7 | 1 | 1 | 1 | 1 | 1 | 3 | 1 | NA | NA | 5 | 1 |
| 10_4134096 | 2 | 0.5 | 4 | 0.25 | NA | NA | NA | NA | 6 | 0.3333 | 1 | 1 | 2 | 0.5 | 2 | 0 | 1 | 1 | 6 | 0.5 |
| 10_4134107 | 1 | 1 | 2 | 1 | NA | NA | NA | NA | 3 | 1 | 1 | 1 | 2 | 1 | 5 | 1 | 1 | 1 | 9 | 1 |
| 10_4136623 | 2 | 0 | 2 | 0 | 2 | 0 | 1 | 1 | 7 | 0.1429 | 1 | 0 | 3 | 0.3333 | 3 | 0.3333 | 1 | 0 | 8 | 0.25 |
| 10_4136627 | 2 | 0.5 | 2 | 0 | 2 | 0 | 1 | 0 | 7 | 0.1429 | 1 | 0 | 3 | 0 | 5 | 0.2 | 1 | 0 | 10 | 0.1 |
| 10_4140484 | 10 | 0.6 | 5 | 0.6 | 4 | 0.5 | 6 | 0.5 | 25 | 0.56 | 7 | 0.5714 | 12 | 0.6667 | 3 | 0.6667 | 10 | 0.5 | 32 | 0.5938 |
| 10_4140486 | 8 | 1 | 3 | 1 | 3 | 1 | 6 | 0.8333 | 20 | 0.95 | 8 | 1 | 11 | 1 | 4 | 1 | 9 | 1 | 32 | 1 |
| 10_4140487 | 9 | 0 | 5 | 0.2 | 4 | 0.25 | 6 | 0 | 24 | 0.0833 | 7 | 0.1429 | 12 | 0.0833 | 4 | 0 | 10 | 0 | 33 | 0.0606 |
| 10_4140508 | 9 | 0.7778 | 5 | 0.8 | 4 | 0.75 | 7 | 1 | 25 | 0.84 | 8 | 0.875 | 12 | 0.8333 | 4 | 1 | 10 | 0.8 | 34 | 0.8529 |

| | | | | | | | | | | | | | | | | | | | | |
|---|---|---|---|---|---|---|---|---|---|---|---|---|---|---|---|---|---|---|---|---|
| 10_414055 | 11 | 0.0909 | 6 | 0 | 5 | 0 | 6 | 0.1667 | 28 | 0.0714 | 8 | 0.125 | 12 | 0.0833 | 6 | 0.1667 | 10 | 0.2 | 36 | 0.1389 |
| 10_414069 | 11 | 0.0909 | 7 | 0.4286 | 7 | 0 | 8 | 0.125 | 33 | 0.1515 | 9 | 0.1111 | 14 | 0.4286 | 4 | 0 | 14 | 0 | 41 | 0.1707 |
| 10_416866 | 1 | 1 | 2 | 0.5 | NA | NA | 1 | 1 | 4 | 0.75 | NA | NA | 2 | 0.5 | 2 | 0.5 | NA | NA | 4 | 0.5 |
| 10_416868 | 1 | 1 | 2 | 0 | NA | NA | 2 | 0.5 | 5 | 0.4 | NA | NA | 2 | 0.5 | 3 | 0.3333 | NA | NA | 5 | 0.4 |
| 10_416869 | 1 | 0 | 3 | 0.6667 | NA | NA | 2 | 0.5 | 6 | 0.5 | NA | NA | 1 | 0 | 4 | 0.75 | NA | NA | 5 | 0.6 |
| 10_416870 | 1 | 1 | 3 | 1 | NA | NA | 2 | 1 | 6 | 1 | NA | NA | 2 | 0.5 | 2 | 1 | NA | NA | 4 | 0.75 |
| 10_416870 | 1 | 1 | 3 | 0 | NA | NA | 1 | 0 | 5 | 0.2 | NA | NA | 2 | 1 | 4 | 0.25 | NA | NA | 6 | 0.5 |
| 10_416871 | NA | NA | 3 | 0.3333 | NA | NA | 2 | 0 | 5 | 0.2 | 1 | 0 | 3 | 0.3333 | 3 | 0.6667 | NA | NA | 7 | 0.4286 |
| 10_416871 | 1 | 1 | 2 | 1 | NA | NA | 2 | 0.5 | 5 | 0.8 | NA | NA | 3 | 1 | 2 | 0.5 | NA | NA | 5 | 0.8 |
| 10_417834 | 7 | 0.5714 | 9 | 0.5556 | 7 | 0.5714 | 12 | 0.3333 | 35 | 0.4857 | 7 | 0.1429 | 20 | 0.35 | 12 | 0.4167 | 4 | 0 | 43 | 0.3023 |
| 10_417836 | 8 | 0.75 | 9 | 0.8889 | 8 | 1 | 11 | 0.8182 | 36 | 0.8611 | 6 | 0.8333 | 19 | 0.9474 | 14 | 0.8571 | 4 | 0.5 | 43 | 0.8605 |
| 10_417836 | 11 | 0.0909 | 7 | 0.1429 | 7 | 0 | 13 | 0.0769 | 38 | 0.0789 | 7 | 0.2857 | 20 | 0.05 | 15 | 0.0667 | 4 | 0 | 46 | 0.087 |
| 10_417836 | 8 | 1 | 9 | 0.8889 | 6 | 0.8333 | 11 | 0.9091 | 34 | 0.9118 | 4 | 1 | 18 | 0.8889 | 12 | 0.8333 | 4 | 1 | 38 | 0.8947 |
| 10_417838 | 9 | 1 | 8 | 0.875 | 6 | 1 | 9 | 0.7778 | 32 | 0.9062 | 5 | 0.8 | 18 | 0.9444 | 11 | 0.7273 | 4 | 0.75 | 38 | 0.8421 |
| 10_417839 | 10 | 0.4 | 10 | 0.1 | 9 | 0.4444 | 13 | 0.3077 | 42 | 0.3095 | 7 | 0.4286 | 19 | 0.4211 | 16 | 0.3125 | 4 | 0.5 | 46 | 0.3913 |
| 10_417840 | 13 | 0.3077 | 10 | 0.5 | 8 | 0.375 | 13 | 0.3846 | 44 | 0.3864 | 6 | 0.1667 | 17 | 0.6471 | 15 | 0.4667 | 3 | 0.6667 | 41 | 0.5122 |
| 10_417842 | 14 | 0.0714 | 10 | 0.1 | 11 | 0.3636 | 13 | 0.3846 | 48 | 0.2292 | 6 | 0.1667 | 19 | 0.2105 | 18 | 0.2778 | 6 | 0.5 | 49 | 0.2653 |
| 10_420580 | 7 | 0.2857 | 6 | 0.1667 | 10 | 0.3 | 16 | 0.4375 | 39 | 0.3333 | NA | NA | 11 | 0.3636 | 10 | 0.2 | 9 | 0.5556 | 30 | 0.3667 |
| 10_420588 | 7 | 0.2857 | 6 | 0 | 9 | 0.2222 | 11 | 0.1818 | 33 | 0.1818 | NA | NA | 9 | 0.2222 | 8 | 0.125 | 2 | 0.5 | 19 | 0.2105 |
| 10_420589 | 7 | 0.1429 | 7 | 0 | 5 | 0.2 | 10 | 0.1 | 29 | 0.1034 | NA | NA | 8 | 0.25 | 7 | 0 | 2 | 0 | 17 | 0.1176 |
| 10_420590 | 6 | 0 | 6 | 0.3333 | 8 | 0.125 | 9 | 0.2222 | 29 | 0.1724 | NA | NA | 6 | 0.1667 | 8 | 0 | 3 | 0 | 17 | 0.0588 |
| 10_420592 | 7 | 0.1429 | 7 | 0.2857 | 9 | 0.2222 | 8 | 0.25 | 31 | 0.2258 | NA | NA | 5 | 0.2 | 8 | 0.25 | 2 | 0 | 15 | 0.2 |
| 10_420592 | 5 | 0.4 | 6 | 0 | 8 | 0 | 9 | 0.1111 | 28 | 0.1071 | NA | NA | 7 | 0.1429 | 6 | 0 | 2 | 0 | 15 | 0.0667 |
| 10_420593 | 6 | 0 | 6 | 0 | 9 | 0 | 9 | 0.2222 | 30 | 0.0667 | NA | NA | 8 | 0.25 | 8 | 0 | 2 | 0 | 18 | 0.1111 |
| 10_420602 | 5 | 0.4 | 5 | 0 | 7 | 0 | 6 | 0 | 23 | 0.087 | NA | NA | 6 | 0.1667 | 4 | 0 | 2 | 0 | 12 | 0.0833 |
| 10_420607 | 5 | 0 | 4 | 0 | 7 | 0.1429 | 4 | 0 | 20 | 0.05 | NA | NA | 6 | 0 | 4 | 0.25 | 2 | 0 | 12 | 0.0833 |
| 10_420621 | 1 | 1 | 2 | 1 | NA | NA | NA | NA | 3 | 1 | NA | NA | 2 | 1 | 1 | 1 | NA | NA | 3 | 1 |
| 10_420621 | 2 | 1 | 1 | 1 | 2 | 1 | NA | NA | 5 | 1 | NA | NA | 3 | 1 | 3 | 1 | NA | NA | 6 | 1 |
| 10_420622 | 1 | 0 | 1 | 1 | 2 | 0.5 | NA | NA | 4 | 0.5 | NA | NA | 3 | 0.3333 | 3 | 0.6667 | NA | NA | 6 | 0.5 |
| 10_420725 | 2 | 0 | 1 | 0 | 2 | 0.5 | NA | NA | 5 | 0.2 | NA | NA | 3 | 0.3333 | 3 | 0 | 1 | 0 | 7 | 0.1429 |
| 10_420726 | 2 | 0.5 | 1 | 1 | 2 | 0.5 | NA | NA | 5 | 0.6 | NA | NA | 1 | 0 | 3 | 0.6667 | 1 | 1 | 5 | 0.6 |
| 10_420726 | 2 | 0 | 1 | 1 | 2 | 1 | NA | NA | 5 | 0.6 | NA | NA | 3 | 0.6667 | 2 | 1 | 1 | 1 | 6 | 0.8333 |
| 10_420726 | 2 | 0.5 | 1 | 1 | 2 | 1 | NA | NA | 5 | 0.8 | NA | NA | 3 | 0.6667 | 2 | 1 | NA | NA | 5 | 0.8 |
| 10_420727 | 2 | 1 | 1 | 1 | 2 | 1 | NA | NA | 5 | 1 | NA | NA | 3 | 0.6667 | NA | NA | NA | NA | 3 | 0.6667 |
| 10_420729 | 2 | 1 | 1 | 1 | 1 | 1 | NA | NA | 4 | 1 | NA | NA | 3 | 1 | 1 | 1 | NA | NA | 4 | 1 |
| 10_420730 | 2 | 0 | 1 | 1 | 1 | 0 | NA | NA | 4 | 0.25 | NA | NA | 3 | 0 | 1 | 1 | NA | NA | 4 | 0.25 |
| 10_421511 | 1 | 1 | 1 | 1 | NA | NA | NA | NA | 2 | 1 | NA | NA | 2 | 1 | NA | NA | NA | NA | 2 | 1 |
| 10_421886 | 6 | 0.5 | 2 | 0 | 1 | 0 | 1 | 1 | 10 | 0.4 | 7 | 0.2857 | 3 | 0 | 3 | 0.6667 | 2 | 0 | 15 | 0.2667 |
| 10_421893 | 6 | 1 | 2 | 1 | 1 | 1 | 1 | 1 | 10 | 1 | 3 | 1 | 2 | 1 | 3 | 1 | 2 | 1 | 10 | 1 |
| 10_421893 | 4 | 1 | 2 | 0.5 | 1 | 1 | 1 | 1 | 8 | 0.875 | 3 | 0.6667 | 2 | 1 | 2 | 0.5 | 2 | 1 | 9 | 0.7778 |
| 10_421894 | 5 | 1 | 1 | 1 | NA | NA | NA | NA | 6 | 1 | 2 | 1 | 2 | 1 | 2 | 0.5 | 1 | 1 | 7 | 0.8571 |
| 10_421895 | 6 | 0.1667 | 1 | 0 | 1 | 0 | 1 | 0 | 9 | 0.1111 | 3 | 0 | 2 | 0.5 | 3 | 0 | 1 | 0 | 9 | 0.1111 |
| 10_424383 | 2 | 0 | 3 | 1 | 5 | 0.4 | 6 | 0.1667 | 16 | 0.375 | 2 | 0.5 | 2 | 1 | 2 | 0 | 3 | 0.3333 | 9 | 0.4444 |
| 10_424383 | 2 | 0 | 3 | 0.3333 | 5 | 0 | 7 | 0 | 17 | 0.0588 | 2 | 0.5 | 3 | 0 | 1 | 0 | 2 | 0 | 8 | 0.125 |
| 10_424386 | 1 | 0 | 3 | 0.3333 | 4 | 0 | 2 | 0.5 | 10 | 0.2 | 2 | 0 | 3 | 0.3333 | 2 | 0 | 3 | 0 | 10 | 0.1 |
| 10_424389 | 2 | 0 | 3 | 0 | 4 | 0.25 | 5 | 0.4 | 14 | 0.2143 | 1 | 0 | 3 | 0.6667 | 1 | 1 | 2 | 0 | 7 | 0.4286 |
| 10_424389 | 1 | 1 | 2 | 0 | 5 | 0.2 | 4 | 0.25 | 12 | 0.25 | 1 | 1 | 2 | 0.5 | 1 | 0 | 2 | 0 | 6 | 0.3333 |
| 10_424392 | 1 | 1 | 2 | 0.5 | 5 | 1 | 5 | 1 | 13 | 0.9231 | 1 | 1 | 3 | 1 | 1 | 1 | 2 | 0.5 | 7 | 0.8571 |
| 10_424392 | NA | NA | 1 | 1 | 3 | 0 | 2 | 0 | 6 | 0.1667 | 1 | 1 | 2 | 0 | NA | NA | 1 | 1 | 4 | 0.5 |
| 10_424392 | 2 | 1 | 2 | 0.5 | 4 | 1 | 4 | 1 | 12 | 0.9167 | NA | NA | 2 | 1 | 1 | 0 | 3 | 0.6667 | 6 | 0.6667 |
| 10_424393 | 2 | 0.5 | 2 | 1 | 3 | 1 | 4 | 0.75 | 11 | 0.8182 | 1 | 1 | 3 | 0.6667 | 1 | 1 | 2 | 0.5 | 7 | 0.7143 |
| 10_424393 | 1 | 1 | 2 | 0.5 | 4 | 0.5 | 3 | 0.6667 | 10 | 0.6 | NA | NA | 2 | 1 | 1 | 0 | 3 | 0.3333 | 6 | 0.5 |
| 10_424591 | NA | NA | 2 | 0.5 | 2 | 1 | 1 | 0 | 5 | 0.6 | NA | NA | 2 | 0.5 | NA | NA | 1 | 1 | 3 | 0.6667 |
| 10_425551 | 4 | 0 | 1 | 0 | 7 | 0.2857 | 4 | 0.25 | 16 | 0.1875 | 2 | 0 | 3 | 0.3333 | 2 | 0 | 2 | 1 | 9 | 0.3333 |
| 10_425552 | 3 | 0.3333 | 1 | 0 | 7 | 0.2857 | 5 | 0.2 | 16 | 0.25 | 2 | 0.5 | 3 | 0 | 2 | 0.5 | 2 | 0.5 | 9 | 0.3333 |
| 10_425765 | 5 | 0.2 | 2 | 0 | 8 | 0 | 7 | 0.2857 | 22 | 0.1364 | 2 | 0.5 | 3 | 0 | 2 | 0 | 2 | 0 | 9 | 0.1111 |
| 10_425766 | 6 | 0.3333 | NA | NA | 5 | 0 | 2 | 0 | 13 | 0.1538 | 1 | 0 | 3 | 0.3333 | 2 | 0 | 2 | 0 | 8 | 0.125 |
| 10_425766 | 4 | 1 | 1 | 1 | 9 | 0.5556 | 6 | 0.6667 | 20 | 0.7 | 2 | 0.5 | 3 | 1 | 2 | 1 | 2 | 1 | 9 | 0.8889 |
| 10_425768 | 5 | 0 | 2 | 0.5 | 5 | 0.4 | 5 | 0.4 | 17 | 0.2941 | 1 | 1 | 3 | 0 | 2 | 1 | 1 | 0 | 7 | 0.4286 |
| 10_425772 | 5 | 0 | 3 | 0.3333 | 7 | 0.4286 | 7 | 0.5714 | 22 | 0.3636 | 2 | 1 | 3 | 0.6667 | 3 | 0 | 4 | 0.5 | 12 | 0.5 |
| 10_427050 | 5 | 0.6 | 6 | 0.6667 | 2 | 0.5 | 4 | 0.5 | 17 | 0.5882 | NA | NA | 4 | 0.75 | 2 | 1 | 1 | 0 | 7 | 0.7143 |
| 10_427059 | 3 | 0 | 4 | 0.25 | 3 | 0 | 3 | 0 | 13 | 0.0769 | 1 | 0 | 5 | 0.2 | 3 | 0 | 3 | 0.3333 | 12 | 0.1667 |
| 10_427062 | 3 | 0 | 4 | 0 | 1 | 0 | 2 | 0.5 | 10 | 0.1 | 2 | 0 | 2 | 0 | 1 | 0 | 3 | 0.3333 | 8 | 0.125 |
| 10_428788 | 1 | 1 | 3 | 1 | 2 | 1 | 3 | 1 | 9 | 1 | 3 | 1 | 2 | 0.5 | 2 | 1 | 2 | 1 | 9 | 0.8889 |
| 10_428790 | 2 | 0 | 3 | 0.3333 | 2 | 1 | 4 | 0.25 | 11 | 0.3636 | 4 | 0.25 | 1 | 0 | 1 | 1 | 2 | 0.5 | 8 | 0.375 |

| | | | | | | | | | | | | | | | | | | | | |
|---|---|---|---|---|---|---|---|---|---|---|---|---|---|---|---|---|---|---|---|---|
| 10_428791 | 2 | 0 | 3 | 1 | 2 | 0.5 | 4 | 0 | 11 | 0.3636 | 4 | 0 | 1 | 0 | 1 | 1 | 2 | 0 | 8 | 0.125 |
| 10_428793 | 2 | 0.5 | 2 | 1 | 2 | 1 | 4 | 0.75 | 10 | 0.8 | 5 | 1 | 2 | 1 | 2 | 1 | 3 | 0.6667 | 12 | 0.9167 |
| 10_428794 | 2 | 0.5 | 1 | 1 | 1 | 1 | 2 | 0.5 | 6 | 0.6667 | 3 | 1 | 2 | 1 | 3 | 1 | 3 | 0.6667 | 11 | 0.9091 |
| 10_428795 | 2 | 0.5 | 1 | 0 | 2 | 1 | 2 | 1 | 7 | 0.7143 | 4 | 1 | 1 | 1 | 2 | 0.5 | 3 | 0.6667 | 10 | 0.8 |
| 10_428795 | 2 | 0 | 1 | 0 | 2 | 0.5 | 2 | 0 | 7 | 0.1429 | 5 | 0 | 1 | 0 | 3 | 0.3333 | 2 | 0 | 11 | 0.0909 |
| 10_428796 | 2 | 0.5 | 1 | 0 | 1 | 0 | 2 | 0.5 | 6 | 0.3333 | 5 | 0.4 | 1 | 0 | 3 | 0.6667 | 2 | 0.5 | 11 | 0.4545 |
| 10_428796 | 2 | 0 | 1 | 1 | 2 | 0 | 2 | 0 | 7 | 0.1429 | 5 | 0.2 | 1 | 0 | 3 | 0 | 3 | 0 | 12 | 0.0833 |
| 10_428799 | 2 | 1 | 1 | 1 | 2 | 1 | 1 | 1 | 6 | 1 | 4 | 1 | 1 | 1 | 1 | 1 | 3 | 1 | 9 | 1 |
| 10_428799 | 3 | 1 | 1 | 1 | 1 | 1 | 1 | 1 | 6 | 1 | 5 | 1 | 1 | 1 | 1 | 1 | 3 | 0.6667 | 10 | 0.9 |
| 10_428800 | 2 | 0 | 1 | 1 | 2 | 0.5 | 2 | 1 | 7 | 0.5714 | 5 | 0.4 | 1 | 0 | 2 | 1 | 3 | 0.3333 | 11 | 0.4545 |
| 10_428800 | 3 | 1 | 1 | 1 | 2 | 1 | 2 | 1 | 8 | 1 | 5 | 1 | 1 | 1 | 2 | 1 | 2 | 1 | 10 | 1 |
| 10_428800 | 2 | 1 | NA | NA | 2 | 1 | 2 | 1 | 6 | 1 | 5 | 0.8 | 1 | 1 | 2 | 0.5 | 4 | 0.75 | 12 | 0.75 |
| 10_428801 | 3 | 1 | 1 | 0 | 1 | 1 | 2 | 0.5 | 7 | 0.7143 | 5 | 0.2 | 1 | 0 | 2 | 1 | 4 | 0.5 | 12 | 0.4167 |
| 10_428802 | 2 | 0.5 | 1 | 1 | 1 | 1 | 2 | 0 | 6 | 0.5 | 5 | 0.4 | 1 | 0 | 2 | 0.5 | 2 | 0.5 | 10 | 0.4 |
| 10_428803 | 2 | 0 | NA | NA | 1 | 1 | 2 | 0 | 5 | 0.2 | 4 | 0.25 | NA | NA | 2 | 0.5 | 4 | 0.25 | 10 | 0.3 |
| 10_428920 | NA | NA | 1 | 1 | NA | NA | NA | NA | 1 | 1 | 1 | 0 | 1 | 1 | NA | NA | NA | NA | 2 | 0.5 |
| 10_428970 | 2 | 0.5 | 1 | 0 | 1 | 1 | 2 | 1 | 6 | 0.6667 | 4 | 0.75 | NA | NA | 2 | 0 | 3 | 0.6667 | 9 | 0.5556 |
| 10_428972 | 2 | 1 | 1 | 0 | 1 | 1 | 1 | 1 | 5 | 0.8 | 3 | 0.3333 | NA | NA | 1 | 1 | 2 | 0.5 | 6 | 0.5 |
| 10_428972 | 2 | 1 | 1 | 1 | 1 | 1 | NA | NA | 4 | 1 | 4 | 0.75 | NA | NA | 2 | 1 | 3 | 1 | 9 | 0.8889 |
| 10_428975 | 2 | 1 | 1 | 1 | 1 | 0 | NA | NA | 4 | 0.75 | 4 | 0.75 | NA | NA | 2 | 0.5 | 2 | 0.5 | 8 | 0.625 |
| 10_428977 | 1 | 1 | 1 | 1 | NA | NA | NA | NA | 2 | 1 | 3 | 0.6667 | NA | NA | 1 | 0 | 2 | 1 | 6 | 0.6667 |
| 10_428978 | 2 | 0.5 | 1 | 1 | 1 | 1 | NA | NA | 4 | 0.75 | 3 | 1 | NA | NA | 2 | 1 | 3 | 1 | 8 | 1 |
| 10_428980 | 1 | 1 | NA | NA | 1 | 1 | NA | NA | 2 | 1 | 2 | 1 | NA | NA | 1 | 1 | 1 | 1 | 4 | 1 |
| 10_428986 | 1 | 1 | 1 | 1 | NA | NA | NA | NA | 2 | 1 | 4 | 1 | NA | NA | 2 | 0.5 | 1 | 0 | 7 | 0.7143 |
| 10_428988 | 1 | 1 | 1 | 1 | NA | NA | NA | NA | 2 | 1 | 4 | 1 | NA | NA | 2 | 1 | NA | NA | 6 | 1 |
| 10_430339 | 1 | 0 | 1 | 1 | NA | NA | NA | NA | 2 | 0.5 | 5 | 0.2 | NA | NA | 2 | 0 | 2 | 0.5 | 9 | 0.2222 |
| 10_430977 | 3 | 0.3333 | 1 | 1 | NA | NA | 1 | 0 | 5 | 0.4 | NA | NA | NA | NA | 3 | 0.3333 | NA | NA | 3 | 0.3333 |
| 10_433159 | 4 | 0.75 | 3 | 0.6667 | 3 | 0.3333 | 2 | 1 | 12 | 0.6667 | NA | NA | 1 | 0 | 2 | 1 | NA | NA | 3 | 0.6667 |
| 10_433160 | 3 | 0.6667 | 3 | 0.3333 | 3 | 0.6667 | 2 | 0.5 | 11 | 0.5455 | NA | NA | 1 | 0 | 3 | 0.6667 | NA | NA | 4 | 0.5 |
| 10_433161 | 3 | 0.3333 | 3 | 0 | 3 | 0 | 2 | 0 | 11 | 0.0909 | NA | NA | 1 | 0 | 4 | 0.25 | NA | NA | 5 | 0.2 |
| 10_433162 | 4 | 0.5 | 3 | 0.3333 | 3 | 1 | 2 | 0.5 | 12 | 0.5833 | 1 | 0 | 1 | 1 | 4 | 0.75 | NA | NA | 6 | 0.6667 |
| 10_433162 | 4 | 1 | 2 | 1 | 3 | 1 | 1 | 1 | 10 | 1 | NA | NA | 1 | 1 | 3 | 1 | NA | NA | 4 | 1 |
| 10_433162 | 3 | 0.6667 | 2 | 0 | 3 | 0.6667 | 2 | 0 | 10 | 0.4 | 1 | 0 | 1 | 0 | 3 | 0.3333 | NA | NA | 5 | 0.2 |
| 10_433163 | 4 | 0.25 | 3 | 0.6667 | 3 | 0 | 2 | 0.5 | 12 | 0.3333 | 1 | 1 | 1 | 0 | 4 | 0.5 | NA | NA | 6 | 0.5 |
| 10_433322 | 28 | 0.3214 | 25 | 0.36 | 21 | 0.2857 | 23 | 0.3913 | 97 | 0.3402 | 24 | 0.2083 | 30 | 0.1667 | 20 | 0.1 | 17 | 0.4706 | 91 | 0.2198 |
| 10_433323 | 42 | 0.2619 | 26 | 0.3462 | 26 | 0.1923 | 24 | 0.2917 | 118 | 0.2712 | 24 | 0.375 | 31 | 0.3871 | 24 | 0.25 | 28 | 0.2143 | 107 | 0.3084 |
| 10_433862 | NA | NA | 1 | 1 | NA | NA | NA | NA | 1 | 1 | NA | NA | NA | NA | 1 | 1 | NA | NA | 1 | 1 |
| 10_433876 | NA | NA | 1 | 1 | NA | NA | NA | NA | 1 | 1 | NA | NA | NA | NA | 1 | 1 | NA | NA | 1 | 1 |
| 10_433881 | NA | NA | 1 | 1 | NA | NA | NA | NA | 1 | 1 | NA | NA | NA | NA | 1 | 1 | NA | NA | 1 | 1 |
| 10_433882 | NA | NA | 1 | 1 | NA | NA | NA | NA | 1 | 1 | NA | NA | NA | NA | 1 | 1 | NA | NA | 1 | 1 |
| 10_434012 | 4 | 0.25 | 1 | 0 | 3 | 0.6667 | NA | NA | 8 | 0.375 | 2 | 0.5 | 2 | 1 | 2 | 0 | 1 | 0 | 7 | 0.4286 |
| 10_434012 | 6 | 0.5 | 7 | 0.7143 | 4 | 0.75 | 3 | 0.3333 | 20 | 0.6 | 5 | 0.4 | 5 | 1 | 9 | 0.6667 | 6 | 0.6667 | 25 | 0.68 |
| 10_434087 | 9 | 0.1111 | 3 | 0.3333 | 4 | 0.25 | 4 | 0.25 | 20 | 0.2 | 3 | 0 | 1 | 0 | 8 | 0.125 | 4 | 0.25 | 16 | 0.125 |
| 10_434088 | 7 | 0.1429 | 5 | 0 | 4 | 0.25 | 3 | 0.3333 | 19 | 0.1579 | 2 | 0.5 | 1 | 0 | 7 | 0 | 4 | 0 | 14 | 0.0714 |
| 10_434089 | 9 | 0.3333 | 4 | 0.25 | 4 | 0 | 3 | 0 | 20 | 0.2 | 4 | 0 | NA | NA | 5 | 0.2 | 4 | 0 | 13 | 0.0769 |
| 10_434090 | 7 | 0 | 4 | 0.25 | 4 | 0 | 4 | 0 | 19 | 0.0526 | 3 | 0 | 1 | 0 | 7 | 0.1429 | 3 | 0 | 14 | 0.0714 |
| 10_434090 | 4 | 0 | 5 | 0.4 | 4 | 0.25 | 2 | 0.5 | 15 | 0.2667 | 4 | 0.25 | 1 | 0 | 6 | 0.1667 | 4 | 0.25 | 15 | 0.2 |
| 10_434090 | 8 | 0.125 | 5 | 0 | 3 | 0 | 3 | 0 | 19 | 0.0526 | 3 | 0.3333 | 1 | 0 | 8 | 0 | 4 | 0.25 | 16 | 0.125 |
| 10_434090 | 7 | 0.4286 | 5 | 0.2 | 4 | 1 | 3 | 0.6667 | 19 | 0.5263 | 4 | 0.75 | 1 | 1 | 7 | 0.4286 | 3 | 1 | 15 | 0.6667 |
| 10_434092 | 1 | 1 | NA | NA | NA | NA | NA | NA | 1 | 1 | NA | NA | NA | NA | NA | NA | 2 | 1 | 2 | 1 |
| 10_434587 | NA | NA | 2 | 1 | 1 | 1 | 1 | 1 | 4 | 1 | NA | NA | NA | NA | 2 | 1 | 2 | 1 | 4 | 1 |
| 10_434590 | NA | NA | 2 | 1 | 1 | 1 | 1 | 1 | 4 | 1 | NA | NA | NA | NA | 3 | 1 | 2 | 1 | 5 | 1 |
| 10_434590 | NA | NA | 2 | 1 | NA | NA | 1 | 0 | 3 | 0.6667 | NA | NA | NA | NA | 3 | 0.6667 | 2 | 0.5 | 5 | 0.6 |
| 10_434593 | NA | NA | 1 | 0 | 1 | 1 | 1 | 0 | 3 | 0.3333 | NA | NA | NA | NA | 3 | 0.3333 | 1 | 0 | 4 | 0.25 |
| 10_434593 | NA | NA | 2 | 0.5 | 1 | 1 | NA | NA | 3 | 0.6667 | NA | NA | NA | NA | 3 | 0.3333 | 2 | 0 | 5 | 0.2 |
| 10_434601 | NA | NA | 2 | 0.5 | 1 | 1 | 1 | 0 | 4 | 0.5 | NA | NA | 1 | 0 | 2 | 0.5 | 2 | 0 | 5 | 0.2 |
| 10_437504 | NA | NA | 1 | 1 | 4 | 1 | 2 | 1 | 7 | 1 | 5 | 1 | 3 | 0.6667 | 1 | 1 | 1 | 0 | 10 | 0.8 |
| 10_437779 | NA | NA | NA | NA | 1 | 1 | 1 | 1 | 2 | 1 | 1 | 1 | NA | NA | NA | NA | NA | NA | 1 | 1 |
| 10_437783 | NA | NA | NA | NA | 1 | 1 | 2 | 0.5 | 3 | 0.6667 | 1 | 1 | NA | NA | NA | NA | NA | NA | 1 | 1 |
| 10_438225 | NA | NA | NA | NA | 2 | 1 | 4 | 1 | 6 | 1 | 1 | 1 | 1 | 1 | NA | NA | 1 | 1 | 3 | 1 |
| 10_438226 | NA | NA | NA | NA | 1 | 1 | 3 | 0.6667 | 4 | 0.75 | 1 | 1 | 1 | 0 | NA | NA | NA | NA | 2 | 0.5 |
| 10_438228 | NA | NA | NA | NA | 1 | 0 | 4 | 0.25 | 5 | 0.2 | 1 | 0 | 1 | 0 | NA | NA | 1 | 1 | 3 | 0.3333 |
| 10_438269 | NA | NA | NA | NA | NA | NA | 1 | 1 | 1 | 1 | NA | NA | NA | NA | 1 | 0 | 1 | 1 | 2 | 0.5 |
| 10_438271 | NA | NA | NA | NA | NA | NA | 1 | 1 | 1 | 1 | NA | NA | NA | NA | 1 | 1 | 1 | 1 | 2 | 1 |
| 10_438272 | NA | NA | NA | NA | NA | NA | 1 | 1 | 1 | 1 | NA | NA | NA | NA | NA | NA | 1 | 1 | 1 | 1 |

| | | | | | | | | | | | | | | | | | | | | |
|---|---|---|---|---|---|---|---|---|---|---|---|---|---|---|---|---|---|---|---|---|
| 10_438272( | NA | NA | NA | NA | NA | NA | 1 | 1 | 1 | 1 | NA | NA | NA | NA | NA | NA | 1 | 1 | 1 | 1 |
| 10_438272( | NA | NA | NA | NA | NA | NA | 1 | 1 | 1 | 1 | NA | NA | NA | NA | NA | NA | 1 | 1 | 1 | 1 |
| 10_441797 | 4 | 0.25 | 2 | 0.5 | 5 | 0 | 5 | 0 | 16 | 0.125 | NA | NA | 7 | 0 | 3 | 0.3333 | 2 | 0 | 12 | 0.0833 |
| 10_442390( | 3 | 0.3333 | 1 | 1 | 3 | 0.6667 | 4 | 0 | 11 | 0.3636 | NA | NA | 6 | 0.3333 | 1 | 1 | 2 | 0.5 | 9 | 0.4444 |
| 10_442390 | 3 | 0.3333 | 2 | 0.5 | 5 | 0 | 5 | 0 | 15 | 0.1333 | 1 | 0 | 5 | 0 | 3 | 0.3333 | 2 | 0 | 11 | 0.0909 |
| 10_442391 | 3 | 0 | 2 | 0 | 5 | 0.2 | 4 | 0 | 14 | 0.0714 | 1 | 0 | 7 | 0.1429 | 3 | 0 | 2 | 0 | 13 | 0.0769 |
| 10_442392 | 4 | 0.25 | 1 | 0 | 5 | 0 | 3 | 0 | 13 | 0.0769 | 1 | 0 | 7 | 0 | 3 | 0 | 2 | 0.5 | 13 | 0.0769 |
| 10_442392 | 4 | 0.5 | 2 | 0.5 | 4 | 0.75 | 5 | 0.2 | 15 | 0.4667 | 1 | 1 | 7 | 0.4286 | 3 | 0.6667 | 2 | 0 | 13 | 0.4615 |
| 10_442394( | 4 | 0.25 | 2 | 0 | 4 | 0 | 5 | 0.4 | 15 | 0.2 | 1 | 0 | 7 | 0 | 3 | 0 | 2 | 0.5 | 13 | 0.0769 |
| 10_442394 | 4 | 0.25 | 2 | 0.5 | 5 | 0.4 | 6 | 0.3333 | 17 | 0.3529 | 1 | 0 | 7 | 0 | 3 | 0.3333 | 2 | 0.5 | 13 | 0.1538 |
| 10_442401( | NA | NA | 2 | 0 | 4 | 0.25 | 3 | 0 | 9 | 0.1111 | NA | NA | 3 | 0 | 2 | 0 | 1 | 1 | 6 | 0.1667 |
| 10_449427 | 1 | 1 | 1 | 1 | NA | NA | 1 | 1 | 3 | 1 | NA | NA | NA | NA | NA | NA | 3 | 1 | 3 | 1 |
| 10_449428 | 1 | 1 | 1 | 1 | NA | NA | 1 | 1 | 3 | 1 | NA | NA | NA | NA | NA | NA | 2 | 1 | 2 | 1 |
| 10_449429( | 2 | 0.5 | 1 | 1 | NA | NA | 1 | 1 | 4 | 0.75 | NA | NA | NA | NA | NA | NA | 2 | 1 | 2 | 1 |
| 10_449429( | 2 | 0.5 | 1 | 0 | NA | NA | 1 | 0 | 4 | 0.25 | NA | NA | NA | NA | NA | NA | 2 | 0.5 | 2 | 0.5 |
| 10_450345 | 1 | 1 | 1 | 1 | NA | NA | NA | NA | 2 | 1 | 3 | 1 | NA | NA | 4 | 0.5 | 2 | 0.5 | 9 | 0.6667 |
| 10_450346( | 2 | 0 | 1 | 1 | 1 | 0 | 1 | 1 | 5 | 0.4 | 3 | 0.3333 | 2 | 0 | 4 | 0 | 2 | 0 | 11 | 0.0909 |
| 10_450347 | 1 | 0 | NA | NA | 1 | 0 | 2 | 0.5 | 4 | 0.25 | 3 | 0.3333 | 2 | 0.5 | 5 | 0.4 | 3 | 0.3333 | 13 | 0.3846 |
| 10_450347( | 2 | 1 | 1 | 1 | 1 | 1 | 2 | 1 | 6 | 1 | 3 | 1 | 2 | 1 | 4 | 1 | 2 | 1 | 11 | 1 |
| 10_450349 | 2 | 0.5 | NA | NA | 1 | 0 | 3 | 0 | 6 | 0.1667 | 3 | 0.3333 | 2 | 0.5 | 4 | 0 | 2 | 0 | 11 | 0.1818 |
| 10_450350 | 1 | 0 | 1 | 1 | 1 | 0 | 2 | 0.5 | 5 | 0.4 | 2 | 0 | 2 | 0 | 3 | 0.3333 | 2 | 0 | 9 | 0.1111 |
| 10_450352( | NA | NA | NA | NA | 1 | 0 | 1 | 1 | 2 | 0.5 | 1 | 1 | NA | NA | NA | NA | NA | NA | 1 | 1 |
| 10_450353( | NA | NA | NA | NA | 1 | 1 | 1 | 1 | 2 | 1 | 1 | 1 | NA | NA | NA | NA | NA | NA | 1 | 1 |
| 10_450356( | NA | NA | NA | NA | 1 | 1 | 1 | 1 | 2 | 1 | 1 | 1 | NA | NA | NA | NA | NA | NA | 1 | 1 |
| 10_452528 | 2 | 0.5 | NA | NA | NA | NA | NA | NA | 2 | 0.5 | NA | NA | 1 | 0 | 2 | 0.5 | NA | NA | 3 | 0.3333 |
| 10_452528 | 2 | 0.5 | NA | NA | NA | NA | NA | NA | 2 | 0.5 | NA | NA | 1 | 1 | 2 | 0 | NA | NA | 3 | 0.3333 |
| 10_452529 | 2 | 0.5 | NA | NA | NA | NA | NA | NA | 2 | 0.5 | NA | NA | 1 | 0 | 2 | 0.5 | NA | NA | 3 | 0.3333 |
| 10_452532 | 1 | 1 | NA | NA | NA | NA | NA | NA | 1 | 1 | NA | NA | 1 | 1 | 2 | 0 | NA | NA | 3 | 0.3333 |
| 10_452533 | 1 | 1 | NA | NA | NA | NA | NA | NA | 1 | 1 | NA | NA | 1 | 0 | 2 | 1 | NA | NA | 3 | 0.6667 |
| 10_452566 | 1 | 1 | NA | NA | NA | NA | NA | NA | 1 | 1 | NA | NA | NA | NA | 1 | 0 | 1 | 1 | 2 | 0.5 |
| 10_455466 | 1 | 0 | 2 | 0.5 | 1 | 0 | 7 | 0 | 11 | 0.0909 | 2 | 0 | NA | NA | 1 | 1 | 3 | 0.6667 | 6 | 0.5 |
| 10_455532 | 4 | 0 | 2 | 0 | 1 | 0 | 8 | 0.125 | 15 | 0.0667 | 2 | 0 | 1 | 0 | 1 | 0 | 5 | 0.2 | 9 | 0.1111 |
| 10_455875 | 4 | 0.25 | 3 | 0.3333 | 2 | 0 | 9 | 0.2222 | 18 | 0.2222 | 4 | 0 | 2 | 0 | 3 | 0.3333 | 5 | 0.2 | 14 | 0.1429 |
| 10_455875 | 4 | 0 | 2 | 0 | 1 | 0 | 9 | 0.1111 | 16 | 0.0625 | 4 | 0 | 2 | 0.5 | 3 | 0.3333 | 6 | 0.1667 | 15 | 0.2 |
| 10_455876 | 4 | 0.5 | 3 | 0 | 2 | 0 | 11 | 0.0909 | 20 | 0.15 | 4 | 0.25 | 2 | 0 | 3 | 0 | 6 | 0 | 15 | 0.0667 |
| 10_455879 | 5 | 0.4 | 4 | 0 | 1 | 1 | 9 | 0 | 19 | 0.1579 | 5 | 0.2 | 2 | 0 | 5 | 0.2 | 5 | 0 | 17 | 0.1176 |
| 10_489699 | NA | NA | 1 | 1 | 1 | 0 | 2 | 1 | 4 | 0.75 | NA | NA | NA | NA | 1 | 1 | 1 | 1 | 2 | 1 |
| 10_489701( | NA | NA | 1 | 1 | 1 | 1 | 2 | 1 | 4 | 1 | NA | NA | 1 | 1 | 1 | 1 | 1 | 1 | 3 | 1 |
| 10_489758 | 1 | 0 | 5 | 0.2 | 8 | 0.125 | 7 | 0 | 21 | 0.0952 | NA | NA | 6 | 0 | 9 | 0 | 4 | 0.25 | 19 | 0.0526 |
| 10_489761 | 3 | 1 | 6 | 1 | 11 | 0.8182 | 11 | 0.8182 | 31 | 0.871 | 2 | 0.5 | 9 | 1 | 10 | 1 | 6 | 1 | 27 | 0.963 |
| 10_489762( | 4 | 0.5 | 5 | 0.4 | 9 | 0.1111 | 10 | 0.7 | 28 | 0.4286 | 2 | 0 | 9 | 0.1111 | 9 | 0.4444 | 5 | 0.4 | 25 | 0.28 |
| 10_489762 | 3 | 0.3333 | 5 | 0.6 | 9 | 0.6667 | 11 | 0.9091 | 28 | 0.7143 | 2 | 1 | 10 | 0.8 | 9 | 0.7778 | 5 | 0.6 | 26 | 0.7692 |
| 10_489762 | 3 | 0 | 4 | 0 | 8 | 0.625 | 11 | 0.4545 | 26 | 0.3846 | 2 | 0.5 | 7 | 0.4286 | 8 | 0.5 | 5 | 0.8 | 22 | 0.5455 |
| 10_489762 | 4 | 0 | 6 | 0.5 | 10 | 0.2 | 11 | 0.5455 | 31 | 0.3548 | 2 | 1 | 9 | 0.2222 | 8 | 0.5 | 5 | 0.2 | 24 | 0.375 |
| 10_489763 | 4 | 0.25 | 6 | 0.3333 | 10 | 0.2 | 10 | 0 | 30 | 0.1667 | 2 | 0 | 9 | 0.2222 | 10 | 0.1 | 6 | 0.1667 | 27 | 0.1481 |
| 10_489766( | 3 | 0.3333 | 6 | 0 | 6 | 0 | 9 | 0.1111 | 24 | 0.0833 | 2 | 0 | 9 | 0.1111 | 8 | 0.125 | 5 | 0.2 | 24 | 0.125 |
| 10_489767 | 4 | 0.5 | 5 | 0 | 6 | 0 | 4 | 0 | 19 | 0.1053 | 1 | 0 | 3 | 0.3333 | 9 | 0.2222 | 3 | 0.3333 | 16 | 0.25 |
| 10_489768 | 3 | 0.3333 | 4 | 0 | 7 | 0.5714 | 9 | 0.3333 | 23 | 0.3478 | 2 | 1 | 6 | 0.3333 | 7 | 0.4286 | 5 | 0.6 | 20 | 0.5 |
| 10_489770 | 5 | 0.8 | 6 | 0.5 | 9 | 0.4444 | 10 | 0.5 | 30 | 0.5333 | NA | NA | 5 | 0.8 | 8 | 0.625 | 6 | 0.5 | 19 | 0.6316 |
| 10_489770( | 5 | 0.8 | 6 | 0.6667 | 8 | 1 | 10 | 0.9 | 29 | 0.8621 | 1 | 1 | 6 | 0.6667 | 8 | 0.875 | 6 | 0.8333 | 21 | 0.8095 |
| 10_489770( | 4 | 0.75 | 4 | 0.75 | 8 | 0.875 | 6 | 0.8333 | 22 | 0.8182 | 1 | 1 | 5 | 0.8 | 6 | 0.8333 | 4 | 1 | 16 | 0.875 |
| 10_489771 | 2 | 1 | 3 | 1 | 6 | 1 | 7 | 0.8571 | 18 | 0.9444 | 1 | 1 | 5 | 1 | 3 | 0.6667 | 3 | 1 | 12 | 0.9167 |
| 10_489773 | 5 | 0 | 6 | 0.1667 | 9 | 0.1111 | 9 | 0 | 29 | 0.069 | 3 | 0.3333 | 7 | 0.4286 | 8 | 0.25 | 6 | 0.1667 | 24 | 0.2917 |
| 10_490088 | 6 | 0.1667 | 5 | 0 | 6 | 0.1667 | 6 | 0.1667 | 23 | 0.1304 | 3 | 0 | 5 | 0 | 6 | 0.1667 | 5 | 0 | 19 | 0.0526 |
| 10_494496 | 4 | 0 | 1 | 0 | 2 | 0 | 3 | 0.3333 | 10 | 0.1 | 1 | 0 | 2 | 0 | 4 | 0 | 4 | 0.25 | 11 | 0.0909 |
| 10_496595 | 3 | 0.3333 | 1 | 1 | 3 | 0.6667 | 3 | 0.6667 | 10 | 0.6 | NA | NA | 1 | 0 | 3 | 0.6667 | 2 | 1 | 6 | 0.6667 |
| 10_496596 | 4 | 0.25 | NA | NA | 1 | 0 | 3 | 0 | 8 | 0.125 | NA | NA | 2 | 0.5 | 2 | 0 | 2 | 0.5 | 6 | 0.3333 |
| 10_504938 | NA | NA | 1 | 1 | NA | NA | 1 | 1 | 2 | 1 | 1 | 1 | NA | NA | NA | NA | NA | NA | 1 | 1 |
| 10_504942 | NA | NA | 1 | 1 | NA | NA | 1 | 1 | 2 | 1 | 1 | 1 | NA | NA | NA | NA | NA | NA | 1 | 1 |
| 10_507265 | NA | NA | 3 | 1 | 4 | 1 | 3 | 0.3333 | 10 | 0.8 | 4 | 0.5 | 2 | 0.5 | 3 | 1 | NA | NA | 9 | 0.6667 |
| 10_507266 | NA | NA | 3 | 1 | 3 | 1 | 3 | 1 | 9 | 1 | 3 | 1 | 4 | 1 | 1 | 1 | NA | NA | 8 | 1 |
| 10_507266 | NA | NA | 3 | 0 | 4 | 0 | 3 | 0.6667 | 10 | 0.2 | 4 | 0 | 4 | 0 | 3 | 0.3333 | NA | NA | 11 | 0.0909 |
| 10_507267 | NA | NA | 3 | 0.3333 | 3 | 0 | 3 | 0 | 9 | 0.1111 | 4 | 0.25 | 4 | 0.25 | 3 | 0.3333 | NA | NA | 11 | 0.2727 |
| 10_507268 | NA | NA | 3 | 0.3333 | 4 | 0.25 | 3 | 0.3333 | 10 | 0.3 | 4 | 0.25 | 3 | 0 | 3 | 0 | NA | NA | 10 | 0.1 |
| 10_513716 | 1 | 1 | NA | NA | NA | NA | NA | NA | 1 | 1 | 1 | 1 | NA | NA | NA | NA | NA | NA | 1 | 1 |

| | | | | | | | | | | | | | | | | | | | | |
|---|---|---|---|---|---|---|---|---|---|---|---|---|---|---|---|---|---|---|---|---|
| 10_513724 | 1 | 1 | NA | NA | NA | NA | NA | NA | 1 | 1 | 2 | 0.5 | NA | NA | NA | NA | NA | NA | 2 | 0.5 |
| 10_521959 | 1 | 1 | 1 | 1 | 1 | 1 | 2 | 1 | 5 | 1 | NA | NA | 1 | 1 | NA | NA | NA | NA | 1 | 1 |
| 10_521959 | NA | NA | 1 | 1 | 1 | 0 | 2 | 0.5 | 4 | 0.5 | NA | NA | 1 | 1 | NA | NA | NA | NA | 1 | 1 |
| 10_521960 | 1 | 1 | 1 | 0 | 1 | 0 | 1 | 0 | 4 | 0.25 | NA | NA | 1 | 1 | NA | NA | NA | NA | 1 | 1 |
| 10_521960 | 1 | 0 | 1 | 0 | 1 | 0 | 1 | 1 | 4 | 0.25 | NA | NA | 1 | 0 | 1 | 1 | NA | NA | 2 | 0.5 |
| 10_521962 | 1 | 1 | 1 | 1 | 1 | 1 | 2 | 1 | 5 | 1 | 1 | 1 | 2 | 1 | 1 | 1 | NA | NA | 4 | 1 |
| 10_521972 | 1 | 1 | NA | NA | NA | NA | 1 | 0 | 2 | 0.5 | 2 | 0 | 2 | 0.5 | 1 | 1 | NA | NA | 5 | 0.4 |
| 10_522139 | 2 | 0 | 2 | 0 | 3 | 0.3333 | 4 | 0 | 11 | 0.0909 | 2 | 0 | 2 | 0 | 2 | 0 | 5 | 0.2 | 11 | 0.0909 |
| 10_522148 | 6 | 0.1667 | 7 | 0.1429 | 5 | 0.2 | 7 | 0 | 25 | 0.12 | 8 | 0.125 | 3 | 0 | 5 | 0.2 | 7 | 0.2857 | 23 | 0.1739 |
| 10_522151 | 7 | 0.7143 | 4 | 0.5 | 2 | 1 | 7 | 0.5714 | 20 | 0.65 | 8 | 0.625 | 3 | 1 | 3 | 1 | 6 | 0.6667 | 20 | 0.75 |
| 10_522154 | 7 | 0.2857 | 6 | 0.3333 | 2 | 0 | 7 | 0.2857 | 22 | 0.2727 | 7 | 0.1429 | 3 | 0 | 4 | 0 | 6 | 0.6667 | 20 | 0.25 |
| 10_522155 | 7 | 0 | 7 | 0.1429 | 3 | 0.3333 | 5 | 0.4 | 22 | 0.1818 | 6 | 0.3333 | 3 | 0.6667 | 3 | 0.3333 | 6 | 0.5 | 18 | 0.4444 |
| 10_522156 | 4 | 0 | 5 | 0 | 4 | 0.25 | 5 | 0.2 | 18 | 0.1111 | 6 | 0.1667 | 2 | 0 | 3 | 0 | 4 | 0.5 | 15 | 0.2 |
| 10_522157 | 6 | 0.5 | 5 | 0.6 | 4 | 0.5 | 5 | 0.8 | 20 | 0.6 | 6 | 0 | 3 | 0.3333 | 4 | 0.25 | 5 | 0.8 | 18 | 0.3333 |
| 10_522157 | 7 | 0 | 5 | 0.6 | 5 | 0.4 | 4 | 0 | 21 | 0.2381 | 6 | 0 | 3 | 0 | 4 | 0 | 4 | 0.5 | 17 | 0.1176 |
| 10_522577 | 3 | 0.6667 | 3 | 0 | 2 | 0.5 | 4 | 0.75 | 12 | 0.5 | 2 | 0 | 1 | 1 | 2 | 0.5 | 3 | 0.3333 | 8 | 0.375 |
| 10_522939 | NA | NA | 2 | 0.5 | 2 | 0 | 6 | 0.1667 | 10 | 0.2 | 2 | 0 | 3 | 0 | 2 | 0 | 5 | 0.2 | 12 | 0.0833 |
| 10_523127 | 7 | 0 | 8 | 0.375 | 6 | 0 | 8 | 0.25 | 29 | 0.1724 | 8 | 0.125 | 6 | 0 | 4 | 0.25 | 11 | 0.0909 | 29 | 0.1034 |
| 10_523128 | 7 | 0.7143 | 10 | 0.3 | 6 | 0.1667 | 8 | 0.5 | 31 | 0.4194 | 9 | 0.4444 | 6 | 0.3333 | 4 | 0.75 | 11 | 0.5455 | 30 | 0.5 |
| 10_523128 | 8 | 0.125 | 9 | 0.4444 | 6 | 0.6667 | 9 | 0.4444 | 32 | 0.4062 | 9 | 0.2222 | 6 | 0 | 5 | 0 | 10 | 0.4 | 30 | 0.2 |
| 10_523130 | 9 | 0 | 9 | 0.1111 | 9 | 0.1111 | 10 | 0 | 37 | 0.0541 | 9 | 0.1111 | 8 | 0.125 | 7 | 0.1429 | 12 | 0.1667 | 36 | 0.1389 |
| 10_523133 | 10 | 0.1 | 12 | 0.1667 | 10 | 0 | 11 | 0 | 43 | 0.0698 | 11 | 0.1818 | 11 | 0.1818 | 10 | 0.1 | 12 | 0.1667 | 44 | 0.1591 |
| 10_523135 | 8 | 0.375 | 14 | 0.2143 | 12 | 0.1667 | 14 | 0.0714 | 48 | 0.1875 | 9 | 0.1111 | 11 | 0.1818 | 12 | 0.0833 | 14 | 0.1429 | 46 | 0.1304 |
| 10_523135 | 10 | 0 | 11 | 0 | 11 | 0.1818 | 15 | 0.0667 | 47 | 0.0638 | 11 | 0.2727 | 12 | 0 | 8 | 0 | 12 | 0.0833 | 43 | 0.093 |
| 10_523137 | 11 | 0.1818 | 15 | 0.1333 | 12 | 0.3333 | 14 | 0.2857 | 52 | 0.2308 | 7 | 0.1429 | 13 | 0.1538 | 13 | 0.1538 | 12 | 0.25 | 45 | 0.1778 |
| 10_523138 | 12 | 0.5833 | 17 | 0.4706 | 13 | 0.6923 | 14 | 0.5 | 56 | 0.5536 | 10 | 0.6 | 12 | 0.5 | 13 | 0.5385 | 13 | 0.8462 | 48 | 0.625 |
| 10_523144 | 11 | 0.6364 | 14 | 0.2857 | 13 | 0.7692 | 15 | 0.2667 | 53 | 0.4717 | 9 | 0.5556 | 15 | 0.4 | 9 | 0.5556 | 13 | 0.4615 | 46 | 0.4783 |
| 10_523145 | 11 | 0.8182 | 14 | 0.5714 | 14 | 0.7857 | 12 | 0.75 | 51 | 0.7255 | 8 | 0.875 | 16 | 0.75 | 8 | 0.625 | 10 | 0.3 | 42 | 0.6429 |
| 10_523145 | 10 | 0.2 | 13 | 0 | 13 | 0.4615 | 15 | 0.2667 | 51 | 0.2353 | 8 | 0.25 | 16 | 0.125 | 6 | 0 | 13 | 0.3846 | 43 | 0.2093 |
| 10_523147 | 10 | 0.1 | 13 | 0.0769 | 12 | 0.1667 | 15 | 0.1333 | 50 | 0.12 | 10 | 0.1 | 14 | 0.1429 | 8 | 0.25 | 13 | 0.2308 | 45 | 0.1778 |
| 10_523147 | 12 | 0.1667 | 14 | 0.1429 | 12 | 0.25 | 17 | 0.2941 | 55 | 0.2182 | 8 | 0 | 15 | 0.2667 | 10 | 0 | 13 | 0.1538 | 46 | 0.1304 |
| 10_523148 | 11 | 0 | 10 | 0 | 14 | 0.1429 | 14 | 0.0714 | 49 | 0.0612 | 8 | 0.125 | 12 | 0.0833 | 10 | 0 | 15 | 0.0667 | 45 | 0.0667 |
| 10_523148 | 12 | 0.5 | 10 | 0.5 | 12 | 0.5 | 17 | 0.6471 | 51 | 0.549 | 9 | 0.7778 | 16 | 0.5 | 9 | 0.3333 | 13 | 0.2308 | 47 | 0.4468 |
| 10_523148 | 12 | 0.0833 | 13 | 0.1538 | 12 | 0.25 | 16 | 0 | 53 | 0.1132 | 12 | 0 | 14 | 0.0714 | 9 | 0 | 14 | 0.2143 | 49 | 0.0816 |
| 10_523149 | 13 | 0.3077 | 12 | 0.1667 | 11 | 0.2727 | 16 | 0.4375 | 52 | 0.3077 | 8 | 0 | 14 | 0.5 | 11 | 0.1818 | 12 | 0.3333 | 45 | 0.2889 |
| 10_523149 | 10 | 0.3 | 14 | 0 | 8 | 0.25 | 15 | 0.0667 | 47 | 0.1277 | 9 | 0.2222 | 14 | 0.1429 | 9 | 0.2222 | 13 | 0.1538 | 45 | 0.1778 |
| 10_523153 | 3 | 0 | 3 | 0 | 3 | 0.3333 | 4 | 0 | 13 | 0.0769 | 4 | 0 | 4 | 0 | 1 | 0 | 5 | 0.8 | 14 | 0.2857 |
| 10_529843 | 7 | 0.2857 | 5 | 0 | 3 | 0.3333 | 4 | 0 | 19 | 0.1579 | 3 | 0.3333 | 7 | 0 | 9 | 0.1111 | 4 | 0 | 23 | 0.087 |
| 10_529843 | 7 | 0.7143 | 4 | 1 | 2 | 1 | 3 | 1 | 16 | 0.875 | 3 | 1 | 4 | 1 | 7 | 0.7143 | 3 | 1 | 17 | 0.8824 |
| 10_529844 | 8 | 0.625 | 6 | 0.1667 | 1 | 0 | 4 | 0.5 | 19 | 0.4211 | 3 | 0.3333 | 7 | 0.8571 | 7 | 0.4286 | 4 | 0 | 21 | 0.4762 |
| 10_529845 | 9 | 0.7778 | 5 | 0.8 | 3 | 1 | 4 | 1 | 21 | 0.8571 | 3 | 1 | 8 | 0.875 | 10 | 0.8 | 3 | 1 | 24 | 0.875 |
| 10_529846 | 9 | 0.2222 | 6 | 0 | 3 | 0.3333 | 4 | 0.5 | 22 | 0.2273 | 3 | 0.3333 | 9 | 0.2222 | 10 | 0 | 4 | 0.5 | 26 | 0.1923 |
| 10_529846 | 8 | 0.25 | 6 | 0 | 3 | 0.3333 | 2 | 0 | 19 | 0.1579 | 3 | 0 | 8 | 0.25 | 8 | 0 | 3 | 0 | 22 | 0.0909 |
| 10_529853 | 10 | 0.1 | 6 | 0 | 4 | 0.25 | 5 | 0 | 25 | 0.08 | 3 | 0 | 9 | 0.2222 | 8 | 0 | 3 | 0 | 23 | 0.087 |
| 10_529864 | 11 | 0.0909 | 9 | 0 | 7 | 0 | 8 | 0.125 | 35 | 0.0571 | 7 | 0 | 11 | 0.0909 | 11 | 0 | 6 | 0.1667 | 35 | 0.0571 |
| 10_529872 | 13 | 0.1538 | 9 | 0.1111 | 9 | 0.1111 | 8 | 0.125 | 39 | 0.1282 | 9 | 0 | 13 | 0.0769 | 10 | 0.1 | 9 | 0.2222 | 41 | 0.0976 |
| 10_531540 | NA | NA | 1 | 1 | 2 | 1 | 1 | 1 | 4 | 1 | 1 | 1 | NA | NA | NA | NA | 1 | 1 | 2 | 1 |
| 10_531542 | NA | NA | 1 | 0 | 2 | 1 | 3 | 0.6667 | 6 | 0.6667 | 1 | 0 | NA | NA | NA | NA | 1 | 1 | 2 | 0.5 |
| 10_531542 | NA | NA | 1 | 1 | 2 | 0.5 | 3 | 0.6667 | 6 | 0.6667 | 1 | 1 | NA | NA | NA | NA | NA | NA | 1 | 1 |
| 10_531543 | NA | NA | 1 | 0 | 1 | 0 | 3 | 0.3333 | 5 | 0.2 | 1 | 1 | NA | NA | NA | NA | 1 | 0 | 2 | 0.5 |
| 10_531575 | NA | NA | 1 | 1 | 2 | 0.5 | 2 | 0.5 | 5 | 0.6 | 1 | 1 | NA | NA | NA | NA | 1 | 0 | 2 | 0.5 |
| 10_532247 | NA | NA | NA | NA | 2 | 1 | NA | NA | 2 | 1 | 1 | 1 | NA | NA | NA | NA | NA | NA | 1 | 1 |
| 10_532247 | NA | NA | 1 | 0 | 2 | 1 | NA | NA | 3 | 0.6667 | 1 | 1 | NA | NA | NA | NA | NA | NA | 1 | 1 |
| 10_532248 | NA | NA | 1 | 1 | 2 | 0.5 | NA | NA | 3 | 0.6667 | 1 | 1 | NA | NA | NA | NA | NA | NA | 1 | 1 |
| 10_532249 | NA | NA | 1 | 1 | 2 | 1 | NA | NA | 3 | 1 | 1 | 1 | NA | NA | NA | NA | NA | NA | 1 | 1 |
| 10_532249 | NA | NA | NA | NA | 2 | 1 | NA | NA | 2 | 1 | 1 | 1 | NA | NA | NA | NA | NA | NA | 1 | 1 |
| 10_532251 | NA | NA | NA | NA | 2 | 1 | NA | NA | 2 | 1 | 1 | 1 | NA | NA | NA | NA | NA | NA | 1 | 1 |
| 10_532251 | NA | NA | NA | NA | 2 | 0.5 | NA | NA | 2 | 0.5 | 1 | 1 | NA | NA | NA | NA | NA | NA | 1 | 1 |
| 10_534732 | 4 | 0 | 5 | 0.4 | NA | NA | 6 | 0 | 15 | 0.1333 | 4 | 0.5 | 2 | 0 | NA | NA | 1 | 0 | 7 | 0.2857 |
| 10_534733 | 6 | 0 | 3 | 0 | 2 | 0 | 7 | 0.1429 | 18 | 0.0556 | 5 | 0 | 1 | 0 | 1 | 0 | 2 | 0.5 | 9 | 0.1111 |
| 10_534839 | 5 | 0 | 5 | 0.4 | 4 | 0 | 8 | 0 | 22 | 0.0909 | 5 | 0.2 | 3 | 0 | 1 | 0 | 2 | 0 | 11 | 0.0909 |
| 10_534839 | 6 | 0.8333 | 4 | 1 | 4 | 1 | 7 | 1 | 21 | 0.9524 | 6 | 1 | 3 | 1 | 1 | 1 | 2 | 1 | 12 | 1 |
| 10_534908 | 1 | 1 | 1 | 0 | 2 | 0 | 1 | 0 | 5 | 0.2 | 1 | 0 | 3 | 0.3333 | 1 | 1 | 2 | 0.5 | 7 | 0.4286 |
| 10_534910 | 1 | 0 | 1 | 1 | 2 | 1 | 1 | 0 | 5 | 0.6 | 1 | 1 | 3 | 0.3333 | 2 | 1 | 2 | 0 | 8 | 0.5 |
| 10_535096 | 7 | 0.1429 | 4 | 0.75 | 3 | 0.3333 | 2 | 0 | 16 | 0.3125 | 1 | 0 | 3 | 0 | 2 | 1 | 5 | 0.6 | 11 | 0.4545 |

| | | | | | | | | | | | | | | | | | | | | |
|---|---|---|---|---|---|---|---|---|---|---|---|---|---|---|---|---|---|---|---|---|
| 10_535097… | 4 | 0.75 | 4 | 1 | 3 | 1 | 2 | 0.5 | 13 | 0.8462 | 1 | 1 | 2 | 1 | 3 | 0.3333 | 5 | 1 | 11 | 0.8182 |
| 10_535097… | 7 | 0 | 4 | 0.5 | 3 | 0 | 2 | 0 | 16 | 0.125 | 1 | 1 | 2 | 0 | 3 | 0 | 5 | 0.4 | 11 | 0.2727 |
| 10_535114… | 4 | 0 | 3 | 0 | 2 | 0.5 | 1 | 0 | 10 | 0.1 | 1 | 0 | 1 | 0 | 2 | 0.5 | 1 | 0 | 5 | 0.2 |
| 10_535512… | 4 | 0 | 2 | 0.5 | 1 | 0 | 1 | 0 | 8 | 0.125 | 1 | 0 | 1 | 0 | 3 | 0.3333 | 2 | 0.5 | 7 | 0.2857 |
| 10_537810… | 1 | 0 | 3 | 0 | 4 | 0 | 2 | 0.5 | 10 | 0.1 | 2 | 0.5 | 2 | 0 | 3 | 0 | 6 | 0 | 13 | 0.0769 |
| 10_537832… | 1 | 0 | 1 | 1 | 2 | 0 | 2 | 0 | 6 | 0.1667 | 2 | 0.5 | 2 | 0 | 3 | 0 | 3 | 0.3333 | 10 | 0.2 |
| 10_537833… | 2 | 0 | NA | NA | 2 | 0 | 3 | 0.3333 | 7 | 0.1429 | 2 | 0.5 | 2 | 0 | 3 | 0 | 4 | 0 | 11 | 0.0909 |
| 10_537835… | 2 | 0 | NA | NA | 1 | 0 | 2 | 0.5 | 5 | 0.2 | 2 | 0.5 | 2 | 0.5 | 2 | 1 | 2 | 0.5 | 8 | 0.625 |
| 10_539505… | 1 | 0 | NA | NA | NA | NA | 2 | 1 | 3 | 0.6667 | NA | NA | 1 | 0 | 1 | 1 | 1 | 1 | 3 | 0.6667 |
| 10_539506… | 2 | 0.5 | NA | NA | NA | NA | 2 | 0.5 | 4 | 0.5 | 1 | 0 | 1 | 1 | 1 | 0 | 2 | 1 | 5 | 0.6 |
| 10_539507… | 2 | 0.5 | NA | NA | NA | NA | 1 | 0 | 3 | 0.3333 | 1 | 1 | 1 | 1 | 1 | 1 | 1 | 0 | 4 | 0.75 |
| 10_539511… | NA | NA | NA | NA | NA | NA | 2 | 0.5 | 2 | 0.5 | 1 | 1 | 1 | 0 | NA | NA | 1 | 1 | 3 | 0.6667 |
| 10_559910… | 1 | 0 | 1 | 0 | NA | NA | 1 | 1 | 3 | 0.3333 | NA | NA | 3 | 0.3333 | 1 | 0 | 3 | 0.3333 | 7 | 0.2857 |
| 10_559911… | 2 | 0 | 1 | 0 | 1 | 1 | 1 | 1 | 5 | 0.4 | NA | NA | 3 | 0.6667 | 1 | 0 | 3 | 0.3333 | 7 | 0.4286 |
| 10_559915… | 3 | 0.6667 | 3 | 0.3333 | 1 | 1 | 1 | 0 | 8 | 0.5 | 1 | 0 | 5 | 0.4 | 1 | 0 | 4 | 0.5 | 11 | 0.3636 |
| 10_559917… | 3 | 0 | 5 | 0.4 | 2 | 0.5 | 1 | 0 | 11 | 0.2727 | 2 | 0.5 | 4 | 0.25 | 1 | 0 | 3 | 0.3333 | 10 | 0.3 |
| 10_559919… | 3 | 0.3333 | 2 | 0.5 | 2 | 0.5 | 2 | 0 | 9 | 0.3333 | 2 | 0 | 5 | 0.8 | 2 | 0 | 5 | 0.6 | 14 | 0.5 |
| 10_559920… | 3 | 0.3333 | 5 | 0.6 | 1 | 0 | 2 | 0.5 | 11 | 0.4545 | 1 | 0 | 6 | 0.5 | 1 | 0 | 2 | 0 | 10 | 0.3 |
| 10_559920… | 4 | 0.75 | 5 | 1 | 2 | 1 | 2 | 1 | 13 | 0.9231 | 1 | 1 | 6 | 0.8333 | 2 | 1 | 4 | 1 | 13 | 0.9231 |
| 10_559921… | 4 | 0.75 | 5 | 0.6 | 1 | 0 | 2 | 0 | 12 | 0.5 | 1 | 0 | 5 | 0.6 | 2 | 0.5 | 4 | 0.75 | 12 | 0.5833 |
| 10_559922… | 4 | 0.25 | 5 | 0.6 | 2 | 1 | 2 | 1 | 13 | 0.6154 | 1 | 1 | 4 | 0.25 | 2 | 0 | 2 | 0.5 | 9 | 0.3333 |
| 10_559923… | 3 | 0 | 4 | 0 | 2 | 0.5 | 2 | 0 | 11 | 0.0909 | 2 | 0 | 6 | 0.1667 | 2 | 0 | 5 | 0 | 15 | 0.0667 |
| 10_559923… | 4 | 1 | 5 | 1 | 2 | 1 | 2 | 1 | 13 | 1 | 2 | 1 | 5 | 1 | 2 | 1 | 5 | 1 | 14 | 1 |
| 10_559927… | 4 | 0.5 | 5 | 0.6 | 2 | 0.5 | 2 | 0 | 13 | 0.4615 | 2 | 0.5 | 5 | 0.2 | 2 | 1 | 4 | 0.25 | 13 | 0.3846 |
| 10_559928… | 3 | 1 | 5 | 1 | 2 | 1 | 2 | 1 | 12 | 1 | 2 | 1 | 6 | 1 | 2 | 1 | 6 | 1 | 16 | 1 |
| 10_559928… | 4 | 0.75 | 5 | 0.8 | 2 | 1 | 1 | 0 | 12 | 0.75 | 2 | 1 | 5 | 1 | 3 | 1 | 5 | 1 | 15 | 1 |
| 10_559928… | 4 | 0.25 | 5 | 0.6 | 1 | 0 | 2 | 0 | 12 | 0.3333 | 2 | 0.5 | 4 | 0 | 3 | 0.3333 | 5 | 0.6 | 14 | 0.3571 |
| 10_559928… | 2 | 0 | 3 | 0 | 1 | 0 | 2 | 0.5 | 8 | 0.125 | 2 | 0.5 | 4 | 0 | 1 | 0 | 5 | 0 | 12 | 0.0833 |
| 10_562637… | 9 | 0.4444 | 12 | 0.1667 | 17 | 0.1765 | 8 | 0.25 | 46 | 0.2391 | 5 | 0.4 | 8 | 0.125 | 6 | 0.6667 | 8 | 0.25 | 27 | 0.3333 |
| 10_562637… | 8 | 1 | 12 | 0.5 | 11 | 0.7273 | 4 | 0.75 | 35 | 0.7143 | 4 | 0.75 | 3 | 1 | 7 | 0.8571 | 8 | 0.625 | 22 | 0.7727 |
| 10_562642… | 11 | 1 | 15 | 0.9333 | 18 | 1 | 9 | 1 | 53 | 0.9811 | 9 | 1 | 13 | 1 | 7 | 1 | 11 | 1 | 40 | 1 |
| 10_562644… | 11 | 0.9091 | 13 | 1 | 19 | 0.9474 | 9 | 1 | 52 | 0.9615 | 11 | 1 | 12 | 0.9167 | 7 | 1 | 11 | 1 | 41 | 0.9756 |
| 10_562645… | 17 | 0.4706 | 17 | 0.2353 | 23 | 0.1739 | 11 | 0 | 68 | 0.2353 | 10 | 0.2 | 12 | 0.1667 | 9 | 0.1111 | 13 | 0.1538 | 44 | 0.1591 |
| 10_562645… | 17 | 0.2353 | 16 | 0.3125 | 21 | 0.381 | 11 | 0.3636 | 65 | 0.3231 | 10 | 0.1 | 10 | 0.4 | 6 | 0 | 11 | 0.0909 | 37 | 0.1622 |
| 10_562645… | 19 | 0.1053 | 16 | 0.3125 | 21 | 0.1905 | 11 | 0.1818 | 67 | 0.194 | 12 | 0.25 | 13 | 0.1538 | 11 | 0.0909 | 14 | 0.0714 | 50 | 0.14 |
| 10_562646… | 14 | 0.9286 | 17 | 0.7647 | 24 | 0.6667 | 14 | 0.7143 | 69 | 0.7536 | 11 | 0.5455 | 10 | 0.8 | 11 | 0.6364 | 12 | 0.6667 | 44 | 0.6591 |
| 10_562646… | 11 | 1 | 8 | 0.625 | 14 | 0.8571 | 7 | 0.7143 | 40 | 0.825 | 7 | 0.8571 | 9 | 0.5556 | 9 | 0.8889 | 5 | 1 | 30 | 0.8 |
| 10_562648… | 16 | 0.25 | 19 | 0.1579 | 20 | 0.05 | 11 | 0.2727 | 66 | 0.1667 | 10 | 0.3 | 11 | 0.0909 | 8 | 0 | 13 | 0.1538 | 42 | 0.1429 |
| 10_562649… | 19 | 1 | 19 | 1 | 27 | 1 | 12 | 1 | 77 | 1 | 12 | 0.9167 | 13 | 0.9231 | 10 | 1 | 18 | 1 | 53 | 0.9623 |
| 10_562653… | 24 | 0.2917 | 23 | 0.3043 | 33 | 0.303 | 17 | 0.1765 | 97 | 0.2784 | 16 | 0.125 | 21 | 0.1429 | 13 | 0.0769 | 20 | 0.2 | 70 | 0.1429 |
| 10_562660… | 29 | 0.1379 | 35 | 0.1714 | 37 | 0.1351 | 19 | 0.1053 | 120 | 0.1417 | 20 | 0.35 | 22 | 0.3182 | 17 | 0.1765 | 23 | 0.1739 | 82 | 0.2561 |
| 10_562660… | 29 | 0.1724 | 34 | 0.1765 | 43 | 0.1395 | 20 | 0.2 | 126 | 0.1667 | 19 | 0.3158 | 22 | 0.2273 | 22 | 0.1818 | 25 | 0.24 | 88 | 0.2386 |
| 10_562662… | 29 | 0.5172 | 26 | 0.6154 | 35 | 0.6857 | 19 | 0.5263 | 109 | 0.5963 | 17 | 0.8235 | 23 | 0.6087 | 18 | 0.8333 | 22 | 0.5455 | 80 | 0.6875 |
| 10_562662… | 32 | 0 | 40 | 0.125 | 42 | 0.0238 | 25 | 0.2 | 139 | 0.0791 | 22 | 0.0909 | 24 | 0.1667 | 23 | 0.1304 | 27 | 0.1111 | 96 | 0.125 |
| 10_562662… | 28 | 0.5714 | 35 | 0.5429 | 40 | 0.4 | 21 | 0.4286 | 124 | 0.4839 | 19 | 0.6842 | 23 | 0.4348 | 19 | 0.4211 | 25 | 0.4 | 86 | 0.4767 |
| 10_562662… | 28 | 0.1071 | 35 | 0.1714 | 38 | 0.1842 | 20 | 0.15 | 121 | 0.157 | 20 | 0.2 | 22 | 0.2727 | 21 | 0.3333 | 24 | 0.2083 | 87 | 0.2529 |
| 10_562663… | 31 | 0.7742 | 37 | 0.7838 | 41 | 0.8293 | 28 | 0.6786 | 137 | 0.7737 | 23 | 0.913 | 24 | 0.8333 | 22 | 0.7273 | 27 | 0.9259 | 96 | 0.8542 |
| 10_562663… | 29 | 0.1724 | 36 | 0.0833 | 42 | 0.0238 | 25 | 0.08 | 132 | 0.0833 | 18 | 0 | 23 | 0.0435 | 23 | 0.0435 | 25 | 0.12 | 89 | 0.0562 |
| 10_573872… | 1 | 1 | NA | NA | 1 | 1 | NA | NA | 2 | 1 | NA | NA | NA | NA | NA | NA | 1 | 1 | 1 | 1 |
| 10_573882… | 6 | 0.1667 | 5 | 0.2 | 4 | 0.25 | 1 | 1 | 16 | 0.25 | 1 | 0 | 2 | 0 | 4 | 0.25 | 3 | 0 | 10 | 0.1 |
| 10_573882… | 4 | 0.75 | 4 | 0.75 | 5 | 0.6 | 2 | 0.5 | 15 | 0.6667 | 1 | 0 | 1 | 1 | 5 | 0.6 | 3 | 0.3333 | 10 | 0.5 |
| 10_573883… | 5 | 0.6 | 4 | 0.75 | 5 | 0.8 | 2 | 1 | 16 | 0.75 | 1 | 0 | 2 | 1 | 5 | 0.6 | 4 | 0.75 | 12 | 0.6667 |
| 10_573930… | 76 | 0.3816 | 107 | 0.4299 | 88 | 0.4091 | 89 | 0.427 | 360 | 0.4139 | 51 | 0.4118 | 65 | 0.2923 | 88 | 0.3864 | 80 | 0.4125 | 284 | 0.3768 |
| 10_573932… | 74 | 0.2703 | 104 | 0.1923 | 78 | 0.2436 | 82 | 0.1585 | 338 | 0.213 | 50 | 0.22 | 65 | 0.2 | 76 | 0.2237 | 76 | 0.1974 | 267 | 0.2097 |
| 10_573932… | NA | NA | NA | NA | 1 | 1 | NA | NA | 1 | 1 | 1 | 1 | NA | NA | NA | NA | NA | NA | 1 | 1 |
| 10_573932… | 88 | 0.3977 | 107 | 0.4019 | 84 | 0.4167 | 83 | 0.4458 | 362 | 0.4144 | 53 | 0.4528 | 64 | 0.4531 | 86 | 0.4186 | 71 | 0.4648 | 274 | 0.4453 |
| 10_576124… | 1 | 0 | 1 | 0 | 2 | 0 | 2 | 0.5 | 6 | 0.1667 | 3 | 0.3333 | 3 | 0 | 3 | 0 | 1 | 0 | 10 | 0.1 |
| 10_576878… | 10 | 0.1 | 5 | 0 | 6 | 0.1667 | 4 | 0 | 25 | 0.08 | 3 | 0.3333 | 5 | 0 | 6 | 0.1667 | 6 | 0.1667 | 20 | 0.15 |
| 10_581056… | 3 | 0.3333 | 1 | 0 | 1 | 0 | NA | NA | 5 | 0.2 | NA | NA | NA | NA | NA | NA | 1 | 1 | 1 | 1 |
| 10_581059… | 2 | 1 | 1 | 1 | NA | NA | NA | NA | 3 | 1 | NA | NA | NA | NA | NA | NA | 3 | 1 | 3 | 1 |
| 10_581063… | 3 | 0.3333 | 2 | 0 | 1 | 0 | NA | NA | 6 | 0.1667 | NA | NA | NA | NA | 1 | 0 | 5 | 0.2 | 6 | 0.1667 |
| 10_581066… | 2 | 1 | 2 | 1 | NA | NA | NA | NA | 4 | 1 | NA | NA | NA | NA | 1 | 1 | 5 | 1 | 6 | 1 |
| 10_581069… | 3 | 0.3333 | 3 | 0.3333 | 2 | 0 | NA | NA | 8 | 0.25 | 1 | 0 | NA | NA | 1 | 0 | 4 | 0.5 | 6 | 0.3333 |
| 10_581072… | 2 | 0 | 3 | 0.3333 | 2 | 0.5 | 1 | 1 | 8 | 0.375 | 1 | 0 | NA | NA | 1 | 0 | 4 | 0.5 | 6 | 0.3333 |
| 10_581072… | 2 | 0 | 3 | 0.3333 | 2 | 0.5 | 1 | 0 | 8 | 0.25 | 1 | 0 | NA | NA | 1 | 1 | 4 | 0.5 | 6 | 0.5 |

| | | | | | | | | | | | | | | | | | | | | |
|---|---|---|---|---|---|---|---|---|---|---|---|---|---|---|---|---|---|---|---|---|
| 10_5810736 | 3 | 0.3333 | 4 | 0.5 | 2 | 0.5 | 1 | 0 | 10 | 0.4 | 1 | 0 | NA | NA | 1 | 0 | 5 | 0.4 | 7 | 0.2857 |
| 10_5813997 | 2 | 0.5 | 7 | 0.2857 | 5 | 0.4 | 5 | 0.4 | 19 | 0.3684 | 1 | 0 | 3 | 0.6667 | 4 | 0.25 | 7 | 0 | 15 | 0.2 |
| 10_5814001 | 5 | 0.2 | 8 | 0.125 | 6 | 0.3333 | 6 | 0.3333 | 25 | 0.24 | 2 | 0 | 3 | 0 | 5 | 0 | 8 | 0.125 | 18 | 0.0556 |
| 10_5814065 | 4 | 0 | 10 | 0.1 | 8 | 0.125 | 7 | 0.1429 | 29 | 0.1034 | 3 | 0.3333 | 3 | 0 | 4 | 0.5 | 8 | 0 | 18 | 0.1667 |
| 10_5831252 | 1 | 1 | 1 | 1 | 1 | 1 | 2 | 1 | 5 | 1 | NA | NA | 1 | 1 | 2 | 1 | 4 | 0.75 | 7 | 0.8571 |
| 10_5831302 | 1 | 0 | 2 | 1 | 1 | 0 | 1 | 0 | 5 | 0.4 | NA | NA | 1 | 1 | 1 | 0 | 4 | 0.5 | 6 | 0.5 |
| 10_5831317 | 1 | 0 | 1 | 1 | 1 | 1 | 2 | 0 | 5 | 0.4 | NA | NA | NA | NA | 1 | 0 | 3 | 0.6667 | 4 | 0.5 |
| 10_5831350 | 1 | 0 | 2 | 0 | 2 | 0.5 | 3 | 0 | 8 | 0.125 | NA | NA | NA | NA | 4 | 0.5 | 3 | 0.3333 | 7 | 0.4286 |
| 10_5831352 | 1 | 1 | 1 | 1 | NA | NA | 3 | 1 | 5 | 1 | NA | NA | 1 | 1 | 3 | 1 | 3 | 1 | 7 | 1 |
| 10_5831396 | 1 | 1 | 3 | 0.6667 | 3 | 0.3333 | 3 | 1 | 10 | 0.7 | NA | NA | 1 | 1 | 1 | 1 | 1 | 1 | 3 | 1 |
| 10_5831396 | 1 | 1 | 1 | 1 | 3 | 0.6667 | 3 | 1 | 8 | 0.875 | NA | NA | NA | NA | 4 | 1 | 3 | 1 | 7 | 1 |
| 10_5831468 | 1 | 0 | 2 | 0 | 3 | 0 | 5 | 0.2 | 11 | 0.0909 | NA | NA | 1 | 0 | 5 | 0.2 | 4 | 0 | 10 | 0.1 |
| 10_5831480 | 1 | 0 | 3 | 0.3333 | 3 | 0.3333 | 5 | 0.8 | 12 | 0.5 | NA | NA | 1 | 0 | 6 | 0.3333 | 4 | 0.5 | 11 | 0.3636 |
| 10_5831481 | 1 | 1 | 3 | 0.6667 | 4 | 0.5 | 5 | 0.2 | 13 | 0.4615 | NA | NA | NA | NA | 6 | 0.6667 | 4 | 0.25 | 10 | 0.5 |
| 10_5831486 | 1 | 0 | 3 | 0 | 4 | 0.25 | 5 | 0.2 | 13 | 0.1538 | NA | NA | 1 | 0 | 6 | 0 | 4 | 0.25 | 11 | 0.0909 |
| 10_5831507 | NA | NA | 3 | 0 | 3 | 0.3333 | 5 | 0.2 | 11 | 0.1818 | NA | NA | 2 | 0 | 4 | 0.25 | 3 | 0 | 9 | 0.1111 |
| 10_5831514 | 1 | 0 | 3 | 0 | 4 | 0 | 5 | 0.2 | 13 | 0.0769 | NA | NA | 1 | 0 | 6 | 0 | 4 | 0.25 | 11 | 0.0909 |
| 10_5831526 | 1 | 0 | 2 | 0.5 | 4 | 0.25 | 4 | 0 | 11 | 0.1818 | NA | NA | 1 | 1 | 6 | 0.3333 | 4 | 0.5 | 11 | 0.4545 |
| 10_5831547 | 1 | 1 | 3 | 0 | 4 | 0 | 5 | 0 | 13 | 0.0769 | NA | NA | 1 | 0 | 7 | 0.2857 | 4 | 0.25 | 12 | 0.25 |
| 10_5831571 | 1 | 1 | 2 | 0.5 | 4 | 1 | 5 | 1 | 12 | 0.9167 | NA | NA | 3 | 1 | 6 | 1 | 4 | 1 | 13 | 1 |
| 10_5831572 | 2 | 0 | 3 | 0 | 5 | 0.2 | 4 | 0.25 | 14 | 0.1429 | NA | NA | 2 | 0 | 6 | 0.1667 | 4 | 0 | 12 | 0.0833 |
| 10_5831580 | 2 | 0 | 3 | 0 | 6 | 0.3333 | 5 | 0.2 | 16 | 0.1875 | NA | NA | 3 | 0 | 7 | 0.4286 | 4 | 0 | 14 | 0.2143 |
| 10_5831591 | 3 | 0 | 3 | 0 | 6 | 0.1667 | 5 | 0.2 | 17 | 0.1176 | NA | NA | 2 | 0.5 | 7 | 0.1429 | 4 | 0 | 13 | 0.1538 |
| 10_5831592 | 3 | 0 | 2 | 0.5 | 6 | 0.3333 | 6 | 0 | 17 | 0.1765 | NA | NA | 3 | 0 | 7 | 0.1429 | 4 | 0 | 14 | 0.0714 |
| 10_5831596 | 3 | 1 | 2 | 0 | 6 | 0.3333 | 6 | 0.3333 | 17 | 0.4118 | NA | NA | 3 | 0.6667 | 6 | 0.5 | 4 | 0.5 | 13 | 0.5385 |
| 10_5831597 | 3 | 0 | 3 | 0.3333 | 7 | 0 | 5 | 0 | 18 | 0.0556 | NA | NA | 3 | 0.3333 | 6 | 0 | 4 | 0 | 13 | 0.0769 |
| 10_5831598 | 2 | 1 | 1 | 0 | 4 | 1 | 5 | 1 | 12 | 0.9167 | NA | NA | 2 | 0.5 | 7 | 0.7143 | 3 | 0.3333 | 12 | 0.5833 |
| 10_5831600 | 3 | 1 | 3 | 1 | 5 | 0.8 | 6 | 1 | 17 | 0.9412 | NA | NA | 3 | 1 | 7 | 1 | 3 | 0.3333 | 13 | 0.8462 |
| 10_5831604 | 2 | 0.5 | 2 | 0 | 6 | 0.5 | 6 | 0.3333 | 16 | 0.375 | NA | NA | 3 | 0.6667 | 8 | 0.25 | 4 | 0 | 15 | 0.2667 |
| 10_5831608 | 3 | 0.3333 | 3 | 0.6667 | 6 | 0.3333 | 6 | 0.3333 | 18 | 0.3889 | NA | NA | 3 | 0.3333 | 6 | 0.1667 | 4 | 0.75 | 13 | 0.3846 |
| 10_5831623 | 4 | 0.75 | 3 | 1 | 6 | 0.8333 | 4 | 1 | 17 | 0.8824 | NA | NA | 3 | 0.3333 | 7 | 0.5714 | 4 | 0.75 | 14 | 0.5714 |
| 10_5831647 | 4 | 0.25 | 2 | 0.5 | 5 | 0.2 | 6 | 0 | 17 | 0.1765 | NA | NA | 3 | 0.3333 | 9 | 0.1111 | 4 | 0.25 | 16 | 0.1875 |
| 10_5831654 | 3 | 1 | 3 | 0 | 7 | 0.1429 | 6 | 0.1667 | 19 | 0.2632 | NA | NA | 3 | 0 | 5 | 0.2 | 4 | 0.75 | 12 | 0.3333 |
| 10_5831660 | 3 | 0.3333 | 2 | 0 | 6 | 0.1667 | 6 | 0.3333 | 17 | 0.2353 | NA | NA | 4 | 0.75 | 8 | 0.25 | 3 | 0.3333 | 15 | 0.4 |
| 10_5831891 | 4 | 0 | 2 | 0 | 5 | 0 | 5 | 0.4 | 16 | 0.125 | NA | NA | 4 | 0.25 | 6 | 0 | 2 | 0 | 12 | 0.0833 |
| 10_5832867 | 5 | 0.2 | 5 | 0.6 | 11 | 0.1818 | 9 | 0.3333 | 30 | 0.3 | 3 | 0 | 7 | 0 | 10 | 0.2 | 6 | 0 | 26 | 0.0769 |
| 10_5832873 | 7 | 0.8571 | 6 | 1 | 11 | 0.6364 | 9 | 0.5556 | 33 | 0.7273 | 3 | 0.3333 | 6 | 0.6667 | 13 | 0.6923 | 4 | 0.25 | 26 | 0.5769 |
| 10_5832952 | 5 | 0.4 | 2 | 1 | 6 | 0.8333 | 4 | 0.5 | 17 | 0.6471 | 2 | 0.5 | 5 | 0.4 | 5 | 0.6 | 1 | 0 | 13 | 0.4615 |
| 10_5832955 | 10 | 0.1 | 5 | 0.2 | 9 | 0.3333 | 9 | 0.1111 | 33 | 0.1818 | 5 | 0.2 | 7 | 0.1429 | 10 | 0.4 | 6 | 0 | 28 | 0.2143 |
| 10_5832966 | 9 | 0.7778 | 5 | 1 | 12 | 1 | 10 | 1 | 36 | 0.9444 | 6 | 0.8333 | 6 | 1 | 13 | 0.9231 | 5 | 0.6 | 30 | 0.8667 |
| 10_5832978 | 10 | 0.4 | 6 | 0.1667 | 14 | 0.1429 | 11 | 0.1818 | 41 | 0.2195 | 6 | 0.1667 | 9 | 0.1111 | 11 | 0.2727 | 5 | 0.4 | 31 | 0.2258 |
| 10_5833014 | 7 | 0.2857 | 3 | 0 | 13 | 0 | 8 | 0.125 | 31 | 0.0968 | 4 | 0.25 | 6 | 0 | 12 | 0.1667 | 6 | 0.1667 | 28 | 0.1429 |
| 10_5897175 | NA | NA | 1 | 1 | NA | NA | NA | NA | 1 | 1 | 1 | 1 | 2 | 1 | NA | NA | NA | NA | 3 | 1 |
| 10_5897199 | NA | NA | 1 | 1 | NA | NA | NA | NA | 1 | 1 | 1 | 0 | 3 | 0.3333 | NA | NA | NA | NA | 4 | 0.25 |
| 10_5897377 | 1 | 0 | 2 | 0 | 1 | 1 | 1 | 0 | 5 | 0.2 | 4 | 0 | 3 | 0 | 1 | 1 | 2 | 0.5 | 10 | 0.2 |
| 10_5897396 | 1 | 1 | 1 | 1 | 1 | 1 | 1 | 1 | 4 | 1 | 3 | 0.6667 | 1 | 0 | 1 | 1 | 2 | 0.5 | 7 | 0.5714 |
| 10_5897415 | 1 | 0 | 3 | 0.3333 | 1 | 0 | 1 | 0 | 6 | 0.1667 | 4 | 0 | 2 | 0.5 | 2 | 0 | 2 | 0 | 10 | 0.1 |
| 10_5897418 | 1 | 0 | 3 | 0.3333 | 1 | 0 | 1 | 0 | 6 | 0.1667 | 3 | 0 | 3 | 0 | 2 | 0 | 2 | 1 | 10 | 0.2 |
| 10_5897446 | 1 | 0 | 3 | 1 | 1 | 0 | 1 | 0 | 6 | 0.5 | 5 | 0.8 | 3 | 1 | 1 | 1 | 2 | 1 | 11 | 0.9091 |
| 10_5905793 | NA | NA | NA | NA | 2 | 0 | 1 | 1 | 3 | 0.3333 | NA | NA | 1 | 1 | NA | NA | NA | NA | 1 | 1 |
| 10_5905802 | NA | NA | NA | NA | 2 | 1 | 1 | 1 | 3 | 1 | NA | NA | 2 | 0.5 | NA | NA | NA | NA | 2 | 0.5 |
| 10_5905818 | NA | NA | NA | NA | 2 | 0 | 2 | 0.5 | 4 | 0.25 | NA | NA | 2 | 0.5 | NA | NA | NA | NA | 2 | 0.5 |
| 10_5905824 | NA | NA | NA | NA | 2 | 1 | 2 | 0.5 | 4 | 0.75 | NA | NA | 2 | 0.5 | NA | NA | NA | NA | 2 | 0.5 |
| 10_5905831 | NA | NA | NA | NA | 1 | 1 | 2 | 0.5 | 3 | 0.6667 | NA | NA | 2 | 1 | NA | NA | NA | NA | 2 | 1 |
| 10_5905837 | NA | NA | NA | NA | 1 | 1 | 1 | 0 | 2 | 0.5 | NA | NA | 2 | 1 | NA | NA | NA | NA | 2 | 1 |
| 10_5905845 | NA | NA | NA | NA | 1 | 1 | 2 | 1 | 3 | 1 | NA | NA | 1 | 1 | NA | NA | NA | NA | 1 | 1 |
| 10_5905846 | NA | NA | NA | NA | 2 | 0.5 | 2 | 0 | 4 | 0.25 | NA | NA | 1 | 1 | NA | NA | NA | NA | 1 | 1 |
| 10_5905866 | NA | NA | NA | NA | 1 | 1 | 1 | 1 | 2 | 1 | NA | NA | 2 | 0.5 | NA | NA | NA | NA | 2 | 0.5 |
| 10_5905882 | NA | NA | NA | NA | 2 | 0.5 | 2 | 0.5 | 4 | 0.5 | NA | NA | 2 | 1 | NA | NA | NA | NA | 2 | 1 |
| 10_5905894 | NA | NA | NA | NA | 1 | 0 | 2 | 1 | 3 | 0.6667 | NA | NA | 2 | 1 | NA | NA | NA | NA | 2 | 1 |
| 10_5905906 | NA | NA | NA | NA | 2 | 1 | 1 | 1 | 3 | 1 | NA | NA | 2 | 1 | NA | NA | NA | NA | 2 | 1 |
| 10_5905910 | NA | NA | 1 | 1 | 1 | 1 | 1 | 1 | 3 | 1 | NA | NA | 1 | 1 | NA | NA | NA | NA | 1 | 1 |
| 10_5905913 | NA | NA | 1 | 0 | 2 | 0.5 | 2 | 0.5 | 5 | 0.4 | NA | NA | 2 | 0.5 | NA | NA | NA | NA | 2 | 0.5 |
| 10_5905916 | NA | NA | 1 | 1 | 2 | 1 | 1 | 1 | 4 | 1 | NA | NA | 2 | 1 | NA | NA | NA | NA | 2 | 1 |
| 10_5905965 | 1 | 1 | 2 | 1 | 2 | 1 | 2 | 0.5 | 7 | 0.8571 | NA | NA | 2 | 0.5 | NA | NA | NA | NA | 2 | 0.5 |

| | | | | | | | | | | | | | | | | | | | | |
|---|---|---|---|---|---|---|---|---|---|---|---|---|---|---|---|---|---|---|---|---|
| 10_590603 | 1 | 0 | 2 | 0.5 | 3 | 0.6667 | 3 | 0 | 9 | 0.3333 | NA | NA | 2 | 0.5 | NA | NA | NA | NA | 2 | 0.5 |
| 10_590620 | 3 | 0 | 3 | 0.6667 | 3 | 0 | 3 | 0 | 12 | 0.1667 | 1 | 0 | 2 | 0.5 | 1 | 0 | NA | NA | 4 | 0.25 |
| 10_592070 | 5 | 0.8 | 8 | 0.75 | 8 | 0.75 | 4 | 0.5 | 25 | 0.72 | 7 | 0.7143 | 5 | 1 | 3 | 0.6667 | 4 | 0.75 | 19 | 0.7895 |
| 10_592073 | 5 | 0 | 5 | 0.2 | 9 | 0.4444 | 5 | 0.4 | 24 | 0.2917 | 6 | 0 | 4 | 0.25 | 4 | 0.25 | 4 | 0 | 18 | 0.1111 |
| 10_592075 | 6 | 0.1667 | 7 | 0.7143 | 12 | 0.5 | 6 | 0.5 | 31 | 0.4839 | 8 | 0.375 | 7 | 0.4286 | 7 | 0.2857 | 6 | 0.3333 | 28 | 0.3571 |
| 10_592076 | 6 | 0.1667 | 6 | 0 | 12 | 0.25 | 5 | 0.4 | 29 | 0.2069 | 8 | 0.125 | 6 | 0.1667 | 6 | 0.5 | 5 | 0 | 25 | 0.2 |
| 10_592321 | 2 | 0 | 3 | 0.3333 | 2 | 0 | 1 | 0 | 8 | 0.125 | 3 | 0 | 4 | 0.25 | 1 | 0 | NA | NA | 8 | 0.125 |
| 10_592322 | 3 | 0.6667 | 2 | 0 | 2 | 0 | 1 | 0 | 8 | 0.25 | 3 | 0.3333 | 5 | 0 | 1 | 1 | NA | NA | 9 | 0.2222 |
| 10_592323 | 2 | 0.5 | 2 | 0 | 1 | 0 | 1 | 1 | 6 | 0.3333 | 2 | 0.5 | 2 | 0 | NA | NA | NA | NA | 4 | 0.25 |
| 10_592323 | 3 | 0.3333 | 3 | 0.6667 | 2 | 1 | 1 | 0 | 9 | 0.5556 | 4 | 0.5 | 5 | 0.6 | 2 | 1 | NA | NA | 11 | 0.6364 |
| 10_592323 | 2 | 1 | 3 | 1 | 2 | 1 | 1 | 1 | 8 | 1 | 4 | 0.75 | 5 | 0.8 | 2 | 0.5 | NA | NA | 11 | 0.7273 |
| 10_592836 | 3 | 0.3333 | 8 | 0.125 | 1 | 1 | 5 | 0.2 | 17 | 0.2353 | 5 | 0.2 | 2 | 0 | 7 | 0.1429 | 5 | 1 | 19 | 0.3684 |
| 10_592846 | 2 | 0 | 6 | 0.1667 | NA | NA | 4 | 0.25 | 12 | 0.1667 | 3 | 0 | 2 | 0 | 4 | 0 | 4 | 0.25 | 13 | 0.0769 |
| 10_593531 | NA | NA | 1 | 1 | NA | NA | NA | NA | 1 | 1 | 1 | 1 | 2 | 0 | NA | NA | NA | NA | 3 | 0.3333 |
| 10_593531 | NA | NA | 1 | 1 | NA | NA | NA | NA | 1 | 1 | 1 | 0 | 2 | 0.5 | NA | NA | NA | NA | 3 | 0.3333 |
| 10_593534 | NA | NA | 1 | 1 | NA | NA | NA | NA | 1 | 1 | NA | NA | 1 | 1 | NA | NA | NA | NA | 1 | 1 |
| 10_596991 | 14 | 0.2857 | 13 | 0.2308 | 7 | 0 | 8 | 0 | 42 | 0.1667 | 12 | 0.1667 | 8 | 0.125 | 14 | 0.1429 | 10 | 0.1 | 44 | 0.1364 |
| 10_596991 | 17 | 0.9412 | 18 | 0.8889 | 8 | 1 | 9 | 0.8889 | 52 | 0.9231 | 10 | 0.8 | 11 | 1 | 15 | 0.8667 | 10 | 0.9 | 46 | 0.8913 |
| 10_596993 | 17 | 0.1765 | 20 | 0.4 | 8 | 0.375 | 13 | 0.1538 | 58 | 0.2759 | 13 | 0.4615 | 10 | 0.2 | 17 | 0.1176 | 8 | 0.5 | 48 | 0.2917 |
| 10_596993 | 18 | 0.1667 | 20 | 0.25 | 9 | 0.1111 | 13 | 0.1538 | 60 | 0.1833 | 13 | 0.0769 | 12 | 0.25 | 19 | 0 | 10 | 0.1 | 54 | 0.0926 |
| 10_596995 | 16 | 0.125 | 17 | 0.0588 | 9 | 0 | 14 | 0.1429 | 56 | 0.0893 | 13 | 0.3077 | 11 | 0 | 18 | 0 | 8 | 0 | 50 | 0.08 |
| 10_596997 | 16 | 0.125 | 16 | 0 | 12 | 0.0833 | 13 | 0.0769 | 57 | 0.0702 | 14 | 0 | 12 | 0.0833 | 18 | 0 | 13 | 0.1538 | 57 | 0.0526 |
| 10_596997 | 17 | 0.2353 | 20 | 0.45 | 11 | 0.4545 | 12 | 0.3333 | 60 | 0.3667 | 15 | 0.3333 | 14 | 0.3571 | 19 | 0.2632 | 13 | 0.4615 | 61 | 0.3443 |
| 10_596998 | 17 | 0.4118 | 18 | 0.4444 | 11 | 0.1818 | 15 | 0.2 | 61 | 0.3279 | 16 | 0.25 | 16 | 0.4375 | 19 | 0.2632 | 15 | 0.4 | 66 | 0.3333 |
| 10_597002 | 1 | 1 | NA | NA | NA | NA | NA | NA | 1 | 1 | 1 | 1 | NA | NA | NA | NA | NA | NA | 1 | 1 |
| 10_597297 | 2 | 0 | NA | NA | 1 | 0 | 2 | 0.5 | 5 | 0.2 | 1 | 1 | NA | NA | NA | NA | 1 | 0 | 2 | 0.5 |
| 10_597297 | 2 | 1 | NA | NA | 1 | 1 | 3 | 1 | 6 | 1 | 1 | 1 | NA | NA | NA | NA | 1 | 1 | 2 | 1 |
| 10_597298 | 2 | 0 | NA | NA | 1 | 1 | 3 | 0 | 6 | 0.1667 | 1 | 0 | NA | NA | NA | NA | 2 | 0.5 | 3 | 0.3333 |
| 10_597299 | 1 | 1 | NA | NA | NA | NA | 2 | 0 | 3 | 0.3333 | NA | NA | NA | NA | NA | NA | 1 | 1 | 1 | 1 |
| 10_597301 | 2 | 1 | NA | NA | 1 | 1 | 2 | 1 | 5 | 1 | NA | NA | NA | NA | NA | NA | 2 | 1 | 2 | 1 |
| 10_597333 | 8 | 1 | 4 | 0.5 | 2 | 1 | 8 | 1 | 22 | 0.9091 | 2 | 1 | 5 | 1 | 7 | 1 | 2 | 1 | 16 | 1 |
| 10_597333 | 8 | 0.125 | 6 | 0.1667 | 3 | 0.3333 | 9 | 0.1111 | 26 | 0.1538 | 2 | 0 | 6 | 0.1667 | 8 | 0 | 3 | 0 | 19 | 0.0526 |
| 10_597334 | 7 | 0.5714 | 5 | 0.4 | 2 | 0.5 | 8 | 0.5 | 22 | 0.5 | 3 | 0.3333 | 6 | 0.1667 | 8 | 0.375 | 3 | 0.6667 | 20 | 0.35 |
| 10_597334 | 8 | 0 | 6 | 0.1667 | 3 | 0 | 9 | 0.1111 | 26 | 0.0769 | 2 | 0 | 7 | 0.1429 | 8 | 0 | 3 | 0.3333 | 20 | 0.1 |
| 10_597334 | 8 | 0.5 | 6 | 0.3333 | 3 | 0.3333 | 8 | 0.375 | 25 | 0.4 | 3 | 0.3333 | 6 | 0.3333 | 6 | 0.5 | 4 | 0.5 | 19 | 0.4211 |
| 10_597336 | 3 | 1 | 5 | 0.4 | 1 | 1 | 4 | 0.75 | 13 | 0.6923 | 3 | 0.3333 | 4 | 0.5 | 1 | 1 | 3 | 0.6667 | 11 | 0.5455 |
| 10_597343 | 9 | 0.4444 | 6 | 0.6667 | 3 | 0 | 7 | 0.2857 | 25 | 0.4 | 5 | 0.4 | 6 | 0.1667 | 7 | 0.8571 | 3 | 0.3333 | 21 | 0.4762 |
| 10_597346 | 8 | 0.375 | 9 | 0.4444 | 3 | 0 | 8 | 0.5 | 28 | 0.3929 | 8 | 0.25 | 6 | 0.1667 | 7 | 0.5714 | 3 | 0.6667 | 24 | 0.375 |
| 10_597862 | 9 | 0.2222 | 6 | 0.1667 | 1 | 0 | 4 | 0 | 20 | 0.15 | 4 | 0.25 | 8 | 0.125 | 5 | 0.4 | 6 | 0.1667 | 23 | 0.2174 |
| 10_597862 | 12 | 0.9167 | 7 | 0.8571 | 3 | 1 | 7 | 1 | 29 | 0.931 | 5 | 0.8 | 8 | 0.875 | 6 | 0.8333 | 7 | 0.8571 | 26 | 0.8462 |
| 10_597862 | 11 | 1 | 5 | 0.6 | 3 | 0.6667 | 7 | 0.7143 | 26 | 0.8077 | 6 | 0.8333 | 5 | 0.8 | 8 | 1 | 7 | 0.8571 | 26 | 0.8846 |
| 10_597863 | 7 | 0.2857 | 7 | 0.1429 | 2 | 0 | 5 | 0.4 | 21 | 0.2381 | 4 | 0 | 8 | 0.375 | 7 | 0.2857 | 7 | 0.1429 | 26 | 0.2308 |
| 10_597863 | 8 | 0 | 4 | 0.5 | 2 | 0 | 6 | 0 | 20 | 0.1 | 5 | 0 | 8 | 0 | 7 | 0.1429 | 9 | 0.3333 | 29 | 0.1379 |
| 10_597865 | 6 | 1 | 6 | 1 | 1 | 1 | 6 | 1 | 19 | 1 | 4 | 1 | 6 | 1 | 4 | 1 | 9 | 1 | 23 | 1 |
| 10_597866 | 7 | 0.4286 | 6 | 0.3333 | 1 | 0 | 5 | 0.2 | 19 | 0.3158 | 2 | 0.5 | 5 | 0.4 | 5 | 0.2 | 9 | 0.4444 | 21 | 0.381 |
| 10_598241 | 8 | 0.5 | 9 | 0.4444 | 12 | 0.4167 | 10 | 0.3 | 39 | 0.4103 | 9 | 0.3333 | 6 | 0.1667 | 8 | 0.25 | 4 | 0.25 | 27 | 0.2593 |
| 10_598242 | 15 | 0 | 12 | 0 | 18 | 0.0556 | 15 | 0.1333 | 60 | 0.05 | 9 | 0 | 11 | 0 | 13 | 0.1538 | 7 | 0 | 40 | 0.05 |
| 10_599422 | 13 | 0.1538 | 10 | 0.3 | 9 | 0 | 7 | 0.1429 | 39 | 0.1538 | 11 | 0 | 18 | 0.0556 | 8 | 0.375 | 12 | 0.1667 | 49 | 0.1224 |
| 10_599671 | 18 | 0 | 20 | 0.05 | 13 | 0.2308 | 22 | 0.2727 | 73 | 0.137 | 12 | 0.1667 | 31 | 0.2258 | 21 | 0.1429 | 19 | 0.1053 | 83 | 0.1687 |
| 10_599673 | 24 | 0.0833 | 27 | 0.1481 | 18 | 0.1667 | 24 | 0.0833 | 93 | 0.1183 | 18 | 0.1667 | 30 | 0.1 | 28 | 0.25 | 22 | 0.2273 | 98 | 0.1837 |
| 10_599673 | 20 | 0.05 | 20 | 0.1 | 13 | 0.0769 | 21 | 0 | 74 | 0.0541 | 15 | 0.2 | 28 | 0.1429 | 23 | 0.3043 | 18 | 0.1667 | 84 | 0.2024 |
| 10_599674 | 18 | 0.1667 | 23 | 0.087 | 13 | 0.3846 | 21 | 0.1905 | 75 | 0.1867 | 11 | 0.0909 | 29 | 0.2759 | 25 | 0.32 | 22 | 0.2727 | 87 | 0.2644 |
| 10_599690 | 42 | 0.119 | 43 | 0.0698 | 37 | 0.0541 | 36 | 0.0833 | 158 | 0.0823 | 30 | 0.0667 | 43 | 0.0465 | 52 | 0.1346 | 39 | 0.2051 | 164 | 0.1159 |
| 10_599690 | 44 | 0.1364 | 49 | 0.0204 | 43 | 0.0233 | 42 | 0.1429 | 178 | 0.0787 | 33 | 0.0909 | 49 | 0.102 | 54 | 0.037 | 39 | 0.1026 | 175 | 0.08 |
| 10_599691 | 38 | 0.7895 | 46 | 0.8696 | 39 | 0.8462 | 39 | 0.7436 | 162 | 0.8148 | 29 | 0.8966 | 46 | 0.8478 | 46 | 0.8696 | 35 | 0.8857 | 156 | 0.8718 |
| 10_599693 | 36 | 0.1389 | 47 | 0.0638 | 36 | 0.1111 | 35 | 0.1143 | 154 | 0.1039 | 29 | 0.1379 | 39 | 0 | 43 | 0.0698 | 40 | 0.025 | 151 | 0.053 |
| 10_599697 | 48 | 0.0208 | 52 | 0.0385 | 42 | 0.0952 | 52 | 0.1154 | 194 | 0.067 | 38 | 0.0526 | 57 | 0.0526 | 59 | 0.0847 | 52 | 0.0769 | 206 | 0.068 |
| 10_600192 | NA | NA | NA | NA | 1 | 1 | NA | NA | 1 | 1 | NA | NA | 1 | 1 | NA | NA | NA | NA | 1 | 1 |
| 10_600193 | NA | NA | NA | NA | 1 | 1 | NA | NA | 1 | 1 | NA | NA | 1 | 1 | NA | NA | NA | NA | 1 | 1 |
| 10_601382 | 207 | 0.2512 | 219 | 0.2922 | 186 | 0.2151 | 176 | 0.2386 | 788 | 0.2513 | 189 | 0.2857 | 157 | 0.2675 | 159 | 0.2642 | 161 | 0.2733 | 666 | 0.2733 |
| 10_602047 | 2 | 0.5 | 5 | 0.4 | 2 | 0 | 1 | 0 | 10 | 0.3 | 1 | 0 | 5 | 0 | 2 | 0.5 | 4 | 0.5 | 12 | 0.25 |
| 10_602047 | NA | NA | 5 | 0.8 | 1 | 0 | 2 | 1 | 8 | 0.75 | NA | NA | 4 | 0.75 | 1 | 1 | 5 | 0.8 | 10 | 0.8 |
| 10_602048 | 2 | 0.5 | 5 | 0.6 | 1 | 0 | 2 | 0 | 10 | 0.4 | NA | NA | 5 | 0.2 | 2 | 0.5 | 5 | 0.4 | 12 | 0.3333 |
| 10_602048 | 2 | 0.5 | 4 | 0.25 | 1 | 0 | 1 | 0 | 8 | 0.25 | NA | NA | 4 | 0 | 2 | 1 | 5 | 0.4 | 11 | 0.3636 |
| 10_602049 | 2 | 1 | 6 | 1 | 2 | 1 | 2 | 1 | 12 | 1 | 1 | 1 | 3 | 1 | 2 | 0.5 | 5 | 0.6 | 11 | 0.7273 |

| | | | | | | | | | | | | | | | | | | | | |
|---|---|---|---|---|---|---|---|---|---|---|---|---|---|---|---|---|---|---|---|---|
| 10_602051 | 2 | 1 | 4 | 1 | 1 | 1 | 2 | 1 | 9 | 1 | 1 | 1 | 4 | 0.5 | 2 | 0.5 | 5 | 0.6 | 12 | 0.5833 |
| 10_602051 | 2 | 0.5 | 6 | 0.8333 | 2 | 1 | 2 | 1 | 12 | 0.8333 | NA | NA | 5 | 0.8 | 2 | 1 | 5 | 1 | 12 | 0.9167 |
| 10_602051 | 1 | 1 | 4 | 1 | 1 | 1 | 2 | 1 | 8 | 1 | 1 | 1 | 3 | 1 | 1 | 1 | 4 | 1 | 9 | 1 |
| 10_602051 | 1 | 0 | 3 | 0 | 2 | 1 | 2 | 0.5 | 8 | 0.375 | 1 | 1 | 3 | 0.6667 | 1 | 1 | 3 | 0.6667 | 8 | 0.75 |
| 10_602054 | 1 | 0 | 5 | 0.4 | 2 | 0.5 | 2 | 1 | 10 | 0.5 | 1 | 0 | 3 | 0.3333 | 3 | 0.3333 | 4 | 0.75 | 11 | 0.4545 |
| 10_602054 | 1 | 1 | 3 | 1 | 1 | 1 | NA | NA | 5 | 1 | 1 | 1 | 2 | 1 | 3 | 1 | 4 | 0.75 | 10 | 0.9 |
| 10_602056 | 1 | 1 | NA | NA | NA | NA | NA | NA | 1 | 1 | NA | NA | NA | NA | 1 | 1 | 2 | 0.5 | 3 | 0.6667 |
| 10_602073 | 2 | 0 | NA | NA | NA | NA | 1 | 1 | 3 | 0.3333 | 1 | 0 | NA | NA | 2 | 0 | 1 | 1 | 4 | 0.25 |
| 10_605518 | 1 | 1 | 1 | 1 | NA | NA | NA | NA | 2 | 1 | 1 | 1 | NA | NA | NA | NA | NA | NA | 1 | 1 |
| 10_605520 | 1 | 1 | 1 | 1 | NA | NA | NA | NA | 2 | 1 | 1 | 1 | NA | NA | NA | NA | NA | NA | 1 | 1 |
| 10_609365 | 3 | 1 | 4 | 0.75 | 2 | 1 | 3 | 0.6667 | 12 | 0.8333 | 3 | 0.3333 | 1 | 1 | 2 | 0.5 | 4 | 0.5 | 10 | 0.5 |
| 10_609365 | 2 | 0.5 | 4 | 0.75 | 2 | 1 | 3 | 1 | 11 | 0.8182 | 3 | 1 | 1 | 1 | 2 | 0.5 | 4 | 0.75 | 10 | 0.8 |
| 10_609365 | 3 | 0 | 4 | 0.5 | 1 | 0 | 3 | 0 | 11 | 0.1818 | 3 | 0.6667 | 1 | 0 | 2 | 0.5 | 4 | 0.5 | 10 | 0.5 |
| 10_609366 | 3 | 0 | 4 | 0.25 | 2 | 0.5 | 3 | 0 | 12 | 0.1667 | 2 | 0.5 | 1 | 0 | 1 | 1 | 4 | 0 | 8 | 0.25 |
| 10_609366 | 3 | 0.3333 | 4 | 0.25 | 1 | 0 | 3 | 0 | 11 | 0.1818 | 2 | 0 | 1 | 0 | 3 | 0 | 3 | 0.3333 | 9 | 0.1111 |
| 10_609367 | 3 | 0.6667 | 2 | 1 | 2 | 0 | 3 | 1 | 10 | 0.7 | 3 | 0.6667 | 1 | 0 | 3 | 0 | 3 | 0.3333 | 10 | 0.3 |
| 10_611568 | NA | NA | NA | NA | 1 | 1 | NA | NA | 1 | 1 | NA | NA | 1 | 0 | 1 | 1 | 1 | 0 | 3 | 0.3333 |
| 10_612645 | 13 | 0.2308 | 14 | 0.2143 | 6 | 0.1667 | 9 | 0.2222 | 42 | 0.2143 | 8 | 0.125 | 2 | 0 | 6 | 0.1667 | 5 | 0 | 21 | 0.0952 |
| 10_612647 | 13 | 0 | 11 | 0.0909 | 5 | 0.2 | 6 | 0.1667 | 35 | 0.0857 | 6 | 0 | 1 | 0 | 5 | 0 | 4 | 0.25 | 16 | 0.0625 |
| 10_612647 | 6 | 0 | 7 | 0 | 3 | 0 | 3 | 0.3333 | 19 | 0.0526 | 4 | 0 | 1 | 0 | 4 | 0.25 | 4 | 0 | 13 | 0.0769 |
| 10_612654 | 6 | 0 | 9 | 0.3333 | 2 | 0.5 | 3 | 0 | 20 | 0.2 | 3 | 0.6667 | NA | NA | 2 | 0 | 4 | 0.5 | 9 | 0.4444 |
| 10_612656 | 8 | 0 | 11 | 0.2727 | 3 | 0 | 6 | 0 | 28 | 0.1071 | 5 | 0 | 2 | 0 | 4 | 0.5 | 4 | 0 | 15 | 0.1333 |
| 10_612657 | 8 | 0.5 | 10 | 0.5 | 2 | 0.5 | 5 | 0.6 | 25 | 0.52 | 4 | 0.25 | 1 | 0 | 3 | 0 | 4 | 0.75 | 12 | 0.3333 |
| 10_612661 | 3 | 0.3333 | 8 | 0.375 | 4 | 0 | 5 | 0.2 | 20 | 0.25 | 5 | 0.2 | 2 | 0.5 | 1 | 0 | 3 | 0 | 11 | 0.1818 |
| 10_612672 | 6 | 0.6667 | 8 | 0.75 | 3 | 1 | 4 | 1 | 21 | 0.8095 | 5 | 0.6 | 1 | 1 | 2 | 1 | 3 | 1 | 11 | 0.8182 |
| 10_612673 | 6 | 0.3333 | 8 | 0.75 | 3 | 0.3333 | 2 | 1 | 19 | 0.5789 | 3 | 0.3333 | 2 | 0.5 | 2 | 0.5 | 2 | 0.5 | 9 | 0.4444 |
| 10_612674 | 4 | 1 | 7 | 0.8571 | 3 | 1 | 2 | 1 | 16 | 0.9375 | 3 | 1 | 2 | 1 | 2 | 1 | 2 | 1 | 9 | 1 |
| 10_612675 | 5 | 0 | 7 | 0.2857 | 3 | 0 | 3 | 0.6667 | 18 | 0.2222 | 4 | 0.5 | 2 | 0.5 | 1 | 0 | 3 | 0.6667 | 10 | 0.5 |
| 10_612677 | 5 | 0.6 | 9 | 0.3333 | 3 | 0.6667 | 2 | 1 | 19 | 0.5263 | 5 | 0.6 | 3 | 0.3333 | 1 | 0 | 4 | 0.75 | 13 | 0.5385 |
| 10_612678 | 3 | 0 | 8 | 0.25 | 4 | 0.25 | 2 | 0.5 | 17 | 0.2353 | 5 | 0 | 3 | 0 | 1 | 0 | 3 | 0.6667 | 12 | 0.1667 |
| 10_612678 | 5 | 0 | 8 | 0 | 4 | 0 | 1 | 1 | 18 | 0.0556 | 5 | 0.2 | 3 | 0 | 1 | 0 | 4 | 0 | 13 | 0.0769 |
| 10_612678 | 4 | 0 | 6 | 0 | 3 | 0.3333 | 1 | 0 | 14 | 0.0714 | 3 | 0 | 2 | 0 | 1 | 0 | 2 | 0.5 | 8 | 0.125 |
| 10_612689 | 2 | 0.5 | 3 | 0 | 1 | 0 | 2 | 0 | 8 | 0.125 | 3 | 0.3333 | 1 | 0 | 1 | 0 | 2 | 0 | 7 | 0.1429 |
| 10_614160 | 1 | 0 | 1 | 1 | 1 | 1 | NA | NA | 3 | 0.6667 | 3 | 1 | 3 | 0.6667 | 1 | 1 | 1 | 0 | 8 | 0.75 |
| 10_614160 | 1 | 0 | 1 | 1 | 1 | 0 | 1 | 0 | 4 | 0.25 | 3 | 0 | 4 | 0.25 | 1 | 1 | NA | NA | 8 | 0.25 |
| 10_614163 | 1 | 0 | 2 | 1 | NA | NA | 2 | 0 | 5 | 0.4 | 3 | 0 | 3 | 0.3333 | 1 | 0 | 1 | 0 | 8 | 0.125 |
| 10_614164 | 1 | 1 | 3 | 1 | 1 | 1 | 2 | 1 | 7 | 1 | 2 | 1 | 4 | 1 | 1 | 1 | 1 | 1 | 8 | 1 |
| 10_614165 | 1 | 0 | NA | NA | NA | NA | 2 | 0.5 | 3 | 0.3333 | 1 | 1 | 3 | 0 | 1 | 0 | 1 | 0 | 6 | 0.1667 |
| 10_614165 | 1 | 1 | 1 | 1 | 1 | 1 | 2 | 1 | 5 | 1 | 2 | 1 | 2 | 1 | 3 | 1 | 1 | 1 | 8 | 1 |
| 10_614166 | 1 | 1 | 2 | 0 | 1 | 1 | 1 | 0 | 5 | 0.4 | 3 | 0.3333 | 3 | 0.3333 | 3 | 0.3333 | 1 | 0 | 10 | 0.3 |
| 10_614166 | 1 | 0 | 2 | 1 | 2 | 0 | 2 | 0 | 7 | 0.2857 | 3 | 0.6667 | 4 | 0.5 | 3 | 0.3333 | NA | NA | 10 | 0.5 |
| 10_614167 | 1 | 1 | 3 | 1 | 2 | 1 | 2 | 1 | 8 | 1 | 3 | 1 | 4 | 1 | 2 | 0.5 | 1 | 1 | 10 | 0.9 |
| 10_614167 | 1 | 0 | 3 | 1 | 2 | 1 | 2 | 0 | 8 | 0.625 | 3 | 0.6667 | 3 | 1 | 3 | 0.3333 | 1 | 0 | 10 | 0.6 |
| 10_614170 | 2 | 0.5 | 2 | 0.5 | 1 | 1 | 2 | 0.5 | 7 | 0.5714 | 4 | 0.25 | 5 | 0.6 | 3 | 0.3333 | 1 | 0 | 13 | 0.3846 |
| 10_614174 | 3 | 0 | 4 | 0 | 2 | 0.5 | 3 | 0 | 12 | 0.0833 | 3 | 0 | 2 | 0.5 | 4 | 0 | 1 | 0 | 10 | 0.1 |
| 10_615097 | 21 | 0.0476 | 22 | 0.0455 | 17 | 0.0588 | 17 | 0.0588 | 77 | 0.0519 | 11 | 0 | 16 | 0.0625 | 17 | 0.0588 | 7 | 0.1429 | 51 | 0.0588 |
| 10_615097 | 23 | 0.0435 | 22 | 0.0455 | 18 | 0.0556 | 17 | 0.0588 | 80 | 0.05 | 11 | 0 | 18 | 0.1111 | 17 | 0.0588 | 7 | 0.1429 | 53 | 0.0755 |
| 10_615097 | 21 | 0.2381 | 21 | 0.1429 | 17 | 0.2353 | 16 | 0.25 | 75 | 0.2133 | 10 | 0.3 | 16 | 0.3125 | 18 | 0.1111 | 8 | 0 | 52 | 0.1923 |
| 10_615271 | 15 | 0.3333 | 9 | 0.3333 | 9 | 0.4444 | 10 | 0.5 | 43 | 0.3953 | 11 | 0.2727 | 13 | 0.6154 | 10 | 0.6 | 14 | 0.4286 | 48 | 0.4792 |
| 10_615271 | 16 | 0.8125 | 11 | 0.5455 | 12 | 0.5833 | 15 | 0.4667 | 54 | 0.6111 | 12 | 0.5833 | 12 | 0.9167 | 7 | 0.5714 | 13 | 0.8462 | 44 | 0.75 |
| 10_615272 | 17 | 0.8235 | 9 | 0.6667 | 10 | 0.5 | 10 | 0.7 | 46 | 0.6957 | 13 | 0.8462 | 9 | 0.6667 | 9 | 0.6667 | 13 | 0.8462 | 44 | 0.7727 |
| 10_615272 | 14 | 0.9286 | 10 | 1 | 11 | 1 | 12 | 0.8333 | 47 | 0.9362 | 13 | 1 | 13 | 1 | 9 | 0.7778 | 15 | 0.9333 | 50 | 0.94 |
| 10_615273 | 13 | 0.2308 | 8 | 0.375 | 8 | 0.125 | 9 | 0.1111 | 38 | 0.2105 | 7 | 0.1429 | 7 | 0.2857 | 7 | 0 | 14 | 0.2143 | 35 | 0.1714 |
| 10_615280 | 17 | 0.2353 | 12 | 0 | 14 | 0.2143 | 15 | 0.1333 | 58 | 0.1552 | 16 | 0.1875 | 16 | 0.1875 | 13 | 0.0769 | 17 | 0.2353 | 62 | 0.1774 |
| 10_615286 | 21 | 0.2381 | 16 | 0.125 | 18 | 0.2222 | 23 | 0.2174 | 78 | 0.2051 | 22 | 0.0909 | 19 | 0.3158 | 15 | 0.2667 | 23 | 0.1739 | 79 | 0.2025 |
| 10_615287 | 21 | 0.4286 | 10 | 0.5 | 16 | 0.25 | 23 | 0.6087 | 70 | 0.4571 | 21 | 0.5714 | 15 | 0.6667 | 12 | 0.3333 | 18 | 0.3889 | 66 | 0.5 |
| 10_615289 | 22 | 0.2273 | 13 | 0.3077 | 20 | 0.3 | 22 | 0.5909 | 77 | 0.3636 | 23 | 0.4783 | 19 | 0.5789 | 17 | 0.4118 | 18 | 0.5 | 77 | 0.4935 |
| 10_615289 | 22 | 0.1818 | 14 | 0.2857 | 18 | 0.1111 | 29 | 0.2069 | 83 | 0.1928 | 23 | 0.087 | 19 | 0.1579 | 21 | 0.1905 | 25 | 0.08 | 88 | 0.125 |
| 10_615316 | 2 | 0.5 | NA | NA | 2 | 1 | 1 | 1 | 5 | 0.8 | NA | NA | 1 | 1 | 1 | 1 | 1 | 0 | 3 | 0.6667 |
| 10_615319 | 3 | 0.3333 | 1 | 0 | 1 | 1 | 3 | 0.3333 | 8 | 0.375 | NA | NA | 4 | 0.5 | 1 | 1 | 2 | 0 | 7 | 0.4286 |
| 10_615320 | 3 | 0.6667 | 2 | 0.5 | 3 | 0.6667 | 3 | 0.6667 | 11 | 0.6364 | NA | NA | 4 | 1 | 2 | 1 | 2 | 0.5 | 8 | 0.875 |
| 10_615323 | 3 | 0.3333 | 1 | 0 | 2 | 0.5 | 3 | 0 | 9 | 0.2222 | NA | NA | 4 | 0 | 1 | 1 | 3 | 0.3333 | 8 | 0.25 |
| 10_615379 | 5 | 0.2 | 3 | 0.3333 | 4 | 0.75 | 3 | 0.6667 | 15 | 0.4667 | NA | NA | 4 | 0.25 | 3 | 0.3333 | 3 | 0.3333 | 10 | 0.3 |
| 10_615380 | 5 | 0.6 | 1 | 0 | 2 | 0.5 | 3 | 0 | 11 | 0.3636 | NA | NA | 4 | 0.5 | 1 | 1 | 2 | 0.5 | 7 | 0.5714 |
| 10_615381 | 5 | 0 | 2 | 0 | 3 | 0.3333 | 4 | 0 | 14 | 0.0714 | NA | NA | 4 | 0 | 3 | 0.3333 | 3 | 0 | 10 | 0.1 |

| | | | | | | | | | | | | | | | | | | | | |
|---|---|---|---|---|---|---|---|---|---|---|---|---|---|---|---|---|---|---|---|---|
| 10_6153824 | 4 | 0.75 | 2 | 0.5 | 4 | 0.5 | 3 | 1 | 13 | 0.6923 | NA | NA | 2 | 0.5 | 3 | 1 | 2 | 0.5 | 7 | 0.7143 |
| 10_6153837 | 4 | 1 | 1 | 1 | 3 | 1 | 2 | 0.5 | 10 | 0.9 | NA | NA | 2 | 0.5 | 2 | 1 | 3 | 1 | 7 | 0.8571 |
| 10_6153847 | 4 | 0 | 2 | 0 | 4 | 0 | 5 | 0.2 | 15 | 0.0667 | 1 | 0 | 1 | 0 | 4 | 0.25 | 3 | 0 | 9 | 0.1111 |
| 10_6157418 | 1 | 1 | 1 | 1 | NA | NA | 2 | 0.5 | 4 | 0.75 | 1 | 0 | NA | NA | 2 | 1 | NA | NA | 3 | 0.6667 |
| 10_6157431 | 1 | 1 | 2 | 0.5 | 1 | 1 | 1 | 0 | 5 | 0.6 | 1 | 1 | NA | NA | 2 | 0 | NA | NA | 3 | 0.3333 |
| 10_6157474 | 6 | 0 | 7 | 0.1429 | 1 | 0 | 3 | 0 | 17 | 0.0588 | 6 | 0.1667 | 1 | 0 | 6 | 0.1667 | 4 | 0 | 17 | 0.1176 |
| 10_6157480 | 5 | 0 | 7 | 0.1429 | NA | NA | 3 | 0.3333 | 15 | 0.1333 | 4 | 0.25 | 1 | 0 | 7 | 0.1429 | 4 | 0 | 16 | 0.125 |
| 10_6157508 | 5 | 0.4 | 2 | 0 | 1 | 1 | 3 | 0.3333 | 11 | 0.3636 | 2 | 0.5 | 1 | 0 | 7 | 0.1429 | 3 | 0.3333 | 13 | 0.2308 |
| 10_6157538 | NA | NA | 1 | 1 | NA | NA | NA | NA | 1 | 1 | 1 | 1 | 1 | 1 | NA | NA | NA | NA | 2 | 1 |
| 10_6157571 | NA | NA | 1 | 1 | NA | NA | NA | NA | 1 | 1 | NA | NA | 1 | 1 | NA | NA | NA | NA | 1 | 1 |
| 10_6161965 | NA | NA | 1 | 0 | 4 | 0.25 | 1 | 0 | 6 | 0.1667 | NA | NA | 1 | 0 | 1 | 0 | 2 | 1 | 4 | 0.5 |
| 10_6161992 | NA | NA | 1 | 0 | 4 | 0.25 | 2 | 0 | 7 | 0.1429 | NA | NA | 2 | 0.5 | 2 | 1 | 3 | 0 | 7 | 0.4286 |
| 10_6162014 | NA | NA | NA | NA | 4 | 0.25 | 1 | 0 | 5 | 0.2 | NA | NA | 1 | 0 | 2 | 0 | 3 | 0.3333 | 6 | 0.1667 |
| 10_6162035 | NA | NA | 1 | 1 | 4 | 1 | 1 | 1 | 6 | 1 | NA | NA | 2 | 1 | 1 | 1 | 2 | 1 | 5 | 1 |
| 10_6162036 | NA | NA | 1 | 1 | 4 | 0 | 2 | 0.5 | 7 | 0.2857 | NA | NA | 2 | 1 | 1 | 1 | 4 | 0 | 7 | 0.4286 |
| 10_6162053 | NA | NA | 1 | 1 | 4 | 1 | 2 | 1 | 7 | 1 | 1 | 1 | 2 | 1 | 3 | 1 | 3 | 0.6667 | 9 | 0.8889 |
| 10_6194429 | 5 | 0 | 8 | 0 | 3 | 0.3333 | 5 | 0.2 | 21 | 0.0952 | 5 | 0 | 7 | 0.1429 | 3 | 0.3333 | 3 | 0 | 18 | 0.1111 |
| 10_6194458 | 5 | 0.4 | 4 | 1 | 1 | 1 | 4 | 0.25 | 14 | 0.5714 | 3 | 0.6667 | 4 | 0.5 | 2 | 1 | 1 | 1 | 10 | 0.7 |
| 10_6194462 | 5 | 0.6 | 4 | 0.75 | NA | NA | 4 | 0.75 | 13 | 0.6923 | 4 | 0.75 | 5 | 0.2 | 2 | 0 | NA | NA | 11 | 0.3636 |
| 10_6202336 | 1 | 1 | 1 | 1 | 3 | 0.3333 | 4 | 1 | 9 | 0.7778 | 2 | 1 | 1 | 0 | 1 | 1 | 3 | 1 | 7 | 0.8571 |
| 10_6202372 | 2 | 0.5 | 1 | 0 | 3 | 0.3333 | 3 | 0 | 9 | 0.2222 | 2 | 0.5 | 1 | 0 | 2 | 0 | 1 | 0 | 6 | 0.1667 |
| 10_6202374 | 1 | 1 | 1 | 0 | 3 | 0.3333 | 4 | 1 | 9 | 0.6667 | 2 | 0.5 | 1 | 1 | 2 | 0.5 | 2 | 0.5 | 7 | 0.5714 |
| 10_6202383 | 2 | 0 | 1 | 0 | 3 | 0 | 3 | 1 | 9 | 0.3333 | 1 | 0 | 1 | 1 | 2 | 0 | 2 | 0 | 6 | 0.1667 |
| 10_6202389 | 2 | 0.5 | 1 | 0 | 2 | 0.5 | 3 | 0.3333 | 8 | 0.375 | 2 | 0 | 1 | 0 | NA | NA | 2 | 0.5 | 5 | 0.2 |
| 10_6210369 | NA | NA | NA | NA | 2 | 0.5 | NA | NA | 2 | 0.5 | NA | NA | 1 | 1 | NA | NA | NA | NA | 1 | 1 |
| 10_6210480 | 2 | 1 | 1 | 1 | 1 | 0 | 1 | 0 | 5 | 0.6 | 1 | 1 | 1 | 1 | NA | NA | NA | NA | 2 | 1 |
| 10_6210507 | 28 | 0.4286 | 24 | 0.2083 | 18 | 0.2778 | 21 | 0.2857 | 91 | 0.3077 | 31 | 0.2903 | 15 | 0.2 | 33 | 0.4242 | 22 | 0.3636 | 101 | 0.3366 |
| 10_6210519 | 32 | 0.2188 | 30 | 0.2 | 21 | 0.3333 | 23 | 0.087 | 106 | 0.2075 | 30 | 0.3 | 17 | 0.2941 | 34 | 0.2647 | 22 | 0.2727 | 103 | 0.2816 |
| 10_6210523 | 30 | 0.9667 | 32 | 1 | 22 | 1 | 22 | 0.9091 | 106 | 0.9717 | 36 | 0.9722 | 18 | 1 | 37 | 1 | 26 | 1 | 117 | 0.9915 |
| 10_6210524 | 35 | 0.2857 | 28 | 0.3214 | 18 | 0.2778 | 21 | 0.0476 | 102 | 0.2451 | 31 | 0.3226 | 14 | 0.2143 | 33 | 0.3333 | 21 | 0.381 | 99 | 0.3232 |
| 10_6210533 | 23 | 1 | 23 | 1 | 18 | 0.9444 | 20 | 0.95 | 84 | 0.9762 | 24 | 1 | 13 | 1 | 26 | 0.8846 | 18 | 1 | 81 | 0.963 |
| 10_6210546 | 28 | 0.1429 | 24 | 0.1667 | 20 | 0.25 | 17 | 0.4118 | 89 | 0.2247 | 30 | 0.1 | 16 | 0.25 | 25 | 0.2 | 22 | 0.1364 | 93 | 0.1613 |
| 10_6210553 | 26 | 0.6154 | 23 | 0.6087 | 22 | 0.3636 | 19 | 0.7895 | 90 | 0.5889 | 31 | 0.5806 | 18 | 0.7222 | 25 | 0.68 | 20 | 0.75 | 94 | 0.6702 |
| 10_6210580 | 29 | 0.1379 | 23 | 0.087 | 18 | 0.3333 | 17 | 0.4118 | 87 | 0.2184 | 24 | 0.125 | 17 | 0.2353 | 23 | 0.0435 | 22 | 0.2727 | 86 | 0.1628 |
| 10_6212213 | 16 | 0.125 | 15 | 0.0667 | 13 | 0.0769 | 9 | 0 | 53 | 0.0755 | 17 | 0.0588 | 8 | 0.125 | 13 | 0.0769 | 11 | 0 | 49 | 0.0612 |
| 10_6226378 | NA | NA | 1 | 1 | NA | NA | NA | NA | 1 | 1 | NA | NA | 1 | 1 | 2 | 0.5 | NA | NA | 3 | 0.6667 |
| 10_6226393 | 1 | 1 | NA | NA | NA | NA | NA | NA | 1 | 1 | NA | NA | 1 | 1 | 2 | 1 | NA | NA | 3 | 1 |
| 10_6226530 | 2 | 0 | 1 | 1 | 1 | 0 | 3 | 0 | 7 | 0.1429 | NA | NA | 2 | 0.5 | 2 | 0 | 5 | 0.2 | 9 | 0.2222 |
| 10_6226531 | 3 | 1 | 1 | 1 | 1 | 1 | 4 | 1 | 9 | 1 | NA | NA | 2 | 1 | 2 | 0.5 | 5 | 0.8 | 9 | 0.7778 |
| 10_6226541 | 3 | 0.6667 | NA | NA | 1 | 0 | 4 | 0.75 | 8 | 0.625 | NA | NA | 2 | 1 | 2 | 0 | 4 | 0.5 | 8 | 0.5 |
| 10_6228674 | NA | NA | NA | NA | 1 | 1 | NA | NA | 1 | 1 | NA | NA | 1 | 1 | NA | NA | NA | NA | 1 | 1 |
| 10_6228675 | NA | NA | NA | NA | 1 | 1 | NA | NA | 1 | 1 | NA | NA | 1 | 1 | NA | NA | NA | NA | 1 | 1 |
| 10_6228702 | NA | NA | NA | NA | 1 | 1 | NA | NA | 1 | 1 | NA | NA | 1 | 1 | NA | NA | NA | NA | 1 | 1 |
| 10_6229176 | 9 | 0.1111 | 12 | 0.1667 | 5 | 0 | 7 | 0 | 33 | 0.0909 | 7 | 0 | 3 | 0.3333 | 2 | 0 | 3 | 0 | 15 | 0.0667 |
| 10_6229223 | 7 | 0.1429 | 11 | 0 | 6 | 0.1667 | 8 | 0 | 32 | 0.0625 | 7 | 0.1429 | 4 | 0.25 | 5 | 0 | 4 | 0 | 20 | 0.1 |
| 10_6229258 | 7 | 0 | 9 | 0.2222 | 4 | 0.25 | 7 | 0.1429 | 27 | 0.1481 | 5 | 0.2 | 4 | 0 | 4 | 0 | 4 | 0 | 17 | 0.0588 |
| 10_6233031 | 5 | 0.2 | 1 | 0 | 3 | 0 | 1 | 0 | 10 | 0.1 | 3 | 0.3333 | 3 | 0 | 3 | 0.3333 | 3 | 0.3333 | 12 | 0.25 |
| 10_6233051 | 8 | 0.875 | 1 | 0 | 5 | 0.6 | 1 | 1 | 15 | 0.7333 | 2 | 0.5 | 4 | 1 | 3 | 0 | 2 | 1 | 11 | 0.6364 |
| 10_6233060 | 7 | 0 | 1 | 0 | 4 | 0.5 | 1 | 0 | 13 | 0.1538 | 3 | 0.3333 | 3 | 0.3333 | 2 | 0 | 2 | 0 | 10 | 0.2 |
| 10_6233084 | 6 | 0.1667 | NA | NA | 4 | 0 | NA | NA | 10 | 0.1 | 2 | 0 | 3 | 0.3333 | 3 | 0 | 2 | 0 | 10 | 0.1 |
| 10_6239493 | 16 | 0.25 | 19 | 0.3158 | 20 | 0.2 | 21 | 0.0476 | 76 | 0.1974 | 9 | 0 | 17 | 0.0588 | 15 | 0.2 | 18 | 0.1111 | 59 | 0.1017 |
| 10_6239507 | 15 | 0.2 | 20 | 0.15 | 20 | 0 | 19 | 0.2105 | 74 | 0.1351 | 7 | 0.1429 | 14 | 0.0714 | 14 | 0.2143 | 15 | 0.1333 | 50 | 0.14 |
| 10_6239508 | 16 | 0.125 | 20 | 0.1 | 21 | 0.0476 | 21 | 0 | 78 | 0.0641 | 8 | 0 | 15 | 0.0667 | 14 | 0 | 17 | 0.1765 | 54 | 0.0741 |
| 10_6239516 | 15 | 0.1333 | 20 | 0.25 | 17 | 0.0588 | 21 | 0.2381 | 73 | 0.1781 | 9 | 0 | 17 | 0.1176 | 14 | 0.0714 | 17 | 0.2353 | 57 | 0.1228 |
| 10_6239519 | 16 | 0.8125 | 19 | 0.7895 | 15 | 0.8667 | 17 | 0.9412 | 67 | 0.8507 | 8 | 1 | 17 | 0.7647 | 13 | 0.9231 | 14 | 1 | 52 | 0.9038 |
| 10_6239527 | 15 | 0.1333 | 14 | 0.0714 | 15 | 0 | 18 | 0.3889 | 62 | 0.1613 | 9 | 0.3333 | 17 | 0.2941 | 14 | 0.1429 | 12 | 0 | 52 | 0.1923 |
| 10_6239528 | 11 | 0.0909 | 18 | 0.1111 | 14 | 0.2143 | 19 | 0.1053 | 62 | 0.129 | 7 | 0.2857 | 14 | 0.0714 | 15 | 0.1333 | 14 | 0.2143 | 50 | 0.16 |
| 10_6239537 | 14 | 0.1429 | 19 | 0.2632 | 18 | 0 | 16 | 0.0625 | 67 | 0.1194 | 8 | 0 | 11 | 0 | 14 | 0.0714 | 14 | 0.1429 | 47 | 0.0638 |
| 10_6239555 | 13 | 0.3846 | 17 | 0.1176 | 13 | 0.0769 | 16 | 0.1875 | 59 | 0.1864 | 6 | 0.1667 | 14 | 0.3571 | 13 | 0.2308 | 14 | 0.2143 | 47 | 0.2553 |
| 10_6241383 | 11 | 0.0909 | 12 | 0.1667 | 11 | 0.1818 | 12 | 0.25 | 46 | 0.1739 | 2 | 0.5 | 12 | 0.25 | 8 | 0.25 | 9 | 0.1111 | 31 | 0.2258 |
| 10_6241617 | 3 | 0 | 1 | 0 | 2 | 0.5 | 2 | 0.5 | 8 | 0.25 | NA | NA | 3 | 0 | 2 | 0 | 3 | 0.3333 | 8 | 0.125 |
| 10_6241800 | 1 | 1 | NA | NA | 1 | 1 | 4 | 0.75 | 6 | 0.8333 | 1 | 1 | 3 | 0.6667 | 2 | 1 | 3 | 1 | 9 | 0.8889 |
| 10_6241811 | NA | NA | NA | NA | 2 | 1 | 4 | 0.75 | 6 | 0.8333 | 1 | 1 | 4 | 1 | 2 | 0.5 | 2 | 1 | 9 | 0.8889 |
| 10_6241822 | 1 | 0 | NA | NA | 2 | 0.5 | 3 | 0 | 6 | 0.1667 | NA | NA | 3 | 0.3333 | 1 | 0 | 2 | 0.5 | 6 | 0.3333 |
| 10_6248283 | 5 | 0.8 | 4 | 0.75 | 3 | 0.6667 | 7 | 0.4286 | 19 | 0.6316 | 1 | 1 | 8 | 0.5 | 6 | 0.5 | 3 | 0 | 18 | 0.4444 |

| | | | | | | | | | | | | | | | | | | | | |
|---|---|---|---|---|---|---|---|---|---|---|---|---|---|---|---|---|---|---|---|---|
| 10_6249513 | 1 | 1 | NA | NA | 1 | 1 | NA | NA | 2 | 1 | 3 | 0.6667 | 1 | 1 | NA | NA | NA | NA | 4 | 0.75 |
| 10_6249514 | 1 | 0 | 1 | 1 | 1 | 0 | NA | NA | 3 | 0.3333 | 2 | 0.5 | 1 | 1 | NA | NA | NA | NA | 3 | 0.6667 |
| 10_6257995 | 2 | 0 | 1 | 0 | NA | NA | 3 | 0.3333 | 6 | 0.1667 | 3 | 0.6667 | 3 | 0.3333 | 6 | 0.5 | 1 | 1 | 13 | 0.5385 |
| 10_6258914 | 6 | 0.8333 | 1 | 1 | 2 | 0 | 1 | 0 | 10 | 0.6 | 2 | 0.5 | 5 | 0.6 | 3 | 0 | 2 | 0 | 12 | 0.3333 |
| 10_6260250 | 3 | 0 | 1 | 1 | 2 | 0 | NA | NA | 6 | 0.1667 | 2 | 0 | 4 | 0 | 3 | 0.3333 | 2 | 0 | 11 | 0.0909 |
| 10_6283875 | 1 | 0 | 1 | 1 | NA | NA | 1 | 0 | 3 | 0.3333 | NA | NA | 1 | 1 | 1 | 0 | 1 | 0 | 3 | 0.3333 |
| 10_6283882 | 1 | 1 | 2 | 0 | NA | NA | 1 | 1 | 4 | 0.5 | NA | NA | 1 | 0 | 1 | 0 | 1 | 1 | 3 | 0.3333 |
| 10_6283902 | 1 | 0 | 1 | 1 | NA | NA | 1 | 1 | 3 | 0.6667 | NA | NA | 2 | 1 | 2 | 0.5 | NA | NA | 4 | 0.75 |
| 10_6284008 | 1 | 0 | 2 | 0.5 | 1 | 1 | 1 | 0 | 5 | 0.4 | 2 | 0 | 2 | 0.5 | 3 | 0.3333 | 1 | 1 | 8 | 0.375 |
| 10_6284751 | 2 | 0.5 | 4 | 0 | 1 | 0 | NA | NA | 7 | 0.1429 | NA | NA | 4 | 0.25 | 4 | 0 | 2 | 0 | 10 | 0.1 |
| 10_6284779 | NA | NA | 4 | 0.75 | NA | NA | NA | NA | 4 | 0.75 | NA | NA | 2 | 1 | 3 | 0.3333 | 2 | 0 | 7 | 0.4286 |
| 10_6285128 | 14 | 0.0714 | 12 | 0.25 | 10 | 0.2 | 16 | 0.0625 | 52 | 0.1346 | 5 | 0 | 11 | 0.0909 | 11 | 0.0909 | 8 | 0.25 | 35 | 0.1143 |
| 10_6285133 | 12 | 0 | 11 | 0.5455 | 10 | 0.3 | 16 | 0.1875 | 49 | 0.2449 | 5 | 0.6 | 10 | 0.3 | 12 | 0.1667 | 8 | 0.125 | 35 | 0.2571 |
| 10_6285137 | 12 | 0 | 12 | 0.1667 | 7 | 0.1429 | 14 | 0.0714 | 45 | 0.0889 | 4 | 0.25 | 9 | 0 | 11 | 0 | 7 | 0.2857 | 31 | 0.0968 |
| 10_6308163 | 1 | 1 | 2 | 0 | NA | NA | 1 | 0 | 4 | 0.25 | 1 | 0 | 6 | 0.1667 | 3 | 0 | NA | NA | 10 | 0.1 |
| 10_6308170 | 1 | 1 | 1 | 1 | 1 | 0 | 1 | 0 | 4 | 0.5 | 1 | 0 | 6 | 0.1667 | 3 | 0.3333 | NA | NA | 10 | 0.2 |
| 10_6308176 | 1 | 1 | 2 | 0 | 1 | 0 | 1 | 0 | 5 | 0.2 | 1 | 0 | 6 | 0.1667 | 3 | 0.3333 | NA | NA | 10 | 0.2 |
| 10_6308180 | 1 | 0 | NA | NA | NA | NA | 1 | 1 | 2 | 0.5 | 1 | 0 | 5 | 0.4 | 3 | 0 | NA | NA | 9 | 0.2222 |
| 10_6308183 | 1 | 1 | 1 | 0 | 1 | 0 | 1 | 0 | 4 | 0.25 | 1 | 0 | 7 | 0.2857 | 2 | 0.5 | NA | NA | 10 | 0.3 |
| 10_6315322 | 3 | 0 | 6 | 0.1667 | 8 | 0 | 3 | 0 | 20 | 0.05 | 1 | 0 | 10 | 0.1 | 6 | 0.1667 | 3 | 0 | 20 | 0.1 |
| 10_6315327 | 3 | 1 | 6 | 0.8333 | 5 | 0.8 | 2 | 1 | 16 | 0.875 | NA | NA | 9 | 0.7778 | 6 | 1 | 2 | 1 | 17 | 0.8824 |
| 10_6315339 | 2 | 0 | 8 | 0.125 | 8 | 0.5 | 3 | 0.3333 | 21 | 0.2857 | 1 | 0 | 10 | 0.1 | 7 | 0.2857 | 4 | 0.25 | 22 | 0.1818 |
| 10_6315525 | 3 | 0 | 4 | 0.25 | 1 | 0 | 5 | 0 | 13 | 0.0769 | 1 | 0 | 2 | 0 | 6 | 0.1667 | 2 | 0 | 11 | 0.0909 |
| 10_6315555 | 1 | 0 | 3 | 0 | 1 | 1 | 5 | 0.2 | 10 | 0.2 | 1 | 0 | 2 | 0.5 | 4 | 0.5 | 1 | 0 | 8 | 0.375 |
| 10_6315629 | 2 | 1 | 4 | 1 | 1 | 1 | 3 | 0.6667 | 10 | 0.9 | 1 | 1 | 2 | 1 | 3 | 1 | 1 | 1 | 7 | 1 |
| 10_6315633 | NA | NA | 3 | 0.6667 | 1 | 1 | 4 | 1 | 8 | 0.875 | NA | NA | 2 | 1 | 6 | 0.8333 | 2 | 0.5 | 10 | 0.8 |
| 10_6315636 | 1 | 0 | 4 | 1 | 1 | 1 | 4 | 1 | 10 | 0.9 | 1 | 1 | 2 | 1 | 4 | 1 | 2 | 1 | 9 | 1 |
| 10_6315642 | 2 | 0 | 4 | 0 | 1 | 0 | 3 | 0.6667 | 10 | 0.2 | NA | NA | 2 | 0 | 6 | 0.1667 | 2 | 0 | 10 | 0.1 |
| 10_6315643 | 2 | 0 | 4 | 0.25 | 1 | 0 | 4 | 0 | 11 | 0.0909 | 1 | 0 | 2 | 0 | 5 | 0.4 | 2 | 0 | 10 | 0.2 |
| 10_6315644 | 2 | 1 | 4 | 0.25 | 1 | 0 | 4 | 0.25 | 11 | 0.3636 | 1 | 0 | 2 | 0 | 5 | 0.4 | 1 | 1 | 9 | 0.3333 |
| 10_6315645 | 2 | 1 | 4 | 0 | 1 | 0 | 4 | 0.25 | 11 | 0.2727 | 1 | 0 | 2 | 0 | 6 | 0.1667 | 2 | 0 | 11 | 0.0909 |
| 10_6315663 | 2 | 0.5 | 2 | 0 | 1 | 1 | 3 | 0.6667 | 8 | 0.5 | NA | NA | 2 | 1 | 5 | 0.6 | 1 | 1 | 8 | 0.75 |
| 10_6315756 | 2 | 0 | 3 | 0 | 1 | 1 | 4 | 0.25 | 10 | 0.2 | 1 | 0 | 2 | 0 | 6 | 0.1667 | 1 | 1 | 10 | 0.2 |
| 10_6315760 | 2 | 0 | 3 | 0.3333 | 1 | 0 | 4 | 0 | 10 | 0.1 | 1 | 0 | 1 | 0 | 6 | 0.3333 | 1 | 0 | 9 | 0.2222 |
| 10_6315766 | 1 | 1 | 2 | 0.5 | 1 | 0 | 4 | 0 | 8 | 0.25 | 1 | 1 | 2 | 0 | 6 | 0.5 | 1 | 0 | 10 | 0.4 |
| 10_6315768 | 2 | 0.5 | 3 | 0 | 1 | 1 | 4 | 0.25 | 10 | 0.3 | 1 | 0 | 2 | 0.5 | 5 | 0.6 | 1 | 0 | 9 | 0.4444 |
| 10_6315779 | 2 | 0.5 | 2 | 0.5 | 1 | 1 | 4 | 0.25 | 9 | 0.4444 | 1 | 0 | 2 | 0.5 | 4 | 0.5 | 1 | 0 | 8 | 0.375 |
| 10_6315780 | 2 | 1 | 1 | 1 | 1 | 0 | 3 | 1 | 7 | 0.8571 | 1 | 0 | 2 | 1 | 3 | 0.6667 | 1 | 1 | 7 | 0.7143 |
| 10_6315799 | 2 | 1 | 3 | 0.6667 | 1 | 1 | 4 | 0 | 10 | 0.5 | 1 | 0 | 2 | 1 | 5 | 0.2 | 1 | 0 | 9 | 0.3333 |
| 10_6374590 | NA | NA | 1 | 1 | NA | NA | NA | NA | 1 | 1 | NA | NA | 1 | 1 | NA | NA | NA | NA | 1 | 1 |
| 10_6376456 | 3 | 0.3333 | NA | NA | 5 | 0.2 | 4 | 0.5 | 12 | 0.3333 | 1 | 0 | 6 | 0.1667 | 4 | 0.5 | 9 | 0.3333 | 20 | 0.3 |
| 10_6376464 | 3 | 0.6667 | NA | NA | 4 | 0.5 | 3 | 1 | 10 | 0.7 | 2 | 0 | 4 | 0.25 | 5 | 0.4 | 5 | 0.4 | 16 | 0.3125 |
| 10_6376492 | 3 | 0.3333 | NA | NA | 4 | 0 | 3 | 0.3333 | 10 | 0.2 | 2 | 0.5 | 8 | 0 | 4 | 0 | 8 | 0.125 | 22 | 0.0909 |
| 10_6376511 | 3 | 0 | 2 | 0 | 11 | 0.1818 | 9 | 0.1111 | 25 | 0.12 | 3 | 0 | 16 | 0.0625 | 12 | 0.0833 | 10 | 0.2 | 41 | 0.0976 |
| 10_6376532 | 5 | 0.4 | 2 | 0 | 10 | 0 | 10 | 0 | 27 | 0.0741 | 2 | 0 | 13 | 0.0769 | 11 | 0.0909 | 9 | 0.2222 | 35 | 0.1143 |
| 10_6376557 | 3 | 0 | 3 | 0 | 10 | 0 | 7 | 0.2857 | 23 | 0.087 | 3 | 0 | 11 | 0.0909 | 12 | 0 | 8 | 0.125 | 34 | 0.0588 |
| 10_6376562 | 6 | 0.1667 | 3 | 0.3333 | 10 | 0.4 | 9 | 0.4444 | 28 | 0.3571 | 4 | 0.25 | 10 | 0.5 | 12 | 0.0833 | 11 | 0.4545 | 37 | 0.3243 |
| 10_6376573 | 9 | 0.2222 | 4 | 0.25 | 5 | 0 | 9 | 0 | 27 | 0.1111 | 5 | 0 | 8 | 0.375 | 13 | 0.2308 | 14 | 0.2143 | 40 | 0.225 |
| 10_6376573 | 7 | 0 | 3 | 0 | 7 | 0.1429 | 10 | 0.1 | 27 | 0.0741 | 4 | 0 | 6 | 0.1667 | 9 | 0.1111 | 11 | 0.1818 | 30 | 0.1333 |
| 10_6376577 | 12 | 0.5833 | 3 | 0 | 9 | 0.4444 | 14 | 0.5 | 38 | 0.4737 | 4 | 0 | 13 | 0.3077 | 16 | 0.375 | 18 | 0.5 | 51 | 0.3725 |
| 10_6376585 | 11 | 0.0909 | 3 | 0 | 9 | 0.1111 | 13 | 0.2308 | 36 | 0.1389 | 5 | 0.2 | 10 | 0.1 | 14 | 0.2143 | 15 | 0.2 | 44 | 0.1818 |
| 10_6376589 | 10 | 1 | 2 | 1 | 9 | 0.8889 | 14 | 0.9286 | 35 | 0.9429 | 3 | 1 | 10 | 0.9 | 14 | 0.8571 | 18 | 1 | 45 | 0.9333 |
| 10_6376590 | 11 | 0.9091 | 3 | 1 | 9 | 1 | 13 | 1 | 36 | 0.9722 | 4 | 1 | 10 | 1 | 15 | 1 | 16 | 1 | 45 | 1 |
| 10_6376596 | 9 | 0.8889 | 3 | 1 | 6 | 0.8333 | 12 | 0.75 | 30 | 0.8333 | 3 | 0.6667 | 9 | 0.7778 | 13 | 0.9231 | 12 | 0.8333 | 37 | 0.8378 |
| 10_6392366 | 11 | 0.6364 | 4 | 0.5 | 9 | 0.2222 | 15 | 0.4 | 39 | 0.4359 | 4 | 0.25 | 10 | 0.1 | 15 | 0.2667 | 16 | 0.25 | 45 | 0.2222 |
| 10_6392371 | 11 | 0 | 4 | 0.25 | 8 | 0 | 10 | 0.1 | 33 | 0.0606 | 5 | 0 | 10 | 0.2 | 13 | 0.1538 | 13 | 0 | 41 | 0.0976 |
| 10_6392393 | 7 | 0.4286 | 3 | 0.6667 | 7 | 0.5714 | 10 | 0.1 | 27 | 0.3704 | 4 | 0 | 10 | 0.6 | 13 | 0.3846 | 15 | 0.6667 | 42 | 0.5 |
| 10_6392394 | 8 | 0.5 | 4 | 0.5 | 9 | 0.5556 | 11 | 0.3636 | 32 | 0.4688 | 5 | 0.6 | 8 | 0.875 | 13 | 0.4615 | 15 | 0.4667 | 41 | 0.561 |
| 10_6392409 | 7 | 0.1429 | 4 | 0 | 4 | 0 | 10 | 0.2 | 25 | 0.12 | 5 | 0 | 7 | 0 | 12 | 0.1667 | 9 | 0 | 33 | 0.0606 |
| 10_6392419 | 8 | 0.25 | 4 | 0.25 | 7 | 0.2857 | 11 | 0.0909 | 30 | 0.2 | 5 | 0.4 | 4 | 0 | 10 | 0.1 | 12 | 0.1667 | 31 | 0.1613 |
| 10_6392428 | 5 | 0.8 | 4 | 0.25 | 4 | 0.25 | 10 | 0.3 | 23 | 0.3913 | 4 | 1 | 5 | 0.8 | 13 | 0.4615 | 11 | 0.5455 | 33 | 0.6061 |
| 10_6392435 | 7 | 0 | 4 | 0.25 | 5 | 0.2 | 11 | 0.3636 | 27 | 0.2222 | 3 | 0 | 6 | 0.5 | 13 | 0 | 12 | 0.1667 | 34 | 0.1471 |
| 10_6392441 | 7 | 0.2857 | 4 | 0.25 | 6 | 0.3333 | 11 | 0.2727 | 28 | 0.2857 | 2 | 0.5 | 6 | 0.3333 | 13 | 0.3846 | 12 | 0.25 | 33 | 0.3333 |
| 10_6392446 | 5 | 1 | 4 | 1 | 7 | 0.7143 | 11 | 1 | 27 | 0.9259 | 3 | 1 | 6 | 1 | 12 | 0.8333 | 10 | 0.8 | 31 | 0.871 |
| 10_6392454 | 5 | 0.4 | 4 | 0.25 | 5 | 0.8 | 11 | 0.3636 | 25 | 0.44 | 2 | 0.5 | 5 | 0.8 | 9 | 0.7778 | 7 | 0.7143 | 23 | 0.7391 |

| | | | | | | | | | | | | | | | | | | | | |
|---|---|---|---|---|---|---|---|---|---|---|---|---|---|---|---|---|---|---|---|---|
| 10_639246 | 6 | 0.3333 | 4 | 0.75 | 4 | 0.5 | 9 | 0.5556 | 23 | 0.5217 | 2 | 0.5 | 5 | 0.6 | 10 | 0.9 | 10 | 0.5 | 27 | 0.6667 |
| 10_639247 | 4 | 0.25 | 1 | 0 | 3 | 0.3333 | 6 | 0.1667 | 14 | 0.2143 | 1 | 0 | 2 | 0 | 7 | 0.1429 | 7 | 0.2857 | 17 | 0.1765 |
| 10_639247 | 5 | 0.4 | 4 | 0.25 | 6 | 0.6667 | 9 | 0.2222 | 24 | 0.375 | 2 | 0 | 4 | 0.5 | 11 | 0.2727 | 8 | 0.375 | 25 | 0.32 |
| 10_639247 | 7 | 0.2857 | 3 | 0 | 6 | 0 | 9 | 0 | 25 | 0.08 | 3 | 0 | 6 | 0.1667 | 12 | 0 | 10 | 0.3 | 31 | 0.129 |
| 10_639248 | 4 | 1 | NA | NA | 5 | 0.6 | 9 | 0.6667 | 18 | 0.7222 | 1 | 1 | 5 | 0.6 | 9 | 0.7778 | 8 | 0.625 | 23 | 0.6957 |
| 10_639249 | 4 | 0.75 | 1 | 1 | 5 | 0.8 | 8 | 0.875 | 18 | 0.8333 | 2 | 1 | 4 | 0.75 | 7 | 0.7143 | 5 | 0.8 | 18 | 0.7778 |
| 10_639250 | 5 | 0.6 | 2 | 0.5 | 6 | 0.6667 | 8 | 0.5 | 21 | 0.5714 | 2 | 1 | 5 | 1 | 11 | 0.6364 | 8 | 0.375 | 26 | 0.6538 |
| 10_639250 | 6 | 0.1667 | 1 | 0 | 4 | 0.25 | 9 | 0.3333 | 20 | 0.25 | 2 | 0.5 | 5 | 0 | 11 | 0.3636 | 6 | 0.3333 | 24 | 0.2917 |
| 10_639257 | 5 | 0.2 | 2 | 0 | 6 | 0.1667 | 7 | 0 | 20 | 0.1 | 3 | 0 | 6 | 0 | 10 | 0.2 | 7 | 0 | 26 | 0.0769 |
| 10_639259 | 3 | 0.3333 | NA | NA | 5 | 0.2 | 7 | 0.2857 | 15 | 0.2667 | 2 | 0 | 3 | 0.3333 | 10 | 0.3 | 7 | 0.5714 | 22 | 0.3636 |
| 10_648478 | NA | NA | 1 | 1 | NA | NA | 1 | 0 | 2 | 0.5 | NA | NA | NA | NA | 1 | 1 | NA | NA | 1 | 1 |
| 10_668452 | 4 | 0 | 3 | 0.3333 | 3 | 0 | 3 | 0 | 13 | 0.0769 | 2 | 0.5 | 3 | 0 | 6 | 0 | 3 | 0 | 14 | 0.0714 |
| 10_668491 | NA | NA | 2 | 0 | 2 | 1 | 2 | 0 | 6 | 0.3333 | NA | NA | NA | NA | 4 | 0.25 | 3 | 0.3333 | 7 | 0.2857 |
| 10_670555 | 2 | 0.5 | 1 | 1 | NA | NA | NA | NA | 3 | 0.6667 | NA | NA | 1 | 0 | 1 | 0 | 2 | 0.5 | 4 | 0.25 |
| 10_670562 | 2 | 1 | 1 | 1 | NA | NA | NA | NA | 3 | 1 | NA | NA | 1 | 1 | 2 | 1 | 2 | 1 | 5 | 1 |
| 10_670606 | 3 | 0.3333 | 1 | 0 | NA | NA | NA | NA | 4 | 0.25 | NA | NA | 1 | 1 | 2 | 1 | 3 | 0.6667 | 6 | 0.8333 |
| 10_670609 | 3 | 0 | 1 | 0 | 1 | 1 | NA | NA | 5 | 0.2 | NA | NA | 1 | 0 | 2 | 0.5 | 3 | 0 | 6 | 0.1667 |
| 10_670611 | 3 | 1 | 1 | 1 | 1 | 0 | NA | NA | 5 | 0.8 | NA | NA | 2 | 0.5 | 2 | 1 | 3 | 0.6667 | 7 | 0.7143 |
| 10_670614 | 3 | 0.6667 | 1 | 1 | 1 | 1 | NA | NA | 5 | 0.8 | NA | NA | 2 | 1 | 2 | 1 | 4 | 0.25 | 8 | 0.625 |
| 10_670616 | 3 | 0.3333 | 1 | 0 | 2 | 0.5 | NA | NA | 6 | 0.3333 | NA | NA | 2 | 1 | 2 | 0.5 | 4 | 0.5 | 8 | 0.625 |
| 10_670617 | 3 | 0.3333 | 1 | 1 | 2 | 0 | NA | NA | 6 | 0.3333 | NA | NA | 2 | 0 | 2 | 0 | 4 | 0.25 | 8 | 0.125 |
| 10_670619 | 3 | 0.6667 | 1 | 0 | 2 | 1 | 1 | 1 | 7 | 0.7143 | NA | NA | 2 | 1 | 2 | 1 | 2 | 0.5 | 6 | 0.8333 |
| 10_670621 | 2 | 0 | 1 | 1 | 2 | 0 | 1 | 0 | 6 | 0.1667 | NA | NA | 2 | 0.5 | 2 | 0 | 4 | 0 | 8 | 0.125 |
| 10_670622 | 3 | 0.3333 | 1 | 0 | 2 | 0 | 1 | 0 | 7 | 0.1429 | NA | NA | 2 | 0 | 2 | 0 | 3 | 0.3333 | 7 | 0.1429 |
| 10_670624 | 2 | 0.5 | NA | NA | 2 | 0 | NA | NA | 4 | 0.25 | NA | NA | 2 | 0 | 1 | 1 | 5 | 0.2 | 8 | 0.25 |
| 10_670625 | 3 | 0.3333 | 1 | 0 | 2 | 0 | 1 | 0 | 7 | 0.1429 | NA | NA | 2 | 0.5 | 2 | 0 | 4 | 0 | 8 | 0.125 |
| 10_670764 | 3 | 0.3333 | 4 | 0 | 3 | 0 | 2 | 0 | 12 | 0.0833 | NA | NA | 4 | 0 | 2 | 0.5 | 6 | 0.1667 | 12 | 0.1667 |
| 10_671027 | 3 | 0.3333 | 9 | 0 | 5 | 0.2 | 6 | 0.1667 | 23 | 0.1304 | 2 | 0.5 | 6 | 0 | 3 | 0.3333 | 3 | 0 | 14 | 0.1429 |
| 10_671027 | 2 | 0 | 10 | 0.1 | 5 | 0.2 | 6 | 0 | 23 | 0.087 | 2 | 0 | 7 | 0 | 3 | 0.3333 | 3 | 0 | 15 | 0.0667 |
| 10_671027 | 2 | 1 | 7 | 0.5714 | 5 | 1 | 5 | 1 | 19 | 0.8421 | 1 | 1 | 6 | 0.6667 | 3 | 0.6667 | 3 | 1 | 13 | 0.7692 |
| 10_671028 | 3 | 0.3333 | 7 | 0.2857 | 3 | 0.3333 | 5 | 0.2 | 18 | 0.2778 | 1 | 1 | 7 | 0.2857 | 3 | 0.3333 | 3 | 0.6667 | 14 | 0.4286 |
| 10_671028 | 3 | 0 | 9 | 0.2222 | 4 | 0.25 | 5 | 0 | 21 | 0.1429 | 2 | 0 | 7 | 0.2857 | 2 | 0 | 2 | 0 | 13 | 0.1538 |
| 10_671029 | 2 | 0.5 | 5 | 0.2 | 4 | 0.25 | 4 | 0.25 | 15 | 0.2667 | 1 | 1 | 5 | 0 | 2 | 0 | 2 | 0.5 | 10 | 0.2 |
| 10_671030 | 2 | 1 | 7 | 1 | 1 | 1 | 3 | 1 | 13 | 1 | 2 | 1 | 5 | 1 | 3 | 1 | 3 | 1 | 13 | 1 |
| 10_671030 | 2 | 0 | 9 | 0.3333 | 5 | 0.2 | 6 | 0 | 22 | 0.1818 | 2 | 1 | 6 | 0.1667 | 3 | 0 | 2 | 0 | 13 | 0.2308 |
| 10_671031 | 3 | 0.3333 | 10 | 0.5 | 5 | 0.2 | 6 | 0.1667 | 24 | 0.3333 | 2 | 0 | 7 | 0.1429 | 3 | 0.3333 | 2 | 0.5 | 14 | 0.2143 |
| 10_671032 | 3 | 0.3333 | 8 | 0.625 | 3 | 0.3333 | 5 | 1 | 19 | 0.6316 | NA | NA | 5 | 0.4 | 2 | 1 | 2 | 0.5 | 9 | 0.5556 |
| 10_671032 | 2 | 1 | 10 | 0.9 | 5 | 1 | 5 | 1 | 22 | 0.9545 | 2 | 1 | 6 | 0.8333 | 3 | 1 | 3 | 1 | 14 | 0.9286 |
| 10_671033 | 3 | 1 | 10 | 1 | 4 | 1 | 5 | 1 | 22 | 1 | 2 | 1 | 7 | 0.8571 | 3 | 1 | 3 | 1 | 15 | 0.9333 |
| 10_671034 | 3 | 1 | 9 | 0.5556 | 3 | 0.6667 | 4 | 0.5 | 19 | 0.6316 | 1 | 1 | 6 | 0.6667 | 2 | 0.5 | 3 | 0.6667 | 12 | 0.6667 |
| 10_671034 | 3 | 0.3333 | 8 | 1 | 5 | 1 | 6 | 1 | 22 | 0.9091 | 1 | 1 | 5 | 1 | 2 | 1 | 2 | 0.5 | 10 | 0.9 |
| 10_671034 | 3 | 0.3333 | 9 | 0.1111 | 5 | 0.8 | 4 | 0.25 | 21 | 0.3333 | 1 | 0 | 6 | 0.5 | 2 | 0.5 | 3 | 0.3333 | 12 | 0.4167 |
| 10_671036 | 3 | 0.6667 | 8 | 0.5 | 4 | 0.5 | 6 | 0.5 | 21 | 0.5238 | 1 | 1 | 4 | 0.25 | 2 | 0.5 | 3 | 0 | 10 | 0.3 |
| 10_673805 | 6 | 0.6667 | NA | NA | 1 | 0 | 5 | 0.6 | 12 | 0.5833 | NA | NA | 1 | 0 | 2 | 1 | 2 | 0.5 | 5 | 0.6 |
| 10_673806 | 2 | 1 | NA | NA | 1 | 0 | 5 | 0.2 | 8 | 0.375 | NA | NA | 1 | 0 | 2 | 0.5 | 2 | 0.5 | 5 | 0.4 |
| 10_673812 | 8 | 0.125 | 9 | 0.1111 | 8 | 0.25 | 8 | 0 | 33 | 0.1212 | 2 | 0 | 2 | 0.5 | 7 | 0 | 7 | 0.1429 | 18 | 0.1111 |
| 10_673816 | 9 | 0.1111 | 9 | 0 | 7 | 0.1429 | 6 | 0 | 31 | 0.0645 | 3 | 0.3333 | 1 | 0 | 7 | 0 | 6 | 0 | 17 | 0.0588 |
| 10_673829 | 2 | 0 | 4 | 0.25 | 1 | 1 | 3 | 0 | 10 | 0.2 | 3 | 0 | NA | NA | 5 | 0.2 | 5 | 0 | 13 | 0.0769 |
| 10_673830 | 7 | 0.2857 | 7 | 0.1429 | 5 | 0.4 | 3 | 0 | 22 | 0.2273 | 3 | 0.3333 | NA | NA | 5 | 0.4 | 7 | 0.1429 | 15 | 0.2667 |
| 10_673834 | 7 | 0.4286 | 4 | 0.5 | 4 | 0.5 | 2 | 1 | 17 | 0.5294 | 4 | 0 | NA | NA | 2 | 0 | 5 | 0.4 | 11 | 0.1818 |
| 10_673835 | 6 | 0 | 6 | 0.1667 | 3 | 0 | 3 | 0 | 18 | 0.0556 | 3 | 0.3333 | 1 | 0 | 4 | 0 | 5 | 0 | 13 | 0.0769 |
| 10_673835 | 7 | 0.8571 | 5 | 0.8 | 4 | 0.75 | 4 | 0.75 | 20 | 0.8 | 3 | 1 | 1 | 1 | 4 | 1 | 6 | 0.6667 | 14 | 0.8571 |
| 10_673836 | 4 | 0 | 2 | 0.5 | 1 | 0 | 1 | 0 | 8 | 0.125 | 2 | 0 | 1 | 0 | 4 | 0.25 | 4 | 0 | 11 | 0.0909 |
| 10_673836 | 7 | 1 | 6 | 1 | 3 | 1 | 4 | 1 | 20 | 1 | 2 | 1 | 1 | 1 | 4 | 1 | 5 | 1 | 12 | 1 |
| 10_673838 | 6 | 1 | 3 | 1 | 2 | 1 | 3 | 1 | 14 | 1 | 3 | 1 | 1 | 1 | 4 | 1 | 5 | 1 | 13 | 1 |
| 10_673839 | 6 | 0.6667 | 5 | 0.6 | 3 | 0.3333 | 2 | 1 | 16 | 0.625 | 3 | 0.6667 | 1 | 1 | 4 | 0.5 | 4 | 0.5 | 12 | 0.5833 |
| 10_673842 | 3 | 0 | 5 | 0.2 | 1 | 0 | 2 | 0.5 | 11 | 0.1818 | 3 | 0 | 1 | 0 | 3 | 0.3333 | 5 | 0.4 | 12 | 0.25 |
| 10_673842 | 6 | 0.8333 | 3 | 0 | 1 | 1 | 1 | 1 | 11 | 0.6364 | 3 | 0.3333 | 1 | 1 | 3 | 1 | 5 | 0.8 | 12 | 0.75 |
| 10_673844 | 4 | 0.25 | 2 | 0.5 | 1 | 0 | 2 | 0.5 | 9 | 0.3333 | 1 | 0 | 1 | 0 | 2 | 0.5 | 4 | 0.25 | 8 | 0.25 |
| 10_677221 | 42 | 0.119 | 57 | 0.0877 | 62 | 0.1129 | 41 | 0.122 | 202 | 0.1089 | 52 | 0.0962 | 44 | 0.0909 | 48 | 0.0417 | 50 | 0.08 | 194 | 0.0773 |
| 10_677222 | 47 | 0.0426 | 58 | 0.0862 | 68 | 0.0294 | 44 | 0.0455 | 217 | 0.0507 | 54 | 0.0741 | 49 | 0.0408 | 53 | 0.1132 | 52 | 0.0385 | 208 | 0.0673 |
| 10_677226 | 37 | 0.2162 | 46 | 0.1087 | 58 | 0.1207 | 41 | 0.2439 | 182 | 0.1648 | 39 | 0.1282 | 35 | 0.1143 | 44 | 0.1591 | 37 | 0.1351 | 155 | 0.1355 |
| 10_677234 | 39 | 0.0256 | 39 | 0.0769 | 57 | 0.1053 | 39 | 0.1026 | 174 | 0.0805 | 41 | 0.0488 | 37 | 0.027 | 38 | 0.0789 | 38 | 0.0526 | 154 | 0.0519 |
| 10_677243 | 23 | 0.1304 | 37 | 0.1351 | 43 | 0.093 | 26 | 0.1538 | 129 | 0.124 | 31 | 0.1613 | 25 | 0.24 | 32 | 0.1875 | 28 | 0.2143 | 116 | 0.1983 |
| 10_677245 | 22 | 0.0455 | 28 | 0 | 36 | 0.0833 | 28 | 0.0714 | 114 | 0.0526 | 27 | 0.037 | 23 | 0 | 24 | 0.0417 | 22 | 0.2727 | 96 | 0.0833 |

| | | | | | | | | | | | | | | | | | | | | |
|---|---|---|---|---|---|---|---|---|---|---|---|---|---|---|---|---|---|---|---|---|
| 10_677247 | 24 | 0.0417 | 29 | 0.069 | 37 | 0.0541 | 26 | 0.1154 | 116 | 0.069 | 20 | 0.05 | 16 | 0.0625 | 27 | 0.1111 | 21 | 0.0952 | 84 | 0.0833 |
| 10_677333 | 27 | 0.0741 | 26 | 0.0385 | 41 | 0.0732 | 28 | 0.1429 | 122 | 0.082 | 25 | 0.2 | 23 | 0.1304 | 25 | 0.08 | 23 | 0.087 | 96 | 0.125 |
| 10_677451 | 20 | 0.75 | 20 | 0.6 | 28 | 0.7143 | 19 | 0.6316 | 87 | 0.6782 | 17 | 0.6471 | 16 | 0.6875 | 15 | 0.8667 | 15 | 0.5333 | 63 | 0.6825 |
| 10_677452 | 21 | 0.0952 | 21 | 0.0476 | 28 | 0.0357 | 18 | 0.0556 | 88 | 0.0568 | 14 | 0.0714 | 18 | 0.0556 | 20 | 0.1 | 17 | 0 | 69 | 0.058 |
| 10_677452 | 19 | 0.0526 | 22 | 0.2273 | 23 | 0.1304 | 18 | 0.0556 | 82 | 0.122 | 16 | 0 | 17 | 0.1176 | 21 | 0.0952 | 14 | 0.0714 | 68 | 0.0735 |
| 10_677453 | 20 | 0.35 | 20 | 0.3 | 24 | 0.25 | 16 | 0.1875 | 80 | 0.275 | 15 | 0.2667 | 17 | 0.1765 | 20 | 0.35 | 16 | 0.4375 | 68 | 0.3088 |
| 10_677454 | 19 | 0.8947 | 17 | 0.9412 | 24 | 0.9167 | 17 | 0.9412 | 77 | 0.9221 | 12 | 0.5833 | 14 | 0.9286 | 15 | 0.8667 | 15 | 0.9333 | 56 | 0.8393 |
| 10_677455 | 20 | 0 | 17 | 0.0588 | 21 | 0.0476 | 13 | 0.1538 | 71 | 0.0563 | 13 | 0.0769 | 16 | 0.0625 | 16 | 0.0625 | 15 | 0.0667 | 60 | 0.0667 |
| 10_677455 | 18 | 0.5556 | 18 | 0.5556 | 22 | 0.5 | 15 | 0.6 | 73 | 0.5479 | 13 | 0.4615 | 14 | 0.7143 | 15 | 0.6667 | 13 | 0.6154 | 55 | 0.6182 |
| 10_677456 | 14 | 0.0714 | 17 | 0.2941 | 23 | 0.1304 | 14 | 0.0714 | 68 | 0.1471 | 14 | 0.2143 | 16 | 0.1875 | 17 | 0.2353 | 15 | 0 | 62 | 0.1613 |
| 10_677457 | 17 | 0.0588 | 15 | 0.1333 | 19 | 0 | 8 | 0.25 | 59 | 0.0847 | 11 | 0 | 14 | 0 | 14 | 0 | 14 | 0.2143 | 53 | 0.0566 |
| 10_678796 | 1 | 1 | NA | NA | NA | NA | NA | NA | 1 | 1 | NA | NA | NA | NA | 1 | 1 | NA | NA | 1 | 1 |
| 10_679287 | 4 | 1 | 7 | 0.7143 | 6 | 1 | 1 | 1 | 18 | 0.8889 | 6 | 0.6667 | 3 | 0.6667 | 1 | 1 | 7 | 0.8571 | 17 | 0.7647 |
| 10_679287 | 3 | 0 | 9 | 0.5556 | 6 | 0.6667 | 1 | 0 | 19 | 0.4737 | 5 | 0.4 | 3 | 0.6667 | 3 | 0.3333 | 7 | 0.4286 | 18 | 0.4444 |
| 10_679288 | 4 | 0.25 | 9 | 0.7778 | 6 | 1 | 1 | 1 | 20 | 0.75 | 6 | 1 | 3 | 0.6667 | 3 | 0.6667 | 7 | 0.2857 | 19 | 0.6316 |
| 10_679288 | 4 | 0 | 9 | 0.2222 | 4 | 0.25 | 1 | 0 | 18 | 0.1667 | 5 | 0.2 | 3 | 0 | 3 | 0 | 7 | 0 | 18 | 0.0556 |
| 10_679291 | 4 | 0 | 9 | 0.2222 | 4 | 0 | 1 | 0 | 18 | 0.1111 | 5 | 0.2 | 3 | 0 | 3 | 0 | 7 | 0 | 18 | 0.0556 |
| 10_679296 | 4 | 0.25 | 7 | 0.2857 | 4 | 0.5 | 1 | 0 | 16 | 0.3125 | 5 | 0.2 | 3 | 0.3333 | 3 | 0 | 6 | 0.1667 | 17 | 0.1765 |
| 10_679297 | 4 | 0 | 6 | 0.1667 | 3 | 0 | 1 | 0 | 14 | 0.0714 | 4 | 0.5 | 3 | 0 | 2 | 0 | 6 | 0.1667 | 15 | 0.2 |
| 10_679314 | 2 | 0 | 5 | 0.2 | 1 | 0 | NA | NA | 8 | 0.125 | 3 | 0 | 3 | 0 | 2 | 1 | 3 | 0 | 11 | 0.1818 |
| 10_679318 | 4 | 0 | 5 | 0.2 | 1 | 1 | 1 | 0 | 11 | 0.1818 | 2 | 0.5 | 3 | 0.6667 | 1 | 1 | 2 | 0 | 8 | 0.5 |
| 10_679319 | 4 | 0.5 | 1 | 1 | 1 | 1 | NA | NA | 6 | 0.6667 | 2 | 0.5 | 3 | 0.3333 | 1 | 1 | 2 | 0.5 | 8 | 0.5 |
| 10_679321 | 4 | 0 | 3 | 0 | 1 | 0 | 1 | 1 | 9 | 0.1111 | 2 | 0 | 3 | 0 | 1 | 0 | 2 | 0.5 | 8 | 0.125 |
| 10_679321 | 4 | 0.5 | 2 | 0.5 | NA | NA | 1 | 0 | 7 | 0.4286 | 2 | 0.5 | 2 | 1 | 1 | 1 | 1 | 1 | 6 | 0.8333 |
| 10_679321 | 4 | 0.75 | 3 | 1 | 1 | 1 | 1 | 0 | 9 | 0.7778 | 2 | 0.5 | 3 | 1 | 1 | 1 | 1 | 1 | 7 | 0.8571 |
| 10_679323 | 2 | 1 | 1 | 1 | NA | NA | NA | NA | 3 | 1 | 2 | 1 | 3 | 1 | 1 | 1 | 2 | 1 | 8 | 1 |
| 10_679324 | 3 | 0 | 3 | 0.6667 | 1 | 0 | 1 | 0 | 8 | 0.25 | 2 | 0 | 3 | 0.3333 | 1 | 0 | 2 | 0 | 8 | 0.125 |
| 10_679324 | 3 | 1 | 2 | 1 | NA | NA | NA | NA | 5 | 1 | 2 | 1 | 2 | 1 | 1 | 1 | 2 | 1 | 7 | 1 |
| 10_679325 | 2 | 0 | 3 | 0.6667 | 1 | 1 | 1 | 0 | 7 | 0.4286 | 2 | 0 | 3 | 0.6667 | 1 | 0 | 2 | 0 | 8 | 0.25 |
| 10_679327 | NA | NA | 2 | 0.5 | 1 | 1 | 1 | 1 | 4 | 0.75 | 2 | 0.5 | 2 | 0.5 | 1 | 0 | 1 | 0 | 6 | 0.3333 |
| 10_679328 | 1 | 1 | 1 | 0 | NA | NA | 1 | 0 | 3 | 0.3333 | 1 | 0 | 1 | 1 | 1 | 0 | 1 | 0 | 4 | 0.25 |
| 10_679329 | NA | NA | 1 | 0 | NA | NA | 1 | 1 | 2 | 0.5 | 2 | 0 | 3 | 0.6667 | 1 | 0 | 1 | 1 | 7 | 0.4286 |
| 10_679329 | NA | NA | 3 | 0.3333 | 1 | 1 | 1 | 0 | 5 | 0.4 | 2 | 0.5 | 3 | 0.6667 | 1 | 0 | 1 | 0 | 7 | 0.4286 |
| 10_679330 | NA | NA | 2 | 0.5 | 1 | 1 | NA | NA | 3 | 0.6667 | 2 | 1 | 3 | 0.6667 | 1 | 1 | 1 | 1 | 7 | 0.8571 |
| 10_679331 | NA | NA | 3 | 0.3333 | 1 | 1 | 1 | 0 | 5 | 0.4 | 2 | 0 | 3 | 0.3333 | 1 | 0 | 1 | 1 | 7 | 0.2857 |
| 10_679334 | NA | NA | 1 | 1 | 1 | 1 | NA | NA | 2 | 1 | 1 | 1 | 3 | 0.3333 | 1 | 1 | 1 | 1 | 6 | 0.6667 |
| 10_679337 | NA | NA | 2 | 1 | 1 | 1 | 1 | 1 | 4 | 1 | 1 | 1 | 2 | 0.5 | NA | NA | 1 | 0 | 4 | 0.5 |
| 10_679338 | NA | NA | 2 | 1 | 1 | 1 | 1 | 1 | 4 | 1 | 2 | 1 | 3 | 1 | NA | NA | 1 | 0 | 6 | 0.8333 |
| 10_679339 | NA | NA | 1 | 1 | 1 | 1 | 1 | 1 | 3 | 1 | 2 | 1 | 1 | 1 | NA | NA | 1 | 1 | 4 | 1 |
| 10_679341 | NA | NA | 2 | 0.5 | 1 | 1 | 1 | 1 | 4 | 0.75 | 2 | 1 | 2 | 1 | NA | NA | 1 | 1 | 5 | 1 |
| 10_679343 | NA | NA | 1 | 0 | 1 | 1 | NA | NA | 2 | 0.5 | 2 | 0.5 | 2 | 0 | NA | NA | 1 | 0 | 5 | 0.2 |
| 10_682669 | NA | NA | NA | NA | NA | NA | 2 | 0.5 | 2 | 0.5 | NA | NA | 1 | 1 | NA | NA | NA | NA | 1 | 1 |
| 10_691060 | NA | NA | NA | NA | 2 | 0.5 | NA | NA | 2 | 0.5 | NA | NA | 1 | 1 | NA | NA | NA | NA | 1 | 1 |
| 10_691064 | NA | NA | NA | NA | 2 | 0.5 | NA | NA | 2 | 0.5 | NA | NA | 1 | 1 | NA | NA | NA | NA | 1 | 1 |
| 10_691262 | NA | NA | NA | NA | 2 | 0.5 | NA | NA | 2 | 0.5 | 1 | 1 | NA | NA | NA | NA | NA | NA | 1 | 1 |
| 10_693756 | NA | NA | 1 | 1 | NA | NA | NA | NA | 1 | 1 | 1 | 1 | NA | NA | 1 | 0 | NA | NA | 2 | 0.5 |
| 10_693758 | NA | NA | 1 | 1 | NA | NA | NA | NA | 1 | 1 | 1 | 1 | NA | NA | NA | NA | NA | NA | 1 | 1 |
| 10_693768 | NA | NA | 1 | 1 | NA | NA | NA | NA | 1 | 1 | NA | NA | NA | NA | 1 | 1 | NA | NA | 1 | 1 |
| 10_693772 | NA | NA | 1 | 1 | NA | NA | NA | NA | 1 | 1 | NA | NA | NA | NA | 1 | 1 | NA | NA | 1 | 1 |
| 10_698222 | 1 | 1 | 1 | 0 | 3 | 1 | 1 | 1 | 6 | 0.8333 | 1 | 0 | 4 | 0.5 | 2 | 1 | 2 | 1 | 9 | 0.6667 |
| 10_698226 | NA | NA | 2 | 0.5 | 3 | 0 | 2 | 0.5 | 7 | 0.2857 | 1 | 1 | 5 | 0.4 | 1 | 0 | 1 | 0 | 8 | 0.375 |
| 10_698227 | 1 | 1 | 3 | 0 | 3 | 0.6667 | 2 | 0.5 | 9 | 0.4444 | 1 | 1 | 6 | 0.5 | 2 | 1 | 2 | 0.5 | 11 | 0.6364 |
| 10_698230 | 1 | 0 | 3 | 0.3333 | 3 | 0.3333 | 2 | 0 | 9 | 0.2222 | 1 | 0 | 6 | 0.5 | 3 | 0.3333 | 3 | 0.3333 | 13 | 0.3846 |
| 10_698230 | 1 | 1 | 3 | 0 | 3 | 0 | 1 | 0 | 8 | 0.125 | 1 | 0 | 4 | 0 | 3 | 0 | 3 | 0.3333 | 11 | 0.0909 |
| 10_698234 | 2 | 0.5 | 6 | 0 | 2 | 0 | 2 | 0 | 12 | 0.0833 | 2 | 0 | 6 | 0 | 3 | 0.3333 | 4 | 0 | 15 | 0.0667 |
| 10_698234 | 2 | 1 | 5 | 0.2 | 3 | 0 | 2 | 0.5 | 12 | 0.3333 | 2 | 0.5 | 4 | 0 | 2 | 0 | 3 | 0.3333 | 11 | 0.1818 |
| 10_698235 | 2 | 0 | 6 | 0.1667 | 4 | 0 | 2 | 0 | 14 | 0.0714 | 2 | 0 | 6 | 0 | 3 | 0 | 4 | 0.25 | 15 | 0.0667 |
| 10_698236 | 2 | 0 | 7 | 0.2857 | 4 | 0 | 2 | 0 | 15 | 0.1333 | 2 | 0 | 6 | 0 | 3 | 0 | 4 | 0.25 | 15 | 0.0667 |
| 10_698237 | NA | NA | 2 | 0.5 | 4 | 1 | 1 | 1 | 7 | 0.8571 | 1 | 0 | 5 | 1 | 2 | 1 | 4 | 1 | 12 | 0.9167 |
| 10_698237 | 2 | 0 | 6 | 0.1667 | 5 | 0 | 2 | 0 | 15 | 0.0667 | 2 | 0 | 4 | 0.25 | 2 | 0 | 4 | 0 | 12 | 0.0833 |
| 10_698239 | 2 | 0.5 | 4 | 0.5 | 5 | 0.4 | 2 | 0.5 | 13 | 0.4615 | 2 | 0 | 6 | 0.6667 | 3 | 0.3333 | 5 | 0.2 | 16 | 0.375 |
| 10_698241 | 3 | 0.3333 | 3 | 0.3333 | 5 | 0.2 | 2 | 1 | 13 | 0.3846 | 3 | 0.6667 | 5 | 0.2 | 3 | 1 | 5 | 0.4 | 16 | 0.5 |
| 10_698244 | 4 | 0.5 | 7 | 0.2857 | 6 | 0.3333 | 2 | 0.5 | 19 | 0.3684 | 2 | 0 | 5 | 0.4 | 3 | 0 | 4 | 0.25 | 14 | 0.2143 |
| 10_698245 | 2 | 0.5 | 4 | 0.5 | 2 | 0.5 | NA | NA | 8 | 0.5 | 3 | 0 | 3 | 0 | 2 | 0.5 | 2 | 0.5 | 10 | 0.2 |
| 10_698247 | 2 | 0 | 2 | 0.5 | 2 | 0 | 1 | 0 | 7 | 0.1429 | 3 | 0 | 3 | 0 | 2 | 0.5 | 2 | 0.5 | 10 | 0.2 |

| | | | | | | | | | | | | | | | | | | | | |
|---|---|---|---|---|---|---|---|---|---|---|---|---|---|---|---|---|---|---|---|---|
| 10_698250 | 1 | 1 | 2 | 1 | 2 | 0 | 1 | 0 | 6 | 0.5 | 3 | 0 | 3 | 0 | 4 | 0 | 3 | 1 | 13 | 0.2308 |
| 10_698251 | 1 | 0 | 3 | 0.3333 | 2 | 0 | 1 | 0 | 7 | 0.1429 | 2 | 0 | 2 | 0.5 | 3 | 0 | 2 | 0 | 9 | 0.1111 |
| 10_698254 | 1 | 0 | 2 | 0 | 3 | 0.3333 | 2 | 1 | 8 | 0.375 | 3 | 0 | 2 | 0 | 3 | 0.6667 | 3 | 0.3333 | 11 | 0.2727 |
| 10_698255 | 1 | 1 | 3 | 0.6667 | 3 | 0.3333 | NA | NA | 7 | 0.5714 | 2 | 1 | 3 | 0.6667 | 5 | 0.8 | 2 | 1 | 12 | 0.8333 |
| 10_698258 | 1 | 1 | 2 | 1 | 4 | 1 | 3 | 1 | 10 | 1 | 3 | 1 | 4 | 1 | 3 | 1 | 4 | 1 | 14 | 1 |
| 10_698259 | 1 | 0 | 3 | 0 | 4 | 0.25 | 2 | 0 | 10 | 0.1 | 3 | 0 | 5 | 0 | 5 | 0.4 | 3 | 0 | 16 | 0.125 |
| 10_698259 | 1 | 0 | 3 | 0.3333 | 5 | 0.4 | 3 | 0.3333 | 12 | 0.3333 | 3 | 0 | 5 | 0 | 5 | 0.2 | 4 | 0.25 | 17 | 0.1176 |
| 10_698259 | 1 | 0 | 2 | 0.5 | 4 | 1 | 3 | 1 | 10 | 0.8 | 3 | 0.6667 | 5 | 0.4 | 5 | 0.6 | 3 | 0.3333 | 16 | 0.5 |
| 10_698259 | 1 | 1 | 3 | 1 | 4 | 1 | 3 | 0.6667 | 11 | 0.9091 | 3 | 1 | 5 | 1 | 5 | 1 | 4 | 1 | 17 | 1 |
| 10_698259 | 1 | 1 | 3 | 1 | 4 | 0.5 | 3 | 0 | 11 | 0.5455 | 3 | 0.6667 | 3 | 0.6667 | 4 | 0.75 | 4 | 0.75 | 14 | 0.7143 |
| 10_698261 | 1 | 0 | 3 | 0 | 6 | 0.6667 | 3 | 0.3333 | 13 | 0.3846 | 2 | 0 | 3 | 0.3333 | 4 | 0.25 | 3 | 0 | 12 | 0.1667 |
| 10_698262 | NA | NA | 3 | 0 | 5 | 0.2 | 4 | 0.25 | 12 | 0.1667 | 3 | 0.3333 | 4 | 0.25 | 4 | 0.5 | 6 | 0 | 17 | 0.2353 |
| 10_698264 | NA | NA | 3 | 0.3333 | 5 | 0 | 2 | 0 | 10 | 0.1 | 4 | 0.25 | 4 | 0.25 | 5 | 0 | 5 | 0.2 | 18 | 0.1667 |
| 10_698266 | 1 | 1 | 1 | 1 | 5 | 0.6 | 4 | 0.25 | 11 | 0.5455 | 3 | 0.6667 | 4 | 0.75 | 4 | 0.5 | 5 | 0.4 | 16 | 0.5625 |
| 10_698271 | 1 | 1 | 3 | 1 | 4 | 0.75 | 3 | 0.6667 | 11 | 0.8182 | 4 | 0.75 | 4 | 1 | 6 | 0.6667 | 5 | 0.8 | 19 | 0.7895 |
| 10_698271 | 1 | 0 | 3 | 0.3333 | 6 | 0 | 4 | 0 | 14 | 0.0714 | 4 | 0.5 | 6 | 0 | 7 | 0.1429 | 6 | 0.1667 | 23 | 0.1739 |
| 10_698272 | 1 | 0 | 3 | 0 | 4 | 0 | 4 | 0.25 | 12 | 0.0833 | 3 | 0 | 6 | 0 | 6 | 0.1667 | 5 | 0 | 20 | 0.05 |
| 10_698273 | 1 | 0 | 3 | 0.3333 | 5 | 0.2 | 4 | 0 | 13 | 0.1538 | 4 | 0 | 6 | 0.3333 | 7 | 0 | 5 | 0 | 22 | 0.0909 |
| 10_698276 | 2 | 0 | 4 | 0 | 6 | 0.1667 | 5 | 0.2 | 17 | 0.1176 | 4 | 0 | 6 | 0 | 8 | 0.125 | 6 | 0.3333 | 24 | 0.125 |
| 10_698279 | 3 | 0 | 4 | 0.25 | 5 | 0.2 | 6 | 0.5 | 18 | 0.2778 | 4 | 0.25 | 7 | 0.2857 | 7 | 0.4286 | 6 | 0.3333 | 24 | 0.3333 |
| 10_698280 | 3 | 0.6667 | 5 | 0.4 | 5 | 0.2 | 7 | 0.5714 | 20 | 0.45 | 5 | 0.4 | 7 | 0.2857 | 7 | 0.5714 | 6 | 0.5 | 25 | 0.44 |
| 10_698280 | 2 | 0 | 4 | 0 | 6 | 0.1667 | 7 | 0.1429 | 19 | 0.1053 | 4 | 0 | 5 | 0 | 7 | 0 | 5 | 0.4 | 21 | 0.0952 |
| 10_698282 | 3 | 0 | 6 | 0.3333 | 7 | 0 | 6 | 0.1667 | 22 | 0.1364 | 5 | 0.2 | 8 | 0.125 | 6 | 0.1667 | 5 | 0.2 | 24 | 0.1667 |
| 10_698283 | 2 | 0 | 1 | 0 | 1 | 0 | 1 | 1 | 5 | 0.2 | 2 | 0 | 4 | 0.25 | 1 | 0 | 1 | 0 | 8 | 0.125 |
| 10_698286 | 2 | 0 | 1 | 0 | 1 | 0 | 2 | 0.5 | 6 | 0.1667 | 1 | 0 | 4 | 0.25 | 1 | 0 | NA | NA | 6 | 0.1667 |
| 10_698288 | 2 | 0 | 1 | 1 | 1 | 0 | 3 | 1 | 7 | 0.5714 | 2 | 0 | 4 | 0.25 | 1 | 0 | 1 | 0 | 8 | 0.125 |
| 10_698289 | 2 | 0.5 | 1 | 0 | 1 | 1 | 3 | 0.3333 | 7 | 0.4286 | 2 | 1 | 4 | 1 | 1 | 1 | 1 | 1 | 8 | 1 |
| 10_698378 | NA | NA | 1 | 0 | 2 | 0.5 | 2 | 0.5 | 5 | 0.4 | 1 | 0 | NA | NA | 1 | 0 | 3 | 0.3333 | 5 | 0.2 |
| 10_698379 | 1 | 0 | 1 | 0 | 2 | 0 | 2 | 0.5 | 6 | 0.1667 | NA | NA | NA | NA | 1 | 0 | 3 | 0.3333 | 4 | 0.25 |
| 10_698379 | 1 | 0 | 1 | 0 | 2 | 0.5 | 1 | 0 | 5 | 0.2 | 1 | 1 | NA | NA | 1 | 0 | 3 | 0.3333 | 5 | 0.4 |
| 10_698380 | 1 | 1 | 1 | 1 | 2 | 0.5 | 2 | 0.5 | 6 | 0.6667 | 1 | 0 | NA | NA | 1 | 0 | 2 | 1 | 4 | 0.5 |
| 10_698382 | NA | NA | 1 | 0 | 1 | 0 | 2 | 1 | 4 | 0.5 | 1 | 1 | NA | NA | 1 | 0 | 2 | 1 | 4 | 0.75 |
| 10_698383 | 1 | 0 | 1 | 0 | 2 | 0.5 | 2 | 1 | 6 | 0.5 | 1 | 0 | NA | NA | 1 | 1 | 3 | 0.6667 | 5 | 0.6 |
| 10_698597 | 14 | 0.3571 | 16 | 0.1875 | 19 | 0.4211 | 18 | 0.2222 | 67 | 0.2985 | 11 | 0.3636 | 15 | 0.2 | 16 | 0.375 | 15 | 0.2 | 57 | 0.2807 |
| 10_698598 | 14 | 0.2143 | 14 | 0.1429 | 19 | 0.2105 | 17 | 0.1765 | 64 | 0.1875 | 8 | 0.375 | 14 | 0.3571 | 15 | 0.1333 | 15 | 0.4 | 52 | 0.3077 |
| 10_698599 | 15 | 0.0667 | 19 | 0.1053 | 21 | 0.1429 | 17 | 0.1176 | 72 | 0.1111 | 10 | 0.1 | 17 | 0 | 17 | 0.0588 | 16 | 0.0625 | 60 | 0.05 |
| 10_698600 | 15 | 0.0667 | 19 | 0.1579 | 21 | 0.0476 | 19 | 0 | 74 | 0.0676 | 12 | 0.1667 | 16 | 0.125 | 16 | 0 | 16 | 0.0625 | 60 | 0.0833 |
| 10_698600 | 15 | 0.8667 | 17 | 0.7059 | 23 | 0.8261 | 18 | 0.6111 | 73 | 0.7534 | 11 | 0.7273 | 18 | 0.7778 | 15 | 0.8 | 15 | 0.8667 | 59 | 0.7966 |
| 10_698600 | 12 | 0.5 | 13 | 0.4615 | 18 | 0.5 | 13 | 0.7692 | 56 | 0.5536 | 6 | 0.5 | 9 | 0.7778 | 10 | 0.4 | 12 | 0.4167 | 37 | 0.5135 |
| 10_698601 | 13 | 0.3077 | 15 | 0.2 | 21 | 0.2857 | 15 | 0.4 | 64 | 0.2969 | 11 | 0.5455 | 17 | 0.1765 | 15 | 0.2667 | 13 | 0.3077 | 56 | 0.3036 |
| 10_698602 | 13 | 0.1538 | 19 | 0.2105 | 20 | 0 | 18 | 0.2222 | 70 | 0.1429 | 9 | 0 | 9 | 0 | 16 | 0.1875 | 16 | 0.0625 | 50 | 0.08 |
| 10_698602 | 14 | 0 | 19 | 0.1053 | 24 | 0.0833 | 20 | 0.05 | 77 | 0.0649 | 13 | 0.0769 | 18 | 0 | 18 | 0.1111 | 18 | 0.0556 | 67 | 0.0597 |
| 10_698602 | 14 | 0.1429 | 17 | 0.0588 | 26 | 0.2308 | 18 | 0.0556 | 75 | 0.1333 | 10 | 0.1 | 18 | 0.3333 | 19 | 0.0526 | 18 | 0.1667 | 65 | 0.1692 |
| 10_698603 | 15 | 0.5333 | 18 | 0.4444 | 24 | 0.4167 | 18 | 0.4444 | 75 | 0.4533 | 13 | 0.5385 | 17 | 0.5294 | 19 | 0.5789 | 18 | 0.6111 | 67 | 0.5672 |
| 10_698603 | 16 | 0.25 | 20 | 0.25 | 27 | 0.1852 | 21 | 0.0952 | 84 | 0.1905 | 15 | 0.1333 | 21 | 0.0952 | 17 | 0.1765 | 19 | 0.0526 | 72 | 0.1111 |
| 10_698603 | 16 | 0.0625 | 19 | 0.0526 | 21 | 0.1429 | 14 | 0.1429 | 70 | 0.1 | 12 | 0.1667 | 15 | 0.0667 | 17 | 0.2353 | 13 | 0.1538 | 57 | 0.1579 |
| 10_698603 | 15 | 0.8667 | 19 | 0.9474 | 27 | 1 | 20 | 0.85 | 81 | 0.9259 | 14 | 0.9286 | 17 | 0.8235 | 16 | 1 | 16 | 1 | 63 | 0.9365 |
| 10_698603 | 13 | 0.9231 | 15 | 0.8 | 22 | 0.7727 | 19 | 0.6842 | 69 | 0.7826 | 13 | 0.9231 | 13 | 0.8462 | 14 | 0.7143 | 13 | 0.5385 | 53 | 0.7547 |
| 10_698604 | 18 | 0.1111 | 19 | 0.0526 | 27 | 0.1111 | 16 | 0.1875 | 80 | 0.1125 | 14 | 0.2143 | 18 | 0.1111 | 19 | 0.2105 | 18 | 0.1667 | 69 | 0.1739 |
| 10_698608 | 22 | 0.8636 | 19 | 0.8947 | 29 | 0.8966 | 18 | 0.7778 | 88 | 0.8636 | 15 | 0.9333 | 21 | 0.7619 | 25 | 0.84 | 17 | 0.9412 | 78 | 0.859 |
| 10_698611 | 24 | 0.0417 | 23 | 0.1304 | 36 | 0.0278 | 28 | 0.0357 | 111 | 0.0541 | 20 | 0.05 | 27 | 0.1481 | 32 | 0.0938 | 24 | 0.1667 | 103 | 0.1165 |
| 10_698627 | 14 | 0.1429 | 15 | 0 | 20 | 0.05 | 8 | 0 | 57 | 0.0526 | 3 | 0 | 9 | 0 | 16 | 0.0625 | 13 | 0.1538 | 41 | 0.0732 |
| 10_698628 | 13 | 0.5385 | 16 | 0.5 | 21 | 0.619 | 10 | 0.5 | 60 | 0.55 | 7 | 0.5714 | 8 | 0.875 | 18 | 0.6111 | 13 | 0.8462 | 46 | 0.7174 |
| 10_698628 | 13 | 0.1538 | 16 | 0.25 | 20 | 0.2 | 9 | 0.2222 | 58 | 0.2069 | 7 | 0.2857 | 9 | 0.4444 | 17 | 0.2353 | 15 | 0.4667 | 48 | 0.3542 |
| 10_698629 | 15 | 0.8 | 15 | 0.7333 | 18 | 0.7778 | 10 | 0.9 | 58 | 0.7931 | 8 | 0.875 | 7 | 1 | 16 | 0.875 | 13 | 0.8462 | 44 | 0.8864 |
| 10_698631 | NA | NA | 2 | 1 | 1 | 0 | NA | NA | 3 | 0.6667 | 2 | 0.5 | 1 | 1 | 1 | 1 | 2 | 1 | 6 | 0.8333 |
| 10_698631 | NA | NA | 2 | 0 | 2 | 0.5 | NA | NA | 4 | 0.25 | 1 | 0 | 1 | 0 | 1 | 0 | 2 | 0.5 | 5 | 0.2 |
| 10_698631 | NA | NA | 1 | 1 | 2 | 1 | NA | NA | 3 | 1 | NA | NA | 1 | 1 | 1 | 0 | 2 | 1 | 4 | 0.75 |
| 10_700251 | 2 | 1 | 1 | 1 | 1 | 1 | 3 | 1 | 7 | 1 | 2 | 1 | 3 | 1 | NA | NA | 1 | 1 | 6 | 1 |
| 10_700255 | 3 | 0 | 4 | 0 | 1 | 0 | 5 | 0.2 | 13 | 0.0769 | 1 | 0 | 2 | 0.5 | NA | NA | 1 | 0 | 4 | 0.25 |
| 10_700258 | 3 | 0 | 3 | 0.3333 | 2 | 0 | 5 | 0.2 | 13 | 0.1538 | 3 | 0.3333 | 4 | 0.25 | NA | NA | 1 | 0 | 8 | 0.25 |
| 10_700259 | 3 | 1 | 5 | 0.2 | 2 | 1 | 7 | 0.1429 | 17 | 0.4118 | 2 | 0 | 4 | 0.25 | NA | NA | 1 | 0 | 7 | 0.1429 |
| 10_700262 | 4 | 0 | 6 | 0.3333 | 2 | 0.5 | 5 | 0.6 | 17 | 0.3529 | 4 | 0.5 | 4 | 0.25 | NA | NA | 1 | 0 | 9 | 0.3333 |
| 10_700264 | 4 | 0.5 | 8 | 0.375 | 2 | 0 | 6 | 0.1667 | 20 | 0.3 | 5 | 0.4 | 5 | 0.4 | NA | NA | 2 | 0.5 | 12 | 0.4167 |
| 10_700267 | 4 | 0.25 | 5 | 0 | 3 | 0 | 6 | 0 | 18 | 0.0556 | 4 | 0 | 4 | 0.5 | 1 | 0 | 1 | 0 | 10 | 0.2 |

| | | | | | | | | | | | | | | | | | | | | |
|---|---|---|---|---|---|---|---|---|---|---|---|---|---|---|---|---|---|---|---|---|
| 10_700267 | 4 | 0.25 | 8 | 0.125 | 3 | 0 | 9 | 0 | 24 | 0.0833 | 5 | 0.2 | 6 | 0.1667 | 1 | 0 | 1 | 0 | 13 | 0.1538 |
| 10_700278 | 3 | 0.6667 | 7 | 0.5714 | 4 | 0.5 | 11 | 0.3636 | 25 | 0.48 | 5 | 0.2 | 6 | 0.8333 | 2 | 0.5 | 4 | 0.25 | 17 | 0.4706 |
| 10_700282 | 6 | 0.3333 | 7 | 0 | 4 | 0.5 | 11 | 0 | 28 | 0.1429 | 6 | 0 | 5 | 0.2 | 4 | 0.25 | 5 | 0.2 | 20 | 0.15 |
| 10_700285 | 6 | 1 | 8 | 0.5 | 4 | 0.75 | 11 | 0.5455 | 29 | 0.6552 | 7 | 0.4286 | 10 | 0.6 | 5 | 0.6 | 4 | 0.25 | 26 | 0.5 |
| 10_700287 | 7 | 0.2857 | 9 | 0.6667 | 5 | 0.6 | 12 | 0.5833 | 33 | 0.5455 | 7 | 0.4286 | 10 | 0.7 | 5 | 0.2 | 8 | 0.5 | 30 | 0.5 |
| 10_700287 | 6 | 1 | 8 | 1 | 4 | 1 | 9 | 1 | 27 | 1 | 6 | 1 | 11 | 1 | 5 | 1 | 5 | 1 | 27 | 1 |
| 10_700287 | 5 | 1 | 9 | 1 | 4 | 1 | 9 | 1 | 27 | 1 | 7 | 1 | 11 | 1 | 4 | 1 | 7 | 1 | 29 | 1 |
| 10_700287 | 5 | 0 | 9 | 0.1111 | 5 | 0 | 11 | 0.0909 | 30 | 0.0667 | 7 | 0.2857 | 11 | 0 | 5 | 0 | 7 | 0 | 30 | 0.0667 |
| 10_700289 | 7 | 0 | 10 | 0.1 | 6 | 0 | 11 | 0.0909 | 34 | 0.0588 | 6 | 0.3333 | 11 | 0.2727 | 5 | 0 | 6 | 0 | 28 | 0.1786 |
| 10_701182 | 1 | 0 | NA | NA | 1 | 0 | 2 | 0.5 | 4 | 0.25 | NA | NA | NA | NA | NA | NA | 1 | 1 | 1 | 1 |
| 10_701184 | 1 | 0 | NA | NA | 1 | 1 | 1 | 1 | 3 | 0.6667 | NA | NA | NA | NA | NA | NA | 1 | 1 | 1 | 1 |
| 10_701185 | NA | NA | NA | NA | 1 | 0 | 2 | 0.5 | 3 | 0.3333 | NA | NA | NA | NA | NA | NA | 1 | 1 | 1 | 1 |
| 10_701189 | 1 | 1 | NA | NA | 1 | 1 | 2 | 1 | 4 | 1 | NA | NA | NA | NA | NA | NA | 1 | 1 | 1 | 1 |
| 10_701189 | 1 | 1 | NA | NA | 1 | 1 | 1 | 1 | 3 | 1 | NA | NA | NA | NA | NA | NA | 1 | 1 | 1 | 1 |
| 10_701200 | 1 | 1 | NA | NA | NA | NA | 2 | 0.5 | 3 | 0.6667 | NA | NA | NA | NA | NA | NA | 1 | 1 | 1 | 1 |
| 10_701205 | 1 | 1 | NA | NA | NA | NA | 1 | 1 | 2 | 1 | NA | NA | NA | NA | NA | NA | 1 | 1 | 1 | 1 |
| 10_701205 | 1 | 1 | NA | NA | NA | NA | 2 | 1 | 3 | 1 | NA | NA | NA | NA | NA | NA | 1 | 1 | 1 | 1 |
| 10_701206 | 1 | 1 | NA | NA | 1 | 1 | 2 | 1 | 4 | 1 | NA | NA | NA | NA | NA | NA | 1 | 1 | 1 | 1 |
| 10_701208 | 1 | 1 | NA | NA | 1 | 1 | 2 | 1 | 4 | 1 | NA | NA | NA | NA | NA | NA | 1 | 1 | 1 | 1 |
| 10_702752 | NA | NA | NA | NA | 1 | 1 | NA | NA | 1 | 1 | NA | NA | 1 | 1 | NA | NA | NA | NA | 1 | 1 |
| 10_703934 | 11 | 0.0909 | 13 | 0.1538 | 8 | 0 | 6 | 0.1667 | 38 | 0.1053 | 6 | 0 | 9 | 0.2222 | 11 | 0.3636 | 7 | 0.2857 | 33 | 0.2424 |
| 10_703953 | 31 | 0.9355 | 34 | 0.8235 | 26 | 0.8462 | 21 | 0.619 | 112 | 0.8214 | 31 | 0.6452 | 38 | 0.8947 | 31 | 0.8065 | 31 | 0.8387 | 131 | 0.8015 |
| 10_703954 | 20 | 0.3 | 23 | 0.1739 | 16 | 0.3125 | 15 | 0.2 | 74 | 0.2432 | 20 | 0.25 | 24 | 0.375 | 25 | 0.36 | 19 | 0.4737 | 88 | 0.3636 |
| 10_703954 | 25 | 0.76 | 28 | 0.8214 | 21 | 0.7619 | 16 | 0.75 | 90 | 0.7778 | 18 | 0.7778 | 29 | 0.7586 | 29 | 0.8276 | 26 | 0.8462 | 102 | 0.8039 |
| 10_703954 | 3 | 1 | 2 | 1 | 3 | 1 | 5 | 0.6 | 13 | 0.8462 | 7 | 0.5714 | 4 | 0.75 | 4 | 0.5 | 4 | 0.75 | 19 | 0.6316 |
| 10_707591 | NA | NA | 1 | 1 | NA | NA | NA | NA | 1 | 1 | NA | NA | 3 | 1 | NA | NA | NA | NA | 3 | 1 |
| 10_710641 | 5 | 0.6 | 4 | 0.75 | 4 | 0.25 | 3 | 1 | 16 | 0.625 | 2 | 0 | 2 | 0.5 | 6 | 0.3333 | 9 | 0.4444 | 19 | 0.3684 |
| 10_710641 | 6 | 1 | 4 | 0.75 | 5 | 0.6 | 4 | 1 | 19 | 0.8421 | 2 | 1 | 3 | 1 | 6 | 0.8333 | 9 | 0.7778 | 20 | 0.85 |
| 10_710641 | 5 | 0.8 | 4 | 0.75 | 5 | 0.8 | 3 | 1 | 17 | 0.8235 | 3 | 0.6667 | 3 | 0.6667 | 7 | 0.8571 | 8 | 0.25 | 21 | 0.5714 |
| 10_710641 | 4 | 0.75 | 2 | 1 | 3 | 0.6667 | 2 | 1 | 11 | 0.8182 | NA | NA | 3 | 1 | 6 | 0.6667 | 6 | 0.6667 | 15 | 0.7333 |
| 10_710642 | 6 | 1 | 3 | 1 | 3 | 1 | 2 | 1 | 14 | 1 | 2 | 1 | 2 | 1 | 4 | 1 | 8 | 0.875 | 16 | 0.9375 |
| 10_710910 | 14 | 0.1429 | 15 | 0.2667 | 9 | 0.3333 | 12 | 0.3333 | 50 | 0.26 | 9 | 0.3333 | 10 | 0.2 | 5 | 0.2 | 10 | 0.1 | 34 | 0.2059 |
| 10_710911 | 13 | 0.1538 | 11 | 0 | 11 | 0.0909 | 12 | 0 | 47 | 0.0638 | 7 | 0.2857 | 7 | 0 | 6 | 0 | 9 | 0.2222 | 29 | 0.1379 |
| 10_710912 | 13 | 0.4615 | 16 | 0.3125 | 12 | 0.4167 | 12 | 0.25 | 53 | 0.3585 | 8 | 0.25 | 9 | 0.4444 | 4 | 0.25 | 10 | 0 | 31 | 0.2258 |
| 10_710913 | 12 | 0.25 | 8 | 0.125 | 11 | 0.0909 | 10 | 0.1 | 41 | 0.1463 | 7 | 0.2857 | 6 | 0.5 | 3 | 0 | 7 | 0 | 23 | 0.2174 |
| 10_710913 | 13 | 0.3846 | 13 | 0.2308 | 12 | 0.5 | 6 | 0.3333 | 44 | 0.3636 | 9 | 0.5556 | 6 | 0.1667 | 5 | 0.6 | 10 | 0.5 | 30 | 0.4667 |
| 10_710914 | 11 | 0.0909 | 14 | 0.3571 | 10 | 0.3 | 7 | 0.1429 | 42 | 0.2381 | 9 | 0.1111 | 7 | 0.1429 | 4 | 0 | 10 | 0.4 | 30 | 0.2 |
| 10_710914 | 12 | 0.75 | 14 | 0.6429 | 11 | 0.6364 | 9 | 0.6667 | 46 | 0.6739 | 8 | 0.375 | 7 | 0.5714 | 3 | 0.6667 | 9 | 0.3333 | 27 | 0.4444 |
| 10_710914 | 12 | 0.0833 | 13 | 0.2308 | 9 | 0.2222 | 8 | 0.125 | 42 | 0.1667 | 6 | 0.1667 | 7 | 0.1429 | 4 | 0.25 | 7 | 0.1429 | 24 | 0.1667 |
| 10_710915 | 11 | 0.3636 | 13 | 0.1538 | 11 | 0.2727 | 9 | 0.3333 | 44 | 0.2727 | 9 | 0.5556 | 8 | 0.25 | 3 | 0 | 10 | 0.1 | 30 | 0.2667 |
| 10_710916 | 12 | 0.5833 | 12 | 0.3333 | 12 | 0.25 | 9 | 0.4444 | 45 | 0.4 | 9 | 0.4444 | 9 | 0.6667 | 3 | 0.6667 | 10 | 0.5 | 31 | 0.5484 |
| 10_711667 | 37 | 0.0811 | 36 | 0.1389 | 23 | 0.0435 | 34 | 0.0882 | 130 | 0.0923 | 28 | 0.1071 | 39 | 0.0256 | 25 | 0.08 | 27 | 0.037 | 119 | 0.0588 |
| 10_711668 | 23 | 1 | 25 | 0.8 | 14 | 1 | 32 | 0.9062 | 94 | 0.9149 | 22 | 0.9545 | 33 | 0.9697 | 17 | 0.9412 | 21 | 0.9048 | 93 | 0.9462 |
| 10_711669 | 35 | 0.0286 | 31 | 0.0323 | 20 | 0.1 | 36 | 0.1111 | 122 | 0.0656 | 23 | 0.087 | 34 | 0.0588 | 24 | 0.0833 | 23 | 0.1304 | 104 | 0.0865 |
| 10_712172 | 1 | 1 | NA | NA | NA | NA | 1 | 1 | 2 | 1 | NA | NA | NA | NA | 1 | 1 | 1 | 1 | 2 | 1 |
| 10_712172 | 2 | 1 | NA | NA | 1 | 1 | 1 | 1 | 4 | 1 | 2 | 1 | 1 | 1 | 2 | 1 | 1 | 1 | 6 | 1 |
| 10_712173 | 2 | 0 | NA | NA | 1 | 1 | 1 | 0 | 4 | 0.25 | 2 | 0.5 | 1 | 0 | 2 | 0 | 1 | 0 | 6 | 0.1667 |
| 10_712174 | 2 | 0.5 | NA | NA | 1 | 1 | 1 | 1 | 4 | 0.75 | 1 | 1 | 1 | 1 | 2 | 1 | 1 | 1 | 5 | 1 |
| 10_712175 | 2 | 1 | NA | NA | 1 | 1 | 1 | 1 | 4 | 1 | 2 | 0.5 | NA | NA | 3 | 1 | 1 | 1 | 6 | 0.8333 |
| 10_712176 | 2 | 1 | NA | NA | 1 | 1 | 1 | 0 | 4 | 0.75 | 2 | 1 | 1 | 1 | 3 | 0.6667 | 1 | 1 | 7 | 0.8571 |
| 10_712180 | 2 | 1 | 1 | 1 | 1 | 0 | 2 | 0 | 6 | 0.5 | 4 | 0.75 | 1 | 0 | 2 | 0.5 | 1 | 0 | 8 | 0.5 |
| 10_712192 | 2 | 1 | NA | NA | 2 | 0.5 | 2 | 0.5 | 6 | 0.6667 | 4 | 1 | 2 | 0.5 | 2 | 1 | 2 | 0.5 | 10 | 0.8 |
| 10_712193 | 1 | 0 | NA | NA | 2 | 0.5 | NA | NA | 3 | 0.3333 | 4 | 0 | 1 | 0 | 3 | 0.3333 | 2 | 0 | 10 | 0.1 |
| 10_712197 | 2 | 0 | 2 | 0.5 | 3 | 1 | 2 | 0 | 9 | 0.4444 | 3 | 0 | 3 | 0.3333 | 3 | 0 | 2 | 0 | 11 | 0.0909 |
| 10_712198 | 2 | 1 | 3 | 0.6667 | 3 | 0.6667 | 1 | 0 | 9 | 0.6667 | 3 | 0.3333 | 2 | 0.5 | 3 | 1 | 2 | 1 | 10 | 0.7 |
| 10_712199 | 2 | 1 | 3 | 1 | 2 | 1 | 2 | 1 | 9 | 1 | 4 | 1 | 3 | 1 | 3 | 1 | 2 | 1 | 12 | 1 |
| 10_712202 | NA | NA | 1 | 0 | NA | NA | 1 | 1 | 2 | 0.5 | 3 | 0.6667 | NA | NA | 2 | 1 | NA | NA | 5 | 0.8 |
| 10_715435 | NA | NA | NA | NA | NA | NA | 1 | 1 | 1 | 1 | 1 | 0 | NA | NA | NA | NA | 1 | 1 | 2 | 0.5 |
| 10_724937 | NA | NA | 1 | 1 | 1 | 1 | 1 | 1 | 3 | 1 | NA | NA | NA | NA | 1 | 1 | NA | NA | 1 | 1 |
| 10_724951 | NA | NA | 1 | 1 | NA | NA | 1 | 0 | 2 | 0.5 | NA | NA | 1 | 1 | 1 | 0 | NA | NA | 2 | 0.5 |
| 10_724952 | NA | NA | 1 | 0 | NA | NA | 1 | 1 | 2 | 0.5 | NA | NA | 1 | 1 | 1 | 0 | NA | NA | 2 | 0.5 |
| 10_725048 | NA | NA | 2 | 0.5 | NA | NA | NA | NA | 2 | 0.5 | NA | NA | NA | NA | 1 | 1 | NA | NA | 1 | 1 |
| 10_725049 | NA | NA | 2 | 1 | NA | NA | NA | NA | 2 | 1 | NA | NA | 1 | 1 | 1 | 1 | NA | NA | 2 | 1 |
| 10_725099 | 93 | 0.1505 | 106 | 0.1038 | 98 | 0.1837 | 115 | 0.113 | 412 | 0.1359 | 87 | 0.0805 | 105 | 0.1524 | 102 | 0.0882 | 83 | 0.2169 | 377 | 0.1326 |
| 10_725103 | 109 | 0.1835 | 106 | 0.2453 | 93 | 0.2258 | 115 | 0.2087 | 423 | 0.2151 | 91 | 0.2857 | 112 | 0.2679 | 107 | 0.2897 | 84 | 0.3214 | 394 | 0.2893 |

| | | | | | | | | | | | | | | | | | | | | |
|---|---|---|---|---|---|---|---|---|---|---|---|---|---|---|---|---|---|---|---|---|
| 10_7251036 | 102 | 0.4412 | 103 | 0.4466 | 94 | 0.4043 | 121 | 0.3388 | 420 | 0.4048 | 82 | 0.4512 | 95 | 0.4316 | 103 | 0.4951 | 72 | 0.4167 | 352 | 0.4517 |
| 10_7251039 | 124 | 0.3387 | 121 | 0.314 | 116 | 0.3534 | 139 | 0.3237 | 500 | 0.332 | 98 | 0.3163 | 127 | 0.3465 | 127 | 0.2913 | 87 | 0.3333 | 439 | 0.3212 |
| 10_7251040 | 94 | 0.3085 | 102 | 0.3725 | 85 | 0.3882 | 109 | 0.4404 | 390 | 0.3795 | 84 | 0.5119 | 110 | 0.4909 | 100 | 0.4 | 71 | 0.4648 | 365 | 0.4658 |
| 10_7251041 | 114 | 0.0965 | 109 | 0.1468 | 93 | 0.129 | 118 | 0.0593 | 434 | 0.106 | 89 | 0.1685 | 95 | 0.1684 | 106 | 0.1509 | 77 | 0.1299 | 367 | 0.1553 |
| 10_7251043 | 96 | 0.0729 | 111 | 0.0721 | 103 | 0.1359 | 120 | 0.15 | 430 | 0.1093 | 91 | 0.1538 | 105 | 0.1143 | 114 | 0.1491 | 76 | 0.2237 | 386 | 0.1554 |
| 10_7251056 | 113 | 0.2035 | 120 | 0.1833 | 114 | 0.1579 | 134 | 0.1642 | 481 | 0.1767 | 99 | 0.1717 | 123 | 0.1789 | 120 | 0.1833 | 87 | 0.1839 | 429 | 0.1795 |
| 10_7251062 | 130 | 0.0923 | 134 | 0.097 | 118 | 0.0847 | 150 | 0.08 | 532 | 0.0883 | 114 | 0.0965 | 136 | 0.0662 | 133 | 0.1579 | 93 | 0.1075 | 476 | 0.1071 |
| 10_7251066 | 131 | 0.0763 | 140 | 0.0857 | 128 | 0.0859 | 145 | 0.0621 | 544 | 0.0772 | 112 | 0.0982 | 134 | 0.0746 | 133 | 0.0827 | 96 | 0.1458 | 475 | 0.0968 |
| 10_7448159 | NA | NA | NA | NA | 1 | 0 | 1 | 1 | 2 | 0.5 | NA | NA | NA | NA | NA | NA | 1 | 1 | 1 | 1 |
| 10_7448161 | NA | NA | NA | NA | 1 | 1 | 1 | 0 | 2 | 0.5 | NA | NA | NA | NA | NA | NA | 1 | 1 | 1 | 1 |
| 10_7448174 | NA | NA | NA | NA | 1 | 1 | 1 | 1 | 2 | 1 | NA | NA | NA | NA | NA | NA | 1 | 1 | 1 | 1 |
| 10_7448189 | NA | NA | NA | NA | 1 | 1 | 1 | 1 | 2 | 1 | NA | NA | NA | NA | NA | NA | 1 | 1 | 1 | 1 |
| 10_7448194 | NA | NA | NA | NA | 1 | 1 | NA | NA | 1 | 1 | NA | NA | NA | NA | NA | NA | 1 | 1 | 1 | 1 |
| 10_7448294 | NA | NA | NA | NA | 1 | 1 | NA | NA | 1 | 1 | NA | NA | NA | NA | NA | NA | 1 | 1 | 1 | 1 |
| 10_7482690 | 3 | 0.3333 | 1 | 1 | 1 | 1 | 8 | 0.75 | 13 | 0.6923 | 1 | 1 | 1 | 1 | 2 | 1 | 2 | 1 | 6 | 1 |
| 10_7482696 | 3 | 0 | 1 | 0 | 1 | 0 | 8 | 0.125 | 13 | 0.0769 | 1 | 0 | 1 | 0 | 2 | 0 | 1 | 1 | 5 | 0.2 |
| 10_7482724 | 4 | 0.75 | 4 | 0.25 | 1 | 1 | 9 | 1 | 18 | 0.7778 | 3 | 0.3333 | 2 | 1 | 4 | 0.75 | 2 | 1 | 11 | 0.7273 |
| 10_7482758 | 2 | 0 | 4 | 0.25 | 1 | 1 | 9 | 0.4444 | 16 | 0.375 | 3 | 0 | 1 | 0 | 3 | 0.3333 | 1 | 0 | 8 | 0.125 |
| 10_7482770 | 3 | 1 | 4 | 0.75 | 1 | 1 | 13 | 1 | 21 | 0.9524 | 4 | 1 | 2 | 0.5 | 4 | 1 | 3 | 1 | 13 | 0.9231 |
| 10_7482778 | 5 | 0.4 | 4 | 0.75 | NA | NA | 14 | 0.7143 | 23 | 0.6522 | 3 | 0.3333 | 2 | 0.5 | 4 | 0.25 | 4 | 0.5 | 13 | 0.3846 |
| 10_7482796 | 4 | 0 | 4 | 0.25 | NA | NA | 14 | 0.0714 | 22 | 0.0909 | 4 | 0 | 2 | 0 | 4 | 0.25 | 3 | 0 | 13 | 0.0769 |
| 10_7485090 | 5 | 0.6 | 8 | 0 | 1 | 0 | 14 | 0.5 | 28 | 0.3571 | 4 | 0.5 | 3 | 0 | 5 | 0.2 | 4 | 0.25 | 16 | 0.25 |
| 10_7485099 | 5 | 0.8 | 7 | 0.4286 | 1 | 1 | 14 | 0.2143 | 27 | 0.4074 | 3 | 0.6667 | 3 | 0 | 6 | 0.1667 | 4 | 0 | 16 | 0.1875 |
| 10_7485101 | 4 | 1 | 2 | 1 | 1 | 1 | 6 | 0.8333 | 13 | 0.9231 | NA | NA | 3 | 0.6667 | 2 | 1 | 4 | 0.75 | 9 | 0.7778 |
| 10_7485145 | 4 | 0.75 | 5 | 0.6 | 2 | 0.5 | 12 | 0.6667 | 23 | 0.6522 | 2 | 1 | 3 | 1 | 4 | 1 | 4 | 0.75 | 13 | 0.9231 |
| 10_7485154 | 5 | 1 | 7 | 1 | 1 | 1 | 12 | 1 | 25 | 1 | 2 | 1 | 3 | 1 | 5 | 1 | 6 | 1 | 16 | 1 |
| 10_7485154 | 6 | 0.6667 | 6 | 1 | 1 | 1 | 14 | 0.9286 | 27 | 0.8889 | 3 | 0.6667 | 3 | 0.3333 | 5 | 1 | 5 | 0.4 | 16 | 0.625 |
| 10_7485162 | 6 | 0.1667 | 7 | 0 | 1 | 0 | 14 | 0.0714 | 28 | 0.0714 | 4 | 0.25 | 3 | 0 | 7 | 0.1429 | 6 | 0.3333 | 20 | 0.2 |
| 10_7485164 | 3 | 1 | 5 | 0.6 | 1 | 1 | 13 | 0.7692 | 22 | 0.7727 | 2 | 1 | 1 | 1 | 4 | 0.75 | 5 | 1 | 12 | 0.9167 |
| 10_7485165 | 6 | 1 | 9 | 0.7778 | 2 | 0 | 14 | 0.7143 | 31 | 0.7419 | 3 | 0.6667 | 3 | 1 | 7 | 1 | 7 | 0.5714 | 20 | 0.8 |
| 10_7485166 | 7 | 0.2857 | 7 | 0 | 1 | 0 | 11 | 0.0909 | 26 | 0.1154 | 5 | 0.2 | 2 | 0 | 7 | 0.1429 | 6 | 0.3333 | 20 | 0.2 |
| 10_7485169 | 5 | 0.2 | 7 | 0 | 1 | 0 | 8 | 0.5 | 21 | 0.2381 | 2 | 0 | 2 | 0 | 5 | 0.6 | 3 | 0 | 12 | 0.25 |
| 10_7485194 | 8 | 0.5 | 9 | 0.1111 | 2 | 0 | 16 | 0.375 | 35 | 0.3143 | 7 | 0.1429 | 7 | 0.4286 | 9 | 0.5556 | 7 | 0.1429 | 30 | 0.3333 |
| 10_7485211 | 6 | 0.5 | 8 | 0.375 | 1 | 0 | 16 | 0.25 | 31 | 0.3226 | 6 | 0.6667 | 7 | 0.5714 | 8 | 0.625 | 8 | 0.25 | 29 | 0.5172 |
| 10_7485245 | 14 | 0.1429 | 14 | 0.1429 | 2 | 0 | 17 | 0.3529 | 47 | 0.2128 | 7 | 0.4286 | 8 | 0.25 | 13 | 0.3846 | 10 | 0.3 | 38 | 0.3421 |
| 10_7485249 | 14 | 0.2143 | 18 | 0 | 3 | 0.3333 | 19 | 0.0526 | 54 | 0.0926 | 8 | 0.125 | 8 | 0.125 | 13 | 0.0769 | 10 | 0.1 | 39 | 0.1026 |
| 10_7485263 | 11 | 0 | 11 | 0.0909 | 1 | 0 | 15 | 0.1333 | 38 | 0.0789 | 5 | 0.2 | 6 | 0.1667 | 11 | 0 | 8 | 0 | 30 | 0.0667 |
| 10_7488859 | 1 | 1 | 1 | 1 | 2 | 0.5 | NA | NA | 4 | 0.75 | 2 | 0 | 3 | 0.3333 | 2 | 0 | 2 | 1 | 9 | 0.3333 |
| 10_7488860 | 1 | 1 | 2 | 1 | 2 | 1 | 3 | 1 | 8 | 1 | 3 | 1 | 4 | 1 | 2 | 0.5 | 2 | 0.5 | 11 | 0.8182 |
| 10_7488872 | 2 | 1 | 2 | 0.5 | 2 | 0.5 | 2 | 0 | 8 | 0.5 | 3 | 0.6667 | 4 | 0 | 2 | 0 | 2 | 1 | 11 | 0.3636 |
| 10_7488874 | 1 | 1 | 2 | 0 | 1 | 0 | 2 | 0 | 6 | 0.1667 | 2 | 0 | 4 | 0.25 | 2 | 0 | 1 | 0 | 9 | 0.1111 |
| 10_7488883 | 1 | 0 | 2 | 0.5 | 2 | 1 | 3 | 1 | 8 | 0.75 | 3 | 0.6667 | 3 | 1 | 2 | 1 | 2 | 0.5 | 10 | 0.8 |
| 10_7488886 | 2 | 1 | 2 | 1 | 1 | 1 | 3 | 1 | 8 | 1 | 3 | 1 | 3 | 1 | 2 | 0.5 | 2 | 1 | 10 | 0.9 |
| 10_7488894 | 2 | 0 | 1 | 1 | 2 | 0.5 | 2 | 0.5 | 7 | 0.4286 | 2 | 0 | 3 | 0.3333 | 2 | 0.5 | 2 | 0 | 9 | 0.2222 |
| 10_7488899 | 1 | 0 | 2 | 0 | 2 | 0.5 | 3 | 0 | 8 | 0.125 | 2 | 0 | 4 | 0.25 | 2 | 0 | 1 | 0 | 9 | 0.1111 |
| 10_7488917 | NA | NA | 2 | 0 | NA | NA | 1 | 1 | 3 | 0.3333 | 2 | 0.5 | 2 | 0 | 1 | 0 | NA | NA | 5 | 0.2 |
| 10_7488920 | 2 | 1 | 1 | 0 | 2 | 1 | 1 | 1 | 6 | 0.8333 | 2 | 0.5 | 3 | 1 | 1 | 0 | 2 | 1 | 8 | 0.75 |
| 10_7488921 | 2 | 0.5 | 2 | 0 | 2 | 0.5 | 3 | 0.3333 | 9 | 0.3333 | 2 | 0 | 4 | 1 | 2 | 0 | 2 | 0.5 | 10 | 0.5 |
| 10_7489020 | 2 | 0 | 3 | 0 | 3 | 0.3333 | 2 | 0 | 10 | 0.1 | 1 | 0 | 5 | 0.6 | 4 | 0.25 | 4 | 0.5 | 14 | 0.4286 |
| 10_7489033 | 1 | 0 | 1 | 0 | 3 | 0.3333 | 1 | 0 | 6 | 0.1667 | 1 | 0 | 4 | 0.5 | 1 | 1 | 2 | 0 | 8 | 0.375 |
| 10_7489426 | 1 | 0 | 2 | 0 | 2 | 0.5 | 4 | 0 | 9 | 0.1111 | 1 | 0 | 3 | 0 | 4 | 0.5 | NA | NA | 8 | 0.25 |
| 10_7489431 | 2 | 0 | NA | NA | 1 | 0 | 4 | 0.5 | 7 | 0.2857 | 1 | 0 | 4 | 0.5 | 4 | 0.25 | NA | NA | 9 | 0.3333 |
| 10_7489464 | 1 | 0 | 2 | 0 | 2 | 0 | 5 | 0.4 | 10 | 0.2 | NA | NA | 2 | 0.5 | 4 | 0 | NA | NA | 6 | 0.1667 |
| 10_7489535 | 1 | 0 | 1 | 0 | 3 | 0 | 5 | 0.4 | 10 | 0.2 | 2 | 0 | 4 | 0.5 | 2 | 0 | NA | NA | 8 | 0.25 |
| 10_7489539 | 1 | 0 | 1 | 0 | 3 | 0.3333 | 4 | 0.75 | 9 | 0.4444 | 2 | 0.5 | 4 | 0.75 | 1 | 0 | NA | NA | 7 | 0.5714 |
| 10_7489548 | 1 | 0 | 1 | 0 | 3 | 0.6667 | 4 | 0 | 9 | 0.2222 | 2 | 0.5 | 5 | 0.4 | 2 | 0.5 | NA | NA | 9 | 0.4444 |
| 10_7489551 | 1 | 0 | 1 | 0 | 3 | 0 | 5 | 0.2 | 10 | 0.1 | 2 | 0 | 3 | 0.3333 | 2 | 0.5 | NA | NA | 7 | 0.2857 |
| 10_7489558 | 1 | 1 | 1 | 1 | 3 | 1 | 4 | 1 | 9 | 1 | 2 | 1 | 4 | 1 | 1 | 1 | NA | NA | 7 | 1 |
| 10_7489562 | NA | NA | 1 | 0 | 3 | 0.3333 | 3 | 0 | 7 | 0.1429 | 2 | 0 | 5 | 0.2 | 2 | 0.5 | NA | NA | 9 | 0.2222 |
| 10_7489567 | NA | NA | 1 | 1 | 2 | 0 | 3 | 0.3333 | 6 | 0.3333 | NA | NA | 5 | 0 | 2 | 0.5 | NA | NA | 7 | 0.1429 |
| 10_7489576 | NA | NA | 1 | 1 | 3 | 1 | 3 | 1 | 7 | 1 | 2 | 0.5 | 4 | 1 | 2 | 1 | NA | NA | 8 | 0.875 |
| 10_7489593 | NA | NA | 1 | 1 | 2 | 1 | 4 | 1 | 7 | 1 | 2 | 1 | 4 | 1 | 2 | 1 | NA | NA | 8 | 1 |
| 10_7489599 | NA | NA | NA | NA | 3 | 0.3333 | 5 | 0 | 8 | 0.125 | 2 | 0 | 3 | 0.3333 | 2 | 0.5 | NA | NA | 7 | 0.2857 |
| 10_7489601 | NA | NA | NA | NA | 2 | 1 | 5 | 1 | 7 | 1 | 2 | 1 | 3 | 1 | 2 | 1 | NA | NA | 7 | 1 |
| 10_7489615 | NA | NA | NA | NA | 2 | 0.5 | 4 | 0.5 | 6 | 0.5 | NA | NA | 3 | 0.3333 | 1 | 1 | NA | NA | 4 | 0.5 |

| | | | | | | | | | | | | | | | | | | | | |
|---|---|---|---|---|---|---|---|---|---|---|---|---|---|---|---|---|---|---|---|---|
| 10_748962 | 1 | 0 | NA | NA | 2 | 0 | 4 | 0.25 | 7 | 0.1429 | NA | NA | 4 | 0.25 | 2 | 0.5 | NA | NA | 6 | 0.3333 |
| 10_748968 | 4 | 0 | 5 | 0.4 | 1 | 1 | 2 | 0 | 12 | 0.25 | 4 | 0 | 2 | 0 | 3 | 0.3333 | 5 | 0.2 | 14 | 0.1429 |
| 10_748968 | 4 | 0.25 | 3 | 0.6667 | 2 | 0 | 1 | 1 | 10 | 0.4 | 2 | 0 | 5 | 0.2 | 5 | 0.4 | 7 | 0.2857 | 19 | 0.2632 |
| 10_748969 | 3 | 0 | 5 | 0 | 3 | 0.3333 | 2 | 0.5 | 13 | 0.1538 | 6 | 0.3333 | 4 | 0 | 6 | 0 | 7 | 0.1429 | 23 | 0.1304 |
| 10_748973 | 5 | 0.2 | 5 | 0 | 4 | 0.25 | 2 | 0 | 16 | 0.125 | 6 | 0.1667 | 6 | 0 | 5 | 0 | 8 | 0.125 | 25 | 0.08 |
| 10_748974 | NA | NA | NA | NA | NA | NA | 1 | 1 | 1 | 1 | NA | NA | 1 | 1 | 1 | 1 | NA | NA | 2 | 1 |
| 10_748975 | NA | NA | NA | NA | NA | NA | 2 | 0.5 | 2 | 0.5 | NA | NA | 1 | 1 | 2 | 1 | NA | NA | 3 | 1 |
| 10_748975 | 6 | 0 | 7 | 0.2857 | 6 | 0 | 4 | 0 | 23 | 0.087 | 4 | 0.25 | 7 | 0.2857 | 8 | 0 | 9 | 0.1111 | 28 | 0.1429 |
| 10_748976 | 5 | 0.6 | 8 | 0.375 | 5 | 0.8 | 4 | 0.5 | 22 | 0.5455 | 7 | 0.4286 | 9 | 0.6667 | 6 | 0.1667 | 9 | 0.4444 | 31 | 0.4516 |
| 10_748978 | 4 | 0.5 | 8 | 0.875 | 7 | 0.5714 | 6 | 0.6667 | 25 | 0.68 | 5 | 0.6 | 9 | 0.5556 | 7 | 0.8571 | 9 | 0.5556 | 30 | 0.6333 |
| 10_748978 | 5 | 1 | 8 | 0.625 | 7 | 0.7143 | 5 | 0.8 | 25 | 0.76 | 5 | 1 | 10 | 0.8 | 7 | 0.8571 | 8 | 0.75 | 30 | 0.8333 |
| 10_748979 | 5 | 0 | 9 | 0.2222 | 7 | 0.1429 | 6 | 0 | 27 | 0.1111 | 4 | 0.25 | 10 | 0.1 | 6 | 0 | 9 | 0.2222 | 29 | 0.1379 |
| 10_750074 | 1 | 1 | NA | NA | NA | NA | 1 | 1 | 2 | 1 | NA | NA | NA | NA | NA | NA | 2 | 0.5 | 2 | 0.5 |
| 10_750075 | 1 | 1 | NA | NA | NA | NA | 1 | 0 | 2 | 0.5 | NA | NA | NA | NA | NA | NA | 2 | 0.5 | 2 | 0.5 |
| 10_750076 | 1 | 1 | NA | NA | NA | NA | 1 | 0 | 2 | 0.5 | NA | NA | NA | NA | NA | NA | 2 | 1 | 2 | 1 |
| 10_750095 | NA | NA | NA | NA | NA | NA | 1 | 1 | 1 | 1 | NA | NA | NA | NA | 1 | 0 | 1 | 1 | 2 | 0.5 |
| 10_750099 | NA | NA | NA | NA | NA | NA | 1 | 1 | 1 | 1 | NA | NA | NA | NA | 1 | 1 | 1 | 1 | 2 | 1 |
| 10_750106 | NA | NA | NA | NA | 1 | 0 | 1 | 1 | 2 | 0.5 | NA | NA | NA | NA | 1 | 0 | 1 | 1 | 2 | 0.5 |
| 10_750181 | 20 | 0.3 | 21 | 0.1429 | 21 | 0.0476 | 19 | 0.0526 | 81 | 0.1358 | 15 | 0 | 26 | 0 | 26 | 0.0769 | 23 | 0.1304 | 90 | 0.0556 |
| 10_750182 | 20 | 0.1 | 22 | 0.0909 | 20 | 0 | 19 | 0.2105 | 81 | 0.0988 | 13 | 0.1538 | 25 | 0.2 | 24 | 0.1667 | 23 | 0.087 | 85 | 0.1529 |
| 10_750182 | 22 | 0.1818 | 20 | 0.25 | 21 | 0.1429 | 20 | 0.2 | 83 | 0.1928 | 15 | 0.1333 | 27 | 0.2593 | 27 | 0.037 | 24 | 0.0833 | 93 | 0.129 |
| 10_750183 | 21 | 0.4762 | 22 | 0.5455 | 21 | 0.4286 | 19 | 0.3684 | 83 | 0.4578 | 15 | 0.5333 | 24 | 0.4583 | 27 | 0.4444 | 24 | 0.5417 | 90 | 0.4889 |
| 10_750183 | 18 | 1 | 19 | 0.9474 | 18 | 0.8889 | 16 | 0.9375 | 71 | 0.9437 | 15 | 0.8667 | 23 | 0.6522 | 31 | 0.9355 | 22 | 0.8636 | 91 | 0.8352 |
| 10_750184 | 17 | 0.4118 | 18 | 0.2778 | 22 | 0.0909 | 19 | 0.3158 | 76 | 0.2632 | 13 | 0.1538 | 26 | 0.3846 | 30 | 0.4 | 23 | 0.3043 | 92 | 0.337 |
| 10_750185 | 20 | 0.35 | 21 | 0.5714 | 20 | 0.35 | 22 | 0.5909 | 83 | 0.4699 | 19 | 0.3684 | 24 | 0.4583 | 31 | 0.5806 | 20 | 0.55 | 94 | 0.5 |
| 10_750185 | 16 | 0.125 | 20 | 0.2 | 14 | 0.2857 | 19 | 0.1579 | 69 | 0.1884 | 18 | 0.0556 | 21 | 0.1429 | 28 | 0.25 | 19 | 0.1053 | 86 | 0.1512 |
| 10_750190 | 10 | 0 | 18 | 0.1111 | 16 | 0.1875 | 17 | 0.1765 | 61 | 0.1311 | 11 | 0.0909 | 17 | 0.2353 | 26 | 0.1154 | 17 | 0.0588 | 71 | 0.1268 |
| 10_750196 | 16 | 0 | 20 | 0.15 | 20 | 0.3 | 24 | 0.0417 | 80 | 0.125 | 15 | 0.1333 | 18 | 0.2222 | 33 | 0.0303 | 19 | 0.0526 | 85 | 0.0941 |
| 10_750197 | 16 | 0.1875 | 20 | 0.1 | 21 | 0.2857 | 22 | 0.1818 | 79 | 0.1899 | 15 | 0 | 21 | 0.1429 | 31 | 0.0645 | 21 | 0.0952 | 88 | 0.0795 |
| 10_750202 | 18 | 0.0556 | 20 | 0.05 | 26 | 0.1154 | 21 | 0.0476 | 85 | 0.0706 | 17 | 0 | 21 | 0.2381 | 33 | 0.0909 | 17 | 0.1176 | 88 | 0.1136 |
| 10_750205 | 14 | 0.4286 | 21 | 0.4762 | 24 | 0.375 | 21 | 0.5238 | 80 | 0.45 | 17 | 0.5882 | 24 | 0.4583 | 32 | 0.375 | 18 | 0.6111 | 91 | 0.4835 |
| 10_750207 | 18 | 0.4444 | 21 | 0.1905 | 26 | 0.3077 | 25 | 0.4 | 90 | 0.3333 | 19 | 0.1579 | 20 | 0.25 | 36 | 0.2778 | 20 | 0.5 | 95 | 0.2947 |
| 10_750811 | NA | NA | 1 | 1 | 1 | 1 | NA | NA | 2 | 1 | NA | NA | NA | NA | NA | NA | 1 | 1 | 1 | 1 |
| 10_750814 | NA | NA | 2 | 1 | NA | NA | NA | NA | 2 | 1 | NA | NA | NA | NA | 1 | 1 | 1 | 1 | 2 | 1 |
| 10_750816 | NA | NA | 3 | 0 | 1 | 1 | NA | NA | 4 | 0.25 | NA | NA | NA | NA | NA | NA | 1 | 1 | 1 | 1 |
| 10_750820 | NA | NA | 2 | 0.5 | 1 | 0 | 1 | 1 | 4 | 0.5 | NA | NA | 1 | 1 | 1 | 1 | 1 | 1 | 3 | 1 |
| 10_750824 | NA | NA | 3 | 0 | 1 | 1 | NA | NA | 4 | 0.25 | NA | NA | 1 | 0 | NA | NA | 1 | 1 | 2 | 0.5 |
| 10_750825 | NA | NA | 2 | 0 | 1 | 1 | NA | NA | 3 | 0.3333 | NA | NA | NA | NA | NA | NA | 1 | 1 | 1 | 1 |
| 10_751457 | 1 | 0 | 5 | 0.4 | 3 | 0.3333 | 2 | 0.5 | 11 | 0.3636 | 1 | 1 | NA | NA | 1 | 0 | NA | NA | 2 | 0.5 |
| 10_751457 | 1 | 0 | 6 | 0 | 4 | 0 | 4 | 0.25 | 15 | 0.0667 | 2 | 0 | 2 | 0.5 | 2 | 0.5 | 2 | 0 | 8 | 0.25 |
| 10_751457 | NA | NA | 5 | 0.4 | 3 | 0.3333 | 4 | 0.25 | 12 | 0.3333 | 1 | 0 | 2 | 0 | 1 | 0 | 2 | 0.5 | 6 | 0.1667 |
| 10_751458 | 1 | 1 | 4 | 1 | 5 | 1 | 5 | 1 | 15 | 1 | 1 | 1 | 2 | 1 | 2 | 1 | 2 | 1 | 7 | 1 |
| 10_751458 | 1 | 1 | 3 | 1 | 3 | 1 | 1 | 1 | 8 | 1 | 1 | 1 | 1 | 1 | NA | NA | NA | NA | 2 | 1 |
| 10_751458 | 1 | 0 | 3 | 1 | 2 | 1 | 2 | 0 | 8 | 0.625 | 2 | 1 | NA | NA | NA | NA | 1 | 1 | 3 | 1 |
| 10_751459 | 1 | 1 | 5 | 0.4 | 4 | 0.25 | 3 | 0.3333 | 13 | 0.3846 | 1 | 0 | NA | NA | NA | NA | 2 | 0.5 | 3 | 0.3333 |
| 10_751460 | 1 | 1 | 7 | 0.2857 | 5 | 0 | 4 | 0.25 | 17 | 0.2353 | 1 | 1 | 2 | 0.5 | 1 | 0 | 2 | 0.5 | 6 | 0.5 |
| 10_751460 | 1 | 0 | 6 | 0.3333 | 5 | 0 | 4 | 0 | 16 | 0.125 | 1 | 0 | NA | NA | 1 | 0 | 2 | 0.5 | 4 | 0.25 |
| 10_751461 | 1 | 0 | 6 | 0 | 5 | 0.2 | 4 | 0 | 16 | 0.0625 | 1 | 0 | 3 | 0 | 2 | 0 | 2 | 0.5 | 8 | 0.125 |
| 10_751463 | 1 | 1 | 6 | 0.6667 | 3 | 0.3333 | 3 | 0.3333 | 13 | 0.5385 | 2 | 0.5 | 3 | 0.6667 | 3 | 0.3333 | 2 | 0.5 | 10 | 0.5 |
| 10_751465 | 1 | 0 | 9 | 0.2222 | 5 | 0.2 | 2 | 0 | 17 | 0.1765 | 2 | 0 | 3 | 0.3333 | 2 | 0 | 2 | 0.5 | 9 | 0.2222 |
| 10_751466 | 1 | 1 | 8 | 0.625 | 5 | 0.8 | 7 | 0.8571 | 21 | 0.7619 | 2 | 0.5 | 5 | 0.6 | 3 | 0.6667 | 2 | 1 | 12 | 0.6667 |
| 10_751467 | 1 | 1 | 9 | 0.1111 | 4 | 0 | 8 | 0 | 22 | 0.0909 | 1 | 0 | 5 | 0 | 3 | 0 | 2 | 0.5 | 11 | 0.0909 |
| 10_751468 | 1 | 0 | 9 | 0.4444 | 5 | 0.2 | 6 | 0.5 | 21 | 0.381 | 2 | 1 | 3 | 0.3333 | 3 | 0.3333 | 2 | 0.5 | 10 | 0.5 |
| 10_751468 | 1 | 0 | 8 | 0 | 4 | 0 | 5 | 0.4 | 18 | 0.1111 | 2 | 1 | 5 | 0 | 2 | 0 | 2 | 0 | 11 | 0.1818 |
| 10_751469 | 1 | 1 | 9 | 1 | 5 | 0.8 | 5 | 1 | 20 | 0.95 | 2 | 1 | 4 | 1 | 3 | 1 | 2 | 1 | 11 | 1 |
| 10_751469 | 1 | 1 | 9 | 0 | 5 | 0 | 7 | 0.2857 | 22 | 0.1364 | 2 | 0 | 4 | 0.25 | 3 | 0 | 1 | 0 | 10 | 0.1 |
| 10_751472 | 1 | 0 | 8 | 0 | 3 | 0.3333 | 4 | 0.25 | 16 | 0.125 | 2 | 0.5 | 4 | 0.5 | 2 | 0 | 1 | 1 | 9 | 0.4444 |
| 10_751472 | 1 | 1 | 10 | 0.2 | 6 | 0.3333 | 8 | 0.125 | 25 | 0.24 | 2 | 0 | 4 | 0.25 | 2 | 0 | 2 | 0 | 10 | 0.1 |
| 10_751616 | NA | NA | NA | NA | NA | NA | 1 | 1 | 1 | 1 | 1 | 1 | NA | NA | NA | NA | NA | NA | 1 | 1 |
| 10_751618 | 8 | 0.875 | 3 | 0.6667 | 7 | 0.8571 | 11 | 0.8182 | 29 | 0.8276 | 5 | 0.8 | 9 | 0.5556 | 20 | 0.85 | 6 | 0.6667 | 40 | 0.75 |
| 10_751689 | 10 | 0.6 | 6 | 0.5 | 9 | 0.4444 | 14 | 0.3571 | 39 | 0.4615 | 4 | 0.5 | 17 | 0.3529 | 22 | 0.4545 | 7 | 0.4286 | 50 | 0.42 |
| 10_751689 | 10 | 0.6 | 4 | 0.25 | 9 | 0.4444 | 13 | 0.3077 | 36 | 0.4167 | 4 | 0.75 | 17 | 0.4706 | 18 | 0.6111 | 6 | 0.3333 | 45 | 0.5333 |
| 10_751690 | 10 | 0 | 5 | 0 | 14 | 0.1429 | 18 | 0.0556 | 47 | 0.0638 | 5 | 0 | 19 | 0.0526 | 25 | 0.04 | 9 | 0.1111 | 58 | 0.0517 |
| 10_751691 | 11 | 0.2727 | 7 | 0.1429 | 15 | 0.0667 | 16 | 0.0625 | 49 | 0.1224 | 5 | 0 | 17 | 0.0588 | 29 | 0.1724 | 11 | 0 | 62 | 0.0968 |
| 10_751691 | 8 | 1 | 7 | 1 | 13 | 1 | 16 | 0.9375 | 44 | 0.9773 | 4 | 1 | 19 | 1 | 24 | 0.9583 | 10 | 1 | 57 | 0.9825 |

| | | | | | | | | | | | | | | | | | | | | |
|---|---|---|---|---|---|---|---|---|---|---|---|---|---|---|---|---|---|---|---|---|
| 10_7516941 | 13 | 0.0769 | 8 | 0.25 | 17 | 0 | 22 | 0.1364 | 60 | 0.1 | 6 | 0.1667 | 23 | 0.0435 | 37 | 0.027 | 14 | 0.2143 | 80 | 0.075 |
| 10_7516943 | 10 | 0.2 | 6 | 0 | 16 | 0.0625 | 15 | 0.0667 | 47 | 0.0851 | 6 | 0 | 17 | 0.1176 | 29 | 0.1379 | 11 | 0.1818 | 63 | 0.127 |
| 10_7516945 | 11 | 0.1818 | 8 | 0.25 | 15 | 0 | 19 | 0.1053 | 53 | 0.1132 | 6 | 0.3333 | 20 | 0.2 | 34 | 0.0588 | 12 | 0.25 | 72 | 0.1528 |
| 10_7516973 | 12 | 0.0833 | 8 | 0 | 13 | 0.0769 | 19 | 0.1053 | 52 | 0.0769 | 3 | 0 | 21 | 0.0952 | 29 | 0.069 | 11 | 0.0909 | 64 | 0.0781 |
| 10_7516977 | 14 | 0.5 | 10 | 0.9 | 20 | 0.85 | 21 | 0.7619 | 65 | 0.7538 | 6 | 0.8333 | 27 | 0.8519 | 35 | 0.7143 | 16 | 0.8125 | 84 | 0.7857 |
| 10_7516980 | 17 | 0.1765 | 10 | 0.1 | 18 | 0.1667 | 24 | 0.0833 | 69 | 0.1304 | 7 | 0 | 28 | 0.1071 | 41 | 0.2683 | 15 | 0.1333 | 91 | 0.1758 |
| 10_7517022 | 18 | 0.2222 | 9 | 0 | 20 | 0.15 | 23 | 0.087 | 70 | 0.1286 | 7 | 0 | 34 | 0.0294 | 41 | 0.1707 | 20 | 0.1 | 102 | 0.098 |
| 10_7517027 | 24 | 0.1667 | 15 | 0.2667 | 24 | 0.25 | 33 | 0.2121 | 96 | 0.2188 | 9 | 0.2222 | 37 | 0.2703 | 58 | 0.2069 | 24 | 0.125 | 128 | 0.2109 |
| 10_7517045 | 30 | 0.0667 | 16 | 0.0625 | 27 | 0.1111 | 37 | 0.0811 | 110 | 0.0818 | 12 | 0.25 | 40 | 0.15 | 59 | 0.0678 | 28 | 0.1071 | 139 | 0.1151 |
| 10_7517046 | 30 | 0.5667 | 15 | 0.6 | 28 | 0.3571 | 38 | 0.5526 | 111 | 0.5135 | 13 | 0.6154 | 38 | 0.5 | 59 | 0.5763 | 25 | 0.4 | 135 | 0.5259 |
| 10_7534324 | NA | NA | 2 | 0 | NA | NA | 2 | 0.5 | 4 | 0.25 | NA | NA | 1 | 1 | 1 | 1 | 2 | 0 | 4 | 0.5 |
| 10_7534351 | NA | NA | 2 | 0 | NA | NA | 1 | 1 | 3 | 0.3333 | NA | NA | 1 | 1 | 2 | 0 | 2 | 0 | 5 | 0.2 |
| 10_7534428 | NA | NA | NA | NA | NA | NA | 1 | 1 | 1 | 1 | NA | NA | 1 | 1 | 2 | 1 | 2 | 1 | 5 | 1 |
| 10_7534443 | NA | NA | NA | NA | NA | NA | 2 | 0.5 | 2 | 0.5 | NA | NA | 1 | 0 | 2 | 0 | 2 | 0.5 | 5 | 0.2 |
| 10_7534457 | NA | NA | NA | NA | NA | NA | 2 | 1 | 2 | 1 | NA | NA | 1 | 1 | 2 | 1 | 2 | 1 | 5 | 1 |
| 10_7545088 | 1 | 1 | NA | NA | NA | NA | 1 | 1 | 2 | 1 | NA | NA | NA | NA | NA | NA | 2 | 1 | 2 | 1 |
| 10_7547280 | 2 | 0.5 | 3 | 0 | NA | NA | NA | NA | 5 | 0.2 | 1 | 0 | 1 | 1 | 1 | 0 | 1 | 0 | 4 | 0.25 |
| 10_7547290 | 2 | 0.5 | 3 | 0.6667 | NA | NA | NA | NA | 5 | 0.6 | 1 | 0 | 1 | 1 | NA | NA | 1 | 1 | 3 | 0.6667 |
| 10_7547304 | NA | NA | 3 | 0.6667 | NA | NA | NA | NA | 3 | 0.6667 | 2 | 0.5 | 1 | 0 | 1 | 1 | 1 | 0 | 5 | 0.4 |
| 10_7548210 | 4 | 0 | 6 | 0.1667 | 14 | 0.2857 | 11 | 0.1818 | 35 | 0.2 | 4 | 0 | 4 | 0 | 8 | 0.125 | 7 | 0.1429 | 23 | 0.087 |
| 10_7549297 | 3 | 0.3333 | 4 | 0 | 5 | 0.4 | 9 | 0.1111 | 21 | 0.1905 | 1 | 1 | 2 | 0 | 5 | 0 | 3 | 0 | 11 | 0.0909 |
| 10_7549304 | 3 | 0.3333 | 3 | 0.3333 | 6 | 0 | 8 | 0 | 20 | 0.1 | 1 | 0 | 2 | 0 | 4 | 0.25 | 3 | 0.3333 | 10 | 0.2 |
| 10_7549306 | 3 | 0.3333 | 4 | 0 | 6 | 0 | 9 | 0.2222 | 22 | 0.1364 | 1 | 0 | 2 | 0 | 4 | 0.25 | 3 | 0 | 10 | 0.1 |
| 10_7549307 | 3 | 0 | 4 | 0.5 | 6 | 0 | 8 | 0 | 21 | 0.0952 | 1 | 0 | 2 | 0.5 | 2 | 0 | 3 | 0.6667 | 8 | 0.375 |
| 10_7549308 | 2 | 0.5 | 4 | 0.5 | 5 | 0.4 | 9 | 0.4444 | 20 | 0.45 | NA | NA | 2 | 0.5 | 2 | 0 | 3 | 0.6667 | 7 | 0.4286 |
| 10_7560728 | 13 | 0.0769 | 9 | 0 | 12 | 0.1667 | 11 | 0 | 45 | 0.0667 | 6 | 0.1667 | 15 | 0.2667 | 9 | 0 | 12 | 0.25 | 42 | 0.1905 |
| 10_7561063 | 2 | 0.5 | 1 | 1 | NA | NA | 2 | 0 | 5 | 0.4 | NA | NA | 3 | 0 | NA | NA | 3 | 1 | 6 | 0.5 |
| 10_7561069 | 2 | 1 | 1 | 0 | 1 | 0 | 1 | 0 | 5 | 0.4 | 1 | 0 | 3 | 0.3333 | NA | NA | 3 | 0.3333 | 7 | 0.2857 |
| 10_7561994 | 3 | 0.3333 | 3 | 0.3333 | NA | NA | 2 | 0.5 | 8 | 0.375 | 1 | 0 | NA | NA | 1 | 0 | 2 | 0.5 | 4 | 0.25 |
| 10_7569588 | 1 | 0 | NA | NA | 2 | 0.5 | 1 | 0 | 4 | 0.25 | NA | NA | 2 | 0.5 | NA | NA | 1 | 1 | 3 | 0.6667 |
| 10_7573131 | NA | NA | 1 | 1 | NA | NA | NA | NA | 1 | 1 | 1 | 1 | 1 | 1 | NA | NA | NA | NA | 2 | 1 |
| 10_7573162 | NA | NA | 1 | 1 | NA | NA | NA | NA | 1 | 1 | 2 | 1 | 1 | 0 | NA | NA | NA | NA | 3 | 0.6667 |
| 10_7573278 | 9 | 0.1111 | 11 | 0 | 12 | 0.1667 | 11 | 0.0909 | 43 | 0.093 | 8 | 0.25 | 9 | 0.1111 | 13 | 0.2308 | 10 | 0.1 | 40 | 0.175 |
| 10_7575751 | NA | NA | NA | NA | 1 | 1 | 1 | 1 | 2 | 1 | NA | NA | NA | NA | 1 | 1 | NA | NA | 1 | 1 |
| 10_7587432 | NA | NA | 1 | 1 | NA | NA | 1 | 1 | 2 | 1 | 2 | 1 | NA | NA | 2 | 1 | NA | NA | 4 | 1 |
| 10_7587477 | NA | NA | 2 | 0.5 | NA | NA | 1 | 0 | 3 | 0.3333 | 2 | 1 | NA | NA | 2 | 0 | NA | NA | 4 | 0.5 |
| 10_7587484 | NA | NA | 1 | 1 | NA | NA | 1 | 0 | 2 | 0.5 | 2 | 0.5 | NA | NA | 2 | 0.5 | 1 | 1 | 5 | 0.6 |
| 10_7587486 | NA | NA | 2 | 0.5 | NA | NA | 1 | 1 | 3 | 0.6667 | 2 | 0.5 | NA | NA | 2 | 0.5 | 1 | 1 | 5 | 0.6 |
| 10_7587490 | NA | NA | 2 | 1 | NA | NA | 1 | 1 | 3 | 1 | 2 | 1 | NA | NA | 2 | 1 | 1 | 1 | 5 | 1 |
| 10_7587495 | NA | NA | NA | NA | NA | NA | 1 | 1 | 1 | 1 | 1 | 1 | NA | NA | 2 | 0.5 | 1 | 1 | 4 | 0.75 |
| 10_7587496 | NA | NA | 2 | 1 | NA | NA | NA | NA | 2 | 1 | 2 | 1 | NA | NA | 2 | 1 | 1 | 1 | 5 | 1 |
| 10_7587520 | 1 | 1 | 2 | 0 | NA | NA | 1 | 0 | 4 | 0.25 | 2 | 0 | NA | NA | 2 | 0 | 1 | 1 | 5 | 0.2 |
| 10_7587527 | 1 | 1 | 2 | 1 | NA | NA | 1 | 1 | 4 | 1 | 2 | 1 | NA | NA | 2 | 1 | 1 | 1 | 5 | 1 |
| 10_7587537 | 1 | 0 | 2 | 0 | NA | NA | 1 | 1 | 4 | 0.25 | 2 | 0 | NA | NA | 2 | 0.5 | 1 | 0 | 5 | 0.2 |
| 10_7587538 | 1 | 1 | 1 | 0 | NA | NA | 1 | 1 | 3 | 0.6667 | 2 | 0.5 | NA | NA | 2 | 1 | 1 | 1 | 5 | 0.8 |
| 10_7587542 | 1 | 0 | 2 | 0 | NA | NA | 1 | 1 | 4 | 0.25 | 2 | 0.5 | NA | NA | 2 | 0 | 1 | 0 | 5 | 0.2 |
| 10_7587553 | 1 | 1 | 2 | 0.5 | NA | NA | 1 | 1 | 4 | 0.75 | 2 | 1 | NA | NA | 2 | 0.5 | 1 | 1 | 5 | 0.8 |
| 10_7587567 | NA | NA | 2 | 0.5 | NA | NA | 1 | 1 | 3 | 0.6667 | 2 | 1 | 1 | 1 | 2 | 0.5 | NA | NA | 5 | 0.8 |
| 10_7587570 | 1 | 1 | 2 | 1 | NA | NA | NA | NA | 3 | 1 | 2 | 1 | 1 | 1 | 2 | 1 | 1 | 1 | 6 | 1 |
| 10_7587575 | 1 | 0 | 3 | 1 | NA | NA | NA | NA | 4 | 0.75 | 2 | 0.5 | 1 | 0 | 1 | 1 | 1 | 1 | 5 | 0.6 |
| 10_7587581 | NA | NA | 2 | 1 | NA | NA | NA | NA | 2 | 1 | 2 | 1 | 1 | 1 | 2 | 1 | 1 | 1 | 6 | 1 |
| 10_7587598 | 1 | 1 | 3 | 1 | NA | NA | 1 | 1 | 5 | 1 | 2 | 1 | 1 | 1 | 1 | 1 | 1 | 1 | 5 | 1 |
| 10_7587621 | 1 | 1 | 4 | 1 | NA | NA | 2 | 0.5 | 7 | 0.8571 | 2 | 1 | 1 | 1 | 2 | 1 | 2 | 1 | 7 | 1 |
| 10_7587624 | 1 | 1 | 4 | 0.5 | NA | NA | 1 | 0 | 6 | 0.5 | 2 | 0.5 | 1 | 0 | 2 | 0 | 2 | 1 | 7 | 0.4286 |
| 10_7587649 | 1 | 1 | 4 | 1 | NA | NA | 2 | 1 | 7 | 1 | 2 | 1 | 1 | 1 | 2 | 1 | 2 | 1 | 7 | 1 |
| 10_7587656 | 1 | 1 | 4 | 1 | NA | NA | 2 | 1 | 7 | 1 | 2 | 1 | 1 | 1 | 2 | 0.5 | 2 | 1 | 7 | 0.8571 |
| 10_7587658 | NA | NA | 4 | 0.25 | NA | NA | 1 | 1 | 5 | 0.4 | 2 | 1 | NA | NA | 2 | 0.5 | 2 | 0.5 | 6 | 0.6667 |
| 10_7587665 | 1 | 1 | 4 | 1 | NA | NA | 2 | 0.5 | 7 | 0.8571 | 1 | 1 | 1 | 1 | 3 | 0.3333 | 2 | 1 | 7 | 0.7143 |
| 10_7587672 | 1 | 1 | 4 | 0.25 | NA | NA | 2 | 0.5 | 7 | 0.4286 | 2 | 0.5 | 1 | 0 | 3 | 0 | 2 | 0.5 | 8 | 0.25 |
| 10_7587675 | 1 | 1 | 4 | 0.25 | NA | NA | 2 | 0.5 | 7 | 0.4286 | 2 | 1 | 1 | 0 | 3 | 1 | 2 | 0.5 | 8 | 0.75 |
| 10_7587677 | 1 | 1 | 4 | 0.75 | NA | NA | 2 | 1 | 7 | 0.8571 | 2 | 1 | NA | NA | 3 | 0 | 1 | 1 | 6 | 0.5 |
| 10_7587734 | 3 | 0.3333 | 6 | 0 | NA | NA | 2 | 0 | 11 | 0.0909 | 2 | 0 | 1 | 0 | 3 | 0.6667 | 2 | 0 | 8 | 0.25 |
| 10_7587763 | 3 | 0.3333 | 6 | 0 | 1 | 0 | 2 | 0 | 12 | 0.0833 | 2 | 0.5 | 1 | 0 | 3 | 0 | 2 | 0 | 8 | 0.125 |
| 10_7587788 | 3 | 0 | 6 | 0 | 1 | 0 | 2 | 0.5 | 12 | 0.0833 | 3 | 0.3333 | 1 | 0 | 6 | 0 | 2 | 0 | 12 | 0.0833 |
| 10_7604314 | 10 | 0.4 | 5 | 0.2 | 10 | 0.3 | 14 | 0.2857 | 39 | 0.3077 | 5 | 0.8 | 6 | 0.8333 | 9 | 0.2222 | 4 | 0.25 | 24 | 0.5 |

| | | | | | | | | | | | | | | | | | | | | |
|---|---|---|---|---|---|---|---|---|---|---|---|---|---|---|---|---|---|---|---|---|
| 10_7604317 | 11 | 0.1818 | 4 | 0 | 10 | 0.1 | 15 | 0.4667 | 40 | 0.25 | 5 | 0.4 | 7 | 0.1429 | 9 | 0.2222 | 4 | 0.5 | 25 | 0.28 |
| 10_7604323 | 11 | 0 | 5 | 0 | 9 | 0.1111 | 16 | 0.125 | 41 | 0.0732 | 5 | 0.2 | 6 | 0.1667 | 7 | 0 | 2 | 0 | 20 | 0.1 |
| 10_7604323 | 10 | 0.7 | 5 | 0.8 | 11 | 0.7273 | 13 | 0.4615 | 39 | 0.641 | 5 | 0.6 | 5 | 0.8 | 9 | 0.7778 | 4 | 0.5 | 23 | 0.6957 |
| 10_7604330 | 11 | 0.1818 | 5 | 0.2 | 9 | 0.1111 | 16 | 0.1875 | 41 | 0.1707 | 5 | 0 | 7 | 0.1429 | 7 | 0.1429 | 4 | 0 | 23 | 0.087 |
| 10_7609595 | 22 | 0 | 19 | 0.1579 | 16 | 0.0625 | 12 | 0 | 69 | 0.058 | 24 | 0.0417 | 15 | 0.1333 | 20 | 0.05 | 12 | 0 | 71 | 0.0563 |
| 10_7609615 | 18 | 0.3333 | 16 | 0.0625 | 17 | 0.1765 | 15 | 0.1333 | 66 | 0.1818 | 18 | 0.2778 | 14 | 0.2143 | 20 | 0.25 | 12 | 0.3333 | 64 | 0.2656 |
| 10_7609641 | 21 | 0.2857 | 18 | 0.3889 | 18 | 0.2222 | 14 | 0.1429 | 71 | 0.2676 | 19 | 0.5789 | 13 | 0.5385 | 19 | 0.1579 | 15 | 0.3333 | 66 | 0.3939 |
| 10_7609656 | 17 | 0.1765 | 14 | 0.5 | 10 | 0.3 | 10 | 0.1 | 51 | 0.2745 | 21 | 0.2381 | 9 | 0.3333 | 16 | 0.25 | 11 | 0.1818 | 57 | 0.2456 |
| 10_7609658 | 22 | 0.1364 | 20 | 0.05 | 16 | 0 | 17 | 0.0588 | 75 | 0.0667 | 23 | 0.087 | 15 | 0.1333 | 21 | 0.0476 | 14 | 0.0714 | 73 | 0.0822 |
| 10_7609664 | 25 | 0.04 | 19 | 0.1579 | 17 | 0.1176 | 16 | 0.125 | 77 | 0.1039 | 20 | 0.2 | 13 | 0.3077 | 19 | 0.0526 | 13 | 0.1538 | 65 | 0.1692 |
| 10_7609686 | 25 | 0.56 | 23 | 0.4783 | 19 | 0.3158 | 16 | 0.6875 | 83 | 0.506 | 23 | 0.7826 | 15 | 0.5333 | 20 | 0.5 | 15 | 0.4 | 73 | 0.5753 |
| 10_7609688 | 22 | 0.2273 | 23 | 0.1739 | 19 | 0.2632 | 16 | 0.1875 | 80 | 0.2125 | 25 | 0.36 | 15 | 0.4 | 21 | 0.1429 | 14 | 0.1429 | 75 | 0.2667 |
| 10_7609980 | 3 | 0 | 4 | 0.25 | 6 | 0 | 4 | 0 | 17 | 0.0588 | 3 | 0 | 5 | 0 | 5 | 0.2 | 5 | 0 | 18 | 0.0556 |
| 10_7609989 | 4 | 0.25 | 3 | 0.6667 | 6 | 0.5 | 4 | 0.5 | 17 | 0.4706 | 3 | 0.3333 | 5 | 0.6 | 4 | 0.5 | 4 | 0.75 | 16 | 0.5625 |
| 10_7610006 | 4 | 0.25 | 4 | 0.25 | 6 | 0.3333 | 4 | 0 | 18 | 0.2222 | NA | NA | 6 | 0.3333 | 6 | 0.1667 | 6 | 0.1667 | 18 | 0.2222 |
| 10_7610042 | 1 | 0 | 2 | 0 | 4 | 0 | 3 | 0.3333 | 10 | 0.1 | 2 | 0 | 4 | 0 | 2 | 0.5 | 4 | 0 | 12 | 0.0833 |
| 10_7610044 | 1 | 0 | 2 | 0 | 3 | 0.3333 | 2 | 0 | 8 | 0.125 | 2 | 0 | 1 | 0 | 1 | 1 | 3 | 0.3333 | 7 | 0.2857 |
| 10_7618721 | 2 | 0 | 1 | 1 | 3 | 0.3333 | 3 | 0 | 9 | 0.2222 | 2 | 1 | 1 | 0 | 2 | 1 | 4 | 0.5 | 9 | 0.6667 |
| 10_7618748 | 1 | 0 | NA | NA | 3 | 0.3333 | 2 | 0.5 | 6 | 0.3333 | 1 | 0 | 1 | 1 | 1 | 0 | 3 | 0.6667 | 6 | 0.5 |
| 10_7618754 | 2 | 0.5 | 1 | 0 | 3 | 0.6667 | 3 | 0 | 9 | 0.3333 | 2 | 0.5 | 1 | 1 | NA | NA | 2 | 0.5 | 5 | 0.6 |
| 10_7620327 | 1 | 0 | 1 | 0 | 2 | 0 | 4 | 0.25 | 8 | 0.125 | 1 | 1 | 2 | 0 | 2 | 0 | 2 | 1 | 7 | 0.4286 |
| 10_7621080 | 1 | 0 | NA | NA | NA | NA | 3 | 0.3333 | 4 | 0.25 | 1 | 1 | 2 | 0.5 | 2 | 0 | 2 | 0 | 7 | 0.2857 |
| 10_7622481 | NA | NA | NA | NA | NA | NA | 2 | 0.5 | 2 | 0.5 | 1 | 1 | 1 | 0 | 1 | 0 | 1 | 0 | 4 | 0.25 |
| 10_7622490 | NA | NA | NA | NA | NA | NA | 2 | 0.5 | 2 | 0.5 | 1 | 0 | NA | NA | 1 | 0 | 1 | 1 | 3 | 0.3333 |
| 10_7622503 | NA | NA | NA | NA | NA | NA | 1 | 1 | 1 | 1 | NA | NA | 1 | 1 | 1 | 1 | 1 | 1 | 3 | 1 |
| 10_7622525 | NA | NA | NA | NA | NA | NA | 1 | 1 | 1 | 1 | 1 | 1 | 1 | 0 | NA | NA | 1 | 1 | 3 | 0.6667 |
| 10_7622808 | NA | NA | NA | NA | NA | NA | 1 | 1 | 1 | 1 | 1 | 0 | NA | NA | NA | NA | 1 | 1 | 2 | 0.5 |
| 10_7622847 | NA | NA | NA | NA | NA | NA | 1 | 1 | 1 | 1 | 1 | 1 | NA | NA | NA | NA | NA | NA | 1 | 1 |
| 10_7622871 | NA | NA | NA | NA | NA | NA | 1 | 1 | 1 | 1 | 1 | 1 | 1 | 1 | NA | NA | NA | NA | 2 | 1 |
| 10_7629671 | NA | NA | NA | NA | 1 | 1 | NA | NA | 1 | 1 | NA | NA | NA | NA | 1 | 1 | NA | NA | 1 | 1 |
| 10_7629823 | NA | NA | NA | NA | 1 | 1 | NA | NA | 1 | 1 | NA | NA | NA | NA | 1 | 1 | NA | NA | 1 | 1 |
| 10_7629846 | NA | NA | NA | NA | 1 | 1 | NA | NA | 1 | 1 | NA | NA | NA | NA | 1 | 1 | NA | NA | 1 | 1 |
| 10_7629904 | NA | NA | NA | NA | 1 | 1 | NA | NA | 1 | 1 | NA | NA | NA | NA | 1 | 1 | NA | NA | 1 | 1 |
| 10_7630600 | 1 | 0 | NA | NA | 1 | 0 | 1 | 1 | 3 | 0.3333 | NA | NA | 1 | 1 | NA | NA | 1 | 0 | 2 | 0.5 |
| 10_7630610 | 1 | 0 | NA | NA | 1 | 1 | 1 | 1 | 3 | 0.6667 | NA | NA | 1 | 1 | NA | NA | 1 | 1 | 2 | 1 |
| 10_7630613 | NA | NA | NA | NA | 1 | 1 | 1 | 0 | 2 | 0.5 | NA | NA | 1 | 1 | NA | NA | 1 | 1 | 2 | 1 |
| 10_7630686 | 1 | 0 | NA | NA | 1 | 0 | 1 | 1 | 3 | 0.3333 | NA | NA | 2 | 0.5 | 1 | 0 | 1 | 0 | 4 | 0.25 |
| 10_7631367 | 1 | 0 | NA | NA | 1 | 1 | 1 | 0 | 3 | 0.3333 | NA | NA | 2 | 0.5 | 1 | 0 | 1 | 0 | 4 | 0.25 |
| 10_7634244 | 4 | 0 | 3 | 0 | 2 | 0 | 9 | 0.1111 | 18 | 0.0556 | 2 | 0.5 | 3 | 0 | 4 | 0 | 2 | 0 | 11 | 0.0909 |
| 10_7634250 | 3 | 0.3333 | 3 | 0.6667 | 1 | 1 | 8 | 0.375 | 15 | 0.4667 | 1 | 0 | 1 | 0 | 5 | 0.2 | 2 | 0 | 9 | 0.1111 |
| 10_7635156 | 7 | 0 | 2 | 0 | 4 | 0 | 10 | 0.2 | 23 | 0.087 | 1 | 0 | 6 | 0 | 5 | 0.2 | 3 | 0.3333 | 15 | 0.1333 |
| 10_7638909 | 1 | 0 | 2 | 0 | 2 | 1 | NA | NA | 5 | 0.4 | 1 | 1 | 2 | 0.5 | 2 | 0.5 | 1 | 0 | 6 | 0.5 |
| 10_7638915 | 1 | 1 | 1 | 0 | 1 | 1 | NA | NA | 3 | 0.6667 | 1 | 0 | 1 | 0 | 1 | 1 | 1 | 1 | 4 | 0.5 |
| 10_7638959 | 2 | 0.5 | 2 | 0 | 2 | 0 | NA | NA | 6 | 0.1667 | 1 | 0 | 3 | 0 | 2 | 0.5 | 1 | 0 | 7 | 0.1429 |
| 10_7639185 | 1 | 0 | 2 | 0 | 2 | 0.5 | NA | NA | 5 | 0.2 | 1 | 0 | 1 | 0 | 4 | 0.25 | NA | NA | 6 | 0.1667 |
| 10_7643191 | 1 | 1 | 1 | 1 | NA | NA | 2 | 0.5 | 4 | 0.75 | NA | NA | 2 | 0.5 | NA | NA | 1 | 1 | 3 | 0.6667 |
| 10_7643210 | 1 | 1 | 1 | 1 | NA | NA | 2 | 0.5 | 4 | 0.75 | NA | NA | 2 | 0.5 | NA | NA | NA | NA | 2 | 0.5 |
| 10_7643216 | 1 | 1 | 1 | 0 | NA | NA | 2 | 1 | 4 | 0.75 | NA | NA | 1 | 1 | NA | NA | 1 | 1 | 2 | 1 |
| 10_7643260 | 1 | 1 | 1 | 1 | NA | NA | NA | NA | 2 | 1 | NA | NA | 1 | 1 | NA | NA | 1 | 1 | 2 | 1 |
| 10_7654480 | 2 | 0 | 3 | 0.3333 | NA | NA | 2 | 0 | 7 | 0.1429 | 1 | 0 | 3 | 0.3333 | 5 | 0.4 | 3 | 0 | 12 | 0.25 |
| 10_7654502 | 4 | 0 | 5 | 0.2 | NA | NA | 3 | 0 | 12 | 0.0833 | 1 | 0 | 4 | 0.25 | 6 | 0 | 3 | 0 | 14 | 0.0714 |
| 10_7654535 | 3 | 0.3333 | 5 | 0.6 | 1 | 0 | 3 | 0.3333 | 12 | 0.4167 | 1 | 0 | 5 | 0 | 6 | 0.1667 | 3 | 1 | 15 | 0.2667 |
| 10_7654537 | 3 | 0.6667 | 4 | 1 | 1 | 0 | 3 | 1 | 11 | 0.8182 | 1 | 0 | 4 | 0.5 | 6 | 0.8333 | 2 | 0.5 | 13 | 0.6154 |
| 10_7654542 | 1 | 0 | 4 | 0.5 | 1 | 1 | 2 | 0.5 | 8 | 0.5 | 1 | 0 | 3 | 0 | 3 | 0.6667 | 1 | 0 | 8 | 0.25 |
| 10_7654550 | 3 | 0.3333 | 3 | 0.6667 | NA | NA | 3 | 1 | 9 | 0.6667 | 1 | 1 | 5 | 0.8 | 5 | 0.8 | 3 | 0.6667 | 14 | 0.7857 |
| 10_7654555 | 3 | 0.3333 | 4 | 0 | 1 | 1 | 3 | 0.3333 | 11 | 0.2727 | 1 | 1 | 5 | 0.8 | 6 | 0.5 | 3 | 1 | 15 | 0.7333 |
| 10_7654571 | 3 | 0 | 5 | 0.6 | 1 | 0 | 3 | 0.3333 | 12 | 0.3333 | 1 | 0 | 3 | 0 | 5 | 0.6 | 3 | 0 | 12 | 0.25 |
| 10_7654576 | 3 | 0 | 4 | 0.25 | 1 | 1 | 2 | 0 | 10 | 0.2 | 1 | 0 | 1 | 1 | 4 | 0.25 | 3 | 0.3333 | 9 | 0.3333 |
| 10_7659774 | 6 | 0.5 | 13 | 0 | 3 | 0.6667 | 6 | 0.3333 | 28 | 0.25 | 12 | 0 | 6 | 0.3333 | 7 | 0.1429 | 5 | 0 | 30 | 0.1 |
| 10_7659795 | 7 | 0.1429 | 12 | 0.0833 | 4 | 0 | 4 | 0 | 27 | 0.0741 | 10 | 0.1 | 4 | 0 | 7 | 0.1429 | 5 | 0 | 26 | 0.0769 |
| 10_7659805 | 7 | 0.1429 | 12 | 0.0833 | 4 | 0 | 5 | 0.2 | 28 | 0.1071 | 11 | 0 | 6 | 0.3333 | 7 | 0.4286 | 5 | 0 | 29 | 0.1724 |
| 10_7659825 | 5 | 0 | 11 | 0.6364 | 4 | 0.5 | 4 | 0.5 | 24 | 0.4583 | 10 | 0.2 | 4 | 0.25 | 6 | 0.3333 | 3 | 0 | 23 | 0.2174 |
| 10_7662123 | NA | NA | NA | NA | 1 | 1 | NA | NA | 1 | 1 | NA | NA | NA | NA | 1 | 1 | NA | NA | 1 | 1 |
| 10_7662183 | NA | NA | NA | NA | 1 | 1 | NA | NA | 1 | 1 | NA | NA | NA | NA | 1 | 1 | NA | NA | 1 | 1 |
| 10_7662312 | NA | NA | NA | NA | 1 | 1 | NA | NA | 1 | 1 | NA | NA | NA | NA | 1 | 1 | NA | NA | 1 | 1 |

| | | | | | | | | | | | | | | | | | | | | |
|---|---|---|---|---|---|---|---|---|---|---|---|---|---|---|---|---|---|---|---|---|
| 10_768779 | NA | NA | 3 | 1 | NA | NA | NA | NA | 3 | 1 | 3 | 1 | NA | NA | NA | NA | 3 | 1 | 6 | 1 |
| 10_768781 | NA | NA | 3 | 0.3333 | 1 | 0 | 2 | 0 | 6 | 0.1667 | 3 | 0.3333 | NA | NA | NA | NA | 3 | 0.3333 | 6 | 0.3333 |
| 10_768781 | NA | NA | 3 | 0.3333 | 1 | 0 | 2 | 0 | 6 | 0.1667 | 2 | 0.5 | NA | NA | NA | NA | 2 | 0 | 4 | 0.25 |
| 10_768782 | NA | NA | 3 | 0.6667 | 1 | 1 | 2 | 0.5 | 6 | 0.6667 | 2 | 0.5 | NA | NA | NA | NA | 3 | 0.3333 | 5 | 0.4 |
| 10_768782 | NA | NA | 1 | 1 | 1 | 1 | 1 | 1 | 3 | 1 | 2 | 1 | NA | NA | NA | NA | 3 | 1 | 5 | 1 |
| 10_768785 | NA | NA | NA | NA | NA | NA | 1 | 1 | 1 | 1 | NA | NA | NA | NA | NA | NA | 2 | 1 | 2 | 1 |
| 10_768786 | NA | NA | 3 | 0.6667 | NA | NA | 1 | 1 | 4 | 0.75 | 3 | 1 | NA | NA | NA | NA | 2 | 0.5 | 5 | 0.8 |
| 10_768787 | NA | NA | 3 | 0 | NA | NA | 2 | 0.5 | 5 | 0.2 | 4 | 0.5 | NA | NA | NA | NA | 3 | 0 | 7 | 0.2857 |
| 10_768857 | 2 | 0 | 2 | 0.5 | 1 | 0 | 2 | 1 | 7 | 0.4286 | 5 | 0.4 | NA | NA | NA | NA | 3 | 0.3333 | 8 | 0.375 |
| 10_768858 | 2 | 0.5 | 4 | 0.5 | 1 | 1 | 2 | 0.5 | 9 | 0.5556 | 4 | 0.25 | NA | NA | NA | NA | 3 | 0.6667 | 7 | 0.4286 |
| 10_771046 | 6 | 0.3333 | 3 | 0.6667 | 3 | 0.6667 | 2 | 0 | 14 | 0.4286 | 4 | 0.5 | 3 | 0.3333 | 2 | 0.5 | 3 | 0.3333 | 12 | 0.4167 |
| 10_771047 | 5 | 0.6 | 4 | 0.5 | 2 | 0 | 2 | 1 | 13 | 0.5385 | 4 | 0.5 | 3 | 0.6667 | 2 | 0.5 | 3 | 0.3333 | 12 | 0.5 |
| 10_771047 | 4 | 0.25 | 4 | 0.25 | 2 | 1 | 2 | 0 | 12 | 0.3333 | 3 | 0.3333 | 3 | 0 | 2 | 0 | 3 | 0 | 11 | 0.0909 |
| 10_771048 | 6 | 0.6667 | 4 | 0.25 | 4 | 0.75 | 1 | 0 | 15 | 0.5333 | 3 | 0.3333 | 2 | 0.5 | 3 | 0.6667 | 3 | 0.3333 | 11 | 0.4545 |
| 10_771050 | 6 | 0.5 | 2 | 1 | 4 | 0.75 | 2 | 0.5 | 14 | 0.6429 | 4 | 0.75 | 4 | 1 | 3 | 1 | 4 | 0.75 | 15 | 0.8667 |
| 10_771050 | 4 | 0.5 | 4 | 0.75 | 4 | 0.25 | 2 | 0 | 14 | 0.4286 | 3 | 0.6667 | 3 | 0.3333 | 4 | 1 | 3 | 0.6667 | 13 | 0.6923 |
| 10_771053 | 5 | 0.2 | 5 | 0.2 | 4 | 0.25 | 3 | 0 | 17 | 0.1765 | 4 | 0.25 | 3 | 0.6667 | 4 | 0 | 4 | 0.25 | 15 | 0.2667 |
| 10_771267 | 23 | 0.1304 | 23 | 0.1739 | 23 | 0.1304 | 13 | 0.1538 | 82 | 0.1463 | 14 | 0 | 11 | 0.3636 | 17 | 0.2353 | 17 | 0.1765 | 59 | 0.1864 |
| 10_771269 | 16 | 0.375 | 20 | 0.35 | 19 | 0.3158 | 10 | 0.2 | 65 | 0.3231 | 9 | 0.3333 | 7 | 0.2857 | 8 | 0.25 | 9 | 0.2222 | 33 | 0.2727 |
| 10_771271 | 11 | 0 | 16 | 0.125 | 14 | 0.0714 | 9 | 0.1111 | 50 | 0.08 | 12 | 0.1667 | 8 | 0.125 | 7 | 0 | 9 | 0 | 36 | 0.0833 |
| 10_771274 | 18 | 0 | 24 | 0.0417 | 18 | 0.0556 | 14 | 0.1429 | 74 | 0.0541 | 11 | 0.0909 | 11 | 0.0909 | 12 | 0.1667 | 10 | 0 | 44 | 0.0909 |
| 10_771304 | 3 | 0.3333 | 4 | 0 | 6 | 0 | 3 | 0.3333 | 16 | 0.125 | 2 | 0.5 | 1 | 0 | 2 | 0 | 2 | 0 | 7 | 0.1429 |
| 10_771305 | 4 | 0 | 6 | 0.8333 | 6 | 0.8333 | 4 | 0.5 | 20 | 0.6 | 3 | 0.6667 | 1 | 1 | 5 | 0.4 | 2 | 1 | 11 | 0.6364 |
| 10_771305 | 3 | 0.6667 | 4 | 0.25 | 3 | 0.3333 | 1 | 0 | 11 | 0.3636 | 1 | 0 | 2 | 0.5 | 3 | 0 | 3 | 0 | 9 | 0.1111 |
| 10_771305 | 4 | 0.25 | 2 | 1 | 4 | 1 | 2 | 1 | 12 | 0.75 | 2 | 0.5 | 2 | 0.5 | 4 | 0.5 | 2 | 0.5 | 10 | 0.5 |
| 10_771305 | 3 | 0.6667 | 4 | 1 | 6 | 0.5 | 2 | 1 | 15 | 0.7333 | 3 | 0.6667 | 2 | 1 | 6 | 0.6667 | 3 | 0.6667 | 14 | 0.7143 |
| 10_771306 | 4 | 0.25 | 4 | 0.25 | 6 | 0.3333 | 2 | 0 | 16 | 0.25 | 2 | 0.5 | 2 | 0 | 5 | 0 | 4 | 0 | 13 | 0.0769 |
| 10_771576 | 1 | 1 | 3 | 0 | 2 | 0 | 1 | 0 | 7 | 0.1429 | 1 | 0 | NA | NA | 4 | 0.5 | 1 | 1 | 6 | 0.5 |
| 10_771578 | 1 | 0 | 3 | 0 | 3 | 0.3333 | 1 | 0 | 8 | 0.125 | 1 | 1 | NA | NA | 2 | 0 | 1 | 0 | 4 | 0.25 |
| 10_771579 | 3 | 0.3333 | 3 | 0.3333 | 3 | 0.3333 | 1 | 1 | 10 | 0.4 | 1 | 0 | NA | NA | 4 | 0.25 | 1 | 0 | 6 | 0.1667 |
| 10_771579 | 1 | 0 | 3 | 0 | 2 | 0.5 | NA | NA | 6 | 0.1667 | 1 | 0 | NA | NA | 3 | 0.3333 | 1 | 0 | 5 | 0.2 |
| 10_771579 | 3 | 1 | 3 | 1 | 3 | 0.3333 | 1 | 1 | 10 | 0.8 | 1 | 1 | NA | NA | 3 | 1 | 1 | 1 | 5 | 1 |
| 10_771581 | 2 | 1 | 3 | 0.3333 | 2 | 0.5 | NA | NA | 7 | 0.5714 | 1 | 1 | NA | NA | 3 | 0.6667 | 1 | 1 | 5 | 0.8 |
| 10_771583 | 1 | 0 | 3 | 0.3333 | 3 | 0.3333 | NA | NA | 7 | 0.2857 | 1 | 1 | NA | NA | 2 | 0.5 | 1 | 1 | 4 | 0.75 |
| 10_772733 | NA | NA | NA | NA | NA | NA | 1 | 1 | 1 | 1 | NA | NA | 1 | 1 | 1 | 1 | NA | NA | 2 | 1 |
| 10_772737 | NA | NA | NA | NA | NA | NA | 1 | 1 | 1 | 1 | NA | NA | 1 | 1 | 1 | 0 | NA | NA | 2 | 0.5 |
| 10_772737 | NA | NA | NA | NA | NA | NA | 1 | 1 | 1 | 1 | NA | NA | 1 | 1 | 1 | 1 | NA | NA | 2 | 1 |
| 10_773181 | NA | NA | 1 | 1 | NA | NA | NA | NA | 1 | 1 | 1 | 1 | NA | NA | NA | NA | NA | NA | 1 | 1 |
| 10_773227 | 6 | 0.1667 | 5 | 0.4 | 5 | 0.2 | 4 | 0 | 20 | 0.2 | 6 | 0.1667 | 2 | 0 | 2 | 0 | 1 | 1 | 11 | 0.1818 |
| 10_773231 | 4 | 0 | 3 | 0.3333 | 5 | 0 | 5 | 0.2 | 17 | 0.1176 | 7 | 0.1429 | 2 | 0 | 2 | 0 | 1 | 0 | 12 | 0.0833 |
| 10_773235 | 5 | 0 | 4 | 0.25 | 4 | 0 | 5 | 0 | 18 | 0.0556 | 6 | 0.5 | 2 | 0.5 | 2 | 0 | 1 | 0 | 11 | 0.3636 |
| 10_773237 | 5 | 0.6 | 3 | 0.6667 | 3 | 0.6667 | 3 | 1 | 14 | 0.7143 | 5 | 0.8 | NA | NA | 1 | 1 | 1 | 1 | 7 | 0.8571 |
| 10_773239 | 4 | 0.5 | 3 | 0.6667 | 3 | 0.6667 | 5 | 0.4 | 15 | 0.5333 | 7 | 0.2857 | NA | NA | 2 | 0 | 1 | 0 | 10 | 0.2 |
| 10_773240 | 4 | 0.75 | 3 | 1 | 3 | 1 | 3 | 0.6667 | 13 | 0.8462 | 6 | 1 | 5 | 0.4 | 5 | 0.8 | 2 | 1 | 18 | 0.7778 |
| 10_773240 | 3 | 0.6667 | 1 | 1 | 4 | 0.25 | 2 | 1 | 10 | 0.6 | 6 | 0.6667 | 4 | 0.5 | 3 | 0.6667 | 2 | 0 | 15 | 0.5333 |
| 10_773240 | 4 | 0.25 | 2 | 0.5 | 4 | 0.5 | 5 | 0.4 | 15 | 0.4 | 7 | 0.4286 | 4 | 0 | 4 | 1 | 3 | 0 | 18 | 0.3889 |
| 10_773241 | 4 | 0.75 | 1 | 1 | 2 | 0 | 4 | 0.75 | 11 | 0.6364 | 7 | 1 | 5 | 0.6 | 4 | 1 | 4 | 0.25 | 20 | 0.75 |
| 10_773241 | 5 | 0.4 | 1 | 0 | 4 | 0.25 | 3 | 0.6667 | 13 | 0.3846 | 7 | 0.4286 | 4 | 0.25 | 4 | 0.5 | 3 | 0.6667 | 18 | 0.4444 |
| 10_774012 | NA | NA | NA | NA | 3 | 0.6667 | 2 | 0 | 5 | 0.4 | NA | NA | NA | NA | 1 | 1 | NA | NA | 1 | 1 |
| 10_774330 | 13 | 0.4615 | 8 | 0.5 | 5 | 0.4 | 17 | 0.4706 | 43 | 0.4651 | 9 | 0.5556 | 10 | 0.4 | 13 | 0.6154 | 5 | 0.4 | 37 | 0.5135 |
| 10_774332 | 10 | 0.5 | 8 | 0.25 | 5 | 0 | 16 | 0.625 | 39 | 0.4359 | 8 | 0.125 | 9 | 0.1111 | 6 | 0.1667 | 4 | 0 | 27 | 0.1111 |
| 10_774336 | 15 | 0.0667 | 10 | 0.1 | 5 | 0 | 18 | 0.1111 | 48 | 0.0833 | 10 | 0 | 9 | 0.1111 | 19 | 0.1053 | 4 | 0 | 42 | 0.0714 |
| 10_774523 | 27 | 0.1852 | 25 | 0.36 | 20 | 0.45 | 23 | 0.2174 | 95 | 0.2947 | 12 | 0.25 | 15 | 0.4 | 21 | 0.1429 | 16 | 0.125 | 64 | 0.2188 |
| 10_774529 | 29 | 0.5517 | 25 | 0.52 | 27 | 0.3704 | 30 | 0.4667 | 111 | 0.4775 | 17 | 0.4118 | 17 | 0.5882 | 29 | 0.5862 | 30 | 0.5 | 93 | 0.5269 |
| 10_774805 | 9 | 0.1111 | 9 | 0.1111 | 9 | 0.1111 | 9 | 0.2222 | 36 | 0.1389 | 4 | 0.25 | 7 | 0.1429 | 11 | 0.1818 | 8 | 0.125 | 30 | 0.1667 |
| 10_774805 | 5 | 0.6 | 8 | 0.875 | 10 | 0.7 | 10 | 0.9 | 33 | 0.7879 | 5 | 0.6 | 8 | 0.75 | 11 | 0.5455 | 5 | 1 | 29 | 0.6897 |
| 10_774807 | 8 | 0.375 | 10 | 0.5 | 10 | 0.6 | 11 | 0.2727 | 39 | 0.4359 | 6 | 0.5 | 8 | 0.375 | 9 | 0.3333 | 9 | 0.6667 | 32 | 0.4688 |
| 10_774807 | 8 | 0.5 | 9 | 0 | 9 | 0.2222 | 10 | 0 | 36 | 0.1667 | 5 | 0.2 | 5 | 0 | 9 | 0.2222 | 7 | 0.2857 | 26 | 0.1923 |
| 10_774810 | 9 | 0 | 10 | 0 | 10 | 0.1 | 11 | 0.0909 | 40 | 0.05 | 5 | 0 | 9 | 0.3333 | 12 | 0.0833 | 10 | 0.2 | 36 | 0.1667 |
| 10_774811 | 8 | 0.375 | 12 | 0.1667 | 9 | 0.1111 | 9 | 0.4444 | 38 | 0.2632 | 5 | 0 | 8 | 0.125 | 10 | 0.2 | 10 | 0.1 | 33 | 0.1212 |
| 10_777357 | NA | NA | NA | NA | 1 | 0 | 1 | 1 | 2 | 0.5 | 2 | 0.5 | NA | NA | NA | NA | 1 | 0 | 3 | 0.3333 |
| 10_777358 | NA | NA | NA | NA | 1 | 1 | 1 | 1 | 2 | 1 | 1 | 1 | NA | NA | NA | NA | 1 | 1 | 2 | 1 |
| 10_777358 | NA | NA | NA | NA | 1 | 0 | 1 | 1 | 2 | 0.5 | 2 | 0.5 | NA | NA | NA | NA | 1 | 1 | 3 | 0.6667 |
| 10_777451 | NA | NA | NA | NA | 1 | 1 | NA | NA | 1 | 1 | 1 | 0 | 1 | 1 | NA | NA | 1 | 1 | 3 | 0.6667 |
| 10_777453 | 2 | 0.5 | NA | NA | 1 | 0 | 2 | 0 | 5 | 0.2 | 3 | 0.6667 | NA | NA | 2 | 0 | 3 | 0.3333 | 8 | 0.375 |

| | | | | | | | | | | | | | | | | | | | | |
|---|---|---|---|---|---|---|---|---|---|---|---|---|---|---|---|---|---|---|---|---|
| 10_777455 | 2 | 0 | 3 | 0.3333 | 2 | 0 | 3 | 0 | 10 | 0.1 | 3 | 0.3333 | 2 | 0.5 | 3 | 0 | 2 | 0 | 10 | 0.2 |
| 10_777458 | 2 | 0 | 2 | 1 | 1 | 0 | 3 | 0.6667 | 8 | 0.5 | 3 | 0.6667 | 2 | 0.5 | 3 | 0.6667 | 1 | 0 | 9 | 0.5556 |
| 10_777459 | 1 | 1 | 1 | 1 | NA | NA | 3 | 1 | 5 | 1 | NA | NA | 2 | 0.5 | 1 | 1 | NA | NA | 3 | 0.6667 |
| 10_777459 | 1 | 0 | 1 | 0 | NA | NA | 3 | 0.6667 | 5 | 0.4 | 1 | 0 | 2 | 0 | 2 | 0.5 | 2 | 0 | 7 | 0.1429 |
| 10_777460 | 1 | 0 | 1 | 1 | NA | NA | 3 | 0.6667 | 5 | 0.6 | NA | NA | 1 | 0 | 2 | 0.5 | 2 | 1 | 5 | 0.6 |
| 10_778156 | NA | NA | 2 | 0.5 | NA | NA | 1 | 1 | 3 | 0.6667 | NA | NA | NA | NA | 1 | 0 | 1 | 1 | 2 | 0.5 |
| 10_778204 | 9 | 0.6667 | 4 | 0.5 | 4 | 0.5 | 4 | 0 | 21 | 0.4762 | 6 | 0.3333 | 6 | 0 | 5 | 0.4 | 4 | 0.25 | 21 | 0.2381 |
| 10_778205 | 7 | 0.8571 | 4 | 0.75 | 4 | 0.75 | 4 | 0.75 | 19 | 0.7895 | 5 | 0.8 | 7 | 1 | 4 | 0.75 | 4 | 0.75 | 20 | 0.85 |
| 10_778208 | 6 | 0.1667 | 2 | 0 | 3 | 0.3333 | 3 | 0 | 14 | 0.1429 | 3 | 0 | 3 | 0 | 2 | 0.5 | 3 | 0 | 11 | 0.0909 |
| 10_778230 | NA | NA | NA | NA | NA | NA | 1 | 1 | 1 | 1 | 1 | 0 | 1 | 1 | 2 | 0 | NA | NA | 4 | 0.25 |
| 10_778239 | 14 | 0.2143 | 14 | 0.2143 | 27 | 0.2222 | 10 | 0.2 | 65 | 0.2154 | 11 | 0.1818 | 20 | 0.25 | 22 | 0.3636 | 22 | 0.1818 | 75 | 0.2533 |
| 10_778240 | 18 | 0.4444 | 13 | 0.6154 | 25 | 0.56 | 13 | 0.7692 | 69 | 0.5797 | 13 | 0.4615 | 16 | 0.6875 | 25 | 0.64 | 19 | 0.4211 | 73 | 0.5616 |
| 10_779137 | 1 | 1 | NA | NA | NA | NA | NA | NA | 1 | 1 | 1 | 1 | NA | NA | 1 | 0 | NA | NA | 2 | 0.5 |
| 10_779139 | 1 | 1 | NA | NA | NA | NA | NA | NA | 1 | 1 | 1 | 1 | NA | NA | 1 | 1 | NA | NA | 2 | 1 |
| 10_779141 | 1 | 1 | NA | NA | NA | NA | NA | NA | 1 | 1 | 1 | 1 | NA | NA | 1 | 1 | NA | NA | 2 | 1 |
| 10_779142 | 1 | 1 | NA | NA | NA | NA | NA | NA | 1 | 1 | 1 | 1 | NA | NA | 1 | 1 | NA | NA | 2 | 1 |
| 10_779143 | 1 | 1 | NA | NA | NA | NA | NA | NA | 1 | 1 | 1 | 1 | NA | NA | 1 | 1 | NA | NA | 2 | 1 |
| 10_779145 | 1 | 1 | NA | NA | NA | NA | NA | NA | 1 | 1 | 1 | 1 | NA | NA | 1 | 1 | NA | NA | 2 | 1 |
| 10_779149 | 1 | 1 | NA | NA | NA | NA | NA | NA | 1 | 1 | 1 | 1 | NA | NA | 1 | 1 | NA | NA | 2 | 1 |
| 10_779937 | 1 | 1 | NA | NA | NA | NA | NA | NA | 1 | 1 | NA | NA | 1 | 1 | NA | NA | NA | NA | 1 | 1 |
| 10_779950 | 1 | 1 | NA | NA | 2 | 1 | 1 | 1 | 4 | 1 | NA | NA | NA | NA | NA | NA | 1 | 1 | 1 | 1 |
| 10_779950 | 1 | 0 | NA | NA | 1 | 0 | 1 | 1 | 3 | 0.3333 | NA | NA | NA | NA | NA | NA | 1 | 1 | 1 | 1 |
| 10_779953 | 1 | 1 | NA | NA | 2 | 1 | 1 | 1 | 4 | 1 | NA | NA | NA | NA | NA | NA | 1 | 1 | 1 | 1 |
| 10_779963 | 1 | 1 | NA | NA | 2 | 0.5 | NA | NA | 3 | 0.6667 | NA | NA | NA | NA | NA | NA | 1 | 1 | 1 | 1 |
| 10_779979 | 11 | 0.0909 | 10 | 0.3 | 12 | 0.1667 | 11 | 0.0909 | 44 | 0.1591 | 10 | 0 | 13 | 0.0769 | 9 | 0.1111 | 8 | 0.125 | 40 | 0.075 |
| 10_779979 | 14 | 0 | 11 | 0.0909 | 14 | 0.0714 | 14 | 0.2143 | 53 | 0.0943 | 12 | 0 | 13 | 0.3077 | 11 | 0.1818 | 13 | 0.0769 | 49 | 0.1429 |
| 10_779981 | 14 | 0.2857 | 13 | 0.2308 | 13 | 0.3846 | 18 | 0.2222 | 58 | 0.2759 | 9 | 0.3333 | 16 | 0.1875 | 11 | 0.5455 | 12 | 0.0833 | 48 | 0.2708 |
| 10_779983 | 13 | 0.5385 | 11 | 0.4545 | 10 | 0.4 | 16 | 0.875 | 50 | 0.6 | 9 | 0.6667 | 13 | 0.6154 | 8 | 0.75 | 12 | 0.5833 | 42 | 0.6429 |
| 10_779983 | 12 | 0.25 | 11 | 0.0909 | 12 | 0 | 18 | 0 | 53 | 0.0755 | 10 | 0.1 | 14 | 0.0714 | 11 | 0.2727 | 11 | 0 | 46 | 0.1087 |
| 10_779983 | 13 | 0.5385 | 9 | 0.4444 | 12 | 0.5833 | 14 | 0.4286 | 48 | 0.5 | 10 | 0.3 | 14 | 0.2857 | 10 | 0.7 | 10 | 0.4 | 44 | 0.4091 |
| 10_779984 | 9 | 0.1111 | 8 | 0 | 11 | 0.1818 | 13 | 0 | 41 | 0.0732 | 8 | 0.25 | 13 | 0.0769 | 10 | 0 | 9 | 0 | 40 | 0.075 |
| 10_779984 | 11 | 0.5455 | 10 | 0 | 10 | 0.1 | 14 | 0.4286 | 45 | 0.2889 | 7 | 0.2857 | 11 | 0.2727 | 9 | 0.3333 | 8 | 0.25 | 35 | 0.2857 |
| 10_780072 | 1 | 1 | 2 | 0.5 | 3 | 1 | 1 | 1 | 7 | 0.8571 | 3 | 0.6667 | 1 | 0 | 2 | 1 | 5 | 0.8 | 11 | 0.7273 |
| 10_780073 | 1 | 0 | 2 | 0.5 | 1 | 0 | 1 | 0 | 5 | 0.2 | 2 | 0 | 2 | 0.5 | 4 | 0.5 | 4 | 0.25 | 12 | 0.3333 |
| 10_780073 | 1 | 1 | 2 | 0 | NA | NA | 1 | 0 | 4 | 0.25 | 3 | 0 | 2 | 0.5 | 4 | 0 | 5 | 0 | 14 | 0.0714 |
| 10_780074 | 1 | 1 | 3 | 1 | 1 | 1 | 1 | 1 | 6 | 1 | 3 | 0.6667 | 2 | 1 | 3 | 1 | 4 | 0.25 | 12 | 0.6667 |
| 10_780075 | 1 | 0 | 2 | 0 | 1 | 0 | 1 | 1 | 5 | 0.2 | 3 | 0.3333 | 2 | 0 | 5 | 0.2 | 5 | 0.4 | 15 | 0.2667 |
| 10_780075 | 1 | 1 | 3 | 1 | 1 | 1 | 1 | 1 | 6 | 1 | 3 | 1 | 2 | 1 | 5 | 1 | 5 | 0.8 | 15 | 0.9333 |
| 10_780075 | 1 | 1 | 2 | 1 | 1 | 0 | 1 | 0 | 5 | 0.6 | 3 | 0.6667 | 1 | 1 | 3 | 1 | 5 | 0.6 | 12 | 0.75 |
| 10_780076 | 1 | 0 | 2 | 0.5 | NA | NA | 1 | 0 | 4 | 0.25 | 3 | 0 | 1 | 0 | 5 | 0.6 | 4 | 0.5 | 13 | 0.3846 |
| 10_780076 | 1 | 1 | 2 | 1 | NA | NA | 1 | 1 | 4 | 1 | 3 | 1 | 2 | 1 | 3 | 1 | 3 | 0.3333 | 11 | 0.8182 |
| 10_780076 | 1 | 1 | 3 | 0.6667 | 1 | 1 | NA | NA | 5 | 0.8 | 3 | 0.6667 | 2 | 1 | 5 | 1 | 2 | 1 | 12 | 0.9167 |
| 10_780077 | 1 | 0 | 3 | 0.3333 | 1 | 0 | 1 | 0 | 6 | 0.1667 | 2 | 0 | 2 | 1 | 5 | 0 | 4 | 0 | 13 | 0.1538 |
| 10_780077 | 1 | 0 | 3 | 0.3333 | 1 | 1 | 1 | 0 | 6 | 0.3333 | 3 | 0.3333 | 1 | 0 | 4 | 0.25 | 4 | 0.25 | 12 | 0.25 |
| 10_780216 | 2 | 0 | 3 | 0.3333 | 5 | 0 | 3 | 0 | 13 | 0.0769 | 1 | 0 | 7 | 0.2857 | 4 | 0 | 3 | 0.3333 | 15 | 0.2 |
| 10_780217 | 2 | 0 | 3 | 0 | 4 | 0.25 | 2 | 0 | 11 | 0.0909 | 1 | 0 | 5 | 0 | 3 | 0.3333 | 4 | 0 | 13 | 0.0769 |
| 10_780217 | 1 | 0 | 2 | 0 | 5 | 0.2 | 2 | 0 | 10 | 0.1 | 1 | 0 | 5 | 0 | 3 | 0 | 3 | 0.3333 | 12 | 0.0833 |
| 10_780228 | 5 | 0.4 | 8 | 0.125 | 15 | 0.3333 | 11 | 0.0909 | 39 | 0.2308 | 13 | 0.0769 | 9 | 0 | 7 | 0.2857 | 12 | 0.5833 | 41 | 0.2439 |
| 10_780228 | 6 | 0.3333 | 8 | 0.625 | 12 | 0.75 | 11 | 0.5455 | 37 | 0.5946 | 13 | 0.3846 | 9 | 0.4444 | 5 | 0.6 | 11 | 0.5455 | 38 | 0.4737 |
| 10_780228 | 5 | 0.2 | 6 | 0.3333 | 10 | 0.2 | 10 | 0.1 | 31 | 0.1935 | 11 | 0.0909 | 7 | 0.2857 | 8 | 0.25 | 10 | 0.6 | 36 | 0.3056 |
| 10_780233 | 7 | 0.2857 | 7 | 0.1429 | 11 | 0.3636 | 6 | 0.3333 | 31 | 0.2903 | 8 | 0.375 | 7 | 0.2857 | 1 | 0 | 8 | 0.75 | 24 | 0.4583 |
| 10_780235 | 5 | 0.2 | 5 | 0.6 | 9 | 0.2222 | 4 | 0 | 23 | 0.2609 | 5 | 0.6 | 5 | 0.2 | 2 | 0 | 4 | 0.25 | 16 | 0.3125 |
| 10_780235 | 7 | 0.7143 | 5 | 0.4 | 11 | 0.9091 | 5 | 0.6 | 28 | 0.7143 | 5 | 0.8 | 5 | 0.8 | 2 | 0.5 | 6 | 0.8333 | 18 | 0.7778 |
| 10_780236 | 6 | 0.5 | 6 | 0.8333 | 10 | 1 | 5 | 1 | 27 | 0.8519 | 8 | 0.5 | 5 | 1 | 2 | 0.5 | 7 | 0.8571 | 22 | 0.7273 |
| 10_780236 | 7 | 0 | 7 | 0.1429 | 11 | 0.0909 | 4 | 0 | 29 | 0.069 | 8 | 0 | 4 | 0.75 | 2 | 1 | 8 | 0.25 | 22 | 0.3182 |
| 10_780238 | 7 | 0 | 5 | 0 | 9 | 0.3333 | 5 | 0.2 | 26 | 0.1538 | 7 | 0.2857 | 2 | 0 | 1 | 0 | 7 | 0.5714 | 17 | 0.3529 |
| 10_780248 | NA | NA | NA | NA | 1 | 1 | NA | NA | 1 | 1 | NA | NA | NA | NA | 1 | 1 | NA | NA | 1 | 1 |
| 10_780253 | NA | NA | NA | NA | 1 | 1 | NA | NA | 1 | 1 | NA | NA | NA | NA | 1 | 1 | NA | NA | 1 | 1 |
| 10_780257 | NA | NA | NA | NA | 1 | 1 | NA | NA | 1 | 1 | NA | NA | NA | NA | 1 | 1 | NA | NA | 1 | 1 |
| 10_780266 | NA | NA | NA | NA | 1 | 1 | NA | NA | 1 | 1 | NA | NA | NA | NA | 1 | 1 | NA | NA | 1 | 1 |
| 10_780292 | 6 | 0 | 5 | 0 | 8 | 0.125 | 5 | 0.4 | 24 | 0.125 | 7 | 0 | 4 | 0 | 1 | 0 | 7 | 0.1429 | 19 | 0.0526 |
| 10_780398 | 2 | 0.5 | 2 | 0 | 1 | 0 | 1 | 0 | 6 | 0.1667 | 3 | 0 | 2 | 0.5 | NA | NA | 3 | 0 | 8 | 0.125 |
| 10_780401 | 3 | 0 | 3 | 0.6667 | 2 | 0 | 3 | 0 | 11 | 0.1818 | 5 | 0.4 | 2 | 0.5 | NA | NA | 2 | 0 | 9 | 0.3333 |
| 10_780586 | 3 | 0 | 1 | 0 | NA | NA | 2 | 0.5 | 6 | 0.1667 | 2 | 0 | 1 | 1 | NA | NA | 1 | 0 | 4 | 0.25 |
| 10_781051 | 12 | 0.3333 | 7 | 0.4286 | 12 | 0.0833 | 8 | 0 | 39 | 0.2051 | 7 | 0.2857 | 9 | 0.2222 | 8 | 0.125 | 12 | 0.25 | 36 | 0.2222 |

| | | | | | | | | | | | | | | | | | | | | |
|---|---|---|---|---|---|---|---|---|---|---|---|---|---|---|---|---|---|---|---|---|
| 10_781052 | 5 | 0.2 | 5 | 0.2 | 4 | 0.25 | 6 | 0.1667 | 20 | 0.2 | 5 | 0.2 | 4 | 0.5 | 3 | 0.6667 | 7 | 0.1429 | 19 | 0.3158 |
| 10_781053 | 10 | 0.9 | 7 | 0.7143 | 12 | 0.9167 | 9 | 0.7778 | 38 | 0.8421 | 6 | 0.8333 | 9 | 1 | 6 | 0.8333 | 11 | 0.9091 | 32 | 0.9062 |
| 10_781054 | 7 | 1 | 7 | 0.8571 | 11 | 1 | 8 | 1 | 33 | 0.9697 | 3 | 1 | 7 | 1 | 6 | 1 | 10 | 0.9 | 26 | 0.9615 |
| 10_781055 | 9 | 0.1111 | 6 | 0.1667 | 11 | 0.0909 | 7 | 0.1429 | 33 | 0.1212 | 5 | 0.2 | 9 | 0 | 5 | 0.4 | 8 | 0.125 | 27 | 0.1481 |
| 10_781055 | 9 | 0.2222 | 7 | 0.4286 | 12 | 0 | 8 | 0.125 | 36 | 0.1667 | 3 | 0 | 5 | 0 | 6 | 0.1667 | 11 | 0.1818 | 25 | 0.12 |
| 10_781055 | 9 | 0 | 7 | 0.1429 | 11 | 0.0909 | 8 | 0.25 | 35 | 0.1143 | 6 | 0.3333 | 9 | 0 | 6 | 0 | 11 | 0.0909 | 32 | 0.0938 |
| 10_781056 | 11 | 0.0909 | 11 | 0.0909 | 10 | 0.1 | 13 | 0.0769 | 45 | 0.0889 | 9 | 0 | 11 | 0.2727 | 6 | 0.3333 | 11 | 0.1818 | 37 | 0.1892 |
| 10_781056 | 12 | 0.3333 | 12 | 0.4167 | 11 | 0.3636 | 13 | 0.3846 | 48 | 0.375 | 7 | 0.1429 | 14 | 0.1429 | 7 | 0.5714 | 12 | 0.5 | 40 | 0.325 |
| 10_781056 | 13 | 0.3846 | 12 | 0.25 | 10 | 0.4 | 11 | 0.5455 | 46 | 0.3913 | 8 | 0.25 | 11 | 0.3636 | 7 | 0.7143 | 12 | 0.5 | 38 | 0.4474 |
| 10_781057 | 13 | 0.2308 | 10 | 0.2 | 10 | 0.1 | 13 | 0.1538 | 46 | 0.1739 | 6 | 0.5 | 14 | 0.2143 | 7 | 0 | 12 | 0 | 39 | 0.1538 |
| 10_781057 | 10 | 0.7 | 11 | 0.6364 | 12 | 0.6667 | 11 | 0.7273 | 44 | 0.6818 | 8 | 0.375 | 11 | 0.8182 | 7 | 0.7143 | 11 | 0.5455 | 37 | 0.6216 |
| 10_781057 | 11 | 0.0909 | 13 | 0.0769 | 9 | 0 | 9 | 0.1111 | 42 | 0.0714 | 7 | 0.2857 | 11 | 0 | 6 | 0.3333 | 11 | 0.2727 | 35 | 0.2 |
| 10_781069 | 7 | 0 | 9 | 0 | 11 | 0.0909 | 6 | 0.1667 | 33 | 0.0606 | 6 | 0 | 8 | 0.125 | 3 | 0 | 8 | 0.125 | 25 | 0.08 |
| 10_781082 | 8 | 0.25 | 4 | 0.25 | 8 | 0.125 | 4 | 0 | 24 | 0.1667 | 5 | 0.2 | 7 | 0.1429 | 8 | 0.25 | 6 | 0.5 | 26 | 0.2692 |
| 10_781086 | 5 | 0 | 4 | 0 | 5 | 0.4 | 4 | 0 | 18 | 0.1111 | 4 | 0.25 | 5 | 0 | 5 | 0 | 6 | 0.1667 | 20 | 0.1 |
| 10_781088 | 4 | 0 | 4 | 0.25 | 6 | 0.1667 | 4 | 0.5 | 18 | 0.2222 | 4 | 0 | 3 | 0 | 3 | 0.3333 | 2 | 0.5 | 12 | 0.1667 |
| 10_781092 | 2 | 0 | 2 | 0 | 2 | 0 | 5 | 0.2 | 11 | 0.0909 | 3 | 0 | 3 | 0.3333 | 7 | 0 | 3 | 0 | 16 | 0.0625 |
| 10_781093 | 4 | 0 | 1 | 0 | 2 | 0 | 4 | 0.25 | 11 | 0.0909 | 4 | 0.25 | 1 | 0 | 5 | 0 | 3 | 0 | 13 | 0.0769 |
| 10_781093 | 4 | 0.25 | 2 | 0.5 | 3 | 0 | 6 | 0 | 15 | 0.1333 | 3 | 0.3333 | 5 | 0 | 6 | 0.5 | 3 | 0.3333 | 17 | 0.2941 |
| 10_781094 | 3 | 0 | 1 | 0 | 2 | 0 | 4 | 0.5 | 10 | 0.2 | 4 | 0.25 | 4 | 0.25 | 6 | 0.1667 | 3 | 0 | 17 | 0.1765 |
| 10_781094 | 4 | 0.5 | 2 | 0.5 | 3 | 0.3333 | 5 | 0.4 | 14 | 0.4286 | 3 | 0 | 3 | 0.3333 | 6 | 0.3333 | 2 | 0 | 14 | 0.2143 |
| 10_781094 | 4 | 0.75 | 2 | 0.5 | 3 | 1 | 5 | 0.4 | 14 | 0.6429 | 2 | 1 | 4 | 1 | 6 | 1 | 3 | 0.6667 | 15 | 0.9333 |
| 10_781094 | 3 | 1 | 2 | 0.5 | 3 | 1 | 6 | 0.5 | 14 | 0.7143 | 4 | 0.5 | 4 | 0.5 | 7 | 0.5714 | 3 | 0.6667 | 18 | 0.5556 |
| 10_782294 | 8 | 0 | 9 | 0 | 11 | 0 | 6 | 0.3333 | 34 | 0.0588 | 6 | 0 | 4 | 0.25 | 12 | 0 | 6 | 0.1667 | 28 | 0.0714 |
| 10_782362 | 48 | 0.0625 | 39 | 0.0513 | 41 | 0.0732 | 47 | 0.0851 | 175 | 0.0686 | 35 | 0.0857 | 36 | 0.1944 | 46 | 0.087 | 40 | 0.075 | 157 | 0.1083 |
| 10_782363 | 60 | 0.1167 | 47 | 0.1915 | 47 | 0.1064 | 51 | 0.1176 | 205 | 0.1317 | 35 | 0.2 | 40 | 0.225 | 53 | 0.1698 | 38 | 0.1316 | 166 | 0.1807 |
| 10_782364 | 48 | 0.0625 | 35 | 0.1143 | 39 | 0.0256 | 43 | 0.0233 | 165 | 0.0545 | 27 | 0.037 | 34 | 0.1765 | 40 | 0.15 | 34 | 0.1176 | 135 | 0.1259 |
| 10_782365 | 56 | 0.2321 | 37 | 0.2432 | 42 | 0.2619 | 51 | 0.1765 | 186 | 0.2258 | 33 | 0.2424 | 39 | 0.2564 | 52 | 0.1923 | 39 | 0.359 | 163 | 0.2577 |
| 10_782410 | 26 | 0.0385 | 16 | 0.0625 | 21 | 0.0476 | 22 | 0.0909 | 85 | 0.0588 | 16 | 0.125 | 12 | 0.0833 | 19 | 0 | 21 | 0.0476 | 68 | 0.0588 |
| 10_782410 | 31 | 0.129 | 16 | 0.25 | 19 | 0.0526 | 30 | 0.0333 | 96 | 0.1042 | 20 | 0.3 | 14 | 0.2143 | 21 | 0.0952 | 21 | 0.0952 | 76 | 0.1711 |
| 10_782634 | 11 | 0.0909 | 6 | 0 | 13 | 0.0769 | 9 | 0.3333 | 39 | 0.1282 | 4 | 0 | 10 | 0.1 | 10 | 0.1 | 14 | 0 | 38 | 0.0526 |
| 10_782634 | 8 | 0.125 | 5 | 0 | 5 | 0 | 9 | 0.2222 | 27 | 0.1111 | 4 | 0.25 | 6 | 0.1667 | 6 | 0 | 10 | 0.1 | 26 | 0.1154 |
| 10_782728 | 39 | 0.0769 | 14 | 0.2143 | 26 | 0.1154 | 38 | 0.0789 | 117 | 0.1026 | 19 | 0 | 42 | 0.0714 | 29 | 0.1724 | 31 | 0.0645 | 121 | 0.0826 |
| 10_782732 | 14 | 0.1429 | 9 | 0 | 16 | 0.1875 | 22 | 0 | 61 | 0.082 | 9 | 0 | 18 | 0.1111 | 20 | 0 | 11 | 0.1818 | 58 | 0.069 |
| 10_782756 | 20 | 0.05 | 7 | 0 | 10 | 0.2 | 18 | 0.0556 | 55 | 0.0727 | 7 | 0.1429 | 13 | 0.1538 | 15 | 0.2667 | 10 | 0.1 | 45 | 0.1778 |
| 10_782760 | 22 | 0.2273 | 9 | 0.2222 | 11 | 0.1818 | 16 | 0 | 58 | 0.1552 | 9 | 0 | 16 | 0.125 | 12 | 0.1667 | 13 | 0.0769 | 50 | 0.1 |
| 10_782762 | 19 | 0.5263 | 10 | 0.6 | 9 | 0.7778 | 14 | 0.5 | 52 | 0.5769 | 6 | 0.5 | 16 | 0.625 | 15 | 0.6667 | 15 | 0.4 | 52 | 0.5577 |
| 10_782763 | 18 | 0.0556 | 9 | 0.2222 | 7 | 0.4286 | 15 | 0.2 | 49 | 0.1837 | 9 | 0.3333 | 18 | 0.1111 | 12 | 0.3333 | 13 | 0.2308 | 52 | 0.2308 |
| 10_782763 | 17 | 0.0588 | 7 | 0.1429 | 6 | 0.1667 | 14 | 0.2857 | 44 | 0.1591 | 8 | 0.375 | 15 | 0.3333 | 14 | 0.3571 | 15 | 0.1333 | 52 | 0.2885 |
| 10_782765 | 18 | 0.2222 | 9 | 0.2222 | 7 | 0.1429 | 15 | 0.2 | 49 | 0.2041 | 10 | 0.1 | 17 | 0.2941 | 14 | 0.1429 | 18 | 0.2222 | 59 | 0.2034 |
| 10_784199 | 3 | 0 | 3 | 0 | 1 | 0 | 5 | 0.2 | 12 | 0.0833 | 1 | 0 | 2 | 0.5 | 7 | 0 | 1 | 0 | 11 | 0.0909 |
| 10_784200 | 3 | 0 | 2 | 0 | 1 | 0 | 3 | 0.3333 | 9 | 0.1111 | NA | NA | 1 | 0 | 6 | 0.1667 | NA | NA | 7 | 0.1429 |
| 10_784200 | 3 | 0.6667 | 1 | 1 | 1 | 1 | 5 | 0.4 | 10 | 0.6 | 1 | 1 | 1 | 0 | 7 | 0.7143 | 1 | 1 | 10 | 0.7 |
| 10_784201 | 2 | 0.5 | 4 | 0.5 | 2 | 1 | 5 | 0.6 | 13 | 0.6154 | 1 | 1 | 2 | 0.5 | 7 | 0.8571 | 1 | 1 | 11 | 0.8182 |
| 10_784203 | 1 | 0 | 6 | 0.1667 | 1 | 0 | 4 | 0.5 | 12 | 0.25 | 2 | 0 | 1 | 0 | 5 | 0.4 | 1 | 0 | 9 | 0.2222 |
| 10_784212 | NA | NA | 1 | 1 | NA | NA | NA | NA | 1 | 1 | NA | NA | NA | NA | NA | NA | 1 | 1 | 1 | 1 |
| 10_784214 | NA | NA | 2 | 0.5 | 1 | 1 | NA | NA | 3 | 0.6667 | NA | NA | NA | NA | NA | NA | 1 | 1 | 1 | 1 |
| 10_784215 | NA | NA | 2 | 0.5 | NA | NA | NA | NA | 2 | 0.5 | NA | NA | NA | NA | NA | NA | 1 | 1 | 1 | 1 |
| 10_784252 | NA | NA | NA | NA | 1 | 1 | NA | NA | 1 | 1 | NA | NA | NA | NA | NA | NA | 1 | 1 | 1 | 1 |
| 10_786317 | 1 | 1 | 1 | 0 | NA | NA | 2 | 0.5 | 4 | 0.5 | NA | NA | 2 | 0.5 | NA | NA | NA | NA | 2 | 0.5 |
| 10_789002 | 4 | 0 | NA | NA | NA | NA | 3 | 0.3333 | 7 | 0.1429 | 4 | 0.25 | 2 | 0.5 | 4 | 0.25 | 2 | 0 | 12 | 0.25 |
| 10_789003 | 3 | 0.3333 | 1 | 0 | 1 | 1 | 3 | 0.3333 | 8 | 0.375 | 1 | 0 | 1 | 1 | 3 | 0 | 2 | 0.5 | 7 | 0.2857 |
| 10_789003 | 3 | 0.3333 | 1 | 0 | 1 | 0 | 2 | 0 | 7 | 0.1429 | 4 | 0 | 2 | 0 | 5 | 0.2 | 3 | 0 | 14 | 0.0714 |
| 10_789004 | 5 | 0.8 | 1 | 0 | 1 | 1 | 3 | 0.3333 | 10 | 0.6 | 3 | 0 | 3 | 0.3333 | 5 | 0.4 | 3 | 0.3333 | 14 | 0.2857 |
| 10_789006 | 4 | 0.5 | 1 | 1 | 1 | 1 | 4 | 0.75 | 10 | 0.7 | 3 | 1 | 2 | 0.5 | 4 | 0.75 | 2 | 1 | 11 | 0.8182 |
| 10_789008 | 4 | 0 | 1 | 1 | 1 | 0 | 4 | 0 | 10 | 0.1 | 2 | 0 | 2 | 0.5 | 6 | 0 | 2 | 0 | 12 | 0.0833 |
| 10_789010 | 3 | 0.3333 | 1 | 1 | 1 | 1 | 4 | 0.25 | 9 | 0.4444 | 2 | 0 | 2 | 1 | 6 | 0.3333 | 2 | 0.5 | 12 | 0.4167 |
| 10_789013 | 2 | 0.5 | 1 | 1 | 1 | 1 | 2 | 0.5 | 6 | 0.6667 | 2 | 1 | 2 | 1 | 4 | 1 | 1 | 1 | 9 | 1 |
| 10_789013 | 3 | 0.6667 | 1 | 1 | 1 | 1 | 3 | 0.6667 | 8 | 0.75 | 2 | 1 | 3 | 1 | 6 | 0.6667 | 2 | 1 | 13 | 0.8462 |
| 10_789016 | 3 | 0 | 1 | 1 | 1 | 0 | 3 | 0.3333 | 8 | 0.25 | 2 | 1 | 2 | 1 | 4 | 1 | 2 | 0.5 | 10 | 0.9 |
| 10_789017 | 3 | 0.3333 | 1 | 1 | 1 | 0 | 3 | 0.6667 | 8 | 0.5 | 2 | 0 | 2 | 0.5 | 4 | 1 | 2 | 0.5 | 10 | 0.6 |
| 10_789017 | 3 | 0.3333 | 1 | 0 | 1 | 0 | 3 | 0 | 8 | 0.125 | 2 | 1 | 2 | 0.5 | 4 | 0.75 | 2 | 1 | 10 | 0.8 |
| 10_789018 | 2 | 1 | NA | NA | 1 | 1 | 3 | 1 | 6 | 1 | 2 | 1 | 2 | 1 | 4 | 0.75 | 2 | 0.5 | 10 | 0.8 |
| 10_789018 | 3 | 1 | 1 | 1 | 1 | 1 | 4 | 1 | 9 | 1 | 2 | 1 | 3 | 1 | 3 | 1 | 2 | 1 | 10 | 1 |
| 10_789019 | 2 | 1 | 1 | 0 | 1 | 0 | 4 | 0 | 8 | 0.25 | 1 | 1 | 3 | 1 | 3 | 0.6667 | 2 | 1 | 9 | 0.8889 |

| | | | | | | | | | | | | | | | | | | | | |
|---|---|---|---|---|---|---|---|---|---|---|---|---|---|---|---|---|---|---|---|---|
| 10_7890194 | 2 | 0 | 1 | 1 | 1 | 1 | 4 | 0.25 | 8 | 0.375 | NA | NA | 2 | 0.5 | 3 | 0.3333 | 2 | 0.5 | 7 | 0.4286 |
| 10_7890198 | 1 | 1 | 1 | 0 | NA | NA | 4 | 0 | 6 | 0.1667 | 2 | 0 | 2 | 0.5 | 4 | 0.25 | 1 | 0 | 9 | 0.2222 |
| 10_7890200 | 3 | 0.3333 | 1 | 0 | NA | NA | 4 | 0 | 8 | 0.125 | 1 | 0 | 3 | 0.3333 | 5 | 0 | 2 | 0 | 11 | 0.0909 |
| 10_7890212 | 3 | 0 | 1 | 0 | NA | NA | 4 | 0.25 | 8 | 0.125 | 2 | 0 | 3 | 0.3333 | 4 | 0.25 | 2 | 0 | 11 | 0.1818 |
| 10_7890242 | 2 | 1 | 1 | 1 | NA | NA | 4 | 1 | 7 | 1 | 1 | 1 | 3 | 0.6667 | 5 | 0.6 | 2 | 1 | 11 | 0.7273 |
| 10_7890250 | 3 | 0.3333 | 1 | 0 | NA | NA | 4 | 0.5 | 8 | 0.375 | 2 | 0.5 | 1 | 1 | 4 | 0.5 | 2 | 0 | 9 | 0.4444 |
| 10_7890255 | 2 | 1 | 1 | 1 | NA | NA | 4 | 1 | 7 | 1 | 2 | 1 | 2 | 1 | 4 | 1 | 1 | 1 | 9 | 1 |
| 10_7890258 | 3 | 0.6667 | 1 | 0 | NA | NA | 4 | 0.5 | 8 | 0.5 | 2 | 0.5 | 2 | 0 | 5 | 0.2 | 1 | 0 | 10 | 0.2 |
| 10_7890262 | 1 | 0 | 1 | 0 | NA | NA | 4 | 0.25 | 6 | 0.1667 | 1 | 0 | 1 | 1 | 3 | 0.6667 | NA | NA | 5 | 0.6 |
| 10_7890272 | 3 | 1 | NA | NA | NA | NA | 3 | 1 | 6 | 1 | 2 | 1 | 1 | 1 | 4 | 1 | 1 | 1 | 8 | 1 |
| 10_7890275 | 3 | 0.6667 | NA | NA | NA | NA | 3 | 0.3333 | 6 | 0.5 | 2 | 1 | 2 | 1 | 4 | 1 | 1 | 1 | 9 | 1 |
| 10_7890280 | 2 | 1 | NA | NA | NA | NA | 3 | 1 | 5 | 1 | 2 | 1 | 2 | 0.5 | 3 | 0.6667 | 1 | 1 | 8 | 0.75 |
| 10_7890284 | 3 | 0.3333 | NA | NA | NA | NA | 3 | 0.3333 | 6 | 0.3333 | 1 | 1 | 2 | 0 | 5 | 0.4 | 1 | 0 | 9 | 0.3333 |
| 10_7890292 | 3 | 1 | NA | NA | NA | NA | 3 | 1 | 6 | 1 | 1 | 1 | 1 | 1 | 5 | 1 | 1 | 1 | 8 | 1 |
| 10_7890300 | 3 | 0.3333 | NA | NA | NA | NA | 3 | 0.3333 | 6 | 0.3333 | 1 | 0 | 2 | 0 | 4 | 0.25 | 1 | 0 | 8 | 0.125 |
| 10_7890313 | 3 | 0 | NA | NA | NA | NA | 2 | 0.5 | 5 | 0.2 | 1 | 0 | 2 | 0 | 5 | 0.2 | NA | NA | 8 | 0.125 |
| 10_7890330 | 2 | 0 | NA | NA | NA | NA | 2 | 0.5 | 4 | 0.25 | 1 | 1 | 2 | 0.5 | 3 | 0.3333 | 1 | 0 | 7 | 0.4286 |
| 10_7890339 | 3 | 0.6667 | NA | NA | NA | NA | 2 | 0.5 | 5 | 0.6 | 1 | 1 | 1 | 1 | 5 | 0.6 | 1 | 1 | 8 | 0.75 |
| 10_7890361 | 3 | 0.6667 | NA | NA | NA | NA | 2 | 0.5 | 5 | 0.6 | 1 | 0 | 1 | 0 | 4 | 0.25 | 1 | 1 | 7 | 0.2857 |
| 10_7890366 | 3 | 0.3333 | NA | NA | NA | NA | 2 | 0 | 5 | 0.2 | 1 | 0 | 2 | 0.5 | 4 | 0.5 | 1 | 0 | 8 | 0.375 |
| 10_7890367 | 3 | 1 | NA | NA | NA | NA | NA | NA | 3 | 1 | 1 | 1 | NA | NA | 5 | 0.8 | NA | NA | 6 | 0.8333 |
| 10_7936265 | NA | NA | NA | NA | NA | NA | 1 | 1 | 1 | 1 | 1 | 0 | 1 | 1 | 3 | 1 | NA | NA | 5 | 0.8 |
| 10_7937612 | 4 | 0 | 3 | 0.3333 | 6 | 0 | 5 | 0.4 | 18 | 0.1667 | 3 | 0.6667 | 5 | 0.2 | 2 | 0 | 6 | 0.3333 | 16 | 0.3125 |
| 10_7937614 | 4 | 0.5 | 3 | 0.3333 | 6 | 0.5 | 4 | 0.5 | 17 | 0.4706 | 2 | 1 | 5 | 0.8 | 2 | 0.5 | 6 | 0.1667 | 15 | 0.5333 |
| 10_7937616 | 3 | 0.6667 | 4 | 0.75 | 5 | 0.8 | 3 | 0 | 15 | 0.6 | 2 | 1 | 5 | 0.6 | 2 | 1 | 3 | 1 | 12 | 0.8333 |
| 10_7937658 | 4 | 1 | 4 | 1 | 5 | 1 | 2 | 1 | 15 | 1 | 2 | 1 | 5 | 1 | 2 | 1 | 5 | 1 | 14 | 1 |
| 10_7937668 | 4 | 0.5 | 3 | 0.6667 | 5 | 1 | 3 | 0.6667 | 15 | 0.7333 | 2 | 1 | 5 | 1 | 2 | 0.5 | 6 | 0.8333 | 15 | 0.8667 |
| 10_7937685 | 4 | 0.5 | 2 | 0 | 5 | 0.2 | 2 | 0 | 13 | 0.2308 | 2 | 0 | 6 | 0.6667 | 1 | 0 | 4 | 0 | 13 | 0.3077 |
| 10_7937689 | 4 | 0 | 2 | 0.5 | 4 | 0 | 3 | 0.3333 | 13 | 0.1538 | 1 | 0 | 4 | 0.25 | 2 | 0.5 | 5 | 0.2 | 12 | 0.25 |
| 10_7937743 | 2 | 0.5 | 3 | 1 | 5 | 1 | 3 | 1 | 13 | 0.9231 | 1 | 1 | 5 | 0.8 | 1 | 1 | 4 | 0.5 | 11 | 0.7273 |
| 10_7937748 | 3 | 0.6667 | 2 | 0 | 4 | 0.75 | 3 | 0.6667 | 12 | 0.5833 | 2 | 1 | 3 | 0.6667 | 1 | 0 | 4 | 0 | 10 | 0.4 |
| 10_7937754 | 3 | 0.6667 | 2 | 0.5 | 4 | 0 | 3 | 0.3333 | 12 | 0.3333 | 2 | 0.5 | 4 | 0.5 | 2 | 0.5 | 5 | 0.2 | 13 | 0.3846 |
| 10_7937773 | 2 | 0 | 3 | 0.6667 | 4 | 0.5 | 1 | 0 | 10 | 0.4 | 2 | 0 | 4 | 0 | 2 | 0.5 | 4 | 0 | 12 | 0.0833 |
| 10_7937775 | 1 | 1 | 2 | 0.5 | 2 | 0.5 | 2 | 0 | 7 | 0.4286 | 2 | 0 | 2 | 0 | 2 | 0 | 4 | 0.5 | 10 | 0.2 |
| 10_7944803 | NA | NA | 2 | 1 | NA | NA | NA | NA | 2 | 1 | NA | NA | NA | NA | NA | NA | 1 | 1 | 1 | 1 |
| 10_7945411 | 3 | 0.6667 | 3 | 1 | 1 | 0 | 2 | 1 | 9 | 0.7778 | 2 | 1 | 4 | 1 | 3 | 1 | NA | NA | 9 | 1 |
| 10_7945412 | 3 | 0.3333 | 4 | 0.25 | 1 | 1 | 1 | 0 | 9 | 0.3333 | 3 | 0 | 4 | 0 | 3 | 0.6667 | NA | NA | 10 | 0.2 |
| 10_7945444 | 4 | 0 | 3 | 0 | 1 | 0 | 2 | 0.5 | 10 | 0.1 | 1 | 1 | 4 | 0.5 | 5 | 0.2 | NA | NA | 10 | 0.4 |
| 10_7945460 | 4 | 0.5 | 2 | 1 | 1 | 1 | 2 | 1 | 9 | 0.7778 | 2 | 0.5 | 3 | 0.6667 | 6 | 0.6667 | NA | NA | 11 | 0.6364 |
| 10_7945470 | 5 | 0.2 | 7 | 0 | 1 | 1 | 1 | 0 | 14 | 0.1429 | 3 | 0 | 3 | 0 | 9 | 0.6667 | 1 | 0 | 16 | 0.375 |
| 10_7945524 | 3 | 0.3333 | 1 | 0 | NA | NA | NA | NA | 4 | 0.25 | 3 | 0 | NA | NA | 2 | 0.5 | 1 | 0 | 6 | 0.1667 |
| 10_7945554 | 5 | 0.2 | 5 | 0.2 | NA | NA | 1 | 0 | 11 | 0.1818 | 3 | 0.3333 | 2 | 0 | 9 | 0.1111 | 1 | 0 | 15 | 0.1333 |
| 10_7945562 | 2 | 0.5 | 5 | 0.2 | 1 | 1 | NA | NA | 8 | 0.375 | 2 | 0.5 | 2 | 0 | 7 | 0 | 1 | 1 | 12 | 0.1667 |
| 10_7945569 | 5 | 0.2 | 4 | 0 | 1 | 0 | NA | NA | 10 | 0.1 | 2 | 0 | 1 | 0 | 8 | 0.125 | 1 | 0 | 12 | 0.0833 |
| 10_7945569 | 4 | 0.5 | 4 | 0.5 | 1 | 0 | NA | NA | 9 | 0.4444 | 1 | 1 | 1 | 1 | 7 | 0.4286 | 1 | 0 | 10 | 0.5 |
| 10_7947331 | 1 | 0 | 1 | 0 | 1 | 1 | NA | NA | 3 | 0.3333 | NA | NA | 1 | 1 | 2 | 0 | 1 | 0 | 4 | 0.25 |
| 10_7947345 | 1 | 1 | 1 | 0 | 1 | 0 | NA | NA | 3 | 0.3333 | NA | NA | 1 | 1 | 2 | 0 | 1 | 0 | 4 | 0.25 |
| 10_7947359 | 1 | 1 | 1 | 0 | 1 | 1 | NA | NA | 3 | 0.6667 | 1 | 1 | 3 | 0.3333 | 2 | 0.5 | 1 | 0 | 7 | 0.4286 |
| 10_7947380 | 2 | 0.5 | 1 | 0 | 2 | 0 | NA | NA | 5 | 0.2 | 2 | 0 | 2 | 0.5 | 3 | 0.6667 | 1 | 0 | 8 | 0.375 |
| 10_7947388 | 2 | 0.5 | 2 | 0.5 | 2 | 0 | NA | NA | 6 | 0.3333 | 2 | 0 | 2 | 0 | 3 | 0.3333 | 1 | 0 | 8 | 0.125 |
| 10_7947389 | 2 | 0.5 | NA | NA | NA | NA | NA | NA | 2 | 0.5 | 2 | 0 | 1 | 1 | 2 | 0 | NA | NA | 5 | 0.2 |
| 10_7947390 | 2 | 1 | 1 | 1 | 2 | 1 | NA | NA | 5 | 1 | 1 | 0 | 3 | 1 | 3 | 0.3333 | 1 | 0 | 8 | 0.5 |
| 10_7947411 | 4 | 0 | 2 | 0.5 | 2 | 0.5 | NA | NA | 8 | 0.25 | 2 | 0 | 3 | 0.3333 | 3 | 0 | 2 | 1 | 10 | 0.3 |
| 10_7947426 | 4 | 0 | 3 | 0 | 4 | 0.25 | 1 | 0 | 12 | 0.0833 | 2 | 0.5 | 3 | 0.3333 | 3 | 0 | 2 | 0 | 10 | 0.2 |
| 10_7947436 | 6 | 0.1667 | 2 | 0 | 3 | 0 | 1 | 0 | 12 | 0.0833 | 2 | 0 | 3 | 0 | 4 | 0 | 2 | 0.5 | 11 | 0.0909 |
| 10_7947437 | 5 | 0.6 | 3 | 0.6667 | 3 | 0 | 1 | 1 | 12 | 0.5 | 2 | 0.5 | 4 | 0.5 | 5 | 0.2 | 2 | 1 | 13 | 0.4615 |
| 10_7947440 | 5 | 0 | 3 | 0.3333 | 3 | 0 | 1 | 0 | 12 | 0.0833 | 2 | 0 | 3 | 0 | 5 | 0.2 | 2 | 0 | 12 | 0.0833 |
| 10_7947443 | 6 | 0.1667 | 2 | 0 | 3 | 0.3333 | 1 | 0 | 12 | 0.1667 | 2 | 0.5 | 4 | 0 | 4 | 0 | 2 | 0.5 | 12 | 0.1667 |
| 10_7947449 | 6 | 0 | 3 | 0.3333 | 3 | 0 | 1 | 0 | 13 | 0.0769 | 2 | 0.5 | 3 | 0.3333 | 5 | 0.4 | 1 | 1 | 11 | 0.4545 |
| 10_7947459 | 7 | 1 | 2 | 1 | 3 | 1 | 1 | 1 | 13 | 1 | 2 | 1 | 4 | 1 | 5 | 1 | 2 | 1 | 13 | 1 |
| 10_7947463 | 4 | 1 | 2 | 0.5 | 3 | 0.3333 | 1 | 1 | 10 | 0.7 | 1 | 1 | 2 | 1 | 5 | 1 | 2 | 0.5 | 10 | 0.9 |
| 10_7947467 | 7 | 0.5714 | 3 | 0.6667 | 3 | 0.6667 | 1 | 1 | 14 | 0.6429 | 2 | 0.5 | 4 | 0 | 4 | 0.25 | 2 | 0.5 | 12 | 0.25 |
| 10_7947477 | 7 | 0.8571 | 1 | 1 | 4 | 1 | 1 | 1 | 13 | 0.9231 | 3 | 1 | 2 | 1 | 5 | 1 | 1 | 1 | 11 | 1 |
| 10_7947485 | 6 | 0 | 3 | 0 | 3 | 0.3333 | 1 | 1 | 13 | 0.1538 | 3 | 0.3333 | 2 | 0.5 | 5 | 0.2 | 1 | 1 | 11 | 0.3636 |
| 10_7947503 | 7 | 0.5714 | 2 | 1 | 4 | 0.5 | 1 | 1 | 14 | 0.6429 | 2 | 1 | 6 | 0.3333 | 5 | 0.4 | 1 | 0 | 14 | 0.4286 |

| | | | | | | | | | | | | | | | | | | | | |
|---|---|---|---|---|---|---|---|---|---|---|---|---|---|---|---|---|---|---|---|---|
| 10_794751 | 6 | 0.3333 | 1 | 0 | 1 | 0 | 1 | 0 | 9 | 0.2222 | 2 | 0 | 4 | 0 | 6 | 0.3333 | 1 | 1 | 13 | 0.2308 |
| 10_794751 | 5 | 0.2 | 3 | 0 | 3 | 0 | 1 | 0 | 12 | 0.0833 | 2 | 0 | 6 | 0 | 7 | 0.1429 | 1 | 1 | 16 | 0.125 |
| 10_795053 | 9 | 0 | 8 | 0 | 10 | 0.1 | 5 | 0.2 | 32 | 0.0625 | 8 | 0.125 | 11 | 0.0909 | 5 | 0.2 | 10 | 0.1 | 34 | 0.1176 |
| 10_795053 | 12 | 0.1667 | 12 | 0.3333 | 15 | 0.3333 | 8 | 0.375 | 47 | 0.2979 | 12 | 0.25 | 11 | 0.4545 | 7 | 0.1429 | 12 | 0.4167 | 42 | 0.3333 |
| 10_795054 | 9 | 0 | 10 | 0.1 | 13 | 0.0769 | 9 | 0.2222 | 41 | 0.0976 | 11 | 0.1818 | 9 | 0 | 7 | 0 | 11 | 0.0909 | 38 | 0.0789 |
| 10_795115 | 14 | 0.6429 | 14 | 0.3571 | 19 | 0.8421 | 8 | 0.625 | 55 | 0.6364 | 14 | 0.6429 | 16 | 0.8125 | 9 | 0.8889 | 12 | 0.75 | 51 | 0.7647 |
| 10_795115 | 13 | 0.9231 | 11 | 0.9091 | 19 | 0.8421 | 7 | 1 | 50 | 0.9 | 13 | 0.6923 | 16 | 0.875 | 6 | 0.8333 | 8 | 0.875 | 43 | 0.814 |
| 10_795115 | NA | NA | 1 | 1 | 1 | 1 | 1 | 1 | 3 | 1 | NA | NA | NA | NA | 1 | 1 | 2 | 1 | 3 | 1 |
| 10_795116 | 11 | 0.1818 | 10 | 0.6 | 12 | 0.25 | 6 | 0.6667 | 39 | 0.3846 | 10 | 0.3 | 12 | 0.25 | 10 | 0.7 | 11 | 0.1818 | 43 | 0.3488 |
| 10_795118 | 16 | 0.75 | 15 | 0.8 | 24 | 0.8333 | 10 | 0.7 | 65 | 0.7846 | 17 | 0.7059 | 16 | 0.9375 | 11 | 1 | 14 | 1 | 58 | 0.8966 |
| 10_795118 | 11 | 0.3636 | 12 | 0.4167 | 20 | 0.4 | 9 | 0.4444 | 52 | 0.4038 | 14 | 0.2143 | 13 | 0.6154 | 9 | 0.2222 | 13 | 0.6154 | 49 | 0.4286 |
| 10_795118 | 14 | 0.2143 | 14 | 0.2857 | 24 | 0.0833 | 11 | 0.0909 | 63 | 0.1587 | 16 | 0.0625 | 15 | 0.0667 | 11 | 0.0909 | 14 | 0.0714 | 56 | 0.0714 |
| 10_795118 | 11 | 1 | 12 | 1 | 17 | 0.9412 | 8 | 0.875 | 48 | 0.9583 | 11 | 0.8182 | 12 | 1 | 8 | 1 | 7 | 0.8571 | 38 | 0.9211 |
| 10_795238 | 8 | 0 | 16 | 0.0625 | 6 | 0.1667 | 8 | 0 | 38 | 0.0526 | 9 | 0 | 8 | 0 | 7 | 0.1429 | 10 | 0.1 | 34 | 0.0588 |
| 10_795239 | 4 | 1 | 12 | 0.8333 | 6 | 0.5 | 3 | 0.6667 | 25 | 0.76 | 9 | 0.8889 | 7 | 1 | 7 | 0.4286 | 7 | 0.7143 | 30 | 0.7667 |
| 10_795239 | 5 | 0.4 | 15 | 0.3333 | 4 | 0.25 | 5 | 0.4 | 29 | 0.3448 | 8 | 0.5 | 4 | 0.75 | 5 | 0 | 7 | 0.1429 | 24 | 0.3333 |
| 10_795239 | 2 | 0.5 | 7 | 0.7143 | 3 | 0.3333 | 6 | 0.6667 | 18 | 0.6111 | 3 | 1 | 7 | 1 | 6 | 0.6667 | 6 | 0.8333 | 22 | 0.8636 |
| 10_795241 | 5 | 0.4 | 16 | 0.375 | 4 | 0.25 | 8 | 0.25 | 33 | 0.3333 | 6 | 0.5 | 7 | 0.4286 | 7 | 0.5714 | 8 | 0.5 | 28 | 0.5 |
| 10_795241 | 6 | 0.8333 | 17 | 0.8824 | 4 | 0.5 | 7 | 0.8571 | 34 | 0.8235 | 8 | 0.25 | 7 | 0.8571 | 8 | 0.875 | 9 | 0.6667 | 32 | 0.6562 |
| 10_795602 | 14 | 0.4286 | 13 | 0.3077 | 15 | 0.2667 | 13 | 0.2308 | 55 | 0.3091 | 18 | 0.3889 | 14 | 0.2143 | 17 | 0.2353 | 8 | 0.375 | 57 | 0.2982 |
| 10_795603 | 12 | 0.6667 | 16 | 0.75 | 14 | 0.9286 | 14 | 0.7857 | 56 | 0.7857 | 15 | 0.6667 | 9 | 0.8889 | 15 | 0.6667 | 6 | 0.8333 | 45 | 0.7333 |
| 10_795603 | 15 | 1 | 18 | 1 | 13 | 1 | 13 | 0.9231 | 59 | 0.9831 | 17 | 0.9412 | 13 | 1 | 18 | 1 | 7 | 1 | 55 | 0.9818 |
| 10_795603 | 14 | 0.5 | 17 | 0.3529 | 17 | 0.2353 | 12 | 0.3333 | 60 | 0.35 | 18 | 0.3889 | 16 | 0.4375 | 17 | 0.1765 | 7 | 0.2857 | 58 | 0.3276 |
| 10_795604 | 15 | 0.0667 | 17 | 0.2353 | 15 | 0.1333 | 14 | 0 | 61 | 0.1148 | 17 | 0.1176 | 15 | 0.1333 | 19 | 0.1579 | 6 | 0 | 57 | 0.1228 |
| 10_795604 | 13 | 0.3846 | 16 | 0.3125 | 16 | 0.5 | 12 | 0.5 | 57 | 0.4211 | 18 | 0.2222 | 12 | 0.75 | 16 | 0.375 | 8 | 0.625 | 54 | 0.4444 |
| 10_795604 | 10 | 0.5 | 11 | 0.7273 | 13 | 0.2308 | 11 | 0.4545 | 45 | 0.4667 | 15 | 0.5333 | 8 | 0.375 | 20 | 0.25 | 6 | 0.3333 | 49 | 0.3673 |
| 10_795710 | 12 | 0.8333 | 20 | 0.85 | 20 | 0.85 | 20 | 0.75 | 72 | 0.8194 | 20 | 0.8 | 12 | 0.9167 | 24 | 0.7083 | 9 | 0.8889 | 65 | 0.8 |
| 10_795712 | 19 | 0.2105 | 21 | 0.0952 | 27 | 0.0741 | 23 | 0.1304 | 90 | 0.1222 | 23 | 0 | 16 | 0.125 | 29 | 0.1034 | 15 | 0.0667 | 83 | 0.0723 |
| 10_795712 | 19 | 0.1579 | 25 | 0.24 | 29 | 0.069 | 25 | 0.08 | 98 | 0.1327 | 30 | 0.1667 | 21 | 0 | 31 | 0.0645 | 14 | 0.1429 | 96 | 0.0938 |
| 10_795712 | 15 | 0.8667 | 16 | 0.8125 | 25 | 0.8 | 20 | 0.85 | 76 | 0.8289 | 29 | 0.8966 | 16 | 0.9375 | 22 | 0.8636 | 13 | 0.7692 | 80 | 0.875 |
| 10_795713 | 19 | 1 | 21 | 1 | 25 | 0.96 | 23 | 1 | 88 | 0.9886 | 22 | 0.9545 | 19 | 0.9474 | 30 | 1 | 16 | 1 | 87 | 0.977 |
| 10_795713 | 16 | 0 | 18 | 0.1111 | 27 | 0.1111 | 23 | 0.1304 | 84 | 0.0952 | 26 | 0.0385 | 18 | 0.1667 | 22 | 0.0455 | 16 | 0.1875 | 82 | 0.0976 |
| 10_795716 | 17 | 0.2353 | 18 | 0.1111 | 27 | 0.1852 | 17 | 0.1176 | 79 | 0.1646 | 23 | 0.1739 | 19 | 0.1053 | 17 | 0.4118 | 16 | 0.1875 | 75 | 0.2133 |
| 10_795716 | 13 | 0.0769 | 20 | 0.2 | 23 | 0.1739 | 22 | 0.0909 | 78 | 0.141 | 22 | 0.0455 | 20 | 0.3 | 25 | 0.16 | 17 | 0.0588 | 84 | 0.1429 |
| 10_795716 | 14 | 0.2143 | 15 | 0.3333 | 28 | 0.4643 | 21 | 0.2857 | 78 | 0.3462 | 26 | 0.2308 | 22 | 0.4545 | 28 | 0.2143 | 14 | 0.2857 | 90 | 0.2889 |
| 10_795749 | 6 | 0.8333 | 5 | 1 | 2 | 1 | 1 | 1 | 14 | 0.9286 | 3 | 1 | 1 | 1 | NA | NA | 1 | 0 | 5 | 0.8 |
| 10_795751 | 5 | 0.8 | 4 | 1 | 2 | 1 | NA | NA | 11 | 0.9091 | 5 | 0.6 | NA | NA | NA | NA | 1 | 0 | 6 | 0.5 |
| 10_795751 | 6 | 0.3333 | 2 | 0.5 | 2 | 0 | 1 | 0 | 11 | 0.2727 | 5 | 0.2 | 1 | 1 | NA | NA | 1 | 0 | 7 | 0.2857 |
| 10_795752 | 4 | 0.75 | 4 | 0.75 | 2 | 1 | NA | NA | 10 | 0.8 | 5 | 0.4 | 1 | 1 | NA | NA | 1 | 1 | 7 | 0.5714 |
| 10_795754 | 5 | 0.2 | 5 | 0.2 | 2 | 0 | NA | NA | 12 | 0.1667 | 4 | 0.5 | 1 | 0 | NA | NA | 1 | 0 | 6 | 0.3333 |
| 10_795754 | 5 | 0.8 | 5 | 0.8 | 2 | 0.5 | NA | NA | 12 | 0.75 | 3 | 1 | 1 | 1 | NA | NA | 1 | 1 | 5 | 1 |
| 10_795755 | 6 | 0.3333 | 5 | 1 | 2 | 0 | NA | NA | 13 | 0.5385 | 4 | 0.25 | 1 | 0 | NA | NA | 1 | 1 | 6 | 0.3333 |
| 10_795792 | 3 | 0.3333 | 2 | 1 | NA | NA | NA | NA | 5 | 0.6 | 2 | 1 | 1 | 0 | NA | NA | 1 | 1 | 4 | 0.75 |
| 10_795944 | 10 | 0 | 18 | 0.1111 | 12 | 0.25 | 10 | 0 | 50 | 0.1 | 10 | 0.2 | 18 | 0.3333 | 12 | 0 | 5 | 0.4 | 45 | 0.2222 |
| 10_795944 | 12 | 0 | 15 | 0.2 | 13 | 0.1538 | 14 | 0.4286 | 54 | 0.2037 | 11 | 0 | 21 | 0.1429 | 13 | 0.2308 | 8 | 0.25 | 53 | 0.1509 |
| 10_795945 | 10 | 0.6 | 17 | 0.4118 | 12 | 0.5 | 12 | 0.6667 | 51 | 0.5294 | 8 | 0.5 | 17 | 0.5882 | 11 | 0.7273 | 7 | 0.5714 | 43 | 0.6047 |
| 10_795947 | 11 | 0.8182 | 21 | 0.7143 | 12 | 0.9167 | 16 | 0.875 | 60 | 0.8167 | 8 | 0.875 | 17 | 0.8824 | 11 | 0.8182 | 8 | 0.875 | 44 | 0.8636 |
| 10_795948 | 11 | 1 | 16 | 0.875 | 14 | 0.9286 | 12 | 0.9167 | 53 | 0.9245 | 10 | 0.8 | 18 | 0.8333 | 13 | 0.7692 | 8 | 0.75 | 49 | 0.7959 |
| 10_796549 | 22 | 0.3636 | 23 | 0.3913 | 32 | 0.4062 | 22 | 0.3636 | 99 | 0.3838 | 26 | 0.3462 | 28 | 0.2143 | 26 | 0.3462 | 24 | 0.2083 | 104 | 0.2788 |
| 10_796550 | 24 | 0.5833 | 21 | 0.4762 | 35 | 0.5429 | 23 | 0.4783 | 103 | 0.5243 | 27 | 0.5556 | 28 | 0.6429 | 30 | 0.6 | 26 | 0.6154 | 111 | 0.6036 |
| 10_796552 | 23 | 0.087 | 25 | 0.28 | 42 | 0.1667 | 23 | 0.087 | 113 | 0.1593 | 32 | 0.1875 | 32 | 0.25 | 31 | 0.1613 | 30 | 0.1333 | 125 | 0.184 |
| 10_796552 | 22 | 0.2727 | 26 | 0.2692 | 36 | 0.5278 | 23 | 0.3478 | 107 | 0.3738 | 28 | 0.3214 | 28 | 0.4286 | 29 | 0.3793 | 29 | 0.3103 | 114 | 0.3596 |
| 10_796558 | 22 | 0.5455 | 25 | 0.52 | 40 | 0.25 | 30 | 0.3333 | 117 | 0.3846 | 35 | 0.2286 | 35 | 0.3429 | 33 | 0.303 | 32 | 0.25 | 135 | 0.2815 |
| 10_796561 | 22 | 0 | 25 | 0.08 | 36 | 0.0278 | 26 | 0.1154 | 109 | 0.055 | 29 | 0.069 | 33 | 0.0909 | 36 | 0.0833 | 30 | 0.0333 | 128 | 0.0703 |
| 10_796563 | 25 | 0.12 | 30 | 0 | 44 | 0.0455 | 36 | 0.1111 | 135 | 0.0667 | 34 | 0.1471 | 31 | 0.1613 | 38 | 0.1053 | 34 | 0.0882 | 137 | 0.1241 |
| 10_796563 | 23 | 0.087 | 26 | 0.0769 | 40 | 0.075 | 33 | 0.0606 | 122 | 0.0738 | 32 | 0.0312 | 36 | 0.0833 | 40 | 0.05 | 29 | 0.1379 | 137 | 0.073 |
| 10_796891 | 1 | 1 | NA | NA | NA | NA | NA | NA | 1 | 1 | NA | NA | 1 | 1 | NA | NA | NA | NA | 1 | 1 |
| 10_796896 | 1 | 1 | NA | NA | NA | NA | NA | NA | 1 | 1 | NA | NA | 1 | 1 | NA | NA | 1 | 1 | 2 | 1 |
| 10_796998 | 1 | 0 | 4 | 0 | 7 | 0 | 4 | 0.25 | 16 | 0.0625 | 5 | 0 | 2 | 0.5 | 7 | 0 | 6 | 0 | 20 | 0.05 |
| 10_797006 | 5 | 0.2 | 7 | 0.1429 | 12 | 0.25 | 14 | 0.1429 | 38 | 0.1842 | 9 | 0.1111 | 4 | 0 | 9 | 0.1111 | 9 | 0.2222 | 31 | 0.129 |
| 10_797006 | 4 | 0 | 8 | 0.125 | 12 | 0.0833 | 14 | 0 | 38 | 0.0526 | 9 | 0.1111 | 6 | 0 | 7 | 0.1429 | 10 | 0.1 | 32 | 0.0938 |
| 10_797011 | 4 | 0.5 | 6 | 0.1667 | 9 | 0.2222 | 13 | 0.2308 | 32 | 0.25 | 9 | 0.3333 | 6 | 0.5 | 6 | 0.3333 | 6 | 0.1667 | 27 | 0.3333 |
| 10_797016 | 2 | 0.5 | 4 | 0 | 8 | 0.25 | 11 | 0.1818 | 25 | 0.2 | 6 | 0 | 6 | 0.1667 | 5 | 0.2 | 5 | 0 | 22 | 0.0909 |
| 10_797017 | 3 | 0 | 5 | 0.4 | 8 | 0 | 12 | 0.25 | 28 | 0.1786 | 8 | 0.375 | 6 | 0.1667 | 5 | 0.2 | 5 | 0.2 | 24 | 0.25 |
| 10_797308 | 11 | 0.0909 | 5 | 0 | 10 | 0 | 6 | 0.1667 | 32 | 0.0625 | 8 | 0.125 | 6 | 0.1667 | 5 | 0 | 6 | 0 | 25 | 0.08 |

| | | | | | | | | | | | | | | | | | | | | |
|---|---|---|---|---|---|---|---|---|---|---|---|---|---|---|---|---|---|---|---|---|
| 10_797310 | 11 | 0.3636 | 5 | 0.2 | 9 | 0.4444 | 4 | 0.5 | 29 | 0.3793 | 10 | 0.5 | 8 | 0.25 | 6 | 0 | 6 | 0.3333 | 30 | 0.3 |
| 10_797310 | 8 | 0.625 | 2 | 0.5 | 6 | 0.3333 | 4 | 0 | 20 | 0.4 | 9 | 0.2222 | 4 | 0.5 | 3 | 0.6667 | 3 | 0.6667 | 19 | 0.4211 |
| 10_797310 | 8 | 0.625 | 5 | 0.4 | 6 | 0.1667 | 4 | 0.25 | 23 | 0.3913 | 7 | 0.1429 | 5 | 0 | 4 | 0.25 | 4 | 0.5 | 20 | 0.2 |
| 10_797390 | 2 | 0.5 | 1 | 0 | 2 | 0 | 1 | 0 | 6 | 0.1667 | 3 | 0.3333 | 3 | 0 | 5 | 0.2 | 1 | 0 | 12 | 0.1667 |
| 10_797463 | 42 | 0.0952 | 39 | 0.1282 | 40 | 0.05 | 40 | 0.025 | 161 | 0.0745 | 37 | 0.1351 | 47 | 0.1064 | 43 | 0.0233 | 27 | 0.0741 | 154 | 0.0844 |
| 10_797463 | 43 | 0.186 | 42 | 0.0952 | 37 | 0.1622 | 37 | 0.2162 | 159 | 0.1635 | 39 | 0.1795 | 44 | 0.1818 | 43 | 0.1163 | 29 | 0.1034 | 155 | 0.1484 |
| 10_797465 | 46 | 0.1087 | 42 | 0.119 | 36 | 0.2222 | 39 | 0.1795 | 163 | 0.1534 | 35 | 0.1714 | 46 | 0.3913 | 43 | 0.186 | 31 | 0.129 | 155 | 0.2323 |
| 10_797465 | 44 | 0.0682 | 43 | 0.1395 | 39 | 0.0513 | 40 | 0.1 | 166 | 0.0904 | 35 | 0.1429 | 49 | 0.1429 | 47 | 0.1064 | 29 | 0.2069 | 160 | 0.1438 |
| 10_797534 | 1 | 1 | NA | NA | 1 | 0 | NA | NA | 2 | 0.5 | NA | NA | 2 | 0.5 | NA | NA | 1 | 1 | 3 | 0.6667 |
| 10_797535 | 1 | 1 | NA | NA | 1 | 1 | NA | NA | 2 | 1 | NA | NA | 2 | 1 | NA | NA | 1 | 0 | 3 | 0.6667 |
| 10_797544 | 1 | 1 | 1 | 0 | 2 | 0 | NA | NA | 4 | 0.25 | NA | NA | 2 | 0.5 | NA | NA | 1 | 0 | 3 | 0.3333 |
| 10_797577 | 1 | 1 | 1 | 1 | 2 | 0.5 | NA | NA | 4 | 0.75 | NA | NA | 2 | 0.5 | NA | NA | 1 | 1 | 3 | 0.6667 |
| 10_797577 | 1 | 0 | 1 | 0 | 2 | 0.5 | NA | NA | 4 | 0.25 | NA | NA | 2 | 0.5 | NA | NA | 2 | 0 | 4 | 0.25 |
| 10_797668 | NA | NA | NA | NA | 1 | 1 | NA | NA | 1 | 1 | NA | NA | NA | NA | 1 | 1 | NA | NA | 1 | 1 |
| 10_797670 | NA | NA | NA | NA | 1 | 1 | NA | NA | 1 | 1 | NA | NA | NA | NA | 1 | 1 | NA | NA | 1 | 1 |
| 10_797888 | NA | NA | NA | NA | 1 | 1 | NA | NA | 1 | 1 | NA | NA | NA | NA | NA | NA | 1 | 1 | 1 | 1 |
| 10_798035 | 11 | 0.4545 | 11 | 0.2727 | 6 | 0 | 8 | 0.5 | 36 | 0.3333 | 3 | 0.6667 | 8 | 0.125 | 3 | 0 | 9 | 0.6667 | 23 | 0.3913 |
| 10_798035 | 7 | 0.5714 | 11 | 0.6364 | 4 | 0.5 | 9 | 0.6667 | 31 | 0.6129 | 4 | 0.75 | 7 | 0.1429 | 3 | 0 | 7 | 0.5714 | 21 | 0.381 |
| 10_798041 | NA | NA | 3 | 0 | 1 | 1 | 3 | 0 | 7 | 0.1429 | 4 | 0.25 | 1 | 0 | NA | NA | 2 | 0.5 | 7 | 0.2857 |
| 10_798048 | 1 | 0 | 4 | 0.5 | 1 | 0 | 2 | 0.5 | 8 | 0.375 | 5 | 0.2 | 1 | 0 | NA | NA | 2 | 0 | 8 | 0.125 |
| 10_798129 | 12 | 0.4167 | 11 | 0.1818 | 8 | 0.5 | 13 | 0.5385 | 44 | 0.4091 | 9 | 0.5556 | 6 | 0.1667 | 11 | 0.5455 | 14 | 0.2857 | 40 | 0.4 |
| 10_798129 | 9 | 0.4444 | 12 | 0 | 12 | 0.3333 | 14 | 0.3571 | 47 | 0.2766 | 14 | 0.3571 | 6 | 0.1667 | 8 | 0.25 | 11 | 0.0909 | 39 | 0.2308 |
| 10_798130 | 13 | 0.8462 | 19 | 0.6842 | 14 | 0.9286 | 16 | 0.8125 | 62 | 0.8065 | 14 | 0.8571 | 7 | 0.8571 | 16 | 0.625 | 15 | 0.8667 | 52 | 0.7885 |
| 10_798132 | 12 | 0.0833 | 18 | 0.1111 | 14 | 0.0714 | 17 | 0.0588 | 61 | 0.082 | 13 | 0.3077 | 8 | 0 | 16 | 0.0625 | 13 | 0.0769 | 50 | 0.12 |
| 10_798132 | 11 | 0.0909 | 19 | 0.1053 | 15 | 0.0667 | 16 | 0.0625 | 61 | 0.082 | 14 | 0.0714 | 11 | 0.0909 | 18 | 0.1111 | 12 | 0.3333 | 55 | 0.1455 |
| 10_798133 | 14 | 0.6429 | 18 | 0.3889 | 15 | 0.7333 | 17 | 0.4118 | 64 | 0.5312 | 15 | 0.7333 | 11 | 0.6364 | 16 | 0.625 | 13 | 0.4615 | 55 | 0.6182 |
| 10_798134 | 10 | 0.1 | 17 | 0.1765 | 10 | 0.2 | 12 | 0.4167 | 49 | 0.2245 | 15 | 0.2 | 7 | 0.1429 | 11 | 0.0909 | 7 | 0.4286 | 40 | 0.2 |
| 10_798309 | 1 | 1 | 3 | 0.6667 | 1 | 0 | NA | NA | 5 | 0.6 | 1 | 0 | NA | NA | 1 | 0 | 2 | 0.5 | 4 | 0.25 |
| 10_798513 | NA | NA | 1 | 1 | 1 | 1 | 1 | 0 | 3 | 0.6667 | 2 | 0 | NA | NA | 2 | 1 | 1 | 1 | 5 | 0.6 |
| 10_798672 | 1 | 1 | NA | NA | NA | NA | NA | NA | 1 | 1 | 2 | 1 | 1 | 1 | NA | NA | 1 | 1 | 4 | 1 |
| 10_798718 | 9 | 0.4444 | 6 | 0.1667 | 11 | 0.3636 | 16 | 0.4375 | 42 | 0.381 | 12 | 0.6667 | 11 | 0.4545 | 10 | 0.7 | 11 | 0.4545 | 44 | 0.5682 |
| 10_798718 | 11 | 0.0909 | 15 | 0.2667 | 15 | 0.0667 | 24 | 0.0833 | 65 | 0.1231 | 13 | 0 | 13 | 0.1538 | 16 | 0.125 | 13 | 0.0769 | 55 | 0.0909 |
| 10_798719 | 9 | 0.5556 | 15 | 0.4667 | 10 | 0.2 | 13 | 0.4615 | 47 | 0.4255 | 7 | 0.1429 | 10 | 0 | 12 | 0.5833 | 7 | 0.4286 | 36 | 0.3056 |
| 10_798719 | 11 | 0.0909 | 14 | 0.0714 | 13 | 0.0769 | 18 | 0.1111 | 56 | 0.0893 | 9 | 0.1111 | 13 | 0 | 13 | 0.2308 | 12 | 0 | 47 | 0.0851 |
| 10_798720 | 11 | 0.0909 | 16 | 0.125 | 12 | 0.0833 | 19 | 0.2105 | 58 | 0.1379 | 13 | 0 | 11 | 0.0909 | 15 | 0.1333 | 9 | 0 | 48 | 0.0625 |
| 10_798721 | 12 | 0 | 19 | 0.1579 | 14 | 0.2143 | 21 | 0.0952 | 66 | 0.1212 | 13 | 0.0769 | 11 | 0.0909 | 16 | 0.1875 | 12 | 0 | 52 | 0.0962 |
| 10_799521 | 4 | 0.25 | 2 | 0 | NA | NA | 2 | 0 | 8 | 0.125 | 5 | 0 | 1 | 0 | 5 | 0.4 | 2 | 0 | 13 | 0.1538 |
| 10_799523 | 3 | 0.3333 | 1 | 1 | 2 | 1 | 2 | 0.5 | 8 | 0.625 | 4 | 0.5 | 5 | 0.8 | 1 | 1 | 1 | 0 | 11 | 0.6364 |
| 10_799523 | 6 | 0.5 | 2 | 1 | 2 | 1 | 1 | 1 | 11 | 0.7273 | 6 | 0.8333 | 5 | 0.8 | 5 | 0.8 | 3 | 0.3333 | 19 | 0.7368 |
| 10_799524 | 7 | 1 | 5 | 0.2 | 4 | 0.5 | 3 | 0.6667 | 19 | 0.6316 | 7 | 0.8571 | 7 | 0.8571 | 4 | 0.75 | 3 | 0.6667 | 21 | 0.8095 |
| 10_799525 | 6 | 0.3333 | 6 | 0.8333 | 5 | 0.2 | 4 | 0.25 | 21 | 0.4286 | 9 | 0.5556 | 8 | 0.25 | 5 | 0.8 | 4 | 0 | 26 | 0.4231 |
| 10_799526 | 5 | 0 | 4 | 0 | 5 | 0 | 4 | 0.25 | 18 | 0.0556 | 5 | 0 | 7 | 0.2857 | 4 | 0 | 3 | 0.3333 | 19 | 0.1579 |
| 10_799526 | 6 | 0.1667 | 5 | 0.6 | 6 | 0.3333 | 6 | 0.3333 | 23 | 0.3478 | 6 | 0.1667 | 9 | 0.2222 | 5 | 0.2 | 5 | 0.8 | 25 | 0.32 |
| 10_799621 | 9 | 0.2222 | 6 | 0 | 7 | 0.1429 | 9 | 0 | 31 | 0.0968 | 9 | 0 | 10 | 0 | 11 | 0.0909 | 5 | 0.2 | 35 | 0.0571 |
| 10_799659 | 15 | 0.0667 | 8 | 0.125 | 11 | 0.0909 | 10 | 0 | 44 | 0.0682 | 10 | 0 | 10 | 0 | 9 | 0.2222 | 7 | 0 | 36 | 0.0556 |
| 10_799663 | 7 | 0.4286 | 5 | 0.4 | 4 | 0.25 | 4 | 0.25 | 20 | 0.35 | 7 | 0.7143 | 3 | 0.6667 | 8 | 0.375 | 6 | 0 | 24 | 0.4167 |
| 10_799665 | 34 | 0.9706 | 46 | 0.9565 | 46 | 0.9348 | 33 | 0.9697 | 159 | 0.956 | 40 | 0.95 | 46 | 0.9783 | 42 | 0.9286 | 40 | 1 | 168 | 0.9643 |
| 10_799665 | 35 | 0.6857 | 43 | 0.6977 | 46 | 0.7609 | 34 | 0.7353 | 158 | 0.7215 | 41 | 0.8049 | 44 | 0.8636 | 42 | 0.8095 | 42 | 0.8095 | 169 | 0.8225 |
| 10_799667 | 31 | 0.9677 | 36 | 0.9722 | 29 | 0.9655 | 33 | 0.9697 | 129 | 0.969 | 35 | 1 | 38 | 0.9737 | 35 | 1 | 31 | 0.9677 | 139 | 0.9856 |
| 10_799668 | 30 | 0.9667 | 42 | 0.9762 | 35 | 0.9714 | 35 | 0.9429 | 142 | 0.9648 | 30 | 0.9667 | 36 | 1 | 32 | 1 | 29 | 1 | 127 | 0.9921 |
| 10_799669 | 26 | 0.5385 | 33 | 0.5152 | 23 | 0.5652 | 18 | 0.3889 | 100 | 0.51 | 31 | 0.5161 | 28 | 0.5357 | 18 | 0.7778 | 25 | 0.48 | 102 | 0.5588 |
| 10_799669 | 33 | 0.697 | 43 | 0.6977 | 38 | 0.7105 | 36 | 0.7778 | 150 | 0.72 | 33 | 0.8182 | 42 | 0.8333 | 36 | 0.75 | 35 | 0.6571 | 146 | 0.7671 |
| 10_799671 | 26 | 0.9231 | 34 | 1 | 30 | 1 | 28 | 0.9643 | 118 | 0.9746 | 24 | 1 | 32 | 1 | 21 | 0.9048 | 30 | 0.9333 | 107 | 0.9626 |
| 10_799672 | 31 | 0.9677 | 37 | 0.973 | 37 | 0.973 | 35 | 0.9714 | 140 | 0.9714 | 32 | 0.9688 | 36 | 1 | 31 | 0.9677 | 35 | 0.9714 | 134 | 0.9776 |
| 10_799673 | 28 | 0.4286 | 34 | 0.4706 | 30 | 0.5 | 27 | 0.4815 | 119 | 0.4706 | 28 | 0.2857 | 31 | 0.3226 | 28 | 0.2857 | 31 | 0.5806 | 118 | 0.3729 |
| 10_799676 | 28 | 0.1071 | 29 | 0.3103 | 34 | 0.1765 | 30 | 0.2333 | 121 | 0.2066 | 29 | 0.3103 | 34 | 0.3824 | 25 | 0.12 | 29 | 0.3448 | 117 | 0.2991 |
| 10_799677 | 18 | 0.9444 | 27 | 1 | 29 | 0.9655 | 19 | 0.8947 | 93 | 0.957 | 21 | 0.9524 | 28 | 0.9643 | 16 | 0.9375 | 23 | 0.9565 | 88 | 0.9545 |
| 10_799678 | 26 | 0.6923 | 26 | 0.5769 | 28 | 0.6071 | 22 | 0.5 | 102 | 0.598 | 21 | 0.381 | 28 | 0.6429 | 15 | 0.8 | 21 | 0.619 | 85 | 0.6 |
| 10_799682 | 4 | 0 | 6 | 0 | 3 | 0 | 2 | 0.5 | 15 | 0.0667 | 6 | 0 | 5 | 0.6 | 4 | 0.25 | 5 | 0.2 | 20 | 0.25 |
| 10_799687 | 3 | 0.6667 | 2 | 0 | 3 | 0.6667 | NA | NA | 8 | 0.5 | 3 | 0.3333 | 3 | 1 | 1 | 0 | 2 | 0.5 | 9 | 0.5556 |
| 10_799687 | 3 | 0 | 3 | 0 | 3 | 0 | 2 | 0.5 | 11 | 0.0909 | 5 | 0 | 3 | 0.3333 | 4 | 0.5 | 3 | 0.3333 | 15 | 0.2667 |
| 10_799781 | 7 | 0 | 4 | 0.25 | 11 | 0.0909 | 9 | 0.1111 | 31 | 0.0968 | 7 | 0 | 9 | 0 | 5 | 0.2 | 6 | 0.1667 | 27 | 0.0741 |
| 10_799858 | NA | NA | 1 | 1 | 3 | 0 | NA | NA | 4 | 0.25 | 2 | 0.5 | NA | NA | NA | NA | NA | NA | 2 | 0.5 |
| 10_799924 | 1 | 1 | 5 | 0.6 | 3 | 0.3333 | 3 | 0.6667 | 12 | 0.5833 | 4 | 0.25 | 2 | 0.5 | 2 | 0 | 1 | 0 | 9 | 0.2222 |
| 10_799925 | 1 | 1 | 6 | 0.3333 | 3 | 0.3333 | 2 | 0.5 | 12 | 0.4167 | 3 | 0.3333 | 3 | 0 | 1 | 0 | NA | NA | 7 | 0.1429 |

| | | | | | | | | | | | | | | | | | | | | |
|---|---|---|---|---|---|---|---|---|---|---|---|---|---|---|---|---|---|---|---|---|
| 10_7999298 | 2 | 0.5 | 5 | 0 | 5 | 0.2 | 4 | 0.25 | 16 | 0.1875 | 4 | 0.5 | 3 | 0.3333 | 2 | 0 | 1 | 0 | 10 | 0.3 |
| 10_7999320 | 1 | 0 | 6 | 0.3333 | 5 | 0.6 | 4 | 0.75 | 16 | 0.5 | 3 | 0.6667 | 3 | 0.3333 | 2 | 0 | 2 | 1 | 10 | 0.5 |
| 10_7999328 | 2 | 0 | 5 | 0.4 | 5 | 0.4 | 3 | 1 | 15 | 0.4667 | 3 | 0.6667 | 3 | 0 | 3 | 0 | 4 | 0.5 | 13 | 0.3077 |
| 10_7999344 | 2 | 1 | 5 | 0.8 | 3 | 0.6667 | 4 | 0.75 | 14 | 0.7857 | 3 | 0.3333 | 3 | 0.6667 | 2 | 1 | 4 | 0.5 | 12 | 0.5833 |
| 10_8000723 | 14 | 0.6429 | 13 | 0.7692 | 7 | 0.4286 | 23 | 0.3913 | 57 | 0.5439 | 9 | 0.5556 | 13 | 0.6154 | 13 | 0.3846 | 7 | 0.4286 | 42 | 0.5 |
| 10_8002481 | NA | NA | 3 | 0.6667 | 1 | 0 | NA | NA | 4 | 0.5 | 1 | 1 | 2 | 0.5 | NA | NA | 2 | 0 | 5 | 0.4 |
| 10_8002487 | NA | NA | 4 | 1 | 1 | 1 | NA | NA | 5 | 1 | 1 | 1 | 1 | 1 | 1 | 0 | 2 | 1 | 5 | 0.8 |
| 10_8002492 | NA | NA | 2 | 0.5 | 1 | 1 | NA | NA | 3 | 0.6667 | 1 | 0 | 1 | 1 | 1 | 0 | 3 | 0.6667 | 6 | 0.5 |
| 10_8002506 | NA | NA | 2 | 1 | 1 | 1 | NA | NA | 3 | 1 | 1 | 1 | 1 | 1 | 1 | 1 | 2 | 1 | 5 | 1 |
| 10_8002513 | NA | NA | 1 | 1 | NA | NA | NA | NA | 1 | 1 | 1 | 1 | NA | NA | NA | NA | 1 | 0 | 2 | 0.5 |
| 10_8002534 | NA | NA | 4 | 1 | 1 | 1 | NA | NA | 5 | 1 | NA | NA | 1 | 0 | 1 | 1 | 2 | 0.5 | 4 | 0.5 |
| 10_8002557 | 1 | 0 | 3 | 0.6667 | 2 | 1 | NA | NA | 6 | 0.6667 | 2 | 1 | 1 | 0 | NA | NA | 3 | 1 | 6 | 0.8333 |
| 10_8002561 | NA | NA | 4 | 1 | 1 | 1 | NA | NA | 5 | 1 | 2 | 1 | 1 | 1 | 1 | 1 | 3 | 1 | 7 | 1 |
| 10_8006846 | 2 | 0.5 | NA | NA | 2 | 1 | 1 | 1 | 5 | 0.8 | NA | NA | 3 | 0.6667 | 1 | 1 | NA | NA | 4 | 0.75 |
| 10_8006858 | 2 | 0.5 | 1 | 0 | 2 | 0.5 | 1 | 1 | 6 | 0.5 | NA | NA | 3 | 1 | 1 | 0 | NA | NA | 4 | 0.75 |
| 10_8006878 | 2 | 0.5 | 1 | 1 | 3 | 1 | 1 | 0 | 7 | 0.7143 | NA | NA | 3 | 0.6667 | 1 | 0 | 1 | 1 | 5 | 0.6 |
| 10_8007955 | 4 | 0.5 | 5 | 0.8 | 4 | 0.25 | 3 | 0 | 16 | 0.4375 | 1 | 0 | 4 | 0.75 | 4 | 0.75 | 1 | 0 | 10 | 0.6 |
| 10_8007982 | 4 | 0.5 | 7 | 0.4286 | 4 | 0.5 | 5 | 0 | 20 | 0.35 | NA | NA | 3 | 0 | 3 | 0.3333 | 1 | 0 | 7 | 0.1429 |
| 10_8008017 | 1 | 1 | 3 | 0.6667 | NA | NA | 1 | 0 | 5 | 0.6 | NA | NA | 1 | 0 | 1 | 1 | NA | NA | 2 | 0.5 |
| 10_8008060 | NA | NA | 3 | 0.3333 | NA | NA | 2 | 0 | 5 | 0.2 | NA | NA | 1 | 1 | 1 | 0 | NA | NA | 2 | 0.5 |
| 10_8008073 | NA | NA | 3 | 0.3333 | NA | NA | 2 | 0 | 5 | 0.2 | NA | NA | 1 | 1 | 2 | 0 | NA | NA | 3 | 0.3333 |
| 10_8008174 | NA | NA | 3 | 0.3333 | NA | NA | 1 | 0 | 4 | 0.25 | NA | NA | 1 | 0 | 2 | 1 | NA | NA | 3 | 0.6667 |
| 10_8012358 | 3 | 0 | 4 | 0.25 | 8 | 0.25 | 5 | 0.6 | 20 | 0.3 | 3 | 0.3333 | 8 | 0.75 | 7 | 0.4286 | 6 | 0.3333 | 24 | 0.5 |
| 10_8012358 | 4 | 0.25 | 4 | 0.25 | 8 | 0.125 | 6 | 0.3333 | 22 | 0.2273 | 4 | 0 | 7 | 0.2857 | 7 | 0.1429 | 8 | 0.25 | 26 | 0.1923 |
| 10_8012361 | 5 | 0.4 | 4 | 0 | 7 | 0.2857 | 7 | 0.2857 | 23 | 0.2609 | 3 | 0 | 6 | 0.1667 | 5 | 0.4 | 2 | 0 | 16 | 0.1875 |
| 10_8012366 | 6 | 0.6667 | 4 | 0.25 | 8 | 0.375 | 7 | 0.7143 | 25 | 0.52 | 4 | 0.5 | 4 | 0.75 | 6 | 0.3333 | 7 | 0.4286 | 21 | 0.4762 |
| 10_8012993 | 180 | 0.1056 | 208 | 0.1923 | 203 | 0.0788 | 227 | 0.163 | 818 | 0.1369 | 159 | 0.1384 | 184 | 0.1196 | 219 | 0.1416 | 184 | 0.125 | 746 | 0.1314 |
| 10_8014720 | 1 | 1 | NA | NA | 1 | 1 | 1 | 1 | 3 | 1 | NA | NA | NA | NA | 3 | 0.6667 | NA | NA | 3 | 0.6667 |
| 10_8014732 | 1 | 1 | NA | NA | 1 | 1 | 1 | 1 | 3 | 1 | NA | NA | NA | NA | 3 | 1 | NA | NA | 3 | 1 |
| 10_8014758 | 2 | 0 | NA | NA | 1 | 0 | 1 | 1 | 4 | 0.25 | NA | NA | NA | NA | 3 | 0.6667 | NA | NA | 3 | 0.6667 |
| 10_8014764 | 2 | 1 | NA | NA | 2 | 1 | 1 | 1 | 5 | 1 | NA | NA | NA | NA | 3 | 1 | NA | NA | 3 | 1 |
| 10_8014791 | 2 | 0.5 | NA | NA | 2 | 0 | 1 | 1 | 5 | 0.4 | NA | NA | NA | NA | 3 | 0.6667 | NA | NA | 3 | 0.6667 |
| 10_8014791 | 1 | 0 | NA | NA | 2 | 1 | 1 | 1 | 4 | 0.75 | NA | NA | NA | NA | 2 | 0.5 | NA | NA | 2 | 0.5 |
| 10_8014795 | 2 | 0.5 | NA | NA | 2 | 0 | 1 | 0 | 5 | 0.2 | NA | NA | NA | NA | 3 | 0.3333 | NA | NA | 3 | 0.3333 |
| 10_8014813 | 2 | 0.5 | 2 | 0.5 | 2 | 0.5 | 1 | 1 | 7 | 0.5714 | NA | NA | NA | NA | 2 | 0.5 | NA | NA | 2 | 0.5 |
| 10_8014827 | 2 | 0 | 1 | 0 | 2 | 0.5 | 1 | 0 | 6 | 0.1667 | NA | NA | NA | NA | 2 | 0.5 | NA | NA | 2 | 0.5 |
| 10_8014872 | 3 | 0.3333 | 2 | 1 | 2 | 0 | 1 | 0 | 8 | 0.375 | NA | NA | NA | NA | 3 | 0.6667 | 1 | 1 | 4 | 0.75 |
| 10_8014900 | 2 | 1 | 3 | 1 | 2 | 1 | 1 | 1 | 8 | 1 | NA | NA | NA | NA | 5 | 1 | 2 | 1 | 7 | 1 |
| 10_8014914 | 3 | 0 | 2 | 0.5 | 2 | 0 | 1 | 0 | 8 | 0.125 | NA | NA | 1 | 1 | 5 | 0 | 2 | 0 | 8 | 0.125 |
| 10_8014915 | 3 | 0.6667 | 3 | 1 | 2 | 1 | 1 | 1 | 9 | 0.8889 | NA | NA | 1 | 1 | 5 | 1 | 2 | 1 | 8 | 1 |
| 10_8014922 | 3 | 0.6667 | 3 | 0.3333 | 2 | 0.5 | 1 | 1 | 9 | 0.5556 | NA | NA | 1 | 1 | 5 | 0.6 | 2 | 0.5 | 8 | 0.625 |
| 10_8014923 | 3 | 0.6667 | 2 | 0.5 | 2 | 0 | NA | NA | 7 | 0.4286 | NA | NA | 1 | 1 | 5 | 0.4 | 2 | 0 | 8 | 0.375 |
| 10_8014925 | 3 | 0.3333 | 2 | 0 | 2 | 0 | 1 | 1 | 8 | 0.25 | NA | NA | NA | NA | 5 | 0.2 | 2 | 0.5 | 7 | 0.2857 |
| 10_8014934 | 2 | 0 | 2 | 0 | 1 | 0 | 1 | 1 | 6 | 0.1667 | NA | NA | 1 | 0 | 5 | 0.2 | 2 | 0 | 8 | 0.125 |
| 10_8014951 | 3 | 0 | 3 | 0.3333 | 2 | 0.5 | 1 | 1 | 9 | 0.3333 | NA | NA | NA | NA | 5 | 0.4 | 2 | 1 | 7 | 0.5714 |
| 10_8014961 | 3 | 0.3333 | 4 | 0.25 | 1 | 1 | 1 | 0 | 9 | 0.3333 | NA | NA | 1 | 0 | 5 | 0 | 2 | 0.5 | 8 | 0.125 |
| 10_8014966 | 1 | 1 | 4 | 0.5 | 1 | 1 | 1 | 1 | 7 | 0.7143 | NA | NA | 1 | 1 | 3 | 1 | 2 | 1 | 6 | 1 |
| 10_8014970 | 2 | 0.5 | 4 | 0 | 2 | 0 | 1 | 0 | 9 | 0.1111 | NA | NA | 2 | 0.5 | 3 | 0 | 2 | 0 | 7 | 0.1429 |
| 10_8014974 | 2 | 0.5 | 4 | 0 | 1 | 0 | NA | NA | 7 | 0.1429 | NA | NA | 1 | 0 | 4 | 0 | 2 | 0.5 | 7 | 0.1429 |
| 10_8014979 | 3 | 0.6667 | 4 | 0.5 | 1 | 1 | 1 | 1 | 9 | 0.6667 | NA | NA | 1 | 1 | 4 | 0.75 | 2 | 1 | 7 | 0.8571 |
| 10_8014998 | 2 | 0.5 | 3 | 0 | 2 | 0 | NA | NA | 7 | 0.1429 | NA | NA | 2 | 0 | 2 | 0 | 2 | 0.5 | 6 | 0.1667 |
| 10_8015032 | 4 | 0.75 | 5 | 0.6 | 2 | 0.5 | 1 | 1 | 12 | 0.6667 | NA | NA | 3 | 0.6667 | 6 | 0.6667 | 3 | 0.3333 | 12 | 0.5833 |
| 10_8015037 | 4 | 0.75 | 5 | 0.4 | 3 | 0.3333 | 1 | 0 | 13 | 0.4615 | NA | NA | 3 | 0 | 6 | 0.6667 | 4 | 0.25 | 13 | 0.3846 |
| 10_8015040 | 4 | 0.5 | 5 | 0.2 | 2 | 0.5 | 1 | 0 | 12 | 0.3333 | NA | NA | 2 | 0.5 | 6 | 0.3333 | 4 | 0.25 | 12 | 0.3333 |
| 10_8015042 | 4 | 1 | 4 | 0.5 | 3 | 0.6667 | 1 | 0 | 12 | 0.6667 | NA | NA | 2 | 1 | 5 | 0.2 | 3 | 0.3333 | 10 | 0.4 |
| 10_8015058 | 4 | 0 | 5 | 0.2 | 2 | 0 | 1 | 0 | 12 | 0.0833 | NA | NA | 4 | 0 | 6 | 0.1667 | 4 | 0 | 14 | 0.0714 |
| 10_8015064 | 4 | 0.75 | 2 | 0.5 | 2 | 1 | NA | NA | 8 | 0.75 | NA | NA | 3 | 0.3333 | 3 | 0.3333 | 3 | 0.6667 | 9 | 0.4444 |
| 10_8015071 | 4 | 0.5 | 5 | 0 | 3 | 0.3333 | 1 | 0 | 13 | 0.2308 | NA | NA | 4 | 0 | 6 | 0 | 4 | 0.25 | 14 | 0.0714 |
| 10_8015093 | 5 | 0 | 5 | 0 | 3 | 0.6667 | 1 | 0 | 14 | 0.1429 | NA | NA | 4 | 0.25 | 6 | 0.3333 | 4 | 0.25 | 14 | 0.2857 |
| 10_8015099 | 5 | 0.4 | 5 | 0.8 | 3 | 1 | 2 | 0.5 | 15 | 0.6667 | NA | NA | 4 | 0.5 | 6 | 0.8333 | 7 | 1 | 17 | 0.8235 |
| 10_8015122 | 4 | 0.25 | 2 | 0.5 | 2 | 0 | 1 | 1 | 9 | 0.3333 | NA | NA | 1 | 0 | 4 | 1 | 5 | 1 | 10 | 0.9 |
| 10_8015148 | 4 | 0.25 | 5 | 0.4 | 5 | 0.4 | 2 | 0 | 16 | 0.3125 | NA | NA | 5 | 0.6 | 6 | 0.3333 | 6 | 0.1667 | 17 | 0.3529 |
| 10_8015154 | 4 | 0 | 4 | 0.25 | 4 | 0 | 1 | 0 | 13 | 0.0769 | NA | NA | 4 | 0 | 5 | 0.4 | 6 | 0.1667 | 15 | 0.2 |
| 10_8015164 | 3 | 1 | 4 | 1 | 7 | 1 | NA | NA | 14 | 1 | NA | NA | 3 | 1 | 6 | 1 | 7 | 1 | 16 | 1 |
| 10_8015180 | 5 | 1 | 6 | 0.8333 | 5 | 1 | 2 | 0.5 | 18 | 0.8889 | NA | NA | 5 | 0.4 | 7 | 0.8571 | 5 | 0.6 | 17 | 0.6471 |

| | | | | | | | | | | | | | | | | | | | | |
|---|---|---|---|---|---|---|---|---|---|---|---|---|---|---|---|---|---|---|---|---|
| 10_801518 | 5 | 1 | 5 | 1 | 5 | 1 | 2 | 1 | 17 | 1 | NA | NA | 3 | 1 | 6 | 1 | 5 | 1 | 14 | 1 |
| 10_801520 | 4 | 0.75 | 4 | 1 | 3 | 1 | 1 | 0 | 12 | 0.8333 | 1 | 1 | 5 | 0.8 | 6 | 0.6667 | 6 | 0.5 | 18 | 0.6667 |
| 10_801521 | 5 | 0.4 | 3 | 0 | 5 | 0.4 | 2 | 0 | 15 | 0.2667 | 1 | 1 | 6 | 0.1667 | 6 | 0.3333 | 8 | 0.25 | 21 | 0.2857 |
| 10_801522 | 2 | 0.5 | 5 | 0.4 | 5 | 0.4 | 2 | 0 | 14 | 0.3571 | 2 | 1 | 5 | 0.4 | 5 | 0.4 | 5 | 0.6 | 17 | 0.5294 |
| 10_801523 | 5 | 0.6 | 7 | 0.7143 | 7 | 0.5714 | 2 | 0 | 21 | 0.5714 | 2 | 0.5 | 6 | 0.3333 | 4 | 0.5 | 7 | 0.5714 | 19 | 0.4737 |
| 10_801523 | 5 | 0.8 | 8 | 0.875 | 7 | 1 | 3 | 0.6667 | 23 | 0.8696 | 2 | 1 | 6 | 1 | 5 | 0.8 | 7 | 0.7143 | 20 | 0.85 |
| 10_801524 | 4 | 0 | 5 | 0 | 5 | 0.2 | 4 | 0 | 18 | 0.0556 | 1 | 0 | 4 | 0 | 5 | 0 | 8 | 0.125 | 18 | 0.0556 |
| 10_801525 | 5 | 0.4 | 6 | 0.5 | 9 | 0.5556 | 4 | 1 | 24 | 0.5833 | 2 | 0.5 | 5 | 0.8 | 8 | 0.625 | 8 | 0.5 | 23 | 0.6087 |
| 10_801525 | 1 | 1 | 3 | 0.6667 | 7 | 0.5714 | 3 | 0.6667 | 14 | 0.6429 | 1 | 0 | 5 | 0.4 | 7 | 0.1429 | 8 | 0.75 | 21 | 0.4286 |
| 10_801526 | 5 | 0.2 | 9 | 0.1111 | 10 | 0.2 | 6 | 0 | 30 | 0.1333 | 3 | 0 | 8 | 0.375 | 10 | 0.1 | 9 | 0 | 30 | 0.1333 |
| 10_801527 | 4 | 0.25 | 7 | 0.4286 | 7 | 0 | 3 | 0.6667 | 21 | 0.2857 | 2 | 0.5 | 5 | 0.4 | 10 | 0.3 | 6 | 0.5 | 23 | 0.3913 |
| 10_801528 | 5 | 0.4 | 9 | 0 | 7 | 0.1429 | 6 | 0 | 27 | 0.1111 | 3 | 0 | 6 | 0.1667 | 9 | 0.1111 | 6 | 0.1667 | 24 | 0.125 |
| 10_801528 | 5 | 0.4 | 9 | 0.3333 | 11 | 0.1818 | 6 | 0.3333 | 31 | 0.2903 | 3 | 0.6667 | 8 | 0.375 | 9 | 0.2222 | 8 | 0.25 | 28 | 0.3214 |
| 10_801534 | 6 | 0.3333 | 10 | 0.1 | 12 | 0.1667 | 6 | 0 | 34 | 0.1471 | 2 | 0 | 7 | 0.1429 | 10 | 0.1 | 7 | 0.1429 | 26 | 0.1154 |
| 10_801535 | 5 | 0.4 | 11 | 0 | 11 | 0 | 7 | 0.1429 | 34 | 0.0882 | 3 | 0.6667 | 8 | 0.125 | 10 | 0.4 | 7 | 0.1429 | 28 | 0.2857 |
| 10_801536 | 6 | 0.1667 | 11 | 0.1818 | 15 | 0.1333 | 6 | 0.1667 | 38 | 0.1579 | 3 | 0.3333 | 8 | 0 | 12 | 0.1667 | 10 | 0.2 | 33 | 0.1515 |
| 10_801537 | 4 | 0 | 11 | 0.1818 | 13 | 0 | 6 | 0 | 34 | 0.0588 | 3 | 0.3333 | 8 | 0 | 9 | 0.1111 | 8 | 0 | 28 | 0.0714 |
| 10_801543 | 5 | 0.4 | 2 | 0.5 | 2 | 0.5 | 6 | 0.3333 | 15 | 0.4 | 2 | 1 | 1 | 0 | 4 | 0.5 | 2 | 0.5 | 9 | 0.5556 |
| 10_801544 | 6 | 0.6667 | NA | NA | 2 | 0.5 | 6 | 0.6667 | 14 | 0.6429 | 2 | 1 | NA | NA | 2 | 1 | 3 | 0.6667 | 7 | 0.8571 |
| 10_801545 | 7 | 0.8571 | 5 | 0.6 | 4 | 0.5 | 6 | 0.8333 | 22 | 0.7273 | 6 | 0.8333 | 1 | 1 | 8 | 0.875 | 7 | 0.8571 | 22 | 0.8636 |
| 10_801546 | 7 | 0.4286 | 4 | 0.25 | 6 | 0.1667 | 6 | 0 | 23 | 0.2174 | 4 | 0.75 | 3 | 0.3333 | 7 | 0.1429 | 6 | 0.3333 | 20 | 0.35 |
| 10_801549 | 6 | 1 | 6 | 1 | 5 | 1 | 5 | 1 | 22 | 1 | 4 | 1 | 2 | 0.5 | 8 | 1 | 8 | 1 | 22 | 0.9545 |
| 10_801549 | 5 | 0 | 5 | 0 | 6 | 0 | 4 | 0.25 | 20 | 0.05 | 4 | 0 | 3 | 0 | 9 | 0.1111 | 8 | 0.25 | 24 | 0.125 |
| 10_801550 | 5 | 1 | 5 | 0.4 | 6 | 0.5 | 4 | 0.25 | 20 | 0.55 | 4 | 0.75 | 3 | 0.3333 | 8 | 0.625 | 6 | 1 | 21 | 0.7143 |
| 10_801550 | 7 | 0.8571 | 5 | 0.8 | 6 | 0.8333 | 5 | 1 | 23 | 0.8696 | 3 | 1 | 1 | 0 | 8 | 1 | 8 | 0.875 | 20 | 0.9 |
| 10_801711 | 2 | 0.5 | 2 | 0 | 1 | 0 | 1 | 0 | 6 | 0.1667 | 1 | 0 | 3 | 0 | 5 | 0 | 2 | 1 | 11 | 0.1818 |
| 10_801711 | 1 | 1 | 2 | 0.5 | 1 | 1 | NA | NA | 4 | 0.75 | 1 | 0 | 5 | 0.6 | 5 | 0.8 | 2 | 0.5 | 13 | 0.6154 |
| 10_801713 | 2 | 0 | 2 | 0.5 | 1 | 1 | NA | NA | 5 | 0.4 | 1 | 0 | 3 | 0.3333 | 3 | 0.6667 | 2 | 0 | 9 | 0.3333 |
| 10_801713 | 2 | 0.5 | 1 | 1 | 1 | 0 | 1 | 0 | 5 | 0.4 | 1 | 1 | 4 | 0.75 | 4 | 0.5 | 2 | 1 | 11 | 0.7273 |
| 10_801714 | 2 | 0.5 | 2 | 0.5 | 1 | 0 | 1 | 1 | 6 | 0.5 | 1 | 1 | 5 | 0.6 | 5 | 1 | 3 | 0.6667 | 14 | 0.7857 |
| 10_801714 | 2 | 0.5 | 1 | 1 | 1 | 0 | 1 | 1 | 5 | 0.6 | 1 | 0 | 4 | 1 | 4 | 0.5 | 3 | 1 | 12 | 0.75 |
| 10_801716 | 2 | 1 | 2 | 1 | 1 | 1 | 1 | 0 | 6 | 0.8333 | 1 | 1 | 5 | 0.6 | 5 | 1 | 3 | 1 | 14 | 0.8571 |
| 10_801718 | 2 | 0.5 | 2 | 0.5 | 1 | 0 | 1 | 0 | 6 | 0.3333 | 1 | 1 | 5 | 0.2 | 5 | 0.4 | 3 | 0 | 14 | 0.2857 |
| 10_801719 | 2 | 0 | 3 | 0.6667 | 1 | 0 | 1 | 0 | 7 | 0.2857 | 1 | 0 | 5 | 0.4 | 5 | 0.4 | 3 | 0 | 14 | 0.2857 |
| 10_801721 | 2 | 0.5 | 4 | 1 | 1 | 1 | 1 | 0 | 8 | 0.75 | 1 | 1 | 4 | 1 | 6 | 1 | 3 | 1 | 14 | 1 |
| 10_8017213 | NA | NA | NA | NA | 1 | 1 | NA | NA | 1 | 1 | NA | NA | NA | NA | 1 | 1 | NA | NA | 1 | 1 |
| 10_802011 | 2 | 0.5 | 4 | 0 | 3 | 0 | 3 | 0 | 12 | 0.0833 | 3 | 0 | 3 | 0.3333 | 3 | 0 | 2 | 0 | 11 | 0.0909 |
| 10_802165 | 1 | 0 | 2 | 0.5 | 1 | 0 | 3 | 0.6667 | 7 | 0.4286 | 1 | 0 | 5 | 0.4 | 2 | 0 | 2 | 0.5 | 10 | 0.3 |
| 10_802168 | 1 | 0 | 1 | 0 | 1 | 1 | 2 | 0 | 5 | 0.2 | 1 | 1 | 6 | 0.5 | 2 | 0.5 | 3 | 0.3333 | 12 | 0.5 |
| 10_802169 | 1 | 1 | 1 | 1 | 1 | 1 | 3 | 0.6667 | 6 | 0.8333 | 1 | 1 | 4 | 1 | 2 | 0.5 | 1 | 1 | 8 | 0.875 |
| 10_802172 | 1 | 1 | NA | NA | NA | NA | 4 | 1 | 5 | 1 | NA | NA | 4 | 1 | 1 | 0 | 1 | 0 | 6 | 0.6667 |
| 10_802174 | 1 | 1 | NA | NA | 1 | 1 | 3 | 1 | 5 | 1 | 1 | 1 | 4 | 1 | 2 | 1 | 1 | 1 | 8 | 1 |
| 10_802176 | 1 | 1 | NA | NA | 1 | 0 | 4 | 0.25 | 6 | 0.3333 | 1 | 0 | 5 | 0.4 | 1 | 0 | 1 | 0 | 8 | 0.25 |
| 10_802176 | 1 | 1 | NA | NA | 1 | 1 | 4 | 0.75 | 6 | 0.8333 | 1 | 1 | 4 | 1 | 1 | 1 | 1 | 1 | 7 | 1 |
| 10_802177 | 1 | 1 | NA | NA | NA | NA | 3 | 1 | 4 | 1 | 1 | 1 | 5 | 1 | 1 | 1 | 1 | 1 | 8 | 1 |
| 10_802178 | 1 | 1 | NA | NA | NA | NA | 2 | 1 | 3 | 1 | 1 | 1 | 4 | 1 | NA | NA | 1 | 0 | 6 | 0.8333 |
| 10_802179 | 1 | 1 | NA | NA | 1 | 1 | 3 | 1 | 5 | 1 | NA | NA | 3 | 1 | 1 | 1 | 1 | 1 | 5 | 1 |
| 10_802180 | 1 | 1 | NA | NA | 1 | 0 | 3 | 0 | 5 | 0.2 | NA | NA | 2 | 0.5 | 1 | 0 | 1 | 0 | 4 | 0.25 |
| 10_802226 | 1 | 0 | NA | NA | NA | NA | 3 | 0.6667 | 4 | 0.5 | NA | NA | 3 | 0.3333 | 1 | 0 | 1 | 0 | 5 | 0.2 |
| 10_802229 | 1 | 0 | NA | NA | NA | NA | 3 | 0.3333 | 4 | 0.25 | NA | NA | 4 | 0.5 | 1 | 0 | 1 | 1 | 6 | 0.5 |
| 10_802229 | 1 | 1 | NA | NA | NA | NA | 2 | 0 | 3 | 0.3333 | NA | NA | 3 | 0.6667 | NA | NA | 1 | 0 | 4 | 0.5 |
| 10_802283 | 20 | 0.45 | 25 | 0.2 | 21 | 0.3333 | 23 | 0.0435 | 89 | 0.2472 | 21 | 0.1905 | 21 | 0.0952 | 24 | 0.125 | 20 | 0.25 | 86 | 0.1628 |
| 10_802284 | 26 | 0.8462 | 22 | 0.7273 | 22 | 0.8182 | 20 | 0.85 | 90 | 0.8111 | 20 | 0.85 | 17 | 0.8235 | 21 | 0.8571 | 15 | 0.8667 | 73 | 0.8493 |
| 10_802285 | 20 | 0.35 | 18 | 0.2222 | 20 | 0.2 | 21 | 0.1429 | 79 | 0.2278 | 20 | 0.3 | 15 | 0.2 | 15 | 0.1333 | 19 | 0.4737 | 69 | 0.2899 |
| 10_802286 | 23 | 0.4783 | 23 | 0.3913 | 22 | 0.3182 | 23 | 0.3913 | 91 | 0.3956 | 19 | 0.2632 | 19 | 0.3158 | 24 | 0.4583 | 17 | 0.1765 | 79 | 0.3165 |
| 10_802287 | 17 | 0.4118 | 19 | 0.2105 | 20 | 0.35 | 17 | 0.3529 | 73 | 0.3288 | 22 | 0.4091 | 14 | 0.2143 | 20 | 0.3 | 13 | 0.5385 | 69 | 0.3623 |
| 10_802287 | 27 | 0.3704 | 24 | 0.25 | 24 | 0.3333 | 21 | 0.5238 | 96 | 0.3646 | 24 | 0.2917 | 21 | 0.5714 | 25 | 0.4 | 19 | 0.7368 | 89 | 0.4831 |
| 10_802288 | 23 | 0.1304 | 21 | 0.1429 | 22 | 0.0455 | 22 | 0.1364 | 88 | 0.1136 | 23 | 0.1304 | 19 | 0.1579 | 22 | 0.0455 | 22 | 0.1818 | 86 | 0.1279 |
| 10_802388 | 67 | 0.0746 | 79 | 0.038 | 69 | 0.0725 | 66 | 0.1061 | 281 | 0.0712 | 75 | 0.0933 | 64 | 0.125 | 68 | 0.1029 | 51 | 0.098 | 258 | 0.1047 |
| 10_802455 | 2 | 0 | 3 | 0 | 2 | 0 | 3 | 0.3333 | 10 | 0.1 | 2 | 0 | 7 | 0.1429 | 1 | 0 | 2 | 0 | 12 | 0.0833 |
| 10_802456 | 2 | 1 | 3 | 0.6667 | 2 | 0.5 | 2 | 1 | 9 | 0.7778 | 3 | 0.6667 | 7 | 0.5714 | 1 | 0 | 1 | 1 | 12 | 0.5833 |
| 10_802457 | 2 | 1 | 2 | 1 | 3 | 0.6667 | 4 | 0.5 | 11 | 0.7273 | 2 | 1 | 6 | 0.6667 | 1 | 1 | 2 | 1 | 11 | 0.8182 |
| 10_802458 | 2 | 0.5 | 3 | 0.3333 | 2 | 1 | 3 | 0.3333 | 10 | 0.5 | 3 | 0.3333 | 6 | 0.1667 | NA | NA | 2 | 1 | 11 | 0.3636 |
| 10_803781 | 5 | 0.4 | 6 | 0.1667 | 2 | 0 | 1 | 0 | 14 | 0.2143 | 3 | 0 | 4 | 0 | 2 | 1 | 3 | 0 | 12 | 0.1667 |
| 10_803851 | 1 | 1 | 4 | 0 | 2 | 0 | 2 | 0 | 9 | 0.1111 | 1 | 1 | NA | NA | 1 | 0 | 1 | 0 | 3 | 0.3333 |

| | | | | | | | | | | | | | | | | | | | | |
|---|---|---|---|---|---|---|---|---|---|---|---|---|---|---|---|---|---|---|---|---|
| 10_8038525 | 3 | 0 | 4 | 0.25 | 2 | 0 | 2 | 0 | 11 | 0.0909 | 2 | 1 | NA | NA | 1 | 0 | 3 | 0.3333 | 6 | 0.5 |
| 10_8038542 | 3 | 0.3333 | 4 | 0.5 | 2 | 0 | 1 | 0 | 10 | 0.3 | 1 | 0 | NA | NA | 1 | 1 | 2 | 0 | 4 | 0.25 |
| 10_8038550 | 2 | 1 | 3 | 0.6667 | 2 | 1 | 1 | 1 | 8 | 0.875 | 2 | 1 | NA | NA | 1 | 0 | 2 | 0.5 | 5 | 0.6 |
| 10_8038553 | 2 | 0.5 | 2 | 1 | NA | NA | 1 | 1 | 5 | 0.8 | 2 | 1 | NA | NA | 1 | 0 | 1 | 0 | 4 | 0.5 |
| 10_8038557 | 2 | 0.5 | 3 | 0.3333 | 1 | 0 | 1 | 0 | 7 | 0.2857 | 1 | 1 | NA | NA | NA | NA | 2 | 0 | 3 | 0.3333 |
| 10_8038569 | 3 | 0.6667 | 2 | 0.5 | 1 | 1 | NA | NA | 6 | 0.6667 | 1 | 1 | NA | NA | 1 | 1 | 1 | 1 | 3 | 1 |
| 10_8038577 | 2 | 1 | NA | NA | NA | NA | NA | NA | 2 | 1 | 3 | 0 | NA | NA | 1 | 0 | 2 | 0.5 | 6 | 0.1667 |
| 10_8038605 | 2 | 0.5 | 3 | 0 | NA | NA | 1 | 0 | 6 | 0.1667 | 3 | 0 | NA | NA | 1 | 1 | 2 | 0 | 6 | 0.1667 |
| 10_8038622 | 3 | 0.6667 | 3 | 0 | NA | NA | 1 | 0 | 7 | 0.2857 | 3 | 0.3333 | NA | NA | 1 | 1 | 2 | 1 | 6 | 0.6667 |
| 10_8038631 | 2 | 1 | 3 | 0.6667 | NA | NA | 1 | 1 | 6 | 0.8333 | 2 | 0.5 | NA | NA | 1 | 1 | 2 | 0.5 | 5 | 0.6 |
| 10_8040198 | 1 | 0 | NA | NA | NA | NA | 1 | 1 | 2 | 0.5 | NA | NA | NA | NA | 1 | 1 | NA | NA | 1 | 1 |
| 10_8040310 | 8 | 0.125 | 9 | 0.1111 | 9 | 0 | 12 | 0.0833 | 38 | 0.0789 | 2 | 0 | 12 | 0.25 | 11 | 0.0909 | 8 | 0.125 | 33 | 0.1515 |
| 10_8040377 | 7 | 0.1429 | 9 | 0 | 4 | 0 | 9 | 0.1111 | 29 | 0.069 | 2 | 0 | 7 | 0.1429 | 8 | 0.125 | 6 | 0.1667 | 23 | 0.1304 |
| 10_8040381 | 7 | 0.4286 | 5 | 0.2 | 2 | 0 | 9 | 0.5556 | 23 | 0.3913 | 1 | 0 | 2 | 0.5 | 7 | 0.2857 | 6 | 0.5 | 16 | 0.375 |
| 10_8040406 | 5 | 0.4 | 4 | 0.25 | 2 | 0 | 6 | 0.5 | 17 | 0.3529 | 1 | 0 | 4 | 0.5 | 3 | 0 | 3 | 0.3333 | 11 | 0.2727 |
| 10_8040417 | 8 | 0 | 5 | 0 | 5 | 0.2 | 7 | 0.2857 | 25 | 0.12 | 2 | 0.5 | 6 | 0 | 7 | 0.4286 | 5 | 0 | 20 | 0.2 |
| 10_8040439 | 7 | 0 | 4 | 0 | 4 | 0.5 | 8 | 0.25 | 23 | 0.1739 | 2 | 0 | 6 | 0.1667 | 5 | 0.4 | 3 | 0 | 16 | 0.1875 |
| 10_8040461 | 5 | 1 | 6 | 1 | 2 | 1 | 6 | 1 | 19 | 1 | 2 | 1 | 6 | 1 | 6 | 1 | 3 | 1 | 17 | 1 |
| 10_8040462 | 7 | 1 | 5 | 1 | 2 | 1 | 6 | 1 | 20 | 1 | 2 | 1 | 6 | 1 | 5 | 1 | 2 | 1 | 15 | 1 |
| 10_8040469 | 5 | 0.6 | 4 | 1 | 3 | 1 | 5 | 0.6 | 17 | 0.7647 | 2 | 1 | 5 | 0.6 | 5 | 1 | 2 | 0.5 | 14 | 0.7857 |
| 10_8041962 | 22 | 0.5455 | 35 | 0.4286 | 41 | 0.2683 | 38 | 0.3421 | 136 | 0.375 | 38 | 0.3421 | 28 | 0.2857 | 29 | 0.2759 | 27 | 0.4815 | 122 | 0.3443 |
| 10_8041965 | 23 | 0.2174 | 35 | 0.0571 | 45 | 0.1778 | 38 | 0.1842 | 141 | 0.156 | 38 | 0.1579 | 29 | 0.1379 | 28 | 0.0714 | 26 | 0.1923 | 121 | 0.1405 |
| 10_8041974 | 25 | 0.24 | 34 | 0.2647 | 41 | 0.2439 | 35 | 0.2571 | 135 | 0.2519 | 34 | 0.2353 | 25 | 0.24 | 29 | 0.2069 | 23 | 0.3043 | 111 | 0.2432 |
| 10_8042276 | 15 | 0.6667 | 24 | 0.6667 | 29 | 0.5172 | 23 | 0.4783 | 91 | 0.5714 | 22 | 0.6364 | 16 | 0.8125 | 18 | 0.5556 | 20 | 0.6 | 76 | 0.6447 |
| 10_8042280 | 11 | 0.0909 | 16 | 0.0625 | 26 | 0.1154 | 17 | 0.1765 | 70 | 0.1143 | 24 | 0 | 14 | 0.2143 | 19 | 0.1053 | 13 | 0 | 70 | 0.0714 |
| 10_8042312 | NA | NA | NA | NA | 1 | 1 | 1 | 0 | 2 | 0.5 | 2 | 0 | 1 | 1 | NA | NA | 2 | 0.5 | 5 | 0.4 |
| 10_8044251 | 1 | 1 | NA | NA | NA | NA | NA | NA | 1 | 1 | NA | NA | NA | NA | NA | NA | 1 | 1 | 1 | 1 |
| 10_8044280 | 1 | 1 | NA | NA | NA | NA | NA | NA | 1 | 1 | NA | NA | NA | NA | NA | NA | 1 | 1 | 1 | 1 |
| 10_8044297 | 1 | 1 | NA | NA | NA | NA | NA | NA | 1 | 1 | 1 | 1 | NA | NA | NA | NA | 1 | 1 | 2 | 1 |
| 10_8044320 | 1 | 1 | NA | NA | NA | NA | NA | NA | 1 | 1 | 1 | 1 | NA | NA | NA | NA | 1 | 0 | 2 | 0.5 |
| 10_8044332 | 1 | 1 | NA | NA | NA | NA | NA | NA | 1 | 1 | 1 | 1 | NA | NA | NA | NA | 1 | 1 | 2 | 1 |
| 10_8044334 | 1 | 1 | NA | NA | NA | NA | NA | NA | 1 | 1 | 1 | 1 | NA | NA | NA | NA | 1 | 1 | 2 | 1 |
| 10_8045090 | 5 | 0 | 8 | 0.25 | 6 | 0.5 | 3 | 0 | 22 | 0.2273 | 7 | 0 | 10 | 0.1 | 4 | 0.25 | 12 | 0.1667 | 33 | 0.1212 |
| 10_8045093 | 3 | 1 | 5 | 0.8 | 3 | 1 | 2 | 1 | 13 | 0.9231 | 6 | 1 | 8 | 1 | 2 | 1 | 8 | 1 | 24 | 1 |
| 10_8045100 | 5 | 0.6 | 8 | 0.625 | 6 | 0.6667 | 4 | 1 | 23 | 0.6957 | 8 | 0.875 | 10 | 0.4 | 4 | 0.5 | 12 | 0.8333 | 34 | 0.6765 |
| 10_8045103 | 5 | 0.4 | 5 | 0.2 | 4 | 0 | 4 | 0 | 18 | 0.1667 | 6 | 0.1667 | 8 | 0.125 | 4 | 0 | 10 | 0.2 | 28 | 0.1429 |
| 10_8045110 | 5 | 0 | 7 | 0 | 6 | 0.1667 | 3 | 0.3333 | 21 | 0.0952 | 8 | 0 | 9 | 0.1111 | 3 | 0 | 12 | 0.0833 | 32 | 0.0625 |
| 10_8045118 | 6 | 0 | 6 | 0.3333 | 6 | 0 | 5 | 0 | 23 | 0.087 | 7 | 0.1429 | 6 | 0 | 3 | 0.3333 | 10 | 0 | 26 | 0.0769 |
| 10_8045122 | 6 | 0.6667 | 8 | 0.625 | 6 | 0.6667 | 5 | 0.8 | 25 | 0.68 | 9 | 0.4444 | 11 | 0.7273 | 4 | 0.75 | 11 | 0.5455 | 35 | 0.6 |
| 10_8045123 | 6 | 0.1667 | 7 | 0.7143 | 6 | 0.5 | 4 | 0.5 | 23 | 0.4783 | 8 | 0.625 | 8 | 0.375 | 4 | 0.5 | 13 | 0.5385 | 33 | 0.5152 |
| 10_8045125 | 6 | 0.3333 | 7 | 0.4286 | 6 | 0.5 | 6 | 0.3333 | 25 | 0.4 | 9 | 0.3333 | 10 | 0.4 | 4 | 0 | 13 | 0.4615 | 36 | 0.3611 |
| 10_8045135 | 4 | 1 | 8 | 1 | 6 | 1 | 5 | 1 | 23 | 1 | 7 | 1 | 11 | 1 | 4 | 1 | 13 | 1 | 35 | 1 |
| 10_8045143 | 7 | 0.5714 | 6 | 0.8333 | 3 | 1 | 5 | 0.6 | 21 | 0.7143 | 6 | 0.6667 | 7 | 0.5714 | 2 | 0.5 | 10 | 0.6 | 25 | 0.6 |
| 10_8047028 | 5 | 0.6 | 13 | 0.4615 | 12 | 0.25 | 9 | 0.3333 | 39 | 0.3846 | 14 | 0.5714 | 5 | 0.4 | 10 | 0.4 | 7 | 0.7143 | 36 | 0.5278 |
| 10_8047595 | 13 | 0.0769 | 20 | 0 | 27 | 0.0741 | 18 | 0.1111 | 78 | 0.0641 | 20 | 0.1 | 9 | 0.3333 | 20 | 0 | 17 | 0.1765 | 66 | 0.1212 |
| 10_8047601 | 11 | 0.2727 | 23 | 0.2609 | 22 | 0.3182 | 19 | 0.1579 | 75 | 0.2533 | 20 | 0.25 | 10 | 0.3 | 17 | 0.2941 | 15 | 0.2667 | 62 | 0.2742 |
| 10_8047604 | 12 | 0.9167 | 24 | 0.9167 | 24 | 0.8333 | 19 | 0.9474 | 79 | 0.8987 | 18 | 1 | 9 | 0.8889 | 19 | 0.8947 | 13 | 0.8462 | 59 | 0.9153 |
| 10_8047631 | 10 | 0.1 | 21 | 0.381 | 24 | 0.25 | 15 | 0.4667 | 70 | 0.3143 | 17 | 0.2941 | 10 | 0.3 | 22 | 0.2727 | 13 | 0.6154 | 62 | 0.3548 |
| 10_8047638 | 11 | 0 | 23 | 0.1739 | 26 | 0.1154 | 16 | 0.125 | 76 | 0.1184 | 17 | 0.0588 | 12 | 0.1667 | 20 | 0.2 | 17 | 0.1176 | 66 | 0.1364 |
| 10_8047645 | 12 | 0.0833 | 25 | 0.2 | 29 | 0.1724 | 17 | 0.1765 | 83 | 0.1687 | 22 | 0.2273 | 14 | 0 | 24 | 0.125 | 17 | 0.1176 | 77 | 0.1299 |
| 10_8047649 | 12 | 0.25 | 19 | 0.3158 | 24 | 0.3333 | 17 | 0.2353 | 72 | 0.2917 | 19 | 0.3684 | 13 | 0.3846 | 18 | 0.5 | 14 | 0.2857 | 64 | 0.3906 |
| 10_8047915 | 11 | 1 | 9 | 1 | 7 | 1 | 17 | 1 | 44 | 1 | 10 | 0.9 | 12 | 0.9167 | 10 | 1 | 5 | 1 | 37 | 0.9459 |
| 10_8047917 | 7 | 0.5714 | 9 | 0.4444 | 6 | 0.3333 | 17 | 0.4706 | 39 | 0.4615 | 9 | 0.3333 | 9 | 0.3333 | 12 | 0.4167 | 6 | 0.6667 | 36 | 0.4167 |
| 10_8047918 | 8 | 0.5 | 9 | 0.1111 | 6 | 0.1667 | 18 | 0.1111 | 41 | 0.1951 | 11 | 0.0909 | 9 | 0.2222 | 11 | 0.3636 | 6 | 0.5 | 37 | 0.2703 |
| 10_8047920 | 7 | 0.1429 | 9 | 0.3333 | 6 | 0 | 17 | 0.2941 | 39 | 0.2308 | 11 | 0.2727 | 12 | 0.0833 | 9 | 0.2222 | 5 | 0.4 | 37 | 0.2162 |
| 10_8050123 | 6 | 0.1667 | 9 | 0.2222 | 3 | 0 | 7 | 0.2857 | 25 | 0.2 | 7 | 0 | 5 | 0.2 | 2 | 0.5 | 6 | 0 | 20 | 0.1 |
| 10_8050149 | 7 | 0.1429 | 6 | 0.3333 | 2 | 0 | 6 | 0.3333 | 21 | 0.2381 | 4 | 0.5 | 5 | 0.8 | 1 | 1 | 6 | 0.6667 | 16 | 0.6875 |
| 10_8050156 | 6 | 0.1667 | 7 | 0.2857 | 2 | 0 | 4 | 0.5 | 19 | 0.2632 | 5 | 0.2 | 3 | 0.3333 | 2 | 0 | 6 | 0.1667 | 16 | 0.1875 |
| 10_8050193 | 6 | 0.1667 | 6 | 0.1667 | 2 | 0 | 2 | 0 | 16 | 0.125 | 6 | 0.3333 | 7 | 0.2857 | 2 | 0 | 5 | 0.2 | 20 | 0.25 |
| 10_8052255 | 1 | 1 | NA | NA | 2 | 0.5 | NA | NA | 3 | 0.6667 | NA | NA | 1 | 0 | 1 | 1 | 1 | 1 | 3 | 0.6667 |
| 10_8052278 | 1 | 1 | NA | NA | 2 | 1 | NA | NA | 3 | 1 | NA | NA | 1 | 0 | NA | NA | 1 | 1 | 2 | 0.5 |
| 10_8052279 | 1 | 1 | NA | NA | 2 | 0 | NA | NA | 3 | 0.3333 | NA | NA | 1 | 0 | NA | NA | 1 | 1 | 2 | 0.5 |
| 10_8052284 | NA | NA | NA | NA | 2 | 1 | NA | NA | 2 | 1 | NA | NA | 1 | 1 | NA | NA | 1 | 1 | 2 | 1 |
| 10_8052723 | NA | NA | NA | NA | 1 | 1 | NA | NA | 1 | 1 | NA | NA | NA | NA | NA | NA | 1 | 1 | 1 | 1 |
| 10_8052728 | NA | NA | NA | NA | 1 | 1 | NA | NA | 1 | 1 | NA | NA | NA | NA | NA | NA | 1 | 1 | 1 | 1 |

| | | | | | | | | | | | | | | | | | | | | |
|---|---|---|---|---|---|---|---|---|---|---|---|---|---|---|---|---|---|---|---|---|
| 10_8054092 | 2 | 0 | 3 | 0 | 4 | 0.25 | 5 | 0 | 14 | 0.0714 | 2 | 0 | 1 | 0 | 3 | 0.3333 | 5 | 0.4 | 11 | 0.2727 |
| 10_8054102 | 2 | 0.5 | 3 | 0.6667 | 3 | 1 | 5 | 0.8 | 13 | 0.7692 | 3 | 0.6667 | 2 | 0.5 | 2 | 1 | 6 | 1 | 13 | 0.8462 |
| 10_8054297 | 16 | 0.3125 | 12 | 0.1667 | 15 | 0.0667 | 15 | 0 | 58 | 0.1379 | 15 | 0.0667 | 19 | 0.2105 | 25 | 0.08 | 14 | 0.0714 | 73 | 0.1096 |
| 10_8054303 | 12 | 0.0833 | 12 | 0.0833 | 16 | 0.0625 | 13 | 0 | 53 | 0.0566 | 13 | 0 | 16 | 0.25 | 23 | 0.087 | 14 | 0.1429 | 66 | 0.1212 |
| 10_8054321 | 13 | 0.4615 | 8 | 0.25 | 11 | 0.5455 | 12 | 0.75 | 44 | 0.5227 | 12 | 0.5 | 15 | 0.4667 | 15 | 0.3333 | 11 | 0.5455 | 53 | 0.4528 |
| 10_8054347 | 13 | 0.8462 | 9 | 0.6667 | 13 | 0.5385 | 11 | 0.8182 | 46 | 0.7174 | 12 | 0.9167 | 13 | 0.7692 | 22 | 0.7273 | 15 | 0.6 | 62 | 0.7419 |
| 10_8055513 | 5 | 0.2 | 3 | 0.3333 | 1 | 0 | 4 | 0 | 13 | 0.1538 | 5 | 0.2 | 6 | 0.1667 | 4 | 0 | 3 | 0 | 18 | 0.1111 |
| 10_8062177 | 1 | 1 | 2 | 0.5 | 1 | 1 | 1 | 1 | 5 | 0.8 | 1 | 0 | 1 | 1 | 1 | 1 | 1 | 0 | 4 | 0.5 |
| 10_8062462 | 2 | 0 | 1 | 0 | 1 | 0 | 1 | 1 | 5 | 0.2 | 1 | 0 | 2 | 0 | 2 | 0 | 2 | 0.5 | 7 | 0.1429 |
| 10_8062475 | 2 | 0 | 2 | 0 | 1 | 1 | 1 | 0 | 6 | 0.1667 | 1 | 0 | 2 | 1 | 2 | 0.5 | 2 | 0.5 | 7 | 0.5714 |
| 10_8062478 | NA | NA | 1 | 1 | NA | NA | NA | NA | 1 | 1 | NA | NA | 3 | 0.3333 | 1 | 0 | 2 | 0.5 | 6 | 0.3333 |
| 10_8062655 | 1 | 0 | 2 | 0.5 | NA | NA | 1 | 1 | 4 | 0.5 | NA | NA | | | 3 | 0.6667 | 2 | 1 | 5 | 0.8 |
| 10_8062681 | 1 | 1 | 2 | 0 | 1 | 0 | 1 | 1 | 5 | 0.4 | NA | NA | NA | NA | 4 | 0.5 | 2 | 1 | 6 | 0.6667 |
| 10_8062686 | 1 | 1 | 3 | 1 | 1 | 0 | 1 | 1 | 6 | 0.8333 | NA | NA | 3 | 0.6667 | 4 | 1 | 2 | 1 | 9 | 0.8889 |
| 10_8062698 | 2 | 0.5 | 2 | 0.5 | 1 | 1 | 1 | 0 | 6 | 0.5 | NA | NA | 1 | 0 | 3 | 0.6667 | 2 | 0.5 | 6 | 0.5 |
| 10_8062704 | 2 | 1 | 2 | 1 | 1 | 1 | 1 | 1 | 6 | 1 | NA | NA | 2 | 1 | 5 | 0.6 | 2 | 0.5 | 9 | 0.6667 |
| 10_8062712 | 1 | 1 | 1 | 1 | NA | NA | 1 | 1 | 3 | 1 | 1 | 1 | 1 | 1 | 4 | 0.5 | 1 | 1 | 7 | 0.7143 |
| 10_8062947 | 2 | 1 | 4 | 0.5 | 1 | 1 | 1 | 0 | 8 | 0.625 | 1 | 1 | 2 | 0.5 | 6 | 0.8333 | 4 | 0.75 | 13 | 0.7692 |
| 10_8062954 | 3 | 0.3333 | 5 | 0.2 | 2 | 0 | 1 | 0 | 11 | 0.1818 | 1 | 0 | 2 | 0 | 9 | 0.2222 | 5 | 0 | 17 | 0.1176 |
| 10_8062962 | 3 | 0 | 5 | 1 | 1 | 0 | 2 | 1 | 11 | 0.6364 | 1 | 1 | 4 | 0 | 6 | 0.5 | 4 | 0.25 | 15 | 0.3333 |
| 10_8062965 | 3 | 0.6667 | 4 | 0.75 | 2 | 1 | 3 | 1 | 12 | 0.8333 | 1 | 1 | 5 | 0.8 | 8 | 0.875 | 3 | 0.6667 | 17 | 0.8235 |
| 10_8062973 | 4 | 1 | 5 | 0.6 | 2 | 1 | 2 | 1 | 13 | 0.8462 | 1 | 1 | 4 | 0.5 | 6 | 1 | 3 | 1 | 14 | 0.8571 |
| 10_8062984 | 4 | 0.5 | 5 | 0.4 | NA | NA | 2 | 1 | 11 | 0.5455 | 1 | 1 | 3 | 0.3333 | 6 | 0.3333 | 3 | 0.3333 | 13 | 0.3846 |
| 10_8063017 | 3 | 0.3333 | 6 | 0 | 1 | 1 | 4 | 0.25 | 14 | 0.2143 | 1 | 0 | 4 | 0 | 7 | 0.1429 | 3 | 0 | 15 | 0.0667 |
| 10_8063021 | 3 | 0 | 6 | 0 | 3 | 0.3333 | 4 | 0 | 16 | 0.0625 | NA | NA | 5 | 0 | 9 | 0.1111 | 4 | 0.25 | 18 | 0.1111 |
| 10_8063103 | 3 | 0.3333 | 6 | 0 | 3 | 0.3333 | 6 | 0 | 18 | 0.1111 | 1 | 0 | 3 | 0.3333 | 10 | 0.1 | 6 | 0 | 20 | 0.1 |
| 10_8063225 | 7 | 0.4286 | 9 | 0.6667 | 9 | 0.5556 | 8 | 0.5 | 33 | 0.5455 | 7 | 0.8571 | 6 | 0.5 | 13 | 0.6923 | 11 | 0.4545 | 37 | 0.6216 |
| 10_8063228 | 8 | 0 | 10 | 0.1 | 9 | 0.2222 | 7 | 0.1429 | 34 | 0.1176 | 10 | 0.1 | 10 | 0.1 | 15 | 0.2667 | 11 | 0.1818 | 46 | 0.1739 |
| 10_8063991 | 6 | 0 | 4 | 0.25 | 9 | 0 | 10 | 0.2 | 29 | 0.1034 | 8 | 0 | 7 | 0.5714 | 9 | 0.2222 | 6 | 0.5 | 30 | 0.3 |
| 10_8064441 | 4 | 0.5 | 2 | 0.5 | 4 | 0.25 | 5 | 0.6 | 15 | 0.4667 | NA | NA | 5 | 0.2 | 2 | 1 | 2 | 0 | 9 | 0.3333 |
| 10_8064450 | 4 | 0.25 | 2 | 0 | 4 | 0 | 6 | 0 | 16 | 0.0625 | NA | NA | 5 | 0.6 | 2 | 0.5 | 2 | 0.5 | 9 | 0.5556 |
| 10_8064451 | 4 | 0.5 | 2 | 0.5 | 5 | 0.4 | 6 | 0.3333 | 17 | 0.4118 | NA | NA | 5 | 0.6 | 2 | 0 | 2 | 0.5 | 9 | 0.4444 |
| 10_8066483 | 107 | 0.0654 | 114 | 0.0526 | 86 | 0.0465 | 114 | 0.0526 | 421 | 0.0546 | 74 | 0.0811 | 103 | 0.0583 | 124 | 0.0887 | 90 | 0.0333 | 391 | 0.0665 |
| 10_8066877 | 12 | 0.1667 | 9 | 0.1111 | 8 | 0.25 | 24 | 0.125 | 53 | 0.1509 | 4 | 0 | 10 | 0.1 | 23 | 0.087 | 16 | 0.0625 | 53 | 0.0755 |
| 10_8066886 | 11 | 1 | 7 | 0.8571 | 9 | 0.7778 | 21 | 0.8095 | 48 | 0.8542 | 1 | 1 | 10 | 0.8 | 20 | 0.55 | 11 | 0.7273 | 42 | 0.6667 |
| 10_8066888 | 13 | 0.1538 | 9 | 0.3333 | 10 | 0.8 | 27 | 0.4074 | 59 | 0.4068 | 4 | 0.25 | 13 | 0.6154 | 24 | 0.375 | 14 | 0.4286 | 55 | 0.4364 |
| 10_8066893 | 13 | 0.2308 | 7 | 0.1429 | 9 | 0.1111 | 25 | 0.12 | 54 | 0.1481 | 2 | 0.5 | 11 | 0.3636 | 23 | 0.087 | 15 | 0.2 | 51 | 0.1961 |
| 10_8066893 | 12 | 0.0833 | 8 | 0.125 | 10 | 0.2 | 25 | 0.08 | 55 | 0.1091 | 3 | 0 | 12 | 0.0833 | 26 | 0.0385 | 15 | 0.0667 | 56 | 0.0536 |
| 10_8066895 | 7 | 0.1429 | 8 | 0.125 | 7 | 0.1429 | 23 | 0.3043 | 45 | 0.2222 | 2 | 0.5 | 11 | 0.1818 | 21 | 0.2381 | 12 | 0.25 | 46 | 0.2391 |
| 10_8066901 | 12 | 0.5 | 7 | 0.8571 | 6 | 0.6667 | 15 | 0.5333 | 40 | 0.6 | 2 | 0.5 | 10 | 0.4 | 19 | 0.5789 | 13 | 0.4615 | 44 | 0.5 |
| 10_8066901 | 11 | 0.2727 | 9 | 0.2222 | 8 | 0 | 23 | 0.087 | 51 | 0.1373 | 2 | 0 | 10 | 0.2 | 20 | 0.05 | 12 | 0.1667 | 44 | 0.1136 |
| 10_8066904 | 12 | 0 | 8 | 0.125 | 10 | 0.3 | 27 | 0.0741 | 57 | 0.1053 | 4 | 0.25 | 13 | 0.0769 | 24 | 0.0833 | 15 | 0.1333 | 56 | 0.1071 |
| 10_8066921 | 10 | 0.3 | 7 | 0.7143 | 8 | 0.75 | 13 | 0.3077 | 38 | 0.4737 | 2 | 0.5 | 10 | 0.4 | 22 | 0.5455 | 8 | 0.625 | 42 | 0.5238 |
| 10_8066924 | 12 | 0.5833 | 9 | 0.5556 | 9 | 0.6667 | 24 | 0.625 | 54 | 0.6111 | 3 | 0.3333 | 9 | 0.6667 | 24 | 0.4167 | 12 | 0.5833 | 48 | 0.5 |
| 10_8066925 | 13 | 0.4615 | 7 | 0.7143 | 9 | 0.5556 | 20 | 0.4 | 49 | 0.4898 | 2 | 1 | 10 | 0.2 | 20 | 0.35 | 10 | 0.5 | 42 | 0.381 |
| 10_8066927 | 12 | 0.0833 | 8 | 0.125 | 9 | 0 | 22 | 0.0909 | 51 | 0.0784 | 2 | 0 | 11 | 0 | 22 | 0.0909 | 12 | 0.1667 | 47 | 0.0851 |
| 10_8066939 | 10 | 1 | 2 | 1 | 8 | 1 | 15 | 1 | 35 | 1 | 1 | 0 | 8 | 1 | 18 | 0.8889 | 8 | 1 | 35 | 0.9143 |
| 10_8066948 | 11 | 0.7273 | 8 | 0.75 | 8 | 0.875 | 20 | 0.75 | 47 | 0.766 | 3 | 1 | 12 | 0.9167 | 19 | 0.9474 | 11 | 0.9091 | 45 | 0.9333 |
| 10_8066956 | 10 | 0.6 | 5 | 0.6 | 7 | 0.8571 | 13 | 0.4615 | 35 | 0.6 | 1 | 0 | 8 | 0.75 | 10 | 0.5 | 8 | 0.375 | 27 | 0.5185 |
| 10_8066969 | 11 | 0.0909 | 8 | 0.25 | 10 | 0.4 | 26 | 0.2308 | 55 | 0.2364 | 2 | 0 | 11 | 0 | 19 | 0.1579 | 10 | 0.2 | 42 | 0.119 |
| 10_8066970 | 12 | 0.9167 | 8 | 0.875 | 7 | 1 | 21 | 0.9524 | 48 | 0.9375 | 2 | 1 | 10 | 1 | 21 | 0.9524 | 13 | 0.9231 | 46 | 0.9565 |
| 10_8066977 | 12 | 0.8333 | 7 | 0.2857 | 8 | 0.75 | 22 | 0.6818 | 49 | 0.6735 | 2 | 0.5 | 11 | 0.5455 | 20 | 0.55 | 10 | 0.7 | 43 | 0.5814 |
| 10_8066978 | 11 | 0.0909 | 6 | 0 | 9 | 0 | 23 | 0.087 | 49 | 0.0612 | 2 | 0 | 11 | 0 | 20 | 0.05 | 11 | 0.2727 | 44 | 0.0909 |
| 10_8066988 | 10 | 0.9 | 5 | 0.8 | 7 | 1 | 16 | 0.9375 | 38 | 0.9211 | 1 | 1 | 8 | 0.5 | 17 | 0.8235 | 9 | 0.7778 | 35 | 0.7429 |
| 10_8067000 | 7 | 1 | 7 | 0.7143 | 7 | 1 | 20 | 0.85 | 41 | 0.878 | 2 | 1 | 8 | 1 | 20 | 0.95 | 11 | 0.9091 | 41 | 0.9512 |
| 10_8067002 | 4 | 1 | 5 | 0.6 | 3 | 1 | 12 | 0.6667 | 24 | 0.75 | 1 | 0 | 8 | 0.375 | 13 | 0.6154 | 6 | 1 | 28 | 0.6071 |
| 10_8067005 | 6 | 0 | 7 | 0.5714 | 6 | 0.1667 | 18 | 0.2222 | 37 | 0.2432 | 2 | 0.5 | 9 | 0.1111 | 19 | 0.2105 | 11 | 0.3636 | 41 | 0.2439 |
| 10_8067011 | 8 | 0.375 | 6 | 0 | 7 | 0 | 18 | 0 | 39 | 0.0769 | 2 | 0 | 8 | 0.125 | 17 | 0.0588 | 8 | 0 | 35 | 0.0571 |
| 10_8067012 | 9 | 0.6667 | 5 | 0.8 | 7 | 1 | 18 | 0.6111 | 39 | 0.7179 | 1 | 1 | 9 | 0.8889 | 16 | 0.875 | 8 | 0.875 | 34 | 0.8824 |
| 10_8067025 | 6 | 0.6667 | 7 | 0.1429 | 7 | 0.4286 | 15 | 0.4667 | 35 | 0.4286 | 1 | 0 | 5 | 0.6 | 14 | 0.5 | 8 | 0.125 | 28 | 0.3929 |
| 10_8067026 | 7 | 0.8571 | 6 | 0.8333 | 5 | 1 | 19 | 0.5789 | 37 | 0.7297 | 2 | 1 | 6 | 0.6667 | 13 | 0.7692 | 8 | 1 | 29 | 0.8276 |
| 10_8067037 | 8 | 0.5 | 4 | 1 | 5 | 0.4 | 12 | 0.4167 | 29 | 0.5172 | 1 | 1 | 6 | 0.6667 | 10 | 0.5 | 8 | 0.5 | 25 | 0.56 |
| 10_8067042 | 8 | 0.625 | 5 | 0.4 | 4 | 0.75 | 14 | 0.6429 | 31 | 0.6129 | 1 | 1 | 7 | 0.8571 | 12 | 0.5 | 7 | 0.7143 | 27 | 0.6667 |
| 10_8067051 | 8 | 1 | 4 | 0.75 | 4 | 1 | 15 | 0.9333 | 31 | 0.9355 | 3 | 1 | 6 | 1 | 14 | 1 | 7 | 0.8571 | 30 | 0.9667 |
| 10_8067052 | 5 | 0.4 | 3 | 0.3333 | 4 | 0.5 | 12 | 0.5833 | 24 | 0.5 | 1 | 1 | 5 | 0.6 | 12 | 0.3333 | 6 | 0.3333 | 24 | 0.4167 |

| | | | | | | | | | | | | | | | | | | | | |
|---|---|---|---|---|---|---|---|---|---|---|---|---|---|---|---|---|---|---|---|---|
| 10_8067082 | 3 | 0.3333 | 2 | 0 | 2 | 0 | 6 | 0 | 13 | 0.0769 | 1 | 0 | 3 | 0.3333 | 7 | 0.2857 | 4 | 0.5 | 15 | 0.3333 |
| 10_8067082 | 9 | 0.1111 | 4 | 0 | 4 | 0 | 16 | 0.125 | 33 | 0.0909 | 2 | 0 | 6 | 0.1667 | 12 | 0.1667 | 6 | 0 | 26 | 0.1154 |
| 10_8067090 | 7 | 0.1429 | 4 | 0 | 2 | 0.5 | 10 | 0 | 23 | 0.087 | 2 | 0.5 | 4 | 0.25 | 8 | 0.125 | 6 | 0 | 20 | 0.15 |
| 10_8067095 | 7 | 0.4286 | 4 | 0.5 | 3 | 0.3333 | 14 | 0.2143 | 28 | 0.3214 | 2 | 0 | 5 | 0.4 | 11 | 0.3636 | 6 | 0.1667 | 24 | 0.2917 |
| 10_8067105 | 7 | 0.7143 | 4 | 0.75 | 4 | 1 | 14 | 1 | 29 | 0.8966 | 2 | 1 | 4 | 1 | 13 | 0.6154 | 7 | 0.8571 | 26 | 0.7692 |
| 10_8067126 | 5 | 0.6 | 3 | 0.6667 | 3 | 1 | 11 | 1 | 22 | 0.8636 | 1 | 1 | 3 | 1 | 10 | 0.9 | 7 | 0.8571 | 21 | 0.9048 |
| 10_8067128 | 5 | 0.2 | 2 | 0.5 | 3 | 0.3333 | 13 | 0.0769 | 23 | 0.1739 | 1 | 0 | 3 | 0.3333 | 8 | 0.125 | 6 | 0.1667 | 18 | 0.1667 |
| 10_8067148 | 6 | 0.5 | 2 | 0 | 2 | 0.5 | 12 | 0 | 22 | 0.1818 | NA | NA | 2 | 0.5 | 11 | 0.1818 | 6 | 0.3333 | 19 | 0.2632 |
| 10_8067153 | 6 | 0.3333 | 2 | 0.5 | 2 | 0.5 | 9 | 0.3333 | 19 | 0.3684 | NA | NA | 4 | 0.5 | 10 | 0.3 | 5 | 0.2 | 19 | 0.3158 |
| 10_8067155 | 5 | 0.2 | 3 | 0 | 2 | 0 | 7 | 0.1429 | 17 | 0.1176 | NA | NA | 3 | 0 | 7 | 0.2857 | 4 | 0.25 | 14 | 0.2143 |
| 10_8067168 | 5 | 0.4 | 1 | 0 | 1 | 1 | 8 | 0 | 15 | 0.2 | NA | NA | 3 | 0 | 10 | 0.2 | 6 | 0.5 | 19 | 0.2632 |
| 10_8067171 | 6 | 0.1667 | 1 | 0 | 2 | 0 | 9 | 0.3333 | 18 | 0.2222 | NA | NA | 2 | 0.5 | 11 | 0.1818 | 6 | 0.3333 | 19 | 0.2632 |
| 10_8067196 | 3 | 0 | 1 | 1 | 1 | 0 | 4 | 0 | 9 | 0.1111 | NA | NA | 1 | 0 | 6 | 0 | 4 | 0.5 | 11 | 0.1818 |
| 10_8068879 | 6 | 0 | 3 | 0.3333 | 5 | 0 | 2 | 0 | 16 | 0.0625 | 2 | 0 | 5 | 0 | 5 | 0.2 | 5 | 0 | 17 | 0.0588 |
| 10_8070293 | 1 | 1 | 2 | 0 | 2 | 0.5 | 1 | 1 | 6 | 0.5 | NA | NA | 1 | 0 | 2 | 0.5 | NA | NA | 3 | 0.3333 |
| 10_8070346 | 2 | 0 | 7 | 0.4286 | 5 | 0.4 | 6 | 1 | 20 | 0.55 | 2 | 0.5 | NA | NA | 5 | 0.8 | 2 | 0.5 | 9 | 0.6667 |
| 10_8070353 | 1 | 0 | 7 | 0.1429 | 4 | 0 | 4 | 0 | 16 | 0.0625 | 3 | 0.3333 | 3 | 0 | 5 | 0 | 3 | 0 | 14 | 0.0714 |
| 10_8070354 | 1 | 1 | 7 | 0.8571 | 4 | 1 | 4 | 0.75 | 16 | 0.875 | 2 | 1 | 2 | 1 | 5 | 0.6 | 3 | 0.3333 | 12 | 0.6667 |
| 10_8070355 | 1 | 0 | 8 | 0.125 | 3 | 0 | 5 | 0 | 17 | 0.0588 | 3 | 0 | 3 | 0.3333 | 4 | 0 | 2 | 0 | 12 | 0.0833 |
| 10_8070355 | 1 | 0 | 9 | 0.4444 | 5 | 0.2 | 6 | 0.8333 | 21 | 0.4762 | 2 | 0.5 | 3 | 0.3333 | 6 | 0.1667 | 4 | 1 | 15 | 0.4667 |
| 10_8070406 | 2 | 1 | 9 | 0.5556 | 3 | 0.3333 | 7 | 0.5714 | 21 | 0.5714 | 3 | 1 | 4 | 0.75 | 6 | 0.8333 | 4 | 0.75 | 17 | 0.8235 |
| 10_8070409 | 2 | 1 | 7 | 1 | 5 | 1 | 6 | 1 | 20 | 1 | 4 | 1 | 5 | 0.8 | 6 | 1 | 4 | 1 | 19 | 0.9474 |
| 10_8070413 | 2 | 1 | 8 | 1 | 4 | 1 | 5 | 1 | 19 | 1 | 4 | 0.75 | 5 | 1 | 6 | 1 | 4 | 1 | 19 | 0.9474 |
| 10_8070421 | 1 | 0 | 7 | 1 | 3 | 0.6667 | 6 | 0.8333 | 17 | 0.8235 | 3 | 1 | 5 | 0.6 | 6 | 0.8333 | 5 | 0.6 | 19 | 0.7368 |
| 10_8070428 | 2 | 0.5 | 5 | 0.4 | 4 | 0.5 | 6 | 0.6667 | 17 | 0.5294 | 4 | 0.75 | 3 | 1 | 5 | 0.8 | 5 | 0.6 | 17 | 0.7647 |
| 10_8073582 | 39 | 0 | 29 | 0.2069 | 25 | 0.16 | 35 | 0.0286 | 128 | 0.0859 | 24 | 0.0417 | 38 | 0.0526 | 35 | 0.1429 | 26 | 0.1538 | 123 | 0.0976 |
| 10_8073595 | 27 | 0.8889 | 15 | 0.8667 | 16 | 1 | 23 | 0.7391 | 81 | 0.8642 | 14 | 0.9286 | 26 | 0.8077 | 27 | 0.8519 | 18 | 1 | 85 | 0.8824 |
| 10_8073808 | 3 | 0.3333 | NA | NA | NA | NA | NA | NA | 3 | 0.3333 | NA | NA | 4 | 0.75 | NA | NA | 3 | 0.3333 | 7 | 0.5714 |
| 10_8073841 | 1 | 1 | NA | NA | 1 | 0 | NA | NA | 2 | 0.5 | NA | NA | 3 | 0.3333 | NA | NA | 2 | 0.5 | 5 | 0.4 |
| 10_8073846 | 1 | 1 | NA | NA | 1 | 0 | NA | NA | 2 | 0.5 | NA | NA | 3 | 0.6667 | NA | NA | 2 | 0.5 | 5 | 0.6 |
| 10_8076763 | 3 | 0 | 4 | 0.25 | 3 | 0 | 3 | 0.6667 | 13 | 0.2308 | 1 | 0 | 1 | 1 | 2 | 0.5 | 1 | 0 | 5 | 0.4 |
| 10_8076774 | 5 | 0.4 | 4 | 0 | 3 | 0 | 3 | 0 | 15 | 0.1333 | 1 | 1 | 1 | 1 | 2 | 0.5 | 1 | 0 | 5 | 0.6 |
| 10_8076775 | 5 | 0.2 | 4 | 0 | 3 | 0.3333 | 3 | 0.6667 | 15 | 0.2667 | 1 | 0 | 1 | 1 | 2 | 0.5 | 2 | 0 | 6 | 0.3333 |
| 10_8076786 | 4 | 1 | 2 | 1 | 2 | 0.5 | 2 | 0.5 | 10 | 0.8 | 1 | 1 | 1 | 1 | 1 | 1 | 1 | 0 | 4 | 0.75 |
| 10_8078532 | 24 | 0.0417 | 24 | 0 | 34 | 0.0588 | 48 | 0.1042 | 130 | 0.0615 | 23 | 0.0435 | 56 | 0.125 | 71 | 0.0986 | 31 | 0.0968 | 181 | 0.0994 |
| 10_8078613 | 40 | 0.1 | 24 | 0.0833 | 58 | 0.0345 | 58 | 0.0517 | 180 | 0.0611 | 24 | 0.125 | 71 | 0.0704 | 88 | 0.0568 | 37 | 0.0811 | 220 | 0.0727 |
| 10_8078960 | 1 | 0 | NA | NA | 1 | 1 | 1 | 0 | 3 | 0.3333 | 1 | 1 | 2 | 0 | 2 | 0 | NA | NA | 5 | 0.2 |
| 10_8078965 | NA | NA | NA | NA | 1 | 0 | 1 | 1 | 2 | 0.5 | 1 | 0 | 1 | 1 | 2 | 0.5 | NA | NA | 4 | 0.5 |
| 10_8083389 | 8 | 0.625 | 6 | 0.1667 | 11 | 0.3636 | 12 | 0.5833 | 37 | 0.4595 | 6 | 0.5 | 13 | 0.3846 | 13 | 0.1538 | 8 | 0.5 | 40 | 0.35 |
| 10_8085617 | 5 | 0 | 21 | 0.1905 | 12 | 0.0833 | 17 | 0.1176 | 55 | 0.1273 | 9 | 0.1111 | 10 | 0 | 9 | 0 | 7 | 0.2857 | 35 | 0.0857 |
| 10_8085655 | 5 | 0 | 15 | 0.1333 | 8 | 0 | 11 | 0.3636 | 39 | 0.1538 | 6 | 0.5 | 8 | 0.125 | 6 | 0.1667 | 5 | 0 | 25 | 0.2 |
| 10_8085656 | 4 | 0.5 | 12 | 0.5833 | 8 | 0.25 | 10 | 0.4 | 34 | 0.4412 | 6 | 0.6667 | 5 | 0.6 | 6 | 0.6667 | 5 | 0.6 | 22 | 0.6364 |
| 10_8085661 | 5 | 1 | 13 | 0.9231 | 8 | 0.875 | 10 | 0.9 | 36 | 0.9167 | 7 | 0.8571 | 7 | 1 | 3 | 1 | 5 | 1 | 22 | 0.9545 |
| 10_8085668 | 4 | 1 | 10 | 0.9 | 8 | 0.75 | 13 | 0.9231 | 35 | 0.8857 | 7 | 1 | 7 | 0.8571 | 4 | 1 | 5 | 1 | 23 | 0.9565 |
| 10_8085679 | 4 | 1 | 12 | 1 | 8 | 1 | 12 | 1 | 36 | 1 | 7 | 0.8571 | 9 | 1 | 3 | 1 | 4 | 1 | 23 | 0.9565 |
| 10_8085682 | 4 | 0.25 | 11 | 0.7273 | 7 | 0.8571 | 10 | 0.9 | 32 | 0.75 | 5 | 0.4 | 9 | 0.5556 | 4 | 0.5 | 3 | 0.3333 | 21 | 0.4762 |
| 10_8085683 | 4 | 0.75 | 12 | 0.9167 | 8 | 0.75 | 11 | 0.9091 | 35 | 0.8571 | 7 | 0.7143 | 9 | 0.8889 | 4 | 0.75 | 4 | 1 | 24 | 0.8333 |
| 10_8085686 | 2 | 0.5 | 11 | 0.8182 | 6 | 0.6667 | 11 | 0.4545 | 30 | 0.6333 | 6 | 0.8333 | 7 | 0.7143 | 3 | 1 | 2 | 0.5 | 18 | 0.7778 |
| 10_8085686 | 4 | 0.25 | 12 | 0.5 | 8 | 0.25 | 12 | 0.0833 | 36 | 0.2778 | 7 | 0.2857 | 9 | 0.1111 | 4 | 0 | 3 | 0 | 23 | 0.1304 |
| 10_8085689 | 3 | 1 | 9 | 1 | 6 | 1 | 9 | 1 | 27 | 1 | 5 | 1 | 7 | 1 | 3 | 1 | 2 | 1 | 17 | 1 |
| 10_8085726 | 2 | 0.5 | 8 | 0.875 | 7 | 1 | 10 | 0.7 | 27 | 0.8148 | 6 | 1 | 4 | 1 | 1 | 1 | 3 | 0.3333 | 14 | 0.8571 |
| 10_8085727 | 3 | 0.6667 | 7 | 0.4286 | 8 | 0.75 | 12 | 0.3333 | 30 | 0.5 | 7 | 0.4286 | 6 | 0.3333 | 1 | 0 | 3 | 0.3333 | 17 | 0.3529 |
| 10_8085778 | 1 | 0 | 2 | 0 | 5 | 0.4 | 8 | 0.25 | 16 | 0.25 | 3 | 0.3333 | 3 | 0.3333 | NA | NA | 2 | 0 | 8 | 0.25 |
| 10_8085791 | 1 | 1 | 5 | 0.2 | 4 | 0 | 6 | 0.1667 | 16 | 0.1875 | 5 | 0.6 | 1 | 0 | NA | NA | 1 | 0 | 7 | 0.4286 |
| 10_8085806 | 1 | 0 | 4 | 0.5 | 4 | 0.25 | 10 | 0.5 | 19 | 0.4211 | 4 | 0.25 | 2 | 0.5 | NA | NA | 2 | 0.5 | 8 | 0.375 |
| 10_8085812 | 1 | 0 | 5 | 0.2 | 4 | 0.25 | 11 | 0.1818 | 21 | 0.1905 | 3 | 0.3333 | 1 | 0 | NA | NA | 1 | 0 | 5 | 0.2 |
| 10_8086461 | 9 | 0.6667 | 6 | 0.1667 | 2 | 0.5 | 7 | 0.7143 | 24 | 0.5417 | 6 | 0.1667 | 8 | 0.375 | 8 | 0.625 | 7 | 0 | 29 | 0.3103 |
| 10_8086522 | 7 | 0.2857 | 4 | 0 | 1 | 1 | 8 | 0.25 | 20 | 0.25 | 6 | 0.1667 | 7 | 0.5714 | 7 | 0 | 3 | 0 | 23 | 0.2174 |
| 10_8086591 | 4 | 0 | 2 | 0 | 1 | 0 | 5 | 0.4 | 12 | 0.1667 | 2 | 0 | 6 | 0 | 5 | 0.6 | 3 | 0 | 16 | 0.1875 |
| 10_8086598 | 8 | 0.25 | 3 | 0 | 1 | 0 | 5 | 0.4 | 17 | 0.2353 | 4 | 0 | 7 | 0 | 4 | 0.25 | 6 | 0.1667 | 21 | 0.0952 |
| 10_8086612 | 7 | 0.1429 | 3 | 0 | NA | NA | 4 | 0.5 | 14 | 0.2143 | 4 | 0 | 8 | 0 | 4 | 0 | 7 | 0.2857 | 23 | 0.087 |
| 10_8086613 | 7 | 0.7143 | 2 | 1 | 1 | 1 | 4 | 0.5 | 14 | 0.7143 | 5 | 0.4 | 7 | 0.4286 | 4 | 1 | 5 | 1 | 21 | 0.6667 |
| 10_8086615 | 6 | 0.1667 | 2 | 0 | NA | NA | 4 | 0.5 | 12 | 0.25 | 5 | 0.2 | 5 | 0.2 | 4 | 0 | 5 | 0 | 19 | 0.1053 |
| 10_8086636 | 8 | 0 | 3 | 0 | 1 | 1 | 4 | 0.5 | 16 | 0.1875 | 4 | 0.25 | 6 | 0.1667 | 4 | 0 | 5 | 0.4 | 19 | 0.2105 |
| 10_8086637 | 8 | 0 | 2 | 0 | 1 | 1 | 3 | 0 | 14 | 0.0714 | 4 | 0.25 | 6 | 0.1667 | 4 | 0 | 5 | 0 | 19 | 0.1053 |

| | | | | | | | | | | | | | | | | | | | | |
|---|---|---|---|---|---|---|---|---|---|---|---|---|---|---|---|---|---|---|---|---|
| 10_808664 | 4 | 0.25 | 2 | 0 | NA | NA | 2 | 0.5 | 8 | 0.25 | 3 | 0 | 4 | 0.25 | 1 | 0 | 4 | 0 | 12 | 0.0833 |
| 10_808665 | 9 | 0 | 3 | 0 | 1 | 1 | 4 | 0.75 | 17 | 0.2353 | 4 | 0 | 7 | 0.1429 | 4 | 0.5 | 5 | 0 | 20 | 0.15 |
| 10_808665 | 9 | 0 | 3 | 0.3333 | 1 | 0 | 4 | 0.25 | 17 | 0.1176 | 4 | 0 | 7 | 0.1429 | 4 | 0 | 5 | 0.2 | 20 | 0.1 |
| 10_808665 | 7 | 0.5714 | 1 | 0 | 1 | 1 | 3 | 0.6667 | 12 | 0.5833 | 4 | 0 | 3 | 0.6667 | 2 | 1 | 3 | 0 | 12 | 0.3333 |
| 10_808666 | 9 | 0.8889 | 4 | 0.75 | 1 | 1 | 7 | 0.8571 | 21 | 0.8571 | 6 | 0.6667 | 6 | 1 | 5 | 1 | 3 | 1 | 20 | 0.9 |
| 10_808667 | 8 | 1 | 4 | 0.75 | 1 | 1 | 5 | 1 | 18 | 0.9444 | 6 | 1 | 6 | 1 | 3 | 1 | 2 | 1 | 17 | 1 |
| 10_808670 | 8 | 0.125 | 4 | 0 | 1 | 0 | 6 | 0.1667 | 19 | 0.1053 | 6 | 0.1667 | 6 | 0 | 4 | 0 | 1 | 0 | 17 | 0.0588 |
| 10_808672 | 6 | 0.8333 | 2 | 0.5 | 1 | 1 | 6 | 0.6667 | 15 | 0.7333 | 6 | 1 | 3 | 0.6667 | 2 | 1 | 1 | 1 | 12 | 0.9167 |
| 10_808673 | 8 | 0.5 | 2 | 0 | 1 | 0 | 5 | 0.2 | 16 | 0.3125 | 6 | 0.5 | 5 | 0.6 | 4 | 0.25 | 1 | 0 | 16 | 0.4375 |
| 10_808800 | 14 | 0.1429 | 23 | 0.087 | 20 | 0.1 | 22 | 0.2273 | 79 | 0.1392 | 15 | 0.0667 | 14 | 0.1429 | 15 | 0.3333 | 19 | 0.0526 | 63 | 0.1429 |
| 10_808800 | 16 | 0.125 | 23 | 0.2174 | 20 | 0.25 | 22 | 0.1364 | 81 | 0.1852 | 15 | 0.3333 | 15 | 0.3333 | 17 | 0.2941 | 16 | 0.3125 | 63 | 0.3175 |
| 10_808802 | 14 | 0.4286 | 19 | 0.5789 | 17 | 0.5294 | 20 | 0.5 | 70 | 0.5143 | 10 | 0.3 | 11 | 0.2727 | 14 | 0.5714 | 17 | 0.2353 | 52 | 0.3462 |
| 10_808803 | 8 | 0 | 19 | 0.2105 | 16 | 0.3125 | 16 | 0.0625 | 59 | 0.1695 | 9 | 0.3333 | 6 | 0.1667 | 15 | 0.2 | 13 | 0.2308 | 43 | 0.2326 |
| 10_808803 | 15 | 0.0667 | 21 | 0.0952 | 19 | 0.2105 | 22 | 0.1818 | 77 | 0.1429 | 12 | 0.3333 | 13 | 0.1538 | 18 | 0.2778 | 18 | 0.1111 | 61 | 0.2131 |
| 10_808803 | 13 | 0.6154 | 19 | 0.8421 | 18 | 0.6667 | 21 | 0.8095 | 71 | 0.7465 | 10 | 0.8 | 12 | 0.8333 | 19 | 0.7368 | 18 | 0.8889 | 59 | 0.8136 |
| 10_808805 | 14 | 0.0714 | 22 | 0.1364 | 19 | 0.1579 | 17 | 0.2941 | 72 | 0.1667 | 11 | 0.0909 | 13 | 0.2308 | 18 | 0.2778 | 18 | 0.1667 | 60 | 0.2 |
| 10_808805 | 10 | 0.7 | 20 | 0.8 | 18 | 0.7778 | 20 | 0.6 | 68 | 0.7206 | 10 | 0.9 | 11 | 0.6364 | 18 | 0.8889 | 18 | 0.8889 | 57 | 0.8421 |
| 10_808807 | 11 | 0.5455 | 18 | 0.6111 | 13 | 0.5385 | 20 | 0.85 | 62 | 0.6613 | 9 | 0.6667 | 11 | 0.4545 | 17 | 0.7059 | 17 | 0.7059 | 54 | 0.6481 |
| 10_809180 | 9 | 0.8889 | 7 | 0.8571 | 5 | 0.8 | 4 | 1 | 25 | 0.88 | 7 | 0.8571 | 5 | 1 | 3 | 0.6667 | 5 | 0.6 | 20 | 0.8 |
| 10_809581 | 1 | 0 | 1 | 0 | NA | NA | 4 | 0.25 | 6 | 0.1667 | 1 | 0 | 1 | 0 | 3 | 0 | 2 | 0.5 | 7 | 0.1429 |
| 10_809595 | 11 | 0.1818 | 16 | 0 | 14 | 0.1429 | 17 | 0.2353 | 58 | 0.1379 | 7 | 0 | 10 | 0.2 | 18 | 0.0556 | 14 | 0.1429 | 49 | 0.102 |
| 10_809596 | 10 | 0.6 | 12 | 0.5833 | 10 | 0.3 | 12 | 0.6667 | 44 | 0.5455 | 2 | 1 | 11 | 0.5455 | 13 | 0.3846 | 11 | 0.6364 | 37 | 0.5405 |
| 10_809596 | 9 | 0.2222 | 16 | 0.25 | 13 | 0.5385 | 17 | 0.5294 | 55 | 0.4 | 8 | 0.375 | 12 | 0.1667 | 17 | 0.3529 | 16 | 0.3125 | 53 | 0.3019 |
| 10_809599 | NA | NA | 5 | 0 | 2 | 0 | 2 | 0.5 | 9 | 0.1111 | 1 | 1 | 2 | 0.5 | 6 | 0.1667 | NA | NA | 9 | 0.3333 |
| 10_809798 | 2 | 0.5 | 4 | 0 | 1 | 0 | 2 | 0.5 | 9 | 0.2222 | 7 | 0.2857 | 6 | 0.5 | 3 | 0.6667 | 3 | 0.3333 | 19 | 0.4211 |
| 10_809800 | 2 | 0 | 3 | 0.3333 | 1 | 0 | 3 | 0 | 9 | 0.1111 | 6 | 0.1667 | 7 | 0 | 2 | 0 | 2 | 0 | 17 | 0.0588 |
| 10_809800 | 2 | 1 | 4 | 1 | 2 | 1 | 3 | 1 | 11 | 1 | 6 | 1 | 7 | 1 | 3 | 1 | 2 | 0.5 | 18 | 0.9444 |
| 10_809804 | 1 | 1 | 2 | 1 | 2 | 1 | 2 | 1 | 7 | 1 | 6 | 0.8333 | 7 | 1 | 2 | 1 | 3 | 1 | 18 | 0.9444 |
| 10_809805 | 1 | 1 | 3 | 0.3333 | 1 | 1 | 4 | 0.25 | 9 | 0.4444 | 7 | 0.1429 | 7 | 0.2857 | 2 | 0.5 | 2 | 0 | 18 | 0.2222 |
| 10_809807 | 2 | 0.5 | 4 | 0.25 | NA | NA | 3 | 0 | 9 | 0.2222 | 8 | 0.75 | 8 | 0.25 | 3 | 1 | 2 | 0.5 | 21 | 0.5714 |
| 10_809808 | 3 | 1 | 5 | 0.6 | 2 | 1 | 5 | 1 | 15 | 0.8667 | 8 | 0.875 | 8 | 0.75 | 3 | 1 | 2 | 1 | 21 | 0.8571 |
| 10_809842 | 3 | 0.3333 | 7 | 0.5714 | 6 | 0.5 | 5 | 0.6 | 21 | 0.5238 | 7 | 0.8571 | 10 | 0.7 | 3 | 0.6667 | 3 | 1 | 23 | 0.7826 |
| 10_809842 | 2 | 0 | 6 | 0.1667 | 6 | 0.1667 | 7 | 0.1429 | 21 | 0.1429 | 7 | 0 | 7 | 0.1429 | 3 | 0 | 3 | 0.3333 | 20 | 0.1 |
| 10_809842 | 3 | 0 | 5 | 0.2 | 3 | 0.3333 | 5 | 0 | 16 | 0.125 | 4 | 0 | 7 | 0.2857 | 3 | 0 | 1 | 0 | 15 | 0.1333 |
| 10_809843 | 2 | 0 | 6 | 0.5 | 5 | 0.4 | 3 | 0.6667 | 16 | 0.4375 | 8 | 0.5 | 9 | 0.4444 | 4 | 0.75 | 2 | 1 | 23 | 0.5652 |
| 10_809843 | 4 | 0.5 | 6 | 0.3333 | 4 | 0.5 | 3 | 0.6667 | 17 | 0.4706 | 8 | 0.75 | 8 | 0.5 | 4 | 0.5 | 3 | 0.6667 | 23 | 0.6087 |
| 10_809845 | 4 | 0 | 8 | 0.375 | 6 | 0.3333 | 6 | 0 | 24 | 0.2083 | 6 | 0.3333 | 8 | 0.25 | 4 | 0 | 1 | 1 | 19 | 0.2632 |
| 10_809863 | 5 | 0.2 | 5 | 0 | 8 | 0.25 | 3 | 0 | 21 | 0.1429 | 7 | 0.1429 | 9 | 0.1111 | 8 | 0.25 | 3 | 0 | 27 | 0.1481 |
| 10_809877 | 7 | 0 | 9 | 0 | 10 | 0.1 | 9 | 0.1111 | 35 | 0.0571 | 9 | 0 | 12 | 0.0833 | 15 | 0.0667 | 4 | 0 | 40 | 0.05 |
| 10_809878 | 8 | 0.125 | 16 | 0.125 | 13 | 0.0769 | 12 | 0.0833 | 49 | 0.102 | 11 | 0 | 17 | 0 | 16 | 0.125 | 6 | 0.1667 | 50 | 0.06 |
| 10_809878 | 9 | 0.1111 | 15 | 0.0667 | 13 | 0.1538 | 11 | 0 | 48 | 0.0833 | 12 | 0.0833 | 17 | 0.1176 | 15 | 0.2 | 5 | 0 | 49 | 0.1224 |
| 10_809892 | 1 | 1 | NA | NA | 2 | 0.5 | NA | NA | 3 | 0.6667 | NA | NA | 1 | 0 | 3 | 0.6667 | NA | NA | 4 | 0.5 |
| 10_809895 | 3 | 0.3333 | 1 | 1 | 3 | 0 | 2 | 0.5 | 9 | 0.3333 | NA | NA | 4 | 0.25 | 3 | 0.3333 | 2 | 0 | 9 | 0.2222 |
| 10_809895 | 4 | 0.75 | 1 | 0 | 3 | 1 | 3 | 1 | 11 | 0.8182 | NA | NA | 4 | 0.75 | 3 | 1 | 3 | 0.6667 | 10 | 0.8 |
| 10_809897 | 3 | 1 | 1 | 1 | 4 | 0.75 | 3 | 0.6667 | 11 | 0.8182 | NA | NA | 2 | 1 | 3 | 1 | 3 | 1 | 8 | 1 |
| 10_809897 | 6 | 0.1667 | 1 | 0 | 2 | 1 | 2 | 0.5 | 11 | 0.3636 | NA | NA | 3 | 0.3333 | 1 | 0 | 3 | 0.6667 | 7 | 0.4286 |
| 10_809898 | 6 | 0 | 1 | 0 | 4 | 0.25 | 2 | 0 | 13 | 0.0769 | NA | NA | 5 | 0.2 | 2 | 0 | 3 | 0 | 10 | 0.1 |
| 10_809898 | 2 | 0.5 | 1 | 1 | 2 | 1 | 1 | 1 | 6 | 0.8333 | NA | NA | 3 | 0.6667 | 2 | 0.5 | 1 | 1 | 6 | 0.6667 |
| 10_809900 | 1 | 0 | 1 | 0 | 1 | 1 | NA | NA | 3 | 0.3333 | NA | NA | 1 | 0 | 1 | 0 | 1 | 1 | 3 | 0.3333 |
| 10_809997 | 2 | 0.5 | 1 | 1 | NA | NA | 1 | 1 | 4 | 0.75 | NA | NA | 1 | 1 | 2 | 0.5 | 1 | 0 | 4 | 0.5 |
| 10_810178 | 127 | 0.0157 | 110 | 0.0727 | 114 | 0.0702 | 121 | 0.0744 | 472 | 0.0572 | 104 | 0.0865 | 104 | 0.0865 | 117 | 0.094 | 101 | 0.0792 | 426 | 0.0869 |
| 10_810179 | 95 | 0.1895 | 87 | 0.1609 | 88 | 0.2273 | 106 | 0.2358 | 376 | 0.2048 | 82 | 0.2561 | 97 | 0.2887 | 94 | 0.1383 | 70 | 0.1714 | 343 | 0.2157 |
| 10_810179 | 124 | 0.2016 | 115 | 0.2174 | 108 | 0.1667 | 117 | 0.1538 | 464 | 0.1853 | 105 | 0.1619 | 117 | 0.2308 | 124 | 0.1935 | 97 | 0.2268 | 443 | 0.2032 |
| 10_810179 | 142 | 0.5211 | 138 | 0.4638 | 121 | 0.4545 | 130 | 0.4769 | 531 | 0.4802 | 113 | 0.5221 | 127 | 0.5354 | 130 | 0.4769 | 118 | 0.5593 | 488 | 0.5225 |
| 10_810180 | 148 | 0.777 | 129 | 0.8372 | 119 | 0.8319 | 137 | 0.8248 | 533 | 0.8161 | 121 | 0.8182 | 138 | 0.8623 | 148 | 0.8514 | 116 | 0.8534 | 523 | 0.847 |
| 10_810262 | 9 | 0 | 9 | 0 | 1 | 0 | 9 | 0.2222 | 28 | 0.0714 | 6 | 0.1667 | 4 | 0 | 6 | 0.1667 | 8 | 0 | 24 | 0.0833 |
| 10_810262 | 5 | 0.2 | 6 | 0 | 3 | 0.3333 | 9 | 0.1111 | 23 | 0.1304 | 6 | 0 | 4 | 0 | 3 | 0 | 8 | 0.25 | 21 | 0.0952 |
| 10_810264 | 9 | 0.1111 | 8 | 0 | 5 | 0.4 | 9 | 0.2222 | 31 | 0.1613 | 6 | 0.3333 | 5 | 0.2 | 7 | 0 | 8 | 0.375 | 26 | 0.2308 |
| 10_810277 | 12 | 0.0833 | 9 | 0.1111 | 5 | 0 | 10 | 0.2 | 36 | 0.1111 | 8 | 0.125 | 6 | 0.3333 | 7 | 0.1429 | 11 | 0.4545 | 32 | 0.2812 |
| 10_810286 | 8 | 0.625 | 8 | 0.125 | 5 | 0 | 10 | 0.2 | 31 | 0.2581 | 8 | 0 | 10 | 0.1 | 6 | 0 | 8 | 0.25 | 32 | 0.0938 |
| 10_810290 | NA | NA | NA | NA | 1 | 1 | NA | NA | 1 | 1 | 1 | 1 | NA | NA | NA | NA | NA | NA | 1 | 1 |
| 10_810290 | NA | NA | NA | NA | 1 | 1 | NA | NA | 1 | 1 | 1 | 1 | NA | NA | NA | NA | NA | NA | 1 | 1 |
| 10_810405 | 12 | 0.25 | 16 | 0.1875 | 18 | 0.2222 | 12 | 0.25 | 58 | 0.2241 | 13 | 0.3846 | 11 | 0.0909 | 13 | 0.1538 | 18 | 0.2778 | 55 | 0.2364 |
| 10_810406 | 9 | 0.2222 | 8 | 0.25 | 7 | 0.1429 | 6 | 0.1667 | 30 | 0.2 | 6 | 0.3333 | 5 | 0.4 | 9 | 0 | 8 | 0.25 | 28 | 0.2143 |
| 10_810407 | 9 | 0 | 8 | 0.125 | 8 | 0.25 | 4 | 0 | 29 | 0.1034 | 4 | 0.25 | 7 | 0 | 9 | 0 | 11 | 0.1818 | 31 | 0.0968 |

| | | | | | | | | | | | | | | | | | | | | |
|---|---|---|---|---|---|---|---|---|---|---|---|---|---|---|---|---|---|---|---|---|
| 10_810408 | 11 | 0.8182 | 14 | 0.7857 | 14 | 0.7143 | 11 | 0.7273 | 50 | 0.76 | 12 | 0.6667 | 13 | 0.6154 | 14 | 0.7857 | 21 | 0.8095 | 60 | 0.7333 |
| 10_810409 | 10 | 0.1 | 11 | 0.3636 | 13 | 0 | 8 | 0 | 42 | 0.119 | 9 | 0.1111 | 8 | 0.25 | 9 | 0.1111 | 18 | 0 | 44 | 0.0909 |
| 10_810415 | 12 | 0.25 | 9 | 0.3333 | 11 | 0.2727 | 9 | 0.4444 | 41 | 0.3171 | 4 | 0 | 10 | 0.2 | 13 | 0.3077 | 18 | 0.2222 | 45 | 0.2222 |
| 10_810417 | 11 | 0.0909 | 8 | 0.25 | 10 | 0.1 | 8 | 0.125 | 37 | 0.1351 | 5 | 0.2 | 10 | 0.2 | 13 | 0.0769 | 19 | 0.1053 | 47 | 0.1277 |
| 10_810418 | 11 | 0.1818 | 9 | 0.1111 | 12 | 0 | 8 | 0 | 40 | 0.075 | 5 | 0 | 10 | 0.1 | 13 | 0 | 18 | 0.1667 | 46 | 0.087 |
| 10_810418 | 12 | 0 | 9 | 0.1111 | 11 | 0.1818 | 7 | 0.1429 | 39 | 0.1026 | 5 | 0.2 | 9 | 0.1111 | 13 | 0 | 18 | 0.0556 | 45 | 0.0667 |
| 10_810419 | 12 | 0.1667 | 10 | 0.1 | 13 | 0.2308 | 8 | 0.25 | 43 | 0.186 | 8 | 0 | 12 | 0.0833 | 17 | 0.0588 | 17 | 0.1176 | 54 | 0.0741 |
| 10_810420 | 16 | 0.8125 | 5 | 1 | 9 | 0.8889 | 4 | 0.75 | 34 | 0.8529 | 7 | 0.8571 | 11 | 0.5455 | 8 | 0.75 | 12 | 1 | 38 | 0.7895 |
| 10_810420 | 16 | 0.8125 | 10 | 0.9 | 9 | 0.6667 | 8 | 0.5 | 43 | 0.7442 | 7 | 0.7143 | 10 | 0.8 | 16 | 0.5625 | 16 | 0.875 | 49 | 0.7347 |
| 10_810421 | 15 | 0.6 | 9 | 0.7778 | 11 | 1 | 8 | 0.75 | 43 | 0.7674 | 9 | 0.5556 | 9 | 0.4444 | 16 | 0.625 | 17 | 0.8235 | 51 | 0.6471 |
| 10_810425 | 15 | 0.2 | 7 | 0.1429 | 9 | 0.3333 | 7 | 0.4286 | 38 | 0.2632 | 6 | 0.1667 | 6 | 0 | 14 | 0.1429 | 10 | 0 | 36 | 0.0833 |
| 10_810760 | NA | NA | NA | NA | NA | NA | 1 | 1 | 1 | 1 | 1 | 1 | NA | NA | NA | NA | NA | NA | 1 | 1 |
| 10_810871 | 4 | 0.25 | 2 | 0.5 | 4 | 0 | 2 | 0 | 12 | 0.1667 | 2 | 0 | 3 | 0.3333 | 2 | 0 | 2 | 0.5 | 9 | 0.2222 |
| 10_810876 | 4 | 0.25 | 6 | 0.1667 | 4 | 0 | 2 | 0 | 16 | 0.125 | 2 | 0.5 | 6 | 0.1667 | 4 | 0 | 3 | 0 | 15 | 0.1333 |
| 10_810879 | 4 | 0.25 | 6 | 0.1667 | 3 | 0 | 2 | 0 | 15 | 0.1333 | 1 | 0 | 5 | 0 | 3 | 0 | 3 | 0.3333 | 12 | 0.0833 |
| 10_810978 | 19 | 0.0526 | 23 | 0.087 | 19 | 0.0526 | 20 | 0.05 | 81 | 0.0617 | 19 | 0.0526 | 17 | 0 | 20 | 0.1 | 14 | 0.1429 | 70 | 0.0714 |
| 10_810979 | 15 | 0 | 16 | 0 | 10 | 0.1 | 21 | 0.1429 | 62 | 0.0645 | 19 | 0.1579 | 18 | 0.1111 | 21 | 0.2381 | 11 | 0 | 69 | 0.1449 |
| 10_810986 | 23 | 0.1304 | 21 | 0 | 17 | 0.1765 | 20 | 0.15 | 81 | 0.1111 | 18 | 0.0556 | 16 | 0.1875 | 15 | 0.2667 | 15 | 0.0667 | 64 | 0.1406 |
| 10_810986 | 23 | 0.2609 | 24 | 0.125 | 25 | 0.12 | 26 | 0.1923 | 98 | 0.1735 | 22 | 0.2727 | 22 | 0.2273 | 25 | 0.16 | 18 | 0.0556 | 87 | 0.1839 |
| 10_810987 | 17 | 0.4706 | 19 | 0.4737 | 14 | 0.2143 | 16 | 0.3125 | 66 | 0.3788 | 16 | 0.5625 | 15 | 0.5333 | 14 | 0.4286 | 12 | 0.3333 | 57 | 0.4737 |
| 10_810987 | 21 | 0.7143 | 24 | 0.4167 | 22 | 0.5 | 23 | 0.3478 | 90 | 0.4889 | 17 | 0.6471 | 20 | 0.45 | 22 | 0.6364 | 15 | 0.4 | 74 | 0.5405 |
| 10_811005 | 5 | 0.8 | 4 | 0.75 | 3 | 0.3333 | 7 | 0.5714 | 19 | 0.6316 | 6 | 0.3333 | 6 | 0.6667 | 11 | 0.5455 | 3 | 0.6667 | 26 | 0.5385 |
| 10_811006 | 5 | 0.6 | 4 | 0 | 5 | 0.6 | 7 | 0.2857 | 21 | 0.381 | 4 | 0.25 | 4 | 0.75 | 6 | 0.3333 | 4 | 0.5 | 18 | 0.4444 |
| 10_811006 | 10 | 0 | 5 | 0 | 6 | 0.1667 | 8 | 0.125 | 29 | 0.069 | 6 | 0 | 6 | 0.1667 | 12 | 0.0833 | 4 | 0 | 28 | 0.0714 |
| 10_811047 | NA | NA | 1 | 1 | 1 | 1 | 1 | 0 | 3 | 0.6667 | 1 | 0 | NA | NA | NA | NA | 2 | 1 | 3 | 0.6667 |
| 10_811049 | NA | NA | 1 | 1 | 1 | 1 | 1 | 0 | 3 | 0.6667 | 1 | 1 | NA | NA | 1 | 0 | 2 | 1 | 4 | 0.75 |
| 10_811089 | 1 | 0 | 5 | 0.4 | 1 | 1 | 1 | 0 | 8 | 0.375 | 4 | 0 | 1 | 0 | 3 | 0.3333 | 2 | 0 | 10 | 0.1 |
| 10_811539 | 18 | 0.0556 | 20 | 0.05 | 11 | 0.0909 | 14 | 0.0714 | 63 | 0.0635 | 15 | 0.2 | 9 | 0.1111 | 12 | 0.0833 | 17 | 0.0588 | 53 | 0.1132 |
| 10_811539 | 13 | 0.7692 | 14 | 0.5 | 9 | 0.7778 | 10 | 0.7 | 46 | 0.6739 | 12 | 0.75 | 7 | 0.5714 | 7 | 0.4286 | 15 | 0.7333 | 41 | 0.6585 |
| 10_811541 | 21 | 0.619 | 23 | 0.5652 | 15 | 0.4 | 13 | 0.3846 | 72 | 0.5139 | 17 | 0.6471 | 10 | 0.6 | 12 | 0.6667 | 19 | 0.3684 | 58 | 0.5517 |
| 10_811541 | 19 | 0.8947 | 22 | 0.8636 | 13 | 0.9231 | 15 | 1 | 69 | 0.913 | 18 | 0.8889 | 10 | 0.9 | 12 | 0.8333 | 19 | 1 | 59 | 0.9153 |
| 10_811542 | 19 | 0.2632 | 20 | 0.6 | 14 | 0.7857 | 14 | 0.6429 | 67 | 0.5522 | 20 | 0.65 | 9 | 0.2222 | 14 | 0.6429 | 20 | 0.55 | 63 | 0.5556 |
| 10_811542 | 12 | 0.6667 | 19 | 0.7368 | 13 | 0.7692 | 10 | 0.6 | 54 | 0.7037 | 16 | 0.75 | 7 | 0.7143 | 11 | 0.6364 | 11 | 0.5455 | 45 | 0.6667 |
| 10_811542 | 16 | 0.9375 | 16 | 0.875 | 14 | 0.9286 | 14 | 0.8571 | 60 | 0.9 | 15 | 1 | 6 | 1 | 10 | 0.9 | 15 | 0.8667 | 46 | 0.9348 |
| 10_811543 | 23 | 0.0435 | 22 | 0.0455 | 22 | 0.0909 | 16 | 0.125 | 83 | 0.0723 | 18 | 0.1111 | 9 | 0 | 15 | 0.2 | 18 | 0 | 60 | 0.0833 |
| 10_811544 | 26 | 0.1923 | 23 | 0.0435 | 21 | 0.0476 | 16 | 0.25 | 86 | 0.1279 | 21 | 0.0952 | 8 | 0 | 15 | 0 | 21 | 0.2381 | 65 | 0.1077 |
| 10_811544 | 13 | 0.2308 | 15 | 0.0667 | 12 | 0 | 9 | 0 | 49 | 0.0816 | 11 | 0 | 6 | 0 | 11 | 0.0909 | 11 | 0.1818 | 39 | 0.0769 |
| 10_811546 | 24 | 0.7083 | 27 | 0.5185 | 17 | 0.6471 | 16 | 0.5 | 84 | 0.5952 | 21 | 0.5714 | 7 | 0.4286 | 17 | 0.6471 | 19 | 0.4737 | 64 | 0.5469 |
| 10_811546 | 25 | 0.28 | 28 | 0.1786 | 20 | 0.2 | 18 | 0.2222 | 91 | 0.2198 | 22 | 0.0909 | 8 | 0.25 | 18 | 0 | 22 | 0.0909 | 70 | 0.0857 |
| 10_811546 | 20 | 0.9 | 26 | 0.9615 | 20 | 0.9 | 15 | 0.8667 | 81 | 0.9136 | 21 | 0.9524 | 9 | 1 | 18 | 1 | 22 | 0.9091 | 70 | 0.9571 |
| 10_811547 | 22 | 0.3636 | 22 | 0.3636 | 15 | 0.6667 | 13 | 0.3846 | 72 | 0.4306 | 19 | 0.2105 | 6 | 0.1667 | 14 | 0.3571 | 19 | 0.2105 | 58 | 0.2414 |
| 10_811547 | 22 | 0.8182 | 27 | 0.8148 | 16 | 0.8125 | 15 | 0.8 | 80 | 0.8125 | 18 | 0.7222 | 12 | 0.5833 | 16 | 0.875 | 18 | 0.7778 | 64 | 0.75 |
| 10_811547 | 27 | 0.3704 | 25 | 0.28 | 17 | 0.4706 | 16 | 0.125 | 85 | 0.3176 | 21 | 0.2381 | 12 | 0.0833 | 20 | 0.25 | 23 | 0.2609 | 76 | 0.2237 |
| 10_811548 | 21 | 0.4286 | 25 | 0.24 | 17 | 0.3529 | 17 | 0.4118 | 80 | 0.35 | 21 | 0.2857 | 11 | 0.0909 | 18 | 0.3889 | 22 | 0.3636 | 72 | 0.3056 |
| 10_811553 | 19 | 0.8947 | 28 | 0.6071 | 18 | 0.6667 | 15 | 0.6 | 80 | 0.6875 | 18 | 0.6667 | 11 | 0.7273 | 16 | 0.6875 | 21 | 0.7619 | 66 | 0.7121 |
| 10_811556 | 32 | 0.1562 | 35 | 0.1429 | 24 | 0.1667 | 26 | 0.0385 | 117 | 0.1282 | 32 | 0.125 | 15 | 0.0667 | 25 | 0.24 | 30 | 0.2 | 102 | 0.1667 |
| 10_811584 | 2 | 0.5 | 1 | 1 | 1 | 1 | NA | NA | 4 | 0.75 | NA | NA | 1 | 1 | 1 | 1 | 1 | 1 | 3 | 1 |
| 10_811590 | 2 | 0.5 | 1 | 0 | 1 | 0 | NA | NA | 4 | 0.25 | 1 | 0 | 3 | 0.6667 | 2 | 0 | 1 | 0 | 7 | 0.2857 |
| 10_811615 | 2 | 0 | 1 | 0 | 5 | 0.2 | 1 | 0 | 9 | 0.1111 | 2 | 0.5 | 4 | 0.25 | 5 | 0 | 2 | 0 | 13 | 0.1538 |
| 10_811616 | 5 | 1 | 2 | 1 | 3 | 1 | 1 | 1 | 11 | 1 | 2 | 1 | 6 | 1 | 5 | 1 | 2 | 1 | 15 | 1 |
| 10_811618 | 1 | 1 | NA | NA | 3 | 0.6667 | NA | NA | 4 | 0.75 | NA | NA | 5 | 1 | 2 | 0.5 | NA | NA | 7 | 0.8571 |
| 10_811618 | 4 | 0 | 2 | 0.5 | 5 | 0.2 | 1 | 0 | 12 | 0.1667 | 3 | 0.6667 | 6 | 0.5 | 5 | 0.4 | 3 | 0 | 17 | 0.4118 |
| 10_811618 | 3 | 1 | 2 | 1 | 4 | 1 | 1 | 1 | 10 | 1 | 3 | 1 | 7 | 1 | 6 | 1 | 3 | 1 | 19 | 1 |
| 10_811619 | 3 | 0.6667 | 2 | 1 | 3 | 1 | NA | NA | 8 | 0.875 | 3 | 0.6667 | 4 | 0.75 | 4 | 0.75 | 1 | 1 | 12 | 0.75 |
| 10_811619 | 3 | 1 | 2 | 1 | 3 | 1 | 1 | 1 | 9 | 1 | 3 | 0.6667 | 6 | 1 | 5 | 0.8 | 4 | 1 | 18 | 0.8889 |
| 10_811620 | 5 | 1 | 1 | 1 | 5 | 1 | 1 | 1 | 12 | 1 | 3 | 1 | 6 | 1 | 6 | 1 | 3 | 1 | 18 | 1 |
| 10_811819 | 3 | 0.6667 | 3 | 0.6667 | NA | NA | 3 | 0 | 9 | 0.4444 | 5 | 0.4 | 3 | 0.3333 | 7 | 0.5714 | 6 | 0.6667 | 21 | 0.5238 |
| 10_811820 | 4 | 0 | 5 | 0.4 | 1 | 0 | 2 | 0 | 12 | 0.1667 | 5 | 0.4 | 3 | 0.3333 | 7 | 0.4286 | 7 | 0.2857 | 22 | 0.3636 |
| 10_811824 | 5 | 0.2 | 3 | 0 | 1 | 0 | 2 | 0 | 11 | 0.0909 | 4 | 0 | 2 | 0 | 7 | 0.1429 | 7 | 0.1429 | 20 | 0.1 |
| 10_811998 | 1 | 1 | 1 | 1 | NA | NA | 1 | 1 | 3 | 1 | 1 | 1 | NA | NA | NA | NA | 1 | 1 | 2 | 1 |
| 10_811998 | 1 | 0 | NA | NA | NA | NA | 1 | 1 | 2 | 0.5 | 1 | 1 | NA | NA | NA | NA | 1 | 1 | 2 | 1 |
| 10_811998 | 1 | 1 | 1 | 1 | NA | NA | 1 | 1 | 3 | 1 | 1 | 1 | NA | NA | NA | NA | NA | NA | 1 | 1 |
| 10_812029 | 9 | 0.1111 | 9 | 0.2222 | 11 | 0 | 11 | 0.4545 | 40 | 0.2 | 5 | 0.4 | 2 | 0 | 15 | 0.1333 | 6 | 0.1667 | 28 | 0.1786 |
| 10_812029 | 10 | 0.2 | 9 | 0.2222 | 11 | 0.1818 | 13 | 0.2308 | 43 | 0.2093 | 6 | 0.1667 | 3 | 0 | 15 | 0.0667 | 8 | 0.375 | 32 | 0.1562 |
| 10_812030 | 9 | 0.1111 | 9 | 0.2222 | 11 | 0.1818 | 11 | 0.0909 | 40 | 0.15 | 7 | 0.1429 | 3 | 0 | 13 | 0.0769 | 7 | 0.2857 | 30 | 0.1333 |

| | | | | | | | | | | | | | | | | | | | | |
|---|---|---|---|---|---|---|---|---|---|---|---|---|---|---|---|---|---|---|---|---|
| 10_8120305 | 9 | 0.1111 | 7 | 0.2857 | 12 | 0.1667 | 13 | 0.2308 | 41 | 0.1951 | 5 | 0.2 | 3 | 0.3333 | 14 | 0.0714 | 6 | 0.1667 | 28 | 0.1429 |
| 10_8120311 | 11 | 0.4545 | 6 | 0.5 | 10 | 0.5 | 11 | 0.7273 | 38 | 0.5526 | 5 | 0.8 | 3 | 0.6667 | 9 | 0.5556 | 8 | 0.5 | 25 | 0.6 |
| 10_8120315 | 11 | 1 | 8 | 1 | 11 | 1 | 12 | 0.9167 | 42 | 0.9762 | 4 | 1 | 3 | 1 | 12 | 0.75 | 8 | 0.75 | 27 | 0.8148 |
| 10_8120316 | 11 | 0.1818 | 7 | 0.2857 | 10 | 0 | 11 | 0 | 39 | 0.1026 | 2 | 0.5 | 3 | 0 | 12 | 0 | 7 | 0.1429 | 24 | 0.0833 |
| 10_8120317 | 8 | 0 | 9 | 0.2222 | 9 | 0.2222 | 12 | 0 | 38 | 0.1053 | 4 | 0 | 2 | 0 | 12 | 0.0833 | 7 | 0.4286 | 25 | 0.16 |
| 10_8122374 | 7 | 0 | 10 | 0 | 15 | 0.1333 | 6 | 0.3333 | 38 | 0.1053 | 11 | 0.2727 | 11 | 0.0909 | 8 | 0.125 | 9 | 0 | 39 | 0.1282 |
| 10_8122378 | 5 | 0.6 | 9 | 0.3333 | 15 | 0.4667 | 3 | 0 | 32 | 0.4062 | 11 | 0.5455 | 10 | 0.4 | 7 | 0.2857 | 8 | 0.25 | 36 | 0.3889 |
| 10_8122682 | NA | NA | 1 | 0 | 1 | 1 | NA | NA | 2 | 0.5 | NA | NA | NA | NA | NA | NA | 1 | 1 | 1 | 1 |
| 10_8122703 | NA | NA | 1 | 1 | 2 | 1 | NA | NA | 3 | 1 | NA | NA | NA | NA | NA | NA | 1 | 1 | 1 | 1 |
| 10_8122706 | 1 | 1 | 1 | 1 | 2 | 0 | NA | NA | 4 | 0.5 | NA | NA | NA | NA | NA | NA | 1 | 1 | 1 | 1 |
| 10_8122727 | 2 | 0.5 | 1 | 0 | 2 | 0 | NA | NA | 5 | 0.2 | NA | NA | NA | NA | NA | NA | 1 | 1 | 1 | 1 |
| 10_8123081 | NA | NA | NA | NA | NA | NA | 1 | 1 | 1 | 1 | NA | NA | 2 | 0.5 | NA | NA | NA | NA | 2 | 0.5 |
| 10_8123361 | 1 | 0 | 2 | 0 | 7 | 0.1429 | 1 | 0 | 11 | 0.0909 | 3 | 0 | 3 | 0 | 3 | 0.3333 | 4 | 0 | 13 | 0.0769 |
| 10_8123493 | 5 | 0.2 | 6 | 0.1667 | 3 | 0 | 3 | 0 | 17 | 0.1176 | 5 | 0.2 | 5 | 0.2 | 5 | 0 | 6 | 0 | 21 | 0.0952 |
| 10_8123570 | 4 | 0 | 2 | 0 | 3 | 0.3333 | 2 | 0 | 11 | 0.0909 | 4 | 0.25 | 3 | 0 | 1 | 0 | 3 | 0.3333 | 11 | 0.1818 |
| 10_8123603 | 3 | 0.3333 | 1 | 0 | 2 | 0.5 | 2 | 0 | 8 | 0.25 | 4 | 0 | 2 | 0 | 2 | 0.5 | 4 | 0 | 12 | 0.0833 |
| 10_8123614 | 2 | 0 | NA | NA | 2 | 0.5 | 2 | 0 | 6 | 0.1667 | 2 | 1 | 1 | 0 | 1 | 0 | 3 | 0 | 7 | 0.2857 |
| 10_8123625 | 5 | 0.8 | 1 | 1 | 3 | 1 | 2 | 1 | 11 | 0.9091 | 3 | 1 | 3 | 1 | 2 | 1 | 4 | 1 | 12 | 1 |
| 10_8123627 | 3 | 0.6667 | 1 | 0 | 3 | 0.6667 | 2 | 0.5 | 9 | 0.5556 | 4 | 0.25 | 2 | 0.5 | 2 | 1 | 4 | 0.25 | 12 | 0.4167 |
| 10_8123633 | 5 | 1 | 1 | 0 | NA | NA | 2 | 0.5 | 8 | 0.75 | 4 | 0 | 3 | 0.6667 | 2 | 0 | 3 | 0.3333 | 12 | 0.25 |
| 10_8123635 | 5 | 0 | 1 | 0 | 2 | 0 | 2 | 0.5 | 10 | 0.1 | 3 | 0.6667 | 2 | 0.5 | 2 | 0 | 3 | 0.3333 | 10 | 0.4 |
| 10_8123635 | 5 | 0.8 | 1 | 1 | 3 | 1 | NA | NA | 9 | 0.8889 | 2 | 0.5 | 2 | 0.5 | 2 | 1 | 3 | 0.3333 | 9 | 0.5556 |
| 10_8123642 | 4 | 0.5 | 1 | 1 | 3 | 1 | 2 | 1 | 10 | 0.8 | 4 | 0.75 | 2 | 0.5 | 1 | 1 | 2 | 1 | 9 | 0.7778 |
| 10_8123643 | 2 | 0.5 | 1 | 0 | 3 | 0 | 2 | 0.5 | 8 | 0.25 | 4 | 0.5 | 2 | 0.5 | 2 | 0 | 2 | 0 | 10 | 0.3 |
| 10_8123671 | 4 | 0 | 1 | 0 | 3 | 0.3333 | 2 | 0 | 10 | 0.1 | 2 | 0 | 3 | 0 | 2 | 0 | 3 | 0.3333 | 10 | 0.1 |
| 10_8123673 | 4 | 0 | 1 | 0 | 3 | 0.6667 | 2 | 0 | 10 | 0.2 | 4 | 0.25 | 3 | 0 | 2 | 0.5 | 3 | 0 | 12 | 0.1667 |
| 10_8123674 | 5 | 0.8 | 1 | 1 | 2 | 0.5 | 2 | 0.5 | 10 | 0.7 | 4 | 1 | 3 | 0.6667 | 2 | 0.5 | 3 | 1 | 12 | 0.8333 |
| 10_8123675 | 5 | 0 | 1 | 0 | 2 | 0 | 2 | 0.5 | 10 | 0.1 | 3 | 0.6667 | 2 | 0 | 2 | 0 | 3 | 0 | 10 | 0.2 |
| 10_8123678 | 4 | 0.25 | 1 | 1 | 2 | 0 | 2 | 0 | 9 | 0.2222 | 4 | 0.5 | 3 | 0.3333 | 2 | 0 | 3 | 0 | 12 | 0.25 |
| 10_8125613 | 3 | 0.3333 | 1 | 0 | 1 | 0 | 2 | 0 | 7 | 0.1429 | 2 | 0.5 | 2 | 0 | 1 | 0 | 3 | 0 | 8 | 0.125 |
| 10_8125627 | 4 | 0 | 1 | 0 | 1 | 1 | 2 | 0 | 8 | 0.125 | 2 | 0.5 | 1 | 0 | 1 | 0 | 3 | 0 | 7 | 0.1429 |
| 10_8125633 | 3 | 0 | 1 | 0 | 1 | 1 | 2 | 0.5 | 7 | 0.2857 | 2 | 0.5 | 2 | 0 | 1 | 0 | 3 | 0 | 8 | 0.125 |
| 10_8125637 | 2 | 0 | NA | NA | 1 | 1 | 2 | 1 | 5 | 0.6 | 2 | 0.5 | 2 | 0 | NA | NA | 3 | 0.6667 | 7 | 0.4286 |
| 10_8125641 | 4 | 0 | 1 | 0 | 1 | 0 | 2 | 0.5 | 8 | 0.125 | 1 | 0 | 2 | 0 | 1 | 1 | 2 | 0 | 6 | 0.1667 |
| 10_8129532 | 29 | 0.2759 | 26 | 0.2692 | 21 | 0.3333 | 27 | 0.1111 | 103 | 0.2427 | 16 | 0.25 | 36 | 0.3333 | 29 | 0.3793 | 28 | 0.25 | 109 | 0.3119 |
| 10_8129569 | 28 | 0.0714 | 27 | 0.1481 | 21 | 0.1905 | 28 | 0.0714 | 104 | 0.1154 | 16 | 0.25 | 34 | 0.2353 | 29 | 0.069 | 26 | 0.1538 | 105 | 0.1714 |
| 10_8129585 | 21 | 0.0476 | 19 | 0.0526 | 15 | 0.1333 | 18 | 0.1667 | 73 | 0.0959 | 12 | 0.1667 | 19 | 0.1579 | 23 | 0.1304 | 18 | 0.1667 | 72 | 0.1528 |
| 10_8129590 | 23 | 0.087 | 23 | 0.2609 | 18 | 0.1667 | 26 | 0.1154 | 90 | 0.1556 | 14 | 0.1429 | 31 | 0.2581 | 26 | 0.1923 | 22 | 0.3182 | 93 | 0.2366 |
| 10_8129639 | 14 | 0.4286 | 17 | 0.0588 | 15 | 0.4667 | 19 | 0.3684 | 65 | 0.3231 | 11 | 0.3636 | 23 | 0.1304 | 22 | 0.2273 | 20 | 0.25 | 76 | 0.2237 |
| 10_8129648 | 18 | 0.3333 | 18 | 0.2222 | 14 | 0.1429 | 18 | 0.3889 | 68 | 0.2794 | 8 | 0.125 | 24 | 0.2917 | 22 | 0.1818 | 22 | 0.4545 | 76 | 0.2895 |
| 10_8129676 | 14 | 0 | 15 | 0.3333 | 11 | 0.0909 | 19 | 0.0526 | 59 | 0.1186 | 10 | 0.1 | 20 | 0.05 | 20 | 0.2 | 13 | 0.0769 | 63 | 0.1111 |
| 10_8129679 | 15 | 0.8667 | 20 | 0.8 | 10 | 0.6 | 21 | 0.5714 | 66 | 0.7121 | 10 | 0.6 | 22 | 0.5455 | 20 | 0.65 | 16 | 0.875 | 68 | 0.6618 |
| 10_8129690 | 8 | 0.25 | 11 | 0.0909 | 9 | 0.1111 | 16 | 0.125 | 44 | 0.1364 | 7 | 0 | 20 | 0.1 | 14 | 0.0714 | 11 | 0.0909 | 52 | 0.0769 |
| 10_8129693 | 14 | 0.5 | 15 | 0.6667 | 12 | 0.5 | 20 | 0.35 | 61 | 0.4918 | 9 | 0.7778 | 24 | 0.5 | 20 | 0.75 | 16 | 0.5 | 69 | 0.6087 |
| 10_8129694 | 15 | 0.1333 | 18 | 0.3333 | 14 | 0.1429 | 21 | 0.1905 | 68 | 0.2059 | 10 | 0.4 | 25 | 0.32 | 21 | 0.0952 | 17 | 0.4118 | 73 | 0.2877 |
| 10_8129695 | 15 | 0.1333 | 19 | 0.1053 | 14 | 0 | 20 | 0.05 | 68 | 0.0735 | 9 | 0.1111 | 26 | 0.0769 | 21 | 0 | 17 | 0.0588 | 73 | 0.0548 |
| 10_8129701 | 11 | 0.0909 | 15 | 0.2 | 8 | 0.125 | 17 | 0.1176 | 51 | 0.1373 | 6 | 0 | 20 | 0 | 17 | 0.2941 | 12 | 0 | 55 | 0.0909 |
| 10_8129730 | 11 | 0 | 12 | 0.25 | 11 | 0 | 14 | 0.0714 | 48 | 0.0833 | 9 | 0.2222 | 21 | 0.1905 | 16 | 0.1875 | 14 | 0.2857 | 60 | 0.2167 |
| 10_8129734 | 7 | 0.7143 | 8 | 0.375 | 6 | 0.5 | 7 | 0.4286 | 28 | 0.5 | 6 | 0.6667 | 13 | 0.6154 | 12 | 0.6667 | 7 | 0.5714 | 38 | 0.6316 |
| 10_8129734 | 8 | 0.75 | 12 | 1 | 9 | 0.8889 | 11 | 0.9091 | 40 | 0.9 | 6 | 1 | 18 | 0.8333 | 14 | 0.8571 | 13 | 0.9231 | 51 | 0.8824 |
| 10_8129740 | 8 | 0.25 | 11 | 0.1818 | 8 | 0 | 11 | 0.0909 | 38 | 0.1316 | 7 | 0.1429 | 14 | 0.0714 | 11 | 0.1818 | 9 | 0 | 41 | 0.0976 |
| 10_8129750 | 11 | 0.3636 | 14 | 0.0714 | 12 | 0.1667 | 14 | 0.0714 | 51 | 0.1569 | 9 | 0.1111 | 18 | 0.2222 | 16 | 0.0625 | 12 | 0.1667 | 55 | 0.1455 |
| 10_8129758 | 8 | 0.75 | 12 | 0.75 | 10 | 0.7 | 8 | 0.5 | 38 | 0.6842 | 8 | 0.75 | 13 | 0.7692 | 13 | 0.4615 | 13 | 0.6923 | 47 | 0.6596 |
| 10_8129761 | 11 | 0.2727 | 11 | 0.2727 | 12 | 0.1667 | 14 | 0.3571 | 48 | 0.2708 | 6 | 0 | 15 | 0.4667 | 17 | 0.3529 | 9 | 0.4444 | 47 | 0.3617 |
| 10_8129773 | 6 | 0 | 8 | 0.125 | 7 | 0 | 11 | 0.0909 | 32 | 0.0625 | 7 | 0 | 12 | 0 | 12 | 0 | 9 | 0.2222 | 40 | 0.05 |
| 10_8132240 | 8 | 0.375 | 7 | 0.2857 | 11 | 0.3636 | 7 | 0.4286 | 33 | 0.3636 | 7 | 0.7143 | 11 | 0.4545 | 7 | 0.5714 | 6 | 0.3333 | 31 | 0.5161 |
| 10_8132269 | 8 | 0 | 6 | 0.1667 | 9 | 0.1111 | 8 | 0 | 31 | 0.0645 | 8 | 0 | 12 | 0 | 7 | 0.1429 | 6 | 0.1667 | 33 | 0.0606 |
| 10_8132277 | 6 | 0 | 1 | 0 | 3 | 0.6667 | 7 | 0 | 17 | 0.1176 | 4 | 0.25 | 4 | 0 | 4 | 0.25 | 3 | 0 | 15 | 0.1333 |
| 10_8132299 | 6 | 0.5 | 5 | 1 | 6 | 0.5 | 6 | 0.3333 | 23 | 0.5652 | 6 | 0.6667 | 11 | 0.4545 | 7 | 0.8571 | 2 | 1 | 26 | 0.6538 |
| 10_8132305 | 5 | 0 | 4 | 0 | 8 | 0.125 | 8 | 0.125 | 25 | 0.08 | 8 | 0.125 | 10 | 0 | 6 | 0.1667 | 4 | 0 | 28 | 0.0714 |
| 10_8132327 | 6 | 0.1667 | 5 | 0 | 10 | 0.4 | 7 | 0.4286 | 28 | 0.2857 | 6 | 0.3333 | 9 | 0.4444 | 4 | 1 | 4 | 0.25 | 23 | 0.4783 |
| 10_8132331 | 5 | 0.2 | 3 | 0 | 7 | 0 | 5 | 0.2 | 20 | 0.1 | 5 | 0 | 8 | 0.125 | 6 | 0.3333 | 4 | 0.5 | 23 | 0.2174 |
| 10_8132340 | 6 | 0.1667 | 5 | 0.2 | 8 | 0 | 8 | 0.25 | 27 | 0.1481 | 7 | 0.4286 | 8 | 0.125 | 6 | 0.1667 | 5 | 0.2 | 26 | 0.2308 |
| 10_8132928 | 3 | 0.3333 | 2 | 0 | 3 | 0.3333 | 2 | 0.5 | 10 | 0.3 | 4 | 0 | 2 | 0.5 | 1 | 0 | 1 | 0 | 8 | 0.125 |
| 10_8133840 | NA | NA | 1 | 1 | NA | NA | NA | NA | 1 | 1 | NA | NA | 1 | 1 | NA | NA | NA | NA | 1 | 1 |

| | | | | | | | | | | | | | | | | | | | | |
|---|---|---|---|---|---|---|---|---|---|---|---|---|---|---|---|---|---|---|---|---|
| 10_813385( | NA | NA | 1 | 1 | NA | NA | NA | NA | 1 | 1 | NA | NA | 1 | 1 | NA | NA | NA | NA | 1 | 1 |
| 10_813386 | NA | NA | 1 | 1 | NA | NA | NA | NA | 1 | 1 | NA | NA | 1 | 1 | NA | NA | 1 | 1 | 2 | 1 |
| 10_813386 | NA | NA | 1 | 1 | NA | NA | NA | NA | 1 | 1 | NA | NA | 1 | 1 | NA | NA | 1 | 1 | 2 | 1 |
| 10_813647! | 18 | 0.1111 | 20 | 0.05 | 18 | 0 | 17 | 0.0588 | 73 | 0.0548 | 13 | 0.1538 | 16 | 0 | 16 | 0.125 | 15 | 0.0667 | 60 | 0.0833 |
| 10_813652: | 14 | 0.2857 | 13 | 0.3846 | 11 | 0.0909 | 17 | 0.2353 | 55 | 0.2545 | 11 | 0.2727 | 15 | 0.2 | 16 | 0.0625 | 12 | 0.4167 | 54 | 0.2222 |
| 10_813654( | 11 | 0.0909 | 11 | 0.1818 | 10 | 0 | 12 | 0.0833 | 44 | 0.0909 | 8 | 0 | 13 | 0.2308 | 11 | 0.1818 | 12 | 0 | 44 | 0.1136 |
| 10_813655! | 12 | 0.1667 | 13 | 0.1538 | 6 | 0 | 15 | 0.2 | 46 | 0.1522 | 6 | 0.1667 | 11 | 0.0909 | 9 | 0 | 14 | 0.1429 | 40 | 0.1 |
| 10_813656! | 13 | 0.0769 | 9 | 0 | 9 | 0 | 13 | 0.1538 | 44 | 0.0682 | 11 | 0.1818 | 9 | 0 | 11 | 0 | 14 | 0.0714 | 45 | 0.0667 |
| 10_813659( | 14 | 0.1429 | 10 | 0.3 | 12 | 0.25 | 12 | 0.1667 | 48 | 0.2083 | 10 | 0.6 | 10 | 0.2 | 10 | 0.2 | 12 | 0.25 | 42 | 0.3095 |
| 10_813662( | 9 | 1 | 6 | 0.6667 | 11 | 0.7273 | 8 | 0.75 | 34 | 0.7941 | 8 | 0.875 | 9 | 0.8889 | 10 | 1 | 6 | 0.8333 | 33 | 0.9091 |
| 10_813664 | 9 | 0.1111 | 5 | 0 | 7 | 0.1429 | 7 | 0.1429 | 28 | 0.1071 | 6 | 0.1667 | 7 | 0.1429 | 7 | 0 | 7 | 0.1429 | 27 | 0.1111 |
| 10_813665( | 7 | 0.2857 | 5 | 0.8 | 6 | 0.8333 | 7 | 0.8571 | 25 | 0.68 | 3 | 1 | 4 | 0.75 | 5 | 0.6 | 6 | 0.8333 | 18 | 0.7778 |
| 10_813666 | 8 | 0.375 | 6 | 0 | 7 | 0.1429 | 8 | 0.125 | 29 | 0.1724 | 7 | 0.2857 | 7 | 0.1429 | 7 | 0.2857 | 6 | 0 | 27 | 0.1852 |
| 10_813667( | 8 | 0 | 3 | 0.6667 | 6 | 0 | 8 | 0 | 25 | 0.08 | 3 | 0 | 5 | 0 | 6 | 0.1667 | 4 | 0.25 | 18 | 0.1111 |
| 10_814014 | 91 | 0.0769 | 113 | 0.0619 | 92 | 0.1196 | 102 | 0.1176 | 398 | 0.093 | 93 | 0.1505 | 101 | 0.1386 | 104 | 0.125 | 73 | 0.1233 | 371 | 0.1348 |
| 10_814019: | 42 | 0.0952 | 33 | 0.0909 | 30 | 0.1667 | 44 | 0.2273 | 149 | 0.1477 | 40 | 0.125 | 41 | 0.1463 | 29 | 0.069 | 23 | 0.1304 | 133 | 0.1203 |
| 10_814251 | NA | NA | 1 | 1 | NA | NA | NA | NA | 1 | 1 | NA | NA | 2 | 0.5 | NA | NA | NA | NA | 2 | 0.5 |
| 10_814989: | 1 | 1 | 2 | 0 | NA | NA | 3 | 0 | 6 | 0.1667 | 2 | 0.5 | NA | NA | 1 | 0 | NA | NA | 3 | 0.3333 |
| 10_814989! | 1 | 1 | 2 | 0 | NA | NA | 3 | 0 | 6 | 0.1667 | 2 | 0.5 | NA | NA | 1 | 0 | NA | NA | 3 | 0.3333 |
| 10_814990: | 1 | 1 | 1 | 1 | NA | NA | 3 | 1 | 5 | 1 | NA | NA | NA | NA | 1 | 1 | NA | NA | 1 | 1 |
| 10_814991( | 1 | 1 | 2 | 1 | NA | NA | 3 | 1 | 6 | 1 | 1 | 1 | NA | NA | 1 | 1 | NA | NA | 2 | 1 |
| 10_814993 | 1 | 1 | 1 | 1 | NA | NA | 4 | 0.75 | 6 | 0.8333 | 2 | 1 | NA | NA | 1 | 1 | NA | NA | 3 | 1 |
| 10_814994 | 1 | 1 | 2 | 1 | NA | NA | 3 | 1 | 6 | 1 | 2 | 1 | NA | NA | 1 | 1 | NA | NA | 3 | 1 |
| 10_814994! | 1 | 0 | 2 | 0 | NA | NA | 3 | 0.3333 | 6 | 0.1667 | 1 | 0 | NA | NA | 1 | 1 | NA | NA | 2 | 0.5 |
| 10_814997! | 1 | 1 | 2 | 0.5 | NA | NA | 3 | 1 | 6 | 0.8333 | 2 | 1 | NA | NA | 1 | 1 | NA | NA | 3 | 1 |
| 10_814998( | 1 | 1 | 2 | 1 | NA | NA | 3 | 1 | 6 | 1 | 1 | 1 | NA | NA | NA | NA | NA | NA | 1 | 1 |
| 10_814999: | 1 | 1 | 2 | 1 | NA | NA | 4 | 0.5 | 7 | 0.7143 | 2 | 0.5 | NA | NA | 1 | 1 | NA | NA | 3 | 0.6667 |
| 10_814999( | 1 | 1 | 2 | 1 | NA | NA | 3 | 1 | 6 | 1 | 2 | 1 | NA | NA | 1 | 1 | NA | NA | 3 | 1 |
| 10_815001( | 1 | 1 | 1 | 1 | NA | NA | 2 | 1 | 4 | 1 | 1 | 1 | NA | NA | 1 | 1 | NA | NA | 2 | 1 |
| 10_815021( | 1 | 1 | 1 | 0 | NA | NA | 2 | 0 | 4 | 0.25 | 1 | 1 | NA | NA | 2 | 0.5 | NA | NA | 3 | 0.6667 |
| 10_815022! | 1 | 1 | 1 | 0 | NA | NA | 2 | 0.5 | 4 | 0.5 | 1 | 1 | NA | NA | 2 | 0 | NA | NA | 3 | 0.3333 |
| 10_815024 | 1 | 1 | NA | NA | NA | NA | 2 | 0 | 3 | 0.3333 | 1 | 1 | NA | NA | 2 | 0.5 | NA | NA | 3 | 0.6667 |
| 10_815559! | NA | NA | 1 | 1 | 1 | 0 | NA | NA | 2 | 0.5 | NA | NA | NA | NA | 1 | 0 | 1 | 1 | 2 | 0.5 |
| 10_815560 | NA | NA | 1 | 1 | 1 | 0 | NA | NA | 2 | 0.5 | NA | NA | NA | NA | 1 | 1 | 1 | 1 | 2 | 1 |
| 10_815562! | NA | NA | NA | NA | 1 | 1 | NA | NA | 1 | 1 | NA | NA | NA | NA | 1 | 1 | 1 | 1 | 2 | 1 |
| 10_815568! | NA | NA | 1 | 1 | 1 | 1 | NA | NA | 2 | 1 | NA | NA | NA | NA | 1 | 1 | 1 | 1 | 2 | 1 |
| 10_815570! | 1 | 1 | 1 | 0 | 1 | 1 | 1 | 1 | 4 | 0.75 | NA | NA | NA | NA | 1 | 1 | 1 | 0 | 2 | 0.5 |
| 10_815570 | 1 | 1 | 1 | 1 | 1 | 1 | 1 | 1 | 4 | 1 | NA | NA | NA | NA | 1 | 1 | 1 | 1 | 2 | 1 |
| 10_815573! | 1 | 1 | NA | NA | 1 | 1 | 1 | 1 | 3 | 1 | NA | NA | NA | NA | 1 | 1 | 1 | 1 | 2 | 1 |
| 10_815575( | 1 | 1 | 1 | 0 | 2 | 0 | NA | NA | 4 | 0.25 | NA | NA | NA | NA | 1 | 1 | 1 | 0 | 2 | 0.5 |
| 10_815575 | 1 | 1 | 1 | 1 | 2 | 0.5 | 1 | 1 | 5 | 0.8 | NA | NA | NA | NA | 1 | 1 | 1 | 1 | 2 | 1 |
| 10_815577 | 1 | 0 | 1 | 1 | 2 | 0 | 1 | 1 | 5 | 0.4 | NA | NA | NA | NA | 1 | 1 | 1 | 0 | 2 | 0.5 |
| 10_815805 | NA | NA | NA | NA | 1 | 1 | NA | NA | 1 | 1 | NA | NA | 1 | 1 | 1 | 1 | 2 | 1 | 4 | 1 |
| 10_815806 | NA | NA | NA | NA | 1 | 1 | NA | NA | 1 | 1 | NA | NA | 1 | 1 | 1 | 1 | NA | NA | 2 | 1 |
| 10_815806: | NA | NA | NA | NA | 1 | 1 | NA | NA | 1 | 1 | NA | NA | 1 | 1 | NA | NA | 1 | 1 | 2 | 1 |
| 10_815811! | NA | NA | NA | NA | 1 | 1 | NA | NA | 1 | 1 | NA | NA | NA | NA | 1 | 1 | 1 | 1 | 2 | 1 |
| 10_815813( | NA | NA | NA | NA | 1 | 1 | NA | NA | 1 | 1 | NA | NA | NA | NA | 1 | 0 | 1 | 1 | 2 | 0.5 |
| 10_815815! | NA | NA | NA | NA | 1 | 1 | NA | NA | 1 | 1 | NA | NA | NA | NA | 1 | 1 | 1 | 1 | 2 | 1 |
| 10_815816 | NA | NA | NA | NA | 1 | 1 | NA | NA | 1 | 1 | NA | NA | NA | NA | 1 | 1 | 1 | 1 | 2 | 1 |
| 10_815819 | NA | NA | NA | NA | 1 | 1 | NA | NA | 1 | 1 | NA | NA | NA | NA | 1 | 1 | NA | NA | 1 | 1 |
| 10_815821: | NA | NA | NA | NA | 1 | 1 | NA | NA | 1 | 1 | NA | NA | NA | NA | 1 | 1 | NA | NA | 1 | 1 |
| 10_816070! | 1 | 1 | NA | NA | NA | NA | NA | NA | 1 | 1 | NA | NA | 1 | 1 | NA | NA | NA | NA | 1 | 1 |
| 10_817151! | NA | NA | 1 | 1 | NA | NA | NA | NA | 1 | 1 | NA | NA | NA | NA | 3 | 1 | 2 | 0.5 | 5 | 0.8 |
| 10_817159! | NA | NA | 1 | 1 | NA | NA | NA | NA | 1 | 1 | NA | NA | NA | NA | 2 | 1 | NA | NA | 2 | 1 |
| 10_817161! | NA | NA | 1 | 1 | NA | NA | NA | NA | 1 | 1 | NA | NA | NA | NA | 3 | 1 | 2 | 0.5 | 5 | 0.8 |
| 10_817164! | 1 | 1 | NA | NA | NA | NA | NA | NA | 1 | 1 | NA | NA | NA | NA | 2 | 1 | 1 | 1 | 3 | 1 |
| 10_817165: | 1 | 0 | 1 | 1 | NA | NA | NA | NA | 2 | 0.5 | NA | NA | NA | NA | 3 | 0.3333 | 1 | 1 | 4 | 0.5 |
| 10_817166! | 1 | 1 | 1 | 1 | NA | NA | NA | NA | 2 | 1 | NA | NA | NA | NA | 3 | 1 | 2 | 1 | 5 | 1 |
| 10_817168! | 1 | 1 | NA | NA | 1 | 1 | NA | NA | 2 | 1 | NA | NA | NA | NA | 3 | 0.3333 | 1 | 1 | 4 | 0.5 |
| 10_817391: | NA | NA | 2 | 0.5 | NA | NA | NA | NA | 2 | 0.5 | NA | NA | 2 | 0.5 | 1 | 0 | NA | NA | 3 | 0.3333 |
| 10_818489! | NA | NA | 1 | 1 | NA | NA | NA | NA | 1 | 1 | 1 | 1 | NA | NA | NA | NA | NA | NA | 1 | 1 |
| 10_818494! | NA | NA | 1 | 1 | NA | NA | NA | NA | 1 | 1 | 1 | 1 | NA | NA | NA | NA | NA | NA | 1 | 1 |
| 10_818496! | NA | NA | 1 | 1 | NA | NA | NA | NA | 1 | 1 | 1 | 1 | 1 | 1 | 1 | 1 | NA | NA | 3 | 1 |
| 10_818499! | 1 | 1 | 1 | 1 | NA | NA | NA | NA | 2 | 1 | 1 | 1 | NA | NA | 1 | 1 | NA | NA | 2 | 1 |
| 10_818500 | NA | NA | 1 | 1 | NA | NA | NA | NA | 1 | 1 | 1 | 0 | 1 | 1 | 1 | 0 | NA | NA | 3 | 0.3333 |

| | | | | | | | | | | | | | | | | | | | | |
|---|---|---|---|---|---|---|---|---|---|---|---|---|---|---|---|---|---|---|---|---|
| 10_818501 | NA | NA | 1 | 1 | NA | NA | NA | NA | 1 | 1 | NA | NA | 1 | 1 | 1 | 1 | NA | NA | 2 | 1 |
| 10_818957 | NA | NA | 2 | 0.5 | 1 | 0 | NA | NA | 3 | 0.3333 | NA | NA | NA | NA | 1 | 1 | 1 | 0 | 2 | 0.5 |
| 10_818960 | NA | NA | 2 | 1 | 1 | 1 | NA | NA | 3 | 1 | NA | NA | NA | NA | NA | NA | 2 | 1 | 2 | 1 |
| 10_818964 | NA | NA | 1 | 1 | 1 | 1 | NA | NA | 2 | 1 | NA | NA | NA | NA | 1 | 1 | 2 | 0.5 | 3 | 0.6667 |
| 10_818964 | NA | NA | 2 | 0.5 | 1 | 1 | NA | NA | 3 | 0.6667 | NA | NA | NA | NA | 1 | 1 | 2 | 1 | 3 | 1 |
| 10_818964 | NA | NA | 2 | 1 | 1 | 1 | NA | NA | 3 | 1 | NA | NA | NA | NA | 1 | 0 | 1 | 1 | 2 | 0.5 |
| 10_818968 | NA | NA | 2 | 1 | 1 | 0 | 1 | 1 | 4 | 0.75 | NA | NA | NA | NA | NA | NA | 2 | 1 | 2 | 1 |
| 10_818972 | NA | NA | 2 | 1 | 1 | 1 | 1 | 0 | 4 | 0.75 | NA | NA | NA | NA | 1 | 1 | 2 | 0.5 | 3 | 0.6667 |
| 10_818998 | NA | NA | 2 | 0.5 | 1 | 0 | NA | NA | 3 | 0.3333 | NA | NA | 1 | 1 | NA | NA | NA | NA | 1 | 1 |
| 10_819448 | NA | NA | NA | NA | NA | NA | 1 | 1 | 1 | 1 | NA | NA | 1 | 1 | NA | NA | 1 | 1 | 2 | 1 |
| 10_819449 | NA | NA | NA | NA | NA | NA | 1 | 1 | 1 | 1 | NA | NA | NA | NA | NA | NA | 1 | 1 | 1 | 1 |
| 10_819450 | NA | NA | NA | NA | NA | NA | 1 | 1 | 1 | 1 | NA | NA | 2 | 1 | NA | NA | 1 | 1 | 3 | 1 |
| 10_819453 | NA | NA | NA | NA | NA | NA | 1 | 1 | 1 | 1 | NA | NA | 2 | 0.5 | NA | NA | 2 | 1 | 4 | 0.75 |
| 10_819454 | NA | NA | NA | NA | NA | NA | 1 | 1 | 1 | 1 | NA | NA | 2 | 1 | NA | NA | 2 | 1 | 4 | 1 |
| 10_819463 | NA | NA | 1 | 1 | NA | NA | 1 | 1 | 2 | 1 | NA | NA | 1 | 1 | NA | NA | 2 | 1 | 3 | 1 |
| 10_819463 | NA | NA | 1 | 1 | NA | NA | 1 | 1 | 2 | 1 | NA | NA | 2 | 1 | NA | NA | 2 | 1 | 4 | 1 |
| 10_819464 | NA | NA | 1 | 1 | NA | NA | 2 | 1 | 3 | 1 | NA | NA | 2 | 1 | NA | NA | 2 | 1 | 4 | 1 |
| 10_819469 | NA | NA | 2 | 0.5 | NA | NA | 3 | 0.3333 | 5 | 0.4 | NA | NA | 1 | 1 | NA | NA | 2 | 0 | 3 | 0.3333 |
| 10_819867 | NA | NA | 1 | 1 | NA | NA | 1 | 0 | 2 | 0.5 | 1 | 1 | NA | NA | NA | NA | NA | NA | 1 | 1 |
| 10_819868 | NA | NA | 1 | 1 | NA | NA | 1 | 0 | 2 | 0.5 | 1 | 1 | NA | NA | NA | NA | NA | NA | 1 | 1 |
| 10_819868 | NA | NA | 1 | 1 | NA | NA | 1 | 1 | 2 | 1 | 1 | 1 | NA | NA | NA | NA | NA | NA | 1 | 1 |
| 10_819869 | NA | NA | 1 | 1 | NA | NA | 1 | 1 | 2 | 1 | 1 | 1 | NA | NA | NA | NA | NA | NA | 1 | 1 |
| 10_819873 | NA | NA | 1 | 1 | NA | NA | NA | NA | 1 | 1 | 1 | 1 | NA | NA | NA | NA | NA | NA | 1 | 1 |
| 10_819874 | NA | NA | 1 | 1 | NA | NA | 1 | 0 | 2 | 0.5 | 1 | 1 | NA | NA | NA | NA | NA | NA | 1 | 1 |
| 10_819879 | NA | NA | 1 | 1 | NA | NA | NA | NA | 1 | 1 | 1 | 1 | NA | NA | NA | NA | NA | NA | 1 | 1 |
| 10_819880 | 1 | 1 | 1 | 1 | NA | NA | NA | NA | 2 | 1 | 1 | 1 | NA | NA | NA | NA | NA | NA | 1 | 1 |
| 10_820609 | 2 | 0.5 | 3 | 0 | 6 | 0.1667 | NA | NA | 11 | 0.1818 | 1 | 1 | 2 | 0 | 2 | 0.5 | 1 | 0 | 6 | 0.3333 |
| 10_820619 | 2 | 0 | 3 | 0.3333 | 6 | 0.1667 | NA | NA | 11 | 0.1818 | NA | NA | 1 | 1 | 2 | 0 | NA | NA | 3 | 0.3333 |
| 10_824850 | 3 | 0.3333 | 5 | 0.4 | 2 | 1 | 8 | 0.5 | 18 | 0.5 | 4 | 1 | 7 | 0.2857 | 4 | 0.25 | 2 | 0.5 | 17 | 0.4706 |
| 10_824850 | 4 | 0 | 5 | 0.2 | 3 | 0.3333 | 8 | 0.5 | 20 | 0.3 | 3 | 0.6667 | 8 | 0.25 | 5 | 0.4 | 2 | 0.5 | 18 | 0.3889 |
| 10_824851 | 4 | 0.25 | 4 | 0.5 | 2 | 0.5 | 7 | 0.7143 | 17 | 0.5294 | 3 | 0.3333 | 9 | 0.3333 | 4 | 0.5 | 3 | 0.3333 | 19 | 0.3684 |
| 10_824852 | 3 | 0.3333 | 3 | 0.3333 | 1 | 1 | 5 | 0.8 | 12 | 0.5833 | 2 | 1 | 8 | 0.5 | 2 | 0.5 | 3 | 1 | 15 | 0.6667 |
| 10_824854 | 5 | 0.2 | 3 | 0 | 2 | 0 | 5 | 0.2 | 15 | 0.1333 | 3 | 0.6667 | 9 | 0.4444 | 6 | 0.3333 | 2 | 0.5 | 20 | 0.45 |
| 10_824856 | 3 | 0 | 3 | 0 | 3 | 0 | 6 | 0.1667 | 15 | 0.0667 | 4 | 0.25 | 6 | 0 | 3 | 0 | 1 | 0 | 14 | 0.0714 |
| 10_824859 | 4 | 0 | 4 | 0 | 3 | 0.3333 | 6 | 0 | 17 | 0.0588 | 4 | 0.25 | 12 | 0.1667 | 5 | 0 | 4 | 0.25 | 25 | 0.16 |
| 10_824872 | NA | NA | NA | NA | NA | NA | 1 | 1 | 1 | 1 | 1 | 1 | NA | NA | 1 | 1 | 2 | 1 | 4 | 1 |
| 10_825749 | 4 | 1 | 3 | 1 | 4 | 1 | NA | NA | 11 | 1 | 3 | 1 | 2 | 1 | 1 | 1 | NA | NA | 6 | 1 |
| 10_825750 | 4 | 0.25 | 5 | 0.4 | 4 | 0.75 | NA | NA | 13 | 0.4615 | 2 | 1 | 2 | 0.5 | 1 | 0 | 1 | 1 | 6 | 0.6667 |
| 10_827320 | 4 | 0.25 | 6 | 0 | 8 | 0.125 | 8 | 0.375 | 26 | 0.1923 | 1 | 0 | 7 | 0.7143 | 9 | 0.3333 | 7 | 0.4286 | 24 | 0.4583 |
| 10_827320 | 5 | 0.6 | 7 | 0.2857 | 8 | 0.125 | 8 | 0.125 | 28 | 0.25 | 1 | 0 | 6 | 0.3333 | 9 | 0.3333 | 6 | 0.5 | 22 | 0.3636 |
| 10_830274 | 1 | 1 | 1 | 0 | NA | NA | NA | NA | 2 | 0.5 | NA | NA | 1 | 1 | NA | NA | 2 | 1 | 3 | 1 |
| 10_830275 | 1 | 1 | NA | NA | NA | NA | NA | NA | 1 | 1 | NA | NA | 1 | 1 | NA | NA | 1 | 1 | 2 | 1 |
| 10_830277 | 2 | 0.5 | 1 | 0 | NA | NA | NA | NA | 3 | 0.3333 | 2 | 0 | 2 | 0 | NA | NA | 2 | 0.5 | 6 | 0.1667 |
| 10_830278 | 2 | 0.5 | 1 | 0 | NA | NA | NA | NA | 3 | 0.3333 | NA | NA | 2 | 0 | NA | NA | 2 | 0.5 | 4 | 0.25 |
| 10_830301 | 4 | 0.5 | 2 | 0.5 | 3 | 0 | 2 | 0 | 11 | 0.2727 | 3 | 0.3333 | 4 | 0 | 2 | 0 | 4 | 0 | 13 | 0.0769 |
| 10_830315 | 7 | 0.4286 | 2 | 0.5 | 4 | 0.25 | 7 | 0 | 20 | 0.25 | 4 | 0 | 8 | 0.5 | 4 | 0.25 | 4 | 0.25 | 20 | 0.3 |
| 10_830691 | NA | NA | 1 | 1 | 2 | 0 | 3 | 0 | 6 | 0.1667 | 1 | 0 | 1 | 0 | 3 | 0.3333 | 1 | 0 | 6 | 0.1667 |
| 10_830694 | NA | NA | NA | NA | 2 | 1 | 3 | 0.6667 | 5 | 0.8 | 1 | 0 | 1 | 1 | 3 | 1 | NA | NA | 5 | 0.8 |
| 10_830694 | NA | NA | 1 | 1 | 2 | 0 | 3 | 0.3333 | 6 | 0.3333 | 1 | 1 | 1 | 1 | 3 | 0.6667 | NA | NA | 5 | 0.8 |
| 10_830695 | NA | NA | 1 | 1 | 1 | 0 | 3 | 0.6667 | 5 | 0.6 | 1 | 1 | 1 | 0 | 2 | 0.5 | NA | NA | 4 | 0.5 |
| 10_831523 | NA | NA | NA | NA | 1 | 1 | 1 | 1 | 2 | 1 | 1 | 1 | 1 | 0 | NA | NA | NA | NA | 2 | 0.5 |
| 10_831523 | NA | NA | NA | NA | 1 | 1 | 1 | 0 | 2 | 0.5 | 1 | 1 | 1 | 0 | NA | NA | NA | NA | 2 | 0.5 |
| 10_832017 | 2 | 0.5 | NA | NA | 2 | 1 | 4 | 0.75 | 8 | 0.75 | NA | NA | 1 | 1 | 2 | 1 | 2 | 1 | 5 | 1 |
| 10_832018 | 2 | 0.5 | NA | NA | 3 | 0.3333 | 4 | 0.5 | 9 | 0.4444 | NA | NA | 2 | 0.5 | 2 | 0.5 | 2 | 0.5 | 6 | 0.5 |
| 10_832019 | 2 | 1 | NA | NA | 3 | 1 | 4 | 1 | 9 | 1 | NA | NA | 2 | 1 | 2 | 1 | 2 | 1 | 6 | 1 |
| 10_832019 | 2 | 0.5 | NA | NA | 3 | 0.3333 | 3 | 0 | 8 | 0.25 | NA | NA | 2 | 0.5 | 2 | 0 | 2 | 0 | 6 | 0.1667 |
| 10_832021 | 3 | 0.3333 | NA | NA | 3 | 0 | 4 | 0.25 | 10 | 0.2 | NA | NA | 2 | 0 | 2 | 0 | 1 | 1 | 5 | 0.2 |
| 10_834301 | 4 | 0.5 | 2 | 0.5 | 5 | 0.6 | 3 | 0.3333 | 14 | 0.5 | 1 | 0 | 1 | 1 | 3 | 0.6667 | NA | NA | 5 | 0.6 |
| 10_834302 | 3 | 0.6667 | 2 | 1 | 4 | 0.75 | 3 | 0.6667 | 12 | 0.75 | 1 | 1 | 3 | 0.6667 | 3 | 1 | NA | NA | 7 | 0.8571 |
| 10_834302 | 4 | 0.75 | 1 | 1 | 5 | 1 | 3 | 1 | 13 | 0.9231 | 1 | 1 | 3 | 1 | 3 | 1 | NA | NA | 7 | 1 |
| 10_834302 | 3 | 0.6667 | 2 | 0 | 2 | 1 | 3 | 0 | 10 | 0.4 | 1 | 1 | 2 | 1 | 1 | 0 | NA | NA | 4 | 0.75 |
| 10_834305 | 5 | 0.4 | 3 | 0.6667 | 5 | 0.8 | 3 | 0.3333 | 16 | 0.5625 | 1 | 1 | 3 | 0.6667 | 5 | 0.4 | 1 | 1 | 10 | 0.6 |
| 10_834305 | 5 | 0.4 | 3 | 0 | 5 | 0.4 | 3 | 0.3333 | 16 | 0.3125 | 1 | 0 | 3 | 0.6667 | 5 | 0 | 1 | 1 | 10 | 0.3 |
| 10_834307 | 4 | 0 | 3 | 0 | 4 | 0 | 3 | 0.3333 | 14 | 0.0714 | 1 | 0 | 2 | 0 | 5 | 0.2 | NA | NA | 8 | 0.125 |
| 10_834316 | 5 | 0 | 4 | 0.25 | 9 | 0.3333 | 3 | 0.3333 | 21 | 0.2381 | 1 | 1 | 3 | 0 | 7 | 0.5714 | 3 | 0 | 14 | 0.3571 |

| | | | | | | | | | | | | | | | | | | | | |
|---|---|---|---|---|---|---|---|---|---|---|---|---|---|---|---|---|---|---|---|---|
| 10_8343616 | 12 | 0.1667 | 5 | 0.2 | 4 | 0.5 | 4 | 0.25 | 25 | 0.24 | 7 | 0.2857 | 8 | 0.375 | 4 | 0 | 8 | 0 | 27 | 0.1852 |
| 10_8343653 | 12 | 0.5 | 6 | 0.3333 | 4 | 0.75 | 5 | 0.2 | 27 | 0.4444 | 6 | 0.5 | 7 | 0.4286 | 5 | 0.4 | 7 | 0.2857 | 25 | 0.4 |
| 10_8343662 | 10 | 0.2 | 6 | 0.1667 | 4 | 0.5 | 3 | 0.6667 | 23 | 0.3043 | 5 | 1 | 8 | 0.25 | 6 | 0.1667 | 5 | 0.2 | 24 | 0.375 |
| 10_8343667 | 6 | 0 | 4 | 0 | 2 | 0.5 | 2 | 0 | 14 | 0.0714 | 4 | 0.25 | 5 | 0 | 5 | 0 | 3 | 0 | 17 | 0.0588 |
| 10_8344854 | NA | NA | 1 | 1 | 1 | 0 | NA | NA | 2 | 0.5 | 1 | 0 | 1 | 1 | NA | NA | NA | NA | 2 | 0.5 |
| 10_8346584 | NA | NA | 2 | 0.5 | NA | NA | NA | NA | 2 | 0.5 | NA | NA | 2 | 0.5 | NA | NA | NA | NA | 2 | 0.5 |
| 10_8359794 | 5 | 1 | NA | NA | 5 | 1 | 5 | 0.8 | 15 | 0.9333 | 3 | 0.6667 | 1 | 1 | 4 | 0.25 | 4 | 0.5 | 12 | 0.5 |
| 10_8359796 | 6 | 0 | NA | NA | 6 | 0.1667 | 5 | 0.2 | 17 | 0.1176 | 3 | 0.6667 | 1 | 0 | 4 | 0 | 4 | 0.25 | 12 | 0.25 |
| 10_8359830 | 2 | 0.5 | NA | NA | 6 | 0 | 2 | 0 | 10 | 0.1 | 3 | 0 | 1 | 0 | 3 | 0 | 3 | 0.3333 | 10 | 0.1 |
| 10_8359832 | 3 | 0 | NA | NA | 5 | 0.2 | 3 | 0.3333 | 11 | 0.1818 | 3 | 0 | 1 | 0 | 4 | 0.25 | 3 | 0 | 11 | 0.0909 |
| 10_8420715 | 1 | 0 | 6 | 0.5 | 2 | 0 | 1 | 0 | 10 | 0.3 | 3 | 0.6667 | 1 | 0 | 3 | 0 | 2 | 0.5 | 9 | 0.3333 |
| 10_8420725 | 2 | 0.5 | 8 | 1 | 2 | 1 | 1 | 1 | 13 | 0.9231 | 2 | 1 | 1 | 1 | 3 | 1 | 2 | 1 | 8 | 1 |
| 10_8420772 | 1 | 0 | 7 | 0.1429 | 2 | 0 | 1 | 0 | 11 | 0.0909 | 3 | 0 | NA | NA | 1 | 0 | 1 | 1 | 5 | 0.2 |
| 10_8420920 | NA | NA | 1 | 1 | 1 | 0 | 1 | 0 | 3 | 0.3333 | 1 | 1 | NA | NA | NA | NA | NA | NA | 1 | 1 |
| 10_8420999 | NA | NA | 1 | 1 | 1 | 1 | 1 | 1 | 3 | 1 | 1 | 1 | NA | NA | NA | NA | NA | NA | 1 | 1 |
| 10_8421015 | NA | NA | 1 | 0 | 1 | 1 | 1 | 1 | 3 | 0.6667 | 1 | 1 | NA | NA | NA | NA | NA | NA | 1 | 1 |
| 10_8421024 | NA | NA | 1 | 1 | 1 | 1 | 1 | 1 | 3 | 1 | 1 | 1 | NA | NA | NA | NA | NA | NA | 1 | 1 |
| 10_8421050 | NA | NA | 1 | 0 | 1 | 1 | 1 | 0 | 3 | 0.3333 | 1 | 1 | NA | NA | NA | NA | NA | NA | 1 | 1 |
| 10_8421060 | NA | NA | 1 | 1 | 1 | 0 | NA | NA | 2 | 0.5 | 1 | 1 | NA | NA | NA | NA | NA | NA | 1 | 1 |
| 10_8421077 | NA | NA | 1 | 1 | 1 | 1 | 1 | 1 | 3 | 1 | 1 | 1 | NA | NA | NA | NA | NA | NA | 1 | 1 |
| 10_8436223 | 1 | 0 | 5 | 0.4 | 1 | 0 | 1 | 0 | 8 | 0.25 | 3 | 0.6667 | 1 | 1 | 3 | 0.6667 | 6 | 0.6667 | 13 | 0.6923 |
| 10_8436279 | 2 | 0 | 3 | 1 | NA | NA | 1 | 0 | 6 | 0.5 | 2 | 0 | 1 | 0 | 1 | 1 | 6 | 0.5 | 10 | 0.4 |
| 10_8436293 | 1 | 1 | 4 | 0.5 | NA | NA | 1 | 1 | 6 | 0.6667 | 3 | 0.6667 | 2 | 0 | 1 | 1 | 5 | 0.6 | 11 | 0.5455 |
| 10_8436318 | 1 | 1 | 4 | 0.75 | NA | NA | NA | NA | 5 | 0.8 | 3 | 0.6667 | 2 | 0 | 1 | 1 | 4 | 0.5 | 10 | 0.5 |
| 10_8436328 | NA | NA | 4 | 0.5 | NA | NA | NA | NA | 4 | 0.5 | 3 | 0 | 1 | 0 | 1 | 1 | 4 | 0 | 9 | 0.1111 |
| 10_8436332 | NA | NA | 3 | 1 | NA | NA | NA | NA | 3 | 1 | 3 | 1 | NA | NA | 1 | 1 | 2 | 1 | 6 | 1 |
| 10_8436342 | NA | NA | 3 | 0.3333 | NA | NA | NA | NA | 3 | 0.3333 | 1 | 0 | 2 | 0 | 1 | 0 | 5 | 0.4 | 9 | 0.2222 |
| 10_8436347 | 1 | 1 | 4 | 1 | NA | NA | NA | NA | 5 | 1 | 2 | 1 | 2 | 0.5 | 1 | 1 | 5 | 1 | 10 | 0.9 |
| 10_8436348 | 1 | 1 | 4 | 1 | NA | NA | NA | NA | 5 | 1 | 2 | 1 | 2 | 1 | 1 | 1 | 5 | 1 | 10 | 1 |
| 10_8436349 | 1 | 0 | 4 | 0.25 | NA | NA | NA | NA | 5 | 0.2 | 2 | 1 | 2 | 1 | 1 | 0 | 5 | 0.2 | 10 | 0.5 |
| 10_8436353 | 1 | 0 | 4 | 0.25 | NA | NA | NA | NA | 5 | 0.2 | 2 | 0 | 2 | 0 | 1 | 0 | 5 | 0.2 | 10 | 0.1 |
| 10_8436354 | 1 | 1 | 4 | 0.75 | NA | NA | NA | NA | 5 | 0.8 | NA | NA | NA | NA | 1 | 1 | 3 | 1 | 4 | 1 |
| 10_8436356 | 1 | 1 | 4 | 1 | NA | NA | NA | NA | 5 | 1 | 2 | 1 | 2 | 1 | 1 | 1 | 5 | 1 | 10 | 1 |
| 10_8436391 | 1 | 1 | 3 | 1 | NA | NA | NA | NA | 4 | 1 | 2 | 1 | 1 | 1 | 1 | 1 | 4 | 1 | 8 | 1 |
| 10_8436393 | 1 | 1 | 2 | 0 | NA | NA | NA | NA | 3 | 0.3333 | 2 | 1 | 1 | 1 | 1 | 1 | 4 | 1 | 8 | 1 |
| 10_8436414 | 1 | 0 | 3 | 0.3333 | NA | NA | NA | NA | 4 | 0.25 | 2 | 0.5 | 1 | 1 | 1 | 1 | 4 | 0.5 | 8 | 0.625 |
| 10_8436420 | NA | NA | 3 | 0.6667 | NA | NA | NA | NA | 3 | 0.6667 | 1 | 0 | 1 | 1 | 1 | 1 | 4 | 0.75 | 7 | 0.7143 |
| 10_8444950 | 3 | 0.6667 | 6 | 0.6667 | 2 | 0.5 | 5 | 1 | 16 | 0.75 | 2 | 1 | 4 | 0.75 | 4 | 1 | 6 | 1 | 16 | 0.9375 |
| 10_8444967 | 2 | 0 | 4 | 0.25 | 3 | 0.3333 | 4 | 0.25 | 13 | 0.2308 | 2 | 0 | 5 | 0.2 | 4 | 0.25 | 5 | 0 | 16 | 0.125 |
| 10_8444985 | 3 | 0.3333 | 4 | 0 | 2 | 0 | 4 | 0.25 | 13 | 0.1538 | 2 | 0 | 5 | 0.2 | 2 | 0 | 6 | 0.1667 | 15 | 0.1333 |
| 10_8456160 | NA | NA | 3 | 0.3333 | NA | NA | 1 | 0 | 4 | 0.25 | 1 | 0 | 1 | 1 | NA | NA | 2 | 0.5 | 4 | 0.5 |
| 10_8456171 | 1 | 1 | 1 | 0 | 2 | 0 | 1 | 0 | 5 | 0.2 | 2 | 0 | 1 | 1 | 1 | 1 | 2 | 0 | 6 | 0.3333 |
| 10_8456193 | 1 | 1 | 2 | 1 | 1 | 1 | 1 | 1 | 5 | 1 | 2 | 1 | 2 | 1 | 2 | 1 | 2 | 1 | 8 | 1 |
| 10_8456202 | 1 | 0 | 2 | 0.5 | 2 | 1 | 1 | 1 | 6 | 0.6667 | 2 | 1 | 1 | 0 | 1 | 1 | 1 | 0 | 5 | 0.6 |
| 10_8456212 | NA | NA | 2 | 1 | 2 | 0 | NA | NA | 4 | 0.5 | 3 | 0.3333 | 1 | 0 | 2 | 0 | 2 | 0 | 8 | 0.125 |
| 10_8456223 | 1 | 0 | 3 | 0.3333 | 2 | 0 | NA | NA | 6 | 0.1667 | 3 | 0.3333 | 2 | 1 | 1 | 1 | 2 | 0.5 | 8 | 0.625 |
| 10_8456258 | 1 | 0 | 3 | 0.3333 | 2 | 0 | NA | NA | 6 | 0.1667 | 3 | 0 | 2 | 0.5 | 2 | 0 | 2 | 0 | 9 | 0.1111 |
| 10_8456270 | 1 | 0 | 3 | 0.3333 | NA | NA | NA | NA | 4 | 0.25 | 3 | 0 | 1 | 1 | 2 | 0.5 | 1 | 0 | 7 | 0.2857 |
| 10_8456294 | 1 | 0 | 3 | 0 | 2 | 0.5 | NA | NA | 6 | 0.1667 | 3 | 0 | 2 | 0.5 | 2 | 1 | 2 | 0.5 | 9 | 0.4444 |
| 10_8457969 | NA | NA | NA | NA | NA | NA | 1 | 1 | 1 | 1 | NA | NA | NA | NA | 1 | 1 | NA | NA | 1 | 1 |
| 10_8457971 | NA | NA | NA | NA | NA | NA | 1 | 1 | 1 | 1 | NA | NA | NA | NA | 1 | 1 | NA | NA | 1 | 1 |
| 10_8457972 | NA | NA | NA | NA | NA | NA | 1 | 1 | 1 | 1 | NA | NA | NA | NA | 1 | 1 | NA | NA | 1 | 1 |
| 10_8457981 | NA | NA | NA | NA | NA | NA | 1 | 1 | 1 | 1 | NA | NA | NA | NA | 1 | 1 | NA | NA | 1 | 1 |
| 10_8458096 | NA | NA | NA | NA | NA | NA | 1 | 1 | 1 | 1 | NA | NA | NA | NA | 1 | 1 | NA | NA | 1 | 1 |
| 10_8458115 | NA | NA | NA | NA | NA | NA | 1 | 1 | 1 | 1 | NA | NA | NA | NA | 1 | 1 | NA | NA | 1 | 1 |
| 10_8474089 | 3 | 0 | 5 | 0.4 | 1 | 1 | 2 | 0 | 11 | 0.2727 | 2 | 0 | 3 | 0.3333 | 1 | 0 | NA | NA | 6 | 0.1667 |
| 10_8474109 | 3 | 0.6667 | 5 | 1 | 1 | 1 | 2 | 1 | 11 | 0.9091 | 2 | 0.5 | 2 | 0.5 | 1 | 1 | NA | NA | 5 | 0.6 |
| 10_8474116 | 3 | 0.3333 | 5 | 0.2 | 1 | 1 | 1 | 1 | 10 | 0.4 | 2 | 0 | 3 | 0 | 1 | 1 | NA | NA | 6 | 0.1667 |
| 10_8474125 | 2 | 0 | 5 | 0.2 | 1 | 0 | 2 | 0.5 | 10 | 0.2 | 2 | 1 | 3 | 0.3333 | 2 | 0 | NA | NA | 7 | 0.4286 |
| 10_8474132 | 3 | 0 | 5 | 0.6 | 1 | 0 | 2 | 0.5 | 11 | 0.3636 | 2 | 0.5 | 2 | 0 | 2 | 0.5 | NA | NA | 6 | 0.3333 |
| 10_8474169 | 3 | 0.3333 | 5 | 0.4 | 1 | 0 | 1 | 1 | 10 | 0.4 | 2 | 0 | 3 | 0.3333 | 2 | 0.5 | NA | NA | 7 | 0.2857 |
| 10_8478355 | 3 | 1 | 4 | 0.5 | 4 | 0.75 | 6 | 1 | 17 | 0.8235 | 4 | 1 | 7 | 0.5714 | 5 | 0.8 | 2 | 0.5 | 18 | 0.7222 |
| 10_8478360 | 4 | 0.75 | 6 | 0.5 | 6 | 0.8333 | 11 | 0.8182 | 27 | 0.7407 | 3 | 0.6667 | 9 | 0.6667 | 5 | 0.8 | 2 | 1 | 19 | 0.7368 |
| 10_8478361 | 4 | 0.25 | 7 | 0.4286 | 4 | 0.25 | 11 | 0.1818 | 26 | 0.2692 | 2 | 0.5 | 9 | 0.2222 | 5 | 0.4 | 2 | 0.5 | 18 | 0.3333 |
| 10_8478363 | 4 | 0.5 | 7 | 0.2857 | 6 | 0.3333 | 9 | 0.2222 | 26 | 0.3077 | 4 | 0.5 | 8 | 0.125 | 6 | 0.1667 | 2 | 0 | 20 | 0.2 |

| | | | | | | | | | | | | | | | | | | | | |
|---|---|---|---|---|---|---|---|---|---|---|---|---|---|---|---|---|---|---|---|---|
| 10_847837( | 4 | 0.75 | 7 | 0.7143 | 7 | 1 | 11 | 1 | 29 | 0.8966 | 3 | 1 | 9 | 0.8889 | 5 | 0.8 | 2 | 1 | 19 | 0.8947 |
| 10_847837! | 4 | 0.25 | 7 | 0.5714 | 7 | 0.1429 | 10 | 0.4 | 28 | 0.3571 | 4 | 0.5 | 9 | 0.5556 | 6 | 0.3333 | 2 | 0 | 21 | 0.4286 |
| 10_847837! | 4 | 0.75 | 6 | 1 | 7 | 0.5714 | 7 | 0.8571 | 24 | 0.7917 | 4 | 1 | 9 | 0.7778 | 6 | 0.6667 | 2 | 0.5 | 21 | 0.7619 |
| 10_847838' | 4 | 0.5 | 4 | 0.75 | 7 | 0.5714 | 11 | 0.5455 | 26 | 0.5769 | 3 | 0.6667 | 9 | 0.7778 | 5 | 0.6 | 1 | 1 | 18 | 0.7222 |
| 10_848509( | 4 | 0.25 | 9 | 0.2222 | 11 | 0.1818 | 11 | 0.2727 | 35 | 0.2286 | 3 | 0.3333 | 11 | 0.0909 | 12 | 0.3333 | 5 | 0 | 31 | 0.1935 |
| 10_848509! | 5 | 0 | 9 | 0 | 8 | 0.125 | 10 | 0.1 | 32 | 0.0625 | 4 | 0 | 11 | 0.1818 | 10 | 0.1 | 5 | 0 | 30 | 0.1 |
| 10_848510‹ | 5 | 0.2 | 11 | 0 | 10 | 0.1 | 12 | 0 | 38 | 0.0526 | 5 | 0 | 12 | 0.25 | 11 | 0.0909 | 4 | 0 | 32 | 0.125 |
| 10_848510‹ | 4 | 0.5 | 9 | 0.3333 | 8 | 0.375 | 12 | 0.5 | 33 | 0.4242 | 5 | 0 | 13 | 0.3077 | 11 | 0.3636 | 5 | 0.4 | 34 | 0.2941 |
| 10_848511 | 3 | 0.6667 | 8 | 0.625 | 11 | 0.4545 | 12 | 0.75 | 34 | 0.6176 | 3 | 0.6667 | 7 | 0.2857 | 10 | 0.4 | 5 | 0.4 | 25 | 0.4 |
| 10_848513( | 5 | 0.2 | 11 | 0.1818 | 12 | 0.1667 | 14 | 0.2857 | 42 | 0.2143 | 6 | 0.1667 | 12 | 0.3333 | 12 | 0.1667 | 2 | 0.5 | 32 | 0.25 |
| 10_848513' | 5 | 0.6 | 8 | 0.75 | 12 | 0.75 | 15 | 0.8 | 40 | 0.75 | 5 | 0.8 | 12 | 0.5 | 13 | 0.4615 | 5 | 0.6 | 35 | 0.5429 |
| 10_848514! | 4 | 0.25 | 10 | 0.1 | 12 | 0.4167 | 12 | 0.3333 | 38 | 0.2895 | 5 | 0 | 11 | 0.2727 | 13 | 0.4615 | 7 | 0.4286 | 36 | 0.3333 |
| 10_848514! | 4 | 0 | 10 | 0 | 12 | 0.0833 | 16 | 0.125 | 42 | 0.0714 | 5 | 0.2 | 12 | 0.25 | 13 | 0.1538 | 6 | 0.1667 | 36 | 0.1944 |
| 10_849503: | 3 | 0.3333 | 6 | 0.1667 | 2 | 1 | 7 | 0.2857 | 18 | 0.3333 | 2 | 1 | 8 | 0 | 4 | 0.25 | 4 | 0 | 18 | 0.1667 |
| 10_849504( | 3 | 0.3333 | 5 | 0.2 | 2 | 0 | 7 | 0.4286 | 17 | 0.2941 | 2 | 0 | 9 | 0.1111 | 5 | 0.2 | 5 | 0.4 | 21 | 0.1905 |
| 10_849504! | 2 | 0 | 4 | 0 | 1 | 0 | 7 | 0.2857 | 14 | 0.1429 | 1 | 0 | 5 | 0.2 | 5 | 0.2 | 5 | 0 | 16 | 0.125 |
| 10_849505‹ | 4 | 0.25 | 6 | 0 | 2 | 0.5 | 7 | 0.2857 | 19 | 0.2105 | 2 | 1 | 9 | 0.3333 | 6 | 0.5 | 4 | 0.25 | 21 | 0.4286 |
| 10_849506( | 3 | 1 | 4 | 1 | 1 | 1 | 7 | 1 | 15 | 1 | 1 | 1 | 7 | 1 | 3 | 1 | NA | NA | 11 | 1 |
| 10_849506: | 3 | 1 | 5 | 0.8 | 2 | 1 | 5 | 1 | 15 | 0.9333 | 2 | 1 | 9 | 0.7778 | 5 | 1 | 5 | 0.8 | 21 | 0.8571 |
| 10_849506‹ | 4 | 0 | 4 | 0.75 | 1 | 0 | 7 | 0.4286 | 16 | 0.375 | 2 | 0 | 5 | 0.2 | 6 | 0 | 4 | 0.75 | 17 | 0.2353 |
| 10_849506( | 4 | 1 | 5 | 0.8 | 1 | 1 | 6 | 1 | 16 | 0.9375 | 1 | 1 | 7 | 1 | 4 | 1 | 5 | 1 | 17 | 1 |
| 10_849508: | 4 | 0 | 5 | 0 | 2 | 0 | 7 | 0.1429 | 18 | 0.0556 | 1 | 0 | 8 | 0.125 | 4 | 0 | 6 | 0.3333 | 19 | 0.1579 |
| 10_849509' | 3 | 0.3333 | 4 | 0.25 | 2 | 0.5 | 6 | 0.5 | 15 | 0.4 | NA | NA | 5 | 0.2 | 5 | 0.4 | 6 | 0.6667 | 16 | 0.4375 |
| 10_849509: | 4 | 0.25 | 4 | 0.5 | 2 | 0 | 4 | 0 | 14 | 0.2143 | NA | NA | 8 | 0.375 | 5 | 0.2 | 5 | 0.6 | 18 | 0.3889 |
| 10_849525( | 4 | 0.25 | 5 | 0.2 | 4 | 0.25 | 7 | 0 | 20 | 0.15 | NA | NA | 9 | 0.3333 | 6 | 0.1667 | 8 | 0.125 | 23 | 0.2174 |
| 10_849525! | 5 | 0 | 6 | 0 | 4 | 0.25 | 9 | 0.1111 | 24 | 0.0833 | 2 | 0.5 | 10 | 0.1 | 5 | 0.2 | 8 | 0.125 | 25 | 0.16 |
| 10_849534: | 7 | 0.2857 | 8 | 0.5 | 4 | 0.5 | 11 | 0.2727 | 30 | 0.3667 | 5 | 0 | 16 | 0.375 | 8 | 0.125 | 9 | 0.4444 | 38 | 0.2895 |
| 10_849553: | 2 | 0 | 4 | 0 | 3 | 0.3333 | 1 | 0 | 10 | 0.1 | 1 | 0 | 2 | 0 | 3 | 0.3333 | 1 | 0 | 7 | 0.1429 |
| 10_849554' | 3 | 0.3333 | 5 | 0.2 | 3 | 0 | 1 | 0 | 12 | 0.1667 | 1 | 0 | 2 | 0.5 | 3 | 0.6667 | 1 | 1 | 7 | 0.5714 |
| 10_849554! | 3 | 0.6667 | 5 | 1 | 2 | 1 | 1 | 1 | 11 | 0.9091 | 1 | 0 | 2 | 1 | 3 | 1 | 1 | 1 | 7 | 0.8571 |
| 10_849554( | 3 | 0.3333 | 5 | 0.4 | 3 | 0 | 1 | 0 | 12 | 0.25 | 1 | 0 | 2 | 0 | 3 | 0.6667 | 1 | 0 | 7 | 0.2857 |
| 10_849555( | 2 | 0 | 4 | 0.5 | 3 | 0 | 1 | 0 | 10 | 0.2 | 1 | 0 | 2 | 0 | 3 | 0.3333 | 1 | 1 | 7 | 0.2857 |
| 10_849555! | 3 | 0.3333 | 3 | 0.6667 | 3 | 0.6667 | 1 | 0 | 10 | 0.5 | 1 | 1 | 3 | 0.6667 | 3 | 0.6667 | 1 | 0 | 8 | 0.625 |
| 10_849557! | 3 | 0.3333 | 4 | 0.75 | 3 | 0.3333 | 1 | 0 | 11 | 0.4545 | 1 | 1 | 3 | 0 | 2 | 1 | 1 | 1 | 7 | 0.5714 |
| 10_849558( | 1 | 0 | 3 | 0.6667 | 1 | 0 | 1 | 0 | 6 | 0.3333 | 1 | 0 | 2 | 0 | 2 | 0.5 | 1 | 0 | 6 | 0.1667 |
| 10_849559( | 2 | 1 | 4 | 0.5 | 2 | 0.5 | 1 | 1 | 9 | 0.6667 | 1 | 0 | 3 | 0 | 1 | 0 | 1 | 1 | 6 | 0.1667 |
| 10_849560 | 1 | 0 | 3 | 0.3333 | 1 | 1 | 1 | 0 | 6 | 0.3333 | 1 | 0 | 2 | 0 | 1 | 0 | 1 | 1 | 5 | 0.2 |
| 10_849561‹ | 2 | 1 | 3 | 1 | 1 | 1 | 1 | 1 | 7 | 1 | 1 | 1 | 1 | 1 | 2 | 1 | 1 | 1 | 5 | 1 |
| 10_849562 | 2 | 0.5 | 3 | 0.3333 | 2 | 0 | NA | NA | 7 | 0.2857 | 1 | 1 | 1 | 1 | 1 | 0 | 1 | 0 | 4 | 0.5 |
| 10_849562! | NA | NA | NA | NA | 1 | 1 | NA | NA | 1 | 1 | NA | NA | 1 | 1 | NA | NA | 1 | 1 | 2 | 1 |
| 10_849563‹ | 1 | 1 | 4 | 1 | 2 | 0.5 | 1 | 1 | 8 | 0.875 | 1 | 0 | NA | NA | 2 | 0.5 | 1 | 1 | 4 | 0.5 |
| 10_849565' | 2 | 1 | 5 | 0.6 | 1 | 0 | 1 | 0 | 9 | 0.5556 | 1 | 0 | 3 | 0.3333 | 2 | 0.5 | NA | NA | 6 | 0.3333 |
| 10_849565! | 1 | 0 | 5 | 0.2 | NA | NA | 1 | 0 | 7 | 0.1429 | 1 | 0 | 3 | 0.3333 | 3 | 0 | 1 | 0 | 8 | 0.125 |
| 10_849566( | 2 | 1 | 5 | 0.6 | 1 | 0 | 1 | 1 | 9 | 0.6667 | 1 | 1 | 3 | 0.6667 | 3 | 0.3333 | 1 | 1 | 8 | 0.625 |
| 10_850203( | 3 | 0.3333 | NA | NA | 3 | 0.3333 | 1 | 0 | 7 | 0.2857 | 2 | 0.5 | 4 | 0.25 | 3 | 0 | 3 | 0 | 12 | 0.1667 |
| 10_852232 | NA | NA | NA | NA | NA | NA | 1 | 1 | 1 | 1 | 2 | 1 | 1 | 0 | NA | NA | NA | NA | 3 | 0.6667 |
| 10_852234: | NA | NA | NA | NA | NA | NA | 1 | 1 | 1 | 1 | 1 | 1 | NA | NA | NA | NA | NA | NA | 1 | 1 |
| 10_852235( | NA | NA | NA | NA | NA | NA | 1 | 1 | 1 | 1 | 2 | 1 | 1 | 0 | NA | NA | NA | NA | 3 | 0.6667 |
| 10_852240( | NA | NA | NA | NA | NA | NA | 1 | 1 | 1 | 1 | 3 | 0.3333 | 1 | 1 | NA | NA | NA | NA | 4 | 0.5 |
| 10_854942! | 1 | 1 | 1 | 1 | 1 | 0 | 2 | 1 | 5 | 0.8 | 1 | 1 | 1 | 1 | 2 | 0 | 1 | 1 | 5 | 0.6 |
| 10_854943( | 1 | 1 | 1 | 1 | NA | NA | 3 | 1 | 5 | 1 | 3 | 1 | 1 | 1 | 2 | 0.5 | 2 | 0.5 | 8 | 0.75 |
| 10_854945( | 1 | 1 | 2 | 1 | 1 | 1 | 4 | 1 | 8 | 1 | 3 | 1 | 1 | 1 | 2 | 1 | 2 | 1 | 8 | 1 |
| 10_854945: | 2 | 1 | 2 | 1 | 1 | 1 | 3 | 1 | 8 | 1 | 4 | 0.75 | 1 | 1 | 2 | 1 | 2 | 1 | 9 | 0.8889 |
| 10_854945! | 1 | 1 | 2 | 1 | 1 | 1 | 4 | 0.5 | 8 | 0.75 | 4 | 0.75 | 1 | 1 | 1 | 1 | 2 | 1 | 8 | 0.875 |
| 10_854946' | 1 | 0 | 2 | 1 | 1 | 1 | 4 | 0.5 | 8 | 0.625 | 4 | 0.5 | 1 | 0 | 1 | 0 | 2 | 0.5 | 8 | 0.375 |
| 10_854947' | 1 | 1 | 1 | 1 | 1 | 1 | 4 | 1 | 7 | 1 | 4 | 1 | 1 | 1 | 2 | 1 | 2 | 1 | 9 | 1 |
| 10_854950! | 2 | 0 | 1 | 0 | 1 | 0 | 4 | 0.25 | 8 | 0.125 | 5 | 0 | 1 | 1 | 2 | 0.5 | 2 | 0.5 | 10 | 0.3 |
| 10_854957: | 2 | 0 | 2 | 0.5 | NA | NA | 5 | 0.2 | 9 | 0.2222 | 6 | 0.1667 | 2 | 0 | 3 | 0.3333 | 2 | 0.5 | 13 | 0.2308 |
| 10_854957! | 3 | 0.6667 | 2 | 0.5 | 2 | 0 | 5 | 0.6 | 12 | 0.5 | 5 | 0.8 | 2 | 0 | 3 | 0.3333 | 2 | 1 | 12 | 0.5833 |
| 10_854959 | 2 | 1 | 3 | 0.3333 | 2 | 0 | 5 | 0.4 | 12 | 0.4167 | 5 | 0.6 | 1 | 0 | 3 | 0.3333 | 2 | 0.5 | 11 | 0.4545 |
| 10_854959! | 3 | 0 | 3 | 1 | 2 | 0 | 4 | 0.5 | 12 | 0.4167 | 7 | 0.1429 | 2 | 0.5 | 3 | 0.6667 | 1 | 0 | 13 | 0.3077 |
| 10_854960! | 1 | 1 | 1 | 1 | NA | NA | 5 | 0.8 | 7 | 0.8571 | 5 | 0.6 | 2 | 0 | 2 | 0 | 2 | 0 | 11 | 0.2727 |
| 10_854960( | 1 | 1 | 2 | 1 | 2 | 0 | 4 | 0.5 | 9 | 0.5556 | 5 | 0.6 | 2 | 0.5 | 4 | 0.25 | 1 | 1 | 12 | 0.5 |
| 10_857240! | 1 | 1 | 2 | 1 | 8 | 0.875 | 3 | 1 | 14 | 0.9286 | 2 | 1 | 3 | 0.6667 | 2 | 1 | 2 | 1 | 9 | 0.8889 |
| 10_857242‹ | 1 | 0 | 4 | 0.5 | 6 | 0.3333 | 2 | 0 | 13 | 0.3077 | 1 | 0 | 4 | 0.5 | 6 | 0.5 | 2 | 0 | 13 | 0.3846 |

| | | | | | | | | | | | | | | | | | | | | |
|---|---|---|---|---|---|---|---|---|---|---|---|---|---|---|---|---|---|---|---|---|
| 10_8572793 | 3 | 0.6667 | 2 | 0.5 | 2 | 1 | 5 | 0.8 | 12 | 0.75 | 2 | 0.5 | 3 | 0.3333 | 3 | 0.6667 | NA | NA | 8 | 0.5 |
| 10_8572798 | 2 | 0 | 2 | 0 | 3 | 0.3333 | 5 | 0.2 | 12 | 0.1667 | 1 | 0 | 4 | 0.25 | 3 | 0 | NA | NA | 8 | 0.125 |
| 10_8572799 | 2 | 0 | 1 | 0 | 3 | 0.3333 | 4 | 0 | 10 | 0.1 | 3 | 0.3333 | 4 | 0.25 | 3 | 0 | NA | NA | 10 | 0.2 |
| 10_8572849 | 1 | 0 | 3 | 0.6667 | 3 | 0 | 4 | 0.25 | 11 | 0.2727 | 3 | 0.3333 | 3 | 0.3333 | 3 | 0.3333 | NA | NA | 9 | 0.3333 |
| 10_8572886 | 1 | 1 | 2 | 1 | 1 | 1 | 2 | 1 | 6 | 1 | 2 | 1 | 3 | 1 | 2 | 1 | NA | NA | 7 | 1 |
| 10_8572891 | 1 | 1 | 2 | 0.5 | 2 | 1 | 4 | 1 | 9 | 0.8889 | 3 | 1 | 4 | 0.75 | 4 | 0.5 | NA | NA | 11 | 0.7273 |
| 10_8572905 | 1 | 1 | 3 | 0.6667 | 1 | 0 | 4 | 0.75 | 9 | 0.6667 | 2 | 0.5 | 3 | 0.3333 | 3 | 1 | NA | NA | 8 | 0.625 |
| 10_8572905 | 1 | 0 | 2 | 0 | 2 | 1 | 4 | 0.5 | 9 | 0.4444 | 2 | 0 | 3 | 0.3333 | 3 | 0.3333 | NA | NA | 8 | 0.25 |
| 10_8572927 | 1 | 1 | 3 | 0 | 2 | 0.5 | 3 | 0.3333 | 9 | 0.3333 | 1 | 0 | 3 | 0 | 3 | 0.3333 | NA | NA | 7 | 0.1429 |
| 10_8574324 | 1 | 0 | 2 | 1 | 1 | 0 | 2 | 0.5 | 6 | 0.5 | 1 | 1 | 2 | 0 | 2 | 0 | NA | NA | 5 | 0.2 |
| 10_8574340 | NA | NA | 1 | 0 | 1 | 0 | 3 | 0.3333 | 5 | 0.2 | 1 | 1 | 2 | 0.5 | 2 | 0 | NA | NA | 5 | 0.4 |
| 10_8574351 | NA | NA | 1 | 0 | 1 | 1 | 4 | 0.75 | 6 | 0.6667 | 1 | 1 | 2 | 1 | 2 | 1 | NA | NA | 5 | 1 |
| 10_8574370 | NA | NA | 1 | 0 | 1 | 1 | 3 | 1 | 5 | 0.8 | 1 | 1 | 2 | 0 | 2 | 0.5 | NA | NA | 5 | 0.4 |
| 10_8574376 | NA | NA | 2 | 0.5 | 1 | 1 | 2 | 1 | 5 | 0.8 | 1 | 1 | 2 | 1 | 1 | 0 | NA | NA | 4 | 0.75 |
| 10_8574378 | NA | NA | 2 | 0.5 | 1 | 0 | 3 | 0.3333 | 6 | 0.3333 | 1 | 1 | 2 | 0 | 2 | 0.5 | NA | NA | 5 | 0.4 |
| 10_8577465 | 15 | 0.4 | 11 | 0.5455 | 11 | 0.3636 | 14 | 0.4286 | 51 | 0.4314 | 10 | 0.4 | 16 | 0.5 | 10 | 0.5 | 16 | 0.4375 | 52 | 0.4615 |
| 10_8577481 | 18 | 0.2222 | 13 | 0.3077 | 17 | 0.2941 | 16 | 0.0625 | 64 | 0.2188 | 9 | 0.2222 | 19 | 0 | 12 | 0 | 13 | 0.1538 | 53 | 0.0755 |
| 10_8577484 | 18 | 0.8889 | 12 | 1 | 14 | 0.8571 | 10 | 0.8 | 54 | 0.8889 | 9 | 0.7778 | 13 | 0.7692 | 13 | 0.9231 | 13 | 0.6154 | 48 | 0.7708 |
| 10_8577486 | 19 | 0.0526 | 10 | 0.2 | 15 | 0.1333 | 15 | 0 | 59 | 0.0847 | 9 | 0 | 18 | 0.0556 | 13 | 0 | 19 | 0.1053 | 59 | 0.0508 |
| 10_8579363 | 5 | 0 | 2 | 0 | 5 | 0.4 | 7 | 0 | 19 | 0.1053 | 2 | 0.5 | 4 | 0 | 6 | 0.1667 | 8 | 0 | 20 | 0.1 |
| 10_8586037 | NA | NA | 5 | 0.4 | 2 | 0.5 | NA | NA | 7 | 0.4286 | 1 | 1 | 1 | 1 | 1 | 1 | 6 | 0.8333 | 9 | 0.8889 |
| 10_8586051 | 2 | 1 | 8 | 0 | 3 | 0.6667 | 1 | 0 | 14 | 0.2857 | 1 | 0 | 1 | 0 | 2 | 0.5 | 5 | 0.2 | 9 | 0.2222 |
| 10_8586056 | 2 | 0 | 6 | 0.1667 | 3 | 0.3333 | 1 | 0 | 12 | 0.1667 | 1 | 1 | 1 | 0 | 2 | 0 | 5 | 0.2 | 9 | 0.2222 |
| 10_8588377 | 4 | 0.25 | 11 | 0.0909 | 6 | 0.1667 | 1 | 0 | 22 | 0.1364 | 1 | 1 | 3 | 0 | 7 | 0 | 5 | 0.4 | 16 | 0.1875 |
| 10_8588387 | 5 | 0.2 | 12 | 0.25 | 5 | 0.4 | 1 | 1 | 23 | 0.3043 | 1 | 0 | 3 | 0.3333 | 6 | 0.5 | 3 | 1 | 13 | 0.5385 |
| 10_8588398 | 4 | 0.5 | 15 | 0.8 | 7 | 0.4286 | 1 | 1 | 27 | 0.6667 | 2 | 0.5 | 5 | 0.6 | 7 | 0.4286 | 4 | 0.75 | 18 | 0.5556 |
| 10_8588399 | 5 | 0 | 13 | 0.1538 | 7 | 0 | 1 | 0 | 26 | 0.0769 | 2 | 0 | 5 | 0 | 6 | 0 | 5 | 0.2 | 18 | 0.0556 |
| 10_8588405 | 5 | 0.2 | 14 | 0.1429 | 7 | 0.2857 | 1 | 0 | 27 | 0.1852 | 2 | 0.5 | 5 | 0 | 7 | 0 | 4 | 0 | 18 | 0.0556 |
| 10_8588409 | 5 | 0.4 | 13 | 0.2308 | 5 | 0.4 | 1 | 0 | 24 | 0.2917 | 2 | 0 | 4 | 0.25 | 7 | 0.2857 | 5 | 0.2 | 18 | 0.2222 |
| 10_8588526 | NA | NA | 2 | 0 | 3 | 0.6667 | 5 | 0 | 10 | 0.2 | 1 | 0 | 1 | 1 | 3 | 0 | 4 | 0 | 9 | 0.1111 |
| 10_8588536 | NA | NA | 1 | 1 | 2 | 1 | 6 | 1 | 9 | 1 | 1 | 0 | 2 | 1 | 3 | 1 | 5 | 1 | 11 | 0.9091 |
| 10_8588564 | NA | NA | 2 | 1 | 2 | 0 | 6 | 0.5 | 10 | 0.5 | 1 | 0 | 2 | 0.5 | 2 | 0 | 4 | 0 | 9 | 0.1111 |
| 10_8588565 | NA | NA | 1 | 0 | 2 | 0 | 4 | 0.5 | 7 | 0.2857 | 1 | 0 | 2 | 0.5 | 1 | 0 | 3 | 0 | 7 | 0.1429 |
| 10_8588568 | NA | NA | 2 | 0 | 2 | 0 | 6 | 0.1667 | 10 | 0.1 | 1 | 0 | 2 | 0.5 | 3 | 0.3333 | 3 | 0.3333 | 9 | 0.3333 |
| 10_8588574 | NA | NA | 1 | 0 | 2 | 1 | 6 | 0.8333 | 9 | 0.7778 | 1 | 1 | 2 | 1 | 3 | 0.6667 | 3 | 0.6667 | 9 | 0.7778 |
| 10_8588587 | NA | NA | 2 | 0.5 | 2 | 0 | 4 | 0 | 8 | 0.125 | 1 | 0 | NA | NA | 3 | 0.3333 | 3 | 0.3333 | 7 | 0.2857 |
| 10_8588604 | NA | NA | 2 | 0.5 | 2 | 0.5 | 4 | 0.5 | 8 | 0.5 | 1 | 0 | NA | NA | 3 | 0.3333 | 3 | 0.6667 | 7 | 0.4286 |
| 10_8588669 | NA | NA | 2 | 0.5 | 2 | 1 | 2 | 0.5 | 6 | 0.6667 | NA | NA | NA | NA | 2 | 0.5 | 2 | 0.5 | 4 | 0.5 |
| 10_8588689 | NA | NA | 2 | 1 | 2 | 1 | 2 | 0.5 | 6 | 0.8333 | NA | NA | NA | NA | 3 | 0.3333 | 1 | 1 | 4 | 0.5 |
| 10_8588692 | NA | NA | 2 | 0.5 | 1 | 1 | 2 | 1 | 5 | 0.8 | NA | NA | NA | NA | 3 | 0.6667 | 2 | 0.5 | 5 | 0.6 |
| 10_8588751 | NA | NA | 1 | 0 | 2 | 1 | 2 | 0.5 | 5 | 0.6 | NA | NA | NA | NA | NA | NA | 2 | 0.5 | 2 | 0.5 |
| 10_8588760 | NA | NA | 2 | 0.5 | 2 | 0.5 | 2 | 0 | 6 | 0.3333 | NA | NA | NA | NA | NA | NA | 2 | 0.5 | 2 | 0.5 |
| 10_8588950 | NA | NA | 1 | 1 | 2 | 0.5 | 1 | 0 | 4 | 0.5 | NA | NA | NA | NA | NA | NA | 2 | 0.5 | 2 | 0.5 |
| 10_8589005 | NA | NA | 1 | 1 | 2 | 1 | 1 | 1 | 4 | 1 | 1 | 1 | NA | NA | 1 | 1 | 3 | 1 | 5 | 1 |
| 10_8589016 | NA | NA | 1 | 0 | 1 | 1 | 1 | 0 | 3 | 0.3333 | 1 | 1 | NA | NA | NA | NA | 1 | 1 | 2 | 1 |
| 10_8589026 | NA | NA | 2 | 0 | 2 | 0.5 | 1 | 0 | 5 | 0.2 | 1 | 0 | NA | NA | 1 | 0 | 3 | 0.3333 | 5 | 0.2 |
| 10_8589030 | NA | NA | 1 | 1 | 2 | 0.5 | NA | NA | 3 | 0.6667 | 1 | 1 | NA | NA | NA | NA | 2 | 0.5 | 3 | 0.6667 |
| 10_8589033 | NA | NA | 2 | 0.5 | 2 | 0.5 | NA | NA | 4 | 0.5 | NA | NA | NA | NA | 1 | 0 | 3 | 0.3333 | 4 | 0.25 |
| 10_8589079 | NA | NA | 1 | 1 | 2 | 0 | 1 | 1 | 4 | 0.5 | 1 | 1 | NA | NA | 1 | 1 | 2 | 0.5 | 4 | 0.75 |
| 10_8589099 | NA | NA | 3 | 0.3333 | 2 | 1 | 1 | 0 | 6 | 0.5 | 1 | 1 | NA | NA | 2 | 0.5 | 2 | 1 | 5 | 0.8 |
| 10_8589103 | NA | NA | 3 | 0.3333 | 1 | 0 | 1 | 0 | 5 | 0.2 | 1 | 1 | NA | NA | 2 | 0.5 | 2 | 0 | 5 | 0.4 |
| 10_8589116 | NA | NA | 3 | 0.3333 | 2 | 1 | 1 | 0 | 6 | 0.5 | 1 | 1 | NA | NA | 2 | 0.5 | 2 | 0 | 5 | 0.4 |
| 10_8589121 | NA | NA | 2 | 0.5 | 2 | 0.5 | NA | NA | 4 | 0.5 | 1 | 1 | NA | NA | 1 | 1 | 1 | 1 | 3 | 1 |
| 10_8589127 | NA | NA | 2 | 0.5 | 2 | 1 | 1 | 1 | 5 | 0.8 | 1 | 1 | NA | NA | 2 | 1 | 1 | 0 | 4 | 0.75 |
| 10_8589135 | NA | NA | 3 | 0 | 2 | 0 | 1 | 1 | 6 | 0.1667 | 1 | 1 | NA | NA | 2 | 0 | 1 | 1 | 4 | 0.5 |
| 10_8589140 | NA | NA | 3 | 0.3333 | 2 | 0 | 1 | 0 | 6 | 0.1667 | 1 | 0 | NA | NA | 2 | 0.5 | 1 | 1 | 4 | 0.5 |
| 10_8613651 | 5 | 0.4 | 13 | 0.0769 | 9 | 0.2222 | 7 | 0.1429 | 34 | 0.1765 | 4 | 0.5 | 10 | 0.2 | 10 | 0 | 8 | 0.25 | 32 | 0.1875 |
| 10_8613664 | 4 | 0.5 | 14 | 0.7857 | 9 | 0.2222 | 8 | 0.625 | 35 | 0.5714 | 3 | 1 | 11 | 0.8182 | 13 | 0.6923 | 7 | 0.8571 | 34 | 0.7941 |
| 10_8618162 | 6 | 0 | 15 | 0.0667 | 9 | 0 | 9 | 0.2222 | 39 | 0.0769 | 2 | 0.5 | 11 | 0.0909 | 13 | 0 | 8 | 0 | 34 | 0.0588 |
| 10_8618188 | 6 | 0.1667 | 15 | 0.1333 | 8 | 0.25 | 11 | 0.2727 | 40 | 0.2 | 2 | 0.5 | 13 | 0.0769 | 12 | 0.25 | 10 | 0.4 | 37 | 0.2432 |
| 10_8618244 | 10 | 0.1 | 15 | 0.1333 | 9 | 0.1111 | 10 | 0.1 | 44 | 0.1136 | 3 | 0.6667 | 12 | 0 | 16 | 0.0625 | 10 | 0.3 | 41 | 0.1463 |
| 10_8618393 | 12 | 0.25 | 14 | 0.2857 | 7 | 0.2857 | 11 | 0.5455 | 44 | 0.3409 | 2 | 0.5 | 16 | 0.125 | 18 | 0.0556 | 12 | 0.25 | 48 | 0.1458 |
| 10_8630314 | NA | NA | 2 | 0.5 | 1 | 1 | 1 | 0 | 4 | 0.5 | 1 | 0 | NA | NA | 1 | 1 | 1 | 1 | 3 | 0.6667 |
| 10_8652689 | 34 | 0.5882 | 51 | 0.5882 | 31 | 0.6774 | 32 | 0.6875 | 148 | 0.6284 | 36 | 0.6389 | 43 | 0.6047 | 36 | 0.6667 | 38 | 0.6842 | 153 | 0.6471 |
| 10_8653773 | 17 | 0.0588 | 29 | 0.2414 | 17 | 0.2353 | 28 | 0.2143 | 91 | 0.1978 | 15 | 0.1333 | 26 | 0.1538 | 20 | 0.25 | 25 | 0.12 | 86 | 0.1628 |

| | | | | | | | | | | | | | | | | | | | | |
|---|---|---|---|---|---|---|---|---|---|---|---|---|---|---|---|---|---|---|---|---|
| 10_8666996 | 2 | 0.5 | 3 | 0.6667 | 1 | 0 | 2 | 0 | 8 | 0.375 | 2 | 1 | 3 | 1 | 1 | 1 | 1 | 0 | 7 | 0.8571 |
| 10_8666999 | 3 | 0.6667 | 2 | 1 | 2 | 0 | 2 | 0.5 | 9 | 0.5556 | NA | NA | 3 | 1 | 1 | 1 | 1 | 0 | 5 | 0.8 |
| 10_8667025 | 2 | 0.5 | 3 | 0.3333 | 2 | 0 | 2 | 0.5 | 9 | 0.3333 | 2 | 1 | 3 | 0.6667 | 1 | 0 | 2 | 1 | 8 | 0.75 |
| 10_8667027 | 2 | 0 | 2 | 0 | 1 | 1 | 1 | 0 | 6 | 0.1667 | 2 | 0 | 2 | 0 | 1 | 0 | 2 | 0.5 | 7 | 0.1429 |
| 10_8667036 | 2 | 1 | 1 | 1 | 2 | 1 | 2 | 1 | 7 | 1 | 1 | 1 | 3 | 1 | 1 | 1 | 3 | 1 | 8 | 1 |
| 10_8667038 | 3 | 0.3333 | 3 | 1 | 2 | 1 | 2 | 0.5 | 10 | 0.7 | 2 | 1 | 3 | 1 | NA | NA | 3 | 0.6667 | 8 | 0.875 |
| 10_8667042 | 3 | 0.6667 | 2 | 0.5 | 2 | 0 | 2 | 0.5 | 9 | 0.4444 | 1 | 0 | 2 | 1 | 1 | 1 | 2 | 0.5 | 6 | 0.6667 |
| 10_8667052 | 3 | 1 | 2 | 1 | 2 | 1 | 3 | 1 | 10 | 1 | 2 | 1 | 3 | 0.6667 | 1 | 1 | 3 | 1 | 9 | 0.8889 |
| 10_8667056 | 2 | 1 | 3 | 1 | 1 | 1 | 2 | 1 | 8 | 1 | 1 | 1 | 2 | 1 | 2 | 1 | 2 | 1 | 7 | 1 |
| 10_8667070 | 3 | 1 | 3 | 1 | 2 | 1 | 4 | 1 | 12 | 1 | 2 | 1 | 3 | 0.6667 | 3 | 1 | 3 | 1 | 11 | 0.9091 |
| 10_8667072 | 3 | 0.6667 | 3 | 0.6667 | 2 | 1 | 4 | 0.75 | 12 | 0.75 | 3 | 1 | 3 | 0.3333 | 3 | 1 | 2 | 1 | 11 | 0.8182 |
| 10_8667075 | 2 | 0 | 3 | 0.3333 | 3 | 0 | 4 | 0.25 | 12 | 0.1667 | 3 | 0 | 3 | 0 | 3 | 0.3333 | 3 | 0 | 12 | 0.0833 |
| 10_8667077 | 2 | 0 | 2 | 0 | 3 | 0.3333 | 3 | 0 | 10 | 0.1 | 3 | 0 | 3 | 0 | 3 | 0.3333 | 3 | 0.3333 | 12 | 0.1667 |
| 10_8667088 | 3 | 0 | 3 | 1 | 2 | 0.5 | 3 | 0.3333 | 11 | 0.4545 | 2 | 0.5 | 3 | 0.6667 | 2 | 0.5 | 4 | 0.25 | 11 | 0.4545 |
| 10_8667091 | 3 | 1 | 4 | 0.75 | 3 | 0.6667 | 4 | 1 | 14 | 0.8571 | 3 | 1 | 3 | 0.3333 | 2 | 0.5 | 4 | 0.5 | 12 | 0.5833 |
| 10_8667103 | 3 | 0 | 6 | 0 | 3 | 0 | 4 | 0.25 | 16 | 0.0625 | 2 | 0 | 3 | 0 | 3 | 0 | 5 | 0.2 | 13 | 0.0769 |
| 10_8667113 | 2 | 0.5 | 6 | 0 | 3 | 0.3333 | 4 | 0.25 | 15 | 0.2 | 3 | 0.3333 | 3 | 0 | 3 | 0 | 3 | 0.3333 | 12 | 0.1667 |
| 10_8667372 | 9 | 0 | 8 | 0.375 | 8 | 0.125 | 9 | 0.2222 | 34 | 0.1765 | 5 | 0.2 | 6 | 0.3333 | 9 | 0.2222 | 8 | 0.125 | 28 | 0.2143 |
| 10_8732702 | 3 | 0 | 2 | 0.5 | 3 | 0.3333 | NA | NA | 8 | 0.25 | NA | NA | NA | NA | 2 | 0.5 | NA | NA | 2 | 0.5 |
| 10_8732736 | 2 | 0.5 | 3 | 0.6667 | 3 | 0.6667 | NA | NA | 8 | 0.625 | NA | NA | NA | NA | 1 | 1 | NA | NA | 1 | 1 |
| 10_8732740 | 1 | 1 | 3 | 0.6667 | 3 | 1 | NA | NA | 7 | 0.8571 | NA | NA | NA | NA | 1 | 1 | NA | NA | 1 | 1 |
| 10_8777178 | NA | NA | 3 | 0.3333 | NA | NA | NA | NA | 3 | 0.3333 | 2 | 0.5 | NA | NA | 1 | 0 | NA | NA | 3 | 0.3333 |
| 10_8777188 | NA | NA | 3 | 0.3333 | NA | NA | NA | NA | 3 | 0.3333 | 2 | 0 | NA | NA | 1 | 1 | NA | NA | 3 | 0.3333 |
| 10_8826955 | 1 | 1 | 2 | 0.5 | 2 | 1 | 3 | 0 | 8 | 0.5 | 2 | 0 | 1 | 1 | 3 | 0.6667 | 3 | 0 | 9 | 0.3333 |
| 10_8826965 | 2 | 0.5 | 2 | 1 | 1 | 1 | 2 | 1 | 7 | 0.8571 | 1 | 0 | 1 | 1 | 1 | 1 | 1 | 1 | 4 | 0.75 |
| 10_8826966 | 2 | 0.5 | 2 | 0 | 1 | 0 | 3 | 0 | 8 | 0.125 | 1 | 0 | 1 | 0 | 3 | 0.3333 | 3 | 0 | 8 | 0.125 |
| 10_8826975 | 2 | 1 | 2 | 1 | 2 | 0.5 | 3 | 1 | 9 | 0.8889 | 2 | 0.5 | NA | NA | 2 | 0.5 | 2 | 0.5 | 6 | 0.5 |
| 10_8828181 | 5 | 0.6 | 5 | 0.2 | 2 | 0 | 5 | 0.4 | 17 | 0.3529 | 4 | 0.25 | 2 | 0.5 | 3 | 0 | 3 | 1 | 12 | 0.4167 |
| 10_8828188 | 4 | 0.5 | 6 | 0.5 | 5 | 0.8 | 5 | 0.4 | 20 | 0.55 | 4 | 0.5 | 2 | 0 | 5 | 0.8 | 5 | 0.6 | 16 | 0.5625 |
| 10_8828193 | 6 | 0.6667 | 7 | 0.7143 | 5 | 0.8 | 7 | 0.8571 | 25 | 0.76 | 4 | 0.25 | 2 | 1 | 5 | 0.6 | 3 | 0.6667 | 14 | 0.5714 |
| 10_8828200 | 5 | 0.2 | 5 | 0.8 | 5 | 0.6 | 7 | 0.4286 | 22 | 0.5 | 4 | 0.25 | 1 | 1 | 5 | 0.6 | 4 | 0.5 | 14 | 0.5 |
| 10_8828218 | 6 | 0 | 7 | 0.1429 | 5 | 0.2 | 7 | 0.1429 | 25 | 0.12 | 4 | 0.25 | 3 | 0 | 5 | 0.2 | 5 | 0.2 | 17 | 0.1765 |
| 10_8828236 | 6 | 0 | 7 | 0 | 4 | 0.25 | 6 | 0.1667 | 23 | 0.087 | 4 | 0 | 2 | 0 | 4 | 0 | 5 | 0.2 | 15 | 0.0667 |
| 10_8828259 | 6 | 0.1667 | 6 | 0.3333 | 5 | 0 | 5 | 0 | 22 | 0.1364 | 4 | 0 | 2 | 0 | 5 | 0.2 | 4 | 0 | 15 | 0.0667 |
| 10_8828260 | 6 | 0.3333 | 7 | 0.7143 | 5 | 0 | 5 | 0 | 23 | 0.3043 | 3 | 0.6667 | 3 | 0 | 6 | 0.5 | 5 | 0.4 | 17 | 0.4118 |
| 10_8828277 | 7 | 0.1429 | 8 | 0.125 | 5 | 0 | 5 | 0 | 25 | 0.08 | 4 | 0 | 4 | 0 | 4 | 0.25 | 5 | 0 | 17 | 0.0588 |
| 10_8830549 | 9 | 0 | 4 | 0 | 3 | 0.3333 | 1 | 0 | 17 | 0.0588 | 2 | 0.5 | 4 | 0 | 2 | 0 | 2 | 0 | 10 | 0.1 |
| 10_8831004 | 3 | 0 | 5 | 0 | 4 | 0.25 | 5 | 0.2 | 17 | 0.1176 | 1 | 0 | 1 | 1 | 2 | 0 | 2 | 0 | 6 | 0.1667 |
| 10_8831091 | 3 | 0 | 4 | 0 | 3 | 0 | 6 | 0.1667 | 16 | 0.0625 | NA | NA | 2 | 0 | 3 | 0.3333 | 1 | 0 | 6 | 0.1667 |
| 10_8831127 | 3 | 0.6667 | 4 | 0.5 | 4 | 0.5 | 6 | 0.6667 | 17 | 0.5882 | 2 | 0.5 | 3 | 0.3333 | 3 | 0.3333 | 2 | 0.5 | 10 | 0.4 |
| 10_8858262 | 2 | 1 | NA | NA | 1 | 0 | NA | NA | 3 | 0.6667 | 3 | 0.3333 | 2 | 0.5 | 4 | 0 | 3 | 1 | 12 | 0.4167 |
| 10_8858278 | 3 | 0.6667 | 1 | 0 | 3 | 1 | 1 | 0 | 8 | 0.625 | 1 | 1 | 1 | 1 | 4 | 0.5 | 3 | 0.6667 | 9 | 0.6667 |
| 10_8878524 | 3 | 0 | 2 | 0 | 2 | 0 | 4 | 0.25 | 11 | 0.0909 | 3 | 0.3333 | 3 | 0.3333 | 1 | 0 | 3 | 0 | 10 | 0.2 |
| 10_8878525 | 4 | 0.5 | 3 | 0.3333 | 1 | 0 | 4 | 0.75 | 12 | 0.5 | 2 | 0 | 3 | 0.6667 | 3 | 0.3333 | 3 | 0.6667 | 11 | 0.4545 |
| 10_8878527 | 4 | 0.5 | 1 | 1 | 1 | 1 | 5 | 0.2 | 11 | 0.4545 | 3 | 0 | 4 | 0 | 3 | 0.6667 | 3 | 0 | 13 | 0.1538 |
| 10_8878535 | 3 | 0 | 3 | 0 | NA | NA | 4 | 0.25 | 10 | 0.1 | 3 | 0 | 4 | 0.25 | 3 | 0 | 3 | 0 | 13 | 0.0769 |
| 10_8878542 | 3 | 0.3333 | 2 | 0 | 1 | 0 | 4 | 0.25 | 10 | 0.2 | 2 | 0 | 4 | 0.25 | 3 | 0 | 3 | 0 | 12 | 0.0833 |
| 10_8881456 | 2 | 0 | 2 | 0 | 1 | 0 | 3 | 0.3333 | 8 | 0.125 | 3 | 0 | 1 | 0 | 2 | 0 | 2 | 0.5 | 8 | 0.125 |
| 10_8881463 | 3 | 0.3333 | 2 | 0 | 1 | 0 | 3 | 0.3333 | 9 | 0.2222 | 3 | 0 | 1 | 0 | 2 | 0.5 | 2 | 0 | 8 | 0.125 |
| 10_8909341 | 2 | 0 | 1 | 0 | 1 | 0 | 2 | 0.5 | 6 | 0.1667 | 1 | 0 | 1 | 1 | 1 | 0 | NA | NA | 3 | 0.3333 |
| 10_8947473 | NA | NA | NA | NA | NA | NA | 3 | 0.6667 | 3 | 0.6667 | NA | NA | NA | NA | 1 | 1 | 1 | 1 | 2 | 1 |
| 10_8947480 | NA | NA | NA | NA | NA | NA | 1 | 1 | 1 | 1 | NA | NA | 1 | 0 | 1 | 0 | 2 | 0.5 | 4 | 0.25 |
| 10_8947487 | NA | NA | NA | NA | NA | NA | 3 | 0.6667 | 3 | 0.6667 | NA | NA | 1 | 1 | 1 | 0 | 1 | 1 | 3 | 0.6667 |
| 10_8947553 | 2 | 0.5 | 4 | 0.25 | 1 | 0 | 4 | 0 | 11 | 0.1818 | 4 | 0 | 1 | 1 | 1 | 0 | 2 | 0.5 | 8 | 0.25 |
| 10_8947558 | 3 | 0 | 3 | 0.3333 | NA | NA | 3 | 0.6667 | 9 | 0.3333 | 3 | 0.3333 | 1 | 1 | 1 | 0 | 3 | 1 | 8 | 0.625 |
| 10_8947587 | 3 | 0.3333 | 2 | 0.5 | 1 | 0 | 2 | 0.5 | 8 | 0.375 | 3 | 0 | 1 | 1 | 1 | 1 | 3 | 0.6667 | 8 | 0.5 |
| 10_8947605 | 3 | 1 | 1 | 1 | NA | NA | 2 | 1 | 6 | 1 | 4 | 1 | 1 | 1 | 1 | 1 | 4 | 0.75 | 10 | 0.9 |
| 10_8947611 | 1 | 1 | 1 | 1 | NA | NA | 2 | 1 | 4 | 1 | 3 | 1 | NA | NA | NA | NA | NA | NA | 3 | 1 |
| 10_8947624 | 3 | 0.6667 | 1 | 1 | NA | NA | 2 | 0.5 | 6 | 0.6667 | 3 | 1 | 1 | 1 | 1 | 0 | 4 | 0.75 | 9 | 0.7778 |
| 10_8947640 | 2 | 1 | 1 | 1 | NA | NA | 1 | 1 | 4 | 1 | 3 | 1 | 1 | 1 | 1 | 1 | 3 | 1 | 8 | 1 |
| 10_8947656 | 2 | 0 | NA | NA | NA | NA | 2 | 0.5 | 4 | 0.25 | 4 | 0.25 | 1 | 0 | 1 | 0 | 3 | 0.6667 | 9 | 0.3333 |
| 10_8947658 | 3 | 0 | 1 | 0 | NA | NA | 2 | 0.5 | 6 | 0.1667 | 4 | 0.5 | 1 | 0 | 1 | 0 | 2 | 0 | 8 | 0.25 |
| 10_8947662 | 3 | 0.6667 | 1 | 1 | NA | NA | 2 | 1 | 6 | 0.8333 | 3 | 1 | 1 | 0 | 1 | 1 | 3 | 1 | 8 | 0.875 |
| 10_8947666 | 2 | 0.5 | 1 | 1 | NA | NA | 2 | 0 | 5 | 0.4 | 3 | 1 | 1 | 1 | NA | NA | 3 | 1 | 7 | 1 |
| 10_8947667 | 3 | 0.3333 | 1 | 1 | NA | NA | 2 | 1 | 6 | 0.6667 | 2 | 1 | 1 | 1 | 1 | 1 | 3 | 1 | 7 | 1 |

| | | | | | | | | | | | | | | | | | | | | |
|---|---|---|---|---|---|---|---|---|---|---|---|---|---|---|---|---|---|---|---|---|
| 10_894767 | 1 | 1 | NA | NA | NA | NA | 1 | 1 | 2 | 1 | 3 | 1 | 1 | 1 | NA | NA | 2 | 1 | 6 | 1 |
| 10_894767 | 2 | 0.5 | NA | NA | NA | NA | 2 | 0.5 | 4 | 0.5 | 4 | 0.75 | 1 | 1 | 1 | 0 | 2 | 0.5 | 8 | 0.625 |
| 10_894768 | 3 | 0 | NA | NA | NA | NA | 2 | 0.5 | 5 | 0.2 | 3 | 0.6667 | 1 | 0 | 1 | 0 | 1 | 0 | 6 | 0.3333 |
| 10_894769 | 3 | 0.3333 | NA | NA | NA | NA | 2 | 0.5 | 5 | 0.4 | 4 | 0.5 | 1 | 0 | 1 | 0 | 2 | 0.5 | 8 | 0.375 |
| 10_895482 | 1 | 0 | 5 | 0.2 | 7 | 0.4286 | 4 | 0 | 17 | 0.2353 | 4 | 0.25 | 5 | 0.4 | 4 | 0.5 | 1 | 0 | 14 | 0.3571 |
| 10_895483 | 1 | 1 | 4 | 0 | 7 | 0 | 3 | 0.3333 | 15 | 0.1333 | 4 | 0 | 5 | 0 | 4 | 0.25 | 1 | 0 | 14 | 0.0714 |
| 10_896406 | NA | NA | 1 | 1 | NA | NA | 4 | 0.25 | 5 | 0.4 | NA | NA | 3 | 0 | 3 | 0.3333 | NA | NA | 6 | 0.1667 |
| 10_896407 | NA | NA | 1 | 0 | 1 | 0 | 5 | 0.4 | 7 | 0.2857 | 1 | 0 | 4 | 0.75 | 4 | 0 | NA | NA | 9 | 0.3333 |
| 10_896407 | NA | NA | 1 | 1 | NA | NA | 4 | 0.5 | 5 | 0.6 | 1 | 1 | 3 | 0.6667 | 4 | 0.75 | NA | NA | 8 | 0.75 |
| 10_896408 | NA | NA | 1 | 0 | 1 | 0 | 5 | 0.2 | 7 | 0.1429 | 1 | 0 | 4 | 0.75 | 3 | 0 | NA | NA | 8 | 0.375 |
| 10_896408 | NA | NA | NA | NA | 1 | 1 | 4 | 0.25 | 5 | 0.4 | 1 | 1 | 4 | 0 | 3 | 0 | NA | NA | 8 | 0.125 |
| 10_896408 | NA | NA | 1 | 1 | 1 | 1 | 3 | 0.6667 | 5 | 0.8 | 1 | 1 | 3 | 0.3333 | 4 | 0.5 | NA | NA | 8 | 0.5 |
| 10_896409 | NA | NA | 1 | 1 | 1 | 0 | 5 | 0.6 | 7 | 0.5714 | NA | NA | 4 | 0.75 | 4 | 0.25 | NA | NA | 8 | 0.5 |
| 10_896410 | NA | NA | 1 | 0 | 1 | 0 | 5 | 0.2 | 7 | 0.1429 | 1 | 0 | 4 | 0 | 6 | 0.1667 | NA | NA | 11 | 0.0909 |
| 10_896410 | 1 | 0 | 1 | 0 | 1 | 0 | 5 | 0.2 | 8 | 0.125 | 1 | 0 | 4 | 0.25 | 6 | 0 | NA | NA | 11 | 0.0909 |
| 10_896411 | 1 | 1 | 1 | 0 | 1 | 0 | 5 | 0 | 8 | 0.125 | 1 | 0 | 3 | 0.3333 | 6 | 0 | NA | NA | 10 | 0.1 |
| 10_896412 | NA | NA | 1 | 0 | NA | NA | 4 | 0.25 | 5 | 0.2 | NA | NA | 2 | 0 | 4 | 0.5 | NA | NA | 6 | 0.3333 |
| 10_896413 | 1 | 1 | 1 | 0 | 1 | 0 | 4 | 0.25 | 7 | 0.2857 | 1 | 0 | 4 | 0.25 | 6 | 0.5 | NA | NA | 11 | 0.3636 |
| 10_896415 | 2 | 0 | 1 | 0 | 1 | 1 | 5 | 0 | 9 | 0.1111 | 1 | 0 | 4 | 0 | 6 | 0.1667 | NA | NA | 11 | 0.0909 |
| 10_896417 | 2 | 0.5 | 1 | 0 | 1 | 0 | 5 | 0.2 | 9 | 0.2222 | 1 | 0 | 4 | 0.25 | 5 | 0 | NA | NA | 10 | 0.1 |
| 10_896417 | 2 | 0.5 | 2 | 1 | 1 | 1 | 5 | 1 | 10 | 0.9 | 1 | 1 | 4 | 0.75 | 5 | 1 | NA | NA | 10 | 0.9 |
| 10_896418 | 2 | 1 | 2 | 0.5 | 1 | 1 | 5 | 0.6 | 10 | 0.7 | 1 | 1 | 4 | 0.75 | 6 | 0.6667 | NA | NA | 11 | 0.7273 |
| 10_896419 | 2 | 0.5 | 2 | 0.5 | 1 | 0 | 4 | 0 | 9 | 0.2222 | NA | NA | 3 | 0.3333 | 5 | 0 | NA | NA | 8 | 0.125 |
| 10_896538 | 5 | 0.4 | 3 | 0 | 2 | 0.5 | 5 | 0 | 15 | 0.2 | 1 | 0 | 5 | 0.8 | 7 | 0.1429 | 3 | 0.6667 | 16 | 0.4375 |
| 10_896540 | 5 | 0.2 | 3 | 0 | 2 | 0 | 6 | 0.1667 | 16 | 0.125 | 1 | 0 | 5 | 0.6 | 7 | 0.1429 | 3 | 0.6667 | 16 | 0.375 |
| 10_896541 | 6 | 0.1667 | 3 | 0 | 2 | 0 | 6 | 0 | 17 | 0.0588 | 1 | 0 | 5 | 0.2 | 7 | 0 | 4 | 0.25 | 17 | 0.1176 |
| 10_896544 | 5 | 0.6 | 1 | 1 | 4 | 0 | 7 | 0.2857 | 17 | 0.3529 | NA | NA | 5 | 0.4 | 9 | 0.4444 | 4 | 0.25 | 18 | 0.3889 |
| 10_896550 | 6 | 0.1667 | 3 | 0 | 3 | 0.6667 | 5 | 0.4 | 17 | 0.2941 | NA | NA | 7 | 0.5714 | 6 | 0.5 | 4 | 0.75 | 17 | 0.5882 |
| 10_896551 | 8 | 0.125 | 3 | 0.3333 | 3 | 0.3333 | 6 | 0 | 20 | 0.15 | NA | NA | 7 | 0 | 8 | 0.5 | 6 | 0.3333 | 21 | 0.2857 |
| 10_896552 | 5 | 0 | 2 | 0 | 3 | 0.3333 | 6 | 0 | 16 | 0.0625 | NA | NA | 4 | 0.25 | 7 | 0.1429 | 3 | 0 | 14 | 0.1429 |
| 10_896554 | 9 | 0.1111 | 3 | 0 | 4 | 0.5 | 7 | 0 | 23 | 0.1304 | NA | NA | 8 | 0 | 10 | 0.2 | 5 | 0 | 23 | 0.087 |
| 10_896555 | 8 | 0.125 | 3 | 0 | 4 | 0.5 | 8 | 0.375 | 23 | 0.2609 | 1 | 0 | 8 | 0.375 | 11 | 0.3636 | 7 | 0.1429 | 27 | 0.2963 |
| 10_896555 | 7 | 0.4286 | 3 | 0 | 4 | 0.75 | 7 | 0.2857 | 21 | 0.381 | 1 | 0 | 7 | 0.2857 | 11 | 0.1818 | 7 | 0.2857 | 26 | 0.2308 |
| 10_896555 | 8 | 1 | 2 | 1 | 4 | 1 | 8 | 0.625 | 22 | 0.8636 | 1 | 0 | 8 | 1 | 7 | 0.8571 | 7 | 1 | 23 | 0.913 |
| 10_908030 | 3 | 0.3333 | 1 | 0 | 2 | 0 | 3 | 0.3333 | 9 | 0.2222 | 3 | 0 | 8 | 0.375 | 5 | 0.4 | 2 | 0 | 18 | 0.2778 |
| 10_908031 | 3 | 1 | 1 | 1 | 2 | 0 | 5 | 0.6 | 11 | 0.6364 | 2 | 0 | 9 | 0.4444 | 8 | 0.625 | 2 | 1 | 21 | 0.5238 |
| 10_908041 | 6 | 0 | 4 | 0 | 2 | 0 | 6 | 0.3333 | 18 | 0.1111 | 4 | 0.25 | 11 | 0.0909 | 11 | 0.3636 | 3 | 0.3333 | 29 | 0.2414 |
| 10_908043 | 5 | 0.4 | 5 | 0.4 | 2 | 0.5 | 5 | 0.8 | 17 | 0.5294 | 4 | 0.25 | 9 | 0.6667 | 10 | 0.3 | 3 | 0.6667 | 26 | 0.4615 |
| 10_908056 | 30 | 0.6333 | 28 | 0.7857 | 22 | 0.6364 | 27 | 0.7778 | 107 | 0.7103 | 18 | 0.5556 | 38 | 0.8158 | 25 | 0.68 | 29 | 0.7586 | 110 | 0.7273 |
| 10_908058 | 33 | 0.0909 | 26 | 0.1538 | 18 | 0 | 31 | 0.1935 | 108 | 0.1204 | 17 | 0.0588 | 30 | 0.1 | 28 | 0.25 | 28 | 0.1071 | 103 | 0.1359 |
| 10_908059 | 35 | 0.4 | 28 | 0.6071 | 21 | 0.5238 | 30 | 0.4667 | 114 | 0.4912 | 20 | 0.6 | 40 | 0.475 | 29 | 0.6897 | 28 | 0.6071 | 117 | 0.5812 |
| 10_908062 | 37 | 0.0541 | 32 | 0.1562 | 28 | 0.0357 | 35 | 0.1143 | 132 | 0.0909 | 20 | 0.1 | 44 | 0.0227 | 38 | 0.1053 | 34 | 0.0294 | 136 | 0.0588 |
| 10_908074 | 28 | 0.1429 | 25 | 0.28 | 22 | 0.2727 | 35 | 0.2 | 110 | 0.2182 | 19 | 0.2105 | 30 | 0.2667 | 25 | 0.4 | 19 | 0.2632 | 93 | 0.2903 |
| 10_908087 | 7 | 0.1429 | 12 | 0.1667 | 2 | 0 | 11 | 0.4545 | 32 | 0.25 | 3 | 0.3333 | 7 | 0 | 6 | 0.1667 | 6 | 0 | 22 | 0.0909 |
| 10_908253 | NA | NA | 2 | 0.5 | 1 | 0 | 2 | 0.5 | 5 | 0.4 | 4 | 0 | 1 | 0 | NA | NA | 2 | 0.5 | 7 | 0.1429 |
| 10_908253 | NA | NA | 2 | 0 | 1 | 0 | 2 | 1 | 5 | 0.4 | 4 | 0 | 1 | 1 | NA | NA | 2 | 0 | 7 | 0.1429 |
| 10_909318 | 1 | 1 | NA | NA | NA | NA | NA | NA | 1 | 1 | NA | NA | NA | NA | NA | NA | 1 | 1 | 1 | 1 |
| 10_909322 | 1 | 1 | NA | NA | NA | NA | NA | NA | 1 | 1 | NA | NA | NA | NA | NA | NA | 1 | 1 | 1 | 1 |
| 10_909323 | 1 | 1 | NA | NA | NA | NA | NA | NA | 1 | 1 | NA | NA | NA | NA | NA | NA | 1 | 1 | 1 | 1 |
| 10_909324 | 1 | 1 | NA | NA | NA | NA | NA | NA | 1 | 1 | NA | NA | NA | NA | NA | NA | 1 | 1 | 1 | 1 |
| 10_909332 | 1 | 1 | NA | NA | NA | NA | 1 | 0 | 2 | 0.5 | NA | NA | 1 | 0 | NA | NA | 1 | 1 | 2 | 0.5 |
| 10_909377 | 3 | 0.6667 | 2 | 0 | 1 | 0 | 3 | 0.3333 | 9 | 0.3333 | NA | NA | 2 | 0 | 1 | 0 | 1 | 1 | 4 | 0.25 |
| 10_909377 | 2 | 1 | 1 | 1 | 1 | 1 | 3 | 0.6667 | 7 | 0.8571 | NA | NA | 2 | 0.5 | 1 | 0 | NA | NA | 3 | 0.3333 |
| 10_909377 | 3 | 1 | 3 | 0.6667 | 1 | 1 | 3 | 1 | 10 | 0.9 | NA | NA | 2 | 1 | 1 | 1 | 2 | 1 | 5 | 1 |
| 10_909381 | 2 | 1 | 3 | 0.6667 | 1 | 1 | 3 | 0.6667 | 9 | 0.7778 | NA | NA | 2 | 1 | 1 | 1 | 2 | 1 | 5 | 1 |
| 10_909382 | 2 | 0 | 3 | 0.3333 | 1 | 0 | 1 | 0 | 7 | 0.1429 | NA | NA | 2 | 0.5 | 1 | 0 | 2 | 1 | 5 | 0.6 |
| 10_909383 | 2 | 1 | 3 | 1 | 2 | 1 | 3 | 0.6667 | 10 | 0.9 | 1 | 0 | 2 | 1 | 1 | 1 | 2 | 1 | 6 | 0.8333 |
| 10_909384 | 2 | 0 | 3 | 0.3333 | 2 | 0 | 3 | 0 | 10 | 0.1 | 1 | 0 | 2 | 0.5 | 1 | 0 | 2 | 0 | 6 | 0.1667 |
| 10_909422 | NA | NA | NA | NA | 1 | 1 | 1 | 1 | 2 | 1 | NA | NA | 1 | 0 | 2 | 0.5 | NA | NA | 3 | 0.3333 |
| 10_909527 | 141 | 0.1702 | 119 | 0.1597 | 115 | 0.2348 | 106 | 0.1415 | 481 | 0.1767 | 98 | 0.1735 | 116 | 0.0776 | 96 | 0.1562 | 99 | 0.1111 | 409 | 0.1271 |
| 10_909528 | 140 | 0.5643 | 136 | 0.6176 | 122 | 0.5492 | 108 | 0.6389 | 506 | 0.5909 | 97 | 0.5464 | 127 | 0.622 | 106 | 0.6321 | 103 | 0.5922 | 433 | 0.6005 |
| 10_909563 | 5 | 0.2 | 1 | 0 | 2 | 0 | 5 | 0 | 13 | 0.0769 | 1 | 1 | 2 | 0 | 2 | 0 | 4 | 0.25 | 9 | 0.2222 |
| 10_909580 | 6 | 0.1667 | 3 | 0 | 6 | 0.1667 | 3 | 0.3333 | 18 | 0.1667 | 1 | 0 | 3 | 0 | 4 | 0.25 | 5 | 0 | 13 | 0.0769 |
| 10_909849 | NA | NA | 5 | 0.2 | 1 | 0 | 2 | 0 | 8 | 0.125 | NA | NA | 2 | 0.5 | 4 | 0 | NA | NA | 6 | 0.1667 |
| 10_909850 | NA | NA | 6 | 0.5 | 4 | 1 | 3 | 0.6667 | 13 | 0.6923 | 1 | 1 | 4 | 0.75 | 4 | 0.75 | 6 | 0.3333 | 15 | 0.6 |

| | | | | | | | | | | | | | | | | | | | | |
|---|---|---|---|---|---|---|---|---|---|---|---|---|---|---|---|---|---|---|---|---|
| 10_909851 | 1 | 1 | 7 | 0.4286 | 4 | 0.75 | 3 | 1 | 15 | 0.6667 | 1 | 1 | 4 | 0.75 | 5 | 0.6 | 7 | 0.2857 | 17 | 0.5294 |
| 10_909851 | 2 | 0 | 5 | 1 | 4 | 0.75 | 3 | 0.6667 | 14 | 0.7143 | 1 | 1 | 3 | 0.3333 | 5 | 1 | 6 | 0.6667 | 15 | 0.7333 |
| 10_909852 | 2 | 0.5 | 7 | 0.8571 | 4 | 0 | 3 | 0.3333 | 16 | 0.5 | 1 | 1 | 4 | 0.5 | 5 | 0.6 | 6 | 0.1667 | 16 | 0.4375 |
| 10_909852 | 2 | 0 | 6 | 0.5 | 4 | 0.5 | 2 | 0.5 | 14 | 0.4286 | 1 | 1 | 3 | 0.3333 | 5 | 0.2 | 4 | 0.25 | 13 | 0.3077 |
| 10_909852 | 2 | 1 | 5 | 0.2 | 1 | 1 | 3 | 0.6667 | 11 | 0.5455 | 1 | 0 | 4 | 0.5 | 2 | 0.5 | 4 | 0.25 | 11 | 0.3636 |
| 10_909852 | 2 | 0.5 | 3 | 1 | 4 | 1 | 1 | 1 | 10 | 0.9 | NA | NA | 3 | 0.6667 | 4 | 0.75 | 4 | 0.75 | 11 | 0.7273 |
| 10_930749 | 33 | 0.1212 | 25 | 0.04 | 29 | 0 | 36 | 0.1111 | 123 | 0.0732 | 21 | 0.0952 | 27 | 0.037 | 37 | 0.0541 | 27 | 0.1852 | 112 | 0.0893 |
| 10_930751 | 13 | 0.3077 | 9 | 0.5556 | 4 | 0.5 | 7 | 0.4286 | 33 | 0.4242 | 6 | 0.1667 | 9 | 0.2222 | 10 | 0.4 | 10 | 0.4 | 35 | 0.3143 |
| 10_930752 | 10 | 0.5 | 6 | 0.3333 | 5 | 0.2 | 6 | 0.6667 | 27 | 0.4444 | 5 | 0.2 | 10 | 0.7 | 10 | 0.5 | 9 | 0.5556 | 34 | 0.5294 |
| 10_930753 | 11 | 0 | 9 | 0.1111 | 5 | 0.4 | 6 | 0.1667 | 31 | 0.129 | 6 | 0.3333 | 11 | 0.1818 | 9 | 0 | 11 | 0.1818 | 37 | 0.1622 |
| 10_930753 | 11 | 0 | 8 | 0.125 | 5 | 0.2 | 5 | 0 | 29 | 0.069 | 7 | 0 | 10 | 0 | 13 | 0.0769 | 10 | 0.1 | 40 | 0.05 |
| 10_930754 | 12 | 0.75 | 7 | 1 | 5 | 0.6 | 7 | 1 | 31 | 0.8387 | 7 | 0.7143 | 11 | 0.9091 | 11 | 1 | 9 | 1 | 38 | 0.9211 |
| 10_930754 | 13 | 0.6154 | 8 | 1 | 4 | 1 | 7 | 0.5714 | 32 | 0.75 | 7 | 0.4286 | 10 | 0.9 | 11 | 0.7273 | 6 | 0.8333 | 34 | 0.7353 |
| 10_930754 | 27 | 0 | 27 | 0.1481 | 36 | 0.1111 | 35 | 0.1143 | 125 | 0.096 | 23 | 0 | 30 | 0.2333 | 36 | 0.0833 | 30 | 0.1 | 119 | 0.1092 |
| 10_930755 | 33 | 0.7879 | 25 | 0.8 | 30 | 0.8 | 32 | 0.875 | 120 | 0.8167 | 22 | 0.7273 | 25 | 0.92 | 38 | 0.9474 | 29 | 0.931 | 114 | 0.8947 |
| 10_930755 | 25 | 0.32 | 24 | 0.4167 | 27 | 0.3704 | 30 | 0.3 | 106 | 0.3491 | 21 | 0.5238 | 24 | 0.4167 | 33 | 0.4242 | 24 | 0.4167 | 102 | 0.4412 |
| 10_930755 | 32 | 0.0312 | 29 | 0.2069 | 37 | 0.1081 | 35 | 0.0857 | 133 | 0.1053 | 23 | 0.0435 | 31 | 0.129 | 43 | 0.0698 | 30 | 0.1333 | 127 | 0.0945 |
| 10_930757 | 33 | 0.8485 | 29 | 0.7586 | 35 | 0.5429 | 35 | 0.6857 | 132 | 0.7045 | 20 | 0.7 | 31 | 0.4839 | 36 | 0.6389 | 24 | 0.5833 | 111 | 0.5946 |
| 10_930764 | 41 | 0.2439 | 36 | 0.2778 | 37 | 0.4054 | 34 | 0.2353 | 148 | 0.2905 | 19 | 0.3158 | 33 | 0.3636 | 37 | 0.1622 | 26 | 0.3077 | 115 | 0.2783 |
| 10_930765 | 45 | 0.4222 | 32 | 0.6562 | 47 | 0.4468 | 35 | 0.5143 | 159 | 0.4969 | 18 | 0.4444 | 38 | 0.5 | 41 | 0.439 | 29 | 0.5172 | 126 | 0.4762 |
| 10_930765 | 47 | 0.1277 | 37 | 0.1622 | 50 | 0.14 | 36 | 0.0556 | 170 | 0.1235 | 23 | 0.1739 | 38 | 0.1053 | 45 | 0.1333 | 35 | 0.2571 | 141 | 0.1631 |
| 10_930766 | 49 | 0.0408 | 37 | 0.1081 | 51 | 0.0392 | 37 | 0.0541 | 174 | 0.0575 | 24 | 0 | 38 | 0.0789 | 43 | 0.093 | 31 | 0.0645 | 136 | 0.0662 |
| 10_930766 | 45 | 0.1556 | 33 | 0.303 | 47 | 0.3404 | 39 | 0.2564 | 164 | 0.2622 | 23 | 0.3043 | 36 | 0.25 | 38 | 0.3421 | 34 | 0.4118 | 131 | 0.3282 |
| 10_930835 | NA | NA | 1 | 1 | NA | NA | NA | NA | 1 | 1 | 1 | 1 | NA | NA | NA | NA | NA | NA | 1 | 1 |
| 10_930837 | NA | NA | 1 | 1 | NA | NA | NA | NA | 1 | 1 | 1 | 1 | NA | NA | NA | NA | NA | NA | 1 | 1 |
| 10_930845 | NA | NA | 1 | 1 | NA | NA | NA | NA | 1 | 1 | 1 | 1 | NA | NA | NA | NA | NA | NA | 1 | 1 |
| 10_930846 | NA | NA | 1 | 1 | NA | NA | NA | NA | 1 | 1 | 1 | 1 | NA | NA | NA | NA | NA | NA | 1 | 1 |
| 10_930852 | 1 | 1 | 1 | 0 | 1 | 0 | NA | NA | 3 | 0.3333 | 2 | 0.5 | 1 | 0 | 1 | 0 | NA | NA | 4 | 0.25 |
| 10_930852 | 1 | 1 | 2 | 0 | 1 | 0 | NA | NA | 4 | 0.25 | 2 | 0.5 | 1 | 0 | 1 | 1 | NA | NA | 4 | 0.5 |
| 10_930853 | 1 | 0 | 1 | 1 | 1 | 0 | NA | NA | 3 | 0.3333 | 2 | 0.5 | 1 | 0 | 1 | 0 | 1 | 1 | 5 | 0.4 |
| 10_930853 | NA | NA | 2 | 0.5 | NA | NA | NA | NA | 2 | 0.5 | 2 | 0.5 | 1 | 0 | 1 | 0 | NA | NA | 4 | 0.25 |
| 10_930857 | 1 | 1 | 2 | 1 | 1 | 0 | 1 | 0 | 5 | 0.6 | 2 | 0.5 | 1 | 0 | 1 | 1 | 1 | 1 | 5 | 0.6 |
| 10_930858 | NA | NA | 2 | 1 | 1 | 0 | 1 | 1 | 4 | 0.75 | 1 | 1 | 1 | 0 | 1 | 0 | NA | NA | 3 | 0.3333 |
| 10_931009 | NA | NA | 2 | 0.5 | 1 | 0 | 1 | 0 | 4 | 0.25 | NA | NA | NA | NA | 1 | 0 | 1 | 1 | 2 | 0.5 |
| 10_931012 | NA | NA | 1 | 1 | NA | NA | 1 | 1 | 2 | 1 | NA | NA | NA | NA | NA | NA | 1 | 1 | 1 | 1 |
| 10_933206 | 5 | 0 | 8 | 0 | 1 | 0 | 5 | 0.2 | 19 | 0.0526 | 10 | 0.2 | 5 | 0.2 | 6 | 0.1667 | 6 | 0 | 27 | 0.1481 |
| 10_934804 | NA | NA | 1 | 1 | 1 | 1 | NA | NA | 2 | 1 | 1 | 1 | NA | NA | 1 | 1 | 2 | 0.5 | 4 | 0.75 |
| 10_934807 | NA | NA | NA | NA | 1 | 1 | NA | NA | 1 | 1 | 1 | 1 | NA | NA | 1 | 1 | 2 | 1 | 4 | 1 |
| 10_934807 | NA | NA | 1 | 1 | NA | NA | NA | NA | 1 | 1 | 1 | 0 | NA | NA | 2 | 1 | 2 | 1 | 5 | 0.8 |
| 10_935817 | NA | NA | 2 | 0.5 | 2 | 1 | NA | NA | 4 | 0.75 | 1 | 1 | NA | NA | 2 | 1 | 2 | 0.5 | 5 | 0.8 |
| 10_935829 | NA | NA | 2 | 0.5 | 2 | 0 | 1 | 0 | 5 | 0.2 | 1 | 0 | NA | NA | 1 | 0 | 2 | 0.5 | 4 | 0.25 |
| 10_936967 | 1 | 1 | 1 | 1 | NA | NA | NA | NA | 2 | 1 | NA | NA | NA | NA | 1 | 0 | 1 | 1 | 2 | 0.5 |
| 10_936968 | 7 | 0 | 10 | 0.4 | 4 | 0 | 6 | 0 | 27 | 0.1481 | 13 | 0 | 11 | 0.0909 | 16 | 0 | 18 | 0.1111 | 58 | 0.0517 |
| 10_936970 | 9 | 0.1111 | 14 | 0.2143 | 6 | 0.3333 | 12 | 0 | 41 | 0.1463 | 17 | 0.2941 | 13 | 0.0769 | 22 | 0.1818 | 22 | 0.1818 | 74 | 0.1892 |
| 10_936971 | 9 | 0.3333 | 14 | 0.3571 | 5 | 0.6 | 12 | 0.25 | 40 | 0.35 | 13 | 0.2308 | 13 | 0.0769 | 21 | 0.2381 | 19 | 0.2632 | 66 | 0.2121 |
| 10_936972 | 5 | 0.2 | 11 | 0 | 5 | 0 | 10 | 0.1 | 31 | 0.0645 | 11 | 0 | 12 | 0.0833 | 16 | 0.0625 | 19 | 0.1579 | 58 | 0.0862 |
| 10_936973 | 8 | 0.125 | 12 | 0.0833 | 6 | 0 | 10 | 0.3 | 36 | 0.1389 | 12 | 0.1667 | 12 | 0 | 17 | 0.1765 | 18 | 0.1111 | 59 | 0.1186 |
| 10_936973 | 7 | 0.2857 | 13 | 0.1538 | 5 | 0.2 | 11 | 0.1818 | 36 | 0.1944 | 12 | 0 | 13 | 0.0769 | 21 | 0.0476 | 20 | 0.2 | 66 | 0.0909 |
| 10_937979 | 2 | 0 | 2 | 0.5 | 1 | 0 | NA | NA | 5 | 0.2 | NA | NA | 2 | 0.5 | 1 | 1 | NA | NA | 3 | 0.6667 |
| 10_937984 | 2 | 0.5 | 1 | 0 | 1 | 1 | 2 | 0 | 6 | 0.3333 | NA | NA | 2 | 1 | 2 | 1 | NA | NA | 4 | 1 |
| 10_937993 | 1 | 1 | 5 | 0 | NA | NA | 1 | 0 | 7 | 0.1429 | NA | NA | 2 | 0.5 | 2 | 0 | NA | NA | 4 | 0.25 |
| 10_937998 | 2 | 0.5 | 3 | 0.3333 | NA | NA | NA | NA | 5 | 0.4 | NA | NA | 1 | 0 | 2 | 0.5 | NA | NA | 3 | 0.3333 |
| 10_937999 | 2 | 0.5 | 4 | 0 | NA | NA | NA | NA | 6 | 0.1667 | NA | NA | 2 | 0 | 2 | 0.5 | NA | NA | 4 | 0.25 |
| 10_938000 | 2 | 1 | 4 | 0.5 | NA | NA | NA | NA | 6 | 0.6667 | NA | NA | 2 | 0 | 2 | 0.5 | NA | NA | 4 | 0.25 |
| 10_938028 | NA | NA | 1 | 1 | NA | NA | NA | NA | 1 | 1 | NA | NA | NA | NA | 1 | 1 | NA | NA | 1 | 1 |
| 10_938049 | 1 | 1 | NA | NA | NA | NA | NA | NA | 1 | 1 | NA | NA | NA | NA | 1 | 1 | NA | NA | 1 | 1 |
| 10_938055 | 6 | 0 | 5 | 0 | 3 | 0 | 6 | 0.3333 | 20 | 0.1 | 1 | 0 | 4 | 0.25 | 5 | 0 | 6 | 0 | 16 | 0.0625 |
| 10_938057 | 4 | 0.25 | 3 | 0 | 2 | 0 | 3 | 0.3333 | 12 | 0.1667 | NA | NA | 3 | 0 | 4 | 0.25 | 5 | 0 | 12 | 0.0833 |
| 10_938057 | 6 | 0 | 4 | 0.25 | 2 | 0 | 8 | 0.125 | 20 | 0.1 | 1 | 0 | 4 | 0.25 | 5 | 0 | 5 | 0.8 | 15 | 0.3333 |
| 10_938058 | 5 | 0.2 | 2 | 0.5 | 2 | 0 | 4 | 0.25 | 13 | 0.2308 | 1 | 0 | 2 | 0 | 4 | 0.5 | 4 | 0.5 | 11 | 0.3636 |
| 10_938058 | 6 | 0.6667 | 5 | 0.6 | 1 | 1 | 8 | 0.25 | 20 | 0.5 | 1 | 1 | 3 | 0.3333 | 7 | 0.7143 | 5 | 0.4 | 16 | 0.5625 |
| 10_938059 | 5 | 1 | 4 | 1 | 1 | 1 | 4 | 1 | 14 | 1 | NA | NA | 4 | 1 | 5 | 0.8 | 5 | 0.4 | 14 | 0.7143 |
| 10_938060 | 5 | 0.2 | 4 | 0 | 2 | 0.5 | 6 | 0.1667 | 17 | 0.1765 | NA | NA | 9 | 0.1111 | 5 | 0 | 6 | 0.1667 | 20 | 0.1 |
| 10_938061 | 5 | 1 | 5 | 1 | 3 | 1 | 8 | 0.75 | 21 | 0.9048 | NA | NA | 7 | 0.7143 | 4 | 0.5 | 4 | 0.75 | 15 | 0.6667 |
| 10_938061 | 4 | 0 | 3 | 0.3333 | 3 | 0 | 6 | 0.1667 | 16 | 0.125 | NA | NA | 7 | 0.2857 | 4 | 0.25 | 4 | 0.25 | 15 | 0.2667 |

| | | | | | | | | | | | | | | | | | | | | |
|---|---|---|---|---|---|---|---|---|---|---|---|---|---|---|---|---|---|---|---|---|
| 10_938062 | 6 | 0.8333 | 5 | 1 | 3 | 1 | 9 | 1 | 23 | 0.9565 | NA | NA | 8 | 1 | 6 | 1 | 6 | 0.8333 | 20 | 0.95 |
| 10_938063 | 6 | 0 | 4 | 0.25 | 2 | 0 | 8 | 0 | 20 | 0.05 | NA | NA | 8 | 0 | 5 | 0 | 3 | 0.3333 | 16 | 0.0625 |
| 10_938063 | 5 | 0.2 | 3 | 0.3333 | 3 | 0 | 8 | 0.125 | 19 | 0.1579 | NA | NA | 7 | 0 | 5 | 0.2 | 2 | 0.5 | 14 | 0.1429 |
| 10_938063 | 7 | 0.5714 | 4 | 0.5 | 3 | 1 | 7 | 0.7143 | 21 | 0.6667 | NA | NA | 6 | 0.5 | 6 | 0.6667 | 4 | 0.75 | 16 | 0.625 |
| 10_938064 | 7 | 1 | 3 | 1 | 3 | 0.6667 | 6 | 0.8333 | 19 | 0.8947 | NA | NA | 8 | 0.75 | 6 | 1 | 4 | 1 | 18 | 0.8889 |
| 10_938064 | 6 | 0.1667 | 3 | 0.6667 | 3 | 0.3333 | 8 | 0 | 20 | 0.2 | NA | NA | 7 | 0.2857 | 6 | 0 | 3 | 0 | 16 | 0.125 |
| 10_938064 | 6 | 0 | 5 | 0.6 | 2 | 0.5 | 9 | 0.1111 | 22 | 0.2273 | NA | NA | 7 | 0 | 6 | 0.5 | 4 | 0.25 | 17 | 0.2353 |
| 10_938065 | 6 | 0 | 5 | 0 | 3 | 0 | 6 | 0.1667 | 20 | 0.05 | NA | NA | 6 | 0 | 4 | 0 | 2 | 0.5 | 12 | 0.0833 |
| 10_938065 | 6 | 0 | 5 | 0.2 | 3 | 0.3333 | 6 | 0.1667 | 20 | 0.15 | NA | NA | 8 | 0.5 | 6 | 0 | 4 | 0.5 | 18 | 0.3333 |
| 10_938328 | 6 | 0.3333 | 1 | 1 | 3 | 0.6667 | 4 | 0.25 | 14 | 0.4286 | 1 | 1 | 4 | 0.5 | 3 | 0.3333 | 4 | 0.5 | 12 | 0.5 |
| 10_938328 | 5 | 0 | 2 | 0.5 | 2 | 0.5 | 2 | 0.5 | 11 | 0.2727 | 1 | 1 | 5 | 0 | 3 | 0.3333 | 4 | 0.25 | 13 | 0.2308 |
| 10_938429 | 4 | 0.25 | 3 | 0 | 3 | 0 | 4 | 0 | 14 | 0.0714 | 1 | 0 | 5 | 0.2 | 3 | 0 | 3 | 0 | 12 | 0.0833 |
| 10_940312 | 2 | 1 | 3 | 1 | NA | NA | NA | NA | 5 | 1 | 2 | 1 | NA | NA | 1 | 0 | 1 | 0 | 4 | 0.5 |
| 10_940312 | 3 | 0 | 4 | 0.5 | 1 | 0 | 1 | 1 | 9 | 0.3333 | 1 | 1 | NA | NA | 2 | 0 | 1 | 0 | 4 | 0.25 |
| 10_940313 | 3 | 1 | 4 | 0.75 | 1 | 1 | NA | NA | 8 | 0.875 | 3 | 0.6667 | NA | NA | 2 | 1 | 1 | 0 | 6 | 0.6667 |
| 10_940566 | 39 | 0.0513 | 47 | 0.1277 | 28 | 0.0357 | 37 | 0.027 | 151 | 0.0662 | 40 | 0.05 | 36 | 0.0556 | 33 | 0.0909 | 42 | 0.0238 | 151 | 0.053 |
| 10_940567 | 38 | 0.2895 | 50 | 0.22 | 29 | 0.3448 | 41 | 0.2439 | 158 | 0.2658 | 42 | 0.2143 | 40 | 0.325 | 39 | 0.2564 | 39 | 0.3333 | 160 | 0.2812 |
| 10_940570 | 5 | 0 | 7 | 0.1429 | 2 | 0.5 | 8 | 0.125 | 22 | 0.1364 | 4 | 0.25 | 9 | 0.2222 | 5 | 0.2 | 5 | 0.2 | 23 | 0.2174 |
| 10_944144 | 2 | 0.5 | 2 | 0.5 | 1 | 1 | NA | NA | 5 | 0.6 | NA | NA | NA | NA | 1 | 1 | 1 | 0 | 2 | 0.5 |
| 10_944146 | 2 | 1 | 2 | 1 | 1 | 1 | NA | NA | 5 | 1 | NA | NA | NA | NA | 1 | 1 | 1 | 0 | 2 | 0.5 |
| 10_944150 | 2 | 0.5 | 2 | 0 | NA | NA | NA | NA | 4 | 0.25 | NA | NA | 1 | 0 | NA | NA | 1 | 1 | 2 | 0.5 |
| 10_944150 | 2 | 0 | 2 | 0.5 | 1 | 1 | NA | NA | 5 | 0.4 | NA | NA | 1 | 0 | 1 | 1 | 1 | 0 | 3 | 0.3333 |
| 10_944228 | 2 | 1 | 3 | 1 | 1 | 1 | NA | NA | 6 | 1 | NA | NA | NA | NA | 1 | 1 | 1 | 1 | 2 | 1 |
| 10_944228 | 2 | 1 | 3 | 1 | 1 | 1 | NA | NA | 6 | 1 | NA | NA | 1 | 1 | NA | NA | 1 | 1 | 2 | 1 |
| 10_944228 | 2 | 1 | 3 | 0.6667 | 1 | 1 | NA | NA | 6 | 0.8333 | NA | NA | 1 | 1 | 1 | 1 | NA | NA | 2 | 1 |
| 10_944230 | 2 | 0 | 3 | 0.6667 | 1 | 1 | NA | NA | 6 | 0.5 | 1 | 1 | 1 | 1 | 2 | 1 | 2 | 0 | 6 | 0.6667 |
| 10_944230 | 1 | 1 | 3 | 1 | 1 | 1 | NA | NA | 5 | 1 | 1 | 1 | 1 | 1 | 2 | 1 | 2 | 1 | 6 | 1 |
| 10_944230 | 2 | 0.5 | 3 | 0.3333 | 1 | 1 | NA | NA | 6 | 0.5 | 1 | 1 | 1 | 1 | 2 | 0.5 | NA | NA | 4 | 0.75 |
| 10_944232 | 2 | 1 | 3 | 0.6667 | 1 | 1 | NA | NA | 6 | 0.8333 | 1 | 0 | 1 | 1 | 2 | 1 | 2 | 1 | 6 | 0.8333 |
| 10_944232 | 2 | 1 | 3 | 0.6667 | 1 | 1 | NA | NA | 6 | 0.8333 | 1 | 0 | 1 | 1 | 2 | 1 | 2 | 1 | 6 | 0.8333 |
| 10_944236 | 3 | 0 | 2 | 0 | 1 | 0 | 1 | 1 | 7 | 0.1429 | 2 | 0 | 1 | 1 | 2 | 0 | 3 | 0 | 8 | 0.125 |
| 10_944238 | 1 | 0 | 3 | 0.3333 | 1 | 0 | NA | NA | 5 | 0.2 | 1 | 0 | 1 | 1 | 1 | 0 | 1 | 0 | 4 | 0.25 |
| 10_944250 | 2 | 0 | 2 | 0 | 1 | 0 | 2 | 0.5 | 7 | 0.1429 | 2 | 0.5 | 1 | 0 | 1 | 0 | 2 | 0.5 | 6 | 0.3333 |
| 10_944256 | 4 | 0.25 | 3 | 0 | 2 | 0 | 3 | 0 | 12 | 0.0833 | 3 | 0 | 2 | 0 | 2 | 0 | 3 | 0.3333 | 10 | 0.1 |
| 10_944257 | 1 | 0 | 4 | 0.25 | 2 | 0 | 2 | 0.5 | 9 | 0.2222 | 4 | 0.25 | NA | NA | 1 | 0 | 3 | 0.3333 | 8 | 0.25 |
| 10_944262 | 3 | 0 | 5 | 0.4 | 5 | 0.2 | 3 | 0.3333 | 16 | 0.25 | 5 | 0.2 | 3 | 0.6667 | 3 | 0 | 5 | 0 | 16 | 0.1875 |
| 10_944266 | 5 | 0.2 | 7 | 0.1429 | 4 | 0.5 | 4 | 0.25 | 20 | 0.25 | 6 | 0 | 6 | 0.3333 | 3 | 0.3333 | 5 | 0 | 20 | 0.15 |
| 10_945076 | 1 | 0 | 1 | 1 | NA | NA | NA | NA | 2 | 0.5 | NA | NA | 2 | 1 | NA | NA | 1 | 1 | 3 | 1 |
| 10_945087 | 1 | 1 | NA | NA | NA | NA | NA | NA | 1 | 1 | NA | NA | 1 | 1 | NA | NA | 1 | 1 | 2 | 1 |
| 10_945088 | 2 | 1 | NA | NA | NA | NA | NA | NA | 2 | 1 | NA | NA | NA | NA | NA | NA | 1 | 1 | 1 | 1 |
| 10_945089 | 2 | 0.5 | NA | NA | NA | NA | NA | NA | 2 | 0.5 | 1 | 1 | 1 | 1 | NA | NA | 1 | 1 | 3 | 1 |
| 10_945090 | 1 | 1 | NA | NA | NA | NA | NA | NA | 1 | 1 | 1 | 0 | NA | NA | NA | NA | 1 | 1 | 2 | 0.5 |
| 10_945090 | 2 | 1 | NA | NA | NA | NA | NA | NA | 2 | 1 | 1 | 1 | 1 | 1 | NA | NA | 1 | 1 | 3 | 1 |
| 10_945090 | 2 | 0.5 | 1 | 0 | NA | NA | NA | NA | 3 | 0.3333 | 1 | 1 | 1 | 0 | NA | NA | 1 | 1 | 3 | 0.6667 |
| 10_945092 | 2 | 1 | NA | NA | NA | NA | NA | NA | 2 | 1 | 1 | 1 | 1 | 1 | NA | NA | 1 | 1 | 3 | 1 |
| 10_945094 | 1 | 1 | NA | NA | NA | NA | NA | NA | 1 | 1 | 1 | 1 | 1 | 0 | NA | NA | 1 | 1 | 3 | 0.6667 |
| 10_945096 | 7 | 0.2857 | 3 | 1 | 2 | 0.5 | 2 | 0 | 14 | 0.4286 | 2 | 0.5 | 4 | 0.75 | 4 | 0.75 | 3 | 0 | 13 | 0.5385 |
| 10_945098 | 6 | 0.1667 | 4 | 0 | 2 | 0 | 2 | 0 | 14 | 0.0714 | 2 | 0.5 | 4 | 0 | 1 | 0 | 2 | 0 | 9 | 0.1111 |
| 10_945106 | 4 | 0 | 4 | 0.25 | 2 | 0 | 2 | 0 | 12 | 0.0833 | 2 | 0 | 5 | 0 | 3 | 0 | 2 | 0.5 | 12 | 0.0833 |
| 10_945128 | 1 | 0 | 3 | 0 | NA | NA | 1 | 1 | 5 | 0.2 | 2 | 0.5 | 1 | 0 | 1 | 0 | NA | NA | 4 | 0.25 |
| 10_945129 | 2 | 1 | 2 | 0 | NA | NA | 2 | 0.5 | 6 | 0.5 | 2 | 0 | 2 | 0.5 | 1 | 0 | 1 | 0 | 6 | 0.1667 |
| 10_945548 | 3 | 0.6667 | NA | NA | NA | NA | 1 | 0 | 4 | 0.5 | 1 | 1 | 4 | 0 | 1 | 0 | 3 | 0 | 9 | 0.1111 |
| 10_945566 | 1 | 1 | NA | NA | NA | NA | NA | NA | 1 | 1 | NA | NA | NA | NA | NA | NA | 2 | 0.5 | 2 | 0.5 |
| 10_945568 | 2 | 0 | NA | NA | 1 | 1 | NA | NA | 3 | 0.3333 | NA | NA | NA | NA | NA | NA | 3 | 0.3333 | 3 | 0.3333 |
| 10_946256 | 4 | 0.25 | NA | NA | 2 | 0 | 4 | 0 | 10 | 0.1 | 3 | 0 | 5 | 0 | 1 | 1 | 4 | 0.25 | 13 | 0.1538 |
| 10_946256 | 3 | 1 | NA | NA | 2 | 0.5 | 4 | 0.5 | 9 | 0.6667 | 4 | 0.75 | 5 | 0.6 | 1 | 1 | 4 | 0.5 | 14 | 0.6429 |
| 10_946258 | 3 | 0.3333 | NA | NA | 4 | 0.25 | 4 | 0.25 | 11 | 0.2727 | 5 | 0.4 | 5 | 0.2 | 1 | 0 | 4 | 0 | 15 | 0.2 |
| 10_946277 | NA | NA | 4 | 0.5 | 1 | 1 | 3 | 0.3333 | 8 | 0.5 | 1 | 0 | 3 | 0 | 1 | 0 | 3 | 0.6667 | 8 | 0.25 |
| 10_950106 | 1 | 1 | 1 | 0 | NA | NA | NA | NA | 2 | 0.5 | NA | NA | NA | NA | 2 | 0.5 | NA | NA | 2 | 0.5 |
| 10_950111 | 3 | 0.6667 | 2 | 0.5 | 1 | 0 | 1 | 1 | 7 | 0.5714 | NA | NA | NA | NA | 2 | 0.5 | NA | NA | 2 | 0.5 |
| 10_950113 | NA | NA | NA | NA | 1 | 1 | NA | NA | 1 | 1 | NA | NA | NA | NA | 1 | 1 | NA | NA | 1 | 1 |
| 10_950113 | 3 | 0 | 1 | 1 | NA | NA | NA | NA | 4 | 0.25 | NA | NA | NA | NA | 2 | 0.5 | NA | NA | 2 | 0.5 |
| 10_950113 | 3 | 0.6667 | 2 | 0.5 | 1 | 0 | 1 | 1 | 7 | 0.5714 | NA | NA | NA | NA | 2 | 1 | NA | NA | 2 | 1 |
| 10_950114 | 2 | 0.5 | 1 | 1 | 1 | 1 | NA | NA | 4 | 0.75 | NA | NA | NA | NA | 2 | 1 | NA | NA | 2 | 1 |
| 10_950117 | 3 | 0.3333 | 1 | 1 | 1 | 0 | 1 | 0 | 6 | 0.3333 | NA | NA | NA | NA | 2 | 1 | NA | NA | 2 | 1 |

| | | | | | | | | | | | | | | | | | | | | |
|---|---|---|---|---|---|---|---|---|---|---|---|---|---|---|---|---|---|---|---|---|
| 10_9515857 | 2 | 0 | 1 | 1 | 1 | 0 | NA | NA | 4 | 0.25 | NA | NA | NA | NA | 1 | 1 | NA | NA | 1 | 1 |
| 10_9515867 | 3 | 1 | 1 | 1 | 1 | 1 | 1 | 1 | 6 | 1 | 1 | 1 | NA | NA | 1 | 1 | NA | NA | 2 | 1 |
| 10_9522368 | 1 | 1 | 1 | 0 | NA | NA | NA | NA | 2 | 0.5 | NA | NA | NA | NA | NA | NA | 1 | 1 | 1 | 1 |
| 10_9524755 | 2 | 1 | 1 | 1 | 1 | 0 | 3 | 0 | 7 | 0.4286 | NA | NA | NA | NA | 1 | 1 | 1 | 0 | 2 | 0.5 |
| 10_9524763 | 2 | 1 | 1 | 1 | 1 | 0 | 3 | 0.6667 | 7 | 0.7143 | NA | NA | NA | NA | 1 | 0 | 1 | 1 | 2 | 0.5 |
| 10_9524773 | 2 | 0.5 | 1 | 0 | 1 | 1 | 3 | 0.6667 | 7 | 0.5714 | NA | NA | NA | NA | 1 | 0 | 1 | 1 | 2 | 0.5 |
| 10_9524845 | 1 | 1 | 1 | 0 | 1 | 1 | 3 | 0.3333 | 6 | 0.5 | NA | NA | NA | NA | NA | NA | 1 | 1 | 1 | 1 |
| 10_9524855 | 2 | 0 | 1 | 0 | 1 | 1 | 3 | 0.6667 | 7 | 0.4286 | NA | NA | NA | NA | NA | NA | 1 | 1 | 1 | 1 |
| 10_9524857 | 2 | 1 | 1 | 1 | 1 | 1 | 3 | 1 | 7 | 1 | NA | NA | NA | NA | NA | NA | 1 | 1 | 1 | 1 |
| 10_9524867 | 2 | 0.5 | NA | NA | 1 | 1 | 2 | 1 | 5 | 0.8 | NA | NA | NA | NA | NA | NA | 1 | 1 | 1 | 1 |
| 10_9524870 | 2 | 1 | NA | NA | 1 | 1 | 3 | 1 | 6 | 1 | NA | NA | NA | NA | NA | NA | 1 | 1 | 1 | 1 |
| 10_9524875 | 2 | 0 | NA | NA | 1 | 0 | 3 | 0.3333 | 6 | 0.1667 | NA | NA | NA | NA | NA | NA | 1 | 1 | 1 | 1 |
| 10_9524887 | 2 | 0 | 1 | 0 | 1 | 0 | 1 | 1 | 5 | 0.2 | NA | NA | NA | NA | NA | NA | 1 | 1 | 1 | 1 |
| 10_9535144 | NA | NA | 2 | 0.5 | 1 | 1 | NA | NA | 3 | 0.6667 | 1 | 0 | 1 | 1 | NA | NA | NA | NA | 2 | 0.5 |
| 10_9537718 | 20 | 0.6 | 32 | 0.5625 | 15 | 0.6667 | 13 | 0.5385 | 80 | 0.5875 | 17 | 0.7647 | 21 | 0.5714 | 15 | 0.4 | 16 | 0.6875 | 69 | 0.6087 |
| 10_9537720 | 16 | 0.1875 | 27 | 0.2222 | 13 | 0.2308 | 11 | 0 | 67 | 0.1791 | 18 | 0.2222 | 19 | 0 | 14 | 0.2143 | 13 | 0.2308 | 64 | 0.1562 |
| 10_9537814 | 18 | 0.2778 | 34 | 0.1765 | 13 | 0.1538 | 7 | 0.2857 | 72 | 0.2083 | 17 | 0.3529 | 17 | 0.0588 | 12 | 0.25 | 16 | 0.25 | 62 | 0.2258 |
| 10_9537825 | 23 | 0.6087 | 35 | 0.6 | 19 | 0.9474 | 11 | 0.5455 | 88 | 0.6705 | 19 | 0.8421 | 21 | 0.7619 | 18 | 0.7778 | 16 | 0.625 | 74 | 0.7568 |
| 10_9537841 | 22 | 0.1364 | 36 | 0.1111 | 17 | 0.0588 | 15 | 0 | 90 | 0.0889 | 22 | 0.1818 | 22 | 0 | 17 | 0.0588 | 15 | 0.2667 | 76 | 0.1184 |
| 10_9537873 | 30 | 0.1333 | 38 | 0.0263 | 19 | 0 | 19 | 0.0526 | 106 | 0.0566 | 24 | 0.125 | 26 | 0.1154 | 16 | 0 | 19 | 0.1579 | 85 | 0.1059 |
| 10_9538059 | 7 | 0.1429 | 16 | 0.125 | 8 | 0.125 | 11 | 0.1818 | 42 | 0.1429 | 7 | 0.5714 | 14 | 0.1429 | 7 | 0.1429 | 9 | 0.1111 | 37 | 0.2162 |
| 10_9538075 | 6 | 0.5 | 14 | 0.6429 | 7 | 0.4286 | 11 | 0.4545 | 38 | 0.5263 | 6 | 0.6667 | 14 | 0.5714 | 5 | 0.6 | 9 | 0.3333 | 34 | 0.5294 |
| 10_9538117 | 8 | 0 | 19 | 0.2632 | 7 | 0.1429 | 10 | 0.1 | 44 | 0.1591 | 5 | 0.4 | 14 | 0.1429 | 9 | 0.3333 | 10 | 0 | 38 | 0.1842 |
| 10_9538309 | 12 | 0.4167 | 13 | 0.1538 | 12 | 0.4167 | 13 | 0.0769 | 50 | 0.26 | 9 | 0 | 13 | 0.2308 | 13 | 0.2308 | 12 | 0.25 | 47 | 0.1915 |
| 10_9538311 | 12 | 0.0833 | 16 | 0 | 16 | 0.125 | 14 | 0 | 58 | 0.0517 | 13 | 0.2308 | 13 | 0 | 11 | 0.1818 | 13 | 0.1538 | 50 | 0.14 |
| 10_9538345 | 5 | 0 | 6 | 0 | 5 | 0.2 | 8 | 0.125 | 24 | 0.0833 | 4 | 0.25 | 7 | 0 | 7 | 0.2857 | 6 | 0.3333 | 24 | 0.2083 |
| 10_9538365 | 11 | 0.2727 | 10 | 0.3 | 14 | 0.3571 | 15 | 0.3333 | 50 | 0.32 | 12 | 0.25 | 13 | 0.5385 | 9 | 0.5556 | 11 | 0.1818 | 45 | 0.3778 |
| 10_9538440 | 6 | 0 | 8 | 0 | 12 | 0.0833 | 9 | 0.1111 | 35 | 0.0571 | 7 | 0.1429 | 9 | 0.1111 | 6 | 0.3333 | 10 | 0 | 32 | 0.125 |
| 10_9538519 | 7 | 0.4286 | 9 | 0.2222 | 9 | 0.3333 | 10 | 0 | 35 | 0.2286 | 7 | 0.2857 | 7 | 0 | 8 | 0.125 | 10 | 0.1 | 32 | 0.125 |
| 10_9587256 | 1 | 0 | 3 | 0.3333 | 1 | 0 | 1 | 0 | 6 | 0.1667 | NA | NA | NA | NA | 4 | 0.25 | 3 | 0 | 7 | 0.1429 |
| 10_9587810 | 3 | 0.3333 | 6 | 0 | 1 | 0 | 1 | 0 | 11 | 0.0909 | NA | NA | NA | NA | 3 | 0 | 3 | 0.3333 | 6 | 0.1667 |
| 10_9587812 | 1 | 1 | 6 | 1 | 1 | 1 | 1 | 1 | 9 | 1 | NA | NA | NA | NA | 5 | 1 | 3 | 1 | 8 | 1 |
| 10_9587821 | 3 | 0 | 6 | 0.3333 | 1 | 0 | 1 | 0 | 11 | 0.1818 | NA | NA | NA | NA | 6 | 0.1667 | 4 | 0 | 10 | 0.1 |
| 10_9588010 | 3 | 0.3333 | 7 | 0 | NA | NA | 1 | 1 | 11 | 0.1818 | NA | NA | 2 | 0 | 10 | 0.2 | 4 | 0.5 | 16 | 0.25 |
| 10_9588040 | 3 | 0 | 7 | 0.1429 | NA | NA | 1 | 0 | 11 | 0.0909 | NA | NA | 2 | 0 | 9 | 0.2222 | 5 | 0 | 16 | 0.125 |
| 10_9588074 | 5 | 0 | 9 | 0.1111 | NA | NA | 1 | 0 | 15 | 0.0667 | NA | NA | 2 | 0 | 8 | 0.125 | 4 | 0 | 14 | 0.0714 |
| 10_9645301 | 4 | 0 | 9 | 0.1111 | 9 | 0.1111 | 5 | 0 | 27 | 0.0741 | 9 | 0.1111 | 11 | 0.2727 | 14 | 0 | 5 | 0.2 | 39 | 0.1282 |
| 10_9645423 | 7 | 0.4286 | 10 | 0.7 | 9 | 0.3333 | 9 | 0.2222 | 35 | 0.4286 | 9 | 0.2222 | 11 | 0.3636 | 14 | 0.7143 | 4 | 0.75 | 38 | 0.5 |
| 10_9645425 | 5 | 1 | 7 | 0.8571 | 9 | 1 | 9 | 0.8889 | 30 | 0.9333 | 10 | 0.9 | 10 | 1 | 15 | 1 | 5 | 1 | 40 | 0.975 |
| 10_9645429 | 3 | 0.3333 | 3 | 1 | 4 | 0.5 | 5 | 0.4 | 15 | 0.5333 | 5 | 0.8 | 4 | 0.5 | 7 | 0.5714 | 5 | 0.8 | 21 | 0.6667 |
| 10_9645442 | 6 | 0 | 10 | 0.1 | 9 | 0.1111 | 10 | 0.2 | 35 | 0.1143 | 10 | 0.1 | 12 | 0 | 17 | 0.1765 | 6 | 0.1667 | 45 | 0.1111 |
| 10_9645444 | 2 | 0.5 | 3 | 1 | 2 | 0.5 | 7 | 0.7143 | 14 | 0.7143 | 5 | 0.8 | 8 | 0.875 | 4 | 1 | 3 | 1 | 20 | 0.9 |
| 10_9645447 | 5 | 0.4 | 7 | 0.4286 | 6 | 0.1667 | 8 | 0.375 | 26 | 0.3462 | 6 | 0.6667 | 12 | 0.5 | 9 | 0.4444 | 4 | 0.5 | 31 | 0.5161 |
| 10_9645449 | 6 | 0.8333 | 4 | 1 | 7 | 0.8571 | 8 | 1 | 25 | 0.92 | 9 | 0.6667 | 10 | 0.9 | 11 | 0.9091 | 3 | 0.6667 | 33 | 0.8182 |
| 10_9645470 | 6 | 0.1667 | 8 | 0.25 | 5 | 0.2 | 5 | 0.4 | 24 | 0.25 | 4 | 0 | 11 | 0.4545 | 15 | 0.0667 | 3 | 0.3333 | 33 | 0.2121 |
| 10_9645533 | 11 | 0 | 12 | 0.1667 | 15 | 0.0667 | 14 | 0.0714 | 52 | 0.0769 | 9 | 0 | 17 | 0.1765 | 25 | 0.08 | 10 | 0.2 | 61 | 0.1148 |
| 10_9734254 | 8 | 0.125 | 3 | 0 | 1 | 0 | 2 | 0 | 14 | 0.0714 | NA | NA | NA | NA | 2 | 0.5 | 2 | 0 | 4 | 0.25 |
| 10_9750966 | 2 | 0 | 1 | 0 | 3 | 0.3333 | 6 | 0.3333 | 12 | 0.25 | NA | NA | 3 | 0.3333 | 4 | 0 | NA | NA | 7 | 0.1429 |
| 10_9751335 | 1 | 1 | NA | NA | NA | NA | 1 | 1 | 2 | 1 | NA | NA | 1 | 0 | NA | NA | 1 | 1 | 2 | 0.5 |
| 10_9751833 | NA | NA | 1 | 1 | NA | NA | NA | NA | 1 | 1 | NA | NA | NA | NA | 3 | 1 | 1 | 1 | 4 | 1 |
| 10_9880474 | 4 | 0.25 | NA | NA | 3 | 0.6667 | 5 | 0.4 | 12 | 0.4167 | 4 | 0.25 | 4 | 0 | 1 | 0 | 3 | 0.3333 | 12 | 0.1667 |
| 10_9880482 | 4 | 0.75 | 1 | 1 | 2 | 1 | 5 | 1 | 12 | 0.9167 | 4 | 0.75 | 3 | 1 | 1 | 1 | 4 | 1 | 12 | 0.9167 |
| 10_9880485 | 4 | 0.75 | 2 | 0 | 3 | 1 | 5 | 0.4 | 14 | 0.5714 | 3 | 0.3333 | 6 | 0.5 | NA | NA | 5 | 0.6 | 14 | 0.5 |
| 10_9881562 | 4 | 0.75 | NA | NA | 4 | 0 | 5 | 0.2 | 13 | 0.3077 | 4 | 0.25 | 6 | 0 | 3 | 0.6667 | 4 | 0.25 | 17 | 0.2353 |
| 10_9885353 | NA | NA | 2 | 0.5 | NA | NA | 2 | 0 | 4 | 0.25 | NA | NA | NA | NA | 1 | 1 | 4 | 0 | 5 | 0.2 |
| 10_9885892 | 23 | 0.2609 | 33 | 0.2727 | 18 | 0.1111 | 20 | 0.15 | 94 | 0.2128 | 13 | 0.2308 | 20 | 0.35 | 35 | 0.2286 | 21 | 0.381 | 89 | 0.2921 |
| 10_9885902 | 25 | 0.12 | 37 | 0.0541 | 29 | 0.069 | 30 | 0.0333 | 121 | 0.0661 | 16 | 0.0625 | 30 | 0.1 | 39 | 0.0513 | 28 | 0.1071 | 113 | 0.0796 |
| 10_9885903 | 25 | 0.72 | 32 | 0.7188 | 21 | 0.7143 | 19 | 0.8421 | 97 | 0.7423 | 13 | 0.7692 | 22 | 0.8636 | 37 | 0.8649 | 20 | 0.8 | 92 | 0.837 |
| 10_9885906 | 25 | 0.36 | 39 | 0.359 | 25 | 0.32 | 30 | 0.2333 | 119 | 0.3193 | 15 | 0.2667 | 31 | 0.4839 | 38 | 0.3947 | 26 | 0.5 | 110 | 0.4273 |
| 10_9885908 | 23 | 1 | 42 | 0.9762 | 28 | 1 | 28 | 1 | 121 | 0.9917 | 15 | 1 | 30 | 1 | 40 | 0.95 | 28 | 1 | 113 | 0.9823 |
| 10_9885909 | 25 | 0.16 | 43 | 0.0465 | 28 | 0.0714 | 30 | 0.1 | 126 | 0.0873 | 17 | 0.1765 | 30 | 0.2333 | 42 | 0.0952 | 29 | 0.1034 | 118 | 0.1441 |
| 10_9885915 | 24 | 0.9583 | 37 | 1 | 28 | 0.9643 | 26 | 0.9231 | 115 | 0.9652 | 15 | 1 | 21 | 0.8571 | 34 | 0.9412 | 25 | 0.92 | 95 | 0.9263 |
| 10_9885919 | 23 | 0.8696 | 34 | 0.8824 | 31 | 0.871 | 29 | 0.931 | 117 | 0.8889 | 17 | 1 | 30 | 0.9667 | 38 | 0.9211 | 24 | 0.9167 | 109 | 0.945 |
| 10_9885920 | 20 | 0.25 | 33 | 0.3636 | 20 | 0.2 | 17 | 0.1765 | 90 | 0.2667 | 10 | 0.5 | 24 | 0.3333 | 28 | 0.3571 | 20 | 0.4 | 82 | 0.378 |
| 10_9885921 | 27 | 0.1111 | 36 | 0.1389 | 31 | 0.0645 | 31 | 0.0968 | 125 | 0.104 | 20 | 0 | 31 | 0.0645 | 40 | 0.05 | 30 | 0.1333 | 121 | 0.0661 |

| | | | | | | | | | | | | | | | | | | | | |
|---|---|---|---|---|---|---|---|---|---|---|---|---|---|---|---|---|---|---|---|---|
| 10_9885978 | 41 | 0.1463 | 62 | 0.2097 | 54 | 0.1111 | 45 | 0.2 | 202 | 0.1683 | 26 | 0.3462 | 46 | 0.1522 | 59 | 0.2203 | 40 | 0.35 | 171 | 0.2515 |
| 10_9886055 | 13 | 0.1538 | 25 | 0.12 | 23 | 0.087 | 18 | 0.0556 | 79 | 0.1013 | 9 | 0.1111 | 24 | 0.0833 | 22 | 0.1364 | 20 | 0.1 | 75 | 0.1067 |
| 10_9886183 | 23 | 0.0435 | 34 | 0.0588 | 33 | 0.0606 | 30 | 0.0667 | 120 | 0.0583 | 17 | 0 | 30 | 0.0667 | 34 | 0.1176 | 40 | 0.025 | 121 | 0.0579 |
| 10_9894516 | NA | NA | 1 | 0 | 1 | 1 | NA | NA | 2 | 0.5 | 1 | 0 | NA | NA | NA | NA | 2 | 0.5 | 3 | 0.3333 |
| 10_9894521 | NA | NA | NA | NA | 1 | 1 | NA | NA | 1 | 1 | 1 | 1 | NA | NA | NA | NA | 2 | 1 | 3 | 1 |
| 10_9894577 | NA | NA | 1 | 1 | 1 | 1 | NA | NA | 2 | 1 | 1 | 0 | NA | NA | NA | NA | 1 | 1 | 2 | 0.5 |
| 10_9894591 | NA | NA | 1 | 1 | 1 | 1 | NA | NA | 2 | 1 | 1 | 1 | NA | NA | NA | NA | 1 | 1 | 2 | 1 |
| 10_9894593 | NA | NA | 1 | 1 | 1 | 1 | NA | NA | 2 | 1 | 1 | 1 | NA | NA | NA | NA | 2 | 1 | 3 | 1 |
| 10_9894608 | NA | NA | 1 | 1 | 1 | 1 | NA | NA | 2 | 1 | 1 | 1 | NA | NA | NA | NA | 2 | 0.5 | 3 | 0.6667 |
| 10_9894628 | NA | NA | 1 | 1 | 1 | 1 | NA | NA | 2 | 1 | 1 | 1 | NA | NA | NA | NA | 2 | 1 | 3 | 1 |
| 10_9894639 | NA | NA | 1 | 1 | 1 | 0 | NA | NA | 2 | 0.5 | 1 | 0 | NA | NA | NA | NA | 2 | 0.5 | 3 | 0.3333 |
| 10_9894687 | NA | NA | 1 | 0 | 1 | 1 | NA | NA | 2 | 0.5 | 1 | 0 | NA | NA | NA | NA | 1 | 1 | 2 | 0.5 |
| 10_9903254 | 1 | 1 | 2 | 0.5 | NA | NA | 1 | 0 | 4 | 0.5 | NA | NA | 1 | 0 | 2 | 0 | 1 | 1 | 4 | 0.25 |
| 10_9903274 | 1 | 1 | 2 | 1 | NA | NA | NA | NA | 3 | 1 | NA | NA | 1 | 1 | 2 | 0.5 | 1 | 1 | 4 | 0.75 |
| 10_9903282 | 1 | 1 | 1 | 0 | NA | NA | 1 | 0 | 3 | 0.3333 | NA | NA | 1 | 1 | 2 | 0.5 | NA | NA | 3 | 0.6667 |
| 10_9909923 | NA | NA | 5 | 0 | 4 | 0.25 | 1 | 0 | 10 | 0.1 | 1 | 0 | 1 | 0 | 2 | 0 | 2 | 0.5 | 6 | 0.1667 |
| 10_9909935 | 1 | 0 | 6 | 0 | 5 | 0.2 | 2 | 0 | 14 | 0.0714 | 1 | 0 | 1 | 0 | 2 | 0.5 | 1 | 0 | 5 | 0.2 |
| 10_9909966 | 1 | 0 | 7 | 0.8571 | 5 | 0.6 | 2 | 0.5 | 15 | 0.6667 | 1 | 1 | 1 | 0 | 2 | 0.5 | 2 | 0.5 | 6 | 0.5 |
| 10_9909980 | 1 | 0 | 5 | 0 | 5 | 0.4 | 2 | 0 | 13 | 0.1538 | 1 | 0 | 1 | 0 | 2 | 0.5 | 3 | 0 | 7 | 0.1429 |
| 10_9909987 | 1 | 0 | 5 | 0 | 4 | 0.25 | 2 | 0 | 12 | 0.0833 | 1 | 0 | 1 | 1 | 3 | 0 | 2 | 0 | 7 | 0.1429 |
| 10_9910052 | 2 | 0.5 | 8 | 0.375 | 4 | 1 | 2 | 1 | 16 | 0.625 | 1 | 1 | 1 | 1 | 4 | 1 | 3 | 1 | 9 | 1 |
| 10_9910054 | 2 | 0.5 | 6 | 0.5 | 3 | 0.6667 | 2 | 1 | 13 | 0.6154 | 1 | 1 | 1 | 1 | 2 | 0 | 3 | 0.6667 | 7 | 0.5714 |
| 10_9910058 | 2 | 0.5 | 9 | 0.8889 | 5 | 1 | 2 | 1 | 18 | 0.8889 | 1 | 1 | 1 | 1 | 5 | 0.8 | 3 | 1 | 10 | 0.9 |
| 10_9910066 | 2 | 0.5 | 9 | 0.6667 | 5 | 1 | 2 | 1 | 18 | 0.7778 | 1 | 1 | NA | NA | 5 | 0.4 | 3 | 1 | 9 | 0.6667 |
| 10_9910187 | 2 | 0.5 | 9 | 0.2222 | 5 | 0.6 | 1 | 0 | 17 | 0.3529 | 1 | 0 | 1 | 0 | 5 | 0 | 2 | 0.5 | 9 | 0.1111 |
| 10_9910204 | 3 | 0 | 10 | 0.1 | 5 | 0 | 2 | 0 | 20 | 0.05 | NA | NA | NA | NA | 5 | 0 | 4 | 0.25 | 9 | 0.1111 |
| 10_9910205 | 3 | 0.3333 | 10 | 0.6 | 6 | 0.6667 | 2 | 0 | 21 | 0.5238 | 1 | 0 | 1 | 1 | 5 | 0.4 | 4 | 1 | 11 | 0.6364 |
| 10_9910205 | 3 | 1 | 8 | 1 | 6 | 0.8333 | 2 | 0.5 | 19 | 0.8947 | 1 | 1 | 1 | 1 | 6 | 0.8333 | 4 | 0.75 | 12 | 0.8333 |
| 10_9910214 | 4 | 0.25 | 7 | 0.8571 | 5 | 0.2 | 2 | 0.5 | 18 | 0.5 | 1 | 0 | NA | NA | 6 | 0.1667 | 4 | 0.75 | 11 | 0.3636 |
| 10_9910236 | 4 | 0.25 | 6 | 0.1667 | 6 | 0.1667 | 2 | 0 | 18 | 0.1667 | 1 | 0 | NA | NA | 6 | 0.1667 | 4 | 0.25 | 11 | 0.1818 |
| 10_9910236 | 4 | 0 | 10 | 0.3 | 5 | 0 | 1 | 0 | 20 | 0.15 | 1 | 0 | 1 | 0 | 6 | 0 | 4 | 0.5 | 12 | 0.1667 |
| 10_9910242 | 4 | 0.5 | 8 | 0.5 | 6 | 0.3333 | 2 | 0 | 20 | 0.4 | 1 | 0 | 1 | 1 | 7 | 0.2857 | 4 | 0.75 | 13 | 0.4615 |
| 10_9910243 | 4 | 0.5 | 9 | 0.4444 | 6 | 0.1667 | 2 | 0.5 | 21 | 0.381 | 1 | 0 | NA | NA | 6 | 0.1667 | 4 | 0.25 | 11 | 0.1818 |
| 10_9910252 | 4 | 0.25 | 9 | 0.1111 | 7 | 0 | 2 | 0.5 | 22 | 0.1364 | 2 | 0.5 | 1 | 0 | 6 | 0.1667 | 4 | 0.25 | 13 | 0.2308 |
| 10_9910255 | 5 | 0.2 | 8 | 0.375 | 8 | 0 | 2 | 0 | 23 | 0.1739 | 3 | 0.3333 | 1 | 0 | 8 | 0.125 | 4 | 0 | 16 | 0.125 |
| 10_9910288 | 5 | 0 | 11 | 0.1818 | 8 | 0.375 | 2 | 0.5 | 26 | 0.2308 | 3 | 0 | 1 | 1 | 6 | 0 | 5 | 0.2 | 15 | 0.1333 |
| 10_9993207 | 3 | 1 | 7 | 1 | 2 | 1 | 4 | 1 | 16 | 1 | NA | NA | 4 | 1 | 4 | 0.75 | 4 | 0.75 | 12 | 0.8333 |
| 10_9993210 | 3 | 1 | 5 | 0.4 | 1 | 1 | 3 | 1 | 12 | 0.75 | NA | NA | 2 | 1 | 4 | 1 | 3 | 1 | 9 | 1 |
| 10_9993211 | 3 | 1 | 7 | 0.5714 | 2 | 1 | 4 | 1 | 16 | 0.8125 | NA | NA | 5 | 0.8 | 4 | 1 | 2 | 0.5 | 11 | 0.8182 |
| 10_9993229 | 3 | 0.6667 | 7 | 0.4286 | 2 | 0 | 4 | 0.5 | 16 | 0.4375 | NA | NA | 6 | 0.5 | 4 | 0.5 | 1 | 0 | 11 | 0.4545 |
| 10_9993230 | 2 | 1 | 6 | 0.1667 | 2 | 1 | 4 | 0.25 | 14 | 0.4286 | NA | NA | 4 | 0.5 | 3 | 0.3333 | 3 | 0 | 10 | 0.3 |
| 10_9993231 | 3 | 0.3333 | 5 | 0.2 | 2 | 1 | 3 | 1 | 13 | 0.5385 | NA | NA | 4 | 0 | 4 | 0.25 | 3 | 0 | 11 | 0.0909 |
| 10_9993232 | 1 | 0 | 5 | 0 | 2 | 0 | 1 | 1 | 9 | 0.1111 | NA | NA | 3 | 0.6667 | 4 | 0.25 | 4 | 0.25 | 11 | 0.3636 |
| 10_9993248 | 3 | 1 | 6 | 1 | NA | NA | 4 | 1 | 13 | 1 | NA | NA | 7 | 1 | 4 | 1 | 3 | 1 | 14 | 1 |
| 10_9993252 | 1 | 1 | 5 | 0.8 | 2 | 0.5 | 2 | 1 | 10 | 0.8 | NA | NA | 6 | 1 | 4 | 1 | 3 | 0.6667 | 13 | 0.9231 |
| 10_9993255 | 3 | 0.6667 | 7 | 0.7143 | 2 | 0 | 4 | 0.5 | 16 | 0.5625 | NA | NA | 8 | 0.375 | 5 | 0.6 | 5 | 0.6 | 18 | 0.5 |
| 10_9993260 | 2 | 0 | 7 | 0 | 2 | 0.5 | 4 | 0.25 | 15 | 0.1333 | NA | NA | 7 | 0 | 5 | 0.2 | 5 | 0 | 17 | 0.0588 |
| 10_9993262 | 2 | 0 | 7 | 0.1429 | 2 | 0 | 4 | 0 | 15 | 0.0667 | NA | NA | 8 | 0.375 | 5 | 0.2 | 3 | 0 | 16 | 0.25 |
| 10_9993266 | 3 | 0 | 6 | 0.1667 | 1 | 0 | 4 | 0.25 | 14 | 0.1429 | NA | NA | 7 | 0.1429 | 5 | 0.2 | 5 | 0 | 17 | 0.1176 |
| 10_9993271 | 3 | 0 | 6 | 0.1667 | 1 | 0 | 4 | 0 | 14 | 0.0714 | NA | NA | 8 | 0.125 | 5 | 0.2 | 5 | 0 | 18 | 0.1111 |
| 10_9993287 | 3 | 1 | 7 | 0.7143 | 2 | 1 | 4 | 1 | 16 | 0.875 | NA | NA | 8 | 0.875 | 3 | 1 | 5 | 0.6 | 16 | 0.8125 |
| 10_9993293 | 3 | 0.3333 | 6 | 0 | 3 | 1 | 4 | 0.5 | 16 | 0.375 | NA | NA | 8 | 0.875 | 4 | 0.5 | 4 | 0.5 | 16 | 0.6875 |
| 10_9993467 | 5 | 0.6 | 8 | 0.125 | 3 | 0 | 4 | 0.25 | 20 | 0.25 | 2 | 0 | 10 | 0.1 | 7 | 0.1429 | 6 | 0 | 25 | 0.08 |
| 10_9993579 | 6 | 0.3333 | 6 | 0.1667 | 4 | 0 | 9 | 0.3333 | 25 | 0.24 | 3 | 0.3333 | 12 | 0 | 7 | 0.1429 | 11 | 0.2727 | 33 | 0.1515 |
| 10_9993598 | 6 | 0.1667 | 8 | 0.25 | 7 | 0.2857 | 9 | 0.3333 | 30 | 0.2667 | 3 | 1 | 11 | 0.2727 | 7 | 0.5714 | 10 | 0.2 | 31 | 0.3871 |
| 10_1002823 | NA | NA | 2 | 1 | NA | NA | 1 | 0 | 3 | 0.6667 | NA | NA | 1 | 1 | NA | NA | 2 | 0 | 3 | 0.3333 |
| 10_1002823 | NA | NA | 3 | 1 | NA | NA | 1 | 1 | 4 | 1 | NA | NA | 1 | 1 | NA | NA | 2 | 1 | 3 | 1 |
| 10_1002823 | NA | NA | 2 | 0.5 | NA | NA | 1 | 0 | 3 | 0.3333 | NA | NA | 1 | 0 | NA | NA | 2 | 1 | 3 | 0.6667 |
| 10_1002825 | NA | NA | 2 | 0.5 | NA | NA | 1 | 1 | 3 | 0.6667 | NA | NA | 1 | 1 | NA | NA | 2 | 0 | 3 | 0.3333 |
| 10_1002825 | 1 | 1 | 3 | 1 | NA | NA | 1 | 0 | 5 | 0.8 | NA | NA | 1 | 1 | NA | NA | 2 | 0 | 3 | 0.3333 |
| 10_1002826 | NA | NA | 2 | 1 | NA | NA | 1 | 1 | 3 | 1 | NA | NA | 1 | 1 | NA | NA | 1 | 1 | 2 | 1 |
| 10_1002828 | 1 | 0 | 2 | 0.5 | NA | NA | 1 | 1 | 4 | 0.5 | NA | NA | 1 | 0 | NA | NA | 1 | 1 | 2 | 0.5 |
| 10_1002829 | NA | NA | 1 | 1 | NA | NA | 1 | 1 | 2 | 1 | NA | NA | 1 | 1 | NA | NA | 1 | 1 | 2 | 1 |
| 10_1002829 | NA | NA | 2 | 0.5 | NA | NA | 1 | 0 | 3 | 0.3333 | NA | NA | 1 | 1 | NA | NA | NA | NA | 1 | 1 |
| 10_1002830 | 1 | 1 | 2 | 0.5 | NA | NA | 1 | 1 | 4 | 0.75 | NA | NA | 1 | 0 | NA | NA | 1 | 1 | 2 | 0.5 |

| | | | | | | | | | | | | | | | | | | | | |
|---|---|---|---|---|---|---|---|---|---|---|---|---|---|---|---|---|---|---|---|---|
| 10_100283 | 1 | 1 | NA | NA | NA | NA | 1 | 1 | 2 | 1 | NA | NA | 1 | 1 | NA | NA | NA | NA | 1 | 1 |
| 10_100283 | NA | NA | NA | NA | NA | NA | 1 | 1 | 1 | 1 | NA | NA | 1 | 1 | NA | NA | NA | NA | 1 | 1 |
| 10_100408 | 2 | 0 | 4 | 0 | 5 | 0 | 2 | 0.5 | 13 | 0.0769 | 2 | 0 | 2 | 0.5 | 9 | 0.1111 | 1 | 0 | 14 | 0.1429 |
| 10_100408 | 1 | 1 | 4 | 0 | 4 | 0 | 3 | 0 | 12 | 0.0833 | 3 | 0 | 2 | 0 | 8 | 0.25 | 1 | 0 | 14 | 0.1429 |
| 10_100408 | NA | NA | NA | NA | 1 | 0 | 1 | 1 | 2 | 0.5 | NA | NA | 1 | 1 | NA | NA | NA | NA | 1 | 1 |
| 10_100408 | 2 | 0.5 | 4 | 0 | 3 | 0.3333 | 4 | 0 | 13 | 0.1538 | 2 | 0 | 1 | 0 | 7 | 0.1429 | 1 | 0 | 11 | 0.0909 |
| 10_100409 | 2 | 0 | 4 | 0.25 | 4 | 0 | 3 | 0 | 13 | 0.0769 | 2 | 0.5 | 1 | 0 | 8 | 0.125 | 1 | 0 | 12 | 0.1667 |
| 10_100409 | 1 | 1 | 3 | 0.3333 | 4 | 0 | 4 | 0 | 12 | 0.1667 | 2 | 0.5 | 1 | 0 | 4 | 0.25 | 1 | 1 | 8 | 0.375 |
| 10_100409 | 1 | 1 | 3 | 0 | 3 | 0 | 2 | 0 | 9 | 0.1111 | 2 | 0.5 | 1 | 0 | 5 | 0 | 1 | 0 | 9 | 0.1111 |
| 10_102224 | NA | NA | NA | NA | 1 | 1 | 1 | 1 | 2 | 1 | 1 | 0 | 1 | 1 | NA | NA | NA | NA | 2 | 0.5 |
| 10_102224 | 1 | 0 | NA | NA | NA | NA | 1 | 1 | 2 | 0.5 | 1 | 1 | NA | NA | NA | NA | NA | NA | 1 | 1 |
| 10_102224 | 1 | 1 | NA | NA | 1 | 0 | 2 | 0 | 4 | 0.25 | 1 | 1 | 1 | 1 | NA | NA | NA | NA | 2 | 1 |
| 10_102224 | 1 | 1 | NA | NA | 2 | 0 | 2 | 1 | 5 | 0.6 | 2 | 0.5 | 1 | 1 | NA | NA | NA | NA | 3 | 0.6667 |
| 10_102224 | 1 | 0 | NA | NA | 2 | 0 | 1 | 1 | 4 | 0.25 | 2 | 0 | 1 | 1 | NA | NA | NA | NA | 3 | 0.3333 |
| 10_102224 | 1 | 0 | NA | NA | 1 | 1 | 2 | 0 | 4 | 0.25 | 2 | 0.5 | NA | NA | NA | NA | NA | NA | 2 | 0.5 |
| 10_102224 | 1 | 0 | NA | NA | 2 | 0.5 | 1 | 0 | 4 | 0.25 | 1 | 1 | 1 | 1 | NA | NA | NA | NA | 2 | 1 |
| 10_102224 | 1 | 0 | NA | NA | 2 | 0.5 | 2 | 0 | 5 | 0.2 | 2 | 0.5 | 1 | 0 | NA | NA | NA | NA | 3 | 0.3333 |
| 10_102224 | 1 | 1 | NA | NA | 1 | 1 | 2 | 1 | 4 | 1 | 2 | 0.5 | 1 | 0 | NA | NA | NA | NA | 3 | 0.3333 |
| 10_102224 | 1 | 1 | NA | NA | 2 | 1 | 2 | 1 | 5 | 1 | 2 | 1 | 1 | 1 | NA | NA | 1 | 1 | 4 | 1 |
| 10_102224 | 1 | 1 | NA | NA | 2 | 0 | 2 | 0.5 | 5 | 0.4 | 2 | 1 | 1 | 0 | NA | NA | 1 | 0 | 4 | 0.5 |
| 10_102224 | 1 | 1 | NA | NA | 2 | 0.5 | 1 | 1 | 4 | 0.75 | 1 | 1 | 1 | 0 | NA | NA | 1 | 0 | 3 | 0.3333 |
| 10_102224 | 1 | 1 | NA | NA | 1 | 1 | 2 | 1 | 4 | 1 | 2 | 1 | 1 | 1 | NA | NA | 1 | 1 | 4 | 1 |
| 10_102224 | 1 | 1 | NA | NA | 2 | 0 | 2 | 0 | 5 | 0.2 | 2 | 0.5 | 1 | 1 | NA | NA | NA | NA | 3 | 0.6667 |
| 10_102224 | 1 | 0 | NA | NA | 2 | 0.5 | 2 | 0 | 5 | 0.2 | 2 | 0.5 | 1 | 1 | NA | NA | 1 | 0 | 4 | 0.5 |
| 10_102225 | NA | NA | NA | NA | 2 | 0.5 | 2 | 0.5 | 4 | 0.5 | 1 | 1 | 1 | 1 | NA | NA | 1 | 1 | 3 | 1 |
| 10_102225 | NA | NA | NA | NA | 3 | 0.3333 | 2 | 1 | 5 | 0.6 | 1 | 1 | 1 | 1 | NA | NA | 1 | 1 | 3 | 1 |
| 10_102225 | 1 | 1 | NA | NA | 4 | 1 | 2 | 0.5 | 7 | 0.8571 | 1 | 1 | 1 | 1 | NA | NA | 1 | 1 | 3 | 1 |
| 10_102225 | 1 | 1 | NA | NA | 3 | 1 | 3 | 1 | 7 | 1 | 1 | 1 | 1 | 1 | NA | NA | 1 | 1 | 3 | 1 |
| 10_102225 | NA | NA | NA | NA | 5 | 0.4 | 2 | 0 | 7 | 0.2857 | 1 | 1 | 1 | 0 | NA | NA | 2 | 0 | 4 | 0.25 |
| 10_102225 | 1 | 0 | NA | NA | 5 | 0.2 | 4 | 0.25 | 10 | 0.2 | NA | NA | 1 | 0 | NA | NA | 1 | 1 | 2 | 0.5 |
| 10_102226 | 1 | 0 | NA | NA | 2 | 0.5 | 2 | 0 | 5 | 0.2 | 1 | 0 | 1 | 1 | NA | NA | NA | NA | 2 | 0.5 |
| 10_102317 | 6 | 0.1667 | 3 | 0 | 9 | 0.1111 | 9 | 0.1111 | 27 | 0.1111 | 3 | 0 | 4 | 0 | 4 | 0 | 4 | 0.25 | 15 | 0.0667 |
| 10_103224 | 5 | 0.2 | 5 | 0 | 1 | 0 | 6 | 0.1667 | 17 | 0.1176 | 1 | 0 | 2 | 0 | 4 | 0 | 7 | 0.1429 | 14 | 0.0714 |
| 10_103229 | 5 | 0 | 5 | 0 | 1 | 0 | 5 | 0.2 | 16 | 0.0625 | 1 | 0 | 2 | 0 | 5 | 0 | 6 | 0.1667 | 14 | 0.0714 |
| 10_103229 | 4 | 0.25 | 4 | 0 | 2 | 0 | 4 | 0 | 14 | 0.0714 | 1 | 0 | 2 | 0 | 3 | 0 | 6 | 0.1667 | 12 | 0.0833 |
| 10_103254 | 6 | 0 | 12 | 0.0833 | 2 | 0 | 5 | 0.2 | 25 | 0.08 | 1 | 0 | 3 | 0.3333 | 3 | 0.3333 | 8 | 0.125 | 15 | 0.2 |
| 10_103254 | 3 | 0.3333 | 6 | 0.3333 | NA | NA | 1 | 1 | 10 | 0.4 | NA | NA | 2 | 0 | 2 | 0.5 | 6 | 0.6667 | 10 | 0.5 |
| 10_103254 | 6 | 0.1667 | 14 | 0.5 | 3 | 0.6667 | 9 | 0.3333 | 32 | 0.4062 | 3 | 0.3333 | 5 | 0.4 | 8 | 0.375 | 10 | 0.4 | 26 | 0.3846 |
| 10_103255 | 7 | 0.4286 | 13 | 0.6154 | 3 | 0.3333 | 7 | 0.4286 | 30 | 0.5 | 4 | 0 | 7 | 0.2857 | 9 | 0.4444 | 10 | 0.5 | 30 | 0.3667 |
| 10_105661 | 1 | 0 | 1 | 1 | NA | NA | NA | NA | 2 | 0.5 | NA | NA | NA | NA | 2 | 1 | NA | NA | 2 | 1 |
| 10_105662 | 1 | 1 | 1 | 0 | NA | NA | NA | NA | 2 | 0.5 | NA | NA | NA | NA | 2 | 1 | NA | NA | 2 | 1 |
| 10_105678 | 6 | 0.3333 | 5 | 0 | 1 | 0 | 6 | 0 | 18 | 0.1111 | 1 | 0 | 5 | 0.2 | 8 | 0.125 | 4 | 0 | 18 | 0.1111 |
| 10_105682 | 10 | 0.3 | 4 | 0.75 | 3 | 0.3333 | 8 | 0.5 | 25 | 0.44 | 1 | 1 | 4 | 0.75 | 10 | 0.5 | 5 | 0.6 | 20 | 0.6 |
| 10_105683 | 8 | 0.625 | 4 | 0.25 | 4 | 0.75 | 7 | 0.8571 | 23 | 0.6522 | 1 | 1 | 5 | 1 | 11 | 0.8182 | 3 | 0.6667 | 20 | 0.85 |
| 10_105683 | 9 | 0.7778 | 4 | 1 | 4 | 1 | 7 | 0.7143 | 24 | 0.8333 | 1 | 1 | 6 | 1 | 10 | 0.8 | 5 | 1 | 22 | 0.9091 |
| 10_105683 | 10 | 0.2 | 5 | 0.6 | 4 | 0 | 8 | 0.75 | 27 | 0.4074 | 1 | 1 | 5 | 0.2 | 11 | 0.5455 | 3 | 0.3333 | 20 | 0.45 |
| 10_105683 | 9 | 0.7778 | 5 | 0.6 | 4 | 0.5 | 7 | 0.7143 | 25 | 0.68 | 1 | 1 | 5 | 0.8 | 11 | 0.8182 | 5 | 0.4 | 22 | 0.7273 |
| 10_105683 | 9 | 0.2222 | 4 | 0.25 | 1 | 1 | 4 | 0.5 | 18 | 0.3333 | 2 | 0.5 | 5 | 0.6 | 9 | 0.6667 | 4 | 0.5 | 20 | 0.6 |
| 10_106306 | 1 | 0 | 1 | 0 | NA | NA | 1 | 1 | 3 | 0.3333 | NA | NA | 1 | 1 | NA | NA | 1 | 0 | 2 | 0.5 |
| 10_106310 | 3 | 0 | 2 | 0.5 | NA | NA | 2 | 0 | 7 | 0.1429 | 1 | 0 | 1 | 0 | NA | NA | 2 | 0.5 | 4 | 0.25 |
| 10_106767 | 4 | 0.75 | 5 | 1 | 3 | 0.3333 | 6 | 1 | 18 | 0.8333 | 3 | 0.6667 | 3 | 1 | 3 | 0.6667 | 7 | 0.8571 | 16 | 0.8125 |
| 10_107256 | 5 | 0.2 | 1 | 0 | NA | NA | 5 | 0.4 | 11 | 0.2727 | 1 | 1 | 6 | 0.3333 | 4 | 0.25 | 5 | 0.2 | 16 | 0.3125 |
| 10_107256 | 4 | 0.25 | 2 | 0 | NA | NA | 5 | 0.4 | 11 | 0.2727 | 2 | 0.5 | 6 | 0.1667 | 5 | 0 | 5 | 0.2 | 18 | 0.1667 |
| 10_107256 | 4 | 0.5 | 3 | 0.3333 | NA | NA | 3 | 0 | 10 | 0.3 | 2 | 0.5 | 6 | 0.1667 | 6 | 0.1667 | 5 | 0 | 19 | 0.1579 |
| 10_107256 | 4 | 0.25 | 2 | 0 | NA | NA | 4 | 0 | 10 | 0.1 | 2 | 0.5 | 4 | 0 | 5 | 0 | 3 | 0.3333 | 14 | 0.1429 |
| 10_107258 | 8 | 0.5 | 7 | 0.2857 | 3 | 0 | 6 | 0.1667 | 24 | 0.2917 | 2 | 0 | 6 | 0.5 | 4 | 0.25 | 6 | 0.1667 | 18 | 0.2778 |
| 10_107258 | 5 | 0.2 | 6 | 0 | 3 | 0 | 6 | 0 | 20 | 0.05 | 2 | 0 | 6 | 0 | 4 | 0 | 7 | 0.1429 | 19 | 0.0526 |
| 10_107997 | NA | NA | NA | NA | NA | NA | 1 | 1 | 1 | 1 | NA | NA | NA | NA | 1 | 1 | NA | NA | 1 | 1 |
| 10_108113 | 2 | 0 | 5 | 0.2 | 3 | 0 | 5 | 0 | 15 | 0.0667 | 2 | 0 | 8 | 0.375 | 6 | 0 | 6 | 0 | 22 | 0.1364 |
| 10_108113 | 2 | 0 | 5 | 0.2 | 5 | 0 | 7 | 0.2857 | 19 | 0.1579 | 2 | 0 | 6 | 0 | 8 | 0.125 | 6 | 0.1667 | 22 | 0.0909 |
| 10_108113 | 2 | 1 | 6 | 0.3333 | 5 | 0.8 | 5 | 0.8 | 18 | 0.6667 | 2 | 1 | 8 | 0.5 | 7 | 0.4286 | 7 | 0.5714 | 24 | 0.5417 |
| 10_108113 | 1 | 0 | 5 | 0.2 | 4 | 0.25 | 6 | 0.1667 | 16 | 0.1875 | 2 | 0.5 | 5 | 0 | 5 | 0 | 7 | 0 | 19 | 0.0526 |
| 10_108113 | 3 | 0.3333 | 5 | 0.6 | 5 | 0.6 | 7 | 0.2857 | 20 | 0.45 | 1 | 1 | 6 | 0.3333 | 6 | 0.5 | 6 | 0.3333 | 19 | 0.4211 |
| 10_108113 | 3 | 0 | 6 | 0.8333 | 5 | 0.4 | 6 | 0.8333 | 20 | 0.6 | 2 | 0 | 8 | 0.625 | 7 | 0.7143 | 7 | 0.7143 | 24 | 0.625 |
| 10_108113 | 3 | 1 | 6 | 0.8333 | 3 | 1 | 5 | 0.4 | 17 | 0.7647 | 2 | 0 | 8 | 0.875 | 5 | 1 | 6 | 0.8333 | 21 | 0.8095 |

| | | | | | | | | | | | | | | | | | | | | |
|---|---|---|---|---|---|---|---|---|---|---|---|---|---|---|---|---|---|---|---|---|
| 10_108113 | 3 | 0 | 8 | 0.125 | 4 | 0.25 | 7 | 0.1429 | 22 | 0.1364 | 2 | 0 | 9 | 0.1111 | 8 | 0.125 | 6 | 0.1667 | 25 | 0.12 |
| 10_108113 | 4 | 0.25 | 8 | 0.25 | 6 | 0.5 | 10 | 0.3 | 28 | 0.3214 | 2 | 0 | 11 | 0.0909 | 9 | 0.1111 | 6 | 0.3333 | 28 | 0.1429 |
| 10_108249 | 1 | 1 | 2 | 0.5 | 1 | 1 | 1 | 1 | 5 | 0.8 | 1 | 1 | NA | NA | NA | NA | NA | NA | 1 | 1 |
| 10_108335 | 54 | 0.1111 | 78 | 0.1026 | 61 | 0.1311 | 58 | 0.0862 | 251 | 0.1076 | 47 | 0.1489 | 49 | 0.2449 | 45 | 0.1778 | 62 | 0.1935 | 203 | 0.1921 |
| 10_108335 | 58 | 0.0517 | 86 | 0.0698 | 72 | 0.125 | 62 | 0.0806 | 278 | 0.0827 | 52 | 0.0769 | 53 | 0.1509 | 51 | 0.098 | 69 | 0.1449 | 225 | 0.12 |
| 10_108336 | 39 | 0.5128 | 45 | 0.6 | 38 | 0.4737 | 40 | 0.25 | 162 | 0.463 | 28 | 0.5357 | 36 | 0.3333 | 26 | 0.4615 | 36 | 0.5556 | 126 | 0.4683 |
| 10_109517 | 6 | 0.1667 | 7 | 0.4286 | 6 | 0 | 2 | 0 | 21 | 0.1905 | 4 | 0.25 | 5 | 0 | 1 | 1 | 6 | 0.1667 | 16 | 0.1875 |
| 10_109517 | 4 | 0.75 | 6 | 0.6667 | 7 | 0.4286 | 2 | 0.5 | 19 | 0.5789 | 4 | 0.75 | 6 | 0.3333 | 1 | 1 | 4 | 0 | 15 | 0.4 |
| 10_109517 | 5 | 0.2 | 5 | 0 | 7 | 0 | 2 | 0 | 19 | 0.0526 | 3 | 0 | 7 | 0 | NA | NA | 4 | 0.25 | 14 | 0.0714 |
| 10_109517 | 6 | 0.1667 | 6 | 0.1667 | 6 | 0 | 2 | 0 | 20 | 0.1 | 4 | 0.25 | 6 | 0.1667 | 1 | 0 | 4 | 0.25 | 15 | 0.2 |
| 10_109518 | 5 | 0 | 5 | 0 | 7 | 0.1429 | 2 | 0 | 19 | 0.0526 | 4 | 0 | 6 | 0.1667 | 1 | 1 | 3 | 0.3333 | 14 | 0.2143 |
| 10_109518 | 3 | 0 | 3 | 0 | 5 | 0.2 | 2 | 0 | 13 | 0.0769 | 4 | 0.5 | 5 | 0 | 1 | 0 | 3 | 0 | 13 | 0.1538 |
| 10_109518 | 6 | 0 | 3 | 0.3333 | 6 | 0 | 2 | 0 | 17 | 0.0588 | 3 | 0 | 6 | 0 | 1 | 0 | 3 | 0.3333 | 13 | 0.0769 |
| 10_109518 | 5 | 0 | 4 | 0.5 | 4 | 0.25 | 2 | 0 | 15 | 0.2 | 3 | 0.3333 | 5 | 0 | 1 | 0 | 3 | 0 | 12 | 0.0833 |
| 10_109519 | 4 | 0 | 3 | 0 | 4 | 0.25 | 2 | 0 | 13 | 0.0769 | 3 | 0 | 6 | 0 | 1 | 0 | 3 | 0.3333 | 13 | 0.0769 |
| 10_109519 | 2 | 0.5 | 2 | 0 | 4 | 0 | 2 | 0 | 10 | 0.1 | 2 | 0 | 5 | 0.2 | 1 | 0 | 1 | 1 | 9 | 0.2222 |
| 10_109519 | 3 | 0 | 2 | 0 | 4 | 0.25 | 2 | 0 | 11 | 0.0909 | 3 | 0 | 5 | 0.2 | 1 | 0 | 1 | 0 | 10 | 0.1 |
| 10_109519 | 3 | 0.6667 | 2 | 0.5 | 2 | 0.5 | 2 | 0 | 9 | 0.4444 | 2 | 0 | 4 | 0.5 | NA | NA | 1 | 1 | 7 | 0.4286 |
| 10_109519 | 2 | 0 | 1 | 1 | 3 | 0.3333 | 1 | 0 | 7 | 0.2857 | 2 | 0 | 4 | 0 | 1 | 0 | 1 | 1 | 8 | 0.125 |
| 10_109519 | 3 | 0.3333 | NA | NA | 3 | 0.3333 | 2 | 0.5 | 8 | 0.375 | 2 | 0.5 | 5 | 0.2 | 1 | 0 | 1 | 0 | 9 | 0.2222 |
| 10_109519 | 2 | 1 | 1 | 1 | 3 | 1 | 2 | 1 | 8 | 1 | 2 | 1 | 3 | 1 | 1 | 1 | 1 | 1 | 7 | 1 |
| 10_110290 | 1 | 1 | 1 | 1 | 1 | 0 | NA | NA | 3 | 0.6667 | 1 | 1 | NA | NA | 1 | 1 | NA | NA | 2 | 1 |
| 10_110778 | 11 | 0.0909 | 14 | 0 | 14 | 0 | 20 | 0.1 | 59 | 0.0508 | 8 | 0.125 | 18 | 0.0556 | 17 | 0.1176 | 6 | 0.1667 | 49 | 0.102 |
| 10_110779 | 8 | 0 | 11 | 0.0909 | 14 | 0.0714 | 16 | 0.0625 | 49 | 0.0612 | 5 | 0 | 13 | 0 | 16 | 0.1875 | 5 | 0 | 39 | 0.0769 |
| 10_110779 | 6 | 0.1667 | 10 | 0.4 | 12 | 0.4167 | 10 | 0.4 | 38 | 0.3684 | 6 | 0.3333 | 9 | 0.4444 | 14 | 0.4286 | 2 | 1 | 31 | 0.4516 |
| 10_110780 | 6 | 0.1667 | 10 | 0.6 | 11 | 0.1818 | 14 | 0.5714 | 41 | 0.4146 | 4 | 0.5 | 8 | 0.625 | 11 | 0.5455 | 5 | 0.2 | 28 | 0.5 |
| 10_110780 | 6 | 0.8333 | 10 | 1 | 7 | 1 | 12 | 1 | 35 | 0.9714 | 2 | 0.5 | 10 | 0.9 | 10 | 0.8 | 4 | 1 | 26 | 0.8462 |
| 10_110780 | 5 | 0.4 | 7 | 0.1429 | 11 | 0.3636 | 15 | 0.4667 | 38 | 0.3684 | 3 | 0.3333 | 9 | 0.5556 | 14 | 0.6429 | 5 | 0.8 | 31 | 0.6129 |
| 10_110780 | 7 | 0.1429 | 11 | 0.0909 | 12 | 0.0833 | 15 | 0.1333 | 45 | 0.1111 | 3 | 0.3333 | 11 | 0.1818 | 14 | 0.1429 | 5 | 0 | 33 | 0.1515 |
| 10_110780 | 7 | 0 | 10 | 0 | 12 | 0.25 | 12 | 0.0833 | 41 | 0.0976 | 4 | 0.25 | 9 | 0 | 13 | 0.0769 | 4 | 0 | 30 | 0.0667 |
| 10_110780 | 6 | 0.5 | 9 | 0.8889 | 11 | 0.5455 | 15 | 0.7333 | 41 | 0.6829 | 3 | 0.3333 | 9 | 0.7778 | 14 | 0.7143 | 4 | 0.25 | 30 | 0.6333 |
| 10_110780 | 5 | 0.6 | 10 | 0.4 | 10 | 0.5 | 14 | 0.5714 | 39 | 0.5128 | 4 | 0.75 | 10 | 0.3 | 14 | 0.5 | 4 | 0.5 | 32 | 0.4688 |
| 10_111129 | 6 | 0.5 | 9 | 0.2222 | 4 | 0.25 | 18 | 0.3889 | 37 | 0.3514 | 6 | 0.1667 | 14 | 0.2857 | 13 | 0.3846 | 11 | 0.1818 | 44 | 0.2727 |
| 10_111129 | 6 | 0.8333 | 9 | 0.8889 | 3 | 1 | 13 | 1 | 31 | 0.9355 | 5 | 0.8 | 15 | 0.8 | 11 | 0.8182 | 14 | 0.8571 | 45 | 0.8222 |
| 10_111129 | 6 | 0.8333 | 10 | 1 | 4 | 1 | 17 | 0.8824 | 37 | 0.9189 | 6 | 0.6667 | 16 | 0.875 | 13 | 1 | 13 | 0.7692 | 48 | 0.8542 |
| 10_111129 | 7 | 0.1429 | 8 | 0 | 3 | 0 | 16 | 0.125 | 34 | 0.0882 | 6 | 0 | 14 | 0.1429 | 11 | 0.2727 | 9 | 0 | 40 | 0.125 |
| 10_111129 | 8 | 0.125 | 6 | 0.1667 | 5 | 0.6 | 14 | 0.0714 | 33 | 0.1818 | 6 | 0.1667 | 14 | 0.2143 | 14 | 0.1429 | 14 | 0.2857 | 48 | 0.2083 |
| 10_111129 | 8 | 0.75 | 9 | 0.7778 | 4 | 1 | 13 | 0.8462 | 34 | 0.8235 | 7 | 0.7143 | 17 | 0.8824 | 13 | 0.8462 | 8 | 0.875 | 45 | 0.8444 |
| 10_111129 | 9 | 0.4444 | 12 | 0.25 | 6 | 0.3333 | 17 | 0.1765 | 44 | 0.2727 | 6 | 0.3333 | 21 | 0.381 | 11 | 0.5455 | 16 | 0.125 | 54 | 0.3333 |
| 10_111130 | 11 | 0.0909 | 9 | 0.1111 | 6 | 0.1667 | 22 | 0.1818 | 48 | 0.1458 | 5 | 0 | 21 | 0.1429 | 17 | 0.2941 | 15 | 0.0667 | 58 | 0.1552 |
| 10_111131 | 10 | 0.1 | 12 | 0.1667 | 5 | 0.2 | 16 | 0.1875 | 43 | 0.1628 | 3 | 0 | 19 | 0.2632 | 18 | 0.2222 | 12 | 0.3333 | 52 | 0.25 |
| 10_111132 | 14 | 0.1429 | 17 | 0.1176 | 14 | 0.1429 | 16 | 0 | 61 | 0.0984 | 7 | 0.2857 | 25 | 0.04 | 30 | 0.2 | 23 | 0.2609 | 85 | 0.1765 |
| 10_111132 | 14 | 0.3571 | 16 | 0.125 | 14 | 0.3571 | 19 | 0.3158 | 63 | 0.2857 | 8 | 0.25 | 29 | 0.3448 | 31 | 0.3871 | 26 | 0.1538 | 94 | 0.2979 |
| 10_111135 | NA | NA | NA | NA | NA | NA | 2 | 0.5 | 2 | 0.5 | 1 | 0 | 1 | 1 | NA | NA | 1 | 0 | 3 | 0.3333 |
| 10_111135 | NA | NA | NA | NA | NA | NA | 1 | 1 | 1 | 1 | 1 | 1 | 1 | 1 | NA | NA | 1 | 1 | 3 | 1 |
| 10_111135 | NA | NA | NA | NA | NA | NA | 2 | 1 | 2 | 1 | 1 | 1 | 1 | 0 | NA | NA | 1 | 0 | 3 | 0.3333 |
| 10_111135 | NA | NA | NA | NA | NA | NA | 2 | 1 | 2 | 1 | 1 | 1 | 1 | 1 | NA | NA | NA | NA | 2 | 1 |
| 10_111135 | NA | NA | NA | NA | NA | NA | 2 | 1 | 2 | 1 | 1 | 1 | 1 | 1 | NA | NA | NA | NA | 2 | 1 |
| 10_111135 | NA | NA | NA | NA | NA | NA | 2 | 0.5 | 2 | 0.5 | 1 | 1 | 1 | 1 | NA | NA | 1 | 0 | 3 | 0.6667 |
| 10_111135 | NA | NA | NA | NA | NA | NA | 1 | 1 | 1 | 1 | 1 | 1 | 1 | 1 | NA | NA | 1 | 0 | 3 | 0.6667 |
| 10_111135 | NA | NA | NA | NA | NA | NA | 2 | 0.5 | 2 | 0.5 | 1 | 1 | 1 | 1 | NA | NA | 1 | 0 | 3 | 0.6667 |
| 10_111136 | NA | NA | NA | NA | NA | NA | 1 | 1 | 1 | 1 | NA | NA | 2 | 1 | NA | NA | 1 | 0 | 3 | 0.6667 |
| 10_111136 | NA | NA | NA | NA | NA | NA | 2 | 1 | 2 | 1 | 1 | 1 | 2 | 1 | NA | NA | 1 | 0 | 4 | 0.75 |
| 10_111136 | NA | NA | NA | NA | NA | NA | 1 | 1 | 1 | 1 | NA | NA | 2 | 1 | NA | NA | 1 | 0 | 3 | 0.6667 |
| 10_111137 | NA | NA | NA | NA | NA | NA | 1 | 1 | 1 | 1 | 1 | 1 | 1 | 1 | NA | NA | 1 | 0 | 3 | 0.6667 |
| 10_111137 | NA | NA | NA | NA | NA | NA | 2 | 1 | 2 | 1 | 1 | 1 | 2 | 0.5 | NA | NA | 1 | 0 | 4 | 0.5 |
| 10_111332 | 5 | 0.2 | 13 | 0.1538 | 8 | 0 | 14 | 0.4286 | 40 | 0.225 | 6 | 0.1667 | 8 | 0.375 | 5 | 0.2 | 7 | 0.2857 | 26 | 0.2692 |
| 10_111336 | NA | NA | 1 | 0 | 1 | 1 | NA | NA | 2 | 0.5 | 1 | 1 | NA | NA | NA | NA | NA | NA | 1 | 1 |
| 10_111337 | NA | NA | 1 | 1 | 1 | 1 | NA | NA | 2 | 1 | 1 | 1 | NA | NA | NA | NA | NA | NA | 1 | 1 |
| 10_111342 | 5 | 0.4 | 8 | 0.25 | 4 | 0.5 | 8 | 0.375 | 25 | 0.36 | 2 | 0.5 | 5 | 0.2 | 4 | 0.25 | 5 | 0 | 16 | 0.1875 |
| 10_111342 | 7 | 0 | 8 | 0.375 | 7 | 0.4286 | 9 | 0.1111 | 31 | 0.2258 | 3 | 0.3333 | 5 | 0.2 | 10 | 0.1 | 7 | 0.1429 | 25 | 0.16 |
| 10_111343 | 6 | 0 | 14 | 0 | 8 | 0.125 | 9 | 0.3333 | 37 | 0.1081 | 4 | 0 | 9 | 0 | 11 | 0.0909 | 6 | 0.1667 | 30 | 0.0667 |
| 10_111344 | 5 | 1 | 13 | 0.6154 | 8 | 0.75 | 10 | 0.4 | 36 | 0.6389 | 4 | 0.75 | 10 | 0.8 | 10 | 0.8 | 9 | 0.5556 | 33 | 0.7273 |
| 10_111344 | 6 | 0.3333 | 15 | 0.1333 | 10 | 0.2 | 9 | 0.2222 | 40 | 0.2 | 4 | 0.25 | 10 | 0.1 | 11 | 0.3636 | 9 | 0.3333 | 34 | 0.2647 |
| 10_111344 | 7 | 0.8571 | 15 | 0.9333 | 12 | 1 | 10 | 1 | 44 | 0.9545 | 3 | 1 | 10 | 1 | 10 | 0.9 | 10 | 0.9 | 33 | 0.9394 |

| | | | | | | | | | | | | | | | | | | | | |
|---|---|---|---|---|---|---|---|---|---|---|---|---|---|---|---|---|---|---|---|---|
| 10_1113441 | 7 | 1 | 14 | 1 | 12 | 0.9167 | 10 | 1 | 43 | 0.9767 | 3 | 1 | 10 | 1 | 11 | 0.9091 | 9 | 0.8889 | 33 | 0.9394 |
| 10_1113442 | 8 | 0 | 14 | 0.1429 | 11 | 0.1818 | 9 | 0.2222 | 42 | 0.1429 | 4 | 0.25 | 8 | 0.25 | 8 | 0.25 | 8 | 0.25 | 28 | 0.25 |
| 10_1113443 | 9 | 0.3333 | 14 | 0.0714 | 12 | 0.1667 | 10 | 0.3 | 45 | 0.2 | 4 | 0.5 | 10 | 0.2 | 10 | 0.3 | 8 | 0.375 | 32 | 0.3125 |
| 10_1113444 | 7 | 0.4286 | 15 | 0.4667 | 11 | 0.2727 | 7 | 0.2857 | 40 | 0.375 | 5 | 0.2 | 8 | 0.375 | 13 | 0.3077 | 8 | 0.125 | 34 | 0.2647 |
| 10_1118206 | 6 | 0.5 | NA | NA | 3 | 0.3333 | 5 | 0.6 | 14 | 0.5 | 1 | 1 | 5 | 0.4 | 4 | 0.25 | 7 | 0.4286 | 17 | 0.4118 |
| 10_1118216 | 11 | 0.1818 | 2 | 0 | 3 | 0 | 5 | 0 | 21 | 0.0952 | 3 | 0.3333 | 3 | 0 | 6 | 0.1667 | 8 | 0 | 20 | 0.1 |
| 10_1118226 | 1 | 1 | 1 | 0 | NA | NA | 1 | 0 | 3 | 0.3333 | NA | NA | 1 | 1 | 1 | 0 | NA | NA | 2 | 0.5 |
| 10_1121077 | NA | NA | 1 | 1 | 3 | 0 | NA | NA | 4 | 0.25 | NA | NA | NA | NA | 1 | 1 | 2 | 0.5 | 3 | 0.6667 |
| 10_1121077 | 2 | 0 | 1 | 1 | 2 | 0 | 1 | 0 | 6 | 0.1667 | NA | NA | 1 | 0 | 1 | 1 | 1 | 0 | 3 | 0.3333 |
| 10_1121081 | 2 | 0 | 1 | 1 | 3 | 0.6667 | NA | NA | 6 | 0.5 | NA | NA | 1 | 0 | 2 | 0.5 | 2 | 0.5 | 5 | 0.4 |
| 10_1121081 | 3 | 0.3333 | 1 | 1 | 3 | 1 | NA | NA | 7 | 0.7143 | NA | NA | 1 | 1 | 2 | 1 | 2 | 0.5 | 5 | 0.8 |
| 10_1121082 | 3 | 0 | 1 | 1 | 3 | 0.3333 | 1 | 0 | 8 | 0.25 | NA | NA | 1 | 0 | 1 | 1 | 2 | 0 | 4 | 0.25 |
| 10_1121082 | 2 | 0 | 2 | 0.5 | 3 | 0.3333 | 1 | 1 | 8 | 0.375 | NA | NA | 1 | 0 | 1 | 1 | 2 | 0.5 | 4 | 0.5 |
| 10_1122917 | 3 | 0.6667 | 1 | 1 | 3 | 0.6667 | 1 | 1 | 8 | 0.75 | NA | NA | 1 | 1 | 1 | 1 | 2 | 0.5 | 4 | 0.75 |
| 10_1122917 | 3 | 1 | 1 | 1 | 2 | 1 | NA | NA | 6 | 1 | NA | NA | 1 | 1 | 2 | 1 | 2 | 1 | 5 | 1 |
| 10_1122918 | 2 | 0.5 | 3 | 1 | 3 | 1 | 1 | 1 | 9 | 0.8889 | NA | NA | 1 | 0 | 1 | 1 | 2 | 1 | 4 | 0.75 |
| 10_1122919 | 3 | 0 | 3 | 0.6667 | 2 | 0.5 | 1 | 1 | 9 | 0.4444 | 1 | 1 | NA | NA | 1 | 0 | 2 | 1 | 4 | 0.75 |
| 10_1122920 | 3 | 0 | 3 | 0 | 3 | 0.6667 | 2 | 0 | 11 | 0.1818 | 1 | 1 | 1 | 0 | 2 | 0 | 2 | 0.5 | 6 | 0.3333 |
| 10_1122921 | 3 | 0.3333 | 2 | 0 | 4 | 0 | 2 | 0 | 11 | 0.0909 | 2 | 0.5 | 1 | 1 | 2 | 1 | 2 | 1 | 7 | 0.8571 |
| 10_1122921 | 3 | 0 | 2 | 0.5 | 3 | 0.3333 | 2 | 0 | 10 | 0.2 | 2 | 0.5 | NA | NA | 1 | 0 | 2 | 0 | 5 | 0.2 |
| 10_1122994 | 5 | 0.2 | 3 | 0.6667 | 3 | 0.6667 | 4 | 0.25 | 15 | 0.4 | 2 | 0 | 1 | 0 | 1 | 0 | 4 | 0.25 | 8 | 0.125 |
| 10_1122995 | 4 | 0.25 | 2 | 1 | 2 | 0.5 | 3 | 0.3333 | 11 | 0.4545 | 2 | 1 | 1 | 0 | 2 | 1 | 4 | 0.25 | 9 | 0.5556 |
| 10_1122995 | 5 | 0 | 2 | 0.5 | 3 | 0.3333 | 4 | 0 | 14 | 0.1429 | 2 | 0 | 1 | 1 | 2 | 0 | 3 | 0 | 8 | 0.125 |
| 10_1122995 | 4 | 0.5 | 3 | 0.3333 | 3 | 0.6667 | 4 | 0.5 | 14 | 0.5 | 2 | 0 | 1 | 0 | 1 | 1 | 4 | 0.5 | 8 | 0.375 |
| 10_1122995 | 5 | 0 | 3 | 0.3333 | 4 | 0 | 3 | 0 | 15 | 0.0667 | 1 | 0 | NA | NA | 2 | 0.5 | 4 | 0 | 7 | 0.1429 |
| 10_1122995 | 5 | 0.4 | 3 | 1 | 3 | 0.3333 | 3 | 0.3333 | 14 | 0.5 | 2 | 0.5 | 1 | 0 | 2 | 1 | 3 | 0.6667 | 8 | 0.625 |
| 10_1122996 | 4 | 0 | 3 | 0.3333 | 3 | 0.3333 | 4 | 0.25 | 14 | 0.2143 | 2 | 0 | 1 | 0 | 2 | 0.5 | 3 | 0.3333 | 8 | 0.25 |
| 10_1123002 | 5 | 0 | 4 | 0 | 3 | 0 | 5 | 0.4 | 17 | 0.1176 | 3 | 0 | 2 | 0 | 2 | 0 | 4 | 0.25 | 11 | 0.0909 |
| 10_1123009 | 4 | 1 | 5 | 0.8 | 4 | 0.5 | 6 | 0.3333 | 19 | 0.6316 | 3 | 1 | 3 | 0.6667 | 3 | 1 | 6 | 0.5 | 15 | 0.7333 |
| 10_1123010 | 3 | 0 | 6 | 0.3333 | 3 | 0.3333 | 6 | 0.3333 | 18 | 0.2778 | 2 | 0 | 2 | 0.5 | 2 | 0 | 6 | 0.1667 | 12 | 0.1667 |
| 10_1123010 | 4 | 0.25 | 4 | 0 | 3 | 0 | 5 | 0.2 | 16 | 0.125 | 3 | 0.3333 | 3 | 0.3333 | 4 | 0 | 6 | 0.3333 | 16 | 0.25 |
| 10_1123010 | 5 | 1 | 4 | 0.75 | 3 | 1 | 5 | 1 | 17 | 0.9412 | 3 | 1 | 3 | 1 | 3 | 0.6667 | 6 | 1 | 15 | 0.9333 |
| 10_1123012 | 5 | 1 | 6 | 1 | 3 | 1 | 6 | 1 | 20 | 1 | 2 | 1 | 2 | 1 | 3 | 1 | 4 | 1 | 11 | 1 |
| 10_1123012 | 5 | 1 | 7 | 1 | 3 | 0.6667 | 6 | 1 | 21 | 0.9524 | 4 | 1 | 2 | 1 | 4 | 1 | 4 | 0.5 | 14 | 0.8571 |
| 10_1123012 | 5 | 0 | 4 | 0.25 | 4 | 0.25 | 6 | 0.1667 | 19 | 0.1579 | 4 | 0 | 3 | 0 | 4 | 0 | 6 | 0.1667 | 17 | 0.0588 |
| 10_1123013 | 4 | 0.75 | 5 | 0.8 | 4 | 1 | 4 | 1 | 17 | 0.8824 | 4 | 1 | 3 | 1 | 4 | 1 | 4 | 0.75 | 15 | 0.9333 |
| 10_1123013 | 5 | 0.4 | 4 | 0.75 | 4 | 0.75 | 5 | 0.4 | 18 | 0.5556 | 4 | 0.5 | 3 | 0.3333 | 3 | 1 | 5 | 0.8 | 15 | 0.6667 |
| 10_1123021 | 6 | 0.3333 | 9 | 0.3333 | 6 | 0.3333 | 5 | 0.2 | 26 | 0.3077 | 4 | 0 | 3 | 0 | 5 | 0.2 | 5 | 0.4 | 17 | 0.1765 |
| 10_1127622 | 13 | 0.1538 | 18 | 0.1111 | 14 | 0.0714 | 10 | 0.1 | 55 | 0.1091 | 10 | 0.1 | 13 | 0.0769 | 12 | 0.0833 | 8 | 0.25 | 43 | 0.1163 |
| 10_1127623 | 11 | 0.4545 | 20 | 0.25 | 12 | 0.3333 | 10 | 0.2 | 53 | 0.3019 | 8 | 0.5 | 12 | 0.25 | 10 | 0.4 | 11 | 0.6364 | 41 | 0.439 |
| 10_1127624 | 10 | 0.3 | 18 | 0.1667 | 14 | 0.3571 | 10 | 0.2 | 52 | 0.25 | 10 | 0.6 | 12 | 0.1667 | 9 | 0.3333 | 10 | 0.3 | 41 | 0.3415 |
| 10_1127630 | 7 | 0.5714 | 13 | 0.3846 | 8 | 0.25 | 6 | 0.1667 | 34 | 0.3529 | 9 | 0.2222 | 7 | 0.1429 | 7 | 0.2857 | 4 | 0 | 27 | 0.1852 |
| 10_1127631 | 5 | 1 | 13 | 0.9231 | 8 | 1 | 7 | 1 | 33 | 0.9697 | 10 | 1 | 9 | 0.8889 | 7 | 1 | 8 | 1 | 34 | 0.9706 |
| 10_1127631 | 5 | 0 | 14 | 0.2143 | 9 | 0.2222 | 7 | 0 | 35 | 0.1429 | 10 | 0 | 9 | 0.2222 | 7 | 0 | 8 | 0.125 | 34 | 0.0882 |
| 10_1127632 | 5 | 0.8 | 10 | 1 | 4 | 1 | 6 | 1 | 25 | 0.96 | 9 | 0.8889 | 7 | 1 | 7 | 1 | 5 | 1 | 28 | 0.9643 |
| 10_1127633 | 6 | 1 | 12 | 1 | 8 | 1 | 7 | 1 | 33 | 1 | 10 | 1 | 7 | 1 | 7 | 1 | 8 | 1 | 32 | 1 |
| 10_1127633 | 5 | 0.4 | 11 | 0.5455 | 7 | 0.4286 | 5 | 0.2 | 28 | 0.4286 | 6 | 0.1667 | 6 | 0.5 | 5 | 0.6 | 7 | 0 | 24 | 0.2917 |
| 10_1127634 | 4 | 1 | 13 | 0.9231 | 9 | 1 | 3 | 1 | 29 | 0.9655 | 10 | 1 | 6 | 1 | 4 | 0.75 | 8 | 1 | 28 | 0.9643 |
| 10_1127634 | 3 | 0.6667 | 11 | 0.3636 | 8 | 0.375 | 7 | 0.4286 | 29 | 0.4138 | 10 | 0.3 | 8 | 0.25 | 6 | 0.3333 | 5 | 0.4 | 29 | 0.3103 |
| 10_1127635 | 6 | 0.6667 | 14 | 0.7857 | 9 | 0.8889 | 6 | 0.8333 | 35 | 0.8 | 9 | 0.8889 | 8 | 0.625 | 6 | 0.6667 | 6 | 0.8333 | 29 | 0.7586 |
| 10_1127636 | 6 | 0 | 12 | 0.25 | 9 | 0 | 7 | 0 | 34 | 0.0882 | 8 | 0.125 | 7 | 0.1429 | 5 | 0 | 7 | 0 | 27 | 0.0741 |
| 10_1127636 | 4 | 0.25 | 9 | 0.6667 | 7 | 0.4286 | 7 | 0.1429 | 27 | 0.4074 | 7 | 0.2857 | 6 | 0.1667 | 5 | 0.4 | 8 | 0.25 | 26 | 0.2692 |
| 10_1127637 | 6 | 0.8333 | 11 | 0.9091 | 9 | 0.7778 | 7 | 0.5714 | 33 | 0.7879 | 10 | 0.8 | 7 | 0.5714 | 5 | 0.8 | 8 | 0.875 | 30 | 0.7667 |
| 10_1127638 | 5 | 0.8 | 11 | 1 | 8 | 1 | 7 | 1 | 31 | 0.9677 | 9 | 1 | 6 | 1 | 6 | 1 | 6 | 1 | 27 | 1 |
| 10_1127639 | NA | NA | 2 | 1 | 7 | 1 | 4 | 0.5 | 13 | 0.8462 | 1 | 1 | 3 | 1 | NA | NA | NA | NA | 4 | 1 |
| 10_1127639 | 5 | 0.6 | 10 | 0.8 | 9 | 0.7778 | 7 | 0.5714 | 31 | 0.7097 | 7 | 0.7143 | 6 | 0.5 | 6 | 1 | 5 | 0.6 | 24 | 0.7083 |
| 10_1127641 | 1 | 0 | 4 | 0.25 | 8 | 0.125 | 3 | 0.3333 | 16 | 0.1875 | 7 | 0.2857 | 1 | 1 | 2 | 0 | 6 | 0.1667 | 16 | 0.25 |
| 10_1127642 | 4 | 1 | 10 | 1 | 7 | 0.8571 | 3 | 1 | 24 | 0.9583 | 6 | 1 | 5 | 1 | 4 | 1 | 3 | 1 | 18 | 1 |
| 10_1127642 | 5 | 0.6 | 10 | 0.3 | 7 | 0.5714 | 3 | 0.6667 | 25 | 0.48 | 9 | 0.4444 | 6 | 0.3333 | 4 | 1 | 6 | 0.5 | 25 | 0.52 |
| 10_1127644 | 5 | 0.4 | 7 | 0.8571 | 5 | 0 | 1 | 1 | 18 | 0.5 | 8 | 0.125 | 5 | 0.6 | 2 | 0 | 5 | 0.2 | 20 | 0.25 |
| 10_1127644 | 4 | 0 | 10 | 0.2 | 5 | 0 | 3 | 0.3333 | 22 | 0.1364 | 8 | 0.375 | 4 | 0.25 | 4 | 0 | 5 | 0.4 | 21 | 0.2857 |
| 10_1127645 | 5 | 0 | 9 | 0.1111 | 6 | 0.1667 | 1 | 0 | 21 | 0.0952 | 9 | 0.1111 | 5 | 0 | 4 | 0.5 | 5 | 0 | 23 | 0.1304 |
| 10_1142356 | NA | NA | 2 | 0 | 1 | 1 | NA | NA | 3 | 0.3333 | NA | NA | NA | NA | 1 | 0 | 1 | 1 | 2 | 0.5 |
| 10_1142359 | NA | NA | 2 | 0.5 | 1 | 1 | 1 | 0 | 4 | 0.5 | 2 | 0 | NA | NA | 2 | 0.5 | 1 | 0 | 5 | 0.2 |
| 10_1142364 | NA | NA | 4 | 0.75 | 1 | 0 | 1 | 0 | 6 | 0.5 | 2 | 0 | NA | NA | 3 | 0.3333 | NA | NA | 5 | 0.2 |

| | | | | | | | | | | | | | | | | | | | | |
|---|---|---|---|---|---|---|---|---|---|---|---|---|---|---|---|---|---|---|---|---|
| 10_114236 | NA | NA | 4 | 0.5 | 1 | 1 | 1 | 0 | 6 | 0.5 | 2 | 1 | NA | NA | 3 | 0.3333 | NA | NA | 5 | 0.6 |
| 10_114236 | NA | NA | 4 | 1 | 1 | 1 | 1 | 1 | 6 | 1 | 1 | 1 | NA | NA | 3 | 1 | NA | NA | 4 | 1 |
| 10_114236 | NA | NA | 4 | 0 | 1 | 0 | 1 | 1 | 6 | 0.1667 | 1 | 0 | NA | NA | 3 | 0.3333 | NA | NA | 4 | 0.25 |
| 10_114237 | NA | NA | 2 | 1 | NA | NA | 1 | 1 | 3 | 1 | 1 | 1 | NA | NA | 3 | 1 | NA | NA | 4 | 1 |
| 10_114237 | NA | NA | 3 | 0.6667 | 1 | 1 | 1 | 1 | 5 | 0.8 | 1 | 1 | NA | NA | 3 | 0.6667 | NA | NA | 4 | 0.75 |
| 10_114237 | NA | NA | 3 | 0.3333 | 1 | 1 | NA | NA | 4 | 0.5 | 1 | 1 | NA | NA | 3 | 0.3333 | NA | NA | 4 | 0.5 |
| 10_114237 | NA | NA | 2 | 1 | 1 | 1 | NA | NA | 3 | 1 | 1 | 1 | NA | NA | 3 | 1 | NA | NA | 4 | 1 |
| 10_114237 | NA | NA | 4 | 0.5 | 1 | 1 | NA | NA | 5 | 0.6 | 1 | 1 | NA | NA | 3 | 0 | NA | NA | 4 | 0.25 |
| 10_114237 | NA | NA | 4 | 0.25 | 1 | 1 | NA | NA | 5 | 0.4 | 1 | 1 | NA | NA | 3 | 0 | NA | NA | 4 | 0.25 |
| 10_114636 | NA | NA | 2 | 0.5 | NA | NA | NA | NA | 2 | 0.5 | NA | NA | NA | NA | 1 | 1 | NA | NA | 1 | 1 |
| 10_115034 | 11 | 0.0909 | 10 | 0 | 2 | 0 | 5 | 0.2 | 28 | 0.0714 | 2 | 0.5 | 4 | 0.5 | 7 | 0.1429 | 5 | 0.2 | 18 | 0.2778 |
| 10_115035 | 9 | 0.5556 | 8 | 0.625 | 1 | 0 | 4 | 0.25 | 22 | 0.5 | 2 | 1 | 4 | 0.75 | 6 | 0.6667 | 5 | 0.6 | 17 | 0.7059 |
| 10_115035 | 9 | 0.3333 | 9 | 0.3333 | 2 | 0 | 4 | 0 | 24 | 0.25 | 2 | 0.5 | 4 | 0 | 5 | 0 | 4 | 0.75 | 15 | 0.2667 |
| 10_115035 | 12 | 0.3333 | 8 | 0.25 | 2 | 0 | 2 | 0 | 24 | 0.25 | 2 | 0.5 | 2 | 0 | 7 | 0 | 5 | 0.2 | 16 | 0.125 |
| 10_115035 | 10 | 0.1 | 8 | 0.25 | 2 | 0.5 | 4 | 0.5 | 24 | 0.25 | 2 | 0.5 | 4 | 0.25 | 7 | 0 | 4 | 0.25 | 17 | 0.1765 |
| 10_115123 | 7 | 0 | 2 | 0 | 5 | 0.4 | 4 | 0 | 18 | 0.1111 | 2 | 0.5 | 3 | 0.3333 | 2 | 0.5 | 3 | 0 | 10 | 0.3 |
| 10_115124 | 5 | 0.4 | 3 | 0 | 5 | 0.2 | 5 | 0.4 | 18 | 0.2778 | 4 | 0 | 6 | 0 | 7 | 0.2857 | 7 | 0.1429 | 24 | 0.125 |
| 10_115125 | 5 | 0 | 2 | 0.5 | 5 | 0 | 3 | 0 | 15 | 0.0667 | 4 | 0 | 5 | 0 | 7 | 0 | 7 | 0.2857 | 23 | 0.087 |
| 10_115125 | 7 | 0 | 9 | 0.3333 | 8 | 0 | 7 | 0 | 31 | 0.0968 | 7 | 0.1429 | 8 | 0.125 | 10 | 0.1 | 10 | 0 | 35 | 0.0857 |
| 10_115126 | 15 | 0.3333 | 13 | 0.3077 | 17 | 0.1765 | 12 | 0.3333 | 57 | 0.2807 | 15 | 0.2667 | 17 | 0.4118 | 13 | 0.5385 | 18 | 0.4444 | 63 | 0.4127 |
| 10_115126 | 13 | 0.0769 | 12 | 0.0833 | 16 | 0.1875 | 11 | 0.2727 | 52 | 0.1538 | 15 | 0.1333 | 18 | 0.2222 | 13 | 0.0769 | 18 | 0.1667 | 64 | 0.1562 |
| 10_115126 | 13 | 0.9231 | 10 | 0.9 | 14 | 0.8571 | 10 | 0.7 | 47 | 0.8511 | 16 | 0.9375 | 14 | 0.9286 | 13 | 1 | 16 | 0.875 | 59 | 0.9322 |
| 10_115151 | 8 | 0.375 | 5 | 0.4 | 11 | 0.4545 | 9 | 0.4444 | 33 | 0.4242 | 5 | 0.8 | 16 | 0.6875 | 9 | 0.3333 | 9 | 0.4444 | 39 | 0.5641 |
| 10_115176 | 2 | 0 | 1 | 0 | 1 | 0 | 1 | 1 | 5 | 0.2 | 1 | 0 | 2 | 0 | 2 | 0 | 2 | 0.5 | 7 | 0.1429 |
| 10_115177 | 4 | 0.25 | 3 | 1 | 1 | 1 | 3 | 0.6667 | 11 | 0.6364 | 1 | 1 | 2 | 0.5 | 5 | 0.4 | 1 | 0 | 9 | 0.4444 |
| 10_115177 | 5 | 0.4 | 3 | 0.6667 | 1 | 0 | 3 | 0 | 12 | 0.3333 | 1 | 0 | 2 | 0 | 4 | 0 | 2 | 0.5 | 9 | 0.1111 |
| 10_115177 | 5 | 0.2 | 4 | 0 | 1 | 1 | 4 | 0.25 | 14 | 0.2143 | 4 | 0 | 2 | 0 | 5 | 0 | 3 | 0.3333 | 14 | 0.0714 |
| 10_115178 | 3 | 0.6667 | 4 | 0.75 | 1 | 1 | 4 | 0.75 | 12 | 0.75 | 3 | 1 | 2 | 0 | 5 | 0.6 | 3 | 1 | 13 | 0.6923 |
| 10_115178 | 3 | 1 | 4 | 0.5 | 1 | 1 | 2 | 0.5 | 10 | 0.7 | 3 | 0.6667 | 2 | 0.5 | 5 | 0.8 | 1 | 1 | 11 | 0.7273 |
| 10_115178 | 3 | 0 | 4 | 0.25 | NA | NA | 4 | 0.25 | 11 | 0.1818 | 4 | 0 | 3 | 0 | 6 | 0.1667 | 1 | 0 | 14 | 0.0714 |
| 10_115178 | 5 | 0.8 | 4 | 0.75 | 2 | 1 | 4 | 1 | 15 | 0.8667 | 6 | 0.5 | 4 | 0.75 | 5 | 0.8 | 3 | 1 | 18 | 0.7222 |
| 10_115178 | 6 | 0.3333 | 4 | 0.25 | 2 | 0.5 | 4 | 0.5 | 16 | 0.375 | 5 | 0.2 | 4 | 0 | 4 | 0 | 2 | 0 | 15 | 0.0667 |
| 10_115178 | 6 | 1 | 4 | 1 | 1 | 0 | 3 | 1 | 14 | 0.9286 | 3 | 1 | 1 | 1 | 4 | 1 | 3 | 1 | 11 | 1 |
| 10_115248 | 1 | 1 | NA | NA | NA | NA | NA | NA | 1 | 1 | NA | NA | NA | NA | 1 | 1 | NA | NA | 1 | 1 |
| 10_115267 | 1 | 0 | 2 | 0.5 | 1 | 0 | 2 | 0 | 6 | 0.1667 | 1 | 0 | 3 | 0 | 4 | 0.25 | 1 | 0 | 9 | 0.1111 |
| 10_115267 | 2 | 0.5 | 1 | 0 | NA | NA | 2 | 1 | 5 | 0.6 | 1 | 1 | 2 | 0.5 | 3 | 0.6667 | 2 | 0.5 | 8 | 0.625 |
| 10_115268 | 1 | 0 | 1 | 1 | 1 | 0 | 2 | 1 | 5 | 0.6 | 2 | 0.5 | 3 | 0.6667 | 3 | 0.6667 | 2 | 0.5 | 10 | 0.6 |
| 10_115268 | 3 | 0.3333 | 2 | 1 | 1 | 0 | 2 | 0 | 8 | 0.375 | 2 | 0 | 3 | 0.3333 | 4 | 0 | 2 | 0 | 11 | 0.0909 |
| 10_115268 | 2 | 0 | 2 | 0 | NA | NA | 2 | 1 | 6 | 0.3333 | 1 | 1 | 1 | 0 | 3 | 0.6667 | 1 | 0 | 6 | 0.5 |
| 10_115268 | 2 | 1 | 3 | 0.6667 | NA | NA | 2 | 1 | 7 | 0.8571 | 1 | 0 | 1 | 1 | 2 | 1 | 1 | 0 | 5 | 0.6 |
| 10_115268 | 3 | 1 | 4 | 0.75 | 1 | 1 | 2 | 0.5 | 10 | 0.8 | 3 | 0.6667 | 3 | 0.6667 | 4 | 0.5 | 3 | 0.6667 | 13 | 0.6154 |
| 10_115287 | NA | NA | 2 | 0.5 | NA | NA | NA | NA | 2 | 0.5 | NA | NA | 1 | 1 | NA | NA | NA | NA | 1 | 1 |
| 10_115287 | NA | NA | 2 | 0.5 | NA | NA | NA | NA | 2 | 0.5 | NA | NA | 1 | 1 | NA | NA | NA | NA | 1 | 1 |
| 10_115288 | NA | NA | 2 | 1 | NA | NA | NA | NA | 2 | 1 | NA | NA | 1 | 1 | NA | NA | NA | NA | 1 | 1 |
| 10_115854 | 3 | 0.3333 | 1 | 0 | 2 | 1 | 2 | 0.5 | 8 | 0.5 | 1 | 1 | 1 | 1 | 4 | 1 | 1 | 1 | 7 | 1 |
| 10_115884 | 4 | 0 | 1 | 0 | 2 | 0 | 2 | 0.5 | 9 | 0.1111 | NA | NA | 2 | 0 | 4 | 0.25 | 1 | 0 | 7 | 0.1429 |
| 10_116109 | 5 | 0.2 | 2 | 0 | 2 | 0.5 | 5 | 0.2 | 14 | 0.2143 | 2 | 0.5 | 4 | 0 | 3 | 0 | 4 | 0.5 | 13 | 0.2308 |
| 10_116109 | 4 | 0.25 | 3 | 0.3333 | 2 | 0 | 5 | 0.2 | 14 | 0.2143 | 1 | 0 | 4 | 0.75 | 3 | 0.3333 | 4 | 0 | 12 | 0.3333 |
| 10_116109 | 6 | 1 | 3 | 1 | 1 | 1 | 4 | 1 | 14 | 1 | 2 | 1 | 4 | 1 | 4 | 1 | 4 | 1 | 14 | 1 |
| 10_116110 | 7 | 0.5714 | 3 | 1 | 2 | 0.5 | 5 | 0.8 | 17 | 0.7059 | 2 | 1 | 5 | 1 | 5 | 1 | 4 | 0.5 | 16 | 0.875 |
| 10_116110 | 6 | 0.6667 | 4 | 0.5 | 2 | 1 | 4 | 1 | 16 | 0.75 | 2 | 0.5 | 5 | 0.8 | 5 | 0.4 | 2 | 1 | 14 | 0.6429 |
| 10_116110 | 6 | 0.1667 | 3 | 0.3333 | 2 | 1 | 5 | 0.2 | 16 | 0.3125 | 1 | 0 | 5 | 0 | 5 | 0.2 | 2 | 0 | 13 | 0.0769 |
| 10_116110 | 8 | 0.75 | 4 | 0.5 | 2 | 0.5 | 4 | 0.5 | 18 | 0.6111 | 2 | 1 | 3 | 0.6667 | 4 | 1 | 4 | 0 | 13 | 0.6154 |
| 10_116110 | 7 | 0 | 3 | 0.3333 | 2 | 0 | 5 | 0 | 17 | 0.0588 | 2 | 0 | 4 | 0.25 | 5 | 0.2 | 4 | 0 | 15 | 0.1333 |
| 10_116110 | 8 | 0.125 | 4 | 0.25 | 2 | 1 | 4 | 0.25 | 18 | 0.2778 | 2 | 0.5 | 3 | 0.3333 | 5 | 0.6 | 4 | 0.25 | 14 | 0.4286 |
| 10_116110 | 9 | 0 | 4 | 0 | 1 | 0 | 5 | 0.2 | 19 | 0.0526 | 2 | 0 | 4 | 0.25 | 4 | 0 | 3 | 0 | 13 | 0.0769 |
| 10_116223 | 2 | 0.5 | NA | NA | 2 | 0 | 1 | 0 | 5 | 0.2 | 1 | 0 | 2 | 0 | 2 | 0.5 | NA | NA | 5 | 0.2 |
| 10_116223 | 1 | 1 | 1 | 1 | 2 | 0.5 | 1 | 1 | 5 | 0.8 | 1 | 1 | 2 | 0.5 | 1 | 1 | 1 | 1 | 5 | 0.8 |
| 10_116223 | 2 | 0.5 | 1 | 1 | 2 | 0.5 | 1 | 1 | 6 | 0.6667 | 1 | 0 | 3 | 0.6667 | 2 | 0.5 | 2 | 0.5 | 8 | 0.5 |
| 10_116223 | 2 | 0.5 | 1 | 0 | 2 | 0 | 1 | 0 | 6 | 0.1667 | 1 | 1 | 3 | 0.3333 | 2 | 1 | 2 | 0.5 | 8 | 0.625 |
| 10_116223 | 2 | 0 | 1 | 0 | 2 | 0.5 | 1 | 0 | 6 | 0.1667 | 1 | 0 | 3 | 0.3333 | 2 | 0 | 2 | 0.5 | 8 | 0.25 |
| 10_116223 | 2 | 0.5 | 1 | 0 | 2 | 0 | 1 | 1 | 6 | 0.3333 | NA | NA | 3 | 0.3333 | 2 | 0 | 1 | 1 | 6 | 0.3333 |
| 10_116223 | 2 | 0.5 | 1 | 0 | 1 | 1 | 1 | 0 | 5 | 0.4 | 1 | 0 | 2 | 0.5 | 2 | 0 | 2 | 1 | 7 | 0.4286 |
| 10_116223 | 1 | 1 | 1 | 0 | 2 | 0 | 1 | 1 | 5 | 0.4 | NA | NA | 2 | 0.5 | 2 | 0.5 | 1 | 0 | 5 | 0.4 |
| 10_116223 | 1 | 1 | 1 | 1 | 2 | 0.5 | 1 | 1 | 5 | 0.8 | 1 | 1 | 3 | 0.6667 | 2 | 1 | 2 | 1 | 8 | 0.875 |

| | | | | | | | | | | | | | | | | | | | | |
|---|---|---|---|---|---|---|---|---|---|---|---|---|---|---|---|---|---|---|---|---|
| 10_1162239 | 2 | 0.5 | 1 | 0 | 2 | 0.5 | 1 | 1 | 6 | 0.5 | 1 | 1 | 3 | 0.6667 | 2 | 0.5 | 2 | 0 | 8 | 0.5 |
| 10_1162239 | 2 | 0.5 | 1 | 0 | 2 | 0 | 1 | 0 | 6 | 0.1667 | 1 | 0 | 3 | 0.3333 | 2 | 0 | 2 | 0.5 | 8 | 0.25 |
| 10_1162239 | 2 | 1 | 1 | 0 | 2 | 0.5 | 1 | 0 | 6 | 0.5 | NA | NA | 3 | 0 | 1 | 0 | 2 | 0.5 | 6 | 0.1667 |
| 10_1162240 | 2 | 0.5 | 1 | 0 | 1 | 1 | 1 | 0 | 5 | 0.4 | 1 | 0 | 3 | 1 | 2 | 0.5 | 2 | 0.5 | 8 | 0.625 |
| 10_1162240 | 1 | 0 | 1 | 0 | 1 | 0 | 1 | 1 | 4 | 0.25 | 1 | 1 | 3 | 0 | 1 | 1 | 1 | 1 | 6 | 0.5 |
| 10_1162507 | 1 | 1 | NA | NA | NA | NA | NA | NA | 1 | 1 | 1 | 0 | 1 | 1 | NA | NA | NA | NA | 2 | 0.5 |
| 10_1162507 | 2 | 1 | NA | NA | NA | NA | NA | NA | 2 | 1 | 1 | 1 | 1 | 1 | 1 | 0 | NA | NA | 3 | 0.6667 |
| 10_1162508 | 2 | 0.5 | NA | NA | NA | NA | NA | NA | 2 | 0.5 | 1 | 1 | 1 | 0 | 1 | 0 | NA | NA | 3 | 0.3333 |
| 10_1162537 | 10 | 0 | 9 | 0.1111 | 6 | 0 | 12 | 0.1667 | 37 | 0.0811 | 9 | 0.3333 | 5 | 0 | 12 | 0.1667 | 1 | 0 | 27 | 0.1852 |
| 10_1162538 | 10 | 0.5 | 11 | 0.6364 | 5 | 0.6 | 7 | 0.4286 | 33 | 0.5455 | 10 | 0.5 | 5 | 0.4 | 15 | 0.5333 | 5 | 0.4 | 35 | 0.4857 |
| 10_1162539 | 11 | 0.2727 | 9 | 0.2222 | 7 | 0.1429 | 12 | 0.3333 | 39 | 0.2564 | 11 | 0.1818 | 5 | 0.4 | 13 | 0.2308 | 5 | 0.2 | 34 | 0.2353 |
| 10_1162539 | 13 | 0.9231 | 13 | 0.6154 | 8 | 0.625 | 15 | 0.6 | 49 | 0.6939 | 12 | 0.5 | 7 | 0.5714 | 15 | 0.6667 | 6 | 0.8333 | 40 | 0.625 |
| 10_1162539 | 11 | 0.7273 | 12 | 0.8333 | 6 | 0.5 | 12 | 0.75 | 41 | 0.7317 | 13 | 0.6923 | 6 | 0.5 | 12 | 0.75 | 5 | 0.6 | 36 | 0.6667 |
| 10_1163108 | 1 | 1 | 2 | 0.5 | NA | NA | NA | NA | 3 | 0.6667 | 1 | 1 | NA | NA | NA | NA | NA | NA | 1 | 1 |
| 10_1163211 | 7 | 0.4286 | 4 | 0 | 8 | 0.375 | 2 | 0 | 21 | 0.2857 | 7 | 0.2857 | 5 | 0.4 | 6 | 0.6667 | 8 | 0.25 | 26 | 0.3846 |
| 10_1163212 | 7 | 0 | 4 | 0.5 | 9 | 0.3333 | 2 | 0 | 22 | 0.2273 | 7 | 0.1429 | 5 | 0.2 | 5 | 0.2 | 11 | 0.0909 | 28 | 0.1429 |
| 10_1163212 | 8 | 0.875 | 6 | 0.3333 | 9 | 0.4444 | 2 | 0 | 25 | 0.52 | 6 | 0.3333 | 5 | 0.6 | 6 | 0.3333 | 11 | 0.3636 | 28 | 0.3929 |
| 10_1163212 | 5 | 0 | 4 | 0.5 | 7 | 0.4286 | 1 | 0 | 17 | 0.2941 | 5 | 0.2 | 4 | 0.5 | 6 | 0.6667 | 7 | 0.5714 | 22 | 0.5 |
| 10_1163213 | 5 | 0.6 | 4 | 0.5 | 10 | 0.4 | 2 | 0 | 21 | 0.4286 | 6 | 0.3333 | 4 | 0.5 | 6 | 0.3333 | 9 | 0.6667 | 25 | 0.48 |
| 10_1163217 | 7 | 0.1429 | 7 | 0.4286 | 9 | 0.6667 | 2 | 0.5 | 25 | 0.44 | 5 | 0.4 | 5 | 1 | 9 | 0.4444 | 12 | 0.4167 | 31 | 0.5161 |
| 10_1163218 | 5 | 0.6 | 4 | 0 | 7 | 0.2857 | 1 | 0 | 17 | 0.2941 | 4 | 0 | 5 | 0.2 | 8 | 0.375 | 12 | 0.1667 | 29 | 0.2069 |
| 10_1163221 | 5 | 0.6 | 6 | 0.3333 | 8 | 0.25 | NA | NA | 19 | 0.3684 | 5 | 0.8 | 3 | 0 | 8 | 0.5 | 11 | 0.2727 | 27 | 0.4074 |
| 10_1167417 | 1 | 0 | 1 | 1 | 3 | 1 | 2 | 1 | 7 | 0.8571 | NA | NA | 1 | 0 | 4 | 1 | 2 | 1 | 7 | 0.8571 |
| 10_1167420 | 1 | 0 | NA | NA | 2 | 0.5 | 2 | 0.5 | 5 | 0.4 | NA | NA | 1 | 0 | 2 | 0.5 | 1 | 0 | 4 | 0.25 |
| 10_1167421 | 1 | 1 | 1 | 1 | 2 | 1 | 3 | 1 | 7 | 1 | NA | NA | 1 | 0 | 4 | 0.5 | 1 | 1 | 6 | 0.5 |
| 10_1167427 | 7 | 0.2857 | 8 | 0.625 | 4 | 0.75 | 10 | 0.6 | 29 | 0.5517 | NA | NA | 7 | 0.4286 | 4 | 0.5 | 7 | 0.4286 | 18 | 0.4444 |
| 10_1167428 | 5 | 0.2 | 6 | 0.3333 | 5 | 0.6 | 11 | 0.5455 | 27 | 0.4444 | NA | NA | 7 | 0.2857 | 4 | 0.25 | 7 | 0.1429 | 18 | 0.2222 |
| 10_1167428 | 4 | 0.25 | 5 | 0.6 | 6 | 0.3333 | 10 | 0.4 | 25 | 0.4 | 1 | 1 | 5 | 0.4 | 2 | 0.5 | 7 | 0.2857 | 15 | 0.4 |
| 10_1167429 | 6 | 0 | 7 | 0.1429 | 4 | 0 | 11 | 0.0909 | 28 | 0.0714 | 1 | 0 | 7 | 0.1429 | 2 | 0 | 7 | 0.2857 | 17 | 0.1765 |
| 10_1167429 | 5 | 0.4 | 5 | 0.4 | 5 | 0.8 | 8 | 0.75 | 23 | 0.6087 | NA | NA | 6 | 0.1667 | 1 | 0 | 7 | 0.7143 | 14 | 0.4286 |
| 10_1167981 | NA | NA | 2 | 1 | NA | NA | NA | NA | 2 | 1 | 2 | 1 | NA | NA | NA | NA | NA | NA | 2 | 1 |
| 10_1167981 | NA | NA | 2 | 1 | NA | NA | NA | NA | 2 | 1 | 2 | 1 | NA | NA | NA | NA | NA | NA | 2 | 1 |
| 10_1167987 | NA | NA | 2 | 1 | NA | NA | NA | NA | 2 | 1 | 2 | 1 | 1 | 0 | NA | NA | NA | NA | 3 | 0.6667 |
| 10_1167988 | NA | NA | 2 | 1 | NA | NA | 1 | 1 | 3 | 1 | 2 | 0.5 | NA | NA | NA | NA | NA | NA | 2 | 0.5 |
| 10_1169065 | 1 | 0 | 2 | 0.5 | 2 | 0.5 | 2 | 0.5 | 7 | 0.4286 | 2 | 0.5 | 4 | 0.25 | 1 | 1 | 3 | 0 | 10 | 0.3 |
| 10_1169066 | 1 | 1 | 4 | 0 | 2 | 0 | 3 | 0 | 10 | 0.1 | 3 | 0.3333 | 7 | 0.1429 | 1 | 0 | 3 | 0 | 14 | 0.1429 |
| 10_1169070 | 1 | 0 | 5 | 0 | 4 | 0 | 3 | 0.3333 | 13 | 0.0769 | 1 | 0 | 7 | 0.1429 | 1 | 0 | 2 | 0 | 11 | 0.0909 |
| 10_1169071 | 1 | 0 | 4 | 0.25 | 4 | 0.5 | 3 | 0.3333 | 12 | 0.3333 | 2 | 0.5 | 6 | 0.1667 | 1 | 0 | 1 | 0 | 10 | 0.2 |
| 10_1169072 | 1 | 0 | 5 | 0.4 | 3 | 0 | 3 | 0.6667 | 12 | 0.3333 | 2 | 0.5 | 7 | 0.2857 | 1 | 0 | 1 | 0 | 11 | 0.2727 |
| 10_1169096 | NA | NA | 1 | 0 | NA | NA | 2 | 1 | 3 | 0.6667 | 2 | 0.5 | 5 | 0.8 | 1 | 1 | NA | NA | 8 | 0.75 |
| 10_1169097 | NA | NA | 3 | 1 | NA | NA | 2 | 1 | 5 | 1 | 2 | 1 | 5 | 1 | 1 | 1 | NA | NA | 8 | 1 |
| 10_1169099 | NA | NA | 3 | 1 | NA | NA | 2 | 1 | 5 | 1 | 2 | 1 | 5 | 1 | 1 | 1 | NA | NA | 8 | 1 |
| 10_1169099 | NA | NA | 2 | 0.5 | NA | NA | 2 | 1 | 4 | 0.75 | 2 | 1 | 3 | 1 | 1 | 1 | NA | NA | 6 | 1 |
| 10_1169100 | NA | NA | 3 | 0.6667 | NA | NA | 2 | 0.5 | 5 | 0.6 | 2 | 0.5 | 5 | 0.4 | NA | NA | NA | NA | 7 | 0.4286 |
| 10_1169100 | NA | NA | 2 | 1 | NA | NA | 1 | 1 | 3 | 1 | 2 | 0.5 | 5 | 1 | NA | NA | NA | NA | 7 | 0.8571 |
| 10_1169100 | NA | NA | 3 | 0.3333 | NA | NA | 2 | 0 | 5 | 0.2 | 1 | 0 | 4 | 0.25 | 1 | 0 | NA | NA | 6 | 0.1667 |
| 10_1169101 | NA | NA | 2 | 0.5 | NA | NA | 2 | 0.5 | 4 | 0.5 | 2 | 1 | 3 | 1 | 1 | 1 | NA | NA | 6 | 1 |
| 10_1169101 | NA | NA | 3 | 0.3333 | NA | NA | 2 | 0 | 5 | 0.2 | 2 | 0.5 | 3 | 0 | 1 | 0 | NA | NA | 6 | 0.1667 |
| 10_1169102 | NA | NA | 3 | 0 | NA | NA | 2 | 0.5 | 5 | 0.2 | 2 | 1 | 3 | 0.6667 | 1 | 0 | NA | NA | 6 | 0.6667 |
| 10_1169102 | NA | NA | 3 | 0.6667 | NA | NA | 2 | 0.5 | 5 | 0.6 | 2 | 0 | 3 | 0.6667 | 1 | 1 | NA | NA | 6 | 0.5 |
| 10_1169102 | NA | NA | 3 | 0 | NA | NA | 2 | 0.5 | 5 | 0.2 | 2 | 0.5 | 4 | 0.25 | 1 | 1 | NA | NA | 7 | 0.4286 |
| 10_1169103 | NA | NA | 3 | 0.3333 | NA | NA | 1 | 1 | 4 | 0.5 | 2 | 0.5 | 4 | 0 | NA | NA | NA | NA | 6 | 0.1667 |
| 10_1169104 | NA | NA | 3 | 1 | NA | NA | NA | NA | 3 | 1 | 1 | 1 | 3 | 1 | NA | NA | NA | NA | 4 | 1 |
| 10_1169105 | NA | NA | 3 | 0.3333 | NA | NA | NA | NA | 3 | 0.3333 | 2 | 1 | 3 | 1 | NA | NA | NA | NA | 5 | 1 |
| 10_1169105 | NA | NA | 2 | 0.5 | NA | NA | NA | NA | 2 | 0.5 | 2 | 0 | 3 | 0.3333 | NA | NA | NA | NA | 5 | 0.2 |
| 10_1170513 | 6 | 0.1667 | 3 | 0 | 6 | 0.1667 | 9 | 0 | 24 | 0.0833 | 6 | 0.3333 | 5 | 0 | 6 | 0 | 6 | 0.3333 | 23 | 0.1739 |
| 10_1170513 | 7 | 0.2857 | 3 | 0 | 8 | 0.5 | 11 | 0.3636 | 29 | 0.3448 | 7 | 0.2857 | 8 | 0.25 | 5 | 0.4 | 8 | 0.125 | 28 | 0.25 |
| 10_1170514 | 7 | 0.8571 | 4 | 0.5 | 9 | 0.8889 | 10 | 0.7 | 30 | 0.7667 | 7 | 0.7143 | 8 | 1 | 7 | 0.5714 | 8 | 0.75 | 30 | 0.7667 |
| 10_1170515 | 7 | 0.5714 | 2 | 1 | 7 | 0.4286 | 9 | 0.5556 | 25 | 0.56 | 5 | 0.6 | 8 | 0.75 | 7 | 0.4286 | 8 | 0.375 | 28 | 0.5357 |
| 10_1170515 | 6 | 0.3333 | 4 | 0.5 | 7 | 0.2857 | 10 | 0.2 | 27 | 0.2963 | 6 | 0.3333 | 6 | 0.6667 | 5 | 0.8 | 6 | 0.5 | 23 | 0.5652 |
| 10_1170516 | 6 | 1 | 2 | 1 | 6 | 1 | 12 | 1 | 26 | 1 | 6 | 1 | 5 | 1 | 5 | 1 | 7 | 1 | 23 | 1 |
| 10_1170516 | 7 | 0.2857 | 4 | 0 | 7 | 0.1429 | 9 | 0.1111 | 27 | 0.1481 | 7 | 0.1429 | 8 | 0 | 5 | 0.4 | 8 | 0.25 | 28 | 0.1786 |
| 10_1171136 | 4 | 0 | 4 | 0 | 1 | 1 | 1 | 0 | 10 | 0.1 | 8 | 0 | 3 | 0 | 3 | 0.3333 | 2 | 0.5 | 16 | 0.125 |
| 10_1171811 | 6 | 0.6667 | 10 | 0.4 | 3 | 1 | 9 | 0.5556 | 28 | 0.5714 | 3 | 0 | 7 | 0.1429 | 3 | 0.6667 | 4 | 0.25 | 17 | 0.2353 |
| 10_1171821 | 4 | 0 | 4 | 0 | 2 | 0 | 5 | 0.2 | 15 | 0.0667 | 2 | 0 | 6 | 0.1667 | 1 | 0 | 1 | 0 | 10 | 0.1 |

| | | | | | | | | | | | | | | | | | | | | |
|---|---|---|---|---|---|---|---|---|---|---|---|---|---|---|---|---|---|---|---|---|
| 10_1171821 | 4 | 0.25 | 5 | 0.6 | 2 | 0 | 6 | 0.3333 | 17 | 0.3529 | 1 | 0 | 6 | 0 | 2 | 0.5 | 4 | 0 | 13 | 0.0769 |
| 10_1171834 | 3 | 0 | 4 | 0 | 1 | 1 | 4 | 0.25 | 12 | 0.1667 | 1 | 0 | 7 | 0.1429 | NA | NA | 3 | 0 | 11 | 0.0909 |
| 10_1171840 | 3 | 0 | 5 | 0 | 1 | 0 | 4 | 0.25 | 13 | 0.0769 | 1 | 0 | 7 | 0.1429 | NA | NA | 3 | 0 | 11 | 0.0909 |
| 10_1171846 | 2 | 0.5 | NA | NA | NA | NA | NA | NA | 2 | 0.5 | NA | NA | 2 | 0 | 1 | 1 | NA | NA | 3 | 0.3333 |
| 10_1171849 | 7 | 0.1429 | 5 | 0 | 4 | 0 | 10 | 0.1 | 26 | 0.0769 | 2 | 0 | 7 | 0.1429 | 7 | 0.2857 | 7 | 0.2857 | 23 | 0.2174 |
| 10_1171852 | 6 | 0.3333 | 8 | 0.125 | 4 | 0 | 8 | 0.5 | 26 | 0.2692 | 1 | 0 | 8 | 0.375 | 2 | 0 | 10 | 0.3 | 21 | 0.2857 |
| 10_1171852 | 7 | 0.7143 | 6 | 0.8333 | 3 | 0.3333 | 8 | 0.75 | 24 | 0.7083 | 1 | 1 | 7 | 0.5714 | 6 | 1 | 8 | 0.75 | 22 | 0.7727 |
| 10_1171897 | NA | NA | 1 | 1 | NA | NA | 2 | 1 | 3 | 1 | NA | NA | NA | NA | 1 | 1 | 1 | 0 | 2 | 0.5 |
| 10_1171970 | 5 | 0.4 | 5 | 0 | 1 | 0 | 5 | 0 | 16 | 0.125 | NA | NA | 3 | 0.3333 | 3 | 0 | 6 | 0.1667 | 12 | 0.1667 |
| 10_1171971 | 5 | 0.4 | 6 | 0.3333 | 2 | 0 | 6 | 0.3333 | 19 | 0.3158 | NA | NA | 4 | 0 | 2 | 0.5 | 5 | 0 | 11 | 0.0909 |
| 10_1171971 | 6 | 0.1667 | 5 | 0 | 3 | 0.3333 | 5 | 0.2 | 19 | 0.1579 | NA | NA | 3 | 0 | 3 | 0 | 7 | 0.1429 | 13 | 0.0769 |
| 10_1175193 | NA | NA | NA | NA | 1 | 1 | NA | NA | 1 | 1 | 1 | 1 | 1 | 1 | 1 | 1 | 1 | 1 | 4 | 1 |
| 10_1175195 | NA | NA | NA | NA | 1 | 1 | NA | NA | 1 | 1 | 1 | 1 | 1 | 1 | 1 | 1 | 1 | 1 | 4 | 1 |
| 10_1175195 | NA | NA | NA | NA | 1 | 1 | NA | NA | 1 | 1 | 1 | 1 | 1 | 0 | 1 | 1 | 1 | 1 | 4 | 0.75 |
| 10_1175198 | NA | NA | NA | NA | 1 | 1 | NA | NA | 1 | 1 | 1 | 1 | 2 | 1 | 1 | 1 | 1 | 1 | 5 | 1 |
| 10_1175199 | NA | NA | NA | NA | 1 | 1 | NA | NA | 1 | 1 | 1 | 1 | 1 | 1 | 1 | 1 | 1 | 1 | 4 | 1 |
| 10_1175201 | NA | NA | NA | NA | 1 | 1 | NA | NA | 1 | 1 | 2 | 0.5 | 2 | 0.5 | 1 | 1 | 1 | 1 | 6 | 0.6667 |
| 10_1175203 | NA | NA | NA | NA | 1 | 1 | NA | NA | 1 | 1 | 2 | 0.5 | 2 | 0 | 1 | 0 | 1 | 0 | 6 | 0.1667 |
| 10_1175204 | 1 | 0 | NA | NA | 2 | 0.5 | NA | NA | 3 | 0.3333 | NA | NA | 2 | 0 | 1 | 1 | 1 | 1 | 4 | 0.5 |
| 10_1175206 | 1 | 1 | NA | NA | 1 | 1 | NA | NA | 2 | 1 | 2 | 1 | NA | NA | 1 | 1 | 1 | 1 | 4 | 1 |
| 10_1175206 | 1 | 1 | NA | NA | 2 | 0.5 | NA | NA | 3 | 0.6667 | 2 | 1 | 1 | 1 | 1 | 1 | 1 | 1 | 5 | 1 |
| 10_1175218 | NA | NA | NA | NA | 1 | 0 | 1 | 1 | 2 | 0.5 | 1 | 1 | 1 | 0 | 1 | 0 | 1 | 1 | 4 | 0.5 |
| 10_1175229 | NA | NA | NA | NA | 1 | 0 | 2 | 1 | 3 | 0.6667 | 1 | 0 | 1 | 1 | 2 | 0.5 | 1 | 0 | 5 | 0.4 |
| 10_1175230 | NA | NA | NA | NA | 2 | 0.5 | 2 | 1 | 4 | 0.75 | 2 | 0 | 1 | 1 | 2 | 0 | 1 | 0 | 6 | 0.1667 |
| 10_1175249 | 4 | 0 | 8 | 0.125 | 9 | 0.1111 | 10 | 0 | 31 | 0.0645 | 6 | 0 | 10 | 0.1 | 8 | 0.25 | 4 | 0 | 28 | 0.1071 |
| 10_1175251 | 2 | 0.5 | 7 | 0.2857 | 8 | 0.5 | 6 | 0.3333 | 23 | 0.3913 | 5 | 0.4 | 7 | 0.4286 | 8 | 0.25 | 3 | 0 | 23 | 0.3043 |
| 10_1175256 | 3 | 0.3333 | 8 | 0.375 | 9 | 0.1111 | 8 | 0.375 | 28 | 0.2857 | 6 | 0.3333 | 9 | 0.2222 | 9 | 0.3333 | 4 | 0 | 28 | 0.25 |
| 10_1175257 | NA | NA | 8 | 0.125 | 7 | 0.2857 | 4 | 0.25 | 19 | 0.2105 | 3 | 0 | 6 | 0.1667 | 7 | 0.1429 | 3 | 0.3333 | 19 | 0.1579 |
| 10_1175257 | 1 | 1 | 8 | 0.125 | 9 | 0.5556 | 8 | 0.375 | 26 | 0.3846 | 6 | 0.3333 | 7 | 0.4286 | 9 | 0.4444 | 4 | 0.5 | 26 | 0.4231 |
| 10_1175258 | 1 | 1 | 6 | 1 | 6 | 0.8333 | 8 | 0.875 | 21 | 0.9048 | 4 | 0.5 | 7 | 0.8571 | 7 | 0.7143 | 4 | 0.75 | 22 | 0.7273 |
| 10_1175260 | 2 | 1 | 8 | 0.75 | 5 | 0.6 | 7 | 1 | 22 | 0.8182 | 3 | 0.6667 | 6 | 0.8333 | 6 | 0.6667 | 4 | 0.75 | 19 | 0.7368 |
| 10_1175261 | 1 | 0 | 8 | 0.25 | 7 | 0 | 8 | 0.125 | 24 | 0.125 | 3 | 0.3333 | 5 | 0 | 4 | 0 | 4 | 0 | 16 | 0.0625 |
| 10_1175262 | 2 | 1 | 6 | 1 | 6 | 0.8333 | 8 | 1 | 22 | 0.9545 | 3 | 0.6667 | 6 | 1 | 6 | 1 | 3 | 1 | 18 | 0.9444 |
| 10_1175262 | 2 | 0.5 | 5 | 1 | 7 | 0.7143 | 6 | 1 | 20 | 0.85 | 4 | 0.75 | 4 | 1 | 5 | 1 | 3 | 0.6667 | 16 | 0.875 |
| 10_1175262 | 2 | 0.5 | 8 | 0.5 | 7 | 0.2857 | 7 | 0.4286 | 24 | 0.4167 | 3 | 1 | 5 | 0.4 | 5 | 0.2 | 2 | 0.5 | 15 | 0.4667 |
| 10_1175263 | 2 | 0 | 8 | 0.25 | 8 | 0.25 | 6 | 0.5 | 24 | 0.2917 | 3 | 0 | 3 | 0.6667 | 7 | 0.1429 | 3 | 0.3333 | 16 | 0.25 |
| 10_1175264 | 1 | 0 | 7 | 0.2857 | 4 | 0.25 | 6 | 0.3333 | 18 | 0.2778 | 4 | 0.25 | 7 | 0.2857 | 6 | 0.1667 | 2 | 0 | 19 | 0.2105 |
| 10_1175265 | 1 | 0 | 7 | 0.2857 | 7 | 0.2857 | 8 | 0.125 | 23 | 0.2174 | 2 | 0.5 | 7 | 0.2857 | 6 | 0 | 4 | 0 | 19 | 0.1579 |
| 10_1175265 | 1 | 0 | 7 | 0.1429 | 8 | 0 | 7 | 0.2857 | 23 | 0.1304 | 4 | 0 | 4 | 0.25 | 6 | 0 | 4 | 0.25 | 18 | 0.1111 |
| 10_1175266 | 1 | 0 | 7 | 0.7143 | 7 | 0.2857 | 7 | 0.1429 | 22 | 0.3636 | 4 | 0.25 | 7 | 0.2857 | 6 | 0.5 | 3 | 0.3333 | 20 | 0.35 |
| 10_1175699 | 3 | 0.3333 | 3 | 0.3333 | 5 | 0.6 | 1 | 1 | 12 | 0.5 | 1 | 1 | 1 | 0 | 2 | 1 | 3 | 0.6667 | 7 | 0.7143 |
| 10_1175699 | 4 | 0 | 3 | 0.3333 | 5 | 0 | 1 | 0 | 13 | 0.0769 | 1 | 0 | 1 | 0 | 2 | 0.5 | 3 | 0 | 7 | 0.1429 |
| 10_1175708 | 3 | 0.6667 | 3 | 0.6667 | 3 | 0.6667 | 1 | 0 | 10 | 0.6 | NA | NA | 1 | 0 | 1 | 1 | 2 | 0.5 | 4 | 0.5 |
| 10_1175712 | 2 | 1 | 2 | 0.5 | 2 | 0 | NA | NA | 6 | 0.5 | NA | NA | 1 | 1 | 1 | 1 | 1 | 1 | 3 | 1 |
| 10_1175712 | 1 | 0 | 2 | 0.5 | 3 | 0 | 1 | 0 | 7 | 0.1429 | NA | NA | 1 | 0 | 1 | 1 | 1 | 0 | 3 | 0.3333 |
| 10_1175713 | 2 | 1 | 2 | 1 | 2 | 1 | 1 | 1 | 7 | 1 | NA | NA | 1 | 1 | 1 | 1 | 1 | 1 | 3 | 1 |
| 10_1175713 | 3 | 1 | 3 | 1 | 3 | 1 | 1 | 1 | 10 | 1 | NA | NA | 1 | 1 | 1 | 1 | 1 | 1 | 3 | 1 |
| 10_1175714 | 3 | 1 | 3 | 0.6667 | 3 | 0.3333 | 1 | 1 | 10 | 0.7 | NA | NA | 1 | 1 | 1 | 0 | 1 | 1 | 3 | 0.6667 |
| 10_1175714 | 3 | 1 | 2 | 0.5 | 2 | 1 | NA | NA | 7 | 0.8571 | NA | NA | 1 | 1 | 1 | 1 | NA | NA | 2 | 1 |
| 10_1175717 | 2 | 1 | 3 | 1 | 2 | 1 | 1 | 1 | 8 | 1 | NA | NA | 1 | 1 | NA | NA | 1 | 1 | 2 | 1 |
| 10_1175717 | 2 | 0.5 | 2 | 1 | 2 | 1 | 1 | 1 | 7 | 0.8571 | NA | NA | 1 | 1 | NA | NA | 1 | 1 | 2 | 1 |
| 10_1175718 | 2 | 1 | 3 | 0.6667 | 1 | 0 | NA | NA | 6 | 0.6667 | NA | NA | 1 | 1 | NA | NA | 1 | 1 | 2 | 1 |
| 10_1175720 | 2 | 1 | 3 | 1 | 3 | 0.6667 | 1 | 1 | 9 | 0.8889 | NA | NA | 1 | 1 | NA | NA | 1 | 0 | 2 | 0.5 |
| 10_1175721 | 2 | 0.5 | 1 | 0 | 1 | 0 | 1 | 0 | 5 | 0.2 | NA | NA | 1 | 1 | NA | NA | 1 | 1 | 2 | 1 |
| 10_1175819 | 2 | 1 | 3 | 1 | 2 | 0.5 | NA | NA | 7 | 0.8571 | NA | NA | 1 | 1 | NA | NA | NA | NA | 1 | 1 |
| 10_1176502 | 4 | 0.25 | 3 | 0.3333 | 3 | 0 | 2 | 0 | 12 | 0.1667 | 2 | 0 | 1 | 0 | 1 | 1 | 2 | 0 | 6 | 0.1667 |
| 10_1176514 | 8 | 0.375 | 11 | 0.0909 | 10 | 0.3 | 11 | 0.2727 | 40 | 0.25 | 6 | 0.3333 | 9 | 0.3333 | 5 | 0.4 | 12 | 0.4167 | 32 | 0.375 |
| 10_1176516 | 7 | 0.8571 | 9 | 0.4444 | 11 | 0.2727 | 11 | 0.5455 | 38 | 0.5 | 5 | 0.6 | 8 | 0.625 | 7 | 0.5714 | 12 | 0.5833 | 32 | 0.5938 |
| 10_1179827 | 1 | 1 | 1 | 1 | NA | NA | NA | NA | 2 | 1 | NA | NA | NA | NA | NA | NA | 1 | 1 | 1 | 1 |
| 10_1186917 | 4 | 0.25 | 3 | 0.6667 | 1 | 1 | 1 | 0 | 9 | 0.4444 | 8 | 0.25 | 1 | 0 | 7 | 0.1429 | 3 | 0.6667 | 19 | 0.2632 |
| 10_1186918 | 4 | 0.25 | 3 | 0.6667 | 1 | 0 | 1 | 1 | 9 | 0.4444 | 4 | 0.5 | 1 | 1 | 4 | 0.25 | 3 | 0.3333 | 12 | 0.4167 |
| 10_1186919 | 2 | 1 | 2 | 0.5 | 1 | 1 | 1 | 1 | 6 | 0.8333 | 5 | 0.6 | 1 | 1 | 6 | 0.8333 | 3 | 1 | 15 | 0.8 |
| 10_1186920 | 2 | 0 | 1 | 0 | NA | NA | 1 | 1 | 4 | 0.25 | 4 | 0.25 | NA | NA | 6 | 0 | 2 | 0 | 12 | 0.0833 |
| 10_1186930 | 1 | 0 | 2 | 0 | 1 | 0 | 1 | 1 | 5 | 0.2 | 7 | 0 | NA | NA | 5 | 0 | 3 | 0.3333 | 15 | 0.0667 |
| 10_1186931 | 1 | 0 | 3 | 0 | 1 | 0 | 1 | 1 | 6 | 0.1667 | 4 | 0.25 | NA | NA | 6 | 0.1667 | 3 | 0 | 13 | 0.1538 |

| | | | | | | | | | | | | | | | | | | | | |
|---|---|---|---|---|---|---|---|---|---|---|---|---|---|---|---|---|---|---|---|---|
| 10_1186933 | NA | NA | 2 | 0.5 | NA | NA | 1 | 0 | 3 | 0.3333 | 6 | 0.1667 | NA | NA | 6 | 0 | 2 | 0 | 14 | 0.0714 |
| 10_1186945 | 1 | 0 | 2 | 0 | 1 | 0 | 1 | 1 | 5 | 0.2 | 4 | 0 | NA | NA | 3 | 0.3333 | 2 | 0 | 9 | 0.1111 |
| 10_1186946 | 1 | 0 | 1 | 1 | 1 | 0 | NA | NA | 3 | 0.3333 | 3 | 0 | NA | NA | 4 | 0.25 | 2 | 0 | 9 | 0.1111 |
| 10_1186954 | NA | NA | 1 | 1 | NA | NA | NA | NA | 1 | 1 | 3 | 1 | NA | NA | 3 | 1 | 2 | 1 | 8 | 1 |
| 10_1186955 | NA | NA | 1 | 1 | NA | NA | NA | NA | 1 | 1 | 3 | 0.6667 | NA | NA | 4 | 1 | 1 | 1 | 8 | 0.875 |
| 10_1186959 | NA | NA | 1 | 1 | NA | NA | NA | NA | 1 | 1 | 3 | 1 | NA | NA | 3 | 0.6667 | 2 | 0.5 | 8 | 0.75 |
| 10_1186960 | NA | NA | 1 | 1 | NA | NA | NA | NA | 1 | 1 | 2 | 0 | NA | NA | 3 | 1 | 1 | 1 | 6 | 0.6667 |
| 10_1186960 | NA | NA | 1 | 1 | NA | NA | NA | NA | 1 | 1 | 3 | 1 | NA | NA | 3 | 1 | 2 | 0.5 | 8 | 0.875 |
| 10_1186962 | NA | NA | 1 | 1 | NA | NA | NA | NA | 1 | 1 | 3 | 1 | NA | NA | 3 | 1 | 1 | 1 | 7 | 1 |
| 10_1186963 | NA | NA | 1 | 1 | NA | NA | NA | NA | 1 | 1 | 3 | 0.6667 | NA | NA | 2 | 0 | 2 | 0.5 | 7 | 0.4286 |
| 10_1186966 | NA | NA | 1 | 1 | NA | NA | NA | NA | 1 | 1 | 1 | 1 | NA | NA | 3 | 0.3333 | 2 | 0.5 | 6 | 0.5 |
| 10_1186967 | NA | NA | 1 | 1 | NA | NA | NA | NA | 1 | 1 | 1 | 1 | NA | NA | 2 | 0 | 2 | 0.5 | 5 | 0.4 |
| 10_1190354 | 5 | 0 | 9 | 0.1111 | 4 | 0.5 | 4 | 0 | 22 | 0.1364 | 2 | 0.5 | 4 | 0.25 | 10 | 0.1 | 4 | 0 | 20 | 0.15 |
| 10_1190369 | 5 | 0.4 | 8 | 0.125 | 6 | 0.1667 | 7 | 0 | 26 | 0.1538 | 2 | 0 | 4 | 0.5 | 5 | 0.2 | 4 | 0 | 15 | 0.2 |
| 10_1190372 | 5 | 0 | 8 | 0.125 | 7 | 0.1429 | 9 | 0.1111 | 29 | 0.1034 | 2 | 0 | 3 | 0.3333 | 6 | 0 | 4 | 0.5 | 15 | 0.2 |
| 10_1190375 | 2 | 0 | 5 | 0.4 | 3 | 0.6667 | 7 | 0.4286 | 17 | 0.4118 | 2 | 0.5 | 2 | 1 | 6 | 0.1667 | 1 | 0 | 11 | 0.3636 |
| 10_1190383 | 6 | 0.5 | 10 | 0.3 | 7 | 0 | 7 | 0 | 30 | 0.2 | 2 | 0 | 3 | 0.3333 | 5 | 0.2 | 4 | 0.25 | 14 | 0.2143 |
| 10_1190386 | 10 | 0.3 | 16 | 0.125 | 14 | 0.1429 | 14 | 0.3571 | 54 | 0.2222 | 10 | 0.4 | 10 | 0.2 | 9 | 0.2222 | 16 | 0.375 | 45 | 0.3111 |
| 10_1190388 | 8 | 0.375 | 15 | 0.3333 | 15 | 0.4 | 13 | 0.4615 | 51 | 0.3922 | 7 | 0.4286 | 8 | 0.625 | 9 | 0.4444 | 11 | 0.5455 | 35 | 0.5143 |
| 10_1190388 | 7 | 1 | 14 | 1 | 15 | 1 | 13 | 0.9231 | 49 | 0.9796 | 8 | 1 | 9 | 0.6667 | 9 | 1 | 14 | 0.9286 | 40 | 0.9 |
| 10_1190466 | 7 | 0.1429 | 9 | 0 | 11 | 0 | 9 | 0.1111 | 36 | 0.0556 | 5 | 0 | 7 | 0.1429 | 6 | 0 | 9 | 0.1111 | 27 | 0.0741 |
| 10_1190469 | 6 | 0.6667 | 13 | 0.9231 | 14 | 0.9286 | 10 | 0.8 | 43 | 0.8605 | 6 | 0.5 | 7 | 0.8571 | 5 | 0.4 | 10 | 0.7 | 28 | 0.6429 |
| 10_1190469 | 8 | 0.75 | 13 | 0.5385 | 14 | 0.5714 | 9 | 0.7778 | 44 | 0.6364 | 6 | 0.5 | 7 | 0.4286 | 7 | 0.7143 | 9 | 0.3333 | 29 | 0.4828 |
| 10_1190470 | 8 | 0 | 13 | 0.1538 | 15 | 0.2 | 10 | 0.2 | 46 | 0.1522 | 6 | 0 | 6 | 0.1667 | 7 | 0.2857 | 10 | 0.1 | 29 | 0.1379 |
| 10_1190470 | 6 | 0 | 10 | 0.2 | 14 | 0.0714 | 8 | 0.125 | 38 | 0.1053 | 5 | 0.4 | 6 | 0 | 7 | 0 | 9 | 0.1111 | 27 | 0.1111 |
| 10_1190471 | 8 | 1 | 11 | 1 | 15 | 1 | 10 | 1 | 44 | 1 | 5 | 1 | 7 | 1 | 6 | 1 | 10 | 1 | 28 | 1 |
| 10_1190471 | 4 | 0.25 | 11 | 0.1818 | 15 | 0.0667 | 6 | 0 | 36 | 0.1111 | 4 | 0 | 6 | 0.1667 | 5 | 0 | 8 | 0.25 | 23 | 0.1304 |
| 10_1190472 | 6 | 0.5 | 12 | 0.6667 | 11 | 0.5455 | 8 | 0.75 | 37 | 0.6216 | 4 | 0.5 | 6 | 0.6667 | 6 | 0.6667 | 8 | 0.5 | 24 | 0.5833 |
| 10_1190472 | 7 | 0.8571 | 9 | 1 | 12 | 0.8333 | 6 | 1 | 34 | 0.9118 | 3 | 1 | 5 | 1 | 4 | 1 | 10 | 1 | 22 | 1 |
| 10_1190472 | 4 | 1 | 9 | 0.8889 | 12 | 1 | 10 | 0.9 | 35 | 0.9429 | 3 | 1 | 6 | 0.8333 | 5 | 1 | 10 | 1 | 24 | 0.9583 |
| 10_1190474 | 4 | 0.25 | 11 | 0.4545 | 10 | 0.2 | 9 | 0.1111 | 34 | 0.2647 | 4 | 0.25 | 5 | 0.4 | 4 | 0.25 | 6 | 0.6667 | 19 | 0.4211 |
| 10_1190474 | 5 | 0.4 | 12 | 0.1667 | 13 | 0.3077 | 9 | 0 | 39 | 0.2051 | 3 | 0 | 7 | 0.1429 | 5 | 0.6 | 9 | 0.1111 | 24 | 0.2083 |
| 10_1190474 | 4 | 0 | 12 | 0 | 12 | 0.1667 | 8 | 0 | 36 | 0.0556 | 3 | 0 | 6 | 0.1667 | 4 | 0 | 7 | 0 | 20 | 0.05 |
| 10_1190476 | 2 | 1 | 11 | 0.2727 | 12 | 0.6667 | 8 | 0.875 | 33 | 0.6061 | 3 | 0.3333 | 6 | 0.6667 | 4 | 0.75 | 6 | 0.8333 | 19 | 0.6842 |
| 10_1190476 | 5 | 0.4 | 9 | 0.4444 | 12 | 0.1667 | 8 | 0 | 34 | 0.2353 | 3 | 0 | 6 | 0.1667 | 3 | 0.3333 | 7 | 0.1429 | 19 | 0.1579 |
| 10_1190477 | 4 | 0.75 | 10 | 0.8 | 11 | 0.9091 | 9 | 0.8889 | 34 | 0.8529 | 2 | 1 | 6 | 0.8333 | 5 | 0.8 | 7 | 1 | 20 | 0.9 |
| 10_1196531 | NA | NA | NA | NA | 1 | 1 | NA | NA | 1 | 1 | NA | NA | NA | NA | NA | NA | 1 | 1 | 1 | 1 |
| 10_1199112 | NA | NA | 2 | 0 | 2 | 0.5 | 1 | 0 | 5 | 0.2 | NA | NA | 1 | 0 | 3 | 0.3333 | NA | NA | 4 | 0.25 |
| 10_1199114 | NA | NA | 2 | 0.5 | 2 | 1 | NA | NA | 4 | 0.75 | NA | NA | NA | NA | 2 | 0.5 | NA | NA | 2 | 0.5 |
| 10_1199202 | NA | NA | 1 | 1 | NA | NA | NA | NA | 1 | 1 | 1 | 1 | NA | NA | 2 | 1 | NA | NA | 3 | 1 |
| 10_1199203 | NA | NA | 1 | 1 | NA | NA | NA | NA | 1 | 1 | 2 | 1 | NA | NA | 1 | 1 | NA | NA | 3 | 1 |
| 10_1199203 | NA | NA | 1 | 1 | NA | NA | NA | NA | 1 | 1 | 2 | 0.5 | NA | NA | 1 | 0 | NA | NA | 3 | 0.3333 |
| 10_1199205 | NA | NA | 1 | 1 | NA | NA | NA | NA | 1 | 1 | 1 | 1 | NA | NA | 1 | 1 | NA | NA | 2 | 1 |
| 10_1199787 | NA | NA | 1 | 1 | NA | NA | NA | NA | 1 | 1 | 1 | 1 | NA | NA | NA | NA | NA | NA | 1 | 1 |
| 10_1199789 | NA | NA | 1 | 1 | NA | NA | NA | NA | 1 | 1 | 1 | 1 | NA | NA | NA | NA | NA | NA | 1 | 1 |
| 10_1199789 | NA | NA | 1 | 1 | NA | NA | NA | NA | 1 | 1 | 1 | 1 | NA | NA | NA | NA | NA | NA | 1 | 1 |
| 10_1200607 | 13 | 0.3077 | 11 | 0.1818 | 14 | 0.3571 | 10 | 0.2 | 48 | 0.2708 | 8 | 0.125 | 11 | 0.1818 | 11 | 0.1818 | 9 | 0.5556 | 39 | 0.2564 |
| 10_1200671 | 10 | 0 | 15 | 0.0667 | 7 | 0.1429 | 9 | 0.1111 | 41 | 0.0732 | 8 | 0.25 | 5 | 0.2 | 13 | 0.0769 | 9 | 0.1111 | 35 | 0.1429 |
| 10_1200672 | 10 | 0.2 | 11 | 0.2727 | 8 | 0.125 | 8 | 0.25 | 37 | 0.2162 | 6 | 0 | 4 | 0.5 | 11 | 0.4545 | 7 | 0.2857 | 28 | 0.3214 |
| 10_1200672 | 10 | 1 | 10 | 1 | 8 | 1 | 8 | 0.875 | 36 | 0.9722 | 8 | 1 | 3 | 1 | 11 | 0.9091 | 5 | 1 | 27 | 0.963 |
| 10_1200674 | 9 | 0 | 16 | 0.125 | 8 | 0.25 | 9 | 0.1111 | 42 | 0.119 | 10 | 0.2 | 4 | 0 | 13 | 0 | 8 | 0.25 | 35 | 0.1143 |
| 10_1207602 | 1 | 0 | 1 | 0 | 3 | 0.3333 | 4 | 0 | 9 | 0.1111 | 3 | 0 | 6 | 0 | 1 | 1 | 6 | 0 | 16 | 0.0625 |
| 10_1207603 | 1 | 0 | 1 | 0 | 3 | 0.3333 | 4 | 0.25 | 9 | 0.2222 | 2 | 0 | 5 | 0 | NA | NA | 5 | 0.2 | 12 | 0.0833 |
| 10_1207609 | 1 | 0 | 1 | 0 | 2 | 1 | 2 | 0.5 | 6 | 0.5 | 1 | 0 | 4 | 0.25 | 1 | 0 | 6 | 0 | 12 | 0.0833 |
| 10_1207609 | 1 | 0 | 1 | 0 | 3 | 0.6667 | 1 | 1 | 6 | 0.5 | 1 | 0 | 2 | 0 | 1 | 1 | 6 | 0.6667 | 10 | 0.5 |
| 10_1207612 | 1 | 1 | 1 | 1 | 3 | 1 | 2 | 1 | 7 | 1 | NA | NA | 3 | 0.6667 | NA | NA | 6 | 1 | 9 | 0.8889 |
| 10_1207613 | 1 | 0 | NA | NA | 2 | 1 | 2 | 0.5 | 5 | 0.6 | 1 | 0 | 4 | 0.5 | 1 | 1 | 7 | 0.7143 | 13 | 0.6154 |
| 10_1207613 | 1 | 1 | 1 | 1 | 2 | 1 | 2 | 0.5 | 6 | 0.8333 | 1 | 1 | 3 | 1 | NA | NA | 2 | 1 | 6 | 1 |
| 10_1208137 | NA | NA | NA | NA | 2 | 0.5 | 1 | 1 | 3 | 0.6667 | NA | NA | NA | NA | 1 | 1 | 2 | 0 | 3 | 0.3333 |
| 10_1208156 | NA | NA | NA | NA | NA | NA | 1 | 1 | 1 | 1 | NA | NA | NA | NA | 1 | 0 | 1 | 1 | 2 | 0.5 |
| 10_1208157 | NA | NA | NA | NA | NA | NA | 1 | 1 | 1 | 1 | NA | NA | NA | NA | 1 | 1 | NA | NA | 1 | 1 |
| 10_1208168 | NA | NA | NA | NA | NA | NA | 1 | 1 | 1 | 1 | NA | NA | NA | NA | 1 | 1 | NA | NA | 1 | 1 |
| 10_1211001 | 7 | 0.1429 | 6 | 0.3333 | 4 | 0 | 8 | 0.5 | 25 | 0.28 | 7 | 0.1429 | 7 | 0.1429 | 8 | 0 | 6 | 0.1667 | 28 | 0.1071 |
| 10_1211002 | 9 | 0.3333 | 11 | 0.4545 | 5 | 0.6 | 8 | 0.25 | 33 | 0.3939 | 8 | 0.75 | 10 | 0.5 | 9 | 0.2222 | 8 | 0.25 | 35 | 0.4286 |
| 10_1211003 | 7 | 0.8571 | 9 | 0.8889 | 5 | 0.8 | 8 | 0.875 | 29 | 0.8621 | 6 | 0.6667 | 10 | 0.8 | 8 | 0.75 | 6 | 0.6667 | 30 | 0.7333 |

| | | | | | | | | | | | | | | | | | | | | |
|---|---|---|---|---|---|---|---|---|---|---|---|---|---|---|---|---|---|---|---|---|
| 10_121100 | 8 | 0.125 | 10 | 0.1 | 5 | 0 | 7 | 0.2857 | 30 | 0.1333 | 8 | 0.125 | 10 | 0 | 8 | 0.125 | 8 | 0 | 34 | 0.0588 |
| 10_121100 | 8 | 0.25 | 11 | 0.3636 | 3 | 0.3333 | 8 | 0.5 | 30 | 0.3667 | 7 | 0.7143 | 10 | 0.6 | 8 | 0.5 | 7 | 0.5714 | 32 | 0.5938 |
| 10_121100 | 8 | 0.625 | 10 | 0.5 | 4 | 0.75 | 7 | 0.8571 | 29 | 0.6552 | 7 | 0.8571 | 10 | 0.7 | 8 | 0.625 | 7 | 0.8571 | 32 | 0.75 |
| 10_121100 | 7 | 0.1429 | 10 | 0.2 | 3 | 0.3333 | 7 | 0.2857 | 27 | 0.2222 | 5 | 0 | 9 | 0.1111 | 6 | 0 | 6 | 0.3333 | 26 | 0.1154 |
| 10_121101 | 5 | 1 | 8 | 0.625 | NA | NA | 7 | 0.8571 | 20 | 0.8 | 6 | 0.5 | 6 | 0.3333 | 4 | 0.75 | 7 | 0.8571 | 23 | 0.6087 |
| 10_121101 | 5 | 0.2 | 9 | 0 | 1 | 1 | 5 | 0.4 | 20 | 0.2 | 5 | 0 | 8 | 0.375 | 3 | 0.6667 | 5 | 0.2 | 21 | 0.2857 |
| 10_121101 | 4 | 0.25 | 9 | 0 | NA | NA | 7 | 0 | 20 | 0.05 | 6 | 0.1667 | 9 | 0.2222 | 5 | 0.2 | 7 | 0 | 27 | 0.1481 |
| 10_121101 | 3 | 0 | 7 | 0.1429 | NA | NA | 6 | 0 | 16 | 0.0625 | 6 | 0.1667 | 5 | 0 | 4 | 0 | 6 | 0.1667 | 21 | 0.0952 |
| 10_121101 | 2 | 1 | 7 | 0.1429 | NA | NA | 4 | 0.25 | 13 | 0.3077 | 2 | 0 | 6 | 0.5 | 5 | 0.6 | 3 | 0 | 16 | 0.375 |
| 10_121101 | 2 | 0 | 9 | 0.1111 | NA | NA | 6 | 0.1667 | 17 | 0.1176 | 6 | 0 | 7 | 0.1429 | 4 | 0.5 | 6 | 0.1667 | 23 | 0.1739 |
| 10_121101 | 3 | 1 | 6 | 1 | NA | NA | 2 | 1 | 11 | 1 | 6 | 0.6667 | 5 | 1 | 4 | 0.5 | 4 | 1 | 19 | 0.7895 |
| 10_121101 | 3 | 1 | 4 | 0.25 | NA | NA | 4 | 0 | 11 | 0.3636 | 3 | 0.3333 | 5 | 0.6 | 3 | 0.6667 | 4 | 0.75 | 15 | 0.6 |
| 10_121101 | 4 | 1 | 8 | 1 | NA | NA | 6 | 1 | 18 | 1 | 4 | 1 | 7 | 1 | 4 | 1 | 4 | 1 | 19 | 1 |
| 10_121101 | 3 | 0 | 7 | 0.1429 | NA | NA | 2 | 0.5 | 12 | 0.1667 | 2 | 0 | 2 | 0 | 5 | 0 | 3 | 0.3333 | 12 | 0.0833 |
| 10_121101 | 5 | 0 | 8 | 0.125 | NA | NA | 6 | 0 | 19 | 0.0526 | 5 | 0 | 7 | 0.2857 | 5 | 0.2 | 5 | 0 | 22 | 0.1364 |
| 10_121101 | 4 | 0.25 | 9 | 0.2222 | NA | NA | 6 | 0.1667 | 19 | 0.2105 | 7 | 0.4286 | 7 | 0.4286 | 5 | 0 | 5 | 0 | 24 | 0.25 |
| 10_121101 | 5 | 1 | 5 | 1 | NA | NA | 7 | 1 | 17 | 1 | 5 | 1 | 6 | 1 | 4 | 1 | 3 | 1 | 18 | 1 |
| 10_121102 | 5 | 1 | 8 | 1 | NA | NA | 7 | 1 | 20 | 1 | 5 | 1 | 6 | 1 | 4 | 1 | 3 | 1 | 18 | 1 |
| 10_121102 | 5 | 0.6 | 9 | 0.3333 | NA | NA | 6 | 0.1667 | 20 | 0.35 | 7 | 0.7143 | 7 | 0.5714 | 5 | 0.4 | 5 | 0.2 | 24 | 0.5 |
| 10_121102 | 4 | 0.75 | 7 | 0.5714 | NA | NA | 7 | 0.5714 | 18 | 0.6111 | 4 | 0.75 | 7 | 0 | 5 | 0.8 | 3 | 0.3333 | 19 | 0.4211 |
| 10_121102 | 2 | 0 | 4 | 0.5 | NA | NA | 5 | 0.4 | 11 | 0.3636 | 4 | 0.25 | 4 | 0.25 | 4 | 0 | 3 | 0 | 15 | 0.1333 |
| 10_121102 | 4 | 0.75 | 8 | 0.5 | NA | NA | 7 | 0.7143 | 19 | 0.6316 | 6 | 0.5 | 6 | 0.1667 | 6 | 0.5 | 5 | 0.6 | 23 | 0.4348 |
| 10_121102 | 4 | 0.5 | 7 | 0.2857 | 1 | 1 | 6 | 0.1667 | 18 | 0.3333 | 3 | 0.3333 | 5 | 0.2 | 4 | 0.25 | 3 | 0.6667 | 15 | 0.3333 |
| 10_121102 | 1 | 1 | 3 | 1 | NA | NA | 6 | 1 | 10 | 1 | 5 | 1 | 4 | 1 | 4 | 1 | 5 | 1 | 18 | 1 |
| 10_121102 | 3 | 0 | 7 | 0.5714 | 1 | 1 | 7 | 0.7143 | 18 | 0.5556 | 6 | 0.8333 | 7 | 0.2857 | 5 | 0.8 | 5 | 0.6 | 23 | 0.6087 |
| 10_121102 | 2 | 1 | 7 | 0.8571 | 1 | 0 | 4 | 1 | 14 | 0.8571 | 3 | 1 | 5 | 1 | 4 | 1 | 5 | 0.8 | 17 | 0.9412 |
| 10_121102 | 2 | 0.5 | 5 | 0.6 | 1 | 1 | 6 | 0.5 | 14 | 0.5714 | 4 | 1 | 6 | 0.8333 | 4 | 0.5 | 4 | 0.5 | 18 | 0.7222 |
| 10_121102 | 3 | 0 | 7 | 0.4286 | 1 | 0 | 3 | 0 | 14 | 0.2143 | 3 | 0.3333 | 5 | 0.6 | 4 | 0.25 | 5 | 0.4 | 17 | 0.4118 |
| 10_121102 | 2 | 1 | 8 | 1 | 1 | 1 | 4 | 1 | 15 | 1 | 4 | 1 | 5 | 1 | 5 | 1 | 1 | 1 | 15 | 1 |
| 10_121102 | 2 | 1 | 5 | 0.8 | 2 | 0 | 5 | 0.8 | 14 | 0.7143 | 5 | 0.6 | 6 | 0.1667 | 4 | 0.75 | 4 | 1 | 19 | 0.5789 |
| 10_121102 | 3 | 0 | 6 | 0.1667 | 1 | 0 | 5 | 0.2 | 15 | 0.1333 | 5 | 0 | 4 | 0 | 4 | 0.5 | 3 | 0.3333 | 16 | 0.1875 |
| 10_121102 | 2 | 0.5 | 7 | 0.4286 | 1 | 0 | 6 | 0.3333 | 16 | 0.375 | 4 | 0.25 | 5 | 0.4 | 5 | 0.4 | 2 | 0 | 16 | 0.3125 |
| 10_121102 | 3 | 1 | 7 | 0.8571 | 1 | 1 | 7 | 1 | 18 | 0.9444 | 5 | 1 | 5 | 1 | 4 | 1 | 4 | 1 | 18 | 1 |
| 10_121103 | 2 | 0 | 7 | 0.1429 | 2 | 0 | 7 | 0 | 18 | 0.0556 | 4 | 0.5 | 6 | 0.1667 | 5 | 0 | 3 | 0.3333 | 18 | 0.2222 |
| 10_121103 | 3 | 1 | 7 | 1 | 2 | 1 | 2 | 0.5 | 14 | 0.9286 | 3 | 1 | 5 | 1 | 5 | 1 | 3 | 0.6667 | 16 | 0.9375 |
| 10_121103 | 2 | 0.5 | 3 | 0.3333 | 1 | 0 | 5 | 0.2 | 11 | 0.2727 | 3 | 0.3333 | 1 | 0 | 2 | 0 | 2 | 0 | 8 | 0.125 |
| 10_121103 | 2 | 0.5 | 8 | 0.5 | 2 | 0.5 | 7 | 0 | 19 | 0.3158 | 3 | 0.3333 | 3 | 0 | 5 | 0.2 | 3 | 0 | 14 | 0.1429 |
| 10_121103 | 2 | 0 | 8 | 0.25 | 2 | 0 | 6 | 0.1667 | 18 | 0.1667 | 3 | 0.3333 | 3 | 0 | 6 | 0.1667 | 2 | 0 | 14 | 0.1429 |
| 10_121103 | 2 | 0 | 6 | 0.3333 | 1 | 0 | 4 | 0.5 | 13 | 0.3077 | 2 | 0 | 2 | 0 | 4 | 0 | 2 | 0.5 | 10 | 0.1 |
| 10_121103 | 2 | 0.5 | 6 | 0.3333 | 2 | 0 | 7 | 0.1429 | 17 | 0.2353 | 3 | 0.6667 | 2 | 0.5 | 5 | 0 | 3 | 0.6667 | 13 | 0.3846 |
| 10_121103 | 2 | 0.5 | 7 | 0.1429 | 1 | 0 | 6 | 0.3333 | 16 | 0.25 | 2 | 0 | 3 | 0 | 5 | 0 | 3 | 0.3333 | 13 | 0.0769 |
| 10_121103 | 2 | 1 | 8 | 0.25 | 1 | 0 | 6 | 0 | 17 | 0.2353 | 1 | 0 | 3 | 0.3333 | 5 | 0.2 | 3 | 0.6667 | 12 | 0.3333 |
| 10_121231 | 2 | 0 | 3 | 0.3333 | 9 | 0.2222 | 5 | 0.4 | 19 | 0.2632 | 4 | 0 | 6 | 0.1667 | 4 | 0.25 | 1 | 0 | 15 | 0.1333 |
| 10_121231 | 1 | 0 | 3 | 0.6667 | 9 | 0.3333 | 6 | 0.3333 | 19 | 0.3684 | 4 | 0.75 | 7 | 0.7143 | 3 | 0.3333 | 1 | 0 | 15 | 0.6 |
| 10_121231 | 2 | 0 | 4 | 0.25 | 10 | 0.5 | 3 | 0.6667 | 19 | 0.4211 | 5 | 0.4 | 5 | 0.4 | 6 | 0.5 | 3 | 0.6667 | 19 | 0.4737 |
| 10_121231 | 3 | 0.6667 | 4 | 1 | 11 | 0.7273 | 7 | 0.8571 | 25 | 0.8 | 6 | 1 | 7 | 0.7143 | 6 | 0.8333 | 2 | 1 | 21 | 0.8571 |
| 10_121231 | 3 | 0.6667 | 4 | 0.5 | 11 | 0.2727 | 7 | 0.2857 | 25 | 0.36 | 6 | 0.1667 | 6 | 0.1667 | 6 | 0.3333 | 3 | 0.3333 | 21 | 0.2381 |
| 10_121231 | 3 | 0 | 4 | 0.25 | 12 | 0 | 8 | 0.125 | 27 | 0.0741 | 7 | 0.1429 | 8 | 0.25 | 6 | 0 | 4 | 0.25 | 25 | 0.16 |
| 10_121232 | 3 | 0 | 5 | 0 | 13 | 0.0769 | 9 | 0.2222 | 30 | 0.1 | 9 | 0 | 9 | 0 | 8 | 0.125 | 3 | 0.3333 | 29 | 0.069 |
| 10_121232 | 3 | 0 | 6 | 0.3333 | 11 | 0.0909 | 10 | 0.4 | 30 | 0.2333 | 9 | 0.2222 | 8 | 0.125 | 9 | 0 | 4 | 0.25 | 30 | 0.1333 |
| 10_121232 | 3 | 0.6667 | 5 | 0.6 | 16 | 0.625 | 10 | 0.7 | 34 | 0.6471 | 10 | 0.5 | 11 | 0.8182 | 11 | 0.6364 | 6 | 0.8333 | 38 | 0.6842 |
| 10_121233 | 4 | 1 | 7 | 0.7143 | 15 | 0.9333 | 11 | 0.8182 | 37 | 0.8649 | 10 | 0.9 | 13 | 1 | 10 | 0.9 | 5 | 0.8 | 38 | 0.9211 |
| 10_121233 | 4 | 0 | 8 | 0.125 | 16 | 0.125 | 10 | 0.1 | 38 | 0.1053 | 10 | 0 | 13 | 0.0769 | 13 | 0.1538 | 6 | 0.1667 | 42 | 0.0952 |
| 10_121246 | 1 | 0 | 2 | 1 | 2 | 0.5 | 2 | 1 | 7 | 0.7143 | 6 | 0.6667 | 2 | 0.5 | 2 | 0.5 | 1 | 1 | 11 | 0.6364 |
| 10_121246 | 2 | 0 | 5 | 0.2 | 2 | 0 | 2 | 0 | 11 | 0.0909 | 5 | 0 | 1 | 0 | 2 | 0.5 | 1 | 0 | 9 | 0.1111 |
| 10_121246 | 2 | 0 | 5 | 0.2 | 4 | 0.5 | 2 | 0.5 | 13 | 0.3077 | 6 | 0.5 | 1 | 0 | 1 | 0 | 1 | 0 | 9 | 0.3333 |
| 10_121246 | 2 | 0 | 5 | 0 | 4 | 0.25 | 2 | 0.5 | 13 | 0.1538 | 7 | 0.2857 | 1 | 0 | 3 | 0.3333 | 1 | 0 | 12 | 0.25 |
| 10_121246 | 2 | 0.5 | 5 | 0 | 4 | 0 | 1 | 0 | 12 | 0.0833 | 7 | 0 | 2 | 0 | 3 | 0.3333 | 1 | 0 | 13 | 0.0769 |
| 10_121247 | 2 | 0 | 4 | 0 | 4 | 0.25 | 1 | 0 | 11 | 0.0909 | 6 | 0.5 | 2 | 0 | 3 | 1 | 1 | 1 | 12 | 0.5833 |
| 10_121247 | 2 | 0 | 4 | 0 | 3 | 0.3333 | 2 | 0.5 | 11 | 0.1818 | 5 | 0.2 | 1 | 1 | 2 | 0 | 1 | 0 | 9 | 0.2222 |
| 10_121247 | 2 | 0 | 5 | 0.2 | 4 | 0 | 2 | 0 | 13 | 0.0769 | 7 | 0.2857 | 2 | 0.5 | 2 | 0 | NA | NA | 11 | 0.2727 |
| 10_121248 | 1 | 0 | 4 | 0.75 | 4 | 0.5 | 2 | 0.5 | 11 | 0.5455 | 5 | 0.6 | 1 | 0 | 3 | 0 | NA | NA | 9 | 0.3333 |
| 10_121248 | 2 | 0.5 | 3 | 0 | 2 | 0 | 2 | 0 | 9 | 0.1111 | 6 | 0.1667 | 2 | 0 | 2 | 0 | NA | NA | 10 | 0.1 |
| 10_121248 | 2 | 1 | 2 | 0.5 | 3 | 0 | 1 | 0 | 8 | 0.375 | 4 | 0.25 | 1 | 0 | 2 | 0 | NA | NA | 7 | 0.1429 |
| 10_121248 | 2 | 1 | 2 | 1 | 3 | 1 | 2 | 1 | 9 | 1 | 5 | 1 | 1 | 1 | 2 | 0.5 | NA | NA | 8 | 0.875 |

| | | | | | | | | | | | | | | | | | | | | |
|---|---|---|---|---|---|---|---|---|---|---|---|---|---|---|---|---|---|---|---|---|
| 10_1212483 | 2 | 0.5 | 2 | 1 | 2 | 0.5 | 2 | 1 | 8 | 0.75 | 4 | 0.5 | 1 | 1 | 1 | 0 | NA | NA | 6 | 0.5 |
| 10_1212483 | 2 | 0 | 2 | 0.5 | 3 | 0.6667 | 2 | 0.5 | 9 | 0.4444 | 2 | 1 | 1 | 0 | 1 | 0 | NA | NA | 4 | 0.5 |
| 10_1212484 | 2 | 0 | 2 | 0 | 3 | 0.3333 | 2 | 0 | 9 | 0.1111 | 5 | 0 | 1 | 1 | 1 | 0 | NA | NA | 7 | 0.1429 |
| 10_1212486 | 2 | 1 | 2 | 1 | 3 | 1 | 2 | 1 | 9 | 1 | 4 | 1 | 1 | 1 | 2 | 1 | NA | NA | 7 | 1 |
| 10_1212486 | 2 | 1 | 2 | 1 | 3 | 1 | 2 | 1 | 9 | 1 | 5 | 0.8 | 1 | 1 | 2 | 1 | NA | NA | 8 | 0.875 |
| 10_1213827 | 2 | 0.5 | 9 | 0.5556 | 4 | 0.5 | 2 | 1 | 17 | 0.5882 | 4 | 1 | 7 | 0.5714 | 5 | 0.6 | 8 | 0.875 | 24 | 0.75 |
| 10_1213828 | 2 | 0.5 | 8 | 0.25 | 5 | 0 | 3 | 0 | 18 | 0.1667 | 4 | 0.25 | 8 | 0 | 7 | 0 | 7 | 0.2857 | 26 | 0.1154 |
| 10_1213830 | 2 | 0 | 7 | 0.2857 | 1 | 0 | 1 | 1 | 11 | 0.2727 | 2 | 0 | 4 | 0.5 | 3 | 0.6667 | 4 | 0.5 | 13 | 0.4615 |
| 10_1214251 | 7 | 0.1429 | 2 | 0 | 3 | 0 | 7 | 0.1429 | 19 | 0.1053 | NA | NA | 5 | 0 | 6 | 0.1667 | 3 | 0 | 14 | 0.0714 |
| 10_1214252 | 6 | 0 | 2 | 0 | 2 | 0 | 5 | 0.2 | 15 | 0.0667 | NA | NA | 4 | 0.25 | 5 | 0 | 3 | 0 | 12 | 0.0833 |
| 10_1214389 | 5 | 0.2 | 5 | 0.2 | 3 | 0.3333 | 7 | 0.1429 | 20 | 0.2 | 1 | 0 | 2 | 0 | 5 | 0.2 | 3 | 0.3333 | 11 | 0.1818 |
| 10_1214391 | 6 | 0.3333 | 3 | 0 | 3 | 0.3333 | 6 | 0.6667 | 18 | 0.3889 | 1 | 1 | 2 | 0.5 | 6 | 0.1667 | 2 | 0 | 11 | 0.2727 |
| 10_1214475 | 3 | 0 | 2 | 0 | NA | NA | 5 | 0.2 | 10 | 0.1 | 1 | 1 | NA | NA | 4 | 0 | NA | NA | 5 | 0.2 |
| 10_1215813 | 1 | 0 | 2 | 0.5 | NA | NA | NA | NA | 3 | 0.3333 | NA | NA | 1 | 1 | NA | NA | NA | NA | 1 | 1 |
| 10_1215865 | 1 | 1 | 1 | 1 | NA | NA | NA | NA | 2 | 1 | NA | NA | 2 | 0.5 | NA | NA | 1 | 1 | 3 | 0.6667 |
| 10_1215867 | 1 | 1 | 1 | 1 | NA | NA | 1 | 1 | 3 | 1 | NA | NA | 3 | 0.6667 | NA | NA | 1 | 1 | 4 | 0.75 |
| 10_1215869 | 1 | 1 | 1 | 1 | NA | NA | 1 | 1 | 3 | 1 | NA | NA | 3 | 1 | NA | NA | NA | NA | 3 | 1 |
| 10_1215871 | 1 | 1 | 1 | 1 | NA | NA | NA | NA | 2 | 1 | NA | NA | 3 | 1 | NA | NA | 1 | 1 | 4 | 1 |
| 10_1215871 | 1 | 1 | 1 | 0 | NA | NA | NA | NA | 2 | 0.5 | NA | NA | 2 | 1 | NA | NA | 1 | 1 | 3 | 1 |
| 10_1216178 | NA | NA | 1 | 0 | 1 | 1 | 2 | 0 | 4 | 0.25 | 2 | 0 | 2 | 0 | 1 | 0 | 4 | 0.25 | 9 | 0.1111 |
| 10_1216201 | NA | NA | NA | NA | 1 | 0 | 2 | 0.5 | 3 | 0.3333 | 2 | 0 | 3 | 0.3333 | 1 | 0 | 3 | 0.3333 | 9 | 0.2222 |
| 10_1216204 | NA | NA | NA | NA | 1 | 0 | 2 | 0.5 | 3 | 0.3333 | 2 | 0 | 5 | 0 | 1 | 0 | 3 | 0.6667 | 11 | 0.1818 |
| 10_1216204 | NA | NA | NA | NA | 1 | 1 | 2 | 0 | 3 | 0.3333 | 1 | 1 | 4 | 0.5 | NA | NA | 3 | 0.3333 | 8 | 0.5 |
| 10_1216205 | NA | NA | NA | NA | 1 | 1 | 2 | 1 | 3 | 1 | 2 | 0.5 | 5 | 1 | 1 | 1 | 3 | 1 | 11 | 0.9091 |
| 10_1216205 | NA | NA | NA | NA | 1 | 1 | 2 | 1 | 3 | 1 | 1 | 1 | 2 | 1 | 1 | 1 | 3 | 1 | 7 | 1 |
| 10_1216206 | NA | NA | NA | NA | 1 | 1 | 2 | 1 | 3 | 1 | 2 | 1 | 4 | 0.75 | 1 | 1 | 2 | 0.5 | 9 | 0.7778 |
| 10_1216206 | NA | NA | NA | NA | NA | NA | 2 | 0.5 | 2 | 0.5 | 2 | 0.5 | 4 | 0.25 | 1 | 0 | 1 | 0 | 8 | 0.25 |
| 10_1216206 | NA | NA | NA | NA | NA | NA | 1 | 1 | 1 | 1 | NA | NA | 3 | 0.6667 | 1 | 1 | 2 | 0.5 | 6 | 0.6667 |
| 10_1216206 | NA | NA | NA | NA | 1 | 1 | 3 | 1 | 4 | 1 | 2 | 1 | 2 | 1 | 1 | 1 | 3 | 0.6667 | 8 | 0.875 |
| 10_1216207 | NA | NA | NA | NA | NA | NA | 3 | 1 | 3 | 1 | 1 | 1 | 4 | 1 | 1 | 0 | 3 | 1 | 9 | 0.8889 |
| 10_1216207 | NA | NA | NA | NA | 1 | 0 | 2 | 0.5 | 3 | 0.3333 | 2 | 0 | 4 | 0.5 | NA | NA | 3 | 0.6667 | 9 | 0.4444 |
| 10_1216207 | NA | NA | NA | NA | NA | NA | 2 | 0.5 | 2 | 0.5 | 2 | 0 | 4 | 0.5 | 1 | 0 | 3 | 0 | 10 | 0.2 |
| 10_1216207 | NA | NA | NA | NA | 1 | 0 | 3 | 0.6667 | 4 | 0.5 | 2 | 0.5 | 5 | 0.2 | 1 | 0 | 2 | 0 | 10 | 0.2 |
| 10_1216210 | NA | NA | NA | NA | 1 | 1 | 2 | 0 | 3 | 0.3333 | 2 | 1 | 4 | 0.25 | 1 | 1 | 3 | 0.6667 | 10 | 0.6 |
| 10_1216210 | NA | NA | NA | NA | 1 | 0 | 2 | 0.5 | 3 | 0.3333 | 2 | 0.5 | 4 | 0.5 | 1 | 0 | 2 | 0.5 | 9 | 0.4444 |
| 10_1216211 | NA | NA | NA | NA | NA | NA | 3 | 1 | 3 | 1 | 1 | 1 | 4 | 0.75 | 1 | 1 | 3 | 1 | 9 | 0.8889 |
| 10_1216211 | NA | NA | NA | NA | 1 | 1 | 2 | 1 | 3 | 1 | 2 | 1 | 4 | 0.5 | 2 | 0.5 | 3 | 1 | 11 | 0.7273 |
| 10_1216213 | NA | NA | NA | NA | 1 | 1 | 3 | 0 | 4 | 0.25 | 2 | 0.5 | 5 | 0.8 | 2 | 1 | 2 | 0.5 | 11 | 0.7273 |
| 10_1216213 | NA | NA | NA | NA | 1 | 0 | 2 | 0.5 | 3 | 0.3333 | 1 | 1 | 3 | 0 | NA | NA | NA | NA | 4 | 0.25 |
| 10_1216214 | NA | NA | NA | NA | 1 | 0 | 1 | 1 | 2 | 0.5 | 2 | 0.5 | 4 | 0.25 | 2 | 0.5 | 2 | 0.5 | 10 | 0.4 |
| 10_1219153 | NA | NA | 1 | 1 | NA | NA | NA | NA | 1 | 1 | NA | NA | NA | NA | NA | NA | 1 | 1 | 1 | 1 |
| 10_1219155 | NA | NA | 1 | 1 | NA | NA | NA | NA | 1 | 1 | NA | NA | NA | NA | NA | NA | 1 | 1 | 1 | 1 |
| 10_1219157 | NA | NA | 1 | 1 | NA | NA | NA | NA | 1 | 1 | NA | NA | NA | NA | NA | NA | 1 | 1 | 1 | 1 |
| 10_1219170 | NA | NA | 1 | 1 | NA | NA | NA | NA | 1 | 1 | NA | NA | 1 | 1 | NA | NA | NA | NA | 1 | 1 |
| 10_1219172 | 1 | 1 | 1 | 0 | 4 | 0.5 | 4 | 0.5 | 10 | 0.5 | 4 | 0.75 | 6 | 0.1667 | NA | NA | 3 | 0.6667 | 13 | 0.4615 |
| 10_1219173 | 1 | 0 | 1 | 0 | 4 | 0 | 6 | 0.3333 | 12 | 0.1667 | 4 | 0 | 5 | 0 | 2 | 0.5 | 3 | 0 | 14 | 0.0714 |
| 10_1219173 | 1 | 0 | 1 | 1 | 2 | 0.5 | 5 | 0.2 | 9 | 0.3333 | 4 | 0.5 | 5 | 0.4 | 1 | 0 | 3 | 0 | 13 | 0.3077 |
| 10_1219260 | NA | NA | 2 | 0 | 4 | 0.5 | 3 | 0 | 9 | 0.2222 | 2 | 0 | 4 | 0.5 | 1 | 0 | 1 | 0 | 8 | 0.25 |
| 10_1225151 | 2 | 0 | 3 | 0 | 1 | 0 | 1 | 1 | 7 | 0.1429 | NA | NA | NA | NA | 1 | 0 | 1 | 1 | 2 | 0.5 |
| 10_1227145 | 4 | 0 | 3 | 0.3333 | 2 | 0 | 4 | 0 | 13 | 0.0769 | NA | NA | NA | NA | 3 | 0 | 1 | 1 | 4 | 0.25 |
| 10_1227772 | 3 | 0.6667 | 1 | 1 | 1 | 0 | 3 | 0.6667 | 8 | 0.625 | NA | NA | 2 | 0.5 | 2 | 0 | 2 | 0.5 | 6 | 0.3333 |
| 10_1227772 | 3 | 0.6667 | 3 | 0 | 2 | 0.5 | 3 | 0.3333 | 11 | 0.3636 | NA | NA | 3 | 0.3333 | 3 | 0.6667 | 2 | 0.5 | 8 | 0.5 |
| 10_1227773 | 4 | 0.25 | 3 | 0.6667 | 1 | 1 | 4 | 0.5 | 12 | 0.5 | NA | NA | 2 | 0.5 | 3 | 0 | 2 | 0.5 | 7 | 0.2857 |
| 10_1227774 | 5 | 0.8 | 2 | 0.5 | 2 | 0.5 | 3 | 0.3333 | 12 | 0.5833 | NA | NA | 3 | 0.6667 | 2 | 0 | 2 | 0.5 | 7 | 0.4286 |
| 10_1227778 | 3 | 0.3333 | 2 | 0.5 | 1 | 0 | 3 | 0 | 9 | 0.2222 | NA | NA | NA | NA | 3 | 0.3333 | 3 | 0 | 6 | 0.1667 |
| 10_1227798 | 3 | 0.3333 | 3 | 0 | 1 | 0 | 5 | 0.4 | 12 | 0.25 | 1 | 1 | 3 | 0.3333 | 2 | 0 | 3 | 0.3333 | 9 | 0.3333 |
| 10_1227804 | 2 | 0.5 | 3 | 0 | 3 | 0.3333 | 7 | 0 | 15 | 0.1333 | 1 | 0 | 4 | 0.25 | 4 | 0 | 5 | 0 | 14 | 0.0714 |
| 10_1227814 | 5 | 0.2 | 5 | 0.4 | 5 | 0 | 8 | 0.25 | 23 | 0.2174 | 2 | 0.5 | 5 | 0 | 6 | 0 | 7 | 0 | 20 | 0.05 |
| 10_1228281 | 5 | 0.2 | 3 | 0 | 2 | 0 | 6 | 0.3333 | 16 | 0.1875 | 2 | 0 | 2 | 0 | 3 | 0 | 5 | 0.2 | 12 | 0.0833 |
| 10_1228292 | 4 | 0.25 | 2 | 0 | 2 | 0 | 4 | 0 | 12 | 0.0833 | 2 | 0.5 | 2 | 0 | 2 | 0 | 6 | 0.1667 | 12 | 0.1667 |
| 10_1228294 | 3 | 0.3333 | 1 | 0 | NA | NA | 4 | 0.25 | 8 | 0.25 | 1 | 0 | 2 | 0.5 | 1 | 0 | 6 | 0 | 10 | 0.1 |
| 10_1228299 | 4 | 0 | 1 | 0 | 2 | 0 | 3 | 0.3333 | 10 | 0.1 | 1 | 0 | 3 | 0 | 3 | 0.3333 | 6 | 0 | 13 | 0.0769 |
| 10_1228304 | 3 | 0.3333 | 1 | 0 | NA | NA | 1 | 0 | 5 | 0.2 | NA | NA | 1 | 1 | 2 | 0.5 | 4 | 0.5 | 7 | 0.5714 |
| 10_1228307 | 3 | 1 | NA | NA | 1 | 1 | 2 | 0.5 | 6 | 0.8333 | 1 | 1 | 3 | 0.6667 | 2 | 1 | 3 | 1 | 9 | 0.8889 |
| 10_1228307 | 2 | 1 | NA | NA | NA | NA | 2 | 0.5 | 4 | 0.75 | NA | NA | 2 | 0 | 2 | 1 | 3 | 0.6667 | 7 | 0.5714 |

| | | | | | | | | | | | | | | | | | | | | |
|---|---|---|---|---|---|---|---|---|---|---|---|---|---|---|---|---|---|---|---|---|
| 10_1228308 | 3 | 0.3333 | 1 | 0 | 1 | 0 | 2 | 0 | 7 | 0.1429 | 1 | 0 | 1 | 0 | 1 | 1 | 3 | 0.3333 | 6 | 0.3333 |
| 10_1228310 | 13 | 0.2308 | 6 | 0.3333 | 3 | 0 | 8 | 0.125 | 30 | 0.2 | 5 | 0.6 | 3 | 0 | 4 | 0.5 | 6 | 0.3333 | 18 | 0.3889 |
| 10_1228311 | 12 | 0.0833 | 5 | 0 | 4 | 0 | 7 | 0.1429 | 28 | 0.0714 | 5 | 0.4 | 4 | 0 | 6 | 0.1667 | 4 | 0.25 | 19 | 0.2105 |
| 10_1228313 | 14 | 0.1429 | 5 | 0.2 | 4 | 0.25 | 7 | 0.2857 | 30 | 0.2 | 4 | 0.25 | 3 | 0 | 5 | 0.4 | 8 | 0.25 | 20 | 0.25 |
| 10_1228313 | 9 | 0.3333 | 4 | 0.5 | 3 | 0.6667 | 9 | 0.4444 | 25 | 0.44 | 3 | 1 | 3 | 0.6667 | 2 | 0.5 | 6 | 0.6667 | 14 | 0.7143 |
| 10_1228313 | 12 | 0.25 | 4 | 0.25 | 2 | 0 | 7 | 0 | 25 | 0.16 | 4 | 0 | 3 | 0.3333 | 5 | 0 | 6 | 0 | 18 | 0.0556 |
| 10_1228316 | 12 | 0.5833 | 5 | 0.4 | 4 | 0.75 | 8 | 0.75 | 29 | 0.6207 | 4 | 0.75 | 2 | 1 | 6 | 0.8333 | 7 | 0.8571 | 19 | 0.8421 |
| 10_1228495 | 5 | 1 | 3 | 0.6667 | 3 | 0 | 3 | 0.6667 | 14 | 0.6429 | 2 | 0.5 | 1 | 1 | 5 | 0.6 | 5 | 0.8 | 13 | 0.6923 |
| 10_1228496 | 6 | 0.3333 | 3 | 0.6667 | 2 | 0 | 2 | 0.5 | 13 | 0.3846 | 1 | 0 | 1 | 0 | 6 | 0.1667 | 3 | 0.6667 | 11 | 0.2727 |
| 10_1228496 | 3 | 0.6667 | 3 | 0 | 3 | 0 | 2 | 0 | 11 | 0.1818 | 2 | 0 | 1 | 0 | 5 | 0.2 | 3 | 0 | 11 | 0.0909 |
| 10_1228496 | 3 | 0.6667 | 3 | 0.6667 | 3 | 0 | 1 | 1 | 10 | 0.5 | 1 | 1 | NA | NA | 4 | 0.75 | 4 | 1 | 9 | 0.8889 |
| 10_1229411 | NA | NA | 3 | 0.3333 | 1 | 0 | 1 | 0 | 5 | 0.2 | 1 | 1 | 2 | 0 | 4 | 0 | 2 | 0 | 9 | 0.1111 |
| 10_1229480 | 5 | 0 | 3 | 0.3333 | 6 | 0 | 5 | 0 | 19 | 0.0526 | 5 | 0.2 | 7 | 0 | 6 | 0 | 2 | 0 | 20 | 0.05 |
| 10_1229492 | 2 | 0 | 3 | 0 | 6 | 0.3333 | 6 | 0 | 17 | 0.1176 | 4 | 0.25 | 7 | 0.1429 | 4 | 0.25 | 5 | 0.2 | 20 | 0.2 |
| 10_1229493 | 2 | 1 | 4 | 0.5 | 6 | 0.8333 | 7 | 0.8571 | 19 | 0.7895 | 5 | 0.4 | 7 | 0.7143 | 5 | 0.8 | 3 | 1 | 20 | 0.7 |
| 10_1229494 | 3 | 1 | 4 | 0.75 | 5 | 0.8 | 8 | 1 | 20 | 0.9 | 2 | 1 | 6 | 0.8333 | 2 | 1 | 5 | 1 | 15 | 0.9333 |
| 10_1229494 | 2 | 0.5 | 5 | 0.2 | 7 | 0 | 9 | 0 | 23 | 0.087 | 3 | 0.3333 | 8 | 0 | 4 | 0 | 5 | 0.2 | 20 | 0.1 |
| 10_1229494 | 3 | 0.3333 | 5 | 0 | 6 | 0.1667 | 9 | 0.3333 | 23 | 0.2174 | 3 | 1 | 8 | 0.625 | 4 | 0 | 5 | 0.4 | 20 | 0.5 |
| 10_1229495 | 3 | 0.6667 | 5 | 0.8 | 6 | 0.6667 | 8 | 0.75 | 22 | 0.7273 | 3 | 1 | 8 | 0.625 | 4 | 0.25 | 4 | 0.75 | 19 | 0.6316 |
| 10_1229495 | 3 | 0.3333 | 4 | 0 | 8 | 0.5 | 10 | 0.3 | 25 | 0.32 | 3 | 0.3333 | 8 | 0.125 | 4 | 0.25 | 5 | 0 | 20 | 0.15 |
| 10_1229495 | 3 | 0 | 5 | 0.4 | 6 | 0 | 10 | 0.2 | 24 | 0.1667 | 3 | 0 | 8 | 0.125 | 4 | 0 | 5 | 0 | 20 | 0.05 |
| 10_1229614 | 2 | 0 | 2 | 0 | 5 | 0 | 6 | 0.1667 | 15 | 0.0667 | 2 | 0 | 4 | 0.25 | 3 | 0 | 5 | 0 | 14 | 0.0714 |
| 10_1229669 | 2 | 0 | 2 | 0 | 3 | 0.3333 | 5 | 0 | 12 | 0.0833 | 2 | 0 | 2 | 0 | 2 | 0 | 5 | 0.2 | 11 | 0.0909 |
| 10_1235748 | 6 | 0 | 10 | 0.2 | 3 | 0.3333 | 4 | 0.25 | 23 | 0.1739 | 3 | 0.3333 | 2 | 0 | 1 | 0 | 3 | 0 | 9 | 0.1111 |
| 10_1235748 | 6 | 0.3333 | 9 | 0.5556 | 3 | 0.6667 | 4 | 0.75 | 22 | 0.5455 | 3 | 1 | 3 | 0.6667 | 1 | 1 | 3 | 0.6667 | 10 | 0.8 |
| 10_1235749 | 5 | 1 | 10 | 0.8 | 3 | 1 | 4 | 1 | 22 | 0.9091 | 3 | 1 | 3 | 0.6667 | 1 | 1 | 4 | 1 | 11 | 0.9091 |
| 10_1235749 | 4 | 0.75 | 8 | 1 | 2 | 1 | 3 | 1 | 17 | 0.9412 | 3 | 1 | 2 | 1 | 1 | 1 | 3 | 1 | 9 | 1 |
| 10_1235750 | 6 | 0.3333 | 10 | 0.2 | 3 | 0.3333 | 4 | 0.25 | 23 | 0.2609 | 4 | 0.25 | 1 | 1 | NA | NA | 4 | 0.25 | 9 | 0.3333 |
| 10_1235751 | 4 | 1 | 8 | 0.75 | 3 | 1 | 4 | 0.5 | 19 | 0.7895 | 4 | 0.75 | 3 | 0.6667 | 1 | 1 | 4 | 0.75 | 12 | 0.75 |
| 10_1235751 | 6 | 0.8333 | 11 | 1 | 3 | 1 | 4 | 1 | 24 | 0.9583 | 3 | 1 | 3 | 0.6667 | 1 | 1 | 4 | 0.75 | 11 | 0.8182 |
| 10_1235752 | 6 | 0.6667 | 10 | 0.5 | 3 | 0.3333 | 4 | 0.5 | 23 | 0.5217 | 3 | 1 | 2 | 0 | 1 | 0 | 4 | 0.25 | 10 | 0.4 |
| 10_1235752 | 7 | 0.5714 | 10 | 0.7 | 3 | 1 | 5 | 0.8 | 25 | 0.72 | 4 | 0.75 | 3 | 0.3333 | 1 | 1 | 4 | 0.75 | 12 | 0.6667 |
| 10_1235752 | 7 | 0 | 10 | 0.2 | 2 | 0 | 3 | 0 | 22 | 0.0909 | 4 | 0.25 | 2 | 0 | 1 | 0 | 3 | 0 | 10 | 0.1 |
| 10_1235763 | 8 | 0 | 8 | 0.25 | 4 | 0 | 4 | 0 | 24 | 0.0833 | 5 | 0.2 | 5 | 0 | 2 | 0 | 4 | 0.25 | 16 | 0.125 |
| 10_1235764 | 7 | 0 | 11 | 0.2727 | 4 | 0.25 | 5 | 0.2 | 27 | 0.1852 | 5 | 0 | 6 | 0 | 2 | 0.5 | 5 | 0.4 | 18 | 0.1667 |
| 10_1235765 | 8 | 0 | 10 | 0.5 | 6 | 0 | 4 | 0.25 | 28 | 0.2143 | 4 | 0.25 | 6 | 0.5 | 1 | 0 | 5 | 0.2 | 16 | 0.3125 |
| 10_1235766 | 7 | 0.4286 | 12 | 0.5833 | 8 | 0.625 | 3 | 1 | 30 | 0.6 | 8 | 0.5 | 6 | 0.3333 | 2 | 0.5 | 4 | 0.75 | 20 | 0.5 |
| 10_1250564 | 3 | 0.3333 | 1 | 0 | 1 | 0 | NA | NA | 5 | 0.2 | NA | NA | NA | NA | 2 | 0.5 | 3 | 0 | 5 | 0.2 |
| 10_1250634 | NA | NA | 1 | 1 | 1 | 1 | 1 | 0 | 3 | 0.6667 | 1 | 1 | NA | NA | 4 | 0.25 | 1 | 1 | 6 | 0.5 |
| 10_1250635 | NA | NA | NA | NA | 1 | 0 | 1 | 1 | 2 | 0.5 | 1 | 1 | NA | NA | 4 | 0.75 | 1 | 0 | 6 | 0.6667 |
| 10_1250640 | NA | NA | NA | NA | 1 | 1 | NA | NA | 1 | 1 | 1 | 0 | NA | NA | 4 | 1 | 1 | 1 | 6 | 0.8333 |
| 10_1267232 | 1 | 0 | 2 | 0 | 2 | 0.5 | 2 | 1 | 7 | 0.4286 | NA | NA | 2 | 0.5 | NA | NA | NA | NA | 2 | 0.5 |
| 10_1267233 | 2 | 0 | 2 | 0 | 3 | 0.6667 | 2 | 0.5 | 9 | 0.3333 | NA | NA | 2 | 0.5 | NA | NA | NA | NA | 2 | 0.5 |
| 10_1267233 | 1 | 1 | 1 | 0 | 2 | 0.5 | 2 | 1 | 6 | 0.6667 | NA | NA | 2 | 1 | NA | NA | NA | NA | 2 | 1 |
| 10_1268323 | 5 | 0.6 | 3 | 0.3333 | 8 | 0.625 | 1 | 0 | 17 | 0.5294 | 8 | 0.375 | 2 | 0.5 | 7 | 0.7143 | 4 | 0.5 | 21 | 0.5238 |
| 10_1268323 | 3 | 0.6667 | 2 | 1 | 6 | 1 | 1 | 1 | 12 | 0.9167 | 8 | 0.75 | 1 | 1 | 5 | 1 | 4 | 0.75 | 18 | 0.8333 |
| 10_1268324 | 5 | 0.4 | 3 | 0.3333 | 8 | 0.125 | 2 | 0 | 18 | 0.2222 | 9 | 0.3333 | 2 | 0.5 | 7 | 0.1429 | 4 | 0 | 22 | 0.2273 |
| 10_1268325 | 5 | 0 | 3 | 0 | 8 | 0.125 | 2 | 0 | 18 | 0.0556 | 8 | 0 | 1 | 0 | 5 | 0.2 | 4 | 0 | 18 | 0.0556 |
| 10_1268326 | 5 | 0.8 | 4 | 0.75 | 8 | 0.75 | 2 | 0.5 | 19 | 0.7368 | 9 | 0.6667 | 2 | 1 | 7 | 0.8571 | 4 | 1 | 22 | 0.8182 |
| 10_1268327 | 3 | 0 | 4 | 0 | 6 | 0.1667 | 1 | 1 | 14 | 0.1429 | 8 | 0.125 | 2 | 0.5 | 7 | 0.1429 | 3 | 0 | 20 | 0.15 |
| 10_1268330 | 5 | 0 | 5 | 0.2 | 6 | 0 | 2 | 0 | 18 | 0.0556 | 7 | 0.1429 | 2 | 0.5 | 8 | 0 | 4 | 0 | 21 | 0.0952 |
| 10_1268332 | 7 | 0 | 5 | 0 | 8 | 0.375 | 2 | 0 | 22 | 0.1364 | 6 | 0.6667 | 2 | 0.5 | 7 | 0.2857 | 4 | 0 | 19 | 0.3684 |
| 10_1268333 | 6 | 0.3333 | 4 | 0.25 | 8 | 0.375 | 3 | 0.6667 | 21 | 0.381 | 7 | 0.4286 | 2 | 1 | 8 | 0.375 | 4 | 0 | 21 | 0.381 |
| 10_1268334 | 7 | 0 | 6 | 0.1667 | 8 | 0.25 | 2 | 0 | 23 | 0.1304 | 8 | 0.375 | 2 | 0 | 9 | 0 | 6 | 0.1667 | 25 | 0.16 |
| 10_1268584 | 5 | 0.2 | 4 | 0.5 | 2 | 0.5 | 4 | 0.75 | 15 | 0.4667 | 6 | 0.6667 | 4 | 0.5 | 4 | 0.75 | 6 | 0.8333 | 20 | 0.7 |
| 10_1268584 | 5 | 0.8 | 4 | 0.5 | 2 | 0.5 | 4 | 0.25 | 15 | 0.5333 | 6 | 0.6667 | 3 | 0.3333 | 4 | 0.5 | 5 | 0.2 | 18 | 0.4444 |
| 10_1268910 | NA | NA | 1 | 0 | 1 | 1 | 2 | 0 | 4 | 0.25 | 1 | 0 | 3 | 0 | 1 | 0 | 1 | 1 | 6 | 0.1667 |
| 10_1268912 | NA | NA | 1 | 1 | 1 | 1 | 1 | 0 | 3 | 0.6667 | NA | NA | 3 | 1 | 1 | 1 | 1 | 1 | 5 | 1 |
| 10_1268916 | NA | NA | 1 | 0 | 1 | 0 | 1 | 1 | 3 | 0.3333 | NA | NA | 3 | 1 | NA | NA | NA | NA | 3 | 1 |
| 10_1268919 | NA | NA | 7 | 0.2857 | 3 | 0 | 4 | 0 | 14 | 0.1429 | 1 | 0 | 7 | 0.5714 | 3 | 0 | 3 | 0 | 14 | 0.2857 |
| 10_1268920 | NA | NA | 5 | 0.2 | 3 | 0.3333 | 4 | 0 | 12 | 0.1667 | 1 | 0 | 6 | 0.1667 | 1 | 0 | 1 | 1 | 9 | 0.2222 |
| 10_1268920 | NA | NA | 5 | 1 | 2 | 1 | 6 | 1 | 13 | 1 | 1 | 1 | 7 | 1 | 3 | 1 | 4 | 1 | 15 | 1 |
| 10_1268921 | 1 | 1 | 6 | 1 | 3 | 1 | 5 | 0.4 | 15 | 0.8 | 1 | 1 | 8 | 1 | 3 | 0.3333 | 4 | 0.5 | 16 | 0.75 |
| 10_1268921 | NA | NA | 6 | 0.1667 | 2 | 0 | 6 | 0.1667 | 14 | 0.1429 | 1 | 0 | 5 | 0 | 3 | 0 | 3 | 0.3333 | 12 | 0.0833 |
| 10_1268922 | NA | NA | 6 | 0.5 | 1 | 0 | 4 | 0.5 | 11 | 0.4545 | NA | NA | 6 | 0.3333 | 2 | 0 | 4 | 0.25 | 12 | 0.25 |

| | | | | | | | | | | | | | | | | | | | | |
|---|---|---|---|---|---|---|---|---|---|---|---|---|---|---|---|---|---|---|---|---|
| 10_126892 | 1 | 0 | 6 | 0.1667 | 3 | 0 | 6 | 0.6667 | 16 | 0.3125 | 1 | 1 | 8 | 0.375 | 3 | 0 | 3 | 0 | 15 | 0.2667 |
| 10_126892 | 1 | 0 | 5 | 0.6 | 3 | 0.3333 | 6 | 1 | 15 | 0.6667 | 1 | 0 | 8 | 0.375 | 3 | 0.6667 | 3 | 0.6667 | 15 | 0.4667 |
| 10_126892 | NA | NA | 5 | 0.8 | 3 | 1 | 5 | 0.8 | 13 | 0.8462 | 1 | 1 | 5 | 1 | 3 | 1 | 4 | 1 | 13 | 1 |
| 10_126892 | 1 | 0 | 7 | 0.2857 | 2 | 0 | 4 | 0.75 | 14 | 0.3571 | NA | NA | 5 | 0.2 | 3 | 0.3333 | 4 | 0 | 12 | 0.1667 |
| 10_126892 | 1 | 1 | 6 | 1 | 3 | 1 | 5 | 1 | 15 | 1 | 1 | 1 | 7 | 1 | 3 | 1 | 4 | 1 | 15 | 1 |
| 10_126892 | 1 | 0 | 5 | 0.6 | 3 | 0.6667 | 5 | 0 | 14 | 0.3571 | 1 | 0 | 7 | 0.1429 | 1 | 0 | 3 | 0 | 12 | 0.0833 |
| 10_126892 | NA | NA | 6 | 0.1667 | 3 | 0.3333 | 2 | 0.5 | 11 | 0.2727 | NA | NA | 6 | 0.6667 | 2 | 0.5 | 2 | 0.5 | 10 | 0.6 |
| 10_126892 | 1 | 1 | 7 | 0.5714 | 3 | 0 | 3 | 0 | 14 | 0.3571 | 1 | 0 | 6 | 0.5 | 3 | 0.3333 | 3 | 1 | 13 | 0.5385 |
| 10_126892 | NA | NA | 4 | 0.25 | 2 | 0 | 3 | 0 | 9 | 0.1111 | NA | NA | 3 | 0 | 1 | 0 | 3 | 0.3333 | 7 | 0.1429 |
| 10_126892 | NA | NA | 4 | 0.5 | 1 | 0 | 4 | 0.25 | 9 | 0.3333 | NA | NA | 4 | 0.25 | 3 | 0 | 3 | 0 | 10 | 0.1 |
| 10_126901 | 2 | 0 | 1 | 0 | NA | NA | 1 | 1 | 4 | 0.25 | NA | NA | 1 | 0 | NA | NA | 2 | 0.5 | 3 | 0.3333 |
| 10_126901 | 2 | 0 | 1 | 0 | NA | NA | 2 | 0.5 | 5 | 0.2 | NA | NA | 1 | 1 | NA | NA | 2 | 0 | 3 | 0.3333 |
| 10_126903 | 9 | 0 | 22 | 0.0909 | 14 | 0.1429 | 9 | 0.1111 | 54 | 0.0926 | 11 | 0.0909 | 11 | 0 | 6 | 0 | 14 | 0.2143 | 42 | 0.0952 |
| 10_126903 | 8 | 0.375 | 19 | 0.5789 | 12 | 0.5833 | 7 | 0.2857 | 46 | 0.5 | 12 | 0.5 | 10 | 0.3 | 6 | 0.6667 | 10 | 0.4 | 38 | 0.4474 |
| 10_126903 | 6 | 0.3333 | 16 | 0.25 | 11 | 0.2727 | 7 | 0 | 40 | 0.225 | 10 | 0.2 | 11 | 0.2727 | 4 | 0.5 | 10 | 0.2 | 35 | 0.2571 |
| 10_126903 | 7 | 0.7143 | 16 | 0.625 | 12 | 0.3333 | 6 | 0.6667 | 41 | 0.561 | 12 | 0.25 | 8 | 0.375 | 4 | 0.5 | 11 | 0.4545 | 35 | 0.3714 |
| 10_126903 | 9 | 0.4444 | 16 | 0.125 | 12 | 0.1667 | 4 | 0.25 | 41 | 0.2195 | 11 | 0.2727 | 7 | 0.4286 | 5 | 0.2 | 12 | 0.0833 | 35 | 0.2286 |
| 10_126915 | 9 | 1 | 16 | 0.875 | 10 | 0.7 | 8 | 0.75 | 43 | 0.8372 | 12 | 0.75 | 15 | 0.7333 | 19 | 0.7895 | 4 | 1 | 50 | 0.78 |
| 10_126916 | 9 | 0.3333 | 23 | 0.2174 | 14 | 0.2143 | 9 | 0.3333 | 55 | 0.2545 | 15 | 0.2 | 16 | 0.25 | 23 | 0.2174 | 5 | 0.2 | 59 | 0.2203 |
| 10_126916 | 8 | 0.25 | 19 | 0.1579 | 14 | 0.0714 | 9 | 0.1111 | 50 | 0.14 | 10 | 0.2 | 12 | 0.0833 | 21 | 0.0952 | 4 | 0 | 47 | 0.1064 |
| 10_126916 | 9 | 1 | 21 | 0.9524 | 13 | 0.6154 | 8 | 0.875 | 51 | 0.8627 | 9 | 0.8889 | 12 | 1 | 20 | 1 | 4 | 0.75 | 45 | 0.9556 |
| 10_126916 | 6 | 0 | 20 | 0.15 | 12 | 0.0833 | 7 | 0.1429 | 45 | 0.1111 | 11 | 0.2727 | 12 | 0 | 21 | 0.0476 | 5 | 0 | 49 | 0.0816 |
| 10_126916 | 9 | 0 | 24 | 0.375 | 17 | 0.1765 | 9 | 0 | 59 | 0.2034 | 12 | 0.0833 | 17 | 0.2941 | 24 | 0.3333 | 5 | 0.2 | 58 | 0.2586 |
| 10_126916 | 10 | 0.9 | 21 | 0.7619 | 11 | 0.9091 | 8 | 0.875 | 50 | 0.84 | 13 | 0.8462 | 14 | 0.7857 | 22 | 0.8636 | 6 | 0.8333 | 55 | 0.8364 |
| 10_126916 | 8 | 1 | 22 | 0.7273 | 15 | 0.7333 | 9 | 0.8889 | 54 | 0.7963 | 13 | 0.6923 | 16 | 0.5 | 23 | 0.7391 | 6 | 0.6667 | 58 | 0.6552 |
| 10_126916 | 9 | 0 | 23 | 0.2174 | 15 | 0.1333 | 9 | 0.1111 | 56 | 0.1429 | 13 | 0.2308 | 17 | 0.0588 | 21 | 0.2857 | 6 | 0.1667 | 57 | 0.193 |
| 10_126916 | 10 | 0 | 19 | 0.1053 | 18 | 0 | 7 | 0.1429 | 54 | 0.0556 | 14 | 0 | 15 | 0.0667 | 26 | 0.0769 | 5 | 0.2 | 60 | 0.0667 |
| 10_126927 | 12 | 0.5 | 27 | 0.3704 | 31 | 0.2581 | 13 | 0.3077 | 83 | 0.3373 | 21 | 0.4762 | 37 | 0.3514 | 31 | 0.2581 | 18 | 0.2222 | 107 | 0.3271 |
| 10_126927 | 15 | 0.0667 | 30 | 0.2333 | 32 | 0.0625 | 12 | 0.25 | 89 | 0.1461 | 24 | 0.125 | 33 | 0.0909 | 31 | 0.1935 | 15 | 0.1333 | 103 | 0.1359 |
| 10_126927 | 13 | 0.1538 | 35 | 0.2571 | 34 | 0.2647 | 15 | 0.4 | 97 | 0.268 | 28 | 0.3214 | 39 | 0.2821 | 42 | 0.381 | 20 | 0.2 | 129 | 0.3101 |
| 10_126927 | 14 | 0.5714 | 37 | 0.7027 | 33 | 0.7576 | 12 | 0.8333 | 96 | 0.7188 | 22 | 0.6818 | 36 | 0.75 | 35 | 0.7143 | 19 | 0.7368 | 112 | 0.7232 |
| 10_126927 | 18 | 0.2778 | 35 | 0.2286 | 39 | 0.2821 | 13 | 0.3846 | 105 | 0.2762 | 25 | 0.28 | 40 | 0.325 | 40 | 0.425 | 20 | 0.45 | 125 | 0.368 |
| 10_126927 | 15 | 0.5333 | 28 | 0.8571 | 32 | 0.5938 | 10 | 0.7 | 85 | 0.6824 | 16 | 0.75 | 24 | 0.7083 | 27 | 0.6667 | 18 | 0.8889 | 85 | 0.7412 |
| 10_126927 | 18 | 0.1111 | 43 | 0.1395 | 41 | 0.0244 | 16 | 0 | 118 | 0.0763 | 31 | 0.0968 | 45 | 0.1778 | 45 | 0.0667 | 22 | 0 | 143 | 0.0979 |
| 10_126927 | NA | NA | NA | NA | 1 | 1 | NA | NA | 1 | 1 | 1 | 0 | 1 | 1 | 1 | 0 | NA | NA | 3 | 0.3333 |
| 10_126928 | 13 | 0.9231 | 36 | 0.9167 | 39 | 0.9487 | 15 | 0.9333 | 103 | 0.932 | 27 | 0.9259 | 39 | 0.9231 | 38 | 0.9211 | 22 | 0.8636 | 126 | 0.9127 |
| 10_126928 | 18 | 0.3333 | 39 | 0.1538 | 37 | 0.1351 | 17 | 0.0588 | 111 | 0.1622 | 24 | 0.375 | 38 | 0.1842 | 47 | 0.2553 | 23 | 0.2609 | 132 | 0.2576 |
| 10_126928 | 22 | 0.0455 | 53 | 0.0943 | 43 | 0.0698 | 19 | 0.1579 | 137 | 0.0876 | 30 | 0.1333 | 45 | 0.1333 | 48 | 0.1667 | 23 | 0.0435 | 146 | 0.1301 |
| 10_126928 | 26 | 0.1923 | 76 | 0.1184 | 66 | 0.1364 | 34 | 0.0882 | 202 | 0.1287 | 46 | 0.1304 | 80 | 0.0875 | 80 | 0.15 | 41 | 0.1707 | 247 | 0.1296 |
| 10_126928 | 35 | 0.1143 | 86 | 0.0814 | 84 | 0.1667 | 37 | 0.1622 | 242 | 0.1281 | 56 | 0.0893 | 96 | 0.1354 | 98 | 0.1939 | 48 | 0.2292 | 298 | 0.1611 |
| 10_126930 | 1 | 1 | 1 | 1 | 2 | 1 | NA | NA | 4 | 1 | NA | NA | 1 | 1 | NA | NA | NA | NA | 1 | 1 |
| 10_126930 | 1 | 1 | 1 | 1 | 2 | 1 | NA | NA | 4 | 1 | NA | NA | NA | NA | NA | NA | 1 | 1 | 1 | 1 |
| 10_126930 | 1 | 0 | 1 | 0 | 2 | 0.5 | NA | NA | 4 | 0.25 | NA | NA | 1 | 0 | NA | NA | 1 | 1 | 2 | 0.5 |
| 10_126931 | NA | NA | NA | NA | 1 | 1 | NA | NA | 1 | 1 | NA | NA | 1 | 0 | NA | NA | 1 | 1 | 2 | 0.5 |
| 10_126931 | 2 | 0 | 4 | 0 | 4 | 0.25 | 7 | 0.5714 | 17 | 0.2941 | 4 | 0.25 | 5 | 0 | 5 | 0.4 | 5 | 0.4 | 19 | 0.2632 |
| 10_126931 | 1 | 0 | 2 | 0.5 | 4 | 0.25 | 4 | 0.25 | 11 | 0.2727 | 4 | 0 | 4 | 0 | 4 | 0.25 | 5 | 0 | 17 | 0.0588 |
| 10_126931 | 5 | 0.2 | 21 | 0.0476 | 7 | 0 | 11 | 0.0909 | 44 | 0.0682 | 11 | 0.1818 | 11 | 0.1818 | 9 | 0.1111 | 7 | 0 | 38 | 0.1316 |
| 10_126931 | 9 | 0.7778 | 21 | 0.7143 | 12 | 0.9167 | 11 | 0.6364 | 53 | 0.7547 | 15 | 0.8 | 13 | 0.8462 | 13 | 0.6923 | 13 | 0.6923 | 54 | 0.7593 |
| 10_126931 | 5 | 0.2 | 14 | 0.1429 | 4 | 0 | 7 | 0 | 30 | 0.1 | 11 | 0.2727 | 9 | 0.3333 | 11 | 0.0909 | 11 | 0.1818 | 42 | 0.2143 |
| 10_126931 | 4 | 0.75 | 8 | 1 | 5 | 1 | 5 | 0.8 | 22 | 0.9091 | 9 | 1 | 8 | 1 | 8 | 1 | 10 | 1 | 35 | 1 |
| 10_126955 | 1 | 0 | 6 | 0.1667 | 2 | 0 | 4 | 0 | 13 | 0.0769 | 5 | 0 | 2 | 0 | 4 | 0.25 | 2 | 0 | 13 | 0.0769 |
| 10_127000 | 1 | 1 | NA | NA | NA | NA | NA | NA | 1 | 1 | NA | NA | 1 | 1 | NA | NA | NA | NA | 1 | 1 |
| 10_127001 | 1 | 1 | NA | NA | NA | NA | NA | NA | 1 | 1 | NA | NA | 1 | 1 | NA | NA | NA | NA | 1 | 1 |
| 10_127001 | 4 | 0 | 7 | 0 | 3 | 0 | 5 | 0.2 | 19 | 0.0526 | 1 | 0 | 2 | 0 | 8 | 0 | 4 | 0.25 | 15 | 0.0667 |
| 10_127003 | 1 | 0 | 2 | 0 | NA | NA | 1 | 1 | 4 | 0.25 | NA | NA | 2 | 0.5 | 2 | 0 | NA | NA | 4 | 0.25 |
| 10_127004 | 1 | 0 | 2 | 0.5 | NA | NA | 2 | 0.5 | 5 | 0.4 | NA | NA | 2 | 1 | 2 | 0.5 | 1 | 0 | 5 | 0.6 |
| 10_127005 | 8 | 0.125 | 18 | 0.1667 | 5 | 0 | 8 | 0.25 | 39 | 0.1538 | 7 | 0 | 14 | 0.1429 | 15 | 0.0667 | 8 | 0 | 44 | 0.0682 |
| 10_127006 | NA | NA | NA | NA | NA | NA | 1 | 1 | 1 | 1 | NA | NA | NA | NA | 1 | 1 | 1 | 0 | 2 | 0.5 |
| 10_127007 | 10 | 0.5 | 16 | 0.3125 | 9 | 0.1111 | 10 | 0.6 | 45 | 0.3778 | 8 | 0.25 | 15 | 0.3333 | 16 | 0.25 | 10 | 0.5 | 49 | 0.3265 |
| 10_127007 | 9 | 0.4444 | 11 | 0.1818 | 7 | 0.1429 | 6 | 0 | 33 | 0.2121 | 4 | 0.25 | 13 | 0.0769 | 14 | 0.0714 | 7 | 0 | 38 | 0.0789 |
| 10_127007 | 10 | 0.1 | 18 | 0.1111 | 9 | 0 | 10 | 0 | 47 | 0.0638 | 8 | 0 | 15 | 0.2 | 17 | 0.1176 | 8 | 0 | 48 | 0.1042 |
| 10_127007 | 8 | 0 | 14 | 0.0714 | 7 | 0.1429 | 8 | 0.25 | 37 | 0.1081 | 8 | 0.375 | 9 | 0.2222 | 11 | 0.1818 | 7 | 0.5714 | 35 | 0.3143 |
| 10_127007 | 7 | 0.4286 | 12 | 0.5833 | 6 | 0.8333 | 8 | 0.5 | 33 | 0.5758 | 7 | 0.4286 | 8 | 0.375 | 11 | 0.2727 | 8 | 0.5 | 34 | 0.3824 |
| 10_127008 | 8 | 1 | 20 | 1 | 11 | 1 | 9 | 1 | 48 | 1 | 11 | 1 | 13 | 0.8462 | 17 | 1 | 11 | 1 | 52 | 0.9615 |
| 10_127008 | 9 | 0.3333 | 19 | 0.1579 | 12 | 0.0833 | 9 | 0.1111 | 49 | 0.1633 | 12 | 0 | 15 | 0.2 | 16 | 0.0625 | 11 | 0.2727 | 54 | 0.1296 |

| | | | | | | | | | | | | | | | | | | | | |
|---|---|---|---|---|---|---|---|---|---|---|---|---|---|---|---|---|---|---|---|---|
| 10_127008( | 10 | 0 | 20 | 0.05 | 8 | 0.25 | 3 | 0 | 41 | 0.0732 | 7 | 0.1429 | 10 | 0 | 17 | 0.1765 | 7 | 0 | 41 | 0.0976 |
| 10_127008( | 11 | 0.4545 | 22 | 0.6818 | 13 | 0.6923 | 8 | 1 | 54 | 0.6852 | 10 | 0.6 | 15 | 0.6667 | 18 | 0.8889 | 9 | 0.6667 | 52 | 0.7308 |
| 10_127008' | 10 | 0.6 | 22 | 0.6818 | 11 | 0.7273 | 6 | 0.8333 | 49 | 0.6939 | 10 | 0.8 | 16 | 0.4375 | 14 | 0.7143 | 9 | 0.8889 | 49 | 0.6735 |
| 10_127008: | NA | NA | 1 | 1 | NA | NA | 1 | 1 | 2 | 1 | NA | NA | NA | NA | 1 | 0 | 2 | 1 | 3 | 0.6667 |
| 10_127008 | NA | NA | 1 | 1 | 1 | 0 | 1 | 0 | 3 | 0.3333 | NA | NA | NA | NA | 2 | 0 | 2 | 0.5 | 4 | 0.25 |
| 10_127018! | 21 | 0.3333 | 16 | 0.4375 | 12 | 0.1667 | 4 | 0.5 | 53 | 0.3396 | 17 | 0.0588 | 6 | 0.3333 | 8 | 0.125 | 14 | 0.3571 | 45 | 0.2 |
| 10_127018! | 18 | 0.8333 | 19 | 0.8421 | 10 | 0.7 | 5 | 0.8 | 52 | 0.8077 | 18 | 0.6667 | 6 | 0.6667 | 8 | 0.75 | 15 | 0.5333 | 47 | 0.6383 |
| 10_127018! | 19 | 0.5263 | 18 | 0.6111 | 10 | 0.4 | 4 | 0.5 | 51 | 0.5294 | 19 | 0.5789 | 5 | 0.6 | 8 | 0.625 | 11 | 0.8182 | 43 | 0.6512 |
| 10_127018: | 10 | 0 | 14 | 0.0714 | 9 | 0.1111 | 3 | 0 | 36 | 0.0556 | 14 | 0 | 5 | 0 | 5 | 0.4 | 11 | 0 | 35 | 0.0571 |
| 10_127024: | 6 | 0 | 6 | 0.3333 | 4 | 0 | 2 | 0.5 | 18 | 0.1667 | 7 | 0 | NA | NA | 4 | 0.5 | 2 | 0 | 13 | 0.1538 |
| 10_127024 | 7 | 0.2857 | 5 | 0 | 3 | 0 | 2 | 0.5 | 17 | 0.1765 | 5 | 0.6 | NA | NA | 3 | 0 | 4 | 0 | 12 | 0.25 |
| 10_127026 | 25 | 0.08 | 25 | 0.12 | 27 | 0.1852 | 30 | 0.2 | 107 | 0.1495 | 18 | 0.1111 | 32 | 0.2188 | 29 | 0.2759 | 27 | 0.1852 | 106 | 0.2075 |
| 10_127026 | 17 | 0.7059 | 22 | 0.8182 | 25 | 0.64 | 24 | 0.75 | 88 | 0.7273 | 15 | 0.7333 | 26 | 0.8077 | 26 | 0.6923 | 25 | 0.8 | 92 | 0.7609 |
| 10_127026! | 19 | 0.2632 | 23 | 0.1304 | 25 | 0.32 | 30 | 0.1667 | 97 | 0.2165 | 16 | 0.125 | 31 | 0.2903 | 28 | 0.1429 | 28 | 0.2857 | 103 | 0.2233 |
| 10_127026! | 21 | 0.1429 | 23 | 0.087 | 23 | 0.1304 | 27 | 0.037 | 94 | 0.0957 | 17 | 0 | 32 | 0.0625 | 31 | 0.0323 | 29 | 0.1034 | 109 | 0.055 |
| 10_127026: | 15 | 0.0667 | 21 | 0 | 19 | 0.1053 | 21 | 0.1429 | 76 | 0.0789 | 14 | 0.3571 | 27 | 0.1111 | 25 | 0.08 | 18 | 0.2222 | 84 | 0.1667 |
| 10_127027: | 15 | 0.2667 | 15 | 0 | 16 | 0.1875 | 20 | 0.15 | 66 | 0.1515 | 14 | 0 | 22 | 0.1364 | 22 | 0.1364 | 22 | 0.0455 | 80 | 0.0875 |
| 10_127029( | 15 | 0 | 14 | 0.0714 | 18 | 0.1111 | 21 | 0.0476 | 68 | 0.0588 | 15 | 0.0667 | 27 | 0.037 | 21 | 0 | 24 | 0.125 | 87 | 0.0575 |
| 10_127029' | 15 | 0.2 | 12 | 0.4167 | 17 | 0.0588 | 19 | 0.1579 | 63 | 0.1905 | 14 | 0.1429 | 25 | 0.2 | 19 | 0.0526 | 25 | 0.16 | 83 | 0.1446 |
| 10_127029' | 16 | 0.0625 | 13 | 0.0769 | 18 | 0.0556 | 22 | 0.0455 | 69 | 0.058 | 15 | 0.0667 | 25 | 0.04 | 21 | 0.3333 | 22 | 0.1364 | 83 | 0.1446 |
| 10_127034: | 3 | 0 | 6 | 0 | 7 | 0.4286 | 2 | 0.5 | 18 | 0.2222 | 5 | 0 | 5 | 0.2 | 7 | 0.2857 | 6 | 0 | 23 | 0.1304 |
| 10_127034: | 3 | 0.6667 | 8 | 0.75 | 7 | 0.8571 | 2 | 1 | 20 | 0.8 | 5 | 1 | 5 | 1 | 7 | 0.8571 | 5 | 0.8 | 22 | 0.9091 |
| 10_127034: | 3 | 0 | 10 | 0.1 | 8 | 0.125 | 2 | 0 | 23 | 0.087 | 5 | 0 | 5 | 0.2 | 8 | 0.125 | 6 | 0.1667 | 24 | 0.125 |
| 10_127034: | 3 | 0.6667 | 5 | 0.8 | 4 | 0.75 | NA | NA | 12 | 0.75 | 2 | 1 | 1 | 1 | 1 | 1 | 3 | 1 | 7 | 1 |
| 10_127034! | 4 | 0.25 | 8 | 0.25 | 6 | 0.1667 | 3 | 0 | 21 | 0.1905 | 4 | 0.75 | 5 | 0 | 7 | 0 | 6 | 0 | 22 | 0.1364 |
| 10_127034! | 4 | 0.25 | 6 | 0.5 | 8 | 0.625 | 2 | 1 | 20 | 0.55 | 5 | 0.8 | 4 | 0.75 | 6 | 0.3333 | 7 | 0.4286 | 22 | 0.5455 |
| 10_127034! | 3 | 0 | 7 | 0 | 7 | 0.1429 | 3 | 0 | 20 | 0.05 | 3 | 0 | 3 | 0 | 6 | 0 | 6 | 0.1667 | 18 | 0.0556 |
| 10_127034{ | 3 | 0.6667 | 7 | 0.4286 | 8 | 0.375 | 3 | 0.6667 | 21 | 0.4762 | 4 | 0.5 | 3 | 0.3333 | 6 | 0 | 5 | 0.4 | 18 | 0.2778 |
| 10_127034( | 3 | 0.3333 | 5 | 0 | 6 | 0.5 | 3 | 0.6667 | 17 | 0.3529 | 4 | 0.5 | 3 | 0.3333 | 6 | 0.6667 | 2 | 0.5 | 15 | 0.5333 |
| 10_127035( | 2 | 0 | 4 | 0.75 | 5 | 0.2 | 2 | 0 | 13 | 0.3077 | 4 | 0.25 | 1 | 0 | 5 | 0.2 | NA | NA | 10 | 0.2 |
| 10_127063: | 70 | 0.1 | 130 | 0.1077 | 120 | 0.1 | 59 | 0.0678 | 379 | 0.0976 | 95 | 0.0947 | 77 | 0.0779 | 71 | 0.1127 | 49 | 0.1224 | 292 | 0.0993 |
| 10_127063! | 66 | 0.1818 | 121 | 0.1322 | 117 | 0.188 | 53 | 0.1321 | 357 | 0.1597 | 86 | 0.093 | 68 | 0.0882 | 66 | 0.0758 | 42 | 0.1429 | 262 | 0.0954 |
| 10_127063( | 72 | 0.1111 | 118 | 0.178 | 117 | 0.1453 | 48 | 0.1875 | 355 | 0.1549 | 90 | 0.1222 | 64 | 0.1406 | 65 | 0.0923 | 46 | 0.1304 | 265 | 0.1208 |
| 10_127063( | 57 | 0.2632 | 98 | 0.2551 | 83 | 0.2892 | 41 | 0.3171 | 279 | 0.276 | 72 | 0.2917 | 55 | 0.2364 | 56 | 0.3214 | 35 | 0.2286 | 218 | 0.2752 |
| 10_127064: | 47 | 0.383 | 106 | 0.4434 | 94 | 0.4255 | 44 | 0.2955 | 291 | 0.4055 | 73 | 0.4658 | 54 | 0.3704 | 45 | 0.3333 | 40 | 0.425 | 212 | 0.4057 |
| 10_127117 | 17 | 0.0588 | 26 | 0.1923 | 18 | 0.1111 | 19 | 0 | 80 | 0.1 | 23 | 0.0435 | 27 | 0.1852 | 29 | 0.1379 | 21 | 0 | 100 | 0.1 |
| 10_127118( | 5 | 0.2 | 6 | 0.5 | 3 | 0.3333 | 3 | 0.3333 | 17 | 0.3529 | 8 | 0.75 | 4 | 0.5 | 4 | 0.5 | 5 | 0.4 | 21 | 0.5714 |
| 10_127118' | 5 | 0 | 8 | 0 | 3 | 0.3333 | 3 | 0 | 19 | 0.0526 | 6 | 0.1667 | 5 | 0.2 | 5 | 0 | 5 | 0 | 21 | 0.0952 |
| 10_127118: | 5 | 0.2 | 6 | 0.3333 | 3 | 0.6667 | 3 | 0 | 17 | 0.2941 | 8 | 0.25 | 4 | 0 | 5 | 0.2 | 5 | 0 | 22 | 0.1364 |
| 10_127119! | 2 | 0 | 5 | 0.2 | 2 | 0 | 2 | 0 | 11 | 0.0909 | 5 | 0 | 2 | 0 | 5 | 0 | 5 | 0.2 | 17 | 0.0588 |
| 10_127131 | 4 | 0 | 5 | 0.2 | 4 | 0 | 5 | 0 | 18 | 0.0556 | 6 | 0 | 9 | 0.1111 | 4 | 0.25 | 6 | 0 | 25 | 0.08 |
| 10_127131! | 4 | 1 | 2 | 0.5 | 4 | 0.75 | 4 | 0.25 | 14 | 0.6429 | 5 | 0.2 | 4 | 1 | 5 | 0.4 | 4 | 0.75 | 18 | 0.5556 |
| 10_127132( | 7 | 0.8571 | 4 | 1 | 4 | 0.75 | 6 | 1 | 21 | 0.9048 | 5 | 1 | 9 | 0.8889 | 5 | 0.8 | 7 | 0.8571 | 26 | 0.8846 |
| 10_127132' | 6 | 0.6667 | 6 | 0.6667 | 10 | 0.6 | 4 | 0.75 | 26 | 0.6538 | 3 | 0.6667 | 3 | 0.6667 | 3 | 0.6667 | 3 | 1 | 12 | 0.75 |
| 10_127132' | 4 | 0 | 5 | 1 | 10 | 0.4 | 3 | 0.6667 | 22 | 0.5 | 4 | 0.25 | 2 | 0.5 | 3 | 0.3333 | 3 | 0.6667 | 12 | 0.4167 |
| 10_127132' | 6 | 0.6667 | 4 | 0.25 | 4 | 0.25 | 5 | 0.8 | 19 | 0.5263 | 5 | 0.4 | 7 | 0.5714 | 5 | 0.4 | 7 | 0.5714 | 24 | 0.5 |
| 10_127334: | 13 | 0 | 12 | 0.25 | 10 | 0.3 | 8 | 0.25 | 43 | 0.186 | 13 | 0.1538 | 9 | 0 | 15 | 0.1333 | 16 | 0.25 | 53 | 0.1509 |
| 10_127334: | 12 | 0.9167 | 15 | 0.8667 | 12 | 0.75 | 14 | 0.7857 | 53 | 0.8302 | 22 | 0.9091 | 10 | 0.8 | 14 | 0.8571 | 18 | 0.9444 | 64 | 0.8906 |
| 10_127334 | 13 | 0.3077 | 16 | 0.1875 | 15 | 0.3333 | 12 | 0.3333 | 56 | 0.2857 | 24 | 0.125 | 11 | 0.3636 | 18 | 0.5 | 18 | 0.1111 | 71 | 0.2535 |
| 10_127334! | 14 | 0.2857 | 16 | 0.375 | 16 | 0.3125 | 12 | 0.4167 | 58 | 0.3448 | 21 | 0.381 | 11 | 0.2727 | 20 | 0.2 | 18 | 0.2778 | 70 | 0.2857 |
| 10_127334( | 14 | 0.5714 | 13 | 0.6154 | 12 | 0.5 | 12 | 0.3333 | 51 | 0.5098 | 19 | 0.5263 | 9 | 0.5556 | 23 | 0.3913 | 17 | 0.6471 | 68 | 0.5147 |
| 10_127334: | 14 | 0.1429 | 15 | 0.2 | 15 | 0 | 12 | 0.0833 | 56 | 0.1071 | 19 | 0.1053 | 10 | 0 | 23 | 0.087 | 18 | 0.0556 | 70 | 0.0714 |
| 10_127334( | 16 | 0.5625 | 19 | 0.6316 | 20 | 0.55 | 16 | 0.5 | 71 | 0.5634 | 22 | 0.5 | 10 | 0.5 | 23 | 0.6957 | 20 | 0.6 | 75 | 0.5867 |
| 10_127335( | 17 | 0.1176 | 18 | 0.1111 | 11 | 0.4545 | 11 | 0.2727 | 57 | 0.2105 | 15 | 0.0667 | 10 | 0 | 15 | 0.5333 | 22 | 0.2727 | 62 | 0.2419 |
| 10_127353: | NA | NA | 1 | 1 | NA | NA | 1 | 0 | 2 | 0.5 | 2 | 0 | 2 | 0.5 | 3 | 0.6667 | NA | NA | 7 | 0.4286 |
| 10_127353( | NA | NA | 5 | 0.4 | 1 | 0 | 2 | 0 | 8 | 0.25 | 3 | 0.6667 | 3 | 0 | 4 | 0 | 1 | 0 | 11 | 0.1818 |
| 10_127361( | 1 | 0 | 2 | 0.5 | 1 | 0 | NA | NA | 4 | 0.25 | 2 | 0 | 2 | 0 | 1 | 1 | 3 | 0 | 8 | 0.125 |
| 10_127361{ | 2 | 0.5 | 3 | 1 | 1 | 0 | NA | NA | 6 | 0.6667 | 2 | 0.5 | 1 | 0 | 1 | 1 | 2 | 0.5 | 6 | 0.5 |
| 10_127362( | 2 | 0 | 2 | 0 | 1 | 1 | NA | NA | 5 | 0.2 | 2 | 0.5 | 1 | 0 | 1 | 0 | 2 | 0.5 | 6 | 0.3333 |
| 10_127362( | 1 | 1 | 2 | 0 | NA | NA | NA | NA | 3 | 0.3333 | 2 | 0 | 1 | 1 | NA | NA | 1 | 0 | 4 | 0.25 |
| 10_127362! | 2 | 0.5 | NA | NA | NA | NA | NA | NA | 2 | 0.5 | NA | NA | 1 | 1 | 1 | 0 | 1 | 1 | 3 | 0.6667 |
| 10_127362! | 2 | 1 | 2 | 1 | NA | NA | NA | NA | 4 | 1 | NA | NA | 1 | 1 | NA | NA | 2 | 0.5 | 3 | 0.6667 |
| 10_127495{ | 2 | 0.5 | NA | NA | 1 | 0 | 1 | 0 | 4 | 0.25 | 1 | 1 | NA | NA | NA | NA | 1 | 1 | 2 | 1 |
| 10_127495( | 2 | 1 | NA | NA | 1 | 1 | 1 | 1 | 4 | 1 | 1 | 1 | 1 | 1 | NA | NA | 1 | 1 | 3 | 1 |
| 10_127495( | 2 | 1 | NA | NA | 1 | 0 | 1 | 1 | 4 | 0.75 | 1 | 1 | 1 | 1 | NA | NA | 1 | 1 | 3 | 1 |

| | | | | | | | | | | | | | | | | | | | | |
|---|---|---|---|---|---|---|---|---|---|---|---|---|---|---|---|---|---|---|---|---|
| 10_127496 | 2 | 0.5 | NA | NA | 1 | 0 | 1 | 0 | 4 | 0.25 | 1 | 1 | 1 | 0 | NA | NA | 1 | 1 | 3 | 0.6667 |
| 10_127497 | 3 | 0.3333 | 5 | 0.6 | 4 | 0.5 | 2 | 0 | 14 | 0.4286 | 6 | 0.8333 | 4 | 0.5 | 6 | 0.5 | 2 | 1 | 18 | 0.6667 |
| 10_127497 | 4 | 0.25 | 7 | 0.1429 | 7 | 0 | 3 | 0 | 21 | 0.0952 | 7 | 0.1429 | 5 | 0.2 | 9 | 0.1111 | 2 | 0.5 | 23 | 0.1739 |
| 10_127497 | 4 | 0.25 | 5 | 0.2 | 7 | 0.1429 | 4 | 0 | 20 | 0.15 | 6 | 0 | 6 | 0 | 10 | 0.2 | NA | NA | 22 | 0.0909 |
| 10_127497 | 3 | 0.6667 | 5 | 0.8 | 7 | 0.7143 | 3 | 1 | 18 | 0.7778 | 6 | 1 | 6 | 0.8333 | 9 | 0.8889 | 1 | 0 | 22 | 0.8636 |
| 10_127497 | 4 | 0.25 | 5 | 0.6 | 10 | 0.1 | 4 | 0.75 | 23 | 0.3478 | 8 | 0 | 5 | 0.4 | 11 | 0.3636 | 2 | 1 | 26 | 0.3077 |
| 10_127497 | 3 | 1 | 5 | 0.8 | 10 | 0.9 | 4 | 1 | 22 | 0.9091 | 7 | 1 | 6 | 1 | 11 | 0.9091 | 2 | 1 | 26 | 0.9615 |
| 10_127500 | 1 | 1 | 1 | 1 | 2 | 1 | 2 | 0.5 | 6 | 0.8333 | 3 | 0.6667 | 1 | 1 | 4 | 1 | NA | NA | 8 | 0.875 |
| 10_127500 | 3 | 1 | 1 | 0 | 2 | 0.5 | 2 | 1 | 8 | 0.75 | 3 | 0.6667 | 1 | 0 | 3 | 0.6667 | NA | NA | 7 | 0.5714 |
| 10_127500 | 3 | 0.3333 | 1 | 0 | 2 | 0 | 2 | 0 | 8 | 0.125 | 3 | 0 | 1 | 0 | 4 | 0.5 | NA | NA | 8 | 0.25 |
| 10_127532 | 1 | 1 | 1 | 1 | NA | NA | NA | NA | 2 | 1 | 1 | 1 | NA | NA | NA | NA | NA | NA | 1 | 1 |
| 10_127534 | 2 | 0.5 | 1 | 1 | 1 | 0 | NA | NA | 4 | 0.5 | 1 | 1 | 2 | 0 | NA | NA | NA | NA | 3 | 0.3333 |
| 10_127535 | 2 | 0.5 | 1 | 1 | NA | NA | NA | NA | 3 | 0.6667 | 1 | 1 | 2 | 0.5 | NA | NA | NA | NA | 3 | 0.6667 |
| 10_127535 | 2 | 0 | 1 | 1 | 1 | 0 | NA | NA | 4 | 0.25 | 1 | 0 | 2 | 1 | NA | NA | NA | NA | 3 | 0.6667 |
| 10_127536 | 1 | 1 | 1 | 1 | 1 | 0 | NA | NA | 3 | 0.6667 | 2 | 1 | NA | NA | NA | NA | NA | NA | 2 | 1 |
| 10_127536 | 2 | 1 | 1 | 0 | 1 | 0 | NA | NA | 4 | 0.5 | 1 | 1 | 2 | 0 | NA | NA | NA | NA | 3 | 0.3333 |
| 10_127578 | 1 | 0 | NA | NA | 1 | 0 | 1 | 1 | 3 | 0.3333 | NA | NA | 1 | 1 | 1 | 0 | NA | NA | 2 | 0.5 |
| 10_127578 | 1 | 1 | NA | NA | NA | NA | 1 | 1 | 2 | 1 | NA | NA | 1 | 1 | 1 | 0 | NA | NA | 2 | 0.5 |
| 10_127579 | 1 | 1 | NA | NA | NA | NA | 1 | 1 | 2 | 1 | NA | NA | 1 | 1 | NA | NA | NA | NA | 1 | 1 |
| 10_127579 | 1 | 1 | NA | NA | 1 | 0 | 1 | 0 | 3 | 0.3333 | NA | NA | 1 | 1 | 1 | 0 | NA | NA | 2 | 0.5 |
| 10_127579 | 1 | 1 | NA | NA | 1 | 1 | 1 | 1 | 3 | 1 | NA | NA | NA | NA | 1 | 1 | NA | NA | 1 | 1 |
| 10_127580 | 1 | 0 | NA | NA | 1 | 0 | 1 | 1 | 3 | 0.3333 | NA | NA | 2 | 0.5 | 1 | 0 | 1 | 0 | 4 | 0.25 |
| 10_127580 | 1 | 1 | NA | NA | 1 | 1 | NA | NA | 2 | 1 | NA | NA | 2 | 0.5 | NA | NA | 1 | 0 | 3 | 0.3333 |
| 10_127580 | 1 | 0 | NA | NA | 2 | 1 | 1 | 0 | 4 | 0.5 | NA | NA | 2 | 0.5 | NA | NA | 1 | 1 | 3 | 0.6667 |
| 10_127580 | 1 | 1 | NA | NA | 1 | 1 | 1 | 1 | 3 | 1 | NA | NA | 1 | 1 | 1 | 1 | 1 | 1 | 3 | 1 |
| 10_127580 | 1 | 0 | NA | NA | 2 | 1 | 1 | 1 | 4 | 0.75 | NA | NA | 2 | 0 | 1 | 0 | 1 | 1 | 4 | 0.25 |
| 10_127582 | 2 | 1 | 1 | 1 | 2 | 1 | 2 | 1 | 7 | 1 | NA | NA | 2 | 0.5 | 2 | 1 | 2 | 0.5 | 6 | 0.6667 |
| 10_127582 | 1 | 1 | 1 | 1 | 2 | 1 | 2 | 1 | 6 | 1 | NA | NA | 2 | 1 | 1 | 1 | 2 | 1 | 5 | 1 |
| 10_127847 | 4 | 0.25 | 2 | 0 | 2 | 0 | 3 | 0 | 11 | 0.0909 | 3 | 0 | 4 | 0 | 2 | 0.5 | 3 | 0 | 12 | 0.0833 |
| 10_127860 | NA | NA | 1 | 1 | NA | NA | NA | NA | 1 | 1 | 1 | 1 | NA | NA | 1 | 0 | NA | NA | 2 | 0.5 |
| 10_127861 | NA | NA | 1 | 1 | NA | NA | NA | NA | 1 | 1 | 1 | 1 | NA | NA | 1 | 1 | NA | NA | 2 | 1 |
| 10_127861 | NA | NA | 1 | 1 | NA | NA | NA | NA | 1 | 1 | 1 | 1 | NA | NA | NA | NA | NA | NA | 1 | 1 |
| 10_127865 | 1 | 0 | 1 | 1 | 1 | 0 | NA | NA | 3 | 0.3333 | 1 | 0 | 2 | 0.5 | 1 | 0 | 2 | 0 | 6 | 0.1667 |
| 10_127883 | NA | NA | 1 | 1 | NA | NA | NA | NA | 1 | 1 | NA | NA | 1 | 1 | NA | NA | NA | NA | 1 | 1 |
| 10_127883 | NA | NA | 1 | 1 | NA | NA | NA | NA | 1 | 1 | NA | NA | 1 | 1 | NA | NA | NA | NA | 1 | 1 |
| 10_127883 | NA | NA | 1 | 0 | NA | NA | NA | NA | 1 | 1 | NA | NA | 1 | 1 | NA | NA | NA | NA | 1 | 1 |
| 10_127883 | NA | NA | 1 | 1 | NA | NA | NA | NA | 1 | 1 | NA | NA | 1 | 1 | NA | NA | NA | NA | 1 | 1 |
| 10_127883 | NA | NA | 1 | 0 | NA | NA | NA | NA | 1 | 1 | NA | NA | 1 | 1 | NA | NA | NA | NA | 1 | 1 |
| 10_127883 | NA | NA | 1 | 1 | NA | NA | NA | NA | 1 | 1 | NA | NA | 1 | 1 | NA | NA | NA | NA | 1 | 1 |
| 10_127884 | 7 | 0.1429 | 4 | 0.25 | 7 | 0.1429 | 6 | 0.1667 | 24 | 0.1667 | 3 | 0 | 4 | 0 | 4 | 0.25 | 9 | 0.1111 | 20 | 0.1 |
| 10_127912 | 8 | 0 | 13 | 0.1538 | 9 | 0.1111 | 15 | 0.1333 | 45 | 0.1111 | 12 | 0.25 | 6 | 0.1667 | 10 | 0.1 | 7 | 0.1429 | 35 | 0.1714 |
| 10_127946 | NA | NA | 2 | 0.5 | 1 | 0 | 2 | 0.5 | 5 | 0.4 | NA | NA | NA | NA | NA | NA | 1 | 1 | 1 | 1 |
| 10_127946 | NA | NA | 2 | 1 | 1 | 1 | 2 | 1 | 5 | 1 | NA | NA | NA | NA | NA | NA | 2 | 1 | 2 | 1 |
| 10_127950 | NA | NA | 1 | 1 | 1 | 0 | NA | NA | 2 | 0.5 | NA | NA | NA | NA | NA | NA | 3 | 0.3333 | 3 | 0.3333 |
| 10_127955 | 1 | 0 | 2 | 0.5 | 1 | 0 | 2 | 0 | 6 | 0.1667 | NA | NA | NA | NA | NA | NA | 3 | 0.3333 | 3 | 0.3333 |
| 10_127962 | 2 | 0.5 | 6 | 0.8333 | 4 | 1 | 3 | 0.6667 | 15 | 0.8 | 1 | 1 | 2 | 1 | 1 | 1 | 5 | 0.8 | 9 | 0.8889 |
| 10_127962 | 2 | 0 | 7 | 0.2857 | 5 | 0.4 | 4 | 0 | 18 | 0.2222 | NA | NA | 2 | 0.5 | 1 | 0 | 5 | 0.2 | 8 | 0.25 |
| 10_127963 | 2 | 1 | 4 | 0.75 | 2 | 1 | 5 | 0.6 | 13 | 0.7692 | NA | NA | 3 | 0.3333 | 1 | 1 | 4 | 0.75 | 8 | 0.625 |
| 10_127964 | 4 | 0.25 | 8 | 0 | 5 | 0.2 | 5 | 0.2 | 22 | 0.1364 | 2 | 0 | 4 | 0 | 2 | 0 | 7 | 0.1429 | 15 | 0.0667 |
| 10_127964 | 5 | 0 | 8 | 0 | 6 | 0.6667 | 5 | 0.4 | 24 | 0.25 | 1 | 1 | 5 | 0.2 | 4 | 0.25 | 7 | 0.1429 | 17 | 0.2353 |
| 10_127964 | 3 | 0.3333 | 7 | 0.4286 | 5 | 0.8 | 5 | 0.6 | 20 | 0.55 | 1 | 0 | 5 | 0.4 | 3 | 0 | 5 | 0 | 14 | 0.1429 |
| 10_127964 | 6 | 0.3333 | 7 | 0.4286 | 4 | 1 | 4 | 0.25 | 21 | 0.4762 | 1 | 1 | 5 | 0.2 | 3 | 0.6667 | 8 | 0.375 | 17 | 0.4118 |
| 10_127964 | 6 | 0 | 8 | 0.125 | 5 | 0.2 | 5 | 0 | 24 | 0.0833 | 2 | 0.5 | 5 | 0.2 | 5 | 0 | 8 | 0 | 20 | 0.1 |
| 10_127967 | 1 | 0 | 2 | 0.5 | 1 | 1 | NA | NA | 4 | 0.5 | NA | NA | NA | NA | 2 | 0.5 | 1 | 0 | 3 | 0.3333 |
| 10_127967 | 1 | 1 | 1 | 1 | NA | NA | NA | NA | 2 | 1 | NA | NA | NA | NA | 2 | 1 | 1 | 0 | 3 | 0.6667 |
| 10_127967 | 1 | 0 | 2 | 0.5 | NA | NA | NA | NA | 3 | 0.3333 | NA | NA | NA | NA | 2 | 0.5 | 1 | 1 | 3 | 0.6667 |
| 10_127969 | 2 | 0 | 1 | 0 | 2 | 1 | 1 | 0 | 6 | 0.3333 | NA | NA | NA | NA | 1 | 1 | 2 | 1 | 3 | 1 |
| 10_128045 | 1 | 0 | 2 | 0 | 4 | 0.25 | 3 | 0.3333 | 10 | 0.2 | 7 | 0.1429 | 1 | 1 | 1 | 0 | 3 | 0.3333 | 12 | 0.25 |
| 10_128045 | 1 | 0 | 2 | 0 | 6 | 0 | 3 | 0.6667 | 12 | 0.1667 | 7 | 0.2857 | 1 | 1 | 1 | 1 | 3 | 0.3333 | 12 | 0.4167 |
| 10_128045 | 1 | 0 | 2 | 0 | 5 | 0.4 | 3 | 0 | 11 | 0.1818 | 7 | 0 | 1 | 0 | NA | NA | 3 | 0.3333 | 11 | 0.0909 |
| 10_128045 | 1 | 0 | 2 | 0 | 4 | 0.25 | 2 | 0 | 9 | 0.1111 | 6 | 0.1667 | 1 | 0 | NA | NA | 3 | 0.3333 | 10 | 0.2 |
| 10_128045 | NA | NA | 1 | 1 | 1 | 0 | NA | NA | 2 | 0.5 | NA | NA | NA | NA | NA | NA | 2 | 0.5 | 2 | 0.5 |
| 10_128046 | NA | NA | NA | NA | 1 | 0 | 1 | 1 | 2 | 0.5 | 1 | 0 | NA | NA | 1 | 1 | 2 | 0 | 4 | 0.25 |
| 10_128046 | NA | NA | NA | NA | 1 | 1 | 1 | 1 | 2 | 1 | 1 | 1 | NA | NA | 1 | 1 | 2 | 1 | 4 | 1 |
| 10_128046 | NA | NA | NA | NA | NA | NA | 1 | 1 | 1 | 1 | 1 | 1 | NA | NA | 1 | 1 | 2 | 0.5 | 4 | 0.75 |

| | | | | | | | | | | | | | | | | | | | | |
|---|---|---|---|---|---|---|---|---|---|---|---|---|---|---|---|---|---|---|---|---|
| 10_128046 | NA | NA | NA | NA | NA | NA | 1 | 1 | 1 | 1 | 1 | 1 | NA | NA | 1 | 1 | 1 | 1 | 3 | 1 |
| 10_128047 | NA | NA | NA | NA | 1 | 1 | 1 | 0 | 2 | 0.5 | 1 | 0 | NA | NA | 1 | 1 | 1 | 1 | 3 | 0.6667 |
| 10_128126 | 1 | 1 | NA | NA | NA | NA | 3 | 0 | 4 | 0.25 | 2 | 1 | NA | NA | NA | NA | 1 | 0 | 3 | 0.6667 |
| 10_128127 | 1 | 1 | 2 | 1 | NA | NA | 3 | 1 | 6 | 1 | 2 | 1 | NA | NA | NA | NA | 1 | 1 | 3 | 1 |
| 10_128127 | 2 | 1 | 2 | 1 | NA | NA | 2 | 1 | 6 | 1 | 2 | 1 | NA | NA | NA | NA | 1 | 1 | 3 | 1 |
| 10_128127 | 2 | 1 | 2 | 1 | NA | NA | 3 | 0.6667 | 7 | 0.8571 | 1 | 1 | NA | NA | NA | NA | 1 | 1 | 2 | 1 |
| 10_128127 | 2 | 1 | 2 | 1 | NA | NA | 3 | 1 | 7 | 1 | 2 | 1 | NA | NA | NA | NA | 1 | 1 | 3 | 1 |
| 10_128127 | 2 | 0 | 3 | 0.6667 | NA | NA | 2 | 0 | 7 | 0.2857 | 2 | 0.5 | NA | NA | NA | NA | 1 | 0 | 3 | 0.3333 |
| 10_128127 | 2 | 0.5 | 2 | 0.5 | NA | NA | 3 | 0.6667 | 7 | 0.5714 | 2 | 1 | NA | NA | NA | NA | 1 | 1 | 3 | 1 |
| 10_128128 | 2 | 0 | 3 | 0 | NA | NA | 3 | 0.3333 | 8 | 0.125 | 2 | 0 | NA | NA | NA | NA | 1 | 1 | 3 | 0.3333 |
| 10_128128 | 2 | 0 | 2 | 0 | 1 | 1 | 3 | 0.6667 | 8 | 0.375 | 1 | 1 | NA | NA | NA | NA | 2 | 0 | 3 | 0.3333 |
| 10_128128 | 2 | 0 | 3 | 0.6667 | NA | NA | 3 | 0 | 8 | 0.25 | 2 | 0.5 | NA | NA | NA | NA | 2 | 0.5 | 4 | 0.5 |
| 10_128128 | 1 | 0 | 3 | 0.3333 | 1 | 0 | 2 | 0.5 | 7 | 0.2857 | 2 | 0.5 | NA | NA | NA | NA | 1 | 1 | 3 | 0.6667 |
| 10_128128 | 3 | 0.3333 | 3 | 0 | 1 | 1 | 2 | 0 | 9 | 0.2222 | 2 | 0.5 | NA | NA | NA | NA | 2 | 0.5 | 4 | 0.5 |
| 10_128154 | 2 | 0.5 | 1 | 1 | 4 | 0 | NA | NA | 7 | 0.2857 | 1 | 1 | NA | NA | NA | NA | 1 | 0 | 2 | 0.5 |
| 10_128156 | 2 | 0 | 1 | 1 | 4 | 0 | 2 | 0.5 | 9 | 0.2222 | 2 | 0.5 | NA | NA | NA | NA | 1 | 0 | 3 | 0.3333 |
| 10_128156 | 2 | 0 | 1 | 0 | 5 | 0.2 | 3 | 0.3333 | 11 | 0.1818 | 3 | 0.3333 | NA | NA | NA | NA | 2 | 0 | 5 | 0.2 |
| 10_128163 | 10 | 0.4 | 6 | 0.6667 | 6 | 0.5 | 16 | 0.5625 | 38 | 0.5263 | 7 | 0.7143 | 6 | 0.8333 | 11 | 0.4545 | 7 | 0.2857 | 31 | 0.5484 |
| 10_128163 | 8 | 0.25 | 5 | 0 | 5 | 0 | 15 | 0.2667 | 33 | 0.1818 | 7 | 0 | 4 | 0.25 | 9 | 0.2222 | 7 | 0.1429 | 27 | 0.1481 |
| 10_128196 | 22 | 0.6818 | 38 | 0.7632 | 24 | 0.9167 | 32 | 0.875 | 116 | 0.8103 | 19 | 0.5263 | 33 | 0.8788 | 26 | 0.6538 | 22 | 0.9091 | 100 | 0.76 |
| 10_128198 | 30 | 0.0667 | 50 | 0.06 | 33 | 0.0909 | 42 | 0.119 | 155 | 0.0839 | 32 | 0.0625 | 46 | 0.0435 | 38 | 0.0526 | 25 | 0.16 | 141 | 0.0709 |
| 10_128199 | 6 | 0.5 | 7 | 0.4286 | 4 | 0.25 | 7 | 0.2857 | 24 | 0.375 | 6 | 0.3333 | 7 | 0.2857 | 4 | 0.25 | 7 | 0.1429 | 24 | 0.25 |
| 10_128229 | NA | NA | 4 | 0.25 | 4 | 0.25 | 2 | 0.5 | 10 | 0.3 | 1 | 1 | NA | NA | 4 | 0.25 | 1 | 0 | 6 | 0.3333 |
| 10_128231 | 4 | 0 | 3 | 0.3333 | 5 | 0.2 | 2 | 0 | 14 | 0.1429 | 6 | 0.1667 | 4 | 0.5 | 4 | 0 | 6 | 0 | 20 | 0.15 |
| 10_128231 | 4 | 0.25 | 4 | 0.75 | 5 | 0.2 | 2 | 0 | 15 | 0.3333 | 6 | 0.3333 | 3 | 0.3333 | 5 | 0.4 | 6 | 0.3333 | 20 | 0.35 |
| 10_128237 | 11 | 0.0909 | 24 | 0.0833 | 12 | 0 | 10 | 0.1 | 57 | 0.0702 | 14 | 0 | 23 | 0.1304 | 14 | 0.0714 | 12 | 0 | 63 | 0.0635 |
| 10_128271 | 3 | 0 | 2 | 0 | 5 | 0 | 4 | 0.5 | 14 | 0.1429 | 3 | 0 | 7 | 0 | 7 | 0.1429 | 3 | 0.3333 | 20 | 0.1 |
| 10_128274 | 9 | 0.3333 | 5 | 0.6 | 9 | 0.4444 | 9 | 0.2222 | 32 | 0.375 | 7 | 0.1429 | 10 | 0.3 | 11 | 0.2727 | 4 | 0.25 | 32 | 0.25 |
| 10_128274 | 9 | 0.1111 | 4 | 0 | 7 | 0.1429 | 9 | 0 | 29 | 0.069 | 7 | 0 | 10 | 0.1 | 11 | 0.0909 | 5 | 0.2 | 33 | 0.0909 |
| 10_128274 | 8 | 0.25 | 5 | 0 | 7 | 0 | 8 | 0 | 28 | 0.0714 | 7 | 0 | 11 | 0.1818 | 11 | 0.0909 | 5 | 0 | 34 | 0.0882 |
| 10_128274 | 9 | 0.2222 | 5 | 0 | 9 | 0.2222 | 9 | 0 | 32 | 0.125 | 5 | 0.2 | 10 | 0.2 | 9 | 0.1111 | 4 | 0 | 28 | 0.1429 |
| 10_128274 | 8 | 0.125 | 4 | 0.25 | 9 | 0.1111 | 9 | 0 | 30 | 0.1 | 5 | 0 | 10 | 0.1 | 10 | 0.3 | 2 | 0 | 27 | 0.1481 |
| 10_128274 | 9 | 0.2222 | 6 | 0 | 9 | 0.1111 | 9 | 0.1111 | 33 | 0.1212 | 7 | 0.1429 | 12 | 0.1667 | 10 | 0.3 | 5 | 0 | 34 | 0.1765 |
| 10_128274 | 9 | 1 | 6 | 1 | 9 | 1 | 9 | 1 | 33 | 1 | 7 | 1 | 11 | 1 | 8 | 1 | 3 | 1 | 29 | 1 |
| 10_128274 | 6 | 0.5 | 8 | 0.375 | 10 | 0.1 | 6 | 0.3333 | 30 | 0.3 | 6 | 0.1667 | 11 | 0.5455 | 9 | 0.4444 | 6 | 0.5 | 32 | 0.4375 |
| 10_128275 | 8 | 0.125 | 10 | 0.4 | 12 | 0.0833 | 7 | 0.1429 | 37 | 0.1892 | 12 | 0.6667 | 14 | 0.4286 | 12 | 0.0833 | 6 | 0.5 | 44 | 0.4091 |
| 10_128275 | 1 | 0 | 4 | 0.25 | 4 | 0 | NA | NA | 9 | 0.1111 | 2 | 0.5 | 3 | 0 | 2 | 0 | NA | NA | 7 | 0.1429 |
| 10_128318 | 8 | 0.125 | 12 | 0 | 9 | 0.1111 | 9 | 0.2222 | 38 | 0.1053 | 10 | 0 | 9 | 0.2222 | 6 | 0 | 5 | 0.2 | 30 | 0.1 |
| 10_128319 | 7 | 0.2857 | 11 | 0.1818 | 12 | 0 | 7 | 0.1429 | 37 | 0.1351 | 10 | 0.1 | 10 | 0.2 | 7 | 0.1429 | 8 | 0 | 35 | 0.1143 |
| 10_128319 | 10 | 0.2 | 11 | 0.0909 | 10 | 0.1 | 11 | 0.2727 | 42 | 0.1667 | 10 | 0 | 8 | 0.25 | 8 | 0 | 7 | 0 | 33 | 0.0606 |
| 10_128323 | 9 | 0.1111 | 9 | 0 | 14 | 0.0714 | 11 | 0.0909 | 43 | 0.0698 | 12 | 0 | 9 | 0.1111 | 11 | 0.0909 | 6 | 0 | 38 | 0.0526 |
| 10_128324 | 15 | 0.0667 | 17 | 0.0588 | 18 | 0.0556 | 13 | 0.0769 | 63 | 0.0635 | 13 | 0.0769 | 11 | 0 | 13 | 0.0769 | 12 | 0.0833 | 49 | 0.0612 |
| 10_128325 | 14 | 0.7857 | 17 | 0.9412 | 23 | 0.913 | 14 | 0.8571 | 68 | 0.8824 | 15 | 0.9333 | 12 | 0.8333 | 16 | 0.875 | 12 | 1 | 55 | 0.9091 |
| 10_128325 | 16 | 0.1875 | 12 | 0.4167 | 21 | 0.381 | 18 | 0.1111 | 67 | 0.2687 | 16 | 0.1875 | 14 | 0.3571 | 15 | 0.4 | 12 | 0.1667 | 57 | 0.2807 |
| 10_128325 | 17 | 0.3529 | 13 | 0.3077 | 23 | 0.3043 | 14 | 0.5 | 67 | 0.3582 | 17 | 0.2941 | 14 | 0.5 | 14 | 0.6429 | 13 | 0.5385 | 58 | 0.4828 |
| 10_128325 | 18 | 0.1111 | 11 | 0.4545 | 21 | 0.2381 | 15 | 0.3333 | 65 | 0.2615 | 17 | 0.4118 | 13 | 0.4615 | 16 | 0.4375 | 9 | 0.2222 | 55 | 0.4 |
| 10_128325 | 16 | 0.1875 | 12 | 0.3333 | 16 | 0.25 | 14 | 0.1429 | 58 | 0.2241 | 12 | 0.3333 | 13 | 0.1538 | 14 | 0.2857 | 12 | 0.25 | 51 | 0.2549 |
| 10_128326 | 2 | 0.5 | 3 | 0.3333 | 2 | 0.5 | 1 | 1 | 8 | 0.5 | NA | NA | 2 | 0.5 | 1 | 0 | 2 | 0.5 | 5 | 0.4 |
| 10_128330 | NA | NA | 3 | 0 | 3 | 0 | 6 | 0.1667 | 12 | 0.0833 | 4 | 0 | 2 | 0 | 3 | 0.3333 | 3 | 0 | 12 | 0.0833 |
| 10_128338 | 3 | 1 | 1 | 1 | 1 | 1 | 1 | 1 | 6 | 1 | NA | NA | NA | NA | 1 | 1 | 1 | 1 | 2 | 1 |
| 10_128338 | 4 | 0.5 | 1 | 1 | 1 | 0 | 2 | 0 | 8 | 0.375 | NA | NA | NA | NA | 2 | 0.5 | 1 | 0 | 3 | 0.3333 |
| 10_128338 | 3 | 1 | 1 | 1 | 1 | 1 | 2 | 1 | 7 | 1 | NA | NA | NA | NA | 2 | 0.5 | 1 | 0 | 3 | 0.3333 |
| 10_128338 | 1 | 0 | 1 | 1 | NA | NA | 1 | 1 | 3 | 0.6667 | NA | NA | NA | NA | 2 | 1 | 1 | 1 | 3 | 1 |
| 10_128338 | 4 | 1 | 1 | 1 | NA | NA | NA | NA | 5 | 1 | NA | NA | NA | NA | 1 | 1 | NA | NA | 1 | 1 |
| 10_128338 | 4 | 1 | 1 | 1 | NA | NA | NA | NA | 5 | 1 | NA | NA | NA | NA | 1 | 1 | NA | NA | 1 | 1 |
| 10_128339 | 4 | 1 | 2 | 1 | NA | NA | NA | NA | 6 | 1 | NA | NA | NA | NA | 1 | 1 | NA | NA | 1 | 1 |
| 10_128379 | 3 | 1 | 3 | 1 | 2 | 1 | 2 | 0 | 10 | 0.8 | NA | NA | 4 | 0.25 | 2 | 0.5 | 1 | 1 | 7 | 0.4286 |
| 10_128379 | 4 | 1 | 3 | 0.6667 | 2 | 0.5 | 1 | 0 | 10 | 0.7 | NA | NA | 4 | 1 | 2 | 1 | 1 | 0 | 7 | 0.8571 |
| 10_128379 | 4 | 0.25 | 2 | 0 | 2 | 0 | 2 | 0.5 | 10 | 0.2 | NA | NA | 4 | 0 | 2 | 0 | 1 | 1 | 7 | 0.1429 |
| 10_128379 | 4 | 1 | 3 | 1 | 2 | 1 | 2 | 0.5 | 11 | 0.9091 | NA | NA | 4 | 1 | 2 | 1 | 1 | 1 | 7 | 1 |
| 10_128379 | 4 | 0.25 | 3 | 0.6667 | 2 | 0.5 | 2 | 0 | 11 | 0.3636 | NA | NA | 4 | 0.5 | 1 | 0 | 1 | 0 | 6 | 0.3333 |
| 10_128380 | 4 | 1 | 3 | 1 | 2 | 1 | 2 | 1 | 11 | 1 | NA | NA | 4 | 1 | 2 | 1 | NA | NA | 6 | 1 |
| 10_128380 | 4 | 0.25 | 3 | 0 | 1 | 0 | 2 | 0 | 10 | 0.1 | NA | NA | 4 | 0.5 | 1 | 0 | 1 | 0 | 6 | 0.3333 |
| 10_128380 | 4 | 1 | 2 | 1 | 1 | 0 | 1 | 1 | 8 | 0.875 | NA | NA | 1 | 1 | 1 | 1 | NA | NA | 2 | 1 |
| 10_128380 | 3 | 1 | 2 | 1 | 2 | 1 | NA | NA | 7 | 1 | NA | NA | 3 | 1 | NA | NA | 1 | 1 | 4 | 1 |

| | | | | | | | | | | | | | | | | | | | | |
|---|---|---|---|---|---|---|---|---|---|---|---|---|---|---|---|---|---|---|---|---|
| 10_1283804 | 3 | 1 | 3 | 1 | 2 | 1 | 1 | 1 | 9 | 1 | NA | NA | 2 | 1 | NA | NA | 1 | 1 | 3 | 1 |
| 10_1283805 | 3 | 0.3333 | 3 | 0.6667 | 2 | 0.5 | 1 | 1 | 9 | 0.5556 | NA | NA | 2 | 0.5 | NA | NA | NA | NA | 2 | 0.5 |
| 10_1283806 | 2 | 1 | 3 | 0.3333 | 2 | 0.5 | 1 | 0 | 8 | 0.5 | NA | NA | 2 | 0.5 | NA | NA | 1 | 1 | 3 | 0.6667 |
| 10_1283806 | 2 | 0 | 2 | 0.5 | 2 | 0 | NA | NA | 6 | 0.1667 | NA | NA | 3 | 0.3333 | NA | NA | 1 | 0 | 4 | 0.25 |
| 10_1283807 | 1 | 1 | 3 | 0 | 1 | 0 | 1 | 1 | 6 | 0.3333 | NA | NA | 2 | 0.5 | NA | NA | 1 | 1 | 3 | 0.6667 |
| 10_1283809 | 2 | 1 | 2 | 1 | 2 | 1 | 1 | 1 | 7 | 1 | NA | NA | 1 | 1 | NA | NA | NA | NA | 1 | 1 |
| 10_1283812 | 2 | 0.5 | 3 | 0.3333 | 2 | 0.5 | 1 | 1 | 8 | 0.5 | NA | NA | 2 | 0.5 | NA | NA | 1 | 1 | 3 | 0.6667 |
| 10_1283812 | 2 | 0.5 | 2 | 0 | 1 | 0 | NA | NA | 5 | 0.2 | NA | NA | 1 | 0 | NA | NA | 1 | 1 | 2 | 0.5 |
| 10_1283840 | 116 | 0.3966 | 110 | 0.3 | 116 | 0.319 | 108 | 0.25 | 450 | 0.3178 | 97 | 0.3608 | 117 | 0.3077 | 130 | 0.3385 | 94 | 0.234 | 438 | 0.3128 |
| 10_1283841 | 138 | 0.2826 | 130 | 0.2692 | 127 | 0.2283 | 128 | 0.3047 | 523 | 0.2715 | 124 | 0.3629 | 130 | 0.3231 | 151 | 0.2715 | 104 | 0.2404 | 509 | 0.3006 |
| 10_1283937 | 2 | 0.5 | 3 | 1 | 4 | 0.75 | 2 | 1 | 11 | 0.8182 | 5 | 0.4 | 4 | 0.25 | 6 | 0.6667 | 8 | 0.75 | 23 | 0.5652 |
| 10_1283937 | NA | NA | 4 | 0.75 | 4 | 0.25 | 3 | 0 | 11 | 0.3636 | 4 | 0.25 | 2 | 0 | 6 | 0.3333 | 5 | 0.6 | 17 | 0.3529 |
| 10_1283943 | 1 | 0 | 4 | 0.5 | 2 | 1 | 2 | 0.5 | 9 | 0.5556 | 3 | 0.3333 | 3 | 0.3333 | 6 | 0.5 | 4 | 0.25 | 16 | 0.375 |
| 10_1283945 | 9 | 0 | 12 | 0 | 8 | 0 | 10 | 0.3 | 39 | 0.0769 | 10 | 0.2 | 15 | 0 | 15 | 0.0667 | 10 | 0 | 50 | 0.06 |
| 10_1283945 | 8 | 0.25 | 9 | 0.4444 | 7 | 0.4286 | 7 | 0 | 31 | 0.2903 | 9 | 0.3333 | 13 | 0.3077 | 16 | 0.3125 | 10 | 0.3 | 48 | 0.3125 |
| 10_1284036 | NA | NA | 2 | 0.5 | 1 | 0 | 1 | 0 | 4 | 0.25 | 1 | 0 | 1 | 0 | 1 | 1 | NA | NA | 3 | 0.3333 |
| 10_1284036 | NA | NA | 2 | 0 | 2 | 0 | 1 | 1 | 5 | 0.2 | NA | NA | 1 | 1 | 1 | 0 | NA | NA | 2 | 0.5 |
| 10_1284036 | NA | NA | 2 | 1 | 2 | 1 | 1 | 0 | 5 | 0.8 | 1 | 0 | 1 | 0 | 1 | 1 | NA | NA | 3 | 0.3333 |
| 10_1284043 | NA | NA | 1 | 0 | 1 | 1 | 1 | 1 | 3 | 0.6667 | NA | NA | 1 | 0 | 1 | 1 | NA | NA | 2 | 0.5 |
| 10_1284047 | NA | NA | 1 | 1 | 2 | 0 | NA | NA | 3 | 0.3333 | NA | NA | 1 | 1 | 1 | 0 | NA | NA | 2 | 0.5 |
| 10_1284052 | NA | NA | 2 | 1 | 2 | 0.5 | NA | NA | 4 | 0.75 | NA | NA | 1 | 0 | 1 | 1 | NA | NA | 2 | 0.5 |
| 10_1284053 | NA | NA | 2 | 0.5 | 1 | 0 | NA | NA | 3 | 0.3333 | NA | NA | 1 | 0 | 1 | 1 | NA | NA | 2 | 0.5 |
| 10_1284053 | NA | NA | 2 | 0.5 | 1 | 1 | NA | NA | 3 | 0.6667 | NA | NA | 1 | 0 | 1 | 1 | NA | NA | 2 | 0.5 |
| 10_1284053 | NA | NA | 2 | 0.5 | 2 | 0.5 | NA | NA | 4 | 0.5 | NA | NA | 1 | 0 | 1 | 1 | NA | NA | 2 | 0.5 |
| 10_1284054 | NA | NA | 2 | 0.5 | 2 | 0 | NA | NA | 4 | 0.25 | NA | NA | 1 | 1 | 1 | 1 | NA | NA | 2 | 1 |
| 10_1284700 | NA | NA | NA | NA | NA | NA | 1 | 1 | 1 | 1 | NA | NA | 1 | 1 | NA | NA | 1 | 0 | 2 | 0.5 |
| 10_1284703 | NA | NA | NA | NA | NA | NA | 1 | 1 | 1 | 1 | NA | NA | 2 | 1 | NA | NA | 1 | 1 | 3 | 1 |
| 10_1284703 | NA | NA | NA | NA | NA | NA | 1 | 1 | 1 | 1 | NA | NA | 2 | 0 | NA | NA | 1 | 1 | 3 | 0.3333 |
| 10_1284703 | NA | NA | NA | NA | NA | NA | 1 | 1 | 1 | 1 | NA | NA | 2 | 0.5 | NA | NA | 1 | 1 | 3 | 0.6667 |
| 10_1285068 | 1 | 0 | 2 | 1 | 2 | 0 | 2 | 0.5 | 7 | 0.4286 | 3 | 1 | 2 | 0.5 | NA | NA | 3 | 0.3333 | 8 | 0.625 |
| 10_1285070 | NA | NA | 2 | 0.5 | 1 | 1 | 1 | 1 | 4 | 0.75 | 1 | 1 | 1 | 0 | NA | NA | 1 | 1 | 3 | 0.6667 |
| 10_1285070 | 1 | 1 | 2 | 1 | 1 | 1 | 2 | 1 | 6 | 1 | 3 | 1 | 2 | 1 | NA | NA | 3 | 1 | 8 | 1 |
| 10_1285070 | 1 | 0 | 2 | 0 | 2 | 0.5 | 2 | 0 | 7 | 0.1429 | 2 | 0.5 | 2 | 0 | NA | NA | 2 | 0.5 | 6 | 0.3333 |
| 10_1285073 | NA | NA | 2 | 0.5 | 2 | 1 | 1 | 1 | 5 | 0.8 | 3 | 0.6667 | NA | NA | NA | NA | 3 | 0.6667 | 6 | 0.6667 |
| 10_1285074 | NA | NA | 1 | 1 | 2 | 1 | NA | NA | 3 | 1 | 2 | 1 | 2 | 1 | NA | NA | 3 | 0.6667 | 7 | 0.8571 |
| 10_1285076 | 1 | 1 | 1 | 1 | NA | NA | NA | NA | 2 | 1 | 1 | 1 | 1 | 1 | NA | NA | 3 | 0 | 5 | 0.4 |
| 10_1285076 | 1 | 1 | 2 | 0 | NA | NA | NA | NA | 3 | 0.3333 | 1 | 1 | 3 | 0.6667 | NA | NA | 3 | 0 | 7 | 0.4286 |
| 10_1285127 | NA | NA | 1 | 1 | NA | NA | NA | NA | 1 | 1 | NA | NA | 1 | 1 | NA | NA | NA | NA | 1 | 1 |
| 10_1285130 | NA | NA | 1 | 1 | NA | NA | NA | NA | 1 | 1 | NA | NA | 1 | 1 | NA | NA | 1 | 1 | 2 | 1 |
| 10_1285136 | 14 | 0.0714 | 18 | 0.3333 | 16 | 0.125 | 12 | 0.1667 | 60 | 0.1833 | 19 | 0.3158 | 13 | 0.2308 | 16 | 0.1875 | 15 | 0.3333 | 63 | 0.2698 |
| 10_1285136 | 16 | 0.25 | 17 | 0.4118 | 18 | 0.2778 | 15 | 0.4 | 66 | 0.3333 | 15 | 0.4 | 14 | 0.2857 | 17 | 0.3529 | 16 | 0.5625 | 62 | 0.4032 |
| 10_1285149 | 8 | 0.125 | 10 | 0.1 | 12 | 0.0833 | 13 | 0 | 43 | 0.0698 | 15 | 0.2 | 12 | 0.25 | 11 | 0.1818 | 9 | 0.4444 | 47 | 0.2553 |
| 10_1285151 | 8 | 0.25 | 10 | 0.2 | 12 | 0.1667 | 10 | 0 | 40 | 0.15 | 10 | 0.2 | 10 | 0.2 | 11 | 0.0909 | 6 | 0 | 37 | 0.1351 |
| 10_1285151 | 8 | 0.625 | 14 | 0.3571 | 13 | 0.3077 | 14 | 0.3571 | 49 | 0.3878 | 14 | 0.2143 | 12 | 0.4167 | 12 | 0.0833 | 10 | 0.4 | 48 | 0.2708 |
| 10_1285560 | 1 | 1 | 2 | 0.5 | 2 | 0 | NA | NA | 5 | 0.4 | 1 | 1 | 3 | 0.3333 | 2 | 1 | NA | NA | 6 | 0.6667 |
| 10_1286016 | 5 | 1 | 3 | 1 | 4 | 1 | 3 | 1 | 15 | 1 | 5 | 1 | 5 | 1 | 1 | 1 | 1 | 1 | 12 | 1 |
| 10_1286019 | 6 | 1 | 3 | 1 | 4 | 1 | 3 | 1 | 16 | 1 | 5 | 0.4 | 5 | 1 | NA | NA | 3 | 1 | 13 | 0.7692 |
| 10_1286019 | 7 | 1 | 2 | 1 | 3 | 1 | 2 | 1 | 14 | 1 | 5 | 1 | 4 | 1 | 1 | 1 | 2 | 1 | 12 | 1 |
| 10_1286019 | 2 | 0.5 | 2 | 0 | 4 | 0.5 | 1 | 1 | 9 | 0.4444 | 2 | 0.5 | 2 | 0 | NA | NA | 2 | 1 | 6 | 0.5 |
| 10_1286020 | 5 | 0 | 3 | 0.3333 | 4 | 0 | 2 | 0 | 14 | 0.0714 | 6 | 0.3333 | 4 | 0 | 1 | 0 | 3 | 0.3333 | 14 | 0.2143 |
| 10_1286021 | 5 | 1 | 3 | 1 | 4 | 1 | 2 | 1 | 14 | 1 | 4 | 1 | 3 | 0.6667 | 1 | 1 | 3 | 1 | 11 | 0.9091 |
| 10_1286023 | 4 | 0 | 2 | 0 | 3 | 0.6667 | 2 | 0 | 11 | 0.1818 | 5 | 0.2 | 4 | 0 | 1 | 0 | NA | NA | 10 | 0.1 |
| 10_1286372 | NA | NA | 3 | 0 | 2 | 0 | 3 | 0.3333 | 8 | 0.125 | 2 | 0 | 1 | 0 | 2 | 0.5 | 2 | 0 | 7 | 0.1429 |
| 10_1286379 | NA | NA | 3 | 0 | 1 | 1 | 3 | 0.3333 | 7 | 0.2857 | 2 | 0.5 | 1 | 0 | 1 | 0 | 2 | 0 | 6 | 0.1667 |
| 10_1286379 | NA | NA | 2 | 0 | NA | NA | 3 | 0.3333 | 5 | 0.2 | 2 | 1 | 1 | 0 | 1 | 0 | 2 | 0.5 | 6 | 0.5 |
| 10_1286380 | NA | NA | 3 | 0.3333 | 2 | 0.5 | 2 | 0 | 7 | 0.2857 | 1 | 1 | 1 | 1 | 1 | 0 | 2 | 1 | 5 | 0.8 |
| 10_1286381 | NA | NA | 3 | 0.3333 | 2 | 0 | 3 | 0 | 8 | 0.125 | 2 | 1 | 1 | 0 | 1 | 0 | 2 | 0.5 | 6 | 0.5 |
| 10_1286382 | NA | NA | 2 | 1 | 2 | 1 | 3 | 0.3333 | 7 | 0.7143 | 2 | 1 | 1 | 1 | 1 | 1 | 2 | 1 | 6 | 1 |
| 10_1286421 | 7 | 0.2857 | 2 | 0 | 1 | 0 | NA | NA | 10 | 0.2 | 3 | 0 | 2 | 0 | 3 | 0 | 2 | 0.5 | 10 | 0.1 |
| 10_1286421 | 4 | 0.25 | 2 | 1 | 1 | 1 | NA | NA | 7 | 0.5714 | 1 | 0 | 1 | 0 | 3 | 1 | 2 | 0.5 | 7 | 0.5714 |
| 10_1286422 | 6 | 0.6667 | 2 | 1 | 1 | 0 | NA | NA | 9 | 0.6667 | 1 | 0 | 2 | 0.5 | 3 | 0.3333 | 2 | 0 | 8 | 0.25 |
| 10_1286422 | 7 | 0.8571 | 2 | 1 | 2 | 1 | NA | NA | 11 | 0.9091 | 3 | 0.6667 | 2 | 1 | 3 | 0.6667 | 2 | 1 | 10 | 0.8 |
| 10_1286423 | 8 | 0.125 | 1 | 0 | 2 | 0 | NA | NA | 11 | 0.0909 | 3 | 0.3333 | 2 | 0 | 4 | 0 | 2 | 0 | 11 | 0.0909 |
| 10_1286423 | 8 | 0.375 | 1 | 0 | 2 | 0 | NA | NA | 11 | 0.2727 | 3 | 0.6667 | 2 | 0 | 4 | 0.25 | 1 | 0 | 10 | 0.3 |
| 10_1286424 | 8 | 0.875 | 1 | 1 | 2 | 1 | NA | NA | 11 | 0.9091 | 3 | 0.6667 | 2 | 0.5 | 3 | 0 | 2 | 0 | 10 | 0.3 |

| | | | | | | | | | | | | | | | | | | | | |
|---|---|---|---|---|---|---|---|---|---|---|---|---|---|---|---|---|---|---|---|---|
| 10_1286425 | 7 | 0.5714 | 2 | 0 | 2 | 1 | NA | NA | 11 | 0.5455 | 2 | 0.5 | 1 | 0 | 4 | 0.5 | 2 | 0 | 9 | 0.3333 |
| 10_1286427 | 9 | 1 | 2 | 1 | 3 | 1 | NA | NA | 14 | 1 | 3 | 1 | 2 | 1 | 4 | 1 | 2 | 1 | 11 | 1 |
| 10_1286428 | 9 | 0.2222 | 1 | 0 | 2 | 1 | NA | NA | 12 | 0.3333 | 3 | 0.6667 | 2 | 0.5 | 4 | 0.75 | 2 | 1 | 11 | 0.7273 |
| 10_1286428 | 8 | 0.375 | 3 | 0.3333 | 2 | 0.5 | NA | NA | 13 | 0.3846 | 3 | 0.3333 | 2 | 0.5 | 4 | 0.25 | 2 | 0 | 11 | 0.2727 |
| 10_1286428 | 8 | 1 | 2 | 1 | 2 | 1 | NA | NA | 12 | 1 | 3 | 1 | 2 | 1 | 5 | 1 | 2 | 1 | 12 | 1 |
| 10_1286429 | 6 | 0 | 1 | 1 | 1 | 0 | 1 | 0 | 9 | 0.1111 | 2 | 0 | 2 | 0 | 5 | 0.2 | 2 | 0 | 11 | 0.0909 |
| 10_1286429 | 8 | 0.875 | 2 | 0.5 | 2 | 1 | 1 | 1 | 13 | 0.8462 | 3 | 1 | 2 | 1 | 5 | 0.8 | 2 | 1 | 12 | 0.9167 |
| 10_1286429 | 7 | 0.5714 | 1 | 0 | 2 | 0 | 1 | 1 | 11 | 0.4545 | 2 | 0.5 | 2 | 0.5 | 6 | 0.5 | 1 | 1 | 11 | 0.5455 |
| 10_1286431 | 7 | 1 | 2 | 0.5 | 2 | 0 | 3 | 0.3333 | 14 | 0.6429 | 3 | 1 | 2 | 0.5 | 6 | 0.6667 | 1 | 1 | 12 | 0.75 |
| 10_1286432 | 8 | 0.25 | 2 | 0.5 | 2 | 0 | 3 | 0.3333 | 15 | 0.2667 | 4 | 0 | 2 | 0.5 | 6 | 0.3333 | 2 | 0.5 | 14 | 0.2857 |
| 10_1286433 | 6 | 0.6667 | 2 | 1 | 1 | 1 | 3 | 1 | 12 | 0.8333 | 3 | 1 | 2 | 1 | 4 | 1 | 1 | 1 | 10 | 1 |
| 10_1286434 | 9 | 0.2222 | 2 | 0 | 2 | 0 | 3 | 0.3333 | 16 | 0.1875 | 4 | 0 | 3 | 0.3333 | 4 | 0.25 | 2 | 0 | 13 | 0.1538 |
| 10_1286531 | 11 | 0.0909 | 3 | 0 | NA | NA | 5 | 0 | 19 | 0.0526 | 4 | 0 | 4 | 0 | 6 | 0.1667 | 3 | 0 | 17 | 0.0588 |
| 10_1286534 | 9 | 0.3333 | 3 | 0.3333 | 2 | 0 | 3 | 0 | 17 | 0.2353 | 3 | 0.3333 | 4 | 0 | 7 | 0.2857 | 3 | 0.6667 | 17 | 0.2941 |
| 10_1286535 | 8 | 0 | 1 | 0 | 2 | 0.5 | 2 | 0 | 13 | 0.0769 | 3 | 0 | 4 | 0.25 | 5 | 0.2 | 2 | 0 | 14 | 0.1429 |
| 10_1287183 | NA | NA | 1 | 1 | 2 | 0.5 | NA | NA | 3 | 0.6667 | NA | NA | NA | NA | 2 | 0.5 | NA | NA | 2 | 0.5 |
| 10_1287190 | NA | NA | 1 | 1 | 2 | 0.5 | NA | NA | 3 | 0.6667 | NA | NA | NA | NA | 2 | 0.5 | 1 | 0 | 3 | 0.3333 |
| 10_1287190 | NA | NA | 1 | 1 | 2 | 0.5 | NA | NA | 3 | 0.6667 | NA | NA | NA | NA | 2 | 1 | 1 | 1 | 3 | 1 |
| 10_1287190 | NA | NA | NA | NA | 2 | 0.5 | NA | NA | 2 | 0.5 | NA | NA | 1 | 1 | 2 | 0.5 | 1 | 1 | 4 | 0.75 |
| 10_1287192 | NA | NA | 2 | 1 | 3 | 1 | NA | NA | 5 | 1 | 1 | 1 | 1 | 1 | 2 | 1 | 1 | 1 | 5 | 1 |
| 10_1287193 | NA | NA | 2 | 0.5 | 1 | 1 | NA | NA | 3 | 0.6667 | NA | NA | 1 | 0 | 2 | 0.5 | 1 | 0 | 4 | 0.25 |
| 10_1287193 | NA | NA | 2 | 1 | 2 | 1 | NA | NA | 4 | 1 | 1 | 1 | 1 | 1 | 1 | 1 | 1 | 1 | 4 | 1 |
| 10_1287194 | NA | NA | 2 | 1 | 2 | 1 | NA | NA | 4 | 1 | NA | NA | 1 | 1 | 2 | 0.5 | 1 | 0 | 4 | 0.5 |
| 10_1287212 | 1 | 1 | NA | NA | NA | NA | NA | NA | 1 | 1 | 1 | 1 | NA | NA | NA | NA | NA | NA | 1 | 1 |
| 10_1287212 | 1 | 1 | NA | NA | NA | NA | NA | NA | 1 | 1 | 1 | 1 | NA | NA | NA | NA | NA | NA | 1 | 1 |
| 10_1287222 | 1 | 1 | NA | NA | NA | NA | NA | NA | 1 | 1 | 1 | 1 | NA | NA | NA | NA | NA | NA | 1 | 1 |
| 10_1304122 | NA | NA | NA | NA | 1 | 1 | NA | NA | 1 | 1 | NA | NA | NA | NA | 1 | 1 | 1 | 1 | 2 | 1 |
| 11_3079043 | 1 | 0 | 1 | 1 | NA | NA | 1 | 0 | 3 | 0.3333 | NA | NA | 1 | 0 | NA | NA | 2 | 0.5 | 3 | 0.3333 |
| 11_3079962 | NA | NA | 1 | 1 | NA | NA | 1 | 1 | 2 | 1 | 1 | 1 | NA | NA | NA | NA | NA | NA | 1 | 1 |
| 11_3080000 | NA | NA | 1 | 0 | NA | NA | 1 | 1 | 2 | 0.5 | NA | NA | NA | NA | NA | NA | 1 | 1 | 1 | 1 |
| 11_3080434 | NA | NA | 2 | 0 | NA | NA | 1 | 1 | 3 | 0.3333 | 1 | 0 | NA | NA | 1 | 1 | 2 | 0 | 4 | 0.25 |
| 11_3080794 | 1 | 1 | 3 | 0 | NA | NA | 1 | 1 | 5 | 0.4 | 1 | 0 | 1 | 0 | 1 | 1 | 3 | 0.3333 | 6 | 0.3333 |
| 11_3080940 | NA | NA | 2 | 0.5 | NA | NA | 1 | 1 | 3 | 0.6667 | 1 | 0 | 1 | 1 | 1 | 0 | 2 | 0.5 | 5 | 0.4 |
| 11_3170383 | 6 | 0.1667 | 3 | 0 | 2 | 0 | 2 | 0 | 13 | 0.0769 | NA | NA | 4 | 0.25 | NA | NA | 2 | 0 | 6 | 0.1667 |
| 11_3194071 | 7 | 0.5714 | 7 | 0.4286 | 6 | 0.8333 | 5 | 0.8 | 25 | 0.64 | 6 | 0.6667 | 9 | 0.2222 | 7 | 0.7143 | 5 | 0.6 | 27 | 0.5185 |
| 11_3194083 | 6 | 0.5 | 8 | 0.25 | 5 | 0.2 | 4 | 0.25 | 23 | 0.3043 | 4 | 0.25 | 7 | 0.2857 | 6 | 0.1667 | 1 | 0 | 18 | 0.2222 |
| 11_3194119 | 5 | 0 | 8 | 0.125 | 7 | 0.2857 | 4 | 0.25 | 24 | 0.1667 | 6 | 0 | 8 | 0.125 | 6 | 0 | 5 | 0.2 | 25 | 0.08 |
| 11_3194143 | 8 | 0.375 | 7 | 0.2857 | 7 | 0 | 4 | 0.25 | 26 | 0.2308 | 6 | 0.5 | 8 | 0.25 | 5 | 0.2 | 6 | 0.1667 | 25 | 0.28 |
| 11_3194537 | 5 | 0.2 | 8 | 0.125 | 6 | 0.3333 | 4 | 0 | 23 | 0.1739 | 8 | 0 | 8 | 0 | 7 | 0.4286 | 4 | 0 | 27 | 0.1111 |
| 11_3198669 | 2 | 0.5 | NA | NA | 2 | 0 | NA | NA | 4 | 0.25 | 1 | 1 | NA | NA | NA | NA | NA | NA | 1 | 1 |
| 11_3199949 | 13 | 0.3846 | 6 | 0.1667 | 10 | 0.3 | 14 | 0.2143 | 43 | 0.2791 | 9 | 0.4444 | 11 | 0.1818 | 11 | 0.3636 | 7 | 0.4286 | 38 | 0.3421 |
| 11_3200078 | 10 | 0.3 | 10 | 0.6 | 12 | 0.6667 | 13 | 0.3846 | 45 | 0.4889 | 11 | 0.2727 | 11 | 0.3636 | 9 | 0.4444 | 11 | 0.3636 | 42 | 0.3571 |
| 11_3200142 | 11 | 0.0909 | 9 | 0.2222 | 13 | 0.3846 | 15 | 0.4 | 48 | 0.2917 | 12 | 0.1667 | 12 | 0.0833 | 9 | 0.2222 | 12 | 0.25 | 45 | 0.1778 |
| 11_3200151 | 16 | 0.5 | 10 | 0.3 | 15 | 0.2 | 17 | 0.6471 | 58 | 0.431 | 14 | 0.5714 | 12 | 0.6667 | 13 | 0.2308 | 12 | 0.1667 | 51 | 0.4118 |
| 11_3200220 | 14 | 0.0714 | 7 | 0.1429 | 15 | 0.2 | 15 | 0.1333 | 51 | 0.1373 | 12 | 0.1667 | 11 | 0.0909 | 9 | 0.2222 | 11 | 0.0909 | 43 | 0.1395 |
| 11_3240570 | NA | NA | NA | NA | NA | NA | 2 | 0.5 | 2 | 0.5 | 1 | 1 | 2 | 1 | NA | NA | NA | NA | 3 | 1 |
| 11_3240969 | NA | NA | NA | NA | NA | NA | 3 | 1 | 3 | 1 | 2 | 0.5 | 3 | 1 | NA | NA | 1 | 0 | 6 | 0.6667 |
| 11_3241151 | NA | NA | NA | NA | 1 | 1 | 1 | 1 | 2 | 1 | 1 | 1 | 3 | 1 | NA | NA | NA | NA | 4 | 1 |
| 11_3241158 | NA | NA | NA | NA | NA | NA | 3 | 0.6667 | 3 | 0.6667 | 2 | 0 | 3 | 0.3333 | NA | NA | 1 | 0 | 6 | 0.1667 |
| 11_3241176 | NA | NA | NA | NA | 1 | 1 | 1 | 1 | 2 | 1 | 2 | 0.5 | 3 | 1 | NA | NA | NA | NA | 5 | 0.8 |
| 11_3241311 | NA | NA | NA | NA | 1 | 1 | 2 | 1 | 3 | 1 | 2 | 0.5 | 3 | 0.6667 | 1 | 1 | 1 | 1 | 7 | 0.7143 |
| 11_3241333 | NA | NA | 1 | 0 | 1 | 0 | 3 | 0.3333 | 5 | 0.2 | 2 | 0 | 3 | 0.6667 | 1 | 0 | 1 | 1 | 7 | 0.4286 |
| 11_3241355 | NA | NA | 1 | 0 | 1 | 0 | 1 | 1 | 3 | 0.3333 | 2 | 0.5 | 3 | 0.6667 | 1 | 1 | NA | NA | 6 | 0.6667 |
| 11_3241443 | NA | NA | 1 | 1 | 1 | 1 | 3 | 1 | 5 | 1 | 1 | 1 | 1 | 1 | 1 | 1 | NA | NA | 3 | 1 |
| 11_3257957 | NA | NA | NA | NA | 1 | 1 | 3 | 0 | 4 | 0.25 | 3 | 0.3333 | 2 | 0.5 | 1 | 0 | 1 | 0 | 7 | 0.2857 |
| 11_3258068 | NA | NA | NA | NA | 1 | 0 | 3 | 0.3333 | 4 | 0.25 | 3 | 0.3333 | 2 | 0 | 2 | 0 | 2 | 0 | 9 | 0.1111 |
| 11_3258101 | NA | NA | NA | NA | NA | NA | 3 | 0.6667 | 3 | 0.6667 | 3 | 0.6667 | 2 | 0.5 | 2 | 1 | 2 | 1 | 9 | 0.7778 |
| 11_3258239 | NA | NA | 1 | 0 | 1 | 0 | 2 | 0.5 | 4 | 0.25 | 1 | 1 | 1 | 0 | 2 | 0.5 | 2 | 0 | 6 | 0.3333 |
| 11_3258555 | 3 | 0.3333 | 2 | 0.5 | 2 | 0.5 | 3 | 0 | 10 | 0.3 | 2 | 0 | 2 | 0.5 | 1 | 0 | 2 | 0 | 7 | 0.1429 |
| 11_3258648 | 3 | 0.3333 | 2 | 0.5 | 1 | 0 | 3 | 0.3333 | 9 | 0.3333 | 2 | 0 | 2 | 0.5 | 3 | 0.3333 | 1 | 0 | 8 | 0.25 |
| 11_3283406 | 6 | 0.1667 | 5 | 0 | 2 | 0 | 4 | 0 | 17 | 0.0588 | 4 | 0 | 3 | 0 | 4 | 0.25 | 5 | 0 | 16 | 0.0625 |
| 11_3291662 | 7 | 0.1429 | 6 | 0.5 | 2 | 0 | 5 | 0 | 20 | 0.2 | 5 | 0.2 | 4 | 0 | 5 | 0 | 5 | 0.2 | 19 | 0.1053 |
| 11_3291757 | 6 | 0.3333 | 7 | 0.2857 | 4 | 0 | 6 | 0 | 23 | 0.1739 | 5 | 0.2 | 4 | 0.5 | 6 | 0 | 4 | 0.25 | 19 | 0.2105 |
| 11_3291770 | 7 | 0.5714 | 6 | 0.3333 | 4 | 0.5 | 5 | 0.2 | 22 | 0.4091 | 3 | 0.6667 | 5 | 1 | 1 | 0 | 5 | 0.6 | 14 | 0.7143 |
| 11_3291972 | 7 | 0.7143 | 8 | 0.875 | 4 | 0.25 | 7 | 0.8571 | 26 | 0.7308 | 8 | 0.875 | 5 | 1 | 6 | 0.5 | 5 | 0.4 | 24 | 0.7083 |

| | | | | | | | | | | | | | | | | | | | | |
|---|---|---|---|---|---|---|---|---|---|---|---|---|---|---|---|---|---|---|---|---|
| 11_3292039 | 7 | 0.2857 | 8 | 0.5 | 4 | 0.25 | 5 | 0.6 | 24 | 0.4167 | 7 | 0.7143 | 4 | 0.75 | 6 | 0.1667 | 5 | 0.6 | 22 | 0.5455 |
| 11_3292121 | 7 | 0.2857 | 8 | 0.375 | 3 | 0 | 7 | 0.1429 | 25 | 0.24 | 6 | 0.3333 | 5 | 0.2 | 5 | 0 | 5 | 0.2 | 21 | 0.1905 |
| 11_3292171 | 8 | 0.25 | 9 | 0.3333 | 5 | 0 | 8 | 0.375 | 30 | 0.2667 | 7 | 0.5714 | 5 | 0.2 | 3 | 0 | 3 | 0.3333 | 18 | 0.3333 |
| 11_3292220 | 7 | 0.1429 | 8 | 0 | 6 | 0 | 8 | 0.375 | 29 | 0.1379 | 7 | 0.1429 | 6 | 0 | 6 | 0 | 5 | 0.2 | 24 | 0.0833 |
| 11_3292404 | 5 | 0.4 | 6 | 0.1667 | 6 | 0.3333 | 8 | 0.625 | 25 | 0.4 | 7 | 0.7143 | 5 | 0.8 | 5 | 0.4 | 7 | 0.4286 | 24 | 0.5833 |
| 11_3292821 | 7 | 0 | 6 | 0.1667 | 7 | 0.1429 | 9 | 0 | 29 | 0.069 | 6 | 0 | 5 | 0.4 | 7 | 0.2857 | 6 | 0 | 24 | 0.1667 |
| 11_3294427 | 1 | 1 | 5 | 0.2 | 2 | 0 | 6 | 0 | 14 | 0.1429 | 2 | 0 | 9 | 0.1111 | 3 | 0 | 4 | 0 | 18 | 0.0556 |
| 11_3294658 | 6 | 0.1667 | 10 | 0.1 | 9 | 0 | 10 | 0.3 | 35 | 0.1429 | 4 | 0 | 13 | 0.0769 | 5 | 0.2 | 7 | 0.2857 | 29 | 0.1379 |
| 11_3294943 | 5 | 0.2 | 9 | 0.6667 | 8 | 0.625 | 11 | 0.3636 | 33 | 0.4848 | 4 | 0.5 | 10 | 0.6 | 4 | 0.5 | 6 | 0.6667 | 24 | 0.5833 |
| 11_3295167 | 6 | 0.8333 | 8 | 0.375 | 8 | 0.25 | 7 | 0.4286 | 29 | 0.4483 | 2 | 1 | 10 | 0.5 | 3 | 0 | 7 | 0.7143 | 22 | 0.5455 |
| 11_3295318 | 6 | 0.3333 | 7 | 0.4286 | 7 | 0.5714 | 8 | 0.625 | 28 | 0.5 | 3 | 0.3333 | 10 | 0.5 | 4 | 0 | 6 | 0.6667 | 23 | 0.4348 |
| 11_3295360 | 4 | 0.5 | 5 | 0 | 6 | 0.3333 | 5 | 0.2 | 20 | 0.25 | 1 | 1 | 7 | 0.2857 | 3 | 0 | 4 | 0.25 | 15 | 0.2667 |
| 11_3295444 | 7 | 0.4286 | 7 | 0.2857 | 9 | 0.2222 | 8 | 0.375 | 31 | 0.3226 | 2 | 0 | 6 | 0.5 | 4 | 0 | 5 | 0.4 | 17 | 0.2941 |
| 11_3295454 | 6 | 0.8333 | 5 | 0.8 | 8 | 0.25 | 8 | 0.625 | 27 | 0.5926 | 1 | 1 | 6 | 0.8333 | 3 | 0.6667 | 4 | 0.5 | 14 | 0.7143 |
| 11_3295661 | 10 | 0.4 | 7 | 0.1429 | 12 | 0.3333 | 10 | 0.2 | 39 | 0.2821 | 3 | 1 | 10 | 0.2 | 6 | 0.1667 | 7 | 0 | 26 | 0.2308 |
| 11_3295714 | 9 | 0.2222 | 6 | 0.1667 | 10 | 0 | 10 | 0 | 35 | 0.0857 | 3 | 0.6667 | 7 | 0.1429 | 7 | 0.4286 | 9 | 0 | 26 | 0.2308 |
| 11_3295753 | 8 | 0.5 | 6 | 0.5 | 10 | 0.6 | 10 | 0.8 | 34 | 0.6176 | 3 | 0.6667 | 11 | 0.5455 | 6 | 0.3333 | 7 | 0.8571 | 27 | 0.5926 |
| 11_3295861 | 9 | 0.2222 | 5 | 0 | 8 | 0.25 | 4 | 0 | 26 | 0.1538 | 3 | 0.3333 | 7 | 0 | 6 | 0.1667 | 6 | 0 | 22 | 0.0909 |
| 11_3308072 | 2 | 0.5 | NA | NA | NA | NA | NA | NA | 2 | 0.5 | NA | NA | NA | NA | NA | NA | 1 | 1 | 1 | 1 |
| 11_3366075 | 7 | 0.2857 | 6 | 0.5 | 7 | 0.4286 | 3 | 0.6667 | 23 | 0.4348 | 3 | 0.6667 | 6 | 0.5 | 5 | 0.4 | 3 | 0.6667 | 17 | 0.5294 |
| 11_3366081 | 7 | 0 | 5 | 0.2 | 6 | 0.3333 | 3 | 0.3333 | 21 | 0.1905 | 5 | 0 | 6 | 0.1667 | 4 | 0.25 | 2 | 1 | 17 | 0.2353 |
| 11_3366190 | 7 | 0.7143 | 5 | 1 | 5 | 1 | 3 | 1 | 20 | 0.9 | 4 | 0.75 | 8 | 0.625 | 4 | 0.25 | 2 | 1 | 18 | 0.6111 |
| 11_3366248 | 4 | 0 | 4 | 0 | 7 | 0.1429 | 1 | 0 | 16 | 0.0625 | 3 | 0 | 4 | 0.25 | 3 | 0 | 2 | 0.5 | 12 | 0.1667 |
| 11_3366691 | 5 | 0.2 | 4 | 0 | 3 | 0 | 2 | 0 | 14 | 0.0714 | 3 | 0 | 5 | 0 | 3 | 0.3333 | 3 | 0.3333 | 14 | 0.1429 |
| 11_3366710 | 6 | 0.8333 | 4 | 0.75 | 3 | 1 | 2 | 0.5 | 15 | 0.8 | 2 | 1 | 5 | 0.4 | 4 | 0.75 | 3 | 1 | 14 | 0.7143 |
| 11_3366749 | 6 | 0.1667 | 5 | 0 | 2 | 0.5 | 1 | 0 | 14 | 0.1429 | 2 | 0.5 | 6 | 0.3333 | 4 | 0.5 | 3 | 0.3333 | 15 | 0.4 |
| 11_3366795 | 4 | 0.5 | 6 | 0.5 | 2 | 0.5 | 1 | 1 | 13 | 0.5385 | 2 | 1 | 5 | 0.4 | 3 | 0.6667 | 1 | 1 | 11 | 0.6364 |
| 11_3366957 | 3 | 1 | 6 | 0.8333 | 3 | 1 | 1 | 0 | 13 | 0.8462 | 2 | 0.5 | 5 | 0.6 | 4 | 1 | 3 | 1 | 14 | 0.7857 |
| 11_3367017 | 5 | 0.6 | 6 | 0.5 | 3 | 0.6667 | 2 | 1 | 16 | 0.625 | 2 | 1 | 5 | 0.8 | 4 | 0.75 | 3 | 0.6667 | 14 | 0.7857 |
| 11_3367167 | 5 | 0.4 | 5 | 0.2 | 3 | 0.3333 | 2 | 1 | 15 | 0.4 | 2 | 0.5 | 3 | 0.3333 | 3 | 0 | 2 | 0.5 | 10 | 0.3 |
| 11_3367219 | 6 | 0.5 | 4 | 0.25 | 3 | 0.6667 | 2 | 0.5 | 15 | 0.4667 | 2 | 0 | 3 | 0.6667 | 4 | 0.5 | 2 | 0.5 | 11 | 0.4545 |
| 11_3367269 | 5 | 0.4 | 7 | 0.4286 | 3 | 1 | 2 | 0.5 | 17 | 0.5294 | 1 | 1 | 4 | 0.25 | 4 | 0.5 | 3 | 0.6667 | 12 | 0.5 |
| 11_3367305 | 4 | 0.75 | 4 | 1 | 3 | 0.6667 | 1 | 1 | 12 | 0.8333 | 2 | 0.5 | 3 | 0.6667 | 4 | 0.75 | 3 | 1 | 12 | 0.75 |
| 11_3367309 | 4 | 0.5 | 3 | 0.3333 | 3 | 0.3333 | 1 | 0 | 11 | 0.3636 | 2 | 0.5 | 4 | 0.75 | 3 | 0.3333 | 2 | 0.5 | 11 | 0.5455 |
| 11_3367327 | 3 | 0.3333 | 2 | 0.5 | 3 | 0.3333 | 2 | 0 | 10 | 0.3 | 3 | 0 | 4 | 0.25 | 4 | 0.25 | 1 | 1 | 12 | 0.25 |
| 11_3367461 | 4 | 1 | 4 | 1 | 3 | 1 | 2 | 1 | 13 | 1 | 2 | 1 | 4 | 0.75 | 4 | 0.75 | 1 | 1 | 11 | 0.8182 |
| 11_3367489 | 3 | 0.6667 | 4 | 0.75 | 3 | 0.3333 | 1 | 0 | 11 | 0.5455 | 3 | 0 | 3 | 0.3333 | 2 | 0.5 | 2 | 0.5 | 10 | 0.3 |
| 11_3367576 | 2 | 0 | 4 | 0.75 | 3 | 1 | 2 | 0.5 | 11 | 0.6364 | 2 | 0.5 | 4 | 1 | 4 | 0.75 | 2 | 1 | 12 | 0.8333 |
| 11_3367636 | 4 | 0.25 | 3 | 0.3333 | 3 | 0.3333 | 2 | 1 | 12 | 0.4167 | 1 | 0 | 4 | 0.25 | 2 | 0 | 2 | 0.5 | 9 | 0.2222 |
| 11_3367656 | 4 | 0.25 | 3 | 0 | 3 | 0.3333 | 2 | 0 | 12 | 0.1667 | 3 | 0.3333 | 4 | 0 | 4 | 0.75 | 2 | 0 | 13 | 0.3077 |
| 11_3367684 | 2 | 1 | 1 | 1 | 1 | 1 | 2 | 1 | 6 | 1 | 2 | 1 | 2 | 1 | 2 | 1 | 1 | 1 | 7 | 1 |
| 11_3367702 | 3 | 0.3333 | 2 | 0 | 2 | 0 | 1 | 1 | 8 | 0.25 | 1 | 0 | 3 | 0.6667 | 3 | 0 | 1 | 1 | 8 | 0.375 |
| 11_3367823 | 4 | 0.25 | 2 | 0 | 2 | 0 | 2 | 0 | 10 | 0.1 | 3 | 0 | 4 | 0 | 2 | 0.5 | 2 | 0 | 11 | 0.0909 |
| 11_3367848 | 3 | 0.3333 | 3 | 0 | 3 | 0 | 2 | 0 | 11 | 0.0909 | 3 | 0 | 5 | 0.2 | 3 | 0 | 2 | 0.5 | 13 | 0.1538 |
| 11_3367864 | 4 | 1 | 3 | 0.6667 | 3 | 1 | 1 | 1 | 11 | 0.9091 | 1 | 1 | 5 | 0.8 | 4 | 1 | 2 | 1 | 12 | 0.9167 |
| 11_3367882 | 3 | 0 | 3 | 0.6667 | 3 | 0.3333 | 2 | 0 | 11 | 0.2727 | 2 | 0 | 5 | 0.2 | 2 | 0.5 | 2 | 0 | 11 | 0.1818 |
| 11_3367952 | 4 | 0.75 | 2 | 1 | 2 | 1 | 2 | 1 | 10 | 0.9 | 2 | 1 | 4 | 0.75 | 4 | 0.75 | 2 | 1 | 12 | 0.8333 |
| 11_3407421 | 1 | 1 | NA | NA | 2 | 0 | 1 | 0 | 4 | 0.25 | NA | NA | NA | NA | NA | NA | 2 | 0.5 | 2 | 0.5 |
| 11_3408019 | NA | NA | 2 | 0.5 | 2 | 0 | 1 | 1 | 5 | 0.4 | NA | NA | NA | NA | 1 | 1 | 1 | 0 | 2 | 0.5 |
| 11_3408178 | 1 | 1 | 2 | 1 | 1 | 1 | NA | NA | 4 | 1 | NA | NA | NA | NA | 1 | 1 | 1 | 1 | 2 | 1 |
| 11_3408188 | 1 | 1 | 2 | 0 | 1 | 0 | 1 | 0 | 5 | 0.2 | NA | NA | NA | NA | 1 | 1 | 2 | 0 | 3 | 0.3333 |
| 11_3408266 | 1 | 1 | 2 | 0 | 1 | 1 | 1 | 0 | 5 | 0.4 | NA | NA | NA | NA | 1 | 1 | 2 | 0.5 | 3 | 0.6667 |
| 11_3444435 | 17 | 0.3529 | 18 | 0.3333 | 3 | 0 | 6 | 0.3333 | 44 | 0.3182 | 7 | 0.4286 | 6 | 0.6667 | 12 | 0.3333 | 5 | 0.2 | 30 | 0.4 |
| 11_3444638 | 14 | 0.0714 | 9 | 0.3333 | 2 | 0 | 4 | 0.25 | 29 | 0.1724 | 5 | 0.2 | 4 | 0 | 7 | 0.1429 | 2 | 0 | 18 | 0.1111 |
| 11_3444843 | 17 | 0.5294 | 19 | 0.6842 | 2 | 0.5 | 6 | 0.6667 | 44 | 0.6136 | 7 | 0.1429 | 4 | 0.25 | 10 | 0.3 | 5 | 0.4 | 26 | 0.2692 |
| 11_3452240 | 6 | 0.8333 | 9 | 0.8889 | NA | NA | 2 | 1 | 17 | 0.8824 | 4 | 1 | 1 | 1 | 5 | 0.8 | 3 | 0.6667 | 13 | 0.8462 |
| 11_3452283 | 8 | 1 | 9 | 1 | NA | NA | 2 | 1 | 19 | 1 | 5 | 1 | NA | NA | 6 | 0.6667 | 4 | 1 | 15 | 0.8667 |
| 11_3452538 | 8 | 0 | 9 | 0.2222 | NA | NA | 2 | 0 | 19 | 0.1053 | 5 | 0 | NA | NA | 4 | 0 | 4 | 0.25 | 13 | 0.0769 |
| 11_3452553 | 8 | 0.5 | 9 | 1 | NA | NA | 2 | 1 | 19 | 0.7895 | 6 | 0.8333 | NA | NA | 5 | 0.8 | 4 | 0.75 | 15 | 0.8 |
| 11_3452721 | 4 | 0.75 | 7 | 0.7143 | NA | NA | 2 | 0.5 | 13 | 0.6923 | 4 | 1 | NA | NA | NA | NA | 3 | 0 | 7 | 0.5714 |
| 11_3453021 | 5 | 0.4 | 7 | 0 | NA | NA | 2 | 0 | 14 | 0.1429 | 6 | 0.3333 | NA | NA | 5 | 0.2 | 3 | 0.3333 | 14 | 0.2857 |
| 11_3589840 | 11 | 0.5455 | 11 | 0.6364 | 8 | 0.625 | 13 | 0.8462 | 43 | 0.6744 | 2 | 0.5 | 8 | 0.625 | 13 | 0.1538 | 10 | 0.6 | 33 | 0.4242 |
| 11_3589911 | 13 | 0.7692 | 10 | 0.6 | 8 | 0.75 | 11 | 0.3636 | 42 | 0.619 | 3 | 0 | 8 | 0.75 | 12 | 0.75 | 11 | 0.6364 | 34 | 0.6471 |
| 11_3590002 | 8 | 0.125 | 9 | 0.2222 | 6 | 0.3333 | 12 | 0.25 | 35 | 0.2286 | 3 | 0.3333 | 6 | 0.3333 | 12 | 0 | 6 | 0.1667 | 27 | 0.1481 |
| 11_3590051 | 13 | 0.0769 | 8 | 0 | 9 | 0.1111 | 12 | 0.1667 | 42 | 0.0952 | 1 | 1 | 7 | 0.2857 | 13 | 0.1538 | 9 | 0.1111 | 30 | 0.2 |

| | | | | | | | | | | | | | | | | | | | | |
|---|---|---|---|---|---|---|---|---|---|---|---|---|---|---|---|---|---|---|---|---|
| 11_359017 | 13 | 0 | 12 | 0.1667 | 9 | 0.3333 | 14 | 0.1429 | 48 | 0.1458 | 3 | 0.6667 | 8 | 0.125 | 12 | 0.0833 | 12 | 0.1667 | 35 | 0.1714 |
| 11_359021 | 10 | 1 | 12 | 1 | 9 | 1 | 13 | 1 | 44 | 1 | 2 | 0.5 | 8 | 1 | 13 | 1 | 10 | 0.9 | 33 | 0.9394 |
| 11_359023 | 7 | 0 | 12 | 0.5 | 7 | 0.1429 | 13 | 0.2308 | 39 | 0.2564 | 2 | 0.5 | 7 | 0.2857 | 11 | 0.0909 | 12 | 0.1667 | 32 | 0.1875 |
| 11_359030 | 11 | 0.1818 | 11 | 0.0909 | 8 | 0 | 12 | 0 | 42 | 0.0714 | 3 | 0 | 5 | 0 | 9 | 0.1111 | 10 | 0.3 | 27 | 0.1481 |
| 11_359041 | 11 | 0.4545 | 11 | 0.7273 | 8 | 0.625 | 11 | 0.5455 | 41 | 0.5854 | 3 | 0.3333 | 8 | 0.75 | 11 | 0.3636 | 12 | 0.4167 | 34 | 0.4706 |
| 11_359044 | 10 | 0.1 | 11 | 0 | 6 | 0 | 12 | 0.1667 | 39 | 0.0769 | 2 | 0 | 6 | 0 | 12 | 0 | 11 | 0.1818 | 31 | 0.0645 |
| 11_359045 | 12 | 0.25 | 11 | 0.1818 | 8 | 0.5 | 10 | 0.7 | 41 | 0.3902 | 3 | 1 | 7 | 0.5714 | 11 | 0.3636 | 11 | 0.2727 | 32 | 0.4375 |
| 11_359050 | 12 | 0.5833 | 8 | 0.375 | 8 | 0 | 10 | 0.2 | 38 | 0.3158 | 3 | 0.6667 | 7 | 0.2857 | 10 | 0.2 | 10 | 0.1 | 30 | 0.2333 |
| 11_359052 | 11 | 0.6364 | 11 | 0.6364 | 7 | 0.5714 | 9 | 0.6667 | 38 | 0.6316 | 3 | 1 | 7 | 0.5714 | 13 | 0.8462 | 12 | 0.6667 | 35 | 0.7429 |
| 11_359053 | 10 | 0.7 | 11 | 0.8182 | 5 | 1 | 10 | 0.6 | 36 | 0.75 | 2 | 0.5 | 6 | 0.8333 | 12 | 0.8333 | 11 | 0.6364 | 31 | 0.7419 |
| 11_359061 | 11 | 0.1818 | 11 | 0.2727 | 7 | 0.2857 | 10 | 0.2 | 39 | 0.2308 | 3 | 0 | 6 | 0 | 11 | 0.4545 | 11 | 0.2727 | 31 | 0.2581 |
| 11_359065 | 11 | 0.4545 | 9 | 0.7778 | 6 | 0.5 | 9 | 0.4444 | 35 | 0.5429 | 1 | 1 | 7 | 0.7143 | 10 | 0.5 | 10 | 0.3 | 28 | 0.5 |
| 11_359071 | 10 | 0.1 | 9 | 0 | 4 | 0.25 | 8 | 0.375 | 31 | 0.1613 | 1 | 0 | 3 | 0.3333 | 9 | 0.3333 | 11 | 0.2727 | 24 | 0.2917 |
| 11_359086 | 11 | 0.3636 | 10 | 0.3 | 5 | 1 | 12 | 0.5 | 38 | 0.4737 | 2 | 0 | 6 | 0.6667 | 12 | 0.5833 | 11 | 0.3636 | 31 | 0.4839 |
| 11_359095 | 12 | 0 | 10 | 0.1 | 6 | 0 | 11 | 0.0909 | 39 | 0.0513 | 2 | 0 | 6 | 0 | 12 | 0.1667 | 11 | 0.0909 | 31 | 0.0968 |
| 11_359097 | 8 | 0.125 | 8 | 0 | 5 | 0.2 | 7 | 0 | 28 | 0.0714 | 2 | 0.5 | 5 | 0.4 | 7 | 0.1429 | 8 | 0.125 | 22 | 0.2273 |
| 11_359097 | 11 | 0.1818 | 10 | 0 | 5 | 0 | 9 | 0 | 35 | 0.0571 | 2 | 0 | 5 | 0 | 10 | 0.1 | 11 | 0.1818 | 28 | 0.1071 |
| 11_359111 | 7 | 0.1429 | 7 | 0 | 4 | 0 | 6 | 0.1667 | 24 | 0.0833 | 1 | 0 | 4 | 0.5 | 8 | 0.125 | 7 | 0 | 20 | 0.15 |
| 11_359111 | NA | NA | 1 | 0 | NA | NA | 1 | 1 | 2 | 0.5 | NA | NA | NA | NA | NA | NA | 1 | 1 | 1 | 1 |
| 11_359112 | 10 | 0.1 | 9 | 0.2222 | 5 | 0 | 9 | 0.2222 | 33 | 0.1515 | 2 | 0.5 | 6 | 0 | 11 | 0.1818 | 10 | 0.4 | 29 | 0.2414 |
| 11_359114 | 10 | 1 | 9 | 0.6667 | 6 | 1 | 8 | 0.75 | 33 | 0.8485 | 2 | 0.5 | 5 | 0.8 | 8 | 1 | 9 | 1 | 24 | 0.9167 |
| 11_359127 | 9 | 0.8889 | 8 | 1 | 3 | 0.6667 | 7 | 0.8571 | 27 | 0.8889 | 2 | 1 | 5 | 0.8 | 9 | 0.8889 | 7 | 0.5714 | 23 | 0.7826 |
| 11_359135 | 8 | 0 | 10 | 0.2 | 5 | 0.4 | 8 | 0.25 | 31 | 0.1935 | 2 | 0.5 | 6 | 0 | 9 | 0.3333 | 8 | 0.125 | 25 | 0.2 |
| 11_359140 | 9 | 0.2222 | 9 | 0 | 5 | 0.6 | 8 | 0.25 | 31 | 0.2258 | 2 | 0 | 6 | 0.1667 | 8 | 0.375 | 9 | 0.1111 | 25 | 0.2 |
| 11_359150 | 5 | 0.2 | 4 | 0 | 4 | 0 | 7 | 0.1429 | 20 | 0.1 | 2 | 0.5 | 6 | 0 | 9 | 0.1111 | 7 | 0 | 24 | 0.0833 |
| 11_359151 | 7 | 0.1429 | 6 | 0.1667 | 3 | 0 | 9 | 0.2222 | 25 | 0.16 | 1 | 0 | 3 | 0.3333 | 6 | 0.1667 | 5 | 0 | 15 | 0.1333 |
| 11_359171 | 9 | 0 | 6 | 0 | 5 | 0 | 9 | 0.2222 | 29 | 0.069 | 1 | 0 | 4 | 0.25 | 8 | 0 | 7 | 0.2857 | 20 | 0.15 |
| 11_359173 | 7 | 0.5714 | 7 | 0.4286 | 5 | 0.4 | 8 | 0.75 | 27 | 0.5556 | NA | NA | 3 | 0.6667 | 7 | 0.5714 | 7 | 0.4286 | 17 | 0.5294 |
| 11_359176 | 9 | 0 | 7 | 0.1429 | 4 | 0.75 | 8 | 0 | 28 | 0.1429 | 1 | 0 | 5 | 0 | 8 | 0.25 | 7 | 0.4286 | 21 | 0.2381 |
| 11_359184 | 7 | 0.5714 | 5 | 0.8 | 4 | 0.75 | 7 | 0.7143 | 23 | 0.6957 | 1 | 0 | 4 | 0.25 | 7 | 0.8571 | 5 | 0.6 | 17 | 0.5882 |
| 11_359194 | 9 | 0.2222 | 5 | 0 | 4 | 0 | 7 | 0.5714 | 25 | 0.24 | 1 | 0 | 4 | 0 | 8 | 0 | 5 | 0.2 | 18 | 0.0556 |
| 11_359200 | 7 | 0.2857 | 4 | 0.5 | 4 | 0.25 | 6 | 0.5 | 21 | 0.381 | 1 | 0 | 5 | 0 | 8 | 0.125 | 6 | 0.3333 | 20 | 0.15 |
| 11_359203 | 5 | 0.4 | 4 | 0.25 | 5 | 0.4 | 9 | 0.3333 | 23 | 0.3478 | NA | NA | 2 | 0 | 5 | 0.4 | 3 | 0 | 10 | 0.2 |
| 11_359203 | 5 | 0.4 | 4 | 0.5 | 3 | 0.6667 | 4 | 1 | 16 | 0.625 | 1 | 1 | 3 | 1 | 4 | 0.75 | 5 | 0.8 | 13 | 0.8462 |
| 11_359221 | 7 | 0.1429 | 4 | 0 | 3 | 0.6667 | 8 | 0.125 | 22 | 0.1818 | 1 | 0 | 5 | 0.4 | 6 | 0.6667 | 7 | 0.2857 | 19 | 0.4211 |
| 11_359233 | 8 | 0.375 | 4 | 0 | 3 | 0.6667 | 7 | 0.5714 | 22 | 0.4091 | 1 | 0 | 5 | 1 | 5 | 0.2 | 5 | 0.4 | 16 | 0.5 |
| 11_359236 | 8 | 0 | 5 | 0 | 4 | 0.25 | 8 | 0.125 | 25 | 0.08 | 1 | 1 | 4 | 0.25 | 6 | 0.1667 | 6 | 0.1667 | 17 | 0.2353 |
| 11_359238 | 5 | 0 | 5 | 0 | 2 | 0 | 7 | 0.1429 | 19 | 0.0526 | 1 | 0 | 5 | 0 | 6 | 0 | 6 | 0.1667 | 18 | 0.0556 |
| 11_359240 | 4 | 0.25 | 4 | 0 | 2 | 0 | 7 | 0 | 17 | 0.0588 | 1 | 0 | 3 | 0 | 7 | 0 | 6 | 0.1667 | 17 | 0.0588 |
| 11_359256 | 7 | 1 | 5 | 1 | 4 | 1 | 5 | 1 | 21 | 1 | 1 | 0 | 4 | 1 | 7 | 1 | 5 | 1 | 17 | 0.9412 |
| 11_359273 | 4 | 0.5 | 3 | 0.3333 | 2 | 0.5 | 4 | 0.5 | 13 | 0.4615 | 1 | 0 | 1 | 0 | 6 | 0.5 | 6 | 0.3333 | 14 | 0.3571 |
| 11_359274 | 5 | 0.2 | 4 | 0 | 4 | 0 | 6 | 0.1667 | 19 | 0.1053 | 1 | 0 | 2 | 0 | 4 | 0.25 | 4 | 0 | 11 | 0.0909 |
| 11_359280 | 6 | 0.8333 | 5 | 0.8 | 4 | 1 | 5 | 1 | 20 | 0.9 | 1 | 1 | 4 | 0.75 | 3 | 1 | 5 | 1 | 13 | 0.9231 |
| 11_359291 | 7 | 0.1429 | 3 | 0 | 2 | 0 | 5 | 0 | 17 | 0.0588 | 1 | 1 | 4 | 0 | 5 | 0 | 4 | 0.25 | 14 | 0.1429 |
| 11_359294 | 7 | 0 | 5 | 0 | 3 | 0 | 4 | 0.25 | 19 | 0.0526 | NA | NA | 3 | 0 | 6 | 0.1667 | 4 | 0 | 13 | 0.0769 |
| 11_359299 | 6 | 1 | 4 | 1 | 3 | 1 | 4 | 0.75 | 17 | 0.9412 | NA | NA | 4 | 0.75 | 6 | 0.8333 | 4 | 0.75 | 14 | 0.7857 |
| 11_359453 | 5 | 0.4 | 2 | 1 | 3 | 0 | 1 | 0 | 11 | 0.3636 | 2 | 1 | 3 | 1 | 4 | 1 | 3 | 0.6667 | 12 | 0.9167 |
| 11_359467 | 3 | 0.6667 | 2 | 1 | 2 | 0.5 | 1 | 0 | 8 | 0.625 | 1 | 0 | 3 | 0.6667 | 4 | 0.25 | 3 | 0.6667 | 11 | 0.4545 |
| 11_359478 | 10 | 0 | 7 | 0.1429 | 5 | 0.2 | 5 | 0.2 | 27 | 0.1111 | 2 | 0 | 6 | 0 | 7 | 0 | 4 | 0.25 | 19 | 0.0526 |
| 11_359488 | 11 | 0.7273 | 5 | 0.2 | 4 | 0.25 | 4 | 0.75 | 24 | 0.5417 | 2 | 1 | 6 | 0.5 | 8 | 0.625 | 5 | 0.8 | 21 | 0.6667 |
| 11_359497 | 11 | 0.0909 | 6 | 0.1667 | 3 | 0 | 6 | 0.1667 | 26 | 0.1154 | 2 | 0.5 | 7 | 0 | 7 | 0 | 5 | 0.2 | 21 | 0.0952 |
| 11_359502 | 7 | 1 | 6 | 1 | 3 | 1 | 4 | 1 | 20 | 1 | 2 | 1 | 6 | 1 | 7 | 0.8571 | 5 | 1 | 20 | 0.95 |
| 11_359505 | 9 | 0.7778 | 4 | 0.75 | 4 | 0.5 | 4 | 0.5 | 21 | 0.6667 | 2 | 1 | 5 | 0.6 | 5 | 0.6 | 6 | 0.6667 | 18 | 0.6667 |
| 11_359508 | 10 | 0.8 | 6 | 0.5 | 4 | 0.75 | 5 | 0.4 | 25 | 0.64 | 1 | 1 | 6 | 0.6667 | 7 | 0.8571 | 5 | 0.6 | 19 | 0.7368 |
| 11_359509 | 10 | 0.2 | 6 | 0.1667 | 4 | 0 | 5 | 0.2 | 25 | 0.16 | 2 | 0 | 6 | 0.1667 | 6 | 0.1667 | 4 | 0 | 18 | 0.1111 |
| 11_359515 | 10 | 0.9 | 6 | 0.8333 | 4 | 1 | 5 | 1 | 25 | 0.92 | 2 | 1 | 5 | 0.8 | 7 | 1 | 6 | 1 | 20 | 0.95 |
| 11_359517 | 9 | 0.8889 | 6 | 0.8333 | 4 | 1 | 6 | 0.5 | 25 | 0.8 | 2 | 1 | 6 | 1 | 7 | 1 | 6 | 0.8333 | 21 | 0.9524 |
| 11_359525 | 8 | 0.25 | 5 | 0.4 | 3 | 0.6667 | 4 | 0.25 | 20 | 0.35 | 2 | 0 | 3 | 0.6667 | 6 | 0.5 | 4 | 0.5 | 15 | 0.4667 |
| 11_359537 | 7 | 0 | 5 | 0 | 3 | 0.3333 | 4 | 0 | 19 | 0.0526 | 2 | 0.5 | 4 | 0 | 6 | 0 | 5 | 0.2 | 17 | 0.1176 |
| 11_359656 | 3 | 1 | 6 | 0.6667 | 4 | 0.25 | 5 | 0.2 | 18 | 0.5 | 2 | 0.5 | 2 | 0.5 | 5 | 0.6 | 4 | 0.25 | 13 | 0.4615 |
| 11_359658 | 5 | 0 | 5 | 0.2 | 3 | 0 | 6 | 0.3333 | 19 | 0.1579 | 2 | 0 | 4 | 0 | 6 | 0.1667 | 4 | 0.25 | 16 | 0.125 |
| 11_359666 | 6 | 0.3333 | 4 | 0 | 3 | 0.3333 | 6 | 0 | 19 | 0.1579 | 1 | 0 | 4 | 0.5 | 4 | 0 | 4 | 0.25 | 13 | 0.2308 |
| 11_359669 | 7 | 1 | 4 | 1 | 3 | 1 | 2 | 1 | 16 | 1 | 1 | 1 | 3 | 0.6667 | 2 | 1 | 3 | 1 | 9 | 0.8889 |
| 11_359673 | 5 | 0.4 | 6 | 0.6667 | 2 | 1 | 3 | 0.3333 | 16 | 0.5625 | 2 | 0.5 | 2 | 1 | 5 | 0.6 | 4 | 0.75 | 13 | 0.6923 |
| 11_359676 | 6 | 0.1667 | 6 | 0.1667 | 2 | 0 | 6 | 0 | 20 | 0.1 | 1 | 1 | 4 | 0.25 | 5 | 0.2 | 4 | 0 | 14 | 0.2143 |

| | | | | | | | | | | | | | | | | | | | | |
|---|---|---|---|---|---|---|---|---|---|---|---|---|---|---|---|---|---|---|---|---|
| 11_3596774 | 5 | 0.2 | 6 | 0.1667 | 3 | 0.6667 | 6 | 0.1667 | 20 | 0.25 | 1 | 1 | 3 | 0.3333 | 4 | 0.25 | 4 | 0.25 | 12 | 0.3333 |
| 11_3596799 | 6 | 1 | 6 | 0.8333 | 3 | 1 | 5 | 1 | 20 | 0.95 | 1 | 1 | 4 | 0.75 | 3 | 0.6667 | 4 | 1 | 12 | 0.8333 |
| 11_3596910 | 4 | 0.25 | 5 | 0.6 | 3 | 0.6667 | 4 | 0.75 | 16 | 0.5625 | 2 | 1 | 3 | 0.6667 | 4 | 0.75 | 4 | 1 | 13 | 0.8462 |
| 11_3596966 | 4 | 0.75 | 5 | 1 | 2 | 1 | 4 | 1 | 15 | 0.9333 | 2 | 0.5 | 2 | 1 | 3 | 1 | 3 | 1 | 10 | 0.9 |
| 11_3596976 | 4 | 1 | 6 | 1 | 2 | 1 | 6 | 1 | 18 | 1 | 2 | 1 | 3 | 1 | 5 | 1 | 4 | 1 | 14 | 1 |
| 11_3597027 | 4 | 0.25 | 4 | 0.5 | 1 | 0 | 6 | 0.3333 | 15 | 0.3333 | 2 | 0.5 | 2 | 1 | 3 | 0.6667 | 4 | 0 | 11 | 0.4545 |
| 11_3597043 | 4 | 0.75 | 6 | 1 | 3 | 1 | 4 | 0.75 | 17 | 0.8824 | 2 | 1 | NA | NA | 1 | 1 | 4 | 0.75 | 7 | 0.8571 |
| 11_3597070 | 5 | 0.2 | 6 | 0.3333 | 3 | 0.6667 | 6 | 0.3333 | 20 | 0.35 | 2 | 0.5 | 3 | 0.6667 | 4 | 0.25 | 4 | 0 | 13 | 0.3077 |
| 11_3597253 | 4 | 0.25 | 3 | 0 | 3 | 0 | 3 | 0 | 13 | 0.0769 | NA | NA | 4 | 0.25 | 4 | 0 | 3 | 0 | 11 | 0.0909 |
| 11_3597350 | 5 | 0 | 3 | 0 | 3 | 0 | 5 | 0.4 | 16 | 0.125 | 2 | 0 | 4 | 0 | 5 | 0.4 | 3 | 0 | 14 | 0.1429 |
| 11_3597445 | 5 | 0.4 | 3 | 1 | 3 | 1 | 5 | 0.6 | 16 | 0.6875 | 2 | 1 | 2 | 0.5 | 3 | 1 | 2 | 1 | 9 | 0.8889 |
| 11_3599600 | 1 | 1 | NA | NA | NA | NA | NA | NA | 1 | 1 | 1 | 1 | 1 | 0 | NA | NA | NA | NA | 2 | 0.5 |
| 11_3599640 | 1 | 1 | NA | NA | NA | NA | NA | NA | 1 | 1 | 1 | 1 | 1 | 0 | NA | NA | NA | NA | 2 | 0.5 |
| 11_3599851 | 2 | 1 | 2 | 0.5 | NA | NA | NA | NA | 4 | 0.75 | 1 | 1 | 1 | 0 | NA | NA | 1 | 1 | 3 | 0.6667 |
| 11_3600027 | 2 | 1 | 2 | 0.5 | NA | NA | NA | NA | 4 | 0.75 | 1 | 1 | NA | NA | NA | NA | 1 | 1 | 2 | 1 |
| 11_3600144 | 2 | 1 | 2 | 1 | NA | NA | NA | NA | 4 | 1 | 2 | 1 | NA | NA | NA | NA | 1 | 1 | 3 | 1 |
| 11_3600360 | 3 | 0.6667 | 2 | 0.5 | NA | NA | NA | NA | 5 | 0.6 | 1 | 1 | 1 | 1 | NA | NA | 1 | 1 | 3 | 1 |
| 11_3633933 | 1 | 1 | 4 | 0.5 | NA | NA | 2 | 0 | 7 | 0.4286 | 2 | 0 | 1 | 1 | 2 | 0.5 | 2 | 0 | 7 | 0.2857 |
| 11_3639099 | 4 | 1 | 5 | 0.6 | NA | NA | 2 | 1 | 11 | 0.8182 | 2 | 1 | 3 | 0.6667 | 3 | 1 | 2 | 1 | 10 | 0.9 |
| 11_3639176 | 4 | 0.5 | 6 | 0.3333 | NA | NA | 2 | 0 | 12 | 0.3333 | 2 | 0 | 4 | 0 | 3 | 0.3333 | 3 | 0 | 12 | 0.0833 |
| 11_3639185 | 4 | 1 | 5 | 1 | NA | NA | 2 | 1 | 11 | 1 | 2 | 1 | 4 | 1 | 3 | 1 | 3 | 1 | 12 | 1 |
| 11_3639212 | 2 | 1 | 4 | 1 | NA | NA | 2 | 1 | 8 | 1 | 2 | 1 | 4 | 0.75 | 2 | 1 | 2 | 0 | 10 | 0.7 |
| 11_3639330 | 2 | 0.5 | 7 | 0.2857 | NA | NA | 2 | 0.5 | 11 | 0.3636 | 3 | 0.3333 | 3 | 0 | 2 | 0 | 4 | 0.25 | 12 | 0.1667 |
| 11_3639425 | 4 | 0.25 | 6 | 0.1667 | NA | NA | 2 | 1 | 12 | 0.3333 | 2 | 0 | 3 | 0.6667 | 3 | 0.3333 | 4 | 0.5 | 12 | 0.4167 |
| 11_3639464 | 5 | 0.8 | 7 | 0.8571 | NA | NA | 2 | 1 | 14 | 0.8571 | 2 | 1 | 3 | 1 | 3 | 1 | 4 | 0.75 | 12 | 0.9167 |
| 11_3639519 | 5 | 0.8 | 7 | 0.8571 | NA | NA | 2 | 1 | 14 | 0.8571 | 2 | 0.5 | 3 | 0.6667 | 3 | 0.6667 | 4 | 0.75 | 12 | 0.6667 |
| 11_3639607 | 5 | 0.2 | 5 | 0 | NA | NA | 2 | 0 | 12 | 0.0833 | 2 | 0.5 | 2 | 0.5 | 2 | 0.5 | 4 | 0 | 10 | 0.3 |
| 11_3639725 | 4 | 0.25 | 6 | 0.5 | NA | NA | 2 | 1 | 12 | 0.5 | 1 | 1 | 2 | 0.5 | 2 | 0.5 | 2 | 0 | 7 | 0.4286 |
| 11_3639775 | 2 | 1 | 2 | 0.5 | NA | NA | 3 | 0.3333 | 7 | 0.5714 | NA | NA | 2 | 0.5 | 1 | 1 | 1 | 1 | 4 | 0.75 |
| 11_3639935 | 3 | 1 | 7 | 1 | NA | NA | 3 | 1 | 13 | 1 | 1 | 1 | 2 | 1 | 3 | 0.6667 | 4 | 0.75 | 10 | 0.8 |
| 11_3639939 | 6 | 0.1667 | 8 | 0.125 | NA | NA | 2 | 0 | 16 | 0.125 | 2 | 0.5 | 2 | 0 | 3 | 0 | 6 | 0.1667 | 13 | 0.1538 |
| 11_3639953 | 6 | 0 | 8 | 0.125 | NA | NA | 3 | 0 | 17 | 0.0588 | 2 | 0 | 2 | 0.5 | 3 | 0 | 6 | 0 | 13 | 0.0769 |
| 11_3640046 | 5 | 0.8 | 5 | 1 | NA | NA | 3 | 1 | 13 | 0.9231 | 1 | 1 | 3 | 1 | 3 | 1 | 4 | 1 | 11 | 1 |
| 11_3640158 | 3 | 1 | 9 | 1 | NA | NA | 3 | 1 | 15 | 1 | 2 | 1 | 3 | 1 | 2 | 1 | 5 | 0.8 | 12 | 0.9167 |
| 11_3640244 | 6 | 0.1667 | 7 | 0.1429 | NA | NA | 1 | 0 | 14 | 0.1429 | 1 | 0 | 3 | 0.3333 | 2 | 0.5 | 3 | 0.3333 | 9 | 0.3333 |
| 11_3654582 | 8 | 0.25 | 6 | 0.1667 | 4 | 0.5 | 3 | 0 | 21 | 0.2381 | 3 | 0 | 1 | 1 | NA | NA | 4 | 0 | 8 | 0.125 |
| 11_3655093 | 8 | 0.375 | 7 | 0.4286 | 5 | 0.6 | 1 | 1 | 21 | 0.4762 | 2 | 0 | 4 | 0.25 | NA | NA | 4 | 0.25 | 10 | 0.2 |
| 11_3813375 | 2 | 0 | 1 | 1 | NA | NA | 1 | 0 | 4 | 0.25 | NA | NA | 1 | 1 | 1 | 1 | NA | NA | 2 | 1 |
| 11_3846993 | 3 | 0.6667 | NA | NA | NA | NA | NA | NA | 3 | 0.6667 | 1 | 1 | 2 | 0.5 | 1 | 1 | NA | NA | 4 | 0.75 |
| 11_3847411 | 2 | 1 | NA | NA | NA | NA | 1 | 1 | 3 | 1 | 1 | 1 | 2 | 1 | 1 | 1 | NA | NA | 4 | 1 |
| 11_3847514 | 2 | 1 | NA | NA | NA | NA | NA | NA | 2 | 1 | 1 | 1 | 2 | 1 | 1 | 1 | NA | NA | 4 | 1 |
| 11_3847616 | 1 | 1 | NA | NA | NA | NA | NA | NA | 1 | 1 | 1 | 1 | 2 | 1 | NA | NA | NA | NA | 3 | 1 |
| 11_3847622 | 3 | 1 | NA | NA | NA | NA | NA | NA | 3 | 1 | 1 | 0 | 1 | 1 | 1 | 1 | NA | NA | 3 | 0.6667 |
| 11_3847804 | 3 | 0.3333 | NA | NA | NA | NA | 1 | 0 | 4 | 0.25 | 1 | 0 | 2 | 0.5 | 1 | 0 | NA | NA | 4 | 0.25 |
| 11_3848153 | 3 | 1 | NA | NA | 1 | 1 | 1 | 1 | 5 | 1 | 1 | 1 | 2 | 1 | 1 | 1 | NA | NA | 4 | 1 |
| 11_3848420 | 3 | 0.3333 | 1 | 0 | 1 | 0 | 1 | 1 | 6 | 0.3333 | 1 | 1 | 2 | 0.5 | 1 | 0 | NA | NA | 4 | 0.5 |
| 11_3848442 | 3 | 0.3333 | 1 | 1 | 1 | 0 | 1 | 0 | 6 | 0.3333 | 1 | 1 | 1 | 0 | 1 | 0 | NA | NA | 3 | 0.3333 |
| 11_3848462 | 3 | 0.6667 | 1 | 1 | 1 | 0 | 1 | 1 | 6 | 0.6667 | 1 | 1 | 2 | 1 | 1 | 0 | NA | NA | 4 | 0.75 |
| 11_3848850 | 3 | 0.3333 | 1 | 0 | NA | NA | 1 | 1 | 5 | 0.4 | 1 | 0 | 1 | 0 | 1 | 0 | 1 | 1 | 4 | 0.25 |
| 11_3851150 | 4 | 0 | 3 | 0.3333 | 3 | 0 | 2 | 0 | 12 | 0.0833 | 1 | 0 | 2 | 0.5 | 4 | 0 | 2 | 0.5 | 9 | 0.2222 |
| 11_3857225 | 2 | 0 | 8 | 0 | 4 | 0.25 | 1 | 1 | 15 | 0.1333 | 2 | 0.5 | 3 | 0 | 3 | 0 | 2 | 0.5 | 10 | 0.2 |
| 11_3857385 | 1 | 0 | 7 | 0.2857 | 4 | 0.25 | 1 | 1 | 13 | 0.3077 | 1 | 0 | 1 | 1 | 2 | 0 | 2 | 0 | 6 | 0.1667 |
| 11_3857397 | 2 | 1 | 6 | 0.8333 | 4 | 1 | 1 | 1 | 13 | 0.9231 | 2 | 1 | 3 | 1 | 3 | 0.6667 | 3 | 0.6667 | 11 | 0.8182 |
| 11_3857493 | 1 | 1 | 8 | 1 | 2 | 1 | NA | NA | 11 | 1 | 2 | 1 | 3 | 1 | 2 | 1 | 3 | 1 | 10 | 1 |
| 11_3857520 | 2 | 1 | 5 | 0.2 | 2 | 1 | NA | NA | 9 | 0.5556 | 2 | 1 | 2 | 1 | 3 | 0.3333 | 3 | 0.6667 | 10 | 0.7 |
| 11_3857573 | 2 | 1 | 4 | 0.5 | 4 | 0.75 | 1 | 1 | 11 | 0.7273 | 2 | 0.5 | 2 | 0 | 3 | 0.3333 | 2 | 0.5 | 9 | 0.3333 |
| 11_3857852 | 2 | 0.5 | 5 | 0.6 | 3 | 0.6667 | 1 | 0 | 11 | 0.5455 | 1 | 1 | 2 | 1 | 2 | 0 | 1 | 0 | 6 | 0.5 |
| 11_3862787 | 2 | 0 | 4 | 0.5 | 4 | 0.25 | 1 | 1 | 11 | 0.3636 | 1 | 0 | 2 | 1 | 2 | 0 | 2 | 1 | 7 | 0.5714 |
| 11_3863081 | 2 | 0 | 4 | 0.75 | 3 | 0.6667 | 1 | 1 | 10 | 0.6 | 1 | 0 | 2 | 1 | 2 | 0.5 | 2 | 0.5 | 7 | 0.5714 |
| 11_3863357 | 2 | 0.5 | 2 | 0 | 4 | 0.25 | 1 | 0 | 9 | 0.2222 | NA | NA | 2 | 1 | 1 | 0 | 2 | 0 | 5 | 0.4 |
| 11_3863478 | 1 | 0 | 2 | 0 | 4 | 0.25 | 1 | 0 | 8 | 0.125 | NA | NA | 2 | 0 | 2 | 0 | 2 | 0.5 | 6 | 0.1667 |
| 11_3863489 | 1 | 0 | 2 | 0.5 | 3 | 0 | 1 | 0 | 7 | 0.1429 | NA | NA | 2 | 0 | 2 | 0 | 2 | 0.5 | 6 | 0.1667 |
| 11_3863789 | 1 | 0 | NA | NA | 2 | 1 | NA | NA | 3 | 0.6667 | NA | NA | 1 | 1 | 1 | 0 | NA | NA | 2 | 0.5 |
| 11_3867331 | 3 | 0.6667 | 3 | 0 | 6 | 0.5 | 5 | 0.4 | 17 | 0.4118 | 2 | 0.5 | 6 | 0.1667 | 1 | 0 | 5 | 0.6 | 14 | 0.3571 |
| 11_3867410 | 4 | 0.25 | 3 | 0 | 5 | 0.2 | 3 | 0 | 15 | 0.1333 | 1 | 0 | 5 | 0 | 1 | 0 | 4 | 0.25 | 11 | 0.0909 |

| | | | | | | | | | | | | | | | | | | | | |
|---|---|---|---|---|---|---|---|---|---|---|---|---|---|---|---|---|---|---|---|---|
| 11_386744 | 4 | 0.25 | NA | NA | 5 | 0.2 | 4 | 0.25 | 13 | 0.2308 | 1 | 1 | 5 | 0.2 | NA | NA | 3 | 0.3333 | 9 | 0.3333 |
| 11_387341 | 4 | 0.25 | 3 | 0 | 7 | 0 | 2 | 0 | 16 | 0.0625 | 2 | 0 | 5 | 0.4 | 1 | 0 | 4 | 0 | 12 | 0.1667 |
| 11_392893 | 7 | 0.7143 | 2 | 0.5 | 8 | 0.75 | 6 | 0.5 | 23 | 0.6522 | 4 | 1 | 5 | 0.6 | 7 | 1 | 3 | 0.6667 | 19 | 0.8421 |
| 11_392905 | 8 | 0.125 | 4 | 0.25 | 8 | 0.375 | 6 | 0 | 26 | 0.1923 | 4 | 0.5 | 5 | 0.2 | 7 | 0.1429 | 3 | 0.3333 | 19 | 0.2632 |
| 11_394817 | NA | NA | 1 | 1 | 1 | 1 | 1 | 1 | 3 | 1 | 3 | 1 | 1 | 1 | NA | NA | 1 | 1 | 5 | 1 |
| 11_394848 | 1 | 1 | 1 | 0 | 2 | 0.5 | 1 | 0 | 5 | 0.4 | 4 | 0 | 2 | 1 | 1 | 0 | 1 | 0 | 8 | 0.25 |
| 11_394860 | 1 | 1 | 1 | 0 | 2 | 1 | 1 | 1 | 5 | 0.8 | 4 | 0.5 | 2 | 0.5 | 1 | 1 | 2 | 1 | 9 | 0.6667 |
| 11_394887 | 1 | 1 | 2 | 1 | 3 | 1 | 1 | 1 | 7 | 1 | 4 | 1 | 2 | 1 | 1 | 1 | 2 | 1 | 9 | 1 |
| 11_394900 | 1 | 0 | 2 | 0 | 3 | 0 | 1 | 1 | 7 | 0.1429 | 4 | 0.5 | 2 | 0.5 | 1 | 0 | 2 | 0 | 9 | 0.3333 |
| 11_394905 | 1 | 1 | 2 | 1 | 2 | 1 | 1 | 1 | 6 | 1 | 3 | 1 | 2 | 1 | 1 | 0 | 2 | 1 | 8 | 0.875 |
| 11_394919 | NA | NA | 1 | 1 | 1 | 1 | 1 | 1 | 3 | 1 | 2 | 1 | 1 | 1 | NA | NA | 1 | 1 | 4 | 1 |
| 11_404188 | 12 | 0.1667 | 16 | 0.125 | 11 | 0.0909 | 6 | 0.1667 | 45 | 0.1333 | 9 | 0 | 7 | 0.2857 | 3 | 0 | 7 | 0.1429 | 26 | 0.1154 |
| 11_404201 | 5 | 0.2 | 10 | 0.3 | 8 | 0.25 | 3 | 0.3333 | 26 | 0.2692 | 7 | 0.4286 | 8 | 0.25 | 4 | 0 | 4 | 0.5 | 23 | 0.3043 |
| 11_404792 | 5 | 0.2 | 6 | 0 | 4 | 0 | 1 | 0 | 16 | 0.0625 | 3 | 0 | 3 | 0.6667 | 2 | 0.5 | 4 | 0 | 12 | 0.25 |
| 11_404819 | 4 | 0.25 | 5 | 0 | 3 | 0.3333 | NA | NA | 12 | 0.1667 | 3 | 0 | 3 | 0 | 1 | 1 | 4 | 0.5 | 11 | 0.2727 |
| 11_404819 | 4 | 0 | 4 | 0.25 | 2 | 0 | NA | NA | 10 | 0.1 | 1 | 0 | 2 | 0 | 2 | 0.5 | 3 | 0.3333 | 8 | 0.25 |
| 11_409057 | 1 | 0 | 2 | 0.5 | 2 | 0 | 3 | 0 | 8 | 0.125 | 4 | 0.75 | 4 | 0.25 | 7 | 0.4286 | NA | NA | 15 | 0.4667 |
| 11_409071 | 2 | 0 | 3 | 0.3333 | 2 | 0 | 4 | 0 | 11 | 0.0909 | 4 | 0.5 | 5 | 0 | 9 | 0.1111 | 1 | 0 | 19 | 0.1579 |
| 11_409076 | NA | NA | 3 | 0.3333 | 2 | 0 | 3 | 0 | 8 | 0.125 | 2 | 0.5 | 4 | 0 | 9 | 0 | 1 | 0 | 16 | 0.0625 |
| 11_409087 | 2 | 0.5 | 2 | 0 | 2 | 0 | 5 | 0.2 | 11 | 0.1818 | 4 | 0.25 | 5 | 0.4 | 10 | 0.1 | 1 | 0 | 20 | 0.2 |
| 11_409091 | 1 | 1 | 3 | 1 | 2 | 0.5 | 5 | 0.6 | 11 | 0.7273 | 3 | 0.6667 | 5 | 1 | 8 | 0.875 | 1 | 1 | 17 | 0.8824 |
| 11_409146 | 2 | 1 | 3 | 0 | 2 | 0.5 | 4 | 0 | 11 | 0.2727 | 3 | 0.3333 | NA | NA | 2 | 0 | NA | NA | 5 | 0.2 |
| 11_413033 | 13 | 0.5385 | 6 | 0.5 | 11 | 0.5455 | 11 | 0.4545 | 41 | 0.5122 | 8 | 0.875 | 12 | 0.75 | 5 | 0.4 | 8 | 0.625 | 33 | 0.697 |
| 11_413045 | 12 | 0.4167 | 6 | 0.1667 | 13 | 0.2308 | 11 | 0.6364 | 42 | 0.381 | 3 | 0.3333 | 13 | 0.5385 | 5 | 0.4 | 7 | 0.7143 | 28 | 0.5357 |
| 11_416501 | 1 | 0 | 5 | 0.4 | 3 | 0.6667 | 3 | 0 | 12 | 0.3333 | 2 | 1 | 2 | 1 | 4 | 0.5 | NA | NA | 8 | 0.75 |
| 11_416505 | 3 | 0.6667 | 4 | 0 | 3 | 0.3333 | 1 | 0 | 11 | 0.2727 | 2 | 0.5 | NA | NA | 3 | 0 | NA | NA | 5 | 0.2 |
| 11_416507 | 3 | 0 | 4 | 0.25 | 4 | 0.25 | 3 | 0.3333 | 14 | 0.2143 | 4 | 0.25 | 2 | 0 | 4 | 0 | NA | NA | 10 | 0.1 |
| 11_416508 | 3 | 0.3333 | 5 | 0.6 | 5 | 0.6 | 3 | 0.3333 | 16 | 0.5 | 5 | 0.8 | 3 | 0.6667 | 4 | 0.75 | NA | NA | 12 | 0.75 |
| 11_416514 | 3 | 1 | 5 | 0.2 | 4 | 0.5 | 3 | 0.3333 | 15 | 0.4667 | 5 | 0.6 | 3 | 0.3333 | 4 | 0.25 | NA | NA | 12 | 0.4167 |
| 11_416520 | 3 | 1 | 4 | 0.5 | 5 | 0.8 | 2 | 1 | 14 | 0.7857 | 3 | 0.6667 | 2 | 1 | 3 | 0.6667 | NA | NA | 8 | 0.75 |
| 11_416655 | 1 | 1 | 1 | 1 | 2 | 0 | NA | NA | 4 | 0.5 | 1 | 1 | 1 | 0 | NA | NA | NA | NA | 2 | 0.5 |
| 11_417197 | 2 | 1 | NA | NA | NA | NA | 1 | 0 | 3 | 0.6667 | 2 | 0 | 1 | 0 | 1 | 1 | NA | NA | 4 | 0.25 |
| 11_417205 | 2 | 0.5 | 1 | 0 | NA | NA | 1 | 0 | 4 | 0.25 | 2 | 0.5 | NA | NA | 1 | 1 | NA | NA | 3 | 0.6667 |
| 11_417210 | 2 | 1 | 1 | 1 | NA | NA | 1 | 1 | 4 | 1 | 2 | 0.5 | 1 | 0 | 1 | 1 | 1 | 1 | 5 | 0.6 |
| 11_443207 | 3 | 0 | 6 | 0 | 8 | 0.125 | 11 | 0.0909 | 28 | 0.0714 | 6 | 0 | 11 | 0.0909 | 9 | 0.1111 | 2 | 0 | 28 | 0.0714 |
| 11_443235 | 2 | 0 | 7 | 0.2857 | 7 | 0 | 9 | 0 | 25 | 0.08 | 5 | 0 | 11 | 0.1818 | 6 | 0 | 3 | 0 | 25 | 0.08 |
| 11_443241 | 3 | 0 | 7 | 0.1429 | 9 | 0.1111 | 9 | 0.1111 | 28 | 0.1071 | 5 | 0 | 11 | 0.2727 | 7 | 0 | 3 | 0.3333 | 26 | 0.1538 |
| 11_443275 | 1 | 1 | 3 | 0.3333 | 4 | 0.75 | 3 | 0.3333 | 11 | 0.5455 | 2 | 0.5 | 7 | 0 | 4 | 0.5 | 2 | 0 | 15 | 0.2 |
| 11_443277 | 3 | 0 | 6 | 0.1667 | 4 | 0.5 | 5 | 0 | 18 | 0.1667 | 4 | 0 | 6 | 0.1667 | 5 | 0 | 2 | 0 | 17 | 0.0588 |
| 11_443280 | 1 | 1 | 8 | 0.5 | 3 | 0.6667 | 5 | 0.8 | 17 | 0.6471 | 3 | 1 | 6 | 0.8333 | 4 | 0.75 | 4 | 1 | 17 | 0.8824 |
| 11_443738 | 3 | 0 | 5 | 0.2 | 4 | 0 | 6 | 0.1667 | 18 | 0.1111 | 5 | 0.4 | 10 | 0.2 | 8 | 0.25 | 3 | 0 | 26 | 0.2308 |
| 11_443749 | 2 | 0 | 1 | 0 | 3 | 0.6667 | 4 | 0 | 10 | 0.2 | 3 | 0.3333 | 8 | 0.5 | 4 | 0.25 | 2 | 1 | 17 | 0.4706 |
| 11_443755 | 2 | 1 | 2 | 0.5 | 4 | 1 | 5 | 1 | 13 | 0.9231 | 4 | 1 | 10 | 1 | 6 | 0.6667 | 3 | 1 | 23 | 0.913 |
| 11_443763 | 3 | 0 | 2 | 0 | 2 | 0 | 4 | 0.25 | 11 | 0.0909 | 4 | 0 | 5 | 0 | 5 | 0.2 | 3 | 0 | 17 | 0.0588 |
| 11_443765 | 3 | 1 | 1 | 1 | 3 | 1 | 6 | 0.8333 | 13 | 0.9231 | 4 | 1 | 10 | 0.9 | 7 | 0.8571 | 3 | 1 | 24 | 0.9167 |
| 11_443775 | 2 | 1 | 2 | 0.5 | 4 | 0.75 | 5 | 1 | 13 | 0.8462 | 4 | 1 | 6 | 0.8333 | 5 | 1 | 2 | 1 | 17 | 0.9412 |
| 11_443777 | 2 | 0.5 | 3 | 0.6667 | 4 | 0.5 | 5 | 0.4 | 14 | 0.5 | 4 | 0.5 | 9 | 0.3333 | 5 | 0.6 | 2 | 0.5 | 20 | 0.45 |
| 11_443782 | 2 | 0.5 | 2 | 0 | 4 | 0.5 | 5 | 0.4 | 13 | 0.3846 | 5 | 0.6 | 9 | 0.6667 | 5 | 0.6 | 2 | 0.5 | 21 | 0.619 |
| 11_443794 | 3 | 0.3333 | 2 | 0 | 4 | 0 | 6 | 0.1667 | 15 | 0.1333 | 3 | 0 | 9 | 0.1111 | 5 | 0.2 | 3 | 0.3333 | 20 | 0.15 |
| 11_443809 | 3 | 0.6667 | 2 | 0 | 1 | 0 | 6 | 0.5 | 12 | 0.4167 | 3 | 0.3333 | 3 | 0.3333 | 4 | 0 | 3 | 0.3333 | 13 | 0.2308 |
| 11_443838 | 1 | 1 | 2 | 0.5 | 1 | 0 | 4 | 0.75 | 8 | 0.625 | 3 | 0.6667 | 7 | 1 | 5 | 0.4 | 1 | 1 | 16 | 0.75 |
| 11_443846 | 2 | 0 | 1 | 0 | 1 | 0 | 4 | 0.75 | 8 | 0.375 | 4 | 0.25 | 7 | 0.7143 | 4 | 0.25 | 2 | 0.5 | 17 | 0.4706 |
| 11_461715 | 1 | 1 | NA | NA | NA | NA | NA | NA | 1 | 1 | NA | NA | 1 | 1 | NA | NA | 1 | 1 | 2 | 1 |
| 11_461735 | 1 | 1 | NA | NA | NA | NA | NA | NA | 1 | 1 | NA | NA | NA | NA | NA | NA | 1 | 1 | 1 | 1 |
| 11_463320 | 2 | 0 | 1 | 0 | 1 | 1 | 2 | 0 | 6 | 0.1667 | 3 | 0.3333 | 2 | 0.5 | 1 | 1 | 3 | 0 | 9 | 0.3333 |
| 11_463332 | 3 | 0.3333 | 2 | 0 | 1 | 0 | 2 | 0 | 8 | 0.125 | 2 | 1 | 2 | 0 | 1 | 0 | 3 | 0.3333 | 8 | 0.375 |
| 11_463339 | 2 | 0.5 | 1 | 1 | 1 | 1 | NA | NA | 4 | 0.75 | 2 | 0.5 | NA | NA | 1 | 1 | 3 | 0.6667 | 6 | 0.6667 |
| 11_463343 | 2 | 0 | 1 | 0 | 1 | 0 | 1 | 1 | 5 | 0.2 | 2 | 0.5 | NA | NA | 1 | 0 | 3 | 0 | 6 | 0.1667 |
| 11_463348 | 1 | 0 | 1 | 0 | 1 | 0 | 1 | 1 | 4 | 0.25 | 2 | 1 | NA | NA | 1 | 1 | 3 | 0.6667 | 6 | 0.8333 |
| 11_463355 | 2 | 1 | 1 | 0 | NA | NA | 1 | 1 | 4 | 0.75 | 2 | 0.5 | NA | NA | NA | NA | 2 | 0 | 4 | 0.25 |
| 11_463364 | 2 | 1 | 1 | 0 | 1 | 0 | 2 | 0.5 | 6 | 0.5 | NA | NA | NA | NA | 1 | 0 | 2 | 0.5 | 3 | 0.3333 |
| 11_463451 | 3 | 0 | 1 | 1 | NA | NA | 4 | 0 | 8 | 0.125 | 2 | 0 | NA | NA | 2 | 0.5 | 3 | 0.3333 | 7 | 0.2857 |
| 11_463455 | 4 | 0.25 | 1 | 0 | 1 | 0 | 3 | 0 | 9 | 0.1111 | 3 | 0.3333 | NA | NA | 2 | 0.5 | 3 | 0.6667 | 8 | 0.5 |
| 11_463475 | 4 | 0 | 1 | 0 | 1 | 0 | 3 | 0.3333 | 9 | 0.1111 | 3 | 0.3333 | NA | NA | 1 | 0 | 1 | 0 | 5 | 0.2 |
| 11_463502 | 4 | 0 | 1 | 0 | 1 | 1 | 2 | 0 | 8 | 0.125 | 2 | 0.5 | NA | NA | 2 | 0.5 | 2 | 0.5 | 6 | 0.5 |

| | | | | | | | | | | | | | | | | | | | | |
|---|---|---|---|---|---|---|---|---|---|---|---|---|---|---|---|---|---|---|---|---|
| 11_463511 | 3 | 0.3333 | NA | NA | 1 | 1 | 3 | 0.3333 | 7 | 0.4286 | 3 | 0.6667 | NA | NA | 3 | 0 | 3 | 0 | 9 | 0.2222 |
| 11_463565 | 4 | 0 | 1 | 1 | 1 | 1 | 5 | 0 | 11 | 0.1818 | 3 | 0.3333 | NA | NA | 3 | 0 | 2 | 0.5 | 8 | 0.25 |
| 11_465203 | 36 | 0.2778 | 51 | 0.4118 | 43 | 0.4419 | 42 | 0.4762 | 172 | 0.407 | 46 | 0.3696 | 38 | 0.3947 | 57 | 0.3158 | 45 | 0.4 | 186 | 0.3656 |
| 11_467246 | 2 | 0.5 | 5 | 0.2 | 5 | 0.4 | 3 | 0.3333 | 15 | 0.3333 | 4 | 0.25 | 4 | 0 | 1 | 0 | 3 | 0 | 12 | 0.0833 |
| 11_468710 | 3 | 0 | 2 | 0.5 | 4 | 0 | 5 | 0.4 | 14 | 0.2143 | 3 | 0 | 5 | 0.6 | 9 | 0.7778 | 6 | 0.1667 | 23 | 0.4783 |
| 11_468744 | 2 | 0 | 2 | 0 | 2 | 0.5 | 4 | 0.25 | 10 | 0.2 | 1 | 0 | 4 | 0 | 9 | 0.3333 | 4 | 0 | 18 | 0.1667 |
| 11_468744 | 4 | 0.5 | 2 | 1 | 2 | 1 | 5 | 0.6 | 13 | 0.6923 | 1 | 1 | 5 | 0.4 | 8 | 0.5 | 4 | 0.75 | 18 | 0.5556 |
| 11_468751 | 2 | 1 | 5 | 0.2 | 5 | 0.2 | 4 | 0.25 | 16 | 0.3125 | 6 | 0.1667 | 4 | 0 | 1 | 0 | 5 | 0.4 | 16 | 0.1875 |
| 11_468765 | 4 | 0 | 2 | 0 | 4 | 0.25 | 5 | 0 | 15 | 0.0667 | 2 | 0 | 6 | 0.5 | 7 | 0.2857 | 5 | 0.2 | 20 | 0.3 |
| 11_468766 | 4 | 0.25 | 2 | 0 | 3 | 0 | 4 | 0.25 | 13 | 0.1538 | 2 | 0 | 6 | 0 | 8 | 0.375 | 5 | 0.4 | 21 | 0.2381 |
| 11_468771 | 3 | 0 | 2 | 0 | 3 | 0 | 4 | 0.25 | 12 | 0.0833 | 1 | 1 | 5 | 0 | 8 | 0.125 | 4 | 0 | 18 | 0.1111 |
| 11_468776 | 2 | 0 | 2 | 0 | 1 | 0 | 5 | 0.2 | 10 | 0.1 | 1 | 0 | 5 | 0 | 10 | 0.1 | 4 | 0 | 20 | 0.05 |
| 11_468804 | 3 | 0 | 2 | 0 | 4 | 0.25 | 5 | 0 | 14 | 0.0714 | 1 | 0 | 6 | 0.1667 | 8 | 0.125 | 5 | 0 | 20 | 0.1 |
| 11_468863 | 2 | 0 | 2 | 0 | 2 | 0 | 5 | 0.2 | 11 | 0.0909 | NA | NA | 6 | 0.3333 | 4 | 0.25 | 1 | 0 | 11 | 0.2727 |
| 11_468871 | 3 | 1 | 2 | 0.5 | 2 | 1 | 5 | 0.6 | 12 | 0.75 | NA | NA | 5 | 0.4 | 4 | 0.25 | 3 | 0 | 12 | 0.25 |
| 11_471603 | 11 | 0.1818 | 7 | 0.1429 | 10 | 0.2 | 6 | 0.6667 | 34 | 0.2647 | 12 | 0.25 | 15 | 0.1333 | 8 | 0.25 | 9 | 0.1111 | 44 | 0.1818 |
| 11_471610 | 12 | 0.25 | 7 | 0.1429 | 10 | 0.1 | 6 | 0.3333 | 35 | 0.2 | 12 | 0 | 15 | 0 | 7 | 0.2857 | 8 | 0.125 | 42 | 0.0714 |
| 11_471688 | 7 | 0.1429 | 4 | 0.25 | 8 | 0.125 | 4 | 0 | 23 | 0.1304 | 7 | 0.1429 | 12 | 0.1667 | 7 | 0 | 7 | 0 | 33 | 0.0909 |
| 11_471774 | 6 | 0.1667 | 3 | 0.6667 | 5 | 0 | 4 | 0.25 | 18 | 0.2222 | 4 | 0 | 9 | 0.2222 | 6 | 0.1667 | 8 | 0.25 | 27 | 0.1852 |
| 11_471812 | 3 | 0.3333 | 1 | 1 | 4 | 0.25 | 3 | 1 | 11 | 0.5455 | 2 | 0.5 | 8 | 0.25 | 3 | 0.3333 | 7 | 0.5714 | 20 | 0.4 |
| 11_471822 | 3 | 0.6667 | 1 | 0 | 3 | 0 | 3 | 0 | 10 | 0.2 | 1 | 0 | 8 | 0 | 7 | 0.2857 | 8 | 0 | 24 | 0.0833 |
| 11_484322 | 8 | 0.25 | 6 | 0.6667 | 8 | 0.625 | 7 | 0.7143 | 29 | 0.5517 | 10 | 0.4 | 5 | 0.4 | 6 | 0.5 | 6 | 0.3333 | 27 | 0.4074 |
| 11_484327 | 8 | 0.125 | 4 | 0.5 | 13 | 0.6154 | 10 | 0.2 | 35 | 0.3714 | 11 | 0.2727 | 5 | 0.2 | 9 | 0.2222 | 8 | 0.625 | 33 | 0.3333 |
| 11_484328 | 11 | 0.1818 | 6 | 0.1667 | 12 | 0.1667 | 13 | 0 | 42 | 0.119 | 12 | 0 | 7 | 0 | 10 | 0.1 | 7 | 0.1429 | 36 | 0.0556 |
| 11_484333 | 8 | 0.5 | 6 | 0.5 | 13 | 0.4615 | 14 | 0.5 | 41 | 0.4878 | 8 | 0.5 | 8 | 0.5 | 11 | 0.3636 | 8 | 0.5 | 35 | 0.4571 |
| 11_484346 | 11 | 0.2727 | 6 | 0.3333 | 13 | 0.3077 | 11 | 0.1818 | 41 | 0.2683 | 12 | 0.25 | 5 | 0.2 | 12 | 0 | 8 | 0.125 | 37 | 0.1351 |
| 11_484357 | 12 | 0.25 | 6 | 0.1667 | 13 | 0.2308 | 12 | 0.1667 | 43 | 0.2093 | 12 | 0.0833 | 8 | 0.125 | 13 | 0.1538 | 8 | 0.125 | 41 | 0.122 |
| 11_484364 | 7 | 0.2857 | 3 | 1 | 9 | 0.2222 | 10 | 0.3 | 29 | 0.3448 | 5 | 0 | 5 | 0.2 | 7 | 0.4286 | 4 | 0 | 21 | 0.1905 |
| 11_486246 | 1 | 0 | 1 | 1 | 1 | 0 | NA | NA | 3 | 0.3333 | NA | NA | NA | NA | 1 | 0 | 1 | 1 | 2 | 0.5 |
| 11_488904 | 4 | 0.75 | 3 | 0 | 5 | 0.4 | 7 | 0.5714 | 19 | 0.4737 | 5 | 0.6 | 6 | 0.6667 | 6 | 0.3333 | 2 | 1 | 19 | 0.5789 |
| 11_488911 | 5 | 1 | 3 | 1 | 7 | 1 | 10 | 1 | 25 | 1 | 8 | 1 | 5 | 1 | 9 | 1 | 3 | 1 | 25 | 1 |
| 11_488931 | 4 | 0 | 4 | 0.25 | 6 | 0.1667 | 8 | 0.5 | 22 | 0.2727 | 6 | 0.1667 | 3 | 0.3333 | 10 | 0.3 | 4 | 0.25 | 23 | 0.2609 |
| 11_489134 | 2 | 0.5 | 3 | 0.3333 | NA | NA | 3 | 0 | 8 | 0.25 | 3 | 0.6667 | 2 | 0 | 1 | 0 | 3 | 0 | 9 | 0.2222 |
| 11_496660 | 6 | 0 | 9 | 0 | 4 | 0.75 | 8 | 0.125 | 27 | 0.1481 | 3 | 0 | 3 | 0 | 5 | 0 | 7 | 0.2857 | 18 | 0.1111 |
| 11_496938 | 2 | 0.5 | 7 | 0 | 2 | 0.5 | 8 | 0 | 19 | 0.1053 | 1 | 0 | 3 | 0.6667 | 3 | 0 | 3 | 0.3333 | 10 | 0.3 |
| 11_499205 | 11 | 0.3636 | 14 | 0.6429 | 13 | 0.1538 | 12 | 0.0833 | 50 | 0.32 | 9 | 0.1111 | 15 | 0.2 | 9 | 0.2222 | 5 | 0.4 | 38 | 0.2105 |
| 11_499210 | 18 | 0.2778 | 21 | 0.3333 | 16 | 0.375 | 20 | 0.3 | 75 | 0.32 | 12 | 0 | 20 | 0.4 | 10 | 0.1 | 12 | 0.0833 | 54 | 0.1852 |
| 11_499220 | 19 | 0.5263 | 17 | 0.5882 | 14 | 0.5714 | 18 | 0.7222 | 68 | 0.6029 | 11 | 0.4545 | 15 | 0.6 | 6 | 0.6667 | 9 | 0.2222 | 41 | 0.4878 |
| 11_499245 | 15 | 0.0667 | 19 | 0.0526 | 20 | 0.1 | 20 | 0.05 | 74 | 0.0676 | 17 | 0.1176 | 14 | 0 | 12 | 0.1667 | 11 | 0.1818 | 54 | 0.1111 |
| 11_500956 | 1 | 0 | NA | NA | 1 | 1 | 2 | 0.5 | 4 | 0.5 | 2 | 0.5 | 2 | 0.5 | 2 | 0 | 5 | 0.4 | 11 | 0.3636 |
| 11_500971 | 2 | 0 | NA | NA | 2 | 1 | 2 | 0 | 6 | 0.3333 | 4 | 0.5 | 2 | 0 | 2 | 0 | 4 | 0.25 | 12 | 0.25 |
| 11_500987 | 9 | 0.4444 | 6 | 0.3333 | 12 | 0.0833 | 6 | 0 | 33 | 0.2121 | 5 | 0 | 2 | 1 | 7 | 0.1429 | 5 | 0.2 | 19 | 0.2105 |
| 11_501007 | 8 | 0.875 | 12 | 1 | 10 | 0.9 | 7 | 0.5714 | 37 | 0.8649 | 7 | 1 | 3 | 0.6667 | 7 | 0.8571 | 6 | 0.8333 | 23 | 0.8696 |
| 11_501010 | 9 | 0.1111 | 12 | 0 | 13 | 0 | 8 | 0.25 | 42 | 0.0714 | 7 | 0.2857 | 4 | 0 | 7 | 0.1429 | 6 | 0.1667 | 24 | 0.1667 |
| 11_501012 | 8 | 1 | 10 | 1 | 14 | 1 | 6 | 1 | 38 | 1 | 7 | 1 | 4 | 1 | 7 | 1 | 4 | 0.75 | 22 | 0.9545 |
| 11_501014 | 11 | 0.6364 | 10 | 0.4 | 14 | 0.6429 | 7 | 0.7143 | 42 | 0.5952 | 6 | 0.8333 | 4 | 0.5 | 6 | 0.8333 | 5 | 1 | 21 | 0.8095 |
| 11_501016 | 11 | 0.9091 | 10 | 1 | 8 | 0.875 | 5 | 1 | 34 | 0.9412 | 5 | 0.8 | 3 | 0.6667 | 3 | 0.6667 | 5 | 1 | 16 | 0.8125 |
| 11_501021 | 11 | 1 | 10 | 1 | 10 | 1 | 8 | 1 | 39 | 1 | 7 | 1 | 3 | 1 | 7 | 1 | 5 | 1 | 22 | 1 |
| 11_501022 | 9 | 0.1111 | 10 | 0.5 | 13 | 0.2308 | 7 | 0 | 39 | 0.2308 | 5 | 0.4 | 3 | 0.3333 | 6 | 0.3333 | 3 | 0 | 17 | 0.2941 |
| 11_501025 | 12 | 0 | 11 | 0.2727 | 12 | 0.1667 | 7 | 0.1429 | 42 | 0.1429 | 7 | 0.1429 | 3 | 0 | 6 | 0.1667 | 5 | 0.2 | 21 | 0.1429 |
| 11_501027 | 10 | 0.4 | 11 | 0.2727 | 11 | 0.5455 | 5 | 0.6 | 37 | 0.4324 | 5 | 0.8 | 1 | 1 | 6 | 0.3333 | 4 | 0.25 | 16 | 0.5 |
| 11_501031 | 9 | 0.3333 | 12 | 0.5833 | 11 | 0.9091 | 7 | 0.4286 | 39 | 0.5897 | 6 | 0.6667 | 4 | 0.25 | 7 | 0.7143 | 5 | 0.8 | 22 | 0.6364 |
| 11_501037 | 3 | 0.3333 | 8 | 0 | 7 | 0.1429 | 7 | 0.1429 | 25 | 0.12 | 5 | 0.2 | 3 | 0 | 3 | 0 | 3 | 0 | 14 | 0.0714 |
| 11_501048 | 13 | 0.1538 | 8 | 0.375 | 9 | 0.1111 | 5 | 0.4 | 35 | 0.2286 | 9 | 0 | 6 | 0 | 5 | 0.2 | 6 | 0.1667 | 26 | 0.0769 |
| 11_501052 | 18 | 0.1667 | 20 | 0.1 | 13 | 0 | 8 | 0 | 59 | 0.0847 | 13 | 0 | 11 | 0.0909 | 5 | 0.2 | 10 | 0 | 39 | 0.0513 |
| 11_501055 | 18 | 0 | 19 | 0.0526 | 13 | 0 | 10 | 0.2 | 60 | 0.05 | 12 | 0.0833 | 11 | 0.0909 | 6 | 0 | 10 | 0.1 | 39 | 0.0769 |
| 11_501065 | 14 | 0.2857 | 14 | 0.1429 | 8 | 0.125 | 7 | 0.2857 | 43 | 0.2093 | 12 | 0.25 | 11 | 0.0909 | 3 | 0.3333 | 9 | 0 | 35 | 0.1429 |
| 11_501066 | 16 | 0.375 | 13 | 0.6923 | 11 | 0.3636 | 8 | 0.5 | 48 | 0.4792 | 11 | 0.8182 | 9 | 0.5556 | 3 | 0.6667 | 8 | 0.75 | 31 | 0.7097 |
| 11_501068 | 16 | 0.0625 | 14 | 0.2857 | 12 | 0 | 9 | 0.1111 | 51 | 0.1176 | 12 | 0.0833 | 11 | 0.0909 | 3 | 0.3333 | 10 | 0.2 | 36 | 0.1389 |
| 11_501078 | 15 | 0.5333 | 16 | 0.875 | 12 | 0.5833 | 7 | 0.7143 | 50 | 0.68 | 13 | 0.4615 | 11 | 0.9091 | 4 | 0.75 | 10 | 0.5 | 38 | 0.6316 |
| 11_501081 | 16 | 0.1875 | 15 | 0.0667 | 12 | 0.25 | 8 | 0 | 51 | 0.1373 | 11 | 0.4545 | 11 | 0.0909 | 5 | 0.2 | 9 | 0 | 36 | 0.1944 |
| 11_501085 | 6 | 0.5 | 4 | 0.25 | 7 | 0.5714 | 2 | 0.5 | 19 | 0.4737 | 6 | 0.5 | 4 | 0.25 | 1 | 1 | 3 | 0.3333 | 14 | 0.4286 |
| 11_501086 | 16 | 0.25 | 13 | 0.1538 | 10 | 0 | 8 | 0.25 | 47 | 0.1702 | 13 | 0.2308 | 11 | 0.2727 | 3 | 0 | 9 | 0.2222 | 36 | 0.2222 |
| 11_501092 | 13 | 0.7692 | 14 | 0.7143 | 11 | 0.6364 | 7 | 0.7143 | 45 | 0.7111 | 13 | 0.6923 | 8 | 0.75 | 4 | 0.75 | 9 | 0.7778 | 34 | 0.7353 |
| 11_501116 | 14 | 0.9286 | 12 | 0.8333 | 10 | 0.9 | 7 | 0.8571 | 43 | 0.8837 | 10 | 1 | 11 | 0.9091 | 3 | 1 | 9 | 0.6667 | 33 | 0.8788 |

| | | | | | | | | | | | | | | | | | | | | |
|---|---|---|---|---|---|---|---|---|---|---|---|---|---|---|---|---|---|---|---|---|
| 11_501147 | 11 | 0.5455 | 10 | 0.4 | 10 | 0.8 | 7 | 0.2857 | 38 | 0.5263 | 9 | 0.3333 | 8 | 0.625 | 3 | 0.3333 | 7 | 0.5714 | 27 | 0.4815 |
| 11_501176 | 10 | 0.1 | 9 | 0.1111 | 8 | 0 | 6 | 0.1667 | 33 | 0.0909 | 9 | 0.2222 | 7 | 0.2857 | 2 | 0 | 7 | 0 | 25 | 0.16 |
| 11_501408 | 11 | 0.3636 | 8 | 0.25 | 9 | 0.1111 | 6 | 0 | 34 | 0.2059 | 10 | 0.2 | 6 | 0.1667 | 2 | 0.5 | 5 | 0 | 23 | 0.1739 |
| 11_501447 | 7 | 0.1429 | 4 | 0.25 | 7 | 0.5714 | 5 | 0.2 | 23 | 0.3043 | 7 | 0 | 6 | 0.1667 | 2 | 0.5 | 6 | 0.3333 | 21 | 0.1905 |
| 11_501452 | 6 | 0.1667 | 2 | 0 | 8 | 0 | 2 | 0 | 18 | 0.0556 | 4 | 0.25 | 4 | 0 | 1 | 0 | 4 | 0 | 13 | 0.0769 |
| 11_501453 | 6 | 0.5 | 2 | 1 | 8 | 0.25 | 2 | 0 | 18 | 0.3889 | 3 | 0.3333 | 4 | 0.5 | NA | NA | 4 | 0.25 | 11 | 0.3636 |
| 11_501461 | 6 | 0.3333 | 2 | 0 | 8 | 0 | NA | NA | 16 | 0.125 | 3 | 0.3333 | 4 | 0.5 | NA | NA | 3 | 0.3333 | 10 | 0.4 |
| 11_501468 | 5 | 0.2 | 1 | 0 | 8 | 0 | 1 | 0 | 15 | 0.0667 | 4 | 0.25 | 4 | 0 | NA | NA | 4 | 0 | 12 | 0.0833 |
| 11_501481 | 3 | 0 | 1 | 0 | 8 | 0.125 | 1 | 0 | 13 | 0.0769 | 4 | 0.25 | 4 | 0 | NA | NA | 1 | 0 | 9 | 0.1111 |
| 11_501484 | 5 | 0.4 | 2 | 0.5 | 9 | 0.4444 | 2 | 0.5 | 18 | 0.4444 | 5 | 0.6 | 3 | 1 | 1 | 0 | 3 | 0.6667 | 12 | 0.6667 |
| 11_502034 | 18 | 0.0556 | 24 | 0.0417 | 16 | 0.0625 | 27 | 0.0741 | 85 | 0.0588 | 16 | 0.125 | 22 | 0.0455 | 26 | 0.1538 | 19 | 0 | 83 | 0.0843 |
| 11_503007 | 2 | 0 | 6 | 0.1667 | 1 | 1 | 4 | 0.25 | 13 | 0.2308 | 2 | 0.5 | 2 | 0 | 3 | 0.3333 | 4 | 0.25 | 11 | 0.2727 |
| 11_503035 | 1 | 0 | 5 | 0.2 | 1 | 0 | 4 | 0 | 11 | 0.0909 | 3 | 0 | 4 | 0.25 | 5 | 0.6 | 4 | 0.75 | 16 | 0.4375 |
| 11_505429 | 1 | 0 | 3 | 0 | 3 | 0.3333 | 3 | 0.6667 | 10 | 0.3 | NA | NA | 2 | 0.5 | 1 | 0 | 1 | 0 | 4 | 0.25 |
| 11_505556 | NA | NA | NA | NA | 2 | 0.5 | 2 | 0.5 | 4 | 0.5 | NA | NA | 3 | 0.3333 | NA | NA | 1 | 1 | 4 | 0.5 |
| 11_505669 | 1 | 0 | 2 | 0 | 2 | 0.5 | 2 | 0.5 | 7 | 0.2857 | 1 | 1 | NA | NA | 1 | 0 | NA | NA | 2 | 0.5 |
| 11_514178 | 2 | 0 | 3 | 0.3333 | 1 | 1 | 2 | 0 | 8 | 0.25 | 6 | 0.3333 | 1 | 1 | 1 | 0 | 2 | 1 | 10 | 0.5 |
| 11_514179 | 3 | 1 | 3 | 1 | 1 | 0 | 2 | 0 | 9 | 0.6667 | 6 | 0 | 1 | 0 | 1 | 1 | 3 | 0.6667 | 11 | 0.2727 |
| 11_514200 | 2 | 0 | 2 | 0 | 1 | 1 | 2 | 0.5 | 7 | 0.2857 | 7 | 0.1429 | 1 | 1 | 1 | 0 | 2 | 0 | 11 | 0.1818 |
| 11_514214 | 2 | 1 | 3 | 0.6667 | 1 | 1 | 2 | 0.5 | 8 | 0.75 | 7 | 0.8571 | 1 | 0 | 1 | 1 | 3 | 1 | 12 | 0.8333 |
| 11_514227 | 3 | 1 | 3 | 0 | NA | NA | 2 | 0 | 8 | 0.375 | 7 | 0.1429 | 1 | 1 | 1 | 1 | 3 | 1 | 12 | 0.5 |
| 11_514232 | 1 | 0 | 2 | 0.5 | 1 | 1 | 2 | 0.5 | 6 | 0.5 | 7 | 0.5714 | 1 | 0 | 1 | 1 | 1 | 1 | 10 | 0.6 |
| 11_514232 | 3 | 0.6667 | 3 | 0.6667 | 1 | 1 | 2 | 0.5 | 9 | 0.6667 | 5 | 0.4 | 1 | 1 | 1 | 1 | 3 | 0.6667 | 10 | 0.6 |
| 11_514239 | 2 | 0.5 | 3 | 1 | 1 | 0 | 2 | 0 | 8 | 0.5 | 7 | 0.2857 | 1 | 0 | 1 | 0 | 3 | 0.6667 | 12 | 0.3333 |
| 11_514252 | 3 | 0 | 2 | 0 | 1 | 1 | 2 | 0 | 8 | 0.125 | 7 | 0 | 1 | 0 | 1 | 0 | 3 | 0.6667 | 12 | 0.1667 |
| 11_514255 | 3 | 1 | 1 | 1 | NA | NA | 1 | 1 | 5 | 1 | 6 | 1 | 1 | 1 | 1 | 1 | 2 | 1 | 10 | 1 |
| 11_514256 | 2 | 0.5 | 2 | 0 | NA | NA | 2 | 0 | 6 | 0.1667 | 5 | 0.2 | 1 | 0 | 1 | 0 | 1 | 0 | 8 | 0.125 |
| 11_514270 | 2 | 0 | 1 | 0 | NA | NA | 1 | 1 | 4 | 0.25 | 7 | 0.2857 | 1 | 0 | 1 | 0 | 2 | 0 | 11 | 0.1818 |
| 11_514343 | 3 | 0.3333 | 1 | 0 | NA | NA | 2 | 0 | 6 | 0.1667 | 6 | 0.1667 | 1 | 0 | 1 | 1 | 2 | 0.5 | 10 | 0.3 |
| 11_514351 | 3 | 0.3333 | NA | NA | NA | NA | 2 | 0 | 5 | 0.2 | 6 | 0.5 | 1 | 0 | 1 | 1 | 2 | 0 | 10 | 0.4 |
| 11_514364 | 3 | 0.3333 | 1 | 0 | NA | NA | 2 | 0 | 6 | 0.1667 | 4 | 0.25 | 1 | 0 | 1 | 0 | 2 | 0 | 8 | 0.125 |
| 11_514460 | 2 | 0.5 | NA | NA | NA | NA | 2 | 0.5 | 4 | 0.5 | 6 | 0.3333 | 1 | 0 | 1 | 1 | 2 | 1 | 10 | 0.5 |
| 11_514472 | 2 | 1 | NA | NA | NA | NA | 1 | 1 | 3 | 1 | 5 | 0.8 | 1 | 1 | 1 | 1 | 1 | 0 | 8 | 0.75 |
| 11_514474 | 2 | 0.5 | NA | NA | NA | NA | 2 | 0 | 4 | 0.25 | 7 | 0.7143 | 1 | 0 | 1 | 0 | 1 | 0 | 10 | 0.5 |
| 11_514495 | 2 | 1 | NA | NA | NA | NA | 2 | 0.5 | 4 | 0.75 | 4 | 0 | NA | NA | 1 | 1 | NA | NA | 5 | 0.2 |
| 11_514505 | 2 | 0 | NA | NA | NA | NA | 2 | 0.5 | 4 | 0.25 | 5 | 0.4 | 1 | 0 | 1 | 0 | 1 | 0 | 8 | 0.25 |
| 11_514508 | 2 | 0.5 | NA | NA | NA | NA | 2 | 0 | 4 | 0.25 | 5 | 0.2 | 1 | 0 | 1 | 0 | 1 | 0 | 8 | 0.125 |
| 11_515186 | 1 | 1 | NA | NA | NA | NA | 1 | 0 | 2 | 0.5 | 5 | 0.4 | 1 | 1 | 1 | 1 | 1 | 0 | 8 | 0.5 |
| 11_522971 | NA | NA | 1 | 0 | 3 | 0.3333 | 1 | 1 | 5 | 0.4 | 2 | 0.5 | NA | NA | NA | NA | NA | NA | 2 | 0.5 |
| 11_522984 | NA | NA | 1 | 0 | 3 | 0.3333 | 1 | 1 | 5 | 0.4 | 2 | 0.5 | NA | NA | NA | NA | NA | NA | 2 | 0.5 |
| 11_523084 | NA | NA | NA | NA | NA | NA | 1 | 1 | 1 | 1 | 2 | 1 | NA | NA | NA | NA | NA | NA | 2 | 1 |
| 11_523105 | NA | NA | NA | NA | 1 | 1 | 1 | 1 | 2 | 1 | 1 | 1 | NA | NA | NA | NA | NA | NA | 1 | 1 |
| 11_544512 | NA | NA | 1 | 1 | NA | NA | NA | NA | 1 | 1 | NA | NA | 1 | 1 | NA | NA | NA | NA | 1 | 1 |
| 11_547471 | 10 | 0 | 21 | 0.0952 | 11 | 0.1818 | 17 | 0.0588 | 59 | 0.0847 | 13 | 0.1538 | 22 | 0.0909 | 19 | 0.1053 | 12 | 0.1667 | 66 | 0.1212 |
| 11_547479 | 9 | 0.4444 | 20 | 0.35 | 11 | 0.1818 | 18 | 0.3333 | 58 | 0.3276 | 13 | 0.0769 | 22 | 0.2273 | 21 | 0.4286 | 12 | 0 | 68 | 0.2206 |
| 11_547483 | 9 | 0 | 19 | 0.1053 | 12 | 0 | 16 | 0.1875 | 56 | 0.0893 | 13 | 0.0769 | 21 | 0.1905 | 20 | 0.15 | 14 | 0.1429 | 68 | 0.1471 |
| 11_547486 | 8 | 1 | 18 | 1 | 13 | 0.9231 | 14 | 0.9286 | 53 | 0.9623 | 12 | 1 | 19 | 1 | 17 | 0.9412 | 11 | 1 | 59 | 0.9831 |
| 11_547492 | 8 | 0 | 21 | 0.0476 | 12 | 0.0833 | 16 | 0.1875 | 57 | 0.0877 | 13 | 0.1538 | 19 | 0.2105 | 19 | 0.1053 | 12 | 0.0833 | 63 | 0.1429 |
| 11_547512 | 9 | 0.8889 | 20 | 0.9 | 13 | 1 | 15 | 0.9333 | 57 | 0.9298 | 13 | 1 | 16 | 0.9375 | 19 | 0.8947 | 8 | 1 | 56 | 0.9464 |
| 11_547521 | 9 | 1 | 22 | 0.8182 | 10 | 1 | 16 | 1 | 57 | 0.9298 | 10 | 0.8 | 16 | 0.9375 | 17 | 0.8235 | 9 | 0.7778 | 52 | 0.8462 |
| 11_547565 | 14 | 0.1429 | 26 | 0.0385 | 15 | 0.0667 | 19 | 0.0526 | 74 | 0.0676 | 14 | 0.0714 | 27 | 0.0741 | 23 | 0.0435 | 15 | 0.0667 | 79 | 0.0633 |
| 11_548116 | 2 | 0.5 | 3 | 0.6667 | NA | NA | 2 | 0 | 7 | 0.4286 | 1 | 1 | 1 | 0 | NA | NA | 1 | 0 | 3 | 0.3333 |
| 11_548117 | 2 | 0 | 2 | 0.5 | NA | NA | 2 | 0.5 | 6 | 0.3333 | 1 | 1 | 1 | 1 | 1 | 1 | 1 | 1 | 4 | 1 |
| 11_548138 | 3 | 0.6667 | 3 | 1 | NA | NA | 2 | 0.5 | 8 | 0.75 | NA | NA | 1 | 1 | 1 | 1 | NA | NA | 2 | 1 |
| 11_548144 | 2 | 0 | 3 | 0.3333 | NA | NA | 2 | 0 | 7 | 0.1429 | NA | NA | 1 | 1 | 1 | 0 | 1 | 0 | 3 | 0.3333 |
| 11_565423 | 1 | 0 | NA | NA | NA | NA | 2 | 0.5 | 3 | 0.3333 | NA | NA | 1 | 1 | 1 | 0 | NA | NA | 2 | 0.5 |
| 11_565448 | 11 | 0.2727 | 13 | 0.0769 | 14 | 0.1429 | 13 | 0.0769 | 51 | 0.1373 | 12 | 0.1667 | 10 | 0.3 | 16 | 0.3125 | 9 | 0.1111 | 47 | 0.234 |
| 11_565453 | 9 | 0.4444 | 14 | 0.5714 | 17 | 0.6471 | 11 | 0.5455 | 51 | 0.5686 | 13 | 0.5385 | 16 | 0.8125 | 17 | 0.7059 | 10 | 0.9 | 56 | 0.7321 |
| 11_565460 | 9 | 0.4444 | 12 | 0.25 | 17 | 0.2941 | 13 | 0.2308 | 51 | 0.2941 | 12 | 0.25 | 13 | 0.2308 | 14 | 0.2143 | 8 | 0.125 | 47 | 0.2128 |
| 11_566507 | 5 | 0 | 9 | 0.3333 | 4 | 0.25 | 13 | 0.3077 | 31 | 0.2581 | 9 | 0 | 7 | 0.1429 | 11 | 0 | 2 | 0.5 | 29 | 0.069 |
| 11_566549 | 5 | 0.2 | 9 | 0.5556 | 4 | 0.25 | 11 | 0.7273 | 29 | 0.5172 | 9 | 0.4444 | 6 | 0.8333 | 12 | 0.25 | 3 | 0.3333 | 30 | 0.4333 |
| 11_566551 | 6 | 0.1667 | 9 | 0.1111 | 3 | 0 | 11 | 0.3636 | 29 | 0.2069 | 9 | 0.1111 | 6 | 0.1667 | 10 | 0.1 | 3 | 0 | 28 | 0.1071 |
| 11_566562 | 4 | 0 | 7 | 0.1429 | 2 | 0.5 | 7 | 0.4286 | 20 | 0.25 | 7 | 0.2857 | 4 | 0.25 | 12 | 0.1667 | 3 | 0.3333 | 26 | 0.2308 |
| 11_566567 | 5 | 0.2 | 8 | 0.25 | 4 | 0 | 9 | 0.3333 | 26 | 0.2308 | 8 | 0.5 | 5 | 0.2 | 11 | 0 | 2 | 0.5 | 26 | 0.2308 |
| 11_566568 | 6 | 0.8333 | 9 | 0.7778 | 4 | 0.5 | 8 | 1 | 27 | 0.8148 | 8 | 0.875 | 3 | 1 | 10 | 0.8 | 2 | 1 | 23 | 0.8696 |

| | | | | | | | | | | | | | | | | | | | | |
|---|---|---|---|---|---|---|---|---|---|---|---|---|---|---|---|---|---|---|---|---|
| 11_5665722 | 1 | 0 | 3 | 0 | 2 | 0 | 5 | 0.4 | 11 | 0.1818 | 6 | 0 | 3 | 0.3333 | 6 | 0 | 2 | 0 | 17 | 0.0588 |
| 11_5665728 | 4 | 0.5 | 6 | 0.1667 | 1 | 0 | 6 | 0.5 | 17 | 0.3529 | 7 | 0.1429 | 2 | 0 | 10 | 0.5 | 2 | 0 | 21 | 0.2857 |
| 11_5665888 | 4 | 1 | 6 | 1 | 3 | 0.6667 | 10 | 1 | 23 | 0.9565 | 7 | 1 | 4 | 1 | 10 | 0.9 | 3 | 0.6667 | 24 | 0.9167 |
| 11_5665965 | 3 | 0.6667 | 7 | 0.5714 | 3 | 0.6667 | 10 | 0.5 | 23 | 0.5652 | 8 | 0.75 | 3 | 0.6667 | 10 | 0.3 | 3 | 0 | 24 | 0.4583 |
| 11_5666004 | 5 | 0.2 | 9 | 0.2222 | 1 | 0 | 10 | 0.4 | 25 | 0.28 | 7 | 0.1429 | 3 | 0.3333 | 10 | 0.1 | 2 | 0.5 | 22 | 0.1818 |
| 11_5666085 | 5 | 1 | 9 | 0.7778 | 3 | 1 | 10 | 1 | 27 | 0.9259 | 7 | 0.8571 | 4 | 1 | 9 | 0.8889 | 3 | 1 | 23 | 0.913 |
| 11_5666159 | 1 | 1 | 6 | 1 | NA | NA | 6 | 1 | 13 | 1 | 5 | 1 | 3 | 1 | 8 | 1 | 2 | 1 | 18 | 1 |
| 11_5666169 | 5 | 0 | 9 | 0.2222 | 3 | 0 | 9 | 0.2222 | 26 | 0.1538 | 8 | 0.25 | 4 | 0.75 | 9 | 0.1111 | 3 | 0.3333 | 24 | 0.2917 |
| 11_5666211 | 5 | 1 | 8 | 1 | 3 | 1 | 7 | 1 | 23 | 1 | 5 | 1 | 4 | 0.75 | 6 | 1 | 3 | 1 | 18 | 0.9444 |
| 11_5666241 | 3 | 0.3333 | 8 | 0.5 | 2 | 0 | 8 | 0.25 | 21 | 0.3333 | 7 | 0 | 4 | 0.25 | 8 | 0.625 | 3 | 0.3333 | 22 | 0.3182 |
| 11_5666510 | 4 | 1 | 7 | 1 | 3 | 1 | 5 | 1 | 19 | 1 | 5 | 1 | 3 | 1 | 4 | 1 | 1 | 1 | 13 | 1 |
| 11_5666664 | 4 | 1 | 6 | 1 | 3 | 1 | 5 | 1 | 18 | 1 | 6 | 1 | 4 | 1 | 7 | 0.8571 | 2 | 1 | 19 | 0.9474 |
| 11_5666681 | 4 | 1 | 6 | 0.6667 | 2 | 0.5 | 3 | 1 | 15 | 0.8 | 5 | 0.6 | 3 | 0.6667 | 3 | 1 | 1 | 0 | 12 | 0.6667 |
| 11_5666687 | 4 | 0.25 | 5 | 0.2 | 3 | 0.6667 | 3 | 0.3333 | 15 | 0.3333 | 6 | 0.3333 | 2 | 0 | 4 | 0.5 | 2 | 0 | 14 | 0.2857 |
| 11_5666704 | 4 | 0.75 | 7 | 0.8571 | 3 | 1 | 5 | 1 | 19 | 0.8947 | 6 | 1 | 4 | 1 | 6 | 1 | 2 | 1 | 18 | 1 |
| 11_5666717 | 2 | 0 | 7 | 0.1429 | 3 | 0.3333 | 4 | 0.25 | 16 | 0.1875 | 5 | 0.4 | 4 | 0.25 | 5 | 0.2 | 2 | 0.5 | 16 | 0.3125 |
| 11_5667045 | 4 | 0 | 6 | 0.3333 | 3 | 0.3333 | 5 | 0.2 | 18 | 0.2222 | 6 | 0.3333 | 3 | 0.6667 | 4 | 0 | 2 | 0 | 15 | 0.2667 |
| 11_5667072 | 4 | 1 | 6 | 1 | 3 | 1 | 5 | 1 | 18 | 1 | 6 | 1 | 3 | 1 | 4 | 1 | 2 | 1 | 15 | 1 |
| 11_5667117 | 3 | 0.3333 | 6 | 0.6667 | 3 | 0.6667 | 5 | 0.8 | 17 | 0.6471 | 6 | 0.5 | 3 | 0.6667 | 4 | 0.75 | 2 | 0.5 | 15 | 0.6 |
| 11_5667198 | 3 | 0 | 7 | 0.8571 | 3 | 0.3333 | 5 | 0 | 18 | 0.3889 | 4 | 0 | 1 | 1 | 3 | 0 | 1 | 0 | 9 | 0.1111 |
| 11_5667382 | 2 | 0 | 6 | 0.3333 | 2 | 0 | 5 | 0.4 | 15 | 0.2667 | 5 | 0 | 1 | 0 | 3 | 1 | 1 | 1 | 10 | 0.4 |
| 11_5667447 | 4 | 0 | 5 | 0.4 | 3 | 0 | 5 | 0.2 | 17 | 0.1765 | 4 | 0.25 | 1 | 0 | 2 | 0.5 | 1 | 0 | 8 | 0.25 |
| 11_5667459 | 3 | 0.3333 | 6 | 0.5 | 3 | 1 | 5 | 0.6 | 17 | 0.5882 | 4 | 0 | 1 | 1 | 2 | 0 | NA | NA | 7 | 0.1429 |
| 11_5667495 | 4 | 0.25 | 6 | 0.5 | 3 | 0 | 5 | 0.8 | 18 | 0.4444 | 5 | 1 | 1 | 0 | 1 | 1 | NA | NA | 7 | 0.8571 |
| 11_5667576 | 4 | 0.25 | 6 | 0.6667 | 3 | 0 | 4 | 0.5 | 17 | 0.4118 | 4 | 0.75 | 1 | 0 | 3 | 0.6667 | NA | NA | 8 | 0.625 |
| 11_5667594 | 4 | 1 | 6 | 1 | 3 | 1 | 4 | 1 | 17 | 1 | 4 | 1 | 1 | 1 | 2 | 1 | NA | NA | 7 | 1 |
| 11_5667708 | 3 | 0.3333 | 6 | 0 | 3 | 0.3333 | 4 | 0 | 16 | 0.125 | 3 | 0 | 1 | 1 | 2 | 0 | NA | NA | 6 | 0.1667 |
| 11_5668128 | 4 | 0.25 | 3 | 0.3333 | 2 | 1 | 3 | 0.6667 | 12 | 0.5 | 3 | 0.6667 | 1 | 0 | 2 | 1 | NA | NA | 6 | 0.6667 |
| 11_5668185 | 3 | 0 | 2 | 0.5 | 2 | 0 | 2 | 0.5 | 9 | 0.2222 | 3 | 0 | 1 | 1 | 2 | 0.5 | NA | NA | 6 | 0.3333 |
| 11_5711982 | 2 | 0.5 | 1 | 0 | 1 | 0 | 2 | 0 | 6 | 0.1667 | 1 | 0 | 1 | 0 | 1 | 1 | NA | NA | 3 | 0.3333 |
| 11_5730299 | 4 | 0.75 | 2 | 0 | 1 | 0 | NA | NA | 7 | 0.4286 | 3 | 0.3333 | NA | NA | NA | NA | NA | NA | 3 | 0.3333 |
| 11_5730318 | 3 | 0 | 2 | 0.5 | 2 | 0.5 | NA | NA | 7 | 0.2857 | 2 | 0.5 | NA | NA | 1 | 0 | 2 | 0.5 | 5 | 0.4 |
| 11_5730330 | 2 | 1 | 2 | 0.5 | 2 | 0.5 | NA | NA | 6 | 0.6667 | 2 | 1 | NA | NA | 1 | 1 | 1 | 1 | 4 | 1 |
| 11_5730372 | 2 | 0 | 2 | 0.5 | 1 | 0 | NA | NA | 5 | 0.2 | 3 | 0.3333 | NA | NA | NA | NA | 2 | 0 | 5 | 0.2 |
| 11_5730434 | 2 | 0 | 1 | 0 | 2 | 0.5 | NA | NA | 5 | 0.2 | 3 | 0 | NA | NA | 1 | 0 | 2 | 0.5 | 6 | 0.1667 |
| 11_5730532 | 1 | 1 | 1 | 0 | NA | NA | NA | NA | 2 | 0.5 | 2 | 1 | NA | NA | 2 | 0.5 | 1 | 1 | 5 | 0.8 |
| 11_5730607 | 1 | 0 | 2 | 0.5 | 2 | 1 | NA | NA | 5 | 0.6 | 3 | 1 | NA | NA | 1 | 1 | 2 | 0.5 | 6 | 0.8333 |
| 11_5730633 | 1 | 0 | 1 | 1 | 1 | 1 | NA | NA | 3 | 0.6667 | 3 | 1 | NA | NA | 1 | 1 | 2 | 1 | 6 | 1 |
| 11_5730688 | 1 | 1 | 1 | 1 | 2 | 1 | NA | NA | 4 | 1 | 2 | 1 | NA | NA | 2 | 1 | 1 | 1 | 5 | 1 |
| 11_5730802 | 1 | 1 | 1 | 0 | 1 | 0 | NA | NA | 3 | 0.3333 | 1 | 1 | NA | NA | 2 | 0 | 1 | 0 | 4 | 0.25 |
| 11_5734038 | 1 | 1 | NA | NA | NA | NA | NA | NA | 1 | 1 | 1 | 1 | NA | NA | NA | NA | 2 | 0 | 3 | 0.3333 |
| 11_5734119 | 1 | 1 | NA | NA | NA | NA | NA | NA | 1 | 1 | 1 | 0 | 1 | 0 | NA | NA | 2 | 0.5 | 4 | 0.25 |
| 11_5734671 | 1 | 1 | 1 | 0 | 1 | 1 | NA | NA | 3 | 0.6667 | 2 | 1 | 1 | 1 | NA | NA | 2 | 0.5 | 5 | 0.8 |
| 11_5750096 | 11 | 1 | 8 | 0.625 | 6 | 1 | 13 | 1 | 38 | 0.9211 | 3 | 1 | 8 | 1 | 12 | 0.8333 | 7 | 1 | 30 | 0.9333 |
| 11_5750108 | 10 | 0.2 | 8 | 0 | 5 | 0.2 | 12 | 0.3333 | 35 | 0.2 | 4 | 0 | 12 | 0.25 | 17 | 0.1176 | 9 | 0.1111 | 42 | 0.1429 |
| 11_5750292 | 10 | 0.4 | 9 | 0.2222 | 6 | 0 | 11 | 0.4545 | 36 | 0.3056 | 6 | 0.3333 | 12 | 0.3333 | 16 | 0.125 | 11 | 0.1818 | 45 | 0.2222 |
| 11_5750313 | 10 | 0.4 | 11 | 0.5455 | 6 | 0.6667 | 12 | 0.6667 | 39 | 0.5641 | 5 | 0.4 | 10 | 0.7 | 17 | 0.4118 | 8 | 0.625 | 40 | 0.525 |
| 11_5750420 | 11 | 0.2727 | 11 | 0.0909 | 6 | 0 | 14 | 0.4286 | 42 | 0.2381 | 6 | 0.3333 | 11 | 0.4545 | 20 | 0.1 | 12 | 0.3333 | 49 | 0.2653 |
| 11_5750939 | 2 | 0.5 | 4 | 0.25 | 1 | 1 | 3 | 0 | 10 | 0.3 | 2 | 0 | 5 | 0.2 | 6 | 0.1667 | 2 | 0.5 | 15 | 0.2 |
| 11_5751084 | NA | NA | NA | NA | NA | NA | 1 | 1 | 1 | 1 | 1 | 1 | NA | NA | NA | NA | NA | NA | 1 | 1 |
| 11_5753121 | 2 | 1 | 1 | 1 | 3 | 0.3333 | 3 | 0 | 9 | 0.4444 | 2 | 0.5 | 1 | 0 | 6 | 0 | 2 | 0.5 | 11 | 0.1818 |
| 11_5753465 | 3 | 0.3333 | 1 | 1 | 2 | 0 | 3 | 0.3333 | 9 | 0.3333 | 3 | 0 | 1 | 1 | 7 | 0.2857 | 2 | 0 | 13 | 0.2308 |
| 11_5778186 | 1 | 0 | 2 | 1 | NA | NA | 1 | 1 | 4 | 0.75 | NA | NA | NA | NA | 2 | 1 | NA | NA | 2 | 1 |
| 11_5778240 | 1 | 0 | 3 | 0.3333 | NA | NA | 2 | 0 | 6 | 0.1667 | NA | NA | NA | NA | 2 | 0.5 | NA | NA | 2 | 0.5 |
| 11_5778369 | 1 | 1 | 3 | 1 | NA | NA | 2 | 1 | 6 | 1 | NA | NA | NA | NA | 2 | 1 | NA | NA | 2 | 1 |
| 11_5778708 | 1 | 1 | 3 | 1 | NA | NA | 1 | 1 | 5 | 1 | NA | NA | NA | NA | 2 | 0.5 | NA | NA | 2 | 0.5 |
| 11_5778800 | 1 | 1 | 3 | 1 | NA | NA | 1 | 1 | 5 | 1 | NA | NA | NA | NA | 2 | 1 | NA | NA | 2 | 1 |
| 11_5778889 | 1 | 1 | 3 | 1 | NA | NA | 1 | 1 | 5 | 1 | NA | NA | NA | NA | 1 | 1 | NA | NA | 1 | 1 |
| 11_5779028 | 1 | 0 | 2 | 1 | NA | NA | 1 | 1 | 4 | 0.75 | NA | NA | NA | NA | 1 | 1 | NA | NA | 1 | 1 |
| 11_5806458 | NA | NA | 1 | 1 | NA | NA | NA | NA | 1 | 1 | 1 | 1 | NA | NA | NA | NA | NA | NA | 1 | 1 |
| 11_5821344 | 5 | 0.2 | 2 | 0.5 | 1 | 1 | NA | NA | 8 | 0.375 | 3 | 1 | NA | NA | 1 | 0 | 2 | 0 | 6 | 0.5 |
| 11_5821679 | 9 | 0.5556 | 3 | 0.6667 | 2 | 0 | NA | NA | 14 | 0.5 | 2 | 0.5 | NA | NA | 1 | 1 | 3 | 0.6667 | 6 | 0.6667 |
| 11_5821727 | 8 | 0.5 | 3 | 0.3333 | 1 | 1 | NA | NA | 12 | 0.5 | 2 | 0.5 | 1 | 1 | 1 | 0 | 3 | 0.3333 | 7 | 0.4286 |
| 11_5821785 | 9 | 0.2222 | 3 | 0.6667 | 2 | 0.5 | NA | NA | 14 | 0.3571 | 3 | 0.3333 | 1 | 0 | 1 | 0 | 3 | 0.3333 | 8 | 0.25 |
| 11_5821833 | 9 | 0.7778 | 2 | 1 | 1 | 1 | NA | NA | 12 | 0.8333 | 2 | 0.5 | 1 | 1 | NA | NA | 3 | 1 | 6 | 0.8333 |

| | | | | | | | | | | | | | | | | | | | | |
|---|---|---|---|---|---|---|---|---|---|---|---|---|---|---|---|---|---|---|---|---|
| 11_5821842 | 9 | 0.5556 | 2 | 1 | 2 | 1 | NA | NA | 13 | 0.6923 | 3 | 0 | 1 | 0 | 1 | 1 | 2 | 0.5 | 7 | 0.2857 |
| 11_5821982 | 12 | 0.9167 | 4 | 1 | 3 | 0.6667 | NA | NA | 19 | 0.8947 | 3 | 1 | 1 | 1 | 1 | 0 | 3 | 0.6667 | 8 | 0.75 |
| 11_5821993 | 11 | 0.4545 | 3 | 0 | 2 | 0.5 | NA | NA | 16 | 0.375 | 3 | 1 | 1 | 0 | 1 | 1 | 1 | 0 | 6 | 0.6667 |
| 11_5905839 | 4 | 0.25 | 1 | 0 | 4 | 0 | 2 | 0 | 11 | 0.0909 | 1 | 0 | NA | NA | 4 | 0.25 | 5 | 0 | 10 | 0.1 |
| 11_5916220 | 10 | 0.4 | 3 | 0.3333 | 9 | 0.3333 | 4 | 0.5 | 26 | 0.3846 | 5 | 0.6 | 4 | 0.5 | 5 | 0.2 | 8 | 0.375 | 22 | 0.4091 |
| 11_5916831 | 2 | 0.5 | 6 | 0 | 7 | 0.1429 | 4 | 0 | 19 | 0.1053 | 8 | 0.375 | 3 | 0 | 6 | 0.1667 | 11 | 0.1818 | 28 | 0.2143 |
| 11_5916942 | 2 | 0 | 6 | 0 | 7 | 0 | 4 | 0.25 | 19 | 0.0526 | 8 | 0.125 | 5 | 0.2 | 6 | 0.1667 | 11 | 0 | 30 | 0.1 |
| 11_5916998 | NA | NA | 1 | 1 | NA | NA | NA | NA | 1 | 1 | NA | NA | NA | NA | 1 | 1 | NA | NA | 1 | 1 |
| 11_5917396 | 1 | 0 | 6 | 0 | 8 | 0 | 3 | 0.3333 | 18 | 0.0556 | 9 | 0.1111 | 2 | 0 | 5 | 0.2 | 11 | 0.0909 | 27 | 0.1111 |
| 11_5919898 | 77 | 0.1169 | 66 | 0.1212 | 60 | 0.1 | 59 | 0.1864 | 262 | 0.1298 | 56 | 0.1786 | 62 | 0.2742 | 82 | 0.0976 | 70 | 0.2286 | 270 | 0.1889 |
| 11_5920022 | 85 | 0.8941 | 77 | 0.8961 | 65 | 0.8923 | 73 | 0.8904 | 300 | 0.8933 | 62 | 0.9355 | 68 | 0.8824 | 78 | 0.8846 | 74 | 0.8784 | 282 | 0.8936 |
| 11_5920044 | 65 | 0.2 | 62 | 0.1774 | 59 | 0.2034 | 62 | 0.2742 | 248 | 0.2137 | 50 | 0.26 | 57 | 0.1579 | 71 | 0.1972 | 52 | 0.25 | 230 | 0.213 |
| 11_5920081 | 92 | 0.837 | 82 | 0.7561 | 91 | 0.7692 | 81 | 0.8025 | 346 | 0.7919 | 63 | 0.8413 | 90 | 0.8556 | 101 | 0.802 | 73 | 0.8904 | 327 | 0.844 |
| 11_5920109 | 97 | 0.732 | 84 | 0.7143 | 88 | 0.6932 | 86 | 0.6395 | 355 | 0.6958 | 61 | 0.6885 | 86 | 0.7442 | 91 | 0.7582 | 79 | 0.7722 | 317 | 0.7445 |
| 11_5920130 | 95 | 0.2316 | 94 | 0.266 | 90 | 0.2556 | 91 | 0.2088 | 370 | 0.2405 | 71 | 0.2394 | 91 | 0.1868 | 102 | 0.1863 | 86 | 0.3023 | 350 | 0.2257 |
| 11_5920171 | 95 | 0.2105 | 81 | 0.1481 | 80 | 0.25 | 89 | 0.191 | 345 | 0.2 | 63 | 0.1746 | 84 | 0.2619 | 89 | 0.2472 | 77 | 0.2597 | 313 | 0.2396 |
| 11_5920186 | 90 | 0.4333 | 90 | 0.5 | 85 | 0.5176 | 79 | 0.3924 | 344 | 0.4622 | 63 | 0.4444 | 79 | 0.4684 | 91 | 0.4835 | 67 | 0.5373 | 300 | 0.4833 |
| 11_5920211 | 97 | 0.0722 | 89 | 0.0674 | 89 | 0.0899 | 90 | 0.0667 | 365 | 0.074 | 68 | 0.0735 | 84 | 0.0357 | 98 | 0.102 | 74 | 0.0946 | 324 | 0.0772 |
| 11_5920337 | 66 | 0.1061 | 61 | 0.0492 | 57 | 0.0526 | 69 | 0.0725 | 253 | 0.0711 | 41 | 0.122 | 70 | 0.0714 | 77 | 0.0649 | 63 | 0.0794 | 251 | 0.0797 |
| 11_5920408 | 68 | 0.0441 | 61 | 0.082 | 65 | 0.0462 | 65 | 0.0308 | 259 | 0.0502 | 55 | 0.0364 | 66 | 0.0758 | 67 | 0.0448 | 55 | 0.0545 | 243 | 0.0535 |
| 11_5920431 | 86 | 0.1512 | 78 | 0.1154 | 75 | 0.1467 | 76 | 0.1316 | 315 | 0.1365 | 68 | 0.1324 | 79 | 0.1266 | 93 | 0.0753 | 70 | 0.1286 | 310 | 0.1129 |
| 11_5920461 | 71 | 0.3944 | 69 | 0.4493 | 61 | 0.3607 | 68 | 0.3382 | 269 | 0.3866 | 57 | 0.4035 | 72 | 0.4028 | 82 | 0.3659 | 54 | 0.463 | 265 | 0.4038 |
| 11_5920514 | 87 | 0.1149 | 72 | 0.1806 | 69 | 0.1449 | 69 | 0.1159 | 297 | 0.138 | 61 | 0.2131 | 72 | 0.2778 | 89 | 0.2247 | 62 | 0.2097 | 284 | 0.2324 |
| 11_5920755 | 74 | 0.0541 | 67 | 0.0746 | 62 | 0.0645 | 61 | 0.0656 | 264 | 0.0644 | 49 | 0.1224 | 56 | 0.0357 | 75 | 0.0667 | 55 | 0.0545 | 235 | 0.0681 |
| 11_5920809 | 85 | 0.3765 | 66 | 0.2576 | 65 | 0.2462 | 64 | 0.3281 | 280 | 0.3071 | 60 | 0.3833 | 56 | 0.3393 | 80 | 0.3125 | 61 | 0.3607 | 257 | 0.3463 |
| 11_5921031 | 74 | 0.7838 | 46 | 0.7826 | 51 | 0.7843 | 46 | 0.7826 | 217 | 0.7834 | 47 | 0.8511 | 49 | 0.7551 | 59 | 0.7966 | 41 | 0.8293 | 196 | 0.8061 |
| 11_5921225 | 75 | 0.2667 | 52 | 0.2115 | 56 | 0.1786 | 57 | 0.1404 | 240 | 0.2042 | 51 | 0.2549 | 42 | 0.2143 | 67 | 0.1493 | 42 | 0.2381 | 202 | 0.2079 |
| 11_5921491 | 72 | 0.1667 | 46 | 0.2391 | 51 | 0.1569 | 50 | 0.1 | 219 | 0.1644 | 42 | 0.119 | 42 | 0.1429 | 60 | 0.15 | 43 | 0.186 | 187 | 0.1497 |
| 11_5921585 | 67 | 0.1045 | 43 | 0.0698 | 55 | 0.1273 | 46 | 0.0652 | 211 | 0.0948 | 38 | 0.1842 | 46 | 0.1087 | 59 | 0.1186 | 40 | 0.1 | 183 | 0.1257 |
| 11_5921696 | 64 | 0.0625 | 45 | 0.0444 | 46 | 0.0217 | 47 | 0.0851 | 202 | 0.0545 | 47 | 0.0426 | 47 | 0.1277 | 57 | 0.0702 | 40 | 0.075 | 191 | 0.0785 |
| 11_5959379 | NA | NA | 1 | 1 | NA | NA | NA | NA | 1 | 1 | NA | NA | NA | NA | 1 | 1 | NA | NA | 1 | 1 |
| 11_5959433 | NA | NA | 1 | 1 | NA | NA | NA | NA | 1 | 1 | NA | NA | NA | NA | 1 | 1 | NA | NA | 1 | 1 |
| 11_5960252 | NA | NA | 3 | 0.3333 | NA | NA | 1 | 0 | 4 | 0.25 | 1 | 0 | 1 | 1 | 1 | 1 | NA | NA | 3 | 0.6667 |
| 11_5960259 | NA | NA | 3 | 0.3333 | NA | NA | 1 | 0 | 4 | 0.25 | 1 | 0 | 1 | 1 | 1 | 0 | NA | NA | 3 | 0.3333 |
| 11_5960281 | NA | NA | 2 | 0 | NA | NA | 1 | 1 | 3 | 0.3333 | 1 | 0 | 1 | 1 | 2 | 0 | NA | NA | 4 | 0.25 |
| 11_6056321 | 12 | 0.0833 | 8 | 0 | 13 | 0.2308 | 19 | 0.0526 | 52 | 0.0962 | 15 | 0.1333 | 12 | 0 | 9 | 0 | 4 | 0 | 40 | 0.05 |
| 11_6056467 | 14 | 0.0714 | 12 | 0 | 17 | 0.1176 | 22 | 0.1364 | 65 | 0.0923 | 11 | 0.0909 | 12 | 0.25 | 11 | 0 | 4 | 0 | 38 | 0.1053 |
| 11_6056488 | 14 | 0.2143 | 12 | 0.1667 | 18 | 0.1667 | 23 | 0.1304 | 67 | 0.1642 | 12 | 0.25 | 16 | 0.25 | 10 | 0 | 7 | 0.4286 | 45 | 0.2222 |
| 11_6056687 | 13 | 0.3846 | 7 | 0.7143 | 15 | 0.8667 | 22 | 0.5 | 57 | 0.5965 | 11 | 0.8182 | 16 | 0.5625 | 12 | 0.5833 | 6 | 0.8333 | 45 | 0.6667 |
| 11_6056770 | 12 | 0.9167 | 6 | 0.8333 | 15 | 0.8667 | 20 | 0.95 | 53 | 0.9057 | 10 | 1 | 12 | 0.75 | 9 | 1 | 4 | 1 | 35 | 0.9143 |
| 11_6056782 | 14 | 0.1429 | 9 | 0 | 17 | 0.1176 | 22 | 0.1364 | 62 | 0.1129 | 11 | 0.0909 | 15 | 0.0667 | 10 | 0.2 | 6 | 0.1667 | 42 | 0.119 |
| 11_6056809 | 13 | 0.1538 | 7 | 0.5714 | 14 | 0.2143 | 21 | 0.1429 | 55 | 0.2182 | 10 | 0.3 | 14 | 0.2143 | 11 | 0.1818 | 6 | 0.1667 | 41 | 0.2195 |
| 11_6056819 | 11 | 0.8182 | 9 | 0.7778 | 15 | 0.8667 | 22 | 0.7727 | 57 | 0.807 | 12 | 0.9167 | 13 | 0.8462 | 11 | 0.9091 | 5 | 1 | 41 | 0.9024 |
| 11_6056946 | 10 | 0.8 | 8 | 0.875 | 13 | 0.8462 | 17 | 0.8235 | 48 | 0.8333 | 9 | 0.7778 | 13 | 0.8462 | 8 | 0.625 | 6 | 0.5 | 36 | 0.7222 |
| 11_6057028 | 10 | 0.6 | 7 | 0.8571 | 12 | 0.5833 | 19 | 0.6842 | 48 | 0.6667 | 9 | 0.6667 | 14 | 0.6429 | 8 | 0.75 | 4 | 1 | 35 | 0.7143 |
| 11_6057048 | 13 | 0 | 8 | 0 | 16 | 0 | 23 | 0.1304 | 60 | 0.05 | 12 | 0.1667 | 15 | 0.0667 | 11 | 0 | 7 | 0 | 45 | 0.0667 |
| 11_6057210 | 12 | 0.0833 | 8 | 0.25 | 15 | 0 | 22 | 0.0909 | 57 | 0.0877 | 12 | 0.0833 | 14 | 0.1429 | 11 | 0.0909 | 7 | 0.1429 | 44 | 0.1136 |
| 11_6057216 | 12 | 0.0833 | 7 | 0 | 15 | 0.2667 | 20 | 0.05 | 54 | 0.1111 | 12 | 0.0833 | 14 | 0 | 12 | 0.25 | 6 | 0 | 44 | 0.0909 |
| 11_6057282 | 10 | 0.5 | 6 | 0.6667 | 9 | 0.7778 | 11 | 0.5455 | 36 | 0.6111 | 10 | 0.7 | 9 | 0.6667 | 8 | 0.5 | 3 | 0 | 30 | 0.5667 |
| 11_6057522 | 11 | 0.0909 | 8 | 0.125 | 14 | 0.1429 | 16 | 0.25 | 49 | 0.1633 | 10 | 0 | 14 | 0.1429 | 8 | 0.125 | 7 | 0.1429 | 39 | 0.1026 |
| 11_6057564 | 11 | 0 | 8 | 0 | 14 | 0.0714 | 15 | 0.1333 | 48 | 0.0625 | 10 | 0.2 | 13 | 0.0769 | 7 | 0.1429 | 7 | 0 | 37 | 0.1081 |
| 11_6057647 | 10 | 0.4 | 8 | 0.25 | 12 | 0.6667 | 17 | 0.4118 | 47 | 0.4468 | 11 | 0.2727 | 11 | 0.1818 | 8 | 0.25 | 6 | 0.5 | 36 | 0.2778 |
| 11_6058004 | 10 | 0 | 6 | 0.1667 | 10 | 0 | 13 | 0.0769 | 39 | 0.0513 | 11 | 0.0909 | 11 | 0.0909 | 7 | 0 | 6 | 0.1667 | 35 | 0.0857 |
| 11_6058170 | 10 | 0.3 | 6 | 0.3333 | 10 | 0.8 | 11 | 0.6364 | 37 | 0.5405 | 8 | 0.625 | 9 | 0.6667 | 5 | 0.8 | 4 | 0.25 | 26 | 0.6154 |
| 11_6122613 | 16 | 0.1875 | 18 | 0.4444 | 8 | 0.25 | 26 | 0.1923 | 68 | 0.2647 | 29 | 0.3103 | 26 | 0.2692 | 11 | 0.3636 | 20 | 0.25 | 86 | 0.2907 |
| 11_6143253 | 6 | 0.1667 | 3 | 0 | 5 | 0 | 2 | 0.5 | 16 | 0.125 | 4 | 0 | 8 | 0.125 | 3 | 0 | 1 | 0 | 16 | 0.0625 |
| 11_6149989 | 4 | 0 | 2 | 0.5 | 2 | 0 | 2 | 0 | 10 | 0.1 | 2 | 0 | 7 | 0 | 2 | 0.5 | NA | NA | 11 | 0.0909 |
| 11_6150061 | 3 | 0.3333 | 3 | 0.3333 | 4 | 0 | 1 | 0 | 11 | 0.1818 | 3 | 0.3333 | 8 | 0.125 | 2 | 0 | 1 | 0 | 14 | 0.1429 |
| 11_6209052 | 3 | 0 | 7 | 0.4286 | 3 | 0.6667 | 4 | 0.5 | 17 | 0.4118 | 1 | 1 | 12 | 0.3333 | 3 | 0.3333 | 5 | 0.4 | 21 | 0.381 |
| 11_6209078 | 1 | 1 | 5 | 0.8 | 3 | 0.6667 | 3 | 0.6667 | 12 | 0.75 | 4 | 0.5 | 8 | 0.625 | 3 | 0.6667 | 3 | 0.6667 | 18 | 0.6111 |
| 11_6209082 | 3 | 0.3333 | 7 | 0.5714 | 4 | 0.75 | 4 | 0.5 | 18 | 0.5556 | 3 | 0.6667 | 5 | 0.4 | 3 | 0.6667 | 7 | 0.7143 | 18 | 0.6111 |
| 11_6209162 | 3 | 0 | 7 | 0.2857 | 5 | 0.2 | 5 | 0.4 | 20 | 0.25 | 3 | 0.6667 | 9 | 0.2222 | 4 | 0.5 | 7 | 0.2857 | 23 | 0.3478 |
| 11_6220600 | 4 | 1 | 5 | 0.2 | NA | NA | 1 | 1 | 10 | 0.6 | 3 | 0.6667 | 6 | 0.3333 | 4 | 0.25 | 4 | 1 | 17 | 0.5294 |
| 11_6221182 | 12 | 0.1667 | 13 | 0 | 6 | 0 | 7 | 0.1429 | 38 | 0.0789 | 11 | 0.1818 | 8 | 0 | 12 | 0.1667 | 12 | 0 | 43 | 0.093 |
| 11_6221551 | 6 | 0 | 6 | 0.3333 | 2 | 0 | 5 | 0.4 | 19 | 0.2105 | 6 | 0.1667 | 6 | 0.1667 | 6 | 0 | 5 | 0 | 23 | 0.087 |

| | | | | | | | | | | | | | | | | | | | | |
|---|---|---|---|---|---|---|---|---|---|---|---|---|---|---|---|---|---|---|---|---|
| 11_6221553 | 1 | 1 | 2 | 0.5 | NA | NA | NA | NA | 3 | 0.6667 | 1 | 1 | NA | NA | NA | NA | 1 | 0 | 2 | 0.5 |
| 11_6246993 | 4 | 0 | 9 | 0.4444 | 5 | 0.6 | 9 | 0.4444 | 27 | 0.4074 | 7 | 0.4286 | 8 | 0.25 | 6 | 0.5 | 6 | 0.5 | 27 | 0.4074 |
| 11_6247032 | 5 | 0.6 | 10 | 0.5 | 8 | 0.75 | 10 | 0.6 | 33 | 0.6061 | 7 | 0.2857 | 6 | 0.6667 | 6 | 0.6667 | 6 | 0.8333 | 25 | 0.6 |
| 11_6247041 | 6 | 0.3333 | 10 | 0.6 | 7 | 0.4286 | 11 | 0.1818 | 34 | 0.3824 | 7 | 0.4286 | 8 | 0.375 | 7 | 0.8571 | 6 | 0.5 | 28 | 0.5357 |
| 11_6247134 | 4 | 0.75 | 9 | 0.4444 | 8 | 0.75 | 10 | 0.5 | 31 | 0.5806 | 4 | 0.5 | 9 | 0.6667 | 5 | 1 | 6 | 0.6667 | 24 | 0.7083 |
| 11_6247144 | 4 | 0.75 | 8 | 0.5 | 6 | 0.5 | 10 | 0.4 | 28 | 0.5 | 2 | 0 | 8 | 0.75 | 6 | 0.8333 | 6 | 0.5 | 22 | 0.6364 |
| 11_6305634 | 16 | 0.125 | 15 | 0.2 | 19 | 0.0526 | 31 | 0.0645 | 81 | 0.0988 | 11 | 0.1818 | 25 | 0.04 | 17 | 0.1176 | 13 | 0 | 66 | 0.0758 |
| 11_6305934 | 25 | 0.16 | 20 | 0.05 | 24 | 0.125 | 35 | 0.0571 | 104 | 0.0962 | 15 | 0.0667 | 32 | 0.125 | 26 | 0.1154 | 17 | 0.2941 | 90 | 0.1444 |
| 11_6306191 | 24 | 0.2083 | 22 | 0.1818 | 26 | 0.4231 | 32 | 0.2812 | 104 | 0.2788 | 19 | 0.3158 | 32 | 0.25 | 26 | 0.3077 | 18 | 0.3333 | 95 | 0.2947 |
| 11_6306536 | 28 | 0.25 | 26 | 0.1538 | 26 | 0.1923 | 34 | 0.1765 | 114 | 0.193 | 26 | 0.3462 | 40 | 0.25 | 30 | 0.2667 | 23 | 0.1304 | 119 | 0.2521 |
| 11_6306689 | 2 | 0.5 | NA | NA | NA | NA | 1 | 0 | 3 | 0.3333 | 1 | 0 | NA | NA | 1 | 0 | 1 | 1 | 3 | 0.3333 |
| 11_6306928 | 2 | 0.5 | NA | NA | NA | NA | 1 | 0 | 3 | 0.3333 | 1 | 1 | NA | NA | 1 | 0 | 1 | 0 | 3 | 0.3333 |
| 11_6307695 | 2 | 0 | NA | NA | NA | NA | 2 | 0.5 | 4 | 0.25 | 1 | 0 | 1 | 0 | 1 | 1 | 2 | 0.5 | 5 | 0.4 |
| 11_6307864 | 3 | 0.3333 | NA | NA | NA | NA | 2 | 0 | 5 | 0.2 | 1 | 0 | 1 | 1 | 1 | 1 | 2 | 0.5 | 5 | 0.6 |
| 11_6355019 | 22 | 0.1818 | 30 | 0.1 | 14 | 0.1429 | 23 | 0.087 | 89 | 0.1236 | 19 | 0 | 25 | 0.08 | 17 | 0.1765 | 21 | 0 | 82 | 0.061 |
| 11_6355198 | 16 | 0.1875 | 30 | 0.2333 | 13 | 0.4615 | 16 | 0.125 | 75 | 0.24 | 23 | 0.5217 | 26 | 0.4615 | 15 | 0.4667 | 16 | 0.25 | 80 | 0.4375 |
| 11_6355250 | 18 | 0.8889 | 31 | 0.8065 | 16 | 0.9375 | 22 | 0.9545 | 87 | 0.8851 | 23 | 0.9565 | 28 | 0.9643 | 19 | 0.9474 | 20 | 0.85 | 90 | 0.9333 |
| 11_6355391 | 21 | 0.381 | 31 | 0.3548 | 21 | 0.2857 | 23 | 0.1739 | 96 | 0.3021 | 25 | 0.16 | 32 | 0.2812 | 24 | 0.25 | 26 | 0.1538 | 107 | 0.215 |
| 11_6355751 | 13 | 0.6154 | 25 | 0.52 | 18 | 0.6667 | 21 | 0.619 | 77 | 0.5974 | 18 | 0.4444 | 27 | 0.4074 | 19 | 0.4211 | 22 | 0.4545 | 86 | 0.4302 |
| 11_6356058 | 15 | 0.1333 | 26 | 0.2692 | 14 | 0.1429 | 25 | 0.16 | 80 | 0.1875 | 16 | 0.25 | 28 | 0.1786 | 22 | 0.0455 | 19 | 0.0526 | 85 | 0.1294 |
| 11_6369601 | 129 | 0.1628 | 212 | 0.1462 | 170 | 0.1706 | 158 | 0.1266 | 669 | 0.151 | 135 | 0.1407 | 150 | 0.18 | 164 | 0.1768 | 140 | 0.2 | 589 | 0.1749 |
| 11_6369620 | 129 | 0.0543 | 243 | 0.107 | 184 | 0.0652 | 163 | 0.0736 | 719 | 0.0793 | 139 | 0.0863 | 158 | 0.0886 | 170 | 0.0765 | 159 | 0.0566 | 626 | 0.0767 |
| 11_6369812 | 36 | 0.1389 | 50 | 0.06 | 53 | 0.0755 | 38 | 0.0526 | 177 | 0.0791 | 37 | 0.0541 | 56 | 0.0893 | 36 | 0.1111 | 40 | 0.05 | 169 | 0.0769 |
| 11_6377978 | 4 | 0.25 | 2 | 0 | 2 | 0.5 | 6 | 0 | 14 | 0.1429 | 2 | 0 | 3 | 0 | 6 | 0 | 1 | 1 | 12 | 0.0833 |
| 11_6422919 | 7 | 0 | 20 | 0.05 | 14 | 0.0714 | 15 | 0.0667 | 56 | 0.0536 | 10 | 0.1 | 6 | 0 | 10 | 0.1 | 11 | 0 | 37 | 0.0541 |
| 11_6422987 | 8 | 0.375 | 18 | 0.1667 | 14 | 0 | 13 | 0.0769 | 53 | 0.1321 | 10 | 0.2 | 7 | 0.7143 | 8 | 0.25 | 13 | 0.0769 | 38 | 0.2632 |
| 11_6423319 | 9 | 0.1111 | 18 | 0.2778 | 12 | 0.25 | 11 | 0.2727 | 50 | 0.24 | 8 | 0.375 | 5 | 0.2 | 6 | 0.3333 | 11 | 0.1818 | 30 | 0.2667 |
| 11_6423476 | 6 | 0.3333 | 16 | 0.0625 | 13 | 0.1538 | 12 | 0.0833 | 47 | 0.1277 | 8 | 0.25 | 5 | 0.2 | 8 | 0 | 16 | 0.0625 | 37 | 0.1081 |
| 11_6423521 | 10 | 0.8 | 16 | 0.75 | 12 | 1 | 13 | 0.7692 | 51 | 0.8235 | 8 | 0.625 | 6 | 0.6667 | 8 | 0.875 | 15 | 0.8 | 37 | 0.7568 |
| 11_6423543 | 10 | 1 | 18 | 1 | 14 | 1 | 14 | 1 | 56 | 1 | 11 | 1 | 6 | 1 | 7 | 1 | 13 | 1 | 37 | 1 |
| 11_6423645 | 7 | 1 | 16 | 1 | 12 | 1 | 15 | 1 | 50 | 1 | 11 | 1 | 6 | 1 | 5 | 1 | 16 | 0.9375 | 38 | 0.9737 |
| 11_6423755 | 6 | 0.5 | 16 | 0.875 | 14 | 0.7857 | 14 | 0.7143 | 50 | 0.76 | 11 | 0.8182 | 6 | 1 | 6 | 0.8333 | 13 | 0.8462 | 36 | 0.8611 |
| 11_6423786 | 7 | 0.2857 | 18 | 0.3889 | 12 | 0.25 | 14 | 0.5714 | 51 | 0.3922 | 10 | 0.5 | 6 | 0.5 | 6 | 0.5 | 13 | 0.1538 | 35 | 0.3714 |
| 11_6423822 | 6 | 1 | 13 | 0.8462 | 9 | 0.8889 | 10 | 0.7 | 38 | 0.8421 | 7 | 0.8571 | 5 | 1 | 6 | 0.8333 | 10 | 1 | 28 | 0.9286 |
| 11_6423840 | 8 | 0.625 | 13 | 0.3846 | 12 | 0.5 | 14 | 0.5 | 47 | 0.4894 | 10 | 0.2 | 4 | 0.75 | 5 | 0.8 | 12 | 0.4167 | 31 | 0.4516 |
| 11_6423845 | 3 | 1 | 14 | 0.3571 | 9 | 0.2222 | 9 | 0.4444 | 35 | 0.4 | 9 | 0.3333 | 5 | 0.8 | 4 | 0.25 | 9 | 0.5556 | 27 | 0.4815 |
| 11_6423846 | 1 | 0 | 1 | 1 | NA | NA | 2 | 0 | 4 | 0.25 | NA | NA | 1 | 1 | NA | NA | NA | NA | 1 | 1 |
| 11_6423908 | 7 | 0.2857 | 15 | 0.2667 | 10 | 0.8 | 12 | 0.5 | 44 | 0.4545 | 8 | 0.625 | 3 | 1 | 4 | 0 | 14 | 0.2143 | 29 | 0.3793 |
| 11_6424032 | 4 | 1 | 11 | 1 | 11 | 1 | 14 | 0.9286 | 40 | 0.975 | 11 | 1 | 5 | 1 | 3 | 1 | 5 | 1 | 24 | 1 |
| 11_6424153 | 6 | 0.3333 | 8 | 0.875 | 11 | 0.9091 | 8 | 0.625 | 33 | 0.7273 | 8 | 0.5 | 4 | 0.75 | 4 | 0.5 | 9 | 0.4444 | 25 | 0.52 |
| 11_6424224 | 7 | 0.7143 | 13 | 0.4615 | 9 | 0.3333 | 12 | 0.3333 | 41 | 0.439 | 10 | 0.5 | 5 | 0.2 | 6 | 0.8333 | 13 | 0.3077 | 34 | 0.4412 |
| 11_6424230 | 6 | 0.3333 | 12 | 0.3333 | 12 | 0.1667 | 12 | 0 | 42 | 0.1905 | 10 | 0.1 | 5 | 0.4 | 4 | 0 | 12 | 0.25 | 31 | 0.1935 |
| 11_6424309 | 5 | 1 | 11 | 0.9091 | 10 | 1 | 6 | 1 | 32 | 0.9688 | 8 | 1 | 4 | 1 | 3 | 1 | 5 | 1 | 20 | 1 |
| 11_6424533 | 6 | 0.5 | 14 | 0.2857 | 11 | 0.2727 | 10 | 0.5 | 41 | 0.3659 | 10 | 0.5 | 4 | 0.75 | 5 | 0.2 | 11 | 0.4545 | 30 | 0.4667 |
| 11_6424559 | 8 | 0.5 | 15 | 0.4667 | 9 | 0.4444 | 10 | 0.7 | 42 | 0.5238 | 11 | 0.5455 | 5 | 0.6 | 5 | 0.6 | 10 | 0.8 | 31 | 0.6452 |
| 11_6424694 | 6 | 0.3333 | 4 | 0.75 | 7 | 0.7143 | 7 | 0.5714 | 24 | 0.5833 | 9 | 0.5556 | 1 | 0 | 5 | 0 | 6 | 0.5 | 21 | 0.381 |
| 11_6424726 | 7 | 0.5714 | 8 | 0.5 | 10 | 0.7 | 8 | 0.75 | 33 | 0.6364 | 10 | 0.5 | 3 | 1 | 4 | 0.75 | 9 | 0.5556 | 26 | 0.6154 |
| 11_6425168 | 5 | 0.4 | 6 | 0.3333 | 5 | 0.4 | 10 | 0.4 | 26 | 0.3846 | 4 | 0.5 | 3 | 1 | 5 | 0.8 | 6 | 0.6667 | 18 | 0.7222 |
| 11_6425450 | 5 | 0 | 4 | 0.25 | 5 | 0 | 7 | 0.2857 | 21 | 0.1429 | 4 | 0.25 | 3 | 0.3333 | 3 | 0 | 3 | 0 | 13 | 0.1538 |
| 11_6425484 | 1 | 1 | NA | NA | NA | NA | 1 | 1 | 2 | 1 | 1 | 1 | 1 | 1 | NA | NA | NA | NA | 2 | 1 |
| 11_6425732 | 3 | 0 | 4 | 0.25 | 3 | 0 | 6 | 0 | 16 | 0.0625 | 3 | 0 | 3 | 0.3333 | NA | NA | 3 | 0 | 9 | 0.1111 |
| 11_6426306 | 1 | 1 | 1 | 0 | 1 | 0 | 1 | 0 | 4 | 0.25 | 1 | 1 | 2 | 0 | NA | NA | NA | NA | 3 | 0.3333 |
| 11_6426314 | NA | NA | 1 | 1 | 1 | 1 | 1 | 0 | 3 | 0.6667 | 1 | 1 | 2 | 1 | NA | NA | NA | NA | 3 | 1 |
| 11_6426420 | 1 | 0 | 1 | 0 | 1 | 1 | 1 | 0 | 4 | 0.25 | NA | NA | 2 | 0.5 | NA | NA | NA | NA | 2 | 0.5 |
| 11_6426552 | 1 | 1 | 1 | 0 | 1 | 1 | 1 | 1 | 4 | 0.75 | 3 | 1 | 2 | 1 | NA | NA | NA | NA | 5 | 1 |
| 11_6427680 | 2 | 0.5 | NA | NA | NA | NA | 1 | 0 | 3 | 0.3333 | 2 | 0 | 1 | 1 | NA | NA | NA | NA | 3 | 0.3333 |
| 11_6427850 | 1 | 1 | NA | NA | NA | NA | 1 | 1 | 2 | 1 | 2 | 0 | 1 | 1 | NA | NA | NA | NA | 3 | 0.3333 |
| 11_6428234 | 2 | 0.5 | NA | NA | NA | NA | 1 | 0 | 3 | 0.3333 | 2 | 0.5 | 1 | 0 | NA | NA | NA | NA | 3 | 0.3333 |
| 11_6428316 | 2 | 0.5 | NA | NA | NA | NA | 1 | 0 | 3 | 0.3333 | 2 | 1 | 1 | 0 | NA | NA | NA | NA | 3 | 0.6667 |
| 11_6429206 | 1 | 1 | NA | NA | NA | NA | NA | NA | 1 | 1 | 2 | 0.5 | NA | NA | NA | NA | NA | NA | 2 | 0.5 |
| 11_6429304 | 1 | 1 | NA | NA | NA | NA | NA | NA | 1 | 1 | 2 | 0.5 | NA | NA | NA | NA | NA | NA | 2 | 0.5 |
| 11_6429479 | 1 | 1 | NA | NA | NA | NA | 1 | 0 | 2 | 0.5 | 2 | 1 | NA | NA | 1 | 0 | NA | NA | 3 | 0.6667 |
| 11_6430694 | NA | NA | NA | NA | NA | NA | 1 | 1 | 1 | 1 | NA | NA | NA | NA | 1 | 1 | NA | NA | 1 | 1 |
| 11_6546108 | 24 | 0.0417 | 20 | 0.15 | 15 | 0.0667 | 19 | 0 | 78 | 0.0641 | 9 | 0 | 21 | 0.0476 | 16 | 0.1875 | 14 | 0.0714 | 60 | 0.0833 |
| 11_6546239 | 26 | 0.0769 | 15 | 0 | 14 | 0 | 19 | 0.1579 | 74 | 0.0676 | 9 | 0.2222 | 25 | 0.2 | 18 | 0.1111 | 16 | 0.25 | 68 | 0.1912 |

| | | | | | | | | | | | | | | | | | | | | |
|---|---|---|---|---|---|---|---|---|---|---|---|---|---|---|---|---|---|---|---|---|
| 11_654629 | 26 | 0.2308 | 21 | 0.3333 | 13 | 0.3077 | 18 | 0.1111 | 78 | 0.2436 | 9 | 0.1111 | 26 | 0.1538 | 22 | 0.1818 | 15 | 0.2 | 72 | 0.1667 |
| 11_654630 | 28 | 0.0714 | 23 | 0.1739 | 17 | 0 | 20 | 0.1 | 88 | 0.0909 | 11 | 0.0909 | 27 | 0.037 | 23 | 0.087 | 16 | 0.125 | 77 | 0.0779 |
| 11_654634 | 23 | 0.4783 | 22 | 0.4091 | 16 | 0.1875 | 17 | 0.4706 | 78 | 0.3974 | 10 | 0.7 | 27 | 0.5556 | 23 | 0.3913 | 19 | 0.3158 | 79 | 0.4684 |
| 11_654635 | 26 | 0.5385 | 22 | 0.5455 | 17 | 0.5882 | 17 | 0.5882 | 82 | 0.561 | 10 | 0.6 | 28 | 0.7143 | 21 | 0.6667 | 17 | 0.7059 | 76 | 0.6842 |
| 11_654637 | 26 | 0.4615 | 19 | 0.6316 | 11 | 0.3636 | 16 | 0.6875 | 72 | 0.5417 | 9 | 0.3333 | 24 | 0.4167 | 19 | 0.5789 | 15 | 0.5333 | 67 | 0.4776 |
| 11_654642 | 23 | 0.087 | 23 | 0.087 | 17 | 0 | 18 | 0.0556 | 81 | 0.0617 | 12 | 0 | 24 | 0.0417 | 19 | 0.2105 | 18 | 0.1111 | 73 | 0.0959 |
| 11_654653 | 23 | 0.913 | 21 | 0.8095 | 13 | 0.8462 | 16 | 0.75 | 73 | 0.8356 | 8 | 0.75 | 20 | 0.85 | 20 | 0.85 | 16 | 0.8125 | 64 | 0.8281 |
| 11_654666 | 22 | 0.7273 | 19 | 0.8421 | 14 | 0.5714 | 15 | 0.6 | 70 | 0.7 | 8 | 0.875 | 29 | 0.7931 | 19 | 0.6842 | 17 | 0.7647 | 73 | 0.7671 |
| 11_654668 | 25 | 0.32 | 20 | 0.45 | 17 | 0.5882 | 18 | 0.3333 | 80 | 0.4125 | 11 | 0.5455 | 29 | 0.3793 | 24 | 0.375 | 19 | 0.3684 | 83 | 0.3976 |
| 11_654670 | 26 | 0.5769 | 21 | 0.7143 | 16 | 0.625 | 18 | 0.5556 | 81 | 0.6173 | 10 | 0.7 | 28 | 0.5357 | 21 | 0.4762 | 16 | 0.6875 | 75 | 0.5733 |
| 11_654944 | 4 | 0 | 3 | 0 | 5 | 0.2 | 5 | 0.2 | 17 | 0.1176 | 1 | 0 | 6 | 0.5 | 4 | 0 | 1 | 0 | 12 | 0.25 |
| 11_654949 | 6 | 0.1667 | 4 | 0 | 5 | 0.2 | 6 | 0.1667 | 21 | 0.1429 | 1 | 1 | 6 | 0.3333 | 4 | 0 | 1 | 0 | 12 | 0.25 |
| 11_654976 | 6 | 0.8333 | 3 | 1 | 4 | 1 | 6 | 0.3333 | 19 | 0.7368 | 1 | 1 | 5 | 0.6 | 4 | 0.25 | 1 | 1 | 11 | 0.5455 |
| 11_654991 | 5 | 0.8 | 3 | 0.6667 | 4 | 1 | 6 | 0.8333 | 18 | 0.8333 | NA | NA | 5 | 0.6 | 2 | 0.5 | 1 | 0 | 8 | 0.5 |
| 11_655002 | 4 | 0.5 | 3 | 0.6667 | 2 | 0 | 5 | 0.4 | 14 | 0.4286 | NA | NA | 3 | 0 | 1 | 0 | 1 | 1 | 5 | 0.2 |
| 11_655009 | 4 | 0 | 3 | 0.6667 | 4 | 0.5 | 6 | 0.8333 | 17 | 0.5294 | NA | NA | 4 | 0.5 | 2 | 1 | 1 | 0 | 7 | 0.5714 |
| 11_655016 | 4 | 0.25 | 2 | 0 | 3 | 0 | 4 | 0.25 | 13 | 0.1538 | NA | NA | 3 | 0.3333 | 1 | 0 | 1 | 0 | 5 | 0.2 |
| 11_655025 | 4 | 0.25 | 3 | 0.3333 | 4 | 0 | 5 | 0.2 | 16 | 0.1875 | NA | NA | 3 | 0.3333 | 1 | 1 | 1 | 0 | 5 | 0.4 |
| 11_655034 | 3 | 0 | 3 | 0.3333 | 4 | 0.25 | 3 | 0 | 13 | 0.1538 | NA | NA | 3 | 0.3333 | NA | NA | NA | NA | 3 | 0.3333 |
| 11_655045 | 4 | 0.75 | 2 | 1 | 3 | 0.3333 | 4 | 0.75 | 13 | 0.6923 | NA | NA | 2 | 1 | 1 | 0 | 1 | 1 | 4 | 0.75 |
| 11_655050 | 4 | 0.75 | 2 | 1 | 4 | 1 | 3 | 1 | 13 | 0.9231 | NA | NA | 4 | 1 | 1 | 1 | 1 | 1 | 6 | 1 |
| 11_655056 | 2 | 0.5 | 2 | 0.5 | 2 | 0.5 | 3 | 0.3333 | 9 | 0.4444 | NA | NA | 4 | 0.5 | 1 | 0 | 1 | 0 | 6 | 0.3333 |
| 11_655062 | 4 | 0.25 | 2 | 0 | 4 | 0.5 | 4 | 0.25 | 14 | 0.2857 | NA | NA | 4 | 1 | 1 | 0 | NA | NA | 5 | 0.8 |
| 11_655080 | 4 | 0 | 3 | 0.3333 | 2 | 0 | 3 | 0.3333 | 12 | 0.1667 | NA | NA | 3 | 0.3333 | 1 | 0 | 1 | 0 | 5 | 0.2 |
| 11_655098 | 3 | 0.3333 | 3 | 0 | 3 | 0.6667 | 4 | 1 | 13 | 0.5385 | NA | NA | 3 | 1 | 1 | 0 | 1 | 0 | 5 | 0.6 |
| 11_655178 | 2 | 0.5 | 1 | 1 | 3 | 0.6667 | 3 | 0.3333 | 9 | 0.5556 | NA | NA | 2 | 0.5 | 1 | 0 | 1 | 1 | 4 | 0.5 |
| 11_655214 | 2 | 0 | 1 | 0 | 3 | 0.3333 | 2 | 0 | 8 | 0.125 | NA | NA | 2 | 0.5 | 1 | 0 | 1 | 0 | 4 | 0.25 |
| 11_655220 | 2 | 0 | 1 | 1 | 3 | 0 | 2 | 0 | 8 | 0.125 | NA | NA | 2 | 0 | 1 | 0 | 1 | 1 | 4 | 0.25 |
| 11_655225 | 2 | 1 | 1 | 1 | 2 | 1 | 2 | 0.5 | 7 | 0.8571 | NA | NA | 2 | 1 | 1 | 1 | 1 | 1 | 4 | 1 |
| 11_655227 | 1 | 0 | 1 | 0 | 2 | 0 | 2 | 0.5 | 6 | 0.1667 | NA | NA | 2 | 0.5 | NA | NA | NA | NA | 2 | 0.5 |
| 11_655236 | 2 | 1 | NA | NA | 2 | 1 | 2 | 0.5 | 6 | 0.8333 | NA | NA | 2 | 1 | 1 | 1 | 1 | 1 | 4 | 1 |
| 11_656571 | 3 | 0 | 5 | 0.2 | 3 | 0.3333 | 8 | 0.125 | 19 | 0.1579 | 4 | 0 | 5 | 0.2 | 4 | 0.25 | 1 | 0 | 14 | 0.1429 |
| 11_656576 | 3 | 0 | 5 | 0 | 2 | 0.5 | 5 | 0.2 | 15 | 0.1333 | 6 | 0.1667 | 3 | 0.3333 | 3 | 0.6667 | 2 | 0.5 | 14 | 0.3571 |
| 11_656621 | 4 | 0 | 5 | 0.2 | 2 | 0 | 4 | 0.25 | 15 | 0.1333 | 4 | 0.25 | 2 | 0 | 3 | 0.3333 | 1 | 0 | 10 | 0.2 |
| 11_656657 | 2 | 1 | 3 | 0 | 2 | 0 | 5 | 0 | 12 | 0.1667 | 4 | 0 | 5 | 0.2 | 3 | 0 | 2 | 0 | 14 | 0.0714 |
| 11_656659 | 4 | 0.25 | 5 | 0.4 | 3 | 0 | 6 | 0 | 18 | 0.1667 | 4 | 0.5 | 7 | 0.2857 | 3 | 0 | 3 | 0.3333 | 17 | 0.2941 |
| 11_656661 | 4 | 0 | 6 | 0.1667 | 3 | 0 | 5 | 0 | 18 | 0.0556 | 5 | 0 | 7 | 0 | 4 | 0 | 3 | 0.3333 | 19 | 0.0526 |
| 11_657087 | NA | NA | 3 | 0 | 1 | 0 | 4 | 0.25 | 8 | 0.125 | 4 | 0.25 | 6 | 0 | 3 | 0 | 2 | 0 | 15 | 0.0667 |
| 11_657400 | NA | NA | 3 | 0.3333 | 1 | 1 | 2 | 1 | 6 | 0.6667 | 1 | 0 | 4 | 1 | 2 | 0.5 | 1 | 0 | 8 | 0.625 |
| 11_657410 | NA | NA | 2 | 0.5 | 1 | 1 | 3 | 0 | 6 | 0.3333 | 2 | 0.5 | 5 | 0.4 | 3 | 0.6667 | 2 | 0 | 12 | 0.4167 |
| 11_657413 | NA | NA | 3 | 0 | 1 | 0 | 3 | 0.3333 | 7 | 0.1429 | 2 | 0.5 | 5 | 0.4 | 3 | 0.3333 | 2 | 0 | 12 | 0.3333 |
| 11_662667 | 2 | 0.5 | NA | NA | NA | NA | 2 | 0 | 4 | 0.25 | 1 | 0 | 4 | 0 | 3 | 0.3333 | 1 | 1 | 9 | 0.2222 |
| 11_662680 | 2 | 0 | NA | NA | NA | NA | 2 | 0.5 | 4 | 0.25 | 1 | 0 | 3 | 0.6667 | 3 | 0 | 1 | 0 | 8 | 0.25 |
| 11_662696 | 2 | 0.5 | NA | NA | NA | NA | 2 | 0 | 4 | 0.25 | NA | NA | 4 | 0.25 | 2 | 0 | 2 | 0 | 8 | 0.125 |
| 11_662719 | 1 | 1 | NA | NA | NA | NA | 3 | 0.6667 | 4 | 0.75 | 1 | 1 | 4 | 1 | 3 | 1 | 1 | 1 | 9 | 1 |
| 11_711952 | NA | NA | 1 | 1 | 1 | 1 | NA | NA | 2 | 1 | 2 | 1 | 2 | 1 | NA | NA | NA | NA | 4 | 1 |
| 11_711979 | NA | NA | 1 | 0 | 1 | 1 | 1 | 1 | 3 | 0.6667 | 2 | 0.5 | 2 | 0 | NA | NA | NA | NA | 4 | 0.25 |
| 11_711991 | NA | NA | 1 | 1 | 1 | 0 | 1 | 1 | 3 | 0.6667 | 2 | 0 | 2 | 0.5 | NA | NA | NA | NA | 4 | 0.25 |
| 11_711995 | NA | NA | 1 | 0 | 1 | 1 | 1 | 1 | 3 | 0.6667 | 2 | 0.5 | 2 | 0.5 | NA | NA | NA | NA | 4 | 0.5 |
| 11_712022 | NA | NA | NA | NA | 1 | 0 | 1 | 1 | 2 | 0.5 | 3 | 0.6667 | 1 | 1 | NA | NA | NA | NA | 4 | 0.75 |
| 11_712034 | NA | NA | 1 | 0 | 1 | 1 | 1 | 0 | 3 | 0.3333 | 3 | 1 | 2 | 0 | NA | NA | NA | NA | 5 | 0.6 |
| 11_712050 | NA | NA | 1 | 0 | 1 | 0 | 1 | 1 | 3 | 0.3333 | 3 | 0.6667 | NA | NA | NA | NA | NA | NA | 3 | 0.6667 |
| 11_712143 | NA | NA | 3 | 0.3333 | 4 | 0 | 1 | 0 | 8 | 0.125 | 5 | 0.4 | 2 | 0 | NA | NA | NA | NA | 7 | 0.2857 |
| 11_712200 | NA | NA | 4 | 0.25 | 4 | 0.25 | 1 | 1 | 9 | 0.3333 | 5 | 0.2 | 2 | 0.5 | NA | NA | NA | NA | 7 | 0.2857 |
| 11_712214 | NA | NA | 5 | 0.2 | 3 | 0.3333 | 1 | 1 | 9 | 0.3333 | 6 | 0.5 | 2 | 0 | NA | NA | NA | NA | 8 | 0.375 |
| 11_712221 | NA | NA | 4 | 0.25 | 3 | 0 | 1 | 0 | 8 | 0.125 | 6 | 0.1667 | 2 | 0 | NA | NA | NA | NA | 8 | 0.125 |
| 11_712222 | NA | NA | 6 | 0 | 4 | 0.25 | 1 | 0 | 11 | 0.0909 | 4 | 0.25 | 2 | 0 | NA | NA | NA | NA | 6 | 0.1667 |
| 11_712231 | NA | NA | 5 | 0.4 | 4 | 0.5 | NA | NA | 9 | 0.4444 | 5 | 0.4 | 1 | 1 | NA | NA | NA | NA | 6 | 0.5 |
| 11_712233 | NA | NA | 4 | 0.5 | 3 | 0.6667 | 1 | 1 | 8 | 0.625 | 5 | 1 | 1 | 1 | NA | NA | NA | NA | 6 | 1 |
| 11_712236 | NA | NA | 6 | 0.5 | 4 | 0.5 | 1 | 1 | 11 | 0.5455 | 5 | 0.8 | 1 | 0 | NA | NA | NA | NA | 6 | 0.6667 |
| 11_712238 | NA | NA | 5 | 0.2 | 4 | 0.75 | 1 | 0 | 10 | 0.4 | 5 | 0.8 | 2 | 0.5 | NA | NA | NA | NA | 7 | 0.7143 |
| 11_712239 | NA | NA | 6 | 0 | 3 | 0 | 1 | 1 | 10 | 0.1 | 6 | 0.1667 | 2 | 0 | NA | NA | NA | NA | 8 | 0.125 |
| 11_712257 | NA | NA | 5 | 0.2 | 4 | 0 | 1 | 1 | 10 | 0.2 | 3 | 0.3333 | 2 | 0 | NA | NA | NA | NA | 5 | 0.2 |
| 11_712268 | NA | NA | 5 | 0.2 | 3 | 0 | 1 | 0 | 9 | 0.1111 | 4 | 0.5 | 2 | 0 | NA | NA | NA | NA | 6 | 0.3333 |
| 11_712270 | NA | NA | 6 | 0.8333 | 4 | 1 | 1 | 1 | 11 | 0.9091 | 5 | 1 | 1 | 1 | NA | NA | NA | NA | 6 | 1 |

| | | | | | | | | | | | | | | | | | | | | |
|---|---|---|---|---|---|---|---|---|---|---|---|---|---|---|---|---|---|---|---|---|
| 11_7123524 | 1 | 0 | 6 | 0.1667 | 5 | 0 | 1 | 0 | 13 | 0.0769 | 5 | 0 | 2 | 0 | NA | NA | 1 | 1 | 8 | 0.125 |
| 11_7123848 | 1 | 0 | 5 | 0.4 | 6 | 0.3333 | NA | NA | 12 | 0.3333 | 10 | 0.5 | 1 | 0 | NA | NA | 1 | 0 | 12 | 0.4167 |
| 11_7123935 | NA | NA | 5 | 0.4 | 4 | 0.25 | NA | NA | 9 | 0.3333 | 4 | 0.25 | 2 | 0 | NA | NA | 1 | 0 | 7 | 0.1429 |
| 11_7128148 | 6 | 0.3333 | 15 | 0.1333 | 17 | 0.1765 | 5 | 0.2 | 43 | 0.186 | 16 | 0.25 | 7 | 0.2857 | 8 | 0.25 | 13 | 0.1538 | 44 | 0.2273 |
| 11_7156263 | 3 | 0.3333 | 4 | 0 | 3 | 0 | 5 | 0.4 | 15 | 0.2 | 1 | 0 | NA | NA | 4 | 0.25 | 2 | 0 | 7 | 0.1429 |
| 11_7156287 | 3 | 0.3333 | 4 | 0.25 | 3 | 0.3333 | 4 | 0 | 14 | 0.2143 | 1 | 0 | NA | NA | 4 | 0.25 | 2 | 0.5 | 7 | 0.2857 |
| 11_7156772 | 3 | 0.3333 | 3 | 0 | 1 | 0 | 3 | 0.6667 | 10 | 0.3 | 1 | 0 | NA | NA | 4 | 0 | 1 | 1 | 6 | 0.1667 |
| 11_7156844 | 2 | 0 | 2 | 0 | 1 | 1 | 2 | 1 | 7 | 0.4286 | 1 | 0 | NA | NA | 4 | 0.25 | 2 | 0 | 7 | 0.1429 |
| 11_7156856 | 3 | 0.3333 | 4 | 0.5 | 1 | 0 | 5 | 0.2 | 13 | 0.3077 | 1 | 0 | NA | NA | 4 | 0 | 2 | 0.5 | 7 | 0.1429 |
| 11_7156894 | 1 | 0 | 3 | 0.6667 | 1 | 1 | 1 | 1 | 6 | 0.6667 | 1 | 0 | NA | NA | 4 | 0.25 | 1 | 1 | 6 | 0.3333 |
| 11_7156935 | 3 | 1 | 2 | 0.5 | NA | NA | 2 | 0.5 | 7 | 0.7143 | 1 | 0 | NA | NA | 4 | 0.75 | 1 | 1 | 6 | 0.6667 |
| 11_7157300 | 3 | 0.3333 | 3 | 0 | NA | NA | 4 | 0 | 10 | 0.1 | 1 | 1 | NA | NA | 3 | 0 | 1 | 0 | 5 | 0.2 |
| 11_7157377 | 3 | 0.6667 | 4 | 0.5 | 1 | 1 | 5 | 0.6 | 13 | 0.6154 | 1 | 0 | NA | NA | 4 | 0.5 | 2 | 0.5 | 7 | 0.4286 |
| 11_8382068 | 11 | 0.0909 | 18 | 0.3333 | 8 | 0.5 | 9 | 0.1111 | 46 | 0.2609 | 9 | 0.1111 | 6 | 0.1667 | 8 | 0.25 | 5 | 0.4 | 28 | 0.2143 |
| 11_8382080 | 13 | 0 | 16 | 0 | 8 | 0.25 | 10 | 0.1 | 47 | 0.0638 | 9 | 0.1111 | 5 | 0 | 8 | 0.125 | 4 | 0 | 26 | 0.0769 |
| 11_8382605 | 8 | 0.125 | 14 | 0.2857 | 3 | 0.3333 | 5 | 0 | 30 | 0.2 | 5 | 0.2 | 4 | 0.5 | 7 | 0.1429 | 3 | 0 | 19 | 0.2105 |
| 11_8382837 | 6 | 0 | 16 | 0.0625 | 4 | 0 | 8 | 0.125 | 34 | 0.0588 | 7 | 0.1429 | 5 | 0.2 | 5 | 0.2 | 3 | 0 | 20 | 0.15 |
| 11_8382987 | 5 | 0 | 9 | 0 | 3 | 0.6667 | 5 | 0 | 22 | 0.0909 | 8 | 0 | 4 | 0.25 | 4 | 0 | 2 | 0.5 | 18 | 0.1111 |
| 11_8383089 | 4 | 0 | 15 | 0.1333 | 4 | 0 | 5 | 0 | 28 | 0.0714 | 7 | 0.1429 | 4 | 0.25 | 3 | 0 | 3 | 0 | 17 | 0.1176 |
| 11_8383297 | 5 | 0.2 | 13 | 0.3077 | 4 | 0.5 | 4 | 0.75 | 26 | 0.3846 | 6 | 0.1667 | 3 | 0 | 3 | 0 | 3 | 0.3333 | 15 | 0.1333 |
| 11_8383346 | 4 | 0.25 | 12 | 0.4167 | 2 | 1 | 4 | 0.25 | 22 | 0.4091 | 6 | 0.5 | 3 | 0.6667 | 4 | 0.75 | 2 | 0.5 | 15 | 0.6 |
| 11_8383437 | 3 | 1 | 12 | 0.8333 | 1 | 0 | 3 | 0.6667 | 19 | 0.7895 | 6 | 1 | 4 | 0.75 | 2 | 1 | 1 | 1 | 13 | 0.9231 |
| 11_8383469 | 5 | 0.2 | 13 | 0.4615 | 2 | 0.5 | 4 | 0.75 | 24 | 0.4583 | 7 | 0.5714 | 4 | 0.25 | 3 | 0.3333 | 3 | 0.3333 | 17 | 0.4118 |
| 11_8383475 | 4 | 0 | 13 | 0.1538 | 2 | 0 | 5 | 0 | 24 | 0.0833 | 8 | 0.125 | 4 | 0.5 | 4 | 0 | 3 | 0 | 19 | 0.1579 |
| 11_8383634 | 2 | 0 | 9 | 0.3333 | 1 | 1 | 5 | 0.4 | 17 | 0.3529 | 3 | 0 | 4 | 0.25 | 3 | 0 | 3 | 0 | 13 | 0.0769 |
| 11_8383661 | 3 | 0.6667 | 13 | 0.3846 | 3 | 0.6667 | 5 | 1 | 24 | 0.5833 | 7 | 0.5714 | 4 | 0.25 | 3 | 0.6667 | 2 | 0.5 | 16 | 0.5 |
| 11_8383672 | 3 | 0.3333 | 11 | 0.2727 | NA | NA | 3 | 0 | 17 | 0.2353 | 7 | 0.1429 | 4 | 0.5 | 2 | 0 | 4 | 0.25 | 17 | 0.2353 |
| 11_8383837 | 2 | 0.5 | 11 | 0.2727 | 3 | 1 | 5 | 0.6 | 21 | 0.4762 | 7 | 0.4286 | 4 | 0 | 4 | 0 | 2 | 0 | 17 | 0.1765 |
| 11_8383889 | 2 | 1 | 11 | 1 | 3 | 1 | 4 | 1 | 20 | 1 | 7 | 0.8571 | 4 | 1 | 3 | 1 | 3 | 1 | 17 | 0.9412 |
| 11_8383927 | 2 | 0.5 | 8 | 0.125 | 2 | 0.5 | 5 | 0.2 | 17 | 0.2353 | 6 | 0.3333 | 2 | 0 | 2 | 0 | 2 | 0 | 12 | 0.1667 |
| 11_8383996 | 3 | 0.3333 | 7 | 0.2857 | 2 | 0.5 | 5 | 0.6 | 17 | 0.4118 | 7 | 0.4286 | 3 | 0.6667 | 4 | 0.75 | 2 | 0.5 | 16 | 0.5625 |
| 11_8384007 | 2 | 1 | 9 | 0.2222 | 2 | 0.5 | 5 | 0.4 | 18 | 0.3889 | 7 | 0.4286 | 3 | 0 | 3 | 0.3333 | 1 | 0 | 14 | 0.2857 |
| 11_8384052 | 2 | 1 | 9 | 0.8889 | 2 | 1 | 5 | 1 | 18 | 0.9444 | 5 | 1 | 4 | 1 | 4 | 1 | 1 | 0 | 14 | 0.9286 |
| 11_8384123 | 2 | 1 | 9 | 1 | 2 | 1 | 5 | 1 | 18 | 1 | 6 | 1 | 4 | 1 | 3 | 0.6667 | 2 | 1 | 15 | 0.9333 |
| 11_8384225 | 2 | 1 | 7 | 1 | 2 | 1 | 4 | 0.75 | 15 | 0.9333 | 6 | 1 | 4 | 0.75 | 3 | 0.6667 | 2 | 0.5 | 15 | 0.8 |
| 11_8384326 | 1 | 0 | 7 | 0.8571 | 1 | 0 | 4 | 0.75 | 13 | 0.6923 | 4 | 0.75 | 3 | 0.6667 | 1 | 1 | 1 | 0 | 9 | 0.6667 |
| 11_8384388 | 3 | 0.3333 | 5 | 0.2 | 1 | 0 | 4 | 0 | 13 | 0.1538 | 4 | 0.25 | 2 | 0 | 3 | 0 | 1 | 0 | 10 | 0.1 |
| 11_8384406 | 2 | 0.5 | 6 | 0.3333 | 1 | 1 | 5 | 0.4 | 14 | 0.4286 | 5 | 0.6 | 2 | 0 | 3 | 1 | NA | NA | 10 | 0.6 |
| 11_8401173 | 3 | 0.3333 | 2 | 0 | NA | NA | 2 | 0 | 7 | 0.1429 | 3 | 0.3333 | NA | NA | 2 | 0 | NA | NA | 5 | 0.2 |
| 11_8401373 | 3 | 1 | 2 | 1 | NA | NA | 3 | 1 | 8 | 1 | 4 | 1 | NA | NA | 2 | 0.5 | 1 | 1 | 7 | 0.8571 |
| 11_8442233 | 1 | 0 | 1 | 1 | NA | NA | NA | NA | 2 | 0.5 | 2 | 0.5 | NA | NA | NA | NA | NA | NA | 2 | 0.5 |
| 11_1106613 | 2 | 0 | 1 | 1 | 2 | 1 | 1 | 1 | 6 | 0.6667 | NA | NA | 1 | 0 | 1 | 1 | 1 | 1 | 3 | 0.6667 |
| 11_1106623 | 1 | 1 | NA | NA | 2 | 1 | 1 | 0 | 4 | 0.75 | NA | NA | 1 | 1 | 2 | 1 | 1 | 1 | 4 | 1 |
| 11_1106631 | 2 | 0 | 1 | 0 | 2 | 0.5 | NA | NA | 5 | 0.2 | NA | NA | 1 | 0 | 2 | 1 | 1 | 0 | 4 | 0.5 |
| 11_1106635 | 2 | 1 | NA | NA | 1 | 1 | 1 | 0 | 4 | 0.75 | NA | NA | 1 | 0 | 1 | 1 | 1 | 1 | 3 | 0.6667 |
| 11_1121152 | 2 | 0.5 | 1 | 0 | 1 | 0 | 1 | 0 | 5 | 0.2 | NA | NA | 1 | 0 | 2 | 1 | 1 | 0 | 4 | 0.5 |
| 11_1121153 | 2 | 0.5 | 1 | 1 | 1 | 1 | NA | NA | 4 | 0.75 | NA | NA | 1 | 1 | 2 | 0.5 | 1 | 0 | 4 | 0.5 |
| 11_1121158 | 2 | 0 | 1 | 1 | 2 | 0 | 1 | 0 | 6 | 0.1667 | NA | NA | 1 | 1 | 2 | 0.5 | 1 | 0 | 4 | 0.5 |
| 11_1121161 | 2 | 0.5 | 1 | 0 | 1 | 1 | 1 | 1 | 5 | 0.6 | NA | NA | 1 | 0 | 2 | 0.5 | 1 | 0 | 4 | 0.25 |
| 11_1121168 | 2 | 0 | 1 | 0 | 2 | 0.5 | 1 | 1 | 6 | 0.3333 | NA | NA | 1 | 1 | 2 | 0 | 1 | 0 | 4 | 0.25 |
| 11_1121171 | 2 | 1 | 1 | 1 | 2 | 1 | 1 | 1 | 6 | 1 | NA | NA | 1 | 1 | 2 | 1 | NA | NA | 3 | 1 |
| 11_1171935 | 1 | 1 | NA | NA | NA | NA | NA | NA | 1 | 1 | NA | NA | 1 | 0 | NA | NA | 1 | 1 | 2 | 0.5 |
| 11_1171941 | 1 | 1 | NA | NA | NA | NA | NA | NA | 1 | 1 | NA | NA | 1 | 1 | NA | NA | 1 | 1 | 2 | 1 |
| 11_1171965 | 1 | 1 | NA | NA | NA | NA | NA | NA | 1 | 1 | NA | NA | 1 | 1 | NA | NA | 1 | 1 | 2 | 1 |
| 11_1188098 | 5 | 0 | 5 | 0 | 2 | 0.5 | 4 | 0.25 | 16 | 0.125 | NA | NA | 6 | 0 | 4 | 0.25 | NA | NA | 10 | 0.1 |
| 11_1188313 | 3 | 1 | 2 | 1 | NA | NA | 4 | 1 | 9 | 1 | NA | NA | 3 | 0.6667 | 2 | 0.5 | 1 | 1 | 6 | 0.6667 |
| 11_1188330 | 3 | 0.6667 | 2 | 1 | NA | NA | 4 | 1 | 9 | 0.8889 | NA | NA | 3 | 1 | 2 | 1 | 1 | 1 | 6 | 1 |
| 11_1188336 | 2 | 1 | 2 | 0.5 | 1 | 1 | 2 | 1 | 7 | 0.8571 | NA | NA | 3 | 1 | 2 | 0.5 | NA | NA | 5 | 0.8 |
| 11_1218714 | 4 | 0 | 5 | 0.2 | 1 | 0 | 9 | 0.1111 | 19 | 0.1053 | 7 | 0.2857 | 4 | 0.25 | 6 | 0.1667 | 4 | 0.25 | 21 | 0.2381 |
| 11_1218733 | 3 | 0 | 5 | 0.2 | 1 | 0 | 7 | 0 | 16 | 0.0625 | 5 | 0.2 | 4 | 0.25 | 5 | 0.4 | 2 | 0 | 16 | 0.25 |
| 11_1218745 | 4 | 1 | 5 | 1 | 1 | 1 | 8 | 1 | 18 | 1 | 7 | 1 | 3 | 1 | 5 | 1 | 3 | 1 | 18 | 1 |
| 11_1218753 | 2 | 0 | 5 | 0.2 | 1 | 0 | 8 | 0.125 | 16 | 0.125 | 7 | 0 | 2 | 0.5 | 6 | 0.1667 | NA | NA | 15 | 0.1333 |
| 11_1218769 | 4 | 1 | 5 | 0.8 | NA | NA | 7 | 0.7143 | 16 | 0.8125 | 6 | 0.6667 | 4 | 0.75 | 4 | 1 | 1 | 1 | 15 | 0.8 |
| 11_1218780 | 3 | 0.6667 | 4 | 1 | 1 | 0 | 4 | 1 | 12 | 0.8333 | 5 | 0.4 | 4 | 0.75 | 3 | 0.6667 | 1 | 1 | 13 | 0.6154 |
| 11_1218784 | 4 | 1 | 4 | 0.75 | 1 | 1 | 8 | 1 | 17 | 0.9412 | 7 | 1 | 4 | 0.75 | 5 | 0.8 | 1 | 1 | 17 | 0.8824 |

| | | | | | | | | | | | | | | | | | | | | |
|---|---|---|---|---|---|---|---|---|---|---|---|---|---|---|---|---|---|---|---|---|
| 11_1218785 | 4 | 0.25 | 4 | 0.25 | 1 | 1 | 8 | 0.625 | 17 | 0.4706 | 7 | 0.4286 | 4 | 0.25 | 5 | 0.4 | 1 | 0 | 17 | 0.3529 |
| 11_1220327 | 3 | 0.3333 | 3 | 0 | 1 | 0 | 6 | 0.1667 | 13 | 0.1538 | 3 | 0.3333 | 2 | 0.5 | 4 | 0 | 1 | 0 | 10 | 0.2 |
| 11_1220329 | 3 | 0.3333 | 3 | 0 | 1 | 0 | 6 | 0.1667 | 13 | 0.1538 | 3 | 0 | 2 | 0 | 4 | 0.25 | NA | NA | 9 | 0.1111 |
| 11_1220334 | 3 | 0 | 3 | 0 | 1 | 0 | 6 | 0.1667 | 13 | 0.0769 | 2 | 0 | 1 | 0 | 4 | 0.25 | NA | NA | 7 | 0.1429 |
| 11_1220338 | 3 | 1 | 3 | 1 | NA | NA | 6 | 0.6667 | 12 | 0.8333 | 2 | 1 | 2 | 1 | 4 | 0.75 | 1 | 1 | 9 | 0.8889 |
| 11_1220342 | 3 | 0.6667 | 3 | 0 | NA | NA | 5 | 0 | 11 | 0.1818 | 3 | 0.3333 | 2 | 0 | 4 | 0 | 1 | 0 | 10 | 0.1 |
| 11_1220350 | 3 | 0 | 3 | 0.3333 | 1 | 1 | 4 | 0.5 | 11 | 0.3636 | 2 | 0.5 | 1 | 0 | 4 | 0.25 | NA | NA | 7 | 0.2857 |
| 11_1220357 | 3 | 0.3333 | 3 | 0 | 1 | 0 | 6 | 0 | 13 | 0.0769 | 3 | 0 | 2 | 0 | 4 | 0.5 | NA | NA | 9 | 0.2222 |
| 11_1220368 | 3 | 0.3333 | 3 | 0.3333 | 1 | 1 | 4 | 0.5 | 11 | 0.4545 | 3 | 0.6667 | 2 | 0 | 3 | 0.3333 | NA | NA | 8 | 0.375 |
| 11_1220378 | 3 | 0.3333 | 3 | 0 | 1 | 0 | 4 | 0 | 11 | 0.0909 | 2 | 0.5 | 1 | 0 | 3 | 0.3333 | 1 | 0 | 7 | 0.2857 |
| 11_1220419 | 2 | 0 | 2 | 0 | NA | NA | 4 | 0.25 | 8 | 0.125 | 3 | 0.3333 | 2 | 0 | 1 | 0 | 1 | 0 | 7 | 0.1429 |
| 11_1220427 | 2 | 0.5 | 3 | 0.3333 | NA | NA | 4 | 0 | 9 | 0.2222 | 3 | 0.6667 | 2 | 0.5 | 1 | 0 | NA | NA | 6 | 0.5 |
| 11_1220437 | 2 | 1 | 3 | 1 | NA | NA | 4 | 1 | 9 | 1 | 2 | 1 | NA | NA | 1 | 1 | NA | NA | 3 | 1 |
| 11_1220469 | 2 | 0 | 2 | 1 | NA | NA | 4 | 0 | 8 | 0.25 | 2 | 0 | 1 | 1 | 1 | 0 | 1 | 1 | 5 | 0.4 |
| 11_1220472 | 2 | 1 | 2 | 1 | NA | NA | 4 | 1 | 8 | 1 | 2 | 1 | 1 | 1 | 1 | 1 | NA | NA | 4 | 1 |
| 11_1221704 | 1 | 0 | 3 | 0 | NA | NA | 3 | 0.6667 | 7 | 0.2857 | 2 | 0 | 1 | 1 | 1 | 0 | NA | NA | 4 | 0.25 |
| 11_1621300 | 11 | 0.1818 | 12 | 0.0833 | 6 | 0.3333 | 3 | 0 | 32 | 0.1562 | 8 | 0 | 7 | 0 | 8 | 0.125 | 11 | 0.2727 | 34 | 0.1176 |
| 11_1623186 | 4 | 1 | 3 | 1 | 2 | 1 | 2 | 0.5 | 11 | 0.9091 | 2 | 0.5 | 3 | 1 | 2 | 1 | 3 | 1 | 10 | 0.9 |
| 11_1623193 | 3 | 1 | 3 | 1 | 2 | 1 | 2 | 0.5 | 10 | 0.9 | 2 | 1 | 2 | 1 | 2 | 1 | 3 | 1 | 9 | 1 |
| 11_1623199 | 13 | 0.4615 | 10 | 0.2 | 8 | 0.375 | 3 | 0.3333 | 34 | 0.3529 | 7 | 0.2857 | 7 | 0.1429 | 8 | 0.125 | 10 | 0.3 | 32 | 0.2188 |
| 11_1623209 | 12 | 0.1667 | 10 | 0.2 | 6 | 0.5 | 5 | 0.2 | 33 | 0.2424 | 7 | 0.2857 | 7 | 0.4286 | 8 | 0.75 | 12 | 0.3333 | 34 | 0.4412 |
| 11_1623217 | 13 | 1 | 12 | 1 | 7 | 0.8571 | 4 | 1 | 36 | 0.9722 | 10 | 1 | 7 | 1 | 10 | 1 | 13 | 1 | 40 | 1 |
| 11_1623219 | 11 | 0.2727 | 9 | 0.3333 | 6 | 0.1667 | 4 | 0.25 | 30 | 0.2667 | 8 | 0.25 | 6 | 0.1667 | 9 | 0.4444 | 13 | 0.3077 | 36 | 0.3056 |
| 11_1623224 | 12 | 1 | 9 | 1 | 7 | 1 | 2 | 1 | 30 | 1 | 10 | 1 | 5 | 1 | 8 | 1 | 13 | 1 | 36 | 1 |
| 11_1623226 | 13 | 0.2308 | 13 | 0.0769 | 7 | 0.4286 | 5 | 0.4 | 38 | 0.2368 | 10 | 0.4 | 6 | 0.3333 | 10 | 0.5 | 13 | 0.3846 | 39 | 0.4103 |
| 11_1623231 | 14 | 0.8571 | 9 | 0.7778 | 5 | 1 | 5 | 1 | 33 | 0.8788 | 9 | 0.8889 | 4 | 1 | 9 | 0.6667 | 12 | 1 | 34 | 0.8824 |
| 11_1623236 | 7 | 0.4286 | 8 | 0.375 | 5 | 0.4 | 5 | 0.2 | 25 | 0.36 | 7 | 0.1429 | 5 | 0.2 | 6 | 0.1667 | 10 | 0.5 | 28 | 0.2857 |
| 11_1623236 | 7 | 1 | 7 | 1 | 6 | 1 | 3 | 1 | 23 | 1 | 5 | 1 | 4 | 0.75 | 6 | 1 | 10 | 1 | 25 | 0.96 |
| 11_1623239 | 12 | 0.1667 | 10 | 0.3 | 7 | 0.5714 | 5 | 0.6 | 34 | 0.3529 | 10 | 0.2 | 5 | 0.4 | 9 | 0.5556 | 12 | 0.4167 | 36 | 0.3889 |
| 11_1623244 | 13 | 0.4615 | 12 | 0.1667 | 8 | 0.625 | 4 | 0.25 | 37 | 0.3784 | 10 | 0.4 | 7 | 0.1429 | 10 | 0.3 | 10 | 0.3 | 37 | 0.2973 |
| 11_1623275 | 11 | 0.3636 | 15 | 0.6 | 9 | 0.6667 | 8 | 0.625 | 43 | 0.5581 | 8 | 0.25 | 5 | 0.4 | 9 | 0.2222 | 12 | 0.75 | 34 | 0.4412 |
| 11_1623312 | 15 | 0.2667 | 13 | 0.2308 | 8 | 0.625 | 9 | 0 | 45 | 0.2667 | 9 | 0.1111 | 9 | 0.1111 | 13 | 0.3077 | 14 | 0.4286 | 45 | 0.2667 |
| 11_1623315 | 16 | 0.25 | 15 | 0.1333 | 9 | 0 | 9 | 0 | 49 | 0.1224 | 9 | 0.1111 | 10 | 0.3 | 12 | 0.1667 | 13 | 0.1538 | 44 | 0.1818 |
| 11_1623319 | 13 | 0.2308 | 15 | 0.1333 | 8 | 0.25 | 6 | 0 | 42 | 0.1667 | 9 | 0 | 10 | 0.1 | 13 | 0.2308 | 15 | 0.4667 | 47 | 0.234 |
| 11_1623322 | 13 | 0.3846 | 15 | 0.3333 | 9 | 0.3333 | 9 | 0.2222 | 46 | 0.3261 | 9 | 0 | 11 | 0.1818 | 11 | 0.2727 | 14 | 0.3571 | 45 | 0.2222 |
| 11_1623333 | 16 | 0.1875 | 14 | 0.3571 | 8 | 0.5 | 9 | 0 | 47 | 0.2553 | 8 | 0 | 9 | 0.1111 | 15 | 0.2 | 13 | 0 | 45 | 0.0889 |
| 11_1623367 | 15 | 0.1333 | 15 | 0.2 | 10 | 0.3 | 9 | 0.1111 | 49 | 0.1837 | 9 | 0.2222 | 7 | 0.1429 | 15 | 0.2667 | 16 | 0.25 | 47 | 0.234 |
| 11_1623675 | 1 | 0 | 3 | 0.3333 | NA | NA | 1 | 0 | 5 | 0.2 | 1 | 0 | 1 | 1 | 1 | 1 | 4 | 0.25 | 7 | 0.4286 |
| 11_1645073 | 1 | 1 | NA | NA | 1 | 0 | 2 | 0 | 4 | 0.25 | NA | NA | 1 | 0 | 2 | 0.5 | 2 | 0 | 5 | 0.2 |
| 11_1645078 | 1 | 1 | 1 | 0 | 1 | 0 | 2 | 1 | 5 | 0.6 | NA | NA | 1 | 0 | 3 | 0.6667 | 4 | 0.5 | 8 | 0.5 |
| 11_1645084 | 1 | 1 | 1 | 0 | 1 | 0 | 2 | 1 | 5 | 0.6 | NA | NA | 2 | 0 | 3 | 0.6667 | 3 | 0.6667 | 8 | 0.5 |
| 11_1645150 | 1 | 1 | 1 | 0 | 1 | 0 | 1 | 0 | 4 | 0.25 | NA | NA | 2 | 0 | 2 | 0.5 | 3 | 0 | 7 | 0.1429 |
| 11_1686158 | 1 | 1 | 1 | 0 | NA | NA | NA | NA | 2 | 0.5 | 1 | 0 | 3 | 0.3333 | NA | NA | NA | NA | 4 | 0.25 |
| 11_1686174 | 1 | 1 | 1 | 1 | NA | NA | NA | NA | 2 | 1 | 2 | 1 | 3 | 1 | NA | NA | NA | NA | 5 | 1 |
| 11_1686189 | 1 | 1 | 1 | 1 | NA | NA | NA | NA | 2 | 1 | 2 | 1 | 3 | 1 | NA | NA | NA | NA | 5 | 1 |
| 11_1686191 | 1 | 1 | 1 | 1 | NA | NA | NA | NA | 2 | 1 | 2 | 0.5 | 3 | 1 | NA | NA | NA | NA | 5 | 0.8 |
| 11_1686197 | 1 | 1 | 1 | 1 | NA | NA | NA | NA | 2 | 1 | 2 | 1 | 3 | 0.6667 | NA | NA | NA | NA | 5 | 0.8 |
| 11_1686207 | 1 | 1 | 1 | 1 | NA | NA | NA | NA | 2 | 1 | 1 | 0 | 3 | 0.6667 | NA | NA | NA | NA | 4 | 0.5 |
| 11_1686211 | 1 | 1 | NA | NA | NA | NA | NA | NA | 1 | 1 | NA | NA | 3 | 0.6667 | NA | NA | NA | NA | 3 | 0.6667 |
| 11_1686217 | 1 | 1 | NA | NA | NA | NA | NA | NA | 1 | 1 | 2 | 0.5 | 3 | 0.6667 | NA | NA | NA | NA | 5 | 0.6 |
| 11_1686219 | 1 | 1 | 1 | 1 | NA | NA | NA | NA | 2 | 1 | 1 | 1 | 3 | 1 | NA | NA | NA | NA | 4 | 1 |
| 11_1692214 | 2 | 0 | 3 | 0.3333 | NA | NA | 2 | 0 | 7 | 0.1429 | 1 | 0 | 2 | 0.5 | 2 | 0.5 | 2 | 0 | 7 | 0.2857 |
| 11_1700218 | 9 | 0 | 13 | 0.2308 | 6 | 0 | 3 | 0.3333 | 31 | 0.129 | 7 | 0.1429 | 6 | 0.1667 | 4 | 0 | 3 | 0.3333 | 20 | 0.15 |
| 11_1702859 | 11 | 0.1818 | 15 | 0.0667 | 5 | 0 | 4 | 0.25 | 35 | 0.1143 | 8 | 0.125 | 7 | 0.2857 | 4 | 0 | 7 | 0.2857 | 26 | 0.1923 |
| 11_1702860 | 7 | 0.7143 | 10 | 0.7 | 4 | 0.5 | 1 | 1 | 22 | 0.6818 | 8 | 0.875 | 5 | 0.6 | 3 | 0.3333 | 6 | 0.6667 | 22 | 0.6818 |
| 11_1702861 | 8 | 0.125 | 11 | 0 | 5 | 0 | 4 | 0.25 | 28 | 0.0714 | 7 | 0 | 6 | 0.3333 | 3 | 0 | 7 | 0.1429 | 23 | 0.1304 |
| 11_1702870 | 13 | 0.1538 | 14 | 0.0714 | 5 | 0.4 | 3 | 0.3333 | 35 | 0.1714 | 8 | 0.25 | 6 | 0 | 3 | 0 | 7 | 0.2857 | 24 | 0.1667 |
| 11_1702886 | 13 | 0.5385 | 14 | 0.2857 | 6 | 0 | 5 | 0 | 38 | 0.2895 | 8 | 0 | 7 | 0.2857 | 3 | 0 | 7 | 0.2857 | 25 | 0.16 |
| 11_1702900 | 12 | 0 | 17 | 0 | 6 | 0.1667 | 5 | 0.2 | 40 | 0.05 | 9 | 0.3333 | 7 | 0 | 4 | 0 | 7 | 0 | 27 | 0.1111 |
| 11_1702925 | 14 | 0.3571 | 15 | 0.3333 | 4 | 0.5 | 4 | 0 | 37 | 0.3243 | 8 | 0.25 | 7 | 0.1429 | 5 | 0.2 | 6 | 0 | 26 | 0.1538 |
| 11_1714830 | 57 | 0.0702 | 41 | 0.1463 | 38 | 0.0789 | 45 | 0.0667 | 181 | 0.0884 | 35 | 0.0286 | 56 | 0.0714 | 45 | 0.0222 | 34 | 0.0882 | 170 | 0.0529 |
| 11_1714846 | 53 | 0.2264 | 36 | 0.3056 | 34 | 0.1471 | 40 | 0.075 | 163 | 0.1902 | 36 | 0.1111 | 51 | 0.3529 | 39 | 0.1795 | 34 | 0.2647 | 160 | 0.2375 |
| 11_1714851 | 54 | 0.1667 | 36 | 0.1667 | 38 | 0.1579 | 43 | 0.186 | 171 | 0.1696 | 40 | 0.15 | 50 | 0.1 | 42 | 0.119 | 32 | 0.125 | 164 | 0.122 |
| 11_1714929 | 76 | 0.3289 | 60 | 0.3667 | 49 | 0.2449 | 57 | 0.2456 | 242 | 0.3017 | 41 | 0.2439 | 64 | 0.3594 | 55 | 0.3273 | 39 | 0.4103 | 199 | 0.3367 |
| 11_1714934 | 78 | 0.1923 | 57 | 0.2281 | 46 | 0.1522 | 53 | 0.1698 | 234 | 0.188 | 47 | 0.1915 | 64 | 0.2188 | 57 | 0.193 | 43 | 0.1628 | 211 | 0.1943 |

| | | | | | | | | | | | | | | | | | | | | |
|---|---|---|---|---|---|---|---|---|---|---|---|---|---|---|---|---|---|---|---|---|
| 11_1715363 | 7 | 0.1429 | 2 | 0.5 | 4 | 0 | 4 | 0 | 17 | 0.1176 | 5 | 0 | 4 | 0 | 8 | 0.25 | 6 | 0 | 23 | 0.087 |
| 11_1715381 | 8 | 0.375 | 2 | 0.5 | 5 | 0.2 | 4 | 0.75 | 19 | 0.4211 | 4 | 0 | 5 | 0.2 | 7 | 0.4286 | 7 | 0.1429 | 23 | 0.2174 |
| 11_1718292 | 2 | 0.5 | 2 | 1 | 1 | 0 | NA | NA | 5 | 0.6 | 1 | 0 | 2 | 1 | 2 | 0 | 2 | 0.5 | 7 | 0.4286 |
| 11_1789013 | 3 | 0 | 2 | 0 | 3 | 0 | 2 | 0.5 | 10 | 0.1 | 2 | 0 | 2 | 0 | 4 | 0.25 | 1 | 0 | 9 | 0.1111 |
| 11_1789017 | 1 | 1 | 2 | 1 | 3 | 1 | 1 | 1 | 7 | 1 | 2 | 0.5 | 2 | 0.5 | 4 | 0.5 | 1 | 1 | 9 | 0.5556 |
| 11_1789568 | 2 | 0.5 | 2 | 0 | 3 | 1 | 1 | 1 | 8 | 0.625 | 2 | 0 | 1 | 0 | 3 | 0.6667 | 1 | 0 | 7 | 0.2857 |
| 11_1789595 | 2 | 0 | 2 | 0.5 | 3 | 0.6667 | 2 | 1 | 9 | 0.5556 | 2 | 0.5 | 1 | 0 | 2 | 1 | 1 | 1 | 6 | 0.6667 |
| 11_1789605 | 2 | 0.5 | 2 | 0 | 3 | 0.3333 | 2 | 0 | 9 | 0.2222 | 2 | 0.5 | 2 | 0.5 | 2 | 0.5 | NA | NA | 6 | 0.5 |
| 11_1789624 | 3 | 0.6667 | 2 | 0.5 | 3 | 0.6667 | 1 | 1 | 9 | 0.6667 | 2 | 1 | 1 | 0 | 2 | 0.5 | 1 | 1 | 6 | 0.6667 |
| 11_1789717 | 1 | 0 | 1 | 0 | 2 | 0.5 | 2 | 0 | 6 | 0.1667 | 1 | 0 | 2 | 0.5 | 2 | 0.5 | 1 | 0 | 6 | 0.3333 |
| 11_1789719 | 2 | 0.5 | 2 | 0 | 2 | 0.5 | 2 | 1 | 8 | 0.5 | 1 | 0 | 2 | 0 | 2 | 0.5 | 1 | 0 | 6 | 0.1667 |
| 11_1789722 | 2 | 0 | 2 | 0.5 | 3 | 0.6667 | 2 | 0 | 9 | 0.3333 | 1 | 1 | 2 | 0.5 | 2 | 0.5 | 1 | 1 | 6 | 0.6667 |
| 11_1789730 | 1 | 0 | 2 | 0 | 3 | 0 | 2 | 0.5 | 8 | 0.125 | 1 | 1 | 2 | 0 | 2 | 0 | 1 | 1 | 6 | 0.3333 |
| 11_1789743 | 2 | 1 | 2 | 1 | 1 | 1 | 2 | 1 | 7 | 1 | NA | NA | 2 | 1 | 2 | 1 | 1 | 1 | 5 | 1 |
| 11_1789754 | 2 | 0 | 2 | 0.5 | 3 | 0 | 2 | 0 | 9 | 0.1111 | NA | NA | 2 | 0 | 2 | 0.5 | 1 | 0 | 5 | 0.2 |
| 11_1883080 | NA | NA | NA | NA | NA | NA | 1 | 1 | 1 | 1 | NA | NA | NA | NA | 1 | 1 | 3 | 0.3333 | 4 | 0.5 |
| 11_1997092 | 4 | 0.25 | 5 | 0 | 3 | 0.3333 | 4 | 0 | 16 | 0.125 | 4 | 0.25 | 5 | 0 | 1 | 0 | 2 | 0 | 12 | 0.0833 |
| 11_1997094 | 5 | 1 | 5 | 1 | 4 | 1 | 5 | 1 | 19 | 1 | 4 | 1 | 5 | 0.8 | 1 | 1 | 2 | 0.5 | 12 | 0.8333 |
| 11_1997101 | 4 | 1 | 4 | 1 | 4 | 0.75 | 2 | 1 | 14 | 0.9286 | 4 | 1 | 5 | 0.8 | 1 | 1 | 1 | 1 | 11 | 0.9091 |
| 11_1997105 | 2 | 0.5 | 6 | 0.1667 | 3 | 0.3333 | 3 | 0 | 14 | 0.2143 | 4 | 0.25 | 3 | 0 | NA | NA | 2 | 0 | 9 | 0.1111 |
| 11_1997119 | 5 | 0.6 | 6 | 0.8333 | 3 | 0.3333 | 6 | 0.6667 | 20 | 0.65 | 3 | 0.3333 | 4 | 0.75 | 2 | 0.5 | 3 | 0.6667 | 12 | 0.5833 |
| 11_1997123 | 5 | 0.6 | 6 | 0.8333 | 4 | 0.75 | 4 | 0.75 | 19 | 0.7368 | 3 | 0.6667 | 5 | 0.4 | 2 | 1 | 3 | 1 | 13 | 0.6923 |
| 11_1997126 | 5 | 0.2 | 6 | 0 | 4 | 0.25 | 4 | 0.25 | 19 | 0.1579 | 4 | 0 | 5 | 0.2 | 2 | 0 | 3 | 0.3333 | 14 | 0.1429 |
| 11_1997127 | 5 | 0 | 6 | 0.1667 | 4 | 0.25 | 5 | 0 | 20 | 0.1 | 4 | 0.25 | 5 | 0.4 | 2 | 0.5 | 3 | 0 | 14 | 0.2857 |
| 11_1997129 | 5 | 0 | 5 | 0 | 4 | 0.25 | 6 | 0 | 20 | 0.05 | 4 | 0 | 5 | 0.2 | 2 | 0 | 3 | 0 | 14 | 0.0714 |
| 11_1997144 | 4 | 0.25 | 6 | 0.3333 | 4 | 0.5 | 5 | 0.6 | 19 | 0.4211 | 4 | 0.5 | 4 | 0.5 | 1 | 0 | 3 | 0.3333 | 12 | 0.4167 |
| 11_1997159 | 4 | 0 | 6 | 0.1667 | 4 | 0 | 4 | 0.5 | 18 | 0.1667 | 4 | 0 | 5 | 0.4 | 2 | 0 | 3 | 0.3333 | 14 | 0.2143 |
| 11_1997168 | 5 | 0.6 | 8 | 0.25 | 4 | 0.25 | 6 | 0.6667 | 23 | 0.4348 | 4 | 0 | 5 | 0.8 | 2 | 0 | 2 | 0.5 | 13 | 0.3846 |
| 11_1997170 | 5 | 0 | 6 | 0 | 4 | 0.25 | 4 | 0 | 19 | 0.0526 | 4 | 0.25 | 5 | 0 | 2 | 0 | 3 | 0 | 14 | 0.0714 |
| 11_1997180 | 4 | 0.25 | 7 | 0 | 3 | 0.3333 | 4 | 0 | 18 | 0.1111 | 4 | 0 | 6 | 0.1667 | 1 | 0 | 2 | 0.5 | 13 | 0.1538 |
| 11_1997315 | 8 | 0.25 | 12 | 0.0833 | 5 | 0 | 7 | 0 | 32 | 0.0938 | 5 | 0 | 5 | 0 | 2 | 0.5 | 4 | 0 | 16 | 0.0625 |
| 11_1997330 | 9 | 0.1111 | 9 | 0.1111 | 4 | 0 | 6 | 0.3333 | 28 | 0.1429 | 7 | 0.2857 | 7 | 0.4286 | 2 | 0 | 5 | 0.2 | 21 | 0.2857 |
| 11_1997334 | 8 | 0 | 11 | 0.0909 | 4 | 0.5 | 5 | 0.2 | 28 | 0.1429 | 5 | 0 | 6 | 0 | 2 | 0 | 3 | 0.3333 | 16 | 0.0625 |
| 11_1997359 | 9 | 0 | 10 | 0.1 | 3 | 0 | 5 | 0.2 | 27 | 0.0741 | 6 | 0.1667 | 8 | 0 | 3 | 0.3333 | 5 | 0.2 | 22 | 0.1364 |
| 11_1997360 | 7 | 0.4286 | 11 | 0.1818 | 6 | 0.3333 | 5 | 0 | 29 | 0.2414 | 7 | 0.1429 | 6 | 0.1667 | 2 | 0 | 5 | 0.2 | 20 | 0.15 |
| 11_2001291 | 46 | 0.3043 | 49 | 0.2857 | 34 | 0.2941 | 43 | 0.3488 | 172 | 0.3081 | 23 | 0.3913 | 30 | 0.3333 | 46 | 0.2609 | 42 | 0.4762 | 141 | 0.3617 |
| 11_2001293 | 48 | 0.1042 | 44 | 0.1818 | 34 | 0.1176 | 42 | 0.0952 | 168 | 0.125 | 21 | 0.0952 | 33 | 0.1818 | 46 | 0.1522 | 42 | 0.119 | 142 | 0.1408 |
| 11_2001398 | 23 | 0.0435 | 34 | 0.1176 | 23 | 0.0435 | 20 | 0 | 100 | 0.06 | 10 | 0.1 | 20 | 0.1 | 32 | 0.0625 | 27 | 0.1481 | 89 | 0.1011 |
| 11_2001458 | 6 | 0.5 | 11 | 0.3636 | 5 | 0.2 | 4 | 0.5 | 26 | 0.3846 | 4 | 0.25 | 3 | 0.6667 | 5 | 0.8 | 6 | 0.6667 | 18 | 0.6111 |
| 11_2001470 | 10 | 0.2 | 9 | 0.1111 | 5 | 0 | 5 | 0.4 | 29 | 0.1724 | 4 | 0 | 2 | 0 | 8 | 0.125 | 4 | 0.25 | 18 | 0.1111 |
| 11_2017471 | 22 | 0.0455 | 27 | 0 | 23 | 0.1304 | 23 | 0.1739 | 95 | 0.0842 | 15 | 0 | 21 | 0.1429 | 32 | 0.125 | 22 | 0.0455 | 90 | 0.0889 |
| 11_2017499 | 30 | 0.2667 | 32 | 0.0625 | 26 | 0.0769 | 28 | 0.1071 | 116 | 0.1293 | 21 | 0.0952 | 25 | 0.12 | 36 | 0.1111 | 29 | 0.1034 | 111 | 0.1081 |
| 11_2017504 | 22 | 0.3636 | 36 | 0.25 | 26 | 0.1923 | 25 | 0.32 | 109 | 0.2752 | 20 | 0.2 | 24 | 0.2083 | 31 | 0.1935 | 26 | 0.2692 | 101 | 0.2178 |
| 11_2017699 | 2 | 0 | 4 | 0 | 1 | 1 | 2 | 0 | 9 | 0.1111 | NA | NA | 2 | 0.5 | 3 | 0.6667 | 1 | 0 | 6 | 0.5 |
| 11_2017923 | 2 | 0 | 4 | 0 | 1 | 0 | 2 | 0.5 | 9 | 0.1111 | NA | NA | 2 | 0 | 3 | 0.3333 | NA | NA | 5 | 0.2 |
| 11_2018858 | 2 | 0.5 | 5 | 0 | 5 | 0 | 2 | 0 | 14 | 0.0714 | NA | NA | 1 | 0 | 3 | 0 | 3 | 0.3333 | 7 | 0.1429 |
| 11_2018938 | 2 | 1 | 5 | 1 | 4 | 0.75 | 2 | 1 | 13 | 0.9231 | NA | NA | 1 | 1 | 3 | 1 | 2 | 1 | 6 | 1 |
| 11_2018944 | 1 | 0 | 5 | 0.2 | 5 | 0 | 2 | 0 | 13 | 0.0769 | NA | NA | 1 | 1 | 3 | 0.3333 | 3 | 0 | 7 | 0.2857 |
| 11_2018947 | 1 | 1 | 2 | 1 | 2 | 1 | NA | NA | 5 | 1 | NA | NA | NA | NA | 2 | 1 | 2 | 1 | 4 | 1 |
| 11_2018952 | 1 | 0 | 4 | 0.5 | 4 | 0.25 | 1 | 0 | 10 | 0.3 | NA | NA | 1 | 0 | 1 | 0 | 3 | 0.3333 | 5 | 0.2 |
| 11_2018960 | 1 | 0 | 4 | 0.25 | 5 | 0 | 2 | 0.5 | 12 | 0.1667 | NA | NA | 1 | 0 | 1 | 0 | 1 | 1 | 3 | 0.3333 |
| 11_2018979 | 1 | 0 | 4 | 0.25 | 4 | 0 | NA | NA | 9 | 0.1111 | NA | NA | 1 | 0 | 1 | 1 | 3 | 0 | 5 | 0.2 |
| 11_2019005 | 1 | 1 | 4 | 1 | 4 | 0.75 | 2 | 1 | 11 | 0.9091 | NA | NA | 1 | 1 | 1 | 1 | 3 | 0.6667 | 5 | 0.8 |
| 11_2019007 | 1 | 0 | 3 | 0 | 5 | 0.4 | 2 | 0 | 11 | 0.1818 | NA | NA | 1 | 1 | 1 | 0 | 3 | 0 | 5 | 0.2 |
| 11_2019018 | NA | NA | 4 | 0 | 5 | 0 | 2 | 0.5 | 11 | 0.0909 | NA | NA | NA | NA | 1 | 1 | 1 | 1 | 2 | 1 |
| 11_2019026 | NA | NA | 2 | 1 | 4 | 0 | 2 | 0.5 | 8 | 0.375 | NA | NA | NA | NA | 1 | 1 | 2 | 1 | 3 | 1 |
| 11_2019037 | NA | NA | 2 | 1 | 4 | 0.5 | 2 | 1 | 8 | 0.75 | NA | NA | NA | NA | NA | NA | 2 | 1 | 2 | 1 |
| 11_2019041 | NA | NA | 1 | 0 | 5 | 0.6 | 2 | 0 | 8 | 0.375 | NA | NA | NA | NA | NA | NA | 2 | 0.5 | 2 | 0.5 |
| 11_2025229 | 8 | 0.125 | 5 | 0.4 | 3 | 0.3333 | 5 | 0.2 | 21 | 0.2381 | 3 | 1 | 2 | 0.5 | 4 | 0.25 | 5 | 0.2 | 14 | 0.4286 |
| 11_2025243 | 8 | 0.125 | 4 | 0.25 | 2 | 0 | 4 | 0 | 18 | 0.1111 | 4 | 0 | 3 | 0.6667 | 3 | 0.3333 | 5 | 0.2 | 15 | 0.2667 |
| 11_2025250 | 4 | 0.5 | 5 | 0.6 | 3 | 1 | 5 | 0.2 | 17 | 0.5294 | 1 | 1 | 3 | 0.6667 | 5 | 0.8 | 4 | 0.5 | 13 | 0.6923 |
| 11_2025251 | 8 | 0.125 | 6 | 0.1667 | 2 | 0.5 | 5 | 0.2 | 21 | 0.1905 | 3 | 0.3333 | 4 | 0.25 | 4 | 0 | 4 | 0.25 | 15 | 0.2 |
| 11_2025278 | 13 | 0.0769 | 3 | 0.3333 | 3 | 0 | 3 | 0.3333 | 22 | 0.1364 | 4 | 0.25 | 3 | 0.6667 | 7 | 0.2857 | 7 | 0.7143 | 21 | 0.4762 |
| 11_2025283 | 11 | 0.2727 | 5 | 0.4 | 2 | 1 | 3 | 0.6667 | 21 | 0.4286 | 4 | 0.75 | 4 | 0.5 | 5 | 0.6 | 4 | 0.75 | 17 | 0.6471 |
| 11_2025291 | 12 | 0.3333 | 4 | 0.5 | 3 | 1 | 2 | 0.5 | 21 | 0.4762 | 3 | 1 | 4 | 0.75 | 7 | 0.4286 | 4 | 0.75 | 18 | 0.6667 |

| | | | | | | | | | | | | | | | | | | | | |
|---|---|---|---|---|---|---|---|---|---|---|---|---|---|---|---|---|---|---|---|---|
| 11_2025303 | 13 | 0.0769 | 3 | 0 | 4 | 0.25 | 3 | 0 | 23 | 0.087 | 4 | 0 | 4 | 0 | 6 | 0.1667 | 6 | 0 | 20 | 0.05 |
| 11_2025311 | 12 | 0.0833 | 4 | 0.25 | 3 | 0 | 3 | 0.3333 | 22 | 0.1364 | 4 | 0.5 | 5 | 0.2 | 7 | 0 | 5 | 0 | 21 | 0.1429 |
| 11_2025315 | 10 | 0.8 | 3 | 0.6667 | 4 | 1 | 3 | 1 | 20 | 0.85 | 2 | 1 | 3 | 1 | 6 | 1 | 6 | 1 | 17 | 1 |
| 11_2025320 | 11 | 0.7273 | 4 | 0.25 | 3 | 1 | 1 | 0 | 19 | 0.6316 | 4 | 0.75 | 4 | 0.75 | 7 | 0.7143 | 5 | 0.6 | 20 | 0.7 |
| 11_2025342 | 10 | 0.7 | 1 | 0 | 3 | 1 | 2 | 0 | 16 | 0.625 | 4 | 0.25 | 5 | 0.2 | 5 | 0.2 | 5 | 0.6 | 19 | 0.3158 |
| 11_2025351 | 13 | 0.8462 | 3 | 0.6667 | 4 | 1 | 3 | 0.6667 | 23 | 0.8261 | 4 | 1 | 5 | 1 | 6 | 1 | 7 | 0.8571 | 22 | 0.9545 |
| 11_2025357 | 14 | 0.7857 | 3 | 0.6667 | 3 | 0.6667 | 3 | 1 | 23 | 0.7826 | 3 | 1 | 5 | 0.6 | 4 | 0.25 | 5 | 0.8 | 17 | 0.6471 |
| 11_2025361 | 13 | 0.2308 | 3 | 0 | 3 | 0.3333 | 2 | 0.5 | 21 | 0.2381 | 4 | 0.25 | 5 | 0.2 | 6 | 0 | 6 | 0.3333 | 21 | 0.1905 |
| 11_2025366 | 14 | 0.1429 | 3 | 0 | 3 | 0.3333 | 3 | 0.6667 | 23 | 0.2174 | 3 | 0 | 5 | 0.4 | 4 | 0.5 | 4 | 0.75 | 16 | 0.4375 |
| 11_2025367 | 11 | 0 | 2 | 0 | 2 | 0 | 3 | 0.3333 | 18 | 0.0556 | 2 | 0.5 | 4 | 0 | 3 | 0 | 4 | 0 | 13 | 0.0769 |
| 11_2025368 | 10 | 0 | 5 | 0 | 4 | 0.5 | 2 | 0 | 21 | 0.0952 | 2 | 0.5 | 6 | 0.3333 | 5 | 0 | 5 | 0.4 | 18 | 0.2778 |
| 11_2025383 | 11 | 0.3636 | 3 | 0 | 4 | 0.5 | 3 | 0 | 21 | 0.2857 | 1 | 0 | 6 | 0.5 | 3 | 0.3333 | 5 | 0.2 | 15 | 0.3333 |
| 11_2025398 | 12 | 0.5833 | 6 | 0.8333 | 4 | 0.75 | 1 | 1 | 23 | 0.6957 | 2 | 0.5 | 5 | 0.8 | 5 | 1 | 7 | 0.7143 | 19 | 0.7895 |
| 11_2025409 | 9 | 0.6667 | 3 | 0.3333 | 3 | 1 | 2 | 0.5 | 17 | 0.6471 | 2 | 0.5 | 5 | 0.6 | 5 | 0.8 | 4 | 0.75 | 16 | 0.6875 |
| 11_2025429 | 12 | 0.8333 | 5 | 1 | 3 | 1 | 1 | 1 | 21 | 0.9048 | 2 | 1 | 3 | 1 | 3 | 1 | 5 | 0.6 | 13 | 0.8462 |
| 11_2028196 | 10 | 0.6 | 4 | 0.25 | 2 | 1 | 2 | 0.5 | 18 | 0.5556 | 2 | 0.5 | 4 | 0.75 | 3 | 0.6667 | 3 | 0.3333 | 12 | 0.5833 |
| 11_2028197 | 9 | 0.1111 | 4 | 0 | 1 | 1 | 2 | 0.5 | 16 | 0.1875 | 2 | 0 | 4 | 0.25 | 3 | 0 | 1 | 0 | 10 | 0.1 |
| 11_2028259 | 4 | 0 | 4 | 0 | 1 | 1 | 2 | 0.5 | 11 | 0.1818 | 1 | 0 | 2 | 0.5 | 3 | 0.3333 | 2 | 0.5 | 8 | 0.375 |
| 11_2059794 | 1 | 1 | NA | NA | 1 | 1 | 1 | 1 | 3 | 1 | NA | NA | NA | NA | 1 | 1 | 1 | 1 | 2 | 1 |
| 11_2059798 | 1 | 1 | NA | NA | 1 | 1 | 1 | 1 | 3 | 1 | NA | NA | 1 | 1 | 2 | 1 | 1 | 1 | 4 | 1 |
| 11_2059820 | 2 | 0.5 | NA | NA | 1 | 0 | NA | NA | 3 | 0.3333 | NA | NA | 1 | 1 | 2 | 0 | 1 | 0 | 4 | 0.25 |
| 11_2059825 | 1 | 1 | NA | NA | 1 | 1 | 1 | 1 | 3 | 1 | NA | NA | 1 | 1 | 1 | 1 | 1 | 1 | 3 | 1 |
| 11_2059828 | 2 | 0.5 | NA | NA | 1 | 1 | 1 | 1 | 4 | 0.75 | NA | NA | 1 | 0 | 1 | 1 | 1 | 1 | 3 | 0.6667 |
| 11_2059844 | 1 | 1 | NA | NA | 1 | 0 | 1 | 1 | 3 | 0.6667 | NA | NA | 1 | 0 | 1 | 0 | 1 | 1 | 3 | 0.3333 |
| 11_2059845 | 1 | 1 | NA | NA | 1 | 0 | NA | NA | 2 | 0.5 | NA | NA | 1 | 1 | 1 | 1 | 1 | 1 | 3 | 1 |
| 11_2059850 | 2 | 0 | NA | NA | 1 | 1 | NA | NA | 3 | 0.3333 | NA | NA | 2 | 0.5 | 2 | 0.5 | 1 | 1 | 5 | 0.6 |
| 11_2059853 | 2 | 0.5 | NA | NA | 1 | 1 | 1 | 1 | 4 | 0.75 | NA | NA | 1 | 0 | 1 | 0 | 1 | 1 | 3 | 0.3333 |
| 11_2059858 | 1 | 0 | NA | NA | 1 | 1 | 1 | 0 | 3 | 0.3333 | NA | NA | 2 | 1 | 1 | 0 | 1 | 0 | 4 | 0.5 |
| 11_2059870 | 1 | 1 | NA | NA | 1 | 0 | NA | NA | 2 | 0.5 | NA | NA | 1 | 1 | 2 | 0.5 | 1 | 0 | 4 | 0.5 |
| 11_2059875 | 2 | 0.5 | NA | NA | 1 | 0 | 1 | 1 | 4 | 0.5 | NA | NA | 2 | 1 | 2 | 1 | 1 | 1 | 5 | 1 |
| 11_2059881 | 2 | 1 | NA | NA | 1 | 1 | 1 | 1 | 4 | 1 | NA | NA | 2 | 1 | 2 | 1 | 1 | 1 | 5 | 1 |
| 11_2059898 | 2 | 1 | NA | NA | 1 | 1 | 1 | 1 | 4 | 1 | NA | NA | 1 | 1 | 1 | 1 | NA | NA | 2 | 1 |
| 11_2059901 | 2 | 1 | NA | NA | 1 | 1 | NA | NA | 3 | 1 | NA | NA | 1 | 1 | 1 | 1 | NA | NA | 2 | 1 |
| 11_2059942 | 3 | 0.6667 | NA | NA | NA | NA | 2 | 0.5 | 5 | 0.6 | NA | NA | 2 | 0.5 | NA | NA | 1 | 1 | 3 | 0.6667 |
| 11_2059975 | 3 | 0 | NA | NA | 1 | 1 | 1 | 0 | 5 | 0.2 | NA | NA | 1 | 1 | NA | NA | 1 | 0 | 2 | 0.5 |
| 11_2060154 | 3 | 0.3333 | NA | NA | 2 | 0 | 2 | 0 | 7 | 0.1429 | 1 | 1 | 3 | 0 | 2 | 0 | 3 | 0 | 9 | 0.1111 |
| 11_2060244 | 4 | 0.25 | 1 | 1 | 2 | 0 | 2 | 0 | 9 | 0.2222 | 1 | 0 | 5 | 0 | 2 | 0 | 3 | 0.3333 | 11 | 0.0909 |
| 11_2060255 | 4 | 0 | 1 | 0 | 2 | 0.5 | 2 | 0 | 9 | 0.1111 | 2 | 0.5 | 5 | 0 | 2 | 0 | 3 | 0 | 12 | 0.0833 |
| 11_2063603 | 2 | 1 | 7 | 0.7143 | 2 | 0 | 14 | 0.5714 | 25 | 0.6 | 7 | 0.7143 | 6 | 0.5 | 11 | 0.5455 | 11 | 0.4545 | 35 | 0.5429 |
| 11_2067578 | 2 | 1 | 2 | 0.5 | 1 | 1 | 5 | 1 | 10 | 0.9 | 2 | 1 | 5 | 1 | 7 | 1 | 8 | 0.875 | 22 | 0.9545 |
| 11_2067579 | 1 | 1 | 2 | 0.5 | NA | NA | 6 | 0.3333 | 9 | 0.4444 | 2 | 0.5 | 3 | 0.6667 | 6 | 0.8333 | 8 | 0.75 | 19 | 0.7368 |
| 11_2067584 | 2 | 0 | 2 | 1 | 1 | 1 | 8 | 0.25 | 13 | 0.3846 | 2 | 0.5 | 4 | 0.5 | 7 | 0.2857 | 8 | 0.625 | 21 | 0.4762 |
| 11_2067588 | 1 | 0 | 2 | 0 | 1 | 1 | 8 | 0.375 | 12 | 0.3333 | 2 | 0 | 1 | 0 | 6 | 0.1667 | 7 | 0.2857 | 16 | 0.1875 |
| 11_2067594 | 2 | 0 | 2 | 0.5 | 1 | 0 | 7 | 0.1429 | 12 | 0.1667 | 1 | 0 | 4 | 0 | 5 | 0.2 | 8 | 0.25 | 18 | 0.1667 |
| 11_2067595 | 2 | 0.5 | 2 | 0 | 1 | 0 | 6 | 0.1667 | 11 | 0.1818 | 2 | 0 | 5 | 0.2 | 5 | 0.6 | 9 | 0.3333 | 21 | 0.3333 |
| 11_2067601 | 1 | 1 | 2 | 0 | 1 | 0 | 6 | 0 | 10 | 0.1 | 2 | 0 | 5 | 0.2 | 5 | 0.2 | 9 | 0.2222 | 21 | 0.1905 |
| 11_2067603 | 1 | 1 | 1 | 0 | 1 | 1 | 7 | 0.2857 | 10 | 0.4 | 2 | 1 | 4 | 0.75 | 3 | 0.3333 | 7 | 0.4286 | 16 | 0.5625 |
| 11_2067604 | 1 | 0 | 2 | 0 | 1 | 0 | 7 | 0.1429 | 11 | 0.0909 | 2 | 0 | 4 | 0 | 4 | 0.25 | 8 | 0.125 | 18 | 0.1111 |
| 11_2067605 | 1 | 0 | 2 | 1 | 1 | 1 | 4 | 0.25 | 8 | 0.5 | 1 | 1 | 4 | 0.75 | 5 | 0.8 | 9 | 0.7778 | 19 | 0.7895 |
| 11_2067610 | 2 | 0 | 1 | 0 | 1 | 0 | 8 | 0.25 | 12 | 0.1667 | 2 | 0 | 4 | 0 | 4 | 0.5 | 8 | 0.25 | 18 | 0.2222 |
| 11_2067611 | 1 | 0 | 2 | 0.5 | NA | NA | 5 | 0.4 | 8 | 0.375 | 2 | 0.5 | 5 | 0.4 | 4 | 0 | 6 | 0 | 17 | 0.1765 |
| 11_2067611 | 2 | 0 | 2 | 0 | 1 | 0 | 5 | 0.4 | 10 | 0.2 | 2 | 0 | 4 | 0 | 3 | 0.3333 | 8 | 0.25 | 17 | 0.1765 |
| 11_2067619 | 1 | 1 | 1 | 0 | 1 | 0 | 8 | 0.125 | 11 | 0.1818 | 1 | 0 | 5 | 0.2 | 5 | 0.2 | 9 | 0 | 20 | 0.1 |
| 11_2067622 | 1 | 1 | 1 | 0 | 1 | 0 | 7 | 0 | 10 | 0.1 | 1 | 0 | 4 | 0 | 5 | 0.2 | 8 | 0.125 | 18 | 0.1111 |
| 11_2067630 | 2 | 0 | 1 | 0 | 1 | 1 | 6 | 0.3333 | 10 | 0.3 | 1 | 1 | 5 | 0.6 | 4 | 0.75 | 9 | 0.2222 | 19 | 0.4737 |
| 11_2067633 | 2 | 0.5 | 1 | 1 | 1 | 0 | 8 | 0.625 | 12 | 0.5833 | 1 | 1 | 5 | 0.6 | 5 | 0.8 | 9 | 0.6667 | 20 | 0.7 |
| 11_2067635 | 2 | 0 | 1 | 0 | 1 | 0 | 7 | 0.4286 | 11 | 0.2727 | NA | NA | 4 | 0 | 4 | 0.5 | 8 | 0.25 | 16 | 0.25 |
| 11_2067640 | 2 | 0.5 | 1 | 0 | 1 | 0 | 6 | 0.1667 | 10 | 0.2 | 1 | 0 | 5 | 0.2 | 4 | 0.25 | 8 | 0.375 | 18 | 0.2778 |
| 11_2067653 | 1 | 0 | 1 | 0 | NA | NA | 5 | 0.4 | 7 | 0.2857 | 1 | 1 | 3 | 0.6667 | 5 | 0.2 | 6 | 0.6667 | 15 | 0.5333 |
| 11_2067659 | 2 | 0 | 1 | 0 | NA | NA | 6 | 0.1667 | 9 | 0.1111 | 1 | 1 | 3 | 0.6667 | 5 | 0 | 6 | 0.1667 | 15 | 0.2667 |
| 11_2067673 | 1 | 0 | 1 | 0 | NA | NA | 5 | 0.2 | 7 | 0.1429 | 1 | 0 | 2 | 0.5 | 4 | 0 | 7 | 0 | 14 | 0.0714 |
| 11_2067677 | 2 | 0 | 1 | 0 | NA | NA | 6 | 0.1667 | 9 | 0.1111 | 1 | 0 | 4 | 0.25 | 4 | 0 | 7 | 0.4286 | 16 | 0.25 |
| 11_2067700 | 1 | 1 | NA | NA | NA | NA | 5 | 0.6 | 6 | 0.6667 | NA | NA | 4 | 1 | 2 | 1 | 4 | 1 | 10 | 1 |
| 11_2067712 | 1 | 1 | NA | NA | NA | NA | 1 | 0 | 2 | 0.5 | NA | NA | 3 | 1 | 1 | 0 | 5 | 0.6 | 9 | 0.6667 |
| 11_2067713 | 2 | 1 | NA | NA | NA | NA | 2 | 1 | 4 | 1 | NA | NA | 4 | 1 | 3 | 1 | 5 | 0.6 | 12 | 0.8333 |

| | | | | | | | | | | | | | | | | | | | | |
|---|---|---|---|---|---|---|---|---|---|---|---|---|---|---|---|---|---|---|---|---|
| 11_206772 | 1 | 1 | NA | NA | NA | NA | 4 | 0 | 5 | 0.2 | NA | NA | 4 | 0 | 3 | 0.3333 | 7 | 0.1429 | 14 | 0.1429 |
| 11_206773 | 1 | 1 | NA | NA | NA | NA | 2 | 0 | 3 | 0.3333 | NA | NA | 4 | 1 | 3 | 0.6667 | 4 | 0.75 | 11 | 0.8182 |
| 11_206774 | 1 | 1 | NA | NA | NA | NA | 4 | 0.75 | 5 | 0.8 | NA | NA | 4 | 0.25 | 3 | 1 | 6 | 0.3333 | 13 | 0.4615 |
| 11_206775 | 2 | 0 | NA | NA | NA | NA | 4 | 0.25 | 6 | 0.1667 | NA | NA | 4 | 0 | 3 | 0.3333 | 5 | 0 | 12 | 0.0833 |
| 11_206920 | NA | NA | NA | NA | 1 | 1 | NA | NA | 1 | 1 | NA | NA | 1 | 1 | NA | NA | NA | NA | 1 | 1 |
| 11_206923 | 1 | 0 | NA | NA | 2 | 1 | NA | NA | 3 | 0.6667 | NA | NA | 1 | 1 | NA | NA | NA | NA | 1 | 1 |
| 11_206925 | 1 | 1 | NA | NA | 1 | 1 | NA | NA | 2 | 1 | 1 | 0 | 1 | 1 | NA | NA | NA | NA | 2 | 0.5 |
| 11_206925 | 1 | 1 | NA | NA | 2 | 0.5 | NA | NA | 3 | 0.6667 | 1 | 0 | 1 | 1 | NA | NA | NA | NA | 2 | 0.5 |
| 11_206925 | 1 | 1 | NA | NA | 2 | 1 | NA | NA | 3 | 1 | 1 | 1 | 1 | 1 | NA | NA | NA | NA | 2 | 1 |
| 11_206926 | 1 | 1 | NA | NA | 2 | 0 | NA | NA | 3 | 0.3333 | NA | NA | 1 | 1 | NA | NA | NA | NA | 1 | 1 |
| 11_206926 | NA | NA | NA | NA | 2 | 0.5 | NA | NA | 2 | 0.5 | 1 | 1 | NA | NA | NA | NA | NA | NA | 1 | 1 |
| 11_206927 | 1 | 1 | NA | NA | 2 | 1 | NA | NA | 3 | 1 | 1 | 1 | 1 | 1 | NA | NA | NA | NA | 2 | 1 |
| 11_207756 | 7 | 0.4286 | 7 | 0.1429 | 8 | 0.125 | 8 | 0 | 30 | 0.1667 | 5 | 0 | 4 | 0.25 | 7 | 0.2857 | 15 | 0.2667 | 31 | 0.2258 |
| 11_207760 | 7 | 0.4286 | 7 | 0.2857 | 9 | 0.3333 | 9 | 0.5556 | 32 | 0.4062 | 4 | 0.5 | 2 | 0 | 6 | 0.3333 | 14 | 0.6429 | 26 | 0.5 |
| 11_207761 | 8 | 0.375 | 5 | 0 | 9 | 0.5556 | 10 | 0.5 | 32 | 0.4062 | 5 | 0.4 | 2 | 0.5 | 6 | 0.6667 | 13 | 0.3077 | 26 | 0.4231 |
| 11_207761 | 9 | 1 | 8 | 0.875 | 9 | 0.7778 | 9 | 0.7778 | 35 | 0.8571 | 5 | 0.6 | 1 | 0 | 4 | 0.5 | 12 | 0.75 | 22 | 0.6364 |
| 11_207762 | 5 | 0.4 | 6 | 0.6667 | 6 | 0.5 | 7 | 0.8571 | 24 | 0.625 | 3 | 0.3333 | NA | NA | 4 | 0.75 | 10 | 0.5 | 17 | 0.5294 |
| 11_207762 | 7 | 0.7143 | 3 | 1 | 5 | 0.2 | 9 | 0.2222 | 24 | 0.4583 | 3 | 0.6667 | 2 | 0.5 | 5 | 0.8 | 10 | 0.2 | 20 | 0.45 |
| 11_207762 | 8 | 0.875 | 8 | 0.875 | 9 | 0.7778 | 9 | 0.7778 | 34 | 0.8235 | 5 | 1 | 2 | 1 | 5 | 0.8 | 12 | 0.6667 | 24 | 0.7917 |
| 11_207763 | 8 | 0.25 | 8 | 0.125 | 9 | 0.2222 | 10 | 0.3 | 35 | 0.2286 | 5 | 0.4 | 2 | 0.5 | 5 | 0 | 12 | 0.3333 | 24 | 0.2917 |
| 11_207763 | 7 | 0.5714 | 7 | 0.4286 | 6 | 0.6667 | 10 | 0.5 | 30 | 0.5333 | 5 | 0.4 | 1 | 0 | 5 | 0.6 | 12 | 0.3333 | 23 | 0.3913 |
| 11_207764 | 8 | 0.125 | 6 | 0 | 8 | 0.25 | 7 | 0.2857 | 29 | 0.1724 | 5 | 0 | 2 | 0.5 | 4 | 0.25 | 12 | 0.0833 | 23 | 0.1304 |
| 11_207764 | 8 | 0.25 | 5 | 0.6 | 8 | 0.5 | 9 | 0.5556 | 30 | 0.4667 | 5 | 0.6 | 2 | 0 | 5 | 0.2 | 11 | 0.5455 | 23 | 0.4348 |
| 11_210980 | 4 | 0.5 | 6 | 0 | 4 | 0 | 9 | 0.2222 | 23 | 0.1739 | 2 | 0 | 13 | 0.1538 | 5 | 0 | 14 | 0.2857 | 34 | 0.1765 |
| 11_210980 | 5 | 0.2 | 7 | 0.2857 | 3 | 0 | 9 | 0.1111 | 24 | 0.1667 | 2 | 0.5 | 11 | 0.3636 | 5 | 0.2 | 12 | 0.3333 | 30 | 0.3333 |
| 11_210981 | 4 | 0 | 5 | 0.4 | 4 | 0.5 | 6 | 0.1667 | 19 | 0.2632 | NA | NA | 9 | 0.3333 | 3 | 0.3333 | 8 | 0.625 | 20 | 0.45 |
| 11_210982 | 6 | 0.3333 | 6 | 0.1667 | 4 | 0 | 8 | 0.375 | 24 | 0.25 | 2 | 0.5 | 10 | 0.2 | 7 | 0.2857 | 11 | 0.3636 | 30 | 0.3 |
| 11_210983 | 9 | 0.4444 | 9 | 0.1111 | 4 | 0.75 | 10 | 0.4 | 32 | 0.375 | 2 | 1 | 15 | 0.3333 | 7 | 0.1429 | 14 | 0.2143 | 38 | 0.2895 |
| 11_210984 | 9 | 0.7778 | 7 | 0.2857 | 4 | 0.75 | 9 | 0.4444 | 29 | 0.5517 | 2 | 1 | 16 | 0.6875 | 8 | 0.625 | 13 | 0.7692 | 39 | 0.7179 |
| 11_210985 | 7 | 1 | 7 | 0.5714 | 4 | 1 | 10 | 0.8 | 28 | 0.8214 | 2 | 1 | 15 | 0.6 | 7 | 0.7143 | 12 | 0.9167 | 36 | 0.75 |
| 11_210985 | 7 | 0 | 4 | 0.25 | 3 | 0 | 3 | 0 | 17 | 0.0588 | NA | NA | 11 | 0.1818 | 7 | 0 | 12 | 0.1667 | 30 | 0.1333 |
| 11_210985 | 8 | 0.5 | 7 | 0.1429 | 3 | 1 | 11 | 0.0909 | 29 | 0.3103 | 2 | 1 | 17 | 0.4118 | 11 | 0.3636 | 14 | 0.7143 | 44 | 0.5227 |
| 11_210987 | 8 | 0.125 | 7 | 0.1429 | 4 | 0 | 9 | 0 | 28 | 0.0714 | 2 | 0 | 17 | 0.0588 | 11 | 0 | 14 | 0.2143 | 44 | 0.0909 |
| 11_210987 | 7 | 0.4286 | 7 | 0.2857 | 3 | 1 | 10 | 0.4 | 27 | 0.4444 | 1 | 0 | 16 | 0.625 | 9 | 0 | 13 | 0.6154 | 39 | 0.4615 |
| 11_210997 | 10 | 0.3 | 11 | 0.0909 | 4 | 0 | 11 | 0.1818 | 36 | 0.1667 | 2 | 0 | 15 | 0.0667 | 10 | 0 | 12 | 0.1667 | 39 | 0.0769 |
| 11_211000 | 10 | 0.1 | 9 | 0.1111 | 4 | 0 | 11 | 0 | 34 | 0.0588 | 2 | 0 | 16 | 0 | 15 | 0.1333 | 12 | 0.3333 | 45 | 0.1333 |
| 11_212146 | 2 | 0 | 1 | 0 | 3 | 0.6667 | 4 | 0.5 | 10 | 0.4 | NA | NA | NA | NA | 1 | 1 | NA | NA | 1 | 1 |
| 11_212147 | 2 | 0 | 1 | 0 | 3 | 0 | 4 | 0.25 | 10 | 0.1 | NA | NA | NA | NA | 1 | 1 | NA | NA | 1 | 1 |
| 11_212147 | 2 | 0.5 | 1 | 0 | 2 | 0 | 3 | 0 | 8 | 0.125 | NA | NA | NA | NA | 1 | 1 | NA | NA | 1 | 1 |
| 11_212277 | 2 | 1 | 2 | 0.5 | 3 | 0 | 4 | 0 | 11 | 0.2727 | NA | NA | NA | NA | 1 | 1 | NA | NA | 1 | 1 |
| 11_212278 | 1 | 0 | 1 | 0 | 3 | 0.3333 | 4 | 1 | 9 | 0.5556 | NA | NA | NA | NA | 1 | 1 | NA | NA | 1 | 1 |
| 11_212279 | 1 | 1 | 2 | 1 | 3 | 0.3333 | 3 | 0.6667 | 9 | 0.6667 | NA | NA | NA | NA | 1 | 1 | NA | NA | 1 | 1 |
| 11_212698 | 2 | 0.5 | 4 | 0 | 7 | 0.1429 | 5 | 0 | 18 | 0.1111 | 1 | 0 | 2 | 0 | 2 | 0.5 | 2 | 0 | 7 | 0.1429 |
| 11_212699 | 2 | 0 | 4 | 0.5 | 8 | 0 | 5 | 0.6 | 19 | 0.2632 | 1 | 0 | 3 | 0.3333 | 1 | 1 | 2 | 0.5 | 7 | 0.4286 |
| 11_212699 | 2 | 1 | 2 | 1 | 6 | 0.5 | 4 | 1 | 14 | 0.7857 | NA | NA | 4 | 0.75 | 2 | 1 | 2 | 0.5 | 8 | 0.75 |
| 11_212700 | 3 | 0.3333 | 3 | 0 | 7 | 0.2857 | 4 | 0.5 | 17 | 0.2941 | 1 | 1 | 2 | 0 | 2 | 0 | 2 | 0.5 | 7 | 0.2857 |
| 11_212708 | 2 | 0 | 4 | 0.25 | 6 | 0 | 5 | 0 | 17 | 0.0588 | 1 | 1 | 3 | 0 | 4 | 0 | 1 | 0 | 9 | 0.1111 |
| 11_212717 | 14 | 0.0714 | 11 | 0 | 9 | 0 | 17 | 0.2941 | 51 | 0.1176 | 13 | 0 | 10 | 0.1 | 14 | 0.0714 | 14 | 0.0714 | 51 | 0.0588 |
| 11_216617 | 1 | 0 | 1 | 1 | 1 | 0 | NA | NA | 3 | 0.3333 | 2 | 1 | NA | NA | 1 | 1 | 1 | 1 | 4 | 1 |
| 11_216619 | 1 | 0 | 1 | 1 | 1 | 0 | NA | NA | 3 | 0.3333 | 2 | 0.5 | NA | NA | 1 | 0 | 1 | 0 | 4 | 0.25 |
| 11_219455 | NA | NA | 1 | 1 | 1 | 1 | NA | NA | 2 | 1 | NA | NA | NA | NA | NA | NA | 1 | 1 | 1 | 1 |
| 11_219456 | NA | NA | 1 | 1 | 1 | 1 | NA | NA | 2 | 1 | NA | NA | NA | NA | NA | NA | 1 | 1 | 1 | 1 |
| 11_219457 | NA | NA | NA | NA | 1 | 1 | NA | NA | 1 | 1 | NA | NA | NA | NA | NA | NA | 1 | 1 | 1 | 1 |
| 11_219458 | NA | NA | 1 | 1 | 1 | 1 | NA | NA | 2 | 1 | NA | NA | NA | NA | NA | NA | 1 | 1 | 1 | 1 |
| 11_219459 | NA | NA | NA | NA | 1 | 1 | NA | NA | 1 | 1 | NA | NA | NA | NA | NA | NA | 1 | 1 | 1 | 1 |
| 11_219462 | NA | NA | 1 | 1 | 1 | 1 | NA | NA | 2 | 1 | NA | NA | NA | NA | NA | NA | 1 | 1 | 1 | 1 |
| 11_219465 | NA | NA | 1 | 1 | 1 | 1 | NA | NA | 2 | 1 | NA | NA | NA | NA | NA | NA | 1 | 1 | 1 | 1 |
| 11_219565 | 4 | 0 | 5 | 0 | 2 | 0.5 | 2 | 0 | 13 | 0.0769 | 2 | 0 | 1 | 1 | 2 | 0.5 | 3 | 0 | 8 | 0.25 |
| 11_219566 | 4 | 0.5 | 5 | 0.8 | 2 | 1 | 2 | 1 | 13 | 0.7692 | 2 | 1 | 1 | 0 | 2 | 0.5 | 3 | 0.6667 | 8 | 0.625 |
| 11_219567 | 4 | 0.25 | 4 | 0.25 | 1 | 0 | 2 | 0 | 11 | 0.1818 | 2 | 0.5 | NA | NA | 1 | 0 | 1 | 0 | 4 | 0.25 |
| 11_219567 | 4 | 0 | 5 | 0 | 2 | 0 | 2 | 0.5 | 13 | 0.0769 | 2 | 0.5 | NA | NA | 2 | 0 | 3 | 0 | 7 | 0.1429 |
| 11_220455 | 2 | 0.5 | 2 | 0.5 | 1 | 0 | NA | NA | 5 | 0.4 | 1 | 1 | NA | NA | NA | NA | 1 | 1 | 2 | 1 |
| 11_220456 | 2 | 1 | 3 | 0.3333 | 1 | 0 | NA | NA | 6 | 0.5 | 1 | 1 | NA | NA | NA | NA | 1 | 0 | 2 | 0.5 |
| 11_220457 | 2 | 0.5 | 3 | 0 | NA | NA | NA | NA | 5 | 0.2 | 1 | 1 | NA | NA | NA | NA | 1 | 0 | 2 | 0.5 |
| 11_220460 | 2 | 0.5 | 3 | 0 | NA | NA | NA | NA | 5 | 0.2 | 1 | 1 | NA | NA | NA | NA | 1 | 1 | 2 | 1 |

| | | | | | | | | | | | | | | | | | | | | |
|---|---|---|---|---|---|---|---|---|---|---|---|---|---|---|---|---|---|---|---|---|
| 11_220461 | 2 | 1 | 3 | 0 | NA | NA | NA | NA | 5 | 0.4 | 1 | 0 | NA | NA | NA | NA | 1 | 1 | 2 | 0.5 |
| 11_222356 | NA | NA | 2 | 1 | NA | NA | NA | NA | 2 | 1 | NA | NA | NA | NA | NA | NA | 1 | 1 | 1 | 1 |
| 11_224685 | NA | NA | NA | NA | 2 | 0.5 | 1 | 1 | 3 | 0.6667 | NA | NA | 1 | 1 | 2 | 1 | NA | NA | 3 | 1 |
| 11_224687 | NA | NA | NA | NA | 2 | 1 | 1 | 1 | 3 | 1 | NA | NA | 2 | 1 | 2 | 0.5 | NA | NA | 4 | 0.75 |
| 11_224687 | NA | NA | NA | NA | 1 | 1 | 1 | 1 | 2 | 1 | NA | NA | 3 | 0.6667 | 2 | 0 | NA | NA | 5 | 0.4 |
| 11_224690 | NA | NA | NA | NA | 2 | 1 | 1 | 1 | 3 | 1 | NA | NA | 3 | 0.6667 | 1 | 1 | NA | NA | 4 | 0.75 |
| 11_224690 | NA | NA | NA | NA | 2 | 1 | NA | NA | 2 | 1 | NA | NA | 2 | 1 | 1 | 1 | NA | NA | 3 | 1 |
| 11_224690 | NA | NA | NA | NA | 2 | 1 | 1 | 1 | 3 | 1 | NA | NA | 3 | 1 | 1 | 1 | NA | NA | 4 | 1 |
| 11_227848 | 14 | 0.2857 | 6 | 0.5 | 8 | 0.5 | 8 | 0.25 | 36 | 0.3611 | 10 | 0.1 | 8 | 0.25 | 10 | 0.1 | 11 | 0.3636 | 39 | 0.2051 |
| 11_227853 | 9 | 0.5556 | 7 | 0.5714 | 6 | 0.5 | 8 | 0.5 | 30 | 0.5333 | 8 | 0.375 | 8 | 0.375 | 11 | 0.3636 | 10 | 0.2 | 37 | 0.3243 |
| 11_227853 | 10 | 0.1 | 8 | 0.5 | 5 | 0.4 | 8 | 0.375 | 31 | 0.3226 | 8 | 0 | 6 | 0.3333 | 10 | 0.4 | 7 | 0 | 31 | 0.1935 |
| 11_227854 | 7 | 0.1429 | 7 | 0.2857 | 6 | 0 | 6 | 0.1667 | 26 | 0.1538 | 7 | 0.1429 | 7 | 0.2857 | 9 | 0.3333 | 9 | 0.1111 | 32 | 0.2188 |
| 11_227859 | 9 | 0.2222 | 7 | 0.4286 | 6 | 0.3333 | 6 | 0.1667 | 28 | 0.2857 | 7 | 0.2857 | 7 | 0.1429 | 8 | 0.75 | 8 | 0.25 | 30 | 0.3667 |
| 11_227860 | 8 | 0.5 | 8 | 0.625 | 6 | 0.6667 | 6 | 0.3333 | 28 | 0.5357 | 5 | 0.8 | 7 | 0.8571 | 8 | 0.5 | 8 | 0.625 | 28 | 0.6786 |
| 11_227861 | 9 | 1 | 8 | 1 | 6 | 1 | 6 | 1 | 29 | 1 | 6 | 1 | 7 | 1 | 9 | 1 | 9 | 1 | 31 | 1 |
| 11_227861 | 9 | 0.3333 | 8 | 0.625 | 5 | 0.2 | 5 | 0.2 | 27 | 0.3704 | 6 | 0.5 | 6 | 0.3333 | 8 | 0.375 | 8 | 0.25 | 28 | 0.3571 |
| 11_227863 | 9 | 0.8889 | 5 | 1 | 3 | 0.6667 | 6 | 0.8333 | 23 | 0.8696 | 4 | 1 | 6 | 0.6667 | 9 | 0.7778 | 9 | 0.8889 | 28 | 0.8214 |
| 11_227863 | 9 | 1 | 5 | 1 | 4 | 1 | 4 | 1 | 22 | 1 | 5 | 1 | 7 | 1 | 9 | 1 | 5 | 1 | 26 | 1 |
| 11_227864 | 7 | 0.1429 | 6 | 0.1667 | 4 | 0 | 3 | 0 | 20 | 0.1 | 6 | 0.1667 | 7 | 0 | 8 | 0 | 7 | 0.1429 | 28 | 0.0714 |
| 11_227865 | 8 | 1 | 6 | 0.8333 | 6 | 1 | 6 | 0.8333 | 26 | 0.9231 | 4 | 0.5 | 7 | 1 | 8 | 0.75 | 7 | 1 | 26 | 0.8462 |
| 11_227865 | 7 | 0.5714 | 6 | 0.6667 | 5 | 0.4 | 6 | 0.6667 | 24 | 0.5833 | 6 | 0.6667 | 7 | 0.4286 | 7 | 0.8571 | 5 | 0.6 | 25 | 0.64 |
| 11_227865 | 8 | 0.375 | 6 | 0.3333 | 6 | 0.5 | 6 | 0.5 | 26 | 0.4231 | 4 | 0.5 | 7 | 0.1429 | 8 | 0.25 | 7 | 0.5714 | 26 | 0.3462 |
| 11_227865 | 8 | 0.125 | 6 | 0 | 6 | 0 | 6 | 0.1667 | 26 | 0.0769 | 5 | 0 | 7 | 0 | 8 | 0 | 7 | 0.2857 | 27 | 0.0741 |
| 11_227866 | 8 | 0.875 | 5 | 1 | 6 | 1 | 6 | 0.8333 | 25 | 0.92 | 5 | 1 | 6 | 1 | 8 | 1 | 6 | 1 | 25 | 1 |
| 11_227866 | 7 | 0.5714 | 5 | 0.4 | 5 | 0.4 | 5 | 0.2 | 22 | 0.4091 | 3 | 0 | 7 | 0.2857 | 6 | 0.5 | 6 | 0.6667 | 22 | 0.4091 |
| 11_227866 | 5 | 0.8 | 4 | 0.75 | 3 | 0.3333 | 4 | 0.5 | 16 | 0.625 | 1 | 1 | 6 | 0.5 | 5 | 0.4 | 4 | 0.5 | 16 | 0.5 |
| 11_227868 | 7 | 1 | 5 | 1 | 5 | 1 | 5 | 1 | 22 | 1 | 5 | 1 | 6 | 1 | 7 | 1 | 7 | 1 | 25 | 1 |
| 11_227868 | 5 | 1 | 5 | 1 | 6 | 1 | 2 | 1 | 18 | 1 | 5 | 1 | 4 | 1 | 6 | 1 | 6 | 0.8333 | 21 | 0.9524 |
| 11_227868 | 8 | 0 | 4 | 0.25 | 6 | 0 | 5 | 0.2 | 23 | 0.087 | 5 | 0 | 6 | 0 | 7 | 0.2857 | 7 | 0.2857 | 25 | 0.16 |
| 11_227869 | 7 | 0.1429 | 5 | 0.2 | 6 | 0.1667 | 6 | 0.1667 | 24 | 0.1667 | 4 | 0 | 5 | 0 | 6 | 0.3333 | 6 | 0 | 21 | 0.0952 |
| 11_227869 | 5 | 0.6 | 4 | 0 | 4 | 0 | 4 | 0 | 17 | 0.1765 | 4 | 0.5 | 3 | 0.3333 | 6 | 0 | 6 | 0 | 19 | 0.1579 |
| 11_227869 | 6 | 0.1667 | 4 | 0.5 | 4 | 0 | 4 | 0.25 | 18 | 0.2222 | 3 | 0 | 5 | 0.2 | 7 | 0.4286 | 5 | 0 | 20 | 0.2 |
| 11_227870 | 4 | 0 | 3 | 0 | 6 | 0.1667 | 5 | 0.2 | 18 | 0.1111 | 3 | 0.3333 | 4 | 0.5 | 7 | 0 | 7 | 0.2857 | 21 | 0.2381 |
| 11_228095 | NA | NA | 1 | 1 | NA | NA | NA | NA | 1 | 1 | NA | NA | 1 | 0 | 1 | 1 | NA | NA | 2 | 0.5 |
| 11_228100 | NA | NA | 1 | 1 | NA | NA | NA | NA | 1 | 1 | 1 | 0 | 1 | 0 | 1 | 1 | NA | NA | 3 | 0.3333 |
| 11_228104 | NA | NA | 1 | 1 | NA | NA | NA | NA | 1 | 1 | 1 | 1 | 1 | 1 | 1 | 1 | NA | NA | 3 | 1 |
| 11_228108 | NA | NA | 1 | 1 | NA | NA | NA | NA | 1 | 1 | NA | NA | NA | NA | 1 | 1 | NA | NA | 1 | 1 |
| 11_228110 | NA | NA | 1 | 1 | NA | NA | NA | NA | 1 | 1 | 1 | 1 | 1 | 0 | 1 | 1 | 1 | 1 | 4 | 0.75 |
| 11_228111 | NA | NA | 1 | 1 | NA | NA | NA | NA | 1 | 1 | NA | NA | NA | NA | NA | NA | 1 | 1 | 1 | 1 |
| 11_228498 | 11 | 0.2727 | 6 | 0 | 9 | 0.2222 | 8 | 0 | 34 | 0.1471 | 2 | 0 | 7 | 0.1429 | 7 | 0 | 5 | 0.2 | 21 | 0.0952 |
| 11_228498 | 13 | 0.4615 | 6 | 0.3333 | 9 | 0.2222 | 6 | 0.5 | 34 | 0.3824 | 2 | 0.5 | 7 | 0.4286 | 6 | 0.1667 | 5 | 0.6 | 20 | 0.4 |
| 11_229223 | 1 | 0 | 5 | 0.4 | NA | NA | 1 | 0 | 7 | 0.2857 | NA | NA | NA | NA | 2 | 1 | NA | NA | 2 | 1 |
| 11_229223 | 3 | 0 | 4 | 0.25 | 1 | 0 | 1 | 0 | 9 | 0.1111 | 2 | 0 | 1 | 1 | NA | NA | NA | NA | 3 | 0.3333 |
| 11_229223 | 3 | 0.3333 | 7 | 0 | 1 | 0 | 2 | 0 | 13 | 0.0769 | 3 | 0.3333 | 1 | 1 | 2 | 0 | 1 | 0 | 7 | 0.2857 |
| 11_229224 | 3 | 0.6667 | 5 | 0.2 | 1 | 0 | 2 | 0 | 11 | 0.2727 | 3 | 0.6667 | 2 | 0.5 | 2 | 0.5 | NA | NA | 7 | 0.5714 |
| 11_229224 | 2 | 0 | 6 | 0.1667 | 1 | 0 | 2 | 0.5 | 11 | 0.1818 | 3 | 0 | 2 | 0.5 | 2 | 0 | NA | NA | 7 | 0.1429 |
| 11_229225 | 3 | 0 | 5 | 0.2 | 1 | 0 | 2 | 0 | 11 | 0.0909 | 2 | 0 | 1 | 1 | 3 | 0 | NA | NA | 6 | 0.1667 |
| 11_229225 | 3 | 0 | 8 | 0.375 | 2 | 0 | 3 | 0 | 16 | 0.1875 | 2 | 0 | 2 | 0.5 | 3 | 0.3333 | 1 | 0 | 8 | 0.25 |
| 11_229226 | 2 | 0 | 5 | 0 | 1 | 0 | 3 | 0.3333 | 11 | 0.0909 | 3 | 0 | 2 | 0 | 3 | 0.3333 | 1 | 0 | 9 | 0.1111 |
| 11_229227 | 3 | 1 | 7 | 0.8571 | 2 | 1 | 3 | 0.3333 | 15 | 0.8 | 3 | 0.6667 | 2 | 0.5 | 2 | 0.5 | 1 | 1 | 8 | 0.625 |
| 11_229227 | 2 | 0 | 9 | 0.1111 | 2 | 0 | 1 | 0 | 14 | 0.0714 | 3 | 0 | 2 | 0.5 | 3 | 0 | 1 | 0 | 9 | 0.1111 |
| 11_229227 | 3 | 0 | 9 | 0 | 2 | 0 | 3 | 0.3333 | 17 | 0.0588 | 2 | 0.5 | 2 | 0 | 3 | 0.3333 | NA | NA | 7 | 0.2857 |
| 11_229228 | 3 | 0.6667 | 8 | 0.625 | 2 | 0 | 3 | 0.6667 | 16 | 0.5625 | 3 | 1 | 2 | 1 | 3 | 0.6667 | 1 | 1 | 9 | 0.8889 |
| 11_229229 | 4 | 0.25 | 8 | 0 | 2 | 0 | 3 | 0 | 17 | 0.0588 | 3 | 0 | 2 | 0 | 4 | 0 | 1 | 1 | 10 | 0.1 |
| 11_229230 | 3 | 0.6667 | 8 | 0.75 | 2 | 0.5 | 2 | 0.5 | 15 | 0.6667 | 3 | 1 | 2 | 0.5 | 2 | 1 | 1 | 0 | 8 | 0.75 |
| 11_229235 | 3 | 0.3333 | 11 | 0.0909 | 4 | 0 | 3 | 0 | 21 | 0.0952 | 5 | 0 | 2 | 0 | 6 | 0.1667 | 4 | 0 | 17 | 0.0588 |
| 11_229236 | 4 | 0 | 10 | 0.1 | 4 | 0.25 | 3 | 0 | 21 | 0.0952 | 4 | 0 | 3 | 0 | 4 | 0 | 5 | 0.2 | 16 | 0.0625 |
| 11_229238 | 4 | 0 | 14 | 0.0714 | 4 | 0 | 5 | 0.4 | 27 | 0.1111 | 6 | 0 | 4 | 0 | 7 | 0.2857 | 5 | 0 | 22 | 0.0909 |
| 11_229241 | 4 | 0.25 | 14 | 0 | 4 | 0.25 | 6 | 0 | 28 | 0.0714 | 6 | 0 | 4 | 0.25 | 8 | 0.25 | 5 | 0 | 23 | 0.1304 |
| 11_229242 | 4 | 0.25 | 13 | 0 | 4 | 0 | 4 | 0.25 | 25 | 0.08 | 5 | 0.2 | 4 | 0.25 | 9 | 0.1111 | 5 | 0 | 23 | 0.1304 |
| 11_229242 | 4 | 1 | 12 | 0.9167 | 4 | 1 | 7 | 1 | 27 | 0.963 | 6 | 1 | 3 | 1 | 7 | 1 | 6 | 1 | 22 | 1 |
| 11_229242 | 4 | 0 | 8 | 0.25 | 3 | 0.3333 | 6 | 0.3333 | 21 | 0.2381 | 6 | 0.3333 | 4 | 0.25 | 3 | 0 | 4 | 0.75 | 17 | 0.3529 |
| 11_229243 | 4 | 0.25 | 14 | 0.0714 | 4 | 0.25 | 7 | 0 | 29 | 0.1034 | 5 | 0.4 | 4 | 0 | 7 | 0 | 6 | 0.1667 | 22 | 0.1364 |
| 11_229245 | 4 | 0.25 | 13 | 0.0769 | 4 | 0.25 | 7 | 0 | 28 | 0.1071 | 8 | 0 | 4 | 0 | 11 | 0.1818 | 6 | 0.1667 | 29 | 0.1034 |
| 11_229245 | 4 | 1 | 12 | 0.25 | 4 | 0.5 | 7 | 0.1429 | 27 | 0.3704 | 7 | 0.7143 | 2 | 0.5 | 10 | 0.6 | 6 | 0.5 | 25 | 0.6 |

| | | | | | | | | | | | | | | | | | | | | |
|---|---|---|---|---|---|---|---|---|---|---|---|---|---|---|---|---|---|---|---|---|
| 11_2292454 | 3 | 1 | 7 | 0.8571 | 1 | 1 | 6 | 0.8333 | 17 | 0.8824 | 3 | 1 | 2 | 1 | 8 | 1 | 3 | 1 | 16 | 1 |
| 11_2292458 | 4 | 0.25 | 6 | 0 | 2 | 0.5 | 4 | 0.5 | 16 | 0.25 | 5 | 0.4 | 1 | 0 | 9 | 0.6667 | 4 | 0.5 | 19 | 0.5263 |
| 11_2292459 | 3 | 0 | 12 | 0.1667 | 4 | 0.25 | 6 | 0.1667 | 25 | 0.16 | 4 | 0.5 | 3 | 0 | 6 | 0.1667 | 5 | 0.2 | 18 | 0.2222 |
| 11_2292461 | 4 | 0.25 | 13 | 0.3846 | 4 | 0 | 7 | 0.1429 | 28 | 0.25 | 6 | 0 | 4 | 0.25 | 8 | 0.625 | 6 | 0.1667 | 24 | 0.2917 |
| 11_2292465 | 3 | 0.6667 | 10 | 1 | 3 | 0.6667 | 7 | 0.8571 | 23 | 0.8696 | 5 | 1 | 3 | 1 | 7 | 1 | 6 | 1 | 21 | 1 |
| 11_2292468 | 3 | 0.6667 | 15 | 0.5333 | 7 | 0.2857 | 5 | 0.4 | 30 | 0.4667 | 8 | 0.375 | 4 | 0.5 | 12 | 0.75 | 6 | 0.8333 | 30 | 0.6333 |
| 11_2295325 | 14 | 0.2143 | 18 | 0 | 15 | 0.0667 | 20 | 0.2 | 67 | 0.1194 | 14 | 0.2857 | 20 | 0.15 | 15 | 0.2 | 13 | 0.2308 | 62 | 0.2097 |
| 11_2297094 | 1 | 1 | NA | NA | NA | NA | NA | NA | 1 | 1 | 1 | 1 | NA | NA | NA | NA | NA | NA | 1 | 1 |
| 11_2320629 | 1 | 0 | 1 | 1 | 4 | 0.25 | 1 | 0 | 7 | 0.2857 | NA | NA | 2 | 0 | 2 | 0.5 | 1 | 0 | 5 | 0.2 |
| 11_2320633 | 1 | 0 | 1 | 0 | 6 | 0.1667 | 2 | 0 | 10 | 0.1 | 1 | 1 | 2 | 0.5 | 2 | 0 | 3 | 0 | 8 | 0.25 |
| 11_2331411 | NA | NA | NA | NA | NA | NA | 2 | 0.5 | 2 | 0.5 | NA | NA | 1 | 1 | NA | NA | NA | NA | 1 | 1 |
| 11_2331439 | NA | NA | NA | NA | NA | NA | 2 | 0.5 | 2 | 0.5 | NA | NA | 1 | 1 | NA | NA | NA | NA | 1 | 1 |
| 11_2340465 | 6 | 0 | 4 | 0 | NA | NA | 2 | 1 | 12 | 0.1667 | 3 | 0 | 3 | 0.3333 | 5 | 0 | 3 | 0 | 14 | 0.0714 |
| 11_2340490 | 2 | 0 | NA | NA | NA | NA | 2 | 0.5 | 4 | 0.25 | NA | NA | NA | NA | 2 | 0.5 | NA | NA | 2 | 0.5 |
| 11_2340641 | 3 | 0.3333 | NA | NA | NA | NA | 2 | 0.5 | 5 | 0.4 | NA | NA | NA | NA | 2 | 1 | 2 | 0 | 4 | 0.5 |
| 11_2341438 | 5 | 0 | 7 | 0 | 2 | 0 | 6 | 0.1667 | 20 | 0.05 | 1 | 0 | 5 | 0 | 4 | 0 | 3 | 0.3333 | 13 | 0.0769 |
| 11_2341444 | 5 | 0.6 | 7 | 0.7143 | 2 | 1 | 5 | 0.2 | 19 | 0.5789 | 2 | 0.5 | 5 | 0.4 | 5 | 0 | 4 | 0.25 | 16 | 0.25 |
| 11_2341446 | 4 | 0.75 | 5 | 0.2 | 1 | 1 | 5 | 0.6 | 15 | 0.5333 | 2 | 0 | 4 | 0.25 | 4 | 0 | 2 | 0.5 | 12 | 0.1667 |
| 11_2341451 | 4 | 0.25 | 7 | 0.1429 | 2 | 0 | 4 | 0 | 17 | 0.1176 | 2 | 0 | 5 | 0 | 3 | 0 | 4 | 0.5 | 14 | 0.1429 |
| 11_2341452 | 4 | 0.25 | 6 | 0.1667 | 1 | 0 | 6 | 0.1667 | 17 | 0.1765 | 2 | 0.5 | 3 | 0.3333 | 4 | 0.25 | 4 | 0.25 | 13 | 0.3077 |
| 11_2341456 | 5 | 0.2 | 5 | 0 | 2 | 0.5 | 6 | 0 | 18 | 0.1111 | 2 | 0 | 4 | 0 | 5 | 0 | 4 | 0.5 | 15 | 0.1333 |
| 11_2341460 | 5 | 0.2 | 7 | 0.7143 | 1 | 0 | 6 | 0.5 | 19 | 0.4737 | 2 | 0.5 | 5 | 0.4 | 4 | 0.5 | 4 | 0.75 | 15 | 0.5333 |
| 11_2341465 | 5 | 0.2 | 5 | 0.2 | 3 | 0 | 5 | 0.4 | 18 | 0.2222 | 2 | 0 | 4 | 0.25 | 2 | 0 | 4 | 0.25 | 12 | 0.1667 |
| 11_2343864 | 4 | 0.5 | 10 | 0 | 8 | 0.125 | 9 | 0 | 31 | 0.0968 | 4 | 0.25 | 6 | 0.1667 | 6 | 0.1667 | 11 | 0.4545 | 27 | 0.2963 |
| 11_2343867 | 5 | 0.2 | 12 | 0.1667 | 6 | 0.3333 | 7 | 0.1429 | 30 | 0.2 | 3 | 0 | 4 | 0 | 5 | 0 | 6 | 0.3333 | 18 | 0.1111 |
| 11_2343867 | 5 | 0.4 | 10 | 0.1 | 7 | 0.2857 | 9 | 0.2222 | 31 | 0.2258 | 5 | 0.6 | 5 | 0.2 | 5 | 0.2 | 8 | 0 | 23 | 0.2174 |
| 11_2343871 | 4 | 0 | 11 | 0.2727 | 9 | 0.3333 | 10 | 0.1 | 34 | 0.2059 | 5 | 0 | 6 | 0.1667 | 6 | 0.3333 | 10 | 0 | 27 | 0.1111 |
| 11_2343872 | 7 | 0.5714 | 12 | 0.6667 | 9 | 0.5556 | 11 | 0.7273 | 39 | 0.641 | 5 | 0.8 | 6 | 0.8333 | 7 | 0.7143 | 11 | 0.8182 | 29 | 0.7931 |
| 11_2343874 | 7 | 0.5714 | 12 | 0.8333 | 9 | 0.4444 | 10 | 0.6 | 38 | 0.6316 | 5 | 0.4 | 6 | 0.5 | 7 | 0.4286 | 11 | 0.5455 | 29 | 0.4828 |
| 11_2343886 | 7 | 0.4286 | 8 | 0.625 | 7 | 0.4286 | 8 | 0.375 | 30 | 0.4667 | 4 | 1 | 5 | 1 | 7 | 0.8571 | 8 | 0.5 | 24 | 0.7917 |
| 11_2343890 | 6 | 0.3333 | 12 | 0.25 | 9 | 0.3333 | 10 | 0.2 | 37 | 0.2703 | 5 | 0 | 5 | 0 | 9 | 0.3333 | 10 | 0.1 | 29 | 0.1379 |
| 11_2343898 | 7 | 0.4286 | 12 | 0.3333 | 8 | 0.5 | 9 | 0.6667 | 36 | 0.4722 | 5 | 0.4 | 6 | 0.8333 | 8 | 0.25 | 10 | 0.4 | 29 | 0.4483 |
| 11_2343903 | 7 | 0.7143 | 10 | 0.6 | 9 | 0.6667 | 10 | 0.5 | 36 | 0.6111 | 6 | 1 | 6 | 0.5 | 9 | 0.6667 | 9 | 0.6667 | 30 | 0.7 |
| 11_2343912 | 5 | 1 | 12 | 0.75 | 8 | 0.625 | 11 | 0.7273 | 36 | 0.75 | 5 | 0.4 | 6 | 0.5 | 8 | 0.375 | 10 | 0.9 | 29 | 0.5862 |
| 11_2343918 | 6 | 0.3333 | 9 | 0.3333 | 9 | 0.1111 | 8 | 0.125 | 32 | 0.2188 | 6 | 0.5 | 5 | 0.4 | 7 | 0.4286 | 9 | 0.4444 | 27 | 0.4444 |
| 11_2343989 | 4 | 0 | 7 | 0.7143 | 5 | 0.4 | 3 | 0.3333 | 19 | 0.4211 | 2 | 0 | 4 | 0.75 | 3 | 0 | 4 | 0.75 | 13 | 0.4615 |
| 11_2343991 | 4 | 0 | 6 | 0 | 5 | 0.4 | 4 | 0 | 19 | 0.1053 | 2 | 0 | 4 | 0.25 | 2 | 0.5 | 4 | 0 | 12 | 0.1667 |
| 11_2344037 | 5 | 0.4 | 6 | 0.1667 | 5 | 0.4 | 3 | 0 | 19 | 0.2632 | 2 | 0.5 | 3 | 0.3333 | 3 | 0.3333 | 5 | 0 | 13 | 0.2308 |
| 11_2344050 | 3 | 0.6667 | 5 | 0.4 | 4 | 0.25 | 2 | 0 | 14 | 0.3571 | 2 | 0 | 3 | 0.6667 | 3 | 1 | 6 | 0.6667 | 14 | 0.6429 |
| 11_2352498 | 2 | 0 | 7 | 0 | 3 | 0 | 6 | 0.1667 | 18 | 0.0556 | 1 | 0 | 3 | 0 | 4 | 0.5 | 2 | 0 | 10 | 0.2 |
| 11_2359899 | 5 | 0 | 1 | 0 | 5 | 0.6 | 6 | 0.3333 | 17 | 0.2941 | 3 | 0.3333 | 5 | 0.2 | 1 | 0 | 2 | 1 | 11 | 0.3636 |
| 11_2359901 | 7 | 0.2857 | 5 | 0.2 | 4 | 0.25 | 8 | 0.125 | 24 | 0.2083 | 3 | 0.3333 | 4 | 0.25 | 2 | 0.5 | 3 | 0.3333 | 12 | 0.3333 |
| 11_2359909 | 8 | 0.75 | 5 | 0.8 | 5 | 0.8 | 6 | 0.6667 | 24 | 0.75 | 4 | 0.5 | 4 | 0.25 | 3 | 1 | 3 | 1 | 14 | 0.6429 |
| 11_2359909 | 8 | 0.25 | 4 | 0.5 | 6 | 0 | 2 | 0 | 20 | 0.2 | 3 | 0.3333 | 4 | 0 | 3 | 0 | 2 | 0.5 | 12 | 0.1667 |
| 11_2359911 | 10 | 0.1 | 5 | 0.2 | 6 | 0 | 7 | 0 | 28 | 0.0714 | 4 | 0.25 | 5 | 0 | 2 | 0 | 4 | 0.25 | 15 | 0.1333 |
| 11_2359926 | 7 | 0.1429 | 5 | 0.2 | 5 | 0 | 6 | 0 | 23 | 0.087 | 4 | 0.25 | 3 | 0 | 3 | 0 | 3 | 0.3333 | 13 | 0.1538 |
| 11_2359935 | 8 | 0 | 4 | 0.25 | 5 | 0.2 | 5 | 0.4 | 22 | 0.1818 | 2 | 0 | 4 | 0.5 | 2 | 0 | 3 | 0.6667 | 11 | 0.3636 |
| 11_2359938 | 7 | 0.2857 | 5 | 0 | 5 | 0 | 8 | 0.25 | 25 | 0.16 | 4 | 0 | 4 | 0 | 2 | 0.5 | 4 | 0.25 | 14 | 0.1429 |
| 11_2359953 | 8 | 0.75 | 6 | 0.8333 | 5 | 0.8 | 7 | 0.7143 | 26 | 0.7692 | 4 | 0.25 | 5 | 0.6 | 1 | 1 | 4 | 1 | 14 | 0.6429 |
| 11_2360553 | 7 | 0 | 4 | 0 | 3 | 0 | 6 | 0.1667 | 20 | 0.05 | 3 | 0 | 3 | 0.3333 | 2 | 0 | 3 | 0.3333 | 11 | 0.1818 |
| 11_2360557 | 7 | 0.7143 | 6 | 1 | 5 | 0.8 | 6 | 0.6667 | 24 | 0.7917 | 3 | 0.6667 | 4 | 0.25 | 2 | 1 | 3 | 0.6667 | 12 | 0.5833 |
| 11_2360559 | 6 | 0.3333 | 6 | 0.8333 | 4 | 0.75 | 8 | 0.625 | 24 | 0.625 | 4 | 0.75 | 3 | 0 | 2 | 0 | 4 | 1 | 13 | 0.5385 |
| 11_2360566 | 6 | 1 | 5 | 1 | 5 | 1 | 7 | 0.8571 | 23 | 0.9565 | 4 | 0.75 | 4 | 1 | 2 | 1 | 4 | 1 | 14 | 0.9286 |
| 11_2360600 | 6 | 0 | 4 | 0 | 4 | 0.25 | 7 | 0.1429 | 21 | 0.0952 | 4 | 0 | 4 | 0 | 2 | 0 | 4 | 0.25 | 14 | 0.0714 |
| 11_2368143 | NA | NA | 4 | 0.25 | NA | NA | 1 | 1 | 5 | 0.4 | 1 | 0 | 2 | 0 | NA | NA | 2 | 0.5 | 5 | 0.2 |
| 11_2381216 | NA | NA | 2 | 0 | 1 | 1 | NA | NA | 3 | 0.3333 | NA | NA | NA | NA | NA | NA | 1 | 1 | 1 | 1 |
| 11_2381361 | 2 | 0.5 | 2 | 0 | 2 | 0 | NA | NA | 6 | 0.1667 | 1 | 0 | 1 | 0 | NA | NA | 1 | 1 | 3 | 0.3333 |
| 11_2381363 | 2 | 0.5 | 2 | 0.5 | 3 | 1 | NA | NA | 7 | 0.7143 | 1 | 0 | 1 | 1 | NA | NA | 1 | 0 | 3 | 0.3333 |
| 11_2381382 | 1 | 0 | 2 | 1 | 3 | 0.3333 | NA | NA | 6 | 0.5 | 1 | 1 | 1 | 0 | NA | NA | 1 | 0 | 3 | 0.3333 |
| 11_2381385 | 3 | 0.6667 | 2 | 0.5 | 3 | 0.3333 | NA | NA | 8 | 0.5 | 1 | 0 | 1 | 1 | NA | NA | 1 | 1 | 3 | 0.6667 |
| 11_2398394 | NA | NA | 1 | 1 | NA | NA | NA | NA | 1 | 1 | NA | NA | 1 | 1 | NA | NA | NA | NA | 1 | 1 |
| 11_2398400 | NA | NA | 1 | 1 | NA | NA | NA | NA | 1 | 1 | NA | NA | 1 | 1 | NA | NA | NA | NA | 1 | 1 |
| 11_2398988 | NA | NA | 1 | 1 | NA | NA | NA | NA | 1 | 1 | 1 | 1 | NA | NA | NA | NA | NA | NA | 1 | 1 |
| 11_2399026 | NA | NA | 1 | 1 | NA | NA | NA | NA | 1 | 1 | 1 | 1 | NA | NA | NA | NA | NA | NA | 1 | 1 |
| 11_2399034 | NA | NA | 1 | 1 | NA | NA | NA | NA | 1 | 1 | 1 | 1 | NA | NA | NA | NA | NA | NA | 1 | 1 |

| | | | | | | | | | | | | | | | | | | | | |
|---|---|---|---|---|---|---|---|---|---|---|---|---|---|---|---|---|---|---|---|---|
| 11_2399051 | NA | NA | 1 | 1 | NA | NA | NA | NA | 1 | 1 | 1 | 1 | NA | NA | NA | NA | NA | NA | 1 | 1 |
| 11_2399076 | NA | NA | 1 | 1 | NA | NA | NA | NA | 1 | 1 | 1 | 1 | NA | NA | NA | NA | NA | NA | 1 | 1 |
| 11_2399100 | NA | NA | 1 | 1 | NA | NA | NA | NA | 1 | 1 | 1 | 1 | NA | NA | NA | NA | NA | NA | 1 | 1 |
| 11_2403544 | 10 | 1 | 7 | 1 | 7 | 1 | 9 | 1 | 33 | 1 | 8 | 1 | 20 | 0.95 | 5 | 0.8 | 9 | 1 | 42 | 0.9524 |
| 11_2403567 | NA | NA | NA | NA | 1 | 1 | NA | NA | 1 | 1 | NA | NA | NA | NA | 1 | 1 | NA | NA | 1 | 1 |
| 11_2411330 | 4 | 0.5 | 2 | 0.5 | 3 | 0.6667 | 2 | 0.5 | 11 | 0.5455 | 1 | 1 | 7 | 0.5714 | NA | NA | 2 | 0.5 | 10 | 0.6 |
| 11_2411335 | 4 | 0.25 | 3 | 0.6667 | 3 | 0.3333 | 1 | 0 | 11 | 0.3636 | 2 | 0.5 | 6 | 0.3333 | 1 | 1 | 2 | 0.5 | 11 | 0.4545 |
| 11_2411347 | 4 | 0.25 | 1 | 0 | 3 | 0 | 1 | 0 | 9 | 0.1111 | 1 | 0 | 6 | 0.1667 | 1 | 0 | 2 | 0 | 10 | 0.1 |
| 11_2411348 | 4 | 0.25 | 3 | 0.3333 | 3 | 0 | 1 | 0 | 11 | 0.1818 | 2 | 0 | 5 | 0.4 | 1 | 0 | 2 | 0 | 10 | 0.2 |
| 11_2411361 | 2 | 0.5 | 2 | 0 | 3 | 1 | 1 | 1 | 8 | 0.625 | 2 | 0.5 | 6 | 0.6667 | 1 | 1 | NA | NA | 9 | 0.6667 |
| 11_2411363 | 4 | 0.25 | 4 | 0 | 3 | 0 | 2 | 0 | 13 | 0.0769 | 3 | 0.3333 | 6 | 0 | 1 | 0 | NA | NA | 10 | 0.1 |
| 11_2411419 | 5 | 0.6 | 7 | 0.8571 | 4 | 0.25 | 2 | 0.5 | 18 | 0.6111 | 1 | 1 | 6 | 0.8333 | 2 | 0.5 | 3 | 0.3333 | 12 | 0.6667 |
| 11_2411423 | 3 | 0 | 4 | 0 | 4 | 0 | 2 | 0.5 | 13 | 0.0769 | 2 | 0 | 5 | 0.2 | 1 | 0 | 1 | 0 | 9 | 0.1111 |
| 11_2411440 | 6 | 1 | 7 | 1 | 7 | 1 | 2 | 1 | 22 | 1 | 3 | 1 | 8 | 1 | 2 | 1 | 4 | 1 | 17 | 1 |
| 11_2411442 | 6 | 0.1667 | 5 | 0.2 | 7 | 0.2857 | 2 | 1 | 20 | 0.3 | 4 | 0.25 | 8 | 0.25 | 2 | 0.5 | 4 | 0.25 | 18 | 0.2778 |
| 11_2411469 | 6 | 0 | 8 | 0.125 | 8 | 0.25 | 2 | 0.5 | 24 | 0.1667 | 6 | 0.3333 | 7 | 0 | 3 | 0.3333 | 5 | 0 | 21 | 0.1429 |
| 11_2411469 | 6 | 0.1667 | 8 | 0.125 | 7 | 0.1429 | 1 | 0 | 22 | 0.1364 | 7 | 0.2857 | 7 | 0.1429 | 3 | 0 | 5 | 0.2 | 22 | 0.1818 |
| 11_2411491 | 1 | 1 | 2 | 1 | 3 | 1 | NA | NA | 6 | 1 | 2 | 1 | NA | NA | 1 | 1 | 1 | 1 | 4 | 1 |
| 11_2411653 | NA | NA | 6 | 0 | 4 | 0.25 | 3 | 0 | 13 | 0.0769 | 2 | 0 | NA | NA | 2 | 0.5 | 2 | 0 | 6 | 0.1667 |
| 11_2411685 | NA | NA | 6 | 0.1667 | 5 | 0 | 3 | 0 | 14 | 0.0714 | 3 | 0 | NA | NA | 2 | 0.5 | 2 | 0 | 7 | 0.1429 |
| 11_2411693 | NA | NA | 4 | 0.25 | 4 | 0 | 3 | 0 | 11 | 0.0909 | 3 | 0 | NA | NA | 2 | 0.5 | 1 | 0 | 6 | 0.1667 |
| 11_2411699 | NA | NA | 5 | 0 | 3 | 0 | 2 | 0.5 | 10 | 0.1 | 2 | 0 | NA | NA | 2 | 0.5 | 2 | 0 | 6 | 0.1667 |
| 11_2411701 | NA | NA | 5 | 0.4 | 5 | 0 | 3 | 0.3333 | 13 | 0.2308 | 3 | 0.3333 | NA | NA | 2 | 1 | 2 | 0.5 | 7 | 0.5714 |
| 11_2411710 | NA | NA | 6 | 0.1667 | 5 | 0 | 3 | 0 | 14 | 0.0714 | 3 | 0.3333 | NA | NA | 2 | 0.5 | 1 | 0 | 6 | 0.3333 |
| 11_2411718 | NA | NA | 4 | 0 | 4 | 0 | 4 | 0.5 | 12 | 0.1667 | 3 | 0.3333 | NA | NA | 3 | 0 | 2 | 0.5 | 8 | 0.25 |
| 11_2412279 | 15 | 0.0667 | 20 | 0.25 | 11 | 0.1818 | 13 | 0.2308 | 59 | 0.1864 | 19 | 0.1053 | 23 | 0.1739 | 10 | 0.1 | 13 | 0 | 65 | 0.1077 |
| 11_2412300 | 17 | 0.3529 | 16 | 0.25 | 5 | 0 | 12 | 0.4167 | 50 | 0.3 | 14 | 0.3571 | 12 | 0.1667 | 6 | 0.3333 | 11 | 0.2727 | 43 | 0.2791 |
| 11_2412303 | 14 | 0.5714 | 16 | 0.4375 | 9 | 0.7778 | 14 | 0.5714 | 53 | 0.566 | 15 | 0.5333 | 20 | 0.45 | 11 | 0.5455 | 11 | 0.6364 | 57 | 0.5263 |
| 11_2555261 | 1 | 1 | NA | NA | NA | NA | 2 | 0 | 3 | 0.3333 | 1 | 1 | 1 | 0 | 2 | 0.5 | 4 | 0.25 | 8 | 0.375 |
| 11_2555273 | 1 | 1 | NA | NA | NA | NA | 2 | 1 | 3 | 1 | 1 | 1 | 1 | 1 | 2 | 1 | 4 | 1 | 8 | 1 |
| 11_2555297 | 1 | 0 | NA | NA | NA | NA | 2 | 0.5 | 3 | 0.3333 | 1 | 1 | 1 | 1 | 2 | 0.5 | 3 | 0 | 7 | 0.4286 |
| 11_2633693 | 1 | 1 | 1 | 1 | NA | NA | NA | NA | 2 | 1 | 1 | 0 | NA | NA | 1 | 1 | NA | NA | 2 | 0.5 |
| 11_2633700 | 1 | 1 | 1 | 1 | NA | NA | NA | NA | 2 | 1 | 1 | 1 | NA | NA | 1 | 1 | NA | NA | 2 | 1 |
| 11_2633711 | 1 | 1 | 2 | 1 | NA | NA | NA | NA | 3 | 1 | 1 | 1 | NA | NA | 1 | 1 | NA | NA | 2 | 1 |
| 11_2642157 | 3 | 0.3333 | 2 | 1 | 1 | 0 | 1 | 1 | 7 | 0.5714 | 1 | 1 | 3 | 0.3333 | 2 | 0.5 | 1 | 1 | 7 | 0.5714 |
| 11_2642159 | 2 | 0 | 1 | 0 | 1 | 0 | 1 | 1 | 5 | 0.2 | 1 | 1 | 3 | 0 | 2 | 0 | 1 | 0 | 7 | 0.1429 |
| 11_2642169 | 3 | 0.6667 | 1 | 1 | 1 | 0 | 1 | 1 | 6 | 0.6667 | 1 | 0 | 2 | 0 | 2 | 0.5 | 1 | 0 | 6 | 0.1667 |
| 11_2833728 | 3 | 0 | 6 | 0.5 | 6 | 0 | 4 | 0 | 19 | 0.1579 | 6 | 0 | 2 | 0.5 | 8 | 0 | 5 | 0.2 | 21 | 0.0952 |
| 11_2833816 | 3 | 0 | 3 | 0 | 5 | 0.2 | 5 | 0 | 16 | 0.0625 | 4 | 0.25 | NA | NA | 9 | 0 | 5 | 0 | 18 | 0.0556 |
| 11_2833877 | 2 | 0 | 2 | 0 | 3 | 0.3333 | 4 | 0 | 11 | 0.0909 | 2 | 0.5 | NA | NA | 4 | 0.25 | 5 | 0.2 | 11 | 0.2727 |
| 11_2833881 | 3 | 0.3333 | 2 | 0 | 4 | 0 | 4 | 0.25 | 13 | 0.1538 | 3 | 0 | NA | NA | 6 | 0.3333 | 6 | 0.1667 | 15 | 0.2 |
| 11_2852667 | 1 | 1 | 3 | 0 | 3 | 0 | 3 | 0.3333 | 10 | 0.2 | 3 | 0.6667 | NA | NA | 2 | 1 | 3 | 1 | 8 | 0.875 |
| 11_2852679 | 1 | 1 | 2 | 1 | 2 | 1 | 3 | 1 | 8 | 1 | 3 | 1 | NA | NA | 3 | 0.6667 | 3 | 1 | 9 | 0.8889 |
| 11_2852682 | 1 | 1 | 3 | 0.3333 | 3 | 0 | 4 | 0.5 | 11 | 0.3636 | 3 | 0 | NA | NA | 4 | 0.25 | 2 | 0.5 | 9 | 0.2222 |
| 11_2852712 | 1 | 1 | 3 | 0.6667 | 2 | 1 | 2 | 1 | 8 | 0.875 | 3 | 1 | NA | NA | 3 | 1 | 2 | 1 | 8 | 1 |
| 11_2852713 | 1 | 0 | 2 | 0 | 2 | 0 | 2 | 1 | 7 | 0.2857 | 3 | 0 | NA | NA | 4 | 0.75 | 1 | 0 | 8 | 0.375 |
| 11_2853364 | 1 | 0 | 1 | 0 | 1 | 0 | 2 | 0.5 | 5 | 0.2 | 2 | 0.5 | NA | NA | 1 | 1 | 1 | 1 | 4 | 0.75 |
| 11_2853378 | 1 | 0 | NA | NA | 1 | 0 | 2 | 0.5 | 4 | 0.25 | 2 | 0.5 | NA | NA | 1 | 1 | 1 | 1 | 4 | 0.75 |
| 11_2881782 | 5 | 0.8 | 2 | 1 | 4 | 0.75 | 3 | 1 | 14 | 0.8571 | 1 | 1 | NA | NA | 2 | 1 | 1 | 1 | 4 | 1 |
| 11_2881792 | 5 | 0 | 2 | 0 | 4 | 0.25 | 3 | 0 | 14 | 0.0714 | 1 | 0 | NA | NA | 2 | 0.5 | 1 | 0 | 4 | 0.25 |
| 11_2912098 | NA | NA | NA | NA | 1 | 1 | NA | NA | 1 | 1 | NA | NA | 1 | 1 | 1 | 1 | 2 | 0.5 | 4 | 0.75 |
| 11_2913806 | 2 | 0.5 | 1 | 0 | 2 | 1 | 2 | 0 | 7 | 0.4286 | 1 | 0 | 1 | 1 | 1 | 0 | 1 | 1 | 4 | 0.5 |
| 11_2913820 | 3 | 0.3333 | 2 | 0 | 2 | 0 | 5 | 0 | 12 | 0.0833 | 1 | 0 | 1 | 0 | 3 | 0.3333 | 2 | 0 | 7 | 0.1429 |
| 11_2913821 | 3 | 0.3333 | 1 | 0 | 2 | 0 | 5 | 0 | 11 | 0.0909 | 1 | 0 | 1 | 0 | 3 | 0.3333 | 2 | 0 | 7 | 0.1429 |
| 11_2913836 | 2 | 0.5 | 3 | 1 | 2 | 0 | 5 | 0.2 | 12 | 0.4167 | 1 | 0 | 1 | 1 | 3 | 0.6667 | 2 | 1 | 7 | 0.7143 |
| 11_2913838 | 1 | 0 | 3 | 0 | 2 | 0.5 | 3 | 0.3333 | 9 | 0.2222 | 1 | 0 | 1 | 0 | 1 | 1 | 2 | 0.5 | 5 | 0.4 |
| 11_2913844 | 2 | 0.5 | 3 | 0 | 2 | 0.5 | 5 | 0 | 12 | 0.1667 | 1 | 0 | 1 | 1 | 2 | 0.5 | 2 | 0.5 | 6 | 0.5 |
| 11_2913847 | 2 | 0 | 4 | 0.25 | 2 | 0.5 | 4 | 0.5 | 12 | 0.3333 | 1 | 0 | 1 | 1 | 3 | 0 | 2 | 0 | 7 | 0.1429 |
| 11_2916629 | NA | NA | 1 | 0 | NA | NA | 3 | 0.6667 | 4 | 0.5 | NA | NA | NA | NA | NA | NA | 1 | 1 | 1 | 1 |
| 11_2916853 | 4 | 0 | 5 | 0 | 3 | 0.3333 | 4 | 0 | 16 | 0.0625 | 2 | 0 | 3 | 0.3333 | 4 | 0 | 5 | 0 | 14 | 0.0714 |
| 11_2916859 | 3 | 0.6667 | 3 | 0.3333 | 2 | 0 | 4 | 0 | 12 | 0.25 | 1 | 0 | 3 | 0 | 4 | 0.25 | 4 | 0.25 | 12 | 0.1667 |
| 11_2916864 | 4 | 0 | 5 | 0.2 | 4 | 0.25 | 4 | 0.25 | 17 | 0.1765 | 1 | 0 | 1 | 0 | 4 | 0.5 | 3 | 0 | 9 | 0.2222 |
| 11_2916867 | 4 | 0.5 | 5 | 0.4 | 5 | 0 | 4 | 0.25 | 18 | 0.2778 | 2 | 1 | 3 | 0.6667 | 4 | 0.75 | 5 | 0.4 | 14 | 0.6429 |
| 11_2916898 | 5 | 0.2 | 5 | 0.2 | 6 | 0 | 4 | 0 | 20 | 0.1 | 4 | 0 | 3 | 0 | 4 | 0 | 4 | 0.25 | 15 | 0.0667 |
| 11_2917177 | 22 | 0.2273 | 9 | 0.6667 | 9 | 0.2222 | 10 | 0.4 | 50 | 0.34 | 8 | 0.375 | 12 | 0.5 | 13 | 0.3846 | 10 | 0.5 | 43 | 0.4419 |

| | | | | | | | | | | | | | | | | | | | | |
|---|---|---|---|---|---|---|---|---|---|---|---|---|---|---|---|---|---|---|---|---|
| 11_291718 | 19 | 0.6316 | 9 | 0.7778 | 5 | 0.4 | 11 | 0.5455 | 44 | 0.6136 | 8 | 0.875 | 12 | 0.5 | 12 | 0.75 | 12 | 0.9167 | 44 | 0.75 |
| 11_291970 | 22 | 0.1364 | 7 | 0.4286 | 13 | 0.3846 | 12 | 0.0833 | 54 | 0.2222 | 6 | 0.5 | 6 | 0.5 | 14 | 0.2143 | 9 | 0.3333 | 35 | 0.3429 |
| 11_294135 | NA | NA | NA | NA | 1 | 0 | 1 | 1 | 2 | 0.5 | NA | NA | NA | NA | 3 | 0.6667 | NA | NA | 3 | 0.6667 |
| 11_294137 | NA | NA | NA | NA | 1 | 1 | 1 | 0 | 2 | 0.5 | NA | NA | 1 | 1 | 2 | 0 | NA | NA | 3 | 0.3333 |
| 11_294138 | NA | NA | NA | NA | 2 | 0 | 1 | 1 | 3 | 0.3333 | NA | NA | 1 | 0 | 3 | 0.3333 | NA | NA | 4 | 0.25 |
| 11_294138 | NA | NA | NA | NA | 2 | 0.5 | 1 | 0 | 3 | 0.3333 | NA | NA | 1 | 0 | 2 | 0.5 | NA | NA | 3 | 0.3333 |
| 11_294517 | NA | NA | NA | NA | 1 | 1 | NA | NA | 1 | 1 | NA | NA | NA | NA | 2 | 0.5 | NA | NA | 2 | 0.5 |
| 11_294521 | NA | NA | NA | NA | 1 | 1 | NA | NA | 1 | 1 | NA | NA | NA | NA | 1 | 1 | NA | NA | 1 | 1 |
| 11_294534 | NA | NA | 2 | 0 | 3 | 0.3333 | 1 | 0 | 6 | 0.1667 | 1 | 0 | 2 | 0 | 3 | 0.3333 | NA | NA | 6 | 0.1667 |
| 11_294539 | NA | NA | 2 | 0.5 | 3 | 0.3333 | 1 | 1 | 6 | 0.5 | 1 | 1 | 1 | 0 | 4 | 0 | NA | NA | 6 | 0.1667 |
| 11_294539 | NA | NA | 3 | 0.6667 | 3 | 0.6667 | 1 | 0 | 7 | 0.5714 | 1 | 1 | 1 | 1 | 4 | 1 | NA | NA | 6 | 1 |
| 11_294540 | NA | NA | 2 | 0 | 3 | 0.6667 | 1 | 0 | 6 | 0.3333 | 1 | 1 | 1 | 0 | 4 | 0.25 | NA | NA | 6 | 0.3333 |
| 11_294540 | NA | NA | 3 | 0.3333 | 3 | 0.6667 | 1 | 0 | 7 | 0.4286 | 1 | 0 | 1 | 0 | 3 | 1 | NA | NA | 5 | 0.6 |
| 11_294541 | NA | NA | 3 | 0 | 2 | 0.5 | 1 | 1 | 6 | 0.3333 | 1 | 0 | 1 | 0 | 4 | 0.25 | NA | NA | 6 | 0.1667 |
| 11_294541 | NA | NA | 3 | 0.6667 | 3 | 0.6667 | 1 | 0 | 7 | 0.5714 | 1 | 0 | 1 | 0 | 3 | 0.6667 | NA | NA | 5 | 0.4 |
| 11_294542 | NA | NA | 3 | 0 | 2 | 0.5 | 1 | 0 | 6 | 0.1667 | 1 | 0 | 1 | 0 | 4 | 0.25 | 1 | 0 | 7 | 0.1429 |
| 11_294577 | NA | NA | 3 | 0 | 5 | 0.2 | 2 | 0 | 10 | 0.1 | 1 | 0 | 2 | 0 | 4 | 0.25 | 1 | 0 | 8 | 0.125 |
| 11_294577 | NA | NA | 3 | 0.3333 | 3 | 0.6667 | 1 | 0 | 7 | 0.4286 | 1 | 1 | 2 | 0 | 4 | 0.5 | 1 | 1 | 8 | 0.5 |
| 11_294577 | NA | NA | 1 | 1 | 4 | 0.25 | 2 | 0 | 7 | 0.2857 | 1 | 1 | 2 | 0 | 3 | 0 | 1 | 1 | 7 | 0.2857 |
| 11_294579 | NA | NA | 2 | 0.5 | 5 | 1 | 2 | 0.5 | 9 | 0.7778 | 1 | 1 | 2 | 1 | 3 | 0.6667 | 1 | 1 | 7 | 0.8571 |
| 11_294579 | NA | NA | 3 | 1 | 5 | 1 | 2 | 1 | 10 | 1 | 1 | 1 | 2 | 1 | 3 | 0.6667 | 1 | 1 | 7 | 0.8571 |
| 11_294579 | NA | NA | 3 | 0.6667 | 5 | 0.8 | 2 | 0.5 | 10 | 0.7 | 1 | 1 | 2 | 0 | 3 | 0.6667 | NA | NA | 6 | 0.5 |
| 11_294597 | NA | NA | 4 | 0 | 6 | 0.1667 | NA | NA | 10 | 0.1 | 2 | 0.5 | 3 | 0 | 1 | 0 | 1 | 1 | 7 | 0.2857 |
| 11_294604 | 1 | 0 | 3 | 0 | 7 | 0.2857 | 1 | 0 | 12 | 0.1667 | 3 | 0 | 4 | 0 | 2 | 0 | 4 | 0.25 | 13 | 0.0769 |
| 11_294607 | 1 | 0 | 1 | 0 | 5 | 0.2 | 1 | 0 | 8 | 0.125 | 3 | 0 | 1 | 1 | NA | NA | 2 | 0 | 6 | 0.1667 |
| 11_294632 | NA | NA | NA | NA | 2 | 0 | 1 | 1 | 3 | 0.3333 | NA | NA | NA | NA | 1 | 1 | NA | NA | 1 | 1 |
| 11_294632 | NA | NA | NA | NA | 2 | 0.5 | 1 | 0 | 3 | 0.3333 | 2 | 1 | 2 | 0 | 1 | 0 | NA | NA | 5 | 0.4 |
| 11_294633 | NA | NA | NA | NA | 2 | 0 | 1 | 1 | 3 | 0.3333 | 2 | 0.5 | 2 | 0.5 | 1 | 0 | NA | NA | 5 | 0.4 |
| 11_294633 | NA | NA | NA | NA | 2 | 0.5 | NA | NA | 2 | 0.5 | NA | NA | 2 | 1 | 1 | 0 | NA | NA | 3 | 0.6667 |
| 11_294633 | NA | NA | NA | NA | 2 | 0.5 | 1 | 1 | 3 | 0.6667 | 1 | 1 | 2 | 1 | 1 | 1 | NA | NA | 4 | 1 |
| 11_294634 | NA | NA | NA | NA | 2 | 1 | 1 | 1 | 3 | 1 | 2 | 0.5 | 1 | 1 | 1 | 0 | NA | NA | 4 | 0.5 |
| 11_294636 | NA | NA | NA | NA | 2 | 0.5 | 1 | 0 | 3 | 0.3333 | 2 | 1 | 1 | 0 | 1 | 1 | NA | NA | 4 | 0.75 |
| 11_294949 | 4 | 1 | 1 | 1 | 3 | 1 | 5 | 0.6 | 13 | 0.8462 | 1 | 1 | 1 | 1 | NA | NA | NA | NA | 2 | 1 |
| 11_296422 | 1 | 1 | NA | NA | NA | NA | NA | NA | 1 | 1 | NA | NA | 1 | 1 | 1 | 1 | NA | NA | 2 | 1 |
| 11_296434 | 7 | 0.4286 | 4 | 0.5 | 6 | 0.5 | 11 | 0.1818 | 28 | 0.3571 | 5 | 0.2 | 8 | 0.375 | 5 | 0.2 | 8 | 0.25 | 26 | 0.2692 |
| 11_296565 | 6 | 0.3333 | 3 | 0 | 6 | 0.3333 | 12 | 0.25 | 27 | 0.2593 | 6 | 0.6667 | 7 | 0.5714 | 5 | 0.4 | 8 | 0.25 | 26 | 0.4615 |
| 11_296566 | 7 | 0.2857 | 4 | 0.25 | 8 | 0.125 | 11 | 0 | 30 | 0.1333 | 8 | 0.125 | 9 | 0.2222 | 5 | 0 | 7 | 0.1429 | 29 | 0.1379 |
| 11_296568 | 9 | 0 | 6 | 0.1667 | 10 | 0.2 | 15 | 0.2 | 40 | 0.15 | 9 | 0.2222 | 10 | 0 | 5 | 0 | 8 | 0.375 | 32 | 0.1562 |
| 11_296568 | 10 | 1 | 5 | 0.8 | 9 | 1 | 12 | 1 | 36 | 0.9722 | 8 | 1 | 10 | 0.9 | 5 | 1 | 6 | 0.8333 | 29 | 0.931 |
| 11_296570 | 10 | 0.3 | 7 | 0.2857 | 9 | 0.6667 | 14 | 0.4286 | 40 | 0.425 | 10 | 0.5 | 8 | 0.5 | 4 | 0.5 | 7 | 0.1429 | 29 | 0.4138 |
| 11_296571 | 10 | 0.3 | 9 | 0.4444 | 11 | 0.3636 | 15 | 0.2667 | 45 | 0.3333 | 10 | 0.3 | 9 | 0 | 5 | 0.2 | 8 | 0.125 | 32 | 0.1562 |
| 11_296573 | 11 | 0.3636 | 10 | 0.2 | 11 | 0.2727 | 12 | 0.3333 | 44 | 0.2955 | 9 | 0.1111 | 9 | 0.3333 | 5 | 0 | 9 | 0.1111 | 32 | 0.1562 |
| 11_296574 | 12 | 0.5 | 10 | 0.3 | 12 | 0.1667 | 14 | 0.2143 | 48 | 0.2917 | 10 | 0.1 | 8 | 0.375 | 5 | 0.2 | 10 | 0.4 | 33 | 0.2727 |
| 11_296575 | 12 | 0.5833 | 9 | 0.5556 | 10 | 0.7 | 15 | 0.5333 | 46 | 0.587 | 10 | 0.6 | 8 | 0.375 | 5 | 0.6 | 7 | 0.7143 | 30 | 0.5667 |
| 11_296577 | 12 | 0.5 | 14 | 0.7143 | 13 | 0.5385 | 15 | 0.4667 | 54 | 0.5556 | 11 | 0.7273 | 11 | 0.5455 | 4 | 0.25 | 8 | 0.5 | 34 | 0.5588 |
| 11_296580 | 13 | 0.3846 | 14 | 0.2857 | 13 | 0.3077 | 16 | 0.4375 | 56 | 0.3571 | 10 | 0.4 | 12 | 0.1667 | 6 | 0.1667 | 8 | 0.25 | 36 | 0.25 |
| 11_296581 | 13 | 0.1538 | 14 | 0.2857 | 14 | 0.0714 | 16 | 0.0625 | 57 | 0.1404 | 8 | 0 | 11 | 0.0909 | 5 | 0 | 10 | 0.2 | 34 | 0.0882 |
| 11_296581 | 11 | 0.7273 | 14 | 0.4286 | 14 | 0.5 | 15 | 0.6 | 54 | 0.5556 | 12 | 0.5 | 10 | 0.3 | 4 | 0 | 9 | 0.1111 | 35 | 0.2857 |
| 11_296582 | 12 | 0 | 15 | 0.1333 | 15 | 0.2667 | 18 | 0.2778 | 60 | 0.1833 | 10 | 0.1 | 10 | 0.1 | 6 | 0 | 10 | 0.2 | 36 | 0.1111 |
| 11_296583 | 9 | 0.3333 | 12 | 0.3333 | 12 | 0.3333 | 16 | 0.25 | 49 | 0.3061 | 9 | 0.1111 | 10 | 0.2 | 5 | 0.2 | 11 | 0.2727 | 35 | 0.2 |
| 11_296583 | 11 | 0.1818 | 14 | 0.1429 | 15 | 0.3333 | 18 | 0.0556 | 58 | 0.1724 | 12 | 0.25 | 11 | 0.0909 | 5 | 0.2 | 10 | 0 | 38 | 0.1316 |
| 11_296585 | 14 | 0.0714 | 19 | 0.2632 | 15 | 0.2 | 21 | 0.1429 | 69 | 0.1739 | 12 | 0.0833 | 13 | 0.1538 | 7 | 0 | 14 | 0.0714 | 46 | 0.087 |
| 11_296587 | 14 | 0.2857 | 20 | 0.25 | 14 | 0.1429 | 18 | 0.1667 | 66 | 0.2121 | 8 | 0 | 10 | 0.1 | 6 | 0.3333 | 14 | 0.0714 | 38 | 0.1053 |
| 11_296880 | 1 | 1 | NA | NA | 1 | 0 | NA | NA | 2 | 0.5 | NA | NA | 2 | 0.5 | NA | NA | NA | NA | 2 | 0.5 |
| 11_296927 | 88 | 0.1818 | 121 | 0.2066 | 81 | 0.2469 | 89 | 0.2584 | 379 | 0.2216 | 83 | 0.1928 | 79 | 0.2532 | 62 | 0.1613 | 80 | 0.225 | 304 | 0.2105 |
| 11_296932 | 2 | 0.5 | 3 | 0 | 4 | 0 | 2 | 0 | 11 | 0.0909 | 1 | 0 | 1 | 0 | 5 | 0 | 1 | 1 | 8 | 0.125 |
| 11_296936 | 1 | 0 | 1 | 0 | 3 | 0.3333 | 1 | 0 | 6 | 0.1667 | 1 | 1 | 1 | 0 | 5 | 0 | 1 | 0 | 8 | 0.125 |
| 11_296944 | NA | NA | 2 | 0.5 | 3 | 0.6667 | 1 | 0 | 6 | 0.5 | 1 | 1 | 1 | 1 | 5 | 0.8 | 1 | 1 | 8 | 0.875 |
| 11_296944 | NA | NA | 1 | 1 | 3 | 0.3333 | 1 | 0 | 5 | 0.4 | 1 | 0 | 1 | 0 | 3 | 0.3333 | NA | NA | 5 | 0.2 |
| 11_296944 | NA | NA | 2 | 0.5 | 3 | 0.3333 | 1 | 1 | 6 | 0.5 | NA | NA | 1 | 0 | 3 | 0.6667 | 1 | 0 | 5 | 0.4 |
| 11_300494 | 50 | 0.24 | 53 | 0.1698 | 42 | 0.1429 | 50 | 0.26 | 195 | 0.2051 | 26 | 0.2692 | 42 | 0.1429 | 31 | 0.1613 | 49 | 0.2245 | 148 | 0.1959 |
| 11_300501 | 67 | 0.3881 | 66 | 0.3333 | 56 | 0.3571 | 62 | 0.3226 | 251 | 0.3506 | 33 | 0.5758 | 67 | 0.4478 | 44 | 0.3409 | 69 | 0.4203 | 213 | 0.4366 |
| 11_300504 | 52 | 0.25 | 51 | 0.098 | 42 | 0 | 50 | 0.12 | 195 | 0.1231 | 27 | 0.1111 | 49 | 0.1429 | 31 | 0.0968 | 55 | 0.1273 | 162 | 0.1235 |
| 11_300590 | NA | NA | NA | NA | 1 | 0 | 1 | 1 | 2 | 0.5 | NA | NA | 2 | 0.5 | 1 | 1 | 2 | 0 | 5 | 0.4 |
| 11_300590 | NA | NA | NA | NA | 1 | 0 | 1 | 1 | 2 | 0.5 | 1 | 1 | 2 | 0.5 | NA | NA | 3 | 0.3333 | 6 | 0.5 |

| | | | | | | | | | | | | | | | | | | | | |
|---|---|---|---|---|---|---|---|---|---|---|---|---|---|---|---|---|---|---|---|---|
| 11_3006774 | 28 | 0.1429 | 28 | 0.0714 | 27 | 0.1481 | 33 | 0.0606 | 116 | 0.1034 | 20 | 0.1 | 28 | 0.0714 | 24 | 0.0833 | 37 | 0.0541 | 109 | 0.0734 |
| 11_3006779 | 29 | 0.6552 | 29 | 0.8621 | 23 | 0.7826 | 29 | 0.5862 | 110 | 0.7182 | 18 | 0.8889 | 25 | 0.56 | 21 | 0.6667 | 35 | 0.7143 | 99 | 0.697 |
| 11_3006790 | 23 | 0.4348 | 18 | 0.6111 | 15 | 0.5333 | 22 | 0.4091 | 78 | 0.4872 | 13 | 0.6923 | 16 | 0.5 | 12 | 0.5833 | 25 | 0.4 | 66 | 0.5152 |
| 11_3006792 | 29 | 0.3448 | 21 | 0.381 | 23 | 0.3913 | 29 | 0.3793 | 102 | 0.3725 | 18 | 0.5556 | 28 | 0.3214 | 22 | 0.5455 | 35 | 0.2857 | 103 | 0.3981 |
| 11_3008695 | 13 | 0.0769 | 8 | 0 | 11 | 0.2727 | 13 | 0.1538 | 45 | 0.1333 | 6 | 0.1667 | 12 | 0.0833 | 6 | 0 | 15 | 0 | 39 | 0.0513 |
| 11_3008699 | 15 | 0 | 11 | 0.0909 | 12 | 0.25 | 17 | 0.3529 | 55 | 0.1818 | 12 | 0.1667 | 14 | 0.2143 | 9 | 0.3333 | 21 | 0.2381 | 56 | 0.2321 |
| 11_3008704 | 17 | 0.0588 | 11 | 0 | 12 | 0.0833 | 18 | 0.1111 | 58 | 0.069 | 12 | 0.1667 | 14 | 0.2143 | 10 | 0 | 21 | 0.0952 | 57 | 0.1228 |
| 11_3008707 | 10 | 0.3 | 9 | 0.2222 | 10 | 0.3 | 17 | 0.2353 | 46 | 0.2609 | 8 | 0.25 | 11 | 0.1818 | 9 | 0 | 18 | 0.3333 | 46 | 0.2174 |
| 11_3008717 | 15 | 0.6 | 10 | 0.5 | 9 | 0.7778 | 18 | 0.3889 | 52 | 0.5385 | 12 | 0.75 | 11 | 0.6364 | 9 | 0.7778 | 17 | 0.5882 | 49 | 0.6735 |
| 11_3008719 | 16 | 0.3125 | 9 | 0.3333 | 11 | 0.0909 | 18 | 0.2222 | 54 | 0.2407 | 11 | 0.0909 | 11 | 0.2727 | 8 | 0.25 | 17 | 0.2941 | 47 | 0.234 |
| 11_3008722 | 14 | 0.2143 | 9 | 0.4444 | 10 | 0.5 | 20 | 0.4 | 53 | 0.3774 | 11 | 0.6364 | 14 | 0.3571 | 9 | 0.7778 | 12 | 0.4167 | 46 | 0.5217 |
| 11_3008742 | 16 | 0.25 | 7 | 0.4286 | 10 | 0.1 | 17 | 0.4118 | 50 | 0.3 | 11 | 0.1818 | 13 | 0.2308 | 9 | 0.4444 | 15 | 0.2667 | 48 | 0.2708 |
| 11_3008885 | 11 | 0.1818 | 4 | 0 | 7 | 0.2857 | 13 | 0 | 35 | 0.1143 | 10 | 0.1 | 11 | 0.1818 | 4 | 0 | 9 | 0.1111 | 34 | 0.1176 |
| 11_3008913 | 8 | 0.125 | 6 | 0.3333 | 6 | 0.1667 | 11 | 0.3636 | 31 | 0.2581 | 7 | 0.4286 | 9 | 0.2222 | 2 | 0.5 | 11 | 0.5455 | 29 | 0.4138 |
| 11_3008925 | 9 | 0.5556 | 6 | 0.1667 | 8 | 0.375 | 11 | 0.3636 | 34 | 0.3824 | 7 | 0.4286 | 10 | 0.3 | 3 | 0.6667 | 11 | 0.2727 | 31 | 0.3548 |
| 11_3078734 | 3 | 0.3333 | 2 | 0 | 2 | 0 | 2 | 0 | 9 | 0.1111 | 3 | 0.3333 | 2 | 0.5 | NA | NA | 1 | 0 | 6 | 0.3333 |
| 11_3082991 | 6 | 0 | 5 | 0 | 4 | 0.25 | 5 | 0 | 20 | 0.05 | 3 | 0 | 2 | 0 | 4 | 0 | 3 | 0.3333 | 12 | 0.0833 |
| 11_3083023 | 7 | 0.2857 | 5 | 0 | 5 | 0.2 | 7 | 0 | 24 | 0.125 | 4 | 0.5 | 2 | 0 | 3 | 0.6667 | 2 | 0 | 11 | 0.3636 |
| 11_3084108 | 2 | 0.5 | NA | NA | 2 | 0.5 | 6 | 0 | 10 | 0.2 | NA | NA | NA | NA | 1 | 1 | 1 | 0 | 2 | 0.5 |
| 11_3084135 | 2 | 1 | NA | NA | 2 | 1 | 6 | 0.8333 | 10 | 0.9 | NA | NA | NA | NA | 3 | 0.6667 | 1 | 0 | 4 | 0.5 |
| 11_3084164 | 2 | 1 | NA | NA | 1 | 1 | 8 | 0.625 | 11 | 0.7273 | NA | NA | NA | NA | 3 | 0.3333 | 2 | 0 | 5 | 0.2 |
| 11_3084172 | 2 | 1 | 1 | 0 | 2 | 1 | 7 | 0.7143 | 12 | 0.75 | NA | NA | NA | NA | 3 | 0.6667 | 2 | 0.5 | 5 | 0.6 |
| 11_3084193 | 2 | 1 | 1 | 1 | 2 | 1 | 8 | 0.625 | 13 | 0.7692 | NA | NA | NA | NA | 2 | 0.5 | 2 | 0.5 | 4 | 0.5 |
| 11_3084194 | 2 | 0.5 | 1 | 0 | 2 | 0 | 8 | 0.375 | 13 | 0.3077 | NA | NA | NA | NA | 3 | 0 | 2 | 0.5 | 5 | 0.2 |
| 11_3084217 | 1 | 1 | NA | NA | NA | NA | 6 | 1 | 7 | 1 | NA | NA | NA | NA | 2 | 1 | NA | NA | 2 | 1 |
| 11_3084223 | 2 | 1 | 1 | 0 | 1 | 1 | 5 | 0.8 | 9 | 0.7778 | NA | NA | NA | NA | 2 | 1 | 2 | 1 | 4 | 1 |
| 11_3084233 | 2 | 0 | 1 | 1 | 2 | 0.5 | 8 | 0.25 | 13 | 0.3077 | NA | NA | NA | NA | 3 | 0.3333 | 2 | 0 | 5 | 0.2 |
| 11_3084237 | 2 | 0.5 | 1 | 1 | 2 | 0.5 | 6 | 0.5 | 11 | 0.5455 | NA | NA | NA | NA | 2 | 1 | 1 | 0 | 3 | 0.6667 |
| 11_3084242 | 1 | 1 | NA | NA | 1 | 1 | 7 | 0.5714 | 9 | 0.6667 | NA | NA | NA | NA | 3 | 0.6667 | 2 | 0 | 5 | 0.4 |
| 11_3084248 | 2 | 1 | 1 | 1 | 2 | 1 | 9 | 0.7778 | 14 | 0.8571 | NA | NA | NA | NA | 1 | 1 | 2 | 1 | 3 | 1 |
| 11_3084253 | 2 | 0 | 1 | 0 | 2 | 1 | 9 | 0.4444 | 14 | 0.4286 | NA | NA | NA | NA | 2 | 0.5 | 2 | 0.5 | 4 | 0.5 |
| 11_3084260 | 1 | 1 | 1 | 1 | 1 | 1 | 7 | 1 | 10 | 1 | NA | NA | NA | NA | 3 | 1 | 2 | 1 | 5 | 1 |
| 11_3084267 | 2 | 0 | 1 | 0 | 2 | 0 | 9 | 0.2222 | 14 | 0.1429 | NA | NA | NA | NA | 3 | 0.3333 | 2 | 0.5 | 5 | 0.4 |
| 11_3084280 | 1 | 1 | 1 | 1 | 2 | 1 | 8 | 1 | 12 | 1 | NA | NA | NA | NA | 3 | 1 | 2 | 1 | 5 | 1 |
| 11_3084281 | 2 | 1 | 1 | 0 | 1 | 1 | 7 | 0.7143 | 11 | 0.7273 | NA | NA | NA | NA | 2 | 1 | 2 | 0 | 4 | 0.5 |
| 11_3084300 | 2 | 0.5 | 1 | 1 | 3 | 0 | 10 | 0.5 | 16 | 0.4375 | NA | NA | NA | NA | 3 | 0.3333 | 3 | 0 | 6 | 0.1667 |
| 11_3085077 | NA | NA | NA | NA | 1 | 1 | NA | NA | 1 | 1 | NA | NA | NA | NA | 1 | 1 | NA | NA | 1 | 1 |
| 11_3085079 | NA | NA | NA | NA | 1 | 1 | NA | NA | 1 | 1 | NA | NA | NA | NA | 1 | 1 | NA | NA | 1 | 1 |
| 11_3088156 | NA | NA | NA | NA | 2 | 0 | 4 | 0.25 | 6 | 0.1667 | 1 | 1 | NA | NA | NA | NA | NA | NA | 1 | 1 |
| 11_3088166 | 2 | 0.5 | 6 | 1 | 6 | 0.3333 | 6 | 0.6667 | 20 | 0.65 | 3 | 1 | 3 | 0.6667 | 6 | 0.8333 | 3 | 1 | 15 | 0.8667 |
| 11_3088169 | 2 | 0 | 6 | 0.3333 | 5 | 0.2 | 6 | 0.1667 | 19 | 0.2105 | 4 | 0 | 2 | 0 | 7 | 0.2857 | 3 | 0.6667 | 16 | 0.25 |
| 11_3088179 | 2 | 0.5 | 4 | 0.75 | 5 | 0.6 | 7 | 0.2857 | 18 | 0.5 | 4 | 0.5 | 3 | 0.6667 | 5 | 0.8 | 4 | 1 | 16 | 0.75 |
| 11_3092728 | 2 | 1 | 3 | 0.6667 | 3 | 1 | 4 | 0.75 | 12 | 0.8333 | NA | NA | 4 | 1 | 4 | 0.5 | 3 | 0.6667 | 11 | 0.7273 |
| 11_3092730 | 2 | 0 | 2 | 0.5 | 1 | 0 | 3 | 0 | 8 | 0.125 | NA | NA | 4 | 0.25 | 4 | 0.25 | 2 | 0.5 | 10 | 0.3 |
| 11_3092737 | 2 | 1 | 3 | 1 | 3 | 0.6667 | 4 | 0.75 | 12 | 0.8333 | 1 | 0 | 3 | 1 | 3 | 1 | 3 | 1 | 10 | 0.9 |
| 11_3092745 | 2 | 0 | 3 | 0 | 3 | 0 | 3 | 0.3333 | 11 | 0.0909 | 1 | 0 | 3 | 0 | 3 | 0 | 3 | 0.3333 | 10 | 0.1 |
| 11_3092747 | 2 | 1 | 1 | 1 | 1 | 1 | 4 | 1 | 8 | 1 | 1 | 0 | 2 | 1 | 3 | 1 | 3 | 1 | 9 | 0.8889 |
| 11_3092753 | 1 | 0 | NA | NA | 2 | 1 | 3 | 0.3333 | 6 | 0.5 | 1 | 0 | 2 | 0.5 | 3 | 0 | 3 | 0 | 9 | 0.1111 |
| 11_3092758 | 2 | 0.5 | 2 | 0 | 2 | 0 | NA | NA | 6 | 0.1667 | 1 | 0 | 3 | 0.6667 | 2 | 0.5 | 2 | 1 | 8 | 0.625 |
| 11_3094837 | 1 | 1 | 4 | 0.25 | NA | NA | NA | NA | 5 | 0.4 | NA | NA | NA | NA | 1 | 0 | 3 | 0.3333 | 4 | 0.25 |
| 11_3105111 | 3 | 0 | 5 | 0.2 | 2 | 0 | 1 | 0 | 11 | 0.0909 | 1 | 0 | 2 | 0 | 2 | 0 | 3 | 0.3333 | 8 | 0.125 |
| 11_3105115 | 3 | 0 | 5 | 0.2 | 1 | 0 | 2 | 0.5 | 11 | 0.1818 | 1 | 1 | 2 | 0 | 2 | 0.5 | 3 | 0.3333 | 8 | 0.375 |
| 11_3105128 | 2 | 0 | 4 | 0.5 | 2 | 0.5 | 2 | 0.5 | 10 | 0.4 | 1 | 0 | 2 | 0.5 | 2 | 1 | 3 | 0.6667 | 8 | 0.625 |
| 11_3131002 | 1 | 1 | NA | NA | NA | NA | NA | NA | 1 | 1 | NA | NA | 1 | 1 | NA | NA | 1 | 1 | 2 | 1 |
| 11_3131008 | 1 | 1 | NA | NA | NA | NA | NA | NA | 1 | 1 | NA | NA | 1 | 1 | 1 | 1 | 1 | 1 | 3 | 1 |
| 11_3131020 | 1 | 1 | NA | NA | NA | NA | NA | NA | 1 | 1 | NA | NA | 1 | 1 | 1 | 1 | NA | NA | 2 | 1 |
| 11_3161423 | 1 | 1 | NA | NA | NA | NA | NA | NA | 1 | 1 | NA | NA | 1 | 1 | 1 | 0 | 1 | 0 | 3 | 0.3333 |
| 11_3161429 | 1 | 1 | NA | NA | NA | NA | NA | NA | 1 | 1 | NA | NA | 1 | 1 | NA | NA | 1 | 0 | 2 | 0.5 |
| 11_3161444 | 1 | 1 | NA | NA | NA | NA | NA | NA | 1 | 1 | NA | NA | 1 | 1 | NA | NA | 1 | 1 | 2 | 1 |
| 11_3161532 | 17 | 0.1765 | 19 | 0.1053 | 17 | 0 | 14 | 0.0714 | 67 | 0.0896 | 11 | 0 | 18 | 0.1111 | 20 | 0 | 13 | 0.1538 | 62 | 0.0645 |
| 11_3161558 | 13 | 0.3077 | 20 | 0.3 | 11 | 0.2727 | 12 | 0.3333 | 56 | 0.3036 | 8 | 0.5 | 21 | 0.2857 | 16 | 0.375 | 9 | 0.3333 | 54 | 0.3519 |
| 11_3161559 | 15 | 0.9333 | 19 | 0.5789 | 15 | 0.6 | 14 | 0.7143 | 63 | 0.6984 | 8 | 0.875 | 21 | 0.8571 | 17 | 0.8235 | 14 | 0.9286 | 60 | 0.8667 |
| 11_3161571 | 13 | 0.3846 | 20 | 0.5 | 14 | 0.2857 | 15 | 0.3333 | 62 | 0.3871 | 8 | 0.5 | 17 | 0.4706 | 15 | 0.4667 | 12 | 0.4167 | 52 | 0.4615 |
| 11_3161575 | 15 | 0.0667 | 20 | 0.1 | 13 | 0 | 17 | 0.1765 | 65 | 0.0923 | 10 | 0.1 | 19 | 0 | 19 | 0 | 11 | 0.1818 | 59 | 0.0508 |
| 11_3161582 | 14 | 0.2857 | 17 | 0.5882 | 16 | 0.375 | 17 | 0.2941 | 64 | 0.3906 | 10 | 0.5 | 19 | 0.4737 | 18 | 0.3333 | 11 | 0.5455 | 58 | 0.4483 |

| | | | | | | | | | | | | | | | | | | | | |
|---|---|---|---|---|---|---|---|---|---|---|---|---|---|---|---|---|---|---|---|---|
| 11_318210 | 2 | 0 | NA | NA | NA | NA | 1 | 1 | 3 | 0.3333 | 2 | 0.5 | NA | NA | 1 | 1 | 1 | 0 | 4 | 0.5 |
| 11_318212 | 2 | 1 | NA | NA | NA | NA | 3 | 0.3333 | 5 | 0.6 | 2 | 1 | NA | NA | 1 | 1 | 1 | 0 | 4 | 0.75 |
| 11_318212 | 2 | 1 | NA | NA | NA | NA | 2 | 1 | 4 | 1 | 1 | 1 | NA | NA | 2 | 1 | 1 | 1 | 4 | 1 |
| 11_318219 | 1 | 1 | NA | NA | 2 | 0.5 | NA | NA | 3 | 0.6667 | 3 | 0.3333 | NA | NA | 2 | 0.5 | 1 | 1 | 6 | 0.5 |
| 11_318220 | 2 | 0 | 1 | 0 | 2 | 0.5 | 2 | 0.5 | 7 | 0.2857 | 4 | 0.25 | NA | NA | 2 | 0 | 1 | 0 | 7 | 0.1429 |
| 11_318226 | 2 | 0.5 | 2 | 0.5 | 2 | 0 | 4 | 0 | 10 | 0.2 | 3 | 0 | NA | NA | 3 | 0 | 4 | 0.25 | 10 | 0.1 |
| 11_318226 | 1 | 1 | 1 | 0 | 2 | 1 | 4 | 0.75 | 8 | 0.75 | 3 | 0.3333 | NA | NA | 2 | 0.5 | 4 | 0.75 | 9 | 0.5556 |
| 11_318226 | 2 | 0 | 2 | 0 | 2 | 0 | 4 | 0.25 | 10 | 0.1 | 4 | 0 | NA | NA | 3 | 0 | 4 | 0.25 | 11 | 0.0909 |
| 11_318227 | 1 | 1 | 2 | 0.5 | 2 | 0.5 | 3 | 0.6667 | 8 | 0.625 | 3 | 1 | NA | NA | 3 | 0.6667 | 4 | 0.75 | 10 | 0.8 |
| 11_318232 | 2 | 1 | 2 | 1 | 2 | 0.5 | 2 | 0.5 | 8 | 0.75 | 4 | 0.5 | NA | NA | 3 | 0.6667 | 4 | 1 | 11 | 0.7273 |
| 11_318815 | 2 | 1 | 4 | 0.5 | 4 | 0.5 | 2 | 0 | 12 | 0.5 | 5 | 0 | 2 | 0 | 4 | 0 | 4 | 0.25 | 15 | 0.0667 |
| 11_318816 | 2 | 0.5 | 5 | 0.4 | 4 | 0.25 | 4 | 0.5 | 15 | 0.4 | 5 | 0 | 1 | 0 | 4 | 0 | 4 | 0.25 | 14 | 0.0714 |
| 11_318847 | 5 | 0.6 | 14 | 0.1429 | 12 | 0.0833 | 12 | 0.25 | 43 | 0.2093 | 12 | 0.1667 | 9 | 0.3333 | 7 | 0.1429 | 9 | 0.2222 | 37 | 0.2162 |
| 11_318847 | 5 | 0.2 | 14 | 0.1429 | 12 | 0.1667 | 12 | 0.1667 | 43 | 0.1628 | 12 | 0.0833 | 10 | 0.4 | 9 | 0.3333 | 9 | 0.2222 | 40 | 0.25 |
| 11_319829 | 2 | 1 | 1 | 1 | 5 | 0.4 | 4 | 0.5 | 12 | 0.5833 | 2 | 1 | 2 | 0.5 | 1 | 0 | 1 | 1 | 6 | 0.6667 |
| 11_321596 | NA | NA | 3 | 0.6667 | NA | NA | 1 | 0 | 4 | 0.5 | NA | NA | NA | NA | NA | NA | 1 | 1 | 1 | 1 |
| 11_321597 | NA | NA | 2 | 0.5 | 2 | 0.5 | 1 | 1 | 5 | 0.6 | 2 | 1 | NA | NA | NA | NA | 1 | 1 | 3 | 1 |
| 11_321597 | NA | NA | 5 | 0.8 | 1 | 1 | 1 | 0 | 7 | 0.7143 | 1 | 1 | NA | NA | NA | NA | 1 | 1 | 2 | 1 |
| 11_321794 | 1 | 1 | NA | NA | NA | NA | 1 | 1 | 2 | 1 | NA | NA | 1 | 1 | NA | NA | 1 | 1 | 2 | 1 |
| 11_321816 | 3 | 1 | 1 | 0 | 2 | 0.5 | 4 | 0.5 | 10 | 0.6 | 2 | 0.5 | 2 | 0.5 | 4 | 0.75 | 3 | 0.6667 | 11 | 0.6364 |
| 11_321820 | 4 | 0.25 | 1 | 0 | 2 | 0 | 3 | 0.6667 | 10 | 0.3 | 1 | 0 | 2 | 0 | 3 | 0.3333 | 2 | 0 | 8 | 0.125 |
| 11_321822 | 2 | 0 | NA | NA | 2 | 0.5 | 2 | 0.5 | 6 | 0.3333 | 1 | 0 | 2 | 0.5 | 1 | 1 | 2 | 0 | 6 | 0.3333 |
| 11_321822 | 1 | 1 | NA | NA | 2 | 0 | 2 | 1 | 5 | 0.6 | 1 | 1 | 2 | 1 | 1 | 0 | 2 | 0.5 | 6 | 0.6667 |
| 11_321822 | 3 | 0.6667 | NA | NA | 2 | 1 | 1 | 1 | 6 | 0.8333 | 1 | 1 | 1 | 1 | 1 | 1 | 2 | 0.5 | 5 | 0.8 |
| 11_321827 | 4 | 0.25 | 6 | 0 | 3 | 0 | 3 | 0 | 16 | 0.0625 | 4 | 0.25 | 4 | 0 | 1 | 0 | 3 | 0 | 12 | 0.0833 |
| 11_321828 | 4 | 0 | 5 | 0.2 | 2 | 0 | 2 | 0.5 | 13 | 0.1538 | 2 | 0.5 | 4 | 0.25 | 1 | 0 | 2 | 0 | 9 | 0.2222 |
| 11_321829 | 3 | 0.3333 | 5 | 0.4 | 2 | 0.5 | 3 | 0.3333 | 13 | 0.3846 | 4 | 0.75 | 5 | 0.6 | 1 | 0 | 3 | 0.3333 | 13 | 0.5385 |
| 11_324494 | NA | NA | 1 | 0 | 2 | 0.5 | 2 | 0.5 | 5 | 0.4 | NA | NA | NA | NA | 2 | 0 | 5 | 0.2 | 7 | 0.1429 |
| 11_324661 | 1 | 1 | 1 | 1 | 2 | 1 | 2 | 1 | 6 | 1 | NA | NA | NA | NA | 3 | 1 | 4 | 1 | 7 | 1 |
| 11_324662 | 1 | 0 | 1 | 1 | 1 | 1 | 2 | 0.5 | 5 | 0.6 | NA | NA | NA | NA | 1 | 0 | 3 | 0.6667 | 4 | 0.5 |
| 11_324662 | 1 | 0 | 2 | 0 | 2 | 0.5 | 1 | 0 | 6 | 0.1667 | NA | NA | NA | NA | 2 | 0.5 | 4 | 0.5 | 6 | 0.5 |
| 11_324662 | 1 | 0 | NA | NA | 2 | 0.5 | 2 | 0 | 5 | 0.2 | NA | NA | NA | NA | 3 | 0.3333 | 3 | 0 | 6 | 0.1667 |
| 11_324664 | 1 | 1 | 1 | 1 | 1 | 1 | NA | NA | 3 | 1 | NA | NA | 2 | 0.5 | 2 | 1 | 4 | 1 | 8 | 0.875 |
| 11_324664 | 1 | 0 | NA | NA | 1 | 1 | 2 | 0.5 | 4 | 0.5 | NA | NA | NA | NA | 3 | 1 | 4 | 0.75 | 7 | 0.8571 |
| 11_324664 | 1 | 1 | 1 | 1 | 1 | 1 | 2 | 1 | 5 | 1 | NA | NA | 2 | 1 | 3 | 1 | 4 | 1 | 9 | 1 |
| 11_324667 | 1 | 0 | 2 | 0.5 | 1 | 1 | 4 | 0 | 8 | 0.25 | NA | NA | 2 | 0.5 | 2 | 0.5 | 3 | 0.3333 | 7 | 0.4286 |
| 11_324668 | 1 | 0 | 1 | 1 | 1 | 0 | 3 | 0 | 6 | 0.1667 | NA | NA | 2 | 0 | 3 | 0.3333 | 3 | 0.3333 | 8 | 0.25 |
| 11_324671 | 1 | 0 | 1 | 0 | 2 | 0.5 | 5 | 0.2 | 9 | 0.2222 | 1 | 1 | 2 | 0 | 2 | 0 | 5 | 0 | 10 | 0.1 |
| 11_326618 | 9 | 0.6667 | 4 | 0.25 | 4 | 0 | 7 | 0.2857 | 24 | 0.375 | 7 | 0.5714 | 7 | 0.1429 | 12 | 0.25 | 5 | 0 | 31 | 0.2581 |
| 11_326619 | 7 | 0.2857 | 4 | 0 | 3 | 0 | 7 | 0 | 21 | 0.0952 | 7 | 0.2857 | 6 | 0.1667 | 10 | 0.1 | 1 | 0 | 24 | 0.1667 |
| 11_326946 | 12 | 1 | 5 | 0.8 | 9 | 1 | 12 | 0.6667 | 38 | 0.8684 | 10 | 0.7 | 8 | 1 | 13 | 1 | 12 | 0.9167 | 43 | 0.907 |
| 11_326946 | 11 | 0.1818 | 7 | 0.4286 | 8 | 0.25 | 14 | 0.2143 | 40 | 0.25 | 10 | 0 | 8 | 0.375 | 13 | 0 | 11 | 0 | 42 | 0.0714 |
| 11_326946 | 13 | 0.8462 | 7 | 1 | 8 | 1 | 13 | 1 | 41 | 0.9512 | 9 | 0.7778 | 10 | 0.8 | 15 | 1 | 11 | 1 | 45 | 0.9111 |
| 11_326947 | 14 | 0.5714 | 9 | 0.1111 | 6 | 0 | 12 | 0.5833 | 41 | 0.3902 | 10 | 0.7 | 8 | 0.375 | 10 | 0.8 | 10 | 0.5 | 38 | 0.6053 |
| 11_326948 | 14 | 1 | 9 | 0.8889 | 7 | 1 | 10 | 1 | 40 | 0.975 | 8 | 1 | 7 | 1 | 15 | 1 | 9 | 1 | 39 | 1 |
| 11_326948 | 12 | 0.0833 | 6 | 0.3333 | 8 | 0.125 | 13 | 0.1538 | 39 | 0.1538 | 9 | 0 | 8 | 0 | 13 | 0.2308 | 10 | 0 | 40 | 0.075 |
| 11_326950 | 12 | 1 | 9 | 1 | 9 | 1 | 15 | 1 | 45 | 1 | 11 | 1 | 9 | 0.8889 | 15 | 1 | 13 | 1 | 48 | 0.9792 |
| 11_326962 | 14 | 0.1429 | 15 | 0.1333 | 13 | 0.1538 | 16 | 0.0625 | 58 | 0.1207 | 15 | 0.3333 | 12 | 0.1667 | 19 | 0.2632 | 15 | 0.4 | 61 | 0.2951 |
| 11_326963 | 17 | 0.2353 | 13 | 0.1538 | 11 | 0.0909 | 13 | 0 | 54 | 0.1296 | 16 | 0.125 | 12 | 0.25 | 19 | 0.0526 | 13 | 0.1538 | 60 | 0.1333 |
| 11_326964 | 20 | 0.6 | 16 | 0.375 | 16 | 0.5625 | 17 | 0.4118 | 69 | 0.4928 | 20 | 0.35 | 13 | 0.6154 | 18 | 0.7778 | 18 | 0.5 | 69 | 0.5507 |
| 11_330671 | NA | NA | NA | NA | 1 | 1 | NA | NA | 1 | 1 | NA | NA | 1 | 1 | NA | NA | 1 | 1 | 2 | 1 |
| 11_330675 | NA | NA | NA | NA | 1 | 1 | NA | NA | 1 | 1 | NA | NA | 1 | 1 | NA | NA | NA | NA | 1 | 1 |
| 11_330676 | NA | NA | NA | NA | 1 | 1 | NA | NA | 1 | 1 | NA | NA | NA | NA | NA | NA | 1 | 1 | 1 | 1 |
| 11_334199 | 1 | 1 | NA | NA | 1 | 1 | 1 | 1 | 3 | 1 | NA | NA | NA | NA | NA | NA | 1 | 1 | 1 | 1 |
| 11_334199 | 1 | 1 | NA | NA | 1 | 1 | 1 | 1 | 3 | 1 | NA | NA | NA | NA | NA | NA | 1 | 1 | 1 | 1 |
| 11_334200 | 1 | 0 | NA | NA | 1 | 1 | 1 | 1 | 3 | 0.6667 | NA | NA | NA | NA | NA | NA | 1 | 1 | 1 | 1 |
| 11_334200 | NA | NA | NA | NA | 1 | 1 | 1 | 0 | 2 | 0.5 | NA | NA | NA | NA | NA | NA | 1 | 1 | 1 | 1 |
| 11_335800 | 9 | 0.6667 | 3 | 0.6667 | 4 | 0.75 | 8 | 0.75 | 24 | 0.7083 | 2 | 1 | 10 | 0.6 | 5 | 0.8 | 7 | 0.7143 | 24 | 0.7083 |
| 11_335804 | 7 | 0 | 5 | 0.4 | 3 | 0 | 7 | 0.4286 | 22 | 0.2273 | 2 | 0 | 10 | 0.5 | 5 | 0.6 | 6 | 0.1667 | 23 | 0.3913 |
| 11_335804 | 4 | 1 | 4 | 0.75 | 3 | 1 | 7 | 1 | 18 | 0.9444 | 1 | 1 | 9 | 0.5556 | 6 | 0.3333 | 5 | 1 | 21 | 0.619 |
| 11_335805 | 6 | 0.8333 | 5 | 0.6 | 3 | 0.3333 | 7 | 0.5714 | 21 | 0.619 | 1 | 1 | 11 | 0.8182 | 6 | 1 | 7 | 0.4286 | 25 | 0.76 |
| 11_335805 | 6 | 0.3333 | 5 | 0.8 | 2 | 0 | 6 | 0 | 19 | 0.3158 | NA | NA | 10 | 0.2 | 6 | 0 | 7 | 0.1429 | 23 | 0.1304 |
| 11_350690 | NA | NA | NA | NA | 1 | 1 | 2 | 0 | 3 | 0.3333 | NA | NA | NA | NA | 2 | 0.5 | NA | NA | 2 | 0.5 |
| 11_350691 | NA | NA | NA | NA | 1 | 1 | 1 | 1 | 2 | 1 | 1 | 1 | NA | NA | 2 | 1 | NA | NA | 3 | 1 |
| 11_350692 | NA | NA | NA | NA | 1 | 1 | 1 | 1 | 2 | 1 | NA | NA | NA | NA | 2 | 1 | NA | NA | 2 | 1 |

| | | | | | | | | | | | | | | | | | | | | |
|---|---|---|---|---|---|---|---|---|---|---|---|---|---|---|---|---|---|---|---|---|
| 11_3506947 | NA | NA | NA | NA | 1 | 1 | 2 | 0.5 | 3 | 0.6667 | 1 | 1 | NA | NA | 2 | 1 | NA | NA | 3 | 1 |
| 11_3506951 | NA | NA | NA | NA | 1 | 1 | 1 | 1 | 2 | 1 | 1 | 1 | 1 | 1 | 2 | 1 | NA | NA | 4 | 1 |
| 11_3506962 | NA | NA | NA | NA | 1 | 1 | 1 | 1 | 2 | 1 | 1 | 1 | 1 | 1 | 2 | 1 | NA | NA | 4 | 1 |
| 11_3559878 | 5 | 0 | 3 | 0 | 2 | 0 | 4 | 0.25 | 14 | 0.0714 | 3 | 0.6667 | 4 | 0 | 5 | 0 | 2 | 0 | 14 | 0.1429 |
| 11_3559893 | 5 | 0.6 | 3 | 1 | 2 | 0.5 | 4 | 1 | 14 | 0.7857 | 3 | 1 | 5 | 0.4 | 7 | 0.4286 | 2 | 0.5 | 17 | 0.5294 |
| 11_3559900 | 5 | 0.4 | 3 | 0.3333 | 2 | 0 | 3 | 0 | 13 | 0.2308 | 3 | 0 | 5 | 0 | 8 | 0 | 2 | 0.5 | 18 | 0.0556 |
| 11_3559904 | 4 | 0 | 3 | 0.3333 | 2 | 1 | 4 | 0.25 | 13 | 0.3077 | 3 | 1 | 5 | 0.2 | 3 | 0.3333 | 2 | 0.5 | 13 | 0.4615 |
| 11_3559913 | 3 | 0.6667 | 3 | 1 | 1 | 0 | 2 | 0.5 | 9 | 0.6667 | 2 | 0.5 | 5 | 0.6 | 5 | 0.4 | 2 | 1 | 14 | 0.5714 |
| 11_3559915 | 4 | 0.75 | 3 | 1 | 2 | 0 | 3 | 0.6667 | 12 | 0.6667 | 2 | 0 | 2 | 0 | 4 | 0.75 | 3 | 1 | 11 | 0.5455 |
| 11_3559921 | 4 | 0.75 | 3 | 1 | 2 | 0.5 | 4 | 0.75 | 13 | 0.7692 | 3 | 1 | 2 | 1 | 7 | 1 | 3 | 1 | 15 | 1 |
| 11_3566693 | 2 | 0.5 | 4 | 0 | 1 | 0 | 2 | 0.5 | 9 | 0.2222 | 2 | 0 | 3 | 0.3333 | 1 | 1 | 1 | 0 | 7 | 0.2857 |
| 11_3566842 | 3 | 0.6667 | 4 | 0.5 | 2 | 0 | 2 | 0.5 | 11 | 0.4545 | 2 | 0 | 3 | 0 | 1 | 1 | NA | NA | 6 | 0.1667 |
| 11_3566848 | 3 | 0 | 4 | 0.25 | 1 | 0 | 3 | 0.3333 | 11 | 0.1818 | 2 | 0 | 3 | 0 | NA | NA | 1 | 1 | 6 | 0.1667 |
| 11_3567644 | 3 | 0.6667 | 4 | 0.25 | 2 | 0 | 3 | 0.6667 | 12 | 0.4167 | 2 | 0 | 3 | 0.3333 | 1 | 0 | 1 | 1 | 7 | 0.2857 |
| 11_3567653 | 3 | 0.3333 | 3 | 1 | 2 | 1 | 2 | 0.5 | 10 | 0.7 | 2 | 0.5 | 3 | 0.3333 | 1 | 1 | 2 | 0.5 | 8 | 0.5 |
| 11_3567722 | 2 | 0 | 4 | 0 | 2 | 0.5 | 4 | 0 | 12 | 0.0833 | 1 | 0 | 3 | 0.3333 | 1 | 0 | 2 | 0.5 | 7 | 0.2857 |
| 11_3567951 | 3 | 0.3333 | 7 | 0.4286 | 1 | 0 | 3 | 0.3333 | 14 | 0.3571 | 2 | 0.5 | 4 | 0 | NA | NA | 5 | 0.2 | 11 | 0.1818 |
| 11_3568048 | 2 | 0 | 8 | 0 | 2 | 0 | 6 | 0.1667 | 18 | 0.0556 | 2 | 0.5 | 7 | 0 | 2 | 0 | 7 | 0 | 18 | 0.0556 |
| 11_3568102 | 3 | 0 | 8 | 0 | 5 | 0.2 | 4 | 0.25 | 20 | 0.1 | 3 | 0.3333 | 7 | 0.2857 | 1 | 0 | 6 | 0.1667 | 17 | 0.2353 |
| 11_3568110 | 4 | 0 | 7 | 0 | 6 | 0.1667 | 5 | 0.4 | 22 | 0.1364 | 3 | 0 | 6 | 0 | 1 | 1 | 7 | 0 | 17 | 0.0588 |
| 11_3568110 | 3 | 0.6667 | 6 | 0.3333 | 4 | 0.25 | 3 | 0.3333 | 16 | 0.375 | 2 | 0 | 6 | 0.1667 | 2 | 0 | 3 | 0 | 13 | 0.0769 |
| 11_3568195 | 5 | 0 | 11 | 0.0909 | 9 | 0.3333 | 6 | 0.1667 | 31 | 0.1613 | 6 | 0.1667 | 7 | 0.1429 | 5 | 0.4 | 7 | 0 | 25 | 0.16 |
| 11_3568827 | 10 | 0.9 | 23 | 0.9565 | 9 | 1 | 11 | 1 | 53 | 0.9623 | 11 | 0.9091 | 15 | 0.9333 | 18 | 0.7222 | 13 | 0.8462 | 57 | 0.8421 |
| 11_3568838 | 13 | 0.0769 | 20 | 0.2 | 9 | 0 | 10 | 0.2 | 52 | 0.1346 | 11 | 0.0909 | 16 | 0.125 | 19 | 0.0526 | 10 | 0.4 | 56 | 0.1429 |
| 11_3568841 | 10 | 0.7 | 19 | 0.4211 | 8 | 0.25 | 6 | 0 | 43 | 0.3953 | 10 | 0.2 | 12 | 0.3333 | 15 | 0.4 | 9 | 0.2222 | 46 | 0.3043 |
| 11_3568918 | 6 | 0.1667 | 13 | 0.2308 | 5 | 0 | 4 | 0 | 28 | 0.1429 | 5 | 0 | 7 | 0.2857 | 9 | 0.5556 | 4 | 0.5 | 25 | 0.36 |
| 11_3569929 | 5 | 0.2 | 7 | 0 | 6 | 0.1667 | 8 | 0.125 | 26 | 0.1154 | 5 | 0 | 6 | 0 | 7 | 0.2857 | 4 | 0 | 22 | 0.0909 |
| 11_3572935 | 9 | 0.4444 | 7 | 0.1429 | 8 | 0.625 | 3 | 0 | 27 | 0.3704 | 4 | 0.25 | 8 | 0.5 | 8 | 0.375 | 8 | 0.375 | 28 | 0.3929 |
| 11_3572965 | 10 | 0 | 8 | 0 | 11 | 0.2727 | 4 | 0 | 33 | 0.0909 | 2 | 0 | 10 | 0.1 | 14 | 0 | 10 | 0.1 | 36 | 0.0556 |
| 11_3572969 | 5 | 0 | 8 | 0.375 | 9 | 0.1111 | 3 | 0 | 25 | 0.16 | 4 | 0.25 | 9 | 0 | 12 | 0.0833 | 8 | 0 | 33 | 0.0606 |
| 11_3572994 | 6 | 0.3333 | 8 | 0.125 | 10 | 0.1 | 4 | 0.25 | 28 | 0.1786 | 4 | 0 | 11 | 0.0909 | 13 | 0.0769 | 8 | 0.125 | 36 | 0.0833 |
| 11_3576683 | 1 | 0 | 3 | 0.6667 | 4 | 0.25 | 1 | 0 | 9 | 0.3333 | 2 | 0 | 4 | 0.5 | 5 | 0.4 | 4 | 0.5 | 15 | 0.4 |
| 11_3576689 | 1 | 0 | 2 | 0 | 4 | 0.25 | 2 | 0.5 | 9 | 0.2222 | 1 | 0 | 4 | 0 | 6 | 0.3333 | 4 | 0 | 15 | 0.1333 |
| 11_3587108 | NA | NA | 3 | 0 | 1 | 0 | 1 | 1 | 5 | 0.2 | NA | NA | 1 | 0 | 2 | 0.5 | 4 | 0.25 | 7 | 0.2857 |
| 11_3587116 | NA | NA | 2 | 0.5 | 3 | 0 | 1 | 0 | 6 | 0.1667 | NA | NA | 1 | 0 | 1 | 1 | 4 | 0 | 6 | 0.1667 |
| 11_3589765 | 1 | 1 | 6 | 0.5 | 3 | 0.6667 | 9 | 0.6667 | 19 | 0.6316 | 2 | 1 | 5 | 0.2 | 2 | 1 | 3 | 1 | 12 | 0.6667 |
| 11_3589777 | 2 | 0.5 | 6 | 0 | 2 | 0 | 4 | 0 | 14 | 0.0714 | 1 | 0 | 4 | 0 | 2 | 0.5 | 3 | 0 | 10 | 0.1 |
| 11_3589778 | 3 | 0.3333 | 6 | 0.1667 | 3 | 0 | 8 | 0 | 20 | 0.1 | 2 | 0 | 5 | 0.2 | 2 | 0 | 2 | 0 | 11 | 0.0909 |
| 11_3589781 | 3 | 0.3333 | 5 | 0.2 | 3 | 0 | 8 | 0 | 19 | 0.1053 | 2 | 0 | 5 | 0.6 | 1 | 0 | 3 | 0 | 11 | 0.2727 |
| 11_3589788 | 3 | 0.3333 | 6 | 0.1667 | 3 | 0.6667 | 6 | 0.1667 | 18 | 0.2778 | 2 | 0 | 5 | 0.2 | 1 | 0 | 3 | 0.3333 | 11 | 0.1818 |
| 11_3589797 | 2 | 0 | 6 | 0.5 | 2 | 1 | 7 | 0.7143 | 17 | 0.5882 | 2 | 0.5 | 5 | 0.4 | NA | NA | 3 | 0.3333 | 10 | 0.4 |
| 11_3589812 | 2 | 0 | 5 | 0 | 2 | 0 | 4 | 0.25 | 13 | 0.0769 | 1 | 0 | 5 | 0.2 | 1 | 0 | 3 | 0 | 10 | 0.1 |
| 11_3589831 | 1 | 0 | 3 | 0.6667 | 2 | 1 | 3 | 0.3333 | 9 | 0.5556 | 1 | 1 | 5 | 0.4 | 1 | 0 | 3 | 1 | 10 | 0.6 |
| 11_3589832 | 1 | 0 | 2 | 0 | 2 | 0.5 | 4 | 0.5 | 9 | 0.3333 | 2 | 0 | 4 | 0.5 | 1 | 1 | 3 | 0 | 10 | 0.3 |
| 11_3589837 | 1 | 1 | 2 | 0.5 | 2 | 1 | 3 | 1 | 8 | 0.875 | 1 | 1 | 4 | 0.75 | 1 | 1 | 1 | 1 | 7 | 0.8571 |
| 11_3589845 | 1 | 1 | 3 | 0 | 2 | 0 | 4 | 0 | 10 | 0.1 | 2 | 0 | 5 | 0.4 | 1 | 0 | 2 | 0.5 | 10 | 0.3 |
| 11_3589846 | 1 | 0 | 3 | 0.3333 | 2 | 0 | 4 | 0.5 | 10 | 0.3 | 2 | 0 | 5 | 0.2 | 1 | 0 | 2 | 1 | 10 | 0.3 |
| 11_3589850 | 1 | 1 | 3 | 0 | 1 | 0 | 4 | 0 | 9 | 0.1111 | 2 | 0.5 | 5 | 0.2 | 1 | 0 | 2 | 0 | 10 | 0.2 |
| 11_3589863 | NA | NA | 3 | 0.3333 | 1 | 1 | 3 | 0 | 7 | 0.2857 | 1 | 0 | 3 | 0.3333 | 1 | 0 | 2 | 0 | 7 | 0.1429 |
| 11_3589867 | 1 | 0 | 3 | 0.3333 | 2 | 0.5 | 4 | 0.25 | 10 | 0.3 | 2 | 0.5 | 5 | 0.6 | 1 | 0 | 2 | 0.5 | 10 | 0.5 |
| 11_3589878 | 1 | 0 | 2 | 0 | 2 | 0.5 | 4 | 0.25 | 9 | 0.2222 | 2 | 0.5 | 5 | 0.2 | 1 | 0 | 2 | 0 | 10 | 0.2 |
| 11_3589888 | 1 | 0 | 2 | 1 | 2 | 0.5 | 3 | 0.6667 | 8 | 0.625 | 1 | 0 | 3 | 0 | 1 | 1 | 2 | 1 | 7 | 0.4286 |
| 11_3589891 | 1 | 1 | 2 | 0 | 2 | 0 | 4 | 0.25 | 9 | 0.2222 | 2 | 0.5 | 5 | 0.2 | 1 | 0 | 2 | 0.5 | 10 | 0.3 |
| 11_3589909 | 1 | 0 | 1 | 0 | 2 | 0.5 | 4 | 0 | 8 | 0.125 | 1 | 0 | 4 | 0 | 1 | 0 | 2 | 0.5 | 8 | 0.125 |
| 11_3589920 | 1 | 0 | 2 | 1 | 2 | 1 | 4 | 1 | 9 | 0.8889 | 1 | 1 | 4 | 1 | 1 | 1 | 2 | 1 | 8 | 1 |
| 11_3589933 | 1 | 0 | 2 | 1 | 1 | 1 | 4 | 0.75 | 8 | 0.75 | 1 | 1 | 4 | 0.75 | 1 | 1 | 2 | 1 | 8 | 0.875 |
| 11_3589938 | 1 | 1 | 1 | 1 | 1 | 0 | 3 | 0 | 6 | 0.3333 | 1 | 0 | 4 | 0 | 1 | 0 | 2 | 0.5 | 8 | 0.125 |
| 11_3589956 | 1 | 1 | 1 | 1 | 1 | 0 | 4 | 0.5 | 7 | 0.5714 | 1 | 1 | 4 | 0.25 | 1 | 1 | 2 | 0 | 8 | 0.375 |
| 11_3591415 | NA | NA | 1 | 1 | 1 | 0 | 3 | 0.3333 | 5 | 0.4 | 1 | 0 | 4 | 0.25 | 1 | 0 | 1 | 1 | 7 | 0.2857 |
| 11_3591435 | NA | NA | NA | NA | NA | NA | 4 | 0.75 | 4 | 0.75 | 1 | 1 | 3 | 0.6667 | 1 | 0 | NA | NA | 5 | 0.6 |
| 11_3591499 | NA | NA | NA | NA | NA | NA | 2 | 1 | 2 | 1 | NA | NA | 2 | 1 | 1 | 1 | 1 | 1 | 4 | 1 |
| 11_3591502 | NA | NA | 1 | 0 | NA | NA | 2 | 0.5 | 3 | 0.3333 | NA | NA | 3 | 0.3333 | NA | NA | 1 | 1 | 4 | 0.5 |
| 11_3591503 | NA | NA | 1 | 0 | 1 | 0 | 2 | 0.5 | 4 | 0.25 | NA | NA | 3 | 0.3333 | 1 | 1 | 1 | 1 | 5 | 0.6 |
| 11_3591534 | NA | NA | 1 | 0 | 1 | 0 | 3 | 1 | 5 | 0.6 | NA | NA | 3 | 0.6667 | 1 | 1 | 1 | 0 | 5 | 0.6 |
| 11_3591553 | NA | NA | 1 | 0 | 1 | 1 | 4 | 0 | 6 | 0.1667 | NA | NA | 3 | 0.3333 | 1 | 0 | 1 | 0 | 5 | 0.2 |

| | | | | | | | | | | | | | | | | | | | | |
|---|---|---|---|---|---|---|---|---|---|---|---|---|---|---|---|---|---|---|---|---|
| 11_3593787 | NA | NA | 1 | 0 | 1 | 0 | 2 | 0.5 | 4 | 0.25 | NA | NA | 3 | 0.6667 | 1 | 1 | NA | NA | 4 | 0.75 |
| 11_3593802 | NA | NA | 1 | 1 | 1 | 0 | 2 | 0.5 | 4 | 0.5 | NA | NA | 3 | 0 | 1 | 1 | NA | NA | 4 | 0.25 |
| 11_3593814 | NA | NA | 1 | 1 | NA | NA | 2 | 0.5 | 3 | 0.6667 | NA | NA | 3 | 0.6667 | 1 | 1 | NA | NA | 4 | 0.75 |
| 11_3595403 | NA | NA | 1 | 1 | 1 | 1 | 2 | 0 | 4 | 0.5 | NA | NA | 2 | 0.5 | NA | NA | NA | NA | 2 | 0.5 |
| 11_4057059 | 19 | 0.0526 | 19 | 0.0526 | 22 | 0.0455 | 19 | 0.1053 | 79 | 0.0633 | 12 | 0.0833 | 26 | 0.0769 | 18 | 0.2222 | 13 | 0.1538 | 69 | 0.1304 |
| 11_4057086 | 16 | 0.3125 | 14 | 0.5 | 19 | 0.3158 | 18 | 0.2222 | 67 | 0.3284 | 10 | 0.1 | 24 | 0.2917 | 19 | 0.2632 | 11 | 0.1818 | 64 | 0.2344 |
| 11_4057091 | 14 | 0.0714 | 11 | 0 | 19 | 0.2105 | 16 | 0.0625 | 60 | 0.1 | 11 | 0.1818 | 23 | 0.1304 | 17 | 0.1765 | 11 | 0 | 62 | 0.129 |
| 11_4062483 | 5 | 0.2 | 4 | 0.25 | 2 | 0 | 2 | 0 | 13 | 0.1538 | 3 | 0 | 1 | 0 | NA | NA | 1 | 1 | 5 | 0.2 |
| 11_4063004 | 5 | 0 | 5 | 0 | 2 | 0 | 7 | 0.2857 | 19 | 0.1053 | 3 | 0 | 2 | 0 | 1 | 1 | 3 | 0 | 9 | 0.1111 |
| 11_4063006 | 5 | 0.2 | 5 | 0.2 | 2 | 0.5 | 6 | 0.1667 | 18 | 0.2222 | 3 | 0 | 2 | 0.5 | 1 | 0 | 3 | 0.3333 | 9 | 0.2222 |
| 11_4063986 | 14 | 0.1429 | 13 | 0.2308 | 12 | 0.0833 | 19 | 0.1579 | 58 | 0.1552 | 11 | 0.3636 | 5 | 0 | 8 | 0.25 | 7 | 0.1429 | 31 | 0.2258 |
| 11_4064037 | 10 | 0.1 | 10 | 0.1 | 10 | 0.1 | 16 | 0.1875 | 46 | 0.1304 | 6 | 0.1667 | 3 | 0.3333 | 9 | 0 | 7 | 0.1429 | 25 | 0.12 |
| 11_4064123 | 10 | 0.1 | 12 | 0.1667 | 10 | 0.3 | 16 | 0 | 48 | 0.125 | 7 | 0.1429 | 2 | 0 | 7 | 0.2857 | 6 | 0.3333 | 22 | 0.2273 |
| 11_4064155 | 11 | 0 | 12 | 0 | 11 | 0.0909 | 15 | 0.2 | 49 | 0.0816 | 7 | 0.1429 | 2 | 0 | 7 | 0 | 6 | 0.3333 | 22 | 0.1364 |
| 11_4064474 | 9 | 0.3333 | 6 | 0.3333 | 8 | 0.25 | 14 | 0.4286 | 37 | 0.3514 | 6 | 0.5 | 2 | 0.5 | 6 | 0.5 | 6 | 0.3333 | 20 | 0.45 |
| 11_4064475 | 9 | 0 | 7 | 0.1429 | 9 | 0.2222 | 13 | 0.1538 | 38 | 0.1316 | 6 | 0 | 2 | 0 | 5 | 0 | 5 | 0.2 | 18 | 0.0556 |
| 11_4139150 | 2 | 1 | 1 | 0 | 2 | 0.5 | NA | NA | 5 | 0.6 | 1 | 0 | 2 | 0.5 | 3 | 0.3333 | 2 | 0.5 | 8 | 0.375 |
| 11_4139160 | 2 | 0.5 | 2 | 0 | 2 | 0.5 | NA | NA | 6 | 0.3333 | 2 | 0.5 | 2 | 0 | 4 | 0 | 2 | 0 | 10 | 0.1 |
| 11_4180106 | 11 | 0.0909 | 24 | 0.1667 | 16 | 0.125 | 11 | 0.2727 | 62 | 0.1613 | 11 | 0.0909 | 14 | 0.2857 | 14 | 0.0714 | 20 | 0.05 | 59 | 0.1186 |
| 11_4180161 | 12 | 0 | 22 | 0.0455 | 12 | 0.1667 | 11 | 0 | 57 | 0.0526 | 6 | 0.3333 | 16 | 0 | 16 | 0 | 21 | 0.0476 | 59 | 0.0508 |
| 11_4180221 | 11 | 0 | 17 | 0.2353 | 14 | 0.1429 | 9 | 0 | 51 | 0.1176 | 5 | 0.2 | 11 | 0 | 14 | 0.1429 | 19 | 0 | 49 | 0.0612 |
| 11_4180322 | 11 | 0.9091 | 19 | 0.8947 | 13 | 1 | 15 | 1 | 58 | 0.9483 | 8 | 1 | 15 | 0.9333 | 15 | 0.9333 | 27 | 0.9259 | 65 | 0.9385 |
| 11_4180330 | 11 | 0.8182 | 21 | 0.9048 | 16 | 0.8125 | 20 | 1 | 68 | 0.8971 | 9 | 1 | 17 | 1 | 14 | 0.7857 | 27 | 1 | 67 | 0.9552 |
| 11_4180337 | 7 | 0.1429 | 15 | 0.2 | 12 | 0.0833 | 14 | 0.0714 | 48 | 0.125 | 7 | 0.1429 | 10 | 0.1 | 12 | 0.0833 | 19 | 0.1579 | 48 | 0.125 |
| 11_4189127 | 1 | 0 | 2 | 0 | 2 | 0 | 3 | 0.6667 | 8 | 0.25 | 3 | 0.3333 | 1 | 0 | 2 | 0 | 2 | 0 | 8 | 0.125 |
| 11_4202236 | 25 | 0.12 | 54 | 0.1296 | 40 | 0.075 | 46 | 0.0652 | 165 | 0.097 | 27 | 0.0741 | 33 | 0.0909 | 35 | 0.1143 | 31 | 0.0645 | 126 | 0.0873 |
| 11_4202387 | 14 | 0.2857 | 37 | 0.1622 | 18 | 0.1667 | 38 | 0.1579 | 107 | 0.1776 | 19 | 0.1053 | 36 | 0.1944 | 25 | 0.2 | 19 | 0.1579 | 99 | 0.1717 |
| 11_4202635 | 10 | 0.7 | 22 | 0.8182 | 16 | 0.9375 | 29 | 0.7931 | 77 | 0.8182 | 18 | 0.7222 | 24 | 0.8333 | 22 | 0.7727 | 16 | 0.875 | 80 | 0.8 |
| 11_4251738 | 1 | 0 | 3 | 0.3333 | 5 | 0 | 6 | 0 | 15 | 0.0667 | NA | NA | 3 | 0 | 2 | 0.5 | NA | NA | 5 | 0.2 |
| 11_4251744 | 3 | 0.6667 | 4 | 1 | 4 | 1 | 7 | 1 | 18 | 0.9444 | NA | NA | 3 | 1 | 3 | 1 | NA | NA | 6 | 1 |
| 11_4251762 | 4 | 1 | 3 | 1 | 4 | 0.75 | 8 | 1 | 19 | 0.9474 | 1 | 1 | 2 | 1 | 3 | 1 | 1 | 1 | 7 | 1 |
| 11_4251771 | 4 | 1 | 3 | 1 | 4 | 1 | 8 | 0.875 | 19 | 0.9474 | 1 | 1 | 2 | 1 | 3 | 1 | NA | NA | 6 | 1 |
| 11_4251774 | 4 | 1 | 4 | 1 | 4 | 1 | 10 | 0.9 | 22 | 0.9545 | 1 | 1 | 3 | 1 | 3 | 1 | 1 | 1 | 8 | 1 |
| 11_4251778 | 3 | 1 | 3 | 1 | 5 | 1 | 10 | 1 | 21 | 1 | 1 | 1 | 2 | 1 | 3 | 0.6667 | 1 | 1 | 7 | 0.8571 |
| 11_4251780 | 3 | 1 | 3 | 0.6667 | 5 | 0.4 | 8 | 0.625 | 19 | 0.6316 | 1 | 1 | 3 | 1 | 3 | 0.6667 | 1 | 1 | 8 | 0.875 |
| 11_4251813 | 4 | 0.5 | 3 | 0 | 6 | 0.3333 | 10 | 0.1 | 23 | 0.2174 | 1 | 0 | 3 | 0.6667 | 2 | 0.5 | 2 | 1 | 8 | 0.625 |
| 11_4251817 | 3 | 0.3333 | 2 | 0 | 5 | 0 | 8 | 0.375 | 18 | 0.2222 | 1 | 0 | 2 | 1 | 3 | 0.3333 | 2 | 0 | 8 | 0.375 |
| 11_4251819 | 4 | 1 | 3 | 1 | 5 | 0.6 | 11 | 0.9091 | 23 | 0.8696 | 1 | 1 | 3 | 1 | 3 | 1 | 2 | 1 | 9 | 1 |
| 11_4251829 | 4 | 0.5 | 4 | 0.25 | 6 | 0.1667 | 9 | 0.3333 | 23 | 0.3043 | 1 | 0 | 3 | 0 | 3 | 0 | 2 | 1 | 9 | 0.2222 |
| 11_4251831 | 3 | 1 | 3 | 0.3333 | 6 | 0.3333 | 10 | 0.6 | 22 | 0.5455 | 1 | 0 | 3 | 0.6667 | 3 | 0.6667 | 2 | 1 | 9 | 0.6667 |
| 11_4251835 | 4 | 0.5 | 1 | 0 | 6 | 0 | 10 | 0.4 | 21 | 0.2857 | 1 | 1 | 3 | 0.6667 | 3 | 0.6667 | 2 | 1 | 9 | 0.7778 |
| 11_4251860 | 3 | 0.6667 | 5 | 0.4 | 6 | 0 | 11 | 0.0909 | 25 | 0.2 | 1 | 0 | 3 | 0.3333 | 4 | 0 | 2 | 0.5 | 10 | 0.2 |
| 11_4251883 | 4 | 0 | 5 | 0.2 | 6 | 0.1667 | 10 | 0.1 | 25 | 0.12 | 1 | 0 | 4 | 0.5 | 3 | 0.6667 | 2 | 0 | 10 | 0.4 |
| 11_4251897 | 4 | 0 | 7 | 0.5714 | 6 | 0.5 | 11 | 0.1818 | 28 | 0.3214 | 1 | 0 | 5 | 0.2 | 6 | 0.5 | 2 | 0.5 | 14 | 0.3571 |
| 11_4251919 | 4 | 0.5 | 6 | 0.1667 | 4 | 0.25 | 10 | 0.3 | 24 | 0.2917 | NA | NA | 6 | 0.1667 | 7 | 0 | 3 | 0.3333 | 16 | 0.125 |
| 11_4251926 | 3 | 0.6667 | 3 | 0.3333 | 6 | 0 | 9 | 0.2222 | 21 | 0.2381 | 1 | 0 | 5 | 0 | 6 | 0.1667 | 2 | 0.5 | 14 | 0.1429 |
| 11_4251939 | 5 | 0 | 5 | 0.2 | 5 | 0 | 13 | 0.0769 | 28 | 0.0714 | 1 | 0 | 7 | 0 | 7 | 0 | 2 | 0.5 | 17 | 0.0588 |
| 11_4252145 | 13 | 0 | 13 | 0.1538 | 7 | 0.1429 | 22 | 0.0909 | 55 | 0.0909 | 4 | 0.5 | 9 | 0 | 14 | 0.0714 | 6 | 0 | 33 | 0.0909 |
| 11_4252321 | 27 | 0.5185 | 20 | 0.5 | 14 | 0.5714 | 28 | 0.3929 | 89 | 0.4831 | 6 | 0.6667 | 13 | 0.3846 | 17 | 0.2941 | 11 | 0.4545 | 47 | 0.4043 |
| 11_4333701 | 8 | 0.625 | 3 | 0.3333 | 9 | 0.7778 | 5 | 0.6 | 25 | 0.64 | 4 | 1 | 8 | 0.625 | 9 | 0.5556 | 5 | 1 | 26 | 0.7308 |
| 11_4333701 | 10 | 0.2 | 2 | 0.5 | 8 | 0.125 | 5 | 0.4 | 25 | 0.24 | 5 | 0.2 | 8 | 0.5 | 11 | 0.3636 | 6 | 0 | 30 | 0.3 |
| 11_4333703 | 8 | 0.625 | 3 | 0.6667 | 7 | 0.8571 | 7 | 0.4286 | 25 | 0.64 | 6 | 0.5 | 8 | 0.625 | 10 | 0.9 | 6 | 0.5 | 30 | 0.6667 |
| 11_4333711 | 9 | 0.2222 | 3 | 0 | 9 | 0.5556 | 7 | 0.5714 | 28 | 0.3929 | 6 | 0 | 8 | 0.375 | 12 | 0.1667 | 4 | 0.5 | 30 | 0.2333 |
| 11_4333723 | 8 | 0.375 | 1 | 0 | 8 | 0.5 | 4 | 0.5 | 21 | 0.4286 | 6 | 0.6667 | 5 | 0.4 | 6 | 0.8333 | 5 | 0.6 | 22 | 0.6364 |
| 11_4333794 | 1 | 1 | 1 | 1 | 3 | 1 | 2 | 0.5 | 7 | 0.8571 | NA | NA | 1 | 1 | 1 | 1 | 1 | 0 | 3 | 0.6667 |
| 11_4333813 | NA | NA | 1 | 1 | 1 | 0 | 2 | 0 | 4 | 0.25 | NA | NA | 1 | 0 | 1 | 1 | 1 | 0 | 3 | 0.3333 |
| 11_4333822 | NA | NA | 1 | 0 | 1 | 1 | 2 | 0.5 | 4 | 0.5 | NA | NA | 1 | 0 | 1 | 1 | 1 | 0 | 3 | 0.3333 |
| 11_4333830 | NA | NA | 1 | 1 | 1 | 1 | 2 | 1 | 4 | 1 | NA | NA | 1 | 1 | 1 | 1 | 1 | 0 | 3 | 0.6667 |
| 11_4333838 | NA | NA | NA | NA | 1 | 1 | 2 | 0 | 3 | 0.3333 | NA | NA | 1 | 0 | 1 | 1 | 1 | 1 | 3 | 0.6667 |
| 11_4345084 | NA | NA | 2 | 1 | NA | NA | NA | NA | 2 | 1 | NA | NA | 1 | 1 | NA | NA | NA | NA | 1 | 1 |
| 11_4345091 | NA | NA | 2 | 0.5 | NA | NA | NA | NA | 2 | 0.5 | NA | NA | 1 | 1 | NA | NA | NA | NA | 1 | 1 |
| 11_4345095 | NA | NA | 2 | 0.5 | NA | NA | NA | NA | 2 | 0.5 | NA | NA | 1 | 1 | NA | NA | NA | NA | 1 | 1 |
| 11_4345469 | NA | NA | 3 | 0.6667 | NA | NA | NA | NA | 3 | 0.6667 | NA | NA | 1 | 1 | NA | NA | NA | NA | 1 | 1 |
| 11_4347682 | NA | NA | NA | NA | 1 | 1 | NA | NA | 1 | 1 | NA | NA | 1 | 1 | NA | NA | NA | NA | 1 | 1 |
| 11_4347709 | NA | NA | 1 | 1 | NA | NA | NA | NA | 1 | 1 | NA | NA | 1 | 1 | NA | NA | NA | NA | 1 | 1 |

| | | | | | | | | | | | | | | | | | | | | |
|---|---|---|---|---|---|---|---|---|---|---|---|---|---|---|---|---|---|---|---|---|
| 11_4347740 | NA | NA | NA | NA | 1 | 1 | NA | NA | 1 | 1 | NA | NA | 1 | 1 | NA | NA | NA | NA | 1 | 1 |
| 11_4355960 | NA | NA | NA | NA | NA | NA | 3 | 0.6667 | 3 | 0.6667 | NA | NA | 1 | 1 | NA | NA | 3 | 0.3333 | 4 | 0.5 |
| 11_4359578 | NA | NA | 1 | 1 | NA | NA | 4 | 0.75 | 5 | 0.8 | NA | NA | 1 | 1 | 1 | 1 | 2 | 1 | 4 | 1 |
| 11_4359584 | NA | NA | 1 | 1 | NA | NA | 2 | 0 | 3 | 0.3333 | NA | NA | NA | NA | 1 | 1 | 2 | 1 | 3 | 1 |
| 11_4359585 | NA | NA | 1 | 1 | NA | NA | 1 | 1 | 2 | 1 | NA | NA | 1 | 0 | 1 | 1 | 3 | 0 | 5 | 0.2 |
| 11_4359623 | NA | NA | 1 | 1 | NA | NA | 2 | 1 | 3 | 1 | 2 | 1 | 1 | 1 | 2 | 1 | 4 | 0.5 | 9 | 0.7778 |
| 11_4359637 | NA | NA | 1 | 1 | NA | NA | NA | NA | 1 | 1 | 1 | 0 | 1 | 1 | 1 | 1 | 1 | 1 | 4 | 0.75 |
| 11_4359640 | NA | NA | 1 | 1 | NA | NA | NA | NA | 1 | 1 | 1 | 1 | 1 | 0 | 1 | 1 | 1 | 1 | 4 | 0.75 |
| 11_4359648 | NA | NA | 1 | 1 | NA | NA | NA | NA | 1 | 1 | 1 | 1 | 1 | 1 | 1 | 1 | 1 | 1 | 4 | 1 |
| 11_4359673 | NA | NA | 1 | 1 | NA | NA | NA | NA | 1 | 1 | 1 | 1 | NA | NA | NA | NA | 1 | 0 | 2 | 0.5 |
| 11_4361125 | 1 | 0 | NA | NA | NA | NA | 3 | 0.3333 | 4 | 0.25 | 2 | 0.5 | NA | NA | 2 | 0.5 | 3 | 0.3333 | 7 | 0.4286 |
| 11_4361248 | 1 | 0 | NA | NA | NA | NA | 3 | 0.3333 | 4 | 0.25 | 2 | 0 | NA | NA | 2 | 0 | 2 | 0.5 | 6 | 0.1667 |
| 11_4361267 | 1 | 1 | NA | NA | NA | NA | 2 | 0.5 | 3 | 0.6667 | 2 | 0.5 | NA | NA | 2 | 0 | 2 | 0 | 6 | 0.1667 |
| 11_4362094 | 4 | 0 | 10 | 0.1 | 3 | 0.3333 | 8 | 0.125 | 25 | 0.12 | 3 | 0.3333 | 8 | 0.125 | 9 | 0.1111 | 2 | 0 | 22 | 0.1364 |
| 11_4362126 | 6 | 0.8333 | 7 | 0.8571 | 2 | 1 | 8 | 0.75 | 23 | 0.8261 | 3 | 1 | 7 | 1 | 9 | 0.7778 | 4 | 0.75 | 23 | 0.8696 |
| 11_4362241 | 1 | 1 | NA | NA | NA | NA | NA | NA | 1 | 1 | NA | NA | 1 | 1 | NA | NA | NA | NA | 1 | 1 |
| 11_4363594 | 4 | 0 | 9 | 0.1111 | 3 | 0 | 9 | 0.2222 | 25 | 0.12 | 5 | 0 | 7 | 0.1429 | 7 | 0.2857 | 4 | 0.25 | 23 | 0.1739 |
| 11_4363604 | 5 | 0.6 | 7 | 0.1429 | 2 | 0 | 7 | 0.2857 | 21 | 0.2857 | 5 | 0 | 5 | 0 | 6 | 0.3333 | 2 | 0 | 18 | 0.1111 |
| 11_4366643 | 2 | 0.5 | 2 | 0 | 2 | 1 | 1 | 1 | 7 | 0.5714 | 2 | 0.5 | NA | NA | NA | NA | 1 | 1 | 3 | 0.6667 |
| 11_4434553 | 5 | 0.2 | 5 | 0.2 | 7 | 0.1429 | 11 | 0.1818 | 28 | 0.1786 | 9 | 0.2222 | 8 | 0.25 | 7 | 0.1429 | 11 | 0.2727 | 35 | 0.2286 |
| 11_4434562 | 6 | 0.1667 | 3 | 0 | 7 | 0.1429 | 10 | 0.1 | 26 | 0.1154 | 6 | 0.1667 | 3 | 0 | 7 | 0.1429 | 9 | 0 | 25 | 0.08 |
| 11_4434577 | 6 | 0.5 | 3 | 0 | 6 | 0.5 | 9 | 0.7778 | 24 | 0.5417 | 8 | 0.875 | 8 | 0.75 | 6 | 0.8333 | 10 | 0.5 | 32 | 0.7188 |
| 11_4434578 | 5 | 0.4 | 5 | 0.2 | 6 | 0.3333 | 10 | 0.3 | 26 | 0.3077 | 7 | 0.1429 | 8 | 0.125 | 6 | 0.3333 | 10 | 0.2 | 31 | 0.1935 |
| 11_4434579 | 5 | 0.4 | 5 | 0.6 | 5 | 0.4 | 9 | 0.4444 | 24 | 0.4583 | 5 | 0.4 | 6 | 0.5 | 7 | 0.1429 | 8 | 0.375 | 26 | 0.3462 |
| 11_4434582 | 5 | 0.4 | 4 | 1 | 7 | 0.8571 | 11 | 0.6364 | 27 | 0.7037 | 6 | 0.5 | 6 | 0.6667 | 7 | 0.5714 | 8 | 0.875 | 27 | 0.6667 |
| 11_4434610 | 3 | 0 | 4 | 0.5 | 6 | 0.8333 | 11 | 0.5455 | 24 | 0.5417 | 8 | 0.875 | 6 | 0.5 | 6 | 0.5 | 9 | 0.6667 | 29 | 0.6552 |
| 11_4434633 | 5 | 0.4 | 4 | 0.5 | 3 | 1 | 9 | 0.6667 | 21 | 0.619 | 6 | 0.5 | 5 | 0.4 | 4 | 0.75 | 8 | 0.5 | 23 | 0.5217 |
| 11_4434642 | 6 | 0.1667 | 3 | 0.3333 | 3 | 0 | 11 | 0 | 23 | 0.087 | 7 | 0.1429 | 5 | 0.2 | 6 | 0 | 10 | 0 | 28 | 0.0714 |
| 11_4440933 | NA | NA | 1 | 1 | NA | NA | 1 | 1 | 2 | 1 | NA | NA | 1 | 1 | NA | NA | NA | NA | 1 | 1 |
| 11_4440940 | NA | NA | 1 | 0 | NA | NA | 1 | 1 | 2 | 0.5 | NA | NA | 1 | 1 | NA | NA | NA | NA | 1 | 1 |
| 11_4440948 | NA | NA | 1 | 0 | NA | NA | 1 | 1 | 2 | 0.5 | NA | NA | 1 | 1 | NA | NA | NA | NA | 1 | 1 |
| 11_4445246 | 9 | 1 | 8 | 0.875 | 3 | 1 | 7 | 0.8571 | 27 | 0.9259 | 9 | 0.8889 | 9 | 1 | 11 | 0.9091 | 8 | 1 | 37 | 0.9459 |
| 11_4445485 | 6 | 0.1667 | 5 | 0 | 3 | 0 | 6 | 0 | 20 | 0.05 | 6 | 0.3333 | 7 | 0 | 9 | 0 | 8 | 0 | 30 | 0.0667 |
| 11_4445487 | 8 | 0.5 | 7 | 0.8571 | 3 | 0.6667 | 7 | 0.8571 | 25 | 0.72 | 9 | 0.7778 | 12 | 1 | 10 | 0.7 | 9 | 0.4444 | 40 | 0.75 |
| 11_4445488 | 9 | 0.3333 | 7 | 0.7143 | 3 | 0 | 6 | 0.3333 | 25 | 0.4 | 7 | 0.4286 | 9 | 0.5556 | 9 | 0.2222 | 10 | 0.5 | 35 | 0.4286 |
| 11_4445492 | 9 | 0.8889 | 6 | 0.6667 | 3 | 0.6667 | 6 | 0.8333 | 24 | 0.7917 | 7 | 0.5714 | 10 | 0.8 | 10 | 0.7 | 8 | 0.875 | 35 | 0.7429 |
| 11_4445495 | 10 | 0.1 | 7 | 0.1429 | 2 | 0 | 7 | 0.1429 | 26 | 0.1154 | 8 | 0.125 | 11 | 0.0909 | 12 | 0.0833 | 10 | 0.1 | 41 | 0.0976 |
| 11_4445497 | 6 | 0.8333 | 4 | 1 | 1 | 1 | 5 | 1 | 16 | 0.9375 | 7 | 0.4286 | 9 | 0.4444 | 6 | 1 | 8 | 0.75 | 30 | 0.6333 |
| 11_4445498 | 8 | 0.375 | 5 | 0.6 | 3 | 0 | 8 | 0.25 | 24 | 0.3333 | 7 | 0.4286 | 12 | 0.25 | 11 | 0.0909 | 9 | 0.4444 | 39 | 0.2821 |
| 11_4445500 | 10 | 1 | 5 | 1 | 3 | 1 | 8 | 0.875 | 26 | 0.9615 | 7 | 0.7143 | 12 | 0.9167 | 11 | 0.9091 | 10 | 0.7 | 40 | 0.825 |
| 11_4445501 | 5 | 0.2 | 6 | 0.3333 | NA | NA | 2 | 0 | 13 | 0.2308 | 3 | 0.3333 | 8 | 0.5 | 5 | 0.4 | 7 | 0.2857 | 23 | 0.3913 |
| 11_4445513 | 10 | 0.9 | 9 | 0.8889 | 3 | 1 | 8 | 1 | 30 | 0.9333 | 10 | 0.9 | 11 | 1 | 10 | 1 | 9 | 0.8889 | 40 | 0.95 |
| 11_4445520 | 9 | 0.7778 | 9 | 0.8889 | 3 | 1 | 7 | 1 | 28 | 0.8929 | 8 | 0.625 | 11 | 0.8182 | 11 | 0.6364 | 9 | 0.8889 | 39 | 0.7436 |
| 11_4445521 | 10 | 0.3 | 8 | 0.125 | 2 | 0 | 8 | 0.125 | 28 | 0.1786 | 9 | 0 | 12 | 0.0833 | 12 | 0.4167 | 9 | 0.2222 | 42 | 0.1905 |
| 11_4445525 | 7 | 0.7143 | 8 | 0.625 | 2 | 1 | 7 | 0.7143 | 24 | 0.7083 | 9 | 0.4444 | 10 | 0.6 | 8 | 0.625 | 11 | 0.2727 | 38 | 0.4737 |
| 11_4445564 | 8 | 0.875 | 9 | 0.7778 | 2 | 1 | 9 | 0.7778 | 28 | 0.8214 | 10 | 0.6 | 12 | 0.8333 | 12 | 0.9167 | 11 | 1 | 45 | 0.8444 |
| 11_4445564 | 8 | 0 | 4 | 0 | NA | NA | 4 | 0.25 | 16 | 0.0625 | 4 | 0.25 | 8 | 0.25 | 8 | 0 | 4 | 0 | 24 | 0.125 |
| 11_4445574 | 13 | 0.0769 | 8 | 0 | 2 | 0 | 11 | 0.0909 | 34 | 0.0588 | 10 | 0.2 | 13 | 0.1538 | 12 | 0.3333 | 14 | 0.0714 | 49 | 0.1837 |
| 11_4445579 | 12 | 0.0833 | 9 | 0.1111 | 2 | 0 | 10 | 0 | 33 | 0.0606 | 10 | 0.1 | 13 | 0.3077 | 13 | 0 | 15 | 0 | 51 | 0.098 |
| 11_4445601 | 16 | 0.375 | 9 | 0.5556 | 5 | 0.4 | 10 | 0.6 | 40 | 0.475 | 10 | 0.6 | 14 | 0.5714 | 14 | 0.5 | 15 | 0.6 | 53 | 0.566 |
| 11_4489800 | 2 | 0.5 | 2 | 0 | NA | NA | 1 | 0 | 5 | 0.2 | NA | NA | NA | NA | 1 | 1 | NA | NA | 1 | 1 |
| 11_4579989 | 4 | 0.25 | 3 | 0.6667 | 2 | 0 | 8 | 0.25 | 17 | 0.2941 | 4 | 0.25 | 4 | 0.5 | 6 | 0.1667 | 4 | 0 | 18 | 0.2222 |
| 11_4579997 | 4 | 0.5 | 4 | 0.5 | 3 | 0.3333 | 9 | 0.4444 | 20 | 0.45 | 3 | 0 | 3 | 0.6667 | 5 | 0 | 10 | 0.4 | 21 | 0.2857 |
| 11_4579998 | 3 | 0.6667 | 3 | 1 | 1 | 0 | 8 | 1 | 15 | 0.8667 | 3 | 0.3333 | 2 | 1 | 7 | 0.7143 | 5 | 1 | 17 | 0.7647 |
| 11_4580006 | 5 | 1 | 4 | 1 | 1 | 1 | 10 | 1 | 20 | 1 | 3 | 1 | 2 | 1 | 7 | 0.7143 | 10 | 0.9 | 22 | 0.8636 |
| 11_4580009 | 5 | 0.4 | 4 | 0.25 | 3 | 0.3333 | 11 | 0.0909 | 23 | 0.2174 | 4 | 0.25 | 4 | 0.25 | 7 | 0.1429 | 11 | 0.0909 | 26 | 0.1538 |
| 11_4580014 | 5 | 0.2 | 4 | 0.75 | 3 | 0 | 9 | 0.1111 | 21 | 0.2381 | 4 | 0 | 4 | 0.25 | 8 | 0.25 | 10 | 0.2 | 26 | 0.1923 |
| 11_4580015 | 3 | 0.3333 | 5 | 0 | 2 | 0 | 9 | 0.3333 | 19 | 0.2105 | 2 | 0 | 3 | 0.3333 | 8 | 0.25 | 9 | 0 | 22 | 0.1364 |
| 11_4580023 | 5 | 0.2 | 5 | 0.4 | 4 | 0 | 8 | 0 | 22 | 0.1364 | 1 | 0 | 2 | 0 | 7 | 0 | 10 | 0.1 | 20 | 0.05 |
| 11_4580038 | 5 | 1 | 5 | 1 | 3 | 0.6667 | 10 | 0.9 | 23 | 0.913 | 3 | 1 | 3 | 1 | 8 | 0.375 | 9 | 0.7778 | 23 | 0.6957 |
| 11_4580045 | 6 | 0.5 | 5 | 0.4 | 4 | 0 | 11 | 0.1818 | 26 | 0.2692 | 4 | 0 | 3 | 0.6667 | 9 | 0 | 8 | 0.25 | 24 | 0.1667 |
| 11_4580065 | 6 | 0.1667 | 4 | 0 | 6 | 0 | 11 | 0.1818 | 27 | 0.1111 | 5 | 0.2 | 4 | 0 | 9 | 0.2222 | 9 | 0 | 27 | 0.1111 |
| 11_4580076 | 5 | 0 | 4 | 0.25 | 7 | 0.5714 | 11 | 0.2727 | 27 | 0.2963 | 6 | 0.1667 | 4 | 0.5 | 9 | 0.2222 | 9 | 0.3333 | 28 | 0.2857 |
| 11_4580078 | 6 | 0.6667 | 4 | 0.75 | 6 | 0.8333 | 7 | 0.7143 | 23 | 0.7391 | 6 | 0.8333 | 5 | 1 | 9 | 0.5556 | 5 | 0.4 | 25 | 0.68 |
| 11_4580082 | 6 | 0 | 5 | 0.2 | 7 | 0.1429 | 11 | 0.1818 | 29 | 0.1379 | 6 | 0 | 5 | 0 | 10 | 0 | 10 | 0.2 | 31 | 0.0645 |

| | | | | | | | | | | | | | | | | | | | | |
|---|---|---|---|---|---|---|---|---|---|---|---|---|---|---|---|---|---|---|---|---|
| 11_4580087 | 6 | 0.1667 | 4 | 0 | 7 | 0 | 11 | 0.0909 | 28 | 0.0714 | 5 | 0.2 | 5 | 0.2 | 10 | 0.1 | 7 | 0 | 27 | 0.1111 |
| 11_4581929 | 1 | 1 | 1 | 0 | 2 | 0 | 2 | 1 | 6 | 0.5 | 2 | 0.5 | NA | NA | 2 | 0.5 | 2 | 0 | 6 | 0.3333 |
| 11_4581938 | 1 | 1 | 1 | 0 | 1 | 0 | 3 | 0 | 6 | 0.1667 | 2 | 0.5 | 1 | 0 | 2 | 0 | 2 | 0 | 7 | 0.1429 |
| 11_4581961 | 1 | 1 | 1 | 0 | 2 | 0 | 3 | 0 | 7 | 0.1429 | 1 | 0 | 1 | 1 | 2 | 0 | 2 | 0 | 6 | 0.1667 |
| 11_4582439 | NA | NA | 1 | 1 | 1 | 0 | 1 | 1 | 3 | 0.6667 | NA | NA | 1 | 0 | 1 | 1 | NA | NA | 2 | 0.5 |
| 11_4582461 | NA | NA | 1 | 1 | 1 | 0 | 1 | 1 | 3 | 0.6667 | 1 | 1 | NA | NA | 1 | 1 | NA | NA | 2 | 1 |
| 11_4582462 | NA | NA | 1 | 1 | 1 | 0 | 1 | 1 | 3 | 0.6667 | 1 | 0 | 1 | 0 | 1 | 1 | NA | NA | 3 | 0.3333 |
| 11_4582471 | NA | NA | NA | NA | 1 | 1 | 1 | 0 | 2 | 0.5 | NA | NA | 1 | 0 | 1 | 1 | NA | NA | 2 | 0.5 |
| 11_4583888 | NA | NA | 6 | 0.1667 | NA | NA | NA | NA | 6 | 0.1667 | 2 | 0 | NA | NA | 2 | 0.5 | 4 | 0 | 8 | 0.125 |
| 11_4583888 | NA | NA | 4 | 0.25 | NA | NA | 1 | 1 | 5 | 0.4 | 2 | 0.5 | NA | NA | 3 | 0.3333 | 3 | 0 | 8 | 0.25 |
| 11_4583899 | 1 | 1 | 6 | 0.1667 | NA | NA | 1 | 0 | 8 | 0.25 | 2 | 0.5 | NA | NA | 2 | 0.5 | 2 | 0 | 6 | 0.3333 |
| 11_4583905 | 1 | 1 | 6 | 0.6667 | NA | NA | 1 | 0 | 8 | 0.625 | 2 | 0.5 | NA | NA | 3 | 0 | 2 | 0 | 7 | 0.1429 |
| 11_4603990 | NA | NA | 2 | 1 | 1 | 1 | NA | NA | 3 | 1 | 1 | 1 | NA | NA | NA | NA | NA | NA | 1 | 1 |
| 11_4604058 | NA | NA | 2 | 1 | NA | NA | NA | NA | 2 | 1 | 1 | 1 | NA | NA | NA | NA | NA | NA | 1 | 1 |
| 11_4604061 | NA | NA | 2 | 0.5 | NA | NA | NA | NA | 2 | 0.5 | 1 | 1 | NA | NA | NA | NA | NA | NA | 1 | 1 |
| 11_4608504 | 43 | 0.0465 | 51 | 0.1176 | 43 | 0.0465 | 47 | 0.0426 | 184 | 0.0652 | 33 | 0.0909 | 43 | 0.0465 | 52 | 0.0577 | 46 | 0.087 | 174 | 0.069 |
| 11_4608703 | 19 | 0.5263 | 30 | 0.5667 | 29 | 0.4828 | 29 | 0.4483 | 107 | 0.5047 | 21 | 0.4286 | 23 | 0.6522 | 32 | 0.5625 | 27 | 0.4444 | 103 | 0.5243 |
| 11_4608704 | 27 | 0.1481 | 33 | 0.0909 | 35 | 0.0286 | 35 | 0.0857 | 130 | 0.0846 | 24 | 0 | 31 | 0.0968 | 39 | 0.0513 | 32 | 0.0938 | 126 | 0.0635 |
| 11_4608719 | 23 | 0.1304 | 34 | 0.1176 | 31 | 0.1935 | 27 | 0.1481 | 115 | 0.1478 | 25 | 0.24 | 23 | 0.1739 | 38 | 0.2105 | 31 | 0.0323 | 117 | 0.1624 |
| 11_4608720 | 27 | 0.2222 | 34 | 0.1765 | 35 | 0.2 | 33 | 0.3333 | 129 | 0.2326 | 26 | 0.2692 | 28 | 0.1786 | 40 | 0.15 | 37 | 0.3243 | 131 | 0.229 |
| 11_4608724 | 17 | 0 | 26 | 0.0385 | 29 | 0.1034 | 28 | 0.1071 | 100 | 0.07 | 24 | 0.1667 | 23 | 0.1739 | 32 | 0.125 | 24 | 0.0833 | 103 | 0.1359 |
| 11_4612855 | NA | NA | 2 | 1 | 2 | 1 | 2 | 1 | 6 | 1 | NA | NA | 2 | 1 | 1 | 1 | 1 | 1 | 4 | 1 |
| 11_4612880 | NA | NA | 2 | 0 | 2 | 0 | 2 | 0.5 | 6 | 0.1667 | NA | NA | 2 | 0 | 1 | 0 | 1 | 1 | 4 | 0.25 |
| 11_4612890 | NA | NA | 2 | 0.5 | 2 | 0 | 2 | 1 | 6 | 0.5 | NA | NA | 2 | 0 | NA | NA | 1 | 1 | 3 | 0.3333 |
| 11_4612904 | NA | NA | 1 | 1 | 2 | 0.5 | 1 | 0 | 4 | 0.5 | NA | NA | 1 | 1 | NA | NA | 1 | 1 | 2 | 1 |
| 11_4612906 | NA | NA | 2 | 0 | 2 | 0 | 1 | 1 | 5 | 0.2 | NA | NA | 1 | 0 | 1 | 1 | 1 | 0 | 3 | 0.3333 |
| 11_4612924 | NA | NA | 2 | 0.5 | 2 | 0 | 2 | 0 | 6 | 0.1667 | 1 | 0 | 2 | 0.5 | 1 | 0 | 1 | 0 | 5 | 0.2 |
| 11_4612931 | NA | NA | 1 | 0 | 2 | 0.5 | 2 | 1 | 5 | 0.6 | 1 | 0 | 2 | 1 | 1 | 0 | 1 | 0 | 5 | 0.4 |
| 11_4612932 | NA | NA | 2 | 0 | 2 | 0 | 3 | 0.6667 | 7 | 0.2857 | 1 | 0 | 1 | 1 | 1 | 0 | 1 | 0 | 4 | 0.25 |
| 11_4612939 | NA | NA | 3 | 0 | 1 | 1 | 3 | 1 | 7 | 0.5714 | 1 | 1 | 2 | 0.5 | 1 | 0 | 1 | 0 | 5 | 0.4 |
| 11_4612944 | NA | NA | 3 | 1 | 2 | 1 | 3 | 1 | 8 | 1 | 1 | 1 | 2 | 1 | 1 | 1 | 1 | 1 | 5 | 1 |
| 11_4612949 | NA | NA | 3 | 0 | 2 | 0.5 | 3 | 0.3333 | 8 | 0.25 | 1 | 0 | 2 | 0.5 | 1 | 0 | 1 | 0 | 5 | 0.2 |
| 11_4612955 | NA | NA | 3 | 0.3333 | 1 | 0 | 3 | 0.6667 | 7 | 0.4286 | NA | NA | 1 | 0 | NA | NA | 1 | 1 | 2 | 0.5 |
| 11_4612971 | NA | NA | 3 | 0.3333 | 2 | 0 | 3 | 0 | 8 | 0.125 | 1 | 0 | 1 | 0 | 1 | 1 | 1 | 0 | 4 | 0.25 |
| 11_4612974 | NA | NA | 1 | 1 | 2 | 0 | 2 | 0.5 | 5 | 0.4 | 1 | 1 | NA | NA | NA | NA | NA | NA | 1 | 1 |
| 11_4612978 | NA | NA | 2 | 0.5 | 2 | 0 | 3 | 0 | 7 | 0.1429 | 1 | 0 | 2 | 0.5 | 1 | 0 | NA | NA | 4 | 0.25 |
| 11_4613015 | NA | NA | 2 | 0.5 | 2 | 1 | 3 | 0.6667 | 7 | 0.7143 | 1 | 0 | 2 | 0.5 | 1 | 1 | 1 | 1 | 5 | 0.6 |
| 11_4613023 | NA | NA | 3 | 0 | 2 | 0 | 3 | 0.3333 | 8 | 0.125 | 1 | 1 | 1 | 0 | NA | NA | 1 | 0 | 3 | 0.3333 |
| 11_4613024 | NA | NA | 3 | 0.3333 | 2 | 0.5 | 3 | 0.3333 | 8 | 0.375 | 1 | 1 | 2 | 1 | 1 | 1 | 1 | 1 | 5 | 1 |
| 11_4613031 | NA | NA | 3 | 0.3333 | 2 | 0.5 | 3 | 0.6667 | 8 | 0.5 | 1 | 0 | 2 | 0.5 | 1 | 0 | 1 | 0 | 5 | 0.2 |
| 11_4613033 | NA | NA | 3 | 0.6667 | 2 | 0 | 3 | 0.3333 | 8 | 0.375 | 1 | 0 | 1 | 0 | 1 | 0 | 1 | 1 | 4 | 0.25 |
| 11_4633152 | 3 | 0.6667 | 2 | 1 | 3 | 0.3333 | 1 | 1 | 9 | 0.6667 | 3 | 1 | 2 | 1 | 2 | 0.5 | 2 | 1 | 9 | 0.8889 |
| 11_4633164 | 3 | 0.6667 | 3 | 1 | 5 | 1 | 2 | 0.5 | 13 | 0.8462 | 2 | 0.5 | 3 | 0.3333 | 3 | 0 | 2 | 0.5 | 10 | 0.3 |
| 11_4633167 | 5 | 0.2 | 3 | 0.3333 | 5 | 0.2 | 2 | 0.5 | 15 | 0.2667 | 3 | 0.3333 | 1 | 0 | 3 | 0.3333 | 2 | 0.5 | 9 | 0.3333 |
| 11_4633168 | 5 | 0.2 | 3 | 0.3333 | 5 | 0.2 | 1 | 0 | 14 | 0.2143 | 3 | 0 | 3 | 0.3333 | 3 | 0 | 3 | 0 | 12 | 0.0833 |
| 11_4633180 | 4 | 1 | 1 | 1 | 5 | 1 | 3 | 1 | 13 | 1 | 3 | 1 | 4 | 1 | 3 | 1 | 3 | 1 | 13 | 1 |
| 11_4633200 | 5 | 0.8 | 3 | 1 | 5 | 0.6 | 2 | 1 | 15 | 0.8 | 4 | 0.75 | 4 | 0.75 | 3 | 1 | 2 | 1 | 13 | 0.8462 |
| 11_4633204 | 6 | 0.6667 | 3 | 1 | 4 | 0.5 | 3 | 0.6667 | 16 | 0.6875 | 4 | 0.5 | 4 | 0.5 | 3 | 0.6667 | 1 | 1 | 12 | 0.5833 |
| 11_4633220 | 9 | 0.8889 | 4 | 1 | 5 | 1 | 3 | 1 | 21 | 0.9524 | 4 | 0.75 | 3 | 1 | 2 | 1 | 2 | 1 | 11 | 0.9091 |
| 11_4633914 | NA | NA | 1 | 1 | NA | NA | NA | NA | 1 | 1 | NA | NA | NA | NA | 1 | 1 | NA | NA | 1 | 1 |
| 11_4869821 | 5 | 1 | 2 | 1 | NA | NA | 2 | 1 | 9 | 1 | 1 | 1 | 2 | 1 | 3 | 1 | 1 | 1 | 7 | 1 |
| 11_4869855 | 2 | 0.5 | 1 | 0 | 1 | 0 | 5 | 0 | 9 | 0.1111 | NA | NA | 1 | 1 | NA | NA | 1 | 1 | 2 | 1 |
| 11_4871640 | NA | NA | 2 | 0.5 | 2 | 0 | 2 | 0 | 6 | 0.1667 | 2 | 0.5 | 2 | 0 | 2 | 0.5 | NA | NA | 6 | 0.3333 |
| 11_4871670 | NA | NA | 1 | 1 | NA | NA | 2 | 0.5 | 3 | 0.6667 | NA | NA | 2 | 0.5 | 2 | 0.5 | NA | NA | 4 | 0.5 |
| 11_4875669 | 2 | 0 | 2 | 0.5 | NA | NA | 1 | 1 | 5 | 0.4 | NA | NA | 1 | 0 | 1 | 0 | 2 | 1 | 4 | 0.5 |
| 11_4875681 | 2 | 1 | 3 | 0.6667 | NA | NA | 1 | 1 | 6 | 0.8333 | NA | NA | 1 | 1 | 1 | 1 | 3 | 1 | 5 | 1 |
| 11_4875687 | 1 | 1 | 2 | 0.5 | NA | NA | 1 | 0 | 4 | 0.5 | NA | NA | NA | NA | 1 | 1 | 1 | 1 | 2 | 1 |
| 11_4875707 | 1 | 1 | 3 | 0 | NA | NA | 1 | 0 | 5 | 0.2 | NA | NA | NA | NA | 1 | 1 | 3 | 0.3333 | 4 | 0.5 |
| 11_4875712 | 1 | 1 | 3 | 0.3333 | NA | NA | 1 | 0 | 5 | 0.4 | NA | NA | 1 | 0 | 1 | 0 | 2 | 0.5 | 4 | 0.25 |
| 11_4875742 | 1 | 1 | 3 | 1 | NA | NA | 1 | 1 | 5 | 1 | NA | NA | 1 | 1 | 1 | 1 | 2 | 1 | 4 | 1 |
| 11_4875746 | 1 | 1 | 3 | 0.3333 | NA | NA | 1 | 0 | 5 | 0.4 | NA | NA | 1 | 0 | 1 | 0 | 2 | 0.5 | 4 | 0.25 |
| 11_4875750 | 1 | 1 | 3 | 1 | NA | NA | 1 | 1 | 5 | 1 | NA | NA | 1 | 1 | 1 | 1 | 2 | 1 | 4 | 1 |
| 11_4875760 | 1 | 1 | 2 | 1 | NA | NA | 1 | 1 | 4 | 1 | NA | NA | NA | NA | 1 | 1 | 2 | 1 | 3 | 1 |
| 11_4875769 | 1 | 0 | 3 | 0.3333 | NA | NA | 1 | 0 | 5 | 0.2 | NA | NA | NA | NA | 1 | 1 | 2 | 1 | 3 | 1 |
| 11_4875767 | NA | NA | 3 | 0 | NA | NA | 1 | 1 | 4 | 0.25 | NA | NA | 1 | 0 | 1 | 1 | 1 | 0 | 3 | 0.3333 |

| | | | | | | | | | | | | | | | | | | | | |
|---|---|---|---|---|---|---|---|---|---|---|---|---|---|---|---|---|---|---|---|---|
| 11_487620 | 1 | 0 | 3 | 0.3333 | NA | NA | 1 | 0 | 5 | 0.2 | NA | NA | NA | NA | 1 | 0 | 2 | 0.5 | 3 | 0.3333 |
| 11_491059 | NA | NA | NA | NA | 2 | 1 | 2 | 1 | 4 | 1 | NA | NA | 1 | 0 | 1 | 1 | 2 | 0.5 | 4 | 0.5 |
| 11_491060 | NA | NA | NA | NA | 3 | 0.6667 | 4 | 0 | 7 | 0.2857 | NA | NA | 1 | 1 | 2 | 0.5 | NA | NA | 3 | 0.6667 |
| 11_491060 | NA | NA | NA | NA | 3 | 0.3333 | 4 | 0.75 | 7 | 0.5714 | NA | NA | 2 | 1 | 2 | 1 | 1 | 1 | 5 | 1 |
| 11_491061 | NA | NA | NA | NA | 3 | 0 | 4 | 0.5 | 7 | 0.2857 | NA | NA | 2 | 0.5 | 2 | 0 | 2 | 0 | 6 | 0.1667 |
| 11_491061 | NA | NA | NA | NA | 2 | 0.5 | 4 | 0.25 | 6 | 0.3333 | NA | NA | 2 | 0.5 | 2 | 0.5 | 2 | 0 | 6 | 0.3333 |
| 11_491062 | NA | NA | NA | NA | 2 | 1 | 4 | 1 | 6 | 1 | NA | NA | 2 | 1 | 1 | 0 | NA | NA | 3 | 0.6667 |
| 11_491063 | NA | NA | NA | NA | 3 | 0 | 3 | 1 | 6 | 0.5 | NA | NA | 2 | 0.5 | 2 | 1 | 2 | 0.5 | 6 | 0.6667 |
| 11_491063 | NA | NA | NA | NA | 2 | 1 | 4 | 1 | 6 | 1 | NA | NA | 1 | 1 | 2 | 1 | 2 | 1 | 5 | 1 |
| 11_491065 | NA | NA | NA | NA | 3 | 1 | 4 | 0.75 | 7 | 0.8571 | NA | NA | 2 | 1 | 2 | 0.5 | 2 | 1 | 6 | 0.8333 |
| 11_491065 | NA | NA | NA | NA | 3 | 0.3333 | 4 | 0.25 | 7 | 0.2857 | NA | NA | 2 | 0.5 | 2 | 0 | 2 | 0.5 | 6 | 0.3333 |
| 11_491066 | NA | NA | NA | NA | 3 | 0.3333 | 4 | 0.25 | 7 | 0.2857 | NA | NA | 2 | 0.5 | 2 | 0 | 2 | 0 | 6 | 0.1667 |
| 11_491068 | NA | NA | NA | NA | 2 | 1 | 2 | 1 | 4 | 1 | NA | NA | 1 | 1 | 2 | 1 | 2 | 1 | 5 | 1 |
| 11_491069 | NA | NA | NA | NA | 3 | 0 | 4 | 0.25 | 7 | 0.1429 | NA | NA | 2 | 0.5 | 2 | 0 | 2 | 0.5 | 6 | 0.3333 |
| 11_491071 | NA | NA | NA | NA | 3 | 1 | 4 | 1 | 7 | 1 | NA | NA | 2 | 1 | 2 | 0.5 | 2 | 1 | 6 | 0.8333 |
| 11_491072 | NA | NA | NA | NA | 3 | 0.3333 | 4 | 0.5 | 7 | 0.4286 | NA | NA | 2 | 0.5 | 2 | 1 | 2 | 1 | 6 | 0.8333 |
| 11_491072 | NA | NA | NA | NA | 3 | 0.6667 | 3 | 0 | 6 | 0.3333 | NA | NA | 1 | 0 | 2 | 0.5 | 2 | 0.5 | 5 | 0.4 |
| 11_491072 | NA | NA | 1 | 0 | 3 | 0 | 4 | 0.5 | 8 | 0.25 | NA | NA | 2 | 1 | 1 | 0 | 2 | 0 | 5 | 0.4 |
| 11_491074 | NA | NA | 1 | 0 | 2 | 0 | 3 | 0.3333 | 6 | 0.1667 | NA | NA | 2 | 0 | 2 | 0.5 | 2 | 0 | 6 | 0.1667 |
| 11_491074 | NA | NA | 2 | 0 | 3 | 0 | 4 | 0.25 | 9 | 0.1111 | NA | NA | 2 | 0.5 | 2 | 0.5 | 2 | 0 | 6 | 0.3333 |
| 11_491075 | NA | NA | 2 | 0.5 | 3 | 0.3333 | 4 | 0.5 | 9 | 0.4444 | NA | NA | 2 | 1 | 2 | 0.5 | 2 | 0.5 | 6 | 0.6667 |
| 11_491076 | NA | NA | 1 | 0 | 3 | 0.6667 | 2 | 1 | 6 | 0.6667 | NA | NA | 2 | 1 | 2 | 1 | 2 | 1 | 6 | 1 |
| 11_491077 | NA | NA | 2 | 0.5 | 1 | 0 | 3 | 0 | 6 | 0.1667 | NA | NA | NA | NA | 1 | 0 | 2 | 0.5 | 3 | 0.3333 |
| 11_491079 | NA | NA | 2 | 0.5 | 3 | 0 | 3 | 0 | 8 | 0.125 | NA | NA | 2 | 0 | 2 | 0.5 | 2 | 0 | 6 | 0.1667 |
| 11_491080 | NA | NA | NA | NA | 2 | 1 | 2 | 1 | 4 | 1 | NA | NA | 2 | 1 | 2 | 1 | 2 | 1 | 6 | 1 |
| 11_491081 | NA | NA | 1 | 1 | 3 | 0.3333 | 3 | 0 | 7 | 0.2857 | NA | NA | 2 | 0.5 | 2 | 0.5 | 2 | 0.5 | 6 | 0.5 |
| 11_491085 | NA | NA | 2 | 0 | 5 | 0.4 | 4 | 0.75 | 11 | 0.4545 | NA | NA | 2 | 0 | 3 | 0.3333 | 2 | 0 | 7 | 0.1429 |
| 11_491086 | NA | NA | 2 | 0 | 3 | 0.6667 | 3 | 0 | 8 | 0.25 | NA | NA | 2 | 0 | 3 | 0.3333 | 2 | 0 | 7 | 0.1429 |
| 11_491087 | NA | NA | 1 | 0 | 2 | 0 | 4 | 0.5 | 7 | 0.2857 | NA | NA | 2 | 0 | 3 | 0.3333 | 1 | 0 | 6 | 0.1667 |
| 11_491087 | NA | NA | 1 | 0 | 4 | 0 | 4 | 0.25 | 9 | 0.1111 | NA | NA | 3 | 0 | 1 | 0 | 2 | 0.5 | 6 | 0.1667 |
| 11_491087 | NA | NA | 2 | 0 | 4 | 0 | 4 | 0.25 | 10 | 0.1 | NA | NA | 3 | 0 | 3 | 0.3333 | 1 | 0 | 7 | 0.1429 |
| 11_491087 | 1 | 1 | 1 | 0 | 4 | 0.25 | 4 | 0 | 10 | 0.2 | NA | NA | 3 | 0 | 3 | 0.6667 | 1 | 1 | 7 | 0.4286 |
| 11_491088 | 1 | 0 | 2 | 0.5 | 5 | 0.6 | 4 | 0.75 | 12 | 0.5833 | NA | NA | 2 | 0 | 3 | 0 | 2 | 0.5 | 7 | 0.1429 |
| 11_491090 | 1 | 0 | 2 | 0 | 5 | 0 | 4 | 0.25 | 12 | 0.0833 | NA | NA | 5 | 0 | 3 | 0.3333 | 3 | 0.3333 | 11 | 0.1818 |
| 11_491091 | 3 | 0 | 2 | 0.5 | 5 | 0.2 | 2 | 0 | 12 | 0.1667 | NA | NA | 4 | 0.5 | 4 | 0.5 | 2 | 1 | 10 | 0.6 |
| 11_491091 | 3 | 0 | 2 | 0 | 6 | 0.1667 | 4 | 0 | 15 | 0.0667 | NA | NA | 5 | 0 | 4 | 0 | 3 | 0.3333 | 12 | 0.0833 |
| 11_491092 | 3 | 0.3333 | 2 | 0.5 | 5 | 0 | 4 | 0.5 | 14 | 0.2857 | NA | NA | 5 | 0.4 | 2 | 0 | 2 | 1 | 9 | 0.4444 |
| 11_491362 | NA | NA | 1 | 1 | 1 | 1 | 2 | 1 | 4 | 1 | 2 | 1 | 1 | 1 | NA | NA | 1 | 1 | 4 | 1 |
| 11_491362 | NA | NA | 1 | 0 | 1 | 1 | 1 | 1 | 3 | 0.6667 | 1 | 0 | 1 | 0 | 1 | 1 | 2 | 0.5 | 5 | 0.4 |
| 11_491363 | NA | NA | 1 | 1 | 1 | 0 | 3 | 0.6667 | 5 | 0.6 | 1 | 1 | 1 | 1 | 1 | 1 | 2 | 1 | 5 | 1 |
| 11_491514 | 1 | 0 | 6 | 0 | 1 | 0 | 7 | 0.1429 | 15 | 0.0667 | 3 | 0 | 4 | 0.25 | 3 | 0 | 3 | 0 | 13 | 0.0769 |
| 11_491515 | 1 | 1 | 4 | 0.25 | 2 | 0.5 | 6 | 0.8333 | 13 | 0.6154 | 3 | 1 | 3 | 0.3333 | 3 | 1 | 3 | 0.3333 | 12 | 0.6667 |
| 11_491516 | NA | NA | 7 | 0.5714 | 2 | 0 | 8 | 0.375 | 17 | 0.4118 | 1 | 0 | 5 | 0.4 | 3 | 1 | 1 | 1 | 10 | 0.6 |
| 11_491517 | 1 | 1 | 3 | 1 | 2 | 1 | 7 | 1 | 13 | 1 | 3 | 1 | 5 | 0.8 | 2 | 1 | 1 | 1 | 11 | 0.9091 |
| 11_491518 | 1 | 0 | 7 | 0 | 3 | 0 | 8 | 0.125 | 19 | 0.0526 | 3 | 0 | 5 | 0 | 4 | 0.25 | 3 | 0 | 15 | 0.0667 |
| 11_491519 | 3 | 1 | 6 | 1 | 3 | 1 | 8 | 1 | 20 | 1 | 1 | 0 | 6 | 1 | 4 | 1 | 3 | 1 | 14 | 0.9286 |
| 11_491519 | 3 | 0 | 6 | 0.5 | 2 | 0 | 7 | 0.4286 | 18 | 0.3333 | 2 | 0 | 7 | 0.1429 | 4 | 0.25 | 2 | 0.5 | 15 | 0.2 |
| 11_491520 | 3 | 0.6667 | 7 | 0.5714 | 3 | 0.3333 | 8 | 0.75 | 21 | 0.619 | 3 | 0 | 7 | 0.5714 | 4 | 0.25 | 3 | 0.3333 | 17 | 0.3529 |
| 11_491521 | 3 | 1 | 7 | 0.7143 | 3 | 0.3333 | 8 | 0.625 | 21 | 0.6667 | 2 | 0 | 6 | 0.6667 | 3 | 0.3333 | 3 | 0.6667 | 14 | 0.5 |
| 11_491522 | 2 | 0 | 5 | 0 | 2 | 0 | 5 | 0.2 | 14 | 0.0714 | 1 | 0 | 4 | 0 | 4 | 0.25 | 2 | 1 | 11 | 0.2727 |
| 11_491522 | 3 | 1 | 6 | 0.3333 | 3 | 0.3333 | 8 | 0.875 | 20 | 0.65 | 2 | 0.5 | 6 | 0.6667 | 4 | 1 | 3 | 1 | 15 | 0.8 |
| 11_491523 | 2 | 0.5 | 6 | 0.1667 | 2 | 0 | 6 | 0.5 | 16 | 0.3125 | NA | NA | 4 | 0.25 | 2 | 0 | 2 | 0 | 8 | 0.125 |
| 11_491523 | 1 | 1 | 5 | 1 | 3 | 0.6667 | 8 | 1 | 17 | 0.9412 | 1 | 0 | 7 | 0.7143 | 2 | 0.5 | 2 | 1 | 12 | 0.6667 |
| 11_491524 | 2 | 0.5 | 5 | 1 | 2 | 0.5 | 8 | 0.75 | 17 | 0.7647 | 1 | 1 | 5 | 1 | 3 | 1 | 2 | 1 | 11 | 1 |
| 11_491525 | 3 | 1 | 5 | 1 | 3 | 0.3333 | 8 | 1 | 19 | 0.8947 | 3 | 1 | 7 | 0.7143 | 3 | 0.6667 | 3 | 0.6667 | 16 | 0.75 |
| 11_491525 | 3 | 0 | 6 | 0.3333 | 4 | 0 | 7 | 0.5714 | 20 | 0.3 | 2 | 0.5 | 7 | 0.2857 | 4 | 0.75 | 4 | 0.75 | 17 | 0.5294 |
| 11_491526 | 4 | 1 | 6 | 1 | 4 | 1 | 10 | 1 | 24 | 1 | 2 | 1 | 8 | 1 | 4 | 1 | 3 | 1 | 17 | 1 |
| 11_491526 | 3 | 0.6667 | 5 | 0.8 | 3 | 0.6667 | 10 | 0.8 | 21 | 0.7619 | 2 | 1 | 8 | 0.75 | 4 | 1 | 3 | 1 | 17 | 0.8824 |
| 11_491528 | 3 | 0.3333 | 8 | 0.5 | 3 | 0.3333 | 11 | 0.4545 | 25 | 0.44 | 2 | 0 | 9 | 0.3333 | 4 | 0 | 4 | 0.75 | 19 | 0.3158 |
| 11_491528 | 3 | 1 | 8 | 1 | 4 | 1 | 12 | 1 | 27 | 1 | 3 | 0.6667 | 9 | 1 | 4 | 1 | 4 | 1 | 20 | 0.95 |
| 11_491533 | 3 | 0 | 7 | 0.4286 | 4 | 0.25 | 11 | 0.2727 | 25 | 0.28 | 5 | 0.6 | 8 | 0.25 | 5 | 0.4 | 4 | 0.25 | 22 | 0.3636 |
| 11_491712 | NA | NA | NA | NA | 1 | 1 | NA | NA | 1 | 1 | NA | NA | NA | NA | 2 | 1 | 1 | 1 | 3 | 1 |
| 11_491712 | NA | NA | NA | NA | 1 | 1 | NA | NA | 1 | 1 | NA | NA | NA | NA | 1 | 1 | 1 | 0 | 2 | 0.5 |
| 11_491714 | NA | NA | NA | NA | 1 | 1 | 1 | 1 | 2 | 1 | NA | NA | NA | NA | 2 | 1 | NA | NA | 2 | 1 |
| 11_491714 | NA | NA | NA | NA | 1 | 1 | 1 | 1 | 2 | 1 | NA | NA | NA | NA | 2 | 1 | 1 | 1 | 3 | 1 |

| | | | | | | | | | | | | | | | | | | | | |
|---|---|---|---|---|---|---|---|---|---|---|---|---|---|---|---|---|---|---|---|---|
| 11_491715 | NA | NA | NA | NA | 1 | 1 | 1 | 1 | 2 | 1 | NA | NA | NA | NA | 1 | 1 | 1 | 1 | 2 | 1 |
| 11_491716 | NA | NA | NA | NA | 1 | 1 | 1 | 1 | 2 | 1 | NA | NA | NA | NA | 2 | 1 | 1 | 1 | 3 | 1 |
| 11_495425 | NA | NA | NA | NA | 1 | 1 | 1 | 0 | 2 | 0.5 | NA | NA | NA | NA | 1 | 1 | NA | NA | 1 | 1 |
| 11_495426 | NA | NA | NA | NA | 1 | 1 | 1 | 1 | 2 | 1 | NA | NA | NA | NA | 1 | 1 | NA | NA | 1 | 1 |
| 11_495429 | NA | NA | NA | NA | 1 | 0 | 1 | 1 | 2 | 0.5 | NA | NA | NA | NA | 1 | 0 | 1 | 1 | 2 | 0.5 |
| 11_495432 | NA | NA | NA | NA | NA | NA | 1 | 1 | 1 | 1 | NA | NA | NA | NA | 1 | 0 | 1 | 1 | 2 | 0.5 |
| 11_495433 | NA | NA | NA | NA | 1 | 1 | 1 | 1 | 2 | 1 | NA | NA | NA | NA | 1 | 1 | NA | NA | 1 | 1 |
| 11_495624 | 7 | 0.4286 | 4 | 0.25 | 2 | 0.5 | 9 | 0.2222 | 22 | 0.3182 | NA | NA | 2 | 0 | 2 | 0.5 | 5 | 0.2 | 9 | 0.2222 |
| 11_495624 | 6 | 0.5 | 5 | 0.8 | 3 | 0.6667 | 8 | 0.625 | 22 | 0.6364 | NA | NA | 1 | 0 | 4 | 0.5 | 3 | 0 | 8 | 0.25 |
| 11_495635 | 5 | 0.4 | 6 | 0 | 3 | 0.3333 | 5 | 0.2 | 19 | 0.2105 | NA | NA | 2 | 0 | 5 | 0 | 4 | 0.5 | 11 | 0.1818 |
| 11_495639 | 5 | 0.2 | 3 | 0 | 3 | 0 | 5 | 0.6 | 16 | 0.25 | NA | NA | 2 | 0.5 | 3 | 0 | 2 | 0 | 7 | 0.1429 |
| 11_495639 | 3 | 0.3333 | 4 | 0 | 3 | 0 | 3 | 0.6667 | 13 | 0.2308 | NA | NA | 2 | 0.5 | 4 | 0 | 2 | 0 | 8 | 0.125 |
| 11_495644 | 6 | 0.1667 | 6 | 0 | 5 | 0.6 | 6 | 0 | 23 | 0.1739 | 4 | 0.5 | 4 | 0.5 | 5 | 0 | 6 | 0.1667 | 19 | 0.2632 |
| 11_495644 | 6 | 0.1667 | 7 | 0.2857 | 4 | 0 | 7 | 0 | 24 | 0.125 | 4 | 0.25 | 4 | 0.25 | 5 | 0 | 3 | 0.3333 | 16 | 0.1875 |
| 11_495652 | 3 | 0.6667 | 6 | 0.1667 | 2 | 0.5 | 5 | 0.8 | 16 | 0.5 | 4 | 0.5 | 5 | 0.2 | 4 | 0.5 | 5 | 0.2 | 18 | 0.3333 |
| 11_495653 | 3 | 0 | 5 | 0 | 2 | 0.5 | 6 | 0 | 16 | 0.0625 | 4 | 0 | 2 | 0.5 | 4 | 0.25 | 4 | 0 | 14 | 0.1429 |
| 11_495656 | 3 | 0.3333 | 4 | 0.25 | 2 | 0 | 6 | 0 | 15 | 0.1333 | 4 | 0 | 5 | 0 | 3 | 0.6667 | 4 | 0 | 16 | 0.125 |
| 11_495658 | 2 | 0.5 | 4 | 0.25 | 2 | 0.5 | 6 | 0.1667 | 14 | 0.2857 | 3 | 0.3333 | 4 | 0.25 | 3 | 0 | 3 | 0 | 13 | 0.1538 |
| 11_495659 | 2 | 0 | 4 | 0 | 2 | 0.5 | 5 | 0 | 13 | 0.0769 | 4 | 0 | 4 | 0 | 3 | 0.3333 | 4 | 0 | 15 | 0.0667 |
| 11_495660 | 3 | 0 | 5 | 0.4 | 2 | 0 | 5 | 0.6 | 15 | 0.3333 | 4 | 0.5 | 4 | 0.75 | 3 | 0 | 4 | 0.5 | 15 | 0.4667 |
| 11_495661 | 2 | 0 | 2 | 0 | 2 | 0.5 | 3 | 0 | 9 | 0.1111 | 3 | 0 | 4 | 0.5 | 2 | 0 | 2 | 0 | 11 | 0.1818 |
| 11_495682 | NA | NA | 5 | 0 | 2 | 0 | 4 | 0.25 | 11 | 0.0909 | 3 | 0.3333 | 4 | 0 | 3 | 0 | 2 | 0 | 12 | 0.0833 |
| 11_496032 | NA | NA | 2 | 0.5 | NA | NA | 1 | 0 | 3 | 0.3333 | 2 | 0.5 | 2 | 0 | 2 | 0 | 2 | 0 | 8 | 0.125 |
| 11_497286 | NA | NA | 2 | 0 | NA | NA | 1 | 1 | 3 | 0.3333 | 2 | 0 | NA | NA | 1 | 1 | 1 | 0 | 4 | 0.25 |
| 11_497292 | NA | NA | 3 | 0.6667 | NA | NA | 1 | 1 | 4 | 0.75 | 2 | 0.5 | NA | NA | 1 | 0 | 2 | 0.5 | 5 | 0.4 |
| 11_497638 | 1 | 0 | NA | NA | NA | NA | 1 | 1 | 2 | 0.5 | 1 | 1 | 1 | 0 | NA | NA | 1 | 0 | 3 | 0.3333 |
| 11_497638 | 1 | 1 | NA | NA | NA | NA | 1 | 1 | 2 | 1 | 1 | 1 | 1 | 0 | NA | NA | NA | NA | 2 | 0.5 |
| 11_497639 | 1 | 1 | NA | NA | NA | NA | NA | NA | 1 | 1 | 1 | 1 | 1 | 0 | NA | NA | NA | NA | 2 | 0.5 |
| 11_497741 | 8 | 0.75 | 8 | 0.25 | 16 | 0.75 | 11 | 0.7273 | 43 | 0.6512 | 8 | 0.375 | 6 | 0.5 | 13 | 0.8462 | 11 | 0.8182 | 38 | 0.6842 |
| 11_497742 | 10 | 0.1 | 10 | 0.1 | 18 | 0.1667 | 11 | 0.0909 | 49 | 0.1224 | 7 | 0.2857 | 9 | 0.1111 | 16 | 0.25 | 13 | 0.2308 | 45 | 0.2222 |
| 11_497742 | 13 | 0.1538 | 12 | 0.25 | 19 | 0.1579 | 13 | 0.5385 | 57 | 0.2632 | 11 | 0.1818 | 11 | 0.3636 | 15 | 0.4 | 13 | 0.3077 | 50 | 0.32 |
| 11_497742 | 13 | 0 | 12 | 0.0833 | 22 | 0.0455 | 13 | 0.0769 | 60 | 0.05 | 12 | 0 | 11 | 0.2727 | 18 | 0.0556 | 16 | 0.0625 | 57 | 0.0877 |
| 11_497744 | 13 | 0.0769 | 12 | 0.25 | 23 | 0.087 | 12 | 0 | 60 | 0.1 | 11 | 0 | 8 | 0.125 | 15 | 0.2 | 15 | 0.1333 | 49 | 0.1224 |
| 11_497756 | 14 | 0 | 16 | 0.125 | 25 | 0.32 | 17 | 0.0588 | 72 | 0.1528 | 16 | 0.25 | 14 | 0.2143 | 21 | 0.0952 | 20 | 0.2 | 71 | 0.1831 |
| 11_497762 | 14 | 0.0714 | 20 | 0 | 26 | 0.1154 | 19 | 0.0526 | 79 | 0.0633 | 17 | 0.1765 | 15 | 0.1333 | 19 | 0 | 20 | 0 | 71 | 0.0704 |
| 11_498705 | 3 | 0.6667 | 3 | 1 | 6 | 0.5 | 2 | 0.5 | 14 | 0.6429 | 4 | 0.75 | 3 | 0.6667 | 3 | 0.6667 | 3 | 0.3333 | 13 | 0.6154 |
| 11_498705 | 3 | 0.6667 | 3 | 0.6667 | 5 | 0.2 | 2 | 0.5 | 13 | 0.4615 | 5 | 0.4 | 2 | 0.5 | 5 | 0.6 | 3 | 0 | 15 | 0.4 |
| 11_498705 | 3 | 0.3333 | 3 | 0.3333 | 6 | 0.1667 | 2 | 0 | 14 | 0.2143 | 4 | 0.5 | 3 | 0.3333 | 5 | 0 | 4 | 0 | 16 | 0.1875 |
| 11_498706 | 3 | 0.3333 | 3 | 1 | 5 | 0.6 | 2 | 0.5 | 13 | 0.6154 | 4 | 0.5 | 3 | 0.3333 | 6 | 1 | 4 | 0.5 | 17 | 0.6471 |
| 11_498706 | 4 | 0.75 | 3 | 1 | 7 | 0.2857 | 2 | 0.5 | 16 | 0.5625 | 4 | 0.5 | 3 | 1 | 5 | 0.4 | 4 | 0.5 | 16 | 0.5625 |
| 11_498708 | 4 | 0.25 | 3 | 0.6667 | 7 | 0.4286 | 2 | 0.5 | 16 | 0.4375 | 4 | 1 | 3 | 0.6667 | 5 | 0.6 | 4 | 0.75 | 16 | 0.75 |
| 11_498710 | 4 | 0 | 2 | 0 | 5 | 0.2 | 2 | 0 | 13 | 0.0769 | 5 | 0 | 3 | 0 | 6 | 0.1667 | 4 | 0 | 18 | 0.0556 |
| 11_500099 | NA | NA | 5 | 0.6 | 4 | 0.75 | 3 | 1 | 12 | 0.75 | 2 | 0.5 | 1 | 1 | 3 | 1 | 1 | 1 | 7 | 0.8571 |
| 11_500099 | NA | NA | 6 | 0 | 4 | 0.25 | 2 | 0 | 12 | 0.0833 | 2 | 0.5 | 1 | 0 | 2 | 0 | 1 | 1 | 6 | 0.3333 |
| 11_500105 | NA | NA | 4 | 0.25 | 1 | 0 | 2 | 0 | 7 | 0.1429 | 2 | 1 | 1 | 1 | 1 | 0 | 1 | 1 | 5 | 0.8 |
| 11_500106 | NA | NA | 4 | 1 | 1 | 0 | NA | NA | 5 | 0.8 | 2 | 1 | 2 | 1 | 2 | 0.5 | 1 | 1 | 7 | 0.8571 |
| 11_500109 | 1 | 1 | 4 | 0.75 | 1 | 0 | 2 | 0.5 | 8 | 0.625 | 2 | 0 | 2 | 0 | 2 | 0.5 | 1 | 0 | 7 | 0.1429 |
| 11_500109 | 1 | 1 | 3 | 1 | 1 | 1 | 1 | 1 | 6 | 1 | 1 | 1 | 1 | 1 | 1 | 1 | 1 | 1 | 4 | 1 |
| 11_500109 | 1 | 0 | 3 | 0.3333 | NA | NA | 2 | 0 | 6 | 0.1667 | 3 | 0.6667 | 2 | 0 | 1 | 0 | 1 | 0 | 7 | 0.2857 |
| 11_500110 | 1 | 1 | 4 | 0.75 | 1 | 1 | 2 | 0 | 8 | 0.625 | 3 | 0.6667 | 2 | 0 | 2 | 0.5 | 1 | 1 | 8 | 0.5 |
| 11_500112 | NA | NA | 4 | 0.5 | 1 | 0 | 2 | 0.5 | 7 | 0.4286 | 2 | 0 | 2 | 1 | 1 | 1 | 1 | 0 | 6 | 0.5 |
| 11_500112 | NA | NA | 5 | 0.2 | 1 | 1 | 2 | 0.5 | 8 | 0.375 | 3 | 0 | 2 | 0.5 | 2 | 0 | 1 | 0 | 8 | 0.125 |
| 11_500114 | 1 | 1 | 4 | 1 | NA | NA | 2 | 1 | 7 | 1 | 4 | 1 | 1 | 1 | 2 | 0.5 | 1 | 0 | 8 | 0.75 |
| 11_500114 | 1 | 0 | 4 | 0.25 | 1 | 0 | 2 | 0 | 8 | 0.125 | 4 | 0.25 | 2 | 0.5 | 2 | 1 | 1 | 0 | 9 | 0.4444 |
| 11_500119 | NA | NA | 4 | 0.25 | 1 | 1 | 4 | 0 | 9 | 0.2222 | 3 | 0.6667 | 2 | 1 | 1 | 1 | 2 | 1 | 8 | 0.875 |
| 11_500121 | NA | NA | 4 | 0.5 | 1 | 0 | 3 | 0.6667 | 8 | 0.5 | 3 | 0.6667 | 2 | 1 | 1 | 1 | 1 | 0 | 7 | 0.7143 |
| 11_500122 | NA | NA | 4 | 0.25 | 1 | 0 | 4 | 0.5 | 9 | 0.3333 | 3 | 0.3333 | 2 | 0 | 1 | 0 | 2 | 0 | 8 | 0.125 |
| 11_500122 | NA | NA | 4 | 0.25 | NA | NA | 3 | 0 | 7 | 0.1429 | 1 | 0 | 2 | 0.5 | 1 | 0 | 1 | 1 | 5 | 0.4 |
| 11_500123 | NA | NA | 4 | 1 | 1 | 1 | 3 | 0.6667 | 8 | 0.875 | 2 | 1 | 2 | 1 | NA | NA | 2 | 1 | 6 | 1 |
| 11_500125 | NA | NA | 4 | 0.5 | 2 | 0.5 | 4 | 0 | 10 | 0.3 | 2 | 0 | 2 | 0 | 1 | 0 | 2 | 1 | 7 | 0.2857 |
| 11_500125 | NA | NA | 5 | 0 | 2 | 0 | 4 | 0.25 | 11 | 0.0909 | 3 | 0.3333 | 1 | 0 | 1 | 1 | 2 | 0 | 7 | 0.2857 |
| 11_500126 | NA | NA | 4 | 1 | 2 | 1 | 2 | 1 | 8 | 1 | 3 | 1 | 2 | 1 | 1 | 1 | 2 | 1 | 8 | 1 |
| 11_500126 | NA | NA | 3 | 0.6667 | NA | NA | 4 | 0.5 | 7 | 0.5714 | 1 | 1 | NA | NA | 1 | 1 | 1 | 1 | 3 | 1 |
| 11_500127 | 1 | 1 | 4 | 1 | 2 | 1 | 2 | 0.5 | 9 | 0.8889 | 2 | 1 | 2 | 1 | 1 | 1 | 2 | 1 | 7 | 1 |
| 11_500128 | 1 | 1 | 4 | 1 | 2 | 1 | 4 | 0.75 | 11 | 0.9091 | 3 | 1 | 2 | 1 | 1 | 1 | 2 | 1 | 8 | 1 |

| | | | | | | | | | | | | | | | | | | | | |
|---|---|---|---|---|---|---|---|---|---|---|---|---|---|---|---|---|---|---|---|---|
| 11_5001292 | NA | NA | 4 | 0.5 | 2 | 0 | 5 | 0.4 | 11 | 0.3636 | 3 | 0.6667 | 1 | 0 | 1 | 1 | 2 | 0.5 | 7 | 0.5714 |
| 11_5001293 | NA | NA | 3 | 0.3333 | 2 | 0 | 3 | 0.3333 | 8 | 0.25 | 1 | 1 | 1 | 0 | 1 | 0 | 1 | 0 | 4 | 0.25 |
| 11_5001295 | 1 | 1 | 4 | 0.75 | 1 | 1 | 5 | 0.8 | 11 | 0.8182 | 2 | 0.5 | 2 | 0.5 | 1 | 1 | 2 | 0.5 | 7 | 0.5714 |
| 11_5001305 | 2 | 1 | 4 | 1 | 2 | 0.5 | 5 | 0.8 | 13 | 0.8462 | 3 | 1 | 2 | 1 | 2 | 1 | 2 | 0.5 | 9 | 0.8889 |
| 11_5001328 | 1 | 0 | 3 | 0.3333 | 2 | 0.5 | 5 | 0.4 | 11 | 0.3636 | 2 | 1 | 2 | 0.5 | 1 | 0 | 2 | 0.5 | 7 | 0.5714 |
| 11_5001430 | 3 | 0 | 3 | 0 | 3 | 0 | 7 | 0.1429 | 16 | 0.0625 | 5 | 0.2 | 2 | 0 | 2 | 0 | 4 | 0 | 13 | 0.0769 |
| 11_5001545 | 5 | 0.2 | 7 | 0.1429 | 6 | 0 | 8 | 0.25 | 26 | 0.1538 | 6 | 0.1667 | 5 | 0 | 2 | 0.5 | 7 | 0.2857 | 20 | 0.2 |
| 11_5001599 | 5 | 0 | 10 | 0 | 7 | 0.2857 | 8 | 0 | 30 | 0.0667 | 5 | 0 | 7 | 0 | 4 | 0 | 8 | 0.375 | 24 | 0.125 |
| 11_5001621 | 2 | 0 | 4 | 0.25 | 7 | 0.2857 | 6 | 0.3333 | 19 | 0.2632 | 5 | 0.4 | 4 | 0.25 | 4 | 0 | 4 | 0.25 | 17 | 0.2353 |
| 11_5003717 | 4 | 0.5 | 5 | 0.4 | 5 | 0 | 2 | 0 | 16 | 0.25 | 1 | 0 | 1 | 0 | 1 | 0 | 3 | 0.6667 | 6 | 0.3333 |
| 11_5005248 | 8 | 0.25 | 12 | 0.0833 | 11 | 0.0909 | 8 | 0.375 | 39 | 0.1795 | 4 | 0 | 9 | 0 | 11 | 0.3636 | 9 | 0.1111 | 33 | 0.1515 |
| 11_5006158 | 11 | 0.5455 | 13 | 0.8462 | 18 | 0.5556 | 11 | 0.4545 | 53 | 0.6038 | 5 | 0.8 | 13 | 0.7692 | 13 | 0.6923 | 10 | 0.7 | 41 | 0.7317 |
| 11_5006175 | 11 | 0.1818 | 12 | 0 | 18 | 0.1111 | 12 | 0.1667 | 53 | 0.1132 | 7 | 0 | 14 | 0.1429 | 12 | 0.0833 | 11 | 0.2727 | 44 | 0.1364 |
| 11_5006198 | 13 | 0.3077 | 12 | 0.25 | 17 | 0.1765 | 14 | 0 | 56 | 0.1786 | 9 | 0.2222 | 18 | 0.0556 | 13 | 0.2308 | 12 | 0.0833 | 52 | 0.1346 |
| 11_5006205 | 10 | 0.6 | 14 | 0.6429 | 19 | 0.6842 | 12 | 0.5833 | 55 | 0.6364 | 9 | 0.6667 | 15 | 0.6 | 14 | 0.8571 | 11 | 0.8182 | 49 | 0.7347 |
| 11_5006206 | 12 | 0.6667 | 15 | 0.7333 | 13 | 0.7692 | 11 | 0.6364 | 51 | 0.7059 | 8 | 0.625 | 15 | 0.7333 | 14 | 0.6429 | 11 | 0.8182 | 48 | 0.7083 |
| 11_5006212 | 12 | 0.3333 | 12 | 0.3333 | 12 | 0.5 | 11 | 0.3636 | 47 | 0.383 | 10 | 0.5 | 17 | 0.2353 | 11 | 0.3636 | 11 | 0.5455 | 49 | 0.3878 |
| 11_5006215 | 13 | 1 | 15 | 0.7333 | 17 | 0.8235 | 12 | 0.9167 | 57 | 0.8596 | 9 | 0.7778 | 19 | 0.8947 | 16 | 0.75 | 14 | 0.8571 | 58 | 0.8276 |
| 11_5006216 | 11 | 0.3636 | 17 | 0.3529 | 18 | 0.1667 | 13 | 0.0769 | 59 | 0.2373 | 11 | 0.3636 | 19 | 0.2105 | 16 | 0.5625 | 13 | 0.1538 | 59 | 0.322 |
| 11_5006223 | 9 | 0.7778 | 11 | 0.7273 | 12 | 0.8333 | 11 | 0.9091 | 43 | 0.814 | 8 | 0.875 | 12 | 0.8333 | 7 | 0.8571 | 6 | 1 | 33 | 0.8788 |
| 11_5006242 | 12 | 0 | 18 | 0 | 18 | 0.1667 | 18 | 0.1111 | 66 | 0.0758 | 11 | 0 | 19 | 0.0526 | 16 | 0 | 14 | 0.1429 | 60 | 0.05 |
| 11_5006254 | 14 | 0.3571 | 17 | 0.2353 | 15 | 0.2667 | 16 | 0.125 | 62 | 0.2419 | 12 | 0.4167 | 17 | 0.1765 | 19 | 0.2632 | 13 | 0.4615 | 61 | 0.3115 |
| 11_5006265 | 12 | 0.75 | 15 | 0.8 | 17 | 0.7647 | 13 | 0.6154 | 57 | 0.7368 | 8 | 0.625 | 13 | 0.6923 | 18 | 0.7778 | 11 | 0.6364 | 50 | 0.7 |
| 11_5006272 | 13 | 0.6923 | 17 | 0.5882 | 19 | 0.5263 | 15 | 0.4 | 64 | 0.5469 | 12 | 0.5833 | 18 | 0.4444 | 19 | 0.6842 | 12 | 0.5 | 61 | 0.5574 |
| 11_5006274 | 11 | 0.0909 | 14 | 0.0714 | 17 | 0.1765 | 17 | 0.1176 | 59 | 0.1186 | 10 | 0.1 | 17 | 0.1765 | 18 | 0.1111 | 13 | 0.0769 | 58 | 0.1207 |
| 11_5006328 | 2 | 0 | 3 | 0.3333 | 5 | 0 | 5 | 0 | 15 | 0.0667 | 3 | 0 | 4 | 0 | 7 | 0.1429 | 1 | 0 | 15 | 0.0667 |
| 11_5006355 | 2 | 0 | 4 | 0 | 2 | 0.5 | 4 | 0.25 | 12 | 0.1667 | 3 | 0 | 3 | 0.3333 | 6 | 0 | 1 | 0 | 13 | 0.0769 |
| 11_5009060 | 2 | 1 | 1 | 1 | NA | NA | 2 | 1 | 5 | 1 | 1 | 1 | 1 | 1 | 2 | 0.5 | NA | NA | 4 | 0.75 |
| 11_5009069 | 2 | 1 | 1 | 1 | NA | NA | 1 | 1 | 4 | 1 | 1 | 0 | 1 | 0 | 2 | 1 | NA | NA | 4 | 0.5 |
| 11_5009073 | 2 | 0.5 | 1 | 0 | NA | NA | 1 | 0 | 4 | 0.25 | 1 | 0 | 1 | 1 | 3 | 0 | NA | NA | 5 | 0.2 |
| 11_5009073 | 2 | 0.5 | 1 | 1 | NA | NA | 2 | 1 | 5 | 0.8 | 1 | 1 | 1 | 0 | 3 | 0.3333 | NA | NA | 5 | 0.4 |
| 11_5009082 | 2 | 0.5 | 2 | 1 | NA | NA | 1 | 1 | 5 | 0.8 | 1 | 1 | 1 | 1 | 5 | 0.8 | NA | NA | 7 | 0.8571 |
| 11_5009089 | 2 | 0.5 | 2 | 0 | NA | NA | 1 | 1 | 5 | 0.4 | 1 | 0 | 1 | 0 | 4 | 0.5 | NA | NA | 6 | 0.3333 |
| 11_5009120 | 72 | 0.0417 | 77 | 0.0909 | 62 | 0.0806 | 60 | 0.1167 | 271 | 0.0812 | 61 | 0.0328 | 60 | 0.05 | 72 | 0.0694 | 51 | 0.098 | 244 | 0.0615 |
| 11_5012572 | 12 | 0.9167 | 6 | 0.8333 | 10 | 0.9 | 11 | 0.9091 | 39 | 0.8974 | 6 | 1 | 14 | 0.7857 | 10 | 0.9 | 10 | 0.9 | 40 | 0.875 |
| 11_5014677 | 2 | 0 | NA | NA | 1 | 0 | 2 | 0.5 | 5 | 0.2 | NA | NA | 3 | 0.6667 | NA | NA | 2 | 0.5 | 5 | 0.6 |
| 11_5014737 | 1 | 0 | NA | NA | 1 | 0 | 2 | 1 | 4 | 0.5 | NA | NA | 3 | 1 | NA | NA | 2 | 1 | 5 | 1 |
| 11_5014745 | 2 | 0.5 | NA | NA | 1 | 0 | 2 | 0 | 5 | 0.2 | NA | NA | 3 | 0.6667 | NA | NA | 2 | 0 | 5 | 0.4 |
| 11_5014760 | 2 | 0.5 | NA | NA | 1 | 0 | 2 | 1 | 5 | 0.6 | NA | NA | 2 | 0.5 | NA | NA | 2 | 1 | 4 | 0.75 |
| 11_5014768 | 2 | 0 | NA | NA | 1 | 1 | 2 | 0 | 5 | 0.2 | NA | NA | 3 | 0.3333 | NA | NA | 2 | 0 | 5 | 0.2 |
| 11_5014802 | 1 | 0 | NA | NA | 1 | 0 | 1 | 1 | 3 | 0.3333 | NA | NA | 2 | 1 | NA | NA | 2 | 0.5 | 4 | 0.75 |
| 11_5014813 | 2 | 0 | NA | NA | 1 | 0 | 2 | 0.5 | 5 | 0.2 | NA | NA | 3 | 0.3333 | NA | NA | 1 | 0 | 4 | 0.25 |
| 11_5014859 | NA | NA | NA | NA | NA | NA | 2 | 0.5 | 2 | 0.5 | NA | NA | 3 | 0.3333 | NA | NA | 2 | 0 | 5 | 0.2 |
| 11_5014872 | 1 | 0 | NA | NA | NA | NA | 2 | 1 | 3 | 0.6667 | NA | NA | 4 | 1 | NA | NA | 1 | 1 | 5 | 1 |
| 11_5014883 | 1 | 1 | NA | NA | NA | NA | 2 | 1 | 3 | 1 | NA | NA | 3 | 0.6667 | NA | NA | 2 | 0.5 | 5 | 0.6 |
| 11_5014899 | 1 | 1 | NA | NA | NA | NA | 2 | 1 | 3 | 1 | NA | NA | 4 | 0.5 | NA | NA | 1 | 0 | 5 | 0.4 |
| 11_5014932 | 1 | 1 | NA | NA | NA | NA | 1 | 1 | 2 | 1 | NA | NA | 1 | 0 | NA | NA | 2 | 0.5 | 3 | 0.3333 |
| 11_5014933 | NA | NA | NA | NA | NA | NA | 2 | 0.5 | 2 | 0.5 | NA | NA | 3 | 1 | NA | NA | 1 | 1 | 4 | 1 |
| 11_5014950 | 1 | 0 | NA | NA | NA | NA | 2 | 1 | 3 | 0.6667 | NA | NA | 4 | 0.25 | NA | NA | 1 | 1 | 5 | 0.4 |
| 11_5014957 | 1 | 1 | NA | NA | NA | NA | 2 | 0.5 | 3 | 0.6667 | NA | NA | 3 | 0.6667 | NA | NA | 2 | 1 | 5 | 0.8 |
| 11_5015658 | NA | NA | NA | NA | NA | NA | 1 | 1 | 1 | 1 | NA | NA | 1 | 1 | NA | NA | NA | NA | 1 | 1 |
| 11_5015662 | 1 | 1 | NA | NA | NA | NA | 2 | 0.5 | 3 | 0.6667 | NA | NA | 3 | 0.6667 | NA | NA | 1 | 1 | 4 | 0.75 |
| 11_5015678 | 1 | 0 | NA | NA | NA | NA | 2 | 0.5 | 3 | 0.3333 | NA | NA | 3 | 0.3333 | NA | NA | 2 | 0.5 | 5 | 0.4 |
| 11_5015711 | 1 | 1 | NA | NA | NA | NA | 2 | 0 | 3 | 0.3333 | NA | NA | 3 | 0.6667 | NA | NA | 2 | 0.5 | 5 | 0.6 |
| 11_5015725 | 1 | 1 | NA | NA | NA | NA | 1 | 1 | 2 | 1 | NA | NA | 2 | 1 | NA | NA | 1 | 1 | 3 | 1 |
| 11_5015726 | 1 | 1 | NA | NA | NA | NA | 2 | 1 | 3 | 1 | NA | NA | 3 | 0.6667 | NA | NA | 2 | 1 | 5 | 0.8 |
| 11_5015733 | 1 | 1 | NA | NA | NA | NA | 2 | 0.5 | 3 | 0.6667 | NA | NA | 2 | 0.5 | NA | NA | 2 | 0.5 | 4 | 0.5 |
| 11_5018601 | 53 | 0.1509 | 46 | 0.1957 | 50 | 0.2 | 53 | 0.2075 | 202 | 0.1881 | 35 | 0.1143 | 44 | 0.0227 | 51 | 0.2549 | 46 | 0.1739 | 176 | 0.1477 |
| 11_5018638 | 43 | 0.1628 | 38 | 0.0263 | 44 | 0.0909 | 49 | 0.1633 | 174 | 0.1149 | 29 | 0.1379 | 39 | 0.0769 | 43 | 0.0698 | 43 | 0.2326 | 154 | 0.1299 |
| 11_5018709 | 37 | 0.4054 | 29 | 0.2414 | 33 | 0.3333 | 41 | 0.2683 | 140 | 0.3143 | 23 | 0.4348 | 36 | 0.3611 | 32 | 0.3125 | 39 | 0.359 | 130 | 0.3615 |
| 11_5027673 | 28 | 0.1071 | 22 | 0.0909 | 21 | 0 | 30 | 0.0667 | 101 | 0.0693 | 13 | 0.2308 | 26 | 0.0769 | 24 | 0.125 | 22 | 0.0455 | 85 | 0.1059 |
| 11_5027683 | 29 | 0.069 | 23 | 0.0435 | 23 | 0.0435 | 30 | 0.0667 | 105 | 0.0571 | 14 | 0.0714 | 27 | 0.037 | 19 | 0.1053 | 18 | 0 | 78 | 0.0513 |
| 11_5027690 | 28 | 0.0714 | 30 | 0.2333 | 24 | 0.125 | 37 | 0.0811 | 119 | 0.1261 | 16 | 0.125 | 32 | 0.0625 | 28 | 0.0357 | 29 | 0.1034 | 105 | 0.0762 |
| 11_5028035 | 3 | 0.6667 | 4 | 0.75 | 1 | 0 | 2 | 0.5 | 10 | 0.6 | 3 | 0 | 2 | 0.5 | 4 | 0.25 | 6 | 0.5 | 15 | 0.3333 |
| 11_5028041 | 4 | 0.25 | 4 | 0.25 | 1 | 0 | 3 | 0.3333 | 12 | 0.25 | 3 | 0.3333 | 2 | 0 | 4 | 0 | 7 | 0.4286 | 16 | 0.25 |

| | | | | | | | | | | | | | | | | | | | | |
|---|---|---|---|---|---|---|---|---|---|---|---|---|---|---|---|---|---|---|---|---|
| 11_5028052 | 3 | 0.3333 | 5 | 0 | 2 | 0 | 3 | 0 | 13 | 0.0769 | 3 | 0 | 2 | 0 | 5 | 0 | 7 | 0.5714 | 17 | 0.2353 |
| 11_5028061 | 2 | 1 | 5 | 0.4 | 2 | 0.5 | 3 | 0.6667 | 12 | 0.5833 | 3 | 0.3333 | 2 | 0.5 | 4 | 0.25 | 7 | 0.5714 | 16 | 0.4375 |
| 11_5068761 | 1 | 0 | 1 | 0 | 2 | 0.5 | 1 | 0 | 5 | 0.2 | NA | NA | 1 | 1 | NA | NA | NA | NA | 1 | 1 |
| 11_5068811 | 2 | 1 | 2 | 1 | 1 | 0 | 1 | 1 | 6 | 0.8333 | NA | NA | 1 | 1 | 1 | 0 | 2 | 1 | 4 | 0.75 |
| 11_5070488 | NA | NA | 3 | 0.6667 | NA | NA | 4 | 0.5 | 7 | 0.5714 | 1 | 1 | 4 | 0.25 | 4 | 0.5 | 5 | 0.6 | 14 | 0.5 |
| 11_5070518 | NA | NA | 3 | 0.3333 | 1 | 0 | 3 | 0.3333 | 7 | 0.2857 | 1 | 0 | 4 | 0.25 | 3 | 0.3333 | 3 | 0.3333 | 11 | 0.2727 |
| 11_5070524 | NA | NA | 2 | 0.5 | 1 | 0 | 4 | 0.25 | 7 | 0.2857 | 1 | 0 | 4 | 0 | 3 | 0 | 4 | 0.25 | 12 | 0.0833 |
| 11_5070525 | NA | NA | 3 | 1 | NA | NA | 4 | 1 | 7 | 1 | 1 | 1 | 4 | 1 | 3 | 1 | 4 | 1 | 12 | 1 |
| 11_5070529 | NA | NA | 2 | 1 | 1 | 1 | 4 | 1 | 7 | 1 | 1 | 1 | 4 | 1 | 3 | 1 | 3 | 1 | 11 | 1 |
| 11_5070541 | NA | NA | 2 | 1 | 1 | 1 | 2 | 1 | 5 | 1 | 1 | 1 | 2 | 1 | 3 | 0.6667 | 2 | 1 | 8 | 0.875 |
| 11_5070547 | NA | NA | 3 | 1 | 1 | 1 | 5 | 1 | 9 | 1 | 1 | 1 | 4 | 1 | 3 | 1 | 4 | 1 | 12 | 1 |
| 11_5070552 | NA | NA | 3 | 0.3333 | 1 | 1 | 5 | 0.4 | 9 | 0.4444 | 1 | 1 | 4 | 0.5 | 3 | 0.3333 | 4 | 0.25 | 12 | 0.4167 |
| 11_5070553 | NA | NA | 3 | 0.3333 | 1 | 0 | 4 | 0 | 8 | 0.125 | 1 | 0 | 4 | 0.25 | 3 | 0 | 3 | 0 | 11 | 0.0909 |
| 11_5070561 | NA | NA | 2 | 0 | 1 | 0 | 2 | 0.5 | 5 | 0.2 | 1 | 0 | 2 | 0 | 2 | 0.5 | 4 | 0.25 | 9 | 0.2222 |
| 11_5070561 | NA | NA | 2 | 0.5 | NA | NA | 1 | 1 | 3 | 0.6667 | 1 | 0 | 1 | 1 | 1 | 0 | 2 | 0 | 5 | 0.2 |
| 11_5070567 | NA | NA | 3 | 0 | 1 | 0 | 5 | 0.6 | 9 | 0.3333 | 1 | 0 | 4 | 0.25 | 3 | 0 | 4 | 0 | 12 | 0.0833 |
| 11_5070575 | NA | NA | 3 | 0.6667 | NA | NA | 3 | 0.3333 | 6 | 0.5 | 1 | 1 | 4 | 0.5 | 3 | 0.6667 | 3 | 0.3333 | 11 | 0.5455 |
| 11_5070581 | NA | NA | 2 | 0.5 | NA | NA | 4 | 0.25 | 6 | 0.3333 | 1 | 1 | 3 | 0 | 3 | 0.3333 | 3 | 0 | 10 | 0.2 |
| 11_5070584 | NA | NA | 3 | 1 | NA | NA | 3 | 0.6667 | 6 | 0.8333 | 1 | 1 | 4 | 1 | 3 | 1 | 3 | 1 | 11 | 1 |
| 11_5070586 | NA | NA | 2 | 0.5 | NA | NA | 3 | 0 | 5 | 0.2 | 1 | 0 | 2 | 0.5 | 3 | 0.3333 | 2 | 0.5 | 8 | 0.375 |
| 11_5070589 | NA | NA | 3 | 0.6667 | NA | NA | 3 | 0.3333 | 6 | 0.5 | 1 | 0 | 4 | 0.25 | 3 | 1 | 3 | 0.3333 | 11 | 0.4545 |
| 11_5070594 | NA | NA | 3 | 0.6667 | NA | NA | 4 | 1 | 7 | 0.8571 | 1 | 1 | 3 | 1 | 2 | 1 | 3 | 1 | 9 | 1 |
| 11_5070594 | NA | NA | 3 | 0.3333 | NA | NA | 1 | 0 | 4 | 0.25 | 1 | 0 | 3 | 0 | 1 | 0 | 3 | 0.3333 | 8 | 0.125 |
| 11_5070603 | NA | NA | 3 | 0.3333 | NA | NA | 4 | 0.25 | 7 | 0.2857 | 1 | 0 | 4 | 0.25 | 3 | 0.3333 | 3 | 0 | 11 | 0.1818 |
| 11_5070643 | NA | NA | 2 | 0.5 | NA | NA | 3 | 1 | 5 | 0.8 | 1 | 1 | 3 | 0.3333 | 2 | 0 | 3 | 0.6667 | 9 | 0.4444 |
| 11_5070668 | NA | NA | 2 | 0.5 | NA | NA | 3 | 0.6667 | 5 | 0.6 | NA | NA | 2 | 0.5 | 3 | 0.6667 | 1 | 1 | 6 | 0.6667 |
| 11_5070675 | NA | NA | 2 | 0.5 | NA | NA | 3 | 0 | 5 | 0.2 | NA | NA | 3 | 1 | 3 | 0.3333 | 3 | 0.3333 | 9 | 0.5556 |
| 11_5076386 | NA | NA | 2 | 1 | NA | NA | NA | NA | 2 | 1 | NA | NA | NA | NA | 1 | 1 | 1 | 0 | 2 | 0.5 |
| 11_5076412 | NA | NA | 1 | 1 | NA | NA | NA | NA | 1 | 1 | NA | NA | NA | NA | 1 | 1 | 1 | 1 | 2 | 1 |
| 11_5079036 | 9 | 0.4444 | 4 | 0.75 | 4 | 0 | 2 | 0.5 | 19 | 0.4211 | 2 | 1 | NA | NA | 2 | 0.5 | 2 | 1 | 6 | 0.8333 |
| 11_5079039 | 9 | 0.5556 | 6 | 0.5 | 5 | 0.4 | 3 | 0.6667 | 23 | 0.5217 | 2 | 1 | NA | NA | 2 | 0 | 2 | 0 | 6 | 0.3333 |
| 11_5079054 | 7 | 0.1429 | 5 | 0.2 | 4 | 0 | 2 | 0 | 18 | 0.1111 | 2 | 0 | NA | NA | 3 | 0.3333 | NA | NA | 5 | 0.2 |
| 11_5079056 | 7 | 0 | 5 | 0.2 | 4 | 0 | 2 | 0 | 18 | 0.0556 | 1 | 1 | NA | NA | 2 | 0 | NA | NA | 3 | 0.3333 |
| 11_5079067 | 8 | 1 | 6 | 1 | 5 | 1 | 2 | 1 | 21 | 1 | 1 | 1 | NA | NA | 2 | 1 | 1 | 1 | 4 | 1 |
| 11_5079068 | 8 | 0.25 | 6 | 0.1667 | 5 | 0 | 3 | 0 | 22 | 0.1364 | 2 | 0 | NA | NA | 2 | 1 | 1 | 0 | 5 | 0.4 |
| 11_5079074 | 7 | 0 | 6 | 0.1667 | 5 | 0.2 | 3 | 0 | 21 | 0.0952 | 2 | 0.5 | NA | NA | 3 | 0.6667 | NA | NA | 5 | 0.6 |
| 11_5079078 | 6 | 1 | 4 | 1 | 5 | 1 | 2 | 1 | 17 | 1 | 1 | 1 | NA | NA | 2 | 1 | 1 | 1 | 4 | 1 |
| 11_5079085 | 8 | 0.75 | 6 | 0.6667 | 4 | 0.5 | 2 | 0 | 20 | 0.6 | 1 | 0 | NA | NA | 2 | 1 | NA | NA | 3 | 0.6667 |
| 11_5079094 | 8 | 0.75 | 4 | 0.75 | 3 | 0 | 3 | 0.3333 | 18 | 0.5556 | 2 | 1 | NA | NA | 2 | 0.5 | NA | NA | 4 | 0.75 |
| 11_5079109 | 7 | 1 | 5 | 1 | 5 | 1 | 3 | 1 | 20 | 1 | NA | NA | NA | NA | 1 | 1 | NA | NA | 1 | 1 |
| 11_5079124 | 7 | 0 | 5 | 0 | 5 | 0.2 | 3 | 0 | 20 | 0.05 | 1 | 1 | NA | NA | 2 | 0 | 1 | 0 | 4 | 0.25 |
| 11_5079134 | 7 | 0.1429 | 4 | 0.25 | 5 | 0.2 | 2 | 0 | 18 | 0.1667 | 1 | 0 | NA | NA | 1 | 1 | NA | NA | 2 | 0.5 |
| 11_5079151 | 6 | 0.5 | 5 | 0.8 | 4 | 0.5 | 2 | 0 | 17 | 0.5294 | 1 | 1 | NA | NA | 1 | 1 | 1 | 1 | 3 | 1 |
| 11_5079153 | 7 | 0.4286 | 3 | 1 | 4 | 1 | 2 | 1 | 16 | 0.75 | 1 | 1 | NA | NA | 1 | 1 | 1 | 1 | 3 | 1 |
| 11_5079159 | 7 | 1 | 5 | 1 | 5 | 1 | 2 | 1 | 19 | 1 | 1 | 1 | NA | NA | 1 | 1 | 1 | 1 | 3 | 1 |
| 11_5079177 | 5 | 0.4 | 3 | 0.3333 | 5 | 0 | 2 | 0 | 15 | 0.2 | 1 | 0 | NA | NA | 1 | 1 | 1 | 0 | 3 | 0.3333 |
| 11_5079183 | 7 | 0.2857 | 5 | 0.6 | 5 | 0 | 2 | 0 | 19 | 0.2632 | 1 | 0 | NA | NA | 1 | 1 | 1 | 1 | 3 | 0.6667 |
| 11_5079188 | 3 | 1 | 3 | 1 | 2 | 1 | 1 | 1 | 9 | 1 | NA | NA | NA | NA | 1 | 1 | NA | NA | 1 | 1 |
| 11_5079203 | 5 | 1 | 5 | 1 | 5 | 1 | 1 | 1 | 16 | 1 | 1 | 1 | NA | NA | NA | NA | NA | NA | 1 | 1 |
| 11_5081300 | 2 | 0.5 | 1 | 1 | NA | NA | 1 | 1 | 4 | 0.75 | NA | NA | 2 | 0.5 | NA | NA | 1 | 1 | 3 | 0.6667 |
| 11_5081304 | 1 | 1 | 1 | 0 | NA | NA | 1 | 0 | 3 | 0.3333 | NA | NA | 2 | 0.5 | 1 | 0 | 1 | 0 | 4 | 0.25 |
| 11_5081306 | 2 | 0.5 | 1 | 1 | NA | NA | 1 | 1 | 4 | 0.75 | NA | NA | 1 | 1 | 1 | 0 | 1 | 1 | 3 | 0.6667 |
| 11_5081336 | 2 | 1 | 1 | 1 | NA | NA | 1 | 1 | 4 | 1 | NA | NA | 2 | 1 | NA | NA | NA | NA | 2 | 1 |
| 11_5081348 | 2 | 0 | 1 | 1 | NA | NA | 1 | 0 | 4 | 0.25 | NA | NA | 2 | 1 | 1 | 1 | NA | NA | 3 | 1 |
| 11_5081368 | 1 | 1 | NA | NA | NA | NA | NA | NA | 1 | 1 | NA | NA | 2 | 0.5 | 1 | 0 | NA | NA | 3 | 0.3333 |
| 11_5081407 | 1 | 1 | 1 | 1 | NA | NA | NA | NA | 2 | 1 | NA | NA | 2 | 1 | 1 | 1 | NA | NA | 3 | 1 |
| 11_5081421 | 2 | 1 | 1 | 1 | NA | NA | NA | NA | 3 | 1 | NA | NA | 1 | 1 | 1 | 1 | NA | NA | 2 | 1 |
| 11_5081422 | 2 | 1 | 1 | 1 | NA | NA | NA | NA | 3 | 1 | NA | NA | 2 | 1 | 1 | 1 | NA | NA | 3 | 1 |
| 11_5081456 | 2 | 1 | 1 | 1 | NA | NA | NA | NA | 3 | 1 | NA | NA | 2 | 1 | 1 | 1 | NA | NA | 3 | 1 |
| 11_5096024 | NA | NA | NA | NA | NA | NA | 2 | 1 | 2 | 1 | NA | NA | 2 | 1 | NA | NA | NA | NA | 2 | 1 |
| 11_5096117 | NA | NA | NA | NA | 1 | 0 | 3 | 1 | 4 | 0.75 | NA | NA | 3 | 0.6667 | NA | NA | NA | NA | 3 | 0.6667 |
| 11_5096144 | NA | NA | NA | NA | 1 | 0 | 2 | 0.5 | 3 | 0.3333 | NA | NA | 3 | 0 | NA | NA | 1 | 1 | 4 | 0.25 |
| 11_5096150 | 1 | 1 | NA | NA | 1 | 1 | 2 | 0.5 | 4 | 0.75 | NA | NA | 3 | 0.6667 | NA | NA | 1 | 1 | 4 | 0.75 |
| 11_5096195 | 1 | 1 | NA | NA | NA | NA | 3 | 0.6667 | 4 | 0.75 | NA | NA | 3 | 1 | NA | NA | 1 | 1 | 4 | 1 |
| 11_5116372 | 1 | 0 | 4 | 0.25 | 8 | 0.125 | 4 | 0 | 17 | 0.1176 | 3 | 0.3333 | 3 | 0 | 1 | 0 | 2 | 0 | 9 | 0.1111 |

| | | | | | | | | | | | | | | | | | | | | |
|---|---|---|---|---|---|---|---|---|---|---|---|---|---|---|---|---|---|---|---|---|
| 11_5116374 | 1 | 0 | 6 | 0.3333 | 8 | 0.25 | 5 | 0.6 | 20 | 0.35 | 4 | 0.25 | 3 | 0 | 1 | 0 | 2 | 0.5 | 10 | 0.2 |
| 11_5117229 | 8 | 0.25 | 9 | 0.1111 | 13 | 0.0769 | 7 | 0.1429 | 37 | 0.1351 | 5 | 0 | 8 | 0.125 | 3 | 0.6667 | 7 | 0.1429 | 23 | 0.1739 |
| 11_5147094 | NA | NA | NA | NA | 3 | 0.6667 | NA | NA | 3 | 0.6667 | NA | NA | NA | NA | NA | NA | 2 | 0.5 | 2 | 0.5 |
| 11_5147097 | NA | NA | NA | NA | 3 | 0.3333 | NA | NA | 3 | 0.3333 | NA | NA | NA | NA | NA | NA | 1 | 1 | 1 | 1 |
| 11_5147103 | NA | NA | NA | NA | 3 | 0.3333 | NA | NA | 3 | 0.3333 | NA | NA | NA | NA | NA | NA | 1 | 1 | 1 | 1 |
| 11_5147117 | NA | NA | NA | NA | 3 | 0.3333 | NA | NA | 3 | 0.3333 | NA | NA | NA | NA | NA | NA | 2 | 0.5 | 2 | 0.5 |
| 11_5147121 | NA | NA | NA | NA | 3 | 0.3333 | NA | NA | 3 | 0.3333 | NA | NA | NA | NA | NA | NA | 2 | 1 | 2 | 1 |
| 11_5147147 | NA | NA | NA | NA | 4 | 1 | NA | NA | 4 | 1 | NA | NA | NA | NA | NA | NA | 1 | 1 | 1 | 1 |
| 11_5147163 | NA | NA | NA | NA | 3 | 0.3333 | NA | NA | 3 | 0.3333 | NA | NA | NA | NA | NA | NA | 1 | 1 | 1 | 1 |
| 11_5147166 | NA | NA | NA | NA | 4 | 0.75 | NA | NA | 4 | 0.75 | NA | NA | 1 | 1 | NA | NA | 2 | 0.5 | 3 | 0.6667 |
| 11_5147174 | 1 | 0 | NA | NA | 3 | 0.6667 | NA | NA | 4 | 0.5 | NA | NA | 1 | 0 | NA | NA | 2 | 0.5 | 3 | 0.3333 |
| 11_5147366 | 1 | 1 | NA | NA | 2 | 0 | NA | NA | 3 | 0.3333 | NA | NA | 1 | 1 | NA | NA | NA | NA | 1 | 1 |
| 11_5147376 | 1 | 0 | NA | NA | 3 | 0.3333 | NA | NA | 4 | 0.25 | NA | NA | 1 | 1 | NA | NA | NA | NA | 1 | 1 |
| 11_5151404 | 11 | 0.9091 | 10 | 0.8 | 9 | 0.7778 | 7 | 0.7143 | 37 | 0.8108 | 9 | 0.6667 | 9 | 0.8889 | 13 | 0.9231 | 12 | 0.9167 | 43 | 0.8605 |
| 11_5151455 | 8 | 0.375 | 11 | 0.0909 | 9 | 0.4444 | 15 | 0.2667 | 43 | 0.2791 | 8 | 0.375 | 6 | 0 | 8 | 0 | 9 | 0 | 31 | 0.0968 |
| 11_5151473 | 9 | 0.7778 | 11 | 0.8182 | 5 | 1 | 14 | 0.7857 | 39 | 0.8205 | 7 | 0.4286 | 6 | 0.6667 | 8 | 0.375 | 9 | 0.8889 | 30 | 0.6 |
| 11_5151484 | 10 | 0 | 11 | 0.2727 | 8 | 0.125 | 15 | 0.4 | 44 | 0.2273 | 7 | 0.1429 | 5 | 0.4 | 8 | 0 | 11 | 0.5455 | 31 | 0.2903 |
| 11_5151485 | 9 | 0.4444 | 9 | 0.4444 | 7 | 0.2857 | 13 | 0.0769 | 38 | 0.2895 | 7 | 0.1429 | 6 | 0.1667 | 8 | 0.5 | 9 | 0.1111 | 30 | 0.2333 |
| 11_5151869 | NA | NA | 1 | 1 | NA | NA | NA | NA | 1 | 1 | NA | NA | NA | NA | 1 | 1 | NA | NA | 1 | 1 |
| 11_5151877 | NA | NA | 1 | 1 | NA | NA | NA | NA | 1 | 1 | NA | NA | NA | NA | 1 | 1 | NA | NA | 1 | 1 |
| 11_5151896 | NA | NA | 1 | 1 | NA | NA | NA | NA | 1 | 1 | NA | NA | NA | NA | 1 | 1 | NA | NA | 1 | 1 |
| 11_5151905 | NA | NA | 1 | 1 | NA | NA | NA | NA | 1 | 1 | NA | NA | NA | NA | 1 | 1 | NA | NA | 1 | 1 |
| 11_5153396 | 3 | 0 | 3 | 0 | 2 | 0.5 | 3 | 0 | 11 | 0.0909 | NA | NA | 1 | 0 | 1 | 0 | 4 | 0.25 | 6 | 0.1667 |
| 11_5153454 | 1 | 1 | 3 | 1 | 1 | 1 | 4 | 1 | 9 | 1 | 1 | 1 | 1 | 0 | 2 | 0.5 | 4 | 1 | 8 | 0.75 |
| 11_5153473 | 2 | 1 | 4 | 1 | 1 | 1 | 4 | 1 | 11 | 1 | 1 | 1 | 1 | 1 | 1 | 1 | 4 | 1 | 7 | 1 |
| 11_5153474 | 2 | 0 | 5 | 0 | 1 | 0 | 4 | 0.25 | 12 | 0.0833 | 2 | 0 | 1 | 1 | 2 | 0 | 4 | 0 | 9 | 0.1111 |
| 11_5153615 | 1 | 0 | 4 | 0.25 | 1 | 0 | 4 | 0.25 | 10 | 0.2 | 1 | 0 | 1 | 1 | 1 | 0 | 4 | 0.25 | 7 | 0.2857 |
| 11_5153628 | 2 | 1 | 4 | 1 | 1 | 1 | 3 | 1 | 10 | 1 | NA | NA | 1 | 1 | 1 | 1 | 5 | 1 | 7 | 1 |
| 11_5153698 | 1 | 1 | 2 | 0 | NA | NA | 1 | 0 | 4 | 0.25 | 1 | 0 | 1 | 1 | NA | NA | 5 | 0 | 7 | 0.1429 |
| 11_5154774 | 1 | 1 | NA | NA | NA | NA | NA | NA | 1 | 1 | 1 | 1 | 1 | 0 | NA | NA | NA | NA | 2 | 0.5 |
| 11_5154818 | 1 | 1 | NA | NA | NA | NA | NA | NA | 1 | 1 | 1 | 1 | 1 | 1 | NA | NA | NA | NA | 2 | 1 |
| 11_5154834 | 1 | 1 | NA | NA | NA | NA | NA | NA | 1 | 1 | 1 | 1 | 1 | 1 | NA | NA | NA | NA | 2 | 1 |
| 11_5157639 | 5 | 0.2 | 7 | 0 | 10 | 0.3 | 4 | 0 | 26 | 0.1538 | 2 | 0 | 7 | 0.1429 | 8 | 0.25 | 4 | 0 | 21 | 0.1429 |
| 11_5157646 | 7 | 0.1429 | 5 | 0.8 | 5 | 0.2 | 3 | 0 | 20 | 0.3 | 5 | 0 | 5 | 0.4 | 5 | 0.6 | 5 | 0.4 | 20 | 0.35 |
| 11_5157654 | 5 | 1 | 4 | 1 | 7 | 0.7143 | 4 | 1 | 20 | 0.9 | 3 | 0.6667 | 4 | 1 | 7 | 0.8571 | 5 | 0.8 | 19 | 0.8421 |
| 11_5157660 | 8 | 0.375 | 7 | 0.2857 | 8 | 0.375 | 2 | 0 | 25 | 0.32 | 5 | 0 | 4 | 0 | 8 | 0.375 | 6 | 0.1667 | 23 | 0.1739 |
| 11_5157670 | 7 | 0.7143 | 7 | 1 | 6 | 0.8333 | 2 | 1 | 22 | 0.8636 | 3 | 1 | 4 | 0.5 | 7 | 1 | 5 | 0.8 | 19 | 0.8421 |
| 11_5157671 | 5 | 1 | 5 | 0.8 | 7 | 1 | NA | NA | 17 | 0.9412 | 3 | 1 | 2 | 1 | 4 | 1 | 5 | 1 | 14 | 1 |
| 11_5157675 | 7 | 0.4286 | 7 | 0.1429 | 8 | 0.125 | 1 | 0 | 23 | 0.2174 | 4 | 0.75 | 4 | 0.5 | 7 | 0.1429 | 5 | 0 | 20 | 0.3 |
| 11_5158348 | 1 | 0 | 6 | 0.1667 | 2 | 0 | 4 | 0 | 13 | 0.0769 | 3 | 0 | 2 | 0.5 | 2 | 0 | 3 | 0 | 10 | 0.1 |
| 11_5158490 | NA | NA | 5 | 0 | 2 | 0.5 | 2 | 0 | 9 | 0.1111 | 1 | 0 | 1 | 0 | 2 | 0 | 1 | 1 | 5 | 0.2 |
| 11_5163448 | NA | NA | 1 | 0 | 1 | 1 | NA | NA | 2 | 0.5 | NA | NA | 1 | 1 | NA | NA | 1 | 1 | 2 | 1 |
| 11_5163467 | NA | NA | 1 | 1 | 1 | 1 | NA | NA | 2 | 1 | NA | NA | 1 | 1 | NA | NA | NA | NA | 1 | 1 |
| 11_5163472 | NA | NA | 1 | 1 | 1 | 1 | NA | NA | 2 | 1 | NA | NA | 1 | 1 | NA | NA | 1 | 1 | 2 | 1 |
| 11_5168234 | 26 | 0.3462 | 18 | 0.3333 | 14 | 0.4286 | 8 | 0.25 | 66 | 0.3485 | 15 | 0.2667 | 11 | 0.0909 | 16 | 0.25 | 23 | 0.3913 | 65 | 0.2769 |
| 11_5170513 | 6 | 0.1667 | 11 | 0.1818 | 8 | 0.125 | 9 | 0 | 34 | 0.1176 | 7 | 0.2857 | 8 | 0.125 | 9 | 0 | 5 | 0.2 | 29 | 0.1379 |
| 11_5170538 | 4 | 0.75 | 10 | 0.9 | 8 | 0.75 | 8 | 0.5 | 30 | 0.7333 | 5 | 0.6 | 7 | 0.7143 | 8 | 0.875 | 4 | 0.75 | 24 | 0.75 |
| 11_5170567 | 5 | 0.4 | 5 | 0.2 | 4 | 0 | 3 | 0.3333 | 17 | 0.2353 | 4 | 0 | 4 | 0.25 | 6 | 0.1667 | 2 | 0.5 | 16 | 0.1875 |
| 11_5170569 | 5 | 0.4 | 6 | 0.1667 | 6 | 0 | 5 | 0.4 | 22 | 0.2273 | 4 | 0.5 | 5 | 0.4 | 5 | 0.4 | 4 | 0.5 | 18 | 0.4444 |
| 11_5170572 | 4 | 0.75 | 5 | 0.2 | 5 | 0.6 | 4 | 0.25 | 18 | 0.4444 | 2 | 0.5 | 2 | 0 | 2 | 0 | 2 | 0.5 | 8 | 0.25 |
| 11_5170577 | 3 | 0.3333 | 6 | 0.1667 | 7 | 0.1429 | 6 | 0.1667 | 22 | 0.1818 | 4 | 0.25 | 6 | 0.5 | 6 | 0.1667 | 3 | 0.6667 | 19 | 0.3684 |
| 11_5170586 | 2 | 1 | 7 | 1 | 6 | 0.8333 | 5 | 0.8 | 20 | 0.9 | 3 | 1 | 4 | 1 | 4 | 1 | 3 | 1 | 14 | 1 |
| 11_5170593 | 4 | 1 | 8 | 0.875 | 6 | 1 | 6 | 1 | 24 | 0.9583 | 4 | 1 | 3 | 1 | 5 | 1 | 3 | 1 | 15 | 1 |
| 11_5170595 | 5 | 0.6 | 8 | 0.875 | 7 | 0.5714 | 5 | 0.6 | 25 | 0.68 | 4 | 0.75 | 3 | 1 | 5 | 0.6 | 3 | 1 | 15 | 0.8 |
| 11_5170597 | 3 | 0.6667 | 6 | 0.8333 | 6 | 0.5 | 4 | 0.5 | 19 | 0.6316 | 4 | 0.5 | 3 | 1 | 5 | 0.6 | 1 | 0 | 13 | 0.6154 |
| 11_5170605 | 5 | 0.8 | 8 | 0.625 | 7 | 0.5714 | 5 | 0.4 | 25 | 0.6 | 1 | 0 | 4 | 1 | 5 | 0.6 | 3 | 1 | 13 | 0.7692 |
| 11_5170611 | 5 | 1 | 7 | 1 | 4 | 1 | 3 | 1 | 19 | 1 | 4 | 1 | 3 | 1 | 3 | 1 | 3 | 1 | 13 | 1 |
| 11_5170615 | 4 | 0.75 | 3 | 0.3333 | 4 | 1 | 3 | 0.6667 | 14 | 0.7143 | 2 | 1 | 3 | 1 | 4 | 0.75 | 3 | 0.3333 | 12 | 0.75 |
| 11_5170615 | 4 | 1 | 6 | 0 | 5 | 0.4 | 2 | 0.5 | 17 | 0.4118 | NA | NA | 2 | 0 | 4 | 0.5 | 2 | 1 | 8 | 0.5 |
| 11_5170627 | 5 | 1 | 7 | 1 | 4 | 1 | 2 | 1 | 18 | 1 | 4 | 1 | 3 | 1 | 5 | 1 | 3 | 1 | 15 | 1 |
| 11_5170631 | 5 | 0.2 | 5 | 0.2 | 4 | 0.25 | 2 | 0 | 16 | 0.1875 | 3 | 0.6667 | 2 | 1 | 5 | 1 | 3 | 0 | 13 | 0.6923 |
| 11_5170633 | 5 | 0.2 | 7 | 0.1429 | 4 | 0.25 | 3 | 0 | 19 | 0.1579 | 3 | 0.6667 | 2 | 0 | 5 | 0 | 3 | 0.3333 | 13 | 0.2308 |
| 11_5170640 | 5 | 0 | 6 | 0 | 4 | 0.25 | 3 | 0 | 18 | 0.0556 | 3 | 0 | 3 | 0.3333 | 4 | 0 | 3 | 0 | 13 | 0.0769 |
| 11_5170642 | 5 | 0.2 | 6 | 0 | 4 | 0.25 | 3 | 0.3333 | 18 | 0.1667 | 2 | 0 | 3 | 0.3333 | 4 | 0.5 | 1 | 0 | 10 | 0.3 |
| 11_5170701 | 4 | 0.25 | 5 | 0 | 2 | 0.5 | 2 | 0 | 13 | 0.1538 | NA | NA | NA | NA | 1 | 0 | 1 | 1 | 2 | 0.5 |

| | | | | | | | | | | | | | | | | | | | | |
|---|---|---|---|---|---|---|---|---|---|---|---|---|---|---|---|---|---|---|---|---|
| 11_5170739 | 1 | 1 | 4 | 0.75 | 2 | 0 | 1 | 1 | 8 | 0.625 | NA | NA | NA | NA | 1 | 0 | 1 | 1 | 2 | 0.5 |
| 11_5170752 | 1 | 1 | 4 | 0 | 3 | 0 | 1 | 0 | 9 | 0.1111 | NA | NA | NA | NA | 2 | 0 | 1 | 1 | 3 | 0.3333 |
| 11_5170755 | 3 | 0.6667 | 2 | 0 | 3 | 0.3333 | NA | NA | 8 | 0.375 | NA | NA | NA | NA | 2 | 1 | 1 | 1 | 3 | 1 |
| 11_5170779 | 2 | 1 | 4 | 1 | 3 | 1 | NA | NA | 9 | 1 | NA | NA | NA | NA | 2 | 1 | NA | NA | 2 | 1 |
| 11_5170790 | 3 | 1 | 3 | 1 | 2 | 1 | 1 | 1 | 9 | 1 | NA | NA | NA | NA | 2 | 1 | NA | NA | 2 | 1 |
| 11_5170823 | 2 | 0.5 | 2 | 1 | 3 | 0.6667 | NA | NA | 7 | 0.7143 | NA | NA | NA | NA | 1 | 1 | NA | NA | 1 | 1 |
| 11_5186676 | 8 | 0 | 6 | 0 | 5 | 0.2 | 8 | 0.25 | 27 | 0.1111 | 7 | 0.2857 | 2 | 0 | 11 | 0.0909 | 6 | 0.3333 | 26 | 0.1923 |
| 11_5186688 | 10 | 0.1 | 6 | 0 | 5 | 0 | 12 | 0.0833 | 33 | 0.0606 | 7 | 0.1429 | 3 | 0 | 11 | 0.0909 | 6 | 0.1667 | 27 | 0.1111 |
| 11_5186695 | 9 | 0.4444 | 6 | 0.3333 | 5 | 0.8 | 6 | 0.8333 | 26 | 0.5769 | 4 | 0.75 | 2 | 0.5 | 7 | 1 | 5 | 0.4 | 18 | 0.7222 |
| 11_5186719 | 8 | 0.125 | 5 | 0 | 7 | 0 | 11 | 0.1818 | 31 | 0.0968 | 6 | 0.1667 | 4 | 0.25 | 12 | 0 | 6 | 0.3333 | 28 | 0.1429 |
| 11_5186771 | 9 | 0 | 6 | 0.1667 | 6 | 0.1667 | 10 | 0.3 | 31 | 0.1613 | 6 | 0.6667 | 4 | 0 | 10 | 0.4 | 6 | 0.3333 | 26 | 0.3846 |
| 11_5186775 | 7 | 0 | 6 | 0.1667 | 9 | 0.3333 | 10 | 0.2 | 32 | 0.1875 | 6 | 0.5 | 4 | 0.75 | 12 | 0.4167 | 5 | 0.2 | 27 | 0.4444 |
| 11_5186792 | 6 | 0 | 3 | 0.3333 | 4 | 0.25 | 7 | 0.4286 | 20 | 0.25 | 6 | 0 | 3 | 0 | 8 | 0.25 | 6 | 0.1667 | 23 | 0.1304 |
| 11_5186794 | 12 | 0.1667 | 5 | 0.2 | 7 | 0 | 8 | 0 | 32 | 0.0938 | 5 | 0.2 | 4 | 0 | 12 | 0.25 | 5 | 0.2 | 26 | 0.1923 |
| 11_5186797 | 8 | 0 | 6 | 0.1667 | 6 | 0.3333 | 10 | 0.1 | 30 | 0.1333 | 7 | 0 | 5 | 0 | 9 | 0.2222 | 5 | 0.4 | 26 | 0.1538 |
| 11_5186806 | 14 | 0 | 6 | 0 | 9 | 0.1111 | 11 | 0.1818 | 40 | 0.075 | 8 | 0.25 | 6 | 0 | 12 | 0.0833 | 6 | 0 | 32 | 0.0938 |
| 11_5187621 | NA | NA | 3 | 0.3333 | 1 | 0 | 1 | 0 | 5 | 0.2 | 1 | 0 | 1 | 0 | 3 | 0.3333 | 2 | 0 | 7 | 0.1429 |
| 11_5187656 | 10 | 0.1 | 24 | 0.0833 | 23 | 0.2174 | 23 | 0.1739 | 80 | 0.15 | 15 | 0.4 | 21 | 0.1905 | 28 | 0.1786 | 12 | 0.3333 | 76 | 0.25 |
| 11_5187689 | 9 | 0.1111 | 22 | 0.0455 | 21 | 0.1429 | 22 | 0.0455 | 74 | 0.0811 | 17 | 0.0588 | 18 | 0.0556 | 20 | 0.1 | 10 | 0.2 | 65 | 0.0923 |
| 11_5187708 | 9 | 0.1111 | 26 | 0.3462 | 24 | 0.375 | 23 | 0.3913 | 82 | 0.3415 | 16 | 0.25 | 15 | 0.4 | 28 | 0.4286 | 13 | 0.2308 | 72 | 0.3472 |
| 11_5187750 | 19 | 0.1053 | 42 | 0.0476 | 32 | 0.125 | 34 | 0.0294 | 127 | 0.0709 | 19 | 0.0526 | 31 | 0.0323 | 39 | 0.0769 | 23 | 0.1739 | 112 | 0.0804 |
| 11_5191801 | 1 | 0 | NA | NA | 1 | 0 | 4 | 0.25 | 6 | 0.1667 | 1 | 0 | 1 | 1 | NA | NA | 4 | 0.5 | 6 | 0.5 |
| 11_5198816 | 1 | 1 | NA | NA | 1 | 1 | NA | NA | 2 | 1 | NA | NA | NA | NA | NA | NA | 1 | 1 | 1 | 1 |
| 11_5198824 | 1 | 1 | NA | NA | 2 | 1 | NA | NA | 3 | 1 | NA | NA | NA | NA | NA | NA | 1 | 1 | 1 | 1 |
| 11_5198839 | NA | NA | NA | NA | 1 | 1 | NA | NA | 1 | 1 | NA | NA | NA | NA | NA | NA | 1 | 1 | 1 | 1 |
| 11_5198870 | NA | NA | NA | NA | 2 | 0.5 | NA | NA | 2 | 0.5 | NA | NA | NA | NA | NA | NA | 1 | 1 | 1 | 1 |
| 11_5200404 | 35 | 0.2571 | 32 | 0.1562 | 40 | 0.15 | 30 | 0.1667 | 137 | 0.1825 | 18 | 0.3333 | 25 | 0.12 | 44 | 0.1591 | 28 | 0.2143 | 115 | 0.1913 |
| 11_5201283 | 42 | 0.119 | 45 | 0.1556 | 50 | 0.2 | 38 | 0.1842 | 175 | 0.1657 | 28 | 0.1429 | 29 | 0.2759 | 53 | 0.1887 | 40 | 0.15 | 150 | 0.1867 |
| 11_5201290 | 47 | 0.234 | 45 | 0.3111 | 52 | 0.3269 | 39 | 0.3333 | 183 | 0.3005 | 29 | 0.2414 | 30 | 0.3667 | 59 | 0.3051 | 38 | 0.2895 | 156 | 0.3013 |
| 11_5201292 | 50 | 0.58 | 43 | 0.5814 | 60 | 0.5167 | 42 | 0.5238 | 195 | 0.5487 | 31 | 0.4839 | 34 | 0.5 | 59 | 0.4915 | 40 | 0.7 | 164 | 0.5427 |
| 11_5201308 | 52 | 0.5385 | 45 | 0.4667 | 60 | 0.55 | 44 | 0.5227 | 201 | 0.5224 | 30 | 0.7 | 35 | 0.6 | 59 | 0.5424 | 43 | 0.6279 | 167 | 0.6048 |
| 11_5201319 | 46 | 0.1087 | 45 | 0.3556 | 55 | 0.2545 | 39 | 0.1026 | 185 | 0.2108 | 30 | 0.2333 | 28 | 0.1786 | 58 | 0.2414 | 43 | 0.3488 | 159 | 0.2579 |
| 11_5201336 | 50 | 0.28 | 46 | 0.2174 | 55 | 0.3091 | 39 | 0.1795 | 190 | 0.2526 | 33 | 0.2121 | 35 | 0.3143 | 61 | 0.2131 | 39 | 0.1282 | 168 | 0.2143 |
| 11_5201343 | 53 | 0.1132 | 43 | 0.186 | 47 | 0.1277 | 42 | 0.0952 | 185 | 0.1297 | 33 | 0.2424 | 33 | 0.1818 | 56 | 0.1429 | 36 | 0.1389 | 158 | 0.1709 |
| 11_5201360 | 12 | 0.25 | 10 | 0.2 | 11 | 0.0909 | 7 | 0.1429 | 40 | 0.175 | 5 | 0 | 6 | 0.1667 | 6 | 0.1667 | 10 | 0.3 | 27 | 0.1852 |
| 11_5201368 | 10 | 0.1 | 9 | 0.2222 | 10 | 0.3 | 9 | 0 | 38 | 0.1579 | 4 | 0 | 7 | 0 | 8 | 0.125 | 11 | 0.2727 | 30 | 0.1333 |
| 11_5201383 | 9 | 0.6667 | 9 | 0.5556 | 7 | 0.7143 | 8 | 0.5 | 33 | 0.6061 | 4 | 0.25 | 9 | 0.4444 | 7 | 0.5714 | 10 | 0.7 | 30 | 0.5333 |
| 11_5201412 | 10 | 0.1 | 11 | 0.4545 | 12 | 0.0833 | 8 | 0.125 | 41 | 0.1951 | 4 | 0.5 | 10 | 0.1 | 7 | 0 | 10 | 0.1 | 31 | 0.129 |
| 11_5201820 | NA | NA | 3 | 0.3333 | 1 | 0 | 3 | 0.6667 | 7 | 0.4286 | 1 | 1 | 1 | 1 | NA | NA | 1 | 0 | 3 | 0.6667 |
| 11_5213705 | 93 | 0.1935 | 81 | 0.2099 | 62 | 0.2258 | 85 | 0.1647 | 321 | 0.1963 | 66 | 0.2273 | 73 | 0.2192 | 65 | 0.1846 | 74 | 0.2838 | 278 | 0.2302 |
| 11_5227933 | 94 | 0.0851 | 82 | 0.1463 | 79 | 0.0759 | 69 | 0.1594 | 324 | 0.1142 | 55 | 0.1091 | 84 | 0.131 | 87 | 0.1494 | 68 | 0.2059 | 294 | 0.1497 |
| 11_5229407 | 8 | 0.125 | 10 | 0.1 | 6 | 0 | 11 | 0.1818 | 35 | 0.1143 | 4 | 0 | 11 | 0 | 6 | 0.3333 | 9 | 0.2222 | 30 | 0.1333 |
| 11_5229413 | 9 | 0.4444 | 10 | 0 | 7 | 0.2857 | 12 | 0.25 | 38 | 0.2368 | 5 | 0 | 10 | 0.2 | 7 | 0.4286 | 9 | 0.1111 | 31 | 0.1935 |
| 11_5229414 | 8 | 0.5 | 10 | 0.4 | 7 | 0.2857 | 12 | 0.3333 | 37 | 0.3784 | 4 | 0.25 | 10 | 0.1 | 7 | 0.2857 | 9 | 0.3333 | 30 | 0.2333 |
| 11_5229417 | 8 | 0.125 | 10 | 0.2 | 4 | 0.25 | 12 | 0.1667 | 34 | 0.1765 | 3 | 0 | 8 | 0.125 | 7 | 0.2857 | 9 | 0.2222 | 27 | 0.1852 |
| 11_5229426 | 9 | 0.7778 | 11 | 0.9091 | 7 | 0.8571 | 12 | 0.9167 | 39 | 0.8718 | 5 | 0.8 | 11 | 0.9091 | 7 | 1 | 7 | 1 | 30 | 0.9333 |
| 11_5229782 | 10 | 0.1 | 11 | 0.0909 | 8 | 0.125 | 12 | 0 | 41 | 0.0732 | 6 | 0.3333 | 11 | 0 | 6 | 0.1667 | 9 | 0.1111 | 32 | 0.125 |
| 11_5229784 | 10 | 0.3 | 12 | 0.1667 | 9 | 0.2222 | 12 | 0 | 43 | 0.1628 | 6 | 0 | 11 | 0.0909 | 7 | 0 | 10 | 0.1 | 34 | 0.0588 |
| 11_5229791 | 9 | 0.3333 | 10 | 0.5 | 9 | 0.4444 | 12 | 0.0833 | 40 | 0.325 | 4 | 0.5 | 11 | 0.1818 | 7 | 0.1429 | 9 | 0.2222 | 31 | 0.2258 |
| 11_5229941 | 10 | 0.3 | 14 | 0.0714 | 10 | 0 | 17 | 0 | 51 | 0.0784 | 9 | 0.1111 | 12 | 0 | 8 | 0.25 | 13 | 0 | 42 | 0.0714 |
| 11_5229946 | 12 | 0.5 | 13 | 0.1538 | 9 | 0.2222 | 16 | 0.3125 | 50 | 0.3 | 7 | 0 | 10 | 0.2 | 9 | 0.1111 | 14 | 0.2143 | 40 | 0.15 |
| 11_5307712 | 1 | 1 | 1 | 1 | NA | NA | 2 | 0 | 4 | 0.5 | 2 | 1 | 2 | 0 | 1 | 1 | NA | NA | 5 | 0.6 |
| 11_5307722 | 1 | 0 | 1 | 1 | NA | NA | 2 | 0 | 4 | 0.25 | 2 | 0 | 2 | 0.5 | 1 | 0 | 1 | 0 | 6 | 0.1667 |
| 11_5307726 | 1 | 0 | 1 | 0 | 1 | 0 | 2 | 1 | 5 | 0.4 | 2 | 0.5 | 2 | 1 | 1 | 1 | NA | NA | 5 | 0.8 |
| 11_5307727 | 1 | 1 | 1 | 1 | NA | NA | 2 | 1 | 4 | 1 | 1 | 1 | 2 | 0.5 | NA | NA | 1 | 1 | 4 | 0.75 |
| 11_5307756 | 1 | 1 | 1 | 1 | 1 | 0 | 2 | 1 | 5 | 0.8 | 2 | 0.5 | 1 | 1 | 1 | 1 | 1 | 1 | 5 | 0.8 |
| 11_5307763 | 1 | 0 | 1 | 0 | 1 | 0 | 1 | 1 | 4 | 0.25 | 2 | 0.5 | 2 | 0.5 | 2 | 0 | 2 | 1 | 8 | 0.5 |
| 11_5307764 | 1 | 1 | 1 | 1 | NA | NA | 1 | 1 | 3 | 1 | 1 | 1 | 1 | 1 | 2 | 1 | 1 | 1 | 5 | 1 |
| 11_5307783 | 1 | 0 | 1 | 1 | 1 | 1 | 1 | 0 | 4 | 0.5 | 2 | 0.5 | 2 | 1 | 2 | 0.5 | 1 | 1 | 7 | 0.7143 |
| 11_5307810 | NA | NA | 1 | 1 | NA | NA | NA | NA | 1 | 1 | 1 | 0 | 1 | 1 | 2 | 0 | 1 | 0 | 5 | 0.2 |
| 11_5307834 | NA | NA | 1 | 1 | NA | NA | NA | NA | 1 | 1 | 2 | 0 | 2 | 0.5 | 2 | 0.5 | 1 | 1 | 7 | 0.4286 |
| 11_5307924 | 1 | 1 | 1 | 0 | 2 | 0 | 2 | 0 | 6 | 0.1667 | 2 | 0.5 | 1 | 1 | 2 | 0.5 | 1 | 1 | 6 | 0.6667 |
| 11_5308087 | 2 | 1 | 5 | 0.4 | 3 | 0.6667 | 2 | 0 | 12 | 0.5 | 3 | 0.6667 | 1 | 0 | 3 | 0 | 1 | 0 | 8 | 0.25 |
| 11_5308096 | 1 | 1 | 4 | 1 | NA | NA | 1 | 0 | 6 | 0.8333 | 2 | 1 | 1 | 1 | 1 | 1 | NA | NA | 4 | 1 |
| 11_5308098 | 2 | 1 | 5 | 1 | 3 | 1 | 2 | 1 | 12 | 1 | 3 | 1 | 1 | 1 | 3 | 1 | 1 | 1 | 8 | 1 |

| | | | | | | | | | | | | | | | | | | | | |
|---|---|---|---|---|---|---|---|---|---|---|---|---|---|---|---|---|---|---|---|---|
| 11_530812 | 3 | 0.6667 | 6 | 0.1667 | 3 | 0.3333 | 2 | 0.5 | 14 | 0.3571 | 3 | 1 | 1 | 0 | 4 | 1 | 2 | 1 | 10 | 0.9 |
| 11_530813 | 3 | 0 | 6 | 0 | 3 | 0 | 3 | 0.3333 | 15 | 0.0667 | 3 | 0.3333 | 1 | 0 | 3 | 0 | 2 | 0 | 9 | 0.1111 |
| 11_530814 | 3 | 1 | 7 | 0.8571 | 4 | 0.75 | 3 | 0.6667 | 17 | 0.8235 | 3 | 1 | 2 | 1 | 4 | 0.75 | 2 | 1 | 11 | 0.9091 |
| 11_530814 | 4 | 0 | 7 | 0 | 3 | 0 | 3 | 0.3333 | 17 | 0.0588 | 3 | 0.3333 | 2 | 0.5 | 4 | 0.25 | 2 | 0 | 11 | 0.2727 |
| 11_530814 | 4 | 0.5 | 7 | 0.4286 | 3 | 0.6667 | 3 | 1 | 17 | 0.5882 | 2 | 0.5 | 2 | 1 | 4 | 0.25 | 1 | 0 | 9 | 0.4444 |
| 11_530817 | 5 | 0.6 | 7 | 0.4286 | 5 | 0.2 | 3 | 0 | 20 | 0.35 | 3 | 0 | 4 | 0.5 | 4 | 0.5 | 3 | 0.3333 | 14 | 0.3571 |
| 11_530817 | 4 | 0.25 | 7 | 0.1429 | 3 | 0.3333 | 3 | 0.3333 | 17 | 0.2353 | 3 | 0 | 4 | 0.5 | 4 | 0 | 3 | 0.3333 | 14 | 0.2143 |
| 11_530817 | 4 | 0.25 | 3 | 0 | 1 | 0 | NA | NA | 8 | 0.125 | 1 | 0 | 1 | 1 | 4 | 0.25 | NA | NA | 6 | 0.3333 |
| 11_530818 | 5 | 1 | 7 | 1 | 3 | 1 | 3 | 1 | 18 | 1 | 3 | 1 | 3 | 1 | 4 | 1 | 3 | 1 | 13 | 1 |
| 11_530819 | 4 | 0.5 | 8 | 0.25 | 3 | 0.6667 | 2 | 0.5 | 17 | 0.4118 | 2 | 1 | 4 | 0.25 | 5 | 0.4 | 1 | 0 | 12 | 0.4167 |
| 11_530819 | 4 | 1 | 7 | 1 | 3 | 1 | 3 | 1 | 17 | 1 | 3 | 1 | 4 | 1 | 5 | 1 | 3 | 1 | 15 | 1 |
| 11_530819 | 4 | 0.75 | 6 | 0.8333 | 3 | 0.6667 | 2 | 0 | 15 | 0.6667 | 2 | 1 | 3 | 1 | 4 | 1 | 2 | 1 | 11 | 1 |
| 11_530821 | 4 | 1 | 7 | 1 | 3 | 1 | 3 | 1 | 17 | 1 | 3 | 1 | 4 | 1 | 5 | 1 | 3 | 1 | 15 | 1 |
| 11_530822 | 4 | 0 | 5 | 0 | 2 | 0.5 | 3 | 0 | 14 | 0.0714 | 3 | 0.3333 | 2 | 0 | 4 | 0 | 3 | 0.3333 | 12 | 0.1667 |
| 11_530824 | 4 | 1 | 7 | 1 | 2 | 0.5 | 3 | 1 | 16 | 0.9375 | 4 | 1 | 3 | 0.6667 | 5 | 1 | 2 | 1 | 14 | 0.9286 |
| 11_530831 | 8 | 0.75 | 8 | 0.375 | 6 | 0.5 | 6 | 0.1667 | 28 | 0.4643 | 7 | 0.2857 | 8 | 0.375 | 6 | 0.1667 | 4 | 0.5 | 25 | 0.32 |
| 11_531509 | 13 | 0 | 13 | 0.0769 | 9 | 0 | 18 | 0.1111 | 53 | 0.0566 | 10 | 0 | 13 | 0.0769 | 6 | 0.1667 | 4 | 0.5 | 33 | 0.1212 |
| 11_531509 | 15 | 0.0667 | 14 | 0.1429 | 8 | 0.125 | 16 | 0.1875 | 53 | 0.1321 | 9 | 0 | 14 | 0.3571 | 7 | 0.2857 | 5 | 0.2 | 35 | 0.2286 |
| 11_531511 | 12 | 0.4167 | 15 | 0.4 | 8 | 0.25 | 14 | 0.2857 | 49 | 0.3469 | 2 | 0.5 | 10 | 0.6 | 5 | 0.6 | 4 | 0.5 | 21 | 0.5714 |
| 11_531523 | 7 | 0.1429 | 8 | 0.25 | 4 | 0.25 | 11 | 0.0909 | 30 | 0.1667 | 4 | 0.25 | 5 | 0.2 | 6 | 0.1667 | 2 | 0 | 17 | 0.1765 |
| 11_531524 | 10 | 0.3 | 16 | 0.4375 | 7 | 0.5714 | 19 | 0.4211 | 52 | 0.4231 | 13 | 0.3846 | 12 | 0.25 | 10 | 0.3 | 14 | 0.2857 | 49 | 0.3061 |
| 11_531524 | 11 | 0.8182 | 17 | 0.8824 | 8 | 0.875 | 19 | 0.8421 | 55 | 0.8545 | 14 | 0.7143 | 14 | 0.7857 | 9 | 0.6667 | 15 | 0.8667 | 52 | 0.7692 |
| 11_531525 | 11 | 0.2727 | 16 | 0.1875 | 7 | 0 | 19 | 0.1579 | 53 | 0.1698 | 14 | 0.0714 | 13 | 0.0769 | 10 | 0.2 | 14 | 0.0714 | 51 | 0.098 |
| 11_532233 | 2 | 0.5 | NA | NA | 1 | 0 | 1 | 0 | 4 | 0.25 | 2 | 0 | 3 | 0.3333 | 4 | 0.5 | 2 | 0 | 11 | 0.2727 |
| 11_532234 | 2 | 0.5 | NA | NA | 1 | 0 | 1 | 0 | 4 | 0.25 | 2 | 0 | 3 | 0.6667 | 3 | 0 | 2 | 0 | 10 | 0.2 |
| 11_532234 | 2 | 0 | NA | NA | 1 | 0 | 1 | 1 | 4 | 0.25 | 2 | 0 | 3 | 0 | 2 | 0.5 | 2 | 1 | 9 | 0.3333 |
| 11_532235 | 1 | 1 | NA | NA | 1 | 1 | 1 | 0 | 3 | 0.6667 | 1 | 0 | 1 | 1 | 2 | 1 | 2 | 1 | 6 | 0.8333 |
| 11_532236 | 2 | 0.5 | 1 | 0 | 1 | 0 | 1 | 0 | 5 | 0.2 | 2 | 0 | 3 | 0.3333 | 2 | 0.5 | 1 | 0 | 8 | 0.25 |
| 11_532237 | 2 | 1 | 1 | 0 | 1 | 0 | 1 | 0 | 5 | 0.4 | 2 | 0 | 3 | 0.3333 | 4 | 0.75 | 1 | 0 | 10 | 0.4 |
| 11_532633 | 1 | 1 | NA | NA | 3 | 1 | 2 | 0.5 | 6 | 0.8333 | 2 | 1 | 2 | 1 | 1 | 1 | 1 | 1 | 6 | 1 |
| 11_532633 | 1 | 0 | NA | NA | 3 | 0.6667 | 2 | 0.5 | 6 | 0.5 | 2 | 0.5 | 3 | 0.3333 | 1 | 1 | 1 | 1 | 7 | 0.5714 |
| 11_532634 | 1 | 0 | NA | NA | 3 | 0.3333 | 2 | 0 | 6 | 0.1667 | 2 | 0 | 3 | 0 | 1 | 0 | 1 | 1 | 7 | 0.1429 |
| 11_532636 | 1 | 0 | NA | NA | 3 | 0 | 2 | 0.5 | 6 | 0.1667 | 2 | 1 | 3 | 0 | 1 | 0 | 1 | 0 | 7 | 0.2857 |
| 11_532636 | 1 | 0 | NA | NA | 3 | 0.6667 | 1 | 1 | 5 | 0.6 | 2 | 1 | 2 | 0.5 | 1 | 1 | 1 | 1 | 6 | 0.8333 |
| 11_532637 | 1 | 0 | NA | NA | 3 | 0.6667 | 2 | 0.5 | 6 | 0.5 | 2 | 1 | 2 | 1 | 1 | 1 | 1 | 1 | 6 | 1 |
| 11_532905 | NA | NA | NA | NA | 2 | 0.5 | 1 | 0 | 3 | 0.3333 | 1 | 1 | 2 | 0 | 1 | 0 | 2 | 0 | 6 | 0.1667 |
| 11_532906 | NA | NA | 2 | 0.5 | 2 | 0.5 | 2 | 0.5 | 6 | 0.5 | 2 | 1 | 2 | 0 | 1 | 1 | 2 | 1 | 7 | 0.7143 |
| 11_532907 | 1 | 0 | 2 | 0 | 4 | 0.25 | 2 | 0 | 9 | 0.1111 | 2 | 0 | 3 | 0 | 1 | 1 | 3 | 0.3333 | 9 | 0.2222 |
| 11_532907 | 2 | 0 | 2 | 0.5 | 4 | 0 | 2 | 0 | 10 | 0.1 | 2 | 0.5 | 3 | 0 | 1 | 0 | 3 | 0.3333 | 9 | 0.2222 |
| 11_532909 | 1 | 1 | 2 | 0 | 4 | 0.5 | 2 | 1 | 9 | 0.5556 | 2 | 0.5 | 3 | 1 | 2 | 0.5 | 4 | 0.5 | 11 | 0.6364 |
| 11_532910 | 2 | 0.5 | 2 | 0 | 3 | 0.3333 | 2 | 0 | 9 | 0.2222 | 2 | 0.5 | 3 | 0.3333 | 1 | 1 | 4 | 0.75 | 10 | 0.6 |
| 11_532910 | 2 | 0 | 2 | 0.5 | 5 | 0.6 | 2 | 1 | 11 | 0.5455 | 2 | 0.5 | 1 | 0 | 2 | 1 | 4 | 0.25 | 9 | 0.4444 |
| 11_532952 | 2 | 0 | 2 | 0 | 4 | 0.25 | 3 | 0 | 11 | 0.0909 | 1 | 0 | 3 | 0 | 2 | 0.5 | 4 | 0 | 10 | 0.1 |
| 11_533063 | 2 | 0 | 3 | 1 | 5 | 0.8 | 4 | 0.75 | 14 | 0.7143 | 2 | 0.5 | 4 | 0.75 | 3 | 1 | 4 | 0.75 | 13 | 0.7692 |
| 11_533063 | 1 | 1 | 2 | 1 | 4 | 0.75 | 3 | 1 | 10 | 0.9 | 3 | 1 | 3 | 1 | 3 | 1 | 2 | 1 | 11 | 1 |
| 11_533068 | 2 | 0.5 | 3 | 0.3333 | 6 | 0.1667 | 4 | 0.25 | 15 | 0.2667 | 2 | 0 | 5 | 0.2 | 3 | 0.3333 | 4 | 0 | 14 | 0.1429 |
| 11_533071 | 3 | 0 | 3 | 0 | 7 | 0.7143 | 4 | 0.25 | 17 | 0.3529 | 3 | 0 | 4 | 0.5 | 3 | 0.6667 | 4 | 0 | 14 | 0.2857 |
| 11_533106 | 14 | 0.0714 | 13 | 0.2308 | 17 | 0.2941 | 16 | 0.1875 | 60 | 0.2 | 10 | 0.2 | 11 | 0.0909 | 14 | 0.4286 | 11 | 0.2727 | 46 | 0.2609 |
| 11_533110 | 10 | 0.3 | 14 | 0.1429 | 22 | 0.1818 | 21 | 0.1905 | 67 | 0.194 | 10 | 0.2 | 17 | 0.1765 | 16 | 0.25 | 12 | 0.25 | 55 | 0.2182 |
| 11_533197 | 61 | 0.6721 | 42 | 0.5476 | 37 | 0.6486 | 47 | 0.5532 | 187 | 0.6096 | 47 | 0.4468 | 53 | 0.5472 | 46 | 0.4348 | 44 | 0.6136 | 190 | 0.5105 |
| 11_533548 | NA | NA | NA | NA | NA | NA | 2 | 1 | 2 | 1 | NA | NA | NA | NA | 1 | 1 | NA | NA | 1 | 1 |
| 11_533548 | NA | NA | NA | NA | NA | NA | 2 | 0.5 | 2 | 0.5 | NA | NA | NA | NA | 1 | 1 | NA | NA | 1 | 1 |
| 11_533566 | NA | NA | NA | NA | 1 | 1 | 2 | 0.5 | 3 | 0.6667 | NA | NA | NA | NA | 1 | 1 | NA | NA | 1 | 1 |
| 11_533566 | NA | NA | NA | NA | 1 | 1 | 2 | 0 | 3 | 0.3333 | NA | NA | NA | NA | 1 | 1 | NA | NA | 1 | 1 |
| 11_533578 | NA | NA | 1 | 0 | 1 | 0 | 2 | 1 | 4 | 0.5 | NA | NA | 1 | 1 | 1 | 1 | NA | NA | 2 | 1 |
| 11_533676 | 24 | 0.2083 | 13 | 0.3077 | 16 | 0.25 | 33 | 0.1515 | 86 | 0.2093 | 22 | 0.0909 | 36 | 0.2778 | 29 | 0.1379 | 27 | 0.2222 | 114 | 0.193 |
| 11_533678 | 12 | 0.1667 | 8 | 0.125 | 12 | 0.1667 | 19 | 0.3158 | 51 | 0.2157 | 13 | 0.0769 | 16 | 0.0625 | 14 | 0.0714 | 18 | 0.2222 | 61 | 0.1148 |
| 11_533678 | 9 | 0.2222 | 7 | 0.2857 | 2 | 0 | 7 | 0 | 25 | 0.16 | 5 | 0 | 10 | 0.1 | 8 | 0.375 | 6 | 0.5 | 29 | 0.2414 |
| 11_533681 | 19 | 0.1579 | 15 | 0.2 | 18 | 0.0556 | 30 | 0.0667 | 82 | 0.1098 | 19 | 0.0526 | 35 | 0.0286 | 29 | 0.0345 | 27 | 0.1852 | 110 | 0.0727 |
| 11_533682 | 19 | 0.1579 | 15 | 0.1333 | 19 | 0.2105 | 29 | 0.2414 | 82 | 0.1951 | 20 | 0.15 | 32 | 0.2188 | 28 | 0.2857 | 24 | 0.2083 | 104 | 0.2212 |
| 11_533686 | 17 | 0.1176 | 16 | 0 | 18 | 0.1111 | 28 | 0.0357 | 79 | 0.0633 | 20 | 0 | 35 | 0.0571 | 24 | 0.0417 | 22 | 0.1364 | 101 | 0.0594 |
| 11_533687 | 15 | 0.7333 | 14 | 0.8571 | 19 | 0.7368 | 25 | 0.64 | 73 | 0.726 | 18 | 0.8333 | 30 | 0.8 | 22 | 0.8182 | 18 | 0.7778 | 88 | 0.8068 |
| 11_533688 | 16 | 0.75 | 14 | 0.6429 | 12 | 0.6667 | 24 | 0.8333 | 66 | 0.7424 | 14 | 0.5714 | 29 | 0.7931 | 19 | 0.6316 | 17 | 0.7647 | 79 | 0.7089 |
| 11_533691 | 14 | 1 | 13 | 1 | 15 | 1 | 19 | 0.9474 | 61 | 0.9836 | 15 | 1 | 28 | 0.9286 | 17 | 1 | 16 | 1 | 76 | 0.9737 |
| 11_533691 | 12 | 0.4167 | 12 | 0.25 | 14 | 0.4286 | 22 | 0.5909 | 60 | 0.45 | 17 | 0.1765 | 27 | 0.2963 | 17 | 0.5294 | 19 | 0.2632 | 80 | 0.3125 |

| | | | | | | | | | | | | | | | | | | | | |
|---|---|---|---|---|---|---|---|---|---|---|---|---|---|---|---|---|---|---|---|---|
| 11_533692( | 13 | 0.6923 | 11 | 0.5455 | 13 | 0.5385 | 21 | 0.2857 | 58 | 0.4828 | 16 | 0.3125 | 24 | 0.4167 | 17 | 0.5882 | 17 | 0.2941 | 74 | 0.4054 |
| 11_5336928 | 11 | 0.5455 | 13 | 0.5385 | 16 | 0.5625 | 20 | 0.3 | 60 | 0.4667 | 19 | 0.3158 | 27 | 0.5926 | 19 | 0.4737 | 16 | 0.625 | 81 | 0.5062 |
| 11_533694' | 14 | 0.3571 | 11 | 0.3636 | 14 | 0.0714 | 23 | 0.1304 | 62 | 0.2097 | 18 | 0.1667 | 29 | 0.2759 | 19 | 0.1053 | 16 | 0.0625 | 82 | 0.1707 |
| 11_533697: | 11 | 0.0909 | 10 | 0.5 | 15 | 0.4 | 19 | 0.2105 | 55 | 0.2909 | 17 | 0.2941 | 24 | 0.2083 | 17 | 0.2941 | 14 | 0.3571 | 72 | 0.2778 |
| 11_533698( | 8 | 0.375 | 11 | 0.1818 | 12 | 0.0833 | 21 | 0.3333 | 52 | 0.25 | 14 | 0.2143 | 24 | 0.0417 | 16 | 0.3125 | 15 | 0.0667 | 69 | 0.1449 |
| 11_533699: | 11 | 0.4545 | 10 | 0.1 | 13 | 0.3077 | 20 | 0.25 | 54 | 0.2778 | 16 | 0.1875 | 23 | 0.3043 | 18 | 0.6111 | 14 | 0.2857 | 71 | 0.3521 |
| 11_5336994 | 9 | 0.7778 | 9 | 0.7778 | 10 | 0.7 | 20 | 0.7 | 48 | 0.7292 | 13 | 0.4615 | 23 | 0.6522 | 16 | 0.9375 | 12 | 0.75 | 64 | 0.7031 |
| 11_533702( | NA | NA | NA | NA | 1 | 1 | NA | NA | 1 | 1 | 2 | 0.5 | NA | NA | 1 | 1 | NA | NA | 3 | 0.6667 |
| 11_533702! | 8 | 0.25 | 10 | 0 | 12 | 0.1667 | 18 | 0 | 48 | 0.0833 | 13 | 0.0769 | 22 | 0 | 19 | 0.1053 | 12 | 0.25 | 66 | 0.0909 |
| 11_533703: | 8 | 0.375 | 11 | 0 | 10 | 0.2 | 17 | 0.3529 | 46 | 0.2391 | 13 | 0.3077 | 24 | 0.125 | 17 | 0.2353 | 13 | 0.3846 | 67 | 0.2388 |
| 11_533703! | 7 | 0.1429 | 11 | 0.1818 | 10 | 0.3 | 16 | 0.125 | 44 | 0.1818 | 13 | 0.1538 | 21 | 0.2857 | 16 | 0.125 | 9 | 0.1111 | 59 | 0.1864 |
| 11_533704( | 8 | 0.125 | 10 | 0.2 | 12 | 0.1667 | 15 | 0.1333 | 45 | 0.1556 | 13 | 0.1538 | 20 | 0.2 | 16 | 0.25 | 12 | 0.3333 | 61 | 0.2295 |
| 11_5337054 | NA | NA | NA | NA | 1 | 1 | NA | NA | 1 | 1 | 2 | 1 | NA | NA | NA | NA | NA | NA | 2 | 1 |
| 11_533707! | NA | NA | NA | NA | 2 | 1 | NA | NA | 2 | 1 | 2 | 1 | NA | NA | 1 | 1 | 1 | 1 | 4 | 1 |
| 11_533708( | NA | NA | NA | NA | 2 | 0.5 | NA | NA | 2 | 0.5 | 2 | 0.5 | NA | NA | NA | NA | 1 | 0 | 3 | 0.3333 |
| 11_533708' | NA | NA | NA | NA | 1 | 1 | NA | NA | 1 | 1 | 2 | 1 | NA | NA | 1 | 1 | NA | NA | 3 | 1 |
| 11_5337084 | 7 | 0.1429 | 6 | 0 | 8 | 0.125 | 10 | 0 | 31 | 0.0645 | 9 | 0 | 18 | 0.0556 | 14 | 0 | 13 | 0.1538 | 54 | 0.0556 |
| 11_533708( | NA | NA | NA | NA | 1 | 1 | NA | NA | 1 | 1 | 2 | 1 | NA | NA | 1 | 1 | 1 | 1 | 4 | 1 |
| 11_533740! | NA | NA | 1 | 0 | 1 | 1 | 1 | 0 | 3 | 0.3333 | 1 | 0 | 1 | 1 | NA | NA | NA | NA | 2 | 0.5 |
| 11_533742: | NA | NA | 1 | 1 | 1 | 0 | 1 | 1 | 3 | 0.6667 | NA | NA | 1 | 1 | NA | NA | NA | NA | 1 | 1 |
| 11_533742! | NA | NA | 1 | 1 | 1 | 1 | 1 | 1 | 3 | 1 | NA | NA | 1 | 1 | NA | NA | NA | NA | 1 | 1 |
| 11_534320: | 1 | 0 | 5 | 0.2 | 4 | 0.5 | 2 | 0.5 | 12 | 0.3333 | 2 | 0 | 1 | 0 | 2 | 0.5 | NA | NA | 5 | 0.2 |
| 11_5343214 | 1 | 1 | 5 | 0.6 | 3 | 0.6667 | 2 | 0.5 | 11 | 0.6364 | 1 | 1 | 1 | 0 | 2 | 0 | NA | NA | 4 | 0.25 |
| 11_5343274 | 2 | 0.5 | 1 | 0 | 2 | 0 | 2 | 0 | 7 | 0.1429 | 1 | 0 | 1 | 0 | 2 | 0.5 | 1 | 0 | 5 | 0.2 |
| 11_534327! | 2 | 0.5 | 2 | 0 | 2 | 0 | 2 | 0.5 | 8 | 0.25 | 1 | 1 | NA | NA | 1 | 0 | 1 | 0 | 3 | 0.3333 |
| 11_534330( | 2 | 0.5 | 2 | 0.5 | 3 | 0.3333 | 2 | 0.5 | 9 | 0.4444 | 1 | 0 | 1 | 1 | 2 | 0 | 1 | 0 | 5 | 0.2 |
| 11_534337! | 3 | 0.3333 | 2 | 0.5 | 4 | 0.25 | 3 | 0.3333 | 12 | 0.3333 | 1 | 0 | 1 | 1 | 2 | 0 | 2 | 0.5 | 6 | 0.3333 |
| 11_534337( | 3 | 0.3333 | 2 | 0.5 | 3 | 1 | 2 | 1 | 10 | 0.7 | 1 | 1 | 1 | 0 | 1 | 0 | 1 | 1 | 4 | 0.5 |
| 11_534339( | 2 | 0 | 3 | 0.3333 | 4 | 0.25 | 3 | 0 | 12 | 0.1667 | 1 | 0 | 1 | 1 | 1 | 0 | 2 | 0.5 | 5 | 0.4 |
| 11_534342( | 3 | 0 | 4 | 0 | 3 | 0 | 4 | 0.25 | 14 | 0.0714 | 1 | 0 | 1 | 0 | 1 | 1 | 1 | 0 | 4 | 0.25 |
| 11_534345' | 2 | 0 | 5 | 0 | 4 | 0.25 | 2 | 0.5 | 13 | 0.1538 | NA | NA | 1 | 1 | 1 | 0 | 2 | 0 | 4 | 0.25 |
| 11_534345( | 2 | 0 | 6 | 0.1667 | 4 | 0.5 | 2 | 0 | 14 | 0.2143 | 2 | 0 | 1 | 0 | 2 | 0 | 1 | 1 | 6 | 0.1667 |
| 11_534346( | 3 | 0 | 6 | 0.1667 | 2 | 0 | 4 | 0.25 | 15 | 0.1333 | 2 | 1 | 1 | 1 | 2 | 0 | 2 | 0 | 7 | 0.4286 |
| 11_534348! | 3 | 0 | 7 | 0.2857 | 3 | 0 | 2 | 0.5 | 15 | 0.2 | 2 | 0 | 2 | 0.5 | 2 | 0 | 2 | 0.5 | 8 | 0.25 |
| 11_534392( | 15 | 0 | 14 | 0.0714 | 13 | 0.2308 | 10 | 0 | 52 | 0.0769 | 7 | 0 | 6 | 0 | 7 | 0.2857 | 25 | 0.12 | 45 | 0.1111 |
| 11_534393: | 16 | 0.25 | 21 | 0.1429 | 12 | 0.1667 | 15 | 0.0667 | 64 | 0.1562 | 8 | 0.25 | 7 | 0 | 10 | 0.1 | 30 | 0.2 | 55 | 0.1636 |
| 11_534393( | 12 | 0.1667 | 21 | 0.1429 | 14 | 0 | 12 | 0 | 59 | 0.0847 | 11 | 0.1818 | 8 | 0.125 | 9 | 0.1111 | 32 | 0.0938 | 60 | 0.1167 |
| 11_534394! | 17 | 0.4706 | 20 | 0.6 | 14 | 0.2143 | 13 | 0.6154 | 64 | 0.4844 | 9 | 0.6667 | 8 | 0.25 | 12 | 0.0833 | 32 | 0.375 | 61 | 0.3443 |
| 11_534395' | 10 | 0.2 | 20 | 0.1 | 13 | 0.2308 | 16 | 0.0625 | 59 | 0.1356 | 9 | 0 | 8 | 0.25 | 12 | 0 | 30 | 0.1667 | 59 | 0.1186 |
| 11_534396' | 15 | 0.1333 | 26 | 0.2308 | 14 | 0 | 17 | 0.1176 | 72 | 0.1389 | 10 | 0.2 | 8 | 0.25 | 12 | 0 | 33 | 0.1515 | 63 | 0.1429 |
| 11_534402: | 21 | 0.1429 | 31 | 0.2903 | 24 | 0.0833 | 18 | 0.2222 | 94 | 0.1915 | 16 | 0.1875 | 13 | 0.3077 | 15 | 0.3333 | 37 | 0.1081 | 81 | 0.1975 |
| 11_534403: | 17 | 0.1176 | 32 | 0.0938 | 20 | 0.05 | 14 | 0.1429 | 83 | 0.0964 | 19 | 0.2105 | 14 | 0.0714 | 12 | 0.0833 | 36 | 0.1111 | 81 | 0.1235 |
| 11_5349194 | 3 | 0 | 10 | 0.1 | 7 | 0 | 6 | 0.1667 | 26 | 0.0769 | 3 | 0.3333 | 6 | 0 | 3 | 0 | 7 | 0.1429 | 19 | 0.1053 |
| 11_534920' | 3 | 0.6667 | 10 | 0.3 | 8 | 0.875 | 6 | 0.3333 | 27 | 0.5185 | 4 | 0.75 | 8 | 0.625 | 3 | 1 | 6 | 0.3333 | 21 | 0.619 |
| 11_5349204 | 3 | 0.3333 | 10 | 0 | 7 | 0.4286 | 6 | 0 | 26 | 0.1538 | 3 | 0 | 7 | 0.1429 | 2 | 1 | 7 | 0 | 19 | 0.1579 |
| 11_534939: | 1 | 0 | 7 | 0.4286 | 8 | 0.625 | 6 | 0.5 | 22 | 0.5 | 4 | 0.75 | 6 | 0.5 | 4 | 0.75 | 3 | 0.3333 | 17 | 0.5882 |
| 11_535406( | NA | NA | 1 | 1 | 1 | 0 | 5 | 0.4 | 7 | 0.4286 | NA | NA | 1 | 0 | NA | NA | 2 | 0.5 | 3 | 0.3333 |
| 11_535411( | NA | NA | 2 | 1 | 1 | 1 | 5 | 1 | 8 | 1 | NA | NA | 1 | 1 | NA | NA | 1 | 1 | 2 | 1 |
| 11_5375604 | 2 | 0.5 | 3 | 0 | 1 | 0 | 4 | 0.25 | 10 | 0.2 | 2 | 0.5 | NA | NA | 1 | 0 | 2 | 0 | 5 | 0.2 |
| 11_537561( | 1 | 1 | 3 | 0.3333 | 1 | 1 | 3 | 0.3333 | 8 | 0.5 | 2 | 0.5 | NA | NA | 1 | 0 | 3 | 0.6667 | 6 | 0.5 |
| 11_537563! | 2 | 1 | 4 | 0 | 1 | 1 | 5 | 0.2 | 12 | 0.3333 | 2 | 1 | NA | NA | NA | NA | 3 | 0.6667 | 5 | 0.8 |
| 11_537566' | 1 | 1 | 3 | 1 | 1 | 1 | 5 | 0.4 | 10 | 0.7 | 2 | 0.5 | NA | NA | NA | NA | 3 | 0.3333 | 5 | 0.4 |
| 11_538740: | NA | NA | 1 | 1 | 2 | 0 | 1 | 1 | 4 | 0.5 | 2 | 0.5 | NA | NA | NA | NA | 4 | 0.5 | 6 | 0.5 |
| 11_538741( | NA | NA | 1 | 1 | 2 | 1 | 1 | 1 | 4 | 1 | 1 | 0 | NA | NA | NA | NA | 4 | 1 | 5 | 0.8 |
| 11_540222( | 1 | 1 | 2 | 0 | 1 | 0 | 5 | 0 | 9 | 0.1111 | 5 | 0 | 2 | 0 | 1 | 1 | 5 | 0 | 13 | 0.0769 |
| 11_540223: | 3 | 0.6667 | 2 | 0 | NA | NA | 5 | 0.2 | 10 | 0.3 | 5 | 0 | 2 | 0 | 1 | 0 | 5 | 0.2 | 13 | 0.0769 |
| 11_542284' | 1 | 1 | NA | NA | NA | NA | NA | NA | 1 | 1 | NA | NA | NA | NA | NA | NA | 1 | 1 | 1 | 1 |
| 11_542520! | 16 | 0.75 | 13 | 1 | 17 | 0.6471 | 10 | 0.9 | 56 | 0.8036 | 15 | 0.8667 | 17 | 0.7059 | 27 | 0.8148 | 25 | 0.76 | 84 | 0.7857 |
| 11_542522! | 16 | 0.4375 | 17 | 0.3529 | 20 | 0.45 | 12 | 0.5 | 65 | 0.4308 | 16 | 0.5625 | 21 | 0.4286 | 23 | 0.4348 | 30 | 0.4667 | 90 | 0.4667 |
| 11_542524: | 1 | 0 | 4 | 0.5 | 2 | 0 | 4 | 0.25 | 11 | 0.2727 | 1 | 0 | 4 | 0.25 | 3 | 0.3333 | 11 | 0.4545 | 19 | 0.3684 |
| 11_542528' | 1 | 0 | 5 | 0.6 | 3 | 0.6667 | 5 | 0.4 | 14 | 0.5 | 1 | 1 | 6 | 0.5 | 3 | 0.6667 | 11 | 0.5455 | 21 | 0.5714 |
| 11_542531: | 2 | 0.5 | 4 | 0.25 | 1 | 0 | 7 | 0.4286 | 14 | 0.3571 | 4 | 0.25 | 6 | 0 | 4 | 0.25 | 12 | 0.4167 | 26 | 0.2692 |
| 11_542533' | 3 | 0 | 5 | 0.2 | 4 | 0 | 8 | 0.125 | 20 | 0.1 | 5 | 0.2 | 6 | 0.3333 | 6 | 0.1667 | 13 | 0.1538 | 30 | 0.2 |
| 11_5429304 | NA | NA | NA | NA | 1 | 1 | NA | NA | 1 | 1 | NA | NA | NA | NA | 3 | 0.6667 | 2 | 0.5 | 5 | 0.6 |
| 11_5429304 | NA | NA | NA | NA | 1 | 1 | NA | NA | 1 | 1 | NA | NA | NA | NA | 3 | 1 | 1 | 1 | 4 | 1 |

| | | | | | | | | | | | | | | | | | | | | |
|---|---|---|---|---|---|---|---|---|---|---|---|---|---|---|---|---|---|---|---|---|
| 11_5429312 | NA | NA | NA | NA | 1 | 1 | NA | NA | 1 | 1 | NA | NA | NA | NA | 3 | 0.3333 | 2 | 0 | 5 | 0.2 |
| 11_5429315 | NA | NA | NA | NA | 1 | 1 | NA | NA | 1 | 1 | NA | NA | NA | NA | 2 | 0.5 | 2 | 0.5 | 4 | 0.5 |
| 11_5439316 | NA | NA | 3 | 0.3333 | 4 | 0.5 | 3 | 0.6667 | 10 | 0.5 | 2 | 1 | 2 | 0.5 | NA | NA | 6 | 0.8333 | 10 | 0.8 |
| 11_5439337 | 1 | 0 | 3 | 0 | 4 | 0 | 3 | 0.3333 | 11 | 0.0909 | 2 | 0.5 | 2 | 0 | NA | NA | 6 | 0 | 10 | 0.1 |
| 11_5439339 | 1 | 0 | 3 | 0 | 3 | 0 | 3 | 0.3333 | 10 | 0.1 | 2 | 0 | 2 | 0 | NA | NA | 6 | 0.1667 | 10 | 0.1 |
| 11_5439357 | 1 | 1 | 4 | 0.5 | 3 | 0.3333 | 3 | 0 | 11 | 0.3636 | 2 | 0 | 2 | 0.5 | NA | NA | 5 | 0.6 | 9 | 0.4444 |
| 11_5439360 | 1 | 0 | 3 | 1 | 4 | 0.75 | 3 | 0 | 11 | 0.5455 | NA | NA | 2 | 0.5 | NA | NA | 5 | 0.6 | 7 | 0.5714 |
| 11_5439386 | NA | NA | 3 | 0.3333 | 2 | 0 | 2 | 0 | 7 | 0.1429 | 2 | 0 | 1 | 1 | NA | NA | 4 | 0 | 7 | 0.1429 |
| 11_5439397 | 1 | 1 | 5 | 0.4 | 4 | 0 | 4 | 0.5 | 14 | 0.3571 | 1 | 0 | 1 | 1 | NA | NA | 6 | 0.1667 | 8 | 0.25 |
| 11_5439398 | 1 | 0 | 4 | 0.5 | 3 | 0 | 4 | 0.25 | 12 | 0.25 | 2 | 0.5 | 1 | 0 | NA | NA | 6 | 0.3333 | 9 | 0.3333 |
| 11_5439410 | 1 | 1 | 5 | 0.8 | 3 | 1 | 4 | 0.75 | 13 | 0.8462 | 2 | 1 | 2 | 1 | NA | NA | 5 | 1 | 9 | 1 |
| 11_5439419 | 2 | 0.5 | 5 | 0.2 | 3 | 0 | 5 | 0 | 15 | 0.1333 | 1 | 0 | 2 | 0 | NA | NA | 6 | 0.1667 | 9 | 0.1111 |
| 11_5439421 | 2 | 1 | 4 | 1 | 4 | 1 | 5 | 1 | 15 | 1 | 2 | 1 | 1 | 1 | NA | NA | 4 | 0.75 | 7 | 0.8571 |
| 11_5439434 | 2 | 1 | 4 | 0.25 | 4 | 0 | 3 | 0 | 13 | 0.2308 | 2 | 0.5 | 3 | 0.3333 | NA | NA | 6 | 0.5 | 11 | 0.4545 |
| 11_5439435 | 2 | 0.5 | 5 | 1 | 3 | 0.6667 | 5 | 0.8 | 15 | 0.8 | 2 | 1 | 3 | 0.6667 | NA | NA | 5 | 0.8 | 10 | 0.8 |
| 11_5440077 | 2 | 0 | 4 | 0.75 | 4 | 0 | 5 | 0.6 | 15 | 0.4 | 2 | 0 | 2 | 0.5 | NA | NA | 5 | 0.2 | 9 | 0.2222 |
| 11_5440657 | 3 | 0 | 5 | 0 | 6 | 0.1667 | 9 | 0.1111 | 23 | 0.087 | 3 | 0.3333 | 3 | 0.3333 | 1 | 0 | 4 | 0 | 11 | 0.1818 |
| 11_5455176 | NA | NA | 1 | 1 | 1 | 1 | 2 | 1 | 4 | 1 | NA | NA | 1 | 1 | 2 | 1 | 1 | 1 | 4 | 1 |
| 11_5455187 | 1 | 1 | 1 | 1 | 1 | 0 | 2 | 0 | 5 | 0.4 | NA | NA | 1 | 0 | 2 | 0.5 | 1 | 0 | 4 | 0.25 |
| 11_5458218 | 3 | 0.6667 | 3 | 0.3333 | 2 | 1 | 2 | 1 | 10 | 0.7 | NA | NA | 1 | 1 | 2 | 0 | 1 | 1 | 4 | 0.5 |
| 11_5458224 | 2 | 0.5 | 3 | 1 | 2 | 1 | 1 | 1 | 8 | 0.875 | NA | NA | 1 | 1 | 3 | 1 | NA | NA | 4 | 1 |
| 11_5458230 | 2 | 0.5 | 3 | 0.6667 | 2 | 0.5 | 2 | 0 | 9 | 0.4444 | NA | NA | 2 | 1 | 3 | 0.3333 | 1 | 1 | 6 | 0.6667 |
| 11_5458234 | 2 | 0.5 | 3 | 0.6667 | 1 | 1 | 1 | 1 | 7 | 0.7143 | NA | NA | 2 | 1 | 2 | 0.5 | 1 | 1 | 5 | 0.8 |
| 11_5458242 | 3 | 0 | 3 | 0.3333 | 1 | 0 | 2 | 0 | 9 | 0.1111 | NA | NA | 2 | 0.5 | 3 | 0.3333 | 1 | 0 | 6 | 0.3333 |
| 11_5458244 | 2 | 0.5 | 1 | 1 | 1 | 1 | 2 | 0 | 6 | 0.5 | NA | NA | 2 | 0.5 | 2 | 1 | 1 | 0 | 5 | 0.6 |
| 11_5458522 | 3 | 0.3333 | 1 | 0 | NA | NA | 1 | 0 | 5 | 0.2 | NA | NA | NA | NA | 2 | 0.5 | 1 | 0 | 3 | 0.3333 |
| 11_5458717 | 3 | 0.3333 | 3 | 0 | 1 | 0 | 2 | 0 | 9 | 0.1111 | 1 | 0 | 3 | 0 | 3 | 0.6667 | 1 | 1 | 8 | 0.375 |
| 11_5458728 | 4 | 0.5 | 3 | 0.3333 | 2 | 1 | 2 | 1 | 11 | 0.6364 | 1 | 1 | 2 | 0.5 | 4 | 1 | 1 | 1 | 8 | 0.875 |
| 11_5458762 | 4 | 0.5 | 3 | 0.3333 | 4 | 0.75 | 2 | 0.5 | 13 | 0.5385 | 1 | 1 | 3 | 0.6667 | 3 | 0.3333 | 2 | 1 | 9 | 0.6667 |
| 11_5458765 | 2 | 0.5 | 2 | 0.5 | 4 | 0.25 | 2 | 0.5 | 10 | 0.4 | 1 | 1 | 3 | 0 | 3 | 0.6667 | 3 | 1 | 10 | 0.6 |
| 11_5458809 | 5 | 0.4 | 4 | 0 | 4 | 0 | 2 | 0 | 15 | 0.1333 | 2 | 0 | 3 | 0 | 2 | 0.5 | 1 | 0 | 8 | 0.125 |
| 11_5458818 | 4 | 1 | 4 | 0 | 3 | 0.3333 | 2 | 0 | 13 | 0.3846 | NA | NA | 2 | 0.5 | 4 | 0.25 | 1 | 1 | 7 | 0.4286 |
| 11_5458819 | 5 | 0 | 3 | 0 | 5 | 0.6 | 2 | 0 | 15 | 0.2 | 1 | 0 | 3 | 0.3333 | 5 | 0.2 | 2 | 0.5 | 11 | 0.2727 |
| 11_5458825 | 4 | 0 | 3 | 0 | 5 | 0.4 | 2 | 0 | 14 | 0.1429 | 2 | 0 | 2 | 0 | 3 | 0.3333 | 2 | 0 | 9 | 0.1111 |
| 11_5458873 | 4 | 0.25 | 5 | 0.2 | 5 | 0.4 | 4 | 0.5 | 18 | 0.3333 | 1 | 0 | 3 | 0 | 6 | 0.5 | 2 | 0 | 12 | 0.25 |
| 11_5458873 | 4 | 0 | 6 | 0.1667 | 5 | 0.4 | 4 | 0.25 | 19 | 0.2105 | 1 | 1 | 3 | 0 | 6 | 0.1667 | 2 | 0 | 12 | 0.1667 |
| 11_5458894 | 4 | 0.25 | 5 | 0 | 5 | 0 | 5 | 0 | 19 | 0.0526 | 2 | 0 | 3 | 0.3333 | 5 | 0 | 3 | 0 | 13 | 0.0769 |
| 11_5458928 | 5 | 0.2 | 6 | 0 | 5 | 0 | 5 | 0.2 | 21 | 0.0952 | 2 | 0 | 1 | 0 | 6 | 0.5 | 2 | 0 | 11 | 0.2727 |
| 11_5461026 | 19 | 0.0526 | 14 | 0.2143 | 10 | 0 | 17 | 0 | 60 | 0.0667 | 11 | 0.1818 | 14 | 0 | 15 | 0.1333 | 8 | 0.125 | 48 | 0.1042 |
| 11_5461034 | 19 | 0.6842 | 12 | 0.4167 | 11 | 0.6364 | 17 | 0.6471 | 59 | 0.6102 | 11 | 0.4545 | 17 | 0.5294 | 14 | 0.5714 | 8 | 0.875 | 50 | 0.58 |
| 11_5463112 | 14 | 0.3571 | 10 | 0.3 | 7 | 0.4286 | 14 | 0.2857 | 45 | 0.3333 | 10 | 0.3 | 12 | 0.25 | 10 | 0.4 | 8 | 0.5 | 40 | 0.35 |
| 11_5463125 | 10 | 0.2 | 8 | 0.25 | 5 | 0 | 13 | 0 | 36 | 0.1111 | 6 | 0 | 7 | 0.1429 | 8 | 0.125 | 3 | 0 | 24 | 0.0833 |
| 11_5473070 | 4 | 0 | 2 | 0 | 6 | 0.1667 | 3 | 0 | 15 | 0.0667 | 1 | 0 | 3 | 0.3333 | 2 | 0 | 2 | 0 | 8 | 0.125 |
| 11_5473087 | 3 | 0 | 3 | 0 | 6 | 0.1667 | 3 | 0 | 15 | 0.0667 | 1 | 0 | 4 | 0.25 | 1 | 0 | 1 | 0 | 7 | 0.1429 |
| 11_5473168 | 4 | 1 | 3 | 1 | 5 | 0.8 | 4 | 0.5 | 16 | 0.8125 | 1 | 1 | 1 | 1 | 2 | 1 | 2 | 0.5 | 6 | 0.8333 |
| 11_5473169 | 3 | 1 | 1 | 0 | 3 | 0 | 2 | 1 | 9 | 0.5556 | 1 | 1 | NA | NA | 2 | 0 | NA | NA | 3 | 0.3333 |
| 11_5473182 | 4 | 1 | 3 | 0.6667 | 6 | 0.5 | 4 | 0.25 | 17 | 0.5882 | 1 | 1 | 1 | 0 | 1 | 0 | 2 | 0 | 5 | 0.2 |
| 11_5473189 | 4 | 0.75 | 3 | 0.6667 | 4 | 1 | 4 | 0.75 | 15 | 0.8 | 1 | 1 | 1 | 1 | 1 | 1 | 2 | 1 | 5 | 1 |
| 11_5476116 | 50 | 0.14 | 59 | 0.1186 | 44 | 0.25 | 53 | 0.1321 | 206 | 0.1553 | 41 | 0.0976 | 43 | 0.2093 | 49 | 0.1429 | 35 | 0.1714 | 168 | 0.1548 |
| 11_5480163 | 4 | 1 | 2 | 1 | 2 | 1 | 5 | 0.6 | 13 | 0.8462 | 3 | 0.6667 | 4 | 0.75 | 2 | 1 | 2 | 0.5 | 11 | 0.7273 |
| 11_5480172 | 5 | 0.2 | 2 | 1 | 2 | 0 | 5 | 0.2 | 14 | 0.2857 | 5 | 0.2 | 4 | 0 | 4 | 0.5 | 2 | 0 | 15 | 0.2 |
| 11_5480173 | 5 | 0.2 | 2 | 1 | 2 | 0.5 | 5 | 0.4 | 14 | 0.4286 | 5 | 0.4 | 4 | 0.5 | 4 | 0.75 | 2 | 0 | 15 | 0.4667 |
| 11_5480192 | 5 | 0.4 | 2 | 1 | 2 | 0.5 | 2 | 0.5 | 11 | 0.5455 | 4 | 0.5 | 4 | 0.25 | 2 | 0 | 2 | 0 | 12 | 0.25 |
| 11_5480197 | 5 | 0.6 | 2 | 0.5 | 2 | 0.5 | 3 | 0.3333 | 12 | 0.5 | 5 | 0.2 | 4 | 0.5 | 4 | 0.25 | 2 | 1 | 15 | 0.4 |
| 11_5480200 | 5 | 1 | 2 | 1 | 2 | 0.5 | 4 | 1 | 13 | 0.9231 | 2 | 0.5 | 3 | 0.6667 | 3 | 1 | 2 | 1 | 10 | 0.8 |
| 11_5480208 | 5 | 0.2 | 2 | 0 | 2 | 0.5 | 4 | 0 | 13 | 0.1538 | 3 | 0 | 4 | 0.25 | 4 | 0 | 2 | 0 | 13 | 0.0769 |
| 11_5480220 | 2 | 0.5 | 2 | 0.5 | 2 | 1 | 3 | 0.6667 | 9 | 0.6667 | NA | NA | 3 | 0.6667 | 2 | 0.5 | 1 | 1 | 6 | 0.6667 |
| 11_5480238 | NA | NA | NA | NA | NA | NA | 3 | 0.6667 | 3 | 0.6667 | 2 | 0.5 | 1 | 0 | 2 | 0 | NA | NA | 5 | 0.2 |
| 11_5487016 | 13 | 0.6923 | 16 | 0.625 | 6 | 0.6667 | 10 | 0.5 | 45 | 0.6222 | 10 | 0.6 | 9 | 0.5556 | 10 | 0.5 | 7 | 0.5714 | 36 | 0.5556 |
| 11_5500044 | 7 | 0 | 10 | 0.4 | 10 | 0.3 | 13 | 0.0769 | 40 | 0.2 | 6 | 0 | 6 | 0 | 7 | 0.1429 | 4 | 0.25 | 23 | 0.087 |
| 11_5500051 | 9 | 0 | 10 | 0.3 | 12 | 0.1667 | 13 | 0.1538 | 44 | 0.1591 | 7 | 0.2857 | 6 | 0.1667 | 6 | 0.3333 | 7 | 0.4286 | 26 | 0.3077 |
| 11_5500065 | 6 | 0.1667 | 7 | 0.4286 | 10 | 0.5 | 8 | 0.375 | 31 | 0.3871 | 4 | 0.75 | 5 | 0.4 | 5 | 0.6 | 5 | 0 | 19 | 0.4211 |
| 11_5500133 | 10 | 0.4 | 9 | 0.3333 | 12 | 0.5 | 11 | 0.1818 | 42 | 0.3571 | 9 | 0.7778 | 9 | 0.3333 | 8 | 0.25 | 11 | 0.3636 | 37 | 0.4324 |
| 11_5500135 | 8 | 0 | 10 | 0.4 | 13 | 0.1538 | 10 | 0.2 | 41 | 0.1951 | 6 | 0.1667 | 8 | 0 | 8 | 0.125 | 10 | 0.1 | 32 | 0.0938 |
| 11_5500168 | 9 | 0.1111 | 11 | 0.0909 | 12 | 0.0833 | 13 | 0.0769 | 45 | 0.0889 | 12 | 0.0833 | 9 | 0.2222 | 11 | 0.0909 | 11 | 0 | 43 | 0.093 |

| | | | | | | | | | | | | | | | | | | | | |
|---|---|---|---|---|---|---|---|---|---|---|---|---|---|---|---|---|---|---|---|---|
| 11_551235 | 3 | 0.6667 | 1 | 0 | 1 | 1 | 1 | 0 | 6 | 0.5 | 2 | 1 | 1 | 0 | 3 | 0.6667 | 3 | 1 | 9 | 0.7778 |
| 11_551236 | 1 | 0 | 2 | 1 | 1 | 1 | 1 | 1 | 5 | 0.8 | 3 | 0.6667 | NA | NA | 3 | 0.3333 | 3 | 0.3333 | 9 | 0.4444 |
| 11_551236 | 3 | 0.6667 | 1 | 0 | NA | NA | NA | NA | 4 | 0.5 | 1 | 1 | 1 | 0 | 1 | 0 | 3 | 0.6667 | 6 | 0.5 |
| 11_551238 | 4 | 0.75 | 2 | 0.5 | 1 | 0 | 2 | 0 | 9 | 0.4444 | 4 | 0.5 | 1 | 0 | 4 | 0.75 | 4 | 0.5 | 13 | 0.5385 |
| 11_551240 | 3 | 0.3333 | 2 | 0 | 1 | 0 | 2 | 0.5 | 8 | 0.25 | 5 | 0 | 2 | 0 | 4 | 0.25 | 4 | 0 | 15 | 0.0667 |
| 11_551241 | 4 | 0.5 | 3 | 0.3333 | 1 | 1 | 1 | 0 | 9 | 0.4444 | 6 | 0.3333 | 2 | 0 | 5 | 0.2 | 4 | 0.25 | 17 | 0.2353 |
| 11_551241 | 3 | 0.3333 | 3 | 0 | 1 | 0 | 1 | 1 | 8 | 0.25 | 3 | 0.3333 | 1 | 1 | 4 | 0.25 | 5 | 0.2 | 13 | 0.3077 |
| 11_551243 | 4 | 0 | 5 | 0.2 | 1 | 0 | 2 | 0.5 | 12 | 0.1667 | 6 | 0.1667 | 2 | 0 | 4 | 0.25 | 5 | 0.4 | 17 | 0.2353 |
| 11_551244 | 5 | 0 | 6 | 0.3333 | 2 | 0.5 | 3 | 0.3333 | 16 | 0.25 | 6 | 0.1667 | 2 | 0 | 5 | 0 | 5 | 0.6 | 18 | 0.2222 |
| 11_551245 | 5 | 0.6 | 4 | 0.75 | 2 | 1 | 2 | 0.5 | 13 | 0.6923 | 5 | 0.4 | 2 | 1 | 5 | 0.8 | 5 | 1 | 17 | 0.7647 |
| 11_551245 | 3 | 0.6667 | 5 | 1 | 2 | 1 | 1 | 1 | 11 | 0.9091 | 6 | 0.5 | 2 | 1 | 4 | 0.5 | 4 | 1 | 16 | 0.6875 |
| 11_551245 | 4 | 0.5 | 4 | 0 | 2 | 0 | 3 | 0.3333 | 13 | 0.2308 | 4 | 0.25 | 2 | 0 | 4 | 0 | 5 | 0.2 | 15 | 0.1333 |
| 11_551246 | 4 | 0.25 | 4 | 0.75 | 1 | 1 | 2 | 0.5 | 11 | 0.5455 | 5 | 0.4 | 2 | 0.5 | 3 | 0.3333 | 3 | 0.6667 | 13 | 0.4615 |
| 11_551247 | 4 | 0.25 | 3 | 0.6667 | 2 | 0.5 | 1 | 0 | 10 | 0.4 | 4 | 0.25 | 1 | 1 | 3 | 0.3333 | 6 | 0.5 | 14 | 0.4286 |
| 11_551248 | 5 | 0.2 | 4 | 0 | 2 | 0.5 | 3 | 0.3333 | 14 | 0.2143 | 6 | 0 | 4 | 0.25 | 5 | 0.2 | 7 | 0.2857 | 22 | 0.1818 |
| 11_551265 | 7 | 0 | 10 | 0.1 | 5 | 0 | 3 | 0.3333 | 25 | 0.08 | 9 | 0.1111 | 3 | 0.3333 | 8 | 0 | 7 | 0.1429 | 27 | 0.1111 |
| 11_551267 | 7 | 0.1429 | 12 | 0.3333 | 5 | 0.2 | 3 | 0 | 27 | 0.2222 | 9 | 0.3333 | 2 | 0.5 | 7 | 0.2857 | 7 | 0.2857 | 25 | 0.32 |
| 11_551269 | 5 | 0.2 | 10 | 0.3 | 3 | 0 | 2 | 0 | 20 | 0.2 | 7 | 0 | 2 | 0.5 | 9 | 0.4444 | 7 | 0.1429 | 25 | 0.24 |
| 11_553694 | 2 | 0.5 | NA | NA | NA | NA | 3 | 1 | 5 | 0.8 | 1 | 0 | 2 | 0.5 | 1 | 1 | 1 | 0 | 5 | 0.4 |
| 11_553903 | 6 | 0.1667 | 8 | 0.125 | 6 | 0 | 9 | 0.2222 | 29 | 0.1379 | 2 | 0 | 10 | 0.2 | 8 | 0.25 | 4 | 0.25 | 24 | 0.2083 |
| 11_553905 | 7 | 0.5714 | 11 | 0.9091 | 6 | 0.8333 | 12 | 0.9167 | 36 | 0.8333 | 4 | 1 | 13 | 0.9231 | 6 | 1 | 7 | 1 | 30 | 0.9667 |
| 11_553907 | 8 | 0.875 | 9 | 0.4444 | 6 | 0.3333 | 8 | 0.75 | 31 | 0.6129 | 4 | 0.5 | 10 | 0.8 | 8 | 0.5 | 2 | 0.5 | 24 | 0.625 |
| 11_553907 | 3 | 1 | 8 | 0.875 | 3 | 0.6667 | 9 | 0.8889 | 23 | 0.8696 | 6 | 1 | 6 | 0.8333 | 6 | 0.8333 | 6 | 0.8333 | 24 | 0.875 |
| 11_553908 | 7 | 0.2857 | 9 | 0.5556 | 4 | 0.25 | 10 | 0.9 | 30 | 0.5667 | 6 | 0.1667 | 12 | 0.8333 | 11 | 0.2727 | 6 | 0.3333 | 35 | 0.4571 |
| 11_553909 | 6 | 0.3333 | 7 | 0.1429 | 1 | 1 | 12 | 0 | 26 | 0.1538 | 6 | 0 | 9 | 0 | 11 | 0.0909 | 5 | 0.2 | 31 | 0.0645 |
| 11_553910 | 6 | 0.8333 | 8 | 0.5 | 6 | 0.5 | 11 | 0.5455 | 31 | 0.5806 | 5 | 0.6 | 10 | 0.7 | 8 | 0.5 | 8 | 0.5 | 31 | 0.5806 |
| 11_553911 | 6 | 0.3333 | 9 | 0.4444 | 6 | 0.3333 | 13 | 0.3077 | 34 | 0.3529 | 5 | 0.2 | 12 | 0.4167 | 11 | 0.5455 | 7 | 0.4286 | 35 | 0.4286 |
| 11_553912 | 6 | 0 | 8 | 0 | 5 | 0.2 | 12 | 0.0833 | 31 | 0.0645 | 5 | 0.2 | 11 | 0 | 11 | 0.0909 | 7 | 0.1429 | 34 | 0.0882 |
| 11_553915 | 3 | 0.6667 | 6 | 0.6667 | 6 | 0.3333 | 5 | 0.6 | 20 | 0.55 | 3 | 0.6667 | 7 | 0.5714 | 6 | 0.8333 | 4 | 0.75 | 20 | 0.7 |
| 11_553916 | 2 | 0 | 6 | 0 | 4 | 0.25 | 5 | 0.4 | 17 | 0.1765 | 3 | 0 | 5 | 0 | 6 | 0 | 2 | 0.5 | 16 | 0.0625 |
| 11_553952 | 3 | 0.3333 | 3 | 0 | 3 | 0 | 2 | 0.5 | 11 | 0.1818 | 2 | 0 | 1 | 0 | 3 | 0 | 4 | 0.25 | 10 | 0.1 |
| 11_553954 | 3 | 0 | 2 | 0 | 1 | 0 | 2 | 0.5 | 8 | 0.125 | 1 | 0 | 2 | 0.5 | 3 | 0 | 4 | 0.25 | 10 | 0.2 |
| 11_570766 | 4 | 0 | 11 | 0.3636 | 8 | 0.125 | 6 | 0 | 29 | 0.1724 | 1 | 0 | 5 | 0 | 9 | 0.2222 | 5 | 0 | 20 | 0.1 |
| 11_572043 | NA | NA | NA | NA | NA | NA | 2 | 0.5 | 2 | 0.5 | NA | NA | 1 | 1 | NA | NA | 1 | 0 | 2 | 0.5 |
| 11_572044 | NA | NA | NA | NA | NA | NA | 2 | 1 | 2 | 1 | NA | NA | 1 | 0 | NA | NA | 1 | 1 | 2 | 0.5 |
| 11_572046 | NA | NA | NA | NA | NA | NA | 2 | 1 | 2 | 1 | NA | NA | 1 | 1 | NA | NA | 1 | 1 | 2 | 1 |
| 11_572075 | NA | NA | 1 | 1 | NA | NA | 3 | 1 | 4 | 1 | NA | NA | 1 | 1 | 1 | 1 | 1 | 1 | 3 | 1 |
| 11_572075 | NA | NA | 1 | 0 | NA | NA | 1 | 1 | 2 | 0.5 | NA | NA | 1 | 1 | 1 | 1 | 1 | 0 | 3 | 0.6667 |
| 11_572180 | 2 | 0 | 3 | 0.3333 | 1 | 0 | 2 | 0 | 8 | 0.125 | 1 | 1 | 1 | 0 | 6 | 0.1667 | 2 | 0.5 | 10 | 0.3 |
| 11_572183 | 16 | 0.3125 | 31 | 0.4516 | 18 | 0.5556 | 24 | 0.4583 | 89 | 0.4494 | 6 | 0.5 | 19 | 0.6316 | 25 | 0.48 | 16 | 0.5 | 66 | 0.5303 |
| 11_572184 | 15 | 0.2 | 31 | 0.3226 | 16 | 0.1875 | 20 | 0.35 | 82 | 0.2805 | 7 | 0.2857 | 14 | 0.2857 | 21 | 0.1905 | 13 | 0.2308 | 55 | 0.2364 |
| 11_572184 | 16 | 0.125 | 35 | 0 | 18 | 0.1667 | 24 | 0.0417 | 93 | 0.0645 | 6 | 0.1667 | 18 | 0.0556 | 27 | 0.1111 | 13 | 0.0769 | 64 | 0.0938 |
| 11_572185 | 16 | 0.0625 | 25 | 0.16 | 16 | 0.125 | 18 | 0.2222 | 75 | 0.1467 | 4 | 0 | 13 | 0.1538 | 26 | 0.0385 | 13 | 0.2308 | 56 | 0.1071 |
| 11_572186 | 14 | 0.2857 | 25 | 0.16 | 11 | 0.3636 | 15 | 0.2667 | 65 | 0.2462 | 4 | 0.25 | 8 | 0.375 | 15 | 0.2 | 10 | 0.1 | 37 | 0.2162 |
| 11_572187 | 19 | 1 | 28 | 0.7143 | 13 | 0.7692 | 23 | 0.8696 | 83 | 0.8313 | 2 | 1 | 14 | 1 | 23 | 0.7826 | 16 | 0.75 | 55 | 0.8364 |
| 11_572188 | 20 | 0.65 | 35 | 0.6857 | 17 | 0.5294 | 22 | 0.5909 | 94 | 0.6277 | 9 | 0.5556 | 19 | 0.4211 | 28 | 0.5 | 14 | 0.7143 | 70 | 0.5286 |
| 11_572190 | 27 | 0.0741 | 36 | 0.1667 | 13 | 0.1538 | 21 | 0.0476 | 97 | 0.1134 | 10 | 0.1 | 20 | 0.15 | 30 | 0.1 | 19 | 0.1053 | 79 | 0.1139 |
| 11_572190 | 23 | 0.0435 | 34 | 0.0882 | 15 | 0.2667 | 20 | 0 | 92 | 0.087 | 9 | 0 | 19 | 0.1053 | 28 | 0.1071 | 19 | 0.0526 | 75 | 0.08 |
| 11_572190 | 24 | 0.5 | 33 | 0.3939 | 12 | 0.5833 | 21 | 0.2857 | 90 | 0.4222 | 9 | 0.3333 | 18 | 0.5 | 31 | 0.3548 | 18 | 0.1667 | 76 | 0.3421 |
| 11_572191 | 25 | 0.08 | 41 | 0.1463 | 16 | 0.0625 | 23 | 0.087 | 105 | 0.1048 | 9 | 0.1111 | 16 | 0.125 | 28 | 0.0714 | 18 | 0 | 71 | 0.0704 |
| 11_572192 | 30 | 0.5333 | 42 | 0.4762 | 16 | 0.6875 | 23 | 0.4783 | 111 | 0.5225 | 9 | 0.6667 | 18 | 0.6111 | 33 | 0.5455 | 17 | 0.3529 | 77 | 0.5325 |
| 11_572195 | 30 | 0.2333 | 42 | 0.4524 | 19 | 0.3684 | 25 | 0.64 | 116 | 0.4224 | 10 | 0.3 | 18 | 0.3333 | 38 | 0.4474 | 24 | 0.4167 | 90 | 0.4 |
| 11_572196 | 24 | 0.3333 | 39 | 0.2821 | 18 | 0.2778 | 26 | 0.2308 | 107 | 0.2804 | 12 | 0.4167 | 20 | 0.4 | 38 | 0.3421 | 26 | 0.3077 | 96 | 0.3542 |
| 11_572198 | 29 | 0.2414 | 46 | 0.1957 | 25 | 0.28 | 29 | 0.1724 | 129 | 0.2171 | 10 | 0.3 | 23 | 0.0435 | 41 | 0.1951 | 24 | 0.0417 | 98 | 0.1327 |
| 11_572202 | 28 | 0.7143 | 41 | 0.7073 | 26 | 0.8846 | 25 | 0.6 | 120 | 0.725 | 13 | 0.8462 | 26 | 0.7308 | 32 | 0.7188 | 24 | 0.5833 | 95 | 0.7053 |
| 11_572202 | 32 | 0.4688 | 51 | 0.2353 | 31 | 0.2258 | 33 | 0.1515 | 147 | 0.2653 | 18 | 0.3889 | 29 | 0.2759 | 40 | 0.3 | 28 | 0.2857 | 115 | 0.3043 |
| 11_572204 | 28 | 0.75 | 40 | 0.725 | 23 | 0.8696 | 28 | 0.8571 | 119 | 0.7899 | 17 | 0.6471 | 29 | 0.7586 | 32 | 0.7188 | 20 | 0.85 | 98 | 0.7449 |
| 11_572205 | 31 | 0.7742 | 53 | 0.717 | 27 | 0.8148 | 31 | 0.7097 | 142 | 0.7465 | 18 | 0.6667 | 33 | 0.8788 | 40 | 0.75 | 30 | 0.6333 | 121 | 0.7438 |
| 11_572205 | 29 | 0.2414 | 47 | 0.3191 | 25 | 0.08 | 31 | 0.0645 | 132 | 0.197 | 15 | 0.2 | 31 | 0.0968 | 39 | 0.2308 | 28 | 0.1429 | 113 | 0.1681 |
| 11_572205 | 25 | 0.08 | 42 | 0.0952 | 27 | 0.1481 | 27 | 0.0741 | 121 | 0.0992 | 17 | 0.0588 | 29 | 0.0345 | 33 | 0.0606 | 24 | 0.0833 | 103 | 0.0583 |
| 11_572206 | 35 | 1 | 57 | 1 | 34 | 1 | 36 | 0.9722 | 162 | 0.9938 | 17 | 0.9412 | 34 | 1 | 49 | 1 | 33 | 1 | 133 | 0.9925 |
| 11_572206 | 32 | 0.125 | 52 | 0.1154 | 35 | 0.0571 | 36 | 0.1944 | 155 | 0.1226 | 18 | 0.0556 | 37 | 0.0811 | 50 | 0.18 | 36 | 0.1389 | 141 | 0.1277 |
| 11_572207 | 36 | 0.1389 | 51 | 0.2157 | 34 | 0.2353 | 39 | 0.2308 | 160 | 0.2062 | 18 | 0.1111 | 36 | 0.1944 | 47 | 0.1277 | 35 | 0.2286 | 136 | 0.1691 |
| 11_572207 | 33 | 0.3939 | 49 | 0.449 | 26 | 0.3846 | 37 | 0.3243 | 145 | 0.3931 | 14 | 0.6429 | 35 | 0.4571 | 39 | 0.3846 | 35 | 0.3143 | 123 | 0.4146 |
| 11_572209 | 15 | 0.0667 | 18 | 0.2222 | 11 | 0.0909 | 12 | 0.1667 | 56 | 0.1429 | 8 | 0.25 | 12 | 0.3333 | 10 | 0.1 | 8 | 0.125 | 38 | 0.2105 |

| | | | | | | | | | | | | | | | | | | | | |
|---|---|---|---|---|---|---|---|---|---|---|---|---|---|---|---|---|---|---|---|---|
| 11_5722094 | 16 | 0.125 | 23 | 0.1739 | 15 | 0.0667 | 18 | 0.1111 | 72 | 0.125 | 9 | 0.1111 | 13 | 0.0769 | 12 | 0.25 | 9 | 0 | 43 | 0.1163 |
| 11_5722106 | 12 | 0.0833 | 22 | 0.2727 | 12 | 0.1667 | 16 | 0.125 | 62 | 0.1774 | 11 | 0.0909 | 12 | 0.25 | 13 | 0.1538 | 9 | 0.2222 | 45 | 0.1778 |
| 11_5737437 | 3 | 0 | 4 | 0.25 | 6 | 0.3333 | 3 | 0 | 16 | 0.1875 | 6 | 0 | 2 | 0 | 4 | 0.25 | 2 | 0 | 14 | 0.0714 |
| 11_5737437 | 3 | 0.3333 | 4 | 0 | 6 | 0.1667 | 3 | 0 | 16 | 0.125 | 6 | 0.5 | 1 | 0 | 4 | 0.25 | 3 | 0.3333 | 14 | 0.3571 |
| 11_5737452 | 2 | 0.5 | 4 | 0.25 | 5 | 0.4 | 2 | 0 | 13 | 0.3077 | 5 | 0.4 | 1 | 1 | 3 | 0 | 2 | 0 | 11 | 0.2727 |
| 11_5737454 | 3 | 0.6667 | 5 | 0.8 | 6 | 0.3333 | 2 | 0.5 | 16 | 0.5625 | 6 | 0.6667 | 1 | 1 | 4 | 0.5 | 1 | 0 | 12 | 0.5833 |
| 11_5737478 | 2 | 0.5 | 4 | 0.25 | 5 | 0.2 | 3 | 0.6667 | 14 | 0.3571 | 3 | 0 | 1 | 0 | 2 | 0.5 | 2 | 0 | 8 | 0.125 |
| 11_5741882 | NA | NA | 1 | 1 | 1 | 1 | NA | NA | 2 | 1 | NA | NA | NA | NA | 3 | 1 | NA | NA | 3 | 1 |
| 11_5741909 | NA | NA | 1 | 1 | 1 | 0 | NA | NA | 2 | 0.5 | NA | NA | NA | NA | 2 | 0.5 | NA | NA | 2 | 0.5 |
| 11_5742035 | NA | NA | 1 | 0 | 1 | 1 | NA | NA | 2 | 0.5 | NA | NA | NA | NA | 3 | 0.3333 | NA | NA | 3 | 0.3333 |
| 11_5742058 | NA | NA | 1 | 0 | 1 | 1 | NA | NA | 2 | 0.5 | NA | NA | NA | NA | 3 | 0.3333 | NA | NA | 3 | 0.3333 |
| 11_5742070 | NA | NA | 1 | 0 | 1 | 1 | NA | NA | 2 | 0.5 | NA | NA | NA | NA | 3 | 1 | NA | NA | 3 | 1 |
| 11_5742073 | NA | NA | 1 | 1 | NA | NA | NA | NA | 1 | 1 | NA | NA | NA | NA | 3 | 1 | NA | NA | 3 | 1 |
| 11_5742082 | NA | NA | 1 | 0 | 1 | 1 | NA | NA | 2 | 0.5 | NA | NA | NA | NA | 3 | 1 | NA | NA | 3 | 1 |
| 11_5742096 | NA | NA | 1 | 0 | 1 | 1 | NA | NA | 2 | 0.5 | 1 | 0 | NA | NA | 3 | 1 | NA | NA | 4 | 0.75 |
| 11_5742098 | NA | NA | 1 | 0 | 1 | 1 | NA | NA | 2 | 0.5 | 1 | 1 | NA | NA | 3 | 1 | NA | NA | 4 | 1 |
| 11_5742099 | NA | NA | 1 | 1 | 1 | 0 | NA | NA | 2 | 0.5 | NA | NA | NA | NA | 3 | 0.6667 | NA | NA | 3 | 0.6667 |
| 11_5742138 | NA | NA | 1 | 0 | 1 | 1 | NA | NA | 2 | 0.5 | 1 | 0 | NA | NA | 3 | 1 | NA | NA | 4 | 0.75 |
| 11_5742200 | NA | NA | NA | NA | 1 | 1 | NA | NA | 1 | 1 | NA | NA | NA | NA | 3 | 1 | NA | NA | 3 | 1 |
| 11_5742207 | NA | NA | 1 | 1 | 1 | 0 | NA | NA | 2 | 0.5 | 2 | 0.5 | NA | NA | 4 | 0.75 | NA | NA | 6 | 0.6667 |
| 11_5742221 | NA | NA | 1 | 0 | 1 | 1 | NA | NA | 2 | 0.5 | 2 | 1 | NA | NA | 3 | 1 | NA | NA | 5 | 1 |
| 11_5742249 | NA | NA | 1 | 0 | 1 | 1 | NA | NA | 2 | 0.5 | NA | NA | NA | NA | 3 | 0.6667 | NA | NA | 3 | 0.6667 |
| 11_5742283 | NA | NA | NA | NA | 1 | 1 | NA | NA | 1 | 1 | 2 | 0.5 | NA | NA | 5 | 0.4 | NA | NA | 7 | 0.4286 |
| 11_5742419 | NA | NA | 2 | 0 | 3 | 0.3333 | 1 | 1 | 6 | 0.3333 | 3 | 0.3333 | 1 | 0 | 4 | 0 | NA | NA | 8 | 0.125 |
| 11_5742420 | NA | NA | 2 | 0.5 | 2 | 0 | 1 | 0 | 5 | 0.2 | 3 | 0.6667 | 1 | 1 | 5 | 0.8 | 1 | 0 | 10 | 0.7 |
| 11_5743534 | NA | NA | 1 | 1 | NA | NA | NA | NA | 1 | 1 | NA | NA | NA | NA | 1 | 1 | NA | NA | 1 | 1 |
| 11_5769250 | 4 | 0 | 3 | 0 | 2 | 0.5 | 4 | 0.25 | 13 | 0.1538 | NA | NA | 4 | 0 | 5 | 0.4 | 5 | 0.2 | 14 | 0.2143 |
| 11_5770094 | 7 | 0.2857 | 6 | 0.3333 | 6 | 0.3333 | 8 | 0.25 | 27 | 0.2963 | 1 | 0 | 9 | 0.3333 | 9 | 0.1111 | 7 | 0.4286 | 26 | 0.2692 |
| 11_5770099 | 8 | 0.625 | 6 | 0.5 | 4 | 1 | 8 | 0.75 | 26 | 0.6923 | NA | NA | 10 | 0.7 | 11 | 0.2727 | 7 | 0.2857 | 28 | 0.4286 |
| 11_5770525 | NA | NA | 5 | 0 | 4 | 0.25 | 6 | 0 | 15 | 0.0667 | 1 | 0 | 6 | 0.1667 | 4 | 0 | 3 | 0 | 14 | 0.0714 |
| 11_5770561 | 1 | 0 | 3 | 0.3333 | 1 | 0 | 8 | 0.125 | 13 | 0.1538 | NA | NA | 5 | 0 | 3 | 0 | 2 | 0.5 | 10 | 0.1 |
| 11_5794069 | 1 | 0 | 1 | 0 | 1 | 1 | NA | NA | 3 | 0.3333 | 3 | 1 | 2 | 0.5 | 2 | 1 | 2 | 0 | 9 | 0.6667 |
| 11_5794955 | 2 | 0.5 | 2 | 0.5 | 1 | 0 | NA | NA | 5 | 0.4 | 3 | 0.6667 | 4 | 0.25 | 5 | 0.2 | 2 | 0 | 14 | 0.2857 |
| 11_5794956 | 2 | 0 | 2 | 0 | 2 | 0.5 | NA | NA | 6 | 0.1667 | 3 | 0.3333 | 4 | 0 | 4 | 0.5 | 3 | 0.3333 | 14 | 0.2857 |
| 11_5794957 | 2 | 0.5 | 1 | 1 | 1 | 0 | NA | NA | 4 | 0.5 | 3 | 0.6667 | 4 | 0.25 | 5 | 0.4 | 3 | 0.3333 | 15 | 0.4 |
| 11_5794967 | 2 | 0.5 | 2 | 0.5 | 1 | 0 | NA | NA | 5 | 0.4 | 3 | 0 | 2 | 0 | 5 | 0.4 | 3 | 0.6667 | 13 | 0.3077 |
| 11_5795021 | 2 | 1 | 3 | 1 | 4 | 1 | NA | NA | 9 | 1 | 4 | 1 | 6 | 1 | 6 | 0.8333 | 3 | 0.3333 | 19 | 0.8421 |
| 11_5795034 | 1 | 0 | NA | NA | 4 | 0.25 | NA | NA | 5 | 0.2 | 5 | 0.2 | 5 | 0.4 | 6 | 0.3333 | 3 | 0.3333 | 19 | 0.3158 |
| 11_5795050 | 1 | 1 | 1 | 1 | 4 | 0.75 | NA | NA | 6 | 0.8333 | 5 | 0.4 | 5 | 0.4 | 8 | 0.5 | 2 | 0.5 | 20 | 0.45 |
| 11_5795057 | 2 | 1 | 2 | 0.5 | 5 | 0.6 | NA | NA | 9 | 0.6667 | 6 | 0.3333 | 7 | 0.2857 | 8 | 0 | 3 | 0.3333 | 24 | 0.2083 |
| 11_5795057 | 2 | 1 | 1 | 1 | 5 | 1 | NA | NA | 8 | 1 | 5 | 1 | 7 | 1 | 7 | 1 | 2 | 1 | 21 | 1 |
| 11_5795058 | 2 | 1 | 3 | 0.3333 | 4 | 0.75 | NA | NA | 9 | 0.6667 | 6 | 0.6667 | 5 | 0.6 | 8 | 0.625 | 2 | 0.5 | 21 | 0.619 |
| 11_5795059 | 2 | 0.5 | 3 | 0 | 3 | 0.3333 | NA | NA | 8 | 0.25 | 5 | 0.2 | 6 | 0.5 | 8 | 0.25 | 2 | 0.5 | 21 | 0.3333 |
| 11_5795067 | 2 | 0 | 2 | 0.5 | 6 | 0.1667 | NA | NA | 10 | 0.2 | 6 | 0.1667 | 5 | 0 | 8 | 0.25 | 3 | 0 | 22 | 0.1364 |
| 11_5795083 | 2 | 1 | 3 | 0.3333 | 9 | 0.6667 | NA | NA | 14 | 0.6429 | 6 | 0.5 | 7 | 0.8571 | 10 | 0.6 | 3 | 0.6667 | 26 | 0.6538 |
| 11_5795127 | 3 | 0 | 3 | 0.3333 | 9 | 0.2222 | NA | NA | 15 | 0.2 | 7 | 0.4286 | 8 | 0.25 | 13 | 0.1538 | 4 | 0.5 | 32 | 0.2812 |
| 11_5795238 | 4 | 0.25 | 4 | 0.25 | 17 | 0.4118 | 2 | 0 | 27 | 0.3333 | 5 | 0.4 | 8 | 0.5 | 5 | 0.2 | 2 | 0.5 | 20 | 0.4 |
| 11_5800827 | 5 | 0.2 | 5 | 0 | 3 | 0.3333 | 6 | 0 | 19 | 0.1053 | 9 | 0.2222 | 7 | 0 | 7 | 0.2857 | 3 | 0 | 26 | 0.1538 |
| 11_5800835 | 4 | 0.25 | 5 | 0.2 | 7 | 0.1429 | 7 | 0.2857 | 23 | 0.2174 | 12 | 0.5 | 5 | 0.6 | 7 | 0.7143 | 2 | 1 | 26 | 0.6154 |
| 11_5800847 | 3 | 0.6667 | 5 | 0.8 | 7 | 0.8571 | 7 | 0.7143 | 22 | 0.7727 | 13 | 0.8462 | 7 | 0.5714 | 9 | 0.8889 | 3 | 1 | 32 | 0.8125 |
| 11_5800861 | 5 | 0.2 | 4 | 0.75 | 7 | 0.4286 | 7 | 0.2857 | 23 | 0.3913 | 12 | 0.5833 | 7 | 0.7143 | 9 | 0.6667 | 3 | 0.6667 | 31 | 0.6452 |
| 11_5800908 | 5 | 0.2 | 1 | 0 | 5 | 0.2 | 4 | 0 | 15 | 0.1333 | 9 | 0.1111 | 9 | 0.1111 | 7 | 0 | 5 | 0 | 30 | 0.0667 |
| 11_5800921 | 6 | 0 | 6 | 0.1667 | 8 | 0 | 6 | 0.1667 | 26 | 0.0769 | 9 | 0 | 10 | 0.1 | 10 | 0.1 | 5 | 0.4 | 34 | 0.1176 |
| 11_5801129 | 1 | 1 | NA | NA | NA | NA | 2 | 0.5 | 3 | 0.6667 | NA | NA | 2 | 0.5 | 3 | 0.6667 | 1 | 1 | 6 | 0.6667 |
| 11_5801148 | 1 | 0 | NA | NA | NA | NA | 2 | 0.5 | 3 | 0.3333 | NA | NA | 2 | 0.5 | 3 | 0.3333 | 1 | 0 | 6 | 0.3333 |
| 11_5801195 | 1 | 0 | 1 | 0 | NA | NA | 1 | 1 | 3 | 0.3333 | NA | NA | 3 | 0.6667 | 3 | 0 | 1 | 1 | 7 | 0.4286 |
| 11_5801201 | 1 | 1 | 2 | 0 | NA | NA | 2 | 1 | 5 | 0.6 | NA | NA | 2 | 0 | 5 | 0.2 | 1 | 0 | 8 | 0.125 |
| 11_5801203 | 1 | 0 | 2 | 0.5 | NA | NA | 2 | 0 | 5 | 0.2 | NA | NA | 3 | 0 | 5 | 0.2 | 1 | 0 | 9 | 0.1111 |
| 11_5801221 | 1 | 1 | 1 | 1 | NA | NA | 1 | 1 | 3 | 1 | NA | NA | 3 | 1 | 5 | 0.8 | 1 | 1 | 9 | 0.8889 |
| 11_5801223 | 1 | 0 | 2 | 0.5 | NA | NA | 1 | 1 | 4 | 0.5 | NA | NA | 3 | 0.6667 | 3 | 1 | NA | NA | 6 | 0.8333 |
| 11_5801225 | 1 | 1 | 2 | 0 | NA | NA | 1 | 1 | 4 | 0.5 | NA | NA | 2 | 0.5 | 4 | 0.25 | 1 | 1 | 7 | 0.4286 |
| 11_5801244 | 1 | 1 | NA | NA | NA | NA | 1 | 0 | 2 | 0.5 | NA | NA | 3 | 0.3333 | 2 | 0.5 | 1 | 1 | 6 | 0.5 |
| 11_5801603 | 1 | 1 | 1 | 0 | NA | NA | 1 | 1 | 3 | 0.6667 | NA | NA | 3 | 0.3333 | 2 | 0 | 1 | 1 | 6 | 0.3333 |
| 11_5807026 | 5 | 0.2 | 5 | 0 | 4 | 0 | 5 | 0.2 | 19 | 0.1053 | 2 | 0 | 7 | 0.4286 | 6 | 0.1667 | 4 | 0.5 | 19 | 0.3158 |
| 11_5809710 | NA | NA | 1 | 1 | NA | NA | NA | NA | 1 | 1 | NA | NA | 2 | 1 | 1 | 1 | NA | NA | 3 | 1 |

| | | | | | | | | | | | | | | | | | | | | |
|---|---|---|---|---|---|---|---|---|---|---|---|---|---|---|---|---|---|---|---|---|
| 11_5809715 | NA | NA | 1 | 1 | NA | NA | NA | NA | 1 | 1 | NA | NA | 2 | 1 | 2 | 1 | NA | NA | 4 | 1 |
| 11_5809787 | NA | NA | 1 | 1 | NA | NA | NA | NA | 1 | 1 | 1 | 1 | 2 | 1 | 2 | 1 | NA | NA | 5 | 1 |
| 11_5809807 | NA | NA | 2 | 1 | NA | NA | NA | NA | 2 | 1 | 2 | 1 | 2 | 1 | 1 | 1 | NA | NA | 5 | 1 |
| 11_5820610 | 2 | 1 | 4 | 1 | 5 | 0.8 | 5 | 0.6 | 16 | 0.8125 | 1 | 1 | 1 | 1 | 5 | 0.8 | 7 | 0.8571 | 14 | 0.8571 |
| 11_5820623 | 3 | 0.6667 | 2 | 1 | 5 | 0.6 | 5 | 0.8 | 15 | 0.7333 | 1 | 1 | 3 | 0.6667 | 7 | 0.5714 | 7 | 0.5714 | 18 | 0.6111 |
| 11_5820652 | 2 | 0.5 | 1 | 0 | 6 | 0.1667 | 4 | 0 | 13 | 0.1538 | 3 | 0 | 2 | 0 | 6 | 0.1667 | 9 | 0.3333 | 20 | 0.2 |
| 11_5820688 | 3 | 0 | 2 | 0.5 | 6 | 0.1667 | 3 | 0 | 14 | 0.1429 | 3 | 0 | 2 | 0 | 7 | 0.5714 | 9 | 0 | 21 | 0.1905 |
| 11_5820713 | 3 | 0 | 1 | 0 | 6 | 0 | 3 | 0.3333 | 13 | 0.0769 | 3 | 0.3333 | 2 | 0.5 | 7 | 0.7143 | 6 | 0 | 18 | 0.3889 |
| 11_5820714 | 3 | 0 | 1 | 1 | 6 | 0 | 4 | 0 | 14 | 0.0714 | 3 | 0.3333 | 2 | 0 | 7 | 0.1429 | 7 | 0 | 19 | 0.1053 |
| 11_5820734 | 3 | 0 | 2 | 0 | 4 | 0.25 | 4 | 0 | 13 | 0.0769 | 3 | 0.3333 | 2 | 0 | 6 | 0.1667 | 8 | 0.125 | 19 | 0.1579 |
| 11_5820756 | 1 | 1 | NA | NA | 4 | 0.75 | 4 | 0.5 | 9 | 0.6667 | 3 | 0.6667 | 1 | 0 | 6 | 0.8333 | 7 | 0.4286 | 17 | 0.5882 |
| 11_5820773 | 3 | 0 | 1 | 0 | 5 | 0 | 4 | 0.75 | 13 | 0.2308 | 2 | 0 | 1 | 0 | 6 | 0.6667 | 7 | 0.4286 | 16 | 0.4375 |
| 11_5820776 | 3 | 0.3333 | 1 | 0 | 5 | 0.4 | 4 | 0 | 13 | 0.2308 | 3 | 0 | 1 | 0 | 6 | 0.6667 | 7 | 0.1429 | 17 | 0.2941 |
| 11_5820811 | 2 | 0 | 1 | 0 | 4 | 0.25 | 4 | 0.5 | 11 | 0.2727 | 3 | 0 | 1 | 0 | 6 | 0.1667 | 6 | 0 | 16 | 0.0625 |
| 11_5820813 | 3 | 1 | NA | NA | 4 | 1 | 3 | 1 | 10 | 1 | 3 | 1 | NA | NA | 4 | 1 | 4 | 0.75 | 11 | 0.9091 |
| 11_5820828 | 3 | 0 | 1 | 0 | 3 | 0.3333 | 3 | 0.3333 | 10 | 0.2 | 2 | 0 | 1 | 0 | 6 | 0.6667 | 5 | 0.6 | 14 | 0.5 |
| 11_5820833 | 1 | 0 | NA | NA | 2 | 0.5 | 2 | 0.5 | 5 | 0.4 | 2 | 0 | 1 | 0 | 4 | 0.5 | 5 | 0 | 12 | 0.1667 |
| 11_5821029 | 1 | 1 | 1 | 1 | 3 | 1 | 3 | 1 | 8 | 1 | 1 | 1 | NA | NA | 4 | 1 | 3 | 0.6667 | 8 | 0.875 |
| 11_5822228 | 1 | 1 | 1 | 1 | 1 | 1 | NA | NA | 3 | 1 | 2 | 1 | NA | NA | NA | NA | NA | NA | 2 | 1 |
| 11_5822231 | 1 | 1 | 1 | 0 | 1 | 1 | NA | NA | 3 | 0.6667 | 2 | 0.5 | NA | NA | NA | NA | NA | NA | 2 | 0.5 |
| 11_5823693 | 3 | 1 | 2 | 0.5 | 3 | 0.6667 | 2 | 1 | 10 | 0.8 | 4 | 0.5 | 2 | 1 | 1 | 1 | NA | NA | 7 | 0.7143 |
| 11_5823695 | 3 | 1 | 2 | 0 | 2 | 0 | 1 | 1 | 8 | 0.5 | 3 | 0.6667 | 2 | 1 | 1 | 1 | NA | NA | 6 | 0.8333 |
| 11_5823703 | 3 | 0.6667 | 2 | 0.5 | 4 | 0.75 | 3 | 1 | 12 | 0.75 | 4 | 0.75 | 1 | 0 | 1 | 0 | NA | NA | 6 | 0.5 |
| 11_5823711 | 3 | 0 | 2 | 0 | 4 | 0.25 | 3 | 0.3333 | 12 | 0.1667 | 5 | 0 | 2 | 0.5 | 2 | 0 | NA | NA | 9 | 0.1111 |
| 11_5823804 | 3 | 0.3333 | 2 | 0 | 5 | 0.2 | 4 | 0.25 | 14 | 0.2143 | 3 | 0 | 2 | 0 | 2 | 0.5 | 1 | 0 | 8 | 0.125 |
| 11_5823820 | 5 | 0 | 2 | 0.5 | 7 | 0 | 6 | 0.1667 | 20 | 0.1 | 4 | 0.5 | 4 | 0 | 4 | 0 | 1 | 0 | 13 | 0.1538 |
| 11_5824637 | NA | NA | 2 | 0.5 | NA | NA | NA | NA | 2 | 0.5 | 1 | 1 | 1 | 0 | NA | NA | NA | NA | 2 | 0.5 |
| 11_5824638 | NA | NA | 2 | 1 | NA | NA | NA | NA | 2 | 1 | 1 | 1 | 1 | 1 | NA | NA | NA | NA | 2 | 1 |
| 11_5874926 | 3 | 1 | NA | NA | 1 | 0 | NA | NA | 4 | 0.75 | NA | NA | 1 | 1 | 1 | 1 | 2 | 1 | 4 | 1 |
| 11_5874972 | 3 | 1 | NA | NA | 1 | 0 | NA | NA | 4 | 0.75 | NA | NA | 1 | 1 | 1 | 1 | 2 | 0 | 4 | 0.5 |
| 11_5875017 | 2 | 1 | NA | NA | 1 | 0 | NA | NA | 3 | 0.6667 | NA | NA | 1 | 0 | 1 | 1 | 2 | 1 | 4 | 0.75 |
| 11_5875023 | 2 | 1 | NA | NA | 1 | 1 | NA | NA | 3 | 1 | NA | NA | 1 | 0 | 1 | 1 | 1 | 1 | 3 | 0.6667 |
| 11_5875042 | 1 | 1 | NA | NA | 1 | 0 | NA | NA | 2 | 0.5 | NA | NA | 1 | 1 | 1 | 0 | 2 | 0.5 | 4 | 0.5 |
| 11_5875051 | 2 | 1 | NA | NA | 1 | 1 | NA | NA | 3 | 1 | NA | NA | 1 | 0 | 1 | 1 | 1 | 1 | 3 | 0.6667 |
| 11_5875097 | 2 | 0.5 | NA | NA | 1 | 0 | NA | NA | 3 | 0.3333 | NA | NA | 1 | 0 | 1 | 1 | 2 | 0.5 | 4 | 0.5 |
| 11_5875124 | 2 | 0.5 | NA | NA | 1 | 0 | NA | NA | 3 | 0.3333 | NA | NA | 1 | 0 | 1 | 1 | 2 | 1 | 4 | 0.75 |
| 11_5875131 | 1 | 1 | NA | NA | 1 | 0 | NA | NA | 2 | 0.5 | NA | NA | 1 | 1 | 1 | 1 | 2 | 1 | 4 | 1 |
| 11_5875133 | 1 | 1 | NA | NA | NA | NA | NA | NA | 1 | 1 | NA | NA | NA | NA | 1 | 1 | 1 | 1 | 2 | 1 |
| 11_5875147 | 1 | 1 | NA | NA | 1 | 1 | NA | NA | 2 | 1 | NA | NA | 1 | 0 | NA | NA | 2 | 1 | 3 | 0.6667 |
| 11_5875189 | 2 | 0.5 | NA | NA | 1 | 1 | NA | NA | 3 | 0.6667 | NA | NA | 1 | 0 | 1 | 1 | 2 | 0.5 | 4 | 0.5 |
| 11_5875261 | 2 | 0.5 | NA | NA | 1 | 0 | 1 | 0 | 4 | 0.25 | NA | NA | 2 | 1 | 2 | 0 | 2 | 1 | 6 | 0.6667 |
| 11_5877086 | NA | NA | NA | NA | 1 | 1 | NA | NA | 1 | 1 | NA | NA | 1 | 1 | NA | NA | NA | NA | 1 | 1 |
| 11_5877091 | 1 | 1 | NA | NA | 2 | 0.5 | NA | NA | 3 | 0.6667 | NA | NA | 1 | 1 | NA | NA | NA | NA | 1 | 1 |
| 11_5877101 | 1 | 1 | NA | NA | 1 | 1 | NA | NA | 2 | 1 | NA | NA | 1 | 1 | 1 | 1 | NA | NA | 2 | 1 |
| 11_5877107 | 2 | 1 | NA | NA | 2 | 1 | NA | NA | 4 | 1 | NA | NA | 1 | 1 | 1 | 1 | NA | NA | 2 | 1 |
| 11_5877108 | 2 | 0.5 | NA | NA | 2 | 0 | NA | NA | 4 | 0.25 | NA | NA | 1 | 0 | 1 | 1 | NA | NA | 2 | 0.5 |
| 11_5877122 | 1 | 1 | NA | NA | 2 | 1 | NA | NA | 3 | 1 | NA | NA | 1 | 1 | 1 | 1 | NA | NA | 2 | 1 |
| 11_5877124 | 2 | 1 | NA | NA | 2 | 0.5 | NA | NA | 4 | 0.75 | NA | NA | 1 | 1 | 1 | 1 | NA | NA | 2 | 1 |
| 11_5877150 | 2 | 0.5 | NA | NA | 2 | 0 | NA | NA | 4 | 0.25 | NA | NA | 1 | 1 | 2 | 0.5 | NA | NA | 3 | 0.6667 |
| 11_5877162 | 2 | 1 | NA | NA | 3 | 0.6667 | NA | NA | 5 | 0.8 | NA | NA | 1 | 0 | 2 | 0.5 | NA | NA | 3 | 0.3333 |
| 11_5877173 | 2 | 0.5 | NA | NA | 3 | 0 | 1 | 1 | 6 | 0.3333 | NA | NA | 1 | 0 | 2 | 0.5 | NA | NA | 3 | 0.3333 |
| 11_5877218 | 2 | 0.5 | NA | NA | 3 | 0 | 1 | 0 | 6 | 0.1667 | NA | NA | 1 | 0 | 2 | 0.5 | NA | NA | 3 | 0.3333 |
| 11_5877219 | 2 | 0.5 | NA | NA | 3 | 0 | 1 | 0 | 6 | 0.1667 | NA | NA | 1 | 0 | 2 | 0.5 | NA | NA | 3 | 0.3333 |
| 11_5877231 | 2 | 0.5 | NA | NA | 3 | 0 | 1 | 0 | 6 | 0.1667 | NA | NA | 1 | 0 | 2 | 0.5 | NA | NA | 3 | 0.3333 |
| 11_5877374 | 2 | 0.5 | NA | NA | 3 | 0.3333 | 3 | 0.6667 | 8 | 0.5 | NA | NA | 1 | 0 | 2 | 0.5 | NA | NA | 3 | 0.3333 |
| 11_5877997 | NA | NA | 3 | 0.6667 | NA | NA | NA | NA | 3 | 0.6667 | NA | NA | 2 | 0.5 | 2 | 0 | NA | NA | 4 | 0.25 |
| 11_5878008 | NA | NA | 3 | 0.3333 | NA | NA | NA | NA | 3 | 0.3333 | NA | NA | 1 | 0 | 3 | 0.3333 | NA | NA | 4 | 0.25 |
| 11_5878018 | NA | NA | 3 | 0.6667 | NA | NA | NA | NA | 3 | 0.6667 | NA | NA | 1 | 1 | 3 | 0.3333 | NA | NA | 4 | 0.5 |
| 11_5878043 | NA | NA | 2 | 0.5 | NA | NA | NA | NA | 2 | 0.5 | NA | NA | 1 | 0 | 2 | 0.5 | NA | NA | 3 | 0.3333 |
| 11_5878049 | NA | NA | 1 | 1 | NA | NA | NA | NA | 1 | 1 | NA | NA | 1 | 1 | 3 | 0.6667 | NA | NA | 4 | 0.75 |
| 11_5878051 | NA | NA | 2 | 1 | NA | NA | NA | NA | 2 | 1 | NA | NA | 1 | 1 | 3 | 1 | NA | NA | 4 | 1 |
| 11_5878105 | NA | NA | 2 | 0.5 | NA | NA | NA | NA | 2 | 0.5 | NA | NA | 1 | 1 | 3 | 0.3333 | NA | NA | 4 | 0.5 |
| 11_5878130 | NA | NA | 1 | 1 | NA | NA | NA | NA | 1 | 1 | NA | NA | NA | NA | 3 | 0.6667 | NA | NA | 3 | 0.6667 |
| 11_5878132 | NA | NA | 2 | 0.5 | NA | NA | NA | NA | 2 | 0.5 | NA | NA | 1 | 0 | 4 | 0.25 | NA | NA | 5 | 0.2 |
| 11_5878138 | NA | NA | 2 | 1 | NA | NA | NA | NA | 2 | 1 | NA | NA | 1 | 1 | 3 | 0.3333 | NA | NA | 4 | 0.5 |

| | | | | | | | | | | | | | | | | | | | | |
|---|---|---|---|---|---|---|---|---|---|---|---|---|---|---|---|---|---|---|---|---|
| 11_5878142 | NA | NA | 2 | 0.5 | NA | NA | NA | NA | 2 | 0.5 | NA | NA | 1 | 0 | 3 | 0.6667 | NA | NA | 4 | 0.5 |
| 11_5878179 | NA | NA | 2 | 1 | NA | NA | NA | NA | 2 | 1 | NA | NA | NA | NA | 1 | 1 | NA | NA | 1 | 1 |
| 11_5878198 | NA | NA | 2 | 1 | NA | NA | NA | NA | 2 | 1 | NA | NA | NA | NA | 3 | 1 | NA | NA | 3 | 1 |
| 11_5878221 | NA | NA | 1 | 1 | NA | NA | NA | NA | 1 | 1 | NA | NA | NA | NA | 2 | 1 | NA | NA | 2 | 1 |
| 11_5882327 | 58 | 0.069 | 63 | 0.2063 | 62 | 0.0968 | 49 | 0.1429 | 232 | 0.1293 | 47 | 0.1489 | 51 | 0.1765 | 69 | 0.1014 | 56 | 0.125 | 223 | 0.1345 |
| 11_5882394 | 46 | 0.2826 | 54 | 0.3704 | 60 | 0.4667 | 48 | 0.3542 | 208 | 0.375 | 45 | 0.4667 | 56 | 0.4643 | 59 | 0.3051 | 55 | 0.4182 | 215 | 0.4093 |
| 11_5882409 | 48 | 0.1667 | 57 | 0.1053 | 66 | 0.1212 | 54 | 0.1296 | 225 | 0.1289 | 43 | 0.1163 | 67 | 0.0896 | 54 | 0.0926 | 61 | 0.1639 | 225 | 0.1156 |
| 11_5883930 | NA | NA | NA | NA | 1 | 1 | 1 | 1 | 2 | 1 | NA | NA | NA | NA | 1 | 0 | 1 | 1 | 2 | 0.5 |
| 11_5883934 | 1 | 1 | NA | NA | 1 | 1 | 1 | 1 | 3 | 1 | NA | NA | NA | NA | 1 | 1 | 1 | 1 | 2 | 1 |
| 11_5883954 | 1 | 0 | NA | NA | 1 | 1 | 1 | 0 | 3 | 0.3333 | NA | NA | NA | NA | 1 | 0 | 1 | 1 | 2 | 0.5 |
| 11_5883995 | 8 | 0.625 | 2 | 0.5 | 4 | 1 | 7 | 0.7143 | 21 | 0.7143 | 6 | 1 | 9 | 0.7778 | 3 | 1 | 3 | 0.6667 | 21 | 0.8571 |
| 11_5884025 | 2 | 1 | NA | NA | 2 | 1 | 3 | 1 | 7 | 1 | 3 | 0.6667 | 5 | 1 | 1 | 1 | 4 | 0.75 | 13 | 0.8462 |
| 11_5884029 | 8 | 0.75 | 2 | 0 | 4 | 0.75 | 10 | 0.8 | 24 | 0.7083 | 7 | 0.5714 | 8 | 0.5 | 5 | 0.8 | 4 | 0.5 | 24 | 0.5833 |
| 11_5884574 | 6 | 0.5 | 6 | 0.3333 | 4 | 0.25 | 10 | 0.3 | 26 | 0.3462 | 5 | 0.4 | 6 | 0.5 | 3 | 0 | 3 | 0.3333 | 17 | 0.3529 |
| 11_5884599 | 5 | 1 | 3 | 1 | 1 | 1 | 5 | 0.4 | 14 | 0.7857 | 3 | 1 | 6 | 0.8333 | 4 | 1 | NA | NA | 13 | 0.9231 |
| 11_5884608 | 5 | 0.8 | 7 | 0.5714 | 4 | 0.5 | 12 | 0.5833 | 28 | 0.6071 | 5 | 0.6 | 4 | 0.5 | 4 | 0.75 | 2 | 0.5 | 15 | 0.6 |
| 11_5884621 | 7 | 0 | 8 | 0.125 | 4 | 0 | 16 | 0.125 | 35 | 0.0857 | 6 | 0 | 5 | 0.2 | 4 | 0.25 | 3 | 0.3333 | 18 | 0.1667 |
| 11_5884624 | 7 | 1 | 7 | 1 | 4 | 1 | 15 | 1 | 33 | 1 | 4 | 1 | 6 | 1 | 4 | 1 | 3 | 1 | 17 | 1 |
| 11_5884631 | 7 | 0.1429 | 8 | 0.25 | 4 | 0 | 16 | 0.0625 | 35 | 0.1143 | 6 | 0 | 6 | 0 | 3 | 0 | 2 | 0.5 | 17 | 0.0588 |
| 11_5884655 | 3 | 0.3333 | 2 | 0.5 | 4 | 0.25 | 9 | 0.1111 | 18 | 0.2222 | 2 | 0.5 | 2 | 0 | NA | NA | NA | NA | 4 | 0.25 |
| 11_5884691 | 3 | 0.6667 | 3 | 0.3333 | 3 | 0.6667 | 7 | 0.4286 | 16 | 0.5 | 2 | 0.5 | 4 | 0.75 | 1 | 1 | NA | NA | 7 | 0.7143 |
| 11_5886208 | NA | NA | NA | NA | 1 | 1 | NA | NA | 1 | 1 | 2 | 0.5 | NA | NA | NA | NA | NA | NA | 2 | 0.5 |
| 11_5886210 | NA | NA | NA | NA | 1 | 1 | NA | NA | 1 | 1 | 2 | 0.5 | NA | NA | NA | NA | NA | NA | 2 | 0.5 |
| 11_5888106 | 2 | 0 | NA | NA | 2 | 0.5 | 3 | 0 | 7 | 0.1429 | 2 | 0.5 | 2 | 0 | 1 | 0 | 5 | 0 | 10 | 0.1 |
| 11_5888123 | 2 | 0.5 | NA | NA | 2 | 0.5 | 3 | 0 | 7 | 0.2857 | 2 | 0.5 | 3 | 0.3333 | 2 | 0 | 4 | 0 | 11 | 0.1818 |
| 11_5888137 | 1 | 1 | NA | NA | 2 | 1 | 3 | 1 | 6 | 1 | 2 | 1 | 3 | 1 | 3 | 1 | 4 | 1 | 12 | 1 |
| 11_5888146 | 2 | 1 | NA | NA | 1 | 0 | 3 | 0.3333 | 6 | 0.5 | 1 | 1 | 2 | 1 | 3 | 0.6667 | 5 | 0.6 | 11 | 0.7273 |
| 11_5888151 | 2 | 0.5 | NA | NA | 2 | 0.5 | 3 | 0.3333 | 7 | 0.4286 | 2 | 0.5 | 3 | 0 | 3 | 0.3333 | 3 | 0.3333 | 11 | 0.2727 |
| 11_5888154 | 2 | 0 | NA | NA | 3 | 0.3333 | 3 | 0.3333 | 8 | 0.25 | 2 | 1 | 2 | 0.5 | 3 | 0 | 4 | 0.25 | 11 | 0.3636 |
| 11_5888162 | 1 | 0 | 1 | 0 | 3 | 0 | 3 | 0.3333 | 8 | 0.125 | 2 | 0.5 | 4 | 0 | 3 | 0 | 4 | 0 | 13 | 0.0769 |
| 11_5888171 | 2 | 0 | 1 | 0 | 1 | 1 | 3 | 0.3333 | 7 | 0.2857 | 2 | 1 | 4 | 0.5 | 2 | 0 | 4 | 0.5 | 12 | 0.5 |
| 11_5888210 | 2 | 0 | 1 | 0 | 6 | 0.1667 | 3 | 0 | 12 | 0.0833 | 2 | 0.5 | 5 | 0.2 | 4 | 0 | 4 | 0.25 | 15 | 0.2 |
| 11_5888213 | 1 | 0 | 1 | 0 | 6 | 0.1667 | 3 | 0 | 11 | 0.0909 | 2 | 0.5 | 5 | 0 | 4 | 0.25 | 4 | 0.75 | 15 | 0.3333 |
| 11_5888220 | 2 | 0 | 1 | 0 | 6 | 0 | 3 | 0.3333 | 12 | 0.0833 | 2 | 0.5 | 4 | 0.25 | 4 | 0.25 | 5 | 0.4 | 15 | 0.3333 |
| 11_5888222 | 2 | 0 | 1 | 0 | 6 | 0.1667 | 3 | 0 | 12 | 0.0833 | 2 | 0.5 | 5 | 0 | 4 | 0.5 | 5 | 0.4 | 16 | 0.3125 |
| 11_5904827 | 2 | 0.5 | 2 | 0.5 | 1 | 1 | 4 | 0.75 | 9 | 0.6667 | NA | NA | NA | NA | 1 | 0 | 2 | 0.5 | 3 | 0.3333 |
| 11_5904830 | 1 | 0 | 2 | 0 | NA | NA | 5 | 0.4 | 8 | 0.25 | NA | NA | NA | NA | NA | NA | 2 | 0.5 | 2 | 0.5 |
| 11_5904855 | 1 | 0 | 2 | 0 | 1 | 0 | 2 | 0.5 | 6 | 0.1667 | NA | NA | 1 | 1 | 1 | 0 | 2 | 0 | 4 | 0.25 |
| 11_5904864 | 1 | 1 | 2 | 0 | 1 | 1 | 3 | 0.3333 | 7 | 0.4286 | NA | NA | 1 | 1 | 1 | 0 | 1 | 1 | 3 | 0.6667 |
| 11_5904869 | 1 | 0 | 2 | 1 | 1 | 1 | 3 | 1 | 7 | 0.8571 | NA | NA | 1 | 1 | 1 | 1 | 2 | 1 | 4 | 1 |
| 11_5904878 | NA | NA | NA | NA | NA | NA | 2 | 1 | 2 | 1 | NA | NA | NA | NA | NA | NA | 1 | 1 | 1 | 1 |
| 11_5904880 | 1 | 1 | 2 | 1 | 1 | 1 | 3 | 0.6667 | 7 | 0.8571 | NA | NA | NA | NA | 1 | 1 | 2 | 0.5 | 3 | 0.6667 |
| 11_5904886 | 1 | 1 | 1 | 1 | NA | NA | 1 | 1 | 3 | 1 | NA | NA | NA | NA | NA | NA | 2 | 0.5 | 2 | 0.5 |
| 11_5904911 | 1 | 1 | 2 | 1 | 1 | 1 | 3 | 1 | 7 | 1 | NA | NA | NA | NA | 1 | 1 | 2 | 1 | 3 | 1 |
| 11_5904913 | 1 | 1 | 2 | 0.5 | 1 | 1 | 3 | 0.6667 | 7 | 0.7143 | NA | NA | NA | NA | NA | NA | 2 | 1 | 2 | 1 |
| 11_5904918 | 1 | 1 | 1 | 1 | 1 | 1 | 3 | 1 | 6 | 1 | NA | NA | NA | NA | NA | NA | 2 | 1 | 2 | 1 |
| 11_5904920 | 1 | 1 | 2 | 1 | 1 | 1 | 3 | 1 | 7 | 1 | NA | NA | NA | NA | NA | NA | 2 | 1 | 2 | 1 |
| 11_5904921 | 1 | 0 | 2 | 0.5 | 1 | 1 | 3 | 1 | 7 | 0.7143 | NA | NA | NA | NA | NA | NA | 2 | 1 | 2 | 1 |
| 11_5904926 | NA | NA | 2 | 1 | 1 | 1 | 3 | 0.3333 | 6 | 0.6667 | NA | NA | NA | NA | NA | NA | 1 | 1 | 1 | 1 |
| 11_5904977 | 1 | 1 | 1 | 0 | NA | NA | 1 | 1 | 3 | 0.6667 | NA | NA | NA | NA | NA | NA | 2 | 1 | 2 | 1 |
| 11_5906663 | 1 | 1 | 1 | 0 | 2 | 0.5 | NA | NA | 4 | 0.5 | 2 | 0.5 | 1 | 1 | NA | NA | NA | NA | 3 | 0.6667 |
| 11_5906671 | 1 | 0 | 1 | 0 | 2 | 0.5 | NA | NA | 4 | 0.25 | 2 | 0.5 | 1 | 0 | NA | NA | NA | NA | 3 | 0.3333 |
| 11_5906680 | 1 | 1 | 1 | 0 | 2 | 0 | NA | NA | 4 | 0.25 | 1 | 0 | 1 | 1 | NA | NA | NA | NA | 2 | 0.5 |
| 11_5906807 | NA | NA | 1 | 1 | 1 | 1 | NA | NA | 2 | 1 | 1 | 1 | NA | NA | NA | NA | NA | NA | 1 | 1 |
| 11_5907472 | 28 | 0.1071 | 30 | 0.0333 | 20 | 0.1 | 25 | 0.16 | 103 | 0.0971 | 22 | 0.1364 | 19 | 0.1053 | 26 | 0.1154 | 21 | 0.0952 | 88 | 0.1136 |
| 11_5907477 | 26 | 0.1923 | 29 | 0.2759 | 19 | 0.2105 | 26 | 0.3462 | 100 | 0.26 | 19 | 0.3158 | 16 | 0.375 | 24 | 0.1667 | 19 | 0.3684 | 78 | 0.2949 |
| 11_5907489 | 34 | 0.6471 | 31 | 0.4839 | 24 | 0.6667 | 31 | 0.6129 | 120 | 0.6 | 23 | 0.6087 | 21 | 0.7143 | 27 | 0.6296 | 25 | 0.52 | 96 | 0.6146 |
| 11_5908916 | 1 | 1 | 2 | 0.5 | 1 | 0 | 1 | 0 | 5 | 0.4 | 3 | 0.3333 | NA | NA | 2 | 1 | 1 | 0 | 6 | 0.5 |
| 11_5908937 | 1 | 1 | 2 | 0.5 | 1 | 1 | NA | NA | 4 | 0.75 | 3 | 0.3333 | NA | NA | 2 | 1 | 1 | 1 | 6 | 0.6667 |
| 11_5908943 | 1 | 1 | 2 | 0 | 1 | 0 | 1 | 0 | 5 | 0.2 | 3 | 0 | NA | NA | 2 | 0 | 2 | 0.5 | 7 | 0.1429 |
| 11_5908966 | NA | NA | 2 | 0.5 | NA | NA | NA | NA | 2 | 0.5 | 2 | 0.5 | NA | NA | 1 | 1 | 2 | 0 | 5 | 0.4 |
| 11_5909060 | 1 | 1 | 1 | 1 | 1 | 1 | NA | NA | 3 | 1 | 1 | 1 | NA | NA | 1 | 1 | 2 | 1 | 4 | 1 |
| 11_5909065 | 1 | 1 | NA | NA | 1 | 1 | NA | NA | 2 | 1 | 1 | 1 | NA | NA | 1 | 1 | 2 | 0.5 | 4 | 0.75 |
| 11_5909074 | NA | NA | 1 | 1 | 1 | 1 | NA | NA | 2 | 1 | NA | NA | NA | NA | 1 | 1 | 2 | 1 | 3 | 1 |
| 11_5909078 | NA | NA | 1 | 1 | 1 | 1 | NA | NA | 2 | 1 | NA | NA | NA | NA | 1 | 1 | 2 | 0.5 | 3 | 0.6667 |

| | | | | | | | | | | | | | | | | | | | | |
|---|---|---|---|---|---|---|---|---|---|---|---|---|---|---|---|---|---|---|---|---|
| 11_590908 | NA | NA | 1 | 1 | 1 | 1 | NA | NA | 2 | 1 | NA | NA | NA | NA | 1 | 0 | 2 | 0.5 | 3 | 0.3333 |
| 11_590909 | NA | NA | 1 | 1 | 1 | 0 | NA | NA | 2 | 0.5 | NA | NA | NA | NA | 1 | 1 | 2 | 0.5 | 3 | 0.6667 |
| 11_590909 | NA | NA | 1 | 1 | 1 | 1 | NA | NA | 2 | 1 | NA | NA | NA | NA | 1 | 0 | 2 | 1 | 3 | 0.6667 |
| 11_590965 | 9 | 0.7778 | 8 | 0.625 | 4 | 0.5 | 12 | 0.4167 | 33 | 0.5758 | 7 | 0.1429 | 6 | 0.5 | 9 | 0.5556 | 4 | 0.25 | 26 | 0.3846 |
| 11_590966 | 9 | 0.5556 | 9 | 0.4444 | 4 | 0.5 | 12 | 0.25 | 34 | 0.4118 | 6 | 0.3333 | 6 | 0.6667 | 9 | 0.4444 | 5 | 0.4 | 26 | 0.4615 |
| 11_590967 | 3 | 0.3333 | 5 | 0.2 | 1 | 0 | 5 | 0 | 14 | 0.1429 | 2 | 0.5 | 4 | 0.25 | 4 | 0.25 | NA | NA | 10 | 0.3 |
| 11_590968 | 3 | 0.3333 | 2 | 0.5 | 2 | 0.5 | 3 | 0 | 10 | 0.3 | 1 | 1 | 3 | 0 | 2 | 0 | 1 | 0 | 7 | 0.1429 |
| 11_591011 | 7 | 0 | 2 | 0 | 7 | 0.1429 | 3 | 0.3333 | 19 | 0.1053 | 1 | 0 | 3 | 0.3333 | 1 | 0 | 3 | 0 | 8 | 0.125 |
| 11_591014 | 7 | 0.1429 | 4 | 0 | 7 | 0.1429 | 4 | 0 | 22 | 0.0909 | 1 | 1 | 3 | 0.3333 | NA | NA | 4 | 0.5 | 8 | 0.5 |
| 11_591023 | 72 | 0.2083 | 73 | 0.1507 | 67 | 0.209 | 55 | 0.0909 | 267 | 0.1685 | 65 | 0.1538 | 66 | 0.1364 | 57 | 0.1228 | 63 | 0.2381 | 251 | 0.1633 |
| 11_591033 | 66 | 0.5909 | 52 | 0.5769 | 50 | 0.52 | 45 | 0.5333 | 213 | 0.5587 | 54 | 0.5556 | 65 | 0.5231 | 47 | 0.4681 | 65 | 0.6615 | 231 | 0.5584 |
| 11_591034 | 68 | 0.0588 | 56 | 0.0357 | 58 | 0.0862 | 46 | 0.087 | 228 | 0.0658 | 62 | 0.0484 | 73 | 0.0685 | 43 | 0.0698 | 69 | 0.0435 | 247 | 0.0567 |
| 11_593194 | 34 | 0.1765 | 54 | 0.0926 | 39 | 0.0256 | 38 | 0.1579 | 165 | 0.1091 | 33 | 0.1515 | 34 | 0.0882 | 31 | 0.2258 | 44 | 0.25 | 142 | 0.1831 |
| 11_593289 | 34 | 0.8824 | 39 | 0.9487 | 36 | 0.9167 | 34 | 0.8235 | 143 | 0.8951 | 31 | 0.871 | 34 | 0.9118 | 28 | 0.9286 | 40 | 0.9 | 133 | 0.9023 |
| 11_593289 | 30 | 0.2 | 35 | 0.2286 | 31 | 0.129 | 32 | 0.1562 | 128 | 0.1797 | 31 | 0.2581 | 31 | 0.0645 | 27 | 0.1852 | 35 | 0.2 | 124 | 0.1774 |
| 11_593290 | 21 | 0.3333 | 25 | 0.48 | 16 | 0.4375 | 19 | 0.5263 | 81 | 0.4444 | 16 | 0.4375 | 20 | 0.7 | 15 | 0.2667 | 18 | 0.4444 | 69 | 0.4783 |
| 11_593291 | 19 | 0.0526 | 22 | 0 | 16 | 0.0625 | 19 | 0.1579 | 76 | 0.0658 | 14 | 0.1429 | 23 | 0.0435 | 20 | 0 | 22 | 0.0909 | 79 | 0.0633 |
| 11_593291 | NA | NA | 5 | 0.2 | 1 | 0 | NA | NA | 6 | 0.1667 | 4 | 0 | 4 | 0.25 | 3 | 0 | 2 | 0 | 13 | 0.0769 |
| 11_593292 | 25 | 0.04 | 38 | 0.1053 | 28 | 0.0357 | 29 | 0.1034 | 120 | 0.075 | 29 | 0.1724 | 33 | 0.0303 | 27 | 0.1852 | 35 | 0.1714 | 124 | 0.1371 |
| 11_593292 | 24 | 0.125 | 30 | 0.2667 | 21 | 0.0476 | 21 | 0.0952 | 96 | 0.1458 | 21 | 0.0476 | 27 | 0.1481 | 17 | 0 | 31 | 0.129 | 96 | 0.0938 |
| 11_593292 | 26 | 0.6923 | 35 | 0.5143 | 22 | 0.6818 | 24 | 0.7083 | 107 | 0.6355 | 24 | 0.5 | 27 | 0.4074 | 23 | 0.6087 | 31 | 0.6774 | 105 | 0.5524 |
| 11_593462 | NA | NA | NA | NA | 3 | 1 | 2 | 1 | 5 | 1 | NA | NA | 2 | 1 | 1 | 1 | 3 | 1 | 6 | 1 |
| 11_593462 | NA | NA | NA | NA | 2 | 0 | 2 | 0.5 | 4 | 0.25 | NA | NA | 2 | 0.5 | NA | NA | 2 | 1 | 4 | 0.75 |
| 11_593463 | NA | NA | NA | NA | 3 | 0.6667 | 2 | 0 | 5 | 0.4 | NA | NA | NA | NA | 1 | 0 | 2 | 1 | 3 | 0.6667 |
| 11_593463 | NA | NA | NA | NA | 3 | 0.6667 | 2 | 0.5 | 5 | 0.6 | NA | NA | 2 | 0.5 | 1 | 1 | 3 | 0.6667 | 6 | 0.6667 |
| 11_593465 | NA | NA | NA | NA | 3 | 0.3333 | 2 | 0 | 5 | 0.2 | NA | NA | 2 | 0 | 1 | 1 | 3 | 0.3333 | 6 | 0.3333 |
| 11_593467 | NA | NA | NA | NA | 2 | 1 | 2 | 1 | 4 | 1 | NA | NA | 2 | 1 | 1 | 1 | 3 | 1 | 6 | 1 |
| 11_593467 | NA | NA | NA | NA | 3 | 0.6667 | 2 | 0.5 | 5 | 0.6 | NA | NA | 2 | 1 | 1 | 1 | 3 | 1 | 6 | 1 |
| 11_593531 | 1 | 1 | 1 | 0 | 1 | 1 | 2 | 1 | 5 | 0.8 | NA | NA | NA | NA | NA | NA | 1 | 1 | 1 | 1 |
| 11_593533 | 1 | 0 | 1 | 0 | 1 | 1 | 2 | 0.5 | 5 | 0.4 | NA | NA | NA | NA | NA | NA | 1 | 1 | 1 | 1 |
| 11_593534 | NA | NA | 1 | 0 | NA | NA | 2 | 0.5 | 3 | 0.3333 | NA | NA | NA | NA | NA | NA | 1 | 1 | 1 | 1 |
| 11_593534 | 1 | 1 | 1 | 1 | NA | NA | 2 | 1 | 4 | 1 | NA | NA | NA | NA | NA | NA | 1 | 1 | 1 | 1 |
| 11_593535 | 2 | 0.5 | NA | NA | 1 | 0 | 1 | 0 | 4 | 0.25 | 1 | 1 | 1 | 0 | 1 | 0 | 1 | 0 | 4 | 0.25 |
| 11_593536 | 1 | 0 | 1 | 0 | 1 | 0 | 2 | 0.5 | 5 | 0.2 | NA | NA | NA | NA | NA | NA | 1 | 1 | 1 | 1 |
| 11_593544 | 1 | 1 | 1 | 1 | NA | NA | 2 | 1 | 4 | 1 | NA | NA | NA | NA | NA | NA | 1 | 1 | 1 | 1 |
| 11_593547 | 1 | 1 | NA | NA | NA | NA | 2 | 1 | 3 | 1 | NA | NA | NA | NA | NA | NA | 1 | 1 | 1 | 1 |
| 11_593549 | 1 | 1 | 1 | 1 | NA | NA | 2 | 1 | 4 | 1 | NA | NA | NA | NA | NA | NA | 1 | 1 | 1 | 1 |
| 11_593549 | 1 | 0 | 1 | 1 | NA | NA | 2 | 1 | 4 | 0.75 | NA | NA | NA | NA | NA | NA | 1 | 1 | 1 | 1 |
| 11_593764 | 2 | 1 | 1 | 1 | 1 | 0 | 2 | 1 | 6 | 0.8333 | NA | NA | 1 | 0 | 3 | 1 | NA | NA | 4 | 0.75 |
| 11_593764 | 1 | 1 | 1 | 1 | 1 | 1 | 1 | 1 | 4 | 1 | NA | NA | 1 | 1 | 3 | 1 | NA | NA | 4 | 1 |
| 11_593770 | 1 | 0 | 1 | 0 | 2 | 1 | 2 | 1 | 6 | 0.6667 | NA | NA | 1 | 1 | 1 | 0 | NA | NA | 2 | 0.5 |
| 11_593780 | 1 | 1 | 1 | 1 | 2 | 1 | 2 | 1 | 6 | 1 | NA | NA | NA | NA | 1 | 1 | NA | NA | 1 | 1 |
| 11_593782 | 1 | 0 | 1 | 0 | 2 | 1 | 2 | 0.5 | 6 | 0.5 | NA | NA | NA | NA | 1 | 1 | NA | NA | 1 | 1 |
| 11_593784 | 1 | 0 | NA | NA | 1 | 0 | 1 | 1 | 3 | 0.3333 | NA | NA | NA | NA | 1 | 1 | NA | NA | 1 | 1 |
| 11_594017 | 2 | 0.5 | 1 | 1 | NA | NA | NA | NA | 3 | 0.6667 | NA | NA | NA | NA | 1 | 0 | 2 | 0.5 | 3 | 0.3333 |
| 11_596319 | 2 | 0 | 4 | 0.25 | 3 | 0 | 7 | 0 | 16 | 0.0625 | 2 | 0 | 5 | 0 | 3 | 0.3333 | 4 | 0 | 14 | 0.0714 |
| 11_596328 | 5 | 0.8 | 3 | 0.3333 | 2 | 0.5 | 7 | 0.2857 | 17 | 0.4706 | 2 | 0.5 | 8 | 0.75 | 2 | 0.5 | 6 | 0.5 | 18 | 0.6111 |
| 11_596434 | 6 | 0 | 3 | 0 | 5 | 0.4 | 8 | 0 | 22 | 0.0909 | 3 | 0.3333 | 8 | 0 | 3 | 0 | 10 | 0.2 | 24 | 0.125 |
| 11_596435 | 7 | 0.5714 | 4 | 0.25 | 6 | 0.3333 | 9 | 0.2222 | 26 | 0.3462 | 3 | 0 | 10 | 0 | 4 | 0 | 11 | 0.2727 | 28 | 0.1071 |
| 11_596436 | 6 | 0.3333 | 4 | 0 | 5 | 0 | 8 | 0.125 | 23 | 0.1304 | 3 | 0.3333 | 10 | 0.2 | 4 | 0.75 | 11 | 0.3636 | 28 | 0.3571 |
| 11_596436 | 6 | 0.3333 | 4 | 0 | 5 | 0.4 | 5 | 0.2 | 20 | 0.25 | 3 | 0 | 10 | 0 | 4 | 0.5 | 9 | 0.4444 | 26 | 0.2308 |
| 11_596438 | 6 | 0.1667 | 4 | 0.25 | 5 | 0.6 | 8 | 0 | 23 | 0.2174 | 3 | 0 | 9 | 0.1111 | 3 | 0.3333 | 12 | 0 | 27 | 0.0741 |
| 11_596483 | 4 | 0.25 | 2 | 0 | NA | NA | 3 | 0 | 9 | 0.1111 | 1 | 0 | 1 | 0 | 4 | 0.25 | 6 | 0.1667 | 12 | 0.1667 |
| 11_596485 | 4 | 0.25 | 3 | 0.6667 | 1 | 1 | 3 | 0.3333 | 11 | 0.4545 | 2 | 1 | 2 | 0 | 3 | 0.6667 | 5 | 0 | 12 | 0.3333 |
| 11_596492 | 5 | 0.2 | 2 | 0 | 2 | 0 | 4 | 0 | 13 | 0.0769 | 3 | 0 | 3 | 0.3333 | 5 | 0.2 | 8 | 0.375 | 19 | 0.2632 |
| 11_596492 | 5 | 0.2 | 2 | 0 | 2 | 0 | 3 | 0 | 12 | 0.0833 | 3 | 0.3333 | 1 | 0 | 5 | 0.2 | 9 | 0 | 18 | 0.1111 |
| 11_596498 | 5 | 0.8 | 1 | 1 | 2 | 0.5 | 4 | 0.75 | 12 | 0.75 | 4 | 0.75 | 4 | 0.5 | 5 | 0.6 | 9 | 0.7778 | 22 | 0.6818 |
| 11_596500 | 3 | 0.3333 | 2 | 0 | 3 | 0.6667 | 3 | 0.3333 | 11 | 0.3636 | 2 | 0 | 3 | 0.3333 | 4 | 0 | 6 | 0.1667 | 15 | 0.1333 |
| 11_596502 | 6 | 0.1667 | 4 | 0 | 4 | 0.25 | 7 | 0.1429 | 21 | 0.1429 | 4 | 0.25 | 7 | 0 | 5 | 0.6 | 12 | 0.25 | 28 | 0.25 |
| 11_596502 | 6 | 0.6667 | 4 | 0.75 | 4 | 0.5 | 6 | 0.3333 | 20 | 0.55 | 4 | 0.75 | 6 | 0 | 4 | 0.5 | 10 | 0.4 | 24 | 0.375 |
| 11_596505 | 6 | 0.5 | 5 | 0.4 | 5 | 0.8 | 4 | 0.5 | 20 | 0.55 | 5 | 0.2 | 9 | 0.2222 | 5 | 0.4 | 13 | 0.3846 | 32 | 0.3125 |
| 11_596507 | 7 | 0.7143 | 6 | 0.8333 | 5 | 0.4 | 6 | 0.3333 | 24 | 0.5833 | 6 | 0.8333 | 8 | 0.25 | 6 | 0.5 | 12 | 0.5 | 32 | 0.5 |
| 11_596509 | 7 | 0 | 5 | 0.2 | 5 | 0 | 7 | 0.2857 | 24 | 0.125 | 8 | 0.25 | 8 | 0.25 | 7 | 0.2857 | 11 | 0 | 34 | 0.1765 |
| 11_596509 | 8 | 0.5 | 6 | 0.1667 | 5 | 0.2 | 7 | 0.5714 | 26 | 0.3846 | 7 | 0.2857 | 8 | 0.625 | 7 | 0.4286 | 13 | 0.3846 | 35 | 0.4286 |
| 11_596510 | 8 | 0.25 | 7 | 0 | 5 | 0 | 7 | 0 | 27 | 0.0741 | 9 | 0.1111 | 10 | 0.1 | 8 | 0.25 | 13 | 0.1538 | 40 | 0.15 |

| | | | | | | | | | | | | | | | | | | | | |
|---|---|---|---|---|---|---|---|---|---|---|---|---|---|---|---|---|---|---|---|---|
| 11_5965159 | 8 | 0.5 | 8 | 0.75 | 5 | 0.6 | 10 | 0.2 | 31 | 0.4839 | 11 | 0.4545 | 12 | 0.1667 | 8 | 0.25 | 14 | 0.4286 | 45 | 0.3333 |
| 11_5965169 | 7 | 0.2857 | 5 | 0 | 5 | 0 | 9 | 0.1111 | 26 | 0.1154 | 10 | 0 | 9 | 0.1111 | 8 | 0.125 | 12 | 0.0833 | 39 | 0.0769 |
| 11_5965177 | 8 | 0.125 | 6 | 0.1667 | 6 | 0 | 9 | 0.1111 | 29 | 0.1034 | 12 | 0.1667 | 13 | 0.0769 | 8 | 0.25 | 13 | 0.1538 | 46 | 0.1522 |
| 11_5965179 | 5 | 0.2 | 6 | 0.3333 | 5 | 0.4 | 7 | 0.2857 | 23 | 0.3043 | 11 | 0.0909 | 10 | 0 | 6 | 0.1667 | 12 | 0.0833 | 39 | 0.0769 |
| 11_5965182 | 8 | 0.625 | 8 | 0.125 | 7 | 0.1429 | 9 | 0.5556 | 32 | 0.375 | 12 | 0.4167 | 13 | 0.4615 | 7 | 0.4286 | 11 | 0.2727 | 43 | 0.3953 |
| 11_5965189 | 9 | 0.5556 | 5 | 0.6 | 7 | 0.4286 | 7 | 0.2857 | 28 | 0.4643 | 10 | 0.5 | 10 | 0.7 | 6 | 0.6667 | 14 | 0.5 | 40 | 0.575 |
| 11_5965213 | 11 | 0.6364 | 7 | 0.8571 | 7 | 0.8571 | 14 | 0.7857 | 39 | 0.7692 | 11 | 0.9091 | 13 | 0.6154 | 7 | 0.5714 | 13 | 0.8462 | 44 | 0.75 |
| 11_5966639 | 21 | 0.0952 | 18 | 0.0556 | 20 | 0 | 33 | 0.0606 | 92 | 0.0543 | 15 | 0.0667 | 27 | 0.1111 | 14 | 0.2143 | 25 | 0.12 | 81 | 0.1235 |
| 11_5966651 | 19 | 0.3158 | 13 | 0.3077 | 13 | 0.4615 | 28 | 0.4286 | 73 | 0.3836 | 9 | 0.3333 | 21 | 0.2857 | 13 | 0.5385 | 21 | 0.4286 | 64 | 0.3906 |
| 11_5966653 | 20 | 0.6 | 14 | 0.4286 | 18 | 0.5 | 34 | 0.5882 | 86 | 0.5465 | 16 | 0.6875 | 27 | 0.5926 | 15 | 0.7333 | 25 | 0.64 | 83 | 0.6506 |
| 11_5966807 | 4 | 0.25 | 1 | 1 | 1 | 0 | 6 | 0.8333 | 12 | 0.5833 | 1 | 0 | 1 | 1 | 5 | 0.4 | 1 | 0 | 8 | 0.375 |
| 11_5966812 | 3 | 0.3333 | 1 | 0 | 1 | 0 | 6 | 0.1667 | 11 | 0.1818 | 1 | 1 | 2 | 1 | 3 | 0 | 1 | 0 | 7 | 0.4286 |
| 11_5968277 | 8 | 0.375 | 5 | 0.4 | 6 | 0.1667 | 7 | 0.4286 | 26 | 0.3462 | 3 | 0 | 7 | 0.2857 | 5 | 0.4 | 7 | 0.1429 | 22 | 0.2273 |
| 11_5968282 | 7 | 0.4286 | 9 | 0.1111 | 6 | 0 | 5 | 0.4 | 27 | 0.2222 | 2 | 0 | 6 | 0.1667 | 6 | 0.1667 | 6 | 0.3333 | 20 | 0.2 |
| 11_5968284 | 8 | 0.625 | 7 | 1 | 7 | 1 | 6 | 0.8333 | 28 | 0.8571 | 3 | 1 | 7 | 0.8571 | 7 | 0.5714 | 7 | 1 | 24 | 0.8333 |
| 11_5968292 | 5 | 0.8 | 9 | 0.6667 | 5 | 0.4 | 7 | 0.8571 | 26 | 0.6923 | 4 | 0.5 | 5 | 0.2 | 4 | 0.25 | 6 | 0.5 | 19 | 0.3684 |
| 11_5968307 | 4 | 0.5 | 8 | 0.125 | 5 | 0.8 | 5 | 0.2 | 22 | 0.3636 | 3 | 0.6667 | 6 | 0.6667 | 5 | 0.4 | 5 | 0.2 | 19 | 0.4737 |
| 11_5968319 | 7 | 0.8571 | 7 | 0.5714 | 5 | 0.6 | 6 | 0.6667 | 25 | 0.68 | 2 | 0.5 | 7 | 0.8571 | 6 | 0.5 | 7 | 0.7143 | 22 | 0.6818 |
| 11_5968321 | 7 | 0.4286 | 7 | 0.2857 | 4 | 0.5 | 7 | 0.4286 | 25 | 0.4 | 4 | 0.5 | 7 | 0.4286 | 5 | 0.4 | 7 | 0.1429 | 23 | 0.3478 |
| 11_5968324 | 7 | 0.2857 | 9 | 0.2222 | 5 | 0.4 | 8 | 0.375 | 29 | 0.3103 | 4 | 0.5 | 7 | 0.2857 | 6 | 0.3333 | 6 | 0.3333 | 23 | 0.3478 |
| 11_5968325 | 4 | 0 | 7 | 0.1429 | 1 | 0 | 8 | 0.25 | 20 | 0.15 | 2 | 0.5 | 7 | 0.1429 | 3 | 0.6667 | 6 | 0 | 18 | 0.2222 |
| 11_5968327 | 6 | 0.1667 | 9 | 0.3333 | 5 | 0.6 | 7 | 0.4286 | 27 | 0.3704 | 3 | 0 | 6 | 0.5 | 6 | 0.1667 | 7 | 0.2857 | 22 | 0.2727 |
| 11_5968331 | 6 | 0.8333 | 8 | 1 | 3 | 0.6667 | 7 | 0.7143 | 24 | 0.8333 | 4 | 0.75 | 7 | 1 | 5 | 0.8 | 6 | 0.6667 | 22 | 0.8182 |
| 11_5968336 | 5 | 0.6 | 4 | 0.75 | 2 | 1 | 7 | 0.8571 | 18 | 0.7778 | 2 | 1 | 5 | 1 | 2 | 0.5 | 5 | 0.4 | 14 | 0.7143 |
| 11_5968338 | 4 | 0.75 | 5 | 0.2 | 4 | 0.5 | 6 | 0.3333 | 19 | 0.4211 | 2 | 0.5 | 5 | 0.4 | 2 | 0.5 | 6 | 0.3333 | 15 | 0.4 |
| 11_5968338 | 1 | 1 | NA | NA | NA | NA | 1 | 1 | 2 | 1 | 1 | 1 | 1 | 0 | NA | NA | 1 | 1 | 3 | 0.6667 |
| 11_5968342 | 5 | 0.2 | 5 | 0.2 | 5 | 0.2 | 6 | 0.6667 | 21 | 0.3333 | 3 | 0.3333 | 6 | 0.5 | 6 | 0.1667 | 6 | 0.3333 | 21 | 0.3333 |
| 11_5968350 | 5 | 0.6 | 9 | 0.4444 | 5 | 0.2 | 6 | 0.5 | 25 | 0.44 | 4 | 1 | 7 | 0.5714 | 6 | 0.5 | 6 | 0.5 | 23 | 0.6087 |
| 11_5970878 | 15 | 0.2 | 11 | 0.1818 | 10 | 0.3 | 14 | 0.1429 | 50 | 0.2 | 6 | 0.1667 | 13 | 0.3077 | 17 | 0.1176 | 10 | 0 | 46 | 0.1522 |
| 11_5973907 | 2 | 0.5 | 5 | 0.8 | 1 | 0 | 3 | 0.3333 | 11 | 0.5455 | 2 | 0.5 | 3 | 0.6667 | NA | NA | 3 | 0 | 8 | 0.375 |
| 11_5976675 | 3 | 0.3333 | 8 | 0.125 | 5 | 0.2 | 7 | 0 | 23 | 0.1304 | 1 | 0 | 5 | 0 | 3 | 0.3333 | 7 | 0 | 16 | 0.0625 |
| 11_5976677 | 2 | 0.5 | 11 | 0.8182 | 4 | 0.75 | 9 | 0.5556 | 26 | 0.6923 | 3 | 1 | 7 | 0.4286 | 3 | 0.6667 | 8 | 1 | 21 | 0.7619 |
| 11_5976682 | 2 | 0 | 7 | 0.5714 | 4 | 0.25 | 8 | 0.5 | 21 | 0.4286 | 1 | 1 | 5 | 0.6 | 4 | 0.25 | 6 | 0.3333 | 16 | 0.4375 |
| 11_5976689 | 2 | 0 | 7 | 0.4286 | 4 | 0 | 8 | 0 | 21 | 0.1429 | 2 | 0.5 | 7 | 0.4286 | 4 | 0.25 | 7 | 0 | 20 | 0.25 |
| 11_5985159 | 5 | 1 | 14 | 0.9286 | 4 | 0.75 | 3 | 1 | 26 | 0.9231 | 8 | 1 | 9 | 1 | 2 | 1 | 8 | 0.75 | 27 | 0.9259 |
| 11_5985167 | 3 | 0.3333 | 13 | 0.6154 | 2 | 0 | 3 | 0 | 21 | 0.4286 | 6 | 0.5 | 9 | 0.5556 | 2 | 1 | 4 | 0 | 21 | 0.4762 |
| 11_5985186 | 5 | 0 | 14 | 0.0714 | 4 | 0.25 | 3 | 0 | 26 | 0.0769 | 9 | 0.2222 | 10 | 0 | 3 | 0.3333 | 7 | 0 | 29 | 0.1034 |
| 11_5985195 | 4 | 0.5 | 6 | 0.5 | 2 | 0.5 | 2 | 0.5 | 14 | 0.5 | 3 | 0.3333 | 4 | 0.25 | 2 | 0.5 | 5 | 0.6 | 14 | 0.4286 |
| 11_5985202 | 5 | 0.4 | 6 | 0.1667 | 1 | 0 | 2 | 0.5 | 14 | 0.2857 | 3 | 0 | 4 | 0 | 3 | 0 | 5 | 0.4 | 15 | 0.1333 |
| 11_5985250 | 6 | 1 | 6 | 1 | 2 | 1 | 3 | 0.6667 | 17 | 0.9412 | 2 | 1 | 4 | 0.75 | 3 | 0.6667 | 6 | 0.8333 | 15 | 0.8 |
| 11_5985255 | 6 | 0.1667 | 7 | 0 | 3 | 0 | 3 | 0 | 19 | 0.0526 | 3 | 0 | 4 | 0 | 4 | 0.25 | 7 | 0 | 18 | 0.0556 |
| 11_5985260 | 6 | 1 | 6 | 1 | 2 | 1 | 3 | 0.6667 | 17 | 0.9412 | 3 | 1 | 4 | 1 | 3 | 1 | 5 | 0.8 | 15 | 0.9333 |
| 11_5985271 | 6 | 0.3333 | 4 | 0.25 | 4 | 0 | 2 | 0 | 16 | 0.1875 | 3 | 0.6667 | 4 | 0 | 4 | 0.25 | 7 | 0.2857 | 18 | 0.2778 |
| 11_5985274 | 6 | 0 | 6 | 0.1667 | 4 | 0 | 3 | 0 | 19 | 0.0526 | 2 | 0.5 | 5 | 0 | 3 | 0 | 7 | 0 | 17 | 0.0588 |
| 11_5985281 | 6 | 1 | 7 | 0.8571 | 4 | 1 | 3 | 0.6667 | 20 | 0.9 | 3 | 1 | 5 | 1 | 3 | 1 | 7 | 1 | 18 | 1 |
| 11_5985282 | 6 | 1 | 6 | 0.8333 | 4 | 0.75 | 1 | 0 | 17 | 0.8235 | 3 | 1 | 2 | 1 | 2 | 1 | 7 | 0.7143 | 14 | 0.8571 |
| 11_5985288 | 6 | 0 | 6 | 0.1667 | 3 | 0.3333 | 3 | 0 | 18 | 0.1111 | 2 | 0.5 | 4 | 0.25 | 4 | 0.5 | 6 | 0 | 16 | 0.25 |
| 11_5985303 | 1 | 1 | 3 | 0 | 1 | 0 | 1 | 0 | 6 | 0.1667 | 2 | 0.5 | 5 | 0.2 | 2 | 0 | 2 | 0 | 11 | 0.1818 |
| 11_5985417 | 2 | 0.5 | 3 | 0 | NA | NA | 5 | 0 | 10 | 0.1 | 4 | 0.5 | 3 | 0 | 3 | 0.3333 | 1 | 0 | 11 | 0.2727 |
| 11_5985422 | 1 | 0 | 3 | 1 | 1 | 1 | 5 | 0.6 | 10 | 0.7 | 3 | 1 | 4 | 0.5 | 2 | 0.5 | 1 | 1 | 10 | 0.7 |
| 11_5985430 | 1 | 1 | 3 | 0.6667 | 1 | 1 | 5 | 0.4 | 10 | 0.6 | 3 | 0.6667 | 4 | 0.5 | 3 | 0.3333 | NA | NA | 10 | 0.5 |
| 11_5985433 | 1 | 1 | 3 | 0.6667 | 1 | 1 | 5 | 0.6 | 10 | 0.7 | 3 | 0.6667 | 4 | 1 | 2 | 1 | 1 | 1 | 10 | 0.9 |
| 11_5985444 | 2 | 0 | 3 | 0.3333 | 1 | 0 | 5 | 0 | 11 | 0.0909 | 3 | 0.3333 | 3 | 0.3333 | 5 | 0 | 1 | 0 | 12 | 0.1667 |
| 11_5985445 | 2 | 0 | 3 | 1 | 1 | 1 | 4 | 0 | 10 | 0.4 | 3 | 0.6667 | 2 | 0.5 | 4 | 1 | 1 | 1 | 10 | 0.8 |
| 11_5985448 | 2 | 0 | 2 | 1 | 1 | 1 | 2 | 1 | 7 | 0.7143 | 3 | 1 | 2 | 1 | 2 | 1 | NA | NA | 7 | 1 |
| 11_5985456 | 2 | 1 | 3 | 1 | 1 | 1 | 4 | 0.75 | 10 | 0.9 | 2 | 1 | 4 | 0.75 | 5 | 1 | 1 | 1 | 12 | 0.9167 |
| 11_5985460 | 1 | 1 | 3 | 1 | 1 | 1 | 5 | 0.8 | 10 | 0.9 | 3 | 1 | 3 | 0.6667 | 5 | 1 | 1 | 1 | 12 | 0.9167 |
| 11_5985466 | 2 | 0 | 3 | 0 | 1 | 1 | 5 | 0.4 | 11 | 0.2727 | 2 | 1 | 4 | 0.75 | 5 | 0.4 | 1 | 1 | 12 | 0.6667 |
| 11_5985503 | 2 | 0.5 | 3 | 0.3333 | 1 | 0 | 5 | 0.8 | 11 | 0.5455 | NA | NA | 3 | 1 | 4 | 0.25 | 1 | 1 | 8 | 0.625 |
| 11_5985558 | 2 | 0.5 | 1 | 1 | 1 | 0 | 5 | 0.2 | 9 | 0.3333 | NA | NA | 2 | 0 | 3 | 0.3333 | 1 | 0 | 6 | 0.1667 |
| 11_6003114 | 3 | 0.3333 | 1 | 0 | 2 | 0 | 2 | 0 | 8 | 0.125 | 1 | 0 | 1 | 1 | 1 | 0 | 2 | 0.5 | 5 | 0.4 |
| 11_6003174 | 2 | 0.5 | 1 | 0 | 1 | 0 | NA | NA | 4 | 0.25 | 1 | 0 | 1 | 1 | 1 | 0 | 2 | 0 | 5 | 0.2 |
| 11_6003187 | 1 | 1 | 1 | 1 | 2 | 1 | 1 | 1 | 5 | 1 | 1 | 1 | 1 | 1 | 2 | 1 | 2 | 1 | 6 | 1 |
| 11_6007608 | 1 | 1 | 1 | 1 | NA | NA | 1 | 1 | 3 | 1 | 1 | 1 | 1 | 1 | NA | NA | NA | NA | 2 | 1 |
| 11_6007642 | 2 | 1 | 2 | 1 | NA | NA | 1 | 1 | 5 | 1 | 1 | 0 | 1 | 1 | NA | NA | NA | NA | 2 | 0.5 |

| | | | | | | | | | | | | | | | | | | | | |
|---|---|---|---|---|---|---|---|---|---|---|---|---|---|---|---|---|---|---|---|---|
| 11_600768( | 1 | 0 | 2 | 0 | NA | NA | 1 | 1 | 4 | 0.25 | 1 | 1 | 2 | 0 | NA | NA | NA | NA | 3 | 0.3333 |
| 11_600768 | 2 | 1 | 2 | 0.5 | NA | NA | 1 | 1 | 5 | 0.8 | 1 | 0 | 2 | 0.5 | NA | NA | NA | NA | 3 | 0.3333 |
| 11_600768! | 1 | 1 | 1 | 0 | NA | NA | 1 | 0 | 3 | 0.3333 | 1 | 0 | 2 | 0.5 | NA | NA | NA | NA | 3 | 0.3333 |
| 11_600769! | 1 | 0 | 1 | 0 | NA | NA | 1 | 1 | 3 | 0.3333 | 1 | 1 | 1 | 0 | NA | NA | NA | NA | 2 | 0.5 |
| 11_600771( | 1 | 1 | 2 | 0 | NA | NA | 1 | 0 | 4 | 0.25 | 1 | 1 | 2 | 0 | NA | NA | NA | NA | 3 | 0.3333 |
| 11_600774: | 2 | 0.5 | 2 | 1 | NA | NA | 3 | 0.6667 | 7 | 0.7143 | 1 | 1 | 1 | 1 | NA | NA | NA | NA | 2 | 1 |
| 11_600777: | 1 | 1 | 2 | 0.5 | NA | NA | 3 | 1 | 6 | 0.8333 | 1 | 1 | 2 | 1 | NA | NA | NA | NA | 3 | 1 |
| 11_600777! | 2 | 0.5 | 2 | 0.5 | NA | NA | 2 | 1 | 6 | 0.6667 | 1 | 1 | 3 | 1 | NA | NA | NA | NA | 4 | 1 |
| 11_600778: | 2 | 1 | 2 | 0.5 | NA | NA | 3 | 1 | 7 | 0.8571 | 1 | 1 | 3 | 1 | NA | NA | NA | NA | 4 | 1 |
| 11_600778! | 1 | 0 | 1 | 1 | NA | NA | 3 | 1 | 5 | 0.8 | 1 | 1 | 2 | 0 | NA | NA | NA | NA | 3 | 0.3333 |
| 11_600779: | 2 | 1 | 2 | 1 | NA | NA | 2 | 1 | 6 | 1 | 1 | 1 | 3 | 1 | NA | NA | NA | NA | 4 | 1 |
| 11_600780: | 1 | 0 | 3 | 0.6667 | 1 | 1 | 3 | 0.6667 | 8 | 0.625 | 1 | 1 | 2 | 1 | NA | NA | NA | NA | 3 | 1 |
| 11_600780( | 2 | 0.5 | 3 | 0.3333 | NA | NA | 3 | 0 | 8 | 0.25 | 1 | 0 | 3 | 0.6667 | NA | NA | NA | NA | 4 | 0.5 |
| 11_600781! | 1 | 0 | 3 | 0.3333 | 1 | 0 | 4 | 0 | 9 | 0.1111 | 1 | 0 | 3 | 0.3333 | NA | NA | NA | NA | 4 | 0.25 |
| 11_600783' | NA | NA | 2 | 0 | 1 | 1 | 4 | 0.75 | 7 | 0.5714 | 1 | 1 | 3 | 0.6667 | NA | NA | NA | NA | 4 | 0.75 |
| 11_600784: | 2 | 1 | 3 | 0 | 1 | 1 | 4 | 0.25 | 10 | 0.4 | 1 | 0 | 3 | 0.6667 | NA | NA | NA | NA | 4 | 0.5 |
| 11_600785' | 1 | 1 | 3 | 0.6667 | 1 | 1 | 3 | 0.6667 | 8 | 0.75 | NA | NA | 2 | 1 | NA | NA | NA | NA | 2 | 1 |
| 11_600785! | 2 | 0.5 | 3 | 0 | 1 | 0 | 4 | 0 | 10 | 0.1 | 1 | 0 | 5 | 0.4 | NA | NA | NA | NA | 6 | 0.3333 |
| 11_600786' | 2 | 0 | 2 | 0 | 1 | 0 | 3 | 0.3333 | 8 | 0.125 | 1 | 0 | 3 | 0.3333 | NA | NA | NA | NA | 4 | 0.25 |
| 11_600789: | 1 | 0 | 3 | 0.3333 | 2 | 0 | 4 | 0 | 10 | 0.1 | 1 | 0 | 5 | 0.2 | NA | NA | NA | NA | 6 | 0.1667 |
| 11_600789! | 2 | 0.5 | 3 | 0.3333 | 2 | 0 | 4 | 0 | 11 | 0.1818 | 1 | 0 | 5 | 0.2 | NA | NA | NA | NA | 6 | 0.1667 |
| 11_600789! | 1 | 1 | 2 | 0.5 | 1 | 1 | 2 | 0 | 6 | 0.5 | 1 | 1 | 4 | 0.75 | NA | NA | NA | NA | 5 | 0.8 |
| 11_600790! | 2 | 0 | 2 | 0 | 2 | 0 | 4 | 0.5 | 10 | 0.2 | 1 | 0 | 4 | 0.5 | NA | NA | NA | NA | 5 | 0.4 |
| 11_600790! | NA | NA | 2 | 0.5 | 2 | 0.5 | 5 | 0.4 | 9 | 0.4444 | 1 | 1 | 4 | 0.75 | NA | NA | NA | NA | 5 | 0.8 |
| 11_600792! | 2 | 0.5 | 3 | 0 | 2 | 0.5 | 4 | 0.5 | 11 | 0.3636 | NA | NA | 5 | 0.4 | NA | NA | 1 | 0 | 6 | 0.3333 |
| 11_600793! | 2 | 0 | 3 | 0.3333 | 1 | 0 | 5 | 0 | 11 | 0.0909 | 1 | 0 | 4 | 0 | NA | NA | 1 | 1 | 6 | 0.1667 |
| 11_600798( | 2 | 0 | 4 | 0.25 | 2 | 0 | 5 | 0.2 | 13 | 0.1538 | 1 | 1 | 4 | 0 | NA | NA | 1 | 0 | 6 | 0.1667 |
| 11_600798! | 1 | 0 | 3 | 0 | 2 | 0.5 | 6 | 0.1667 | 12 | 0.1667 | 1 | 0 | 5 | 0.2 | NA | NA | 1 | 0 | 7 | 0.1429 |
| 11_600800: | 1 | 0 | 3 | 0 | 2 | 0.5 | 4 | 0.5 | 10 | 0.3 | 1 | 0 | 4 | 0.25 | NA | NA | NA | NA | 5 | 0.2 |
| 11_600801( | 2 | 0.5 | 4 | 0.5 | 2 | 0 | 5 | 0.2 | 13 | 0.3077 | 1 | 1 | 4 | 0.25 | NA | NA | 1 | 1 | 6 | 0.5 |
| 11_600802( | 2 | 0 | 5 | 0.4 | 1 | 0 | 4 | 0.25 | 12 | 0.25 | 1 | 0 | 4 | 0.5 | NA | NA | 1 | 0 | 6 | 0.3333 |
| 11_600802( | 2 | 0.5 | 3 | 0.3333 | 1 | 1 | 5 | 0.6 | 11 | 0.5455 | NA | NA | 5 | 0.6 | NA | NA | 1 | 1 | 6 | 0.6667 |
| 11_600803. | 1 | 0 | 5 | 0 | 2 | 0 | 4 | 0.25 | 12 | 0.0833 | 1 | 0 | 5 | 0.2 | NA | NA | NA | NA | 6 | 0.1667 |
| 11_600809: | NA | NA | 1 | 1 | NA | NA | NA | NA | 1 | 1 | NA | NA | 2 | 1 | NA | NA | 1 | 1 | 3 | 1 |
| 11_600809. | 3 | 0 | 8 | 0 | 2 | 0 | 5 | 0.4 | 18 | 0.1111 | 2 | 0 | 6 | 0.3333 | NA | NA | 1 | 0 | 9 | 0.2222 |
| 11_600809( | 2 | 0.5 | 5 | 0.8 | 1 | 0 | 4 | 0.5 | 12 | 0.5833 | NA | NA | 5 | 1 | NA | NA | 2 | 0 | 7 | 0.7143 |
| 11_600810! | 1 | 0 | 4 | 0.75 | 1 | 1 | 4 | 1 | 10 | 0.8 | 1 | 0 | 3 | 1 | 1 | 1 | NA | NA | 5 | 0.8 |
| 11_600811. | 3 | 1 | 8 | 0.875 | 2 | 1 | 5 | 1 | 18 | 0.9444 | 2 | 1 | 7 | 1 | 2 | 1 | 3 | 1 | 14 | 1 |
| 11_600812! | 2 | 0 | 9 | 0.1111 | 2 | 0 | 5 | 0 | 18 | 0.0556 | 2 | 0 | 6 | 0.1667 | 1 | 0 | 1 | 0 | 10 | 0.1 |
| 11_600813: | 1 | 0 | 6 | 0.8333 | 2 | 0 | 6 | 0.5 | 15 | 0.5333 | 2 | 0.5 | 6 | 0.1667 | 3 | 0.3333 | 2 | 0 | 13 | 0.2308 |
| 11_600813. | 1 | 1 | 8 | 0.125 | 2 | 0 | 4 | 0.25 | 15 | 0.2 | 2 | 0 | 4 | 0.5 | 2 | 0 | 2 | 0 | 10 | 0.2 |
| 11_600813: | 2 | 0.5 | 7 | 0.7143 | 2 | 0.5 | 5 | 0.4 | 16 | 0.5625 | 2 | 0.5 | 8 | 0.25 | 3 | 1 | 2 | 0.5 | 15 | 0.4667 |
| 11_600814. | 1 | 0 | 9 | 0.7778 | 3 | 0.3333 | 5 | 0 | 18 | 0.4444 | 2 | 0 | 8 | 0.375 | 3 | 0.6667 | 2 | 0 | 15 | 0.3333 |
| 11_600891! | 5 | 0.2 | 7 | 0.1429 | 2 | 0.5 | 3 | 0.3333 | 17 | 0.2353 | 5 | 0 | 8 | 0.375 | 1 | 0 | 1 | 0 | 15 | 0.2 |
| 11_600893! | 6 | 0.5 | 13 | 0.0769 | 3 | 0 | 8 | 0.375 | 30 | 0.2333 | 4 | 0 | 9 | 0 | 5 | 0.4 | 4 | 0 | 22 | 0.0909 |
| 11_600894: | 6 | 0.5 | 13 | 0.7692 | 3 | 0.3333 | 6 | 1 | 28 | 0.7143 | 3 | 0.6667 | 9 | 0.7778 | 4 | 0.75 | 3 | 1 | 19 | 0.7895 |
| 11_600894! | 6 | 0.1667 | 14 | 0.0714 | 3 | 0 | 7 | 0.2857 | 30 | 0.1333 | 5 | 0 | 11 | 0 | 5 | 0.2 | 5 | 0.2 | 26 | 0.0769 |
| 11_600906: | 9 | 0.1111 | 10 | 0.2 | 7 | 0.1429 | 12 | 0.0833 | 38 | 0.1316 | 6 | 0.1667 | 9 | 0.4444 | 11 | 0.2727 | 15 | 0.2 | 41 | 0.2683 |
| 11_600907: | 9 | 0.2222 | 9 | 0.3333 | 6 | 0.3333 | 9 | 0 | 33 | 0.2121 | 4 | 0.25 | 8 | 0.375 | 6 | 0 | 10 | 0.2 | 28 | 0.2143 |
| 11_600908( | 5 | 0 | 7 | 0 | 5 | 0.2 | 11 | 0.1818 | 28 | 0.1071 | 6 | 0.1667 | 8 | 0 | 7 | 0 | 11 | 0.2727 | 32 | 0.125 |
| 11_601186! | 24 | 0.0833 | 21 | 0.0476 | 15 | 0 | 15 | 0.2 | 75 | 0.08 | 10 | 0.4 | 13 | 0.1538 | 11 | 0.3636 | 12 | 0.0833 | 46 | 0.2391 |
| 11_601188' | 24 | 0.2917 | 19 | 0.3684 | 14 | 0.5714 | 16 | 0.25 | 73 | 0.3562 | 9 | 0.6667 | 11 | 0 | 13 | 0.4615 | 12 | 0.25 | 45 | 0.3333 |
| 11_601188: | 21 | 0.0952 | 17 | 0.1765 | 10 | 0.1 | 11 | 0.0909 | 59 | 0.1186 | 7 | 0.1429 | 13 | 0.1538 | 12 | 0.3333 | 12 | 0.0833 | 44 | 0.1818 |
| 11_601188! | 25 | 0 | 21 | 0.0952 | 15 | 0.0667 | 14 | 0.1429 | 75 | 0.0667 | 9 | 0.1111 | 14 | 0 | 12 | 0.25 | 13 | 0 | 48 | 0.0833 |
| 11_601190: | 22 | 0.1364 | 17 | 0.1765 | 12 | 0.25 | 11 | 0.2727 | 62 | 0.1935 | 6 | 0 | 10 | 0.2 | 10 | 0.2 | 11 | 0.0909 | 37 | 0.1351 |
| 11_601190! | 23 | 0.0435 | 18 | 0.1111 | 12 | 0.0833 | 9 | 0 | 62 | 0.0645 | 6 | 0 | 10 | 0.1 | 9 | 0 | 11 | 0.0909 | 36 | 0.0556 |
| 11_601191: | 22 | 0.0455 | 15 | 0.0667 | 10 | 0.1 | 10 | 0.4 | 57 | 0.1228 | 8 | 0 | 7 | 0.2857 | 12 | 0.5 | 12 | 0 | 39 | 0.2051 |
| 11_601197! | 17 | 0.1176 | 15 | 0.3333 | 10 | 0.1 | 10 | 0.2 | 52 | 0.1923 | 7 | 0.5714 | 7 | 0.1429 | 4 | 0.5 | 9 | 0 | 27 | 0.2593 |
| 11_601204( | 14 | 0.1429 | 7 | 0.2857 | 8 | 0.125 | 9 | 0.1111 | 38 | 0.1579 | 6 | 0.3333 | 4 | 0.25 | 2 | 0 | 3 | 0.3333 | 15 | 0.2667 |
| 11_601205' | 13 | 0.7692 | 9 | 0.6667 | 6 | 0.8333 | 5 | 0.2 | 33 | 0.6667 | 6 | 1 | 3 | 0.3333 | 2 | 0 | 3 | 1 | 14 | 0.7143 |
| 11_601205: | 12 | 0.0833 | 6 | 0.3333 | 8 | 0.125 | 8 | 0.125 | 34 | 0.1471 | 4 | 0.25 | 3 | 0 | 2 | 0 | 2 | 0 | 11 | 0.0909 |
| 11_601206: | 13 | 0.6923 | 9 | 1 | 7 | 0.8571 | 5 | 0.8 | 34 | 0.8235 | 6 | 0.8333 | 4 | 1 | 2 | 0.5 | 2 | 1 | 14 | 0.8571 |
| 11_601207: | 12 | 0.0833 | 9 | 0.1111 | 7 | 0 | 6 | 0 | 34 | 0.0588 | 6 | 0.1667 | 4 | 0 | 2 | 0 | 2 | 0 | 14 | 0.0714 |
| 11_601208( | 11 | 0.5455 | 9 | 0.6667 | 6 | 0.5 | 7 | 0.7143 | 33 | 0.6061 | 6 | 0.8333 | 3 | 0.6667 | 1 | 1 | 1 | 1 | 11 | 0.8182 |
| 11_601208: | 7 | 0.5714 | 9 | 0.7778 | 7 | 0.4286 | 6 | 0.1667 | 29 | 0.5172 | 5 | 1 | 3 | 0.6667 | 2 | 0.5 | 1 | 1 | 11 | 0.8182 |

| | | | | | | | | | | | | | | | | | | | | |
|---|---|---|---|---|---|---|---|---|---|---|---|---|---|---|---|---|---|---|---|---|
| 11_6028545 | 1 | 0 | 1 | 1 | NA | NA | 1 | 1 | 3 | 0.6667 | 1 | 0 | 1 | 0 | 1 | 1 | 1 | 0 | 4 | 0.25 |
| 11_6028584 | 1 | 0 | 1 | 0 | NA | NA | 1 | 1 | 3 | 0.3333 | 1 | 0 | 1 | 0 | 1 | 1 | 1 | 0 | 4 | 0.25 |
| 11_6028650 | 2 | 0.5 | 1 | 0 | NA | NA | 1 | 0 | 4 | 0.25 | 1 | 0 | 1 | 0 | 1 | 1 | 1 | 0 | 4 | 0.25 |
| 11_6028657 | 2 | 0.5 | 1 | 0 | NA | NA | 1 | 1 | 4 | 0.5 | 1 | 1 | 1 | 1 | NA | NA | 1 | 0 | 3 | 0.6667 |
| 11_6029160 | 5 | 0.2 | 2 | 0 | 3 | 0 | 5 | 0.2 | 15 | 0.1333 | 3 | 0 | 3 | 0 | 3 | 0 | 1 | 1 | 10 | 0.1 |
| 11_6029191 | 5 | 0.4 | 3 | 0.3333 | 4 | 0.25 | 6 | 0.5 | 18 | 0.3889 | 3 | 0.6667 | 3 | 1 | 3 | 0.6667 | 3 | 0.3333 | 12 | 0.6667 |
| 11_6029203 | 6 | 0 | 2 | 0 | 4 | 0.25 | 5 | 0 | 17 | 0.0588 | 4 | 0 | 3 | 0 | 3 | 0.3333 | 3 | 0 | 13 | 0.0769 |
| 11_6029215 | 5 | 0.4 | 2 | 0.5 | 4 | 0.75 | 5 | 0.6 | 16 | 0.5625 | 4 | 0.25 | 3 | 0.3333 | 3 | 0.6667 | 3 | 0.3333 | 13 | 0.3846 |
| 11_6033288 | 4 | 0.5 | 3 | 0 | 6 | 0.5 | 3 | 0 | 16 | 0.3125 | 3 | 0 | 5 | 0.2 | 1 | 1 | 3 | 0.3333 | 12 | 0.25 |
| 11_6033289 | 2 | 0.5 | 3 | 0.3333 | 5 | 1 | 2 | 0.5 | 12 | 0.6667 | 1 | 1 | 5 | 0.8 | 2 | 0.5 | 1 | 1 | 9 | 0.7778 |
| 11_6033290 | 2 | 1 | 4 | 0.5 | 5 | 0.8 | 3 | 1 | 14 | 0.7857 | 3 | 1 | 5 | 0.8 | 2 | 0 | 3 | 0.6667 | 13 | 0.6923 |
| 11_6033294 | 4 | 0 | 3 | 0 | 7 | 0.1429 | 3 | 0 | 17 | 0.0588 | 4 | 0 | 5 | 0 | 2 | 0 | 3 | 0.3333 | 14 | 0.0714 |
| 11_6033298 | 3 | 0 | 2 | 0.5 | 7 | 0.4286 | 3 | 0 | 15 | 0.2667 | 4 | 0.75 | 6 | 0 | 2 | 0 | 3 | 0 | 15 | 0.2 |
| 11_6033311 | 3 | 0.3333 | 2 | 0.5 | 8 | 0.5 | 2 | 0.5 | 15 | 0.4667 | 5 | 0.4 | 5 | 0.2 | 2 | 0 | 2 | 0.5 | 14 | 0.2857 |
| 11_6033339 | 4 | 0 | 3 | 0 | 10 | 0.1 | 3 | 0 | 20 | 0.05 | 5 | 0.2 | 6 | 0.1667 | 3 | 0 | 5 | 0.4 | 19 | 0.2105 |
| 11_6033345 | 3 | 0 | 3 | 0 | 10 | 0 | 4 | 0.5 | 20 | 0.1 | 4 | 0.25 | 6 | 0.1667 | 3 | 0 | 5 | 0.2 | 18 | 0.1667 |
| 11_6033357 | 4 | 0 | 3 | 0 | 8 | 0.375 | 4 | 0.25 | 19 | 0.2105 | 4 | 0.25 | 6 | 0.3333 | 2 | 0 | 3 | 0 | 15 | 0.2 |
| 11_6033364 | 4 | 0 | 3 | 0.3333 | 8 | 0.125 | 5 | 0 | 20 | 0.1 | 4 | 0.25 | 6 | 0.1667 | 3 | 0.3333 | 4 | 0 | 17 | 0.1765 |
| 11_6033372 | 5 | 0.2 | 3 | 0.6667 | 9 | 0.1111 | 3 | 0.3333 | 20 | 0.25 | 5 | 0.6 | 7 | 0 | 2 | 0 | 4 | 0 | 18 | 0.1667 |
| 11_6033383 | 6 | 0.6667 | 3 | 0.3333 | 7 | 0.5714 | 3 | 0 | 19 | 0.4737 | 5 | 0.6 | 6 | 0.8333 | 3 | 0.6667 | 3 | 0.6667 | 17 | 0.7059 |
| 11_6033394 | 6 | 0.3333 | 3 | 0 | 9 | 0 | 5 | 0.2 | 23 | 0.1304 | 5 | 0 | 7 | 0.2857 | 3 | 0 | 4 | 0 | 19 | 0.1053 |
| 11_6033411 | 6 | 0.5 | 3 | 0.6667 | 10 | 0.5 | 4 | 0.25 | 23 | 0.4783 | 4 | 0.25 | 8 | 0.375 | 4 | 0.5 | 3 | 0.3333 | 19 | 0.3684 |
| 11_6033474 | 8 | 0.25 | 4 | 0 | 10 | 0.1 | 7 | 0.2857 | 29 | 0.1724 | 7 | 0.7143 | 8 | 0.25 | 4 | 0.25 | 5 | 0.4 | 24 | 0.4167 |
| 11_6035910 | 7 | 0.1429 | 10 | 0.1 | 5 | 0.4 | 5 | 0 | 27 | 0.1481 | 6 | 0.1667 | 4 | 0 | 4 | 0 | 3 | 0 | 17 | 0.0588 |
| 11_6035914 | 7 | 0.5714 | 9 | 0.3333 | 7 | 0.8571 | 8 | 0.625 | 31 | 0.5806 | 7 | 0.7143 | 5 | 0.6 | 7 | 0.7143 | 5 | 0.2 | 24 | 0.5833 |
| 11_6042808 | 1 | 1 | NA | NA | 1 | 1 | NA | NA | 2 | 1 | NA | NA | 2 | 1 | NA | NA | NA | NA | 2 | 1 |
| 11_6042817 | 1 | 1 | NA | NA | 1 | 0 | NA | NA | 2 | 0.5 | NA | NA | 3 | 0.6667 | NA | NA | NA | NA | 3 | 0.6667 |
| 11_6042843 | 1 | 1 | NA | NA | 1 | 1 | NA | NA | 2 | 1 | NA | NA | 1 | 1 | NA | NA | NA | NA | 1 | 1 |
| 11_6042846 | 1 | 1 | NA | NA | NA | NA | NA | NA | 1 | 1 | NA | NA | 3 | 1 | NA | NA | NA | NA | 3 | 1 |
| 11_6042915 | 4 | 0.25 | NA | NA | NA | NA | NA | NA | 4 | 0.25 | 1 | 1 | NA | NA | 1 | 0 | NA | NA | 2 | 0.5 |
| 11_6042974 | 5 | 0.4 | 1 | 1 | 4 | 0 | 1 | 0 | 11 | 0.2727 | 2 | 0.5 | NA | NA | 1 | 0 | 1 | 1 | 4 | 0.5 |
| 11_6043001 | 4 | 1 | NA | NA | 3 | 0.3333 | 1 | 1 | 8 | 0.75 | 2 | 1 | 1 | 1 | 1 | 0 | 1 | 1 | 5 | 0.8 |
| 11_6044515 | 5 | 1 | NA | NA | 3 | 1 | 1 | 1 | 9 | 1 | 2 | 1 | 1 | 1 | 2 | 0.5 | 1 | 1 | 6 | 0.8333 |
| 11_6044526 | 5 | 1 | NA | NA | 5 | 1 | 1 | 1 | 11 | 1 | 3 | 1 | 2 | 1 | 3 | 1 | 1 | 1 | 9 | 1 |
| 11_6044548 | 4 | 0.25 | NA | NA | 5 | 0.4 | 1 | 0 | 10 | 0.3 | 4 | 0.25 | 2 | 0 | 4 | 1 | 1 | 0 | 11 | 0.4545 |
| 11_6044596 | 5 | 0 | NA | NA | 4 | 0.25 | 2 | 0 | 11 | 0.0909 | 4 | 0.25 | 2 | 0 | 4 | 0 | 1 | 0 | 11 | 0.0909 |
| 11_6044605 | 6 | 0 | 1 | 0 | 5 | 0.2 | 3 | 0 | 15 | 0.0667 | 4 | 0 | 2 | 0 | 5 | 0.2 | 1 | 0 | 12 | 0.0833 |
| 11_6044639 | 6 | 0 | 2 | 0 | 4 | 0 | 6 | 0.1667 | 18 | 0.0556 | 5 | 0 | 2 | 0 | 9 | 0.3333 | 1 | 0 | 17 | 0.1765 |
| 11_6044877 | 6 | 0 | 7 | 0.1429 | 9 | 0.1111 | 4 | 0.25 | 26 | 0.1154 | 2 | 0 | 5 | 0.4 | 2 | 0 | 3 | 0.3333 | 12 | 0.25 |
| 11_6044885 | 4 | 0.25 | 7 | 0.1429 | 9 | 0.1111 | 6 | 0 | 26 | 0.1154 | 3 | 0 | 5 | 0.2 | 3 | 0 | 2 | 0 | 13 | 0.0769 |
| 11_6044915 | 7 | 0.1429 | 7 | 0.2857 | 7 | 0.4286 | 6 | 0.3333 | 27 | 0.2963 | 3 | 0.3333 | 3 | 0.3333 | 2 | 0 | 3 | 0 | 11 | 0.1818 |
| 11_6060396 | 10 | 0.6 | 7 | 0.4286 | 1 | 0 | 5 | 0.8 | 23 | 0.5652 | 7 | 0.4286 | 4 | 0.75 | 2 | 1 | 3 | 0.6667 | 16 | 0.625 |
| 11_6060433 | 9 | 0.8889 | 11 | 0.8182 | 2 | 1 | 7 | 0.8571 | 29 | 0.8621 | 5 | 0.6 | 4 | 1 | 3 | 0.6667 | 3 | 0.6667 | 15 | 0.7333 |
| 11_6060436 | 10 | 0.6 | 11 | 0.4545 | 2 | 0.5 | 9 | 0.3333 | 32 | 0.4688 | 4 | 0.5 | 4 | 0.75 | 2 | 0 | 3 | 0.3333 | 13 | 0.4615 |
| 11_6060438 | 11 | 0.2727 | 11 | 0.0909 | 2 | 0 | 8 | 0.125 | 32 | 0.1562 | 5 | 0 | 4 | 0 | 2 | 0.5 | 2 | 0 | 13 | 0.0769 |
| 11_6060440 | 10 | 0.9 | 10 | 0.6 | 2 | 0.5 | 7 | 0.5714 | 29 | 0.6897 | 5 | 0.2 | 4 | 0.5 | 3 | 0.3333 | 3 | 0.6667 | 15 | 0.4 |
| 11_6060460 | 8 | 0.5 | 10 | 0.1 | 2 | 0 | 8 | 0.25 | 28 | 0.25 | 6 | 0.1667 | 3 | 0.3333 | 2 | 0 | 3 | 0 | 14 | 0.1429 |
| 11_6060473 | 7 | 0.5714 | 7 | 0.5714 | 1 | 1 | 7 | 0.5714 | 22 | 0.5909 | 5 | 0.4 | 3 | 0.6667 | 1 | 0 | 2 | 0.5 | 11 | 0.4545 |
| 11_6060481 | 11 | 0.0909 | 9 | 0.2222 | 2 | 0.5 | 8 | 0.125 | 30 | 0.1667 | 7 | 0.4286 | 4 | 0 | 3 | 0.3333 | 3 | 0.3333 | 17 | 0.2941 |
| 11_6062194 | 1 | 0 | 2 | 0.5 | NA | NA | 1 | 1 | 4 | 0.5 | NA | NA | 1 | 0 | 2 | 0.5 | 1 | 1 | 4 | 0.5 |
| 11_6062199 | 2 | 1 | 3 | 1 | NA | NA | 3 | 1 | 8 | 1 | 2 | 0.5 | 1 | 1 | 2 | 1 | 1 | 1 | 6 | 0.8333 |
| 11_6062207 | NA | NA | 3 | 0.6667 | NA | NA | 3 | 1 | 6 | 0.8333 | 1 | 1 | 1 | 1 | 2 | 1 | 1 | 1 | 5 | 1 |
| 11_6062211 | 2 | 1 | 4 | 1 | NA | NA | 3 | 1 | 9 | 1 | 2 | 1 | 1 | 1 | 2 | 1 | 1 | 1 | 6 | 1 |
| 11_6062215 | 2 | 1 | 4 | 1 | NA | NA | 3 | 0.6667 | 9 | 0.8889 | 2 | 0 | 2 | 0.5 | NA | NA | 1 | 1 | 5 | 0.4 |
| 11_6062244 | 2 | 1 | 3 | 0.6667 | 2 | 1 | 3 | 0.3333 | 10 | 0.7 | 2 | 1 | 2 | 0.5 | 2 | 0.5 | NA | NA | 6 | 0.6667 |
| 11_6062260 | 1 | 1 | 2 | 1 | 2 | 1 | 3 | 1 | 8 | 1 | 2 | 1 | 1 | 0 | 2 | 1 | 1 | 1 | 6 | 0.8333 |
| 11_6062270 | 2 | 0.5 | 2 | 1 | 2 | 0.5 | 2 | 1 | 8 | 0.75 | 2 | 0.5 | 1 | 0 | 1 | 0 | NA | NA | 4 | 0.25 |
| 11_6062272 | 1 | 0 | 4 | 0.25 | 2 | 0 | 2 | 0 | 9 | 0.1111 | 2 | 0 | 2 | 0 | 2 | 0 | 1 | 1 | 7 | 0.1429 |
| 11_6062280 | 2 | 1 | 5 | 1 | 2 | 1 | 3 | 1 | 12 | 1 | 2 | 1 | 2 | 1 | 2 | 1 | 1 | 1 | 7 | 1 |
| 11_6062281 | 2 | 0 | 4 | 0.5 | 1 | 0 | 3 | 0 | 10 | 0.2 | 1 | 0 | 2 | 0 | 2 | 0.5 | 1 | 0 | 6 | 0.1667 |
| 11_6062286 | 2 | 1 | 5 | 1 | 2 | 1 | 3 | 1 | 12 | 1 | 2 | 1 | 1 | 1 | 2 | 0 | 1 | 1 | 6 | 0.6667 |
| 11_6062297 | 1 | 0 | 6 | 0.3333 | 2 | 0.5 | 3 | 1 | 12 | 0.5 | 2 | 0.5 | 1 | 0 | 2 | 0.5 | 1 | 1 | 6 | 0.5 |
| 11_6062307 | 2 | 0 | 5 | 0.4 | 2 | 1 | 3 | 0.3333 | 12 | 0.4167 | 1 | 1 | 2 | 0 | 2 | 1 | NA | NA | 5 | 0.6 |
| 11_6062311 | 1 | 1 | 5 | 1 | 1 | 1 | 3 | 1 | 10 | 1 | NA | NA | NA | NA | 2 | 0.5 | 1 | 1 | 3 | 0.6667 |
| 11_6062316 | 2 | 0 | 6 | 0.6667 | 2 | 0 | 3 | 0.6667 | 13 | 0.4615 | 2 | 0.5 | 2 | 0.5 | 2 | 0.5 | 1 | 1 | 7 | 0.5714 |

| | | | | | | | | | | | | | | | | | | | | |
|---|---|---|---|---|---|---|---|---|---|---|---|---|---|---|---|---|---|---|---|---|
| 11_606232 | 3 | 0.6667 | 5 | 0.2 | 2 | 0.5 | 1 | 1 | 11 | 0.4545 | 1 | 1 | 2 | 0.5 | 2 | 0 | 1 | 1 | 6 | 0.5 |
| 11_606234 | 2 | 0 | 5 | 0.6 | 2 | 0.5 | 3 | 0.6667 | 12 | 0.5 | 2 | 0 | 1 | 0 | 2 | 0 | 2 | 1 | 7 | 0.2857 |
| 11_606690 | 1 | 1 | NA | NA | 1 | 1 | NA | NA | 2 | 1 | NA | NA | 1 | 1 | NA | NA | NA | NA | 1 | 1 |
| 11_606702 | 2 | 0.5 | NA | NA | 1 | 1 | NA | NA | 3 | 0.6667 | NA | NA | 1 | 0 | 1 | 1 | NA | NA | 2 | 0.5 |
| 11_606705 | 2 | 0.5 | NA | NA | 1 | 0 | NA | NA | 3 | 0.3333 | NA | NA | 1 | 1 | 1 | 0 | NA | NA | 2 | 0.5 |
| 11_606706 | 1 | 1 | NA | NA | 1 | 1 | NA | NA | 2 | 1 | NA | NA | 1 | 0 | 1 | 1 | NA | NA | 2 | 0.5 |
| 11_606710 | 1 | 1 | NA | NA | 1 | 1 | NA | NA | 2 | 1 | NA | NA | NA | NA | 1 | 1 | NA | NA | 1 | 1 |
| 11_606710 | 3 | 1 | NA | NA | NA | NA | NA | NA | 3 | 1 | NA | NA | 1 | 1 | 1 | 0 | NA | NA | 2 | 0.5 |
| 11_606746 | 2 | 0.5 | NA | NA | 1 | 1 | 1 | 0 | 4 | 0.5 | NA | NA | 1 | 0 | 1 | 1 | NA | NA | 2 | 0.5 |
| 11_606748 | 3 | 1 | NA | NA | 2 | 0.5 | 1 | 0 | 6 | 0.6667 | NA | NA | 1 | 0 | 1 | 1 | NA | NA | 2 | 0.5 |
| 11_606749 | 2 | 1 | NA | NA | 1 | 1 | 1 | 1 | 4 | 1 | NA | NA | 1 | 1 | 1 | 1 | NA | NA | 2 | 1 |
| 11_606750 | 3 | 0 | NA | NA | 2 | 0.5 | 1 | 0 | 6 | 0.1667 | NA | NA | 1 | 1 | 1 | 0 | NA | NA | 2 | 0.5 |
| 11_606750 | 2 | 1 | NA | NA | 2 | 1 | NA | NA | 4 | 1 | NA | NA | 1 | 1 | 1 | 1 | NA | NA | 2 | 1 |
| 11_606750 | 3 | 0.3333 | NA | NA | 2 | 0 | 1 | 0 | 6 | 0.1667 | NA | NA | 1 | 1 | 1 | 1 | NA | NA | 2 | 1 |
| 11_606752 | 2 | 1 | 1 | 1 | 1 | 1 | NA | NA | 4 | 1 | NA | NA | 1 | 1 | NA | NA | NA | NA | 1 | 1 |
| 11_606753 | 3 | 0.3333 | 1 | 1 | 2 | 0.5 | 1 | 1 | 7 | 0.5714 | NA | NA | 1 | 1 | 1 | 1 | NA | NA | 2 | 1 |
| 11_606755 | 2 | 1 | 1 | 0 | 2 | 0 | NA | NA | 5 | 0.4 | NA | NA | 1 | 1 | 1 | 1 | NA | NA | 2 | 1 |
| 11_606755 | 3 | 1 | 1 | 1 | 2 | 1 | 1 | 1 | 7 | 1 | NA | NA | NA | NA | 1 | 1 | NA | NA | 1 | 1 |
| 11_606758 | 2 | 0.5 | 1 | 0 | 2 | 0 | 1 | 0 | 6 | 0.1667 | NA | NA | 1 | 0 | 1 | 1 | NA | NA | 2 | 0.5 |
| 11_606758 | 3 | 1 | 2 | 1 | 2 | 1 | 1 | 1 | 8 | 1 | NA | NA | 1 | 1 | 1 | 1 | NA | NA | 2 | 1 |
| 11_606761 | 3 | 0.6667 | 2 | 1 | 2 | 0 | 1 | 0 | 8 | 0.5 | NA | NA | 1 | 0 | 1 | 1 | NA | NA | 2 | 0.5 |
| 11_606761 | 3 | 0.3333 | 2 | 1 | 2 | 0 | 1 | 1 | 8 | 0.5 | NA | NA | 1 | 0 | 1 | 1 | NA | NA | 2 | 0.5 |
| 11_606763 | 4 | 0.25 | 1 | 1 | 3 | 0 | 1 | 0 | 9 | 0.2222 | NA | NA | 1 | 0 | 1 | 1 | NA | NA | 2 | 0.5 |
| 11_606764 | 4 | 0.5 | 2 | 1 | 3 | 0.6667 | 1 | 1 | 10 | 0.7 | NA | NA | 1 | 1 | NA | NA | NA | NA | 1 | 1 |
| 11_606767 | 4 | 0.25 | 2 | 1 | 3 | 0.3333 | 1 | 1 | 10 | 0.5 | NA | NA | NA | NA | 1 | 1 | NA | NA | 1 | 1 |
| 11_606768 | 3 | 0.6667 | NA | NA | 3 | 0.6667 | 1 | 0 | 7 | 0.5714 | NA | NA | 1 | 0 | 1 | 1 | NA | NA | 2 | 0.5 |
| 11_606769 | 4 | 0.5 | 2 | 0.5 | 2 | 0 | 1 | 0 | 9 | 0.3333 | NA | NA | 1 | 0 | 1 | 1 | NA | NA | 2 | 0.5 |
| 11_606769 | 3 | 0.3333 | 1 | 0 | 3 | 0 | 1 | 0 | 8 | 0.125 | NA | NA | 1 | 0 | 1 | 1 | NA | NA | 2 | 0.5 |
| 11_606769 | 2 | 0 | 2 | 0.5 | 2 | 0 | 1 | 0 | 7 | 0.1429 | NA | NA | 1 | 0 | 1 | 1 | NA | NA | 2 | 0.5 |
| 11_606790 | 3 | 0.3333 | 2 | 0.5 | 6 | 0.1667 | 2 | 0 | 13 | 0.2308 | 2 | 0 | 3 | 0.3333 | 1 | 0 | NA | NA | 6 | 0.1667 |
| 11_607486 | 6 | 0.8333 | 8 | 0.75 | 3 | 0.6667 | 7 | 0.7143 | 24 | 0.75 | 5 | 0.4 | 5 | 1 | 6 | 0.8333 | 2 | 0.5 | 18 | 0.7222 |
| 11_607486 | 5 | 0.2 | 7 | 0 | 2 | 0 | 5 | 0.2 | 19 | 0.1053 | 5 | 0 | 5 | 0.2 | 3 | 0.6667 | 2 | 0.5 | 15 | 0.2667 |
| 11_607489 | 5 | 0.2 | 9 | 0.1111 | 2 | 1 | 6 | 0.3333 | 22 | 0.2727 | 6 | 0.1667 | 5 | 0 | 7 | 0.1429 | 1 | 0 | 19 | 0.1053 |
| 11_607553 | 11 | 0.7273 | 6 | 0.5 | 4 | 0 | 4 | 1 | 25 | 0.6 | 3 | 0.6667 | 9 | 0.6667 | 11 | 0.5455 | 6 | 0.5 | 29 | 0.5862 |
| 11_607554 | 9 | 0 | 8 | 0.5 | 4 | 0 | 4 | 0.5 | 25 | 0.24 | 4 | 0.5 | 11 | 0.7273 | 13 | 0.3077 | 5 | 0.4 | 33 | 0.4848 |
| 11_607556 | 11 | 1 | 7 | 1 | 5 | 1 | 5 | 0.8 | 28 | 0.9643 | 6 | 0.8333 | 11 | 1 | 12 | 1 | 5 | 1 | 34 | 0.9706 |
| 11_607556 | 9 | 0.1111 | 5 | 0.6 | 5 | 0.2 | 5 | 0.4 | 24 | 0.2917 | 6 | 0.3333 | 8 | 0.375 | 10 | 0.1 | 4 | 0.25 | 28 | 0.25 |
| 11_607557 | 10 | 1 | 7 | 1 | 4 | 1 | 4 | 0.75 | 25 | 0.96 | 5 | 1 | 9 | 1 | 11 | 1 | 3 | 1 | 28 | 1 |
| 11_607558 | 8 | 1 | 4 | 1 | 4 | 1 | 5 | 1 | 21 | 1 | 5 | 1 | 9 | 1 | 9 | 1 | 4 | 1 | 27 | 1 |
| 11_607558 | 9 | 0.2222 | 7 | 0.1429 | 5 | 0.2 | 5 | 0 | 26 | 0.1538 | 6 | 0 | 11 | 0.1818 | 9 | 0 | 4 | 0 | 30 | 0.0667 |
| 11_607946 | NA | NA | 1 | 1 | NA | NA | NA | NA | 1 | 1 | NA | NA | 1 | 1 | NA | NA | 2 | 0 | 3 | 0.3333 |
| 11_607950 | NA | NA | 1 | 1 | NA | NA | NA | NA | 1 | 1 | NA | NA | 2 | 0 | NA | NA | 2 | 0.5 | 4 | 0.25 |
| 11_607960 | NA | NA | 1 | 1 | NA | NA | NA | NA | 1 | 1 | NA | NA | 1 | 1 | NA | NA | 1 | 1 | 2 | 1 |
| 11_607961 | NA | NA | 1 | 1 | NA | NA | NA | NA | 1 | 1 | NA | NA | 1 | 1 | NA | NA | 1 | 1 | 2 | 1 |
| 11_607964 | NA | NA | 1 | 1 | NA | NA | NA | NA | 1 | 1 | NA | NA | 1 | 1 | NA | NA | NA | NA | 1 | 1 |
| 11_608426 | NA | NA | 1 | 0 | 2 | 0 | 2 | 0.5 | 5 | 0.2 | 2 | 0.5 | 5 | 0.2 | 2 | 0.5 | 1 | 0 | 10 | 0.3 |
| 11_608430 | NA | NA | NA | NA | 1 | 1 | 2 | 1 | 3 | 1 | 2 | 1 | 3 | 0.6667 | 2 | 1 | 2 | 1 | 9 | 0.8889 |
| 11_608431 | NA | NA | 1 | 0 | 2 | 0.5 | 2 | 1 | 5 | 0.6 | 2 | 0.5 | 5 | 0.4 | 2 | 0.5 | 2 | 0 | 11 | 0.3636 |
| 11_608431 | NA | NA | 1 | 1 | 2 | 0 | 2 | 0.5 | 5 | 0.4 | 2 | 0 | 4 | 0.5 | 2 | 0 | 1 | 0 | 9 | 0.2222 |
| 11_608431 | NA | NA | 1 | 1 | 1 | 1 | 2 | 0.5 | 4 | 0.75 | 2 | 0.5 | 4 | 0.75 | 2 | 0.5 | 2 | 1 | 10 | 0.7 |
| 11_608432 | NA | NA | 1 | 1 | 2 | 0.5 | NA | NA | 3 | 0.6667 | 2 | 0 | 4 | 0.5 | 2 | 0 | 2 | 1 | 10 | 0.4 |
| 11_608434 | NA | NA | 2 | 0 | 2 | 0.5 | 2 | 0.5 | 6 | 0.3333 | 2 | 0 | 4 | 0.5 | 2 | 0 | 2 | 1 | 10 | 0.4 |
| 11_609602 | 1 | 1 | 1 | 1 | NA | NA | 1 | 1 | 3 | 1 | 1 | 0 | NA | NA | 1 | 1 | NA | NA | 2 | 0.5 |
| 11_609603 | 1 | 1 | 1 | 1 | 1 | 0 | 1 | 1 | 4 | 0.75 | NA | NA | 1 | 1 | 1 | 0 | NA | NA | 2 | 0.5 |
| 11_609604 | 1 | 1 | 1 | 1 | 1 | 1 | 1 | 1 | 4 | 1 | 1 | 1 | 1 | 1 | 1 | 1 | NA | NA | 3 | 1 |
| 11_609605 | 1 | 1 | 1 | 1 | 1 | 1 | 1 | 1 | 4 | 1 | 1 | 1 | 1 | 1 | 1 | 1 | NA | NA | 3 | 1 |
| 11_609605 | 1 | 1 | 1 | 1 | 1 | 1 | 1 | 1 | 4 | 1 | 1 | 1 | 1 | 1 | 1 | 1 | NA | NA | 3 | 1 |
| 11_609608 | 1 | 1 | 1 | 1 | 1 | 1 | 1 | 0 | 4 | 0.75 | 1 | 0 | 1 | 1 | 1 | 1 | NA | NA | 3 | 0.6667 |
| 11_609608 | 1 | 0 | 1 | 0 | 1 | 1 | 1 | 1 | 4 | 0.5 | 1 | 1 | 1 | 0 | 1 | 0 | NA | NA | 3 | 0.3333 |
| 11_609610 | 1 | 1 | 1 | 1 | 1 | 0 | 1 | 1 | 4 | 0.75 | 1 | 0 | 1 | 1 | 1 | 0 | 1 | 1 | 4 | 0.5 |
| 11_609612 | 1 | 1 | 1 | 1 | 1 | 1 | 1 | 1 | 4 | 1 | 1 | 1 | 1 | 1 | 1 | 1 | NA | NA | 3 | 1 |
| 11_609612 | 1 | 1 | 1 | 1 | 1 | 1 | 1 | 1 | 4 | 1 | 1 | 1 | NA | NA | 1 | 1 | NA | NA | 2 | 1 |
| 11_609613 | 1 | 1 | NA | NA | NA | NA | 1 | 1 | 2 | 1 | 1 | 1 | 1 | 1 | 1 | 1 | 1 | 1 | 4 | 1 |
| 11_609614 | NA | NA | 1 | 1 | 1 | 1 | NA | NA | 2 | 1 | 1 | 1 | 1 | 1 | 1 | 1 | 1 | 1 | 4 | 1 |
| 11_609615 | 1 | 0 | 1 | 1 | NA | NA | NA | NA | 2 | 0.5 | 1 | 1 | 1 | 1 | 1 | 1 | 1 | 1 | 4 | 1 |

| | | | | | | | | | | | | | | | | | | | | |
|---|---|---|---|---|---|---|---|---|---|---|---|---|---|---|---|---|---|---|---|---|
| 11_6096155 | 1 | 1 | 1 | 1 | 1 | 1 | 1 | 1 | 4 | 1 | 1 | 1 | 1 | 0 | 1 | 1 | 1 | 1 | 4 | 0.75 |
| 11_6096164 | 1 | 1 | 1 | 1 | 1 | 1 | 1 | 1 | 4 | 1 | 2 | 1 | 1 | 1 | 2 | 1 | 1 | 1 | 6 | 1 |
| 11_6096167 | 1 | 1 | NA | NA | NA | NA | 1 | 0 | 2 | 0.5 | 2 | 0.5 | 1 | 1 | 2 | 0 | 1 | 1 | 6 | 0.5 |
| 11_6096171 | 1 | 0 | 1 | 1 | 1 | 0 | 1 | 0 | 4 | 0.25 | 2 | 0.5 | 1 | 1 | 2 | 0.5 | 1 | 0 | 6 | 0.5 |
| 11_6096175 | 1 | 1 | 1 | 1 | 1 | 1 | 1 | 0 | 4 | 0.75 | 2 | 0.5 | 1 | 1 | 2 | 1 | 2 | 1 | 7 | 0.8571 |
| 11_6103848 | 1 | 1 | NA | NA | NA | NA | NA | NA | 1 | 1 | NA | NA | NA | NA | NA | NA | 1 | 1 | 1 | 1 |
| 11_6104338 | 13 | 0 | 20 | 0.1 | 17 | 0.2353 | 23 | 0.1304 | 73 | 0.1233 | 16 | 0.0625 | 29 | 0.1034 | 16 | 0.25 | 21 | 0.0952 | 82 | 0.122 |
| 11_6104340 | 13 | 0.0769 | 15 | 0.2 | 16 | 0.0625 | 19 | 0 | 63 | 0.0794 | 15 | 0.0667 | 26 | 0.0385 | 13 | 0 | 17 | 0.1176 | 71 | 0.0563 |
| 11_6104358 | 2 | 0 | 2 | 0 | 7 | 0.1429 | 8 | 0 | 19 | 0.0526 | 1 | 0 | 7 | 0 | 3 | 0 | 7 | 0.1429 | 18 | 0.0556 |
| 11_6104364 | 14 | 0.2143 | 14 | 0.0714 | 14 | 0.2143 | 18 | 0.2222 | 60 | 0.1833 | 15 | 0.1333 | 24 | 0.2917 | 11 | 0.2727 | 21 | 0.2857 | 71 | 0.2535 |
| 11_6104366 | 12 | 0.3333 | 10 | 0.4 | 14 | 0.2857 | 15 | 0.3333 | 51 | 0.3333 | 13 | 0.2308 | 22 | 0 | 11 | 0.1818 | 15 | 0.4 | 61 | 0.1803 |
| 11_6113581 | 8 | 0 | 8 | 0.125 | 5 | 0.2 | 6 | 0 | 27 | 0.0741 | 3 | 0 | 7 | 0.1429 | 5 | 0 | 2 | 0 | 17 | 0.0588 |
| 11_6113621 | 6 | 0.1667 | 5 | 0 | 4 | 0.25 | 6 | 0.1667 | 21 | 0.1429 | 3 | 0.3333 | 7 | 0 | 3 | 0 | 2 | 0 | 15 | 0.0667 |
| 11_6113644 | 6 | 1 | 6 | 0.8333 | 3 | 1 | 5 | 0.6 | 20 | 0.85 | 3 | 0.6667 | 6 | 1 | 4 | 0.5 | 1 | 1 | 14 | 0.7857 |
| 11_6113658 | 5 | 0 | 5 | 0 | 3 | 0 | 6 | 0.1667 | 19 | 0.0526 | 3 | 0.3333 | 6 | 0.1667 | 4 | 0 | NA | NA | 13 | 0.1538 |
| 11_6113672 | 4 | 0.25 | 4 | 0 | 3 | 0.6667 | 6 | 0 | 17 | 0.1765 | 2 | 0 | 5 | 0.4 | 4 | 0.25 | NA | NA | 11 | 0.2727 |
| 11_6113693 | 5 | 0 | 3 | 0.3333 | 2 | 0.5 | 4 | 0.25 | 14 | 0.2143 | 1 | 0 | 4 | 0.25 | 3 | 0.3333 | 1 | 1 | 9 | 0.3333 |
| 11_6113699 | 5 | 0 | 3 | 0 | 2 | 0.5 | 6 | 0.1667 | 16 | 0.125 | 3 | 0.6667 | 6 | 0.3333 | 2 | 0 | 1 | 1 | 12 | 0.4167 |
| 11_6115306 | 4 | 0.25 | 5 | 0.2 | 3 | 0.3333 | 5 | 0.2 | 17 | 0.2353 | NA | NA | 5 | 0 | 2 | 0.5 | 1 | 1 | 8 | 0.25 |
| 11_6129952 | NA | NA | NA | NA | 1 | 1 | NA | NA | 1 | 1 | NA | NA | NA | NA | 1 | 1 | NA | NA | 1 | 1 |
| 11_6130216 | NA | NA | NA | NA | 1 | 1 | NA | NA | 1 | 1 | NA | NA | NA | NA | 1 | 1 | NA | NA | 1 | 1 |
| 11_6133935 | 18 | 0.6111 | 18 | 0.3889 | 16 | 0.75 | 15 | 0.4667 | 67 | 0.5522 | 11 | 0.2727 | 15 | 0.6 | 14 | 0.5714 | 13 | 0.4615 | 53 | 0.4906 |
| 11_6133984 | 15 | 0.1333 | 15 | 0.4 | 14 | 0.4286 | 9 | 0.3333 | 53 | 0.3208 | 7 | 0.7143 | 12 | 0.4167 | 11 | 0.1818 | 7 | 0.7143 | 37 | 0.4595 |
| 11_6134017 | 14 | 0.2143 | 13 | 0.6154 | 12 | 0.3333 | 8 | 0.625 | 47 | 0.4255 | 4 | 0.5 | 10 | 0.6 | 9 | 0.6667 | 6 | 0.5 | 29 | 0.5862 |
| 11_6134018 | 12 | 0.3333 | 14 | 0.5 | 10 | 0.2 | 7 | 0.2857 | 43 | 0.3488 | 5 | 0.4 | 9 | 0.5556 | 9 | 0.2222 | 7 | 0.4286 | 30 | 0.4 |
| 11_6134051 | 9 | 0.8889 | 7 | 1 | 8 | 0.875 | 5 | 1 | 29 | 0.931 | 3 | 1 | 6 | 1 | 6 | 1 | 4 | 1 | 19 | 1 |
| 11_6138109 | 4 | 0.5 | 5 | 0.4 | 2 | 0 | NA | NA | 11 | 0.3636 | 1 | 1 | NA | NA | 2 | 0 | NA | NA | 3 | 0.3333 |
| 11_6138112 | 3 | 0 | 6 | 0.5 | 3 | 0.3333 | NA | NA | 12 | 0.3333 | 1 | 1 | NA | NA | 2 | 0.5 | NA | NA | 3 | 0.6667 |
| 11_6138113 | 4 | 0.5 | 4 | 0.5 | 2 | 0.5 | NA | NA | 10 | 0.5 | 1 | 0 | NA | NA | 2 | 1 | NA | NA | 3 | 0.6667 |
| 11_6138162 | 3 | 0 | 6 | 0.1667 | 3 | 0 | NA | NA | 12 | 0.0833 | 1 | 1 | NA | NA | 2 | 0.5 | NA | NA | 3 | 0.6667 |
| 11_6138175 | 1 | 1 | 6 | 0.8333 | 2 | 0.5 | NA | NA | 9 | 0.7778 | 1 | 1 | NA | NA | 1 | 1 | NA | NA | 2 | 1 |
| 11_6138182 | 3 | 1 | 6 | 0.6667 | 1 | 1 | NA | NA | 10 | 0.8 | 1 | 1 | NA | NA | NA | NA | NA | NA | 1 | 1 |
| 11_6138198 | 3 | 0.3333 | 4 | 0 | 3 | 0.3333 | NA | NA | 10 | 0.2 | 1 | 1 | NA | NA | 1 | 0 | NA | NA | 2 | 0.5 |
| 11_6138207 | 3 | 0.3333 | 5 | 0.2 | 2 | 0 | NA | NA | 10 | 0.2 | 1 | 1 | NA | NA | 1 | 0 | NA | NA | 2 | 0.5 |
| 11_6138225 | 2 | 0.5 | 2 | 1 | 2 | 1 | NA | NA | 6 | 0.8333 | 1 | 1 | NA | NA | 1 | 1 | NA | NA | 2 | 1 |
| 11_6138233 | 3 | 1 | 4 | 1 | 1 | 1 | NA | NA | 8 | 1 | 1 | 1 | NA | NA | 1 | 1 | NA | NA | 2 | 1 |
| 11_6140364 | 4 | 0.25 | 1 | 0 | 3 | 0 | 2 | 0 | 10 | 0.1 | 2 | 0 | 4 | 0 | 3 | 0.3333 | 4 | 0 | 13 | 0.0769 |
| 11_6140368 | 5 | 0.4 | 1 | 0 | 3 | 0 | 3 | 0 | 12 | 0.1667 | 2 | 0 | 4 | 0 | 3 | 0.3333 | 4 | 0 | 13 | 0.0769 |
| 11_6140838 | 13 | 0.0769 | 10 | 0.2 | 10 | 0.4 | 6 | 0.3333 | 39 | 0.2308 | 4 | 0.25 | 10 | 0.1 | 9 | 0.2222 | 10 | 0.2 | 33 | 0.1818 |
| 11_6140853 | 10 | 0.1 | 13 | 0 | 13 | 0.1538 | 7 | 0 | 43 | 0.0698 | 6 | 0.1667 | 14 | 0 | 15 | 0 | 13 | 0.3077 | 48 | 0.1042 |
| 11_6140862 | 11 | 0.4545 | 14 | 0.4286 | 12 | 0.4167 | 4 | 0.75 | 41 | 0.4634 | 6 | 0.5 | 12 | 0.5833 | 17 | 0.3529 | 14 | 0.5 | 49 | 0.4694 |
| 11_6140874 | 13 | 0.7692 | 12 | 0.75 | 9 | 0.8889 | 5 | 0.8 | 39 | 0.7949 | 6 | 0.8333 | 13 | 0.6923 | 15 | 0.4667 | 14 | 0.7857 | 48 | 0.6667 |
| 11_6140879 | 11 | 0.1818 | 8 | 0.125 | 11 | 0.1818 | 4 | 0.5 | 34 | 0.2059 | 4 | 0 | 11 | 0.4545 | 12 | 0.25 | 13 | 0.2308 | 40 | 0.275 |
| 11_6140916 | 11 | 0 | 11 | 0.0909 | 10 | 0.3 | 5 | 0 | 37 | 0.1081 | 4 | 0 | 12 | 0.0833 | 15 | 0.2 | 13 | 0.0769 | 44 | 0.1136 |
| 11_6140985 | 13 | 0.0769 | 10 | 0.1 | 12 | 0.0833 | 3 | 0.3333 | 38 | 0.1053 | 4 | 0 | 11 | 0.1818 | 9 | 0.1111 | 11 | 0.1818 | 35 | 0.1429 |
| 11_6140993 | 5 | 0.2 | 8 | 0 | 6 | 0 | 1 | 0 | 20 | 0.05 | 2 | 0 | 7 | 0 | 5 | 0.4 | 6 | 0.1667 | 20 | 0.15 |
| 11_6141018 | 10 | 0.1 | 8 | 0 | 12 | 0.1667 | 3 | 0.3333 | 33 | 0.1212 | 4 | 0.75 | 11 | 0.1818 | 9 | 0.1111 | 11 | 0.1818 | 35 | 0.2286 |
| 11_6141031 | 7 | 0.1429 | 9 | 0.1111 | 9 | 0.1111 | NA | NA | 25 | 0.12 | 3 | 0.3333 | 8 | 0 | 8 | 0.25 | 9 | 0.1111 | 28 | 0.1429 |
| 11_6143721 | 1 | 0 | 4 | 0.25 | 3 | 0 | NA | NA | 8 | 0.125 | 1 | 0 | 4 | 0 | 6 | 0.3333 | 7 | 0 | 18 | 0.1111 |
| 11_6143724 | 1 | 1 | 3 | 1 | 2 | 1 | NA | NA | 6 | 1 | 1 | 0 | 4 | 0.75 | 5 | 1 | 6 | 0.8333 | 16 | 0.8125 |
| 11_6143725 | 1 | 0 | 3 | 0.6667 | 2 | 1 | NA | NA | 6 | 0.6667 | 2 | 1 | 4 | 0.75 | 4 | 0.75 | 3 | 1 | 13 | 0.8462 |
| 11_6143727 | 1 | 0 | 4 | 0.5 | 2 | 0 | NA | NA | 7 | 0.2857 | 2 | 0 | 3 | 0.3333 | 5 | 0.4 | 5 | 0.2 | 15 | 0.2667 |
| 11_6143732 | 1 | 1 | 2 | 0.5 | 1 | 0 | NA | NA | 4 | 0.5 | 1 | 1 | 4 | 1 | 3 | 0.6667 | 5 | 0.2 | 13 | 0.6154 |
| 11_6143736 | 1 | 1 | 4 | 0.75 | 3 | 0.3333 | NA | NA | 8 | 0.625 | 1 | 0 | 3 | 1 | 4 | 0.75 | 7 | 1 | 15 | 0.8667 |
| 11_6143745 | 1 | 1 | 4 | 0.25 | 2 | 1 | NA | NA | 7 | 0.5714 | 1 | 0 | 2 | 0.5 | 5 | 0.6 | 5 | 0.6 | 13 | 0.5385 |
| 11_6143752 | NA | NA | 1 | 0 | 2 | 0.5 | NA | NA | 3 | 0.3333 | NA | NA | 2 | 1 | 4 | 0.5 | 4 | 0.5 | 10 | 0.6 |
| 11_6143753 | 1 | 0 | 3 | 0.6667 | 2 | 0 | NA | NA | 6 | 0.3333 | NA | NA | 3 | 0 | 4 | 0 | 3 | 0.3333 | 10 | 0.1 |
| 11_6143762 | 1 | 0 | 3 | 0.6667 | 3 | 0.6667 | NA | NA | 7 | 0.5714 | 1 | 1 | 3 | 0.3333 | 4 | 0 | 4 | 0 | 12 | 0.1667 |
| 11_6143769 | 1 | 0 | 3 | 0.3333 | 3 | 0.6667 | NA | NA | 7 | 0.4286 | 2 | 0 | 2 | 1 | 3 | 0.3333 | 4 | 0.25 | 11 | 0.3636 |
| 11_6162097 | 8 | 0.25 | 10 | 0.1 | 6 | 0 | 13 | 0.0769 | 37 | 0.1081 | 10 | 0.3 | 9 | 0 | 11 | 0.0909 | 6 | 0 | 36 | 0.1111 |
| 11_6166788 | 9 | 0.2222 | 5 | 0 | 12 | 0.3333 | 9 | 0.3333 | 35 | 0.2571 | 5 | 0.2 | 10 | 0.3 | 9 | 0.3333 | 9 | 0.2222 | 33 | 0.2727 |
| 11_6166795 | 11 | 0.9091 | 8 | 1 | 15 | 0.9333 | 10 | 0.9 | 44 | 0.9318 | 5 | 1 | 10 | 1 | 11 | 0.8182 | 9 | 0.8889 | 35 | 0.9143 |
| 11_6166797 | 8 | 0 | 6 | 0 | 9 | 0.1111 | 10 | 0.1 | 33 | 0.0606 | 4 | 0.25 | 10 | 0.1 | 9 | 0.2222 | 8 | 0.125 | 31 | 0.1613 |
| 11_6166800 | 10 | 0.6 | 6 | 0.5 | 10 | 0.1 | 11 | 0.4545 | 37 | 0.4054 | 4 | 0.5 | 10 | 0.2 | 7 | 0.4286 | 9 | 0.2222 | 30 | 0.3 |
| 11_6166805 | 11 | 0.9091 | 8 | 1 | 14 | 1 | 11 | 0.9091 | 44 | 0.9545 | 5 | 1 | 12 | 0.9167 | 10 | 0.9 | 9 | 0.8889 | 36 | 0.9167 |

| | | | | | | | | | | | | | | | | | | | | |
|---|---|---|---|---|---|---|---|---|---|---|---|---|---|---|---|---|---|---|---|---|
| 11_616680 | 9 | 0.5556 | 8 | 0.875 | 13 | 0.6923 | 10 | 0.7 | 40 | 0.7 | 5 | 0.6 | 11 | 0.3636 | 10 | 0.7 | 8 | 0.75 | 34 | 0.5882 |
| 11_616683 | 9 | 0.8889 | 5 | 0.8 | 7 | 0.8571 | 6 | 0.6667 | 27 | 0.8148 | 1 | 1 | 9 | 0.7778 | 7 | 0.8571 | 7 | 0.8571 | 24 | 0.8333 |
| 11_616683 | 7 | 0.5714 | 6 | 0.5 | 12 | 0.5833 | 11 | 0.3636 | 36 | 0.5 | 5 | 0.2 | 10 | 0.6 | 10 | 0.6 | 7 | 0.4286 | 32 | 0.5 |
| 11_616683 | 7 | 0.1429 | 7 | 0.2857 | 12 | 0.3333 | 11 | 0 | 37 | 0.1892 | 4 | 0.5 | 9 | 0.3333 | 9 | 0.1111 | 6 | 0.1667 | 28 | 0.25 |
| 11_616684 | 5 | 0.2 | 8 | 0.375 | 8 | 0.25 | 8 | 0.125 | 29 | 0.2414 | 3 | 0.3333 | 7 | 0.2857 | 9 | 0.3333 | 4 | 0 | 23 | 0.2609 |
| 11_616684 | 6 | 0.3333 | 8 | 0.5 | 12 | 0.4167 | 10 | 0.2 | 36 | 0.3611 | 5 | 0.2 | 9 | 0.3333 | 11 | 0.3636 | 7 | 0.4286 | 32 | 0.3438 |
| 11_616685 | 8 | 0.125 | 8 | 0.125 | 14 | 0 | 10 | 0 | 40 | 0.05 | 5 | 0 | 10 | 0 | 11 | 0.0909 | 8 | 0.125 | 34 | 0.0588 |
| 11_616685 | 8 | 0.5 | 8 | 0.25 | 11 | 0.6364 | 10 | 0.7 | 37 | 0.5405 | 5 | 0.6 | 8 | 0.75 | 11 | 0.7273 | 8 | 0.5 | 32 | 0.6562 |
| 11_616685 | 4 | 0.25 | 6 | 0 | 8 | 0.125 | 8 | 0.125 | 26 | 0.1154 | 4 | 0 | 10 | 0.3 | 10 | 0.1 | 7 | 0 | 31 | 0.129 |
| 11_616685 | 7 | 1 | 7 | 1 | 12 | 0.9167 | 10 | 0.9 | 36 | 0.9444 | 3 | 1 | 10 | 0.9 | 10 | 1 | 9 | 1 | 32 | 0.9688 |
| 11_617635 | 2 | 0 | 1 | 0 | 1 | 0 | 2 | 0.5 | 6 | 0.1667 | NA | NA | 4 | 0.25 | 1 | 0 | 3 | 0 | 8 | 0.125 |
| 11_618058 | 1 | 0 | 2 | 0 | 1 | 1 | 1 | 0 | 5 | 0.2 | NA | NA | 3 | 0 | 1 | 0 | 4 | 0.5 | 8 | 0.25 |
| 11_618059 | 2 | 0.5 | 2 | 0 | 1 | 0 | 1 | 0 | 6 | 0.1667 | NA | NA | 3 | 0.3333 | NA | NA | 3 | 0.6667 | 6 | 0.5 |
| 11_618060 | 1 | 1 | 1 | 0 | NA | NA | 1 | 1 | 3 | 0.6667 | NA | NA | 2 | 1 | NA | NA | 3 | 0.6667 | 5 | 0.8 |
| 11_618060 | 1 | 1 | 2 | 1 | 1 | 0 | NA | NA | 4 | 0.75 | NA | NA | 3 | 0.6667 | 1 | 0 | 3 | 1 | 7 | 0.7143 |
| 11_618061 | 2 | 0.5 | 2 | 0 | 1 | 1 | 1 | 0 | 6 | 0.3333 | NA | NA | 2 | 0.5 | NA | NA | 2 | 0 | 4 | 0.25 |
| 11_618061 | 1 | 1 | 2 | 0 | 1 | 0 | NA | NA | 4 | 0.25 | NA | NA | 3 | 0 | 1 | 0 | 3 | 0.6667 | 7 | 0.2857 |
| 11_618061 | 1 | 1 | 2 | 0 | 1 | 1 | 1 | 0 | 5 | 0.4 | NA | NA | 2 | 1 | 1 | 1 | 3 | 0 | 6 | 0.5 |
| 11_618062 | 1 | 0 | 2 | 0.5 | 1 | 0 | 1 | 0 | 5 | 0.2 | NA | NA | 2 | 0 | 1 | 0 | 1 | 1 | 4 | 0.25 |
| 11_618069 | 2 | 0 | 2 | 0.5 | NA | NA | 1 | 0 | 5 | 0.2 | NA | NA | 2 | 0.5 | NA | NA | 2 | 0.5 | 4 | 0.5 |
| 11_620087 | NA | NA | NA | NA | 1 | 0 | 3 | 0.3333 | 4 | 0.25 | 1 | 0 | 1 | 0 | 2 | 0.5 | NA | NA | 4 | 0.25 |
| 11_620087 | NA | NA | NA | NA | NA | NA | 3 | 0.6667 | 3 | 0.6667 | 1 | 0 | 1 | 1 | 2 | 0 | NA | NA | 4 | 0.25 |
| 11_620090 | NA | NA | NA | NA | 2 | 0 | 3 | 0.3333 | 5 | 0.2 | 1 | 1 | 1 | 0 | 2 | 0.5 | NA | NA | 4 | 0.5 |
| 11_620090 | NA | NA | NA | NA | 2 | 0.5 | 2 | 0.5 | 4 | 0.5 | 1 | 0 | 1 | 1 | 3 | 0 | NA | NA | 5 | 0.2 |
| 11_620097 | NA | NA | NA | NA | 3 | 0.3333 | 4 | 0 | 7 | 0.1429 | 1 | 0 | 2 | 0.5 | 3 | 0 | 2 | 0.5 | 8 | 0.25 |
| 11_620098 | NA | NA | NA | NA | 3 | 0.3333 | 3 | 0.3333 | 6 | 0.3333 | 1 | 1 | 2 | 0 | 3 | 0 | 2 | 0 | 8 | 0.125 |
| 11_620099 | NA | NA | 1 | 0 | 3 | 0.3333 | 2 | 0.5 | 6 | 0.3333 | 1 | 1 | 2 | 0.5 | 3 | 0.3333 | 2 | 0 | 8 | 0.375 |
| 11_620289 | NA | NA | 2 | 1 | 1 | 1 | 4 | 1 | 7 | 1 | NA | NA | 1 | 1 | 3 | 0.6667 | 2 | 1 | 6 | 0.8333 |
| 11_620298 | NA | NA | 2 | 1 | 2 | 1 | 5 | 1 | 9 | 1 | 1 | 1 | 2 | 0.5 | 3 | 1 | 2 | 0.5 | 8 | 0.75 |
| 11_620298 | NA | NA | 2 | 1 | 3 | 0.6667 | 4 | 0.5 | 9 | 0.6667 | 1 | 0 | 2 | 0.5 | 3 | 1 | 1 | 0 | 7 | 0.5714 |
| 11_620299 | NA | NA | 2 | 0 | 3 | 0 | 5 | 0.2 | 10 | 0.1 | 1 | 0 | 2 | 0 | 3 | 0 | 2 | 0.5 | 8 | 0.125 |
| 11_620300 | NA | NA | 2 | 0 | 3 | 0 | 5 | 0.2 | 10 | 0.1 | 1 | 1 | 2 | 0 | 3 | 0 | 2 | 0 | 8 | 0.125 |
| 11_620301 | NA | NA | 2 | 0 | 3 | 0.3333 | 5 | 0.8 | 10 | 0.5 | 1 | 1 | 4 | 0.5 | 3 | 0.6667 | 2 | 0.5 | 10 | 0.6 |
| 11_620302 | NA | NA | 2 | 0.5 | 1 | 0 | 5 | 0.4 | 8 | 0.375 | NA | NA | 4 | 0.75 | 2 | 0.5 | 2 | 0.5 | 8 | 0.625 |
| 11_620302 | NA | NA | 2 | 0.5 | 3 | 0.6667 | 5 | 0.4 | 10 | 0.5 | NA | NA | 4 | 0.5 | 3 | 0.6667 | 2 | 0 | 9 | 0.4444 |
| 11_620303 | NA | NA | 2 | 1 | 3 | 0.3333 | 4 | 0.5 | 9 | 0.5556 | 1 | 0 | 3 | 0.3333 | 3 | 0.6667 | 2 | 0.5 | 9 | 0.4444 |
| 11_620303 | NA | NA | 2 | 0 | 3 | 0.3333 | 5 | 0.2 | 10 | 0.2 | 1 | 0 | 3 | 0.3333 | 3 | 0 | 2 | 0 | 9 | 0.1111 |
| 11_620303 | NA | NA | 2 | 0 | 3 | 0.3333 | 5 | 0 | 10 | 0.1 | 1 | 0 | 4 | 0.25 | 3 | 0 | 2 | 0 | 10 | 0.1 |
| 11_621103 | NA | NA | 2 | 0.5 | 4 | 0.75 | 1 | 1 | 7 | 0.7143 | 4 | 0.75 | 1 | 0 | 2 | 1 | 1 | 0 | 8 | 0.625 |
| 11_621108 | NA | NA | 2 | 0 | 3 | 0.3333 | 1 | 1 | 6 | 0.3333 | 5 | 0.6 | 1 | 1 | 2 | 0.5 | NA | NA | 8 | 0.625 |
| 11_621110 | NA | NA | 2 | 0 | 4 | 0.25 | NA | NA | 6 | 0.1667 | 5 | 0 | 1 | 0 | 2 | 0.5 | NA | NA | 8 | 0.125 |
| 11_621116 | NA | NA | 1 | 0 | 5 | 0.2 | NA | NA | 6 | 0.1667 | 3 | 0.3333 | 2 | 0.5 | 1 | 0 | NA | NA | 6 | 0.3333 |
| 11_621117 | NA | NA | 2 | 0 | 5 | 0.2 | NA | NA | 7 | 0.1429 | 3 | 0.6667 | 2 | 1 | 2 | 0 | 1 | 0 | 8 | 0.5 |
| 11_621120 | NA | NA | 2 | 1 | 4 | 0.25 | NA | NA | 6 | 0.5 | 2 | 0 | 1 | 0 | 2 | 0.5 | 1 | 1 | 6 | 0.3333 |
| 11_621121 | NA | NA | 1 | 0 | 5 | 0.6 | NA | NA | 6 | 0.5 | 2 | 1 | NA | NA | 1 | 0 | 1 | 0 | 4 | 0.5 |
| 11_621129 | 2 | 0 | 4 | 0.25 | 7 | 0 | 2 | 0 | 15 | 0.0667 | 5 | 0 | 3 | 0 | 2 | 0 | 2 | 0.5 | 12 | 0.0833 |
| 11_621129 | 2 | 0 | 5 | 0.2 | 7 | 0 | 2 | 0 | 16 | 0.0625 | 5 | 0.2 | 3 | 0 | 2 | 0 | 2 | 0 | 12 | 0.0833 |
| 11_621133 | 4 | 0 | 5 | 0 | 9 | 0.2222 | 2 | 0 | 20 | 0.1 | 5 | 0 | 3 | 0.3333 | 2 | 0 | 1 | 0 | 11 | 0.0909 |
| 11_621134 | 4 | 0 | 6 | 0.5 | 7 | 0.2857 | 2 | 0.5 | 19 | 0.3158 | 5 | 0.2 | 4 | 0 | 2 | 0 | 2 | 0.5 | 13 | 0.1538 |
| 11_621136 | 5 | 0 | 6 | 0.1667 | 7 | 0.2857 | 4 | 0.25 | 22 | 0.1818 | 6 | 0.3333 | 3 | 0 | 3 | 0 | 1 | 0 | 13 | 0.1538 |
| 11_621563 | NA | NA | 1 | 1 | NA | NA | NA | NA | 1 | 1 | 1 | 1 | NA | NA | NA | NA | NA | NA | 1 | 1 |
| 11_621565 | NA | NA | 1 | 1 | NA | NA | NA | NA | 1 | 1 | 1 | 1 | NA | NA | NA | NA | NA | NA | 1 | 1 |
| 11_621568 | NA | NA | 1 | 1 | NA | NA | NA | NA | 1 | 1 | 1 | 1 | NA | NA | NA | NA | NA | NA | 1 | 1 |
| 11_621568 | NA | NA | 1 | 1 | NA | NA | NA | NA | 1 | 1 | 2 | 1 | NA | NA | NA | NA | NA | NA | 2 | 1 |
| 11_621582 | NA | NA | 4 | 1 | 6 | 1 | 2 | 0 | 12 | 0.8333 | 1 | 1 | 1 | 0 | 2 | 1 | 2 | 1 | 6 | 0.8333 |
| 11_621582 | 1 | 0 | 6 | 0.1667 | 7 | 0 | 1 | 0 | 15 | 0.0667 | 2 | 1 | 1 | 0 | 2 | 0.5 | 3 | 0 | 8 | 0.375 |
| 11_621583 | 1 | 0 | 6 | 0.5 | 5 | 0.2 | 2 | 0 | 14 | 0.2857 | 2 | 0 | 1 | 0 | 2 | 0 | 3 | 0.6667 | 8 | 0.25 |
| 11_622049 | 3 | 0.3333 | 10 | 0.5 | 8 | 0.125 | 5 | 0.4 | 26 | 0.3462 | 5 | 0.4 | 5 | 0.2 | 6 | 0.6667 | 6 | 0.5 | 22 | 0.4545 |
| 11_622049 | 3 | 0.6667 | 9 | 0.5556 | 7 | 0.1429 | 8 | 0.375 | 27 | 0.4074 | 6 | 0.6667 | 3 | 0.3333 | 5 | 0.2 | 6 | 0.3333 | 20 | 0.4 |
| 11_622050 | 2 | 1 | 7 | 0.2857 | 8 | 0.5 | 7 | 0.5714 | 24 | 0.5 | 4 | 0 | 3 | 0.3333 | 4 | 0.75 | 4 | 0.25 | 15 | 0.3333 |
| 11_622050 | 3 | 0.3333 | 10 | 0.4 | 9 | 0.5556 | 8 | 0.75 | 30 | 0.5333 | 6 | 1 | 5 | 0.8 | 6 | 0.8333 | 6 | 1 | 23 | 0.913 |
| 11_622051 | 4 | 0.25 | 9 | 0.6667 | 9 | 0.2222 | 7 | 0.7143 | 29 | 0.4828 | 6 | 0.5 | 5 | 0.4 | 6 | 0.6667 | 7 | 0.8571 | 24 | 0.625 |
| 11_622053 | 4 | 0.5 | 10 | 0.4 | 10 | 0.3 | 7 | 0.7143 | 31 | 0.4516 | 7 | 0.8571 | 3 | 0.3333 | 5 | 1 | 6 | 1 | 21 | 0.8571 |
| 11_622053 | 4 | 0.25 | 11 | 0.0909 | 10 | 0.3 | 7 | 0.7143 | 32 | 0.3125 | 7 | 0.2857 | 5 | 0.2 | 5 | 0.6 | 7 | 0.7143 | 24 | 0.4583 |
| 11_622055 | 4 | 0.25 | 11 | 0.1818 | 8 | 0 | 9 | 0.1111 | 32 | 0.125 | 7 | 0.4286 | 5 | 0.2 | 5 | 0 | 7 | 0.5714 | 24 | 0.3333 |

| | | | | | | | | | | | | | | | | | | | | |
|---|---|---|---|---|---|---|---|---|---|---|---|---|---|---|---|---|---|---|---|---|
| 11_6220560 | 4 | 0.5 | 11 | 0.5455 | 10 | 0.5 | 8 | 0.5 | 33 | 0.5152 | 7 | 0.7143 | 4 | 0.5 | 6 | 0.8333 | 6 | 0.8333 | 23 | 0.7391 |
| 11_6220601 | 4 | 0.25 | 10 | 0.2 | 10 | 0.3 | 7 | 0.1429 | 31 | 0.2258 | 4 | 0 | 1 | 0 | 4 | 0.25 | 6 | 0.3333 | 15 | 0.2 |
| 11_6220606 | 3 | 0 | 13 | 0.1538 | 10 | 0 | 9 | 0 | 35 | 0.0571 | 5 | 0 | 3 | 0 | 5 | 0.2 | 6 | 0.1667 | 19 | 0.1053 |
| 11_6220684 | 4 | 0.25 | 2 | 0.5 | 4 | 0 | 3 | 0.3333 | 13 | 0.2308 | NA | NA | 1 | 0 | 4 | 0.25 | NA | NA | 5 | 0.2 |
| 11_6220692 | 4 | 0 | 5 | 0.2 | 7 | 0.1429 | 4 | 0 | 20 | 0.1 | 2 | 0 | 2 | 0 | 5 | 0.2 | 2 | 0 | 11 | 0.0909 |
| 11_6220758 | 4 | 0.5 | 4 | 0.25 | 2 | 0.5 | 3 | 0.6667 | 13 | 0.4615 | NA | NA | 3 | 0.6667 | 4 | 0 | 3 | 0.3333 | 10 | 0.3 |
| 11_6220767 | 3 | 0 | 4 | 0 | 2 | 0 | 3 | 0.3333 | 12 | 0.0833 | NA | NA | 1 | 0 | 4 | 0.25 | 3 | 0 | 8 | 0.125 |
| 11_6220841 | 9 | 0.7778 | 10 | 0.6 | 9 | 0.6667 | 8 | 0.375 | 36 | 0.6111 | 5 | 1 | 12 | 0.6667 | 14 | 0.6429 | 14 | 0.6429 | 45 | 0.6889 |
| 11_6220845 | 7 | 0 | 8 | 0.125 | 6 | 0.3333 | 9 | 0 | 30 | 0.1 | 4 | 0 | 11 | 0.1818 | 6 | 0 | 12 | 0 | 33 | 0.0606 |
| 11_6220848 | 7 | 0.4286 | 6 | 0.3333 | 2 | 1 | 6 | 0.3333 | 21 | 0.4286 | NA | NA | 6 | 0.3333 | 6 | 0.5 | 10 | 0.4 | 22 | 0.4091 |
| 11_6220848 | 1 | 1 | NA | NA | NA | NA | NA | NA | 1 | 1 | NA | NA | NA | NA | 1 | 1 | 1 | 0 | 2 | 0.5 |
| 11_6220849 | 8 | 0 | 9 | 0 | 5 | 0.4 | 5 | 0 | 27 | 0.0741 | 5 | 0 | 12 | 0.0833 | 9 | 0.1111 | 10 | 0 | 36 | 0.0556 |
| 11_6220857 | 11 | 0.9091 | 11 | 0.7273 | 8 | 0.625 | 8 | 1 | 38 | 0.8158 | 5 | 1 | 12 | 0.9167 | 11 | 1 | 17 | 0.8824 | 45 | 0.9333 |
| 11_6220869 | 11 | 0.7273 | 9 | 0.5556 | 8 | 0.375 | 8 | 0.625 | 36 | 0.5833 | 6 | 0.6667 | 12 | 0.5 | 6 | 0.5 | 14 | 0.5714 | 38 | 0.5526 |
| 11_6221191 | NA | NA | NA | NA | 1 | 1 | NA | NA | 1 | 1 | NA | NA | 1 | 1 | NA | NA | 4 | 0.75 | 5 | 0.8 |
| 11_6221195 | NA | NA | NA | NA | 1 | 1 | NA | NA | 1 | 1 | NA | NA | 1 | 0 | NA | NA | 5 | 0.4 | 6 | 0.3333 |
| 11_6221210 | NA | NA | NA | NA | 1 | 1 | NA | NA | 1 | 1 | NA | NA | 3 | 0.3333 | NA | NA | 4 | 1 | 7 | 0.7143 |
| 11_6221659 | 1 | 0 | 1 | 0 | 1 | 0 | 2 | 1 | 5 | 0.4 | NA | NA | 2 | 0 | 2 | 1 | NA | NA | 4 | 0.5 |
| 11_6221661 | 1 | 0 | 1 | 0 | 1 | 1 | 3 | 0 | 6 | 0.1667 | NA | NA | 2 | 0 | 2 | 0.5 | NA | NA | 4 | 0.25 |
| 11_6221757 | 1 | 1 | NA | NA | NA | NA | NA | NA | 1 | 1 | NA | NA | 1 | 1 | NA | NA | NA | NA | 1 | 1 |
| 11_6222450 | 5 | 0.6 | 5 | 0.4 | 4 | 0.5 | 3 | 0 | 17 | 0.4118 | 5 | 0.6 | 6 | 0 | 6 | 0.1667 | 14 | 0.2857 | 31 | 0.2581 |
| 11_6222460 | 5 | 0 | 5 | 0 | 4 | 0.25 | 3 | 0 | 17 | 0.0588 | 4 | 0.25 | 6 | 0.1667 | 5 | 0.4 | 13 | 0.2308 | 28 | 0.25 |
| 11_6226411 | 1 | 0 | 2 | 0.5 | 1 | 1 | NA | NA | 4 | 0.5 | 2 | 0.5 | 2 | 0 | 2 | 0 | 1 | 0 | 7 | 0.1429 |
| 11_6235019 | NA | NA | NA | NA | 1 | 1 | NA | NA | 1 | 1 | 1 | 1 | NA | NA | NA | NA | 1 | 1 | 2 | 1 |
| 11_6240396 | NA | NA | 2 | 1 | 2 | 1 | 1 | 1 | 5 | 1 | NA | NA | NA | NA | 2 | 0.5 | NA | NA | 2 | 0.5 |
| 11_6242992 | 5 | 0.2 | 2 | 0.5 | NA | NA | 1 | 0 | 8 | 0.25 | 3 | 0.3333 | 4 | 0 | 4 | 0 | 2 | 0.5 | 13 | 0.1538 |
| 11_6243341 | NA | NA | 1 | 1 | NA | NA | 1 | 1 | 2 | 1 | NA | NA | 1 | 1 | NA | NA | NA | NA | 1 | 1 |
| 11_6249570 | NA | NA | NA | NA | NA | NA | 1 | 1 | 1 | 1 | 1 | 1 | NA | NA | NA | NA | NA | NA | 1 | 1 |
| 11_6249571 | 1 | 0 | 1 | 0 | 2 | 0 | 2 | 0.5 | 6 | 0.1667 | NA | NA | 2 | 0.5 | NA | NA | 3 | 0.3333 | 5 | 0.4 |
| 11_6249575 | 2 | 0.5 | 1 | 1 | 2 | 0.5 | 2 | 1 | 7 | 0.7143 | 1 | 1 | 2 | 1 | 1 | 1 | 4 | 0.75 | 8 | 0.875 |
| 11_6249594 | 2 | 1 | NA | NA | 2 | 0.5 | 2 | 0.5 | 6 | 0.6667 | 1 | 1 | 2 | 0.5 | NA | NA | 4 | 0.75 | 7 | 0.7143 |
| 11_6249597 | 1 | 1 | NA | NA | 2 | 0 | 2 | 0 | 5 | 0.2 | 1 | 0 | 2 | 0.5 | NA | NA | 4 | 0.5 | 7 | 0.4286 |
| 11_6249674 | 1 | 1 | 1 | 1 | 2 | 1 | 2 | 1 | 6 | 1 | NA | NA | 2 | 1 | 1 | 1 | 2 | 1 | 5 | 1 |
| 11_6249681 | 1 | 1 | 1 | 1 | 2 | 1 | 1 | 0 | 5 | 0.8 | NA | NA | 2 | 0.5 | 1 | 1 | 2 | 0.5 | 5 | 0.6 |
| 11_6249695 | NA | NA | 1 | 1 | 2 | 0 | 1 | 0 | 4 | 0.25 | NA | NA | 2 | 0.5 | 1 | 1 | 2 | 0 | 5 | 0.4 |
| 11_6249699 | 1 | 0 | 1 | 0 | 2 | 0 | 1 | 1 | 5 | 0.2 | NA | NA | 2 | 0.5 | 1 | 0 | 2 | 0 | 5 | 0.2 |
| 11_6255556 | 3 | 0.6667 | 3 | 0.3333 | 1 | 1 | 1 | 0 | 8 | 0.5 | 4 | 0.25 | 4 | 0.25 | 2 | 0.5 | 1 | 1 | 11 | 0.3636 |
| 11_6255565 | 3 | 0.3333 | 2 | 0 | 1 | 0 | 1 | 0 | 7 | 0.1429 | 4 | 0.5 | 6 | 0.3333 | 1 | 1 | 1 | 0 | 12 | 0.4167 |
| 11_6255575 | 3 | 0 | 3 | 0.3333 | 1 | 0 | 1 | 0 | 8 | 0.125 | 4 | 0 | 5 | 0 | 2 | 0 | 1 | 1 | 12 | 0.0833 |
| 11_6255579 | 2 | 0 | 3 | 0.3333 | 1 | 0 | 1 | 0 | 7 | 0.1429 | 4 | 0 | 5 | 0.2 | 2 | 0.5 | 1 | 0 | 12 | 0.1667 |
| 11_6255585 | 2 | 1 | 2 | 1 | 1 | 1 | 1 | 1 | 6 | 1 | 4 | 1 | 5 | 0.8 | 2 | 1 | NA | NA | 11 | 0.9091 |
| 11_6255585 | 2 | 1 | 3 | 0.3333 | 1 | 1 | 1 | 1 | 7 | 0.7143 | 4 | 0.25 | 6 | 0.5 | 2 | 1 | 1 | 1 | 13 | 0.5385 |
| 11_6255587 | 3 | 0 | 2 | 1 | 2 | 0.5 | 1 | 1 | 8 | 0.5 | 4 | 0.5 | 5 | 0.2 | 2 | 0.5 | 1 | 0 | 12 | 0.3333 |
| 11_6255589 | 2 | 0.5 | 3 | 0 | 1 | 0 | 1 | 0 | 7 | 0.1429 | 3 | 0.3333 | 4 | 1 | 2 | 0 | 1 | 0 | 10 | 0.5 |
| 11_6255607 | 2 | 1 | 1 | 1 | NA | NA | 1 | 1 | 4 | 1 | 1 | 1 | 3 | 1 | 1 | 1 | NA | NA | 5 | 1 |
| 11_6255615 | 2 | 1 | 1 | 0 | NA | NA | 1 | 0 | 4 | 0.5 | 1 | 1 | 4 | 1 | NA | NA | NA | NA | 5 | 1 |
| 11_6255624 | 2 | 0.5 | 1 | 0 | 1 | 1 | NA | NA | 4 | 0.5 | 1 | 0 | 3 | 1 | 1 | 0 | NA | NA | 5 | 0.6 |
| 11_6255638 | 2 | 1 | 1 | 0 | 1 | 1 | 1 | 1 | 5 | 0.8 | 1 | 0 | 4 | 1 | 1 | 1 | NA | NA | 6 | 0.8333 |
| 11_6255643 | 1 | 1 | NA | NA | 1 | 1 | 1 | 1 | 3 | 1 | 1 | 0 | 2 | 1 | 1 | 1 | NA | NA | 4 | 0.75 |
| 11_6255645 | 2 | 0.5 | 1 | 0 | 1 | 0 | 1 | 0 | 5 | 0.2 | 1 | 0 | 4 | 0.5 | 1 | 0 | NA | NA | 6 | 0.3333 |
| 11_6255651 | 2 | 0.5 | 1 | 0 | 2 | 0.5 | 1 | 1 | 6 | 0.5 | 1 | 1 | 3 | 1 | 1 | 1 | NA | NA | 5 | 1 |
| 11_6255658 | 3 | 0.6667 | 3 | 0.3333 | 4 | 0.5 | 2 | 1 | 12 | 0.5833 | 4 | 0.25 | 6 | 0.6667 | 2 | 1 | 1 | 1 | 13 | 0.6154 |
| 11_6255686 | 4 | 1 | 3 | 1 | 4 | 1 | 3 | 1 | 14 | 1 | 4 | 1 | 2 | 1 | 2 | 1 | 1 | 1 | 9 | 1 |
| 11_6255689 | 4 | 0 | 3 | 0.3333 | 2 | 0 | 2 | 0.5 | 11 | 0.1818 | 3 | 0 | 4 | 0.25 | NA | NA | 1 | 0 | 8 | 0.125 |
| 11_6255692 | 3 | 0.3333 | 2 | 1 | 3 | 0.3333 | 3 | 0.3333 | 11 | 0.4545 | 2 | 0 | 6 | 0.1667 | 2 | 1 | 1 | 0 | 11 | 0.2727 |
| 11_6255698 | 3 | 0 | 3 | 0 | 4 | 0.25 | 3 | 0 | 13 | 0.0769 | 4 | 0 | 7 | 0.1429 | 2 | 0.5 | 1 | 0 | 14 | 0.1429 |
| 11_6260374 | 4 | 0.25 | 1 | 0 | 2 | 0 | 1 | 0 | 8 | 0.125 | NA | NA | 1 | 1 | 1 | 0 | NA | NA | 2 | 0.5 |
| 11_6260397 | 4 | 0 | 1 | 0 | 1 | 0 | 1 | 1 | 7 | 0.1429 | NA | NA | 1 | 1 | 1 | 0 | NA | NA | 2 | 0.5 |
| 11_6260437 | 3 | 0.6667 | 1 | 1 | 1 | 1 | 1 | 1 | 6 | 0.8333 | NA | NA | 1 | 1 | 1 | 1 | NA | NA | 2 | 1 |
| 11_6260461 | 3 | 1 | 1 | 1 | 1 | 1 | NA | NA | 5 | 1 | NA | NA | 1 | 1 | 1 | 1 | NA | NA | 2 | 1 |
| 11_6260462 | 3 | 0.3333 | 1 | 0 | NA | NA | NA | NA | 4 | 0.25 | NA | NA | 1 | 1 | 1 | 0 | NA | NA | 2 | 0.5 |
| 11_6260476 | 2 | 0.5 | 1 | 1 | 1 | 0 | NA | NA | 4 | 0.5 | NA | NA | 1 | 1 | 1 | 1 | NA | NA | 2 | 1 |
| 11_6260482 | 2 | 1 | 1 | 1 | 1 | 1 | NA | NA | 4 | 1 | NA | NA | 1 | 1 | NA | NA | NA | NA | 1 | 1 |
| 11_6260590 | 2 | 1 | 1 | 0 | NA | NA | NA | NA | 3 | 0.6667 | NA | NA | NA | NA | 1 | 1 | NA | NA | 1 | 1 |
| 11_6260595 | 2 | 1 | 1 | 1 | NA | NA | NA | NA | 3 | 1 | NA | NA | NA | NA | 1 | 1 | NA | NA | 1 | 1 |

| | | | | | | | | | | | | | | | | | | | | |
|---|---|---|---|---|---|---|---|---|---|---|---|---|---|---|---|---|---|---|---|---|
| 11_626191 | 1 | 1 | 1 | 1 | NA | NA | NA | NA | 2 | 1 | NA | NA | 1 | 1 | NA | NA | NA | NA | 1 | 1 |
| 11_626193 | 1 | 1 | 1 | 0 | NA | NA | NA | NA | 2 | 0.5 | NA | NA | 1 | 1 | NA | NA | NA | NA | 1 | 1 |
| 11_626194 | 1 | 1 | NA | NA | NA | NA | NA | NA | 1 | 1 | NA | NA | 1 | 1 | 1 | 1 | NA | NA | 2 | 1 |
| 11_626699 | NA | NA | NA | NA | NA | NA | 2 | 0.5 | 2 | 0.5 | 1 | 0 | 1 | 1 | NA | NA | 1 | 0 | 3 | 0.3333 |
| 11_626701 | NA | NA | NA | NA | NA | NA | 3 | 0.3333 | 3 | 0.3333 | 1 | 0 | 1 | 0 | 1 | 1 | NA | NA | 3 | 0.3333 |
| 11_626704 | NA | NA | 1 | 1 | NA | NA | 3 | 0.3333 | 4 | 0.5 | 1 | 0 | 1 | 0 | 1 | 0 | 1 | 1 | 4 | 0.25 |
| 11_626705 | 1 | 1 | 2 | 1 | 1 | 1 | 4 | 0.75 | 8 | 0.875 | NA | NA | 1 | 1 | 2 | 1 | 1 | 1 | 4 | 1 |
| 11_626706 | 1 | 1 | 2 | 0.5 | 2 | 1 | 3 | 0.6667 | 8 | 0.75 | 1 | 1 | NA | NA | 1 | 1 | 1 | 1 | 3 | 1 |
| 11_626708 | 2 | 1 | 2 | 0.5 | 2 | 0.5 | 2 | 0.5 | 8 | 0.625 | 1 | 1 | 1 | 0 | 2 | 0.5 | 1 | 1 | 5 | 0.6 |
| 11_626709 | 2 | 0.5 | 2 | 0 | 2 | 0.5 | 2 | 0 | 8 | 0.25 | 1 | 1 | 1 | 0 | 2 | 0.5 | 1 | 1 | 5 | 0.6 |
| 11_626709 | 2 | 1 | 2 | 0.5 | 2 | 0.5 | 2 | 0.5 | 8 | 0.625 | 1 | 1 | 1 | 1 | 2 | 0.5 | 1 | 0 | 5 | 0.6 |
| 11_626713 | 2 | 0.5 | 2 | 1 | 2 | 0.5 | 1 | 0 | 7 | 0.5714 | 1 | 0 | 1 | 0 | 2 | 1 | 1 | 0 | 5 | 0.4 |
| 11_626713 | 2 | 1 | 2 | 0 | 2 | 1 | 2 | 1 | 8 | 0.75 | 1 | 1 | 1 | 1 | 2 | 1 | 1 | 1 | 5 | 1 |
| 11_626714 | 1 | 1 | 2 | 0.5 | 2 | 1 | 1 | 1 | 6 | 0.8333 | 1 | 1 | 1 | 0 | 2 | 1 | 1 | 1 | 5 | 0.8 |
| 11_626716 | 2 | 1 | 1 | 1 | 2 | 1 | 2 | 0.5 | 7 | 0.8571 | 1 | 1 | NA | NA | 2 | 0 | 1 | 1 | 4 | 0.5 |
| 11_626717 | NA | NA | 1 | 1 | NA | NA | NA | NA | 1 | 1 | 1 | 1 | NA | NA | NA | NA | 1 | 1 | 2 | 1 |
| 11_630502 | 4 | 0 | 10 | 0.1 | 10 | 0.3 | 3 | 0 | 27 | 0.1481 | 5 | 0.2 | 2 | 0.5 | 4 | 0 | 3 | 0.3333 | 14 | 0.2143 |
| 11_638535 | 2 | 0 | 3 | 0 | 2 | 0.5 | 6 | 0 | 13 | 0.0769 | 4 | 0 | 4 | 0 | 3 | 0.3333 | 6 | 0.1667 | 17 | 0.1176 |
| 11_638536 | 1 | 1 | 1 | 0 | 2 | 0 | 4 | 0 | 8 | 0.125 | 2 | 0.5 | 4 | 0.25 | 1 | 1 | 5 | 0.6 | 12 | 0.5 |
| 11_638537 | 2 | 1 | 3 | 0.3333 | 1 | 0 | 5 | 0.8 | 11 | 0.6364 | 3 | 1 | 3 | 0.6667 | 1 | 1 | 4 | 0.75 | 11 | 0.8182 |
| 11_638539 | 2 | 0 | 2 | 0.5 | 1 | 0 | 4 | 0.75 | 9 | 0.4444 | 4 | 0.75 | 4 | 0.5 | 1 | 1 | 4 | 1 | 13 | 0.7692 |
| 11_638539 | 2 | 0 | 3 | 0 | 1 | 0 | 5 | 0.2 | 11 | 0.0909 | 3 | 0.3333 | 3 | 0 | 2 | 0 | 2 | 0 | 10 | 0.1 |
| 11_638547 | 2 | 0.5 | 2 | 1 | 1 | 1 | 3 | 1 | 8 | 0.875 | 1 | 1 | 1 | 1 | 1 | 1 | 3 | 0.3333 | 6 | 0.6667 |
| 11_638548 | 1 | 1 | 2 | 1 | 1 | 1 | 3 | 1 | 7 | 1 | 2 | 1 | 1 | 1 | NA | NA | 2 | 0.5 | 5 | 0.8 |
| 11_638549 | 2 | 0.5 | 2 | 0 | NA | NA | 3 | 0.3333 | 7 | 0.2857 | 1 | 0 | NA | NA | NA | NA | 3 | 0.6667 | 4 | 0.5 |
| 11_638550 | 2 | 1 | 1 | 1 | 1 | 1 | 3 | 1 | 7 | 1 | 1 | 1 | NA | NA | NA | NA | 3 | 1 | 4 | 1 |
| 11_638551 | 2 | 0 | 1 | 1 | NA | NA | 2 | 0.5 | 5 | 0.4 | 2 | 0.5 | 1 | 1 | NA | NA | 3 | 0.3333 | 6 | 0.5 |
| 11_639735 | 4 | 0 | 2 | 1 | 2 | 0.5 | 1 | 1 | 9 | 0.4444 | 5 | 0.2 | 3 | 0.3333 | 4 | 0.25 | 2 | 0.5 | 14 | 0.2857 |
| 11_639736 | 3 | 0.3333 | 1 | 0 | 1 | 0 | 1 | 0 | 6 | 0.1667 | 3 | 0 | NA | NA | 4 | 0.25 | 1 | 0 | 8 | 0.125 |
| 11_639737 | 2 | 0 | 1 | 1 | 2 | 0.5 | 1 | 0 | 6 | 0.3333 | 5 | 0.4 | 3 | 0 | 4 | 0.25 | 1 | 0 | 13 | 0.2308 |
| 11_639738 | 4 | 0.5 | 2 | 0 | 2 | 0.5 | NA | NA | 8 | 0.375 | 4 | 0.25 | 2 | 0.5 | 5 | 0 | 2 | 0 | 13 | 0.1538 |
| 11_639738 | 4 | 0.75 | 2 | 1 | 2 | 0 | 1 | 0 | 9 | 0.5556 | 3 | 0.3333 | 3 | 0 | 3 | 0.3333 | 1 | 0 | 10 | 0.2 |
| 11_643242 | NA | NA | 1 | 1 | NA | NA | 1 | 0 | 2 | 0.5 | 1 | 1 | 1 | 0 | 1 | 0 | NA | NA | 3 | 0.3333 |
| 11_643245 | NA | NA | 1 | 1 | NA | NA | 1 | 0 | 2 | 0.5 | NA | NA | 1 | 1 | 1 | 1 | NA | NA | 2 | 1 |
| 11_643261 | NA | NA | 1 | 0 | NA | NA | 1 | 1 | 2 | 0.5 | 1 | 1 | 1 | 1 | 1 | 1 | NA | NA | 3 | 1 |
| 11_643263 | NA | NA | 1 | 1 | NA | NA | 1 | 1 | 2 | 1 | 1 | 1 | NA | NA | 1 | 0 | NA | NA | 2 | 0.5 |
| 11_643264 | NA | NA | 1 | 1 | NA | NA | NA | NA | 1 | 1 | 1 | 1 | 1 | 1 | 1 | 1 | NA | NA | 3 | 1 |
| 11_643267 | NA | NA | 1 | 1 | NA | NA | NA | NA | 1 | 1 | NA | NA | 1 | 1 | 1 | 1 | NA | NA | 2 | 1 |
| 11_643267 | NA | NA | 1 | 1 | NA | NA | 1 | 0 | 2 | 0.5 | NA | NA | 1 | 1 | 1 | 0 | NA | NA | 2 | 0.5 |
| 11_644112 | 1 | 0 | 3 | 0.6667 | NA | NA | NA | NA | 4 | 0.5 | NA | NA | 1 | 1 | NA | NA | NA | NA | 1 | 1 |
| 11_644115 | 1 | 1 | 2 | 1 | NA | NA | NA | NA | 3 | 1 | NA | NA | 1 | 1 | NA | NA | NA | NA | 1 | 1 |
| 11_644115 | 1 | 1 | 2 | 1 | NA | NA | NA | NA | 3 | 1 | NA | NA | 1 | 1 | NA | NA | NA | NA | 1 | 1 |
| 11_644119 | 1 | 0 | 3 | 0.3333 | NA | NA | NA | NA | 4 | 0.25 | NA | NA | 1 | 1 | NA | NA | NA | NA | 1 | 1 |
| 11_644119 | 1 | 1 | 2 | 1 | NA | NA | NA | NA | 3 | 1 | NA | NA | 1 | 1 | NA | NA | NA | NA | 1 | 1 |
| 11_644134 | 1 | 0 | 2 | 0.5 | NA | NA | NA | NA | 3 | 0.3333 | NA | NA | 2 | 0.5 | NA | NA | NA | NA | 2 | 0.5 |
| 11_648920 | 2 | 0.5 | 4 | 1 | 4 | 0.75 | 3 | 0.6667 | 13 | 0.7692 | 4 | 0.75 | NA | NA | 7 | 0.8571 | 2 | 1 | 13 | 0.8462 |
| 11_648923 | 2 | 1 | 6 | 0.6667 | 4 | 1 | 4 | 0.75 | 16 | 0.8125 | 4 | 0.25 | NA | NA | 8 | 0.625 | 3 | 1 | 15 | 0.6 |
| 11_648923 | 2 | 0.5 | 3 | 1 | 5 | 0.6 | 4 | 1 | 14 | 0.7857 | 5 | 0.4 | NA | NA | 8 | 1 | 3 | 1 | 16 | 0.8125 |
| 11_648924 | 2 | 0.5 | 4 | 0.25 | 4 | 0 | 4 | 0 | 14 | 0.1429 | 4 | 0.25 | NA | NA | 7 | 0 | 4 | 0 | 15 | 0.0667 |
| 11_648925 | 1 | 1 | 1 | 1 | NA | NA | NA | NA | 2 | 1 | 1 | 1 | NA | NA | 1 | 1 | NA | NA | 2 | 1 |
| 11_648927 | 2 | 0.5 | 1 | 1 | 1 | 1 | 1 | 0 | 5 | 0.6 | 1 | 1 | NA | NA | 3 | 0.6667 | NA | NA | 4 | 0.75 |
| 11_648927 | 2 | 0 | 1 | 1 | 1 | 1 | 1 | 0 | 5 | 0.4 | 1 | 1 | NA | NA | 3 | 1 | NA | NA | 4 | 1 |
| 11_648935 | 12 | 0 | 17 | 0.1765 | 10 | 0.1 | 12 | 0 | 51 | 0.0784 | 11 | 0 | 14 | 0.0714 | 15 | 0.1333 | 6 | 0.1667 | 46 | 0.087 |
| 11_648935 | 13 | 0.5385 | 17 | 0.4118 | 8 | 0.5 | 11 | 0.4545 | 49 | 0.4694 | 9 | 0.3333 | 14 | 0.3571 | 18 | 0.5 | 6 | 0.5 | 47 | 0.4255 |
| 11_648938 | 15 | 0.2667 | 20 | 0.2 | 7 | 0.1429 | 14 | 0.1429 | 56 | 0.1964 | 12 | 0.1667 | 13 | 0.0769 | 11 | 0.2727 | 7 | 0.1429 | 43 | 0.1628 |
| 11_648940 | 15 | 0.8667 | 20 | 0.9 | 7 | 0.8571 | 14 | 0.8571 | 56 | 0.875 | 11 | 0.9091 | 11 | 0.7273 | 12 | 0.8333 | 6 | 0.8333 | 40 | 0.825 |
| 11_648941 | 12 | 0.0833 | 17 | 0.0588 | 6 | 0.3333 | 12 | 0.0833 | 47 | 0.1064 | 12 | 0 | 10 | 0.1 | 10 | 0 | 4 | 0.25 | 36 | 0.0556 |
| 11_648991 | 12 | 1 | 16 | 0.5625 | 4 | 0.75 | 10 | 0.5 | 42 | 0.6905 | 11 | 0.9091 | 12 | 0.75 | 10 | 0.7 | 3 | 0.3333 | 36 | 0.75 |
| 11_655794 | 15 | 0.1333 | 21 | 0 | 17 | 0.2353 | 11 | 0.1818 | 64 | 0.125 | 14 | 0.0714 | 14 | 0.1429 | 22 | 0.0909 | 18 | 0.1111 | 68 | 0.1029 |
| 11_655795 | 11 | 0.0909 | 22 | 0.0909 | 13 | 0 | 9 | 0 | 55 | 0.0545 | 15 | 0.1333 | 12 | 0 | 21 | 0.1905 | 17 | 0 | 65 | 0.0923 |
| 11_655796 | 9 | 0.4444 | 20 | 0.15 | 19 | 0.2632 | 10 | 0.3 | 58 | 0.2586 | 13 | 0.2308 | 11 | 0.3636 | 18 | 0.2778 | 13 | 0.3077 | 55 | 0.2909 |
| 11_655797 | 14 | 0.2857 | 20 | 0.25 | 18 | 0.2222 | 9 | 0.3333 | 61 | 0.2623 | 15 | 0.4 | 12 | 0.25 | 18 | 0.2222 | 17 | 0.1765 | 62 | 0.2581 |
| 11_655806 | 9 | 0.6667 | 12 | 0.8333 | 16 | 0.6875 | 8 | 0.625 | 45 | 0.7111 | 10 | 0.5 | 11 | 0.7273 | 11 | 0.8182 | 15 | 0.8 | 47 | 0.7234 |
| 11_655807 | 9 | 0.6667 | 13 | 0.8462 | 14 | 0.8571 | 6 | 0.8333 | 42 | 0.8095 | 11 | 0.7273 | 10 | 0.8 | 15 | 0.8667 | 10 | 0.8 | 46 | 0.8043 |
| 11_655808 | 9 | 0.4444 | 12 | 0.5 | 14 | 0.5714 | 10 | 0.4 | 45 | 0.4889 | 7 | 0.5714 | 11 | 0.6364 | 15 | 0.8 | 10 | 0.5 | 43 | 0.6512 |

| | | | | | | | | | | | | | | | | | | | | |
|---|---|---|---|---|---|---|---|---|---|---|---|---|---|---|---|---|---|---|---|---|
| 11_655808 | 11 | 0 | 14 | 0.2143 | 15 | 0.1333 | 10 | 0.1 | 50 | 0.12 | 10 | 0.1 | 11 | 0.1818 | 16 | 0.1875 | 15 | 0.0667 | 52 | 0.1346 |
| 11_655809 | 11 | 0.1818 | 15 | 0.0667 | 10 | 0 | 7 | 0 | 43 | 0.0698 | 8 | 0.125 | 10 | 0.1 | 17 | 0.1176 | 12 | 0.1667 | 47 | 0.1277 |
| 11_655811 | 10 | 0 | 15 | 0.1333 | 15 | 0.1333 | 7 | 0 | 47 | 0.0851 | 9 | 0.2222 | 11 | 0 | 16 | 0.0625 | 14 | 0.0714 | 50 | 0.08 |
| 11_655811 | 8 | 0.125 | 14 | 0.1429 | 13 | 0.3846 | 9 | 0.3333 | 44 | 0.25 | 10 | 0.4 | 11 | 0.1818 | 16 | 0.3125 | 12 | 0.4167 | 49 | 0.3265 |
| 11_655812 | 7 | 0.7143 | 12 | 0.75 | 13 | 0.7692 | 9 | 0.6667 | 41 | 0.7317 | 8 | 0.875 | 10 | 0.6 | 14 | 0.7143 | 11 | 0.9091 | 43 | 0.7674 |
| 11_655812 | 7 | 1 | 12 | 0.9167 | 11 | 0.6364 | 8 | 0.75 | 38 | 0.8158 | 10 | 0.9 | 11 | 0.9091 | 16 | 1 | 13 | 0.9231 | 50 | 0.94 |
| 11_655812 | 5 | 0.2 | 11 | 0.8182 | 8 | 0.5 | 5 | 0.4 | 29 | 0.5517 | 8 | 0.5 | 8 | 0.5 | 11 | 0.8182 | 6 | 0.5 | 33 | 0.6061 |
| 11_655813 | 7 | 0.4286 | 15 | 0.2 | 15 | 0.2 | 7 | 0.1429 | 44 | 0.2273 | 8 | 0.375 | 11 | 0.2727 | 15 | 0.2667 | 15 | 0.4667 | 49 | 0.3469 |
| 11_655813 | 9 | 0.7778 | 11 | 0.7273 | 13 | 0.8462 | 5 | 0.6 | 38 | 0.7632 | 8 | 0.5 | 9 | 0.7778 | 13 | 0.6923 | 12 | 0.75 | 42 | 0.6905 |
| 11_655813 | 8 | 0.75 | 13 | 0.9231 | 11 | 0.8182 | 5 | 1 | 37 | 0.8649 | 6 | 1 | 11 | 0.9091 | 13 | 0.9231 | 11 | 0.8182 | 41 | 0.9024 |
| 11_655814 | 6 | 0.8333 | 13 | 0.9231 | 14 | 0.6429 | 8 | 0.75 | 41 | 0.7805 | 8 | 0.625 | 10 | 0.7 | 14 | 0.9286 | 14 | 0.9286 | 46 | 0.8261 |
| 11_655814 | 5 | 1 | 9 | 0.7778 | 10 | 0.8 | 5 | 0.6 | 29 | 0.7931 | 4 | 1 | 8 | 0.625 | 13 | 0.8462 | 9 | 0.5556 | 34 | 0.7353 |
| 11_657190 | 2 | 0.5 | 1 | 1 | 1 | 1 | NA | NA | 4 | 0.75 | 1 | 0 | 1 | 1 | 2 | 0.5 | 1 | 0 | 5 | 0.4 |
| 11_657191 | 1 | 0 | 1 | 1 | 1 | 0 | NA | NA | 3 | 0.3333 | NA | NA | 1 | 1 | 2 | 0.5 | 1 | 1 | 4 | 0.75 |
| 11_657192 | 2 | 1 | 1 | 1 | 1 | 1 | NA | NA | 4 | 1 | 1 | 1 | NA | NA | 3 | 1 | 1 | 1 | 5 | 1 |
| 11_657193 | 2 | 1 | 1 | 0 | NA | NA | NA | NA | 3 | 0.6667 | 1 | 1 | 1 | 1 | 2 | 1 | 1 | 0 | 5 | 0.8 |
| 11_657193 | 1 | 1 | 1 | 1 | 1 | 1 | NA | NA | 3 | 1 | 1 | 1 | 1 | 1 | 3 | 1 | 1 | 1 | 6 | 1 |
| 11_657194 | 2 | 1 | 1 | 1 | 1 | 1 | NA | NA | 4 | 1 | 1 | 1 | 1 | 1 | 3 | 0.3333 | 1 | 1 | 6 | 0.6667 |
| 11_657195 | 2 | 0.5 | 1 | 1 | 1 | 1 | NA | NA | 4 | 0.75 | 1 | 1 | 1 | 1 | 3 | 0.6667 | 1 | 1 | 6 | 0.8333 |
| 11_657195 | 2 | 0.5 | 1 | 0 | 1 | 0 | NA | NA | 4 | 0.25 | 1 | 0 | 1 | 1 | 3 | 0 | 2 | 0 | 7 | 0.1429 |
| 11_657197 | 2 | 0.5 | 1 | 0 | 1 | 0 | NA | NA | 4 | 0.25 | 1 | 1 | 1 | 0 | 2 | 0 | 2 | 1 | 6 | 0.5 |
| 11_661029 | NA | NA | 1 | 1 | NA | NA | 1 | 1 | 2 | 1 | 1 | 0 | 2 | 1 | 2 | 1 | 1 | 1 | 6 | 0.8333 |
| 11_661032 | 1 | 0 | 2 | 0.5 | 1 | 0 | 1 | 0 | 5 | 0.2 | 1 | 0 | 2 | 0 | 1 | 1 | NA | NA | 4 | 0.25 |
| 11_661034 | 1 | 1 | 1 | 1 | 1 | 1 | 1 | 1 | 4 | 1 | 1 | 1 | 2 | 1 | 2 | 1 | 1 | 1 | 6 | 1 |
| 11_661034 | 1 | 1 | 2 | 0 | 1 | 0 | 1 | 0 | 5 | 0.2 | 1 | 1 | 2 | 0 | 2 | 0 | 1 | 0 | 6 | 0.1667 |
| 11_661049 | 1 | 1 | 2 | 0 | 1 | 0 | 1 | 1 | 5 | 0.4 | NA | NA | 1 | 1 | 1 | 0 | 1 | 0 | 3 | 0.3333 |
| 11_661069 | 1 | 1 | 1 | 0 | NA | NA | 1 | 1 | 3 | 0.6667 | NA | NA | NA | NA | NA | NA | 1 | 1 | 1 | 1 |
| 11_669290 | 2 | 0 | 3 | 0.3333 | 1 | 1 | 3 | 0 | 9 | 0.2222 | 4 | 0.25 | 2 | 0 | 1 | 0 | 3 | 0.3333 | 10 | 0.2 |
| 11_669290 | 2 | 0 | 3 | 0.3333 | 1 | 0 | 1 | 1 | 7 | 0.2857 | 2 | 0 | 2 | 1 | 1 | 0 | 3 | 1 | 8 | 0.625 |
| 11_669311 | 2 | 0.5 | 2 | 1 | 1 | 1 | 2 | 0.5 | 7 | 0.7143 | 4 | 0.5 | 2 | 1 | 2 | 0.5 | 3 | 0.3333 | 11 | 0.5455 |
| 11_669324 | 2 | 0.5 | 2 | 1 | NA | NA | 1 | 1 | 5 | 0.8 | 4 | 1 | 2 | 0.5 | 2 | 1 | 3 | 1 | 11 | 0.9091 |
| 11_669325 | 2 | 0.5 | 1 | 1 | NA | NA | 1 | 1 | 4 | 0.75 | 4 | 1 | 2 | 1 | 2 | 1 | 3 | 0.6667 | 11 | 0.9091 |
| 11_669326 | 2 | 1 | 1 | 1 | NA | NA | 1 | 1 | 4 | 1 | NA | NA | 2 | 1 | 2 | 0.5 | 2 | 0.5 | 6 | 0.6667 |
| 11_669326 | 1 | 1 | 2 | 0.5 | NA | NA | NA | NA | 3 | 0.6667 | 4 | 1 | 2 | 1 | 2 | 1 | 3 | 1 | 11 | 1 |
| 11_669547 | 1 | 1 | NA | NA | 1 | 1 | 1 | 0 | 3 | 0.6667 | NA | NA | 1 | 1 | NA | NA | 1 | 0 | 2 | 0.5 |
| 11_669549 | 1 | 1 | 1 | 0 | NA | NA | 1 | 1 | 3 | 0.6667 | 1 | 1 | 1 | 1 | NA | NA | 2 | 1 | 4 | 1 |
| 11_675742 | NA | NA | 7 | 0.1429 | 5 | 0.2 | 4 | 0 | 16 | 0.125 | 7 | 0.1429 | 11 | 0.0909 | 7 | 0.2857 | 3 | 0 | 28 | 0.1429 |
| 11_675742 | NA | NA | 8 | 0.875 | 4 | 0.5 | 4 | 0.5 | 16 | 0.6875 | 10 | 0.7 | 11 | 1 | 7 | 0.8571 | 5 | 0.8 | 33 | 0.8485 |
| 11_675744 | NA | NA | 10 | 0.3 | 5 | 0.2 | 5 | 0 | 20 | 0.2 | 11 | 0.1818 | 11 | 0.2727 | 6 | 0.3333 | 6 | 0.6667 | 34 | 0.3235 |
| 11_675744 | 1 | 0 | 9 | 0.1111 | 5 | 0.2 | 5 | 0 | 20 | 0.1 | 12 | 0.0833 | 9 | 0.1111 | 7 | 0.2857 | 6 | 0.3333 | 34 | 0.1765 |
| 11_675746 | 1 | 0 | 6 | 0.3333 | 5 | 0.6 | 5 | 0.4 | 17 | 0.4118 | 12 | 0.1667 | 9 | 0.3333 | 5 | 0.6 | 6 | 0.8333 | 32 | 0.4062 |
| 11_675757 | 1 | 0 | 8 | 0.25 | 8 | 0 | 3 | 0.3333 | 20 | 0.15 | 12 | 0.0833 | 14 | 0.0714 | 7 | 0 | 10 | 0.3 | 43 | 0.1163 |
| 11_675762 | 3 | 0 | 10 | 0.1 | 8 | 0.25 | 7 | 0 | 28 | 0.1071 | 9 | 0.1111 | 13 | 0 | 9 | 0.2222 | 7 | 0 | 38 | 0.0789 |
| 11_675887 | NA | NA | NA | NA | NA | NA | 1 | 1 | 1 | 1 | NA | NA | NA | NA | NA | NA | 1 | 1 | 1 | 1 |
| 11_678641 | 3 | 0 | 7 | 0 | 6 | 0 | 3 | 0.3333 | 19 | 0.0526 | 3 | 0 | 3 | 0 | 1 | 0 | 3 | 0.3333 | 10 | 0.1 |
| 11_681007 | 2 | 0.5 | 3 | 0 | NA | NA | 9 | 0.1111 | 14 | 0.1429 | 5 | 0 | 3 | 0 | 2 | 0.5 | 4 | 0.25 | 14 | 0.1429 |
| 11_681037 | NA | NA | NA | NA | NA | NA | 2 | 1 | 2 | 1 | 1 | 1 | NA | NA | NA | NA | NA | NA | 1 | 1 |
| 11_683877 | 1 | 1 | NA | NA | NA | NA | NA | NA | 1 | 1 | NA | NA | 1 | 0 | NA | NA | 2 | 0.5 | 3 | 0.3333 |
| 11_683881 | 1 | 1 | NA | NA | NA | NA | NA | NA | 1 | 1 | NA | NA | 1 | 1 | NA | NA | 1 | 1 | 2 | 1 |
| 11_683881 | 1 | 1 | NA | NA | NA | NA | NA | NA | 1 | 1 | NA | NA | 1 | 0 | NA | NA | 2 | 0.5 | 3 | 0.3333 |
| 11_683883 | 1 | 1 | NA | NA | NA | NA | NA | NA | 1 | 1 | NA | NA | 1 | 0 | NA | NA | 1 | 1 | 2 | 0.5 |
| 11_687059 | 10 | 0.1 | 5 | 0.2 | 7 | 0.4286 | 10 | 0.6 | 32 | 0.3438 | 4 | 0.25 | 7 | 0.1429 | 8 | 0.375 | 2 | 0.5 | 21 | 0.2857 |
| 11_687060 | 11 | 0.5455 | 5 | 0.8 | 8 | 0.625 | 14 | 0.7857 | 38 | 0.6842 | 6 | 1 | 7 | 0.4286 | 9 | 0.6667 | 2 | 0.5 | 24 | 0.6667 |
| 11_687060 | 9 | 0.2222 | 6 | 0.5 | 9 | 0.4444 | 14 | 0.2143 | 38 | 0.3158 | 7 | 0.5714 | 7 | 0.4286 | 10 | 0.4 | 1 | 0 | 25 | 0.44 |
| 11_687061 | 13 | 0.0769 | 6 | 0.3333 | 11 | 0.0909 | 12 | 0.4167 | 42 | 0.2143 | 7 | 0 | 7 | 0.5714 | 9 | 0.1111 | 2 | 0 | 25 | 0.2 |
| 11_687061 | 13 | 0.5385 | 7 | 0.5714 | 11 | 0.4545 | 13 | 0.3846 | 44 | 0.4773 | 8 | 0.375 | 6 | 0.5 | 9 | 0.4444 | 5 | 0.2 | 28 | 0.3929 |
| 11_687061 | 13 | 0.2308 | 7 | 0.4286 | 11 | 0.0909 | 14 | 0.2857 | 45 | 0.2444 | 9 | 0.1111 | 8 | 0 | 9 | 0.2222 | 5 | 0.2 | 31 | 0.129 |
| 11_687062 | 12 | 0 | 7 | 0 | 10 | 0.1 | 13 | 0.2308 | 42 | 0.0952 | 7 | 0.1429 | 7 | 0 | 9 | 0.2222 | 3 | 0 | 26 | 0.1154 |
| 11_687062 | 13 | 0.4615 | 9 | 0.2222 | 11 | 0.2727 | 15 | 0.4 | 48 | 0.3542 | 9 | 0.4444 | 7 | 0.5714 | 6 | 0.6667 | 5 | 0.2 | 27 | 0.4815 |
| 11_687063 | 15 | 0.1333 | 9 | 0.1111 | 10 | 0.3 | 13 | 0 | 47 | 0.1277 | 9 | 0.2222 | 7 | 0.1429 | 9 | 0 | 7 | 0.1429 | 32 | 0.125 |
| 11_687065 | 16 | 0.0625 | 11 | 0.0909 | 19 | 0.2105 | 20 | 0.15 | 66 | 0.1364 | 11 | 0.0909 | 15 | 0.0667 | 13 | 0.0769 | 11 | 0.2727 | 50 | 0.12 |
| 11_687069 | 19 | 0.3158 | 12 | 0.3333 | 13 | 0.3846 | 18 | 0.1667 | 62 | 0.2903 | 11 | 0.1818 | 16 | 0.4375 | 17 | 0.2353 | 10 | 0.4 | 54 | 0.3148 |
| 11_687069 | 21 | 0.1429 | 12 | 0 | 12 | 0.0833 | 23 | 0.0435 | 68 | 0.0735 | 11 | 0 | 13 | 0.1538 | 13 | 0.0769 | 10 | 0.1 | 47 | 0.0851 |
| 11_687071 | 22 | 0.4091 | 20 | 0.35 | 25 | 0.32 | 25 | 0.32 | 92 | 0.3478 | 21 | 0.2857 | 16 | 0.1875 | 20 | 0.4 | 12 | 0.5 | 69 | 0.3333 |
| 11_687072 | 20 | 0.15 | 22 | 0.0909 | 21 | 0 | 26 | 0.0385 | 89 | 0.0674 | 20 | 0.1 | 18 | 0.1667 | 18 | 0.0556 | 13 | 0 | 69 | 0.087 |

| | | | | | | | | | | | | | | | | | | | | |
|---|---|---|---|---|---|---|---|---|---|---|---|---|---|---|---|---|---|---|---|---|
| 11_687130 | 14 | 0.2143 | 12 | 0.3333 | 12 | 0.1667 | 18 | 0.1667 | 56 | 0.2143 | 9 | 0.2222 | 14 | 0.4286 | 15 | 0.2 | 12 | 0.5833 | 50 | 0.36 |
| 11_687132 | 12 | 0.9167 | 14 | 1 | 8 | 0.875 | 19 | 0.7895 | 53 | 0.8868 | 12 | 0.9167 | 14 | 0.9286 | 19 | 0.6842 | 12 | 0.5833 | 57 | 0.7719 |
| 11_687953 | 88 | 0.0795 | 125 | 0.208 | 115 | 0.1652 | 131 | 0.0992 | 459 | 0.1416 | 124 | 0.1774 | 111 | 0.1622 | 129 | 0.1783 | 109 | 0.1193 | 473 | 0.1607 |
| 11_687970 | 2 | 0 | 3 | 0.3333 | 2 | 0.5 | 1 | 0 | 8 | 0.25 | NA | NA | 1 | 0 | NA | NA | 1 | 1 | 2 | 0.5 |
| 11_687975 | 4 | 0.25 | 3 | 0.3333 | 2 | 0 | 3 | 0.3333 | 12 | 0.25 | NA | NA | 1 | 0 | NA | NA | 2 | 1 | 3 | 0.6667 |
| 11_687976 | 2 | 1 | 4 | 1 | 2 | 1 | 2 | 1 | 10 | 1 | NA | NA | NA | NA | NA | NA | 1 | 1 | 1 | 1 |
| 11_688185 | 1 | 1 | NA | NA | NA | NA | NA | NA | 1 | 1 | NA | NA | NA | NA | 2 | 1 | NA | NA | 2 | 1 |
| 11_688340 | 1 | 1 | 5 | 0.2 | 2 | 0.5 | 7 | 0.1429 | 15 | 0.2667 | 3 | 0.6667 | 2 | 0 | 5 | 0.6 | 4 | 0.75 | 14 | 0.5714 |
| 11_688341 | 1 | 1 | 5 | 0.6 | 2 | 1 | 6 | 1 | 14 | 0.8571 | 3 | 1 | 4 | 1 | 6 | 1 | 3 | 1 | 16 | 1 |
| 11_688341 | 1 | 0 | 4 | 0.75 | 2 | 0 | 7 | 0.7143 | 14 | 0.5714 | 3 | 0.6667 | 3 | 1 | 6 | 0.6667 | 4 | 0.75 | 16 | 0.75 |
| 11_688341 | 1 | 1 | 5 | 0.8 | 2 | 1 | 6 | 0.6667 | 14 | 0.7857 | 3 | 1 | 4 | 1 | 6 | 0.8333 | 4 | 1 | 17 | 0.9412 |
| 11_688342 | 1 | 1 | 5 | 1 | NA | NA | 5 | 0.8 | 11 | 0.9091 | 2 | 1 | 3 | 1 | 4 | 1 | 2 | 1 | 11 | 1 |
| 11_688342 | 1 | 0 | 5 | 0 | 2 | 0 | 7 | 0.1429 | 15 | 0.0667 | 2 | 0.5 | 1 | 0 | 6 | 0.5 | 4 | 0.75 | 13 | 0.5385 |
| 11_688345 | 1 | 0 | 1 | 0 | 1 | 0 | 4 | 0.5 | 7 | 0.2857 | 2 | 0.5 | 2 | 1 | 2 | 0 | 4 | 0.25 | 10 | 0.4 |
| 11_688345 | 1 | 1 | 1 | 0 | 1 | 1 | 5 | 1 | 8 | 0.875 | 1 | 1 | 2 | 1 | 2 | 0.5 | 4 | 0.75 | 9 | 0.7778 |
| 11_688346 | 1 | 0 | 1 | 0 | NA | NA | 5 | 0.6 | 7 | 0.4286 | 2 | 0 | 2 | 0.5 | 2 | 0 | 4 | 0.5 | 10 | 0.3 |
| 11_688346 | 1 | 1 | 1 | 1 | 1 | 1 | 4 | 1 | 7 | 1 | 1 | 1 | 2 | 1 | 2 | 1 | 4 | 1 | 9 | 1 |
| 11_688351 | 1 | 0 | 1 | 0 | 2 | 0 | 5 | 0.4 | 9 | 0.2222 | 2 | 0.5 | 2 | 0.5 | 1 | 0 | 3 | 0.6667 | 8 | 0.5 |
| 11_688351 | 1 | 0 | 1 | 0 | 2 | 0.5 | 4 | 0 | 8 | 0.125 | 2 | 0 | 2 | 0.5 | 1 | 0 | 2 | 0 | 7 | 0.1429 |
| 11_688355 | 1 | 0 | 2 | 0 | 2 | 0 | 6 | 0.3333 | 11 | 0.1818 | 2 | 0.5 | 2 | 0 | 2 | 0.5 | 3 | 0 | 9 | 0.2222 |
| 11_688373 | 1 | 0 | 2 | 0.5 | 1 | 0 | 7 | 0 | 11 | 0.0909 | 2 | 0 | 2 | 0.5 | 4 | 0 | 3 | 0 | 11 | 0.0909 |
| 11_688634 | 6 | 0.3333 | 1 | 0 | 1 | 0 | 2 | 0 | 10 | 0.2 | 1 | 1 | 5 | 0 | 1 | 0 | 3 | 0 | 10 | 0.1 |
| 11_688645 | 5 | 0.2 | 1 | 0 | 1 | 0 | 2 | 0 | 9 | 0.1111 | 1 | 0 | 4 | 0 | NA | NA | 3 | 0.3333 | 8 | 0.125 |
| 11_688768 | 1 | 0 | 3 | 0 | 2 | 0 | 4 | 0.25 | 10 | 0.1 | NA | NA | 2 | 0 | 2 | 0.5 | 1 | 1 | 5 | 0.4 |
| 11_688775 | NA | NA | 3 | 0.6667 | NA | NA | 4 | 0.5 | 7 | 0.5714 | NA | NA | 1 | 0 | 1 | 1 | 3 | 0 | 5 | 0.2 |
| 11_688777 | 2 | 0 | 3 | 0.3333 | NA | NA | 4 | 0 | 9 | 0.1111 | NA | NA | 2 | 0.5 | NA | NA | 3 | 0 | 5 | 0.2 |
| 11_688785 | NA | NA | 2 | 0.5 | NA | NA | 4 | 0.75 | 6 | 0.6667 | NA | NA | 2 | 1 | NA | NA | 3 | 0.3333 | 5 | 0.6 |
| 11_688786 | NA | NA | 2 | 0.5 | NA | NA | 4 | 0.75 | 6 | 0.6667 | NA | NA | NA | NA | NA | NA | 3 | 0.3333 | 3 | 0.3333 |
| 11_688787 | NA | NA | 2 | 0.5 | NA | NA | 2 | 0 | 4 | 0.25 | NA | NA | 1 | 0 | NA | NA | 2 | 0.5 | 3 | 0.3333 |
| 11_688788 | NA | NA | 2 | 0.5 | NA | NA | 1 | 1 | 3 | 0.6667 | NA | NA | 2 | 1 | NA | NA | 3 | 0.3333 | 5 | 0.6 |
| 11_689170 | 3 | 0 | NA | NA | 1 | 0 | 1 | 1 | 5 | 0.2 | 1 | 0 | NA | NA | 2 | 0.5 | NA | NA | 3 | 0.3333 |
| 11_689270 | 6 | 0.1667 | 3 | 0 | 5 | 0.4 | 3 | 0 | 17 | 0.1765 | 6 | 0.1667 | 2 | 0 | 5 | 0 | 2 | 0 | 15 | 0.0667 |
| 11_689272 | 2 | 0 | 1 | 0 | 4 | 0.25 | 3 | 0 | 10 | 0.1 | 5 | 0.2 | 2 | 0 | 4 | 0 | 2 | 0 | 13 | 0.0769 |
| 11_689508 | 3 | 0 | 2 | 0.5 | 2 | 0 | 1 | 0 | 8 | 0.125 | 4 | 0 | 3 | 0.3333 | 4 | 0.5 | 4 | 0.25 | 15 | 0.2667 |
| 11_689508 | 3 | 0.3333 | 3 | 1 | NA | NA | NA | NA | 6 | 0.6667 | 3 | 0.3333 | 3 | 0.6667 | 2 | 0.5 | 3 | 0.6667 | 11 | 0.5455 |
| 11_689511 | 3 | 0.6667 | 3 | 1 | 2 | 0.5 | 1 | 1 | 9 | 0.7778 | 4 | 1 | 3 | 0.6667 | 1 | 1 | 3 | 0.6667 | 11 | 0.8182 |
| 11_689515 | 4 | 0 | 3 | 1 | 4 | 0 | 1 | 0 | 12 | 0.25 | 3 | 0.3333 | 5 | 0 | 4 | 0 | 4 | 0 | 16 | 0.0625 |
| 11_689515 | 4 | 1 | 3 | 1 | 4 | 1 | 2 | 1 | 13 | 1 | 4 | 1 | 6 | 1 | 4 | 1 | 4 | 1 | 18 | 1 |
| 11_689515 | 3 | 1 | 3 | 1 | 4 | 0.75 | 2 | 1 | 12 | 0.9167 | 5 | 1 | 5 | 1 | 3 | 1 | 2 | 1 | 15 | 1 |
| 11_689517 | 5 | 1 | 4 | 0.75 | 3 | 0.3333 | 3 | 0.6667 | 15 | 0.7333 | 4 | 1 | 6 | 0.3333 | 6 | 0.5 | 4 | 0.75 | 20 | 0.6 |
| 11_689517 | 4 | 0.25 | 4 | 0 | 4 | 0 | 1 | 0 | 13 | 0.0769 | 4 | 0.25 | 7 | 0 | 6 | 0 | 3 | 0 | 20 | 0.05 |
| 11_689517 | 4 | 0 | 4 | 0 | 4 | 0.25 | 2 | 0.5 | 14 | 0.1429 | 6 | 0.3333 | 6 | 0.1667 | 3 | 0.6667 | 3 | 0.6667 | 18 | 0.3889 |
| 11_689520 | 6 | 0 | 4 | 0.5 | 4 | 0 | 6 | 0.3333 | 20 | 0.2 | 6 | 0.1667 | 7 | 0 | 6 | 0.1667 | 4 | 0.25 | 23 | 0.1304 |
| 11_689521 | 5 | 0.8 | 3 | 1 | 5 | 0.8 | 7 | 0.7143 | 20 | 0.8 | 5 | 0.8 | 7 | 0.4286 | 5 | 0.2 | 5 | 0.6 | 22 | 0.5 |
| 11_689827 | 163 | 0.0613 | 190 | 0.0895 | 157 | 0.1019 | 144 | 0.1181 | 654 | 0.0917 | 163 | 0.0859 | 144 | 0.0625 | 119 | 0.1429 | 151 | 0.1192 | 577 | 0.1005 |
| 11_689964 | 3 | 0.3333 | 6 | 0 | 4 | 0 | 1 | 0 | 14 | 0.0714 | 6 | 0.1667 | 8 | 0 | 3 | 0 | 4 | 0.25 | 21 | 0.0952 |
| 11_689968 | 1 | 1 | 2 | 0.5 | 1 | 1 | NA | NA | 4 | 0.75 | 3 | 1 | 2 | 0 | NA | NA | 1 | 1 | 6 | 0.6667 |
| 11_689999 | 3 | 0 | 6 | 0.1667 | 4 | 0 | 6 | 0.1667 | 19 | 0.1053 | 9 | 0.2222 | 14 | 0.2857 | 8 | 0.25 | 4 | 0.25 | 35 | 0.2571 |
| 11_690001 | 5 | 0.6 | 8 | 0.625 | 6 | 0.6667 | 9 | 0.6667 | 28 | 0.6429 | 8 | 0.875 | 16 | 0.5625 | 9 | 0.6667 | 4 | 0.5 | 37 | 0.6486 |
| 11_690002 | 5 | 0.4 | 10 | 0.4 | 7 | 0.2857 | 6 | 0.3333 | 28 | 0.3571 | 5 | 0.2 | 14 | 0.1429 | 6 | 0 | 5 | 0.6 | 30 | 0.2 |
| 11_692164 | 6 | 0.1667 | 7 | 0.1429 | 7 | 0.2857 | 7 | 0.1429 | 27 | 0.1852 | 6 | 0.3333 | 3 | 0 | 10 | 0.2 | 7 | 0.2857 | 26 | 0.2308 |
| 11_692233 | 3 | 0.3333 | 5 | 0 | 3 | 0 | 3 | 0.3333 | 14 | 0.1429 | 1 | 1 | 2 | 0 | 2 | 0.5 | NA | NA | 5 | 0.4 |
| 11_692237 | 2 | 0.5 | 4 | 0.75 | 5 | 0.6 | 4 | 0.5 | 15 | 0.6 | 3 | 0.6667 | 1 | 1 | 2 | 0.5 | NA | NA | 6 | 0.6667 |
| 11_692320 | 16 | 0.4375 | 23 | 0.2174 | 14 | 0.5714 | 32 | 0.375 | 85 | 0.3765 | 8 | 0.75 | 20 | 0.55 | 11 | 0.2727 | 14 | 0.5 | 53 | 0.5094 |
| 11_692321 | 16 | 0.0625 | 24 | 0.0417 | 13 | 0.0769 | 29 | 0.1379 | 82 | 0.0854 | 9 | 0.1111 | 19 | 0.0526 | 13 | 0.1538 | 14 | 0.1429 | 55 | 0.1091 |
| 11_692321 | 16 | 0.6875 | 22 | 0.7273 | 13 | 0.6154 | 26 | 0.6923 | 77 | 0.6883 | 9 | 0.6667 | 22 | 0.8636 | 13 | 0.6154 | 13 | 0.7692 | 57 | 0.7544 |
| 11_692322 | 14 | 0.7143 | 25 | 0.76 | 13 | 0.9231 | 29 | 0.5862 | 81 | 0.716 | 8 | 0.75 | 20 | 0.85 | 14 | 0.7857 | 13 | 0.6154 | 55 | 0.7636 |
| 11_692324 | 15 | 0.2667 | 22 | 0.2273 | 15 | 0.2667 | 30 | 0.2333 | 82 | 0.2439 | 9 | 0.1111 | 19 | 0.1579 | 11 | 0.2727 | 10 | 0.1 | 49 | 0.1633 |
| 11_692327 | 12 | 0.0833 | 23 | 0.087 | 14 | 0.0714 | 35 | 0.0857 | 84 | 0.0833 | 10 | 0.1 | 21 | 0.2381 | 17 | 0.0588 | 15 | 0.0667 | 63 | 0.127 |
| 11_692330 | 19 | 0.0526 | 30 | 0.1 | 19 | 0.1579 | 37 | 0.0811 | 105 | 0.0952 | 13 | 0.1538 | 26 | 0.0385 | 21 | 0.1429 | 16 | 0.0625 | 76 | 0.0921 |
| 11_692345 | 30 | 0.1 | 27 | 0.0741 | 29 | 0.0345 | 17 | 0.1765 | 103 | 0.0874 | 34 | 0.0588 | 25 | 0.12 | 30 | 0.1 | 15 | 0.1333 | 104 | 0.0962 |
| 11_692347 | 26 | 0.1154 | 24 | 0.125 | 29 | 0.1379 | 17 | 0 | 96 | 0.1042 | 35 | 0.0857 | 22 | 0.1364 | 20 | 0.1 | 15 | 0.0667 | 92 | 0.0978 |
| 11_692349 | 19 | 0.5263 | 26 | 0.2308 | 22 | 0.3636 | 14 | 0.1429 | 81 | 0.321 | 26 | 0.1538 | 19 | 0.3158 | 17 | 0.2941 | 10 | 0.4 | 72 | 0.2639 |
| 11_692819 | 6 | 0 | 5 | 0.2 | 3 | 0 | 6 | 0.3333 | 20 | 0.15 | 1 | 0 | 5 | 0.2 | 2 | 0 | 7 | 0.1429 | 15 | 0.1333 |
| 11_692823 | 4 | 0 | 7 | 0.1429 | 4 | 0 | 4 | 0 | 19 | 0.0526 | 1 | 0 | 6 | 0 | 3 | 0.3333 | 7 | 0 | 17 | 0.0588 |

| | | | | | | | | | | | | | | | | | | | | |
|---|---|---|---|---|---|---|---|---|---|---|---|---|---|---|---|---|---|---|---|---|
| 11_692825 | 6 | 0 | 6 | 0 | 4 | 0 | 5 | 0.4 | 21 | 0.0952 | 1 | 0 | 5 | 0.2 | 2 | 0 | 8 | 0 | 16 | 0.0625 |
| 11_692832 | 3 | 0 | 4 | 0 | 2 | 0 | 3 | 0.3333 | 12 | 0.0833 | 1 | 0 | 1 | 0 | 2 | 0 | 3 | 0.3333 | 7 | 0.1429 |
| 11_6928328 | 3 | 0.3333 | 5 | 0.2 | 2 | 0.5 | 3 | 0.3333 | 13 | 0.3077 | 1 | 0 | 1 | 0 | 2 | 0 | 4 | 0.5 | 8 | 0.25 |
| 11_692841 | 1 | 1 | 3 | 0 | 1 | 0 | 2 | 0 | 7 | 0.1429 | 1 | 0 | NA | NA | 1 | 0 | 4 | 0.25 | 6 | 0.1667 |
| 11_692844 | 1 | 0 | 2 | 0.5 | 1 | 0 | 2 | 0.5 | 6 | 0.3333 | NA | NA | NA | NA | 1 | 1 | 2 | 0.5 | 3 | 0.6667 |
| 11_692845 | 1 | 0 | 2 | 0.5 | 1 | 1 | 2 | 0.5 | 6 | 0.5 | 1 | 0 | NA | NA | NA | NA | 4 | 0.25 | 5 | 0.2 |
| 11_6928572 | NA | NA | NA | NA | NA | NA | 1 | 1 | 1 | 1 | 1 | 0 | NA | NA | 1 | 1 | 3 | 0.6667 | 5 | 0.6 |
| 11_692881 | 2 | 0.5 | NA | NA | NA | NA | NA | NA | 2 | 0.5 | NA | NA | NA | NA | 1 | 1 | NA | NA | 1 | 1 |
| 11_692881 | 2 | 0.5 | 1 | 0 | NA | NA | NA | NA | 3 | 0.3333 | NA | NA | NA | NA | 1 | 1 | 1 | 0 | 2 | 0.5 |
| 11_692882 | 2 | 0 | 1 | 1 | NA | NA | NA | NA | 3 | 0.3333 | NA | NA | NA | NA | 1 | 1 | 1 | 0 | 2 | 0.5 |
| 11_692883 | 2 | 1 | 1 | 1 | NA | NA | NA | NA | 3 | 1 | NA | NA | NA | NA | 1 | 1 | 1 | 1 | 2 | 1 |
| 11_692952 | 1 | 1 | NA | NA | 1 | 1 | 1 | 1 | 3 | 1 | NA | NA | NA | NA | 2 | 1 | 2 | 0.5 | 4 | 0.75 |
| 11_692954 | 1 | 1 | NA | NA | NA | NA | 1 | 1 | 2 | 1 | NA | NA | NA | NA | 3 | 1 | 1 | 0 | 4 | 0.75 |
| 11_692954 | NA | NA | NA | NA | 1 | 1 | 1 | 1 | 2 | 1 | NA | NA | NA | NA | 3 | 1 | NA | NA | 3 | 1 |
| 11_692954 | 1 | 1 | NA | NA | 1 | 0 | 1 | 0 | 3 | 0.3333 | NA | NA | NA | NA | 3 | 0.6667 | NA | NA | 3 | 0.6667 |
| 11_692954 | NA | NA | NA | NA | 1 | 0 | 1 | 1 | 2 | 0.5 | NA | NA | NA | NA | 2 | 0.5 | 1 | 0 | 3 | 0.3333 |
| 11_692957 | 1 | 0 | NA | NA | NA | NA | 1 | 1 | 2 | 0.5 | NA | NA | NA | NA | 2 | 1 | 1 | 1 | 3 | 1 |
| 11_692958 | 1 | 1 | NA | NA | 1 | 1 | NA | NA | 2 | 1 | NA | NA | NA | NA | 1 | 1 | 1 | 1 | 2 | 1 |
| 11_692960 | 1 | 0 | NA | NA | 1 | 1 | 1 | 0 | 3 | 0.3333 | NA | NA | NA | NA | 1 | 1 | NA | NA | 1 | 1 |
| 11_692968 | NA | NA | NA | NA | 1 | 0 | 1 | 1 | 2 | 0.5 | NA | NA | NA | NA | 2 | 0.5 | 1 | 1 | 3 | 0.6667 |
| 11_694452 | 16 | 0.4375 | 16 | 0.375 | 11 | 0.6364 | 14 | 0.6429 | 57 | 0.5088 | 7 | 0.8571 | 18 | 0.4444 | 14 | 0.5 | 11 | 0.5455 | 50 | 0.54 |
| 11_694455 | 9 | 0.4444 | 11 | 0.1818 | 8 | 0.5 | 17 | 0.3529 | 45 | 0.3556 | 8 | 0.625 | 20 | 0.2 | 12 | 0.4167 | 9 | 0.4444 | 49 | 0.3673 |
| 11_694455 | 14 | 0.0714 | 10 | 0 | 10 | 0.1 | 15 | 0.1333 | 49 | 0.0816 | 11 | 0.0909 | 20 | 0 | 14 | 0.0714 | 12 | 0.0833 | 57 | 0.0526 |
| 11_694457 | 11 | 0.0909 | 13 | 0 | 10 | 0 | 18 | 0.2222 | 52 | 0.0962 | 12 | 0.1667 | 21 | 0.0476 | 10 | 0.1 | 10 | 0.2 | 53 | 0.1132 |
| 11_694458 | 10 | 0.1 | 11 | 0.0909 | 8 | 0.25 | 19 | 0.1053 | 48 | 0.125 | 9 | 0.1111 | 20 | 0.05 | 12 | 0 | 11 | 0.0909 | 52 | 0.0577 |
| 11_694458 | 15 | 0.5333 | 13 | 0.1538 | 9 | 0.2222 | 19 | 0.3158 | 56 | 0.3214 | 11 | 0.3636 | 20 | 0.3 | 13 | 0.1538 | 10 | 0.3 | 54 | 0.2778 |
| 11_694458 | 12 | 0.9167 | 13 | 1 | 7 | 0.8571 | 14 | 0.9286 | 46 | 0.9348 | 11 | 1 | 18 | 0.8333 | 11 | 1 | 10 | 1 | 50 | 0.94 |
| 11_694458 | 12 | 0.25 | 11 | 0.2727 | 7 | 0 | 14 | 0.3571 | 44 | 0.25 | 8 | 0.125 | 18 | 0.1111 | 11 | 0.1818 | 11 | 0 | 48 | 0.1042 |
| 11_694460 | 13 | 0 | 11 | 0.0909 | 6 | 0.1667 | 12 | 0.0833 | 42 | 0.0714 | 7 | 0 | 15 | 0 | 7 | 0.1429 | 9 | 0.1111 | 38 | 0.0526 |
| 11_694506 | 4 | 0.75 | 2 | 0.5 | 2 | 0.5 | 10 | 0.6 | 18 | 0.6111 | 3 | 0.6667 | 7 | 0.7143 | 7 | 1 | 2 | 1 | 19 | 0.8421 |
| 11_694526 | 2 | 0 | 3 | 0.3333 | 1 | 0 | NA | NA | 6 | 0.1667 | 1 | 1 | 1 | 0 | 3 | 0 | 1 | 0 | 6 | 0.1667 |
| 11_694696 | NA | NA | 3 | 1 | 2 | 0.5 | NA | NA | 5 | 0.8 | 2 | 0.5 | 1 | 1 | 2 | 0.5 | 1 | 1 | 6 | 0.6667 |
| 11_694697 | 1 | 1 | 2 | 0.5 | 2 | 0.5 | NA | NA | 5 | 0.6 | 2 | 1 | 1 | 1 | 2 | 0 | NA | NA | 5 | 0.6 |
| 11_694918 | 51 | 0.0588 | 59 | 0.1017 | 45 | 0.0889 | 39 | 0.1026 | 194 | 0.0876 | 44 | 0.1136 | 40 | 0.125 | 39 | 0.1282 | 43 | 0.0698 | 166 | 0.1084 |
| 11_694960 | 34 | 0.0882 | 41 | 0.0732 | 30 | 0.0333 | 34 | 0.0588 | 139 | 0.0647 | 26 | 0.0385 | 27 | 0.0741 | 32 | 0.0312 | 33 | 0.1515 | 118 | 0.0763 |
| 11_694962 | 29 | 0.1034 | 39 | 0.0513 | 29 | 0.1379 | 27 | 0.0741 | 124 | 0.0887 | 26 | 0.0769 | 25 | 0.12 | 29 | 0.1034 | 28 | 0.1071 | 108 | 0.1019 |
| 11_694963 | 29 | 0.1034 | 40 | 0.15 | 29 | 0.069 | 30 | 0 | 128 | 0.0859 | 25 | 0.08 | 25 | 0.04 | 29 | 0.1379 | 27 | 0 | 106 | 0.066 |
| 11_695138 | NA | NA | 3 | 0.3333 | NA | NA | NA | NA | 3 | 0.3333 | NA | NA | NA | NA | NA | NA | 1 | 1 | 1 | 1 |
| 11_695145 | NA | NA | 5 | 0.2 | NA | NA | NA | NA | 5 | 0.2 | 1 | 0 | 2 | 0.5 | NA | NA | 2 | 0.5 | 5 | 0.4 |
| 11_695147 | NA | NA | 2 | 0.5 | NA | NA | NA | NA | 2 | 0.5 | 1 | 0 | 2 | 0.5 | NA | NA | 1 | 1 | 4 | 0.5 |
| 11_695149 | NA | NA | 3 | 0.6667 | NA | NA | NA | NA | 3 | 0.6667 | 1 | 1 | 2 | 0.5 | NA | NA | 1 | 1 | 4 | 0.75 |
| 11_695149 | NA | NA | 3 | 0.3333 | NA | NA | NA | NA | 3 | 0.3333 | 1 | 1 | 2 | 0.5 | NA | NA | NA | NA | 3 | 0.6667 |
| 11_695149 | NA | NA | 2 | 0.5 | NA | NA | NA | NA | 2 | 0.5 | 1 | 1 | 2 | 1 | NA | NA | 1 | 1 | 4 | 1 |
| 11_695424 | 3 | 0.3333 | 10 | 0 | 5 | 0 | 2 | 0 | 20 | 0.05 | 5 | 0 | 5 | 0 | 7 | 0 | 3 | 0.3333 | 20 | 0.05 |
| 11_695437 | 4 | 0.25 | 17 | 0.2941 | 8 | 0 | 4 | 0.25 | 33 | 0.2121 | 8 | 0 | 7 | 0 | 9 | 0.2222 | 3 | 0.3333 | 27 | 0.1111 |
| 11_695477 | 10 | 0.2 | 9 | 0.2222 | 8 | 0 | 10 | 0.2 | 37 | 0.1622 | 5 | 0.2 | 11 | 0.0909 | 4 | 0 | 9 | 0.2222 | 29 | 0.1379 |
| 11_695477 | 8 | 0.125 | 13 | 0.1538 | 9 | 0 | 10 | 0.1 | 40 | 0.1 | 5 | 0.2 | 9 | 0 | 4 | 0 | 10 | 0.3 | 28 | 0.1429 |
| 11_695480 | 7 | 0 | 7 | 0.2857 | 6 | 0 | 7 | 0 | 27 | 0.0741 | 1 | 0 | 9 | 0.1111 | 2 | 0 | 5 | 0.2 | 17 | 0.1176 |
| 11_695551 | 4 | 0.25 | 2 | 0 | 2 | 0 | 2 | 0 | 10 | 0.1 | 3 | 0 | 3 | 0 | 5 | 0.2 | 1 | 0 | 12 | 0.0833 |
| 11_695555 | 6 | 0.1667 | 4 | 0 | 2 | 0 | 3 | 0.3333 | 15 | 0.1333 | 6 | 0.1667 | 4 | 0.25 | 6 | 0 | 1 | 0 | 17 | 0.1176 |
| 11_695579 | 39 | 0.0769 | 32 | 0.125 | 38 | 0.0263 | 40 | 0.025 | 149 | 0.0604 | 20 | 0.15 | 39 | 0.0256 | 32 | 0.0938 | 37 | 0.0541 | 128 | 0.0703 |
| 11_695582 | 46 | 0.1304 | 37 | 0.1081 | 40 | 0.1 | 50 | 0.04 | 173 | 0.0925 | 29 | 0.069 | 43 | 0.1628 | 45 | 0.1111 | 46 | 0.1522 | 163 | 0.1288 |
| 11_695642 | 6 | 0.3333 | 10 | 0.1 | 6 | 0.5 | 7 | 0.2857 | 29 | 0.2759 | 7 | 0.2857 | 11 | 0 | 6 | 0.1667 | 10 | 0.4 | 34 | 0.2059 |
| 11_695643 | 5 | 0 | 10 | 0.2 | 6 | 0.1667 | 8 | 0.25 | 29 | 0.1724 | 6 | 0.1667 | 8 | 0.125 | 6 | 0.1667 | 8 | 0.5 | 28 | 0.25 |
| 11_695710 | 6 | 0.1667 | 6 | 0.3333 | 4 | 0.5 | 7 | 0.1429 | 23 | 0.2609 | 4 | 0.75 | 6 | 0.1667 | 3 | 0.3333 | 7 | 0.1429 | 20 | 0.3 |
| 11_695711 | 3 | 0.3333 | 5 | 0 | 2 | 0 | 2 | 0 | 12 | 0.0833 | 3 | 0 | 4 | 0 | 2 | 0.5 | 6 | 0.1667 | 15 | 0.1333 |
| 11_695712 | 4 | 0 | 6 | 0.1667 | 5 | 0 | 5 | 0 | 20 | 0.05 | 5 | 0.2 | 5 | 0 | 2 | 0 | 6 | 0.1667 | 18 | 0.1111 |
| 11_695716 | 2 | 0 | 6 | 0.3333 | 3 | 0.6667 | 3 | 0.3333 | 14 | 0.3571 | 4 | 0.25 | 3 | 0.3333 | 1 | 0 | 5 | 0.4 | 13 | 0.3077 |
| 11_695773 | 2 | 0 | 3 | 0.3333 | 2 | 0 | 5 | 0 | 12 | 0.0833 | 4 | 0 | 2 | 0.5 | 1 | 0 | 2 | 0 | 9 | 0.1111 |
| 11_695773 | 1 | 0 | 3 | 0.3333 | 2 | 0 | 7 | 0.4286 | 13 | 0.3077 | 4 | 0.25 | 3 | 0.3333 | 1 | 0 | 2 | 0.5 | 10 | 0.3 |
| 11_695776 | 1 | 1 | 3 | 0.3333 | 2 | 1 | 4 | 0.5 | 10 | 0.6 | 5 | 0.8 | 1 | 1 | 1 | 1 | 1 | 1 | 8 | 0.875 |
| 11_696249 | 1 | 0 | 12 | 0 | 9 | 0.1111 | 8 | 0.125 | 30 | 0.0667 | 4 | 0.25 | 12 | 0 | 6 | 0 | 6 | 0.1667 | 28 | 0.0714 |
| 11_696250 | 1 | 0 | 8 | 0.375 | 8 | 0.375 | 4 | 0.25 | 21 | 0.3333 | 4 | 0.5 | 10 | 0.4 | 4 | 0.75 | 5 | 0.4 | 23 | 0.4783 |
| 11_696672 | NA | NA | 4 | 0.5 | 1 | 0 | 1 | 0 | 6 | 0.3333 | 2 | 0 | 5 | 0.6 | NA | NA | 1 | 0 | 8 | 0.375 |
| 11_696679 | NA | NA | 1 | 1 | NA | NA | NA | NA | 1 | 1 | NA | NA | 1 | 1 | NA | NA | NA | NA | 1 | 1 |

| | | | | | | | | | | | | | | | | | | | | |
|---|---|---|---|---|---|---|---|---|---|---|---|---|---|---|---|---|---|---|---|---|
| 11_696689 | NA | NA | 4 | 1 | 6 | 1 | 2 | 1 | 12 | 1 | 3 | 1 | 6 | 1 | 1 | 1 | 1 | 1 | 11 | 1 |
| 11_696689 | NA | NA | 4 | 0.5 | 6 | 0.1667 | 4 | 0.5 | 14 | 0.3571 | 3 | 0.3333 | 7 | 0.4286 | 1 | 1 | 1 | 0 | 12 | 0.4167 |
| 11_696689 | NA | NA | 2 | 0 | 6 | 0.1667 | 4 | 0 | 12 | 0.0833 | 2 | 0 | 5 | 0.2 | 1 | 0 | 1 | 0 | 9 | 0.1111 |
| 11_696689 | NA | NA | 3 | 0.6667 | 5 | 0.8 | 2 | 1 | 10 | 0.8 | 3 | 1 | 5 | 1 | 1 | 1 | 2 | 1 | 11 | 1 |
| 11_696690 | NA | NA | 5 | 0.2 | 6 | 0.5 | 4 | 0.25 | 15 | 0.3333 | 3 | 0 | 8 | 0.25 | NA | NA | 1 | 0 | 12 | 0.1667 |
| 11_696692 | NA | NA | 5 | 0.8 | 5 | 0.8 | 3 | 0.3333 | 13 | 0.6923 | 2 | 0 | 7 | 0.7143 | 1 | 1 | 1 | 1 | 11 | 0.6364 |
| 11_696693 | NA | NA | 6 | 1 | 7 | 1 | 3 | 1 | 16 | 1 | 3 | 1 | 8 | 1 | 1 | 1 | 2 | 1 | 14 | 1 |
| 11_696694 | 1 | 1 | 5 | 1 | 6 | 1 | 3 | 1 | 15 | 1 | 3 | 0.6667 | 7 | 1 | 1 | 1 | 2 | 1 | 13 | 0.9231 |
| 11_696695 | 1 | 0 | 3 | 0 | 5 | 0.4 | 3 | 0.3333 | 12 | 0.25 | 3 | 0 | 6 | 0.1667 | 2 | 0 | 1 | 0 | 12 | 0.0833 |
| 11_696697 | 1 | 1 | 6 | 0.6667 | 4 | 1 | 4 | 0.75 | 15 | 0.8 | 3 | 0.3333 | 6 | 0.6667 | 2 | 0.5 | 2 | 0.5 | 13 | 0.5385 |
| 11_696699 | 1 | 1 | 7 | 0.4286 | 7 | 0.8571 | 4 | 0.5 | 19 | 0.6316 | 4 | 0.75 | 9 | 0.6667 | 2 | 0.5 | 3 | 0.3333 | 18 | 0.6111 |
| 11_696726 | 5 | 0 | 8 | 0 | 13 | 0.1538 | 4 | 0 | 30 | 0.0667 | 4 | 0.25 | 12 | 0.25 | 7 | 0.1429 | 2 | 0 | 25 | 0.2 |
| 11_697140 | 38 | 0.2632 | 30 | 0.2667 | 30 | 0.2667 | 35 | 0.4571 | 133 | 0.3158 | 21 | 0.1429 | 23 | 0.2609 | 24 | 0.2917 | 18 | 0.2778 | 86 | 0.2442 |
| 11_697150 | 25 | 0.2 | 24 | 0.125 | 18 | 0.2222 | 24 | 0.2917 | 91 | 0.2088 | 11 | 0.4545 | 13 | 0.2308 | 17 | 0.1176 | 12 | 0.1667 | 53 | 0.2264 |
| 11_697150 | 24 | 0.3333 | 23 | 0.2609 | 19 | 0.3158 | 20 | 0.25 | 86 | 0.2907 | 11 | 0.3636 | 10 | 0.2 | 15 | 0.3333 | 11 | 0.1818 | 47 | 0.2766 |
| 11_697155 | 16 | 0.125 | 16 | 0.125 | 12 | 0.1667 | 14 | 0.0714 | 58 | 0.1207 | 11 | 0.1818 | 10 | 0.2 | 11 | 0.1818 | 8 | 0 | 40 | 0.15 |
| 11_697157 | 15 | 0.2 | 14 | 0 | 9 | 0.2222 | 11 | 0 | 49 | 0.102 | 7 | 0.1429 | 8 | 0 | 11 | 0.1818 | 8 | 0 | 34 | 0.0882 |
| 11_697157 | 14 | 0.4286 | 14 | 0.4286 | 11 | 0.3636 | 15 | 0.6667 | 54 | 0.4815 | 8 | 0.75 | 7 | 0.2857 | 12 | 0.5 | 9 | 0.3333 | 36 | 0.4722 |
| 11_697158 | 10 | 1 | 11 | 0.7273 | 5 | 1 | 7 | 0.7143 | 33 | 0.8485 | 1 | 1 | 5 | 0.8 | 8 | 0.875 | 3 | 0.6667 | 17 | 0.8235 |
| 11_697158 | 14 | 0.7857 | 16 | 0.75 | 12 | 0.8333 | 14 | 0.8571 | 56 | 0.8036 | 9 | 1 | 8 | 0.875 | 13 | 0.6923 | 8 | 0.875 | 38 | 0.8421 |
| 11_697159 | 12 | 0.9167 | 15 | 0.8 | 13 | 0.9231 | 9 | 0.7778 | 49 | 0.8571 | 7 | 1 | 4 | 0.75 | 10 | 0.8 | 7 | 0.7143 | 28 | 0.8214 |
| 11_697159 | 14 | 0.9286 | 17 | 0.5882 | 12 | 0.6667 | 11 | 0.6364 | 54 | 0.7037 | 7 | 0.4286 | 9 | 0.4444 | 11 | 0.6364 | 5 | 0.4 | 32 | 0.5 |
| 11_697159 | 10 | 1 | 16 | 1 | 8 | 1 | 9 | 1 | 43 | 1 | 4 | 1 | 8 | 0.875 | 8 | 0.75 | 6 | 0.8333 | 26 | 0.8462 |
| 11_697162 | 11 | 0.1818 | 11 | 0.4545 | 8 | 0.375 | 9 | 0 | 39 | 0.2564 | 5 | 0.4 | 7 | 0.4286 | 11 | 0.2727 | 7 | 0.4286 | 30 | 0.3667 |
| 11_697164 | 11 | 0.2727 | 5 | 0.2 | 7 | 0.1429 | 5 | 0.2 | 28 | 0.2143 | 3 | 0.3333 | 5 | 0.2 | 9 | 0.1111 | 4 | 0 | 21 | 0.1429 |
| 11_697165 | 8 | 0.875 | 5 | 1 | 6 | 1 | 8 | 0.875 | 27 | 0.9259 | 1 | 1 | 4 | 1 | 6 | 0.6667 | 3 | 1 | 14 | 0.8571 |
| 11_697165 | 9 | 0.7778 | 3 | 1 | 7 | 1 | 9 | 1 | 28 | 0.9286 | 2 | 1 | 7 | 0.8571 | 10 | 0.9 | 4 | 1 | 23 | 0.913 |
| 11_697165 | 10 | 0.8 | 5 | 0.8 | 7 | 0.8571 | 8 | 0.75 | 30 | 0.8 | 3 | 1 | 6 | 0.5 | 8 | 0.75 | 4 | 0.75 | 21 | 0.7143 |
| 11_697165 | 6 | 0 | 4 | 0 | 4 | 0.5 | 5 | 0 | 19 | 0.1053 | 1 | 0 | 2 | 0 | 6 | 0.1667 | 2 | 0.5 | 11 | 0.1818 |
| 11_697175 | 7 | 0.1429 | 4 | 0.5 | 5 | 0.4 | 3 | 0.3333 | 19 | 0.3158 | 2 | 0 | 6 | 0.1667 | 6 | 0 | 2 | 0 | 16 | 0.0625 |
| 11_697178 | 4 | 0.75 | 5 | 0.8 | 5 | 0.8 | 6 | 0.6667 | 20 | 0.75 | 3 | 1 | 6 | 0.6667 | 7 | 0.4286 | 2 | 1 | 18 | 0.6667 |
| 11_697178 | 5 | 1 | 5 | 1 | 6 | 1 | 4 | 0.75 | 20 | 0.95 | 3 | 0.6667 | 4 | 1 | 8 | 0.75 | 3 | 1 | 18 | 0.8333 |
| 11_697179 | 5 | 0 | 5 | 0.2 | 6 | 0.3333 | 4 | 0.5 | 20 | 0.25 | 2 | 0.5 | 5 | 0 | 8 | 0.5 | 3 | 0.6667 | 18 | 0.3889 |
| 11_697180 | 5 | 0.2 | 5 | 0.8 | 6 | 0.3333 | 5 | 0.6 | 21 | 0.4762 | 3 | 0 | 6 | 0.3333 | 7 | 0.1429 | 2 | 0.5 | 18 | 0.2222 |
| 11_697254 | 4 | 0.25 | 4 | 0.25 | 4 | 0 | 5 | 0 | 17 | 0.1176 | 2 | 0 | 4 | 0.25 | 5 | 0 | 1 | 1 | 12 | 0.1667 |
| 11_697282 | 2 | 0.5 | 3 | 0.3333 | 4 | 0.75 | 5 | 0.8 | 14 | 0.6429 | 2 | 0.5 | 1 | 0 | 4 | 0.75 | 1 | 1 | 8 | 0.625 |
| 11_697284 | 2 | 0 | 4 | 0 | 3 | 0 | 3 | 0.3333 | 12 | 0.0833 | 1 | 1 | 2 | 0 | 5 | 0.2 | 1 | 0 | 9 | 0.2222 |
| 11_697302 | 27 | 0.0741 | 16 | 0.25 | 18 | 0.1667 | 21 | 0.1905 | 82 | 0.1585 | 17 | 0.0588 | 15 | 0.0667 | 18 | 0.3333 | 13 | 0.0769 | 63 | 0.1429 |
| 11_697413 | 1 | 1 | NA | NA | 1 | 0 | 1 | 0 | 3 | 0.3333 | 1 | 1 | 1 | 0 | 1 | 0 | 1 | 0 | 4 | 0.25 |
| 11_697699 | NA | NA | 1 | 0 | 2 | 0.5 | NA | NA | 3 | 0.3333 | NA | NA | NA | NA | 1 | 1 | NA | NA | 1 | 1 |
| 11_697701 | NA | NA | 1 | 0 | 2 | 1 | NA | NA | 3 | 0.6667 | NA | NA | NA | NA | 1 | 1 | NA | NA | 1 | 1 |
| 11_697704 | NA | NA | 1 | 1 | 2 | 1 | NA | NA | 3 | 1 | NA | NA | NA | NA | 1 | 1 | NA | NA | 1 | 1 |
| 11_697704 | NA | NA | 1 | 1 | 2 | 0.5 | NA | NA | 3 | 0.6667 | NA | NA | NA | NA | 1 | 1 | NA | NA | 1 | 1 |
| 11_697705 | NA | NA | 1 | 0 | 2 | 1 | NA | NA | 3 | 0.6667 | NA | NA | NA | NA | 1 | 1 | NA | NA | 1 | 1 |
| 11_697705 | NA | NA | 1 | 0 | 2 | 0.5 | NA | NA | 3 | 0.3333 | NA | NA | NA | NA | 1 | 1 | NA | NA | 1 | 1 |
| 11_697709 | NA | NA | 1 | 1 | NA | NA | NA | NA | 1 | 1 | NA | NA | NA | NA | 1 | 1 | NA | NA | 1 | 1 |
| 11_698021 | 2 | 0 | 3 | 0.3333 | 3 | 0 | 3 | 0.3333 | 11 | 0.1818 | 4 | 0.25 | 9 | 0.1111 | 7 | 0 | 8 | 0.125 | 28 | 0.1071 |
| 11_698040 | 5 | 0.4 | 5 | 0 | 5 | 0.2 | 5 | 0.2 | 20 | 0.2 | 4 | 0 | 9 | 0.1111 | 8 | 0.125 | 10 | 0.2 | 31 | 0.129 |
| 11_698042 | 4 | 1 | 6 | 0.8333 | 5 | 1 | 6 | 0.6667 | 21 | 0.8571 | 3 | 0.6667 | 10 | 0.8 | 8 | 0.75 | 9 | 1 | 30 | 0.8333 |
| 11_698042 | 4 | 0.25 | 6 | 0.3333 | 6 | 0 | 4 | 0.5 | 20 | 0.25 | 3 | 0.3333 | 7 | 0.4286 | 7 | 0.4286 | 8 | 0.625 | 25 | 0.48 |
| 11_698043 | 2 | 0 | 3 | 0.3333 | 8 | 0.125 | 7 | 0.4286 | 20 | 0.25 | 4 | 0 | 4 | 0.5 | 5 | 0.6 | 8 | 0.125 | 21 | 0.2857 |
| 11_698712 | 4 | 0.5 | 2 | 0 | 1 | 1 | 4 | 0.25 | 11 | 0.3636 | 3 | 0.6667 | 1 | 1 | 1 | 0 | 1 | 1 | 6 | 0.6667 |
| 11_698809 | 15 | 0.1333 | 13 | 0.2308 | 16 | 0.125 | 17 | 0 | 61 | 0.1148 | 12 | 0.0833 | 16 | 0.1875 | 16 | 0.125 | 27 | 0.0741 | 71 | 0.1127 |
| 11_698809 | 12 | 0.5 | 9 | 0.6667 | 14 | 0.6429 | 15 | 0.4667 | 50 | 0.56 | 9 | 0.5556 | 11 | 0.5455 | 14 | 0.7143 | 15 | 0.8667 | 49 | 0.6939 |
| 11_698810 | 16 | 0.5 | 13 | 0.6923 | 20 | 0.4 | 19 | 0.5789 | 68 | 0.5294 | 14 | 0.4286 | 18 | 0.6111 | 17 | 0.4706 | 28 | 0.6071 | 77 | 0.5455 |
| 11_698811 | 14 | 0.3571 | 10 | 0.4 | 17 | 0.3529 | 18 | 0.5 | 59 | 0.4068 | 15 | 0.4667 | 16 | 0.5 | 16 | 0.125 | 25 | 0.44 | 72 | 0.3889 |
| 11_698812 | 13 | 0.4615 | 11 | 0.4545 | 13 | 0.3846 | 15 | 0.4 | 52 | 0.4231 | 9 | 0.3333 | 14 | 0.5 | 12 | 0.4167 | 23 | 0.4348 | 58 | 0.431 |
| 11_698812 | 12 | 0.4167 | 10 | 0.1 | 9 | 0.3333 | 13 | 0.2308 | 44 | 0.2727 | 10 | 0.1 | 13 | 0.2308 | 9 | 0.2222 | 21 | 0.1905 | 53 | 0.1887 |
| 11_698892 | 2 | 0 | 4 | 0.25 | 1 | 0 | 2 | 0 | 9 | 0.1111 | 2 | 0 | 2 | 0.5 | 2 | 0 | 2 | 0.5 | 8 | 0.25 |
| 11_698892 | 2 | 0 | 4 | 0.25 | 1 | 0 | 2 | 0 | 9 | 0.1111 | 3 | 0.3333 | 2 | 0 | 2 | 0.5 | 3 | 0 | 10 | 0.2 |
| 11_698892 | 2 | 0.5 | 4 | 0 | 1 | 0 | 1 | 0 | 8 | 0.125 | 3 | 0.3333 | 2 | 0.5 | 1 | 1 | 3 | 0.3333 | 9 | 0.4444 |
| 11_698893 | 3 | 0.6667 | 5 | 0.6 | 3 | 0.6667 | 3 | 1 | 14 | 0.7143 | 3 | 1 | 2 | 1 | 1 | 1 | 6 | 0.8333 | 12 | 0.9167 |
| 11_698895 | 3 | 0.3333 | 2 | 0.5 | 2 | 1 | 4 | 0.5 | 11 | 0.5455 | 5 | 0.6 | 2 | 0 | 2 | 0.5 | 5 | 0.4 | 14 | 0.4286 |
| 11_698895 | 1 | 0 | 4 | 1 | 2 | 0.5 | 4 | 0.5 | 11 | 0.6364 | 4 | 0.5 | 2 | 0 | 1 | 0 | 5 | 0 | 12 | 0.1667 |
| 11_698897 | 3 | 0.6667 | 4 | 0.25 | 3 | 0.6667 | 5 | 0.4 | 15 | 0.4667 | 5 | 0.4 | 2 | 0 | 2 | 1 | 5 | 0.4 | 14 | 0.4286 |

| | | | | | | | | | | | | | | | | | | | | |
|---|---|---|---|---|---|---|---|---|---|---|---|---|---|---|---|---|---|---|---|---|
| 11_6988975 | 3 | 0.3333 | 3 | 0 | 2 | 0 | 2 | 0 | 10 | 0.1 | 4 | 0 | NA | NA | NA | NA | 4 | 0.25 | 8 | 0.125 |
| 11_6988985 | 3 | 0 | 5 | 0.2 | 2 | 0 | 4 | 0.25 | 14 | 0.1429 | 6 | 0.1667 | 3 | 0 | 3 | 0 | 7 | 0.1429 | 19 | 0.1053 |
| 11_6989022 | 3 | 1 | 2 | 0 | 3 | 0.6667 | 6 | 0.3333 | 14 | 0.5 | 5 | 0.4 | 3 | 0.3333 | 2 | 0 | 6 | 0.3333 | 16 | 0.3125 |
| 11_6989029 | 2 | 1 | 5 | 0.8 | 2 | 0 | 6 | 0.6667 | 15 | 0.6667 | 6 | 0.6667 | 3 | 0.6667 | 3 | 1 | 9 | 0.6667 | 21 | 0.7143 |
| 11_6989030 | 4 | 0.25 | 5 | 0.4 | 2 | 1 | 4 | 0.5 | 15 | 0.4667 | 6 | 0.5 | 3 | 0.6667 | 3 | 0.3333 | 8 | 0.375 | 20 | 0.45 |
| 11_6989031 | 4 | 0.75 | 4 | 1 | NA | NA | 6 | 1 | 14 | 0.9286 | 2 | 0.5 | 2 | 0.5 | 2 | 0.5 | 8 | 0.875 | 14 | 0.7143 |
| 11_6989044 | 3 | 1 | 4 | 1 | 2 | 1 | 6 | 0.8333 | 15 | 0.9333 | 6 | 1 | 2 | 1 | 6 | 1 | 8 | 1 | 22 | 1 |
| 11_6990065 | 1 | 0 | 4 | 0 | NA | NA | 4 | 0.25 | 9 | 0.1111 | 3 | 0 | 1 | 0 | NA | NA | 2 | 0.5 | 6 | 0.1667 |
| 11_6993551 | 25 | 0.6 | 44 | 0.6591 | 47 | 0.5532 | 23 | 0.4783 | 139 | 0.5827 | 31 | 0.6129 | 53 | 0.5094 | 54 | 0.4259 | 36 | 0.5833 | 174 | 0.5172 |
| 11_6993554 | 25 | 0.28 | 43 | 0.3023 | 40 | 0.325 | 21 | 0.1905 | 129 | 0.2868 | 27 | 0.3333 | 46 | 0.2609 | 43 | 0.3721 | 31 | 0.3226 | 147 | 0.3197 |
| 11_6993558 | 25 | 0.68 | 42 | 0.7619 | 47 | 0.8298 | 24 | 0.7917 | 138 | 0.7754 | 25 | 0.96 | 51 | 0.7059 | 50 | 0.76 | 33 | 0.8788 | 159 | 0.7987 |
| 11_6993564 | 30 | 0.0667 | 46 | 0.0652 | 47 | 0.1277 | 21 | 0.1429 | 144 | 0.0972 | 32 | 0.2188 | 58 | 0.1034 | 53 | 0.1509 | 37 | 0.1351 | 180 | 0.1444 |
| 11_6993578 | 29 | 0.0345 | 46 | 0.0652 | 47 | 0.0426 | 20 | 0.1 | 142 | 0.0563 | 32 | 0 | 59 | 0.0847 | 53 | 0.0377 | 34 | 0.0882 | 178 | 0.0562 |
| 11_6993668 | NA | NA | 2 | 0 | NA | NA | 3 | 0.3333 | 5 | 0.2 | 2 | 1 | 3 | 0.6667 | 1 | 0 | 1 | 0 | 7 | 0.5714 |
| 11_6993689 | NA | NA | 3 | 0 | NA | NA | 3 | 0.3333 | 6 | 0.1667 | 2 | 0 | 3 | 0.3333 | 1 | 0 | 1 | 1 | 7 | 0.2857 |
| 11_6993755 | 1 | 0 | 3 | 0 | NA | NA | 4 | 0.25 | 8 | 0.125 | 2 | 0 | 2 | 0.5 | 1 | 0 | 1 | 0 | 6 | 0.1667 |
| 11_6993786 | 2 | 0 | 3 | 0.3333 | NA | NA | 1 | 0 | 6 | 0.1667 | 3 | 0.6667 | NA | NA | NA | NA | NA | NA | 3 | 0.6667 |
| 11_7012607 | 9 | 0.7778 | 11 | 0.8182 | 6 | 0.6667 | 2 | 1 | 28 | 0.7857 | 7 | 0.7143 | 10 | 0.8 | 14 | 0.7857 | 7 | 1 | 38 | 0.8158 |
| 11_7012616 | 6 | 0.1667 | 10 | 0.4 | 6 | 0.5 | 3 | 0.3333 | 25 | 0.36 | 4 | 0 | 9 | 0.2222 | 14 | 0.2143 | 4 | 0.5 | 31 | 0.2258 |
| 11_7012617 | 8 | 0.375 | 10 | 0.3 | 7 | 0.2857 | 2 | 0.5 | 27 | 0.3333 | 7 | 0.1429 | 9 | 0.1111 | 12 | 0.25 | 7 | 0.1429 | 35 | 0.1714 |
| 11_7012897 | 59 | 0.1525 | 56 | 0.0179 | 43 | 0.1163 | 54 | 0.1111 | 212 | 0.0991 | 46 | 0.0435 | 51 | 0.0392 | 78 | 0.1282 | 36 | 0.1667 | 211 | 0.0948 |
| 11_7012906 | 59 | 0.2542 | 65 | 0.2615 | 63 | 0.2698 | 63 | 0.2857 | 250 | 0.268 | 65 | 0.2769 | 57 | 0.2632 | 88 | 0.3409 | 42 | 0.3333 | 252 | 0.3056 |
| 11_7012910 | 58 | 0.3793 | 60 | 0.2333 | 59 | 0.3051 | 61 | 0.3115 | 238 | 0.3067 | 59 | 0.1695 | 59 | 0.3051 | 87 | 0.2989 | 43 | 0.2326 | 248 | 0.2581 |
| 11_7028481 | 4 | 0.75 | 2 | 1 | 1 | 1 | 1 | 1 | 8 | 0.875 | 4 | 1 | 6 | 0.8333 | 2 | 0.5 | 5 | 1 | 17 | 0.8824 |
| 11_7028490 | 4 | 0.5 | 2 | 0.5 | 1 | 1 | 1 | 0 | 8 | 0.5 | 4 | 0.5 | 6 | 0.8333 | 2 | 1 | 4 | 0.5 | 16 | 0.6875 |
| 11_7028524 | 4 | 0 | 2 | 0 | 1 | 1 | 2 | 0 | 9 | 0.1111 | 4 | 0.25 | 3 | 0 | 2 | 0 | 5 | 0 | 14 | 0.0714 |
| 11_7028525 | 3 | 0 | 2 | 0.5 | 1 | 0 | 1 | 0 | 7 | 0.1429 | 4 | 0.25 | 4 | 0.5 | 2 | 0.5 | 4 | 0 | 14 | 0.2857 |
| 11_7028535 | 3 | 0.6667 | 2 | 0 | 1 | 0 | 1 | 1 | 7 | 0.4286 | 4 | 0.75 | 3 | 0.6667 | 2 | 1 | 4 | 1 | 13 | 0.8462 |
| 11_7028549 | 4 | 0.25 | 1 | 0 | 1 | 0 | 2 | 0 | 8 | 0.125 | 4 | 0.25 | 3 | 0 | 2 | 0 | 4 | 0 | 13 | 0.0769 |
| 11_7028558 | 3 | 1 | 1 | 0 | 1 | 1 | 1 | 1 | 6 | 0.8333 | 3 | 0.6667 | 4 | 0.75 | 2 | 0.5 | 5 | 0.8 | 14 | 0.7143 |
| 11_7028574 | 3 | 0 | 1 | 0 | 1 | 0 | 1 | 1 | 6 | 0.1667 | 3 | 0.3333 | 3 | 0 | 2 | 0.5 | 5 | 0.2 | 13 | 0.2308 |
| 11_7033102 | 8 | 0.25 | 9 | 0.1111 | 3 | 0.3333 | 3 | 0 | 23 | 0.1739 | 5 | 0 | 7 | 0.1429 | 4 | 0 | 6 | 0.5 | 22 | 0.1818 |
| 11_7033106 | 8 | 0.125 | 9 | 0 | 5 | 0.2 | 3 | 0 | 25 | 0.08 | 6 | 0.1667 | 8 | 0 | 5 | 0 | 6 | 0.1667 | 25 | 0.08 |
| 11_7033111 | 7 | 0 | 9 | 0.1111 | 6 | 0.1667 | 3 | 0.3333 | 25 | 0.12 | 5 | 0 | 7 | 0.1429 | 3 | 0 | 6 | 0.1667 | 21 | 0.0952 |
| 11_7033120 | 7 | 0 | 10 | 0.3 | 6 | 0 | 3 | 0.3333 | 26 | 0.1538 | 5 | 0 | 8 | 0.25 | 4 | 0.25 | 5 | 0.2 | 22 | 0.1818 |
| 11_7033142 | 7 | 0.7143 | 10 | 0.6 | 4 | 0.75 | 4 | 0.25 | 25 | 0.6 | 6 | 0.6667 | 7 | 0.2857 | 4 | 0.5 | 6 | 1 | 23 | 0.6087 |
| 11_7034293 | 1 | 0 | 1 | 1 | NA | NA | 1 | 1 | 3 | 0.6667 | NA | NA | NA | NA | 2 | 1 | NA | NA | 2 | 1 |
| 11_7034397 | 1 | 0 | 4 | 0.5 | NA | NA | 2 | 0.5 | 7 | 0.4286 | 1 | 1 | 1 | 0 | 6 | 0.6667 | 1 | 1 | 9 | 0.6667 |
| 11_7034406 | 2 | 1 | 3 | 0.6667 | NA | NA | NA | NA | 5 | 0.8 | 1 | 0 | 3 | 0.3333 | 6 | 0.3333 | 1 | 1 | 11 | 0.3636 |
| 11_7034410 | 2 | 0.5 | 5 | 0.2 | NA | NA | 2 | 0 | 9 | 0.2222 | 1 | 1 | 3 | 0 | 5 | 0 | 1 | 1 | 10 | 0.2 |
| 11_7034415 | 1 | 0 | 5 | 0.2 | NA | NA | 2 | 0.5 | 8 | 0.25 | 1 | 0 | 3 | 0 | 4 | 0.25 | 1 | 0 | 9 | 0.1111 |
| 11_7034432 | 1 | 1 | 4 | 0.25 | NA | NA | 2 | 0 | 7 | 0.2857 | 1 | 0 | 3 | 0.3333 | 5 | 0 | 1 | 0 | 10 | 0.1 |
| 11_7043346 | 2 | 0 | 3 | 0 | 1 | 0 | 1 | 1 | 7 | 0.1429 | NA | NA | 2 | 0 | 2 | 0.5 | NA | NA | 4 | 0.25 |
| 11_7044838 | 2 | 0 | 6 | 0.3333 | 2 | 0 | 2 | 0.5 | 12 | 0.25 | 1 | 0 | 1 | 1 | 4 | 0.25 | 2 | 0.5 | 8 | 0.375 |
| 11_7044849 | 2 | 0 | 6 | 0.1667 | 3 | 0 | 2 | 0 | 13 | 0.0769 | 1 | 0 | 1 | 0 | 4 | 0.25 | 2 | 0 | 8 | 0.125 |
| 11_7044851 | 2 | 0 | 6 | 0.6667 | 3 | 0.3333 | 3 | 0.6667 | 14 | 0.5 | 1 | 0 | 2 | 0 | 4 | 0.25 | 2 | 0 | 9 | 0.1111 |
| 11_7044853 | 3 | 0.6667 | 5 | 0.6 | 3 | 0.3333 | 2 | 1 | 13 | 0.6154 | 1 | 1 | 2 | 1 | 4 | 0.5 | 2 | 1 | 9 | 0.7778 |
| 11_7044867 | 3 | 0 | 5 | 0 | 3 | 0 | 3 | 0.3333 | 14 | 0.0714 | 1 | 0 | 2 | 0 | 4 | 0.25 | 2 | 0 | 9 | 0.1111 |
| 11_7044870 | 2 | 0.5 | 6 | 0.5 | 2 | 0.5 | 3 | 0.3333 | 13 | 0.4615 | 1 | 0 | 2 | 1 | 3 | 0.3333 | 3 | 1 | 9 | 0.6667 |
| 11_7050453 | 27 | 0.1852 | 27 | 0.1481 | 27 | 0.1481 | 32 | 0.1875 | 113 | 0.1681 | 17 | 0.1176 | 36 | 0.0556 | 33 | 0.0909 | 31 | 0.2903 | 117 | 0.1368 |
| 11_7050457 | 29 | 0.1379 | 33 | 0.1212 | 32 | 0.0625 | 34 | 0.2059 | 128 | 0.1328 | 17 | 0.1176 | 39 | 0.1538 | 37 | 0.0541 | 28 | 0.1071 | 121 | 0.1074 |
| 11_7050473 | 30 | 0.1333 | 37 | 0.1351 | 34 | 0.1176 | 34 | 0.0588 | 135 | 0.1111 | 18 | 0.1667 | 44 | 0.1136 | 36 | 0.1111 | 25 | 0.16 | 123 | 0.1301 |
| 11_7050480 | 30 | 0.3667 | 37 | 0.5676 | 36 | 0.6111 | 37 | 0.5135 | 140 | 0.5214 | 18 | 0.5556 | 41 | 0.6098 | 34 | 0.5 | 25 | 0.48 | 118 | 0.5424 |
| 11_7050494 | 21 | 0.3333 | 21 | 0.4286 | 28 | 0.4286 | 19 | 0.4737 | 89 | 0.4157 | 9 | 0.2222 | 33 | 0.5152 | 20 | 0.5 | 19 | 0.3158 | 81 | 0.4321 |
| 11_7053507 | 24 | 0.2083 | 18 | 0 | 17 | 0.1765 | 15 | 0.0667 | 74 | 0.1216 | 13 | 0.1538 | 13 | 0 | 17 | 0.2353 | 12 | 0 | 55 | 0.1091 |
| 11_7053517 | 30 | 0.9 | 21 | 0.9524 | 18 | 1 | 20 | 0.85 | 89 | 0.9213 | 13 | 0.7692 | 19 | 0.8947 | 19 | 0.8947 | 15 | 0.6667 | 66 | 0.8182 |
| 11_7053534 | 31 | 0.1613 | 20 | 0.1 | 21 | 0.0952 | 21 | 0.3333 | 93 | 0.172 | 15 | 0.2 | 21 | 0.1905 | 23 | 0.1739 | 18 | 0.2778 | 77 | 0.2078 |
| 11_7053544 | 33 | 0.6364 | 20 | 0.6 | 20 | 0.8 | 23 | 0.5652 | 96 | 0.6458 | 14 | 0.3571 | 21 | 0.4762 | 20 | 0.8 | 16 | 0.5625 | 71 | 0.5634 |
| 11_7053551 | 33 | 0.4848 | 21 | 0.3333 | 22 | 0.4545 | 22 | 0.4545 | 98 | 0.4388 | 15 | 0.2667 | 23 | 0.3913 | 22 | 0.3636 | 17 | 0.2353 | 77 | 0.3247 |
| 11_7053557 | 31 | 0.5484 | 18 | 0.0556 | 23 | 0.3478 | 19 | 0.2632 | 91 | 0.3407 | 12 | 0.3333 | 22 | 0.2727 | 20 | 0.25 | 17 | 0.2941 | 71 | 0.2817 |
| 11_7053817 | 2 | 0.5 | 5 | 0.6 | 6 | 0.3333 | 2 | 1 | 15 | 0.5333 | 3 | 0 | 13 | 0.3846 | 9 | 0.3333 | 5 | 0.2 | 30 | 0.3 |
| 11_7053835 | 2 | 0 | 7 | 0.2857 | 5 | 0 | 2 | 0 | 16 | 0.125 | 4 | 0.25 | 9 | 0 | 7 | 0 | 6 | 0.3333 | 26 | 0.1154 |
| 11_7054191 | 7 | 0.4286 | 13 | 0.0769 | 13 | 0.3846 | 8 | 0.125 | 41 | 0.2439 | 5 | 0 | 23 | 0.0435 | 12 | 0 | 8 | 0.375 | 48 | 0.0833 |
| 11_7054208 | 7 | 0 | 11 | 0 | 12 | 0.0833 | 10 | 0.1 | 40 | 0.05 | 9 | 0.2222 | 19 | 0.2632 | 10 | 0 | 6 | 0 | 44 | 0.1591 |
| 11_7054212 | 8 | 0.125 | 14 | 0 | 15 | 0.2 | 9 | 0 | 46 | 0.087 | 7 | 0 | 18 | 0.0556 | 11 | 0.1818 | 8 | 0.25 | 44 | 0.1136 |

| | | | | | | | | | | | | | | | | | | | | |
|---|---|---|---|---|---|---|---|---|---|---|---|---|---|---|---|---|---|---|---|---|
| 11_7054522 | 33 | 0.1818 | 33 | 0.2121 | 29 | 0.2414 | 39 | 0.1026 | 134 | 0.1791 | 45 | 0.2222 | 26 | 0.0769 | 29 | 0.1379 | 32 | 0.1562 | 132 | 0.1591 |
| 11_7055488 | 12 | 0.0833 | 12 | 0.1667 | 14 | 0 | 15 | 0.0667 | 53 | 0.0755 | 5 | 0.2 | 15 | 0.1333 | 10 | 0 | 13 | 0.3077 | 43 | 0.1628 |
| 11_7056045 | 13 | 0 | 23 | 0.2174 | 19 | 0 | 16 | 0.125 | 71 | 0.0986 | 21 | 0.0476 | 11 | 0.0909 | 19 | 0.1053 | 16 | 0.25 | 67 | 0.1194 |
| 11_7056054 | 9 | 0 | 22 | 0.0909 | 14 | 0.2143 | 12 | 0 | 57 | 0.0877 | 13 | 0.0769 | 8 | 0 | 16 | 0.125 | 10 | 0.1 | 47 | 0.0851 |
| 11_7056060 | 17 | 0.1176 | 22 | 0.0909 | 21 | 0.1429 | 16 | 0.3125 | 76 | 0.1579 | 22 | 0.1364 | 12 | 0.3333 | 17 | 0.2941 | 14 | 0.0714 | 65 | 0.2 |
| 11_7056071 | 15 | 0.0667 | 25 | 0.12 | 23 | 0 | 18 | 0.1111 | 81 | 0.0741 | 21 | 0.1429 | 14 | 0.1429 | 16 | 0.3125 | 16 | 0.1875 | 67 | 0.194 |
| 11_7056072 | 15 | 0.4 | 26 | 0.5385 | 19 | 0.4737 | 17 | 0.2941 | 77 | 0.4416 | 22 | 0.5455 | 14 | 0.5714 | 17 | 0.4706 | 17 | 0.5294 | 70 | 0.5286 |
| 11_7056073 | 16 | 0.0625 | 26 | 0.0769 | 22 | 0.0909 | 18 | 0 | 82 | 0.061 | 24 | 0.125 | 12 | 0.0833 | 18 | 0 | 12 | 0 | 66 | 0.0606 |
| 11_7056095 | 13 | 0.3846 | 18 | 0.6667 | 14 | 0.5714 | 14 | 0.7857 | 59 | 0.6102 | 8 | 1 | 9 | 0.8889 | 13 | 0.7692 | 12 | 0.5833 | 42 | 0.7857 |
| 11_7056103 | 14 | 0 | 21 | 0.2381 | 20 | 0.15 | 16 | 0.0625 | 71 | 0.1268 | 16 | 0.1875 | 12 | 0.25 | 15 | 0.2667 | 14 | 0.3571 | 57 | 0.2632 |
| 11_7056489 | 8 | 0.375 | 17 | 0.7059 | 16 | 0.625 | 14 | 0.6429 | 55 | 0.6182 | 9 | 0.5556 | 10 | 0.4 | 13 | 0.6154 | 8 | 1 | 40 | 0.625 |
| 11_7056558 | 8 | 0 | 10 | 0 | 9 | 0.2222 | 13 | 0.2308 | 40 | 0.125 | 9 | 0.2222 | 8 | 0 | 10 | 0.1 | 7 | 0.2857 | 34 | 0.1471 |
| 11_7057193 | 7 | 0.4286 | 10 | 0.7 | 7 | 0.4286 | 10 | 0.4 | 34 | 0.5 | 6 | 0.6667 | 4 | 0.25 | 7 | 0.4286 | 4 | 1 | 21 | 0.5714 |
| 11_7057212 | 6 | 0.1667 | 8 | 0.125 | 7 | 0 | 9 | 0 | 30 | 0.0667 | 4 | 0.5 | 5 | 0.4 | 7 | 0.2857 | 5 | 0.2 | 21 | 0.3333 |
| 11_7057230 | 5 | 1 | 8 | 0.625 | 7 | 0.5714 | 10 | 0.6 | 30 | 0.6667 | 4 | 0.75 | 7 | 1 | 7 | 0.7143 | 5 | 1 | 23 | 0.8696 |
| 11_7061719 | 3 | 0 | 3 | 0 | 7 | 0.1429 | 6 | 0 | 19 | 0.0526 | 6 | 0.1667 | 2 | 0 | 3 | 0 | 1 | 0 | 12 | 0.0833 |
| 11_7061925 | 3 | 1 | 4 | 1 | 8 | 0.75 | 6 | 1 | 21 | 0.9048 | 3 | 1 | 1 | 1 | 2 | 0.5 | 2 | 1 | 8 | 0.875 |
| 11_7061928 | 3 | 0.3333 | 2 | 0.5 | 7 | 0.5714 | 4 | 0.5 | 16 | 0.5 | 4 | 0.25 | 1 | 0 | 2 | 0 | 2 | 1 | 9 | 0.3333 |
| 11_7062119 | 2 | 0 | 3 | 0 | 1 | 0 | 6 | 0.1667 | 12 | 0.0833 | 4 | 0 | 1 | 1 | 2 | 0 | 2 | 0 | 9 | 0.1111 |
| 11_7066218 | 2 | 1 | 2 | 1 | 1 | 1 | 1 | 1 | 6 | 1 | 1 | 1 | 1 | 1 | 3 | 1 | 2 | 1 | 7 | 1 |
| 11_7066224 | 2 | 1 | 1 | 1 | 1 | 1 | 1 | 1 | 5 | 1 | 1 | 1 | 1 | 0 | 4 | 0.75 | 2 | 1 | 8 | 0.75 |
| 11_7066248 | 1 | 0 | 2 | 1 | 1 | 1 | 1 | 1 | 5 | 0.8 | 1 | 1 | 1 | 1 | 4 | 1 | 2 | 1 | 8 | 1 |
| 11_7066256 | 2 | 0.5 | 2 | 0.5 | 1 | 0 | 1 | 1 | 6 | 0.5 | 1 | 1 | 1 | 1 | 6 | 0.5 | 2 | 0.5 | 10 | 0.6 |
| 11_7066269 | 2 | 0 | 2 | 0 | 1 | 0 | 1 | 1 | 6 | 0.1667 | NA | NA | 1 | 0 | 6 | 0.5 | 2 | 0.5 | 9 | 0.4444 |
| 11_7066273 | 2 | 0.5 | 1 | 1 | 1 | 0 | 1 | 0 | 5 | 0.4 | NA | NA | NA | NA | 6 | 0.1667 | 2 | 1 | 8 | 0.375 |
| 11_7066307 | 2 | 1 | 1 | 1 | 1 | 0 | NA | NA | 4 | 0.75 | 1 | 1 | 1 | 1 | 5 | 0.6 | 1 | 1 | 8 | 0.75 |
| 11_7066308 | 2 | 0 | 2 | 0 | NA | NA | 1 | 1 | 5 | 0.2 | 1 | 1 | 1 | 0 | 6 | 0.1667 | 2 | 0 | 10 | 0.2 |
| 11_7066317 | 2 | 0.5 | 2 | 0.5 | 1 | 0 | 1 | 0 | 6 | 0.3333 | 1 | 0 | 1 | 0 | 5 | 0 | 2 | 0.5 | 9 | 0.1111 |
| 11_7066324 | 2 | 0.5 | 2 | 0.5 | 1 | 0 | 1 | 1 | 6 | 0.5 | 1 | 1 | 1 | 1 | 5 | 0.4 | 2 | 0.5 | 9 | 0.5556 |
| 11_7066331 | 1 | 1 | 1 | 0 | 1 | 0 | 1 | 0 | 4 | 0.25 | 1 | 0 | 1 | 0 | 6 | 0 | 2 | 0.5 | 10 | 0.1 |
| 11_7066336 | 1 | 0 | 2 | 0 | 1 | 1 | 1 | 0 | 5 | 0.2 | 1 | 1 | 1 | 0 | 5 | 0 | 2 | 0 | 9 | 0.1111 |
| 11_7066357 | 2 | 1 | 2 | 0.5 | 1 | 0 | 1 | 1 | 6 | 0.6667 | NA | NA | 1 | 0 | 6 | 0.3333 | 2 | 0.5 | 9 | 0.3333 |
| 11_7066362 | 1 | 1 | 2 | 0 | 1 | 0 | 1 | 0 | 5 | 0.2 | NA | NA | NA | NA | 3 | 0.3333 | 2 | 0.5 | 5 | 0.4 |
| 11_7066369 | 2 | 1 | 2 | 1 | 1 | 1 | 1 | 1 | 6 | 1 | 1 | 1 | 1 | 1 | 5 | 1 | 3 | 1 | 10 | 1 |
| 11_7066472 | NA | NA | 1 | 1 | NA | NA | NA | NA | 1 | 1 | NA | NA | NA | NA | 1 | 0 | 1 | 1 | 2 | 0.5 |
| 11_7080822 | 6 | 0.8333 | 4 | 0.5 | 9 | 0.4444 | 11 | 0.7273 | 30 | 0.6333 | 4 | 0.25 | 7 | 0.2857 | 6 | 0.3333 | 10 | 0.5 | 27 | 0.3704 |
| 11_7080824 | 5 | 0.6 | 6 | 0.6667 | 9 | 0.7778 | 11 | 0.7273 | 31 | 0.7097 | 4 | 0.75 | 6 | 0.8333 | 8 | 0.875 | 10 | 0.6 | 28 | 0.75 |
| 11_7080827 | 7 | 0.1429 | 6 | 0 | 10 | 0.1 | 11 | 0 | 34 | 0.0588 | 4 | 0.5 | 6 | 0 | 8 | 0 | 10 | 0 | 28 | 0.0714 |
| 11_7080836 | 5 | 0 | 4 | 0 | 9 | 0.1111 | 10 | 0.1 | 28 | 0.0714 | 3 | 0 | 4 | 0 | 8 | 0.125 | 7 | 0.5714 | 22 | 0.2273 |
| 11_7080839 | 5 | 0.8 | 5 | 1 | 8 | 1 | 10 | 0.7 | 28 | 0.8571 | 4 | 1 | 6 | 0.5 | 6 | 0.8333 | 10 | 0.9 | 26 | 0.8077 |
| 11_7083127 | 11 | 0 | 7 | 0 | 8 | 0.125 | 12 | 0.0833 | 38 | 0.0526 | 7 | 0 | 8 | 0 | 12 | 0.1667 | 11 | 0.1818 | 38 | 0.1053 |
| 11_7083128 | 10 | 0.6 | 8 | 0.5 | 10 | 0.5 | 13 | 0.6154 | 41 | 0.561 | 6 | 0.8333 | 10 | 0.5 | 11 | 0.5455 | 10 | 0.5 | 37 | 0.5676 |
| 11_7083130 | 12 | 0.1667 | 8 | 0.5 | 11 | 0.2727 | 14 | 0.2857 | 45 | 0.2889 | 7 | 0 | 9 | 0 | 11 | 0.1818 | 11 | 0.2727 | 38 | 0.1316 |
| 11_7083135 | 10 | 0.1 | 8 | 0 | 10 | 0.2 | 12 | 0.25 | 40 | 0.15 | 6 | 0 | 8 | 0.125 | 11 | 0 | 10 | 0.4 | 35 | 0.1429 |
| 11_7083138 | 8 | 0 | 6 | 0 | 10 | 0.3 | 12 | 0.1667 | 36 | 0.1389 | 6 | 0 | 8 | 0 | 9 | 0.3333 | 7 | 0.1429 | 30 | 0.1333 |
| 11_7083141 | 11 | 0 | 9 | 0 | 11 | 0.0909 | 15 | 0.1333 | 46 | 0.0652 | 8 | 0.125 | 10 | 0 | 12 | 0.0833 | 12 | 0.0833 | 42 | 0.0714 |
| 11_7083145 | 10 | 0.6 | 5 | 0.6 | 10 | 0.7 | 12 | 0.4167 | 37 | 0.5676 | 6 | 0.6667 | 7 | 0.2857 | 10 | 0.2 | 6 | 0.1667 | 29 | 0.3103 |
| 11_7083148 | 9 | 0.2222 | 8 | 0.375 | 11 | 0.2727 | 13 | 0.2308 | 41 | 0.2683 | 6 | 0.1667 | 7 | 0.1429 | 10 | 0.4 | 12 | 0.5 | 35 | 0.3429 |
| 11_7083149 | 11 | 0.5455 | 7 | 0.8571 | 8 | 0.875 | 12 | 0.5833 | 38 | 0.6842 | 7 | 0.8571 | 7 | 0.4286 | 6 | 0.8333 | 12 | 0.5 | 32 | 0.625 |
| 11_7083150 | 11 | 1 | 8 | 1 | 7 | 1 | 15 | 0.9333 | 41 | 0.9756 | 7 | 1 | 6 | 1 | 11 | 1 | 11 | 1 | 35 | 1 |
| 11_7083158 | 11 | 0.2727 | 7 | 0.2857 | 11 | 0.1818 | 16 | 0.25 | 45 | 0.2444 | 7 | 0.4286 | 7 | 0.2857 | 10 | 0.3 | 12 | 0.1667 | 36 | 0.2778 |
| 11_7083187 | 12 | 0.25 | 6 | 0.6667 | 10 | 0.2 | 12 | 0.1667 | 40 | 0.275 | 6 | 0.6667 | 8 | 0 | 11 | 0.2727 | 10 | 0.2 | 35 | 0.2571 |
| 11_7083188 | 12 | 0 | 8 | 0 | 10 | 0.2 | 12 | 0.1667 | 42 | 0.0952 | 10 | 0.1 | 10 | 0 | 11 | 0.0909 | 11 | 0.1818 | 42 | 0.0952 |
| 11_7083190 | 12 | 0.0833 | 6 | 0.3333 | 10 | 0.2 | 11 | 0 | 39 | 0.1282 | 9 | 0.2222 | 9 | 0 | 10 | 0 | 11 | 0.1818 | 39 | 0.1026 |
| 11_7083201 | 8 | 0.125 | 6 | 0.3333 | 6 | 0.3333 | 10 | 0 | 30 | 0.1667 | 5 | 0 | 5 | 0 | 7 | 0.2857 | 4 | 0.25 | 21 | 0.1429 |
| 11_7083260 | 7 | 0.5714 | 7 | 0.4286 | 9 | 0.2222 | 11 | 0.0909 | 34 | 0.2941 | 6 | 0.5 | 6 | 0.6667 | 8 | 0.125 | 9 | 0 | 29 | 0.2759 |
| 11_7083296 | 5 | 0 | 7 | 0.2857 | 7 | 0.1429 | 5 | 0 | 24 | 0.125 | 6 | 0 | 5 | 0.2 | 10 | 0.3 | 9 | 0.2222 | 30 | 0.2 |
| 11_7083298 | 5 | 0.2 | 6 | 0.5 | 9 | 0 | 10 | 0.1 | 30 | 0.1667 | 6 | 0.5 | 6 | 0 | 9 | 0 | 7 | 0 | 28 | 0.1071 |
| 11_7083314 | 6 | 0 | 8 | 0 | 9 | 0.1111 | 8 | 0.125 | 31 | 0.0645 | 6 | 0.1667 | 5 | 0.2 | 11 | 0 | 11 | 0.1818 | 33 | 0.1212 |
| 11_7083348 | 6 | 0 | 8 | 0.125 | 8 | 0 | 9 | 0.1111 | 31 | 0.0645 | 6 | 0.1667 | 4 | 0 | 11 | 0 | 10 | 0.1 | 31 | 0.0645 |
| 11_7083367 | 6 | 0 | 9 | 0.3333 | 11 | 0.0909 | 10 | 0.1 | 36 | 0.1389 | 8 | 0 | 6 | 0 | 11 | 0.0909 | 10 | 0.2 | 35 | 0.0857 |
| 11_7083422 | NA | NA | 1 | 0 | NA | NA | 1 | 1 | 2 | 0.5 | NA | NA | 1 | 1 | 1 | 0 | 1 | 0 | 3 | 0.3333 |
| 11_7086870 | 12 | 0.5 | 13 | 0.4615 | 8 | 0.25 | 9 | 0.3333 | 42 | 0.4048 | 14 | 0.7143 | 23 | 0.5217 | 11 | 0.3636 | 12 | 0.4167 | 60 | 0.5167 |
| 11_7086891 | 14 | 0.8571 | 8 | 1 | 9 | 0.7778 | 10 | 0.9 | 41 | 0.878 | 12 | 1 | 22 | 0.8636 | 16 | 0.875 | 11 | 0.8182 | 61 | 0.8852 |
| 11_7086891 | 13 | 0.3846 | 11 | 0.3636 | 8 | 0 | 6 | 0.3333 | 38 | 0.2895 | 13 | 0.3846 | 22 | 0.2273 | 15 | 0.2667 | 11 | 0.0909 | 61 | 0.2459 |

| | | | | | | | | | | | | | | | | | | | | |
|---|---|---|---|---|---|---|---|---|---|---|---|---|---|---|---|---|---|---|---|---|
| 11_7087508 | 4 | 0.25 | 1 | 1 | 1 | 1 | 1 | 1 | 7 | 0.5714 | 2 | 1 | 4 | 0.25 | 3 | 0.3333 | 5 | 0.6 | 14 | 0.5 |
| 11_7087512 | 5 | 0.8 | 1 | 0 | 1 | 1 | 1 | 1 | 8 | 0.75 | 4 | 1 | 4 | 0.25 | 4 | 0.5 | 4 | 1 | 16 | 0.6875 |
| 11_7087515 | 5 | 1 | 1 | 1 | 1 | 1 | 2 | 1 | 9 | 1 | 3 | 0.6667 | 3 | 1 | 4 | 1 | 5 | 0.8 | 15 | 0.8667 |
| 11_7087518 | 5 | 1 | 1 | 1 | NA | NA | 2 | 1 | 8 | 1 | 3 | 0.6667 | 3 | 1 | 4 | 0.25 | 2 | 0.5 | 12 | 0.5833 |
| 11_7087519 | 4 | 0.25 | 1 | 0 | NA | NA | 1 | 1 | 6 | 0.3333 | 4 | 0.25 | 4 | 0.25 | 4 | 0.25 | 3 | 0 | 15 | 0.2 |
| 11_7087523 | 4 | 1 | 1 | 1 | 1 | 1 | 1 | 1 | 7 | 1 | 2 | 1 | 3 | 1 | 2 | 1 | 3 | 0.6667 | 10 | 0.9 |
| 11_7087532 | 4 | 1 | NA | NA | 1 | 1 | 1 | 1 | 6 | 1 | 2 | 1 | 1 | 0 | 2 | 0.5 | 4 | 0.75 | 9 | 0.6667 |
| 11_7087542 | 2 | 0.5 | NA | NA | 1 | 1 | NA | NA | 3 | 0.6667 | 2 | 1 | 3 | 1 | 1 | 1 | 5 | 0.8 | 11 | 0.9091 |
| 11_7087545 | 5 | 0.4 | 1 | 0 | 1 | 1 | NA | NA | 7 | 0.4286 | 3 | 0 | 3 | 0.6667 | 4 | 0 | 5 | 0.8 | 15 | 0.4 |
| 11_7087556 | 5 | 0.8 | NA | NA | 1 | 1 | NA | NA | 6 | 0.8333 | 2 | 1 | 3 | 0.3333 | 4 | 1 | 3 | 0.6667 | 12 | 0.75 |
| 11_7087580 | 4 | 0.75 | NA | NA | NA | NA | 1 | 1 | 5 | 0.8 | 3 | 0.3333 | 4 | 0.5 | 3 | 0.3333 | 4 | 0.75 | 14 | 0.5 |
| 11_7087601 | 4 | 0.25 | 1 | 0 | 1 | 0 | 1 | 0 | 7 | 0.1429 | 2 | 0.5 | 3 | 0 | 4 | 0 | 4 | 0 | 13 | 0.0769 |
| 11_7087754 | 3 | 0 | 1 | 0 | 2 | 0 | 1 | 1 | 7 | 0.1429 | 1 | 0 | 3 | 0.3333 | 3 | 0.3333 | 1 | 0 | 8 | 0.25 |
| 11_7087760 | 2 | 1 | 1 | 0 | 1 | 1 | 2 | 0.5 | 6 | 0.6667 | 1 | 1 | 4 | 0.75 | 3 | 0.3333 | 2 | 1 | 10 | 0.7 |
| 11_7087761 | 2 | 1 | 1 | 0 | 2 | 1 | 2 | 0 | 7 | 0.5714 | 1 | 0 | 3 | 0.6667 | 3 | 0.6667 | 2 | 0 | 9 | 0.4444 |
| 11_7087763 | 2 | 0.5 | 1 | 0 | 2 | 1 | 2 | 1 | 7 | 0.7143 | 1 | 0 | 4 | 0.75 | 2 | 0 | 2 | 1 | 9 | 0.5556 |
| 11_7087765 | 2 | 0 | 1 | 0 | 2 | 1 | 2 | 1 | 7 | 0.5714 | 1 | 1 | 2 | 0 | 1 | 1 | 2 | 0 | 6 | 0.3333 |
| 11_7087769 | 2 | 1 | 1 | 0 | 2 | 0.5 | 2 | 1 | 7 | 0.7143 | 1 | 1 | 4 | 1 | 3 | 0.6667 | 2 | 0.5 | 10 | 0.8 |
| 11_7087773 | 2 | 0 | 1 | 0 | 2 | 0 | 2 | 0.5 | 7 | 0.1429 | 1 | 0 | 3 | 0 | 2 | 0 | 2 | 1 | 8 | 0.25 |
| 11_7087774 | 2 | 1 | 1 | 1 | 2 | 1 | 3 | 1 | 8 | 1 | 1 | 1 | 4 | 0.75 | 2 | 1 | 2 | 1 | 9 | 0.8889 |
| 11_7087782 | 2 | 0 | 1 | 0 | 2 | 0.5 | 2 | 0 | 7 | 0.1429 | 1 | 1 | 4 | 0.75 | 2 | 0.5 | 2 | 0.5 | 9 | 0.6667 |
| 11_7087785 | 2 | 0 | 1 | 1 | 2 | 0 | 3 | 0.3333 | 8 | 0.25 | 1 | 0 | 4 | 0 | 1 | 0 | 2 | 0.5 | 8 | 0.125 |
| 11_7087786 | 2 | 0 | 1 | 0 | 2 | 1 | 3 | 0.3333 | 8 | 0.375 | 1 | 0 | 4 | 0.25 | 2 | 0 | 2 | 0 | 9 | 0.1111 |
| 11_7087794 | 2 | 1 | 1 | 1 | 1 | 1 | 2 | 0 | 6 | 0.6667 | NA | NA | 4 | 1 | 2 | 0 | 1 | 1 | 7 | 0.7143 |
| 11_7087813 | 2 | 0.5 | 1 | 0 | 2 | 0.5 | 2 | 0.5 | 7 | 0.4286 | NA | NA | 4 | 0.5 | 1 | 1 | 2 | 0 | 7 | 0.4286 |
| 11_7087816 | 2 | 0 | 1 | 1 | NA | NA | 2 | 0.5 | 5 | 0.4 | NA | NA | 3 | 0 | 2 | 0 | 1 | 1 | 6 | 0.1667 |
| 11_7089062 | 2 | 0 | 1 | 1 | NA | NA | NA | NA | 3 | 0.3333 | NA | NA | 2 | 0 | 2 | 0 | 2 | 0.5 | 6 | 0.1667 |
| 11_7090077 | 3 | 0.3333 | 4 | 0.25 | 3 | 0.3333 | 3 | 0.3333 | 13 | 0.3077 | 3 | 0 | 3 | 0 | 4 | 0.75 | 1 | 0 | 11 | 0.2727 |
| 11_7090078 | 4 | 0 | 5 | 0.4 | 3 | 0 | 4 | 0.25 | 16 | 0.1875 | 3 | 0 | 5 | 0.2 | 3 | 0 | 2 | 0 | 13 | 0.0769 |
| 11_7090091 | 3 | 0.3333 | 3 | 0.3333 | 3 | 0 | 4 | 0.5 | 13 | 0.3077 | 2 | 0 | 1 | 0 | 2 | 1 | 3 | 0.6667 | 8 | 0.5 |
| 11_7090098 | 3 | 0 | 2 | 0 | 4 | 0 | 3 | 0.3333 | 12 | 0.0833 | 2 | 0 | 3 | 0 | 5 | 0.4 | 4 | 0.5 | 14 | 0.2857 |
| 11_7090099 | 4 | 0.25 | 3 | 0.3333 | 4 | 0.75 | 4 | 0.5 | 15 | 0.4667 | 3 | 0 | 5 | 0.4 | 5 | 0.2 | 3 | 0.3333 | 16 | 0.25 |
| 11_7090105 | 3 | 1 | 3 | 0.6667 | 4 | 1 | 4 | 1 | 14 | 0.9286 | 1 | 1 | 5 | 1 | 3 | 1 | 4 | 0.75 | 13 | 0.9231 |
| 11_7090111 | 3 | 1 | 3 | 1 | 4 | 1 | 4 | 1 | 14 | 1 | 3 | 0.6667 | 4 | 1 | 5 | 1 | 4 | 0.75 | 16 | 0.875 |
| 11_7091084 | 2 | 0 | 2 | 0.5 | 2 | 0 | 3 | 0.3333 | 9 | 0.2222 | 1 | 0 | 1 | 0 | NA | NA | 1 | 1 | 3 | 0.3333 |
| 11_7091091 | 2 | 0 | 3 | 0.3333 | 1 | 1 | 3 | 0.3333 | 9 | 0.3333 | 2 | 0.5 | 1 | 0 | NA | NA | 1 | 0 | 4 | 0.25 |
| 11_7091098 | 2 | 0 | 2 | 0.5 | 2 | 0.5 | 3 | 0 | 9 | 0.2222 | 2 | 0 | 2 | 0.5 | NA | NA | 2 | 0 | 6 | 0.1667 |
| 11_7091602 | 2 | 0.5 | 2 | 0.5 | 3 | 0.3333 | 4 | 0.25 | 11 | 0.3636 | 1 | 1 | 3 | 0.3333 | NA | NA | 2 | 0.5 | 6 | 0.5 |
| 11_7091611 | 2 | 1 | 2 | 1 | 3 | 0.6667 | 4 | 0.75 | 11 | 0.8182 | 2 | 0.5 | 3 | 1 | NA | NA | 2 | 1 | 7 | 0.8571 |
| 11_7091612 | 2 | 0.5 | 1 | 1 | 2 | 1 | 4 | 0.5 | 9 | 0.6667 | 2 | 0 | 2 | 0.5 | NA | NA | 1 | 0 | 5 | 0.2 |
| 11_7091614 | 2 | 0.5 | 3 | 0.3333 | 3 | 0.3333 | 4 | 0.25 | 12 | 0.3333 | 2 | 0.5 | 3 | 0 | NA | NA | 2 | 0 | 7 | 0.1429 |
| 11_7091641 | 2 | 0.5 | 2 | 0.5 | 3 | 1 | 4 | 1 | 11 | 0.8182 | 2 | 0.5 | 3 | 1 | NA | NA | 1 | 1 | 6 | 0.8333 |
| 11_7091643 | 2 | 1 | 3 | 1 | 4 | 1 | 4 | 0.75 | 13 | 0.9231 | 2 | 0.5 | 3 | 0.6667 | NA | NA | 1 | 1 | 6 | 0.6667 |
| 11_7091644 | 1 | 0 | 2 | 0.5 | 3 | 0.6667 | 3 | 0.6667 | 9 | 0.5556 | 1 | 0 | 2 | 1 | NA | NA | 1 | 0 | 4 | 0.5 |
| 11_7091648 | 1 | 1 | 3 | 0.3333 | 3 | 0.3333 | 3 | 0 | 10 | 0.3 | 1 | 0 | 3 | 0 | NA | NA | 2 | 0.5 | 6 | 0.1667 |
| 11_7091654 | 2 | 1 | 3 | 0.6667 | 4 | 0.5 | 4 | 0.75 | 13 | 0.6923 | 2 | 1 | 3 | 1 | NA | NA | 1 | 1 | 6 | 1 |
| 11_7091657 | 2 | 0 | 3 | 0.3333 | 3 | 0.6667 | 4 | 0.25 | 12 | 0.3333 | 2 | 0.5 | 3 | 0.3333 | NA | NA | 2 | 1 | 7 | 0.5714 |
| 11_7091678 | 3 | 0 | 3 | 0.3333 | 2 | 0 | 4 | 0 | 12 | 0.0833 | 3 | 0 | 3 | 0.3333 | 1 | 0 | 2 | 0 | 9 | 0.1111 |
| 11_7091688 | 1 | 0 | 3 | 0.3333 | 3 | 0.6667 | 3 | 0 | 10 | 0.3 | 3 | 0.3333 | 1 | 0 | 2 | 0 | 2 | 0 | 8 | 0.125 |
| 11_7091786 | 5 | 0 | 2 | 0 | 4 | 0.5 | 4 | 0 | 15 | 0.1333 | 3 | 0.3333 | 3 | 0.3333 | 2 | 0 | 3 | 0.6667 | 11 | 0.3636 |
| 11_7091787 | 5 | 0.2 | 4 | 0.25 | 5 | 0.4 | 5 | 0.6 | 19 | 0.3684 | 5 | 0.4 | 3 | 0 | 2 | 0.5 | 3 | 0.3333 | 13 | 0.3077 |
| 11_7091807 | 7 | 0.4286 | 3 | 0 | 6 | 0.1667 | 5 | 0 | 21 | 0.1905 | 4 | 0.5 | 2 | 0 | 2 | 0 | 2 | 0 | 10 | 0.2 |
| 11_7092291 | 52 | 0.1154 | 58 | 0.2069 | 44 | 0.2045 | 57 | 0.193 | 211 | 0.1801 | 38 | 0.1842 | 57 | 0.193 | 64 | 0.2656 | 39 | 0.1538 | 198 | 0.2071 |
| 11_7092294 | 55 | 0.2182 | 62 | 0.1774 | 50 | 0.3 | 66 | 0.303 | 233 | 0.2489 | 39 | 0.2821 | 56 | 0.1964 | 67 | 0.3134 | 35 | 0.4571 | 197 | 0.2995 |
| 11_7092298 | 61 | 0.8689 | 72 | 0.8611 | 54 | 0.8519 | 72 | 0.875 | 259 | 0.8649 | 43 | 0.9535 | 70 | 0.9714 | 70 | 0.9714 | 49 | 0.9388 | 232 | 0.9612 |
| 11_7092298 | 63 | 0.3492 | 65 | 0.4923 | 55 | 0.5455 | 64 | 0.375 | 247 | 0.4372 | 42 | 0.2857 | 67 | 0.5224 | 71 | 0.4648 | 48 | 0.5 | 228 | 0.4561 |
| 11_7092304 | 71 | 0.1549 | 73 | 0.1644 | 57 | 0.1228 | 75 | 0.16 | 276 | 0.1522 | 47 | 0.1702 | 68 | 0.1029 | 77 | 0.1169 | 46 | 0.1957 | 238 | 0.1387 |
| 11_7092306 | 63 | 0.9048 | 66 | 0.9545 | 53 | 0.9623 | 76 | 0.9211 | 258 | 0.9341 | 43 | 1 | 63 | 0.9524 | 75 | 0.9867 | 42 | 0.9762 | 223 | 0.9776 |
| 11_7092310 | 59 | 0.339 | 62 | 0.4194 | 49 | 0.4694 | 64 | 0.4844 | 234 | 0.4274 | 41 | 0.5366 | 60 | 0.5 | 68 | 0.4706 | 39 | 0.4103 | 208 | 0.4808 |
| 11_7092317 | 62 | 0.6452 | 55 | 0.7636 | 51 | 0.7255 | 66 | 0.7879 | 234 | 0.7308 | 40 | 0.775 | 54 | 0.8333 | 66 | 0.7727 | 42 | 0.8095 | 202 | 0.797 |
| 11_7092337 | 60 | 0.3167 | 60 | 0.1833 | 51 | 0.2353 | 70 | 0.2143 | 241 | 0.2365 | 40 | 0.25 | 65 | 0.2 | 67 | 0.2687 | 37 | 0.1892 | 209 | 0.2297 |
| 11_7092371 | 43 | 0.1163 | 56 | 0.1071 | 38 | 0.0526 | 51 | 0.0392 | 188 | 0.0798 | 25 | 0.24 | 50 | 0.08 | 53 | 0.0943 | 34 | 0.0882 | 162 | 0.1111 |
| 11_7092427 | 43 | 0.0698 | 43 | 0.0465 | 36 | 0.1667 | 52 | 0.0577 | 174 | 0.0805 | 24 | 0.1667 | 48 | 0.0833 | 50 | 0.08 | 31 | 0.129 | 153 | 0.1046 |
| 11_7165051 | 4 | 0.25 | 2 | 0.5 | 2 | 0 | 2 | 0 | 10 | 0.2 | 2 | 0 | 2 | 0.5 | 1 | 0 | 2 | 0 | 7 | 0.1429 |
| 11_7165053 | 3 | 0.3333 | 3 | 0.3333 | 3 | 1 | 2 | 0.5 | 11 | 0.5455 | 2 | 0 | 2 | 0 | 1 | 1 | 2 | 0.5 | 7 | 0.2857 |

| | | | | | | | | | | | | | | | | | | | | |
|---|---|---|---|---|---|---|---|---|---|---|---|---|---|---|---|---|---|---|---|---|
| 11_716506 | 3 | 0 | 4 | 0.25 | 4 | 0.25 | 2 | 1 | 13 | 0.3077 | 2 | 0 | 2 | 1 | 1 | 0 | 3 | 0.3333 | 8 | 0.375 |
| 11_716509 | 2 | 0.5 | 2 | 0 | 4 | 0.5 | 2 | 0.5 | 10 | 0.4 | 3 | 0.6667 | 2 | 1 | 2 | 1 | 3 | 1 | 10 | 0.9 |
| 11_716795 | 3 | 0 | 8 | 0.125 | 15 | 0.1333 | 7 | 0 | 33 | 0.0909 | 5 | 0 | 5 | 0.2 | 6 | 0 | 4 | 0.25 | 20 | 0.1 |
| 11_716797 | 3 | 1 | 8 | 0.875 | 13 | 0.8462 | 6 | 0.8333 | 30 | 0.8667 | 5 | 0.8 | 5 | 1 | 6 | 1 | 6 | 0.8333 | 22 | 0.9091 |
| 11_716797 | 5 | 0.8 | 6 | 1 | 14 | 0.9286 | 7 | 0.7143 | 32 | 0.875 | 5 | 0.8 | 4 | 1 | 6 | 1 | 4 | 0.75 | 19 | 0.8947 |
| 11_716797 | 4 | 0.5 | 8 | 0.5 | 15 | 0.3333 | 6 | 0.3333 | 33 | 0.3939 | 6 | 0.5 | 3 | 0 | 4 | 0.25 | 6 | 0.5 | 19 | 0.3684 |
| 11_716799 | 5 | 0.2 | 7 | 0.2857 | 9 | 0.5556 | 6 | 0.1667 | 27 | 0.3333 | 4 | 0.5 | 4 | 0.25 | 4 | 0.5 | 4 | 0 | 16 | 0.3125 |
| 11_716799 | 5 | 0.8 | 8 | 0.875 | 15 | 0.9333 | 7 | 1 | 35 | 0.9143 | 6 | 0.8333 | 6 | 0.8333 | 4 | 1 | 7 | 0.8571 | 23 | 0.8696 |
| 11_716800 | 5 | 0.4 | 9 | 0.3333 | 15 | 0.3333 | 6 | 0.3333 | 35 | 0.3429 | 6 | 0.5 | 5 | 0.4 | 6 | 0.6667 | 5 | 0.2 | 22 | 0.4545 |
| 11_716800 | 5 | 0.2 | 7 | 0.5714 | 15 | 0.4 | 8 | 0.625 | 35 | 0.4571 | 6 | 0.5 | 6 | 0.6667 | 5 | 0.4 | 6 | 0.6667 | 23 | 0.5652 |
| 11_716801 | 4 | 0 | 7 | 0.1429 | 13 | 0.1538 | 4 | 0 | 28 | 0.1071 | 6 | 0 | 6 | 0.3333 | 6 | 0 | 6 | 0 | 24 | 0.0833 |
| 11_716802 | 5 | 0.2 | 6 | 0.3333 | 12 | 0.5 | 8 | 0.625 | 31 | 0.4516 | 8 | 0.125 | 5 | 1 | 6 | 0.8333 | 6 | 0.3333 | 25 | 0.52 |
| 11_716803 | 5 | 0.2 | 10 | 0.7 | 15 | 0.4 | 8 | 0.375 | 38 | 0.4474 | 7 | 0.4286 | 6 | 0.8333 | 5 | 0.6 | 8 | 0.5 | 26 | 0.5769 |
| 11_716803 | 3 | 0.3333 | 9 | 0.1111 | 16 | 0.0625 | 9 | 0.6667 | 37 | 0.2432 | 5 | 0.6 | 4 | 0.5 | 5 | 0.2 | 8 | 0.375 | 22 | 0.4091 |
| 11_719390 | 2 | 0 | 4 | 0 | 2 | 0 | 6 | 0.3333 | 14 | 0.1429 | 3 | 0 | 5 | 0 | 4 | 0 | 7 | 0.4286 | 19 | 0.1579 |
| 11_719393 | 2 | 0.5 | 5 | 0.2 | 3 | 0 | 5 | 0 | 15 | 0.1333 | 2 | 0 | 3 | 0 | 5 | 0.2 | 6 | 0 | 16 | 0.0625 |
| 11_719420 | 2 | 0 | 3 | 0.3333 | 1 | 0 | 1 | 0 | 7 | 0.1429 | NA | NA | 3 | 0.3333 | 1 | 0 | 1 | 0 | 5 | 0.2 |
| 11_719420 | 1 | 0 | 3 | 0 | NA | NA | 1 | 1 | 5 | 0.2 | NA | NA | 3 | 0 | 1 | 0 | 1 | 1 | 5 | 0.2 |
| 11_720450 | 2 | 0.5 | 1 | 1 | 2 | 0.5 | 2 | 1 | 7 | 0.7143 | NA | NA | 2 | 0.5 | 3 | 0.6667 | 1 | 1 | 6 | 0.6667 |
| 11_720450 | 2 | 0 | 2 | 0.5 | 3 | 0.3333 | 3 | 0.6667 | 10 | 0.4 | NA | NA | 2 | 0.5 | 3 | 0 | 1 | 1 | 6 | 0.3333 |
| 11_720452 | 1 | 1 | 1 | 1 | 4 | 1 | 2 | 0.5 | 8 | 0.875 | NA | NA | 1 | 1 | 2 | 1 | 1 | 1 | 4 | 1 |
| 11_720453 | 2 | 0 | 1 | 1 | 4 | 0.25 | 3 | 0.3333 | 10 | 0.3 | NA | NA | 1 | 1 | 1 | 0 | 1 | 1 | 3 | 0.6667 |
| 11_720453 | 2 | 1 | 1 | 1 | 3 | 1 | 3 | 0.6667 | 9 | 0.8889 | NA | NA | 1 | 1 | 3 | 1 | 1 | 1 | 5 | 1 |
| 11_720455 | 2 | 1 | 1 | 1 | 3 | 0.6667 | 3 | 0.6667 | 9 | 0.7778 | NA | NA | 1 | 1 | 2 | 1 | 1 | 1 | 4 | 1 |
| 11_721003 | 18 | 0.5 | 7 | 0.1429 | 12 | 0.5 | 8 | 0.75 | 45 | 0.4889 | 5 | 0.4 | 13 | 0.5385 | 6 | 0.6667 | 4 | 0.75 | 28 | 0.5714 |
| 11_721004 | 15 | 0.0667 | 4 | 0 | 9 | 0.2222 | 7 | 0.5714 | 35 | 0.2 | 4 | 0.5 | 10 | 0.1 | 5 | 0.2 | 5 | 0 | 24 | 0.1667 |
| 11_721026 | 6 | 0.1667 | 3 | 0.3333 | 4 | 0.25 | 5 | 0.4 | 18 | 0.2778 | 3 | 0 | 5 | 0.2 | 7 | 0.5714 | 3 | 0.3333 | 18 | 0.3333 |
| 11_721333 | NA | NA | NA | NA | NA | NA | 2 | 0.5 | 2 | 0.5 | NA | NA | 1 | 1 | 2 | 1 | NA | NA | 3 | 1 |
| 11_721338 | NA | NA | NA | NA | NA | NA | 2 | 0.5 | 2 | 0.5 | NA | NA | NA | NA | 1 | 1 | NA | NA | 1 | 1 |
| 11_721995 | NA | NA | 2 | 1 | 1 | 0 | 2 | 1 | 5 | 0.8 | 1 | 1 | NA | NA | NA | NA | NA | NA | 1 | 1 |
| 11_723418 | 4 | 0 | 5 | 0.2 | 2 | 0 | 8 | 0 | 19 | 0.0526 | NA | NA | 10 | 0.1 | 3 | 0 | 5 | 0 | 18 | 0.0556 |
| 11_723418 | 4 | 0 | 5 | 0.2 | 2 | 0 | 9 | 0 | 20 | 0.05 | NA | NA | 10 | 0.1 | 3 | 0 | 5 | 0 | 18 | 0.0556 |
| 11_723423 | 4 | 0.5 | 7 | 0.4286 | 2 | 0.5 | 10 | 0.5 | 23 | 0.4783 | 2 | 1 | 10 | 0.8 | 5 | 0.8 | 6 | 0.5 | 23 | 0.7391 |
| 11_723425 | 7 | 0 | 4 | 0.5 | 7 | 0.4286 | 2 | 0 | 20 | 0.25 | 9 | 0.1111 | 2 | 0 | 2 | 0 | 4 | 0 | 17 | 0.0588 |
| 11_723430 | 2 | 0 | 3 | 0.3333 | 3 | 0 | 1 | 0 | 9 | 0.1111 | 3 | 0 | NA | NA | 2 | 0.5 | 1 | 0 | 6 | 0.1667 |
| 11_723430 | 4 | 0 | 5 | 0.8 | 8 | 0.125 | 4 | 0.25 | 21 | 0.2857 | 9 | 0.2222 | 2 | 1 | 2 | 0.5 | 4 | 0.5 | 17 | 0.4118 |
| 11_723432 | 5 | 0 | 3 | 0.3333 | 6 | 0 | 3 | 0.3333 | 17 | 0.1176 | 6 | 0 | 2 | 0 | 2 | 0 | 4 | 0.25 | 14 | 0.0714 |
| 11_723433 | 2 | 0.5 | 2 | 0 | 5 | 0.2 | 2 | 0.5 | 11 | 0.2727 | 8 | 0.5 | 1 | 0 | 2 | 0 | 3 | 0 | 14 | 0.2857 |
| 11_723434 | 3 | 0 | 4 | 0 | 4 | 0.25 | 1 | 0 | 12 | 0.0833 | 5 | 0.2 | 1 | 0 | 1 | 1 | 4 | 0 | 11 | 0.1818 |
| 11_723434 | 3 | 0 | 5 | 0.2 | 7 | 0.1429 | 3 | 0 | 18 | 0.1111 | 8 | 0.125 | 1 | 0 | 2 | 0 | 4 | 0.25 | 15 | 0.1333 |
| 11_723434 | 5 | 0 | 5 | 0.2 | 7 | 0.1429 | 4 | 0.25 | 21 | 0.1429 | 10 | 0.2 | 2 | 0 | 2 | 0 | 4 | 0 | 18 | 0.1111 |
| 11_723435 | 5 | 0 | 5 | 0 | 6 | 0.1667 | 4 | 0.25 | 20 | 0.1 | 9 | 0.1111 | 2 | 0 | 2 | 0 | 3 | 0 | 16 | 0.0625 |
| 11_723437 | 3 | 0.6667 | 5 | 0.8 | 5 | 0.2 | 2 | 0.5 | 15 | 0.5333 | 6 | 0.1667 | 2 | 1 | 2 | 0 | 2 | 0.5 | 12 | 0.3333 |
| 11_723437 | 3 | 1 | 5 | 0.6 | 5 | 0.8 | 3 | 1 | 16 | 0.8125 | 2 | 1 | NA | NA | 2 | 1 | NA | NA | 4 | 1 |
| 11_723438 | 5 | 0.8 | 4 | 0.75 | 7 | 0.8571 | 4 | 0.75 | 20 | 0.8 | 6 | 0.5 | 2 | 1 | 2 | 1 | 3 | 0.3333 | 13 | 0.6154 |
| 11_723438 | 4 | 0 | 3 | 0 | 7 | 0.1429 | 4 | 0 | 18 | 0.0556 | 6 | 0.5 | 2 | 0 | 1 | 0 | 1 | 0 | 10 | 0.3 |
| 11_723439 | 3 | 0.3333 | 4 | 0 | 6 | 0.1667 | 2 | 0 | 15 | 0.1333 | 5 | 0.2 | 2 | 0 | 2 | 0 | 2 | 0 | 11 | 0.0909 |
| 11_725744 | 11 | 0.8182 | 5 | 0.4 | 6 | 0.8333 | 6 | 0.6667 | 28 | 0.7143 | 4 | 0.75 | 9 | 0.3333 | 8 | 0.75 | 4 | 1 | 25 | 0.64 |
| 11_725744 | 12 | 0.25 | 8 | 0.625 | 6 | 0.3333 | 5 | 0.6 | 31 | 0.4194 | 5 | 0.2 | 10 | 0.4 | 10 | 0.1 | 5 | 0.4 | 30 | 0.2667 |
| 11_725745 | 11 | 1 | 6 | 0.8333 | 5 | 0.8 | 3 | 0.6667 | 25 | 0.88 | 5 | 1 | 7 | 1 | 8 | 0.75 | 6 | 1 | 26 | 0.9231 |
| 11_725745 | 12 | 0.1667 | 7 | 0 | 6 | 0.1667 | 6 | 0.3333 | 31 | 0.1613 | 7 | 0.1429 | 9 | 0.1111 | 10 | 0.1 | 5 | 0 | 31 | 0.0968 |
| 11_725746 | 13 | 0.1538 | 8 | 0.125 | 5 | 0 | 6 | 0.3333 | 32 | 0.1562 | 7 | 0.1429 | 10 | 0.2 | 9 | 0.2222 | 6 | 0 | 32 | 0.1562 |
| 11_726584 | 2 | 0.5 | 1 | 1 | 1 | 1 | 2 | 0.5 | 6 | 0.6667 | NA | NA | 2 | 0 | 1 | 1 | 1 | 1 | 4 | 0.5 |
| 11_726586 | 2 | 1 | NA | NA | 1 | 1 | 3 | 1 | 6 | 1 | NA | NA | 2 | 1 | 1 | 1 | 2 | 1 | 5 | 1 |
| 11_726586 | 2 | 1 | 1 | 1 | 2 | 1 | 3 | 0.6667 | 8 | 0.875 | 2 | 1 | 2 | 1 | 1 | 1 | 3 | 1 | 8 | 1 |
| 11_726586 | 2 | 0.5 | 1 | 0 | NA | NA | 3 | 0.3333 | 6 | 0.3333 | 1 | 0 | 2 | 1 | 1 | 1 | 3 | 0 | 7 | 0.4286 |
| 11_726587 | 2 | 1 | 1 | 1 | 2 | 1 | 3 | 1 | 8 | 1 | 2 | 1 | 3 | 1 | 2 | 1 | 3 | 1 | 10 | 1 |
| 11_726587 | 2 | 0.5 | 1 | 0 | 2 | 0 | 3 | 0.3333 | 8 | 0.25 | 2 | 0 | 3 | 0.3333 | 2 | 1 | 3 | 0.6667 | 10 | 0.5 |
| 11_726588 | 1 | 0 | 1 | 0 | 1 | 0 | 3 | 0.3333 | 6 | 0.1667 | 1 | 0 | 2 | 0 | 2 | 0 | 2 | 0.5 | 7 | 0.1429 |
| 11_726592 | 2 | 0.5 | 1 | 0 | 3 | 1 | 4 | 0.5 | 10 | 0.6 | 2 | 0 | 5 | 0.6 | 2 | 0.5 | 3 | 1 | 12 | 0.5833 |
| 11_726593 | 2 | 1 | 1 | 1 | 3 | 1 | 2 | 1 | 8 | 1 | 2 | 1 | 5 | 0.6 | 2 | 1 | 6 | 0.5 | 15 | 0.6667 |
| 11_726593 | 2 | 0.5 | 1 | 0 | 3 | 0.3333 | 3 | 0.3333 | 9 | 0.3333 | 1 | 0 | 5 | 0 | 2 | 0 | 4 | 0.75 | 12 | 0.25 |
| 11_726594 | 2 | 1 | 1 | 1 | 3 | 1 | 4 | 0.5 | 10 | 0.8 | 2 | 0.5 | 4 | 0.75 | 2 | 1 | 6 | 0.5 | 14 | 0.6429 |
| 11_726597 | 2 | 0.5 | NA | NA | 3 | 0 | 4 | 0 | 9 | 0.1111 | 2 | 0 | 4 | 0 | 3 | 0.3333 | 4 | 0.25 | 13 | 0.1538 |
| 11_726597 | 2 | 0.5 | 1 | 0 | 3 | 0 | 4 | 0.25 | 10 | 0.2 | 2 | 0 | 5 | 0.4 | 3 | 0.3333 | 4 | 0.25 | 14 | 0.2857 |

| | | | | | | | | | | | | | | | | | | | | |
|---|---|---|---|---|---|---|---|---|---|---|---|---|---|---|---|---|---|---|---|---|
| 11_726598 | 1 | 1 | 1 | 0 | 2 | 0.5 | 3 | 0.6667 | 7 | 0.5714 | 2 | 0 | 4 | 0.5 | 1 | 0 | 4 | 0.25 | 11 | 0.2727 |
| 11_726598 | 2 | 0.5 | 1 | 0 | 3 | 0 | 4 | 0.25 | 10 | 0.2 | 1 | 0 | 4 | 0.25 | 3 | 0 | 3 | 0.6667 | 11 | 0.2727 |
| 11_726599 | 2 | 0.5 | 1 | 0 | 1 | 0 | 4 | 0.75 | 8 | 0.5 | 2 | 1 | 5 | 0.6 | 3 | 0.6667 | 5 | 0.4 | 15 | 0.6 |
| 11_726610 | 3 | 0 | 1 | 0 | 4 | 0 | 5 | 0.2 | 13 | 0.0769 | 2 | 0 | 5 | 0.2 | 2 | 0 | 4 | 0 | 13 | 0.0769 |
| 11_726615 | 3 | 0 | 1 | 0 | 4 | 0 | 5 | 0.4 | 13 | 0.1538 | 3 | 0.3333 | 6 | 0 | 3 | 0.3333 | 4 | 0 | 16 | 0.125 |
| 11_726616 | 3 | 0.3333 | 1 | 0 | 4 | 0 | 4 | 0 | 12 | 0.0833 | 4 | 0.25 | 5 | 0 | 3 | 0 | 3 | 0 | 15 | 0.0667 |
| 11_726617 | 3 | 0.3333 | 1 | 0 | 4 | 0 | 5 | 0.2 | 13 | 0.1538 | 4 | 0.25 | 6 | 0 | 3 | 0.3333 | 5 | 0.2 | 18 | 0.1667 |
| 11_726617 | 3 | 0.3333 | 1 | 1 | 4 | 0.25 | 5 | 0.2 | 13 | 0.3077 | 3 | 0.6667 | 5 | 0.4 | 2 | 0.5 | 1 | 0 | 11 | 0.4545 |
| 11_726618 | 4 | 0.25 | 1 | 0 | 3 | 0 | 5 | 0.4 | 13 | 0.2308 | 4 | 0.25 | 6 | 0.1667 | 4 | 0.25 | 4 | 0.25 | 18 | 0.2222 |
| 11_726619 | 4 | 0.25 | 1 | 0 | 4 | 0 | 6 | 0 | 15 | 0.0667 | 4 | 0.25 | 6 | 0 | 4 | 0.25 | 5 | 0 | 19 | 0.1053 |
| 11_726688 | NA | NA | 3 | 0 | 2 | 0.5 | 4 | 0 | 9 | 0.1111 | 2 | 0 | 4 | 0.25 | NA | NA | NA | NA | 6 | 0.1667 |
| 11_726689 | 1 | 1 | 4 | 0.25 | 3 | 0.3333 | 4 | 0 | 12 | 0.25 | 1 | 0 | 3 | 0.3333 | NA | NA | NA | NA | 4 | 0.25 |
| 11_726690 | 1 | 1 | 2 | 0.5 | 3 | 0.6667 | 3 | 0 | 9 | 0.4444 | 2 | 0.5 | 3 | 0 | NA | NA | NA | NA | 5 | 0.2 |
| 11_726690 | 1 | 1 | 2 | 1 | 3 | 0.6667 | 3 | 0.6667 | 9 | 0.7778 | 2 | 1 | 3 | 1 | NA | NA | NA | NA | 5 | 1 |
| 11_726691 | NA | NA | 3 | 0.3333 | 2 | 0.5 | 2 | 0.5 | 7 | 0.4286 | 2 | 0 | 4 | 0.5 | NA | NA | NA | NA | 6 | 0.3333 |
| 11_726691 | 1 | 1 | 4 | 0.75 | 3 | 0 | 3 | 0.6667 | 11 | 0.5455 | 2 | 0 | 5 | 0.4 | NA | NA | NA | NA | 7 | 0.2857 |
| 11_726691 | 1 | 1 | 4 | 1 | 3 | 1 | 3 | 1 | 11 | 1 | 2 | 1 | 4 | 1 | NA | NA | NA | NA | 6 | 1 |
| 11_726692 | NA | NA | 3 | 0.6667 | 3 | 0.6667 | 3 | 1 | 9 | 0.7778 | 2 | 1 | 5 | 0.6 | NA | NA | NA | NA | 7 | 0.7143 |
| 11_726692 | 1 | 1 | 4 | 1 | 3 | 1 | 3 | 1 | 11 | 1 | 2 | 1 | 4 | 1 | NA | NA | NA | NA | 6 | 1 |
| 11_726693 | NA | NA | 4 | 0 | 3 | 1 | 1 | 0 | 8 | 0.375 | 2 | 1 | 4 | 1 | NA | NA | NA | NA | 6 | 1 |
| 11_726693 | 1 | 1 | 4 | 0.5 | 3 | 0.3333 | 4 | 0.5 | 12 | 0.5 | 2 | 0.5 | 4 | 0.25 | NA | NA | NA | NA | 6 | 0.3333 |
| 11_726694 | 1 | 1 | 4 | 1 | 3 | 1 | 4 | 1 | 12 | 1 | 2 | 1 | 4 | 1 | NA | NA | NA | NA | 6 | 1 |
| 11_726696 | 1 | 1 | 4 | 0.75 | 2 | 1 | 4 | 0 | 11 | 0.5455 | 1 | 0 | 3 | 0.6667 | NA | NA | NA | NA | 4 | 0.5 |
| 11_726696 | 1 | 0 | 4 | 0.25 | 1 | 1 | 3 | 0.6667 | 9 | 0.4444 | 2 | 0.5 | 2 | 0 | NA | NA | NA | NA | 4 | 0.25 |
| 11_726865 | NA | NA | 2 | 1 | 3 | 0.6667 | 2 | 1 | 7 | 0.8571 | 2 | 1 | 2 | 1 | NA | NA | NA | NA | 4 | 1 |
| 11_728040 | 1 | 0 | 1 | 1 | 1 | 0 | 4 | 0.75 | 7 | 0.5714 | NA | NA | NA | NA | NA | NA | 2 | 0.5 | 2 | 0.5 |
| 11_728155 | 2 | 1 | NA | NA | 2 | 1 | 6 | 0.8333 | 10 | 0.9 | 1 | 1 | 2 | 1 | 1 | 1 | 4 | 1 | 8 | 1 |
| 11_728157 | 2 | 0 | 2 | 0.5 | 2 | 1 | 7 | 0.8571 | 13 | 0.6923 | 1 | 1 | 3 | 1 | 1 | 1 | 4 | 0.75 | 9 | 0.8889 |
| 11_728159 | 2 | 0.5 | 1 | 0 | 2 | 0.5 | 7 | 0.8571 | 12 | 0.6667 | NA | NA | 2 | 0.5 | 1 | 1 | 3 | 0.3333 | 6 | 0.5 |
| 11_728159 | 2 | 0 | 1 | 1 | 2 | 0 | 6 | 0.1667 | 11 | 0.1818 | 1 | 1 | 2 | 0 | 1 | 0 | 4 | 0 | 8 | 0.125 |
| 11_728162 | 2 | 0 | 2 | 0 | 1 | 1 | 7 | 0.1429 | 12 | 0.1667 | 1 | 1 | 3 | 0 | NA | NA | 4 | 0.25 | 8 | 0.25 |
| 11_728162 | 1 | 0 | 3 | 0 | 1 | 0 | 7 | 0.1429 | 12 | 0.0833 | NA | NA | 3 | 0 | 1 | 1 | 5 | 0.2 | 9 | 0.2222 |
| 11_728163 | 2 | 0 | 2 | 0.5 | 2 | 0.5 | 6 | 0.3333 | 12 | 0.3333 | 1 | 0 | 2 | 0 | 1 | 0 | 5 | 0.4 | 9 | 0.2222 |
| 11_728165 | 1 | 0 | 1 | 0 | 2 | 1 | 5 | 0.8 | 9 | 0.6667 | 1 | 1 | 3 | 0.3333 | 1 | 1 | 6 | 0.3333 | 11 | 0.4545 |
| 11_728166 | 2 | 0.5 | 3 | 0 | 2 | 0 | 6 | 0 | 13 | 0.0769 | 1 | 0 | 4 | 0 | 1 | 0 | 6 | 0.1667 | 12 | 0.0833 |
| 11_728167 | 2 | 0.5 | 2 | 0 | 2 | 0 | 6 | 0.3333 | 12 | 0.25 | NA | NA | 4 | 0.5 | 1 | 1 | 5 | 0.4 | 10 | 0.5 |
| 11_728172 | 2 | 0 | 2 | 0 | 1 | 1 | 7 | 0.1429 | 12 | 0.1667 | 1 | 1 | 5 | 0.2 | 1 | 0 | 7 | 0.1429 | 14 | 0.2143 |
| 11_728172 | 1 | 1 | 2 | 0.5 | 2 | 1 | 5 | 0.4 | 10 | 0.6 | 1 | 1 | 3 | 0.3333 | 1 | 0 | 3 | 0.3333 | 8 | 0.375 |
| 11_728173 | 1 | 1 | 3 | 0.6667 | 2 | 1 | 7 | 1 | 13 | 0.9231 | 1 | 1 | 5 | 0.4 | 1 | 1 | 6 | 0.8333 | 13 | 0.6923 |
| 11_728174 | 1 | 0 | 3 | 0.3333 | 1 | 1 | 6 | 0 | 11 | 0.1818 | 1 | 1 | 2 | 0 | 1 | 0 | 7 | 0.2857 | 11 | 0.2727 |
| 11_728176 | 1 | 0 | 3 | 0.3333 | 2 | 0.5 | 7 | 0.2857 | 13 | 0.3077 | 1 | 0 | 5 | 0.4 | 2 | 0.5 | 7 | 0 | 15 | 0.2 |
| 11_728176 | 1 | 1 | 3 | 0 | 1 | 1 | 7 | 0 | 12 | 0.1667 | 1 | 0 | 4 | 0 | 2 | 0.5 | 8 | 0.375 | 15 | 0.2667 |
| 11_728176 | 2 | 0 | 3 | 0 | 3 | 0.3333 | 7 | 0.2857 | 15 | 0.2 | 1 | 0 | 5 | 0 | 2 | 0 | 7 | 0.1429 | 15 | 0.0667 |
| 11_728176 | 2 | 0 | 3 | 0 | 3 | 0 | 7 | 0.1429 | 15 | 0.0667 | 1 | 1 | 4 | 0.25 | 2 | 0.5 | 7 | 0.5714 | 14 | 0.5 |
| 11_728177 | 1 | 0 | 5 | 0.2 | 4 | 0.5 | 6 | 0.1667 | 16 | 0.25 | 2 | 0 | 5 | 0.2 | 2 | 0.5 | 7 | 0.2857 | 16 | 0.25 |
| 11_728177 | 2 | 0 | 3 | 0.3333 | 4 | 0 | 7 | 0.2857 | 16 | 0.1875 | 2 | 0.5 | 5 | 0 | 2 | 0.5 | 8 | 0.125 | 17 | 0.1765 |
| 11_729312 | 5 | 0 | 8 | 0.25 | 4 | 0.25 | 8 | 0.25 | 25 | 0.2 | NA | NA | 4 | 0 | 6 | 0.3333 | 7 | 0 | 17 | 0.1176 |
| 11_729312 | 7 | 0.5714 | 7 | 0.8571 | 4 | 0.5 | 9 | 0.4444 | 27 | 0.5926 | 1 | 1 | 5 | 0.2 | 5 | 0.4 | 5 | 0 | 16 | 0.25 |
| 11_729313 | 7 | 0 | 11 | 0.4545 | 5 | 0.2 | 8 | 0.125 | 31 | 0.2258 | 2 | 0.5 | 6 | 0.1667 | 6 | 0.3333 | 7 | 0.2857 | 21 | 0.2857 |
| 11_729314 | 8 | 0.375 | 9 | 0.3333 | 4 | 0.5 | 9 | 0.4444 | 30 | 0.4 | 2 | 0.5 | 6 | 0 | 8 | 0 | 7 | 0.1429 | 23 | 0.087 |
| 11_729316 | 7 | 0.2857 | 11 | 0.2727 | 4 | 0 | 9 | 0 | 31 | 0.1613 | 2 | 0 | 4 | 0.5 | 8 | 0 | 7 | 0.1429 | 21 | 0.1429 |
| 11_729317 | 9 | 0.7778 | 12 | 0.8333 | 6 | 1 | 13 | 0.9231 | 40 | 0.875 | 3 | 0.6667 | 6 | 0.6667 | 7 | 0.5714 | 9 | 0.6667 | 25 | 0.64 |
| 11_729317 | 7 | 0.1429 | 9 | 0.2222 | 4 | 0 | 11 | 0 | 31 | 0.0968 | 3 | 0 | 5 | 0.4 | 6 | 0.1667 | 9 | 0.1111 | 23 | 0.1739 |
| 11_729317 | 8 | 1 | 12 | 0.9167 | 5 | 1 | 12 | 1 | 37 | 0.973 | 3 | 1 | 5 | 0.8 | 8 | 0.875 | 9 | 1 | 25 | 0.92 |
| 11_729319 | 8 | 0.75 | 11 | 0.5455 | 4 | 0.75 | 14 | 0.5714 | 37 | 0.6216 | 4 | 1 | 8 | 0.5 | 7 | 0.7143 | 8 | 0.625 | 27 | 0.6667 |
| 11_729320 | 9 | 0.2222 | 9 | 0.3333 | 6 | 0.3333 | 13 | 0.1538 | 37 | 0.2432 | 5 | 0.2 | 5 | 0.2 | 6 | 0.1667 | 8 | 0.125 | 24 | 0.1667 |
| 11_729321 | 12 | 0.25 | 10 | 0.4 | 7 | 0.8571 | 14 | 0.2857 | 43 | 0.3953 | 4 | 0 | 9 | 0.6667 | 9 | 0.2222 | 10 | 0.2 | 32 | 0.3125 |
| 11_729321 | 11 | 0.8182 | 10 | 1 | 7 | 1 | 10 | 0.8 | 38 | 0.8947 | 5 | 1 | 9 | 0.7778 | 7 | 1 | 7 | 1 | 28 | 0.9286 |
| 11_729321 | 9 | 0.1111 | 9 | 0.1111 | 5 | 0.2 | 12 | 0.0833 | 35 | 0.1143 | 4 | 0 | 9 | 0.2222 | 10 | 0 | 9 | 0 | 32 | 0.0625 |
| 11_729322 | 13 | 0.3077 | 11 | 0.2727 | 7 | 0.4286 | 14 | 0.3571 | 45 | 0.3333 | 6 | 0.6667 | 9 | 0.3333 | 11 | 0.4545 | 10 | 0.1 | 36 | 0.3611 |
| 11_729323 | 8 | 0.125 | 10 | 0.3 | 6 | 0.8333 | 11 | 0.2727 | 35 | 0.3429 | 5 | 0.6 | 7 | 0.5714 | 9 | 0.6667 | 10 | 0.1 | 31 | 0.4516 |
| 11_729700 | 4 | 0.75 | 2 | 0.5 | 1 | 1 | 2 | 0.5 | 9 | 0.6667 | 2 | 0.5 | 1 | 0 | NA | NA | 1 | 0 | 4 | 0.25 |
| 11_729702 | 3 | 0 | 4 | 0 | 1 | 0 | 3 | 0.3333 | 11 | 0.0909 | 2 | 0.5 | 1 | 1 | NA | NA | 3 | 0.3333 | 6 | 0.5 |
| 11_729703 | 4 | 0.5 | 4 | 0.25 | 1 | 0 | 2 | 1 | 11 | 0.4545 | 3 | 0 | 1 | 1 | NA | NA | 3 | 0.6667 | 7 | 0.4286 |
| 11_729704 | 2 | 0 | 1 | 1 | NA | NA | 1 | 1 | 4 | 0.5 | 1 | 1 | 1 | 1 | NA | NA | 3 | 1 | 5 | 1 |

| | | | | | | | | | | | | | | | | | | | | |
|---|---|---|---|---|---|---|---|---|---|---|---|---|---|---|---|---|---|---|---|---|
| 11_7297064 | 4 | 0 | 3 | 0 | 1 | 0 | 3 | 0.3333 | 11 | 0.0909 | 3 | 0.3333 | 2 | 1 | NA | NA | 3 | 0.3333 | 8 | 0.5 |
| 11_7306648 | 4 | 0.5 | 3 | 0 | 5 | 0.2 | 2 | 0 | 14 | 0.2143 | 4 | 0.5 | 3 | 0.3333 | 6 | 0.5 | 5 | 0.4 | 18 | 0.4444 |
| 11_7306690 | 3 | 0 | 5 | 0.4 | 5 | 0 | 3 | 0.6667 | 16 | 0.25 | 4 | 0.25 | 4 | 0.5 | 6 | 0.3333 | 5 | 0.4 | 19 | 0.3684 |
| 11_7306690 | NA | NA | NA | NA | 1 | 1 | NA | NA | 1 | 1 | NA | NA | NA | NA | 1 | 1 | NA | NA | 1 | 1 |
| 11_7306713 | 8 | 0.125 | 10 | 0.2 | 9 | 0.3333 | 4 | 0.25 | 31 | 0.2258 | 13 | 0.2308 | 16 | 0.125 | 14 | 0.1429 | 11 | 0.1818 | 54 | 0.1667 |
| 11_7306718 | 8 | 0.375 | 10 | 0.3 | 10 | 0.5 | 4 | 0.5 | 32 | 0.4062 | 10 | 0.3 | 14 | 0.7143 | 11 | 0.4545 | 11 | 0.4545 | 46 | 0.5 |
| 11_7306722 | 8 | 0.875 | 9 | 0.7778 | 9 | 0.6667 | 4 | 0.5 | 30 | 0.7333 | 11 | 0.9091 | 14 | 0.8571 | 12 | 0.75 | 10 | 1 | 47 | 0.8723 |
| 11_7306728 | 6 | 0 | 8 | 0.375 | 8 | 0.125 | 4 | 0.25 | 26 | 0.1923 | 9 | 0.1111 | 10 | 0 | 11 | 0.0909 | 11 | 0.0909 | 41 | 0.0732 |
| 11_7306730 | 8 | 0.375 | 8 | 0.125 | 9 | 0.3333 | 4 | 0.25 | 29 | 0.2759 | 11 | 0.2727 | 13 | 0.1538 | 12 | 0.4167 | 11 | 0.3636 | 47 | 0.2979 |
| 11_7306743 | 6 | 0 | 10 | 0 | 10 | 0.2 | 4 | 0 | 30 | 0.0667 | 12 | 0.1667 | 9 | 0.1111 | 12 | 0.0833 | 11 | 0 | 44 | 0.0909 |
| 11_7307471 | 2 | 0.5 | 1 | 0 | 2 | 0 | NA | NA | 5 | 0.2 | NA | NA | 2 | 0 | 1 | 0 | 2 | 1 | 5 | 0.4 |
| 11_7307508 | 1 | 1 | 1 | 0 | 2 | 0 | NA | NA | 4 | 0.25 | NA | NA | 1 | 1 | 1 | 0 | 1 | 0 | 3 | 0.3333 |
| 11_7307516 | 1 | 1 | 1 | 1 | 2 | 1 | NA | NA | 4 | 1 | NA | NA | 1 | 1 | 1 | 1 | 2 | 1 | 4 | 1 |
| 11_7307520 | 1 | 1 | NA | NA | 2 | 1 | NA | NA | 3 | 1 | NA | NA | 2 | 1 | 1 | 1 | 2 | 1 | 5 | 1 |
| 11_7307523 | 1 | 1 | NA | NA | 1 | 0 | NA | NA | 2 | 0.5 | NA | NA | 2 | 0.5 | NA | NA | 2 | 0.5 | 4 | 0.5 |
| 11_7307551 | 1 | 0 | 1 | 1 | 2 | 1 | NA | NA | 4 | 0.75 | NA | NA | 3 | 1 | 1 | 0 | 2 | 1 | 6 | 0.8333 |
| 11_7307554 | 1 | 0 | NA | NA | 2 | 1 | NA | NA | 3 | 0.6667 | NA | NA | 2 | 0.5 | 1 | 1 | 2 | 0.5 | 5 | 0.6 |
| 11_7307564 | 1 | 1 | 1 | 0 | 2 | 0.5 | NA | NA | 4 | 0.5 | NA | NA | 2 | 0.5 | 1 | 0 | 1 | 1 | 4 | 0.5 |
| 11_7307573 | 1 | 1 | NA | NA | 2 | 0.5 | NA | NA | 3 | 0.6667 | NA | NA | 3 | 1 | 1 | 0 | 2 | 1 | 6 | 0.8333 |
| 11_7307583 | 1 | 0 | NA | NA | 2 | 0.5 | NA | NA | 3 | 0.3333 | NA | NA | 2 | 0.5 | 1 | 0 | 2 | 0 | 5 | 0.2 |
| 11_7311475 | NA | NA | NA | NA | 1 | 1 | NA | NA | 1 | 1 | 1 | 1 | NA | NA | 1 | 1 | NA | NA | 2 | 1 |
| 11_7311488 | NA | NA | NA | NA | 1 | 1 | 1 | 1 | 2 | 1 | 1 | 1 | NA | NA | 1 | 1 | 1 | 1 | 3 | 1 |
| 11_7311492 | NA | NA | NA | NA | NA | NA | 1 | 1 | 1 | 1 | NA | NA | NA | NA | NA | NA | 1 | 1 | 1 | 1 |
| 11_7311520 | NA | NA | NA | NA | 1 | 1 | 1 | 1 | 2 | 1 | 1 | 1 | NA | NA | 1 | 1 | 1 | 1 | 3 | 1 |
| 11_7319621 | 2 | 1 | NA | NA | 1 | 1 | 2 | 0.5 | 5 | 0.8 | NA | NA | 1 | 1 | 1 | 1 | 2 | 1 | 4 | 1 |
| 11_7319631 | 3 | 0.3333 | NA | NA | 4 | 0 | 2 | 0.5 | 9 | 0.2222 | 1 | 0 | 1 | 1 | 2 | 0 | 3 | 0 | 7 | 0.1429 |
| 11_7319638 | 3 | 0.3333 | NA | NA | 4 | 0.5 | 2 | 0.5 | 9 | 0.4444 | 1 | 1 | 1 | 0 | 2 | 0.5 | 2 | 0.5 | 6 | 0.5 |
| 11_7321524 | 2 | 0.5 | NA | NA | 3 | 1 | NA | NA | 5 | 0.8 | NA | NA | 1 | 1 | 2 | 1 | 3 | 0.3333 | 6 | 0.6667 |
| 11_7321537 | 1 | 0 | NA | NA | 3 | 0.3333 | NA | NA | 4 | 0.25 | 1 | 0 | 1 | 0 | 2 | 1 | 2 | 0.5 | 6 | 0.5 |
| 11_7427576 | 8 | 0.25 | 12 | 0.25 | 7 | 0 | 9 | 0.1111 | 36 | 0.1667 | 3 | 0.6667 | 11 | 0.1818 | 5 | 0.2 | 7 | 0.2857 | 26 | 0.2692 |
| 11_7427585 | 8 | 0.5 | 11 | 0.1818 | 8 | 0 | 10 | 0.5 | 37 | 0.2973 | 4 | 0.5 | 10 | 0.4 | 5 | 0.6 | 7 | 0.7143 | 26 | 0.5385 |
| 11_7427792 | 4 | 0.25 | 8 | 0 | 5 | 0 | 3 | 0 | 20 | 0.05 | 3 | 0.3333 | 5 | 0 | 2 | 0 | 3 | 0 | 13 | 0.0769 |
| 11_7427801 | 3 | 0 | 5 | 0.4 | 2 | 1 | 2 | 0 | 12 | 0.3333 | 2 | 1 | 3 | 0.6667 | 1 | 1 | 3 | 0.6667 | 9 | 0.7778 |
| 11_7427806 | 3 | 0.3333 | 6 | 0.1667 | 1 | 1 | 1 | 0 | 11 | 0.2727 | 2 | 0.5 | 4 | 0 | 1 | 0 | 2 | 0.5 | 9 | 0.2222 |
| 11_7452905 | 2 | 1 | 1 | 1 | NA | NA | 1 | 1 | 4 | 1 | 1 | 0 | 3 | 1 | 1 | 1 | NA | NA | 5 | 0.8 |
| 11_7452916 | 2 | 1 | 1 | 1 | NA | NA | 1 | 1 | 4 | 1 | 1 | 1 | 3 | 1 | 1 | 1 | NA | NA | 5 | 1 |
| 11_7452919 | 1 | 1 | 1 | 1 | NA | NA | NA | NA | 2 | 1 | 1 | 0 | 2 | 0.5 | 2 | 0 | NA | NA | 5 | 0.2 |
| 11_7452928 | 2 | 0 | 1 | 1 | NA | NA | 1 | 0 | 4 | 0.25 | NA | NA | 3 | 0.3333 | 2 | 0.5 | NA | NA | 5 | 0.4 |
| 11_7452960 | 2 | 1 | 2 | 1 | NA | NA | 3 | 1 | 7 | 1 | 2 | 1 | 5 | 1 | 1 | 1 | NA | NA | 8 | 1 |
| 11_7452961 | 2 | 0.5 | 3 | 0 | NA | NA | 3 | 0 | 8 | 0.125 | 2 | 0.5 | 5 | 0 | 2 | 0 | NA | NA | 9 | 0.1111 |
| 11_7452974 | 1 | 1 | 2 | 0 | NA | NA | 3 | 0 | 6 | 0.1667 | 1 | 0 | 4 | 0.25 | 1 | 0 | NA | NA | 6 | 0.1667 |
| 11_7453105 | 4 | 0 | 4 | 0.25 | 1 | 0 | 4 | 0 | 13 | 0.0769 | 3 | 0.6667 | 5 | 0 | 4 | 0 | NA | NA | 12 | 0.1667 |
| 11_7453174 | 4 | 0 | 4 | 0.25 | 2 | 0 | 4 | 0.25 | 14 | 0.1429 | 5 | 0 | 7 | 0.2857 | 4 | 0.25 | NA | NA | 16 | 0.1875 |
| 11_7453180 | 4 | 0.5 | 5 | 0 | 2 | 0 | 4 | 0 | 15 | 0.1333 | 5 | 0.2 | 7 | 0.1429 | 4 | 0.25 | NA | NA | 16 | 0.1875 |
| 11_7453182 | 4 | 0.25 | 6 | 0.6667 | 1 | 0 | 4 | 0.25 | 15 | 0.4 | 5 | 0.4 | 5 | 0.2 | 4 | 0.5 | NA | NA | 14 | 0.3571 |
| 11_7453188 | 3 | 0 | 6 | 0 | 2 | 0.5 | 5 | 0.2 | 16 | 0.125 | 5 | 0.2 | 7 | 0.1429 | 3 | 0 | NA | NA | 15 | 0.1333 |
| 11_7453192 | 5 | 0.2 | 5 | 0.2 | 1 | 0 | 3 | 0.3333 | 14 | 0.2143 | 5 | 0.2 | 7 | 0.1429 | 3 | 0.3333 | NA | NA | 15 | 0.2 |
| 11_7455238 | 2 | 1 | NA | NA | 1 | 0 | 2 | 0.5 | 5 | 0.6 | 1 | 1 | 1 | 1 | NA | NA | 3 | 0.6667 | 5 | 0.8 |
| 11_7455246 | 2 | 1 | NA | NA | 1 | 1 | 2 | 1 | 5 | 1 | 1 | 0 | 2 | 1 | NA | NA | 2 | 1 | 5 | 0.8 |
| 11_7455258 | 2 | 1 | NA | NA | 1 | 0 | 2 | 0.5 | 5 | 0.6 | 1 | 0 | 2 | 1 | 1 | 0 | 4 | 0.5 | 8 | 0.5 |
| 11_7455605 | 2 | 0 | 1 | 0 | 1 | 1 | 4 | 0.5 | 8 | 0.375 | 1 | 1 | 2 | 0 | 1 | 0 | 4 | 0 | 8 | 0.125 |
| 11_7456024 | 5 | 0 | 4 | 0 | 5 | 0.2 | 6 | 0.3333 | 20 | 0.15 | 5 | 0.2 | 6 | 0.3333 | 5 | 0.4 | 4 | 0 | 20 | 0.25 |
| 11_7456039 | 7 | 0.2857 | 5 | 0 | 6 | 0.5 | 5 | 0 | 23 | 0.2174 | 6 | 0.6667 | 3 | 0.3333 | 5 | 0.4 | 5 | 0.2 | 19 | 0.4211 |
| 11_7456045 | 5 | 0.8 | 3 | 0.6667 | 4 | 0.75 | 4 | 0.75 | 16 | 0.75 | 5 | 1 | 4 | 1 | 3 | 1 | 6 | 0.6667 | 18 | 0.8889 |
| 11_7456154 | 11 | 0.2727 | 11 | 0.3636 | 8 | 0.375 | 10 | 0.2 | 40 | 0.3 | 10 | 0.3 | 8 | 0.25 | 9 | 0.1111 | 8 | 0.375 | 35 | 0.2571 |
| 11_7456506 | 37 | 0.0811 | 54 | 0.1667 | 33 | 0.0909 | 50 | 0.18 | 174 | 0.1379 | 26 | 0.1538 | 34 | 0.0588 | 47 | 0.0213 | 39 | 0.0769 | 146 | 0.0685 |
| 11_7456519 | 35 | 0.3143 | 42 | 0.4048 | 28 | 0.3571 | 48 | 0.5833 | 153 | 0.4314 | 17 | 0.3529 | 27 | 0.3704 | 42 | 0.3095 | 35 | 0.4 | 121 | 0.3554 |
| 11_7456542 | 34 | 0.1765 | 39 | 0.2051 | 25 | 0.16 | 38 | 0.2632 | 136 | 0.2059 | 15 | 0.4667 | 25 | 0.28 | 34 | 0.2059 | 27 | 0.2593 | 101 | 0.2772 |
| 11_7456595 | 29 | 0.0345 | 41 | 0.0732 | 30 | 0.0667 | 37 | 0.1081 | 137 | 0.073 | 17 | 0 | 29 | 0.0345 | 33 | 0.0606 | 28 | 0.1071 | 107 | 0.0561 |
| 11_7456598 | 29 | 0.0345 | 39 | 0.1538 | 32 | 0.125 | 37 | 0.027 | 137 | 0.0876 | 17 | 0.0588 | 27 | 0.1111 | 32 | 0.125 | 28 | 0.1429 | 104 | 0.1154 |
| 11_7456752 | 13 | 0.2308 | 15 | 0.0667 | 15 | 0.1333 | 18 | 0.1667 | 61 | 0.1475 | 10 | 0.2 | 15 | 0 | 15 | 0.2 | 18 | 0.1667 | 58 | 0.1379 |
| 11_7456771 | 17 | 0.1176 | 19 | 0.2632 | 18 | 0.1667 | 22 | 0.0909 | 76 | 0.1579 | 8 | 0.25 | 16 | 0.125 | 15 | 0.3333 | 20 | 0.3 | 59 | 0.2542 |
| 11_7459020 | 16 | 0.125 | 14 | 0.1429 | 10 | 0.1 | 15 | 0.0667 | 55 | 0.1091 | 9 | 0 | 9 | 0.1111 | 8 | 0 | 15 | 0.2 | 41 | 0.0976 |
| 11_7470804 | 1 | 0 | 3 | 0 | 1 | 1 | NA | NA | 5 | 0.2 | NA | NA | 3 | 0 | 3 | 0 | 2 | 0.5 | 8 | 0.125 |
| 11_7470829 | 1 | 0 | 3 | 0.3333 | 1 | 0 | NA | NA | 5 | 0.2 | NA | NA | 2 | 0 | 2 | 0.5 | 2 | 0.5 | 6 | 0.3333 |

| | | | | | | | | | | | | | | | | | | | | |
|---|---|---|---|---|---|---|---|---|---|---|---|---|---|---|---|---|---|---|---|---|
| 11_747086 | 2 | 0 | 5 | 0.2 | 3 | 0 | NA | NA | 10 | 0.1 | 3 | 0 | 4 | 0 | 4 | 0.25 | 4 | 0 | 15 | 0.0667 |
| 11_747189 | 2 | 1 | 1 | 0 | NA | NA | 1 | 0 | 4 | 0.5 | NA | NA | 1 | 0 | 1 | 1 | NA | NA | 2 | 0.5 |
| 11_747362 | 2 | 0.5 | 6 | 0.3333 | 2 | 0 | 2 | 0 | 12 | 0.25 | NA | NA | 4 | 0 | 5 | 0 | 3 | 0.3333 | 12 | 0.0833 |
| 11_747402 | 6 | 0.6667 | 13 | 0.6923 | 3 | 0.6667 | 7 | 0.2857 | 29 | 0.5862 | 5 | 0.6 | 9 | 0.4444 | 9 | 0.4444 | 8 | 0 | 31 | 0.3548 |
| 11_747408 | 4 | 0.25 | 10 | 0.2 | 2 | 0.5 | 5 | 0 | 21 | 0.1905 | 5 | 0.4 | 5 | 0 | 6 | 0.3333 | 7 | 0 | 23 | 0.1739 |
| 11_747410 | 5 | 0.2 | 9 | 0.1111 | 2 | 0 | 5 | 0 | 21 | 0.0952 | 5 | 0.2 | 4 | 0.25 | 6 | 0.1667 | 6 | 0.3333 | 21 | 0.2381 |
| 11_747412 | 6 | 0.8333 | 7 | 0.2857 | 2 | 1 | 7 | 0.7143 | 22 | 0.6364 | 4 | 0.75 | 5 | 0.6 | 8 | 0.875 | 5 | 0.4 | 22 | 0.6818 |
| 11_747414 | 4 | 0.75 | 7 | 0.2857 | 1 | 1 | 5 | 0.4 | 17 | 0.4706 | 2 | 0.5 | 5 | 0.2 | 8 | 0.125 | 6 | 0.5 | 21 | 0.2857 |
| 11_747419 | 4 | 0 | 8 | 0.125 | 2 | 0 | 7 | 0.1429 | 21 | 0.0952 | 2 | 0.5 | 4 | 0 | 6 | 0 | 5 | 0 | 17 | 0.0588 |
| 11_747452 | 4 | 0 | 6 | 0.3333 | 1 | 0 | 4 | 0.75 | 15 | 0.3333 | 3 | 0 | 4 | 0.25 | 6 | 0.1667 | 2 | 0 | 15 | 0.1333 |
| 11_747911 | 1 | 0 | 1 | 1 | 1 | 0 | 1 | 1 | 4 | 0.5 | NA | NA | 5 | 0.2 | 2 | 0 | 2 | 0 | 9 | 0.1111 |
| 11_747912 | 1 | 0 | 1 | 1 | 1 | 0 | 1 | 0 | 4 | 0.25 | NA | NA | 5 | 0.4 | 2 | 0 | 2 | 0.5 | 9 | 0.3333 |
| 11_747912 | 1 | 1 | 1 | 1 | 1 | 1 | 1 | 1 | 4 | 1 | NA | NA | 5 | 0.4 | 1 | 1 | 2 | 1 | 8 | 0.625 |
| 11_747913 | NA | NA | 1 | 0 | NA | NA | 1 | 1 | 2 | 0.5 | NA | NA | 5 | 0 | 2 | 0 | 2 | 0.5 | 9 | 0.1111 |
| 11_747953 | 9 | 0.1111 | 6 | 0 | 9 | 0 | 12 | 0.0833 | 36 | 0.0556 | 2 | 0 | 9 | 0.1111 | 9 | 0.1111 | 6 | 0 | 26 | 0.0769 |
| 11_747958 | 7 | 0 | 5 | 0.2 | 9 | 0 | 9 | 0.1111 | 30 | 0.0667 | 4 | 0 | 8 | 0 | 8 | 0.125 | 5 | 0.2 | 25 | 0.08 |
| 11_747959 | 9 | 0.3333 | 5 | 0.2 | 9 | 0.2222 | 10 | 0.2 | 33 | 0.2424 | 4 | 0.5 | 8 | 0 | 8 | 0 | 5 | 0.2 | 25 | 0.12 |
| 11_747960 | 9 | 0 | 5 | 0 | 7 | 0 | 10 | 0.2 | 31 | 0.0645 | 4 | 0.25 | 8 | 0.25 | 9 | 0 | 5 | 0.2 | 26 | 0.1538 |
| 11_747966 | 7 | 0.7143 | 3 | 1 | 8 | 0.5 | 9 | 0.8889 | 27 | 0.7407 | 1 | 1 | 5 | 1 | 8 | 0.875 | 5 | 0.6 | 19 | 0.8421 |
| 11_747967 | 9 | 0.1111 | 3 | 0.3333 | 9 | 0 | 10 | 0.1 | 31 | 0.0968 | 3 | 0 | 5 | 0 | 9 | 0.1111 | 5 | 0.2 | 22 | 0.0909 |
| 11_747969 | 8 | 0.625 | 1 | 1 | 5 | 0.4 | 9 | 1 | 23 | 0.7391 | 3 | 0.6667 | 4 | 0.75 | 7 | 0.5714 | 5 | 0.8 | 19 | 0.6842 |
| 11_747969 | 7 | 0.5714 | 3 | 0.3333 | 3 | 0.3333 | 8 | 0 | 21 | 0.2857 | 2 | 1 | 7 | 0.1429 | 4 | 0.5 | 3 | 0.3333 | 16 | 0.375 |
| 11_747969 | 9 | 0.4444 | 2 | 0 | 6 | 0 | 10 | 0 | 27 | 0.1481 | 3 | 0.3333 | 7 | 0.1429 | 7 | 0.2857 | 3 | 0.3333 | 20 | 0.25 |
| 11_747971 | 8 | 0.125 | 3 | 0.3333 | 5 | 0 | 8 | 0.125 | 24 | 0.125 | 2 | 0 | 6 | 0 | 7 | 0.1429 | 3 | 0 | 18 | 0.0556 |
| 11_747991 | 2 | 1 | 2 | 0.5 | 1 | 0 | 3 | 0.3333 | 8 | 0.5 | 1 | 1 | 4 | 1 | 4 | 1 | 2 | 0.5 | 11 | 0.9091 |
| 11_747992 | 2 | 1 | 3 | 1 | 1 | 1 | 2 | 1 | 8 | 1 | 1 | 0 | 5 | 1 | 4 | 1 | 3 | 0.6667 | 13 | 0.8462 |
| 11_747993 | 5 | 0.2 | 9 | 0.1111 | 5 | 0.2 | 10 | 0.3 | 29 | 0.2069 | 3 | 0 | 8 | 0.125 | 7 | 0.2857 | 8 | 0.625 | 26 | 0.3077 |
| 11_747994 | 2 | 0.5 | 3 | 0 | 2 | 0 | 2 | 0.5 | 9 | 0.2222 | 1 | 0 | 5 | 0 | 5 | 0.2 | 2 | 0 | 13 | 0.0769 |
| 11_747995 | 1 | 1 | 3 | 0.6667 | 3 | 0.3333 | 3 | 0.6667 | 10 | 0.6 | 1 | 0 | 5 | 0.6 | 4 | 0.75 | 1 | 1 | 11 | 0.6364 |
| 11_748104 | 2 | 0 | 2 | 0.5 | NA | NA | NA | NA | 4 | 0.25 | 1 | 1 | 1 | 1 | 3 | 0.3333 | 3 | 0.3333 | 8 | 0.5 |
| 11_748210 | 2 | 0 | 1 | 1 | 1 | 1 | NA | NA | 4 | 0.5 | NA | NA | NA | NA | 1 | 1 | NA | NA | 1 | 1 |
| 11_748210 | 2 | 0.5 | 1 | 0 | 2 | 0.5 | NA | NA | 5 | 0.4 | 1 | 1 | NA | NA | 1 | 1 | NA | NA | 2 | 1 |
| 11_748211 | 1 | 1 | 1 | 1 | 2 | 1 | NA | NA | 4 | 1 | 1 | 1 | NA | NA | 1 | 1 | NA | NA | 2 | 1 |
| 11_748213 | 2 | 0.5 | 1 | 0 | 1 | 1 | NA | NA | 4 | 0.5 | 1 | 1 | NA | NA | 1 | 1 | NA | NA | 2 | 1 |
| 11_750537 | 2 | 0.5 | 3 | 0.3333 | 2 | 0 | 1 | 1 | 8 | 0.375 | 2 | 0.5 | NA | NA | 2 | 0.5 | 1 | 0 | 5 | 0.4 |
| 11_750539 | 2 | 0.5 | 3 | 0 | 2 | 0 | 1 | 0 | 8 | 0.125 | 1 | 0 | NA | NA | 2 | 0.5 | 1 | 0 | 4 | 0.25 |
| 11_750539 | 2 | 0.5 | 3 | 0 | 3 | 0.3333 | 1 | 0 | 9 | 0.2222 | 2 | 0.5 | NA | NA | 2 | 0 | 1 | 0 | 5 | 0.2 |
| 11_750550 | 2 | 0 | 5 | 0.4 | 4 | 0.25 | 1 | 0 | 12 | 0.25 | 3 | 0.3333 | 2 | 0 | 5 | 0 | 1 | 0 | 11 | 0.0909 |
| 11_750563 | 1 | 0 | 2 | 0.5 | NA | NA | NA | NA | 3 | 0.3333 | 1 | 1 | NA | NA | NA | NA | NA | NA | 1 | 1 |
| 11_750564 | 1 | 1 | 1 | 1 | NA | NA | NA | NA | 2 | 1 | 1 | 1 | NA | NA | NA | NA | NA | NA | 1 | 1 |
| 11_750565 | 1 | 1 | 2 | 0.5 | NA | NA | NA | NA | 3 | 0.6667 | 1 | 1 | NA | NA | NA | NA | NA | NA | 1 | 1 |
| 11_750637 | 2 | 0 | NA | NA | NA | NA | 1 | 1 | 3 | 0.3333 | 1 | 0 | 1 | 1 | 2 | 0 | NA | NA | 4 | 0.25 |
| 11_750638 | 2 | 0.5 | NA | NA | NA | NA | NA | NA | 2 | 0.5 | 1 | 0 | 1 | 1 | 2 | 0 | NA | NA | 4 | 0.25 |
| 11_750638 | 2 | 0.5 | NA | NA | NA | NA | 1 | 0 | 3 | 0.3333 | 1 | 0 | 1 | 1 | 2 | 0 | NA | NA | 4 | 0.25 |
| 11_750639 | 2 | 1 | NA | NA | 1 | 1 | 1 | 1 | 4 | 1 | 1 | 1 | 1 | 1 | 1 | 1 | NA | NA | 3 | 1 |
| 11_750639 | 2 | 0.5 | NA | NA | 1 | 1 | NA | NA | 3 | 0.6667 | 1 | 1 | 1 | 1 | 1 | 1 | NA | NA | 3 | 1 |
| 11_750640 | 1 | 1 | NA | NA | 1 | 1 | 1 | 1 | 3 | 1 | 1 | 1 | 1 | 1 | 1 | 1 | NA | NA | 3 | 1 |
| 11_750641 | 2 | 1 | NA | NA | 1 | 1 | 1 | 1 | 4 | 1 | 1 | 1 | 1 | 1 | 2 | 1 | NA | NA | 4 | 1 |
| 11_750642 | 2 | 1 | NA | NA | NA | NA | NA | NA | 2 | 1 | 1 | 1 | NA | NA | 1 | 1 | NA | NA | 2 | 1 |
| 11_750644 | 2 | 0.5 | NA | NA | 1 | 0 | 1 | 0 | 4 | 0.25 | NA | NA | 1 | 1 | 2 | 1 | NA | NA | 3 | 1 |
| 11_750647 | 2 | 1 | 1 | 1 | 2 | 1 | 2 | 1 | 7 | 1 | 2 | 1 | 1 | 1 | 4 | 1 | NA | NA | 7 | 1 |
| 11_750648 | 2 | 0.5 | 1 | 0 | 2 | 0.5 | 2 | 0 | 7 | 0.2857 | 1 | 0 | 1 | 0 | 3 | 0.3333 | NA | NA | 5 | 0.2 |
| 11_750650 | 2 | 0.5 | 1 | 0 | 3 | 0.3333 | 2 | 0 | 8 | 0.25 | 2 | 0.5 | NA | NA | 4 | 0 | NA | NA | 6 | 0.1667 |
| 11_750652 | 2 | 0.5 | 1 | 0 | 3 | 0.3333 | 1 | 1 | 7 | 0.4286 | 2 | 1 | 1 | 1 | 4 | 0.5 | 1 | 1 | 8 | 0.75 |
| 11_750655 | 3 | 0.3333 | 1 | 0 | 3 | 0 | 1 | 0 | 8 | 0.125 | 1 | 0 | 1 | 1 | 4 | 0.25 | 1 | 1 | 7 | 0.4286 |
| 11_750670 | 1 | 0 | 4 | 0.25 | 1 | 0 | 4 | 0 | 10 | 0.1 | 3 | 0 | 2 | 0 | 3 | 0 | 6 | 0.5 | 14 | 0.2143 |
| 11_750684 | 2 | 1 | 3 | 0.6667 | 1 | 0 | 3 | 0.3333 | 9 | 0.5556 | 1 | 0 | 1 | 1 | 2 | 0 | 5 | 0.6 | 9 | 0.4444 |
| 11_750685 | 2 | 0 | 4 | 0.25 | 7 | 0.4286 | 5 | 0.2 | 18 | 0.2778 | 2 | 0 | 6 | 0.5 | 5 | 0.8 | 7 | 0.5714 | 20 | 0.55 |
| 11_750685 | 2 | 1 | 4 | 0.25 | 8 | 0.125 | 5 | 0.2 | 19 | 0.2632 | 4 | 0.25 | 6 | 0.3333 | 5 | 0.2 | 7 | 0.4286 | 22 | 0.3182 |
| 11_750686 | 2 | 0.5 | 3 | 0.6667 | 7 | 1 | 5 | 0.4 | 17 | 0.7059 | 2 | 0 | 7 | 0.8571 | 5 | 0.4 | 7 | 0.8571 | 21 | 0.6667 |
| 11_750686 | 2 | 0 | 4 | 0.25 | 7 | 0 | 5 | 0 | 18 | 0.0556 | 1 | 0 | 5 | 0.2 | 5 | 0.2 | 8 | 0 | 19 | 0.1053 |
| 11_750687 | 2 | 1 | 4 | 0.25 | 8 | 0.5 | 6 | 0.8333 | 20 | 0.6 | 2 | 1 | 6 | 0.8333 | 4 | 0.25 | 8 | 0.875 | 20 | 0.75 |
| 11_750687 | 2 | 0 | 3 | 0.6667 | 8 | 0.25 | 5 | 0.6 | 18 | 0.3889 | 2 | 0 | 8 | 0.625 | 5 | 0.4 | 8 | 0.75 | 23 | 0.5652 |
| 11_750689 | 1 | 1 | 3 | 1 | 8 | 0.25 | 4 | 1 | 16 | 0.625 | 3 | 0.3333 | 7 | 0.4286 | 5 | 0.2 | 7 | 0.7143 | 22 | 0.4545 |
| 11_750690 | 1 | 1 | 1 | 1 | 5 | 1 | 5 | 1 | 12 | 1 | 2 | 1 | 6 | 1 | 4 | 1 | 5 | 1 | 17 | 1 |

| | | | | | | | | | | | | | | | | | | | | |
|---|---|---|---|---|---|---|---|---|---|---|---|---|---|---|---|---|---|---|---|---|
| 11_7515565 | 3 | 0.3333 | 1 | 1 | 3 | 0.3333 | 2 | 0 | 9 | 0.3333 | 3 | 1 | 2 | 0.5 | 4 | 0.25 | 2 | 0 | 11 | 0.4545 |
| 11_7515567 | 3 | 0.6667 | 1 | 0 | 3 | 0.3333 | 2 | 1 | 9 | 0.5556 | 2 | 1 | 3 | 0 | 3 | 0 | 2 | 0 | 10 | 0.2 |
| 11_7515573 | 3 | 0.3333 | 3 | 0.3333 | 3 | 0 | 2 | 0.5 | 11 | 0.2727 | 3 | 0.3333 | 3 | 0.3333 | 4 | 0.75 | 2 | 0.5 | 12 | 0.5 |
| 11_7515575 | 3 | 0.3333 | 3 | 0 | 3 | 0 | 2 | 0 | 11 | 0.0909 | 3 | 0.3333 | 2 | 0.5 | 4 | 0 | 1 | 0 | 10 | 0.2 |
| 11_7515597 | 2 | 1 | 3 | 1 | 3 | 1 | 1 | 1 | 9 | 1 | 1 | 1 | 4 | 1 | 3 | 1 | 2 | 1 | 10 | 1 |
| 11_7515604 | 3 | 0 | 3 | 0 | 3 | 0.3333 | 3 | 0 | 12 | 0.0833 | 3 | 0 | 4 | 0.25 | 4 | 0 | 2 | 0 | 13 | 0.0769 |
| 11_7515608 | 3 | 0.3333 | 3 | 0.6667 | 3 | 0.3333 | 3 | 0.3333 | 12 | 0.4167 | 3 | 0.3333 | 4 | 0.5 | 3 | 0.6667 | 2 | 1 | 12 | 0.5833 |
| 11_7515616 | 2 | 0.5 | 3 | 0.3333 | 3 | 0 | 3 | 0 | 11 | 0.1818 | 2 | 0.5 | 3 | 0 | 3 | 0.3333 | 2 | 0 | 10 | 0.2 |
| 11_7515628 | 2 | 1 | 4 | 0.75 | 3 | 0.6667 | 3 | 0.6667 | 12 | 0.75 | 3 | 0.3333 | 3 | 0.6667 | 3 | 0.6667 | 2 | 0.5 | 11 | 0.5455 |
| 11_7515634 | 3 | 1 | 4 | 1 | 2 | 1 | 2 | 1 | 11 | 1 | 2 | 1 | 4 | 1 | 4 | 1 | 2 | 1 | 12 | 1 |
| 11_7515646 | 4 | 0 | 5 | 0.4 | 3 | 0 | 3 | 0.3333 | 15 | 0.2 | 3 | 0.3333 | 5 | 0.2 | 4 | 0 | 2 | 0.5 | 14 | 0.2143 |
| 11_7515647 | 4 | 0.25 | 5 | 0.2 | 3 | 0 | 2 | 0 | 14 | 0.1429 | 3 | 0 | 4 | 0 | 4 | 0.25 | 3 | 0 | 14 | 0.0714 |
| 11_7515658 | 3 | 0.6667 | 4 | 0.5 | 3 | 0.3333 | 2 | 0.5 | 12 | 0.5 | 3 | 0.6667 | 2 | 0.5 | 4 | 0.75 | 3 | 1 | 12 | 0.75 |
| 11_7515664 | 3 | 1 | 5 | 0.4 | 3 | 0.6667 | 2 | 0.5 | 13 | 0.6154 | 3 | 0.3333 | 4 | 1 | 3 | 1 | 2 | 0 | 12 | 0.6667 |
| 11_7515670 | 2 | 0 | 5 | 0 | 2 | 0.5 | 2 | 0.5 | 11 | 0.1818 | 3 | 0 | 4 | 0.25 | 4 | 0.75 | 3 | 0 | 14 | 0.2857 |
| 11_7515671 | 4 | 0.75 | 4 | 1 | 4 | 1 | 4 | 1 | 16 | 0.9375 | 3 | 1 | 5 | 1 | 4 | 0.75 | 3 | 1 | 15 | 0.9333 |
| 11_7515680 | 5 | 0.2 | 5 | 0.2 | 3 | 0.3333 | 4 | 0.25 | 17 | 0.2353 | 3 | 0 | 4 | 0.25 | 2 | 0 | 3 | 0.3333 | 12 | 0.1667 |
| 11_7515691 | 4 | 0.5 | 6 | 0.1667 | 4 | 0.25 | 3 | 0.3333 | 17 | 0.2941 | 3 | 1 | 5 | 0.6 | 4 | 0.5 | 3 | 0.3333 | 15 | 0.6 |
| 11_7515695 | 2 | 0 | 6 | 0 | 3 | 0.3333 | 4 | 0.5 | 15 | 0.2 | 2 | 0.5 | 2 | 0.5 | 3 | 0.6667 | 3 | 0.3333 | 10 | 0.5 |
| 11_7515706 | 5 | 1 | 7 | 0.8571 | 4 | 0.75 | 5 | 0.8 | 21 | 0.8571 | 3 | 1 | 4 | 1 | 4 | 0.75 | 3 | 1 | 14 | 0.9286 |
| 11_7515715 | 5 | 1 | 5 | 0.8 | 5 | 0.6 | 4 | 1 | 19 | 0.8421 | 3 | 1 | 5 | 0.8 | 5 | 0.8 | 3 | 0.3333 | 16 | 0.75 |
| 11_7515721 | 5 | 0.8 | 7 | 0.8571 | 5 | 1 | 5 | 0.8 | 22 | 0.8636 | 3 | 0.6667 | 5 | 1 | 5 | 1 | 2 | 1 | 15 | 0.9333 |
| 11_7515727 | 5 | 1 | 7 | 0.4286 | 6 | 1 | 5 | 0.8 | 23 | 0.7826 | 3 | 1 | 5 | 0.6 | 4 | 0.75 | 3 | 1 | 15 | 0.8 |
| 11_7515730 | 5 | 1 | 6 | 1 | 7 | 1 | 3 | 1 | 21 | 1 | 3 | 1 | 4 | 1 | 5 | 1 | 2 | 1 | 14 | 1 |
| 11_7515731 | 4 | 0.5 | 5 | 0 | 3 | 0 | 3 | 0.6667 | 15 | 0.2667 | 2 | 0 | 2 | 1 | 4 | 0.5 | 1 | 1 | 9 | 0.5556 |
| 11_7515753 | 5 | 1 | 7 | 1 | 7 | 1 | 3 | 1 | 22 | 1 | 2 | 1 | 5 | 1 | 5 | 1 | 2 | 1 | 14 | 1 |
| 11_7515768 | 7 | 0 | 5 | 0.2 | 8 | 0.125 | 6 | 0 | 26 | 0.0769 | 3 | 0 | 4 | 0.75 | 6 | 0.1667 | 3 | 0.3333 | 16 | 0.3125 |
| 11_7515771 | 7 | 0.2857 | 6 | 0 | 8 | 0 | 7 | 0 | 28 | 0.0714 | 3 | 0 | 6 | 0.1667 | 6 | 0.3333 | 3 | 0 | 18 | 0.1667 |
| 11_7515774 | 6 | 0.8333 | 8 | 1 | 7 | 1 | 5 | 0.8 | 26 | 0.9231 | 3 | 1 | 5 | 1 | 5 | 1 | 3 | 1 | 16 | 1 |
| 11_7515778 | 6 | 0.1667 | 7 | 0.2857 | 8 | 0.5 | 6 | 0.1667 | 27 | 0.2963 | 3 | 0.3333 | 4 | 0.25 | 4 | 0 | 3 | 0.3333 | 14 | 0.2143 |
| 11_7515813 | 6 | 0.1667 | 8 | 0 | 8 | 0.125 | 5 | 0 | 27 | 0.0741 | 6 | 0 | 3 | 0 | 7 | 0.2857 | 3 | 0 | 19 | 0.1053 |
| 11_7519194 | 13 | 0.1538 | 8 | 0 | 3 | 0.3333 | 5 | 0.2 | 29 | 0.1379 | 7 | 0 | 5 | 0 | 8 | 0.125 | 6 | 0.5 | 26 | 0.1538 |
| 11_7519197 | 11 | 0.3636 | 8 | 0.375 | 2 | 0.5 | 5 | 0.6 | 26 | 0.4231 | 4 | 0.25 | 5 | 0.4 | 8 | 0.5 | 5 | 0.4 | 22 | 0.4091 |
| 11_7519199 | 11 | 0.7273 | 7 | 0.5714 | 4 | 0.5 | 5 | 0.2 | 27 | 0.5556 | 8 | 0.875 | 4 | 1 | 7 | 0.8571 | 5 | 0.6 | 24 | 0.8333 |
| 11_7519205 | 13 | 0.1538 | 6 | 0.3333 | 3 | 0.3333 | 5 | 0 | 27 | 0.1852 | 7 | 0 | 5 | 0.2 | 8 | 0.25 | 4 | 0 | 24 | 0.125 |
| 11_7519217 | 13 | 0 | 8 | 0.125 | 4 | 0.25 | 5 | 0.4 | 30 | 0.1333 | 7 | 0.2857 | 4 | 0.25 | 8 | 0 | 6 | 0.1667 | 25 | 0.16 |
| 11_7528128 | NA | NA | 1 | 1 | NA | NA | NA | NA | 1 | 1 | 1 | 1 | 3 | 0.6667 | NA | NA | NA | NA | 4 | 0.75 |
| 11_7528151 | NA | NA | 1 | 1 | NA | NA | NA | NA | 1 | 1 | 2 | 0.5 | 3 | 0.6667 | NA | NA | NA | NA | 5 | 0.6 |
| 11_7528159 | NA | NA | 1 | 1 | NA | NA | NA | NA | 1 | 1 | 1 | 1 | 3 | 1 | NA | NA | NA | NA | 4 | 1 |
| 11_7528197 | NA | NA | 1 | 1 | NA | NA | NA | NA | 1 | 1 | 2 | 1 | 2 | 0.5 | NA | NA | NA | NA | 4 | 0.75 |
| 11_7529571 | NA | NA | 1 | 1 | 1 | 1 | 1 | 0 | 3 | 0.6667 | 1 | 1 | 3 | 0.3333 | 2 | 1 | 1 | 1 | 7 | 0.7143 |
| 11_7529583 | NA | NA | 1 | 1 | 2 | 0.5 | 1 | 1 | 4 | 0.75 | 3 | 1 | 3 | 0.3333 | 2 | 1 | NA | NA | 8 | 0.75 |
| 11_7529589 | NA | NA | 1 | 0 | 2 | 0.5 | 1 | 1 | 4 | 0.5 | 3 | 0 | 3 | 0.3333 | 2 | 0.5 | 1 | 0 | 9 | 0.2222 |
| 11_7529596 | NA | NA | 1 | 1 | 2 | 1 | 1 | 1 | 4 | 1 | 3 | 1 | 3 | 1 | 1 | 1 | 1 | 1 | 8 | 1 |
| 11_7529641 | NA | NA | 1 | 1 | 2 | 0 | 2 | 0 | 5 | 0.2 | 2 | 0 | 2 | 0 | 2 | 1 | 1 | 0 | 7 | 0.2857 |
| 11_7530101 | 1 | 1 | 2 | 0.5 | 3 | 0 | 2 | 0.5 | 8 | 0.375 | NA | NA | 1 | 1 | 3 | 0.6667 | 3 | 0.3333 | 7 | 0.5714 |
| 11_7530101 | 1 | 1 | 2 | 0 | 2 | 0 | 1 | 1 | 6 | 0.3333 | 1 | 1 | 1 | 0 | 2 | 0 | 3 | 0.3333 | 7 | 0.2857 |
| 11_7530105 | 1 | 1 | 2 | 0 | 2 | 0 | 1 | 0 | 6 | 0.1667 | 1 | 1 | 1 | 0 | 3 | 0 | 2 | 0 | 7 | 0.1429 |
| 11_7533216 | 2 | 0 | 3 | 0.3333 | 2 | 0 | 4 | 0.25 | 11 | 0.1818 | 2 | 0 | 2 | 0 | 6 | 0.1667 | 3 | 0 | 13 | 0.0769 |
| 11_7533219 | 2 | 0.5 | 2 | 0.5 | 2 | 0.5 | 4 | 0.25 | 10 | 0.4 | 1 | 1 | 2 | 0.5 | 6 | 0.5 | 2 | 0.5 | 11 | 0.5455 |
| 11_7533232 | 2 | 0 | 2 | 0.5 | 1 | 0 | 3 | 0 | 8 | 0.125 | 2 | 0 | 1 | 1 | 4 | 0.25 | 2 | 0 | 9 | 0.2222 |
| 11_7533241 | 2 | 0.5 | 2 | 0 | 1 | 0 | 4 | 0.25 | 9 | 0.2222 | 2 | 0 | 1 | 1 | 5 | 0.4 | 2 | 0 | 10 | 0.3 |
| 11_7533262 | 2 | 1 | 3 | 1 | 2 | 0.5 | 3 | 1 | 10 | 0.9 | NA | NA | 2 | 1 | 5 | 1 | 2 | 1 | 9 | 1 |
| 11_7533268 | 2 | 0.5 | 3 | 1 | 1 | 0 | 3 | 1 | 9 | 0.7778 | NA | NA | 2 | 1 | 5 | 0.8 | 1 | 1 | 8 | 0.875 |
| 11_7533635 | 2 | 0.5 | 3 | 0 | 1 | 0 | 2 | 0 | 8 | 0.125 | NA | NA | 1 | 0 | 3 | 0 | 1 | 1 | 5 | 0.2 |
| 11_7533639 | 2 | 0.5 | 3 | 0.6667 | 1 | 0 | 2 | 0.5 | 8 | 0.5 | NA | NA | 1 | 0 | 3 | 1 | 2 | 1 | 6 | 0.8333 |
| 11_7533640 | 2 | 0.5 | 3 | 0 | 1 | 0 | 3 | 0.6667 | 9 | 0.3333 | NA | NA | NA | NA | 4 | 0.25 | 2 | 0.5 | 6 | 0.3333 |
| 11_7533656 | 1 | 0 | 3 | 0.6667 | 1 | 1 | 2 | 0 | 7 | 0.4286 | NA | NA | NA | NA | 4 | 0.5 | 1 | 0 | 5 | 0.4 |
| 11_7534184 | 2 | 0.5 | NA | NA | 1 | 0 | 1 | 1 | 4 | 0.5 | NA | NA | NA | NA | 4 | 0.5 | 1 | 1 | 5 | 0.6 |
| 11_7534188 | 2 | 0.5 | NA | NA | 1 | 1 | NA | NA | 3 | 0.6667 | NA | NA | NA | NA | 3 | 0.3333 | 1 | 0 | 4 | 0.25 |
| 11_7534201 | 2 | 1 | NA | NA | 1 | 0 | 1 | 1 | 4 | 0.75 | NA | NA | NA | NA | 3 | 1 | NA | NA | 3 | 1 |
| 11_7534209 | 2 | 1 | NA | NA | 1 | 1 | 1 | 1 | 4 | 1 | NA | NA | NA | NA | 3 | 1 | 1 | 1 | 4 | 1 |
| 11_7534219 | 2 | 1 | NA | NA | 1 | 0 | 1 | 0 | 4 | 0.5 | NA | NA | NA | NA | 2 | 0.5 | 1 | 0 | 3 | 0.3333 |
| 11_7534278 | 2 | 0.5 | NA | NA | NA | NA | NA | NA | 2 | 0.5 | NA | NA | NA | NA | 2 | 0.5 | NA | NA | 2 | 0.5 |
| 11_7534284 | 2 | 1 | NA | NA | 1 | 0 | NA | NA | 3 | 0.6667 | NA | NA | NA | NA | 2 | 0.5 | NA | NA | 2 | 0.5 |

| | | | | | | | | | | | | | | | | | | | | |
|---|---|---|---|---|---|---|---|---|---|---|---|---|---|---|---|---|---|---|---|---|
| 11_753431 | 2 | 0.5 | NA | NA | NA | NA | NA | NA | 2 | 0.5 | NA | NA | NA | NA | 1 | 1 | NA | NA | 1 | 1 |
| 11_753433 | 2 | 1 | NA | NA | 1 | 1 | NA | NA | 3 | 1 | NA | NA | NA | NA | 1 | 1 | NA | NA | 1 | 1 |
| 11_753434 | 1 | 1 | NA | NA | 1 | 1 | NA | NA | 2 | 1 | NA | NA | NA | NA | 1 | 1 | NA | NA | 1 | 1 |
| 11_753434 | 1 | 1 | NA | NA | 1 | 0 | NA | NA | 2 | 0.5 | NA | NA | NA | NA | 1 | 1 | NA | NA | 1 | 1 |
| 11_753440 | 1 | 1 | NA | NA | 1 | 0 | NA | NA | 2 | 0.5 | NA | NA | NA | NA | 1 | 1 | NA | NA | 1 | 1 |
| 11_753562 | 2 | 0 | 5 | 0.2 | 1 | 0 | 3 | 0.3333 | 11 | 0.1818 | 5 | 0 | NA | NA | 5 | 0.4 | 4 | 0 | 14 | 0.1429 |
| 11_753562 | 2 | 0.5 | 5 | 1 | 1 | 1 | 3 | 1 | 11 | 0.9091 | 4 | 1 | NA | NA | 5 | 0.8 | 5 | 0.8 | 14 | 0.8571 |
| 11_753565 | 2 | 0 | 3 | 0.3333 | 1 | 0 | 1 | 0 | 7 | 0.1429 | 2 | 0 | NA | NA | 5 | 0.4 | 4 | 0 | 11 | 0.1818 |
| 11_753565 | 2 | 0 | 5 | 0 | 1 | 1 | 3 | 0.3333 | 11 | 0.1818 | 4 | 0.5 | 1 | 0 | 3 | 0.3333 | 4 | 0.5 | 12 | 0.4167 |
| 11_753566 | 2 | 0.5 | 4 | 0 | 1 | 0 | 2 | 0.5 | 9 | 0.2222 | 4 | 0 | 1 | 0 | 5 | 0.2 | 6 | 0.3333 | 16 | 0.1875 |
| 11_753566 | 1 | 0 | 3 | 0.3333 | NA | NA | 1 | 0 | 5 | 0.2 | 3 | 0 | 1 | 0 | 4 | 0 | 3 | 0.3333 | 11 | 0.0909 |
| 11_753567 | 2 | 0.5 | 6 | 0.8333 | 3 | 1 | 3 | 1 | 14 | 0.8571 | 4 | 1 | 1 | 1 | 4 | 0.75 | 6 | 0.6667 | 15 | 0.8 |
| 11_753569 | 3 | 0 | 7 | 0.2857 | 3 | 0.6667 | 3 | 0 | 16 | 0.25 | 5 | 0.6 | 1 | 1 | 5 | 0.6 | 5 | 0.6 | 16 | 0.625 |
| 11_753571 | 3 | 0 | 7 | 0 | 4 | 0.25 | 3 | 0 | 17 | 0.0588 | 5 | 0 | 1 | 0 | 5 | 0 | 7 | 0.4286 | 18 | 0.1667 |
| 11_753922 | 7 | 0 | 18 | 0.1111 | 8 | 0 | 5 | 0 | 38 | 0.0526 | 13 | 0.1538 | 8 | 0 | 9 | 0.1111 | 11 | 0 | 41 | 0.0732 |
| 11_754039 | NA | NA | NA | NA | NA | NA | 1 | 1 | 1 | 1 | NA | NA | NA | NA | NA | NA | 1 | 1 | 1 | 1 |
| 11_754168 | 1 | 1 | 1 | 1 | NA | NA | NA | NA | 2 | 1 | 2 | 1 | NA | NA | NA | NA | NA | NA | 2 | 1 |
| 11_754223 | 1 | 1 | 1 | 1 | NA | NA | NA | NA | 2 | 1 | NA | NA | NA | NA | NA | NA | 1 | 1 | 1 | 1 |
| 11_754633 | 2 | 1 | NA | NA | NA | NA | 1 | 1 | 3 | 1 | 3 | 0.6667 | 2 | 0.5 | 2 | 0.5 | 1 | 1 | 8 | 0.625 |
| 11_754633 | 1 | 0 | 1 | 1 | NA | NA | 1 | 1 | 3 | 0.6667 | 3 | 0 | 1 | 0 | 2 | 0.5 | 1 | 0 | 7 | 0.1429 |
| 11_754633 | 2 | 0.5 | 3 | 0 | NA | NA | NA | NA | 5 | 0.2 | 3 | 1 | 2 | 0.5 | 2 | 0.5 | 1 | 1 | 8 | 0.75 |
| 11_754634 | 2 | 1 | 3 | 0.3333 | NA | NA | 1 | 1 | 6 | 0.6667 | 2 | 1 | 1 | 1 | 3 | 1 | 2 | 1 | 8 | 1 |
| 11_754634 | 2 | 0.5 | 2 | 0.5 | NA | NA | 1 | 1 | 5 | 0.6 | 3 | 1 | 3 | 0.6667 | 3 | 1 | 2 | 0.5 | 11 | 0.8182 |
| 11_754635 | 2 | 0 | 2 | 0.5 | NA | NA | 1 | 0 | 5 | 0.2 | 3 | 0 | 1 | 1 | 3 | 0.3333 | 2 | 0 | 9 | 0.2222 |
| 11_754636 | 2 | 0.5 | 3 | 0.3333 | NA | NA | 1 | 1 | 6 | 0.5 | 3 | 0.3333 | 3 | 0.6667 | 4 | 0.75 | 2 | 0.5 | 12 | 0.5833 |
| 11_754637 | 1 | 1 | 3 | 0.3333 | NA | NA | 1 | 1 | 5 | 0.6 | 3 | 0.6667 | 3 | 0.3333 | 5 | 0.6 | 2 | 0.5 | 13 | 0.5385 |
| 11_754638 | 2 | 0.5 | 3 | 0.3333 | NA | NA | 2 | 0.5 | 7 | 0.4286 | 3 | 0.3333 | 3 | 0.6667 | 5 | 0.6 | 2 | 0.5 | 13 | 0.5385 |
| 11_754639 | 2 | 0.5 | 2 | 0.5 | NA | NA | 2 | 0 | 6 | 0.3333 | 3 | 0.3333 | 2 | 1 | 5 | 0.6 | 1 | 1 | 11 | 0.6364 |
| 11_754640 | 1 | 1 | 3 | 0 | NA | NA | 2 | 0 | 6 | 0.1667 | 3 | 1 | 3 | 1 | 4 | 0.5 | 2 | 0 | 12 | 0.6667 |
| 11_754640 | 1 | 1 | 2 | 0 | NA | NA | 1 | 0 | 4 | 0.25 | 3 | 0.3333 | 3 | 0.3333 | 4 | 0 | 2 | 0.5 | 12 | 0.25 |
| 11_754644 | 2 | 0.5 | 3 | 0 | NA | NA | 2 | 0 | 7 | 0.1429 | 4 | 0 | 4 | 0.25 | 4 | 0 | 3 | 0 | 15 | 0.0667 |
| 11_754664 | 5 | 0 | 4 | 0 | 1 | 0 | 4 | 0.25 | 14 | 0.0714 | NA | NA | 1 | 0 | 2 | 0 | 3 | 1 | 6 | 0.5 |
| 11_754667 | 3 | 0 | 4 | 0 | 1 | 0 | 3 | 0.3333 | 11 | 0.0909 | NA | NA | 1 | 0 | 3 | 0.6667 | 2 | 0 | 6 | 0.3333 |
| 11_754668 | 4 | 0 | 3 | 0 | 1 | 0 | 4 | 0.5 | 12 | 0.1667 | NA | NA | 1 | 0 | 3 | 0.3333 | 3 | 0.3333 | 7 | 0.2857 |
| 11_754677 | 5 | 0 | 4 | 0 | 3 | 0 | 5 | 0.2 | 17 | 0.0588 | 2 | 0.5 | 2 | 0 | 3 | 0 | 2 | 0 | 9 | 0.1111 |
| 11_755179 | 15 | 0.3333 | 14 | 0.0714 | 12 | 0.4167 | 8 | 0.125 | 49 | 0.2449 | 5 | 0.2 | 7 | 0.2857 | 7 | 0.2857 | 10 | 0.3 | 29 | 0.2759 |
| 11_755283 | 2 | 0 | 1 | 0 | 2 | 0.5 | 3 | 0.6667 | 8 | 0.375 | NA | NA | 2 | 0 | 2 | 0.5 | 4 | 0 | 8 | 0.125 |
| 11_755289 | 2 | 0.5 | 2 | 0 | 2 | 1 | 3 | 0.3333 | 9 | 0.4444 | NA | NA | 2 | 0.5 | 3 | 0 | 4 | 0.5 | 9 | 0.3333 |
| 11_755362 | 2 | 0 | 1 | 1 | 2 | 0.5 | 3 | 0 | 8 | 0.25 | 1 | 0 | 3 | 1 | 2 | 0 | 3 | 0.3333 | 9 | 0.4444 |
| 11_755363 | 3 | 0 | 1 | 0 | 3 | 0 | 5 | 0.2 | 12 | 0.0833 | 1 | 0 | 2 | 0.5 | 2 | 0 | 5 | 0 | 10 | 0.1 |
| 11_755364 | 4 | 0 | 1 | 0 | 3 | 0.3333 | 5 | 0.2 | 13 | 0.1538 | 1 | 0 | 2 | 0.5 | 3 | 0 | 4 | 0.25 | 10 | 0.2 |
| 11_755376 | 3 | 0 | 2 | 0 | 2 | 0.5 | 5 | 0 | 12 | 0.0833 | 1 | 0 | 2 | 0.5 | 4 | 0 | 4 | 0 | 11 | 0.0909 |
| 11_755377 | 4 | 0 | 2 | 0 | 3 | 0 | 5 | 0.2 | 14 | 0.0714 | 1 | 0 | 2 | 0.5 | 5 | 0.2 | 4 | 0 | 12 | 0.1667 |
| 11_755377 | 2 | 0 | 2 | 0 | 2 | 1 | 5 | 0.6 | 11 | 0.4545 | 1 | 1 | 3 | 0 | 5 | 0 | 2 | 0.5 | 11 | 0.1818 |
| 11_755380 | 2 | 0 | 2 | 0 | 3 | 0 | 5 | 0.2 | 12 | 0.0833 | 1 | 0 | 2 | 0 | 5 | 0.2 | 3 | 0 | 11 | 0.0909 |
| 11_755386 | 5 | 0 | 3 | 0.3333 | 3 | 0 | 6 | 0.1667 | 17 | 0.1176 | 3 | 0 | 4 | 0 | 5 | 0 | 3 | 0.3333 | 15 | 0.0667 |
| 11_755387 | 4 | 0 | 3 | 0.3333 | 2 | 0 | 5 | 0 | 14 | 0.0714 | 3 | 0 | 4 | 0.25 | 4 | 0 | 3 | 0 | 14 | 0.0714 |
| 11_755390 | 5 | 0.6 | 3 | 1 | 2 | 1 | 6 | 0.1667 | 16 | 0.5625 | 2 | 0.5 | 5 | 0.6 | 2 | 1 | 2 | 0.5 | 11 | 0.6364 |
| 11_755392 | 5 | 0.2 | 3 | 0.3333 | 3 | 0.3333 | 4 | 0 | 15 | 0.2 | 1 | 0 | 3 | 0.6667 | 3 | 0 | 3 | 0 | 10 | 0.2 |
| 11_755393 | 5 | 0.2 | 3 | 0.3333 | 3 | 0.3333 | 6 | 0.1667 | 17 | 0.2353 | 2 | 0 | 5 | 0.4 | 4 | 0.25 | 3 | 0 | 14 | 0.2143 |
| 11_755393 | 4 | 0.25 | 3 | 0.3333 | 3 | 0 | 5 | 0.2 | 15 | 0.2 | 2 | 0 | 4 | 1 | 3 | 0.3333 | 2 | 0.5 | 11 | 0.5455 |
| 11_755393 | 4 | 0.75 | NA | NA | 2 | 0.5 | 6 | 0 | 12 | 0.3333 | 3 | 0 | 5 | 0.4 | 3 | 0 | 3 | 0.3333 | 14 | 0.2143 |
| 11_755393 | 5 | 0.4 | 1 | 0 | 3 | 0.3333 | 6 | 0.5 | 15 | 0.4 | 3 | 0 | 5 | 0.4 | 4 | 0.25 | 2 | 0.5 | 14 | 0.2857 |
| 11_755394 | 4 | 0.75 | 2 | 0 | 3 | 0.6667 | 4 | 0.5 | 13 | 0.5385 | 3 | 1 | 2 | 1 | 3 | 1 | 3 | 1 | 11 | 1 |
| 11_755638 | 2 | 1 | NA | NA | NA | NA | 1 | 0 | 3 | 0.6667 | 2 | 0 | 2 | 0.5 | 1 | 0 | 1 | 1 | 6 | 0.3333 |
| 11_755703 | 10 | 0.4 | 5 | 0.4 | 3 | 0.3333 | 10 | 0.5 | 28 | 0.4286 | 4 | 0.5 | 7 | 0 | 8 | 0.375 | 12 | 0.3333 | 31 | 0.2903 |
| 11_755703 | 8 | 0.75 | 4 | 0.75 | 3 | 0.6667 | 8 | 0.5 | 23 | 0.6522 | 3 | 1 | 7 | 0.7143 | 8 | 0.875 | 12 | 1 | 30 | 0.9 |
| 11_755831 | 8 | 1 | 5 | 1 | 3 | 1 | 13 | 0.8462 | 29 | 0.931 | 4 | 1 | 7 | 1 | 8 | 1 | 7 | 1 | 26 | 1 |
| 11_755831 | 7 | 0 | 7 | 0.2857 | 2 | 0 | 12 | 0.1667 | 28 | 0.1429 | 6 | 0 | 7 | 0.1429 | 9 | 0.2222 | 13 | 0.2308 | 35 | 0.1714 |
| 11_755832 | 8 | 1 | 7 | 1 | 1 | 1 | 10 | 0.9 | 26 | 0.9615 | 5 | 0.4 | 10 | 0.9 | 8 | 1 | 15 | 0.8667 | 38 | 0.8421 |
| 11_755832 | 10 | 0.3 | 5 | 0 | 2 | 0 | 14 | 0.0714 | 31 | 0.129 | 8 | 0 | 9 | 0.4444 | 11 | 0.1818 | 15 | 0 | 43 | 0.1395 |
| 11_755833 | 7 | 0.1429 | 6 | 0.3333 | 2 | 0.5 | 13 | 0.5385 | 28 | 0.3929 | 7 | 0.4286 | 10 | 0.3 | 11 | 0.4545 | 14 | 0.5 | 42 | 0.4286 |
| 11_755833 | 10 | 0.3 | 9 | 0.3333 | 3 | 0 | 14 | 0.2857 | 36 | 0.2778 | 8 | 0.375 | 10 | 0.2 | 11 | 0.3636 | 15 | 0.2 | 44 | 0.2727 |
| 11_755835 | 5 | 0 | 2 | 0 | 1 | 0 | 9 | 0.2222 | 17 | 0.1176 | 3 | 0 | 7 | 0.4286 | 3 | 0.3333 | 6 | 0.1667 | 19 | 0.2632 |
| 11_755943 | 10 | 0.4 | 7 | 0.7143 | 3 | 1 | 18 | 0.5 | 38 | 0.5526 | 7 | 0.5714 | 11 | 0.3636 | 12 | 0.5833 | 20 | 0.45 | 50 | 0.48 |

| | | | | | | | | | | | | | | | | | | | | |
|---|---|---|---|---|---|---|---|---|---|---|---|---|---|---|---|---|---|---|---|---|
| 11_755943 | 8 | 0.25 | 8 | 0.25 | 3 | 0 | 14 | 0.2143 | 33 | 0.2121 | 7 | 0.1429 | 9 | 0.4444 | 10 | 0.3 | 17 | 0.1176 | 43 | 0.2326 |
| 11_755944 | 9 | 0.6667 | 5 | 0.6 | 2 | 1 | 15 | 0.7333 | 31 | 0.7097 | 5 | 1 | 10 | 0.5 | 10 | 0.7 | 15 | 0.9333 | 40 | 0.775 |
| 11_755944 | 6 | 0.3333 | 4 | 0 | 2 | 0.5 | 14 | 0.2857 | 26 | 0.2692 | 8 | 0.25 | 9 | 0 | 8 | 0 | 12 | 0.25 | 37 | 0.1351 |
| 11_755947 | 11 | 0.9091 | 7 | 1 | 3 | 1 | 18 | 0.8889 | 39 | 0.9231 | 8 | 0.625 | 13 | 0.6923 | 12 | 0.6667 | 21 | 0.7143 | 54 | 0.6852 |
| 11_755948 | 10 | 1 | 7 | 0.8571 | 3 | 1 | 18 | 0.8889 | 38 | 0.9211 | 7 | 0.7143 | 12 | 0.8333 | 12 | 0.8333 | 20 | 0.95 | 51 | 0.8627 |
| 11_755948 | 10 | 0.5 | 8 | 0.25 | 4 | 0.75 | 16 | 0.3125 | 38 | 0.3947 | 9 | 0.2222 | 11 | 0.0909 | 11 | 0.2727 | 20 | 0.4 | 51 | 0.2745 |
| 11_755948 | 9 | 0.2222 | 8 | 0.25 | 3 | 0.3333 | 15 | 0.1333 | 35 | 0.2 | 7 | 0.1429 | 9 | 0.2222 | 8 | 0.25 | 15 | 0.1333 | 39 | 0.1795 |
| 11_755948 | 8 | 0.375 | 6 | 0.3333 | 3 | 1 | 13 | 0.2308 | 30 | 0.3667 | 7 | 0.4286 | 12 | 0.0833 | 11 | 0.2727 | 18 | 0.2778 | 48 | 0.25 |
| 11_755951 | 9 | 0.2222 | 7 | 0.4286 | 2 | 0 | 15 | 0.1333 | 33 | 0.2121 | 9 | 0.2222 | 11 | 0.0909 | 11 | 0.1818 | 20 | 0.15 | 51 | 0.1569 |
| 11_755951 | 10 | 0.9 | 8 | 0.5 | 6 | 0.5 | 14 | 0.5 | 38 | 0.6053 | 7 | 0.4286 | 11 | 0.6364 | 12 | 0.5833 | 15 | 0.7333 | 45 | 0.6222 |
| 11_755952 | 13 | 0.0769 | 9 | 0.1111 | 6 | 0 | 20 | 0.1 | 48 | 0.0833 | 9 | 0.1111 | 15 | 0.2 | 13 | 0.1538 | 21 | 0.2381 | 58 | 0.1897 |
| 11_755954 | 12 | 0.0833 | 8 | 0.125 | 6 | 0.1667 | 17 | 0.0588 | 43 | 0.093 | 8 | 0 | 16 | 0.1875 | 13 | 0 | 19 | 0 | 56 | 0.0536 |
| 11_755959 | 13 | 0.0769 | 12 | 0.1667 | 7 | 0 | 21 | 0.1429 | 53 | 0.1132 | 13 | 0.2308 | 18 | 0.1111 | 13 | 0.2308 | 18 | 0.2778 | 62 | 0.2097 |
| 11_755961 | 15 | 0.1333 | 11 | 0.0909 | 10 | 0 | 23 | 0.087 | 59 | 0.0847 | 13 | 0 | 20 | 0.1 | 14 | 0.1429 | 22 | 0.0909 | 69 | 0.087 |
| 11_755963 | 14 | 0.5714 | 11 | 0.6364 | 11 | 0.5455 | 20 | 0.45 | 56 | 0.5357 | 13 | 0.1538 | 19 | 0.6316 | 13 | 0.3077 | 20 | 0.55 | 65 | 0.4462 |
| 11_755964 | 10 | 0.8 | 7 | 0.8571 | 9 | 1 | 13 | 0.8462 | 39 | 0.8718 | 5 | 0.6 | 12 | 1 | 5 | 1 | 12 | 0.9167 | 34 | 0.9118 |
| 11_755964 | 15 | 0.4667 | 10 | 0.3 | 12 | 0.25 | 23 | 0.4348 | 60 | 0.3833 | 14 | 0.2143 | 20 | 0.2 | 13 | 0.2308 | 20 | 0.4 | 67 | 0.2687 |
| 11_755964 | 12 | 0.9167 | 11 | 0.8182 | 13 | 0.9231 | 21 | 1 | 57 | 0.9298 | 14 | 0.8571 | 21 | 1 | 13 | 1 | 23 | 0.9565 | 71 | 0.9577 |
| 11_755965 | 18 | 0.2778 | 13 | 0.1538 | 13 | 0 | 22 | 0.0455 | 66 | 0.1212 | 16 | 0 | 20 | 0.05 | 11 | 0.2727 | 25 | 0.08 | 72 | 0.0833 |
| 11_756555 | 110 | 0.0545 | 138 | 0.1087 | 117 | 0.1282 | 111 | 0.1171 | 476 | 0.1029 | 98 | 0.1429 | 102 | 0.1078 | 133 | 0.1729 | 96 | 0.0938 | 429 | 0.1329 |
| 11_756557 | 122 | 0.459 | 148 | 0.4932 | 128 | 0.4141 | 116 | 0.4741 | 514 | 0.4611 | 101 | 0.396 | 110 | 0.4818 | 140 | 0.4714 | 103 | 0.5728 | 454 | 0.4802 |
| 11_756560 | 129 | 0.0543 | 155 | 0.071 | 129 | 0.0388 | 125 | 0.08 | 538 | 0.0613 | 101 | 0.0891 | 113 | 0.0531 | 140 | 0.0786 | 110 | 0.0636 | 464 | 0.0711 |
| 11_756600 | 7 | 1 | 7 | 0.5714 | 9 | 0.8889 | 15 | 0.9333 | 38 | 0.8684 | 8 | 0.875 | 17 | 0.9412 | 18 | 1 | 8 | 0.875 | 51 | 0.9412 |
| 11_756601 | 9 | 1 | 7 | 1 | 11 | 0.9091 | 14 | 1 | 41 | 0.9756 | 7 | 0.8571 | 13 | 1 | 14 | 1 | 6 | 0.8333 | 40 | 0.95 |
| 11_756602 | 6 | 0.6667 | 6 | 0.3333 | 8 | 0.125 | 10 | 0.3 | 30 | 0.3333 | 3 | 0.6667 | 8 | 0.125 | 14 | 0.5714 | 5 | 0.4 | 30 | 0.4333 |
| 11_756605 | 5 | 0.4 | 6 | 0.5 | 8 | 0.375 | 15 | 0.1333 | 34 | 0.2941 | 6 | 0.1667 | 17 | 0.4118 | 16 | 0.375 | 7 | 0.5714 | 46 | 0.3913 |
| 11_756606 | 4 | 0 | 2 | 0 | 3 | 0 | 8 | 0.25 | 17 | 0.1176 | 3 | 0 | 10 | 0.3 | 13 | 0.2308 | 3 | 0.3333 | 29 | 0.2414 |
| 11_756606 | 5 | 0.6 | 5 | 0.4 | 6 | 0.1667 | 13 | 0.4615 | 29 | 0.4138 | 7 | 0.4286 | 17 | 0.4118 | 17 | 0.2353 | 8 | 0.5 | 49 | 0.3673 |
| 11_756609 | 4 | 0.25 | 7 | 0.1429 | 5 | 0.2 | 15 | 0.2 | 31 | 0.1935 | 8 | 0.125 | 14 | 0.0714 | 13 | 0.1538 | 7 | 0.1429 | 42 | 0.119 |
| 11_756611 | 4 | 0.25 | 8 | 0.25 | 5 | 0 | 12 | 0.25 | 29 | 0.2069 | 8 | 0.125 | 15 | 0.2 | 13 | 0.3077 | 8 | 0.375 | 44 | 0.25 |
| 11_756613 | 4 | 0 | 8 | 0.125 | 5 | 0 | 11 | 0.0909 | 28 | 0.0714 | 8 | 0.125 | 15 | 0.2667 | 13 | 0.1538 | 7 | 0 | 43 | 0.1628 |
| 11_756614 | 4 | 0 | 6 | 0 | 4 | 0.25 | 10 | 0.2 | 24 | 0.125 | 7 | 0.1429 | 14 | 0.0714 | 10 | 0 | 8 | 0.125 | 39 | 0.0769 |
| 11_756621 | 2 | 0 | 3 | 0 | 5 | 0 | 7 | 0.1429 | 17 | 0.0588 | 3 | 0.3333 | 13 | 0 | 11 | 0.0909 | 6 | 0.1667 | 33 | 0.0909 |
| 11_756623 | 2 | 1 | 2 | 0.5 | 4 | 0.75 | 7 | 0.4286 | 15 | 0.6 | 3 | 0.3333 | 10 | 0.7 | 8 | 0.25 | 7 | 0.7143 | 28 | 0.5357 |
| 11_756623 | 2 | 0 | 3 | 0.3333 | 5 | 0.4 | 8 | 0 | 18 | 0.1667 | 2 | 0 | 11 | 0.1818 | 9 | 0.5556 | 7 | 0.4286 | 29 | 0.3448 |
| 11_756623 | 2 | 1 | 3 | 0.6667 | 3 | 0.6667 | 7 | 0.4286 | 15 | 0.6 | NA | NA | 12 | 0.5 | 7 | 0.4286 | 6 | 0.8333 | 25 | 0.56 |
| 11_756623 | 2 | 1 | 3 | 0.3333 | 3 | 0.6667 | 8 | 0.375 | 16 | 0.5 | 2 | 1 | 12 | 0.5833 | 9 | 0.8889 | 6 | 0.8333 | 29 | 0.7586 |
| 11_756624 | NA | NA | NA | NA | 1 | 0 | 7 | 0.2857 | 8 | 0.25 | 2 | 0 | 7 | 0.2857 | 5 | 0.4 | 1 | 0 | 15 | 0.2667 |
| 11_756624 | 2 | 0 | 3 | 0 | 4 | 0.25 | 8 | 0.375 | 17 | 0.2353 | 4 | 0 | 11 | 0.1818 | 9 | 0.4444 | 5 | 0.4 | 29 | 0.2759 |
| 11_756624 | 2 | 1 | 3 | 0.6667 | 4 | 0.75 | 6 | 1 | 15 | 0.8667 | 4 | 0.75 | 9 | 0.7778 | 8 | 0.875 | 5 | 0.6 | 26 | 0.7692 |
| 11_756625 | 1 | 0 | 3 | 1 | 4 | 1 | 6 | 1 | 14 | 0.9286 | 4 | 1 | 10 | 0.8 | 9 | 0.5556 | 6 | 0.6667 | 29 | 0.7241 |
| 11_756626 | NA | NA | 3 | 0.3333 | 2 | 1 | 2 | 1 | 7 | 0.7143 | 2 | 0 | 9 | 0.8889 | 5 | 0.2 | 3 | 0.6667 | 19 | 0.5789 |
| 11_756626 | NA | NA | 2 | 1 | 3 | 1 | 4 | 0.25 | 9 | 0.6667 | 3 | 0.6667 | 8 | 0.625 | 5 | 1 | 3 | 0.6667 | 19 | 0.7368 |
| 11_756627 | NA | NA | 3 | 0.3333 | 4 | 0.5 | 4 | 0.25 | 11 | 0.3636 | 3 | 0.3333 | 10 | 0.5 | 9 | 0.7778 | 4 | 0.5 | 26 | 0.5769 |
| 11_756628 | 1 | 0 | 2 | 0.5 | 4 | 0.5 | 5 | 0 | 12 | 0.25 | 2 | 0 | 9 | 0.1111 | 5 | 0 | 3 | 0.3333 | 19 | 0.1053 |
| 11_757228 | NA | NA | NA | NA | 3 | 0.6667 | NA | NA | 3 | 0.6667 | NA | NA | NA | NA | 1 | 1 | NA | NA | 1 | 1 |
| 11_758939 | NA | NA | 1 | 1 | NA | NA | NA | NA | 1 | 1 | NA | NA | 1 | 1 | NA | NA | NA | NA | 1 | 1 |
| 11_759101 | 3 | 1 | 1 | 1 | 3 | 1 | 1 | 1 | 8 | 1 | 1 | 1 | NA | NA | 2 | 1 | 3 | 1 | 6 | 1 |
| 11_759102 | 3 | 0.6667 | 1 | 0 | 3 | 0 | 1 | 0 | 8 | 0.25 | 1 | 1 | NA | NA | 1 | 0 | 2 | 0.5 | 4 | 0.5 |
| 11_759104 | 4 | 0.5 | 1 | 0 | 2 | 0.5 | 1 | 1 | 8 | 0.5 | 1 | 0 | NA | NA | 2 | 0 | 2 | 0.5 | 5 | 0.2 |
| 11_759109 | 3 | 0.6667 | 1 | 1 | 2 | 0 | 1 | 1 | 7 | 0.5714 | 1 | 1 | NA | NA | 1 | 1 | 1 | 0 | 3 | 0.6667 |
| 11_759110 | 3 | 1 | NA | NA | 2 | 0.5 | 1 | 1 | 6 | 0.8333 | 1 | 1 | NA | NA | 2 | 1 | 3 | 0.6667 | 6 | 0.8333 |
| 11_759110 | 3 | 1 | NA | NA | 2 | 0.5 | 1 | 1 | 6 | 0.8333 | 1 | 1 | NA | NA | 2 | 1 | 3 | 1 | 6 | 1 |
| 11_759116 | 2 | 0.5 | 1 | 1 | NA | NA | 1 | 1 | 4 | 0.75 | NA | NA | NA | NA | 1 | 0 | 3 | 0.3333 | 4 | 0.25 |
| 11_759116 | 3 | 0.6667 | 1 | 1 | NA | NA | 1 | 1 | 5 | 0.8 | NA | NA | NA | NA | 2 | 0 | 2 | 1 | 4 | 0.5 |
| 11_759123 | 2 | 0.5 | NA | NA | NA | NA | NA | NA | 2 | 0.5 | NA | NA | NA | NA | 1 | 0 | 3 | 0.3333 | 4 | 0.25 |
| 11_759123 | 1 | 1 | NA | NA | NA | NA | NA | NA | 1 | 1 | NA | NA | NA | NA | 1 | 1 | 2 | 1 | 3 | 1 |
| 11_759126 | 1 | 1 | NA | NA | NA | NA | NA | NA | 1 | 1 | 1 | 1 | NA | NA | 1 | 1 | 3 | 1 | 5 | 1 |
| 11_759127 | 1 | 1 | NA | NA | NA | NA | NA | NA | 1 | 1 | 1 | 1 | NA | NA | 1 | 0 | 2 | 1 | 4 | 0.75 |
| 11_759349 | 1 | 0 | NA | NA | 2 | 0.5 | NA | NA | 3 | 0.3333 | NA | NA | 1 | 1 | 2 | 0.5 | NA | NA | 3 | 0.6667 |
| 11_759351 | 1 | 1 | NA | NA | 2 | 0 | NA | NA | 3 | 0.3333 | 1 | 0 | NA | NA | 4 | 0.5 | NA | NA | 5 | 0.4 |
| 11_759351 | 1 | 0 | NA | NA | 2 | 1 | NA | NA | 3 | 0.6667 | 1 | 0 | 1 | 1 | 4 | 0.5 | NA | NA | 6 | 0.5 |
| 11_759364 | NA | NA | NA | NA | 1 | 1 | NA | NA | 1 | 1 | 1 | 0 | NA | NA | 4 | 1 | NA | NA | 5 | 0.8 |
| 11_759367 | NA | NA | NA | NA | 1 | 1 | NA | NA | 1 | 1 | 1 | 1 | NA | NA | 2 | 1 | NA | NA | 3 | 1 |
| 11_759371 | 7 | 0.2857 | 3 | 0.6667 | 8 | 0.25 | 5 | 0 | 23 | 0.2609 | 4 | 0.5 | 3 | 0 | 3 | 0.3333 | 10 | 0.2 | 20 | 0.25 |

| | | | | | | | | | | | | | | | | | | | | |
|---|---|---|---|---|---|---|---|---|---|---|---|---|---|---|---|---|---|---|---|---|
| 11_7593724 | 7 | 1 | 7 | 0.8571 | 7 | 0.8571 | 5 | 1 | 26 | 0.9231 | 4 | 1 | 3 | 1 | 6 | 1 | 8 | 1 | 21 | 1 |
| 11_7602703 | 4 | 0.25 | 3 | 0 | 4 | 0 | 3 | 0 | 14 | 0.0714 | 2 | 0 | 2 | 0 | 1 | 0 | 6 | 0.1667 | 11 | 0.0909 |
| 11_7609905 | 1 | 0 | 1 | 1 | 2 | 1 | NA | NA | 4 | 0.75 | NA | NA | 1 | 1 | 1 | 0 | NA | NA | 2 | 0.5 |
| 11_7609980 | 1 | 1 | NA | NA | 1 | 0 | 2 | 0 | 4 | 0.25 | 1 | 0 | 1 | 1 | 1 | 1 | 2 | 0 | 5 | 0.4 |
| 11_7610008 | 1 | 1 | 1 | 1 | 2 | 0.5 | 2 | 0.5 | 6 | 0.6667 | NA | NA | 2 | 1 | 1 | 0 | 1 | 1 | 4 | 0.75 |
| 11_7610009 | 1 | 1 | 1 | 1 | 1 | 1 | 2 | 1 | 5 | 1 | 1 | 0 | 2 | 0.5 | 1 | 0 | 1 | 1 | 5 | 0.4 |
| 11_7610163 | 2 | 0.5 | NA | NA | 1 | 1 | NA | NA | 3 | 0.6667 | NA | NA | 1 | 1 | 2 | 0.5 | 2 | 0.5 | 5 | 0.6 |
| 11_7610165 | 1 | 0 | 2 | 0.5 | NA | NA | NA | NA | 3 | 0.3333 | NA | NA | 1 | 1 | NA | NA | NA | NA | 1 | 1 |
| 11_7610172 | 1 | 1 | NA | NA | 1 | 1 | NA | NA | 2 | 1 | NA | NA | 3 | 1 | 2 | 1 | 3 | 1 | 8 | 1 |
| 11_7610186 | 1 | 1 | NA | NA | 1 | 1 | NA | NA | 2 | 1 | NA | NA | 3 | 0.6667 | 2 | 1 | 3 | 1 | 8 | 0.875 |
| 11_7610189 | NA | NA | 2 | 0.5 | NA | NA | NA | NA | 2 | 0.5 | NA | NA | 1 | 1 | NA | NA | NA | NA | 1 | 1 |
| 11_7610192 | 1 | 0 | 1 | 1 | NA | NA | NA | NA | 2 | 0.5 | NA | NA | 1 | 1 | NA | NA | NA | NA | 1 | 1 |
| 11_7610207 | 1 | 1 | 2 | 0.5 | NA | NA | NA | NA | 3 | 0.6667 | NA | NA | 1 | 1 | NA | NA | NA | NA | 1 | 1 |
| 11_7610210 | 1 | 1 | 2 | 0.5 | NA | NA | NA | NA | 3 | 0.6667 | NA | NA | 1 | 1 | NA | NA | NA | NA | 1 | 1 |
| 11_7610221 | 1 | 1 | NA | NA | 1 | 1 | NA | NA | 2 | 1 | NA | NA | 2 | 1 | 2 | 1 | 2 | 1 | 6 | 1 |
| 11_7610228 | 1 | 0 | 2 | 0.5 | NA | NA | NA | NA | 3 | 0.3333 | NA | NA | 1 | 1 | NA | NA | NA | NA | 1 | 1 |
| 11_7610243 | NA | NA | NA | NA | 1 | 1 | NA | NA | 1 | 1 | NA | NA | 2 | 1 | 1 | 1 | 1 | 1 | 4 | 1 |
| 11_7610245 | 1 | 1 | NA | NA | NA | NA | NA | NA | 1 | 1 | NA | NA | 2 | 1 | 2 | 1 | 3 | 1 | 7 | 1 |
| 11_7610255 | 1 | 1 | NA | NA | NA | NA | NA | NA | 1 | 1 | NA | NA | NA | NA | 2 | 0.5 | 2 | 0.5 | 4 | 0.5 |
| 11_7610262 | 1 | 1 | 2 | 0 | NA | NA | NA | NA | 3 | 0.3333 | 1 | 1 | 2 | 1 | NA | NA | NA | NA | 3 | 1 |
| 11_7610289 | 1 | 1 | 1 | 0 | NA | NA | NA | NA | 2 | 0.5 | 1 | 0 | 3 | 0.3333 | NA | NA | NA | NA | 4 | 0.25 |
| 11_7610291 | 1 | 0 | 2 | 0.5 | NA | NA | NA | NA | 3 | 0.3333 | 1 | 1 | 3 | 0 | NA | NA | NA | NA | 4 | 0.25 |
| 11_7610299 | 1 | 0 | 2 | 0.5 | NA | NA | NA | NA | 3 | 0.3333 | 1 | 0 | 2 | 0.5 | NA | NA | NA | NA | 3 | 0.3333 |
| 11_7610323 | 1 | 1 | 2 | 1 | NA | NA | NA | NA | 3 | 1 | 1 | 0 | 3 | 0.6667 | NA | NA | NA | NA | 4 | 0.5 |
| 11_7610326 | 1 | 1 | 2 | 1 | NA | NA | NA | NA | 3 | 1 | 1 | 1 | 2 | 1 | NA | NA | NA | NA | 3 | 1 |
| 11_7610339 | 1 | 1 | 2 | 0.5 | NA | NA | 1 | 1 | 4 | 0.75 | 1 | 1 | 1 | 0 | NA | NA | NA | NA | 2 | 0.5 |
| 11_7610340 | 2 | 1 | NA | NA | NA | NA | NA | NA | 2 | 1 | NA | NA | 2 | 0.5 | 1 | 0 | 3 | 0.6667 | 6 | 0.5 |
| 11_7610342 | 1 | 1 | 1 | 1 | NA | NA | 1 | 0 | 3 | 0.6667 | 1 | 1 | 3 | 0.6667 | NA | NA | NA | NA | 4 | 0.75 |
| 11_7610360 | 2 | 1 | NA | NA | NA | NA | NA | NA | 2 | 1 | NA | NA | 2 | 0 | 1 | 1 | 3 | 0 | 6 | 0.1667 |
| 11_7610368 | 2 | 1 | NA | NA | NA | NA | NA | NA | 2 | 1 | NA | NA | 2 | 0 | 1 | 1 | 3 | 0.3333 | 6 | 0.3333 |
| 11_7610377 | 2 | 0.5 | NA | NA | NA | NA | NA | NA | 2 | 0.5 | NA | NA | NA | NA | 1 | 0 | 3 | 0.6667 | 4 | 0.5 |
| 11_7610382 | 2 | 1 | NA | NA | NA | NA | NA | NA | 2 | 1 | NA | NA | 1 | 1 | 1 | 1 | 3 | 0.6667 | 5 | 0.8 |
| 11_7610406 | 1 | 0 | 2 | 0.5 | NA | NA | 1 | 1 | 4 | 0.5 | NA | NA | 2 | 0.5 | NA | NA | NA | NA | 2 | 0.5 |
| 11_7610414 | 1 | 1 | 2 | 1 | NA | NA | 1 | 0 | 4 | 0.75 | 1 | 1 | 1 | 1 | NA | NA | NA | NA | 2 | 1 |
| 11_7610431 | 1 | 1 | NA | NA | NA | NA | NA | NA | 1 | 1 | NA | NA | 1 | 1 | 1 | 1 | 2 | 1 | 4 | 1 |
| 11_7610439 | 2 | 1 | NA | NA | NA | NA | NA | NA | 2 | 1 | NA | NA | 1 | 1 | 1 | 1 | 2 | 1 | 4 | 1 |
| 11_7610440 | 1 | 1 | 2 | 1 | NA | NA | 1 | 1 | 4 | 1 | 2 | 0.5 | 2 | 0.5 | NA | NA | NA | NA | 4 | 0.5 |
| 11_7610442 | 1 | 1 | 2 | 1 | NA | NA | 1 | 1 | 4 | 1 | 2 | 1 | 2 | 1 | NA | NA | NA | NA | 4 | 1 |
| 11_7610465 | 1 | 1 | 2 | 0.5 | NA | NA | 1 | 0 | 4 | 0.5 | 2 | 0.5 | 3 | 0 | NA | NA | NA | NA | 5 | 0.2 |
| 11_7610496 | 1 | 1 | 2 | 0.5 | NA | NA | 1 | 1 | 4 | 0.75 | 2 | 1 | 2 | 1 | NA | NA | NA | NA | 4 | 1 |
| 11_7610512 | 1 | 1 | 1 | 0 | 1 | 1 | 1 | 1 | 4 | 0.75 | 2 | 1 | 1 | 1 | NA | NA | NA | NA | 3 | 1 |
| 11_7610516 | 1 | 0 | 2 | 0.5 | 1 | 1 | 1 | 1 | 5 | 0.6 | 2 | 1 | 2 | 1 | NA | NA | NA | NA | 4 | 1 |
| 11_7610542 | 1 | 1 | 2 | 1 | 2 | 1 | 1 | 1 | 6 | 1 | 2 | 0.5 | 1 | 1 | NA | NA | NA | NA | 3 | 0.6667 |
| 11_7610545 | 1 | 0 | 2 | 0.5 | 2 | 0 | 1 | 0 | 6 | 0.1667 | 1 | 1 | 2 | 0.5 | NA | NA | NA | NA | 3 | 0.6667 |
| 11_7610555 | NA | NA | 1 | 1 | NA | NA | NA | NA | 1 | 1 | 2 | 0.5 | 2 | 1 | NA | NA | NA | NA | 4 | 0.75 |
| 11_7611173 | 6 | 0.1667 | 9 | 0.2222 | 5 | 0.4 | 4 | 0.25 | 24 | 0.25 | 5 | 0.4 | 5 | 0.8 | 14 | 0.1429 | 3 | 0 | 27 | 0.2963 |
| 11_7611184 | 2 | 0.5 | 7 | 0 | 5 | 0 | 1 | 0 | 15 | 0.0667 | 5 | 0.2 | 3 | 0.3333 | 14 | 0 | 3 | 0 | 25 | 0.08 |
| 11_7613399 | 1 | 0 | 1 | 1 | 2 | 0.5 | 1 | 0 | 5 | 0.4 | 1 | 0 | 1 | 0 | 1 | 0 | 1 | 1 | 4 | 0.25 |
| 11_7613419 | NA | NA | 1 | 0 | 2 | 1 | NA | NA | 3 | 0.6667 | 1 | 1 | 1 | 1 | 1 | 1 | 1 | 0 | 4 | 0.75 |
| 11_7614099 | NA | NA | NA | NA | 1 | 0 | 4 | 0.5 | 5 | 0.4 | 2 | 0 | 4 | 0.25 | 4 | 0.25 | 1 | 0 | 11 | 0.1818 |
| 11_7614096 | NA | NA | NA | NA | 1 | 0 | 4 | 0.25 | 5 | 0.2 | 1 | 1 | 3 | 0 | 4 | 0 | 1 | 0 | 9 | 0.1111 |
| 11_7614098 | NA | NA | NA | NA | 2 | 1 | 4 | 1 | 6 | 1 | 2 | 1 | 4 | 1 | 3 | 1 | 1 | 1 | 10 | 1 |
| 11_7614102 | NA | NA | NA | NA | 1 | 1 | 3 | 1 | 4 | 1 | 3 | 1 | 3 | 0.6667 | 4 | 1 | 1 | 1 | 11 | 0.9091 |
| 11_7614111 | NA | NA | NA | NA | 2 | 1 | 4 | 1 | 6 | 1 | 3 | 0.6667 | 4 | 0.75 | 3 | 1 | 1 | 1 | 11 | 0.8182 |
| 11_7614114 | NA | NA | NA | NA | 1 | 1 | 3 | 0.6667 | 4 | 0.75 | 1 | 1 | 3 | 1 | 1 | 0 | NA | NA | 5 | 0.8 |
| 11_7614115 | NA | NA | NA | NA | NA | NA | 3 | 1 | 3 | 1 | 1 | 1 | 2 | 0.5 | 4 | 0 | NA | NA | 7 | 0.2857 |
| 11_7614122 | NA | NA | NA | NA | 1 | 0 | 4 | 0.5 | 5 | 0.4 | 3 | 0.3333 | 4 | 0.25 | 4 | 0 | 1 | 1 | 12 | 0.25 |
| 11_7614123 | NA | NA | NA | NA | 1 | 0 | 4 | 0.75 | 5 | 0.6 | 2 | 0.5 | 4 | 0.25 | 4 | 0.5 | 1 | 0 | 11 | 0.3636 |
| 11_7614884 | 5 | 0.2 | 3 | 0 | 2 | 1 | 3 | 0.3333 | 13 | 0.3077 | 1 | 1 | 2 | 0.5 | 1 | 1 | 3 | 0.6667 | 7 | 0.7143 |
| 11_7614886 | 5 | 0 | 5 | 0 | 2 | 0 | 5 | 0.2 | 17 | 0.0588 | NA | NA | 1 | 1 | 2 | 0.5 | 6 | 0.1667 | 9 | 0.3333 |
| 11_7614901 | 7 | 0.5714 | 4 | 0.5 | 1 | 1 | 5 | 0.2 | 17 | 0.4706 | 1 | 0 | 1 | 0 | 2 | 1 | 5 | 0.6 | 9 | 0.5556 |
| 11_7614905 | 7 | 0.5714 | 2 | 1 | 1 | 1 | 5 | 0.8 | 15 | 0.7333 | 1 | 1 | NA | NA | 1 | 1 | 5 | 0.6 | 7 | 0.7143 |
| 11_7614907 | 6 | 0 | 3 | 0.6667 | 1 | 1 | 3 | 0.6667 | 13 | 0.3846 | 1 | 1 | 1 | 1 | 1 | 1 | 6 | 0.8333 | 9 | 0.8889 |
| 11_7614914 | 5 | 0.6 | 3 | 0 | 1 | 1 | 4 | 0 | 13 | 0.3077 | 1 | 0 | 1 | 0 | 2 | 0 | 2 | 0.5 | 6 | 0.1667 |
| 11_7614932 | 7 | 0.8571 | 3 | 1 | 1 | 1 | 3 | 1 | 14 | 0.9286 | 1 | 1 | 1 | 1 | 2 | 0.5 | 4 | 1 | 8 | 0.875 |

| | | | | | | | | | | | | | | | | | | | | |
|---|---|---|---|---|---|---|---|---|---|---|---|---|---|---|---|---|---|---|---|---|
| 11_761493 | 7 | 0.2857 | 4 | 0 | 1 | 1 | 4 | 0.5 | 16 | 0.3125 | 1 | 1 | 2 | 0 | 2 | 0.5 | 4 | 0.25 | 9 | 0.3333 |
| 11_761493 | 7 | 0.4286 | 4 | 0.5 | 1 | 0 | 4 | 0.25 | 16 | 0.375 | 1 | 1 | 1 | 0 | 2 | 0 | 4 | 0.5 | 8 | 0.375 |
| 11_761494 | 1 | 1 | NA | NA | NA | NA | NA | NA | 1 | 1 | NA | NA | NA | NA | NA | NA | 1 | 1 | 1 | 1 |
| 11_761494 | 5 | 0.4 | 4 | 1 | 1 | 0 | 4 | 0.75 | 14 | 0.6429 | NA | NA | 2 | 1 | 1 | 0 | 4 | 0.25 | 7 | 0.4286 |
| 11_761494 | 5 | 0.6 | 4 | 0.25 | 1 | 1 | 4 | 0.25 | 14 | 0.4286 | 1 | 0 | 2 | 1 | 2 | 0.5 | 4 | 0.5 | 9 | 0.5556 |
| 11_763050 | 7 | 1 | 3 | 1 | 3 | 0.6667 | 4 | 1 | 17 | 0.9412 | 6 | 1 | 5 | 1 | 4 | 1 | 4 | 1 | 19 | 1 |
| 11_763051 | 8 | 0.25 | 5 | 0.2 | 5 | 0 | 4 | 0 | 22 | 0.1364 | 4 | 0 | 6 | 0 | 4 | 0.25 | 3 | 0 | 17 | 0.0588 |
| 11_763052 | 7 | 1 | 4 | 1 | 6 | 1 | 5 | 1 | 22 | 1 | 6 | 1 | 8 | 1 | 7 | 1 | 5 | 1 | 26 | 1 |
| 11_763058 | 3 | 0 | 2 | 1 | 2 | 0.5 | 3 | 0 | 10 | 0.3 | NA | NA | 3 | 0.6667 | 3 | 0 | 1 | 0 | 7 | 0.2857 |
| 11_763062 | 2 | 0 | 2 | 0 | 2 | 0.5 | 2 | 0 | 8 | 0.125 | 2 | 0 | 3 | 0 | 3 | 0.3333 | 3 | 0 | 11 | 0.0909 |
| 11_763092 | 25 | 0.12 | 30 | 0.1333 | 29 | 0.069 | 32 | 0.125 | 116 | 0.1121 | 20 | 0.1 | 25 | 0.08 | 33 | 0.0606 | 41 | 0.122 | 119 | 0.0924 |
| 11_763095 | 25 | 0.48 | 29 | 0.6207 | 30 | 0.4 | 27 | 0.5185 | 111 | 0.5045 | 20 | 0.6 | 24 | 0.625 | 30 | 0.7333 | 38 | 0.5 | 112 | 0.6071 |
| 11_763096 | 24 | 0.0417 | 28 | 0.0357 | 29 | 0.069 | 28 | 0.1786 | 109 | 0.0826 | 16 | 0.125 | 26 | 0.1154 | 31 | 0.0968 | 37 | 0.0541 | 110 | 0.0909 |
| 11_763097 | 17 | 0.0588 | 18 | 0.0556 | 25 | 0.08 | 24 | 0.0417 | 84 | 0.0595 | 15 | 0.2 | 19 | 0.0526 | 24 | 0.0417 | 30 | 0.0333 | 88 | 0.0682 |
| 11_763097 | 19 | 0.3158 | 21 | 0.4286 | 24 | 0.4583 | 20 | 0.5 | 84 | 0.4286 | 15 | 0.4667 | 17 | 0.5294 | 21 | 0.5238 | 29 | 0.5172 | 82 | 0.5122 |
| 11_763098 | 23 | 0.7826 | 23 | 0.8696 | 28 | 0.7857 | 27 | 0.8519 | 101 | 0.8218 | 18 | 0.6111 | 21 | 0.8095 | 29 | 0.8621 | 38 | 0.7368 | 106 | 0.7642 |
| 11_763098 | 19 | 0.5789 | 24 | 0.875 | 26 | 0.6538 | 23 | 0.7391 | 92 | 0.7174 | 14 | 0.8571 | 20 | 0.85 | 23 | 0.8261 | 28 | 0.75 | 85 | 0.8118 |
| 11_763099 | 21 | 0.0952 | 23 | 0 | 27 | 0.037 | 23 | 0.1304 | 94 | 0.0638 | 20 | 0.05 | 23 | 0 | 28 | 0.0357 | 36 | 0.1389 | 107 | 0.0654 |
| 11_766175 | 6 | 0.1667 | 8 | 0.25 | 5 | 0 | 4 | 0 | 23 | 0.1304 | 5 | 0.4 | 6 | 0.1667 | 4 | 0.5 | 4 | 0 | 19 | 0.2632 |
| 11_766180 | 5 | 0 | 7 | 0.1429 | 4 | 0 | 2 | 0 | 18 | 0.0556 | 4 | 0 | 5 | 0 | 4 | 0.25 | 3 | 0 | 16 | 0.0625 |
| 11_766182 | 3 | 0 | 8 | 0 | 4 | 0 | 4 | 0.25 | 19 | 0.0526 | 6 | 0.3333 | 4 | 0 | 3 | 0.3333 | 2 | 0 | 15 | 0.2 |
| 11_766183 | 4 | 0 | 6 | 0.1667 | 4 | 0.25 | 3 | 0.3333 | 17 | 0.1765 | 5 | 0.4 | 4 | 0 | 3 | 0 | 3 | 0.3333 | 15 | 0.2 |
| 11_766183 | 5 | 0.2 | 8 | 0 | 3 | 0 | 3 | 0.3333 | 19 | 0.1053 | 6 | 0.1667 | 4 | 0 | 4 | 0 | 3 | 0 | 17 | 0.0588 |
| 11_766185 | 3 | 0.6667 | 7 | 0 | 3 | 0 | 3 | 0.6667 | 16 | 0.25 | 5 | 0.4 | 5 | 0.2 | 2 | 0.5 | 3 | 0 | 15 | 0.2667 |
| 11_766196 | 5 | 0.2 | 7 | 0 | 3 | 0 | 4 | 0 | 19 | 0.0526 | 5 | 0.6 | 4 | 0.25 | 2 | 0.5 | 2 | 0 | 13 | 0.3846 |
| 11_766199 | 2 | 0 | 7 | 0.1429 | 3 | 0 | 4 | 0 | 16 | 0.0625 | 3 | 0.3333 | 4 | 0 | 2 | 0 | 2 | 0 | 11 | 0.0909 |
| 11_766205 | 1 | 1 | 6 | 0.5 | 3 | 0.6667 | 3 | 0.6667 | 13 | 0.6154 | 4 | 0.5 | 4 | 0.5 | 1 | 0 | 1 | 1 | 10 | 0.5 |
| 11_766206 | 2 | 1 | 6 | 0.8333 | 3 | 1 | 3 | 1 | 14 | 0.9286 | 2 | 0.5 | 3 | 1 | 2 | 1 | 2 | 0.5 | 9 | 0.7778 |
| 11_766206 | 1 | 0 | 6 | 0.6667 | 3 | 0.6667 | 3 | 0.6667 | 13 | 0.6154 | 4 | 0.75 | 1 | 0 | 2 | 1 | 1 | 1 | 8 | 0.75 |
| 11_766208 | 2 | 0.5 | 5 | 0.2 | 3 | 0 | 2 | 0.5 | 12 | 0.25 | 4 | 0.5 | 3 | 0 | 2 | 0.5 | 2 | 0 | 11 | 0.2727 |
| 11_766208 | 1 | 1 | 4 | 0 | 2 | 0.5 | 1 | 0 | 8 | 0.25 | 4 | 0.5 | 4 | 0 | 2 | 0.5 | 2 | 0 | 12 | 0.25 |
| 11_766208 | 2 | 1 | 6 | 1 | 3 | 1 | 3 | 1 | 14 | 1 | 4 | 0.75 | 3 | 1 | 2 | 1 | 2 | 1 | 11 | 0.9091 |
| 11_766209 | 2 | 1 | 6 | 1 | 3 | 0.6667 | 3 | 1 | 14 | 0.9286 | 4 | 1 | 4 | 1 | 2 | 1 | 2 | 1 | 12 | 1 |
| 11_766693 | 3 | 0.3333 | 4 | 0.5 | 2 | 0.5 | 3 | 0.3333 | 12 | 0.4167 | 2 | 0.5 | 7 | 0.7143 | 1 | 0 | 5 | 0.2 | 15 | 0.4667 |
| 11_766703 | 4 | 0.75 | 5 | 0 | 3 | 0.3333 | 9 | 0.3333 | 21 | 0.3333 | 4 | 0 | 11 | 0.2727 | 3 | 0.3333 | 7 | 0.4286 | 25 | 0.28 |
| 11_766703 | 6 | 0.8333 | 5 | 1 | 5 | 1 | 9 | 0.8889 | 25 | 0.92 | 6 | 0.8333 | 15 | 0.9333 | 3 | 1 | 8 | 1 | 32 | 0.9375 |
| 11_766703 | 6 | 0 | 2 | 0 | 5 | 0.2 | 7 | 0.1429 | 20 | 0.1 | 4 | 0 | 8 | 0.125 | 2 | 0 | 5 | 0 | 19 | 0.0526 |
| 11_766704 | 5 | 0.8 | 4 | 1 | 6 | 1 | 7 | 0.8571 | 22 | 0.9091 | 5 | 1 | 11 | 0.8182 | 1 | 1 | 8 | 1 | 25 | 0.92 |
| 11_766705 | 3 | 0 | 5 | 0 | 5 | 0 | 7 | 0.1429 | 20 | 0.05 | 5 | 0.4 | 12 | 0.0833 | 3 | 0.3333 | 7 | 0.1429 | 27 | 0.1852 |
| 11_766706 | 5 | 1 | 4 | 1 | 4 | 1 | 8 | 0.5 | 21 | 0.8095 | 6 | 0.6667 | 13 | 0.6923 | 3 | 0.6667 | 7 | 1 | 29 | 0.7586 |
| 11_766707 | 4 | 0.5 | 5 | 0.6 | 4 | 0.75 | 8 | 0.25 | 21 | 0.4762 | 4 | 0.5 | 12 | 0.5 | 2 | 0.5 | 6 | 0.5 | 24 | 0.5 |
| 11_766708 | 4 | 0.5 | 4 | 0.5 | 5 | 0.8 | 8 | 0.625 | 21 | 0.619 | 6 | 0.8333 | 12 | 0.4167 | 2 | 1 | 5 | 1 | 25 | 0.68 |
| 11_766708 | 5 | 0.2 | 2 | 0 | 3 | 0.3333 | 6 | 0.5 | 16 | 0.3125 | 4 | 0 | 10 | 0.3 | 2 | 0.5 | 5 | 0.2 | 21 | 0.2381 |
| 11_766708 | 5 | 0.8 | 4 | 0.5 | 4 | 1 | 5 | 0.8 | 18 | 0.7778 | 5 | 0.8 | 13 | 0.6154 | 2 | 1 | 5 | 1 | 25 | 0.76 |
| 11_766710 | 2 | 0.5 | 4 | 1 | 2 | 1 | 6 | 1 | 14 | 0.9286 | 5 | 1 | 11 | 0.5455 | 2 | 1 | 6 | 0.8333 | 24 | 0.75 |
| 11_766711 | 3 | 0.6667 | 4 | 0.25 | 4 | 0 | 6 | 0.3333 | 17 | 0.2941 | 5 | 0.4 | 10 | 0.2 | 2 | 0.5 | 6 | 0.1667 | 23 | 0.2609 |
| 11_766717 | 3 | 0 | 3 | 0 | 2 | 0 | 5 | 0.2 | 13 | 0.0769 | 4 | 0.75 | 8 | 0.25 | NA | NA | 3 | 0 | 15 | 0.3333 |
| 11_766717 | 2 | 1 | 3 | 0.6667 | 3 | 1 | 5 | 1 | 13 | 0.9231 | 5 | 1 | 8 | 0.875 | 2 | 1 | 5 | 1 | 20 | 0.95 |
| 11_766720 | 3 | 0.3333 | 3 | 0 | 3 | 0 | 6 | 0 | 15 | 0.0667 | 6 | 0 | 10 | 0.2 | 1 | 0 | 6 | 0 | 23 | 0.087 |
| 11_766720 | 3 | 0 | 3 | 0.3333 | 2 | 0 | 5 | 0 | 13 | 0.0769 | 5 | 0 | 9 | 0.1111 | NA | NA | 4 | 0.25 | 18 | 0.1111 |
| 11_766723 | 3 | 0.6667 | 3 | 0.3333 | 3 | 0.3333 | 3 | 0 | 12 | 0.3333 | 6 | 0.8333 | 7 | 0.2857 | 1 | 1 | 6 | 0.3333 | 20 | 0.5 |
| 11_766725 | 2 | 0 | 2 | 0.5 | 2 | 0.5 | 5 | 0.2 | 11 | 0.2727 | 4 | 0.25 | 7 | 0.5714 | 1 | 1 | 5 | 0 | 17 | 0.3529 |
| 11_766725 | 3 | 0.3333 | 2 | 0 | 3 | 0.6667 | 5 | 0.2 | 13 | 0.3077 | 5 | 0.4 | 8 | 0.375 | 1 | 0 | 6 | 0.3333 | 20 | 0.35 |
| 11_766725 | 2 | 0 | 3 | 0 | 3 | 0.3333 | 6 | 0.1667 | 14 | 0.1429 | 5 | 0.2 | 8 | 0 | 1 | 0 | 4 | 0 | 18 | 0.0556 |
| 11_766726 | 2 | 1 | 3 | 0.3333 | 3 | 0 | 6 | 0 | 14 | 0.2143 | 5 | 0 | 10 | 0.1 | 1 | 0 | 5 | 0.2 | 21 | 0.0952 |
| 11_766726 | 3 | 0.3333 | 2 | 0 | 1 | 0 | 6 | 0.1667 | 12 | 0.1667 | 4 | 0 | 9 | 0.1111 | 1 | 0 | 6 | 0.1667 | 20 | 0.1 |
| 11_766727 | 3 | 0.3333 | 2 | 0 | 2 | 0 | 6 | 0.3333 | 13 | 0.2308 | 4 | 0.75 | 11 | 0.1818 | 1 | 0 | 5 | 0.2 | 21 | 0.2857 |
| 11_766727 | 2 | 1 | 3 | 0.3333 | 2 | 1 | 5 | 1 | 12 | 0.8333 | 5 | 1 | 7 | 0.7143 | 1 | 1 | 4 | 1 | 17 | 0.8824 |
| 11_766728 | 1 | 1 | 3 | 0.6667 | 2 | 0.5 | 3 | 0.3333 | 9 | 0.5556 | 4 | 1 | 8 | 0.625 | 1 | 1 | 5 | 0.8 | 18 | 0.7778 |
| 11_766729 | 1 | 0 | 1 | 0 | 1 | 0 | 3 | 0.3333 | 6 | 0.1667 | 3 | 0.3333 | 6 | 0.5 | 1 | 0 | 6 | 0.6667 | 16 | 0.5 |
| 11_766730 | NA | NA | 3 | 0 | 2 | 0 | 5 | 0.2 | 10 | 0.1 | 3 | 0 | 6 | 0.1667 | 1 | 0 | 5 | 0 | 15 | 0.0667 |
| 11_766730 | NA | NA | 2 | 0 | 2 | 0 | 5 | 0.2 | 9 | 0.1111 | 4 | 0.75 | 4 | 0.25 | 1 | 0 | 6 | 0.3333 | 15 | 0.4 |
| 11_766730 | NA | NA | 3 | 1 | 2 | 1 | 5 | 1 | 10 | 1 | 3 | 0.6667 | 7 | 0.7143 | 1 | 1 | 6 | 1 | 17 | 0.8235 |
| 11_766884 | NA | NA | 3 | 0 | 2 | 0.5 | 4 | 0 | 9 | 0.1111 | NA | NA | 5 | 0.2 | 1 | 0 | 3 | 0 | 9 | 0.1111 |
| 11_766884 | NA | NA | 3 | 0 | 2 | 0.5 | 4 | 0.25 | 9 | 0.2222 | 1 | 0 | 6 | 0 | 1 | 0 | 4 | 0.5 | 12 | 0.1667 |

| | | | | | | | | | | | | | | | | | | | | |
|---|---|---|---|---|---|---|---|---|---|---|---|---|---|---|---|---|---|---|---|---|
| 11_766884( | NA | NA | 3 | 0.6667 | 2 | 0 | 4 | 0.25 | 9 | 0.3333 | 1 | 0 | 6 | 0 | 1 | 0 | 4 | 0.5 | 12 | 0.1667 |
| 11_767033( | NA | NA | 1 | 0 | 2 | 0 | 1 | 1 | 4 | 0.25 | 1 | 1 | 3 | 0 | NA | NA | 2 | 0.5 | 6 | 0.3333 |
| 11_767313 | NA | NA | 1 | 1 | 2 | 1 | 1 | 1 | 4 | 1 | NA | NA | 1 | 1 | NA | NA | 1 | 0 | 2 | 0.5 |
| 11_768774( | 8 | 0.25 | 9 | 0.6667 | 4 | 0.75 | 8 | 0.875 | 29 | 0.6207 | 4 | 1 | 9 | 0.3333 | 5 | 0.8 | 3 | 0.6667 | 21 | 0.619 |
| 11_768775 | 9 | 0.4444 | 6 | 0.6667 | 2 | 0.5 | 5 | 0.2 | 22 | 0.4545 | 4 | 0.5 | 10 | 0.4 | 7 | 0.2857 | 3 | 0.3333 | 24 | 0.375 |
| 11_768775: | 8 | 0.625 | 7 | 0.8571 | 6 | 1 | 8 | 0.875 | 29 | 0.8276 | 5 | 0.4 | 10 | 0.3 | 7 | 0.5714 | 1 | 0 | 23 | 0.3913 |
| 11_768775 | 10 | 0.8 | 9 | 0.7778 | 6 | 0.5 | 10 | 0.9 | 35 | 0.7714 | 5 | 0.8 | 10 | 0.6 | 9 | 0.7778 | 4 | 0.75 | 28 | 0.7143 |
| 11_768775 | 9 | 0.2222 | 9 | 0 | 5 | 0 | 10 | 0.2 | 33 | 0.1212 | 4 | 0.25 | 11 | 0 | 9 | 0 | 4 | 0.5 | 28 | 0.1071 |
| 11_768778( | 10 | 0.2 | 9 | 0 | 6 | 0.1667 | 7 | 0 | 32 | 0.0938 | 8 | 0.25 | 10 | 0 | 6 | 0.1667 | 3 | 0.3333 | 27 | 0.1481 |
| 11_769361 | 2 | 1 | NA | NA | 2 | 0 | NA | NA | 4 | 0.5 | 1 | 1 | NA | NA | NA | NA | NA | NA | 1 | 1 |
| 11_769365 | 2 | 1 | 2 | 1 | 2 | 1 | 1 | 1 | 7 | 1 | 2 | 1 | 3 | 1 | 2 | 1 | 3 | 1 | 10 | 1 |
| 11_769365: | 2 | 1 | 2 | 0.5 | 5 | 0.2 | 1 | 1 | 10 | 0.5 | 2 | 0 | 2 | 0.5 | 1 | 1 | 3 | 0 | 8 | 0.25 |
| 11_769365( | 2 | 1 | 3 | 1 | 4 | 1 | 1 | 1 | 10 | 1 | 1 | 1 | 2 | 1 | 2 | 1 | 3 | 1 | 8 | 1 |
| 11_769365 | 2 | 1 | 3 | 1 | 5 | 1 | 1 | 1 | 11 | 1 | 1 | 1 | 2 | 1 | 2 | 1 | 3 | 1 | 8 | 1 |
| 11_769366: | 2 | 1 | 3 | 1 | 4 | 0.75 | 1 | 1 | 10 | 0.9 | 2 | 1 | 3 | 1 | 2 | 1 | 3 | 1 | 10 | 1 |
| 11_769366 | 1 | 0 | 2 | 0 | 4 | 0.5 | 1 | 0 | 8 | 0.25 | 2 | 0.5 | 2 | 1 | 2 | 0 | 3 | 0.3333 | 9 | 0.4444 |
| 11_769367 | NA | NA | 2 | 1 | 4 | 1 | 1 | 1 | 7 | 1 | 1 | 1 | 2 | 1 | 1 | 0 | 3 | 1 | 7 | 0.8571 |
| 11_769367 | 1 | 1 | 3 | 1 | 4 | 1 | 1 | 0 | 9 | 0.8889 | 1 | 0 | 2 | 0 | 2 | 1 | 2 | 1 | 7 | 0.5714 |
| 11_769369 | 2 | 0.5 | 2 | 0 | 5 | 0.6 | 1 | 0 | 10 | 0.4 | 1 | 0 | 2 | 0.5 | 1 | 0 | 2 | 0.5 | 6 | 0.3333 |
| 11_769370 | 1 | 0 | 1 | 1 | 3 | 0.3333 | 1 | 0 | 6 | 0.3333 | 1 | 0 | 2 | 0 | 2 | 0 | 2 | 0.5 | 7 | 0.1429 |
| 11_769372 | 2 | 0 | 3 | 0.3333 | 1 | 0 | 1 | 0 | 7 | 0.1429 | 1 | 0 | 2 | 0 | 1 | 0 | 2 | 0.5 | 6 | 0.1667 |
| 11_773458: | NA | NA | NA | NA | NA | NA | 1 | 1 | 1 | 1 | 1 | 1 | 1 | 1 | 2 | 1 | 2 | 1 | 6 | 1 |
| 11_773458 | NA | NA | NA | NA | NA | NA | 1 | 1 | 1 | 1 | 1 | 0 | 1 | 1 | 2 | 0.5 | 2 | 1 | 6 | 0.6667 |
| 11_773461: | NA | NA | NA | NA | NA | NA | 1 | 1 | 1 | 1 | 1 | 1 | 1 | 0 | 2 | 0 | 2 | 1 | 6 | 0.5 |
| 11_773462: | NA | NA | NA | NA | NA | NA | 1 | 1 | 1 | 1 | NA | NA | NA | NA | NA | NA | 1 | 1 | 1 | 1 |
| 11_773471 | NA | NA | NA | NA | 1 | 0 | 1 | 1 | 2 | 0.5 | 1 | 0 | 1 | 1 | 2 | 1 | 2 | 1 | 6 | 0.8333 |
| 11_773487: | 2 | 0.5 | NA | NA | 2 | 0 | 1 | 1 | 5 | 0.4 | 1 | 1 | NA | NA | 3 | 0.3333 | 2 | 1 | 6 | 0.6667 |
| 11_773487 | 3 | 0.6667 | NA | NA | 2 | 0.5 | 1 | 1 | 6 | 0.6667 | 1 | 0 | 1 | 1 | 3 | 0 | 2 | 1 | 7 | 0.4286 |
| 11_773488: | 3 | 0.3333 | NA | NA | 2 | 0 | NA | NA | 5 | 0.2 | 1 | 0 | 2 | 0.5 | 4 | 0 | 2 | 1 | 9 | 0.3333 |
| 11_773491 | 4 | 0.5 | NA | NA | 3 | 0.3333 | 3 | 0.3333 | 10 | 0.4 | 1 | 0 | 2 | 1 | 5 | 0 | 1 | 0 | 9 | 0.2222 |
| 11_773501 | 4 | 1 | 2 | 1 | 4 | 0.25 | 3 | 1 | 13 | 0.7692 | 1 | 0 | 1 | 0 | 5 | 0.4 | 2 | 1 | 9 | 0.4444 |
| 11_773502: | 5 | 0.2 | NA | NA | 3 | 0.3333 | 3 | 0.3333 | 11 | 0.2727 | 1 | 0 | 1 | 0 | 5 | 0.6 | 2 | 0.5 | 9 | 0.4444 |
| 11_773503: | 4 | 0.5 | 2 | 1 | 4 | 0.25 | 2 | 0 | 12 | 0.4167 | 2 | 0 | 2 | 0.5 | 5 | 0.4 | 3 | 0.6667 | 12 | 0.4167 |
| 11_773504: | 5 | 0.2 | 2 | 0 | 4 | 0 | 3 | 0 | 14 | 0.0714 | 3 | 0 | 2 | 0 | 3 | 0.3333 | 3 | 0 | 11 | 0.0909 |
| 11_773504 | 5 | 0.4 | 2 | 0 | 4 | 0 | 3 | 0.3333 | 14 | 0.2143 | 3 | 0.6667 | 2 | 0 | 5 | 0 | 3 | 0.3333 | 13 | 0.2308 |
| 11_773504( | 5 | 0.4 | 2 | 0 | 4 | 0 | 3 | 0 | 14 | 0.1429 | 3 | 0 | 2 | 0 | 4 | 0 | 3 | 0.3333 | 12 | 0.0833 |
| 11_773507( | 5 | 0.2 | 2 | 0 | 3 | 0 | 2 | 0 | 12 | 0.0833 | 3 | 0.3333 | 2 | 0 | 5 | 0.2 | 3 | 1 | 13 | 0.3846 |
| 11_773507 | 4 | 0 | 2 | 0.5 | 3 | 0.3333 | 2 | 0.5 | 11 | 0.2727 | 3 | 0.6667 | 1 | 1 | 4 | 0.75 | 2 | 0 | 10 | 0.6 |
| 11_773508 | 5 | 0 | 2 | 0.5 | 5 | 0.2 | 3 | 0 | 15 | 0.1333 | 3 | 0.3333 | 2 | 0 | 5 | 0.2 | 3 | 0.3333 | 13 | 0.2308 |
| 11_773603: | 1 | 0 | 2 | 0.5 | 1 | 1 | NA | NA | 4 | 0.5 | 2 | 0.5 | NA | NA | NA | NA | 1 | 0 | 3 | 0.3333 |
| 11_773604 | 1 | 1 | 3 | 0.6667 | 1 | 1 | NA | NA | 5 | 0.8 | 1 | 0 | NA | NA | NA | NA | 2 | 1 | 3 | 0.6667 |
| 11_773604( | 1 | 0 | 2 | 1 | 1 | 1 | NA | NA | 4 | 0.75 | 1 | 0 | NA | NA | NA | NA | 2 | 0.5 | 3 | 0.3333 |
| 11_773605 | 1 | 0 | 3 | 0.6667 | 1 | 1 | NA | NA | 5 | 0.6 | 2 | 0.5 | NA | NA | NA | NA | 1 | 1 | 3 | 0.6667 |
| 11_773611 | 1 | 0 | 5 | 0.2 | NA | NA | NA | NA | 6 | 0.1667 | 2 | 0 | NA | NA | 1 | 0 | 2 | 0.5 | 5 | 0.2 |
| 11_773611 | 2 | 1 | 7 | 0.4286 | 1 | 1 | NA | NA | 10 | 0.6 | 1 | 1 | NA | NA | 1 | 0 | 1 | 1 | 3 | 0.6667 |
| 11_773613 | 5 | 0.4 | 11 | 0.1818 | 10 | 0.1 | 13 | 0.0769 | 39 | 0.1538 | 5 | 0 | 8 | 0 | 14 | 0.0714 | 13 | 0.2308 | 40 | 0.1 |
| 11_773620 | 2 | 1 | 8 | 0.75 | 11 | 0.8182 | 14 | 0.7143 | 35 | 0.7714 | 5 | 0.8 | 8 | 0.625 | 10 | 0.8 | 12 | 0.6667 | 35 | 0.7143 |
| 11_773622 | 3 | 0.3333 | 7 | 0 | 9 | 0.1111 | 14 | 0 | 33 | 0.0606 | 4 | 0.25 | 8 | 0 | 10 | 0 | 10 | 0.2 | 32 | 0.0938 |
| 11_773624( | 2 | 0 | 8 | 0.25 | 7 | 0.2857 | 11 | 0.1818 | 28 | 0.2143 | 4 | 0 | 8 | 0.375 | 9 | 0.2222 | 10 | 0.1 | 31 | 0.1935 |
| 11_773624: | 1 | 1 | 5 | 0 | 8 | 0.125 | 9 | 0 | 23 | 0.087 | 2 | 0.5 | 5 | 0 | 10 | 0.1 | 7 | 0 | 24 | 0.0833 |
| 11_773627 | NA | NA | 3 | 0 | 3 | 0 | 4 | 0.5 | 10 | 0.2 | 1 | 0 | NA | NA | 2 | 0 | 6 | 0.5 | 9 | 0.3333 |
| 11_773957( | 14 | 0.1429 | 31 | 0 | 20 | 0.1 | 16 | 0.0625 | 81 | 0.0617 | 19 | 0.1053 | 16 | 0.0625 | 14 | 0.1429 | 19 | 0.1053 | 68 | 0.1029 |
| 11_773973 | 18 | 0.1667 | 41 | 0.122 | 21 | 0 | 17 | 0 | 97 | 0.0825 | 29 | 0.0345 | 24 | 0.125 | 21 | 0.0476 | 22 | 0.0909 | 96 | 0.0729 |
| 11_773974( | 20 | 0.3 | 34 | 0.2059 | 18 | 0.1667 | 14 | 0.2143 | 86 | 0.2209 | 22 | 0.2727 | 27 | 0.2593 | 18 | 0.3889 | 24 | 0.0833 | 91 | 0.2418 |
| 11_773974 | 18 | 0.1667 | 34 | 0.1765 | 16 | 0.125 | 20 | 0.2 | 88 | 0.1705 | 21 | 0.1429 | 24 | 0.125 | 26 | 0.1154 | 26 | 0.0385 | 97 | 0.1031 |
| 11_773976 | 23 | 0.4348 | 39 | 0.641 | 23 | 0.3478 | 21 | 0.5238 | 106 | 0.5094 | 27 | 0.5556 | 26 | 0.5 | 20 | 0.6 | 26 | 0.4231 | 99 | 0.5152 |
| 11_773977( | 22 | 0.0455 | 38 | 0.1053 | 23 | 0.087 | 22 | 0.0455 | 105 | 0.0762 | 30 | 0.1 | 23 | 0.2174 | 28 | 0.1429 | 25 | 0.08 | 106 | 0.1321 |
| 11_773982 | 31 | 0.1613 | 55 | 0.0364 | 29 | 0.0345 | 23 | 0.1304 | 138 | 0.0797 | 32 | 0.0625 | 28 | 0.0714 | 31 | 0.0645 | 34 | 0.1471 | 125 | 0.088 |
| 11_773983( | 27 | 0.1111 | 47 | 0.0851 | 29 | 0.1034 | 21 | 0.0476 | 124 | 0.0887 | 33 | 0.1212 | 31 | 0.0323 | 25 | 0.08 | 27 | 0.037 | 116 | 0.069 |
| 11_773983( | 32 | 0.375 | 50 | 0.34 | 30 | 0.3 | 26 | 0.1923 | 138 | 0.3116 | 32 | 0.2188 | 31 | 0.0645 | 29 | 0.2759 | 35 | 0.3143 | 127 | 0.2205 |
| 11_773995 | 1 | 1 | 1 | 0 | 1 | 0 | 2 | 0.5 | 5 | 0.4 | NA | NA | NA | NA | NA | NA | 2 | 0.5 | 2 | 0.5 |
| 11_773995 | 1 | 1 | NA | NA | NA | NA | NA | NA | 1 | 1 | NA | NA | NA | NA | NA | NA | 1 | 1 | 1 | 1 |
| 11_773996: | 1 | 1 | NA | NA | NA | NA | 1 | 0 | 2 | 0.5 | 1 | 1 | 1 | 1 | NA | NA | 1 | 1 | 3 | 1 |
| 11_773998: | 1 | 1 | NA | NA | NA | NA | 1 | 1 | 2 | 1 | NA | NA | NA | NA | NA | NA | 1 | 1 | 1 | 1 |
| 11_774064: | NA | NA | 1 | 1 | NA | NA | NA | NA | 1 | 1 | 1 | 1 | NA | NA | NA | NA | NA | NA | 1 | 1 |

| | | | | | | | | | | | | | | | | | | | | |
|---|---|---|---|---|---|---|---|---|---|---|---|---|---|---|---|---|---|---|---|---|
| 11_774095( | NA | NA | 1 | 1 | NA | NA | NA | NA | 1 | 1 | 1 | 1 | NA | NA | NA | NA | NA | NA | 1 | 1 |
| 11_774095! | NA | NA | 1 | 1 | NA | NA | NA | NA | 1 | 1 | 1 | 1 | NA | NA | NA | NA | NA | NA | 1 | 1 |
| 11_774096! | NA | NA | 1 | 1 | NA | NA | NA | NA | 1 | 1 | 1 | 1 | NA | NA | NA | NA | NA | NA | 1 | 1 |
| 11_774100: | NA | NA | 1 | 1 | NA | NA | NA | NA | 1 | 1 | 1 | 1 | NA | NA | NA | NA | NA | NA | 1 | 1 |
| 11_774226( | 5 | 0.2 | 6 | 0.1667 | 9 | 0.1111 | 6 | 0 | 26 | 0.1154 | 5 | 0.2 | 7 | 0.1429 | 7 | 0 | 6 | 0.1667 | 25 | 0.12 |
| 11_774238 | 3 | 0 | 5 | 0.4 | 11 | 0.1818 | 5 | 0 | 24 | 0.1667 | 4 | 0 | 7 | 0.2857 | 5 | 0.4 | 4 | 0 | 20 | 0.2 |
| 11_774239( | 1 | 0 | 4 | 0.25 | 10 | 0 | 4 | 0.5 | 19 | 0.1579 | 4 | 0 | 6 | 0 | 4 | 0 | 3 | 0.3333 | 17 | 0.0588 |
| 11_774247: | 3 | 0 | 4 | 0 | 6 | 0.1667 | 2 | 0 | 15 | 0.0667 | 3 | 0 | 5 | 0 | 1 | 1 | 4 | 0 | 13 | 0.0769 |
| 11_774286 | 1 | 0 | 1 | 0 | 2 | 0.5 | NA | NA | 4 | 0.25 | 1 | 1 | 3 | 0 | NA | NA | 1 | 1 | 5 | 0.4 |
| 11_774287 | NA | NA | 2 | 1 | 1 | 0 | NA | NA | 3 | 0.6667 | 1 | 0 | 3 | 0 | NA | NA | 2 | 1 | 6 | 0.3333 |
| 11_774289 | 1 | 0 | 2 | 0.5 | 2 | 0.5 | NA | NA | 5 | 0.4 | 1 | 0 | 1 | 0 | NA | NA | 1 | 1 | 3 | 0.3333 |
| 11_774290( | 1 | 1 | 2 | 0 | 2 | 1 | NA | NA | 5 | 0.6 | 1 | 0 | 1 | 1 | NA | NA | 2 | 1 | 4 | 0.75 |
| 11_774325 | 1 | 1 | 2 | 0 | 2 | 0 | NA | NA | 5 | 0.2 | 1 | 1 | 1 | 0 | NA | NA | 1 | 0 | 3 | 0.3333 |
| 11_774325: | 1 | 1 | 2 | 0 | 2 | 0 | NA | NA | 5 | 0.2 | 1 | 1 | 1 | 0 | NA | NA | 1 | 0 | 3 | 0.3333 |
| 11_775825 | 1 | 1 | NA | NA | 1 | 1 | 2 | 0 | 4 | 0.5 | NA | NA | NA | NA | 2 | 0 | 1 | 1 | 3 | 0.3333 |
| 11_775828: | 1 | 0 | NA | NA | 1 | 0 | 2 | 0.5 | 4 | 0.25 | 1 | 0 | 1 | 1 | 2 | 0 | NA | NA | 4 | 0.25 |
| 11_775828 | 1 | 0 | NA | NA | 1 | 0 | 1 | 1 | 3 | 0.3333 | NA | NA | 1 | 0 | 2 | 1 | 1 | 1 | 4 | 0.75 |
| 11_775830 | 1 | 1 | NA | NA | 1 | 1 | 1 | 1 | 3 | 1 | 1 | 1 | 2 | 1 | 2 | 1 | 1 | 1 | 6 | 1 |
| 11_775831: | 1 | 0 | NA | NA | 1 | 0 | 2 | 1 | 4 | 0.5 | 1 | 1 | 1 | 1 | 2 | 1 | 1 | 1 | 5 | 1 |
| 11_775833 | 1 | 1 | NA | NA | 1 | 1 | NA | NA | 2 | 1 | 1 | 1 | 2 | 1 | 2 | 1 | 1 | 0 | 6 | 0.8333 |
| 11_775834: | 1 | 1 | NA | NA | 1 | 1 | 2 | 1 | 4 | 1 | 1 | 0 | 2 | 1 | 1 | 1 | 1 | 1 | 5 | 0.8 |
| 11_775836 | 1 | 1 | NA | NA | 1 | 1 | 2 | 1 | 4 | 1 | 1 | 0 | 2 | 1 | 2 | 1 | 2 | 1 | 7 | 0.8571 |
| 11_775836 | 1 | 0 | NA | NA | NA | NA | 2 | 0.5 | 3 | 0.3333 | 1 | 0 | 2 | 0 | 2 | 0 | 2 | 0.5 | 7 | 0.1429 |
| 11_775837 | 1 | 1 | NA | NA | 1 | 1 | 2 | 0.5 | 4 | 0.75 | 1 | 0 | 1 | 1 | 2 | 1 | 2 | 0.5 | 6 | 0.6667 |
| 11_775837 | 1 | 1 | NA | NA | 1 | 1 | 1 | 1 | 3 | 1 | 1 | 1 | 1 | 1 | 2 | 1 | 2 | 1 | 6 | 1 |
| 11_775838: | NA | NA | NA | NA | 1 | 1 | 2 | 1 | 3 | 1 | 1 | 1 | 2 | 1 | 2 | 0.5 | 1 | 1 | 6 | 0.8333 |
| 11_775838 | 1 | 1 | NA | NA | 1 | 0 | 2 | 0 | 4 | 0.25 | 1 | 0 | 2 | 1 | 2 | 0 | 2 | 0 | 7 | 0.2857 |
| 11_775839: | 1 | 0 | NA | NA | 1 | 0 | 1 | 1 | 3 | 0.3333 | 1 | 0 | 2 | 0.5 | 2 | 1 | 2 | 0.5 | 7 | 0.5714 |
| 11_775839 | 1 | 1 | NA | NA | 1 | 1 | 1 | 1 | 3 | 1 | 1 | 1 | 1 | 1 | 1 | 1 | NA | NA | 3 | 1 |
| 11_776005 | 1 | 0 | NA | NA | 1 | 0 | 2 | 0.5 | 4 | 0.25 | 2 | 0.5 | 2 | 0.5 | 1 | 0 | 2 | 0.5 | 7 | 0.4286 |
| 11_776006: | 1 | 1 | NA | NA | 1 | 1 | 2 | 1 | 4 | 1 | 2 | 0.5 | 2 | 0.5 | 2 | 0.5 | 2 | 0.5 | 8 | 0.5 |
| 11_776007 | 1 | 1 | NA | NA | 1 | 0 | 2 | 0.5 | 4 | 0.5 | 1 | 0 | 1 | 1 | 3 | 0.6667 | 2 | 0.5 | 7 | 0.5714 |
| 11_776007 | 1 | 0 | NA | NA | 1 | 1 | 2 | 0.5 | 4 | 0.5 | 2 | 0.5 | 2 | 1 | 3 | 1 | 2 | 1 | 9 | 0.8889 |
| 11_776008 | 2 | 0 | 1 | 0 | NA | NA | 2 | 0.5 | 5 | 0.2 | 2 | 1 | 3 | 0 | 3 | 0.6667 | 2 | 0.5 | 10 | 0.5 |
| 11_776013 | 3 | 0.3333 | 2 | 0 | 1 | 1 | 4 | 0.75 | 10 | 0.5 | 4 | 0.25 | 4 | 0.5 | 4 | 0.25 | 4 | 0.25 | 16 | 0.3125 |
| 11_776034 | 1 | 0 | 4 | 0.25 | 1 | 0 | 2 | 1 | 8 | 0.375 | 5 | 0.8 | 2 | 0 | 4 | 0.25 | 3 | 0 | 14 | 0.3571 |
| 11_776034: | 4 | 0.25 | 5 | 0 | 1 | 0 | 4 | 0.25 | 14 | 0.1429 | 7 | 0.1429 | 2 | 0 | 7 | 0 | 2 | 0 | 18 | 0.0556 |
| 11_776034 | 3 | 0 | 5 | 0.4 | 1 | 0 | 4 | 0 | 13 | 0.1538 | 6 | 0.1667 | 2 | 0 | 8 | 0.125 | 2 | 0 | 18 | 0.1111 |
| 11_776047 | 2 | 0 | 4 | 0.25 | 1 | 0 | 2 | 0 | 9 | 0.1111 | 3 | 0.6667 | NA | NA | 3 | 0 | NA | NA | 6 | 0.3333 |
| 11_776062: | 2 | 0 | 6 | 0 | 2 | 0 | 4 | 0.25 | 14 | 0.0714 | 3 | 0 | 2 | 0 | 5 | 0.2 | 1 | 0 | 11 | 0.0909 |
| 11_776076: | 1 | 0 | 4 | 0 | 3 | 0.3333 | 1 | 0 | 9 | 0.1111 | 4 | 0.25 | 2 | 0 | 3 | 0 | 2 | 0 | 11 | 0.0909 |
| 11_776085 | 1 | 0 | 4 | 0 | 5 | 0.2 | 3 | 0 | 13 | 0.0769 | 6 | 0.1667 | 3 | 0.3333 | 4 | 0 | 3 | 0 | 16 | 0.125 |
| 11_776684 | 1 | 0 | NA | NA | NA | NA | 1 | 1 | 2 | 0.5 | NA | NA | NA | NA | NA | NA | 1 | 1 | 1 | 1 |
| 11_776685: | 1 | 0 | NA | NA | NA | NA | 1 | 1 | 2 | 0.5 | NA | NA | NA | NA | NA | NA | 1 | 1 | 1 | 1 |
| 11_777398 | 1 | 1 | NA | NA | NA | NA | NA | NA | 1 | 1 | NA | NA | 1 | 1 | NA | NA | NA | NA | 1 | 1 |
| 11_777501 | 1 | 1 | NA | NA | 1 | 1 | NA | NA | 2 | 1 | 1 | 1 | 1 | 0 | NA | NA | NA | NA | 2 | 0.5 |
| 11_777504 | 1 | 1 | NA | NA | 1 | 1 | NA | NA | 2 | 1 | 1 | 1 | NA | NA | NA | NA | NA | NA | 1 | 1 |
| 11_777556 | 7 | 0.2857 | 7 | 0.1429 | 10 | 0.2 | 14 | 0.3571 | 38 | 0.2632 | 5 | 0.2 | 16 | 0.1875 | 8 | 0.125 | 14 | 0.0714 | 43 | 0.1395 |
| 11_777557: | 5 | 1 | 6 | 0.5 | 8 | 0.25 | 15 | 0.4667 | 34 | 0.5 | 4 | 1 | 15 | 0.5333 | 6 | 0.6667 | 13 | 0.3846 | 38 | 0.5526 |
| 11_777557 | 7 | 0.8571 | 7 | 1 | 8 | 0.875 | 12 | 0.9167 | 34 | 0.9118 | 4 | 1 | 15 | 0.9333 | 7 | 1 | 11 | 1 | 37 | 0.973 |
| 11_777559 | 2 | 0.5 | 8 | 0.25 | 4 | 0.25 | 7 | 0.7143 | 21 | 0.4286 | 6 | 0.6667 | 16 | 0.375 | 6 | 0.8333 | 9 | 0.5556 | 37 | 0.5405 |
| 11_777559 | 6 | 0.3333 | 8 | 0.5 | 9 | 0.6667 | 14 | 0.7143 | 37 | 0.5946 | 7 | 0.4286 | 20 | 0.45 | 8 | 0.375 | 15 | 0.5333 | 50 | 0.46 |
| 11_777564 | 9 | 0.2222 | 10 | 0.3 | 12 | 0.0833 | 20 | 0.2 | 51 | 0.1961 | 9 | 0.2222 | 18 | 0.2222 | 13 | 0.3077 | 17 | 0.2941 | 57 | 0.2632 |
| 11_777566 | 10 | 0.1 | 15 | 0.2667 | 14 | 0.0714 | 22 | 0 | 61 | 0.0984 | 12 | 0.0833 | 20 | 0.15 | 14 | 0 | 19 | 0.0526 | 65 | 0.0769 |
| 11_777567 | 10 | 0 | 14 | 0.2143 | 14 | 0.2143 | 22 | 0.1364 | 60 | 0.15 | 11 | 0.0909 | 20 | 0.1 | 12 | 0.0833 | 19 | 0.1053 | 62 | 0.0968 |
| 11_778442 | 10 | 0.1 | 11 | 0.1818 | 10 | 0.1 | 15 | 0.2 | 46 | 0.1522 | 12 | 0 | 9 | 0.2222 | 11 | 0.1818 | 7 | 0 | 39 | 0.1026 |
| 11_778444 | 11 | 0.6364 | 12 | 0.8333 | 14 | 0.8571 | 18 | 0.5 | 55 | 0.6909 | 12 | 0.5 | 11 | 0.6364 | 13 | 0.8462 | 11 | 0.7273 | 47 | 0.6809 |
| 11_778445 | 11 | 0.0909 | 14 | 0.4286 | 13 | 0.3077 | 18 | 0.0556 | 56 | 0.2143 | 12 | 0.5 | 10 | 0.1 | 13 | 0.0769 | 12 | 0.3333 | 47 | 0.2553 |
| 11_778445 | 13 | 0 | 15 | 0.2 | 15 | 0 | 22 | 0.0909 | 65 | 0.0769 | 13 | 0.1538 | 11 | 0.2727 | 16 | 0.1875 | 14 | 0 | 54 | 0.1481 |
| 11_778446 | 14 | 0.5 | 12 | 0.3333 | 15 | 0.4 | 20 | 0.4 | 61 | 0.4098 | 11 | 0.3636 | 9 | 0.4444 | 14 | 0.5 | 14 | 0.4286 | 48 | 0.4375 |
| 11_778447 | 13 | 0.8462 | 9 | 0.8889 | 13 | 0.9231 | 24 | 0.7917 | 59 | 0.8475 | 10 | 0.9 | 9 | 1 | 16 | 0.9375 | 15 | 1 | 50 | 0.96 |
| 11_778447 | 10 | 0.4 | 11 | 0.8182 | 12 | 0.75 | 23 | 0.4783 | 56 | 0.5893 | 7 | 0.2857 | 11 | 0.5455 | 11 | 0.6364 | 13 | 0.6154 | 42 | 0.5476 |
| 11_778447 | 15 | 0.2667 | 12 | 0.25 | 15 | 0.3333 | 23 | 0.3043 | 65 | 0.2923 | 11 | 0.2727 | 9 | 0.3333 | 10 | 0.4 | 15 | 0.4 | 45 | 0.3556 |
| 11_778449 | 1 | 0 | 2 | 0.5 | NA | NA | 1 | 1 | 4 | 0.5 | NA | NA | 1 | 0 | NA | NA | 1 | 1 | 2 | 0.5 |
| 11_778456 | 14 | 0.7857 | 14 | 0.7857 | 15 | 0.6 | 29 | 0.4828 | 72 | 0.625 | 13 | 0.6923 | 13 | 0.3846 | 19 | 0.7368 | 16 | 0.75 | 61 | 0.6557 |

| | | | | | | | | | | | | | | | | | | | | |
|---|---|---|---|---|---|---|---|---|---|---|---|---|---|---|---|---|---|---|---|---|
| 11_778646 | 24 | 0.4583 | 22 | 0.1818 | 25 | 0.36 | 39 | 0.2308 | 110 | 0.3 | 17 | 0.4118 | 17 | 0.2941 | 28 | 0.1071 | 25 | 0.32 | 87 | 0.2644 |
| 11_778690 | 40 | 0.2 | 37 | 0.0541 | 42 | 0.1429 | 59 | 0.1525 | 178 | 0.1404 | 30 | 0.1 | 43 | 0.2093 | 56 | 0.2143 | 38 | 0.0789 | 167 | 0.1617 |
| 11_778691 | 43 | 0.2558 | 35 | 0.4 | 40 | 0.325 | 59 | 0.3729 | 177 | 0.339 | 28 | 0.25 | 42 | 0.2857 | 51 | 0.3333 | 37 | 0.3784 | 158 | 0.3165 |
| 11_778691 | 34 | 0.0882 | 30 | 0.0333 | 37 | 0 | 50 | 0.1 | 151 | 0.0596 | 26 | 0.0385 | 34 | 0.0588 | 44 | 0.0455 | 31 | 0.0645 | 135 | 0.0519 |
| 11_778692 | 44 | 0.6591 | 40 | 0.7 | 40 | 0.75 | 65 | 0.7077 | 189 | 0.7037 | 29 | 0.7241 | 45 | 0.6667 | 57 | 0.614 | 42 | 0.6429 | 173 | 0.6532 |
| 11_778692 | 43 | 0.0233 | 36 | 0.1944 | 43 | 0.093 | 57 | 0.0877 | 179 | 0.095 | 31 | 0.0968 | 41 | 0.1463 | 58 | 0.0862 | 42 | 0.0476 | 172 | 0.093 |
| 11_778693 | 46 | 0.6304 | 37 | 0.6486 | 38 | 0.7105 | 61 | 0.5574 | 182 | 0.6264 | 30 | 0.7 | 39 | 0.6667 | 54 | 0.6667 | 36 | 0.5556 | 159 | 0.6478 |
| 11_778694 | 46 | 0.587 | 42 | 0.5238 | 46 | 0.413 | 66 | 0.6061 | 200 | 0.54 | 31 | 0.7742 | 46 | 0.5217 | 63 | 0.5238 | 47 | 0.4255 | 187 | 0.5401 |
| 11_778695 | 46 | 0.1304 | 43 | 0.1163 | 45 | 0.0222 | 63 | 0.0952 | 197 | 0.0914 | 32 | 0 | 45 | 0.2 | 60 | 0.0833 | 44 | 0.1364 | 181 | 0.1105 |
| 11_778697 | 59 | 0.0678 | 51 | 0.1961 | 55 | 0.0909 | 80 | 0.05 | 245 | 0.0939 | 36 | 0.1389 | 49 | 0.0816 | 71 | 0.0845 | 56 | 0.1607 | 212 | 0.1132 |
| 11_778697 | 58 | 0.069 | 49 | 0.0612 | 53 | 0.0566 | 72 | 0.0833 | 232 | 0.069 | 37 | 0.0811 | 50 | 0.06 | 71 | 0.0563 | 56 | 0.1071 | 214 | 0.0748 |
| 11_778721 | 1 | 1 | NA | NA | NA | NA | NA | NA | 1 | 1 | 1 | 1 | NA | NA | NA | NA | NA | NA | 1 | 1 |
| 11_778722 | 1 | 1 | NA | NA | NA | NA | NA | NA | 1 | 1 | 1 | 1 | NA | NA | NA | NA | NA | NA | 1 | 1 |
| 11_778724 | 1 | 1 | NA | NA | 1 | 0 | NA | NA | 2 | 0.5 | 2 | 0.5 | NA | NA | NA | NA | NA | NA | 2 | 0.5 |
| 11_778726 | 1 | 1 | NA | NA | 1 | 1 | NA | NA | 2 | 1 | 2 | 0.5 | NA | NA | NA | NA | NA | NA | 2 | 0.5 |
| 11_778726 | 1 | 1 | NA | NA | NA | NA | NA | NA | 1 | 1 | 2 | 0.5 | NA | NA | NA | NA | NA | NA | 2 | 0.5 |
| 11_778730 | 1 | 1 | NA | NA | NA | NA | NA | NA | 1 | 1 | 2 | 1 | NA | NA | NA | NA | NA | NA | 2 | 1 |
| 11_778733 | 1 | 1 | NA | NA | NA | NA | NA | NA | 1 | 1 | 1 | 1 | NA | NA | NA | NA | NA | NA | 1 | 1 |
| 11_778739 | NA | NA | 1 | 1 | 1 | 0 | NA | NA | 2 | 0.5 | NA | NA | 1 | 0 | NA | NA | 1 | 1 | 2 | 0.5 |
| 11_778743 | 1 | 1 | NA | NA | NA | NA | NA | NA | 1 | 1 | 1 | 1 | NA | NA | NA | NA | NA | NA | 1 | 1 |
| 11_778745 | 1 | 1 | NA | NA | NA | NA | NA | NA | 1 | 1 | 1 | 1 | NA | NA | NA | NA | NA | NA | 1 | 1 |
| 11_778749 | 1 | 1 | NA | NA | NA | NA | NA | NA | 1 | 1 | 1 | 1 | NA | NA | NA | NA | NA | NA | 1 | 1 |
| 11_779002 | NA | NA | 2 | 0 | 2 | 0.5 | NA | NA | 4 | 0.25 | 1 | 1 | 2 | 0.5 | 3 | 0 | 2 | 0 | 8 | 0.25 |
| 11_779002 | NA | NA | 2 | 0 | 2 | 0.5 | 1 | 0 | 5 | 0.2 | 1 | 1 | 4 | 0.5 | 4 | 0.25 | 3 | 0.6667 | 12 | 0.5 |
| 11_779005 | NA | NA | 2 | 0.5 | 2 | 1 | 1 | 0 | 5 | 0.6 | 1 | 1 | 4 | 0.75 | 3 | 1 | 3 | 0.6667 | 11 | 0.8182 |
| 11_779005 | NA | NA | 2 | 1 | 2 | 0.5 | 1 | 0 | 5 | 0.6 | 1 | 1 | 4 | 0.5 | 2 | 0.5 | 2 | 0 | 9 | 0.4444 |
| 11_779139 | NA | NA | 3 | 0.6667 | 3 | 1 | 1 | 1 | 7 | 0.8571 | 2 | 1 | 3 | 1 | 6 | 0.8333 | 2 | 1 | 13 | 0.9231 |
| 11_779139 | NA | NA | 3 | 0.3333 | 2 | 0 | 1 | 1 | 6 | 0.3333 | 2 | 0.5 | 4 | 0.25 | 6 | 0.1667 | 2 | 0 | 14 | 0.2143 |
| 11_779139 | NA | NA | 3 | 0.6667 | 2 | 0 | 1 | 1 | 6 | 0.5 | 2 | 1 | 3 | 0.3333 | 7 | 0.2857 | 3 | 0.6667 | 15 | 0.4667 |
| 11_779141 | NA | NA | 2 | 1 | 2 | 1 | 1 | 1 | 5 | 1 | 2 | 1 | 3 | 1 | 6 | 1 | 2 | 1 | 13 | 1 |
| 11_779141 | NA | NA | 2 | 0.5 | 2 | 0.5 | 1 | 1 | 5 | 0.6 | 2 | 0 | 3 | 0 | 8 | 0.375 | 2 | 0 | 15 | 0.2 |
| 11_779144 | NA | NA | 4 | 1 | 3 | 1 | 3 | 1 | 10 | 1 | 2 | 1 | 5 | 0.8 | 6 | 0.6667 | 3 | 1 | 16 | 0.8125 |
| 11_779145 | NA | NA | 4 | 0.25 | 2 | 0 | 3 | 0.6667 | 9 | 0.3333 | 2 | 0 | 5 | 0.2 | 6 | 0.1667 | 3 | 0 | 16 | 0.125 |
| 11_779168 | 1 | 1 | 2 | 0 | 2 | 0 | 3 | 0 | 8 | 0.125 | 2 | 0 | 5 | 0.4 | 8 | 0.125 | 3 | 0.3333 | 18 | 0.2222 |
| 11_779204 | 3 | 0.3333 | 7 | 0.2857 | 3 | 0.3333 | 1 | 0 | 14 | 0.2857 | 1 | 1 | 6 | 0.1667 | 6 | 0.3333 | 1 | 0 | 14 | 0.2857 |
| 11_779205 | 4 | 0.5 | 7 | 0.7143 | 3 | 0.3333 | 1 | 0 | 15 | 0.5333 | 1 | 1 | 5 | 0.8 | 6 | 0.5 | 3 | 0.6667 | 15 | 0.6667 |
| 11_779205 | 3 | 0 | 5 | 0 | 3 | 0.3333 | NA | NA | 11 | 0.0909 | 1 | 1 | 1 | 0 | 4 | 0.25 | 2 | 0 | 8 | 0.25 |
| 11_779206 | 3 | 0.3333 | 6 | 0.3333 | NA | NA | 1 | 1 | 10 | 0.4 | 1 | 1 | 6 | 0.1667 | 5 | 0.4 | 1 | 0 | 13 | 0.3077 |
| 11_779206 | 3 | 0.6667 | 6 | 0.8333 | 3 | 0.6667 | 1 | 1 | 13 | 0.7692 | 1 | 1 | 6 | 0.6667 | 7 | 0.7143 | 2 | 0.5 | 16 | 0.6875 |
| 11_779207 | 4 | 0 | 6 | 0.5 | 2 | 0.5 | 2 | 0.5 | 14 | 0.3571 | 1 | 1 | 6 | 0.5 | 7 | 0.8571 | 3 | 0 | 17 | 0.5882 |
| 11_779208 | 5 | 0 | 7 | 0.2857 | 1 | 0 | 2 | 0 | 15 | 0.1333 | NA | NA | 7 | 0.1429 | 8 | 0.25 | 4 | 0.25 | 19 | 0.2105 |
| 11_779209 | 3 | 0 | 4 | 0.25 | 2 | 0.5 | NA | NA | 9 | 0.2222 | 1 | 0 | 3 | 0.6667 | 5 | 0.2 | 2 | 1 | 11 | 0.4545 |
| 11_779210 | 1 | 1 | 5 | 0.6 | 3 | 0.3333 | NA | NA | 9 | 0.5556 | 1 | 0 | 4 | 0.5 | 6 | 0.1667 | 3 | 0.6667 | 14 | 0.3571 |
| 11_779252 | 1 | 1 | NA | NA | NA | NA | NA | NA | 1 | 1 | 2 | 1 | NA | NA | 1 | 0 | NA | NA | 3 | 0.6667 |
| 11_779280 | 4 | 1 | NA | NA | 1 | 1 | NA | NA | 5 | 1 | 1 | 1 | 2 | 1 | 1 | 1 | NA | NA | 4 | 1 |
| 11_779280 | 4 | 0.75 | 2 | 0.5 | 1 | 0 | NA | NA | 7 | 0.5714 | 2 | 0.5 | 2 | 1 | 1 | 1 | 1 | 1 | 6 | 0.8333 |
| 11_779281 | 4 | 0.25 | 2 | 0 | 1 | 0 | NA | NA | 7 | 0.1429 | 1 | 0 | 2 | 0.5 | 1 | 0 | 1 | 1 | 5 | 0.4 |
| 11_779283 | 3 | 0.6667 | 2 | 1 | 1 | 1 | NA | NA | 6 | 0.8333 | 2 | 1 | 2 | 1 | 1 | 1 | 1 | 1 | 6 | 1 |
| 11_779284 | 4 | 0.5 | 2 | 0.5 | 1 | 0 | NA | NA | 7 | 0.4286 | 2 | 0.5 | 2 | 0 | 1 | 0 | 1 | 1 | 6 | 0.3333 |
| 11_779284 | 2 | 0 | 1 | 1 | 1 | 0 | NA | NA | 4 | 0.25 | 2 | 0.5 | 2 | 0.5 | 1 | 0 | 1 | 1 | 6 | 0.5 |
| 11_779285 | 3 | 0.6667 | 2 | 0.5 | 2 | 0.5 | NA | NA | 7 | 0.5714 | 2 | 0 | 2 | 0.5 | 1 | 0 | 1 | 1 | 6 | 0.3333 |
| 11_779499 | 8 | 0 | 13 | 0 | 6 | 0.3333 | 5 | 0 | 32 | 0.0625 | 8 | 0.375 | 7 | 0 | 8 | 0 | 4 | 0.25 | 27 | 0.1481 |
| 11_779501 | 8 | 0.125 | 13 | 0.0769 | 6 | 0.1667 | 7 | 0.2857 | 34 | 0.1471 | 8 | 0.125 | 7 | 0.1429 | 8 | 0.125 | 5 | 0.2 | 28 | 0.1429 |
| 11_779501 | 7 | 0.1429 | 14 | 0.0714 | 6 | 0.1667 | 6 | 0.1667 | 33 | 0.1212 | 8 | 0.125 | 7 | 0.1429 | 8 | 0 | 5 | 0 | 28 | 0.0714 |
| 11_779647 | 2 | 0 | 3 | 0 | 2 | 0 | 6 | 0.1667 | 13 | 0.0769 | 4 | 0.5 | 2 | 0 | 2 | 0.5 | 1 | 1 | 9 | 0.4444 |
| 11_779647 | 1 | 1 | 3 | 1 | 1 | 1 | 6 | 1 | 11 | 1 | 3 | 1 | 1 | 1 | 2 | 1 | 2 | 1 | 8 | 1 |
| 11_779647 | 2 | 0 | 3 | 0.3333 | 2 | 1 | 7 | 1 | 14 | 0.7143 | 3 | 0.6667 | 2 | 1 | 2 | 0.5 | 2 | 0.5 | 9 | 0.6667 |
| 11_779649 | 2 | 0 | 3 | 1 | 1 | 0 | 4 | 0.25 | 10 | 0.4 | 5 | 0.4 | 1 | 1 | 2 | 0 | 2 | 0 | 10 | 0.3 |
| 11_780445 | 3 | 0.3333 | 1 | 1 | 2 | 0 | 2 | 0 | 8 | 0.25 | 2 | 0.5 | 1 | 0 | 1 | 0 | 1 | 0 | 5 | 0.2 |
| 11_780446 | 2 | 1 | 1 | 1 | 1 | 1 | 1 | 0 | 5 | 0.8 | 2 | 0.5 | 1 | 0 | 2 | 0.5 | 1 | 1 | 6 | 0.5 |
| 11_780447 | 3 | 1 | 1 | 1 | 3 | 1 | 2 | 1 | 9 | 1 | 2 | 1 | 1 | 1 | 2 | 1 | 1 | 1 | 6 | 1 |
| 11_780448 | 3 | 0 | 1 | 0 | 2 | 0 | 2 | 0.5 | 8 | 0.125 | 2 | 0 | 1 | 0 | 2 | 0.5 | 1 | 0 | 6 | 0.1667 |
| 11_780449 | 3 | 0.3333 | 1 | 0 | 4 | 0.25 | 2 | 0 | 10 | 0.2 | 3 | 0.3333 | 1 | 0 | 3 | 0.6667 | NA | NA | 7 | 0.4286 |
| 11_780449 | 1 | 0 | 1 | 0 | 4 | 0.25 | 2 | 0.5 | 8 | 0.25 | 2 | 0.5 | 1 | 1 | 3 | 0 | NA | NA | 6 | 0.3333 |
| 11_780451 | 4 | 0.25 | 1 | 1 | 5 | 0.2 | 2 | 0.5 | 12 | 0.3333 | 3 | 0.3333 | 1 | 1 | 2 | 0.5 | 1 | 0 | 7 | 0.4286 |

| | | | | | | | | | | | | | | | | | | | | |
|---|---|---|---|---|---|---|---|---|---|---|---|---|---|---|---|---|---|---|---|---|
| 11_780456 | 4 | 1 | 1 | 1 | 4 | 1 | 2 | 1 | 11 | 1 | 3 | 0.6667 | 2 | 1 | 3 | 1 | 1 | 0 | 9 | 0.7778 |
| 11_780458 | 3 | 0 | 2 | 0.5 | 4 | 0.25 | 3 | 0.6667 | 12 | 0.3333 | 2 | 0.5 | 2 | 0 | 3 | 0 | 1 | 0 | 8 | 0.125 |
| 11_780458 | 4 | 0 | 3 | 0 | 4 | 0.25 | 3 | 0 | 14 | 0.0714 | 4 | 0.25 | 2 | 0 | 2 | 0 | 1 | 0 | 9 | 0.1111 |
| 11_780459 | 4 | 1 | 3 | 1 | 4 | 1 | 3 | 1 | 14 | 1 | 3 | 0.6667 | 2 | 1 | 3 | 1 | 1 | 1 | 9 | 0.8889 |
| 11_780459 | 4 | 0.5 | 2 | 0.5 | 5 | 0.2 | 3 | 0 | 14 | 0.2857 | 4 | 0 | 1 | 0 | 3 | 0.3333 | 1 | 0 | 9 | 0.1111 |
| 11_780464 | 4 | 0 | 3 | 0.3333 | 6 | 0.1667 | 4 | 0.25 | 17 | 0.1765 | 2 | 0 | 2 | 0 | 3 | 0 | 1 | 1 | 8 | 0.125 |
| 11_780467 | 3 | 0.3333 | 4 | 0.25 | 6 | 0.1667 | 4 | 0 | 17 | 0.1765 | 5 | 0.2 | 3 | 0 | 3 | 0 | 1 | 0 | 12 | 0.0833 |
| 11_780472 | 3 | 0 | 4 | 0.5 | 6 | 0 | 4 | 0 | 17 | 0.1176 | 4 | 0 | 3 | 0 | 3 | 0.3333 | NA | NA | 10 | 0.1 |
| 11_780472 | 5 | 0 | 4 | 0.25 | 3 | 0.3333 | 4 | 0 | 16 | 0.125 | 4 | 0.25 | 2 | 0 | 4 | 0.25 | NA | NA | 10 | 0.2 |
| 11_780474 | 5 | 0 | 3 | 0 | 6 | 0 | 5 | 0.2 | 19 | 0.0526 | 3 | 0.6667 | 3 | 0 | 5 | 0 | NA | NA | 11 | 0.1818 |
| 11_780504 | 1 | 0 | 5 | 1 | 3 | 1 | 2 | 1 | 11 | 0.9091 | 1 | 1 | 1 | 1 | NA | NA | 1 | 1 | 3 | 1 |
| 11_780506 | 1 | 1 | 4 | 0 | 4 | 0 | 2 | 0.5 | 11 | 0.1818 | 1 | 0 | 1 | 0 | NA | NA | 1 | 1 | 3 | 0.3333 |
| 11_780507 | 1 | 1 | 5 | 0.6 | 4 | 1 | 2 | 1 | 12 | 0.8333 | 1 | 1 | 1 | 1 | NA | NA | 1 | 1 | 3 | 1 |
| 11_780507 | 1 | 1 | 5 | 1 | 2 | 1 | 2 | 1 | 10 | 1 | 1 | 1 | 1 | 1 | NA | NA | 1 | 1 | 3 | 1 |
| 11_780508 | 1 | 0 | 5 | 0.2 | 4 | 0.25 | 2 | 0.5 | 12 | 0.25 | 1 | 0 | 1 | 0 | NA | NA | 1 | 1 | 3 | 0.3333 |
| 11_780508 | 1 | 0 | 5 | 0.2 | 4 | 0.25 | 2 | 0.5 | 12 | 0.25 | 1 | 0 | 1 | 1 | NA | NA | 1 | 0 | 3 | 0.3333 |
| 11_780509 | 1 | 0 | 5 | 0.8 | 2 | 1 | 2 | 0.5 | 10 | 0.7 | NA | NA | NA | NA | NA | NA | 1 | 1 | 1 | 1 |
| 11_780509 | 1 | 1 | 4 | 0.5 | 4 | 0.5 | 2 | 0.5 | 11 | 0.5455 | 1 | 0 | 1 | 0 | NA | NA | 1 | 1 | 3 | 0.3333 |
| 11_780509 | 1 | 0 | 5 | 0.2 | 4 | 0.25 | 2 | 0 | 12 | 0.1667 | 1 | 0 | 1 | 1 | NA | NA | 1 | 0 | 3 | 0.3333 |
| 11_780510 | 1 | 1 | 4 | 0.25 | 4 | 0.75 | 1 | 0 | 10 | 0.5 | 1 | 1 | 1 | 1 | NA | NA | 1 | 1 | 3 | 1 |
| 11_780696 | 3 | 0 | 4 | 0 | 1 | 1 | 1 | 0 | 9 | 0.1111 | 1 | 1 | 2 | 0 | 2 | 0 | 3 | 0 | 8 | 0.125 |
| 11_780710 | 3 | 1 | 7 | 0.8571 | 4 | 1 | 2 | 1 | 16 | 0.9375 | 3 | 1 | 3 | 1 | 4 | 1 | 4 | 0.75 | 14 | 0.9286 |
| 11_780710 | 3 | 1 | 5 | 0.8 | 4 | 0.75 | 2 | 1 | 14 | 0.8571 | 3 | 0.6667 | 3 | 1 | 4 | 1 | 4 | 1 | 14 | 0.9286 |
| 11_780711 | 3 | 0.3333 | 4 | 0.25 | 3 | 0 | 1 | 0 | 11 | 0.1818 | 3 | 0 | 3 | 0.3333 | 4 | 0.25 | 4 | 0.25 | 14 | 0.2143 |
| 11_780711 | 2 | 1 | 7 | 0.7143 | 4 | 0.25 | 1 | 0 | 14 | 0.5714 | 3 | 0.6667 | 3 | 0.3333 | 3 | 0.6667 | 3 | 0.3333 | 12 | 0.5 |
| 11_780711 | 3 | 0 | 6 | 0.1667 | 4 | 0 | 2 | 0 | 15 | 0.0667 | 3 | 0.3333 | 4 | 0 | 3 | 0 | 4 | 0 | 14 | 0.0714 |
| 11_780711 | 3 | 0.6667 | 7 | 1 | 3 | 0.6667 | 2 | 1 | 15 | 0.8667 | 3 | 1 | 4 | 1 | 4 | 0.5 | 3 | 1 | 14 | 0.8571 |
| 11_780712 | 2 | 0 | 6 | 0.5 | 3 | 0 | 1 | 0 | 12 | 0.25 | 2 | 0 | 3 | 0 | 3 | 0 | 4 | 0.5 | 12 | 0.1667 |
| 11_780713 | 1 | 0 | 4 | 0.5 | 1 | 0 | 2 | 0 | 8 | 0.25 | 3 | 0 | 1 | 0 | 2 | 0 | 3 | 0.3333 | 9 | 0.1111 |
| 11_780718 | 2 | 0.5 | 2 | 0.5 | 1 | 0 | 1 | 0 | 6 | 0.3333 | NA | NA | NA | NA | 1 | 1 | 2 | 0 | 3 | 0.3333 |
| 11_780722 | 2 | 0.5 | NA | NA | 1 | 0 | 1 | 0 | 4 | 0.25 | NA | NA | NA | NA | 2 | 0.5 | 2 | 0.5 | 4 | 0.5 |
| 11_780728 | 2 | 0 | 2 | 0 | 1 | 1 | 2 | 0.5 | 7 | 0.2857 | NA | NA | 1 | 0 | 3 | 0 | 2 | 0.5 | 6 | 0.1667 |
| 11_780732 | 2 | 0 | 2 | 0.5 | 1 | 0 | 1 | 0 | 6 | 0.1667 | NA | NA | NA | NA | 1 | 1 | 2 | 0 | 3 | 0.3333 |
| 11_780741 | NA | NA | 1 | 1 | 1 | 1 | 1 | 1 | 3 | 1 | 3 | 0.6667 | 1 | 1 | 1 | 1 | NA | NA | 5 | 0.8 |
| 11_780828 | 4 | 0.25 | 1 | 0 | 3 | 0 | 1 | 1 | 9 | 0.2222 | 7 | 0.1429 | 1 | 0 | 4 | 0 | 1 | 0 | 13 | 0.0769 |
| 11_780828 | 4 | 0.5 | 1 | 0 | 3 | 0.3333 | 1 | 0 | 9 | 0.3333 | 7 | 0.2857 | 1 | 1 | 4 | 0.5 | 1 | 0 | 13 | 0.3846 |
| 11_780976 | 3 | 1 | NA | NA | 1 | 1 | 4 | 0.5 | 8 | 0.75 | 2 | 0 | 3 | 0.3333 | 2 | 1 | NA | NA | 7 | 0.4286 |
| 11_780977 | 2 | 0 | NA | NA | NA | NA | 1 | 1 | 3 | 0.3333 | 1 | 0 | 1 | 1 | 2 | 0.5 | NA | NA | 4 | 0.5 |
| 11_780978 | 11 | 0.0909 | 8 | 0.125 | 7 | 0 | 11 | 0.1818 | 37 | 0.1081 | 9 | 0 | 9 | 0.1111 | 7 | 0.1429 | 6 | 0.3333 | 31 | 0.129 |
| 11_780979 | 16 | 0.3125 | 10 | 0.6 | 11 | 0.7273 | 17 | 0.3529 | 54 | 0.463 | 10 | 0.3 | 11 | 0.6364 | 11 | 0.7273 | 10 | 0.6 | 42 | 0.5714 |
| 11_780980 | 18 | 0.0556 | 12 | 0.1667 | 11 | 0.0909 | 13 | 0 | 54 | 0.0741 | 12 | 0.0833 | 13 | 0.0769 | 12 | 0.1667 | 11 | 0.0909 | 48 | 0.1042 |
| 11_780983 | 15 | 0.9333 | 11 | 0.9091 | 13 | 1 | 14 | 0.8571 | 53 | 0.9245 | 14 | 0.8571 | 12 | 1 | 13 | 0.9231 | 10 | 0.8 | 49 | 0.898 |
| 11_781364 | 2 | 0.5 | 1 | 0 | NA | NA | NA | NA | 3 | 0.3333 | 1 | 0 | 1 | 1 | 1 | 0 | 1 | 0 | 4 | 0.25 |
| 11_781457 | 8 | 0.5 | 10 | 0.3 | 12 | 0.75 | 11 | 0.3636 | 41 | 0.4878 | 11 | 0.6364 | 11 | 0.0909 | 6 | 0.6667 | 10 | 0.6 | 38 | 0.4737 |
| 11_781457 | 10 | 0.6 | 11 | 0.9091 | 11 | 0.7273 | 13 | 0.6923 | 45 | 0.7333 | 13 | 0.6154 | 10 | 0.8 | 8 | 0.75 | 9 | 1 | 40 | 0.775 |
| 11_781457 | 11 | 0.0909 | 8 | 0.125 | 11 | 0.2727 | 14 | 0.0714 | 44 | 0.1364 | 11 | 0 | 10 | 0.2 | 8 | 0.25 | 9 | 0 | 38 | 0.1053 |
| 11_781458 | 10 | 0.6 | 10 | 0.6 | 14 | 0.5714 | 13 | 0.3846 | 47 | 0.5319 | 12 | 0.3333 | 11 | 0.4545 | 7 | 0.5714 | 7 | 0.4286 | 37 | 0.4324 |
| 11_781459 | 12 | 0.6667 | 9 | 0.7778 | 14 | 0.7143 | 13 | 0.6923 | 48 | 0.7083 | 12 | 0.5833 | 10 | 0.8 | 8 | 0.875 | 8 | 0.625 | 38 | 0.7105 |
| 11_781460 | 10 | 0.1 | 11 | 0.1818 | 14 | 0.2857 | 14 | 0.1429 | 49 | 0.1837 | 14 | 0.0714 | 12 | 0.0833 | 10 | 0.3 | 8 | 0.375 | 44 | 0.1818 |
| 11_781460 | 10 | 0.1 | 11 | 0.2727 | 14 | 0 | 16 | 0.25 | 51 | 0.1569 | 15 | 0 | 12 | 0.0833 | 8 | 0.125 | 10 | 0.1 | 45 | 0.0667 |
| 11_781461 | 11 | 1 | 11 | 0.9091 | 10 | 1 | 15 | 0.9333 | 47 | 0.9574 | 13 | 1 | 13 | 0.9231 | 10 | 1 | 10 | 1 | 46 | 0.9783 |
| 11_781462 | 11 | 0.0909 | 10 | 0.2 | 10 | 0.3 | 12 | 0 | 43 | 0.1395 | 14 | 0.2143 | 12 | 0.1667 | 6 | 0 | 8 | 0.25 | 40 | 0.175 |
| 11_781463 | 1 | 0 | 1 | 0 | 1 | 1 | 4 | 0 | 7 | 0.1429 | 3 | 0.3333 | 5 | 0 | 1 | 1 | NA | NA | 9 | 0.2222 |
| 11_781603 | 4 | 0.25 | 4 | 0.5 | 3 | 0.3333 | 6 | 0.1667 | 17 | 0.2941 | 2 | 0 | 8 | 0 | 3 | 0.3333 | 3 | 0.3333 | 16 | 0.125 |
| 11_781623 | 3 | 1 | 3 | 0.3333 | 4 | 0.25 | 6 | 0.5 | 16 | 0.5 | 2 | 0 | 6 | 0.6667 | 3 | 0.6667 | 3 | 0.3333 | 14 | 0.5 |
| 11_781626 | 3 | 0 | 4 | 0.25 | 3 | 1 | 5 | 0.2 | 15 | 0.3333 | 2 | 0 | 7 | 0.2857 | 3 | 0 | 3 | 0.3333 | 15 | 0.2 |
| 11_781626 | 3 | 1 | 4 | 1 | 1 | 1 | 2 | 1 | 10 | 1 | 1 | 1 | 6 | 1 | 2 | 1 | 1 | 1 | 10 | 1 |
| 11_781627 | 4 | 1 | 5 | 0.8 | 3 | 1 | 5 | 0.6 | 17 | 0.8235 | 1 | 1 | 7 | 0.5714 | 2 | 1 | 2 | 0.5 | 12 | 0.6667 |
| 11_781628 | 4 | 0.25 | 5 | 0.2 | 3 | 0 | 5 | 0 | 17 | 0.1176 | 2 | 0 | 8 | 0 | 3 | 0 | 4 | 0.25 | 17 | 0.0588 |
| 11_781628 | 4 | 1 | 5 | 1 | 3 | 1 | 6 | 0.8333 | 18 | 0.9444 | 3 | 1 | 8 | 1 | 3 | 0.6667 | 4 | 1 | 18 | 0.9444 |
| 11_781629 | 2 | 0.5 | 3 | 0 | 1 | 0 | 6 | 0.1667 | 12 | 0.1667 | 3 | 0 | 8 | 0.625 | 2 | 0.5 | 4 | 0.5 | 17 | 0.4706 |
| 11_781631 | 4 | 0.75 | 4 | 1 | 3 | 1 | 7 | 0.8571 | 18 | 0.8889 | 3 | 0.3333 | 8 | 0.75 | 3 | 0.3333 | 5 | 0.4 | 19 | 0.5263 |
| 11_781636 | 5 | 0.2 | 4 | 0.5 | 4 | 0.25 | 7 | 0.5714 | 20 | 0.4 | 3 | 0 | 9 | 0.1111 | 4 | 0.25 | 5 | 0 | 21 | 0.0952 |
| 11_781636 | 5 | 0 | 5 | 0 | 4 | 0.25 | 7 | 0.1429 | 21 | 0.0952 | 3 | 0.3333 | 9 | 0 | 3 | 0 | 4 | 0 | 19 | 0.0526 |
| 11_781639 | 3 | 0 | 3 | 0 | 3 | 0.3333 | 4 | 0.25 | 13 | 0.1538 | 2 | 0 | 9 | 0 | 5 | 0.2 | 5 | 0.2 | 21 | 0.0952 |

| | | | | | | | | | | | | | | | | | | | | |
|---|---|---|---|---|---|---|---|---|---|---|---|---|---|---|---|---|---|---|---|---|
| 11_7816399 | 5 | 0 | 3 | 0.6667 | 4 | 0.25 | 8 | 0 | 20 | 0.15 | 2 | 0 | 7 | 0.1429 | 5 | 0 | 4 | 0 | 18 | 0.0556 |
| 11_7816794 | 9 | 0.2222 | 5 | 0 | 5 | 0 | 7 | 0 | 26 | 0.0769 | 3 | 0.3333 | 5 | 0.2 | 6 | 0 | 5 | 0 | 19 | 0.1053 |
| 11_7816795 | 8 | 0.5 | 8 | 0.5 | 5 | 0 | 9 | 0.2222 | 30 | 0.3333 | 4 | 0.5 | 10 | 0.4 | 5 | 0.2 | 6 | 0.1667 | 25 | 0.32 |
| 11_7818084 | 16 | 0.8125 | 15 | 0.8667 | 14 | 0.4286 | 14 | 0.3571 | 59 | 0.6271 | 12 | 0.5833 | 15 | 0.6 | 12 | 0.6667 | 7 | 0.7143 | 46 | 0.6304 |
| 11_7818086 | 18 | 0.8333 | 16 | 0.6875 | 14 | 0.7143 | 14 | 0.5714 | 62 | 0.7097 | 12 | 0.75 | 17 | 1 | 13 | 0.6923 | 9 | 0.7778 | 51 | 0.8235 |
| 11_7818092 | 17 | 0.4118 | 13 | 0.5385 | 12 | 0.5 | 15 | 0.3333 | 57 | 0.4386 | 13 | 0.2308 | 16 | 0.3125 | 14 | 0.5 | 7 | 0.1429 | 50 | 0.32 |
| 11_7818092 | 16 | 0.375 | 13 | 0.5385 | 11 | 0.5455 | 16 | 0.6875 | 56 | 0.5357 | 10 | 0.6 | 16 | 0.6875 | 13 | 0.3077 | 8 | 0.5 | 47 | 0.5319 |
| 11_7818096 | 12 | 0.4167 | 11 | 0.1818 | 9 | 0.1111 | 11 | 0.2727 | 43 | 0.2558 | 14 | 0.2857 | 13 | 0.3077 | 12 | 0.25 | 10 | 0.3 | 49 | 0.2857 |
| 11_7821741 | 103 | 0.0777 | 150 | 0.06 | 154 | 0.0974 | 97 | 0.0825 | 504 | 0.0794 | 141 | 0.0638 | 166 | 0.0723 | 137 | 0.0803 | 105 | 0.0381 | 549 | 0.0656 |
| 11_7821984 | 5 | 0.2 | 3 | 0 | 2 | 0.5 | 3 | 0 | 13 | 0.1538 | 4 | 0.25 | 2 | 0 | 6 | 0 | 5 | 0 | 17 | 0.0588 |
| 11_7823283 | 4 | 0.25 | 6 | 0.6667 | 2 | 0.5 | 3 | 0 | 15 | 0.4 | 2 | 0 | 5 | 0.6 | 7 | 0.7143 | 9 | 0.3333 | 23 | 0.4783 |
| 11_7823306 | 6 | 0.3333 | 9 | 0.5556 | 5 | 0.8 | 7 | 0.7143 | 27 | 0.5926 | 6 | 0.6667 | 7 | 0.5714 | 4 | 0.5 | 8 | 0.375 | 25 | 0.52 |
| 11_7823309 | 3 | 0 | 9 | 0.3333 | 5 | 0.4 | 5 | 0 | 22 | 0.2273 | 3 | 0 | 5 | 0 | 5 | 0.4 | 12 | 0.1667 | 25 | 0.16 |
| 11_7823338 | 8 | 0 | 11 | 0.1818 | 7 | 0 | 10 | 0 | 36 | 0.0556 | 6 | 0.1667 | 6 | 0.5 | 9 | 0.2222 | 12 | 0.0833 | 33 | 0.2121 |
| 11_7823340 | 8 | 0.5 | 11 | 0.3636 | 7 | 0 | 10 | 0.3 | 36 | 0.3056 | 5 | 0.4 | 8 | 0 | 9 | 0.3333 | 12 | 0.0833 | 34 | 0.1765 |
| 11_7823357 | 6 | 0 | 10 | 0.2 | 4 | 0 | 9 | 0 | 29 | 0.069 | 6 | 0.1667 | 7 | 0 | 7 | 0 | 10 | 0.2 | 30 | 0.1 |
| 11_7823956 | 4 | 0 | 2 | 0.5 | 5 | 0 | 4 | 0 | 15 | 0.0667 | 5 | 0 | 2 | 0 | 1 | 0 | 3 | 0.3333 | 11 | 0.0909 |
| 11_7823994 | 5 | 0 | 2 | 0 | 4 | 0 | 5 | 0.2 | 16 | 0.0625 | 7 | 0 | 2 | 0.5 | 1 | 0 | 3 | 0 | 13 | 0.0769 |
| 11_7835723 | NA | NA | 1 | 1 | 3 | 0.3333 | NA | NA | 4 | 0.5 | 1 | 0 | 4 | 0.5 | 2 | 0.5 | 1 | 0 | 8 | 0.375 |
| 11_7835752 | NA | NA | 1 | 1 | 2 | 0.5 | NA | NA | 3 | 0.6667 | 1 | 0 | 4 | 1 | 2 | 0 | 1 | 0 | 8 | 0.5 |
| 11_7835784 | NA | NA | 2 | 0.5 | 2 | 1 | NA | NA | 4 | 0.75 | 1 | 1 | 4 | 1 | 2 | 1 | 1 | 1 | 8 | 1 |
| 11_7836148 | NA | NA | 1 | 0 | 1 | 0 | 2 | 0.5 | 4 | 0.25 | 2 | 0 | 1 | 1 | 1 | 0 | NA | NA | 4 | 0.25 |
| 11_7836428 | 2 | 0 | 2 | 0.5 | 2 | 0.5 | 3 | 0.6667 | 9 | 0.4444 | 2 | 0.5 | 5 | 0.2 | 3 | 0 | 2 | 0.5 | 12 | 0.25 |
| 11_7836454 | 1 | 0 | 2 | 0.5 | 1 | 0 | 3 | 0.3333 | 7 | 0.2857 | 2 | 1 | 4 | 0.25 | 3 | 0 | 2 | 0 | 11 | 0.2727 |
| 11_7836517 | 1 | 0 | 1 | 1 | 2 | 0.5 | 2 | 0 | 6 | 0.3333 | 2 | 1 | 2 | 0 | 3 | 0 | 1 | 0 | 8 | 0.25 |
| 11_7836517 | 2 | 0 | 1 | 1 | 2 | 0 | 2 | 0 | 7 | 0.1429 | 2 | 1 | 3 | 0 | 3 | 0.3333 | 1 | 0 | 9 | 0.3333 |
| 11_7836554 | NA | NA | 1 | 1 | NA | NA | 1 | 0 | 2 | 0.5 | 2 | 0.5 | 3 | 0 | 2 | 0 | 1 | 0 | 8 | 0.125 |
| 11_7836562 | 1 | 0 | 1 | 1 | 1 | 0 | 1 | 0 | 4 | 0.25 | 2 | 1 | 3 | 0.6667 | 2 | 0.5 | 1 | 0 | 8 | 0.625 |
| 11_7836564 | 2 | 0 | 1 | 0 | 1 | 1 | 1 | 0 | 5 | 0.2 | 1 | 1 | 3 | 0 | 1 | 0 | 1 | 0 | 6 | 0.1667 |
| 11_7839269 | 19 | 0.4737 | 18 | 0.3889 | 14 | 0.3571 | 16 | 0.3125 | 67 | 0.3881 | 12 | 0.25 | 8 | 0.625 | 11 | 0.2727 | 5 | 0 | 36 | 0.3056 |
| 11_7839300 | 21 | 0.0476 | 18 | 0.1111 | 13 | 0 | 16 | 0.1875 | 68 | 0.0882 | 12 | 0.0833 | 9 | 0 | 10 | 0.1 | 4 | 0 | 35 | 0.0571 |
| 11_7839820 | 6 | 0.5 | 2 | 0 | 5 | 0.6 | 6 | 0.6667 | 19 | 0.5263 | 6 | 0.3333 | 6 | 0.5 | 5 | 0.6 | 4 | 0.75 | 21 | 0.5238 |
| 11_7839832 | 7 | 0.5714 | NA | NA | 5 | 0.6 | 10 | 0.7 | 22 | 0.6364 | 7 | 0.4286 | 9 | 0.6667 | 5 | 0.8 | 6 | 0.6667 | 27 | 0.6296 |
| 11_7839847 | 7 | 0.8571 | 1 | 0 | 6 | 0 | 8 | 0 | 22 | 0.2727 | 5 | 0.2 | 9 | 0.3333 | 6 | 0.8333 | 4 | 0.75 | 24 | 0.5 |
| 11_7839855 | 7 | 0 | 2 | 0 | 8 | 0.125 | 10 | 0.1 | 27 | 0.0741 | 8 | 0.125 | 10 | 0 | 6 | 0.1667 | 5 | 0.2 | 29 | 0.1034 |
| 11_7839858 | 6 | 0 | 2 | 0 | 7 | 0.1429 | 9 | 0.1111 | 24 | 0.0833 | 8 | 0.125 | 10 | 0.1 | 5 | 0.2 | 6 | 0 | 29 | 0.1034 |
| 11_7841279 | 9 | 0.4444 | 17 | 0.5294 | 13 | 0.3846 | 12 | 0.4167 | 51 | 0.451 | 15 | 0.5333 | 7 | 0.5714 | 9 | 0.4444 | 6 | 0.8333 | 37 | 0.5676 |
| 11_7841287 | 8 | 0 | 19 | 0.1053 | 15 | 0.1333 | 13 | 0.2308 | 55 | 0.1273 | 11 | 0.0909 | 6 | 0 | 9 | 0.1111 | 8 | 0.625 | 34 | 0.2059 |
| 11_7841290 | 8 | 0.125 | 17 | 0.0588 | 14 | 0.1429 | 13 | 0.0769 | 52 | 0.0962 | 15 | 0.2 | 7 | 0.2857 | 10 | 0.1 | 7 | 0.1429 | 39 | 0.1795 |
| 11_7841293 | 12 | 0.1667 | 17 | 0.1176 | 17 | 0.0588 | 15 | 0.2 | 61 | 0.1311 | 16 | 0.125 | 8 | 0 | 10 | 0.1 | 7 | 0.1429 | 41 | 0.0976 |
| 11_7841305 | 9 | 0.4444 | 17 | 0.3529 | 13 | 0.1538 | 9 | 0.2222 | 48 | 0.2917 | 12 | 0.3333 | 7 | 0.2857 | 10 | 0.2 | 8 | 0.25 | 37 | 0.2703 |
| 11_7841313 | 6 | 0.5 | 15 | 0.2 | 11 | 0.2727 | 10 | 0.1 | 42 | 0.2381 | 7 | 0.4286 | 8 | 0.125 | 8 | 0 | 5 | 0.4 | 28 | 0.2143 |
| 11_7841316 | 6 | 0.5 | 10 | 0.4 | 9 | 0.5556 | 6 | 0.6667 | 31 | 0.5161 | 7 | 0.1429 | 4 | 0 | 7 | 0.4286 | 6 | 0.8333 | 24 | 0.375 |
| 11_7841322 | 8 | 0.375 | 16 | 0.6875 | 10 | 0.5 | 9 | 0.6667 | 43 | 0.5814 | 8 | 0.5 | 6 | 0.6667 | 10 | 0.4 | 6 | 0.8333 | 30 | 0.5667 |
| 11_7841331 | 8 | 0.125 | 18 | 0.5556 | 14 | 0.5 | 9 | 0.5556 | 49 | 0.4694 | 13 | 0.3077 | 8 | 0.25 | 9 | 0.2222 | 8 | 0.375 | 38 | 0.2895 |
| 11_7841346 | 9 | 0.5556 | 16 | 0.75 | 12 | 0.4167 | 8 | 0.75 | 45 | 0.6222 | 11 | 0.7273 | 9 | 0.3333 | 10 | 0.5 | 6 | 0.5 | 36 | 0.5278 |
| 11_7841386 | 4 | 0.25 | 3 | 0 | 3 | 0 | 5 | 0.2 | 15 | 0.1333 | 8 | 0.125 | 1 | 0 | 7 | 0.1429 | 3 | 0 | 19 | 0.1053 |
| 11_7841387 | 4 | 0.25 | 2 | 0 | 3 | 0 | 3 | 0.3333 | 12 | 0.1667 | 7 | 0.2857 | NA | NA | 5 | 0 | 3 | 0 | 15 | 0.1333 |
| 11_7841394 | 4 | 0.25 | 3 | 0 | 3 | 0.3333 | 5 | 0 | 15 | 0.1333 | 9 | 0.1111 | NA | NA | 7 | 0.4286 | 4 | 0.25 | 20 | 0.25 |
| 11_7841402 | 4 | 0.75 | 2 | 0 | 2 | 0.5 | 4 | 0.5 | 12 | 0.5 | 10 | 0.6 | NA | NA | 3 | 0.3333 | 4 | 0.5 | 17 | 0.5294 |
| 11_7841407 | 3 | 0.3333 | 1 | 0 | 2 | 0 | 5 | 0 | 11 | 0.0909 | 10 | 0.2 | NA | NA | 5 | 0.2 | 2 | 0 | 17 | 0.1765 |
| 11_7841474 | 4 | 0.5 | 2 | 0.5 | 1 | 1 | 2 | 1 | 9 | 0.6667 | 7 | 0.5714 | NA | NA | 5 | 0.8 | 3 | 1 | 15 | 0.7333 |
| 11_7841484 | 3 | 0 | 1 | 0 | 1 | 0 | 1 | 1 | 6 | 0.1667 | 7 | 0.1429 | 1 | 0 | 5 | 0.2 | 4 | 0 | 17 | 0.1176 |
| 11_7841486 | 4 | 0 | 2 | 0.5 | 1 | 0 | 2 | 0 | 9 | 0.1111 | 8 | 0 | NA | NA | 4 | 0 | 5 | 0.2 | 17 | 0.0588 |
| 11_7841572 | 2 | 0 | 1 | 0 | 1 | 1 | 2 | 0 | 6 | 0.1667 | 3 | 0 | 1 | 0 | 1 | 0 | 3 | 0.3333 | 8 | 0.125 |
| 11_7841573 | 4 | 0 | 3 | 0 | 1 | 0 | 2 | 0.5 | 10 | 0.1 | 7 | 0.1429 | 2 | 0 | 1 | 1 | 5 | 0 | 15 | 0.1333 |
| 11_7841577 | 4 | 0.25 | 2 | 0 | 1 | 0 | 1 | 0 | 8 | 0.125 | 6 | 0 | 1 | 0 | 1 | 1 | 5 | 0.2 | 13 | 0.1538 |
| 11_7841612 | 3 | 0 | 3 | 0 | 1 | 1 | 1 | 0 | 8 | 0.125 | 4 | 0 | 2 | 0 | NA | NA | 4 | 0.25 | 10 | 0.1 |
| 11_7841613 | 1 | 0 | 3 | 0.3333 | 1 | 0 | 1 | 1 | 6 | 0.3333 | 3 | 0.3333 | 1 | 0 | NA | NA | 3 | 0.6667 | 7 | 0.4286 |
| 11_7841621 | 2 | 0 | 3 | 0 | 1 | 1 | 1 | 0 | 7 | 0.1429 | 5 | 0 | 1 | 0 | NA | NA | 4 | 0.75 | 10 | 0.3 |
| 11_7841622 | 2 | 0.5 | 3 | 0 | 1 | 1 | NA | NA | 6 | 0.3333 | 5 | 0.4 | 1 | 0 | NA | NA | 2 | 0 | 8 | 0.25 |
| 11_7841629 | NA | NA | 2 | 0.5 | NA | NA | 2 | 0.5 | 4 | 0.5 | 5 | 0.2 | 1 | 0 | NA | NA | 2 | 0 | 8 | 0.125 |
| 11_7841851 | 1 | 0 | 2 | 0.5 | 1 | 0 | NA | NA | 4 | 0.25 | 6 | 0 | 1 | 0 | NA | NA | 4 | 0.25 | 11 | 0.0909 |
| 11_7842728 | 6 | 0.1667 | 7 | 0.2857 | 3 | 0 | 3 | 0 | 19 | 0.1579 | 1 | 0 | 2 | 0.5 | 3 | 0 | 3 | 0 | 9 | 0.1111 |
| 11_7843226 | 6 | 0.6667 | 10 | 0.5 | 6 | 0.5 | 10 | 0.5 | 32 | 0.5312 | 3 | 0.6667 | 10 | 0.8 | 6 | 1 | 3 | 1 | 22 | 0.8636 |

| | | | | | | | | | | | | | | | | | | | | |
|---|---|---|---|---|---|---|---|---|---|---|---|---|---|---|---|---|---|---|---|---|
| 11_7843279 | 4 | 0.25 | 3 | 0.6667 | 1 | 1 | 2 | 0.5 | 10 | 0.5 | NA | NA | 3 | 0.6667 | 1 | 0 | 1 | 1 | 5 | 0.6 |
| 11_7843280 | 3 | 0 | 3 | 0.3333 | 2 | 0 | 2 | 0 | 10 | 0.1 | NA | NA | 3 | 0.3333 | 1 | 0 | 2 | 0 | 6 | 0.1667 |
| 11_7843323 | 3 | 1 | 3 | 0.6667 | 1 | 1 | 2 | 0.5 | 9 | 0.7778 | NA | NA | NA | NA | 1 | 0 | 2 | 0.5 | 3 | 0.3333 |
| 11_7843331 | 4 | 0.25 | 3 | 0.6667 | 1 | 1 | 2 | 0 | 10 | 0.4 | NA | NA | 1 | 1 | 1 | 0 | 2 | 0 | 4 | 0.25 |
| 11_7843350 | 2 | 1 | 2 | 0.5 | 1 | 0 | 2 | 0.5 | 7 | 0.5714 | NA | NA | 1 | 1 | NA | NA | 2 | 0 | 3 | 0.3333 |
| 11_7845841 | 14 | 0.4286 | 7 | 0.4286 | 5 | 0.6 | 8 | 0.625 | 34 | 0.5 | 8 | 0.375 | 6 | 0.3333 | 11 | 0.4545 | 5 | 0.6 | 30 | 0.4333 |
| 11_7845855 | 14 | 0.2857 | 7 | 0.1429 | 5 | 0 | 8 | 0 | 34 | 0.1471 | 9 | 0 | 6 | 0.1667 | 8 | 0.125 | 6 | 0 | 29 | 0.069 |
| 11_7845859 | 14 | 0.1429 | 5 | 0.2 | 4 | 0.25 | 6 | 0.3333 | 29 | 0.2069 | 11 | 0 | 5 | 0 | 11 | 0.1818 | 5 | 0.4 | 32 | 0.125 |
| 11_7845868 | 14 | 0.3571 | 6 | 0.1667 | 3 | 0.3333 | 5 | 0 | 28 | 0.25 | 10 | 0 | 5 | 0.4 | 11 | 0.3636 | 7 | 0 | 33 | 0.1818 |
| 11_7845886 | 12 | 0.0833 | 6 | 0.1667 | 4 | 0 | 2 | 0 | 24 | 0.0833 | 9 | 0 | 6 | 0 | 9 | 0.1111 | 6 | 0.3333 | 30 | 0.1 |
| 11_7858668 | 3 | 0.6667 | 2 | 0 | 2 | 0 | 1 | 0 | 8 | 0.25 | 4 | 0 | 3 | 0.3333 | 3 | 0.6667 | 2 | 0 | 12 | 0.25 |
| 11_7864142 | 5 | 0.4 | NA | NA | 3 | 0.3333 | NA | NA | 8 | 0.375 | 4 | 0.75 | 5 | 1 | 2 | 0.5 | NA | NA | 11 | 0.8182 |
| 11_7864146 | 7 | 0.8571 | NA | NA | 4 | 0.5 | 1 | 1 | 12 | 0.75 | 5 | 0.8 | 7 | 0.7143 | 5 | 1 | 1 | 1 | 18 | 0.8333 |
| 11_7864153 | 6 | 1 | NA | NA | 4 | 0.75 | 1 | 1 | 11 | 0.9091 | 5 | 1 | 6 | 0.6667 | 2 | 1 | 1 | 1 | 14 | 0.8571 |
| 11_7864160 | 6 | 0.6667 | 1 | 0 | 4 | 0.5 | NA | NA | 11 | 0.5455 | 5 | 0.6 | 7 | 0.7143 | 5 | 0.6 | 1 | 1 | 18 | 0.6667 |
| 11_7864164 | 4 | 1 | 1 | 1 | 4 | 1 | 1 | 1 | 10 | 1 | 3 | 1 | 6 | 1 | 4 | 1 | 1 | 1 | 14 | 1 |
| 11_7864171 | 6 | 0 | 1 | 1 | 4 | 0.25 | 1 | 1 | 12 | 0.25 | 4 | 0 | 7 | 0 | 6 | 0.1667 | 1 | 0 | 18 | 0.0556 |
| 11_7864171 | 5 | 0.8 | NA | NA | 3 | 0.3333 | 1 | 0 | 9 | 0.5556 | 5 | 0.8 | 5 | 1 | 4 | 0.75 | 1 | 1 | 15 | 0.8667 |
| 11_7864179 | 5 | 0.6 | NA | NA | 4 | 0.75 | 1 | 1 | 10 | 0.7 | 3 | 1 | 2 | 1 | 4 | 0.75 | 1 | 1 | 10 | 0.9 |
| 11_7864181 | 7 | 0 | 1 | 0 | 4 | 0.25 | 1 | 0 | 13 | 0.0769 | 6 | 0.1667 | 7 | 0.2857 | 5 | 0.2 | 1 | 0 | 19 | 0.2105 |
| 11_7864199 | 7 | 0.1429 | 1 | 0 | 4 | 0 | 1 | 0 | 13 | 0.0769 | 5 | 0 | 7 | 0.2857 | 5 | 0.2 | 1 | 0 | 18 | 0.1667 |
| 11_7864202 | 5 | 0.6 | NA | NA | 3 | 0.6667 | 1 | 0 | 9 | 0.5556 | 4 | 0.5 | 7 | 0.7143 | 3 | 1 | 1 | 1 | 15 | 0.7333 |
| 11_7864212 | 7 | 0.1429 | 1 | 0 | 4 | 0.25 | 1 | 0 | 13 | 0.1538 | 6 | 0 | 7 | 0 | 6 | 0.1667 | 1 | 0 | 20 | 0.05 |
| 11_7873009 | 1 | 0 | 1 | 0 | 1 | 0 | 2 | 0.5 | 5 | 0.2 | NA | NA | 1 | 0 | 1 | 0 | 1 | 1 | 3 | 0.3333 |
| 11_7873009 | 1 | 0 | NA | NA | 1 | 1 | 2 | 1 | 4 | 0.75 | NA | NA | 1 | 1 | NA | NA | 1 | 1 | 2 | 1 |
| 11_7873458 | 1 | 1 | NA | NA | 1 | 0 | 2 | 0 | 4 | 0.25 | 2 | 0.5 | 2 | 0 | NA | NA | 1 | 0 | 5 | 0.2 |
| 11_7873468 | NA | NA | NA | NA | 1 | 1 | 2 | 1 | 3 | 1 | 2 | 1 | 2 | 0 | NA | NA | NA | NA | 4 | 0.5 |
| 11_7873474 | 1 | 0 | NA | NA | NA | NA | 1 | 1 | 2 | 0.5 | 1 | 1 | NA | NA | NA | NA | NA | NA | 1 | 1 |
| 11_7873550 | 1 | 1 | NA | NA | 1 | 1 | NA | NA | 2 | 1 | 1 | 1 | 1 | 1 | 1 | 1 | NA | NA | 3 | 1 |
| 11_7873553 | 1 | 1 | NA | NA | 1 | 1 | 1 | 1 | 3 | 1 | 2 | 1 | NA | NA | 1 | 0 | NA | NA | 3 | 0.6667 |
| 11_7873566 | 1 | 0 | NA | NA | 1 | 1 | 1 | 1 | 3 | 0.6667 | 2 | 1 | 2 | 1 | NA | NA | NA | NA | 4 | 1 |
| 11_7873595 | 1 | 1 | NA | NA | 1 | 1 | NA | NA | 2 | 1 | 2 | 1 | 2 | 0.5 | 1 | 0 | NA | NA | 5 | 0.6 |
| 11_7873709 | 1 | 1 | NA | NA | 1 | 1 | 1 | 0 | 3 | 0.6667 | 2 | 0.5 | 2 | 1 | 2 | 1 | NA | NA | 6 | 0.8333 |
| 11_7873714 | 1 | 0 | NA | NA | 1 | 1 | 1 | 0 | 3 | 0.3333 | 1 | 1 | 3 | 0.6667 | 2 | 1 | NA | NA | 6 | 0.8333 |
| 11_7873731 | 1 | 1 | NA | NA | NA | NA | 1 | 1 | 2 | 1 | 2 | 1 | 3 | 1 | 2 | 1 | NA | NA | 7 | 1 |
| 11_7873758 | 2 | 0 | 1 | 1 | 1 | 0 | 1 | 0 | 5 | 0.2 | 1 | 0 | 2 | 0 | 2 | 0.5 | NA | NA | 5 | 0.2 |
| 11_7890492 | 9 | 0.4444 | 10 | 0.4 | 2 | 0.5 | 4 | 0.25 | 25 | 0.4 | 4 | 0.5 | 4 | 0 | 3 | 1 | 3 | 0.6667 | 14 | 0.5 |
| 11_7890556 | 9 | 0 | 10 | 0 | 2 | 1 | 2 | 0 | 23 | 0.087 | 2 | 0.5 | 3 | 0 | 1 | 0 | 3 | 0.3333 | 9 | 0.2222 |
| 11_7890588 | 6 | 0.3333 | 7 | 0.2857 | 3 | 0.3333 | 2 | 0.5 | 18 | 0.3333 | 1 | 0 | 2 | 0 | 2 | 0 | 2 | 0.5 | 7 | 0.1429 |
| 11_7890645 | 4 | 0.5 | 5 | 0.4 | 1 | 1 | 1 | 1 | 11 | 0.5455 | 2 | 0.5 | 2 | 1 | 1 | 1 | 2 | 1 | 7 | 0.8571 |
| 11_7890690 | 4 | 1 | 4 | 1 | 1 | 1 | 1 | 1 | 10 | 1 | 1 | 1 | 1 | 1 | NA | NA | 1 | 1 | 3 | 1 |
| 11_7890704 | 7 | 0.2857 | 6 | 0.3333 | 1 | 0 | 2 | 0 | 16 | 0.25 | 2 | 0 | 2 | 0 | NA | NA | 2 | 0.5 | 6 | 0.1667 |
| 11_7890714 | 2 | 1 | 6 | 1 | 1 | 1 | 1 | 1 | 10 | 1 | 1 | 1 | 2 | 1 | NA | NA | 1 | 1 | 4 | 1 |
| 11_7890714 | 4 | 0 | 5 | 0 | 1 | 0 | 1 | 1 | 11 | 0.0909 | 1 | 0 | NA | NA | NA | NA | 1 | 1 | 2 | 0.5 |
| 11_7890721 | 5 | 0.4 | 5 | 0.2 | 1 | 0 | 1 | 0 | 12 | 0.25 | 1 | 0 | 2 | 1 | NA | NA | 1 | 0 | 4 | 0.5 |
| 11_7890729 | 5 | 0.4 | 6 | 0.1667 | 1 | 0 | 2 | 0.5 | 14 | 0.2857 | 1 | 0 | 2 | 0.5 | NA | NA | 1 | 0 | 4 | 0.25 |
| 11_7890769 | 5 | 0.4 | 3 | 0 | 1 | 0 | 2 | 0.5 | 11 | 0.2727 | 1 | 1 | 1 | 0 | NA | NA | NA | NA | 2 | 0.5 |
| 11_7890775 | 5 | 0.6 | 3 | 0.3333 | NA | NA | 2 | 1 | 10 | 0.6 | 2 | 1 | 1 | 1 | NA | NA | NA | NA | 3 | 1 |
| 11_7890776 | 5 | 0.2 | 3 | 0 | 1 | 0 | 2 | 0 | 11 | 0.0909 | 2 | 0 | 1 | 1 | NA | NA | NA | NA | 3 | 0.3333 |
| 11_7890796 | 4 | 0.25 | 3 | 0.3333 | NA | NA | 2 | 0.5 | 9 | 0.3333 | 3 | 0.3333 | 1 | 1 | NA | NA | NA | NA | 4 | 0.5 |
| 11_7890808 | 3 | 0.3333 | 3 | 0.3333 | NA | NA | 3 | 1 | 9 | 0.5556 | 3 | 0.3333 | 1 | 0 | NA | NA | NA | NA | 4 | 0.25 |
| 11_7891848 | 3 | 0.3333 | 1 | 0 | NA | NA | 1 | 0 | 5 | 0.2 | 2 | 0.5 | 1 | 1 | NA | NA | 1 | 0 | 4 | 0.5 |
| 11_7893590 | 2 | 0.5 | 1 | 0 | NA | NA | NA | NA | 3 | 0.3333 | 1 | 1 | NA | NA | NA | NA | 1 | 0 | 2 | 0.5 |
| 11_7893602 | 2 | 1 | 1 | 1 | NA | NA | NA | NA | 3 | 1 | 1 | 1 | NA | NA | NA | NA | NA | NA | 1 | 1 |
| 11_7893607 | 2 | 0.5 | NA | NA | NA | NA | NA | NA | 2 | 0.5 | 1 | 1 | NA | NA | NA | NA | NA | NA | 1 | 1 |
| 11_7913091 | 3 | 0.3333 | 4 | 0 | 2 | 0 | 7 | 0 | 16 | 0.0625 | 1 | 0 | 2 | 0 | 7 | 0 | 5 | 0.2 | 15 | 0.0667 |
| 11_7913097 | 4 | 0.75 | 2 | 0.5 | 3 | 1 | 6 | 0.8333 | 15 | 0.8 | 2 | 1 | 2 | 0.5 | 6 | 0.6667 | 2 | 1 | 12 | 0.75 |
| 11_7913112 | 4 | 0 | 4 | 0.25 | 3 | 0.3333 | 7 | 0.2857 | 18 | 0.2222 | 2 | 0.5 | 1 | 0 | 5 | 0.2 | 3 | 0 | 11 | 0.1818 |
| 11_7913117 | 4 | 0.25 | 2 | 0 | 2 | 0.5 | 4 | 0.25 | 12 | 0.25 | 2 | 0 | 1 | 0 | 4 | 0.25 | 2 | 0.5 | 9 | 0.2222 |
| 11_7933468 | NA | NA | 1 | 0 | 2 | 0 | 1 | 1 | 4 | 0.25 | 1 | 1 | NA | NA | NA | NA | 3 | 0 | 4 | 0.25 |
| 11_7939230 | 15 | 0 | 21 | 0.0952 | 14 | 0 | 24 | 0.125 | 74 | 0.0676 | 21 | 0.0952 | 18 | 0.0556 | 28 | 0.1429 | 18 | 0.1111 | 85 | 0.1059 |
| 11_7939239 | 19 | 0.2105 | 22 | 0.2273 | 19 | 0.0526 | 24 | 0.4167 | 84 | 0.2381 | 21 | 0.2857 | 19 | 0.1053 | 30 | 0.2 | 17 | 0.2941 | 87 | 0.2184 |
| 11_7939249 | 15 | 0.2 | 20 | 0 | 20 | 0.25 | 22 | 0.1364 | 77 | 0.1429 | 21 | 0.0476 | 20 | 0.15 | 25 | 0.16 | 16 | 0.0625 | 82 | 0.1098 |
| 11_7939261 | 12 | 0.1667 | 22 | 0.0909 | 16 | 0 | 22 | 0.2273 | 72 | 0.125 | 21 | 0.1429 | 16 | 0.0625 | 22 | 0.0455 | 13 | 0.0769 | 72 | 0.0833 |
| 11_7939268 | 11 | 0.1818 | 13 | 0.3846 | 13 | 0.3077 | 12 | 0.1667 | 49 | 0.2653 | 14 | 0.3571 | 10 | 0.4 | 22 | 0.3636 | 11 | 0.2727 | 57 | 0.3509 |

| | | | | | | | | | | | | | | | | | | | | |
|---|---|---|---|---|---|---|---|---|---|---|---|---|---|---|---|---|---|---|---|---|
| 11_7939297 | 10 | 0.1 | 9 | 0.1111 | 8 | 0.125 | 10 | 0.1 | 37 | 0.1081 | 13 | 0.0769 | 8 | 0.125 | 18 | 0.1111 | 12 | 0.0833 | 51 | 0.098 |
| 11_7939298 | 14 | 0.5 | 21 | 0.1905 | 15 | 0.4667 | 20 | 0.6 | 70 | 0.4286 | 15 | 0.4667 | 15 | 0.5333 | 23 | 0.4348 | 16 | 0.25 | 69 | 0.4203 |
| 11_7939299 | 13 | 0 | 21 | 0.1429 | 16 | 0.25 | 19 | 0.0526 | 69 | 0.1159 | 18 | 0 | 16 | 0.125 | 26 | 0.1538 | 17 | 0.1765 | 77 | 0.1169 |
| 11_7939305 | 12 | 0 | 14 | 0.2143 | 14 | 0.0714 | 15 | 0.1333 | 55 | 0.1091 | 15 | 0.2 | 14 | 0.1429 | 26 | 0.1923 | 17 | 0.0588 | 72 | 0.1528 |
| 11_7939309 | 15 | 0.8667 | 20 | 0.75 | 16 | 0.6875 | 19 | 0.7368 | 70 | 0.7571 | 18 | 0.7222 | 15 | 0.8667 | 27 | 0.7778 | 17 | 0.7647 | 77 | 0.7792 |
| 11_7939314 | 9 | 0.2222 | 15 | 0.2 | 10 | 0.2 | 11 | 0.1818 | 45 | 0.2 | 15 | 0 | 14 | 0.2143 | 18 | 0.3889 | 16 | 0.125 | 63 | 0.1905 |
| 11_7939314 | NA | NA | NA | NA | 1 | 1 | 2 | 0 | 3 | 0.3333 | 1 | 1 | NA | NA | 2 | 0.5 | NA | NA | 3 | 0.6667 |
| 11_7939318 | 10 | 0.1 | 12 | 0 | 12 | 0.1667 | 13 | 0.0769 | 47 | 0.0851 | 17 | 0.1765 | 11 | 0 | 21 | 0.0952 | 11 | 0 | 60 | 0.0833 |
| 11_7939322 | 9 | 0.2222 | 16 | 0.4375 | 14 | 0.2143 | 18 | 0.3889 | 57 | 0.3333 | 16 | 0.3125 | 15 | 0.2667 | 21 | 0.381 | 15 | 0.1333 | 67 | 0.2836 |
| 11_7939329 | 14 | 0.8571 | 19 | 0.9474 | 16 | 0.875 | 19 | 1 | 68 | 0.9265 | 15 | 1 | 15 | 0.8667 | 26 | 0.8846 | 18 | 0.8889 | 74 | 0.9054 |
| 11_7939331 | 12 | 0 | 17 | 0.2353 | 11 | 0 | 14 | 0.1429 | 54 | 0.1111 | 10 | 0.2 | 14 | 0.0714 | 20 | 0 | 12 | 0 | 56 | 0.0536 |
| 11_7939331 | 13 | 0.0769 | 18 | 0.1667 | 14 | 0.1429 | 19 | 0.2105 | 64 | 0.1562 | 16 | 0.125 | 15 | 0.0667 | 21 | 0.1429 | 18 | 0.1111 | 70 | 0.1143 |
| 11_7939333 | 11 | 0.7273 | 19 | 0.3158 | 16 | 0.75 | 17 | 0.7059 | 63 | 0.6032 | 15 | 0.4667 | 16 | 0.6875 | 22 | 0.6818 | 16 | 0.5625 | 69 | 0.6087 |
| 11_7939337 | 11 | 0.9091 | 16 | 0.9375 | 13 | 0.9231 | 18 | 1 | 58 | 0.9483 | 15 | 0.9333 | 15 | 0.9333 | 19 | 1 | 16 | 1 | 65 | 0.9692 |
| 11_7939347 | 12 | 0.1667 | 14 | 0.1429 | 14 | 0.2857 | 16 | 0 | 56 | 0.1429 | 13 | 0 | 15 | 0.1333 | 21 | 0.2381 | 18 | 0.1667 | 67 | 0.1493 |
| 11_7939351 | 6 | 0.1667 | 17 | 0.1176 | 11 | 0.0909 | 15 | 0.0667 | 49 | 0.102 | 11 | 0.0909 | 8 | 0.625 | 20 | 0.1 | 12 | 0.1667 | 51 | 0.1961 |
| 11_7939354 | 9 | 0.2222 | 13 | 0.1538 | 13 | 0.0769 | 14 | 0.1429 | 49 | 0.1429 | 11 | 0.0909 | 14 | 0.2857 | 20 | 0.2 | 16 | 0.125 | 61 | 0.1803 |
| 11_7939360 | 9 | 0.2222 | 15 | 0.0667 | 10 | 0 | 14 | 0.1429 | 48 | 0.1042 | 14 | 0.0714 | 12 | 0 | 17 | 0.1176 | 13 | 0 | 56 | 0.0536 |
| 11_7939368 | 11 | 0.5455 | 18 | 0.2778 | 13 | 0.6154 | 17 | 0.5882 | 59 | 0.4915 | 12 | 0.5 | 13 | 0.5385 | 19 | 0.4211 | 15 | 0.2667 | 59 | 0.4237 |
| 11_7939369 | 9 | 0.7778 | 18 | 0.8333 | 7 | 0.5714 | 15 | 0.8 | 49 | 0.7755 | 13 | 0.7692 | 13 | 0.7692 | 20 | 0.9 | 15 | 0.6667 | 61 | 0.7869 |
| 11_7939371 | 11 | 0.1818 | 15 | 0 | 15 | 0 | 18 | 0.1111 | 59 | 0.0678 | 13 | 0 | 15 | 0.1333 | 18 | 0.1667 | 16 | 0.1875 | 62 | 0.129 |
| 11_7939372 | 10 | 1 | 15 | 0.9333 | 15 | 0.9333 | 18 | 0.8333 | 58 | 0.9138 | 11 | 0.9091 | 13 | 1 | 18 | 0.9444 | 16 | 0.9375 | 58 | 0.9483 |
| 11_7941739 | NA | NA | 9 | 0.2222 | 7 | 0.1429 | 1 | 0 | 17 | 0.1765 | 2 | 0 | 6 | 0.1667 | 4 | 0 | 1 | 0 | 13 | 0.0769 |
| 11_7941745 | 1 | 0 | 9 | 0 | 8 | 0.25 | 1 | 0 | 19 | 0.1053 | 3 | 0.3333 | 5 | 0.6 | 4 | 0.25 | 1 | 0 | 13 | 0.3846 |
| 11_7941764 | 2 | 0 | 8 | 0.125 | 9 | 0.3333 | 1 | 0 | 20 | 0.2 | 3 | 0 | 5 | 0.2 | 3 | 0 | 1 | 1 | 12 | 0.1667 |
| 11_7941795 | 2 | 1 | 7 | 1 | 7 | 0.8571 | 1 | 1 | 17 | 0.9412 | 1 | 1 | 4 | 1 | 1 | 1 | NA | NA | 6 | 1 |
| 11_7941805 | 2 | 1 | 6 | 1 | 5 | 1 | NA | NA | 13 | 1 | 1 | 1 | 4 | 1 | 2 | 1 | 1 | 1 | 8 | 1 |
| 11_7941806 | 2 | 1 | 8 | 0.875 | 7 | 1 | 1 | 1 | 18 | 0.9444 | 1 | 1 | 5 | 1 | 3 | 1 | 1 | 1 | 10 | 1 |
| 11_7941809 | NA | NA | 6 | 0.1667 | 8 | 0.5 | 1 | 1 | 15 | 0.4 | 1 | 0 | 5 | 1 | 3 | 0.6667 | 1 | 1 | 10 | 0.8 |
| 11_7941831 | 2 | 0.5 | 6 | 0.5 | 5 | 0 | NA | NA | 13 | 0.3077 | 1 | 1 | 3 | 0.6667 | 2 | 0 | NA | NA | 6 | 0.5 |
| 11_7941844 | 2 | 0.5 | 7 | 0.4286 | 7 | 0.1429 | 1 | 0 | 17 | 0.2941 | 1 | 0 | 3 | 0.3333 | 2 | 0.5 | 1 | 1 | 7 | 0.4286 |
| 11_7941859 | 2 | 0.5 | 7 | 0.2857 | 5 | 0.4 | 1 | 0 | 15 | 0.3333 | 1 | 1 | 4 | 0.5 | 2 | 0.5 | 1 | 0 | 8 | 0.5 |
| 11_7941896 | NA | NA | 1 | 1 | 1 | 1 | NA | NA | 2 | 1 | NA | NA | 1 | 1 | NA | NA | NA | NA | 1 | 1 |
| 11_7942652 | NA | NA | 2 | 0.5 | 4 | 0 | 4 | 0.5 | 10 | 0.3 | 1 | 0 | NA | NA | NA | NA | 4 | 0.25 | 5 | 0.2 |
| 11_7942656 | NA | NA | 2 | 0.5 | 4 | 0.25 | 4 | 0.5 | 10 | 0.4 | 1 | 0 | NA | NA | NA | NA | 3 | 0.6667 | 4 | 0.5 |
| 11_7942663 | NA | NA | 1 | 0 | 3 | 0.3333 | 1 | 0 | 5 | 0.2 | NA | NA | NA | NA | NA | NA | 2 | 0.5 | 2 | 0.5 |
| 11_7945575 | 2 | 1 | 6 | 0.1667 | 6 | 0 | 5 | 0.2 | 19 | 0.2105 | 1 | 0 | 3 | 0 | 2 | 0.5 | 6 | 0 | 12 | 0.0833 |
| 11_7946913 | 2 | 1 | 7 | 0.2857 | 6 | 0.5 | 6 | 0.5 | 21 | 0.4762 | 3 | 0.3333 | 3 | 0.3333 | 5 | 0.4 | 7 | 0.4286 | 18 | 0.3889 |
| 11_7946927 | 1 | 1 | 5 | 0.4 | 6 | 0.1667 | 5 | 0.6 | 17 | 0.4118 | 3 | 0.3333 | 2 | 1 | 3 | 0.3333 | 6 | 0.5 | 14 | 0.5 |
| 11_7946943 | 2 | 1 | 5 | 0.8 | 6 | 0.8333 | 5 | 0.8 | 18 | 0.8333 | 2 | 1 | 3 | 1 | 3 | 1 | 4 | 1 | 12 | 1 |
| 11_7946944 | 1 | 1 | 5 | 0.6 | 4 | 0.5 | 4 | 1 | 14 | 0.7143 | 1 | 0 | 2 | 1 | 5 | 0.8 | 5 | 0.8 | 13 | 0.7692 |
| 11_7946946 | 2 | 0.5 | 7 | 0.2857 | 6 | 0 | 6 | 0 | 21 | 0.1429 | 2 | 0 | 3 | 0.3333 | 4 | 0.25 | 7 | 0 | 16 | 0.125 |
| 11_7946948 | 1 | 1 | 6 | 0.8333 | 6 | 1 | 6 | 1 | 19 | 0.9474 | 3 | 1 | 2 | 1 | 3 | 1 | 7 | 1 | 15 | 1 |
| 11_7946949 | 2 | 0 | 6 | 0.1667 | 4 | 0 | 5 | 0 | 17 | 0.0588 | 3 | 0 | 3 | 0.6667 | 4 | 0.25 | 7 | 0.1429 | 17 | 0.2353 |
| 11_7946966 | NA | NA | 6 | 0.8333 | 5 | 0.4 | 4 | 0.75 | 15 | 0.6667 | 3 | 0.3333 | 3 | 1 | 2 | 1 | 4 | 1 | 12 | 0.8333 |
| 11_7946970 | 1 | 0 | 4 | 0 | 4 | 0 | 5 | 0.2 | 14 | 0.0714 | 3 | 0 | 3 | 0 | 5 | 0 | 5 | 0.2 | 16 | 0.0625 |
| 11_7946974 | 2 | 1 | 7 | 0 | 5 | 0.2 | 6 | 0.6667 | 20 | 0.35 | 2 | 0.5 | 5 | 0.4 | 5 | 0.4 | 8 | 0.625 | 20 | 0.5 |
| 11_7947008 | 2 | 0.5 | 8 | 0 | 6 | 0 | 7 | 0.1429 | 23 | 0.087 | 3 | 0 | 5 | 0.2 | 6 | 0.1667 | 9 | 0 | 23 | 0.087 |
| 11_7947041 | 3 | 0.3333 | 8 | 0.5 | 8 | 0.375 | 10 | 0.3 | 29 | 0.3793 | 4 | 0 | 7 | 0.5714 | 7 | 0.7143 | 9 | 0.1111 | 27 | 0.3704 |
| 11_7947147 | 3 | 0 | 14 | 0 | 9 | 0.1111 | 11 | 0.0909 | 37 | 0.0541 | 5 | 0 | 7 | 0.1429 | 9 | 0.1111 | 9 | 0.2222 | 30 | 0.1333 |
| 11_7947167 | 2 | 0 | 9 | 0.4444 | 10 | 0.1 | 10 | 0.1 | 31 | 0.1935 | 6 | 0.1667 | 6 | 0.5 | 4 | 0 | 8 | 0.125 | 24 | 0.2083 |
| 11_7947191 | 5 | 0.4 | 11 | 0.2727 | 14 | 0.6429 | 14 | 0.6429 | 44 | 0.5227 | 5 | 0.4 | 6 | 0.6667 | 8 | 0.875 | 10 | 0.6 | 29 | 0.6552 |
| 11_7947229 | 7 | 0.2857 | 15 | 0.0667 | 13 | 0.0769 | 14 | 0 | 49 | 0.0816 | 7 | 0.2857 | 7 | 0.1429 | 11 | 0 | 9 | 0.1111 | 34 | 0.1176 |
| 11_7958849 | 18 | 0.1667 | 11 | 0 | 17 | 0.0588 | 15 | 0.0667 | 61 | 0.082 | 17 | 0.0588 | 13 | 0 | 13 | 0.2308 | 14 | 0 | 57 | 0.0702 |
| 11_7958851 | 18 | 0.4444 | 11 | 0.4545 | 26 | 0.3846 | 25 | 0.48 | 80 | 0.4375 | 25 | 0.48 | 16 | 0.625 | 13 | 0.2308 | 16 | 0 | 70 | 0.3571 |
| 11_7958873 | 16 | 0.1875 | 13 | 0 | 23 | 0.3478 | 20 | 0.15 | 72 | 0.1944 | 22 | 0.1818 | 14 | 0.3571 | 11 | 0.0909 | 16 | 0.0625 | 63 | 0.1746 |
| 11_7982492 | 4 | 0 | 3 | 0 | 4 | 0.25 | 2 | 0 | 13 | 0.0769 | NA | NA | 2 | 0.5 | 1 | 0 | 4 | 0.5 | 7 | 0.4286 |
| 11_7982518 | 4 | 0 | 3 | 0 | 5 | 0.2 | 3 | 0 | 15 | 0.0667 | NA | NA | 2 | 0 | 1 | 0 | 2 | 0.5 | 5 | 0.2 |
| 11_7982596 | 3 | 0 | 2 | 0 | 3 | 0.3333 | 1 | 0 | 9 | 0.1111 | NA | NA | 2 | 0 | 2 | 0.5 | 6 | 0.3333 | 10 | 0.3 |
| 11_7982605 | 3 | 0 | 1 | 0 | 2 | 0.5 | 1 | 0 | 7 | 0.1429 | NA | NA | 2 | 0 | 1 | 0 | 6 | 0.3333 | 9 | 0.2222 |
| 11_7982610 | 2 | 0 | 1 | 0 | 3 | 0.6667 | 1 | 0 | 7 | 0.2857 | NA | NA | 2 | 0.5 | 2 | 1 | 6 | 1 | 10 | 0.9 |
| 11_7982637 | 3 | 0.6667 | 2 | 0.5 | 3 | 0.3333 | 1 | 1 | 9 | 0.5556 | NA | NA | 2 | 1 | 2 | 1 | 5 | 1 | 9 | 1 |
| 11_7982641 | 3 | 0.6667 | 3 | 0.3333 | 3 | 0.6667 | 1 | 0 | 10 | 0.5 | NA | NA | 2 | 1 | 2 | 1 | 4 | 1 | 8 | 1 |
| 11_7982687 | 3 | 0 | 2 | 0 | 2 | 0.5 | 2 | 0 | 9 | 0.1111 | NA | NA | 2 | 0.5 | 3 | 0 | 7 | 0.2857 | 12 | 0.25 |
| 11_7992272 | 2 | 1 | 3 | 0.6667 | 4 | 1 | 3 | 0.6667 | 12 | 0.8333 | 2 | 0.5 | 4 | 1 | 3 | 0.6667 | NA | NA | 9 | 0.7778 |

| | | | | | | | | | | | | | | | | | | | | |
|---|---|---|---|---|---|---|---|---|---|---|---|---|---|---|---|---|---|---|---|---|
| 11_799227 | 2 | 0 | 3 | 0.3333 | 5 | 0.6 | 2 | 0.5 | 12 | 0.4167 | 2 | 0.5 | 5 | 0.6 | 3 | 1 | 1 | 0 | 11 | 0.6364 |
| 11_799229 | 2 | 0.5 | 3 | 0 | 7 | 0.1429 | 1 | 1 | 13 | 0.2308 | 1 | 0 | 4 | 0.25 | 2 | 0 | 1 | 0 | 8 | 0.125 |
| 11_799229 | 2 | 0 | 3 | 0 | 6 | 0.1667 | 2 | 0 | 13 | 0.0769 | 2 | 0 | 5 | 0 | 2 | 0.5 | NA | NA | 9 | 0.1111 |
| 11_799229 | 2 | 0 | 3 | 0.3333 | 4 | 0.5 | 2 | 0.5 | 11 | 0.3636 | 2 | 0.5 | 6 | 0.6667 | 3 | 0.6667 | 1 | 0 | 12 | 0.5833 |
| 11_799230 | 1 | 1 | 3 | 0.6667 | 6 | 0.6667 | 2 | 0 | 12 | 0.5833 | 1 | 1 | 6 | 0.5 | 3 | 0.6667 | 1 | 1 | 11 | 0.6364 |
| 11_799230 | 2 | 1 | 3 | 0.3333 | 6 | 0.6667 | 3 | 0.3333 | 14 | 0.5714 | 2 | 1 | 5 | 0.8 | 3 | 0.6667 | 1 | 0 | 11 | 0.7273 |
| 11_799230 | 2 | 0.5 | 3 | 0.3333 | 5 | 0.2 | 3 | 0.6667 | 13 | 0.3846 | 2 | 0 | 4 | 0.25 | 2 | 1 | NA | NA | 8 | 0.375 |
| 11_799233 | 3 | 0.3333 | 3 | 0 | 7 | 0 | 4 | 0 | 17 | 0.0588 | 2 | 0 | 6 | 0.1667 | 4 | 0.25 | 2 | 0 | 14 | 0.1429 |
| 11_799381 | 2 | 0 | 3 | 0.3333 | NA | NA | 2 | 0.5 | 7 | 0.2857 | NA | NA | 3 | 0.6667 | 3 | 0.3333 | 4 | 0.25 | 10 | 0.4 |
| 11_799383 | 2 | 1 | 4 | 0.5 | NA | NA | 3 | 0 | 9 | 0.4444 | NA | NA | 3 | 0.3333 | 3 | 0 | 5 | 0.2 | 11 | 0.1818 |
| 11_799384 | 1 | 1 | 4 | 0.25 | NA | NA | 4 | 0 | 9 | 0.2222 | NA | NA | 3 | 0 | 3 | 0.6667 | 5 | 0.2 | 11 | 0.2727 |
| 11_799384 | 1 | 0 | 4 | 0.25 | NA | NA | 4 | 0 | 9 | 0.1111 | NA | NA | 3 | 0.3333 | 3 | 0 | 5 | 0.2 | 11 | 0.1818 |
| 11_800276 | 1 | 1 | 1 | 1 | NA | NA | 3 | 1 | 5 | 1 | 2 | 0.5 | 2 | 1 | 1 | 0 | 3 | 0.6667 | 8 | 0.625 |
| 11_800279 | NA | NA | NA | NA | NA | NA | 3 | 0.6667 | 3 | 0.6667 | 2 | 0 | 2 | 0.5 | NA | NA | 3 | 0 | 7 | 0.1429 |
| 11_800280 | 1 | 1 | NA | NA | NA | NA | 3 | 0.6667 | 4 | 0.75 | 2 | 1 | 2 | 1 | NA | NA | 3 | 1 | 7 | 1 |
| 11_800292 | NA | NA | NA | NA | 2 | 0.5 | NA | NA | 2 | 0.5 | NA | NA | 1 | 1 | NA | NA | 1 | 1 | 2 | 1 |
| 11_800306 | 1 | 1 | NA | NA | NA | NA | 2 | 1 | 3 | 1 | 2 | 0.5 | 1 | 1 | NA | NA | 2 | 0 | 5 | 0.4 |
| 11_800308 | 1 | 1 | NA | NA | NA | NA | 1 | 1 | 2 | 1 | 2 | 1 | 3 | 1 | NA | NA | 2 | 0.5 | 7 | 0.8571 |
| 11_800309 | 1 | 1 | NA | NA | NA | NA | 1 | 1 | 2 | 1 | 2 | 1 | 2 | 1 | NA | NA | 3 | 0.6667 | 7 | 0.8571 |
| 11_800309 | 1 | 1 | NA | NA | NA | NA | 1 | 0 | 2 | 0.5 | 2 | 1 | 3 | 0.6667 | NA | NA | 3 | 0 | 8 | 0.5 |
| 11_800683 | NA | NA | 1 | 1 | NA | NA | NA | NA | 1 | 1 | NA | NA | NA | NA | NA | NA | 1 | 1 | 1 | 1 |
| 11_800688 | NA | NA | 1 | 1 | NA | NA | NA | NA | 1 | 1 | NA | NA | NA | NA | NA | NA | 1 | 1 | 1 | 1 |
| 11_800695 | NA | NA | 1 | 1 | NA | NA | NA | NA | 1 | 1 | 1 | 0 | 1 | 1 | 1 | 0 | 1 | 1 | 4 | 0.5 |
| 11_800696 | NA | NA | 1 | 0 | 1 | 0 | 1 | 1 | 3 | 0.3333 | 1 | 0 | 1 | 1 | 1 | 0 | 1 | 1 | 4 | 0.5 |
| 11_800705 | NA | NA | 1 | 1 | 1 | 1 | 1 | 1 | 3 | 1 | 1 | 1 | 1 | 1 | 2 | 1 | 1 | 1 | 5 | 1 |
| 11_800895 | NA | NA | 1 | 1 | NA | NA | NA | NA | 1 | 1 | 1 | 0 | NA | NA | NA | NA | 1 | 1 | 2 | 0.5 |
| 11_800896 | NA | NA | 1 | 1 | NA | NA | NA | NA | 1 | 1 | 1 | 1 | 1 | 1 | NA | NA | 2 | 1 | 4 | 1 |
| 11_800903 | NA | NA | 1 | 1 | NA | NA | NA | NA | 1 | 1 | 1 | 0 | 1 | 1 | NA | NA | 2 | 0 | 4 | 0.25 |
| 11_802444 | 1 | 1 | 2 | 0 | 2 | 0 | 2 | 0 | 7 | 0.1429 | 2 | 0 | 1 | 0 | 1 | 1 | 2 | 0 | 6 | 0.1667 |
| 11_802444 | 1 | 1 | 2 | 0.5 | 2 | 0.5 | 2 | 0 | 7 | 0.4286 | 2 | 1 | 1 | 1 | NA | NA | 2 | 1 | 5 | 1 |
| 11_802445 | 1 | 1 | 2 | 0.5 | 2 | 0 | 1 | 0 | 6 | 0.3333 | 2 | 0.5 | 1 | 1 | 1 | 0 | 2 | 0.5 | 6 | 0.5 |
| 11_802446 | NA | NA | 2 | 1 | 2 | 1 | 2 | 1 | 6 | 1 | 2 | 1 | 1 | 1 | 1 | 1 | 3 | 0.6667 | 7 | 0.8571 |
| 11_802448 | 1 | 1 | 2 | 0 | 2 | 0 | 2 | 0 | 7 | 0.1429 | 2 | 0 | 1 | 0 | 1 | 1 | 3 | 0 | 7 | 0.1429 |
| 11_802448 | NA | NA | 1 | 1 | 2 | 0 | 2 | 0 | 5 | 0.2 | 1 | 0 | 1 | 0 | 1 | 0 | 2 | 0.5 | 5 | 0.2 |
| 11_802452 | 1 | 1 | 3 | 0.6667 | 2 | 0 | 1 | 1 | 7 | 0.5714 | 4 | 0.25 | 3 | 0.6667 | 2 | 1 | 2 | 1 | 11 | 0.6364 |
| 11_802455 | 1 | 0 | 2 | 1 | 1 | 0 | 1 | 1 | 5 | 0.6 | 4 | 0.25 | 3 | 0 | 3 | 0.3333 | 3 | 0.3333 | 13 | 0.2308 |
| 11_802458 | 2 | 0 | 4 | 0.25 | 2 | 0 | 2 | 0 | 10 | 0.1 | 4 | 0 | 4 | 0 | 2 | 0 | 4 | 0.25 | 14 | 0.0714 |
| 11_802463 | 4 | 0.25 | 3 | 0.3333 | 2 | 0 | 2 | 0 | 11 | 0.1818 | 4 | 0.25 | 3 | 0 | 2 | 0 | 3 | 0 | 12 | 0.0833 |
| 11_802466 | 4 | 0.5 | 6 | 0.5 | 2 | 0 | 2 | 0 | 14 | 0.3571 | 4 | 0.5 | 3 | 0.6667 | 2 | 0 | 4 | 0.25 | 13 | 0.3846 |
| 11_802545 | 2 | 0 | NA | NA | 2 | 0.5 | 3 | 0 | 7 | 0.1429 | 2 | 0 | 1 | 1 | 2 | 0.5 | NA | NA | 5 | 0.4 |
| 11_802547 | 2 | 1 | NA | NA | 1 | 1 | 3 | 0.6667 | 6 | 0.8333 | 2 | 1 | 1 | 1 | 2 | 1 | NA | NA | 5 | 1 |
| 11_802589 | 2 | 0.5 | NA | NA | 2 | 0 | 3 | 0 | 7 | 0.1429 | 2 | 0 | 1 | 0 | 2 | 0.5 | NA | NA | 5 | 0.2 |
| 11_802862 | 9 | 0.5556 | 14 | 0.7143 | 9 | 0.5556 | 9 | 0.7778 | 41 | 0.6585 | 12 | 0.6667 | 11 | 0.2727 | 14 | 0.6429 | 13 | 0.3846 | 50 | 0.5 |
| 11_802862 | 12 | 0.3333 | 10 | 0.4 | 8 | 0.25 | 8 | 0.125 | 38 | 0.2895 | 9 | 0.2222 | 15 | 0.2667 | 14 | 0.0714 | 12 | 0.3333 | 50 | 0.22 |
| 11_802864 | 14 | 0.1429 | 13 | 0.1538 | 12 | 0.1667 | 12 | 0.1667 | 51 | 0.1569 | 17 | 0.1765 | 18 | 0.2222 | 16 | 0.125 | 14 | 0.1429 | 65 | 0.1692 |
| 11_802864 | 13 | 0.7692 | 17 | 0.9412 | 12 | 0.6667 | 13 | 0.8462 | 55 | 0.8182 | 15 | 0.9333 | 17 | 0.5294 | 15 | 0.7333 | 16 | 0.75 | 63 | 0.7302 |
| 11_802865 | 11 | 0.4545 | 16 | 0.4375 | 10 | 0.4 | 14 | 0.2857 | 51 | 0.3922 | 16 | 0.5625 | 17 | 0.2353 | 16 | 0.375 | 16 | 0.4375 | 65 | 0.4 |
| 11_802865 | 15 | 1 | 16 | 0.9375 | 12 | 0.9167 | 14 | 1 | 57 | 0.9649 | 17 | 1 | 18 | 0.9444 | 17 | 0.9412 | 14 | 1 | 66 | 0.9697 |
| 11_802865 | 13 | 0.3846 | 18 | 0.2778 | 10 | 0.4 | 13 | 0.3077 | 54 | 0.3333 | 16 | 0.3125 | 18 | 0.3889 | 15 | 0.1333 | 18 | 0.3333 | 67 | 0.2985 |
| 11_802867 | 9 | 0 | 12 | 0.0833 | 9 | 0.1111 | 9 | 0.2222 | 39 | 0.1026 | 13 | 0.1538 | 15 | 0.1333 | 14 | 0.1429 | 10 | 0.2 | 52 | 0.1538 |
| 11_803622 | 6 | 0 | 5 | 0.2 | 5 | 0.2 | 8 | 0 | 24 | 0.0833 | 3 | 0 | 5 | 0 | 2 | 1 | 8 | 0.25 | 18 | 0.2222 |
| 11_803623 | 7 | 0.1429 | 10 | 0.3 | 5 | 0.2 | 10 | 0.1 | 32 | 0.1875 | 6 | 0 | 5 | 0.2 | 7 | 0.1429 | 7 | 0.2857 | 25 | 0.16 |
| 11_803625 | 7 | 0 | 9 | 0 | 5 | 0.4 | 11 | 0 | 32 | 0.0625 | 5 | 0.2 | 6 | 0.1667 | 8 | 0.125 | 9 | 0.1111 | 28 | 0.1429 |
| 11_803629 | 7 | 0.8571 | 10 | 0.8 | 3 | 1 | 7 | 1 | 27 | 0.8889 | 6 | 0.8333 | 5 | 1 | 8 | 0.75 | 10 | 0.8 | 29 | 0.8276 |
| 11_803630 | 5 | 0.4 | 9 | 0.1111 | 4 | 0 | 7 | 0 | 25 | 0.12 | 6 | 0.1667 | 5 | 0.2 | 8 | 0 | 10 | 0.1 | 29 | 0.1034 |
| 11_803682 | 9 | 0.2222 | 13 | 0.2308 | 8 | 0.25 | 10 | 0.3 | 40 | 0.25 | 5 | 0.4 | 6 | 0.1667 | 4 | 0 | 6 | 0.6667 | 21 | 0.3333 |
| 11_803684 | 7 | 0 | 16 | 0.1875 | 6 | 0 | 10 | 0.1 | 39 | 0.1026 | 8 | 0.25 | 9 | 0 | 6 | 0 | 6 | 0 | 29 | 0.069 |
| 11_803687 | 7 | 0 | 18 | 0 | 10 | 0 | 12 | 0.25 | 47 | 0.0638 | 10 | 0.2 | 11 | 0.0909 | 7 | 0 | 6 | 0.1667 | 34 | 0.1176 |
| 11_803688 | 9 | 0.4444 | 20 | 0.15 | 10 | 0.2 | 13 | 0.2308 | 52 | 0.2308 | 10 | 0.4 | 13 | 0.2308 | 5 | 0.6 | 10 | 0 | 38 | 0.2632 |
| 11_803688 | 8 | 0.625 | 11 | 0.2727 | 5 | 0 | 9 | 0.4444 | 33 | 0.3636 | 6 | 0.1667 | 11 | 0.4545 | 6 | 0 | 7 | 0.4286 | 30 | 0.3 |
| 11_803689 | 10 | 0.2 | 22 | 0.1364 | 9 | 0 | 17 | 0.0588 | 58 | 0.1034 | 11 | 0.1818 | 13 | 0.0769 | 8 | 0 | 10 | 0.2 | 42 | 0.119 |
| 11_803691 | 12 | 0.75 | 23 | 0.6522 | 12 | 0.75 | 19 | 0.7368 | 66 | 0.7121 | 13 | 0.8462 | 13 | 0.8462 | 11 | 0.5455 | 12 | 0.75 | 49 | 0.7551 |
| 11_803691 | 12 | 0.1667 | 24 | 0.0833 | 13 | 0 | 19 | 0 | 68 | 0.0588 | 14 | 0.2857 | 16 | 0.375 | 12 | 0.0833 | 11 | 0.1818 | 53 | 0.2453 |
| 11_803692 | 10 | 0.2 | 21 | 0.0952 | 14 | 0.0714 | 19 | 0.1579 | 64 | 0.125 | 14 | 0.0714 | 16 | 0.0625 | 12 | 0.25 | 12 | 0.1667 | 54 | 0.1296 |
| 11_803694 | 12 | 1 | 21 | 1 | 15 | 0.8667 | 20 | 1 | 68 | 0.9706 | 10 | 1 | 14 | 1 | 9 | 1 | 11 | 1 | 44 | 1 |

| | | | | | | | | | | | | | | | | | | | | |
|---|---|---|---|---|---|---|---|---|---|---|---|---|---|---|---|---|---|---|---|---|
| 11_8036942 | 13 | 1 | 24 | 0.9167 | 15 | 1 | 23 | 0.9565 | 75 | 0.96 | 11 | 1 | 14 | 1 | 13 | 1 | 12 | 1 | 50 | 1 |
| 11_8036943 | 12 | 1 | 24 | 1 | 14 | 0.8571 | 21 | 0.9524 | 71 | 0.9577 | 9 | 0.8889 | 16 | 1 | 10 | 1 | 13 | 0.7692 | 48 | 0.9167 |
| 11_8036943 | 10 | 0.6 | 22 | 0.4545 | 15 | 0.6667 | 22 | 0.4545 | 69 | 0.5217 | 14 | 0.1429 | 16 | 0.3125 | 12 | 0.5 | 12 | 0.25 | 54 | 0.2963 |
| 11_8036950 | 12 | 1 | 20 | 1 | 11 | 1 | 20 | 1 | 63 | 1 | 11 | 1 | 14 | 1 | 9 | 1 | 15 | 0.9333 | 49 | 0.9796 |
| 11_8036951 | 15 | 1 | 26 | 0.9615 | 16 | 0.9375 | 23 | 1 | 80 | 0.975 | 13 | 1 | 16 | 1 | 15 | 1 | 15 | 1 | 59 | 1 |
| 11_8036951 | 13 | 0.0769 | 22 | 0.0909 | 14 | 0.0714 | 20 | 0 | 69 | 0.058 | 10 | 0 | 17 | 0.0588 | 13 | 0.0769 | 12 | 0.0833 | 52 | 0.0577 |
| 11_8036967 | 13 | 0.0769 | 30 | 0.1 | 16 | 0.0625 | 23 | 0.1739 | 82 | 0.1098 | 13 | 0.1538 | 16 | 0.125 | 15 | 0.0667 | 14 | 0.1429 | 58 | 0.1207 |
| 11_8036974 | 16 | 1 | 27 | 0.9259 | 20 | 1 | 24 | 1 | 87 | 0.977 | 14 | 1 | 18 | 1 | 16 | 1 | 15 | 1 | 63 | 1 |
| 11_8036975 | 11 | 1 | 23 | 1 | 14 | 0.9286 | 15 | 1 | 63 | 0.9841 | 12 | 1 | 12 | 1 | 11 | 1 | 12 | 1 | 47 | 1 |
| 11_8036982 | 17 | 0.1765 | 28 | 0.25 | 20 | 0.25 | 22 | 0.3182 | 87 | 0.2529 | 14 | 0.4286 | 20 | 0.35 | 15 | 0.3333 | 18 | 0.2222 | 67 | 0.3284 |
| 11_8037028 | 17 | 0.2941 | 33 | 0.2424 | 23 | 0.3478 | 23 | 0.1739 | 96 | 0.2604 | 14 | 0.2143 | 16 | 0.4375 | 16 | 0.375 | 13 | 0.5385 | 59 | 0.3898 |
| 11_8037033 | 23 | 0.2609 | 34 | 0.2353 | 24 | 0.25 | 23 | 0.2609 | 104 | 0.25 | 15 | 0.2667 | 17 | 0.3529 | 16 | 0.25 | 16 | 0.3125 | 64 | 0.2969 |
| 11_8037041 | 21 | 0.3333 | 33 | 0.3636 | 20 | 0.35 | 25 | 0.4 | 99 | 0.3636 | 14 | 0.3571 | 15 | 0.5333 | 17 | 0.2353 | 16 | 0.3125 | 62 | 0.3548 |
| 11_8037044 | 25 | 0.04 | 38 | 0.1316 | 26 | 0.1538 | 30 | 0.1667 | 119 | 0.1261 | 18 | 0.1111 | 17 | 0.1176 | 18 | 0.0556 | 18 | 0.1111 | 71 | 0.0986 |
| 11_8037066 | 30 | 0.0333 | 44 | 0.1364 | 34 | 0.0882 | 31 | 0.0968 | 139 | 0.0935 | 22 | 0 | 25 | 0.16 | 23 | 0.3043 | 23 | 0.1739 | 93 | 0.1613 |
| 11_8037080 | 30 | 0.5667 | 47 | 0.6596 | 33 | 0.6061 | 32 | 0.5625 | 142 | 0.6056 | 19 | 0.4737 | 21 | 0.7619 | 26 | 0.5769 | 24 | 0.625 | 90 | 0.6111 |
| 11_8037081 | 26 | 0.4231 | 41 | 0.6585 | 29 | 0.6207 | 30 | 0.6 | 126 | 0.5873 | 16 | 0.6875 | 25 | 0.72 | 24 | 0.5417 | 24 | 0.75 | 89 | 0.6742 |
| 11_8037099 | 26 | 0 | 47 | 0.0851 | 33 | 0 | 31 | 0.0968 | 137 | 0.0511 | 20 | 0.15 | 28 | 0.2143 | 27 | 0.037 | 26 | 0.0385 | 101 | 0.1089 |
| 11_8037110 | 29 | 0.931 | 45 | 0.8667 | 33 | 0.8485 | 33 | 0.9394 | 140 | 0.8929 | 21 | 0.9524 | 27 | 0.8889 | 25 | 0.76 | 24 | 0.9583 | 97 | 0.8866 |
| 11_8037113 | 26 | 0.1154 | 34 | 0.0882 | 26 | 0.1154 | 31 | 0.0968 | 117 | 0.1026 | 17 | 0.0588 | 19 | 0.1579 | 20 | 0.05 | 23 | 0.1304 | 79 | 0.1013 |
| 11_8037114 | 22 | 0.2273 | 39 | 0.3846 | 24 | 0.5 | 27 | 0.2222 | 112 | 0.3393 | 15 | 0.4 | 18 | 0.3333 | 16 | 0.25 | 22 | 0.3636 | 71 | 0.338 |
| 11_8037127 | 31 | 0.4839 | 46 | 0.5217 | 32 | 0.5 | 29 | 0.3103 | 138 | 0.4638 | 19 | 0.3684 | 25 | 0.52 | 23 | 0.4348 | 26 | 0.4615 | 93 | 0.4516 |
| 11_8037130 | 34 | 0.2059 | 50 | 0.2 | 37 | 0.2162 | 38 | 0.1842 | 159 | 0.2013 | 23 | 0.2174 | 30 | 0.1667 | 29 | 0.2759 | 29 | 0.3793 | 111 | 0.2613 |
| 11_8037132 | 34 | 0.3235 | 52 | 0.3462 | 37 | 0.1351 | 38 | 0.2105 | 161 | 0.2609 | 22 | 0.2273 | 30 | 0.2667 | 30 | 0.2 | 32 | 0.3125 | 114 | 0.2544 |
| 11_8037133 | 25 | 0.12 | 43 | 0.093 | 25 | 0.12 | 26 | 0.1538 | 119 | 0.1176 | 15 | 0.1333 | 25 | 0.04 | 18 | 0.1111 | 26 | 0.2692 | 84 | 0.1429 |
| 11_8037320 | NA | NA | 1 | 1 | NA | NA | NA | NA | 1 | 1 | 1 | 1 | NA | NA | 1 | 0 | 1 | 0 | 3 | 0.3333 |
| 11_8071253 | 2 | 1 | 2 | 0 | 4 | 0.25 | NA | NA | 8 | 0.375 | 4 | 0.25 | 2 | 0.5 | 3 | 0.6667 | 2 | 1 | 11 | 0.5455 |
| 11_8071259 | 3 | 0.6667 | 2 | 0.5 | 4 | 0.25 | NA | NA | 9 | 0.4444 | 5 | 0.8 | 1 | 1 | 1 | 1 | 1 | 1 | 8 | 0.875 |
| 11_8077133 | 2 | 0 | 7 | 0.1429 | 7 | 0.1429 | 3 | 0 | 19 | 0.1053 | 6 | 0 | 4 | 0 | 2 | 0 | 8 | 0.125 | 20 | 0.05 |
| 11_8077142 | 1 | 1 | 4 | 0.75 | 5 | 1 | 2 | 0.5 | 12 | 0.8333 | 4 | 0.75 | 4 | 1 | 1 | 1 | 3 | 0.6667 | 12 | 0.8333 |
| 11_8077144 | 1 | 0 | 6 | 0.5 | 6 | 0.5 | 4 | 0.5 | 17 | 0.4706 | 5 | 0.2 | 3 | 0.3333 | 2 | 0.5 | 7 | 0.4286 | 17 | 0.3529 |
| 11_8077149 | 2 | 0.5 | 7 | 0.2857 | 7 | 0.1429 | 3 | 0 | 19 | 0.2105 | 5 | 0.4 | 5 | 0 | 2 | 0 | 9 | 0 | 21 | 0.0952 |
| 11_8077159 | 2 | 0.5 | 6 | 0.5 | 6 | 0.5 | 4 | 0.75 | 18 | 0.5556 | 5 | 0.8 | 5 | 0.4 | 2 | 0 | 9 | 0.3333 | 21 | 0.4286 |
| 11_8077164 | 1 | 0 | 6 | 0.1667 | 6 | 0.3333 | 3 | 0 | 16 | 0.1875 | 1 | 0 | 4 | 0.5 | 2 | 0.5 | 9 | 0.3333 | 16 | 0.375 |
| 11_8077165 | 2 | 0.5 | 6 | 0.8333 | 7 | 0.5714 | 3 | 0.3333 | 18 | 0.6111 | 3 | 1 | 5 | 0.8 | 2 | 0.5 | 8 | 0.625 | 18 | 0.7222 |
| 11_8077167 | 2 | 0 | 7 | 0.2857 | 6 | 0.5 | 4 | 0.5 | 19 | 0.3684 | 4 | 0.5 | 5 | 0.2 | 2 | 0 | 10 | 0.1 | 21 | 0.1905 |
| 11_8077174 | 2 | 0.5 | 6 | 0.5 | 6 | 0.8333 | 3 | 0.6667 | 17 | 0.6471 | 3 | 0.6667 | 4 | 0.75 | 2 | 1 | 9 | 0.7778 | 18 | 0.7778 |
| 11_8077178 | 1 | 0 | 7 | 0.4286 | 7 | 0.1429 | 4 | 0 | 19 | 0.2105 | 3 | 0 | 3 | 0.3333 | 2 | 0.5 | 8 | 0.125 | 16 | 0.1875 |
| 11_8077182 | NA | NA | 2 | 0.5 | 2 | 0.5 | 2 | 0 | 6 | 0.3333 | 3 | 0.6667 | 3 | 0 | 1 | 0 | 7 | 0.4286 | 14 | 0.3571 |
| 11_8077182 | 1 | 0 | 5 | 0.6 | 6 | 0 | 4 | 0.25 | 16 | 0.25 | 4 | 0 | 4 | 0.25 | 2 | 0.5 | 7 | 0 | 17 | 0.1176 |
| 11_8077185 | 1 | 0 | 7 | 0.2857 | 7 | 0.2857 | 3 | 0 | 18 | 0.2222 | 3 | 0.3333 | 2 | 0 | 1 | 0 | 6 | 0.1667 | 12 | 0.1667 |
| 11_8104264 | 1 | 0 | 4 | 0.25 | 3 | 0.3333 | NA | NA | 8 | 0.25 | 2 | 0.5 | 1 | 0 | NA | NA | 3 | 0 | 6 | 0.1667 |
| 11_8104281 | 1 | 0 | 1 | 0 | 3 | 1 | NA | NA | 5 | 0.6 | 1 | 1 | 1 | 1 | NA | NA | 1 | 1 | 3 | 1 |
| 11_8104335 | NA | NA | 1 | 1 | 1 | 1 | NA | NA | 2 | 1 | 1 | 1 | NA | NA | NA | NA | NA | NA | 1 | 1 |
| 11_8104358 | NA | NA | 1 | 1 | 2 | 0.5 | NA | NA | 3 | 0.6667 | 1 | 1 | NA | NA | NA | NA | NA | NA | 1 | 1 |
| 11_8104377 | NA | NA | 1 | 1 | 2 | 1 | NA | NA | 3 | 1 | 1 | 1 | NA | NA | NA | NA | NA | NA | 1 | 1 |
| 11_8104402 | NA | NA | NA | NA | 2 | 0.5 | NA | NA | 2 | 0.5 | 1 | 1 | NA | NA | NA | NA | NA | NA | 1 | 1 |
| 11_8227865 | NA | NA | NA | NA | NA | NA | 1 | 1 | 1 | 1 | 2 | 1 | NA | NA | NA | NA | NA | NA | 2 | 1 |
| 11_8227868 | NA | NA | NA | NA | NA | NA | 2 | 0.5 | 2 | 0.5 | 2 | 0.5 | NA | NA | NA | NA | NA | NA | 2 | 0.5 |
| 11_8227878 | NA | NA | NA | NA | NA | NA | 2 | 0.5 | 2 | 0.5 | 2 | 1 | NA | NA | NA | NA | NA | NA | 2 | 1 |
| 11_8227916 | NA | NA | NA | NA | NA | NA | 2 | 0.5 | 2 | 0.5 | 1 | 1 | NA | NA | NA | NA | NA | NA | 1 | 1 |
| 11_8232534 | NA | NA | NA | NA | NA | NA | 2 | 0.5 | 2 | 0.5 | 2 | 0.5 | NA | NA | 1 | 1 | NA | NA | 3 | 0.6667 |
| 11_8232536 | NA | NA | NA | NA | NA | NA | 2 | 0.5 | 2 | 0.5 | 1 | 0 | NA | NA | 1 | 1 | NA | NA | 2 | 0.5 |
| 11_8232565 | NA | NA | NA | NA | NA | NA | 2 | 0.5 | 2 | 0.5 | 3 | 0.6667 | NA | NA | 1 | 0 | NA | NA | 4 | 0.5 |
| 11_8233516 | NA | NA | NA | NA | NA | NA | 7 | 0.7143 | 7 | 0.7143 | 3 | 0.3333 | 2 | 1 | 1 | 0 | 1 | 1 | 7 | 0.5714 |
| 11_8233552 | NA | NA | 1 | 1 | NA | NA | 7 | 1 | 8 | 1 | 3 | 1 | 1 | 1 | 1 | 1 | 1 | 1 | 6 | 1 |
| 11_8233557 | NA | NA | 1 | 1 | NA | NA | 5 | 0.8 | 6 | 0.8333 | 4 | 0.75 | 2 | 1 | 2 | 0.5 | 1 | 1 | 9 | 0.7778 |
| 11_8233571 | NA | NA | 1 | 1 | NA | NA | 8 | 1 | 9 | 1 | 5 | 1 | 1 | 1 | 2 | 1 | 1 | 1 | 9 | 1 |
| 11_8233591 | NA | NA | 1 | 1 | NA | NA | 7 | 0.8571 | 8 | 0.875 | 5 | 0.8 | 2 | 0.5 | 2 | 1 | 1 | 1 | 10 | 0.8 |
| 11_8233592 | NA | NA | 1 | 0 | NA | NA | 8 | 0.5 | 9 | 0.4444 | 4 | 0.25 | 2 | 0.5 | 2 | 0 | 1 | 0 | 9 | 0.2222 |
| 11_8233600 | NA | NA | NA | NA | NA | NA | 7 | 0.5714 | 7 | 0.5714 | 4 | 0.5 | 1 | 1 | 2 | 1 | 1 | 1 | 8 | 0.75 |
| 11_8233604 | NA | NA | NA | NA | NA | NA | 8 | 0.875 | 8 | 0.875 | 4 | 1 | 2 | 0.5 | 2 | 1 | 1 | 1 | 9 | 0.8889 |
| 11_8233610 | NA | NA | 1 | 0 | NA | NA | 8 | 0.5 | 9 | 0.4444 | 2 | 0.5 | 2 | 0.5 | 3 | 0 | 1 | 1 | 8 | 0.375 |
| 11_8233613 | NA | NA | 1 | 1 | NA | NA | 6 | 0.8333 | 7 | 0.8571 | 4 | 1 | 2 | 1 | 3 | 1 | 1 | 1 | 10 | 1 |
| 11_8233615 | NA | NA | 1 | 1 | NA | NA | 6 | 0.8333 | 7 | 0.8571 | 5 | 0.8 | 1 | 1 | 3 | 1 | 1 | 1 | 10 | 0.9 |

| | | | | | | | | | | | | | | | | | | | | |
|---|---|---|---|---|---|---|---|---|---|---|---|---|---|---|---|---|---|---|---|---|
| 11_8233619 | NA | NA | 1 | 0 | NA | NA | 6 | 0.3333 | 7 | 0.2857 | 5 | 0.2 | 2 | 0.5 | 4 | 0.25 | NA | NA | 11 | 0.2727 |
| 11_8233621 | NA | NA | 1 | 0 | NA | NA | 8 | 0.625 | 9 | 0.5556 | 5 | 0.6 | 2 | 0.5 | 4 | 0.25 | 1 | 0 | 12 | 0.4167 |
| 11_8233623 | NA | NA | 1 | 1 | NA | NA | 7 | 1 | 8 | 1 | 5 | 1 | 2 | 1 | 4 | 1 | 1 | 1 | 12 | 1 |
| 11_8233656 | 1 | 1 | 2 | 1 | NA | NA | 8 | 0.875 | 11 | 0.9091 | 5 | 1 | 2 | 1 | 4 | 1 | 1 | 1 | 12 | 1 |
| 11_8233665 | 1 | 0 | 1 | 0 | NA | NA | 8 | 0.125 | 10 | 0.1 | 2 | 0.5 | 1 | 0 | 4 | 0.25 | 1 | 1 | 8 | 0.375 |
| 11_8233667 | 1 | 0 | 1 | 0 | NA | NA | 4 | 0.25 | 6 | 0.1667 | 4 | 0.25 | 1 | 0 | 4 | 0 | 2 | 0 | 11 | 0.0909 |
| 11_8233671 | 1 | 0 | NA | NA | NA | NA | 8 | 0.125 | 9 | 0.1111 | 3 | 0 | 2 | 0 | 5 | 0.2 | 2 | 0.5 | 12 | 0.1667 |
| 11_8233714 | 1 | 1 | 2 | 0 | NA | NA | 6 | 0.3333 | 9 | 0.3333 | 1 | 0 | 1 | 0 | 3 | 0.6667 | 2 | 0.5 | 7 | 0.4286 |
| 11_8265563 | 1 | 0 | NA | NA | NA | NA | 2 | 0.5 | 3 | 0.3333 | 1 | 0 | 3 | 0.3333 | 1 | 0 | 3 | 0.6667 | 8 | 0.375 |
| 11_8265570 | 2 | 0.5 | 1 | 0 | NA | NA | 2 | 0.5 | 5 | 0.4 | 1 | 0 | 3 | 0 | NA | NA | 4 | 0.25 | 8 | 0.125 |
| 11_8265574 | 2 | 0 | 1 | 0 | NA | NA | 2 | 0.5 | 5 | 0.2 | NA | NA | 3 | 0.6667 | 1 | 0 | 4 | 0.25 | 8 | 0.375 |
| 11_8265598 | 1 | 1 | NA | NA | NA | NA | 3 | 1 | 4 | 1 | 1 | 1 | 2 | 0.5 | 2 | 1 | 4 | 1 | 9 | 0.8889 |
| 11_8265616 | 2 | 0.5 | NA | NA | NA | NA | 3 | 1 | 5 | 0.8 | 1 | 0 | 2 | 1 | 2 | 1 | 3 | 1 | 8 | 0.875 |
| 11_8265619 | 2 | 1 | NA | NA | NA | NA | 3 | 0.6667 | 5 | 0.8 | 1 | 1 | 3 | 0.6667 | 2 | 0.5 | 1 | 1 | 7 | 0.7143 |
| 11_8265621 | 2 | 0.5 | 1 | 1 | NA | NA | 1 | 1 | 4 | 0.75 | 1 | 1 | 3 | 0.6667 | 2 | 1 | 2 | 1 | 8 | 0.875 |
| 11_8265626 | 2 | 1 | NA | NA | NA | NA | 3 | 1 | 5 | 1 | NA | NA | 3 | 1 | 2 | 1 | 4 | 1 | 9 | 1 |
| 11_8265664 | 2 | 0.5 | 1 | 0 | NA | NA | 3 | 0 | 6 | 0.1667 | NA | NA | 4 | 0.5 | 3 | 0 | 3 | 0 | 10 | 0.2 |
| 11_8267423 | NA | NA | NA | NA | NA | NA | 2 | 0.5 | 2 | 0.5 | NA | NA | NA | NA | NA | NA | 1 | 1 | 1 | 1 |
| 11_8267441 | NA | NA | NA | NA | NA | NA | 2 | 1 | 2 | 1 | NA | NA | NA | NA | NA | NA | 1 | 1 | 1 | 1 |
| 11_8267446 | NA | NA | NA | NA | NA | NA | 2 | 1 | 2 | 1 | NA | NA | NA | NA | NA | NA | 1 | 1 | 1 | 1 |
| 11_8269226 | 1 | 1 | NA | NA | NA | NA | 3 | 1 | 4 | 1 | NA | NA | NA | NA | 2 | 1 | 1 | 1 | 3 | 1 |
| 11_8269234 | 1 | 1 | NA | NA | 1 | 1 | 4 | 1 | 6 | 1 | NA | NA | NA | NA | 3 | 1 | 2 | 0.5 | 5 | 0.8 |
| 11_8269249 | NA | NA | NA | NA | 1 | 1 | 4 | 0.5 | 5 | 0.6 | NA | NA | NA | NA | 3 | 0.6667 | 2 | 0.5 | 5 | 0.6 |
| 11_8269251 | 1 | 1 | NA | NA | 1 | 1 | 4 | 1 | 6 | 1 | NA | NA | NA | NA | 3 | 0.6667 | 1 | 1 | 4 | 0.75 |
| 11_8269257 | 1 | 0 | NA | NA | 1 | 1 | 3 | 1 | 5 | 0.8 | NA | NA | NA | NA | 4 | 1 | 2 | 1 | 6 | 1 |
| 11_8269258 | 1 | 0 | NA | NA | 1 | 1 | 4 | 0.25 | 6 | 0.3333 | NA | NA | NA | NA | 3 | 0.3333 | 2 | 0.5 | 5 | 0.4 |
| 11_8269283 | 1 | 1 | NA | NA | NA | NA | 3 | 0.3333 | 4 | 0.5 | 1 | 0 | NA | NA | 5 | 0.4 | 2 | 0.5 | 8 | 0.375 |
| 11_8269302 | NA | NA | NA | NA | 1 | 1 | 1 | 1 | 2 | 1 | NA | NA | NA | NA | 1 | 1 | 1 | 0 | 2 | 0.5 |
| 11_8269321 | 1 | 0 | NA | NA | 1 | 0 | 3 | 0.3333 | 5 | 0.2 | NA | NA | NA | NA | 3 | 0.3333 | 2 | 0 | 5 | 0.2 |
| 11_8269323 | 1 | 1 | NA | NA | 1 | 0 | 3 | 0.3333 | 5 | 0.4 | NA | NA | NA | NA | 2 | 0.5 | 2 | 1 | 4 | 0.75 |
| 11_8269349 | NA | NA | NA | NA | 1 | 1 | 2 | 1 | 3 | 1 | NA | NA | NA | NA | 3 | 0.6667 | 2 | 1 | 5 | 0.8 |
| 11_8269350 | NA | NA | NA | NA | 1 | 1 | 2 | 1 | 3 | 1 | NA | NA | NA | NA | 3 | 0.3333 | 2 | 1 | 5 | 0.6 |
| 11_8269361 | NA | NA | NA | NA | 1 | 1 | NA | NA | 1 | 1 | NA | NA | NA | NA | 2 | 0.5 | 2 | 0.5 | 4 | 0.5 |
| 11_8269463 | NA | NA | NA | NA | 2 | 0.5 | 2 | 0.5 | 4 | 0.5 | NA | NA | 1 | 1 | 3 | 0 | 2 | 0.5 | 6 | 0.3333 |
| 11_8269465 | NA | NA | NA | NA | 2 | 0.5 | 2 | 0 | 4 | 0.25 | NA | NA | 1 | 0 | 5 | 0.2 | 2 | 0 | 8 | 0.125 |
| 11_8269480 | NA | NA | NA | NA | 2 | 0.5 | 2 | 0.5 | 4 | 0.5 | NA | NA | 1 | 0 | 5 | 0.4 | 3 | 1 | 9 | 0.5556 |
| 11_8269671 | 1 | 0 | 1 | 1 | NA | NA | 2 | 0.5 | 4 | 0.5 | 1 | 0 | NA | NA | 1 | 0 | 3 | 0.3333 | 5 | 0.2 |
| 11_8269680 | 1 | 1 | 1 | 1 | NA | NA | 2 | 0.5 | 4 | 0.75 | 1 | 0 | NA | NA | 2 | 1 | 3 | 0.6667 | 6 | 0.6667 |
| 11_8269695 | NA | NA | 1 | 1 | NA | NA | 1 | 0 | 2 | 0.5 | 1 | 0 | NA | NA | 1 | 0 | 2 | 0.5 | 4 | 0.25 |
| 11_8269696 | 1 | 0 | 1 | 1 | NA | NA | 2 | 0.5 | 4 | 0.5 | NA | NA | NA | NA | NA | NA | 3 | 1 | 3 | 1 |
| 11_8270338 | 1 | 1 | NA | NA | NA | NA | 2 | 1 | 3 | 1 | NA | NA | NA | NA | 1 | 1 | 3 | 0.6667 | 4 | 0.75 |
| 11_8270338 | 1 | 1 | 1 | 1 | NA | NA | 1 | 0 | 3 | 0.6667 | NA | NA | NA | NA | 1 | 1 | 3 | 0.6667 | 4 | 0.75 |
| 11_8270348 | NA | NA | 1 | 1 | NA | NA | 2 | 0.5 | 3 | 0.6667 | NA | NA | NA | NA | 1 | 1 | 3 | 1 | 4 | 1 |
| 11_8276957 | 3 | 0 | 2 | 0 | 1 | 0 | 5 | 0.2 | 11 | 0.0909 | NA | NA | 5 | 0.2 | NA | NA | 2 | 0.5 | 7 | 0.2857 |
| 11_8276968 | 3 | 0 | 1 | 0 | 1 | 1 | 5 | 0.2 | 10 | 0.2 | NA | NA | 5 | 0.2 | NA | NA | 1 | 0 | 6 | 0.1667 |
| 11_8276974 | 1 | 0 | NA | NA | NA | NA | 4 | 0.25 | 5 | 0.2 | NA | NA | 5 | 0.6 | NA | NA | 1 | 0 | 6 | 0.5 |
| 11_8276980 | 3 | 0 | 1 | 0 | 1 | 0 | 4 | 0.25 | 9 | 0.1111 | NA | NA | 4 | 0.25 | NA | NA | 1 | 0 | 5 | 0.2 |
| 11_8276981 | NA | NA | 1 | 0 | 1 | 0 | 2 | 0.5 | 4 | 0.25 | NA | NA | 3 | 0.6667 | NA | NA | 1 | 0 | 4 | 0.5 |
| 11_8276982 | 3 | 0 | 1 | 0 | 1 | 1 | 4 | 0 | 9 | 0.1111 | NA | NA | 5 | 0.6 | NA | NA | 2 | 0.5 | 7 | 0.5714 |
| 11_8278809 | 1 | 1 | NA | NA | NA | NA | NA | NA | 1 | 1 | NA | NA | NA | NA | 1 | 1 | NA | NA | 1 | 1 |
| 11_8278818 | 1 | 1 | NA | NA | NA | NA | NA | NA | 1 | 1 | NA | NA | NA | NA | 1 | 1 | NA | NA | 1 | 1 |
| 11_8278859 | 1 | 1 | NA | NA | NA | NA | NA | NA | 1 | 1 | NA | NA | NA | NA | 1 | 1 | NA | NA | 1 | 1 |
| 11_8289318 | NA | NA | 1 | 1 | NA | NA | NA | NA | 1 | 1 | 1 | 1 | 1 | 1 | NA | NA | NA | NA | 2 | 1 |
| 11_8289323 | NA | NA | 1 | 1 | NA | NA | NA | NA | 1 | 1 | 1 | 1 | 1 | 1 | NA | NA | NA | NA | 2 | 1 |
| 11_8289354 | NA | NA | 1 | 1 | NA | NA | NA | NA | 1 | 1 | 1 | 1 | 1 | 1 | NA | NA | NA | NA | 2 | 1 |
| 11_8289360 | NA | NA | 1 | 1 | NA | NA | NA | NA | 1 | 1 | NA | NA | 1 | 1 | NA | NA | NA | NA | 1 | 1 |
| 11_8289366 | NA | NA | 1 | 1 | NA | NA | NA | NA | 1 | 1 | 1 | 1 | 1 | 1 | NA | NA | NA | NA | 2 | 1 |
| 11_8289406 | NA | NA | 1 | 1 | NA | NA | NA | NA | 1 | 1 | NA | NA | 1 | 1 | NA | NA | NA | NA | 1 | 1 |
| 11_8289452 | NA | NA | 1 | 1 | NA | NA | NA | NA | 1 | 1 | NA | NA | 1 | 1 | NA | NA | NA | NA | 1 | 1 |
| 11_8289502 | NA | NA | 1 | 1 | NA | NA | NA | NA | 1 | 1 | NA | NA | 1 | 1 | NA | NA | NA | NA | 1 | 1 |
| 11_8289515 | NA | NA | 1 | 1 | NA | NA | NA | NA | 1 | 1 | NA | NA | 1 | 1 | NA | NA | NA | NA | 1 | 1 |
| 11_8289535 | NA | NA | 1 | 1 | NA | NA | NA | NA | 1 | 1 | NA | NA | 1 | 1 | NA | NA | NA | NA | 1 | 1 |
| 11_8289553 | NA | NA | 1 | 1 | NA | NA | NA | NA | 1 | 1 | NA | NA | 1 | 1 | NA | NA | NA | NA | 1 | 1 |
| 11_8290767 | NA | NA | 1 | 1 | NA | NA | NA | NA | 1 | 1 | NA | NA | 1 | 1 | NA | NA | NA | NA | 1 | 1 |
| 11_8290774 | NA | NA | 1 | 1 | NA | NA | NA | NA | 1 | 1 | NA | NA | 1 | 1 | NA | NA | NA | NA | 1 | 1 |

| | | | | | | | | | | | | | | | | | | | | |
|---|---|---|---|---|---|---|---|---|---|---|---|---|---|---|---|---|---|---|---|---|
| 11_829083 | NA | NA | 1 | 1 | NA | NA | NA | NA | 1 | 1 | NA | NA | 1 | 1 | NA | NA | NA | NA | 1 | 1 |
| 11_829084 | NA | NA | 1 | 1 | NA | NA | NA | NA | 1 | 1 | NA | NA | 1 | 1 | NA | NA | NA | NA | 1 | 1 |
| 11_831863 | 1 | 0 | 1 | 0 | NA | NA | 2 | 0.5 | 4 | 0.25 | NA | NA | NA | NA | 1 | 1 | NA | NA | 1 | 1 |
| 11_831871 | 2 | 0 | 1 | 1 | NA | NA | 1 | 1 | 4 | 0.5 | 1 | 1 | NA | NA | 2 | 0.5 | NA | NA | 3 | 0.6667 |
| 11_831883 | 2 | 0.5 | 1 | 1 | NA | NA | 1 | 0 | 4 | 0.5 | 1 | 0 | NA | NA | 2 | 0.5 | NA | NA | 3 | 0.3333 |
| 11_831887 | 2 | 1 | NA | NA | NA | NA | NA | NA | 2 | 1 | NA | NA | NA | NA | 2 | 1 | NA | NA | 2 | 1 |
| 11_832935 | 11 | 0.5455 | 12 | 0.3333 | 13 | 0.1538 | 7 | 0 | 43 | 0.2791 | 13 | 0.3077 | 17 | 0.1765 | 8 | 0 | 10 | 0.5 | 48 | 0.25 |
| 11_832941 | 13 | 0.0769 | 8 | 0.125 | 10 | 0 | 8 | 0.25 | 39 | 0.1026 | 11 | 0 | 10 | 0.1 | 5 | 0.2 | 7 | 0.2857 | 33 | 0.1212 |
| 11_832948 | 26 | 0.2692 | 20 | 0.3 | 18 | 0.3333 | 13 | 0.0769 | 77 | 0.2597 | 24 | 0.25 | 19 | 0.3158 | 12 | 0.0833 | 18 | 0.3333 | 73 | 0.2603 |
| 11_832948 | 22 | 0.0455 | 20 | 0.2 | 17 | 0.1176 | 13 | 0.1538 | 72 | 0.125 | 23 | 0.0435 | 17 | 0.2353 | 12 | 0 | 18 | 0.1667 | 70 | 0.1143 |
| 11_832949 | 27 | 0.2593 | 23 | 0.3478 | 18 | 0.2222 | 14 | 0.2143 | 82 | 0.2683 | 26 | 0.2692 | 20 | 0.4 | 12 | 0.0833 | 18 | 0.2778 | 76 | 0.2763 |
| 11_832955 | 30 | 0.7333 | 22 | 0.5909 | 28 | 0.5714 | 15 | 0.8 | 95 | 0.6632 | 25 | 0.68 | 24 | 0.625 | 12 | 0.3333 | 21 | 0.4762 | 82 | 0.561 |
| 11_832955 | 27 | 0.5926 | 25 | 0.4 | 29 | 0.3793 | 14 | 0.3571 | 95 | 0.4421 | 25 | 0.56 | 23 | 0.3478 | 13 | 0.3077 | 20 | 0.6 | 81 | 0.4691 |
| 11_833097 | 24 | 0.4167 | 32 | 0.5 | 31 | 0.2581 | 25 | 0.2 | 112 | 0.3482 | 18 | 0.5556 | 34 | 0.4412 | 33 | 0.4242 | 25 | 0.32 | 110 | 0.4273 |
| 11_833097 | 17 | 0.5882 | 19 | 0.5263 | 18 | 0.4444 | 15 | 0.3333 | 69 | 0.4783 | 13 | 0.3846 | 22 | 0.4545 | 23 | 0.6957 | 17 | 0.6471 | 75 | 0.56 |
| 11_833105 | 30 | 0.2333 | 33 | 0.4242 | 33 | 0.3939 | 29 | 0.3103 | 125 | 0.344 | 22 | 0.4091 | 36 | 0.3333 | 34 | 0.3529 | 35 | 0.1714 | 127 | 0.3071 |
| 11_833106 | 37 | 0.027 | 44 | 0.0909 | 44 | 0.1136 | 41 | 0.1463 | 166 | 0.0964 | 34 | 0.0294 | 44 | 0.1136 | 51 | 0.0784 | 36 | 0.1111 | 165 | 0.0848 |
| 11_833110 | 31 | 0.129 | 36 | 0.0278 | 43 | 0.1395 | 34 | 0.0588 | 144 | 0.0903 | 27 | 0.2593 | 41 | 0.0244 | 45 | 0 | 36 | 0.1667 | 149 | 0.094 |
| 11_833121 | NA | NA | NA | NA | NA | NA | 1 | 1 | 1 | 1 | NA | NA | NA | NA | 1 | 1 | NA | NA | 1 | 1 |
| 11_833126 | NA | NA | NA | NA | 1 | 1 | NA | NA | 1 | 1 | NA | NA | NA | NA | 1 | 1 | NA | NA | 1 | 1 |
| 11_833973 | 3 | 0 | 5 | 0.4 | 1 | 1 | 5 | 0 | 14 | 0.2143 | 3 | 0.6667 | 2 | 0.5 | 2 | 1 | 4 | 0.5 | 11 | 0.6364 |
| 11_834656 | 1 | 1 | 1 | 0 | NA | NA | NA | NA | 2 | 0.5 | 1 | 0 | 1 | 0 | NA | NA | 2 | 0.5 | 4 | 0.25 |
| 11_834795 | NA | NA | NA | NA | 2 | 0.5 | 2 | 1 | 4 | 0.75 | NA | NA | 1 | 1 | NA | NA | NA | NA | 1 | 1 |
| 11_836720 | NA | NA | 1 | 1 | 1 | 0 | 2 | 0.5 | 4 | 0.5 | NA | NA | 2 | 0 | 3 | 0 | 1 | 1 | 6 | 0.1667 |
| 11_836722 | NA | NA | 1 | 1 | 3 | 0.6667 | NA | NA | 4 | 0.75 | NA | NA | 1 | 1 | 3 | 1 | 2 | 1 | 6 | 1 |
| 11_836722 | NA | NA | 1 | 1 | 2 | 1 | 1 | 1 | 4 | 1 | NA | NA | 2 | 0.5 | 3 | 0.3333 | 2 | 0.5 | 7 | 0.4286 |
| 11_836724 | NA | NA | 1 | 1 | 3 | 0.3333 | 2 | 0.5 | 6 | 0.5 | NA | NA | 3 | 0.6667 | 2 | 1 | 2 | 0.5 | 7 | 0.7143 |
| 11_836725 | NA | NA | NA | NA | 3 | 1 | 2 | 0 | 5 | 0.6 | NA | NA | 3 | 0.6667 | 2 | 1 | 1 | 0 | 6 | 0.6667 |
| 11_836738 | NA | NA | NA | NA | 1 | 1 | 1 | 1 | 2 | 1 | NA | NA | 1 | 0 | NA | NA | 2 | 0.5 | 3 | 0.3333 |
| 11_836739 | NA | NA | 1 | 0 | 2 | 1 | 1 | 0 | 4 | 0.5 | NA | NA | 1 | 1 | NA | NA | 3 | 0 | 4 | 0.25 |
| 11_836741 | 1 | 0 | 1 | 0 | 1 | 0 | 1 | 1 | 4 | 0.25 | NA | NA | 1 | 1 | NA | NA | 3 | 0 | 4 | 0.25 |
| 11_838527 | 2 | 0 | 7 | 0.1429 | 1 | 0 | 2 | 0 | 12 | 0.0833 | 3 | 0.3333 | NA | NA | 1 | 0 | NA | NA | 4 | 0.25 |
| 11_838527 | 3 | 1 | 7 | 0.8571 | 1 | 1 | 2 | 0.5 | 13 | 0.8462 | 3 | 1 | 1 | 1 | 3 | 1 | NA | NA | 7 | 1 |
| 11_838529 | 3 | 0.6667 | 10 | 0.4 | 1 | 0 | 2 | 1 | 16 | 0.5 | 2 | 1 | 1 | 0 | 3 | 0.6667 | 1 | 1 | 7 | 0.7143 |
| 11_838529 | 2 | 0 | 9 | 0.2222 | 1 | 0 | 2 | 0 | 14 | 0.1429 | 3 | 0 | 1 | 0 | 3 | 0 | 1 | 1 | 8 | 0.125 |
| 11_838533 | 1 | 1 | 8 | 0.125 | NA | NA | 2 | 0 | 11 | 0.1818 | 3 | 0 | 1 | 1 | 1 | 0 | 1 | 0 | 6 | 0.1667 |
| 11_838534 | 1 | 0 | 4 | 0.5 | NA | NA | 2 | 0.5 | 7 | 0.4286 | 3 | 0 | 1 | 1 | 1 | 0 | 1 | 1 | 6 | 0.3333 |
| 11_838534 | 1 | 0 | 8 | 0.25 | NA | NA | 1 | 0 | 10 | 0.2 | 3 | 0 | 1 | 1 | 1 | 0 | NA | NA | 5 | 0.2 |
| 11_838535 | 1 | 0 | 7 | 0.2857 | NA | NA | 2 | 0.5 | 10 | 0.3 | 2 | 0.5 | 1 | 0 | 1 | 0 | 1 | 0 | 5 | 0.2 |
| 11_838536 | 1 | 0 | 7 | 0.2857 | NA | NA | 2 | 0 | 10 | 0.2 | 2 | 0 | 1 | 0 | 1 | 0 | 1 | 1 | 5 | 0.2 |
| 11_838995 | 3 | 0.3333 | 3 | 0.6667 | 3 | 0.3333 | 4 | 1 | 13 | 0.6154 | 2 | 1 | 8 | 1 | 2 | 1 | 5 | 0.8 | 17 | 0.9412 |
| 11_838995 | 3 | 0.3333 | 3 | 0.3333 | 3 | 0.6667 | 5 | 0.2 | 14 | 0.3571 | 2 | 0 | 9 | 0.2222 | 3 | 0 | 5 | 0 | 19 | 0.1053 |
| 11_838996 | 2 | 1 | 3 | 0.3333 | 2 | 0 | 5 | 1 | 12 | 0.6667 | 2 | 0.5 | 8 | 0.625 | 4 | 0.5 | 4 | 0.75 | 18 | 0.6111 |
| 11_838997 | 2 | 0.5 | 4 | 0.5 | 2 | 0 | 4 | 0.5 | 12 | 0.4167 | 2 | 0.5 | 8 | 0.625 | 3 | 0.3333 | 4 | 1 | 17 | 0.6471 |
| 11_838999 | 3 | 0.3333 | 4 | 0.25 | 3 | 0 | 4 | 0.5 | 14 | 0.2857 | 2 | 0.5 | 9 | 0.3333 | 4 | 0 | 4 | 0.5 | 19 | 0.3158 |
| 11_839000 | 3 | 0 | 4 | 0 | 2 | 0.5 | 4 | 0.25 | 13 | 0.1538 | 2 | 0.5 | 7 | 0.4286 | 2 | 0.5 | 4 | 0.25 | 15 | 0.4 |
| 11_839000 | 2 | 0 | 5 | 0.4 | 3 | 0 | 5 | 0.6 | 15 | 0.3333 | 2 | 0 | 10 | 0.4 | 3 | 0.3333 | 5 | 0.6 | 20 | 0.4 |
| 11_839001 | 3 | 0.3333 | 5 | 0 | 3 | 0.6667 | 5 | 0.2 | 16 | 0.25 | 1 | 1 | 9 | 0.1111 | 4 | 0 | 5 | 0.2 | 19 | 0.1579 |
| 11_839001 | 3 | 0.3333 | 5 | 0.2 | 3 | 0.6667 | 5 | 0.4 | 16 | 0.375 | 1 | 0 | 10 | 0.4 | 4 | 0.25 | 5 | 0.4 | 20 | 0.35 |
| 11_839002 | 3 | 1 | 4 | 1 | 2 | 0.5 | 5 | 0.8 | 14 | 0.8571 | 2 | 1 | 8 | 1 | 4 | 1 | 5 | 0.8 | 19 | 0.9474 |
| 11_839002 | 3 | 0 | 5 | 0.4 | 3 | 0 | 4 | 0 | 15 | 0.1333 | 1 | 1 | 9 | 0.2222 | 3 | 0.3333 | 5 | 0.2 | 18 | 0.2778 |
| 11_839004 | 1 | 1 | 1 | 1 | 1 | 1 | 3 | 1 | 6 | 1 | 1 | 1 | 6 | 1 | NA | NA | 4 | 1 | 11 | 1 |
| 11_839004 | 1 | 0 | 1 | 0 | 1 | 1 | 4 | 0 | 7 | 0.1429 | 1 | 1 | 4 | 0.25 | NA | NA | 4 | 0.5 | 9 | 0.4444 |
| 11_839008 | 1 | 0 | 1 | 0 | 1 | 0 | 4 | 0.25 | 7 | 0.1429 | 1 | 0 | 5 | 0.2 | NA | NA | 3 | 0 | 9 | 0.1111 |
| 11_839017 | 1 | 0 | 1 | 1 | 2 | 0.5 | 3 | 0.3333 | 7 | 0.4286 | 1 | 1 | 9 | 0.8889 | NA | NA | 4 | 1 | 14 | 0.9286 |
| 11_839019 | 2 | 0 | 3 | 0 | 2 | 0.5 | 5 | 0.2 | 12 | 0.1667 | 1 | 0 | 9 | 0.1111 | NA | NA | 4 | 0.5 | 14 | 0.2143 |
| 11_839022 | 2 | 0 | 3 | 0 | 4 | 0.25 | 5 | 0 | 14 | 0.0714 | 2 | 0.5 | 11 | 0 | NA | NA | 3 | 0.3333 | 16 | 0.125 |
| 11_839026 | 3 | 0 | 3 | 0.3333 | 3 | 0.3333 | 5 | 0.4 | 14 | 0.2857 | 3 | 0 | 10 | 0.1 | NA | NA | 2 | 0 | 15 | 0.0667 |
| 11_839104 | 3 | 0 | 3 | 0 | 2 | 0 | 2 | 0.5 | 10 | 0.1 | 1 | 0 | 5 | 0.2 | 4 | 0.5 | 2 | 0 | 12 | 0.25 |
| 11_839104 | 3 | 0 | 4 | 0.25 | NA | NA | 2 | 0 | 9 | 0.1111 | 1 | 0 | 4 | 0 | 4 | 0.5 | 2 | 0 | 11 | 0.1818 |
| 11_839105 | 2 | 0 | 4 | 0.75 | 2 | 0.5 | 2 | 0 | 10 | 0.4 | NA | NA | 1 | 0 | 5 | 0.6 | 1 | 1 | 7 | 0.5714 |
| 11_839317 | 2 | 0 | 6 | 0.5 | 5 | 0.4 | 2 | 0.5 | 15 | 0.4 | 2 | 0.5 | 5 | 0.8 | 5 | 0.6 | 2 | 0 | 14 | 0.5714 |
| 11_839317 | 3 | 0.6667 | 6 | 0.8333 | 2 | 1 | 2 | 1 | 13 | 0.8462 | 1 | 1 | 6 | 1 | 5 | 1 | 4 | 1 | 16 | 1 |
| 11_839318 | 2 | 0 | 8 | 0.125 | 4 | 0 | 2 | 0 | 16 | 0.0625 | 2 | 0 | 6 | 0 | 6 | 0.1667 | 4 | 0 | 18 | 0.0556 |
| 11_839319 | 1 | 1 | 6 | 0.8333 | 6 | 0.6667 | 1 | 1 | 14 | 0.7857 | 2 | 0.5 | 6 | 0.6667 | 3 | 0.6667 | 4 | 0.75 | 15 | 0.6667 |

| | | | | | | | | | | | | | | | | | | | | |
|---|---|---|---|---|---|---|---|---|---|---|---|---|---|---|---|---|---|---|---|---|
| 11_8393197 | 1 | 1 | 5 | 1 | 3 | 1 | 2 | 1 | 11 | 1 | 1 | 1 | 6 | 1 | 5 | 0.8 | 4 | 0.75 | 16 | 0.875 |
| 11_8393233 | NA | NA | 8 | 0.25 | 5 | 0.2 | 1 | 0 | 14 | 0.2143 | 2 | 0.5 | 5 | 0.4 | 5 | 0.4 | 3 | 0 | 15 | 0.3333 |
| 11_8393236 | NA | NA | 7 | 0.1429 | 4 | 0 | 1 | 0 | 12 | 0.0833 | 2 | 0 | 7 | 0.1429 | 5 | 0.2 | 3 | 0.3333 | 17 | 0.1765 |
| 11_8393245 | NA | NA | 5 | 1 | 1 | 1 | NA | NA | 6 | 1 | 1 | 1 | 4 | 1 | 1 | 1 | 2 | 1 | 8 | 1 |
| 11_8393247 | NA | NA | 4 | 0.75 | NA | NA | NA | NA | 4 | 0.75 | 2 | 1 | 6 | 0.6667 | 1 | 0 | 2 | 0.5 | 11 | 0.6364 |
| 11_8393251 | NA | NA | 5 | 0.2 | 1 | 0 | NA | NA | 6 | 0.1667 | 1 | 1 | 4 | 0 | 2 | 0 | 3 | 0.6667 | 10 | 0.3 |
| 11_8393267 | NA | NA | 4 | 0.5 | NA | NA | NA | NA | 4 | 0.5 | 1 | 0 | 3 | 0.6667 | 2 | 1 | 3 | 0.6667 | 9 | 0.6667 |
| 11_8393271 | NA | NA | 6 | 0.3333 | 1 | 0 | NA | NA | 7 | 0.2857 | 1 | 0 | 5 | 0.4 | 3 | 0 | 2 | 1 | 11 | 0.3636 |
| 11_8393283 | NA | NA | 4 | 0.75 | 1 | 1 | NA | NA | 5 | 0.8 | 2 | 0.5 | 4 | 0.25 | 3 | 0 | 3 | 0 | 12 | 0.1667 |
| 11_8393286 | NA | NA | 7 | 0.2857 | 1 | 1 | NA | NA | 8 | 0.375 | 2 | 0 | 5 | 0.2 | 3 | 0 | 4 | 0 | 14 | 0.0714 |
| 11_8393292 | NA | NA | 4 | 0.75 | 1 | 0 | NA | NA | 5 | 0.6 | 1 | 1 | 2 | 0.5 | 1 | 1 | 3 | 0 | 7 | 0.4286 |
| 11_8393294 | NA | NA | 4 | 0.75 | 1 | 1 | NA | NA | 5 | 0.8 | 1 | 1 | 3 | 0.6667 | NA | NA | 3 | 0.6667 | 7 | 0.7143 |
| 11_8393297 | NA | NA | 4 | 0.5 | 1 | 0 | NA | NA | 5 | 0.4 | NA | NA | 2 | 0.5 | 1 | 0 | 3 | 0.3333 | 6 | 0.3333 |
| 11_8393301 | NA | NA | 4 | 0.75 | 1 | 1 | NA | NA | 5 | 0.8 | 1 | 1 | 3 | 1 | 1 | 1 | 1 | 1 | 6 | 1 |
| 11_8393424 | 2 | 0 | 5 | 0.4 | 2 | 0.5 | 4 | 0.5 | 13 | 0.3846 | 1 | 0 | 6 | 0.1667 | 1 | 0 | 2 | 0 | 10 | 0.1 |
| 11_8393485 | 1 | 0 | 8 | 0 | 1 | 1 | 4 | 0 | 14 | 0.0714 | NA | NA | 6 | 0.1667 | 1 | 0 | 4 | 0 | 11 | 0.0909 |
| 11_8393919 | 11 | 0.1818 | 4 | 0.25 | 4 | 0 | 10 | 0 | 29 | 0.1034 | 6 | 0 | 12 | 0 | 10 | 0.1 | 8 | 0.125 | 36 | 0.0556 |
| 11_8393951 | 10 | 0.1 | 3 | 0 | 3 | 0 | 9 | 0.3333 | 25 | 0.16 | 6 | 0 | 10 | 0.3 | 8 | 0 | 7 | 0.2857 | 31 | 0.1613 |
| 11_8393956 | 9 | 1 | 3 | 0.3333 | 3 | 0.3333 | 9 | 0.6667 | 24 | 0.7083 | 5 | 0.2 | 10 | 0.7 | 7 | 0.2857 | 7 | 0.7143 | 29 | 0.5172 |
| 11_8393977 | 9 | 1 | 2 | 1 | 2 | 1 | 5 | 1 | 18 | 1 | 6 | 1 | 8 | 1 | 6 | 1 | 3 | 1 | 23 | 1 |
| 11_8393980 | 7 | 0.5714 | 1 | 1 | 1 | 1 | 8 | 0.5 | 17 | 0.5882 | 6 | 0.5 | 8 | 0.25 | 7 | 0.8571 | 5 | 0.4 | 26 | 0.5 |
| 11_8394000 | 9 | 1 | 2 | 1 | 2 | 1 | 8 | 1 | 21 | 1 | 5 | 1 | 11 | 1 | 9 | 0.8889 | 5 | 1 | 30 | 0.9667 |
| 11_8396841 | 1 | 1 | NA | NA | NA | NA | NA | NA | 1 | 1 | 1 | 1 | 1 | 1 | NA | NA | NA | NA | 2 | 1 |
| 11_8396947 | 1 | 1 | NA | NA | NA | NA | NA | NA | 1 | 1 | NA | NA | 1 | 1 | NA | NA | NA | NA | 1 | 1 |
| 11_8396988 | 4 | 0 | 2 | 0 | 4 | 0.25 | 2 | 0 | 12 | 0.0833 | 2 | 0 | 7 | 0.1429 | 4 | 0 | 2 | 0 | 15 | 0.0667 |
| 11_8397011 | 5 | 0.2 | 1 | 0 | 5 | 0.4 | 1 | 0 | 12 | 0.25 | 3 | 0.3333 | 8 | 0.125 | 4 | 0.25 | 3 | 0.3333 | 18 | 0.2222 |
| 11_8397015 | 5 | 0 | 1 | 0 | 5 | 0.2 | 1 | 0 | 12 | 0.0833 | 3 | 0 | 7 | 0.1429 | 4 | 0 | 3 | 0 | 17 | 0.0588 |
| 11_8397031 | 5 | 1 | NA | NA | 5 | 0.6 | 1 | 0 | 11 | 0.7273 | 3 | 0.6667 | 6 | 0.6667 | 4 | 0.75 | 3 | 1 | 16 | 0.75 |
| 11_8397034 | 5 | 0.4 | 1 | 1 | 5 | 0.8 | 1 | 0 | 12 | 0.5833 | 3 | 0 | 6 | 0.3333 | 4 | 0.25 | 3 | 0 | 16 | 0.1875 |
| 11_8397037 | 2 | 0.5 | 1 | 0 | 4 | 0.25 | 1 | 0 | 8 | 0.25 | 2 | 1 | 5 | 0.2 | 2 | 0 | 1 | 0 | 10 | 0.3 |
| 11_8397041 | 5 | 0.4 | 1 | 0 | 5 | 0.4 | 1 | 1 | 12 | 0.4167 | 2 | 0.5 | 6 | 0.5 | 2 | 0.5 | 3 | 0.3333 | 13 | 0.4615 |
| 11_8397045 | 3 | 0.3333 | 1 | 0 | 5 | 0 | 1 | 0 | 10 | 0.1 | 3 | 0 | 6 | 0.1667 | 4 | 0 | 2 | 0 | 15 | 0.0667 |
| 11_8397051 | 5 | 0.8 | 1 | 0 | 4 | 0 | NA | NA | 10 | 0.4 | 3 | 0.6667 | 6 | 0 | 4 | 0 | 3 | 0.6667 | 16 | 0.25 |
| 11_8429057 | 4 | 0 | 6 | 0 | 3 | 0.3333 | 2 | 0.5 | 15 | 0.1333 | 4 | 0.25 | 3 | 0 | 2 | 0 | 2 | 0 | 11 | 0.0909 |
| 11_8429061 | 5 | 1 | 9 | 1 | 3 | 0.6667 | 5 | 1 | 22 | 0.9545 | 5 | 0.6 | 2 | 1 | 1 | 1 | 2 | 1 | 10 | 0.8 |
| 11_8429075 | 4 | 1 | 8 | 1 | 2 | 1 | 3 | 1 | 17 | 1 | 4 | 1 | 3 | 1 | 2 | 1 | 1 | 1 | 10 | 1 |
| 11_8429080 | 5 | 0.2 | 6 | 0.6667 | 1 | 0 | 4 | 0.5 | 16 | 0.4375 | 5 | 0.6 | 1 | 1 | 2 | 1 | 2 | 0.5 | 10 | 0.7 |
| 11_8429081 | 7 | 0.7143 | 7 | 1 | 4 | 0.5 | 4 | 0.75 | 22 | 0.7727 | 4 | 0.75 | 3 | 0.3333 | 1 | 1 | 2 | 0.5 | 10 | 0.6 |
| 11_8429085 | 4 | 0.5 | 7 | 0.2857 | 4 | 1 | 4 | 0.5 | 19 | 0.5263 | 4 | 1 | 2 | 0.5 | 1 | 1 | 2 | 1 | 9 | 0.8889 |
| 11_8429098 | 6 | 0.5 | 7 | 0.8571 | 4 | 1 | 3 | 0.6667 | 20 | 0.75 | 3 | 1 | 3 | 0.6667 | 1 | 1 | 2 | 1 | 9 | 0.8889 |
| 11_8429122 | 6 | 0 | 8 | 0.125 | 5 | 0.2 | 5 | 0.4 | 24 | 0.1667 | 5 | 0.2 | 4 | 0 | 3 | 1 | 2 | 0 | 14 | 0.2857 |
| 11_8429146 | 10 | 0.3 | 9 | 0.5556 | 6 | 0.3333 | 4 | 0 | 29 | 0.3448 | 2 | 0 | 3 | 0.3333 | 2 | 0 | 1 | 0 | 8 | 0.125 |
| 11_8429156 | 9 | 0.5556 | 6 | 0.8333 | 4 | 0.75 | 3 | 0.3333 | 22 | 0.6364 | 4 | 0.25 | 2 | 0 | 2 | 0 | 3 | 0.6667 | 11 | 0.2727 |
| 11_8429172 | 8 | 0.25 | 6 | 0 | 3 | 0.6667 | 4 | 0 | 21 | 0.1905 | 5 | 0.2 | 3 | 0 | 3 | 0 | 3 | 0.3333 | 14 | 0.1429 |
| 11_8429187 | 9 | 0.8889 | 10 | 0.9 | 5 | 1 | 5 | 0.6 | 29 | 0.8621 | 5 | 0.6 | 5 | 0.6 | 2 | 0.5 | 3 | 0.3333 | 15 | 0.5333 |
| 11_8429190 | 7 | 0.8571 | 9 | 0.6667 | 5 | 0.2 | 4 | 0.75 | 25 | 0.64 | 5 | 0.2 | 5 | 0.6 | 3 | 0.6667 | 3 | 0.6667 | 16 | 0.5 |
| 11_8429191 | 8 | 1 | 9 | 1 | 5 | 1 | 5 | 1 | 27 | 1 | 6 | 1 | 5 | 1 | 3 | 1 | 2 | 0.5 | 16 | 0.9375 |
| 11_8429196 | 8 | 0.125 | 6 | 0.5 | 3 | 0 | 4 | 0.25 | 21 | 0.2381 | 4 | 0.75 | 3 | 0 | 1 | 0 | 3 | 0 | 11 | 0.2727 |
| 11_8429219 | 12 | 0.25 | 10 | 0.1 | 6 | 0.3333 | 3 | 0 | 31 | 0.1935 | 6 | 0.1667 | 4 | 0.5 | 3 | 0.3333 | 3 | 0.3333 | 16 | 0.3125 |
| 11_8429231 | 10 | 0.3 | 10 | 0.4 | 5 | 0.2 | 4 | 0 | 29 | 0.2759 | 6 | 0.5 | 5 | 0.8 | 3 | 0.6667 | 3 | 1 | 17 | 0.7059 |
| 11_8429232 | 11 | 0.6364 | 10 | 0.9 | 6 | 1 | 5 | 0.8 | 32 | 0.8125 | 6 | 0.8333 | 5 | 0.8 | 3 | 1 | 2 | 1 | 16 | 0.875 |
| 11_8429239 | 9 | 0.4444 | 10 | 0.8 | 4 | 0.75 | 5 | 0.6 | 28 | 0.6429 | 5 | 0.6 | 5 | 0.4 | 3 | 1 | 2 | 1 | 15 | 0.6667 |
| 11_8472334 | 1 | 0 | 4 | 0 | 1 | 0 | 2 | 0.5 | 8 | 0.125 | 3 | 0.6667 | 5 | 0.2 | 3 | 0 | 2 | 0 | 13 | 0.2308 |
| 11_8472357 | 10 | 0 | 8 | 0.125 | 7 | 0.1429 | 13 | 0.0769 | 38 | 0.0789 | 5 | 0.4 | 16 | 0.0625 | 11 | 0.0909 | 19 | 0.1053 | 51 | 0.1176 |
| 11_8472375 | 7 | 0.2857 | 8 | 0.75 | 7 | 0.5714 | 9 | 0.2222 | 31 | 0.4516 | 6 | 0.6667 | 15 | 0.4 | 11 | 0.7273 | 18 | 0.5 | 50 | 0.54 |
| 11_8472376 | 9 | 0.3333 | 8 | 0.125 | 7 | 0.4286 | 9 | 0.3333 | 33 | 0.303 | 6 | 0.3333 | 14 | 0.0714 | 6 | 0.3333 | 14 | 0.1429 | 40 | 0.175 |
| 11_8476211 | 2 | 0.5 | NA | NA | 1 | 1 | NA | NA | 3 | 0.6667 | NA | NA | 1 | 0 | NA | NA | 1 | 1 | 2 | 0.5 |
| 11_8476218 | 3 | 1 | NA | NA | 1 | 1 | NA | NA | 4 | 1 | NA | NA | 1 | 1 | NA | NA | 1 | 1 | 2 | 1 |
| 11_8476221 | 2 | 0.5 | NA | NA | 1 | 0 | NA | NA | 3 | 0.3333 | NA | NA | 1 | 1 | NA | NA | 1 | 0 | 2 | 0.5 |
| 11_8476224 | 2 | 1 | NA | NA | 1 | 1 | NA | NA | 3 | 1 | NA | NA | 1 | 0 | 1 | 1 | 1 | 1 | 3 | 0.6667 |
| 11_8480142 | 1 | 0 | 1 | 0 | 1 | 1 | NA | NA | 3 | 0.3333 | 1 | 1 | NA | NA | 1 | 0 | NA | NA | 2 | 0.5 |
| 11_8480178 | 1 | 0 | 1 | 1 | 1 | 0 | NA | NA | 3 | 0.3333 | 1 | 0 | NA | NA | 1 | 1 | NA | NA | 2 | 0.5 |
| 11_8480207 | 9 | 0.1111 | 7 | 0.2857 | 2 | 0 | 3 | 0.3333 | 21 | 0.1905 | 6 | 0.3333 | 10 | 0.2 | 6 | 0.1667 | 3 | 0 | 25 | 0.2 |
| 11_8480209 | 7 | 0.5714 | 10 | 0.3 | 2 | 0.5 | 3 | 0.3333 | 22 | 0.4091 | 8 | 0.25 | 10 | 0.3 | 5 | 0.2 | 3 | 0 | 26 | 0.2308 |
| 11_8480211 | 6 | 1 | 7 | 0.5714 | 2 | 1 | 2 | 0.5 | 17 | 0.7647 | 5 | 1 | 10 | 0.9 | 4 | 0.5 | 1 | 1 | 20 | 0.85 |

| | | | | | | | | | | | | | | | | | | | | |
|---|---|---|---|---|---|---|---|---|---|---|---|---|---|---|---|---|---|---|---|---|
| 11_8480229 | 9 | 0.7778 | 9 | 0.7778 | 2 | 1 | 1 | 1 | 21 | 0.8095 | 6 | 1 | 9 | 1 | 6 | 0.8333 | 2 | 0.5 | 23 | 0.913 |
| 11_8480233 | 9 | 0.3333 | 8 | 0.5 | 1 | 0 | 1 | 0 | 19 | 0.3684 | 7 | 0.7143 | 9 | 0.5556 | 6 | 0.1667 | 2 | 0.5 | 24 | 0.5 |
| 11_8485667 | 2 | 0.5 | 1 | 0 | 1 | 0 | 2 | 0 | 6 | 0.1667 | 4 | 0.25 | 1 | 0 | 3 | 0 | 5 | 0.2 | 13 | 0.1538 |
| 11_8485671 | 3 | 1 | 1 | 1 | 2 | 0 | 2 | 0.5 | 8 | 0.625 | 3 | 1 | NA | NA | 4 | 0.75 | 5 | 0.6 | 12 | 0.75 |
| 11_8485673 | 2 | 0.5 | 1 | 0 | 1 | 1 | 2 | 0 | 6 | 0.3333 | 3 | 0.6667 | 1 | 0 | 4 | 0 | 5 | 0 | 13 | 0.1538 |
| 11_8487719 | 5 | 0 | 7 | 0.1429 | 2 | 0.5 | 8 | 0.375 | 22 | 0.2273 | 4 | 0.25 | 8 | 0.125 | 8 | 0.125 | 5 | 0 | 25 | 0.12 |
| 11_8487734 | 4 | 0.75 | 7 | 0.5714 | 1 | 0 | 8 | 0.75 | 20 | 0.65 | 4 | 0.25 | 9 | 0.6667 | 8 | 0.375 | 5 | 0.4 | 26 | 0.4615 |
| 11_8487938 | 5 | 0.6 | 5 | 0.6 | 1 | 1 | 6 | 0.8333 | 17 | 0.7059 | 3 | 0.3333 | 7 | 0.8571 | 6 | 0.6667 | 3 | 1 | 19 | 0.7368 |
| 11_8487949 | 5 | 0 | 5 | 0 | 1 | 0 | 7 | 0.1429 | 18 | 0.0556 | 1 | 0 | 8 | 0.125 | 5 | 0 | 3 | 0 | 17 | 0.0588 |
| 11_8487950 | 5 | 1 | 5 | 0.8 | 1 | 1 | 8 | 0.375 | 19 | 0.6842 | 3 | 0.6667 | 7 | 0.8571 | 6 | 0.6667 | 4 | 0.75 | 20 | 0.75 |
| 11_8488345 | 5 | 0.2 | 5 | 0.2 | 1 | 0 | 6 | 0.3333 | 17 | 0.2353 | 3 | 0 | 7 | 0.1429 | 5 | 0 | 4 | 0 | 19 | 0.0526 |
| 11_8491303 | 3 | 0 | 3 | 0 | 1 | 0 | 3 | 0.3333 | 10 | 0.1 | 2 | 0 | 6 | 0.1667 | 3 | 0 | 3 | 0 | 14 | 0.0714 |
| 11_8507919 | 4 | 0 | 13 | 0.1538 | 3 | 0 | 5 | 0 | 25 | 0.08 | 2 | 0 | 1 | 0 | 10 | 0.1 | 6 | 0.1667 | 19 | 0.1053 |
| 11_8508057 | 7 | 0.1429 | 9 | 0 | 2 | 0.5 | 4 | 0 | 22 | 0.0909 | 2 | 0 | 3 | 0 | 4 | 0.25 | 7 | 0.1429 | 16 | 0.125 |
| 11_8508072 | 6 | 0 | 11 | 0.2727 | 2 | 0 | 3 | 0.3333 | 22 | 0.1818 | 1 | 1 | 2 | 0 | 4 | 0 | 7 | 0.1429 | 14 | 0.1429 |
| 11_8508132 | 5 | 0.2 | 9 | 0.1111 | 2 | 0 | 3 | 0.3333 | 19 | 0.1579 | 1 | 0 | 1 | 0 | 3 | 0.3333 | 6 | 0.1667 | 11 | 0.1818 |
| 11_8508162 | 5 | 0.2 | 6 | 0.3333 | 1 | 1 | 3 | 0.6667 | 15 | 0.4 | NA | NA | 1 | 1 | 2 | 0.5 | 2 | 1 | 5 | 0.8 |
| 11_8508169 | 5 | 0.2 | 7 | 0.4286 | 1 | 0 | 3 | 0.3333 | 16 | 0.3125 | 1 | 0 | 2 | 0 | 2 | 0 | 5 | 0.4 | 10 | 0.2 |
| 11_8508173 | 4 | 0 | 8 | 0.125 | 1 | 0 | 3 | 0 | 16 | 0.0625 | 1 | 0 | 2 | 0 | 2 | 0.5 | 5 | 0 | 10 | 0.1 |
| 11_8508176 | 5 | 0.2 | 4 | 0.75 | 1 | 0 | 3 | 0.6667 | 13 | 0.4615 | 1 | 0 | 2 | 0 | 2 | 0 | 5 | 0.4 | 10 | 0.2 |
| 11_8508191 | 5 | 0.4 | 6 | 0.8333 | NA | NA | 2 | 0.5 | 13 | 0.6154 | NA | NA | 1 | 0 | 2 | 1 | 5 | 0.6 | 8 | 0.625 |
| 11_8508237 | 4 | 0 | 6 | 0 | 3 | 0 | 6 | 0.1667 | 19 | 0.0526 | 2 | 0.5 | 4 | 0 | 4 | 0 | 5 | 0.2 | 15 | 0.1333 |
| 11_8508246 | 4 | 0 | 5 | 0.2 | 1 | 1 | 6 | 0.3333 | 16 | 0.25 | 3 | 0 | 4 | 0.25 | 3 | 0 | 5 | 0 | 15 | 0.0667 |
| 11_8508248 | 2 | 0.5 | 1 | 1 | 1 | 1 | 2 | 1 | 6 | 0.8333 | 2 | 1 | 1 | 0 | 1 | 0 | NA | NA | 4 | 0.5 |
| 11_8508266 | 5 | 0.8 | 4 | 0.25 | 3 | 0 | 4 | 0 | 16 | 0.3125 | 2 | 0.5 | 4 | 0 | 3 | 0.3333 | 3 | 0.3333 | 12 | 0.25 |
| 11_8508269 | 4 | 1 | 3 | 0.6667 | 2 | 1 | 5 | 0.2 | 14 | 0.6429 | 3 | 0.3333 | 4 | 0.25 | 2 | 1 | 4 | 0.75 | 13 | 0.5385 |
| 11_8512621 | NA | NA | 1 | 1 | NA | NA | NA | NA | 1 | 1 | 1 | 1 | NA | NA | NA | NA | NA | NA | 1 | 1 |
| 11_8512622 | NA | NA | 1 | 1 | NA | NA | NA | NA | 1 | 1 | 1 | 1 | NA | NA | NA | NA | NA | NA | 1 | 1 |
| 11_8520233 | 1 | 0 | NA | NA | 1 | 0 | 2 | 0.5 | 4 | 0.25 | NA | NA | 1 | 1 | 1 | 0 | NA | NA | 2 | 0.5 |
| 11_8523650 | 3 | 1 | NA | NA | 1 | 0 | 4 | 0.25 | 8 | 0.5 | 2 | 0.5 | 1 | 0 | 1 | 0 | 1 | 1 | 5 | 0.4 |
| 11_8523658 | 2 | 0 | 1 | 1 | 1 | 1 | 4 | 0.25 | 8 | 0.375 | 3 | 0.3333 | 1 | 0 | 1 | 0 | 1 | 0 | 6 | 0.1667 |
| 11_8523661 | 3 | 1 | 1 | 1 | 1 | 1 | 4 | 1 | 9 | 1 | 2 | 1 | 1 | 0 | 2 | 0.5 | 1 | 1 | 6 | 0.6667 |
| 11_8523661 | 3 | 1 | 1 | 1 | 1 | 1 | 4 | 1 | 9 | 1 | 3 | 1 | 1 | 1 | 2 | 0.5 | 1 | 1 | 7 | 0.8571 |
| 11_8523672 | 2 | 1 | 1 | 0 | 1 | 1 | 4 | 1 | 8 | 0.875 | 3 | 1 | 1 | 1 | 2 | 1 | 1 | 1 | 7 | 1 |
| 11_8523673 | 2 | 0 | NA | NA | NA | NA | 3 | 0.6667 | 5 | 0.4 | 1 | 0 | NA | NA | 1 | 0 | 1 | 1 | 3 | 0.3333 |
| 11_8523681 | 3 | 1 | 1 | 1 | 1 | 1 | 4 | 1 | 9 | 1 | 3 | 1 | 1 | 1 | 2 | 1 | 1 | 1 | 7 | 1 |
| 11_8523695 | 3 | 0.3333 | 1 | 0 | 1 | 1 | 4 | 0.25 | 9 | 0.3333 | 3 | 0.3333 | 1 | 0 | 2 | 0 | 1 | 0 | 7 | 0.1429 |
| 11_8523708 | 2 | 0.5 | 1 | 0 | NA | NA | 3 | 1 | 6 | 0.6667 | 3 | 0.6667 | 1 | 0 | 3 | 0.6667 | 1 | 1 | 8 | 0.625 |
| 11_8571622 | NA | NA | 1 | 0 | 2 | 0.5 | NA | NA | 3 | 0.3333 | 3 | 0.3333 | 1 | 0 | 1 | 1 | 1 | 1 | 6 | 0.5 |
| 11_8571644 | 1 | 1 | 1 | 1 | 3 | 0.6667 | 1 | 0 | 6 | 0.6667 | 4 | 0.5 | 2 | 0.5 | 2 | 0 | 2 | 0 | 10 | 0.3 |
| 11_8609226 | NA | NA | 2 | 0.5 | 1 | 0 | 2 | 0.5 | 5 | 0.4 | NA | NA | NA | NA | NA | NA | 1 | 1 | 1 | 1 |
| 11_8609785 | NA | NA | 2 | 0.5 | NA | NA | 2 | 0 | 4 | 0.25 | NA | NA | NA | NA | NA | NA | 1 | 1 | 1 | 1 |
| 11_8609833 | NA | NA | 3 | 0.6667 | NA | NA | 2 | 1 | 5 | 0.8 | NA | NA | NA | NA | NA | NA | 1 | 1 | 1 | 1 |
| 11_8609840 | NA | NA | 2 | 1 | NA | NA | 2 | 1 | 4 | 1 | NA | NA | NA | NA | NA | NA | 1 | 1 | 1 | 1 |
| 11_8609852 | NA | NA | 2 | 0.5 | 1 | 1 | 2 | 0 | 5 | 0.4 | NA | NA | NA | NA | NA | NA | 1 | 1 | 1 | 1 |
| 11_8609859 | NA | NA | 2 | 1 | 1 | 1 | 2 | 1 | 5 | 1 | NA | NA | NA | NA | NA | NA | 1 | 1 | 1 | 1 |
| 11_8609872 | NA | NA | 1 | 1 | 2 | 1 | NA | NA | 3 | 1 | NA | NA | NA | NA | NA | NA | 1 | 1 | 1 | 1 |
| 11_8609895 | NA | NA | 4 | 1 | 2 | 1 | 2 | 1 | 8 | 1 | NA | NA | NA | NA | 1 | 1 | 1 | 1 | 2 | 1 |
| 11_8609905 | NA | NA | 3 | 0.6667 | 1 | 1 | 2 | 0.5 | 6 | 0.6667 | NA | NA | NA | NA | NA | NA | 1 | 1 | 1 | 1 |
| 11_8609916 | NA | NA | 3 | 0.3333 | 2 | 0.5 | 2 | 0.5 | 7 | 0.4286 | NA | NA | NA | NA | 1 | 0 | 1 | 1 | 2 | 0.5 |
| 11_8610004 | NA | NA | 4 | 0.75 | 1 | 1 | 2 | 0 | 7 | 0.5714 | NA | NA | NA | NA | 1 | 0 | 1 | 1 | 2 | 0.5 |
| 11_8610027 | NA | NA | 4 | 0.75 | 2 | 1 | 2 | 0.5 | 8 | 0.75 | NA | NA | NA | NA | NA | NA | 2 | 0.5 | 2 | 0.5 |
| 11_8610340 | 1 | 0 | 1 | 0 | 4 | 0.25 | 3 | 0.3333 | 9 | 0.2222 | 1 | 0 | NA | NA | 1 | 1 | 1 | 0 | 3 | 0.3333 |
| 11_8610341 | 1 | 0 | 1 | 1 | 4 | 0.5 | 3 | 1 | 9 | 0.6667 | 1 | 1 | NA | NA | 1 | 0 | 1 | 1 | 3 | 0.6667 |
| 11_8613523 | 1 | 0 | NA | NA | 2 | 0.5 | 1 | 1 | 4 | 0.5 | NA | NA | 1 | 1 | NA | NA | NA | NA | 1 | 1 |
| 11_8615800 | 11 | 0.1818 | 17 | 0.1176 | 4 | 0 | 14 | 0.2143 | 46 | 0.1522 | 4 | 0.25 | 7 | 0.1429 | 7 | 0 | 7 | 0.2857 | 25 | 0.16 |
| 11_8615827 | 3 | 0.3333 | 8 | 0 | 1 | 0 | 8 | 0 | 20 | 0.05 | 4 | 0 | 3 | 0 | 4 | 0.25 | 4 | 0 | 15 | 0.0667 |
| 11_8615902 | 5 | 0.4 | 9 | 0.1111 | 3 | 0 | 8 | 0.125 | 25 | 0.16 | 4 | 0.25 | 4 | 0 | 7 | 0.1429 | 4 | 0 | 19 | 0.1053 |
| 11_8616011 | 4 | 0.25 | 7 | 0 | 1 | 0 | 5 | 0.2 | 17 | 0.1176 | 4 | 0 | 5 | 0 | 5 | 0 | 4 | 0.25 | 18 | 0.0556 |
| 11_8616161 | 3 | 0.3333 | 8 | 0.125 | NA | NA | 4 | 0.25 | 15 | 0.2 | 3 | 0.6667 | 1 | 0 | 3 | 0.3333 | 3 | 0.3333 | 10 | 0.4 |
| 11_8616164 | 3 | 0.3333 | 7 | 0.5714 | NA | NA | 4 | 0.75 | 14 | 0.5714 | 3 | 0 | 1 | 1 | 3 | 0.3333 | 3 | 1 | 10 | 0.5 |
| 11_8616167 | 3 | 0.6667 | 6 | 0.8333 | NA | NA | 4 | 1 | 13 | 0.8462 | 3 | 0.3333 | NA | NA | 2 | 0.5 | 3 | 0.6667 | 8 | 0.5 |
| 11_8616188 | 3 | 0.3333 | 6 | 0 | NA | NA | 3 | 0 | 12 | 0.0833 | 2 | 0.5 | 1 | 0 | 3 | 0 | 1 | 0 | 7 | 0.1429 |
| 11_8617601 | 2 | 0 | 4 | 0.5 | NA | NA | 3 | 0 | 9 | 0.2222 | 2 | 0 | 1 | 1 | 1 | 1 | 1 | 0 | 5 | 0.4 |
| 11_8617741 | 2 | 1 | 2 | 0.5 | NA | NA | 3 | 0.3333 | 7 | 0.5714 | 1 | 1 | 1 | 1 | 1 | 1 | 2 | 0 | 5 | 0.6 |

| | | | | | | | | | | | | | | | | | | | | |
|---|---|---|---|---|---|---|---|---|---|---|---|---|---|---|---|---|---|---|---|---|
| 11_8617755 | 2 | 0 | 3 | 0 | NA | NA | 3 | 0.3333 | 8 | 0.125 | 1 | 0 | 1 | 0 | 1 | 1 | 2 | 0 | 5 | 0.2 |
| 11_8618921 | 1 | 1 | 3 | 0.3333 | NA | NA | 1 | 1 | 5 | 0.6 | NA | NA | 1 | 1 | 1 | 0 | NA | NA | 2 | 0.5 |
| 11_8618923 | 1 | 1 | 2 | 1 | NA | NA | 2 | 1 | 5 | 1 | NA | NA | NA | NA | 1 | 1 | NA | NA | 1 | 1 |
| 11_8618945 | NA | NA | 2 | 1 | NA | NA | NA | NA | 2 | 1 | NA | NA | 1 | 1 | NA | NA | NA | NA | 1 | 1 |
| 11_8618947 | NA | NA | 3 | 0.6667 | NA | NA | 1 | 1 | 4 | 0.75 | NA | NA | 1 | 1 | 1 | 1 | NA | NA | 2 | 1 |
| 11_8618965 | 1 | 1 | 3 | 1 | NA | NA | NA | NA | 4 | 1 | NA | NA | 1 | 0 | 1 | 1 | NA | NA | 2 | 0.5 |
| 11_8618973 | 1 | 1 | 3 | 0 | NA | NA | 1 | 0 | 5 | 0.2 | NA | NA | 1 | 0 | 1 | 1 | NA | NA | 2 | 0.5 |
| 11_8618979 | 1 | 0 | 3 | 0.3333 | NA | NA | 1 | 0 | 5 | 0.2 | NA | NA | 1 | 1 | 1 | 0 | NA | NA | 2 | 0.5 |
| 11_861898 | 1 | 0 | 3 | 0.6667 | NA | NA | 1 | 1 | 5 | 0.6 | NA | NA | 1 | 0 | 1 | 1 | NA | NA | 2 | 0.5 |
| 11_862105 | NA | NA | 3 | 1 | NA | NA | 1 | 1 | 4 | 1 | NA | NA | 1 | 1 | 1 | 1 | NA | NA | 2 | 1 |
| 11_8621058 | NA | NA | 3 | 1 | NA | NA | 1 | 1 | 4 | 1 | NA | NA | 1 | 1 | 1 | 0 | NA | NA | 2 | 0.5 |
| 11_862106 | NA | NA | 3 | 0.3333 | NA | NA | 1 | 0 | 4 | 0.25 | NA | NA | NA | NA | 1 | 1 | NA | NA | 1 | 1 |
| 11_8621065 | NA | NA | 3 | 0.3333 | NA | NA | 1 | 1 | 4 | 0.5 | NA | NA | 1 | 1 | 1 | 0 | NA | NA | 2 | 0.5 |
| 11_8621067 | NA | NA | 3 | 0.3333 | NA | NA | 1 | 1 | 4 | 0.5 | NA | NA | 1 | 0 | 1 | 1 | NA | NA | 2 | 0.5 |
| 11_863767 | NA | NA | 2 | 1 | NA | NA | 1 | 0 | 3 | 0.6667 | NA | NA | NA | NA | 2 | 1 | NA | NA | 2 | 1 |
| 11_863770 | 3 | 0.3333 | 6 | 0.1667 | NA | NA | 3 | 0 | 12 | 0.1667 | 3 | 0 | NA | NA | 3 | 0.3333 | NA | NA | 6 | 0.1667 |
| 11_8637707 | 3 | 1 | 6 | 0.3333 | 1 | 0 | 3 | 1 | 13 | 0.6154 | 3 | 0.6667 | NA | NA | 3 | 0.3333 | NA | NA | 6 | 0.5 |
| 11_863772 | 4 | 0.5 | 6 | 0.1667 | 1 | 0 | 2 | 0 | 13 | 0.2308 | 3 | 0 | 1 | 0 | 2 | 0.5 | NA | NA | 6 | 0.1667 |
| 11_8638671 | 6 | 0.3333 | 7 | 0.4286 | 1 | 0 | 3 | 0.3333 | 17 | 0.3529 | 2 | 0 | 1 | 0 | 6 | 0.5 | NA | NA | 9 | 0.3333 |
| 11_8639011 | 4 | 0 | 4 | 0.25 | 7 | 0.1429 | 1 | 0 | 16 | 0.125 | 1 | 0 | 2 | 0 | 3 | 0.3333 | 2 | 0 | 8 | 0.125 |
| 11_8639099 | 4 | 0.5 | 2 | 0 | 4 | 0.25 | 1 | 0 | 11 | 0.2727 | 2 | 0.5 | 2 | 0 | 3 | 0 | 2 | 0.5 | 9 | 0.2222 |
| 11_8639118 | 3 | 0 | 2 | 0 | 5 | 0.2 | 1 | 0 | 11 | 0.0909 | 2 | 0 | 3 | 0 | 4 | 0 | 2 | 0.5 | 11 | 0.0909 |
| 11_8639257 | 1 | 1 | 1 | 0 | 3 | 0.6667 | NA | NA | 5 | 0.6 | 1 | 0 | 4 | 0 | NA | NA | 2 | 0.5 | 7 | 0.1429 |
| 11_8639268 | 1 | 0 | 1 | 1 | 2 | 0 | NA | NA | 4 | 0.25 | 1 | 0 | 3 | 0 | 1 | 0 | 3 | 0.3333 | 8 | 0.125 |
| 11_8639282 | 1 | 1 | 1 | 0 | 2 | 0.5 | NA | NA | 4 | 0.5 | 1 | 0 | 2 | 0.5 | 1 | 0 | 2 | 0 | 6 | 0.1667 |
| 11_8639294 | 1 | 1 | 1 | 0 | 4 | 0.25 | 1 | 0 | 7 | 0.2857 | 1 | 1 | 4 | 0.25 | NA | NA | 3 | 0 | 8 | 0.25 |
| 11_8639305 | 1 | 1 | 1 | 1 | 6 | 0.6667 | 2 | 0.5 | 10 | 0.7 | 1 | 1 | 3 | 0.6667 | NA | NA | 2 | 1 | 6 | 0.8333 |
| 11_863930 | 2 | 0.5 | 1 | 0 | 4 | 0.25 | 3 | 1 | 10 | 0.5 | 1 | 1 | 7 | 0.1429 | NA | NA | 3 | 0.6667 | 11 | 0.3636 |
| 11_8639309 | 2 | 0 | 1 | 1 | 6 | 0.3333 | 3 | 0.3333 | 12 | 0.3333 | 1 | 1 | 7 | 0.5714 | NA | NA | 3 | 0.6667 | 11 | 0.6364 |
| 11_8639311 | 2 | 0.5 | 1 | 1 | 6 | 0.8333 | 3 | 1 | 12 | 0.8333 | 1 | 1 | 7 | 0.7143 | NA | NA | 2 | 1 | 10 | 0.8 |
| 11_8639323 | 2 | 0 | 1 | 0 | 4 | 1 | 2 | 0 | 9 | 0.4444 | NA | NA | 5 | 0.6 | NA | NA | 2 | 0.5 | 7 | 0.5714 |
| 11_8639323 | 1 | 1 | 1 | 1 | 4 | 0.25 | 3 | 0.3333 | 9 | 0.4444 | 1 | 0 | 6 | 0.1667 | NA | NA | 2 | 0.5 | 9 | 0.2222 |
| 11_8639329 | 1 | 0 | 1 | 1 | 4 | 0.5 | 2 | 0.5 | 8 | 0.5 | 1 | 1 | 5 | 0.2 | NA | NA | 2 | 1 | 8 | 0.5 |
| 11_863933 | 1 | 0 | 1 | 1 | 5 | 0.2 | 2 | 0.5 | 9 | 0.3333 | 1 | 0 | 7 | 0.2857 | NA | NA | 1 | 0 | 9 | 0.2222 |
| 11_8639333 | NA | NA | 1 | 1 | 3 | 0.6667 | 2 | 1 | 6 | 0.8333 | 1 | 1 | 5 | 0.6 | NA | NA | 2 | 1 | 8 | 0.75 |
| 11_8639335 | 1 | 0 | 1 | 0 | 4 | 0 | 3 | 0.3333 | 9 | 0.1111 | 1 | 0 | 7 | 0.1429 | NA | NA | 2 | 0 | 10 | 0.1 |
| 11_8639339 | 1 | 0 | 1 | 0 | 4 | 0.5 | 2 | 0.5 | 8 | 0.375 | NA | NA | 6 | 0.1667 | NA | NA | 2 | 0.5 | 8 | 0.25 |
| 11_8639354 | 1 | 0 | 1 | 1 | 5 | 0.2 | 3 | 0 | 10 | 0.2 | NA | NA | 7 | 0.1429 | NA | NA | 2 | 0 | 9 | 0.1111 |
| 11_8647535 | NA | NA | 1 | 0 | NA | NA | 2 | 0.5 | 3 | 0.3333 | NA | NA | NA | NA | NA | NA | 1 | 1 | 1 | 1 |
| 11_8647615 | 15 | 0.2 | 19 | 0.1053 | 20 | 0.25 | 12 | 0.25 | 66 | 0.197 | 15 | 0.0667 | 16 | 0.1875 | 17 | 0.1765 | 13 | 0.2308 | 61 | 0.1639 |
| 11_8658051 | 3 | 0.3333 | 5 | 0 | NA | NA | 2 | 0.5 | 10 | 0.2 | 4 | 0.25 | NA | NA | 4 | 0.25 | 3 | 0.3333 | 11 | 0.2727 |
| 11_8658053 | 3 | 0 | 6 | 0.1667 | 1 | 0 | 2 | 1 | 12 | 0.25 | 3 | 0.6667 | 1 | 0 | 4 | 0.5 | 3 | 0 | 11 | 0.3636 |
| 11_8658054 | 3 | 0.6667 | 6 | 0.6667 | 1 | 1 | 2 | 1 | 12 | 0.75 | 4 | 0.5 | 1 | 0 | 5 | 0.8 | 3 | 1 | 13 | 0.6923 |
| 11_8658055 | 2 | 0.5 | 6 | 0.3333 | 1 | 0 | 1 | 0 | 10 | 0.3 | 4 | 0.5 | 1 | 0 | 4 | 0.75 | 4 | 1 | 13 | 0.6923 |
| 11_865806 | 1 | 0 | 4 | 1 | NA | NA | 1 | 1 | 6 | 0.8333 | 2 | 1 | NA | NA | 3 | 1 | 2 | 1 | 7 | 1 |
| 11_8658061 | 3 | 0.3333 | 4 | 0.25 | NA | NA | 2 | 0 | 9 | 0.2222 | 4 | 0.25 | 1 | 0 | 5 | 0.4 | 4 | 0 | 14 | 0.2143 |
| 11_8658079 | 3 | 0 | 5 | 0 | 1 | 0 | 4 | 0.25 | 13 | 0.0769 | 3 | 0.3333 | 1 | 0 | 2 | 0.5 | 3 | 0.3333 | 9 | 0.3333 |
| 11_865808 | 3 | 1 | 6 | 1 | NA | NA | 4 | 1 | 13 | 1 | 3 | 1 | 1 | 0 | 4 | 1 | 4 | 1 | 12 | 0.9167 |
| 11_865808 | 2 | 0 | 5 | 0 | NA | NA | 2 | 0.5 | 9 | 0.1111 | 3 | 1 | 1 | 1 | 4 | 0.25 | 2 | 0.5 | 10 | 0.6 |
| 11_8658085 | 3 | 0.6667 | 3 | 0.3333 | NA | NA | 4 | 0.75 | 10 | 0.6 | 3 | 0.6667 | 1 | 1 | 3 | 0.3333 | 4 | 0.25 | 11 | 0.4545 |
| 11_8658089 | 3 | 0.3333 | 6 | 0.3333 | NA | NA | 5 | 0.2 | 14 | 0.2857 | 5 | 0.6 | 1 | 0 | 5 | 0.4 | 5 | 0.4 | 16 | 0.4375 |
| 11_8658092 | 4 | 0.5 | 5 | 0.2 | NA | NA | 2 | 0 | 11 | 0.2727 | 3 | 0.3333 | NA | NA | 3 | 0 | 4 | 0.5 | 10 | 0.3 |
| 11_8658347 | 1 | 0 | 1 | 0 | 1 | 1 | 5 | 0.2 | 8 | 0.25 | 1 | 1 | NA | NA | NA | NA | 1 | 0 | 2 | 0.5 |
| 11_8658369 | 3 | 0.3333 | 2 | 0.5 | 1 | 1 | 7 | 0.4286 | 13 | 0.4615 | 2 | 1 | NA | NA | NA | NA | 1 | 0 | 3 | 0.6667 |
| 11_8658374 | 2 | 0 | 1 | 1 | 1 | 1 | 6 | 0 | 10 | 0.2 | 2 | 1 | NA | NA | NA | NA | 1 | 0 | 3 | 0.6667 |
| 11_865839 | 2 | 0 | 2 | 0 | 1 | 0 | 7 | 0.1429 | 12 | 0.0833 | 2 | 0.5 | NA | NA | NA | NA | 1 | 0 | 3 | 0.3333 |
| 11_8658549 | 48 | 0.125 | 62 | 0.129 | 47 | 0.1064 | 43 | 0.1163 | 200 | 0.12 | 25 | 0.16 | 27 | 0.1852 | 54 | 0.0741 | 41 | 0.1463 | 147 | 0.1293 |
| 11_8658594 | 65 | 0.1231 | 66 | 0.1364 | 63 | 0.127 | 45 | 0.0889 | 239 | 0.1213 | 35 | 0.0571 | 30 | 0.1 | 59 | 0.1017 | 44 | 0.0455 | 168 | 0.0774 |
| 11_8658607 | 63 | 0.0159 | 66 | 0.1061 | 60 | 0.0667 | 43 | 0.0233 | 232 | 0.056 | 32 | 0.1562 | 35 | 0.0571 | 58 | 0.0517 | 45 | 0.0444 | 170 | 0.0706 |
| 11_8658614 | 60 | 0.4333 | 67 | 0.2687 | 58 | 0.3448 | 40 | 0.3 | 225 | 0.3378 | 34 | 0.3824 | 33 | 0.3636 | 60 | 0.3833 | 42 | 0.4524 | 169 | 0.3964 |
| 11_8658623 | 60 | 0.0333 | 59 | 0.0847 | 52 | 0.0769 | 33 | 0.0909 | 204 | 0.0686 | 29 | 0.069 | 32 | 0.0625 | 45 | 0.0444 | 39 | 0.0513 | 145 | 0.0552 |
| 11_8658625 | 58 | 0.4138 | 61 | 0.3279 | 52 | 0.4038 | 34 | 0.2941 | 205 | 0.3659 | 30 | 0.5333 | 31 | 0.3226 | 55 | 0.4 | 36 | 0.3889 | 152 | 0.4079 |
| 11_8658627 | 54 | 0.7407 | 56 | 0.625 | 48 | 0.6667 | 32 | 0.8438 | 190 | 0.7053 | 30 | 0.7 | 29 | 0.8621 | 49 | 0.6531 | 39 | 0.7179 | 147 | 0.7211 |
| 11_8666077 | 1 | 0 | NA | NA | 9 | 0.5556 | 9 | 0.4444 | 19 | 0.4737 | 1 | 0 | 5 | 0.2 | 6 | 0.1667 | 5 | 0.4 | 17 | 0.2353 |
| 11_8666094 | 1 | 1 | 1 | 1 | 10 | 0.9 | 10 | 0.9 | 22 | 0.9091 | 1 | 1 | 4 | 0.75 | 5 | 0.6 | 6 | 1 | 16 | 0.8125 |

| | | | | | | | | | | | | | | | | | | | | |
|---|---|---|---|---|---|---|---|---|---|---|---|---|---|---|---|---|---|---|---|---|
| 11_8666097 | 1 | 0 | 1 | 0 | 10 | 0.1 | 10 | 0.1 | 22 | 0.0909 | 1 | 0 | 5 | 0.2 | 8 | 0.125 | 6 | 0 | 20 | 0.1 |
| 11_8666195 | 1 | 0 | 1 | 1 | 8 | 0.25 | 10 | 0.1 | 20 | 0.2 | 1 | 0 | 5 | 0 | 8 | 0.25 | 5 | 0.2 | 19 | 0.1579 |
| 11_8666197 | NA | NA | 1 | 0 | 9 | 0.2222 | 9 | 0.1111 | 19 | 0.1579 | 1 | 0 | 4 | 0 | 6 | 0 | 5 | 0.4 | 16 | 0.125 |
| 11_8670302 | 1 | 1 | NA | NA | NA | NA | 1 | 1 | 2 | 1 | 2 | 1 | NA | NA | NA | NA | 1 | 1 | 3 | 1 |
| 11_8670311 | 1 | 1 | NA | NA | NA | NA | 1 | 1 | 2 | 1 | 1 | 1 | NA | NA | NA | NA | 1 | 1 | 2 | 1 |
| 11_8670385 | 1 | 0 | NA | NA | NA | NA | 1 | 1 | 2 | 0.5 | 1 | 1 | NA | NA | NA | NA | 1 | 1 | 2 | 1 |
| 11_8670490 | 1 | 1 | NA | NA | NA | NA | 1 | 1 | 2 | 1 | 1 | 1 | NA | NA | NA | NA | 1 | 1 | 2 | 1 |
| 11_8682754 | 9 | 0 | 11 | 0.1818 | 4 | 0.25 | 12 | 0.1667 | 36 | 0.1389 | 8 | 0.125 | 16 | 0.25 | 16 | 0.0625 | 14 | 0.2143 | 54 | 0.1667 |
| 11_8682812 | 6 | 0 | 9 | 0 | 3 | 0.3333 | 8 | 0.125 | 26 | 0.0769 | 5 | 0 | 13 | 0.1538 | 17 | 0 | 10 | 0.1 | 45 | 0.0667 |
| 11_8683078 | NA | NA | 1 | 1 | 1 | 0 | 2 | 0 | 4 | 0.25 | 1 | 0 | 4 | 0.25 | 5 | 0.2 | 5 | 0 | 15 | 0.1333 |
| 11_8683093 | 4 | 0.5 | 3 | 0 | 1 | 0 | 2 | 0 | 10 | 0.2 | 1 | 0 | 4 | 0.5 | 5 | 0.2 | 5 | 0 | 15 | 0.2 |
| 11_8683095 | 3 | 0 | 3 | 0.3333 | 1 | 0 | 1 | 0 | 8 | 0.125 | 2 | 0 | 4 | 0.25 | 6 | 0.1667 | 6 | 0 | 18 | 0.1111 |
| 11_8683115 | 4 | 0 | 2 | 0.5 | 1 | 0 | 1 | 0 | 8 | 0.125 | 1 | 0 | 4 | 0.25 | 4 | 0 | 3 | 0.3333 | 12 | 0.1667 |
| 11_8683126 | 3 | 0 | 2 | 0.5 | 1 | 0 | 1 | 0 | 7 | 0.1429 | 2 | 0 | 5 | 0.2 | 4 | 0 | 3 | 0 | 14 | 0.0714 |
| 11_8683134 | 3 | 0.3333 | 2 | 0.5 | 1 | 1 | 1 | 1 | 7 | 0.5714 | 1 | 0 | 5 | 0.8 | 5 | 0.2 | 3 | 0.6667 | 14 | 0.5 |
| 11_8683137 | 2 | 0 | 1 | 0 | 1 | 1 | 1 | 0 | 5 | 0.2 | 2 | 0 | 4 | 0 | 4 | 0.25 | 1 | 0 | 11 | 0.0909 |
| 11_8683142 | 3 | 0.6667 | 2 | 0.5 | 1 | 1 | 1 | 1 | 7 | 0.7143 | 2 | 0.5 | 4 | 0.75 | 3 | 0.6667 | 3 | 0.3333 | 12 | 0.5833 |
| 11_8683144 | 3 | 1 | 1 | 1 | 1 | 1 | 1 | 0 | 6 | 0.8333 | 1 | 1 | 5 | 1 | 3 | 0.3333 | 3 | 0.3333 | 12 | 0.6667 |
| 11_8692604 | 6 | 0.3333 | 3 | 0 | 4 | 0.5 | 6 | 0.3333 | 19 | 0.3158 | 6 | 0.5 | 5 | 0.2 | 5 | 0.4 | 4 | 0 | 20 | 0.3 |
| 11_8696864 | NA | NA | 2 | 0 | 1 | 0 | 1 | 1 | 4 | 0.25 | 2 | 0 | 1 | 0 | 2 | 0.5 | 3 | 0 | 8 | 0.125 |
| 11_8696886 | 1 | 1 | 3 | 0 | 2 | 1 | 5 | 0.2 | 11 | 0.3636 | 3 | 0.3333 | 1 | 0 | 2 | 0 | 3 | 0.6667 | 9 | 0.3333 |
| 11_8696887 | 1 | 1 | 5 | 0.4 | 2 | 1 | 4 | 0.75 | 12 | 0.6667 | 1 | 0 | 1 | 1 | 1 | 1 | 3 | 0.3333 | 6 | 0.5 |
| 11_8697111 | 1 | 1 | 4 | 0.5 | 2 | 0 | 4 | 0.25 | 11 | 0.3636 | 2 | 0 | 1 | 0 | 2 | 0 | 3 | 0.3333 | 8 | 0.125 |
| 11_8697114 | 1 | 1 | 4 | 0.5 | 2 | 1 | 4 | 0 | 11 | 0.4545 | 1 | 1 | 1 | 0 | 2 | 1 | 3 | 0.3333 | 7 | 0.5714 |
| 11_8697121 | NA | NA | 4 | 0.75 | 2 | 1 | 3 | 0.3333 | 9 | 0.6667 | NA | NA | 1 | 0 | 2 | 0.5 | 2 | 1 | 5 | 0.6 |
| 11_8697132 | NA | NA | 3 | 0.3333 | 2 | 1 | 1 | 1 | 6 | 0.6667 | 2 | 0.5 | 1 | 1 | 2 | 1 | 3 | 0.6667 | 8 | 0.75 |
| 11_8697136 | 1 | 1 | 3 | 0.3333 | 2 | 0 | 2 | 0 | 8 | 0.25 | 2 | 0 | 1 | 0 | 2 | 0 | 2 | 1 | 7 | 0.2857 |
| 11_8697137 | 1 | 1 | 2 | 1 | 2 | 1 | 3 | 1 | 8 | 1 | 2 | 0.5 | 1 | 1 | 1 | 1 | 3 | 1 | 7 | 0.8571 |
| 11_8697141 | 1 | 1 | 1 | 0 | 1 | 1 | 3 | 0.3333 | 6 | 0.5 | 1 | 0 | 1 | 0 | 1 | 0 | 3 | 0.3333 | 6 | 0.1667 |
| 11_8697591 | 1 | 0 | 2 | 0.5 | 1 | 0 | NA | NA | 4 | 0.25 | 2 | 0.5 | 1 | 1 | 1 | 1 | 2 | 0.5 | 6 | 0.6667 |
| 11_8697618 | NA | NA | 1 | 1 | 1 | 1 | NA | NA | 2 | 1 | 2 | 1 | 1 | 1 | 1 | 1 | 2 | 1 | 6 | 1 |
| 11_8697625 | 1 | 1 | NA | NA | 1 | 1 | 1 | 0 | 3 | 0.6667 | 1 | 1 | 1 | 1 | 1 | 1 | 2 | 1 | 5 | 1 |
| 11_8697645 | NA | NA | 1 | 0 | 1 | 1 | 2 | 0 | 4 | 0.25 | 2 | 0 | 1 | 1 | 1 | 0 | 2 | 0 | 6 | 0.1667 |
| 11_8697678 | 1 | 0 | 2 | 1 | 1 | 0 | 2 | 0 | 6 | 0.3333 | 2 | 0.5 | NA | NA | 1 | 0 | 2 | 0 | 5 | 0.2 |
| 11_8697682 | 1 | 1 | 2 | 0.5 | 1 | 0 | 2 | 0 | 6 | 0.3333 | 2 | 0.5 | NA | NA | 1 | 0 | 1 | 1 | 4 | 0.5 |
| 11_8697688 | 1 | 1 | 2 | 1 | 1 | 1 | 2 | 0.5 | 6 | 0.8333 | 1 | 1 | 1 | 1 | 1 | 1 | 2 | 1 | 5 | 1 |
| 11_8701282 | 2 | 0.5 | NA | NA | NA | NA | NA | NA | 2 | 0.5 | 1 | 1 | 2 | 1 | 3 | 1 | NA | NA | 6 | 1 |
| 11_8701298 | 2 | 1 | NA | NA | 1 | 1 | 1 | 1 | 4 | 1 | 1 | 1 | 2 | 1 | 3 | 0.6667 | NA | NA | 6 | 0.8333 |
| 11_8701528 | 2 | 0 | 2 | 1 | 1 | 0 | 2 | 0 | 7 | 0.2857 | 1 | 0 | 2 | 0.5 | 3 | 0 | 1 | 1 | 7 | 0.2857 |
| 11_8710925 | 11 | 0.1818 | 12 | 0 | 13 | 0.0769 | 10 | 0.4 | 46 | 0.1522 | 11 | 0.0909 | 18 | 0.1667 | 11 | 0.0909 | 18 | 0.1111 | 58 | 0.1207 |
| 11_8711791 | 11 | 0.6364 | 36 | 0.5833 | 21 | 0.9048 | 1 | 1 | 69 | 0.6957 | 14 | 0.6429 | 7 | 0.4286 | 8 | 0.5 | 9 | 0.5556 | 38 | 0.5526 |
| 11_8711814 | 12 | 0.3333 | 36 | 0.2778 | 16 | 0.375 | 2 | 0.5 | 66 | 0.3182 | 10 | 0.2 | 5 | 0.4 | 7 | 0.1429 | 10 | 0.6 | 32 | 0.3438 |
| 11_8711817 | 12 | 0.1667 | 37 | 0.027 | 19 | 0.0526 | 2 | 0 | 70 | 0.0571 | 13 | 0.2308 | 7 | 0.1429 | 8 | 0.125 | 8 | 0.375 | 36 | 0.2222 |
| 11_8711846 | 11 | 0.1818 | 36 | 0.0833 | 15 | 0.1333 | 2 | 0 | 64 | 0.1094 | 13 | 0.2308 | 6 | 0 | 7 | 0 | 8 | 0 | 34 | 0.0882 |
| 11_8711850 | 12 | 0.25 | 27 | 0.1852 | 15 | 0.3333 | 3 | 0.3333 | 57 | 0.2456 | 13 | 0.1538 | 4 | 0 | 7 | 0.1429 | 7 | 0 | 31 | 0.0968 |
| 11_8711863 | 12 | 0.4167 | 34 | 0.2059 | 15 | 0.4 | 1 | 0 | 62 | 0.2903 | 14 | 0.2857 | 4 | 0 | 5 | 0.6 | 8 | 0.125 | 31 | 0.2581 |
| 11_8712130 | 6 | 0.3333 | 16 | 0.1875 | 8 | 0.25 | 2 | 0 | 32 | 0.2188 | 3 | 0 | 3 | 0.3333 | 2 | 0 | 3 | 0.3333 | 11 | 0.1818 |
| 11_8712170 | 5 | 0.4 | 17 | 0.1765 | 8 | 0.25 | 3 | 0.3333 | 33 | 0.2424 | 2 | 0 | 3 | 0.3333 | 2 | 0.5 | 3 | 0.3333 | 10 | 0.3 |
| 11_8712173 | 6 | 0 | 17 | 0.1765 | 8 | 0 | 2 | 0 | 33 | 0.0909 | 2 | 0 | 4 | 0 | 2 | 0.5 | 3 | 0 | 11 | 0.0909 |
| 11_8712200 | 3 | 0.6667 | 13 | 0.5385 | 5 | 0.8 | 1 | 1 | 22 | 0.6364 | 1 | 1 | 2 | 0.5 | 1 | 1 | NA | NA | 4 | 0.75 |
| 11_8712226 | 5 | 0.2 | 16 | 0.0625 | 7 | 0 | 3 | 0 | 31 | 0.0645 | 2 | 0.5 | 4 | 0.25 | 2 | 0.5 | 3 | 0.3333 | 11 | 0.3636 |
| 11_8712230 | 3 | 0.6667 | 14 | 0.4286 | 7 | 0.4286 | 1 | 0 | 25 | 0.44 | 2 | 0.5 | 3 | 0.6667 | 1 | 0 | 2 | 0.5 | 8 | 0.5 |
| 11_8712240 | 4 | 0.25 | 12 | 0.4167 | 7 | 0.2857 | 3 | 0.3333 | 26 | 0.3462 | 2 | 0.5 | 4 | 0.25 | 2 | 0.5 | 2 | 0.5 | 10 | 0.4 |
| 11_8712244 | 4 | 0.5 | 11 | 0.6364 | 7 | 1 | 3 | 0.6667 | 25 | 0.72 | 1 | 1 | 2 | 1 | 2 | 0.5 | 2 | 0.5 | 7 | 0.7143 |
| 11_8712245 | 2 | 0 | 8 | 0.625 | 7 | 0.5714 | 1 | 1 | 18 | 0.5556 | 1 | 1 | 3 | 1 | 2 | 0.5 | 2 | 0.5 | 8 | 0.75 |
| 11_8712249 | 3 | 0 | 12 | 0.25 | 6 | 0.1667 | 2 | 0 | 23 | 0.1739 | 1 | 1 | 3 | 0.6667 | 2 | 0 | 2 | 0.5 | 8 | 0.5 |
| 11_8712249 | 4 | 0.5 | 11 | 0.9091 | 7 | 0.8571 | 3 | 0.6667 | 25 | 0.8 | 2 | 0.5 | 4 | 0.5 | 2 | 0.5 | 3 | 0.6667 | 11 | 0.5455 |
| 11_8712258 | 3 | 0 | 7 | 0.1429 | 3 | 0 | 2 | 0 | 15 | 0.0667 | 2 | 0 | 3 | 0.3333 | 2 | 0 | 2 | 0.5 | 9 | 0.2222 |
| 11_8712268 | 4 | 0.75 | 9 | 0.4444 | 4 | 0.5 | 3 | 0.6667 | 20 | 0.55 | 2 | 1 | 3 | 0 | 2 | 0.5 | 2 | 0 | 9 | 0.3333 |
| 11_8726974 | 1 | 1 | 1 | 0 | 1 | 0 | NA | NA | 3 | 0.3333 | 1 | 1 | NA | NA | NA | NA | NA | NA | 1 | 1 |
| 11_8726978 | 1 | 0 | 2 | 0 | 1 | 1 | NA | NA | 4 | 0.25 | 1 | 1 | NA | NA | NA | NA | NA | NA | 1 | 1 |
| 11_8748107 | 2 | 0.5 | NA | NA | 1 | 0 | NA | NA | 3 | 0.3333 | 1 | 1 | NA | NA | NA | NA | NA | NA | 1 | 1 |
| 11_8748121 | 19 | 0.3158 | 32 | 0.2812 | 43 | 0.2093 | 30 | 0.0667 | 124 | 0.2097 | 28 | 0.1071 | 18 | 0.1111 | 26 | 0.0769 | 13 | 0.2308 | 85 | 0.1176 |
| 11_8748133 | 17 | 0.0588 | 31 | 0.2581 | 40 | 0.15 | 33 | 0.1212 | 121 | 0.157 | 27 | 0.1481 | 17 | 0.1176 | 27 | 0.1111 | 13 | 0.3077 | 84 | 0.1548 |
| 11_8750191 | 3 | 0.6667 | 2 | 1 | 2 | 1 | 1 | 1 | 8 | 0.875 | 1 | 1 | NA | NA | 6 | 0.5 | 3 | 0.6667 | 10 | 0.6 |

| | | | | | | | | | | | | | | | | | | | | |
|---|---|---|---|---|---|---|---|---|---|---|---|---|---|---|---|---|---|---|---|---|
| 11_875019 | 2 | 1 | 1 | 0 | 2 | 1 | 1 | 1 | 6 | 0.8333 | 2 | 1 | NA | NA | 6 | 1 | 3 | 0.6667 | 11 | 0.9091 |
| 11_875019 | 2 | 0.5 | 2 | 0.5 | 2 | 1 | 2 | 0.5 | 8 | 0.625 | 3 | 1 | NA | NA | 7 | 0.4286 | 3 | 0.6667 | 13 | 0.6154 |
| 11_875021 | 3 | 0.3333 | 1 | 0 | 3 | 0 | 1 | 0 | 8 | 0.125 | 3 | 0 | 1 | 0 | 9 | 0.1111 | 2 | 0 | 15 | 0.0667 |
| 11_875021 | 2 | 1 | 2 | 1 | 3 | 0.6667 | 2 | 0.5 | 9 | 0.7778 | 1 | 1 | 1 | 1 | 8 | 0.875 | 2 | 0.5 | 12 | 0.8333 |
| 11_875021 | 2 | 0.5 | 2 | 0 | 2 | 0.5 | 2 | 0.5 | 8 | 0.375 | 1 | 0 | NA | NA | 9 | 0.2222 | 3 | 0.3333 | 13 | 0.2308 |
| 11_875022 | 3 | 0.6667 | 2 | 0 | 3 | 0 | 3 | 0.3333 | 11 | 0.2727 | 3 | 0 | 1 | 0 | 10 | 0.1 | 3 | 0.6667 | 17 | 0.1765 |
| 11_875023 | 4 | 0 | 2 | 0 | 3 | 0 | 2 | 0.5 | 11 | 0.0909 | 3 | 0 | 1 | 0 | 9 | 0.2222 | 3 | 0 | 16 | 0.125 |
| 11_875026 | 4 | 0.75 | 2 | 0.5 | 4 | 0.75 | 4 | 0.5 | 14 | 0.6429 | 4 | 1 | NA | NA | 8 | 0.875 | 3 | 1 | 15 | 0.9333 |
| 11_875026 | 5 | 0.2 | 2 | 0 | 4 | 0 | 4 | 0 | 15 | 0.0667 | 5 | 0.2 | 1 | 0 | 9 | 0.2222 | 3 | 0 | 18 | 0.1667 |
| 11_875026 | 4 | 0.25 | 2 | 0 | 5 | 0 | 2 | 0 | 13 | 0.0769 | 4 | 0.5 | 2 | 0 | 10 | 0 | 2 | 0.5 | 18 | 0.1667 |
| 11_875028 | 4 | 1 | 2 | 1 | 5 | 0.8 | 4 | 0.75 | 15 | 0.8667 | 4 | 1 | 2 | 1 | 9 | 1 | 5 | 1 | 20 | 1 |
| 11_875029 | 5 | 1 | 2 | 1 | 4 | 1 | 4 | 1 | 15 | 1 | 5 | 1 | 2 | 1 | 9 | 0.7778 | 5 | 0.6 | 21 | 0.8095 |
| 11_875050 | 4 | 0 | 2 | 0.5 | 4 | 0.75 | 4 | 0 | 14 | 0.2857 | 3 | 0.6667 | 3 | 0.3333 | 7 | 0.4286 | 4 | 0 | 17 | 0.3529 |
| 11_875050 | 5 | 0.2 | 3 | 0 | 5 | 0 | 3 | 0.3333 | 16 | 0.125 | 3 | 0 | 4 | 0.25 | 10 | 0.5 | 4 | 0.25 | 21 | 0.3333 |
| 11_875051 | 5 | 0 | 4 | 0.5 | 5 | 0.2 | 5 | 0 | 19 | 0.1579 | 4 | 0.5 | 4 | 0.25 | 9 | 0.1111 | 5 | 0.4 | 22 | 0.2727 |
| 11_875055 | 7 | 0.1429 | 1 | 1 | 7 | 0.2857 | 2 | 0.5 | 17 | 0.2941 | 5 | 0.4 | 2 | 0.5 | 10 | 0.5 | 5 | 0.4 | 22 | 0.4545 |
| 11_875056 | 5 | 0.2 | 2 | 1 | 7 | 0.4286 | 5 | 0.6 | 19 | 0.4737 | 7 | 0.8571 | 4 | 0.25 | 11 | 0.4545 | 7 | 0.5714 | 29 | 0.5517 |
| 11_875056 | 6 | 0 | 2 | 0 | 7 | 0.1429 | 5 | 0.2 | 20 | 0.1 | 6 | 0 | 4 | 0 | 11 | 0.1818 | 7 | 0 | 28 | 0.0714 |
| 11_875057 | 7 | 0.2857 | 3 | 0.6667 | 4 | 0.25 | 3 | 0.3333 | 17 | 0.3529 | 6 | 0.1667 | 3 | 0 | 11 | 0.6364 | 7 | 0.4286 | 27 | 0.4074 |
| 11_875057 | 6 | 0.3333 | 3 | 1 | 6 | 0.5 | 5 | 0.2 | 20 | 0.45 | 6 | 0.5 | 3 | 0.3333 | 11 | 0.6364 | 6 | 0.6667 | 26 | 0.5769 |
| 11_875059 | 7 | 0 | 3 | 0 | 7 | 0.1429 | 8 | 0.125 | 25 | 0.08 | 5 | 0.4 | 4 | 0.25 | 12 | 0 | 6 | 0.1667 | 27 | 0.1481 |
| 11_876219 | 1 | 1 | NA | NA | NA | NA | NA | NA | 1 | 1 | NA | NA | NA | NA | 1 | 1 | NA | NA | 1 | 1 |
| 11_876224 | 1 | 1 | NA | NA | NA | NA | NA | NA | 1 | 1 | NA | NA | NA | NA | 1 | 1 | NA | NA | 1 | 1 |
| 11_876225 | 1 | 1 | NA | NA | NA | NA | NA | NA | 1 | 1 | NA | NA | NA | NA | 1 | 1 | NA | NA | 1 | 1 |
| 11_876228 | 1 | 1 | NA | NA | NA | NA | NA | NA | 1 | 1 | NA | NA | NA | NA | 1 | 1 | NA | NA | 1 | 1 |
| 11_876229 | 1 | 1 | NA | NA | NA | NA | NA | NA | 1 | 1 | NA | NA | NA | NA | 1 | 1 | NA | NA | 1 | 1 |
| 11_876341 | 17 | 0.0588 | 10 | 0.1 | 9 | 0 | 15 | 0.0667 | 51 | 0.0588 | 11 | 0.1818 | 14 | 0 | 16 | 0 | 12 | 0.1667 | 53 | 0.0755 |
| 11_876341 | 14 | 0.1429 | 4 | 0 | 6 | 0.1667 | 12 | 0.1667 | 36 | 0.1389 | 8 | 0.375 | 9 | 0 | 9 | 0.1111 | 8 | 0 | 34 | 0.1176 |
| 11_876342 | 16 | 0.125 | 11 | 0 | 11 | 0 | 14 | 0.2143 | 52 | 0.0962 | 11 | 0.0909 | 13 | 0.1538 | 15 | 0 | 13 | 0.0769 | 52 | 0.0769 |
| 11_876342 | 17 | 0.1765 | 11 | 0 | 11 | 0.1818 | 14 | 0.0714 | 53 | 0.1132 | 12 | 0.1667 | 13 | 0.2308 | 15 | 0.1333 | 11 | 0 | 51 | 0.1373 |
| 11_876344 | 12 | 1 | 9 | 0.7778 | 7 | 1 | 11 | 1 | 39 | 0.9487 | 11 | 1 | 13 | 1 | 13 | 0.9231 | 12 | 0.8333 | 49 | 0.9388 |
| 11_876346 | NA | NA | 1 | 1 | NA | NA | NA | NA | 1 | 1 | 1 | 1 | NA | NA | NA | NA | NA | NA | 1 | 1 |
| 11_876521 | 3 | 0.3333 | 1 | 0 | 1 | 0 | NA | NA | 5 | 0.2 | 2 | 1 | NA | NA | 2 | 0.5 | 1 | 1 | 5 | 0.8 |
| 11_876523 | 2 | 0.5 | 1 | 1 | 2 | 0.5 | NA | NA | 5 | 0.6 | 2 | 0.5 | 1 | 0 | NA | NA | 1 | 1 | 4 | 0.5 |
| 11_876655 | 3 | 1 | 2 | 0.5 | 6 | 0.8333 | 3 | 1 | 14 | 0.8571 | 1 | 0 | 1 | 1 | 4 | 0.75 | 2 | 0.5 | 8 | 0.625 |
| 11_876656 | 3 | 1 | 1 | 1 | 8 | 0.625 | 3 | 0.3333 | 15 | 0.6667 | 2 | 0.5 | 1 | 1 | 5 | 1 | NA | NA | 8 | 0.875 |
| 11_876657 | 2 | 0 | 2 | 0 | 7 | 0 | 1 | 1 | 12 | 0.0833 | 1 | 1 | 1 | 0 | 4 | 0 | 1 | 0 | 7 | 0.1429 |
| 11_876657 | 4 | 1 | 1 | 1 | 6 | 1 | 3 | 0.6667 | 14 | 0.9286 | 2 | 1 | 1 | 1 | 5 | 1 | 1 | 1 | 9 | 1 |
| 11_876658 | 3 | 0.6667 | 2 | 1 | 7 | 1 | 3 | 0.3333 | 15 | 0.8 | 1 | 1 | 1 | 1 | 6 | 1 | 2 | 1 | 10 | 1 |
| 11_876674 | 4 | 0.5 | 2 | 0.5 | 6 | 0.1667 | 5 | 0 | 17 | 0.2353 | 2 | 0 | 4 | 0 | 5 | 0.4 | 2 | 0 | 13 | 0.1538 |
| 11_876687 | 4 | 0.25 | 4 | 0.75 | 6 | 0.5 | 3 | 0 | 17 | 0.4118 | 2 | 1 | 6 | 0.1667 | 4 | 0.5 | 4 | 0.5 | 16 | 0.4375 |
| 11_876688 | 5 | 0.2 | 5 | 0.2 | 9 | 0.3333 | 5 | 0 | 24 | 0.2083 | 2 | 0.5 | 7 | 0.2857 | 7 | 0.2857 | 4 | 0 | 20 | 0.25 |
| 11_876689 | 5 | 0.4 | 5 | 0.2 | 8 | 0 | 6 | 0.1667 | 24 | 0.1667 | 2 | 0 | 6 | 0.1667 | 7 | 0 | 5 | 0 | 20 | 0.05 |
| 11_876689 | 5 | 0.4 | 4 | 0.25 | 7 | 0.5714 | 4 | 0.25 | 20 | 0.4 | 1 | 1 | 5 | 0.4 | 5 | 0.2 | 4 | 1 | 15 | 0.5333 |
| 11_876690 | 4 | 0.25 | 6 | 0.8333 | 6 | 0.5 | 4 | 0.5 | 20 | 0.55 | 1 | 1 | 5 | 0.6 | 5 | 0.6 | 2 | 0 | 13 | 0.5385 |
| 11_876757 | 7 | 0.2857 | 10 | 0.1 | 8 | 0.25 | 8 | 0 | 33 | 0.1515 | 3 | 0 | 7 | 0 | 9 | 0.1111 | 7 | 0.1429 | 26 | 0.0769 |
| 11_876757 | 10 | 0.4 | 11 | 0.1818 | 9 | 0.3333 | 12 | 0.0833 | 42 | 0.2381 | 5 | 0 | 8 | 0.125 | 11 | 0.1818 | 9 | 0.2222 | 33 | 0.1515 |
| 11_876757 | 8 | 0 | 11 | 0.0909 | 7 | 0.2857 | 9 | 0 | 35 | 0.0857 | 4 | 0 | 8 | 0 | 10 | 0.3 | 7 | 0 | 29 | 0.1034 |
| 11_876758 | 8 | 0.375 | 10 | 0.5 | 9 | 0.3333 | 10 | 0.2 | 37 | 0.3514 | 5 | 0.6 | 8 | 0.625 | 9 | 0.3333 | 7 | 0.4286 | 29 | 0.4828 |
| 11_876758 | 7 | 0.2857 | 12 | 0.5833 | 8 | 0.625 | 10 | 0.2 | 37 | 0.4324 | 5 | 0.4 | 8 | 0.875 | 10 | 0.6 | 6 | 0.5 | 29 | 0.6207 |
| 11_876759 | 7 | 0.1429 | 12 | 0.5833 | 9 | 0.3333 | 11 | 0.1818 | 39 | 0.3333 | 5 | 0.6 | 9 | 0.3333 | 12 | 0.5 | 10 | 0.4 | 36 | 0.4444 |
| 11_876761 | 4 | 0.5 | 5 | 0.4 | 5 | 1 | 2 | 1 | 16 | 0.6875 | 3 | 0.6667 | 6 | 0.6667 | 5 | 0.8 | 2 | 0 | 16 | 0.625 |
| 11_877539 | NA | NA | 2 | 0.5 | NA | NA | NA | NA | 2 | 0.5 | NA | NA | 1 | 1 | NA | NA | 1 | 1 | 2 | 1 |
| 11_877540 | NA | NA | 2 | 0.5 | NA | NA | NA | NA | 2 | 0.5 | NA | NA | 1 | 1 | NA | NA | 1 | 1 | 2 | 1 |
| 11_878721 | 9 | 0.5556 | 5 | 0.6 | 7 | 0.8571 | 5 | 0.8 | 26 | 0.6923 | 7 | 0.7143 | 7 | 1 | 7 | 0.8571 | 5 | 0.8 | 26 | 0.8462 |
| 11_878721 | 8 | 0.25 | 3 | 0 | 7 | 0.5714 | 3 | 0.3333 | 21 | 0.3333 | 7 | 0.5714 | 7 | 0.2857 | 7 | 0.1429 | 4 | 0.25 | 25 | 0.32 |
| 11_879379 | NA | NA | NA | NA | NA | NA | 1 | 1 | 1 | 1 | NA | NA | NA | NA | NA | NA | 1 | 1 | 1 | 1 |
| 11_879383 | NA | NA | NA | NA | NA | NA | 1 | 1 | 1 | 1 | NA | NA | NA | NA | NA | NA | 1 | 1 | 1 | 1 |
| 11_879397 | 3 | 0 | 2 | 0.5 | 2 | 0.5 | 2 | 0 | 9 | 0.2222 | NA | NA | 2 | 0 | 2 | 0.5 | 1 | 1 | 5 | 0.4 |
| 11_879404 | 10 | 0 | 15 | 0.1333 | 17 | 0.0588 | 8 | 0.125 | 50 | 0.08 | 11 | 0 | 19 | 0 | 16 | 0.1875 | 15 | 0.0667 | 61 | 0.0656 |
| 11_879419 | 15 | 0.0667 | 12 | 0.0833 | 15 | 0.0667 | 17 | 0 | 59 | 0.0508 | 12 | 0.1667 | 27 | 0.0741 | 16 | 0 | 19 | 0 | 74 | 0.0541 |
| 11_879421 | 15 | 0.1333 | 15 | 0.2667 | 20 | 0.15 | 19 | 0.1579 | 69 | 0.1739 | 12 | 0.1667 | 27 | 0.1111 | 18 | 0.0556 | 21 | 0.0952 | 78 | 0.1026 |
| 11_879442 | 5 | 0.2 | 3 | 0 | 5 | 0 | 7 | 0 | 20 | 0.05 | 1 | 0 | 8 | 0.125 | 8 | 0 | 6 | 0.1667 | 23 | 0.087 |
| 11_879585 | NA | NA | NA | NA | NA | NA | 1 | 1 | 1 | 1 | 1 | 1 | 1 | 0 | NA | NA | NA | NA | 2 | 0.5 |
| 11_879640 | 3 | 0.6667 | 2 | 0.5 | 2 | 0.5 | 1 | 1 | 8 | 0.625 | 1 | 0 | 4 | 0.5 | 1 | 1 | 1 | 0 | 7 | 0.4286 |

| | | | | | | | | | | | | | | | | | | | | |
|---|---|---|---|---|---|---|---|---|---|---|---|---|---|---|---|---|---|---|---|---|
| 11_879640 | 3 | 0.3333 | 2 | 0 | 3 | 0.3333 | 1 | 0 | 9 | 0.2222 | 2 | 0 | 4 | 0.5 | 1 | 0 | NA | NA | 7 | 0.2857 |
| 11_879641 | 3 | 0.6667 | 2 | 0.5 | 2 | 0.5 | 1 | 1 | 8 | 0.625 | NA | NA | 4 | 0.5 | 1 | 1 | NA | NA | 5 | 0.6 |
| 11_879642 | 3 | 0 | 2 | 0.5 | 3 | 0.3333 | 1 | 0 | 9 | 0.2222 | 2 | 0.5 | 3 | 0.3333 | 1 | 0 | 1 | 0 | 7 | 0.2857 |
| 11_879642 | 2 | 1 | 2 | 1 | 3 | 0.6667 | 1 | 1 | 8 | 0.875 | 2 | 0.5 | 3 | 0.3333 | 1 | 1 | 1 | 0 | 7 | 0.4286 |
| 11_879722 | 4 | 0 | 3 | 0.3333 | 3 | 0 | 2 | 0 | 12 | 0.0833 | 2 | 0 | 3 | 0.3333 | 2 | 0 | 3 | 0 | 10 | 0.1 |
| 11_879723 | 5 | 0.2 | 3 | 0 | 3 | 0.3333 | 2 | 0.5 | 13 | 0.2308 | 1 | 0 | 3 | 0 | 2 | 0.5 | 1 | 0 | 7 | 0.1429 |
| 11_879737 | 4 | 0.25 | 2 | 0 | 1 | 1 | 1 | 0 | 8 | 0.25 | NA | NA | 3 | 0.6667 | 2 | 0.5 | NA | NA | 5 | 0.6 |
| 11_879739 | 6 | 0.1667 | 4 | 0 | 3 | 0.3333 | 2 | 0 | 15 | 0.1333 | 3 | 0 | 5 | 0.2 | 3 | 0 | 4 | 0 | 15 | 0.0667 |
| 11_879739 | 6 | 0.5 | 4 | 0.25 | 3 | 1 | 1 | 1 | 14 | 0.5714 | 3 | 0.3333 | 5 | 0.4 | 3 | 0.6667 | 3 | 0.3333 | 14 | 0.4286 |
| 11_879741 | 7 | 0.1429 | 4 | 0 | 4 | 0 | 3 | 0 | 18 | 0.0556 | 3 | 0 | 5 | 0.4 | 3 | 0 | 4 | 0 | 15 | 0.1333 |
| 11_879743 | 5 | 0 | 5 | 0 | 3 | 0 | 3 | 0.3333 | 16 | 0.0625 | 4 | 0.25 | 3 | 0 | 2 | 0 | 3 | 0 | 12 | 0.0833 |
| 11_880607 | 1 | 0 | NA | NA | 2 | 0.5 | 1 | 0 | 4 | 0.25 | 1 | 0 | NA | NA | 1 | 1 | 1 | 0 | 3 | 0.3333 |
| 11_880608 | 1 | 0 | 1 | 0 | 2 | 1 | 1 | 1 | 5 | 0.6 | 1 | 0 | NA | NA | NA | NA | 1 | 1 | 2 | 0.5 |
| 11_880835 | 1 | 0 | 2 | 0.5 | 1 | 0 | 3 | 0.6667 | 7 | 0.4286 | 3 | 0.6667 | NA | NA | NA | NA | 2 | 0.5 | 5 | 0.6 |
| 11_880835 | 2 | 1 | 1 | 0 | 1 | 1 | 3 | 0.6667 | 7 | 0.7143 | 3 | 0.6667 | NA | NA | 1 | 1 | 2 | 1 | 6 | 0.8333 |
| 11_880837 | 2 | 1 | 2 | 0.5 | 2 | 0.5 | 4 | 0.25 | 10 | 0.5 | 3 | 1 | NA | NA | 1 | 0 | 2 | 0 | 6 | 0.5 |
| 11_880839 | 1 | 1 | 2 | 1 | 2 | 1 | 4 | 1 | 9 | 1 | 2 | 1 | NA | NA | 2 | 1 | 2 | 1 | 6 | 1 |
| 11_880839 | 1 | 1 | 2 | 0.5 | 2 | 1 | 5 | 0.4 | 10 | 0.6 | 2 | 0.5 | NA | NA | 2 | 0 | 2 | 0 | 6 | 0.1667 |
| 11_880839 | 1 | 1 | NA | NA | 1 | 1 | 2 | 1 | 4 | 1 | 2 | 1 | NA | NA | 2 | 0.5 | 2 | 0.5 | 6 | 0.6667 |
| 11_880839 | 1 | 1 | 2 | 0 | 2 | 0.5 | 5 | 0.8 | 10 | 0.6 | 3 | 1 | NA | NA | 2 | 0.5 | 2 | 0 | 7 | 0.5714 |
| 11_880842 | 1 | 1 | 3 | 0 | 1 | 1 | 2 | 0.5 | 7 | 0.4286 | 1 | 0 | NA | NA | 2 | 1 | 2 | 0 | 5 | 0.4 |
| 11_880843 | 1 | 0 | 2 | 0 | 2 | 0.5 | 3 | 0.6667 | 8 | 0.375 | 3 | 0.3333 | NA | NA | 2 | 0.5 | 1 | 1 | 6 | 0.5 |
| 11_880843 | 2 | 0.5 | 2 | 0 | 2 | 0.5 | 5 | 0.2 | 11 | 0.2727 | 3 | 0.6667 | NA | NA | 2 | 0 | 2 | 0 | 7 | 0.2857 |
| 11_880844 | NA | NA | 2 | 0 | 2 | 0.5 | 2 | 0 | 6 | 0.1667 | 2 | 0 | NA | NA | 2 | 0 | 1 | 1 | 5 | 0.2 |
| 11_880845 | 2 | 0.5 | 2 | 0 | 2 | 0 | 4 | 0.25 | 10 | 0.2 | 3 | 1 | NA | NA | 2 | 0.5 | 2 | 0.5 | 7 | 0.7143 |
| 11_880849 | 2 | 0 | 2 | 0 | 3 | 0.3333 | 2 | 0 | 9 | 0.1111 | 3 | 0.3333 | NA | NA | 2 | 0 | 2 | 0 | 7 | 0.1429 |
| 11_881016 | 16 | 0.625 | 6 | 0.6667 | 8 | 0.875 | 8 | 0.625 | 38 | 0.6842 | 10 | 1 | 5 | 1 | 9 | 0.8889 | 8 | 1 | 32 | 0.9688 |
| 11_881017 | 16 | 0.6875 | 7 | 0.7143 | 9 | 0.7778 | 12 | 0.75 | 44 | 0.7273 | 12 | 1 | 5 | 0.6 | 9 | 0.7778 | 8 | 0.875 | 34 | 0.8529 |
| 11_881018 | 11 | 0.4545 | 6 | 0.6667 | 5 | 1 | 10 | 0.6 | 32 | 0.625 | 7 | 0.8571 | 2 | 0 | 7 | 0.8571 | 7 | 0.7143 | 23 | 0.7391 |
| 11_881018 | 9 | 0.7778 | 4 | 0.5 | 6 | 0.5 | 8 | 0.75 | 27 | 0.6667 | 5 | 1 | 4 | 0.5 | 7 | 0.7143 | 4 | 0.75 | 20 | 0.75 |
| 11_881021 | 13 | 0.6154 | 6 | 0.1667 | 5 | 0.4 | 16 | 0.5 | 40 | 0.475 | 8 | 0.75 | 6 | 0.8333 | 6 | 0.3333 | 7 | 0.5714 | 27 | 0.6296 |
| 11_881021 | 9 | 0.6667 | 5 | 0.6 | 7 | 0.7143 | 14 | 0.8571 | 35 | 0.7429 | 7 | 1 | 6 | 1 | 6 | 0.6667 | 5 | 1 | 24 | 0.9167 |
| 11_882305 | 10 | 0.1 | 5 | 0.2 | 10 | 0.2 | 12 | 0.0833 | 37 | 0.1351 | 7 | 0.2857 | 8 | 0.125 | 12 | 0 | 7 | 0 | 34 | 0.0882 |
| 11_882305 | 9 | 0 | 4 | 0 | 9 | 0.2222 | 10 | 0.1 | 32 | 0.0938 | 10 | 0 | 8 | 0 | 11 | 0.1818 | 9 | 0.1111 | 38 | 0.0789 |
| 11_882307 | 9 | 0.3333 | 2 | 0 | 8 | 0 | 8 | 0 | 27 | 0.1111 | 6 | 0 | 8 | 0 | 9 | 0.1111 | 8 | 0.125 | 31 | 0.0645 |
| 11_882331 | 3 | 0 | 3 | 0.3333 | 1 | 0 | 5 | 0 | 12 | 0.0833 | 4 | 0 | 1 | 0 | 5 | 0.2 | 3 | 0 | 13 | 0.0769 |
| 11_882349 | 2 | 0 | 1 | 0 | NA | NA | 5 | 0.2 | 8 | 0.125 | 2 | 0 | 3 | 0.3333 | 1 | 0 | 1 | 0 | 7 | 0.1429 |
| 11_882349 | 2 | 0 | 1 | 0 | NA | NA | 5 | 0.6 | 8 | 0.375 | 2 | 0.5 | 3 | 0 | 1 | 1 | 1 | 0 | 7 | 0.2857 |
| 11_886192 | NA | NA | 1 | 0 | NA | NA | 1 | 1 | 2 | 0.5 | 1 | 1 | NA | NA | 1 | 0 | NA | NA | 2 | 0.5 |
| 11_886193 | NA | NA | 1 | 0 | NA | NA | 1 | 1 | 2 | 0.5 | 1 | 0 | NA | NA | 1 | 1 | NA | NA | 2 | 0.5 |
| 11_887218 | 9 | 0.3333 | 7 | 0 | 4 | 0 | 11 | 0.1818 | 31 | 0.1613 | 9 | 0.1111 | 6 | 0 | 6 | 0.1667 | 7 | 0 | 28 | 0.0714 |
| 11_887221 | 7 | 0.1429 | 7 | 0.1429 | 3 | 0 | 12 | 0 | 29 | 0.069 | 8 | 0.125 | 6 | 0 | 6 | 0 | 5 | 0.2 | 25 | 0.08 |
| 11_887222 | 6 | 0.3333 | 5 | 0.6 | 3 | 0.3333 | 8 | 0.375 | 22 | 0.4091 | 6 | 0.1667 | 5 | 0.6 | 6 | 0.5 | 5 | 0.4 | 22 | 0.4091 |
| 11_887225 | 6 | 0 | 5 | 0.4 | 3 | 0 | 10 | 0.2 | 24 | 0.1667 | 6 | 0 | 7 | 0.1429 | 6 | 0.1667 | 6 | 0.3333 | 25 | 0.16 |
| 11_887351 | 3 | 0.3333 | 2 | 0.5 | 2 | 1 | 4 | 0.25 | 11 | 0.4545 | 3 | 0.6667 | 2 | 0.5 | 4 | 0.5 | 3 | 0 | 12 | 0.4167 |
| 11_887351 | 1 | 0 | 2 | 0.5 | 1 | 1 | 4 | 0 | 8 | 0.25 | 3 | 0.3333 | 1 | 0 | 4 | 0.5 | 3 | 0.3333 | 11 | 0.3636 |
| 11_887351 | 4 | 0.25 | 2 | 0 | 2 | 0 | 5 | 0.2 | 13 | 0.1538 | 4 | 0.25 | 2 | 0 | 4 | 0 | 3 | 0.3333 | 13 | 0.1538 |
| 11_887352 | 4 | 0.25 | 1 | 0 | 2 | 0.5 | 4 | 0.25 | 11 | 0.2727 | 4 | 0.25 | 2 | 0 | 4 | 0.5 | 3 | 0.3333 | 13 | 0.3077 |
| 11_887354 | 1 | 0 | 1 | 0 | 2 | 1 | 3 | 1 | 7 | 0.7143 | 4 | 0.5 | NA | NA | 4 | 0.75 | 3 | 0.6667 | 11 | 0.6364 |
| 11_887354 | 4 | 0.5 | 2 | 0.5 | 2 | 1 | 4 | 0.25 | 12 | 0.5 | 4 | 0.25 | 2 | 0 | 4 | 0 | 3 | 0.3333 | 13 | 0.1538 |
| 11_887354 | 4 | 0 | 2 | 0 | 2 | 0.5 | 4 | 0.25 | 12 | 0.1667 | 3 | 0.6667 | 2 | 0 | 4 | 0.5 | 3 | 1 | 12 | 0.5833 |
| 11_887355 | 4 | 0.5 | 2 | 0 | 2 | 0 | 3 | 0.3333 | 11 | 0.2727 | 2 | 0.5 | 1 | 0 | 3 | 0.3333 | 2 | 0.5 | 8 | 0.375 |
| 11_887356 | 2 | 0 | 2 | 0 | 1 | 0 | 4 | 0.25 | 9 | 0.1111 | 3 | 0 | 1 | 1 | 4 | 0 | 3 | 0 | 11 | 0.0909 |
| 11_887359 | 3 | 0.3333 | 1 | 1 | 2 | 0.5 | 2 | 0 | 8 | 0.375 | 3 | 1 | 1 | 0 | 4 | 0.75 | 3 | 0.6667 | 11 | 0.7273 |
| 11_887360 | 2 | 0.5 | 1 | 0 | 2 | 1 | 2 | 0.5 | 7 | 0.5714 | 2 | 0.5 | 1 | 0 | 4 | 0.5 | 2 | 1 | 9 | 0.5556 |
| 11_887360 | 2 | 1 | 1 | 0 | 1 | 0 | 2 | 0.5 | 6 | 0.5 | 2 | 1 | NA | NA | 3 | 0.3333 | 1 | 1 | 6 | 0.6667 |
| 11_887360 | 3 | 0.6667 | 1 | 0 | NA | NA | 2 | 1 | 6 | 0.6667 | 2 | 1 | 1 | 1 | 4 | 0.75 | 2 | 0 | 9 | 0.6667 |
| 11_888149 | 2 | 0 | 3 | 0 | 3 | 0.3333 | 2 | 0 | 10 | 0.1 | 6 | 0.3333 | 1 | 0 | 3 | 0 | 4 | 0 | 14 | 0.1429 |
| 11_888152 | 3 | 0.3333 | 3 | 0.3333 | 1 | 1 | NA | NA | 7 | 0.4286 | 5 | 0 | 1 | 1 | 1 | 0 | 4 | 0.25 | 11 | 0.1818 |
| 11_888153 | 3 | 0.3333 | 4 | 0.75 | 2 | 0.5 | 2 | 1 | 11 | 0.6364 | 4 | 0.75 | 2 | 0.5 | 1 | 1 | 3 | 0.6667 | 10 | 0.7 |
| 11_888719 | 2 | 0 | 2 | 0.5 | 1 | 0 | NA | NA | 5 | 0.2 | 3 | 0.3333 | 2 | 0.5 | 2 | 0 | 2 | 0.5 | 9 | 0.3333 |
| 11_888720 | 2 | 0 | 2 | 0.5 | NA | NA | NA | NA | 4 | 0.25 | 4 | 0.25 | 2 | 0.5 | 2 | 0 | 2 | 0 | 10 | 0.2 |
| 11_888722 | 2 | 0.5 | 1 | 0 | 1 | 1 | NA | NA | 4 | 0.5 | 3 | 0.3333 | 2 | 0.5 | 1 | 1 | 2 | 0 | 8 | 0.375 |
| 11_888724 | 2 | 1 | 1 | 1 | NA | NA | 1 | 0 | 4 | 0.75 | 4 | 0.25 | 2 | 1 | 2 | 1 | 2 | 1 | 10 | 0.7 |
| 11_888725 | 2 | 0 | 2 | 1 | 1 | 0 | 1 | 1 | 6 | 0.5 | 4 | 0.5 | 2 | 0.5 | 1 | 0 | 2 | 0 | 9 | 0.3333 |

| | | | | | | | | | | | | | | | | | | | | |
|---|---|---|---|---|---|---|---|---|---|---|---|---|---|---|---|---|---|---|---|---|
| 11_8887259 | 2 | 0 | 2 | 0.5 | 1 | 0 | 1 | 0 | 6 | 0.1667 | 4 | 0 | 4 | 0 | 2 | 0.5 | 2 | 0 | 12 | 0.0833 |
| 11_8887263 | NA | NA | 2 | 1 | 1 | 0 | 1 | 0 | 4 | 0.5 | 2 | 0.5 | 2 | 0.5 | 1 | 0 | 2 | 0 | 7 | 0.2857 |
| 11_8887272 | 3 | 1 | 2 | 1 | 1 | 1 | 1 | 1 | 7 | 1 | 4 | 1 | 4 | 1 | 2 | 1 | 2 | 1 | 12 | 1 |
| 11_8887283 | 2 | 1 | 1 | 1 | 1 | 1 | 1 | 1 | 5 | 1 | 3 | 1 | 4 | 1 | 2 | 0.5 | 2 | 1 | 11 | 0.9091 |
| 11_8887285 | 2 | 1 | 2 | 1 | 1 | 1 | 1 | 1 | 6 | 1 | 3 | 1 | 3 | 1 | 1 | 1 | 2 | 1 | 9 | 1 |
| 11_8887314 | 2 | 0 | 3 | 0 | 1 | 0 | 1 | 1 | 7 | 0.1429 | 3 | 0.3333 | 4 | 0 | 2 | 0.5 | 1 | 0 | 10 | 0.2 |
| 11_8888162 | 1 | 0 | 3 | 0.6667 | 1 | 0 | NA | NA | 5 | 0.4 | 5 | 0 | 4 | 0.75 | 3 | 0.3333 | 2 | 0.5 | 14 | 0.3571 |
| 11_8888167 | 2 | 0.5 | 3 | 0.3333 | 1 | 1 | 1 | 0 | 7 | 0.4286 | 4 | 0.75 | 5 | 0 | 2 | 0 | 2 | 0.5 | 13 | 0.3077 |
| 11_8888235 | 2 | 0 | 4 | 0.25 | NA | NA | 1 | 0 | 7 | 0.1429 | 5 | 0 | 5 | 0.2 | 4 | 0.25 | 2 | 0 | 16 | 0.125 |
| 11_8890995 | 1 | 1 | 1 | 0 | 1 | 0 | NA | NA | 3 | 0.3333 | 1 | 0 | NA | NA | NA | NA | 2 | 0.5 | 3 | 0.3333 |
| 11_8892640 | 5 | 0 | 1 | 0 | 4 | 0.5 | 3 | 0 | 13 | 0.1538 | NA | NA | 3 | 0.3333 | 2 | 0 | 5 | 0 | 10 | 0.1 |
| 11_8893040 | 5 | 0.2 | 2 | 0 | 7 | 0.1429 | 2 | 0 | 16 | 0.125 | 2 | 0.5 | 1 | 0 | 4 | 0.25 | 6 | 0.3333 | 13 | 0.3077 |
| 11_8893048 | 4 | 0.25 | 2 | 0 | 7 | 0.1429 | 4 | 0 | 17 | 0.1176 | 4 | 0.25 | 1 | 1 | 5 | 0 | 4 | 0.5 | 14 | 0.2857 |
| 11_8893068 | 7 | 0.5714 | 2 | 1 | 8 | 0.875 | 5 | 0.8 | 22 | 0.7727 | 5 | 0.2 | 1 | 1 | 5 | 0.6 | 6 | 0.8333 | 17 | 0.5882 |
| 11_8893077 | 6 | 0.1667 | 2 | 0.5 | 8 | 0.25 | 5 | 0.4 | 21 | 0.2857 | 4 | 0.25 | 1 | 0 | 4 | 0.25 | 5 | 0 | 14 | 0.1429 |
| 11_8893078 | 7 | 0 | 2 | 0 | 7 | 0 | 5 | 0.4 | 21 | 0.0952 | 5 | 0 | 1 | 0 | 5 | 0.2 | 6 | 0.1667 | 17 | 0.1176 |
| 11_8893083 | 4 | 0.75 | NA | NA | 2 | 1 | 2 | 1 | 8 | 0.875 | 5 | 1 | NA | NA | 1 | 0 | 5 | 0.6 | 11 | 0.7273 |
| 11_8893083 | 1 | 0 | 2 | 0 | 3 | 1 | 1 | 0 | 7 | 0.4286 | NA | NA | 1 | 1 | NA | NA | 1 | 1 | 2 | 1 |
| 11_8893091 | 6 | 0 | 1 | 0 | 6 | 0 | 5 | 0.2 | 18 | 0.0556 | 6 | 0 | 1 | 0 | 4 | 0.25 | 5 | 0.2 | 16 | 0.125 |
| 11_8893100 | 6 | 0.8333 | 2 | 0.5 | 8 | 1 | 5 | 1 | 21 | 0.9048 | 3 | 0.6667 | 1 | 1 | 4 | 1 | 6 | 1 | 14 | 0.9286 |
| 11_8893101 | 7 | 0.8571 | 1 | 1 | 5 | 0.4 | 3 | 0.3333 | 16 | 0.625 | 3 | 1 | NA | NA | 4 | 0.5 | 5 | 0.6 | 12 | 0.6667 |
| 11_8893113 | 7 | 1 | 2 | 1 | 6 | 1 | 4 | 1 | 19 | 1 | 3 | 1 | 1 | 0 | 4 | 1 | 5 | 1 | 13 | 0.9231 |
| 11_8893120 | 5 | 1 | 2 | 1 | 5 | 1 | 4 | 1 | 16 | 1 | 3 | 1 | 1 | 1 | 3 | 1 | 4 | 1 | 11 | 1 |
| 11_8893122 | 6 | 1 | NA | NA | 4 | 0.75 | 4 | 1 | 14 | 0.9286 | 4 | 1 | 1 | 1 | 3 | 1 | 4 | 1 | 12 | 1 |
| 11_8893124 | 6 | 0.1667 | 1 | 0 | 7 | 0.1429 | 5 | 0.4 | 19 | 0.2105 | 4 | 0.25 | 1 | 1 | 4 | 0 | 6 | 0.3333 | 15 | 0.2667 |
| 11_8893128 | 7 | 0.8571 | 2 | 1 | 7 | 1 | 4 | 1 | 20 | 0.95 | 5 | 1 | 1 | 1 | 4 | 1 | 6 | 1 | 16 | 1 |
| 11_8893134 | 5 | 0.2 | 2 | 0 | 5 | 0.2 | 5 | 0.4 | 17 | 0.2353 | 4 | 0.75 | 1 | 0 | 3 | 1 | 5 | 0 | 13 | 0.4615 |
| 11_8893135 | 7 | 0.4286 | 2 | 0 | 6 | 0.1667 | 4 | 0.25 | 19 | 0.2632 | 3 | 0.6667 | NA | NA | 3 | 0.6667 | 5 | 0.6 | 11 | 0.6364 |
| 11_8893142 | 6 | 0.3333 | 2 | 0.5 | 7 | 0.1429 | 4 | 0.25 | 19 | 0.2632 | 4 | 1 | 1 | 1 | 4 | 0.5 | 6 | 1 | 15 | 0.8667 |
| 11_8893143 | 7 | 0.8571 | 1 | 1 | 6 | 1 | 3 | 0.6667 | 17 | 0.8824 | 3 | 1 | 1 | 1 | 2 | 0.5 | 6 | 0.6667 | 12 | 0.75 |
| 11_8893153 | 5 | 0 | 2 | 0.5 | 5 | 0.2 | 5 | 0.2 | 17 | 0.1765 | 3 | 0 | NA | NA | 4 | 0.25 | 5 | 0 | 12 | 0.0833 |
| 11_8894153 | 1 | 0 | 1 | 0 | 2 | 0 | 2 | 1 | 6 | 0.3333 | 1 | 0 | NA | NA | 3 | 0.3333 | 2 | 0 | 6 | 0.1667 |
| 11_8919169 | 2 | 0.5 | 2 | 0.5 | 1 | 0 | NA | NA | 5 | 0.4 | 3 | 0.6667 | 2 | 0.5 | NA | NA | 2 | 1 | 7 | 0.7143 |
| 11_8919203 | 2 | 1 | 2 | 1 | NA | NA | NA | NA | 4 | 1 | 2 | 1 | 2 | 1 | NA | NA | 2 | 1 | 6 | 1 |
| 11_8919214 | NA | NA | 1 | 1 | NA | NA | NA | NA | 1 | 1 | NA | NA | NA | NA | NA | NA | 1 | 1 | 1 | 1 |
| 11_8919239 | 2 | 1 | 3 | 0 | NA | NA | NA | NA | 5 | 0.4 | 2 | 0 | 2 | 0 | NA | NA | 3 | 0.3333 | 7 | 0.1429 |
| 11_8919253 | 1 | 0 | 2 | 0.5 | NA | NA | NA | NA | 3 | 0.3333 | 1 | 0 | 2 | 0.5 | NA | NA | 2 | 0.5 | 5 | 0.4 |
| 11_8919254 | 2 | 1 | 3 | 0.6667 | NA | NA | NA | NA | 5 | 0.8 | 1 | 0 | 1 | 0 | NA | NA | 3 | 1 | 5 | 0.6 |
| 11_8919256 | 1 | 1 | 3 | 0 | NA | NA | NA | NA | 4 | 0.25 | NA | NA | 1 | 1 | NA | NA | 2 | 0 | 3 | 0.3333 |
| 11_8919259 | 1 | 0 | 2 | 1 | NA | NA | NA | NA | 3 | 0.6667 | 1 | 1 | 2 | 1 | NA | NA | 3 | 0.6667 | 6 | 0.8333 |
| 11_8919291 | 1 | 1 | 1 | 1 | NA | NA | NA | NA | 2 | 1 | 1 | 1 | 2 | 1 | NA | NA | 2 | 1 | 5 | 1 |
| 11_8987436 | 1 | 0 | NA | NA | 2 | 0.5 | NA | NA | 3 | 0.3333 | 1 | 1 | NA | NA | NA | NA | NA | NA | 1 | 1 |
| 11_8992623 | 1 | 0 | 4 | 0.25 | 2 | 0 | 1 | 0 | 8 | 0.125 | 5 | 0 | 5 | 0.2 | 1 | 0 | 2 | 0 | 13 | 0.0769 |
| 11_8992632 | 1 | 1 | 2 | 1 | 2 | 1 | 1 | 1 | 6 | 1 | 5 | 1 | 4 | 0.75 | NA | NA | 2 | 1 | 11 | 0.9091 |
| 11_8992646 | 1 | 1 | 4 | 1 | 2 | 1 | 1 | 1 | 8 | 1 | 5 | 0.6 | 6 | 1 | 1 | 0 | 2 | 1 | 14 | 0.7857 |
| 11_8992651 | 1 | 1 | 4 | 0.25 | 2 | 0 | 2 | 0 | 9 | 0.2222 | 5 | 0.4 | 5 | 0.2 | 1 | 0 | 2 | 0 | 13 | 0.2308 |
| 11_8992658 | 1 | 1 | 4 | 0.75 | 2 | 1 | 1 | 1 | 8 | 0.875 | 3 | 1 | 5 | 1 | 1 | 0 | 2 | 1 | 11 | 0.9091 |
| 11_8992663 | NA | NA | 3 | 0.6667 | 2 | 0.5 | 2 | 1 | 7 | 0.7143 | 4 | 0.75 | 4 | 0.75 | NA | NA | 1 | 1 | 9 | 0.7778 |
| 11_8992668 | 1 | 1 | 4 | 1 | 2 | 1 | 1 | 0 | 8 | 0.875 | 5 | 0.6 | 5 | 0.8 | NA | NA | 1 | 0 | 11 | 0.6364 |
| 11_8992672 | 1 | 1 | 3 | 1 | 2 | 0.5 | 1 | 1 | 7 | 0.8571 | 5 | 1 | 3 | 1 | 1 | 1 | 1 | 1 | 10 | 1 |
| 11_8992676 | NA | NA | 4 | 0 | 3 | 0 | 2 | 0.5 | 9 | 0.1111 | 6 | 0.1667 | 3 | 0 | 1 | 0 | 2 | 0 | 12 | 0.0833 |
| 11_8992691 | 1 | 0 | 4 | 0.25 | 3 | 0.3333 | 2 | 0.5 | 10 | 0.3 | 7 | 0.4286 | 5 | 0.4 | 1 | 0 | 2 | 0.5 | 15 | 0.4 |
| 11_8992715 | 2 | 1 | 2 | 0 | 3 | 0.3333 | 3 | 1 | 10 | 0.6 | 5 | 0.8 | 5 | 0.8 | 2 | 0.5 | 2 | 1 | 14 | 0.7857 |
| 11_8992722 | 1 | 0 | 4 | 0 | 3 | 0 | 2 | 0.5 | 10 | 0.1 | 7 | 0.2857 | 6 | 0 | 3 | 0.3333 | 2 | 0 | 18 | 0.1667 |
| 11_9014177 | 3 | 1 | 1 | 1 | 1 | 0 | NA | NA | 5 | 0.8 | 4 | 0 | 3 | 0.3333 | 2 | 0.5 | 2 | 0.5 | 11 | 0.2727 |
| 11_9014182 | 2 | 1 | 2 | 1 | 1 | 1 | NA | NA | 5 | 1 | 4 | 1 | 3 | 1 | 2 | 0.5 | 2 | 1 | 11 | 0.9091 |
| 11_9016767 | 30 | 0.3333 | 15 | 0.2 | 16 | 0.1875 | 22 | 0.3636 | 83 | 0.2892 | 19 | 0.6842 | 19 | 0.3684 | 29 | 0.5172 | 17 | 0.2353 | 84 | 0.4643 |
| 11_9016776 | 28 | 0.0357 | 17 | 0 | 16 | 0.0625 | 18 | 0.1111 | 79 | 0.0506 | 19 | 0.1053 | 16 | 0.0625 | 28 | 0.0714 | 20 | 0 | 83 | 0.0602 |
| 11_9016793 | 38 | 0.1842 | 19 | 0.1579 | 31 | 0.0645 | 31 | 0.0323 | 119 | 0.1092 | 28 | 0.25 | 25 | 0.24 | 35 | 0.1714 | 24 | 0.1667 | 112 | 0.2054 |
| 11_9022766 | 13 | 0.4615 | 11 | 0.3636 | 10 | 0.5 | 7 | 0.4286 | 41 | 0.439 | 12 | 0.4167 | 18 | 0.5 | 12 | 0.5 | 6 | 0.5 | 48 | 0.4792 |
| 11_9022822 | 6 | 0.1667 | 8 | 0.25 | 8 | 0 | 7 | 0.1429 | 29 | 0.1379 | 6 | 0.1667 | 11 | 0.0909 | 8 | 0 | 4 | 0.25 | 29 | 0.1034 |
| 11_9022906 | 5 | 0.2 | 5 | 0.2 | 4 | 0 | 7 | 0.2857 | 21 | 0.1905 | 9 | 0.2222 | 8 | 0 | 10 | 0.1 | 4 | 0 | 31 | 0.0968 |
| 11_9022915 | 4 | 0 | 8 | 0.125 | 6 | 0.1667 | 7 | 0.1429 | 25 | 0.12 | 8 | 0.375 | 9 | 0 | 9 | 0.3333 | 4 | 0.25 | 30 | 0.2333 |
| 11_9023014 | 16 | 0 | 13 | 0.0769 | 11 | 0.0909 | 11 | 0.0909 | 51 | 0.0588 | 14 | 0 | 21 | 0.0476 | 21 | 0.0476 | 12 | 0.1667 | 68 | 0.0588 |
| 11_9023018 | 15 | 0.0667 | 12 | 0 | 8 | 0.125 | 11 | 0.0909 | 46 | 0.0652 | 12 | 0.0833 | 19 | 0.1053 | 21 | 0.1429 | 10 | 0.1 | 62 | 0.1129 |

| | | | | | | | | | | | | | | | | | | | | |
|---|---|---|---|---|---|---|---|---|---|---|---|---|---|---|---|---|---|---|---|---|
| 11_902302 | 16 | 0.25 | 12 | 0.5 | 9 | 0.4444 | 11 | 0.3636 | 48 | 0.375 | 15 | 0.4 | 19 | 0.3158 | 22 | 0.3636 | 11 | 0.4545 | 67 | 0.3731 |
| 11_902314 | 7 | 0 | 6 | 0 | 7 | 0.2857 | 8 | 0 | 28 | 0.0714 | 11 | 0.0909 | 15 | 0.0667 | 22 | 0.0455 | 10 | 0.1 | 58 | 0.069 |
| 11_902315 | 9 | 0.7778 | 9 | 0.8889 | 7 | 0.8571 | 10 | 1 | 35 | 0.8857 | 11 | 0.8182 | 16 | 0.9375 | 22 | 1 | 10 | 1 | 59 | 0.9492 |
| 11_902316 | 5 | 0.2 | 5 | 0.4 | 8 | 0.25 | 7 | 0 | 25 | 0.2 | 8 | 0.375 | 9 | 0.2222 | 15 | 0.0667 | 9 | 0.4444 | 41 | 0.2439 |
| 11_902316 | 7 | 0 | 4 | 0 | 8 | 0.25 | 8 | 0 | 27 | 0.0741 | 8 | 0.25 | 10 | 0.2 | 16 | 0.1875 | 8 | 0.125 | 42 | 0.1905 |
| 11_902788 | 7 | 1 | 6 | 0.8333 | 8 | 1 | 6 | 0.8333 | 27 | 0.9259 | 6 | 1 | 12 | 0.8333 | 17 | 0.8235 | 7 | 1 | 42 | 0.881 |
| 11_902788 | 7 | 0.2857 | 5 | 0.2 | 7 | 0.1429 | 4 | 0.25 | 23 | 0.2174 | 6 | 0.3333 | 11 | 0.3636 | 17 | 0.2941 | 7 | 0.2857 | 41 | 0.3171 |
| 11_902788 | 7 | 0.2857 | 6 | 0.1667 | 8 | 0 | 5 | 0.2 | 26 | 0.1538 | 4 | 0.25 | 11 | 0.1818 | 17 | 0.0588 | 6 | 0 | 38 | 0.1053 |
| 11_902812 | 4 | 0.5 | 5 | 0.2 | 7 | 0.1429 | 2 | 0 | 18 | 0.2222 | 6 | 0.3333 | 9 | 0.2222 | 12 | 0.5 | 3 | 0.3333 | 30 | 0.3667 |
| 11_902814 | 3 | 0 | 5 | 0 | 5 | 0.6 | 2 | 0 | 15 | 0.2 | 4 | 0 | 7 | 0.1429 | 10 | 0.2 | 4 | 0 | 25 | 0.12 |
| 11_902817 | NA | NA | 1 | 1 | 1 | 0 | 2 | 0 | 4 | 0.25 | 2 | 0.5 | 2 | 0.5 | 3 | 0.3333 | 1 | 0 | 8 | 0.375 |
| 11_903676 | 1 | 0 | 1 | 1 | 1 | 0 | 1 | 0 | 4 | 0.25 | 1 | 0 | NA | NA | 1 | 1 | NA | NA | 2 | 0.5 |
| 11_903698 | 1 | 0 | 1 | 0 | 1 | 0 | 1 | 1 | 4 | 0.25 | 1 | 0 | NA | NA | 1 | 1 | NA | NA | 2 | 0.5 |
| 11_905290 | 2 | 0.5 | 4 | 0.5 | 4 | 0.25 | 2 | 0.5 | 12 | 0.4167 | 3 | 0 | 1 | 1 | 2 | 0.5 | NA | NA | 6 | 0.3333 |
| 11_905355 | 4 | 0.25 | 4 | 0 | 6 | 0.1667 | 3 | 0 | 17 | 0.1176 | 2 | 0 | 3 | 0.3333 | 3 | 0 | NA | NA | 8 | 0.125 |
| 11_905356 | 4 | 0.25 | 3 | 0.6667 | 6 | 0 | 2 | 0 | 15 | 0.2 | 3 | 0.3333 | 2 | 0 | 4 | 0 | 1 | 0 | 10 | 0.1 |
| 11_905369 | 2 | 0.5 | 1 | 1 | 1 | 1 | NA | NA | 4 | 0.75 | NA | NA | 2 | 0.5 | 2 | 0.5 | 2 | 0.5 | 6 | 0.5 |
| 11_905384 | 2 | 1 | NA | NA | NA | NA | 1 | 1 | 3 | 1 | NA | NA | NA | NA | 2 | 1 | NA | NA | 2 | 1 |
| 11_905385 | 2 | 0.5 | 1 | 0 | NA | NA | 1 | 0 | 4 | 0.25 | NA | NA | NA | NA | 2 | 0.5 | NA | NA | 2 | 0.5 |
| 11_905386 | 2 | 1 | 1 | 0 | NA | NA | 1 | 1 | 4 | 0.75 | NA | NA | NA | NA | 1 | 0 | 1 | 1 | 2 | 0.5 |
| 11_929903 | 1 | 1 | 1 | 1 | NA | NA | NA | NA | 2 | 1 | NA | NA | NA | NA | NA | NA | 2 | 0.5 | 2 | 0.5 |
| 11_929904 | 2 | 0 | 1 | 1 | NA | NA | 1 | 0 | 4 | 0.25 | NA | NA | NA | NA | NA | NA | 3 | 0.3333 | 3 | 0.3333 |
| 11_929906 | 2 | 0.5 | NA | NA | NA | NA | 1 | 0 | 3 | 0.3333 | NA | NA | NA | NA | NA | NA | 4 | 0.5 | 4 | 0.5 |
| 11_929908 | 2 | 0.5 | NA | NA | NA | NA | 1 | 0 | 3 | 0.3333 | NA | NA | NA | NA | NA | NA | 5 | 0.6 | 5 | 0.6 |
| 11_934916 | 1 | 0 | 4 | 0.25 | 2 | 0 | 3 | 0 | 10 | 0.1 | 2 | 0 | 4 | 0.5 | 1 | 0 | 4 | 0 | 11 | 0.1818 |
| 11_934916 | 3 | 0 | 4 | 0.25 | 2 | 0 | 4 | 0 | 13 | 0.0769 | 3 | 0 | 4 | 0.25 | 1 | 1 | 6 | 0.3333 | 14 | 0.2857 |
| 11_934917 | 3 | 0 | 4 | 0.5 | 2 | 0 | 3 | 0.3333 | 12 | 0.25 | 4 | 0 | 5 | 0 | 1 | 0 | 7 | 0.2857 | 17 | 0.1176 |
| 11_934918 | 4 | 0.5 | 5 | 1 | 2 | 1 | 3 | 0.6667 | 14 | 0.7857 | 4 | 0.5 | 6 | 0.5 | 1 | 1 | 9 | 0.5556 | 20 | 0.55 |
| 11_934918 | 3 | 0.6667 | 5 | 0.4 | 1 | 1 | 2 | 0.5 | 11 | 0.5455 | 4 | 0.25 | 6 | 0.3333 | 2 | 1 | 8 | 0.625 | 20 | 0.5 |
| 11_938347 | NA | NA | 2 | 0.5 | 1 | 0 | NA | NA | 3 | 0.3333 | NA | NA | 2 | 0.5 | NA | NA | 2 | 0 | 4 | 0.25 |
| 11_938348 | NA | NA | 2 | 1 | 1 | 1 | NA | NA | 3 | 1 | NA | NA | 2 | 0.5 | NA | NA | 2 | 0 | 4 | 0.25 |
| 11_938350 | NA | NA | 2 | 1 | 2 | 0.5 | NA | NA | 4 | 0.75 | NA | NA | 3 | 0.6667 | NA | NA | 2 | 0 | 5 | 0.4 |
| 11_938366 | NA | NA | 1 | 1 | 4 | 0.25 | NA | NA | 5 | 0.4 | 1 | 0 | 2 | 0.5 | 1 | 0 | 4 | 0 | 8 | 0.125 |
| 11_938371 | NA | NA | 3 | 0.6667 | 4 | 0 | 1 | 1 | 8 | 0.375 | 1 | 0 | 2 | 0.5 | 1 | 0 | 6 | 0.1667 | 10 | 0.2 |
| 11_938371 | NA | NA | 3 | 0 | 4 | 0.25 | NA | NA | 7 | 0.1429 | NA | NA | 2 | 0 | 1 | 0 | 6 | 0.3333 | 9 | 0.2222 |
| 11_938406 | 27 | 0.1852 | 27 | 0.0741 | 23 | 0.2174 | 38 | 0.1316 | 115 | 0.1478 | 26 | 0.1154 | 37 | 0.0811 | 22 | 0.1818 | 31 | 0.1613 | 116 | 0.1293 |
| 11_938406 | 31 | 0.2258 | 34 | 0.2647 | 30 | 0.2667 | 38 | 0.2632 | 133 | 0.2556 | 33 | 0.3333 | 38 | 0.1842 | 24 | 0.25 | 33 | 0.1515 | 128 | 0.2266 |
| 11_938407 | 17 | 0.1765 | 18 | 0.0556 | 10 | 0 | 21 | 0.0476 | 66 | 0.0758 | 13 | 0.1538 | 16 | 0.0625 | 9 | 0.2222 | 14 | 0 | 52 | 0.0962 |
| 11_938499 | 21 | 0.1905 | 23 | 0.4348 | 16 | 0.3125 | 21 | 0.1905 | 81 | 0.284 | 14 | 0.4286 | 11 | 0.2727 | 16 | 0.25 | 16 | 0.0625 | 57 | 0.2456 |
| 11_938501 | 63 | 0.2381 | 55 | 0.2364 | 33 | 0.3333 | 40 | 0.2 | 191 | 0.2461 | 44 | 0.2045 | 44 | 0.1591 | 42 | 0.1667 | 45 | 0.2 | 175 | 0.1829 |
| 11_938502 | 59 | 0.1186 | 54 | 0.037 | 34 | 0.1176 | 38 | 0.0526 | 185 | 0.0811 | 43 | 0.1163 | 32 | 0.0625 | 39 | 0.0256 | 42 | 0.0714 | 156 | 0.0705 |
| 11_939351 | NA | NA | 1 | 1 | 3 | 0.3333 | NA | NA | 4 | 0.5 | 1 | 0 | 1 | 0 | 3 | 0.3333 | 1 | 1 | 6 | 0.3333 |
| 11_940137 | 10 | 0.3 | 11 | 0.0909 | 11 | 0.2727 | 12 | 0.3333 | 44 | 0.25 | 5 | 0.2 | 9 | 0.3333 | 16 | 0.375 | 7 | 0.7143 | 37 | 0.4054 |
| 11_940140 | 9 | 0.1111 | 17 | 0.1176 | 16 | 0 | 12 | 0.0833 | 54 | 0.0741 | 10 | 0.1 | 13 | 0 | 14 | 0.1429 | 7 | 0 | 44 | 0.0682 |
| 11_940140 | 10 | 0.3 | 17 | 0.2353 | 16 | 0.1875 | 14 | 0.2143 | 57 | 0.2281 | 10 | 0.5 | 16 | 0 | 20 | 0.25 | 9 | 0.3333 | 55 | 0.2364 |
| 11_940150 | 15 | 0 | 17 | 0.1176 | 20 | 0.1 | 16 | 0 | 68 | 0.0588 | 12 | 0.1667 | 19 | 0.0526 | 21 | 0.1429 | 15 | 0.0667 | 67 | 0.1045 |
| 11_940164 | 19 | 0.0526 | 28 | 0.0714 | 28 | 0.0714 | 20 | 0.15 | 95 | 0.0842 | 20 | 0.1 | 28 | 0.1429 | 32 | 0.0625 | 23 | 0.2174 | 103 | 0.1262 |
| 11_940164 | 23 | 0.7391 | 25 | 0.68 | 26 | 0.8077 | 20 | 0.65 | 94 | 0.7234 | 16 | 0.75 | 26 | 0.6923 | 28 | 0.8571 | 25 | 0.72 | 95 | 0.7579 |
| 11_940165 | 22 | 0.9091 | 29 | 0.6897 | 32 | 0.7812 | 25 | 0.92 | 108 | 0.8148 | 20 | 0.8 | 28 | 0.8214 | 30 | 0.8333 | 27 | 0.7037 | 105 | 0.7905 |
| 11_941043 | 3 | 1 | 6 | 0.6667 | 4 | 1 | 5 | 0.6 | 18 | 0.7778 | 2 | 0.5 | 4 | 1 | 8 | 1 | 4 | 1 | 18 | 0.9444 |
| 11_941044 | 4 | 0.25 | 9 | 0.1111 | 5 | 0.2 | 5 | 0.2 | 23 | 0.1739 | 1 | 1 | 6 | 0.1667 | 9 | 0.1111 | 3 | 0 | 19 | 0.1579 |
| 11_941045 | 6 | 0.6667 | 9 | 0.6667 | 5 | 1 | 6 | 0.8333 | 26 | 0.7692 | 4 | 1 | 6 | 1 | 8 | 1 | 6 | 0.8333 | 24 | 0.9583 |
| 11_941046 | 4 | 1 | 8 | 1 | 5 | 1 | 6 | 1 | 23 | 1 | 4 | 1 | 6 | 1 | 8 | 1 | 5 | 1 | 23 | 1 |
| 11_941047 | 6 | 0.8333 | 8 | 0.625 | 5 | 1 | 7 | 0.1429 | 26 | 0.6154 | 4 | 0.5 | 5 | 0.8 | 9 | 0.5556 | 5 | 0.8 | 23 | 0.6522 |
| 11_941048 | 5 | 0.2 | 4 | 0.25 | 2 | 0.5 | 2 | 0.5 | 13 | 0.3077 | 4 | 0.5 | 4 | 0.25 | 4 | 0 | 6 | 0.6667 | 18 | 0.3889 |
| 11_941050 | 5 | 0.8 | 10 | 0.9 | 6 | 0.6667 | 9 | 0.8889 | 30 | 0.8333 | 5 | 0.6 | 7 | 0.7143 | 10 | 0.9 | 9 | 0.7778 | 31 | 0.7742 |
| 11_941055 | 8 | 0.25 | 11 | 0.2727 | 9 | 0.2222 | 12 | 0.25 | 40 | 0.25 | 6 | 0 | 6 | 0.3333 | 8 | 0 | 9 | 0.1111 | 29 | 0.1034 |
| 11_941056 | 6 | 0 | 8 | 0 | 4 | 0 | 6 | 0.3333 | 24 | 0.0833 | 5 | 0 | 5 | 0.4 | 5 | 0 | 2 | 0 | 17 | 0.1176 |
| 11_941056 | 7 | 0.2857 | 10 | 0.4 | 10 | 0.8 | 10 | 0.4 | 37 | 0.4865 | 6 | 0.1667 | 6 | 0.3333 | 9 | 0.5556 | 6 | 0.6667 | 27 | 0.4444 |
| 11_941057 | 11 | 0.5455 | 11 | 0.5455 | 10 | 0.6 | 13 | 0.5385 | 45 | 0.5556 | 7 | 0.5714 | 8 | 0.5 | 10 | 0.7 | 7 | 0.4286 | 32 | 0.5625 |
| 11_941058 | 13 | 0.3846 | 11 | 0.2727 | 9 | 0.1111 | 14 | 0.2143 | 47 | 0.2553 | 6 | 0.1667 | 9 | 0.4444 | 13 | 0.1538 | 7 | 0.1429 | 35 | 0.2286 |
| 11_941060 | 13 | 0.0769 | 11 | 0 | 11 | 0.0909 | 15 | 0.0667 | 50 | 0.06 | 11 | 0 | 10 | 0 | 15 | 0.1333 | 9 | 0.2222 | 45 | 0.0889 |
| 11_941788 | 1 | 1 | 6 | 0.3333 | 2 | 1 | NA | NA | 9 | 0.5556 | NA | NA | 2 | 1 | 3 | 0.6667 | 2 | 1 | 7 | 0.8571 |
| 11_941789 | 1 | 1 | 5 | 0.4 | 1 | 0 | NA | NA | 7 | 0.4286 | NA | NA | NA | NA | 1 | 1 | 1 | 0 | 2 | 0.5 |
| 11_941789 | 1 | 1 | 5 | 0.2 | 2 | 1 | NA | NA | 8 | 0.5 | NA | NA | 2 | 0.5 | 3 | 0.3333 | 2 | 1 | 7 | 0.5714 |

| | | | | | | | | | | | | | | | | | | | | |
|---|---|---|---|---|---|---|---|---|---|---|---|---|---|---|---|---|---|---|---|---|
| 11_941790 | 1 | 1 | 5 | 0.2 | 1 | 1 | NA | NA | 7 | 0.4286 | NA | NA | 2 | 0.5 | 3 | 0.6667 | 1 | 0 | 6 | 0.5 |
| 11_941791 | 1 | 0 | 5 | 0 | 2 | 0.5 | NA | NA | 8 | 0.125 | NA | NA | 2 | 0.5 | 3 | 0.3333 | 2 | 0 | 7 | 0.2857 |
| 11_941792 | NA | NA | 1 | 0 | 2 | 0.5 | NA | NA | 3 | 0.3333 | NA | NA | NA | NA | 2 | 1 | 1 | 1 | 3 | 1 |
| 11_941792 | 1 | 1 | 1 | 1 | NA | NA | NA | NA | 2 | 1 | NA | NA | 2 | 1 | NA | NA | NA | NA | 2 | 1 |
| 11_941820 | 10 | 0.9 | 13 | 0.7692 | 9 | 0.8889 | 16 | 0.875 | 48 | 0.8542 | 8 | 0.75 | 11 | 0.6364 | 16 | 0.8125 | 16 | 0.875 | 51 | 0.7843 |
| 11_941821 | 9 | 0.8889 | 12 | 0.5833 | 11 | 0.5455 | 18 | 0.7222 | 50 | 0.68 | 7 | 0.7143 | 10 | 0.8 | 17 | 0.7059 | 14 | 0.5 | 48 | 0.6667 |
| 11_941822 | 9 | 0.8889 | 11 | 0.6364 | 11 | 0.8182 | 18 | 0.8889 | 49 | 0.8163 | 9 | 0.7778 | 8 | 0.875 | 18 | 0.9444 | 14 | 0.7857 | 49 | 0.8571 |
| 11_942304 | 12 | 0.6667 | 13 | 0.4615 | 13 | 0.3077 | 12 | 0.0833 | 50 | 0.38 | 19 | 0.1579 | 18 | 0.5 | 9 | 0.3333 | 11 | 0.3636 | 57 | 0.3333 |
| 11_942304 | 14 | 0.2857 | 14 | 0.1429 | 12 | 0.4167 | 11 | 0.2727 | 51 | 0.2745 | 17 | 0.2941 | 20 | 0.5 | 9 | 0.2222 | 10 | 0.5 | 56 | 0.3929 |
| 11_942304 | 14 | 0.9286 | 11 | 0.2727 | 12 | 0.5833 | 10 | 0.6 | 47 | 0.617 | 18 | 0.6667 | 22 | 0.7273 | 9 | 0.5556 | 11 | 0.5455 | 60 | 0.65 |
| 11_942305 | 12 | 0.0833 | 13 | 0.1538 | 14 | 0.1429 | 8 | 0.25 | 47 | 0.1489 | 14 | 0.0714 | 20 | 0.25 | 9 | 0.1111 | 9 | 0.1111 | 52 | 0.1538 |
| 11_942305 | 17 | 0.7059 | 15 | 0.6 | 14 | 0.7143 | 10 | 0.6 | 56 | 0.6607 | 17 | 0.5882 | 22 | 0.5909 | 9 | 0.4444 | 12 | 0.5833 | 60 | 0.5667 |
| 11_942305 | 16 | 0.25 | 13 | 0.1538 | 12 | 0.3333 | 9 | 0 | 50 | 0.2 | 17 | 0.1765 | 19 | 0.1579 | 11 | 0.0909 | 13 | 0.0769 | 60 | 0.1333 |
| 11_942305 | 16 | 0.5625 | 13 | 0.5385 | 13 | 0.7692 | 11 | 0.4545 | 53 | 0.5849 | 17 | 0.5882 | 21 | 0.7143 | 10 | 0.6 | 13 | 0.3846 | 61 | 0.5902 |
| 11_942307 | 15 | 0.6 | 15 | 0.4 | 9 | 0.4444 | 8 | 0.75 | 47 | 0.5319 | 18 | 0.6667 | 20 | 0.65 | 10 | 0.4 | 11 | 0.3636 | 59 | 0.5593 |
| 11_942307 | 4 | 0.25 | 11 | 0.2727 | 7 | 0.1429 | 6 | 0.5 | 28 | 0.2857 | 8 | 0 | 14 | 0.2857 | 3 | 0 | 6 | 0.1667 | 31 | 0.1613 |
| 11_942308 | 10 | 0.2 | 5 | 0 | 8 | 0.25 | 5 | 0.2 | 28 | 0.1786 | 12 | 0.1667 | 16 | 0.125 | 4 | 0 | 6 | 0 | 38 | 0.1053 |
| 11_942309 | 9 | 0.8889 | 10 | 1 | 11 | 0.7273 | 7 | 1 | 37 | 0.8919 | 12 | 1 | 18 | 0.8333 | 6 | 0.6667 | 10 | 0.7 | 46 | 0.8261 |
| 11_942309 | 10 | 0.1 | 12 | 0.1667 | 10 | 0.1 | 8 | 0.125 | 40 | 0.125 | 14 | 0.0714 | 19 | 0.1053 | 7 | 0 | 10 | 0 | 50 | 0.06 |
| 11_942309 | 7 | 0.5714 | 11 | 0.8182 | 9 | 0.6667 | 6 | 0.8333 | 33 | 0.7273 | 10 | 0.7 | 16 | 0.6875 | 4 | 0.5 | 8 | 0.5 | 38 | 0.6316 |
| 11_942313 | 11 | 0.6364 | 11 | 0.5455 | 12 | 0.6667 | 7 | 0.5714 | 41 | 0.6098 | 14 | 0.7143 | 20 | 0.7 | 7 | 0.4286 | 11 | 0.5455 | 52 | 0.6346 |
| 11_942994 | 1 | 0 | 4 | 0.25 | NA | NA | NA | NA | 5 | 0.2 | 1 | 0 | NA | NA | 1 | 0 | 2 | 0.5 | 4 | 0.25 |
| 11_942995 | 1 | 1 | 4 | 0.75 | NA | NA | NA | NA | 5 | 0.8 | 1 | 0 | NA | NA | 1 | 0 | 2 | 1 | 4 | 0.5 |
| 11_942996 | 1 | 0 | 3 | 0.3333 | NA | NA | NA | NA | 4 | 0.25 | 1 | 0 | NA | NA | 1 | 0 | 2 | 0.5 | 4 | 0.25 |
| 11_942997 | 2 | 0 | 5 | 0.4 | 4 | 0 | 2 | 0.5 | 13 | 0.2308 | 4 | 0.25 | NA | NA | 1 | 0 | 3 | 0.3333 | 8 | 0.25 |
| 11_942998 | 2 | 0.5 | 6 | 1 | 3 | 0.6667 | 3 | 0.6667 | 14 | 0.7857 | 3 | 1 | NA | NA | 2 | 1 | 2 | 1 | 7 | 1 |
| 11_942999 | 2 | 0.5 | 5 | 0.2 | 4 | 0.25 | 4 | 0 | 15 | 0.2 | 3 | 0.3333 | NA | NA | 2 | 0 | 3 | 0.3333 | 8 | 0.25 |
| 11_943000 | 1 | 0 | 5 | 0.8 | 4 | 0.5 | 4 | 1 | 14 | 0.7143 | 3 | 0.3333 | NA | NA | 1 | 0 | 3 | 1 | 7 | 0.5714 |
| 11_943001 | 2 | 0 | 5 | 0.4 | 3 | 0.3333 | 3 | 0 | 13 | 0.2308 | 2 | 0 | NA | NA | 2 | 0 | 3 | 0.3333 | 7 | 0.1429 |
| 11_943001 | 1 | 0 | 3 | 1 | 3 | 1 | 2 | 0.5 | 9 | 0.7778 | 1 | 0 | NA | NA | 2 | 0.5 | 2 | 1 | 5 | 0.6 |
| 11_943001 | 1 | 1 | 1 | 1 | NA | NA | NA | NA | 2 | 1 | NA | NA | NA | NA | NA | NA | 1 | 1 | 1 | 1 |
| 11_943002 | 2 | 0.5 | 3 | 0.6667 | 3 | 0 | 3 | 0 | 11 | 0.2727 | 1 | 0 | NA | NA | 2 | 0.5 | 3 | 0.6667 | 6 | 0.5 |
| 11_943003 | 2 | 0 | 4 | 0 | 4 | 0 | 3 | 0.3333 | 13 | 0.0769 | 2 | 0 | NA | NA | 2 | 0.5 | 2 | 0 | 6 | 0.1667 |
| 11_943003 | 2 | 0.5 | 3 | 0.6667 | 3 | 0 | 2 | 0.5 | 10 | 0.4 | 2 | 0.5 | NA | NA | 2 | 0.5 | 2 | 0.5 | 6 | 0.5 |
| 11_943003 | 2 | 0.5 | 3 | 1 | 4 | 1 | 3 | 1 | 12 | 0.9167 | 2 | 1 | NA | NA | 1 | 1 | 3 | 1 | 6 | 1 |
| 11_943004 | 2 | 0 | 3 | 0 | 3 | 0.3333 | 3 | 0 | 11 | 0.0909 | 2 | 0 | NA | NA | 2 | 0 | 3 | 0.3333 | 7 | 0.1429 |
| 11_943500 | NA | NA | 2 | 1 | NA | NA | 1 | 1 | 3 | 1 | NA | NA | NA | NA | 1 | 1 | 1 | 1 | 2 | 1 |
| 11_943502 | NA | NA | 2 | 0.5 | NA | NA | 1 | 0 | 3 | 0.3333 | NA | NA | NA | NA | 1 | 1 | 1 | 0 | 2 | 0.5 |
| 11_943646 | NA | NA | 1 | 1 | NA | NA | NA | NA | 1 | 1 | NA | NA | NA | NA | NA | NA | 1 | 1 | 1 | 1 |
| 11_944333 | 1 | 1 | NA | NA | NA | NA | NA | NA | 1 | 1 | 1 | 1 | NA | NA | NA | NA | NA | NA | 1 | 1 |
| 11_944487 | 3 | 0 | 2 | 0 | 2 | 0 | 3 | 0.3333 | 10 | 0.1 | 1 | 0 | 1 | 0 | 2 | 0 | 3 | 0.3333 | 7 | 0.1429 |
| 11_945344 | 12 | 0.75 | 9 | 0.7778 | 14 | 0.7857 | 6 | 0.6667 | 41 | 0.7561 | 9 | 0.6667 | 7 | 0.7143 | 5 | 0.6 | 8 | 1 | 29 | 0.7586 |
| 11_945344 | 8 | 0.125 | 11 | 0.0909 | 12 | 0.0833 | 6 | 0 | 37 | 0.0811 | 8 | 0 | 8 | 0.125 | 5 | 0.2 | 7 | 0 | 28 | 0.0714 |
| 11_945345 | 9 | 0.5556 | 11 | 0.6364 | 10 | 0.6 | 8 | 0.375 | 38 | 0.5526 | 6 | 0.6667 | 8 | 0.75 | 6 | 0.5 | 7 | 0.4286 | 27 | 0.5926 |
| 11_945356 | 16 | 0.1875 | 17 | 0.0588 | 15 | 0.2 | 10 | 0.2 | 58 | 0.1552 | 14 | 0.1429 | 7 | 0.2857 | 11 | 0.0909 | 12 | 0 | 44 | 0.1136 |
| 11_945505 | 18 | 0.5 | 4 | 0.5 | 11 | 0.6364 | 9 | 0.6667 | 42 | 0.5714 | 9 | 0.5556 | 7 | 0.8571 | 20 | 0.7 | 12 | 0.9167 | 48 | 0.75 |
| 11_945506 | 19 | 0.8421 | 6 | 0.8333 | 11 | 0.7273 | 10 | 0.7 | 46 | 0.7826 | 7 | 0.4286 | 9 | 0.7778 | 18 | 0.6111 | 11 | 0.9091 | 45 | 0.6889 |
| 11_945506 | 16 | 0.4375 | 6 | 0.3333 | 13 | 0.1538 | 10 | 0.6 | 45 | 0.3778 | 11 | 0.7273 | 9 | 0.4444 | 21 | 0.5714 | 11 | 0.4545 | 52 | 0.5577 |
| 11_945507 | 15 | 0.1333 | 3 | 0.3333 | 7 | 0.2857 | 6 | 0.1667 | 31 | 0.1935 | 6 | 0.1667 | 8 | 0 | 14 | 0.0714 | 10 | 0 | 38 | 0.0526 |
| 11_945507 | 18 | 0.0556 | 6 | 0.1667 | 12 | 0.1667 | 8 | 0.25 | 44 | 0.1364 | 12 | 0.1667 | 7 | 0 | 22 | 0.0455 | 11 | 0 | 52 | 0.0577 |
| 11_945526 | 3 | 0.3333 | NA | NA | 1 | 0 | 1 | 0 | 5 | 0.2 | 2 | 0.5 | NA | NA | 3 | 0.3333 | NA | NA | 5 | 0.4 |
| 11_945549 | 3 | 0.6667 | 5 | 1 | NA | NA | 1 | 1 | 9 | 0.8889 | 2 | 1 | NA | NA | NA | NA | 1 | 1 | 3 | 1 |
| 11_945549 | 3 | 0.6667 | 5 | 0.8 | 1 | 1 | 2 | 1 | 11 | 0.8182 | 2 | 1 | NA | NA | NA | NA | 1 | 0 | 3 | 0.6667 |
| 11_945549 | 3 | 1 | 1 | 1 | 1 | 1 | 1 | 0 | 6 | 0.8333 | 2 | 1 | NA | NA | NA | NA | 1 | 1 | 3 | 1 |
| 11_945552 | 1 | 0 | 5 | 0.2 | 1 | 0 | 2 | 0.5 | 9 | 0.2222 | 1 | 1 | NA | NA | NA | NA | 1 | 0 | 2 | 0.5 |
| 11_945553 | 1 | 1 | 4 | 0.25 | 1 | 0 | 2 | 0 | 8 | 0.25 | 2 | 1 | NA | NA | NA | NA | 1 | 0 | 3 | 0.6667 |
| 11_945554 | 1 | 1 | 3 | 1 | 1 | 1 | 2 | 1 | 7 | 1 | 2 | 1 | NA | NA | NA | NA | NA | NA | 2 | 1 |
| 11_945555 | 1 | 1 | 4 | 1 | 1 | 1 | 2 | 1 | 8 | 1 | 1 | 1 | NA | NA | NA | NA | 1 | 1 | 2 | 1 |
| 11_945555 | 1 | 1 | 5 | 1 | 1 | 1 | 2 | 1 | 9 | 1 | 1 | 1 | NA | NA | NA | NA | 1 | 1 | 2 | 1 |
| 11_945571 | 1 | 0 | 3 | 0.3333 | 1 | 0 | 1 | 0 | 6 | 0.1667 | 2 | 0.5 | NA | NA | NA | NA | 1 | 0 | 3 | 0.3333 |
| 11_945585 | 1 | 1 | 5 | 0 | 1 | 0 | NA | NA | 7 | 0.1429 | 2 | 0.5 | NA | NA | NA | NA | NA | NA | 2 | 0.5 |
| 11_945612 | NA | NA | 3 | 0.3333 | 1 | 0 | NA | NA | 4 | 0.25 | 1 | 1 | NA | NA | NA | NA | NA | NA | 1 | 1 |
| 11_948822 | 17 | 0.7647 | 15 | 0.9333 | 9 | 0.7778 | 13 | 0.6923 | 54 | 0.7963 | 15 | 0.6667 | 11 | 1 | 17 | 1 | 9 | 0.7778 | 52 | 0.8654 |
| 11_948829 | 29 | 0.0345 | 26 | 0.2308 | 14 | 0.2857 | 24 | 0.0417 | 93 | 0.129 | 23 | 0.2609 | 23 | 0.087 | 34 | 0.0294 | 17 | 0.0588 | 97 | 0.1031 |
| 11_948829 | 21 | 0.3333 | 20 | 0.25 | 11 | 0.3636 | 19 | 0.3158 | 71 | 0.3099 | 16 | 0.375 | 19 | 0.4211 | 27 | 0.4074 | 11 | 0.3636 | 73 | 0.3973 |

| | | | | | | | | | | | | | | | | | | | | |
|---|---|---|---|---|---|---|---|---|---|---|---|---|---|---|---|---|---|---|---|---|
| 11_9488303 | 28 | 0.0357 | 25 | 0.08 | 14 | 0 | 19 | 0.1053 | 86 | 0.0581 | 21 | 0.0952 | 22 | 0.1364 | 28 | 0.0357 | 17 | 0.1765 | 88 | 0.1023 |
| 11_9488347 | 26 | 0.7308 | 36 | 0.6667 | 14 | 0.7857 | 24 | 0.7083 | 100 | 0.71 | 21 | 0.7143 | 20 | 0.75 | 29 | 0.7931 | 17 | 0.5294 | 87 | 0.7126 |
| 11_9488349 | 33 | 0.1818 | 34 | 0.2647 | 13 | 0.2308 | 26 | 0.3077 | 106 | 0.2453 | 24 | 0.1667 | 20 | 0.35 | 33 | 0.1515 | 26 | 0.2692 | 103 | 0.2233 |
| 11_9488353 | 37 | 0.3784 | 37 | 0.5405 | 22 | 0.4091 | 27 | 0.5926 | 123 | 0.4797 | 27 | 0.4074 | 28 | 0.3929 | 35 | 0.5429 | 27 | 0.5185 | 117 | 0.4701 |
| 11_9488354 | 41 | 0.8293 | 36 | 0.6667 | 23 | 0.5217 | 33 | 0.7273 | 133 | 0.7068 | 27 | 0.5926 | 30 | 0.6333 | 38 | 0.6842 | 29 | 0.6897 | 124 | 0.6532 |
| 11_9488357 | 33 | 0.7879 | 32 | 0.9062 | 18 | 0.8889 | 24 | 0.9583 | 107 | 0.8785 | 24 | 0.9583 | 12 | 0.75 | 38 | 0.9474 | 25 | 1 | 99 | 0.9394 |
| 11_9488358 | 36 | 0.8889 | 33 | 0.9091 | 20 | 0.65 | 27 | 0.8148 | 116 | 0.8362 | 27 | 0.8519 | 27 | 0.8148 | 38 | 0.7895 | 27 | 0.8889 | 119 | 0.8319 |
| 11_9489620 | 67 | 0.2836 | 82 | 0.3902 | 48 | 0.3542 | 58 | 0.3103 | 255 | 0.3373 | 50 | 0.26 | 55 | 0.4727 | 75 | 0.3067 | 65 | 0.4154 | 245 | 0.3633 |
| 11_9489633 | 77 | 0.1818 | 94 | 0.1277 | 52 | 0.0769 | 68 | 0.0882 | 291 | 0.1237 | 54 | 0.1667 | 63 | 0.1429 | 84 | 0.1071 | 73 | 0.137 | 274 | 0.135 |
| 11_9489646 | 75 | 0.3467 | 86 | 0.3953 | 55 | 0.3818 | 63 | 0.2857 | 279 | 0.3548 | 52 | 0.4423 | 64 | 0.4531 | 85 | 0.2824 | 73 | 0.3836 | 274 | 0.3796 |
| 11_9489647 | 83 | 0.1205 | 97 | 0.0722 | 56 | 0.0893 | 68 | 0.1029 | 304 | 0.0954 | 55 | 0.1455 | 70 | 0.0714 | 91 | 0.1099 | 79 | 0.1266 | 295 | 0.1119 |
| 11_9489649 | 76 | 0.3026 | 86 | 0.3605 | 51 | 0.2941 | 63 | 0.4286 | 276 | 0.3478 | 52 | 0.2115 | 68 | 0.3235 | 75 | 0.3733 | 74 | 0.3108 | 269 | 0.3123 |
| 11_9489652 | 55 | 0.0909 | 73 | 0.2055 | 38 | 0.1053 | 50 | 0.18 | 216 | 0.1528 | 39 | 0.1282 | 47 | 0.1702 | 71 | 0.1831 | 53 | 0.1509 | 210 | 0.1619 |
| 11_9489661 | 84 | 0.1548 | 95 | 0.2105 | 59 | 0.1017 | 72 | 0.0972 | 310 | 0.1484 | 56 | 0.2143 | 71 | 0.1408 | 94 | 0.1489 | 76 | 0.0789 | 297 | 0.1414 |
| 11_9491210 | 6 | 0 | 4 | 0.25 | 2 | 0 | 6 | 0 | 18 | 0.0556 | 3 | 0 | 6 | 0 | 4 | 0 | 3 | 0.3333 | 16 | 0.0625 |
| 11_9491568 | 4 | 1 | 6 | 0.8333 | 6 | 0.8333 | 8 | 1 | 24 | 0.9167 | 4 | 1 | 5 | 1 | 4 | 0.75 | 5 | 1 | 18 | 0.9444 |
| 11_9491574 | 5 | 0.2 | 3 | 0.3333 | 6 | 0.1667 | 9 | 0.5556 | 23 | 0.3478 | 3 | 0 | 5 | 0.2 | 2 | 0 | 5 | 0.4 | 15 | 0.2 |
| 11_9491644 | 5 | 0 | 5 | 0.4 | 7 | 0.2857 | 8 | 0 | 25 | 0.16 | 5 | 0.2 | 5 | 0.2 | 4 | 0.25 | 4 | 0.25 | 18 | 0.2222 |
| 11_9491665 | 7 | 0.1429 | 3 | 0 | 8 | 0.125 | 12 | 0.1667 | 30 | 0.1333 | 6 | 0.1667 | 5 | 0 | 3 | 0 | 6 | 0.1667 | 20 | 0.1 |
| 11_9491771 | 21 | 0.0476 | 17 | 0.1176 | 9 | 0.1111 | 14 | 0.0714 | 61 | 0.082 | 11 | 0.1818 | 14 | 0.0714 | 8 | 0.125 | 13 | 0.2308 | 46 | 0.1522 |
| 11_9493555 | 3 | 0 | 2 | 0 | 3 | 0.6667 | 1 | 0 | 9 | 0.2222 | 1 | 1 | 2 | 0.5 | NA | NA | 2 | 0.5 | 5 | 0.6 |
| 11_9493568 | 3 | 0.3333 | 1 | 0 | 3 | 0.3333 | 1 | 1 | 8 | 0.375 | 1 | 0 | 3 | 0.6667 | NA | NA | 2 | 0.5 | 6 | 0.5 |
| 11_9493570 | 2 | 0 | NA | NA | 3 | 0.3333 | 1 | 0 | 6 | 0.1667 | 1 | 0 | 3 | 0 | NA | NA | 2 | 0.5 | 6 | 0.1667 |
| 11_9493575 | 3 | 1 | 1 | 1 | 3 | 1 | 1 | 1 | 8 | 1 | 1 | 1 | 2 | 1 | NA | NA | 2 | 1 | 5 | 1 |
| 11_9493579 | 3 | 0 | 1 | 0 | 2 | 0.5 | 1 | 1 | 7 | 0.2857 | NA | NA | 2 | 0 | NA | NA | 2 | 0.5 | 4 | 0.25 |
| 11_9493589 | 3 | 1 | NA | NA | 3 | 1 | 1 | 0 | 7 | 0.8571 | 1 | 1 | 2 | 1 | NA | NA | 2 | 0.5 | 5 | 0.8 |
| 11_9493622 | 3 | 1 | 1 | 1 | 1 | 1 | 1 | 1 | 6 | 1 | NA | NA | 2 | 1 | NA | NA | 2 | 0.5 | 4 | 0.75 |
| 11_9493626 | 3 | 0 | 1 | 1 | 3 | 0.3333 | 1 | 0 | 8 | 0.25 | NA | NA | 3 | 0.3333 | NA | NA | 2 | 0 | 5 | 0.2 |
| 11_9493637 | 3 | 0.6667 | 1 | 0 | 1 | 0 | 1 | 1 | 6 | 0.5 | NA | NA | 1 | 1 | NA | NA | 1 | 0 | 2 | 0.5 |
| 11_9516593 | 4 | 0 | 5 | 0.4 | 7 | 0.1429 | 3 | 0.3333 | 19 | 0.2105 | 5 | 0.2 | 7 | 0 | 5 | 0 | 5 | 0.4 | 22 | 0.1364 |
| 11_9516607 | 14 | 0 | 10 | 0 | 10 | 0.1 | 5 | 0.2 | 39 | 0.0513 | 9 | 0 | 13 | 0.0769 | 10 | 0.3 | 8 | 0 | 40 | 0.1 |
| 11_9516643 | 11 | 0.2727 | 6 | 0.6667 | 7 | 0.4286 | 6 | 0.3333 | 30 | 0.4 | 8 | 0.625 | 11 | 0.5455 | 9 | 0.5556 | 6 | 0.6667 | 34 | 0.5882 |
| 11_9516645 | 12 | 0 | 8 | 0.25 | 9 | 0.1111 | 6 | 0 | 35 | 0.0857 | 6 | 0 | 13 | 0.1538 | 9 | 0 | 6 | 0 | 34 | 0.0588 |
| 11_9516663 | 9 | 0.3333 | 4 | 0.25 | 8 | 0.25 | 4 | 0.5 | 25 | 0.32 | 3 | 0.3333 | 12 | 0.0833 | 6 | 0.3333 | 5 | 0.4 | 26 | 0.2308 |
| 11_9519090 | 7 | 0 | 4 | 0.25 | 4 | 0 | 4 | 0 | 19 | 0.0526 | 3 | 0 | 8 | 0 | 8 | 0.125 | 1 | 1 | 20 | 0.1 |
| 11_9519091 | 7 | 0.1429 | 5 | 0 | 4 | 0 | 4 | 0 | 20 | 0.05 | 2 | 0 | 7 | 0.2857 | 8 | 0.125 | 1 | 0 | 18 | 0.1667 |
| 11_9527216 | 3 | 0 | 1 | 0 | 2 | 0.5 | 1 | 0 | 7 | 0.1429 | 2 | 0.5 | 6 | 0.1667 | 3 | 0.3333 | 1 | 0 | 12 | 0.25 |
| 11_9528243 | 6 | 0.8333 | 8 | 0.75 | 11 | 0.8182 | 3 | 0.6667 | 28 | 0.7857 | 4 | 1 | 4 | 1 | 6 | 0.6667 | 6 | 0.5 | 20 | 0.75 |
| 11_9538268 | 15 | 0.3333 | 22 | 0.3636 | 15 | 0.5333 | 16 | 0.3125 | 68 | 0.3824 | 15 | 0.2667 | 12 | 0.4167 | 18 | 0.3889 | 18 | 0.3889 | 63 | 0.3651 |
| 11_9538273 | 15 | 0.0667 | 19 | 0.2105 | 14 | 0.2143 | 17 | 0.3529 | 65 | 0.2154 | 14 | 0.5714 | 12 | 0.3333 | 16 | 0.25 | 13 | 0.4615 | 55 | 0.4 |
| 11_9538274 | 16 | 0.125 | 21 | 0.0476 | 15 | 0 | 18 | 0.0556 | 70 | 0.0571 | 14 | 0 | 13 | 0 | 20 | 0.1 | 18 | 0.1111 | 65 | 0.0615 |
| 11_9538278 | 12 | 0.75 | 20 | 0.9 | 16 | 0.875 | 18 | 0.9444 | 66 | 0.8788 | 16 | 0.8125 | 12 | 1 | 16 | 0.8125 | 17 | 0.9412 | 61 | 0.8852 |
| 11_9538280 | 13 | 0.7692 | 19 | 0.6842 | 14 | 0.7857 | 15 | 0.8 | 61 | 0.7541 | 14 | 0.7143 | 12 | 0.75 | 17 | 0.8235 | 13 | 0.7692 | 56 | 0.7679 |
| 11_9538282 | 12 | 1 | 9 | 0.8889 | 10 | 0.8 | 10 | 0.9 | 41 | 0.9024 | 12 | 1 | 10 | 0.9 | 13 | 0.6923 | 8 | 0.875 | 43 | 0.8605 |
| 11_9538283 | 9 | 1 | 14 | 1 | 14 | 0.9286 | 17 | 0.9412 | 54 | 0.963 | 13 | 1 | 11 | 0.9091 | 16 | 1 | 13 | 0.9231 | 53 | 0.9623 |
| 11_9538285 | 16 | 0.4375 | 21 | 0.5238 | 18 | 0.3889 | 20 | 0.5 | 75 | 0.4667 | 18 | 0.5 | 12 | 0.3333 | 18 | 0.7222 | 17 | 0.5882 | 65 | 0.5538 |
| 11_9538292 | 17 | 0.3529 | 17 | 0.2353 | 18 | 0.2778 | 18 | 0.1667 | 70 | 0.2571 | 17 | 0.2941 | 12 | 0.3333 | 16 | 0.5 | 16 | 0.4375 | 61 | 0.3934 |
| 11_9538299 | 16 | 0.375 | 19 | 0.2632 | 17 | 0.2941 | 19 | 0.4211 | 71 | 0.338 | 18 | 0.2778 | 12 | 0.25 | 19 | 0.4211 | 17 | 0.1765 | 66 | 0.2879 |
| 11_9538305 | 19 | 0.9474 | 19 | 0.7895 | 18 | 0.9444 | 20 | 1 | 76 | 0.9211 | 18 | 0.7778 | 14 | 1 | 21 | 0.9048 | 17 | 0.9412 | 70 | 0.9 |
| 11_9538319 | 21 | 0.0952 | 24 | 0.1667 | 17 | 0.2353 | 16 | 0.125 | 78 | 0.1538 | 17 | 0.1765 | 13 | 0.0769 | 24 | 0.125 | 18 | 0.1111 | 72 | 0.125 |
| 11_9538337 | 17 | 0.5294 | 22 | 0.3636 | 15 | 0.6 | 15 | 0.7333 | 69 | 0.5362 | 14 | 0.7857 | 15 | 0.8 | 23 | 0.7391 | 19 | 1 | 71 | 0.831 |
| 11_9538341 | 17 | 0.2353 | 21 | 0.5714 | 18 | 0.3889 | 20 | 0.25 | 76 | 0.3684 | 18 | 0.2778 | 14 | 0.5714 | 19 | 0.2632 | 23 | 0.3043 | 74 | 0.3378 |
| 11_9539081 | 1 | 1 | NA | NA | NA | NA | NA | NA | 1 | 1 | NA | NA | NA | NA | 1 | 0 | 1 | 1 | 2 | 0.5 |
| 11_9540080 | 1 | 0 | 6 | 0.3333 | NA | NA | 1 | 0 | 8 | 0.25 | 5 | 0.2 | 1 | 1 | 5 | 0.6 | 2 | 0.5 | 13 | 0.4615 |
| 11_9540130 | 2 | 0 | 4 | 0.5 | 1 | 0 | 1 | 1 | 8 | 0.375 | 3 | 0 | 1 | 0 | 2 | 0 | 3 | 0.3333 | 9 | 0.1111 |
| 11_9540158 | 2 | 0 | 2 | 1 | NA | NA | 1 | 1 | 5 | 0.6 | 4 | 0.5 | 2 | 0 | 1 | 0 | 1 | 1 | 8 | 0.375 |
| 11_9540160 | 2 | 1 | 4 | 1 | 1 | 1 | 1 | 1 | 8 | 1 | 3 | 1 | 2 | 1 | 2 | 1 | 1 | 1 | 8 | 1 |
| 11_9540172 | 1 | 1 | 4 | 1 | 1 | 1 | 1 | 1 | 7 | 1 | 4 | 1 | 1 | 1 | 1 | 1 | 1 | 1 | 7 | 1 |
| 11_9540202 | NA | NA | 4 | 0.25 | NA | NA | 1 | 0 | 5 | 0.2 | 4 | 0 | 1 | 0 | 2 | 0 | 2 | 0.5 | 9 | 0.1111 |
| 11_9540204 | NA | NA | 2 | 0.5 | NA | NA | 1 | 0 | 3 | 0.3333 | 3 | 0.6667 | 1 | 0 | 2 | 0 | 1 | 0 | 7 | 0.2857 |
| 11_9540206 | 1 | 0 | 4 | 0.5 | 1 | 0 | 1 | 0 | 7 | 0.2857 | 4 | 0.25 | 1 | 1 | 2 | 0 | 1 | 1 | 8 | 0.375 |
| 11_9545993 | 1 | 1 | NA | NA | NA | NA | 1 | 1 | 2 | 1 | NA | NA | 1 | 1 | 1 | 1 | NA | NA | 2 | 1 |
| 11_9545998 | 1 | 1 | NA | NA | NA | NA | 1 | 1 | 2 | 1 | NA | NA | 1 | 1 | 1 | 1 | NA | NA | 2 | 1 |
| 11_9546007 | 1 | 1 | NA | NA | NA | NA | 1 | 0 | 2 | 0.5 | NA | NA | 1 | 0 | 1 | 1 | NA | NA | 2 | 0.5 |
| 11_9546019 | 1 | 0 | NA | NA | NA | NA | 1 | 1 | 2 | 0.5 | NA | NA | 1 | 1 | 1 | 1 | NA | NA | 2 | 1 |

| | | | | | | | | | | | | | | | | | | | | |
|---|---|---|---|---|---|---|---|---|---|---|---|---|---|---|---|---|---|---|---|---|
| 11_9546022 | 1 | 0 | NA | NA | NA | NA | 1 | 1 | 2 | 0.5 | NA | NA | 1 | 1 | 1 | 1 | NA | NA | 2 | 1 |
| 11_9546045 | 1 | 1 | NA | NA | NA | NA | 1 | 1 | 2 | 1 | NA | NA | 1 | 0 | 1 | 1 | NA | NA | 2 | 0.5 |
| 11_9546124 | 1 | 1 | NA | NA | NA | NA | 1 | 0 | 2 | 0.5 | NA | NA | 1 | 1 | NA | NA | NA | NA | 1 | 1 |
| 11_9546126 | 1 | 0 | NA | NA | NA | NA | 1 | 1 | 2 | 0.5 | NA | NA | 1 | 1 | NA | NA | NA | NA | 1 | 1 |
| 11_9546127 | 1 | 1 | NA | NA | NA | NA | 1 | 0 | 2 | 0.5 | NA | NA | 1 | 1 | NA | NA | NA | NA | 1 | 1 |
| 11_9546129 | NA | NA | NA | NA | NA | NA | 1 | 1 | 1 | 1 | NA | NA | 1 | 1 | NA | NA | NA | NA | 1 | 1 |
| 11_9557065 | 9 | 0.3333 | 15 | 0.1333 | 10 | 0.3 | 23 | 0.1739 | 57 | 0.2105 | 11 | 0.4545 | 11 | 0.1818 | 14 | 0.3571 | 14 | 0.3571 | 50 | 0.34 |
| 11_9557093 | 4 | 1 | 6 | 0.3333 | 9 | 0.6667 | 10 | 0.8 | 29 | 0.6897 | 5 | 0.6 | 5 | 0.8 | 10 | 0.3 | 8 | 0.375 | 28 | 0.4643 |
| 11_9557106 | 4 | 0.5 | 6 | 0.3333 | 7 | 0.4286 | 10 | 0.6 | 27 | 0.4815 | 5 | 0.6 | 4 | 0.75 | 10 | 0.7 | 8 | 0.5 | 27 | 0.6296 |
| 11_9557112 | 4 | 0 | 6 | 0.1667 | 8 | 0.125 | 6 | 0.1667 | 24 | 0.125 | 3 | 0 | 5 | 0.4 | 11 | 0.0909 | 8 | 0.125 | 27 | 0.1481 |
| 11_9557112 | 4 | 0.75 | 5 | 0.6 | 9 | 1 | 6 | 0.6667 | 24 | 0.7917 | 3 | 1 | 4 | 0.75 | 8 | 1 | 8 | 1 | 23 | 0.9565 |
| 11_9557130 | 5 | 1 | 8 | 0.75 | 7 | 0.7143 | 11 | 0.8182 | 31 | 0.8065 | 5 | 0.4 | 3 | 1 | 13 | 0.7692 | 8 | 0.875 | 29 | 0.7586 |
| 11_9557131 | 3 | 0 | 5 | 0 | 5 | 0.2 | 10 | 0.1 | 23 | 0.087 | 3 | 0 | 5 | 0.2 | 10 | 0.2 | 7 | 0.1429 | 25 | 0.16 |
| 11_9557132 | 5 | 1 | 8 | 1 | 8 | 0.75 | 9 | 1 | 30 | 0.9333 | 4 | 1 | 4 | 1 | 12 | 0.8333 | 9 | 1 | 29 | 0.931 |
| 11_9557135 | 6 | 0.1667 | 7 | 0.2857 | 9 | 0.2222 | 11 | 0.0909 | 33 | 0.1818 | 5 | 0 | 4 | 0.25 | 11 | 0.0909 | 10 | 0 | 30 | 0.0667 |
| 11_9557145 | 7 | 1 | 8 | 0.75 | 7 | 0.8571 | 11 | 0.6364 | 33 | 0.7879 | 5 | 0.8 | 5 | 0.6 | 13 | 0.7692 | 10 | 0.9 | 33 | 0.7879 |
| 11_9559131 | NA | NA | 1 | 0 | 2 | 0 | 2 | 0.5 | 5 | 0.2 | 2 | 0 | 2 | 0 | 1 | 1 | 1 | 0 | 6 | 0.1667 |
| 11_9564054 | NA | NA | NA | NA | NA | NA | 1 | 1 | 1 | 1 | 1 | 0 | NA | NA | NA | NA | 1 | 1 | 2 | 0.5 |
| 11_9564095 | NA | NA | NA | NA | NA | NA | 1 | 1 | 1 | 1 | 1 | 1 | NA | NA | NA | NA | NA | NA | 1 | 1 |
| 11_9565482 | NA | NA | 1 | 1 | NA | NA | 1 | 0 | 2 | 0.5 | 1 | 1 | NA | NA | NA | NA | 1 | 1 | 2 | 1 |
| 11_9565500 | NA | NA | 1 | 1 | NA | NA | 2 | 0 | 3 | 0.3333 | 1 | 1 | NA | NA | NA | NA | 1 | 0 | 2 | 0.5 |
| 11_9565509 | NA | NA | 1 | 0 | NA | NA | 2 | 1 | 3 | 0.6667 | 1 | 1 | NA | NA | NA | NA | NA | NA | 1 | 1 |
| 11_9565511 | NA | NA | 1 | 1 | NA | NA | 2 | 1 | 3 | 1 | 1 | 1 | NA | NA | NA | NA | 1 | 0 | 2 | 0.5 |
| 11_9565990 | NA | NA | 5 | 0.2 | 1 | 0 | 2 | 0 | 8 | 0.125 | 2 | 0 | 2 | 0 | 4 | 0 | 1 | 1 | 9 | 0.1111 |
| 11_9566079 | NA | NA | 4 | 0.25 | 1 | 0 | 2 | 0.5 | 7 | 0.2857 | 1 | 0 | 4 | 0.25 | 3 | 0 | 1 | 0 | 9 | 0.1111 |
| 11_9572159 | 4 | 0.5 | 11 | 0.0909 | 7 | 0.2857 | 3 | 0.3333 | 25 | 0.24 | 7 | 0.2857 | 2 | 0 | 6 | 0.1667 | 3 | 0.3333 | 18 | 0.2222 |
| 11_9572179 | 3 | 1 | 10 | 0.7 | 8 | 1 | 3 | 1 | 24 | 0.875 | 6 | 1 | 2 | 1 | 5 | 0.8 | 3 | 1 | 16 | 0.9375 |
| 11_9572195 | NA | NA | 10 | 1 | 5 | 1 | 3 | 1 | 18 | 1 | 5 | 1 | 2 | 1 | 4 | 1 | 4 | 1 | 15 | 1 |
| 11_9572208 | 2 | 0 | 6 | 0.1667 | 1 | 0 | NA | NA | 9 | 0.1111 | 4 | 0 | 2 | 0 | 5 | 0.2 | 2 | 0 | 13 | 0.0769 |
| 11_9572216 | 2 | 0.5 | 12 | 0.3333 | 9 | 0.5556 | 3 | 1 | 26 | 0.5 | 5 | 0.2 | 2 | 0 | 6 | 0.5 | 2 | 0 | 15 | 0.2667 |
| 11_9572218 | 1 | 1 | 16 | 0.875 | 8 | 1 | 2 | 1 | 27 | 0.9259 | 8 | 0.875 | 2 | 1 | 6 | 0.8333 | 4 | 1 | 20 | 0.9 |
| 11_9572239 | 2 | 1 | 14 | 0.8571 | 6 | 0.8333 | 1 | 1 | 23 | 0.8696 | 3 | 1 | 2 | 1 | 4 | 0.75 | 3 | 0.6667 | 12 | 0.8333 |
| 11_9572249 | 5 | 1 | 18 | 0.9444 | 9 | 1 | 3 | 1 | 35 | 0.9714 | 10 | 1 | 3 | 1 | 6 | 1 | 4 | 1 | 23 | 1 |
| 11_9572281 | 5 | 0.8 | 19 | 0.6842 | 10 | 0.6 | 3 | 0.6667 | 37 | 0.6757 | 9 | 0.7778 | 4 | 0.5 | 6 | 1 | 4 | 0.75 | 23 | 0.7826 |
| 11_9572322 | NA | NA | 1 | 1 | 1 | 0 | NA | NA | 2 | 0.5 | NA | NA | 1 | 0 | 1 | 1 | 1 | 0 | 3 | 0.3333 |
| 11_9572334 | 1 | 1 | NA | NA | 1 | 0 | 1 | 0 | 3 | 0.3333 | NA | NA | 1 | 1 | 1 | 0 | NA | NA | 2 | 0.5 |
| 11_9572341 | NA | NA | NA | NA | 1 | 1 | 1 | 1 | 2 | 1 | NA | NA | 1 | 1 | 1 | 1 | 1 | 1 | 3 | 1 |
| 11_9572344 | 1 | 0 | 1 | 1 | 1 | 1 | 1 | 1 | 4 | 0.75 | NA | NA | NA | NA | 1 | 1 | 1 | 1 | 2 | 1 |
| 11_9572356 | 1 | 1 | 1 | 1 | 1 | 0 | 1 | 0 | 4 | 0.5 | NA | NA | 1 | 0 | 1 | 1 | 1 | 0 | 3 | 0.3333 |
| 11_9592605 | 1 | 1 | NA | NA | NA | NA | NA | NA | 1 | 1 | 1 | 1 | NA | NA | NA | NA | NA | NA | 1 | 1 |
| 11_9593815 | 13 | 0 | 14 | 0.1429 | 18 | 0 | 14 | 0.0714 | 59 | 0.0508 | 15 | 0.2 | 17 | 0.1176 | 15 | 0.0667 | 11 | 0.0909 | 58 | 0.1207 |
| 11_9593822 | 6 | 0.1667 | 11 | 0 | 10 | 0 | 10 | 0.2 | 37 | 0.0811 | 13 | 0.1538 | 18 | 0.1111 | 12 | 0.25 | 9 | 0.1111 | 52 | 0.1538 |
| 11_9593860 | NA | NA | 1 | 1 | NA | NA | NA | NA | 1 | 1 | 1 | 1 | 1 | 0 | NA | NA | 2 | 0 | 4 | 0.25 |
| 11_9594109 | 7 | 0.1429 | 5 | 0 | 5 | 0.2 | 5 | 0 | 22 | 0.0909 | 10 | 0.2 | 8 | 0.125 | 11 | 0.1818 | 5 | 0 | 34 | 0.1471 |
| 11_9594110 | 6 | 0 | 5 | 0.4 | 6 | 0.3333 | 4 | 0 | 21 | 0.1905 | 11 | 0.1818 | 8 | 0 | 10 | 0.1 | 4 | 0 | 33 | 0.0909 |
| 11_9594112 | 6 | 0.6667 | 5 | 1 | 7 | 0.7143 | 4 | 0.75 | 22 | 0.7727 | 11 | 0.9091 | 8 | 0.875 | 11 | 0.8182 | 5 | 1 | 35 | 0.8857 |
| 11_9596374 | 3 | 0.3333 | 4 | 0.5 | 3 | 0 | 1 | 0 | 11 | 0.2727 | NA | NA | NA | NA | 2 | 0.5 | 3 | 0.3333 | 5 | 0.4 |
| 11_9596397 | 61 | 0.1148 | 68 | 0.0588 | 44 | 0.0455 | 49 | 0.0816 | 222 | 0.0766 | 35 | 0.0857 | 48 | 0.1458 | 75 | 0.1067 | 51 | 0.098 | 209 | 0.11 |
| 11_9665458 | NA | NA | 1 | 1 | 2 | 1 | NA | NA | 3 | 1 | 1 | 1 | NA | NA | 1 | 1 | NA | NA | 2 | 1 |
| 11_9665484 | NA | NA | NA | NA | 2 | 0.5 | NA | NA | 2 | 0.5 | NA | NA | NA | NA | 1 | 1 | NA | NA | 1 | 1 |
| 11_9665508 | NA | NA | 1 | 0 | 1 | 1 | NA | NA | 2 | 0.5 | NA | NA | 1 | 1 | NA | NA | NA | NA | 1 | 1 |
| 11_9665511 | NA | NA | 1 | 1 | 1 | 1 | NA | NA | 2 | 1 | NA | NA | 1 | 1 | NA | NA | NA | NA | 1 | 1 |
| 11_9665528 | NA | NA | 1 | 1 | 1 | 1 | NA | NA | 2 | 1 | NA | NA | 1 | 1 | NA | NA | NA | NA | 1 | 1 |
| 11_9665847 | 1 | 0 | NA | NA | 1 | 1 | 1 | 0 | 3 | 0.3333 | 1 | 0 | 2 | 0 | 2 | 0.5 | 1 | 0 | 6 | 0.1667 |
| 11_9665858 | 1 | 1 | NA | NA | 1 | 0 | NA | NA | 2 | 0.5 | 1 | 0 | 1 | 0 | 1 | 1 | NA | NA | 3 | 0.3333 |
| 11_9665930 | 4 | 0 | 4 | 0.25 | NA | NA | 5 | 0.2 | 13 | 0.1538 | 3 | 0 | 1 | 1 | 11 | 0 | 2 | 0 | 17 | 0.0588 |
| 11_9665933 | 4 | 0.25 | 2 | 1 | NA | NA | 4 | 0.25 | 10 | 0.4 | 3 | 0 | 1 | 1 | 9 | 0.5556 | 2 | 0.5 | 15 | 0.4667 |
| 11_9665939 | 4 | 0.5 | 3 | 0.6667 | NA | NA | 3 | 0.6667 | 10 | 0.6 | 2 | 0.5 | 1 | 1 | 9 | 0.6667 | 3 | 0.3333 | 15 | 0.6 |
| 11_9665952 | 4 | 0.75 | 3 | 0.3333 | NA | NA | 4 | 0.75 | 11 | 0.6364 | 2 | 1 | 1 | 1 | 5 | 0.2 | 1 | 1 | 9 | 0.5556 |
| 11_9665996 | 4 | 0.75 | 3 | 0.6667 | NA | NA | 3 | 0.3333 | 10 | 0.6 | 2 | 0.5 | 1 | 0 | 7 | 0.7143 | 1 | 0 | 11 | 0.5455 |
| 11_9666005 | 4 | 1 | 3 | 0.3333 | NA | NA | 2 | 1 | 9 | 0.7778 | 3 | 1 | NA | NA | 8 | 0.875 | 1 | 1 | 12 | 0.9167 |
| 11_9669818 | 5 | 0.4 | 7 | 0.1429 | 5 | 0 | 2 | 0 | 19 | 0.1579 | 3 | 0 | NA | NA | 12 | 0.0833 | 3 | 0 | 18 | 0.0556 |
| 11_9669819 | 4 | 0.5 | 8 | 0.25 | 4 | 0.75 | 2 | 0.5 | 18 | 0.4444 | 3 | 0.3333 | NA | NA | 12 | 0.25 | 5 | 0.4 | 20 | 0.3 |
| 11_9670884 | 34 | 0.0294 | 26 | 0.0769 | 26 | 0 | 37 | 0.2432 | 123 | 0.0976 | 19 | 0.2105 | 42 | 0.1429 | 25 | 0.08 | 19 | 0.2105 | 105 | 0.1524 |
| 11_9670917 | 25 | 0.2 | 25 | 0 | 20 | 0 | 31 | 0.0968 | 101 | 0.0792 | 20 | 0.05 | 30 | 0.0667 | 18 | 0.1111 | 13 | 0.3077 | 81 | 0.1111 |

| | | | | | | | | | | | | | | | | | | | | |
|---|---|---|---|---|---|---|---|---|---|---|---|---|---|---|---|---|---|---|---|---|
| 11_967095( | 24 | 0.0833 | 22 | 0.1818 | 15 | 0.1333 | 26 | 0.1923 | 87 | 0.1494 | 16 | 0.1875 | 26 | 0.1154 | 18 | 0.1667 | 12 | 0.0833 | 72 | 0.1389 |
| 11_967098( | 19 | 0.3684 | 18 | 0.3333 | 17 | 0.2353 | 26 | 0.3846 | 80 | 0.3375 | 13 | 0.3077 | 23 | 0.4348 | 17 | 0.5294 | 11 | 0.3636 | 64 | 0.4219 |
| 11_967098( | 20 | 0 | 18 | 0.1111 | 19 | 0.1579 | 26 | 0.0769 | 83 | 0.0843 | 14 | 0.2857 | 21 | 0.0476 | 17 | 0.1176 | 14 | 0.0714 | 66 | 0.1212 |
| 11_967098ℰ | 15 | 0.8667 | 10 | 0.5 | 12 | 0.6667 | 18 | 0.5 | 55 | 0.6364 | 14 | 0.6429 | 14 | 0.2857 | 8 | 0.5 | 12 | 0.4167 | 48 | 0.4583 |
| 11_967099; | 17 | 1 | 12 | 1 | 13 | 1 | 21 | 1 | 63 | 1 | 15 | 1 | 15 | 1 | 13 | 1 | 11 | 1 | 54 | 1 |
| 11_967099; | 11 | 0 | 8 | 0.125 | 11 | 0.1818 | 14 | 0 | 44 | 0.0682 | 9 | 0 | 14 | 0.2143 | 13 | 0 | 10 | 0 | 46 | 0.0652 |
| 11_967099⁄ | 18 | 0.8333 | 14 | 0.9286 | 13 | 0.7692 | 23 | 0.8261 | 68 | 0.8382 | 14 | 0.8571 | 18 | 0.7778 | 14 | 0.5714 | 11 | 0.7273 | 57 | 0.7368 |
| 11_967099( | 14 | 0.7857 | 7 | 0.8571 | 10 | 0.8 | 17 | 0.5882 | 48 | 0.7292 | 10 | 0.4 | 15 | 0.6667 | 11 | 0.4545 | 7 | 0.8571 | 43 | 0.5814 |
| 11_967099( | NA | NA | 1 | 0 | NA | NA | 1 | 1 | 2 | 0.5 | NA | NA | 1 | 1 | NA | NA | 1 | 1 | 2 | 1 |
| 11_967099( | 14 | 0 | 13 | 0.0769 | 10 | 0.3 | 19 | 0.1053 | 56 | 0.1071 | 15 | 0.1333 | 15 | 0.0667 | 13 | 0.1538 | 10 | 0.3 | 53 | 0.1509 |
| 11_967100' | 16 | 0.8125 | 10 | 0.9 | 13 | 0.7692 | 17 | 1 | 56 | 0.875 | 13 | 1 | 14 | 0.9286 | 13 | 1 | 9 | 1 | 49 | 0.9796 |
| 11_967102! | 15 | 1 | 9 | 0.8889 | 13 | 0.6154 | 15 | 0.8667 | 52 | 0.8462 | 12 | 0.9167 | 15 | 0.9333 | 12 | 0.9167 | 9 | 0.8889 | 48 | 0.9167 |
| 11_967102! | 11 | 0.3636 | 10 | 0.4 | 12 | 0.1667 | 18 | 0.2222 | 51 | 0.2745 | 12 | 0.25 | 14 | 0.5 | 13 | 0.0769 | 11 | 0 | 50 | 0.22 |
| 11_967103' | 15 | 0.0667 | 9 | 0 | 12 | 0.25 | 17 | 0.1765 | 53 | 0.1321 | 13 | 0.2308 | 10 | 0.2 | 8 | 0.25 | 9 | 0.2222 | 40 | 0.225 |
| 11_967104! | 11 | 0.1818 | 8 | 0 | 10 | 0.1 | 16 | 0.0625 | 45 | 0.0889 | 10 | 0.2 | 8 | 0 | 9 | 0.1111 | 8 | 0 | 35 | 0.0857 |
| 11_967105( | 12 | 0.5833 | 5 | 0.6 | 9 | 0.5556 | 14 | 0.4286 | 40 | 0.525 | 11 | 0.7273 | 10 | 0.4 | 8 | 0.625 | 9 | 0.5556 | 38 | 0.5789 |
| 11_967107; | 11 | 0.0909 | 8 | 0 | 9 | 0.1111 | 16 | 0.0625 | 44 | 0.0682 | 11 | 0.0909 | 12 | 0.0833 | 9 | 0.2222 | 8 | 0.125 | 40 | 0.125 |
| 11_967107; | 7 | 0.2857 | 7 | 0.2857 | 10 | 0.5 | 13 | 0.9231 | 37 | 0.5676 | 9 | 0.5556 | 11 | 0.6364 | 8 | 0.5 | 5 | 0.6 | 33 | 0.5758 |
| 11_967108; | 10 | 0.9 | 7 | 1 | 11 | 0.9091 | 17 | 0.8824 | 45 | 0.9111 | 11 | 1 | 11 | 0.9091 | 7 | 1 | 8 | 1 | 37 | 0.973 |
| 11_967108ℰ | 8 | 0 | 7 | 0.2857 | 10 | 0.1 | 14 | 0.1429 | 39 | 0.1282 | 11 | 0.0909 | 10 | 0 | 5 | 0 | 6 | 0.1667 | 32 | 0.0625 |
| 11_967108! | 9 | 0.4444 | 6 | 0.1667 | 11 | 0.3636 | 14 | 0.6429 | 40 | 0.45 | 9 | 0.2222 | 12 | 0.5 | 7 | 0.2857 | 8 | 0 | 36 | 0.2778 |
| 11_967109! | 10 | 0.8 | 6 | 1 | 8 | 0.875 | 15 | 0.8667 | 39 | 0.8718 | 9 | 0.7778 | 8 | 0.625 | 7 | 0.8571 | 7 | 1 | 31 | 0.8065 |
| 11_967110; | 9 | 0.1111 | 5 | 0 | 10 | 0 | 13 | 0.2308 | 37 | 0.1081 | 8 | 0 | 10 | 0.1 | 7 | 0.1429 | 6 | 0.5 | 31 | 0.1613 |
| 11_967111! | 6 | 1 | 5 | 1 | 8 | 0.875 | 10 | 1 | 29 | 0.9655 | 8 | 1 | 5 | 1 | 5 | 1 | 6 | 0.8333 | 24 | 0.9583 |
| 11_967180; | 2 | 0 | 3 | 0 | 5 | 0 | 5 | 0.2 | 15 | 0.0667 | 2 | 0 | 4 | 0.25 | NA | NA | 2 | 0 | 8 | 0.125 |
| 11_967183' | 4 | 1 | 3 | 1 | 6 | 1 | 5 | 1 | 18 | 1 | 2 | 1 | 4 | 0.75 | NA | NA | 1 | 1 | 7 | 0.8571 |
| 11_967183' | 6 | 0.6667 | 3 | 0 | 7 | 0.2857 | 5 | 0.4 | 21 | 0.381 | 3 | 0.3333 | 4 | 0 | NA | NA | 1 | 0 | 8 | 0.125 |
| 11_967185( | 6 | 1 | 3 | 1 | 5 | 0.4 | 5 | 1 | 19 | 0.8421 | 2 | 1 | 4 | 1 | NA | NA | 1 | 1 | 7 | 1 |
| 11_967187! | 4 | 0.75 | 3 | 1 | 6 | 0.8333 | 3 | 1 | 16 | 0.875 | 1 | 1 | 2 | 1 | NA | NA | 1 | 1 | 4 | 1 |
| 11_967188⁄ | 4 | 0.75 | 3 | 1 | 4 | 1 | 2 | 1 | 13 | 0.9231 | 1 | 1 | 1 | 1 | NA | NA | 1 | 1 | 3 | 1 |
| 11_967518⁄ | 2 | 0 | 1 | 0 | 1 | 1 | NA | NA | 4 | 0.25 | NA | NA | 1 | 1 | NA | NA | 1 | 0 | 2 | 0.5 |
| 11_967986⁄ | 3 | 0.3333 | 5 | 0.6 | 4 | 0.5 | 2 | 0 | 14 | 0.4286 | 3 | 0.6667 | 3 | 0.3333 | 4 | 0.5 | 1 | 0 | 11 | 0.4545 |
| 11_967987ℰ | 6 | 0.3333 | 6 | 0.6667 | 5 | 0.8 | 4 | 0.75 | 21 | 0.619 | 2 | 0 | 3 | 0.6667 | 7 | 0.4286 | 3 | 0 | 15 | 0.3333 |
| 11_967988; | 4 | 0 | 6 | 0 | 4 | 0.25 | 2 | 0 | 16 | 0.0625 | 2 | 0 | 2 | 0.5 | 4 | 0 | 2 | 0 | 10 | 0.1 |
| 11_967988! | 5 | 0 | 4 | 0 | 6 | 0.1667 | 2 | 0 | 17 | 0.0588 | 3 | 0 | 2 | 0 | 7 | 0 | 3 | 0.3333 | 15 | 0.0667 |
| 11_968032' | 3 | 0 | 4 | 0.25 | 5 | 0 | 3 | 0 | 15 | 0.0667 | 3 | 0.3333 | 3 | 0 | 1 | 0 | 2 | 0 | 9 | 0.1111 |
| 11_968288' | 20 | 0.55 | 8 | 0.125 | 8 | 0.125 | 11 | 0.1818 | 47 | 0.3191 | 4 | 0.75 | 6 | 0.1667 | 13 | 0.5385 | 6 | 0.1667 | 29 | 0.4138 |
| 11_968288ℰ | 12 | 0.1667 | 7 | 0.1429 | 7 | 0 | 9 | 0.1111 | 35 | 0.1143 | 4 | 0.5 | 7 | 0.1429 | 8 | 0 | 4 | 0.25 | 23 | 0.1739 |
| 11_968292ℰ | 12 | 0.0833 | 5 | 0.2 | 7 | 0 | 8 | 0.125 | 32 | 0.0938 | 4 | 0 | 6 | 0.1667 | 6 | 0.1667 | 2 | 0 | 18 | 0.1111 |
| 11_968297! | 14 | 0.2857 | 4 | 0 | 4 | 0.5 | 5 | 0 | 27 | 0.2222 | 2 | 0 | 7 | 0.1429 | 4 | 0.75 | 3 | 0.3333 | 16 | 0.3125 |
| 11_968444; | 2 | 0.5 | 1 | 0 | 4 | 0.25 | 1 | 0 | 8 | 0.25 | 1 | 1 | NA | NA | 5 | 0.2 | 1 | 1 | 7 | 0.4286 |
| 11_968450; | 2 | 0.5 | NA | NA | 4 | 0.25 | NA | NA | 6 | 0.3333 | 1 | 0 | NA | NA | 5 | 0.6 | 1 | 1 | 7 | 0.5714 |
| 11_968451! | 2 | 0.5 | NA | NA | 4 | 0.5 | NA | NA | 6 | 0.5 | 1 | 0 | NA | NA | 5 | 0.4 | 1 | 1 | 7 | 0.4286 |
| 11_968545! | NA | NA | NA | NA | NA | NA | 1 | 1 | 1 | 1 | NA | NA | NA | NA | 1 | 1 | NA | NA | 1 | 1 |
| 11_968550( | NA | NA | NA | NA | NA | NA | 1 | 1 | 1 | 1 | NA | NA | NA | NA | 1 | 1 | NA | NA | 1 | 1 |
| 11_969353; | 8 | 0.125 | 12 | 0.0833 | 11 | 0 | 4 | 0 | 35 | 0.0571 | 5 | 0 | 4 | 0 | 7 | 0.1429 | 4 | 0 | 20 | 0.05 |
| 11_969389⁄ | 9 | 0.1111 | 14 | 0.1429 | 14 | 0.2143 | 5 | 0 | 42 | 0.1429 | 5 | 0 | 11 | 0.2727 | 7 | 0.2857 | 4 | 0 | 27 | 0.1852 |
| 11_969392; | 11 | 0.1818 | 16 | 0.125 | 13 | 0.2308 | 5 | 0.2 | 45 | 0.1778 | 6 | 0.3333 | 11 | 0.2727 | 7 | 0 | 5 | 0.4 | 29 | 0.2414 |
| 11_969393( | 5 | 0.6 | 14 | 0.1429 | 13 | 0.3077 | 4 | 1 | 36 | 0.3611 | 6 | 0.3333 | 8 | 0.25 | 6 | 0 | 5 | 0.4 | 25 | 0.24 |
| 11_969393⁄ | 11 | 0.1818 | 15 | 0.2 | 16 | 0.3125 | 5 | 0.4 | 47 | 0.2553 | 6 | 0.1667 | 12 | 0.4167 | 8 | 0.125 | 5 | 0.4 | 31 | 0.2903 |
| 11_969394( | 8 | 0.25 | 13 | 0 | 13 | 0.4615 | 5 | 0.4 | 39 | 0.2564 | 4 | 0.25 | 8 | 0 | 8 | 0.5 | 5 | 0.2 | 25 | 0.24 |
| 11_969394! | 10 | 0.5 | 15 | 0.5333 | 17 | 0.6471 | 5 | 0.6 | 47 | 0.5745 | 5 | 0.4 | 12 | 0.6667 | 9 | 0.7778 | 6 | 0.6667 | 32 | 0.6562 |
| 11_969395; | 11 | 0.0909 | 16 | 0.0625 | 14 | 0.1429 | 5 | 0 | 46 | 0.087 | 4 | 0.25 | 10 | 0.2 | 10 | 0 | 5 | 0.2 | 29 | 0.1379 |
| 11_969395⁄ | 10 | 0.1 | 17 | 0.1176 | 15 | 0.2 | 5 | 0 | 47 | 0.1277 | 5 | 0.2 | 10 | 0 | 9 | 0.2222 | 4 | 0 | 28 | 0.1071 |
| 11_969399( | 11 | 0.0909 | 10 | 0.2 | 11 | 0 | 3 | 0 | 35 | 0.0857 | 6 | 0 | 13 | 0.1538 | 7 | 0.1429 | 5 | 0.2 | 31 | 0.129 |
| 11_969589; | 1 | 1 | NA | NA | NA | NA | NA | NA | 1 | 1 | NA | NA | 4 | 0.5 | NA | NA | NA | NA | 4 | 0.5 |
| 11_969592; | 1 | 1 | 1 | 0 | NA | NA | NA | NA | 2 | 0.5 | NA | NA | 2 | 0.5 | NA | NA | 1 | 0 | 3 | 0.3333 |
| 11_970271; | 4 | 0 | 9 | 0.1111 | 12 | 0 | 4 | 0.25 | 29 | 0.069 | 3 | 0 | 3 | 0.6667 | 13 | 0 | 1 | 0 | 20 | 0.1 |
| 11_970271; | 5 | 0 | 8 | 0.25 | 12 | 0.1667 | 5 | 0.2 | 30 | 0.1667 | 4 | 0.5 | 3 | 0.6667 | 12 | 0 | 1 | 0 | 20 | 0.2 |
| 11_970284( | 2 | 0 | 6 | 0.3333 | 6 | 0.1667 | 3 | 0.6667 | 17 | 0.2941 | 1 | 0 | 3 | 0.6667 | 4 | 0.25 | 1 | 0 | 9 | 0.3333 |
| 11_970508; | 8 | 0.125 | 12 | 0.0833 | 8 | 0 | 5 | 0 | 33 | 0.0606 | 9 | 0.1111 | 11 | 0.0909 | 11 | 0 | 8 | 0 | 39 | 0.0513 |
| 11_970509; | 9 | 0 | 13 | 0.0769 | 8 | 0.5 | 5 | 0.2 | 35 | 0.1714 | 9 | 0.1111 | 12 | 0.1667 | 11 | 0.1818 | 9 | 0.1111 | 41 | 0.1463 |
| 11_970531⁄ | 10 | 0 | 9 | 0.1111 | 8 | 0.125 | 11 | 0.0909 | 38 | 0.0789 | 6 | 0 | 10 | 0.1 | 5 | 0 | 11 | 0.0909 | 32 | 0.0625 |
| 11_970531( | 8 | 0.125 | 10 | 0.2 | 9 | 0.1111 | 5 | 0.4 | 32 | 0.1875 | 6 | 0.1667 | 8 | 0.375 | 5 | 0.4 | 10 | 0.1 | 29 | 0.2414 |
| 11_970792ℰ | NA | NA | 1 | 1 | 1 | 1 | 1 | 1 | 3 | 1 | NA | NA | 1 | 1 | NA | NA | 1 | 1 | 2 | 1 |

| | | | | | | | | | | | | | | | | | | | | |
|---|---|---|---|---|---|---|---|---|---|---|---|---|---|---|---|---|---|---|---|---|
| 11_970794 | NA | NA | 1 | 1 | NA | NA | 1 | 0 | 2 | 0.5 | NA | NA | 1 | 1 | NA | NA | 1 | 1 | 2 | 1 |
| 11_970795 | NA | NA | 1 | 1 | 1 | 0 | 2 | 0 | 4 | 0.25 | NA | NA | 1 | 1 | NA | NA | 1 | 1 | 2 | 1 |
| 11_970796 | NA | NA | NA | NA | 1 | 1 | 2 | 1 | 3 | 1 | NA | NA | 1 | 1 | NA | NA | 1 | 1 | 2 | 1 |
| 11_970796 | NA | NA | 1 | 1 | 1 | 0 | 1 | 0 | 3 | 0.3333 | NA | NA | 1 | 1 | NA | NA | 1 | 0 | 2 | 0.5 |
| 11_970969 | 5 | 0.2 | 10 | 0.1 | 11 | 0.0909 | 4 | 0 | 30 | 0.1 | 5 | 0 | 5 | 0.2 | 13 | 0.1538 | 3 | 0 | 26 | 0.1154 |
| 11_970976 | 8 | 0 | 16 | 0.125 | 13 | 0.0769 | 6 | 0.1667 | 43 | 0.093 | 11 | 0.0909 | 10 | 0 | 17 | 0.1765 | 9 | 0.1111 | 47 | 0.1064 |
| 11_970978 | 8 | 0.375 | 13 | 0.3846 | 10 | 0.3 | 6 | 0.8333 | 37 | 0.4324 | 8 | 0.25 | 8 | 0.125 | 16 | 0.375 | 5 | 0.4 | 37 | 0.2973 |
| 11_970978 | 8 | 0 | 11 | 0.1818 | 12 | 0 | 6 | 0.1667 | 37 | 0.0811 | 8 | 0.125 | 6 | 0 | 14 | 0 | 7 | 0.1429 | 35 | 0.0571 |
| 11_972193 | 7 | 0.4286 | 7 | 0.1429 | 5 | 0 | 6 | 0 | 25 | 0.16 | 7 | 0 | 8 | 0.375 | 7 | 0 | 9 | 0 | 31 | 0.0968 |
| 11_972194 | 7 | 0 | 8 | 0.125 | 5 | 0.2 | 6 | 0.3333 | 26 | 0.1538 | 9 | 0.1111 | 8 | 0.25 | 9 | 0.1111 | 9 | 0.1111 | 35 | 0.1429 |
| 11_972195 | 9 | 0 | 8 | 0.125 | 5 | 0.2 | 7 | 0.1429 | 29 | 0.1034 | 9 | 0 | 8 | 0 | 9 | 0.1111 | 10 | 0.2 | 36 | 0.0833 |
| 11_972195 | 9 | 0 | 6 | 0.3333 | 5 | 0.2 | 6 | 0.1667 | 26 | 0.1538 | 9 | 0.1111 | 8 | 0.125 | 7 | 0 | 9 | 0.1111 | 33 | 0.0909 |
| 11_972195 | 8 | 0.375 | 6 | 0.1667 | 6 | 0.5 | 5 | 0.2 | 25 | 0.32 | 8 | 0.5 | 6 | 0.1667 | 7 | 0.2857 | 8 | 0.25 | 29 | 0.3103 |
| 11_972195 | 8 | 1 | 6 | 1 | 4 | 1 | 4 | 1 | 22 | 1 | 9 | 1 | 7 | 1 | 6 | 0.8333 | 8 | 0.875 | 30 | 0.9333 |
| 11_972198 | 9 | 0.2222 | 8 | 0 | 6 | 0 | 4 | 0 | 27 | 0.0741 | 10 | 0 | 7 | 0.1429 | 7 | 0.2857 | 10 | 0 | 34 | 0.0882 |
| 11_972199 | 1 | 1 | 2 | 1 | 1 | 1 | 1 | 1 | 5 | 1 | 1 | 1 | NA | NA | 1 | 0 | 2 | 0.5 | 4 | 0.5 |
| 11_972199 | 2 | 0.5 | 1 | 0 | 1 | 1 | NA | NA | 4 | 0.5 | NA | NA | NA | NA | 1 | 0 | 1 | 1 | 2 | 0.5 |
| 11_972200 | 2 | 0.5 | 2 | 0.5 | 1 | 1 | 2 | 1 | 7 | 0.7143 | 1 | 1 | NA | NA | 1 | 1 | NA | NA | 2 | 1 |
| 11_972201 | 2 | 0.5 | 2 | 0.5 | 1 | 1 | 2 | 0 | 7 | 0.4286 | 1 | 0 | NA | NA | 1 | 0 | 1 | 1 | 3 | 0.3333 |
| 11_972202 | 2 | 0.5 | 2 | 0 | 1 | 1 | 1 | 1 | 6 | 0.5 | NA | NA | NA | NA | 1 | 1 | 1 | 0 | 2 | 0.5 |
| 11_972202 | 2 | 0 | 2 | 0.5 | NA | NA | 2 | 0 | 6 | 0.1667 | 1 | 0 | NA | NA | 1 | 0 | 1 | 1 | 3 | 0.3333 |
| 11_972203 | 2 | 1 | 2 | 0.5 | 1 | 1 | 2 | 0.5 | 7 | 0.7143 | 1 | 1 | NA | NA | 1 | 1 | 1 | 1 | 3 | 1 |
| 11_972204 | 2 | 1 | 2 | 1 | 1 | 1 | 2 | 1 | 7 | 1 | 1 | 1 | NA | NA | 1 | 1 | 1 | 0 | 3 | 0.6667 |
| 11_972204 | 2 | 1 | 2 | 0 | 1 | 0 | 1 | 1 | 6 | 0.5 | 1 | 0 | NA | NA | 1 | 1 | 1 | 0 | 3 | 0.3333 |
| 11_972536 | 2 | 0.5 | NA | NA | NA | NA | NA | NA | 2 | 0.5 | 2 | 0.5 | NA | NA | NA | NA | NA | NA | 2 | 0.5 |
| 11_972538 | 1 | 0 | NA | NA | NA | NA | 1 | 1 | 2 | 0.5 | 2 | 0.5 | NA | NA | NA | NA | NA | NA | 2 | 0.5 |
| 11_972539 | 1 | 1 | NA | NA | NA | NA | NA | NA | 1 | 1 | 2 | 1 | NA | NA | NA | NA | NA | NA | 2 | 1 |
| 11_972843 | 2 | 1 | 1 | 1 | NA | NA | 1 | 1 | 4 | 1 | 1 | 1 | NA | NA | NA | NA | NA | NA | 1 | 1 |
| 11_973425 | 1 | 1 | 2 | 1 | NA | NA | 1 | 0 | 4 | 0.75 | 3 | 0 | 1 | 0 | 1 | 0 | 1 | 1 | 6 | 0.1667 |
| 11_973426 | 1 | 1 | 3 | 0.3333 | 1 | 0 | 2 | 0.5 | 7 | 0.4286 | 4 | 0.25 | 1 | 0 | 1 | 0 | 1 | 1 | 7 | 0.2857 |
| 11_973429 | NA | NA | 2 | 1 | 1 | 1 | 2 | 1 | 5 | 1 | 4 | 0.75 | 1 | 1 | 1 | 1 | 2 | 1 | 8 | 0.875 |
| 11_973430 | 1 | 1 | 3 | 0.6667 | 1 | 0 | 2 | 1 | 7 | 0.7143 | 4 | 0.75 | 1 | 0 | 1 | 1 | 1 | 1 | 7 | 0.7143 |
| 11_973430 | 1 | 1 | 3 | 1 | NA | NA | 2 | 1 | 6 | 1 | 2 | 1 | 1 | 1 | 1 | 1 | 2 | 1 | 6 | 1 |
| 11_973430 | 1 | 0 | 3 | 0.3333 | 1 | 0 | 1 | 0 | 6 | 0.1667 | 4 | 0.5 | 1 | 0 | 1 | 0 | 1 | 1 | 7 | 0.4286 |
| 11_973911 | 3 | 0 | 7 | 0.5714 | 5 | 0 | 3 | 0 | 18 | 0.2222 | 6 | 0.1667 | 2 | 0 | 4 | 0 | 3 | 0 | 15 | 0.0667 |
| 11_973912 | 3 | 0.6667 | 7 | 0.8571 | 3 | 1 | 3 | 1 | 16 | 0.875 | 5 | 0.6 | 2 | 1 | 4 | 0.5 | 3 | 1 | 14 | 0.7143 |
| 11_973913 | 2 | 1 | 4 | 1 | 3 | 1 | 3 | 0.6667 | 12 | 0.9167 | 5 | 1 | 2 | 1 | 3 | 1 | 3 | 1 | 13 | 1 |
| 11_973918 | 4 | 1 | 6 | 1 | 5 | 1 | 4 | 1 | 19 | 1 | 7 | 1 | 3 | 1 | 6 | 1 | 2 | 1 | 18 | 1 |
| 11_973918 | 4 | 0.25 | 3 | 0.3333 | 7 | 0.2857 | 4 | 0.5 | 18 | 0.3333 | 7 | 0.5714 | 4 | 0.75 | 6 | 0 | 4 | 0.25 | 21 | 0.381 |
| 11_973925 | 6 | 0.1667 | 12 | 0.5 | 8 | 0.5 | 5 | 0.4 | 31 | 0.4194 | 7 | 0.2857 | 9 | 0.4444 | 11 | 0.3636 | 10 | 0.5 | 37 | 0.4054 |
| 11_973927 | 6 | 0.5 | 13 | 0.7692 | 6 | 0.8333 | 6 | 0.8333 | 31 | 0.7419 | 10 | 0.4 | 7 | 0.4286 | 7 | 0.4286 | 8 | 0.125 | 32 | 0.3438 |
| 11_973927 | 7 | 0.4286 | 13 | 0.2308 | 8 | 0.375 | 5 | 0 | 33 | 0.2727 | 11 | 0.2727 | 8 | 0.125 | 12 | 0.25 | 10 | 0.3 | 41 | 0.2439 |
| 11_974016 | 4 | 0 | 8 | 0.25 | 11 | 0.0909 | 3 | 0 | 26 | 0.1154 | 9 | 0.1111 | 6 | 0.1667 | 9 | 0 | 5 | 0 | 29 | 0.069 |
| 11_974019 | 3 | 0 | 15 | 0.2 | 9 | 0.2222 | 8 | 0 | 35 | 0.1429 | 11 | 0.0909 | 7 | 0.1429 | 7 | 0.2857 | 7 | 0 | 32 | 0.125 |
| 11_974026 | 2 | 0 | 10 | 0.1 | 2 | 0 | 3 | 0 | 17 | 0.0588 | 6 | 0.1667 | 4 | 0 | 4 | 0 | 2 | 0 | 16 | 0.0625 |
| 11_974027 | 3 | 1 | 10 | 0.6 | 3 | 1 | 3 | 0.3333 | 19 | 0.6842 | 8 | 0.625 | 4 | 0.75 | 3 | 0.3333 | 4 | 0.25 | 19 | 0.5263 |
| 11_974027 | 2 | 1 | 10 | 0 | 3 | 0.3333 | 5 | 0.2 | 20 | 0.2 | 8 | 0 | 5 | 0.4 | 3 | 0.6667 | 5 | 0 | 21 | 0.1905 |
| 11_974028 | 3 | 1 | 9 | 0.8889 | 6 | 1 | 3 | 1 | 21 | 0.9524 | 7 | 1 | 5 | 0.4 | 3 | 0.6667 | 4 | 0.75 | 19 | 0.7368 |
| 11_974028 | 3 | 0.6667 | 10 | 0.4 | 5 | 0.4 | 5 | 0.4 | 23 | 0.4348 | 6 | 0.1667 | 5 | 0.4 | 1 | 1 | 5 | 0.2 | 17 | 0.2941 |
| 11_974028 | NA | NA | 6 | 1 | 3 | 1 | 4 | 1 | 13 | 1 | 7 | 1 | 4 | 1 | NA | NA | 2 | 0.5 | 13 | 0.9231 |
| 11_974028 | 3 | 1 | 9 | 0.8889 | 3 | 1 | 3 | 0.6667 | 18 | 0.8889 | 9 | 0.8889 | 4 | 0.75 | 2 | 0 | 3 | 0.6667 | 18 | 0.7222 |
| 11_974029 | 3 | 0.3333 | 11 | 0.0909 | 5 | 0 | 5 | 0.6 | 24 | 0.2083 | 8 | 0.125 | 7 | 0.1429 | 3 | 0 | 5 | 0 | 23 | 0.087 |
| 11_974029 | 3 | 0 | 9 | 0.4444 | 6 | 0 | 4 | 0.25 | 22 | 0.2273 | 7 | 0.2857 | 6 | 0.1667 | 3 | 0 | 4 | 0.5 | 20 | 0.25 |
| 11_974029 | 1 | 0 | 5 | 0.2 | 3 | 0.3333 | 4 | 0 | 13 | 0.1538 | 7 | 0.2857 | 5 | 0.2 | 2 | 0 | 4 | 0 | 18 | 0.1667 |
| 11_974035 | 2 | 1 | 3 | 0.3333 | 1 | 1 | 1 | 0 | 7 | 0.5714 | 2 | 0.5 | NA | NA | 1 | 0 | 1 | 0 | 4 | 0.25 |
| 11_974037 | 1 | 1 | 1 | 0 | 1 | 1 | NA | NA | 3 | 0.6667 | NA | NA | NA | NA | 1 | 1 | 1 | 0 | 2 | 0.5 |
| 11_974076 | 2 | 1 | 1 | 1 | NA | NA | 1 | 0 | 4 | 0.75 | NA | NA | 1 | 1 | NA | NA | 2 | 1 | 3 | 1 |
| 11_974102 | 1 | 1 | NA | NA | NA | NA | 1 | 1 | 2 | 1 | 1 | 1 | 1 | 1 | NA | NA | 2 | 0.5 | 4 | 0.75 |
| 11_974106 | NA | NA | NA | NA | NA | NA | 1 | 1 | 1 | 1 | 1 | 1 | NA | NA | NA | NA | 2 | 0 | 3 | 0.3333 |
| 11_974106 | NA | NA | NA | NA | NA | NA | 1 | 1 | 1 | 1 | 1 | 1 | 1 | 1 | NA | NA | 2 | 0 | 4 | 0.5 |
| 11_974106 | 1 | 1 | NA | NA | NA | NA | 1 | 0 | 2 | 0.5 | 1 | 0 | 1 | 1 | NA | NA | 2 | 0.5 | 4 | 0.5 |
| 11_974108 | 1 | 1 | NA | NA | NA | NA | 1 | 1 | 2 | 1 | NA | NA | 1 | 1 | NA | NA | 2 | 1 | 3 | 1 |
| 11_974108 | 1 | 1 | NA | NA | NA | NA | 1 | 1 | 2 | 1 | 1 | 1 | 1 | 1 | NA | NA | 2 | 0 | 4 | 0.5 |
| 11_974108 | 1 | 0 | NA | NA | NA | NA | 1 | 1 | 2 | 0.5 | 1 | 0 | 1 | 1 | NA | NA | 2 | 0.5 | 4 | 0.5 |
| 11_974162 | 3 | 0.3333 | 4 | 0.25 | 2 | 0 | 2 | 0.5 | 11 | 0.2727 | 7 | 0 | 3 | 0 | NA | NA | 3 | 0.3333 | 13 | 0.0769 |

| | | | | | | | | | | | | | | | | | | | | |
|---|---|---|---|---|---|---|---|---|---|---|---|---|---|---|---|---|---|---|---|---|
| 11_974256 | 4 | 0.75 | 3 | 1 | NA | NA | 2 | 1 | 9 | 0.8889 | 8 | 0.625 | 5 | 1 | 1 | 1 | 2 | 1 | 16 | 0.8125 |
| 11_974258 | 4 | 0.25 | 2 | 0.5 | 1 | 0 | 2 | 0 | 9 | 0.2222 | 7 | 0.1429 | 5 | 0.4 | 1 | 0 | 1 | 0 | 14 | 0.2143 |
| 11_974259 | 4 | 0.75 | 2 | 0.5 | NA | NA | 1 | 0 | 7 | 0.5714 | 7 | 0.8571 | 3 | 0.6667 | 1 | 0 | 2 | 0.5 | 13 | 0.6923 |
| 11_974427 | 3 | 0.6667 | 3 | 0.3333 | 1 | 1 | NA | NA | 7 | 0.5714 | 4 | 0 | 4 | 0.25 | 1 | 0 | 1 | 0 | 10 | 0.1 |
| 11_975182 | 8 | 0.25 | 4 | 0 | 6 | 0.1667 | 6 | 0.1667 | 24 | 0.1667 | 7 | 0.1429 | 1 | 0 | 4 | 0 | 4 | 0.25 | 16 | 0.125 |
| 11_975184 | 8 | 0.625 | 4 | 0.5 | 5 | 0 | 7 | 0.7143 | 24 | 0.5 | 8 | 0.5 | 4 | 0.5 | 6 | 0.3333 | 2 | 0.5 | 20 | 0.45 |
| 11_975186 | 6 | 1 | 5 | 1 | 5 | 1 | 8 | 0.875 | 24 | 0.9583 | 6 | 0.8333 | 4 | 1 | 6 | 1 | 3 | 0.6667 | 19 | 0.8947 |
| 11_975186 | 5 | 1 | 5 | 1 | 4 | 0.75 | 8 | 0.75 | 22 | 0.8636 | 7 | 1 | 4 | 1 | 6 | 0.8333 | 2 | 1 | 19 | 0.9474 |
| 11_975186 | 5 | 1 | 3 | 0.6667 | 4 | 1 | 6 | 0.8333 | 18 | 0.8889 | 6 | 0.6667 | 4 | 1 | 4 | 0.75 | 3 | 1 | 17 | 0.8235 |
| 11_975187 | 6 | 0.8333 | 4 | 1 | 4 | 0.5 | 5 | 1 | 19 | 0.8421 | 7 | 0.8571 | 3 | 0.6667 | 4 | 0.75 | 3 | 1 | 17 | 0.8235 |
| 11_975188 | 5 | 0.6 | 4 | 1 | 5 | 0.6 | 6 | 0.5 | 20 | 0.65 | 5 | 0.4 | 4 | 0.75 | NA | NA | 3 | 0.3333 | 12 | 0.5 |
| 11_975188 | 7 | 0.8571 | 4 | 1 | 5 | 1 | 6 | 0.8333 | 22 | 0.9091 | 7 | 1 | 4 | 1 | 4 | 1 | 3 | 1 | 18 | 1 |
| 11_975189 | 3 | 1 | 1 | 1 | 3 | 0.6667 | 5 | 0.8 | 12 | 0.8333 | 3 | 1 | 3 | 1 | 3 | 1 | 2 | 1 | 11 | 1 |
| 11_975189 | 4 | 0 | 1 | 0 | 4 | 0.25 | 6 | 0.1667 | 15 | 0.1333 | 4 | 0.25 | 2 | 0 | 4 | 0.25 | 3 | 0.6667 | 13 | 0.3077 |
| 11_975189 | 4 | 0.25 | 4 | 0.25 | 5 | 0.4 | 4 | 0.5 | 17 | 0.3529 | 4 | 0.5 | 3 | 0.6667 | 3 | 0.3333 | 3 | 0.3333 | 13 | 0.4615 |
| 11_975190 | 6 | 0 | 4 | 0.5 | 4 | 0.5 | 5 | 0.6 | 19 | 0.3684 | 6 | 0.8333 | 4 | 0.5 | 4 | 0.75 | 3 | 1 | 17 | 0.7647 |
| 11_975190 | 5 | 0.4 | 4 | 0.25 | 3 | 0.3333 | 6 | 0.1667 | 18 | 0.2778 | 6 | 0.3333 | 4 | 0.5 | 4 | 0.5 | 4 | 0.25 | 18 | 0.3889 |
| 11_975191 | 3 | 0 | 5 | 0.2 | 4 | 0.25 | 6 | 0 | 18 | 0.1111 | 3 | 0.3333 | 2 | 0 | 2 | 0 | 4 | 0 | 11 | 0.0909 |
| 11_975193 | 4 | 0.75 | 3 | 0 | 3 | 1 | 5 | 0.8 | 15 | 0.6667 | 5 | 0.8 | 4 | 0.5 | 3 | 0.3333 | 4 | 0.75 | 16 | 0.625 |
| 11_975193 | 4 | 0.25 | 4 | 0 | 3 | 0.3333 | 4 | 0 | 15 | 0.1333 | 5 | 0.2 | 4 | 0.25 | 3 | 0 | 2 | 0 | 14 | 0.1429 |
| 11_975196 | 2 | 0.5 | 4 | 0.5 | 3 | 1 | 4 | 0.5 | 13 | 0.6154 | 6 | 0.5 | 4 | 0.75 | 3 | 0.6667 | 4 | 1 | 17 | 0.7059 |
| 11_975198 | 1 | 0 | 3 | 0 | 1 | 1 | 4 | 0 | 9 | 0.1111 | 4 | 0 | 3 | 0.3333 | 3 | 0 | 2 | 0 | 12 | 0.0833 |
| 11_975201 | 2 | 0.5 | 5 | 0.4 | 1 | 0 | 4 | 0 | 12 | 0.25 | 6 | 0.3333 | 4 | 0.75 | 3 | 0.3333 | 3 | 0.6667 | 16 | 0.5 |
| 11_975201 | 2 | 0 | 4 | 0.25 | 2 | 0 | 2 | 0 | 10 | 0.1 | 3 | 0.6667 | 2 | 0 | 2 | 0 | 2 | 0 | 9 | 0.2222 |
| 11_975352 | 2 | 0 | 1 | 0 | NA | NA | 2 | 0.5 | 5 | 0.2 | 1 | 0 | NA | NA | 4 | 0.25 | 1 | 0 | 6 | 0.1667 |
| 11_975646 | 1 | 0 | NA | NA | 2 | 1 | 1 | 0 | 4 | 0.5 | NA | NA | NA | NA | 1 | 1 | NA | NA | 1 | 1 |
| 11_975724 | 1 | 0 | 4 | 1 | 3 | 1 | 3 | 0.6667 | 11 | 0.8182 | 1 | 1 | 2 | 0 | 2 | 1 | 1 | 0 | 6 | 0.5 |
| 11_975725 | 1 | 1 | 5 | 1 | 3 | 0.6667 | 3 | 0.3333 | 12 | 0.75 | 1 | 1 | 2 | 1 | 4 | 1 | 2 | 1 | 9 | 1 |
| 11_975727 | 1 | 0 | 4 | 0.25 | 2 | 0.5 | 3 | 0 | 10 | 0.2 | 2 | 1 | 1 | 1 | 3 | 0 | 2 | 0.5 | 8 | 0.5 |
| 11_975728 | NA | NA | 3 | 0.3333 | 3 | 0.3333 | 3 | 0.3333 | 9 | 0.3333 | 2 | 0 | 2 | 1 | 2 | 1 | 2 | 0 | 8 | 0.5 |
| 11_975729 | 1 | 0 | 6 | 0 | 3 | 0.3333 | 1 | 0 | 11 | 0.0909 | 1 | 0 | 1 | 0 | 3 | 0.3333 | 2 | 0 | 7 | 0.1429 |
| 11_975741 | 1 | 0 | 6 | 0.5 | 5 | 0.2 | 7 | 0.2857 | 19 | 0.3158 | 2 | 0 | 5 | 0.4 | 2 | 0 | 2 | 0 | 11 | 0.1818 |
| 11_975759 | 1 | 0 | 13 | 0.0769 | 10 | 0.1 | 10 | 0.1 | 34 | 0.0882 | 5 | 0.2 | 8 | 0.125 | 9 | 0.4444 | 8 | 0.125 | 30 | 0.2333 |
| 11_975759 | 4 | 0.5 | 15 | 0.4667 | 12 | 0.5833 | 11 | 0.2727 | 42 | 0.4524 | 7 | 0.2857 | 11 | 0.5455 | 9 | 0.3333 | 8 | 0.25 | 35 | 0.3714 |
| 11_975808 | 2 | 0.5 | 3 | 0.3333 | 6 | 0.1667 | 5 | 0.2 | 16 | 0.25 | 2 | 0.5 | 7 | 0.5714 | 5 | 0.4 | 4 | 0.25 | 18 | 0.4444 |
| 11_975808 | 2 | 0.5 | 2 | 0.5 | 7 | 0.5714 | 5 | 0.6 | 16 | 0.5625 | 1 | 1 | 6 | 0.1667 | 4 | 0 | 5 | 0.6 | 16 | 0.3125 |
| 11_975808 | NA | NA | 3 | 0.3333 | 6 | 0.6667 | 3 | 1 | 12 | 0.6667 | 1 | 1 | 6 | 0.1667 | 4 | 0 | 4 | 0.5 | 15 | 0.2667 |
| 11_976078 | 11 | 0.9091 | 10 | 1 | 5 | 1 | 11 | 0.8182 | 37 | 0.9189 | 3 | 0.6667 | 5 | 0.8 | 10 | 1 | 7 | 0.8571 | 25 | 0.88 |
| 11_976078 | 9 | 0.7778 | 9 | 0.7778 | 7 | 1 | 10 | 0.9 | 35 | 0.8571 | 4 | 1 | 5 | 1 | 7 | 0.8571 | 7 | 1 | 23 | 0.9565 |
| 11_976079 | 10 | 0.7 | 8 | 0.875 | 7 | 0.8571 | 9 | 0.7778 | 34 | 0.7941 | 4 | 0.75 | 6 | 0.6667 | 10 | 0.9 | 6 | 0.8333 | 26 | 0.8077 |
| 11_976079 | 9 | 0.4444 | 7 | 0.5714 | 7 | 0.8571 | 7 | 0.8571 | 30 | 0.6667 | 4 | 0.5 | 5 | 0.8 | 9 | 0.7778 | 6 | 0.5 | 24 | 0.6667 |
| 11_976080 | 5 | 0.6 | 7 | 0.5714 | 2 | 1 | 4 | 0.5 | 18 | 0.6111 | 4 | 0.75 | 3 | 0 | 9 | 0.5556 | 5 | 0.8 | 21 | 0.5714 |
| 11_976083 | 5 | 0.2 | 7 | 0 | 2 | 0 | 4 | 0 | 18 | 0.0556 | 2 | 0 | 3 | 0 | 5 | 0 | 5 | 0.2 | 15 | 0.0667 |
| 11_976084 | 7 | 1 | 5 | 1 | 5 | 1 | 6 | 0.8333 | 23 | 0.9565 | 3 | 1 | 5 | 1 | 5 | 1 | 7 | 0.8571 | 20 | 0.95 |
| 11_976086 | 6 | 1 | 7 | 0.7143 | 6 | 1 | 7 | 1 | 26 | 0.9231 | 4 | 1 | 3 | 1 | 7 | 0.5714 | 6 | 0.6667 | 20 | 0.75 |
| 11_976086 | 4 | 0.5 | 7 | 0.2857 | 5 | 0.6 | 7 | 0.5714 | 23 | 0.4783 | 4 | 0.5 | 1 | 1 | 7 | 0.7143 | 5 | 1 | 17 | 0.7647 |
| 11_976087 | 5 | 0.4 | 7 | 0.2857 | 5 | 0.4 | 7 | 0.2857 | 24 | 0.3333 | 3 | 0.3333 | 3 | 0.3333 | 6 | 0.1667 | 4 | 0.5 | 16 | 0.3125 |
| 11_976087 | 7 | 0.2857 | 6 | 0.3333 | 5 | 0 | 6 | 0.3333 | 24 | 0.25 | 3 | 0 | 3 | 0 | 5 | 0.4 | 6 | 0.1667 | 17 | 0.1765 |
| 11_976087 | 5 | 0.2 | 1 | 0 | 1 | 0 | 2 | 0 | 9 | 0.1111 | 3 | 0.3333 | 2 | 0 | 6 | 0 | 5 | 0 | 16 | 0.0625 |
| 11_976088 | 6 | 0.8333 | 6 | 0.6667 | 5 | 0.4 | 7 | 0.8571 | 24 | 0.7083 | 3 | 1 | 3 | 0.3333 | 8 | 0.625 | 6 | 0.5 | 20 | 0.6 |
| 11_976088 | 6 | 1 | 4 | 1 | 4 | 1 | 2 | 1 | 16 | 1 | 3 | 1 | 2 | 1 | 7 | 1 | 4 | 1 | 16 | 1 |
| 11_976088 | 6 | 0.5 | 4 | 0.5 | 3 | 0.6667 | 5 | 0.6 | 18 | 0.5556 | 2 | 0.5 | 3 | 0.3333 | 4 | 0.25 | 6 | 0.3333 | 15 | 0.3333 |
| 11_976090 | 6 | 0 | 2 | 0 | 4 | 0.25 | 4 | 0 | 16 | 0.0625 | 3 | 0 | 3 | 0.3333 | 5 | 0 | 5 | 0 | 16 | 0.0625 |
| 11_976090 | 7 | 0.4286 | 3 | 0 | 5 | 0 | 7 | 0.1429 | 22 | 0.1818 | 3 | 0 | 3 | 0 | 4 | 0.5 | 4 | 0.5 | 14 | 0.2857 |
| 11_976090 | 6 | 0 | 3 | 0 | 3 | 0 | 5 | 0.4 | 17 | 0.1176 | 2 | 0 | 3 | 0.3333 | 5 | 0.2 | 4 | 0 | 14 | 0.1429 |
| 11_976091 | 6 | 1 | 3 | 1 | 4 | 1 | 6 | 1 | 19 | 1 | 3 | 0.6667 | 3 | 1 | 3 | 1 | 5 | 1 | 14 | 0.9286 |
| 11_976093 | 3 | 1 | 3 | 1 | 3 | 0.6667 | 6 | 0.8333 | 15 | 0.8667 | 3 | 1 | 3 | 1 | 4 | 1 | 3 | 0.6667 | 13 | 0.9231 |
| 11_976095 | 2 | 1 | 3 | 0.3333 | 2 | 0.5 | 5 | 0.4 | 12 | 0.5 | 3 | 0.6667 | 2 | 1 | 4 | 0.5 | 4 | 0.75 | 13 | 0.6923 |
| 11_976096 | 2 | 1 | 2 | 0.5 | 1 | 0 | 1 | 1 | 6 | 0.6667 | 2 | 0.5 | NA | NA | 3 | 0.6667 | 2 | 1 | 7 | 0.7143 |
| 11_976096 | 2 | 0.5 | 2 | 0 | 1 | 0 | 1 | 1 | 6 | 0.3333 | 3 | 0 | 1 | 1 | 4 | 0 | 3 | 0.3333 | 11 | 0.1818 |
| 11_976377 | 5 | 0.6 | 4 | 0.75 | 2 | 0.5 | NA | NA | 11 | 0.6364 | 2 | 0.5 | 1 | 1 | 2 | 0.5 | NA | NA | 5 | 0.6 |
| 11_976378 | 5 | 0.8 | 4 | 1 | 3 | 1 | NA | NA | 12 | 0.9167 | 1 | 0 | NA | NA | 3 | 1 | NA | NA | 4 | 0.75 |
| 11_976378 | 3 | 0.3333 | 3 | 1 | 3 | 0 | NA | NA | 9 | 0.4444 | 2 | 1 | 1 | 1 | 3 | 0.3333 | NA | NA | 6 | 0.6667 |
| 11_976378 | 5 | 0 | 2 | 0.5 | 3 | 0 | NA | NA | 10 | 0.1 | 2 | 0 | NA | NA | 3 | 0.3333 | NA | NA | 5 | 0.2 |
| 11_976379 | 4 | 0 | 2 | 0.5 | 2 | 0.5 | NA | NA | 8 | 0.25 | 2 | 0 | NA | NA | 2 | 0.5 | NA | NA | 4 | 0.25 |

| | | | | | | | | | | | | | | | | | | | | |
|---|---|---|---|---|---|---|---|---|---|---|---|---|---|---|---|---|---|---|---|---|
| 11_9763798 | 3 | 1 | 3 | 1 | 2 | 1 | NA | NA | 8 | 1 | 2 | 1 | NA | NA | 3 | 1 | NA | NA | 5 | 1 |
| 11_9763804 | 5 | 0.2 | 4 | 0.75 | 3 | 0.6667 | NA | NA | 12 | 0.5 | 2 | 1 | NA | NA | 3 | 0.3333 | NA | NA | 5 | 0.6 |
| 11_9763813 | 5 | 0.2 | 3 | 1 | 2 | 1 | NA | NA | 10 | 0.6 | 2 | 0 | NA | NA | 2 | 0.5 | NA | NA | 4 | 0.25 |
| 11_9763817 | 3 | 0.3333 | 4 | 0 | 2 | 0 | NA | NA | 9 | 0.1111 | 2 | 0 | NA | NA | 3 | 0.3333 | NA | NA | 5 | 0.2 |
| 11_9763819 | 4 | 0 | 4 | 0.25 | 3 | 0 | NA | NA | 11 | 0.0909 | 2 | 0.5 | NA | NA | 3 | 0 | NA | NA | 5 | 0.2 |
| 11_9765926 | 1 | 0 | 1 | 1 | 3 | 0.3333 | 1 | 0 | 6 | 0.3333 | 2 | 0.5 | 3 | 0 | 1 | 0 | 2 | 0.5 | 8 | 0.25 |
| 11_9772701 | 8 | 0.25 | 2 | 0 | 9 | 0 | 8 | 0.125 | 27 | 0.1111 | 15 | 0.1333 | 1 | 0 | 3 | 0 | 6 | 0.1667 | 25 | 0.12 |
| 11_9772727 | 9 | 0.3333 | 2 | 0 | 8 | 0.375 | 7 | 0.5714 | 26 | 0.3846 | 12 | 0.0833 | 4 | 0 | 4 | 0 | 6 | 0.3333 | 26 | 0.1154 |
| 11_9772746 | 8 | 0 | 4 | 0.25 | 11 | 0.0909 | 9 | 0 | 32 | 0.0625 | 16 | 0.125 | 5 | 0 | 4 | 0 | 7 | 0.2857 | 32 | 0.125 |
| 11_9772779 | 8 | 0.125 | 4 | 0.25 | 10 | 0.1 | 8 | 0.125 | 30 | 0.1333 | 16 | 0.3125 | 5 | 0 | 6 | 0 | 10 | 0.3 | 37 | 0.2162 |
| 11_9772815 | 8 | 0.5 | 6 | 0.5 | 13 | 0.3846 | 10 | 0.7 | 37 | 0.5135 | 13 | 0.5385 | 5 | 0.2 | 4 | 0.5 | 10 | 0.3 | 32 | 0.4062 |
| 11_9772845 | 11 | 0.0909 | 7 | 0.1429 | 15 | 0.4 | 9 | 0.1111 | 42 | 0.2143 | 15 | 0.6 | 4 | 0 | 4 | 0.25 | 11 | 0.4545 | 34 | 0.4412 |
| 11_9772859 | 12 | 0.1667 | 7 | 0.1429 | 16 | 0.0625 | 13 | 0.0769 | 48 | 0.1042 | 18 | 0.1111 | 5 | 0 | 7 | 0 | 11 | 0.2727 | 41 | 0.122 |
| 11_9772913 | 13 | 0.0769 | 8 | 0 | 20 | 0.05 | 15 | 0.2 | 56 | 0.0893 | 24 | 0.1667 | 8 | 0 | 9 | 0.1111 | 15 | 0.1333 | 56 | 0.125 |
| 11_9773314 | 6 | 0 | 7 | 0.2857 | 7 | 0.2857 | 3 | 0.3333 | 23 | 0.2174 | 4 | 0 | 4 | 0.25 | 5 | 0.2 | 2 | 0 | 15 | 0.1333 |
| 11_9773321 | 5 | 0.2 | 7 | 0.8571 | 7 | 0.8571 | 5 | 0.4 | 24 | 0.625 | 5 | 0.4 | 4 | 0.75 | 6 | 0.8333 | 4 | 0.5 | 19 | 0.6316 |
| 11_9773323 | 6 | 0 | 6 | 0.1667 | 7 | 0.1429 | 4 | 0 | 23 | 0.087 | 5 | 0 | 3 | 0 | 7 | 0.1429 | 3 | 0 | 18 | 0.0556 |
| 11_9773327 | 4 | 0.5 | 7 | 0.8571 | 7 | 0.4286 | 3 | 0.3333 | 21 | 0.5714 | 4 | 0.5 | 4 | 0.75 | 5 | 0.4 | 4 | 0.75 | 17 | 0.5882 |
| 11_9773381 | 7 | 0.2857 | 7 | 0.2857 | 9 | 0.3333 | 6 | 0.3333 | 29 | 0.3103 | 5 | 0 | 3 | 0.3333 | 6 | 0.1667 | 3 | 0 | 17 | 0.1176 |
| 11_9773829 | 1 | 1 | 2 | 1 | 4 | 1 | 3 | 1 | 10 | 1 | 3 | 0.6667 | NA | NA | 4 | 1 | 1 | 1 | 8 | 0.875 |
| 11_9781413 | NA | NA | 2 | 1 | 4 | 1 | NA | NA | 6 | 1 | 1 | 1 | NA | NA | 1 | 1 | NA | NA | 2 | 1 |
| 11_9781426 | NA | NA | 2 | 1 | 3 | 0.3333 | NA | NA | 5 | 0.6 | 1 | 1 | NA | NA | NA | NA | NA | NA | 1 | 1 |
| 11_9781480 | 1 | 1 | 2 | 0.5 | 2 | 1 | NA | NA | 5 | 0.8 | 1 | 1 | NA | NA | NA | NA | NA | NA | 1 | 1 |
| 11_9781527 | 1 | 0 | 2 | 0.5 | 2 | 0 | NA | NA | 5 | 0.2 | 1 | 1 | 1 | 0 | NA | NA | NA | NA | 2 | 0.5 |
| 11_9781529 | 1 | 0 | 2 | 0.5 | 2 | 0 | NA | NA | 5 | 0.2 | 1 | 1 | 1 | 0 | NA | NA | NA | NA | 2 | 0.5 |
| 11_9788068 | 1 | 0 | 2 | 0.5 | NA | NA | NA | NA | 3 | 0.3333 | NA | NA | 2 | 1 | NA | NA | NA | NA | 2 | 1 |
| 11_9788097 | 1 | 1 | NA | NA | NA | NA | NA | NA | 1 | 1 | NA | NA | 2 | 0.5 | NA | NA | NA | NA | 2 | 0.5 |
| 11_9789281 | 12 | 0.5 | 9 | 0.5556 | 12 | 0.6667 | 15 | 0.5333 | 48 | 0.5625 | 12 | 0.6667 | 18 | 0.5556 | 16 | 0.625 | 21 | 0.619 | 67 | 0.6119 |
| 11_9789290 | 6 | 0.1667 | 4 | 0.5 | 1 | 1 | 7 | 0.2857 | 18 | 0.3333 | 2 | 0.5 | 6 | 0.5 | 7 | 0.4286 | 7 | 0.4286 | 22 | 0.4545 |
| 11_9789290 | 2 | 0 | 1 | 1 | 4 | 0.25 | 4 | 0.75 | 11 | 0.4545 | 2 | 1 | 4 | 0 | 1 | 0 | 1 | 0 | 8 | 0.25 |
| 11_9789386 | 8 | 0 | 4 | 0 | 5 | 0.2 | 8 | 0.125 | 25 | 0.08 | 6 | 0.1667 | 7 | 0.1429 | 9 | 0.2222 | 8 | 0.25 | 30 | 0.2 |
| 11_9789395 | 4 | 0 | 1 | 1 | 2 | 0 | 4 | 0 | 11 | 0.0909 | 4 | 0.25 | 6 | 0.1667 | 4 | 0.25 | 8 | 0 | 22 | 0.1364 |
| 11_9789395 | 4 | 0.75 | 3 | 1 | 5 | 0.8 | 6 | 0.6667 | 18 | 0.7778 | 5 | 0.8 | 6 | 0.6667 | 4 | 1 | 8 | 0.875 | 23 | 0.8261 |
| 11_9789397 | 7 | 0.1429 | 4 | 0 | 4 | 0 | 8 | 0.125 | 23 | 0.087 | 5 | 0.2 | 7 | 0.1429 | 5 | 0.2 | 8 | 0 | 25 | 0.12 |
| 11_9789402 | 8 | 0.125 | 4 | 0 | 3 | 0.6667 | 7 | 0.1429 | 22 | 0.1818 | 4 | 0.5 | 7 | 0 | 6 | 0 | 7 | 0 | 24 | 0.0833 |
| 11_9789410 | 6 | 0.3333 | 3 | 0 | 1 | 1 | 3 | 0 | 13 | 0.2308 | 2 | 0.5 | 4 | 0 | 4 | 0 | 4 | 0.25 | 14 | 0.1429 |
| 11_9789415 | 8 | 0.25 | 4 | 0 | 2 | 0 | 3 | 0 | 17 | 0.1176 | 3 | 0.3333 | 6 | 0.1667 | 4 | 0.25 | 4 | 0 | 17 | 0.1765 |
| 11_9789418 | 7 | 0.1429 | 3 | 0 | 3 | 0 | 6 | 0 | 19 | 0.0526 | 4 | 0.25 | 7 | 0.1429 | 5 | 0 | 5 | 0.2 | 21 | 0.1429 |
| 11_9789565 | NA | NA | 1 | 1 | NA | NA | NA | NA | 1 | 1 | NA | NA | 1 | 0 | NA | NA | 1 | 1 | 2 | 0.5 |
| 11_9789567 | NA | NA | 1 | 1 | NA | NA | NA | NA | 1 | 1 | NA | NA | 1 | 1 | 1 | 0 | 1 | 0 | 3 | 0.3333 |
| 11_9792765 | 8 | 0.25 | 9 | 0.4444 | 7 | 0 | 10 | 0.1 | 34 | 0.2059 | 9 | 0.1111 | 10 | 0.3 | 6 | 0.1667 | 10 | 0.2 | 35 | 0.2 |
| 11_9792832 | 9 | 0.2222 | 6 | 0.3333 | 6 | 0.1667 | 8 | 0.125 | 29 | 0.2069 | 6 | 0.1667 | 5 | 0.4 | 5 | 0.2 | 9 | 0.2222 | 25 | 0.24 |
| 11_9793455 | 4 | 0 | 4 | 0 | 1 | 0 | 5 | 0.2 | 14 | 0.0714 | 4 | 0 | 3 | 0 | 1 | 0 | 3 | 0.3333 | 11 | 0.0909 |
| 11_9797360 | 6 | 0.5 | 5 | 0.8 | 3 | 0.3333 | 2 | 1 | 16 | 0.625 | 3 | 0 | 3 | 0.3333 | 2 | 0 | 2 | 0.5 | 10 | 0.2 |
| 11_9797365 | 5 | 0 | 2 | 0.5 | 3 | 0 | 3 | 0.3333 | 13 | 0.1538 | NA | NA | 3 | 0.3333 | 1 | 0 | 1 | 1 | 5 | 0.4 |
| 11_9797373 | 4 | 0 | 5 | 0.4 | 1 | 0 | 2 | 0 | 12 | 0.1667 | 2 | 0.5 | 5 | 0.6 | 1 | 1 | 3 | 0.6667 | 11 | 0.6364 |
| 11_9797386 | 5 | 0.2 | 3 | 0.3333 | 1 | 0 | 2 | 0 | 11 | 0.1818 | 3 | 0 | 4 | 0.25 | 1 | 0 | 2 | 0.5 | 10 | 0.2 |
| 11_9797474 | 2 | 0.5 | 2 | 0.5 | NA | NA | 2 | 0 | 6 | 0.3333 | 3 | 0 | 4 | 0.25 | 1 | 0 | 2 | 0 | 10 | 0.1 |
| 11_9798037 | NA | NA | 1 | 1 | NA | NA | NA | NA | 1 | 1 | 2 | 1 | 1 | 0 | 1 | 0 | NA | NA | 4 | 0.5 |
| 11_9798039 | NA | NA | 1 | 1 | NA | NA | NA | NA | 1 | 1 | 2 | 1 | 1 | 1 | NA | NA | NA | NA | 3 | 1 |
| 11_9804370 | 2 | 0.5 | 1 | 0 | 3 | 0.3333 | 3 | 0 | 9 | 0.2222 | 3 | 0 | 2 | 1 | 3 | 0.3333 | 3 | 0 | 11 | 0.2727 |
| 11_9804386 | 2 | 0.5 | 1 | 0 | 2 | 0.5 | 2 | 0.5 | 7 | 0.4286 | 2 | 0.5 | 2 | 0 | 2 | 0.5 | 2 | 0.5 | 8 | 0.375 |
| 11_9804845 | NA | NA | NA | NA | NA | NA | 2 | 0.5 | 2 | 0.5 | 1 | 1 | 1 | 0 | NA | NA | NA | NA | 2 | 0.5 |
| 11_9804863 | NA | NA | NA | NA | NA | NA | 2 | 0.5 | 2 | 0.5 | 1 | 0 | 1 | 1 | NA | NA | NA | NA | 2 | 0.5 |
| 11_9809477 | NA | NA | 2 | 0.5 | NA | NA | 1 | 0 | 3 | 0.3333 | 1 | 1 | 1 | 1 | NA | NA | NA | NA | 2 | 1 |
| 11_9809492 | NA | NA | 2 | 0 | NA | NA | 1 | 1 | 3 | 0.3333 | 1 | 1 | 1 | 0 | NA | NA | NA | NA | 2 | 0.5 |
| 11_9810151 | NA | NA | 2 | 1 | NA | NA | 1 | 1 | 3 | 1 | 1 | 1 | 2 | 1 | NA | NA | NA | NA | 3 | 1 |
| 11_9810190 | NA | NA | 2 | 0.5 | NA | NA | 1 | 0 | 3 | 0.3333 | 1 | 1 | 2 | 0 | NA | NA | NA | NA | 3 | 0.3333 |
| 11_9813621 | NA | NA | 2 | 0 | NA | NA | 2 | 0.5 | 4 | 0.25 | 2 | 0.5 | 2 | 0.5 | NA | NA | 1 | 1 | 5 | 0.6 |
| 11_9813624 | 1 | 0 | 2 | 0.5 | NA | NA | 2 | 1 | 5 | 0.6 | 2 | 1 | 2 | 1 | NA | NA | 1 | 1 | 5 | 1 |
| 11_9814072 | 1 | 1 | 2 | 0 | NA | NA | 2 | 0 | 5 | 0.2 | 2 | 0 | 3 | 0.3333 | NA | NA | 1 | 0 | 6 | 0.1667 |
| 11_9814075 | NA | NA | 1 | 1 | NA | NA | 1 | 1 | 2 | 1 | 2 | 1 | 1 | 1 | NA | NA | 1 | 1 | 4 | 1 |
| 11_9814080 | 1 | 1 | 1 | 1 | NA | NA | 2 | 0 | 4 | 0.5 | 2 | 0 | 3 | 0 | NA | NA | 1 | 1 | 6 | 0.1667 |
| 11_9814086 | 1 | 1 | 1 | 0 | NA | NA | 1 | 0 | 3 | 0.3333 | 2 | 0 | 3 | 0.3333 | NA | NA | NA | NA | 5 | 0.2 |
| 11_9814087 | NA | NA | 2 | 0.5 | NA | NA | 2 | 1 | 4 | 0.75 | 2 | 0.5 | 2 | 1 | NA | NA | 1 | 0 | 5 | 0.6 |

| | | | | | | | | | | | | | | | | | | | | |
|---|---|---|---|---|---|---|---|---|---|---|---|---|---|---|---|---|---|---|---|---|
| 11_9814106 | NA | NA | NA | NA | NA | NA | 1 | 1 | 1 | 1 | 1 | 0 | 3 | 0.6667 | NA | NA | 1 | 1 | 5 | 0.6 |
| 11_9814108 | 1 | 0 | 2 | 0.5 | NA | NA | 1 | 0 | 4 | 0.25 | 2 | 0 | 2 | 0.5 | NA | NA | 1 | 0 | 5 | 0.2 |
| 11_9814122 | 1 | 0 | 2 | 0 | NA | NA | 1 | 1 | 4 | 0.25 | 2 | 0.5 | 3 | 0.3333 | NA | NA | 1 | 0 | 6 | 0.3333 |
| 11_9814127 | 1 | 1 | 1 | 1 | NA | NA | 1 | 0 | 3 | 0.6667 | 2 | 0.5 | 3 | 0.3333 | 1 | 1 | 1 | 0 | 7 | 0.4286 |
| 11_9814153 | 1 | 0 | 2 | 0.5 | NA | NA | 1 | 1 | 4 | 0.5 | 2 | 0.5 | 2 | 0 | 1 | 0 | 2 | 0 | 7 | 0.1429 |
| 11_9814174 | 2 | 0.5 | 2 | 1 | NA | NA | 1 | 1 | 5 | 0.8 | 2 | 0 | 3 | 0.6667 | 1 | 0 | 2 | 1 | 8 | 0.5 |
| 11_9821666 | 6 | 0.1667 | 7 | 0 | 5 | 0 | 1 | 0 | 19 | 0.0526 | 16 | 0.0625 | 3 | 0 | NA | NA | 4 | 0.25 | 23 | 0.087 |
| 11_9821704 | 3 | 0.6667 | 3 | 0.3333 | 3 | 0.6667 | NA | NA | 9 | 0.5556 | 9 | 0.1111 | 2 | 0 | NA | NA | 3 | 0.3333 | 14 | 0.1429 |
| 11_9821717 | 4 | 0.75 | 5 | 0.4 | 4 | 0.5 | NA | NA | 13 | 0.5385 | 10 | 0.3 | 2 | 0 | NA | NA | 3 | 0 | 15 | 0.2 |
| 11_9821720 | 4 | 0.5 | 5 | 0.2 | 5 | 0.4 | NA | NA | 14 | 0.3571 | 11 | 0.3636 | 3 | 0.6667 | NA | NA | 4 | 0.75 | 18 | 0.5 |
| 11_9821726 | 3 | 0.6667 | 4 | 0.75 | 2 | 1 | 1 | 1 | 10 | 0.8 | 7 | 1 | 3 | 0.6667 | NA | NA | 4 | 1 | 14 | 0.9286 |
| 11_9821730 | 5 | 0.2 | 5 | 0 | 2 | 0 | 1 | 0 | 13 | 0.0769 | 11 | 0.1818 | 3 | 0 | NA | NA | 4 | 0.25 | 18 | 0.1667 |
| 11_9821772 | 4 | 1 | 4 | 0.25 | 2 | 1 | 1 | 0 | 11 | 0.6364 | 7 | 0.5714 | 2 | 0 | NA | NA | 3 | 0.6667 | 12 | 0.5 |
| 11_9821781 | 2 | 0.5 | 5 | 0.2 | 1 | 0 | 1 | 0 | 9 | 0.2222 | 9 | 0.1111 | 3 | 0 | NA | NA | 2 | 0 | 14 | 0.0714 |
| 11_9821801 | 3 | 0 | 1 | 1 | 1 | 0 | NA | NA | 5 | 0.2 | 4 | 0.25 | 1 | 0 | NA | NA | NA | NA | 5 | 0.2 |
| 11_9821821 | 2 | 1 | 2 | 1 | 2 | 1 | 1 | 1 | 7 | 1 | 9 | 0.8889 | 3 | 1 | NA | NA | 2 | 1 | 14 | 0.9286 |
| 11_9821829 | 2 | 1 | 3 | 1 | 1 | 1 | 1 | 1 | 7 | 1 | 7 | 0.8571 | 3 | 1 | NA | NA | 2 | 1 | 12 | 0.9167 |
| 11_9821836 | 3 | 0.6667 | 2 | 1 | 3 | 0.6667 | 1 | 1 | 9 | 0.7778 | 7 | 0.5714 | 3 | 0 | NA | NA | 2 | 0.5 | 12 | 0.4167 |
| 11_9821852 | 1 | 1 | 2 | 0.5 | 2 | 1 | 1 | 1 | 6 | 0.8333 | 3 | 1 | 2 | 0.5 | NA | NA | 1 | 1 | 6 | 0.8333 |
| 11_9821858 | 3 | 1 | 2 | 1 | 1 | 0 | 1 | 1 | 7 | 0.8571 | 5 | 1 | 3 | 1 | NA | NA | 1 | 1 | 9 | 1 |
| 11_9821865 | 2 | 1 | 3 | 0.3333 | 2 | 0.5 | 1 | 0 | 8 | 0.5 | 5 | 0.2 | 3 | 0.3333 | NA | NA | 1 | 0 | 9 | 0.2222 |
| 11_9821873 | 3 | 0.6667 | 2 | 0.5 | 2 | 0 | 1 | 0 | 8 | 0.375 | 4 | 0.25 | 2 | 0 | NA | NA | 1 | 0 | 7 | 0.1429 |
| 11_9821888 | 3 | 0.3333 | 2 | 0.5 | 3 | 0.3333 | NA | NA | 8 | 0.375 | 6 | 0.1667 | 3 | 1 | NA | NA | 1 | 1 | 10 | 0.5 |
| 11_9823932 | 1 | 0 | 2 | 0 | 1 | 0 | 1 | 1 | 5 | 0.2 | NA | NA | 5 | 0.2 | 2 | 0.5 | 1 | 0 | 8 | 0.25 |
| 11_9824843 | 219 | 0.0548 | 103 | 0.068 | 158 | 0.0886 | 240 | 0.0875 | 720 | 0.075 | 95 | 0.0737 | 269 | 0.0483 | 285 | 0.0596 | 147 | 0.0476 | 796 | 0.0553 |
| 11_9827145 | 11 | 0.3636 | 5 | 0.2 | 6 | 0.6667 | 1 | 1 | 23 | 0.4348 | NA | NA | 4 | 0 | 6 | 0.5 | 3 | 0.6667 | 13 | 0.3846 |
| 11_9827148 | 11 | 0.7273 | 6 | 0.6667 | 6 | 0.6667 | 2 | 1 | 25 | 0.72 | 1 | 1 | 6 | 0.8333 | 6 | 0.5 | 3 | 0.6667 | 16 | 0.6875 |
| 11_9827158 | 9 | 0.6667 | 6 | 1 | 5 | 0.6 | 1 | 1 | 21 | 0.7619 | NA | NA | 5 | 0.8 | 6 | 0.8333 | 3 | 1 | 14 | 0.8571 |
| 11_9827162 | 10 | 0.5 | 6 | 0.3333 | 4 | 0.25 | 2 | 0 | 22 | 0.3636 | 1 | 0 | 6 | 0.1667 | 5 | 0.4 | 1 | 0 | 13 | 0.2308 |
| 11_9827165 | 11 | 0.3636 | 6 | 0.5 | 6 | 0.6667 | 2 | 1 | 25 | 0.52 | NA | NA | 6 | 0.6667 | 5 | 0.2 | 3 | 0.6667 | 14 | 0.5 |
| 11_9827168 | 10 | 0.3 | 5 | 0 | 6 | 0 | 2 | 0.5 | 23 | 0.1739 | 1 | 0 | 5 | 0.2 | 6 | 0 | 2 | 0 | 14 | 0.0714 |
| 11_9827169 | 10 | 0.3 | 6 | 0.1667 | 4 | 0 | 2 | 0.5 | 22 | 0.2273 | NA | NA | 6 | 0.1667 | 5 | 0.2 | 3 | 0.3333 | 14 | 0.2143 |
| 11_9827998 | 1 | 0 | 2 | 0 | NA | NA | 5 | 0.4 | 8 | 0.25 | NA | NA | 1 | 1 | 2 | 0.5 | 1 | 0 | 4 | 0.5 |
| 11_9832724 | NA | NA | 1 | 1 | NA | NA | NA | NA | 1 | 1 | NA | NA | NA | NA | 1 | 1 | NA | NA | 1 | 1 |
| 11_9832769 | NA | NA | 1 | 1 | NA | NA | NA | NA | 1 | 1 | NA | NA | NA | NA | 1 | 1 | NA | NA | 1 | 1 |
| 11_9832787 | NA | NA | 1 | 1 | NA | NA | NA | NA | 1 | 1 | NA | NA | NA | NA | 1 | 1 | NA | NA | 1 | 1 |
| 11_9832874 | 24 | 0.0833 | 15 | 0.1333 | 11 | 0.0909 | 13 | 0.3077 | 63 | 0.1429 | 13 | 0.0769 | 17 | 0.1176 | 16 | 0 | 14 | 0.2143 | 60 | 0.1 |
| 11_9837548 | NA | NA | NA | NA | 1 | 1 | NA | NA | 1 | 1 | NA | NA | NA | NA | NA | NA | 1 | 1 | 1 | 1 |
| 11_9847284 | 5 | 0 | 6 | 0 | 5 | 0 | 3 | 0.3333 | 19 | 0.0526 | 3 | 0 | 7 | 0 | 5 | 0.2 | 3 | 0 | 18 | 0.0556 |
| 11_9847345 | 1 | 1 | 1 | 0 | NA | NA | 1 | 1 | 3 | 0.6667 | 1 | 0 | 2 | 0 | 2 | 0 | 1 | 1 | 6 | 0.1667 |
| 11_9857374 | 9 | 0.2222 | 7 | 0.2857 | 11 | 0 | 7 | 0 | 34 | 0.1176 | 10 | 0.3 | 6 | 0 | 10 | 0.2 | 10 | 0.3 | 36 | 0.2222 |
| 11_9858645 | 15 | 0.0667 | 21 | 0.0952 | 22 | 0.1818 | 10 | 0 | 68 | 0.1029 | 15 | 0.1333 | 16 | 0.0625 | 18 | 0.2778 | 25 | 0.08 | 74 | 0.1351 |
| 11_9858653 | 15 | 0.0667 | 20 | 0.35 | 22 | 0.2727 | 11 | 0.6364 | 68 | 0.3088 | 14 | 0.2857 | 14 | 0.2857 | 19 | 0.4211 | 22 | 0.4091 | 69 | 0.3623 |
| 11_9858669 | 15 | 0.2 | 19 | 0.0526 | 15 | 0.2667 | 9 | 0 | 58 | 0.1379 | 14 | 0 | 15 | 0.0667 | 16 | 0.1875 | 19 | 0.0526 | 64 | 0.0781 |
| 11_9858670 | 15 | 0.1333 | 22 | 0.0909 | 15 | 0 | 8 | 0.125 | 60 | 0.0833 | 13 | 0 | 17 | 0.0588 | 20 | 0.15 | 19 | 0.0526 | 69 | 0.0725 |
| 11_9858672 | 13 | 0 | 21 | 0.1905 | 12 | 0.1667 | 6 | 0.1667 | 52 | 0.1346 | 10 | 0.5 | 12 | 0.25 | 17 | 0.2353 | 18 | 0.0556 | 57 | 0.2281 |
| 11_9859552 | 12 | 0.25 | 15 | 0 | 13 | 0.3846 | 8 | 0 | 48 | 0.1667 | 12 | 0.0833 | 12 | 0 | 17 | 0.1765 | 8 | 0.25 | 49 | 0.1224 |
| 11_9859557 | 11 | 0.4545 | 13 | 0.3077 | 14 | 0.3571 | 7 | 0.2857 | 45 | 0.3556 | 10 | 0.7 | 12 | 0.3333 | 16 | 0.3125 | 8 | 0.375 | 46 | 0.413 |
| 11_9863225 | 1 | 0 | 4 | 0 | 1 | 1 | NA | NA | 6 | 0.1667 | 1 | 0 | 5 | 0 | 1 | 1 | 1 | 0 | 8 | 0.125 |
| 11_9865530 | NA | NA | 3 | 0.6667 | 1 | 0 | 2 | 0.5 | 6 | 0.5 | 1 | 0 | NA | NA | 2 | 1 | 2 | 0.5 | 5 | 0.6 |
| 11_9865554 | NA | NA | 5 | 0.2 | 2 | 0.5 | 2 | 0.5 | 9 | 0.3333 | 2 | 0 | NA | NA | 2 | 1 | 1 | 1 | 5 | 0.6 |
| 11_9865677 | 1 | 0 | 7 | 0.1429 | 3 | 0 | 2 | 0.5 | 13 | 0.1538 | 2 | 0 | NA | NA | 4 | 0.25 | 4 | 0.25 | 10 | 0.2 |
| 11_9865900 | 6 | 0.8333 | 7 | 0.5714 | 10 | 0.8 | 15 | 0.6667 | 38 | 0.7105 | 1 | 0 | 13 | 0.7692 | 15 | 0.6667 | 9 | 0.5556 | 48 | 0.6042 |
| 11_9865900 | 6 | 0.8333 | 7 | 0.5714 | 10 | 0.8 | 15 | 0.6667 | 38 | 0.7105 | 11 | 0.3636 | 13 | 0.7692 | 15 | 0.6667 | 9 | 0.5556 | 48 | 0.6042 |
| 11_9865906 | 45 | 0.0444 | 86 | 0.1047 | 47 | 0.1064 | 64 | 0.1094 | 242 | 0.095 | 80 | 0.0875 | 79 | 0.0759 | 74 | 0.1081 | 51 | 0.0588 | 284 | 0.0845 |
| 11_9865930 | 56 | 0.2143 | 94 | 0.1383 | 65 | 0.2923 | 74 | 0.2027 | 289 | 0.2042 | 83 | 0.2892 | 94 | 0.2553 | 88 | 0.1818 | 60 | 0.2167 | 325 | 0.2369 |
| 11_9865932 | 63 | 0.0635 | 101 | 0.1287 | 70 | 0.0571 | 79 | 0.0633 | 313 | 0.0831 | 94 | 0.1383 | 95 | 0.1053 | 100 | 0.09 | 67 | 0.0896 | 356 | 0.1067 |
| 11_9866154 | 6 | 0.1667 | 7 | 0 | 6 | 0 | 15 | 0.1333 | 34 | 0.0882 | 5 | 0 | 12 | 0.0833 | 15 | 0.1333 | 7 | 0 | 39 | 0.0769 |
| 11_9866568 | 4 | 0.5 | 3 | 0 | 3 | 0 | 8 | 0.5 | 18 | 0.3333 | 2 | 0 | 6 | 0.6667 | 8 | 0.375 | 4 | 0.25 | 20 | 0.4 |
| 11_9866583 | 4 | 0.5 | 3 | 0.6667 | 4 | 0 | 8 | 0.25 | 19 | 0.3158 | 2 | 0 | 11 | 0.3636 | 7 | 0.1429 | 5 | 0 | 25 | 0.2 |
| 11_9866594 | 3 | 0.3333 | 4 | 0.25 | 4 | 0.5 | 7 | 0.7143 | 18 | 0.5 | 2 | 0.5 | 9 | 0.5556 | 7 | 0.8571 | 5 | 0.8 | 23 | 0.6957 |
| 11_9868665 | 5 | 0.2 | NA | NA | 5 | 0.2 | 7 | 0.1429 | 17 | 0.1765 | 6 | 0 | 4 | 0.25 | 4 | 0 | 5 | 0 | 19 | 0.0526 |
| 11_9868672 | 2 | 0 | 1 | 1 | 3 | 0.6667 | 8 | 0.875 | 14 | 0.7143 | 5 | 0.8 | 3 | 1 | 3 | 0.6667 | 2 | 1 | 13 | 0.8462 |
| 11_9868712 | 2 | 0 | 3 | 0 | 3 | 0.3333 | 5 | 0.2 | 13 | 0.1538 | 5 | 0 | 2 | 0.5 | 4 | 0.25 | 3 | 0 | 14 | 0.1429 |
| 11_9868716 | 4 | 0.75 | 4 | 0.25 | 5 | 0.4 | 10 | 0.4 | 23 | 0.4348 | 8 | 0.5 | 5 | 0.6 | 5 | 0.4 | 8 | 0.375 | 26 | 0.4615 |

| | | | | | | | | | | | | | | | | | | | | |
|---|---|---|---|---|---|---|---|---|---|---|---|---|---|---|---|---|---|---|---|---|
| 11_9868723 | 6 | 0.1667 | 3 | 0.3333 | 5 | 0 | 10 | 0.2 | 24 | 0.1667 | 8 | 0.125 | 3 | 0.3333 | 6 | 0.1667 | 10 | 0.4 | 27 | 0.2593 |
| 11_9868735 | 8 | 0.25 | 6 | 0.5 | 6 | 0.1667 | 9 | 0.2222 | 29 | 0.2759 | 8 | 0.25 | 4 | 0.5 | 8 | 0.375 | 12 | 0.25 | 32 | 0.3125 |
| 11_9869018 | 8 | 0.25 | 10 | 0.7 | 10 | 0.2 | 15 | 0.3333 | 43 | 0.3721 | 10 | 0.1 | 5 | 0 | 9 | 0 | 11 | 0.2727 | 35 | 0.1143 |
| 11_9869085 | 8 | 0.5 | 11 | 0.5455 | 10 | 0.8 | 15 | 0.3333 | 44 | 0.5227 | 10 | 0.4 | 6 | 0.1667 | 10 | 0.5 | 12 | 0.5 | 38 | 0.4211 |
| 11_9869090 | 6 | 0.6667 | 6 | 0.8333 | 7 | 0.8571 | 10 | 0.7 | 29 | 0.7586 | 8 | 0.625 | 5 | 1 | 6 | 0.8333 | 11 | 0.8182 | 30 | 0.8 |
| 11_9869095 | 9 | 1 | 9 | 1 | 11 | 0.9091 | 10 | 0.9 | 39 | 0.9487 | 7 | 0.8571 | 5 | 1 | 9 | 0.8889 | 10 | 1 | 31 | 0.9355 |
| 11_9869103 | 7 | 0.4286 | 9 | 0.6667 | 10 | 0.4 | 11 | 0.6364 | 37 | 0.5405 | 9 | 0.5556 | 5 | 0.2 | 10 | 0.3 | 12 | 0.5 | 36 | 0.4167 |
| 11_9869105 | 9 | 0.2222 | 8 | 0.25 | 7 | 0 | 14 | 0.2143 | 38 | 0.1842 | 10 | 0.5 | 6 | 0.5 | 10 | 0.3 | 9 | 0.3333 | 35 | 0.4 |
| 11_9869119 | 10 | 0.8 | 9 | 0.5556 | 11 | 1 | 13 | 0.5385 | 43 | 0.7209 | 12 | 0.8333 | 9 | 0.6667 | 11 | 0.6364 | 13 | 1 | 45 | 0.8 |
| 11_9869122 | 6 | 0.6667 | 7 | 0.5714 | 4 | 0.25 | 3 | 0.6667 | 20 | 0.55 | 5 | 0.8 | 3 | 0 | 2 | 1 | 6 | 0.6667 | 16 | 0.625 |
| 11_9869134 | 6 | 0.3333 | 7 | 0.7143 | 3 | 0.6667 | 1 | 0 | 17 | 0.5294 | 7 | 0.5714 | 2 | 1 | 3 | 0.6667 | 8 | 0.375 | 20 | 0.55 |
| 11_9869135 | 6 | 0.6667 | 6 | 0.3333 | 3 | 0.3333 | 2 | 0 | 17 | 0.4118 | 7 | 0.5714 | 4 | 0.5 | 2 | 0.5 | 8 | 0.5 | 21 | 0.5238 |
| 11_9869146 | 7 | 1 | 4 | 1 | 2 | 1 | 3 | 0.6667 | 16 | 0.9375 | 6 | 0.8333 | 2 | 1 | 2 | 1 | 6 | 1 | 16 | 0.9375 |
| 11_9869154 | 7 | 0.1429 | 6 | 0.5 | 3 | 0.3333 | 2 | 1 | 18 | 0.3889 | 6 | 0 | 1 | 0 | 3 | 0.6667 | 7 | 0.2857 | 17 | 0.2353 |
| 11_9869188 | NA | NA | 1 | 1 | NA | NA | NA | NA | 1 | 1 | 1 | 1 | 1 | 1 | NA | NA | NA | NA | 2 | 1 |
| 11_9869632 | 4 | 0.25 | 7 | 0 | 9 | 0 | 12 | 0.0833 | 32 | 0.0625 | 6 | 0 | 9 | 0 | 10 | 0.1 | 5 | 0.2 | 30 | 0.0667 |
| 11_9869717 | 7 | 0 | 12 | 0.0833 | 10 | 0.2 | 12 | 0 | 41 | 0.0732 | 5 | 0.2 | 10 | 0.1 | 10 | 0.2 | 4 | 0.5 | 29 | 0.2069 |
| 11_9869824 | 10 | 0.1 | 13 | 0.3077 | 15 | 0 | 13 | 0.0769 | 51 | 0.1176 | 8 | 0 | 13 | 0.0769 | 13 | 0.0769 | 9 | 0.2222 | 43 | 0.093 |
| 11_9871364 | 2 | 1 | 2 | 0.5 | 3 | 0.6667 | 5 | 0.6 | 12 | 0.6667 | 4 | 1 | NA | NA | 3 | 0.3333 | 3 | 1 | 10 | 0.8 |
| 11_9871378 | 2 | 0.5 | 2 | 1 | 3 | 0.3333 | 5 | 0.6 | 12 | 0.5833 | 4 | 0.5 | 1 | 1 | 3 | 0.6667 | 3 | 0.3333 | 11 | 0.5455 |
| 11_9871398 | 2 | 0 | 2 | 0 | 3 | 0.3333 | 4 | 0.25 | 11 | 0.1818 | 5 | 0 | 1 | 0 | 4 | 0.25 | 4 | 0 | 14 | 0.0714 |
| 11_9871399 | 2 | 0.5 | 1 | 1 | 3 | 0.6667 | 5 | 0 | 11 | 0.3636 | 5 | 0.4 | 1 | 0 | 4 | 0.75 | 4 | 0.5 | 14 | 0.5 |
| 11_9871409 | 1 | 1 | 2 | 0 | 2 | 0 | 3 | 0.3333 | 8 | 0.25 | 3 | 0.6667 | 1 | 0 | 3 | 0.3333 | 3 | 0.3333 | 10 | 0.4 |
| 11_9871410 | 2 | 0 | 2 | 0 | 2 | 0.5 | 4 | 0 | 10 | 0.1 | 4 | 0.5 | 1 | 0 | 3 | 0.3333 | 4 | 0.75 | 12 | 0.5 |
| 11_9871412 | 1 | 0 | 2 | 0.5 | 4 | 0 | 5 | 0.4 | 12 | 0.25 | 5 | 0.2 | 1 | 0 | 4 | 0.25 | 4 | 0.25 | 14 | 0.2143 |
| 11_9871413 | 2 | 0.5 | 2 | 0.5 | 4 | 0.25 | 4 | 0.5 | 12 | 0.4167 | 5 | 0.6 | 1 | 1 | 3 | 1 | 4 | 0 | 13 | 0.5385 |
| 11_9872327 | 4 | 0.5 | 4 | 0 | 5 | 0.2 | 10 | 0.6 | 23 | 0.3913 | 7 | 0.1429 | 3 | 0.6667 | 4 | 0.5 | 3 | 0.6667 | 17 | 0.4118 |
| 11_9872329 | 4 | 0.75 | 5 | 1 | 7 | 0.7143 | 10 | 0.9 | 26 | 0.8462 | 7 | 0.7143 | 3 | 1 | 6 | 0.5 | 5 | 0.8 | 21 | 0.7143 |
| 11_9872329 | 4 | 0 | 4 | 0 | 3 | 0 | 8 | 0.125 | 19 | 0.0526 | 4 | 0.25 | 2 | 0.5 | 4 | 0.25 | NA | NA | 10 | 0.3 |
| 11_9872334 | 3 | 1 | 4 | 0.25 | 6 | 0.6667 | 10 | 0.9 | 23 | 0.7391 | 7 | 0.7143 | 2 | 0.5 | 5 | 0.8 | 5 | 1 | 19 | 0.7895 |
| 11_9872449 | 10 | 0 | 8 | 0 | 6 | 0 | 10 | 0.2 | 34 | 0.0588 | 8 | 0.125 | 4 | 0.25 | 6 | 0.3333 | 7 | 0 | 25 | 0.16 |
| 11_9872781 | 1 | 1 | NA | NA | NA | NA | 4 | 0.5 | 5 | 0.6 | 3 | 0.6667 | NA | NA | NA | NA | 2 | 1 | 5 | 0.8 |
| 11_9872793 | 1 | 0 | 1 | 1 | NA | NA | 3 | 0.6667 | 5 | 0.6 | 5 | 0.6 | NA | NA | 1 | 0 | 3 | 0 | 9 | 0.3333 |
| 11_9872804 | 3 | 0 | 2 | 0.5 | NA | NA | 4 | 0 | 9 | 0.1111 | 5 | 0 | 1 | 0 | 1 | 0 | 3 | 0.3333 | 10 | 0.1 |
| 11_9872812 | 4 | 0 | 3 | 0.3333 | 1 | 0 | 2 | 0 | 10 | 0.1 | 4 | 0 | NA | NA | 1 | 1 | 2 | 0.5 | 7 | 0.2857 |
| 11_9874210 | 8 | 0.5 | 11 | 0.1818 | 10 | 0.4 | 8 | 0.5 | 37 | 0.3784 | 10 | 0.4 | 8 | 0.5 | 3 | 0.3333 | 11 | 0.3636 | 32 | 0.4062 |
| 11_9874217 | 9 | 0.7778 | 13 | 0.5385 | 11 | 0.3636 | 9 | 0.5556 | 42 | 0.5476 | 12 | 0.8333 | 10 | 0.6 | 3 | 0.3333 | 12 | 0.6667 | 37 | 0.6757 |
| 11_9874266 | 10 | 0.2 | 14 | 0.2143 | 12 | 0.3333 | 9 | 0.1111 | 45 | 0.2222 | 11 | 0.2727 | 8 | 0.375 | 3 | 0 | 13 | 0.2308 | 35 | 0.2571 |
| 11_9874326 | 10 | 0.1 | 19 | 0.1579 | 17 | 0 | 11 | 0.3636 | 57 | 0.1404 | 18 | 0.1667 | 11 | 0.0909 | 9 | 0.3333 | 17 | 0.1765 | 55 | 0.1818 |
| 11_9874336 | 11 | 0 | 20 | 0.15 | 19 | 0.2632 | 16 | 0.3125 | 66 | 0.197 | 21 | 0.3333 | 13 | 0.2308 | 10 | 0.4 | 19 | 0.3158 | 63 | 0.3175 |
| 11_9878303 | 4 | 0 | 6 | 0.1667 | 1 | 0 | 1 | 0 | 12 | 0.0833 | 4 | 0 | 2 | 0.5 | 3 | 0 | NA | NA | 9 | 0.1111 |
| 11_9878304 | 4 | 0 | 6 | 0.1667 | 1 | 0 | 1 | 0 | 12 | 0.0833 | 4 | 0.25 | 2 | 0 | 3 | 0 | NA | NA | 9 | 0.1111 |
| 11_9878324 | 4 | 0.5 | 6 | 0.3333 | NA | NA | NA | NA | 10 | 0.4 | 4 | 0.25 | 2 | 0.5 | 3 | 0.3333 | NA | NA | 9 | 0.3333 |
| 11_9878328 | 3 | 0.3333 | 4 | 0 | 1 | 0 | NA | NA | 8 | 0.125 | 2 | 0 | 2 | 0.5 | 2 | 0 | NA | NA | 6 | 0.1667 |
| 11_9878330 | 3 | 0.6667 | 6 | 0.8333 | 1 | 0 | 1 | 0 | 11 | 0.6364 | 3 | 0.6667 | 1 | 1 | 3 | 0.6667 | NA | NA | 7 | 0.7143 |
| 11_9878336 | 3 | 0.6667 | 6 | 0.5 | 1 | 0 | 1 | 1 | 11 | 0.5455 | 4 | 1 | 2 | 0 | 3 | 0 | NA | NA | 9 | 0.4444 |
| 11_9878341 | 4 | 0.75 | 7 | 0.8571 | NA | NA | 1 | 0 | 12 | 0.75 | 4 | 0.75 | 2 | 0.5 | 1 | 1 | NA | NA | 7 | 0.7143 |
| 11_9879054 | 3 | 0 | 7 | 0.4286 | 1 | 1 | 2 | 0 | 13 | 0.3077 | 4 | 0.25 | 2 | 0.5 | 2 | 0 | NA | NA | 8 | 0.25 |
| 11_9879067 | 4 | 1 | 8 | 0.875 | 1 | 1 | 2 | 1 | 15 | 0.9333 | 4 | 1 | 1 | 1 | 1 | 1 | NA | NA | 6 | 1 |
| 11_9879069 | 3 | 1 | 8 | 1 | 1 | 1 | 2 | 1 | 14 | 1 | 3 | 1 | 1 | 1 | 2 | 1 | NA | NA | 6 | 1 |
| 11_9879082 | 4 | 1 | 5 | 1 | 1 | 1 | 1 | 1 | 11 | 1 | 3 | 1 | 2 | 1 | 3 | 1 | NA | NA | 8 | 1 |
| 11_9879219 | 6 | 0.1667 | 5 | 0.2 | 4 | 0 | 6 | 0 | 21 | 0.0952 | 3 | 0 | 3 | 0 | 7 | 0.1429 | NA | NA | 13 | 0.0769 |
| 11_9879237 | 6 | 0.5 | 7 | 0.5714 | 4 | 0.25 | 5 | 0.8 | 22 | 0.5455 | 4 | 0.75 | 3 | 0.6667 | 8 | 0.25 | 1 | 1 | 16 | 0.5 |
| 11_9879241 | 6 | 0.6667 | 7 | 0.2857 | 4 | 0.25 | 6 | 0.3333 | 23 | 0.3913 | 5 | 0.6 | 3 | 0.6667 | 8 | 0.25 | 1 | 1 | 17 | 0.4706 |
| 11_9879282 | 8 | 0.25 | 8 | 0 | 4 | 0 | 5 | 0.4 | 25 | 0.16 | 7 | 0.1429 | 6 | 0.5 | 10 | 0 | 1 | 1 | 24 | 0.2083 |
| 11_9879635 | NA | NA | NA | NA | NA | NA | 1 | 1 | 1 | 1 | NA | NA | 1 | 0 | NA | NA | 1 | 1 | 2 | 0.5 |
| 11_9886433 | 1 | 1 | 3 | 0.6667 | 1 | 1 | 1 | 0 | 6 | 0.6667 | 1 | 1 | 1 | 0 | NA | NA | 1 | 0 | 3 | 0.3333 |
| 11_9886448 | 2 | 0 | 1 | 0 | 1 | 1 | 1 | 0 | 5 | 0.2 | 1 | 1 | 1 | 0 | NA | NA | NA | NA | 2 | 0.5 |
| 11_9886460 | 2 | 0.5 | 3 | 0.3333 | NA | NA | 1 | 0 | 6 | 0.3333 | 1 | 1 | NA | NA | NA | NA | NA | NA | 1 | 1 |
| 11_9886461 | 2 | 1 | 1 | 1 | 1 | 0 | NA | NA | 4 | 0.75 | 1 | 1 | 1 | 0 | NA | NA | 1 | 1 | 3 | 0.6667 |
| 11_9886464 | NA | NA | 1 | 1 | 1 | 1 | 1 | 1 | 3 | 1 | 1 | 1 | 1 | 1 | NA | NA | 1 | 1 | 3 | 1 |
| 11_9886477 | 2 | 0 | 3 | 0 | NA | NA | 1 | 1 | 6 | 0.1667 | 1 | 0 | 1 | 1 | NA | NA | 1 | 0 | 3 | 0.3333 |
| 11_9894229 | 11 | 0.0909 | 11 | 0.0909 | 7 | 0 | 5 | 0 | 34 | 0.0588 | 9 | 0 | 5 | 0.2 | 4 | 0 | 6 | 0.1667 | 24 | 0.0833 |
| 11_9894237 | 4 | 0.25 | 6 | 0.1667 | 7 | 0.2857 | 3 | 0 | 20 | 0.2 | 5 | 0.4 | 3 | 0.6667 | 3 | 0.6667 | 4 | 0.25 | 15 | 0.4667 |
| 11_9894242 | 8 | 0.125 | 9 | 0 | 6 | 0 | 3 | 0.3333 | 26 | 0.0769 | 9 | 0.1111 | 4 | 0 | 3 | 0.3333 | 4 | 0.25 | 20 | 0.15 |

| | | | | | | | | | | | | | | | | | | | | |
|---|---|---|---|---|---|---|---|---|---|---|---|---|---|---|---|---|---|---|---|---|
| 11_991001 | 3 | 0 | 9 | 0.2222 | 5 | 0.6 | 7 | 0.1429 | 24 | 0.25 | NA | NA | 3 | 0.3333 | 5 | 0.2 | 4 | 0.25 | 12 | 0.25 |
| 11_991001 | 2 | 0 | 8 | 0.875 | 4 | 0.5 | 7 | 0.7143 | 21 | 0.6667 | 1 | 1 | 3 | 1 | 5 | 0.4 | 5 | 0.8 | 14 | 0.7143 |
| 11_991006 | 3 | 0.3333 | 7 | 0.1429 | 3 | 0.3333 | 4 | 0.25 | 17 | 0.2353 | 1 | 0 | 2 | 0 | 2 | 0 | 4 | 0.25 | 9 | 0.1111 |
| 11_991006 | 3 | 0 | 8 | 0 | 3 | 0 | 4 | 0.25 | 18 | 0.0556 | 1 | 1 | 1 | 0 | 3 | 0 | 3 | 0 | 8 | 0.125 |
| 11_991009 | 4 | 0.75 | 6 | 0.1667 | 4 | 0.25 | 6 | 0 | 20 | 0.25 | 2 | 0 | 6 | 0.1667 | 2 | 0 | 6 | 0 | 16 | 0.0625 |
| 11_991009 | 5 | 0 | 10 | 0.2 | 6 | 0.3333 | 7 | 0.1429 | 28 | 0.1786 | 2 | 0 | 6 | 0 | 4 | 0.25 | 7 | 0.1429 | 19 | 0.1053 |
| 11_991237 | 3 | 0 | 4 | 0 | 4 | 0 | 3 | 0.3333 | 14 | 0.0714 | 2 | 0 | 6 | 0.1667 | 4 | 0 | 5 | 0 | 17 | 0.0588 |
| 11_991241 | 3 | 0.3333 | 3 | 0 | 4 | 0 | 3 | 0 | 13 | 0.0769 | 2 | 0 | 4 | 0 | 5 | 0.2 | 4 | 0 | 15 | 0.0667 |
| 11_991248 | 3 | 0 | 3 | 0.6667 | 4 | 0.25 | 2 | 0.5 | 12 | 0.3333 | 2 | 0.5 | 5 | 0.4 | 5 | 0 | 4 | 0 | 16 | 0.1875 |
| 11_991249 | 2 | 0 | 2 | 1 | 3 | 0.3333 | 2 | 0 | 9 | 0.3333 | 1 | 0 | 4 | 0.5 | 6 | 0.1667 | 2 | 0 | 13 | 0.2308 |
| 11_991251 | 3 | 0.6667 | 3 | 1 | 5 | 0.6 | 3 | 0.3333 | 14 | 0.6429 | 2 | 0.5 | 4 | 0.5 | 6 | 1 | 4 | 0.5 | 16 | 0.6875 |
| 11_991251 | 2 | 0 | 3 | 0.3333 | 4 | 0 | 3 | 0 | 12 | 0.0833 | 2 | 0 | 4 | 0.25 | 6 | 0.3333 | 4 | 0.5 | 16 | 0.3125 |
| 11_991252 | 2 | 0 | 2 | 0 | 5 | 0.4 | 3 | 0 | 12 | 0.1667 | 2 | 0 | 5 | 0.2 | 6 | 0 | 4 | 0 | 17 | 0.0588 |
| 11_991254 | 3 | 0.3333 | 3 | 0.3333 | 5 | 0 | 3 | 0.3333 | 14 | 0.2143 | 2 | 0.5 | 5 | 0 | 3 | 0 | 3 | 0 | 13 | 0.0769 |
| 11_991257 | 4 | 0 | 3 | 0.3333 | 5 | 0 | 4 | 0.25 | 16 | 0.125 | 2 | 0 | 5 | 0.2 | 4 | 0.25 | 2 | 0 | 13 | 0.1538 |
| 11_991258 | 3 | 0.3333 | 3 | 0.3333 | 5 | 0.2 | 4 | 0 | 15 | 0.2 | 2 | 0 | 5 | 0.2 | 5 | 0.2 | 2 | 0.5 | 14 | 0.2143 |
| 11_992763 | 4 | 0.25 | 5 | 0.2 | 6 | 0.3333 | 3 | 0.6667 | 18 | 0.3333 | 6 | 0.1667 | 6 | 0.1667 | 2 | 0.5 | 5 | 0 | 19 | 0.1579 |
| 11_992771 | NA | NA | 1 | 0 | 1 | 0 | 2 | 0.5 | 4 | 0.25 | NA | NA | 5 | 0.4 | NA | NA | NA | NA | 5 | 0.4 |
| 11_993130 | NA | NA | 1 | 1 | NA | NA | NA | NA | 1 | 1 | NA | NA | 1 | 1 | 1 | 0 | 1 | 0 | 3 | 0.3333 |
| 11_993130 | NA | NA | 1 | 1 | NA | NA | NA | NA | 1 | 1 | NA | NA | 1 | 1 | 1 | 0 | NA | NA | 2 | 0.5 |
| 11_993135 | NA | NA | 1 | 1 | NA | NA | NA | NA | 1 | 1 | NA | NA | 1 | 1 | NA | NA | 1 | 0 | 2 | 0.5 |
| 11_993136 | NA | NA | 1 | 1 | NA | NA | NA | NA | 1 | 1 | NA | NA | 1 | 1 | NA | NA | 1 | 1 | 2 | 1 |
| 11_993145 | NA | NA | NA | NA | 2 | 0.5 | NA | NA | 2 | 0.5 | NA | NA | 1 | 0 | 1 | 1 | 2 | 0.5 | 4 | 0.5 |
| 11_993146 | 1 | 1 | NA | NA | 1 | 1 | NA | NA | 2 | 1 | NA | NA | 1 | 1 | NA | NA | 2 | 1 | 3 | 1 |
| 11_993148 | 1 | 1 | NA | NA | 2 | 1 | NA | NA | 3 | 1 | NA | NA | 1 | 1 | NA | NA | 2 | 1 | 3 | 1 |
| 11_993149 | 1 | 1 | NA | NA | 2 | 1 | NA | NA | 3 | 1 | NA | NA | 1 | 1 | NA | NA | 1 | 1 | 2 | 1 |
| 11_993150 | 1 | 1 | NA | NA | 1 | 1 | NA | NA | 2 | 1 | NA | NA | 1 | 1 | NA | NA | 2 | 1 | 3 | 1 |
| 11_993152 | 1 | 1 | NA | NA | 2 | 1 | NA | NA | 3 | 1 | NA | NA | 1 | 1 | NA | NA | 2 | 1 | 3 | 1 |
| 11_993152 | 1 | 1 | NA | NA | 1 | 1 | NA | NA | 2 | 1 | NA | NA | 1 | 1 | NA | NA | 1 | 1 | 2 | 1 |
| 11_993153 | 1 | 1 | NA | NA | 2 | 0.5 | NA | NA | 3 | 0.6667 | NA | NA | 1 | 1 | NA | NA | 2 | 0.5 | 3 | 0.6667 |
| 11_993153 | NA | NA | NA | NA | 2 | 0.5 | NA | NA | 2 | 0.5 | NA | NA | 1 | 1 | NA | NA | 1 | 1 | 2 | 1 |
| 11_993153 | 1 | 1 | NA | NA | 2 | 1 | NA | NA | 3 | 1 | NA | NA | NA | NA | NA | NA | 2 | 0.5 | 2 | 0.5 |
| 11_993153 | 1 | 1 | NA | NA | 2 | 1 | NA | NA | 3 | 1 | NA | NA | 1 | 1 | NA | NA | 1 | 1 | 2 | 1 |
| 11_993153 | NA | NA | NA | NA | 1 | 1 | NA | NA | 1 | 1 | NA | NA | 1 | 1 | NA | NA | 2 | 0.5 | 3 | 0.6667 |
| 11_993155 | 1 | 0 | 1 | 1 | 1 | 1 | NA | NA | 3 | 0.6667 | NA | NA | 1 | 0 | NA | NA | 2 | 1 | 3 | 0.6667 |
| 11_993159 | 1 | 0 | 2 | 0.5 | 2 | 0 | 1 | 0 | 6 | 0.1667 | NA | NA | 1 | 0 | 1 | 0 | 3 | 0.6667 | 5 | 0.4 |
| 11_993289 | NA | NA | NA | NA | 1 | 1 | 1 | 1 | 2 | 1 | NA | NA | NA | NA | 1 | 1 | NA | NA | 1 | 1 |
| 11_100212 | 120 | 0.0583 | 121 | 0.0413 | 126 | 0.0873 | 134 | 0.0597 | 501 | 0.0619 | 103 | 0.0583 | 162 | 0.0494 | 145 | 0.0483 | 148 | 0.0541 | 558 | 0.052 |
| 11_100212 | 67 | 0.2985 | 47 | 0.1702 | 48 | 0.3333 | 72 | 0.3889 | 234 | 0.3077 | 42 | 0.381 | 60 | 0.3667 | 55 | 0.2545 | 67 | 0.4776 | 224 | 0.375 |
| 11_100238 | 12 | 0 | 17 | 0.1176 | 14 | 0.1429 | 15 | 0.0667 | 58 | 0.0862 | 14 | 0 | 13 | 0.2308 | 25 | 0.08 | 15 | 0.2667 | 67 | 0.1343 |
| 11_100238 | 11 | 0.0909 | 14 | 0 | 17 | 0.1176 | 12 | 0.25 | 54 | 0.1111 | 15 | 0 | 12 | 0.0833 | 23 | 0.087 | 15 | 0.1333 | 65 | 0.0769 |
| 11_100238 | 10 | 0 | 13 | 0.0769 | 17 | 0.1176 | 13 | 0.1538 | 53 | 0.0943 | 14 | 0 | 13 | 0.0769 | 25 | 0.12 | 15 | 0.2 | 67 | 0.1045 |
| 11_100238 | 11 | 0 | 13 | 0.0769 | 19 | 0.1053 | 12 | 0.1667 | 55 | 0.0909 | 11 | 0.0909 | 14 | 0.0714 | 25 | 0.12 | 14 | 0.2143 | 64 | 0.125 |
| 11_100238 | 11 | 0.5455 | 13 | 0.5385 | 16 | 0.4375 | 11 | 0.7273 | 51 | 0.549 | 10 | 0.6 | 12 | 0.6667 | 22 | 0.5909 | 9 | 0.7778 | 53 | 0.6415 |
| 11_100239 | 6 | 0 | 4 | 0 | 12 | 0.1667 | 8 | 0.125 | 30 | 0.1 | 7 | 0.1429 | 7 | 0 | 11 | 0.0909 | 7 | 0.1429 | 32 | 0.0938 |
| 11_100239 | 11 | 0.0909 | 14 | 0 | 14 | 0.1429 | 11 | 0.1818 | 50 | 0.1 | 9 | 0.1111 | 10 | 0.1 | 21 | 0.0952 | 8 | 0.25 | 48 | 0.125 |
| 11_100239 | 9 | 0.1111 | 10 | 0 | 6 | 0 | 6 | 0.1667 | 31 | 0.0645 | 6 | 0 | 6 | 0.1667 | 11 | 0.0909 | 4 | 0 | 27 | 0.0741 |
| 11_100239 | 8 | 0.25 | 11 | 0.0909 | 9 | 0.2222 | 10 | 0.1 | 38 | 0.1579 | 6 | 0 | 7 | 0.1429 | 15 | 0.0667 | 5 | 0 | 33 | 0.0606 |
| 11_100239 | 5 | 0.2 | 6 | 0.1667 | 7 | 0.2857 | 4 | 0 | 22 | 0.1818 | 4 | 0.25 | 9 | 0 | 12 | 0.0833 | 5 | 0.2 | 30 | 0.1 |
| 11_100239 | 8 | 0.75 | 7 | 0.8571 | 6 | 0.6667 | 5 | 1 | 26 | 0.8077 | 3 | 1 | 9 | 0.7778 | 9 | 0.7778 | 3 | 1 | 24 | 0.8333 |
| 11_100240 | 5 | 0 | 5 | 0 | 5 | 0 | 5 | 0.2 | 20 | 0.05 | 4 | 0 | 8 | 0.25 | 8 | 0 | 3 | 0 | 23 | 0.087 |
| 11_100240 | 8 | 0.625 | 6 | 0.6667 | 7 | 0.4286 | 8 | 0.625 | 29 | 0.5862 | 4 | 0.25 | 6 | 0.3333 | 10 | 0.7 | 4 | 0.5 | 24 | 0.5 |
| 11_100240 | 5 | 0.4 | 6 | 0.1667 | 3 | 0.3333 | 8 | 0.125 | 22 | 0.2273 | 4 | 0 | 5 | 0.4 | 8 | 0.5 | 3 | 0.3333 | 20 | 0.35 |
| 11_100252 | 3 | 0 | 13 | 0.2308 | 8 | 0 | 7 | 0.1429 | 31 | 0.129 | 4 | 0 | 9 | 0.3333 | 4 | 0 | 5 | 0.2 | 22 | 0.1818 |
| 11_100261 | 3 | 0.6667 | 10 | 0 | 4 | 0 | 6 | 0.3333 | 23 | 0.1739 | 4 | 0 | 1 | 0 | 12 | 0 | 7 | 0.2857 | 24 | 0.0833 |
| 11_100262 | 2 | 0 | 11 | 0.0909 | 4 | 0.5 | 7 | 0.1429 | 24 | 0.1667 | 3 | 0 | 3 | 0 | 12 | 0.4167 | 7 | 0.2857 | 25 | 0.28 |
| 11_100268 | 3 | 0.3333 | 7 | 0.1429 | 1 | 0 | 5 | 0 | 16 | 0.125 | 2 | 0.5 | 3 | 0.3333 | 10 | 0 | 3 | 0.6667 | 18 | 0.2222 |
| 11_100273 | 3 | 0 | 5 | 0.2 | 1 | 0 | 2 | 0 | 11 | 0.0909 | 1 | 0 | 1 | 0 | 7 | 0.1429 | 2 | 0 | 11 | 0.0909 |
| 11_100299 | 2 | 0.5 | 4 | 0 | 1 | 0 | 2 | 0 | 9 | 0.1111 | 1 | 1 | 1 | 0 | NA | NA | 1 | 0 | 3 | 0.3333 |
| 11_100303 | NA | NA | 2 | 0 | 1 | 1 | 2 | 1 | 5 | 0.6 | 1 | 0 | 1 | 1 | NA | NA | NA | NA | 2 | 0.5 |
| 11_100304 | NA | NA | 3 | 0 | 1 | 1 | 2 | 0.5 | 6 | 0.3333 | 1 | 1 | 1 | 0 | NA | NA | 1 | 0 | 3 | 0.3333 |
| 11_100304 | NA | NA | 2 | 0.5 | 1 | 1 | 2 | 0 | 5 | 0.4 | 1 | 0 | 1 | 1 | NA | NA | NA | NA | 2 | 0.5 |
| 11_100315 | 5 | 0.2 | NA | NA | 1 | 1 | NA | NA | 6 | 0.3333 | 1 | 0 | NA | NA | 2 | 0.5 | 2 | 0 | 5 | 0.2 |
| 11_100320 | 3 | 0.3333 | 5 | 0.2 | 3 | 0.3333 | 8 | 0.375 | 19 | 0.3158 | 2 | 0.5 | 2 | 0.5 | 3 | 0.6667 | 2 | 1 | 9 | 0.6667 |
| 11_100320 | 4 | 0.25 | 8 | 0 | 2 | 0 | 6 | 0.1667 | 20 | 0.1 | 2 | 0 | 1 | 0 | 6 | 0.1667 | 1 | 0 | 10 | 0.1 |

| | | | | | | | | | | | | | | | | | | | | |
|---|---|---|---|---|---|---|---|---|---|---|---|---|---|---|---|---|---|---|---|---|
| 11_1003206 | 3 | 0 | 5 | 0.2 | 4 | 0 | 7 | 0.1429 | 19 | 0.1053 | 2 | 0.5 | 2 | 0 | 6 | 0.6667 | 2 | 0 | 12 | 0.4167 |
| 11_1003207 | 3 | 0.3333 | 7 | 0.1429 | 5 | 0 | 8 | 0 | 23 | 0.087 | 3 | 0 | 3 | 0 | 6 | 0.1667 | 2 | 0.5 | 14 | 0.1429 |
| 11_1003483 | 13 | 0.3077 | 18 | 0.3889 | 6 | 0.5 | 25 | 0.2 | 62 | 0.3065 | 11 | 0.5455 | 14 | 0.3571 | 10 | 0.5 | 13 | 0.3846 | 48 | 0.4375 |
| 11_1003528 | 4 | 0.25 | 2 | 0 | 2 | 0 | 4 | 0 | 12 | 0.0833 | 1 | 1 | 2 | 0 | 1 | 0 | 5 | 0 | 9 | 0.1111 |
| 11_1003533 | 2 | 0.5 | 2 | 0.5 | 2 | 0 | 5 | 0.2 | 11 | 0.2727 | 1 | 0 | 3 | 0 | NA | NA | 4 | 0.5 | 8 | 0.25 |
| 11_1003535 | 3 | 1 | 1 | 1 | 1 | 1 | 4 | 1 | 9 | 1 | 1 | 1 | 3 | 1 | 2 | 1 | 4 | 1 | 10 | 1 |
| 11_1003536 | 3 | 0.3333 | 2 | 0.5 | 1 | 1 | 4 | 0.5 | 10 | 0.5 | 1 | 0 | 3 | 0 | 2 | 0.5 | 5 | 0.8 | 11 | 0.4545 |
| 11_1003536 | 3 | 1 | 2 | 1 | 1 | 1 | 4 | 1 | 10 | 1 | 1 | 1 | 3 | 1 | 2 | 1 | 5 | 1 | 11 | 1 |
| 11_1003536 | 2 | 1 | 2 | 1 | 1 | 1 | 2 | 1 | 7 | 1 | 1 | 1 | 1 | 1 | 1 | 1 | 3 | 0.3333 | 6 | 0.6667 |
| 11_1003537 | 3 | 1 | 2 | 0.5 | 1 | 1 | 4 | 0.25 | 10 | 0.6 | 1 | 0 | 2 | 0.5 | 2 | 0.5 | 5 | 0.4 | 10 | 0.4 |
| 11_1003539 | 1 | 0 | NA | NA | 1 | 1 | NA | NA | 2 | 0.5 | NA | NA | 2 | 0.5 | NA | NA | 2 | 0 | 4 | 0.25 |
| 11_1003539 | 2 | 0.5 | NA | NA | NA | NA | 2 | 0 | 4 | 0.25 | 1 | 0 | 2 | 1 | 2 | 1 | 4 | 0.25 | 9 | 0.5556 |
| 11_1003541 | 1 | 1 | 1 | 0 | 1 | 1 | 2 | 0.5 | 5 | 0.6 | NA | NA | 3 | 0.3333 | 1 | 0 | 2 | 0 | 6 | 0.1667 |
| 11_1003541 | 1 | 0 | 1 | 1 | 1 | 1 | 4 | 0.5 | 7 | 0.5714 | 1 | 0 | 2 | 0.5 | 2 | 0.5 | 2 | 0 | 7 | 0.2857 |
| 11_1003542 | 3 | 0.6667 | NA | NA | 1 | 1 | 4 | 0.25 | 8 | 0.5 | 1 | 1 | 3 | 0.6667 | 2 | 1 | 4 | 0.5 | 10 | 0.7 |
| 11_1003542 | 2 | 1 | NA | NA | NA | NA | 3 | 1 | 5 | 1 | 1 | 0 | 3 | 1 | 1 | 1 | 1 | 1 | 6 | 0.8333 |
| 11_1003544 | 1 | 1 | NA | NA | 1 | 1 | 2 | 1 | 4 | 1 | 1 | 1 | 2 | 1 | 1 | 1 | 3 | 1 | 7 | 1 |
| 11_1003544 | 2 | 0.5 | NA | NA | 1 | 0 | 3 | 1 | 6 | 0.6667 | 1 | 0 | 3 | 0 | 2 | 0.5 | 3 | 0.3333 | 9 | 0.2222 |
| 11_1003545 | 2 | 1 | NA | NA | NA | NA | 3 | 0 | 5 | 0.4 | 1 | 0 | 3 | 1 | 1 | 1 | 4 | 0.75 | 9 | 0.7778 |
| 11_1003546 | 2 | 0.5 | NA | NA | NA | NA | 3 | 1 | 5 | 0.8 | 1 | 1 | 3 | 0 | 2 | 1 | 4 | 1 | 10 | 0.7 |
| 11_1003547 | 2 | 1 | NA | NA | 1 | 1 | 3 | 0.6667 | 6 | 0.8333 | NA | NA | 3 | 0.3333 | 2 | 1 | 4 | 0.75 | 9 | 0.6667 |
| 11_1003550 | 2 | 1 | NA | NA | 1 | 1 | 3 | 1 | 6 | 1 | 1 | 1 | 3 | 0.3333 | 2 | 0.5 | 4 | 0.5 | 10 | 0.5 |
| 11_1003550 | 2 | 1 | NA | NA | 1 | 1 | 2 | 0.5 | 5 | 0.8 | 1 | 1 | 3 | 1 | 2 | 1 | 4 | 0.75 | 10 | 0.9 |
| 11_1003680 | 9 | 0 | 15 | 0.2667 | 9 | 0 | 16 | 0.125 | 49 | 0.1224 | 9 | 0.2222 | 16 | 0 | 17 | 0.0588 | 16 | 0.0625 | 58 | 0.069 |
| 11_1004624 | NA | NA | NA | NA | 1 | 1 | NA | NA | 1 | 1 | NA | NA | NA | NA | NA | NA | 1 | 1 | 1 | 1 |
| 11_1004626 | NA | NA | NA | NA | 1 | 1 | NA | NA | 1 | 1 | NA | NA | NA | NA | NA | NA | 1 | 1 | 1 | 1 |
| 11_1004669 | 8 | 0.125 | 7 | 0.2857 | 5 | 0 | 12 | 0.0833 | 32 | 0.125 | 9 | 0.2222 | 3 | 0 | 6 | 0 | 2 | 0 | 20 | 0.1 |
| 11_1004669 | 7 | 0.1429 | 7 | 0.5714 | 7 | 0.7143 | 11 | 0.5455 | 32 | 0.5 | 6 | 0.5 | 3 | 0.3333 | 4 | 0.5 | 3 | 0.6667 | 16 | 0.5 |
| 11_1004673 | 22 | 0.0455 | 27 | 0.1111 | 21 | 0.0476 | 24 | 0.0833 | 94 | 0.0745 | 18 | 0.1111 | 14 | 0.2857 | 18 | 0 | 10 | 0 | 60 | 0.1 |
| 11_1004675 | 22 | 0.2727 | 31 | 0.4194 | 25 | 0.16 | 24 | 0.4167 | 102 | 0.3235 | 26 | 0.3462 | 16 | 0.1875 | 21 | 0.2381 | 10 | 0.2 | 73 | 0.2603 |
| 11_1004676 | 27 | 0.963 | 29 | 0.8966 | 29 | 0.8276 | 27 | 0.963 | 112 | 0.9107 | 27 | 0.8889 | 15 | 0.8667 | 22 | 0.8182 | 9 | 0.8889 | 73 | 0.863 |
| 11_1004713 | 25 | 0.08 | 36 | 0.0833 | 29 | 0.1379 | 30 | 0.2 | 120 | 0.125 | 24 | 0.125 | 17 | 0.0588 | 24 | 0.1667 | 12 | 0.0833 | 77 | 0.1169 |
| 11_1004718 | 29 | 0.1379 | 41 | 0.0976 | 39 | 0.0256 | 26 | 0.0385 | 135 | 0.0741 | 33 | 0.0606 | 19 | 0 | 31 | 0.0645 | 15 | 0.1333 | 98 | 0.0612 |
| 11_1004719 | 36 | 0.2778 | 41 | 0.2683 | 42 | 0.1667 | 31 | 0.1613 | 150 | 0.22 | 33 | 0.3333 | 16 | 0.4375 | 32 | 0.25 | 16 | 0.25 | 97 | 0.3093 |
| 11_1004724 | 37 | 0.1622 | 45 | 0.2222 | 47 | 0.3404 | 43 | 0.2326 | 172 | 0.2442 | 34 | 0.2647 | 22 | 0.1818 | 37 | 0.2162 | 14 | 0.2143 | 107 | 0.2243 |
| 11_1004725 | 35 | 0.1429 | 44 | 0.1136 | 49 | 0.1224 | 42 | 0.119 | 170 | 0.1235 | 38 | 0.2895 | 19 | 0.1053 | 37 | 0.0811 | 22 | 0.1364 | 116 | 0.1638 |
| 11_1004738 | 6 | 0.3333 | 12 | 0.25 | 14 | 0.1429 | 3 | 0.6667 | 35 | 0.2571 | 7 | 0.4286 | 10 | 0.1 | 10 | 0.1 | 8 | 0.125 | 35 | 0.1714 |
| 11_1004879 | NA | NA | NA | NA | 3 | 0.3333 | NA | NA | 3 | 0.3333 | NA | NA | 1 | 1 | NA | NA | 1 | 0 | 2 | 0.5 |
| 11_1005180 | 1 | 0 | 1 | 0 | 1 | 0 | 6 | 0.5 | 9 | 0.3333 | 1 | 1 | 4 | 0.25 | 3 | 0.6667 | NA | NA | 8 | 0.5 |
| 11_1005859 | 1 | 1 | 1 | 1 | NA | NA | NA | NA | 2 | 1 | NA | NA | NA | NA | 1 | 1 | NA | NA | 1 | 1 |
| 11_1005956 | 9 | 0 | 9 | 0.1111 | 13 | 0.0769 | 13 | 0.1538 | 44 | 0.0909 | 4 | 0 | 7 | 0 | 11 | 0.1818 | 5 | 0 | 27 | 0.0741 |
| 11_1005957 | 9 | 0.2222 | 6 | 0.3333 | 12 | 0.25 | 13 | 0.0769 | 40 | 0.2 | 4 | 0.25 | 7 | 0 | 11 | 0.3636 | 5 | 0.2 | 27 | 0.2222 |
| 11_1005957 | 8 | 0.5 | 8 | 0.5 | 12 | 0.4167 | 12 | 0.75 | 40 | 0.55 | 4 | 0.5 | 7 | 0.4286 | 9 | 0.6667 | 5 | 0.6 | 25 | 0.56 |
| 11_1005958 | 8 | 1 | 7 | 1 | 9 | 0.7778 | 10 | 1 | 34 | 0.9412 | 3 | 1 | 5 | 0.8 | 7 | 1 | 4 | 1 | 19 | 0.9474 |
| 11_1005959 | 9 | 0.8889 | 8 | 1 | 11 | 0.8182 | 10 | 0.8 | 38 | 0.8684 | 4 | 1 | 7 | 1 | 9 | 0.8889 | 4 | 1 | 24 | 0.9583 |
| 11_1005959 | 10 | 0.1 | 9 | 0.1111 | 13 | 0.1538 | 10 | 0.4 | 42 | 0.1905 | 3 | 1 | 7 | 0 | 10 | 0.1 | 5 | 0.6 | 25 | 0.28 |
| 11_1005959 | 8 | 0.625 | 6 | 0.8333 | 10 | 0.6 | 10 | 0.6 | 34 | 0.6471 | 4 | 1 | 4 | 0.25 | 8 | 1 | 2 | 0.5 | 18 | 0.7778 |
| 11_1006058 | 3 | 0 | 3 | 0.3333 | 8 | 0.5 | 1 | 1 | 15 | 0.4 | 1 | 0 | 1 | 0 | 3 | 0.3333 | 1 | 0 | 6 | 0.1667 |
| 11_1006065 | 27 | 0 | 25 | 0.2 | 18 | 0.1667 | 18 | 0.1111 | 88 | 0.1136 | 17 | 0 | 12 | 0.1667 | 34 | 0.0588 | 21 | 0.0476 | 84 | 0.0595 |
| 11_1006321 | 2 | 0 | NA | NA | 3 | 0.3333 | 2 | 0.5 | 7 | 0.2857 | 2 | 0 | 2 | 0.5 | 2 | 0 | NA | NA | 6 | 0.1667 |
| 11_1006322 | 1 | 1 | 1 | 1 | 4 | 1 | 2 | 1 | 8 | 1 | 2 | 1 | 2 | 1 | 4 | 1 | 1 | 1 | 9 | 1 |
| 11_1006323 | 2 | 0.5 | 1 | 0 | 4 | 0.75 | 2 | 1 | 9 | 0.6667 | 2 | 0 | 1 | 0 | 4 | 0.5 | 1 | 1 | 8 | 0.375 |
| 11_1006325 | 2 | 0 | 1 | 0 | 4 | 0.25 | 2 | 0 | 9 | 0.1111 | 2 | 0 | 2 | 0 | 4 | 0.25 | 1 | 0 | 9 | 0.1111 |
| 11_1006325 | 2 | 1 | 1 | 0 | 3 | 0.3333 | 2 | 0.5 | 8 | 0.5 | 2 | 0 | 2 | 0.5 | 4 | 0.25 | NA | NA | 8 | 0.25 |
| 11_1006326 | 2 | 0.5 | 1 | 0 | 2 | 0 | 2 | 1 | 7 | 0.4286 | 1 | 1 | 1 | 0 | 4 | 0.75 | 1 | 1 | 7 | 0.7143 |
| 11_1006574 | NA | NA | NA | NA | 3 | 0.6667 | 1 | 1 | 4 | 0.75 | NA | NA | 1 | 0 | 1 | 1 | NA | NA | 2 | 0.5 |
| 11_1006575 | NA | NA | NA | NA | 3 | 0.6667 | 1 | 1 | 4 | 0.75 | NA | NA | 1 | 1 | 1 | 0 | NA | NA | 2 | 0.5 |
| 11_1006735 | 2 | 0.5 | NA | NA | 2 | 0 | NA | NA | 4 | 0.25 | NA | NA | NA | NA | 3 | 0.3333 | NA | NA | 3 | 0.3333 |
| 11_1006739 | 1 | 1 | NA | NA | 1 | 1 | NA | NA | 2 | 1 | NA | NA | NA | NA | 3 | 0.6667 | NA | NA | 3 | 0.6667 |
| 11_1007781 | 3 | 0 | 2 | 0 | 5 | 0.4 | 5 | 0 | 15 | 0.1333 | 2 | 0 | 6 | 0 | NA | NA | 8 | 0.125 | 16 | 0.0625 |
| 11_1008479 | 3 | 0.3333 | 5 | 0.6 | 2 | 0 | 3 | 0.3333 | 13 | 0.3846 | NA | NA | 4 | 0.25 | NA | NA | NA | NA | 4 | 0.25 |
| 11_1008482 | 3 | 0 | 2 | 0 | 1 | 0 | 2 | 0.5 | 8 | 0.125 | NA | NA | 3 | 0.3333 | NA | NA | NA | NA | 3 | 0.3333 |
| 11_1008491 | 1 | 1 | 3 | 0.6667 | 1 | 1 | 2 | 1 | 7 | 0.8571 | NA | NA | 1 | 1 | NA | NA | NA | NA | 1 | 1 |
| 11_1009119 | 13 | 0.1538 | 16 | 0 | 5 | 0 | 15 | 0.0667 | 49 | 0.0612 | 8 | 0.125 | 29 | 0.0345 | 19 | 0.0526 | 19 | 0.1053 | 75 | 0.0667 |
| 11_1009531 | 28 | 0.0714 | 33 | 0.0606 | 21 | 0.0476 | 33 | 0.0606 | 115 | 0.0609 | 16 | 0.25 | 40 | 0.15 | 30 | 0.0667 | 28 | 0.1429 | 114 | 0.1404 |

| | | | | | | | | | | | | | | | | | | | | |
|---|---|---|---|---|---|---|---|---|---|---|---|---|---|---|---|---|---|---|---|---|
| 11_100953 | 22 | 0.0909 | 38 | 0.1579 | 27 | 0.2222 | 32 | 0.1875 | 119 | 0.1681 | 16 | 0.1875 | 34 | 0.0294 | 31 | 0.0968 | 27 | 0.1111 | 108 | 0.0926 |
| 11_100953 | 35 | 0.4 | 41 | 0.2195 | 33 | 0.2727 | 46 | 0.2826 | 155 | 0.2903 | 22 | 0.3182 | 46 | 0.1522 | 37 | 0.1351 | 36 | 0.2222 | 141 | 0.1915 |
| 11_100967 | 3 | 0 | 1 | 0 | NA | NA | 1 | 1 | 5 | 0.2 | 2 | 1 | 1 | 1 | 2 | 0 | 1 | 1 | 6 | 0.6667 |
| 11_100967 | 2 | 0 | 1 | 0 | NA | NA | 2 | 0.5 | 5 | 0.2 | 1 | 0 | 1 | 0 | 2 | 0.5 | 1 | 1 | 5 | 0.4 |
| 11_100967 | 2 | 1 | 1 | 1 | NA | NA | 2 | 0.5 | 5 | 0.8 | 2 | 1 | NA | NA | 1 | 1 | 1 | 1 | 4 | 1 |
| 11_100967 | 3 | 0.3333 | 2 | 0 | NA | NA | 2 | 0 | 7 | 0.1429 | 3 | 0.3333 | 2 | 0 | 2 | 0 | 1 | 0 | 8 | 0.125 |
| 11_100967 | 3 | 1 | 2 | 0.5 | NA | NA | 2 | 1 | 7 | 0.8571 | 3 | 0.6667 | 2 | 0.5 | 2 | 1 | 1 | 1 | 8 | 0.75 |
| 11_100967 | 3 | 1 | 2 | 1 | NA | NA | 2 | 1 | 7 | 1 | 2 | 0.5 | 2 | 1 | 2 | 0.5 | 1 | 0 | 7 | 0.5714 |
| 11_100967 | 1 | 1 | 2 | 0.5 | NA | NA | NA | NA | 3 | 0.6667 | 2 | 1 | 1 | 0 | 2 | 0.5 | 1 | 1 | 6 | 0.6667 |
| 11_100968 | 2 | 1 | 2 | 1 | NA | NA | 1 | 1 | 5 | 1 | 3 | 1 | 2 | 1 | 2 | 1 | 1 | 1 | 8 | 1 |
| 11_100968 | 3 | 1 | 2 | 0.5 | NA | NA | 2 | 1 | 7 | 0.8571 | 3 | 0.3333 | 3 | 1 | 2 | 0.5 | 1 | 0 | 9 | 0.5556 |
| 11_100968 | 3 | 0.3333 | 2 | 0 | NA | NA | 3 | 0 | 8 | 0.125 | 3 | 0.3333 | 2 | 0 | 1 | 0 | 1 | 1 | 7 | 0.2857 |
| 11_100968 | 1 | 1 | 2 | 0 | NA | NA | 3 | 0.6667 | 6 | 0.5 | 3 | 0.3333 | 2 | 0 | 2 | 0 | 1 | 0 | 8 | 0.125 |
| 11_100968 | 3 | 0 | 2 | 0.5 | NA | NA | 3 | 0.3333 | 8 | 0.25 | 3 | 0 | 3 | 0.3333 | 1 | 0 | 1 | 0 | 8 | 0.125 |
| 11_100968 | 3 | 0 | 2 | 0 | NA | NA | 3 | 0.3333 | 8 | 0.125 | 3 | 0 | 3 | 0.3333 | 2 | 0 | 1 | 0 | 9 | 0.1111 |
| 11_100968 | 3 | 0.3333 | 2 | 0 | NA | NA | 2 | 0 | 7 | 0.1429 | 2 | 0 | 3 | 0.3333 | 2 | 0 | 1 | 1 | 8 | 0.25 |
| 11_100968 | 2 | 0 | 2 | 0.5 | NA | NA | 3 | 0.3333 | 7 | 0.2857 | 3 | 0.3333 | 3 | 0.3333 | 2 | 1 | 1 | 0 | 9 | 0.4444 |
| 11_100973 | 1 | 0 | 2 | 0.5 | NA | NA | NA | NA | 3 | 0.3333 | NA | NA | 2 | 0.5 | NA | NA | NA | NA | 2 | 0.5 |
| 11_100973 | NA | NA | 1 | 0 | 2 | 0.5 | NA | NA | 3 | 0.3333 | NA | NA | 2 | 0.5 | 1 | 1 | 1 | 0 | 4 | 0.5 |
| 11_100973 | 2 | 0 | 2 | 0.5 | 1 | 0 | NA | NA | 5 | 0.2 | 1 | 0 | 4 | 0.25 | 2 | 0 | NA | NA | 7 | 0.1429 |
| 11_100974 | 2 | 1 | 2 | 1 | 2 | 1 | NA | NA | 6 | 1 | 1 | 0 | 4 | 1 | 3 | 1 | 1 | 1 | 9 | 0.8889 |
| 11_100974 | 2 | 1 | 2 | 1 | 1 | 1 | NA | NA | 5 | 1 | 1 | 0 | 3 | 1 | 2 | 1 | 1 | 1 | 7 | 0.8571 |
| 11_100974 | 2 | 0 | 2 | 0.5 | 2 | 0 | NA | NA | 6 | 0.1667 | 1 | 0 | 4 | 0.25 | 4 | 0 | 1 | 0 | 10 | 0.1 |
| 11_100975 | 2 | 0 | 2 | 0.5 | 2 | 0 | NA | NA | 6 | 0.1667 | 2 | 0 | 4 | 0.25 | 3 | 0 | 1 | 0 | 10 | 0.1 |
| 11_100976 | 1 | 0 | 3 | 0 | 4 | 0.25 | NA | NA | 8 | 0.125 | 2 | 0 | 3 | 0.3333 | 4 | 0.5 | NA | NA | 9 | 0.3333 |
| 11_100977 | 2 | 0 | 3 | 0.3333 | 5 | 0.2 | NA | NA | 10 | 0.2 | 1 | 0 | 3 | 0.3333 | 5 | 0.2 | 1 | 0 | 10 | 0.2 |
| 11_100977 | 2 | 0 | 3 | 0 | 2 | 0.5 | 1 | 0 | 8 | 0.125 | 2 | 0 | 4 | 0 | 5 | 0.2 | 2 | 0 | 13 | 0.0769 |
| 11_100977 | 1 | 1 | 3 | 0.6667 | 4 | 0.5 | 2 | 0 | 10 | 0.5 | 1 | 1 | 4 | 0.25 | 4 | 0.5 | 2 | 0.5 | 11 | 0.4545 |
| 11_100982 | 7 | 0.2857 | 9 | 0 | 6 | 0 | 2 | 0 | 24 | 0.0833 | 6 | 0.1667 | 2 | 1 | 4 | 0.25 | 1 | 0 | 13 | 0.3077 |
| 11_100982 | 6 | 0.1667 | 7 | 0 | 4 | 0.25 | 4 | 0.5 | 21 | 0.1905 | 4 | 0 | 3 | 0.3333 | 4 | 0.75 | 1 | 0 | 12 | 0.3333 |
| 11_100988 | 3 | 0 | 2 | 0 | 3 | 0.3333 | 4 | 0 | 12 | 0.0833 | 2 | 0 | NA | NA | 2 | 0 | 7 | 0.2857 | 11 | 0.1818 |
| 11_100988 | 4 | 0 | 4 | 0.5 | 3 | 0 | 2 | 0 | 13 | 0.1538 | 1 | 1 | 2 | 0 | 3 | 0 | 8 | 0 | 14 | 0.0714 |
| 11_100988 | 5 | 1 | 3 | 1 | 3 | 0.6667 | 3 | 0.6667 | 14 | 0.8571 | 2 | 1 | 2 | 0.5 | 1 | 1 | 7 | 0.8571 | 12 | 0.8333 |
| 11_100989 | 4 | 0.5 | 4 | 0.5 | 3 | 0 | 3 | 0.6667 | 14 | 0.4286 | 2 | 1 | 1 | 1 | 3 | 0.3333 | 9 | 0.7778 | 15 | 0.7333 |
| 11_100989 | 3 | 1 | 3 | 0.6667 | 3 | 1 | 2 | 0.5 | 11 | 0.8182 | 2 | 0.5 | 2 | 0.5 | 3 | 1 | 8 | 0.875 | 15 | 0.8 |
| 11_100989 | 3 | 1 | 4 | 1 | 1 | 1 | 3 | 0.3333 | 11 | 0.8182 | 2 | 0.5 | 1 | 1 | 2 | 1 | 8 | 1 | 13 | 0.9231 |
| 11_100989 | 3 | 0.3333 | 4 | 0.25 | 3 | 0.3333 | 3 | 0 | 13 | 0.2308 | 2 | 0 | 1 | 0 | 2 | 0.5 | 8 | 0.125 | 13 | 0.1538 |
| 11_100989 | 1 | 0 | 4 | 0.25 | 2 | 0 | NA | NA | 7 | 0.1429 | NA | NA | 1 | 0 | 1 | 0 | 5 | 0.4 | 7 | 0.2857 |
| 11_101011 | 8 | 0.125 | 5 | 0 | 4 | 0 | 4 | 0.25 | 21 | 0.0952 | 3 | 0 | 2 | 0.5 | 3 | 0 | 5 | 0.2 | 13 | 0.1538 |
| 11_101016 | 8 | 0.25 | 8 | 0 | 7 | 0.1429 | 4 | 0.5 | 27 | 0.1852 | 7 | 0.4286 | 4 | 0 | 6 | 0.1667 | 6 | 0.5 | 23 | 0.3043 |
| 11_101016 | 8 | 0.25 | 10 | 0.1 | 9 | 0.2222 | 3 | 0 | 30 | 0.1667 | 6 | 0.1667 | 4 | 0.25 | 5 | 0 | 7 | 0 | 22 | 0.0909 |
| 11_101017 | 7 | 0 | 8 | 0.125 | 6 | 0.3333 | 1 | 0 | 22 | 0.1364 | 4 | 0 | 3 | 0 | 5 | 0.2 | 6 | 0.1667 | 18 | 0.1111 |
| 11_101017 | 8 | 0.375 | 10 | 0.1 | 8 | 0.375 | 4 | 0.5 | 30 | 0.3 | 7 | 0.5714 | 4 | 0.75 | 6 | 0.1667 | 8 | 0.125 | 25 | 0.36 |
| 11_101017 | 8 | 0.125 | 10 | 0.6 | 6 | 0.1667 | 4 | 0 | 28 | 0.2857 | 5 | 0.4 | 3 | 0.3333 | 6 | 0.5 | 8 | 0 | 22 | 0.2727 |
| 11_101052 | 6 | 0.1667 | 9 | 0.1111 | 8 | 0 | 6 | 0 | 29 | 0.069 | 5 | 0.2 | 3 | 0 | 9 | 0.1111 | 9 | 0 | 26 | 0.0769 |
| 11_101052 | 4 | 0.25 | 7 | 0.7143 | 8 | 0.5 | 6 | 0.3333 | 25 | 0.48 | 6 | 0.3333 | 3 | 0 | 8 | 0.25 | 8 | 0.5 | 25 | 0.32 |
| 11_101052 | 7 | 0.1429 | 8 | 0.125 | 10 | 0.4 | 6 | 0.1667 | 31 | 0.2258 | 5 | 0.2 | 3 | 0 | 11 | 0.4545 | 10 | 0.4 | 29 | 0.3448 |
| 11_101053 | 1 | 1 | 1 | 0 | NA | NA | NA | NA | 2 | 0.5 | 3 | 0.6667 | 1 | 0 | 1 | 0 | 1 | 0 | 6 | 0.3333 |
| 11_101053 | 2 | 1 | 1 | 1 | NA | NA | 2 | 0.5 | 5 | 0.8 | 3 | 0.6667 | 1 | 1 | 1 | 1 | 1 | 1 | 6 | 0.8333 |
| 11_101053 | 2 | 1 | 1 | 1 | NA | NA | 2 | 0.5 | 5 | 0.8 | 4 | 0.25 | 1 | 1 | 1 | 1 | 1 | 1 | 7 | 0.5714 |
| 11_101053 | 2 | 1 | 1 | 1 | NA | NA | 2 | 0.5 | 5 | 0.8 | 4 | 0.75 | 1 | 1 | 1 | 1 | 1 | 1 | 7 | 0.8571 |
| 11_101053 | 2 | 0 | 1 | 1 | NA | NA | 1 | 0 | 4 | 0.25 | 4 | 0.5 | 1 | 0 | NA | NA | NA | NA | 5 | 0.4 |
| 11_101053 | 1 | 0 | 1 | 0 | NA | NA | 1 | 1 | 3 | 0.3333 | 1 | 0 | NA | NA | 1 | 1 | 1 | 0 | 3 | 0.3333 |
| 11_101061 | 1 | 0 | 1 | 1 | NA | NA | 1 | 0 | 3 | 0.3333 | 1 | 1 | 1 | 1 | NA | NA | 1 | 0 | 3 | 0.6667 |
| 11_101069 | NA | NA | 2 | 0 | 2 | 0.5 | 1 | 0 | 5 | 0.2 | 3 | 0 | 1 | 0 | 2 | 0 | 3 | 0.3333 | 9 | 0.1111 |
| 11_101080 | 13 | 0.4615 | 12 | 0.3333 | 14 | 0.2857 | 4 | 0.5 | 43 | 0.3721 | 9 | 0.4444 | 15 | 0.2 | 11 | 0.1818 | 7 | 0.1429 | 42 | 0.2381 |
| 11_101080 | 12 | 0.0833 | 8 | 0.375 | 10 | 0.5 | 2 | 0 | 32 | 0.2812 | 7 | 0 | 10 | 0.3 | 11 | 0.2727 | 5 | 0.4 | 33 | 0.2424 |
| 11_101080 | 15 | 0.2667 | 13 | 0.0769 | 13 | 0.1538 | 4 | 0 | 45 | 0.1556 | 9 | 0.3333 | 16 | 0.375 | 20 | 0.15 | 7 | 0.2857 | 52 | 0.2692 |
| 11_101080 | 12 | 0.0833 | 13 | 0.0769 | 14 | 0.1429 | 5 | 0.2 | 44 | 0.1136 | 8 | 0 | 14 | 0.1429 | 15 | 0.0667 | 5 | 0.2 | 42 | 0.0952 |
| 11_101080 | 9 | 0 | 9 | 0.1111 | 6 | 0.1667 | 2 | 0 | 26 | 0.0769 | 6 | 0.1667 | 11 | 0.0909 | 13 | 0.1538 | 6 | 0 | 36 | 0.1111 |
| 11_101081 | 16 | 0 | 15 | 0.0667 | 17 | 0.1176 | 6 | 0 | 54 | 0.0556 | 10 | 0 | 15 | 0.0667 | 20 | 0.1 | 9 | 0 | 54 | 0.0556 |
| 11_101081 | 15 | 0.0667 | 14 | 0 | 20 | 0.05 | 9 | 0.1111 | 58 | 0.0517 | 9 | 0.1111 | 20 | 0.05 | 19 | 0 | 9 | 0.1111 | 57 | 0.0526 |
| 11_101081 | 14 | 0.0714 | 14 | 0.0714 | 17 | 0.1176 | 6 | 0 | 51 | 0.0784 | 9 | 0.1111 | 15 | 0.0667 | 19 | 0.0526 | 8 | 0.125 | 51 | 0.0784 |
| 11_101082 | 6 | 0.5 | 6 | 0 | 4 | 0.5 | 4 | 0.5 | 20 | 0.35 | 1 | 1 | 6 | 0 | 7 | 0.2857 | 2 | 1 | 16 | 0.3125 |
| 11_101083 | 4 | 0.25 | 5 | 0.4 | 1 | 0 | 2 | 0.5 | 12 | 0.3333 | NA | NA | 4 | 0.75 | 5 | 0.6 | 1 | 1 | 10 | 0.7 |

| | | | | | | | | | | | | | | | | | | | | |
|---|---|---|---|---|---|---|---|---|---|---|---|---|---|---|---|---|---|---|---|---|
| 11_101089 | 3 | 0.3333 | 2 | 0 | 1 | 0 | 3 | 0 | 9 | 0.1111 | 1 | 0 | 2 | 0.5 | 2 | 0.5 | 5 | 0.6 | 10 | 0.5 |
| 11_101090 | 3 | 0.3333 | 2 | 0.5 | 1 | 0 | 3 | 0 | 9 | 0.2222 | 1 | 0 | 1 | 1 | 2 | 0.5 | 5 | 0.6 | 9 | 0.5556 |
| 11_101090 | 7 | 0.2857 | 2 | 0.5 | 4 | 0 | 4 | 0 | 17 | 0.1765 | 1 | 1 | 5 | 0 | 6 | 0.1667 | 5 | 0.4 | 17 | 0.2353 |
| 11_101138 | NA | NA | NA | NA | 1 | 0 | 1 | 1 | 2 | 0.5 | 1 | 1 | 1 | 0 | NA | NA | NA | NA | 2 | 0.5 |
| 11_101138 | 3 | 0 | 4 | 0 | 6 | 0.1667 | 6 | 0.3333 | 19 | 0.1579 | 9 | 0 | 11 | 0.2727 | 7 | 0.1429 | 5 | 0 | 32 | 0.125 |
| 11_101138 | 2 | 0 | 4 | 0.5 | 3 | 0.3333 | 7 | 0.2857 | 16 | 0.3125 | 7 | 0.4286 | 4 | 0.5 | 4 | 0 | 3 | 0 | 18 | 0.2778 |
| 11_101149 | 11 | 0.3636 | 13 | 0 | 7 | 0.2857 | 10 | 0 | 41 | 0.1463 | 9 | 0 | 20 | 0.1 | 5 | 0.2 | 9 | 0.1111 | 43 | 0.093 |
| 11_101149 | 10 | 0.2 | 13 | 0.0769 | 6 | 0 | 9 | 0 | 38 | 0.0789 | 9 | 0.1111 | 18 | 0.1111 | 5 | 0 | 7 | 0.2857 | 39 | 0.1282 |
| 11_101154 | 1 | 1 | NA | NA | 2 | 0 | NA | NA | 3 | 0.3333 | NA | NA | 1 | 1 | NA | NA | NA | NA | 1 | 1 |
| 11_101166 | NA | NA | NA | NA | 3 | 0 | 2 | 0.5 | 5 | 0.2 | 1 | 1 | 1 | 1 | NA | NA | 1 | 0 | 3 | 0.6667 |
| 11_101167 | NA | NA | NA | NA | NA | NA | 2 | 0.5 | 2 | 0.5 | NA | NA | NA | NA | 1 | 1 | NA | NA | 1 | 1 |
| 11_101168 | NA | NA | 1 | 1 | NA | NA | NA | NA | 1 | 1 | NA | NA | NA | NA | 1 | 1 | NA | NA | 1 | 1 |
| 11_101168 | 20 | 0.1 | 17 | 0.0588 | 22 | 0.0455 | 10 | 0 | 69 | 0.058 | 10 | 0.1 | 19 | 0.0526 | 20 | 0 | 15 | 0.1333 | 64 | 0.0625 |
| 11_101169 | 21 | 1 | 13 | 1 | 18 | 1 | 12 | 0.9167 | 64 | 0.9844 | 13 | 1 | 20 | 1 | 21 | 0.9048 | 16 | 1 | 70 | 0.9714 |
| 11_101169 | 22 | 1 | 14 | 0.9286 | 19 | 0.8947 | 11 | 0.9091 | 66 | 0.9394 | 14 | 1 | 20 | 1 | 22 | 0.9091 | 16 | 1 | 72 | 0.9722 |
| 11_101169 | 18 | 0.1667 | 15 | 0.0667 | 17 | 0.0588 | 9 | 0.1111 | 59 | 0.1017 | 11 | 0 | 20 | 0.05 | 21 | 0.0952 | 15 | 0.1333 | 67 | 0.0746 |
| 11_101169 | 20 | 0.55 | 16 | 0.625 | 15 | 0.7333 | 10 | 0.6 | 61 | 0.623 | 14 | 0.6429 | 20 | 0.5 | 19 | 0.6842 | 13 | 0.8462 | 66 | 0.6515 |
| 11_101169 | 20 | 0 | 16 | 0.125 | 14 | 0.0714 | 6 | 0 | 56 | 0.0536 | 14 | 0.2143 | 19 | 0 | 19 | 0.1053 | 13 | 0.2308 | 65 | 0.1231 |
| 11_101169 | 23 | 0.0435 | 15 | 0.2 | 19 | 0.1053 | 10 | 0.1 | 67 | 0.1045 | 14 | 0.1429 | 22 | 0.1364 | 22 | 0.0455 | 15 | 0.0667 | 73 | 0.0959 |
| 11_101179 | 10 | 0 | 11 | 0.1818 | 13 | 0.1538 | 13 | 0 | 47 | 0.0851 | 12 | 0 | 16 | 0.125 | 10 | 0 | 12 | 0.1667 | 50 | 0.08 |
| 11_101179 | 6 | 0.1667 | 14 | 0.2857 | 13 | 0.1538 | 12 | 0.1667 | 45 | 0.2 | 13 | 0.2308 | 15 | 0.1333 | 10 | 0.1 | 11 | 0.3636 | 49 | 0.2041 |
| 11_101212 | 9 | 0 | 9 | 0.2222 | 9 | 0.1111 | 11 | 0.0909 | 38 | 0.1053 | 7 | 0.1429 | 11 | 0.1818 | 10 | 0.1 | 8 | 0.125 | 36 | 0.1389 |
| 11_101212 | 8 | 0.25 | 10 | 0.6 | 13 | 0.6923 | 12 | 0.5 | 43 | 0.5349 | 8 | 0.75 | 11 | 0.6364 | 10 | 0.5 | 10 | 0.6 | 39 | 0.6154 |
| 11_101297 | 12 | 0.1667 | 7 | 0.1429 | 8 | 0 | 7 | 0 | 34 | 0.0882 | 15 | 0.0667 | 8 | 0 | 10 | 0.1 | 13 | 0.1538 | 46 | 0.087 |
| 11_101297 | 15 | 0.0667 | 6 | 0.3333 | 9 | 0.4444 | 9 | 0.1111 | 39 | 0.2051 | 14 | 0.2143 | 9 | 0 | 9 | 0.5556 | 11 | 0.3636 | 43 | 0.2791 |
| 11_101309 | 2 | 0.5 | NA | NA | 1 | 0 | 4 | 0 | 7 | 0.1429 | NA | NA | 1 | 1 | NA | NA | NA | NA | 1 | 1 |
| 11_101316 | NA | NA | 1 | 1 | 1 | 0 | 1 | 0 | 3 | 0.3333 | NA | NA | NA | NA | 2 | 0 | 3 | 0.3333 | 5 | 0.2 |
| 11_101324 | NA | NA | 5 | 0.8 | 1 | 1 | 3 | 1 | 9 | 0.8889 | 1 | 1 | 2 | 1 | 3 | 1 | 3 | 0.6667 | 9 | 0.8889 |
| 11_101324 | NA | NA | 6 | 0.6667 | 1 | 1 | 3 | 1 | 10 | 0.8 | 1 | 0 | 3 | 1 | 4 | 0.5 | 4 | 1 | 12 | 0.75 |
| 11_101324 | NA | NA | 5 | 0.4 | NA | NA | 4 | 0 | 9 | 0.2222 | NA | NA | 5 | 0.2 | 4 | 0 | 2 | 0.5 | 11 | 0.1818 |
| 11_101324 | NA | NA | 5 | 1 | NA | NA | 4 | 0.75 | 9 | 0.8889 | 1 | 1 | 5 | 0.8 | 7 | 0.8571 | 4 | 0.75 | 17 | 0.8235 |
| 11_101324 | NA | NA | 5 | 0.8 | NA | NA | 4 | 1 | 9 | 0.8889 | 1 | 1 | 5 | 1 | 6 | 1 | 4 | 1 | 16 | 1 |
| 11_101324 | NA | NA | 5 | 0.2 | NA | NA | 3 | 0.3333 | 8 | 0.25 | 2 | 0 | 3 | 0.3333 | 7 | 0.1429 | 2 | 0 | 14 | 0.1429 |
| 11_101324 | 1 | 0 | 5 | 0.2 | NA | NA | 4 | 0 | 10 | 0.1 | 2 | 0 | 3 | 0 | 6 | 0.1667 | 4 | 0 | 15 | 0.0667 |
| 11_101324 | 1 | 0 | 3 | 0 | 1 | 0 | 4 | 0.25 | 9 | 0.1111 | 2 | 0 | 2 | 0 | 5 | 0.2 | 4 | 0.25 | 13 | 0.1538 |
| 11_101324 | 1 | 0 | 3 | 1 | 1 | 1 | 3 | 1 | 8 | 0.875 | 1 | 1 | 4 | 1 | 6 | 0.6667 | 3 | 0.6667 | 14 | 0.7857 |
| 11_101325 | 2 | 0.5 | 4 | 0.25 | 1 | 0 | 2 | 0 | 9 | 0.2222 | 3 | 0.6667 | 4 | 0.25 | 6 | 0.5 | 4 | 0.25 | 17 | 0.4118 |
| 11_101325 | 3 | 0.3333 | 5 | 0 | 1 | 0 | 4 | 0.25 | 13 | 0.1538 | 2 | 0 | 5 | 0 | 5 | 0.4 | 5 | 0 | 17 | 0.1176 |
| 11_101325 | NA | NA | 5 | 0 | 2 | 0 | 5 | 0.2 | 12 | 0.0833 | 4 | 0.25 | 4 | 0 | 7 | 0 | 3 | 0 | 18 | 0.0556 |
| 11_101325 | 2 | 1 | 6 | 0.3333 | 1 | 1 | 4 | 0.5 | 13 | 0.5385 | 3 | 0.3333 | 4 | 0.25 | 5 | 0 | 4 | 0.5 | 16 | 0.25 |
| 11_101325 | 3 | 0.3333 | 7 | 0.1429 | 2 | 0 | 4 | 0.5 | 16 | 0.25 | 2 | 0 | 4 | 0.25 | 6 | 0.1667 | 4 | 0 | 16 | 0.125 |
| 11_101325 | 3 | 1 | 5 | 0.6 | 2 | 1 | 5 | 0.8 | 15 | 0.8 | 3 | 1 | 2 | 1 | 4 | 1 | 3 | 1 | 12 | 1 |
| 11_101326 | 3 | 1 | 7 | 0.7143 | 2 | 1 | 6 | 1 | 18 | 0.8889 | 4 | 1 | 5 | 1 | 9 | 1 | 5 | 1 | 23 | 1 |
| 11_101326 | 3 | 1 | 8 | 0.25 | 2 | 1 | 5 | 0.8 | 18 | 0.6111 | 4 | 0.25 | 5 | 0.4 | 8 | 0.625 | 5 | 0.4 | 22 | 0.4545 |
| 11_101326 | 3 | 0 | 7 | 0.1429 | 2 | 0 | 5 | 0.2 | 17 | 0.1176 | 3 | 0 | 5 | 0.2 | 8 | 0.25 | 5 | 0 | 21 | 0.1429 |
| 11_101326 | 3 | 1 | 4 | 0.75 | NA | NA | 4 | 0.75 | 11 | 0.8182 | 4 | 0.5 | 4 | 0.75 | 5 | 0 | 3 | 0.6667 | 16 | 0.4375 |
| 11_101326 | 3 | 0 | 7 | 0.1429 | 2 | 0 | 6 | 0.1667 | 18 | 0.1111 | 4 | 0 | 4 | 0 | 9 | 0.1111 | 2 | 0.5 | 19 | 0.1053 |
| 11_101326 | 3 | 0.6667 | 8 | 0.75 | NA | NA | 4 | 1 | 15 | 0.8 | 4 | 1 | 6 | 1 | 6 | 1 | 5 | 1 | 21 | 1 |
| 11_101326 | 2 | 0 | 4 | 0.25 | 2 | 0 | 5 | 0 | 13 | 0.0769 | 2 | 0 | 6 | 0.3333 | 4 | 0.75 | 3 | 0 | 15 | 0.3333 |
| 11_101326 | 1 | 0 | 4 | 0.25 | 2 | 0.5 | 4 | 0.5 | 11 | 0.3636 | 3 | 0.3333 | 7 | 0.2857 | 8 | 0.25 | 4 | 0 | 22 | 0.2273 |
| 11_101350 | 6 | 0.3333 | 7 | 0.2857 | 3 | 0.6667 | 7 | 0.1429 | 23 | 0.3043 | 9 | 0.4444 | 3 | 0.3333 | 7 | 0.2857 | 5 | 0.2 | 24 | 0.3333 |
| 11_101362 | 15 | 0.0667 | 30 | 0.1 | 26 | 0 | 24 | 0.0833 | 95 | 0.0632 | 17 | 0.0588 | 24 | 0.0833 | 26 | 0 | 11 | 0.2727 | 78 | 0.0769 |
| 11_101362 | 18 | 0.8333 | 27 | 0.8148 | 20 | 0.75 | 22 | 0.8636 | 87 | 0.8161 | 15 | 0.8667 | 24 | 0.9167 | 22 | 0.7727 | 11 | 0.6364 | 72 | 0.8194 |
| 11_101362 | 18 | 0.1667 | 18 | 0 | 18 | 0.1111 | 17 | 0.1765 | 71 | 0.1127 | 9 | 0.2222 | 20 | 0.3 | 13 | 0.0769 | 10 | 0.2 | 52 | 0.2115 |
| 11_101362 | 22 | 0.1364 | 30 | 0.0667 | 28 | 0.0357 | 29 | 0.1034 | 109 | 0.0826 | 23 | 0.0435 | 29 | 0.069 | 26 | 0.0385 | 11 | 0.1818 | 89 | 0.0674 |
| 11_101466 | 9 | 0.3333 | 10 | 0.2 | 11 | 0.1818 | 10 | 0.4 | 40 | 0.275 | 8 | 0.5 | 2 | 1 | 8 | 0.125 | 8 | 0.25 | 26 | 0.3462 |
| 11_101466 | 8 | 0 | 10 | 0 | 14 | 0 | 8 | 0.25 | 40 | 0.05 | 8 | 0 | 2 | 0 | 8 | 0 | 11 | 0.1818 | 29 | 0.069 |
| 11_101468 | 1 | 0 | 1 | 1 | NA | NA | NA | NA | 2 | 0.5 | 2 | 0.5 | NA | NA | NA | NA | NA | NA | 2 | 0.5 |
| 11_101471 | 10 | 0.9 | 8 | 1 | 12 | 1 | 9 | 0.7778 | 39 | 0.9231 | 6 | 0.8333 | 6 | 1 | 8 | 1 | 11 | 0.7273 | 31 | 0.871 |
| 11_101471 | 8 | 0.125 | 12 | 0.1667 | 14 | 0.2857 | 5 | 0.2 | 39 | 0.2051 | 7 | 0.1429 | 7 | 0 | 8 | 0.5 | 12 | 0 | 34 | 0.1471 |
| 11_101471 | 9 | 0.5556 | 12 | 0.1667 | 16 | 0.25 | 12 | 0.3333 | 49 | 0.3061 | 5 | 0.2 | 5 | 0.6 | 9 | 0.3333 | 11 | 0.0909 | 30 | 0.2667 |
| 11_101471 | 10 | 0.1 | 10 | 0 | 13 | 0.3846 | 6 | 0 | 39 | 0.1538 | 5 | 0 | 6 | 0 | 7 | 0.1429 | 7 | 0.1429 | 25 | 0.08 |
| 11_101471 | 11 | 0.3636 | 12 | 0.0833 | 13 | 0 | 11 | 0.0909 | 47 | 0.1277 | 7 | 0.2857 | 7 | 0.1429 | 9 | 0.3333 | 13 | 0.0769 | 36 | 0.1944 |
| 11_101471 | 10 | 0.8 | 14 | 0.7143 | 15 | 0.6 | 13 | 0.6923 | 52 | 0.6923 | 9 | 0.7778 | 9 | 0.6667 | 7 | 0.8571 | 13 | 0.6923 | 38 | 0.7368 |
| 11_101472 | 10 | 0.1 | 17 | 0 | 16 | 0.125 | 17 | 0.0588 | 60 | 0.0667 | 8 | 0 | 8 | 0.25 | 13 | 0.1538 | 14 | 0 | 43 | 0.093 |

| | | | | | | | | | | | | | | | | | | | | |
|---|---|---|---|---|---|---|---|---|---|---|---|---|---|---|---|---|---|---|---|---|
| 11_101472 | 13 | 0.0769 | 19 | 0 | 15 | 0.1333 | 18 | 0.0556 | 65 | 0.0615 | 10 | 0.1 | 9 | 0.1111 | 11 | 0.0909 | 16 | 0.125 | 46 | 0.1087 |
| 11_101557 | 6 | 0.3333 | 4 | 0.5 | 4 | 0 | 7 | 0.1429 | 21 | 0.2381 | 10 | 0.2 | 10 | 0.2 | 4 | 0.75 | 2 | 0.5 | 26 | 0.3077 |
| 11_101557 | 7 | 0.4286 | 5 | 0.2 | 2 | 0.5 | 7 | 0.4286 | 21 | 0.381 | 9 | 1 | 9 | 0.3333 | 7 | 0.2857 | 2 | 1 | 27 | 0.5926 |
| 11_101557 | 9 | 1 | 5 | 1 | 4 | 1 | 8 | 1 | 26 | 1 | 9 | 1 | 10 | 0.9 | 8 | 0.875 | 3 | 1 | 30 | 0.9333 |
| 11_101557 | 10 | 0.9 | 4 | 1 | 3 | 1 | 9 | 1 | 26 | 0.9615 | 8 | 1 | 11 | 1 | 8 | 0.875 | 4 | 1 | 31 | 0.9677 |
| 11_101557 | 10 | 1 | 3 | 1 | 4 | 1 | 8 | 1 | 25 | 1 | 9 | 0.8889 | 13 | 1 | 8 | 1 | 3 | 1 | 33 | 0.9697 |
| 11_101557 | 7 | 0.4286 | 6 | 1 | 2 | 0.5 | 7 | 0.4286 | 22 | 0.5909 | 7 | 0.8571 | 10 | 0.4 | 9 | 0.4444 | 3 | 0.6667 | 29 | 0.5517 |
| 11_101558 | 9 | 0.6667 | 5 | 0.8 | 4 | 1 | 8 | 0.625 | 26 | 0.7308 | 11 | 0.4545 | 10 | 0.6 | 6 | 0.6667 | 4 | 0.75 | 31 | 0.5806 |
| 11_101558 | 10 | 0.2 | 6 | 0 | 5 | 0 | 7 | 0 | 28 | 0.0714 | 9 | 0 | 12 | 0.1667 | 8 | 0.125 | 5 | 0.4 | 34 | 0.1471 |
| 11_101558 | 9 | 0.8889 | 5 | 1 | 6 | 1 | 10 | 1 | 30 | 0.9667 | 8 | 1 | 10 | 1 | 11 | 1 | 6 | 1 | 35 | 1 |
| 11_101558 | 10 | 0.1 | 6 | 0 | 6 | 0 | 11 | 0.0909 | 33 | 0.0606 | 11 | 0.0909 | 13 | 0.0769 | 11 | 0 | 4 | 0 | 39 | 0.0513 |
| 11_101629 | 1 | 1 | 1 | 0 | 1 | 1 | 2 | 0 | 5 | 0.4 | NA | NA | 1 | 0 | 3 | 0.6667 | 1 | 1 | 5 | 0.6 |
| 11_101629 | 1 | 1 | 1 | 0 | NA | NA | 2 | 0 | 4 | 0.25 | 1 | 1 | 1 | 0 | 2 | 0 | 1 | 1 | 5 | 0.4 |
| 11_101657 | 4 | 0 | 5 | 0 | 2 | 0.5 | 3 | 0 | 14 | 0.0714 | 1 | 0 | 4 | 0 | 6 | 0 | 3 | 0.3333 | 14 | 0.0714 |
| 11_101657 | 3 | 1 | 6 | 1 | 1 | 1 | 4 | 0.75 | 14 | 0.9286 | 1 | 0 | 2 | 1 | 5 | 0.8 | 1 | 1 | 9 | 0.7778 |
| 11_101657 | 4 | 0.75 | 7 | 0.8571 | 2 | 1 | 1 | 1 | 14 | 0.8571 | 1 | 1 | 5 | 0.8 | 7 | 0.8571 | 3 | 0.6667 | 16 | 0.8125 |
| 11_101657 | 3 | 0 | 7 | 0.4286 | 2 | 0 | 4 | 0.25 | 16 | 0.25 | 1 | 0 | 5 | 0.8 | 6 | 0.3333 | 3 | 0.3333 | 15 | 0.4667 |
| 11_101657 | 3 | 0.6667 | 5 | 0 | 1 | 0 | 4 | 0.25 | 13 | 0.2308 | 1 | 0 | 5 | 0.4 | 6 | 0.1667 | 3 | 0.3333 | 15 | 0.2667 |
| 11_101657 | 3 | 0 | 7 | 0.1429 | 2 | 0.5 | 4 | 0 | 16 | 0.125 | 1 | 0 | 3 | 0 | 6 | 0.1667 | 2 | 0 | 12 | 0.0833 |
| 11_101657 | 2 | 0.5 | 6 | 0.3333 | 1 | 0 | 3 | 0.3333 | 12 | 0.3333 | 1 | 0 | 5 | 0.2 | 4 | 0.25 | 2 | 0.5 | 12 | 0.25 |
| 11_101658 | 2 | 0.5 | 4 | 0.5 | 2 | 1 | 4 | 0.5 | 12 | 0.5833 | 1 | 1 | 5 | 0.6 | 3 | 0.6667 | 3 | 0.3333 | 12 | 0.5833 |
| 11_101658 | 3 | 0.3333 | 3 | 1 | 2 | 0.5 | 3 | 0.3333 | 11 | 0.5455 | 1 | 0 | 4 | 0.25 | 5 | 0.4 | 1 | 0 | 11 | 0.2727 |
| 11_101658 | 3 | 0.6667 | 6 | 0.1667 | 3 | 0.3333 | 2 | 0 | 14 | 0.2857 | 1 | 0 | 6 | 0.3333 | 7 | 0 | 3 | 0 | 17 | 0.1176 |
| 11_101658 | 3 | 0 | 6 | 0.3333 | 3 | 0 | 3 | 0 | 15 | 0.1333 | 1 | 0 | 7 | 0.2857 | 7 | 0.1429 | 3 | 0 | 18 | 0.1667 |
| 11_101658 | 1 | 1 | 1 | 0 | 1 | 1 | 2 | 0 | 5 | 0.4 | 1 | 0 | 3 | 0 | 2 | 0.5 | 1 | 0 | 7 | 0.1429 |
| 11_101659 | 2 | 1 | 1 | 1 | 1 | 1 | 2 | 0.5 | 6 | 0.8333 | 1 | 1 | 3 | 0.3333 | 3 | 1 | 1 | 0 | 8 | 0.625 |
| 11_101659 | NA | NA | NA | NA | NA | NA | 2 | 0.5 | 2 | 0.5 | 1 | 0 | 3 | 0.3333 | 4 | 0.25 | 2 | 0.5 | 10 | 0.3 |
| 11_101659 | 1 | 1 | 1 | 1 | 1 | 1 | 1 | 1 | 4 | 1 | 1 | 1 | 3 | 0.6667 | 2 | 1 | 2 | 0 | 8 | 0.625 |
| 11_101660 | NA | NA | 2 | 0.5 | NA | NA | NA | NA | 2 | 0.5 | NA | NA | 1 | 1 | NA | NA | 2 | 1 | 3 | 1 |
| 11_101660 | NA | NA | 2 | 1 | NA | NA | NA | NA | 2 | 1 | NA | NA | 1 | 0 | NA | NA | 2 | 0.5 | 3 | 0.3333 |
| 11_101661 | NA | NA | 2 | 0.5 | NA | NA | NA | NA | 2 | 0.5 | NA | NA | 1 | 1 | NA | NA | 2 | 1 | 3 | 1 |
| 11_101661 | NA | NA | 2 | 0.5 | NA | NA | NA | NA | 2 | 0.5 | NA | NA | 1 | 0 | NA | NA | 2 | 1 | 3 | 0.6667 |
| 11_101862 | 15 | 0.0667 | 15 | 0.1333 | 21 | 0.0476 | 11 | 0 | 62 | 0.0645 | 10 | 0.1 | 7 | 0.1429 | 13 | 0.0769 | 14 | 0 | 44 | 0.0682 |
| 11_101862 | 13 | 0.0769 | 12 | 0.0833 | 15 | 0.2667 | 8 | 0 | 48 | 0.125 | 7 | 0 | 12 | 0.1667 | 8 | 0.5 | 8 | 0.125 | 35 | 0.2 |
| 11_101863 | 15 | 0.0667 | 15 | 0.0667 | 18 | 0 | 8 | 0.125 | 56 | 0.0536 | 11 | 0.1818 | 8 | 0 | 10 | 0.1 | 10 | 0.1 | 39 | 0.1026 |
| 11_101863 | 11 | 0 | 14 | 0.0714 | 16 | 0.1875 | 7 | 0.1429 | 48 | 0.1042 | 9 | 0 | 10 | 0.1 | 8 | 0 | 11 | 0.0909 | 38 | 0.0526 |
| 11_101863 | 7 | 0.1429 | 12 | 0 | 12 | 0.1667 | 6 | 0.1667 | 37 | 0.1081 | 7 | 0.1429 | 10 | 0.1 | 6 | 0.1667 | 8 | 0 | 31 | 0.0968 |
| 11_101864 | 8 | 0.25 | 12 | 0.25 | 14 | 0.2143 | 6 | 0 | 40 | 0.2 | 7 | 0 | 10 | 0.3 | 6 | 0 | 6 | 0 | 29 | 0.1034 |
| 11_101864 | 7 | 0.8571 | 11 | 1 | 11 | 1 | 5 | 1 | 34 | 0.9706 | 7 | 1 | 11 | 0.9091 | 6 | 1 | 7 | 1 | 31 | 0.9677 |
| 11_101864 | 3 | 0.3333 | 1 | 1 | 9 | 0.1111 | 4 | 0.25 | 17 | 0.2353 | 4 | 0.5 | 7 | 0.2857 | 4 | 0 | 6 | 0 | 21 | 0.1905 |
| 11_101864 | 8 | 0.75 | 13 | 0.8462 | 13 | 0.9231 | 6 | 0.8333 | 40 | 0.85 | 7 | 0.8571 | 12 | 0.9167 | 5 | 1 | 9 | 1 | 33 | 0.9394 |
| 11_101864 | 3 | 1 | 9 | 1 | 11 | 1 | 2 | 1 | 25 | 1 | 4 | 1 | 8 | 1 | 3 | 1 | 7 | 1 | 22 | 1 |
| 11_101864 | 7 | 0 | 10 | 0 | 14 | 0.1429 | 6 | 0 | 37 | 0.0541 | 5 | 0 | 6 | 0 | 6 | 0.1667 | 5 | 0.2 | 22 | 0.0909 |
| 11_101864 | 6 | 0.1667 | 10 | 0.6 | 14 | 0.3571 | 5 | 0.4 | 35 | 0.4 | 5 | 0.2 | 11 | 0.5455 | 6 | 0.3333 | 9 | 0.4444 | 31 | 0.4194 |
| 11_101864 | 6 | 0.6667 | 8 | 0.375 | 11 | 0.6364 | 5 | 0.8 | 30 | 0.6 | 6 | 0.8333 | 9 | 0.8889 | 6 | 1 | 7 | 0.7143 | 28 | 0.8571 |
| 11_101864 | 7 | 0.2857 | 8 | 0.75 | 13 | 0.2308 | 5 | 0.8 | 33 | 0.4545 | 7 | 0.5714 | 11 | 0.2727 | 5 | 0.6 | 7 | 0.4286 | 30 | 0.4333 |
| 11_101864 | 7 | 0 | 10 | 0.1 | 13 | 0.4615 | 5 | 0.2 | 35 | 0.2286 | 5 | 0 | 10 | 0.5 | 5 | 0 | 8 | 0.25 | 28 | 0.25 |
| 11_101864 | 7 | 0.7143 | 12 | 0.8333 | 12 | 0.5833 | 5 | 0.8 | 36 | 0.7222 | 5 | 0.8 | 12 | 0.6667 | 5 | 0.6 | 5 | 0.8 | 27 | 0.7037 |
| 11_101864 | 7 | 0 | 12 | 0.0833 | 11 | 0.0909 | 5 | 0.4 | 35 | 0.1143 | 6 | 0.1667 | 11 | 0.4545 | 5 | 0 | 8 | 0.125 | 30 | 0.2333 |
| 11_101864 | 6 | 0.8333 | 10 | 0.5 | 10 | 0.4 | 5 | 0.6 | 31 | 0.5484 | 5 | 0.6 | 9 | 0.4444 | 4 | 0.5 | 8 | 0.75 | 26 | 0.5769 |
| 11_101865 | 8 | 0.25 | 8 | 0.5 | 11 | 0.4545 | 5 | 0.4 | 32 | 0.4062 | 2 | 0.5 | 11 | 0.2727 | 6 | 0.5 | 6 | 0.3333 | 25 | 0.36 |
| 11_101865 | 8 | 0 | 11 | 0.0909 | 6 | 0 | 2 | 0.5 | 27 | 0.0741 | 3 | 0 | 8 | 0 | 4 | 0 | 4 | 0.25 | 19 | 0.0526 |
| 11_101865 | 8 | 0.25 | 10 | 0.2 | 8 | 0.375 | 3 | 0.3333 | 29 | 0.2759 | 2 | 0 | 9 | 0.4444 | 3 | 0.3333 | 5 | 0.6 | 19 | 0.4211 |
| 11_101865 | 7 | 0.4286 | 7 | 0.1429 | 7 | 0.2857 | 3 | 0.3333 | 24 | 0.2917 | 2 | 0.5 | 6 | 0.1667 | 4 | 0.25 | 5 | 0.6 | 17 | 0.3529 |
| 11_101885 | NA | NA | 1 | 1 | NA | NA | NA | NA | 1 | 1 | NA | NA | NA | NA | 1 | 1 | NA | NA | 1 | 1 |
| 11_101890 | 1 | 0 | 4 | 0.25 | 3 | 0.6667 | 2 | 0.5 | 10 | 0.4 | 3 | 0.6667 | 2 | 0.5 | 4 | 0.25 | 3 | 0.3333 | 12 | 0.4167 |
| 11_101891 | 11 | 0.5455 | 7 | 0.5714 | 6 | 0.6667 | 4 | 0.5 | 28 | 0.5714 | 5 | 0.6 | 5 | 0.8 | 9 | 0.8889 | 2 | 1 | 21 | 0.8095 |
| 11_101891 | 10 | 0.2 | 7 | 0.1429 | 6 | 0 | 3 | 0 | 26 | 0.1154 | 3 | 0 | 5 | 0 | 7 | 0.2857 | 4 | 0 | 19 | 0.1053 |
| 11_101891 | 10 | 0.9 | 6 | 1 | 4 | 1 | 4 | 0.75 | 24 | 0.9167 | 3 | 1 | 5 | 0.8 | 9 | 0.8889 | 5 | 1 | 22 | 0.9091 |
| 11_101891 | 6 | 0.3333 | 5 | 0.4 | 6 | 0.3333 | 2 | 0.5 | 19 | 0.3684 | 3 | 0 | 5 | 0.6 | 7 | 0.1429 | 3 | 0 | 18 | 0.2222 |
| 11_101948 | 14 | 0.0714 | 21 | 0.0952 | 10 | 0.3 | 20 | 0.1 | 65 | 0.1231 | 14 | 0.0714 | 20 | 0.1 | 17 | 0.1176 | 10 | 0.2 | 61 | 0.1148 |
| 11_101948 | 8 | 0.25 | 14 | 0.0714 | 8 | 0 | 14 | 0 | 44 | 0.0682 | 7 | 0.1429 | 11 | 0 | 12 | 0.0833 | 8 | 0 | 38 | 0.0526 |
| 11_101949 | 9 | 0.5556 | 13 | 0.1538 | 7 | 0.1429 | 13 | 0.3077 | 42 | 0.2857 | 7 | 0.1429 | 14 | 0.2857 | 11 | 0 | 6 | 0 | 38 | 0.1316 |
| 11_101949 | 6 | 0.1667 | 11 | 0.3636 | 6 | 0 | 7 | 0.2857 | 30 | 0.2333 | 3 | 0.6667 | 10 | 0.1 | 8 | 0.5 | 4 | 0.25 | 25 | 0.32 |
| 11_101949 | 8 | 0.25 | 9 | 0.4444 | 5 | 0.2 | 12 | 0.4167 | 34 | 0.3529 | 6 | 0.1667 | 9 | 0.1111 | 9 | 0.3333 | 4 | 0 | 28 | 0.1786 |

| | | | | | | | | | | | | | | | | | | | | |
|---|---|---|---|---|---|---|---|---|---|---|---|---|---|---|---|---|---|---|---|---|
| 11_1019500 | 7 | 1 | 12 | 0.9167 | 7 | 0.5714 | 13 | 0.9231 | 39 | 0.8718 | 6 | 0.8333 | 8 | 0.875 | 10 | 0.9 | 6 | 1 | 30 | 0.9 |
| 11_1019501 | 8 | 1 | 13 | 0.9231 | 6 | 0.8333 | 13 | 0.8462 | 40 | 0.9 | 5 | 1 | 11 | 0.7273 | 12 | 0.8333 | 5 | 1 | 33 | 0.8485 |
| 11_1019502 | 7 | 0 | 11 | 0.0909 | 7 | 0 | 9 | 0.1111 | 34 | 0.0588 | 6 | 0.1667 | 9 | 0.1111 | 10 | 0 | 5 | 0 | 30 | 0.0667 |
| 11_1020537 | 6 | 0.3333 | 6 | 0.3333 | 7 | 0.4286 | 2 | 0 | 21 | 0.3333 | 7 | 0.2857 | 3 | 0 | 6 | 0.1667 | 5 | 0.4 | 21 | 0.2381 |
| 11_1020547 | 3 | 0 | 5 | 0.2 | 10 | 0.1 | 4 | 0 | 22 | 0.0909 | 6 | 0.1667 | 4 | 0 | 7 | 0.4286 | 5 | 0 | 22 | 0.1818 |
| 11_1020548 | 1 | 0 | 5 | 0 | 7 | 0 | 2 | 0.5 | 15 | 0.0667 | 6 | 0.1667 | 4 | 0 | 6 | 0 | 5 | 0.2 | 21 | 0.0952 |
| 11_1020840 | 4 | 0 | 5 | 0 | 6 | 0 | 8 | 0.25 | 23 | 0.087 | 5 | 0 | 5 | 0.2 | 6 | 0 | 2 | 0 | 18 | 0.0556 |
| 11_1020843 | 7 | 0.4286 | 5 | 0 | 6 | 0 | 10 | 0.2 | 28 | 0.1786 | 7 | 0.2857 | 7 | 0.2857 | 7 | 0.2857 | 6 | 0.5 | 27 | 0.3333 |
| 11_1020846 | 7 | 0.1429 | 5 | 0 | 6 | 0.1667 | 8 | 0.125 | 26 | 0.1154 | 7 | 0.1429 | 8 | 0 | 5 | 0 | 8 | 0.125 | 28 | 0.0714 |
| 11_1020847 | 8 | 0.125 | 6 | 0.3333 | 7 | 0 | 12 | 0.5833 | 33 | 0.303 | 8 | 0.375 | 10 | 0.5 | 7 | 0.7143 | 9 | 0.5556 | 34 | 0.5294 |
| 11_1021155 | 1 | 0 | 4 | 0 | 1 | 0 | 2 | 0.5 | 8 | 0.125 | 6 | 0.1667 | 5 | 0 | 6 | 0.1667 | 6 | 0.1667 | 23 | 0.1304 |
| 11_1021690 | NA | NA | NA | NA | 1 | 0 | 1 | 1 | 2 | 0.5 | NA | NA | 1 | 1 | NA | NA | 2 | 0 | 3 | 0.3333 |
| 11_1021692 | NA | NA | NA | NA | 1 | 0 | 1 | 1 | 2 | 0.5 | NA | NA | 1 | 1 | 1 | 1 | 2 | 0.5 | 4 | 0.75 |
| 11_1021699 | NA | NA | NA | NA | 1 | 0 | 1 | 1 | 2 | 0.5 | NA | NA | 1 | 0 | 2 | 1 | 1 | 1 | 4 | 0.75 |
| 11_1021782 | 3 | 0.3333 | 5 | 0 | 7 | 0.1429 | 13 | 0.1538 | 28 | 0.1429 | 6 | 0.1667 | 5 | 0.2 | 5 | 0 | 6 | 0.3333 | 22 | 0.1818 |
| 11_1021784 | 3 | 0 | 6 | 0.1667 | 6 | 0.3333 | 10 | 0.2 | 25 | 0.2 | 6 | 0 | 3 | 0.3333 | 3 | 0 | 4 | 0.5 | 16 | 0.1875 |
| 11_1021785 | 2 | 0.5 | 6 | 0.5 | 5 | 0.6 | 9 | 0.5556 | 22 | 0.5455 | 6 | 0.3333 | 2 | 0 | 4 | 0.25 | 5 | 0.2 | 17 | 0.2353 |
| 11_1021789 | 3 | 0.3333 | 9 | 0.3333 | 6 | 0 | 11 | 0.2727 | 29 | 0.2414 | 4 | 0.5 | 4 | 0.25 | 7 | 0.1429 | 5 | 0.2 | 20 | 0.25 |
| 11_1021791 | 3 | 1 | 8 | 0.5 | 8 | 0.625 | 10 | 0.7 | 29 | 0.6552 | 4 | 0.5 | 2 | 0.5 | 6 | 0.3333 | 5 | 0.8 | 17 | 0.5294 |
| 11_1021792 | 4 | 1 | 9 | 0.6667 | 12 | 0.6667 | 13 | 0.8462 | 38 | 0.7632 | 8 | 0.875 | 7 | 0.4286 | 10 | 0.7 | 8 | 0.625 | 33 | 0.6667 |
| 11_1021793 | 3 | 1 | 8 | 0.75 | 9 | 0.8889 | 13 | 0.7692 | 33 | 0.8182 | 5 | 0.8 | 6 | 0.6667 | 9 | 0.5556 | 9 | 1 | 29 | 0.7586 |
| 11_1021796 | 7 | 0.4286 | 11 | 0.2727 | 14 | 0.2143 | 14 | 0.2143 | 46 | 0.2609 | 8 | 0.375 | 7 | 0.1429 | 9 | 0.1111 | 10 | 0.2 | 34 | 0.2059 |
| 11_1021799 | 7 | 0.2857 | 10 | 0.1 | 11 | 0.1818 | 15 | 0 | 43 | 0.1163 | 10 | 0.2 | 6 | 0 | 10 | 0 | 11 | 0.2727 | 37 | 0.1351 |
| 11_1021810 | 14 | 0.2143 | 19 | 0.2632 | 15 | 0.2 | 10 | 0.2 | 58 | 0.2241 | 11 | 0.2727 | 13 | 0.0769 | 15 | 0.2 | 17 | 0.2941 | 56 | 0.2143 |
| 11_1021813 | 14 | 0.1429 | 20 | 0.05 | 14 | 0 | 13 | 0.0769 | 61 | 0.0656 | 10 | 0.1 | 17 | 0.0588 | 16 | 0 | 14 | 0.0714 | 57 | 0.0526 |
| 11_1021818 | 13 | 0.0769 | 21 | 0.0476 | 19 | 0.1579 | 12 | 0 | 65 | 0.0769 | 8 | 0.125 | 11 | 0 | 14 | 0 | 15 | 0.1333 | 48 | 0.0625 |
| 11_1021821 | 9 | 0.7778 | 11 | 0.7273 | 10 | 0.9 | 6 | 0.8333 | 36 | 0.8056 | 7 | 1 | 3 | 1 | 4 | 1 | 8 | 0.625 | 22 | 0.8636 |
| 11_1021821 | 10 | 0.8 | 11 | 0.5455 | 8 | 0.375 | 7 | 0.1429 | 36 | 0.5 | 8 | 0.625 | 5 | 0.2 | 8 | 0.25 | 9 | 0.5556 | 30 | 0.4333 |
| 11_1021822 | 13 | 0.7692 | 14 | 0.8571 | 14 | 0.9286 | 9 | 0.5556 | 50 | 0.8 | 8 | 0.875 | 8 | 0.875 | 9 | 0.6667 | 11 | 1 | 36 | 0.8611 |
| 11_1021822 | 12 | 0.5 | 13 | 0.7692 | 12 | 0.8333 | 8 | 0.25 | 45 | 0.6222 | 7 | 0.7143 | 10 | 0.8 | 9 | 0.4444 | 11 | 0.4545 | 37 | 0.5946 |
| 11_1021822 | 14 | 0.7143 | 15 | 0.6 | 18 | 0.7222 | 11 | 0.6364 | 58 | 0.6724 | 8 | 0.875 | 10 | 0.7 | 11 | 0.6364 | 16 | 0.9375 | 45 | 0.8 |
| 11_1021824 | 8 | 0.875 | 10 | 0.8 | 10 | 0.7 | 8 | 1 | 36 | 0.8333 | 7 | 0.5714 | 9 | 0.7778 | 6 | 1 | 7 | 1 | 29 | 0.8276 |
| 11_1021824 | 11 | 0.8182 | 11 | 0.4545 | 15 | 0.5333 | 9 | 0.7778 | 46 | 0.6304 | 5 | 0.2 | 8 | 0.5 | 9 | 0.5556 | 14 | 0.5714 | 36 | 0.5 |
| 11_1021825 | 8 | 0.625 | 11 | 0.5455 | 12 | 0.3333 | 6 | 0.3333 | 37 | 0.4595 | 5 | 0.8 | 9 | 0.2222 | 6 | 0.5 | 15 | 0.6667 | 35 | 0.5429 |
| 11_1021959 | NA | NA | 1 | 1 | 1 | 0 | 1 | 1 | 3 | 0.6667 | NA | NA | NA | NA | 2 | 0.5 | 1 | 0 | 3 | 0.3333 |
| 11_1021959 | NA | NA | 1 | 0 | 1 | 1 | 1 | 0 | 3 | 0.3333 | NA | NA | 2 | 0 | 2 | 0.5 | NA | NA | 4 | 0.25 |
| 11_1021969 | 17 | 0.3529 | 8 | 0.125 | 13 | 0.0769 | 9 | 0.1111 | 47 | 0.1915 | 8 | 0.375 | 14 | 0.2143 | 10 | 0 | 11 | 0.4545 | 43 | 0.2558 |
| 11_1021971 | 19 | 0.9474 | 9 | 1 | 16 | 0.9375 | 11 | 0.9091 | 55 | 0.9455 | 11 | 0.9091 | 13 | 1 | 12 | 0.8333 | 14 | 0.9286 | 50 | 0.92 |
| 11_1021971 | 15 | 0.8667 | 10 | 0.6 | 18 | 0.6667 | 10 | 0.7 | 53 | 0.717 | 8 | 0.75 | 11 | 0.7273 | 10 | 0.6 | 15 | 0.8667 | 44 | 0.75 |
| 11_1021972 | 18 | 0.1111 | 9 | 0 | 19 | 0.1579 | 12 | 0.1667 | 58 | 0.1207 | 11 | 0.0909 | 13 | 0.0769 | 12 | 0.1667 | 15 | 0.0667 | 51 | 0.098 |
| 11_1021972 | 17 | 0.0588 | 7 | 0 | 17 | 0.1765 | 11 | 0 | 52 | 0.0769 | 12 | 0.0833 | 11 | 0 | 14 | 0.1429 | 14 | 0.0714 | 51 | 0.0784 |
| 11_1021972 | 12 | 0.25 | 6 | 0 | 11 | 0.0909 | 7 | 0.1429 | 36 | 0.1389 | 8 | 0 | 7 | 0.2857 | 9 | 0.2222 | 7 | 0.2857 | 31 | 0.1935 |
| 11_1021974 | 12 | 0.3333 | 10 | 0.2 | 17 | 0.2353 | 10 | 0.4 | 49 | 0.2857 | 10 | 0.3 | 12 | 0.25 | 10 | 0.5 | 15 | 0.3333 | 47 | 0.3404 |
| 11_1022078 | 1 | 0 | 1 | 0 | NA | NA | 2 | 1 | 4 | 0.5 | NA | NA | 1 | 0 | NA | NA | 1 | 1 | 2 | 0.5 |
| 11_1022081 | 1 | 1 | 1 | 1 | 1 | 0 | 1 | 0 | 4 | 0.5 | NA | NA | 1 | 0 | NA | NA | 1 | 1 | 2 | 0.5 |
| 11_1022093 | 2 | 0.5 | 3 | 0.3333 | 1 | 0 | 2 | 0 | 8 | 0.25 | 1 | 1 | 2 | 0 | 2 | 0.5 | 2 | 0.5 | 7 | 0.4286 |
| 11_1022094 | 1 | 1 | 2 | 0 | 2 | 0 | 2 | 0 | 7 | 0.1429 | 1 | 1 | 2 | 0 | 2 | 0 | 2 | 0 | 7 | 0.1429 |
| 11_1022922 | 14 | 0.1429 | 6 | 0.1667 | 13 | 0 | 9 | 0 | 42 | 0.0714 | 13 | 0.1538 | 8 | 0 | 12 | 0.0833 | 8 | 0.125 | 41 | 0.0976 |
| 11_1022924 | 12 | 0.0833 | 10 | 0.1 | 13 | 0.2308 | 8 | 0.25 | 43 | 0.1628 | 11 | 0.0909 | 8 | 0 | 14 | 0 | 9 | 0.2222 | 42 | 0.0714 |
| 11_1022929 | 19 | 0.5789 | 23 | 0.6522 | 34 | 0.4706 | 18 | 0.3889 | 94 | 0.5213 | 19 | 0.6316 | 25 | 0.44 | 36 | 0.5833 | 28 | 0.4643 | 108 | 0.5278 |
| 11_1022931 | 21 | 0.1429 | 22 | 0.2727 | 39 | 0.3846 | 19 | 0.5263 | 101 | 0.3366 | 21 | 0.1429 | 19 | 0.3684 | 36 | 0.3889 | 26 | 0.2692 | 102 | 0.3039 |
| 11_1022931 | 21 | 0.1429 | 19 | 0.2632 | 40 | 0.05 | 18 | 0.2778 | 98 | 0.1531 | 18 | 0.2778 | 26 | 0.1154 | 35 | 0 | 28 | 0.1429 | 107 | 0.1121 |
| 11_1022933 | 14 | 0.6429 | 20 | 0.25 | 31 | 0.2581 | 16 | 0.625 | 81 | 0.3951 | 20 | 0.3 | 24 | 0.5 | 31 | 0.3226 | 24 | 0.3333 | 99 | 0.3636 |
| 11_1022942 | 12 | 0.0833 | 14 | 0 | 14 | 0.2143 | 15 | 0.2667 | 55 | 0.1455 | 11 | 0.1818 | 16 | 0.0625 | 16 | 0.0625 | 19 | 0.2632 | 62 | 0.1452 |
| 11_1023272 | 7 | 0.1429 | 15 | 0.0667 | 12 | 0 | 5 | 0.2 | 39 | 0.0769 | 7 | 0.1429 | 9 | 0 | 8 | 0 | 10 | 0.2 | 34 | 0.0882 |
| 11_1023273 | 7 | 0.2857 | 16 | 0.25 | 13 | 0.3077 | 5 | 0.2 | 41 | 0.2683 | 10 | 0.5 | 7 | 0.2857 | 9 | 0.3333 | 11 | 0.0909 | 37 | 0.2973 |
| 11_1023273 | 7 | 0.1429 | 12 | 0.1667 | 13 | 0 | 5 | 0 | 37 | 0.0811 | 5 | 0 | 10 | 0.4 | 8 | 0.25 | 11 | 0 | 34 | 0.1765 |
| 11_1023275 | 6 | 0.1667 | 14 | 0.1429 | 13 | 0.3846 | 7 | 0 | 40 | 0.2 | 13 | 0.0769 | 6 | 0.1667 | 8 | 0.375 | 10 | 0 | 37 | 0.1351 |
| 11_1023279 | 9 | 0.4444 | 6 | 0.6667 | 8 | 0.125 | 11 | 0.4545 | 34 | 0.4118 | 8 | 0.5 | 9 | 0.4444 | 14 | 0.5714 | 8 | 0.375 | 39 | 0.4872 |
| 11_1023281 | 8 | 0.125 | 5 | 0.4 | 7 | 0.1429 | 9 | 0 | 29 | 0.1379 | 6 | 0.3333 | 8 | 0.375 | 10 | 0.2 | 8 | 0.125 | 32 | 0.25 |
| 11_1023284 | 4 | 1 | 3 | 1 | 4 | 1 | 7 | 0.8571 | 18 | 0.9444 | 3 | 1 | 6 | 1 | 6 | 1 | 6 | 1 | 21 | 1 |
| 11_1023285 | 7 | 0.7143 | 4 | 0.75 | 6 | 0.5 | 7 | 0.2857 | 24 | 0.5417 | 2 | 0 | 5 | 1 | 7 | 0.5714 | 7 | 0.8571 | 21 | 0.7143 |
| 11_1023287 | 6 | 1 | 5 | 1 | 4 | 1 | 8 | 0.875 | 23 | 0.9565 | 2 | 1 | 7 | 1 | 7 | 1 | 4 | 0.75 | 20 | 0.95 |
| 11_1023288 | 5 | 0.6 | 5 | 0.6 | 5 | 0.6 | 8 | 0.625 | 23 | 0.6087 | 2 | 0.5 | 6 | 0.6667 | 7 | 0.7143 | 5 | 0.6 | 20 | 0.65 |
| 11_1023291 | 6 | 0.8333 | 5 | 1 | 4 | 1 | 8 | 0.875 | 23 | 0.913 | 2 | 0.5 | 7 | 1 | 7 | 1 | 5 | 0.8 | 21 | 0.9048 |

| | | | | | | | | | | | | | | | | | | | | |
|---|---|---|---|---|---|---|---|---|---|---|---|---|---|---|---|---|---|---|---|---|
| 11_1023292 | 5 | 0 | 5 | 0.2 | 4 | 0 | 10 | 0.1 | 24 | 0.0833 | 3 | 0.3333 | 6 | 0 | 7 | 0.1429 | 5 | 0 | 21 | 0.0952 |
| 11_1023295 | 5 | 0.6 | 4 | 0 | 3 | 0.3333 | 9 | 0.4444 | 21 | 0.381 | 2 | 0 | 6 | 0.3333 | 6 | 0.3333 | 2 | 0 | 16 | 0.25 |
| 11_1023677 | 1 | 0 | 1 | 0 | 4 | 0 | 2 | 0.5 | 8 | 0.125 | 4 | 0.25 | 3 | 0 | 3 | 0 | 3 | 0 | 13 | 0.0769 |
| 11_1023677 | NA | NA | 1 | 1 | 2 | 0.5 | 1 | 0 | 4 | 0.5 | 3 | 0.3333 | 3 | 0.6667 | 2 | 0.5 | 2 | 0.5 | 10 | 0.5 |
| 11_1023678 | 1 | 1 | 2 | 1 | 2 | 1 | 3 | 0.6667 | 8 | 0.875 | 2 | 1 | 1 | 0 | 2 | 1 | 2 | 0.5 | 7 | 0.7143 |
| 11_1023679 | 1 | 1 | 2 | 0.5 | 4 | 0.25 | 4 | 0.25 | 11 | 0.3636 | 3 | 0.3333 | 3 | 0 | 2 | 0 | 2 | 0.5 | 10 | 0.2 |
| 11_1023682 | NA | NA | 3 | 0.3333 | 4 | 0 | 3 | 0.3333 | 10 | 0.2 | 5 | 0.2 | 3 | 0.3333 | 2 | 1 | NA | NA | 10 | 0.4 |
| 11_1023685 | 1 | 0 | 4 | 0 | 3 | 0.3333 | 3 | 0 | 11 | 0.0909 | 2 | 0.5 | 3 | 0 | 2 | 0 | 2 | 0 | 9 | 0.1111 |
| 11_1023687 | 1 | 0 | 5 | 0.4 | 4 | 0.25 | 4 | 0.25 | 14 | 0.2857 | 5 | 0.2 | 3 | 0.3333 | 2 | 0.5 | 1 | 0 | 11 | 0.2727 |
| 11_1023690 | 1 | 1 | 5 | 0.6 | 4 | 0.75 | 3 | 1 | 13 | 0.7692 | 5 | 0.2 | 4 | 0.5 | 2 | 0.5 | 1 | 0 | 12 | 0.3333 |
| 11_1023696 | 1 | 1 | 4 | 1 | 3 | 0.6667 | 3 | 0.6667 | 11 | 0.8182 | 3 | 1 | 3 | 1 | 2 | 1 | 1 | 1 | 9 | 1 |
| 11_1023696 | 1 | 0 | 5 | 0.2 | 3 | 0 | 3 | 0 | 12 | 0.0833 | 1 | 1 | 1 | 0 | 2 | 0 | 1 | 0 | 5 | 0.2 |
| 11_1023697 | 1 | 1 | 5 | 0.4 | 3 | 0.3333 | 4 | 0.75 | 13 | 0.5385 | 2 | 0 | 2 | 0.5 | 2 | 0.5 | 1 | 1 | 7 | 0.4286 |
| 11_1023698 | NA | NA | 3 | 1 | 2 | 1 | 2 | 1 | 7 | 1 | 2 | 1 | 2 | 0.5 | 2 | 1 | 1 | 1 | 7 | 0.8571 |
| 11_1023699 | 1 | 0 | 3 | 0 | 3 | 0 | 4 | 0.25 | 11 | 0.0909 | 1 | 0 | NA | NA | 1 | 0 | 1 | 1 | 3 | 0.3333 |
| 11_1023700 | 1 | 0 | 3 | 0.6667 | 3 | 0.6667 | 3 | 0.3333 | 10 | 0.5 | 1 | 0 | 2 | 0.5 | 1 | 0 | NA | NA | 4 | 0.25 |
| 11_1023702 | 1 | 1 | 3 | 1 | 2 | 0.5 | 3 | 0 | 9 | 0.5556 | 2 | 0 | 2 | 0.5 | 1 | 0 | NA | NA | 5 | 0.2 |
| 11_1023702 | 1 | 1 | 2 | 0.5 | 2 | 0.5 | 4 | 0.75 | 9 | 0.6667 | 2 | 0.5 | 2 | 1 | 1 | 1 | NA | NA | 5 | 0.8 |
| 11_1023707 | 1 | 1 | 4 | 0 | 1 | 0 | 4 | 0 | 10 | 0.1 | 2 | 0.5 | 2 | 0 | 1 | 1 | NA | NA | 5 | 0.4 |
| 11_1023708 | NA | NA | 4 | 0.75 | NA | NA | 3 | 0.6667 | 7 | 0.7143 | 2 | 0 | 2 | 0.5 | 1 | 0 | NA | NA | 5 | 0.2 |
| 11_1023720 | NA | NA | 2 | 1 | NA | NA | 1 | 1 | 3 | 1 | 1 | 1 | 1 | 1 | NA | NA | NA | NA | 2 | 1 |
| 11_1023720 | NA | NA | 1 | 0 | NA | NA | 2 | 0.5 | 3 | 0.3333 | 1 | 1 | 1 | 0 | NA | NA | NA | NA | 2 | 0.5 |
| 11_1023725 | NA | NA | NA | NA | NA | NA | 1 | 1 | 1 | 1 | 1 | 1 | NA | NA | NA | NA | NA | NA | 1 | 1 |
| 11_1023726 | NA | NA | 1 | 0 | NA | NA | 3 | 0.3333 | 4 | 0.25 | 1 | 1 | NA | NA | NA | NA | NA | NA | 1 | 1 |
| 11_1023727 | NA | NA | 1 | 1 | NA | NA | 2 | 1 | 3 | 1 | 1 | 1 | NA | NA | NA | NA | NA | NA | 1 | 1 |
| 11_1024672 | 1 | 0 | 6 | 0.1667 | 7 | 0 | 2 | 0 | 16 | 0.0625 | 2 | 0 | 2 | 0 | 4 | 0.25 | 5 | 0 | 13 | 0.0769 |
| 11_1024957 | 4 | 0.25 | 3 | 0.3333 | 2 | 0 | 5 | 0.2 | 14 | 0.2143 | 1 | 1 | 1 | 0 | 1 | 0 | 2 | 0 | 5 | 0.2 |
| 11_1024959 | 4 | 1 | 4 | 0.75 | 2 | 0.5 | 5 | 0.8 | 15 | 0.8 | 1 | 1 | 2 | 0.5 | 1 | 1 | 2 | 1 | 6 | 0.8333 |
| 11_1024963 | 4 | 1 | 2 | 1 | 3 | 1 | 5 | 1 | 14 | 1 | 1 | 1 | 1 | 1 | 5 | 1 | 2 | 1 | 9 | 1 |
| 11_1024968 | 5 | 0.8 | 4 | 0.5 | 4 | 0.5 | 4 | 1 | 17 | 0.7059 | 1 | 1 | 3 | 0.3333 | 6 | 0.5 | 2 | 1 | 12 | 0.5833 |
| 11_1024972 | 6 | 0.3333 | 3 | 0.3333 | 3 | 0 | 6 | 0.1667 | 18 | 0.2222 | 1 | 0 | 5 | 0 | 6 | 0.1667 | 4 | 0 | 16 | 0.0625 |
| 11_1024973 | 5 | 1 | 4 | 0.75 | 4 | 1 | 5 | 1 | 18 | 0.9444 | 1 | 1 | 5 | 1 | 5 | 1 | 4 | 1 | 15 | 1 |
| 11_1024973 | 6 | 1 | 3 | 1 | 3 | 1 | 5 | 1 | 17 | 1 | NA | NA | 6 | 1 | 5 | 1 | 4 | 1 | 15 | 1 |
| 11_1024974 | 5 | 1 | 2 | 1 | 3 | 1 | 3 | 1 | 13 | 1 | 1 | 1 | 3 | 1 | 3 | 1 | 2 | 1 | 9 | 1 |
| 11_1024974 | 5 | 0.4 | 2 | 0 | 3 | 0 | 3 | 0 | 13 | 0.1538 | NA | NA | 3 | 0 | 3 | 0.3333 | 2 | 0 | 8 | 0.125 |
| 11_1024984 | 5 | 0.8 | 1 | 1 | 6 | 0.5 | 4 | 0.75 | 16 | 0.6875 | 2 | 1 | 3 | 0.6667 | 2 | 0 | 4 | 0.5 | 11 | 0.5455 |
| 11_1025757 | 2 | 0 | 4 | 0.5 | 2 | 0 | 1 | 0 | 9 | 0.2222 | 2 | 0 | 4 | 0 | 1 | 0 | 2 | 0.5 | 9 | 0.1111 |
| 11_1025764 | 2 | 0 | 4 | 0 | 3 | 0.3333 | 1 | 0 | 10 | 0.1 | 3 | 0 | 5 | 0.2 | 1 | 0 | 2 | 0.5 | 11 | 0.1818 |
| 11_1025770 | 1 | 1 | 3 | 0.6667 | NA | NA | 1 | 1 | 5 | 0.8 | 2 | 1 | 4 | 0.25 | NA | NA | 2 | 1 | 8 | 0.625 |
| 11_1025771 | 1 | 0 | 3 | 0.3333 | NA | NA | 1 | 0 | 5 | 0.2 | 2 | 0 | 4 | 0.5 | NA | NA | 2 | 1 | 8 | 0.5 |
| 11_1025772 | 1 | 0 | 3 | 0.3333 | NA | NA | 1 | 0 | 5 | 0.2 | 2 | 0 | 4 | 0.25 | NA | NA | 2 | 0 | 8 | 0.125 |
| 11_1025772 | 1 | 0 | 2 | 0.5 | 1 | 0 | NA | NA | 4 | 0.25 | 2 | 0.5 | 1 | 0 | NA | NA | 1 | 1 | 4 | 0.5 |
| 11_1026690 | 4 | 0 | 5 | 0.2 | 3 | 0 | 4 | 0 | 16 | 0.0625 | 5 | 0.2 | 5 | 0 | 2 | 0 | 5 | 0 | 17 | 0.0588 |
| 11_1026691 | 5 | 0.4 | 6 | 0.5 | 3 | 1 | 4 | 0.5 | 18 | 0.5556 | 7 | 0.5714 | 5 | 0.6 | 3 | 0.3333 | 5 | 0.6 | 20 | 0.55 |
| 11_1026692 | 5 | 0.8 | 4 | 0.75 | 2 | 0 | 3 | 0.3333 | 14 | 0.5714 | 5 | 0.8 | 6 | 0.6667 | 1 | 0 | 6 | 0.6667 | 18 | 0.6667 |
| 11_1026692 | 4 | 0.25 | 7 | 0.2857 | 3 | 0.3333 | 4 | 0 | 18 | 0.2222 | 4 | 0.25 | 5 | 0 | 1 | 1 | 6 | 0 | 16 | 0.125 |
| 11_1026693 | 3 | 0 | 7 | 0.2857 | 4 | 0.5 | 4 | 0 | 18 | 0.2222 | 7 | 0.1429 | 3 | 0 | 3 | 0 | 6 | 0.1667 | 19 | 0.1053 |
| 11_1026697 | 6 | 0 | 7 | 0.2857 | 4 | 0 | 8 | 0.125 | 25 | 0.12 | 9 | 0.2222 | 5 | 0.2 | 2 | 0.5 | 6 | 0 | 22 | 0.1818 |
| 11_1026699 | 5 | 0.2 | 4 | 0.5 | 3 | 0.3333 | 8 | 0.25 | 20 | 0.3 | 5 | 0.4 | 3 | 0 | 2 | 0.5 | 4 | 0 | 14 | 0.2143 |
| 11_1026704 | 6 | 0.5 | 7 | 1 | 4 | 0.25 | 8 | 0.25 | 25 | 0.52 | 4 | 0.5 | 5 | 0.8 | 3 | 0.3333 | 4 | 0.5 | 16 | 0.5625 |
| 11_1026706 | 6 | 0.1667 | 8 | 0.25 | 6 | 0.3333 | 9 | 0.2222 | 29 | 0.2414 | 6 | 0.1667 | 5 | 0.2 | 5 | 0.2 | 5 | 0.2 | 21 | 0.1905 |
| 11_1026708 | 7 | 0 | 7 | 0 | 5 | 0 | 10 | 0.4 | 29 | 0.1379 | 5 | 0.4 | 4 | 0.25 | 5 | 0.2 | 5 | 0 | 19 | 0.2105 |
| 11_1026904 | NA | NA | 1 | 1 | 1 | 0 | 1 | 1 | 3 | 0.6667 | NA | NA | 1 | 1 | NA | NA | NA | NA | 1 | 1 |
| 11_1026905 | NA | NA | 1 | 1 | NA | NA | 1 | 0 | 2 | 0.5 | NA | NA | 1 | 0 | NA | NA | 1 | 1 | 2 | 0.5 |
| 11_1026905 | NA | NA | 1 | 1 | NA | NA | 1 | 0 | 2 | 0.5 | NA | NA | 1 | 0 | NA | NA | 1 | 1 | 2 | 0.5 |
| 11_1026905 | NA | NA | 1 | 1 | 1 | 1 | 1 | 0 | 3 | 0.6667 | NA | NA | 1 | 1 | NA | NA | NA | NA | 1 | 1 |
| 11_1026906 | NA | NA | 1 | 1 | 1 | 1 | 1 | 1 | 3 | 1 | NA | NA | 1 | 1 | NA | NA | 1 | 1 | 2 | 1 |
| 11_1026907 | NA | NA | 1 | 1 | 1 | 0 | 1 | 1 | 3 | 0.6667 | NA | NA | 1 | 1 | NA | NA | NA | NA | 1 | 1 |
| 11_1026909 | NA | NA | NA | NA | 1 | 1 | NA | NA | 1 | 1 | NA | NA | 1 | 1 | NA | NA | NA | NA | 1 | 1 |
| 11_1026909 | NA | NA | NA | NA | 1 | 1 | 1 | 1 | 2 | 1 | NA | NA | 1 | 1 | NA | NA | NA | NA | 1 | 1 |
| 11_1026912 | NA | NA | NA | NA | 1 | 1 | 1 | 1 | 2 | 1 | NA | NA | 1 | 1 | NA | NA | NA | NA | 1 | 1 |
| 11_1026912 | NA | NA | NA | NA | 1 | 1 | 1 | 1 | 2 | 1 | NA | NA | 1 | 1 | NA | NA | NA | NA | 1 | 1 |
| 11_1026913 | NA | NA | NA | NA | 1 | 0 | 1 | 1 | 2 | 0.5 | NA | NA | 1 | 1 | NA | NA | NA | NA | 1 | 1 |
| 11_1026913 | NA | NA | NA | NA | 1 | 1 | 1 | 1 | 2 | 1 | NA | NA | 1 | 1 | NA | NA | NA | NA | 1 | 1 |
| 11_1027293 | 13 | 0.4615 | 10 | 0.6 | 6 | 0.5 | 10 | 0.6 | 39 | 0.5385 | 13 | 0.3846 | 9 | 0.5556 | 3 | 0.6667 | 7 | 0.5714 | 32 | 0.5 |

| | | | | | | | | | | | | | | | | | | | | |
|---|---|---|---|---|---|---|---|---|---|---|---|---|---|---|---|---|---|---|---|---|
| 11_1027293 | 16 | 0.5625 | 10 | 0.4 | 11 | 0.6364 | 9 | 0.6667 | 46 | 0.5652 | 16 | 0.5625 | 12 | 0.5 | 5 | 0.6 | 8 | 0.375 | 41 | 0.5122 |
| 11_1027295 | 12 | 0.5833 | 9 | 0.7778 | 10 | 0.3 | 6 | 0.8333 | 37 | 0.5946 | 12 | 0.5833 | 9 | 0.3333 | 5 | 0.2 | 5 | 0.2 | 31 | 0.3871 |
| 11_1027295 | 15 | 0.1333 | 11 | 0.0909 | 9 | 0.1111 | 8 | 0 | 43 | 0.093 | 15 | 0.0667 | 8 | 0.125 | 4 | 0 | 6 | 0.1667 | 33 | 0.0909 |
| 11_1027308 | NA | NA | NA | NA | NA | NA | 1 | 1 | 1 | 1 | 2 | 0.5 | NA | NA | NA | NA | NA | NA | 2 | 0.5 |
| 11_1027314 | NA | NA | NA | NA | NA | NA | 1 | 1 | 1 | 1 | 2 | 0.5 | NA | NA | NA | NA | NA | NA | 2 | 0.5 |
| 11_1027723 | NA | NA | 2 | 0.5 | 1 | 1 | 1 | 0 | 4 | 0.5 | NA | NA | NA | NA | 1 | 1 | NA | NA | 1 | 1 |
| 11_1027726 | NA | NA | 1 | 0 | 1 | 1 | NA | NA | 2 | 0.5 | NA | NA | NA | NA | 1 | 1 | NA | NA | 1 | 1 |
| 11_1027748 | NA | NA | NA | NA | 1 | 1 | 1 | 1 | 2 | 1 | NA | NA | 1 | 0 | 1 | 1 | NA | NA | 2 | 0.5 |
| 11_1027750 | NA | NA | 1 | 1 | 1 | 1 | 1 | 1 | 3 | 1 | NA | NA | NA | NA | 1 | 1 | NA | NA | 1 | 1 |
| 11_1027751 | NA | NA | 2 | 1 | 1 | 1 | 1 | 1 | 4 | 1 | NA | NA | 1 | 1 | 1 | 1 | NA | NA | 2 | 1 |
| 11_1027751 | NA | NA | 2 | 0.5 | NA | NA | 1 | 1 | 3 | 0.6667 | NA | NA | 1 | 1 | 1 | 1 | NA | NA | 2 | 1 |
| 11_1027784 | 9 | 0.1111 | 25 | 0.2 | 15 | 0.0667 | 7 | 0 | 56 | 0.125 | 38 | 0.2105 | 7 | 0 | 9 | 0.1111 | 14 | 0.2857 | 68 | 0.1912 |
| 11_1027785 | 7 | 0 | 27 | 0.0741 | 14 | 0.0714 | 9 | 0 | 57 | 0.0526 | 32 | 0.0312 | 9 | 0.1111 | 11 | 0.0909 | 15 | 0.0667 | 67 | 0.0597 |
| 11_1027786 | 7 | 0.1429 | 27 | 0.1852 | 13 | 0.0769 | 8 | 0.125 | 55 | 0.1455 | 35 | 0.1714 | 9 | 0.2222 | 10 | 0.3 | 14 | 0.2143 | 68 | 0.2059 |
| 11_1027787 | 10 | 0.4 | 23 | 0.5652 | 13 | 0.1538 | 8 | 0.125 | 54 | 0.3704 | 35 | 0.4571 | 7 | 0.4286 | 9 | 0.2222 | 14 | 0.2143 | 65 | 0.3692 |
| 11_1027795 | 8 | 0 | 18 | 0.1111 | 12 | 0.1667 | 8 | 0 | 46 | 0.087 | 26 | 0.0385 | 6 | 0 | 9 | 0 | 14 | 0.1429 | 55 | 0.0545 |
| 11_1027819 | 1 | 1 | NA | NA | 2 | 0 | NA | NA | 3 | 0.3333 | 2 | 0.5 | 1 | 0 | 2 | 0 | 2 | 0 | 7 | 0.1429 |
| 11_1027878 | 7 | 0 | 8 | 0.25 | 8 | 0.375 | 5 | 0 | 28 | 0.1786 | 15 | 0 | 3 | 0 | 3 | 0 | 7 | 0.4286 | 28 | 0.1071 |
| 11_1028302 | 5 | 0.8 | 2 | 1 | 3 | 1 | 3 | 1 | 13 | 0.9231 | 4 | 1 | 2 | 0.5 | 4 | 0.75 | 5 | 1 | 15 | 0.8667 |
| 11_1028305 | 5 | 1 | 2 | 1 | 1 | 0 | 1 | 0 | 9 | 0.7778 | 3 | 1 | 1 | 1 | 2 | 0.5 | 5 | 1 | 11 | 0.9091 |
| 11_1028850 | 10 | 0.2 | 11 | 0.0909 | 8 | 0.125 | 8 | 0 | 37 | 0.1081 | 7 | 0 | 7 | 0.4286 | 9 | 0 | 10 | 0 | 33 | 0.0909 |
| 11_1028853 | 8 | 0.375 | 10 | 0 | 9 | 0.2222 | 7 | 0.1429 | 34 | 0.1765 | 7 | 0 | 6 | 0.1667 | 8 | 0.125 | 9 | 0 | 30 | 0.0667 |
| 11_1028853 | 7 | 0.1429 | 10 | 0 | 8 | 0.25 | 5 | 0.2 | 30 | 0.1333 | 5 | 0 | 6 | 0.1667 | 6 | 0 | 10 | 0.1 | 27 | 0.0741 |
| 11_1028855 | 8 | 0.625 | 11 | 1 | 9 | 0.5556 | 7 | 0.5714 | 35 | 0.7143 | 6 | 0.8333 | 7 | 0.5714 | 6 | 0.6667 | 10 | 0.4 | 29 | 0.5862 |
| 11_1028856 | 5 | 0 | 8 | 0.25 | 6 | 0.1667 | 7 | 0 | 26 | 0.1154 | 5 | 0.2 | 5 | 0.4 | 4 | 0 | 9 | 0 | 23 | 0.1304 |
| 11_1028858 | 4 | 0.25 | 6 | 0 | 7 | 0.1429 | 4 | 0.25 | 21 | 0.1429 | 4 | 0.25 | 3 | 0 | 6 | 0 | 6 | 0 | 19 | 0.0526 |
| 11_1028859 | 5 | 0.4 | 3 | 0 | 6 | 0 | 4 | 0.5 | 18 | 0.2222 | 3 | 0 | 4 | 0.25 | 7 | 0 | 6 | 0.1667 | 20 | 0.1 |
| 11_1028903 | 1 | 1 | NA | NA | NA | NA | NA | NA | 1 | 1 | 1 | 1 | NA | NA | NA | NA | NA | NA | 1 | 1 |
| 11_1028907 | 1 | 1 | NA | NA | NA | NA | NA | NA | 1 | 1 | 1 | 1 | NA | NA | NA | NA | NA | NA | 1 | 1 |
| 11_1028913 | 1 | 1 | NA | NA | NA | NA | NA | NA | 1 | 1 | 1 | 1 | NA | NA | NA | NA | NA | NA | 1 | 1 |
| 11_1029567 | 11 | 0.7273 | 13 | 0.6154 | 6 | 0.8333 | 8 | 0.625 | 38 | 0.6842 | 8 | 0.75 | 12 | 0.75 | 9 | 1 | 9 | 0.8889 | 38 | 0.8421 |
| 11_1029567 | 13 | 0.4615 | 17 | 0.8235 | 7 | 0.7143 | 8 | 0.5 | 45 | 0.6444 | 8 | 0.375 | 10 | 0.8 | 10 | 0.5 | 13 | 0.4615 | 41 | 0.5366 |
| 11_1029568 | 14 | 0.0714 | 19 | 0.1053 | 8 | 0.125 | 9 | 0 | 50 | 0.08 | 11 | 0.0909 | 11 | 0.1818 | 9 | 0.1111 | 15 | 0 | 46 | 0.087 |
| 11_1029569 | 4 | 1 | 8 | 0.5 | 2 | 0.5 | 6 | 0.5 | 20 | 0.6 | 7 | 0.4286 | 8 | 0.375 | 6 | 0.6667 | 7 | 0.8571 | 28 | 0.5714 |
| 11_1029570 | 4 | 0 | 8 | 0.75 | 2 | 1 | 6 | 0.1667 | 20 | 0.45 | 8 | 0.5 | 8 | 0.25 | 6 | 0 | 7 | 0.2857 | 29 | 0.2759 |
| 11_1029571 | 3 | 0 | 7 | 0.1429 | 2 | 0 | 5 | 0 | 17 | 0.0588 | 5 | 0.2 | 9 | 0 | 3 | 0.3333 | 5 | 0.4 | 22 | 0.1818 |
| 11_1029574 | 4 | 0.25 | 10 | 0.3 | 3 | 0.3333 | 8 | 0.5 | 25 | 0.36 | 7 | 0.2857 | 11 | 0.0909 | 5 | 0.4 | 7 | 0.5714 | 30 | 0.3 |
| 11_1029574 | 4 | 0 | 11 | 0.4545 | 3 | 0.3333 | 7 | 0.1429 | 25 | 0.28 | 7 | 0.2857 | 11 | 0.1818 | 5 | 0.8 | 7 | 0.2857 | 30 | 0.3333 |
| 11_1029574 | 4 | 0.25 | 10 | 0.2 | 2 | 0 | 8 | 0 | 24 | 0.125 | 6 | 0.3333 | 9 | 0.2222 | 5 | 0.4 | 7 | 0.5714 | 27 | 0.3704 |
| 11_1029579 | 4 | 0 | 11 | 0.1818 | 4 | 0.25 | 4 | 0 | 23 | 0.1304 | 7 | 0.4286 | 9 | 0 | 4 | 0.25 | 5 | 0.6 | 25 | 0.28 |
| 11_1029581 | 5 | 0.4 | 10 | 0.4 | 4 | 0.5 | 7 | 0.1429 | 26 | 0.3462 | 3 | 0.6667 | 11 | 0.3636 | 5 | 0.4 | 7 | 0.4286 | 26 | 0.4231 |
| 11_1029581 | 5 | 0.4 | 13 | 0 | 5 | 0 | 9 | 0.1111 | 32 | 0.0938 | 7 | 0.2857 | 12 | 0.0833 | 5 | 0 | 9 | 0 | 33 | 0.0909 |
| 11_1029582 | 5 | 0 | 12 | 0.1667 | 5 | 0.6 | 7 | 0 | 29 | 0.1724 | 7 | 0.5714 | 12 | 0.25 | 6 | 0.3333 | 10 | 0.3 | 35 | 0.3429 |
| 11_1029582 | 5 | 0 | 11 | 0.1818 | 3 | 0.3333 | 7 | 0 | 26 | 0.1154 | 5 | 0.2 | 10 | 0.1 | 6 | 0.1667 | 10 | 0.2 | 31 | 0.1613 |
| 11_1029582 | 5 | 0.2 | 13 | 0.3077 | 5 | 0.6 | 9 | 0.1111 | 32 | 0.2812 | 8 | 0 | 13 | 0 | 6 | 0.3333 | 8 | 0.25 | 35 | 0.1143 |
| 11_1029583 | 5 | 0.2 | 13 | 0.0769 | 6 | 0.1667 | 7 | 0.1429 | 31 | 0.129 | 7 | 0.1429 | 13 | 0.1538 | 6 | 0 | 10 | 0 | 36 | 0.0833 |
| 11_1029583 | 5 | 0 | 12 | 0.3333 | 6 | 0.3333 | 9 | 0.1111 | 32 | 0.2188 | 4 | 0.5 | 13 | 0.3077 | 7 | 0.4286 | 10 | 0.5 | 34 | 0.4118 |
| 11_1029584 | 7 | 0.1429 | 12 | 0.25 | 6 | 0.5 | 9 | 0.3333 | 34 | 0.2941 | 8 | 0.5 | 13 | 0.1538 | 8 | 0.5 | 10 | 0.4 | 39 | 0.359 |
| 11_1029584 | 6 | 1 | 12 | 0.75 | 6 | 1 | 7 | 0.5714 | 31 | 0.8065 | 7 | 0.8571 | 12 | 0.9167 | 6 | 0.6667 | 8 | 1 | 33 | 0.8788 |
| 11_1029586 | 7 | 0.2857 | 12 | 0.3333 | 6 | 0.1667 | 8 | 0.25 | 33 | 0.2727 | 7 | 0.4286 | 13 | 0.3077 | 6 | 0.1667 | 11 | 0.6364 | 37 | 0.4054 |
| 11_1029586 | 6 | 0 | 8 | 0.25 | 3 | 0 | 9 | 0 | 26 | 0.0769 | 6 | 0.1667 | 6 | 0 | 2 | 0 | 9 | 0.1111 | 23 | 0.087 |
| 11_1029589 | 8 | 0.25 | 10 | 0.1 | 5 | 0.4 | 10 | 0.3 | 33 | 0.2424 | 6 | 0 | 10 | 0.1 | 7 | 0.2857 | 8 | 0.5 | 31 | 0.2258 |
| 11_1029590 | 7 | 0.4286 | 10 | 0.3 | 7 | 0.2857 | 12 | 0.6667 | 36 | 0.4444 | 7 | 0.1429 | 12 | 0.1667 | 9 | 0.3333 | 11 | 0.2727 | 39 | 0.2308 |
| 11_1029594 | 8 | 1 | 13 | 0.9231 | 7 | 1 | 12 | 0.8333 | 40 | 0.925 | 10 | 0.9 | 12 | 0.9167 | 11 | 0.8182 | 12 | 0.9167 | 45 | 0.8889 |
| 11_1029612 | 1 | 1 | NA | NA | NA | NA | 1 | 1 | 2 | 1 | 2 | 1 | 1 | 1 | NA | NA | NA | NA | 3 | 1 |
| 11_1029613 | 1 | 1 | NA | NA | NA | NA | 3 | 0.3333 | 4 | 0.5 | 2 | 0 | 2 | 0.5 | NA | NA | NA | NA | 4 | 0.25 |
| 11_1029936 | 1 | 1 | 2 | 0 | 3 | 0 | 1 | 1 | 7 | 0.2857 | NA | NA | 1 | 0 | 2 | 0.5 | 1 | 0 | 4 | 0.25 |
| 11_1029942 | NA | NA | 3 | 0.3333 | NA | NA | 1 | 0 | 4 | 0.25 | 1 | 1 | NA | NA | 1 | 1 | NA | NA | 2 | 1 |
| 11_1030023 | 7 | 0.1429 | 6 | 0.1667 | 7 | 0 | 4 | 0 | 24 | 0.0833 | 3 | 0 | 6 | 0.1667 | 5 | 0 | 3 | 0 | 17 | 0.0588 |
| 11_1030024 | 3 | 0 | 6 | 0 | 4 | 0 | 4 | 0.25 | 17 | 0.0588 | 2 | 0 | 6 | 0 | 2 | 0 | 4 | 0.25 | 14 | 0.0714 |
| 11_1030027 | 2 | 0 | 3 | 0 | 3 | 0.3333 | NA | NA | 8 | 0.125 | 2 | 0 | 5 | 0.2 | 4 | 0.25 | 1 | 0 | 12 | 0.1667 |
| 11_1030027 | 6 | 0.1667 | 5 | 0 | 7 | 0 | 4 | 0.25 | 22 | 0.0909 | 4 | 0 | 8 | 0.125 | 6 | 0.1667 | 4 | 0.25 | 22 | 0.1364 |
| 11_1030028 | 5 | 0.2 | 6 | 0.5 | 7 | 1 | 3 | 0.6667 | 21 | 0.619 | 4 | 0.75 | 7 | 0.5714 | 6 | 0.8333 | 5 | 0.6 | 22 | 0.6818 |
| 11_1030028 | 7 | 0.1429 | 5 | 0.2 | 6 | 0.3333 | 4 | 0.5 | 22 | 0.2727 | 5 | 0.4 | 7 | 0.2857 | 6 | 0.1667 | 5 | 0.4 | 23 | 0.3043 |
| 11_1030028 | 7 | 0.1429 | 6 | 0 | 8 | 0.25 | 4 | 0.5 | 25 | 0.2 | 5 | 0.2 | 8 | 0.375 | 5 | 0.2 | 5 | 0 | 23 | 0.2174 |

| | | | | | | | | | | | | | | | | | | | | |
|---|---|---|---|---|---|---|---|---|---|---|---|---|---|---|---|---|---|---|---|---|
| 11_103002 | 7 | 0.1429 | 6 | 0.3333 | 8 | 0.5 | 3 | 0 | 24 | 0.2917 | 6 | 0.1667 | 8 | 0.125 | 6 | 0.1667 | 4 | 0 | 24 | 0.125 |
| 11_103006 | NA | NA | NA | NA | NA | NA | 1 | 1 | 1 | 1 | NA | NA | NA | NA | NA | NA | 1 | 1 | 1 | 1 |
| 11_103007 | 2 | 0.5 | 2 | 1 | 2 | 1 | 3 | 1 | 9 | 0.8889 | 1 | 1 | 2 | 1 | 2 | 1 | 2 | 1 | 7 | 1 |
| 11_103007 | 3 | 1 | 3 | 1 | 3 | 1 | 1 | 1 | 10 | 1 | 2 | 0.5 | 2 | 1 | NA | NA | 4 | 1 | 8 | 0.875 |
| 11_103007 | 3 | 0 | 4 | 0.25 | 3 | 0 | 3 | 0.3333 | 13 | 0.1538 | 1 | 1 | 3 | 0.3333 | 3 | 0.6667 | 3 | 0.6667 | 10 | 0.6 |
| 11_103007 | 2 | 0.5 | 4 | 1 | 3 | 0.3333 | 3 | 0.6667 | 12 | 0.6667 | 1 | 0 | 3 | 0.6667 | 3 | 1 | 3 | 0 | 10 | 0.5 |
| 11_103007 | 1 | 0 | 2 | 0.5 | 2 | 0 | 1 | 0 | 6 | 0.1667 | NA | NA | 4 | 0 | 3 | 0.3333 | 4 | 0.5 | 11 | 0.2727 |
| 11_103007 | 5 | 0.6 | 5 | 0.6 | 4 | 1 | 3 | 0.6667 | 17 | 0.7059 | 2 | 0 | 3 | 0.6667 | 4 | 0.75 | 4 | 0 | 13 | 0.3846 |
| 11_103007 | 4 | 1 | 4 | 1 | 3 | 0.6667 | 3 | 0.6667 | 14 | 0.8571 | 1 | 1 | 1 | 0 | 4 | 0.5 | 4 | 0.75 | 10 | 0.6 |
| 11_103007 | 4 | 0.25 | 3 | 0.3333 | 3 | 0.3333 | 3 | 0.3333 | 13 | 0.3077 | 2 | 0 | 4 | 0.75 | 4 | 0.25 | 4 | 0 | 14 | 0.2857 |
| 11_103007 | 7 | 0.2857 | 4 | 0.25 | 3 | 0.3333 | 3 | 0.3333 | 17 | 0.2941 | 1 | 1 | 5 | 0.4 | 5 | 0.6 | 4 | 0.25 | 15 | 0.4667 |
| 11_103008 | 8 | 0.875 | 6 | 0.6667 | 4 | 0.75 | 4 | 0.75 | 22 | 0.7727 | 2 | 0 | 5 | 0.4 | 6 | 0.5 | 4 | 0.5 | 17 | 0.4118 |
| 11_103008 | 9 | 0.3333 | 6 | 0.1667 | 4 | 0 | 5 | 0 | 24 | 0.1667 | 3 | 0 | 6 | 0 | 5 | 0 | 3 | 0.3333 | 17 | 0.0588 |
| 11_103008 | 8 | 0.75 | 6 | 0.6667 | 3 | 0.6667 | 5 | 0.8 | 22 | 0.7273 | 4 | 1 | 5 | 0.8 | 5 | 0.8 | 3 | 0.6667 | 17 | 0.8235 |
| 11_103008 | 8 | 0 | 6 | 0.1667 | 3 | 0 | 4 | 0.25 | 21 | 0.0952 | 5 | 0 | 4 | 0 | 6 | 0.1667 | 4 | 0 | 19 | 0.0526 |
| 11_103008 | 5 | 0.2 | 5 | 0.2 | 4 | 0 | 4 | 0.25 | 18 | 0.1667 | 6 | 0.3333 | 5 | 0.2 | 7 | 0 | 4 | 0 | 22 | 0.1364 |
| 11_103008 | 7 | 0.8571 | 6 | 1 | 4 | 1 | 5 | 0.8 | 22 | 0.9091 | 4 | 0.75 | 6 | 0.8333 | 7 | 0.8571 | 5 | 0.8 | 22 | 0.8182 |
| 11_103008 | 8 | 0.5 | 6 | 1 | 3 | 0.6667 | 3 | 1 | 20 | 0.75 | 5 | 1 | 3 | 0.6667 | 4 | 0.75 | 3 | 1 | 15 | 0.8667 |
| 11_103008 | 6 | 0.8333 | 4 | 0.75 | 2 | 1 | 5 | 1 | 17 | 0.8824 | 2 | 0.5 | 3 | 0.6667 | 6 | 1 | 4 | 1 | 15 | 0.8667 |
| 11_103008 | 8 | 0.875 | 6 | 0.8333 | 6 | 0.1667 | 5 | 0.6 | 25 | 0.64 | 5 | 0.6 | 6 | 0.5 | 6 | 0.8333 | 5 | 0.4 | 22 | 0.5909 |
| 11_103008 | 9 | 0.7778 | 7 | 0.5714 | 7 | 0.7143 | 5 | 0.4 | 28 | 0.6429 | 4 | 0.5 | 7 | 0.8571 | 8 | 0.75 | 4 | 0.75 | 23 | 0.7391 |
| 11_103009 | 1 | 1 | 2 | 0 | 1 | 0 | NA | NA | 4 | 0.25 | NA | NA | 1 | 1 | 1 | 1 | NA | NA | 2 | 1 |
| 11_103010 | 2 | 0 | 3 | 0.3333 | 2 | 1 | 1 | 0 | 8 | 0.375 | 1 | 1 | 2 | 0.5 | 3 | 0 | 1 | 1 | 7 | 0.4286 |
| 11_103010 | 2 | 0 | 1 | 0 | 2 | 0.5 | 1 | 0 | 6 | 0.1667 | 1 | 0 | 1 | 0 | 3 | 0.3333 | 2 | 0 | 7 | 0.1429 |
| 11_103029 | NA | NA | 1 | 1 | NA | NA | 1 | 1 | 2 | 1 | 3 | 0.6667 | 3 | 1 | 3 | 1 | NA | NA | 9 | 0.8889 |
| 11_103029 | 1 | 1 | 1 | 1 | NA | NA | 1 | 1 | 3 | 1 | 2 | 0.5 | 3 | 1 | 2 | 1 | NA | NA | 7 | 0.8571 |
| 11_103029 | 1 | 0 | 1 | 1 | NA | NA | 1 | 1 | 3 | 0.6667 | 1 | 1 | 3 | 1 | 3 | 0.6667 | NA | NA | 7 | 0.8571 |
| 11_103029 | 1 | 1 | 1 | 0 | NA | NA | 1 | 0 | 3 | 0.3333 | 4 | 0.5 | 1 | 0 | 3 | 0.6667 | NA | NA | 8 | 0.5 |
| 11_103029 | 1 | 1 | 1 | 1 | 1 | 0 | 1 | 1 | 4 | 0.75 | 3 | 0.6667 | 3 | 0.3333 | 3 | 0.3333 | NA | NA | 9 | 0.4444 |
| 11_103029 | 1 | 1 | 1 | 1 | NA | NA | 1 | 1 | 3 | 1 | 3 | 1 | 3 | 1 | 3 | 1 | NA | NA | 9 | 1 |
| 11_103029 | 1 | 0 | 1 | 0 | 1 | 0 | 1 | 1 | 4 | 0.25 | 3 | 0 | 2 | 1 | 2 | 0 | NA | NA | 7 | 0.2857 |
| 11_103030 | 1 | 1 | NA | NA | 1 | 1 | 1 | 0 | 3 | 0.6667 | 2 | 0 | 3 | 1 | 2 | 0.5 | NA | NA | 7 | 0.5714 |
| 11_103030 | 1 | 1 | NA | NA | NA | NA | 1 | 1 | 2 | 1 | 3 | 1 | 2 | 1 | 1 | 1 | NA | NA | 6 | 1 |
| 11_103030 | 1 | 1 | NA | NA | NA | NA | NA | NA | 1 | 1 | 3 | 1 | 2 | 1 | 1 | 1 | NA | NA | 6 | 1 |
| 11_103030 | 1 | 0 | NA | NA | 1 | 1 | 2 | 0.5 | 4 | 0.5 | 1 | 0 | 3 | 0.3333 | 3 | 0 | NA | NA | 7 | 0.1429 |
| 11_103030 | 2 | 0.5 | NA | NA | NA | NA | 2 | 1 | 4 | 0.75 | 1 | 1 | 2 | 0.5 | 3 | 0.6667 | NA | NA | 6 | 0.6667 |
| 11_103089 | 5 | 0 | 9 | 0.1111 | 14 | 0 | 3 | 0.3333 | 31 | 0.0645 | 9 | 0.2222 | 6 | 0.3333 | 4 | 0.25 | 12 | 0.0833 | 31 | 0.1935 |
| 11_103089 | 6 | 0.5 | 13 | 0.0769 | 14 | 0.2857 | 2 | 0 | 35 | 0.2286 | 11 | 0.3636 | 9 | 0.2222 | 4 | 0.25 | 13 | 0.3846 | 37 | 0.3243 |
| 11_103222 | NA | NA | NA | NA | NA | NA | 1 | 1 | 1 | 1 | 1 | 1 | NA | NA | NA | NA | NA | NA | 1 | 1 |
| 11_103223 | 2 | 0 | NA | NA | NA | NA | 4 | 0.25 | 6 | 0.1667 | NA | NA | NA | NA | 1 | 0 | 3 | 0.3333 | 4 | 0.25 |
| 11_103223 | 2 | 0 | NA | NA | NA | NA | 4 | 0.25 | 6 | 0.1667 | NA | NA | 1 | 0 | 2 | 0 | 3 | 0.3333 | 6 | 0.1667 |
| 11_103223 | 2 | 0.5 | 1 | 0 | 1 | 0 | 3 | 0 | 7 | 0.1429 | NA | NA | 1 | 1 | 1 | 0 | 2 | 0 | 4 | 0.25 |
| 11_103223 | 2 | 0.5 | 1 | 0 | NA | NA | 3 | 0.3333 | 6 | 0.3333 | NA | NA | 1 | 0 | 1 | 0 | 1 | 1 | 3 | 0.3333 |
| 11_103223 | 2 | 1 | NA | NA | NA | NA | 3 | 0.6667 | 5 | 0.8 | NA | NA | 1 | 0 | 1 | 1 | 2 | 0.5 | 4 | 0.5 |
| 11_103223 | 2 | 0.5 | NA | NA | NA | NA | 3 | 0.3333 | 5 | 0.4 | NA | NA | NA | NA | NA | NA | 1 | 1 | 1 | 1 |
| 11_103251 | 1 | 1 | 1 | 0 | NA | NA | NA | NA | 2 | 0.5 | 1 | 1 | 2 | 0.5 | NA | NA | NA | NA | 3 | 0.6667 |
| 11_103251 | 1 | 1 | 1 | 1 | NA | NA | 1 | 0 | 3 | 0.6667 | 1 | 1 | 2 | 0.5 | NA | NA | NA | NA | 3 | 0.6667 |
| 11_103256 | 3 | 0.3333 | 5 | 0.6 | 3 | 1 | 3 | 0.6667 | 14 | 0.6429 | NA | NA | 1 | 0 | 2 | 1 | 2 | 1 | 5 | 0.8 |
| 11_103256 | 2 | 0 | 4 | 0.25 | 2 | 0 | 3 | 0.3333 | 11 | 0.1818 | NA | NA | 1 | 0 | 2 | 0 | 2 | 0.5 | 5 | 0.2 |
| 11_103257 | 1 | 1 | 2 | 0.5 | 3 | 0.6667 | NA | NA | 6 | 0.6667 | NA | NA | 1 | 0 | 2 | 0.5 | 1 | 0 | 4 | 0.25 |
| 11_103257 | 1 | 1 | 2 | 0.5 | 2 | 0.5 | 1 | 1 | 6 | 0.6667 | NA | NA | 1 | 0 | 2 | 0.5 | 2 | 0.5 | 5 | 0.4 |
| 11_103267 | NA | NA | 1 | 0 | 2 | 0.5 | 1 | 1 | 4 | 0.5 | NA | NA | 1 | 1 | 2 | 0 | NA | NA | 3 | 0.3333 |
| 11_103267 | NA | NA | 1 | 1 | 2 | 0.5 | NA | NA | 3 | 0.6667 | NA | NA | 1 | 1 | 2 | 0.5 | NA | NA | 3 | 0.6667 |
| 11_103267 | NA | NA | NA | NA | 2 | 0.5 | 1 | 1 | 3 | 0.6667 | NA | NA | 1 | 1 | 2 | 0.5 | NA | NA | 3 | 0.6667 |
| 11_103267 | NA | NA | NA | NA | 3 | 0.3333 | 1 | 1 | 4 | 0.5 | NA | NA | 1 | 1 | 1 | 0 | NA | NA | 2 | 0.5 |
| 11_103268 | NA | NA | NA | NA | 3 | 0.6667 | NA | NA | 3 | 0.6667 | NA | NA | 1 | 0 | 2 | 1 | NA | NA | 3 | 0.6667 |
| 11_103285 | NA | NA | NA | NA | 3 | 0.6667 | 1 | 0 | 4 | 0.5 | NA | NA | NA | NA | 1 | 1 | NA | NA | 1 | 1 |
| 11_103285 | NA | NA | NA | NA | 3 | 0.6667 | 1 | 1 | 4 | 0.75 | NA | NA | NA | NA | 1 | 1 | NA | NA | 1 | 1 |
| 11_103285 | NA | NA | NA | NA | 3 | 0.3333 | 1 | 0 | 4 | 0.25 | NA | NA | NA | NA | 2 | 0.5 | NA | NA | 2 | 0.5 |
| 11_103285 | NA | NA | NA | NA | 3 | 1 | 1 | 1 | 4 | 1 | NA | NA | NA | NA | 2 | 1 | NA | NA | 2 | 1 |
| 11_103540 | 13 | 0.0769 | 12 | 0.25 | 8 | 0.25 | 6 | 0.1667 | 39 | 0.1795 | 9 | 0.1111 | 7 | 0.4286 | 11 | 0.0909 | 7 | 0.2857 | 34 | 0.2059 |
| 11_103540 | 16 | 0.375 | 15 | 0.4667 | 11 | 0.3636 | 8 | 0.25 | 50 | 0.38 | 11 | 0.1818 | 10 | 0.1 | 15 | 0.2 | 12 | 0.3333 | 48 | 0.2083 |
| 11_103540 | NA | NA | NA | NA | 1 | 1 | NA | NA | 1 | 1 | NA | NA | NA | NA | 1 | 1 | NA | NA | 1 | 1 |
| 11_103540 | 20 | 1 | 17 | 1 | 11 | 1 | 8 | 0.875 | 56 | 0.9821 | 11 | 1 | 9 | 0.8889 | 13 | 1 | 10 | 0.9 | 43 | 0.9535 |
| 11_103541 | 18 | 0.2222 | 19 | 0.1053 | 14 | 0.2857 | 10 | 0.2 | 61 | 0.1967 | 15 | 0.2 | 10 | 0.1 | 14 | 0.0714 | 15 | 0.3333 | 54 | 0.1852 |

| | | | | | | | | | | | | | | | | | | | | |
|---|---|---|---|---|---|---|---|---|---|---|---|---|---|---|---|---|---|---|---|---|
| 11_103541 | 17 | 0.2941 | 14 | 0.1429 | 14 | 0.2143 | 8 | 0.125 | 53 | 0.2075 | 13 | 0.3077 | 9 | 0.4444 | 14 | 0.2143 | 14 | 0.2857 | 50 | 0.3 |
| 11_103541 | 21 | 0.7143 | 17 | 0.7059 | 12 | 0.9167 | 9 | 0.7778 | 59 | 0.7627 | 15 | 0.7333 | 6 | 0.8333 | 14 | 0.5714 | 13 | 0.7692 | 48 | 0.7083 |
| 11_103541 | 19 | 1 | 17 | 1 | 14 | 1 | 9 | 1 | 59 | 1 | 13 | 1 | 6 | 0.8333 | 12 | 1 | 11 | 1 | 42 | 0.9762 |
| 11_103541 | 20 | 0.25 | 17 | 0.2353 | 15 | 0.3333 | 10 | 0.1 | 62 | 0.2419 | 15 | 0.2 | 10 | 0.4 | 15 | 0.0667 | 16 | 0.25 | 56 | 0.2143 |
| 11_103541 | 17 | 0.2353 | 14 | 0 | 14 | 0.1429 | 8 | 0.25 | 53 | 0.1509 | 12 | 0.25 | 11 | 0 | 10 | 0.5 | 14 | 0.2143 | 47 | 0.234 |
| 11_103568 | 9 | 0.2222 | 9 | 0.3333 | 6 | 0.1667 | 9 | 0.2222 | 33 | 0.2424 | 4 | 0 | 7 | 0.1429 | 10 | 0.2 | 4 | 0.5 | 25 | 0.2 |
| 11_103568 | 12 | 0 | 10 | 0.2 | 6 | 0 | 9 | 0.1111 | 37 | 0.0811 | 5 | 0.2 | 6 | 0 | 11 | 0.0909 | 3 | 0.3333 | 25 | 0.12 |
| 11_103569 | 7 | 0.8571 | 5 | 0.6 | 4 | 0.25 | 7 | 0.4286 | 23 | 0.5652 | 2 | 0.5 | 3 | 1 | 7 | 0.5714 | 1 | 0 | 13 | 0.6154 |
| 11_103569 | 7 | 0.7143 | 7 | 0.7143 | 4 | 0.75 | 7 | 1 | 25 | 0.8 | 4 | 1 | 5 | 0.8 | 7 | 0.7143 | 1 | 1 | 17 | 0.8235 |
| 11_103569 | 5 | 0.2 | 5 | 0.2 | 3 | 0.3333 | 8 | 0.25 | 21 | 0.2381 | 4 | 0.25 | 4 | 0 | 6 | 0.5 | NA | NA | 14 | 0.2857 |
| 11_103569 | 7 | 1 | 7 | 1 | 6 | 0.5 | 7 | 1 | 27 | 0.8889 | 5 | 1 | 4 | 0.5 | 6 | 1 | 1 | 1 | 16 | 0.875 |
| 11_103569 | 5 | 0.2 | 5 | 0 | 6 | 0.1667 | 9 | 0.2222 | 25 | 0.16 | 5 | 0 | 3 | 0 | 7 | 0.1429 | 1 | 0 | 16 | 0.0625 |
| 11_103570 | 3 | 1 | 6 | 1 | 7 | 0.5714 | 8 | 0.875 | 24 | 0.8333 | 4 | 0.75 | 4 | 0.75 | 3 | 1 | 1 | 1 | 12 | 0.8333 |
| 11_103570 | 7 | 0.1429 | 6 | 0 | 6 | 0 | 9 | 0.1111 | 28 | 0.0714 | 5 | 0 | 3 | 0.3333 | 6 | 0 | 1 | 0 | 15 | 0.0667 |
| 11_103613 | NA | NA | NA | NA | NA | NA | 1 | 1 | 1 | 1 | NA | NA | 1 | 1 | NA | NA | NA | NA | 1 | 1 |
| 11_103614 | 1 | 0 | NA | NA | NA | NA | 1 | 1 | 2 | 0.5 | NA | NA | 1 | 1 | NA | NA | NA | NA | 1 | 1 |
| 11_103614 | 1 | 1 | NA | NA | NA | NA | 1 | 0 | 2 | 0.5 | NA | NA | 2 | 0.5 | NA | NA | NA | NA | 2 | 0.5 |
| 11_103614 | 1 | 1 | NA | NA | NA | NA | 1 | 0 | 2 | 0.5 | NA | NA | 2 | 0.5 | NA | NA | NA | NA | 2 | 0.5 |
| 11_103615 | 1 | 0 | NA | NA | NA | NA | 1 | 1 | 2 | 0.5 | NA | NA | 2 | 0.5 | NA | NA | 1 | 1 | 3 | 0.6667 |
| 11_103615 | 1 | 1 | NA | NA | NA | NA | NA | NA | 1 | 1 | NA | NA | 2 | 0 | NA | NA | 1 | 1 | 3 | 0.3333 |
| 11_103615 | 1 | 1 | NA | NA | NA | NA | 1 | 1 | 2 | 1 | NA | NA | 2 | 1 | NA | NA | 1 | 1 | 3 | 1 |
| 11_103616 | 1 | 1 | NA | NA | NA | NA | 1 | 1 | 2 | 1 | NA | NA | 1 | 1 | NA | NA | 1 | 1 | 2 | 1 |
| 11_103616 | 1 | 1 | 1 | 0 | NA | NA | 1 | 0 | 3 | 0.3333 | NA | NA | 2 | 0.5 | NA | NA | 1 | 0 | 3 | 0.3333 |
| 11_103616 | NA | NA | 2 | 0 | NA | NA | 1 | 1 | 3 | 0.3333 | NA | NA | 3 | 0.6667 | NA | NA | 1 | 0 | 4 | 0.5 |
| 11_103713 | 61 | 0.7869 | 76 | 0.7632 | 59 | 0.8305 | 55 | 0.8182 | 251 | 0.7968 | 51 | 0.8039 | 51 | 0.7255 | 62 | 0.7903 | 55 | 0.7455 | 219 | 0.7671 |
| 11_103713 | 62 | 0.0806 | 84 | 0.0714 | 61 | 0.0656 | 58 | 0.1897 | 265 | 0.0981 | 51 | 0.1176 | 50 | 0.24 | 62 | 0.129 | 63 | 0.0952 | 226 | 0.1416 |
| 11_103713 | 48 | 0.1458 | 59 | 0.0678 | 39 | 0.0513 | 34 | 0.1176 | 180 | 0.0944 | 40 | 0.05 | 37 | 0.027 | 42 | 0.0476 | 45 | 0.1333 | 164 | 0.0671 |
| 11_103714 | 63 | 0.3492 | 78 | 0.2821 | 59 | 0.3559 | 56 | 0.3036 | 256 | 0.3203 | 50 | 0.28 | 43 | 0.2791 | 60 | 0.2 | 64 | 0.3125 | 217 | 0.2673 |
| 11_103874 | 10 | 0.1 | 7 | 0 | 3 | 0 | 7 | 0.2857 | 27 | 0.1111 | 4 | 0.25 | 3 | 0 | 8 | 0.25 | 8 | 0.375 | 23 | 0.2609 |
| 11_103875 | 11 | 0.0909 | 9 | 0.2222 | 3 | 0 | 6 | 0.1667 | 29 | 0.1379 | 5 | 0 | 5 | 0 | 8 | 0 | 10 | 0.2 | 28 | 0.0714 |
| 11_104065 | 1 | 1 | 4 | 0.25 | 1 | 0 | 4 | 0.5 | 10 | 0.4 | NA | NA | 1 | 1 | 2 | 0 | NA | NA | 3 | 0.3333 |
| 11_104065 | 1 | 0 | 4 | 0 | 2 | 0.5 | 4 | 0.75 | 11 | 0.3636 | NA | NA | 1 | 1 | 2 | 0 | NA | NA | 3 | 0.3333 |
| 11_104065 | 1 | 0 | 3 | 1 | 2 | 0.5 | 3 | 0.3333 | 9 | 0.5556 | NA | NA | NA | NA | 1 | 1 | NA | NA | 1 | 1 |
| 11_104066 | NA | NA | 4 | 0.5 | 2 | 0.5 | 5 | 0 | 11 | 0.2727 | NA | NA | 2 | 0.5 | 1 | 0 | NA | NA | 3 | 0.3333 |
| 11_104066 | 1 | 0 | 4 | 0.25 | 2 | 0.5 | 6 | 0.5 | 13 | 0.3846 | NA | NA | 3 | 0.3333 | 2 | 0 | NA | NA | 5 | 0.2 |
| 11_104066 | 1 | 0 | 4 | 0.5 | 2 | 0.5 | 5 | 0.6 | 12 | 0.5 | NA | NA | 2 | 0.5 | 2 | 0.5 | 1 | 1 | 5 | 0.6 |
| 11_104218 | 2 | 1 | 5 | 0.8 | 7 | 0.7143 | 10 | 0.6 | 24 | 0.7083 | 5 | 0.8 | 10 | 0.5 | 8 | 0.375 | 4 | 0.75 | 27 | 0.5556 |
| 11_104219 | 3 | 0.6667 | 7 | 0.4286 | 13 | 0.5385 | 12 | 0.25 | 35 | 0.4286 | 9 | 0.1111 | 11 | 0.3636 | 8 | 0.375 | 6 | 0.3333 | 34 | 0.2941 |
| 11_104219 | 4 | 0.25 | 6 | 0.5 | 14 | 0.3571 | 15 | 0.4 | 39 | 0.3846 | 11 | 0.2727 | 14 | 0.5 | 7 | 0.1429 | 5 | 0 | 37 | 0.2973 |
| 11_104220 | 4 | 0.25 | 8 | 0.375 | 13 | 0.5385 | 14 | 0.5 | 39 | 0.4615 | 12 | 0.4167 | 13 | 0.3846 | 9 | 0.2222 | 4 | 0.5 | 38 | 0.3684 |
| 11_104220 | 5 | 0.2 | 17 | 0.2353 | 16 | 0.3125 | 21 | 0.2857 | 59 | 0.2712 | 13 | 0.1538 | 14 | 0.3571 | 13 | 0.3077 | 6 | 0.3333 | 46 | 0.2826 |
| 11_104221 | 5 | 0.2 | 18 | 0.5 | 17 | 0.2353 | 19 | 0.2105 | 59 | 0.3051 | 16 | 0.4375 | 16 | 0.4375 | 12 | 0.6667 | 7 | 0.4286 | 51 | 0.4902 |
| 11_104221 | 4 | 0.25 | 18 | 0.1667 | 15 | 0.1333 | 22 | 0.0909 | 59 | 0.1356 | 13 | 0.2308 | 13 | 0 | 14 | 0.1429 | 7 | 0.1429 | 47 | 0.1277 |
| 11_104221 | 4 | 0.5 | 18 | 0.2778 | 16 | 0.4375 | 21 | 0.2857 | 59 | 0.339 | 15 | 0.3333 | 14 | 0.5714 | 15 | 0.4 | 6 | 0.5 | 50 | 0.44 |
| 11_104221 | 5 | 0.4 | 15 | 0.0667 | 14 | 0.1429 | 16 | 0.3125 | 50 | 0.2 | 13 | 0.2308 | 14 | 0.5714 | 13 | 0.2308 | 7 | 0.2857 | 47 | 0.3404 |
| 11_104221 | 4 | 0 | 15 | 0.2667 | 10 | 0.3 | 19 | 0.3158 | 48 | 0.2708 | 14 | 0.3571 | 13 | 0.6923 | 13 | 0.3846 | 7 | 0.2857 | 47 | 0.4468 |
| 11_104221 | 5 | 0 | 17 | 0.1765 | 16 | 0.125 | 22 | 0.1818 | 60 | 0.15 | 15 | 0.0667 | 15 | 0.3333 | 14 | 0 | 8 | 0.125 | 52 | 0.1346 |
| 11_104221 | 5 | 0 | 16 | 0.375 | 15 | 0.0667 | 20 | 0.05 | 56 | 0.1429 | 17 | 0.1176 | 14 | 0.2857 | 15 | 0.1333 | 8 | 0.375 | 54 | 0.2037 |
| 11_104221 | 5 | 0.2 | 18 | 0.2778 | 15 | 0.2 | 23 | 0.3043 | 61 | 0.2623 | 16 | 0.0625 | 13 | 0.3077 | 14 | 0.2857 | 7 | 0 | 50 | 0.18 |
| 11_104221 | 5 | 0.6 | 18 | 0.5556 | 16 | 0.5 | 20 | 0.65 | 59 | 0.5763 | 14 | 0.7143 | 14 | 0.5 | 12 | 0.5 | 9 | 0.7778 | 49 | 0.6122 |
| 11_104221 | 5 | 0 | 20 | 0.1 | 15 | 0.0667 | 16 | 0.125 | 56 | 0.0893 | 15 | 0.1333 | 10 | 0.1 | 15 | 0 | 10 | 0 | 50 | 0.06 |
| 11_104221 | 5 | 0 | 22 | 0.0909 | 16 | 0.0625 | 22 | 0.1364 | 65 | 0.0923 | 16 | 0 | 17 | 0.0588 | 16 | 0 | 7 | 0.2857 | 56 | 0.0536 |
| 11_104221 | 4 | 0.25 | 20 | 0.25 | 18 | 0.2778 | 23 | 0.1304 | 65 | 0.2154 | 16 | 0.1875 | 18 | 0.2778 | 15 | 0.3333 | 8 | 0.375 | 57 | 0.2807 |
| 11_104221 | 4 | 0.25 | 16 | 0.1875 | 11 | 0.3636 | 16 | 0.1875 | 47 | 0.234 | 15 | 0.0667 | 12 | 0.3333 | 16 | 0.25 | 8 | 0.25 | 51 | 0.2157 |
| 11_104222 | 6 | 0 | 37 | 0.1081 | 23 | 0 | 29 | 0.0345 | 95 | 0.0526 | 24 | 0.0833 | 23 | 0.0435 | 25 | 0.12 | 20 | 0 | 92 | 0.0652 |
| 11_104222 | 8 | 0.375 | 25 | 0.4 | 22 | 0.5 | 22 | 0.4545 | 77 | 0.4416 | 25 | 0.52 | 19 | 0.5263 | 21 | 0.619 | 19 | 0.4211 | 84 | 0.5238 |
| 11_104225 | 2 | 0.5 | 12 | 0.3333 | 7 | 0.4286 | 3 | 0.3333 | 24 | 0.375 | 6 | 0.1667 | 5 | 0.2 | 3 | 0.3333 | 5 | 0.6 | 19 | 0.3158 |
| 11_104225 | 2 | 1 | 9 | 0.7778 | 7 | 0.5714 | 5 | 0.6 | 23 | 0.6957 | 4 | 0.5 | 4 | 0.5 | 3 | 0.3333 | 5 | 1 | 16 | 0.625 |
| 11_104225 | 2 | 0 | 11 | 0.5455 | 7 | 0.4286 | 4 | 0.25 | 24 | 0.4167 | 6 | 0.6667 | 5 | 0.2 | 3 | 0.3333 | 6 | 0.3333 | 20 | 0.4 |
| 11_104225 | 1 | 0 | 4 | 0.5 | 4 | 0.25 | 2 | 0 | 11 | 0.2727 | 3 | 0.3333 | 1 | 1 | 1 | 0 | 4 | 0.5 | 9 | 0.4444 |
| 11_104314 | NA | NA | 5 | 0.4 | 2 | 0.5 | 1 | 0 | 8 | 0.375 | 3 | 0 | 2 | 0 | 3 | 0.6667 | 4 | 0 | 12 | 0.1667 |
| 11_104314 | NA | NA | 7 | 0.1429 | 2 | 1 | 1 | 0 | 10 | 0.3 | 3 | 0 | 1 | 1 | 3 | 0.3333 | 4 | 0 | 11 | 0.1818 |
| 11_104314 | NA | NA | 7 | 0.8571 | 2 | 1 | NA | NA | 9 | 0.8889 | 3 | 1 | 1 | 1 | 3 | 0.3333 | 4 | 0.75 | 11 | 0.7273 |
| 11_104315 | NA | NA | 8 | 0.125 | 1 | 0 | NA | NA | 9 | 0.1111 | 3 | 0 | 1 | 0 | 1 | 1 | 3 | 0.3333 | 8 | 0.25 |
| 11_104315 | NA | NA | 5 | 0.6 | 2 | 0.5 | NA | NA | 7 | 0.5714 | 3 | 0.6667 | NA | NA | 1 | 1 | 4 | 0.75 | 8 | 0.75 |

| | | | | | | | | | | | | | | | | | | | | |
|---|---|---|---|---|---|---|---|---|---|---|---|---|---|---|---|---|---|---|---|---|
| 11_1043152 | NA | NA | 7 | 0.7143 | 1 | 1 | NA | NA | 8 | 0.75 | 3 | 0.6667 | NA | NA | NA | NA | 2 | 1 | 5 | 0.8 |
| 11_1043152 | NA | NA | 6 | 0.8333 | 2 | 1 | NA | NA | 8 | 0.875 | 3 | 0 | NA | NA | NA | NA | 2 | 1 | 5 | 0.4 |
| 11_1043153 | NA | NA | 7 | 0.5714 | 2 | 0.5 | NA | NA | 9 | 0.5556 | 3 | 0.3333 | NA | NA | NA | NA | 3 | 0.6667 | 6 | 0.5 |
| 11_1043154 | NA | NA | 6 | 0.8333 | 2 | 1 | NA | NA | 8 | 0.875 | 3 | 1 | NA | NA | NA | NA | 3 | 1 | 6 | 1 |
| 11_1043154 | NA | NA | 7 | 0.8571 | 2 | 1 | NA | NA | 9 | 0.8889 | 3 | 1 | NA | NA | NA | NA | 3 | 1 | 6 | 1 |
| 11_1043155 | NA | NA | 7 | 0.4286 | 2 | 0.5 | NA | NA | 9 | 0.4444 | 2 | 0.5 | 1 | 0 | NA | NA | 3 | 0.6667 | 6 | 0.5 |
| 11_1043156 | NA | NA | 4 | 0.5 | 1 | 1 | NA | NA | 5 | 0.6 | 3 | 0.6667 | NA | NA | NA | NA | 1 | 1 | 4 | 0.75 |
| 11_1043157 | NA | NA | 4 | 0.5 | NA | NA | NA | NA | 4 | 0.5 | 2 | 1 | 1 | 1 | NA | NA | 1 | 1 | 4 | 1 |
| 11_1043158 | NA | NA | 6 | 0.1667 | 2 | 0 | NA | NA | 8 | 0.125 | 3 | 0.3333 | NA | NA | NA | NA | 3 | 0 | 6 | 0.1667 |
| 11_1043952 | 2 | 0.5 | 2 | 0 | 1 | 0 | 4 | 0.5 | 9 | 0.3333 | NA | NA | 1 | 1 | 3 | 0 | 5 | 0.4 | 9 | 0.3333 |
| 11_1043953 | 1 | 1 | 1 | 0 | 1 | 0 | 4 | 0 | 7 | 0.1429 | 1 | 0 | 1 | 0 | 3 | 0.3333 | 4 | 0 | 9 | 0.1111 |
| 11_1043955 | 1 | 1 | 2 | 0 | 1 | 1 | 3 | 0.6667 | 7 | 0.5714 | 1 | 1 | 1 | 0 | 3 | 0.3333 | 4 | 0.25 | 9 | 0.3333 |
| 11_1043957 | 1 | 1 | 2 | 0 | 1 | 1 | 4 | 0 | 8 | 0.25 | 1 | 0 | 1 | 0 | 4 | 0.25 | 4 | 0 | 10 | 0.1 |
| 11_1043958 | 1 | 1 | 2 | 1 | 1 | 1 | 1 | 1 | 5 | 1 | 1 | 1 | 1 | 1 | 3 | 1 | 3 | 1 | 8 | 1 |
| 11_1043959 | 1 | 1 | 2 | 1 | 1 | 1 | 1 | 0 | 5 | 0.8 | 1 | 0 | 2 | 0.5 | 3 | 0.6667 | 3 | 1 | 9 | 0.6667 |
| 11_1043959 | 1 | 1 | 3 | 0.3333 | 1 | 0 | 2 | 0 | 7 | 0.2857 | 1 | 0 | 1 | 0 | 3 | 0.6667 | 3 | 0 | 8 | 0.25 |
| 11_1043960 | NA | NA | 3 | 0.6667 | 1 | 0 | 1 | 0 | 5 | 0.4 | 1 | 0 | 2 | 0.5 | 3 | 0 | 3 | 0.3333 | 9 | 0.2222 |
| 11_1043961 | 1 | 1 | 3 | 0.3333 | 1 | 1 | 2 | 1 | 7 | 0.7143 | 1 | 0 | 2 | 1 | 4 | 0.5 | 4 | 1 | 11 | 0.7273 |
| 11_1043962 | 1 | 1 | 3 | 0.6667 | 1 | 1 | NA | NA | 5 | 0.8 | NA | NA | 2 | 0.5 | 2 | 1 | 3 | 1 | 7 | 0.8571 |
| 11_1043962 | 1 | 1 | 3 | 0.3333 | 1 | 1 | 2 | 1 | 7 | 0.7143 | 1 | 1 | 2 | 0.5 | 4 | 0.75 | 4 | 0.75 | 11 | 0.7273 |
| 11_1043962 | 1 | 1 | 3 | 0 | 1 | 1 | 1 | 1 | 6 | 0.5 | 1 | 0 | 2 | 0.5 | 3 | 0 | 3 | 0 | 9 | 0.1111 |
| 11_1043962 | 1 | 0 | 3 | 0.6667 | 1 | 0 | 2 | 1 | 7 | 0.5714 | 1 | 1 | 2 | 1 | 3 | 1 | 3 | 0.6667 | 9 | 0.8889 |
| 11_1043964 | NA | NA | 2 | 0.5 | NA | NA | 2 | 1 | 4 | 0.75 | 1 | 1 | 2 | 0.5 | 2 | 1 | 4 | 0.5 | 9 | 0.6667 |
| 11_1043964 | 1 | 0 | 2 | 0 | 1 | 1 | 2 | 0.5 | 6 | 0.3333 | 1 | 0 | 1 | 1 | 2 | 0 | 4 | 0.5 | 8 | 0.375 |
| 11_1050315 | 1 | 1 | 1 | 0 | NA | NA | NA | NA | 2 | 0.5 | NA | NA | 4 | 0.5 | NA | NA | NA | NA | 4 | 0.5 |
| 11_1050707 | NA | NA | NA | NA | 2 | 1 | NA | NA | 2 | 1 | 2 | 0.5 | NA | NA | NA | NA | NA | NA | 2 | 0.5 |
| 11_1050709 | NA | NA | NA | NA | 2 | 1 | NA | NA | 2 | 1 | 2 | 1 | NA | NA | NA | NA | NA | NA | 2 | 1 |
| 11_1050711 | NA | NA | NA | NA | 2 | 0.5 | NA | NA | 2 | 0.5 | 2 | 1 | NA | NA | NA | NA | NA | NA | 2 | 1 |
| 11_1050712 | NA | NA | NA | NA | 2 | 0.5 | NA | NA | 2 | 0.5 | 2 | 0.5 | NA | NA | NA | NA | NA | NA | 2 | 0.5 |
| 11_1050713 | NA | NA | NA | NA | 2 | 1 | NA | NA | 2 | 1 | 1 | 1 | NA | NA | NA | NA | NA | NA | 1 | 1 |
| 11_1051668 | NA | NA | 1 | 1 | 1 | 0 | NA | NA | 2 | 0.5 | NA | NA | 1 | 1 | NA | NA | NA | NA | 1 | 1 |
| 11_1051721 | 4 | 0.75 | 8 | 0.5 | 5 | 0.6 | 1 | 0 | 18 | 0.5556 | 1 | 1 | 3 | 1 | 7 | 0.7143 | 3 | 1 | 14 | 0.8571 |
| 11_1051721 | 4 | 1 | 7 | 1 | 7 | 1 | 3 | 0.6667 | 21 | 0.9524 | 1 | 1 | 4 | 1 | 6 | 0.8333 | 3 | 1 | 14 | 0.9286 |
| 11_1051721 | 4 | 0 | 8 | 0.5 | 8 | 0 | 3 | 0.3333 | 23 | 0.2174 | NA | NA | 4 | 0.25 | 7 | 0.4286 | 1 | 1 | 12 | 0.4167 |
| 11_1051723 | 4 | 0 | 7 | 0.2857 | 7 | 0.4286 | 3 | 0.6667 | 21 | 0.3333 | 1 | 0 | 4 | 0.5 | 7 | 0.1429 | 3 | 0.3333 | 15 | 0.2667 |
| 11_1051723 | 3 | 0 | 7 | 0.2857 | 7 | 0.4286 | 2 | 0 | 19 | 0.2632 | 1 | 0 | 4 | 0.5 | 8 | 0.5 | 2 | 0.5 | 15 | 0.4667 |
| 11_1051724 | 3 | 0.6667 | 7 | 0.5714 | 9 | 0.4444 | 3 | 0 | 22 | 0.4545 | 1 | 0 | 4 | 0.25 | 6 | 0.3333 | 1 | 1 | 12 | 0.3333 |
| 11_1051724 | 4 | 1 | 6 | 1 | 8 | 0.75 | 3 | 1 | 21 | 0.9048 | 1 | 1 | 3 | 1 | 7 | 1 | 2 | 1 | 13 | 1 |
| 11_1051724 | 4 | 1 | 9 | 0.2222 | 8 | 0.375 | 1 | 0 | 22 | 0.4091 | 1 | 1 | 4 | 0.5 | 8 | 0.75 | 2 | 0 | 15 | 0.6 |
| 11_1051724 | 4 | 0.25 | 6 | 0 | 8 | 0.125 | 3 | 0 | 21 | 0.0952 | 1 | 0 | 4 | 0 | 8 | 0.125 | 1 | 0 | 14 | 0.0714 |
| 11_1058127 | NA | NA | 2 | 1 | 2 | 1 | 2 | 0.5 | 6 | 0.8333 | 4 | 0.75 | 1 | 1 | 2 | 1 | 1 | 1 | 8 | 0.875 |
| 11_1058128 | NA | NA | 2 | 0 | 1 | 0 | 2 | 1 | 5 | 0.4 | 3 | 0.3333 | 2 | 0 | 2 | 0 | NA | NA | 7 | 0.1429 |
| 11_1058129 | NA | NA | 2 | 1 | 2 | 1 | 3 | 1 | 7 | 1 | 4 | 1 | 2 | 0.5 | 2 | 1 | 1 | 1 | 9 | 0.8889 |
| 11_1058130 | NA | NA | 2 | 0 | 2 | 1 | 1 | 1 | 5 | 0.6 | 3 | 1 | 2 | 0.5 | 1 | 1 | NA | NA | 6 | 0.8333 |
| 11_1058137 | 1 | 0 | 3 | 0 | 2 | 0.5 | 3 | 0 | 9 | 0.1111 | 5 | 0 | 2 | 1 | 2 | 0 | 1 | 0 | 10 | 0.2 |
| 11_1058138 | 1 | 1 | 4 | 0.5 | 3 | 0.3333 | 5 | 0.4 | 13 | 0.4615 | 5 | 0 | 2 | 0.5 | 2 | 0.5 | 2 | 0 | 11 | 0.1818 |
| 11_1058139 | NA | NA | 3 | 0.3333 | 3 | 0.3333 | 5 | 0.4 | 11 | 0.3636 | 5 | 0.2 | 2 | 0.5 | 1 | 0 | 1 | 0 | 9 | 0.2222 |
| 11_1058140 | 1 | 1 | 4 | 1 | 3 | 1 | 4 | 0.75 | 12 | 0.9167 | 5 | 1 | 2 | 1 | 1 | 1 | 2 | 1 | 10 | 1 |
| 11_1058141 | 1 | 0 | 3 | 0 | 2 | 0.5 | 3 | 0 | 9 | 0.1111 | 3 | 0.3333 | 1 | 1 | 2 | 0 | 1 | 0 | 7 | 0.2857 |
| 11_1058141 | 1 | 1 | 4 | 1 | 3 | 1 | 5 | 1 | 13 | 1 | 5 | 0.8 | 2 | 1 | 2 | 0.5 | 1 | 1 | 10 | 0.8 |
| 11_1058141 | 1 | 1 | 2 | 0.5 | 3 | 0.6667 | 4 | 1 | 10 | 0.8 | 5 | 0.8 | 2 | 1 | 2 | 0.5 | 2 | 0.5 | 11 | 0.7273 |
| 11_1058144 | 2 | 1 | 4 | 1 | 3 | 1 | 5 | 1 | 14 | 1 | 5 | 1 | 2 | 1 | 2 | 1 | 1 | 1 | 10 | 1 |
| 11_1058144 | 2 | 1 | 3 | 1 | 3 | 1 | 5 | 1 | 13 | 1 | 5 | 0.4 | 2 | 1 | 2 | 1 | 2 | 1 | 11 | 0.7273 |
| 11_1058144 | 2 | 1 | 4 | 1 | 2 | 1 | 5 | 1 | 13 | 1 | 4 | 1 | 1 | 1 | 3 | 0.6667 | 2 | 1 | 10 | 0.9 |
| 11_1058145 | 1 | 1 | 4 | 0.75 | 3 | 0.6667 | 4 | 1 | 12 | 0.8333 | 5 | 0.8 | 2 | 1 | 3 | 1 | 2 | 1 | 12 | 0.9167 |
| 11_1058145 | 2 | 1 | 3 | 1 | 3 | 0.6667 | 5 | 1 | 13 | 0.9231 | 5 | 0.8 | 1 | 1 | 3 | 1 | NA | NA | 9 | 0.8889 |
| 11_1058145 | 2 | 0.5 | 4 | 0.25 | 3 | 0 | 5 | 0 | 14 | 0.1429 | 5 | 0.2 | 2 | 0.5 | 3 | 0 | 2 | 0 | 12 | 0.1667 |
| 11_1058146 | 2 | 0.5 | 3 | 0 | 2 | 0 | 5 | 0 | 12 | 0.0833 | 5 | 0 | 1 | 0 | 3 | 0.3333 | 1 | 0 | 10 | 0.1 |
| 11_1058146 | 2 | 1 | 3 | 0.3333 | 3 | 0.6667 | 5 | 0.4 | 13 | 0.5385 | 5 | 0.2 | 3 | 0.3333 | 3 | 0.6667 | 1 | 0 | 12 | 0.3333 |
| 11_1058147 | 2 | 1 | 3 | 1 | 3 | 1 | 4 | 1 | 12 | 1 | 5 | 0.6 | 3 | 1 | 3 | 0.6667 | 2 | 1 | 13 | 0.7692 |
| 11_1058148 | 2 | 0 | 3 | 0.3333 | 3 | 0 | 6 | 0 | 14 | 0.0714 | 4 | 0.25 | 3 | 0 | 3 | 0 | 2 | 0 | 12 | 0.0833 |
| 11_1058148 | 2 | 0 | 4 | 0 | 3 | 0 | 5 | 0.4 | 14 | 0.1429 | 4 | 0.25 | 3 | 0.3333 | 3 | 0 | 2 | 0 | 12 | 0.1667 |
| 11_1058151 | 2 | 0 | 3 | 0 | 3 | 0.3333 | 5 | 0 | 13 | 0.0769 | 4 | 0.25 | 3 | 0 | 4 | 0 | 1 | 0 | 12 | 0.0833 |
| 11_1058157 | 4 | 0.5 | 4 | 0.25 | 4 | 0.75 | 7 | 0.1429 | 19 | 0.3684 | 5 | 0.2 | 3 | 0 | 5 | 0.2 | 3 | 0 | 16 | 0.125 |
| 11_1058163 | 5 | 0.2 | 6 | 0 | 6 | 0 | 7 | 0.1429 | 24 | 0.0833 | 6 | 0.1667 | 3 | 0 | 7 | 0 | 3 | 0.3333 | 19 | 0.1053 |

| | | | | | | | | | | | | | | | | | | | | |
|---|---|---|---|---|---|---|---|---|---|---|---|---|---|---|---|---|---|---|---|---|
| 11_105817( | 1 | 0 | 5 | 0.4 | 5 | 0 | 4 | 0 | 15 | 0.1333 | 6 | 0 | 2 | 0 | 7 | 0.1429 | 3 | 0 | 18 | 0.0556 |
| 11_1058174 | 4 | 0 | 8 | 0 | 8 | 0 | 10 | 0.2 | 30 | 0.0667 | 6 | 0.1667 | 3 | 0 | 7 | 0.1429 | 5 | 0 | 21 | 0.0952 |
| 11_105817! | 6 | 0 | 11 | 0 | 10 | 0.3 | 9 | 0 | 36 | 0.0833 | 8 | 0.125 | 4 | 0.25 | 10 | 0 | 5 | 0.2 | 27 | 0.1111 |
| 11_105818: | 8 | 0.125 | 8 | 0.375 | 9 | 0.2222 | 10 | 0.2 | 35 | 0.2286 | 7 | 0 | 4 | 0 | 10 | 0 | 7 | 0.2857 | 28 | 0.0714 |
| 11_105818: | 7 | 0 | 12 | 0.3333 | 11 | 0.4545 | 9 | 0.2222 | 39 | 0.2821 | 9 | 0.2222 | 7 | 0.1429 | 10 | 0.4 | 7 | 0.2857 | 33 | 0.2727 |
| 11_105818! | 8 | 0.125 | 15 | 0.1333 | 9 | 0.2222 | 15 | 0.0667 | 47 | 0.1277 | 10 | 0.2 | 9 | 0 | 11 | 0.0909 | 9 | 0.1111 | 39 | 0.1026 |
| 11_105818( | 8 | 0 | 15 | 0.2667 | 10 | 0.5 | 15 | 0.0667 | 48 | 0.2083 | 10 | 0.1 | 9 | 0.3333 | 12 | 0.0833 | 10 | 0.4 | 41 | 0.2195 |
| 11_105818( | 5 | 0 | 11 | 0.2727 | 7 | 0.2857 | 9 | 0.1111 | 32 | 0.1875 | 8 | 0.5 | 6 | 0 | 5 | 0.2 | 6 | 0 | 25 | 0.2 |
| 11_105818 | 7 | 0.2857 | 13 | 0.3846 | 7 | 0.1429 | 12 | 0.3333 | 39 | 0.3077 | 9 | 0.4444 | 9 | 0.2222 | 10 | 0.2 | 8 | 0.25 | 36 | 0.2778 |
| 11_105818 | 8 | 0.25 | 10 | 0.6 | 10 | 0.4 | 9 | 0.2222 | 37 | 0.3784 | 8 | 0.125 | 8 | 0.5 | 9 | 0.1111 | 8 | 0.375 | 33 | 0.2727 |
| 11_105818! | 10 | 0.6 | 15 | 0.3333 | 11 | 0.4545 | 13 | 0.4615 | 49 | 0.449 | 12 | 0.5833 | 10 | 0.4 | 13 | 0.4615 | 9 | 0.7778 | 44 | 0.5455 |
| 11_105819( | 7 | 0 | 15 | 0.0667 | 11 | 0.1818 | 12 | 0.0833 | 45 | 0.0889 | 10 | 0.1 | 9 | 0.1111 | 12 | 0.0833 | 11 | 0.2727 | 42 | 0.1429 |
| 11_105819' | 8 | 0.125 | 12 | 0.0833 | 7 | 0.1429 | 16 | 0.0625 | 43 | 0.093 | 7 | 0.1429 | 9 | 0.1111 | 13 | 0.0769 | 9 | 0.1111 | 38 | 0.1053 |
| 11_105819: | 10 | 0.6 | 19 | 0.5789 | 11 | 0.4545 | 16 | 0.5625 | 56 | 0.5536 | 12 | 0.6667 | 11 | 0.6364 | 14 | 0.6429 | 11 | 0.5455 | 48 | 0.625 |
| 11_105819! | 14 | 0.6429 | 22 | 0.6364 | 13 | 0.6154 | 16 | 0.625 | 65 | 0.6308 | 14 | 0.6429 | 14 | 0.7857 | 17 | 0.7647 | 14 | 0.5714 | 59 | 0.6949 |
| 11_105819! | 9 | 0.2222 | 19 | 0.3158 | 9 | 0 | 13 | 0.3846 | 50 | 0.26 | 11 | 0.2727 | 12 | 0.25 | 16 | 0.3125 | 12 | 0.0833 | 51 | 0.2353 |
| 11_105819! | 13 | 0.0769 | 22 | 0.0455 | 15 | 0.1333 | 19 | 0.2105 | 69 | 0.1159 | 12 | 0.0833 | 13 | 0.0769 | 17 | 0.1176 | 16 | 0.1875 | 58 | 0.1207 |
| 11_105819! | 11 | 0.0909 | 18 | 0.1111 | 12 | 0.25 | 17 | 0.2353 | 58 | 0.1724 | 12 | 0.1667 | 14 | 0.1429 | 15 | 0 | 14 | 0.0714 | 55 | 0.0909 |
| 11_105820( | 11 | 0.2727 | 21 | 0.1905 | 12 | 0 | 19 | 0.1579 | 63 | 0.1587 | 14 | 0.2143 | 12 | 0.0833 | 17 | 0.1176 | 16 | 0.125 | 59 | 0.1356 |
| 11_105825: | 23 | 0.0435 | 7 | 0 | 5 | 0.2 | 4 | 0 | 39 | 0.0513 | 8 | 0.25 | 12 | 0.0833 | 6 | 0 | 8 | 0.125 | 34 | 0.1176 |
| 11_105825! | 18 | 0.3889 | 7 | 0.1429 | 2 | 0 | 6 | 0.3333 | 33 | 0.303 | 7 | 0 | 8 | 0.125 | 5 | 0 | 6 | 0.1667 | 26 | 0.0769 |
| 11_105826: | 16 | 0.25 | 6 | 0.6667 | 2 | 0 | 4 | 0.25 | 28 | 0.3214 | 6 | 0.1667 | 8 | 0.125 | 5 | 0.6 | 4 | 0.25 | 23 | 0.2609 |
| 11_105880: | 1 | 1 | NA | NA | NA | NA | 1 | 0 | 2 | 0.5 | 1 | 1 | 1 | 1 | NA | NA | NA | NA | 2 | 1 |
| 11_105918! | NA | NA | 1 | 1 | NA | NA | NA | NA | 1 | 1 | 2 | 0.5 | 1 | 0 | NA | NA | NA | NA | 3 | 0.3333 |
| 11_105918! | NA | NA | 1 | 1 | NA | NA | NA | NA | 1 | 1 | 1 | 1 | NA | NA | NA | NA | NA | NA | 1 | 1 |
| 11_105923! | 1 | 0 | 1 | 0 | 2 | 1 | 3 | 0.6667 | 7 | 0.5714 | NA | NA | NA | NA | 1 | 1 | 3 | 0.3333 | 4 | 0.5 |
| 11_105928( | 4 | 0 | 5 | 0 | 2 | 0 | 5 | 0.2 | 16 | 0.0625 | 1 | 0 | 1 | 0 | 3 | 0.3333 | 4 | 0 | 9 | 0.1111 |
| 11_105945! | 6 | 0.6667 | 8 | 0.875 | 1 | 0 | 4 | 0.25 | 19 | 0.6316 | 2 | 1 | 6 | 0.8333 | 5 | 1 | 6 | 0.6667 | 19 | 0.8421 |
| 11_105949 | 29 | 0.2069 | 19 | 0.2632 | 10 | 0.3 | 12 | 0.1667 | 70 | 0.2286 | 8 | 0.625 | 14 | 0.4286 | 20 | 0.55 | 21 | 0.381 | 63 | 0.4762 |
| 11_105949 | 31 | 0.0645 | 21 | 0.0476 | 10 | 0.1 | 13 | 0.0769 | 75 | 0.0667 | 9 | 0 | 11 | 0.2727 | 22 | 0 | 20 | 0.1 | 62 | 0.0806 |
| 11_105949! | 33 | 0.5455 | 16 | 0.6875 | 10 | 0.5 | 11 | 0.6364 | 70 | 0.5857 | 8 | 0.75 | 12 | 0.5 | 22 | 0.7273 | 20 | 0.85 | 62 | 0.7258 |
| 11_105949! | 29 | 0.2759 | 19 | 0.1053 | 11 | 0.2727 | 11 | 0.2727 | 70 | 0.2286 | 10 | 0.1 | 12 | 0.1667 | 20 | 0.2 | 24 | 0.1667 | 66 | 0.1667 |
| 11_105957: | NA | NA | 1 | 1 | 2 | 0 | 1 | 0 | 4 | 0.25 | 1 | 0 | NA | NA | 1 | 1 | NA | NA | 2 | 0.5 |
| 11_105963! | 1 | 0 | 1 | 1 | 2 | 0.5 | NA | NA | 4 | 0.5 | 1 | 0 | NA | NA | 1 | 1 | NA | NA | 2 | 0.5 |
| 11_105964' | 2 | 0.5 | 1 | 0 | 2 | 0.5 | NA | NA | 5 | 0.4 | 1 | 1 | NA | NA | 1 | 1 | NA | NA | 2 | 1 |
| 11_105964' | NA | NA | 1 | 1 | 2 | 0.5 | 1 | 1 | 4 | 0.75 | 1 | 1 | NA | NA | 1 | 1 | NA | NA | 2 | 1 |
| 11_105964: | 2 | 1 | 1 | 1 | 2 | 1 | 1 | 1 | 6 | 1 | NA | NA | NA | NA | 1 | 1 | NA | NA | 1 | 1 |
| 11_105964: | 2 | 1 | 1 | 1 | 2 | 1 | 1 | 1 | 6 | 1 | 1 | 1 | NA | NA | 1 | 1 | NA | NA | 2 | 1 |
| 11_105964 | 1 | 1 | NA | NA | 2 | 0.5 | NA | NA | 3 | 0.6667 | NA | NA | NA | NA | 1 | 1 | NA | NA | 1 | 1 |
| 11_105964 | 1 | 1 | 1 | 1 | 2 | 0.5 | NA | NA | 4 | 0.75 | NA | NA | NA | NA | 1 | 1 | NA | NA | 1 | 1 |
| 11_105965' | NA | NA | NA | NA | 2 | 0.5 | NA | NA | 2 | 0.5 | 1 | 0 | NA | NA | 1 | 1 | NA | NA | 2 | 0.5 |
| 11_106044! | NA | NA | 2 | 0.5 | 4 | 1 | 2 | 0.5 | 8 | 0.75 | NA | NA | 3 | 1 | 1 | 0 | 3 | 0.6667 | 7 | 0.7143 |
| 11_106045' | NA | NA | 2 | 1 | 3 | 1 | 4 | 1 | 9 | 1 | NA | NA | 3 | 1 | 2 | 1 | 3 | 1 | 8 | 1 |
| 11_106045: | NA | NA | 2 | 1 | 2 | 1 | 4 | 0.5 | 8 | 0.75 | NA | NA | 3 | 0.6667 | 2 | 1 | 3 | 1 | 8 | 0.875 |
| 11_106045: | NA | NA | 2 | 0.5 | 4 | 0.5 | 4 | 0.5 | 10 | 0.5 | NA | NA | 3 | 0 | 2 | 0 | 3 | 0.6667 | 8 | 0.25 |
| 11_106045: | NA | NA | 2 | 1 | 3 | 1 | 4 | 0.25 | 9 | 0.6667 | NA | NA | 3 | 1 | 1 | 1 | 2 | 1 | 6 | 1 |
| 11_106045: | NA | NA | 1 | 1 | 4 | 1 | 2 | 1 | 7 | 1 | NA | NA | 3 | 1 | 2 | 1 | 4 | 1 | 9 | 1 |
| 11_106045 | NA | NA | 2 | 0.5 | 2 | 0 | 3 | 0 | 7 | 0.1429 | NA | NA | 3 | 0 | 1 | 0 | 5 | 0.2 | 9 | 0.1111 |
| 11_106045 | NA | NA | 2 | 0 | 4 | 0.25 | 2 | 0.5 | 8 | 0.25 | NA | NA | 3 | 0 | 2 | 0.5 | 5 | 0.6 | 10 | 0.4 |
| 11_106045 | NA | NA | 2 | 1 | 4 | 1 | 1 | 1 | 7 | 1 | 1 | 1 | 3 | 0.3333 | 1 | 1 | 4 | 1 | 9 | 0.7778 |
| 11_106046! | NA | NA | 3 | 0.6667 | 4 | 0.25 | 3 | 0 | 10 | 0.3 | 1 | 1 | 3 | 0.6667 | 3 | 0.3333 | 5 | 0.6 | 12 | 0.5833 |
| 11_106047( | 3 | 0.6667 | 1 | 0 | 2 | 0 | NA | NA | 6 | 0.3333 | 1 | 1 | NA | NA | NA | NA | 1 | 1 | 2 | 1 |
| 11_106047 | 3 | 0.3333 | 1 | 0 | 2 | 0 | 1 | 0 | 7 | 0.1429 | 1 | 1 | NA | NA | NA | NA | 1 | 0 | 2 | 0.5 |
| 11_106047! | 1 | 0 | NA | NA | 2 | 0 | 1 | 1 | 4 | 0.25 | 1 | 0 | NA | NA | NA | NA | 1 | 1 | 2 | 0.5 |
| 11_106047! | 3 | 0.6667 | 1 | 0 | 2 | 0.5 | 1 | 0 | 7 | 0.4286 | 1 | 0 | NA | NA | NA | NA | 1 | 1 | 2 | 0.5 |
| 11_106048' | 2 | 0.5 | 1 | 0 | 1 | 1 | NA | NA | 4 | 0.5 | 1 | 1 | NA | NA | NA | NA | 1 | 1 | 2 | 1 |
| 11_106048: | 3 | 0.3333 | 1 | 0 | 2 | 0.5 | NA | NA | 6 | 0.3333 | 1 | 0 | NA | NA | NA | NA | 1 | 1 | 2 | 0.5 |
| 11_106049( | 28 | 0.2857 | 19 | 0.1053 | 14 | 0.2857 | 22 | 0.2727 | 83 | 0.241 | 19 | 0.0526 | 29 | 0.3103 | 25 | 0.16 | 20 | 0.35 | 93 | 0.2258 |
| 11_106054: | 7 | 0.1429 | 2 | 0.5 | 5 | 0.2 | NA | NA | 14 | 0.2143 | 2 | 0 | 2 | 0.5 | 4 | 0.25 | 2 | 0 | 10 | 0.2 |
| 11_106054! | 7 | 0.1429 | 1 | 0 | 6 | 0.3333 | 1 | 0 | 15 | 0.2 | NA | NA | 4 | 0.25 | 4 | 0.5 | 4 | 0.25 | 12 | 0.3333 |
| 11_106054( | 7 | 0.2857 | 1 | 0 | 4 | 0.75 | NA | NA | 12 | 0.4167 | 1 | 0 | 4 | 0.5 | 4 | 0.5 | 4 | 0 | 13 | 0.3077 |
| 11_106054 | 7 | 0.5714 | 1 | 0 | 5 | 0.8 | NA | NA | 13 | 0.6154 | 1 | 0 | 4 | 0 | 3 | 0.6667 | 4 | 0.5 | 12 | 0.3333 |
| 11_106054 | 6 | 0.8333 | 1 | 1 | 4 | 1 | NA | NA | 11 | 0.9091 | NA | NA | 2 | 0.5 | 3 | 1 | 4 | 0.5 | 9 | 0.6667 |
| 11_106054 | 7 | 0.7143 | 1 | 0 | 5 | 0.2 | 1 | 0 | 14 | 0.4286 | 1 | 0 | 4 | 0.5 | 2 | 0.5 | 4 | 0.75 | 11 | 0.5455 |
| 11_106088! | NA | NA | 2 | 0.5 | 2 | 0.5 | 1 | 0 | 5 | 0.4 | NA | NA | 2 | 0.5 | NA | NA | 2 | 0 | 4 | 0.25 |

| | | | | | | | | | | | | | | | | | | | | |
|---|---|---|---|---|---|---|---|---|---|---|---|---|---|---|---|---|---|---|---|---|
| 11_106091 | 6 | 0 | 10 | 0.4 | 6 | 0.1667 | 2 | 0 | 24 | 0.2083 | 1 | 0 | 9 | 0.2222 | 7 | 0.2857 | 8 | 0.5 | 25 | 0.32 |
| 11_106091 | 6 | 0.1667 | 11 | 0.0909 | 6 | 0 | 1 | 0 | 24 | 0.0833 | 1 | 1 | 8 | 0 | 9 | 0 | 11 | 0.0909 | 29 | 0.069 |
| 11_106138 | NA | NA | 1 | 1 | 1 | 1 | 1 | 1 | 3 | 1 | NA | NA | NA | NA | 1 | 1 | NA | NA | 1 | 1 |
| 11_106138 | NA | NA | 1 | 1 | 1 | 1 | 1 | 0 | 3 | 0.6667 | NA | NA | NA | NA | 1 | 1 | NA | NA | 1 | 1 |
| 11_106138 | 8 | 0.875 | 5 | 0.8 | 3 | 1 | 7 | 0.8571 | 23 | 0.8696 | 1 | 1 | 1 | 0 | 2 | 1 | 2 | 1 | 6 | 0.8333 |
| 11_106138 | NA | NA | 1 | 1 | 1 | 1 | 1 | 0 | 3 | 0.6667 | NA | NA | NA | NA | 1 | 1 | NA | NA | 1 | 1 |
| 11_106138 | 6 | 0.3333 | 3 | 0.3333 | 2 | 0.5 | 6 | 0.1667 | 17 | 0.2941 | 2 | 0.5 | 2 | 0 | 2 | 0.5 | 3 | 0 | 9 | 0.2222 |
| 11_106138 | 9 | 0.1111 | 5 | 0 | 9 | 0 | 9 | 0.2222 | 32 | 0.0938 | 2 | 0 | 3 | 0 | 2 | 0 | 4 | 0.25 | 11 | 0.0909 |
| 11_106138 | 10 | 0.4 | 5 | 0.4 | 8 | 0.625 | 10 | 0.4 | 33 | 0.4545 | 3 | 0.6667 | 4 | 0.5 | 3 | 0.3333 | 3 | 0 | 13 | 0.3846 |
| 11_106138 | 8 | 0.625 | 4 | 0.75 | 6 | 1 | 6 | 0.8333 | 24 | 0.7917 | 3 | 0.6667 | 4 | 0.5 | 2 | 1 | 4 | 0.75 | 13 | 0.6923 |
| 11_106138 | 9 | 0.3333 | 3 | 0 | 6 | 0.6667 | 8 | 0.625 | 26 | 0.4615 | 1 | 1 | 4 | 0.5 | 1 | 1 | 4 | 0.25 | 10 | 0.5 |
| 11_106138 | 10 | 0.1 | 5 | 0 | 6 | 0 | 8 | 0.125 | 29 | 0.069 | 3 | 0.6667 | 3 | 0 | 3 | 0.3333 | 4 | 0 | 13 | 0.2308 |
| 11_106139 | 8 | 0.25 | 4 | 0.25 | 10 | 0.6 | 8 | 0.375 | 30 | 0.4 | 3 | 0.3333 | 3 | 0.3333 | 6 | 0.3333 | 4 | 0.75 | 16 | 0.4375 |
| 11_106139 | 10 | 0.3 | 5 | 0.6 | 8 | 0.375 | 10 | 0.5 | 33 | 0.4242 | 2 | 1 | 5 | 0.2 | 7 | 0.2857 | 3 | 0.3333 | 17 | 0.3529 |
| 11_106139 | NA | NA | 1 | 0 | 1 | 1 | 1 | 0 | 3 | 0.3333 | NA | NA | NA | NA | 1 | 1 | NA | NA | 1 | 1 |
| 11_106139 | 14 | 0.2143 | 6 | 0.1667 | 11 | 0 | 12 | 0.0833 | 43 | 0.1163 | 3 | 0 | 6 | 0.1667 | 7 | 0.1429 | 4 | 0 | 20 | 0.1 |
| 11_106139 | 15 | 0.5333 | 6 | 0.3333 | 11 | 0.3636 | 11 | 0.5455 | 43 | 0.4651 | 4 | 0.25 | 6 | 0.1667 | 7 | 0.1429 | 5 | 0.4 | 22 | 0.2273 |
| 11_106139 | 9 | 0 | 3 | 0.3333 | 7 | 0.1429 | 12 | 0 | 31 | 0.0645 | 4 | 0 | 7 | 0.2857 | 8 | 0 | 5 | 0 | 24 | 0.0833 |
| 11_106140 | 3 | 1 | 3 | 1 | 1 | 1 | 5 | 1 | 12 | 1 | 4 | 0.75 | 3 | 0.6667 | 2 | 0.5 | 2 | 0.5 | 11 | 0.6364 |
| 11_106140 | 3 | 0.6667 | 4 | 0.5 | 1 | 1 | 5 | 1 | 13 | 0.7692 | 3 | 1 | 2 | 1 | 5 | 0.8 | 2 | 1 | 12 | 0.9167 |
| 11_106140 | 3 | 1 | 4 | 1 | 1 | 0 | 6 | 0.8333 | 14 | 0.8571 | 3 | 0.6667 | 4 | 0.75 | 3 | 0.3333 | 1 | 1 | 11 | 0.6364 |
| 11_106140 | 3 | 0.3333 | 4 | 0 | 1 | 0 | 5 | 0 | 13 | 0.0769 | 4 | 0 | 3 | 0 | 3 | 0.3333 | 2 | 0 | 12 | 0.0833 |
| 11_106140 | 2 | 1 | 4 | 0.5 | NA | NA | 6 | 0.3333 | 12 | 0.5 | 4 | 0.25 | 4 | 0.5 | 3 | 0.3333 | 2 | 0 | 13 | 0.3077 |
| 11_106140 | 3 | 0 | 4 | 0 | 1 | 0 | 6 | 0.1667 | 14 | 0.0714 | 4 | 0.5 | 3 | 0.6667 | 4 | 0 | 1 | 0 | 12 | 0.3333 |
| 11_106163 | 1 | 1 | NA | NA | 1 | 0 | 1 | 1 | 3 | 0.6667 | NA | NA | NA | NA | 1 | 1 | NA | NA | 1 | 1 |
| 11_106393 | 4 | 0.5 | 10 | 0.3 | 8 | 0.25 | 9 | 0.3333 | 31 | 0.3226 | 6 | 0.5 | 5 | 0.6 | 5 | 0.6 | 13 | 0.5385 | 29 | 0.5517 |
| 11_106394 | 4 | 0 | 7 | 0.1429 | 6 | 0.1667 | 8 | 0.25 | 25 | 0.16 | 6 | 0.1667 | 3 | 0.3333 | 1 | 0 | 13 | 0.0769 | 23 | 0.1304 |
| 11_106394 | 6 | 0.1667 | 4 | 0 | 5 | 0.2 | 8 | 0.125 | 23 | 0.1304 | 5 | 0.4 | 2 | 0 | 2 | 0 | 8 | 0.25 | 17 | 0.2353 |
| 11_106394 | 5 | 0.8 | 5 | 0.8 | 6 | 0.8333 | 10 | 0.8 | 26 | 0.8077 | 5 | 0.8 | 3 | 1 | 1 | 1 | 12 | 0.9167 | 21 | 0.9048 |
| 11_106394 | 5 | 0 | 6 | 0.6667 | 6 | 0.1667 | 10 | 0.3 | 27 | 0.2963 | 5 | 0.4 | 3 | 0.3333 | 2 | 0 | 11 | 0.4545 | 21 | 0.381 |
| 11_106435 | 1 | 0 | 4 | 0 | 5 | 0 | 7 | 0.1429 | 17 | 0.0588 | 4 | 0 | 2 | 0 | 1 | 0 | 11 | 0.0909 | 18 | 0.0556 |
| 11_106437 | 3 | 0 | 4 | 0.25 | 3 | 0.3333 | 7 | 0.2857 | 17 | 0.2353 | 2 | 0.5 | 2 | 0 | NA | NA | 7 | 0 | 11 | 0.0909 |
| 11_106439 | 3 | 0 | 4 | 0 | 3 | 0.3333 | 6 | 0 | 16 | 0.0625 | 3 | 0.3333 | 2 | 0 | NA | NA | 10 | 0.1 | 15 | 0.1333 |
| 11_106458 | 2 | 0 | 4 | 0.75 | 3 | 0.6667 | 6 | 0.5 | 15 | 0.5333 | 2 | 0 | 2 | 0.5 | NA | NA | 9 | 0.4444 | 13 | 0.3846 |
| 11_106458 | 2 | 0.5 | 4 | 0.25 | 3 | 0.3333 | 5 | 0.4 | 14 | 0.3571 | 2 | 0.5 | 2 | 0 | NA | NA | 9 | 0.3333 | 13 | 0.3077 |
| 11_106458 | 1 | 1 | 2 | 1 | 2 | 0 | 6 | 0 | 11 | 0.2727 | 1 | 0 | NA | NA | NA | NA | 8 | 0.375 | 9 | 0.3333 |
| 11_106458 | 2 | 1 | 3 | 1 | 3 | 1 | 6 | 1 | 14 | 1 | 2 | 1 | 2 | 0.5 | NA | NA | 7 | 1 | 11 | 0.9091 |
| 11_106458 | 2 | 0.5 | 2 | 1 | 1 | 1 | 7 | 0.8571 | 12 | 0.8333 | 1 | 0 | 2 | 1 | NA | NA | 7 | 1 | 10 | 0.9 |
| 11_106458 | 2 | 1 | 3 | 1 | 2 | 0 | 6 | 0.6667 | 13 | 0.6923 | 2 | 0.5 | 2 | 0 | NA | NA | 8 | 0.625 | 12 | 0.5 |
| 11_106458 | 1 | 0 | 2 | 0 | 2 | 0 | 5 | 0.2 | 10 | 0.1 | 1 | 0 | 2 | 0.5 | NA | NA | 8 | 0.25 | 11 | 0.2727 |
| 11_106458 | 2 | 0 | 3 | 0 | 2 | 0.5 | 6 | 0 | 13 | 0.0769 | 1 | 0 | 2 | 0 | NA | NA | 7 | 0.1429 | 10 | 0.1 |
| 11_106458 | 2 | 0 | 3 | 0.3333 | 2 | 0 | 6 | 0.1667 | 13 | 0.1538 | 1 | 1 | 2 | 0 | NA | NA | 8 | 0.375 | 11 | 0.3636 |
| 11_106459 | 1 | 1 | 3 | 1 | 1 | 1 | 4 | 0.75 | 9 | 0.8889 | NA | NA | NA | NA | NA | NA | 4 | 0.75 | 4 | 0.75 |
| 11_106459 | 1 | 1 | 3 | 1 | 1 | 1 | 3 | 0.6667 | 8 | 0.875 | 1 | 0 | 2 | 0.5 | NA | NA | 3 | 0.6667 | 6 | 0.5 |
| 11_106545 | 1 | 1 | 4 | 1 | 2 | 1 | NA | NA | 7 | 1 | 2 | 1 | NA | NA | NA | NA | NA | NA | 2 | 1 |
| 11_106579 | 1 | 0 | 3 | 0 | 2 | 0.5 | NA | NA | 6 | 0.1667 | 1 | 1 | NA | NA | NA | NA | NA | NA | 1 | 1 |
| 11_106671 | 50 | 0.22 | 60 | 0.1667 | 54 | 0.1111 | 70 | 0.2 | 234 | 0.1752 | 30 | 0.2 | 44 | 0.0682 | 55 | 0.2 | 49 | 0.1837 | 178 | 0.1629 |
| 11_106672 | 90 | 0.1 | 97 | 0.1134 | 79 | 0.0759 | 108 | 0.1019 | 374 | 0.0989 | 51 | 0.0588 | 87 | 0.092 | 94 | 0.1702 | 83 | 0.0723 | 315 | 0.1048 |
| 11_106672 | 86 | 0.2674 | 90 | 0.3 | 76 | 0.2632 | 102 | 0.2843 | 354 | 0.2797 | 50 | 0.2 | 84 | 0.3452 | 90 | 0.2778 | 75 | 0.32 | 299 | 0.2943 |
| 11_106672 | 79 | 0.1772 | 89 | 0.1124 | 79 | 0.2785 | 102 | 0.1569 | 349 | 0.1777 | 44 | 0.2045 | 80 | 0.2125 | 84 | 0.1786 | 71 | 0.2817 | 279 | 0.2186 |
| 11_106672 | 78 | 0.0513 | 80 | 0.0875 | 71 | 0.0563 | 93 | 0.043 | 322 | 0.059 | 42 | 0.0476 | 67 | 0.1194 | 77 | 0.026 | 67 | 0.0746 | 253 | 0.0672 |
| 11_106673 | 70 | 0.1857 | 76 | 0.2105 | 68 | 0.2647 | 91 | 0.2747 | 305 | 0.2361 | 44 | 0.1818 | 65 | 0.3692 | 75 | 0.2533 | 65 | 0.2462 | 249 | 0.2691 |
| 11_106673 | 22 | 0.1364 | 18 | 0.1667 | 24 | 0.0417 | 27 | 0.1852 | 91 | 0.1319 | 15 | 0.1333 | 21 | 0.0952 | 31 | 0.1613 | 17 | 0.3529 | 84 | 0.1786 |
| 11_106673 | 23 | 0.1304 | 17 | 0.1176 | 22 | 0.0455 | 29 | 0.1724 | 91 | 0.1209 | 12 | 0 | 20 | 0 | 28 | 0.1429 | 14 | 0.0714 | 74 | 0.0676 |
| 11_106674 | 20 | 0.15 | 16 | 0.125 | 24 | 0.0833 | 26 | 0.1538 | 86 | 0.1279 | 12 | 0.0833 | 19 | 0.1579 | 26 | 0.0769 | 12 | 0.1667 | 69 | 0.1159 |
| 11_106674 | 19 | 0.1579 | 14 | 0.2143 | 22 | 0.1818 | 19 | 0.0526 | 74 | 0.1486 | 6 | 0 | 18 | 0.0556 | 24 | 0.1667 | 10 | 0.4 | 58 | 0.1552 |
| 11_106674 | 18 | 0.1111 | 12 | 0.4167 | 20 | 0.15 | 25 | 0.32 | 75 | 0.24 | 8 | 0 | 18 | 0.1667 | 23 | 0.2174 | 13 | 0.3077 | 62 | 0.1935 |
| 11_106680 | 1 | 1 | NA | NA | 1 | 1 | 1 | 1 | 3 | 1 | NA | NA | 1 | 1 | NA | NA | NA | NA | 1 | 1 |
| 11_106680 | 1 | 0 | NA | NA | 1 | 1 | 1 | 1 | 3 | 0.6667 | NA | NA | 2 | 1 | NA | NA | NA | NA | 2 | 1 |
| 11_106680 | 1 | 1 | NA | NA | 1 | 1 | NA | NA | 2 | 1 | NA | NA | 2 | 1 | NA | NA | NA | NA | 2 | 1 |
| 11_106709 | 1 | 0 | 4 | 0.25 | NA | NA | NA | NA | 5 | 0.2 | NA | NA | 2 | 0 | 2 | 0 | 2 | 1 | 6 | 0.3333 |
| 11_106712 | 5 | 0 | 4 | 0 | 3 | 0.3333 | 4 | 0 | 16 | 0.0625 | 1 | 0 | 3 | 0 | 3 | 0.3333 | 4 | 0 | 11 | 0.0909 |
| 11_106713 | 8 | 0.375 | 1 | 0 | 3 | 0.3333 | 5 | 0 | 17 | 0.2353 | 1 | 0 | 6 | 0.1667 | 4 | 0.25 | 4 | 0 | 15 | 0.1333 |
| 11_106713 | 8 | 0.25 | 2 | 1 | 3 | 0.3333 | 4 | 0 | 17 | 0.2941 | 1 | 0 | 6 | 0.1667 | 4 | 0.25 | 4 | 0.75 | 15 | 0.3333 |
| 11_106713 | 8 | 0.25 | 2 | 0 | 3 | 0.3333 | 5 | 0.2 | 18 | 0.2222 | 1 | 0 | 5 | 0.2 | 3 | 0 | 4 | 0.5 | 13 | 0.2308 |

| | | | | | | | | | | | | | | | | | | | | |
|---|---|---|---|---|---|---|---|---|---|---|---|---|---|---|---|---|---|---|---|---|
| 11_1067133 | 7 | 1 | 2 | 1 | 3 | 0.6667 | 4 | 0.75 | 16 | 0.875 | 1 | 1 | 4 | 1 | 3 | 1 | 4 | 0.75 | 12 | 0.9167 |
| 11_1067133 | 7 | 1 | 2 | 1 | 3 | 1 | 3 | 0.6667 | 15 | 0.9333 | 1 | 1 | 5 | 0.8 | 2 | 1 | 4 | 0.75 | 12 | 0.8333 |
| 11_1067134 | 4 | 0.5 | 2 | 0 | 2 | 0.5 | 3 | 0 | 11 | 0.2727 | 1 | 0 | 5 | 0.2 | 3 | 0.6667 | 3 | 0 | 12 | 0.25 |
| 11_1067134 | 7 | 0.7143 | 2 | 0.5 | 3 | 1 | 2 | 0.5 | 14 | 0.7143 | 1 | 1 | 4 | 0.75 | 3 | 0.6667 | 4 | 0.75 | 12 | 0.75 |
| 11_1067135 | 7 | 0.2857 | 2 | 0.5 | 2 | 0 | 3 | 0 | 14 | 0.2143 | 1 | 0 | 4 | 0.25 | 3 | 0.3333 | 2 | 0 | 10 | 0.2 |
| 11_1068796 | 4 | 0 | 3 | 0 | 5 | 0 | 2 | 0.5 | 14 | 0.0714 | 3 | 0 | 5 | 0.4 | 4 | 0.25 | 3 | 0.6667 | 15 | 0.3333 |
| 11_1069206 | 2 | 1 | NA | NA | 1 | 0 | NA | NA | 3 | 0.6667 | NA | NA | 1 | 1 | NA | NA | NA | NA | 1 | 1 |
| 11_1069244 | 1 | 0 | 3 | 0.3333 | NA | NA | 1 | 0 | 5 | 0.2 | 1 | 1 | 1 | 0 | NA | NA | 1 | 0 | 3 | 0.3333 |
| 11_1069244 | 1 | 0 | 3 | 0.3333 | NA | NA | 1 | 0 | 5 | 0.2 | 1 | 0 | 1 | 0 | NA | NA | 1 | 1 | 3 | 0.3333 |
| 11_1069259 | 6 | 0.3333 | 13 | 0.3846 | 3 | 0.3333 | 7 | 0.4286 | 29 | 0.3793 | 4 | 0.25 | 7 | 0.4286 | 10 | 0.4 | 3 | 0.3333 | 24 | 0.375 |
| 11_1069259 | 6 | 1 | 10 | 1 | 2 | 0.5 | 8 | 0.75 | 26 | 0.8846 | 4 | 1 | 7 | 1 | 8 | 0.75 | 3 | 0.6667 | 22 | 0.8636 |
| 11_1069262 | 5 | 0.8 | 9 | 0.5556 | 4 | 0.5 | 5 | 0.4 | 23 | 0.5652 | 1 | 1 | 7 | 0.4286 | 7 | 0.2857 | 3 | 0.6667 | 18 | 0.4444 |
| 11_1069263 | 5 | 0.2 | 11 | 0.2727 | 3 | 0 | 6 | 0.6667 | 25 | 0.32 | 2 | 0 | 7 | 0.5714 | 8 | 0.125 | 4 | 0.5 | 21 | 0.3333 |
| 11_1069264 | 4 | 0.25 | 10 | 0.1 | 2 | 0 | 6 | 0.3333 | 22 | 0.1818 | 2 | 0 | 7 | 0.1429 | 8 | 0.25 | 3 | 0.3333 | 20 | 0.2 |
| 11_1069265 | 5 | 0 | 11 | 0.0909 | 2 | 0 | 6 | 0.1667 | 24 | 0.0833 | 2 | 0.5 | 6 | 0.1667 | 6 | 0 | 4 | 0.25 | 18 | 0.1667 |
| 11_1069265 | 5 | 0.4 | 7 | 0.8571 | 3 | 0.6667 | 4 | 0.75 | 19 | 0.6842 | 2 | 1 | 5 | 1 | 3 | 0 | 4 | 1 | 14 | 0.7857 |
| 11_1069379 | 3 | 0.3333 | 7 | 0 | 2 | 0 | 5 | 0.2 | 17 | 0.1176 | 2 | 0.5 | 3 | 0 | 6 | 0 | 2 | 0.5 | 13 | 0.1538 |
| 11_1069458 | 1 | 0 | 3 | 0.6667 | 1 | 0 | 4 | 0.5 | 9 | 0.4444 | 1 | 0 | 2 | 0.5 | 3 | 0.6667 | 1 | 1 | 7 | 0.5714 |
| 11_1069638 | 1 | 0 | 1 | 1 | 1 | 1 | 2 | 0.5 | 5 | 0.6 | NA | NA | 3 | 1 | 3 | 0.6667 | 1 | 1 | 7 | 0.8571 |
| 11_1069640 | 1 | 1 | NA | NA | 1 | 1 | 1 | 1 | 3 | 1 | NA | NA | 1 | 0 | 3 | 0.6667 | 1 | 0 | 5 | 0.4 |
| 11_1069641 | NA | NA | 1 | 1 | 2 | 0 | 1 | 1 | 4 | 0.5 | NA | NA | 2 | 0.5 | 2 | 1 | 1 | 0 | 5 | 0.6 |
| 11_1069642 | 1 | 0 | 1 | 0 | 2 | 0 | 2 | 0.5 | 6 | 0.1667 | NA | NA | 3 | 0 | 2 | 0.5 | 1 | 0 | 6 | 0.1667 |
| 11_1069646 | 1 | 0 | 1 | 1 | 2 | 0 | 2 | 0 | 6 | 0.1667 | NA | NA | 2 | 0.5 | 2 | 0.5 | 1 | 1 | 5 | 0.6 |
| 11_1069649 | 1 | 0 | 1 | 0 | 2 | 0.5 | 2 | 0 | 6 | 0.1667 | NA | NA | 2 | 1 | 3 | 0 | 1 | 1 | 6 | 0.5 |
| 11_1069649 | 1 | 1 | 1 | 1 | 1 | 0 | 2 | 0 | 5 | 0.4 | NA | NA | 2 | 0.5 | 3 | 0 | 1 | 1 | 6 | 0.3333 |
| 11_1069649 | NA | NA | 1 | 1 | 2 | 0.5 | 2 | 0 | 5 | 0.4 | NA | NA | 1 | 1 | 3 | 0.6667 | 1 | 1 | 5 | 0.8 |
| 11_1069651 | 1 | 0 | 1 | 0 | 2 | 0.5 | 2 | 0 | 6 | 0.1667 | NA | NA | 1 | 0 | 3 | 0.3333 | 1 | 1 | 5 | 0.4 |
| 11_1069652 | 1 | 1 | 1 | 0 | 1 | 0 | 1 | 0 | 4 | 0.25 | NA | NA | 2 | 0 | 3 | 0.6667 | 1 | 0 | 6 | 0.3333 |
| 11_1069654 | 1 | 1 | 1 | 1 | 1 | 1 | 1 | 0 | 4 | 0.75 | NA | NA | 1 | 1 | 2 | 0 | NA | NA | 3 | 0.3333 |
| 11_1070017 | NA | NA | 1 | 0 | 1 | 1 | 1 | 0 | 3 | 0.3333 | NA | NA | 1 | 0 | 2 | 0 | 1 | 1 | 4 | 0.25 |
| 11_1070017 | NA | NA | NA | NA | 1 | 1 | 1 | 1 | 2 | 1 | NA | NA | 1 | 1 | 1 | 1 | 1 | 0 | 3 | 0.6667 |
| 11_1070018 | NA | NA | 1 | 1 | 1 | 1 | 1 | 0 | 3 | 0.6667 | NA | NA | 1 | 0 | 2 | 0.5 | 1 | 0 | 4 | 0.25 |
| 11_1070020 | NA | NA | 1 | 0 | 1 | 1 | 1 | 1 | 3 | 0.6667 | NA | NA | 1 | 1 | 1 | 0 | 1 | 1 | 3 | 0.6667 |
| 11_1070021 | NA | NA | NA | NA | 1 | 0 | 1 | 1 | 2 | 0.5 | NA | NA | 1 | 0 | 1 | 0 | 1 | 1 | 3 | 0.3333 |
| 11_1070021 | NA | NA | 1 | 1 | 1 | 1 | 1 | 0 | 3 | 0.6667 | NA | NA | 1 | 0 | 1 | 1 | 1 | 1 | 3 | 0.6667 |
| 11_1070022 | NA | NA | 1 | 1 | 1 | 1 | 1 | 1 | 3 | 1 | NA | NA | 1 | 1 | 2 | 0.5 | 1 | 1 | 4 | 0.75 |
| 11_1070023 | NA | NA | 1 | 0 | 1 | 1 | 1 | 1 | 3 | 0.6667 | NA | NA | 1 | 1 | 2 | 1 | 1 | 1 | 4 | 1 |
| 11_1070229 | NA | NA | 1 | 0 | NA | NA | 1 | 1 | 2 | 0.5 | NA | NA | 1 | 1 | NA | NA | 1 | 1 | 2 | 1 |
| 11_1070575 | 4 | 1 | NA | NA | 2 | 1 | NA | NA | 6 | 1 | NA | NA | 2 | 1 | 1 | 1 | NA | NA | 3 | 1 |
| 11_1070577 | 4 | 0.75 | NA | NA | 3 | 0.6667 | 4 | 0.5 | 11 | 0.6364 | 1 | 0 | 6 | 0.6667 | 2 | 0.5 | 3 | 0.6667 | 12 | 0.5833 |
| 11_1070579 | 5 | 0.2 | NA | NA | 2 | 0 | 2 | 0 | 9 | 0.1111 | 2 | 0 | 9 | 0 | NA | NA | 5 | 0.2 | 16 | 0.0625 |
| 11_1070581 | 5 | 0.4 | NA | NA | 3 | 0 | 4 | 0 | 12 | 0.1667 | 2 | 0 | 9 | 0.2222 | 2 | 0 | 4 | 0 | 17 | 0.1176 |
| 11_1070581 | 3 | 0.3333 | NA | NA | 3 | 0 | 3 | 0 | 9 | 0.1111 | 2 | 0 | 9 | 0.2222 | 2 | 0 | 4 | 0 | 17 | 0.1176 |
| 11_1070583 | 6 | 0.5 | NA | NA | 2 | 0 | 3 | 0.6667 | 11 | 0.4545 | 2 | 1 | 7 | 0.4286 | 1 | 1 | 4 | 0.5 | 14 | 0.5714 |
| 11_1070583 | 5 | 0.8 | NA | NA | 3 | 0.3333 | 4 | 0.5 | 12 | 0.5833 | 1 | 0 | 10 | 0.4 | 1 | 1 | 4 | 0.75 | 16 | 0.5 |
| 11_1070583 | 5 | 0.8 | NA | NA | 3 | 0.3333 | 4 | 0.75 | 12 | 0.6667 | 3 | 0.6667 | 8 | 0.5 | 1 | 1 | 4 | 0.75 | 16 | 0.625 |
| 11_1070583 | 5 | 0.4 | NA | NA | 2 | 0 | 4 | 0 | 11 | 0.1818 | 2 | 0.5 | 8 | 0.375 | 1 | 0 | 4 | 0 | 15 | 0.2667 |
| 11_1070583 | 6 | 0.5 | NA | NA | 2 | 0 | 4 | 0.25 | 12 | 0.3333 | 3 | 0.3333 | 8 | 0.375 | 1 | 1 | 4 | 0.25 | 16 | 0.375 |
| 11_1070988 | 2 | 1 | 7 | 0 | 5 | 0 | 2 | 0 | 16 | 0.125 | 3 | 0 | 5 | 0.2 | 5 | 0.2 | 5 | 0.2 | 18 | 0.1667 |
| 11_1070990 | 1 | 0 | 5 | 0 | 5 | 0 | 2 | 0.5 | 13 | 0.0769 | 1 | 0 | 5 | 0.4 | 1 | 0 | 4 | 0 | 11 | 0.1818 |
| 11_1070991 | 1 | 0 | 4 | 0 | 6 | 0.3333 | 1 | 0 | 12 | 0.1667 | 2 | 0 | 4 | 0.25 | 2 | 0 | 4 | 0.25 | 12 | 0.1667 |
| 11_1070995 | 2 | 0.5 | 3 | 0 | 5 | 0 | 3 | 0 | 13 | 0.0769 | 2 | 0 | 5 | 0.4 | 1 | 0 | 4 | 0.25 | 12 | 0.25 |
| 11_1070996 | 2 | 1 | 2 | 0 | 4 | 0 | 3 | 0 | 11 | 0.1818 | 2 | 0 | 6 | 0.1667 | 1 | 0 | 4 | 0 | 13 | 0.0769 |
| 11_1070998 | 2 | 0.5 | 3 | 0 | 5 | 0.2 | 3 | 0.3333 | 13 | 0.2308 | 3 | 0 | 5 | 0.2 | 2 | 0 | 5 | 0 | 15 | 0.0667 |
| 11_1071006 | 1 | 1 | 2 | 0.5 | 4 | 0.5 | 2 | 0.5 | 9 | 0.5556 | 3 | 0 | 4 | 0.75 | 4 | 0.75 | 2 | 1 | 13 | 0.6154 |
| 11_1071012 | 1 | 0 | 1 | 1 | 5 | 0 | 3 | 0 | 10 | 0.1 | 2 | 0 | 3 | 0.3333 | 3 | 0 | 2 | 0 | 10 | 0.1 |
| 11_1071017 | 1 | 0 | 1 | 0 | 2 | 0.5 | 3 | 0.3333 | 7 | 0.2857 | 1 | 1 | 2 | 0 | 3 | 0 | 3 | 0.3333 | 9 | 0.2222 |
| 11_1071024 | 3 | 0.6667 | 1 | 1 | 3 | 0.3333 | 1 | 1 | 8 | 0.625 | 2 | 1 | 3 | 0.3333 | 4 | 0.5 | 2 | 1 | 11 | 0.6364 |
| 11_1071026 | 4 | 0.5 | NA | NA | 2 | 0 | 3 | 0.6667 | 9 | 0.4444 | 3 | 0 | 4 | 0 | 4 | 0.5 | 3 | 0.3333 | 14 | 0.2143 |
| 11_1071028 | 4 | 0 | 1 | 1 | 2 | 0.5 | 2 | 0.5 | 9 | 0.3333 | 2 | 0.5 | 4 | 0.5 | 4 | 0.5 | 3 | 0.6667 | 13 | 0.5385 |
| 11_1071030 | 5 | 0.4 | 1 | 0 | 2 | 0.5 | 3 | 0.6667 | 11 | 0.4545 | 2 | 0.5 | 2 | 0 | 5 | 0 | 3 | 0.3333 | 12 | 0.1667 |
| 11_1071030 | 5 | 1 | 1 | 1 | 2 | 1 | 3 | 1 | 11 | 1 | 2 | 1 | 3 | 1 | 5 | 0.8 | 3 | 0.6667 | 13 | 0.8462 |
| 11_1071031 | 3 | 0 | 1 | 1 | 1 | 1 | 1 | 0 | 6 | 0.3333 | 2 | 1 | 1 | 0 | 5 | 0 | 3 | 0.3333 | 11 | 0.2727 |
| 11_1071031 | 5 | 0.4 | 1 | 1 | 2 | 0.5 | 2 | 0.5 | 10 | 0.5 | 1 | 1 | 4 | 0.25 | 4 | 0.5 | 2 | 0.5 | 11 | 0.4545 |
| 11_1073613 | 1 | 0 | NA | NA | 2 | 0 | 1 | 1 | 4 | 0.25 | 1 | 0 | NA | NA | 1 | 1 | NA | NA | 2 | 0.5 |

| | | | | | | | | | | | | | | | | | | | | |
|---|---|---|---|---|---|---|---|---|---|---|---|---|---|---|---|---|---|---|---|---|
| 11_1073884 | 16 | 0.25 | 10 | 0.3 | 6 | 0.8333 | 19 | 0.4211 | 51 | 0.3922 | 8 | 0.125 | 14 | 0.3571 | 13 | 0.3846 | 17 | 0.4706 | 52 | 0.3654 |
| 11_1073884 | 17 | 0.3529 | 11 | 0.1818 | 6 | 0.1667 | 20 | 0.05 | 54 | 0.1852 | 7 | 0.1429 | 14 | 0.0714 | 13 | 0.1538 | 16 | 0.125 | 50 | 0.12 |
| 11_1073886 | 13 | 0.6154 | 12 | 0.6667 | 7 | 0.5714 | 21 | 0.619 | 53 | 0.6226 | 7 | 0.4286 | 15 | 0.7333 | 12 | 0.3333 | 16 | 0.625 | 50 | 0.56 |
| 11_1075635 | NA | NA | 1 | 1 | NA | NA | NA | NA | 1 | 1 | NA | NA | 1 | 1 | 2 | 1 | NA | NA | 3 | 1 |
| 11_1075636 | NA | NA | 1 | 1 | NA | NA | 1 | 0 | 2 | 0.5 | NA | NA | 1 | 1 | 1 | 1 | NA | NA | 2 | 1 |
| 11_1075842 | 5 | 0.6 | 2 | 0.5 | 1 | 1 | 6 | 0.5 | 14 | 0.5714 | 1 | 0 | NA | NA | 3 | 1 | 6 | 0.6667 | 10 | 0.7 |
| 11_1075844 | 7 | 0.1429 | 3 | 0 | 3 | 0 | 8 | 0.5 | 21 | 0.2381 | NA | NA | 1 | 0 | 4 | 0.5 | 7 | 0 | 12 | 0.1667 |
| 11_1076233 | 3 | 1 | 4 | 1 | 9 | 1 | 6 | 0.8333 | 22 | 0.9545 | 3 | 1 | 13 | 0.9231 | 11 | 0.9091 | 4 | 0.75 | 31 | 0.9032 |
| 11_1076233 | 5 | 0.8 | 3 | 1 | 7 | 0.8571 | 4 | 1 | 19 | 0.8947 | 2 | 1 | 13 | 1 | 7 | 0.8571 | 3 | 1 | 25 | 0.96 |
| 11_1076234 | 5 | 0 | 4 | 0.25 | 8 | 0.25 | 7 | 0.5714 | 24 | 0.2917 | 4 | 0 | 15 | 0.2667 | 10 | 0.3 | 4 | 0 | 33 | 0.2121 |
| 11_1076234 | 5 | 0.6 | 4 | 0.25 | 9 | 0.2222 | 6 | 0.3333 | 24 | 0.3333 | 3 | 0.3333 | 13 | 0.2308 | 7 | 0.1429 | 4 | 0.25 | 27 | 0.2222 |
| 11_1076235 | 4 | 0.25 | 5 | 0.2 | 5 | 0 | 6 | 0.1667 | 20 | 0.15 | 4 | 0.25 | 14 | 0.2857 | 8 | 0.375 | 3 | 0.6667 | 29 | 0.3448 |
| 11_1076235 | 4 | 0.25 | 5 | 0 | 8 | 0.125 | 5 | 0.2 | 22 | 0.1364 | 5 | 0 | 15 | 0.2667 | 8 | 0.25 | 5 | 0.2 | 33 | 0.2121 |
| 11_1076238 | 4 | 0.25 | 2 | 1 | 6 | 0.1667 | 5 | 0.2 | 17 | 0.2941 | 3 | 0.3333 | 13 | 0.3077 | 6 | 0.1667 | 4 | 0.25 | 26 | 0.2692 |
| 11_1076238 | 3 | 0 | 2 | 0.5 | 7 | 0 | 7 | 0.2857 | 19 | 0.1579 | 4 | 0.25 | 13 | 0.1538 | 6 | 0.3333 | 4 | 0.25 | 27 | 0.2222 |
| 11_1076239 | 3 | 0.3333 | 2 | 0 | 8 | 0.25 | 7 | 0.1429 | 20 | 0.2 | 4 | 0 | 16 | 0.125 | 6 | 0.3333 | 5 | 0 | 31 | 0.129 |
| 11_1076241 | 3 | 0.3333 | 2 | 0 | 5 | 0.6 | 3 | 0 | 13 | 0.3077 | 3 | 0 | 7 | 0.4286 | 6 | 0.3333 | 3 | 0.6667 | 19 | 0.3684 |
| 11_1076241 | 2 | 0.5 | 1 | 1 | 5 | 0.8 | 6 | 0.6667 | 14 | 0.7143 | 2 | 0 | 8 | 0.5 | 5 | 0.6 | 1 | 0 | 16 | 0.4375 |
| 11_1076241 | 3 | 0.3333 | 3 | 0.3333 | 6 | 0.8333 | 6 | 0.5 | 18 | 0.5556 | 2 | 1 | 13 | 0.6154 | 7 | 1 | 3 | 1 | 25 | 0.8 |
| 11_1076242 | 2 | 0 | 3 | 0.3333 | 4 | 0 | 5 | 0.2 | 14 | 0.1429 | 4 | 0.25 | 14 | 0.2857 | 7 | 0.2857 | 1 | 0 | 26 | 0.2692 |
| 11_1076242 | 2 | 1 | 2 | 1 | 7 | 0.5714 | 6 | 1 | 17 | 0.8235 | 4 | 0.75 | 15 | 0.8667 | 7 | 0.8571 | 2 | 1 | 28 | 0.8571 |
| 11_1076243 | 3 | 0.6667 | 3 | 0.3333 | 7 | 0.2857 | 7 | 0.2857 | 20 | 0.35 | 3 | 0.3333 | 14 | 0.2857 | 8 | 0.625 | 2 | 1 | 27 | 0.4444 |
| 11_1076244 | 2 | 0.5 | 3 | 1 | 6 | 1 | 7 | 1 | 18 | 0.9444 | 4 | 1 | 13 | 1 | 8 | 1 | 2 | 1 | 27 | 1 |
| 11_1076248 | 1 | 1 | 1 | 1 | 6 | 0.5 | 4 | 0.5 | 12 | 0.5833 | 2 | 0 | 11 | 1 | 4 | 0.5 | 1 | 0 | 18 | 0.7222 |
| 11_1076249 | 1 | 1 | 1 | 0 | 3 | 0.6667 | 4 | 0.75 | 9 | 0.6667 | NA | NA | 9 | 0.5556 | 6 | 0.3333 | 2 | 0.5 | 17 | 0.4706 |
| 11_1076249 | 2 | 0.5 | 3 | 1 | 7 | 0.8571 | 4 | 1 | 16 | 0.875 | 2 | 1 | 13 | 0.8462 | 7 | 1 | 2 | 0.5 | 24 | 0.875 |
| 11_1076252 | 2 | 0 | 3 | 0.6667 | 6 | 0.3333 | 5 | 0.4 | 16 | 0.375 | 3 | 0 | 11 | 0.5455 | 7 | 0.5714 | 1 | 1 | 22 | 0.5 |
| 11_1076253 | 2 | 0 | 3 | 0 | 6 | 0.5 | 5 | 0.8 | 16 | 0.4375 | 2 | 0.5 | 12 | 0.5 | 6 | 0.3333 | 2 | 1 | 22 | 0.5 |
| 11_1076254 | 1 | 0 | 3 | 0 | 4 | 0 | 5 | 0.2 | 13 | 0.0769 | 2 | 0 | 13 | 0.3077 | 5 | 0.2 | 2 | 0.5 | 22 | 0.2727 |
| 11_1076255 | 1 | 1 | 1 | 1 | 4 | 0.25 | 3 | 0.3333 | 9 | 0.4444 | 1 | 1 | 7 | 0.2857 | 5 | 0.4 | 1 | 0 | 14 | 0.3571 |
| 11_1076256 | 2 | 0 | 2 | 0.5 | 4 | 0.25 | 4 | 0.5 | 12 | 0.3333 | 2 | 0.5 | 11 | 0.1818 | 6 | 0.3333 | 2 | 0 | 21 | 0.2381 |
| 11_1076259 | 1 | 0 | 1 | 1 | 2 | 0 | 3 | 0 | 7 | 0.1429 | 1 | 0 | 9 | 0.1111 | 6 | 0 | 3 | 0 | 19 | 0.0526 |
| 11_1076260 | 2 | 1 | 1 | 0 | 2 | 0 | 3 | 0 | 8 | 0.25 | 3 | 0 | 11 | 0.0909 | 5 | 0.2 | 3 | 0.3333 | 22 | 0.1364 |
| 11_1077657 | 5 | 0 | 7 | 0.1429 | 3 | 0 | 1 | 0 | 16 | 0.0625 | 1 | 0 | 5 | 0.2 | 3 | 0 | 1 | 0 | 10 | 0.1 |
| 11_1077660 | 5 | 0.6 | 5 | 0.6 | 4 | 0.25 | 1 | 1 | 15 | 0.5333 | 1 | 0 | 5 | 1 | 4 | 0.75 | 2 | 1 | 12 | 0.8333 |
| 11_1077665 | 6 | 0.1667 | 3 | 0.6667 | 3 | 0 | 1 | 0 | 13 | 0.2308 | 1 | 0 | 2 | 0.5 | 3 | 0 | NA | NA | 6 | 0.1667 |
| 11_1077666 | 5 | 1 | 3 | 1 | 4 | 1 | 1 | 1 | 13 | 1 | 1 | 1 | 2 | 0.5 | 1 | 1 | 1 | 1 | 5 | 0.8 |
| 11_1077667 | 6 | 0.8333 | 2 | 1 | 3 | 1 | 1 | 1 | 12 | 0.9167 | NA | NA | 3 | 1 | 3 | 1 | 2 | 1 | 8 | 1 |
| 11_1077668 | 4 | 0.25 | NA | NA | 3 | 0.3333 | 1 | 1 | 8 | 0.375 | 1 | 0 | 3 | 0.6667 | 3 | 0 | 1 | 0 | 8 | 0.25 |
| 11_1077669 | 6 | 0.8333 | 2 | 0.5 | 3 | 0.3333 | NA | NA | 11 | 0.6364 | NA | NA | 3 | 0.6667 | 3 | 0.3333 | 1 | 1 | 7 | 0.5714 |
| 11_1077670 | 6 | 0.1667 | 2 | 0 | 4 | 0 | NA | NA | 12 | 0.0833 | NA | NA | 3 | 0 | 4 | 0.25 | 1 | 0 | 8 | 0.125 |
| 11_1077673 | 6 | 0.5 | 2 | 1 | 4 | 0.5 | NA | NA | 12 | 0.5833 | NA | NA | 3 | 0.3333 | 4 | 0.5 | 1 | 0 | 8 | 0.375 |
| 11_1077673 | 6 | 0 | 2 | 0.5 | 4 | 0 | NA | NA | 12 | 0.0833 | NA | NA | 3 | 0 | 4 | 0 | 1 | 1 | 8 | 0.125 |
| 11_1077674 | 6 | 1 | 2 | 0.5 | 2 | 1 | NA | NA | 10 | 0.9 | NA | NA | 2 | 1 | 1 | 1 | 1 | 1 | 4 | 1 |
| 11_1077676 | 4 | 0 | 1 | 1 | 3 | 0 | NA | NA | 8 | 0.125 | NA | NA | 3 | 1 | 4 | 0.5 | 2 | 1 | 9 | 0.7778 |
| 11_1077676 | 5 | 1 | 1 | 1 | 3 | 1 | NA | NA | 9 | 1 | NA | NA | 3 | 0.6667 | 4 | 1 | 2 | 1 | 9 | 0.8889 |
| 11_1077676 | 6 | 1 | 2 | 1 | 1 | 1 | NA | NA | 9 | 1 | NA | NA | 3 | 1 | 3 | 1 | 2 | 1 | 8 | 1 |
| 11_1077677 | 5 | 1 | 2 | 1 | 4 | 1 | NA | NA | 11 | 1 | NA | NA | 3 | 1 | 3 | 1 | 1 | 1 | 7 | 1 |
| 11_1077678 | 6 | 0.5 | 2 | 0.5 | 4 | 0.5 | NA | NA | 12 | 0.5 | NA | NA | 3 | 1 | 3 | 0.3333 | 2 | 0 | 8 | 0.5 |
| 11_1077681 | 5 | 0.6 | 2 | 0.5 | 2 | 0 | NA | NA | 9 | 0.4444 | NA | NA | 3 | 1 | 4 | 0.75 | 2 | 1 | 9 | 0.8889 |
| 11_1077681 | 5 | 1 | 1 | 1 | 4 | 1 | NA | NA | 10 | 1 | NA | NA | 3 | 1 | 2 | 1 | NA | NA | 5 | 1 |
| 11_1077682 | 5 | 0.6 | 2 | 0 | 3 | 0.6667 | NA | NA | 10 | 0.5 | NA | NA | 2 | 1 | 3 | 0.3333 | 1 | 0 | 6 | 0.5 |
| 11_1077682 | 2 | 0 | 1 | 0 | 3 | 0.3333 | NA | NA | 6 | 0.1667 | NA | NA | 3 | 0.6667 | 2 | 0 | NA | NA | 5 | 0.4 |
| 11_1077682 | 5 | 0.2 | 1 | 0 | 3 | 0 | NA | NA | 9 | 0.1111 | NA | NA | 3 | 0 | 3 | 0.6667 | 1 | 0 | 7 | 0.2857 |
| 11_1077683 | 4 | 0 | 2 | 0.5 | 2 | 0 | NA | NA | 8 | 0.125 | NA | NA | 3 | 0.3333 | 3 | 0.3333 | NA | NA | 6 | 0.3333 |
| 11_1078245 | 13 | 0.1538 | 26 | 0.1154 | 20 | 0.15 | 8 | 0.125 | 67 | 0.1343 | 15 | 0.0667 | 6 | 0 | 11 | 0.3636 | 20 | 0.1 | 52 | 0.1346 |
| 11_1078254 | 14 | 0.0714 | 17 | 0 | 15 | 0.1333 | 6 | 0.3333 | 52 | 0.0962 | 10 | 0.1 | 5 | 0.6 | 8 | 0.125 | 19 | 0.1053 | 42 | 0.1667 |
| 11_1078255 | 13 | 0.2308 | 19 | 0.1053 | 14 | 0.1429 | 6 | 0.3333 | 52 | 0.1731 | 10 | 0 | 6 | 0.3333 | 9 | 0 | 20 | 0.1 | 45 | 0.0889 |
| 11_1078258 | 5 | 0 | 12 | 0 | 8 | 0.375 | 5 | 0 | 30 | 0.1 | 9 | 0.1111 | 4 | 0 | 5 | 0.2 | 10 | 0 | 28 | 0.0714 |
| 11_1078258 | 11 | 0 | 14 | 0.3571 | 14 | 0.1429 | 6 | 0.3333 | 45 | 0.2 | 9 | 0 | 5 | 0.2 | 6 | 0.3333 | 18 | 0.1111 | 38 | 0.1316 |
| 11_1078259 | 7 | 0 | 14 | 0.0714 | 12 | 0.0833 | 4 | 0.5 | 37 | 0.1081 | 4 | 0 | 5 | 0.2 | 5 | 0 | 18 | 0.0556 | 32 | 0.0625 |
| 11_1078263 | 5 | 0.6 | 13 | 0.3846 | 11 | 0.3636 | 6 | 0.8333 | 35 | 0.4857 | 8 | 0.25 | 5 | 0.4 | 8 | 0.5 | 15 | 0.5333 | 36 | 0.4444 |
| 11_1078272 | 4 | 0 | 9 | 0.1111 | 6 | 0.3333 | 6 | 0 | 25 | 0.12 | 4 | 0.25 | 2 | 0 | 2 | 0.5 | 11 | 0.2727 | 19 | 0.2632 |
| 11_1078273 | 3 | 0 | 7 | 0.4286 | 8 | 0.375 | 4 | 0.25 | 22 | 0.3182 | 3 | 0.3333 | 3 | 0.3333 | 2 | 0 | 9 | 0.2222 | 17 | 0.2353 |
| 11_1078277 | 4 | 0 | 7 | 0.4286 | 6 | 0.1667 | 6 | 0.1667 | 23 | 0.2174 | 3 | 0 | 2 | 0 | 2 | 0.5 | 6 | 0.1667 | 13 | 0.1538 |

| | | | | | | | | | | | | | | | | | | | | |
|---|---|---|---|---|---|---|---|---|---|---|---|---|---|---|---|---|---|---|---|---|
| 11_1078277 | 4 | 0.25 | 7 | 0.4286 | 8 | 0 | 6 | 0 | 25 | 0.16 | 3 | 0.3333 | 3 | 0.3333 | 2 | 0.5 | 6 | 0 | 14 | 0.2143 |
| 11_1078278 | 4 | 0.25 | 6 | 0.5 | 8 | 0.25 | 6 | 0.6667 | 24 | 0.4167 | 4 | 0.75 | 3 | 0.3333 | 2 | 0.5 | 6 | 0.3333 | 15 | 0.4667 |
| 11_1078278 | 4 | 0.5 | 6 | 0 | 5 | 0.2 | 4 | 0.25 | 19 | 0.2105 | 4 | 0 | 2 | 0 | NA | NA | 7 | 0.1429 | 13 | 0.0769 |
| 11_1078279 | 3 | 0.6667 | 4 | 1 | 7 | 0.7143 | 5 | 0.8 | 19 | 0.7895 | 3 | 0.6667 | 2 | 1 | 1 | 1 | 7 | 0.7143 | 13 | 0.7692 |
| 11_1078282 | 4 | 0.25 | 5 | 0.4 | 5 | 0.2 | 5 | 0.2 | 19 | 0.2632 | 4 | 0 | 2 | 0.5 | 1 | 0 | 6 | 0.1667 | 13 | 0.1538 |
| 11_1078283 | 4 | 0.25 | 7 | 0.2857 | 5 | 0.2 | 3 | 0.3333 | 19 | 0.2632 | 3 | 1 | 3 | 0.6667 | 1 | 1 | 6 | 0.5 | 13 | 0.6923 |
| 11_1078283 | 4 | 0 | 6 | 0.1667 | 7 | 0.2857 | 4 | 0.25 | 21 | 0.1905 | 3 | 0 | 3 | 0.3333 | 1 | 1 | 6 | 0 | 13 | 0.1538 |
| 11_1078284 | 4 | 0.25 | 6 | 0.6667 | 6 | 0.3333 | 4 | 0.5 | 20 | 0.45 | 3 | 0 | 2 | 0 | 1 | 1 | 7 | 0.2857 | 13 | 0.2308 |
| 11_1078296 | 7 | 0.8571 | 7 | 0.8571 | 6 | 1 | 8 | 0.75 | 28 | 0.8571 | 6 | 0.8333 | 6 | 0.8333 | 3 | 0.3333 | 10 | 0.7 | 25 | 0.72 |
| 11_1078298 | 5 | 0 | 5 | 0 | 6 | 0.3333 | 7 | 0 | 23 | 0.087 | 5 | 0.2 | 3 | 0 | 2 | 0.5 | 10 | 0 | 20 | 0.1 |
| 11_1078300 | 4 | 0.25 | 5 | 0.2 | 6 | 0.3333 | 7 | 0.2857 | 22 | 0.2727 | 6 | 0.3333 | 4 | 0.25 | 3 | 0.6667 | 10 | 0.3 | 23 | 0.3478 |
| 11_1078302 | 4 | 0 | 5 | 0.4 | 6 | 0.1667 | 8 | 0.25 | 23 | 0.2174 | 5 | 0.2 | 3 | 0.3333 | 2 | 0.5 | 10 | 0.3 | 20 | 0.3 |
| 11_1078304 | 4 | 0.25 | 6 | 0 | 5 | 0 | 7 | 0.2857 | 22 | 0.1364 | 4 | 0.25 | 3 | 0 | 3 | 0.6667 | 9 | 0.1111 | 19 | 0.2105 |
| 11_1087616 | NA | NA | 1 | 1 | NA | NA | NA | NA | 1 | 1 | NA | NA | 1 | 1 | NA | NA | NA | NA | 1 | 1 |
| 11_1087621 | NA | NA | 1 | 1 | NA | NA | NA | NA | 1 | 1 | NA | NA | 1 | 1 | NA | NA | NA | NA | 1 | 1 |
| 11_1087622 | NA | NA | 1 | 1 | NA | NA | NA | NA | 1 | 1 | NA | NA | 1 | 1 | NA | NA | NA | NA | 1 | 1 |
| 11_1088404 | 1 | 1 | NA | NA | 4 | 0.5 | 2 | 0.5 | 7 | 0.5714 | NA | NA | NA | NA | 1 | 1 | NA | NA | 1 | 1 |
| 11_1088405 | NA | NA | NA | NA | 4 | 0.75 | 2 | 1 | 6 | 0.8333 | NA | NA | 1 | 1 | NA | NA | NA | NA | 1 | 1 |
| 11_1088405 | 1 | 1 | NA | NA | 4 | 0 | 1 | 0 | 6 | 0.1667 | NA | NA | 2 | 0.5 | 1 | 0 | NA | NA | 3 | 0.3333 |
| 11_1088406 | NA | NA | NA | NA | 4 | 0 | 3 | 0.3333 | 7 | 0.1429 | NA | NA | 1 | 1 | NA | NA | NA | NA | 1 | 1 |
| 11_1088413 | 1 | 0 | NA | NA | 4 | 0.5 | 3 | 0 | 8 | 0.25 | 2 | 0 | 1 | 1 | NA | NA | NA | NA | 3 | 0.3333 |
| 11_1088414 | 2 | 0 | NA | NA | 5 | 0.2 | 4 | 0 | 11 | 0.0909 | 2 | 0.5 | 2 | 0 | NA | NA | NA | NA | 4 | 0.25 |
| 11_1091161 | 1 | 1 | 1 | 1 | NA | NA | 1 | 0 | 3 | 0.6667 | 1 | 0 | 6 | 0.5 | 3 | 0.3333 | 2 | 0.5 | 12 | 0.4167 |
| 11_1091163 | 26 | 0.4615 | 21 | 0.4762 | 25 | 0.44 | 23 | 0.5217 | 95 | 0.4737 | 13 | 0.4615 | 45 | 0.4 | 56 | 0.4464 | 23 | 0.4348 | 137 | 0.4307 |
| 11_1091165 | 27 | 0.1481 | 22 | 0.1818 | 29 | 0.2069 | 33 | 0.2121 | 111 | 0.1892 | 14 | 0.0714 | 56 | 0.1964 | 68 | 0.2059 | 30 | 0.1 | 168 | 0.1726 |
| 11_1091182 | 20 | 0.5 | 8 | 0.5 | 23 | 0.3913 | 23 | 0.3043 | 74 | 0.4054 | 9 | 0.1111 | 38 | 0.3947 | 47 | 0.4468 | 22 | 0.4091 | 116 | 0.3966 |
| 11_1091194 | 1 | 1 | NA | NA | 1 | 0 | 3 | 0.6667 | 5 | 0.6 | 1 | 0 | 4 | 0.25 | 3 | 0.3333 | 2 | 0.5 | 10 | 0.3 |
| 11_1091209 | 1 | 0 | NA | NA | 3 | 0 | 6 | 0.1667 | 10 | 0.1 | 2 | 0 | 3 | 0.3333 | 5 | 0 | 2 | 0 | 12 | 0.0833 |
| 11_1092868 | 2 | 0.5 | 1 | 1 | 1 | 0 | 1 | 0 | 5 | 0.4 | 1 | 0 | NA | NA | 2 | 0.5 | 2 | 0 | 5 | 0.2 |
| 11_1092869 | 2 | 0.5 | 1 | 0 | 1 | 0 | 1 | 0 | 5 | 0.2 | 2 | 0.5 | NA | NA | 1 | 1 | 5 | 0.8 | 8 | 0.75 |
| 11_1092869 | 2 | 1 | 1 | 0 | 1 | 0 | NA | NA | 4 | 0.5 | 2 | 1 | NA | NA | 1 | 1 | 4 | 0.75 | 7 | 0.8571 |
| 11_1092870 | 2 | 1 | NA | NA | 1 | 1 | 1 | 1 | 4 | 1 | 1 | 1 | NA | NA | 1 | 1 | 5 | 0.8 | 7 | 0.8571 |
| 11_1092870 | 2 | 1 | 1 | 1 | 1 | 0 | 1 | 1 | 5 | 0.8 | 2 | 1 | NA | NA | 2 | 1 | 5 | 1 | 9 | 1 |
| 11_1092871 | 2 | 0.5 | 2 | 0.5 | 1 | 0 | 1 | 0 | 6 | 0.3333 | 2 | 0 | NA | NA | 2 | 0 | 5 | 0.2 | 9 | 0.1111 |
| 11_1092873 | 2 | 1 | 2 | 1 | 1 | 1 | 1 | 0 | 6 | 0.8333 | 1 | 0 | NA | NA | 2 | 0 | 5 | 0.6 | 8 | 0.375 |
| 11_1092881 | 2 | 0.5 | 2 | 0 | 1 | 0 | 1 | 0 | 6 | 0.1667 | 2 | 0 | 1 | 0 | 2 | 0.5 | 5 | 0.4 | 10 | 0.3 |
| 11_1092883 | 1 | 0 | 2 | 0.5 | 1 | 0 | 1 | 0 | 5 | 0.2 | 4 | 0.25 | 1 | 0 | 2 | 0 | 4 | 0 | 11 | 0.0909 |
| 11_1092887 | 2 | 0.5 | 5 | 0 | 2 | 0 | 1 | 0 | 10 | 0.1 | 6 | 0 | 1 | 0 | 3 | 0.3333 | 6 | 0 | 16 | 0.0625 |
| 11_1092901 | 3 | 0 | 4 | 0 | 2 | 0 | 4 | 0.25 | 13 | 0.0769 | 10 | 0 | 3 | 0 | 2 | 0.5 | 5 | 0.2 | 20 | 0.1 |
| 11_1092907 | 4 | 0 | 6 | 0 | 2 | 0.5 | 4 | 0 | 16 | 0.0625 | 10 | 0.1 | 3 | 0 | 5 | 0 | 7 | 0.1429 | 25 | 0.08 |
| 11_1092917 | 5 | 0 | 6 | 0.1667 | 3 | 0 | 8 | 0.125 | 22 | 0.0909 | 10 | 0.1 | 4 | 0.25 | 6 | 0 | 8 | 0.125 | 28 | 0.1071 |
| 11_1093254 | 4 | 0.25 | 3 | 0.3333 | 4 | 0.25 | 3 | 0 | 14 | 0.2143 | 2 | 0 | 2 | 1 | 5 | 0 | 4 | 0.25 | 13 | 0.2308 |
| 11_1093275 | 9 | 1 | 3 | 1 | 5 | 0.8 | 6 | 0.8333 | 23 | 0.913 | 3 | 0.6667 | 5 | 0.6 | 6 | 1 | 6 | 0.8333 | 20 | 0.8 |
| 11_1093275 | 6 | 0.3333 | 3 | 0 | 5 | 0.2 | 6 | 0 | 20 | 0.15 | 3 | 0 | 5 | 0.2 | 5 | 0 | 5 | 0 | 18 | 0.0556 |
| 11_1093283 | 6 | 0.1667 | 5 | 0 | 3 | 0 | 2 | 0 | 16 | 0.0625 | 6 | 0.1667 | 3 | 0 | 3 | 0 | 2 | 0 | 14 | 0.0714 |
| 11_1093285 | 3 | 0 | 5 | 0.2 | 4 | 0.5 | 2 | 0 | 14 | 0.2143 | 6 | 0.5 | 3 | 0.3333 | 3 | 0.3333 | 2 | 1 | 14 | 0.5 |
| 11_1093285 | 4 | 1 | 4 | 0.75 | 4 | 0.75 | 2 | 0.5 | 14 | 0.7857 | 5 | 0.6 | 3 | 1 | 2 | 1 | 1 | 1 | 11 | 0.8182 |
| 11_1093458 | 1 | 1 | NA | NA | NA | NA | NA | NA | 1 | 1 | NA | NA | 1 | 1 | NA | NA | NA | NA | 1 | 1 |
| 11_1093807 | NA | NA | NA | NA | 1 | 1 | NA | NA | 1 | 1 | NA | NA | NA | NA | 1 | 1 | NA | NA | 1 | 1 |
| 11_1093810 | NA | NA | NA | NA | 1 | 1 | NA | NA | 1 | 1 | NA | NA | NA | NA | 1 | 1 | NA | NA | 1 | 1 |
| 11_1093811 | NA | NA | NA | NA | 1 | 1 | NA | NA | 1 | 1 | NA | NA | NA | NA | 1 | 1 | NA | NA | 1 | 1 |
| 11_1093811 | NA | NA | 1 | 1 | 1 | 0 | NA | NA | 2 | 0.5 | NA | NA | NA | NA | 1 | 1 | NA | NA | 1 | 1 |
| 11_1094634 | NA | NA | 4 | 0.5 | 1 | 1 | 2 | 0.5 | 7 | 0.5714 | 1 | 1 | 1 | 1 | 2 | 0.5 | 1 | 1 | 5 | 0.8 |
| 11_1094635 | NA | NA | 3 | 0 | 2 | 0 | 2 | 1 | 7 | 0.2857 | 2 | 0 | 1 | 0 | 1 | 0 | 1 | 1 | 5 | 0.2 |
| 11_1094636 | NA | NA | 3 | 0.3333 | 2 | 0 | NA | NA | 5 | 0.2 | 1 | 0 | 1 | 1 | 2 | 0.5 | NA | NA | 4 | 0.5 |
| 11_1095447 | 33 | 0.1515 | 32 | 0.0625 | 23 | 0.087 | 42 | 0.0714 | 130 | 0.0923 | 26 | 0.1538 | 23 | 0.0435 | 23 | 0 | 20 | 0.1 | 92 | 0.0761 |
| 11_1095584 | 54 | 0.3148 | 65 | 0.2462 | 63 | 0.2222 | 66 | 0.2576 | 248 | 0.2581 | 41 | 0.2195 | 49 | 0.2449 | 52 | 0.3077 | 40 | 0.125 | 182 | 0.2308 |
| 11_1095586 | 63 | 0.4762 | 63 | 0.6825 | 57 | 0.4912 | 69 | 0.4493 | 252 | 0.5238 | 44 | 0.5455 | 42 | 0.5 | 43 | 0.5581 | 40 | 0.55 | 169 | 0.5385 |
| 11_1095602 | 81 | 0.284 | 80 | 0.225 | 80 | 0.2875 | 78 | 0.2564 | 319 | 0.2633 | 61 | 0.2131 | 68 | 0.1618 | 62 | 0.3065 | 50 | 0.28 | 241 | 0.2365 |
| 11_1095637 | 4 | 0.5 | 3 | 0.6667 | 4 | 0.25 | 1 | 1 | 12 | 0.5 | 2 | 1 | 3 | 0.3333 | 3 | 0.3333 | 1 | 0 | 9 | 0.4444 |
| 11_1095638 | 3 | 0.6667 | 5 | 0 | 3 | 0.3333 | 1 | 0 | 12 | 0.25 | 2 | 0 | 3 | 0.3333 | 2 | 0.5 | 1 | 0 | 8 | 0.25 |
| 11_1095638 | 4 | 0.5 | 4 | 0.5 | 4 | 0.75 | 1 | 1 | 13 | 0.6154 | 1 | 1 | 3 | 1 | 3 | 1 | 1 | 1 | 8 | 1 |
| 11_1095639 | 2 | 0 | 5 | 0 | 3 | 1 | 1 | 0 | 11 | 0.2727 | 3 | 0.3333 | 3 | 0 | 3 | 0 | 1 | 0 | 10 | 0.1 |
| 11_1095641 | 3 | 1 | 4 | 1 | 3 | 0.3333 | 1 | 0 | 11 | 0.7273 | 2 | 1 | 3 | 1 | 3 | 0.6667 | 1 | 1 | 9 | 0.8889 |
| 11_1095642 | 3 | 0.3333 | 4 | 0.75 | 3 | 0.3333 | 1 | 1 | 11 | 0.5455 | 1 | 0 | 3 | 0.3333 | 3 | 0.3333 | 1 | 1 | 8 | 0.375 |

| | | | | | | | | | | | | | | | | | | | | |
|---|---|---|---|---|---|---|---|---|---|---|---|---|---|---|---|---|---|---|---|---|
| 11_1096127 | 1 | 1 | 3 | 0.3333 | NA | NA | NA | NA | 4 | 0.5 | 1 | 1 | NA | NA | NA | NA | NA | NA | 1 | 1 |
| 11_1098242 | NA | NA | 1 | 0 | NA | NA | 2 | 0.5 | 3 | 0.3333 | NA | NA | 1 | 0 | 1 | 1 | NA | NA | 2 | 0.5 |
| 11_1098251 | NA | NA | 1 | 1 | 1 | 0 | 3 | 0.3333 | 5 | 0.4 | NA | NA | NA | NA | 1 | 1 | NA | NA | 1 | 1 |
| 11_1098253 | NA | NA | 1 | 1 | 1 | 1 | 3 | 1 | 5 | 1 | NA | NA | NA | NA | 1 | 1 | NA | NA | 1 | 1 |
| 11_1099918 | 3 | 0.3333 | 1 | 0 | 2 | 0.5 | NA | NA | 6 | 0.3333 | NA | NA | NA | NA | NA | NA | 2 | 0.5 | 2 | 0.5 |
| 11_1099918 | 3 | 0 | 1 | 0 | 2 | 1 | NA | NA | 6 | 0.3333 | NA | NA | NA | NA | NA | NA | 2 | 0.5 | 2 | 0.5 |
| 11_1099921 | 3 | 1 | 1 | 0 | 1 | 0 | NA | NA | 5 | 0.6 | NA | NA | NA | NA | NA | NA | 2 | 0.5 | 2 | 0.5 |
| 11_1099925 | 2 | 0.5 | 1 | 1 | 2 | 1 | NA | NA | 5 | 0.8 | NA | NA | 1 | 1 | NA | NA | 2 | 1 | 3 | 1 |
| 11_1099927 | 2 | 0.5 | 1 | 0 | 2 | 1 | NA | NA | 5 | 0.6 | NA | NA | 1 | 0 | NA | NA | 2 | 0.5 | 3 | 0.3333 |
| 11_1101607 | 3 | 0 | 5 | 0.2 | 6 | 0 | 3 | 0 | 17 | 0.0588 | 1 | 0 | 2 | 0 | 2 | 0.5 | 4 | 0.25 | 9 | 0.2222 |
| 11_1104037 | 1 | 1 | NA | NA | 1 | 1 | 2 | 0.5 | 4 | 0.75 | NA | NA | NA | NA | 1 | 1 | NA | NA | 1 | 1 |
| 11_1104043 | 1 | 0 | NA | NA | 1 | 1 | 2 | 0.5 | 4 | 0.5 | NA | NA | 1 | 1 | 1 | 1 | NA | NA | 2 | 1 |
| 11_1109152 | 1 | 1 | 1 | 0 | NA | NA | NA | NA | 2 | 0.5 | 1 | 0 | 2 | 0 | 2 | 0 | 1 | 1 | 6 | 0.1667 |
| 11_1109153 | 1 | 1 | NA | NA | NA | NA | NA | NA | 1 | 1 | NA | NA | 1 | 0 | 1 | 1 | 2 | 1 | 4 | 0.75 |
| 11_1109156 | 1 | 0 | 1 | 0 | 1 | 1 | NA | NA | 3 | 0.3333 | 1 | 0 | 2 | 0.5 | 2 | 0 | 3 | 0 | 8 | 0.125 |
| 11_1109157 | 1 | 1 | 1 | 0 | 1 | 0 | NA | NA | 3 | 0.3333 | 1 | 0 | 2 | 0 | 1 | 0 | 2 | 0.5 | 6 | 0.1667 |
| 11_1109159 | 1 | 0 | 1 | 0 | 1 | 1 | NA | NA | 3 | 0.3333 | 1 | 1 | 2 | 0 | 1 | 1 | 2 | 0 | 6 | 0.3333 |
| 11_1109160 | 1 | 1 | 1 | 1 | 1 | 1 | NA | NA | 3 | 1 | 1 | 1 | 2 | 1 | 2 | 1 | 2 | 1 | 7 | 1 |
| 11_1109161 | 1 | 0 | 1 | 1 | 2 | 0 | NA | NA | 4 | 0.25 | 1 | 0 | 2 | 0.5 | 1 | 0 | 3 | 0 | 7 | 0.1429 |
| 11_1109161 | 1 | 1 | 1 | 0 | 3 | 1 | NA | NA | 5 | 0.8 | 1 | 1 | 2 | 1 | 1 | 0 | 3 | 0.6667 | 7 | 0.7143 |
| 11_1109162 | 1 | 0 | 1 | 0 | 2 | 0.5 | NA | NA | 4 | 0.25 | 1 | 0 | 1 | 1 | 1 | 1 | 3 | 0.3333 | 6 | 0.5 |
| 11_1109163 | 1 | 1 | 1 | 1 | 2 | 0.5 | NA | NA | 4 | 0.75 | 1 | 0 | 1 | 1 | 1 | 1 | 2 | 1 | 5 | 0.8 |
| 11_1109163 | NA | NA | 1 | 1 | 2 | 1 | NA | NA | 3 | 1 | 1 | 1 | 2 | 0 | 1 | 0 | 3 | 1 | 7 | 0.5714 |
| 11_1109163 | 1 | 1 | 1 | 0 | 3 | 0 | NA | NA | 5 | 0.2 | 1 | 0 | 2 | 0 | 2 | 1 | 3 | 0.3333 | 8 | 0.375 |
| 11_1109165 | 1 | 1 | 1 | 0 | 2 | 1 | NA | NA | 4 | 0.75 | 1 | 1 | 2 | 0.5 | 1 | 1 | 1 | 1 | 5 | 0.8 |
| 11_1109166 | 1 | 1 | 1 | 1 | 3 | 0.6667 | NA | NA | 5 | 0.8 | 1 | 1 | 2 | 0.5 | 2 | 0.5 | 3 | 0.6667 | 8 | 0.625 |
| 11_1109169 | 1 | 1 | 1 | 0 | 2 | 0 | NA | NA | 4 | 0.25 | 1 | 0 | 2 | 0.5 | 2 | 0 | 3 | 0.6667 | 8 | 0.375 |
| 11_1109172 | 1 | 1 | 1 | 1 | 3 | 1 | NA | NA | 5 | 1 | 1 | 1 | 1 | 1 | 2 | 0.5 | 3 | 0.6667 | 7 | 0.7143 |
| 11_1109173 | 1 | 0 | 1 | 0 | 3 | 0.3333 | NA | NA | 5 | 0.2 | 1 | 0 | 1 | 0 | 3 | 0.3333 | 4 | 0.75 | 9 | 0.4444 |
| 11_1109174 | 1 | 0 | 1 | 1 | 1 | 0 | NA | NA | 3 | 0.3333 | NA | NA | 1 | 0 | 3 | 0 | 4 | 0.5 | 8 | 0.25 |
| 11_1109175 | 1 | 0 | 1 | 1 | 3 | 1 | 1 | 1 | 6 | 0.8333 | 1 | 1 | 1 | 0 | 3 | 0 | 3 | 0.3333 | 8 | 0.25 |
| 11_1109177 | 1 | 0 | 1 | 1 | 3 | 0.6667 | 1 | 0 | 6 | 0.5 | 1 | 0 | 1 | 0 | 1 | 0 | 2 | 0.5 | 5 | 0.2 |
| 11_1109182 | 1 | 1 | 1 | 0 | 3 | 0 | 1 | 0 | 6 | 0.1667 | 3 | 0.3333 | 1 | 0 | 2 | 0 | 3 | 0.3333 | 9 | 0.2222 |
| 11_1109187 | 2 | 0.5 | 1 | 0 | 2 | 0 | 2 | 0 | 7 | 0.1429 | 3 | 0 | 2 | 0 | NA | NA | 3 | 0.3333 | 8 | 0.125 |
| 11_1109625 | NA | NA | NA | NA | 1 | 1 | 1 | 0 | 2 | 0.5 | NA | NA | 2 | 0.5 | 1 | 1 | NA | NA | 3 | 0.6667 |
| 11_1109625 | NA | NA | 2 | 0.5 | 1 | 1 | 1 | 1 | 4 | 0.75 | NA | NA | 1 | 1 | NA | NA | NA | NA | 1 | 1 |
| 11_1109625 | NA | NA | 2 | 1 | 2 | 1 | NA | NA | 4 | 1 | NA | NA | 2 | 1 | 1 | 1 | NA | NA | 3 | 1 |
| 11_1109628 | NA | NA | 1 | 0 | 1 | 1 | 1 | 1 | 3 | 0.6667 | NA | NA | 1 | 1 | NA | NA | NA | NA | 1 | 1 |
| 11_1109631 | NA | NA | 2 | 1 | 2 | 0.5 | 1 | 1 | 5 | 0.8 | NA | NA | 2 | 0.5 | 1 | 1 | NA | NA | 3 | 0.6667 |
| 11_1109633 | NA | NA | 2 | 0 | 2 | 1 | 2 | 0.5 | 6 | 0.5 | NA | NA | 2 | 0 | 1 | 1 | NA | NA | 3 | 0.3333 |
| 11_1109633 | NA | NA | 2 | 0.5 | 2 | 0 | 2 | 1 | 6 | 0.5 | NA | NA | 2 | 1 | 1 | 1 | NA | NA | 3 | 1 |
| 11_1109634 | NA | NA | 2 | 0 | 2 | 0.5 | 2 | 0 | 6 | 0.1667 | NA | NA | 1 | 0 | 1 | 1 | NA | NA | 2 | 0.5 |
| 11_1109634 | NA | NA | 2 | 0.5 | 2 | 1 | 2 | 1 | 6 | 0.8333 | NA | NA | 2 | 1 | 1 | 1 | NA | NA | 3 | 1 |
| 11_1109634 | NA | NA | 2 | 0.5 | 2 | 0.5 | 2 | 1 | 6 | 0.6667 | NA | NA | 2 | 0.5 | 1 | 1 | NA | NA | 3 | 0.6667 |
| 11_1109636 | NA | NA | 2 | 0.5 | 2 | 0.5 | 2 | 0.5 | 6 | 0.5 | NA | NA | 1 | 1 | 1 | 1 | NA | NA | 2 | 1 |
| 11_1109636 | NA | NA | 2 | 0.5 | 2 | 1 | 2 | 0.5 | 6 | 0.6667 | NA | NA | 2 | 1 | 1 | 1 | NA | NA | 3 | 1 |
| 11_1109636 | NA | NA | 2 | 0.5 | 2 | 1 | 2 | 1 | 6 | 0.8333 | NA | NA | 1 | 1 | 1 | 1 | NA | NA | 2 | 1 |
| 11_1109638 | NA | NA | 2 | 0 | 1 | 1 | 1 | 1 | 4 | 0.5 | NA | NA | 2 | 0.5 | 1 | 1 | NA | NA | 3 | 0.6667 |
| 11_1109639 | NA | NA | 2 | 0.5 | 1 | 0 | 3 | 0.6667 | 6 | 0.5 | NA | NA | 1 | 1 | 1 | 1 | NA | NA | 2 | 1 |
| 11_1109639 | NA | NA | 2 | 0.5 | 2 | 0.5 | 3 | 0 | 7 | 0.2857 | NA | NA | 2 | 1 | 1 | 0 | NA | NA | 3 | 0.6667 |
| 11_1109640 | NA | NA | 2 | 1 | 1 | 1 | 3 | 1 | 6 | 1 | NA | NA | 2 | 0.5 | 1 | 1 | NA | NA | 3 | 0.6667 |
| 11_1109640 | NA | NA | 1 | 0 | NA | NA | 3 | 0.6667 | 4 | 0.5 | NA | NA | 2 | 0 | 1 | 1 | NA | NA | 3 | 0.3333 |
| 11_1109640 | NA | NA | 2 | 0 | 2 | 0.5 | 3 | 0.6667 | 7 | 0.4286 | NA | NA | 2 | 0.5 | 1 | 0 | NA | NA | 3 | 0.3333 |
| 11_1109643 | NA | NA | 1 | 1 | 2 | 0.5 | 3 | 0.3333 | 6 | 0.5 | NA | NA | 2 | 0 | 2 | 0.5 | NA | NA | 4 | 0.25 |
| 11_1109655 | 1 | 1 | 3 | 0 | 4 | 0 | 4 | 0 | 12 | 0.0833 | 1 | 0 | 3 | 0.3333 | 2 | 0 | 1 | 0 | 7 | 0.1429 |
| 11_1109659 | 1 | 0 | 3 | 0 | 5 | 0 | 4 | 0.25 | 13 | 0.0769 | 1 | 1 | 3 | 0 | 2 | 0.5 | 2 | 0 | 8 | 0.25 |
| 11_1109660 | NA | NA | 3 | 0 | 6 | 0.1667 | 4 | 0.5 | 13 | 0.2308 | 1 | 0 | 2 | 0.5 | 1 | 0 | 2 | 0 | 6 | 0.1667 |
| 11_1109661 | 2 | 0 | 3 | 0.3333 | 4 | 0.75 | 4 | 0.5 | 13 | 0.4615 | 1 | 0 | 1 | 0 | 2 | 0.5 | 3 | 0 | 7 | 0.1429 |
| 11_1109662 | 2 | 0.5 | 2 | 0 | 6 | 0.6667 | 4 | 0.25 | 14 | 0.4286 | 1 | 0 | 3 | 0.3333 | 2 | 1 | 3 | 0.3333 | 9 | 0.4444 |
| 11_1109666 | 2 | 0 | 3 | 0.3333 | 7 | 0 | 4 | 0 | 16 | 0.0625 | 1 | 0 | 3 | 0.3333 | 3 | 0.3333 | 4 | 0 | 11 | 0.1818 |
| 11_1126731 | 1 | 1 | 1 | 1 | NA | NA | NA | NA | 2 | 1 | 1 | 0 | NA | NA | 1 | 0 | 1 | 1 | 3 | 0.3333 |
| 11_1126747 | 2 | 0 | 1 | 0 | 2 | 0.5 | NA | NA | 5 | 0.2 | 3 | 0.3333 | 1 | 0 | 1 | 0 | 1 | 0 | 6 | 0.1667 |
| 11_1126756 | 2 | 0 | 1 | 0 | 2 | 0.5 | 1 | 0 | 6 | 0.1667 | 1 | 0 | 1 | 0 | 1 | 0 | 2 | 0.5 | 5 | 0.2 |
| 11_1126761 | 3 | 0.3333 | 3 | 0 | 2 | 0 | 1 | 0 | 9 | 0.1111 | 3 | 0 | 1 | 0 | 1 | 0 | 3 | 1 | 8 | 0.375 |
| 11_1126762 | 3 | 0.3333 | 2 | 0.5 | 2 | 0 | 1 | 0 | 8 | 0.25 | 2 | 0 | 1 | 0 | 1 | 0 | 2 | 0.5 | 6 | 0.1667 |

| | | | | | | | | | | | | | | | | | | | | |
|---|---|---|---|---|---|---|---|---|---|---|---|---|---|---|---|---|---|---|---|---|
| 11_1126764 | 3 | 1 | 2 | 1 | 2 | 1 | NA | NA | 7 | 1 | 4 | 1 | 1 | 1 | 1 | 1 | 3 | 1 | 9 | 1 |
| 11_1126764 | 3 | 0 | 2 | 0 | 2 | 0.5 | 1 | 1 | 8 | 0.25 | 3 | 0.3333 | 1 | 0 | 2 | 0 | 3 | 0.3333 | 9 | 0.2222 |
| 11_1126765 | 3 | 1 | 2 | 1 | 1 | 1 | 1 | 1 | 7 | 1 | 5 | 1 | NA | NA | 2 | 1 | 3 | 1 | 10 | 1 |
| 11_1126765 | 3 | 1 | 2 | 1 | 2 | 1 | 1 | 1 | 8 | 1 | 6 | 1 | 1 | 1 | 2 | 1 | 3 | 1 | 12 | 1 |
| 11_1126766 | 2 | 1 | 1 | 1 | 2 | 1 | 1 | 1 | 6 | 1 | 5 | 0.8 | 1 | 1 | 1 | 1 | 3 | 1 | 10 | 0.9 |
| 11_1126766 | 3 | 1 | 1 | 1 | 2 | 1 | NA | NA | 6 | 1 | 5 | 0.8 | NA | NA | 2 | 1 | 3 | 1 | 10 | 0.9 |
| 11_1126767 | 3 | 1 | 2 | 1 | 2 | 1 | 1 | 1 | 8 | 1 | 6 | 1 | 1 | 1 | 2 | 1 | 3 | 1 | 12 | 1 |
| 11_1126767 | 2 | 0.5 | 3 | 0.3333 | 2 | 0.5 | 1 | 0 | 8 | 0.375 | 5 | 0 | 1 | 0 | 2 | 0 | 3 | 0.3333 | 11 | 0.0909 |
| 11_1126779 | 2 | 0 | 4 | 0 | 3 | 0.3333 | 3 | 0 | 12 | 0.0833 | 8 | 0.25 | 2 | 0 | 4 | 0 | 2 | 0.5 | 16 | 0.1875 |
| 11_1126780 | 3 | 0.3333 | 5 | 0.4 | 4 | 0.5 | 4 | 0.5 | 16 | 0.4375 | 5 | 0.8 | 1 | 1 | 4 | 0.75 | 4 | 0 | 14 | 0.5714 |
| 11_1126781 | 1 | 0 | 6 | 0 | 6 | 0.3333 | 3 | 0 | 16 | 0.125 | 8 | 0.25 | 2 | 0 | 4 | 0 | 3 | 0.3333 | 17 | 0.1765 |
| 11_1129348 | 1 | 1 | 1 | 0 | 4 | 0.25 | 2 | 0 | 8 | 0.25 | NA | NA | 4 | 0.75 | 4 | 0.25 | 3 | 0 | 11 | 0.3636 |
| 11_1129348 | 1 | 0 | 1 | 1 | 3 | 0 | 1 | 1 | 6 | 0.3333 | NA | NA | 3 | 0.3333 | 3 | 0.6667 | 3 | 0.3333 | 9 | 0.4444 |
| 11_1129349 | 1 | 1 | 1 | 1 | 3 | 1 | 2 | 1 | 7 | 1 | NA | NA | 4 | 1 | 4 | 1 | 2 | 1 | 10 | 1 |
| 11_1129349 | NA | NA | 1 | 0 | 2 | 0.5 | NA | NA | 3 | 0.3333 | 2 | 0 | 4 | 0.5 | 4 | 0.25 | 2 | 0 | 12 | 0.25 |
| 11_1129349 | 1 | 1 | 1 | 1 | 3 | 1 | 1 | 1 | 6 | 1 | 2 | 1 | 3 | 1 | 3 | 1 | 3 | 1 | 11 | 1 |
| 11_1129351 | 1 | 1 | 1 | 1 | 2 | 0.5 | 1 | 1 | 5 | 0.8 | 2 | 0.5 | 3 | 0.3333 | 4 | 0.5 | 3 | 1 | 12 | 0.5833 |
| 11_1129352 | NA | NA | 1 | 1 | 2 | 1 | 1 | 1 | 4 | 1 | 2 | 1 | 4 | 1 | 3 | 1 | 3 | 1 | 12 | 1 |
| 11_1129352 | NA | NA | 1 | 0 | 2 | 0 | 1 | 1 | 4 | 0.25 | 2 | 0 | 4 | 0 | 3 | 0.3333 | 4 | 0.25 | 13 | 0.1538 |
| 11_1129354 | NA | NA | 1 | 1 | 2 | 0 | 1 | 1 | 4 | 0.5 | 1 | 0 | 4 | 0.25 | 3 | 0.3333 | 3 | 1 | 11 | 0.4545 |
| 11_1129355 | NA | NA | NA | NA | 1 | 1 | 1 | 1 | 2 | 1 | 2 | 1 | 4 | 0.5 | 4 | 0.75 | 3 | 0.3333 | 13 | 0.6154 |
| 11_1129357 | NA | NA | NA | NA | 1 | 0 | 1 | 1 | 2 | 0.5 | 1 | 1 | 3 | 0.6667 | 4 | 1 | 3 | 0.6667 | 11 | 0.8182 |
| 11_1129357 | NA | NA | 1 | 0 | 2 | 1 | 1 | 0 | 4 | 0.5 | 2 | 0 | 3 | 0.3333 | 3 | 0.3333 | 3 | 0.6667 | 11 | 0.3636 |
| 11_1129358 | NA | NA | 1 | 1 | 1 | 1 | NA | NA | 2 | 1 | 2 | 1 | 4 | 0.75 | 3 | 1 | 3 | 1 | 12 | 0.9167 |
| 11_1129358 | NA | NA | 1 | 1 | 2 | 0 | 1 | 1 | 4 | 0.5 | 2 | 0.5 | 2 | 0 | 4 | 0.5 | 2 | 0 | 10 | 0.3 |
| 11_1129358 | NA | NA | 1 | 1 | 1 | 1 | 1 | 0 | 3 | 0.6667 | 2 | 0.5 | 4 | 1 | 4 | 0.75 | 3 | 0.3333 | 13 | 0.6923 |
| 11_1129359 | NA | NA | 1 | 0 | 2 | 0.5 | NA | NA | 3 | 0.3333 | 2 | 0 | 4 | 0 | 4 | 0 | 3 | 0.3333 | 13 | 0.0769 |
| 11_1130922 | NA | NA | NA | NA | 2 | 1 | NA | NA | 2 | 1 | 1 | 0 | 1 | 1 | 3 | 0.6667 | NA | NA | 5 | 0.6 |
| 11_1131371 | NA | NA | 2 | 0.5 | 2 | 0 | 1 | 0 | 5 | 0.2 | NA | NA | 4 | 0.25 | NA | NA | 2 | 0.5 | 6 | 0.3333 |
| 11_1135153 | NA | NA | 3 | 1 | 1 | 1 | NA | NA | 4 | 1 | NA | NA | NA | NA | 1 | 1 | 1 | 1 | 2 | 1 |
| 11_1135154 | NA | NA | 3 | 1 | NA | NA | NA | NA | 3 | 1 | NA | NA | 1 | 0 | 1 | 1 | 1 | 0 | 3 | 0.3333 |
| 11_1135155 | NA | NA | 3 | 0.3333 | 1 | 0 | NA | NA | 4 | 0.25 | NA | NA | 1 | 1 | 1 | 1 | 1 | 1 | 3 | 1 |
| 11_1135159 | NA | NA | 2 | 1 | 2 | 1 | NA | NA | 4 | 1 | NA | NA | 1 | 1 | 1 | 1 | 2 | 1 | 4 | 1 |
| 11_1135161 | NA | NA | 3 | 0.3333 | 3 | 0 | NA | NA | 6 | 0.1667 | NA | NA | 1 | 0 | 1 | 1 | 2 | 0.5 | 4 | 0.5 |
| 11_1135165 | NA | NA | 2 | 0.5 | 4 | 0.5 | NA | NA | 6 | 0.5 | NA | NA | 1 | 0 | 1 | 1 | 2 | 0 | 4 | 0.25 |
| 11_1135166 | NA | NA | 3 | 1 | 4 | 1 | 1 | 1 | 8 | 1 | 1 | 1 | 2 | 1 | 1 | 1 | 2 | 1 | 6 | 1 |
| 11_1135428 | 2 | 0.5 | 3 | 0 | NA | NA | 4 | 0 | 9 | 0.1111 | 2 | 0.5 | 6 | 0.1667 | NA | NA | 2 | 0 | 10 | 0.2 |
| 11_1135431 | 2 | 0 | 4 | 0.25 | NA | NA | 5 | 0 | 11 | 0.0909 | 1 | 0 | 6 | 0.1667 | NA | NA | 2 | 0 | 9 | 0.1111 |
| 11_1135469 | 2 | 0.5 | 3 | 0.3333 | 2 | 1 | 6 | 0.5 | 13 | 0.5385 | 3 | 1 | 8 | 0.5 | 1 | 0 | 1 | 1 | 13 | 0.6154 |
| 11_1135469 | 2 | 0 | 4 | 0 | 2 | 0 | 3 | 0.3333 | 11 | 0.0909 | 3 | 0 | 8 | 0.125 | 1 | 1 | NA | NA | 12 | 0.1667 |
| 11_1135470 | 3 | 0 | 4 | 0 | 2 | 0.5 | 4 | 0 | 13 | 0.0769 | 3 | 0 | 7 | 0.1429 | 1 | 0 | 2 | 0 | 13 | 0.0769 |
| 11_1135470 | 3 | 1 | 4 | 1 | 2 | 1 | 3 | 1 | 12 | 1 | 2 | 1 | 6 | 1 | 1 | 1 | 2 | 1 | 11 | 1 |
| 11_1135470 | 2 | 0.5 | 2 | 0 | 2 | 1 | 4 | 0.75 | 10 | 0.6 | 3 | 0.6667 | 8 | 0.375 | 1 | 1 | 3 | 0.3333 | 15 | 0.4667 |
| 11_1135471 | 2 | 0.5 | 2 | 0 | 2 | 0.5 | 5 | 0 | 11 | 0.1818 | 3 | 0.6667 | 5 | 0 | 1 | 1 | 1 | 0 | 10 | 0.3 |
| 11_1135472 | 3 | 0.3333 | 4 | 0.25 | 2 | 0.5 | 5 | 0 | 14 | 0.2143 | 3 | 0 | 8 | 0 | 1 | 0 | 3 | 0.3333 | 15 | 0.0667 |
| 11_1135473 | 2 | 1 | 4 | 1 | 2 | 1 | 4 | 1 | 12 | 1 | 3 | 1 | 8 | 1 | NA | NA | 2 | 1 | 13 | 1 |
| 11_1135479 | 3 | 0 | 4 | 0.25 | 2 | 1 | 8 | 0.25 | 17 | 0.2941 | 1 | 1 | 7 | 0.4286 | 1 | 1 | 6 | 0.3333 | 15 | 0.4667 |
| 11_1135525 | 2 | 1 | 4 | 0.75 | 3 | 0.6667 | 3 | 1 | 12 | 0.8333 | 2 | 1 | 6 | 0.8333 | 3 | 0.6667 | 3 | 1 | 14 | 0.8571 |
| 11_1135525 | 2 | 0 | 4 | 0.25 | 3 | 0 | 2 | 0 | 11 | 0.0909 | 2 | 0 | 2 | 0.5 | 3 | 0 | 2 | 0 | 9 | 0.1111 |
| 11_1135526 | 1 | 1 | 1 | 0 | 2 | 0.5 | 1 | 1 | 5 | 0.6 | NA | NA | 4 | 1 | 2 | 1 | 2 | 1 | 8 | 1 |
| 11_1135528 | 2 | 0.5 | 3 | 0 | 2 | 0.5 | 1 | 0 | 8 | 0.25 | NA | NA | 4 | 0.25 | 1 | 0 | 1 | 0 | 6 | 0.1667 |
| 11_1135528 | NA | NA | 1 | 0 | NA | NA | 6 | 0.3333 | 7 | 0.2857 | NA | NA | NA | NA | NA | NA | 1 | 1 | 1 | 1 |
| 11_1135541 | 5 | 1 | 7 | 0.8571 | 5 | 0.8 | 5 | 1 | 22 | 0.9091 | 3 | 0.3333 | 5 | 1 | 5 | 1 | 7 | 1 | 20 | 0.9 |
| 11_1135787 | NA | NA | NA | NA | 1 | 1 | NA | NA | 1 | 1 | NA | NA | NA | NA | 2 | 0.5 | NA | NA | 2 | 0.5 |
| 11_1135791 | NA | NA | NA | NA | 1 | 1 | NA | NA | 1 | 1 | NA | NA | NA | NA | 2 | 0.5 | NA | NA | 2 | 0.5 |
| 11_1135792 | NA | NA | NA | NA | 1 | 1 | NA | NA | 1 | 1 | NA | NA | NA | NA | 1 | 1 | NA | NA | 1 | 1 |
| 11_1135794 | NA | NA | NA | NA | 1 | 1 | NA | NA | 1 | 1 | NA | NA | NA | NA | 1 | 1 | NA | NA | 1 | 1 |
| 11_1135795 | NA | NA | NA | NA | 1 | 1 | NA | NA | 1 | 1 | NA | NA | NA | NA | 1 | 1 | NA | NA | 1 | 1 |
| 11_1136178 | 3 | 0.3333 | 3 | 0.3333 | 3 | 0 | 11 | 0.3636 | 20 | 0.3 | 3 | 0.3333 | 1 | 1 | 2 | 0 | 5 | 0.2 | 11 | 0.2727 |
| 11_1136183 | 2 | 0 | 3 | 0.3333 | 4 | 0.75 | 9 | 0.6667 | 18 | 0.5556 | 3 | 0.3333 | 1 | 1 | 2 | 0.5 | 4 | 1 | 10 | 0.7 |
| 11_1136185 | 1 | 0 | 3 | 0.6667 | NA | NA | 6 | 0.6667 | 10 | 0.6 | 1 | 1 | 1 | 0 | NA | NA | 3 | 1 | 5 | 0.8 |
| 11_1136185 | 2 | 1 | 4 | 1 | 3 | 1 | 12 | 1 | 21 | 1 | 2 | 1 | 1 | 1 | 2 | 1 | 2 | 1 | 7 | 1 |
| 11_1136187 | 1 | 0 | 4 | 0.75 | 3 | 1 | 9 | 0.6667 | 17 | 0.7059 | 2 | 0 | 1 | 0 | 2 | 1 | 3 | 1 | 8 | 0.625 |
| 11_1136189 | 1 | 1 | 3 | 1 | 3 | 1 | 10 | 1 | 17 | 1 | NA | NA | NA | NA | 2 | 1 | 2 | 1 | 4 | 1 |
| 11_1136191 | 1 | 1 | 3 | 1 | 4 | 0.75 | 9 | 0.8889 | 17 | 0.8824 | NA | NA | 1 | 1 | 2 | 1 | 3 | 1 | 6 | 1 |

| | | | | | | | | | | | | | | | | | | | | |
|---|---|---|---|---|---|---|---|---|---|---|---|---|---|---|---|---|---|---|---|---|
| 11_113619 | 1 | 1 | 3 | 1 | 4 | 1 | 9 | 1 | 17 | 1 | NA | NA | 1 | 1 | 2 | 1 | 3 | 1 | 6 | 1 |
| 11_113619 | 1 | 0 | 1 | 0 | 2 | 0.5 | 10 | 0.1 | 14 | 0.1429 | NA | NA | NA | NA | 1 | 0 | 2 | 0.5 | 3 | 0.3333 |
| 11_113619 | 1 | 1 | 1 | 0 | 4 | 0.75 | 5 | 0.8 | 11 | 0.7273 | NA | NA | 1 | 1 | 2 | 0 | 3 | 1 | 6 | 0.6667 |
| 11_113619 | 1 | 0 | 2 | 1 | 3 | 0.6667 | 9 | 0.7778 | 15 | 0.7333 | NA | NA | 1 | 1 | 2 | 0 | 3 | 0 | 6 | 0.1667 |
| 11_113619 | 1 | 0 | 2 | 0 | 3 | 0.3333 | 9 | 0 | 15 | 0.0667 | NA | NA | 1 | 0 | 2 | 0.5 | 2 | 0 | 5 | 0.2 |
| 11_113619 | 1 | 0 | 3 | 1 | 3 | 0.6667 | 9 | 0.3333 | 16 | 0.5 | NA | NA | 1 | 0 | 2 | 0.5 | 1 | 1 | 4 | 0.5 |
| 11_113619 | 1 | 1 | 3 | 1 | 3 | 1 | 8 | 0.875 | 15 | 0.9333 | NA | NA | NA | NA | 2 | 0.5 | 1 | 1 | 3 | 0.6667 |
| 11_113619 | 1 | 1 | 2 | 1 | 3 | 1 | 7 | 1 | 13 | 1 | NA | NA | 1 | 1 | 2 | 1 | 1 | 1 | 4 | 1 |
| 11_113619 | 1 | 0 | 3 | 0.3333 | 2 | 0 | 9 | 0.2222 | 15 | 0.2 | NA | NA | 1 | 0 | 2 | 0.5 | 2 | 0 | 5 | 0.2 |
| 11_113620 | 1 | 0 | 2 | 1 | 2 | 1 | 7 | 0.4286 | 12 | 0.5833 | NA | NA | 1 | 1 | 1 | 0 | 2 | 1 | 4 | 0.75 |
| 11_113620 | 1 | 0 | 2 | 0.5 | 2 | 0 | 7 | 0.4286 | 12 | 0.3333 | NA | NA | 1 | 0 | 2 | 0 | 2 | 0.5 | 5 | 0.2 |
| 11_113620 | 1 | 0 | 2 | 0.5 | 2 | 0 | 9 | 0.2222 | 14 | 0.2143 | NA | NA | 1 | 0 | 1 | 1 | 1 | 0 | 3 | 0.3333 |
| 11_113620 | 1 | 1 | 2 | 0 | 1 | 0 | 9 | 0.1111 | 13 | 0.1538 | NA | NA | 1 | 0 | 1 | 0 | 2 | 0.5 | 4 | 0.25 |
| 11_113620 | 1 | 1 | 2 | 0.5 | 2 | 0.5 | 6 | 0.3333 | 11 | 0.4545 | NA | NA | 1 | 1 | 1 | 0 | NA | NA | 2 | 0.5 |
| 11_113620 | 1 | 1 | 1 | 1 | 2 | 0 | 4 | 0.75 | 8 | 0.625 | NA | NA | 1 | 1 | NA | NA | 1 | 1 | 2 | 1 |
| 11_113620 | NA | NA | 2 | 0.5 | 2 | 0.5 | 9 | 0.2222 | 13 | 0.3077 | NA | NA | 1 | 0 | 2 | 0 | 2 | 0.5 | 5 | 0.2 |
| 11_113669 | NA | NA | NA | NA | 2 | 0.5 | NA | NA | 2 | 0.5 | 2 | 0.5 | NA | NA | NA | NA | 1 | 0 | 3 | 0.3333 |
| 11_114610 | 3 | 0.3333 | 8 | 0.125 | 4 | 0.25 | 3 | 0 | 18 | 0.1667 | 1 | 0 | 2 | 0.5 | 1 | 1 | 2 | 0 | 6 | 0.3333 |
| 11_114610 | 2 | 1 | 8 | 0.625 | 4 | 0.25 | 3 | 0.3333 | 17 | 0.5294 | 3 | 0.6667 | 2 | 1 | 1 | 1 | 2 | 0.5 | 8 | 0.75 |
| 11_114610 | 3 | 0 | 7 | 0.1429 | 4 | 0 | 3 | 0.6667 | 17 | 0.1765 | 2 | 0 | 2 | 0 | 1 | 0 | 2 | 0.5 | 7 | 0.1429 |
| 11_114610 | 3 | 0.3333 | 7 | 0.8571 | 3 | 1 | 4 | 1 | 17 | 0.8235 | 3 | 0.3333 | 2 | 0 | 1 | 1 | 2 | 0 | 8 | 0.25 |
| 11_114610 | 3 | 0.3333 | 8 | 0.875 | 5 | 0.8 | 4 | 1 | 20 | 0.8 | 3 | 0.3333 | 2 | 0.5 | 1 | 1 | 2 | 0.5 | 8 | 0.5 |
| 11_114610 | 4 | 0 | 8 | 0.25 | 4 | 0.5 | 3 | 0.3333 | 19 | 0.2632 | 1 | 0 | 2 | 0 | 1 | 1 | 1 | 0 | 5 | 0.2 |
| 11_114610 | 3 | 0 | 8 | 0.125 | 3 | 0.3333 | 2 | 0.5 | 16 | 0.1875 | 3 | 0 | 2 | 0 | 1 | 1 | 2 | 0 | 8 | 0.125 |
| 11_114610 | 4 | 0 | 8 | 0.25 | 3 | 0 | 2 | 0 | 17 | 0.1176 | 3 | 0.3333 | 2 | 0 | 1 | 0 | 1 | 0 | 7 | 0.1429 |
| 11_114611 | 6 | 0 | 7 | 0.4286 | 7 | 0 | 3 | 0 | 23 | 0.1304 | 4 | 0.25 | 2 | 0 | 1 | 0 | 3 | 0 | 10 | 0.1 |
| 11_114611 | 6 | 0 | 8 | 0 | 6 | 0.1667 | 3 | 0.3333 | 23 | 0.087 | 5 | 0.2 | 1 | 0 | 1 | 0 | 3 | 0 | 10 | 0.1 |
| 11_114611 | 6 | 0.8333 | 9 | 0.7778 | 7 | 0.8571 | 3 | 0.6667 | 25 | 0.8 | 4 | 1 | 2 | 1 | 1 | 1 | 3 | 0.6667 | 10 | 0.9 |
| 11_114682 | 7 | 0.8571 | 6 | 0.6667 | 7 | 1 | 7 | 0.8571 | 27 | 0.8519 | 6 | 0.8333 | 3 | 1 | 4 | 0.75 | 5 | 0.6 | 18 | 0.7778 |
| 11_114682 | 6 | 1 | 4 | 1 | 4 | 0.75 | 5 | 1 | 19 | 0.9474 | 3 | 1 | 2 | 1 | 2 | 1 | 5 | 1 | 12 | 1 |
| 11_114682 | 6 | 0.8333 | 6 | 1 | 7 | 0.8571 | 8 | 1 | 27 | 0.9259 | 6 | 0.8333 | 3 | 1 | 5 | 1 | 4 | 0.75 | 18 | 0.8889 |
| 11_114682 | 5 | 0 | 4 | 0.25 | 5 | 0.4 | 7 | 0.5714 | 21 | 0.3333 | 3 | 1 | 2 | 1 | 3 | 0 | 4 | 0.75 | 12 | 0.6667 |
| 11_114682 | 7 | 0.7143 | 6 | 0.3333 | 7 | 0.2857 | 7 | 0.7143 | 27 | 0.5185 | 4 | 1 | 3 | 0.6667 | 5 | 0.6 | 5 | 0.8 | 17 | 0.7647 |
| 11_114683 | 8 | 1 | 4 | 1 | 6 | 1 | 7 | 1 | 25 | 1 | 4 | 0.5 | 1 | 1 | 4 | 1 | 5 | 0.8 | 14 | 0.7857 |
| 11_114683 | 5 | 0.2 | 5 | 0 | 6 | 0.3333 | 6 | 0.1667 | 22 | 0.1818 | 5 | 0.2 | 2 | 0 | 4 | 0.25 | 5 | 0.4 | 16 | 0.25 |
| 11_114683 | 7 | 0.1429 | 5 | 0.4 | 6 | 0.3333 | 6 | 0 | 24 | 0.2083 | 5 | 0.8 | 3 | 0 | 4 | 0 | 5 | 0 | 17 | 0.2353 |
| 11_114684 | 7 | 0 | 5 | 0.2 | 6 | 0.1667 | 6 | 0.1667 | 24 | 0.125 | 6 | 0.3333 | 2 | 0.5 | 4 | 0 | 3 | 0 | 15 | 0.2 |
| 11_114688 | NA | NA | 1 | 0 | 1 | 1 | NA | NA | 2 | 0.5 | 1 | 0 | NA | NA | 1 | 1 | NA | NA | 2 | 0.5 |
| 11_114762 | NA | NA | 2 | 1 | NA | NA | NA | NA | 2 | 1 | NA | NA | NA | NA | NA | NA | 1 | 1 | 1 | 1 |
| 11_114762 | NA | NA | 1 | 1 | NA | NA | NA | NA | 1 | 1 | NA | NA | NA | NA | NA | NA | 1 | 1 | 1 | 1 |
| 11_114762 | NA | NA | 2 | 0.5 | NA | NA | NA | NA | 2 | 0.5 | NA | NA | NA | NA | NA | NA | 1 | 1 | 1 | 1 |
| 11_115071 | 6 | 0.3333 | 5 | 0 | 11 | 0.0909 | 13 | 0.1538 | 35 | 0.1429 | 13 | 0.3077 | 9 | 0.3333 | 12 | 0.25 | 7 | 0.2857 | 41 | 0.2927 |
| 11_115071 | 5 | 0.6 | 4 | 0.5 | 11 | 0.4545 | 14 | 0.2857 | 34 | 0.4118 | 11 | 0.3636 | 8 | 0.25 | 10 | 0.4 | 8 | 0.375 | 37 | 0.3514 |
| 11_115071 | 6 | 0.6667 | 5 | 0.4 | 12 | 0.0833 | 13 | 0.2308 | 36 | 0.2778 | 15 | 0.4667 | 10 | 0.1 | 13 | 0.3846 | 8 | 0.375 | 46 | 0.3478 |
| 11_115071 | 6 | 0.6667 | 4 | 0.5 | 12 | 0.6667 | 12 | 0.5 | 34 | 0.5882 | 17 | 0.6471 | 9 | 0.3333 | 12 | 0.6667 | 8 | 0.375 | 46 | 0.5435 |
| 11_115071 | 8 | 0.5 | 4 | 0.5 | 11 | 0.5455 | 8 | 0.125 | 31 | 0.4194 | 14 | 0.2143 | 8 | 0.5 | 9 | 0.3333 | 7 | 0.1429 | 38 | 0.2895 |
| 11_115073 | 2 | 0 | 1 | 0 | 1 | 0 | 3 | 0.3333 | 7 | 0.1429 | 4 | 0.25 | 2 | 0 | 6 | 0 | 6 | 0.1667 | 18 | 0.1111 |
| 11_115073 | 1 | 1 | 2 | 0 | 1 | 0 | 3 | 0.6667 | 7 | 0.4286 | 5 | 0.6 | 2 | 1 | 3 | 0.3333 | 5 | 0.6 | 15 | 0.6 |
| 11_115134 | 2 | 1 | 3 | 0.6667 | NA | NA | NA | NA | 5 | 0.8 | 1 | 1 | NA | NA | 1 | 1 | 1 | 0 | 3 | 0.6667 |
| 11_115134 | 1 | 1 | 4 | 0.25 | NA | NA | NA | NA | 5 | 0.4 | 2 | 0.5 | NA | NA | 1 | 0 | 1 | 0 | 4 | 0.25 |
| 11_115134 | 1 | 1 | 4 | 1 | NA | NA | NA | NA | 5 | 1 | 2 | 1 | NA | NA | 1 | 1 | 2 | 1 | 5 | 1 |
| 11_115134 | 1 | 1 | 4 | 0.25 | 1 | 0 | NA | NA | 6 | 0.3333 | 2 | 0.5 | NA | NA | 1 | 1 | 2 | 1 | 5 | 0.8 |
| 11_115134 | 1 | 0 | 4 | 0.25 | 1 | 1 | NA | NA | 6 | 0.3333 | 2 | 0.5 | NA | NA | 1 | 0 | 2 | 0.5 | 5 | 0.4 |
| 11_115134 | 1 | 1 | 4 | 1 | NA | NA | NA | NA | 5 | 1 | 1 | 1 | NA | NA | 1 | 1 | 2 | 1 | 4 | 1 |
| 11_115135 | 2 | 0.5 | 3 | 0.6667 | 1 | 0 | NA | NA | 6 | 0.5 | 2 | 0.5 | NA | NA | 1 | 0 | 2 | 0.5 | 5 | 0.4 |
| 11_115135 | 1 | 0 | 4 | 0.25 | 1 | 0 | NA | NA | 6 | 0.1667 | 2 | 0.5 | NA | NA | NA | NA | 1 | 0 | 3 | 0.3333 |
| 11_115135 | 1 | 1 | 4 | 0 | 1 | 0 | 1 | 0 | 7 | 0.1429 | 1 | 1 | NA | NA | 1 | 0 | 2 | 0 | 4 | 0.25 |
| 11_115135 | 1 | 1 | 3 | 1 | NA | NA | 1 | 1 | 5 | 1 | 2 | 1 | NA | NA | 1 | 0 | 2 | 0.5 | 5 | 0.6 |
| 11_115135 | 1 | 1 | 1 | 1 | 2 | 1 | 1 | 0 | 5 | 0.8 | 1 | 1 | NA | NA | NA | NA | 1 | 1 | 2 | 1 |
| 11_115135 | 1 | 0 | 4 | 0.5 | 2 | 1 | 1 | 0 | 8 | 0.5 | 2 | 0.5 | NA | NA | 1 | 0 | 3 | 0.3333 | 6 | 0.3333 |
| 11_115135 | 2 | 1 | 4 | 1 | 3 | 1 | NA | NA | 9 | 1 | 2 | 1 | NA | NA | 2 | 1 | 3 | 1 | 7 | 1 |
| 11_115136 | 2 | 0 | 4 | 0 | 3 | 0.3333 | 1 | 0 | 10 | 0.1 | 2 | 0 | NA | NA | 2 | 0 | 3 | 0.6667 | 7 | 0.2857 |
| 11_115136 | 1 | 1 | 2 | 1 | 2 | 1 | 1 | 1 | 6 | 1 | 2 | 1 | NA | NA | 1 | 1 | NA | NA | 3 | 1 |
| 11_115136 | NA | NA | 1 | 1 | 1 | 1 | NA | NA | 2 | 1 | NA | NA | NA | NA | NA | NA | 1 | 1 | 1 | 1 |
| 11_115136 | 2 | 0.5 | 4 | 0 | 2 | 0 | 1 | 0 | 9 | 0.1111 | 2 | 0.5 | NA | NA | 3 | 0.3333 | 3 | 0 | 8 | 0.25 |

| | | | | | | | | | | | | | | | | | | | | |
|---|---|---|---|---|---|---|---|---|---|---|---|---|---|---|---|---|---|---|---|---|
| 11_115136 | 1 | 1 | 3 | 1 | 2 | 1 | 1 | 1 | 7 | 1 | 1 | 1 | NA | NA | 2 | 1 | 3 | 1 | 6 | 1 |
| 11_115136 | 3 | 0 | 4 | 0.25 | 2 | 0 | 1 | 0 | 10 | 0.1 | 2 | 0.5 | NA | NA | 3 | 0.3333 | 3 | 0 | 8 | 0.25 |
| 11_115136 | 3 | 0.3333 | 4 | 0.25 | 2 | 0 | 1 | 0 | 10 | 0.2 | 2 | 0.5 | NA | NA | 2 | 0.5 | 3 | 0 | 7 | 0.2857 |
| 11_115136 | 2 | 0 | 4 | 0.25 | 3 | 0 | 1 | 0 | 10 | 0.1 | 2 | 1 | 1 | 1 | 3 | 0.3333 | 2 | 0.5 | 8 | 0.625 |
| 11_115136 | 2 | 0.5 | 4 | 0.25 | 3 | 0.3333 | 1 | 0 | 10 | 0.3 | 2 | 0.5 | NA | NA | 4 | 0.25 | 2 | 0 | 8 | 0.25 |
| 11_115136 | 2 | 0 | 4 | 0.25 | 3 | 0 | 1 | 0 | 10 | 0.1 | 2 | 0.5 | NA | NA | 4 | 0.5 | 3 | 0 | 9 | 0.3333 |
| 11_115136 | 1 | 0 | 2 | 0.5 | 1 | 0 | 1 | 1 | 5 | 0.4 | 2 | 0.5 | NA | NA | 1 | 1 | 2 | 1 | 5 | 0.8 |
| 11_115137 | 1 | 0 | 2 | 0.5 | NA | NA | 3 | 0 | 6 | 0.1667 | 4 | 0.25 | NA | NA | 2 | 0 | 3 | 0.3333 | 9 | 0.2222 |
| 11_115140 | 2 | 0.5 | 11 | 0 | 7 | 0.1429 | 7 | 0 | 27 | 0.0741 | 1 | 0 | 7 | 0.4286 | 11 | 0.2727 | 5 | 0.4 | 24 | 0.3333 |
| 11_115140 | 4 | 0 | 12 | 0.0833 | 7 | 0.1429 | 6 | 0 | 29 | 0.069 | 1 | 0 | 7 | 0 | 13 | 0.0769 | 6 | 0.1667 | 27 | 0.0741 |
| 11_115140 | 5 | 0.4 | 9 | 0.2222 | 5 | 0 | 6 | 0.5 | 25 | 0.28 | 1 | 1 | 7 | 0.4286 | 10 | 0.1 | 2 | 0 | 20 | 0.25 |
| 11_115140 | 5 | 0.4 | 8 | 0.625 | 5 | 0.6 | 6 | 0.5 | 24 | 0.5417 | 2 | 0.5 | 5 | 0.2 | 8 | 0.5 | 5 | 0 | 20 | 0.3 |
| 11_115140 | 3 | 0 | 8 | 0.25 | 5 | 0.4 | 6 | 0.1667 | 22 | 0.2273 | 1 | 0 | 5 | 0.4 | 10 | 0.2 | 6 | 0.5 | 22 | 0.3182 |
| 11_115140 | 5 | 0.6 | 10 | 0.3 | 5 | 0.4 | 5 | 0.4 | 25 | 0.4 | 2 | 0.5 | 7 | 0.2857 | 10 | 0.4 | 6 | 0.6667 | 25 | 0.44 |
| 11_115140 | 4 | 0 | 11 | 0.4545 | 6 | 0.3333 | 6 | 0.1667 | 27 | 0.2963 | 1 | 0 | 7 | 0 | 11 | 0.2727 | 6 | 0 | 25 | 0.12 |
| 11_115140 | 5 | 0.4 | 11 | 0.8182 | 5 | 0.4 | 6 | 0.3333 | 27 | 0.5556 | 2 | 0.5 | 7 | 0.4286 | 11 | 0.5455 | 6 | 0.6667 | 26 | 0.5385 |
| 11_115141 | 3 | 0 | 8 | 0.25 | 3 | 0 | 7 | 0.2857 | 21 | 0.1905 | 2 | 0 | 7 | 0 | 7 | 0.1429 | 5 | 0.4 | 21 | 0.1429 |
| 11_115141 | 4 | 0.25 | 7 | 0.2857 | 3 | 0 | 4 | 0.5 | 18 | 0.2778 | 2 | 1 | 5 | 0.6 | 9 | 0.5556 | 4 | 0.25 | 20 | 0.55 |
| 11_115141 | 5 | 0 | 7 | 0 | 2 | 0 | 6 | 0.1667 | 20 | 0.05 | 2 | 0 | 6 | 0.3333 | 8 | 0 | 3 | 0 | 19 | 0.1053 |
| 11_115148 | 2 | 1 | 4 | 0.5 | 3 | 0 | 2 | 0.5 | 11 | 0.4545 | 2 | 0 | 4 | 0.25 | 4 | 0.75 | 1 | 1 | 11 | 0.4545 |
| 11_115151 | NA | NA | 2 | 0 | 2 | 0 | 2 | 0.5 | 6 | 0.1667 | 2 | 0.5 | 1 | 0 | 1 | 0 | NA | NA | 4 | 0.25 |
| 11_115158 | NA | NA | 1 | 1 | 3 | 0.6667 | 3 | 0.6667 | 7 | 0.7143 | 1 | 1 | 2 | 1 | 1 | 1 | 2 | 1 | 6 | 1 |
| 11_115158 | NA | NA | 2 | 0.5 | 4 | 0 | 3 | 0 | 9 | 0.1111 | 1 | 0 | 2 | 0.5 | 1 | 1 | 3 | 0.3333 | 7 | 0.4286 |
| 11_115159 | NA | NA | 1 | 1 | 4 | 1 | 3 | 1 | 8 | 1 | 1 | 0 | 2 | 1 | 1 | 1 | 3 | 0.6667 | 7 | 0.7143 |
| 11_115159 | NA | NA | NA | NA | 3 | 0.6667 | 1 | 0 | 4 | 0.5 | 1 | 0 | 2 | 0 | NA | NA | 2 | 0.5 | 5 | 0.2 |
| 11_115238 | 14 | 0.1429 | 3 | 0 | 11 | 0.4545 | 10 | 0.4 | 38 | 0.2895 | 8 | 0.375 | 8 | 0.125 | 11 | 0.4545 | 10 | 0.1 | 37 | 0.2703 |
| 11_115238 | 12 | 0.0833 | 2 | 0 | 11 | 0.0909 | 11 | 0.0909 | 36 | 0.0833 | 8 | 0 | 9 | 0 | 12 | 0.4167 | 8 | 0.125 | 37 | 0.1622 |
| 11_115284 | 1 | 0 | 4 | 0.25 | 1 | 0 | 2 | 0.5 | 8 | 0.25 | 3 | 0.3333 | NA | NA | 2 | 0 | NA | NA | 5 | 0.2 |
| 11_115284 | 1 | 1 | 2 | 0.5 | 1 | 1 | 2 | 0.5 | 6 | 0.6667 | 3 | 0.6667 | NA | NA | 2 | 0.5 | 1 | 1 | 6 | 0.6667 |
| 11_115284 | 1 | 0 | 4 | 0.25 | 2 | 0 | 2 | 0.5 | 9 | 0.2222 | 4 | 0.5 | NA | NA | 2 | 0 | 1 | 0 | 7 | 0.2857 |
| 11_115284 | 1 | 0 | 4 | 0.75 | 2 | 0 | NA | NA | 7 | 0.4286 | 4 | 0.75 | NA | NA | 1 | 1 | 1 | 1 | 6 | 0.8333 |
| 11_115284 | 1 | 0 | 4 | 0 | 2 | 0.5 | 2 | 0 | 9 | 0.1111 | 4 | 0 | NA | NA | 2 | 0 | 1 | 1 | 7 | 0.1429 |
| 11_115284 | 1 | 1 | 4 | 0.5 | 2 | 0.5 | NA | NA | 7 | 0.5714 | 3 | 0.6667 | NA | NA | NA | NA | NA | NA | 3 | 0.6667 |
| 11_115284 | 1 | 1 | 5 | 0.8 | 1 | 0 | NA | NA | 7 | 0.7143 | 4 | 1 | NA | NA | 1 | 1 | NA | NA | 5 | 1 |
| 11_115285 | 1 | 1 | 4 | 1 | 2 | 1 | 2 | 0.5 | 9 | 0.8889 | 3 | 1 | NA | NA | 2 | 1 | 1 | 1 | 6 | 1 |
| 11_115285 | 1 | 0 | 3 | 0.3333 | 2 | 0 | 2 | 0.5 | 8 | 0.25 | 3 | 0.6667 | NA | NA | 2 | 0.5 | 1 | 0 | 6 | 0.5 |
| 11_115285 | 2 | 0 | 5 | 0.2 | 2 | 0.5 | 2 | 1 | 11 | 0.3636 | 4 | 0.5 | 1 | 1 | 2 | 0.5 | 1 | 0 | 8 | 0.5 |
| 11_115286 | 2 | 0.5 | 3 | 0.3333 | 3 | 0 | 1 | 0 | 9 | 0.2222 | 4 | 0.25 | 1 | 0 | 2 | 0.5 | NA | NA | 7 | 0.2857 |
| 11_115286 | 2 | 0 | 4 | 0.25 | 2 | 0 | 2 | 0 | 10 | 0.1 | 4 | 0 | 1 | 0 | 2 | 0.5 | 1 | 0 | 8 | 0.125 |
| 11_115286 | 3 | 1 | 4 | 0.5 | 3 | 0 | 3 | 0.6667 | 13 | 0.5385 | 5 | 0 | 1 | 1 | 2 | 1 | NA | NA | 8 | 0.375 |
| 11_115286 | 3 | 0 | 3 | 0 | 3 | 0.3333 | 3 | 0 | 12 | 0.0833 | 5 | 0.4 | 1 | 0 | 1 | 0 | 1 | 0 | 8 | 0.25 |
| 11_115286 | 3 | 0 | 4 | 0 | 3 | 0 | 2 | 0.5 | 12 | 0.0833 | 5 | 0.2 | 2 | 0 | 2 | 0 | 1 | 1 | 10 | 0.2 |
| 11_115286 | 3 | 0.3333 | 5 | 0.8 | 3 | 0.6667 | 3 | 0.6667 | 14 | 0.6429 | 3 | 0.6667 | 1 | 0 | 2 | 1 | 1 | 1 | 7 | 0.7143 |
| 11_115286 | 1 | 1 | 5 | 0.8 | 2 | 1 | 3 | 0.6667 | 11 | 0.8182 | 4 | 1 | 2 | 1 | 2 | 1 | 1 | 1 | 9 | 1 |
| 11_115286 | 3 | 0.6667 | 4 | 0.75 | 2 | 0 | 3 | 0.3333 | 12 | 0.5 | 2 | 0 | 1 | 0 | 1 | 1 | 1 | 0 | 5 | 0.2 |
| 11_115300 | NA | NA | 3 | 0 | 1 | 1 | NA | NA | 4 | 0.25 | 2 | 1 | NA | NA | 1 | 0 | NA | NA | 3 | 0.6667 |
| 11_115301 | 6 | 0 | 10 | 0.1 | 7 | 0 | 9 | 0.1111 | 32 | 0.0625 | 10 | 0.2 | 8 | 0.125 | 7 | 0.1429 | 8 | 0 | 33 | 0.1212 |
| 11_115301 | 6 | 0.1667 | 10 | 0.1 | 8 | 0.25 | 9 | 0 | 33 | 0.1212 | 10 | 0.1 | 8 | 0 | 7 | 0.2857 | 8 | 0 | 33 | 0.0909 |
| 11_115301 | 6 | 0.5 | 9 | 0.4444 | 6 | 0.5 | 5 | 0.6 | 26 | 0.5 | 8 | 0.75 | 8 | 0.25 | 8 | 0.5 | 7 | 0.5714 | 31 | 0.5161 |
| 11_115301 | 5 | 0.4 | 9 | 0.3333 | 9 | 0.2222 | 8 | 0.25 | 31 | 0.2903 | 8 | 0.25 | 10 | 0.2 | 6 | 0.5 | 7 | 0.4286 | 31 | 0.3226 |
| 11_115321 | 3 | 0 | 5 | 0.2 | 10 | 0.1 | 4 | 0 | 22 | 0.0909 | 5 | 0 | 4 | 0 | 6 | 0.3333 | 4 | 0 | 19 | 0.1053 |
| 11_115321 | 4 | 0.75 | 6 | 0.1667 | 9 | 0 | 5 | 0.2 | 24 | 0.2083 | 6 | 0 | 5 | 0.2 | 3 | 0 | 5 | 0 | 19 | 0.0526 |
| 11_115321 | 3 | 0.3333 | 4 | 0 | 7 | 0.4286 | 4 | 0.5 | 18 | 0.3333 | 5 | 0.2 | 5 | 0.4 | 4 | 0.25 | 3 | 0 | 17 | 0.2353 |
| 11_115321 | 4 | 0.5 | 5 | 0.4 | 7 | 0.5714 | 6 | 0.6667 | 22 | 0.5455 | 9 | 0.7778 | 6 | 0.8333 | 6 | 1 | 5 | 1 | 26 | 0.8846 |
| 11_115321 | 3 | 0.6667 | 5 | 0.2 | 5 | 0.6 | 5 | 0.4 | 18 | 0.4444 | 6 | 0.5 | 6 | 0.8333 | 5 | 0.6 | 5 | 0.4 | 22 | 0.5909 |
| 11_115321 | 4 | 0.25 | 5 | 0 | 10 | 0.2 | 6 | 0.1667 | 25 | 0.16 | 7 | 0.2857 | 6 | 0.3333 | 5 | 0.2 | 3 | 0.3333 | 21 | 0.2857 |
| 11_115321 | 5 | 0.4 | 6 | 0.3333 | 10 | 0.4 | 6 | 0.3333 | 27 | 0.3704 | 9 | 0.6667 | 6 | 0.8333 | 6 | 0.3333 | 5 | 0.4 | 26 | 0.5769 |
| 11_115321 | 3 | 0.3333 | 3 | 0.3333 | 4 | 0.25 | 4 | 0 | 14 | 0.2143 | 5 | 0.4 | 5 | 0.6 | 3 | 0 | 5 | 0.2 | 18 | 0.3333 |
| 11_115321 | 4 | 1 | 4 | 0 | 9 | 0.4444 | 7 | 0.5714 | 24 | 0.5 | 8 | 0.5 | 3 | 1 | 6 | 0.8333 | 5 | 0.4 | 22 | 0.6364 |
| 11_115321 | 5 | 0.8 | 4 | 1 | 12 | 0.8333 | 8 | 0.75 | 29 | 0.8276 | 8 | 0.75 | 6 | 0.8333 | 7 | 0.7143 | 4 | 1 | 25 | 0.8 |
| 11_115321 | 5 | 0.6 | 4 | 0.25 | 12 | 0.3333 | 6 | 1 | 27 | 0.5185 | 11 | 0.2727 | 6 | 0.5 | 6 | 0.3333 | 5 | 0.2 | 28 | 0.3214 |
| 11_115321 | 3 | 0 | 6 | 0.1667 | 10 | 0.3 | 5 | 0.2 | 24 | 0.2083 | 11 | 0.0909 | 7 | 0.2857 | 4 | 0.25 | 4 | 0 | 26 | 0.1538 |
| 11_115321 | 3 | 1 | 6 | 0.6667 | 11 | 1 | 9 | 1 | 29 | 0.931 | 11 | 0.9091 | 6 | 1 | 7 | 0.8571 | 4 | 1 | 28 | 0.9286 |
| 11_115322 | 1 | 0 | 1 | 0 | 2 | 0.5 | 2 | 0.5 | 6 | 0.3333 | NA | NA | 3 | 1 | NA | NA | NA | NA | 3 | 1 |
| 11_115367 | 1 | 1 | NA | NA | NA | NA | NA | NA | 1 | 1 | NA | NA | 2 | 0.5 | NA | NA | NA | NA | 2 | 0.5 |

| | | | | | | | | | | | | | | | | | | | | |
|---|---|---|---|---|---|---|---|---|---|---|---|---|---|---|---|---|---|---|---|---|
| 11_1153798 | 1 | 1 | 1 | 1 | NA | NA | NA | NA | 2 | 1 | 1 | 1 | 2 | 1 | NA | NA | NA | NA | 3 | 1 |
| 11_1153799 | 1 | 1 | 2 | 1 | NA | NA | 1 | 1 | 4 | 1 | 1 | 1 | 2 | 1 | 1 | 1 | NA | NA | 4 | 1 |
| 11_1153801 | 1 | 1 | 2 | 1 | NA | NA | NA | NA | 3 | 1 | 1 | 1 | 2 | 1 | 1 | 1 | NA | NA | 4 | 1 |
| 11_1153801 | 1 | 0 | 2 | 0.5 | NA | NA | NA | NA | 3 | 0.3333 | 1 | 1 | 2 | 0 | NA | NA | NA | NA | 3 | 0.3333 |
| 11_1153801 | 1 | 0 | 2 | 0 | NA | NA | 1 | 1 | 4 | 0.25 | 1 | 0 | 2 | 0.5 | NA | NA | NA | NA | 3 | 0.3333 |
| 11_1153803 | 1 | 0 | 2 | 1 | NA | NA | 1 | 0 | 4 | 0.5 | 1 | 0 | 2 | 0.5 | 1 | 1 | NA | NA | 4 | 0.5 |
| 11_1153805 | 1 | 1 | 1 | 1 | NA | NA | 1 | 0 | 3 | 0.6667 | 1 | 1 | 2 | 1 | 1 | 1 | NA | NA | 4 | 1 |
| 11_1153805 | 1 | 1 | 2 | 0.5 | NA | NA | 2 | 0.5 | 5 | 0.6 | 1 | 1 | 2 | 1 | 1 | 1 | 1 | 0 | 5 | 0.8 |
| 11_1153806 | 1 | 0 | 2 | 1 | NA | NA | 1 | 0 | 4 | 0.5 | 1 | 0 | 2 | 0.5 | 1 | 1 | 1 | 1 | 5 | 0.6 |
| 11_1153807 | NA | NA | 2 | 0.5 | NA | NA | 1 | 0 | 3 | 0.3333 | 1 | 0 | 2 | 0.5 | 1 | 1 | 1 | 0 | 5 | 0.4 |
| 11_1153807 | NA | NA | 2 | 0.5 | NA | NA | 2 | 0.5 | 4 | 0.5 | 1 | 1 | 1 | 1 | 1 | 0 | NA | NA | 3 | 0.6667 |
| 11_1153808 | 1 | 1 | NA | NA | NA | NA | 2 | 1 | 3 | 1 | 1 | 1 | 2 | 1 | 1 | 1 | 1 | 0 | 5 | 0.8 |
| 11_1153808 | 1 | 0 | 1 | 1 | NA | NA | 2 | 0 | 4 | 0.25 | 1 | 1 | 2 | 0 | NA | NA | 1 | 0 | 4 | 0.25 |
| 11_1153811 | 1 | 0 | 2 | 0.5 | NA | NA | 2 | 0.5 | 5 | 0.4 | 1 | 1 | 2 | 1 | 1 | 1 | 1 | 1 | 5 | 1 |
| 11_1153813 | 5 | 0.6 | 8 | 0.625 | 8 | 0.625 | 5 | 0.8 | 26 | 0.6538 | 8 | 0.75 | 2 | 1 | 8 | 0.75 | 2 | 1 | 20 | 0.8 |
| 11_1153813 | 3 | 0 | 6 | 0.3333 | 7 | 0.2857 | 4 | 0 | 20 | 0.2 | 8 | 0.125 | 1 | 0 | 5 | 0 | 2 | 0.5 | 16 | 0.125 |
| 11_1153813 | 6 | 0.5 | 9 | 0.2222 | 12 | 0.3333 | 5 | 0.2 | 32 | 0.3125 | 9 | 0.2222 | 3 | 0 | 8 | 0.25 | 3 | 0.3333 | 23 | 0.2174 |
| 11_1153813 | 5 | 0.2 | 9 | 0.2222 | 12 | 0.1667 | 6 | 0 | 32 | 0.1562 | 8 | 0.125 | 3 | 0.3333 | 7 | 0 | 1 | 0 | 19 | 0.1053 |
| 11_1153814 | 6 | 0.1667 | 10 | 0.2 | 11 | 0 | 5 | 0 | 32 | 0.0938 | 8 | 0.25 | 3 | 0 | 8 | 0 | 3 | 0 | 22 | 0.0909 |
| 11_1153814 | 6 | 0.1667 | 3 | 0.3333 | 7 | 0.1429 | 6 | 0.1667 | 22 | 0.1818 | 9 | 0.2222 | NA | NA | 5 | 0.2 | 2 | 0 | 16 | 0.1875 |
| 11_1153814 | 6 | 0.3333 | 10 | 0.3 | 11 | 0.0909 | 6 | 0.1667 | 33 | 0.2121 | 8 | 0.25 | 3 | 0.6667 | 8 | 0.375 | 3 | 0 | 22 | 0.3182 |
| 11_1153817 | 5 | 0 | 8 | 0.25 | 11 | 0 | 5 | 0 | 29 | 0.069 | 7 | 0.1429 | 2 | 0.5 | 6 | 0 | 2 | 0 | 17 | 0.1176 |
| 11_1153882 | 11 | 0.8182 | 13 | 0.8462 | 9 | 0.4444 | 9 | 0.8889 | 42 | 0.7619 | 9 | 0.8889 | 13 | 0.7692 | 9 | 0.7778 | 10 | 0.9 | 41 | 0.8293 |
| 11_1153883 | 11 | 0.8182 | 13 | 0.6923 | 6 | 0.6667 | 7 | 0.7143 | 37 | 0.7297 | 9 | 0.8889 | 13 | 0.6923 | 7 | 0.5714 | 10 | 0.7 | 39 | 0.7179 |
| 11_1153883 | 7 | 0.1429 | 12 | 0.5 | 7 | 0.4286 | 8 | 0.75 | 34 | 0.4706 | 7 | 0.1429 | 9 | 0.3333 | 7 | 0.1429 | 8 | 0.125 | 31 | 0.1935 |
| 11_1153886 | 10 | 0.1 | 12 | 0.1667 | 7 | 0.4286 | 8 | 0.25 | 37 | 0.2162 | 9 | 0.2222 | 12 | 0.4167 | 6 | 0.3333 | 9 | 0.2222 | 36 | 0.3056 |
| 11_1153887 | 12 | 1 | 13 | 1 | 8 | 0.875 | 7 | 0.8571 | 40 | 0.95 | 9 | 0.7778 | 13 | 0.8462 | 8 | 0.75 | 10 | 0.8 | 40 | 0.8 |
| 11_1153887 | 11 | 0.2727 | 14 | 0.3571 | 8 | 0.125 | 8 | 0.375 | 41 | 0.2927 | 9 | 0.2222 | 14 | 0.2143 | 8 | 0.25 | 9 | 0.2222 | 40 | 0.225 |
| 11_1153889 | 16 | 0.125 | 16 | 0.0625 | 10 | 0 | 11 | 0.0909 | 53 | 0.0755 | 11 | 0.0909 | 15 | 0.0667 | 5 | 0.2 | 10 | 0.1 | 41 | 0.0976 |
| 11_1154140 | 47 | 0.0426 | 42 | 0.0238 | 35 | 0.1429 | 35 | 0 | 159 | 0.0503 | 31 | 0.0645 | 38 | 0.1053 | 37 | 0.0541 | 37 | 0.1351 | 143 | 0.0909 |
| 11_1154752 | 4 | 0.75 | 6 | 0.8333 | 2 | 1 | 5 | 0.6 | 17 | 0.7647 | 5 | 0.6 | 13 | 0.4615 | 11 | 0.7273 | 9 | 0.7778 | 38 | 0.6316 |
| 11_1154753 | 4 | 0 | 6 | 0 | 2 | 0 | 5 | 0.2 | 17 | 0.0588 | 5 | 0 | 14 | 0 | 13 | 0.1538 | 9 | 0.1111 | 41 | 0.0732 |
| 11_1154754 | 4 | 0.25 | 6 | 0 | 2 | 0 | 5 | 0 | 17 | 0.0588 | 3 | 0 | 14 | 0.0714 | 11 | 0 | 10 | 0.2 | 38 | 0.0789 |
| 11_1154755 | 4 | 0.25 | 6 | 0 | 2 | 0.5 | 3 | 0 | 15 | 0.1333 | 4 | 0 | 14 | 0.0714 | 12 | 0.0833 | 9 | 0.3333 | 39 | 0.1282 |
| 11_1154756 | 3 | 0 | 6 | 0.3333 | 1 | 0 | 3 | 0.6667 | 13 | 0.3077 | 3 | 0.6667 | 13 | 0.3077 | 10 | 0.4 | 8 | 0.375 | 34 | 0.3824 |
| 11_1154756 | 3 | 0 | 5 | 0 | 2 | 0 | 4 | 0.25 | 14 | 0.0714 | 3 | 0 | 13 | 0.0769 | 11 | 0.0909 | 7 | 0.1429 | 34 | 0.0882 |
| 11_1154757 | 4 | 0.5 | 6 | 0 | 2 | 0.5 | 2 | 0 | 14 | 0.2143 | 3 | 0.6667 | 12 | 0.25 | 10 | 0.2 | 10 | 0 | 35 | 0.2 |
| 11_1154759 | 3 | 0.3333 | 4 | 0.25 | 2 | 0 | 2 | 0 | 11 | 0.1818 | 2 | 0 | 11 | 0.1818 | 11 | 0 | 11 | 0 | 35 | 0.0571 |
| 11_1154759 | 4 | 0.25 | 5 | 0.2 | 2 | 0 | 1 | 0 | 12 | 0.1667 | 2 | 0.5 | 11 | 0.3636 | 10 | 0.2 | 9 | 0.1111 | 32 | 0.25 |
| 11_1154760 | 3 | 0.6667 | 6 | 0.3333 | 2 | 1 | 1 | 1 | 12 | 0.5833 | 2 | 0.5 | 12 | 0.5 | 10 | 0.3 | 8 | 0.625 | 32 | 0.4688 |
| 11_1154760 | 2 | 0.5 | 1 | 0 | NA | NA | 3 | 0 | 6 | 0.1667 | NA | NA | 6 | 0.1667 | 6 | 0 | 5 | 0 | 17 | 0.0588 |
| 11_1154761 | 4 | 0.25 | 4 | 0.5 | 2 | 0 | 3 | 0.3333 | 13 | 0.3077 | 2 | 0.5 | 10 | 0 | 11 | 0.4545 | 10 | 0.3 | 33 | 0.2727 |
| 11_1154761 | 4 | 0.75 | 3 | 0.6667 | 2 | 0.5 | 3 | 0.6667 | 12 | 0.6667 | 1 | 1 | 9 | 0.6667 | 7 | 1 | 8 | 0.75 | 25 | 0.8 |
| 11_1154762 | 3 | 0.6667 | 4 | 0.75 | 2 | 0.5 | 2 | 1 | 11 | 0.7273 | 2 | 0.5 | 9 | 0.6667 | 9 | 0.5556 | 9 | 0.5556 | 29 | 0.5862 |
| 11_1154763 | 3 | 0 | 4 | 0.25 | 2 | 0.5 | 3 | 0 | 12 | 0.1667 | 2 | 1 | 9 | 0.1111 | 11 | 0.1818 | 8 | 0.25 | 30 | 0.2333 |
| 11_1154763 | 3 | 0.3333 | 1 | 0 | 2 | 0.5 | 3 | 0.3333 | 9 | 0.3333 | 2 | 1 | 8 | 0.375 | 10 | 0.6 | 9 | 0.7778 | 29 | 0.6207 |
| 11_1154764 | 3 | 0.3333 | 4 | 0.25 | 2 | 0.5 | 2 | 1 | 11 | 0.4545 | 2 | 0.5 | 8 | 0.625 | 9 | 0.6667 | 8 | 0.75 | 27 | 0.6667 |
| 11_1154766 | 3 | 0.3333 | 4 | 0 | 1 | 1 | NA | NA | 8 | 0.25 | 2 | 0 | 6 | 0.1667 | 11 | 0.0909 | 7 | 0 | 26 | 0.0769 |
| 11_1154953 | 1 | 0 | NA | NA | NA | NA | 1 | 1 | 2 | 0.5 | NA | NA | 1 | 1 | NA | NA | NA | NA | 1 | 1 |
| 11_1154977 | 9 | 0.2222 | 9 | 0.1111 | 11 | 0.1818 | 9 | 0 | 38 | 0.1316 | 6 | 0.3333 | 10 | 0.1 | 17 | 0.1765 | 8 | 0.5 | 41 | 0.2439 |
| 11_1154977 | 8 | 0.125 | 10 | 0.2 | 12 | 0.1667 | 9 | 0.2222 | 39 | 0.1795 | 6 | 0 | 12 | 0.1667 | 17 | 0.3529 | 9 | 0.3333 | 44 | 0.25 |
| 11_1154978 | 10 | 0.9 | 11 | 0.9091 | 12 | 0.8333 | 9 | 0.8889 | 42 | 0.881 | 6 | 1 | 10 | 0.9 | 16 | 0.8125 | 9 | 0.8889 | 41 | 0.878 |
| 11_1154979 | 9 | 0.8889 | 10 | 0.9 | 9 | 0.7778 | 10 | 0.9 | 38 | 0.8684 | 5 | 1 | 11 | 0.7273 | 18 | 0.9444 | 7 | 0.7143 | 41 | 0.8537 |
| 11_1154980 | 9 | 0.2222 | 10 | 0.2 | 9 | 0.1111 | 8 | 0.125 | 36 | 0.1667 | 6 | 0 | 11 | 0 | 18 | 0.1111 | 9 | 0.1111 | 44 | 0.0682 |
| 11_1154980 | 9 | 0.1111 | 9 | 0 | 12 | 0.1667 | 7 | 0 | 37 | 0.0811 | 6 | 0 | 10 | 0 | 18 | 0.1111 | 10 | 0.2 | 44 | 0.0909 |
| 11_1154980 | 11 | 0.0909 | 10 | 0.5 | 10 | 0.3 | 10 | 0.3 | 41 | 0.2927 | 6 | 0.1667 | 12 | 0.25 | 18 | 0.0556 | 9 | 0.6667 | 45 | 0.2444 |
| 11_1154982 | 2 | 0 | 5 | 0.2 | 3 | 0 | 2 | 0 | 12 | 0.0833 | 3 | 0.3333 | 1 | 0 | 6 | 0 | 2 | 0 | 12 | 0.0833 |
| 11_1154982 | 2 | 1 | 6 | 0.5 | 3 | 1 | 3 | 0.3333 | 14 | 0.6429 | 2 | 1 | 1 | 1 | 6 | 0.8333 | 2 | 0.5 | 11 | 0.8182 |
| 11_1154983 | 2 | 0.5 | 6 | 0.1667 | 3 | 0 | 2 | 0 | 13 | 0.1538 | 2 | 0 | 1 | 0 | 5 | 0.4 | 1 | 0 | 9 | 0.2222 |
| 11_1154984 | NA | NA | 5 | 0.6 | 2 | 0 | 2 | 0.5 | 9 | 0.4444 | 3 | 0 | 1 | 1 | 3 | 0.3333 | 2 | 0.5 | 9 | 0.3333 |
| 11_1154984 | 1 | 0 | 6 | 0.3333 | 3 | 1 | 2 | 0 | 12 | 0.4167 | 3 | 0 | 1 | 1 | 4 | 0 | 1 | 1 | 9 | 0.2222 |
| 11_1154986 | NA | NA | 1 | 1 | NA | NA | NA | NA | 1 | 1 | NA | NA | NA | NA | NA | NA | 2 | 1 | 2 | 1 |
| 11_1154988 | 4 | 0.25 | 1 | 0 | 3 | 0.3333 | 2 | 1 | 10 | 0.4 | 1 | 0 | 1 | 1 | 1 | 1 | NA | NA | 3 | 0.6667 |
| 11_1154989 | 3 | 1 | 1 | 1 | 2 | 1 | 2 | 0.5 | 8 | 0.875 | 1 | 0 | 2 | 1 | NA | NA | NA | NA | 3 | 0.6667 |
| 11_1154990 | 5 | 0.2 | 3 | 0 | 3 | 0.3333 | 2 | 0 | 13 | 0.1538 | 1 | 0 | 2 | 0.5 | 2 | 1 | NA | NA | 5 | 0.6 |

| | | | | | | | | | | | | | | | | | | | | |
|---|---|---|---|---|---|---|---|---|---|---|---|---|---|---|---|---|---|---|---|---|
| 11_115499( | 5 | 0.2 | 3 | 0.6667 | 3 | 0 | 3 | 0.6667 | 14 | 0.3571 | 1 | 0 | 2 | 0 | 1 | 1 | NA | NA | 4 | 0.25 |
| 11_115499( | 5 | 1 | 2 | 1 | 3 | 0.6667 | 3 | 0.3333 | 13 | 0.7692 | NA | NA | 3 | 0.6667 | 1 | 1 | NA | NA | 4 | 0.75 |
| 11_115499' | 4 | 0.5 | 3 | 0.3333 | 3 | 0.6667 | 2 | 1 | 12 | 0.5833 | 1 | 1 | 3 | 0.6667 | 2 | 0.5 | NA | NA | 6 | 0.6667 |
| 11_115499: | 5 | 0.2 | 2 | 0 | 2 | 0 | 3 | 0 | 12 | 0.0833 | 1 | 0 | 3 | 0 | 2 | 0.5 | NA | NA | 6 | 0.1667 |
| 11_115505: | 5 | 0.4 | 2 | 0.5 | 1 | 0 | 6 | 0.3333 | 14 | 0.3571 | 3 | 0.3333 | 4 | 0.25 | 1 | 0 | 6 | 0.1667 | 14 | 0.2143 |
| 11_115505: | 4 | 0.25 | 4 | 0.25 | 1 | 0 | 8 | 0.5 | 17 | 0.3529 | 4 | 0.5 | 5 | 0.4 | 4 | 0 | 6 | 0.5 | 19 | 0.3684 |
| 11_115505: | 5 | 0 | 3 | 0 | 1 | 0 | 7 | 0.1429 | 16 | 0.0625 | 5 | 0.2 | 4 | 0 | 4 | 0 | 5 | 0 | 18 | 0.0556 |
| 11_115505( | 5 | 0.4 | 8 | 0.375 | 8 | 0.25 | 6 | 0.5 | 27 | 0.3704 | 9 | 0.4444 | 7 | 0.4286 | 7 | 0.2857 | 8 | 0.375 | 31 | 0.3871 |
| 11_115505! | 9 | 0.1111 | 7 | 0 | 6 | 0 | 9 | 0.2222 | 31 | 0.0968 | 8 | 0.25 | 7 | 0 | 7 | 0 | 7 | 0 | 29 | 0.069 |
| 11_115505! | 9 | 0.6667 | 6 | 0.6667 | 6 | 0.5 | 9 | 0.3333 | 30 | 0.5333 | 8 | 0.75 | 6 | 0.6667 | 7 | 0.8571 | 8 | 0.375 | 29 | 0.6552 |
| 11_115506' | 4 | 0.5 | 6 | 0.6667 | 5 | 0.4 | 1 | 1 | 16 | 0.5625 | 4 | 1 | 6 | 0.6667 | 3 | 0 | 5 | 1 | 18 | 0.7222 |
| 11_115506' | 8 | 0.375 | 7 | 0.4286 | 7 | 0.2857 | 9 | 0.6667 | 31 | 0.4516 | 6 | 0.3333 | 6 | 0.8333 | 6 | 0.5 | 8 | 0.5 | 26 | 0.5385 |
| 11_115535' | 24 | 0.6667 | 29 | 0.6207 | 18 | 0.6667 | 23 | 0.6087 | 94 | 0.6383 | 18 | 0.7778 | 20 | 0.55 | 23 | 0.5652 | 22 | 0.5909 | 83 | 0.6145 |
| 11_115535' | 20 | 0.55 | 30 | 0.3667 | 18 | 0.5556 | 20 | 0.2 | 88 | 0.4091 | 15 | 0.4667 | 21 | 0.381 | 24 | 0.2917 | 21 | 0.4286 | 81 | 0.3827 |
| 11_115536: | 18 | 0.2778 | 24 | 0.4583 | 13 | 0.3846 | 13 | 0.3846 | 68 | 0.3824 | 7 | 0.4286 | 13 | 0.6154 | 13 | 0.6923 | 22 | 0.3636 | 55 | 0.5091 |
| 11_115536‹ | 14 | 0.0714 | 22 | 0.0455 | 15 | 0.1333 | 15 | 0.2 | 66 | 0.1061 | 10 | 0.3 | 12 | 0.1667 | 16 | 0.25 | 16 | 0.3125 | 54 | 0.2593 |
| 11_115536! | 18 | 0.1667 | 25 | 0.2 | 17 | 0.2941 | 17 | 0.1765 | 77 | 0.2078 | 12 | 0.4167 | 17 | 0.1176 | 20 | 0.2 | 20 | 0.2 | 69 | 0.2174 |
| 11_115536 | 18 | 0.1111 | 21 | 0.0476 | 18 | 0.1111 | 15 | 0.1333 | 72 | 0.0972 | 13 | 0 | 14 | 0.0714 | 18 | 0.1667 | 16 | 0.125 | 61 | 0.0984 |
| 11_115548( | 4 | 0.75 | 3 | 0.3333 | 1 | 0 | 1 | 0 | 9 | 0.4444 | 3 | 1 | 4 | 0.25 | 5 | 0.2 | 3 | 1 | 15 | 0.5333 |
| 11_115548( | 4 | 0.25 | 4 | 0 | 1 | 0 | 1 | 0 | 10 | 0.1 | 5 | 0 | 4 | 0 | 5 | 0 | 3 | 0.3333 | 17 | 0.0588 |
| 11_115664! | 1 | 1 | NA | NA | NA | NA | NA | NA | 1 | 1 | NA | NA | NA | NA | 1 | 1 | NA | NA | 1 | 1 |
| 11_115664! | 1 | 1 | NA | NA | NA | NA | NA | NA | 1 | 1 | NA | NA | NA | NA | 1 | 1 | NA | NA | 1 | 1 |
| 11_115664( | 1 | 1 | NA | NA | NA | NA | NA | NA | 1 | 1 | NA | NA | NA | NA | 1 | 1 | NA | NA | 1 | 1 |
| 11_115683! | 4 | 0 | 5 | 0 | 1 | 1 | 4 | 0 | 14 | 0.0714 | 3 | 0 | 2 | 0.5 | 1 | 0 | 4 | 0 | 10 | 0.1 |
| 11_115689' | 6 | 0.1667 | 11 | 0.0909 | 2 | 0.5 | 6 | 0 | 25 | 0.12 | 7 | 0 | 4 | 0 | 5 | 0.2 | 7 | 0.2857 | 23 | 0.1304 |
| 11_115689 | 7 | 0.1429 | 11 | 0.0909 | 2 | 0.5 | 9 | 0.3333 | 29 | 0.2069 | 10 | 0.2 | 6 | 0 | 5 | 0.6 | 7 | 0.1429 | 28 | 0.2143 |
| 11_115690: | 2 | 0 | 4 | 0.25 | 2 | 0 | NA | NA | 8 | 0.125 | 2 | 0 | 1 | 0 | 1 | 1 | 1 | 0 | 5 | 0.2 |
| 11_115690! | NA | NA | NA | NA | 3 | 0.3333 | NA | NA | 3 | 0.3333 | NA | NA | 1 | 0 | 1 | 1 | NA | NA | 2 | 0.5 |
| 11_115690! | NA | NA | 3 | 0.3333 | 2 | 0 | NA | NA | 5 | 0.2 | 1 | 0 | 1 | 0 | 2 | 0.5 | NA | NA | 4 | 0.25 |
| 11_115690( | 1 | 1 | 3 | 0.3333 | 2 | 0.5 | NA | NA | 6 | 0.5 | 3 | 0.3333 | 2 | 1 | NA | NA | 1 | 0 | 6 | 0.5 |
| 11_115690( | 1 | 1 | 4 | 0.5 | 2 | 1 | NA | NA | 7 | 0.7143 | 1 | 1 | 2 | 1 | 1 | 1 | 1 | 1 | 5 | 1 |
| 11_115690 | 2 | 0.5 | 3 | 0 | 3 | 0.3333 | NA | NA | 8 | 0.25 | 1 | 1 | 2 | 0.5 | 2 | 0 | 1 | 0 | 6 | 0.3333 |
| 11_115690 | 1 | 0 | 3 | 0.6667 | 3 | 1 | NA | NA | 7 | 0.7143 | 1 | 1 | 2 | 1 | 2 | 0.5 | 1 | 0 | 6 | 0.6667 |
| 11_115692! | NA | NA | 2 | 0.5 | 1 | 0 | NA | NA | 3 | 0.3333 | 1 | 1 | 1 | 0 | NA | NA | 1 | 0 | 3 | 0.3333 |
| 11_115692! | 1 | 0 | 2 | 1 | 3 | 0.3333 | NA | NA | 6 | 0.5 | 1 | 0 | 2 | 1 | NA | NA | 1 | 0 | 4 | 0.5 |
| 11_115702( | NA | NA | 1 | 1 | NA | NA | NA | NA | 1 | 1 | NA | NA | 1 | 1 | NA | NA | NA | NA | 1 | 1 |
| 11_115702! | NA | NA | 2 | 0.5 | NA | NA | NA | NA | 2 | 0.5 | NA | NA | 1 | 1 | NA | NA | NA | NA | 1 | 1 |
| 11_115711: | 3 | 0.3333 | NA | NA | 3 | 0.6667 | NA | NA | 6 | 0.5 | NA | NA | NA | NA | NA | NA | 1 | 1 | 1 | 1 |
| 11_115711‹ | 4 | 0.25 | NA | NA | 2 | 0.5 | NA | NA | 6 | 0.3333 | NA | NA | NA | NA | NA | NA | 1 | 1 | 1 | 1 |
| 11_115713! | 4 | 0.25 | NA | NA | 1 | 1 | NA | NA | 5 | 0.4 | 1 | 0 | NA | NA | NA | NA | 1 | 1 | 2 | 0.5 |
| 11_115713! | 1 | 0 | NA | NA | 2 | 1 | NA | NA | 3 | 0.6667 | 1 | 1 | NA | NA | NA | NA | 1 | 0 | 2 | 0.5 |
| 11_115713( | 5 | 0.6 | NA | NA | 2 | 0.5 | NA | NA | 7 | 0.5714 | NA | NA | NA | NA | NA | NA | 1 | 1 | 1 | 1 |
| 11_115713 | 4 | 1 | NA | NA | 3 | 0.6667 | NA | NA | 7 | 0.8571 | 1 | 0 | NA | NA | NA | NA | 1 | 1 | 2 | 0.5 |
| 11_115713 | 4 | 0.5 | NA | NA | 2 | 1 | NA | NA | 6 | 0.6667 | 1 | 1 | NA | NA | NA | NA | 1 | 1 | 2 | 1 |
| 11_115785( | 2 | 0 | 2 | 0.5 | 3 | 1 | NA | NA | 7 | 0.5714 | NA | NA | NA | NA | 6 | 0.6667 | 1 | 1 | 7 | 0.7143 |
| 11_115785' | 2 | 0.5 | 2 | 0 | 3 | 0 | NA | NA | 7 | 0.1429 | NA | NA | NA | NA | 6 | 0.1667 | 1 | 1 | 7 | 0.2857 |
| 11_115785: | 2 | 0.5 | 2 | 0 | 3 | 0 | NA | NA | 7 | 0.1429 | NA | NA | NA | NA | 5 | 0.2 | 1 | 0 | 6 | 0.1667 |
| 11_115818 | 2 | 0.5 | 1 | 1 | 3 | 0.6667 | NA | NA | 6 | 0.6667 | NA | NA | NA | NA | 2 | 0.5 | NA | NA | 2 | 0.5 |
| 11_115818 | 1 | 0 | 1 | 1 | 1 | 0 | NA | NA | 3 | 0.3333 | NA | NA | NA | NA | 2 | 0.5 | NA | NA | 2 | 0.5 |
| 11_115818 | 1 | 0 | 1 | 0 | 3 | 0.3333 | NA | NA | 5 | 0.2 | NA | NA | NA | NA | 2 | 0.5 | NA | NA | 2 | 0.5 |
| 11_115855( | 9 | 0.1111 | 6 | 0.1667 | 4 | 0 | 3 | 0.3333 | 22 | 0.1364 | 7 | 0 | 5 | 0.2 | 3 | 0 | 3 | 0 | 18 | 0.0556 |
| 11_115855: | 11 | 0.5455 | 5 | 0.4 | 7 | 0.5714 | 3 | 0.3333 | 26 | 0.5 | 9 | 0.6667 | 5 | 0.4 | 3 | 0.3333 | 3 | 1 | 20 | 0.6 |
| 11_115855: | 11 | 0.3636 | 5 | 0.6 | 7 | 0.4286 | 3 | 0.3333 | 26 | 0.4231 | 9 | 0.5556 | 3 | 0.6667 | 3 | 0.6667 | 3 | 0.3333 | 18 | 0.5556 |
| 11_115855: | 11 | 0.9091 | 6 | 0.8333 | 6 | 1 | 2 | 1 | 25 | 0.92 | 9 | 0.7778 | 5 | 1 | 3 | 0.6667 | 3 | 1 | 20 | 0.85 |
| 11_115855: | 11 | 0.3636 | 6 | 0.1667 | 9 | 0.3333 | 2 | 0 | 28 | 0.2857 | 8 | 0.125 | 5 | 0.2 | 3 | 0.6667 | 3 | 0 | 19 | 0.2105 |
| 11_115855: | 8 | 0.75 | 4 | 0.75 | 7 | 0.7143 | NA | NA | 19 | 0.7368 | 5 | 1 | 4 | 0.75 | 1 | 1 | 3 | 1 | 13 | 0.9231 |
| 11_115855‹ | 9 | 0.1111 | 2 | 0 | 6 | 0.1667 | 1 | 0 | 18 | 0.1111 | 5 | 0 | 3 | 0.3333 | 3 | 0.3333 | 2 | 0 | 13 | 0.1538 |
| 11_115855‹ | 5 | 1 | 2 | 1 | 4 | 1 | 1 | 1 | 12 | 1 | 4 | 1 | NA | NA | 2 | 1 | 3 | 1 | 9 | 1 |
| 11_115855! | 3 | 0.6667 | 2 | 1 | 2 | 0.5 | 1 | 1 | 8 | 0.75 | 3 | 1 | 1 | 1 | 2 | 1 | 2 | 1 | 8 | 1 |
| 11_115855! | 4 | 0 | 1 | 0 | 4 | 0.25 | NA | NA | 9 | 0.1111 | 3 | 0 | NA | NA | 1 | 1 | 3 | 0.3333 | 7 | 0.2857 |
| 11_115856' | 7 | 0.1429 | 3 | 0 | 6 | 0 | 1 | 0 | 17 | 0.0588 | 3 | 0.3333 | 2 | 0 | 2 | 0 | 4 | 0 | 11 | 0.0909 |
| 11_115856' | 7 | 0.1429 | 2 | 0.5 | 6 | 0.1667 | 1 | 0 | 16 | 0.1875 | 4 | 0.25 | 3 | 0 | 1 | 0 | 4 | 0.25 | 12 | 0.1667 |
| 11_115856: | 5 | 1 | 3 | 0 | 5 | 0.6 | 1 | 1 | 14 | 0.6429 | 2 | 0.5 | 4 | 0.5 | 2 | 1 | 4 | 0.5 | 12 | 0.5833 |
| 11_115856 | 10 | 0.7 | 1 | 0 | 6 | 0.3333 | 2 | 1 | 19 | 0.5789 | 1 | 0 | 4 | 0.5 | NA | NA | 6 | 1 | 11 | 0.7273 |
| 11_115856 | 10 | 0.4 | 3 | 0.3333 | 7 | 0.2857 | 4 | 1 | 24 | 0.4583 | 2 | 0 | 4 | 0.5 | 2 | 0 | 7 | 0.4286 | 15 | 0.3333 |

| | | | | | | | | | | | | | | | | | | | | |
|---|---|---|---|---|---|---|---|---|---|---|---|---|---|---|---|---|---|---|---|---|
| 11_1158568 | 9 | 0.2222 | 3 | 0.3333 | 7 | 0 | 4 | 0 | 23 | 0.1304 | 2 | 0 | 4 | 0.5 | 2 | 0 | 7 | 0.2857 | 15 | 0.2667 |
| 11_1158570 | 9 | 0.7778 | 2 | 1 | 6 | 0.8333 | 4 | 0.75 | 21 | 0.8095 | 2 | 1 | 3 | 0.6667 | 3 | 0.3333 | 6 | 0.5 | 14 | 0.5714 |
| 11_1158574 | 4 | 0 | 2 | 0.5 | 4 | 0 | 3 | 0 | 13 | 0.0769 | 2 | 0 | 2 | 0.5 | 3 | 0 | 5 | 0 | 12 | 0.0833 |
| 11_1159140 | 44 | 0.0227 | 44 | 0.0682 | 29 | 0.1379 | 38 | 0.0263 | 155 | 0.0581 | 30 | 0.0667 | 42 | 0.0476 | 41 | 0.0732 | 40 | 0.05 | 153 | 0.0588 |
| 11_1159508 | NA | NA | 2 | 0 | 1 | 1 | 1 | 1 | 4 | 0.5 | NA | NA | 2 | 0.5 | 1 | 0 | 2 | 0.5 | 5 | 0.4 |
| 11_1159696 | 76 | 0.0658 | 60 | 0.1333 | 59 | 0.1017 | 71 | 0.1268 | 266 | 0.1053 | 49 | 0.1224 | 50 | 0.12 | 62 | 0.1129 | 55 | 0.1091 | 216 | 0.1157 |
| 11_1159698 | 70 | 0.1857 | 60 | 0.0667 | 58 | 0.1552 | 64 | 0.125 | 252 | 0.1349 | 46 | 0.1087 | 50 | 0.14 | 66 | 0.0303 | 53 | 0.1132 | 215 | 0.093 |
| 11_1159700 | 59 | 0.2881 | 47 | 0.2553 | 51 | 0.3137 | 60 | 0.2833 | 217 | 0.2857 | 43 | 0.2093 | 39 | 0.2564 | 54 | 0.2593 | 45 | 0.2889 | 181 | 0.2541 |
| 11_1159966 | 2 | 1 | NA | NA | NA | NA | NA | NA | 2 | 1 | 1 | 1 | NA | NA | NA | NA | 1 | 1 | 2 | 1 |
| 11_1159967 | 2 | 1 | NA | NA | NA | NA | NA | NA | 2 | 1 | 2 | 1 | 1 | 1 | NA | NA | 1 | 1 | 4 | 1 |
| 11_1159969 | 2 | 0.5 | NA | NA | NA | NA | NA | NA | 2 | 0.5 | 2 | 0.5 | 1 | 0 | NA | NA | 1 | 0 | 4 | 0.25 |
| 11_1160004 | 1 | 1 | NA | NA | NA | NA | NA | NA | 1 | 1 | NA | NA | 1 | 0 | NA | NA | 1 | 1 | 2 | 0.5 |
| 11_1160228 | 8 | 0.25 | 11 | 0.4545 | 8 | 0.625 | 6 | 0.1667 | 33 | 0.3939 | 6 | 0.3333 | 10 | 0.5 | 12 | 0.4167 | 4 | 0.75 | 32 | 0.4688 |
| 11_1160228 | 7 | 0 | 8 | 0.125 | 7 | 0.1429 | 5 | 0 | 27 | 0.0741 | 5 | 0 | 11 | 0.4545 | 9 | 0.2222 | 4 | 0.25 | 29 | 0.2759 |
| 11_1160334 | 9 | 0.2222 | 7 | 0 | 5 | 0.2 | 5 | 0 | 26 | 0.1154 | 3 | 0 | 7 | 0 | 4 | 0 | 5 | 0.2 | 19 | 0.0526 |
| 11_1160340 | 8 | 0.375 | 6 | 0.5 | 5 | 0.4 | 3 | 0.3333 | 22 | 0.4091 | 3 | 1 | 4 | 0.75 | 2 | 0.5 | 5 | 0.6 | 14 | 0.7143 |
| 11_1160628 | 18 | 0.2778 | 25 | 0.16 | 22 | 0.0909 | 24 | 0.0833 | 89 | 0.1461 | 20 | 0.1 | 25 | 0.08 | 23 | 0.3043 | 16 | 0.125 | 84 | 0.1548 |
| 11_1160630 | 22 | 0.1818 | 24 | 0.0833 | 21 | 0.1905 | 28 | 0.1429 | 95 | 0.1474 | 20 | 0 | 21 | 0.1429 | 22 | 0.2727 | 19 | 0.3158 | 82 | 0.1829 |
| 11_1160634 | 20 | 0.15 | 18 | 0 | 16 | 0 | 23 | 0.0435 | 77 | 0.0519 | 16 | 0.0625 | 16 | 0.0625 | 16 | 0 | 17 | 0.1765 | 65 | 0.0769 |
| 11_1160635 | 20 | 0.05 | 18 | 0.0556 | 12 | 0.1667 | 20 | 0.15 | 70 | 0.1 | 22 | 0.0909 | 18 | 0.0556 | 18 | 0.1667 | 20 | 0.1 | 78 | 0.1026 |
| 11_1160636 | 18 | 0.1111 | 17 | 0.0588 | 14 | 0.2143 | 20 | 0 | 69 | 0.087 | 20 | 0.15 | 15 | 0.0667 | 16 | 0.125 | 15 | 0.2667 | 66 | 0.1515 |
| 11_1160639 | 10 | 0.5 | 13 | 0.5385 | 7 | 0.1429 | 16 | 0.5 | 46 | 0.4565 | 10 | 0.3 | 12 | 0.4167 | 10 | 0.4 | 15 | 0.4667 | 47 | 0.4043 |
| 11_1160640 | 11 | 0.1818 | 17 | 0.4118 | 12 | 0.4167 | 14 | 0.2857 | 54 | 0.3333 | 15 | 0.4667 | 18 | 0.5556 | 15 | 0.3333 | 17 | 0.4706 | 65 | 0.4615 |
| 11_1160641 | 9 | 0.2222 | 11 | 0.3636 | 11 | 0.1818 | 15 | 0.0667 | 46 | 0.1957 | 13 | 0.0769 | 18 | 0.0556 | 15 | 0.3333 | 17 | 0.1176 | 63 | 0.1429 |
| 11_1160642 | 13 | 0.0769 | 16 | 0.0625 | 14 | 0.3571 | 15 | 0.1333 | 58 | 0.1552 | 16 | 0.0625 | 20 | 0.1 | 17 | 0.1765 | 18 | 0 | 71 | 0.0845 |
| 11_1160642 | 12 | 0.4167 | 14 | 0.2857 | 14 | 0.2857 | 14 | 0.1429 | 54 | 0.2778 | 14 | 0.2143 | 18 | 0.3333 | 17 | 0.1765 | 18 | 0.2222 | 67 | 0.2388 |
| 11_1160643 | 11 | 0.1818 | 15 | 0.0667 | 14 | 0.1429 | 15 | 0.0667 | 55 | 0.1091 | 14 | 0.0714 | 18 | 0 | 13 | 0.1538 | 16 | 0.1875 | 61 | 0.0984 |
| 11_1161058 | 1 | 1 | 8 | 0.75 | 6 | 0.6667 | 4 | 1 | 19 | 0.7895 | 3 | 0.6667 | 4 | 0.75 | 7 | 0.5714 | 3 | 0.6667 | 17 | 0.6471 |
| 11_1161118 | 1 | 1 | 2 | 0.5 | NA | NA | 1 | 1 | 4 | 0.75 | 1 | 1 | 4 | 0.75 | NA | NA | 1 | 0 | 6 | 0.6667 |
| 11_1161119 | 2 | 1 | 2 | 0.5 | NA | NA | 1 | 0 | 5 | 0.6 | 1 | 0 | 5 | 0.2 | NA | NA | 1 | 0 | 7 | 0.1429 |
| 11_1161120 | 2 | 1 | 1 | 1 | NA | NA | 1 | 1 | 4 | 1 | 1 | 1 | 5 | 0.8 | NA | NA | 1 | 1 | 7 | 0.8571 |
| 11_1161121 | 2 | 0.5 | 1 | 0 | NA | NA | NA | NA | 3 | 0.3333 | 1 | 0 | 3 | 0.3333 | NA | NA | NA | NA | 4 | 0.25 |
| 11_1161136 | NA | NA | NA | NA | 1 | 1 | NA | NA | 1 | 1 | NA | NA | NA | NA | 1 | 1 | NA | NA | 1 | 1 |
| 11_1161146 | NA | NA | NA | NA | 1 | 1 | NA | NA | 1 | 1 | 1 | 1 | NA | NA | 1 | 1 | NA | NA | 2 | 1 |
| 11_1161150 | NA | NA | NA | NA | 1 | 1 | NA | NA | 1 | 1 | 1 | 1 | NA | NA | NA | NA | NA | NA | 1 | 1 |
| 11_1161150 | NA | NA | NA | NA | 1 | 1 | NA | NA | 1 | 1 | 1 | 1 | NA | NA | NA | NA | NA | NA | 1 | 1 |
| 11_1161154 | NA | NA | NA | NA | 1 | 1 | NA | NA | 1 | 1 | 1 | 1 | NA | NA | NA | NA | NA | NA | 1 | 1 |
| 11_1161154 | NA | NA | NA | NA | 1 | 1 | NA | NA | 1 | 1 | 1 | 1 | NA | NA | 1 | 0 | NA | NA | 2 | 0.5 |
| 11_1161156 | NA | NA | NA | NA | 1 | 1 | NA | NA | 1 | 1 | 1 | 1 | NA | NA | 1 | 1 | NA | NA | 2 | 1 |
| 11_1161157 | NA | NA | NA | NA | 1 | 1 | NA | NA | 1 | 1 | 1 | 1 | NA | NA | 1 | 1 | NA | NA | 2 | 1 |
| 11_1161244 | NA | NA | NA | NA | 2 | 0.5 | NA | NA | 2 | 0.5 | 1 | 0 | NA | NA | 2 | 1 | NA | NA | 3 | 0.6667 |
| 11_1161245 | 1 | 1 | NA | NA | 1 | 1 | NA | NA | 2 | 1 | 1 | 0 | NA | NA | 2 | 1 | NA | NA | 3 | 0.6667 |
| 11_1161249 | 1 | 1 | NA | NA | 1 | 1 | NA | NA | 2 | 1 | NA | NA | NA | NA | 2 | 0.5 | NA | NA | 2 | 0.5 |
| 11_1161258 | 1 | 0 | NA | NA | 1 | 1 | NA | NA | 2 | 0.5 | NA | NA | NA | NA | 2 | 0.5 | NA | NA | 2 | 0.5 |
| 11_1161259 | 1 | 0 | NA | NA | 2 | 0.5 | NA | NA | 3 | 0.3333 | NA | NA | NA | NA | 2 | 0.5 | NA | NA | 2 | 0.5 |
| 11_1161260 | 1 | 0 | NA | NA | 2 | 0.5 | NA | NA | 3 | 0.3333 | NA | NA | NA | NA | 2 | 1 | NA | NA | 2 | 1 |
| 11_1161278 | 1 | 0 | NA | NA | 2 | 0.5 | NA | NA | 3 | 0.3333 | NA | NA | NA | NA | 1 | 1 | NA | NA | 1 | 1 |
| 11_1161291 | NA | NA | NA | NA | 2 | 1 | 1 | 1 | 3 | 1 | NA | NA | NA | NA | 1 | 1 | NA | NA | 1 | 1 |
| 11_1161361 | 12 | 0 | 11 | 0 | 13 | 0.2308 | 13 | 0.1538 | 49 | 0.102 | 14 | 0.1429 | 12 | 0.0833 | 8 | 0.25 | 14 | 0 | 48 | 0.1042 |
| 11_1161363 | 11 | 0.2727 | 10 | 0.3 | 11 | 0 | 12 | 0.25 | 44 | 0.2045 | 13 | 0.1538 | 12 | 0.1667 | 7 | 0.4286 | 9 | 0.1111 | 41 | 0.1951 |
| 11_1161363 | 10 | 0 | 13 | 0.0769 | 12 | 0.0833 | 13 | 0.0769 | 48 | 0.0625 | 13 | 0.0769 | 11 | 0.0909 | 7 | 0 | 12 | 0.1667 | 43 | 0.093 |
| 11_1161647 | 6 | 0.1667 | 4 | 0 | 6 | 0 | 7 | 0.1429 | 23 | 0.087 | 3 | 0 | 6 | 0.1667 | 2 | 0 | 1 | 0 | 12 | 0.0833 |
| 11_1161795 | 1 | 1 | 2 | 0 | NA | NA | 1 | 0 | 4 | 0.25 | NA | NA | 1 | 1 | 1 | 1 | 2 | 1 | 4 | 1 |
| 11_1161798 | 1 | 0 | 2 | 0 | NA | NA | 1 | 1 | 4 | 0.25 | NA | NA | 1 | 0 | 1 | 0 | 2 | 0.5 | 4 | 0.25 |
| 11_1161799 | 7 | 0.7143 | 3 | 0.6667 | 5 | 0.8 | 5 | 0.4 | 20 | 0.65 | 11 | 0.7273 | 5 | 0.6 | 4 | 0.5 | 11 | 0.8182 | 31 | 0.7097 |
| 11_1162221 | 3 | 0.3333 | NA | NA | 1 | 0 | NA | NA | 4 | 0.25 | NA | NA | 1 | 1 | NA | NA | NA | NA | 1 | 1 |
| 11_1162224 | 2 | 0.5 | NA | NA | 1 | 0 | NA | NA | 3 | 0.3333 | NA | NA | 1 | 1 | NA | NA | NA | NA | 1 | 1 |
| 11_1162225 | 2 | 0.5 | NA | NA | 1 | 0 | NA | NA | 3 | 0.3333 | NA | NA | 1 | 1 | NA | NA | NA | NA | 1 | 1 |
| 11_1162225 | 1 | 1 | NA | NA | 1 | 0 | NA | NA | 2 | 0.5 | NA | NA | 1 | 1 | NA | NA | NA | NA | 1 | 1 |
| 11_1162226 | 2 | 1 | NA | NA | 1 | 1 | NA | NA | 3 | 1 | NA | NA | 1 | 1 | NA | NA | NA | NA | 1 | 1 |
| 11_1162250 | 1 | 1 | NA | NA | 1 | 1 | NA | NA | 2 | 1 | NA | NA | 1 | 1 | NA | NA | NA | NA | 1 | 1 |
| 11_1162254 | 2 | 1 | NA | NA | 1 | 1 | NA | NA | 3 | 1 | NA | NA | 1 | 1 | NA | NA | NA | NA | 1 | 1 |
| 11_1162257 | 1 | 1 | NA | NA | NA | NA | NA | NA | 1 | 1 | NA | NA | 1 | 1 | NA | NA | NA | NA | 1 | 1 |
| 11_1162265 | 1 | 1 | NA | NA | NA | NA | NA | NA | 1 | 1 | NA | NA | 1 | 1 | NA | NA | NA | NA | 1 | 1 |
| 11_1162274 | 25 | 0.2 | 41 | 0.1463 | 15 | 0.1333 | 16 | 0.1875 | 97 | 0.1649 | 13 | 0.3077 | 25 | 0.12 | 17 | 0.2353 | 19 | 0 | 74 | 0.1486 |

| | | | | | | | | | | | | | | | | | | | | |
|---|---|---|---|---|---|---|---|---|---|---|---|---|---|---|---|---|---|---|---|---|
| 11_1162283 | 18 | 0.5556 | 27 | 0.4815 | 14 | 0.4286 | 16 | 0.3125 | 75 | 0.4533 | 10 | 0.4 | 18 | 0.3889 | 12 | 0.4167 | 15 | 0.6667 | 55 | 0.4727 |
| 11_1162287 | 17 | 0 | 27 | 0.0741 | 11 | 0.0909 | 16 | 0.0625 | 71 | 0.0563 | 11 | 0.1818 | 17 | 0.1176 | 9 | 0.1111 | 11 | 0.0909 | 48 | 0.125 |
| 11_1162288 | 12 | 0.25 | 29 | 0.1379 | 11 | 0.2727 | 14 | 0.2857 | 66 | 0.2121 | 8 | 0.375 | 18 | 0.2778 | 10 | 0.1 | 11 | 0.2727 | 47 | 0.2553 |
| 11_1162290 | 12 | 1 | 20 | 1 | 8 | 0.75 | 12 | 1 | 52 | 0.9615 | 8 | 0.875 | 13 | 0.9231 | 8 | 0.875 | 7 | 1 | 36 | 0.9167 |
| 11_1162290 | 11 | 0 | 23 | 0.1304 | 11 | 0.0909 | 15 | 0 | 60 | 0.0667 | 9 | 0.1111 | 15 | 0.3333 | 9 | 0 | 7 | 0.1429 | 40 | 0.175 |
| 11_1162291 | 7 | 0.7143 | 19 | 0.6842 | 9 | 0.8889 | 12 | 0.75 | 47 | 0.7447 | 9 | 0.7778 | 15 | 0.7333 | 8 | 0.75 | 6 | 0.8333 | 38 | 0.7632 |
| 11_1162291 | 7 | 1 | 18 | 0.9444 | 7 | 0.8571 | 11 | 0.9091 | 43 | 0.9302 | 8 | 1 | 14 | 1 | 8 | 1 | 9 | 1 | 39 | 1 |
| 11_1162291 | 12 | 0.0833 | 20 | 0.2 | 10 | 0.2 | 12 | 0.0833 | 54 | 0.1481 | 7 | 0.1429 | 14 | 0 | 8 | 0.125 | 6 | 0 | 35 | 0.0571 |
| 11_1162291 | 14 | 0.0714 | 24 | 0.0833 | 13 | 0.0769 | 14 | 0.2857 | 65 | 0.1231 | 11 | 0.1818 | 15 | 0.2 | 7 | 0 | 9 | 0.3333 | 42 | 0.1905 |
| 11_1162291 | 14 | 0.6429 | 18 | 0.8889 | 11 | 0.8182 | 12 | 0.8333 | 55 | 0.8 | 6 | 0.6667 | 12 | 0.75 | 5 | 1 | 9 | 0.8889 | 32 | 0.8125 |
| 11_1162291 | 18 | 1 | 25 | 1 | 14 | 1 | 20 | 1 | 77 | 1 | 11 | 1 | 17 | 0.8824 | 11 | 1 | 12 | 1 | 51 | 0.9608 |
| 11_1162292 | 15 | 0.5333 | 24 | 0.5833 | 13 | 0.6923 | 19 | 0.5263 | 71 | 0.5775 | 9 | 0.5556 | 18 | 0.6667 | 8 | 0.75 | 9 | 0.6667 | 44 | 0.6591 |
| 11_1162292 | 16 | 0.125 | 24 | 0.0417 | 13 | 0.1538 | 18 | 0.0556 | 71 | 0.0845 | 10 | 0.1 | 18 | 0.1111 | 10 | 0.2 | 10 | 0.1 | 48 | 0.125 |
| 11_1162293 | 16 | 0.625 | 23 | 0.6522 | 15 | 0.5333 | 15 | 0.4667 | 69 | 0.5797 | 9 | 0.3333 | 19 | 0.5263 | 11 | 0.3636 | 10 | 0.5 | 49 | 0.449 |
| 11_1162294 | 15 | 0.9333 | 17 | 0.8824 | 11 | 0.9091 | 11 | 1 | 54 | 0.9259 | 9 | 1 | 15 | 1 | 6 | 1 | 7 | 0.8571 | 37 | 0.973 |
| 11_1162294 | 16 | 0.25 | 16 | 0.125 | 14 | 0.2143 | 18 | 0.3889 | 64 | 0.25 | 8 | 0.375 | 14 | 0.4286 | 7 | 0.2857 | 9 | 0.4444 | 38 | 0.3947 |
| 11_1162294 | 18 | 0.1111 | 22 | 0.2273 | 11 | 0.1818 | 18 | 0.2222 | 69 | 0.1884 | 10 | 0.3 | 11 | 0.2727 | 11 | 0.2727 | 9 | 0.1111 | 41 | 0.2439 |
| 11_1162295 | 15 | 0.1333 | 15 | 0.0667 | 12 | 0 | 14 | 0.0714 | 56 | 0.0714 | 8 | 0.25 | 9 | 0.3333 | 7 | 0 | 4 | 0 | 28 | 0.1786 |
| 11_1162349 | 1 | 1 | NA | NA | NA | NA | NA | NA | 1 | 1 | NA | NA | 1 | 1 | NA | NA | 1 | 1 | 2 | 1 |
| 11_1162350 | 15 | 0.1333 | 17 | 0 | 10 | 0.1 | 13 | 0 | 55 | 0.0545 | 4 | 0 | 14 | 0.0714 | 10 | 0.1 | 6 | 0.1667 | 34 | 0.0882 |
| 11_1162807 | NA | NA | NA | NA | 1 | 1 | NA | NA | 1 | 1 | NA | NA | NA | NA | 1 | 1 | 1 | 1 | 2 | 1 |
| 11_1163291 | 6 | 0.5 | 12 | 0.75 | 5 | 0.8 | 5 | 0.2 | 28 | 0.6071 | 4 | 0.25 | 8 | 0.625 | 7 | 0.2857 | 5 | 0.8 | 24 | 0.5 |
| 11_1163291 | 7 | 0 | 9 | 0.2222 | 5 | 0.4 | 5 | 0.2 | 26 | 0.1923 | 3 | 0.3333 | 8 | 0.125 | 6 | 0.3333 | 7 | 0.1429 | 24 | 0.2083 |
| 11_1163296 | 6 | 1 | 9 | 1 | 5 | 0.8 | 5 | 0.8 | 25 | 0.92 | 6 | 0.8333 | 6 | 1 | 5 | 0.6 | 3 | 1 | 20 | 0.85 |
| 11_1163296 | 5 | 0 | 10 | 0.4 | 5 | 0.2 | 5 | 0.2 | 25 | 0.24 | 7 | 0.2857 | 9 | 0.5556 | 7 | 0 | 3 | 0.3333 | 26 | 0.3077 |
| 11_1163297 | 6 | 0.6667 | 10 | 0.3 | 7 | 0.1429 | 5 | 0 | 28 | 0.2857 | 7 | 0 | 10 | 0.2 | 7 | 0 | 4 | 0 | 28 | 0.0714 |
| 11_1163298 | 3 | 0 | 9 | 0.1111 | 2 | 0 | 4 | 0.25 | 18 | 0.1111 | 7 | 0.1429 | 5 | 0.4 | 4 | 0 | 4 | 0 | 20 | 0.15 |
| 11_1163618 | 6 | 0 | 6 | 0.3333 | 2 | 0.5 | 6 | 0.3333 | 20 | 0.25 | 11 | 0.2727 | 4 | 0.75 | 12 | 0.25 | 7 | 0 | 34 | 0.2647 |
| 11_1163621 | 5 | 0.4 | 8 | 0.25 | 2 | 0 | 5 | 0.2 | 20 | 0.25 | 10 | 0.2 | 4 | 0 | 10 | 0.4 | 5 | 0.2 | 29 | 0.2414 |
| 11_1163621 | 6 | 0.1667 | 6 | 0 | 1 | 0 | 5 | 0.2 | 18 | 0.1111 | 7 | 0 | 4 | 0.25 | 9 | 0.1111 | 4 | 0.25 | 24 | 0.125 |
| 11_1163623 | 7 | 0.1429 | 6 | 0.3333 | NA | NA | 7 | 0.1429 | 20 | 0.2 | 9 | 0.2222 | 6 | 0 | 10 | 0.6 | 5 | 0.4 | 30 | 0.3333 |
| 11_1164265 | 1 | 1 | NA | NA | NA | NA | NA | NA | 1 | 1 | NA | NA | 2 | 0.5 | NA | NA | NA | NA | 2 | 0.5 |
| 11_1164267 | 1 | 1 | NA | NA | NA | NA | NA | NA | 1 | 1 | NA | NA | 1 | 1 | NA | NA | NA | NA | 1 | 1 |
| 11_1164268 | 1 | 1 | NA | NA | NA | NA | NA | NA | 1 | 1 | 1 | 1 | 2 | 0.5 | NA | NA | NA | NA | 3 | 0.6667 |
| 11_1164269 | 1 | 1 | NA | NA | NA | NA | NA | NA | 1 | 1 | NA | NA | 2 | 1 | NA | NA | NA | NA | 2 | 1 |
| 11_1164271 | 1 | 1 | NA | NA | NA | NA | NA | NA | 1 | 1 | NA | NA | 2 | 0.5 | NA | NA | NA | NA | 2 | 0.5 |
| 11_1164274 | 1 | 1 | NA | NA | NA | NA | NA | NA | 1 | 1 | 1 | 0 | 2 | 0.5 | NA | NA | NA | NA | 3 | 0.3333 |
| 11_1164286 | 1 | 1 | 11 | 0 | 5 | 0 | 3 | 0 | 20 | 0.05 | 4 | 0 | 6 | 0 | 2 | 0 | 5 | 0.2 | 17 | 0.0588 |
| 11_1164287 | 2 | 0 | 9 | 0.2222 | 2 | 0 | 6 | 0.3333 | 19 | 0.2105 | 5 | 0 | 4 | 0.25 | 2 | 0 | 3 | 0 | 14 | 0.0714 |
| 11_1164288 | 2 | 0.5 | 11 | 0.1818 | 7 | 0.2857 | 6 | 0.3333 | 26 | 0.2692 | 6 | 0.1667 | 7 | 0 | 5 | 0.2 | 8 | 0.25 | 26 | 0.1538 |
| 11_1164288 | 2 | 0 | 13 | 0.2308 | 7 | 0 | 9 | 0.3333 | 31 | 0.1935 | 5 | 0.2 | 8 | 0.25 | 3 | 0 | 7 | 0.1429 | 23 | 0.1739 |
| 11_1164290 | 2 | 0.5 | 9 | 0.6667 | 8 | 0.75 | 5 | 1 | 24 | 0.75 | 4 | 0.5 | 3 | 1 | 4 | 0.5 | 6 | 0.8333 | 17 | 0.7059 |
| 11_1164292 | 2 | 0.5 | 11 | 0.4545 | 6 | 0.3333 | 7 | 0.4286 | 26 | 0.4231 | 6 | 0.5 | 6 | 0.1667 | 5 | 0.2 | 8 | 0.5 | 25 | 0.36 |
| 11_1164293 | 2 | 0 | 8 | 0.25 | 7 | 0.2857 | 7 | 0.1429 | 24 | 0.2083 | 6 | 0 | 6 | 0 | 4 | 0.25 | 7 | 0.1429 | 23 | 0.087 |
| 11_1164293 | 2 | 0.5 | 8 | 0.5 | 5 | 0.6 | 6 | 0.8333 | 21 | 0.619 | 6 | 0.5 | 6 | 0.6667 | 3 | 1 | 7 | 0.7143 | 22 | 0.6818 |
| 11_1164298 | 2 | 1 | 8 | 0.875 | 7 | 0.8571 | 6 | 0.8333 | 23 | 0.8696 | 5 | 1 | 6 | 0.8333 | 4 | 0.75 | 9 | 1 | 24 | 0.9167 |
| 11_1164300 | NA | NA | 10 | 0.7 | 6 | 0.8333 | 5 | 0.8 | 21 | 0.7619 | 6 | 0.6667 | 6 | 0.8333 | 4 | 0.75 | 6 | 1 | 22 | 0.8182 |
| 11_1164301 | 1 | 0 | 10 | 0.2 | 7 | 0.1429 | 5 | 0 | 23 | 0.1304 | 6 | 0.1667 | 6 | 0.1667 | 4 | 0.25 | 8 | 0.25 | 24 | 0.2083 |
| 11_1164301 | 1 | 0 | 6 | 0.5 | 6 | 0.8333 | 3 | 0.6667 | 16 | 0.625 | 4 | 0.5 | 5 | 0.6 | 4 | 0.75 | 6 | 0.5 | 19 | 0.5789 |
| 11_1164302 | 1 | 1 | 8 | 0.375 | 7 | 0.5714 | 5 | 0.2 | 21 | 0.4286 | 5 | 0.4 | 6 | 0.6667 | 4 | 0.5 | 7 | 0.8571 | 22 | 0.6364 |
| 11_1164302 | 1 | 0 | 8 | 1 | 5 | 0.8 | 4 | 0.75 | 18 | 0.8333 | 4 | 1 | 4 | 1 | 4 | 1 | 4 | 1 | 16 | 1 |
| 11_1164302 | 1 | 1 | 6 | 0.6667 | 5 | 0.6 | 6 | 0.8333 | 18 | 0.7222 | 3 | 0.6667 | 4 | 1 | 3 | 0.6667 | 1 | 1 | 11 | 0.8182 |
| 11_1164304 | 1 | 0 | 7 | 0.2857 | 4 | 0.25 | 5 | 0.2 | 17 | 0.2353 | 5 | 0.2 | 5 | 0 | 4 | 0.25 | 6 | 0.1667 | 20 | 0.15 |
| 11_1164305 | 1 | 0 | 7 | 0.1429 | 5 | 0 | 4 | 0 | 17 | 0.0588 | 5 | 0 | 5 | 0 | 4 | 0.25 | 6 | 0 | 20 | 0.05 |
| 11_1164324 | NA | NA | 4 | 0 | 3 | 0.3333 | 2 | 0 | 9 | 0.1111 | 3 | 0 | 3 | 0.6667 | 3 | 0 | 2 | 0.5 | 11 | 0.2727 |
| 11_1165370 | 11 | 0.0909 | 9 | 0.2222 | 12 | 0.0833 | 6 | 0 | 38 | 0.1053 | 8 | 0.125 | 11 | 0 | 7 | 0.2857 | 10 | 0.1 | 36 | 0.1111 |
| 11_1165374 | 10 | 0 | 7 | 0.1429 | 12 | 0.3333 | 6 | 0 | 35 | 0.1429 | 5 | 0.2 | 10 | 0.2 | 4 | 0.25 | 8 | 0 | 27 | 0.1481 |
| 11_1165375 | 8 | 0.5 | 6 | 0.1667 | 7 | 0.1429 | 5 | 0.4 | 26 | 0.3077 | 4 | 0.25 | 7 | 0 | 2 | 0 | 5 | 0 | 18 | 0.0556 |
| 11_1165675 | 1 | 1 | NA | NA | 1 | 1 | NA | NA | 2 | 1 | 1 | 0 | 1 | 1 | 2 | 0 | NA | NA | 4 | 0.25 |
| 11_1165681 | 1 | 1 | NA | NA | NA | NA | NA | NA | 1 | 1 | 1 | 0 | 1 | 1 | NA | NA | NA | NA | 2 | 0.5 |
| 11_1166951 | 1 | 1 | NA | NA | 4 | 0.75 | 1 | 1 | 6 | 0.8333 | NA | NA | 1 | 1 | 2 | 0.5 | 1 | 1 | 4 | 0.75 |
| 11_1166952 | 2 | 0.5 | 1 | 0 | 3 | 1 | 1 | 1 | 7 | 0.7143 | 1 | 1 | 1 | 1 | 2 | 1 | 1 | 0 | 5 | 0.8 |
| 11_1166952 | 2 | 1 | 1 | 1 | 4 | 1 | 1 | 1 | 8 | 1 | 2 | 1 | 2 | 1 | 1 | 1 | 2 | 0.5 | 7 | 0.8571 |
| 11_1166953 | 1 | 1 | NA | NA | 3 | 1 | 1 | 1 | 5 | 1 | 2 | 1 | 2 | 0 | 2 | 1 | 2 | 1 | 8 | 0.75 |
| 11_1167283 | 4 | 0.5 | 2 | 1 | 2 | 0.5 | 4 | 0.75 | 12 | 0.6667 | 4 | 1 | 6 | 0.8333 | 5 | 0.8 | 5 | 0.8 | 20 | 0.85 |

| | | | | | | | | | | | | | | | | | | | | |
|---|---|---|---|---|---|---|---|---|---|---|---|---|---|---|---|---|---|---|---|---|
| 11_116728 | 4 | 0 | 2 | 0 | 1 | 0 | 4 | 0.25 | 11 | 0.0909 | 3 | 0 | 6 | 0.1667 | 5 | 0 | 3 | 0 | 17 | 0.0588 |
| 11_116728 | 4 | 0.5 | 2 | 0.5 | 2 | 0 | 4 | 0.25 | 12 | 0.3333 | 4 | 0 | 6 | 0.3333 | 4 | 0 | 3 | 0 | 17 | 0.1176 |
| 11_116729 | 5 | 0.4 | 2 | 0.5 | 1 | 0 | 4 | 0.75 | 12 | 0.5 | 4 | 0.5 | 6 | 1 | 3 | 0.3333 | 2 | 0 | 15 | 0.6 |
| 11_116729 | NA | NA | NA | NA | NA | NA | 2 | 0.5 | 2 | 0.5 | NA | NA | 1 | 1 | NA | NA | NA | NA | 1 | 1 |
| 11_116731 | 4 | 0.5 | 2 | 0.5 | 2 | 0 | 2 | 0 | 10 | 0.3 | 4 | 0 | 4 | 0.25 | 4 | 0.25 | 3 | 0 | 15 | 0.1333 |
| 11_116731 | 4 | 0.5 | 2 | 0.5 | 2 | 1 | 3 | 0.3333 | 11 | 0.5455 | 4 | 0.5 | 4 | 0.5 | 4 | 0 | 3 | 0 | 14 | 0.2857 |
| 11_116732 | 4 | 0.25 | 2 | 0 | 1 | 0 | 3 | 0 | 10 | 0.1 | 4 | 0 | 4 | 0.25 | 4 | 0 | 2 | 0 | 14 | 0.0714 |
| 11_116732 | 4 | 0.25 | 2 | 0 | 1 | 0 | 3 | 0 | 10 | 0.1 | 3 | 0 | 4 | 0.25 | 3 | 0 | 2 | 0 | 12 | 0.0833 |
| 11_116733 | 4 | 0.5 | 3 | 0.6667 | 2 | 0.5 | 2 | 1 | 11 | 0.6364 | 2 | 0.5 | 3 | 1 | 2 | 0.5 | 1 | 0 | 8 | 0.625 |
| 11_116734 | 4 | 0 | 1 | 0 | 7 | 0.1429 | 1 | 0 | 13 | 0.0769 | 2 | 0 | 1 | 0 | 1 | 0 | 3 | 0.3333 | 7 | 0.1429 |
| 11_116739 | 6 | 0.5 | 5 | 0.6 | 9 | 0.1111 | 4 | 0.25 | 24 | 0.3333 | 3 | 0.3333 | 5 | 0.2 | 1 | 0 | 5 | 0.6 | 14 | 0.3571 |
| 11_116739 | 7 | 0.2857 | 4 | 0.25 | 11 | 0.0909 | 5 | 0.2 | 27 | 0.1852 | 4 | 0.75 | 6 | 0.3333 | 2 | 1 | 5 | 0.8 | 17 | 0.6471 |
| 11_116740 | 8 | 0.75 | 6 | 0.8333 | 10 | 0.8 | 4 | 0.75 | 28 | 0.7857 | 6 | 0.8333 | 5 | 1 | 1 | 1 | 5 | 1 | 17 | 0.9412 |
| 11_116740 | 6 | 0.8333 | 7 | 0.7143 | 12 | 0.8333 | 5 | 0.6 | 30 | 0.7667 | 6 | 0.8333 | 5 | 0.6 | 2 | 1 | 5 | 0.6 | 18 | 0.7222 |
| 11_116740 | 28 | 0.4286 | 19 | 0.4211 | 24 | 0.25 | 28 | 0.25 | 99 | 0.3333 | 24 | 0.2083 | 29 | 0.3793 | 30 | 0.3 | 22 | 0.4091 | 105 | 0.3238 |
| 11_116741 | 27 | 0.1852 | 19 | 0 | 22 | 0.0909 | 29 | 0 | 97 | 0.0722 | 20 | 0.1 | 24 | 0.1667 | 26 | 0.0769 | 20 | 0 | 90 | 0.0889 |
| 11_116741 | 25 | 0.2 | 20 | 0.3 | 20 | 0.25 | 28 | 0.25 | 93 | 0.2473 | 18 | 0.2222 | 25 | 0.44 | 25 | 0.32 | 19 | 0.3158 | 87 | 0.3333 |
| 11_116741 | 27 | 0.0741 | 21 | 0.0952 | 22 | 0.0909 | 30 | 0.0333 | 100 | 0.07 | 19 | 0.0526 | 25 | 0.12 | 24 | 0.25 | 19 | 0.1053 | 87 | 0.1379 |
| 11_116741 | 22 | 0.0455 | 17 | 0.1176 | 14 | 0.0714 | 30 | 0.0333 | 83 | 0.0602 | 19 | 0 | 22 | 0.1364 | 22 | 0.0455 | 22 | 0.0455 | 85 | 0.0588 |
| 11_116741 | 21 | 0.0952 | 17 | 0.1765 | 22 | 0.1364 | 23 | 0.1739 | 83 | 0.1446 | 20 | 0.3 | 21 | 0.2857 | 22 | 0.2727 | 23 | 0.3043 | 86 | 0.2907 |
| 11_116742 | 7 | 0.1429 | 4 | 0 | 4 | 0.5 | 3 | 0.6667 | 18 | 0.2778 | 4 | 0 | 8 | 0.625 | 3 | 0.3333 | 2 | 0 | 17 | 0.3529 |
| 11_116743 | 15 | 0.2667 | 13 | 0.2308 | 12 | 0.25 | 17 | 0.2941 | 57 | 0.2632 | 18 | 0.1667 | 16 | 0.25 | 17 | 0.2353 | 19 | 0.2105 | 70 | 0.2143 |
| 11_116744 | 2 | 0 | 1 | 1 | 3 | 0 | 3 | 0.3333 | 9 | 0.2222 | 3 | 0 | 1 | 1 | 4 | 0 | 2 | 0 | 10 | 0.1 |
| 11_116744 | 2 | 0.5 | 1 | 0 | 3 | 0.3333 | 4 | 0 | 10 | 0.2 | 4 | 0.25 | 4 | 0 | 4 | 0 | 2 | 0 | 14 | 0.0714 |
| 11_116764 | 4 | 0.25 | NA | NA | 4 | 0 | 5 | 0.6 | 13 | 0.3077 | 6 | 0.1667 | 6 | 0.1667 | 4 | 0 | 3 | 0.3333 | 19 | 0.1579 |
| 11_116764 | 3 | 0.6667 | NA | NA | 4 | 0.25 | 3 | 0.6667 | 10 | 0.5 | 6 | 0.3333 | 6 | 0.5 | 4 | 0.25 | 1 | 1 | 17 | 0.4118 |
| 11_116765 | 4 | 0.75 | NA | NA | 5 | 0.8 | 5 | 1 | 14 | 0.8571 | 7 | 0.2857 | 7 | 0.5714 | 4 | 0.25 | 3 | 0 | 21 | 0.3333 |
| 11_116765 | 2 | 0 | NA | NA | 5 | 0.4 | 5 | 0.8 | 12 | 0.5 | 6 | 0.1667 | 6 | 0.8333 | 4 | 0.5 | 4 | 0.75 | 20 | 0.55 |
| 11_116765 | 3 | 1 | NA | NA | 5 | 1 | 5 | 1 | 13 | 1 | 5 | 0.4 | 6 | 0.8333 | 4 | 0.25 | 5 | 0.8 | 20 | 0.6 |
| 11_116765 | 3 | 0.3333 | NA | NA | 3 | 0.3333 | 5 | 0.4 | 11 | 0.3636 | 7 | 0 | 3 | 0.3333 | 1 | 1 | 3 | 0 | 14 | 0.1429 |
| 11_116765 | 1 | 1 | NA | NA | NA | NA | NA | NA | 1 | 1 | 1 | 1 | NA | NA | NA | NA | 1 | 1 | 2 | 1 |
| 11_116869 | 5 | 0.2 | 4 | 0.25 | 10 | 0.1 | 12 | 0.1667 | 31 | 0.1613 | 1 | 0 | 4 | 0 | 5 | 0 | 5 | 0.2 | 15 | 0.0667 |
| 11_116869 | 3 | 0 | 3 | 0 | 8 | 0.5 | 10 | 0.3 | 24 | 0.2917 | 1 | 0 | 6 | 0 | 3 | 0 | 5 | 0.2 | 15 | 0.0667 |
| 11_116876 | 5 | 0.4 | 4 | 0.25 | 6 | 0.5 | 11 | 0.3636 | 26 | 0.3846 | NA | NA | 7 | 0.7143 | 3 | 0.3333 | 5 | 0.6 | 15 | 0.6 |
| 11_116876 | 5 | 0 | 4 | 0 | 10 | 0.1 | 13 | 0.0769 | 32 | 0.0625 | 1 | 0 | 6 | 0.1667 | 5 | 0 | 5 | 0 | 17 | 0.0588 |
| 11_116876 | 6 | 0.6667 | 5 | 0.6 | 9 | 0.4444 | 14 | 0.8571 | 34 | 0.6765 | 2 | 0.5 | 6 | 0.5 | 5 | 1 | 9 | 0.7778 | 22 | 0.7273 |
| 11_116876 | 6 | 0.3333 | 3 | 0.3333 | 9 | 0.4444 | 12 | 0.4167 | 30 | 0.4 | 2 | 0.5 | 7 | 0.1429 | 5 | 0.8 | 8 | 0.125 | 22 | 0.3182 |
| 11_116876 | 6 | 0.3333 | 5 | 0.6 | 9 | 0.2222 | 12 | 0.5833 | 32 | 0.4375 | 2 | 0 | 8 | 0.375 | 5 | 0.6 | 11 | 0.3636 | 26 | 0.3846 |
| 11_116877 | 7 | 0.2857 | 5 | 0 | 6 | 0.1667 | 11 | 0.1818 | 29 | 0.1724 | 2 | 0 | 4 | 0 | 5 | 0.2 | 8 | 0.125 | 19 | 0.1053 |
| 11_116877 | 6 | 0.1667 | 6 | 0 | 8 | 0 | 16 | 0.125 | 36 | 0.0833 | 4 | 0 | 9 | 0.1111 | 4 | 0 | 9 | 0.1111 | 26 | 0.0769 |
| 11_116877 | 6 | 0.5 | 5 | 0.2 | 8 | 0.125 | 17 | 0.4706 | 36 | 0.3611 | 4 | 0.75 | 8 | 0 | 5 | 0.6 | 11 | 0.5455 | 28 | 0.4286 |
| 11_116877 | 7 | 0.1429 | 6 | 0.3333 | 10 | 0 | 17 | 0 | 40 | 0.075 | 4 | 0 | 9 | 0 | 6 | 0.1667 | 11 | 0.1818 | 30 | 0.1 |
| 11_116877 | 9 | 0.1111 | 6 | 0.1667 | 10 | 0.1 | 17 | 0.1765 | 42 | 0.1429 | 3 | 0 | 8 | 0.25 | 5 | 0.2 | 11 | 0.1818 | 27 | 0.1852 |
| 11_116877 | 9 | 0.7778 | 5 | 0.8 | 9 | 0.3333 | 15 | 0.2667 | 38 | 0.4737 | 4 | 0.25 | 8 | 0.375 | 5 | 0.6 | 10 | 0.3 | 27 | 0.3704 |
| 11_116877 | 6 | 0.1667 | 4 | 0.75 | 8 | 0.25 | 17 | 0.3529 | 35 | 0.3429 | 2 | 0.5 | 8 | 0.125 | 6 | 0 | 11 | 0.1818 | 27 | 0.1481 |
| 11_116877 | 9 | 0 | 6 | 0.1667 | 10 | 0.1 | 17 | 0.1176 | 42 | 0.0952 | 4 | 0 | 8 | 0.125 | 6 | 0 | 11 | 0.1818 | 29 | 0.1034 |
| 11_116877 | 8 | 0.625 | 3 | 0.6667 | 9 | 0.4444 | 15 | 0.7333 | 35 | 0.6286 | 3 | 0.6667 | 9 | 0.7778 | 6 | 0.6667 | 11 | 0.7273 | 29 | 0.7241 |
| 11_117047 | 13 | 0.3077 | 7 | 0.1429 | 2 | 0 | 5 | 0.2 | 27 | 0.2222 | 9 | 0 | 10 | 0.3 | 12 | 0.1667 | 9 | 0.1111 | 40 | 0.15 |
| 11_117048 | 11 | 0.8182 | 9 | 0.4444 | 4 | 0.75 | 7 | 0.8571 | 31 | 0.7097 | 8 | 0.625 | 9 | 0.7778 | 11 | 0.8182 | 11 | 0.7273 | 39 | 0.7436 |
| 11_117048 | 13 | 0.0769 | 8 | 0 | 4 | 0 | 7 | 0.1429 | 32 | 0.0625 | 8 | 0.25 | 11 | 0.3636 | 11 | 0 | 10 | 0.2 | 40 | 0.2 |
| 11_117048 | 13 | 0.3846 | 9 | 0.3333 | 4 | 0.25 | 7 | 0 | 33 | 0.2727 | 9 | 0.2222 | 12 | 0.5 | 13 | 0.3846 | 10 | 0.4 | 44 | 0.3864 |
| 11_117048 | 13 | 0.6923 | 10 | 0.5 | 4 | 0.75 | 6 | 0.5 | 33 | 0.6061 | 10 | 0.7 | 12 | 0.4167 | 12 | 0.5833 | 8 | 0.625 | 42 | 0.5714 |
| 11_117048 | 11 | 0.8182 | 10 | 0.8 | 3 | 1 | 7 | 0.8571 | 31 | 0.8387 | 10 | 0.9 | 11 | 0.9091 | 14 | 0.6429 | 10 | 0.7 | 45 | 0.7778 |
| 11_117251 | 2 | 0.5 | 5 | 0.4 | 3 | 0.3333 | 1 | 0 | 11 | 0.3636 | 6 | 0 | 3 | 0.3333 | NA | NA | 1 | 0 | 10 | 0.1 |
| 11_117252 | 2 | 0.5 | 10 | 0.5 | 3 | 0 | 1 | 1 | 16 | 0.4375 | 8 | 0.125 | 5 | 0 | 1 | 0 | 2 | 0 | 16 | 0.0625 |
| 11_117252 | 2 | 0.5 | 10 | 0 | 4 | 0.5 | NA | NA | 16 | 0.1875 | 10 | 0.1 | 5 | 0.2 | NA | NA | 2 | 0 | 17 | 0.1176 |
| 11_117252 | 2 | 1 | 6 | 0.1667 | 4 | 0 | NA | NA | 12 | 0.25 | 8 | 0.25 | 3 | 0 | 1 | 0 | 2 | 0 | 14 | 0.1429 |
| 11_117252 | 2 | 1 | 7 | 0.8571 | 4 | 1 | NA | NA | 13 | 0.9231 | 9 | 0.8889 | 5 | 0.8 | 1 | 1 | 4 | 1 | 19 | 0.8947 |
| 11_117252 | 2 | 0.5 | 11 | 0.0909 | 4 | 0 | NA | NA | 17 | 0.1176 | 10 | 0.1 | 6 | 0 | 1 | 0 | 4 | 0.25 | 21 | 0.0952 |
| 11_117252 | 2 | 0 | 11 | 0 | 4 | 0 | 2 | 0.5 | 19 | 0.0526 | 10 | 0 | 6 | 0.1667 | 1 | 1 | 4 | 0.25 | 21 | 0.1429 |
| 11_117252 | 1 | 0 | 8 | 0.25 | 3 | 0.3333 | 2 | 0 | 14 | 0.2143 | 9 | 0.1111 | 7 | 0.1429 | 1 | 0 | 4 | 0.5 | 21 | 0.1905 |
| 11_117503 | 2 | 1 | 1 | 1 | NA | NA | NA | NA | 3 | 1 | NA | NA | 1 | 1 | NA | NA | NA | NA | 1 | 1 |
| 11_117612 | 1 | 1 | NA | NA | NA | NA | NA | NA | 1 | 1 | 1 | 1 | NA | NA | NA | NA | NA | NA | 1 | 1 |
| 11_117612 | 1 | 1 | NA | NA | NA | NA | NA | NA | 1 | 1 | 1 | 1 | NA | NA | NA | NA | NA | NA | 1 | 1 |
| 11_117612 | 1 | 1 | NA | NA | NA | NA | NA | NA | 1 | 1 | 1 | 1 | NA | NA | NA | NA | NA | NA | 1 | 1 |

| | | | | | | | | | | | | | | | | | | | | |
|---|---|---|---|---|---|---|---|---|---|---|---|---|---|---|---|---|---|---|---|---|
| 11_1176128 | 1 | 1 | NA | NA | NA | NA | NA | NA | 1 | 1 | 1 | 1 | NA | NA | NA | NA | NA | NA | 1 | 1 |
| 11_1176134 | 2 | 1 | NA | NA | NA | NA | NA | NA | 2 | 1 | 1 | 1 | NA | NA | NA | NA | NA | NA | 1 | 1 |
| 11_1176134 | 2 | 0.5 | NA | NA | NA | NA | NA | NA | 2 | 0.5 | 1 | 1 | NA | NA | NA | NA | NA | NA | 1 | 1 |
| 11_1176136 | 2 | 1 | NA | NA | NA | NA | NA | NA | 2 | 1 | 1 | 1 | NA | NA | NA | NA | NA | NA | 1 | 1 |
| 11_1176136 | 2 | 1 | NA | NA | NA | NA | NA | NA | 2 | 1 | 1 | 1 | NA | NA | NA | NA | NA | NA | 1 | 1 |
| 11_1176139 | 2 | 1 | NA | NA | NA | NA | NA | NA | 2 | 1 | 1 | 1 | NA | NA | NA | NA | NA | NA | 1 | 1 |
| 11_1176142 | 2 | 0 | NA | NA | 1 | 1 | NA | NA | 3 | 0.3333 | 1 | 1 | NA | NA | NA | NA | NA | NA | 1 | 1 |
| 11_1176516 | 2 | 1 | NA | NA | 3 | 0 | 3 | 0.3333 | 8 | 0.375 | NA | NA | 1 | 0 | 3 | 0 | 4 | 0.25 | 8 | 0.125 |
| 11_1176517 | 3 | 0.6667 | NA | NA | 4 | 0.5 | 3 | 0.3333 | 10 | 0.5 | 1 | 0 | 1 | 1 | 3 | 0 | 5 | 0.6 | 10 | 0.4 |
| 11_1176517 | 2 | 0 | NA | NA | 3 | 0.6667 | 2 | 0 | 7 | 0.2857 | 1 | 1 | 1 | 0 | 3 | 0 | 3 | 0.6667 | 8 | 0.375 |
| 11_1176518 | 3 | 1 | NA | NA | 3 | 1 | 2 | 1 | 8 | 1 | 1 | 1 | 1 | 1 | 2 | 1 | 5 | 1 | 9 | 1 |
| 11_1176518 | 3 | 0 | NA | NA | 3 | 0.6667 | 3 | 0 | 9 | 0.2222 | 1 | 1 | 1 | 0 | 2 | 0 | 3 | 0.3333 | 7 | 0.2857 |
| 11_1176518 | 3 | 0.3333 | NA | NA | 4 | 0.5 | 2 | 0 | 9 | 0.3333 | 1 | 0 | 1 | 1 | 4 | 0.25 | 5 | 0.6 | 11 | 0.4545 |
| 11_1176522 | 2 | 0 | 1 | 0 | 6 | 0 | 3 | 0.3333 | 12 | 0.0833 | 3 | 0.3333 | 1 | 0 | 4 | 0.25 | 5 | 0 | 13 | 0.1538 |
| 11_1176527 | 4 | 0.5 | 1 | 1 | 7 | 0.7143 | 6 | 0.5 | 18 | 0.6111 | 1 | 0 | 1 | 1 | 5 | 0.8 | 7 | 0.8571 | 14 | 0.7857 |
| 11_1176528 | 4 | 0.75 | 2 | 1 | 7 | 0.8571 | 6 | 0.8333 | 19 | 0.8421 | 3 | 1 | 1 | 0 | 5 | 0.8 | 6 | 0.6667 | 15 | 0.7333 |
| 11_1176528 | 3 | 0.3333 | 2 | 0 | 6 | 0 | 4 | 0.25 | 15 | 0.1333 | 2 | 0 | 1 | 1 | 3 | 0.3333 | 8 | 0.375 | 14 | 0.3571 |
| 11_1176528 | 3 | 0.6667 | 2 | 1 | 5 | 1 | 3 | 1 | 13 | 0.9231 | 3 | 1 | NA | NA | 3 | 1 | 6 | 1 | 12 | 1 |
| 11_1176529 | 3 | 1 | 2 | 1 | 5 | 1 | 2 | 0.5 | 12 | 0.9167 | 3 | 1 | 2 | 0.5 | 3 | 0.6667 | 7 | 1 | 15 | 0.8667 |
| 11_1176529 | 2 | 0.5 | 3 | 0.3333 | 6 | 0.1667 | 3 | 0 | 14 | 0.2143 | 3 | 0.3333 | 2 | 0 | 4 | 0 | 6 | 0.1667 | 15 | 0.1333 |
| 11_1176530 | 5 | 0.6 | 3 | 1 | 7 | 0.8571 | 2 | 1 | 17 | 0.8235 | 1 | 0 | 2 | 0.5 | 5 | 0.6 | 6 | 0.3333 | 14 | 0.4286 |
| 11_1176530 | 5 | 0.2 | 4 | 0.5 | 7 | 0.1429 | 4 | 0 | 20 | 0.2 | 3 | 0 | 1 | 0 | 5 | 0.4 | 8 | 0.5 | 17 | 0.3529 |
| 11_1176531 | 5 | 0.6 | 3 | 1 | 7 | 0.7143 | 4 | 0.75 | 19 | 0.7368 | 3 | 0.6667 | 3 | 0.3333 | 5 | 0.8 | 6 | 0.8333 | 17 | 0.7059 |
| 11_1176532 | 5 | 0.4 | 3 | 0.6667 | 7 | 0.4286 | 4 | 0.5 | 19 | 0.4737 | 3 | 0.3333 | 3 | 0 | 5 | 0.6 | 8 | 0.375 | 19 | 0.3684 |
| 11_1176532 | 4 | 0.5 | 3 | 1 | 8 | 0.75 | 4 | 1 | 19 | 0.7895 | 3 | 1 | 3 | 0.6667 | 5 | 1 | 6 | 0.8333 | 17 | 0.8824 |
| 11_1176535 | 5 | 0.8 | 4 | 1 | 9 | 1 | 3 | 1 | 21 | 0.9524 | 2 | 0.5 | 3 | 0.3333 | 3 | 1 | 8 | 1 | 16 | 0.8125 |
| 11_1176536 | 4 | 0 | 3 | 0.3333 | 7 | 0.2857 | 2 | 0.5 | 16 | 0.25 | NA | NA | 2 | 0 | 5 | 0.4 | 5 | 0.2 | 12 | 0.25 |
| 11_1176538 | 5 | 0.2 | 3 | 0 | 10 | 0.6 | 4 | 0 | 22 | 0.3182 | 2 | 0.5 | 3 | 0.3333 | 5 | 0.6 | 7 | 0.1429 | 17 | 0.3529 |
| 11_1176538 | 5 | 0.4 | 3 | 0.6667 | 8 | 0.625 | 4 | 0.5 | 20 | 0.55 | 2 | 1 | 3 | 0.3333 | 4 | 1 | 7 | 0 | 16 | 0.4375 |
| 11_1176540 | 6 | 0 | 3 | 0 | 10 | 0 | 3 | 0.6667 | 22 | 0.0909 | 3 | 0 | 3 | 0 | 5 | 0.2 | 7 | 0 | 18 | 0.0556 |
| 11_1176541 | 5 | 0 | 1 | 0 | 7 | 0 | 4 | 0.25 | 17 | 0.0588 | 3 | 0 | 2 | 0 | 5 | 0 | 8 | 0.125 | 18 | 0.0556 |
| 11_1176542 | 4 | 0 | 3 | 0 | 10 | 0.1 | 4 | 0.25 | 21 | 0.0952 | 3 | 0 | 4 | 0.25 | 6 | 0 | 8 | 0.125 | 21 | 0.0952 |
| 11_1177043 | NA | NA | 2 | 1 | 1 | 1 | 1 | 1 | 4 | 1 | 1 | 1 | 2 | 1 | 2 | 0.5 | 1 | 1 | 6 | 0.8333 |
| 11_1177232 | NA | NA | 1 | 0 | 1 | 0 | 1 | 1 | 3 | 0.3333 | 1 | 1 | NA | NA | NA | NA | 1 | 1 | 2 | 1 |
| 11_1177296 | NA | NA | 2 | 1 | NA | NA | 1 | 1 | 3 | 1 | 1 | 1 | NA | NA | NA | NA | 1 | 1 | 2 | 1 |
| 11_1177298 | NA | NA | NA | NA | 1 | 0 | 1 | 1 | 2 | 0.5 | 2 | 0.5 | NA | NA | NA | NA | 1 | 1 | 3 | 0.6667 |
| 11_1177299 | NA | NA | 2 | 0 | 1 | 1 | 1 | 0 | 4 | 0.25 | 2 | 0.5 | NA | NA | NA | NA | NA | NA | 2 | 0.5 |
| 11_1177299 | NA | NA | 2 | 0.5 | 1 | 1 | 1 | 1 | 4 | 0.75 | 2 | 1 | NA | NA | NA | NA | 1 | 1 | 3 | 1 |
| 11_1177301 | NA | NA | 2 | 0 | 1 | 1 | 1 | 1 | 4 | 0.5 | 2 | 0 | NA | NA | NA | NA | 1 | 1 | 3 | 0.3333 |
| 11_1177302 | NA | NA | 2 | 1 | 1 | 1 | 1 | 1 | 4 | 1 | 1 | 0 | NA | NA | NA | NA | 1 | 1 | 2 | 0.5 |
| 11_1177304 | NA | NA | 2 | 1 | 1 | 0 | 1 | 0 | 4 | 0.5 | 2 | 0 | NA | NA | NA | NA | 1 | 1 | 3 | 0.3333 |
| 11_1177306 | NA | NA | 1 | 1 | NA | NA | 1 | 1 | 2 | 1 | 2 | 0 | NA | NA | NA | NA | 1 | 1 | 3 | 0.3333 |
| 11_1177588 | NA | NA | 1 | 1 | 1 | 1 | NA | NA | 2 | 1 | 1 | 1 | NA | NA | NA | NA | NA | NA | 1 | 1 |
| 11_1178960 | NA | NA | NA | NA | 1 | 1 | NA | NA | 1 | 1 | NA | NA | 1 | 1 | 1 | 1 | NA | NA | 2 | 1 |
| 11_1178962 | 7 | 0.2857 | 8 | 0.125 | 7 | 0 | 4 | 0 | 26 | 0.1154 | 5 | 0.2 | 8 | 0.25 | 7 | 0.1429 | 7 | 0.2857 | 27 | 0.2222 |
| 11_1178965 | 7 | 0.8571 | 8 | 0.625 | 11 | 0.7273 | 4 | 1 | 30 | 0.7667 | 7 | 0.7143 | 8 | 0.75 | 6 | 0.8333 | 8 | 0.875 | 29 | 0.7931 |
| 11_1179144 | 10 | 0.2 | 8 | 0.25 | 12 | 0.0833 | 5 | 0.2 | 35 | 0.1714 | 7 | 0.2857 | 11 | 0.1818 | 9 | 0.2222 | 12 | 0.0833 | 39 | 0.1795 |
| 11_1179144 | 10 | 0.1 | 8 | 0 | 11 | 0.2727 | 5 | 0.2 | 34 | 0.1471 | 9 | 0.4444 | 12 | 0 | 8 | 0.25 | 14 | 0.2857 | 43 | 0.2326 |
| 11_1179145 | 10 | 0.6 | 7 | 0.2857 | 11 | 0.8182 | 6 | 0.1667 | 34 | 0.5294 | 10 | 0.3 | 13 | 0.3077 | 8 | 0.375 | 13 | 0.4615 | 44 | 0.3636 |
| 11_1179145 | 11 | 0.0909 | 7 | 0 | 13 | 0.0769 | 6 | 0.1667 | 37 | 0.0811 | 9 | 0.2222 | 13 | 0.2308 | 8 | 0.125 | 14 | 0.1429 | 44 | 0.1818 |
| 11_1180553 | 6 | 0.3333 | 4 | 0.25 | 12 | 0.1667 | 5 | 0 | 27 | 0.1852 | 10 | 0.3 | 10 | 0.1 | 9 | 0 | 6 | 0 | 35 | 0.1143 |
| 11_1180556 | 4 | 0 | 3 | 0 | 12 | 0.1667 | 9 | 0.1111 | 28 | 0.1071 | 8 | 0 | 9 | 0.1111 | 8 | 0.125 | 5 | 0.2 | 30 | 0.1 |
| 11_1180559 | 7 | 0 | 1 | 0 | 11 | 0.1818 | 3 | 0.6667 | 22 | 0.1818 | 4 | 0 | 4 | 0.25 | 5 | 0.4 | 6 | 0 | 19 | 0.1579 |
| 11_1180562 | 7 | 0 | 1 | 0 | 13 | 0.1538 | 7 | 0.1429 | 28 | 0.1071 | 6 | 0 | 7 | 0.1429 | 9 | 0.1111 | 5 | 0 | 27 | 0.0741 |
| 11_1180567 | 6 | 0.6667 | 2 | 1 | 11 | 0.8182 | 8 | 0.625 | 27 | 0.7407 | 6 | 0.8333 | 6 | 0.5 | 6 | 0.5 | 4 | 0.5 | 22 | 0.5909 |
| 11_1180568 | 3 | 0.3333 | 1 | 0 | 11 | 0.2727 | 6 | 0.6667 | 21 | 0.381 | 4 | 0.5 | 5 | 0.6 | 6 | 0.1667 | 3 | 0.6667 | 18 | 0.4444 |
| 11_1181515 | 1 | 0 | 1 | 0 | 3 | 0.3333 | 1 | 0 | 6 | 0.1667 | 3 | 0 | 3 | 0 | 2 | 0.5 | 3 | 0 | 11 | 0.0909 |
| 11_1181515 | 1 | 1 | 1 | 1 | 3 | 1 | 1 | 1 | 6 | 1 | 2 | 1 | 3 | 1 | NA | NA | 2 | 0.5 | 7 | 0.8571 |
| 11_1181515 | 1 | 1 | 2 | 0.5 | 2 | 0 | 1 | 0 | 6 | 0.3333 | 3 | 0 | 3 | 0 | 2 | 0.5 | 3 | 0 | 11 | 0.0909 |
| 11_1181516 | 1 | 0 | 2 | 0 | 2 | 0 | 1 | 1 | 6 | 0.1667 | 2 | 0 | 2 | 0 | 2 | 0 | 3 | 0.3333 | 9 | 0.1111 |
| 11_1181518 | 1 | 0 | NA | NA | 3 | 0.3333 | 3 | 0.3333 | 7 | 0.2857 | 2 | 0 | 3 | 0.6667 | 1 | 0 | 2 | 0.5 | 8 | 0.375 |
| 11_1181519 | 1 | 0 | 1 | 0 | 3 | 0.3333 | 3 | 1 | 8 | 0.5 | 3 | 0.3333 | 2 | 0.5 | 2 | 0.5 | 4 | 0.75 | 11 | 0.5455 |
| 11_1181519 | 1 | 0 | 2 | 0 | 3 | 0 | 3 | 0.6667 | 9 | 0.2222 | 2 | 1 | 2 | 0.5 | NA | NA | 2 | 0 | 6 | 0.5 |
| 11_1181519 | 1 | 1 | NA | NA | 1 | 1 | 3 | 0.6667 | 5 | 0.8 | 2 | 1 | 1 | 1 | NA | NA | 1 | 1 | 4 | 1 |
| 11_1181520 | NA | NA | 2 | 0.5 | 1 | 0 | 1 | 0 | 4 | 0.25 | 3 | 0.6667 | 1 | 1 | 1 | 0 | 2 | 0 | 7 | 0.4286 |

| | | | | | | | | | | | | | | | | | | | | |
|---|---|---|---|---|---|---|---|---|---|---|---|---|---|---|---|---|---|---|---|---|
| 11_1181520 | 1 | 1 | 2 | 0 | 2 | 0 | 1 | 0 | 6 | 0.1667 | 3 | 0.3333 | 2 | 0 | 2 | 0 | 3 | 0 | 10 | 0.1 |
| 11_1181521 | 1 | 1 | 2 | 1 | 2 | 0.5 | 2 | 1 | 7 | 0.8571 | 2 | 1 | 1 | 0 | 1 | 1 | 3 | 0.6667 | 7 | 0.7143 |
| 11_1181521 | 1 | 1 | 2 | 0 | 3 | 0.3333 | 2 | 0.5 | 8 | 0.375 | 2 | 1 | 2 | 0.5 | 2 | 0 | 3 | 0 | 9 | 0.3333 |
| 11_1181521 | 1 | 0 | 1 | 1 | 3 | 0.3333 | 2 | 1 | 7 | 0.5714 | 3 | 0 | 2 | 0.5 | 1 | 0 | 3 | 0.3333 | 9 | 0.2222 |
| 11_1181521 | NA | NA | 2 | 0.5 | 2 | 0.5 | 2 | 0 | 6 | 0.3333 | 3 | 0.3333 | 1 | 0 | 2 | 0.5 | 3 | 0.3333 | 9 | 0.3333 |
| 11_1181556 | NA | NA | NA | NA | 2 | 0.5 | 1 | 0 | 3 | 0.3333 | 1 | 0 | 1 | 1 | 1 | 1 | 2 | 1 | 5 | 0.8 |
| 11_1181558 | 1 | 0 | 1 | 1 | 3 | 1 | 1 | 0 | 6 | 0.6667 | 3 | 0.6667 | 1 | 1 | 2 | 0.5 | 3 | 0.6667 | 9 | 0.6667 |
| 11_1181559 | NA | NA | 1 | 1 | 3 | 1 | 1 | 1 | 5 | 1 | 2 | 0.5 | 1 | 1 | 1 | 1 | 3 | 1 | 7 | 0.8571 |
| 11_1181920 | 23 | 0.2174 | 25 | 0.16 | 23 | 0.2174 | 21 | 0.0952 | 92 | 0.1739 | 22 | 0.0455 | 36 | 0.1111 | 28 | 0.1071 | 22 | 0.0455 | 108 | 0.0833 |
| 11_1181923 | 32 | 0.125 | 37 | 0.1081 | 27 | 0.1852 | 30 | 0.1333 | 126 | 0.1349 | 32 | 0.0312 | 47 | 0.1489 | 32 | 0.1562 | 31 | 0.129 | 142 | 0.1197 |
| 11_1181923 | 36 | 0.1944 | 35 | 0.0571 | 27 | 0.037 | 32 | 0.0312 | 130 | 0.0846 | 31 | 0.0645 | 52 | 0.1731 | 33 | 0.1818 | 34 | 0.0882 | 150 | 0.1333 |
| 11_1181929 | 32 | 0.125 | 34 | 0.0294 | 24 | 0 | 29 | 0.069 | 119 | 0.0588 | 29 | 0.0345 | 52 | 0.0769 | 35 | 0.0571 | 27 | 0.037 | 143 | 0.0559 |
| 11_1181939 | 18 | 0.2222 | 20 | 0.2 | 15 | 0.2667 | 15 | 0.3333 | 68 | 0.25 | 19 | 0.2105 | 37 | 0.5946 | 25 | 0.4 | 15 | 0.2667 | 96 | 0.4167 |
| 11_1181941 | 20 | 0.3 | 21 | 0.3333 | 14 | 0.2143 | 16 | 0.0625 | 71 | 0.2394 | 22 | 0.1818 | 39 | 0.1795 | 25 | 0.12 | 20 | 0.25 | 106 | 0.1792 |
| 11_1181942 | 20 | 0.15 | 19 | 0 | 16 | 0 | 18 | 0.0556 | 73 | 0.0548 | 20 | 0.15 | 39 | 0.0513 | 20 | 0.1 | 21 | 0.0952 | 100 | 0.09 |
| 11_1181944 | 21 | 0.3333 | 17 | 0.4706 | 16 | 0.5625 | 20 | 0.4 | 74 | 0.4324 | 18 | 0.3889 | 42 | 0.4048 | 22 | 0.5455 | 23 | 0.5652 | 105 | 0.4667 |
| 11_1181945 | 26 | 0.1538 | 18 | 0.0556 | 14 | 0.1429 | 18 | 0.0556 | 76 | 0.1053 | 18 | 0.0556 | 36 | 0.1944 | 24 | 0.1667 | 22 | 0.2273 | 100 | 0.17 |
| 11_1181946 | 19 | 0.4211 | 19 | 0.4211 | 17 | 0.1176 | 20 | 0.3 | 75 | 0.32 | 18 | 0.4444 | 33 | 0.4242 | 21 | 0.4286 | 22 | 0.1818 | 94 | 0.3723 |
| 11_1182837 | 2 | 0.5 | 3 | 1 | 1 | 1 | 1 | 1 | 7 | 0.8571 | 1 | 1 | 3 | 1 | 4 | 0.5 | 3 | 0.6667 | 11 | 0.7273 |
| 11_1182837 | 2 | 1 | 3 | 1 | 1 | 1 | NA | NA | 6 | 1 | 1 | 1 | 2 | 1 | 1 | 1 | 3 | 0.6667 | 7 | 0.8571 |
| 11_1182838 | 1 | 0 | 3 | 0 | 1 | 0 | 1 | 1 | 6 | 0.1667 | NA | NA | 3 | 0 | 2 | 0.5 | 1 | 0 | 6 | 0.1667 |
| 11_1182839 | 2 | 0 | 3 | 0.6667 | 1 | 0 | NA | NA | 6 | 0.3333 | NA | NA | 4 | 0.25 | 4 | 0.5 | 4 | 0.25 | 12 | 0.3333 |
| 11_1182839 | 2 | 1 | 2 | 1 | NA | NA | 1 | 1 | 5 | 1 | NA | NA | 2 | 1 | 3 | 1 | 4 | 0.5 | 9 | 0.7778 |
| 11_1182840 | 2 | 0.5 | 2 | 0 | NA | NA | 1 | 0 | 5 | 0.2 | NA | NA | 4 | 0.75 | 3 | 0.6667 | 3 | 0.3333 | 10 | 0.6 |
| 11_1182841 | 2 | 0.5 | 3 | 1 | NA | NA | NA | NA | 5 | 0.8 | NA | NA | 4 | 0.5 | 3 | 0.6667 | 4 | 0 | 11 | 0.3636 |
| 11_1182841 | 2 | 0 | 3 | 0.3333 | 1 | 0 | NA | NA | 6 | 0.1667 | NA | NA | 4 | 0.5 | 4 | 0.25 | 3 | 0.3333 | 11 | 0.3636 |
| 11_1182843 | 1 | 0 | 3 | 0.3333 | 1 | 0 | NA | NA | 5 | 0.2 | NA | NA | 3 | 0.3333 | 3 | 0.3333 | 3 | 0.3333 | 9 | 0.3333 |
| 11_1182847 | 1 | 1 | 3 | 0.6667 | 1 | 0 | NA | NA | 5 | 0.6 | NA | NA | 3 | 0 | 3 | 0.6667 | 3 | 0.6667 | 9 | 0.4444 |
| 11_1182847 | 1 | 1 | 3 | 1 | 1 | 1 | NA | NA | 5 | 1 | NA | NA | 3 | 1 | 3 | 1 | 3 | 1 | 9 | 1 |
| 11_1182847 | NA | NA | 2 | 0.5 | NA | NA | NA | NA | 2 | 0.5 | NA | NA | 4 | 1 | 3 | 0.6667 | 3 | 0.3333 | 10 | 0.7 |
| 11_1182848 | 1 | 1 | 3 | 0.3333 | NA | NA | NA | NA | 4 | 0.5 | NA | NA | 4 | 1 | 3 | 0.6667 | 3 | 1 | 10 | 0.9 |
| 11_1182974 | 7 | 0.1429 | 4 | 0.25 | 3 | 0.3333 | 12 | 0.3333 | 26 | 0.2692 | 4 | 0.25 | 8 | 0.125 | 6 | 0.1667 | 1 | 0 | 19 | 0.1579 |
| 11_1182976 | 6 | 0.1667 | 4 | 0.25 | 3 | 0.3333 | 10 | 0.2 | 23 | 0.2174 | 4 | 0 | 8 | 0.25 | 6 | 0 | 1 | 0 | 19 | 0.1053 |
| 11_1182977 | 4 | 0.25 | 3 | 0.3333 | 1 | 1 | 12 | 0.75 | 20 | 0.6 | 4 | 0.5 | 7 | 0.4286 | 5 | 0 | 1 | 0 | 17 | 0.2941 |
| 11_1182978 | 4 | 0.25 | 4 | 0 | 2 | 0 | 9 | 0.3333 | 19 | 0.2105 | 3 | 0 | 8 | 0 | 5 | 0.4 | 1 | 0 | 17 | 0.1176 |
| 11_1182980 | 4 | 0 | 3 | 0 | 2 | 0 | 7 | 0.1429 | 16 | 0.0625 | 1 | 0 | 7 | 0.2857 | 5 | 0 | 1 | 0 | 14 | 0.1429 |
| 11_1182983 | 3 | 0.6667 | 3 | 1 | 1 | 1 | 4 | 0.5 | 11 | 0.7273 | 2 | 1 | 5 | 1 | 3 | 0.3333 | 1 | 1 | 11 | 0.8182 |
| 11_1182984 | 4 | 0.75 | 2 | 1 | NA | NA | 4 | 1 | 10 | 0.9 | 2 | 1 | 7 | 1 | 4 | 1 | 1 | 1 | 14 | 1 |
| 11_1182985 | 5 | 0.4 | 2 | 0 | 1 | 1 | 3 | 0 | 11 | 0.2727 | 2 | 0.5 | 6 | 0.1667 | 3 | 0 | 1 | 0 | 12 | 0.1667 |
| 11_1182986 | 2 | 0.5 | 1 | 1 | 1 | 1 | 4 | 1 | 8 | 0.875 | 1 | 1 | 5 | 1 | 3 | 1 | NA | NA | 9 | 1 |
| 11_1182987 | 3 | 0 | 2 | 0 | 1 | 1 | 3 | 0.3333 | 9 | 0.2222 | 2 | 1 | 5 | 0.2 | 2 | 0 | 1 | 0 | 10 | 0.3 |
| 11_1182987 | 4 | 0.75 | 2 | 0.5 | 1 | 1 | 5 | 0.4 | 12 | 0.5833 | 1 | 1 | 5 | 0.6 | 4 | 0 | 1 | 0 | 11 | 0.3636 |
| 11_1182987 | 4 | 0.5 | 1 | 1 | 1 | 1 | 3 | 0.3333 | 9 | 0.5556 | 1 | 0 | 4 | 0.25 | 4 | 0 | 1 | 0 | 10 | 0.1 |
| 11_1183018 | 2 | 1 | 1 | 1 | NA | NA | 4 | 0.5 | 7 | 0.7143 | 1 | 0 | 3 | 0 | 1 | 1 | NA | NA | 5 | 0.2 |
| 11_1183018 | NA | NA | 1 | 1 | 1 | 1 | 3 | 0 | 5 | 0.4 | 1 | 1 | 5 | 0.4 | 1 | 1 | NA | NA | 7 | 0.5714 |
| 11_1183020 | NA | NA | 1 | 1 | 1 | 1 | 4 | 0.75 | 6 | 0.8333 | NA | NA | 4 | 1 | NA | NA | NA | NA | 4 | 1 |
| 11_1183022 | 2 | 0.5 | NA | NA | 1 | 0 | 3 | 0.3333 | 6 | 0.3333 | 1 | 0 | 3 | 0.3333 | 1 | 1 | NA | NA | 5 | 0.4 |
| 11_1183086 | 1 | 1 | NA | NA | NA | NA | NA | NA | 1 | 1 | 1 | 0 | 1 | 1 | NA | NA | NA | NA | 2 | 0.5 |
| 11_1183391 | NA | NA | 1 | 1 | NA | NA | NA | NA | 1 | 1 | NA | NA | 1 | 1 | NA | NA | NA | NA | 1 | 1 |
| 11_1183414 | NA | NA | 1 | 1 | NA | NA | NA | NA | 1 | 1 | NA | NA | 1 | 0 | 1 | 1 | NA | NA | 2 | 0.5 |
| 11_1183418 | NA | NA | 1 | 1 | NA | NA | NA | NA | 1 | 1 | NA | NA | 1 | 1 | 1 | 1 | NA | NA | 2 | 1 |
| 11_1183418 | NA | NA | 1 | 1 | NA | NA | NA | NA | 1 | 1 | NA | NA | 1 | 1 | 1 | 1 | NA | NA | 2 | 1 |
| 11_1183419 | NA | NA | 1 | 1 | NA | NA | NA | NA | 1 | 1 | NA | NA | 1 | 1 | 1 | 1 | NA | NA | 2 | 1 |
| 11_1183419 | NA | NA | 1 | 1 | NA | NA | NA | NA | 1 | 1 | NA | NA | 1 | 1 | NA | NA | NA | NA | 1 | 1 |
| 11_1183420 | NA | NA | 1 | 1 | NA | NA | NA | NA | 1 | 1 | NA | NA | 1 | 1 | NA | NA | NA | NA | 1 | 1 |
| 11_1183422 | NA | NA | 1 | 1 | NA | NA | NA | NA | 1 | 1 | NA | NA | 1 | 1 | 1 | 0 | NA | NA | 2 | 0.5 |
| 11_1183422 | NA | NA | 1 | 1 | NA | NA | NA | NA | 1 | 1 | NA | NA | 1 | 1 | 1 | 1 | NA | NA | 2 | 1 |
| 11_1183781 | 2 | 1 | 2 | 1 | NA | NA | NA | NA | 4 | 1 | 1 | 1 | NA | NA | NA | NA | 1 | 1 | 2 | 1 |
| 11_1183781 | 1 | 1 | 2 | 0.5 | NA | NA | NA | NA | 3 | 0.6667 | 1 | 1 | NA | NA | NA | NA | 1 | 1 | 2 | 1 |
| 11_1183783 | 1 | 1 | 2 | 0 | NA | NA | NA | NA | 3 | 0.3333 | 1 | 1 | NA | NA | NA | NA | 1 | 1 | 2 | 1 |
| 11_1183784 | 2 | 1 | 1 | 1 | 1 | 1 | NA | NA | 4 | 1 | 1 | 1 | NA | NA | NA | NA | 1 | 1 | 2 | 1 |
| 11_1183784 | 2 | 0 | 2 | 0 | 1 | 1 | NA | NA | 5 | 0.2 | 1 | 0 | NA | NA | NA | NA | 1 | 1 | 2 | 0.5 |
| 11_1183807 | 29 | 0.1724 | 42 | 0.4286 | 36 | 0.3056 | 31 | 0.3548 | 138 | 0.3261 | 21 | 0.4286 | 31 | 0.2903 | 25 | 0.28 | 26 | 0.3462 | 103 | 0.3301 |
| 11_1183807 | 19 | 0.1579 | 31 | 0.3871 | 33 | 0.4848 | 23 | 0.3913 | 106 | 0.3774 | 16 | 0.625 | 24 | 0.5 | 22 | 0.3636 | 16 | 0.375 | 78 | 0.4615 |
| 11_1183811 | 26 | 0.1154 | 32 | 0.1875 | 28 | 0.25 | 23 | 0.1739 | 109 | 0.1835 | 19 | 0.1579 | 26 | 0.1538 | 24 | 0.1667 | 17 | 0.2353 | 86 | 0.1744 |

| | | | | | | | | | | | | | | | | | | | | |
|---|---|---|---|---|---|---|---|---|---|---|---|---|---|---|---|---|---|---|---|---|
| 11_1183812 | 23 | 0 | 33 | 0.1515 | 26 | 0.1538 | 23 | 0.0435 | 105 | 0.0952 | 18 | 0 | 21 | 0 | 22 | 0.0909 | 17 | 0.1176 | 78 | 0.0513 |
| 11_1183813 | 20 | 0.8 | 25 | 0.68 | 14 | 0.7857 | 14 | 0.7143 | 73 | 0.7397 | 9 | 0.7778 | 13 | 0.7692 | 15 | 0.8667 | 9 | 1 | 46 | 0.8478 |
| 11_1183814 | 9 | 0.5556 | 13 | 0.1538 | 10 | 0.6 | 14 | 0.2857 | 46 | 0.3696 | 9 | 0.4444 | 13 | 0.0769 | 13 | 0.1538 | 5 | 0.2 | 40 | 0.2 |
| 11_1183816 | 24 | 0.1667 | 28 | 0.1071 | 20 | 0.1 | 22 | 0.1818 | 94 | 0.1383 | 15 | 0.0667 | 22 | 0.1818 | 19 | 0.2105 | 10 | 0.3 | 66 | 0.1818 |
| 11_1183817 | 24 | 0.0833 | 23 | 0.087 | 17 | 0.0588 | 17 | 0 | 81 | 0.0617 | 15 | 0.1333 | 21 | 0.0476 | 16 | 0.0625 | 10 | 0 | 62 | 0.0645 |
| 11_1183818 | 20 | 0.25 | 24 | 0.0833 | 20 | 0.1 | 22 | 0.1818 | 86 | 0.1512 | 13 | 0.0769 | 21 | 0.1429 | 15 | 0.1333 | 9 | 0.2222 | 58 | 0.1379 |
| 11_1183820 | 20 | 0.05 | 23 | 0.0435 | 18 | 0.0556 | 14 | 0.0714 | 75 | 0.0533 | 12 | 0.0833 | 18 | 0.1111 | 14 | 0 | 7 | 0 | 51 | 0.0588 |
| 11_1183828 | 2 | 0 | 2 | 0.5 | NA | NA | 1 | 1 | 5 | 0.4 | 1 | 1 | NA | NA | 1 | 0 | NA | NA | 2 | 0.5 |
| 11_1183832 | 2 | 0 | 2 | 0 | NA | NA | 1 | 1 | 5 | 0.2 | 1 | 1 | 1 | 0 | 1 | 0 | NA | NA | 3 | 0.3333 |
| 11_1189294 | 1 | 1 | 2 | 0 | 2 | 0 | 1 | 1 | 6 | 0.3333 | NA | NA | 2 | 0 | 2 | 0.5 | NA | NA | 4 | 0.25 |
| 11_1189295 | 1 | 1 | 2 | 1 | 2 | 0.5 | NA | NA | 5 | 0.8 | NA | NA | 1 | 1 | 1 | 1 | NA | NA | 2 | 1 |
| 11_1189296 | NA | NA | 2 | 0.5 | 2 | 0 | 2 | 1 | 6 | 0.5 | NA | NA | 1 | 1 | 2 | 0.5 | NA | NA | 3 | 0.6667 |
| 11_1189297 | 1 | 1 | 3 | 1 | 2 | 1 | 2 | 1 | 8 | 1 | NA | NA | 1 | 1 | 2 | 1 | NA | NA | 3 | 1 |
| 11_1189297 | NA | NA | 3 | 0.6667 | 2 | 0 | 2 | 0.5 | 7 | 0.4286 | NA | NA | 1 | 1 | 2 | 0.5 | NA | NA | 3 | 0.6667 |
| 11_1189298 | 1 | 1 | 3 | 1 | 2 | 1 | 2 | 1 | 8 | 1 | NA | NA | 1 | 1 | 2 | 1 | NA | NA | 3 | 1 |
| 11_1189299 | NA | NA | 2 | 1 | 2 | 1 | 2 | 1 | 6 | 1 | NA | NA | 1 | 1 | 2 | 1 | NA | NA | 3 | 1 |
| 11_1189299 | 1 | 0 | 3 | 0.3333 | 2 | 0 | 2 | 0.5 | 8 | 0.25 | NA | NA | NA | NA | 2 | 1 | NA | NA | 2 | 1 |
| 11_1189299 | 1 | 1 | 3 | 0 | 2 | 0 | 1 | 1 | 7 | 0.2857 | NA | NA | 1 | 0 | 2 | 0.5 | NA | NA | 3 | 0.3333 |
| 11_1189300 | 1 | 1 | 2 | 0.5 | 2 | 1 | 2 | 0.5 | 7 | 0.7143 | NA | NA | NA | NA | 2 | 1 | NA | NA | 2 | 1 |
| 11_1189710 | 18 | 0.0556 | 12 | 0.1667 | 12 | 0 | 14 | 0.1429 | 56 | 0.0893 | 6 | 0.1667 | 12 | 0.1667 | 16 | 0.0625 | 10 | 0 | 44 | 0.0909 |
| 11_1189711 | 19 | 0.1053 | 11 | 0.1818 | 11 | 0.1818 | 12 | 0.5833 | 53 | 0.2453 | 7 | 0.4286 | 14 | 0.3571 | 14 | 0.3571 | 10 | 0.2 | 45 | 0.3333 |
| 11_1189712 | 20 | 0.35 | 13 | 0.3846 | 13 | 0.3077 | 15 | 0.1333 | 61 | 0.2951 | 5 | 0 | 14 | 0.2857 | 12 | 0.6667 | 9 | 0.2222 | 40 | 0.35 |
| 11_1189714 | 20 | 0.15 | 11 | 0.3636 | 12 | 0.5 | 13 | 0.3077 | 56 | 0.3036 | 6 | 0.3333 | 13 | 0.2308 | 15 | 0.4 | 9 | 0.3333 | 43 | 0.3256 |
| 11_1189714 | 19 | 0.1579 | 8 | 0.125 | 10 | 0.1 | 11 | 0.2727 | 48 | 0.1667 | 5 | 0.2 | 13 | 0.0769 | 13 | 0.1538 | 9 | 0.2222 | 40 | 0.15 |
| 11_1189715 | 12 | 1 | 6 | 1 | 5 | 1 | 11 | 0.9091 | 34 | 0.9706 | 2 | 1 | 7 | 0.8571 | 12 | 1 | 5 | 1 | 26 | 0.9615 |
| 11_1189715 | 15 | 0.4 | 9 | 0.2222 | 10 | 0.5 | 10 | 0.4 | 44 | 0.3864 | 5 | 0.2 | 12 | 0.6667 | 14 | 0.7143 | 8 | 0.5 | 39 | 0.5897 |
| 11_1189716 | 15 | 0.9333 | 11 | 1 | 7 | 1 | 13 | 1 | 46 | 0.9783 | 5 | 1 | 11 | 1 | 15 | 1 | 8 | 1 | 39 | 1 |
| 11_1189717 | 7 | 1 | 5 | 1 | 6 | 0.8333 | 6 | 1 | 24 | 0.9583 | 2 | 1 | 10 | 1 | 11 | 1 | 8 | 0.875 | 31 | 0.9677 |
| 11_1189719 | 9 | 1 | 9 | 1 | 9 | 1 | 10 | 1 | 37 | 1 | 5 | 1 | 9 | 0.8889 | 13 | 1 | 8 | 1 | 35 | 0.9714 |
| 11_1189719 | 12 | 0.25 | 10 | 0.2 | 9 | 0.2222 | 13 | 0.1538 | 44 | 0.2045 | 5 | 0.2 | 10 | 0.1 | 14 | 0.2857 | 9 | 0.3333 | 38 | 0.2368 |
| 11_1189719 | 12 | 0.8333 | 9 | 0.7778 | 8 | 1 | 12 | 0.8333 | 41 | 0.8537 | 4 | 0.5 | 10 | 0.8 | 13 | 0.8462 | 7 | 0.4286 | 34 | 0.7059 |
| 11_1189721 | 9 | 0.2222 | 10 | 0 | 7 | 0.4286 | 11 | 0.1818 | 37 | 0.1892 | 3 | 0.3333 | 10 | 0.1 | 12 | 0.1667 | 8 | 0.125 | 33 | 0.1515 |
| 11_1189722 | 9 | 0.3333 | 11 | 0.0909 | 9 | 0.3333 | 12 | 0.1667 | 41 | 0.2195 | 4 | 0.25 | 13 | 0.3077 | 12 | 0.25 | 10 | 0.4 | 39 | 0.3077 |
| 11_1189723 | 9 | 0.3333 | 9 | 0.5556 | 9 | 0.5556 | 12 | 0.4167 | 39 | 0.4615 | 5 | 0.4 | 11 | 0.4545 | 12 | 0.6667 | 9 | 0.6667 | 37 | 0.5676 |
| 11_1189724 | 11 | 0.9091 | 8 | 1 | 9 | 0.8889 | 10 | 1 | 38 | 0.9474 | 4 | 1 | 11 | 1 | 11 | 1 | 8 | 0.875 | 34 | 0.9706 |
| 11_1189726 | 10 | 0.1 | 7 | 0.1429 | 8 | 0 | 10 | 0.2 | 35 | 0.1143 | 5 | 0.4 | 11 | 0.1818 | 9 | 0.1111 | 9 | 0.2222 | 34 | 0.2059 |
| 11_1189727 | 6 | 0.1667 | 6 | 0 | 6 | 0 | 8 | 0.125 | 26 | 0.0769 | 5 | 0.2 | 9 | 0.1111 | 7 | 0 | 9 | 0.1111 | 30 | 0.1 |
| 11_1190339 | 15 | 0.1333 | 8 | 0.125 | 10 | 0.2 | 5 | 0.4 | 38 | 0.1842 | 4 | 0.25 | 10 | 0 | 17 | 0.0588 | 12 | 0.25 | 43 | 0.1163 |
| 11_1190340 | 12 | 0 | 7 | 0 | 8 | 0.25 | 5 | 0.2 | 32 | 0.0938 | 3 | 0.3333 | 11 | 0.2727 | 17 | 0 | 9 | 0 | 40 | 0.1 |
| 11_1190341 | 15 | 0 | 8 | 0.125 | 7 | 0.1429 | 6 | 0 | 36 | 0.0556 | 5 | 0.4 | 11 | 0 | 16 | 0.1875 | 12 | 0 | 44 | 0.1136 |
| 11_1190342 | 13 | 0.6154 | 7 | 0.7143 | 8 | 0.25 | 4 | 0.25 | 32 | 0.5 | 4 | 0.75 | 13 | 0.4615 | 14 | 0.5 | 13 | 0.3077 | 44 | 0.4545 |
| 11_1190345 | 13 | 0.5385 | 6 | 0.5 | 9 | 0.7778 | 5 | 0.8 | 33 | 0.6364 | 5 | 0.6 | 9 | 0.4444 | 13 | 0.5385 | 13 | 0.3077 | 40 | 0.45 |
| 11_1190345 | 6 | 0.3333 | 5 | 0.4 | 6 | 0.5 | 5 | 0.2 | 22 | 0.3636 | 4 | 0.75 | 9 | 0.4444 | 11 | 0.5455 | 8 | 0.75 | 32 | 0.5938 |
| 11_1190346 | 11 | 0.0909 | 5 | 0.4 | 7 | 0.2857 | 4 | 0.25 | 27 | 0.2222 | 5 | 0.2 | 9 | 0.2222 | 15 | 0.0667 | 9 | 0.1111 | 38 | 0.1316 |
| 11_1190348 | 10 | 0.1 | 4 | 0 | 9 | 0 | 4 | 0.25 | 27 | 0.0741 | 4 | 0.25 | 9 | 0 | 13 | 0 | 13 | 0.1538 | 39 | 0.0769 |
| 11_1190362 | 5 | 0 | 4 | 0 | 5 | 0 | 3 | 0.3333 | 17 | 0.0588 | 3 | 0.3333 | 2 | 0 | 8 | 0 | 3 | 0 | 16 | 0.0625 |
| 11_1190382 | 1 | 1 | NA | NA | NA | NA | NA | NA | 1 | 1 | NA | NA | NA | NA | 1 | 1 | NA | NA | 1 | 1 |
| 11_1190385 | 1 | 1 | NA | NA | NA | NA | NA | NA | 1 | 1 | NA | NA | NA | NA | 2 | 0.5 | NA | NA | 2 | 0.5 |
| 11_1191609 | 8 | 0.125 | 10 | 0.4 | 8 | 0.25 | 15 | 0.6667 | 41 | 0.4146 | 8 | 0.5 | 24 | 0.4167 | 11 | 0.4545 | 11 | 0.6364 | 54 | 0.4815 |
| 11_1191611 | 18 | 0.1667 | 11 | 0.0909 | 9 | 0.1111 | 22 | 0.3636 | 60 | 0.2167 | 17 | 0.4118 | 33 | 0.1818 | 17 | 0.1176 | 17 | 0.1176 | 84 | 0.2024 |
| 11_1191611 | 16 | 0.375 | 12 | 0.6667 | 9 | 0.3333 | 25 | 0.48 | 62 | 0.4677 | 15 | 0.6 | 30 | 0.4 | 16 | 0.4375 | 18 | 0.5 | 79 | 0.4684 |
| 11_1191612 | 20 | 0.85 | 16 | 0.5625 | 9 | 0.7778 | 25 | 0.64 | 70 | 0.7 | 15 | 0.4667 | 30 | 0.4 | 17 | 0.6471 | 21 | 0.7143 | 83 | 0.5422 |
| 11_1191759 | 56 | 0.125 | 28 | 0.1071 | 37 | 0.1892 | 51 | 0.0784 | 172 | 0.1221 | 38 | 0.0789 | 63 | 0.0794 | 44 | 0.0909 | 42 | 0.119 | 187 | 0.0909 |
| 11_1191762 | 54 | 0.037 | 29 | 0.1034 | 36 | 0.0556 | 54 | 0.0556 | 173 | 0.0578 | 43 | 0.0698 | 69 | 0.0145 | 51 | 0.0392 | 48 | 0.125 | 211 | 0.0569 |
| 11_1191792 | 12 | 0.5833 | 8 | 0.375 | 11 | 0.7273 | 2 | 0.5 | 33 | 0.5758 | 5 | 0.4 | 11 | 0.6364 | 9 | 0.4444 | 7 | 0.7143 | 32 | 0.5625 |
| 11_1191793 | 17 | 0.2353 | 13 | 0.5385 | 13 | 0.3846 | 3 | 0.6667 | 46 | 0.3913 | 9 | 0.3333 | 9 | 0.4444 | 13 | 0.1538 | 9 | 0.5556 | 40 | 0.35 |
| 11_1192346 | 1 | 1 | 1 | 1 | NA | NA | NA | NA | 2 | 1 | 1 | 1 | 2 | 1 | NA | NA | NA | NA | 3 | 1 |
| 11_1192348 | 1 | 0 | 1 | 1 | NA | NA | NA | NA | 2 | 0.5 | 1 | 1 | 2 | 0 | NA | NA | NA | NA | 3 | 0.3333 |
| 11_1192348 | NA | NA | 1 | 1 | NA | NA | NA | NA | 1 | 1 | 1 | 1 | 2 | 1 | NA | NA | NA | NA | 3 | 1 |
| 11_1192349 | 1 | 0 | 1 | 1 | NA | NA | NA | NA | 2 | 0.5 | 1 | 0 | 2 | 0.5 | NA | NA | NA | NA | 3 | 0.3333 |
| 11_1192349 | 1 | 0 | 1 | 1 | NA | NA | NA | NA | 2 | 0.5 | 1 | 1 | 1 | 1 | NA | NA | NA | NA | 2 | 1 |
| 11_1193772 | NA | NA | NA | NA | NA | NA | 2 | 0.5 | 2 | 0.5 | NA | NA | 1 | 1 | NA | NA | 1 | 0 | 2 | 0.5 |
| 11_1193772 | 1 | 1 | NA | NA | NA | NA | 2 | 0 | 3 | 0.3333 | NA | NA | NA | NA | NA | NA | 1 | 1 | 1 | 1 |
| 11_1193983 | NA | NA | NA | NA | 2 | 1 | 1 | 1 | 3 | 1 | NA | NA | 1 | 1 | 2 | 1 | NA | NA | 3 | 1 |
| 11_1193984 | NA | NA | 2 | 1 | 3 | 1 | 1 | 1 | 6 | 1 | NA | NA | 1 | 1 | 2 | 1 | NA | NA | 3 | 1 |

| | | | | | | | | | | | | | | | | | | | | |
|---|---|---|---|---|---|---|---|---|---|---|---|---|---|---|---|---|---|---|---|---|
| 11_1193984 | NA | NA | 2 | 1 | 2 | 1 | 1 | 1 | 5 | 1 | NA | NA | NA | NA | 2 | 1 | NA | NA | 2 | 1 |
| 11_1193985 | NA | NA | 1 | 1 | 2 | 1 | 1 | 1 | 4 | 1 | NA | NA | 1 | 0 | 2 | 1 | NA | NA | 3 | 0.6667 |
| 11_1193986 | NA | NA | 1 | 1 | NA | NA | NA | NA | 1 | 1 | NA | NA | 1 | 1 | 1 | 1 | NA | NA | 2 | 1 |
| 11_1193987 | NA | NA | 1 | 1 | 1 | 1 | NA | NA | 2 | 1 | NA | NA | 1 | 1 | 1 | 0 | NA | NA | 2 | 0.5 |
| 11_1193990 | NA | NA | 1 | 1 | 1 | 1 | NA | NA | 2 | 1 | NA | NA | 1 | 1 | 1 | 0 | NA | NA | 2 | 0.5 |
| 11_1194001 | 2 | 0 | NA | NA | 4 | 0.5 | NA | NA | 6 | 0.3333 | NA | NA | NA | NA | 3 | 0.3333 | 1 | 1 | 4 | 0.5 |
| 11_1194019 | 2 | 0 | 2 | 1 | 1 | 0 | NA | NA | 5 | 0.4 | 1 | 0 | 1 | 1 | 3 | 0 | NA | NA | 5 | 0.2 |
| 11_1194299 | 20 | 0.4 | 32 | 0.3438 | 15 | 0.2667 | 10 | 0.3 | 77 | 0.3377 | 21 | 0.2857 | 10 | 0.6 | 15 | 0.2667 | 9 | 0.4444 | 55 | 0.3636 |
| 11_1194304 | 11 | 0.2727 | 18 | 0.0556 | 7 | 0 | 7 | 0 | 43 | 0.093 | 19 | 0.0526 | 8 | 0.25 | 12 | 0.0833 | 7 | 0 | 46 | 0.087 |
| 11_1194309 | 9 | 0 | 15 | 0.1333 | 6 | 0.3333 | 6 | 0.1667 | 36 | 0.1389 | 11 | 0 | 7 | 0.4286 | 11 | 0.4545 | 8 | 0.125 | 37 | 0.2432 |
| 11_1194310 | 11 | 0.0909 | 15 | 0 | 6 | 0.1667 | 6 | 0 | 38 | 0.0526 | 10 | 0.1 | 8 | 0.125 | 11 | 0 | 7 | 0 | 36 | 0.0556 |
| 11_1194312 | 10 | 0 | 14 | 0.1429 | 6 | 0 | 4 | 0 | 34 | 0.0588 | 11 | 0 | 8 | 0 | 10 | 0.1 | 6 | 0.1667 | 35 | 0.0571 |
| 11_1194319 | 7 | 0.2857 | 13 | 0.3077 | 3 | 1 | 3 | 0 | 26 | 0.3462 | 7 | 0.4286 | 3 | 0 | 5 | 0 | 4 | 0 | 19 | 0.1579 |
| 11_1194324 | 6 | 0.1667 | 11 | 0.0909 | 3 | 0.3333 | 3 | 0 | 23 | 0.1304 | 9 | 0.1111 | 6 | 0 | 6 | 0 | 5 | 0.2 | 26 | 0.0769 |
| 11_1194325 | 4 | 0 | 11 | 0.1818 | 3 | 0.3333 | 3 | 0 | 21 | 0.1429 | 9 | 0.2222 | 5 | 0.4 | 5 | 0.2 | 4 | 0.25 | 23 | 0.2609 |
| 11_1194325 | 6 | 0.1667 | 11 | 0.7273 | 3 | 0.3333 | 3 | 0 | 23 | 0.4348 | 8 | 0.5 | 5 | 0.2 | 5 | 0.8 | 4 | 0 | 22 | 0.4091 |
| 11_1194331 | 5 | 0.6 | 6 | 0.8333 | 3 | 0.6667 | 1 | 0 | 15 | 0.6667 | 5 | 0.4 | 4 | 0.25 | 2 | 0 | 1 | 0 | 12 | 0.25 |
| 11_1194332 | 3 | 0.3333 | 7 | 0.1429 | 3 | 0.3333 | 2 | 0.5 | 15 | 0.2667 | 7 | 0.1429 | 2 | 0.5 | 3 | 0.3333 | 2 | 0.5 | 14 | 0.2857 |
| 11_1194333 | 4 | 1 | 7 | 0.8571 | 2 | 0.5 | 3 | 0.6667 | 16 | 0.8125 | 8 | 1 | 2 | 0.5 | 2 | 1 | 2 | 1 | 14 | 0.9286 |
| 11_1194334 | 6 | 0.1667 | 6 | 0.1667 | 2 | 0 | 2 | 0 | 16 | 0.125 | 8 | 0.125 | 2 | 0 | 1 | 0 | 2 | 0 | 13 | 0.0769 |
| 11_1194335 | 4 | 0.5 | 6 | 0.3333 | 2 | 0.5 | 2 | 0.5 | 14 | 0.4286 | 8 | 0.5 | 3 | 0.3333 | 2 | 0 | 1 | 0 | 14 | 0.3571 |
| 11_1194354 | 5 | 0.2 | 6 | 0.1667 | 3 | 0.3333 | 2 | 0 | 16 | 0.1875 | 7 | 0 | 1 | 0 | 2 | 0.5 | 1 | 0 | 11 | 0.0909 |
| 11_1194381 | 5 | 0.4 | 3 | 0.3333 | 2 | 0 | 1 | 0 | 11 | 0.2727 | 6 | 0 | 1 | 0 | 1 | 0 | 1 | 1 | 9 | 0.1111 |
| 11_1194399 | NA | NA | 1 | 1 | NA | NA | NA | NA | 1 | 1 | 1 | 1 | NA | NA | NA | NA | NA | NA | 1 | 1 |
| 11_1194946 | NA | NA | NA | NA | 1 | 0 | 1 | 1 | 2 | 0.5 | NA | NA | NA | NA | 1 | 1 | NA | NA | 1 | 1 |
| 11_1197882 | 5 | 0.2 | 8 | 0 | 3 | 0 | 4 | 0 | 20 | 0.05 | 5 | 0 | 1 | 0 | 4 | 0.25 | 4 | 0.25 | 14 | 0.1429 |
| 11_1197883 | 3 | 0 | 6 | 0.3333 | 4 | 0 | 3 | 0 | 16 | 0.125 | 4 | 0.25 | NA | NA | 4 | 0.25 | 3 | 0 | 11 | 0.1818 |
| 11_1197885 | 6 | 0 | 8 | 0 | 4 | 0 | 6 | 0.3333 | 24 | 0.0833 | 5 | 0.4 | 2 | 0.5 | 3 | 0.3333 | 6 | 0.3333 | 16 | 0.375 |
| 11_1197886 | 6 | 0.3333 | 10 | 0.5 | 3 | 0.3333 | 6 | 0.3333 | 25 | 0.4 | 4 | 0.75 | 3 | 0 | 3 | 0.3333 | 6 | 0.8333 | 16 | 0.5625 |
| 11_1197892 | 5 | 0 | 13 | 0.1538 | 2 | 0 | 7 | 0.1429 | 27 | 0.1111 | 7 | 0.1429 | 2 | 0 | 2 | 0.5 | 6 | 0.5 | 17 | 0.2941 |
| 11_1197893 | 8 | 0.375 | 14 | 0.2857 | 5 | 0.2 | 9 | 0.2222 | 36 | 0.2778 | 7 | 0.1429 | 3 | 0 | 5 | 0.2 | 8 | 0.25 | 23 | 0.1739 |
| 11_1197894 | 6 | 0.6667 | 13 | 0.2308 | 5 | 0.6 | 7 | 0.1429 | 31 | 0.3548 | 8 | 0.625 | 3 | 0.3333 | 4 | 0.5 | 8 | 0.375 | 23 | 0.4783 |
| 11_1197894 | 7 | 0.1429 | 13 | 0.0769 | 6 | 0.5 | 9 | 0.2222 | 35 | 0.2 | 8 | 0.5 | 2 | 0 | 6 | 0.1667 | 5 | 0.2 | 21 | 0.2857 |
| 11_1197895 | 8 | 0.5 | 14 | 0.2857 | 6 | 0.5 | 9 | 0.1111 | 37 | 0.3243 | 8 | 0.125 | 3 | 0.3333 | 6 | 0.5 | 7 | 0.2857 | 24 | 0.2917 |
| 11_1197895 | 6 | 0.3333 | 14 | 0.1429 | 5 | 0.2 | 9 | 0.3333 | 34 | 0.2353 | 8 | 0.375 | 3 | 0 | 6 | 0.5 | 8 | 0.125 | 25 | 0.28 |
| 11_1197896 | 7 | 0.2857 | 13 | 0.2308 | 6 | 0.3333 | 9 | 0 | 35 | 0.2 | 5 | 0.4 | 3 | 0.3333 | 5 | 0 | 8 | 0.25 | 21 | 0.2381 |
| 11_1197896 | 7 | 0.5714 | 14 | 0.3571 | 6 | 0.1667 | 8 | 0.5 | 35 | 0.4 | 8 | 0.25 | 2 | 0.5 | 6 | 0.5 | 7 | 0.1429 | 23 | 0.3043 |
| 11_1197896 | 7 | 0.4286 | 14 | 0.0714 | 5 | 0.6 | 9 | 0.2222 | 35 | 0.2571 | 7 | 0.4286 | 1 | 0 | 5 | 0.8 | 8 | 0.5 | 21 | 0.5238 |
| 11_1197896 | 8 | 0.125 | 14 | 0.3571 | 6 | 0.5 | 8 | 0 | 36 | 0.25 | 7 | 0.2857 | 3 | 0 | 6 | 0.5 | 7 | 0.5714 | 23 | 0.3913 |
| 11_1197899 | 7 | 0.5714 | 12 | 0.5 | 4 | 0.5 | 7 | 0.4286 | 30 | 0.5 | 5 | 0.6 | 1 | 0 | 4 | 0.25 | 8 | 0.625 | 18 | 0.5 |
| 11_1197900 | 7 | 0 | 14 | 0.1429 | 7 | 0.1429 | 12 | 0.1667 | 40 | 0.125 | 8 | 0.125 | 4 | 0 | 7 | 0 | 8 | 0.125 | 27 | 0.0741 |
| 11_1197901 | 6 | 0.6667 | 14 | 0.9286 | 6 | 0.6667 | 10 | 0.7 | 36 | 0.7778 | 10 | 0.9 | 5 | 1 | 6 | 1 | 8 | 0.875 | 29 | 0.931 |
| 11_1197901 | 7 | 0.1429 | 15 | 0.0667 | 7 | 0 | 10 | 0 | 39 | 0.0513 | 8 | 0.375 | 5 | 0.2 | 7 | 0 | 9 | 0.2222 | 29 | 0.2069 |
| 11_1197901 | 6 | 0.3333 | 11 | 0.1818 | 7 | 0.4286 | 11 | 0.1818 | 35 | 0.2571 | 10 | 0.3 | 4 | 0.5 | 6 | 0.1667 | 7 | 0.2857 | 27 | 0.2963 |
| 11_1198095 | 2 | 1 | 3 | 0 | 2 | 0.5 | 1 | 0 | 8 | 0.375 | 4 | 0.25 | 2 | 0.5 | 6 | 0.5 | 2 | 0 | 14 | 0.3571 |
| 11_1198095 | 2 | 0 | 4 | 0 | 1 | 1 | 1 | 0 | 8 | 0.125 | 3 | 0 | 1 | 0 | 4 | 0.25 | 1 | 0 | 9 | 0.1111 |
| 11_1198096 | 2 | 1 | 2 | 1 | NA | NA | 1 | 1 | 5 | 1 | 4 | 0.75 | 2 | 0.5 | 4 | 0.75 | 1 | 1 | 11 | 0.7273 |
| 11_1198100 | 2 | 0.5 | 5 | 0.6 | 2 | 0.5 | 1 | 1 | 10 | 0.6 | 3 | 1 | 2 | 0.5 | 4 | 0.75 | 1 | 0 | 10 | 0.7 |
| 11_1198101 | 2 | 0 | 5 | 0.6 | 2 | 0.5 | 1 | 0 | 10 | 0.4 | 3 | 1 | 3 | 0.6667 | 4 | 0 | 2 | 0.5 | 12 | 0.5 |
| 11_1198101 | 2 | 1 | 4 | 0.75 | 3 | 1 | 1 | 1 | 10 | 0.9 | 4 | 1 | 3 | 1 | 6 | 0.5 | 2 | 1 | 15 | 0.8 |
| 11_1198101 | 2 | 1 | 2 | 1 | 2 | 0.5 | 1 | 0 | 7 | 0.7143 | 4 | 0.5 | 3 | 0.3333 | 5 | 0.2 | 2 | 0.5 | 14 | 0.3571 |
| 11_1198102 | NA | NA | 4 | 1 | 1 | 1 | 1 | 1 | 6 | 1 | 5 | 0.8 | 2 | 1 | 5 | 0.8 | 2 | 1 | 14 | 0.8571 |
| 11_1198265 | NA | NA | NA | NA | 1 | 1 | NA | NA | 1 | 1 | NA | NA | NA | NA | 1 | 1 | NA | NA | 1 | 1 |
| 11_1198265 | NA | NA | NA | NA | 1 | 1 | 2 | 0 | 3 | 0.3333 | NA | NA | 1 | 0 | 2 | 0.5 | NA | NA | 3 | 0.3333 |
| 11_1198267 | NA | NA | NA | NA | 2 | 1 | 1 | 0 | 3 | 0.6667 | NA | NA | 1 | 0 | 1 | 1 | NA | NA | 2 | 0.5 |
| 11_1198267 | NA | NA | NA | NA | 2 | 0.5 | NA | NA | 2 | 0.5 | NA | NA | 1 | 0 | 2 | 0.5 | NA | NA | 3 | 0.3333 |
| 11_1198920 | 31 | 0.129 | 45 | 0.1778 | 29 | 0.1034 | 44 | 0.2273 | 149 | 0.1678 | 20 | 0.05 | 46 | 0.1304 | 36 | 0.1944 | 26 | 0.1923 | 128 | 0.1484 |
| 11_1198923 | 37 | 0.8649 | 48 | 0.8125 | 28 | 0.8929 | 51 | 0.7255 | 164 | 0.811 | 26 | 0.8462 | 50 | 0.86 | 36 | 0.8056 | 32 | 0.8125 | 144 | 0.8333 |
| 11_1198923 | 39 | 0.641 | 53 | 0.7547 | 30 | 0.7333 | 53 | 0.7736 | 175 | 0.7314 | 28 | 0.6786 | 54 | 0.5926 | 43 | 0.5581 | 34 | 0.4706 | 159 | 0.5723 |
| 11_1198924 | 17 | 0.1765 | 38 | 0.2368 | 18 | 0.1667 | 17 | 0.1176 | 90 | 0.1889 | 14 | 0.0714 | 33 | 0.2727 | 30 | 0.1 | 22 | 0.1818 | 99 | 0.1717 |
| 11_1198924 | 40 | 0.1 | 54 | 0.1111 | 32 | 0.1875 | 54 | 0.0926 | 180 | 0.1167 | 25 | 0.08 | 56 | 0.0893 | 45 | 0.0444 | 35 | 0.1143 | 161 | 0.0807 |
| 11_1198927 | 38 | 0.1053 | 45 | 0.1778 | 28 | 0.1071 | 41 | 0.122 | 152 | 0.1316 | 24 | 0.0833 | 53 | 0.1509 | 43 | 0.093 | 39 | 0.0769 | 159 | 0.1069 |
| 11_1198929 | 43 | 0.5116 | 61 | 0.5246 | 44 | 0.5455 | 59 | 0.5932 | 207 | 0.5459 | 30 | 0.6333 | 69 | 0.4783 | 50 | 0.36 | 44 | 0.5455 | 193 | 0.487 |
| 11_1198931 | 44 | 0.0455 | 61 | 0.1311 | 47 | 0.1489 | 60 | 0.2 | 212 | 0.1368 | 28 | 0.2143 | 67 | 0.209 | 50 | 0.18 | 44 | 0.2727 | 189 | 0.2169 |
| 11_1198932 | 48 | 0.125 | 65 | 0.1538 | 51 | 0.0588 | 65 | 0.1846 | 229 | 0.1354 | 31 | 0.129 | 75 | 0.08 | 55 | 0.0727 | 45 | 0.1111 | 206 | 0.0922 |

| | | | | | | | | | | | | | | | | | | | | |
|---|---|---|---|---|---|---|---|---|---|---|---|---|---|---|---|---|---|---|---|---|
| 11_1198932 | 51 | 0.6471 | 62 | 0.7581 | 51 | 0.7059 | 63 | 0.6667 | 227 | 0.696 | 34 | 0.7059 | 70 | 0.5714 | 54 | 0.537 | 46 | 0.5652 | 204 | 0.5833 |
| 11_1198933 | 41 | 0.0976 | 50 | 0.02 | 39 | 0.1538 | 51 | 0.0784 | 181 | 0.0829 | 24 | 0 | 53 | 0.0566 | 45 | 0.0444 | 37 | 0.0811 | 159 | 0.0503 |
| 11_1198974 | 9 | 0.5556 | 11 | 0.4545 | 13 | 0.3077 | 12 | 0.5 | 45 | 0.4444 | 6 | 0.5 | 10 | 0.5 | 9 | 0.3333 | 6 | 0.1667 | 31 | 0.3871 |
| 11_1198974 | 6 | 0.3333 | 10 | 0.1 | 11 | 0.0909 | 8 | 0 | 35 | 0.1143 | 5 | 0 | 10 | 0 | 8 | 0.125 | 5 | 0.2 | 28 | 0.0714 |
| 11_1198974 | 4 | 0 | 8 | 0.25 | 12 | 0.0833 | 8 | 0.375 | 32 | 0.1875 | 6 | 0.1667 | 9 | 0.1111 | 10 | 0.1 | 5 | 0 | 30 | 0.1 |
| 11_1198975 | 6 | 0.6667 | 11 | 0.6364 | 11 | 0.3636 | 11 | 0.4545 | 39 | 0.5128 | 7 | 0.5714 | 12 | 0.4167 | 8 | 1 | 6 | 0.3333 | 33 | 0.5758 |
| 11_1198975 | 8 | 0.125 | 11 | 0.1818 | 14 | 0.1429 | 12 | 0.1667 | 45 | 0.1556 | 6 | 0.1667 | 9 | 0.1111 | 8 | 0 | 5 | 0.2 | 28 | 0.1071 |
| 11_1198982 | 14 | 0 | 22 | 0.0909 | 21 | 0.0476 | 22 | 0.0455 | 79 | 0.0506 | 28 | 0.1071 | 25 | 0.12 | 18 | 0.0556 | 15 | 0 | 86 | 0.0814 |
| 11_1198986 | 11 | 0.0909 | 13 | 0.1538 | 14 | 0.1429 | 16 | 0.125 | 54 | 0.1296 | 16 | 0.0625 | 21 | 0.1429 | 15 | 0.1333 | 8 | 0 | 60 | 0.1 |
| 11_1198990 | 12 | 0 | 18 | 0.1111 | 15 | 0.3333 | 18 | 0 | 63 | 0.1111 | 22 | 0.0455 | 18 | 0.2778 | 17 | 0 | 14 | 0.2143 | 71 | 0.1268 |
| 11_1198991 | 15 | 0.2 | 17 | 0 | 14 | 0.0714 | 17 | 0.1176 | 63 | 0.0952 | 22 | 0.0455 | 17 | 0.1176 | 20 | 0.1 | 14 | 0.2143 | 73 | 0.1096 |
| 11_1198991 | 15 | 0.2667 | 17 | 0.1176 | 13 | 0.2308 | 14 | 0.2143 | 59 | 0.2034 | 22 | 0.2273 | 11 | 0.1818 | 16 | 0.0625 | 12 | 0.3333 | 61 | 0.1967 |
| 11_1198997 | 3 | 0.3333 | 5 | 0 | 6 | 0 | 8 | 0.125 | 22 | 0.0909 | 7 | 0 | 7 | 0.2857 | 8 | 0 | 4 | 0.25 | 26 | 0.1154 |
| 11_1198998 | 3 | 0 | 5 | 0.2 | 5 | 0.2 | 8 | 0 | 21 | 0.0952 | 5 | 0.2 | 8 | 0.25 | 8 | 0 | 3 | 0.3333 | 24 | 0.1667 |
| 11_1198999 | 2 | 0 | 5 | 0 | 5 | 0 | 7 | 0.1429 | 19 | 0.0526 | 7 | 0 | 8 | 0.125 | 7 | 0 | 3 | 0.6667 | 25 | 0.12 |
| 11_1199000 | 1 | 0 | 5 | 0.4 | 5 | 0.4 | 4 | 0.5 | 15 | 0.4 | 7 | 0.2857 | 5 | 0.4 | 5 | 0.4 | 3 | 0.3333 | 20 | 0.35 |
| 11_1199010 | 12 | 0.5 | 15 | 0.9333 | 13 | 0.4615 | 13 | 0.4615 | 53 | 0.6038 | 16 | 0.8125 | 13 | 0.7692 | 11 | 0.4545 | 9 | 0.4444 | 49 | 0.6531 |
| 11_1199013 | 13 | 0.2308 | 9 | 0.2222 | 11 | 0.3636 | 13 | 0.3846 | 46 | 0.3043 | 12 | 0.4167 | 13 | 0.5385 | 12 | 0.25 | 8 | 0.375 | 45 | 0.4 |
| 11_1199013 | 14 | 0.5714 | 12 | 0.5833 | 8 | 0.5 | 10 | 0.4 | 44 | 0.5227 | 10 | 0.7 | 14 | 0.5 | 9 | 0.6667 | 9 | 0.6667 | 42 | 0.619 |
| 11_1199015 | 8 | 0.5 | 13 | 0.6923 | 8 | 1 | 12 | 0.5 | 41 | 0.6585 | 10 | 0.9 | 12 | 0.75 | 10 | 0.8 | 7 | 0.5714 | 39 | 0.7692 |
| 11_1199015 | 12 | 0.8333 | 9 | 0.8889 | 10 | 0.8 | 10 | 0.6 | 41 | 0.7805 | 10 | 0.7 | 12 | 0.4167 | 11 | 0.6364 | 8 | 0.875 | 41 | 0.6341 |
| 11_1199016 | 11 | 0.5455 | 10 | 0.8 | 8 | 0.75 | 9 | 0.6667 | 38 | 0.6842 | 13 | 0.7692 | 13 | 0.6923 | 8 | 0.625 | 9 | 0.8889 | 43 | 0.7442 |
| 11_1199016 | 14 | 0.9286 | 11 | 0.9091 | 9 | 1 | 10 | 1 | 44 | 0.9545 | 14 | 1 | 13 | 0.9231 | 10 | 0.9 | 9 | 1 | 46 | 0.9565 |
| 11_1199016 | 15 | 0.0667 | 13 | 0.1538 | 11 | 0 | 12 | 0 | 51 | 0.0588 | 13 | 0 | 15 | 0.1333 | 10 | 0 | 8 | 0.125 | 46 | 0.0652 |
| 11_1199020 | 6 | 0.1667 | 9 | 0.1111 | 8 | 0 | 10 | 0 | 33 | 0.0606 | 9 | 0.1111 | 9 | 0 | 8 | 0.125 | 8 | 0 | 34 | 0.0588 |
| 11_1199020 | 7 | 0.5714 | 9 | 0.7778 | 5 | 0.6 | 10 | 0.8 | 31 | 0.7097 | 9 | 1 | 6 | 0.5 | 7 | 0.8571 | 8 | 0.875 | 30 | 0.8333 |
| 11_1199022 | 9 | 0.1111 | 8 | 0.25 | 6 | 0.5 | 6 | 0.1667 | 29 | 0.2414 | 8 | 0.125 | 6 | 0 | 8 | 0 | 5 | 0.2 | 27 | 0.0741 |
| 11_1199023 | 11 | 0.4545 | 7 | 0.7143 | 4 | 1 | 9 | 0.5556 | 31 | 0.6129 | 13 | 0.7692 | 8 | 0.75 | 7 | 0.7143 | 6 | 0.6667 | 34 | 0.7353 |
| 11_1199024 | 5 | 0.6 | 4 | 1 | 1 | 1 | 1 | 1 | 11 | 0.8182 | 5 | 1 | 2 | 1 | 4 | 1 | 4 | 1 | 15 | 1 |
| 11_1199024 | 7 | 0.1429 | 6 | 0.3333 | 2 | 0 | 3 | 0 | 18 | 0.1667 | 7 | 0.1429 | 6 | 0.3333 | 6 | 0 | 3 | 0 | 22 | 0.1364 |
| 11_1199025 | 9 | 0.1111 | 10 | 0.2 | 7 | 0 | 9 | 0.2222 | 35 | 0.1429 | 12 | 0 | 8 | 0.25 | 11 | 0 | 7 | 0.1429 | 38 | 0.0789 |
| 11_1199026 | 7 | 0.2857 | 7 | 0.7143 | 3 | 1 | 6 | 0.6667 | 23 | 0.6087 | 8 | 0.625 | 7 | 0.4286 | 4 | 0.75 | 3 | 0.6667 | 22 | 0.5909 |
| 11_1199026 | 8 | 1 | 10 | 0.8 | 6 | 1 | 9 | 1 | 33 | 0.9394 | 11 | 1 | 10 | 0.9 | 10 | 1 | 7 | 1 | 38 | 0.9737 |
| 11_1199026 | 8 | 0.625 | 6 | 0.8333 | 4 | 1 | 9 | 0.3333 | 27 | 0.6296 | 10 | 0.7 | 8 | 0.75 | 6 | 0.8333 | 5 | 0.8 | 29 | 0.7586 |
| 11_1199027 | 8 | 0.75 | 10 | 0.7 | 5 | 0.8 | 7 | 0.5714 | 30 | 0.7 | 11 | 0.7273 | 9 | 0.5556 | 8 | 0.875 | 8 | 0.625 | 36 | 0.6944 |
| 11_1199028 | 6 | 0.1667 | 4 | 0 | 4 | 0.25 | 6 | 0.1667 | 20 | 0.15 | 9 | 0.2222 | 6 | 0.3333 | 7 | 0.1429 | 6 | 0 | 28 | 0.1786 |
| 11_1199029 | 6 | 0.8333 | 9 | 0.8889 | 4 | 1 | 9 | 0.7778 | 28 | 0.8571 | 11 | 0.9091 | 9 | 0.8889 | 8 | 0.875 | 7 | 1 | 35 | 0.9143 |
| 11_1199817 | 2 | 0 | 3 | 0.3333 | NA | NA | 1 | 0 | 6 | 0.1667 | 3 | 0.6667 | NA | NA | 2 | 0 | 3 | 0 | 8 | 0.25 |
| 11_1199817 | 3 | 1 | 3 | 1 | NA | NA | 1 | 1 | 7 | 1 | 2 | 0.5 | 2 | 1 | 1 | 1 | 2 | 1 | 7 | 0.8571 |
| 11_1199818 | 3 | 1 | 3 | 0.6667 | NA | NA | 1 | 1 | 7 | 0.8571 | 1 | 0 | 2 | 1 | NA | NA | 3 | 0.6667 | 6 | 0.6667 |
| 11_1199818 | 3 | 0 | 3 | 0.3333 | NA | NA | NA | NA | 6 | 0.1667 | 2 | 0 | 2 | 0.5 | 1 | 0 | 2 | 0.5 | 7 | 0.2857 |
| 11_1199818 | 2 | 0 | 3 | 0 | NA | NA | 1 | 1 | 6 | 0.1667 | 1 | 0 | 2 | 0.5 | NA | NA | 3 | 0 | 6 | 0.1667 |
| 11_1199820 | 3 | 0.6667 | 3 | 0.6667 | NA | NA | 1 | 1 | 7 | 0.7143 | 2 | 1 | 2 | 1 | 2 | 1 | 2 | 1 | 8 | 1 |
| 11_1199820 | 3 | 1 | 2 | 1 | NA | NA | 1 | 1 | 6 | 1 | 2 | 0.5 | 1 | 1 | 1 | 0 | 2 | 1 | 6 | 0.6667 |
| 11_1199821 | 1 | 0 | 2 | 1 | NA | NA | 1 | 0 | 4 | 0.5 | 1 | 0 | 1 | 0 | 1 | 0 | 1 | 1 | 4 | 0.25 |
| 11_1199821 | 2 | 0 | 3 | 0 | NA | NA | 1 | 1 | 6 | 0.1667 | 1 | 1 | 2 | 0.5 | 1 | 0 | 1 | 0 | 5 | 0.4 |
| 11_1199822 | 3 | 0.6667 | 3 | 0 | NA | NA | 1 | 0 | 7 | 0.2857 | NA | NA | 2 | 0.5 | 1 | 0 | 2 | 1 | 5 | 0.6 |
| 11_1199823 | 3 | 1 | 3 | 1 | NA | NA | 1 | 1 | 7 | 1 | 1 | 1 | NA | NA | 1 | 1 | 2 | 1 | 4 | 1 |
| 11_1199824 | 3 | 0.3333 | 3 | 0.6667 | NA | NA | 1 | 1 | 7 | 0.5714 | 1 | 1 | 1 | 1 | 1 | 0 | 3 | 0.3333 | 6 | 0.5 |
| 11_1199825 | 3 | 0 | 2 | 0 | NA | NA | 1 | 1 | 6 | 0.1667 | 1 | 1 | 1 | 0 | 1 | 0 | 3 | 0 | 6 | 0.1667 |
| 11_1199825 | 2 | 1 | 2 | 1 | NA | NA | 1 | 1 | 5 | 1 | 1 | 1 | NA | NA | 2 | 0.5 | 3 | 0.6667 | 6 | 0.6667 |
| 11_1199825 | 3 | 0.3333 | 3 | 0 | NA | NA | 1 | 0 | 7 | 0.1429 | 1 | 0 | 1 | 0 | 1 | 0 | 3 | 0.3333 | 6 | 0.1667 |
| 11_1199912 | 4 | 0 | 1 | 0 | 5 | 0.4 | 5 | 0 | 15 | 0.1333 | 6 | 0.5 | 9 | 0.1111 | 6 | 0.1667 | 8 | 0 | 29 | 0.1724 |
| 11_1199912 | 1 | 1 | NA | NA | NA | NA | 2 | 0.5 | 3 | 0.6667 | 2 | 1 | NA | NA | NA | NA | NA | NA | 2 | 1 |
| 11_1199954 | 6 | 0.3333 | 7 | 0.5714 | 10 | 0.3 | 3 | 0.6667 | 26 | 0.4231 | 7 | 0.5714 | 12 | 0.4167 | 4 | 0.75 | 6 | 0.3333 | 29 | 0.4828 |
| 11_1199954 | 6 | 0.1667 | 5 | 0 | 11 | 0.0909 | 4 | 0.5 | 26 | 0.1538 | 6 | 0.1667 | 11 | 0.0909 | 3 | 0 | 5 | 0 | 25 | 0.08 |
| 11_1199955 | 4 | 0.5 | 6 | 0.6667 | 11 | 0.6364 | 4 | 0.75 | 25 | 0.64 | 5 | 0.6 | 10 | 0.8 | 4 | 0.75 | 4 | 0.5 | 23 | 0.6957 |
| 11_1199957 | 5 | 1 | 6 | 1 | 11 | 0.7273 | 3 | 0.6667 | 25 | 0.84 | 6 | 0.8333 | 12 | 0.75 | 4 | 1 | 5 | 1 | 27 | 0.8519 |
| 11_1199957 | 4 | 0 | 6 | 0.1667 | 8 | 0.125 | 3 | 0 | 21 | 0.0952 | 5 | 0.2 | 10 | 0.1 | 2 | 0 | 5 | 0.2 | 22 | 0.1364 |
| 11_1199959 | 5 | 0.6 | 6 | 0.3333 | 8 | 0.25 | 4 | 0.5 | 23 | 0.3913 | 6 | 0 | 12 | 0.4167 | 4 | 0.75 | 4 | 0.5 | 26 | 0.3846 |
| 11_1199960 | 3 | 0 | 4 | 0.25 | 9 | 0.2222 | 4 | 0 | 20 | 0.15 | 3 | 0 | 7 | 0.1429 | 2 | 0 | 5 | 0 | 17 | 0.0588 |
| 11_1199961 | 3 | 0 | 6 | 0 | 10 | 0 | 4 | 0.5 | 23 | 0.087 | 6 | 0 | 10 | 0.1 | 3 | 0.3333 | 5 | 0 | 24 | 0.0833 |
| 11_1199961 | 4 | 0.25 | 6 | 0.1667 | 10 | 0.2 | 4 | 0 | 24 | 0.1667 | 6 | 0.1667 | 10 | 0.1 | 3 | 0 | 5 | 0.6 | 24 | 0.2083 |
| 11_1199963 | 4 | 0.75 | 5 | 1 | 10 | 1 | 4 | 1 | 23 | 0.9565 | 6 | 0.8333 | 7 | 0.7143 | 3 | 0.6667 | 5 | 0.6 | 21 | 0.7143 |
| 11_1199964 | 3 | 0.6667 | 3 | 0 | 9 | 0.2222 | 4 | 0 | 19 | 0.2105 | 5 | 0 | 6 | 0.5 | 2 | 0 | 5 | 0.4 | 18 | 0.2778 |

| | | | | | | | | | | | | | | | | | | | | |
|---|---|---|---|---|---|---|---|---|---|---|---|---|---|---|---|---|---|---|---|---|
| 11_1199965 | 3 | 0.3333 | 4 | 0.5 | 10 | 0.8 | 3 | 0.6667 | 20 | 0.65 | 5 | 0.8 | 7 | 0.4286 | 2 | 1 | 5 | 0.6 | 19 | 0.6316 |
| 11_1199965 | 3 | 0 | 4 | 0 | 9 | 0.2222 | 2 | 0 | 18 | 0.1111 | 4 | 0 | 5 | 0 | 2 | 0 | 4 | 0.25 | 15 | 0.0667 |
| 11_1199966 | 2 | 1 | 4 | 0 | 8 | 0.375 | 3 | 0.6667 | 17 | 0.4118 | 5 | 0.2 | 6 | 0.3333 | 3 | 0.3333 | 4 | 0.5 | 18 | 0.3333 |
| 11_1200009 | 2 | 0 | 2 | 1 | 9 | 0.5556 | 3 | 0.6667 | 16 | 0.5625 | 3 | 0.6667 | 3 | 0.3333 | NA | NA | 2 | 0.5 | 8 | 0.5 |
| 11_1200783 | 1 | 0 | 1 | 1 | NA | NA | NA | NA | 2 | 0.5 | NA | NA | NA | NA | 1 | 1 | NA | NA | 1 | 1 |
| 11_1200805 | 1 | 0 | 1 | 1 | NA | NA | 1 | 1 | 3 | 0.6667 | 1 | 1 | NA | NA | 1 | 1 | 1 | 1 | 3 | 1 |
| 11_1201268 | 1 | 1 | NA | NA | NA | NA | NA | NA | 1 | 1 | 1 | 0 | 1 | 1 | NA | NA | NA | NA | 2 | 0.5 |
| 11_1201807 | 1 | 0 | 1 | 0 | 2 | 0 | 2 | 0.5 | 6 | 0.1667 | NA | NA | NA | NA | 1 | 1 | NA | NA | 1 | 1 |
| 11_1201815 | 2 | 0 | 1 | 0 | 2 | 0.5 | 2 | 0 | 7 | 0.1429 | NA | NA | NA | NA | 2 | 0.5 | NA | NA | 2 | 0.5 |
| 11_1201828 | 2 | 0.5 | 4 | 0.5 | 2 | 0.5 | 3 | 0.3333 | 11 | 0.4545 | NA | NA | NA | NA | 4 | 0.25 | NA | NA | 4 | 0.25 |
| 11_1202370 | 245 | 0.049 | 229 | 0.0611 | 205 | 0.0878 | 254 | 0.0433 | 933 | 0.0589 | 210 | 0.0952 | 214 | 0.0561 | 183 | 0.0656 | 182 | 0.0604 | 789 | 0.0697 |
| 11_1202371 | 244 | 0.0984 | 227 | 0.0925 | 203 | 0.1133 | 253 | 0.1107 | 927 | 0.1036 | 205 | 0.1561 | 210 | 0.119 | 185 | 0.1027 | 184 | 0.1522 | 784 | 0.1327 |
| 11_1202372 | 195 | 0.7282 | 170 | 0.7647 | 155 | 0.7548 | 192 | 0.7396 | 712 | 0.7458 | 162 | 0.784 | 160 | 0.7 | 146 | 0.7123 | 154 | 0.7078 | 622 | 0.7267 |
| 11_1202380 | 173 | 0.8266 | 160 | 0.8562 | 140 | 0.7929 | 203 | 0.8177 | 676 | 0.824 | 149 | 0.8591 | 160 | 0.8062 | 132 | 0.8636 | 145 | 0.8345 | 586 | 0.8396 |
| 11_1202419 | 3 | 0 | 2 | 0.5 | 3 | 0 | 1 | 0 | 9 | 0.1111 | 1 | 0 | 3 | 0.3333 | NA | NA | 1 | 1 | 5 | 0.4 |
| 11_1202606 | NA | NA | 1 | 1 | 2 | 0 | 1 | 1 | 4 | 0.5 | 2 | 1 | NA | NA | 2 | 0.5 | NA | NA | 4 | 0.75 |
| 11_1202649 | NA | NA | 1 | 0 | 1 | 0 | 1 | 1 | 3 | 0.3333 | NA | NA | NA | NA | 1 | 1 | NA | NA | 1 | 1 |
| 11_1202651 | NA | NA | 1 | 1 | 1 | 1 | 1 | 0 | 3 | 0.6667 | 1 | 0 | NA | NA | 2 | 1 | NA | NA | 3 | 0.6667 |
| 11_1202652 | NA | NA | 1 | 1 | 2 | 1 | 1 | 1 | 4 | 1 | 1 | 1 | NA | NA | 2 | 1 | NA | NA | 3 | 1 |
| 11_1202676 | NA | NA | 1 | 0 | 2 | 0.5 | 1 | 0 | 4 | 0.25 | 1 | 1 | NA | NA | 2 | 0.5 | NA | NA | 3 | 0.6667 |
| 11_1202678 | NA | NA | NA | NA | 2 | 1 | 1 | 1 | 3 | 1 | 1 | 0 | NA | NA | 1 | 1 | NA | NA | 2 | 0.5 |
| 11_1202678 | NA | NA | 1 | 1 | 2 | 1 | 1 | 1 | 4 | 1 | 1 | 1 | NA | NA | 1 | 1 | NA | NA | 2 | 1 |
| 11_1202681 | NA | NA | 1 | 0 | 2 | 0.5 | 1 | 1 | 4 | 0.5 | NA | NA | NA | NA | 1 | 1 | NA | NA | 1 | 1 |
| 11_1202711 | 1 | 1 | 1 | 1 | NA | NA | NA | NA | 2 | 1 | 1 | 1 | 2 | 0.5 | 2 | 0.5 | 2 | 1 | 7 | 0.7143 |
| 11_1202720 | 4 | 0.5 | 5 | 0.4 | 9 | 0.4444 | 4 | 0.25 | 22 | 0.4091 | 6 | 0.5 | 7 | 0.4286 | 7 | 0.2857 | 5 | 0.2 | 25 | 0.36 |
| 11_1202722 | 1 | 1 | 5 | 0.2 | 8 | 0.25 | 3 | 0 | 17 | 0.2353 | 4 | 0.25 | 6 | 0 | 6 | 0.1667 | 4 | 0.5 | 20 | 0.2 |
| 11_1202725 | 2 | 0 | 4 | 0.25 | 9 | 0.4444 | 1 | 0 | 16 | 0.3125 | 5 | 0 | 5 | 0 | 5 | 0.4 | 3 | 0.3333 | 18 | 0.1667 |
| 11_1202730 | 2 | 1 | 5 | 1 | 7 | 0.8571 | 3 | 1 | 17 | 0.9412 | 3 | 1 | 3 | 1 | 4 | 0.75 | 4 | 1 | 14 | 0.9286 |
| 11_1202730 | 2 | 0 | 4 | 0 | 8 | 0.125 | 3 | 0 | 17 | 0.0588 | 4 | 0.25 | 3 | 0 | 5 | 0.2 | 4 | 0.25 | 16 | 0.1875 |
| 11_1202731 | 1 | 0 | 3 | 0.3333 | 5 | 0.2 | 2 | 0 | 11 | 0.1818 | 2 | 0 | 3 | 0.3333 | 2 | 0 | 3 | 0.3333 | 10 | 0.2 |
| 11_1202731 | 2 | 0.5 | 5 | 0 | 6 | 0 | 3 | 0 | 16 | 0.0625 | 4 | 0 | 4 | 0.25 | 4 | 0 | 3 | 0 | 15 | 0.0667 |
| 11_1202732 | 2 | 1 | 3 | 0.6667 | 5 | 1 | 3 | 1 | 13 | 0.9231 | 2 | 1 | 4 | 0.5 | 3 | 0.3333 | 4 | 1 | 13 | 0.6923 |
| 11_1202733 | 2 | 0.5 | 4 | 0.5 | 4 | 0.5 | 3 | 1 | 13 | 0.6154 | 3 | 0.3333 | 3 | 0.3333 | 3 | 0.6667 | 4 | 0.25 | 13 | 0.3846 |
| 11_1202733 | 2 | 0 | 3 | 0.3333 | 6 | 0.3333 | 3 | 0 | 14 | 0.2143 | 2 | 0 | 4 | 0 | 5 | 0 | 2 | 0.5 | 13 | 0.0769 |
| 11_1203475 | 2 | 1 | 1 | 1 | 2 | 1 | 1 | 1 | 6 | 1 | NA | NA | NA | NA | 1 | 1 | 1 | 1 | 2 | 1 |
| 11_1203475 | 2 | 0.5 | 1 | 1 | 2 | 1 | 1 | 0 | 6 | 0.6667 | NA | NA | NA | NA | 2 | 0.5 | 1 | 1 | 3 | 0.6667 |
| 11_1203536 | 3 | 0.6667 | NA | NA | 1 | 0 | 1 | 1 | 5 | 0.6 | 2 | 0.5 | 1 | 1 | 1 | 1 | 3 | 0.3333 | 7 | 0.5714 |
| 11_1203544 | 2 | 0 | 1 | 0 | 2 | 0.5 | 2 | 0.5 | 7 | 0.2857 | 1 | 0 | 3 | 0 | 4 | 0 | 3 | 0.3333 | 11 | 0.0909 |
| 11_1203552 | 2 | 1 | NA | NA | 2 | 0 | 1 | 1 | 5 | 0.6 | 1 | 1 | 1 | 0 | 4 | 0 | NA | NA | 6 | 0.1667 |
| 11_1203552 | 2 | 0 | 1 | 0 | 1 | 1 | 1 | 0 | 5 | 0.2 | 1 | 0 | 1 | 1 | 4 | 0.5 | NA | NA | 6 | 0.5 |
| 11_1203558 | 4 | 0.5 | 4 | 0.25 | 2 | 0 | 2 | 0.5 | 12 | 0.3333 | 3 | 0.3333 | 3 | 0.6667 | 2 | 0.5 | 3 | 0.3333 | 11 | 0.4545 |
| 11_1203560 | 4 | 0 | 3 | 0.3333 | 2 | 1 | 2 | 0.5 | 11 | 0.3636 | 2 | 0 | 5 | 0.2 | 2 | 0 | 2 | 0.5 | 11 | 0.1818 |
| 11_1203561 | 4 | 0 | 2 | 0 | 2 | 0.5 | 1 | 0 | 9 | 0.1111 | 2 | 0 | 3 | 0 | 2 | 0.5 | 1 | 0 | 8 | 0.125 |
| 11_1203561 | 4 | 0.75 | 2 | 1 | 2 | 1 | 1 | 1 | 9 | 0.8889 | 2 | 1 | 5 | 0.8 | 2 | 1 | 1 | 1 | 10 | 0.9 |
| 11_1203562 | 4 | 1 | 1 | 1 | 2 | 1 | 1 | 1 | 8 | 1 | 2 | 1 | 4 | 1 | 2 | 1 | 1 | 1 | 9 | 1 |
| 11_1203562 | 3 | 0 | 2 | 1 | 2 | 0.5 | 1 | 0 | 8 | 0.375 | 1 | 1 | 3 | 0 | 2 | 0 | 1 | 0 | 7 | 0.1429 |
| 11_1203562 | 3 | 0 | 2 | 0.5 | 2 | 0.5 | 1 | 0 | 8 | 0.25 | 1 | 0 | 4 | 0 | 2 | 0.5 | 1 | 0 | 8 | 0.125 |
| 11_1203563 | 5 | 0 | 3 | 0 | 5 | 0.2 | 1 | 0 | 14 | 0.0714 | 2 | 0 | 6 | 0 | 2 | 0.5 | 1 | 0 | 11 | 0.0909 |
| 11_1203739 | NA | NA | NA | NA | 1 | 0 | 2 | 1 | 3 | 0.6667 | NA | NA | NA | NA | 2 | 0.5 | NA | NA | 2 | 0.5 |
| 11_1203746 | 5 | 0.4 | 9 | 0.1111 | 10 | 0.3 | 3 | 0.3333 | 27 | 0.2593 | 8 | 0.25 | 9 | 0.2222 | 5 | 0.4 | 5 | 0.2 | 27 | 0.2593 |
| 11_1203746 | 7 | 0.2857 | 8 | 0.25 | 8 | 0.125 | 2 | 1 | 25 | 0.28 | 4 | 0.25 | 6 | 0.1667 | 5 | 0.2 | 3 | 0.3333 | 18 | 0.2222 |
| 11_1203792 | 29 | 0.8621 | 16 | 0.8125 | 11 | 0.8182 | 14 | 1 | 70 | 0.8714 | 12 | 0.9167 | 19 | 0.9474 | 18 | 0.7778 | 20 | 0.85 | 69 | 0.8696 |
| 11_1203792 | 26 | 0.9231 | 15 | 0.9333 | 13 | 0.9231 | 14 | 1 | 68 | 0.9412 | 11 | 0.9091 | 18 | 1 | 16 | 0.875 | 19 | 0.9474 | 64 | 0.9375 |
| 11_1203793 | 25 | 0.2 | 14 | 0.2857 | 14 | 0.3571 | 14 | 0.2143 | 67 | 0.2537 | 10 | 0.2 | 17 | 0.1176 | 17 | 0.2353 | 18 | 0.1667 | 62 | 0.1774 |
| 11_1203794 | 24 | 0.75 | 12 | 0.8333 | 13 | 0.7692 | 12 | 1 | 61 | 0.8197 | 9 | 0.6667 | 19 | 0.6842 | 16 | 0.625 | 17 | 0.8235 | 61 | 0.7049 |
| 11_1203794 | 25 | 0.12 | 13 | 0.4615 | 15 | 0.2 | 14 | 0.6429 | 67 | 0.3134 | 13 | 0.6154 | 17 | 0.5882 | 19 | 0.2105 | 19 | 0.5263 | 68 | 0.4706 |
| 11_1203795 | 12 | 0.1667 | 4 | 0 | 5 | 0.2 | 2 | 0 | 23 | 0.1304 | 3 | 0 | 4 | 0.25 | 2 | 0 | 8 | 0 | 17 | 0.0588 |
| 11_1203890 | 3 | 0.6667 | 3 | 0.3333 | 1 | 0 | 2 | 0 | 9 | 0.3333 | 3 | 0.3333 | 1 | 0 | 4 | 0.25 | 4 | 0 | 12 | 0.1667 |
| 11_1203893 | 3 | 0.3333 | 3 | 0 | NA | NA | 3 | 0.3333 | 9 | 0.2222 | 3 | 0.3333 | 1 | 0 | 3 | 1 | 6 | 0.1667 | 13 | 0.3846 |
| 11_1203894 | 3 | 0.3333 | 5 | 0 | NA | NA | 3 | 0.6667 | 11 | 0.2727 | 3 | 0.6667 | 1 | 0 | 4 | 0.75 | 5 | 0 | 13 | 0.3846 |
| 11_1203894 | 2 | 0 | 4 | 0.25 | NA | NA | 2 | 0 | 8 | 0.125 | 2 | 0.5 | 1 | 0 | 3 | 0.3333 | 5 | 0 | 11 | 0.1818 |
| 11_1203895 | 3 | 0 | 5 | 0 | NA | NA | 3 | 0.3333 | 11 | 0.0909 | 3 | 0 | 1 | 0 | 3 | 0.3333 | 5 | 0 | 12 | 0.0833 |
| 11_1203896 | 2 | 1 | 3 | 1 | NA | NA | 3 | 0.3333 | 8 | 0.75 | 2 | 1 | 1 | 0 | 4 | 1 | 6 | 0.6667 | 13 | 0.7692 |
| 11_1203897 | 3 | 0.3333 | 6 | 0.3333 | NA | NA | 4 | 0.75 | 13 | 0.4615 | 2 | 0.5 | 1 | 1 | 4 | 0.5 | 6 | 0.1667 | 13 | 0.3846 |
| 11_1203900 | 1 | 1 | NA | NA | NA | NA | NA | NA | 1 | 1 | NA | NA | 1 | 0 | 1 | 1 | NA | NA | 2 | 0.5 |

| | | | | | | | | | | | | | | | | | | | | |
|---|---|---|---|---|---|---|---|---|---|---|---|---|---|---|---|---|---|---|---|---|
| 11_120433 | 11 | 0.6364 | 16 | 0.5625 | 14 | 0.5 | 9 | 0.5556 | 50 | 0.56 | 12 | 0.6667 | 12 | 0.5833 | 15 | 0.5333 | 13 | 0.3077 | 52 | 0.5192 |
| 11_120434 | 6 | 0 | 11 | 0.0909 | 11 | 0.0909 | 12 | 0 | 40 | 0.05 | 8 | 0.125 | 7 | 0.1429 | 10 | 0.2 | 10 | 0 | 35 | 0.1143 |
| 11_120434 | 12 | 0.25 | 16 | 0 | 18 | 0.2222 | 14 | 0.0714 | 60 | 0.1333 | 10 | 0.2 | 14 | 0.2143 | 16 | 0 | 16 | 0.1875 | 56 | 0.1429 |
| 11_120434 | 11 | 0.3636 | 15 | 0.2 | 17 | 0.2941 | 12 | 0.5 | 55 | 0.3273 | 10 | 0.3 | 12 | 0.4167 | 15 | 0.4 | 14 | 0.5 | 51 | 0.4118 |
| 11_120434 | 14 | 0.1429 | 16 | 0.0625 | 19 | 0.1053 | 13 | 0 | 62 | 0.0806 | 13 | 0.3077 | 14 | 0.0714 | 16 | 0 | 16 | 0 | 59 | 0.0847 |
| 11_120434 | 13 | 0.0769 | 16 | 0.1875 | 19 | 0.1053 | 13 | 0 | 61 | 0.0984 | 12 | 0.0833 | 13 | 0.1538 | 14 | 0 | 15 | 0.1333 | 54 | 0.0926 |
| 11_120451 | 19 | 0.3684 | 18 | 0.4444 | 13 | 0.7692 | 17 | 0.5882 | 67 | 0.5224 | 16 | 0.5625 | 22 | 0.7273 | 11 | 0.6364 | 13 | 0.6923 | 62 | 0.6613 |
| 11_120451 | 17 | 0.0588 | 17 | 0.0588 | 14 | 0.0714 | 14 | 0.2857 | 62 | 0.1129 | 14 | 0.2143 | 20 | 0.1 | 12 | 0.1667 | 14 | 0.0714 | 60 | 0.1333 |
| 11_120451 | 17 | 0.0588 | 16 | 0.25 | 13 | 0.0769 | 13 | 0.0769 | 59 | 0.1186 | 14 | 0.1429 | 20 | 0.1 | 11 | 0.0909 | 12 | 0.1667 | 57 | 0.1228 |
| 11_120451 | 19 | 0.2632 | 16 | 0.3125 | 12 | 0.1667 | 15 | 0.2 | 62 | 0.2419 | 15 | 0 | 22 | 0.2727 | 10 | 0.3 | 11 | 0.0909 | 58 | 0.1724 |
| 11_120451 | 15 | 0.0667 | 13 | 0.0769 | 12 | 0.3333 | 15 | 0.2 | 55 | 0.1636 | 11 | 0.2727 | 18 | 0.1667 | 11 | 0 | 10 | 0.2 | 50 | 0.16 |
| 11_120451 | 19 | 0.2105 | 16 | 0.3125 | 11 | 0.0909 | 12 | 0.1667 | 58 | 0.2069 | 15 | 0.3333 | 20 | 0.5 | 9 | 0.4444 | 12 | 0.1667 | 56 | 0.375 |
| 11_120452 | 13 | 0.1538 | 16 | 0.25 | 10 | 0.2 | 12 | 0.4167 | 51 | 0.2549 | 17 | 0.4118 | 21 | 0.381 | 9 | 0.2222 | 10 | 0.1 | 57 | 0.3158 |
| 11_120453 | 15 | 0.2 | 14 | 0.0714 | 11 | 0.0909 | 12 | 0 | 52 | 0.0962 | 8 | 0.25 | 19 | 0.1579 | 9 | 0 | 10 | 0.3 | 46 | 0.1739 |
| 11_120469 | 42 | 0.0952 | 46 | 0.1522 | 49 | 0.1224 | 41 | 0.0976 | 178 | 0.118 | 36 | 0.0556 | 40 | 0.05 | 42 | 0.1429 | 37 | 0.027 | 155 | 0.071 |
| 11_120469 | 49 | 0.0612 | 47 | 0.1064 | 51 | 0.098 | 44 | 0.1364 | 191 | 0.0995 | 39 | 0.0769 | 45 | 0.1778 | 42 | 0.0952 | 42 | 0.119 | 168 | 0.119 |
| 11_120470 | 31 | 0.0323 | 32 | 0.1875 | 29 | 0.069 | 26 | 0.0769 | 118 | 0.0932 | 27 | 0.1111 | 30 | 0.2 | 25 | 0.2 | 25 | 0.2 | 107 | 0.1776 |
| 11_120470 | 36 | 0.5 | 35 | 0.6 | 38 | 0.6053 | 31 | 0.5161 | 140 | 0.5571 | 27 | 0.6296 | 30 | 0.6667 | 26 | 0.4231 | 30 | 0.5333 | 113 | 0.5664 |
| 11_120496 | 16 | 0.0625 | 17 | 0.1176 | 16 | 0.125 | 17 | 0.0588 | 66 | 0.0909 | 12 | 0.25 | 27 | 0.037 | 21 | 0.0476 | 14 | 0 | 74 | 0.0676 |
| 11_120496 | 19 | 0.0526 | 17 | 0.1765 | 19 | 0.0526 | 19 | 0.1053 | 74 | 0.0946 | 12 | 0.0833 | 25 | 0.24 | 24 | 0.125 | 15 | 0.2667 | 76 | 0.1842 |
| 11_120496 | 17 | 0.1176 | 13 | 0.0769 | 15 | 0 | 16 | 0.0625 | 61 | 0.0656 | 10 | 0.1 | 21 | 0.1905 | 19 | 0.1053 | 12 | 0 | 62 | 0.1129 |
| 11_120500 | 7 | 1 | 15 | 0.7333 | 8 | 1 | 12 | 0.8333 | 42 | 0.8571 | 7 | 0.8571 | 7 | 0.8571 | 11 | 0.7273 | 4 | 1 | 29 | 0.8276 |
| 11_120501 | 4 | 0.25 | 13 | 0.0769 | 6 | 0.3333 | 6 | 0.6667 | 29 | 0.2759 | 7 | 0.1429 | 6 | 0.3333 | 10 | 0.4 | 3 | 0.3333 | 26 | 0.3077 |
| 11_120501 | 7 | 0.1429 | 15 | 0.2667 | 7 | 0 | 11 | 0.4545 | 40 | 0.25 | 8 | 0.25 | 7 | 0.2857 | 11 | 0.1818 | 5 | 0.4 | 31 | 0.2581 |
| 11_120501 | 7 | 0.2857 | 15 | 0.1333 | 7 | 0 | 11 | 0 | 40 | 0.1 | 8 | 0.375 | 5 | 0 | 12 | 0.1667 | 4 | 0.25 | 29 | 0.2069 |
| 11_120501 | 5 | 1 | 10 | 0.6 | 4 | 0.5 | 8 | 0.625 | 27 | 0.6667 | 6 | 0.1667 | 6 | 0.6667 | 10 | 0.4 | 3 | 0 | 25 | 0.36 |
| 11_120501 | 1 | 0 | 8 | 0.75 | 5 | 0.6 | 9 | 0.6667 | 23 | 0.6522 | 6 | 0.6667 | 5 | 0.2 | 8 | 0.375 | 4 | 0.5 | 23 | 0.4348 |
| 11_120501 | 5 | 1 | 13 | 1 | 6 | 1 | 11 | 1 | 35 | 1 | 6 | 0.8333 | 7 | 0.8571 | 8 | 0.75 | 5 | 0.8 | 26 | 0.8077 |
| 11_120501 | 4 | 1 | 9 | 0.7778 | 3 | 1 | 7 | 0.8571 | 23 | 0.8696 | 5 | 0.6 | 6 | 0.8333 | 7 | 0.4286 | 1 | 1 | 19 | 0.6316 |
| 11_120502 | 1 | 0 | 4 | 0.25 | 2 | 0 | 2 | 0 | 9 | 0.1111 | NA | NA | 1 | 1 | 1 | 0 | NA | NA | 2 | 0.5 |
| 11_120502 | 4 | 0 | 6 | 0.1667 | 4 | 0.25 | 10 | 0 | 24 | 0.0833 | 7 | 0.1429 | 4 | 0.25 | 8 | 0 | 1 | 0 | 20 | 0.1 |
| 11_120509 | 6 | 0.8333 | 6 | 1 | 4 | 0.75 | 3 | 0.3333 | 19 | 0.7895 | 1 | 0 | 4 | 0.75 | 3 | 1 | 2 | 0.5 | 10 | 0.7 |
| 11_120509 | 5 | 0.2 | 6 | 0.3333 | 4 | 0.75 | 3 | 0.6667 | 18 | 0.4444 | 1 | 0 | 4 | 0 | 3 | 0.3333 | 3 | 0.3333 | 11 | 0.1818 |
| 11_120509 | 4 | 0.75 | 6 | 0.6667 | 3 | 1 | 4 | 1 | 17 | 0.8235 | 1 | 1 | 4 | 0.75 | 4 | 1 | 1 | 1 | 10 | 0.9 |
| 11_120516 | 8 | 0.375 | 8 | 0.5 | 4 | 0.75 | 4 | 0.5 | 24 | 0.5 | 3 | 0.6667 | 6 | 0.3333 | 4 | 1 | 6 | 0.6667 | 19 | 0.6316 |
| 11_120516 | 4 | 0 | 9 | 0.2222 | 4 | 0 | 2 | 0 | 19 | 0.1053 | 3 | 0.3333 | 6 | 0 | 4 | 0 | 5 | 0 | 18 | 0.0556 |
| 11_120517 | 4 | 0 | 7 | 0 | 6 | 0 | 2 | 0.5 | 19 | 0.0526 | 4 | 0 | 5 | 0 | 4 | 0 | 5 | 0.2 | 18 | 0.0556 |
| 11_120517 | 3 | 0.3333 | 9 | 0.1111 | 4 | 0 | 3 | 0.3333 | 19 | 0.1579 | 4 | 0 | 5 | 0 | 4 | 0.25 | 4 | 0 | 17 | 0.0588 |
| 11_120517 | 2 | 1 | 8 | 0.25 | 5 | 0.4 | 3 | 1 | 18 | 0.5 | 1 | 1 | 5 | 0.6 | 4 | 1 | 4 | 0.5 | 14 | 0.7143 |
| 11_120519 | 2 | 0 | 2 | 0 | 3 | 0 | 2 | 0.5 | 9 | 0.1111 | 4 | 0 | 3 | 0 | 3 | 0.3333 | 2 | 0 | 12 | 0.0833 |
| 11_120520 | 3 | 0.6667 | 1 | 0 | 2 | 1 | NA | NA | 6 | 0.6667 | 2 | 1 | 2 | 0.5 | 3 | 1 | 3 | 0.3333 | 10 | 0.7 |
| 11_120520 | 3 | 0.6667 | 1 | 0 | 2 | 0 | 1 | 0 | 7 | 0.2857 | 2 | 0.5 | 1 | 0 | 3 | 1 | 2 | 0.5 | 8 | 0.625 |
| 11_120520 | 3 | 1 | 1 | 1 | 2 | 1 | 1 | 1 | 7 | 1 | 1 | 1 | 1 | 1 | 3 | 1 | 3 | 1 | 8 | 1 |
| 11_120523 | NA | NA | 1 | 1 | NA | NA | 1 | 0 | 2 | 0.5 | NA | NA | NA | NA | 1 | 1 | 1 | 1 | 2 | 1 |
| 11_120524 | NA | NA | 1 | 1 | NA | NA | 1 | 1 | 2 | 1 | NA | NA | NA | NA | 1 | 1 | 1 | 1 | 2 | 1 |
| 11_120524 | NA | NA | 2 | 0.5 | NA | NA | 1 | 0 | 3 | 0.3333 | NA | NA | NA | NA | 2 | 0.5 | 1 | 1 | 3 | 0.6667 |
| 11_120524 | NA | NA | 2 | 1 | NA | NA | 1 | 1 | 3 | 1 | 1 | 1 | NA | NA | 2 | 0.5 | 1 | 0 | 4 | 0.5 |
| 11_120524 | NA | NA | 2 | 1 | NA | NA | 1 | 1 | 3 | 1 | 1 | 0 | NA | NA | 1 | 1 | 1 | 0 | 3 | 0.3333 |
| 11_120524 | NA | NA | 2 | 0.5 | NA | NA | 1 | 1 | 3 | 0.6667 | 1 | 1 | NA | NA | 1 | 1 | 1 | 1 | 3 | 1 |
| 11_120525 | NA | NA | 1 | 1 | NA | NA | 1 | 1 | 2 | 1 | 1 | 1 | NA | NA | 2 | 1 | 1 | 1 | 4 | 1 |
| 11_120525 | NA | NA | 2 | 0.5 | NA | NA | 1 | 1 | 3 | 0.6667 | 1 | 1 | NA | NA | 2 | 0.5 | 1 | 0 | 4 | 0.5 |
| 11_120536 | 2 | 1 | 3 | 1 | 3 | 1 | 2 | 1 | 10 | 1 | 1 | 1 | 2 | 1 | 2 | 1 | NA | NA | 5 | 1 |
| 11_120537 | 3 | 1 | 4 | 0.75 | 3 | 1 | 2 | 1 | 12 | 0.9167 | 1 | 0 | 1 | 1 | 2 | 1 | 1 | 1 | 5 | 0.8 |
| 11_120537 | 2 | 1 | 3 | 1 | 2 | 1 | 2 | 1 | 9 | 1 | 1 | 1 | 1 | 1 | 2 | 1 | 1 | 1 | 5 | 1 |
| 11_120537 | 3 | 0.6667 | 4 | 0.75 | 3 | 1 | 2 | 1 | 12 | 0.8333 | 1 | 1 | 2 | 0.5 | 2 | 0.5 | 1 | 0 | 6 | 0.5 |
| 11_120537 | 2 | 1 | 3 | 1 | 3 | 1 | 2 | 1 | 10 | 1 | 1 | 1 | 1 | 1 | 2 | 1 | 1 | 1 | 5 | 1 |
| 11_120537 | 1 | 1 | 4 | 0.25 | 2 | 1 | NA | NA | 7 | 0.5714 | 1 | 1 | NA | NA | 1 | 1 | NA | NA | 2 | 1 |
| 11_120537 | 1 | 1 | 4 | 0 | 2 | 0.5 | 1 | 1 | 8 | 0.375 | 1 | 1 | NA | NA | NA | NA | NA | NA | 1 | 1 |
| 11_120537 | NA | NA | 3 | 0.3333 | 2 | 0.5 | 2 | 0.5 | 7 | 0.4286 | 1 | 1 | NA | NA | NA | NA | NA | NA | 1 | 1 |
| 11_120538 | NA | NA | 4 | 0.75 | 2 | 1 | 2 | 1 | 8 | 0.875 | 1 | 1 | NA | NA | NA | NA | NA | NA | 1 | 1 |
| 11_120538 | NA | NA | 2 | 1 | 1 | 1 | 1 | 1 | 4 | 1 | 1 | 1 | NA | NA | NA | NA | NA | NA | 1 | 1 |
| 11_120579 | 4 | 0 | 2 | 0 | 6 | 0.3333 | 4 | 0.25 | 16 | 0.1875 | 4 | 0 | 2 | 0 | 2 | 0 | 3 | 0.3333 | 11 | 0.0909 |
| 11_120579 | 5 | 0.2 | 2 | 0.5 | 7 | 0 | 3 | 0 | 17 | 0.1176 | 6 | 0 | 2 | 0 | 2 | 0 | 5 | 0.2 | 15 | 0.0667 |
| 11_120609 | 1 | 0 | 2 | 0.5 | NA | NA | 1 | 1 | 4 | 0.5 | 1 | 0 | NA | NA | 5 | 0.6 | 2 | 0 | 8 | 0.375 |
| 11_120611 | 3 | 0.6667 | 2 | 1 | 1 | 0 | 3 | 0.3333 | 9 | 0.5556 | 1 | 0 | 1 | 0 | 6 | 0.3333 | 2 | 0 | 10 | 0.2 |

| | | | | | | | | | | | | | | | | | | | | |
|---|---|---|---|---|---|---|---|---|---|---|---|---|---|---|---|---|---|---|---|---|
| 11_1206118 | 2 | 0 | 2 | 0 | 1 | 0 | 2 | 0.5 | 7 | 0.1429 | 1 | 0 | NA | NA | 5 | 0.2 | NA | NA | 6 | 0.1667 |
| 11_1206119 | 2 | 1 | 2 | 0.5 | 1 | 1 | 3 | 0 | 8 | 0.5 | 1 | 1 | 1 | 1 | 8 | 0.25 | 2 | 0.5 | 12 | 0.4167 |
| 11_1206145 | 3 | 0 | 5 | 0.2 | 3 | 0 | NA | NA | 11 | 0.0909 | 4 | 0 | 3 | 0 | 4 | 0.25 | 1 | 0 | 12 | 0.0833 |
| 11_1206158 | NA | NA | NA | NA | NA | NA | 1 | 1 | 1 | 1 | 1 | 1 | NA | NA | NA | NA | 1 | 0 | 2 | 0.5 |
| 11_1206162 | NA | NA | 1 | 0 | 1 | 0 | 2 | 1 | 4 | 0.5 | 6 | 0 | 4 | 0.25 | 2 | 0 | 7 | 0.1429 | 19 | 0.1053 |
| 11_1206717 | 9 | 0.4444 | 7 | 0.1429 | 17 | 0.2941 | 11 | 0.2727 | 44 | 0.2955 | 9 | 0 | 13 | 0.3846 | 7 | 0.4286 | 7 | 0.4286 | 36 | 0.3056 |
| 11_1206718 | 10 | 0.7 | 12 | 0.9167 | 19 | 0.7368 | 11 | 0.5455 | 52 | 0.7308 | 10 | 0.9 | 13 | 0.6154 | 8 | 1 | 7 | 0.7143 | 38 | 0.7895 |
| 11_1206718 | 9 | 0 | 11 | 0.0909 | 18 | 0 | 12 | 0.1667 | 50 | 0.06 | 9 | 0.1111 | 13 | 0 | 7 | 0.1429 | 6 | 0.1667 | 35 | 0.0857 |
| 11_1206718 | 10 | 0 | 12 | 0.0833 | 20 | 0.1 | 12 | 0.0833 | 54 | 0.0741 | 10 | 0 | 13 | 0.0769 | 7 | 0.1429 | 6 | 0.1667 | 36 | 0.0833 |
| 11_1206718 | 10 | 0.2 | 12 | 0.0833 | 18 | 0.0556 | 12 | 0.0833 | 52 | 0.0962 | 10 | 0.1 | 14 | 0.1429 | 6 | 0.1667 | 7 | 0.1429 | 37 | 0.1351 |
| 11_1206718 | 8 | 1 | 9 | 1 | 18 | 0.6111 | 11 | 0.9091 | 46 | 0.8261 | 9 | 0.7778 | 11 | 0.9091 | 5 | 0.8 | 6 | 1 | 31 | 0.871 |
| 11_1206721 | 8 | 0.25 | 12 | 0.25 | 17 | 0.1765 | 11 | 0.2727 | 48 | 0.2292 | 10 | 0.4 | 14 | 0.2857 | 7 | 0.1429 | 7 | 0.1429 | 38 | 0.2632 |
| 11_1206722 | 8 | 0.125 | 12 | 0.5 | 15 | 0.2 | 13 | 0.0769 | 48 | 0.2292 | 9 | 0.1111 | 14 | 0.1429 | 6 | 0.1667 | 6 | 0.1667 | 35 | 0.1429 |
| 11_1206722 | 5 | 0.2 | 9 | 0 | 12 | 0.0833 | 11 | 0.0909 | 37 | 0.0811 | 9 | 0.1111 | 10 | 0 | 4 | 0.25 | 5 | 0 | 28 | 0.0714 |
| 11_1206723 | 7 | 0.1429 | 9 | 0.4444 | 15 | 0.2 | 12 | 0.3333 | 43 | 0.2791 | 9 | 0.1111 | 13 | 0.1538 | 5 | 0.2 | 6 | 0.3333 | 33 | 0.1818 |
| 11_1206723 | 7 | 0 | 9 | 0.2222 | 9 | 0 | 12 | 0.0833 | 37 | 0.0811 | 7 | 0 | 8 | 0.125 | 5 | 0.2 | 6 | 0.1667 | 26 | 0.1154 |
| 11_1206723 | 7 | 0.4286 | 10 | 0.6 | 16 | 0.375 | 11 | 0.1818 | 44 | 0.3864 | 8 | 0.125 | 12 | 0.5 | 6 | 0.3333 | 6 | 0.3333 | 32 | 0.3438 |
| 11_1206724 | 6 | 0 | 8 | 0 | 14 | 0 | 13 | 0.3077 | 41 | 0.0976 | 8 | 0 | 11 | 0.2727 | 5 | 0.2 | 6 | 0.1667 | 30 | 0.1667 |
| 11_1206774 | 15 | 0 | 16 | 0 | 15 | 0.1333 | 22 | 0.0909 | 68 | 0.0588 | 11 | 0.1818 | 10 | 0 | 16 | 0.125 | 12 | 0.25 | 49 | 0.1429 |
| 11_1206787 | NA | NA | NA | NA | NA | NA | 1 | 1 | 1 | 1 | NA | NA | 1 | 1 | NA | NA | NA | NA | 1 | 1 |
| 11_1206796 | 21 | 0.4762 | 21 | 0.4762 | 24 | 0.5417 | 27 | 0.4444 | 93 | 0.4839 | 17 | 0.4706 | 17 | 0.4118 | 21 | 0.5714 | 20 | 0.3 | 75 | 0.44 |
| 11_1206797 | 23 | 0.3043 | 22 | 0.0455 | 23 | 0.2174 | 28 | 0.1429 | 96 | 0.1771 | 17 | 0.2353 | 16 | 0 | 24 | 0.0417 | 17 | 0.1765 | 74 | 0.1081 |
| 11_1206798 | 21 | 0.0476 | 22 | 0.0455 | 25 | 0.12 | 27 | 0.0741 | 95 | 0.0737 | 17 | 0.0588 | 16 | 0.0625 | 26 | 0.0385 | 20 | 0.05 | 79 | 0.0506 |
| 11_1206800 | 2 | 1 | 3 | 1 | 2 | 1 | 4 | 0.5 | 11 | 0.8182 | NA | NA | 3 | 1 | 5 | 0.6 | 4 | 0.5 | 12 | 0.6667 |
| 11_1206801 | 4 | 0.75 | 3 | 0.6667 | 1 | 1 | 5 | 0.8 | 13 | 0.7692 | NA | NA | 3 | 0.6667 | 4 | 0.75 | 2 | 0.5 | 9 | 0.6667 |
| 11_1206803 | 4 | 0.25 | 4 | 0.25 | 2 | 0 | 4 | 0.25 | 14 | 0.2143 | NA | NA | 3 | 0.3333 | 5 | 0.2 | 2 | 0.5 | 10 | 0.3 |
| 11_1206973 | 1 | 0 | 1 | 0 | 1 | 1 | 1 | 1 | 4 | 0.5 | NA | NA | 1 | 0 | 1 | 1 | NA | NA | 2 | 0.5 |
| 11_1206979 | 19 | 0.5789 | 27 | 0.5926 | 23 | 0.7391 | 21 | 0.8095 | 90 | 0.6778 | 12 | 0.5833 | 23 | 0.6522 | 13 | 0.8462 | 23 | 0.6522 | 71 | 0.6761 |
| 11_1206980 | 18 | 0.1111 | 27 | 0.1481 | 24 | 0.2083 | 20 | 0.3 | 89 | 0.191 | 12 | 0.0833 | 23 | 0.087 | 18 | 0.2778 | 23 | 0.1739 | 76 | 0.1579 |
| 11_1206980 | 17 | 0.1765 | 21 | 0.2857 | 23 | 0.1739 | 19 | 0.1579 | 80 | 0.2 | 9 | 0.4444 | 24 | 0.0833 | 18 | 0.2778 | 17 | 0.1176 | 68 | 0.1912 |
| 11_1206981 | 20 | 0.1 | 25 | 0 | 26 | 0.0769 | 22 | 0.1364 | 93 | 0.0753 | 10 | 0 | 27 | 0.037 | 20 | 0.1 | 23 | 0.087 | 80 | 0.0625 |
| 11_1206982 | 18 | 0.1111 | 16 | 0.125 | 22 | 0.1364 | 16 | 0.25 | 72 | 0.1528 | 9 | 0.1111 | 21 | 0.1429 | 16 | 0.125 | 19 | 0.0526 | 65 | 0.1077 |
| 11_1206985 | 16 | 0.125 | 15 | 0.1333 | 18 | 0.0556 | 16 | 0 | 65 | 0.0769 | 8 | 0.125 | 18 | 0 | 10 | 0.1 | 16 | 0.1875 | 52 | 0.0962 |
| 11_1206986 | 13 | 0 | 20 | 0.1 | 15 | 0.0667 | 13 | 0.0769 | 61 | 0.0656 | 5 | 0 | 18 | 0.1667 | 9 | 0.1111 | 17 | 0.0588 | 49 | 0.102 |
| 11_1207045 | 6 | 0.3333 | 7 | 0.2857 | 7 | 0 | 6 | 0.1667 | 26 | 0.1923 | 1 | 0 | 6 | 0 | 4 | 0.25 | 2 | 0 | 13 | 0.0769 |
| 11_1207046 | 5 | 0.4 | 6 | 0.1667 | 5 | 0.2 | 5 | 0 | 21 | 0.1905 | 3 | 0.3333 | 5 | 0.2 | 1 | 0 | 1 | 0 | 10 | 0.2 |
| 11_1207061 | 5 | 0.2 | 4 | 0.5 | 4 | 0.25 | 4 | 0.25 | 17 | 0.2941 | 2 | 0 | 3 | 0 | 2 | 0.5 | 2 | 0 | 9 | 0.1111 |
| 11_1207062 | 5 | 0 | 3 | 0 | 4 | 0.25 | 5 | 0 | 17 | 0.0588 | 1 | 1 | 4 | 0.5 | 2 | 0.5 | 2 | 0 | 9 | 0.4444 |
| 11_1207097 | 1 | 0 | 3 | 0 | 2 | 0 | 1 | 1 | 7 | 0.1429 | 1 | 0 | 3 | 0.3333 | NA | NA | 1 | 0 | 5 | 0.2 |
| 11_1207175 | 1 | 1 | NA | NA | 2 | 0 | 3 | 0 | 6 | 0.1667 | NA | NA | NA | NA | 2 | 0.5 | 2 | 0 | 4 | 0.25 |
| 11_1207175 | 1 | 1 | 1 | 0 | 2 | 1 | 2 | 1 | 6 | 0.8333 | NA | NA | NA | NA | 2 | 1 | 2 | 1 | 4 | 1 |
| 11_1208474 | 1 | 1 | NA | NA | NA | NA | NA | NA | 1 | 1 | NA | NA | NA | NA | 1 | 1 | 1 | 1 | 2 | 1 |
| 11_1208475 | 1 | 1 | NA | NA | NA | NA | NA | NA | 1 | 1 | 1 | 1 | NA | NA | 1 | 1 | 1 | 1 | 3 | 1 |
| 11_1208488 | 1 | 1 | NA | NA | NA | NA | 1 | 1 | 2 | 1 | 1 | 1 | NA | NA | 1 | 1 | 1 | 1 | 3 | 1 |
| 11_1208490 | 1 | 1 | NA | NA | NA | NA | 2 | 1 | 3 | 1 | 1 | 1 | NA | NA | 1 | 1 | 1 | 1 | 3 | 1 |
| 11_1208528 | 10 | 0 | 14 | 0 | 8 | 0.375 | 12 | 0 | 44 | 0.0682 | 12 | 0.0833 | 14 | 0.0714 | 10 | 0.1 | 9 | 0.1111 | 45 | 0.0889 |
| 11_1208529 | 10 | 0.2 | 14 | 0 | 9 | 0.1111 | 8 | 0.125 | 41 | 0.0976 | 15 | 0.2667 | 12 | 0.25 | 11 | 0.2727 | 9 | 0.2222 | 47 | 0.2553 |
| 11_1208530 | 9 | 0.2222 | 14 | 0.2143 | 10 | 0 | 7 | 0.2857 | 40 | 0.175 | 13 | 0.0769 | 11 | 0.1818 | 9 | 0.1111 | 13 | 0.2308 | 46 | 0.1522 |
| 11_1208534 | 8 | 0 | 10 | 0.2 | 8 | 0.25 | 4 | 0.25 | 30 | 0.1667 | 11 | 0.1818 | 9 | 0.2222 | 4 | 0 | 8 | 0.125 | 32 | 0.1562 |
| 11_1208535 | 8 | 0.375 | 12 | 0.1667 | 9 | 0 | 5 | 0.4 | 34 | 0.2059 | 8 | 0.375 | 6 | 0.3333 | 4 | 0.5 | 9 | 0.3333 | 27 | 0.3704 |
| 11_1208535 | 7 | 0 | 11 | 0 | 9 | 0.2222 | 5 | 0.2 | 32 | 0.0938 | 9 | 0 | 9 | 0.2222 | 4 | 0 | 10 | 0 | 32 | 0.0625 |
| 11_1208536 | 9 | 0.2222 | 11 | 0 | 8 | 0 | 5 | 0.2 | 33 | 0.0909 | 8 | 0.25 | 9 | 0 | 6 | 0.3333 | 11 | 0.1818 | 34 | 0.1765 |
| 11_1208543 | 4 | 0.25 | 9 | 0.2222 | 7 | 0.4286 | 1 | 0 | 21 | 0.2857 | 7 | 0 | 6 | 0.3333 | 6 | 0.3333 | 6 | 0.1667 | 25 | 0.2 |
| 11_1208544 | 5 | 0.4 | 9 | 0 | 7 | 0.1429 | 3 | 0 | 24 | 0.125 | 6 | 0.1667 | 7 | 0 | 6 | 0.1667 | 5 | 0.2 | 24 | 0.125 |
| 11_1208545 | 2 | 1 | 4 | 1 | 5 | 1 | 3 | 1 | 14 | 1 | 5 | 0.4 | 3 | 1 | 4 | 1 | 2 | 1 | 14 | 0.7857 |
| 11_1208545 | 3 | 0.3333 | 5 | 0.4 | 5 | 0.8 | 4 | 0.25 | 17 | 0.4706 | 5 | 0.2 | 5 | 0.4 | 3 | 0.6667 | 4 | 0.5 | 17 | 0.4118 |
| 11_1208546 | 3 | 0 | 8 | 0.125 | 5 | 0 | 3 | 0 | 19 | 0.0526 | 5 | 0 | 4 | 0 | 4 | 0.25 | 5 | 0 | 18 | 0.0556 |
| 11_1208546 | 1 | 1 | 3 | 1 | 3 | 0.3333 | 4 | 0.25 | 11 | 0.5455 | 5 | 0.4 | 4 | 0.25 | 4 | 0.25 | 4 | 0.5 | 17 | 0.3529 |
| 11_1210957 | NA | NA | 1 | 0 | NA | NA | 1 | 1 | 2 | 0.5 | NA | NA | 1 | 1 | NA | NA | NA | NA | 1 | 1 |
| 11_1210960 | NA | NA | 1 | 1 | NA | NA | NA | NA | 1 | 1 | NA | NA | 1 | 1 | NA | NA | NA | NA | 1 | 1 |
| 11_1210966 | NA | NA | 1 | 1 | NA | NA | NA | NA | 1 | 1 | NA | NA | 2 | 0.5 | 1 | 1 | NA | NA | 3 | 0.6667 |
| 11_1210968 | NA | NA | 1 | 1 | NA | NA | NA | NA | 1 | 1 | NA | NA | 2 | 0.5 | 1 | 1 | NA | NA | 3 | 0.6667 |
| 11_1210969 | NA | NA | 1 | 1 | NA | NA | NA | NA | 1 | 1 | NA | NA | 2 | 0.5 | 1 | 1 | NA | NA | 3 | 0.6667 |
| 11_1210971 | NA | NA | 1 | 1 | NA | NA | NA | NA | 1 | 1 | NA | NA | 1 | 1 | 1 | 1 | NA | NA | 2 | 1 |
| 11_1210972 | NA | NA | 1 | 1 | NA | NA | 1 | 1 | 2 | 1 | NA | NA | 3 | 1 | 1 | 1 | NA | NA | 4 | 1 |

| | | | | | | | | | | | | | | | | | | | | |
|---|---|---|---|---|---|---|---|---|---|---|---|---|---|---|---|---|---|---|---|---|
| 11_1210974 | NA | NA | 1 | 1 | NA | NA | NA | NA | 1 | 1 | NA | NA | 3 | 1 | 1 | 1 | NA | NA | 4 | 1 |
| 11_1210976 | NA | NA | NA | NA | NA | NA | 1 | 1 | 1 | 1 | NA | NA | 4 | 0.75 | 1 | 1 | NA | NA | 5 | 0.8 |
| 11_1210977 | NA | NA | NA | NA | NA | NA | 1 | 1 | 1 | 1 | NA | NA | 4 | 0.75 | 1 | 1 | NA | NA | 5 | 0.8 |
| 11_1210977 | NA | NA | NA | NA | NA | NA | 1 | 1 | 1 | 1 | NA | NA | 4 | 0.25 | 1 | 0 | NA | NA | 5 | 0.2 |
| 11_1211117 | 4 | 0.5 | 4 | 0.5 | 3 | 0.3333 | NA | NA | 11 | 0.4545 | 1 | 0 | 1 | 1 | 1 | 0 | 1 | 1 | 4 | 0.5 |
| 11_1211131 | 4 | 0.5 | 5 | 0.2 | 2 | 0 | 1 | 1 | 12 | 0.3333 | NA | NA | 2 | 0 | 1 | 0 | 2 | 0.5 | 5 | 0.2 |
| 11_1211132 | 3 | 0 | 5 | 0 | 4 | 0.25 | 1 | 0 | 13 | 0.0769 | 2 | 0 | 3 | 0 | 2 | 0.5 | 1 | 0 | 8 | 0.125 |
| 11_1211132 | 3 | 0 | 6 | 0.3333 | 4 | 0 | 2 | 1 | 15 | 0.2667 | 2 | 0.5 | 4 | 0.25 | 1 | 0 | 1 | 1 | 8 | 0.375 |
| 11_1211133 | 4 | 0.25 | 6 | 0.1667 | 4 | 0.25 | 2 | 0.5 | 16 | 0.25 | 3 | 0.3333 | 4 | 0.5 | 2 | 0.5 | 2 | 0.5 | 11 | 0.4545 |
| 11_1211133 | 3 | 1 | 4 | 0.75 | 4 | 1 | 3 | 1 | 14 | 0.9286 | 1 | 1 | 4 | 0.75 | 2 | 1 | 1 | 1 | 8 | 0.875 |
| 11_1211137 | 4 | 0.25 | 1 | 0 | 2 | 0 | 1 | 0 | 8 | 0.125 | 1 | 0 | 1 | 0 | NA | NA | 1 | 1 | 3 | 0.3333 |
| 11_1211137 | 2 | 0.5 | 1 | 0 | 2 | 0 | 1 | 1 | 6 | 0.3333 | 2 | 0.5 | 3 | 0.6667 | 1 | 1 | 2 | 1 | 8 | 0.75 |
| 11_1211137 | 4 | 0.25 | 1 | 1 | 2 | 0.5 | 2 | 0 | 9 | 0.3333 | 2 | 0.5 | 3 | 0.3333 | 1 | 0 | 2 | 0.5 | 8 | 0.375 |
| 11_1211138 | 4 | 0.25 | 1 | 1 | 2 | 0 | 2 | 0.5 | 9 | 0.3333 | 1 | 1 | 3 | 0.3333 | NA | NA | 2 | 0 | 6 | 0.3333 |
| 11_1211139 | 3 | 1 | NA | NA | 1 | 1 | 2 | 1 | 6 | 1 | 2 | 1 | 2 | 1 | 1 | 1 | 2 | 1 | 7 | 1 |
| 11_1211140 | 3 | 0.6667 | NA | NA | NA | NA | 2 | 1 | 5 | 0.8 | 1 | 1 | 1 | 1 | NA | NA | 2 | 0.5 | 4 | 0.75 |
| 11_1211141 | 4 | 0.25 | NA | NA | 1 | 0 | 2 | 0 | 7 | 0.1429 | 2 | 0 | 2 | 0.5 | 1 | 1 | 2 | 0 | 7 | 0.2857 |
| 11_1211141 | 4 | 0.25 | NA | NA | 1 | 0 | 1 | 0 | 6 | 0.1667 | 2 | 0 | 2 | 0.5 | 1 | 0 | 2 | 0 | 7 | 0.1429 |
| 11_1211141 | 4 | 0.25 | NA | NA | 1 | 0 | 2 | 0.5 | 7 | 0.2857 | 2 | 0 | 2 | 0.5 | 1 | 0 | 2 | 0 | 7 | 0.1429 |
| 11_1211143 | 4 | 0.25 | NA | NA | 1 | 0 | 2 | 0 | 7 | 0.1429 | NA | NA | NA | NA | 1 | 1 | 2 | 0 | 3 | 0.3333 |
| 11_1211143 | 4 | 0.5 | NA | NA | 1 | 0 | 2 | 0 | 7 | 0.2857 | 1 | 1 | 1 | 0 | 1 | 0 | 2 | 1 | 5 | 0.6 |
| 11_1211144 | 3 | 1 | NA | NA | 1 | 1 | 2 | 1 | 6 | 1 | NA | NA | 2 | 1 | 1 | 1 | 1 | 1 | 4 | 1 |
| 11_1211147 | 1 | 1 | NA | NA | 1 | 1 | 2 | 0.5 | 4 | 0.75 | 1 | 1 | NA | NA | 1 | 1 | 2 | 1 | 4 | 1 |
| 11_1211149 | 2 | 1 | NA | NA | 1 | 0 | 2 | 1 | 5 | 0.8 | 1 | 1 | 2 | 1 | 1 | 1 | 2 | 0.5 | 6 | 0.8333 |
| 11_1211437 | 6 | 0.6667 | 3 | 0.6667 | 5 | 0.8 | 2 | 0 | 16 | 0.625 | 7 | 0.7143 | 8 | 0.5 | 3 | 1 | 5 | 0.8 | 23 | 0.6957 |
| 11_1211438 | 8 | 0.25 | 7 | 0.5714 | 6 | 0.3333 | 3 | 0.6667 | 24 | 0.4167 | 5 | 0.6 | 7 | 0.1429 | 3 | 0.3333 | 4 | 0 | 19 | 0.2632 |
| 11_1211439 | 4 | 0.25 | 6 | 0.3333 | 6 | 0.1667 | 3 | 0 | 19 | 0.2105 | 2 | 0 | 5 | 0.2 | 3 | 0 | 4 | 0 | 14 | 0.0714 |
| 11_1211445 | 7 | 0.5714 | 5 | 0.2 | 5 | 0.6 | 3 | 0.6667 | 20 | 0.5 | 7 | 0.2857 | 8 | 0.375 | 4 | 0.25 | 3 | 0.3333 | 22 | 0.3182 |
| 11_1211446 | 8 | 0.125 | 7 | 0 | 7 | 0.1429 | 1 | 0 | 23 | 0.087 | 8 | 0.125 | 9 | 0.2222 | 4 | 0 | 3 | 0 | 24 | 0.125 |
| 11_1211895 | 4 | 0 | 4 | 0 | 4 | 0.25 | 5 | 0 | 17 | 0.0588 | 3 | 0 | 4 | 0 | 2 | 0 | 4 | 0.25 | 13 | 0.0769 |
| 11_1211895 | 5 | 0 | 5 | 0 | 3 | 0 | 5 | 0.2 | 18 | 0.0556 | 4 | 0.25 | 4 | 0 | 2 | 0 | 4 | 0 | 14 | 0.0714 |
| 11_1211905 | 3 | 0 | 6 | 0 | 3 | 0.6667 | 3 | 0 | 15 | 0.1333 | 3 | 0.3333 | 3 | 0 | 2 | 0 | 4 | 0.25 | 12 | 0.1667 |
| 11_1211977 | 4 | 1 | 8 | 0.625 | 4 | 1 | 6 | 1 | 22 | 0.8636 | 4 | 0.25 | 7 | 0.8571 | 1 | 1 | 5 | 0.8 | 17 | 0.7059 |
| 11_1211978 | 5 | 0.2 | 7 | 0.1429 | 6 | 0.1667 | 7 | 0.2857 | 25 | 0.2 | 4 | 0 | 7 | 0.4286 | 2 | 0 | 7 | 0 | 20 | 0.15 |
| 11_1211978 | 4 | 0 | 6 | 0.1667 | 6 | 0.1667 | 4 | 0 | 20 | 0.1 | 3 | 0.3333 | 5 | 0.2 | 2 | 0.5 | 5 | 0 | 15 | 0.2 |
| 11_1211979 | 5 | 1 | 7 | 1 | 5 | 0.8 | 6 | 1 | 23 | 0.9565 | 3 | 1 | 8 | 0.875 | 2 | 1 | 5 | 1 | 18 | 0.9444 |
| 11_1211979 | 4 | 0.25 | 5 | 0.2 | 6 | 0 | 3 | 0 | 18 | 0.1111 | 3 | 0 | 4 | 0 | 1 | 0 | 5 | 0.2 | 13 | 0.0769 |
| 11_1211979 | 5 | 0.8 | 7 | 0.5714 | 7 | 0.7143 | 5 | 1 | 24 | 0.75 | 6 | 0.6667 | 8 | 0.875 | 1 | 0 | 5 | 0.4 | 20 | 0.65 |
| 11_1211979 | 5 | 0.6 | 3 | 1 | 8 | 0.75 | 8 | 0.75 | 24 | 0.75 | 6 | 0.8333 | 8 | 0.5 | 2 | 0.5 | 6 | 0.5 | 22 | 0.5909 |
| 11_1211979 | 5 | 0.8 | 6 | 1 | 8 | 0.875 | 7 | 0.8571 | 26 | 0.8846 | 6 | 0.8333 | 8 | 1 | 2 | 1 | 6 | 0.8333 | 22 | 0.9091 |
| 11_1211980 | 3 | 0.3333 | 6 | 0.3333 | 3 | 0 | 5 | 0.4 | 17 | 0.2941 | 4 | 0.25 | 3 | 0 | NA | NA | 3 | 0.3333 | 10 | 0.2 |
| 11_1211980 | 3 | 0.3333 | 7 | 0.1429 | 6 | 0.1667 | 6 | 0 | 22 | 0.1364 | 6 | 0.3333 | 7 | 0 | 2 | 0 | 6 | 0.5 | 21 | 0.2381 |
| 11_1211981 | 5 | 1 | 8 | 1 | 9 | 1 | 8 | 1 | 30 | 1 | 6 | 1 | 7 | 1 | 2 | 1 | 7 | 1 | 22 | 1 |
| 11_1211981 | 5 | 0.6 | 6 | 0.8333 | 7 | 0.8571 | 4 | 1 | 22 | 0.8182 | 6 | 0.6667 | 5 | 1 | 1 | 1 | 7 | 0.7143 | 19 | 0.7895 |
| 11_1211982 | 5 | 0.8 | 8 | 0.625 | 8 | 0.625 | 8 | 0.5 | 29 | 0.6207 | 6 | 0.6667 | 8 | 0.75 | 2 | 0 | 7 | 0.5714 | 23 | 0.6087 |
| 11_1211982 | 5 | 0.6 | 9 | 0.6667 | 10 | 1 | 7 | 0.4286 | 31 | 0.7097 | 6 | 0.3333 | 8 | 0.5 | 2 | 1 | 7 | 0.7143 | 23 | 0.5652 |
| 11_1211982 | 4 | 0.5 | 9 | 0.8889 | 9 | 0.5556 | 7 | 0.4286 | 29 | 0.6207 | 6 | 0.8333 | 8 | 0.375 | 2 | 0.5 | 6 | 0.5 | 22 | 0.5455 |
| 11_1211983 | 7 | 0.4286 | 5 | 0.2 | 10 | 0.5 | 7 | 0.1429 | 29 | 0.3448 | 3 | 0 | 8 | 0.25 | 2 | 0.5 | 4 | 0.5 | 17 | 0.2941 |
| 11_1211984 | 7 | 0 | 10 | 0.1 | 9 | 0.1111 | 9 | 0 | 35 | 0.0571 | 8 | 0.125 | 11 | 0.1818 | 2 | 0 | 7 | 0.1429 | 28 | 0.1429 |
| 11_1211985 | 7 | 0.1429 | 8 | 0.125 | 9 | 0.3333 | 9 | 0.3333 | 33 | 0.2424 | 6 | 0.5 | 9 | 0.3333 | 2 | 0 | 7 | 0.1429 | 24 | 0.2917 |
| 11_1211985 | 7 | 0 | 10 | 0.3 | 9 | 0.2222 | 10 | 0.4 | 36 | 0.25 | 8 | 0.125 | 12 | 0.0833 | 2 | 0.5 | 7 | 0.2857 | 29 | 0.1724 |
| 11_1211988 | 9 | 0 | 8 | 0 | 10 | 0.1 | 11 | 0.1818 | 38 | 0.0789 | 9 | 0 | 12 | 0.0833 | 3 | 0 | 4 | 0.25 | 28 | 0.0714 |
| 11_1211996 | 3 | 0.3333 | 3 | 0.3333 | 7 | 0.1429 | 7 | 0.1429 | 20 | 0.2 | 6 | 0 | 7 | 0.1429 | 3 | 0 | 6 | 0.1667 | 22 | 0.0909 |
| 11_1211997 | 1 | 0 | 3 | 0 | 6 | 0.3333 | 6 | 0.5 | 16 | 0.3125 | 6 | 0.3333 | 5 | 0.4 | 3 | 0 | 4 | 0 | 18 | 0.2222 |
| 11_1211997 | 4 | 0 | 3 | 0.3333 | 9 | 0.3333 | 6 | 0 | 22 | 0.1818 | 6 | 0.1667 | 6 | 0 | 4 | 0 | 5 | 0.2 | 21 | 0.0952 |
| 11_1212005 | 5 | 0 | 3 | 0 | 10 | 0.2 | 9 | 0.2222 | 27 | 0.1481 | 8 | 0.125 | 7 | 0.4286 | 5 | 0 | 6 | 0.1667 | 26 | 0.1923 |
| 11_1212188 | 9 | 0.1111 | 7 | 0.2857 | 6 | 0 | 5 | 0 | 27 | 0.1111 | 9 | 0.2222 | 10 | 0 | 12 | 0.1667 | 6 | 0.3333 | 37 | 0.1622 |
| 11_1212189 | 8 | 0.375 | 7 | 0.1429 | 3 | 0.3333 | 5 | 0 | 23 | 0.2174 | 8 | 0.125 | 8 | 0.25 | 8 | 0.25 | 5 | 0 | 29 | 0.1724 |
| 11_1212190 | 7 | 0.8571 | 7 | 0.5714 | 5 | 1 | 4 | 1 | 23 | 0.8261 | 8 | 0.625 | 9 | 0.7778 | 7 | 0.8571 | 5 | 0.8 | 29 | 0.7586 |
| 11_1212191 | 7 | 0.8571 | 8 | 0.75 | 5 | 0.6 | 2 | 0 | 22 | 0.6818 | 6 | 0.5 | 9 | 0.5556 | 9 | 0.8889 | 6 | 0.8333 | 30 | 0.7 |
| 11_1212191 | 8 | 0.25 | 6 | 0 | 4 | 0.25 | 2 | 0 | 20 | 0.15 | 7 | 0.4286 | 9 | 0.1111 | 11 | 0.0909 | 6 | 0.5 | 33 | 0.2424 |
| 11_1212469 | 24 | 0.0833 | 9 | 0.2222 | 12 | 0.1667 | 16 | 0 | 61 | 0.0984 | 16 | 0 | 23 | 0.1304 | 17 | 0.0588 | 23 | 0.1739 | 79 | 0.1013 |
| 11_1212474 | 25 | 0.04 | 7 | 0 | 11 | 0.0909 | 13 | 0.1538 | 56 | 0.0714 | 11 | 0.0909 | 17 | 0.1176 | 14 | 0.0714 | 19 | 0.1053 | 61 | 0.0984 |
| 11_1212475 | 22 | 0.2273 | 7 | 0 | 8 | 0 | 11 | 0.0909 | 48 | 0.125 | 9 | 0 | 15 | 0.2667 | 12 | 0.25 | 16 | 0.0625 | 52 | 0.1538 |
| 11_1212475 | 22 | 0.5909 | 7 | 0.5714 | 9 | 0.5556 | 13 | 0.6154 | 51 | 0.5882 | 11 | 0.4545 | 16 | 0.6875 | 13 | 0.7692 | 17 | 0.5882 | 57 | 0.6316 |

| | | | | | | | | | | | | | | | | | | | | |
|---|---|---|---|---|---|---|---|---|---|---|---|---|---|---|---|---|---|---|---|---|
| 11_1212476 | 23 | 0.4348 | 8 | 0.375 | 8 | 0.25 | 12 | 0.4167 | 51 | 0.3922 | 9 | 0.3333 | 16 | 0.375 | 15 | 0.4 | 20 | 0.2 | 60 | 0.3167 |
| 11_1212476 | 25 | 0.24 | 7 | 0.4286 | 9 | 0.2222 | 12 | 0.1667 | 53 | 0.2453 | 9 | 0.3333 | 19 | 0.1579 | 16 | 0.1875 | 17 | 0.1176 | 61 | 0.1803 |
| 11_1212476 | 26 | 0.8462 | 7 | 0.7143 | 9 | 0.7778 | 14 | 0.7857 | 56 | 0.8036 | 11 | 0.4545 | 19 | 0.7895 | 16 | 0.625 | 20 | 0.7 | 66 | 0.6667 |
| 11_1212477 | 23 | 0.6957 | 6 | 0.6667 | 8 | 0.625 | 10 | 0.8 | 47 | 0.7021 | 8 | 0.75 | 18 | 0.6111 | 16 | 0.5 | 20 | 0.65 | 62 | 0.6129 |
| 11_1212478 | 21 | 0.381 | 8 | 0 | 8 | 0 | 11 | 0.2727 | 48 | 0.2292 | 7 | 0.1429 | 18 | 0.2778 | 15 | 0.1333 | 17 | 0.2353 | 57 | 0.2105 |
| 11_1212478 | 21 | 0.8095 | 8 | 0.625 | 7 | 0.8571 | 12 | 0.75 | 48 | 0.7708 | 9 | 0.8889 | 19 | 0.7368 | 15 | 0.8 | 17 | 0.8824 | 60 | 0.8167 |
| 11_1212479 | 21 | 0.0952 | 8 | 0 | 7 | 0 | 14 | 0.0714 | 50 | 0.06 | 9 | 0 | 16 | 0.0625 | 15 | 0.0667 | 15 | 0.0667 | 55 | 0.0545 |
| 11_1213203 | 2 | 1 | 3 | 0.6667 | 6 | 1 | 3 | 1 | 14 | 0.9286 | 1 | 1 | 2 | 0.5 | 3 | 1 | 3 | 1 | 9 | 0.8889 |
| 11_1213205 | 1 | 0 | 2 | 0 | 2 | 0.5 | 2 | 0.5 | 7 | 0.2857 | 1 | 0 | 1 | 0 | 2 | 0.5 | 4 | 0.25 | 8 | 0.25 |
| 11_1213206 | 2 | 0 | 4 | 0 | 8 | 0.125 | 3 | 0 | 17 | 0.0588 | 2 | 0 | 2 | 0 | 4 | 0.25 | 5 | 0 | 13 | 0.0769 |
| 11_1213215 | 4 | 0 | 6 | 0 | 9 | 0.2222 | 1 | 0 | 20 | 0.1 | 2 | 0.5 | 4 | 0.25 | 5 | 0.2 | 6 | 0.3333 | 17 | 0.2941 |
| 11_1213410 | 10 | 0.1 | 11 | 0.3636 | 10 | 0.3 | 6 | 0 | 37 | 0.2162 | 8 | 0.375 | 7 | 0.4286 | 4 | 0.25 | 5 | 0.2 | 24 | 0.3333 |
| 11_1213412 | 8 | 0.125 | 12 | 0.25 | 11 | 0.2727 | 6 | 0 | 37 | 0.1892 | 8 | 0.125 | 6 | 0.1667 | 5 | 0 | 6 | 0.6667 | 25 | 0.24 |
| 11_1215078 | 5 | 0.2 | 6 | 0.1667 | 2 | 0 | 2 | 0 | 15 | 0.1333 | 4 | 0.25 | 4 | 0.5 | 3 | 0 | 5 | 0.2 | 16 | 0.25 |
| 11_1215079 | 4 | 0.75 | 3 | 0.6667 | 2 | 1 | 2 | 1 | 11 | 0.8182 | 3 | 1 | 2 | 0.5 | 3 | 0.6667 | 6 | 0.6667 | 14 | 0.7143 |
| 11_1215986 | 1 | 1 | NA | NA | NA | NA | NA | NA | 1 | 1 | 1 | 1 | NA | NA | NA | NA | NA | NA | 1 | 1 |
| 11_1215987 | 1 | 1 | 1 | 0 | NA | NA | NA | NA | 2 | 0.5 | 1 | 1 | NA | NA | NA | NA | NA | NA | 1 | 1 |
| 11_1215987 | 1 | 1 | 1 | 0 | NA | NA | NA | NA | 2 | 0.5 | 1 | 1 | NA | NA | NA | NA | NA | NA | 1 | 1 |
| 11_1215987 | 1 | 1 | NA | NA | NA | NA | NA | NA | 1 | 1 | 1 | 1 | NA | NA | NA | NA | NA | NA | 1 | 1 |
| 11_1216068 | 1 | 1 | NA | NA | NA | NA | NA | NA | 1 | 1 | 1 | 1 | NA | NA | NA | NA | NA | NA | 1 | 1 |
| 12_3283627 | 1 | 1 | NA | NA | NA | NA | 2 | 0.5 | 3 | 0.6667 | NA | NA | NA | NA | NA | NA | 1 | 1 | 1 | 1 |
| 12_3284127 | NA | NA | NA | NA | NA | NA | 1 | 1 | 1 | 1 | NA | NA | NA | NA | NA | NA | 1 | 1 | 1 | 1 |
| 12_3284219 | NA | NA | NA | NA | NA | NA | 1 | 1 | 1 | 1 | NA | NA | NA | NA | NA | NA | 1 | 1 | 1 | 1 |
| 12_3284449 | NA | NA | NA | NA | NA | NA | 1 | 1 | 1 | 1 | NA | NA | NA | NA | NA | NA | 1 | 1 | 1 | 1 |
| 12_3284518 | NA | NA | NA | NA | NA | NA | 1 | 1 | 1 | 1 | NA | NA | NA | NA | NA | NA | 1 | 1 | 1 | 1 |
| 12_3284787 | NA | NA | NA | NA | NA | NA | 1 | 1 | 1 | 1 | NA | NA | NA | NA | NA | NA | 1 | 1 | 1 | 1 |
| 12_3284870 | NA | NA | NA | NA | NA | NA | 1 | 1 | 1 | 1 | NA | NA | NA | NA | NA | NA | 1 | 1 | 1 | 1 |
| 12_3284945 | NA | NA | NA | NA | NA | NA | 1 | 1 | 1 | 1 | NA | NA | NA | NA | NA | NA | 1 | 1 | 1 | 1 |
| 12_3285432 | NA | NA | NA | NA | NA | NA | 1 | 1 | 1 | 1 | NA | NA | NA | NA | NA | NA | 1 | 1 | 1 | 1 |
| 12_3292935 | NA | NA | 1 | 0 | 5 | 0.2 | 2 | 0 | 8 | 0.125 | 1 | 1 | 1 | 0 | 2 | 0 | 2 | 0.5 | 6 | 0.3333 |
| 12_3299315 | 9 | 0.1111 | 20 | 0.05 | 11 | 0.1818 | 10 | 0.2 | 50 | 0.12 | 9 | 0 | 12 | 0.0833 | 5 | 0 | 9 | 0.1111 | 35 | 0.0571 |
| 12_3299625 | 13 | 0.2308 | 24 | 0.625 | 12 | 0.0833 | 16 | 0.5625 | 65 | 0.4308 | 12 | 0.4167 | 18 | 0.5556 | 7 | 0.7143 | 9 | 0.6667 | 46 | 0.5652 |
| 12_3300022 | 1 | 0 | NA | NA | 1 | 1 | 1 | 1 | 3 | 0.6667 | NA | NA | NA | NA | 1 | 1 | NA | NA | 1 | 1 |
| 12_3300237 | 1 | 1 | NA | NA | NA | NA | NA | NA | 1 | 1 | NA | NA | NA | NA | 1 | 1 | NA | NA | 1 | 1 |
| 12_3359369 | 10 | 0 | 16 | 0.25 | 8 | 0.125 | 12 | 0.0833 | 46 | 0.1304 | 9 | 0.2222 | 12 | 0.1667 | 4 | 0 | 8 | 0.25 | 33 | 0.1818 |
| 12_3415490 | 3 | 0 | 2 | 0 | 2 | 0 | 3 | 0.3333 | 10 | 0.1 | 1 | 1 | 3 | 0 | 1 | 0 | 4 | 0.25 | 9 | 0.2222 |
| 12_3415544 | 3 | 0 | 3 | 0.3333 | 1 | 0 | 3 | 0 | 10 | 0.1 | 1 | 0 | 2 | 0 | 1 | 0 | 3 | 0.6667 | 7 | 0.2857 |
| 12_3415772 | 3 | 0.3333 | 4 | 0 | 2 | 0 | 5 | 0.4 | 14 | 0.2143 | NA | NA | 3 | 0 | 2 | 0 | 3 | 0.3333 | 8 | 0.125 |
| 12_3415811 | 2 | 0.5 | 4 | 0 | 4 | 0.25 | 4 | 0 | 14 | 0.1429 | 1 | 0 | 2 | 1 | 2 | 1 | 5 | 0 | 10 | 0.4 |
| 12_3415968 | 4 | 0.75 | 3 | 0.3333 | 4 | 0.75 | 4 | 0.5 | 15 | 0.6 | 1 | 1 | 3 | 0.3333 | 2 | 0.5 | 5 | 0.6 | 11 | 0.5455 |
| 12_3416094 | 4 | 0 | 4 | 0 | 3 | 0 | 7 | 0.4286 | 18 | 0.1667 | 1 | 0 | 3 | 0.3333 | 2 | 0 | 5 | 0 | 11 | 0.0909 |
| 12_3416107 | 4 | 0.75 | 4 | 0.5 | 4 | 0.25 | 7 | 0.5714 | 19 | 0.5263 | 1 | 1 | 3 | 0.3333 | 2 | 0 | 5 | 0.2 | 11 | 0.2727 |
| 12_3416228 | 4 | 0.75 | 4 | 0.75 | 5 | 0.6 | 7 | 0.8571 | 20 | 0.75 | NA | NA | 3 | 1 | 2 | 1 | 5 | 0.4 | 10 | 0.7 |
| 12_3416275 | 4 | 0 | 4 | 0 | 5 | 0 | 6 | 0.1667 | 19 | 0.0526 | 1 | 0 | 3 | 0.3333 | 2 | 0 | 5 | 0.2 | 11 | 0.1818 |
| 12_3416308 | 3 | 0 | 4 | 0.25 | 3 | 0.3333 | 6 | 0.3333 | 16 | 0.25 | 1 | 1 | 3 | 0.3333 | 2 | 0.5 | 5 | 0 | 11 | 0.2727 |
| 12_3416516 | 3 | 0.3333 | 4 | 0 | 4 | 0 | 7 | 0 | 18 | 0.0556 | 1 | 1 | 3 | 0.3333 | 2 | 0 | 6 | 0 | 12 | 0.1667 |
| 12_3416567 | 4 | 1 | 4 | 0.25 | 4 | 0.25 | 7 | 0.4286 | 19 | 0.4737 | 1 | 0 | 4 | 0.5 | 2 | 1 | 4 | 0.5 | 11 | 0.5455 |
| 12_3416736 | 2 | 0.5 | 4 | 0.5 | 4 | 0.75 | 5 | 0.6 | 15 | 0.6 | 2 | 1 | 3 | 1 | 2 | 1 | 6 | 0.8333 | 13 | 0.9231 |
| 12_3416942 | 5 | 0.2 | 4 | 0 | 5 | 0.4 | 6 | 0.5 | 20 | 0.3 | 2 | 1 | 5 | 0.6 | 3 | 0.6667 | 5 | 0.4 | 15 | 0.6 |
| 12_3416969 | 3 | 0.3333 | 5 | 0.4 | 4 | 0.25 | 9 | 0.4444 | 21 | 0.381 | 1 | 0 | 5 | 0.6 | 2 | 0 | 5 | 0.6 | 13 | 0.4615 |
| 12_3417116 | 6 | 0.1667 | 7 | 0 | 4 | 0 | 8 | 0.25 | 25 | 0.12 | 4 | 0.25 | 6 | 0.8333 | 6 | 0.6667 | 5 | 0.4 | 21 | 0.5714 |
| 12_3453175 | 9 | 0.1111 | 9 | 0.1111 | 11 | 0.0909 | 17 | 0.2353 | 46 | 0.1522 | 10 | 0.1 | 13 | 0.1538 | 13 | 0.2308 | 12 | 0.3333 | 48 | 0.2083 |
| 12_3453332 | 11 | 0 | 9 | 0 | 12 | 0.25 | 18 | 0.3333 | 50 | 0.18 | 7 | 0 | 13 | 0.3077 | 13 | 0.2308 | 11 | 0.1818 | 44 | 0.2045 |
| 12_3453394 | 12 | 0.75 | 9 | 0.6667 | 6 | 0.5 | 15 | 0.8667 | 42 | 0.7381 | 8 | 0.5 | 13 | 0.8462 | 13 | 0.5385 | 9 | 0.7778 | 43 | 0.6744 |
| 12_3453427 | 11 | 0.0909 | 9 | 0.1111 | 10 | 0.1 | 15 | 0.2667 | 45 | 0.1556 | 10 | 0.1 | 11 | 0.5455 | 15 | 0.1333 | 9 | 0.4444 | 45 | 0.2889 |
| 12_3453459 | 8 | 0.875 | 7 | 0.8571 | 11 | 0.8182 | 14 | 0.7857 | 40 | 0.825 | 8 | 0.625 | 12 | 1 | 10 | 0.9 | 8 | 1 | 38 | 0.8947 |
| 12_3453476 | 11 | 0 | 10 | 0.1 | 11 | 0.0909 | 17 | 0.0588 | 49 | 0.0612 | 10 | 0 | 11 | 0.1818 | 11 | 0 | 11 | 0.1818 | 43 | 0.093 |
| 12_3453507 | 12 | 0 | 9 | 0.2222 | 11 | 0 | 17 | 0.1176 | 49 | 0.0816 | 9 | 0 | 14 | 0 | 16 | 0.125 | 12 | 0.1667 | 51 | 0.0784 |
| 12_3455257 | 1 | 0 | 1 | 0 | 1 | 1 | NA | NA | 3 | 0.3333 | 2 | 0.5 | NA | NA | NA | NA | 1 | 0 | 3 | 0.3333 |
| 12_3455560 | NA | NA | 2 | 0 | 1 | 1 | NA | NA | 3 | 0.3333 | 1 | 0 | NA | NA | NA | NA | 1 | 1 | 2 | 0.5 |
| 12_3455583 | 1 | 0 | 2 | 1 | 1 | 1 | 1 | 1 | 5 | 0.8 | 2 | 0 | 1 | 1 | NA | NA | NA | NA | 3 | 0.3333 |
| 12_3455630 | 1 | 0 | 1 | 0 | 1 | 1 | 1 | 1 | 4 | 0.5 | 2 | 0.5 | 1 | 0 | NA | NA | 1 | 0 | 4 | 0.25 |
| 12_3455688 | 4 | 0.5 | 3 | 0.6667 | 3 | 0.3333 | 7 | 0.7143 | 17 | 0.5882 | 6 | 0.3333 | 3 | 0.3333 | 2 | 1 | 3 | 1 | 14 | 0.5714 |
| 12_3455832 | 5 | 0.2 | 3 | 0 | 5 | 0.2 | 7 | 0.4286 | 20 | 0.25 | 8 | 0.25 | 2 | 0 | 3 | 0 | 5 | 0.2 | 18 | 0.1667 |
| 12_3457723 | NA | NA | 1 | 0 | 2 | 0.5 | 1 | 0 | 4 | 0.25 | 2 | 0.5 | NA | NA | NA | NA | NA | NA | 2 | 0.5 |

| | | | | | | | | | | | | | | | | | | | | |
|---|---|---|---|---|---|---|---|---|---|---|---|---|---|---|---|---|---|---|---|---|
| 12_355019 | NA | NA | 1 | 1 | NA | NA | 1 | 0 | 2 | 0.5 | 1 | 1 | 1 | 0 | NA | NA | NA | NA | 2 | 0.5 |
| 12_355021 | NA | NA | 1 | 1 | NA | NA | 2 | 0.5 | 3 | 0.6667 | 1 | 0 | 1 | 1 | NA | NA | NA | NA | 2 | 0.5 |
| 12_355030 | NA | NA | 1 | 1 | NA | NA | 1 | 0 | 2 | 0.5 | 1 | 1 | 1 | 0 | NA | NA | NA | NA | 2 | 0.5 |
| 12_355034 | NA | NA | NA | NA | NA | NA | 2 | 0.5 | 2 | 0.5 | NA | NA | 1 | 1 | NA | NA | NA | NA | 1 | 1 |
| 12_355044 | NA | NA | NA | NA | NA | NA | 2 | 0.5 | 2 | 0.5 | 1 | 1 | 2 | 0 | NA | NA | NA | NA | 3 | 0.3333 |
| 12_355088 | NA | NA | NA | NA | NA | NA | 1 | 1 | 1 | 1 | 1 | 0 | 2 | 0.5 | NA | NA | NA | NA | 3 | 0.3333 |
| 12_355090 | NA | NA | 1 | 1 | NA | NA | 2 | 1 | 3 | 1 | 1 | 1 | 2 | 1 | NA | NA | NA | NA | 3 | 1 |
| 12_355128 | NA | NA | 1 | 1 | NA | NA | 2 | 1 | 3 | 1 | 1 | 1 | 1 | 1 | NA | NA | NA | NA | 2 | 1 |
| 12_355131 | NA | NA | 2 | 1 | NA | NA | 1 | 1 | 3 | 1 | 1 | 1 | 2 | 1 | NA | NA | NA | NA | 3 | 1 |
| 12_355142 | NA | NA | 2 | 1 | NA | NA | 2 | 1 | 4 | 1 | 1 | 1 | 2 | 1 | NA | NA | NA | NA | 3 | 1 |
| 12_355150 | NA | NA | 2 | 1 | NA | NA | 2 | 1 | 4 | 1 | 1 | 1 | 1 | 1 | NA | NA | NA | NA | 2 | 1 |
| 12_355176 | NA | NA | 2 | 1 | NA | NA | 2 | 1 | 4 | 1 | 1 | 1 | 2 | 1 | NA | NA | NA | NA | 3 | 1 |
| 12_355188 | NA | NA | 1 | 1 | NA | NA | 2 | 0.5 | 3 | 0.6667 | NA | NA | 2 | 0.5 | NA | NA | NA | NA | 2 | 0.5 |
| 12_355190 | NA | NA | 2 | 0.5 | NA | NA | 2 | 0 | 4 | 0.25 | 1 | 0 | 2 | 0.5 | NA | NA | NA | NA | 3 | 0.3333 |
| 12_355207 | NA | NA | NA | NA | NA | NA | 1 | 1 | 1 | 1 | 1 | 1 | 2 | 0 | NA | NA | NA | NA | 3 | 0.3333 |
| 12_355208 | NA | NA | 1 | 1 | NA | NA | NA | NA | 1 | 1 | 1 | 1 | 2 | 1 | NA | NA | NA | NA | 3 | 1 |
| 12_355215 | NA | NA | 1 | 1 | NA | NA | 1 | 1 | 2 | 1 | 1 | 1 | 1 | 1 | NA | NA | NA | NA | 2 | 1 |
| 12_355222 | NA | NA | 2 | 1 | NA | NA | 2 | 1 | 4 | 1 | 1 | 1 | 2 | 1 | NA | NA | NA | NA | 3 | 1 |
| 12_355224 | NA | NA | 2 | 0 | NA | NA | 1 | 1 | 3 | 0.3333 | 1 | 1 | 1 | 0 | NA | NA | NA | NA | 2 | 0.5 |
| 12_355234 | NA | NA | 2 | 0.5 | NA | NA | 2 | 0.5 | 4 | 0.5 | 1 | 1 | 2 | 0 | NA | NA | NA | NA | 3 | 0.3333 |
| 12_355237 | NA | NA | 2 | 1 | NA | NA | 1 | 1 | 3 | 1 | 1 | 1 | NA | NA | NA | NA | NA | NA | 1 | 1 |
| 12_355247 | NA | NA | NA | NA | NA | NA | 2 | 1 | 2 | 1 | 1 | 1 | 2 | 0.5 | NA | NA | NA | NA | 3 | 0.6667 |
| 12_355255 | NA | NA | 1 | 1 | NA | NA | 1 | 0 | 2 | 0.5 | 1 | 1 | 2 | 0.5 | NA | NA | NA | NA | 3 | 0.6667 |
| 12_355302 | NA | NA | 1 | 1 | NA | NA | 1 | 0 | 2 | 0.5 | 1 | 0 | 1 | 1 | NA | NA | NA | NA | 2 | 0.5 |
| 12_355558 | 2 | 0 | 2 | 0 | 2 | 0 | 2 | 1 | 8 | 0.25 | 1 | 1 | 3 | 0 | 5 | 0 | NA | NA | 9 | 0.1111 |
| 12_355613 | 2 | 0.5 | 2 | 0 | 3 | 0 | 2 | 0 | 9 | 0.1111 | 1 | 0 | 3 | 0 | 5 | 0.2 | NA | NA | 9 | 0.1111 |
| 12_355638 | 4 | 0.75 | 2 | 0 | 3 | 0.3333 | 2 | 0 | 11 | 0.3636 | 1 | 0 | 3 | 0 | 5 | 0.2 | NA | NA | 9 | 0.1111 |
| 12_355660 | 4 | 0.5 | 3 | 0 | 3 | 0 | 2 | 0 | 12 | 0.1667 | 1 | 0 | 3 | 0 | 5 | 0.2 | NA | NA | 9 | 0.1111 |
| 12_361798 | NA | NA | NA | NA | NA | NA | 2 | 0.5 | 2 | 0.5 | NA | NA | NA | NA | NA | NA | 1 | 1 | 1 | 1 |
| 12_361802 | NA | NA | NA | NA | NA | NA | 2 | 0.5 | 2 | 0.5 | NA | NA | NA | NA | NA | NA | 1 | 1 | 1 | 1 |
| 12_361810 | NA | NA | NA | NA | NA | NA | 1 | 1 | 1 | 1 | NA | NA | NA | NA | NA | NA | 1 | 1 | 1 | 1 |
| 12_361821 | NA | NA | NA | NA | NA | NA | 1 | 1 | 1 | 1 | NA | NA | NA | NA | NA | NA | 1 | 1 | 1 | 1 |
| 12_361838 | NA | NA | NA | NA | NA | NA | 2 | 1 | 2 | 1 | NA | NA | NA | NA | NA | NA | 1 | 1 | 1 | 1 |
| 12_361934 | 1 | 0 | NA | NA | NA | NA | 2 | 0.5 | 3 | 0.3333 | 1 | 0 | NA | NA | NA | NA | 1 | 1 | 2 | 0.5 |
| 12_362062 | 1 | 1 | NA | NA | 2 | 1 | 1 | 1 | 4 | 1 | 2 | 1 | 1 | 1 | NA | NA | 3 | 1 | 6 | 1 |
| 12_362064 | 2 | 0 | 1 | 0 | 2 | 0 | 2 | 0.5 | 7 | 0.1429 | 1 | 1 | 1 | 0 | NA | NA | 4 | 0.25 | 6 | 0.3333 |
| 12_362069 | 2 | 0.5 | 1 | 1 | 2 | 1 | 3 | 0.6667 | 8 | 0.75 | 1 | 0 | 1 | 1 | NA | NA | 4 | 0.75 | 6 | 0.6667 |
| 12_362071 | 2 | 0.5 | 1 | 0 | 2 | 0 | 3 | 0.6667 | 8 | 0.375 | 1 | 0 | 1 | 1 | NA | NA | 4 | 0.25 | 6 | 0.3333 |
| 12_362088 | 1 | 1 | 1 | 0 | 2 | 0.5 | 2 | 0.5 | 6 | 0.5 | 2 | 0.5 | 1 | 1 | NA | NA | 4 | 0.25 | 7 | 0.4286 |
| 12_362094 | 2 | 0.5 | 1 | 0 | 1 | 1 | 2 | 0.5 | 6 | 0.5 | 1 | 0 | 2 | 0 | NA | NA | 4 | 0.25 | 7 | 0.1429 |
| 12_362095 | 2 | 1 | 1 | 0 | 1 | 0 | 2 | 0.5 | 6 | 0.5 | 2 | 1 | 1 | 0 | NA | NA | 4 | 0 | 7 | 0.2857 |
| 12_362160 | 1 | 1 | 1 | 0 | 1 | 1 | 1 | 1 | 4 | 0.75 | 2 | 1 | 1 | 1 | NA | NA | 2 | 0.5 | 5 | 0.8 |
| 12_362170 | 1 | 0 | NA | NA | 1 | 1 | 1 | 1 | 3 | 0.6667 | 1 | 0 | 1 | 0 | NA | NA | 2 | 0.5 | 4 | 0.25 |
| 12_362172 | 2 | 0 | NA | NA | 1 | 0 | 1 | 1 | 4 | 0.25 | 2 | 0 | 1 | 1 | NA | NA | 3 | 0 | 6 | 0.1667 |
| 12_362189 | 1 | 1 | 2 | 1 | 1 | 0 | 1 | 1 | 5 | 0.8 | NA | NA | 1 | 1 | NA | NA | 3 | 1 | 4 | 1 |
| 12_362193 | 1 | 1 | 1 | 1 | 1 | 1 | 1 | 1 | 4 | 1 | NA | NA | 1 | 1 | NA | NA | 2 | 1 | 3 | 1 |
| 12_362207 | 1 | 1 | 1 | 0 | 1 | 1 | 1 | 1 | 4 | 0.75 | 1 | 1 | 1 | 1 | NA | NA | 2 | 0.5 | 4 | 0.75 |
| 12_362213 | 1 | 1 | NA | NA | NA | NA | 1 | 1 | 2 | 1 | NA | NA | NA | NA | NA | NA | 2 | 0.5 | 2 | 0.5 |
| 12_362220 | 1 | 1 | 2 | 0.5 | NA | NA | 1 | 0 | 4 | 0.5 | NA | NA | 1 | 0 | NA | NA | 2 | 0.5 | 3 | 0.3333 |
| 12_362225 | NA | NA | 2 | 1 | 1 | 1 | NA | NA | 3 | 1 | NA | NA | 1 | 1 | NA | NA | 2 | 0.5 | 3 | 0.6667 |
| 12_382435 | NA | NA | NA | NA | 1 | 1 | NA | NA | 1 | 1 | NA | NA | 2 | 0.5 | 1 | 0 | NA | NA | 3 | 0.3333 |
| 12_382452 | NA | NA | NA | NA | 1 | 1 | NA | NA | 1 | 1 | NA | NA | 2 | 0.5 | 1 | 0 | NA | NA | 3 | 0.3333 |
| 12_383127 | 9 | 0.4444 | 10 | 0.4 | 8 | 0.75 | 9 | 0.1111 | 36 | 0.4167 | 8 | 0.25 | 10 | 0.6 | 6 | 0.6667 | 4 | 0.75 | 28 | 0.5357 |
| 12_383138 | 8 | 0.25 | 9 | 0.1111 | 7 | 0 | 8 | 0.5 | 32 | 0.2188 | 6 | 0.1667 | 12 | 0.1667 | 10 | 0.4 | 7 | 0.5714 | 35 | 0.3143 |
| 12_383168 | 1 | 0 | NA | NA | 2 | 0.5 | 1 | 0 | 4 | 0.25 | NA | NA | 2 | 0.5 | NA | NA | NA | NA | 2 | 0.5 |
| 12_383173 | 1 | 0 | NA | NA | 2 | 1 | 1 | 0 | 4 | 0.5 | NA | NA | 2 | 0 | 1 | 1 | NA | NA | 3 | 0.3333 |
| 12_383214 | 2 | 0 | 1 | 0 | 4 | 0.25 | 1 | 0 | 8 | 0.125 | NA | NA | 2 | 0 | 2 | 0.5 | 1 | 0 | 5 | 0.2 |
| 12_383231 | 1 | 0 | 1 | 0 | 4 | 0.25 | 1 | 0 | 7 | 0.1429 | NA | NA | 2 | 0 | 2 | 0.5 | 1 | 1 | 5 | 0.4 |
| 12_383294 | 1 | 0 | NA | NA | 3 | 1 | 1 | 1 | 5 | 0.8 | NA | NA | 3 | 1 | 1 | 1 | NA | NA | 4 | 1 |
| 12_383299 | 2 | 0.5 | 1 | 1 | 2 | 1 | 1 | 1 | 6 | 0.8333 | NA | NA | 3 | 0.3333 | 1 | 0 | 1 | 0 | 5 | 0.2 |
| 12_383311 | 2 | 0.5 | 1 | 0 | 3 | 1 | 1 | 1 | 7 | 0.7143 | NA | NA | 3 | 1 | 2 | 0 | 1 | 0 | 6 | 0.5 |
| 12_383315 | 2 | 0.5 | 1 | 0 | 3 | 0 | 1 | 0 | 7 | 0.1429 | NA | NA | 3 | 0.3333 | 2 | 0 | 1 | 0 | 6 | 0.1667 |
| 12_383324 | NA | NA | 1 | 1 | 3 | 1 | 1 | 1 | 5 | 1 | NA | NA | 3 | 1 | 2 | 1 | 1 | 1 | 6 | 1 |
| 12_383353 | 1 | 0 | 1 | 1 | 3 | 1 | 1 | 1 | 6 | 0.8333 | NA | NA | 2 | 1 | 2 | 1 | NA | NA | 4 | 1 |
| 12_383369 | 1 | 1 | 1 | 0 | 3 | 0.6667 | 1 | 0 | 6 | 0.5 | NA | NA | 3 | 0.6667 | 2 | 0 | NA | NA | 5 | 0.4 |

| | | | | | | | | | | | | | | | | | | | | |
|---|---|---|---|---|---|---|---|---|---|---|---|---|---|---|---|---|---|---|---|---|
| 12_395766 | 2 | 0.5 | 1 | 1 | 1 | 1 | NA | NA | 4 | 0.75 | 1 | 1 | 1 | 0 | 1 | 0 | 1 | 0 | 4 | 0.25 |
| 12_395775 | 2 | 0.5 | 1 | 1 | 1 | 1 | NA | NA | 4 | 0.75 | 1 | 1 | 1 | 1 | 1 | 1 | 1 | 1 | 4 | 1 |
| 12_395776 | 2 | 0.5 | NA | NA | 1 | 0 | NA | NA | 3 | 0.3333 | 1 | 0 | 1 | 0 | NA | NA | 1 | 1 | 3 | 0.3333 |
| 12_395779 | 1 | 0 | 1 | 1 | 1 | 0 | NA | NA | 3 | 0.3333 | 1 | 1 | 1 | 0 | NA | NA | 1 | 0 | 3 | 0.3333 |
| 12_395803 | 2 | 0 | 1 | 1 | 1 | 1 | NA | NA | 4 | 0.5 | 1 | 0 | 1 | 0 | 1 | 1 | 1 | 1 | 4 | 0.5 |
| 12_396346 | 6 | 0.8333 | 2 | 1 | 3 | 0.3333 | 5 | 0.4 | 16 | 0.625 | 2 | 0.5 | 6 | 0.5 | 4 | 0.25 | 4 | 0.25 | 16 | 0.375 |
| 12_396381 | 5 | 0.2 | 3 | 0.3333 | 3 | 0 | 4 | 0.25 | 15 | 0.2 | 3 | 0.6667 | 4 | 0 | 7 | 0 | 3 | 0.3333 | 17 | 0.1765 |
| 12_396412 | 4 | 0.25 | 2 | 0.5 | 4 | 0.5 | 5 | 0.6 | 15 | 0.4667 | 3 | 1 | 3 | 0 | 7 | 0.5714 | 2 | 0.5 | 15 | 0.5333 |
| 12_396420 | 5 | 0.2 | 3 | 0.3333 | 4 | 0 | 5 | 0 | 17 | 0.1176 | 3 | 0 | 4 | 0 | 7 | 0.2857 | 3 | 0 | 17 | 0.1176 |
| 12_401289 | 12 | 0.0833 | 16 | 0.25 | 14 | 0.3571 | 16 | 0.3125 | 58 | 0.2586 | 9 | 0.4444 | 7 | 0.1429 | 17 | 0.3529 | 15 | 0.2 | 48 | 0.2917 |
| 12_401290 | 11 | 0.6364 | 11 | 0.4545 | 13 | 0.6154 | 13 | 0.4615 | 48 | 0.5417 | 10 | 0.7 | 5 | 0.6 | 14 | 0.5714 | 14 | 0.5714 | 43 | 0.6047 |
| 12_401301 | 12 | 0.4167 | 14 | 0.6429 | 14 | 0.6429 | 12 | 0.5833 | 52 | 0.5769 | 12 | 0.75 | 8 | 0.5 | 17 | 0.7059 | 16 | 0.5625 | 53 | 0.6415 |
| 12_401318 | 11 | 0.1818 | 13 | 0.0769 | 12 | 0.0833 | 12 | 0 | 48 | 0.0833 | 13 | 0.2308 | 7 | 0.1429 | 17 | 0 | 15 | 0.2667 | 52 | 0.1538 |
| 12_401336 | 12 | 0.1667 | 13 | 0 | 12 | 0.1667 | 10 | 0.2 | 47 | 0.1277 | 10 | 0.2 | 7 | 0.1429 | 17 | 0.1176 | 15 | 0.0667 | 49 | 0.1224 |
| 12_401529 | 9 | 0 | 4 | 0 | 5 | 0.2 | 6 | 0.1667 | 24 | 0.0833 | 5 | 0 | 5 | 0 | 9 | 0.1111 | 10 | 0.1 | 29 | 0.069 |
| 12_401590 | 6 | 0 | 2 | 0 | 4 | 0.5 | 3 | 0.3333 | 15 | 0.2 | 2 | 0.5 | 1 | 1 | 3 | 0.6667 | 3 | 0.6667 | 9 | 0.6667 |
| 12_401592 | 9 | 0.6667 | 2 | 1 | 3 | 0.6667 | 3 | 1 | 17 | 0.7647 | 3 | 0.6667 | 1 | 1 | 3 | 1 | 3 | 1 | 10 | 0.9 |
| 12_401618 | 8 | 0.75 | 2 | 1 | 3 | 1 | 4 | 0.75 | 17 | 0.8235 | 2 | 1 | 1 | 1 | 3 | 1 | 4 | 0.75 | 10 | 0.9 |
| 12_401620 | 6 | 0.5 | 2 | 1 | 3 | 1 | 3 | 0.6667 | 14 | 0.7143 | 2 | 0.5 | 1 | 1 | 3 | 1 | 4 | 0.75 | 10 | 0.8 |
| 12_401633 | 6 | 0.8333 | 2 | 1 | 3 | 1 | 3 | 1 | 14 | 0.9286 | 1 | 1 | 1 | 1 | 3 | 1 | 2 | 1 | 7 | 1 |
| 12_401634 | 7 | 0.7143 | 2 | 1 | 3 | 1 | 3 | 0.6667 | 15 | 0.8 | NA | NA | 1 | 1 | 2 | 1 | 4 | 1 | 7 | 1 |
| 12_401639 | 7 | 0.1429 | 2 | 0 | 3 | 0.3333 | 4 | 0 | 16 | 0.125 | 1 | 1 | 1 | 0 | 2 | 0 | 3 | 0.3333 | 7 | 0.2857 |
| 12_401641 | 6 | 0.5 | 2 | 0.5 | 3 | 0 | 2 | 0 | 13 | 0.3077 | 1 | 1 | 1 | 0 | 3 | 0 | 3 | 0.6667 | 8 | 0.375 |
| 12_401644 | 7 | 0.1429 | 2 | 0.5 | 3 | 0.6667 | 4 | 0.25 | 16 | 0.3125 | 1 | 0 | NA | NA | 3 | 0.6667 | 4 | 0.75 | 8 | 0.625 |
| 12_4143070 | NA | NA | 1 | 1 | NA | NA | NA | NA | 1 | 1 | NA | NA | 1 | 1 | NA | NA | NA | NA | 1 | 1 |
| 12_415685 | 4 | 0 | 5 | 0 | 5 | 0.2 | 4 | 0.25 | 18 | 0.1111 | 5 | 0.2 | 8 | 0 | 4 | 0 | 9 | 0.1111 | 26 | 0.0769 |
| 12_415697 | 2 | 0 | 4 | 0 | 5 | 0.4 | 5 | 0.4 | 16 | 0.25 | 5 | 0.4 | 6 | 0.3333 | 3 | 0 | 8 | 0.375 | 22 | 0.3182 |
| 12_415808 | 5 | 0 | 8 | 0.5 | 7 | 0 | 7 | 0.2857 | 27 | 0.2222 | 5 | 0 | 10 | 0.4 | 4 | 0 | 7 | 0.1429 | 26 | 0.1923 |
| 12_416025 | 6 | 0.3333 | 9 | 0.2222 | 12 | 0.1667 | 17 | 0.2353 | 44 | 0.2273 | 13 | 0.1538 | 11 | 0.1818 | 8 | 0.125 | 13 | 0.3077 | 45 | 0.2 |
| 12_416039 | 8 | 0.625 | 8 | 0.375 | 14 | 0.2143 | 18 | 0.2222 | 48 | 0.3125 | 14 | 0.4286 | 11 | 0.3636 | 9 | 0.7778 | 12 | 0.25 | 46 | 0.4348 |
| 12_416391 | NA | NA | 1 | 1 | NA | NA | NA | NA | 1 | 1 | NA | NA | NA | NA | NA | NA | 1 | 1 | 1 | 1 |
| 12_418442 | NA | NA | 2 | 0.5 | NA | NA | 1 | 1 | 3 | 0.6667 | NA | NA | 1 | 1 | 1 | 0 | 1 | 1 | 3 | 0.6667 |
| 12_418445 | NA | NA | 2 | 0 | 1 | 0 | 2 | 1 | 5 | 0.4 | NA | NA | NA | NA | 1 | 1 | 1 | 1 | 2 | 1 |
| 12_418455 | NA | NA | 2 | 0 | NA | NA | 2 | 0.5 | 4 | 0.25 | NA | NA | 1 | 0 | 1 | 0 | 3 | 0.3333 | 5 | 0.2 |
| 12_418459 | NA | NA | 2 | 1 | NA | NA | 2 | 0.5 | 4 | 0.75 | NA | NA | NA | NA | 1 | 1 | 3 | 1 | 4 | 1 |
| 12_418467 | NA | NA | 2 | 1 | NA | NA | 1 | 0 | 3 | 0.6667 | 1 | 1 | 1 | 1 | 1 | 1 | 3 | 1 | 6 | 1 |
| 12_418479 | NA | NA | 2 | 1 | NA | NA | 3 | 1 | 5 | 1 | NA | NA | 1 | 0 | 1 | 1 | 1 | 1 | 3 | 0.6667 |
| 12_418482 | NA | NA | 2 | 1 | NA | NA | 3 | 1 | 5 | 1 | 1 | 0 | 1 | 0 | 1 | 1 | 2 | 1 | 5 | 0.6 |
| 12_426155 | 3 | 0 | 7 | 0.1429 | 8 | 0.25 | 9 | 0 | 27 | 0.1111 | 4 | 0 | 2 | 0 | 5 | 0 | 5 | 0.2 | 16 | 0.0625 |
| 12_426277 | 3 | 0.3333 | 7 | 0.1429 | 7 | 0 | 8 | 0 | 25 | 0.08 | 4 | 0 | 3 | 0 | 5 | 0 | 4 | 0.25 | 16 | 0.0625 |
| 12_428653 | 2 | 0.5 | 2 | 1 | 2 | 1 | 4 | 1 | 10 | 0.9 | 2 | 0.5 | 5 | 0.6 | 3 | 1 | 4 | 0.75 | 14 | 0.7143 |
| 12_428655 | 3 | 0 | 2 | 0 | 2 | 0.5 | 5 | 0.4 | 12 | 0.25 | 2 | 0.5 | 4 | 0.75 | 3 | 0.6667 | 2 | 0.5 | 11 | 0.6364 |
| 12_428681 | NA | NA | NA | NA | NA | NA | 1 | 1 | 1 | 1 | 1 | 1 | 2 | 0 | NA | NA | NA | NA | 3 | 0.3333 |
| 12_428805 | NA | NA | NA | NA | NA | NA | 1 | 1 | 1 | 1 | 1 | 1 | 2 | 1 | NA | NA | 1 | 1 | 4 | 1 |
| 12_428834 | NA | NA | NA | NA | NA | NA | 1 | 1 | 1 | 1 | NA | NA | 2 | 0.5 | NA | NA | 1 | 1 | 3 | 0.6667 |
| 12_429759 | 13 | 0.1538 | 13 | 0 | 16 | 0 | 8 | 0.125 | 50 | 0.06 | 5 | 0 | 7 | 0.1429 | 10 | 0.1 | 7 | 0 | 29 | 0.069 |
| 12_429760 | 14 | 0.6429 | 16 | 0.375 | 15 | 0.5333 | 11 | 0.3636 | 56 | 0.4821 | 6 | 0.8333 | 8 | 0.25 | 11 | 0.2727 | 9 | 0.6667 | 34 | 0.4706 |
| 12_429767 | 13 | 0.7143 | 15 | 0 | 14 | 0.0714 | 11 | 0.2727 | 53 | 0.1132 | 7 | 0 | 7 | 0 | 13 | 0.0769 | 9 | 0.2222 | 36 | 0.0833 |
| 12_429781 | 13 | 0.9231 | 14 | 0.7857 | 15 | 0.9333 | 10 | 0.7 | 52 | 0.8462 | 5 | 0.8 | 5 | 0.6 | 11 | 0.7273 | 8 | 0.75 | 29 | 0.7241 |
| 12_429783 | 13 | 0.2308 | 14 | 0.6429 | 13 | 0.3846 | 11 | 0.2727 | 51 | 0.3922 | 5 | 0.4 | 7 | 0.1429 | 8 | 0.375 | 8 | 0.625 | 28 | 0.3929 |
| 12_429784 | 15 | 0.4 | 14 | 0.2857 | 15 | 0.2667 | 11 | 0.2727 | 55 | 0.3091 | 5 | 0.2 | 7 | 0.2857 | 10 | 0.3 | 7 | 0.4286 | 29 | 0.3103 |
| 12_429799 | 12 | 0.4167 | 12 | 0.5 | 12 | 0.5833 | 10 | 0.6 | 46 | 0.5217 | 5 | 0.4 | 5 | 0.6 | 9 | 0.5556 | 6 | 0 | 25 | 0.4 |
| 12_429807 | 11 | 0.9091 | 6 | 0.8333 | 11 | 1 | 8 | 0.75 | 36 | 0.8889 | 4 | 1 | 5 | 1 | 8 | 0.875 | 6 | 1 | 23 | 0.9565 |
| 12_429819 | 12 | 0.3333 | 8 | 0.375 | 11 | 0.5455 | 9 | 0.4444 | 40 | 0.425 | 4 | 0 | 6 | 0.1667 | 7 | 0.4286 | 6 | 0.6667 | 23 | 0.3478 |
| 12_429819 | 11 | 0.5455 | 10 | 0.4 | 11 | 0.4545 | 5 | 0.6 | 37 | 0.4865 | 2 | 0.5 | 7 | 0.8571 | 8 | 0.5 | 4 | 0.5 | 21 | 0.619 |
| 12_429834 | 9 | 0.3333 | 9 | 0.1111 | 10 | 0.5 | 9 | 0.5556 | 37 | 0.3784 | 2 | 1 | 5 | 0.8 | 9 | 0.4444 | 6 | 0.5 | 22 | 0.5909 |
| 12_429886 | 10 | 0.6 | 12 | 0.5833 | 12 | 0.8333 | 8 | 0.25 | 42 | 0.5952 | 3 | 0 | 3 | 1 | 8 | 0.625 | 7 | 0.5714 | 21 | 0.5714 |
| 12_429932 | 9 | 1 | 10 | 1 | 9 | 1 | 5 | 1 | 33 | 1 | 3 | 1 | 3 | 1 | 5 | 0.8 | 6 | 1 | 17 | 0.9412 |
| 12_429933 | 8 | 0.25 | 10 | 0.3 | 8 | 0.25 | 3 | 0 | 29 | 0.2414 | 2 | 0.5 | 4 | 1 | 7 | 0.2857 | 6 | 0.1667 | 19 | 0.4211 |
| 12_429954 | 8 | 0.875 | 11 | 0.8182 | 9 | 0.7778 | 4 | 0.75 | 32 | 0.8125 | 3 | 1 | 3 | 1 | 6 | 0.5 | 5 | 0.4 | 17 | 0.6471 |
| 12_429958 | 7 | 0.7143 | 9 | 0.6667 | 6 | 0.8333 | 4 | 0.75 | 26 | 0.7308 | 2 | 0.5 | 3 | 1 | 5 | 0.6 | 3 | 0.6667 | 13 | 0.6923 |
| 12_431766 | 5 | 0.2 | 5 | 0.2 | 4 | 0 | 2 | 0 | 16 | 0.125 | 2 | 0 | 1 | 0 | 3 | 0 | 2 | 0.5 | 8 | 0.125 |
| 12_431784 | 4 | 0.25 | 6 | 0 | 8 | 0.125 | 3 | 0.3333 | 21 | 0.1429 | 1 | 0 | 1 | 1 | 4 | 0.25 | 2 | 0 | 8 | 0.25 |
| 12_431786 | 6 | 0.1667 | 6 | 0.6667 | 6 | 0 | 3 | 1 | 21 | 0.381 | 2 | 0 | 1 | 1 | 4 | 0.5 | 1 | 1 | 8 | 0.5 |
| 12_434512 | 6 | 0.3333 | 5 | 0.4 | 4 | 1 | NA | NA | 15 | 0.5333 | 1 | 0 | 1 | 0 | 2 | 0 | 2 | 1 | 6 | 0.3333 |

| | | | | | | | | | | | | | | | | | | | | |
|---|---|---|---|---|---|---|---|---|---|---|---|---|---|---|---|---|---|---|---|---|
| 12_4345132 | 5 | 0.2 | 5 | 0.2 | 4 | 0 | NA | NA | 14 | 0.1429 | 2 | 0 | 2 | 0 | 4 | 0 | 1 | 1 | 9 | 0.1111 |
| 12_4345159 | 6 | 0.8333 | 4 | 0.75 | 3 | 0.6667 | NA | NA | 13 | 0.7692 | 2 | 1 | 2 | 0.5 | 4 | 0.25 | 2 | 0.5 | 10 | 0.5 |
| 12_4345196 | 6 | 0.8333 | 5 | 0 | 4 | 0.5 | NA | NA | 15 | 0.4667 | 2 | 0.5 | 2 | 1 | 4 | 0.75 | 3 | 0.6667 | 11 | 0.7273 |
| 12_4345284 | 3 | 0 | 5 | 0.2 | 3 | 0.3333 | NA | NA | 11 | 0.1818 | 2 | 0.5 | 2 | 0.5 | 3 | 0 | 1 | 0 | 8 | 0.25 |
| 12_4345382 | 5 | 0.2 | 4 | 0 | 3 | 0 | NA | NA | 12 | 0.0833 | 2 | 0.5 | 1 | 1 | 4 | 0 | 2 | 0.5 | 9 | 0.3333 |
| 12_4345466 | 6 | 0.1667 | 4 | 0.25 | 3 | 0 | NA | NA | 13 | 0.1538 | 2 | 0 | NA | NA | 3 | 0.3333 | 2 | 0 | 7 | 0.1429 |
| 12_4345480 | 5 | 0.8 | 4 | 0.75 | 2 | 0.5 | NA | NA | 11 | 0.7273 | 2 | 1 | 1 | 0 | 4 | 1 | 1 | 1 | 8 | 0.875 |
| 12_4345501 | 6 | 0.5 | 4 | 0.5 | 3 | 0.6667 | NA | NA | 13 | 0.5385 | 2 | 1 | 1 | 1 | 4 | 0 | 2 | 0.5 | 9 | 0.4444 |
| 12_4345508 | 6 | 0 | 3 | 0.6667 | 3 | 0.3333 | NA | NA | 12 | 0.25 | 2 | 0.5 | NA | NA | 4 | 0.25 | 2 | 0 | 8 | 0.25 |
| 12_4345702 | 5 | 0.2 | 3 | 0.3333 | 2 | 0 | NA | NA | 10 | 0.2 | 1 | 0 | 2 | 0 | 3 | 0.3333 | 2 | 0 | 8 | 0.125 |
| 12_4345969 | 5 | 0.4 | 3 | 0.6667 | 3 | 0.6667 | NA | NA | 11 | 0.5455 | 1 | 0 | 2 | 0 | 4 | 0.25 | 1 | 0 | 8 | 0.125 |
| 12_4346024 | 4 | 0.5 | 3 | 0.3333 | 3 | 0.6667 | NA | NA | 10 | 0.5 | NA | NA | 2 | 0 | 3 | 0.3333 | 2 | 0.5 | 7 | 0.2857 |
| 12_4346045 | 2 | 0 | 3 | 0.3333 | 2 | 0 | NA | NA | 7 | 0.1429 | 1 | 1 | 2 | 0 | 2 | 0.5 | 1 | 0 | 6 | 0.3333 |
| 12_4642014 | NA | NA | 1 | 0 | 1 | 1 | NA | NA | 2 | 0.5 | 1 | 0 | NA | NA | NA | NA | 1 | 1 | 2 | 0.5 |
| 12_4642184 | NA | NA | 1 | 1 | 1 | 1 | NA | NA | 2 | 1 | 1 | 1 | NA | NA | NA | NA | 1 | 1 | 2 | 1 |
| 12_4642264 | NA | NA | 1 | 1 | 1 | 1 | NA | NA | 2 | 1 | 1 | 0 | NA | NA | NA | NA | 1 | 1 | 2 | 0.5 |
| 12_4642326 | NA | NA | 1 | 1 | 1 | 1 | NA | NA | 2 | 1 | 1 | 1 | NA | NA | NA | NA | 1 | 1 | 2 | 1 |
| 12_4642533 | NA | NA | 1 | 1 | 1 | 1 | NA | NA | 2 | 1 | 1 | 1 | NA | NA | NA | NA | 1 | 1 | 2 | 1 |
| 12_4642683 | NA | NA | 1 | 1 | NA | NA | NA | NA | 1 | 1 | 1 | 1 | NA | NA | NA | NA | 1 | 0 | 2 | 0.5 |
| 12_4727076 | 2 | 0 | NA | NA | 4 | 0.5 | 2 | 0 | 8 | 0.25 | 1 | 1 | 2 | 0 | 1 | 0 | 4 | 0 | 8 | 0.125 |
| 12_4728828 | 5 | 0.2 | 2 | 0 | 5 | 0.2 | 2 | 0 | 14 | 0.1429 | 1 | 0 | 2 | 0 | 3 | 0.3333 | 4 | 0 | 10 | 0.1 |
| 12_4763186 | 2 | 0 | NA | NA | 1 | 0 | 3 | 0.3333 | 6 | 0.1667 | 2 | 0 | 3 | 1 | 1 | 1 | 1 | 0 | 7 | 0.5714 |
| 12_4763250 | 4 | 0.75 | 4 | 1 | 1 | 0 | 3 | 0.3333 | 12 | 0.6667 | 3 | 0.6667 | 2 | 0.5 | 1 | 0 | 2 | 1 | 8 | 0.625 |
| 12_4763270 | 4 | 0.25 | 4 | 0.25 | 1 | 0 | 3 | 0 | 12 | 0.1667 | 3 | 0.3333 | 2 | 1 | 1 | 0 | 4 | 0 | 10 | 0.3 |
| 12_4819244 | 1 | 1 | 1 | 0 | 1 | 0 | 3 | 0.6667 | 6 | 0.5 | 2 | 0 | NA | NA | 1 | 1 | NA | NA | 3 | 0.3333 |
| 12_4819370 | 1 | 1 | 2 | 0.5 | 2 | 0 | 3 | 1 | 8 | 0.625 | 2 | 1 | 2 | 0 | NA | NA | 1 | 0 | 5 | 0.4 |
| 12_4819455 | 1 | 0 | 2 | 0.5 | 3 | 0 | 3 | 0 | 9 | 0.1111 | 2 | 0 | 2 | 0 | 2 | 0.5 | 1 | 0 | 7 | 0.1429 |
| 12_4819577 | 1 | 1 | 2 | 0 | 1 | 0 | 3 | 0 | 7 | 0.1429 | 2 | 0 | 2 | 0 | 2 | 0.5 | 1 | 0 | 7 | 0.1429 |
| 12_4819769 | 1 | 1 | 2 | 1 | 2 | 1 | 2 | 0.5 | 7 | 0.8571 | 3 | 1 | 3 | 1 | 3 | 0.3333 | 1 | 1 | 10 | 0.8 |
| 12_4819819 | 1 | 1 | 2 | 0 | 3 | 0 | 1 | 1 | 7 | 0.2857 | 3 | 0.3333 | 2 | 0 | 3 | 0 | 1 | 0 | 9 | 0.1111 |
| 12_4819826 | 1 | 0 | 1 | 1 | 3 | 0 | 2 | 0 | 7 | 0.1429 | 3 | 0 | 3 | 0 | 3 | 0.3333 | NA | NA | 9 | 0.1111 |
| 12_4819865 | 1 | 0 | 2 | 0.5 | 3 | 0 | 3 | 0.3333 | 9 | 0.2222 | 3 | 0 | 4 | 0 | 3 | 0.3333 | 1 | 0 | 11 | 0.0909 |
| 12_4820023 | 1 | 1 | 2 | 0.5 | 2 | 0.5 | 3 | 0.3333 | 8 | 0.5 | 3 | 0.3333 | 3 | 1 | 2 | 1 | 1 | 0 | 9 | 0.6667 |
| 12_4820056 | 1 | 0 | 2 | 0.5 | 3 | 0 | 3 | 0 | 9 | 0.1111 | 3 | 0.6667 | 3 | 1 | 2 | 1 | 1 | 0 | 9 | 0.7778 |
| 12_4820169 | NA | NA | NA | NA | 1 | 1 | 2 | 0 | 3 | 0.3333 | 3 | 1 | 2 | 0.5 | 1 | 0 | NA | NA | 6 | 0.6667 |
| 12_4820208 | 1 | 0 | 1 | 0 | 3 | 0.3333 | 3 | 0.3333 | 8 | 0.25 | 3 | 0.3333 | 4 | 0.5 | 4 | 0.5 | 1 | 0 | 12 | 0.4167 |
| 12_4828318 | 1 | 0 | 1 | 1 | 1 | 1 | 2 | 0.5 | 5 | 0.6 | 2 | 1 | 4 | 1 | 3 | 0.6667 | 1 | 1 | 10 | 0.9 |
| 12_4828376 | NA | NA | 1 | 0 | 3 | 0.3333 | 3 | 0 | 7 | 0.1429 | 3 | 0 | 2 | 0.5 | 4 | 0.25 | 1 | 0 | 10 | 0.2 |
| 12_4867651 | 13 | 0.1538 | 16 | 0 | 7 | 0.1429 | 14 | 0 | 50 | 0.06 | 12 | 0 | 16 | 0 | 17 | 0.1176 | 12 | 0.0833 | 57 | 0.0526 |
| 12_4883388 | 3 | 0 | 6 | 0.1667 | 2 | 0 | 3 | 0 | 14 | 0.0714 | 3 | 0.3333 | 2 | 0 | 1 | 0 | 1 | 0 | 7 | 0.1429 |
| 12_4906898 | 1 | 1 | 1 | 0 | NA | NA | 1 | 0 | 3 | 0.3333 | NA | NA | NA | NA | 1 | 1 | NA | NA | 1 | 1 |
| 12_4908143 | 1 | 1 | NA | NA | 2 | 0.5 | NA | NA | 3 | 0.6667 | 1 | 1 | NA | NA | NA | NA | NA | NA | 1 | 1 |
| 12_4908156 | 1 | 1 | NA | NA | 1 | 1 | NA | NA | 2 | 1 | 1 | 1 | NA | NA | NA | NA | NA | NA | 1 | 1 |
| 12_4908187 | 1 | 1 | NA | NA | 2 | 1 | NA | NA | 3 | 1 | 1 | 1 | NA | NA | NA | NA | NA | NA | 1 | 1 |
| 12_4928470 | 5 | 0.2 | 5 | 0.4 | 7 | 0.4286 | 3 | 0.6667 | 20 | 0.4 | 2 | 0.5 | 3 | 0.3333 | 5 | 0.8 | 8 | 0 | 18 | 0.3333 |
| 12_4928541 | 6 | 0.5 | 7 | 0 | 7 | 0.1429 | 5 | 0.2 | 25 | 0.2 | 3 | 0.6667 | 3 | 0 | 6 | 0.1667 | 7 | 0 | 19 | 0.1579 |
| 12_4929969 | 5 | 0.2 | 6 | 0 | 6 | 0.1667 | 5 | 0 | 22 | 0.0909 | 3 | 0.3333 | 1 | 0 | 3 | 0 | 6 | 0.1667 | 13 | 0.1538 |
| 12_4930562 | 3 | 1 | 6 | 1 | 6 | 1 | 6 | 0.8333 | 21 | 0.9524 | 3 | 1 | NA | NA | 4 | 1 | 4 | 1 | 11 | 1 |
| 12_4930618 | 5 | 0.8 | 5 | 1 | 7 | 0.4286 | 6 | 0.8333 | 23 | 0.7391 | 3 | 1 | NA | NA | 4 | 1 | 6 | 0.8333 | 13 | 0.9231 |
| 12_5081601 | 1 | 1 | 1 | 0 | NA | NA | NA | NA | 2 | 0.5 | 1 | 1 | NA | NA | NA | NA | NA | NA | 1 | 1 |
| 12_5081742 | 1 | 0 | 1 | 1 | NA | NA | NA | NA | 2 | 0.5 | 1 | 1 | NA | NA | NA | NA | NA | NA | 1 | 1 |
| 12_5094018 | NA | NA | 1 | 1 | NA | NA | 1 | 0 | 2 | 0.5 | 1 | 0 | NA | NA | 1 | 1 | NA | NA | 2 | 0.5 |
| 12_5094048 | 1 | 1 | NA | NA | NA | NA | 1 | 1 | 2 | 1 | 2 | 1 | NA | NA | NA | NA | NA | NA | 2 | 1 |
| 12_5094121 | 1 | 1 | 1 | 1 | NA | NA | NA | NA | 2 | 1 | 1 | 1 | NA | NA | 1 | 1 | NA | NA | 2 | 1 |
| 12_5094320 | 1 | 0 | 1 | 0 | NA | NA | 1 | 1 | 3 | 0.3333 | 2 | 1 | NA | NA | 1 | 0 | NA | NA | 3 | 0.6667 |
| 12_5094400 | 1 | 1 | 1 | 0 | NA | NA | 1 | 0 | 3 | 0.3333 | 1 | 0 | NA | NA | 1 | 1 | NA | NA | 2 | 0.5 |
| 12_5094419 | 1 | 0 | 1 | 1 | NA | NA | 1 | 1 | 3 | 0.6667 | 1 | 1 | NA | NA | 1 | 0 | NA | NA | 2 | 0.5 |
| 12_5094429 | 1 | 1 | 1 | 1 | NA | NA | 1 | 1 | 3 | 1 | 1 | 1 | NA | NA | 1 | 1 | NA | NA | 2 | 1 |
| 12_5095531 | 1 | 1 | 2 | 0 | NA | NA | 1 | 0 | 4 | 0.25 | 1 | 1 | NA | NA | 1 | 0 | NA | NA | 2 | 0.5 |
| 12_5097147 | 3 | 0 | 3 | 0 | 1 | 1 | 1 | 0 | 8 | 0.125 | 2 | 0 | NA | NA | 2 | 0 | 2 | 0.5 | 6 | 0.1667 |
| 12_5127618 | 1 | 1 | 2 | 0.5 | 5 | 1 | 1 | 1 | 9 | 0.8889 | NA | NA | 1 | 1 | 1 | 1 | 3 | 1 | 5 | 1 |
| 12_5127739 | 1 | 0 | 4 | 0 | 5 | 0.2 | 3 | 0 | 13 | 0.0769 | NA | NA | 1 | 0 | 3 | 0 | 3 | 0.3333 | 7 | 0.1429 |
| 12_5128075 | 1 | 0 | 4 | 0.5 | 3 | 0 | 2 | 1 | 10 | 0.4 | NA | NA | 1 | 0 | 1 | 0 | 2 | 0.5 | 4 | 0.25 |
| 12_5129250 | 1 | 0 | 9 | 0.2222 | 10 | 0 | 6 | 0 | 26 | 0.0769 | 1 | 0 | 5 | 0 | 8 | 0.25 | 5 | 0 | 19 | 0.1053 |
| 12_5129261 | 1 | 0 | 9 | 0.3333 | 10 | 0.3 | 6 | 0.1667 | 26 | 0.2692 | 1 | 0 | 5 | 0 | 8 | 0.125 | 5 | 0 | 19 | 0.0526 |

| | | | | | | | | | | | | | | | | | | | | |
|---|---|---|---|---|---|---|---|---|---|---|---|---|---|---|---|---|---|---|---|---|
| 12_512935 | 1 | 1 | 7 | 0.7143 | 9 | 1 | 5 | 1 | 22 | 0.9091 | 1 | 1 | 4 | 1 | 7 | 1 | 4 | 1 | 16 | 1 |
| 12_512958 | NA | NA | 8 | 0.625 | 8 | 0.875 | 3 | 0.3333 | 19 | 0.6842 | 1 | 1 | 4 | 0.75 | 8 | 0.75 | 4 | 1 | 17 | 0.8235 |
| 12_512959 | NA | NA | 8 | 0.125 | 6 | 0 | 2 | 0 | 16 | 0.0625 | 1 | 0 | 4 | 0.25 | 6 | 0.1667 | 4 | 0 | 15 | 0.1333 |
| 12_512960 | NA | NA | 9 | 0.1111 | 9 | 0 | 3 | 0.3333 | 21 | 0.0952 | 1 | 0 | 4 | 0.25 | 7 | 0 | 4 | 0 | 16 | 0.0625 |
| 12_513060 | NA | NA | 4 | 0.75 | 3 | 1 | NA | NA | 7 | 0.8571 | NA | NA | 1 | 1 | 2 | 0 | 1 | 1 | 4 | 0.5 |
| 12_513103 | NA | NA | NA | NA | 4 | 0.5 | NA | NA | 4 | 0.5 | 1 | 1 | 2 | 0.5 | 3 | 0.6667 | 1 | 0 | 7 | 0.5714 |
| 12_513111 | NA | NA | 1 | 1 | 2 | 0.5 | NA | NA | 3 | 0.6667 | 1 | 0 | 1 | 1 | 4 | 0.5 | 1 | 1 | 7 | 0.5714 |
| 12_513122 | NA | NA | 1 | 1 | 4 | 1 | NA | NA | 5 | 1 | 1 | 1 | 2 | 1 | 2 | 1 | 1 | 1 | 6 | 1 |
| 12_542606 | NA | NA | 1 | 1 | 2 | 0 | 1 | 1 | 4 | 0.5 | NA | NA | 1 | 0 | 1 | 1 | 2 | 1 | 4 | 0.75 |
| 12_542627 | NA | NA | 1 | 1 | 2 | 1 | 1 | 1 | 4 | 1 | NA | NA | 1 | 0 | 1 | 1 | 1 | 1 | 3 | 0.6667 |
| 12_542628 | NA | NA | 1 | 1 | 3 | 0 | 2 | 0 | 6 | 0.1667 | NA | NA | 1 | 0 | 1 | 0 | 2 | 0.5 | 4 | 0.25 |
| 12_542641 | NA | NA | 1 | 0 | 2 | 0.5 | 2 | 0.5 | 5 | 0.4 | NA | NA | NA | NA | NA | NA | 2 | 1 | 2 | 1 |
| 12_546495 | NA | NA | 1 | 1 | 3 | 1 | 2 | 1 | 6 | 1 | NA | NA | NA | NA | 1 | 1 | 1 | 1 | 2 | 1 |
| 12_546505 | NA | NA | 1 | 1 | 3 | 0.3333 | 2 | 0.5 | 6 | 0.5 | NA | NA | 1 | 0 | 2 | 0.5 | 1 | 1 | 4 | 0.5 |
| 12_546514 | NA | NA | 2 | 0 | 4 | 0.25 | 2 | 0 | 8 | 0.125 | NA | NA | 1 | 0 | 2 | 0.5 | 1 | 1 | 4 | 0.5 |
| 12_546516 | NA | NA | 1 | 0 | 4 | 0.5 | 2 | 0 | 7 | 0.2857 | 1 | 0 | 1 | 0 | 2 | 0 | 1 | 1 | 5 | 0.2 |
| 12_546518 | NA | NA | 2 | 0 | 4 | 0.25 | 2 | 0 | 8 | 0.125 | NA | NA | 1 | 0 | 1 | 1 | 1 | 1 | 3 | 0.6667 |
| 12_546521 | NA | NA | 2 | 1 | 2 | 1 | 3 | 1 | 7 | 1 | NA | NA | NA | NA | 1 | 1 | NA | NA | 1 | 1 |
| 12_546546 | NA | NA | 2 | 0.5 | 3 | 0.6667 | 3 | 0 | 8 | 0.375 | NA | NA | 1 | 1 | 1 | 1 | 1 | 1 | 3 | 1 |
| 12_546552 | NA | NA | 2 | 0 | 4 | 0.25 | 3 | 0 | 9 | 0.1111 | NA | NA | 1 | 0 | 2 | 0.5 | 1 | 1 | 4 | 0.5 |
| 12_546646 | NA | NA | 1 | 1 | NA | NA | NA | NA | 1 | 1 | NA | NA | NA | NA | 1 | 1 | 1 | 1 | 2 | 1 |
| 12_546656 | NA | NA | 1 | 1 | NA | NA | NA | NA | 1 | 1 | NA | NA | NA | NA | 1 | 1 | 1 | 1 | 2 | 1 |
| 12_791144 | 2 | 1 | NA | NA | NA | NA | NA | NA | 2 | 1 | NA | NA | NA | NA | 1 | 1 | 1 | 0 | 2 | 0.5 |
| 12_791150 | 3 | 0.3333 | NA | NA | 1 | 0 | NA | NA | 4 | 0.25 | NA | NA | 1 | 0 | 2 | 1 | 1 | 0 | 4 | 0.5 |
| 12_791152 | 3 | 0.6667 | NA | NA | 1 | 1 | NA | NA | 4 | 0.75 | NA | NA | 1 | 0 | 2 | 1 | 1 | 1 | 4 | 0.75 |
| 12_833407 | 3 | 0 | 4 | 0.25 | 6 | 0 | 4 | 0.25 | 17 | 0.1176 | 2 | 0 | 3 | 0 | 3 | 0.3333 | 1 | 0 | 9 | 0.1111 |
| 12_833413 | 2 | 0.5 | 1 | 1 | 3 | 0.3333 | 4 | 0.75 | 10 | 0.6 | 2 | 0 | 2 | 0.5 | 3 | 0 | 2 | 0 | 9 | 0.1111 |
| 12_833439 | 5 | 0.4 | 5 | 0.6 | 5 | 0 | 4 | 0 | 19 | 0.2632 | 2 | 0 | 3 | 0.3333 | 4 | 0.5 | 2 | 0.5 | 11 | 0.3636 |
| 12_833443 | 4 | 0.25 | 4 | 0 | 5 | 0.2 | 3 | 0.3333 | 16 | 0.1875 | 2 | 0.5 | 3 | 0.3333 | 4 | 0.25 | 2 | 0 | 11 | 0.2727 |
| 12_833453 | 4 | 0.5 | 5 | 0 | 4 | 0 | 4 | 0 | 17 | 0.1176 | NA | NA | 3 | 0.3333 | 4 | 0.25 | 2 | 0 | 9 | 0.2222 |
| 12_833463 | 5 | 0 | 2 | 0 | 5 | 0.2 | 3 | 0 | 15 | 0.0667 | 1 | 0 | 2 | 0 | 3 | 0 | 2 | 0.5 | 8 | 0.125 |
| 12_833480 | 6 | 0.5 | 5 | 0.4 | 6 | 0.5 | 5 | 1 | 22 | 0.5909 | 3 | 0.6667 | 3 | 1 | 3 | 1 | 2 | 0.5 | 11 | 0.8182 |
| 12_833485 | 2 | 0.5 | 4 | 0.5 | 4 | 0 | 2 | 0 | 12 | 0.25 | 1 | 0 | 5 | 0.2 | 4 | 0.25 | 2 | 0 | 12 | 0.1667 |
| 12_833486 | 4 | 1 | 5 | 0.8 | 6 | 1 | 4 | 1 | 19 | 0.9474 | 2 | 1 | 5 | 0.8 | 4 | 1 | 2 | 1 | 13 | 0.9231 |
| 12_833488 | 5 | 0.6 | 4 | 0.5 | 6 | 0 | 4 | 0 | 19 | 0.2632 | 3 | 0 | 2 | 0.5 | 5 | 0.2 | 2 | 0.5 | 12 | 0.25 |
| 12_833495 | 6 | 0.1667 | 4 | 0 | 6 | 0 | 4 | 0.25 | 20 | 0.1 | NA | NA | 4 | 0 | 5 | 0.4 | 1 | 0 | 10 | 0.2 |
| 12_833498 | 4 | 1 | 6 | 0.8333 | 5 | 1 | 4 | 1 | 19 | 0.9474 | 2 | 0.5 | 4 | 1 | 5 | 1 | 2 | 1 | 13 | 0.9231 |
| 12_833510 | 6 | 0.6667 | 6 | 0.5 | 7 | 0.7143 | 3 | 0.3333 | 22 | 0.5909 | 3 | 0.6667 | 4 | 0.5 | 4 | 0.75 | 2 | 1 | 13 | 0.6923 |
| 12_833512 | 3 | 1 | 4 | 0.75 | 5 | 1 | 4 | 1 | 16 | 0.9375 | 3 | 0.3333 | 4 | 0.75 | 3 | 0.3333 | 1 | 1 | 11 | 0.5455 |
| 12_833517 | 5 | 0.4 | 5 | 0 | 7 | 0 | 3 | 0 | 20 | 0.1 | 3 | 0 | 3 | 0.3333 | 4 | 0.25 | 2 | 0.5 | 12 | 0.25 |
| 12_833518 | 3 | 0.3333 | 4 | 0.25 | 7 | 0.2857 | 4 | 0.25 | 18 | 0.2778 | 2 | 0 | 4 | 0.25 | 4 | 0 | 1 | 0 | 11 | 0.0909 |
| 12_839202 | 1 | 1 | NA | NA | NA | NA | 2 | 1 | 3 | 1 | 2 | 1 | 3 | 1 | NA | NA | NA | NA | 5 | 1 |
| 12_839208 | 1 | 0 | 1 | 1 | NA | NA | 2 | 0 | 4 | 0.25 | 2 | 0.5 | 2 | 0 | NA | NA | NA | NA | 4 | 0.25 |
| 12_839214 | NA | NA | 1 | 1 | NA | NA | NA | NA | 1 | 1 | 2 | 0.5 | 1 | 1 | NA | NA | NA | NA | 3 | 0.6667 |
| 12_839217 | 1 | 0 | 1 | 1 | NA | NA | NA | NA | 2 | 0.5 | 2 | 0.5 | 3 | 0.6667 | NA | NA | NA | NA | 5 | 0.6 |
| 12_839223 | 1 | 1 | 1 | 1 | NA | NA | 1 | 1 | 3 | 1 | 1 | 0 | 3 | 1 | NA | NA | NA | NA | 4 | 0.75 |
| 12_839234 | 1 | 0 | 1 | 0 | NA | NA | 1 | 1 | 3 | 0.3333 | 2 | 0 | 3 | 1 | 1 | 0 | NA | NA | 6 | 0.5 |
| 12_839236 | 1 | 0 | 1 | 1 | NA | NA | 2 | 0.5 | 4 | 0.5 | 1 | 0 | 1 | 1 | 1 | 1 | NA | NA | 3 | 0.6667 |
| 12_839366 | NA | NA | 2 | 0 | NA | NA | 2 | 0.5 | 4 | 0.25 | 1 | 0 | 1 | 1 | 1 | 0 | NA | NA | 3 | 0.3333 |
| 12_839490 | NA | NA | NA | NA | NA | NA | 1 | 1 | 1 | 1 | NA | NA | NA | NA | NA | NA | 1 | 1 | 1 | 1 |
| 12_839558 | NA | NA | NA | NA | NA | NA | 1 | 1 | 1 | 1 | NA | NA | NA | NA | NA | NA | 1 | 1 | 1 | 1 |
| 12_854806 | 79 | 0.1899 | 67 | 0.1045 | 78 | 0.1282 | 64 | 0.0781 | 288 | 0.1285 | 58 | 0.2069 | 83 | 0.1687 | 92 | 0.1087 | 48 | 0.1042 | 281 | 0.1459 |
| 12_854824 | 76 | 0.9737 | 65 | 0.9538 | 74 | 0.973 | 55 | 0.9455 | 270 | 0.963 | 57 | 0.9825 | 80 | 0.975 | 94 | 0.9894 | 50 | 0.92 | 281 | 0.9715 |
| 12_854825 | 55 | 0.6182 | 34 | 0.7647 | 48 | 0.6667 | 45 | 0.6444 | 182 | 0.6648 | 36 | 0.7222 | 59 | 0.678 | 70 | 0.7286 | 37 | 0.6216 | 202 | 0.6931 |
| 12_854828 | 68 | 0.9265 | 56 | 0.9464 | 65 | 0.9846 | 47 | 0.9149 | 236 | 0.9449 | 50 | 0.9 | 66 | 0.9091 | 82 | 0.8415 | 46 | 0.8913 | 244 | 0.8811 |
| 12_854836 | 69 | 0.1159 | 57 | 0.0877 | 67 | 0.0597 | 50 | 0.06 | 243 | 0.0823 | 52 | 0.0577 | 72 | 0.1111 | 81 | 0.0494 | 44 | 0.0682 | 249 | 0.0723 |
| 12_854839 | 64 | 0.4688 | 56 | 0.3393 | 63 | 0.4444 | 49 | 0.6327 | 232 | 0.4655 | 49 | 0.4286 | 74 | 0.4595 | 73 | 0.5068 | 47 | 0.5957 | 243 | 0.4938 |
| 12_854840 | 72 | 0.0417 | 60 | 0.0833 | 71 | 0.0563 | 52 | 0.0769 | 255 | 0.0627 | 54 | 0.0741 | 77 | 0.0779 | 88 | 0.1136 | 50 | 0.12 | 269 | 0.0967 |
| 12_854843 | 61 | 0.9016 | 61 | 0.918 | 55 | 0.9091 | 47 | 0.8936 | 224 | 0.9062 | 46 | 0.8696 | 74 | 0.7973 | 73 | 0.8356 | 43 | 0.7907 | 236 | 0.822 |
| 12_854844 | 50 | 0.84 | 40 | 0.875 | 52 | 0.7885 | 34 | 0.8824 | 176 | 0.8409 | 39 | 0.9487 | 52 | 0.7692 | 63 | 0.7302 | 38 | 0.8421 | 192 | 0.8073 |
| 12_854860 | 66 | 0.2121 | 53 | 0.1698 | 50 | 0.26 | 49 | 0.1429 | 218 | 0.1972 | 46 | 0.1304 | 64 | 0.1875 | 72 | 0.2361 | 39 | 0.1795 | 221 | 0.19 |
| 12_854893 | 52 | 0.75 | 39 | 0.5897 | 43 | 0.6047 | 31 | 0.6452 | 165 | 0.6545 | 39 | 0.6154 | 45 | 0.4222 | 60 | 0.65 | 34 | 0.7353 | 178 | 0.6011 |
| 12_854894 | 60 | 0.8833 | 52 | 0.8846 | 50 | 0.94 | 43 | 0.814 | 205 | 0.8829 | 43 | 0.8605 | 65 | 0.8615 | 72 | 0.875 | 39 | 0.9231 | 219 | 0.8767 |
| 12_854916 | 50 | 0.24 | 43 | 0.2791 | 45 | 0.1778 | 38 | 0.2632 | 176 | 0.2386 | 36 | 0.2778 | 59 | 0.1017 | 57 | 0.193 | 37 | 0.2432 | 189 | 0.1905 |
| 12_854927 | 59 | 0.2542 | 47 | 0.2766 | 47 | 0.2979 | 40 | 0.25 | 193 | 0.2694 | 35 | 0.1714 | 62 | 0.2097 | 64 | 0.3594 | 42 | 0.3333 | 203 | 0.2759 |

| | | | | | | | | | | | | | | | | | | | | |
|---|---|---|---|---|---|---|---|---|---|---|---|---|---|---|---|---|---|---|---|---|
| 12_8549364 | 52 | 0.9615 | 46 | 0.9348 | 43 | 0.9535 | 40 | 0.975 | 181 | 0.9558 | 37 | 0.9189 | 58 | 0.9828 | 65 | 0.9538 | 39 | 1 | 199 | 0.9648 |
| 12_8549373 | 36 | 0.3056 | 32 | 0.375 | 24 | 0.2083 | 30 | 0.1333 | 122 | 0.2623 | 23 | 0.1739 | 38 | 0.1579 | 44 | 0.2955 | 27 | 0.1111 | 132 | 0.197 |
| 12_8549406 | 51 | 0.1569 | 45 | 0.0889 | 46 | 0.087 | 36 | 0.0556 | 178 | 0.1011 | 29 | 0.069 | 59 | 0.0847 | 55 | 0.1455 | 34 | 0.0294 | 177 | 0.0904 |
| 12_8549439 | 49 | 0.449 | 47 | 0.2766 | 44 | 0.4318 | 36 | 0.3889 | 176 | 0.3864 | 35 | 0.4857 | 60 | 0.3167 | 65 | 0.3538 | 40 | 0.35 | 200 | 0.365 |
| 12_8549457 | 40 | 0.1 | 37 | 0.1622 | 34 | 0.0588 | 31 | 0.0645 | 142 | 0.0986 | 25 | 0.04 | 48 | 0.0833 | 49 | 0.0816 | 36 | 0.0278 | 158 | 0.0633 |
| 12_8549511 | 48 | 0.1042 | 40 | 0.05 | 41 | 0.0732 | 33 | 0.0606 | 162 | 0.0741 | 31 | 0.0645 | 56 | 0.1607 | 55 | 0.1273 | 33 | 0.0909 | 175 | 0.12 |
| 12_8636228 | 1 | 1 | NA | NA | NA | NA | NA | NA | 1 | 1 | NA | NA | 1 | 1 | NA | NA | NA | NA | 1 | 1 |
| 12_8636707 | 1 | 1 | NA | NA | NA | NA | NA | NA | 1 | 1 | NA | NA | 1 | 1 | NA | NA | NA | NA | 1 | 1 |
| 12_8636891 | 1 | 1 | NA | NA | NA | NA | NA | NA | 1 | 1 | NA | NA | 1 | 1 | NA | NA | NA | NA | 1 | 1 |
| 12_8779055 | 6 | 0 | 8 | 0.25 | 4 | 0.5 | 10 | 0.2 | 28 | 0.2143 | 5 | 0.2 | 7 | 0.1429 | 9 | 0.2222 | 6 | 0.5 | 27 | 0.2593 |
| 12_8799699 | 7 | 0 | 14 | 0.2143 | 7 | 0 | 8 | 0.125 | 36 | 0.1111 | 14 | 0.1429 | 7 | 0 | 11 | 0.0909 | 5 | 0 | 37 | 0.0811 |
| 12_8799723 | 11 | 0.0909 | 17 | 0.4706 | 10 | 0.3 | 13 | 0.2308 | 51 | 0.2941 | 12 | 0.1667 | 9 | 0 | 13 | 0.3077 | 9 | 0.1111 | 43 | 0.1628 |
| 12_8799836 | 11 | 0.0909 | 17 | 0.2941 | 11 | 0.1818 | 11 | 0.3636 | 50 | 0.24 | 12 | 0.4167 | 11 | 0 | 10 | 0 | 12 | 0.25 | 45 | 0.1778 |
| 12_8801185 | 17 | 0.4118 | 18 | 0.7222 | 14 | 0.4286 | 19 | 0.3684 | 68 | 0.4853 | 13 | 0.3846 | 14 | 0.4286 | 13 | 0.5385 | 15 | 0.5333 | 55 | 0.4727 |
| 12_8801332 | 12 | 0 | 20 | 0 | 9 | 0.2222 | 11 | 0.1818 | 52 | 0.0769 | 6 | 0 | 9 | 0.2222 | 6 | 0 | 10 | 0.2 | 31 | 0.129 |
| 12_8988340 | 35 | 0.2286 | 37 | 0.5405 | 33 | 0.2424 | 22 | 0.3636 | 127 | 0.3465 | 35 | 0.1714 | 34 | 0.4706 | 25 | 0.24 | 23 | 0.3478 | 117 | 0.3077 |
| 12_8988381 | 42 | 0.0952 | 38 | 0.0789 | 39 | 0.0513 | 23 | 0.087 | 142 | 0.0775 | 36 | 0.0833 | 37 | 0.1081 | 28 | 0.1786 | 26 | 0.0769 | 127 | 0.1102 |
| 12_9078167 | 5 | 0 | 8 | 0.25 | 5 | 0.6 | 5 | 0.6 | 23 | 0.3478 | 4 | 0.25 | 3 | 0 | 3 | 0 | 4 | 0.75 | 14 | 0.2857 |
| 12_9078430 | 6 | 0.1667 | 3 | 0.3333 | 3 | 0 | 7 | 0.2857 | 19 | 0.2105 | 5 | 0 | 3 | 0 | 2 | 0.5 | 4 | 0 | 14 | 0.0714 |
| 12_9078756 | 5 | 0 | 3 | 0.3333 | 3 | 0 | 6 | 0.1667 | 17 | 0.1176 | 5 | 0.2 | 3 | 0 | 3 | 0 | 3 | 0 | 14 | 0.0714 |
| 12_9085998 | 2 | 0.5 | 2 | 1 | 2 | 0.5 | NA | NA | 6 | 0.6667 | 1 | 0 | 1 | 0 | 3 | 1 | 3 | 0.6667 | 8 | 0.625 |
| 12_1044117 | 6 | 0.5 | 10 | 0.2 | 4 | 0 | 5 | 0 | 25 | 0.2 | 3 | 0 | 8 | 0.125 | 3 | 0 | 2 | 0 | 16 | 0.0625 |
| 12_1044121 | 5 | 0.6 | 6 | 0.6667 | 3 | 0.3333 | 4 | 0.25 | 18 | 0.5 | 4 | 0.25 | 4 | 0.75 | 5 | 0.6 | 2 | 0 | 15 | 0.4667 |
| 12_1044467 | 5 | 1 | 8 | 0.75 | 6 | 1 | 3 | 0.6667 | 22 | 0.8636 | 5 | 0.4 | 7 | 0.7143 | 6 | 0.6667 | 2 | 0.5 | 20 | 0.6 |
| 12_1044475 | 4 | 0.5 | 9 | 0.4444 | 6 | 0.3333 | 5 | 0.4 | 24 | 0.4167 | 5 | 0.2 | 8 | 0.75 | 6 | 0.6667 | 2 | 0.5 | 21 | 0.5714 |
| 12_1044476 | 4 | 0.25 | 7 | 0.5714 | 4 | 0.5 | 6 | 0.6667 | 21 | 0.5238 | 1 | 1 | 4 | 0.75 | 6 | 0.1667 | 2 | 0 | 13 | 0.3846 |
| 12_1044481 | 2 | 0 | 7 | 0.1429 | 5 | 0 | 6 | 0.1667 | 20 | 0.1 | 3 | 0 | 7 | 0.1429 | 5 | 0 | 2 | 0 | 17 | 0.0588 |
| 12_1044486 | 1 | 1 | 7 | 0.7143 | 3 | 0.6667 | 4 | 0.5 | 15 | 0.6667 | 4 | 0.75 | 5 | 1 | 6 | 1 | 3 | 1 | 18 | 0.9444 |
| 12_1044497 | 2 | 0 | 8 | 0.125 | 4 | 0.25 | 5 | 0 | 19 | 0.1053 | 4 | 0 | 6 | 0 | 7 | 0.1429 | 2 | 0 | 19 | 0.0526 |
| 12_1044511 | 3 | 0 | 9 | 0.4444 | 6 | 0.6667 | 6 | 0.3333 | 24 | 0.4167 | 4 | 0 | 8 | 0.125 | 7 | 0.5714 | 3 | 1 | 22 | 0.3636 |
| 12_1044517 | 3 | 0.3333 | 10 | 0.6 | 8 | 0.875 | 7 | 0.2857 | 28 | 0.5714 | 5 | 0.8 | 8 | 0.5 | 6 | 0.6667 | 3 | 1 | 22 | 0.6818 |
| 12_1044528 | 4 | 0.25 | 10 | 0.4 | 8 | 0.625 | 6 | 0.3333 | 28 | 0.4286 | 4 | 0.5 | 7 | 0.5714 | 7 | 0.7143 | 2 | 1 | 20 | 0.65 |
| 12_1094878 | 6 | 0 | 4 | 0 | 4 | 0.25 | NA | NA | 14 | 0.0714 | 2 | 0 | 2 | 0 | 6 | 0 | 3 | 0.6667 | 13 | 0.1538 |
| 12_1114109 | 1 | 1 | 4 | 1 | 1 | 1 | NA | NA | 6 | 1 | NA | NA | NA | NA | NA | NA | 1 | 1 | 1 | 1 |
| 12_1114123 | 1 | 1 | 4 | 1 | 1 | 1 | NA | NA | 6 | 1 | NA | NA | NA | NA | NA | NA | 1 | 1 | 1 | 1 |
| 12_1134158 | 2 | 0 | 2 | 0.5 | NA | NA | 2 | 0 | 6 | 0.1667 | 1 | 0 | 1 | 0 | 3 | 0.3333 | 1 | 0 | 6 | 0.1667 |
| 12_1136793 | 3 | 0.3333 | 2 | 0.5 | 3 | 0.6667 | 2 | 0.5 | 10 | 0.5 | 1 | 1 | 1 | 0 | 3 | 0.6667 | 5 | 1 | 10 | 0.8 |
| 12_1137615 | 61 | 0.918 | 86 | 0.9302 | 57 | 0.8772 | 59 | 0.9322 | 263 | 0.9163 | 54 | 0.9259 | 32 | 0.8125 | 51 | 0.9412 | 57 | 0.8947 | 194 | 0.9021 |
| 12_1137617 | 57 | 0.6491 | 81 | 0.716 | 56 | 0.5893 | 54 | 0.6852 | 248 | 0.6653 | 49 | 0.7755 | 39 | 0.7692 | 47 | 0.6383 | 58 | 0.6897 | 193 | 0.715 |
| 12_1137628 | 52 | 0.9423 | 87 | 0.931 | 62 | 0.9677 | 59 | 1 | 260 | 0.9577 | 52 | 0.9808 | 34 | 1 | 50 | 0.96 | 67 | 0.9851 | 203 | 0.9803 |
| 12_1137632 | 56 | 0.1964 | 91 | 0.0549 | 63 | 0.0794 | 59 | 0.0508 | 269 | 0.0892 | 52 | 0.0385 | 37 | 0.1081 | 54 | 0.2037 | 60 | 0.1333 | 203 | 0.1232 |
| 12_1148640 | 47 | 0.0851 | 77 | 0.0649 | 52 | 0.0769 | 52 | 0.0962 | 228 | 0.0789 | 45 | 0.0889 | 30 | 0.0667 | 42 | 0.0714 | 53 | 0.1321 | 170 | 0.0941 |
| 12_1242294 | 14 | 1 | 10 | 1 | 12 | 1 | 24 | 0.9167 | 60 | 0.9667 | 11 | 1 | 25 | 0.88 | 17 | 1 | 20 | 0.95 | 73 | 0.9452 |
| 12_1242295 | 14 | 0.5 | 10 | 0.4 | 10 | 0.3 | 21 | 0.381 | 55 | 0.4 | 9 | 0.4444 | 24 | 0.2083 | 13 | 0.3077 | 15 | 0.5333 | 61 | 0.3443 |
| 12_1242298 | 14 | 0.3571 | 11 | 0.2727 | 9 | 0.5556 | 24 | 0.3333 | 58 | 0.3621 | 8 | 0.375 | 22 | 0.4091 | 16 | 0.75 | 16 | 0.5625 | 62 | 0.5323 |
| 12_1242349 | 11 | 0.0909 | 7 | 0.4286 | 9 | 0 | 13 | 0.1538 | 40 | 0.15 | 7 | 0.1429 | 14 | 0.2143 | 11 | 0.4545 | 13 | 0.2308 | 45 | 0.2667 |
| 12_1242352 | 11 | 0 | 8 | 0.25 | 10 | 0.1 | 16 | 0.0625 | 45 | 0.0889 | 6 | 0.3333 | 17 | 0 | 17 | 0.0588 | 15 | 0.0667 | 55 | 0.0727 |
| 12_1242360 | 11 | 0.4545 | 8 | 0.5 | 8 | 0.375 | 15 | 0.5333 | 42 | 0.4762 | 5 | 0.4 | 16 | 0.375 | 17 | 0.3529 | 18 | 0.5 | 56 | 0.4107 |
| 12_1242363 | 11 | 0 | 8 | 0 | 11 | 0.0909 | 17 | 0.1176 | 47 | 0.0638 | 7 | 0 | 18 | 0 | 17 | 0.1176 | 18 | 0.1111 | 60 | 0.0667 |
| 12_1242365 | 11 | 0.0909 | 8 | 0.625 | 11 | 0.0909 | 17 | 0.2941 | 47 | 0.2553 | 7 | 0.4286 | 16 | 0.125 | 16 | 0.375 | 19 | 0.4211 | 58 | 0.3276 |
| 12_1242375 | 12 | 0.5 | 8 | 0.875 | 11 | 0.8182 | 16 | 0.8125 | 47 | 0.7447 | 6 | 1 | 19 | 0.6842 | 16 | 0.75 | 17 | 0.8824 | 58 | 0.7931 |
| 12_1242423 | 13 | 0 | 7 | 0.4286 | 9 | 0.3333 | 19 | 0.4737 | 48 | 0.3125 | 7 | 0.7143 | 19 | 0.2105 | 15 | 0.4667 | 18 | 0.3889 | 59 | 0.3898 |
| 12_1242426 | 14 | 0 | 9 | 0.2222 | 11 | 0.0909 | 20 | 0 | 54 | 0.0556 | 5 | 0.4 | 21 | 0.0952 | 17 | 0.0588 | 18 | 0.1111 | 61 | 0.1148 |
| 12_1242590 | 17 | 0.1176 | 14 | 0 | 16 | 0.0625 | 26 | 0.2308 | 73 | 0.1233 | 10 | 0 | 27 | 0.1481 | 20 | 0.05 | 21 | 0.1429 | 78 | 0.1026 |
| 12_1242654 | 4 | 0 | 4 | 0.25 | 3 | 0 | 3 | 0 | 14 | 0.0714 | NA | NA | 6 | 0 | 4 | 0.25 | NA | NA | 10 | 0.1 |
| 12_1242790 | 2 | 0 | 3 | 0 | 2 | 0 | 4 | 0.25 | 11 | 0.0909 | 2 | 0 | 2 | 0.5 | 3 | 0 | NA | NA | 7 | 0.1429 |
| 12_1242830 | 3 | 0 | 5 | 0 | 3 | 0 | 4 | 0.25 | 15 | 0.0667 | 2 | 0 | 1 | 0 | 5 | 0.2 | 1 | 0 | 9 | 0.1111 |
| 12_1242913 | 4 | 0 | 7 | 0.1429 | 2 | 0.5 | 5 | 0.2 | 18 | 0.1667 | 3 | 0.3333 | 1 | 0 | 6 | 0.1667 | 1 | 0 | 11 | 0.1818 |
| 12_1242942 | 3 | 0 | 5 | 0.4 | 1 | 0 | 2 | 0 | 11 | 0.1818 | 1 | 1 | 1 | 0 | 4 | 0 | 1 | 0 | 7 | 0.1429 |
| 12_1243073 | 1 | 0 | 2 | 0 | 2 | 0.5 | 3 | 0 | 8 | 0.125 | 1 | 1 | 2 | 0.5 | 3 | 0.3333 | 2 | 0.5 | 8 | 0.5 |
| 12_1298653 | NA | NA | NA | NA | 3 | 0.3333 | NA | NA | 3 | 0.3333 | 2 | 1 | NA | NA | 1 | 0 | 1 | 0 | 4 | 0.5 |
| 12_1298670 | NA | NA | NA | NA | 3 | 0.3333 | NA | NA | 3 | 0.3333 | 2 | 1 | NA | NA | 1 | 1 | 2 | 1 | 5 | 1 |
| 12_1298683 | NA | NA | NA | NA | 1 | 1 | NA | NA | 1 | 1 | NA | NA | NA | NA | NA | NA | 1 | 1 | 1 | 1 |
| 12_1298699 | NA | NA | NA | NA | 2 | 1 | NA | NA | 2 | 1 | 2 | 1 | NA | NA | 1 | 1 | NA | NA | 3 | 1 |
| 12_1298701 | NA | NA | NA | NA | 3 | 1 | NA | NA | 3 | 1 | 2 | 1 | NA | NA | 1 | 1 | 1 | 1 | 4 | 1 |

| | | | | | | | | | | | | | | | | | | | | |
|---|---|---|---|---|---|---|---|---|---|---|---|---|---|---|---|---|---|---|---|---|
| 12_129870( | NA | NA | NA | NA | 2 | 0.5 | NA | NA | 2 | 0.5 | 2 | 0.5 | NA | NA | 1 | 1 | 1 | 0 | 4 | 0.5 |
| 12_129871ɛ | NA | NA | NA | NA | 3 | 0.6667 | NA | NA | 3 | 0.6667 | 2 | 1 | NA | NA | 1 | 0 | 1 | 0 | 4 | 0.5 |
| 12_129872( | NA | NA | NA | NA | 2 | 0.5 | NA | NA | 2 | 0.5 | 2 | 0 | NA | NA | 1 | 1 | 1 | 0 | 4 | 0.25 |
| 12_129900! | NA | NA | NA | NA | 2 | 1 | NA | NA | 2 | 1 | 1 | 1 | NA | NA | 1 | 1 | 1 | 1 | 3 | 1 |
| 12_136335: | 5 | 0.4 | 5 | 0.4 | 3 | 0.6667 | 4 | 0.5 | 17 | 0.4706 | 5 | 0.4 | 5 | 0.6 | 5 | 1 | 4 | 0.5 | 19 | 0.6316 |
| 12_136336( | 6 | 0 | 4 | 0.25 | 3 | 0 | 3 | 0.3333 | 16 | 0.125 | 4 | 0.25 | 6 | 0.5 | 5 | 0.2 | 4 | 0.25 | 19 | 0.3158 |
| 12_136336: | 5 | 0.2 | 4 | 0 | 2 | 0 | 4 | 0.5 | 15 | 0.2 | 3 | 0.3333 | 6 | 0.1667 | 2 | 0.5 | 2 | 0.5 | 13 | 0.3077 |
| 12_136338( | 2 | 0.5 | 1 | 0 | 1 | 0 | 4 | 0 | 8 | 0.125 | 5 | 0.2 | 3 | 0 | 2 | 0 | 3 | 0 | 13 | 0.0769 |
| 12_141971( | NA | NA | 2 | 1 | NA | NA | 1 | 0 | 3 | 0.6667 | NA | NA | 1 | 0 | 2 | 0.5 | NA | NA | 3 | 0.3333 |
| 12_141975· | NA | NA | NA | NA | NA | NA | 1 | 1 | 1 | 1 | NA | NA | 1 | 1 | 2 | 0.5 | NA | NA | 3 | 0.6667 |
| 12_141975ɛ | NA | NA | 1 | 1 | NA | NA | NA | NA | 1 | 1 | NA | NA | 1 | 0 | 2 | 1 | NA | NA | 3 | 0.6667 |
| 12_141977! | 17 | 0.2353 | 14 | 0.3571 | 10 | 0.3 | 10 | 0.1 | 51 | 0.2549 | 7 | 0.4286 | 8 | 0.125 | 10 | 0.5 | 15 | 0.2667 | 40 | 0.325 |
| 12_141978: | 14 | 0.5 | 15 | 0.2 | 8 | 0.375 | 10 | 0.2 | 47 | 0.3191 | 8 | 0.25 | 9 | 0.3333 | 10 | 0 | 17 | 0.3529 | 44 | 0.25 |
| 12_141980ɛ | 7 | 0.1429 | 14 | 0.0714 | 5 | 0 | 7 | 0.2857 | 33 | 0.1212 | 4 | 0.25 | 6 | 0 | 10 | 0.1 | 8 | 0 | 28 | 0.0714 |
| 12_141990( | 12 | 0.1667 | 13 | 0.3077 | 4 | 0 | 7 | 0.1429 | 36 | 0.1944 | 3 | 0.3333 | 4 | 0.5 | 8 | 0.125 | 10 | 0.1 | 25 | 0.2 |
| 12_141990: | 13 | 0.4615 | 9 | 0 | 7 | 0 | 7 | 0.1429 | 36 | 0.1944 | 5 | 0.6 | 2 | 0 | 4 | 0.75 | 8 | 0.25 | 19 | 0.4211 |
| 12_141992· | 13 | 0.3077 | 11 | 0.1818 | 6 | 0.1667 | 8 | 0.25 | 38 | 0.2368 | 4 | 0.25 | 5 | 0.2 | 6 | 0.1667 | 11 | 0 | 26 | 0.1154 |
| 12_141993· | 12 | 0.9167 | 11 | 0.8182 | 7 | 0.5714 | 6 | 0.3333 | 36 | 0.7222 | 5 | 0.8 | 5 | 0.8 | 6 | 1 | 10 | 0.6 | 26 | 0.7692 |
| 12_141993· | 10 | 0.4 | 9 | 0.2222 | 5 | 0.6 | 8 | 0.25 | 32 | 0.3438 | 7 | 0.5714 | 5 | 0.2 | 3 | 0.3333 | 11 | 0.2727 | 26 | 0.3462 |
| 12_141994: | 11 | 0.1818 | 10 | 0.2 | 5 | 0.6 | 8 | 0.25 | 34 | 0.2647 | 5 | 0.2 | 4 | 0 | 6 | 0.3333 | 11 | 0.0909 | 26 | 0.1538 |
| 12_141995· | 11 | 0.3636 | 9 | 0.4444 | 6 | 0.3333 | 8 | 0.25 | 34 | 0.3529 | 6 | 0.5 | 4 | 0.25 | 4 | 0.5 | 10 | 0.6 | 24 | 0.5 |
| 12_141995: | 10 | 0.9 | 9 | 1 | 4 | 0.75 | 7 | 0.8571 | 30 | 0.9 | 4 | 1 | 5 | 0.4 | 4 | 1 | 10 | 1 | 23 | 0.8696 |
| 12_141996: | 9 | 0.3333 | 8 | 0 | 2 | 0 | 7 | 0 | 26 | 0.1154 | 4 | 0 | 3 | 0 | 4 | 0.25 | 9 | 0.1111 | 20 | 0.1 |
| 12_141997: | 4 | 1 | 4 | 0.75 | 1 | 1 | 4 | 0.75 | 13 | 0.8462 | 4 | 1 | 2 | 0.5 | 4 | 0.75 | 4 | 1 | 14 | 0.8571 |
| 12_141998( | 9 | 1 | 9 | 1 | 3 | 1 | 7 | 1 | 28 | 1 | 5 | 1 | 5 | 1 | 5 | 1 | 8 | 1 | 23 | 1 |
| 12_141999: | 10 | 0.9 | 8 | 0.875 | 4 | 0.75 | 7 | 0.4286 | 29 | 0.7586 | 5 | 0.4 | 3 | 0.6667 | 5 | 1 | 7 | 0.8571 | 20 | 0.75 |
| 12_142000! | 6 | 0.6667 | 7 | 0.8571 | 4 | 0.75 | 7 | 0.7143 | 24 | 0.75 | 5 | 0.8 | 3 | 0.3333 | 3 | 1 | 6 | 0.5 | 17 | 0.6471 |
| 12_142002: | 4 | 0.75 | 5 | 0.8 | 4 | 0.75 | 5 | 1 | 18 | 0.8333 | 5 | 1 | 4 | 0.75 | 4 | 0.75 | 5 | 1 | 18 | 0.8889 |
| 12_142004: | 3 | 0.3333 | 5 | 0.6 | 4 | 1 | 5 | 0.4 | 17 | 0.5882 | 5 | 0.6 | 3 | 0.3333 | 4 | 0.5 | 4 | 0.25 | 16 | 0.4375 |
| 12_142005: | 5 | 1 | 5 | 0.4 | 3 | 0.6667 | 4 | 0.75 | 17 | 0.7059 | 4 | 1 | 3 | 0.3333 | 5 | 0.8 | 4 | 0.75 | 16 | 0.75 |
| 12_142005! | 5 | 0.2 | 5 | 0.2 | 3 | 0 | 5 | 0.2 | 18 | 0.1667 | 4 | 0.5 | 4 | 0 | 4 | 0 | 6 | 0.3333 | 18 | 0.2222 |
| 12_142006( | 5 | 0 | 4 | 0 | 3 | 0.3333 | 6 | 0.3333 | 18 | 0.1667 | 4 | 0.25 | 4 | 0.25 | 4 | 0 | 6 | 0.1667 | 18 | 0.1667 |
| 12_158418· | 2 | 0 | 1 | 1 | 4 | 0 | 2 | 0.5 | 9 | 0.2222 | 3 | 0 | 3 | 0 | 6 | 0.1667 | 1 | 0 | 13 | 0.0769 |
| 12_158418ɛ | 1 | 1 | 2 | 1 | 6 | 0.8333 | 2 | 1 | 11 | 0.9091 | 3 | 1 | 3 | 1 | 5 | 1 | 2 | 1 | 13 | 1 |
| 12_158420· | 1 | 0 | 2 | 0 | 6 | 0.3333 | 2 | 0 | 11 | 0.1818 | 3 | 0 | 3 | 0.3333 | 5 | 0 | 1 | 0 | 12 | 0.0833 |
| 12_158420( | 1 | 1 | 2 | 1 | 5 | 0.8 | 2 | 0.5 | 10 | 0.8 | 3 | 0.6667 | 2 | 0.5 | 5 | 0.8 | 2 | 1 | 12 | 0.75 |
| 12_158421: | 1 | 1 | 2 | 1 | 4 | 0.5 | 2 | 0.5 | 9 | 0.6667 | 2 | 1 | 3 | 0.6667 | 5 | 1 | 1 | 1 | 11 | 0.9091 |
| 12_158435· | 1 | 1 | 1 | 1 | 3 | 1 | NA | NA | 5 | 1 | 3 | 1 | 1 | 1 | 3 | 1 | 1 | 1 | 8 | 1 |
| 12_158435! | 1 | 1 | NA | NA | 3 | 1 | NA | NA | 4 | 1 | 3 | 1 | 2 | 1 | 2 | 1 | 1 | 1 | 8 | 1 |
| 12_158436: | 1 | 1 | 1 | 1 | 3 | 0.3333 | NA | NA | 5 | 0.6 | 3 | 1 | 2 | 1 | 3 | 1 | NA | NA | 8 | 1 |
| 12_158436: | 1 | 0 | 1 | 1 | 3 | 0.6667 | NA | NA | 5 | 0.6 | 3 | 0.6667 | 2 | 0.5 | 2 | 0.5 | 2 | 0 | 9 | 0.4444 |
| 12_158437: | NA | NA | 1 | 1 | 3 | 1 | NA | NA | 4 | 1 | 2 | 1 | 2 | 1 | 2 | 0.5 | 2 | 1 | 8 | 0.875 |
| 12_158438· | 1 | 0 | 1 | 1 | 3 | 1 | NA | NA | 5 | 0.8 | 3 | 1 | 2 | 1 | 1 | 0 | 2 | 0.5 | 8 | 0.75 |
| 12_158438: | 1 | 1 | 1 | 1 | 2 | 0 | NA | NA | 4 | 0.5 | 2 | 0 | 2 | 0.5 | 3 | 0.6667 | NA | NA | 7 | 0.4286 |
| 12_158653· | NA | NA | 1 | 0 | 3 | 0.3333 | NA | NA | 4 | 0.25 | 1 | 0 | 2 | 0.5 | 2 | 0.5 | NA | NA | 5 | 0.4 |
| 12_158653! | NA | NA | 1 | 0 | 2 | 0.5 | NA | NA | 3 | 0.3333 | 1 | 1 | 2 | 0 | 3 | 0.6667 | 2 | 0 | 8 | 0.375 |
| 12_158654! | NA | NA | NA | NA | 2 | 1 | NA | NA | 2 | 1 | 2 | 0.5 | 2 | 0.5 | 1 | 1 | 1 | 0 | 6 | 0.5 |
| 12_158656! | NA | NA | 1 | 0 | 1 | 1 | NA | NA | 2 | 0.5 | 2 | 0.5 | 1 | 1 | 1 | 0 | NA | NA | 4 | 0.5 |
| 12_158657· | NA | NA | 1 | 1 | 1 | 1 | NA | NA | 2 | 1 | 2 | 0.5 | 2 | 0.5 | 2 | 0 | 1 | 0 | 7 | 0.2857 |
| 12_158661: | NA | NA | 1 | 1 | 1 | 1 | NA | NA | 2 | 1 | 2 | 0 | NA | NA | 2 | 0 | 1 | 1 | 5 | 0.2 |
| 12_165866( | 3 | 0.3333 | 1 | 0 | 2 | 0 | 7 | 0.1429 | 13 | 0.1538 | 5 | 0 | 4 | 0 | 3 | 0.6667 | 1 | 0 | 13 | 0.1538 |
| 12_165867: | 3 | 0.6667 | 1 | 1 | 2 | 1 | 5 | 0.6 | 11 | 0.7273 | 5 | 0.6 | 4 | 0.75 | 3 | 0.3333 | 1 | 1 | 13 | 0.6154 |
| 12_165870· | 2 | 1 | 1 | 1 | 2 | 0.5 | 5 | 0.2 | 10 | 0.5 | 4 | 0.25 | 2 | 0.5 | 1 | 1 | 1 | 0 | 8 | 0.375 |
| 12_165875! | 5 | 0.2 | 1 | 1 | 1 | 0 | 6 | 0.1667 | 13 | 0.2308 | 4 | 0.25 | 2 | 0 | 2 | 0.5 | 1 | 0 | 9 | 0.2222 |
| 12_165875! | 5 | 0 | 1 | 1 | 1 | 0 | 6 | 0.1667 | 13 | 0.1538 | 4 | 0.25 | 2 | 0.5 | 1 | 1 | 1 | 0 | 8 | 0.375 |
| 12_165878! | 4 | 0.25 | NA | NA | 1 | 0 | 7 | 0.1429 | 12 | 0.1667 | 4 | 0 | 2 | 1 | 2 | 0.5 | 1 | 0 | 9 | 0.3333 |
| 12_165879ɛ | 5 | 0 | 1 | 0 | 1 | 0 | 7 | 0.4286 | 14 | 0.2143 | 3 | 0 | 3 | 0.3333 | 2 | 0 | 1 | 0 | 9 | 0.1111 |
| 12_165880· | 4 | 0.5 | 1 | 1 | 1 | 0 | 4 | 0.5 | 10 | 0.5 | 3 | 0.3333 | 2 | 1 | 2 | 1 | NA | NA | 7 | 0.7143 |
| 12_165883ɛ | 4 | 1 | 1 | 1 | 1 | 1 | 4 | 0.75 | 10 | 0.9 | 2 | 1 | 1 | 1 | 2 | 1 | NA | NA | 5 | 1 |
| 12_165884: | 4 | 1 | 1 | 0 | 1 | 1 | 4 | 0.5 | 10 | 0.7 | 2 | 0.5 | 2 | 1 | 2 | 1 | NA | NA | 6 | 0.8333 |
| 12_165884ɛ | 4 | 0 | NA | NA | NA | NA | 4 | 0.25 | 8 | 0.125 | 2 | 0.5 | 2 | 0 | 2 | 0 | NA | NA | 6 | 0.1667 |
| 12_165884! | 5 | 0.2 | 1 | 0 | 1 | 1 | 4 | 0.25 | 11 | 0.2727 | 2 | 1 | 2 | 0.5 | 2 | 0 | NA | NA | 6 | 0.5 |
| 12_166155( | 1 | 1 | NA | NA | NA | NA | 1 | 0 | 2 | 0.5 | 1 | 0 | 2 | 0.5 | 2 | 0 | NA | NA | 5 | 0.2 |
| 12_167035: | 1 | 0 | 3 | 0 | 2 | 0 | 5 | 0.2 | 11 | 0.0909 | 4 | 0.25 | 3 | 0.6667 | 6 | 0.1667 | 1 | 0 | 14 | 0.2857 |
| 12_167040: | 27 | 0.037 | 24 | 0.0833 | 22 | 0.0455 | 21 | 0.0952 | 94 | 0.0638 | 16 | 0.125 | 15 | 0.0667 | 22 | 0 | 18 | 0.0556 | 71 | 0.0563 |

| | | | | | | | | | | | | | | | | | | | | |
|---|---|---|---|---|---|---|---|---|---|---|---|---|---|---|---|---|---|---|---|---|
| 12_1670768 | NA | NA | 4 | 0.25 | 1 | 1 | NA | NA | 5 | 0.4 | NA | NA | 1 | 0 | 1 | 1 | NA | NA | 2 | 0.5 |
| 12_1670997 | 39 | 0.1026 | 16 | 0.125 | 20 | 0 | 20 | 0.2 | 95 | 0.1053 | 21 | 0.1905 | 15 | 0 | 23 | 0.0435 | 24 | 0.1667 | 83 | 0.1084 |
| 12_1670999 | 46 | 0.5217 | 23 | 0.5217 | 23 | 0.6522 | 26 | 0.7308 | 118 | 0.5932 | 24 | 0.5833 | 20 | 0.7 | 30 | 0.6 | 28 | 0.4643 | 102 | 0.5784 |
| 12_1671004 | 40 | 0.375 | 24 | 0.5 | 24 | 0.4583 | 28 | 0.3571 | 116 | 0.4138 | 24 | 0.4583 | 17 | 0.4706 | 29 | 0.3448 | 22 | 0.4545 | 92 | 0.4239 |
| 12_1687124 | 1 | 1 | 1 | 1 | 1 | 0 | 1 | 1 | 4 | 0.75 | 1 | 1 | 1 | 1 | 1 | 1 | NA | NA | 3 | 1 |
| 12_1687555 | 7 | 0.2857 | 5 | 0.8 | 3 | 0.6667 | 2 | 1 | 17 | 0.5882 | 3 | 0.6667 | 8 | 0.5 | 2 | 0.5 | 4 | 0.75 | 17 | 0.5882 |
| 12_1687564 | 9 | 0 | 4 | 0.25 | 4 | 0 | 1 | 0 | 18 | 0.0556 | 1 | 0 | 7 | 0 | 5 | 0.2 | 4 | 0.25 | 17 | 0.1176 |
| 12_1687567 | 9 | 0.4444 | 6 | 0.3333 | 3 | 0.6667 | 2 | 0.5 | 20 | 0.45 | 2 | 0 | 8 | 0.125 | 5 | 0.4 | 3 | 0 | 18 | 0.1667 |
| 12_1687571 | 9 | 0.6667 | 3 | 0.6667 | 4 | 0.5 | 1 | 1 | 17 | 0.6471 | 3 | 0 | 8 | 0.625 | 3 | 0.6667 | 4 | 0 | 18 | 0.3889 |
| 12_1687572 | 9 | 0.4444 | 4 | 0.5 | 2 | 1 | 3 | 0.6667 | 18 | 0.5556 | 2 | 0 | 6 | 0.6667 | 4 | 0.75 | 4 | 0.75 | 16 | 0.625 |
| 12_1687573 | 6 | 0.1667 | 4 | 0 | 3 | 0 | 2 | 0 | 15 | 0.0667 | 3 | 0 | 4 | 0.25 | 2 | 0 | 4 | 0.25 | 13 | 0.1538 |
| 12_1687574 | 8 | 1 | 6 | 1 | 3 | 1 | 2 | 1 | 19 | 1 | 3 | 1 | 8 | 1 | 5 | 1 | 4 | 0.75 | 20 | 0.95 |
| 12_1687589 | 9 | 0.5556 | 6 | 0.6667 | 4 | 0.25 | 3 | 1 | 22 | 0.5909 | 3 | 0.3333 | 5 | 0.6 | 5 | 0.2 | 4 | 0.25 | 17 | 0.3529 |
| 12_1687607 | 7 | 0.5714 | 5 | 0.4 | 4 | 0.75 | 2 | 0 | 18 | 0.5 | 3 | 0 | 8 | 0.5 | 4 | 0.25 | 3 | 0.3333 | 18 | 0.3333 |
| 12_1687622 | 8 | 0.375 | 6 | 0.6667 | 3 | 0.6667 | 1 | 0 | 18 | 0.5 | 2 | 0 | 7 | 0.2857 | 5 | 0.2 | 5 | 0.2 | 19 | 0.2105 |
| 12_1687650 | 9 | 0.2222 | 8 | 0.25 | 5 | 0 | 4 | 0.25 | 26 | 0.1923 | 3 | 0 | 8 | 0.125 | 5 | 0.2 | 6 | 0.1667 | 22 | 0.1364 |
| 12_1699291 | 2 | 0.5 | 7 | 0.1429 | 5 | 0.2 | 3 | 0 | 17 | 0.1765 | NA | NA | 2 | 0.5 | 3 | 0 | 6 | 0 | 11 | 0.0909 |
| 12_1702763 | 11 | 0.1818 | 13 | 0.3077 | 9 | 0.2222 | 9 | 0.1111 | 42 | 0.2143 | 2 | 0.5 | 7 | 0 | 11 | 0.4545 | 11 | 0.2727 | 31 | 0.2903 |
| 12_1703469 | 11 | 0.7273 | 17 | 0.7647 | 9 | 1 | 11 | 0.6364 | 48 | 0.7708 | 3 | 1 | 7 | 0.4286 | 13 | 0.6154 | 11 | 0.7273 | 34 | 0.6471 |
| 12_1703471 | 11 | 0.2727 | 18 | 0.1667 | 10 | 0.4 | 9 | 0.3333 | 48 | 0.2708 | 3 | 0.3333 | 7 | 0.1429 | 13 | 0.2308 | 13 | 0.1538 | 36 | 0.1944 |
| 12_1703474 | 13 | 0.4615 | 18 | 0.5556 | 8 | 0.875 | 12 | 0.5833 | 51 | 0.5882 | 3 | 0.3333 | 7 | 0.4286 | 12 | 0.5 | 12 | 0.5 | 34 | 0.4706 |
| 12_1703484 | 11 | 0.0909 | 12 | 0.1667 | 7 | 0.2857 | 7 | 0.2857 | 37 | 0.1892 | 2 | 0.5 | 7 | 0 | 11 | 0.2727 | 10 | 0.3 | 30 | 0.2333 |
| 12_1703489 | 13 | 0.3077 | 20 | 0.35 | 9 | 0.2222 | 11 | 0.4545 | 53 | 0.3396 | 3 | 0.6667 | 8 | 0.375 | 13 | 0.3846 | 13 | 0.3077 | 37 | 0.3784 |
| 12_1703492 | 12 | 1 | 16 | 1 | 8 | 1 | 12 | 0.75 | 48 | 0.9375 | 3 | 1 | 7 | 0.8571 | 12 | 1 | 13 | 1 | 35 | 0.9714 |
| 12_1703502 | 14 | 0.2143 | 20 | 0.3 | 10 | 0 | 12 | 0.25 | 56 | 0.2143 | 3 | 0 | 9 | 0.1111 | 13 | 0.2308 | 12 | 0.0833 | 37 | 0.1351 |
| 12_1703504 | 11 | 0.1818 | 17 | 0.2353 | 8 | 0 | 6 | 0.1667 | 42 | 0.1667 | 2 | 0 | 8 | 0.125 | 7 | 0.2857 | 7 | 0 | 24 | 0.125 |
| 12_1703568 | 16 | 0.0625 | 21 | 0.0952 | 12 | 0.0833 | 14 | 0.1429 | 63 | 0.0952 | 6 | 0.1667 | 11 | 0.1818 | 14 | 0 | 15 | 0.2 | 46 | 0.1304 |
| 12_1703759 | 23 | 0 | 20 | 0.05 | 14 | 0.2857 | 15 | 0.0667 | 72 | 0.0833 | 6 | 0 | 16 | 0.125 | 18 | 0.1111 | 13 | 0 | 53 | 0.0755 |
| 12_1703762 | 22 | 0.0909 | 21 | 0.0476 | 17 | 0.0588 | 17 | 0 | 77 | 0.0519 | 7 | 0 | 17 | 0.0588 | 20 | 0.05 | 13 | 0.1538 | 57 | 0.0702 |
| 12_1705542 | NA | NA | NA | NA | 1 | 1 | NA | NA | 1 | 1 | 1 | 1 | NA | NA | NA | NA | NA | NA | 1 | 1 |
| 12_1722225 | 15 | 0 | 21 | 0.1429 | 13 | 0.1538 | 14 | 0.0714 | 63 | 0.0952 | 19 | 0.1579 | 7 | 0 | 7 | 0 | 13 | 0 | 46 | 0.0652 |
| 12_1722228 | 12 | 0.1667 | 17 | 0.1176 | 12 | 0.1667 | 12 | 0.1667 | 53 | 0.1509 | 14 | 0.3571 | 5 | 0.2 | 7 | 0 | 13 | 0.3077 | 39 | 0.2564 |
| 12_1722265 | 12 | 0 | 19 | 0.1053 | 8 | 0 | 12 | 0.0833 | 51 | 0.0588 | 15 | 0.1333 | 5 | 0 | 5 | 0 | 12 | 0.0833 | 37 | 0.0811 |
| 12_1722404 | 7 | 0 | 6 | 0 | 1 | 1 | 6 | 0 | 20 | 0.05 | 7 | 0.1429 | 1 | 0 | 2 | 0 | 6 | 0 | 16 | 0.0625 |
| 12_1722437 | 5 | 0 | 7 | 0.1429 | 1 | 0 | 5 | 0.2 | 18 | 0.1111 | 3 | 0.3333 | 2 | 0 | 2 | 0 | 7 | 0.2857 | 14 | 0.2143 |
| 12_1722452 | 6 | 0.1667 | 5 | 0 | 2 | 0 | 5 | 0.2 | 18 | 0.1111 | 5 | 0.2 | 2 | 0 | 2 | 0.5 | 7 | 0 | 16 | 0.125 |
| 12_1722455 | 7 | 0.4286 | 7 | 0.2857 | 3 | 0.3333 | 5 | 0.4 | 22 | 0.3636 | 6 | 0 | 2 | 0 | 3 | 0.6667 | 5 | 0.2 | 16 | 0.1875 |
| 12_1722456 | 7 | 0.4286 | 9 | 0.3333 | 3 | 0.6667 | 5 | 0.4 | 24 | 0.4167 | 6 | 0.3333 | 2 | 0.5 | 3 | 0 | 6 | 0.3333 | 17 | 0.2941 |
| 12_1722466 | 6 | 1 | 6 | 0.8333 | 2 | 1 | 5 | 1 | 19 | 0.9474 | 7 | 1 | 2 | 1 | 2 | 1 | 4 | 1 | 15 | 1 |
| 12_1722467 | 6 | 1 | 9 | 1 | 3 | 1 | 3 | 1 | 21 | 1 | 5 | 1 | 2 | 1 | 3 | 1 | 5 | 1 | 15 | 1 |
| 12_1722469 | 5 | 0.8 | 6 | 0.8333 | 3 | 1 | 3 | 0.6667 | 17 | 0.8235 | 5 | 1 | 2 | 1 | 2 | 1 | 6 | 1 | 15 | 1 |
| 12_1722472 | 7 | 0.5714 | 9 | 0.3333 | 3 | 0 | 5 | 0.8 | 24 | 0.4583 | 8 | 0.375 | 2 | 0 | 3 | 0.6667 | 6 | 0.6667 | 19 | 0.4737 |
| 12_1722483 | 6 | 0.8333 | 8 | 0.75 | 3 | 0.6667 | 5 | 0.6 | 22 | 0.7273 | 7 | 0.7143 | 1 | 1 | 2 | 1 | 6 | 1 | 16 | 0.875 |
| 12_1722504 | 4 | 0 | 6 | 0 | 3 | 0 | 4 | 0.25 | 17 | 0.0588 | 4 | 0 | 1 | 0 | 1 | 0 | 4 | 0.25 | 10 | 0.1 |
| 12_1722513 | 2 | 0 | 6 | 0.5 | 3 | 0.6667 | 3 | 0.6667 | 14 | 0.5 | 4 | 0.5 | 1 | 1 | 1 | 0 | 5 | 0.2 | 11 | 0.3636 |
| 12_1722521 | 3 | 1 | 5 | 0.6 | 2 | 0.5 | 4 | 0.5 | 14 | 0.6429 | 4 | 0.75 | 1 | 1 | 1 | 0 | 5 | 1 | 11 | 0.8182 |
| 12_1722525 | 2 | 0.5 | 6 | 0 | 2 | 0.5 | 3 | 0.3333 | 13 | 0.2308 | 5 | 0 | 1 | 0 | 1 | 0 | 5 | 0.2 | 12 | 0.0833 |
| 12_1722541 | 2 | 0.5 | 4 | 0 | 2 | 1 | 3 | 0 | 11 | 0.2727 | 6 | 0 | NA | NA | 1 | 0 | 4 | 0.75 | 11 | 0.2727 |
| 12_1722555 | 1 | 0 | 6 | 0.1667 | 2 | 0.5 | 1 | 1 | 10 | 0.3 | 4 | 0.5 | 1 | 0 | 1 | 1 | 3 | 0.3333 | 9 | 0.4444 |
| 12_1722581 | 1 | 0 | 5 | 0.4 | 2 | 0.5 | 2 | 0 | 10 | 0.3 | 4 | 0.25 | 1 | 0 | 1 | 0 | 1 | 1 | 7 | 0.2857 |
| 12_1722584 | 1 | 1 | 5 | 0.8 | 2 | 1 | 2 | 0.5 | 10 | 0.8 | 4 | 0.25 | 1 | 1 | 1 | 1 | 1 | 1 | 7 | 0.5714 |
| 12_1733422 | 19 | 0 | 28 | 0.1429 | 14 | 0 | 8 | 0 | 69 | 0.058 | 13 | 0.1538 | 18 | 0.0556 | 21 | 0 | 24 | 0.2917 | 76 | 0.1316 |
| 12_1733425 | 19 | 0.0526 | 30 | 0.0667 | 13 | 0 | 11 | 0.0909 | 73 | 0.0548 | 13 | 0.1538 | 18 | 0.1111 | 22 | 0.0909 | 21 | 0.1429 | 74 | 0.1216 |
| 12_1733436 | 4 | 0.25 | 12 | 0.4167 | 6 | 0.1667 | 4 | 0.5 | 26 | 0.3462 | 7 | 0 | 8 | 0.625 | 8 | 0.5 | 9 | 0.3333 | 32 | 0.375 |
| 12_1733472 | 2 | 0.5 | 2 | 0.5 | NA | NA | 1 | 0 | 5 | 0.4 | 5 | 0 | 1 | 1 | 4 | 0.25 | 4 | 0 | 14 | 0.1429 |
| 12_1747953 | 6 | 0.1667 | 3 | 0 | 1 | 0 | 1 | 1 | 11 | 0.1818 | 2 | 1 | 3 | 0.6667 | NA | NA | 5 | 0.2 | 10 | 0.5 |
| 12_1747995 | 8 | 0.125 | 2 | 0.5 | 1 | 0 | 1 | 0 | 12 | 0.1667 | 2 | 0 | 4 | 0.25 | NA | NA | 5 | 0.2 | 11 | 0.1818 |
| 12_1760128 | 1 | 0 | 2 | 0.5 | 1 | 0 | 3 | 0.3333 | 7 | 0.2857 | 1 | 0 | NA | NA | NA | NA | 1 | 1 | 2 | 0.5 |
| 12_1783628 | 13 | 0.5385 | 6 | 0 | 7 | 0.2857 | 3 | 0.3333 | 29 | 0.3448 | 13 | 0.4615 | 3 | 0 | 8 | 0.625 | 6 | 0.1667 | 30 | 0.4 |
| 12_1783630 | 15 | 0.0667 | 10 | 0.2 | 8 | 0 | 4 | 0 | 37 | 0.0811 | 15 | 0 | 5 | 0 | 8 | 0.25 | 8 | 0.125 | 36 | 0.0833 |
| 12_1784122 | 15 | 1 | 12 | 1 | 9 | 1 | 5 | 1 | 41 | 1 | 17 | 1 | 5 | 1 | 10 | 1 | 9 | 0.7778 | 41 | 0.9512 |
| 12_1784131 | 14 | 0.2143 | 12 | 0.1667 | 11 | 0.1818 | 5 | 0.2 | 42 | 0.1905 | 18 | 0.1667 | 7 | 0.1429 | 11 | 0.1818 | 10 | 0.2 | 46 | 0.1739 |
| 12_1784145 | 14 | 0.8571 | 12 | 0.3333 | 10 | 0.7 | 5 | 1 | 41 | 0.6829 | 19 | 0.5263 | 7 | 1 | 9 | 0.5556 | 10 | 0.8 | 45 | 0.6667 |
| 12_1784147 | 13 | 0.9231 | 13 | 0.6923 | 11 | 0.9091 | 5 | 1 | 42 | 0.8571 | 18 | 0.9444 | 7 | 0.8571 | 12 | 1 | 9 | 1 | 46 | 0.9565 |
| 12_1784152 | 14 | 0.3571 | 13 | 0.2308 | 11 | 0 | 5 | 0.4 | 43 | 0.2326 | 18 | 0.2778 | 5 | 0.2 | 9 | 0 | 9 | 0.2222 | 41 | 0.1951 |

| | | | | | | | | | | | | | | | | | | | | |
|---|---|---|---|---|---|---|---|---|---|---|---|---|---|---|---|---|---|---|---|---|
| 12_1784167 | 15 | 0.2667 | 16 | 0.3125 | 12 | 0.1667 | 5 | 0.4 | 48 | 0.2708 | 18 | 0.5556 | 6 | 0.3333 | 12 | 0.4167 | 10 | 0.3 | 46 | 0.4348 |
| 12_1797909 | 1 | 1 | 1 | 0 | 1 | 0 | NA | NA | 3 | 0.3333 | NA | NA | 1 | 1 | 1 | 1 | NA | NA | 2 | 1 |
| 12_1797913 | 2 | 0.5 | NA | NA | 1 | 0 | NA | NA | 3 | 0.3333 | NA | NA | 1 | 0 | 1 | 1 | NA | NA | 2 | 0.5 |
| 12_1797920 | 2 | 0.5 | 1 | 1 | 1 | 1 | NA | NA | 4 | 0.75 | NA | NA | NA | NA | 1 | 1 | NA | NA | 1 | 1 |
| 12_1798401 | 1 | 1 | NA | NA | NA | NA | NA | NA | 1 | 1 | NA | NA | 1 | 0 | NA | NA | 1 | 1 | 2 | 0.5 |
| 12_1798410 | 1 | 1 | NA | NA | NA | NA | NA | NA | 1 | 1 | NA | NA | 1 | 0 | NA | NA | 1 | 1 | 2 | 0.5 |
| 12_1798503 | 1 | 1 | NA | NA | NA | NA | NA | NA | 1 | 1 | NA | NA | 1 | 1 | NA | NA | 1 | 1 | 2 | 1 |
| 12_2072022 | NA | NA | NA | NA | 1 | 1 | NA | NA | 1 | 1 | 1 | 1 | NA | NA | NA | NA | NA | NA | 1 | 1 |
| 12_2092382 | 2 | 0.5 | 4 | 1 | 1 | 1 | 1 | 1 | 8 | 0.875 | 2 | 1 | NA | NA | NA | NA | NA | NA | 2 | 1 |
| 12_2092384 | 1 | 0 | 3 | 0.3333 | 1 | 1 | 1 | 1 | 6 | 0.5 | 3 | 0.6667 | NA | NA | NA | NA | NA | NA | 3 | 0.6667 |
| 12_2092675 | 2 | 0 | 2 | 0.5 | NA | NA | 1 | 0 | 5 | 0.2 | 3 | 0.6667 | NA | NA | NA | NA | NA | NA | 3 | 0.6667 |
| 12_2092693 | 1 | 1 | 3 | 1 | NA | NA | 1 | 1 | 5 | 1 | 2 | 1 | NA | NA | NA | NA | NA | NA | 2 | 1 |
| 12_2092694 | 1 | 1 | 3 | 0.6667 | NA | NA | 1 | 1 | 5 | 0.8 | 2 | 1 | NA | NA | NA | NA | NA | NA | 2 | 1 |
| 12_2092696 | 1 | 1 | 1 | 1 | NA | NA | 1 | 1 | 3 | 1 | 2 | 1 | NA | NA | NA | NA | NA | NA | 2 | 1 |
| 12_2109839 | NA | NA | NA | NA | NA | NA | 2 | 0.5 | 2 | 0.5 | NA | NA | 1 | 0 | 1 | 1 | 1 | 0 | 3 | 0.3333 |
| 12_2109856 | 1 | 0 | NA | NA | 1 | 0 | 2 | 0.5 | 4 | 0.25 | 1 | 0 | 1 | 0 | 1 | 1 | 1 | 0 | 4 | 0.25 |
| 12_2130960 | NA | NA | NA | NA | 1 | 1 | NA | NA | 1 | 1 | NA | NA | 1 | 1 | NA | NA | 1 | 1 | 2 | 1 |
| 12_2131029 | NA | NA | NA | NA | 1 | 1 | NA | NA | 1 | 1 | NA | NA | 1 | 1 | NA | NA | 1 | 1 | 2 | 1 |
| 12_2131787 | 5 | 0.2 | 2 | 0.5 | NA | NA | 1 | 0 | 8 | 0.25 | NA | NA | 2 | 0.5 | 6 | 0.5 | 4 | 0 | 12 | 0.3333 |
| 12_2131986 | 6 | 0.6667 | 4 | 0.25 | 1 | 0 | 1 | 0 | 12 | 0.4167 | 1 | 0 | 2 | 0.5 | 8 | 0.375 | 6 | 0.1667 | 17 | 0.2941 |
| 12_2132035 | 4 | 0.5 | NA | NA | NA | NA | 2 | 0.5 | 6 | 0.5 | 2 | 0.5 | 1 | 0 | 2 | 0 | NA | NA | 5 | 0.2 |
| 12_2132082 | 9 | 0.8889 | 2 | 1 | 3 | 0.3333 | 8 | 0.875 | 22 | 0.8182 | 4 | 1 | 7 | 0.8571 | 9 | 0.8889 | 3 | 1 | 23 | 0.913 |
| 12_2136071 | NA | NA | 1 | 0 | 1 | 0 | 3 | 0.3333 | 5 | 0.2 | NA | NA | NA | NA | 1 | 0 | 2 | 0.5 | 3 | 0.3333 |
| 12_2136470 | 5 | 0.2 | 1 | 0 | 6 | 0.1667 | 11 | 0 | 23 | 0.087 | 3 | 0 | 4 | 0.25 | 2 | 0 | 3 | 0 | 12 | 0.0833 |
| 12_2136472 | 8 | 0 | 6 | 0.3333 | 7 | 0.2857 | 15 | 0.3333 | 36 | 0.25 | 5 | 0.6 | 11 | 0.2727 | 7 | 0.4286 | 9 | 0.2222 | 32 | 0.3438 |
| 12_2136477 | 9 | 0 | 7 | 0 | 8 | 0.125 | 16 | 0.0625 | 40 | 0.05 | 5 | 0.2 | 12 | 0.1667 | 7 | 0 | 10 | 0 | 34 | 0.0882 |
| 12_2137346 | 4 | 0 | 6 | 0 | 4 | 0.25 | 1 | 0 | 15 | 0.0667 | 3 | 0 | 4 | 0.25 | 4 | 0.25 | 2 | 0 | 13 | 0.1538 |
| 12_2137403 | 2 | 0.5 | 1 | 1 | NA | NA | 1 | 0 | 4 | 0.5 | 1 | 0 | 1 | 1 | 1 | 0 | 2 | 0.5 | 5 | 0.4 |
| 12_2137424 | 1 | 0 | 1 | 0 | NA | NA | 1 | 1 | 3 | 0.3333 | 1 | 1 | 2 | 0 | 3 | 0.3333 | 2 | 0 | 8 | 0.25 |
| 12_2137433 | 2 | 0.5 | NA | NA | NA | NA | 1 | 1 | 3 | 0.6667 | 1 | 0 | 1 | 0 | 2 | 0 | 2 | 0.5 | 6 | 0.1667 |
| 12_2137437 | 2 | 1 | 1 | 1 | NA | NA | NA | NA | 3 | 1 | 1 | 1 | 2 | 1 | 3 | 0.6667 | 2 | 1 | 8 | 0.875 |
| 12_2137446 | 3 | 0.3333 | 1 | 0 | NA | NA | 1 | 1 | 5 | 0.4 | 1 | 0 | 2 | 0 | 3 | 0 | 2 | 0.5 | 8 | 0.125 |
| 12_2137466 | 2 | 0 | NA | NA | NA | NA | 1 | 1 | 3 | 0.3333 | 1 | 0 | 2 | 0.5 | 3 | 0 | 2 | 0 | 8 | 0.125 |
| 12_2137468 | NA | NA | 1 | 1 | NA | NA | 1 | 1 | 2 | 1 | 1 | 0 | 1 | 0 | 3 | 0 | 2 | 0.5 | 7 | 0.1429 |
| 12_2137525 | 2 | 0.5 | 1 | 0 | NA | NA | 1 | 0 | 4 | 0.25 | NA | NA | 2 | 0 | 2 | 0.5 | 1 | 0 | 5 | 0.2 |
| 12_2137527 | 1 | 1 | NA | NA | NA | NA | NA | NA | 1 | 1 | 1 | 0 | 2 | 0.5 | 1 | 0 | NA | NA | 4 | 0.25 |
| 12_2137528 | 3 | 1 | 1 | 1 | NA | NA | 1 | 1 | 5 | 1 | 1 | 1 | 2 | 0.5 | 1 | 0 | 1 | 0 | 5 | 0.4 |
| 12_2137541 | 2 | 1 | 1 | 0 | NA | NA | 1 | 0 | 4 | 0.5 | NA | NA | 2 | 0.5 | 2 | 0.5 | 1 | 0 | 5 | 0.4 |
| 12_2144103 | 75 | 0.2133 | 89 | 0.2697 | 71 | 0.2394 | 76 | 0.2895 | 311 | 0.254 | 66 | 0.3333 | 61 | 0.3279 | 79 | 0.2911 | 80 | 0.3 | 286 | 0.3112 |
| 12_2146685 | 57 | 0.2281 | 64 | 0.1562 | 58 | 0.1379 | 63 | 0.2222 | 242 | 0.186 | 52 | 0.1731 | 46 | 0.2391 | 60 | 0.2333 | 65 | 0.2154 | 223 | 0.2152 |
| 12_2160448 | 3 | 0.3333 | 3 | 0 | 1 | 0 | 2 | 0 | 9 | 0.1111 | 2 | 0 | 2 | 0.5 | 1 | 0 | 3 | 0 | 8 | 0.125 |
| 12_2160449 | 3 | 0 | 3 | 0.3333 | 1 | 1 | 2 | 0.5 | 9 | 0.3333 | 3 | 0.3333 | 2 | 0.5 | 1 | 0 | 3 | 0 | 9 | 0.2222 |
| 12_2161126 | NA | NA | 1 | 1 | NA | NA | NA | NA | 1 | 1 | NA | NA | NA | NA | NA | NA | 1 | 1 | 1 | 1 |
| 12_2161566 | NA | NA | NA | NA | NA | NA | 1 | 1 | 1 | 1 | NA | NA | NA | NA | NA | NA | 1 | 1 | 1 | 1 |
| 12_2300444 | 1 | 0 | NA | NA | NA | NA | 1 | 1 | 2 | 0.5 | NA | NA | 1 | 1 | NA | NA | 2 | 0.5 | 3 | 0.6667 |
| 12_2300445 | 1 | 0 | NA | NA | NA | NA | 1 | 1 | 2 | 0.5 | NA | NA | 1 | 0 | NA | NA | 2 | 0.5 | 3 | 0.3333 |
| 12_2300861 | 1 | 1 | NA | NA | NA | NA | NA | NA | 1 | 1 | NA | NA | NA | NA | NA | NA | 1 | 1 | 1 | 1 |
| 12_2300952 | 1 | 1 | NA | NA | NA | NA | NA | NA | 1 | 1 | NA | NA | NA | NA | NA | NA | 1 | 1 | 1 | 1 |
| 12_2300974 | 1 | 1 | NA | NA | NA | NA | NA | NA | 1 | 1 | NA | NA | NA | NA | NA | NA | 1 | 1 | 1 | 1 |
| 12_2301009 | 1 | 1 | NA | NA | NA | NA | 1 | 1 | 2 | 1 | NA | NA | 1 | 1 | NA | NA | 1 | 1 | 2 | 1 |
| 12_2301010 | 1 | 1 | NA | NA | NA | NA | 1 | 0 | 2 | 0.5 | NA | NA | 1 | 1 | NA | NA | NA | NA | 1 | 1 |
| 12_2301012 | 1 | 1 | NA | NA | NA | NA | 1 | 1 | 2 | 1 | NA | NA | 1 | 1 | NA | NA | 1 | 1 | 2 | 1 |
| 12_2409383 | 1 | 0 | 1 | 1 | NA | NA | NA | NA | 2 | 0.5 | NA | NA | NA | NA | NA | NA | 1 | 1 | 1 | 1 |
| 12_2409428 | NA | NA | 1 | 1 | NA | NA | NA | NA | 1 | 1 | NA | NA | NA | NA | NA | NA | 1 | 1 | 1 | 1 |
| 12_2409434 | 1 | 1 | 1 | 1 | NA | NA | NA | NA | 2 | 1 | NA | NA | NA | NA | NA | NA | 1 | 1 | 1 | 1 |
| 12_2409440 | 1 | 1 | 1 | 1 | NA | NA | NA | NA | 2 | 1 | NA | NA | NA | NA | NA | NA | 1 | 1 | 1 | 1 |
| 12_2409472 | NA | NA | 1 | 1 | NA | NA | NA | NA | 1 | 1 | NA | NA | NA | NA | NA | NA | 1 | 1 | 1 | 1 |
| 12_2422603 | 3 | 0.3333 | NA | NA | 1 | 0 | NA | NA | 4 | 0.25 | NA | NA | 1 | 1 | NA | NA | NA | NA | 1 | 1 |
| 12_2422609 | 3 | 0.6667 | NA | NA | NA | NA | NA | NA | 3 | 0.6667 | NA | NA | 1 | 1 | NA | NA | NA | NA | 1 | 1 |
| 12_2422634 | 3 | 0.3333 | NA | NA | 1 | 0 | NA | NA | 4 | 0.25 | NA | NA | 1 | 1 | NA | NA | NA | NA | 1 | 1 |
| 12_2426365 | NA | NA | NA | NA | NA | NA | 1 | 1 | 1 | 1 | NA | NA | NA | NA | 1 | 1 | NA | NA | 1 | 1 |
| 12_2460829 | NA | NA | 3 | 0.6667 | 4 | 0.25 | 3 | 0 | 10 | 0.3 | 2 | 0 | 6 | 0.3333 | 2 | 0.5 | 4 | 0.75 | 14 | 0.4286 |
| 12_2466619 | NA | NA | 1 | 1 | 1 | 1 | 1 | 1 | 3 | 1 | 1 | 1 | 1 | 1 | NA | NA | NA | NA | 2 | 1 |
| 12_2466685 | NA | NA | NA | NA | 1 | 1 | 1 | 1 | 2 | 1 | 1 | 0 | 1 | 1 | NA | NA | NA | NA | 2 | 0.5 |
| 12_2466691 | NA | NA | NA | NA | 1 | 0 | 1 | 1 | 2 | 0.5 | 1 | 1 | 1 | 0 | NA | NA | NA | NA | 2 | 0.5 |

| | | | | | | | | | | | | | | | | | | | | |
|---|---|---|---|---|---|---|---|---|---|---|---|---|---|---|---|---|---|---|---|---|
| 12_247051 | NA | NA | NA | NA | 1 | 0 | 1 | 1 | 2 | 0.5 | NA | NA | NA | NA | NA | NA | 1 | 1 | 1 | 1 |
| 12_247053 | NA | NA | NA | NA | 1 | 1 | 1 | 1 | 2 | 1 | NA | NA | 1 | 1 | NA | NA | 1 | 1 | 2 | 1 |
| 12_247054 | NA | NA | NA | NA | 1 | 1 | 1 | 1 | 2 | 1 | NA | NA | 1 | 1 | NA | NA | 1 | 1 | 2 | 1 |
| 12_247055 | NA | NA | NA | NA | 1 | 1 | 1 | 1 | 2 | 1 | NA | NA | 1 | 1 | NA | NA | 1 | 0 | 2 | 0.5 |
| 12_247056 | NA | NA | NA | NA | 1 | 0 | 1 | 1 | 2 | 0.5 | NA | NA | NA | NA | NA | NA | 1 | 1 | 1 | 1 |
| 12_247102 | NA | NA | NA | NA | 1 | 1 | 1 | 1 | 2 | 1 | NA | NA | 1 | 1 | NA | NA | NA | NA | 1 | 1 |
| 12_247113 | NA | NA | NA | NA | 1 | 1 | 1 | 1 | 2 | 1 | NA | NA | 2 | 0.5 | NA | NA | 1 | 0 | 3 | 0.3333 |
| 12_247115 | NA | NA | NA | NA | 1 | 0 | 1 | 1 | 2 | 0.5 | NA | NA | 2 | 1 | NA | NA | 1 | 0 | 3 | 0.6667 |
| 12_247127 | NA | NA | NA | NA | 1 | 1 | 1 | 0 | 2 | 0.5 | NA | NA | 1 | 1 | NA | NA | NA | NA | 1 | 1 |
| 12_247159 | 5 | 1 | NA | NA | 2 | 1 | 5 | 0.6 | 12 | 0.8333 | NA | NA | 1 | 1 | 2 | 1 | 3 | 1 | 6 | 1 |
| 12_247160 | 4 | 1 | NA | NA | 2 | 1 | 3 | 1 | 9 | 1 | NA | NA | 1 | 1 | 2 | 1 | 2 | 1 | 5 | 1 |
| 12_247160 | 4 | 0.25 | NA | NA | 2 | 0.5 | 4 | 0.25 | 10 | 0.3 | NA | NA | NA | NA | 1 | 0 | 2 | 0.5 | 3 | 0.3333 |
| 12_247160 | 4 | 1 | NA | NA | 2 | 1 | 3 | 1 | 9 | 1 | NA | NA | 1 | 1 | 1 | 1 | 1 | 0 | 3 | 0.6667 |
| 12_247166 | 2 | 0 | NA | NA | 1 | 1 | 3 | 0.3333 | 6 | 0.3333 | NA | NA | 1 | 0 | 1 | 0 | 3 | 0.3333 | 5 | 0.2 |
| 12_248459 | 1 | 1 | 1 | 1 | 1 | 0 | NA | NA | 3 | 0.6667 | NA | NA | 2 | 1 | 3 | 0 | 1 | 1 | 6 | 0.5 |
| 12_248462 | 1 | 1 | 1 | 1 | 1 | 1 | NA | NA | 3 | 1 | NA | NA | 1 | 1 | 2 | 1 | 1 | 1 | 4 | 1 |
| 12_248464 | 1 | 1 | NA | NA | 1 | 1 | NA | NA | 2 | 1 | NA | NA | 2 | 1 | 2 | 1 | 1 | 1 | 5 | 1 |
| 12_248465 | 1 | 1 | NA | NA | 1 | 0 | NA | NA | 2 | 0.5 | NA | NA | 1 | 1 | 2 | 0.5 | 1 | 0 | 4 | 0.5 |
| 12_249965 | 3 | 0 | NA | NA | 5 | 0 | 1 | 1 | 9 | 0.1111 | 1 | 0 | NA | NA | 2 | 0.5 | 2 | 0 | 5 | 0.2 |
| 12_250090 | 3 | 1 | 1 | 1 | 6 | 0.6667 | 4 | 0.25 | 14 | 0.6429 | 3 | 0.3333 | NA | NA | 3 | 1 | 4 | 0.5 | 10 | 0.6 |
| 12_250090 | 3 | 1 | 2 | 0 | 6 | 0.6667 | 4 | 0.5 | 15 | 0.6 | 1 | 1 | 1 | 0 | 3 | 0.6667 | 3 | 1 | 8 | 0.75 |
| 12_250090 | 3 | 0 | 2 | 0 | 7 | 0.1429 | 4 | 0.5 | 16 | 0.1875 | 2 | 0.5 | 1 | 0 | 3 | 0 | 3 | 0 | 9 | 0.1111 |
| 12_250091 | 2 | 0 | 2 | 0.5 | 5 | 0.4 | 4 | 0.5 | 13 | 0.3846 | 3 | 0.6667 | 1 | 0 | 3 | 0.3333 | 4 | 0 | 11 | 0.2727 |
| 12_250091 | 4 | 0.25 | 2 | 0.5 | 6 | 0.3333 | 3 | 0 | 15 | 0.2667 | 3 | 0.3333 | 1 | 0 | 3 | 0.3333 | 5 | 0.6 | 12 | 0.4167 |
| 12_250091 | 4 | 0.75 | 2 | 0.5 | 5 | 0.8 | 3 | 0.3333 | 14 | 0.6429 | 2 | 1 | 1 | 1 | 3 | 0.6667 | 5 | 0.8 | 11 | 0.8182 |
| 12_250091 | 4 | 0.25 | 2 | 0 | 4 | 0 | 3 | 0 | 13 | 0.0769 | 3 | 0 | 1 | 0 | 2 | 0 | 4 | 0.25 | 10 | 0.1 |
| 12_250092 | 4 | 1 | 2 | 1 | 5 | 0.8 | 3 | 1 | 14 | 0.9286 | 3 | 1 | 1 | 1 | 4 | 1 | 5 | 0.8 | 13 | 0.9231 |
| 12_250092 | 2 | 0.5 | 1 | 0 | 5 | 0 | 2 | 0 | 10 | 0.1 | 2 | 0.5 | 1 | 0 | 1 | 0 | 2 | 0 | 6 | 0.1667 |
| 12_250097 | 3 | 0.3333 | 1 | 0 | 5 | 0 | 5 | 0 | 14 | 0.0714 | 4 | 0.5 | NA | NA | 1 | 0 | 4 | 0 | 9 | 0.2222 |
| 12_250100 | 2 | 1 | 4 | 0.75 | 6 | 0.5 | 5 | 0.2 | 17 | 0.5294 | 4 | 0.5 | NA | NA | 1 | 0 | 3 | 0.3333 | 8 | 0.375 |
| 12_250104 | 1 | 1 | 3 | 0 | 5 | 0 | 2 | 0 | 11 | 0.0909 | 2 | 0 | 1 | 1 | NA | NA | 3 | 0 | 6 | 0.1667 |
| 12_250127 | 2 | 0 | 4 | 0.25 | 7 | 0.1429 | 5 | 0 | 18 | 0.1111 | 3 | 0 | 2 | 0 | 1 | 1 | 3 | 0.3333 | 9 | 0.2222 |
| 12_250128 | 1 | 0 | 3 | 0 | 7 | 0 | 5 | 0.2 | 16 | 0.0625 | 3 | 0.3333 | 2 | 0 | 1 | 0 | 2 | 0 | 8 | 0.125 |
| 12_250129 | 2 | 0.5 | 4 | 0 | 7 | 0 | 5 | 0.2 | 18 | 0.1111 | 3 | 0.3333 | 2 | 0 | 1 | 0 | 3 | 0.3333 | 9 | 0.2222 |
| 12_250130 | 2 | 0 | 3 | 0 | 7 | 0 | 5 | 0.2 | 17 | 0.0588 | 3 | 0.3333 | 2 | 0 | NA | NA | 3 | 0 | 8 | 0.125 |
| 12_250132 | 2 | 0 | 5 | 0.2 | 6 | 0 | 5 | 0 | 18 | 0.0556 | 3 | 0.6667 | 2 | 0 | 1 | 0 | 3 | 0 | 9 | 0.2222 |
| 12_250135 | 3 | 0.3333 | 3 | 1 | 4 | 0.75 | 4 | 0.75 | 14 | 0.7143 | 3 | 1 | 1 | 1 | 1 | 0 | 2 | 1 | 7 | 0.8571 |
| 12_250140 | 2 | 0.5 | 2 | 0 | 6 | 0 | 3 | 0.3333 | 13 | 0.1538 | 2 | 0 | NA | NA | NA | NA | 1 | 1 | 3 | 0.3333 |
| 12_250586 | 4 | 0 | 6 | 0.1667 | 4 | 0 | 4 | 0.25 | 18 | 0.1111 | 3 | 0 | 3 | 0 | 4 | 0.25 | 4 | 0.25 | 14 | 0.1429 |
| 12_250602 | 1 | 0 | 5 | 0.2 | 4 | 0 | 3 | 0 | 13 | 0.0769 | 4 | 0.25 | 3 | 0 | 4 | 0 | 5 | 0 | 16 | 0.0625 |
| 12_250610 | 4 | 0.25 | 6 | 0.3333 | 6 | 0.3333 | 6 | 0 | 22 | 0.2273 | 6 | 0.1667 | 2 | 0 | 5 | 0 | 4 | 0.25 | 17 | 0.1176 |
| 12_251066 | 5 | 0.2 | 4 | 0.25 | 6 | 0.1667 | 6 | 0 | 21 | 0.1429 | 8 | 0.125 | 5 | 0.2 | 13 | 0.3077 | 6 | 0 | 32 | 0.1875 |
| 12_251070 | 5 | 0 | 8 | 0.25 | 9 | 0.3333 | 10 | 0.1 | 32 | 0.1875 | 7 | 0 | 5 | 0.4 | 12 | 0.0833 | 11 | 0.1818 | 35 | 0.1429 |
| 12_251071 | 6 | 0.3333 | 8 | 0 | 10 | 0.3 | 11 | 0 | 35 | 0.1429 | 6 | 0.3333 | 5 | 0.4 | 16 | 0.1875 | 10 | 0.1 | 37 | 0.2162 |
| 12_251072 | 7 | 0.5714 | 8 | 0.5 | 8 | 0.75 | 9 | 0.3333 | 32 | 0.5312 | 6 | 0.3333 | 5 | 0.6 | 16 | 0.5 | 9 | 0.7778 | 36 | 0.5556 |
| 12_251075 | 3 | 0.3333 | 5 | 0.4 | 6 | 0.1667 | 10 | 0.3 | 24 | 0.2917 | 7 | 0.2857 | 4 | 0 | 12 | 0.1667 | 9 | 0.3333 | 32 | 0.2188 |
| 12_251075 | 5 | 1 | 9 | 0.8889 | 10 | 0.8 | 11 | 1 | 35 | 0.9143 | 7 | 1 | 7 | 0.7143 | 17 | 0.7647 | 10 | 0.6 | 41 | 0.7561 |
| 12_251075 | 7 | 0 | 7 | 0 | 9 | 0.1111 | 8 | 0.125 | 31 | 0.0645 | 6 | 0 | 6 | 0 | 14 | 0.1429 | 7 | 0 | 33 | 0.0606 |
| 12_251076 | 5 | 1 | 6 | 0.6667 | 12 | 0.9167 | 11 | 1 | 34 | 0.9118 | 6 | 1 | 7 | 1 | 17 | 0.9412 | 12 | 1 | 42 | 0.9762 |
| 12_251078 | 7 | 0.1429 | 9 | 0 | 12 | 0.25 | 12 | 0.1667 | 40 | 0.15 | 8 | 0.375 | 8 | 0.25 | 20 | 0.25 | 14 | 0.1429 | 50 | 0.24 |
| 12_251079 | 8 | 0 | 12 | 0 | 14 | 0.0714 | 14 | 0.2857 | 48 | 0.1042 | 10 | 0 | 7 | 0 | 21 | 0.1905 | 15 | 0.1333 | 53 | 0.1132 |
| 12_251079 | 6 | 0.5 | 13 | 0.2308 | 14 | 0.5714 | 12 | 0.3333 | 45 | 0.4 | 7 | 0.2857 | 6 | 0.5 | 19 | 0.1579 | 13 | 0.5385 | 45 | 0.3333 |
| 12_251081 | 8 | 0 | 15 | 0.2667 | 13 | 0.3077 | 15 | 0 | 51 | 0.1569 | 8 | 0.375 | 10 | 0.4 | 20 | 0.15 | 13 | 0.1538 | 51 | 0.2353 |
| 12_251090 | 13 | 0.0769 | 18 | 0.2222 | 21 | 0.1429 | 19 | 0 | 71 | 0.1127 | 14 | 0.0714 | 16 | 0.125 | 21 | 0.1905 | 22 | 0.1364 | 73 | 0.137 |
| 12_251091 | 14 | 0.2143 | 20 | 0.1 | 22 | 0.1364 | 19 | 0.2105 | 75 | 0.16 | 13 | 0.0769 | 18 | 0.2778 | 22 | 0.1364 | 21 | 0.0476 | 74 | 0.1351 |
| 12_251094 | 14 | 0.0714 | 19 | 0.2105 | 27 | 0.1111 | 20 | 0.15 | 80 | 0.1375 | 15 | 0.2 | 17 | 0.1765 | 24 | 0.125 | 21 | 0.2381 | 77 | 0.1818 |
| 12_251114 | NA | NA | 1 | 1 | NA | NA | 3 | 0 | 4 | 0.25 | NA | NA | 2 | 0.5 | NA | NA | 1 | 0 | 3 | 0.3333 |
| 12_251122 | NA | NA | 2 | 0.5 | NA | NA | 3 | 0 | 5 | 0.2 | NA | NA | 2 | 0 | NA | NA | 2 | 0.5 | 4 | 0.25 |
| 12_251125 | NA | NA | 2 | 0 | NA | NA | 4 | 0.25 | 6 | 0.1667 | NA | NA | 2 | 0.5 | NA | NA | 2 | 0.5 | 4 | 0.5 |
| 12_251440 | 31 | 0.5484 | 45 | 0.6222 | 43 | 0.5814 | 26 | 0.5769 | 145 | 0.5862 | 25 | 0.4 | 23 | 0.6087 | 16 | 0.4375 | 28 | 0.6071 | 92 | 0.5217 |
| 12_251442 | 31 | 0.1935 | 55 | 0.3455 | 48 | 0.25 | 30 | 0.1333 | 164 | 0.25 | 29 | 0.1724 | 29 | 0.2414 | 16 | 0.3125 | 32 | 0.2188 | 106 | 0.2264 |
| 12_251442 | 32 | 0.25 | 46 | 0.3696 | 47 | 0.2979 | 28 | 0.4286 | 153 | 0.3333 | 26 | 0.3462 | 28 | 0.3929 | 14 | 0.3571 | 27 | 0.4074 | 95 | 0.3789 |
| 12_251443 | 26 | 0.8462 | 48 | 0.9583 | 45 | 0.9556 | 30 | 0.9333 | 149 | 0.9329 | 25 | 1 | 27 | 0.9259 | 13 | 1 | 30 | 1 | 95 | 0.9789 |
| 12_251443 | 26 | 0.6154 | 47 | 0.5745 | 44 | 0.6591 | 29 | 0.5517 | 146 | 0.6027 | 26 | 0.6154 | 29 | 0.6207 | 15 | 0.8 | 27 | 0.7037 | 97 | 0.6701 |
| 12_251444 | 26 | 0.6154 | 44 | 0.5909 | 45 | 0.5333 | 28 | 0.6429 | 143 | 0.5874 | 26 | 0.6538 | 28 | 0.7143 | 15 | 0.5333 | 28 | 0.6071 | 97 | 0.6392 |

| | | | | | | | | | | | | | | | | | | | | |
|---|---|---|---|---|---|---|---|---|---|---|---|---|---|---|---|---|---|---|---|---|
| 12_2514449 | 26 | 0.4231 | 43 | 0.2558 | 44 | 0.2727 | 28 | 0.25 | 141 | 0.2908 | 26 | 0.3462 | 27 | 0.5556 | 14 | 0.0714 | 23 | 0.3043 | 90 | 0.3556 |
| 12_2514456 | 25 | 0.16 | 38 | 0.0789 | 39 | 0.0256 | 25 | 0.08 | 127 | 0.0787 | 26 | 0.0385 | 28 | 0.0714 | 10 | 0.2 | 24 | 0.0417 | 88 | 0.0682 |
| 12_2635820 | 1 | 0 | 2 | 0 | 2 | 0.5 | 5 | 0 | 10 | 0.1 | 1 | 0 | 3 | 0.3333 | 2 | 0 | 1 | 0 | 7 | 0.1429 |
| 12_2636015 | NA | NA | 1 | 0 | 1 | 0 | 4 | 0.5 | 6 | 0.3333 | NA | NA | 2 | 0 | 1 | 1 | NA | NA | 3 | 0.3333 |
| 12_2636025 | NA | NA | 1 | 1 | 1 | 0 | 4 | 0.75 | 6 | 0.6667 | 1 | 0 | 1 | 0 | 1 | 1 | NA | NA | 3 | 0.3333 |
| 12_2651991 | NA | NA | 1 | 1 | 1 | 0 | 1 | 1 | 3 | 0.6667 | NA | NA | NA | NA | 1 | 1 | NA | NA | 1 | 1 |
| 12_2652713 | 1 | 0 | 5 | 0.2 | 4 | 0 | 3 | 0 | 13 | 0.0769 | 2 | 0 | 2 | 0 | 3 | 0 | 3 | 0.3333 | 10 | 0.1 |
| 12_2652808 | NA | NA | 1 | 0 | 3 | 0.3333 | 1 | 0 | 5 | 0.2 | 1 | 0 | 2 | 0 | 1 | 0 | 1 | 1 | 5 | 0.2 |
| 12_2652825 | 1 | 0 | 5 | 1 | 4 | 0.5 | 2 | 1 | 12 | 0.75 | NA | NA | 1 | 1 | 1 | 0 | 3 | 1 | 5 | 0.8 |
| 12_2652825 | NA | NA | 3 | 1 | 5 | 1 | 3 | 1 | 11 | 1 | 2 | 1 | 2 | 1 | 3 | 1 | 3 | 1 | 10 | 1 |
| 12_2652828 | NA | NA | 3 | 1 | 3 | 0.6667 | 3 | 0.6667 | 9 | 0.7778 | 2 | 1 | 1 | 1 | 1 | 1 | 2 | 0 | 6 | 0.6667 |
| 12_2652828 | NA | NA | 2 | 1 | 4 | 0.75 | 3 | 1 | 9 | 0.8889 | 2 | 0.5 | 1 | 1 | 1 | 1 | 3 | 1 | 7 | 0.8571 |
| 12_2652839 | 2 | 1 | 4 | 1 | 5 | 1 | 3 | 1 | 14 | 1 | 3 | 1 | 2 | 1 | 3 | 1 | 3 | 1 | 11 | 1 |
| 12_2652849 | 2 | 1 | 4 | 1 | 5 | 1 | 3 | 1 | 14 | 1 | 3 | 1 | 2 | 1 | 2 | 1 | 3 | 1 | 10 | 1 |
| 12_2652852 | 1 | 1 | 5 | 1 | 5 | 0.8 | 3 | 1 | 14 | 0.9286 | 3 | 1 | 2 | 1 | 2 | 1 | 4 | 1 | 11 | 1 |
| 12_2652859 | 2 | 0 | 3 | 0.6667 | 4 | 0.25 | 3 | 0.3333 | 12 | 0.3333 | 3 | 0 | 1 | 0 | 3 | 0.3333 | 3 | 0.6667 | 10 | 0.3 |
| 12_2652860 | 2 | 1 | 3 | 1 | 3 | 1 | 1 | 1 | 9 | 1 | 2 | 1 | 2 | 1 | 1 | 1 | 4 | 1 | 9 | 1 |
| 12_2652865 | 3 | 0.6667 | 5 | 1 | 5 | 0.4 | 2 | 1 | 15 | 0.7333 | 2 | 0.5 | 2 | 1 | 1 | 0 | 3 | 1 | 8 | 0.75 |
| 12_2652924 | 2 | 0 | 4 | 0.25 | 5 | 0 | 3 | 0 | 14 | 0.0714 | 3 | 0 | 2 | 0 | 2 | 0 | 4 | 0.25 | 11 | 0.0909 |
| 12_2652960 | 2 | 0.5 | 3 | 0 | 6 | 0 | 3 | 0 | 14 | 0.0714 | 3 | 0 | 1 | 0 | 2 | 0.5 | 4 | 0.25 | 10 | 0.2 |
| 12_2719090 | 1 | 0 | NA | NA | 1 | 1 | NA | NA | 2 | 0.5 | 1 | 1 | NA | NA | NA | NA | 1 | 0 | 2 | 0.5 |
| 12_2719109 | 1 | 1 | 4 | 0.75 | 2 | 1 | 1 | 1 | 8 | 0.875 | 2 | 1 | 1 | 1 | 1 | 1 | 1 | 0 | 5 | 0.8 |
| 12_2719115 | 2 | 0.5 | 4 | 0 | 2 | 0.5 | 1 | 0 | 9 | 0.2222 | 1 | 1 | NA | NA | 1 | 0 | 1 | 0 | 3 | 0.3333 |
| 12_2719133 | 3 | 0.6667 | 3 | 0 | 2 | 0 | NA | NA | 8 | 0.25 | 2 | 1 | 1 | 1 | 1 | 0 | 1 | 0 | 5 | 0.6 |
| 12_2719144 | 1 | 1 | 3 | 1 | 1 | 1 | 1 | 1 | 6 | 1 | 2 | 1 | 1 | 1 | 1 | 1 | 1 | 1 | 5 | 1 |
| 12_2719151 | 1 | 0 | 3 | 0.6667 | 1 | 0 | 1 | 0 | 6 | 0.3333 | 2 | 0 | 1 | 0 | NA | NA | 1 | 1 | 4 | 0.25 |
| 12_2719165 | 1 | 1 | 3 | 1 | 1 | 1 | 1 | 1 | 6 | 1 | NA | NA | 1 | 1 | NA | NA | 1 | 1 | 2 | 1 |
| 12_2719168 | NA | NA | 3 | 1 | 1 | 1 | 1 | 1 | 5 | 1 | 2 | 1 | 1 | 1 | NA | NA | 1 | 1 | 4 | 1 |
| 12_2719174 | 1 | 1 | 3 | 1 | 2 | 1 | 1 | 1 | 7 | 1 | 2 | 1 | 1 | 1 | 1 | 1 | 1 | 1 | 5 | 1 |
| 12_2719181 | 1 | 1 | 3 | 0.3333 | 2 | 0 | 1 | 0 | 7 | 0.2857 | 2 | 0.5 | 1 | 0 | 1 | 0 | 1 | 1 | 5 | 0.4 |
| 12_2719194 | 1 | 1 | 2 | 1 | 2 | 0.5 | 1 | 1 | 6 | 0.8333 | 2 | 1 | 1 | 1 | NA | NA | 1 | 1 | 4 | 1 |
| 12_2719195 | 1 | 0 | 2 | 1 | 1 | 1 | 1 | 0 | 5 | 0.6 | 2 | 1 | 1 | 1 | 1 | 1 | 1 | 1 | 5 | 1 |
| 12_2719196 | 1 | 1 | 3 | 1 | 2 | 1 | 1 | 1 | 7 | 1 | 2 | 1 | 1 | 1 | 1 | 1 | 1 | 1 | 5 | 1 |
| 12_2738501 | 3 | 1 | 4 | 0.25 | 3 | 0.6667 | 2 | 0.5 | 12 | 0.5833 | 3 | 0.6667 | 2 | 0 | 3 | 0 | 4 | 1 | 12 | 0.5 |
| 12_2738503 | 2 | 0 | 4 | 0 | 3 | 0.3333 | 3 | 0 | 12 | 0.0833 | 3 | 0 | 4 | 0 | 4 | 0 | 5 | 0.2 | 16 | 0.0625 |
| 12_2738519 | 3 | 0 | 4 | 0 | 3 | 0.3333 | 3 | 0 | 13 | 0.0769 | 3 | 0 | 4 | 0.25 | 4 | 0 | 5 | 0 | 16 | 0.0625 |
| 12_2738558 | 3 | 0 | 6 | 0 | 3 | 0.3333 | 4 | 0.25 | 16 | 0.125 | 3 | 0.3333 | 5 | 0 | 2 | 0 | 6 | 0 | 16 | 0.0625 |
| 12_2738594 | 3 | 0.3333 | 7 | 0.5714 | 5 | 0.6 | 4 | 0.75 | 19 | 0.5789 | 2 | 1 | 7 | 0.7143 | 3 | 0.3333 | 6 | 1 | 18 | 0.7778 |
| 12_2738603 | 4 | 0.5 | 7 | 0.1429 | 6 | 0.1667 | 5 | 0 | 22 | 0.1818 | 4 | 0 | 6 | 0.1667 | 2 | 0.5 | 4 | 0.25 | 16 | 0.1875 |
| 12_2738723 | 4 | 0 | 4 | 0 | 4 | 0 | 4 | 0.25 | 16 | 0.0625 | 4 | 0 | 3 | 0 | 5 | 0 | 5 | 0.2 | 17 | 0.0588 |
| 12_2738756 | 3 | 0 | 1 | 1 | 3 | 0.3333 | 3 | 0 | 10 | 0.2 | 2 | 0 | NA | NA | 3 | 0 | 3 | 0.3333 | 8 | 0.125 |
| 12_2739053 | NA | NA | 1 | 0 | 1 | 1 | 1 | 1 | 3 | 0.6667 | NA | NA | NA | NA | NA | NA | 4 | 0.75 | 4 | 0.75 |
| 12_2739067 | NA | NA | 2 | 1 | 1 | 1 | 1 | 1 | 4 | 1 | 1 | 0 | NA | NA | NA | NA | 2 | 0.5 | 3 | 0.3333 |
| 12_2739070 | NA | NA | 2 | 1 | 1 | 1 | NA | NA | 3 | 1 | 1 | 0 | NA | NA | NA | NA | 4 | 0.5 | 5 | 0.4 |
| 12_2739077 | NA | NA | 2 | 0 | 1 | 1 | 1 | 0 | 4 | 0.25 | 1 | 0 | NA | NA | NA | NA | 4 | 0.25 | 5 | 0.2 |
| 12_2739124 | NA | NA | 1 | 1 | 1 | 0 | 1 | 0 | 3 | 0.3333 | 1 | 1 | NA | NA | NA | NA | 2 | 0.5 | 3 | 0.6667 |
| 12_2739126 | NA | NA | NA | NA | 1 | 1 | 1 | 1 | 2 | 1 | 1 | 1 | NA | NA | NA | NA | NA | NA | 1 | 1 |
| 12_2739135 | NA | NA | 1 | 1 | 1 | 1 | 1 | 1 | 3 | 1 | 1 | 1 | NA | NA | NA | NA | 2 | 1 | 3 | 1 |
| 12_2739249 | NA | NA | NA | NA | 1 | 1 | NA | NA | 1 | 1 | NA | NA | NA | NA | NA | NA | 1 | 1 | 1 | 1 |
| 12_2868101 | 1 | 1 | 2 | 1 | 1 | 1 | 1 | 0 | 5 | 0.8 | NA | NA | 2 | 0.5 | NA | NA | 1 | 1 | 3 | 0.6667 |
| 12_2870915 | 1 | 0 | 1 | 0 | NA | NA | 4 | 0.25 | 6 | 0.1667 | 1 | 0 | 2 | 0 | 3 | 0.3333 | 1 | 1 | 7 | 0.2857 |
| 12_2870940 | NA | NA | 1 | 1 | 2 | 0.5 | 3 | 0 | 6 | 0.3333 | 1 | 0 | 1 | 1 | 2 | 1 | NA | NA | 4 | 0.75 |
| 12_2873107 | 2 | 0.5 | 7 | 0 | 2 | 0 | 9 | 0 | 20 | 0.05 | 4 | 0.25 | 2 | 0 | 1 | 0 | 3 | 0.6667 | 10 | 0.3 |
| 12_2874082 | 8 | 0.125 | 10 | 0.1 | 12 | 0.0833 | 6 | 0.1667 | 36 | 0.1111 | 5 | 0 | 6 | 0.1667 | 3 | 0.6667 | 8 | 0 | 22 | 0.1364 |
| 12_2874111 | 9 | 0.5556 | 9 | 0.5556 | 13 | 0.4615 | 7 | 0.8571 | 38 | 0.5789 | 8 | 0.75 | 5 | 0.6 | 2 | 0 | 11 | 0.6364 | 26 | 0.6154 |
| 12_2874128 | 9 | 0.5556 | 6 | 1 | 13 | 0.2308 | 5 | 1 | 33 | 0.5758 | 7 | 0.8571 | 4 | 1 | 3 | 0 | 11 | 0.4545 | 25 | 0.6 |
| 12_2874134 | 9 | 0.5556 | 11 | 0.4545 | 11 | 0.4545 | 5 | 0.8 | 36 | 0.5278 | 7 | 0.5714 | 5 | 0.4 | 2 | 0.5 | 10 | 0.6 | 24 | 0.5417 |
| 12_2874150 | 10 | 0.4 | 9 | 0.2222 | 11 | 0.2727 | 6 | 0.3333 | 36 | 0.3056 | 8 | 0.125 | 5 | 0.2 | 3 | 0.6667 | 8 | 0.25 | 24 | 0.25 |
| 12_2876053 | NA | NA | NA | NA | NA | NA | 1 | 1 | 1 | 1 | NA | NA | NA | NA | NA | NA | 1 | 1 | 1 | 1 |
| 12_2876073 | NA | NA | NA | NA | NA | NA | 1 | 1 | 1 | 1 | NA | NA | NA | NA | NA | NA | 1 | 1 | 1 | 1 |
| 12_2879643 | 5 | 0.8 | 6 | 0.8333 | 9 | 0.7778 | 4 | 0.75 | 24 | 0.7917 | 4 | 0.75 | 7 | 0.7143 | 2 | 1 | 8 | 1 | 21 | 0.8571 |
| 12_2879671 | 7 | 0.7143 | 5 | 0.2 | 8 | 0.75 | 3 | 0 | 23 | 0.5217 | 5 | 0.4 | 6 | 0.6667 | 1 | 0 | 8 | 0.375 | 20 | 0.45 |
| 12_2879700 | 7 | 0.1429 | 5 | 0.4 | 7 | 0 | 3 | 0 | 22 | 0.1364 | 6 | 0 | 5 | 0.2 | 1 | 1 | 7 | 0.5714 | 19 | 0.3158 |
| 12_2879716 | 6 | 1 | 6 | 1 | 6 | 1 | 3 | 0.6667 | 21 | 0.9524 | 6 | 1 | 4 | 1 | 1 | 0 | 4 | 0.75 | 15 | 0.8667 |
| 12_2879740 | 5 | 0 | 5 | 0.2 | 6 | 0.1667 | 1 | 1 | 17 | 0.1765 | 4 | 0.25 | 4 | 0 | 1 | 0 | 6 | 0 | 15 | 0.0667 |

| | | | | | | | | | | | | | | | | | | | | |
|---|---|---|---|---|---|---|---|---|---|---|---|---|---|---|---|---|---|---|---|---|
| 12_289169 | 7 | 0.2857 | 8 | 0.625 | 5 | 0.4 | 7 | 1 | 27 | 0.5926 | 4 | 0.5 | 8 | 0.5 | 5 | 0.8 | 2 | 1 | 19 | 0.6316 |
| 12_289170 | 8 | 0.25 | 9 | 0.5556 | 5 | 0.2 | 9 | 0.4444 | 31 | 0.3871 | 5 | 0.2 | 7 | 0.4286 | 5 | 0 | 2 | 0 | 19 | 0.2105 |
| 12_289171 | 5 | 0.6 | 8 | 0 | 7 | 0.2857 | 8 | 0.375 | 28 | 0.2857 | 3 | 0 | 8 | 0.625 | 1 | 0 | 3 | 0 | 15 | 0.3333 |
| 12_289172 | 8 | 0.875 | 8 | 0.75 | 5 | 1 | 9 | 0.8889 | 30 | 0.8667 | 5 | 1 | 10 | 0.7 | 5 | 0.8 | 4 | 0.5 | 24 | 0.75 |
| 12_289172 | 8 | 0.25 | 8 | 0.125 | 6 | 0.3333 | 7 | 0 | 29 | 0.1724 | 5 | 0.2 | 10 | 0.2 | 4 | 0 | 2 | 0.5 | 21 | 0.1905 |
| 12_298769 | 3 | 1 | 2 | 0.5 | 2 | 1 | 4 | 0.75 | 11 | 0.8182 | 2 | 1 | 3 | 0.6667 | 5 | 0.6 | 2 | 1 | 12 | 0.75 |
| 12_298770 | 4 | 0.25 | 1 | 1 | 6 | 0.3333 | 3 | 0.6667 | 14 | 0.4286 | 2 | 0.5 | 3 | 0.3333 | 6 | 0.8333 | 2 | 0.5 | 13 | 0.6154 |
| 12_298770 | 3 | 0.6667 | 2 | 0 | 5 | 0.4 | 5 | 0.2 | 15 | 0.3333 | 2 | 0 | 3 | 1 | 6 | 0.3333 | 2 | 0 | 13 | 0.3846 |
| 12_298771 | 4 | 0.5 | 1 | 0 | 5 | 0.8 | 3 | 0.6667 | 13 | 0.6154 | 2 | 0.5 | 3 | 0.6667 | 5 | 1 | 2 | 1 | 12 | 0.8333 |
| 12_298771 | 4 | 0.5 | 2 | 0 | 6 | 0.5 | 5 | 0.6 | 17 | 0.4706 | 1 | 1 | 3 | 0 | 6 | 0.5 | 2 | 0.5 | 12 | 0.4167 |
| 12_298771 | 3 | 0.3333 | 2 | 0.5 | 6 | 0.1667 | 5 | 0 | 16 | 0.1875 | 2 | 0 | 3 | 0 | 5 | 0 | 2 | 0.5 | 12 | 0.0833 |
| 12_298772 | 4 | 0 | 2 | 0.5 | 6 | 0 | 4 | 0.25 | 16 | 0.125 | 2 | 0 | 3 | 0 | 7 | 0.1429 | 2 | 0 | 14 | 0.0714 |
| 12_298772 | 4 | 0.5 | 2 | 0 | 5 | 1 | 3 | 0.6667 | 14 | 0.6429 | 2 | 1 | 3 | 1 | 4 | 0.75 | 2 | 1 | 11 | 0.9091 |
| 12_298772 | 3 | 1 | 2 | 0.5 | 6 | 0.6667 | 5 | 0.8 | 16 | 0.75 | 2 | 1 | 3 | 1 | 3 | 1 | 2 | 1 | 10 | 1 |
| 12_298774 | 2 | 0 | 4 | 0 | 6 | 0.1667 | 6 | 0 | 18 | 0.0556 | 3 | 0.3333 | 4 | 0 | 6 | 0.1667 | 4 | 0 | 17 | 0.1176 |
| 12_298776 | 3 | 0.6667 | 1 | 1 | 3 | 0.6667 | 2 | 1 | 9 | 0.7778 | 1 | 0 | 2 | 0 | 1 | 1 | 3 | 1 | 7 | 0.5714 |
| 12_298776 | 4 | 0 | 5 | 0.2 | 8 | 0.125 | 6 | 0 | 23 | 0.087 | 3 | 0 | 4 | 0 | 6 | 0.1667 | 4 | 0 | 17 | 0.0588 |
| 12_298776 | 4 | 0.25 | 4 | 0 | 8 | 0.625 | 7 | 0.5714 | 23 | 0.4348 | 4 | 0.25 | 4 | 0.25 | 6 | 0.5 | 4 | 0.25 | 18 | 0.3333 |
| 12_299171 | NA | NA | 1 | 1 | NA | NA | NA | NA | 1 | 1 | NA | NA | NA | NA | NA | NA | 1 | 1 | 1 | 1 |
| 12_299704 | 12 | 0.4167 | 8 | 0.25 | 14 | 0.2857 | 11 | 0.0909 | 45 | 0.2667 | 8 | 0.25 | 5 | 0.4 | 12 | 0.25 | 11 | 0.7273 | 36 | 0.4167 |
| 12_299705 | 11 | 0 | 8 | 0.375 | 12 | 0.0833 | 11 | 0.0909 | 42 | 0.119 | 5 | 0.6 | 2 | 0 | 10 | 0.1 | 10 | 0.1 | 27 | 0.1852 |
| 12_299708 | 13 | 0.3077 | 7 | 0.1429 | 15 | 0.2 | 15 | 0.0667 | 50 | 0.18 | 8 | 0.125 | 6 | 0.1667 | 9 | 0.3333 | 11 | 0.1818 | 34 | 0.2059 |
| 12_299708 | 13 | 0.2308 | 6 | 0.3333 | 15 | 0.2 | 15 | 0.4667 | 49 | 0.3061 | 6 | 0.6667 | 6 | 0 | 10 | 0.1 | 10 | 0.4 | 32 | 0.2812 |
| 12_299709 | 14 | 0.2857 | 7 | 0.2857 | 16 | 0.3125 | 15 | 0 | 52 | 0.2115 | 8 | 0.125 | 6 | 0 | 13 | 0.2308 | 11 | 0.3636 | 38 | 0.2105 |
| 12_299712 | 11 | 0.0909 | 10 | 0 | 16 | 0.1875 | 15 | 0.0667 | 52 | 0.0962 | 8 | 0.125 | 9 | 0.3333 | 9 | 0.1111 | 11 | 0.1818 | 37 | 0.1892 |
| 12_299712 | 17 | 0.8824 | 8 | 0.625 | 17 | 0.8235 | 16 | 0.75 | 58 | 0.7931 | 11 | 0.8182 | 10 | 1 | 11 | 0.7273 | 13 | 0.8462 | 45 | 0.8444 |
| 12_299712 | 15 | 0.2 | 10 | 0.2 | 17 | 0 | 16 | 0 | 58 | 0.0862 | 11 | 0 | 11 | 0.1818 | 13 | 0 | 12 | 0.0833 | 47 | 0.0638 |
| 12_299712 | 14 | 0.1429 | 11 | 0.1818 | 19 | 0.2105 | 17 | 0.1176 | 61 | 0.1639 | 11 | 0.0909 | 8 | 0.125 | 12 | 0.1667 | 14 | 0.2143 | 45 | 0.1556 |
| 12_299716 | 11 | 0.2727 | 12 | 0.0833 | 12 | 0.1667 | 16 | 0.125 | 51 | 0.1569 | 11 | 0.0909 | 7 | 0.1429 | 12 | 0.0833 | 8 | 0.375 | 38 | 0.1579 |
| 12_299716 | 5 | 0 | 7 | 0.1429 | 13 | 0.1538 | 12 | 0.0833 | 37 | 0.1081 | 7 | 0 | 2 | 0.5 | 10 | 0.1 | 6 | 0 | 25 | 0.08 |
| 12_299717 | 19 | 0.3684 | 12 | 0.25 | 18 | 0.2778 | 19 | 0.2632 | 68 | 0.2941 | 11 | 0.1818 | 10 | 0.3 | 15 | 0.0667 | 14 | 0.4286 | 50 | 0.24 |
| 12_299717 | 12 | 0.0833 | 10 | 0 | 16 | 0.125 | 16 | 0 | 54 | 0.0556 | 12 | 0.0833 | 9 | 0.1111 | 16 | 0 | 13 | 0.0769 | 50 | 0.06 |
| 12_299717 | 16 | 0.6875 | 13 | 0.5385 | 17 | 0.5294 | 18 | 0.4444 | 64 | 0.5469 | 12 | 0.5833 | 10 | 0.8 | 15 | 0.4667 | 10 | 0.5 | 47 | 0.5745 |
| 12_299718 | 15 | 0.2667 | 12 | 0 | 16 | 0.25 | 18 | 0.2222 | 61 | 0.1967 | 10 | 0.3 | 8 | 0.375 | 12 | 0.1667 | 13 | 0.3846 | 43 | 0.3023 |
| 12_299721 | 17 | 0.1765 | 15 | 0 | 21 | 0.0952 | 18 | 0.0556 | 71 | 0.0845 | 12 | 0.25 | 13 | 0.1538 | 15 | 0.1333 | 11 | 0.0909 | 51 | 0.1569 |
| 12_299724 | 19 | 0.3684 | 18 | 0.3889 | 24 | 0.2917 | 23 | 0.3478 | 84 | 0.3452 | 12 | 0.3333 | 15 | 0.3333 | 17 | 0.1765 | 15 | 0.4 | 59 | 0.3051 |
| 12_300522 | NA | NA | NA | NA | 2 | 1 | 2 | 1 | 4 | 1 | 1 | 0 | 2 | 1 | 1 | 0 | 2 | 1 | 6 | 0.6667 |
| 12_300523 | NA | NA | 1 | 1 | 3 | 0.3333 | 2 | 1 | 6 | 0.6667 | 1 | 1 | NA | NA | 1 | 0 | 2 | 0.5 | 4 | 0.5 |
| 12_300524 | NA | NA | 1 | 1 | 3 | 1 | 1 | 1 | 5 | 1 | 1 | 1 | 1 | 1 | NA | NA | 1 | 1 | 3 | 1 |
| 12_300525 | NA | NA | 1 | 1 | 3 | 0.6667 | 2 | 1 | 6 | 0.8333 | 1 | 1 | 2 | 0.5 | 1 | 0 | 2 | 0.5 | 6 | 0.5 |
| 12_300528 | NA | NA | 1 | 1 | 3 | 1 | 2 | 1 | 6 | 1 | 1 | 1 | 2 | 1 | 1 | 1 | 3 | 1 | 7 | 1 |
| 12_300530 | NA | NA | 1 | 1 | 3 | 0.6667 | 2 | 1 | 6 | 0.8333 | 1 | 1 | 2 | 0.5 | 1 | 1 | 4 | 0.75 | 8 | 0.75 |
| 12_300656 | 2 | 1 | 1 | 0 | 4 | 0 | 2 | 0.5 | 9 | 0.3333 | 3 | 0 | 2 | 0 | 1 | 0 | 4 | 0.25 | 10 | 0.1 |
| 12_300656 | 2 | 0.5 | 1 | 0 | 4 | 0.25 | 2 | 0 | 9 | 0.2222 | 3 | 0 | 2 | 0 | 1 | 0 | 4 | 0.25 | 10 | 0.1 |
| 12_300656 | 2 | 0 | 1 | 0 | 4 | 0 | 2 | 0.5 | 9 | 0.1111 | 3 | 0.3333 | 2 | 0 | 1 | 1 | 3 | 0 | 9 | 0.2222 |
| 12_300661 | 4 | 0.25 | 1 | 1 | 4 | 0.25 | 3 | 1 | 12 | 0.5 | 4 | 0.25 | 2 | 0 | 1 | 0 | 3 | 0.3333 | 10 | 0.2 |
| 12_300673 | 4 | 0 | 2 | 1 | 4 | 0 | 4 | 0.25 | 14 | 0.2143 | 3 | 0 | 2 | 0 | 3 | 0 | 5 | 0.2 | 13 | 0.0769 |
| 12_306387 | 6 | 0.1667 | 5 | 0 | 1 | 0 | 1 | 0 | 13 | 0.0769 | 3 | 0 | 4 | 0.25 | 5 | 0.4 | 4 | 0 | 16 | 0.1875 |
| 12_306388 | 4 | 0.25 | 4 | 0 | 1 | 0 | 1 | 0 | 10 | 0.1 | 3 | 0 | 2 | 0 | 5 | 0.2 | 4 | 0 | 14 | 0.0714 |
| 12_306389 | 5 | 0.6 | 5 | 0.2 | 1 | 1 | 1 | 0 | 12 | 0.4167 | 2 | 0 | 4 | 0.25 | 5 | 0.2 | 5 | 0.4 | 16 | 0.25 |
| 12_306390 | 6 | 0.5 | 6 | 0.1667 | 1 | 1 | 1 | 0 | 14 | 0.3571 | 3 | 0.3333 | 3 | 0 | 4 | 0 | 5 | 0.2 | 15 | 0.1333 |
| 12_306391 | 7 | 0.1429 | 6 | 0.3333 | 1 | 1 | 1 | 0 | 15 | 0.2667 | 2 | 0 | 4 | 0 | 5 | 0.2 | 5 | 0 | 16 | 0.0625 |
| 12_306392 | 6 | 0.1667 | 4 | 0 | 1 | 0 | 1 | 0 | 12 | 0.0833 | 3 | 0 | 4 | 0.25 | 4 | 0 | 2 | 0 | 13 | 0.0769 |
| 12_306395 | 7 | 0.1429 | 6 | 0 | 1 | 0 | 1 | 0 | 15 | 0.0667 | 2 | 0 | 4 | 0.25 | 5 | 0 | 5 | 0 | 16 | 0.0625 |
| 12_306403 | 7 | 0.2857 | 6 | 0.1667 | 1 | 0 | 1 | 0 | 15 | 0.2 | 3 | 0.3333 | 2 | 0 | 5 | 0.2 | 4 | 0.25 | 14 | 0.2143 |
| 12_306406 | 8 | 0 | 5 | 0.2 | 1 | 0 | 1 | 0 | 15 | 0.0667 | 3 | 0.3333 | 5 | 0 | 7 | 0 | 3 | 0 | 18 | 0.0556 |
| 12_306407 | 8 | 0.125 | 5 | 0 | 1 | 0 | 2 | 0 | 16 | 0.0625 | 3 | 0 | 4 | 0 | 5 | 0 | 5 | 0.2 | 17 | 0.0588 |
| 12_306409 | 7 | 0.2857 | 4 | 0 | 1 | 0 | 2 | 0 | 14 | 0.1429 | 4 | 0 | 6 | 0 | 7 | 0.1429 | 6 | 0.5 | 23 | 0.1739 |
| 12_306410 | 6 | 0.1667 | 7 | 0.1429 | 2 | 0 | 2 | 0.5 | 17 | 0.1765 | 3 | 0.6667 | 7 | 0 | 7 | 0 | 7 | 0.4286 | 24 | 0.2083 |
| 12_309344 | NA | NA | 1 | 1 | NA | NA | NA | NA | 1 | 1 | NA | NA | NA | NA | 1 | 1 | NA | NA | 1 | 1 |
| 12_309347 | NA | NA | NA | NA | 1 | 1 | NA | NA | 1 | 1 | NA | NA | NA | NA | 1 | 1 | NA | NA | 1 | 1 |
| 12_309452 | 6 | 0.3333 | 9 | 0.2222 | 9 | 0.1111 | 8 | 0.375 | 32 | 0.25 | 5 | 0.2 | 12 | 0.3333 | 16 | 0.1875 | 12 | 0.3333 | 45 | 0.2667 |
| 12_309453 | 8 | 0 | 9 | 0 | 8 | 0.125 | 9 | 0.2222 | 34 | 0.0882 | 8 | 0 | 13 | 0.1538 | 15 | 0 | 11 | 0.0909 | 47 | 0.0638 |
| 12_313685 | 2 | 0 | 1 | 1 | 1 | 1 | NA | NA | 4 | 0.5 | 1 | 1 | 1 | 1 | 2 | 0.5 | 3 | 0.3333 | 7 | 0.5714 |
| 12_313794 | 2 | 0 | 2 | 1 | 1 | 1 | 1 | 0 | 6 | 0.5 | 2 | 1 | 4 | 0.75 | 3 | 1 | 2 | 0.5 | 11 | 0.8182 |

| | | | | | | | | | | | | | | | | | | | | |
|---|---|---|---|---|---|---|---|---|---|---|---|---|---|---|---|---|---|---|---|---|
| 12_313816 | 16 | 0 | 27 | 0.1852 | 20 | 0 | 17 | 0 | 80 | 0.0625 | 14 | 0.1429 | 14 | 0 | 18 | 0.1111 | 9 | 0.1111 | 55 | 0.0909 |
| 12_313818 | 16 | 0.1875 | 24 | 0.375 | 25 | 0.32 | 15 | 0.3333 | 80 | 0.3125 | 17 | 0.4118 | 16 | 0.3125 | 21 | 0.3333 | 10 | 0.2 | 64 | 0.3281 |
| 12_313820 | 14 | 0.1429 | 16 | 0.4375 | 19 | 0.3158 | 12 | 0.3333 | 61 | 0.3115 | 14 | 0.4286 | 15 | 0.2667 | 17 | 0.1765 | 7 | 0.1429 | 53 | 0.2642 |
| 12_313820 | 17 | 0.2941 | 19 | 0.3158 | 25 | 0.48 | 14 | 0.2143 | 75 | 0.3467 | 17 | 0.3529 | 15 | 0.4667 | 17 | 0.4706 | 10 | 0.3 | 59 | 0.4068 |
| 12_313821 | 11 | 0.0909 | 14 | 0.0714 | 18 | 0.1111 | 10 | 0.2 | 53 | 0.1132 | 19 | 0.1053 | 13 | 0.0769 | 14 | 0.1429 | 6 | 0.1667 | 52 | 0.1154 |
| 12_315364 | 4 | 0.25 | NA | NA | NA | NA | 1 | 1 | 5 | 0.4 | NA | NA | 2 | 0.5 | NA | NA | 2 | 0.5 | 4 | 0.5 |
| 12_315367 | 5 | 0 | NA | NA | 2 | 0 | 1 | 1 | 8 | 0.125 | NA | NA | 2 | 0.5 | NA | NA | 2 | 0 | 4 | 0.25 |
| 12_315372 | 4 | 1 | NA | NA | 3 | 0.6667 | NA | NA | 7 | 0.8571 | NA | NA | 4 | 1 | NA | NA | 2 | 1 | 6 | 1 |
| 12_315372 | 3 | 0.3333 | NA | NA | 2 | 0 | NA | NA | 5 | 0.2 | NA | NA | 4 | 0.25 | NA | NA | 2 | 0.5 | 6 | 0.3333 |
| 12_315372 | 4 | 1 | NA | NA | 4 | 0.5 | NA | NA | 8 | 0.75 | NA | NA | 5 | 0.8 | NA | NA | 1 | 1 | 6 | 0.8333 |
| 12_315372 | 4 | 0.25 | NA | NA | 2 | 0.5 | NA | NA | 6 | 0.3333 | NA | NA | 5 | 0.4 | NA | NA | 2 | 0 | 7 | 0.2857 |
| 12_315373 | 4 | 0.5 | NA | NA | 2 | 0.5 | NA | NA | 6 | 0.5 | NA | NA | 4 | 0.25 | NA | NA | 2 | 0.5 | 6 | 0.3333 |
| 12_315374 | 3 | 1 | NA | NA | 3 | 0.6667 | NA | NA | 6 | 0.8333 | NA | NA | 5 | 1 | NA | NA | 2 | 1 | 7 | 1 |
| 12_315374 | 3 | 0.3333 | NA | NA | 1 | 0 | NA | NA | 4 | 0.25 | NA | NA | 4 | 0 | NA | NA | 2 | 0.5 | 6 | 0.1667 |
| 12_315375 | 2 | 1 | NA | NA | 1 | 1 | NA | NA | 3 | 1 | NA | NA | 2 | 0 | NA | NA | 1 | 1 | 3 | 0.3333 |
| 12_315375 | 3 | 0.3333 | NA | NA | 2 | 0 | NA | NA | 5 | 0.2 | NA | NA | 4 | 0.25 | NA | NA | 1 | 0 | 5 | 0.2 |
| 12_315375 | 3 | 0.6667 | NA | NA | 3 | 0.3333 | NA | NA | 6 | 0.5 | NA | NA | 4 | 0.75 | NA | NA | 2 | 0.5 | 6 | 0.6667 |
| 12_315376 | 3 | 0.6667 | NA | NA | 3 | 0 | NA | NA | 6 | 0.3333 | NA | NA | 5 | 0.2 | NA | NA | 1 | 0 | 6 | 0.1667 |
| 12_315378 | 3 | 0.3333 | NA | NA | 2 | 0 | NA | NA | 5 | 0.2 | NA | NA | 3 | 0 | NA | NA | 1 | 1 | 4 | 0.25 |
| 12_315382 | 2 | 1 | NA | NA | 2 | 1 | NA | NA | 4 | 1 | NA | NA | 3 | 0.3333 | NA | NA | 2 | 1 | 5 | 0.6 |
| 12_316459 | 5 | 0 | 7 | 0 | 6 | 0.1667 | 10 | 0.2 | 28 | 0.1071 | 3 | 0 | 5 | 0.2 | 5 | 0 | 12 | 0.0833 | 25 | 0.08 |
| 12_316462 | 4 | 0.25 | 7 | 0 | 6 | 0 | 9 | 0.2222 | 26 | 0.1154 | 3 | 0.3333 | 4 | 0 | 5 | 0.6 | 14 | 0.0714 | 26 | 0.1923 |
| 12_316901 | 1 | 0 | 2 | 1 | NA | NA | 1 | 0 | 4 | 0.5 | 2 | 0 | NA | NA | 2 | 1 | NA | NA | 4 | 0.5 |
| 12_316903 | 1 | 0 | 1 | 1 | NA | NA | NA | NA | 2 | 0.5 | 2 | 0 | NA | NA | 2 | 0.5 | NA | NA | 4 | 0.25 |
| 12_316904 | 1 | 1 | 1 | 1 | NA | NA | 1 | 1 | 3 | 1 | 2 | 1 | NA | NA | 1 | 1 | NA | NA | 3 | 1 |
| 12_3169045 | NA | NA | NA | NA | NA | NA | 1 | 1 | 1 | 1 | 1 | 1 | NA | NA | NA | NA | NA | NA | 1 | 1 |
| 12_316904 | 1 | 1 | 1 | 0 | NA | NA | 1 | 1 | 3 | 0.6667 | 2 | 1 | NA | NA | 2 | 1 | NA | NA | 4 | 1 |
| 12_3169071 | NA | NA | 1 | 1 | NA | NA | 1 | 1 | 2 | 1 | 2 | 0.5 | NA | NA | 2 | 0.5 | NA | NA | 4 | 0.5 |
| 12_3169072 | NA | NA | 1 | 1 | NA | NA | 1 | 1 | 2 | 1 | 2 | 0.5 | NA | NA | 1 | 1 | NA | NA | 3 | 0.6667 |
| 12_3169086 | NA | NA | 1 | 1 | NA | NA | 1 | 1 | 2 | 1 | 1 | 1 | NA | NA | 2 | 0.5 | NA | NA | 3 | 0.6667 |
| 12_317578 | 2 | 0 | 4 | 0.25 | 1 | 0 | 2 | 0 | 9 | 0.1111 | 2 | 0.5 | 2 | 0 | 2 | 0 | 1 | 0 | 7 | 0.1429 |
| 12_317587 | 7 | 0.1429 | 11 | 0.1818 | 3 | 0 | 3 | 0.3333 | 24 | 0.1667 | 1 | 0 | 2 | 0 | 4 | 0 | 4 | 0.25 | 11 | 0.0909 |
| 12_317589 | 4 | 0.25 | 8 | 0.25 | 3 | 0 | 3 | 0.3333 | 18 | 0.2222 | 1 | 0 | 2 | 0.5 | 4 | 0 | 3 | 0 | 10 | 0.1 |
| 12_317589 | 5 | 0.8 | 11 | 1 | 3 | 0.6667 | 3 | 1 | 22 | 0.9091 | 1 | 1 | 2 | 1 | 4 | 0.75 | 3 | 1 | 10 | 0.9 |
| 12_317590 | 4 | 1 | 10 | 1 | 3 | 1 | 3 | 1 | 20 | 1 | 1 | 1 | 2 | 1 | 4 | 1 | 3 | 1 | 10 | 1 |
| 12_317591 | 3 | 1 | 10 | 0.6 | 3 | 0.6667 | 1 | 1 | 17 | 0.7059 | 1 | 1 | 2 | 1 | 4 | 1 | 2 | 1 | 9 | 1 |
| 12_317592 | 3 | 0.3333 | 7 | 0.7143 | 2 | 0.5 | 2 | 1 | 14 | 0.6429 | 1 | 0 | 2 | 0 | 4 | 0.25 | 1 | 1 | 8 | 0.25 |
| 12_317592 | 4 | 0.25 | 10 | 0 | 2 | 0 | 2 | 0 | 18 | 0.0556 | 1 | 0 | 2 | 0.5 | 2 | 0 | 2 | 0 | 7 | 0.1429 |
| 12_317592 | 2 | 1 | 10 | 1 | 3 | 1 | 2 | 1 | 17 | 1 | 1 | 1 | 2 | 1 | 2 | 1 | 2 | 1 | 7 | 1 |
| 12_317592 | 4 | 0.25 | 8 | 0.125 | 3 | 0.3333 | 2 | 0.5 | 17 | 0.2353 | 1 | 1 | 2 | 0 | 2 | 1 | 1 | 0 | 6 | 0.5 |
| 12_317594 | 4 | 0.25 | 9 | 0.2222 | 3 | 0.6667 | 1 | 0 | 17 | 0.2941 | 1 | 1 | 1 | 0 | 2 | 0.5 | 2 | 0.5 | 6 | 0.5 |
| 12_317595 | 3 | 0 | 9 | 0 | 2 | 1 | 2 | 0.5 | 16 | 0.1875 | 1 | 0 | 1 | 0 | 2 | 0.5 | 2 | 0 | 6 | 0.1667 |
| 12_317595 | 3 | 0.6667 | 5 | 0.2 | 3 | 0.3333 | 1 | 1 | 12 | 0.4167 | 1 | 1 | 1 | 1 | 2 | 1 | 1 | 0 | 5 | 0.8 |
| 12_317595 | 4 | 0 | 9 | 0 | 3 | 0.3333 | 2 | 0 | 18 | 0.0556 | 1 | 0 | NA | NA | 2 | 0.5 | 2 | 1 | 5 | 0.6 |
| 12_317595 | 4 | 0.25 | 8 | 0.375 | 3 | 0.6667 | 2 | 0 | 17 | 0.3529 | 1 | 0 | 1 | 0 | 2 | 0 | 2 | 0.5 | 6 | 0.1667 |
| 12_317595 | 3 | 1 | 6 | 1 | 2 | 1 | 2 | 1 | 13 | 1 | 1 | 1 | NA | NA | 2 | 1 | 2 | 1 | 5 | 1 |
| 12_317597 | 2 | 0.5 | 5 | 0.2 | 3 | 0 | 2 | 0 | 12 | 0.1667 | 1 | 0 | 1 | 0 | 1 | 0 | 2 | 0.5 | 5 | 0.2 |
| 12_317597 | 3 | 1 | 7 | 1 | 3 | 1 | 1 | 1 | 14 | 1 | 1 | 1 | 1 | 1 | 2 | 0 | 1 | 1 | 5 | 0.6 |
| 12_317597 | 3 | 0.3333 | 7 | 0.4286 | 3 | 0.3333 | 2 | 0.5 | 15 | 0.4 | 1 | 1 | 1 | 0 | 1 | 0 | 2 | 0 | 5 | 0.2 |
| 12_317620 | 1 | 0 | 7 | 0 | 1 | 0 | 1 | 1 | 10 | 0.1 | 1 | 0 | NA | NA | 1 | 1 | 2 | 0 | 4 | 0.25 |
| 12_3176207 | NA | NA | 7 | 0.2857 | 1 | 0 | NA | NA | 8 | 0.25 | 1 | 0 | NA | NA | 1 | 1 | 2 | 0 | 4 | 0.25 |
| 12_317622 | 1 | 1 | 6 | 1 | NA | NA | 1 | 1 | 8 | 1 | 1 | 1 | NA | NA | 1 | 1 | 2 | 1 | 4 | 1 |
| 12_3176234 | NA | NA | 7 | 0.5714 | NA | NA | 1 | 0 | 8 | 0.5 | 1 | 1 | NA | NA | 1 | 1 | 1 | 1 | 3 | 1 |
| 12_317623 | 1 | 1 | 7 | 0.4286 | NA | NA | 1 | 0 | 9 | 0.4444 | 1 | 1 | NA | NA | 1 | 1 | 2 | 0.5 | 4 | 0.75 |
| 12_317624 | 1 | 1 | 9 | 0.1111 | NA | NA | 1 | 1 | 11 | 0.2727 | 1 | 1 | NA | NA | 1 | 1 | 2 | 0 | 4 | 0.5 |
| 12_3176247 | NA | NA | 8 | 0.125 | NA | NA | 1 | 0 | 9 | 0.1111 | 1 | 0 | NA | NA | 1 | 0 | 2 | 0.5 | 4 | 0.25 |
| 12_319760 | 1 | 0 | 3 | 0 | 1 | 0 | 2 | 0.5 | 7 | 0.1429 | 2 | 0 | NA | NA | 1 | 0 | 1 | 1 | 4 | 0.25 |
| 12_319767 | 3 | 0 | 5 | 0.2 | 6 | 0 | 6 | 0 | 20 | 0.05 | 3 | 0 | 6 | 0.1667 | 5 | 0 | 5 | 0.4 | 19 | 0.1579 |
| 12_319767 | 1 | 0 | 3 | 0.3333 | 1 | 0 | 3 | 0.3333 | 8 | 0.25 | 2 | 0 | 2 | 0.5 | 1 | 0 | 1 | 0 | 6 | 0.1667 |
| 12_319768 | 1 | 1 | 3 | 1 | 1 | 0 | 3 | 0.3333 | 8 | 0.625 | 2 | 0.5 | 2 | 0.5 | 1 | 1 | 1 | 1 | 6 | 0.6667 |
| 12_319768 | 1 | 0 | 4 | 0.5 | 5 | 0.6 | 4 | 0.25 | 14 | 0.4286 | 3 | 0.3333 | 6 | 0.5 | 3 | 0 | 3 | 0.6667 | 15 | 0.4 |
| 12_319768 | 1 | 1 | 2 | 0.5 | 1 | 0 | 2 | 0 | 6 | 0.3333 | NA | NA | 1 | 0 | NA | NA | 1 | 1 | 2 | 0.5 |
| 12_319771 | 3 | 0.3333 | 2 | 0 | 5 | 0 | 5 | 0 | 15 | 0.0667 | 3 | 0.3333 | 5 | 0 | 3 | 0 | 4 | 0 | 15 | 0.0667 |
| 12_319774 | 3 | 0 | 2 | 0.5 | 3 | 0 | 6 | 0.3333 | 14 | 0.2143 | 3 | 0 | 5 | 0.2 | 3 | 0 | 3 | 0 | 14 | 0.0714 |
| 12_319785 | 2 | 0 | 2 | 0.5 | 1 | 0 | 5 | 0 | 10 | 0.1 | 2 | 0.5 | 2 | 0 | 1 | 0 | NA | NA | 5 | 0.2 |

| | | | | | | | | | | | | | | | | | | | | |
|---|---|---|---|---|---|---|---|---|---|---|---|---|---|---|---|---|---|---|---|---|
| 12_3197855 | 2 | 0.5 | 2 | 0 | 1 | 0 | 5 | 0 | 10 | 0.1 | 2 | 0.5 | 2 | 0 | 1 | 0 | NA | NA | 5 | 0.2 |
| 12_3198079 | 3 | 0.3333 | 2 | 1 | 4 | 0.5 | 5 | 0.8 | 14 | 0.6429 | 3 | 0.6667 | 5 | 0.6 | 2 | 0 | 3 | 0.6667 | 13 | 0.5385 |
| 12_3198098 | 2 | 0 | 3 | 0 | 4 | 0.25 | 5 | 0.2 | 14 | 0.1429 | 2 | 0 | 5 | 0.2 | 1 | 0 | 3 | 0 | 11 | 0.0909 |
| 12_3198741 | 3 | 0.3333 | NA | NA | NA | NA | 1 | 0 | 4 | 0.25 | NA | NA | NA | NA | 2 | 0 | 2 | 0.5 | 4 | 0.25 |
| 12_3200902 | 17 | 0 | 22 | 0.0455 | 20 | 0.15 | 29 | 0.069 | 88 | 0.0682 | 10 | 0.1 | 20 | 0.15 | 28 | 0.1429 | 10 | 0 | 68 | 0.1176 |
| 12_3201013 | 13 | 0 | 11 | 0.0909 | 17 | 0.1176 | 26 | 0.2308 | 67 | 0.1343 | 7 | 0.1429 | 14 | 0.0714 | 23 | 0.087 | 8 | 0 | 52 | 0.0769 |
| 12_3201015 | 14 | 0.2143 | 15 | 0.1333 | 18 | 0.1111 | 25 | 0.12 | 72 | 0.1389 | 6 | 0.5 | 16 | 0.0625 | 26 | 0.1923 | 10 | 0.1 | 58 | 0.1724 |
| 12_3201048 | 14 | 0.2143 | 15 | 0.0667 | 17 | 0.1176 | 27 | 0.1111 | 73 | 0.1233 | 7 | 0 | 12 | 0.0833 | 24 | 0.125 | 8 | 0.125 | 51 | 0.098 |
| 12_3211091 | 6 | 0.6667 | 12 | 0.6667 | 12 | 0.5833 | 15 | 0.7333 | 45 | 0.6667 | 3 | 0.6667 | 4 | 0.75 | 19 | 0.6842 | 5 | 0.6 | 31 | 0.6774 |
| 12_3242938 | NA | NA | 5 | 0.4 | 1 | 1 | 1 | 0 | 7 | 0.4286 | 1 | 1 | 2 | 0.5 | 1 | 0 | 2 | 0 | 6 | 0.3333 |
| 12_3242959 | 1 | 0 | 4 | 0 | 1 | 0 | 1 | 1 | 7 | 0.1429 | 1 | 1 | 1 | 1 | 1 | 1 | 2 | 1 | 5 | 1 |
| 12_3242959 | 1 | 0 | 5 | 0.2 | 1 | 0 | 1 | 1 | 8 | 0.25 | 2 | 1 | 2 | 1 | 1 | 0 | 2 | 0 | 7 | 0.5714 |
| 12_3242970 | 1 | 1 | 5 | 0.8 | 1 | 1 | 1 | 1 | 8 | 0.875 | 2 | 1 | 2 | 1 | 1 | 1 | 2 | 1 | 7 | 1 |
| 12_3242973 | 1 | 0 | 5 | 0.2 | 1 | 0 | NA | NA | 7 | 0.1429 | 2 | 0.5 | 2 | 0 | 2 | 0.5 | NA | NA | 6 | 0.3333 |
| 12_3242976 | NA | NA | 1 | 1 | 1 | 0 | 1 | 0 | 3 | 0.3333 | NA | NA | 1 | 1 | 1 | 1 | NA | NA | 2 | 1 |
| 12_3242980 | 1 | 0 | 5 | 0.4 | 1 | 1 | 1 | 1 | 8 | 0.5 | 2 | 1 | 2 | 0.5 | 2 | 1 | 1 | 1 | 7 | 0.8571 |
| 12_3242984 | 1 | 0 | 5 | 0.4 | 1 | 0 | 1 | 1 | 8 | 0.375 | 2 | 0.5 | 2 | 0 | 2 | 0 | 2 | 0 | 8 | 0.125 |
| 12_3242993 | 1 | 1 | 5 | 1 | 1 | 1 | NA | NA | 7 | 1 | 1 | 1 | 2 | 1 | 1 | 1 | 2 | 1 | 6 | 1 |
| 12_3242996 | 1 | 1 | 5 | 0.2 | 1 | 1 | 1 | 1 | 8 | 0.5 | NA | NA | 2 | 0.5 | 2 | 0.5 | 2 | 1 | 6 | 0.6667 |
| 12_3243006 | 1 | 1 | 4 | 0.75 | NA | NA | NA | NA | 5 | 0.8 | 1 | 1 | 2 | 1 | NA | NA | NA | NA | 3 | 1 |
| 12_3243008 | 1 | 1 | 4 | 0.75 | 1 | 0 | 1 | 1 | 7 | 0.7143 | 1 | 1 | 1 | 1 | 1 | 0 | 2 | 0.5 | 5 | 0.6 |
| 12_3243022 | 1 | 1 | 3 | 0.6667 | 1 | 1 | NA | NA | 5 | 0.8 | 2 | 1 | 1 | 1 | 2 | 1 | 1 | 1 | 6 | 1 |
| 12_3243041 | 2 | 0 | 6 | 0.1667 | 1 | 0 | 1 | 1 | 10 | 0.2 | 2 | 0 | 2 | 0.5 | 3 | 0 | 1 | 0 | 8 | 0.125 |
| 12_3243051 | 2 | 0.5 | 4 | 0.5 | 2 | 0.5 | 1 | 0 | 9 | 0.4444 | 2 | 0.5 | 1 | 1 | 2 | 0.5 | 1 | 0 | 6 | 0.5 |
| 12_3243063 | 2 | 0.5 | 5 | 1 | 1 | 0 | 1 | 1 | 9 | 0.7778 | 2 | 0.5 | 1 | 1 | 2 | 0.5 | 1 | 0 | 6 | 0.5 |
| 12_3243105 | 2 | 0.5 | 4 | 0 | 2 | 0 | 1 | 0 | 9 | 0.1111 | 2 | 0.5 | 1 | 0 | 2 | 0 | 2 | 0 | 7 | 0.1429 |
| 12_3243116 | 2 | 0.5 | 3 | 0 | 2 | 0 | NA | NA | 7 | 0.1429 | 2 | 0 | NA | NA | 2 | 0.5 | 2 | 0 | 6 | 0.1667 |
| 12_3243213 | 3 | 0.3333 | 3 | 0 | 2 | 0 | NA | NA | 8 | 0.125 | 2 | 0.5 | NA | NA | 1 | 0 | 2 | 0 | 5 | 0.2 |
| 12_3243214 | 4 | 0.25 | 4 | 0.25 | 3 | 0 | 1 | 0 | 12 | 0.1667 | 3 | 0.3333 | 1 | 0 | 2 | 0 | 2 | 0 | 8 | 0.125 |
| 12_3243254 | 4 | 0.25 | 3 | 0 | 4 | 0.25 | 1 | 0 | 12 | 0.1667 | 3 | 0.6667 | 1 | 0 | 2 | 0 | 2 | 0 | 8 | 0.25 |
| 12_3290068 | 9 | 0.1111 | 4 | 0 | 4 | 0.25 | 4 | 0 | 21 | 0.0952 | NA | NA | 2 | 1 | 7 | 0 | 5 | 0 | 14 | 0.1429 |
| 12_3316184 | NA | NA | NA | NA | 1 | 1 | NA | NA | 1 | 1 | NA | NA | 1 | 1 | NA | NA | NA | NA | 1 | 1 |
| 12_3316193 | NA | NA | 1 | 1 | 1 | 1 | NA | NA | 2 | 1 | NA | NA | 1 | 1 | NA | NA | NA | NA | 1 | 1 |
| 12_3316196 | NA | NA | 1 | 1 | 1 | 1 | NA | NA | 2 | 1 | NA | NA | 1 | 1 | NA | NA | NA | NA | 1 | 1 |
| 12_3316199 | NA | NA | NA | NA | 1 | 1 | NA | NA | 1 | 1 | NA | NA | 1 | 1 | NA | NA | NA | NA | 1 | 1 |
| 12_3316271 | NA | NA | 1 | 1 | 1 | 1 | NA | NA | 2 | 1 | NA | NA | 1 | 1 | NA | NA | NA | NA | 1 | 1 |
| 12_3316274 | NA | NA | 1 | 0 | 1 | 1 | NA | NA | 2 | 0.5 | NA | NA | 1 | 1 | NA | NA | NA | NA | 1 | 1 |
| 12_3316276 | NA | NA | 1 | 1 | 1 | 0 | NA | NA | 2 | 0.5 | NA | NA | 1 | 1 | NA | NA | NA | NA | 1 | 1 |
| 12_3316290 | NA | NA | 1 | 1 | 1 | 1 | NA | NA | 2 | 1 | NA | NA | 1 | 1 | NA | NA | NA | NA | 1 | 1 |
| 12_3316309 | NA | NA | 1 | 1 | NA | NA | NA | NA | 1 | 1 | NA | NA | 1 | 1 | NA | NA | NA | NA | 1 | 1 |
| 12_3316310 | NA | NA | 1 | 1 | NA | NA | NA | NA | 1 | 1 | NA | NA | 1 | 1 | NA | NA | NA | NA | 1 | 1 |
| 12_3317542 | 1 | 1 | NA | NA | NA | NA | NA | NA | 1 | 1 | 1 | 1 | 1 | 1 | NA | NA | NA | NA | 2 | 1 |
| 12_3317560 | 1 | 1 | NA | NA | NA | NA | NA | NA | 1 | 1 | NA | NA | 1 | 1 | NA | NA | NA | NA | 1 | 1 |
| 12_3317582 | 1 | 1 | NA | NA | NA | NA | NA | NA | 1 | 1 | 1 | 0 | 1 | 1 | NA | NA | NA | NA | 2 | 0.5 |
| 12_3318317 | 4 | 0 | 3 | 0.3333 | 1 | 0 | NA | NA | 8 | 0.125 | 5 | 0.4 | 1 | 0 | 2 | 0.5 | 1 | 0 | 9 | 0.3333 |
| 12_3318326 | 3 | 0.3333 | 2 | 0.5 | NA | NA | NA | NA | 5 | 0.4 | 4 | 0.5 | 1 | 0 | 2 | 0 | 1 | 0 | 8 | 0.25 |
| 12_3318367 | NA | NA | 4 | 0.5 | 1 | 1 | NA | NA | 5 | 0.6 | 4 | 0.75 | 1 | 1 | 2 | 0 | 1 | 1 | 8 | 0.625 |
| 12_3318382 | NA | NA | 4 | 0.5 | 1 | 0 | NA | NA | 5 | 0.4 | 2 | 0.5 | NA | NA | 2 | 0 | 1 | 0 | 5 | 0.2 |
| 12_3318418 | NA | NA | 3 | 0.3333 | NA | NA | NA | NA | 3 | 0.3333 | 3 | 0.3333 | 1 | 0 | 2 | 0 | 1 | 1 | 7 | 0.2857 |
| 12_3318471 | NA | NA | 4 | 0.5 | NA | NA | NA | NA | 4 | 0.5 | 1 | 1 | 1 | 1 | 2 | 0 | 1 | 1 | 5 | 0.6 |
| 12_3318476 | NA | NA | 4 | 1 | NA | NA | NA | NA | 4 | 1 | 3 | 1 | 1 | 1 | 2 | 1 | 1 | 1 | 7 | 1 |
| 12_3318484 | NA | NA | 5 | 0.2 | NA | NA | NA | NA | 5 | 0.2 | 3 | 0 | 2 | 0.5 | 1 | 1 | 1 | 0 | 7 | 0.2857 |
| 12_3318487 | NA | NA | 5 | 0.8 | 1 | 1 | NA | NA | 6 | 0.8333 | 3 | 1 | 1 | 0 | 1 | 0 | 1 | 1 | 6 | 0.6667 |
| 12_3318494 | NA | NA | 5 | 0.4 | 1 | 0 | NA | NA | 6 | 0.3333 | 3 | 0.6667 | 2 | 0 | 1 | 1 | 1 | 0 | 7 | 0.4286 |
| 12_3318497 | NA | NA | 4 | 0 | 1 | 1 | NA | NA | 5 | 0.2 | 2 | 0.5 | 1 | 0 | 1 | 0 | 1 | 1 | 5 | 0.4 |
| 12_3318503 | NA | NA | 5 | 1 | 1 | 1 | NA | NA | 6 | 1 | 3 | 1 | 2 | 1 | 1 | 1 | 1 | 1 | 7 | 1 |
| 12_3318506 | NA | NA | 3 | 1 | 1 | 1 | NA | NA | 4 | 1 | 2 | 1 | 2 | 0.5 | 1 | 1 | 1 | 1 | 6 | 0.8333 |
| 12_3318509 | NA | NA | 3 | 0.3333 | 1 | 1 | NA | NA | 4 | 0.5 | 3 | 0.3333 | 1 | 1 | NA | NA | 1 | 0 | 5 | 0.4 |
| 12_3318511 | NA | NA | 4 | 0.75 | 1 | 1 | NA | NA | 5 | 0.8 | 2 | 0.5 | 2 | 0.5 | 1 | 0 | 1 | 1 | 6 | 0.5 |
| 12_3318528 | NA | NA | 4 | 0.25 | 1 | 1 | NA | NA | 5 | 0.4 | 2 | 0 | 2 | 0.5 | NA | NA | 1 | 0 | 5 | 0.2 |
| 12_3318537 | NA | NA | 4 | 0.5 | 1 | 1 | NA | NA | 5 | 0.6 | 1 | 1 | 2 | 0.5 | 1 | 1 | NA | NA | 4 | 0.75 |
| 12_3318542 | NA | NA | 4 | 0.75 | 1 | 1 | NA | NA | 5 | 0.8 | 1 | 1 | 2 | 1 | 1 | 1 | NA | NA | 4 | 1 |
| 12_3318549 | NA | NA | 2 | 1 | NA | NA | NA | NA | 2 | 1 | NA | NA | 2 | 1 | NA | NA | NA | NA | 2 | 1 |
| 12_3319726 | NA | NA | 1 | 1 | NA | NA | NA | NA | 1 | 1 | 2 | 0.5 | NA | NA | 1 | 0 | NA | NA | 3 | 0.3333 |
| 12_3329998 | 2 | 0.5 | 5 | 0.2 | 3 | 0 | 1 | 1 | 11 | 0.2727 | 3 | 0.6667 | 3 | 0.3333 | 4 | 1 | 1 | 1 | 11 | 0.7273 |

| | | | | | | | | | | | | | | | | | | | | |
|---|---|---|---|---|---|---|---|---|---|---|---|---|---|---|---|---|---|---|---|---|
| 12_334172 | NA | NA | 1 | 0 | 1 | 1 | NA | NA | 2 | 0.5 | NA | NA | 1 | 1 | NA | NA | NA | NA | 1 | 1 |
| 12_334175 | NA | NA | 1 | 1 | NA | NA | NA | NA | 1 | 1 | NA | NA | NA | NA | 1 | 1 | NA | NA | 1 | 1 |
| 12_334176 | NA | NA | 1 | 0 | 1 | 1 | NA | NA | 2 | 0.5 | NA | NA | 1 | 1 | 1 | 1 | NA | NA | 2 | 1 |
| 12_334250 | 2 | 0.5 | 1 | 0 | 1 | 0 | 1 | 0 | 5 | 0.2 | NA | NA | 1 | 0 | 2 | 0.5 | 1 | 0 | 4 | 0.25 |
| 12_334252 | 2 | 0.5 | 1 | 1 | 1 | 1 | 1 | 0 | 5 | 0.6 | NA | NA | 1 | 0 | 2 | 1 | NA | NA | 3 | 0.6667 |
| 12_334253 | 2 | 0.5 | 1 | 1 | 1 | 1 | 1 | 1 | 5 | 0.8 | 1 | 1 | 2 | 1 | 2 | 1 | NA | NA | 5 | 1 |
| 12_334254 | 2 | 0.5 | 2 | 0 | 1 | 1 | 1 | 0 | 6 | 0.3333 | 1 | 0 | 2 | 0.5 | 1 | 1 | 1 | 1 | 5 | 0.6 |
| 12_334256 | 2 | 0 | 2 | 0.5 | 1 | 0 | 1 | 0 | 6 | 0.1667 | 1 | 0 | 2 | 0.5 | 2 | 0 | 1 | 0 | 6 | 0.1667 |
| 12_334257 | 3 | 0.3333 | 2 | 0.5 | 2 | 0.5 | 1 | 0 | 8 | 0.375 | 1 | 0 | 2 | 0 | 1 | 1 | 1 | 0 | 5 | 0.2 |
| 12_334878 | 2 | 1 | NA | NA | NA | NA | 3 | 0.6667 | 5 | 0.8 | NA | NA | 1 | 1 | NA | NA | NA | NA | 1 | 1 |
| 12_334878 | 2 | 0.5 | NA | NA | NA | NA | 3 | 0.6667 | 5 | 0.6 | 1 | 0 | 1 | 1 | NA | NA | NA | NA | 2 | 0.5 |
| 12_334880 | 2 | 0.5 | NA | NA | NA | NA | 3 | 0.3333 | 5 | 0.4 | 1 | 0 | 1 | 1 | 1 | 1 | NA | NA | 3 | 0.6667 |
| 12_334881 | 2 | 1 | NA | NA | NA | NA | 3 | 1 | 5 | 1 | 1 | 1 | 1 | 1 | 1 | 1 | NA | NA | 3 | 1 |
| 12_334883 | 2 | 0.5 | NA | NA | NA | NA | 3 | 0.3333 | 5 | 0.4 | 1 | 1 | 1 | 1 | 1 | 1 | NA | NA | 3 | 1 |
| 12_334897 | NA | NA | NA | NA | NA | NA | 1 | 1 | 1 | 1 | 1 | 1 | NA | NA | NA | NA | NA | NA | 1 | 1 |
| 12_344232 | 2 | 0.5 | 2 | 0.5 | 1 | 0 | NA | NA | 5 | 0.4 | NA | NA | NA | NA | 2 | 0.5 | NA | NA | 2 | 0.5 |
| 12_350971 | NA | NA | NA | NA | NA | NA | 1 | 1 | 1 | 1 | NA | NA | NA | NA | 1 | 1 | NA | NA | 1 | 1 |
| 12_350978 | NA | NA | NA | NA | NA | NA | 1 | 1 | 1 | 1 | NA | NA | NA | NA | 1 | 1 | NA | NA | 1 | 1 |
| 12_351942 | 1 | 0 | 5 | 0.2 | 3 | 0.3333 | 4 | 0.25 | 13 | 0.2308 | 2 | 0 | 3 | 0.6667 | 5 | 0.8 | 4 | 0.25 | 14 | 0.5 |
| 12_351942 | 1 | 1 | 5 | 1 | 3 | 0.6667 | 3 | 1 | 12 | 0.9167 | 2 | 1 | 3 | 1 | 5 | 0.8 | 3 | 1 | 13 | 0.9231 |
| 12_351942 | 1 | 0 | 5 | 0.2 | 3 | 0.6667 | 2 | 0.5 | 11 | 0.3636 | 2 | 0.5 | 2 | 0.5 | 3 | 0.3333 | 5 | 0.6 | 12 | 0.5 |
| 12_351942 | 1 | 1 | 5 | 0 | 2 | 0 | 4 | 0.25 | 12 | 0.1667 | 2 | 0 | 2 | 0 | 5 | 0 | 2 | 0.5 | 11 | 0.0909 |
| 12_351942 | 1 | 0 | 3 | 0.3333 | 1 | 0 | 3 | 0.6667 | 8 | 0.375 | 1 | 1 | 3 | 0.6667 | 5 | 1 | 3 | 0.6667 | 12 | 0.8333 |
| 12_351943 | 1 | 0 | 3 | 0.3333 | 2 | 0 | 3 | 0 | 9 | 0.1111 | 2 | 0.5 | 2 | 0.5 | 5 | 0.6 | 5 | 0.4 | 14 | 0.5 |
| 12_351943 | 1 | 0 | 4 | 1 | 3 | 1 | 4 | 1 | 12 | 0.9167 | 2 | 1 | 3 | 1 | 4 | 1 | 3 | 1 | 12 | 1 |
| 12_351943 | 1 | 1 | 2 | 1 | 3 | 0.6667 | 2 | 1 | 8 | 0.875 | 2 | 1 | 1 | 1 | 3 | 1 | 2 | 1 | 8 | 1 |
| 12_351944 | 1 | 0 | 4 | 0.25 | 3 | 0.3333 | 4 | 0 | 12 | 0.1667 | 3 | 0 | 3 | 0 | 5 | 0 | 5 | 0.2 | 16 | 0.0625 |
| 12_351944 | 1 | 1 | 5 | 0.8 | 3 | 0.6667 | 3 | 1 | 12 | 0.8333 | 2 | 1 | 2 | 1 | 5 | 0.8 | 5 | 1 | 14 | 0.9286 |
| 12_351951 | 3 | 0 | 8 | 0.125 | 3 | 0 | 3 | 0 | 17 | 0.0588 | 3 | 0 | 4 | 0.25 | 8 | 0.125 | 4 | 0 | 19 | 0.1053 |
| 12_351953 | 4 | 0 | 9 | 0.1111 | 3 | 0.3333 | 4 | 0 | 20 | 0.1 | 3 | 0.3333 | 5 | 0 | 7 | 0 | 4 | 0 | 19 | 0.0526 |
| 12_351962 | 5 | 0 | 7 | 0 | 3 | 0 | 4 | 0.25 | 19 | 0.0526 | 2 | 0 | 3 | 0.3333 | 9 | 0 | 4 | 0 | 18 | 0.0556 |
| 12_351963 | 3 | 0.6667 | 6 | 0 | 1 | 0 | 2 | 0.5 | 12 | 0.25 | 2 | 0 | 2 | 0.5 | 9 | 0.3333 | 3 | 0.3333 | 16 | 0.3125 |
| 12_351971 | 5 | 0.2 | 10 | 0.4 | 6 | 0.1667 | 6 | 0.5 | 27 | 0.3333 | 4 | 0 | 3 | 0.3333 | 10 | 0.3 | 4 | 0.5 | 21 | 0.2857 |
| 12_355505 | 1 | 1 | 1 | 1 | NA | NA | 1 | 1 | 3 | 1 | 2 | 0 | NA | NA | 1 | 1 | 1 | 0 | 4 | 0.25 |
| 12_355539 | 1 | 1 | NA | NA | NA | NA | NA | NA | 1 | 1 | NA | NA | NA | NA | 1 | 1 | 1 | 0 | 2 | 0.5 |
| 12_355539 | 1 | 0 | 1 | 1 | NA | NA | NA | NA | 2 | 0.5 | 1 | 1 | NA | NA | NA | NA | 1 | 0 | 2 | 0.5 |
| 12_355540 | 1 | 1 | 1 | 1 | NA | NA | NA | NA | 2 | 1 | 1 | 1 | NA | NA | NA | NA | NA | NA | 1 | 1 |
| 12_355540 | 1 | 1 | NA | NA | NA | NA | NA | NA | 1 | 1 | 1 | 1 | NA | NA | 1 | 1 | NA | NA | 2 | 1 |
| 12_355540 | 1 | 0 | 1 | 1 | NA | NA | NA | NA | 2 | 0.5 | 1 | 1 | NA | NA | 1 | 1 | 1 | 1 | 3 | 1 |
| 12_355545 | 1 | 1 | NA | NA | NA | NA | NA | NA | 1 | 1 | 1 | 0 | NA | NA | NA | NA | 1 | 1 | 2 | 0.5 |
| 12_360652 | 40 | 0.175 | 26 | 0.3077 | 40 | 0.2 | 31 | 0.2903 | 137 | 0.2336 | 21 | 0.0952 | 29 | 0.1724 | 35 | 0.3143 | 41 | 0.2195 | 126 | 0.2143 |
| 12_361420 | 4 | 0.25 | 7 | 0.7143 | 5 | 0.6 | 9 | 0.5556 | 25 | 0.56 | 7 | 0.7143 | 8 | 0.625 | 7 | 0.7143 | 8 | 0.875 | 30 | 0.7333 |
| 12_361516 | NA | NA | NA | NA | 1 | 1 | NA | NA | 1 | 1 | NA | NA | NA | NA | 1 | 1 | NA | NA | 1 | 1 |
| 12_362463 | 7 | 0.4286 | 5 | 0.4 | 2 | 0 | 4 | 0.25 | 18 | 0.3333 | 6 | 0.3333 | 6 | 0 | 4 | 0.25 | 4 | 0.5 | 20 | 0.25 |
| 12_362464 | 10 | 0.8 | 5 | 0.6 | 4 | 0.75 | 5 | 0.8 | 24 | 0.75 | 8 | 0.75 | 8 | 0.625 | 8 | 0.625 | 5 | 1 | 29 | 0.7241 |
| 12_362468 | 10 | 0 | 5 | 0.2 | 6 | 0.1667 | 6 | 0 | 27 | 0.0741 | 7 | 0.1429 | 6 | 0.1667 | 6 | 0 | 5 | 0 | 24 | 0.0833 |
| 12_386806 | 14 | 0.0714 | 13 | 0.0769 | 9 | 0 | 11 | 0.1818 | 47 | 0.0851 | 3 | 0 | 14 | 0.0714 | 15 | 0.0667 | 8 | 0 | 40 | 0.05 |
| 12_386807 | 13 | 0.2308 | 14 | 0.4286 | 8 | 0.375 | 10 | 0.3 | 45 | 0.3333 | 4 | 0.25 | 11 | 0.3636 | 11 | 0.1818 | 5 | 0 | 31 | 0.2258 |
| 12_386809 | 17 | 0.0588 | 15 | 0 | 6 | 0.1667 | 10 | 0.4 | 48 | 0.125 | 4 | 0 | 14 | 0.1429 | 11 | 0.0909 | 7 | 0 | 36 | 0.0833 |
| 12_386837 | 31 | 0.1613 | 29 | 0.0345 | 24 | 0 | 39 | 0.0769 | 123 | 0.0732 | 16 | 0 | 25 | 0.08 | 29 | 0.0345 | 19 | 0.1053 | 89 | 0.0562 |
| 12_389156 | 1 | 0 | 11 | 0.2727 | 3 | 0.3333 | 3 | 0.3333 | 18 | 0.2778 | 3 | 0.3333 | 1 | 1 | 2 | 0 | 4 | 0.25 | 10 | 0.3 |
| 12_389159 | 2 | 1 | 12 | 1 | 3 | 1 | 4 | 1 | 21 | 1 | 3 | 1 | 1 | 1 | 1 | 1 | 5 | 0.8 | 10 | 0.9 |
| 12_389159 | 2 | 0 | 12 | 0.0833 | 3 | 0.3333 | 4 | 0 | 21 | 0.0952 | 3 | 0.3333 | 1 | 0 | 2 | 0 | 5 | 0 | 11 | 0.0909 |
| 12_389164 | 2 | 0.5 | 11 | 0.1818 | 3 | 0 | 3 | 0.3333 | 19 | 0.2105 | 1 | 0 | 2 | 0.5 | 2 | 0 | 3 | 0.6667 | 8 | 0.375 |
| 12_389171 | 2 | 0 | 13 | 0.0769 | 4 | 0.25 | 7 | 0 | 26 | 0.0769 | 4 | 0 | 3 | 0 | 4 | 0 | 6 | 0.1667 | 17 | 0.0588 |
| 12_389171 | 2 | 0 | 13 | 0.7692 | 4 | 0.5 | 7 | 0.2857 | 26 | 0.5385 | 4 | 0.25 | 2 | 1 | 4 | 0.75 | 4 | 0.75 | 14 | 0.6429 |
| 12_389180 | NA | NA | 6 | 0.3333 | 1 | 0 | NA | NA | 7 | 0.2857 | 1 | 0 | 2 | 0.5 | 1 | 0 | 2 | 0 | 6 | 0.1667 |
| 12_400847 | NA | NA | 5 | 0.8 | NA | NA | 2 | 0 | 7 | 0.5714 | 1 | 1 | 7 | 0.4286 | 4 | 0.25 | 5 | 0.6 | 17 | 0.4706 |
| 12_400858 | 6 | 0.6667 | 7 | 0.2857 | 1 | 0 | 8 | 0.75 | 22 | 0.5455 | 5 | 0.2 | 9 | 0.5556 | 6 | 0 | 6 | 0.5 | 26 | 0.3462 |
| 12_400858 | 7 | 0.1429 | 8 | 0.25 | 4 | 0.25 | 7 | 0.4286 | 26 | 0.2692 | 5 | 0.4 | 9 | 0 | 6 | 0 | 8 | 0 | 28 | 0.0714 |
| 12_400860 | 5 | 0.8 | 4 | 0.75 | 1 | 0 | 6 | 0.8333 | 16 | 0.75 | 3 | 1 | 5 | 1 | 4 | 0.75 | 5 | 1 | 17 | 0.9412 |
| 12_400862 | 7 | 0.4286 | 7 | 0 | 2 | 0.5 | 6 | 0 | 22 | 0.1818 | 5 | 0.4 | 8 | 0.75 | 6 | 0 | 5 | 0.2 | 24 | 0.375 |
| 12_400862 | 5 | 1 | 8 | 1 | 2 | 1 | 7 | 1 | 22 | 1 | 6 | 1 | 9 | 1 | 4 | 1 | 7 | 0.8571 | 26 | 0.9615 |
| 12_400862 | 6 | 0.8333 | 6 | 0.8333 | 2 | 1 | 6 | 0.8333 | 20 | 0.85 | 3 | 1 | 9 | 0.8889 | 6 | 0.6667 | 7 | 1 | 25 | 0.88 |
| 12_400863 | 6 | 0.8333 | 7 | 1 | 3 | 1 | 5 | 0.8 | 21 | 0.9048 | 5 | 1 | 8 | 1 | 5 | 1 | 5 | 1 | 23 | 1 |

| | | | | | | | | | | | | | | | | | | | | |
|---|---|---|---|---|---|---|---|---|---|---|---|---|---|---|---|---|---|---|---|---|
| 12_4008633 | 7 | 0.1429 | 7 | 0.2857 | 4 | 0.5 | 6 | 0.3333 | 24 | 0.2917 | 6 | 0.3333 | 8 | 0.25 | 4 | 0 | 7 | 0.1429 | 25 | 0.2 |
| 12_4008640 | 3 | 1 | 5 | 0.8 | 3 | 1 | 6 | 1 | 17 | 0.9412 | 4 | 1 | 3 | 1 | 3 | 0.6667 | 6 | 1 | 16 | 0.9375 |
| 12_4008640 | 3 | 1 | 4 | 0.5 | 2 | 0.5 | 3 | 0.3333 | 12 | 0.5833 | 5 | 0.6 | 7 | 0.8571 | 4 | 0.5 | 5 | 0.4 | 21 | 0.619 |
| 12_4008644 | 6 | 1 | 6 | 0.8333 | 3 | 1 | 6 | 1 | 21 | 0.9524 | 6 | 1 | 7 | 1 | 4 | 1 | 7 | 1 | 24 | 1 |
| 12_4008656 | 5 | 0.2 | 6 | 0.5 | 3 | 0.6667 | 5 | 0.4 | 19 | 0.4211 | 4 | 0.5 | 7 | 0.5714 | 5 | 0.6 | 7 | 0.7143 | 23 | 0.6087 |
| 12_4008664 | 5 | 0 | 8 | 0.25 | 3 | 0.3333 | 7 | 0.2857 | 23 | 0.2174 | 6 | 0.6667 | 7 | 0.2857 | 5 | 0.2 | 7 | 0.5714 | 25 | 0.44 |
| 12_4008666 | 6 | 0.8333 | 8 | 1 | 3 | 1 | 6 | 0.8333 | 23 | 0.913 | 6 | 0.8333 | 7 | 0.8571 | 5 | 1 | 7 | 1 | 25 | 0.92 |
| 12_4008673 | 6 | 0.8333 | 6 | 0.8333 | 3 | 1 | 7 | 0.7143 | 22 | 0.8182 | 6 | 0.8333 | 6 | 0.6667 | 3 | 1 | 6 | 0.8333 | 21 | 0.8095 |
| 12_4026299 | 5 | 0 | 3 | 0 | 5 | 0 | 3 | 0.3333 | 16 | 0.0625 | 4 | 0.25 | 1 | 0 | 3 | 0 | 4 | 0 | 12 | 0.0833 |
| 12_4026305 | 4 | 0 | 5 | 0 | 7 | 0 | 4 | 0.25 | 20 | 0.05 | 5 | 0.2 | 1 | 0 | 4 | 0 | 4 | 0 | 14 | 0.0714 |
| 12_4027332 | NA | NA | NA | NA | 1 | 1 | NA | NA | 1 | 1 | NA | NA | NA | NA | 1 | 1 | NA | NA | 1 | 1 |
| 12_4027369 | NA | NA | NA | NA | 1 | 1 | NA | NA | 1 | 1 | NA | NA | NA | NA | 1 | 1 | NA | NA | 1 | 1 |
| 12_4027371 | NA | NA | NA | NA | 1 | 1 | NA | NA | 1 | 1 | NA | NA | NA | NA | 1 | 1 | NA | NA | 1 | 1 |
| 12_4027468 | NA | NA | NA | NA | 1 | 1 | NA | NA | 1 | 1 | NA | NA | NA | NA | 2 | 1 | NA | NA | 2 | 1 |
| 12_4027480 | NA | NA | NA | NA | 1 | 1 | NA | NA | 1 | 1 | NA | NA | NA | NA | 2 | 1 | NA | NA | 2 | 1 |
| 12_4027483 | NA | NA | NA | NA | 1 | 1 | NA | NA | 1 | 1 | NA | NA | 1 | 1 | 2 | 0.5 | NA | NA | 3 | 0.6667 |
| 12_4036550 | 1 | 1 | 1 | 1 | 2 | 0 | NA | NA | 4 | 0.5 | NA | NA | NA | NA | NA | NA | 1 | 1 | 1 | 1 |
| 12_4036554 | 1 | 0 | 1 | 1 | 3 | 0.3333 | NA | NA | 5 | 0.4 | NA | NA | NA | NA | NA | NA | 1 | 1 | 1 | 1 |
| 12_4036555 | NA | NA | NA | NA | 3 | 1 | 1 | 1 | 4 | 1 | NA | NA | 1 | 1 | NA | NA | 1 | 1 | 2 | 1 |
| 12_4036578 | 2 | 0 | 3 | 0 | 6 | 0.1667 | 3 | 0.3333 | 14 | 0.1429 | NA | NA | 4 | 0 | 1 | 0 | 3 | 0.3333 | 8 | 0.125 |
| 12_4036715 | 3 | 0 | 2 | 0 | 6 | 0.1667 | 3 | 0 | 14 | 0.0714 | NA | NA | 4 | 0.25 | 1 | 0 | 2 | 0 | 7 | 0.1429 |
| 12_4089425 | 1 | 0 | 3 | 0.3333 | 9 | 0.7778 | 7 | 0.8571 | 20 | 0.7 | NA | NA | 4 | 0.5 | 1 | 0 | 6 | 0.6667 | 11 | 0.5455 |
| 12_4089436 | NA | NA | 1 | 1 | 7 | 0.8571 | 4 | 1 | 12 | 0.9167 | NA | NA | 1 | 0 | NA | NA | 2 | 0.5 | 3 | 0.3333 |
| 12_4089443 | 1 | 0 | 5 | 0 | 8 | 0 | 8 | 0.25 | 22 | 0.0909 | NA | NA | 5 | 0.2 | 1 | 0 | 7 | 0 | 13 | 0.0769 |
| 12_4089448 | 1 | 1 | 5 | 0.8 | 5 | 0.6 | 6 | 0.8333 | 17 | 0.7647 | NA | NA | 4 | 0.5 | 1 | 1 | 8 | 0.875 | 13 | 0.7692 |
| 12_4089460 | 1 | 0 | 5 | 0.4 | 10 | 0.4 | 8 | 0.25 | 24 | 0.3333 | 1 | 1 | 6 | 0.3333 | 1 | 0 | 7 | 0.4286 | 15 | 0.4 |
| 12_4089461 | 1 | 1 | 5 | 1 | 8 | 1 | 7 | 0.7143 | 21 | 0.9048 | 1 | 1 | 4 | 1 | 1 | 1 | 8 | 0.625 | 14 | 0.7857 |
| 12_4089491 | 1 | 0 | 5 | 0.4 | 10 | 0.1 | 9 | 0 | 25 | 0.12 | 1 | 1 | 5 | 0 | 1 | 0 | 6 | 0 | 13 | 0.0769 |
| 12_4150175 | 8 | 0.25 | 8 | 0.125 | 6 | 0 | 12 | 0.25 | 34 | 0.1765 | 6 | 0 | 21 | 0.1429 | 17 | 0.1765 | 16 | 0.25 | 60 | 0.1667 |
| 12_4150179 | 5 | 0 | 6 | 0.6667 | 5 | 0.4 | 11 | 0.4545 | 27 | 0.4074 | 4 | 0.5 | 16 | 0.375 | 13 | 0.4615 | 13 | 0.5385 | 46 | 0.4565 |
| 12_4150182 | 7 | 0.2857 | 7 | 0.4286 | 5 | 0.4 | 13 | 0.2308 | 32 | 0.3125 | 5 | 0 | 22 | 0.1364 | 17 | 0.1176 | 16 | 0.25 | 60 | 0.15 |
| 12_4150183 | 7 | 0.2857 | 7 | 0 | 6 | 0 | 14 | 0 | 34 | 0.0588 | 5 | 0 | 21 | 0.0476 | 17 | 0.0588 | 16 | 0.0625 | 59 | 0.0508 |
| 12_4150189 | 7 | 0.7143 | 7 | 0.7143 | 5 | 0.6 | 10 | 0.7 | 29 | 0.6897 | 4 | 0.75 | 20 | 0.6 | 16 | 0.8125 | 15 | 0.8 | 55 | 0.7273 |
| 12_4150193 | 6 | 0.3333 | 5 | 0 | 3 | 1 | 8 | 0.75 | 22 | 0.5 | 4 | 0.25 | 19 | 0.5789 | 11 | 0.8182 | 11 | 0.5455 | 45 | 0.6 |
| 12_4150194 | 6 | 1 | 4 | 0.75 | 3 | 1 | 9 | 0.8889 | 22 | 0.9091 | 4 | 1 | 15 | 1 | 15 | 0.9333 | 12 | 0.9167 | 46 | 0.9565 |
| 12_4150196 | 6 | 0.5 | 6 | 0.1667 | 5 | 0.2 | 10 | 0.2 | 27 | 0.2593 | 3 | 0.3333 | 16 | 0.1875 | 16 | 0.25 | 10 | 0.3 | 45 | 0.2444 |
| 12_4150197 | 4 | 1 | 4 | 1 | 2 | 1 | 11 | 0.8182 | 21 | 0.9048 | 3 | 1 | 20 | 0.8 | 15 | 0.8667 | 14 | 0.8571 | 52 | 0.8462 |
| 12_4150199 | 6 | 0.1667 | 6 | 0.1667 | 4 | 0 | 12 | 0.3333 | 28 | 0.2143 | 4 | 0.25 | 16 | 0.125 | 15 | 0.0667 | 13 | 0.2308 | 48 | 0.1458 |
| 12_4150202 | 6 | 1 | 5 | 1 | 2 | 1 | 9 | 1 | 22 | 1 | 2 | 1 | 15 | 0.9333 | 16 | 1 | 14 | 1 | 47 | 0.9787 |
| 12_4150218 | 7 | 0.1429 | 4 | 0.25 | 5 | 0 | 8 | 0 | 24 | 0.0833 | 3 | 0.3333 | 16 | 0.125 | 15 | 0 | 12 | 0.0833 | 46 | 0.087 |
| 12_4150250 | 5 | 0.2 | 5 | 0 | 3 | 0 | 11 | 0.2727 | 24 | 0.1667 | 2 | 0 | 17 | 0.0588 | 12 | 0.0833 | 13 | 0.0769 | 44 | 0.0682 |
| 12_4150263 | 4 | 1 | 5 | 0.6 | 4 | 0.25 | 10 | 0.4 | 23 | 0.5217 | 2 | 0.5 | 17 | 0.6471 | 16 | 0.375 | 9 | 0.5556 | 44 | 0.5227 |
| 12_4150263 | 5 | 0.2 | 3 | 0 | 4 | 0.5 | 7 | 0.5714 | 19 | 0.3684 | 1 | 0 | 16 | 0.25 | 14 | 0.3571 | 7 | 0.5714 | 38 | 0.3421 |
| 12_4150265 | 5 | 0.8 | 5 | 0.4 | 4 | 1 | 10 | 0.4 | 24 | 0.5833 | 2 | 0.5 | 20 | 0.65 | 16 | 0.75 | 10 | 0.9 | 48 | 0.7292 |
| 12_4150292 | 3 | 1 | 3 | 1 | 2 | 0.5 | 8 | 0.75 | 16 | 0.8125 | 1 | 1 | 14 | 0.7857 | 9 | 0.8889 | 10 | 0.8 | 34 | 0.8235 |
| 12_4150312 | 4 | 0.25 | 3 | 1 | 4 | 0.75 | 8 | 0.625 | 19 | 0.6316 | 1 | 0 | 17 | 0.5294 | 12 | 0.5833 | 8 | 0.625 | 38 | 0.5526 |
| 12_4150314 | 4 | 0.5 | 3 | 0.3333 | 4 | 0 | 8 | 0.125 | 19 | 0.2105 | 1 | 0 | 17 | 0.4118 | 12 | 0.0833 | 8 | 0.375 | 38 | 0.2895 |
| 12_4150317 | 2 | 0 | 1 | 0 | 2 | 0 | 6 | 0.1667 | 11 | 0.0909 | NA | NA | 9 | 0.2222 | 9 | 0.1111 | 5 | 0.2 | 23 | 0.1739 |
| 12_4150324 | 3 | 0 | 3 | 0 | 4 | 0 | 8 | 0.125 | 18 | 0.0556 | 1 | 0 | 17 | 0.0588 | 11 | 0 | 10 | 0.1 | 39 | 0.0513 |
| 12_4150332 | 4 | 0.25 | 3 | 0 | 4 | 0.25 | 8 | 0 | 19 | 0.1053 | 1 | 0 | 16 | 0.125 | 11 | 0.6364 | 8 | 0.375 | 36 | 0.3333 |
| 12_4150341 | 1 | 0 | 3 | 0 | 2 | 0.5 | 6 | 0 | 12 | 0.0833 | 1 | 0 | 11 | 0.2727 | 9 | 0 | 7 | 0.5714 | 28 | 0.25 |
| 12_4150342 | 3 | 1 | 2 | 1 | 1 | 1 | 6 | 0.8333 | 12 | 0.9167 | 1 | 0 | 16 | 0.8125 | 9 | 0.5556 | 7 | 0.8571 | 33 | 0.7273 |
| 12_4150344 | 4 | 0 | 3 | 0 | 3 | 0 | 6 | 0.1667 | 16 | 0.0625 | 1 | 0 | 16 | 0.0625 | 10 | 0.3 | 7 | 0.1429 | 34 | 0.1471 |
| 12_4150360 | 3 | 0.3333 | 3 | 0 | 3 | 0 | 7 | 0 | 16 | 0.0625 | 1 | 0 | 14 | 0.2857 | 11 | 0.0909 | 9 | 0 | 35 | 0.1429 |
| 12_4150369 | 1 | 1 | 2 | 1 | 3 | 1 | 5 | 1 | 11 | 1 | 1 | 1 | 13 | 1 | 9 | 1 | 6 | 1 | 29 | 1 |
| 12_4150370 | 3 | 1 | 3 | 1 | 3 | 0.6667 | 7 | 1 | 16 | 0.9375 | 1 | 0 | 16 | 0.875 | 11 | 0.9091 | 9 | 0.8889 | 37 | 0.8649 |
| 12_4150379 | 3 | 0.6667 | 3 | 0.6667 | 3 | 0.3333 | 7 | 0.4286 | 16 | 0.5 | 1 | 0 | 15 | 0.4667 | 10 | 0.7 | 8 | 0.625 | 34 | 0.5588 |
| 12_4150388 | 2 | 1 | 2 | 0.5 | 2 | 0.5 | 6 | 0.6667 | 12 | 0.6667 | NA | NA | 16 | 0.3125 | 10 | 0.3 | 8 | 0.375 | 34 | 0.3235 |
| 12_4150390 | 2 | 1 | 3 | 1 | 3 | 1 | 5 | 1 | 13 | 1 | 1 | 1 | 15 | 1 | 9 | 0.8889 | 6 | 1 | 31 | 0.9677 |
| 12_4150402 | 3 | 1 | 3 | 1 | 2 | 0.5 | 7 | 1 | 15 | 0.9333 | NA | NA | 16 | 1 | 10 | 0.9 | 9 | 1 | 35 | 0.9714 |
| 12_4150403 | 3 | 0.3333 | 2 | 0 | 3 | 0 | 5 | 0 | 13 | 0.0769 | 1 | 0 | 15 | 0.1333 | 9 | 0 | 9 | 0.1111 | 34 | 0.0882 |
| 12_4150403 | 3 | 0 | 3 | 0.3333 | 3 | 0 | 6 | 0 | 15 | 0.0667 | 1 | 0 | 15 | 0.0667 | 10 | 0.1 | 8 | 0 | 34 | 0.0588 |
| 12_4150405 | 3 | 0 | 3 | 0.3333 | 3 | 0.6667 | 7 | 0.4286 | 16 | 0.375 | 1 | 0 | 14 | 0.2857 | 10 | 0.4 | 8 | 0.125 | 33 | 0.2727 |
| 12_4150406 | 2 | 0 | 3 | 0 | 1 | 0 | 5 | 0.6 | 11 | 0.2727 | 1 | 0 | 10 | 0.4 | 8 | 0.5 | 8 | 0.625 | 27 | 0.4815 |
| 12_4150408 | 3 | 0.6667 | 3 | 0.3333 | 2 | 0.5 | 7 | 0.1429 | 15 | 0.3333 | NA | NA | 16 | 0.25 | 11 | 0.3636 | 9 | 0.6667 | 36 | 0.3889 |

| | | | | | | | | | | | | | | | | | | | | |
|---|---|---|---|---|---|---|---|---|---|---|---|---|---|---|---|---|---|---|---|---|
| 12_4150415 | 3 | 1 | 3 | 1 | 2 | 1 | 4 | 1 | 12 | 1 | 1 | 1 | 13 | 1 | 9 | 1 | 9 | 0.8889 | 32 | 0.9688 |
| 12_4150430 | 1 | 1 | 3 | 0.3333 | 3 | 0.3333 | 5 | 0.8 | 12 | 0.5833 | NA | NA | 12 | 0.75 | 6 | 0.6667 | 7 | 0.8571 | 25 | 0.76 |
| 12_4150433 | 3 | 0 | 3 | 0.3333 | 3 | 0 | 5 | 0.2 | 14 | 0.1429 | 1 | 0 | 12 | 0.1667 | 8 | 0 | 9 | 0 | 30 | 0.0667 |
| 12_4150436 | 3 | 0.6667 | 2 | 0 | 2 | 0.5 | 5 | 0.4 | 12 | 0.4167 | NA | NA | 11 | 0.5455 | 8 | 0.5 | 6 | 0.6667 | 25 | 0.56 |
| 12_4150437 | 3 | 0.3333 | 1 | 0 | 3 | 0.6667 | 5 | 0.6 | 12 | 0.5 | NA | NA | 13 | 0.7692 | 9 | 0.6667 | 8 | 0.875 | 30 | 0.7667 |
| 12_4150441 | 3 | 0 | 3 | 0.3333 | 3 | 0 | 5 | 0 | 14 | 0.0714 | 1 | 0 | 13 | 0.0769 | 7 | 0 | 9 | 0.1111 | 30 | 0.0667 |
| 12_4150455 | 3 | 0 | 1 | 0 | 2 | 0 | 5 | 0.2 | 11 | 0.0909 | 1 | 1 | 13 | 0 | 9 | 0.1111 | 8 | 0.125 | 31 | 0.0968 |
| 12_4153574 | 3 | 0.3333 | 3 | 0 | 2 | 0 | 6 | 0.1667 | 14 | 0.1429 | NA | NA | 12 | 0.0833 | 10 | 0.3 | 8 | 0.25 | 30 | 0.2 |
| 12_4425237 | NA | NA | 3 | 0 | 3 | 0.3333 | 5 | 0.2 | 11 | 0.1818 | 3 | 0 | 1 | 1 | 2 | 0.5 | 2 | 0.5 | 8 | 0.375 |
| 12_4425241 | 1 | 0 | 4 | 0 | 3 | 0.3333 | 6 | 0 | 14 | 0.0714 | 3 | 0.3333 | 4 | 0 | 2 | 0.5 | 2 | 0 | 11 | 0.1818 |
| 12_4425243 | 1 | 0 | 4 | 0 | 4 | 0.25 | 4 | 0 | 13 | 0.0769 | 3 | 0 | 4 | 0.25 | 2 | 0 | 2 | 0 | 11 | 0.0909 |
| 12_4425335 | 6 | 0 | 6 | 0.3333 | 9 | 0.2222 | 11 | 0 | 32 | 0.125 | 9 | 0.1111 | 8 | 0.375 | 6 | 0.1667 | 8 | 0.25 | 31 | 0.2258 |
| 12_4425359 | 7 | 0 | 8 | 0.25 | 11 | 0 | 8 | 0 | 34 | 0.0588 | 10 | 0.3 | 8 | 0.125 | 6 | 0 | 9 | 0.1111 | 33 | 0.1515 |
| 12_4425365 | 7 | 0.1429 | 8 | 0.125 | 12 | 0 | 9 | 0.1111 | 36 | 0.0833 | 9 | 0 | 7 | 0.1429 | 7 | 0 | 8 | 0.125 | 31 | 0.0645 |
| 12_4425374 | 8 | 1 | 8 | 0.875 | 12 | 1 | 10 | 0.8 | 38 | 0.9211 | 10 | 0.6 | 6 | 0.8333 | 8 | 0.5 | 7 | 0.7143 | 31 | 0.6452 |
| 12_4425379 | 8 | 1 | 8 | 1 | 11 | 0.9091 | 10 | 0.9 | 37 | 0.9459 | 10 | 1 | 7 | 0.8571 | 7 | 0.8571 | 7 | 1 | 31 | 0.9355 |
| 12_4425383 | 6 | 0.8333 | 4 | 1 | 11 | 1 | 6 | 0.8333 | 27 | 0.9259 | 9 | 1 | 7 | 1 | 3 | 1 | 7 | 1 | 26 | 1 |
| 12_4425401 | 6 | 0 | 8 | 0.25 | 12 | 0.0833 | 8 | 0 | 34 | 0.0882 | 8 | 0.125 | 6 | 0 | 5 | 0.4 | 7 | 0 | 26 | 0.1154 |
| 12_4425402 | 6 | 0 | 9 | 0.2222 | 10 | 0 | 9 | 0 | 34 | 0.0588 | 10 | 0.1 | 7 | 0 | 5 | 0 | 7 | 0.1429 | 29 | 0.069 |
| 12_4425404 | 8 | 1 | 8 | 1 | 11 | 1 | 10 | 1 | 37 | 1 | 10 | 0.9 | 6 | 1 | 5 | 1 | 8 | 1 | 29 | 0.9655 |
| 12_4425405 | 8 | 0.5 | 7 | 0.7143 | 12 | 0.6667 | 11 | 0.4545 | 38 | 0.5789 | 10 | 0.7 | 5 | 0.4 | 5 | 0.4 | 6 | 0.1667 | 26 | 0.4615 |
| 12_4425408 | 4 | 0.75 | 6 | 0.6667 | 7 | 0.7143 | 7 | 0.4286 | 24 | 0.625 | 4 | 0.75 | 4 | 0.25 | 2 | 0.5 | 7 | 0.2857 | 17 | 0.4118 |
| 12_4425410 | 5 | 0.4 | 9 | 0.3333 | 8 | 0.75 | 8 | 0.5 | 30 | 0.5 | 9 | 0.6667 | 4 | 0.75 | 5 | 0.4 | 7 | 0.5714 | 25 | 0.6 |
| 12_4425413 | 7 | 0.2857 | 8 | 0.375 | 11 | 0.6364 | 11 | 0.4545 | 37 | 0.4595 | 10 | 0.4 | 5 | 0.8 | 5 | 0.6 | 8 | 0.375 | 28 | 0.5 |
| 12_4431602 | 4 | 0.5 | 7 | 0 | 9 | 0 | 2 | 0 | 22 | 0.0909 | 3 | 0.3333 | 5 | 0.2 | 1 | 0 | 9 | 0.1111 | 18 | 0.1667 |
| 12_4432946 | 5 | 0.6 | 9 | 0.7778 | 8 | 0.5 | 2 | 0.5 | 24 | 0.625 | 5 | 0.2 | 3 | 0.6667 | 2 | 1 | 9 | 0.3333 | 19 | 0.4211 |
| 12_4432957 | 5 | 0 | 9 | 0 | 9 | 0.3333 | 3 | 0 | 26 | 0.1154 | 6 | 0 | 4 | 0 | 2 | 0 | 8 | 0.25 | 20 | 0.1 |
| 12_4432969 | 5 | 0 | 9 | 0.3333 | 10 | 0 | 4 | 0.5 | 28 | 0.1786 | 6 | 0.6667 | 5 | 0.4 | 2 | 0 | 8 | 0.125 | 21 | 0.3333 |
| 12_4432972 | 4 | 1 | 8 | 0.875 | 8 | 1 | 4 | 1 | 24 | 0.9583 | 6 | 1 | 5 | 1 | 2 | 0.5 | 7 | 0.8571 | 20 | 0.9 |
| 12_4432991 | 6 | 0 | 8 | 0.125 | 9 | 0.1111 | 3 | 0 | 26 | 0.0769 | 6 | 0.3333 | 5 | 0.2 | 2 | 0 | 10 | 0.3 | 23 | 0.2609 |
| 12_4433023 | 6 | 0.3333 | 5 | 0 | 10 | 0.2 | 4 | 0.25 | 25 | 0.2 | 6 | 0.1667 | 5 | 0 | 3 | 0 | 11 | 0.0909 | 25 | 0.08 |
| 12_4433035 | 7 | 0.8571 | 9 | 0.7778 | 10 | 1 | 4 | 0.75 | 30 | 0.8667 | 6 | 0.5 | 5 | 0.8 | 1 | 0 | 11 | 0.9091 | 23 | 0.7391 |
| 12_4435814 | 10 | 0.1 | 11 | 0 | 11 | 0 | 6 | 0.1667 | 38 | 0.0526 | 7 | 0.1429 | 6 | 0 | 7 | 0 | 14 | 0.0714 | 34 | 0.0588 |
| 12_4435821 | 7 | 0.2857 | 7 | 0.1429 | 6 | 0.1667 | 5 | 0.4 | 25 | 0.24 | 4 | 0 | 5 | 0.2 | 7 | 0.1429 | 8 | 0.125 | 24 | 0.125 |
| 12_4435834 | 11 | 0.5455 | 10 | 0.4 | 10 | 0.3 | 5 | 0.4 | 36 | 0.4167 | 6 | 0.3333 | 5 | 0.2 | 7 | 0.5714 | 12 | 0.5 | 30 | 0.4333 |
| 12_4435846 | 10 | 0 | 13 | 0 | 11 | 0.0909 | 5 | 0.4 | 39 | 0.0769 | 8 | 0 | 6 | 0.3333 | 8 | 0.125 | 15 | 0 | 37 | 0.0811 |
| 12_4435889 | 7 | 0.4286 | 9 | 0.4444 | 9 | 0.3333 | 5 | 0.4 | 30 | 0.4 | 6 | 0.8333 | 5 | 0.6 | 7 | 0.4286 | 9 | 0.5556 | 27 | 0.5926 |
| 12_4462345 | 10 | 0 | 22 | 0.0909 | 14 | 0.0714 | 11 | 0.0909 | 57 | 0.0702 | 5 | 0.2 | 9 | 0 | 17 | 0.1176 | 11 | 0 | 42 | 0.0714 |
| 12_4464518 | 13 | 0.7692 | 25 | 0.72 | 22 | 0.5909 | 14 | 0.5 | 74 | 0.6486 | 6 | 0.5 | 11 | 0.4545 | 19 | 0.6842 | 13 | 0.5385 | 49 | 0.5714 |
| 12_4464522 | 13 | 0.3846 | 23 | 0.4348 | 21 | 0.381 | 14 | 0.2857 | 71 | 0.3803 | 6 | 0.6667 | 12 | 0.4167 | 16 | 0.1875 | 12 | 0.5833 | 46 | 0.413 |
| 12_4464535 | 10 | 1 | 16 | 0.9375 | 12 | 1 | 10 | 1 | 48 | 0.9792 | 5 | 1 | 7 | 0.8571 | 12 | 0.9167 | 7 | 0.8571 | 31 | 0.9032 |
| 12_4464542 | NA | NA | 1 | 1 | NA | NA | NA | NA | 1 | 1 | NA | NA | NA | NA | NA | NA | 1 | 1 | 1 | 1 |
| 12_4464544 | 15 | 0.9333 | 25 | 0.76 | 24 | 0.9167 | 14 | 1 | 78 | 0.8846 | 7 | 0.7143 | 13 | 0.8462 | 21 | 0.9524 | 15 | 1 | 56 | 0.9107 |
| 12_4464550 | 9 | 0.1111 | 20 | 0.1 | 18 | 0.1111 | 14 | 0.0714 | 61 | 0.0984 | 6 | 0.3333 | 13 | 0.3846 | 17 | 0.1765 | 11 | 0.0909 | 47 | 0.234 |
| 12_4464551 | 14 | 0.2143 | 22 | 0.3636 | 22 | 0.2727 | 14 | 0.2143 | 72 | 0.2778 | 7 | 0.4286 | 12 | 0.25 | 20 | 0.15 | 13 | 0.0769 | 52 | 0.1923 |
| 12_4464553 | 11 | 0.9091 | 24 | 0.875 | 20 | 0.85 | 12 | 0.9167 | 67 | 0.8806 | 7 | 1 | 10 | 0.8 | 13 | 0.8462 | 12 | 0.8333 | 42 | 0.8571 |
| 12_4464559 | 10 | 0.8 | 19 | 0.8421 | 20 | 0.55 | 14 | 0.7143 | 63 | 0.7143 | 8 | 0.625 | 12 | 0.75 | 15 | 0.8 | 15 | 0.6667 | 50 | 0.72 |
| 12_4464569 | 9 | 0.1111 | 13 | 0.2308 | 7 | 0.1429 | 5 | 0 | 34 | 0.1471 | 2 | 0.5 | 7 | 0.2857 | 7 | 0.1429 | 3 | 0.3333 | 19 | 0.2632 |
| 12_4464571 | 9 | 0.4444 | 13 | 0.6154 | 7 | 0.5714 | 4 | 0.5 | 33 | 0.5455 | 2 | 0.5 | 4 | 0.25 | 8 | 0.25 | 3 | 1 | 17 | 0.4118 |
| 12_4464583 | 6 | 0 | 12 | 0.0833 | 5 | 0.2 | 2 | 0 | 25 | 0.08 | 2 | 0 | 5 | 0 | 5 | 0.2 | 3 | 0 | 15 | 0.0667 |
| 12_4464631 | 6 | 0.3333 | 12 | 0.3333 | 6 | 0.1667 | 3 | 0 | 27 | 0.2593 | 6 | 0.1667 | 4 | 0.25 | 3 | 0 | 3 | 0.6667 | 16 | 0.25 |
| 12_4464762 | 2 | 0.5 | 5 | 0 | NA | NA | 2 | 0 | 9 | 0.1111 | 2 | 0 | 3 | 0 | 1 | 0 | 2 | 0.5 | 8 | 0.125 |
| 12_4464772 | 2 | 0 | 9 | 0.2222 | NA | NA | 3 | 0 | 14 | 0.1429 | 2 | 0.5 | 3 | 0 | 1 | 0 | 2 | 0 | 8 | 0.125 |
| 12_4464847 | 2 | 0 | 12 | 0.0833 | NA | NA | 3 | 0 | 17 | 0.0588 | 3 | 0 | 4 | 0 | 3 | 0 | 2 | 0.5 | 12 | 0.0833 |
| 12_4464855 | 2 | 1 | 11 | 0.7273 | NA | NA | 4 | 0.25 | 17 | 0.6471 | 3 | 0.6667 | 4 | 0 | 2 | 1 | 2 | 1 | 11 | 0.5455 |
| 12_4464858 | 2 | 0.5 | 12 | 0.1667 | NA | NA | 4 | 0 | 18 | 0.1667 | 3 | 0.3333 | 3 | 0 | 3 | 0 | 1 | 0 | 10 | 0.1 |
| 12_4465175 | 7 | 0.2857 | 18 | 0.1111 | 5 | 0 | 8 | 0 | 38 | 0.1053 | 6 | 0 | 9 | 0.1111 | 8 | 0.125 | 3 | 0.3333 | 26 | 0.1154 |
| 12_4489978 | 6 | 0.5 | 2 | 0.5 | 1 | 1 | 5 | 1 | 14 | 0.7143 | 2 | 0 | 8 | 0.5 | 3 | 0.6667 | 1 | 0 | 14 | 0.4286 |
| 12_4490241 | 1 | 0 | 2 | 1 | NA | NA | 5 | 0.6 | 8 | 0.625 | 2 | 0 | 4 | 0.5 | 2 | 0 | NA | NA | 8 | 0.25 |
| 12_4674493 | 4 | 0.75 | 6 | 1 | 3 | 1 | 4 | 0.75 | 17 | 0.8824 | 3 | 0.3333 | 7 | 1 | 7 | 0.8571 | 4 | 1 | 21 | 0.8571 |
| 12_4674497 | 7 | 0.2857 | 4 | 0.25 | 8 | 0 | 7 | 0 | 26 | 0.1154 | 2 | 0.5 | 10 | 0.2 | 3 | 0 | 6 | 0.1667 | 21 | 0.1905 |
| 12_4674504 | 6 | 0.6667 | 5 | 0.4 | 8 | 0.375 | 7 | 0.4286 | 26 | 0.4615 | 2 | 0.5 | 10 | 0.2 | 5 | 1 | 7 | 0.2857 | 24 | 0.4167 |
| 12_4674505 | 7 | 0.8571 | 5 | 1 | 9 | 1 | 6 | 1 | 27 | 0.963 | 4 | 1 | 8 | 1 | 5 | 1 | 6 | 1 | 23 | 1 |
| 12_4674509 | 7 | 1 | 5 | 1 | 7 | 1 | 7 | 1 | 26 | 1 | 4 | 1 | 10 | 1 | 5 | 1 | 6 | 1 | 25 | 1 |
| 12_4674510 | 5 | 0.4 | 5 | 0.4 | 6 | 0.5 | 7 | 0.5714 | 23 | 0.4783 | 4 | 0.25 | 7 | 0.4286 | 5 | 0.4 | 6 | 0.5 | 22 | 0.4091 |

| | | | | | | | | | | | | | | | | | | | | |
|---|---|---|---|---|---|---|---|---|---|---|---|---|---|---|---|---|---|---|---|---|
| 12_4674512 | 6 | 1 | 2 | 1 | 7 | 1 | 5 | 0.8 | 20 | 0.95 | 4 | 1 | 7 | 1 | 4 | 1 | 4 | 1 | 19 | 1 |
| 12_4674514 | 3 | 1 | 5 | 1 | 6 | 0.8333 | 6 | 0.8333 | 20 | 0.9 | 4 | 1 | 8 | 1 | 4 | 1 | 5 | 1 | 21 | 1 |
| 12_4674519 | 5 | 1 | 4 | 0.5 | 6 | 0.3333 | 5 | 0.8 | 20 | 0.65 | 3 | 1 | 5 | 0.8 | 1 | 1 | 3 | 0.3333 | 12 | 0.75 |
| 12_4674534 | 6 | 0 | 3 | 0 | 6 | 0.3333 | 5 | 0 | 20 | 0.1 | 3 | 0 | 5 | 0 | 4 | 0 | 5 | 0.2 | 17 | 0.0588 |
| 12_4674535 | 6 | 1 | 3 | 1 | 6 | 1 | 7 | 1 | 22 | 1 | 3 | 1 | 9 | 1 | 4 | 1 | 5 | 1 | 21 | 1 |
| 12_4674598 | 4 | 0 | 4 | 0.25 | 7 | 0 | 5 | 0 | 20 | 0.05 | 3 | 0 | 4 | 0.25 | 4 | 0 | 5 | 0 | 16 | 0.0625 |
| 12_4674692 | 4 | 0.25 | 3 | 1 | 6 | 0.6667 | 4 | 0.25 | 17 | 0.5294 | 3 | 0.6667 | 6 | 0.5 | 5 | 0.6 | 3 | 0.3333 | 17 | 0.5294 |
| 12_4674698 | 4 | 0 | 3 | 0 | 7 | 0.1429 | 4 | 0.25 | 18 | 0.1111 | 3 | 0.3333 | 4 | 0.5 | 5 | 0.2 | 4 | 0.25 | 16 | 0.3125 |
| 12_4674703 | 4 | 0.75 | 3 | 1 | 4 | 1 | 4 | 0.25 | 15 | 0.7333 | 2 | 0.5 | 5 | 1 | 5 | 0.8 | 3 | 0.3333 | 15 | 0.7333 |
| 12_4674713 | 4 | 1 | 3 | 1 | 6 | 0.8333 | 4 | 1 | 17 | 0.9412 | 3 | 0.6667 | 5 | 1 | 5 | 1 | 3 | 0.6667 | 16 | 0.875 |
| 12_4674740 | 4 | 1 | 3 | 1 | 5 | 0.8 | 4 | 1 | 16 | 0.9375 | 3 | 0.6667 | 5 | 0.8 | 4 | 0.75 | 3 | 1 | 15 | 0.8 |
| 12_4674750 | 3 | 0.3333 | 3 | 0.6667 | 5 | 0.6 | 3 | 0.6667 | 14 | 0.5714 | 3 | 0.3333 | 3 | 0.6667 | 4 | 0.75 | 3 | 0 | 13 | 0.4615 |
| 12_4674757 | 4 | 1 | 3 | 0.6667 | 5 | 0.8 | 4 | 0.5 | 16 | 0.75 | 3 | 1 | 4 | 1 | 4 | 1 | 3 | 0.6667 | 14 | 0.9286 |
| 12_4674759 | 3 | 0.3333 | 2 | 0 | 5 | 0 | 3 | 0 | 13 | 0.0769 | 3 | 0.6667 | 5 | 0 | 5 | 0 | 3 | 0 | 16 | 0.125 |
| 12_4674804 | 1 | 0 | 1 | 1 | 1 | 0 | 1 | 0 | 4 | 0.25 | 1 | 1 | 1 | 0 | NA | NA | NA | NA | 2 | 0.5 |
| 12_4686554 | 1 | 0 | 4 | 0 | 1 | 0 | 2 | 0.5 | 8 | 0.125 | 2 | 0.5 | 1 | 0 | 4 | 0 | 1 | 0 | 8 | 0.125 |
| 12_4686568 | 1 | 0 | 2 | 0 | 2 | 0 | 2 | 0.5 | 7 | 0.1429 | 2 | 0.5 | 2 | 0 | 5 | 0 | 2 | 0 | 11 | 0.0909 |
| 12_4941578 | NA | NA | NA | NA | NA | NA | 1 | 1 | 1 | 1 | NA | NA | NA | NA | 1 | 1 | NA | NA | 1 | 1 |
| 12_4942963 | 2 | 0 | 2 | 0.5 | NA | NA | 1 | 1 | 5 | 0.4 | NA | NA | 2 | 0.5 | 1 | 0 | NA | NA | 3 | 0.3333 |
| 12_4943001 | 2 | 1 | 2 | 1 | NA | NA | 1 | 1 | 5 | 1 | 2 | 1 | 2 | 0.5 | 2 | 0.5 | NA | NA | 6 | 0.6667 |
| 12_4943127 | 8 | 0.125 | 8 | 0.375 | 5 | 0.4 | 14 | 0.2857 | 35 | 0.2857 | 5 | 0.4 | 12 | 0.4167 | 13 | 0.1538 | 7 | 0.1429 | 37 | 0.2703 |
| 12_4943188 | 17 | 0.1765 | 15 | 0.4667 | 14 | 0.2857 | 22 | 0.3636 | 68 | 0.3235 | 7 | 0.4286 | 18 | 0.2778 | 16 | 0.5 | 11 | 0.3636 | 52 | 0.3846 |
| 12_4943192 | 20 | 0.3 | 18 | 0.5 | 16 | 0.0625 | 22 | 0.5 | 76 | 0.3553 | 9 | 0.2222 | 23 | 0.4783 | 18 | 0.3889 | 15 | 0.3333 | 65 | 0.3846 |
| 12_4943214 | 19 | 0.7368 | 18 | 0.8889 | 13 | 1 | 22 | 0.9545 | 72 | 0.8889 | 8 | 1 | 19 | 1 | 17 | 0.6471 | 12 | 1 | 56 | 0.8929 |
| 12_4943223 | 16 | 0.375 | 18 | 0.4444 | 11 | 0.5455 | 19 | 0.4211 | 64 | 0.4375 | 10 | 0.5 | 19 | 0.6316 | 15 | 0.6 | 14 | 0.5 | 58 | 0.569 |
| 12_4943228 | 20 | 0.9 | 15 | 0.9333 | 13 | 0.6154 | 20 | 0.9 | 68 | 0.8529 | 11 | 1 | 18 | 0.8333 | 20 | 0.9 | 13 | 0.8462 | 62 | 0.8871 |
| 12_4943228 | 21 | 0.2381 | 16 | 0.25 | 16 | 0.4375 | 19 | 0.5263 | 72 | 0.3611 | 12 | 0.3333 | 19 | 0.4211 | 18 | 0.2222 | 15 | 0.2667 | 64 | 0.3125 |
| 12_4943233 | 21 | 0.4286 | 18 | 0.6667 | 15 | 0.4667 | 24 | 0.4167 | 78 | 0.4872 | 11 | 0.5455 | 21 | 0.4286 | 19 | 0.5789 | 18 | 0.7222 | 69 | 0.5652 |
| 12_4943243 | 22 | 0.6364 | 18 | 1 | 16 | 0.8125 | 26 | 0.9231 | 82 | 0.8415 | 10 | 0.9 | 23 | 0.913 | 22 | 1 | 16 | 0.9375 | 71 | 0.9437 |
| 12_4943248 | 23 | 0.2174 | 20 | 0.15 | 17 | 0.1765 | 25 | 0.24 | 85 | 0.2 | 14 | 0.0714 | 25 | 0.12 | 21 | 0.2381 | 18 | 0.2222 | 78 | 0.1667 |
| 12_4943252 | 24 | 0.375 | 17 | 0.4706 | 16 | 0.5 | 17 | 0.5294 | 74 | 0.4595 | 12 | 0.5 | 23 | 0.5217 | 22 | 0.4091 | 14 | 0.5 | 71 | 0.4789 |
| 12_4943261 | 22 | 0.4545 | 20 | 0.7 | 14 | 0.5714 | 29 | 0.4483 | 85 | 0.5294 | 13 | 0.3846 | 25 | 0.36 | 23 | 0.4348 | 18 | 0.6111 | 79 | 0.443 |
| 12_4943264 | 18 | 0.9444 | 17 | 1 | 16 | 1 | 19 | 1 | 70 | 0.9857 | 13 | 1 | 20 | 1 | 20 | 0.95 | 16 | 1 | 69 | 0.9855 |
| 12_4943265 | 22 | 1 | 17 | 1 | 19 | 1 | 29 | 1 | 87 | 1 | 14 | 1 | 26 | 1 | 25 | 1 | 18 | 1 | 83 | 1 |
| 12_4943271 | 13 | 0.1538 | 7 | 0.1429 | 10 | 0.3 | 18 | 0.0556 | 48 | 0.1458 | 11 | 0.1818 | 16 | 0.0625 | 21 | 0.0476 | 13 | 0.0769 | 61 | 0.082 |
| 12_4943272 | 20 | 0.25 | 17 | 0.2941 | 19 | 0.3684 | 26 | 0.4615 | 82 | 0.3537 | 15 | 0.4667 | 20 | 0.25 | 26 | 0.5 | 17 | 0.4118 | 78 | 0.4103 |
| 12_4943273 | 16 | 0.75 | 15 | 0.8667 | 12 | 0.75 | 18 | 0.7222 | 61 | 0.7705 | 11 | 0.6364 | 21 | 0.8095 | 17 | 0.7647 | 17 | 0.7647 | 66 | 0.7576 |
| 12_4943296 | 25 | 0.52 | 21 | 0.2857 | 17 | 0.5882 | 27 | 0.5185 | 90 | 0.4778 | 14 | 0.4286 | 28 | 0.3214 | 25 | 0.44 | 18 | 0.5 | 85 | 0.4118 |
| 12_4943319 | 25 | 0.48 | 22 | 0.4545 | 18 | 0.5556 | 28 | 0.5357 | 93 | 0.5054 | 13 | 0.7692 | 28 | 0.3571 | 22 | 0.5455 | 21 | 0.5714 | 84 | 0.5238 |
| 12_4943376 | 3 | 0.6667 | 1 | 0 | NA | NA | 3 | 0.6667 | 7 | 0.5714 | 1 | 1 | 4 | 0.25 | 3 | 0 | NA | NA | 8 | 0.25 |
| 12_5038827 | 2 | 0 | 2 | 0.5 | 3 | 0 | 3 | 0 | 10 | 0.1 | 1 | 0 | NA | NA | 4 | 0.25 | NA | NA | 5 | 0.2 |
| 12_5038900 | 1 | 0 | 1 | 0 | 3 | 0 | 1 | 1 | 6 | 0.1667 | 1 | 0 | 1 | 0 | 4 | 0.25 | NA | NA | 6 | 0.1667 |
| 12_5041143 | NA | NA | 1 | 0 | 2 | 0.5 | 1 | 1 | 4 | 0.5 | 1 | 1 | NA | NA | 2 | 0.5 | NA | NA | 3 | 0.6667 |
| 12_5142345 | NA | NA | 2 | 0.5 | 1 | 0 | 2 | 0 | 5 | 0.2 | 2 | 0.5 | NA | NA | 2 | 0 | NA | NA | 4 | 0.25 |
| 12_5165028 | 1 | 1 | 2 | 0.5 | 1 | 1 | 1 | 1 | 5 | 0.8 | 2 | 0.5 | NA | NA | NA | NA | NA | NA | 2 | 0.5 |
| 12_5165179 | 2 | 1 | 3 | 0.6667 | 1 | 0 | 1 | 1 | 7 | 0.7143 | 2 | 1 | NA | NA | NA | NA | NA | NA | 2 | 1 |
| 12_5165568 | 14 | 0.0714 | 11 | 0.2727 | 12 | 0.1667 | 15 | 0.3333 | 52 | 0.2115 | 4 | 0 | 8 | 0.375 | 23 | 0.1739 | 9 | 0.1111 | 44 | 0.1818 |
| 12_5165588 | 10 | 0.1 | 9 | 0.1111 | 8 | 0 | 14 | 0.1429 | 41 | 0.0976 | 1 | 0 | 8 | 0.25 | 19 | 0.0526 | 6 | 0 | 34 | 0.0882 |
| 12_5165606 | 12 | 0.5 | 9 | 0.4444 | 10 | 0.1 | 15 | 0.2 | 46 | 0.3043 | 2 | 0 | 8 | 0.375 | 17 | 0.2353 | 7 | 0.2857 | 34 | 0.2647 |
| 12_5165611 | 12 | 0.5 | 13 | 0.3077 | 10 | 0.3 | 16 | 0.3125 | 51 | 0.3529 | 2 | 0 | 8 | 0.625 | 19 | 0.5789 | 8 | 0.375 | 37 | 0.5135 |
| 12_5165656 | 6 | 0.5 | 8 | 0.375 | 5 | 0.2 | 11 | 0.0909 | 30 | 0.2667 | 1 | 0 | 6 | 0.3333 | 12 | 0.3333 | 7 | 0.4286 | 26 | 0.3462 |
| 12_5165657 | 9 | 0.2222 | 10 | 0.3 | 10 | 0 | 16 | 0.125 | 45 | 0.1556 | 2 | 0 | 6 | 0.1667 | 16 | 0.1875 | 7 | 0 | 31 | 0.129 |
| 12_5165664 | 10 | 0.4 | 10 | 0.5 | 11 | 0.6364 | 16 | 0.5 | 47 | 0.5106 | 2 | 1 | 8 | 0.375 | 12 | 0.3333 | 8 | 0.625 | 30 | 0.4667 |
| 12_5165667 | 10 | 0.1 | 9 | 0.3333 | 11 | 0.0909 | 17 | 0.3529 | 47 | 0.234 | 2 | 1 | 8 | 0.375 | 17 | 0.1765 | 9 | 0.2222 | 36 | 0.2778 |
| 12_5165669 | 11 | 0.2727 | 10 | 0.2 | 10 | 0.2 | 13 | 0.3846 | 44 | 0.2727 | 1 | 0 | 6 | 0.1667 | 15 | 0 | 8 | 0.375 | 30 | 0.1333 |
| 12_5165670 | 8 | 0.125 | 8 | 0 | 6 | 0 | 14 | 0.2857 | 36 | 0.1389 | 2 | 0.5 | 6 | 0 | 12 | 0 | 6 | 0.1667 | 26 | 0.0769 |
| 12_5165687 | 8 | 0.625 | 9 | 0.8889 | 9 | 0.5556 | 14 | 0.5714 | 40 | 0.65 | 2 | 0.5 | 5 | 0.8 | 11 | 0.4545 | 7 | 0.8571 | 25 | 0.64 |
| 12_5165702 | 6 | 0.1667 | 6 | 0 | 7 | 0.1429 | 13 | 0.3846 | 32 | 0.2188 | 2 | 0 | 6 | 0 | 15 | 0.2 | 8 | 0 | 31 | 0.0968 |
| 12_5169955 | 3 | 0 | 2 | 0 | 4 | 0.25 | 8 | 0.125 | 17 | 0.1176 | 1 | 1 | 3 | 0 | 6 | 0 | 2 | 0 | 12 | 0.0833 |
| 12_5173841 | 2 | 0 | 2 | 0.5 | 2 | 0 | 8 | 0.25 | 14 | 0.2143 | NA | NA | 3 | 0.3333 | 7 | 0.1429 | 2 | 0 | 12 | 0.1667 |
| 12_5178481 | 2 | 0 | 1 | 1 | 1 | 0 | 1 | 0 | 5 | 0.2 | NA | NA | 2 | 0 | NA | NA | 1 | 1 | 3 | 0.3333 |
| 12_5178482 | 2 | 1 | 1 | 1 | 1 | 1 | 1 | 1 | 5 | 1 | NA | NA | 2 | 1 | NA | NA | 1 | 1 | 3 | 1 |
| 12_5178518 | 2 | 0 | 1 | 1 | 1 | 0 | 1 | 1 | 5 | 0.4 | NA | NA | 3 | 0.3333 | NA | NA | 1 | 1 | 4 | 0.5 |
| 12_5179105 | 5 | 0.2 | 6 | 0.5 | 2 | 0 | 6 | 0.1667 | 19 | 0.2632 | 7 | 0.1429 | 8 | 0.25 | 11 | 0.1818 | 7 | 0.1429 | 33 | 0.1818 |
| 12_5179107 | 5 | 0.6 | 7 | 0.4286 | 3 | 0.6667 | 6 | 0.8333 | 21 | 0.619 | 7 | 0.5714 | 9 | 0.4444 | 10 | 0.6 | 6 | 0.5 | 32 | 0.5312 |

| | | | | | | | | | | | | | | | | | | | | |
|---|---|---|---|---|---|---|---|---|---|---|---|---|---|---|---|---|---|---|---|---|
| 12_5179110 | 6 | 0.5 | 7 | 0.7143 | 3 | 0.6667 | 7 | 0.7143 | 23 | 0.6522 | 7 | 0.7143 | 9 | 0.6667 | 11 | 0.8182 | 11 | 0.4545 | 38 | 0.6579 |
| 12_5179115 | 6 | 0 | 7 | 0 | 3 | 0 | 7 | 0.2857 | 23 | 0.087 | 8 | 0 | 9 | 0.2222 | 12 | 0.25 | 11 | 0.0909 | 40 | 0.15 |
| 12_5179118 | 7 | 0 | 7 | 0.2857 | 3 | 0 | 7 | 0.5714 | 24 | 0.25 | 6 | 0.3333 | 10 | 0.2 | 10 | 0 | 10 | 0.2 | 36 | 0.1667 |
| 12_5179122 | 6 | 0.1667 | 5 | 1 | 2 | 0.5 | 5 | 0.6 | 18 | 0.5556 | 6 | 0.8333 | 10 | 0.5 | 11 | 0.5455 | 7 | 0.7143 | 34 | 0.6176 |
| 12_5179128 | 6 | 0.5 | 5 | 0.2 | 3 | 0.3333 | 6 | 0.5 | 20 | 0.4 | 3 | 0.6667 | 8 | 0.375 | 12 | 0.25 | 10 | 0.3 | 33 | 0.3333 |
| 12_5179303 | NA | NA | 1 | 1 | NA | NA | NA | NA | 1 | 1 | NA | NA | 1 | 1 | NA | NA | 1 | 1 | 2 | 1 |
| 12_5179346 | NA | NA | 1 | 1 | NA | NA | NA | NA | 1 | 1 | NA | NA | 1 | 1 | NA | NA | NA | NA | 1 | 1 |
| 12_5179531 | 5 | 0.4 | 5 | 0.8 | 2 | 0.5 | 5 | 0 | 17 | 0.4118 | 6 | 0.3333 | 4 | 0.75 | 10 | 0.5 | 10 | 0.5 | 30 | 0.5 |
| 12_5179536 | 5 | 0.4 | 6 | 0.3333 | 1 | 0 | 2 | 0.5 | 14 | 0.3571 | 6 | 0.5 | 6 | 0.3333 | 9 | 0.4444 | 9 | 0.4444 | 30 | 0.4333 |
| 12_5179552 | 5 | 0.2 | 5 | 0.6 | 2 | 0.5 | 5 | 0.6 | 17 | 0.4706 | 5 | 0.8 | 6 | 0.5 | 10 | 0.5 | 11 | 0.7273 | 32 | 0.625 |
| 12_5179562 | 5 | 0.8 | 4 | 1 | 2 | 1 | 4 | 1 | 15 | 0.9333 | 4 | 0.75 | 4 | 1 | 9 | 1 | 11 | 1 | 28 | 0.9643 |
| 12_5179563 | 5 | 0.8 | 4 | 0.5 | 1 | 1 | 5 | 0.6 | 15 | 0.6667 | 4 | 0.75 | 3 | 0.6667 | 5 | 0.6 | 10 | 0.8 | 22 | 0.7273 |
| 12_5179564 | 5 | 0.4 | 5 | 0.4 | 2 | 0.5 | 5 | 0.6 | 17 | 0.4706 | 5 | 0.6 | 3 | 1 | 8 | 0.625 | 8 | 0.625 | 24 | 0.6667 |
| 12_5179571 | 5 | 0 | 5 | 0.2 | 2 | 0 | 5 | 0 | 17 | 0.0588 | 4 | 0 | 4 | 0 | 8 | 0.125 | 10 | 0.2 | 26 | 0.1154 |
| 12_5179573 | 2 | 0 | 5 | 0 | NA | NA | 3 | 0.3333 | 10 | 0.1 | 5 | 0.2 | 2 | 0.5 | 7 | 0.4286 | 8 | 0.375 | 22 | 0.3636 |
| 12_5181539 | 2 | 0.5 | 1 | 0 | NA | NA | 2 | 0 | 5 | 0.2 | 2 | 1 | 4 | 0.5 | 6 | 0.3333 | 5 | 0.6 | 17 | 0.5294 |
| 12_5181544 | 3 | 0 | 1 | 0 | NA | NA | 2 | 0.5 | 6 | 0.1667 | 2 | 0.5 | 4 | 0 | 5 | 0.4 | 4 | 0 | 15 | 0.2 |
| 12_5181552 | 2 | 1 | 1 | 0 | NA | NA | 2 | 0.5 | 5 | 0.6 | 2 | 0.5 | 4 | 1 | 5 | 1 | 4 | 0.75 | 15 | 0.8667 |
| 12_5181579 | 3 | 1 | 1 | 1 | NA | NA | 2 | 0 | 6 | 0.6667 | 2 | 0.5 | 4 | 0.25 | 4 | 0 | 3 | 0 | 13 | 0.1538 |
| 12_5181583 | 2 | 0 | 1 | 0 | NA | NA | 2 | 0.5 | 5 | 0.2 | 2 | 0 | 2 | 0 | 4 | 0 | 3 | 0.3333 | 11 | 0.0909 |
| 12_5192277 | 2 | 0 | 1 | 1 | NA | NA | 2 | 0 | 5 | 0.2 | 2 | 0 | NA | NA | 2 | 0 | 1 | 1 | 5 | 0.2 |
| 12_5192373 | 1 | 1 | 1 | 1 | 1 | 1 | 2 | 0.5 | 5 | 0.8 | 2 | 0.5 | NA | NA | 1 | 1 | 1 | 1 | 4 | 0.75 |
| 12_5192390 | 1 | 0 | NA | NA | NA | NA | 1 | 1 | 2 | 0.5 | 1 | 1 | NA | NA | 1 | 1 | 1 | 1 | 3 | 1 |
| 12_5192396 | 2 | 0.5 | NA | NA | 1 | 1 | 2 | 0.5 | 5 | 0.6 | 1 | 1 | NA | NA | 1 | 1 | 1 | 0 | 3 | 0.6667 |
| 12_5192402 | NA | NA | NA | NA | 1 | 0 | 2 | 0.5 | 3 | 0.3333 | NA | NA | NA | NA | 1 | 1 | 1 | 1 | 2 | 1 |
| 12_5192403 | 2 | 0.5 | NA | NA | NA | NA | 2 | 1 | 4 | 0.75 | 1 | 1 | NA | NA | 1 | 1 | 1 | 1 | 3 | 1 |
| 12_5192410 | 2 | 0.5 | NA | NA | 1 | 0 | 2 | 0.5 | 5 | 0.4 | 1 | 1 | NA | NA | 1 | 1 | 1 | 0 | 3 | 0.6667 |
| 12_5203988 | NA | NA | 1 | 1 | 1 | 1 | NA | NA | 2 | 1 | NA | NA | 2 | 0.5 | 1 | 1 | NA | NA | 3 | 0.6667 |
| 12_5204047 | NA | NA | 1 | 0 | 1 | 1 | NA | NA | 2 | 0.5 | NA | NA | 2 | 1 | NA | NA | NA | NA | 2 | 1 |
| 12_5204165 | NA | NA | 1 | 0 | 1 | 1 | NA | NA | 2 | 0.5 | NA | NA | 2 | 0.5 | NA | NA | NA | NA | 2 | 0.5 |
| 12_5204618 | 3 | 0 | 7 | 0.1429 | 2 | 0 | 3 | 0 | 15 | 0.0667 | 1 | 1 | 2 | 0 | 1 | 1 | 3 | 0.6667 | 7 | 0.5714 |
| 12_5204619 | 3 | 0 | 7 | 0.4286 | 2 | 0 | 3 | 0.3333 | 15 | 0.2667 | 1 | 0 | 2 | 0 | 2 | 0 | 3 | 0.3333 | 8 | 0.125 |
| 12_5204628 | 3 | 1 | 6 | 0 | 1 | 0 | 3 | 0.6667 | 13 | 0.3846 | 1 | 0 | 1 | 1 | 2 | 0.5 | 3 | 0.3333 | 7 | 0.4286 |
| 12_5204635 | 3 | 1 | 7 | 0.8571 | 1 | 1 | 3 | 1 | 14 | 0.9286 | 1 | 1 | 1 | 1 | 2 | 1 | 3 | 0.6667 | 7 | 0.8571 |
| 12_5204648 | 3 | 1 | 7 | 0.8571 | NA | NA | 2 | 0.5 | 12 | 0.8333 | 1 | 0 | 2 | 1 | 1 | 1 | 3 | 1 | 7 | 0.8571 |
| 12_5235759 | NA | NA | 2 | 1 | 1 | 0 | 2 | 1 | 5 | 0.8 | NA | NA | NA | NA | NA | NA | 1 | 1 | 1 | 1 |
| 12_5256301 | 3 | 1 | 4 | 0 | NA | NA | 1 | 0 | 8 | 0.375 | NA | NA | 1 | 1 | 4 | 0.25 | 2 | 0.5 | 7 | 0.4286 |
| 12_5256308 | 4 | 1 | 4 | 1 | 3 | 1 | 2 | 1 | 13 | 1 | 1 | 1 | 1 | 1 | 4 | 0.75 | 2 | 1 | 8 | 0.875 |
| 12_5256309 | 3 | 1 | 4 | 1 | 3 | 0.6667 | 2 | 1 | 12 | 0.9167 | 1 | 0 | 1 | 1 | 4 | 1 | 2 | 1 | 8 | 0.875 |
| 12_5256311 | 5 | 0.2 | 4 | 0.25 | 3 | 0 | 2 | 0.5 | 14 | 0.2143 | 1 | 1 | 1 | 1 | 4 | 0.5 | 2 | 0 | 8 | 0.5 |
| 12_5256321 | 4 | 0.75 | 4 | 1 | 3 | 0.6667 | 2 | 1 | 13 | 0.8462 | 1 | 1 | 1 | 1 | 3 | 1 | 2 | 1 | 7 | 1 |
| 12_5256333 | 6 | 0.6667 | 4 | 0.5 | 2 | 0.5 | 2 | 1 | 14 | 0.6429 | 1 | 1 | 1 | 0 | 4 | 0.5 | 2 | 0.5 | 8 | 0.5 |
| 12_5256345 | 7 | 0.2857 | 4 | 0.25 | 3 | 0 | 2 | 0 | 16 | 0.1875 | 1 | 0 | 1 | 0 | 3 | 0.3333 | 2 | 0.5 | 7 | 0.2857 |
| 12_5256373 | 4 | 0.5 | 5 | 0 | 2 | 0 | 1 | 0 | 12 | 0.1667 | 1 | 0 | 1 | 0 | 4 | 0 | 2 | 0.5 | 8 | 0.125 |
| 12_5256386 | 6 | 0 | 5 | 0.2 | 3 | 0 | 2 | 0 | 16 | 0.0625 | 2 | 0 | 1 | 1 | 4 | 0 | 2 | 0 | 9 | 0.1111 |
| 12_5256387 | 6 | 1 | 4 | 0.75 | 3 | 1 | 1 | 1 | 14 | 0.9286 | 1 | 1 | 1 | 1 | 4 | 0.75 | 2 | 1 | 8 | 0.875 |
| 12_5256390 | 6 | 0.5 | 4 | 0.25 | 3 | 0.6667 | 1 | 0 | 14 | 0.4286 | 1 | 1 | 1 | 0 | 4 | 0.5 | 2 | 0 | 8 | 0.375 |
| 12_5256410 | 6 | 0 | 5 | 0.2 | 3 | 0 | 2 | 0 | 16 | 0.0625 | 2 | 0.5 | 1 | 0 | 2 | 0.5 | 1 | 1 | 6 | 0.5 |
| 12_5256419 | 6 | 0 | 5 | 0 | 3 | 0 | 2 | 0.5 | 16 | 0.0625 | 2 | 0 | 1 | 1 | 3 | 0 | 2 | 0 | 8 | 0.125 |
| 12_5256420 | 4 | 0 | 4 | 0 | 3 | 0.3333 | 2 | 0 | 13 | 0.0769 | 2 | 0 | NA | NA | 4 | 0.25 | 2 | 0 | 8 | 0.125 |
| 12_5256452 | 6 | 0.8333 | 5 | 0.8 | 3 | 1 | 2 | 0.5 | 16 | 0.8125 | 4 | 1 | NA | NA | 5 | 1 | 3 | 1 | 12 | 1 |
| 12_5256460 | 6 | 0.1667 | 5 | 0 | 3 | 0 | 2 | 0 | 16 | 0.0625 | 4 | 0 | 1 | 0 | 5 | 0.2 | 3 | 0 | 13 | 0.0769 |
| 12_5256462 | 4 | 0 | 5 | 0 | 3 | 0 | 3 | 0.3333 | 15 | 0.0667 | 4 | 0 | 1 | 0 | 5 | 0.2 | 3 | 0 | 13 | 0.0769 |
| 12_5256472 | 5 | 0.6 | 5 | 0 | 3 | 0.3333 | 2 | 0 | 15 | 0.2667 | 4 | 0 | 1 | 0 | 5 | 0.2 | 3 | 0.6667 | 13 | 0.2308 |
| 12_5256487 | 6 | 1 | 5 | 1 | 3 | 0.3333 | 3 | 1 | 17 | 0.8824 | 4 | 1 | 1 | 0 | 5 | 0.8 | 2 | 1 | 12 | 0.8333 |
| 12_5256492 | 3 | 0.3333 | 4 | 0.25 | 1 | 1 | 3 | 0.3333 | 11 | 0.3636 | 2 | 0.5 | 1 | 1 | 4 | 0 | 1 | 0 | 8 | 0.25 |
| 12_5256497 | 5 | 0.8 | 5 | 1 | 3 | 0.3333 | 3 | 0.6667 | 16 | 0.75 | 3 | 0.3333 | 1 | 1 | 5 | 1 | 3 | 0.6667 | 12 | 0.75 |
| 12_5256536 | 7 | 0 | 5 | 0 | 4 | 0.25 | 3 | 0 | 19 | 0.0526 | 4 | 0 | 1 | 1 | 5 | 0 | 6 | 0 | 16 | 0.0625 |
| 12_5256543 | 6 | 0.1667 | 5 | 0.4 | 4 | 0.25 | 3 | 0 | 18 | 0.2222 | 4 | 0.25 | 1 | 1 | 4 | 0.5 | 3 | 0.3333 | 12 | 0.4167 |
| 12_5256546 | 7 | 0.2857 | 5 | 0.4 | 2 | 0 | 2 | 0.5 | 16 | 0.3125 | 2 | 1 | 1 | 1 | 5 | 0.6 | 6 | 0.5 | 14 | 0.6429 |
| 12_5256556 | 6 | 1 | 5 | 1 | 3 | 1 | 2 | 1 | 16 | 1 | 3 | 1 | NA | NA | 4 | 1 | 6 | 0.8333 | 13 | 0.9231 |
| 12_5256573 | 7 | 0.5714 | 6 | 0.5 | 3 | 0.3333 | 5 | 0.6 | 21 | 0.5238 | 3 | 0.3333 | 1 | 0 | 5 | 0.4 | 6 | 0.5 | 15 | 0.4 |
| 12_5256594 | 8 | 0.875 | 7 | 0.8571 | 6 | 0.5 | 5 | 0.8 | 26 | 0.7692 | 4 | 1 | 1 | 1 | 5 | 0.8 | 8 | 0.625 | 18 | 0.7778 |
| 12_5256618 | 7 | 0.4286 | 7 | 0.4286 | 5 | 0.4 | 4 | 0.5 | 23 | 0.4348 | 4 | 0.75 | 1 | 0 | 5 | 0.4 | 9 | 0.5556 | 19 | 0.5263 |
| 12_5256625 | 6 | 0 | 5 | 0.2 | 4 | 0.25 | NA | NA | 15 | 0.1333 | 1 | 1 | 1 | 1 | 5 | 1 | 6 | 0.5 | 13 | 0.7692 |

| | | | | | | | | | | | | | | | | | | | | |
|---|---|---|---|---|---|---|---|---|---|---|---|---|---|---|---|---|---|---|---|---|
| 12_525662 | 8 | 0 | 8 | 0.25 | 5 | 0 | 4 | 0 | 25 | 0.08 | 3 | 0 | 1 | 0 | 5 | 0 | 10 | 0.1 | 19 | 0.0526 |
| 12_525664 | 7 | 0.2857 | 8 | 0.25 | 3 | 0.3333 | 4 | 0.5 | 22 | 0.3182 | 4 | 0 | 1 | 0 | 5 | 0.2 | 10 | 0.1 | 20 | 0.1 |
| 12_525665 | 8 | 0.375 | 7 | 0.5714 | 4 | 0.75 | 4 | 0.75 | 23 | 0.5652 | 1 | 1 | NA | NA | 5 | 0.6 | 9 | 0.5556 | 15 | 0.6 |
| 12_525665 | 9 | 0 | 8 | 0.125 | 5 | 0.2 | 6 | 0 | 28 | 0.0714 | 3 | 0 | 2 | 0 | 5 | 0 | 10 | 0.3 | 20 | 0.15 |
| 12_526138 | 11 | 0.1818 | 9 | 0.3333 | 11 | 0 | 10 | 0.1 | 41 | 0.1463 | 5 | 0.2 | 6 | 0.1667 | 9 | 0.1111 | 12 | 0.0833 | 32 | 0.125 |
| 12_526139 | 9 | 0.3333 | 9 | 0.3333 | 10 | 0.2 | 8 | 0 | 36 | 0.2222 | 5 | 0.4 | 6 | 0.3333 | 5 | 0.4 | 11 | 0.2727 | 27 | 0.3333 |
| 12_526139 | 10 | 0.2 | 9 | 0.4444 | 9 | 0.4444 | 8 | 0.375 | 36 | 0.3611 | 4 | 0 | 5 | 0.4 | 7 | 0.4286 | 12 | 0.1667 | 28 | 0.25 |
| 12_526139 | 9 | 0.2222 | 8 | 0.125 | 10 | 0.3 | 9 | 0.4444 | 36 | 0.2778 | 4 | 0.25 | 7 | 0.1429 | 9 | 0.2222 | 13 | 0.5385 | 33 | 0.3333 |
| 12_526140 | 10 | 0.8 | 9 | 0.7778 | 10 | 0.8 | 8 | 0.875 | 37 | 0.8108 | 4 | 0.75 | 5 | 1 | 8 | 0.75 | 11 | 0.9091 | 28 | 0.8571 |
| 12_526140 | 8 | 0.25 | 10 | 0.2 | 10 | 0.2 | 9 | 0 | 37 | 0.1622 | 4 | 0 | 7 | 0.2857 | 9 | 0.1111 | 13 | 0 | 33 | 0.0909 |
| 12_526141 | 9 | 0.2222 | 10 | 0.4 | 11 | 0.6364 | 9 | 0.5556 | 39 | 0.4615 | 5 | 0.6 | 7 | 0.2857 | 10 | 0.5 | 12 | 0.25 | 34 | 0.3824 |
| 12_526142 | 11 | 0.2727 | 8 | 0.25 | 8 | 0 | 8 | 0.125 | 35 | 0.1714 | 4 | 0 | 5 | 0 | 10 | 0 | 11 | 0.2727 | 30 | 0.1 |
| 12_526142 | 10 | 0.2 | 8 | 0.25 | 10 | 0 | 7 | 0.1429 | 35 | 0.1429 | 3 | 0 | 4 | 0.25 | 10 | 0 | 10 | 0.2 | 27 | 0.1111 |
| 12_526159 | 7 | 0.1429 | 7 | 0.1429 | 7 | 0 | 10 | 0 | 31 | 0.0645 | 4 | 0.25 | 6 | 0 | 9 | 0.2222 | 6 | 0 | 25 | 0.12 |
| 12_529331 | NA | NA | 2 | 0.5 | NA | NA | 1 | 0 | 3 | 0.3333 | NA | NA | 2 | 0.5 | 2 | 1 | NA | NA | 4 | 0.75 |
| 12_529332 | 1 | 1 | 2 | 0.5 | 2 | 0 | NA | NA | 5 | 0.4 | NA | NA | NA | NA | 3 | 0.3333 | NA | NA | 3 | 0.3333 |
| 12_529333 | NA | NA | 2 | 0.5 | 1 | 1 | 2 | 0.5 | 5 | 0.6 | NA | NA | 2 | 1 | 3 | 0.6667 | NA | NA | 5 | 0.8 |
| 12_529336 | 1 | 1 | 2 | 1 | 2 | 1 | 1 | 1 | 6 | 1 | NA | NA | 1 | 1 | 3 | 1 | 1 | 0 | 5 | 0.8 |
| 12_529337 | 1 | 1 | 3 | 0.3333 | 2 | 0.5 | 2 | 1 | 8 | 0.625 | NA | NA | 2 | 0.5 | 3 | 0.3333 | 1 | 0 | 6 | 0.3333 |
| 12_529339 | 1 | 1 | 4 | 0.5 | 2 | 0.5 | 2 | 0.5 | 9 | 0.5556 | NA | NA | 4 | 0.25 | 2 | 0 | 1 | 0 | 7 | 0.1429 |
| 12_529340 | 1 | 0 | 4 | 0 | 2 | 0 | 2 | 0.5 | 9 | 0.1111 | NA | NA | 4 | 0.25 | 4 | 0 | 1 | 0 | 9 | 0.1111 |
| 12_529344 | 1 | 1 | 3 | 0.3333 | 2 | 1 | 2 | 1 | 8 | 0.75 | NA | NA | 4 | 0.25 | 4 | 0.25 | 1 | 0 | 9 | 0.2222 |
| 12_529345 | 1 | 1 | 4 | 0.5 | 3 | 0.3333 | 2 | 0.5 | 10 | 0.5 | NA | NA | 5 | 0.4 | 4 | 0.25 | 1 | 1 | 10 | 0.4 |
| 12_529345 | 1 | 1 | 4 | 0.5 | 3 | 0 | 2 | 0 | 10 | 0.3 | NA | NA | 5 | 0.4 | 4 | 0.75 | 1 | 0 | 10 | 0.5 |
| 12_529346 | 1 | 0 | 4 | 0 | 2 | 0.5 | 2 | 0.5 | 9 | 0.2222 | NA | NA | 5 | 0.2 | 2 | 0 | NA | NA | 7 | 0.1429 |
| 12_529346 | 1 | 1 | 3 | 1 | 3 | 1 | 2 | 1 | 9 | 1 | NA | NA | 2 | 1 | 4 | 1 | 1 | 1 | 7 | 1 |
| 12_529347 | 1 | 0 | 4 | 0 | 3 | 0.3333 | 2 | 0 | 10 | 0.1 | NA | NA | 6 | 0.1667 | 4 | 0 | 1 | 0 | 11 | 0.0909 |
| 12_531862 | 1 | 0 | 6 | 0.1667 | 5 | 0 | 3 | 0 | 15 | 0.0667 | NA | NA | 7 | 0.1429 | 5 | 0 | 2 | 0 | 14 | 0.0714 |
| 12_531863 | 1 | 0 | 6 | 0.3333 | 7 | 0.5714 | 3 | 0.3333 | 17 | 0.4118 | NA | NA | 7 | 0.2857 | 6 | 0.5 | 2 | 0 | 15 | 0.3333 |
| 12_531863 | NA | NA | 4 | 0.75 | 5 | 0.4 | 3 | 0 | 12 | 0.4167 | NA | NA | 4 | 0.5 | 6 | 0.5 | 2 | 0 | 12 | 0.4167 |
| 12_531864 | 1 | 1 | 6 | 0.8333 | 7 | 0.1429 | 3 | 0 | 17 | 0.4118 | NA | NA | 7 | 0.2857 | 5 | 0.2 | 2 | 0.5 | 14 | 0.2857 |
| 12_531866 | 2 | 0 | 5 | 0 | 6 | 0.3333 | 3 | 0.3333 | 16 | 0.1875 | NA | NA | 6 | 0.1667 | 6 | 0.5 | 2 | 0.5 | 14 | 0.3571 |
| 12_531867 | 2 | 0 | 5 | 0.2 | 8 | 0.25 | 3 | 0 | 18 | 0.1667 | NA | NA | 6 | 0.5 | 6 | 0 | 2 | 0 | 14 | 0.2143 |
| 12_531869 | 2 | 1 | 6 | 1 | 7 | 1 | 3 | 1 | 18 | 1 | NA | NA | 7 | 1 | 6 | 1 | 2 | 1 | 15 | 1 |
| 12_531869 | 1 | 0 | 4 | 0 | 7 | 0.1429 | 2 | 0 | 14 | 0.0714 | NA | NA | 5 | 0.2 | 7 | 0 | 2 | 0 | 14 | 0.0714 |
| 12_531870 | 2 | 0 | 6 | 0.1667 | 8 | 0.125 | 3 | 0.3333 | 19 | 0.1579 | 1 | 0 | 6 | 0.3333 | 5 | 0 | 3 | 0 | 15 | 0.1333 |
| 12_531871 | 1 | 0 | 5 | 0.2 | 7 | 0 | 2 | 0 | 15 | 0.0667 | 1 | 1 | 7 | 0 | 7 | 0.2857 | 3 | 0.3333 | 18 | 0.2222 |
| 12_531874 | 1 | 1 | 6 | 1 | 7 | 0.8571 | 3 | 0.6667 | 17 | 0.8824 | 2 | 0 | 7 | 0.4286 | 8 | 0.5 | 5 | 0.2 | 22 | 0.3636 |
| 12_531874 | 2 | 0.5 | 6 | 0.6667 | 7 | 1 | 3 | 1 | 18 | 0.8333 | 2 | 1 | 5 | 0.6 | 8 | 0.75 | 5 | 1 | 20 | 0.8 |
| 12_531874 | 1 | 0 | 5 | 0 | 5 | 0 | 2 | 0.5 | 13 | 0.0769 | 2 | 0 | 5 | 0.2 | 6 | 0 | 3 | 0.3333 | 16 | 0.125 |
| 12_531876 | 2 | 0.5 | 5 | 0.2 | 9 | 0.1111 | 2 | 0 | 18 | 0.1667 | 1 | 0 | 7 | 0.2857 | 6 | 0 | 7 | 0.5714 | 21 | 0.2857 |
| 12_531876 | 2 | 0 | 6 | 0.5 | 9 | 0.4444 | 2 | 0 | 19 | 0.3684 | 2 | 0 | 7 | 0.7143 | 8 | 0.625 | 7 | 0.7143 | 24 | 0.625 |
| 12_531879 | 2 | 0 | 7 | 0.1429 | 5 | 0.4 | 3 | 0 | 17 | 0.1765 | 2 | 0 | 5 | 0 | 9 | 0.5556 | 6 | 0.1667 | 22 | 0.2727 |
| 12_531879 | 2 | 0.5 | 3 | 0.3333 | 7 | 0.5714 | 3 | 0.6667 | 15 | 0.5333 | 1 | 0 | 9 | 0.3333 | 8 | 0.25 | 6 | 0.3333 | 24 | 0.2917 |
| 12_531880 | 2 | 0 | 6 | 0.1667 | 9 | 0.1111 | 3 | 0 | 20 | 0.1 | 2 | 0 | 7 | 0 | 10 | 0.1 | 8 | 0.25 | 27 | 0.1111 |
| 12_531881 | 2 | 0 | 7 | 0.7143 | 7 | 0.7143 | 3 | 1 | 19 | 0.6842 | 3 | 0 | 6 | 0.6667 | 8 | 0.375 | 7 | 0.2857 | 24 | 0.375 |
| 12_531881 | 2 | 0 | 6 | 0.1667 | 8 | 0.125 | 1 | 0 | 17 | 0.1176 | 3 | 0 | 8 | 0.25 | 8 | 0.125 | 7 | 0.2857 | 26 | 0.1923 |
| 12_531885 | 2 | 0 | 8 | 0.5 | 10 | 0.4 | 3 | 0.3333 | 23 | 0.3913 | 2 | 0 | 9 | 0.5556 | 9 | 0.3333 | 7 | 0.4286 | 27 | 0.4074 |
| 12_531885 | 3 | 0.6667 | 6 | 0.5 | 11 | 0.0909 | 2 | 0 | 22 | 0.2727 | 3 | 0.6667 | 8 | 0.5 | 9 | 0.3333 | 7 | 0.5714 | 27 | 0.4815 |
| 12_531885 | 3 | 1 | 8 | 0.875 | 11 | 0.9091 | 3 | 1 | 25 | 0.92 | 3 | 1 | 7 | 1 | 8 | 1 | 7 | 1 | 25 | 1 |
| 12_531887 | 3 | 0.3333 | 8 | 0.5 | 11 | 0.0909 | 2 | 1 | 24 | 0.3333 | 3 | 0 | 8 | 0.25 | 10 | 0.1 | 7 | 0.1429 | 28 | 0.1429 |
| 12_531887 | 3 | 0 | 8 | 0.125 | 10 | 0.1 | 3 | 0 | 24 | 0.0833 | 2 | 0 | 8 | 0.125 | 9 | 0 | 7 | 0.2857 | 26 | 0.1154 |
| 12_531889 | 3 | 0.3333 | 8 | 0.875 | 8 | 0.375 | 3 | 0.6667 | 22 | 0.5909 | 2 | 0.5 | 6 | 1 | 6 | 0.8333 | 7 | 0.5714 | 21 | 0.7619 |
| 12_531889 | 4 | 0.75 | 8 | 0.625 | 11 | 0.1818 | 3 | 0.3333 | 26 | 0.4231 | 3 | 0 | 8 | 0.375 | 9 | 0.5556 | 8 | 0.375 | 28 | 0.3929 |
| 12_531890 | 4 | 0.75 | 9 | 0.4444 | 11 | 0.3636 | 1 | 1 | 25 | 0.48 | 3 | 0.6667 | 7 | 0.4286 | 8 | 0.625 | 6 | 0.1667 | 24 | 0.4583 |
| 12_531891 | 4 | 0.75 | 8 | 1 | 9 | 0.8889 | 3 | 1 | 24 | 0.9167 | 3 | 1 | 8 | 0.625 | 10 | 0.9 | 8 | 0.875 | 29 | 0.8276 |
| 12_531894 | 6 | 0 | 8 | 0.25 | 13 | 0.2308 | 4 | 0.5 | 31 | 0.2258 | 4 | 0.25 | 8 | 0.125 | 11 | 0.2727 | 9 | 0.3333 | 32 | 0.25 |
| 12_531946 | 6 | 0 | 10 | 0.1 | 14 | 0.0714 | 5 | 0.2 | 35 | 0.0857 | 4 | 0 | 10 | 0.3 | 12 | 0 | 9 | 0.1111 | 35 | 0.1143 |
| 12_531948 | 6 | 0.8333 | 11 | 0.3636 | 16 | 0.75 | 5 | 0.4 | 38 | 0.6053 | 4 | 0 | 9 | 0.6667 | 13 | 0.6923 | 11 | 0.8182 | 37 | 0.6486 |
| 12_531950 | 6 | 0.5 | 12 | 0.25 | 14 | 0.4286 | 6 | 0.1667 | 38 | 0.3421 | 5 | 0.2 | 11 | 0.1818 | 14 | 0.5 | 12 | 0.5 | 42 | 0.381 |
| 12_531951 | 6 | 0 | 12 | 0 | 14 | 0.1429 | 5 | 0 | 37 | 0.0541 | 5 | 0 | 11 | 0.1818 | 14 | 0.1429 | 14 | 0.1429 | 44 | 0.1364 |
| 12_531970 | 7 | 0 | 9 | 0 | 10 | 0.2 | 8 | 0.125 | 34 | 0.0882 | 6 | 0.1667 | 12 | 0 | 12 | 0.0833 | 10 | 0.1 | 40 | 0.075 |
| 12_531970 | 3 | 0 | 6 | 0.3333 | 7 | 0.5714 | 2 | 0.5 | 18 | 0.3889 | 5 | 0.4 | 11 | 0.2727 | 12 | 0.25 | 11 | 0.1818 | 39 | 0.2564 |
| 12_531976 | 9 | 0.1111 | 9 | 0 | 10 | 0.1 | 6 | 0 | 34 | 0.0588 | 6 | 0 | 17 | 0.0588 | 11 | 0.0909 | 14 | 0.0714 | 48 | 0.0625 |
| 12_531976 | 10 | 0.1 | 10 | 0.2 | 12 | 0 | 9 | 0.1111 | 41 | 0.0976 | 6 | 0.1667 | 19 | 0.1053 | 14 | 0.2143 | 13 | 0.0769 | 52 | 0.1346 |

| | | | | | | | | | | | | | | | | | | | | |
|---|---|---|---|---|---|---|---|---|---|---|---|---|---|---|---|---|---|---|---|---|
| 12_5320079 | 1 | 0 | NA | NA | 5 | 0.2 | 4 | 0.25 | 10 | 0.2 | 1 | 1 | 2 | 0.5 | 2 | 1 | 8 | 0.375 | 13 | 0.5385 |
| 12_5320119 | 2 | 0 | 1 | 0 | 5 | 0.4 | 4 | 0.5 | 12 | 0.3333 | 1 | 1 | 2 | 0.5 | 2 | 1 | 8 | 0.125 | 13 | 0.3846 |
| 12_5320125 | 2 | 0.5 | 1 | 0 | 4 | 0 | 4 | 0 | 11 | 0.0909 | 1 | 0 | 1 | 0 | 2 | 0.5 | 8 | 0 | 12 | 0.0833 |
| 12_5320168 | 2 | 0.5 | 1 | 1 | 3 | 0.3333 | 3 | 0.6667 | 9 | 0.5556 | 1 | 0 | 2 | 1 | 2 | 1 | 7 | 0.8571 | 12 | 0.8333 |
| 12_5320193 | 2 | 0.5 | 1 | 1 | 4 | 0.75 | 4 | 0.25 | 11 | 0.5455 | 1 | 0 | 2 | 0.5 | 3 | 1 | 9 | 0.7778 | 15 | 0.7333 |
| 12_5320200 | 3 | 0 | 1 | 0 | 5 | 0.2 | 4 | 0 | 13 | 0.0769 | 1 | 0 | 2 | 0.5 | 3 | 0.6667 | 8 | 0 | 14 | 0.2143 |
| 12_5411542 | NA | NA | NA | NA | 1 | 1 | NA | NA | 1 | 1 | 1 | 1 | NA | NA | NA | NA | 1 | 0 | 2 | 0.5 |
| 12_5411570 | NA | NA | NA | NA | 1 | 1 | NA | NA | 1 | 1 | 1 | 1 | NA | NA | NA | NA | 1 | 1 | 2 | 1 |
| 12_5411577 | NA | NA | NA | NA | 1 | 1 | NA | NA | 1 | 1 | 1 | 1 | NA | NA | NA | NA | 1 | 1 | 2 | 1 |
| 12_5411585 | NA | NA | NA | NA | 1 | 1 | NA | NA | 1 | 1 | 1 | 1 | NA | NA | NA | NA | 1 | 1 | 2 | 1 |
| 12_5411600 | NA | NA | NA | NA | 1 | 1 | NA | NA | 1 | 1 | 1 | 1 | NA | NA | NA | NA | 1 | 1 | 2 | 1 |
| 12_5411623 | NA | NA | NA | NA | 1 | 1 | NA | NA | 1 | 1 | 1 | 1 | NA | NA | NA | NA | 1 | 0 | 2 | 0.5 |
| 12_5411824 | 1 | 1 | NA | NA | 1 | 1 | 1 | 0 | 3 | 0.6667 | 2 | 1 | 1 | 1 | NA | NA | 1 | 0 | 4 | 0.75 |
| 12_5411874 | NA | NA | NA | NA | 1 | 1 | 1 | 1 | 2 | 1 | 2 | 1 | 2 | 0.5 | NA | NA | 1 | 0 | 5 | 0.6 |
| 12_5422607 | 5 | 0.2 | 2 | 0 | NA | NA | NA | NA | 7 | 0.1429 | NA | NA | NA | NA | 2 | 0.5 | 2 | 0 | 4 | 0.25 |
| 12_5422628 | 3 | 0.3333 | 2 | 0 | NA | NA | NA | NA | 5 | 0.2 | NA | NA | NA | NA | 2 | 0.5 | 2 | 0.5 | 4 | 0.5 |
| 12_5422645 | 4 | 0 | 2 | 0 | NA | NA | 1 | 1 | 7 | 0.1429 | NA | NA | NA | NA | 2 | 0.5 | 2 | 0 | 4 | 0.25 |
| 12_5422649 | 4 | 0 | 3 | 0.3333 | NA | NA | 1 | 1 | 8 | 0.25 | NA | NA | NA | NA | 2 | 0.5 | 2 | 0.5 | 4 | 0.5 |
| 12_5422694 | 4 | 0.25 | 3 | 0 | NA | NA | 1 | 1 | 8 | 0.25 | NA | NA | NA | NA | 1 | 0 | 2 | 0.5 | 3 | 0.3333 |
| 12_5422705 | 2 | 0 | 3 | 0.3333 | NA | NA | 1 | 0 | 6 | 0.1667 | NA | NA | NA | NA | 1 | 0 | 2 | 0.5 | 3 | 0.3333 |
| 12_5422713 | 4 | 1 | 3 | 1 | NA | NA | 1 | 1 | 8 | 1 | NA | NA | NA | NA | 1 | 1 | 2 | 1 | 3 | 1 |
| 12_5422722 | 3 | 0.6667 | 3 | 1 | NA | NA | NA | NA | 6 | 0.8333 | NA | NA | NA | NA | NA | NA | 1 | 1 | 1 | 1 |
| 12_5422724 | 4 | 0.25 | 4 | 0 | NA | NA | 1 | 0 | 9 | 0.1111 | NA | NA | NA | NA | 1 | 1 | 2 | 0 | 3 | 0.3333 |
| 12_5422726 | 4 | 1 | 3 | 1 | NA | NA | 1 | 1 | 8 | 1 | NA | NA | NA | NA | 1 | 1 | 2 | 1 | 3 | 1 |
| 12_5469253 | 10 | 0.3 | 7 | 0.5714 | 10 | 0.8 | 4 | 0.75 | 31 | 0.5806 | 7 | 0.1429 | 13 | 0.5385 | 9 | 0.4444 | 9 | 0.6667 | 38 | 0.4737 |
| 12_5469308 | 1 | 0 | 2 | 0 | NA | NA | 2 | 0.5 | 5 | 0.2 | 2 | 0.5 | 4 | 0.25 | 3 | 0.3333 | NA | NA | 9 | 0.3333 |
| 12_5469308 | 2 | 0 | 1 | 0 | 3 | 0.6667 | NA | NA | 6 | 0.3333 | NA | NA | 1 | 0 | 2 | 0 | 3 | 0.6667 | 6 | 0.3333 |
| 12_5469309 | 8 | 1 | 6 | 1 | 9 | 0.8889 | 3 | 1 | 26 | 0.9615 | 5 | 1 | 7 | 1 | 6 | 0.8333 | 5 | 1 | 23 | 0.9565 |
| 12_5471959 | NA | NA | 3 | 0 | 1 | 0 | 2 | 0.5 | 6 | 0.1667 | 2 | 0 | 1 | 0 | 2 | 0.5 | NA | NA | 5 | 0.2 |
| 12_5472032 | 2 | 0.5 | 7 | 0.5714 | 5 | 0.2 | 2 | 0.5 | 16 | 0.4375 | 2 | 0.5 | 3 | 0.3333 | 4 | 0.25 | 3 | 1 | 12 | 0.5 |
| 12_5479268 | 1 | 1 | 2 | 1 | 1 | 1 | NA | NA | 4 | 1 | NA | NA | 1 | 1 | NA | NA | NA | NA | 1 | 1 |
| 12_5479324 | 15 | 0.2667 | 9 | 0.2222 | 5 | 0.4 | 13 | 0.0769 | 42 | 0.2143 | 7 | 0.5714 | 11 | 0.1818 | 6 | 0.3333 | 2 | 1 | 26 | 0.3846 |
| 12_5479365 | 16 | 0.1875 | 7 | 0 | 3 | 0.6667 | 10 | 0.5 | 36 | 0.2778 | 7 | 0.1429 | 9 | 0.2222 | 6 | 0.3333 | 2 | 0.5 | 24 | 0.25 |
| 12_5490602 | 1 | 0 | 7 | 0.1429 | 7 | 0.4286 | 1 | 1 | 16 | 0.3125 | 1 | 0 | 7 | 0.4286 | 2 | 0.5 | 5 | 0.6 | 15 | 0.4667 |
| 12_5490640 | 1 | 0 | 6 | 0 | 5 | 0.2 | 1 | 0 | 13 | 0.0769 | 2 | 0 | 5 | 0.2 | 2 | 0 | 4 | 0 | 13 | 0.0769 |
| 12_5490646 | 1 | 0 | 7 | 0.2857 | 5 | 0.4 | 1 | 1 | 14 | 0.3571 | 1 | 1 | 4 | 0.25 | 1 | 1 | 4 | 0.25 | 10 | 0.4 |
| 12_5490768 | 6 | 0 | 24 | 0.0833 | 13 | 0.0769 | 17 | 0.1765 | 60 | 0.1 | 10 | 0.2 | 19 | 0.1053 | 7 | 0 | 14 | 0.0714 | 50 | 0.1 |
| 12_5490774 | 5 | 0.2 | 25 | 0.64 | 16 | 0.1875 | 13 | 0.5385 | 59 | 0.4576 | 11 | 0.5455 | 18 | 0.3889 | 10 | 0.4 | 17 | 0.5294 | 56 | 0.4643 |
| 12_5490795 | 6 | 0 | 16 | 0.125 | 16 | 0 | 14 | 0.0714 | 52 | 0.0577 | 10 | 0 | 15 | 0.2 | 11 | 0.0909 | 15 | 0.2 | 51 | 0.1373 |
| 12_5494067 | NA | NA | 1 | 1 | 1 | 1 | 1 | 1 | 3 | 1 | NA | NA | 1 | 1 | 1 | 1 | 1 | 0 | 3 | 0.6667 |
| 12_5494103 | 1 | 1 | NA | NA | 1 | 0 | 1 | 0 | 3 | 0.3333 | NA | NA | NA | NA | 1 | 1 | 1 | 1 | 2 | 1 |
| 12_5494531 | 1 | 1 | NA | NA | 1 | 0 | NA | NA | 2 | 0.5 | NA | NA | NA | NA | 1 | 1 | 1 | 0 | 2 | 0.5 |
| 12_5515969 | 9 | 0.3333 | 7 | 0.2857 | 13 | 0.4615 | 7 | 0.1429 | 36 | 0.3333 | 5 | 0.4 | 5 | 0.6 | 6 | 0.3333 | 13 | 0.6154 | 29 | 0.5172 |
| 12_5518571 | 3 | 0 | 5 | 0 | 2 | 0 | 4 | 0.75 | 14 | 0.2143 | 5 | 0 | 9 | 0.2222 | 5 | 0.4 | 2 | 0.5 | 21 | 0.2381 |
| 12_5518577 | 3 | 1 | 5 | 0.8 | 2 | 1 | 3 | 0.3333 | 13 | 0.7692 | 5 | 0.4 | 9 | 0.5556 | 5 | 1 | 1 | 0 | 20 | 0.6 |
| 12_5518583 | 2 | 1 | 2 | 1 | 4 | 0.75 | 3 | 0.3333 | 11 | 0.7273 | 3 | 0.3333 | 8 | 0.75 | 4 | 0.5 | 1 | 1 | 16 | 0.625 |
| 12_5518605 | 2 | 0 | NA | NA | 4 | 0.5 | 4 | 0.25 | 10 | 0.3 | 3 | 0.3333 | 9 | 0.4444 | 4 | 0.75 | 1 | 1 | 17 | 0.5294 |
| 12_5518607 | 2 | 0 | 2 | 0.5 | 2 | 0 | 4 | 0 | 10 | 0.1 | NA | NA | 7 | 0.1429 | 4 | 0.5 | 1 | 0 | 12 | 0.25 |
| 12_5518865 | NA | NA | NA | NA | NA | NA | 2 | 1 | 2 | 1 | 1 | 0 | 1 | 0 | 1 | 1 | 2 | 0.5 | 5 | 0.4 |
| 12_5518874 | NA | NA | NA | NA | 3 | 0.3333 | NA | NA | 3 | 0.3333 | NA | NA | NA | NA | NA | NA | 1 | 1 | 1 | 1 |
| 12_5518874 | NA | NA | NA | NA | NA | NA | 2 | 1 | 2 | 1 | NA | NA | 1 | 1 | 1 | 1 | 2 | 1 | 4 | 1 |
| 12_5518890 | NA | NA | NA | NA | NA | NA | 1 | 1 | 1 | 1 | NA | NA | NA | NA | NA | NA | 3 | 0.3333 | 3 | 0.3333 |
| 12_5523475 | 10 | 0.4 | 9 | 0.6667 | 7 | 0.5714 | 1 | 0 | 27 | 0.5185 | 4 | 0.25 | 8 | 0.375 | 12 | 0.5 | 10 | 0.2 | 34 | 0.3529 |
| 12_5526077 | 5 | 0.4 | 4 | 0.25 | 5 | 0.6 | 1 | 0 | 15 | 0.4 | 5 | 0.2 | 4 | 0 | 4 | 0 | 4 | 0 | 17 | 0.0588 |
| 12_5526083 | 3 | 0.6667 | 5 | 0.8 | 4 | 0.5 | 1 | 1 | 13 | 0.6923 | 5 | 0.8 | 4 | 0.5 | 3 | 0.3333 | 3 | 0.6667 | 15 | 0.6 |
| 12_5526090 | 5 | 0.8 | 4 | 1 | 5 | 0.6 | 1 | 1 | 15 | 0.8 | 5 | 0.8 | 4 | 0.5 | 3 | 1 | 4 | 0.75 | 16 | 0.75 |
| 12_5526111 | 2 | 0.5 | 6 | 0.1667 | 2 | 0.5 | 1 | 0 | 11 | 0.2727 | 6 | 0.1667 | 4 | 0.25 | 3 | 0.6667 | 4 | 0.25 | 17 | 0.2941 |
| 12_5526113 | 4 | 0.25 | 6 | 0.3333 | 4 | 0.25 | 2 | 0.5 | 16 | 0.3125 | 6 | 0.3333 | 4 | 0.75 | 2 | 0.5 | 4 | 0.5 | 16 | 0.5 |
| 12_5526126 | 4 | 0 | 6 | 0.3333 | 6 | 0 | 2 | 0 | 18 | 0.1111 | 5 | 0 | 4 | 0 | 3 | 0.6667 | 4 | 0 | 16 | 0.125 |
| 12_5526352 | NA | NA | 2 | 0.5 | 1 | 0 | NA | NA | 3 | 0.3333 | 2 | 0.5 | NA | NA | 2 | 0 | NA | NA | 4 | 0.25 |
| 12_5526358 | NA | NA | 2 | 0.5 | 1 | 0 | NA | NA | 3 | 0.3333 | 2 | 0 | NA | NA | 2 | 0.5 | NA | NA | 4 | 0.25 |
| 12_5526371 | NA | NA | NA | NA | NA | NA | 1 | 1 | 1 | 1 | NA | NA | 2 | 1 | NA | NA | 2 | 1 | 4 | 1 |
| 12_5526380 | NA | NA | 1 | 1 | NA | NA | NA | NA | 1 | 1 | 1 | 1 | NA | NA | 1 | 1 | NA | NA | 2 | 1 |
| 12_5526381 | NA | NA | NA | NA | NA | NA | 1 | 1 | 1 | 1 | NA | NA | 3 | 0.6667 | NA | NA | 2 | 1 | 5 | 0.8 |
| 12_5526390 | NA | NA | NA | NA | NA | NA | 1 | 1 | 1 | 1 | NA | NA | 2 | 0 | NA | NA | 2 | 0.5 | 4 | 0.25 |

| | | | | | | | | | | | | | | | | | | | | |
|---|---|---|---|---|---|---|---|---|---|---|---|---|---|---|---|---|---|---|---|---|
| 12_553090 | 4 | 0 | 5 | 0.2 | 4 | 0.5 | 3 | 0.6667 | 16 | 0.3125 | 3 | 0.3333 | 8 | 0.25 | 4 | 0.5 | 5 | 0.8 | 20 | 0.45 |
| 12_553255 | NA | NA | 1 | 1 | NA | NA | NA | NA | 1 | 1 | NA | NA | 1 | 1 | 2 | 1 | 1 | 0 | 4 | 0.75 |
| 12_553282 | 2 | 0 | NA | NA | NA | NA | 1 | 1 | 3 | 0.3333 | NA | NA | 1 | 0 | 2 | 0.5 | NA | NA | 3 | 0.3333 |
| 12_553282 | 2 | 0.5 | NA | NA | NA | NA | 1 | 0 | 3 | 0.3333 | NA | NA | 1 | 0 | 2 | 0.5 | NA | NA | 3 | 0.3333 |
| 12_553283 | 5 | 0.8 | 4 | 1 | 4 | 1 | 5 | 1 | 18 | 0.9444 | 2 | 0.5 | 7 | 0.7143 | 3 | 1 | 3 | 0.6667 | 15 | 0.7333 |
| 12_553284 | 5 | 0.4 | 6 | 0.3333 | 3 | 0 | 4 | 0.5 | 18 | 0.3333 | 2 | 0 | 5 | 0 | 3 | 0.3333 | 3 | 0.3333 | 13 | 0.1538 |
| 12_553284 | 5 | 0 | 6 | 0.1667 | 4 | 0 | 4 | 0 | 19 | 0.0526 | 2 | 0 | 7 | 0 | 3 | 0.6667 | 3 | 0.3333 | 15 | 0.2 |
| 12_553501 | 1 | 1 | NA | NA | 1 | 0 | NA | NA | 2 | 0.5 | NA | NA | NA | NA | 2 | 0 | 2 | 0.5 | 4 | 0.25 |
| 12_553502 | 1 | 0 | NA | NA | 1 | 1 | NA | NA | 2 | 0.5 | NA | NA | NA | NA | 2 | 0.5 | 2 | 0.5 | 4 | 0.5 |
| 12_553502 | 1 | 1 | NA | NA | NA | NA | NA | NA | 1 | 1 | NA | NA | NA | NA | 2 | 1 | 2 | 0.5 | 4 | 0.75 |
| 12_553508 | 2 | 0.5 | NA | NA | 1 | 0 | NA | NA | 3 | 0.3333 | NA | NA | NA | NA | 2 | 0 | 2 | 0.5 | 4 | 0.25 |
| 12_553514 | 1 | 1 | NA | NA | 1 | 1 | NA | NA | 2 | 1 | NA | NA | NA | NA | 2 | 1 | 1 | 1 | 3 | 1 |
| 12_553514 | 1 | 1 | NA | NA | 1 | 1 | NA | NA | 2 | 1 | NA | NA | NA | NA | 1 | 1 | 1 | 1 | 2 | 1 |
| 12_553515 | 2 | 0.5 | NA | NA | 1 | 0 | NA | NA | 3 | 0.3333 | NA | NA | NA | NA | 2 | 0 | 2 | 0.5 | 4 | 0.25 |
| 12_553515 | 2 | 0.5 | NA | NA | 1 | 0 | NA | NA | 3 | 0.3333 | NA | NA | NA | NA | 2 | 1 | 2 | 0.5 | 4 | 0.75 |
| 12_554650 | 32 | 0.1562 | 36 | 0.1111 | 17 | 0.0588 | 22 | 0 | 107 | 0.0935 | 20 | 0.1 | 30 | 0.0333 | 30 | 0.2 | 26 | 0.1154 | 106 | 0.1132 |
| 12_554652 | 27 | 0.1111 | 33 | 0.1212 | 17 | 0 | 24 | 0 | 101 | 0.0693 | 20 | 0.1 | 31 | 0.0323 | 28 | 0.1429 | 25 | 0 | 104 | 0.0673 |
| 12_555366 | 11 | 0.5455 | 4 | 0.25 | 4 | 0.75 | 8 | 0.625 | 27 | 0.5556 | 9 | 0.6667 | 13 | 0.6923 | 10 | 0.3 | 6 | 0.5 | 38 | 0.5526 |
| 12_556505 | 5 | 0 | 10 | 0 | 11 | 0.0909 | 11 | 0.0909 | 37 | 0.0541 | 10 | 0.1 | 7 | 0.1429 | 10 | 0.2 | 7 | 0 | 34 | 0.1176 |
| 12_556507 | 8 | 0.75 | 11 | 0.7273 | 8 | 0.875 | 10 | 0.8 | 37 | 0.7838 | 11 | 0.9091 | 8 | 0.875 | 12 | 0.8333 | 7 | 0.7143 | 38 | 0.8421 |
| 12_556507 | 8 | 0.75 | 10 | 0.7 | 10 | 0.9 | 9 | 1 | 37 | 0.8378 | 10 | 0.9 | 8 | 0.875 | 13 | 0.7692 | 6 | 0.8333 | 37 | 0.8378 |
| 12_556509 | 5 | 0.2 | 11 | 0 | 9 | 0.1111 | 10 | 0.1 | 35 | 0.0857 | 9 | 0.1111 | 8 | 0 | 13 | 0 | 6 | 0.1667 | 36 | 0.0556 |
| 12_557059 | 1 | 0 | 5 | 0.2 | 2 | 0 | 4 | 0 | 12 | 0.0833 | 6 | 0 | 4 | 0 | 5 | 0.2 | 3 | 0 | 18 | 0.0556 |
| 12_557233 | 1 | 0 | 6 | 0.1667 | 5 | 0.2 | 3 | 0.3333 | 15 | 0.2 | 9 | 0.2222 | 4 | 0 | 7 | 0.2857 | 6 | 0 | 26 | 0.1538 |
| 12_557235 | 1 | 0 | 6 | 0.1667 | 6 | 0.1667 | 5 | 0.2 | 18 | 0.1667 | 7 | 0.5714 | 4 | 0.25 | 5 | 0.2 | 7 | 0.2857 | 23 | 0.3478 |
| 12_557236 | 1 | 1 | 7 | 0 | 6 | 0.3333 | 4 | 0 | 18 | 0.1667 | 7 | 0 | 4 | 0.25 | 7 | 0 | 7 | 0.2857 | 25 | 0.12 |
| 12_557236 | 1 | 1 | 7 | 0.5714 | 5 | 0.4 | 5 | 0.4 | 18 | 0.5 | 8 | 0.75 | 4 | 0.75 | 7 | 0.5714 | 5 | 0.6 | 24 | 0.6667 |
| 12_557238 | 1 | 1 | 5 | 0.6 | 4 | 1 | 5 | 0.6 | 15 | 0.7333 | 7 | 0.8571 | 4 | 1 | 6 | 0.6667 | 7 | 1 | 24 | 0.875 |
| 12_557863 | 1 | 0 | 4 | 0.25 | 1 | 0 | 1 | 0 | 7 | 0.1429 | 3 | 0 | 1 | 0 | 5 | 0.2 | 2 | 0 | 11 | 0.0909 |
| 12_559150 | 2 | 0 | 3 | 0 | 4 | 0 | 12 | 0.1667 | 21 | 0.0952 | 2 | 0.5 | 6 | 0 | 7 | 0 | 9 | 0.1111 | 24 | 0.0833 |
| 12_559151 | 2 | 1 | 9 | 0.6667 | 5 | 0.8 | 12 | 0.5833 | 28 | 0.6786 | 3 | 1 | 9 | 1 | 9 | 0.7778 | 9 | 0.6667 | 30 | 0.8333 |
| 12_559152 | 2 | 0 | 8 | 0.375 | 5 | 0 | 12 | 0.1667 | 27 | 0.1852 | 3 | 0.3333 | 6 | 0.3333 | 6 | 0.1667 | 6 | 0.3333 | 21 | 0.2857 |
| 12_559153 | 2 | 1 | 5 | 0.8 | 2 | 1 | 11 | 0.7273 | 20 | 0.8 | 3 | 0.6667 | 7 | 0.7143 | 5 | 1 | 7 | 1 | 22 | 0.8636 |
| 12_559154 | 2 | 1 | 8 | 0.75 | 2 | 0.5 | 12 | 0.6667 | 24 | 0.7083 | 3 | 1 | 7 | 0.4286 | 6 | 0.6667 | 8 | 0.875 | 24 | 0.7083 |
| 12_559155 | 2 | 1 | 9 | 0.3333 | 3 | 0 | 11 | 0.4545 | 25 | 0.4 | 3 | 0.3333 | 8 | 0.375 | 8 | 0.5 | 10 | 0.3 | 29 | 0.3793 |
| 12_559156 | 2 | 0.5 | 10 | 0.3 | 1 | 0 | 13 | 0.0769 | 26 | 0.1923 | 3 | 0.6667 | 7 | 0 | 8 | 0.25 | 9 | 0.3333 | 27 | 0.2593 |
| 12_559158 | 2 | 0.5 | 11 | 0.3636 | 3 | 0.3333 | 11 | 0.2727 | 27 | 0.3333 | 3 | 0.3333 | 7 | 0.4286 | 7 | 0.4286 | 7 | 0.4286 | 24 | 0.4167 |
| 12_559160 | 3 | 0 | 10 | 0.2 | 3 | 0 | 13 | 0.3846 | 29 | 0.2414 | 5 | 0.4 | 9 | 0 | 7 | 0.4286 | 10 | 0.2 | 31 | 0.2258 |
| 12_559160 | 3 | 0 | 11 | 0.0909 | 2 | 0 | 12 | 0.3333 | 28 | 0.1786 | 4 | 0 | 9 | 0.3333 | 4 | 0 | 10 | 0.3 | 27 | 0.2222 |
| 12_559162 | 4 | 0 | 13 | 0 | 4 | 0.25 | 14 | 0.0714 | 35 | 0.0571 | 7 | 0.1429 | 9 | 0.1111 | 9 | 0.1111 | 12 | 0 | 37 | 0.0811 |
| 12_559163 | 4 | 0 | 13 | 0.2308 | 4 | 0.5 | 14 | 0.3571 | 35 | 0.2857 | 5 | 0.2 | 9 | 0.2222 | 9 | 0.1111 | 11 | 0.0909 | 34 | 0.1471 |
| 12_559163 | 4 | 0 | 13 | 0.3846 | 6 | 0.1667 | 18 | 0.1111 | 41 | 0.1951 | 8 | 0 | 10 | 0.2 | 11 | 0.3636 | 12 | 0.1667 | 41 | 0.1951 |
| 12_563752 | 6 | 0.6667 | 5 | 0.8 | 2 | 1 | 6 | 0.6667 | 19 | 0.7368 | 1 | 0 | 4 | 0.5 | 4 | 0.75 | 4 | 0 | 13 | 0.3846 |
| 12_563752 | 6 | 0.3333 | 4 | 0.75 | 2 | 0.5 | 6 | 0.1667 | 18 | 0.3889 | 1 | 1 | 4 | 0.25 | 4 | 0.5 | 3 | 0.3333 | 12 | 0.4167 |
| 12_563752 | 4 | 0.25 | 5 | 0 | 2 | 0.5 | 6 | 0.1667 | 17 | 0.1765 | 1 | 0 | 4 | 0 | 4 | 0.25 | 3 | 0.6667 | 12 | 0.25 |
| 12_563753 | 6 | 0.1667 | 5 | 0 | 2 | 0 | 5 | 0.4 | 18 | 0.1667 | 1 | 0 | 3 | 0.3333 | 3 | 0.3333 | 3 | 0 | 10 | 0.2 |
| 12_563754 | 5 | 0 | 5 | 0 | 2 | 0 | 5 | 0.2 | 17 | 0.0588 | 1 | 0 | 3 | 0 | 2 | 0.5 | 3 | 0 | 9 | 0.1111 |
| 12_563755 | 5 | 0.2 | 4 | 0 | 2 | 0 | 4 | 0.25 | 15 | 0.1333 | 1 | 0 | 3 | 0 | 3 | 0.3333 | 3 | 0 | 10 | 0.1 |
| 12_563864 | 1 | 0 | 1 | 1 | NA | NA | 1 | 0 | 3 | 0.3333 | NA | NA | 2 | 1 | 2 | 0.5 | 2 | 0.5 | 6 | 0.6667 |
| 12_563872 | 2 | 0.5 | NA | NA | 2 | 0.5 | 2 | 0 | 6 | 0.3333 | 1 | 0 | 2 | 0.5 | 3 | 0.6667 | 2 | 0.5 | 8 | 0.5 |
| 12_565789 | NA | NA | NA | NA | NA | NA | 2 | 0.5 | 2 | 0.5 | NA | NA | 1 | 1 | 2 | 0.5 | 1 | 0 | 4 | 0.5 |
| 12_565792 | NA | NA | NA | NA | NA | NA | 2 | 0.5 | 2 | 0.5 | NA | NA | 1 | 1 | 2 | 0.5 | 1 | 1 | 4 | 0.75 |
| 12_565793 | NA | NA | NA | NA | NA | NA | 2 | 1 | 2 | 1 | NA | NA | 1 | 1 | 2 | 0.5 | 1 | 1 | 4 | 0.75 |
| 12_565798 | NA | NA | NA | NA | NA | NA | 2 | 1 | 2 | 1 | NA | NA | 1 | 0 | 2 | 1 | 1 | 1 | 4 | 0.75 |
| 12_565798 | NA | NA | NA | NA | NA | NA | 1 | 1 | 1 | 1 | NA | NA | 1 | 1 | 2 | 1 | 1 | 1 | 4 | 1 |
| 12_565798 | NA | NA | NA | NA | NA | NA | 1 | 1 | 1 | 1 | NA | NA | 1 | 1 | 1 | 0 | 1 | 1 | 3 | 0.6667 |
| 12_565832 | NA | NA | NA | NA | NA | NA | 1 | 1 | 1 | 1 | NA | NA | NA | NA | 2 | 0.5 | NA | NA | 2 | 0.5 |
| 12_572852 | 2 | 1 | NA | NA | 1 | 0 | NA | NA | 3 | 0.6667 | NA | NA | 2 | 1 | 2 | 1 | NA | NA | 4 | 1 |
| 12_572854 | 2 | 1 | NA | NA | 1 | 1 | 1 | 1 | 4 | 1 | NA | NA | 1 | 1 | 2 | 1 | NA | NA | 3 | 1 |
| 12_572856 | 1 | 1 | 1 | 1 | 1 | 1 | 1 | 1 | 4 | 1 | NA | NA | 2 | 1 | 2 | 1 | 1 | 1 | 5 | 1 |
| 12_572857 | 1 | 1 | 1 | 1 | 1 | 1 | 1 | 1 | 4 | 1 | NA | NA | 2 | 1 | 2 | 1 | 1 | 1 | 5 | 1 |
| 12_572858 | 2 | 0.5 | 1 | 1 | 1 | 1 | 1 | 1 | 5 | 0.8 | NA | NA | 2 | 0.5 | 2 | 1 | 1 | 0 | 5 | 0.6 |
| 12_572859 | 1 | 1 | 1 | 1 | NA | NA | NA | NA | 2 | 1 | NA | NA | NA | NA | 1 | 1 | NA | NA | 1 | 1 |
| 12_572860 | 2 | 0 | 1 | 1 | 1 | 1 | 1 | 0 | 5 | 0.4 | NA | NA | 2 | 0.5 | 2 | 0.5 | NA | NA | 4 | 0.5 |
| 12_572867 | 3 | 0.3333 | 1 | 1 | 2 | 0 | 2 | 1 | 8 | 0.5 | NA | NA | 2 | 0 | 1 | 0 | 1 | 1 | 4 | 0.25 |

| | | | | | | | | | | | | | | | | | | | | |
|---|---|---|---|---|---|---|---|---|---|---|---|---|---|---|---|---|---|---|---|---|
| 12_5825959 | NA | NA | 1 | 1 | NA | NA | 2 | 1 | 3 | 1 | NA | NA | NA | NA | 2 | 1 | NA | NA | 2 | 1 |
| 12_5825960 | NA | NA | 1 | 0 | NA | NA | 1 | 1 | 2 | 0.5 | NA | NA | NA | NA | 2 | 1 | NA | NA | 2 | 1 |
| 12_5825982 | NA | NA | 1 | 1 | NA | NA | 2 | 1 | 3 | 1 | NA | NA | NA | NA | 2 | 1 | NA | NA | 2 | 1 |
| 12_5825983 | NA | NA | 1 | 1 | NA | NA | 2 | 0.5 | 3 | 0.6667 | NA | NA | NA | NA | 1 | 1 | 1 | 1 | 2 | 1 |
| 12_5826036 | 2 | 1 | 1 | 1 | NA | NA | 2 | 0.5 | 5 | 0.8 | 1 | 1 | NA | NA | 3 | 1 | 1 | 1 | 5 | 1 |
| 12_5826047 | 1 | 1 | 2 | 0.5 | NA | NA | 1 | 1 | 4 | 0.75 | 1 | 0 | NA | NA | 3 | 1 | 1 | 0 | 5 | 0.6 |
| 12_5826068 | 2 | 1 | 2 | 0.5 | 3 | 0.6667 | 2 | 0.5 | 9 | 0.6667 | 3 | 0.3333 | NA | NA | 2 | 1 | 1 | 1 | 6 | 0.6667 |
| 12_5826070 | 1 | 1 | 2 | 1 | 3 | 0.6667 | 3 | 0.3333 | 9 | 0.6667 | 2 | 0.5 | NA | NA | 1 | 1 | NA | NA | 3 | 0.6667 |
| 12_5826073 | 1 | 0 | 1 | 1 | 3 | 0.6667 | 1 | 0 | 6 | 0.5 | 3 | 0.3333 | NA | NA | 3 | 0.3333 | NA | NA | 6 | 0.3333 |
| 12_5826074 | 1 | 1 | 1 | 1 | 3 | 0.6667 | 2 | 1 | 7 | 0.8571 | 1 | 1 | NA | NA | 3 | 0.6667 | NA | NA | 4 | 0.75 |
| 12_5826083 | 2 | 0.5 | 2 | 0.5 | 3 | 0.3333 | 3 | 0.3333 | 10 | 0.4 | 3 | 0 | NA | NA | 3 | 0.3333 | 1 | 0 | 7 | 0.1429 |
| 12_5826212 | 1 | 0 | 3 | 0 | 3 | 0 | 3 | 0.6667 | 10 | 0.2 | 4 | 0 | NA | NA | 4 | 0.25 | 2 | 0 | 10 | 0.1 |
| 12_5826218 | 1 | 1 | 3 | 0.3333 | 3 | 0.3333 | 2 | 0.5 | 9 | 0.4444 | 4 | 0.25 | NA | NA | 3 | 0 | 2 | 0 | 9 | 0.1111 |
| 12_5826231 | 2 | 0.5 | 3 | 0 | 3 | 0 | 3 | 0 | 11 | 0.0909 | 4 | 0 | NA | NA | 4 | 0.25 | 2 | 0 | 10 | 0.1 |
| 12_5826234 | 2 | 0 | 3 | 0.3333 | 3 | 0 | 3 | 0 | 11 | 0.0909 | 3 | 0 | NA | NA | 4 | 0 | 2 | 0.5 | 9 | 0.1111 |
| 12_5826258 | 2 | 0.5 | 3 | 0.6667 | 3 | 0.3333 | 2 | 0.5 | 10 | 0.5 | 2 | 0 | NA | NA | 4 | 0.25 | 2 | 0 | 8 | 0.125 |
| 12_5826273 | 2 | 0.5 | 3 | 0 | 2 | 0 | 3 | 0 | 10 | 0.1 | 4 | 0 | NA | NA | 5 | 0.2 | 2 | 0 | 11 | 0.0909 |
| 12_5908963 | NA | NA | 1 | 1 | NA | NA | NA | NA | 1 | 1 | NA | NA | 2 | 0.5 | 2 | 1 | NA | NA | 4 | 0.75 |
| 12_5909025 | 24 | 0.0833 | 24 | 0 | 13 | 0.0769 | 18 | 0.1111 | 79 | 0.0633 | 14 | 0.0714 | 11 | 0.0909 | 15 | 0 | 22 | 0.0909 | 62 | 0.0645 |
| 12_5909028 | 23 | 0.2609 | 23 | 0.4348 | 11 | 0.3636 | 16 | 0.3125 | 73 | 0.3425 | 14 | 0.3571 | 10 | 0.2 | 14 | 0.1429 | 23 | 0.3913 | 61 | 0.2951 |
| 12_5909028 | 21 | 0.4762 | 24 | 0.25 | 14 | 0.5 | 18 | 0.2778 | 77 | 0.3636 | 13 | 0.1538 | 16 | 0.25 | 16 | 0.3125 | 24 | 0.3333 | 69 | 0.2754 |
| 12_5911895 | 2 | 0 | 11 | 0 | 5 | 0.2 | 6 | 0.1667 | 24 | 0.0833 | 1 | 0 | 3 | 0.3333 | 8 | 0.25 | 5 | 0 | 17 | 0.1765 |
| 12_5911936 | 2 | 0 | 10 | 0.2 | 7 | 0.2857 | 7 | 0 | 26 | 0.1538 | 1 | 1 | 8 | 0.125 | 8 | 0.125 | 5 | 0 | 22 | 0.1364 |
| 12_5911939 | 2 | 0 | 12 | 0.0833 | 6 | 0.1667 | 7 | 0 | 27 | 0.0741 | NA | NA | 7 | 0.1429 | 5 | 0 | 4 | 0 | 16 | 0.0625 |
| 12_5911942 | 3 | 0 | 11 | 0.1818 | 7 | 0.4286 | 7 | 0.1429 | 28 | 0.2143 | 1 | 1 | 8 | 0.375 | 7 | 0.1429 | 5 | 0 | 21 | 0.2381 |
| 12_5911956 | 3 | 0 | 13 | 0.0769 | 7 | 0.2857 | 7 | 0 | 30 | 0.1 | 1 | 1 | 5 | 0.2 | 8 | 0.25 | 5 | 0 | 19 | 0.2105 |
| 12_5911972 | 3 | 0 | 14 | 0.5714 | 7 | 0.5714 | 5 | 0.6 | 29 | 0.5172 | 1 | 1 | 8 | 0.625 | 7 | 0.7143 | 6 | 0.6667 | 22 | 0.6818 |
| 12_5924787 | 1 | 0 | 6 | 0.3333 | 2 | 0 | 8 | 0.375 | 17 | 0.2941 | 2 | 0.5 | 2 | 0 | 1 | 0 | 7 | 0.1429 | 12 | 0.1667 |
| 12_5924803 | NA | NA | 6 | 0 | 3 | 0.3333 | 7 | 0 | 16 | 0.0625 | 2 | 0.5 | 2 | 0 | 2 | 0 | 8 | 0.125 | 14 | 0.1429 |
| 12_5924895 | NA | NA | 7 | 0 | 2 | 0 | 7 | 0.1429 | 16 | 0.0625 | 2 | 0 | 2 | 0.5 | 1 | 0 | 6 | 0 | 11 | 0.0909 |
| 12_5924920 | NA | NA | 5 | 0.2 | 2 | 0 | 5 | 0.2 | 12 | 0.1667 | 2 | 0 | 1 | 0 | 2 | 0.5 | 6 | 0 | 11 | 0.0909 |
| 12_5924927 | NA | NA | 5 | 1 | 2 | 1 | 3 | 1 | 10 | 1 | 2 | 1 | 1 | 1 | 2 | 1 | 5 | 0.8 | 10 | 0.9 |
| 12_5924935 | NA | NA | 4 | 0.75 | 1 | 0 | 3 | 0 | 8 | 0.375 | 2 | 1 | 1 | 0 | 2 | 0.5 | 6 | 0.3333 | 11 | 0.4545 |
| 12_5925134 | NA | NA | 4 | 0.25 | 2 | 0 | 3 | 0.6667 | 9 | 0.3333 | 2 | 0.5 | 1 | 0 | 2 | 0 | 5 | 0.4 | 10 | 0.3 |
| 12_5925143 | NA | NA | 4 | 0 | 2 | 0 | 3 | 0.3333 | 9 | 0.1111 | 2 | 0 | 1 | 0 | 2 | 0 | 6 | 0.1667 | 11 | 0.0909 |
| 12_5925154 | NA | NA | 4 | 1 | 2 | 1 | 4 | 1 | 10 | 1 | 2 | 1 | NA | NA | 1 | 1 | 5 | 0.8 | 8 | 0.875 |
| 12_5925159 | NA | NA | 4 | 0.25 | 2 | 1 | 3 | 0.6667 | 9 | 0.5556 | 2 | 1 | 1 | 1 | 2 | 1 | 5 | 0.8 | 10 | 0.9 |
| 12_5925163 | NA | NA | 4 | 0.25 | 2 | 0.5 | 4 | 0.5 | 10 | 0.4 | 2 | 1 | 1 | 0 | 2 | 0 | 6 | 0.1667 | 11 | 0.2727 |
| 12_6157030 | 2 | 1 | 1 | 1 | 1 | 1 | 1 | 1 | 5 | 1 | NA | NA | 3 | 1 | 1 | 1 | 1 | 1 | 5 | 1 |
| 12_6157047 | 2 | 0.5 | 2 | 0.5 | 1 | 0 | 1 | 0 | 6 | 0.3333 | NA | NA | 2 | 1 | 2 | 0 | 3 | 0 | 7 | 0.2857 |
| 12_6157079 | 3 | 0.3333 | 2 | 0 | 1 | 0 | 2 | 0 | 8 | 0.125 | NA | NA | 3 | 0 | 2 | 0 | 3 | 0.6667 | 8 | 0.25 |
| 12_6157088 | 3 | 1 | 2 | 1 | 1 | 1 | 2 | 0.5 | 8 | 0.875 | NA | NA | 4 | 0.75 | 2 | 1 | 2 | 0.5 | 8 | 0.75 |
| 12_6157091 | 3 | 0.3333 | 2 | 0.5 | 1 | 1 | 2 | 0.5 | 8 | 0.5 | NA | NA | 4 | 0.25 | 1 | 1 | 1 | 0 | 6 | 0.3333 |
| 12_6157097 | 3 | 0 | 2 | 0.5 | 1 | 0 | 2 | 0 | 8 | 0.125 | NA | NA | 4 | 0 | 2 | 0 | 3 | 0.3333 | 9 | 0.1111 |
| 12_6157102 | 3 | 0.3333 | NA | NA | 1 | 1 | 2 | 1 | 6 | 0.6667 | NA | NA | 2 | 0.5 | 2 | 0.5 | 2 | 0.5 | 6 | 0.5 |
| 12_6157107 | 2 | 1 | 2 | 1 | 1 | 0 | 2 | 1 | 7 | 0.8571 | NA | NA | 2 | 1 | 2 | 1 | 3 | 1 | 7 | 1 |
| 12_6157113 | 3 | 0.6667 | 2 | 0 | 1 | 1 | 2 | 0.5 | 8 | 0.5 | NA | NA | 4 | 0.5 | 1 | 1 | 3 | 0 | 8 | 0.375 |
| 12_6157122 | 2 | 0.5 | 2 | 0 | 1 | 0 | 1 | 0 | 6 | 0.1667 | NA | NA | 4 | 0.75 | 2 | 0.5 | 2 | 1 | 8 | 0.75 |
| 12_6157123 | 1 | 1 | 2 | 0.5 | 1 | 1 | 2 | 1 | 6 | 0.8333 | NA | NA | 3 | 0.3333 | 1 | 1 | 1 | 0 | 5 | 0.4 |
| 12_6157146 | 3 | 0.3333 | 3 | 0.3333 | 1 | 0 | 2 | 0 | 9 | 0.2222 | NA | NA | 5 | 0.2 | 3 | 0.3333 | 4 | 0.25 | 12 | 0.25 |
| 12_6188637 | 4 | 1 | 1 | 1 | 1 | 0 | 2 | 1 | 8 | 0.875 | 1 | 0 | 8 | 0.75 | 3 | 0.6667 | 5 | 0.8 | 17 | 0.7059 |
| 12_6188639 | 6 | 0.3333 | 3 | 0 | 1 | 0 | 2 | 0.5 | 12 | 0.25 | 1 | 0 | 8 | 0.25 | 3 | 0 | 4 | 0.5 | 16 | 0.25 |
| 12_6188644 | 6 | 0.3333 | 3 | 0.6667 | 1 | 1 | 2 | 1 | 12 | 0.5833 | NA | NA | 8 | 0.625 | 3 | 1 | 4 | 1 | 15 | 0.8 |
| 12_6188645 | 6 | 0.6667 | 3 | 0.6667 | NA | NA | 2 | 0 | 11 | 0.5455 | 1 | 1 | 8 | 0.375 | 2 | 0.5 | 4 | 0.25 | 15 | 0.4 |
| 12_6188654 | 3 | 0.6667 | 3 | 0.6667 | 1 | 0 | 2 | 0.5 | 9 | 0.5556 | 1 | 0 | 7 | 0.8571 | 2 | 0 | 5 | 0.8 | 15 | 0.6667 |
| 12_6188660 | 2 | 0 | 3 | 0.3333 | 1 | 0 | 1 | 0 | 7 | 0.1429 | 1 | 0 | 4 | 0 | 4 | 0.25 | 2 | 1 | 11 | 0.2727 |
| 12_6188661 | 5 | 0.6 | 2 | 0 | 1 | 1 | 2 | 0.5 | 10 | 0.5 | NA | NA | 7 | 0.5714 | 3 | 1 | 5 | 1 | 15 | 0.8 |
| 12_6188667 | 6 | 0.1667 | 3 | 1 | 1 | 1 | 3 | 0.6667 | 13 | 0.5385 | 1 | 0 | 8 | 0.625 | 4 | 0.75 | 5 | 0.6 | 18 | 0.6111 |
| 12_6188680 | 6 | 0.1667 | 1 | 0 | 1 | 0 | 3 | 0 | 11 | 0.0909 | 1 | 0 | 8 | 0.375 | 4 | 0 | 5 | 0.2 | 18 | 0.2222 |
| 12_6188683 | 4 | 1 | 2 | 1 | NA | NA | 2 | 0.5 | 8 | 0.875 | 1 | 1 | 5 | 1 | 3 | 0.6667 | 3 | 0.6667 | 12 | 0.8333 |
| 12_6188702 | 4 | 0.5 | 3 | 0.3333 | 1 | 0 | 1 | 0 | 9 | 0.3333 | 1 | 0 | 5 | 0.2 | 1 | 0 | 5 | 0.2 | 12 | 0.1667 |
| 12_6188715 | 6 | 0.1667 | 2 | 0 | 1 | 0 | 5 | 0.2 | 14 | 0.1429 | 1 | 0 | 6 | 0.1667 | 5 | 0 | 4 | 0.25 | 16 | 0.125 |
| 12_6188723 | 5 | 1 | 4 | 1 | 1 | 1 | 5 | 1 | 15 | 1 | 1 | 1 | 8 | 0.875 | 5 | 1 | 4 | 1 | 18 | 0.9444 |
| 12_6188725 | 3 | 0 | 3 | 0.3333 | 1 | 0 | 4 | 0 | 11 | 0.0909 | NA | NA | 4 | 0 | 4 | 0 | 4 | 0.25 | 12 | 0.0833 |
| 12_6188734 | 3 | 0.3333 | 3 | 0 | 1 | 1 | 4 | 0 | 11 | 0.1818 | 1 | 0 | 6 | 0.3333 | 4 | 0.25 | 5 | 0 | 16 | 0.1875 |

| | | | | | | | | | | | | | | | | | | | | |
|---|---|---|---|---|---|---|---|---|---|---|---|---|---|---|---|---|---|---|---|---|
| 12_6189015 | 6 | 0.8333 | 4 | 1 | 1 | 1 | 4 | 0.75 | 15 | 0.8667 | 2 | 0.5 | 6 | 0.6667 | 5 | 0.8 | 4 | 1 | 17 | 0.7647 |
| 12_6189016 | 4 | 0.25 | 3 | 0.3333 | 1 | 0 | 4 | 0.5 | 12 | 0.3333 | NA | NA | 8 | 0.625 | 5 | 0.4 | 3 | 0.6667 | 16 | 0.5625 |
| 12_6189024 | 3 | 0 | 2 | 0.5 | NA | NA | 3 | 0.6667 | 8 | 0.375 | 2 | 0.5 | 6 | 0.1667 | 6 | 0.3333 | 4 | 0.25 | 18 | 0.2778 |
| 12_6189037 | 7 | 0.1429 | 5 | 0 | 1 | 0 | 6 | 0 | 19 | 0.0526 | 2 | 0.5 | 8 | 0.125 | 6 | 0.3333 | 4 | 0 | 20 | 0.2 |
| 12_6189042 | 4 | 1 | 3 | 1 | 1 | 0 | 3 | 1 | 11 | 0.9091 | 2 | 1 | 7 | 1 | 7 | 0.7143 | 4 | 1 | 20 | 0.9 |
| 12_6189053 | 7 | 0.8571 | 5 | 0.8 | 1 | 1 | 6 | 0.8333 | 19 | 0.8421 | 2 | 1 | 10 | 0.7 | 7 | 0.8571 | 6 | 0.8333 | 25 | 0.8 |
| 12_6189055 | 7 | 0.1429 | 5 | 0.2 | 1 | 0 | 5 | 0.2 | 18 | 0.1667 | 2 | 0 | 10 | 0 | 8 | 0.25 | 5 | 0 | 25 | 0.08 |
| 12_6189057 | 7 | 1 | 5 | 1 | NA | NA | 5 | 1 | 17 | 1 | 2 | 0.5 | 10 | 1 | 8 | 0.875 | 6 | 0.6667 | 26 | 0.8462 |
| 12_6189061 | 5 | 1 | 3 | 0.6667 | NA | NA | 6 | 1 | 14 | 0.9286 | 2 | 1 | 8 | 0.875 | 6 | 0.8333 | 6 | 0.8333 | 22 | 0.8636 |
| 12_6189065 | 7 | 1 | 5 | 1 | 1 | 1 | 6 | 1 | 19 | 1 | 2 | 1 | 9 | 1 | 7 | 0.8571 | 6 | 1 | 24 | 0.9583 |
| 12_6189071 | 6 | 0.3333 | 4 | 0.75 | 1 | 0 | 5 | 0.2 | 16 | 0.375 | 1 | 0 | 7 | 0.1429 | 7 | 0.1429 | 5 | 0.2 | 20 | 0.15 |
| 12_6189072 | 6 | 0.5 | 4 | 0.5 | 1 | 1 | 5 | 0 | 16 | 0.375 | 1 | 0 | 9 | 0.2222 | 6 | 0.5 | 4 | 0.25 | 20 | 0.3 |
| 12_6189094 | 1 | 1 | 2 | 0.5 | NA | NA | NA | NA | 3 | 0.6667 | NA | NA | 1 | 1 | NA | NA | NA | NA | 1 | 1 |
| 12_6189138 | 1 | 0 | 2 | 0.5 | NA | NA | NA | NA | 3 | 0.3333 | NA | NA | 2 | 0.5 | NA | NA | NA | NA | 2 | 0.5 |
| 12_6190436 | 2 | 0.5 | 3 | 0.6667 | 1 | 0 | 4 | 0 | 10 | 0.3 | 2 | 0 | 8 | 0 | 6 | 0.1667 | 6 | 0.1667 | 22 | 0.0909 |
| 12_6190443 | 6 | 0.8333 | 3 | 1 | 1 | 1 | 6 | 0.1667 | 16 | 0.625 | 4 | 0.25 | 10 | 0.6 | 9 | 0.4444 | 7 | 0.4286 | 30 | 0.4667 |
| 12_6190465 | 6 | 0.3333 | 2 | 1 | 2 | 0.5 | 7 | 0.1429 | 17 | 0.3529 | 4 | 0.25 | 12 | 0.3333 | 8 | 0.625 | 7 | 0.5714 | 31 | 0.4516 |
| 12_6496773 | NA | NA | 1 | 0 | 1 | 1 | 3 | 1 | 5 | 0.8 | 1 | 0 | 1 | 1 | 2 | 0.5 | NA | NA | 4 | 0.5 |
| 12_6496780 | NA | NA | 1 | 1 | 1 | 1 | 3 | 1 | 5 | 1 | 1 | 1 | 1 | 1 | 1 | 1 | NA | NA | 3 | 1 |
| 12_6496791 | NA | NA | 1 | 0 | 1 | 1 | 3 | 0 | 5 | 0.2 | 1 | 1 | 1 | 1 | 2 | 0.5 | NA | NA | 4 | 0.75 |
| 12_6497100 | NA | NA | 1 | 1 | 1 | 0 | 3 | 0.3333 | 5 | 0.4 | 1 | 1 | 1 | 1 | 2 | 0.5 | NA | NA | 4 | 0.75 |
| 12_6497112 | NA | NA | 1 | 1 | 1 | 0 | 2 | 1 | 4 | 0.75 | 1 | 1 | 1 | 1 | 1 | 0 | NA | NA | 3 | 0.6667 |
| 12_6497121 | NA | NA | 1 | 1 | 1 | 0 | 2 | 0.5 | 4 | 0.5 | 1 | 1 | 1 | 0 | 1 | 0 | NA | NA | 3 | 0.3333 |
| 12_6498997 | 1 | 0 | 1 | 0 | 1 | 1 | 2 | 0 | 5 | 0.2 | NA | NA | 1 | 0 | 1 | 1 | 1 | 1 | 3 | 0.6667 |
| 12_6499001 | 1 | 0 | 1 | 1 | 1 | 0 | 1 | 0 | 4 | 0.25 | NA | NA | 1 | 0 | 1 | 1 | 1 | 0 | 3 | 0.3333 |
| 12_6499008 | 1 | 1 | NA | NA | 1 | 1 | 1 | 1 | 3 | 1 | NA | NA | 1 | 1 | 1 | 1 | 1 | 1 | 3 | 1 |
| 12_6499011 | 1 | 1 | NA | NA | NA | NA | 1 | 1 | 2 | 1 | NA | NA | 1 | 0 | 1 | 1 | 1 | 0 | 3 | 0.3333 |
| 12_6499016 | 1 | 1 | 1 | 0 | 1 | 0 | 1 | 0 | 4 | 0.25 | NA | NA | 1 | 0 | 1 | 1 | 1 | 0 | 3 | 0.3333 |
| 12_6499021 | 1 | 1 | NA | NA | 1 | 1 | 1 | 1 | 3 | 1 | NA | NA | 1 | 1 | 1 | 1 | NA | NA | 2 | 1 |
| 12_6499023 | 1 | 1 | 1 | 1 | 1 | 1 | 1 | 0 | 4 | 0.75 | NA | NA | 1 | 1 | 1 | 0 | NA | NA | 2 | 0.5 |
| 12_6499839 | NA | NA | 1 | 1 | 1 | 0 | 1 | 1 | 3 | 0.6667 | NA | NA | 1 | 1 | 1 | 1 | NA | NA | 2 | 1 |
| 12_6499845 | NA | NA | 1 | 0 | 1 | 1 | 2 | 0.5 | 4 | 0.5 | NA | NA | 1 | 1 | 1 | 1 | NA | NA | 2 | 1 |
| 12_6500363 | NA | NA | NA | NA | 1 | 1 | 1 | 1 | 2 | 1 | NA | NA | 1 | 1 | 1 | 1 | NA | NA | 2 | 1 |
| 12_6500369 | NA | NA | NA | NA | 1 | 1 | 2 | 0 | 3 | 0.3333 | NA | NA | 1 | 1 | 1 | 0 | NA | NA | 2 | 0.5 |
| 12_6500371 | NA | NA | NA | NA | 1 | 1 | 2 | 1 | 3 | 1 | NA | NA | 1 | 1 | 1 | 1 | NA | NA | 2 | 1 |
| 12_6500384 | NA | NA | NA | NA | NA | NA | 1 | 1 | 1 | 1 | NA | NA | NA | NA | 1 | 1 | NA | NA | 1 | 1 |
| 12_6500392 | NA | NA | NA | NA | 1 | 1 | 2 | 1 | 3 | 1 | NA | NA | NA | NA | 1 | 1 | NA | NA | 1 | 1 |
| 12_6500407 | NA | NA | NA | NA | 1 | 1 | 2 | 1 | 3 | 1 | NA | NA | 1 | 1 | 1 | 1 | NA | NA | 2 | 1 |
| 12_6500420 | NA | NA | NA | NA | 1 | 1 | 2 | 0.5 | 3 | 0.6667 | NA | NA | 1 | 1 | NA | NA | NA | NA | 1 | 1 |
| 12_6500441 | NA | NA | NA | NA | 1 | 1 | 1 | 1 | 2 | 1 | NA | NA | 1 | 1 | NA | NA | NA | NA | 1 | 1 |
| 12_6501284 | NA | NA | 2 | 1 | 1 | 0 | 1 | 1 | 4 | 0.75 | NA | NA | 1 | 1 | NA | NA | NA | NA | 1 | 1 |
| 12_6501289 | NA | NA | 2 | 0.5 | 1 | 0 | 1 | 0 | 4 | 0.25 | NA | NA | 1 | 1 | NA | NA | NA | NA | 1 | 1 |
| 12_6501385 | 1 | 1 | 3 | 1 | 1 | 1 | 1 | 1 | 6 | 1 | NA | NA | 1 | 1 | NA | NA | 1 | 1 | 2 | 1 |
| 12_6501422 | 2 | 1 | 3 | 1 | 1 | 1 | 1 | 1 | 7 | 1 | NA | NA | NA | NA | NA | NA | 1 | 1 | 1 | 1 |
| 12_6501427 | 2 | 0.5 | 4 | 0.5 | 1 | 1 | 2 | 0 | 9 | 0.4444 | NA | NA | 1 | 1 | NA | NA | 1 | 0 | 2 | 0.5 |
| 12_6501447 | 1 | 1 | 3 | 1 | 1 | 1 | 3 | 0.6667 | 8 | 0.875 | NA | NA | 1 | 1 | NA | NA | 1 | 1 | 2 | 1 |
| 12_6501457 | NA | NA | 2 | 0.5 | 1 | 1 | 2 | 1 | 5 | 0.8 | NA | NA | 1 | 1 | NA | NA | NA | NA | 1 | 1 |
| 12_6501460 | 2 | 1 | 4 | 1 | 1 | 1 | 2 | 1 | 9 | 1 | NA | NA | 1 | 1 | NA | NA | 1 | 1 | 2 | 1 |
| 12_6502759 | 4 | 0.5 | 4 | 0.25 | 1 | 0 | 6 | 0 | 15 | 0.2 | NA | NA | 1 | 0 | 1 | 1 | NA | NA | 2 | 0.5 |
| 12_6506845 | 5 | 0 | 6 | 0 | 4 | 0 | 9 | 0.2222 | 24 | 0.0833 | 3 | 0 | 2 | 0.5 | 3 | 0 | 2 | 0 | 10 | 0.1 |
| 12_6506849 | 7 | 0 | 6 | 0.1667 | 5 | 0.2 | 11 | 0.3636 | 29 | 0.2069 | 5 | 0.2 | 2 | 0 | 3 | 0 | 2 | 0 | 12 | 0.0833 |
| 12_6506850 | 3 | 0.3333 | 7 | 0.2857 | 4 | 0 | 7 | 0.1429 | 21 | 0.1905 | 4 | 0 | 2 | 0.5 | 3 | 0 | 1 | 1 | 10 | 0.2 |
| 12_6508929 | 2 | 0 | 1 | 0 | 4 | 0 | 1 | 1 | 8 | 0.125 | 2 | 0 | 2 | 0.5 | 1 | 0 | 2 | 0.5 | 7 | 0.2857 |
| 12_6510822 | 2 | 0.5 | 3 | 0 | 5 | 0.8 | 2 | 0.5 | 12 | 0.5 | 5 | 0.4 | 3 | 0 | 3 | 0.3333 | 4 | 0.25 | 15 | 0.2667 |
| 12_6510874 | 3 | 0.3333 | 4 | 0.25 | 6 | 0.3333 | 7 | 0.2857 | 20 | 0.3 | 6 | 0.1667 | 2 | 0.5 | 4 | 1 | 4 | 0 | 16 | 0.375 |
| 12_6510988 | 1 | 0 | 3 | 0 | 3 | 0 | 5 | 0.2 | 12 | 0.0833 | 4 | 0 | 1 | 0 | 2 | 0 | 6 | 0.1667 | 13 | 0.0769 |
| 12_6512062 | 22 | 0 | 21 | 0 | 21 | 0.1429 | 26 | 0.1154 | 90 | 0.0667 | 21 | 0.0952 | 30 | 0.1333 | 16 | 0.0625 | 22 | 0.1364 | 89 | 0.1124 |
| 12_6651389 | 2 | 0.5 | 3 | 0 | 3 | 0 | NA | NA | 8 | 0.125 | 2 | 0 | 3 | 0.3333 | 3 | 0 | 1 | 0 | 9 | 0.1111 |
| 12_6651423 | 3 | 0.3333 | 3 | 0 | 3 | 0 | NA | NA | 9 | 0.1111 | 1 | 0 | 3 | 0.3333 | 3 | 0 | 1 | 1 | 8 | 0.25 |
| 12_6651436 | 3 | 0.3333 | 3 | 0 | 4 | 0 | NA | NA | 10 | 0.1 | 1 | 0 | 4 | 0.25 | 3 | 0 | 1 | 0 | 9 | 0.1111 |
| 12_6651439 | 1 | 1 | 3 | 0 | 4 | 0.5 | NA | NA | 8 | 0.375 | 1 | 0 | 4 | 0 | 3 | 0.3333 | 1 | 0 | 9 | 0.1111 |
| 12_6651449 | 2 | 0.5 | 3 | 0 | 4 | 0.5 | NA | NA | 9 | 0.3333 | 1 | 1 | 4 | 0.5 | 3 | 0.3333 | 1 | 1 | 9 | 0.5556 |
| 12_6651464 | 2 | 1 | 3 | 0 | 4 | 0.25 | NA | NA | 9 | 0.3333 | 1 | 0 | 4 | 0.5 | 3 | 0.6667 | 1 | 1 | 9 | 0.5556 |
| 12_6651660 | 3 | 0.6667 | NA | NA | 2 | 0.5 | 3 | 0.6667 | 8 | 0.625 | 1 | 1 | NA | NA | 3 | 0.6667 | 1 | 1 | 5 | 0.8 |
| 12_6651673 | 3 | 0 | NA | NA | 2 | 0 | 3 | 0.3333 | 8 | 0.125 | NA | NA | 1 | 1 | 3 | 0.3333 | 1 | 1 | 5 | 0.6 |

| | | | | | | | | | | | | | | | | | | | | |
|---|---|---|---|---|---|---|---|---|---|---|---|---|---|---|---|---|---|---|---|---|
| 12_6651683 | 3 | 0.6667 | NA | NA | 2 | 0 | 3 | 1 | 8 | 0.625 | 1 | 1 | NA | NA | 3 | 0.6667 | 1 | 1 | 5 | 0.8 |
| 12_6651690 | 3 | 0.6667 | NA | NA | 2 | 0.5 | 3 | 0.6667 | 8 | 0.625 | 1 | 1 | 1 | 0 | 3 | 1 | 1 | 1 | 6 | 0.8333 |
| 12_6651691 | 2 | 1 | NA | NA | NA | NA | 3 | 0.3333 | 5 | 0.6 | NA | NA | NA | NA | 2 | 1 | 1 | 1 | 3 | 1 |
| 12_6651711 | 3 | 0 | NA | NA | 1 | 0 | 3 | 0.3333 | 7 | 0.1429 | 1 | 0 | 1 | 0 | 3 | 0.3333 | 1 | 0 | 6 | 0.1667 |
| 12_6651721 | 3 | 1 | NA | NA | 2 | 0.5 | 3 | 0.6667 | 8 | 0.75 | 1 | 0 | 1 | 0 | 3 | 1 | 1 | 1 | 6 | 0.6667 |
| 12_6651729 | 3 | 0.6667 | NA | NA | 1 | 1 | 2 | 0.5 | 6 | 0.6667 | 1 | 1 | 1 | 1 | 3 | 1 | NA | NA | 5 | 1 |
| 12_6651784 | 2 | 0 | NA | NA | 4 | 0.25 | 2 | 0.5 | 8 | 0.25 | 1 | 0 | NA | NA | 3 | 0.3333 | 3 | 0 | 7 | 0.1429 |
| 12_6651800 | 2 | 1 | NA | NA | 3 | 1 | 2 | 1 | 7 | 1 | 1 | 1 | NA | NA | 2 | 1 | 2 | 1 | 5 | 1 |
| 12_6651810 | 1 | 1 | 1 | 1 | 4 | 0.5 | 3 | 0.6667 | 9 | 0.6667 | NA | NA | NA | NA | 4 | 0.5 | 3 | 0.3333 | 7 | 0.4286 |
| 12_6651811 | 2 | 0.5 | 1 | 1 | 4 | 1 | 2 | 1 | 9 | 0.8889 | NA | NA | NA | NA | 3 | 1 | 2 | 1 | 5 | 1 |
| 12_6651815 | 1 | 0 | NA | NA | 4 | 0 | 2 | 0.5 | 7 | 0.1429 | NA | NA | NA | NA | 2 | 0.5 | 2 | 0.5 | 4 | 0.5 |
| 12_6651817 | 1 | 1 | NA | NA | 3 | 0.6667 | 2 | 1 | 6 | 0.8333 | NA | NA | NA | NA | 3 | 1 | 3 | 1 | 6 | 1 |
| 12_6651818 | 2 | 0.5 | 1 | 0 | 2 | 1 | 1 | 0 | 6 | 0.5 | NA | NA | NA | NA | 2 | 1 | 3 | 1 | 5 | 1 |
| 12_6651822 | 1 | 1 | 1 | 1 | 3 | 1 | 3 | 1 | 8 | 1 | NA | NA | NA | NA | 2 | 1 | 2 | 1 | 4 | 1 |
| 12_6667679 | 1 | 0 | 1 | 0 | 3 | 0 | 3 | 0.3333 | 8 | 0.125 | NA | NA | NA | NA | 1 | 1 | 2 | 1 | 3 | 1 |
| 12_6667682 | 1 | 0 | 1 | 0 | 3 | 0.3333 | 2 | 1 | 7 | 0.4286 | NA | NA | NA | NA | 1 | 1 | 2 | 1 | 3 | 1 |
| 12_6670196 | 1 | 1 | 1 | 1 | 2 | 1 | 3 | 0.3333 | 7 | 0.7143 | NA | NA | NA | NA | 1 | 0 | 2 | 0.5 | 3 | 0.3333 |
| 12_6670198 | 1 | 1 | NA | NA | 2 | 0 | 2 | 0 | 5 | 0.2 | NA | NA | NA | NA | 1 | 0 | 2 | 0.5 | 3 | 0.3333 |
| 12_6670199 | 1 | 0 | 1 | 0 | 1 | 0 | 3 | 0.3333 | 6 | 0.1667 | NA | NA | NA | NA | 1 | 0 | 2 | 0.5 | 3 | 0.3333 |
| 12_6670202 | 1 | 0 | 1 | 1 | NA | NA | 3 | 0 | 5 | 0.2 | NA | NA | NA | NA | NA | NA | 2 | 0.5 | 2 | 0.5 |
| 12_6673614 | 1 | 0 | NA | NA | 2 | 0 | 2 | 0.5 | 5 | 0.2 | NA | NA | NA | NA | 1 | 1 | 1 | 0 | 2 | 0.5 |
| 12_6726900 | NA | NA | 1 | 1 | NA | NA | 1 | 0 | 2 | 0.5 | NA | NA | NA | NA | 1 | 1 | NA | NA | 1 | 1 |
| 12_6726967 | NA | NA | 1 | 1 | NA | NA | 1 | 1 | 2 | 1 | NA | NA | NA | NA | 2 | 0.5 | NA | NA | 2 | 0.5 |
| 12_6726971 | NA | NA | 1 | 1 | NA | NA | 1 | 1 | 2 | 1 | NA | NA | NA | NA | 1 | 1 | NA | NA | 1 | 1 |
| 12_6922958 | 33 | 0.0606 | 37 | 0.1351 | 22 | 0.0455 | 31 | 0.0968 | 123 | 0.0894 | 22 | 0.1364 | 23 | 0.1304 | 20 | 0.05 | 20 | 0 | 85 | 0.0824 |
| 12_6922963 | 32 | 0.5312 | 39 | 0.4359 | 23 | 0.4348 | 28 | 0.4286 | 122 | 0.459 | 22 | 0.4545 | 22 | 0.3182 | 21 | 0.2857 | 21 | 0.6667 | 86 | 0.4302 |
| 12_6922988 | 31 | 0.9355 | 32 | 0.75 | 23 | 0.7826 | 33 | 0.8182 | 119 | 0.8235 | 21 | 0.7143 | 23 | 0.6522 | 17 | 0.7647 | 23 | 0.7391 | 84 | 0.7143 |
| 12_6923166 | 2 | 0 | 4 | 0 | 5 | 0.4 | 5 | 0.6 | 16 | 0.3125 | 3 | 0.3333 | 2 | 0.5 | 5 | 0.6 | 1 | 0 | 11 | 0.4545 |
| 12_6923170 | 1 | 0 | 4 | 0.25 | 4 | 0.25 | 6 | 0.1667 | 15 | 0.2 | 4 | 0.5 | 4 | 0.25 | 3 | 0.6667 | 1 | 0 | 12 | 0.4167 |
| 12_6923181 | 2 | 1 | 5 | 0.8 | 4 | 1 | 7 | 0.8571 | 18 | 0.8889 | 4 | 1 | 4 | 1 | 5 | 1 | 1 | 1 | 14 | 1 |
| 12_6923187 | 1 | 0 | 4 | 0.25 | 5 | 0 | 6 | 0.1667 | 16 | 0.125 | 3 | 0 | 4 | 0.25 | 4 | 0.25 | 1 | 0 | 12 | 0.1667 |
| 12_6923198 | 2 | 0 | 6 | 0.1667 | 5 | 0.2 | 7 | 0.2857 | 20 | 0.2 | 4 | 0.25 | 4 | 0 | 5 | 0.2 | 1 | 0 | 14 | 0.1429 |
| 12_6923200 | 2 | 0.5 | 5 | 0.8 | 5 | 0.2 | 7 | 0.4286 | 19 | 0.4737 | 4 | 0.5 | 4 | 1 | 4 | 0.5 | 1 | 1 | 13 | 0.6923 |
| 12_6923210 | 1 | 0 | 4 | 0 | 6 | 0.3333 | 6 | 0.3333 | 17 | 0.2353 | 3 | 0.3333 | 4 | 0 | 4 | 0 | 1 | 0 | 12 | 0.0833 |
| 12_6923219 | 2 | 1 | 5 | 0.4 | 6 | 0.1667 | 9 | 0.3333 | 22 | 0.3636 | 4 | 0.25 | 4 | 0.75 | 4 | 0.25 | 2 | 0 | 14 | 0.3571 |
| 12_6923221 | 2 | 0.5 | 4 | 0.75 | 4 | 1 | 5 | 1 | 15 | 0.8667 | 1 | 1 | 3 | 1 | 3 | 0.6667 | 2 | 1 | 9 | 0.8889 |
| 12_6923224 | 2 | 0.5 | 3 | 0 | 4 | 0 | 5 | 0.4 | 14 | 0.2143 | 2 | 0 | 2 | 0 | 4 | 0.5 | 1 | 0 | 9 | 0.2222 |
| 12_6923228 | 2 | 0.5 | 5 | 0.6 | 6 | 0.5 | 9 | 0.4444 | 22 | 0.5 | 4 | 0 | 3 | 0.6667 | 4 | 0.5 | 2 | 0.5 | 13 | 0.3846 |
| 12_6923232 | 2 | 0 | 6 | 0.1667 | 6 | 0 | 9 | 0.1111 | 23 | 0.087 | 2 | 0 | 4 | 0 | 4 | 0.25 | 2 | 0 | 12 | 0.0833 |
| 12_6923234 | 2 | 0 | 5 | 0.2 | 5 | 0.6 | 8 | 0.375 | 20 | 0.35 | 4 | 0 | 3 | 0 | 5 | 0.2 | 2 | 0 | 14 | 0.0714 |
| 12_6923241 | 3 | 1 | 6 | 1 | 5 | 1 | 8 | 1 | 22 | 1 | 4 | 1 | 4 | 1 | 5 | 1 | 3 | 1 | 16 | 1 |
| 12_6923244 | 2 | 0 | 3 | 0 | 5 | 0 | 5 | 0.4 | 15 | 0.1333 | 4 | 0 | 4 | 0 | 3 | 0.3333 | 2 | 0 | 13 | 0.0769 |
| 12_6923245 | 2 | 1 | 4 | 1 | 5 | 1 | 6 | 1 | 17 | 1 | 4 | 1 | 4 | 1 | 3 | 1 | 2 | 1 | 13 | 1 |
| 12_6923245 | 2 | 1 | 6 | 1 | 5 | 1 | 8 | 1 | 21 | 1 | 5 | 1 | 3 | 1 | 4 | 1 | 2 | 1 | 14 | 1 |
| 12_6923247 | 3 | 0.6667 | 4 | 1 | 6 | 0.3333 | 8 | 0.5 | 21 | 0.5714 | 3 | 0.6667 | 4 | 0.75 | 2 | 1 | 4 | 0.75 | 13 | 0.7692 |
| 12_6923254 | 4 | 0.5 | 6 | 0.8333 | 5 | 0.6 | 7 | 0.8571 | 22 | 0.7273 | 5 | 0.4 | 4 | 0.75 | 3 | 0.6667 | 4 | 1 | 16 | 0.6875 |
| 12_6923262 | 3 | 0.3333 | 3 | 0 | 4 | 0 | 7 | 0 | 17 | 0.0588 | 5 | 0 | 3 | 0 | 3 | 0.3333 | 3 | 0 | 14 | 0.0714 |
| 12_6923271 | 3 | 0 | 5 | 0.2 | 5 | 0 | 9 | 0.1111 | 22 | 0.0909 | 5 | 0.2 | 5 | 0 | 4 | 0 | 4 | 0.25 | 18 | 0.1111 |
| 12_6923272 | 4 | 0.75 | 5 | 0.6 | 7 | 1 | 6 | 0.8333 | 22 | 0.8182 | 3 | 1 | 5 | 0.8 | 3 | 0.3333 | 2 | 1 | 13 | 0.7692 |
| 12_6923279 | 3 | 0 | 5 | 0.2 | 5 | 0.2 | 9 | 0.1111 | 22 | 0.1364 | 4 | 0.25 | 5 | 0 | 5 | 0 | 3 | 0 | 17 | 0.0588 |
| 12_6923292 | 4 | 0 | 5 | 0.2 | 6 | 0.1667 | 8 | 0.25 | 23 | 0.1739 | 5 | 0.2 | 4 | 0 | 6 | 0 | 3 | 0 | 18 | 0.0556 |
| 12_6923296 | 4 | 0.25 | 4 | 0.5 | 4 | 0.25 | 5 | 0 | 17 | 0.2353 | 3 | 0.3333 | 5 | 0.2 | 5 | 0.2 | 3 | 0 | 16 | 0.1875 |
| 12_6923666 | 3 | 0 | NA | NA | 2 | 0 | 1 | 1 | 6 | 0.1667 | 2 | 0 | 4 | 0.25 | 1 | 0 | 1 | 0 | 8 | 0.125 |
| 12_6923677 | 3 | 0 | NA | NA | 2 | 0 | 2 | 0.5 | 7 | 0.1429 | 2 | 0 | 4 | 0.25 | NA | NA | 1 | 0 | 7 | 0.1429 |
| 12_6923677 | 3 | 0.3333 | NA | NA | 2 | 0.5 | 2 | 1 | 7 | 0.5714 | 2 | 0.5 | 3 | 0.3333 | 1 | 1 | 1 | 0 | 7 | 0.4286 |
| 12_6923679 | 3 | 0.6667 | NA | NA | 2 | 0 | 1 | 0 | 6 | 0.3333 | 2 | 1 | 4 | 1 | 1 | 1 | 1 | 0 | 8 | 0.875 |
| 12_6924331 | 1 | 1 | NA | NA | 2 | 1 | 3 | 0.6667 | 6 | 0.8333 | 2 | 0.5 | 2 | 0.5 | 1 | 0 | 1 | 1 | 6 | 0.5 |
| 12_6924337 | 2 | 0.5 | NA | NA | 2 | 0 | 3 | 0 | 7 | 0.1429 | 2 | 0 | 2 | 0.5 | 1 | 0 | 1 | 1 | 6 | 0.3333 |
| 12_6924343 | 2 | 0 | NA | NA | 2 | 0 | 4 | 0.25 | 8 | 0.125 | 2 | 0.5 | 1 | 0 | 1 | 0 | 1 | 1 | 5 | 0.4 |
| 12_6924347 | 1 | 0 | NA | NA | 2 | 0.5 | 3 | 0.6667 | 6 | 0.5 | 2 | 1 | 2 | 1 | 1 | 1 | 1 | 0 | 6 | 0.8333 |
| 12_6924373 | 1 | 1 | NA | NA | 2 | 1 | 2 | 0.5 | 5 | 0.8 | 2 | 0.5 | NA | NA | 1 | 1 | NA | NA | 3 | 0.6667 |
| 12_6924377 | 1 | 1 | NA | NA | 2 | 0 | 3 | 0 | 6 | 0.1667 | 2 | 0.5 | 2 | 0 | 1 | 0 | 1 | 1 | 6 | 0.3333 |
| 12_6924453 | 1 | 0 | 3 | 0 | 1 | 1 | 1 | 0 | 6 | 0.1667 | NA | NA | 3 | 0.3333 | 2 | 0 | 1 | 1 | 6 | 0.3333 |
| 12_6924470 | 2 | 0.5 | 3 | 1 | 4 | 0.75 | 2 | 1 | 11 | 0.8182 | 1 | 1 | 5 | 0.6 | 2 | 0 | 1 | 1 | 9 | 0.5556 |
| 12_6924485 | 3 | 0 | 3 | 0.3333 | 5 | 0 | 2 | 0 | 13 | 0.0769 | 1 | 0 | 3 | 0.3333 | 2 | 0 | 1 | 0 | 7 | 0.1429 |

| | | | | | | | | | | | | | | | | | | | | |
|---|---|---|---|---|---|---|---|---|---|---|---|---|---|---|---|---|---|---|---|---|
| 12_692449 | 1 | 0 | 3 | 0 | 3 | 0 | 2 | 0.5 | 9 | 0.1111 | 1 | 1 | 5 | 0.4 | 2 | 0 | 1 | 0 | 9 | 0.3333 |
| 12_692449 | 2 | 0.5 | 2 | 1 | 5 | 0.2 | 2 | 1 | 11 | 0.5455 | 1 | 0 | 3 | 1 | 2 | 1 | 1 | 1 | 7 | 0.8571 |
| 12_692450 | 3 | 0.3333 | 3 | 1 | 4 | 0 | 1 | 0 | 11 | 0.3636 | 1 | 0 | 5 | 0 | 3 | 0 | 1 | 1 | 10 | 0.1 |
| 12_692451 | 2 | 0.5 | 2 | 1 | 3 | 0.3333 | 2 | 1 | 9 | 0.6667 | NA | NA | 4 | 1 | 3 | 1 | 1 | 1 | 8 | 1 |
| 12_692453 | 1 | 0 | 3 | 0 | 4 | 0.25 | 2 | 0 | 10 | 0.1 | NA | NA | 3 | 0.3333 | 1 | 0 | 1 | 0 | 5 | 0.2 |
| 12_692461 | 2 | 0 | 4 | 0.25 | 5 | 0 | 2 | 0 | 13 | 0.0769 | 1 | 0 | 4 | 0.25 | 2 | 0 | 2 | 0.5 | 9 | 0.2222 |
| 12_692462 | 4 | 0 | 3 | 0.3333 | 3 | 0.3333 | 2 | 0 | 12 | 0.1667 | 1 | 0 | 4 | 0.5 | 3 | 0 | 1 | 0 | 9 | 0.2222 |
| 12_692463 | 2 | 0 | 3 | 0.6667 | 2 | 1 | 3 | 0.3333 | 10 | 0.5 | 1 | 0 | 4 | 0.5 | NA | NA | 2 | 0.5 | 7 | 0.4286 |
| 12_692464 | 2 | 0.5 | 4 | 0.25 | 2 | 0 | 2 | 0.5 | 10 | 0.3 | 1 | 0 | 4 | 0.25 | NA | NA | 1 | 0 | 6 | 0.1667 |
| 12_692465 | 2 | 0.5 | 2 | 0.5 | 3 | 0 | 2 | 1 | 9 | 0.4444 | 1 | 1 | 3 | 1 | 1 | 1 | 1 | 1 | 6 | 1 |
| 12_693300 | 2 | 0.5 | 1 | 1 | 1 | 1 | 1 | 1 | 5 | 0.8 | 1 | 1 | 2 | 1 | 2 | 1 | NA | NA | 5 | 1 |
| 12_693301 | 2 | 0.5 | 1 | 0 | 1 | 0 | 1 | 1 | 5 | 0.4 | 1 | 0 | 2 | 1 | 3 | 1 | NA | NA | 6 | 0.8333 |
| 12_693312 | 3 | 0 | 1 | 0 | 1 | 1 | 1 | 0 | 6 | 0.1667 | 2 | 0 | 4 | 0.75 | 4 | 0 | 1 | 0 | 11 | 0.2727 |
| 12_693574 | 8 | 0.125 | 9 | 0.5556 | 2 | 0.5 | 16 | 0.25 | 35 | 0.3143 | 9 | 0.2222 | 6 | 0.1667 | 9 | 0.3333 | 10 | 0.3 | 34 | 0.2647 |
| 12_693577 | 9 | 0 | 9 | 0.1111 | 3 | 0.3333 | 14 | 0.2143 | 35 | 0.1429 | 11 | 0 | 4 | 0 | 11 | 0.0909 | 10 | 0.1 | 36 | 0.0556 |
| 12_693586 | 7 | 0.5714 | 5 | 0.2 | 7 | 0.8571 | 6 | 0.8333 | 25 | 0.64 | 2 | 0 | 2 | 0 | 1 | 1 | 7 | 0.7143 | 12 | 0.5 |
| 12_693586 | 2 | 0 | 3 | 0.3333 | 4 | 0 | 6 | 0 | 15 | 0.0667 | NA | NA | 2 | 0 | 1 | 0 | 4 | 0.5 | 7 | 0.2857 |
| 12_693587 | 7 | 0.5714 | 5 | 0.4 | 7 | 0.4286 | 4 | 0.5 | 23 | 0.4783 | 1 | 1 | 2 | 0 | 1 | 0 | 10 | 0.4 | 14 | 0.3571 |
| 12_693690 | 5 | 0.2 | 6 | 0.3333 | 5 | 0 | 2 | 0 | 18 | 0.1667 | NA | NA | 1 | 0 | NA | NA | 5 | 0.4 | 6 | 0.3333 |
| 12_693690 | 5 | 0 | 5 | 0.2 | 5 | 0.2 | 1 | 0 | 16 | 0.125 | NA | NA | 1 | 0 | NA | NA | 4 | 0.25 | 5 | 0.2 |
| 12_693690 | 5 | 0 | 5 | 0.4 | 6 | 0.3333 | 2 | 0 | 18 | 0.2222 | NA | NA | 1 | 0 | NA | NA | 7 | 0.4286 | 8 | 0.375 |
| 12_693691 | NA | NA | 1 | 0 | 2 | 0.5 | NA | NA | 3 | 0.3333 | NA | NA | 1 | 0 | NA | NA | 1 | 1 | 2 | 0.5 |
| 12_694195 | 5 | 0.6 | 4 | 0.25 | 6 | 0.1667 | 5 | 0.4 | 20 | 0.35 | 2 | 0.5 | 3 | 0.3333 | 9 | 0.3333 | 4 | 0.25 | 18 | 0.3333 |
| 12_694197 | 5 | 0.4 | 3 | 0.3333 | 5 | 0.2 | 5 | 0.8 | 18 | 0.4444 | 3 | 0.6667 | 2 | 1 | 9 | 0.5556 | 4 | 1 | 18 | 0.7222 |
| 12_694197 | 6 | 0 | 3 | 0.3333 | 4 | 0.5 | 4 | 0.75 | 17 | 0.3529 | 4 | 0.25 | 3 | 0.3333 | 9 | 0.3333 | 3 | 0 | 19 | 0.2632 |
| 12_694198 | 6 | 0.5 | 3 | 0.6667 | 6 | 0.6667 | 3 | 0.6667 | 18 | 0.6111 | 4 | 0.5 | 4 | 0 | 9 | 0.4444 | 4 | 0.75 | 21 | 0.4286 |
| 12_694198 | 4 | 0 | 4 | 0.25 | 4 | 0.25 | 4 | 0.25 | 16 | 0.1875 | 4 | 0.25 | 3 | 0.6667 | 11 | 0.0909 | 4 | 0 | 22 | 0.1818 |
| 12_694199 | 6 | 0.5 | 3 | 1 | 4 | 0.75 | 5 | 0.2 | 18 | 0.5556 | 4 | 0.5 | 4 | 0.25 | 11 | 0.6364 | 5 | 0.8 | 24 | 0.5833 |
| 12_694199 | 5 | 0.6 | 4 | 0.5 | 4 | 0.25 | 4 | 0.75 | 17 | 0.5294 | 7 | 0.1429 | 3 | 0 | 9 | 0.8889 | 3 | 1 | 22 | 0.5455 |
| 12_694200 | 6 | 0.6667 | 1 | 0 | 6 | 0.6667 | 4 | 0.75 | 17 | 0.6471 | 8 | 0.5 | 3 | 0.6667 | 9 | 0.6667 | 5 | 0.6 | 25 | 0.6 |
| 12_694200 | 7 | 0.8571 | 3 | 1 | 5 | 0.8 | 4 | 0.25 | 19 | 0.7368 | 8 | 0.75 | 2 | 1 | 7 | 0.5714 | 5 | 0.6 | 22 | 0.6818 |
| 12_694200 | 7 | 1 | 4 | 1 | 6 | 1 | 4 | 1 | 21 | 1 | 6 | 1 | 3 | 1 | 10 | 1 | 5 | 1 | 24 | 1 |
| 12_694200 | 6 | 0 | 3 | 0 | 3 | 0 | 4 | 0.5 | 16 | 0.125 | 7 | 0.2857 | 1 | 0 | 4 | 0 | 3 | 0 | 15 | 0.1333 |
| 12_694202 | 7 | 0.8571 | 2 | 1 | 5 | 1 | 4 | 0.75 | 18 | 0.8889 | 6 | 1 | 1 | 1 | 9 | 1 | 3 | 1 | 19 | 1 |
| 12_694203 | 7 | 0.4286 | 5 | 0 | 7 | 0 | 4 | 0 | 23 | 0.1304 | 8 | 0.125 | 6 | 0.1667 | 11 | 0.1818 | 5 | 0 | 30 | 0.1333 |
| 12_694203 | 8 | 0.375 | 6 | 0.1667 | 6 | 0.1667 | 4 | 0.25 | 24 | 0.25 | 8 | 0.125 | 5 | 0 | 11 | 0.1818 | 6 | 0.5 | 30 | 0.2 |
| 12_696245 | 1 | 1 | 3 | 1 | 6 | 0.8333 | 2 | 0.5 | 12 | 0.8333 | 1 | 1 | 1 | 1 | NA | NA | 1 | 0 | 3 | 0.6667 |
| 12_696293 | 3 | 0 | 6 | 0.3333 | 6 | 0.3333 | 2 | 0 | 17 | 0.2353 | 1 | 1 | 1 | 0 | NA | NA | 2 | 0 | 4 | 0.25 |
| 12_696313 | 6 | 0.5 | 8 | 0.25 | 2 | 0 | 5 | 0.4 | 21 | 0.3333 | 2 | 0.5 | 4 | 0.5 | 4 | 0.25 | 3 | 0.3333 | 13 | 0.3846 |
| 12_696314 | 6 | 0 | 8 | 0.25 | 3 | 0 | 5 | 0 | 22 | 0.0909 | 3 | 0 | 5 | 0 | 3 | 0 | 3 | 0.3333 | 14 | 0.0714 |
| 12_696314 | 6 | 0.6667 | 8 | 0.75 | 3 | 0.6667 | 5 | 0.8 | 22 | 0.7273 | 3 | 1 | 5 | 0.6 | 4 | 0.75 | 3 | 1 | 15 | 0.8 |
| 12_696318 | 7 | 0.2857 | 9 | 0 | 2 | 0 | 6 | 0.1667 | 24 | 0.125 | 3 | 0 | 4 | 0.25 | 3 | 0.3333 | 3 | 0 | 13 | 0.1538 |
| 12_696318 | 7 | 0.1429 | 9 | 0.1111 | 2 | 0 | 6 | 0 | 24 | 0.0833 | 3 | 0 | 4 | 0.25 | 3 | 0.3333 | 3 | 0 | 13 | 0.1538 |
| 12_696319 | 4 | 1 | 5 | 1 | 2 | 1 | 4 | 1 | 15 | 1 | 3 | 1 | 3 | 1 | 3 | 1 | 2 | 1 | 11 | 1 |
| 12_696319 | 7 | 1 | 5 | 1 | 2 | 0.5 | 5 | 1 | 19 | 0.9474 | 1 | 1 | 2 | 1 | 1 | 0 | 1 | 1 | 5 | 0.8 |
| 12_696321 | 5 | 0.2 | 7 | 0 | 2 | 0 | 6 | 0.3333 | 20 | 0.15 | 3 | 0 | 5 | 0.4 | 3 | 0 | 3 | 0 | 14 | 0.1429 |
| 12_696321 | 6 | 0.3333 | 7 | 0.2857 | 2 | 0.5 | 6 | 0 | 21 | 0.2381 | 3 | 0.6667 | 5 | 0 | 3 | 0 | 3 | 0 | 14 | 0.1429 |
| 12_696322 | 4 | 1 | 4 | 1 | 1 | 1 | 6 | 1 | 15 | 1 | NA | NA | 4 | 1 | 3 | 1 | 3 | 1 | 10 | 1 |
| 12_696322 | 3 | 0 | 7 | 0.5714 | 2 | 0.5 | 6 | 0.5 | 18 | 0.4444 | NA | NA | 4 | 0.75 | 1 | 0 | 3 | 0.6667 | 8 | 0.625 |
| 12_696322 | 3 | 0.3333 | 7 | 0.5714 | 2 | 0.5 | 6 | 0.3333 | 18 | 0.4444 | 2 | 0.5 | 5 | 0.2 | 2 | 0 | 3 | 0.6667 | 12 | 0.3333 |
| 12_696637 | 4 | 0.5 | 4 | 0.25 | 1 | 0 | 3 | 0.6667 | 12 | 0.4167 | 2 | 1 | 1 | 1 | NA | NA | 2 | 0.5 | 5 | 0.8 |
| 12_696638 | 4 | 1 | 4 | 0.75 | 1 | 0 | 2 | 0.5 | 11 | 0.7273 | 3 | 1 | 2 | 1 | NA | NA | 2 | 1 | 7 | 1 |
| 12_696638 | 3 | 0 | 2 | 1 | NA | NA | 3 | 0 | 8 | 0.25 | 3 | 0.3333 | 1 | 0 | NA | NA | 1 | 1 | 5 | 0.4 |
| 12_696639 | 3 | 0 | 4 | 0.25 | 1 | 0 | 3 | 0 | 11 | 0.0909 | 1 | 1 | 2 | 0 | NA | NA | 2 | 0 | 5 | 0.2 |
| 12_696641 | 2 | 0.5 | 3 | 0.3333 | NA | NA | 2 | 1 | 7 | 0.5714 | 1 | 0 | 2 | 0 | NA | NA | 1 | 1 | 4 | 0.25 |
| 12_696642 | 2 | 0 | 3 | 0 | 1 | 0 | 3 | 0.3333 | 9 | 0.1111 | 2 | 0 | 2 | 0 | NA | NA | 1 | 1 | 5 | 0.2 |
| 12_697383 | 2 | 0 | 6 | 0 | 4 | 0.5 | 2 | 0 | 14 | 0.1429 | 2 | 0.5 | 2 | 0 | 5 | 0.4 | NA | NA | 9 | 0.3333 |
| 12_697385 | 2 | 0.5 | 6 | 0 | 3 | 0.3333 | 2 | 0.5 | 13 | 0.2308 | 2 | 0.5 | 2 | 0 | 4 | 0.25 | NA | NA | 8 | 0.25 |
| 12_697387 | 2 | 0 | 6 | 0.3333 | 2 | 0.5 | NA | NA | 10 | 0.3 | 2 | 1 | 1 | 0 | 4 | 1 | NA | NA | 7 | 0.8571 |
| 12_697391 | 2 | 0.5 | 4 | 0.25 | 3 | 0.6667 | 1 | 0 | 10 | 0.4 | 2 | 1 | NA | NA | 4 | 0.25 | NA | NA | 6 | 0.5 |
| 12_697905 | 5 | 0.2 | 2 | 0 | 3 | 0 | 2 | 0 | 12 | 0.0833 | 2 | 0 | 3 | 0.3333 | 5 | 0 | 3 | 0 | 13 | 0.0769 |
| 12_697906 | 3 | 0.3333 | NA | NA | 1 | 1 | 1 | 1 | 5 | 0.6 | 2 | 1 | 2 | 1 | 4 | 1 | 3 | 0.6667 | 11 | 0.9091 |
| 12_697907 | 5 | 0.2 | 2 | 0.5 | 1 | 0 | NA | NA | 8 | 0.25 | 3 | 0 | 3 | 0.3333 | 5 | 0 | 4 | 0 | 15 | 0.0667 |
| 12_697913 | 4 | 0.25 | 3 | 0 | 1 | 0 | 1 | 0 | 9 | 0.1111 | 3 | 0 | 2 | 0 | 5 | 0.2 | 4 | 0 | 14 | 0.0714 |
| 12_697930 | 1 | 1 | 1 | 0 | 1 | 1 | 2 | 1 | 5 | 0.8 | NA | NA | 3 | 0.6667 | 2 | 0.5 | NA | NA | 5 | 0.6 |

| | | | | | | | | | | | | | | | | | | | | |
|---|---|---|---|---|---|---|---|---|---|---|---|---|---|---|---|---|---|---|---|---|
| 12_6979378 | 2 | 0.5 | 2 | 0 | 1 | 0 | 1 | 0 | 6 | 0.1667 | NA | NA | 4 | 0.25 | 1 | 0 | NA | NA | 5 | 0.2 |
| 12_6979382 | 2 | 0.5 | 1 | 1 | 1 | 1 | 1 | 1 | 5 | 0.8 | NA | NA | 2 | 1 | 2 | 1 | 1 | 0 | 5 | 0.8 |
| 12_6979383 | NA | NA | 1 | 1 | 1 | 1 | 1 | 1 | 3 | 1 | NA | NA | 3 | 1 | 2 | 0.5 | 1 | 0 | 6 | 0.6667 |
| 12_6979395 | 2 | 0 | 1 | 1 | 1 | 1 | 1 | 1 | 5 | 0.6 | NA | NA | 2 | 0 | 1 | 1 | 1 | 0 | 4 | 0.25 |
| 12_6979413 | 2 | 0.5 | NA | NA | 1 | 1 | 2 | 0.5 | 5 | 0.6 | NA | NA | 3 | 0.6667 | 1 | 1 | NA | NA | 4 | 0.75 |
| 12_6980339 | 1 | 1 | 1 | 0 | 1 | 0 | 2 | 0 | 5 | 0.2 | NA | NA | 2 | 0 | 1 | 1 | 1 | 0 | 4 | 0.25 |
| 12_6985190 | 4 | 0.75 | 3 | 0.3333 | 3 | 0.3333 | 2 | 0 | 12 | 0.4167 | 4 | 0.25 | 8 | 0.25 | 1 | 0 | 4 | 0.5 | 17 | 0.2941 |
| 12_7000597 | 18 | 0.8333 | 9 | 1 | 6 | 1 | 12 | 0.9167 | 45 | 0.9111 | 9 | 1 | 16 | 0.6875 | 22 | 0.9545 | 16 | 0.75 | 63 | 0.8413 |
| 12_7000600 | 18 | 0.2222 | 10 | 0.1 | 7 | 0.1429 | 17 | 0.1176 | 52 | 0.1538 | 8 | 0.125 | 18 | 0.2778 | 17 | 0.2353 | 16 | 0.1875 | 59 | 0.2203 |
| 12_7000608 | 19 | 0.4211 | 12 | 0.3333 | 7 | 0.4286 | 17 | 0.5294 | 55 | 0.4364 | 9 | 0.4444 | 21 | 0.2381 | 20 | 0.35 | 17 | 0.2353 | 67 | 0.2985 |
| 12_7000610 | 20 | 0.65 | 10 | 0.4 | 5 | 0.8 | 15 | 0.8 | 50 | 0.66 | 6 | 0.8333 | 14 | 0.7857 | 20 | 0.55 | 14 | 0.5714 | 54 | 0.6481 |
| 12_7000614 | 16 | 0.9375 | 8 | 0.625 | 7 | 1 | 16 | 0.75 | 47 | 0.8298 | 8 | 0.875 | 18 | 0.7222 | 21 | 0.8571 | 18 | 0.8889 | 65 | 0.8308 |
| 12_7001015 | 15 | 0.1333 | 11 | 0.0909 | 9 | 0 | 16 | 0 | 51 | 0.0588 | 9 | 0 | 18 | 0.0556 | 26 | 0.0385 | 20 | 0.15 | 73 | 0.0685 |
| 12_7001022 | 21 | 0.4762 | 14 | 0.5 | 8 | 0.5 | 17 | 0.7059 | 60 | 0.55 | 9 | 0.7778 | 21 | 0.6667 | 26 | 0.5 | 22 | 0.5909 | 78 | 0.6026 |
| 12_7001023 | 19 | 0.7895 | 11 | 0.6364 | 9 | 0.7778 | 12 | 0.5833 | 51 | 0.7059 | 11 | 0.6364 | 21 | 0.7619 | 25 | 0.76 | 21 | 0.8571 | 78 | 0.7692 |
| 12_7001232 | 1 | 0 | NA | NA | 1 | 0 | 5 | 0.2 | 7 | 0.1429 | 3 | 0.6667 | 1 | 1 | 3 | 0 | NA | NA | 7 | 0.4286 |
| 12_7001246 | 1 | 0 | NA | NA | 1 | 1 | 5 | 0.4 | 7 | 0.4286 | 4 | 0.5 | NA | NA | 3 | 0.3333 | NA | NA | 7 | 0.4286 |
| 12_7001260 | 1 | 0 | NA | NA | 1 | 1 | 3 | 1 | 5 | 0.8 | 4 | 0.75 | 1 | 1 | 1 | 1 | NA | NA | 6 | 0.8333 |
| 12_7001280 | 1 | 0 | NA | NA | 1 | 0 | 4 | 0.5 | 6 | 0.3333 | 3 | 1 | 1 | 0 | 1 | 0 | NA | NA | 5 | 0.6 |
| 12_7001592 | NA | NA | 1 | 1 | NA | NA | NA | NA | 1 | 1 | NA | NA | NA | NA | 1 | 1 | NA | NA | 1 | 1 |
| 12_7001636 | 1 | 1 | 1 | 1 | NA | NA | NA | NA | 2 | 1 | NA | NA | NA | NA | 1 | 1 | 1 | 0 | 2 | 0.5 |
| 12_7002284 | 3 | 0 | NA | NA | 1 | 1 | 3 | 0.3333 | 7 | 0.2857 | 3 | 0.6667 | 1 | 0 | 3 | 0.6667 | NA | NA | 7 | 0.5714 |
| 12_7002292 | 1 | 0 | NA | NA | NA | NA | 2 | 0.5 | 3 | 0.3333 | 3 | 0 | 1 | 0 | 3 | 0.6667 | NA | NA | 7 | 0.2857 |
| 12_7002292 | 2 | 1 | NA | NA | 1 | 1 | 3 | 0.6667 | 6 | 0.8333 | 4 | 0.75 | 1 | 1 | 3 | 1 | NA | NA | 8 | 0.875 |
| 12_7003109 | 2 | 1 | 1 | 0 | 1 | 0 | 2 | 0.5 | 6 | 0.5 | 2 | 1 | 1 | 1 | 2 | 1 | NA | NA | 5 | 1 |
| 12_7003110 | NA | NA | NA | NA | 1 | 1 | 1 | 0 | 2 | 0.5 | 3 | 0.3333 | 1 | 0 | 2 | 0 | NA | NA | 6 | 0.1667 |
| 12_7003113 | 3 | 0.6667 | 1 | 1 | 1 | 0 | 3 | 1 | 8 | 0.75 | 4 | 0.25 | 1 | 1 | 2 | 0.5 | NA | NA | 7 | 0.4286 |
| 12_7003124 | 2 | 1 | 1 | 1 | 1 | 1 | 3 | 1 | 7 | 1 | 4 | 1 | 1 | 1 | 2 | 1 | NA | NA | 7 | 1 |
| 12_7003127 | 1 | 0 | 1 | 1 | 1 | 0 | 3 | 0.6667 | 6 | 0.5 | 4 | 0.75 | 1 | 0 | 2 | 0 | NA | NA | 7 | 0.4286 |
| 12_7003129 | 2 | 0.5 | 1 | 1 | 1 | 0 | 2 | 1 | 6 | 0.6667 | 4 | 1 | 1 | 0 | NA | NA | NA | NA | 5 | 0.8 |
| 12_7003137 | 1 | 0 | 1 | 0 | 1 | 1 | 3 | 0.3333 | 6 | 0.3333 | 4 | 0.5 | NA | NA | NA | NA | NA | NA | 4 | 0.5 |
| 12_7005851 | 5 | 0.2 | 1 | 1 | 3 | 0 | NA | NA | 9 | 0.2222 | 2 | 0.5 | 4 | 0.25 | 4 | 0.5 | 2 | 0 | 12 | 0.3333 |
| 12_7005963 | 4 | 0 | NA | NA | 3 | 0.3333 | 3 | 0 | 10 | 0.1 | 1 | 0 | 4 | 0.25 | 2 | 0 | 1 | 0 | 8 | 0.125 |
| 12_7005987 | 3 | 0.3333 | NA | NA | 3 | 0 | 2 | 0 | 8 | 0.125 | 1 | 0 | 1 | 0 | 2 | 0.5 | 1 | 0 | 5 | 0.2 |
| 12_7006010 | 4 | 0.25 | NA | NA | 2 | 0 | 3 | 0.3333 | 9 | 0.2222 | NA | NA | 4 | 0.25 | 2 | 0 | 1 | 1 | 7 | 0.2857 |
| 12_7006011 | 4 | 0.25 | NA | NA | 1 | 0 | 3 | 0.3333 | 8 | 0.25 | NA | NA | 4 | 0.25 | 2 | 0 | 1 | 0 | 7 | 0.1429 |
| 12_7006015 | 2 | 0.5 | NA | NA | 1 | 0 | 2 | 0 | 5 | 0.2 | NA | NA | 3 | 0 | 2 | 0.5 | 1 | 0 | 6 | 0.1667 |
| 12_7006029 | 1 | 1 | NA | NA | 2 | 0 | 3 | 0 | 6 | 0.1667 | NA | NA | 2 | 1 | 1 | 1 | 1 | 0 | 4 | 0.75 |
| 12_7006040 | 3 | 0.3333 | NA | NA | 2 | 0 | 3 | 0.3333 | 8 | 0.25 | NA | NA | 1 | 1 | 1 | 1 | NA | NA | 2 | 1 |
| 12_7006045 | 3 | 1 | NA | NA | 2 | 1 | 3 | 1 | 8 | 1 | NA | NA | 1 | 1 | 1 | 1 | 1 | 1 | 3 | 1 |
| 12_7006059 | 2 | 1 | NA | NA | 2 | 1 | 2 | 1 | 6 | 1 | NA | NA | 3 | 1 | 1 | 1 | 1 | 1 | 5 | 1 |
| 12_7006061 | 2 | 0.5 | NA | NA | NA | NA | 2 | 1 | 4 | 0.75 | NA | NA | 3 | 0.3333 | 1 | 0 | 1 | 1 | 5 | 0.4 |
| 12_7006062 | 2 | 1 | NA | NA | 2 | 1 | 2 | 1 | 6 | 1 | NA | NA | 2 | 1 | 1 | 1 | 1 | 0 | 4 | 0.75 |
| 12_7006077 | 1 | 1 | NA | NA | 1 | 1 | 2 | 1 | 4 | 1 | NA | NA | 3 | 1 | 1 | 1 | 1 | 1 | 5 | 1 |
| 12_7006078 | 3 | 1 | NA | NA | 1 | 1 | 2 | 1 | 6 | 1 | NA | NA | 1 | 1 | 1 | 1 | 1 | 1 | 3 | 1 |
| 12_7006081 | 2 | 1 | NA | NA | 1 | 1 | NA | NA | 3 | 1 | NA | NA | 2 | 1 | 1 | 1 | 1 | 1 | 4 | 1 |
| 12_7006083 | 3 | 1 | NA | NA | 1 | 1 | 3 | 1 | 7 | 1 | NA | NA | 2 | 1 | 1 | 1 | 1 | 1 | 4 | 1 |
| 12_7006092 | 3 | 1 | NA | NA | NA | NA | 3 | 1 | 6 | 1 | NA | NA | 2 | 1 | 1 | 0 | 1 | 1 | 4 | 0.75 |
| 12_7006097 | 3 | 1 | NA | NA | 1 | 1 | 3 | 0.3333 | 7 | 0.7143 | NA | NA | 3 | 0.6667 | 1 | 1 | 1 | 1 | 5 | 0.8 |
| 12_7006106 | 3 | 0 | NA | NA | 1 | 0 | 3 | 0.3333 | 7 | 0.1429 | NA | NA | 2 | 0.5 | 1 | 0 | 1 | 0 | 4 | 0.25 |
| 12_7006112 | 1 | 0 | NA | NA | 1 | 1 | 2 | 0 | 4 | 0.25 | NA | NA | 2 | 0 | 1 | 1 | 1 | 0 | 4 | 0.25 |
| 12_7006120 | 2 | 0.5 | NA | NA | NA | NA | 2 | 0.5 | 4 | 0.5 | NA | NA | 3 | 0.3333 | 1 | 0 | 1 | 1 | 5 | 0.4 |
| 12_7006123 | 2 | 1 | NA | NA | NA | NA | 2 | 1 | 4 | 1 | NA | NA | 3 | 1 | 1 | 1 | 1 | 1 | 5 | 1 |
| 12_7006143 | 1 | 1 | NA | NA | NA | NA | 2 | 0 | 3 | 0.3333 | NA | NA | 3 | 1 | NA | NA | 1 | 1 | 4 | 1 |
| 12_7006146 | 2 | 0.5 | NA | NA | NA | NA | 2 | 0.5 | 4 | 0.5 | NA | NA | 2 | 0.5 | NA | NA | 1 | 0 | 3 | 0.3333 |
| 12_7006154 | 1 | 1 | NA | NA | NA | NA | NA | NA | 1 | 1 | NA | NA | 1 | 1 | NA | NA | 1 | 1 | 2 | 1 |
| 12_7006577 | 2 | 0.5 | 2 | 0 | 2 | 0 | 3 | 0.3333 | 9 | 0.2222 | NA | NA | 2 | 0 | 2 | 0.5 | 1 | 0 | 5 | 0.2 |
| 12_7006587 | 2 | 0.5 | 2 | 0 | 2 | 0.5 | 2 | 0 | 8 | 0.25 | NA | NA | 2 | 0 | 2 | 0.5 | 1 | 0 | 5 | 0.2 |
| 12_7009205 | NA | NA | 1 | 1 | NA | NA | 1 | 0 | 2 | 0.5 | NA | NA | NA | NA | NA | NA | 1 | 1 | 1 | 1 |
| 12_7009207 | NA | NA | 1 | 1 | 1 | 1 | 1 | 0 | 3 | 0.6667 | NA | NA | NA | NA | NA | NA | 1 | 1 | 1 | 1 |
| 12_7015867 | 2 | 0.5 | NA | NA | NA | NA | 1 | 1 | 3 | 0.6667 | NA | NA | NA | NA | NA | NA | 1 | 1 | 1 | 1 |
| 12_7015909 | 1 | 1 | NA | NA | NA | NA | 1 | 1 | 2 | 1 | NA | NA | NA | NA | NA | NA | 1 | 1 | 1 | 1 |
| 12_7015919 | 2 | 1 | NA | NA | NA | NA | 1 | 1 | 3 | 1 | NA | NA | NA | NA | NA | NA | 1 | 1 | 1 | 1 |
| 12_7015922 | 2 | 1 | NA | NA | NA | NA | 1 | 1 | 3 | 1 | NA | NA | NA | NA | NA | NA | 1 | 1 | 1 | 1 |
| 12_7029153 | 28 | 0.2143 | 23 | 0.1304 | 16 | 0.1875 | 14 | 0.1429 | 81 | 0.1728 | 19 | 0.1579 | 10 | 0 | 23 | 0.087 | 18 | 0.1111 | 70 | 0.1 |

| | | | | | | | | | | | | | | | | | | | | |
|---|---|---|---|---|---|---|---|---|---|---|---|---|---|---|---|---|---|---|---|---|
| 12_702919 | 23 | 0.0435 | 22 | 0.0455 | 15 | 0.1333 | 16 | 0 | 76 | 0.0526 | 17 | 0.1176 | 10 | 0.1 | 23 | 0.087 | 21 | 0.0476 | 71 | 0.0845 |
| 12_702924 | 12 | 0.0833 | 14 | 0.1429 | 6 | 0.1667 | 10 | 0 | 42 | 0.0952 | 11 | 0.0909 | 8 | 0 | 10 | 0.1 | 11 | 0 | 40 | 0.05 |
| 12_702924 | 16 | 0 | 23 | 0.1304 | 10 | 0.1 | 14 | 0 | 63 | 0.0635 | 16 | 0.0625 | 11 | 0 | 18 | 0.0556 | 19 | 0.1053 | 64 | 0.0625 |
| 12_702925 | 17 | 0.1765 | 23 | 0.1739 | 10 | 0.1 | 14 | 0.2857 | 64 | 0.1875 | 17 | 0.1765 | 10 | 0 | 16 | 0.4375 | 18 | 0.2778 | 61 | 0.2459 |
| 12_702927 | 13 | 0.2308 | 21 | 0.1429 | 9 | 0.1111 | 13 | 0.2308 | 56 | 0.1786 | 14 | 0.0714 | 6 | 0 | 13 | 0.0769 | 16 | 0.0625 | 49 | 0.0612 |
| 12_702927 | 15 | 0 | 19 | 0.1579 | 8 | 0 | 13 | 0.0769 | 55 | 0.0727 | 17 | 0.0588 | 9 | 0 | 17 | 0.1176 | 18 | 0.0556 | 61 | 0.0656 |
| 12_702928 | 15 | 0.0667 | 24 | 0.0833 | 8 | 0.25 | 14 | 0.0714 | 61 | 0.0984 | 17 | 0.1765 | 9 | 0 | 17 | 0 | 18 | 0.0556 | 61 | 0.0656 |
| 12_702930 | 10 | 0.6 | 10 | 0.2 | 5 | 0 | 9 | 0.2222 | 34 | 0.2941 | 10 | 0.4 | 3 | 0 | 11 | 0.2727 | 13 | 0.3077 | 37 | 0.2973 |
| 12_702933 | 2 | 0 | 5 | 0 | 2 | 0.5 | 2 | 0 | 11 | 0.0909 | 2 | 0 | 1 | 0 | 1 | 0 | 3 | 0.3333 | 7 | 0.1429 |
| 12_702977 | 7 | 0.2857 | 3 | 0.3333 | 3 | 0 | 2 | 0 | 15 | 0.2 | 4 | 0.25 | 1 | 0 | 1 | 0 | 5 | 0 | 11 | 0.0909 |
| 12_703006 | 2 | 0.5 | 2 | 0.5 | 1 | 0 | NA | NA | 5 | 0.4 | 1 | 1 | NA | NA | NA | NA | NA | NA | 1 | 1 |
| 12_703008 | 2 | 0.5 | 2 | 0 | 1 | 0 | NA | NA | 5 | 0.2 | 1 | 1 | NA | NA | NA | NA | NA | NA | 1 | 1 |
| 12_703131 | 5 | 0.6 | 5 | 0.2 | 1 | 0 | 4 | 0.25 | 15 | 0.3333 | 6 | 0.5 | 4 | 0.25 | 3 | 0 | 2 | 0 | 15 | 0.2667 |
| 12_703131 | 3 | 1 | 4 | 0.5 | 1 | 0 | 2 | 0.5 | 10 | 0.6 | 3 | 0.3333 | 3 | 0 | 3 | 0 | 3 | 0 | 12 | 0.0833 |
| 12_703131 | 4 | 0 | 5 | 0.4 | 1 | 1 | 4 | 0 | 14 | 0.2143 | 6 | 0.3333 | 4 | 0 | 2 | 0 | 3 | 0.3333 | 15 | 0.2 |
| 12_703131 | 4 | 0.25 | 5 | 0.4 | 1 | 0 | 4 | 0.25 | 14 | 0.2857 | 5 | 0 | 3 | 0.6667 | 2 | 0 | 1 | 0 | 11 | 0.1818 |
| 12_703132 | 4 | 0.25 | 4 | 0 | NA | NA | 3 | 0 | 11 | 0.0909 | 6 | 0.1667 | 4 | 0 | 3 | 0 | 3 | 0 | 16 | 0.0625 |
| 12_703133 | 3 | 1 | 5 | 0.2 | NA | NA | 2 | 0 | 10 | 0.4 | 6 | 0.1667 | 3 | 0 | 3 | 0 | 3 | 0.3333 | 15 | 0.1333 |
| 12_703138 | 3 | 0.3333 | 3 | 0 | NA | NA | 2 | 0.5 | 8 | 0.25 | 3 | 0.6667 | 3 | 0 | 1 | 0 | 1 | 0 | 8 | 0.25 |
| 12_703139 | 3 | 1 | 4 | 0.75 | 1 | 0 | 2 | 0.5 | 10 | 0.7 | 4 | 0.5 | 2 | 0.5 | 1 | 1 | 2 | 1 | 9 | 0.6667 |
| 12_703141 | 3 | 1 | 4 | 0.5 | 1 | 0 | 2 | 0.5 | 10 | 0.6 | 3 | 1 | 2 | 0 | 2 | 1 | 2 | 0 | 9 | 0.5556 |
| 12_703142 | 2 | 0.5 | 3 | 1 | 1 | 0 | 2 | 1 | 8 | 0.75 | 4 | 0.5 | 2 | 1 | 2 | 1 | 2 | 0 | 10 | 0.6 |
| 12_703143 | 2 | 1 | 4 | 1 | 1 | 1 | 3 | 1 | 10 | 1 | 3 | 0.6667 | 2 | 0.5 | 2 | 0.5 | 2 | 0 | 9 | 0.4444 |
| 12_703143 | 2 | 0.5 | 4 | 0.25 | 1 | 0 | 2 | 0 | 9 | 0.2222 | 3 | 0.3333 | 2 | 0.5 | 2 | 1 | 3 | 0 | 10 | 0.4 |
| 12_703144 | 2 | 1 | 4 | 0 | NA | NA | 2 | 0.5 | 8 | 0.375 | 2 | 0.5 | 2 | 0 | 2 | 0.5 | 2 | 0.5 | 8 | 0.375 |
| 12_703146 | 1 | 1 | 4 | 0.25 | NA | NA | 3 | 0.3333 | 8 | 0.375 | 3 | 0 | 2 | 0 | 2 | 0.5 | 3 | 0 | 10 | 0.1 |
| 12_703147 | 1 | 1 | 4 | 0.5 | 1 | 0 | 3 | 0.6667 | 9 | 0.5556 | 2 | 1 | 1 | 0 | 2 | 0.5 | 2 | 0 | 7 | 0.4286 |
| 12_703933 | 7 | 0.2857 | 5 | 0.2 | 1 | 0 | 2 | 1 | 15 | 0.3333 | 7 | 0.1429 | 7 | 0.1429 | 3 | 0.6667 | 2 | 0 | 19 | 0.2105 |
| 12_703935 | 7 | 0.1429 | 5 | 0 | 2 | 0 | 1 | 0 | 15 | 0.0667 | 7 | 0.1429 | 6 | 0 | 2 | 0 | 3 | 0 | 18 | 0.0556 |
| 12_703937 | 6 | 0 | 4 | 0.25 | 2 | 0 | 2 | 0 | 14 | 0.0714 | 6 | 0.1667 | 8 | 0.125 | 3 | 0 | 2 | 0 | 19 | 0.1053 |
| 12_703937 | 6 | 0.1667 | 4 | 0 | 2 | 0 | 2 | 0 | 14 | 0.0714 | 6 | 0.1667 | 8 | 0.25 | 3 | 0 | 4 | 0.25 | 21 | 0.1905 |
| 12_703937 | 5 | 0.6 | 4 | 0 | 2 | 0 | 2 | 0 | 13 | 0.2308 | 5 | 0.2 | 8 | 0.25 | 2 | 0 | 4 | 0 | 19 | 0.1579 |
| 12_705129 | 6 | 0.1667 | 9 | 0.3333 | 5 | 0.4 | 3 | 0.3333 | 23 | 0.3043 | 4 | 0.5 | 6 | 0 | 3 | 0.3333 | 1 | 0 | 14 | 0.2143 |
| 12_705129 | 6 | 0.3333 | 10 | 0.1 | 5 | 0.6 | 3 | 0.3333 | 24 | 0.2917 | 3 | 0.3333 | 5 | 0.4 | 3 | 0.3333 | NA | NA | 11 | 0.3636 |
| 12_705129 | 6 | 0.1667 | 10 | 0.1 | 6 | 0 | 3 | 0.3333 | 25 | 0.12 | 4 | 0.25 | 7 | 0.1429 | 3 | 0.3333 | 1 | 0 | 15 | 0.2 |
| 12_705130 | 7 | 0.2857 | 10 | 0.3 | 2 | 0.5 | 2 | 0 | 21 | 0.2857 | 4 | 0.25 | 5 | 0 | 1 | 0 | 1 | 0 | 11 | 0.0909 |
| 12_705137 | 8 | 0 | 14 | 0.1429 | 5 | 0.2 | 3 | 0 | 30 | 0.1 | 3 | 0 | 7 | 0 | 3 | 0.3333 | 2 | 0 | 15 | 0.0667 |
| 12_705140 | 14 | 0.1429 | 14 | 0.2857 | 4 | 0 | 7 | 0.2857 | 39 | 0.2051 | 4 | 0 | 7 | 0 | 5 | 0.6 | 4 | 0 | 20 | 0.15 |
| 12_705141 | 13 | 0.1538 | 17 | 0.0588 | 4 | 0 | 9 | 0.1111 | 43 | 0.093 | 5 | 0 | 9 | 0.2222 | 7 | 0 | 4 | 0.25 | 25 | 0.12 |
| 12_709462 | 3 | 0.3333 | 2 | 0 | 1 | 0 | 2 | 0 | 8 | 0.125 | 1 | 0 | 2 | 0 | 3 | 0.3333 | 3 | 0 | 9 | 0.1111 |
| 12_709462 | 3 | 0 | 3 | 0.3333 | 1 | 1 | 1 | 1 | 8 | 0.375 | 1 | 0 | 3 | 0.6667 | 2 | 0 | 3 | 0.6667 | 9 | 0.4444 |
| 12_709463 | 2 | 0.5 | 2 | 1 | 1 | 0 | NA | NA | 5 | 0.6 | 1 | 1 | 1 | 1 | NA | NA | NA | NA | 2 | 1 |
| 12_709464 | 1 | 1 | 3 | 0.6667 | 1 | 1 | 2 | 1 | 7 | 0.8571 | NA | NA | 1 | 1 | 1 | 1 | 2 | 0 | 4 | 0.5 |
| 12_709465 | 3 | 0.3333 | 3 | 0 | 1 | 0 | 1 | 0 | 8 | 0.125 | 2 | 0 | 3 | 0.3333 | 2 | 0 | 3 | 0 | 10 | 0.1 |
| 12_709465 | 2 | 1 | 4 | 1 | 1 | 1 | 2 | 1 | 9 | 1 | 3 | 1 | 2 | 1 | 2 | 1 | 3 | 1 | 10 | 1 |
| 12_709467 | 3 | 1 | 5 | 0 | 1 | 1 | NA | NA | 9 | 0.4444 | 2 | 0 | 3 | 0 | 3 | 0 | 3 | 0.3333 | 11 | 0.0909 |
| 12_709467 | 3 | 0 | 4 | 0.25 | 1 | 0 | 2 | 0 | 10 | 0.1 | 2 | 0 | 3 | 0.3333 | 3 | 0 | 3 | 0 | 11 | 0.0909 |
| 12_709469 | 3 | 0 | 4 | 0.25 | 1 | 0 | 1 | 0 | 9 | 0.1111 | 1 | 0 | 3 | 0.3333 | 1 | 0 | 2 | 0.5 | 7 | 0.2857 |
| 12_709470 | 1 | 0 | 3 | 0.3333 | 1 | 0 | 2 | 0 | 7 | 0.1429 | 2 | 0.5 | 2 | 0.5 | 3 | 0.3333 | 4 | 0.25 | 11 | 0.3636 |
| 12_709473 | 3 | 0.3333 | 4 | 0 | 1 | 0 | 2 | 0 | 10 | 0.1 | 2 | 0 | 3 | 0 | 3 | 0.3333 | 4 | 0 | 12 | 0.0833 |
| 12_709486 | 2 | 0 | 5 | 0 | 1 | 1 | 2 | 0 | 10 | 0.1 | 2 | 0.5 | 2 | 0 | 3 | 0 | 4 | 0 | 11 | 0.0909 |
| 12_710086 | 18 | 0.1111 | 12 | 0.1667 | 11 | 0.1818 | 17 | 0.0588 | 58 | 0.1207 | 16 | 0.0625 | 12 | 0.1667 | 13 | 0 | 17 | 0.1176 | 58 | 0.0862 |
| 12_710097 | 4 | 0 | 5 | 0 | 3 | 0 | 5 | 0.2 | 17 | 0.0588 | 2 | 0 | 5 | 0 | 3 | 0 | 7 | 0.1429 | 17 | 0.0588 |
| 12_710107 | 9 | 0.1111 | 8 | 0 | 5 | 0.2 | 4 | 0.25 | 26 | 0.1154 | 4 | 0.25 | 8 | 0.125 | 4 | 0.25 | 12 | 0.25 | 28 | 0.2143 |
| 12_710111 | 12 | 0.0833 | 13 | 0.1538 | 7 | 0.1429 | 5 | 0.2 | 37 | 0.1351 | 5 | 0.2 | 9 | 0.1111 | 4 | 0.25 | 15 | 0 | 33 | 0.0909 |
| 12_710112 | 14 | 0.0714 | 10 | 0.2 | 7 | 0.1429 | 7 | 0.4286 | 38 | 0.1842 | 5 | 0 | 7 | 0.2857 | 5 | 0.4 | 15 | 0.2 | 32 | 0.2188 |
| 12_710412 | 23 | 0.1304 | 25 | 0.4 | 25 | 0.44 | 15 | 0.1333 | 88 | 0.2955 | 12 | 0.1667 | 18 | 0.1667 | 27 | 0.1852 | 19 | 0.1053 | 76 | 0.1579 |
| 12_710422 | NA | NA | 4 | 0 | 5 | 0.2 | NA | NA | 9 | 0.1111 | 5 | 0 | 4 | 0 | 1 | 1 | 4 | 0 | 14 | 0.0714 |
| 12_710423 | NA | NA | 3 | 0.3333 | 7 | 0.1429 | 1 | 0 | 11 | 0.1818 | 7 | 0.1429 | 4 | 0 | 1 | 0 | 4 | 0.25 | 16 | 0.125 |
| 12_710424 | NA | NA | 1 | 0 | 6 | 0.1667 | 1 | 0 | 8 | 0.125 | 6 | 0 | 3 | 0 | 2 | 0 | 5 | 0.2 | 16 | 0.0625 |
| 12_710424 | NA | NA | 2 | 0.5 | 5 | 0 | 1 | 0 | 8 | 0.125 | 5 | 0.2 | 4 | 0 | 4 | 0 | 3 | 0.3333 | 16 | 0.125 |
| 12_710427 | NA | NA | NA | NA | 1 | 1 | NA | NA | 1 | 1 | NA | NA | 1 | 0 | NA | NA | 1 | 1 | 2 | 0.5 |
| 12_710570 | 1 | 0 | 2 | 0 | 1 | 1 | 3 | 0 | 7 | 0.1429 | 1 | 0 | NA | NA | 3 | 0.3333 | NA | NA | 4 | 0.25 |
| 12_710570 | 1 | 0 | 2 | 0 | 1 | 0 | 3 | 0.3333 | 7 | 0.1429 | 1 | 0 | NA | NA | 3 | 0.3333 | NA | NA | 4 | 0.25 |
| 12_710571 | 1 | 0 | 2 | 0.5 | NA | NA | 3 | 0 | 6 | 0.1667 | 1 | 0 | NA | NA | 3 | 0.3333 | NA | NA | 4 | 0.25 |

| | | | | | | | | | | | | | | | | | | | | |
|---|---|---|---|---|---|---|---|---|---|---|---|---|---|---|---|---|---|---|---|---|
| 12_7112188 | 2 | 0 | 1 | 1 | 1 | 0 | 1 | 0 | 5 | 0.2 | 1 | 0 | 3 | 0.6667 | 2 | 0 | 1 | 1 | 7 | 0.4286 |
| 12_7112203 | 2 | 0 | 1 | 0 | 1 | 1 | 2 | 0 | 6 | 0.1667 | 1 | 0 | 3 | 0 | 3 | 0.3333 | 2 | 0 | 9 | 0.1111 |
| 12_7112221 | 2 | 0 | 1 | 0 | 1 | 1 | 2 | 1 | 6 | 0.5 | 1 | 0 | 3 | 0.3333 | 1 | 0 | 1 | 1 | 6 | 0.3333 |
| 12_7112240 | NA | NA | 1 | 0 | 1 | 1 | 2 | 1 | 4 | 0.75 | NA | NA | 3 | 0.6667 | 3 | 0.3333 | 2 | 0.5 | 8 | 0.5 |
| 12_7112254 | 2 | 0 | 1 | 1 | NA | NA | 2 | 0 | 5 | 0.2 | 2 | 0 | 3 | 0.3333 | 3 | 0.3333 | 2 | 0 | 10 | 0.2 |
| 12_7114438 | 1 | 1 | 1 | 0 | 1 | 1 | 3 | 0 | 6 | 0.3333 | 2 | 0 | 3 | 0 | 4 | 0 | 1 | 1 | 10 | 0.1 |
| 12_7114441 | 1 | 1 | 1 | 1 | 2 | 1 | 3 | 1 | 7 | 1 | 2 | 0.5 | 3 | 1 | 2 | 1 | 1 | 1 | 8 | 0.875 |
| 12_7114504 | 2 | 0.5 | 2 | 0 | 4 | 0.25 | 5 | 0 | 13 | 0.1538 | 3 | 0 | 3 | 0 | 6 | 0 | 2 | 0.5 | 14 | 0.0714 |
| 12_7114511 | NA | NA | 2 | 0 | 4 | 0.25 | 5 | 0 | 11 | 0.0909 | 1 | 0 | 3 | 0 | 6 | 0.3333 | 2 | 0 | 12 | 0.1667 |
| 12_7117003 | 10 | 0.5 | 10 | 0.3 | 6 | 0.1667 | 5 | 0.4 | 31 | 0.3548 | 5 | 0.6 | 4 | 0.25 | 4 | 0.25 | 6 | 0.1667 | 19 | 0.3158 |
| 12_7117017 | 9 | 0.4444 | 10 | 0.2 | 5 | 0 | 5 | 0.4 | 29 | 0.2759 | 5 | 0.2 | 3 | 0.6667 | 2 | 0.5 | 8 | 0.375 | 18 | 0.3889 |
| 12_7117020 | 9 | 0 | 10 | 0 | 7 | 0.2857 | 5 | 0 | 31 | 0.0645 | 5 | 0.2 | 4 | 0.25 | 3 | 0 | 9 | 0.1111 | 21 | 0.1429 |
| 12_7117030 | 10 | 1 | 11 | 0.9091 | 7 | 0.8571 | 5 | 1 | 33 | 0.9394 | 4 | 1 | 2 | 1 | 3 | 1 | 8 | 0.875 | 17 | 0.9412 |
| 12_7117044 | 10 | 0.1 | 10 | 0.2 | 6 | 0 | 5 | 0.4 | 31 | 0.1613 | 5 | 0.2 | 4 | 0.25 | 3 | 0 | 8 | 0 | 20 | 0.1 |
| 12_7117167 | 1 | 1 | NA | NA | NA | NA | NA | NA | 1 | 1 | 2 | 0.5 | NA | NA | NA | NA | 1 | 0 | 3 | 0.3333 |
| 12_7117227 | 8 | 0.25 | 9 | 0 | 5 | 0.8 | 5 | 0.2 | 27 | 0.2593 | 4 | 0.25 | 4 | 0.25 | 3 | 0 | 6 | 0.1667 | 17 | 0.1765 |
| 12_7136420 | NA | NA | 1 | 1 | NA | NA | NA | NA | 1 | 1 | 1 | 0 | 2 | 1 | NA | NA | NA | NA | 3 | 0.6667 |
| 12_7136437 | NA | NA | 1 | 1 | NA | NA | NA | NA | 1 | 1 | 1 | 1 | 2 | 1 | NA | NA | NA | NA | 3 | 1 |
| 12_7136552 | NA | NA | 2 | 1 | NA | NA | NA | NA | 2 | 1 | 1 | 1 | 3 | 0.6667 | NA | NA | NA | NA | 4 | 0.75 |
| 12_7136565 | NA | NA | 2 | 1 | NA | NA | NA | NA | 2 | 1 | 1 | 1 | 3 | 1 | NA | NA | NA | NA | 4 | 1 |
| 12_7136657 | NA | NA | 2 | 0.5 | NA | NA | NA | NA | 2 | 0.5 | 2 | 0 | 2 | 0.5 | NA | NA | NA | NA | 4 | 0.25 |
| 12_7203657 | 3 | 0 | 5 | 0 | 1 | 0 | 4 | 0.25 | 13 | 0.0769 | 2 | 0 | 2 | 0 | 1 | 0 | 3 | 0.3333 | 8 | 0.125 |
| 12_7203658 | 2 | 0 | 4 | 0.5 | 1 | 0 | 4 | 0.25 | 11 | 0.2727 | 2 | 0 | 2 | 0.5 | 2 | 0 | 4 | 0.25 | 10 | 0.2 |
| 12_7203662 | 1 | 0 | 4 | 0.25 | 1 | 1 | 3 | 0.6667 | 9 | 0.4444 | 2 | 0.5 | 1 | 1 | 2 | 0 | 3 | 1 | 8 | 0.625 |
| 12_7203665 | 2 | 0 | 3 | 0 | 1 | 1 | 4 | 0 | 10 | 0.1 | 2 | 0.5 | 1 | 1 | 2 | 0 | 2 | 0.5 | 7 | 0.4286 |
| 12_7203671 | 2 | 0.5 | 3 | 1 | 1 | 1 | NA | NA | 6 | 0.8333 | 2 | 1 | 2 | 1 | 2 | 1 | 2 | 0.5 | 8 | 0.875 |
| 12_7203690 | 3 | 0.3333 | 3 | 0.6667 | 1 | 0 | 3 | 0 | 10 | 0.3 | 3 | 0 | 2 | 0.5 | 2 | 0 | 5 | 0.4 | 12 | 0.25 |
| 12_7203719 | 2 | 0.5 | 6 | 0.5 | 1 | 1 | 2 | 0.5 | 11 | 0.5455 | 3 | 0.3333 | 3 | 0.3333 | 2 | 0.5 | 4 | 0.5 | 12 | 0.4167 |
| 12_7203749 | 3 | 0 | 6 | 0.1667 | 1 | 1 | 4 | 0 | 14 | 0.1429 | 3 | 0 | 6 | 0.1667 | 2 | 0 | 5 | 0 | 16 | 0.0625 |
| 12_7203752 | 3 | 0.3333 | 6 | 0.1667 | 1 | 0 | 4 | 0.25 | 14 | 0.2143 | 3 | 0.3333 | 6 | 0 | 2 | 0 | 6 | 0 | 17 | 0.0588 |
| 12_7203754 | 3 | 0 | 6 | 0.1667 | 1 | 0 | 4 | 0 | 14 | 0.0714 | 3 | 0 | 4 | 0.25 | 2 | 0 | 5 | 0.2 | 14 | 0.1429 |
| 12_7203756 | 2 | 0 | 7 | 0.1429 | 1 | 0 | 4 | 0.25 | 14 | 0.1429 | 2 | 0 | 6 | 0.3333 | 2 | 0 | 6 | 0 | 16 | 0.125 |
| 12_7203765 | 3 | 0.6667 | 7 | 0.7143 | 1 | 0 | 4 | 0.75 | 15 | 0.6667 | 3 | 0 | 6 | 0.6667 | 2 | 0.5 | 4 | 0.5 | 15 | 0.4667 |
| 12_7203766 | 3 | 0.6667 | 6 | 0.3333 | 1 | 1 | 4 | 1 | 14 | 0.6429 | 3 | 0.3333 | 6 | 0.5 | 2 | 0.5 | 5 | 0.6 | 16 | 0.5 |
| 12_7203791 | 3 | 0 | 6 | 0.1667 | 1 | 0 | 5 | 0.2 | 15 | 0.1333 | 3 | 0 | 5 | 0.4 | 2 | 0.5 | 4 | 0 | 14 | 0.2143 |
| 12_7203864 | 3 | 0.3333 | 8 | 0.125 | 2 | 0.5 | 3 | 0.3333 | 16 | 0.25 | 3 | 0 | 5 | 0.2 | 3 | 0 | 4 | 0.25 | 15 | 0.1333 |
| 12_7203877 | 3 | 1 | 8 | 0.875 | 4 | 0.25 | 4 | 0.75 | 19 | 0.7368 | 3 | 0.3333 | 7 | 0.7143 | 4 | 0.5 | 6 | 0.6667 | 20 | 0.6 |
| 12_7211490 | 4 | 0.25 | 2 | 0 | 1 | 0 | 1 | 0 | 8 | 0.125 | 3 | 0 | 1 | 0 | 1 | 0 | 1 | 1 | 6 | 0.1667 |
| 12_7211499 | 4 | 0.5 | 1 | 0 | 1 | 0 | NA | NA | 6 | 0.3333 | 3 | 0.3333 | 1 | 0 | 2 | 1 | NA | NA | 6 | 0.5 |
| 12_7211577 | 3 | 0.3333 | 1 | 1 | 1 | 0 | NA | NA | 5 | 0.4 | 1 | 0 | 1 | 1 | 2 | 0 | 1 | 1 | 5 | 0.4 |
| 12_7211597 | 2 | 0.5 | 1 | 1 | NA | NA | NA | NA | 3 | 0.6667 | NA | NA | NA | NA | 1 | 1 | 1 | 1 | 2 | 1 |
| 12_7211604 | 3 | 0.6667 | NA | NA | 1 | 0 | NA | NA | 4 | 0.5 | 1 | 1 | 1 | 1 | 1 | 1 | 1 | 1 | 4 | 1 |
| 12_7211604 | 2 | 1 | 1 | 1 | 1 | 1 | NA | NA | 4 | 1 | 1 | 1 | 1 | 1 | 1 | 1 | NA | NA | 3 | 1 |
| 12_7211610 | 1 | 1 | 1 | 1 | NA | NA | NA | NA | 2 | 1 | 1 | 0 | NA | NA | 1 | 0 | 1 | 1 | 3 | 0.3333 |
| 12_7211622 | 2 | 0.5 | 2 | 1 | 1 | 0 | NA | NA | 5 | 0.6 | 1 | 1 | NA | NA | NA | NA | 1 | 1 | 2 | 1 |
| 12_7211630 | 1 | 1 | NA | NA | 1 | 1 | NA | NA | 2 | 1 | 1 | 0 | NA | NA | 1 | 0 | 1 | 1 | 3 | 0.3333 |
| 12_7211633 | 1 | 0 | NA | NA | 1 | 1 | NA | NA | 2 | 0.5 | 1 | 0 | NA | NA | 1 | 1 | 1 | 0 | 3 | 0.3333 |
| 12_7211637 | 1 | 1 | NA | NA | 1 | 0 | NA | NA | 2 | 0.5 | 1 | 0 | NA | NA | NA | NA | 1 | 1 | 2 | 0.5 |
| 12_7211640 | 1 | 1 | NA | NA | NA | NA | NA | NA | 1 | 1 | NA | NA | NA | NA | 1 | 1 | 1 | 1 | 2 | 1 |
| 12_7211642 | 1 | 0 | NA | NA | 1 | 1 | NA | NA | 2 | 0.5 | NA | NA | NA | NA | 2 | 0.5 | 1 | 0 | 3 | 0.3333 |
| 12_7211645 | 1 | 1 | NA | NA | 1 | 1 | NA | NA | 2 | 1 | 1 | 1 | NA | NA | 1 | 0 | 1 | 1 | 3 | 0.6667 |
| 12_7211665 | 1 | 1 | NA | NA | 1 | 1 | NA | NA | 2 | 1 | 1 | 1 | NA | NA | 2 | 1 | 1 | 1 | 4 | 1 |
| 12_7211703 | 1 | 1 | NA | NA | NA | NA | NA | NA | 1 | 1 | 1 | 1 | NA | NA | 2 | 1 | 1 | 1 | 4 | 1 |
| 12_7211708 | 1 | 0 | NA | NA | 1 | 1 | NA | NA | 2 | 0.5 | NA | NA | NA | NA | 2 | 0.5 | NA | NA | 2 | 0.5 |
| 12_7211714 | 1 | 0 | NA | NA | 1 | 1 | NA | NA | 2 | 0.5 | 1 | 0 | NA | NA | 2 | 0.5 | NA | NA | 3 | 0.3333 |
| 12_7211735 | 1 | 1 | NA | NA | 1 | 1 | NA | NA | 2 | 1 | 1 | 1 | NA | NA | 2 | 1 | NA | NA | 3 | 1 |
| 12_7211741 | NA | NA | NA | NA | 1 | 1 | NA | NA | 1 | 1 | 1 | 1 | NA | NA | 2 | 1 | NA | NA | 3 | 1 |
| 12_7214741 | 8 | 0 | 12 | 0.1667 | 8 | 0 | 10 | 0 | 38 | 0.0526 | 3 | 0 | 11 | 0.0909 | 10 | 0.1 | 7 | 0 | 31 | 0.0645 |
| 12_7214748 | 8 | 0 | 9 | 0 | 8 | 0.125 | 9 | 0.2222 | 34 | 0.0882 | 2 | 0 | 10 | 0.2 | 10 | 0 | 6 | 0 | 28 | 0.0714 |
| 12_7214759 | 8 | 0.875 | 11 | 1 | 10 | 0.8 | 12 | 0.8333 | 41 | 0.878 | 4 | 1 | 11 | 0.9091 | 11 | 0.8182 | 7 | 1 | 33 | 0.9091 |
| 12_7214777 | 10 | 0.5 | 14 | 0.7143 | 9 | 0.5556 | 9 | 0.7778 | 42 | 0.6429 | 3 | 0.6667 | 10 | 0.8 | 10 | 0.6 | 7 | 0.8571 | 30 | 0.7333 |
| 12_7214779 | 9 | 0 | 11 | 0 | 8 | 0.25 | 9 | 0 | 37 | 0.0541 | 5 | 0 | 11 | 0.2727 | 10 | 0.1 | 7 | 0 | 33 | 0.1212 |
| 12_7214793 | 8 | 0 | 12 | 0.0833 | 8 | 0.125 | 9 | 0.1111 | 37 | 0.0811 | 3 | 0 | 11 | 0.1818 | 11 | 0.1818 | 7 | 0 | 32 | 0.125 |
| 12_7214795 | 7 | 0.4286 | 10 | 0.1 | 9 | 0.4444 | 7 | 0.7143 | 33 | 0.3939 | 5 | 0.6 | 11 | 0.5455 | 8 | 0.75 | 6 | 0.5 | 30 | 0.6 |
| 12_7214814 | 6 | 0.3333 | 13 | 0.1538 | 8 | 0.125 | 7 | 0.2857 | 34 | 0.2059 | 5 | 0 | 9 | 0.1111 | 12 | 0 | 7 | 0.1429 | 33 | 0.0606 |

| | | | | | | | | | | | | | | | | | | | | |
|---|---|---|---|---|---|---|---|---|---|---|---|---|---|---|---|---|---|---|---|---|
| 12_722585 | 219 | 0.0548 | 258 | 0.0504 | 181 | 0.0994 | 204 | 0.0539 | 862 | 0.0626 | 179 | 0.067 | 212 | 0.1179 | 179 | 0.0782 | 214 | 0.1028 | 784 | 0.0931 |
| 12_723504 | 24 | 0.0417 | 31 | 0.129 | 32 | 0.1562 | 29 | 0.069 | 116 | 0.1034 | 20 | 0.15 | 31 | 0.2258 | 28 | 0.1071 | 31 | 0.129 | 110 | 0.1545 |
| 12_723507 | 33 | 0.303 | 39 | 0.2308 | 33 | 0.4242 | 36 | 0.3333 | 141 | 0.3191 | 25 | 0.36 | 39 | 0.2821 | 32 | 0.2812 | 37 | 0.3514 | 133 | 0.3158 |
| 12_723508 | 27 | 0.3333 | 34 | 0.4118 | 33 | 0.1818 | 32 | 0.4688 | 126 | 0.3492 | 18 | 0.3333 | 29 | 0.4483 | 30 | 0.2333 | 30 | 0.3333 | 107 | 0.3364 |
| 12_723509 | 30 | 0.1 | 39 | 0.0769 | 34 | 0.0882 | 34 | 0.0882 | 137 | 0.0876 | 21 | 0.0476 | 37 | 0.0811 | 30 | 0.0667 | 37 | 0.0811 | 125 | 0.072 |
| 12_723509 | 29 | 0.3448 | 36 | 0.1111 | 29 | 0.1724 | 33 | 0.2424 | 127 | 0.2126 | 21 | 0.1429 | 36 | 0.25 | 28 | 0.3571 | 35 | 0.3143 | 120 | 0.275 |
| 12_723550 | 23 | 0.3043 | 24 | 0.4167 | 21 | 0.619 | 26 | 0.3077 | 94 | 0.4043 | 17 | 0.2941 | 31 | 0.2581 | 22 | 0.3636 | 27 | 0.3333 | 97 | 0.3093 |
| 12_723550 | 19 | 0.1579 | 30 | 0.1333 | 21 | 0.2381 | 31 | 0.2258 | 101 | 0.1881 | 15 | 0.4 | 32 | 0.2188 | 25 | 0.36 | 25 | 0.32 | 97 | 0.3093 |
| 12_723551 | 22 | 0.0909 | 29 | 0.3448 | 22 | 0.3182 | 32 | 0.2188 | 105 | 0.2476 | 17 | 0.2353 | 34 | 0.1176 | 25 | 0.28 | 28 | 0.1429 | 104 | 0.1827 |
| 12_723552 | 19 | 0.4211 | 30 | 0.4667 | 22 | 0.6818 | 27 | 0.4074 | 98 | 0.4898 | 16 | 0.3125 | 28 | 0.4643 | 22 | 0.5455 | 26 | 0.5769 | 92 | 0.4891 |
| 12_723552 | 16 | 0.9375 | 22 | 0.9091 | 14 | 0.8571 | 26 | 0.8462 | 78 | 0.8846 | 9 | 0.8889 | 21 | 0.8571 | 16 | 0.6875 | 19 | 0.7895 | 65 | 0.8 |
| 12_723552 | 23 | 0.4348 | 27 | 0.3704 | 22 | 0.1818 | 32 | 0.1562 | 104 | 0.2788 | 15 | 0.3333 | 28 | 0.3214 | 21 | 0.3333 | 27 | 0.2963 | 91 | 0.3187 |
| 12_723555 | 18 | 0.1667 | 25 | 0.08 | 17 | 0.1765 | 27 | 0.037 | 87 | 0.1034 | 15 | 0.0667 | 29 | 0.1379 | 19 | 0 | 23 | 0.0435 | 86 | 0.0698 |
| 12_723555 | 19 | 0.5789 | 23 | 0.4783 | 20 | 0.3 | 24 | 0.5417 | 86 | 0.4767 | 12 | 0.5 | 28 | 0.5714 | 18 | 0.3889 | 17 | 0.5882 | 75 | 0.52 |
| 12_724558 | 1 | 1 | NA | NA | NA | NA | NA | NA | 1 | 1 | NA | NA | NA | NA | NA | NA | 1 | 1 | 1 | 1 |
| 12_724559 | 1 | 1 | NA | NA | NA | NA | NA | NA | 1 | 1 | NA | NA | NA | NA | NA | NA | 1 | 1 | 1 | 1 |
| 12_724560 | 1 | 1 | NA | NA | NA | NA | NA | NA | 1 | 1 | NA | NA | NA | NA | NA | NA | 1 | 1 | 1 | 1 |
| 12_724562 | 1 | 1 | NA | NA | NA | NA | NA | NA | 1 | 1 | NA | NA | NA | NA | NA | NA | 1 | 1 | 1 | 1 |
| 12_724565 | 1 | 1 | NA | NA | NA | NA | NA | NA | 1 | 1 | NA | NA | NA | NA | NA | NA | 1 | 1 | 1 | 1 |
| 12_725881 | 1 | 0 | 2 | 0.5 | 1 | 0 | NA | NA | 4 | 0.25 | 3 | 0 | 1 | 0 | NA | NA | 3 | 0.3333 | 7 | 0.1429 |
| 12_725882 | 1 | 1 | 2 | 0.5 | 1 | 1 | NA | NA | 4 | 0.75 | 2 | 0.5 | 1 | 1 | NA | NA | 3 | 0.6667 | 6 | 0.6667 |
| 12_725882 | 1 | 1 | 2 | 1 | 1 | 1 | NA | NA | 4 | 1 | 1 | 1 | 1 | 1 | NA | NA | 2 | 0.5 | 4 | 0.75 |
| 12_725884 | 1 | 1 | 2 | 1 | NA | NA | 1 | 0 | 4 | 0.75 | 3 | 1 | 1 | 1 | NA | NA | 3 | 1 | 7 | 1 |
| 12_725884 | NA | NA | 2 | 1 | 1 | 1 | 1 | 0 | 4 | 0.75 | 3 | 1 | 1 | 1 | NA | NA | 3 | 1 | 7 | 1 |
| 12_725885 | 1 | 0 | 2 | 0 | 1 | 1 | 2 | 0 | 6 | 0.1667 | 3 | 0 | 1 | 1 | NA | NA | 3 | 0.6667 | 7 | 0.4286 |
| 12_725886 | 1 | 1 | 1 | 0 | NA | NA | 3 | 0.6667 | 5 | 0.6 | 2 | 0.5 | 1 | 0 | NA | NA | 2 | 0 | 5 | 0.2 |
| 12_725888 | 1 | 0 | 2 | 0 | 1 | 0 | 1 | 1 | 5 | 0.2 | 2 | 0.5 | 1 | 0 | NA | NA | 1 | 0 | 4 | 0.25 |
| 12_725963 | NA | NA | NA | NA | 1 | 1 | NA | NA | 1 | 1 | NA | NA | 1 | 0 | 1 | 1 | 2 | 1 | 4 | 0.75 |
| 12_725964 | NA | NA | NA | NA | 2 | 0.5 | NA | NA | 2 | 0.5 | NA | NA | 1 | 1 | 1 | 0 | 2 | 0.5 | 4 | 0.5 |
| 12_725971 | NA | NA | NA | NA | 1 | 1 | NA | NA | 1 | 1 | NA | NA | 1 | 1 | 1 | 1 | 2 | 0.5 | 4 | 0.75 |
| 12_725972 | NA | NA | NA | NA | 2 | 0.5 | NA | NA | 2 | 0.5 | NA | NA | 1 | 1 | 1 | 1 | 2 | 1 | 4 | 1 |
| 12_725973 | NA | NA | NA | NA | 1 | 1 | NA | NA | 1 | 1 | NA | NA | 1 | 1 | 1 | 1 | 2 | 1 | 4 | 1 |
| 12_725982 | 1 | 1 | NA | NA | 2 | 0.5 | NA | NA | 3 | 0.6667 | NA | NA | 1 | 0 | NA | NA | 3 | 0.3333 | 4 | 0.25 |
| 12_726027 | NA | NA | 1 | 0 | 1 | 1 | 1 | 1 | 3 | 0.6667 | 2 | 0 | NA | NA | NA | NA | 1 | 1 | 3 | 0.3333 |
| 12_726029 | NA | NA | 2 | 0 | 1 | 1 | 2 | 0.5 | 5 | 0.4 | 3 | 1 | NA | NA | NA | NA | 1 | 0 | 4 | 0.75 |
| 12_726036 | NA | NA | 3 | 0 | 1 | 0 | 2 | 0.5 | 6 | 0.1667 | 3 | 0 | NA | NA | NA | NA | 1 | 1 | 4 | 0.25 |
| 12_726137 | NA | NA | 2 | 0.5 | NA | NA | NA | NA | 2 | 0.5 | 3 | 0.3333 | NA | NA | NA | NA | 1 | 0 | 4 | 0.25 |
| 12_726204 | 1 | 0 | 5 | 0.4 | 2 | 0.5 | 1 | 0 | 9 | 0.3333 | 3 | 0.6667 | NA | NA | NA | NA | 1 | 0 | 4 | 0.5 |
| 12_726205 | 1 | 0 | 3 | 0.3333 | 1 | 1 | 2 | 0 | 7 | 0.2857 | 2 | 0.5 | NA | NA | NA | NA | 1 | 0 | 3 | 0.3333 |
| 12_726206 | 1 | 0 | 5 | 0.2 | 2 | 0.5 | 3 | 0.6667 | 11 | 0.3636 | 3 | 0.3333 | NA | NA | NA | NA | 1 | 1 | 4 | 0.5 |
| 12_726206 | 1 | 1 | 4 | 0.75 | 1 | 1 | 3 | 0.6667 | 9 | 0.7778 | 3 | 0.6667 | NA | NA | NA | NA | 1 | 1 | 4 | 0.75 |
| 12_726208 | 1 | 1 | 3 | 1 | 1 | 1 | 4 | 0.75 | 9 | 0.8889 | 1 | 1 | 1 | 1 | NA | NA | 1 | 1 | 3 | 1 |
| 12_726208 | 1 | 0 | 4 | 0.25 | 2 | 0 | 1 | 0 | 8 | 0.125 | 2 | 0 | 2 | 0 | NA | NA | 1 | 1 | 5 | 0.2 |
| 12_726210 | 1 | 1 | 5 | 0.4 | 2 | 1 | 3 | 0.6667 | 11 | 0.6364 | 3 | 0.3333 | 2 | 1 | NA | NA | 1 | 1 | 6 | 0.6667 |
| 12_726211 | 1 | 1 | 5 | 1 | 2 | 0.5 | 3 | 1 | 11 | 0.9091 | 3 | 0.6667 | 2 | 0.5 | NA | NA | 1 | 1 | 6 | 0.6667 |
| 12_726211 | 1 | 0 | 5 | 0.4 | 2 | 0.5 | 1 | 0 | 9 | 0.3333 | 3 | 0.3333 | 1 | 1 | NA | NA | 1 | 0 | 5 | 0.4 |
| 12_726261 | 4 | 0 | 3 | 0.3333 | 2 | 1 | 3 | 0.3333 | 12 | 0.3333 | 2 | 0 | 2 | 0.5 | NA | NA | 1 | 0 | 5 | 0.2 |
| 12_726264 | 4 | 0.25 | 4 | 0.5 | NA | NA | 1 | 1 | 9 | 0.4444 | 3 | 0.3333 | 2 | 1 | NA | NA | 2 | 0 | 7 | 0.4286 |
| 12_726264 | 2 | 0.5 | 4 | 0.75 | 2 | 0.5 | 2 | 0 | 10 | 0.5 | 1 | 1 | 1 | 1 | NA | NA | 2 | 0 | 4 | 0.5 |
| 12_728306 | 15 | 1 | 13 | 1 | 19 | 0.9474 | 8 | 1 | 55 | 0.9818 | 10 | 0.9 | 10 | 1 | 17 | 1 | 5 | 1 | 42 | 0.9762 |
| 12_728306 | 13 | 0.1538 | 14 | 0.2857 | 17 | 0.1176 | 8 | 0.25 | 52 | 0.1923 | 12 | 0.25 | 10 | 0.1 | 17 | 0.1765 | 7 | 0 | 46 | 0.1522 |
| 12_728307 | 15 | 0 | 14 | 0.1429 | 18 | 0.0556 | 9 | 0 | 56 | 0.0536 | 12 | 0 | 8 | 0 | 16 | 0.125 | 9 | 0.1111 | 45 | 0.0667 |
| 12_728308 | 14 | 0.2143 | 12 | 0.4167 | 16 | 0.5 | 10 | 0.5 | 52 | 0.4038 | 12 | 0.5833 | 11 | 0.4545 | 15 | 0.4 | 9 | 0.4444 | 47 | 0.4681 |
| 12_728309 | 10 | 1 | 9 | 0.7778 | 11 | 0.9091 | 5 | 0.4 | 35 | 0.8286 | 8 | 0.625 | 7 | 0.7143 | 8 | 0.875 | 5 | 1 | 28 | 0.7857 |
| 12_728309 | 16 | 0.4375 | 12 | 0.6667 | 18 | 0.6667 | 10 | 0.7 | 56 | 0.6071 | 13 | 0.6154 | 10 | 0.6 | 15 | 0.6 | 7 | 0.5714 | 45 | 0.6 |
| 12_728309 | 19 | 0 | 13 | 0.0769 | 19 | 0.1053 | 14 | 0.0714 | 65 | 0.0615 | 13 | 0.1538 | 13 | 0 | 18 | 0.1667 | 9 | 0 | 53 | 0.0943 |
| 12_728312 | 20 | 0.2 | 14 | 0.1429 | 20 | 0.3 | 14 | 0.2857 | 68 | 0.2353 | 14 | 0.5 | 15 | 0.2667 | 17 | 0.1176 | 8 | 0.25 | 54 | 0.2778 |
| 12_728337 | NA | NA | 1 | 1 | 5 | 0.4 | NA | NA | 6 | 0.5 | 2 | 0.5 | 3 | 0.6667 | 3 | 0.6667 | NA | NA | 8 | 0.625 |
| 12_728352 | NA | NA | NA | NA | 6 | 0.1667 | NA | NA | 6 | 0.1667 | 2 | 0 | 4 | 0 | 2 | 0 | 1 | 1 | 9 | 0.1111 |
| 12_728405 | 13 | 0.1538 | 16 | 0.0625 | 8 | 0.25 | 11 | 0.1818 | 48 | 0.1458 | 10 | 0 | 8 | 0.25 | 13 | 0.2308 | 6 | 0 | 37 | 0.1351 |
| 12_728406 | 22 | 0.1364 | 15 | 0.2 | 16 | 0.0625 | 19 | 0.1053 | 72 | 0.125 | 15 | 0.2 | 13 | 0 | 18 | 0.0556 | 9 | 0.2222 | 55 | 0.1091 |
| 12_728406 | 22 | 0.3636 | 17 | 0.1765 | 17 | 0.0588 | 20 | 0.15 | 76 | 0.1974 | 14 | 0.2857 | 16 | 0.1875 | 20 | 0.15 | 10 | 0.2 | 60 | 0.2 |
| 12_728408 | 23 | 0.5652 | 18 | 0.4444 | 11 | 0.4545 | 18 | 0.5 | 70 | 0.5 | 16 | 0.5625 | 13 | 0.5385 | 18 | 0.5 | 9 | 0.4444 | 56 | 0.5179 |
| 12_728409 | 21 | 0.7619 | 16 | 0.6875 | 14 | 0.8571 | 21 | 0.7143 | 72 | 0.75 | 18 | 0.8889 | 10 | 0.7 | 14 | 0.7857 | 11 | 0.5455 | 53 | 0.7547 |
| 12_728410 | 25 | 0.32 | 20 | 0.2 | 16 | 0.25 | 25 | 0.16 | 86 | 0.2326 | 20 | 0.25 | 15 | 0.0667 | 21 | 0.1905 | 10 | 0.3 | 66 | 0.197 |

| | | | | | | | | | | | | | | | | | | | | |
|---|---|---|---|---|---|---|---|---|---|---|---|---|---|---|---|---|---|---|---|---|
| 12_7284103 | 23 | 0.1739 | 22 | 0.1818 | 13 | 0 | 25 | 0.32 | 83 | 0.1928 | 18 | 0.1111 | 14 | 0.3571 | 20 | 0.35 | 10 | 0.2 | 62 | 0.2581 |
| 12_7284261 | 2 | 0.5 | 3 | 0.3333 | 2 | 0 | 2 | 0 | 9 | 0.2222 | 1 | 0 | 1 | 0 | 2 | 1 | 1 | 0 | 5 | 0.4 |
| 12_7284391 | 2 | 0 | 4 | 0 | 3 | 0.3333 | 2 | 0 | 11 | 0.0909 | 2 | 0 | 3 | 0.3333 | 3 | 0 | 1 | 0 | 9 | 0.1111 |
| 12_7331945 | 5 | 0.2 | 5 | 0 | 4 | 0 | 3 | 0 | 17 | 0.0588 | 1 | 1 | 1 | 0 | 5 | 0 | 1 | 0 | 8 | 0.125 |
| 12_7331953 | 6 | 0 | 5 | 0.2 | 5 | 0 | 3 | 0 | 19 | 0.0526 | 1 | 0 | 1 | 0 | 4 | 0.25 | 2 | 0 | 8 | 0.125 |
| 12_7332806 | 2 | 1 | NA | NA | 1 | 1 | 3 | 1 | 6 | 1 | 1 | 1 | 1 | 1 | 1 | 1 | NA | NA | 3 | 1 |
| 12_7332815 | 2 | 0.5 | NA | NA | 1 | 0 | 3 | 0.3333 | 6 | 0.3333 | 1 | 1 | 2 | 0.5 | 1 | 1 | NA | NA | 4 | 0.75 |
| 12_7332820 | 2 | 1 | NA | NA | 1 | 0 | 2 | 0 | 5 | 0.4 | 1 | 1 | 2 | 0.5 | 1 | 1 | NA | NA | 4 | 0.75 |
| 12_7332826 | 2 | 1 | NA | NA | NA | NA | 1 | 1 | 3 | 1 | 1 | 1 | 2 | 1 | 1 | 0 | NA | NA | 4 | 0.75 |
| 12_7332834 | 2 | 0 | NA | NA | 1 | 0 | 3 | 0.3333 | 6 | 0.1667 | 1 | 1 | 3 | 0 | 1 | 1 | NA | NA | 5 | 0.4 |
| 12_7333155 | 2 | 0 | NA | NA | 1 | 1 | 3 | 0.3333 | 6 | 0.3333 | 1 | 1 | 2 | 0.5 | 1 | 1 | NA | NA | 4 | 0.75 |
| 12_7333159 | 2 | 1 | NA | NA | 1 | 1 | 2 | 0.5 | 5 | 0.8 | 1 | 1 | 2 | 1 | 1 | 1 | NA | NA | 4 | 1 |
| 12_7333161 | 2 | 0.5 | NA | NA | 1 | 0 | 3 | 0.3333 | 6 | 0.3333 | 1 | 0 | 1 | 1 | 1 | 1 | NA | NA | 3 | 0.6667 |
| 12_7333165 | 2 | 0.5 | NA | NA | 1 | 0 | 3 | 0.3333 | 6 | 0.3333 | 1 | 1 | 1 | 0 | 1 | 1 | NA | NA | 3 | 0.6667 |
| 12_7333182 | 2 | 0 | NA | NA | 1 | 0 | 3 | 0.3333 | 6 | 0.1667 | NA | NA | 1 | 1 | 1 | 0 | NA | NA | 2 | 0.5 |
| 12_7333190 | 1 | 1 | NA | NA | 1 | 1 | 3 | 1 | 5 | 1 | 1 | 1 | 2 | 1 | 1 | 1 | NA | NA | 4 | 1 |
| 12_7333201 | 2 | 0 | NA | NA | 1 | 0 | 3 | 0.3333 | 6 | 0.1667 | 1 | 0 | 2 | 0.5 | 1 | 0 | NA | NA | 4 | 0.25 |
| 12_7363186 | 1 | 1 | 1 | 0 | 1 | 0 | 2 | 1 | 5 | 0.6 | NA | NA | 3 | 1 | 2 | 0 | 1 | 0 | 6 | 0.5 |
| 12_7380640 | 3 | 0 | 2 | 0 | 6 | 0 | 6 | 0.3333 | 17 | 0.1176 | 2 | 0.5 | 10 | 0 | 4 | 0.25 | 4 | 0 | 20 | 0.1 |
| 12_7382376 | 2 | 0.5 | 3 | 0.3333 | 8 | 0.75 | 6 | 0.6667 | 19 | 0.6316 | 2 | 1 | 10 | 0.7 | 4 | 0.25 | 4 | 0.75 | 20 | 0.65 |
| 12_7382379 | 2 | 0.5 | 3 | 0.3333 | 8 | 0.375 | 6 | 1 | 19 | 0.5789 | 1 | 0 | 10 | 0.5 | 4 | 0.25 | 3 | 0.6667 | 18 | 0.4444 |
| 12_7382385 | 1 | 0 | 2 | 0 | 8 | 0 | 5 | 0.4 | 16 | 0.125 | 2 | 0.5 | 7 | 0 | 4 | 0 | 4 | 0.5 | 17 | 0.1765 |
| 12_7382391 | 2 | 0 | 3 | 0 | 7 | 0.1429 | 4 | 0 | 16 | 0.0625 | 3 | 0 | 8 | 0.125 | 3 | 0 | 4 | 0.25 | 18 | 0.1111 |
| 12_7382410 | 3 | 1 | 2 | 0.5 | 6 | 0.8333 | 6 | 0.8333 | 17 | 0.8235 | 3 | 0.3333 | 8 | 0.875 | 3 | 0.6667 | 4 | 0.75 | 18 | 0.7222 |
| 12_7382419 | 3 | 1 | 3 | 0.6667 | 7 | 0.7143 | 8 | 0.5 | 21 | 0.6667 | 2 | 0.5 | 10 | 0.3 | 4 | 0.25 | 4 | 0.5 | 20 | 0.35 |
| 12_7382433 | 1 | 0 | 1 | 1 | 7 | 0.4286 | 8 | 0.5 | 17 | 0.4706 | 3 | 0.3333 | 8 | 0.25 | 4 | 0 | 1 | 0 | 16 | 0.1875 |
| 12_7382439 | 3 | 0 | 3 | 0 | 10 | 0.5 | 10 | 0.3 | 26 | 0.3077 | 3 | 0.6667 | 10 | 0.2 | 3 | 0 | 4 | 0.5 | 20 | 0.3 |
| 12_7382467 | 2 | 1 | 3 | 0.6667 | 9 | 1 | 9 | 0.8889 | 23 | 0.913 | 3 | 1 | 11 | 0.9091 | 4 | 1 | 2 | 1 | 20 | 0.95 |
| 12_7382473 | 2 | 1 | 2 | 0 | 8 | 0.5 | 10 | 0.8 | 22 | 0.6364 | 3 | 0.6667 | 10 | 0.8 | 4 | 0.25 | 4 | 0.5 | 21 | 0.619 |
| 12_7382475 | 1 | 0 | 3 | 0.3333 | 10 | 0.3 | 6 | 0.3333 | 20 | 0.3 | 2 | 0 | 10 | 0.1 | 5 | 0 | 4 | 0.5 | 21 | 0.1429 |
| 12_7382480 | 2 | 1 | 2 | 0.5 | 9 | 0.3333 | 12 | 0.3333 | 25 | 0.4 | 3 | 0.3333 | 11 | 0.4545 | 5 | 0.2 | 4 | 0.25 | 23 | 0.3478 |
| 12_7395496 | 4 | 0.25 | 5 | 0.2 | 1 | 0 | 7 | 0.4286 | 17 | 0.2941 | 1 | 0 | 4 | 0.5 | 4 | 0.5 | NA | NA | 9 | 0.4444 |
| 12_7398732 | 8 | 0.125 | 8 | 0.125 | 2 | 0.5 | 10 | 0.3 | 28 | 0.2143 | 6 | 0.5 | 12 | 0.3333 | 9 | 0.4444 | 4 | 0.25 | 31 | 0.3871 |
| 12_7398873 | 5 | 0 | 7 | 0.2857 | 3 | 0 | 9 | 0.1111 | 24 | 0.125 | 5 | 0 | 11 | 0.1818 | 8 | 0 | 2 | 0 | 26 | 0.0769 |
| 12_7398895 | 7 | 0.2857 | 9 | 0.3333 | 4 | 0 | 11 | 0.3636 | 31 | 0.2903 | 8 | 0.5 | 12 | 0.6667 | 9 | 0.6667 | 6 | 0.5 | 35 | 0.6 |
| 12_7398906 | 7 | 0.1429 | 7 | 0.2857 | 5 | 0 | 8 | 0 | 27 | 0.1111 | 7 | 0.1429 | 12 | 0 | 7 | 0 | 5 | 0.4 | 31 | 0.0968 |
| 12_7399256 | 9 | 0 | 6 | 0.3333 | 4 | 0 | 11 | 0.1818 | 30 | 0.1333 | 7 | 0.1429 | 13 | 0.0769 | 9 | 0 | 4 | 0 | 33 | 0.0606 |
| 12_7399335 | 17 | 0.0588 | 13 | 0.0769 | 8 | 0.125 | 15 | 0.1333 | 53 | 0.0943 | 9 | 0.2222 | 15 | 0.1333 | 10 | 0.2 | 10 | 0.2 | 44 | 0.1818 |
| 12_7401138 | 3 | 0 | 1 | 1 | 2 | 0.5 | 2 | 0 | 8 | 0.25 | 2 | 0 | 2 | 0 | 1 | 1 | 3 | 0 | 8 | 0.125 |
| 12_7403075 | 5 | 0.2 | 4 | 0.25 | 3 | 0.3333 | 2 | 0 | 14 | 0.2143 | 3 | 0.6667 | 5 | 0 | 2 | 0 | 3 | 0 | 13 | 0.1538 |
| 12_7403561 | 3 | 0 | 5 | 0.2 | 5 | 0 | 2 | 0.5 | 15 | 0.1333 | NA | NA | 3 | 0 | 1 | 0 | 1 | 1 | 5 | 0.2 |
| 12_7420887 | 1 | 0 | 5 | 0 | 3 | 0.3333 | 3 | 0 | 12 | 0.0833 | NA | NA | 1 | 0 | 4 | 0.25 | 4 | 0 | 9 | 0.1111 |
| 12_7420891 | 1 | 0 | 4 | 0.25 | 3 | 0.3333 | 4 | 0.25 | 12 | 0.25 | NA | NA | 1 | 0 | 4 | 0.25 | 3 | 0.3333 | 8 | 0.25 |
| 12_7420892 | 1 | 0 | 5 | 0.8 | 3 | 0.3333 | 2 | 0.5 | 11 | 0.5455 | NA | NA | 1 | 1 | 4 | 1 | 4 | 0.25 | 9 | 0.6667 |
| 12_7420904 | 1 | 1 | 4 | 0.75 | 3 | 0.6667 | 4 | 1 | 12 | 0.8333 | NA | NA | 1 | 1 | 5 | 0.8 | 4 | 1 | 10 | 0.9 |
| 12_7420911 | 1 | 1 | 4 | 0.75 | 3 | 1 | 4 | 1 | 12 | 0.9167 | NA | NA | 1 | 1 | 5 | 1 | 3 | 1 | 9 | 1 |
| 12_7420917 | 1 | 1 | 2 | 0 | 2 | 0.5 | 4 | 0.5 | 9 | 0.4444 | NA | NA | 1 | 1 | 6 | 0.6667 | 4 | 0.5 | 11 | 0.6364 |
| 12_7420930 | 1 | 1 | 4 | 1 | 3 | 0.6667 | 4 | 0.75 | 12 | 0.8333 | NA | NA | 1 | 1 | 6 | 0.6667 | 2 | 1 | 9 | 0.7778 |
| 12_7420936 | 1 | 0 | 5 | 0.2 | 3 | 0 | 4 | 0 | 13 | 0.0769 | NA | NA | 1 | 0 | 6 | 0.1667 | 3 | 0.3333 | 10 | 0.2 |
| 12_7420940 | 1 | 0 | 4 | 0.25 | 3 | 0.6667 | 4 | 0.25 | 12 | 0.3333 | NA | NA | 1 | 0 | 5 | 0 | 4 | 0.25 | 10 | 0.1 |
| 12_7420944 | 1 | 0 | 5 | 0.8 | 4 | 0.5 | 2 | 0 | 12 | 0.5 | NA | NA | 1 | 1 | 6 | 0.3333 | 4 | 0.5 | 11 | 0.4545 |
| 12_7428173 | 3 | 0.3333 | 9 | 0.4444 | 4 | 0 | 7 | 0.4286 | 23 | 0.3478 | 3 | 0.3333 | 5 | 0.4 | 7 | 0.5714 | 8 | 0.375 | 23 | 0.4348 |
| 12_7428505 | 3 | 0 | 10 | 0.1 | 5 | 0.4 | 7 | 0 | 25 | 0.12 | 4 | 0 | 4 | 0.25 | 7 | 0.4286 | 8 | 0 | 23 | 0.1739 |
| 12_7431351 | 3 | 1 | 11 | 0.6364 | 6 | 0.6667 | 9 | 0.6667 | 29 | 0.6897 | 5 | 0.6 | 4 | 0.75 | 8 | 0.75 | 8 | 0.875 | 25 | 0.76 |
| 12_7431363 | 3 | 0 | 9 | 0.1111 | 4 | 0 | 6 | 0.1667 | 22 | 0.0909 | 1 | 0 | 3 | 0 | 7 | 0.2857 | 8 | 0.25 | 19 | 0.2105 |
| 12_7431365 | 2 | 0 | 9 | 0.3333 | 5 | 0.4 | 7 | 0.4286 | 23 | 0.3478 | 5 | 0.4 | 5 | 0.6 | 8 | 0.75 | 11 | 0.4545 | 29 | 0.5517 |
| 12_7431369 | 3 | 0 | 11 | 0 | 6 | 0 | 9 | 0.2222 | 29 | 0.069 | 5 | 0.2 | 6 | 0 | 8 | 0.125 | 11 | 0 | 30 | 0.0667 |
| 12_7431381 | 1 | 0 | 7 | 0.5714 | 6 | 0.5 | 4 | 0.75 | 18 | 0.5556 | 4 | 0.5 | 3 | 0.3333 | 3 | 1 | 8 | 0.5 | 18 | 0.5556 |
| 12_7431381 | 2 | 0 | 9 | 0 | 6 | 0 | 9 | 0.2222 | 26 | 0.0769 | 6 | 0 | 6 | 0.1667 | 9 | 0.1111 | 9 | 0.1111 | 30 | 0.1 |
| 12_7431384 | 2 | 1 | 12 | 0.9167 | 6 | 1 | 10 | 1 | 30 | 0.9667 | 6 | 0.8333 | 6 | 1 | 8 | 1 | 10 | 1 | 30 | 0.9667 |
| 12_7431408 | 3 | 0.3333 | 13 | 0.4615 | 6 | 0.5 | 9 | 0.3333 | 31 | 0.4194 | 6 | 0.3333 | 7 | 0.2857 | 9 | 0.4444 | 10 | 0.4 | 32 | 0.375 |
| 12_7431436 | 4 | 0.25 | 13 | 0.3077 | 3 | 0 | 10 | 0.5 | 30 | 0.3333 | 4 | 0.25 | 7 | 0.4286 | 9 | 0.3333 | 7 | 0.4286 | 27 | 0.3704 |
| 12_7434235 | 6 | 0.5 | 8 | 0.75 | 2 | 1 | 9 | 0.5556 | 25 | 0.64 | 6 | 0.1667 | 6 | 0.5 | 4 | 1 | 9 | 0.5556 | 25 | 0.52 |
| 12_7434243 | 3 | 0 | 7 | 0 | 1 | 0 | 9 | 0.2222 | 20 | 0.1 | 4 | 0.25 | 6 | 0 | 5 | 0.2 | 7 | 0 | 22 | 0.0909 |
| 12_7434251 | 4 | 1 | 8 | 0.875 | 2 | 1 | 8 | 1 | 22 | 0.9545 | 5 | 1 | 6 | 0.8333 | 5 | 1 | 8 | 0.875 | 24 | 0.9167 |

| | | | | | | | | | | | | | | | | | | | | |
|---|---|---|---|---|---|---|---|---|---|---|---|---|---|---|---|---|---|---|---|---|
| 12_743427 | 3 | 0.3333 | 10 | 0.5 | 2 | 1 | 8 | 0.375 | 23 | 0.4783 | 5 | 0.4 | 4 | 0 | 5 | 0.4 | 7 | 0.2857 | 21 | 0.2857 |
| 12_743427 | 5 | 0.8 | 9 | 0.6667 | 1 | 0 | 4 | 0.75 | 19 | 0.6842 | 4 | 0.75 | 4 | 0.5 | 2 | 1 | 5 | 0.4 | 15 | 0.6 |
| 12_743428 | 4 | 0.25 | 10 | 0 | 1 | 0 | 5 | 0 | 20 | 0.05 | 4 | 0 | 4 | 0 | 5 | 0.2 | 9 | 0.1111 | 22 | 0.0909 |
| 12_743429 | 4 | 0.5 | 9 | 0.3333 | 2 | 0.5 | 4 | 0.5 | 19 | 0.4211 | 4 | 0.5 | 3 | 0.3333 | 5 | 0.6 | 9 | 0.2222 | 21 | 0.381 |
| 12_743455 | 2 | 0 | 4 | 0.25 | 1 | 0 | 1 | 0 | 8 | 0.125 | 1 | 0 | 2 | 0 | 1 | 0 | 2 | 0.5 | 6 | 0.1667 |
| 12_743461 | 2 | 0.5 | 7 | 0.4286 | 1 | 0 | 3 | 0 | 13 | 0.3077 | 1 | 0 | 2 | 0 | 1 | 0 | 3 | 0.3333 | 7 | 0.1429 |
| 12_743463 | 2 | 0.5 | 7 | 0 | 1 | 0 | 3 | 0 | 13 | 0.0769 | 1 | 1 | 2 | 0 | 2 | 0 | 4 | 0.5 | 9 | 0.3333 |
| 12_743470 | 1 | 0 | 6 | 0.1667 | 2 | 0 | 2 | 0.5 | 11 | 0.1818 | 1 | 0 | 3 | 0.3333 | 2 | 0 | 5 | 0 | 11 | 0.0909 |
| 12_743475 | 2 | 0.5 | 5 | 0 | 3 | 0.3333 | 3 | 0.3333 | 13 | 0.2308 | 2 | 0.5 | 4 | 0.25 | 3 | 0 | 6 | 0.5 | 15 | 0.3333 |
| 12_749343 | 5 | 0 | 8 | 0.125 | 12 | 0.0833 | 6 | 0.1667 | 31 | 0.0968 | 4 | 0 | 2 | 0.5 | 7 | 0.1429 | 6 | 0 | 19 | 0.1053 |
| 12_749353 | 4 | 0 | 5 | 0.2 | 7 | 0 | 6 | 0.1667 | 22 | 0.0909 | 2 | 0.5 | 1 | 0 | 6 | 0 | 4 | 0 | 13 | 0.0769 |
| 12_749438 | 1 | 0 | 2 | 0.5 | NA | NA | NA | NA | 3 | 0.3333 | NA | NA | 2 | 0.5 | 1 | 0 | 1 | 1 | 4 | 0.5 |
| 12_749441 | 1 | 1 | 2 | 1 | NA | NA | NA | NA | 3 | 1 | 2 | 1 | 2 | 1 | 2 | 0.5 | 1 | 1 | 7 | 0.8571 |
| 12_749441 | 1 | 0 | 1 | 1 | NA | NA | NA | NA | 2 | 0.5 | 2 | 0 | 2 | 0.5 | 2 | 0.5 | 1 | 1 | 7 | 0.4286 |
| 12_749444 | NA | NA | 2 | 0.5 | NA | NA | NA | NA | 2 | 0.5 | NA | NA | NA | NA | 2 | 0.5 | 1 | 0 | 3 | 0.3333 |
| 12_749444 | 1 | 1 | 2 | 1 | NA | NA | NA | NA | 3 | 1 | 2 | 0.5 | 1 | 0 | 1 | 1 | 1 | 1 | 5 | 0.6 |
| 12_749445 | 1 | 1 | 2 | 1 | NA | NA | NA | NA | 3 | 1 | 2 | 1 | NA | NA | 1 | 1 | NA | NA | 3 | 1 |
| 12_749446 | NA | NA | 3 | 1 | NA | NA | NA | NA | 3 | 1 | 2 | 0 | 1 | 0 | 1 | 1 | 1 | 0 | 5 | 0.2 |
| 12_749446 | 1 | 1 | 2 | 1 | NA | NA | NA | NA | 3 | 1 | 1 | 1 | 1 | 0 | 2 | 1 | 1 | 0 | 5 | 0.6 |
| 12_749446 | 1 | 0 | 2 | 1 | NA | NA | NA | NA | 3 | 0.6667 | 1 | 1 | NA | NA | 2 | 1 | 1 | 1 | 4 | 1 |
| 12_749447 | 1 | 1 | 3 | 0.3333 | NA | NA | NA | NA | 4 | 0.5 | 1 | 1 | 1 | 1 | 2 | 1 | 1 | 1 | 5 | 1 |
| 12_749447 | 1 | 0 | 3 | 0.3333 | NA | NA | NA | NA | 4 | 0.25 | 2 | 0.5 | 1 | 1 | 2 | 1 | 1 | 0 | 6 | 0.6667 |
| 12_749447 | 1 | 0 | 2 | 1 | NA | NA | NA | NA | 3 | 0.6667 | 2 | 0 | 1 | 0 | 1 | 0 | 1 | 1 | 5 | 0.2 |
| 12_749448 | 1 | 1 | 3 | 0 | NA | NA | NA | NA | 4 | 0.25 | 2 | 0.5 | 1 | 1 | 1 | 0 | 1 | 1 | 5 | 0.6 |
| 12_749448 | 1 | 0 | 2 | 0.5 | NA | NA | NA | NA | 3 | 0.3333 | 2 | 1 | 1 | 0 | 2 | 1 | NA | NA | 5 | 0.8 |
| 12_749448 | 1 | 1 | 1 | 1 | NA | NA | NA | NA | 2 | 1 | NA | NA | 1 | 0 | 1 | 1 | 1 | 1 | 3 | 0.6667 |
| 12_749450 | 1 | 1 | NA | NA | NA | NA | NA | NA | 1 | 1 | 2 | 1 | 1 | 0 | 2 | 0 | 1 | 1 | 6 | 0.5 |
| 12_751343 | 1 | 1 | 2 | 1 | NA | NA | NA | NA | 3 | 1 | 2 | 1 | NA | NA | 1 | 1 | NA | NA | 3 | 1 |
| 12_751343 | 1 | 0 | 3 | 0.6667 | NA | NA | NA | NA | 4 | 0.5 | 2 | 0.5 | NA | NA | NA | NA | NA | NA | 2 | 0.5 |
| 12_751343 | 1 | 0 | 2 | 0.5 | NA | NA | NA | NA | 3 | 0.3333 | 2 | 1 | NA | NA | 1 | 0 | NA | NA | 3 | 0.6667 |
| 12_751610 | 1 | 0 | 3 | 0.6667 | NA | NA | NA | NA | 4 | 0.5 | 2 | 0.5 | NA | NA | 1 | 1 | NA | NA | 3 | 0.6667 |
| 12_751611 | 1 | 1 | 2 | 1 | NA | NA | NA | NA | 3 | 1 | 2 | 1 | NA | NA | 1 | 1 | NA | NA | 3 | 1 |
| 12_751611 | 1 | 1 | 3 | 1 | NA | NA | NA | NA | 4 | 1 | 2 | 1 | NA | NA | NA | NA | NA | NA | 2 | 1 |
| 12_751611 | 1 | 0 | 2 | 0.5 | NA | NA | NA | NA | 3 | 0.3333 | 2 | 0.5 | NA | NA | NA | NA | NA | NA | 2 | 0.5 |
| 12_752238 | 1 | 1 | NA | NA | NA | NA | NA | NA | 1 | 1 | 1 | 1 | NA | NA | NA | NA | NA | NA | 1 | 1 |
| 12_754473 | 1 | 1 | NA | NA | NA | NA | NA | NA | 1 | 1 | NA | NA | 1 | 1 | NA | NA | NA | NA | 1 | 1 |
| 12_754476 | 1 | 1 | NA | NA | NA | NA | NA | NA | 1 | 1 | NA | NA | 1 | 1 | NA | NA | NA | NA | 1 | 1 |
| 12_754477 | 1 | 1 | NA | NA | NA | NA | NA | NA | 1 | 1 | NA | NA | 1 | 1 | NA | NA | NA | NA | 1 | 1 |
| 12_754965 | 4 | 0 | 2 | 0 | 2 | 0 | 6 | 0.5 | 14 | 0.2143 | 1 | 0 | 5 | 0.2 | 1 | 0 | 1 | 0 | 8 | 0.125 |
| 12_754971 | 6 | 0.5 | 6 | 1 | 6 | 0.5 | 8 | 0.75 | 26 | 0.6923 | 2 | 1 | 7 | 0.8571 | 5 | 0.8 | 6 | 0.6667 | 20 | 0.8 |
| 12_754977 | 6 | 0 | 7 | 0.1429 | 6 | 0.1667 | 10 | 0.1 | 29 | 0.1034 | 2 | 0 | 11 | 0.2727 | 5 | 0 | 7 | 0 | 25 | 0.12 |
| 12_754981 | 5 | 0 | 2 | 0 | 4 | 0.25 | 8 | 0.125 | 19 | 0.1053 | 2 | 0 | 7 | 0.1429 | 5 | 0 | 3 | 0 | 17 | 0.0588 |
| 12_754995 | 9 | 0.3333 | 13 | 0.2308 | 7 | 0.2857 | 11 | 0.4545 | 40 | 0.325 | 3 | 0 | 12 | 0.1667 | 5 | 0.4 | 10 | 0.3 | 30 | 0.2333 |
| 12_754996 | 10 | 0 | 13 | 0 | 8 | 0.25 | 12 | 0.1667 | 43 | 0.093 | 5 | 0 | 15 | 0 | 5 | 0 | 10 | 0.2 | 35 | 0.0571 |
| 12_754998 | 6 | 0.3333 | 10 | 0.3 | 6 | 0.1667 | 10 | 0.5 | 32 | 0.3438 | 5 | 0.4 | 13 | 0.1538 | 2 | 0 | 9 | 0 | 29 | 0.1379 |
| 12_755002 | 9 | 0.4444 | 9 | 0.6667 | 7 | 0.2857 | 10 | 0.8 | 35 | 0.5714 | 5 | 0.2 | 10 | 0.3 | 4 | 0.25 | 8 | 0.25 | 27 | 0.2593 |
| 12_755002 | 9 | 1 | 10 | 0.9 | 8 | 1 | 10 | 1 | 37 | 0.973 | 3 | 1 | 10 | 0.8 | 4 | 1 | 7 | 0.8571 | 24 | 0.875 |
| 12_755003 | 8 | 0.5 | 8 | 0.5 | 7 | 0.4286 | 9 | 0.5556 | 32 | 0.5 | 5 | 0.4 | 10 | 0.7 | 2 | 1 | 7 | 0.5714 | 24 | 0.625 |
| 12_755003 | 8 | 0.75 | 10 | 0.8 | 7 | 0.7143 | 11 | 0.8182 | 36 | 0.7778 | 5 | 0.8 | 9 | 0.8889 | 3 | 0.6667 | 8 | 0.75 | 25 | 0.8 |
| 12_755623 | 7 | 0 | 7 | 0 | 5 | 0 | 8 | 0.25 | 27 | 0.0741 | 3 | 0 | 8 | 0 | 4 | 0 | 4 | 0.25 | 19 | 0.0526 |
| 12_755624 | 6 | 0.5 | 5 | 0.2 | 3 | 0 | 6 | 0.8333 | 20 | 0.45 | 3 | 1 | 6 | 0.1667 | 3 | 0.6667 | 4 | 0.5 | 16 | 0.5 |
| 12_755624 | 4 | 0.25 | 5 | 0 | 5 | 0.8 | 7 | 0.7143 | 21 | 0.4762 | 3 | 0.6667 | 8 | 0.25 | 4 | 0 | 4 | 0.5 | 19 | 0.3158 |
| 12_755624 | 7 | 0.5714 | 5 | 0 | 5 | 0.4 | 7 | 0.7143 | 24 | 0.4583 | 4 | 0.5 | 7 | 0.1429 | 3 | 1 | 4 | 0.25 | 18 | 0.3889 |
| 12_755625 | 6 | 0.1667 | 5 | 0.2 | 5 | 0 | 8 | 0.125 | 24 | 0.125 | 3 | 0.3333 | 8 | 0 | 3 | 0 | 4 | 0.25 | 18 | 0.1111 |
| 12_756425 | 6 | 0.1667 | 2 | 0 | 3 | 0 | 6 | 0 | 17 | 0.0588 | 1 | 0 | 4 | 0.25 | 2 | 0 | 2 | 0 | 9 | 0.1111 |
| 12_757623 | 4 | 0 | 3 | 0 | 7 | 0.2857 | 4 | 0.25 | 18 | 0.1667 | 4 | 0 | 7 | 0.2857 | 9 | 0.1111 | 7 | 0.1429 | 27 | 0.1481 |
| 12_757630 | 3 | 0.3333 | 3 | 0.3333 | 6 | 0.5 | 5 | 0.2 | 17 | 0.3529 | 2 | 0 | 7 | 0.1429 | 7 | 0.1429 | 9 | 0.1111 | 25 | 0.12 |
| 12_757631 | 1 | 0 | 2 | 0 | 7 | 0.7143 | 3 | 1 | 13 | 0.6154 | 2 | 0.5 | 6 | 0.6667 | 5 | 0.6 | 7 | 0.1429 | 20 | 0.45 |
| 12_757631 | 2 | 0.5 | 2 | 0.5 | 7 | 0.8571 | 5 | 0.6 | 16 | 0.6875 | 2 | 0 | 7 | 0.7143 | 6 | 0.1667 | 8 | 0.625 | 23 | 0.4783 |
| 12_757633 | 4 | 0.75 | 2 | 1 | 5 | 0.8 | 5 | 0.6 | 16 | 0.75 | 2 | 0 | 7 | 0.4286 | 6 | 0.5 | 8 | 0.5 | 23 | 0.4348 |
| 12_757818 | 3 | 1 | 1 | 0 | 4 | 0.75 | 5 | 0.4 | 13 | 0.6154 | NA | NA | 7 | 0.2857 | 4 | 0.25 | 5 | 0.4 | 16 | 0.3125 |
| 12_757819 | 1 | 0 | 1 | 1 | 5 | 0.6 | 4 | 0.25 | 11 | 0.4545 | 1 | 1 | 2 | 0 | 3 | 0 | 4 | 0 | 10 | 0.1 |
| 12_757821 | 2 | 0.5 | 1 | 0 | 6 | 0.3333 | 4 | 0.25 | 13 | 0.3077 | 1 | 1 | 4 | 0.5 | 6 | 0.3333 | 5 | 0.2 | 16 | 0.375 |
| 12_761569 | NA | NA | 1 | 0 | 2 | 0.5 | NA | NA | 3 | 0.3333 | 1 | 0 | 1 | 1 | 5 | 0.6 | NA | NA | 7 | 0.5714 |
| 12_761569 | NA | NA | 2 | 0.5 | NA | NA | NA | NA | 2 | 0.5 | 1 | 1 | 1 | 0 | 5 | 0.8 | NA | NA | 7 | 0.7143 |

| | | | | | | | | | | | | | | | | | | | | |
|---|---|---|---|---|---|---|---|---|---|---|---|---|---|---|---|---|---|---|---|---|
| 12_761569 | NA | NA | 2 | 0.5 | 1 | 0 | NA | NA | 3 | 0.3333 | 1 | 1 | 1 | 1 | 4 | 0.75 | NA | NA | 6 | 0.8333 |
| 12_761570 | NA | NA | 2 | 0.5 | 2 | 1 | NA | NA | 4 | 0.75 | 1 | 1 | 1 | 1 | 4 | 0.5 | NA | NA | 6 | 0.6667 |
| 12_763745 | 2 | 1 | NA | NA | NA | NA | 2 | 1 | 4 | 1 | NA | NA | 2 | 1 | 4 | 1 | 3 | 0.6667 | 9 | 0.8889 |
| 12_763746 | 2 | 0 | NA | NA | 2 | 0.5 | 3 | 0.6667 | 7 | 0.4286 | NA | NA | 1 | 1 | 4 | 0 | 3 | 0 | 8 | 0.125 |
| 12_763747 | 2 | 0.5 | NA | NA | 2 | 0.5 | 3 | 0.3333 | 7 | 0.4286 | NA | NA | 1 | 0 | 5 | 0.4 | 3 | 1 | 9 | 0.5556 |
| 12_763749 | 2 | 0 | 1 | 1 | 2 | 0.5 | 2 | 0.5 | 7 | 0.4286 | NA | NA | 2 | 0.5 | 4 | 0.25 | 2 | 0.5 | 8 | 0.375 |
| 12_763752 | 2 | 0.5 | 1 | 0 | 3 | 1 | 3 | 1 | 9 | 0.7778 | 1 | 1 | 1 | 1 | 5 | 1 | 3 | 0.6667 | 10 | 0.9 |
| 12_763752 | 2 | 0 | 1 | 0 | 3 | 0.3333 | 3 | 0 | 9 | 0.1111 | 1 | 1 | 2 | 0 | 6 | 0 | 3 | 0 | 12 | 0.0833 |
| 12_763753 | 2 | 0 | 1 | 0 | 2 | 0.5 | 5 | 0 | 10 | 0.1 | 1 | 0 | 2 | 0 | 6 | 0.1667 | 3 | 0 | 12 | 0.0833 |
| 12_763754 | 1 | 1 | 1 | 1 | 3 | 0.3333 | 4 | 0.75 | 9 | 0.6667 | NA | NA | 2 | 1 | 5 | 1 | 2 | 0 | 9 | 0.7778 |
| 12_763754 | 2 | 1 | 1 | 1 | 3 | 1 | 4 | 0.25 | 10 | 0.7 | 1 | 0 | 1 | 1 | 6 | 0.8333 | 2 | 1 | 10 | 0.8 |
| 12_763765 | 6 | 0.3333 | 2 | 1 | 4 | 1 | 7 | 0.8571 | 19 | 0.7368 | 2 | 1 | 3 | 1 | 6 | 0.6667 | 4 | 1 | 15 | 0.8667 |
| 12_763766 | 8 | 0.75 | 5 | 0.4 | 4 | 0.75 | 5 | 1 | 22 | 0.7273 | 2 | 1 | 3 | 0.6667 | 10 | 0.7 | 4 | 0.75 | 19 | 0.7368 |
| 12_763767 | 8 | 1 | 5 | 0.6 | 3 | 0.3333 | 6 | 0.8333 | 22 | 0.7727 | 2 | 1 | 2 | 1 | 7 | 0.5714 | 3 | 0.6667 | 14 | 0.7143 |
| 12_763767 | 6 | 0.5 | 4 | 0.75 | 2 | 0.5 | 5 | 0.6 | 17 | 0.5882 | 2 | 1 | 3 | 0.3333 | 8 | 0.625 | 4 | 0.25 | 17 | 0.5294 |
| 12_763767 | 7 | 0.1429 | 5 | 0 | 4 | 0.25 | 8 | 0.375 | 24 | 0.2083 | 2 | 0.5 | 3 | 0.3333 | 8 | 0.125 | 4 | 0.25 | 17 | 0.2353 |
| 12_763767 | 8 | 0.375 | 5 | 0.6 | 3 | 0.3333 | 7 | 0.5714 | 23 | 0.4783 | 2 | 0.5 | 3 | 0.6667 | 7 | 0.4286 | 4 | 1 | 16 | 0.625 |
| 12_763768 | 8 | 1 | 5 | 1 | 4 | 1 | 8 | 1 | 25 | 1 | 2 | 1 | 3 | 0.6667 | 10 | 1 | 4 | 1 | 19 | 0.9474 |
| 12_763768 | 7 | 0.1429 | 5 | 0.2 | 4 | 0.25 | 11 | 0.3636 | 27 | 0.2593 | 2 | 0 | 2 | 0.5 | 11 | 0.0909 | 4 | 0 | 19 | 0.1053 |
| 12_763770 | 6 | 1 | 5 | 1 | 2 | 1 | 10 | 0.9 | 23 | 0.9565 | 2 | 1 | 2 | 0.5 | 9 | 0.7778 | 4 | 1 | 17 | 0.8235 |
| 12_763770 | 5 | 1 | 4 | 1 | 2 | 1 | 10 | 1 | 21 | 1 | 1 | 1 | 1 | 1 | 9 | 1 | 3 | 1 | 14 | 1 |
| 12_763770 | 5 | 0.8 | 4 | 1 | 2 | 1 | 9 | 1 | 20 | 0.95 | 3 | 1 | 2 | 1 | 10 | 0.9 | 2 | 1 | 17 | 0.9412 |
| 12_763771 | 5 | 0.4 | 5 | 0.4 | 3 | 0 | 10 | 0.1 | 23 | 0.2174 | 3 | 0 | 2 | 0 | 9 | 0.4444 | 5 | 0.4 | 19 | 0.3158 |
| 12_763773 | 7 | 0.4286 | 6 | 0.5 | 4 | 0.5 | 14 | 0.5 | 31 | 0.4839 | 4 | 0.5 | 6 | 0.5 | 12 | 0.1667 | 6 | 0.1667 | 28 | 0.2857 |
| 12_763777 | 7 | 0.4286 | 7 | 0.4286 | 5 | 0.4 | 12 | 0.6667 | 31 | 0.5161 | 4 | 0.5 | 8 | 0.625 | 9 | 0.5556 | 4 | 0.25 | 25 | 0.52 |
| 12_763781 | 8 | 0.125 | 5 | 0.4 | 3 | 0.3333 | 13 | 0.3077 | 29 | 0.2759 | 4 | 0.25 | 5 | 0.2 | 11 | 0.2727 | 4 | 0.25 | 24 | 0.25 |
| 12_763785 | 8 | 0 | 6 | 0.1667 | 2 | 0.5 | 13 | 0.0769 | 29 | 0.1034 | 3 | 0.3333 | 9 | 0.1111 | 4 | 0.5 | 4 | 0.25 | 20 | 0.25 |
| 12_763798 | 10 | 0 | 9 | 0 | 5 | 0 | 16 | 0.125 | 40 | 0.05 | 4 | 0 | 15 | 0.1333 | 10 | 0 | 9 | 0 | 38 | 0.0526 |
| 12_763810 | 1 | 0 | 1 | 1 | NA | NA | 5 | 0 | 7 | 0.1429 | NA | NA | 3 | 0.3333 | NA | NA | 2 | 0 | 5 | 0.2 |
| 12_763811 | 1 | 0 | NA | NA | NA | NA | 3 | 0.6667 | 4 | 0.5 | NA | NA | 2 | 1 | NA | NA | NA | NA | 2 | 1 |
| 12_763818 | 1 | 0 | 3 | 0 | 1 | 0 | 5 | 0.2 | 10 | 0.1 | NA | NA | 4 | 0 | NA | NA | 2 | 0.5 | 6 | 0.1667 |
| 12_763818 | 1 | 0 | 3 | 0 | 1 | 1 | 5 | 0.6 | 10 | 0.4 | NA | NA | 2 | 0 | NA | NA | 2 | 0.5 | 4 | 0.25 |
| 12_763827 | 1 | 0 | 3 | 0 | 1 | 0 | 4 | 0.75 | 9 | 0.3333 | 1 | 0 | 2 | 0 | 3 | 0.3333 | 2 | 0.5 | 8 | 0.25 |
| 12_763828 | 1 | 1 | 2 | 0.5 | 1 | 1 | 2 | 1 | 6 | 0.8333 | 1 | 0 | 1 | 1 | 3 | 0.6667 | 4 | 0.25 | 9 | 0.4444 |
| 12_763830 | 1 | 0 | 2 | 1 | 1 | 1 | 4 | 1 | 8 | 0.875 | 1 | 1 | 1 | 1 | 3 | 1 | 4 | 0.75 | 9 | 0.8889 |
| 12_763832 | 1 | 0 | 2 | 0.5 | 1 | 0 | 4 | 0.75 | 8 | 0.5 | 1 | 0 | 1 | 0 | 3 | 0 | 3 | 0.3333 | 8 | 0.125 |
| 12_763832 | 1 | 1 | 3 | 1 | 2 | 1 | 4 | 1 | 10 | 1 | 1 | 1 | 2 | 1 | 2 | 1 | 3 | 0.6667 | 8 | 0.875 |
| 12_763833 | NA | NA | 3 | 0.6667 | 2 | 1 | 3 | 1 | 8 | 0.875 | 1 | 0 | 2 | 1 | 1 | 1 | 4 | 0.25 | 8 | 0.5 |
| 12_763834 | 1 | 0 | 3 | 0.6667 | 2 | 0.5 | 4 | 0.5 | 10 | 0.5 | 1 | 0 | 2 | 0.5 | 3 | 0 | 3 | 0 | 9 | 0.1111 |
| 12_763835 | 1 | 0 | 2 | 0 | 1 | 1 | 1 | 1 | 5 | 0.4 | NA | NA | 2 | 0.5 | 2 | 0 | 4 | 0 | 8 | 0.125 |
| 12_763835 | 1 | 1 | 3 | 0 | 2 | 0.5 | 4 | 0.25 | 10 | 0.3 | 1 | 0 | 1 | 1 | 3 | 0.3333 | 4 | 0 | 9 | 0.2222 |
| 12_763837 | 1 | 0 | 2 | 0 | 2 | 0 | 4 | 0.25 | 9 | 0.1111 | 1 | 0 | 2 | 0.5 | 3 | 0.3333 | 4 | 0 | 10 | 0.2 |
| 12_763839 | 2 | 0.5 | 3 | 0 | 2 | 0 | 3 | 0.3333 | 10 | 0.2 | 1 | 0 | 2 | 0 | 4 | 0.25 | 4 | 0 | 11 | 0.0909 |
| 12_763839 | 2 | 0.5 | 3 | 0.6667 | 2 | 0 | 4 | 0.25 | 11 | 0.3636 | 1 | 0 | 2 | 1 | 3 | 0.3333 | 5 | 0 | 11 | 0.2727 |
| 12_763839 | 2 | 0.5 | 3 | 0.6667 | 2 | 1 | 3 | 0.3333 | 10 | 0.6 | 1 | 1 | 2 | 1 | 3 | 1 | 5 | 0.8 | 11 | 0.9091 |
| 12_763915 | NA | NA | NA | NA | NA | NA | 1 | 1 | 1 | 1 | 2 | 1 | 1 | 0 | 1 | 1 | NA | NA | 4 | 0.75 |
| 12_763958 | 2 | 1 | 2 | 1 | 2 | 0.5 | 1 | 0 | 7 | 0.7143 | 3 | 0.6667 | 1 | 0 | 3 | 0.6667 | NA | NA | 7 | 0.5714 |
| 12_763959 | 2 | 0.5 | 2 | 1 | 2 | 0.5 | 1 | 1 | 7 | 0.7143 | 4 | 0.5 | NA | NA | 4 | 0 | NA | NA | 8 | 0.25 |
| 12_763959 | 2 | 0.5 | 1 | 1 | NA | NA | 1 | 1 | 4 | 0.75 | 2 | 0.5 | 1 | 0 | 4 | 0 | NA | NA | 7 | 0.1429 |
| 12_763959 | NA | NA | 2 | 1 | 1 | 0 | 1 | 1 | 4 | 0.75 | 2 | 0.5 | 1 | 1 | 3 | 0.6667 | NA | NA | 6 | 0.6667 |
| 12_763960 | 2 | 0 | 2 | 0.5 | 2 | 0.5 | 1 | 0 | 7 | 0.2857 | 4 | 0.5 | 1 | 0 | 4 | 0 | NA | NA | 9 | 0.2222 |
| 12_763961 | 2 | 0 | 2 | 1 | NA | NA | NA | NA | 4 | 0.5 | 3 | 1 | NA | NA | 2 | 0.5 | NA | NA | 5 | 0.8 |
| 12_763962 | 2 | 1 | 1 | 0 | NA | NA | 1 | 0 | 4 | 0.5 | 3 | 0.6667 | 1 | 1 | 3 | 0.6667 | NA | NA | 7 | 0.7143 |
| 12_763963 | 2 | 0.5 | 1 | 0 | NA | NA | 1 | 1 | 4 | 0.5 | 4 | 0.25 | 1 | 1 | 1 | 1 | NA | NA | 6 | 0.5 |
| 12_763963 | 2 | 0.5 | 2 | 0 | NA | NA | 1 | 0 | 5 | 0.2 | 3 | 0.3333 | 1 | 0 | 4 | 0.5 | NA | NA | 8 | 0.375 |
| 12_763965 | 2 | 0 | NA | NA | NA | NA | 1 | 1 | 3 | 0.3333 | 1 | 0 | NA | NA | 3 | 0.3333 | NA | NA | 4 | 0.25 |
| 12_763965 | 1 | 0 | 2 | 1 | NA | NA | 1 | 1 | 4 | 0.75 | 3 | 0.6667 | 1 | 1 | 4 | 0.75 | NA | NA | 8 | 0.75 |
| 12_763966 | 2 | 1 | 2 | 0 | NA | NA | 1 | 0 | 5 | 0.4 | 3 | 0.3333 | 1 | 0 | 4 | 0.5 | NA | NA | 8 | 0.375 |
| 12_763967 | 2 | 0 | 2 | 0.5 | NA | NA | 1 | 0 | 5 | 0.2 | 3 | 0 | 1 | 0 | 3 | 0.3333 | NA | NA | 7 | 0.1429 |
| 12_763968 | 2 | 0.5 | 2 | 0.5 | NA | NA | NA | NA | 4 | 0.5 | 3 | 0.6667 | 1 | 0 | 4 | 0 | NA | NA | 8 | 0.25 |
| 12_763968 | 2 | 1 | 2 | 1 | NA | NA | 1 | 0 | 5 | 0.8 | 2 | 0.5 | 1 | 1 | 4 | 1 | NA | NA | 7 | 0.8571 |
| 12_764160 | NA | NA | NA | NA | NA | NA | 1 | 1 | 1 | 1 | 1 | 1 | NA | NA | 1 | 0 | NA | NA | 2 | 0.5 |
| 12_764323 | 2 | 1 | 1 | 1 | 1 | 1 | NA | NA | 4 | 1 | 1 | 1 | 1 | 1 | NA | NA | 3 | 1 | 5 | 1 |
| 12_764323 | 1 | 0 | 1 | 1 | 2 | 1 | NA | NA | 4 | 0.75 | 1 | 1 | 1 | 1 | NA | NA | 3 | 1 | 5 | 1 |
| 12_764327 | 2 | 0.5 | 1 | 0 | 2 | 0.5 | NA | NA | 5 | 0.4 | 1 | 0 | NA | NA | NA | NA | 3 | 0.6667 | 4 | 0.5 |

| | | | | | | | | | | | | | | | | | | | | |
|---|---|---|---|---|---|---|---|---|---|---|---|---|---|---|---|---|---|---|---|---|
| 12_764327 | 1 | 1 | NA | NA | 2 | 1 | NA | NA | 3 | 1 | 1 | 0 | NA | NA | NA | NA | 2 | 0.5 | 3 | 0.3333 |
| 12_764330 | 2 | 0 | 1 | 0 | 1 | 1 | NA | NA | 4 | 0.25 | 1 | 0 | 1 | 1 | NA | NA | 3 | 0.6667 | 5 | 0.6 |
| 12_764332 | 2 | 1 | 2 | 1 | 1 | 1 | NA | NA | 5 | 1 | 1 | 1 | 1 | 1 | NA | NA | 3 | 1 | 5 | 1 |
| 12_764333 | 2 | 0.5 | 2 | 1 | 2 | 0.5 | NA | NA | 6 | 0.6667 | 1 | 0 | NA | NA | NA | NA | 3 | 1 | 4 | 0.75 |
| 12_764333 | 2 | 1 | 2 | 1 | 2 | 1 | NA | NA | 6 | 1 | 1 | 1 | 1 | 0 | NA | NA | 2 | 1 | 4 | 0.75 |
| 12_764334 | 2 | 1 | 2 | 0.5 | 2 | 1 | NA | NA | 6 | 0.8333 | 1 | 1 | 1 | 1 | NA | NA | 3 | 1 | 5 | 1 |
| 12_764335 | 3 | 0.6667 | 2 | 0.5 | 2 | 1 | NA | NA | 7 | 0.7143 | 1 | 1 | NA | NA | NA | NA | 2 | 1 | 3 | 1 |
| 12_764335 | 3 | 1 | 2 | 1 | 2 | 1 | NA | NA | 7 | 1 | 1 | 1 | 1 | 1 | NA | NA | 3 | 1 | 5 | 1 |
| 12_764336 | 2 | 1 | 2 | 1 | 1 | 1 | NA | NA | 5 | 1 | 1 | 1 | NA | NA | NA | NA | 2 | 0.5 | 3 | 0.6667 |
| 12_764336 | 3 | 1 | 2 | 0 | 2 | 0 | NA | NA | 7 | 0.4286 | NA | NA | 1 | 1 | NA | NA | 2 | 0 | 3 | 0.3333 |
| 12_764337 | 1 | 0 | 2 | 0.5 | 2 | 0.5 | NA | NA | 5 | 0.4 | 1 | 0 | 1 | 0 | NA | NA | 3 | 0.3333 | 5 | 0.2 |
| 12_764338 | 2 | 1 | 2 | 0.5 | 2 | 0 | NA | NA | 6 | 0.5 | 1 | 0 | 1 | 0 | NA | NA | 3 | 0.6667 | 5 | 0.4 |
| 12_764340 | 2 | 0 | 2 | 1 | 1 | 0 | NA | NA | 5 | 0.4 | NA | NA | NA | NA | NA | NA | 2 | 1 | 2 | 1 |
| 12_764341 | 2 | 0 | 2 | 0.5 | 2 | 0 | NA | NA | 6 | 0.1667 | 1 | 0 | NA | NA | NA | NA | 3 | 0.6667 | 4 | 0.5 |
| 12_764342 | 2 | 0.5 | 2 | 0.5 | 2 | 0 | NA | NA | 6 | 0.3333 | NA | NA | 1 | 0 | NA | NA | 2 | 1 | 3 | 0.6667 |
| 12_764351 | 4 | 0.25 | 2 | 0 | 2 | 0.5 | NA | NA | 8 | 0.25 | NA | NA | 1 | 0 | 2 | 0 | 1 | 1 | 4 | 0.25 |
| 12_764417 | 1 | 1 | 1 | 1 | NA | NA | 1 | 1 | 3 | 1 | NA | NA | 1 | 1 | NA | NA | NA | NA | 1 | 1 |
| 12_764419 | 1 | 1 | 1 | 0 | NA | NA | 1 | 0 | 3 | 0.3333 | NA | NA | 1 | 1 | NA | NA | NA | NA | 1 | 1 |
| 12_764419 | 1 | 1 | 1 | 0 | NA | NA | 1 | 1 | 3 | 0.6667 | NA | NA | 1 | 1 | NA | NA | NA | NA | 1 | 1 |
| 12_764420 | 1 | 1 | 1 | 1 | NA | NA | 1 | 1 | 3 | 1 | NA | NA | 1 | 1 | NA | NA | NA | NA | 1 | 1 |
| 12_764520 | 6 | 1 | 5 | 0.8 | 4 | 0.75 | 2 | 1 | 17 | 0.8824 | 3 | 0.6667 | 3 | 1 | 4 | 1 | 9 | 1 | 19 | 0.9474 |
| 12_764522 | 7 | 0.1429 | 4 | 0 | 6 | 0.1667 | 1 | 0 | 18 | 0.1111 | 4 | 0 | 3 | 0.3333 | 5 | 0.4 | 10 | 0.1 | 22 | 0.1818 |
| 12_764525 | 6 | 0 | 5 | 0.2 | 7 | 0 | 2 | 0 | 20 | 0.05 | 4 | 0 | NA | NA | 5 | 0.2 | 8 | 0 | 17 | 0.0588 |
| 12_764526 | 6 | 1 | 8 | 0.875 | 7 | 1 | 4 | 1 | 25 | 0.96 | 4 | 1 | 2 | 1 | 3 | 1 | 9 | 0.7778 | 18 | 0.8889 |
| 12_764529 | 8 | 0.5 | 9 | 0.3333 | 9 | 0.4444 | 5 | 0.2 | 31 | 0.3871 | 6 | 0.5 | 3 | 0.6667 | 5 | 0.4 | 9 | 0.3333 | 23 | 0.4348 |
| 12_764530 | 7 | 0.2857 | 8 | 0.125 | 9 | 0.1111 | 4 | 0.25 | 28 | 0.1786 | 6 | 0 | 3 | 0.3333 | 4 | 0.25 | 10 | 0.4 | 23 | 0.2609 |
| 12_764530 | 9 | 0.2222 | 8 | 0.375 | 9 | 0.3333 | 4 | 0.25 | 30 | 0.3 | 4 | 0.5 | 3 | 0.3333 | 5 | 0.6 | 8 | 0.125 | 20 | 0.35 |
| 12_764531 | 8 | 0.125 | 5 | 0.6 | 8 | 0.25 | 3 | 0.3333 | 24 | 0.2917 | 5 | 0.4 | 2 | 0.5 | 3 | 0 | 10 | 0.5 | 20 | 0.4 |
| 12_764532 | 8 | 0.875 | 8 | 0.875 | 9 | 1 | 4 | 1 | 29 | 0.931 | 6 | 1 | 3 | 1 | 4 | 1 | 9 | 1 | 22 | 1 |
| 12_764950 | NA | NA | NA | NA | NA | NA | 1 | 1 | 1 | 1 | 1 | 1 | NA | NA | NA | NA | NA | NA | 1 | 1 |
| 12_764951 | NA | NA | NA | NA | 1 | 1 | 1 | 0 | 2 | 0.5 | 1 | 1 | NA | NA | NA | NA | NA | NA | 1 | 1 |
| 12_764967 | NA | NA | NA | NA | 1 | 1 | NA | NA | 1 | 1 | 1 | 1 | NA | NA | NA | NA | NA | NA | 1 | 1 |
| 12_764972 | NA | NA | NA | NA | 2 | 1 | NA | NA | 2 | 1 | 2 | 1 | NA | NA | NA | NA | NA | NA | 2 | 1 |
| 12_766227 | 1 | 1 | 1 | 1 | 1 | 1 | 1 | 0 | 4 | 0.75 | 1 | 0 | 1 | 0 | 1 | 1 | NA | NA | 3 | 0.3333 |
| 12_766230 | 1 | 1 | 1 | 1 | 1 | 0 | 3 | 0.6667 | 6 | 0.6667 | 2 | 0.5 | 1 | 0 | 1 | 1 | 1 | 0 | 5 | 0.4 |
| 12_766230 | 1 | 0 | 1 | 1 | 1 | 1 | 2 | 0 | 5 | 0.4 | 2 | 0.5 | NA | NA | 1 | 1 | 1 | 0 | 4 | 0.5 |
| 12_766231 | 1 | 0 | NA | NA | 1 | 0 | 1 | 1 | 3 | 0.3333 | 2 | 0.5 | 1 | 0 | 1 | 1 | 1 | 1 | 5 | 0.6 |
| 12_766232 | 1 | 0 | 1 | 0 | 1 | 0 | 1 | 1 | 4 | 0.25 | 2 | 0 | 1 | 0 | 1 | 1 | 1 | 0 | 5 | 0.2 |
| 12_766232 | 2 | 0.5 | 1 | 1 | 1 | 0 | 2 | 0.5 | 6 | 0.5 | 1 | 0 | 1 | 0 | 1 | 1 | 1 | 0 | 4 | 0.25 |
| 12_766232 | 2 | 0 | 1 | 0 | 1 | 1 | 2 | 0 | 6 | 0.1667 | 2 | 0.5 | 1 | 1 | 1 | 0 | 1 | 0 | 5 | 0.4 |
| 12_766234 | 2 | 1 | 1 | 1 | 1 | 0 | 3 | 0.6667 | 7 | 0.7143 | 2 | 1 | 1 | 1 | 1 | 1 | 2 | 1 | 6 | 1 |
| 12_766235 | 2 | 0 | 1 | 0 | 1 | 0 | 3 | 0.6667 | 7 | 0.2857 | 2 | 1 | 1 | 1 | 1 | 1 | 1 | 1 | 5 | 1 |
| 12_766235 | 2 | 0.5 | 1 | 0 | 1 | 0 | 2 | 0 | 6 | 0.1667 | 2 | 0 | 1 | 0 | 1 | 0 | 2 | 0.5 | 6 | 0.1667 |
| 12_766236 | 2 | 1 | 1 | 1 | 1 | 1 | 2 | 0.5 | 6 | 0.8333 | 3 | 1 | 1 | 1 | 1 | 1 | 1 | 0 | 6 | 0.8333 |
| 12_766238 | 2 | 1 | 1 | 1 | 1 | 1 | 2 | 0.5 | 6 | 0.8333 | NA | NA | 1 | 0 | NA | NA | 1 | 1 | 2 | 0.5 |
| 12_766247 | 1 | 0 | 1 | 1 | 1 | 1 | 1 | 0 | 4 | 0.5 | 3 | 1 | 1 | 1 | 1 | 1 | NA | NA | 5 | 1 |
| 12_766247 | 3 | 0.3333 | 1 | 0 | 1 | 0 | 3 | 0.3333 | 8 | 0.25 | 2 | 0.5 | 1 | 0 | 1 | 0 | 2 | 0 | 6 | 0.1667 |
| 12_766248 | 3 | 0.6667 | 1 | 0 | 1 | 1 | 2 | 0.5 | 7 | 0.5714 | 2 | 1 | 1 | 0 | 1 | 0 | NA | NA | 4 | 0.5 |
| 12_766250 | 3 | 1 | 1 | 1 | 1 | 1 | 3 | 0 | 8 | 0.625 | 3 | 1 | 1 | 1 | NA | NA | 1 | 1 | 5 | 1 |
| 12_766250 | 3 | 0.6667 | 1 | 1 | 2 | 0.5 | 2 | 0.5 | 8 | 0.625 | 3 | 0.3333 | 1 | 0 | 1 | 1 | 2 | 0.5 | 7 | 0.4286 |
| 12_766250 | 2 | 0 | 1 | 0 | NA | NA | 2 | 1 | 5 | 0.4 | 2 | 0.5 | NA | NA | 1 | 1 | 2 | 0 | 5 | 0.4 |
| 12_766251 | 3 | 0.6667 | 1 | 0 | 2 | 1 | 3 | 1 | 9 | 0.7778 | 3 | 0.6667 | 1 | 0 | 1 | 1 | 2 | 1 | 7 | 0.7143 |
| 12_766251 | 2 | 0 | 1 | 1 | 2 | 0.5 | 2 | 0 | 7 | 0.2857 | 3 | 0 | NA | NA | 1 | 0 | 1 | 1 | 5 | 0.2 |
| 12_766251 | 3 | 0.3333 | 1 | 1 | 2 | 0 | 3 | 0 | 9 | 0.2222 | 2 | 0.5 | 1 | 0 | 1 | 1 | 2 | 0 | 6 | 0.3333 |
| 12_766251 | 3 | 1 | 1 | 1 | 2 | 1 | 2 | 1 | 8 | 1 | 3 | 0.3333 | NA | NA | 1 | 1 | 1 | 1 | 5 | 0.6 |
| 12_766251 | 2 | 0 | 1 | 0 | 3 | 0.3333 | 3 | 0.3333 | 9 | 0.2222 | 2 | 1 | 1 | 1 | 1 | 1 | 2 | 0.5 | 6 | 0.8333 |
| 12_766252 | 3 | 0.6667 | 1 | 1 | 3 | 1 | 1 | 1 | 8 | 0.875 | 3 | 0.6667 | 1 | 1 | 1 | 1 | 2 | 1 | 7 | 0.8571 |
| 12_766253 | 2 | 1 | 1 | 1 | 3 | 1 | 3 | 0.6667 | 9 | 0.8889 | 2 | 1 | 1 | 1 | NA | NA | 1 | 1 | 4 | 1 |
| 12_766253 | 3 | 0.6667 | 1 | 0 | 3 | 0.3333 | 3 | 0.3333 | 10 | 0.4 | 3 | 0 | 1 | 0 | 1 | 1 | 2 | 0 | 7 | 0.1429 |
| 12_766253 | 2 | 0.5 | NA | NA | 2 | 1 | 3 | 0.3333 | 7 | 0.5714 | 3 | 0.6667 | 1 | 0 | 1 | 1 | 1 | 1 | 6 | 0.6667 |
| 12_766255 | 2 | 1 | 2 | 0 | 3 | 0 | 4 | 0.25 | 11 | 0.2727 | 3 | 0.3333 | 1 | 0 | 1 | 1 | 2 | 0 | 7 | 0.2857 |
| 12_766257 | 3 | 0.3333 | 2 | 1 | 3 | 0.6667 | 5 | 0.2 | 13 | 0.4615 | 3 | 0.6667 | 1 | 0 | 1 | 1 | 1 | 1 | 6 | 0.6667 |
| 12_766274 | 13 | 0.2308 | 10 | 0 | 9 | 0.2222 | 9 | 0 | 41 | 0.122 | 6 | 0.1667 | 14 | 0.1429 | 11 | 0 | 9 | 0.3333 | 40 | 0.15 |
| 12_766279 | 18 | 0.1667 | 15 | 0.1333 | 11 | 0.0909 | 14 | 0.2143 | 58 | 0.1552 | 9 | 0 | 15 | 0.0667 | 14 | 0.0714 | 16 | 0.1875 | 54 | 0.0926 |
| 12_766279 | 15 | 0.3333 | 10 | 0.3 | 8 | 0.5 | 7 | 0.1429 | 40 | 0.325 | 6 | 0.3333 | 9 | 0.5556 | 13 | 0.3846 | 13 | 0.6923 | 41 | 0.5122 |

| | | | | | | | | | | | | | | | | | | | | |
|---|---|---|---|---|---|---|---|---|---|---|---|---|---|---|---|---|---|---|---|---|
| 12_766294 | 14 | 0.0714 | 9 | 0.1111 | 7 | 0.2857 | 8 | 0 | 38 | 0.1053 | 4 | 0.25 | 9 | 0.1111 | 8 | 0.125 | 8 | 0.125 | 29 | 0.1379 |
| 12_766297 | 11 | 0.8182 | 8 | 1 | 6 | 0.5 | 8 | 0.5 | 33 | 0.7273 | 5 | 0.4 | 8 | 0.5 | 7 | 0.5714 | 8 | 1 | 28 | 0.6429 |
| 12_766299 | 11 | 0.4545 | 7 | 0.2857 | 4 | 0.5 | 8 | 0.25 | 30 | 0.3667 | 5 | 1 | 9 | 0.5556 | 8 | 0.625 | 9 | 0.4444 | 31 | 0.6129 |
| 12_766299 | 11 | 0.1818 | 8 | 0.5 | 4 | 0.25 | 7 | 0.7143 | 30 | 0.4 | 5 | 0.8 | 9 | 0.3333 | 9 | 0.4444 | 8 | 0.625 | 31 | 0.5161 |
| 12_766503 | 6 | 0 | 4 | 0 | 1 | 0 | 8 | 0.25 | 19 | 0.1053 | 2 | 0.5 | 4 | 0 | 2 | 0 | 4 | 0 | 12 | 0.0833 |
| 12_766503 | 4 | 0.75 | 4 | 1 | 1 | 0 | 6 | 0.6667 | 15 | 0.7333 | 2 | 1 | 3 | 1 | 3 | 0.3333 | 4 | 0.75 | 12 | 0.75 |
| 12_766594 | 3 | 0.3333 | 2 | 0 | NA | NA | 1 | 0 | 6 | 0.1667 | NA | NA | 4 | 0.75 | 2 | 0 | 2 | 0 | 8 | 0.375 |
| 12_766596 | 2 | 0 | 2 | 1 | NA | NA | 1 | 0 | 5 | 0.4 | NA | NA | 4 | 0.75 | 2 | 0.5 | 2 | 1 | 8 | 0.75 |
| 12_766599 | 2 | 0.5 | 2 | 1 | NA | NA | 1 | 0 | 5 | 0.6 | NA | NA | 4 | 0.5 | 2 | 0 | 2 | 0.5 | 8 | 0.375 |
| 12_766599 | 2 | 1 | 1 | 1 | NA | NA | 1 | 1 | 4 | 1 | NA | NA | 4 | 0.75 | 1 | 0 | 2 | 1 | 7 | 0.7143 |
| 12_766599 | 2 | 0 | 2 | 0 | NA | NA | 1 | 1 | 5 | 0.2 | NA | NA | 1 | 0 | 2 | 0.5 | 2 | 0 | 5 | 0.2 |
| 12_766675 | 2 | 1 | 1 | 1 | NA | NA | 1 | 1 | 4 | 1 | NA | NA | 3 | 0.6667 | 1 | 1 | 1 | 1 | 5 | 0.8 |
| 12_766679 | 2 | 0.5 | 1 | 0 | NA | NA | 1 | 0 | 4 | 0.25 | NA | NA | 3 | 0 | 1 | 1 | 1 | 0 | 5 | 0.2 |
| 12_766681 | 2 | 0 | 1 | 1 | NA | NA | 1 | 0 | 4 | 0.25 | NA | NA | 2 | 0.5 | NA | NA | NA | NA | 2 | 0.5 |
| 12_768297 | 11 | 0.1818 | 10 | 0 | 9 | 0 | 9 | 0.1111 | 39 | 0.0769 | 13 | 0.2308 | 15 | 0.1333 | 8 | 0 | 8 | 0.375 | 44 | 0.1818 |
| 12_768449 | 23 | 0.3043 | 19 | 0.3158 | 14 | 0.2143 | 21 | 0.1429 | 77 | 0.2468 | 11 | 0.3636 | 21 | 0.3333 | 30 | 0.1667 | 17 | 0.3529 | 79 | 0.2785 |
| 12_768451 | 30 | 0.4 | 25 | 0.28 | 22 | 0.3636 | 27 | 0.3333 | 104 | 0.3462 | 14 | 0.3571 | 29 | 0.3793 | 33 | 0.4848 | 23 | 0.4348 | 99 | 0.4242 |
| 12_768452 | 24 | 0.0417 | 21 | 0.1429 | 15 | 0.1333 | 19 | 0.1053 | 79 | 0.1013 | 12 | 0.1667 | 24 | 0.125 | 27 | 0.1852 | 17 | 0.1176 | 80 | 0.15 |
| 12_768452 | 22 | 0.0909 | 16 | 0.0625 | 13 | 0.0769 | 22 | 0 | 73 | 0.0548 | 10 | 0.1 | 28 | 0.1429 | 24 | 0.0417 | 20 | 0 | 82 | 0.0732 |
| 12_768462 | 16 | 0.125 | 12 | 0.0833 | 15 | 0 | 17 | 0.1176 | 60 | 0.0833 | 9 | 0.2222 | 16 | 0.0625 | 18 | 0.1667 | 17 | 0 | 60 | 0.1 |
| 12_768463 | 20 | 0.3 | 11 | 0.4545 | 11 | 0.2727 | 14 | 0.4286 | 56 | 0.3571 | 8 | 0.5 | 18 | 0.5556 | 21 | 0.5238 | 14 | 0.3571 | 61 | 0.4918 |
| 12_768464 | 18 | 0.1667 | 12 | 0.1667 | 14 | 0.1429 | 13 | 0.2308 | 57 | 0.1754 | 7 | 0.2857 | 16 | 0.3125 | 19 | 0.3158 | 15 | 0.1333 | 57 | 0.2632 |
| 12_768465 | 20 | 0.15 | 13 | 0.2308 | 15 | 0.1333 | 14 | 0.0714 | 62 | 0.1452 | 7 | 0.4286 | 18 | 0.2222 | 18 | 0 | 15 | 0 | 58 | 0.1207 |
| 12_768469 | 17 | 0.1176 | 9 | 0.3333 | 12 | 0.25 | 8 | 0.25 | 46 | 0.2174 | 5 | 0.2 | 14 | 0.5 | 17 | 0.2941 | 7 | 0 | 43 | 0.3023 |
| 12_769671 | 3 | 0.3333 | 5 | 0 | 4 | 0.75 | 4 | 0.75 | 16 | 0.4375 | 4 | 1 | 3 | 0.6667 | 5 | 0.8 | 4 | 0.75 | 16 | 0.8125 |
| 12_769674 | 5 | 0 | 6 | 0 | 6 | 0.3333 | 3 | 0.3333 | 20 | 0.15 | 5 | 0 | 2 | 0 | 5 | 0.4 | 6 | 0 | 18 | 0.1111 |
| 12_769843 | 8 | 0.25 | 7 | 0.2857 | 9 | 0.3333 | 5 | 0 | 29 | 0.2414 | 11 | 0.0909 | 6 | 0.1667 | 11 | 0.0909 | 11 | 0.1818 | 39 | 0.1282 |
| 12_769847 | 5 | 0.8 | 5 | 0.8 | 8 | 0.625 | 4 | 0.75 | 22 | 0.7273 | 7 | 0.8571 | 5 | 0.8 | 12 | 0.8333 | 11 | 0.9091 | 35 | 0.8571 |
| 12_769848 | 4 | 0.25 | 5 | 0.6 | 11 | 0.6364 | 4 | 0.5 | 24 | 0.5417 | 7 | 0.7143 | 6 | 1 | 10 | 0.6 | 8 | 0.75 | 31 | 0.7419 |
| 12_769849 | 5 | 1 | 6 | 1 | 6 | 1 | 3 | 1 | 20 | 1 | 4 | 1 | 8 | 0.875 | 10 | 0.8 | 6 | 0.8333 | 28 | 0.8571 |
| 12_769849 | 2 | 1 | 6 | 0.8333 | 10 | 0.9 | 3 | 1 | 21 | 0.9048 | 9 | 0.8889 | 5 | 0.8 | 12 | 0.8333 | 7 | 1 | 33 | 0.8788 |
| 12_769849 | 3 | 0.6667 | 6 | 0.8333 | 9 | 1 | 5 | 1 | 23 | 0.913 | 10 | 1 | 6 | 1 | 14 | 1 | 10 | 1 | 40 | 1 |
| 12_769850 | 2 | 0 | 5 | 0.4 | 9 | 0.8889 | 3 | 0 | 19 | 0.5263 | 9 | 0.2222 | 8 | 0.25 | 12 | 0.4167 | 10 | 0.4 | 39 | 0.3333 |
| 12_769851 | 4 | 0.5 | 7 | 0.8571 | 11 | 0.5455 | 5 | 0.8 | 27 | 0.6667 | 10 | 1 | 8 | 0.375 | 12 | 0.5 | 9 | 0.6667 | 39 | 0.641 |
| 12_769852 | 3 | 0.6667 | 6 | 0.5 | 8 | 0.875 | 5 | 1 | 22 | 0.7727 | 10 | 0.9 | 8 | 0.75 | 13 | 0.9231 | 10 | 0.8 | 41 | 0.8537 |
| 12_769853 | 4 | 0.25 | 5 | 1 | 4 | 0.75 | 4 | 0.75 | 17 | 0.7059 | 8 | 0.5 | 4 | 0.75 | 7 | 0.8571 | 6 | 0.3333 | 25 | 0.6 |
| 12_769855 | 1 | 1 | NA | NA | 1 | 1 | 2 | 0 | 4 | 0.5 | 2 | 0.5 | 2 | 0 | 1 | 1 | 2 | 0.5 | 7 | 0.4286 |
| 12_769858 | 1 | 1 | NA | NA | 1 | 0 | 2 | 0 | 4 | 0.25 | 2 | 0.5 | 2 | 0.5 | NA | NA | 2 | 0.5 | 6 | 0.5 |
| 12_773788 | 2 | 0.5 | 4 | 0.25 | 3 | 0.3333 | 2 | 0 | 11 | 0.2727 | 2 | 0 | 5 | 0.2 | 1 | 0 | 5 | 0 | 13 | 0.0769 |
| 12_773788 | 2 | 0 | 5 | 0 | 3 | 0.3333 | 2 | 0 | 12 | 0.0833 | 2 | 0 | 5 | 0.2 | 1 | 0 | 5 | 0.2 | 13 | 0.1538 |
| 12_773789 | 2 | 1 | 5 | 1 | 2 | 1 | 1 | 1 | 10 | 1 | 2 | 1 | 5 | 1 | 1 | 1 | 5 | 1 | 13 | 1 |
| 12_773789 | 2 | 0.5 | 3 | 1 | 3 | 0.6667 | 1 | 1 | 9 | 0.7778 | 2 | 1 | 6 | 1 | 1 | 0 | 5 | 0.8 | 14 | 0.8571 |
| 12_773790 | 2 | 0 | 5 | 0.4 | 4 | 0.25 | 2 | 0 | 13 | 0.2308 | 2 | 0 | 5 | 0 | 1 | 1 | 5 | 0.2 | 13 | 0.1538 |
| 12_775218 | 5 | 0 | 7 | 0 | 6 | 0 | 2 | 0.5 | 20 | 0.05 | 2 | 0 | 5 | 0.4 | 3 | 0 | 8 | 0.125 | 18 | 0.1667 |
| 12_775219 | 4 | 0.5 | 1 | 0 | 2 | 0.5 | 1 | 1 | 8 | 0.5 | NA | NA | 5 | 0.6 | 3 | 0.3333 | 6 | 0.3333 | 14 | 0.4286 |
| 12_775219 | 7 | 0.1429 | 7 | 0.1429 | 6 | 0.3333 | 3 | 0 | 23 | 0.1739 | 3 | 0 | 8 | 0 | 4 | 0.25 | 8 | 0.25 | 23 | 0.1304 |
| 12_775220 | 7 | 0.1429 | 6 | 0.3333 | 4 | 0 | 3 | 0 | 20 | 0.15 | 1 | 0 | 6 | 0.1667 | 4 | 0.25 | 8 | 0.125 | 19 | 0.1579 |
| 12_775220 | 4 | 0.25 | 5 | 0 | 4 | 0 | 3 | 0 | 16 | 0.0625 | 1 | 0 | 7 | 0 | 5 | 0 | 5 | 0.2 | 18 | 0.0556 |
| 12_775220 | 3 | 0.3333 | 4 | 0.25 | 4 | 0 | 2 | 0 | 13 | 0.1538 | 2 | 0 | 5 | 0.4 | 3 | 0 | 5 | 0 | 15 | 0.1333 |
| 12_775221 | 3 | 0.3333 | 1 | 0 | 4 | 0.25 | 2 | 0.5 | 10 | 0.3 | 2 | 0 | 7 | 0.5714 | 2 | 0.5 | 8 | 0 | 19 | 0.2632 |
| 12_775221 | 7 | 0.1429 | 5 | 0 | 7 | 0.2857 | 4 | 0.25 | 23 | 0.1739 | 2 | 0 | 7 | 0.1429 | 3 | 0 | 10 | 0.1 | 22 | 0.0909 |
| 12_775221 | 4 | 0 | 6 | 0.3333 | 7 | 0.2857 | 4 | 0.5 | 21 | 0.2857 | 3 | 0 | 8 | 0.5 | 5 | 0.2 | 8 | 0.375 | 24 | 0.3333 |
| 12_775223 | 4 | 0 | 4 | 0 | 7 | 0.1429 | 2 | 0 | 17 | 0.0588 | 2 | 0.5 | 5 | 0.4 | 3 | 0 | 7 | 0 | 17 | 0.1765 |
| 12_775223 | 7 | 0.4286 | 5 | 0.4 | 8 | 0.125 | 4 | 0.5 | 24 | 0.3333 | 4 | 0 | 8 | 0.125 | 5 | 0.4 | 9 | 0 | 26 | 0.1154 |
| 12_775223 | 8 | 0.125 | 6 | 0.5 | 8 | 0.125 | 5 | 0.2 | 27 | 0.2222 | 4 | 0 | 6 | 0 | 5 | 0.2 | 10 | 0.2 | 25 | 0.12 |
| 12_775223 | 7 | 0.5714 | 4 | 1 | 8 | 0.875 | 5 | 0.6 | 24 | 0.75 | 3 | 0.3333 | 7 | 0.5714 | 4 | 0.75 | 10 | 1 | 24 | 0.75 |
| 12_775224 | 9 | 0.4444 | 8 | 0.875 | 9 | 0.8889 | 5 | 0.4 | 31 | 0.6774 | 3 | 0.6667 | 9 | 0.8889 | 4 | 0.5 | 10 | 0.8 | 26 | 0.7692 |
| 12_775224 | 8 | 0.5 | 8 | 0.875 | 9 | 0.6667 | 6 | 0.5 | 31 | 0.6452 | 4 | 0.75 | 9 | 0.6667 | 6 | 0.6667 | 10 | 0.4 | 29 | 0.5862 |
| 12_775224 | 9 | 0.4444 | 7 | 0.4286 | 9 | 0.2222 | 3 | 1 | 28 | 0.4286 | 3 | 0.3333 | 7 | 0.2857 | 6 | 0.5 | 8 | 0.625 | 24 | 0.4583 |
| 12_775226 | 5 | 0.2 | 6 | 0.1667 | 5 | 0 | 5 | 0 | 21 | 0.0952 | 3 | 0 | 5 | 0 | 5 | 0.6 | 8 | 0.375 | 21 | 0.2857 |
| 12_775227 | 8 | 0.875 | 6 | 1 | 9 | 0.7778 | 4 | 0.75 | 27 | 0.8519 | 5 | 0.8 | 8 | 1 | 4 | 0.75 | 6 | 1 | 23 | 0.913 |
| 12_782724 | NA | NA | 1 | 1 | 2 | 0 | 1 | 0 | 4 | 0.25 | 2 | 0 | 1 | 0 | 1 | 1 | NA | NA | 4 | 0.25 |
| 12_782726 | 1 | 0 | NA | NA | 2 | 0 | 1 | 1 | 4 | 0.25 | 2 | 0 | 1 | 1 | 1 | 0 | NA | NA | 4 | 0.25 |
| 12_782727 | 1 | 0 | 1 | 1 | 2 | 0 | 1 | 0 | 5 | 0.2 | 1 | 0 | 2 | 0.5 | 1 | 0 | NA | NA | 4 | 0.25 |
| 12_782728 | 1 | 0 | 1 | 0 | 2 | 0.5 | 1 | 0 | 5 | 0.2 | 1 | 0 | 2 | 0 | 1 | 1 | NA | NA | 4 | 0.25 |

| | | | | | | | | | | | | | | | | | | | | |
|---|---|---|---|---|---|---|---|---|---|---|---|---|---|---|---|---|---|---|---|---|
| 12_782730 | 1 | 1 | 1 | 1 | 2 | 1 | 1 | 1 | 5 | 1 | 1 | 1 | 2 | 1 | 1 | 1 | NA | NA | 4 | 1 |
| 12_782731 | NA | NA | NA | NA | 1 | 1 | NA | NA | 1 | 1 | NA | NA | 1 | 1 | 1 | 1 | NA | NA | 2 | 1 |
| 12_782731 | 1 | 1 | NA | NA | 1 | 1 | 1 | 1 | 3 | 1 | NA | NA | 1 | 1 | 1 | 1 | NA | NA | 2 | 1 |
| 12_785389 | 5 | 0.2 | 3 | 0 | 3 | 0 | 4 | 0.5 | 15 | 0.0667 | 1 | 0 | 9 | 0.1111 | 4 | 0.25 | 2 | 0 | 16 | 0.125 |
| 12_787307 | 6 | 0.5 | 4 | 0.5 | 5 | 0.4 | 8 | 0.5 | 23 | 0.4783 | 5 | 0.4 | 14 | 0.4286 | 6 | 0.3333 | 4 | 0.75 | 29 | 0.4483 |
| 12_787308 | 5 | 0.4 | 6 | 0.5 | 4 | 0.25 | 7 | 0.2857 | 22 | 0.3636 | 5 | 0.8 | 14 | 0.3571 | 8 | 0.375 | 6 | 0.3333 | 33 | 0.4242 |
| 12_787312 | 8 | 0.25 | 5 | 0.2 | 5 | 0.2 | 8 | 0.375 | 26 | 0.2692 | 3 | 0 | 16 | 0.125 | 8 | 0.125 | 6 | 0 | 33 | 0.0909 |
| 12_787313 | 8 | 0.125 | 6 | 0.3333 | 6 | 0 | 9 | 0.1111 | 29 | 0.1379 | 5 | 0 | 13 | 0.0769 | 9 | 0.2222 | 7 | 0 | 34 | 0.0882 |
| 12_787313 | 7 | 0.2857 | 2 | 0.5 | 7 | 0 | 9 | 0 | 25 | 0.12 | 6 | 0.3333 | 16 | 0.1875 | 10 | 0.3 | 7 | 0.2857 | 39 | 0.2564 |
| 12_788440 | NA | NA | 3 | 0 | 1 | 1 | 5 | 0.2 | 9 | 0.2222 | NA | NA | 1 | 1 | 3 | 0.3333 | NA | NA | 4 | 0.5 |
| 12_788440 | NA | NA | 3 | 1 | 1 | 1 | 5 | 0.4 | 9 | 0.6667 | NA | NA | 1 | 1 | 2 | 1 | NA | NA | 3 | 1 |
| 12_788842 | 1 | 0 | 4 | 0.25 | 1 | 0 | 6 | 0.1667 | 12 | 0.1667 | NA | NA | 3 | 0.3333 | 5 | 0 | 2 | 0 | 10 | 0.1 |
| 12_788843 | 1 | 0 | 3 | 0.6667 | 1 | 0 | 5 | 0.6 | 10 | 0.5 | NA | NA | 1 | 1 | 4 | 0.25 | 2 | 1 | 7 | 0.5714 |
| 12_788843 | 1 | 1 | 1 | 0 | 1 | 0 | 6 | 0 | 9 | 0.1111 | NA | NA | 3 | 0.3333 | 4 | 0.25 | 2 | 1 | 9 | 0.4444 |
| 12_788844 | 1 | 1 | 4 | 1 | 1 | 1 | 7 | 1 | 13 | 1 | NA | NA | 3 | 0.6667 | 4 | 1 | 2 | 0.5 | 9 | 0.7778 |
| 12_788844 | 1 | 1 | 4 | 1 | 1 | 1 | 6 | 1 | 12 | 1 | NA | NA | 3 | 1 | 2 | 1 | 2 | 1 | 7 | 1 |
| 12_788845 | 1 | 0 | 4 | 0.5 | 1 | 0 | 7 | 0.4286 | 13 | 0.3846 | NA | NA | 2 | 0.5 | 4 | 0.5 | 2 | 0 | 8 | 0.375 |
| 12_788845 | 1 | 0 | 4 | 0 | 1 | 0 | 7 | 0.1429 | 13 | 0.0769 | NA | NA | 2 | 0 | 2 | 0.5 | 1 | 0 | 5 | 0.2 |
| 12_788901 | 44 | 0.3864 | 60 | 0.3167 | 44 | 0.2727 | 42 | 0.2619 | 190 | 0.3105 | 23 | 0.1739 | 37 | 0.2973 | 53 | 0.3208 | 34 | 0.2941 | 147 | 0.2857 |
| 12_788902 | 44 | 0.0682 | 57 | 0.0877 | 41 | 0.0488 | 42 | 0.0714 | 184 | 0.0707 | 26 | 0.1154 | 36 | 0.0556 | 57 | 0.0175 | 35 | 0.0857 | 154 | 0.0584 |
| 12_788903 | 34 | 0.2059 | 48 | 0.1458 | 35 | 0.1143 | 37 | 0.0541 | 154 | 0.1299 | 21 | 0.1429 | 33 | 0.0606 | 49 | 0.102 | 27 | 0 | 130 | 0.0769 |
| 12_789317 | 13 | 0.0769 | 8 | 0.375 | 10 | 0.3 | 7 | 0.1429 | 38 | 0.2105 | 9 | 0.2222 | 10 | 0.4 | 6 | 0.3333 | 9 | 0 | 34 | 0.2353 |
| 12_789319 | 13 | 0 | 10 | 0.2 | 11 | 0.2727 | 7 | 0 | 41 | 0.122 | 9 | 0.1111 | 13 | 0.2308 | 10 | 0 | 11 | 0 | 43 | 0.093 |
| 12_789320 | 12 | 0.3333 | 9 | 0.3333 | 11 | 0.2727 | 7 | 0.4286 | 39 | 0.3333 | 9 | 0.4444 | 13 | 0.3077 | 9 | 0.4444 | 10 | 0.3 | 41 | 0.3659 |
| 12_789322 | NA | NA | 1 | 0 | 2 | 0.5 | NA | NA | 3 | 0.3333 | 1 | 0 | 2 | 0 | 1 | 1 | NA | NA | 4 | 0.25 |
| 12_790086 | 2 | 0 | 4 | 0.25 | 5 | 0.2 | 1 | 1 | 12 | 0.25 | 2 | 0.5 | 3 | 0 | 1 | 0 | 2 | 0 | 8 | 0.125 |
| 12_790087 | 3 | 1 | 4 | 0.5 | 5 | 0.2 | 1 | 0 | 13 | 0.4615 | 3 | 0.6667 | 4 | 0.25 | 2 | 1 | 2 | 0 | 11 | 0.4545 |
| 12_790099 | 3 | 0 | 3 | 0 | 3 | 0.3333 | 1 | 0 | 10 | 0.1 | 2 | 0 | 2 | 0 | 2 | 0.5 | 2 | 0 | 8 | 0.125 |
| 12_790106 | 3 | 0 | 2 | 0 | 4 | 0.5 | NA | NA | 9 | 0.2222 | 2 | 0 | 2 | 0 | NA | NA | 2 | 0.5 | 6 | 0.1667 |
| 12_790110 | 1 | 0 | NA | NA | 1 | 1 | NA | NA | 2 | 0.5 | 1 | 1 | NA | NA | NA | NA | NA | NA | 1 | 1 |
| 12_790111 | 2 | 1 | 2 | 1 | 2 | 1 | NA | NA | 6 | 1 | 2 | 1 | 2 | 1 | NA | NA | 1 | 1 | 5 | 1 |
| 12_790111 | 1 | 1 | 2 | 0 | 2 | 1 | NA | NA | 5 | 0.6 | 2 | 1 | 2 | 1 | NA | NA | 1 | 1 | 5 | 1 |
| 12_790111 | 2 | 1 | 2 | 1 | 2 | 0.5 | NA | NA | 6 | 0.8333 | 2 | 1 | 1 | 1 | NA | NA | 1 | 1 | 4 | 1 |
| 12_790111 | 2 | 1 | 2 | 1 | 2 | 1 | NA | NA | 6 | 1 | 2 | 1 | 2 | 1 | NA | NA | 1 | 1 | 5 | 1 |
| 12_790111 | 1 | 1 | 1 | 0 | 2 | 1 | NA | NA | 4 | 0.75 | 2 | 1 | 2 | 1 | NA | NA | 1 | 1 | 5 | 1 |
| 12_790112 | 1 | 0 | 2 | 0.5 | 1 | 0 | NA | NA | 4 | 0.25 | 2 | 1 | 2 | 0.5 | NA | NA | 1 | 1 | 5 | 0.8 |
| 12_790112 | 1 | 1 | 1 | 1 | 1 | 1 | NA | NA | 3 | 1 | 2 | 1 | 2 | 1 | NA | NA | 1 | 1 | 5 | 1 |
| 12_790113 | 2 | 0.5 | 2 | 1 | 2 | 0.5 | NA | NA | 6 | 0.6667 | 2 | 1 | 1 | 1 | NA | NA | 1 | 1 | 4 | 1 |
| 12_790113 | 1 | 1 | 2 | 1 | NA | NA | NA | NA | 3 | 1 | 1 | 1 | NA | NA | NA | NA | NA | NA | 1 | 1 |
| 12_790113 | 2 | 1 | 2 | 1 | 2 | 0.5 | NA | NA | 6 | 0.8333 | 2 | 1 | 2 | 1 | NA | NA | 1 | 0 | 5 | 0.8 |
| 12_790114 | 1 | 0 | 2 | 1 | 2 | 0.5 | NA | NA | 5 | 0.6 | 2 | 1 | 2 | 0.5 | NA | NA | 1 | 0 | 5 | 0.6 |
| 12_790116 | 2 | 1 | 2 | 1 | 2 | 0.5 | NA | NA | 6 | 0.8333 | NA | NA | 2 | 1 | NA | NA | 1 | 1 | 3 | 1 |
| 12_790116 | 2 | 1 | 2 | 1 | 1 | 1 | NA | NA | 5 | 1 | 2 | 1 | 2 | 1 | NA | NA | 1 | 1 | 5 | 1 |
| 12_791263 | 1 | 0 | 2 | 0 | 1 | 0 | 1 | 1 | 5 | 0.2 | 2 | 0 | NA | NA | NA | NA | 1 | 1 | 3 | 0.3333 |
| 12_791268 | 1 | 0 | 1 | 1 | NA | NA | 1 | 1 | 3 | 0.6667 | 3 | 1 | NA | NA | 1 | 0 | NA | NA | 4 | 0.75 |
| 12_791291 | 1 | 0 | 1 | 1 | 1 | 1 | 1 | 1 | 4 | 0.75 | 1 | 1 | 1 | 0 | 1 | 1 | NA | NA | 3 | 0.6667 |
| 12_791291 | NA | NA | 1 | 1 | 1 | 1 | 1 | 1 | 3 | 1 | 1 | 0 | 1 | 0 | 1 | 1 | NA | NA | 3 | 0.3333 |
| 12_791882 | NA | NA | NA | NA | 2 | 0.5 | NA | NA | 2 | 0.5 | NA | NA | NA | NA | NA | NA | 1 | 1 | 1 | 1 |
| 12_792027 | 2 | 0.5 | NA | NA | 2 | 0 | NA | NA | 4 | 0.25 | 1 | 0 | 4 | 0 | 1 | 0 | 2 | 1 | 8 | 0.25 |
| 12_792028 | 2 | 0.5 | NA | NA | NA | NA | NA | NA | 2 | 0.5 | 1 | 1 | 4 | 1 | 1 | 1 | 2 | 1 | 8 | 1 |
| 12_792031 | 9 | 0.7778 | 15 | 0.7333 | 13 | 0.8462 | 13 | 0.7692 | 50 | 0.78 | 10 | 0.9 | 10 | 0.7 | 12 | 0.6667 | 7 | 1 | 39 | 0.7949 |
| 12_792032 | 9 | 0.3333 | 15 | 0.0667 | 19 | 0.1053 | 15 | 0.2 | 58 | 0.1552 | 15 | 0.0667 | 13 | 0.0769 | 14 | 0.1429 | 13 | 0.2308 | 55 | 0.1273 |
| 12_792032 | 9 | 0.1111 | 15 | 0.2667 | 16 | 0.125 | 11 | 0.1818 | 51 | 0.1765 | 13 | 0.1538 | 7 | 0.1429 | 11 | 0.0909 | 9 | 0.2222 | 40 | 0.15 |
| 12_792220 | 2 | 0 | 2 | 0.5 | 1 | 0 | NA | NA | 5 | 0.2 | 2 | 0 | 4 | 0 | 2 | 0.5 | NA | NA | 8 | 0.125 |
| 12_792224 | 3 | 0.6667 | 5 | 1 | 2 | 1 | 3 | 1 | 13 | 0.9231 | 2 | 0.5 | 4 | 0.75 | 1 | 1 | 2 | 0.5 | 9 | 0.6667 |
| 12_792225 | 2 | 0.5 | 3 | 0.3333 | 2 | 0 | 3 | 0 | 10 | 0.2 | 2 | 0 | 3 | 0.3333 | 1 | 0 | 1 | 1 | 7 | 0.2857 |
| 12_792225 | 2 | 0.5 | 5 | 1 | 2 | 1 | 3 | 0.6667 | 12 | 0.8333 | 2 | 1 | 2 | 0.5 | 1 | 1 | 2 | 0.5 | 7 | 0.7143 |
| 12_792226 | 2 | 0 | 4 | 0.25 | 1 | 0 | 3 | 0 | 10 | 0.1 | 2 | 1 | 2 | 0.5 | 1 | 0 | 1 | 0 | 6 | 0.5 |
| 12_792226 | 2 | 1 | 4 | 0.75 | 1 | 1 | 3 | 1 | 10 | 0.9 | 3 | 1 | 3 | 1 | 1 | 1 | 1 | 1 | 8 | 1 |
| 12_792227 | 2 | 0 | 4 | 0 | 1 | 0 | 4 | 0.25 | 11 | 0.0909 | 2 | 0 | 4 | 0.25 | 1 | 1 | 1 | 1 | 8 | 0.375 |
| 12_792227 | 2 | 1 | 4 | 1 | NA | NA | 4 | 1 | 10 | 1 | 3 | 1 | 3 | 1 | 1 | 1 | 2 | 1 | 9 | 1 |
| 12_792228 | 2 | 0 | 4 | 0.5 | 1 | 0 | 4 | 0.25 | 11 | 0.2727 | 3 | 0.3333 | 4 | 0 | 1 | 0 | 2 | 0 | 10 | 0.1 |
| 12_792229 | 2 | 0.5 | 4 | 1 | 1 | 1 | 4 | 1 | 11 | 0.9091 | 3 | 1 | 3 | 1 | 1 | 1 | 1 | 1 | 8 | 1 |
| 12_792688 | 2 | 1 | NA | NA | NA | NA | NA | NA | 2 | 1 | NA | NA | 1 | 1 | NA | NA | NA | NA | 1 | 1 |
| 12_792792 | 3 | 0 | 1 | 0 | 2 | 0 | 1 | 1 | 7 | 0.1429 | 2 | 0.5 | 1 | 0 | 4 | 0 | 2 | 0.5 | 9 | 0.2222 |

| | | | | | | | | | | | | | | | | | | | | |
|---|---|---|---|---|---|---|---|---|---|---|---|---|---|---|---|---|---|---|---|---|
| 12_792795 | 3 | 0.3333 | NA | NA | 2 | 0 | 2 | 0 | 7 | 0.1429 | 2 | 0.5 | 1 | 0 | 1 | 0 | 2 | 1 | 6 | 0.5 |
| 12_792796 | 3 | 0.3333 | 1 | 0 | 2 | 0 | 2 | 0 | 8 | 0.125 | 2 | 0.5 | 2 | 0 | 2 | 0 | 2 | 0 | 8 | 0.125 |
| 12_792797 | 3 | 0.3333 | 1 | 1 | 1 | 0 | 2 | 0 | 7 | 0.2857 | 2 | 0 | 2 | 0 | 2 | 0 | 2 | 0.5 | 8 | 0.125 |
| 12_792804 | 2 | 0 | 1 | 1 | 1 | 0 | 2 | 0 | 6 | 0.1667 | 1 | 0 | 2 | 0 | 2 | 0 | 2 | 0.5 | 7 | 0.1429 |
| 12_792804 | 3 | 1 | NA | NA | 1 | 0 | 1 | 1 | 5 | 0.8 | 1 | 1 | 2 | 0.5 | 2 | 1 | 2 | 0.5 | 7 | 0.7143 |
| 12_792805 | 3 | 0.3333 | NA | NA | 2 | 0.5 | 2 | 1 | 7 | 0.5714 | 1 | 1 | 1 | 0 | 2 | 1 | 2 | 1 | 6 | 0.8333 |
| 12_792806 | 1 | 1 | NA | NA | 1 | 1 | 2 | 1 | 4 | 1 | 1 | 1 | 1 | 1 | 2 | 1 | 1 | 1 | 5 | 1 |
| 12_792808 | 1 | 0 | NA | NA | 2 | 0 | 2 | 0.5 | 5 | 0.2 | 1 | 0 | 1 | 1 | 2 | 0 | 1 | 1 | 5 | 0.4 |
| 12_801547 | 3 | 0 | 6 | 0 | 3 | 0.3333 | 5 | 0.2 | 17 | 0.1176 | 4 | 0.25 | 4 | 0 | 5 | 0.2 | 6 | 0 | 19 | 0.1053 |
| 12_801548 | 3 | 0.3333 | 6 | 0 | 3 | 0.3333 | 5 | 0.2 | 17 | 0.1765 | 4 | 0.25 | 4 | 0.25 | 4 | 0 | 7 | 0.1429 | 19 | 0.1579 |
| 12_801549 | 2 | 1 | 6 | 0.8333 | 2 | 0.5 | 4 | 0.75 | 14 | 0.7857 | 3 | 0.6667 | 4 | 0.25 | 5 | 0.6 | 6 | 0.8333 | 18 | 0.6111 |
| 12_801555 | 3 | 0 | 6 | 0.1667 | 2 | 0.5 | 3 | 1 | 14 | 0.3571 | 4 | 0 | 2 | 0.5 | 5 | 0.6 | 5 | 0.4 | 16 | 0.375 |
| 12_801555 | 3 | 0.3333 | 5 | 0.8 | 2 | 1 | 4 | 0.5 | 14 | 0.6429 | 4 | 0.5 | 2 | 0.5 | 5 | 0.2 | 6 | 0.6667 | 17 | 0.4706 |
| 12_801556 | 3 | 0 | 5 | 0.4 | 2 | 0 | 4 | 0 | 14 | 0.1429 | 4 | 0 | 1 | 0 | 5 | 0 | 5 | 0.2 | 15 | 0.0667 |
| 12_801557 | 3 | 0.3333 | 4 | 0.75 | 2 | 0 | 3 | 0.6667 | 12 | 0.5 | 3 | 0 | 1 | 0 | 5 | 0.4 | 4 | 0.5 | 13 | 0.3077 |
| 12_801561 | 2 | 0.5 | 5 | 0 | 2 | 0 | 1 | 0 | 10 | 0.1 | 3 | 0 | NA | NA | 3 | 0.3333 | 4 | 0 | 10 | 0.1 |
| 12_801564 | NA | NA | 5 | 0.2 | 1 | 0 | 1 | 0 | 7 | 0.1429 | 2 | 0 | NA | NA | 3 | 0 | 6 | 0.1667 | 11 | 0.0909 |
| 12_801565 | 2 | 0.5 | 6 | 0.5 | NA | NA | NA | NA | 8 | 0.5 | 3 | 0.6667 | 1 | 1 | 2 | 1 | 5 | 0.6 | 11 | 0.7273 |
| 12_801566 | 2 | 0 | 6 | 0.1667 | 1 | 0 | 1 | 0 | 10 | 0.1 | 3 | 0 | 1 | 0 | 4 | 0 | 6 | 0.1667 | 14 | 0.0714 |
| 12_801567 | 2 | 0.5 | 4 | 0.25 | 1 | 0 | 1 | 0 | 8 | 0.25 | NA | NA | 1 | 0 | 4 | 0.25 | 2 | 0 | 7 | 0.1429 |
| 12_801569 | 2 | 1 | 5 | 0.4 | 1 | 1 | 1 | 0 | 9 | 0.5556 | 2 | 1 | 1 | 0 | 3 | 0.6667 | 2 | 1 | 8 | 0.75 |
| 12_801572 | NA | NA | 3 | 0.3333 | 1 | 1 | 1 | 1 | 5 | 0.6 | 2 | 0.5 | NA | NA | 3 | 1 | 2 | 0 | 7 | 0.5714 |
| 12_801716 | 1 | 1 | NA | NA | NA | NA | 1 | 1 | 2 | 1 | 1 | 1 | 1 | 0 | NA | NA | NA | NA | 2 | 0.5 |
| 12_802149 | 15 | 0.3333 | 17 | 0.4118 | 11 | 0.0909 | 22 | 0.1364 | 65 | 0.2462 | 13 | 0 | 11 | 0.1818 | 7 | 0.2857 | 8 | 0.125 | 39 | 0.1282 |
| 12_802162 | NA | NA | 3 | 0.3333 | 2 | 0 | 8 | 0.375 | 13 | 0.3077 | 1 | 0 | NA | NA | 2 | 0 | 2 | 0.5 | 5 | 0.2 |
| 12_802168 | NA | NA | 3 | 0 | 1 | 0 | 6 | 0.5 | 10 | 0.3 | 2 | 0.5 | NA | NA | 2 | 0 | 2 | 0 | 6 | 0.1667 |
| 12_802170 | 14 | 0.1429 | 13 | 0.0769 | 10 | 0.3 | 18 | 0.2778 | 55 | 0.2 | 10 | 0.2 | 7 | 0 | 6 | 0.1667 | 9 | 0.3333 | 32 | 0.1875 |
| 12_802171 | NA | NA | 3 | 0.3333 | NA | NA | 1 | 0 | 4 | 0.25 | 1 | 1 | NA | NA | 1 | 0 | 2 | 0 | 4 | 0.25 |
| 12_802172 | NA | NA | 3 | 0.3333 | NA | NA | 3 | 0 | 6 | 0.1667 | 2 | 0 | NA | NA | 1 | 0 | 2 | 0.5 | 5 | 0.2 |
| 12_803068 | 4 | 0.25 | 4 | 0 | 3 | 0 | 4 | 0 | 15 | 0.0667 | 7 | 0.1429 | 4 | 0.25 | NA | NA | 1 | 0 | 12 | 0.1667 |
| 12_803836 | 7 | 0 | 6 | 0.3333 | 7 | 0.1429 | 4 | 0.25 | 24 | 0.1667 | 1 | 0 | 4 | 0.25 | 4 | 0.5 | 4 | 0 | 13 | 0.2308 |
| 12_803836 | 5 | 0.6 | 5 | 0.6 | 6 | 0.8333 | 3 | 0.6667 | 19 | 0.6842 | 1 | 0 | 5 | 0.4 | 4 | 0.75 | 3 | 1 | 13 | 0.6154 |
| 12_803839 | 6 | 0.1667 | 5 | 0 | 4 | 0 | 2 | 0 | 17 | 0.0588 | 1 | 0 | 3 | 0 | 1 | 1 | 2 | 0 | 7 | 0.1429 |
| 12_803839 | 4 | 0 | 5 | 0.4 | 2 | 0.5 | 2 | 0.5 | 13 | 0.3077 | 1 | 0 | 4 | 0.25 | 1 | 0 | 2 | 0 | 8 | 0.125 |
| 12_803840 | 6 | 0.1667 | 5 | 0.2 | 3 | 0 | 3 | 0 | 17 | 0.1176 | 1 | 0 | 5 | 0.2 | 2 | 0 | 3 | 0 | 11 | 0.0909 |
| 12_803845 | 5 | 0.2 | 5 | 0 | 2 | 0 | 1 | 0 | 13 | 0.0769 | 1 | 0 | 4 | 0.25 | 2 | 0 | 3 | 0 | 10 | 0.1 |
| 12_803850 | 4 | 0.75 | 2 | 0 | 1 | 1 | 2 | 0.5 | 9 | 0.5556 | NA | NA | 2 | 1 | 2 | 1 | 1 | 1 | 5 | 1 |
| 12_803852 | 3 | 0.3333 | 3 | 0.3333 | 1 | 0 | 1 | 0 | 8 | 0.25 | NA | NA | NA | NA | 4 | 0.5 | 2 | 0.5 | 6 | 0.5 |
| 12_803852 | 2 | 0 | 3 | 0.6667 | 2 | 1 | 1 | 0 | 8 | 0.5 | NA | NA | 1 | 1 | 4 | 0.25 | 1 | 0 | 6 | 0.3333 |
| 12_803852 | 1 | 1 | NA | NA | NA | NA | NA | NA | 1 | 1 | NA | NA | NA | NA | NA | NA | 1 | 1 | 1 | 1 |
| 12_803853 | 3 | 1 | 2 | 0 | 2 | 1 | NA | NA | 7 | 0.7143 | NA | NA | 1 | 1 | 1 | 0 | NA | NA | 2 | 0.5 |
| 12_803854 | 4 | 0.25 | 3 | 0 | 2 | 0 | 1 | 1 | 10 | 0.2 | NA | NA | 1 | 0 | 6 | 0.5 | 2 | 0 | 9 | 0.3333 |
| 12_803854 | 4 | 0.5 | 3 | 1 | 2 | 1 | 1 | 1 | 10 | 0.8 | NA | NA | 1 | 1 | 5 | 1 | 2 | 1 | 8 | 1 |
| 12_803855 | 3 | 0.3333 | 3 | 0 | 2 | 0 | 1 | 0 | 9 | 0.1111 | NA | NA | 1 | 1 | 5 | 0.2 | 1 | 0 | 7 | 0.2857 |
| 12_803855 | 4 | 0.75 | 3 | 0.3333 | 2 | 0.5 | NA | NA | 9 | 0.5556 | NA | NA | 1 | 1 | 5 | 0.6 | 1 | 0 | 7 | 0.5714 |
| 12_803855 | 3 | 0.3333 | 3 | 0.6667 | 1 | 0 | 1 | 0 | 8 | 0.375 | NA | NA | 1 | 1 | 4 | 1 | 2 | 1 | 7 | 1 |
| 12_803855 | 4 | 0.25 | 3 | 0.6667 | 2 | 0.5 | 1 | 0 | 10 | 0.4 | NA | NA | 1 | 0 | 3 | 0.3333 | 2 | 0 | 6 | 0.1667 |
| 12_803856 | 4 | 0.25 | 2 | 0.5 | 2 | 0 | 1 | 0 | 9 | 0.2222 | NA | NA | NA | NA | 5 | 0.6 | 2 | 0 | 7 | 0.4286 |
| 12_803858 | 2 | 1 | 2 | 1 | 2 | 0.5 | 1 | 0 | 7 | 0.7143 | NA | NA | 1 | 1 | 3 | 0.6667 | NA | NA | 4 | 0.75 |
| 12_803858 | 4 | 1 | 2 | 1 | 2 | 1 | 1 | 1 | 9 | 1 | NA | NA | 1 | 1 | 3 | 1 | 2 | 1 | 6 | 1 |
| 12_803859 | 2 | 0.5 | 2 | 1 | 1 | 1 | NA | NA | 5 | 0.8 | NA | NA | 1 | 1 | 3 | 0.6667 | 2 | 0.5 | 6 | 0.6667 |
| 12_803859 | 3 | 0.3333 | 2 | 1 | 1 | 0 | 1 | 0 | 7 | 0.4286 | NA | NA | 1 | 1 | 3 | 0.6667 | 2 | 0.5 | 6 | 0.6667 |
| 12_803862 | 3 | 0.3333 | 2 | 0 | 1 | 0 | 1 | 0 | 7 | 0.1429 | NA | NA | 1 | 0 | 2 | 0.5 | 1 | 0 | 4 | 0.25 |
| 12_803864 | 3 | 0.3333 | 1 | 1 | 1 | 1 | 1 | 0 | 6 | 0.5 | NA | NA | 1 | 0 | 1 | 1 | 1 | 0 | 3 | 0.3333 |
| 12_803866 | 3 | 0.3333 | 1 | 0 | 1 | 0 | 1 | 0 | 6 | 0.1667 | NA | NA | 1 | 1 | 2 | 0 | NA | NA | 3 | 0.3333 |
| 12_803867 | 3 | 0.3333 | NA | NA | 1 | 1 | 1 | 1 | 5 | 0.6 | NA | NA | 1 | 1 | 2 | 0.5 | NA | NA | 3 | 0.6667 |
| 12_803868 | 3 | 0.6667 | NA | NA | 1 | 1 | 1 | 0 | 5 | 0.6 | NA | NA | NA | NA | 2 | 0.5 | NA | NA | 2 | 0.5 |
| 12_803868 | 3 | 0.3333 | NA | NA | 1 | 0 | 1 | 0 | 5 | 0.2 | NA | NA | NA | NA | 1 | 1 | NA | NA | 1 | 1 |
| 12_804824 | 1 | 1 | 3 | 0 | 1 | 1 | NA | NA | 5 | 0.4 | NA | NA | 1 | 0 | 2 | 0.5 | NA | NA | 3 | 0.3333 |
| 12_805362 | 4 | 0.25 | 2 | 0 | 2 | 0 | NA | NA | 8 | 0.125 | 3 | 0.6667 | 4 | 0.5 | 7 | 0.2857 | 4 | 0 | 18 | 0.3333 |
| 12_805363 | 5 | 0.6 | 2 | 1 | 3 | 0.3333 | 1 | 0 | 11 | 0.5455 | 2 | 0 | 4 | 0.75 | 6 | 0.6667 | 5 | 1 | 17 | 0.7059 |
| 12_805364 | 5 | 0.6 | 2 | 0 | 4 | 0.25 | 1 | 0 | 12 | 0.3333 | 3 | 0 | 3 | 0.6667 | 7 | 0.2857 | 5 | 0.2 | 18 | 0.2778 |
| 12_805378 | NA | NA | NA | NA | 1 | 1 | NA | NA | 1 | 1 | 1 | 1 | NA | NA | 4 | 0 | 1 | 0 | 6 | 0.1667 |
| 12_805393 | 6 | 0 | 6 | 0.5 | 4 | 0 | 2 | 0 | 18 | 0.1667 | 3 | 0.3333 | 4 | 0.25 | 8 | 0.25 | 7 | 0.1429 | 22 | 0.2273 |
| 12_805395 | 6 | 0 | 7 | 0.1429 | 5 | 0 | 2 | 0 | 20 | 0.05 | 4 | 0.5 | 3 | 0 | 9 | 0.1111 | 5 | 0 | 21 | 0.1429 |

| | | | | | | | | | | | | | | | | | | | | |
|---|---|---|---|---|---|---|---|---|---|---|---|---|---|---|---|---|---|---|---|---|
| 12_805436 | 7 | 0.8571 | 8 | 0.875 | 6 | 0.8333 | 2 | 0.5 | 23 | 0.8261 | 4 | 0.75 | 4 | 1 | 8 | 0.625 | 7 | 0.5714 | 23 | 0.6957 |
| 12_805441 | 5 | 0.4 | 8 | 0.125 | 5 | 0 | 2 | 0 | 20 | 0.15 | 5 | 0 | 4 | 0.5 | 8 | 0 | 6 | 0.3333 | 23 | 0.1739 |
| 12_805446 | 8 | 0.5 | 7 | 0.2857 | 7 | 0.2857 | 2 | 0 | 24 | 0.3333 | 7 | 0.1429 | 5 | 0.2 | 11 | 0.0909 | 6 | 0 | 29 | 0.1034 |
| 12_806486 | 5 | 0.6 | 4 | 0.25 | 3 | 0 | 4 | 0.75 | 16 | 0.4375 | 12 | 0.25 | 2 | 1 | 13 | 0.4615 | 7 | 0.2857 | 34 | 0.3824 |
| 12_806486 | 2 | 0 | 4 | 0.25 | 2 | 0 | 5 | 0.6 | 13 | 0.3077 | 6 | 0.3333 | 3 | 0.6667 | 8 | 0.375 | 7 | 0.1429 | 24 | 0.3333 |
| 12_806488 | 8 | 0 | 6 | 0.1667 | 7 | 0.1429 | 5 | 0 | 26 | 0.0769 | 12 | 0.0833 | 6 | 0 | 19 | 0.2105 | 9 | 0 | 46 | 0.1087 |
| 12_806492 | 10 | 0.1 | 8 | 0 | 8 | 0 | 6 | 0.1667 | 32 | 0.0625 | 14 | 0.0714 | 7 | 0 | 23 | 0.1739 | 8 | 0 | 52 | 0.0962 |
| 12_806494 | 9 | 0.1111 | 8 | 0.125 | 7 | 0.1429 | 6 | 0 | 30 | 0.1 | 8 | 0.25 | 3 | 0 | 23 | 0.3043 | 10 | 0.3 | 44 | 0.2727 |
| 12_806495 | 9 | 0 | 9 | 0 | 9 | 0.2222 | 8 | 0 | 35 | 0.0571 | 15 | 0.1333 | 6 | 0.1667 | 27 | 0.037 | 12 | 0 | 60 | 0.0667 |
| 12_806503 | 11 | 0.0909 | 9 | 0.2222 | 10 | 0.1 | 11 | 0.1818 | 41 | 0.1463 | 17 | 0.3529 | 13 | 0.0769 | 28 | 0.1429 | 13 | 0.0769 | 71 | 0.169 |
| 12_806504 | 12 | 0.3333 | 10 | 0.3 | 11 | 0.0909 | 8 | 0.75 | 41 | 0.3415 | 16 | 0.4375 | 14 | 0.3571 | 23 | 0.2609 | 13 | 0.2308 | 66 | 0.3182 |
| 12_806504 | 15 | 0.4 | 10 | 0.2 | 12 | 0.4167 | 10 | 0.5 | 47 | 0.383 | 20 | 0.4 | 12 | 0.0833 | 31 | 0.129 | 17 | 0.1765 | 80 | 0.2 |
| 12_806809 | 18 | 0.0556 | 10 | 0.1 | 12 | 0 | 17 | 0.0588 | 57 | 0.0526 | 11 | 0 | 9 | 0.2222 | 16 | 0 | 13 | 0.0769 | 49 | 0.0612 |
| 12_806811 | 19 | 0.1579 | 11 | 0.1818 | 9 | 0 | 17 | 0.0588 | 56 | 0.1071 | 7 | 0.1429 | 13 | 0 | 18 | 0.2222 | 11 | 0.2727 | 49 | 0.1633 |
| 12_806912 | 1 | 1 | 1 | 1 | NA | NA | 2 | 0.5 | 4 | 0.75 | 1 | 1 | NA | NA | NA | NA | 1 | 0 | 2 | 0.5 |
| 12_806916 | 1 | 1 | 1 | 0 | NA | NA | 2 | 0.5 | 4 | 0.5 | 1 | 1 | NA | NA | NA | NA | 1 | 0 | 2 | 0.5 |
| 12_807263 | 2 | 1 | 3 | 1 | 2 | 1 | 3 | 1 | 10 | 1 | 1 | 1 | NA | NA | 1 | 1 | 1 | 1 | 3 | 1 |
| 12_807264 | 2 | 1 | 2 | 0.5 | 2 | 1 | 3 | 0.6667 | 9 | 0.7778 | 1 | 0 | NA | NA | 1 | 1 | 1 | 1 | 3 | 0.6667 |
| 12_807266 | 1 | 1 | 3 | 1 | 1 | 1 | 2 | 1 | 7 | 1 | 1 | 1 | NA | NA | 1 | 0 | 2 | 1 | 4 | 0.75 |
| 12_807266 | 2 | 1 | 3 | 1 | 2 | 1 | 3 | 1 | 10 | 1 | 1 | 0 | NA | NA | 1 | 1 | 1 | 1 | 3 | 0.6667 |
| 12_807270 | 2 | 0 | 2 | 0.5 | 1 | 0 | 3 | 0.3333 | 8 | 0.25 | 2 | 0 | NA | NA | 1 | 1 | 2 | 0 | 5 | 0.2 |
| 12_807286 | 5 | 0.2 | 2 | 0 | 1 | 0 | 2 | 0 | 10 | 0.1 | 3 | 0.3333 | NA | NA | 1 | 0 | 2 | 0 | 6 | 0.1667 |
| 12_807295 | 5 | 0.4 | 5 | 0.4 | 2 | 0 | 3 | 0.3333 | 15 | 0.3333 | 5 | 0.4 | 2 | 0 | 2 | 0.5 | 2 | 0.5 | 11 | 0.3636 |
| 12_807296 | 7 | 0.2857 | 5 | 0.2 | 1 | 0 | 3 | 0 | 16 | 0.1875 | 4 | 0.5 | 2 | 0 | 2 | 0 | 2 | 0 | 10 | 0.2 |
| 12_807296 | 7 | 0.1429 | 5 | 0 | 1 | 0 | 2 | 0 | 15 | 0.0667 | 5 | 0.2 | 2 | 0 | 1 | 0 | 2 | 0.5 | 10 | 0.2 |
| 12_807297 | 6 | 0.1667 | 5 | 0.2 | 1 | 0 | 2 | 0 | 14 | 0.1429 | 4 | 0 | 2 | 0.5 | 3 | 0 | 2 | 0 | 11 | 0.0909 |
| 12_807297 | 6 | 0.1667 | 5 | 0 | 2 | 0 | 3 | 0 | 16 | 0.0625 | 5 | 0.2 | 2 | 0 | 2 | 0 | 2 | 0 | 11 | 0.0909 |
| 12_807298 | 7 | 0.7143 | 5 | 0.4 | 1 | 0 | 3 | 0.3333 | 16 | 0.5 | 5 | 0.6 | 2 | 0 | 3 | 0.3333 | 2 | 0.5 | 12 | 0.4167 |
| 12_807298 | 5 | 0.8 | 4 | 1 | 2 | 0.5 | 3 | 1 | 14 | 0.8571 | 4 | 0.75 | 2 | 0 | 2 | 0 | 1 | 0 | 9 | 0.3333 |
| 12_808010 | 1 | 0 | 3 | 0 | 1 | 1 | NA | NA | 5 | 0.2 | NA | NA | NA | NA | 4 | 0.25 | 2 | 0 | 6 | 0.1667 |
| 12_808011 | 1 | 1 | 3 | 0.6667 | NA | NA | NA | NA | 4 | 0.75 | NA | NA | NA | NA | 4 | 0.5 | NA | NA | 4 | 0.5 |
| 12_808012 | 1 | 1 | 4 | 1 | NA | NA | NA | NA | 5 | 1 | NA | NA | NA | NA | 3 | 1 | 2 | 1 | 5 | 1 |
| 12_808013 | 1 | 1 | 4 | 0.75 | NA | NA | NA | NA | 5 | 0.8 | NA | NA | NA | NA | 3 | 0.6667 | 2 | 1 | 5 | 0.8 |
| 12_808015 | 1 | 0 | 4 | 0.5 | 1 | 0 | NA | NA | 6 | 0.3333 | NA | NA | NA | NA | 3 | 0.6667 | 2 | 0.5 | 5 | 0.6 |
| 12_808018 | 1 | 1 | 1 | 0 | NA | NA | NA | NA | 2 | 0.5 | NA | NA | NA | NA | 2 | 0 | 1 | 1 | 3 | 0.3333 |
| 12_808019 | 1 | 1 | 1 | 1 | NA | NA | NA | NA | 2 | 1 | NA | NA | NA | NA | 1 | 1 | 2 | 1 | 3 | 1 |
| 12_808019 | 1 | 0 | 2 | 0.5 | NA | NA | NA | NA | 3 | 0.3333 | NA | NA | NA | NA | 3 | 0.3333 | 1 | 1 | 4 | 0.5 |
| 12_808031 | 1 | 1 | 1 | 0 | 2 | 1 | 1 | 0 | 5 | 0.6 | NA | NA | NA | NA | 3 | 0 | 1 | 1 | 4 | 0.25 |
| 12_808036 | 1 | 1 | NA | NA | 2 | 1 | 1 | 1 | 4 | 1 | NA | NA | NA | NA | 3 | 1 | NA | NA | 3 | 1 |
| 12_808036 | 1 | 1 | NA | NA | 1 | 1 | 1 | 0 | 3 | 0.6667 | NA | NA | NA | NA | 2 | 1 | NA | NA | 2 | 1 |
| 12_808037 | 1 | 1 | NA | NA | 2 | 1 | 1 | 1 | 4 | 1 | NA | NA | NA | NA | 3 | 1 | NA | NA | 3 | 1 |
| 12_808037 | 1 | 0 | NA | NA | 2 | 0.5 | 1 | 0 | 4 | 0.25 | NA | NA | NA | NA | 3 | 0.6667 | NA | NA | 3 | 0.6667 |
| 12_808039 | 1 | 1 | NA | NA | 1 | 1 | 1 | 1 | 3 | 1 | NA | NA | NA | NA | 3 | 1 | NA | NA | 3 | 1 |
| 12_808040 | 1 | 1 | NA | NA | 2 | 0 | 1 | 0 | 4 | 0.25 | NA | NA | NA | NA | 3 | 0.3333 | NA | NA | 3 | 0.3333 |
| 12_808040 | 1 | 1 | NA | NA | 2 | 0.5 | 1 | 1 | 4 | 0.75 | NA | NA | NA | NA | 2 | 0.5 | NA | NA | 2 | 0.5 |
| 12_808040 | NA | NA | NA | NA | 1 | 1 | NA | NA | 1 | 1 | NA | NA | NA | NA | 3 | 1 | NA | NA | 3 | 1 |
| 12_808041 | 1 | 1 | NA | NA | 2 | 0.5 | 1 | 1 | 4 | 0.75 | NA | NA | NA | NA | 3 | 0.3333 | NA | NA | 3 | 0.3333 |
| 12_808041 | 1 | 1 | NA | NA | 2 | 0.5 | 1 | 0 | 4 | 0.5 | NA | NA | NA | NA | 3 | 0.3333 | NA | NA | 3 | 0.3333 |
| 12_808056 | 1 | 1 | NA | NA | NA | NA | NA | NA | 1 | 1 | NA | NA | NA | NA | 2 | 0.5 | NA | NA | 2 | 0.5 |
| 12_808056 | 1 | 1 | NA | NA | 1 | 1 | NA | NA | 2 | 1 | NA | NA | NA | NA | 2 | 1 | NA | NA | 2 | 1 |
| 12_809212 | 3 | 0.6667 | 1 | 1 | 2 | 1 | 1 | 1 | 7 | 0.8571 | 1 | 1 | 1 | 1 | 1 | 0 | 2 | 0.5 | 5 | 0.6 |
| 12_809212 | 4 | 0.25 | 1 | 1 | 2 | 0 | 1 | 1 | 8 | 0.375 | 1 | 1 | 1 | 0 | 1 | 0 | 3 | 0 | 6 | 0.1667 |
| 12_809240 | 5 | 0 | 1 | 1 | 2 | 0.5 | 1 | 0 | 9 | 0.2222 | 1 | 0 | 1 | 0 | 1 | 1 | 3 | 0.6667 | 6 | 0.5 |
| 12_809240 | 5 | 0.4 | 1 | 1 | 1 | 0 | 1 | 0 | 8 | 0.375 | 1 | 1 | 1 | 0 | 1 | 0 | 4 | 0.25 | 7 | 0.2857 |
| 12_809243 | 4 | 0.25 | 2 | 0 | 2 | 0 | NA | NA | 8 | 0.125 | 3 | 0.3333 | 1 | 0 | 1 | 0 | 3 | 0.3333 | 8 | 0.25 |
| 12_809244 | 5 | 0.2 | 2 | 0 | 2 | 0.5 | NA | NA | 9 | 0.2222 | 2 | 0.5 | NA | NA | 1 | 0 | 3 | 0 | 6 | 0.1667 |
| 12_809244 | 4 | 0.5 | 3 | 0.3333 | 2 | 0 | NA | NA | 9 | 0.3333 | 3 | 0.6667 | 1 | 1 | 1 | 0 | 4 | 0.5 | 9 | 0.5556 |
| 12_809247 | 3 | 0.6667 | 3 | 0.6667 | 1 | 0 | 1 | 1 | 8 | 0.625 | 3 | 1 | 1 | 1 | 1 | 0 | 3 | 0.3333 | 8 | 0.625 |
| 12_809262 | 6 | 0 | 7 | 0.1429 | 3 | 0 | 3 | 0 | 19 | 0.0526 | 4 | 0.25 | 2 | 0 | 2 | 0.5 | 4 | 0 | 12 | 0.1667 |
| 12_809262 | 6 | 0 | 8 | 0 | 3 | 0 | 3 | 0.3333 | 20 | 0.05 | 6 | 0.1667 | 2 | 0.5 | 2 | 0 | 5 | 0 | 15 | 0.1333 |
| 12_809263 | 5 | 0 | 6 | 0.1667 | 3 | 0 | 3 | 0 | 17 | 0.0588 | 6 | 0 | 2 | 0 | 2 | 0 | 5 | 0.2 | 15 | 0.0667 |
| 12_809264 | 6 | 0.1667 | 8 | 0.625 | 3 | 0.3333 | 3 | 0.3333 | 20 | 0.4 | 6 | 0.6667 | 3 | 0.3333 | 2 | 0.5 | 5 | 0 | 16 | 0.375 |
| 12_809266 | 5 | 0.2 | 5 | 0.4 | 1 | 0 | 2 | 0 | 13 | 0.2308 | 4 | 0 | 2 | 0.5 | 2 | 0 | 5 | 0 | 13 | 0.0769 |
| 12_809274 | 7 | 0.4286 | 8 | 0.125 | 3 | 0.6667 | 3 | 0 | 21 | 0.2857 | 10 | 0.1 | 4 | 0.5 | 2 | 0 | 7 | 0.1429 | 23 | 0.1739 |
| 12_809967 | 13 | 0.3846 | 16 | 0.125 | 18 | 0.4444 | 18 | 0.5 | 65 | 0.3692 | 12 | 0.3333 | 14 | 0.5 | 12 | 0.0833 | 20 | 0.3 | 58 | 0.3103 |

| | | | | | | | | | | | | | | | | | | | | |
|---|---|---|---|---|---|---|---|---|---|---|---|---|---|---|---|---|---|---|---|---|
| 12_8099675 | 10 | 0.7 | 18 | 0.8889 | 18 | 0.8889 | 20 | 0.9 | 66 | 0.8636 | 11 | 0.9091 | 13 | 0.9231 | 15 | 0.7333 | 21 | 1 | 60 | 0.9 |
| 12_8099676 | 2 | 0 | 9 | 0.2222 | 8 | 0.375 | 5 | 0.2 | 24 | 0.25 | 2 | 0.5 | 2 | 0 | 5 | 0.2 | 14 | 0.2143 | 23 | 0.2174 |
| 12_8099689 | 11 | 0.1818 | 13 | 0.1538 | 11 | 0.0909 | 12 | 0 | 47 | 0.1064 | 6 | 0 | 15 | 0.1333 | 13 | 0 | 17 | 0.1176 | 51 | 0.0784 |
| 12_8099696 | 17 | 0.1176 | 21 | 0.0476 | 28 | 0.0714 | 19 | 0 | 85 | 0.0588 | 13 | 0.1538 | 21 | 0.0476 | 20 | 0.1 | 28 | 0.1429 | 82 | 0.1098 |
| 12_8099704 | 20 | 0.05 | 22 | 0.0455 | 28 | 0.0357 | 20 | 0.15 | 90 | 0.0667 | 14 | 0.2143 | 20 | 0.05 | 25 | 0 | 29 | 0.0345 | 88 | 0.0568 |
| 12_8099712 | 18 | 0.5 | 19 | 0.3684 | 24 | 0.4583 | 18 | 0.6111 | 79 | 0.481 | 15 | 0.6 | 19 | 0.5263 | 15 | 0.3333 | 25 | 0.4 | 74 | 0.4595 |
| 12_8123180 | 1 | 1 | 4 | 0.75 | 3 | 1 | 1 | 1 | 9 | 0.8889 | 2 | 1 | NA | NA | NA | NA | NA | NA | 2 | 1 |
| 12_8123210 | 2 | 0 | 4 | 0.25 | 3 | 0.3333 | 1 | 1 | 10 | 0.3 | 2 | 0.5 | NA | NA | NA | NA | NA | NA | 2 | 0.5 |
| 12_8123221 | 2 | 0 | 4 | 0.75 | 3 | 0 | 1 | 0 | 10 | 0.3 | 2 | 0.5 | NA | NA | NA | NA | NA | NA | 2 | 0.5 |
| 12_8123240 | 1 | 0 | 3 | 1 | 3 | 0.6667 | 1 | 1 | 8 | 0.75 | 1 | 1 | NA | NA | NA | NA | NA | NA | 1 | 1 |
| 12_8123264 | 2 | 0.5 | 4 | 1 | 3 | 0.6667 | 1 | 1 | 10 | 0.8 | 2 | 1 | NA | NA | NA | NA | NA | NA | 2 | 1 |
| 12_8123276 | 2 | 0.5 | 3 | 0.3333 | 3 | 0.3333 | NA | NA | 8 | 0.375 | 1 | 1 | NA | NA | NA | NA | NA | NA | 1 | 1 |
| 12_8124504 | 3 | 0.6667 | 3 | 0.6667 | NA | NA | 2 | 0.5 | 8 | 0.625 | 3 | 0.6667 | 2 | 0.5 | 3 | 0.6667 | 3 | 1 | 11 | 0.7273 |
| 12_8124512 | 2 | 0.5 | 2 | 0.5 | NA | NA | 1 | 0 | 5 | 0.4 | 3 | 0 | 1 | 0 | 3 | 0.3333 | 2 | 0.5 | 9 | 0.2222 |
| 12_8124547 | 2 | 1 | 3 | 0.6667 | NA | NA | 1 | 1 | 6 | 0.8333 | 3 | 1 | 1 | 1 | 3 | 1 | 2 | 1 | 9 | 1 |
| 12_8124552 | 2 | 1 | 3 | 1 | NA | NA | 2 | 1 | 7 | 1 | 2 | 1 | NA | NA | 2 | 1 | 3 | 1 | 7 | 1 |
| 12_8124570 | 3 | 0.6667 | 3 | 0.3333 | NA | NA | 2 | 0 | 8 | 0.375 | 2 | 0.5 | 1 | 1 | 1 | 0 | 3 | 0.3333 | 7 | 0.4286 |
| 12_8124571 | 3 | 0 | 2 | 0.5 | NA | NA | 2 | 0 | 7 | 0.1429 | 3 | 0 | 1 | 0 | 1 | 1 | 2 | 0 | 7 | 0.1429 |
| 12_8124573 | 1 | 1 | 3 | 1 | NA | NA | 2 | 0.5 | 6 | 0.8333 | 3 | 1 | NA | NA | 1 | 1 | 3 | 1 | 7 | 1 |
| 12_8124584 | 1 | 1 | 2 | 0.5 | NA | NA | 1 | 1 | 4 | 0.75 | 3 | 1 | 1 | 1 | NA | NA | 2 | 1 | 6 | 1 |
| 12_8124589 | 2 | 0.5 | 3 | 0.6667 | NA | NA | 1 | 0 | 6 | 0.5 | 3 | 0.3333 | 1 | 0 | NA | NA | 3 | 0.3333 | 7 | 0.2857 |
| 12_8124594 | 2 | 1 | 3 | 0.6667 | NA | NA | 2 | 1 | 7 | 0.8571 | 1 | 1 | 1 | 1 | NA | NA | 2 | 1 | 4 | 1 |
| 12_8124608 | 1 | 1 | 2 | 0.5 | NA | NA | NA | NA | 3 | 0.6667 | 1 | 1 | 1 | 1 | 1 | 1 | 1 | 0 | 4 | 0.75 |
| 12_8124609 | 2 | 0.5 | 2 | 0 | NA | NA | 1 | 1 | 5 | 0.4 | 3 | 0.3333 | 1 | 0 | 1 | 0 | 2 | 0 | 7 | 0.1429 |
| 12_8124614 | 2 | 0.5 | 2 | 0.5 | NA | NA | 1 | 0 | 5 | 0.4 | 3 | 1 | 1 | 0 | 1 | 0 | 2 | 0 | 7 | 0.4286 |
| 12_8124622 | 1 | 1 | 1 | 1 | NA | NA | 1 | 1 | 3 | 1 | 2 | 0.5 | 1 | 1 | 1 | 1 | 2 | 1 | 6 | 0.8333 |
| 12_8124649 | 2 | 1 | 2 | 0.5 | NA | NA | 1 | 1 | 5 | 0.8 | 3 | 0.6667 | 1 | 1 | 1 | 1 | 1 | 1 | 6 | 0.8333 |
| 12_8136108 | 2 | 0.5 | 1 | 1 | NA | NA | 1 | 0 | 4 | 0.5 | 3 | 0.3333 | 1 | 0 | 1 | 1 | NA | NA | 5 | 0.4 |
| 12_8136110 | 1 | 1 | 1 | 0 | NA | NA | 1 | 0 | 3 | 0.3333 | 3 | 0.3333 | NA | NA | 1 | 1 | NA | NA | 4 | 0.5 |
| 12_8136111 | 2 | 1 | 1 | 1 | NA | NA | NA | NA | 3 | 1 | 1 | 1 | 1 | 1 | 1 | 0 | NA | NA | 3 | 0.6667 |
| 12_8136115 | 2 | 0.5 | 1 | 1 | NA | NA | 1 | 1 | 4 | 0.75 | 2 | 0.5 | 1 | 1 | 1 | 1 | NA | NA | 4 | 0.75 |
| 12_8136138 | 1 | 1 | 1 | 1 | NA | NA | 1 | 1 | 3 | 1 | 2 | 1 | 1 | 0 | 1 | 1 | NA | NA | 4 | 0.75 |
| 12_8136152 | 1 | 1 | NA | NA | NA | NA | 1 | 1 | 2 | 1 | 2 | 0.5 | 1 | 1 | 1 | 1 | NA | NA | 4 | 0.75 |
| 12_8136158 | 2 | 1 | NA | NA | NA | NA | 1 | 1 | 3 | 1 | 1 | 1 | 1 | 1 | 1 | 1 | NA | NA | 3 | 1 |
| 12_8136164 | 2 | 0.5 | NA | NA | NA | NA | 1 | 1 | 3 | 0.6667 | 2 | 0.5 | 1 | 1 | 1 | 1 | NA | NA | 4 | 0.75 |
| 12_8136182 | 2 | 1 | NA | NA | NA | NA | 1 | 1 | 3 | 1 | 2 | 0.5 | 1 | 0 | 1 | 1 | NA | NA | 4 | 0.5 |
| 12_8136191 | 1 | 1 | NA | NA | NA | NA | 1 | 1 | 2 | 1 | 2 | 1 | 1 | 1 | NA | NA | NA | NA | 3 | 1 |
| 12_8136232 | 1 | 0 | NA | NA | NA | NA | 1 | 1 | 2 | 0.5 | 2 | 0.5 | 1 | 0 | 1 | 1 | NA | NA | 4 | 0.5 |
| 12_8136242 | 1 | 1 | NA | NA | NA | NA | 1 | 0 | 2 | 0.5 | 2 | 0.5 | NA | NA | 1 | 1 | NA | NA | 3 | 0.6667 |
| 12_8136266 | 1 | 1 | NA | NA | NA | NA | 1 | 1 | 2 | 1 | 2 | 1 | 1 | 1 | 1 | 1 | NA | NA | 4 | 1 |
| 12_8136271 | 1 | 1 | NA | NA | NA | NA | 1 | 1 | 2 | 1 | 2 | 0.5 | 1 | 1 | 1 | 1 | NA | NA | 4 | 0.75 |
| 12_8146676 | NA | NA | NA | NA | NA | NA | 1 | 1 | 1 | 1 | NA | NA | NA | NA | 1 | 1 | NA | NA | 1 | 1 |
| 12_8151911 | 2 | 0 | 3 | 0 | 3 | 0 | 3 | 0.3333 | 11 | 0.0909 | 3 | 0.3333 | 1 | 0 | 3 | 0 | 1 | 0 | 8 | 0.125 |
| 12_8154815 | 4 | 0.25 | 5 | 0.2 | 2 | 0.5 | 2 | 0.5 | 13 | 0.3077 | 1 | 0 | 2 | 0.5 | NA | NA | 1 | 0 | 4 | 0.25 |
| 12_8154854 | 6 | 0.5 | 7 | 0.4286 | 3 | 0 | 4 | 0.75 | 20 | 0.45 | 3 | 0.6667 | 7 | 0.7143 | 1 | 1 | 3 | 0.6667 | 14 | 0.7143 |
| 12_8154869 | 5 | 0.8 | 7 | 0.7143 | 3 | 0.6667 | 4 | 1 | 19 | 0.7895 | 3 | 0.6667 | 7 | 0.8571 | 1 | 1 | 2 | 1 | 13 | 0.8462 |
| 12_8154876 | 6 | 1 | 7 | 1 | 3 | 0.6667 | 4 | 1 | 20 | 0.95 | 2 | 0.5 | 5 | 1 | 1 | 1 | 3 | 1 | 11 | 0.9091 |
| 12_8160153 | 1 | 1 | NA | NA | NA | NA | NA | NA | 1 | 1 | NA | NA | NA | NA | NA | NA | 1 | 1 | 1 | 1 |
| 12_8160165 | 1 | 1 | NA | NA | NA | NA | NA | NA | 1 | 1 | NA | NA | NA | NA | NA | NA | 1 | 1 | 1 | 1 |
| 12_8160175 | 1 | 1 | NA | NA | NA | NA | NA | NA | 1 | 1 | NA | NA | NA | NA | NA | NA | 1 | 1 | 1 | 1 |
| 12_8160206 | 1 | 1 | NA | NA | NA | NA | NA | NA | 1 | 1 | NA | NA | NA | NA | NA | NA | 1 | 1 | 1 | 1 |
| 12_8160207 | 1 | 1 | NA | NA | NA | NA | NA | NA | 1 | 1 | NA | NA | NA | NA | NA | NA | 1 | 1 | 1 | 1 |
| 12_8162041 | 1 | 1 | 3 | 0.6667 | NA | NA | NA | NA | 4 | 0.75 | 3 | 0 | 2 | 0.5 | 3 | 0.3333 | NA | NA | 8 | 0.25 |
| 12_8162065 | 1 | 0 | 4 | 0.5 | NA | NA | 1 | 0 | 6 | 0.3333 | 3 | 0 | 5 | 0.2 | 3 | 0.6667 | 1 | 0 | 12 | 0.25 |
| 12_8171041 | 4 | 0.75 | 1 | 1 | 1 | 1 | NA | NA | 6 | 0.8333 | 1 | 1 | 1 | 0 | 3 | 1 | NA | NA | 5 | 0.8 |
| 12_8171054 | 4 | 0 | 1 | 0 | 2 | 0.5 | NA | NA | 7 | 0.1429 | NA | NA | 3 | 0.6667 | 4 | 0.25 | 1 | 1 | 8 | 0.5 |
| 12_8171057 | 3 | 0.6667 | 1 | 1 | 2 | 1 | NA | NA | 6 | 0.8333 | 1 | 1 | 3 | 0.6667 | 4 | 1 | 1 | 1 | 9 | 0.8889 |
| 12_8171060 | 3 | 0.3333 | 2 | 1 | 2 | 1 | NA | NA | 7 | 0.7143 | 1 | 1 | 2 | 0.5 | 5 | 0.8 | 1 | 1 | 9 | 0.7778 |
| 12_8171062 | 3 | 0.3333 | 2 | 0.5 | 2 | 1 | NA | NA | 7 | 0.5714 | NA | NA | 2 | 0.5 | 2 | 1 | 1 | 1 | 5 | 0.8 |
| 12_8171070 | 3 | 0.6667 | 2 | 0.5 | 2 | 0 | NA | NA | 7 | 0.4286 | 1 | 0 | 2 | 0.5 | 5 | 0.4 | 1 | 0 | 9 | 0.3333 |
| 12_8171074 | 4 | 0.5 | 2 | 0.5 | 2 | 1 | NA | NA | 8 | 0.625 | 1 | 0 | 2 | 0 | 5 | 0.6 | 1 | 1 | 9 | 0.4444 |
| 12_8171077 | 3 | 0.6667 | 1 | 0 | 2 | 0 | NA | NA | 6 | 0.3333 | 1 | 0 | 1 | 0 | 4 | 0.5 | 1 | 0 | 7 | 0.2857 |
| 12_8171084 | 4 | 0 | 2 | 0.5 | 1 | 1 | NA | NA | 7 | 0.2857 | 1 | 0 | 2 | 0.5 | 7 | 0 | 1 | 1 | 11 | 0.1818 |
| 12_8171107 | 4 | 0 | 4 | 0 | 2 | 0.5 | 1 | 0 | 11 | 0.0909 | 1 | 0 | 5 | 0.2 | 6 | 0 | 1 | 0 | 13 | 0.0769 |
| 12_8171136 | 6 | 0.1667 | 3 | 0.3333 | 1 | 0 | 1 | 1 | 11 | 0.2727 | 1 | 0 | 5 | 0.2 | 7 | 0.2857 | 2 | 0 | 15 | 0.2 |

| | | | | | | | | | | | | | | | | | | | | |
|---|---|---|---|---|---|---|---|---|---|---|---|---|---|---|---|---|---|---|---|---|
| 12_817128 | 4 | 0 | 3 | 0.3333 | 3 | 0 | NA | NA | 10 | 0.1 | 2 | 0 | 4 | 0 | 3 | 0.3333 | 1 | 0 | 10 | 0.1 |
| 12_817130 | 3 | 0.6667 | 3 | 1 | 2 | 0.5 | 1 | 1 | 9 | 0.7778 | 1 | 1 | 4 | 0.75 | 3 | 1 | 1 | 1 | 9 | 0.8889 |
| 12_817130 | 3 | 0.3333 | 1 | 1 | 3 | 0.6667 | 1 | 1 | 8 | 0.625 | 2 | 0 | 3 | 1 | 3 | 0.6667 | NA | NA | 8 | 0.625 |
| 12_817131 | 3 | 0.3333 | 1 | 0 | 3 | 0.3333 | 1 | 1 | 8 | 0.375 | 2 | 1 | 4 | 1 | 3 | 0.6667 | 1 | 0 | 10 | 0.8 |
| 12_817131 | 3 | 0.3333 | 2 | 0 | 2 | 0.5 | 1 | 0 | 8 | 0.25 | 1 | 0 | 4 | 0.5 | 2 | 0.5 | 1 | 0 | 8 | 0.375 |
| 12_817132 | 4 | 0.5 | 3 | 0 | 3 | 0.6667 | 1 | 1 | 11 | 0.4545 | 1 | 1 | 4 | 0.25 | 3 | 0 | 1 | 0 | 9 | 0.2222 |
| 12_817134 | 3 | 0 | 5 | 0.2 | 3 | 0 | 2 | 0 | 13 | 0.0769 | 2 | 0.5 | 4 | 0.25 | 3 | 0 | 1 | 0 | 10 | 0.2 |
| 12_817136 | 4 | 0.25 | 4 | 1 | 2 | 0.5 | 2 | 1 | 12 | 0.6667 | 2 | 0 | 3 | 0.3333 | 3 | 0.6667 | 1 | 1 | 9 | 0.4444 |
| 12_817139 | 4 | 0.25 | 3 | 1 | 2 | 0.5 | 2 | 0.5 | 11 | 0.5455 | 2 | 1 | 3 | 0.6667 | 3 | 0.6667 | 3 | 0.6667 | 11 | 0.7273 |
| 12_817617 | 2 | 0 | 7 | 0.1429 | 8 | 0.25 | 1 | 0 | 18 | 0.1667 | 2 | 0.5 | 4 | 0.25 | 6 | 0.1667 | 1 | 0 | 13 | 0.2308 |
| 12_817622 | 1 | 0 | 6 | 0 | 8 | 0.25 | 1 | 0 | 16 | 0.125 | 3 | 0 | 3 | 0 | 6 | 0.5 | 1 | 0 | 13 | 0.2308 |
| 12_817627 | 2 | 0.5 | 5 | 0 | 8 | 0 | 1 | 0 | 16 | 0.0625 | 3 | 0 | 4 | 0.25 | 4 | 0 | 1 | 0 | 12 | 0.0833 |
| 12_817643 | NA | NA | 3 | 0 | 3 | 0.3333 | NA | NA | 6 | 0.1667 | 1 | 0 | 1 | 0 | 2 | 0.5 | 1 | 0 | 5 | 0.2 |
| 12_817680 | NA | NA | 2 | 0.5 | 1 | 1 | NA | NA | 3 | 0.6667 | 1 | 1 | NA | NA | 2 | 0 | 1 | 1 | 4 | 0.5 |
| 12_817685 | NA | NA | NA | NA | NA | NA | 1 | 1 | 1 | 1 | 1 | 0 | NA | NA | 2 | 0.5 | NA | NA | 3 | 0.3333 |
| 12_817689 | 1 | 1 | 1 | 1 | 1 | 1 | 1 | 1 | 4 | 1 | 1 | 1 | NA | NA | 3 | 1 | NA | NA | 4 | 1 |
| 12_817689 | 1 | 1 | 1 | 1 | 2 | 1 | 1 | 1 | 5 | 1 | 1 | 1 | NA | NA | 2 | 0.5 | NA | NA | 3 | 0.6667 |
| 12_817689 | 1 | 1 | 1 | 0 | 2 | 0.5 | 1 | 1 | 5 | 0.6 | 1 | 0 | NA | NA | 3 | 0.3333 | NA | NA | 4 | 0.25 |
| 12_817689 | 1 | 1 | 1 | 1 | 1 | 0 | 1 | 1 | 4 | 0.75 | 1 | 1 | NA | NA | 1 | 1 | NA | NA | 2 | 1 |
| 12_817690 | 1 | 1 | 1 | 0 | 1 | 1 | 1 | 1 | 4 | 0.75 | 1 | 1 | NA | NA | 1 | 1 | NA | NA | 2 | 1 |
| 12_817691 | 1 | 0 | 1 | 0 | 1 | 0 | 1 | 1 | 4 | 0.25 | 1 | 1 | NA | NA | NA | NA | NA | NA | 1 | 1 |
| 12_817709 | 1 | 1 | 1 | 1 | 2 | 1 | 1 | 0 | 5 | 0.8 | 1 | 1 | NA | NA | 2 | 1 | NA | NA | 3 | 1 |
| 12_817709 | 1 | 0 | 1 | 0 | 1 | 1 | 1 | 0 | 4 | 0.25 | NA | NA | NA | NA | 1 | 1 | NA | NA | 1 | 1 |
| 12_819645 | NA | NA | NA | NA | 1 | 1 | NA | NA | 1 | 1 | NA | NA | NA | NA | NA | NA | 1 | 1 | 1 | 1 |
| 12_819647 | NA | NA | NA | NA | 1 | 1 | NA | NA | 1 | 1 | NA | NA | NA | NA | NA | NA | 1 | 1 | 1 | 1 |
| 12_819649 | NA | NA | NA | NA | 1 | 1 | NA | NA | 1 | 1 | NA | NA | 1 | 1 | NA | NA | 1 | 0 | 2 | 0.5 |
| 12_819655 | NA | NA | NA | NA | 2 | 1 | NA | NA | 2 | 1 | NA | NA | NA | NA | 1 | 1 | 1 | 0 | 2 | 0.5 |
| 12_819657 | NA | NA | NA | NA | 2 | 0.5 | NA | NA | 2 | 0.5 | NA | NA | NA | NA | 1 | 0 | 1 | 1 | 2 | 0.5 |
| 12_819658 | NA | NA | NA | NA | 2 | 1 | NA | NA | 2 | 1 | NA | NA | NA | NA | 1 | 1 | 1 | 0 | 2 | 0.5 |
| 12_819658 | NA | NA | NA | NA | 2 | 1 | NA | NA | 2 | 1 | NA | NA | NA | NA | 1 | 1 | 1 | 1 | 2 | 1 |
| 12_819659 | NA | NA | NA | NA | 1 | 1 | NA | NA | 1 | 1 | NA | NA | NA | NA | 1 | 0 | 1 | 1 | 2 | 0.5 |
| 12_820428 | 2 | 0.5 | 1 | 1 | 6 | 0.8333 | 4 | 0.5 | 13 | 0.6923 | 1 | 1 | 2 | 1 | NA | NA | 1 | 1 | 4 | 1 |
| 12_820429 | 3 | 1 | 1 | 1 | 5 | 1 | 3 | 0.6667 | 12 | 0.9167 | 3 | 1 | 3 | 0.6667 | NA | NA | 2 | 1 | 8 | 0.875 |
| 12_820430 | 3 | 0.6667 | 3 | 1 | 6 | 0.5 | 5 | 0.4 | 17 | 0.5882 | 3 | 1 | 4 | 1 | NA | NA | 2 | 1 | 9 | 1 |
| 12_820430 | 3 | 0 | 2 | 0 | 6 | 0.1667 | 5 | 0 | 16 | 0.0625 | 3 | 0 | 4 | 0.25 | NA | NA | 2 | 0 | 9 | 0.1111 |
| 12_820432 | 2 | 1 | 1 | 1 | 3 | 1 | 3 | 1 | 9 | 1 | 2 | 1 | 4 | 1 | NA | NA | 2 | 1 | 8 | 1 |
| 12_820432 | 4 | 0.5 | 1 | 0 | 5 | 0.2 | 5 | 0.2 | 15 | 0.2667 | 3 | 0.3333 | 4 | 0 | NA | NA | 2 | 0.5 | 9 | 0.2222 |
| 12_820433 | 3 | 0 | 2 | 0 | 5 | 0 | 5 | 0.4 | 15 | 0.1333 | 3 | 0 | 4 | 0.25 | NA | NA | 2 | 0.5 | 9 | 0.2222 |
| 12_820434 | 3 | 0.6667 | 1 | 0 | 6 | 0.5 | 4 | 0.25 | 14 | 0.4286 | 3 | 0.3333 | 4 | 0.5 | NA | NA | 1 | 1 | 8 | 0.5 |
| 12_820434 | 6 | 0.5 | 1 | 0 | 7 | 0.2857 | 5 | 0.2 | 19 | 0.3158 | 1 | 0 | 1 | 0 | NA | NA | 2 | 0.5 | 4 | 0.25 |
| 12_820434 | 4 | 0.25 | 1 | 0 | 6 | 0 | 5 | 0.2 | 16 | 0.125 | 2 | 0.5 | 3 | 0 | NA | NA | 1 | 1 | 6 | 0.3333 |
| 12_820435 | 1 | 1 | 2 | 0 | 5 | 0.6 | 5 | 0.8 | 13 | 0.6154 | 2 | 0 | 2 | 0 | NA | NA | 1 | 1 | 5 | 0.2 |
| 12_820436 | 4 | 0.75 | 2 | 1 | 6 | 0.8333 | 4 | 0.75 | 16 | 0.8125 | 1 | 1 | 2 | 0.5 | NA | NA | 1 | 1 | 4 | 0.75 |
| 12_820436 | 4 | 0 | 1 | 1 | 6 | 0.1667 | 6 | 0.6667 | 17 | 0.3529 | 2 | 0.5 | 3 | 0 | 1 | 0 | 1 | 1 | 7 | 0.2857 |
| 12_820437 | 5 | 0.2 | 1 | 0 | 5 | 0.2 | 3 | 0 | 14 | 0.1429 | 2 | 0.5 | 2 | 0 | 1 | 0 | 2 | 0 | 7 | 0.1429 |
| 12_820438 | 3 | 1 | 1 | 0 | 2 | 0.5 | 2 | 0.5 | 8 | 0.625 | NA | NA | 2 | 0.5 | NA | NA | 1 | 1 | 3 | 0.6667 |
| 12_823874 | 3 | 0 | 2 | 0 | 3 | 0.3333 | 9 | 0.1111 | 17 | 0.1176 | 3 | 0.3333 | 8 | 0 | 6 | 0.5 | 2 | 0.5 | 19 | 0.2632 |
| 12_823875 | 2 | 0.5 | 5 | 0.8 | 4 | 0.5 | 7 | 0.4286 | 18 | 0.5556 | 5 | 0.8 | 9 | 0.6667 | 6 | 0.1667 | 2 | 0.5 | 22 | 0.5455 |
| 12_823875 | 3 | 1 | 4 | 0.5 | 5 | 0.8 | 10 | 0.5 | 22 | 0.6364 | 5 | 0.6 | 7 | 0.5714 | 7 | 0.8571 | 2 | 0.5 | 21 | 0.6667 |
| 12_823878 | 3 | 0 | 6 | 0 | 6 | 0 | 11 | 0.3636 | 26 | 0.1538 | 4 | 0.25 | 9 | 0 | 7 | 0.1429 | 3 | 0.3333 | 23 | 0.1304 |
| 12_823881 | 2 | 0 | 7 | 0.1429 | 6 | 0.1667 | 11 | 0.1818 | 26 | 0.1538 | 5 | 0.2 | 8 | 0 | 8 | 0.375 | 5 | 0 | 26 | 0.1538 |
| 12_823882 | 3 | 0 | 6 | 0.3333 | 6 | 0.1667 | 13 | 0.3846 | 28 | 0.2857 | 8 | 0.25 | 9 | 0.3333 | 10 | 0.5 | 6 | 0.1667 | 33 | 0.3333 |
| 12_823885 | 2 | 0.5 | 5 | 0.6 | 5 | 0 | 10 | 0.4 | 22 | 0.3636 | 8 | 0.375 | 8 | 0.5 | 6 | 0.3333 | 3 | 0.3333 | 25 | 0.4 |
| 12_823886 | 6 | 0.1667 | 8 | 0.125 | 11 | 0.0909 | 11 | 0.2727 | 36 | 0.1667 | 10 | 0.2 | 10 | 0.1 | 12 | 0.0833 | 7 | 0.1429 | 39 | 0.1282 |
| 12_823887 | 5 | 0.8 | 7 | 0.4286 | 7 | 0.5714 | 11 | 0.3636 | 30 | 0.5 | 7 | 0.7143 | 8 | 0.375 | 11 | 0.6364 | 4 | 0 | 30 | 0.5 |
| 12_823888 | 4 | 0.5 | 5 | 0.6 | 10 | 0.2 | 9 | 0.2222 | 28 | 0.3214 | 8 | 0.125 | 8 | 0.125 | 9 | 0.1111 | 5 | 0.6 | 30 | 0.2 |
| 12_823888 | 4 | 0.25 | 6 | 0.3333 | 8 | 0.875 | 11 | 0.6364 | 29 | 0.5862 | 7 | 0.5714 | 7 | 0.2857 | 10 | 0.7 | 7 | 0.8571 | 31 | 0.6129 |
| 12_823888 | 5 | 0 | 10 | 0.3 | 9 | 0.1111 | 10 | 0.2 | 34 | 0.1765 | 8 | 0.125 | 11 | 0.1818 | 11 | 0.1818 | 7 | 0 | 37 | 0.1351 |
| 12_823889 | 4 | 0.25 | 10 | 0.3 | 8 | 0.25 | 12 | 0.0833 | 34 | 0.2059 | 10 | 0.4 | 10 | 0.5 | 12 | 0.4167 | 7 | 0.4286 | 39 | 0.4359 |
| 12_823889 | 6 | 0 | 8 | 0.125 | 11 | 0.1818 | 14 | 0.2143 | 39 | 0.1538 | 9 | 0.2222 | 12 | 0.0833 | 14 | 0.2143 | 7 | 0.4286 | 42 | 0.2143 |
| 12_823890 | 3 | 0.6667 | 8 | 0.75 | 9 | 0.6667 | 12 | 0.6667 | 32 | 0.6875 | 10 | 0.6 | 11 | 1 | 10 | 1 | 6 | 0.6667 | 37 | 0.8378 |
| 12_823891 | 4 | 0.5 | 9 | 0.3333 | 8 | 0.625 | 11 | 0.2727 | 32 | 0.4062 | 5 | 0.4 | 8 | 0.125 | 9 | 0.1111 | 6 | 0.3333 | 28 | 0.2143 |
| 12_824430 | 6 | 0 | 11 | 0 | 12 | 0.0833 | 16 | 0.25 | 45 | 0.1111 | 9 | 0 | 14 | 0.0714 | 9 | 0.1111 | 6 | 0.1667 | 38 | 0.0789 |
| 12_824430 | 10 | 0.8 | 15 | 0.6667 | 15 | 0.8 | 18 | 0.8889 | 58 | 0.7931 | 10 | 0.5 | 17 | 0.7647 | 15 | 0.8667 | 6 | 0.6667 | 48 | 0.7292 |
| 12_824431 | 10 | 0.1 | 18 | 0.0556 | 18 | 0.0556 | 21 | 0.1429 | 67 | 0.0896 | 12 | 0.1667 | 17 | 0.1176 | 15 | 0.0667 | 10 | 0.1 | 54 | 0.1111 |

| | | | | | | | | | | | | | | | | | | | | |
|---|---|---|---|---|---|---|---|---|---|---|---|---|---|---|---|---|---|---|---|---|
| 12_824433 | 9 | 0.1111 | 15 | 0.0667 | 18 | 0.2778 | 17 | 0.2941 | 59 | 0.2034 | 10 | 0.6 | 13 | 0.3077 | 13 | 0.3846 | 7 | 0.2857 | 43 | 0.3953 |
| 12_824434 | 8 | 0.625 | 13 | 0.4615 | 18 | 0.2222 | 21 | 0.2381 | 60 | 0.3333 | 13 | 0.4615 | 16 | 0.4375 | 15 | 0.5333 | 9 | 0.4444 | 53 | 0.4717 |
| 12_824720 | 12 | 0.3333 | 19 | 0.1579 | 23 | 0.3478 | 35 | 0.3143 | 89 | 0.2921 | 17 | 0.2941 | 22 | 0.1818 | 25 | 0.44 | 21 | 0.2857 | 85 | 0.3059 |
| 12_824720 | 16 | 0.5625 | 17 | 0.4706 | 23 | 0.6522 | 40 | 0.625 | 96 | 0.5938 | 16 | 0.5 | 25 | 0.6 | 26 | 0.5385 | 22 | 0.7273 | 89 | 0.5955 |
| 12_824721 | 11 | 0.2727 | 15 | 0.4 | 16 | 0.3125 | 31 | 0.3871 | 73 | 0.3562 | 13 | 0.3077 | 21 | 0.4286 | 21 | 0.1429 | 16 | 0.1875 | 71 | 0.2676 |
| 12_824721 | 17 | 0 | 16 | 0.125 | 21 | 0.0952 | 34 | 0.0294 | 88 | 0.0568 | 15 | 0.0667 | 23 | 0.1739 | 25 | 0.04 | 25 | 0.08 | 88 | 0.0909 |
| 12_824796 | 13 | 0.1538 | 19 | 0.3684 | 22 | 0.2273 | 35 | 0.3714 | 89 | 0.3034 | 17 | 0.4118 | 23 | 0.1304 | 22 | 0.1364 | 20 | 0.45 | 82 | 0.2683 |
| 12_824968 | 15 | 0.0667 | 22 | 0.0909 | 27 | 0.1481 | 40 | 0.075 | 104 | 0.0962 | 16 | 0.3125 | 27 | 0 | 27 | 0.1111 | 26 | 0.1538 | 96 | 0.125 |
| 12_824968 | 23 | 0 | 24 | 0.125 | 31 | 0.1935 | 41 | 0.2195 | 119 | 0.1513 | 20 | 0.2 | 36 | 0.1667 | 31 | 0.0968 | 28 | 0.1071 | 115 | 0.1391 |
| 12_824969 | 19 | 0.1579 | 27 | 0.2222 | 31 | 0.1613 | 45 | 0.1778 | 122 | 0.1803 | 21 | 0.3333 | 32 | 0.125 | 32 | 0.0625 | 26 | 0.3077 | 111 | 0.1892 |
| 12_824978 | 24 | 0.5417 | 24 | 0.625 | 31 | 0.4839 | 36 | 0.6389 | 115 | 0.5739 | 19 | 0.6842 | 29 | 0.6207 | 27 | 0.5926 | 28 | 0.5 | 103 | 0.5922 |
| 12_832050 | NA | NA | NA | NA | 1 | 1 | NA | NA | 1 | 1 | 2 | 0.5 | 1 | 0 | NA | NA | NA | NA | 3 | 0.3333 |
| 12_832050 | NA | NA | 1 | 1 | 1 | 1 | NA | NA | 2 | 1 | 2 | 1 | 2 | 1 | NA | NA | NA | NA | 4 | 1 |
| 12_832083 | 1 | 1 | 3 | 0.6667 | 2 | 0 | NA | NA | 6 | 0.5 | 1 | 0 | NA | NA | 2 | 0.5 | NA | NA | 3 | 0.3333 |
| 12_832521 | NA | NA | 1 | 1 | NA | NA | 1 | 1 | 2 | 1 | NA | NA | 1 | 1 | NA | NA | NA | NA | 1 | 1 |
| 12_833072 | 1 | 1 | NA | NA | NA | NA | NA | NA | 1 | 1 | NA | NA | NA | NA | 1 | 1 | NA | NA | 1 | 1 |
| 12_833072 | 1 | 1 | NA | NA | NA | NA | NA | NA | 1 | 1 | NA | NA | NA | NA | 1 | 1 | NA | NA | 1 | 1 |
| 12_835883 | 4 | 0.25 | 2 | 0.5 | 5 | 0.4 | 5 | 0.2 | 16 | 0.3125 | NA | NA | 2 | 0 | 3 | 0.3333 | 3 | 0.3333 | 8 | 0.25 |
| 12_835883 | 2 | 1 | 2 | 1 | 3 | 0.6667 | 3 | 1 | 10 | 0.9 | NA | NA | 2 | 0 | 3 | 1 | 2 | 1 | 7 | 0.7143 |
| 12_835933 | 1 | 1 | 2 | 0 | 2 | 0 | 2 | 0 | 7 | 0.1429 | 3 | 0.3333 | 2 | 0 | 2 | 0 | 3 | 0 | 10 | 0.1 |
| 12_835934 | 1 | 1 | 1 | 0 | 1 | 0 | 2 | 0.5 | 5 | 0.4 | 4 | 0.5 | 2 | 0.5 | 2 | 0 | 3 | 0.3333 | 11 | 0.3636 |
| 12_835935 | 1 | 1 | 1 | 0 | 1 | 1 | 1 | 0 | 4 | 0.5 | NA | NA | 1 | 0 | 2 | 0 | 2 | 0.5 | 5 | 0.2 |
| 12_835935 | 1 | 0 | 2 | 0.5 | 2 | 0.5 | 1 | 0 | 6 | 0.3333 | 3 | 0.3333 | 2 | 0 | 2 | 0 | 2 | 0.5 | 9 | 0.2222 |
| 12_835937 | 6 | 0.1667 | 8 | 0.25 | 5 | 0.4 | 6 | 0.1667 | 25 | 0.24 | 5 | 0.4 | 5 | 0.2 | 6 | 0.3333 | 3 | 0 | 19 | 0.2632 |
| 12_835943 | 2 | 0.5 | 8 | 0.125 | 3 | 0 | 4 | 0.25 | 17 | 0.1765 | 4 | 0.5 | 4 | 0.25 | 5 | 0.2 | 3 | 0.3333 | 16 | 0.3125 |
| 12_835992 | 1 | 1 | 8 | 0.125 | 5 | 0 | 5 | 0 | 19 | 0.1053 | 4 | 0.25 | 4 | 0 | 4 | 0.25 | 2 | 0 | 14 | 0.1429 |
| 12_836214 | 1 | 0 | 4 | 0.75 | 2 | 0 | 5 | 0.2 | 12 | 0.3333 | 3 | 0.6667 | 2 | 1 | 3 | 0.3333 | 1 | 1 | 9 | 0.6667 |
| 12_836215 | 1 | 0 | 6 | 0.6667 | 2 | 0.5 | 3 | 0 | 12 | 0.4167 | 3 | 0 | 1 | 1 | 3 | 0.6667 | NA | NA | 7 | 0.4286 |
| 12_836215 | 1 | 1 | 3 | 0.6667 | 2 | 0.5 | 4 | 1 | 10 | 0.8 | 2 | 0.5 | 2 | 1 | 1 | 1 | 1 | 1 | 6 | 0.8333 |
| 12_836216 | 1 | 0 | 5 | 0.4 | 2 | 0 | 4 | 0.5 | 12 | 0.3333 | 3 | 0 | 2 | 1 | 3 | 0.3333 | 1 | 1 | 9 | 0.4444 |
| 12_836216 | 1 | 0 | 5 | 0 | 2 | 0.5 | 4 | 0 | 12 | 0.0833 | 3 | 0.3333 | 3 | 0 | 3 | 0.3333 | 1 | 0 | 10 | 0.2 |
| 12_836216 | 1 | 1 | 5 | 1 | 2 | 1 | 4 | 1 | 12 | 1 | 3 | 1 | 2 | 1 | 3 | 1 | NA | NA | 8 | 1 |
| 12_836216 | 1 | 1 | 3 | 1 | 2 | 0 | 3 | 0 | 9 | 0.4444 | 2 | 0 | 3 | 1 | 3 | 0.6667 | 1 | 0 | 9 | 0.5556 |
| 12_836218 | 1 | 1 | 3 | 1 | 1 | 1 | 2 | 1 | 7 | 1 | 2 | 1 | 3 | 1 | 2 | 0.5 | 1 | 1 | 8 | 0.875 |
| 12_836218 | NA | NA | 4 | 0.25 | 1 | 0 | 2 | 0 | 7 | 0.1429 | 1 | 1 | 3 | 0 | 2 | 0 | 1 | 0 | 7 | 0.1429 |
| 12_836413 | NA | NA | 3 | 1 | 2 | 0.5 | 2 | 0.5 | 7 | 0.7143 | 1 | 0 | 2 | 0.5 | NA | NA | NA | NA | 3 | 0.3333 |
| 12_836414 | 1 | 1 | 2 | 0.5 | 2 | 0 | 2 | 1 | 7 | 0.5714 | 1 | 0 | 3 | 0.6667 | 1 | 0 | 1 | 1 | 6 | 0.5 |
| 12_836415 | NA | NA | 2 | 1 | 2 | 1 | 2 | 1 | 6 | 1 | 1 | 1 | 2 | 0.5 | 1 | 1 | 1 | 1 | 5 | 0.8 |
| 12_836416 | NA | NA | 1 | 1 | 1 | 1 | 1 | 1 | 3 | 1 | 1 | 1 | 1 | 1 | 1 | 1 | 1 | 1 | 4 | 1 |
| 12_836418 | 1 | 0 | 1 | 1 | NA | NA | 2 | 0.5 | 4 | 0.5 | 1 | 1 | 1 | 1 | 1 | 1 | 1 | 1 | 4 | 1 |
| 12_836418 | 1 | 0 | 2 | 1 | 2 | 1 | 2 | 1 | 7 | 0.8571 | 1 | 0 | 2 | 1 | 1 | 1 | NA | NA | 4 | 0.75 |
| 12_836418 | NA | NA | 2 | 1 | 1 | 1 | 2 | 0.5 | 5 | 0.8 | 1 | 1 | 2 | 1 | 1 | 1 | NA | NA | 4 | 1 |
| 12_836419 | 1 | 1 | 2 | 1 | 2 | 0.5 | 2 | 1 | 7 | 0.8571 | 1 | 0 | 2 | 1 | 1 | 1 | NA | NA | 4 | 0.75 |
| 12_837219 | 4 | 0.25 | 9 | 0.3333 | 8 | 0.625 | 2 | 0.5 | 23 | 0.4348 | 2 | 0.5 | 12 | 0.3333 | 7 | 0.2857 | 7 | 0.2857 | 28 | 0.3214 |
| 12_837283 | 5 | 0.6 | 11 | 0.5455 | 12 | 0.5 | 3 | 0.6667 | 31 | 0.5484 | 2 | 1 | 14 | 0.2143 | 10 | 0.7 | 7 | 0.2857 | 33 | 0.4242 |
| 12_837284 | 5 | 0.8 | 10 | 0.8 | 12 | 0.75 | 2 | 0.5 | 29 | 0.7586 | 2 | 1 | 12 | 0.9167 | 10 | 0.6 | 8 | 0.875 | 32 | 0.8125 |
| 12_837284 | 6 | 0.1667 | 9 | 0.4444 | 13 | 0.3846 | 2 | 0 | 30 | 0.3333 | 2 | 0 | 14 | 0.2143 | 9 | 0.2222 | 7 | 0.2857 | 32 | 0.2188 |
| 12_837290 | 3 | 0.3333 | 8 | 0.625 | 13 | 0.3077 | 3 | 0 | 27 | 0.3704 | 2 | 0.5 | 9 | 0.1111 | 7 | 0.5714 | 5 | 0 | 23 | 0.2609 |
| 12_837295 | 3 | 0.3333 | 10 | 0.5 | 14 | 0.5 | 6 | 0.5 | 33 | 0.4848 | 2 | 0.5 | 11 | 0.0909 | 7 | 0.4286 | 6 | 0.5 | 26 | 0.3077 |
| 12_837296 | 4 | 0 | 11 | 0.1818 | 15 | 0.1333 | 8 | 0.25 | 38 | 0.1579 | 3 | 0 | 11 | 0.0909 | 9 | 0.2222 | 8 | 0.125 | 31 | 0.129 |
| 12_837808 | 5 | 0.8 | 7 | 0.7143 | 8 | 0.875 | NA | NA | 20 | 0.8 | 5 | 0.8 | 8 | 1 | 3 | 1 | 7 | 1 | 23 | 0.9565 |
| 12_837808 | 5 | 0.4 | 7 | 0.1429 | 7 | 0.5714 | 1 | 0 | 20 | 0.35 | 3 | 0 | 9 | 0 | 2 | 0.5 | 7 | 0.7143 | 21 | 0.2857 |
| 12_837808 | 5 | 0.2 | 6 | 0 | 5 | 0.2 | 1 | 0 | 17 | 0.1176 | 2 | 0 | 6 | 0 | 3 | 0.3333 | 7 | 0.1429 | 18 | 0.1111 |
| 12_837809 | 6 | 0.1667 | 6 | 0 | 7 | 0 | 1 | 0 | 20 | 0.05 | 4 | 0 | 9 | 0.1111 | 3 | 0 | 7 | 0.2857 | 23 | 0.1304 |
| 12_837809 | 6 | 0.3333 | 7 | 0.7143 | 8 | 0.5 | 1 | 1 | 22 | 0.5455 | 5 | 0 | 9 | 0.3333 | 3 | 0.3333 | 6 | 0.3333 | 23 | 0.2609 |
| 12_837809 | 5 | 0.6 | 7 | 0.8571 | 8 | 0.5 | 1 | 1 | 21 | 0.6667 | 5 | 0.8 | 8 | 0.625 | 3 | 1 | 5 | 0.6 | 21 | 0.7143 |
| 12_837809 | 6 | 0.3333 | 6 | 0.6667 | 7 | 0.5714 | 1 | 0 | 20 | 0.5 | 5 | 0.6 | 9 | 0.2222 | 3 | 0 | 7 | 0.4286 | 24 | 0.3333 |
| 12_837809 | 5 | 0.6 | 6 | 0.8333 | 6 | 0.8333 | 1 | 1 | 18 | 0.7778 | 5 | 0.8 | 8 | 0.875 | 2 | 1 | 6 | 0.8333 | 21 | 0.8571 |
| 12_837810 | 6 | 1 | 5 | 0.6 | 5 | 0.8 | 1 | 1 | 17 | 0.8235 | 3 | 0.6667 | 5 | 1 | 3 | 1 | 4 | 0.75 | 15 | 0.8667 |
| 12_837810 | 6 | 0.3333 | 6 | 0.3333 | 8 | 0.375 | 1 | 1 | 21 | 0.381 | 4 | 0.75 | 10 | 0.4 | 3 | 0.3333 | 7 | 0.4286 | 24 | 0.4583 |
| 12_837811 | 6 | 0.1667 | 5 | 0.4 | 8 | 0.25 | 1 | 0 | 20 | 0.25 | 4 | 0.25 | 8 | 0.125 | 3 | 0.3333 | 8 | 0 | 23 | 0.1304 |
| 12_837811 | 5 | 1 | 8 | 0.5 | 6 | 0.6667 | NA | NA | 19 | 0.6842 | 5 | 0.4 | 9 | 0.7778 | 3 | 0.6667 | 8 | 0.875 | 25 | 0.72 |
| 12_837811 | 6 | 0 | 8 | 0 | 6 | 0.3333 | 1 | 0 | 21 | 0.0952 | 5 | 0 | 10 | 0.1 | 3 | 0.3333 | 7 | 0 | 25 | 0.08 |
| 12_837811 | 4 | 1 | 6 | 1 | 6 | 1 | 1 | 1 | 17 | 1 | 2 | 1 | 7 | 0.8571 | 3 | 1 | 7 | 1 | 19 | 0.9474 |
| 12_837812 | 6 | 0.1667 | 8 | 0 | 7 | 0.2857 | 1 | 0 | 22 | 0.1364 | 6 | 0 | 9 | 0.1111 | 2 | 0 | 8 | 0.125 | 25 | 0.08 |

| | | | | | | | | | | | | | | | | | | | | |
|---|---|---|---|---|---|---|---|---|---|---|---|---|---|---|---|---|---|---|---|---|
| 12_8378127 | 4 | 0.5 | 7 | 0.2857 | 7 | 0 | 2 | 0.5 | 20 | 0.25 | 4 | 0 | 9 | 0.1111 | 3 | 0 | 7 | 0.1429 | 23 | 0.087 |
| 12_8378136 | 6 | 1 | 7 | 1 | 7 | 1 | 2 | 1 | 22 | 1 | 6 | 0.6667 | 8 | 0.75 | 3 | 1 | 8 | 0.75 | 25 | 0.76 |
| 12_8378140 | 5 | 0.8 | 8 | 0.625 | 8 | 0.375 | 2 | 0.5 | 23 | 0.5652 | 5 | 0.8 | 8 | 0.5 | 3 | 0.6667 | 7 | 0.5714 | 23 | 0.6087 |
| 12_8378143 | 5 | 0.6 | 5 | 0.6 | 6 | 0.3333 | 1 | 0 | 17 | 0.4706 | 4 | 0.25 | 5 | 0.2 | 3 | 0.6667 | 7 | 0.2857 | 19 | 0.3158 |
| 12_8378158 | 6 | 1 | 9 | 0.8889 | 7 | 0.7143 | 2 | 1 | 24 | 0.875 | 7 | 0.5714 | 8 | 0.875 | 2 | 1 | 9 | 0.8889 | 26 | 0.8077 |
| 12_8378233 | 2 | 0.5 | 2 | 0 | 1 | 1 | NA | NA | 5 | 0.4 | NA | NA | 1 | 1 | NA | NA | 2 | 0.5 | 3 | 0.6667 |
| 12_8378485 | 2 | 0 | 4 | 0.5 | 2 | 0.5 | 1 | 0 | 9 | 0.3333 | NA | NA | 2 | 0.5 | 4 | 0.25 | 3 | 0 | 9 | 0.2222 |
| 12_8378510 | 2 | 0 | 5 | 0.2 | 3 | 0 | 2 | 0 | 12 | 0.0833 | NA | NA | 2 | 0 | 4 | 0 | 3 | 0.6667 | 9 | 0.2222 |
| 12_8383863 | NA | NA | NA | NA | NA | NA | 1 | 1 | 1 | 1 | NA | NA | 1 | 1 | NA | NA | NA | NA | 1 | 1 |
| 12_8384202 | 3 | 0 | 6 | 0 | 4 | 0.25 | 3 | 0 | 16 | 0.0625 | 4 | 0.25 | 3 | 0 | 2 | 0 | 1 | 0 | 10 | 0.1 |
| 12_8384214 | 2 | 0 | 6 | 0 | 4 | 0.25 | 4 | 0 | 16 | 0.0625 | 6 | 0.1667 | 4 | 0.25 | 2 | 0 | 2 | 0 | 14 | 0.1429 |
| 12_8384248 | 1 | 0 | 5 | 0 | 3 | 0.3333 | 4 | 0.25 | 13 | 0.1538 | 5 | 0 | 4 | 0.25 | 2 | 0 | 2 | 0 | 13 | 0.0769 |
| 12_8384271 | 2 | 0.5 | 5 | 0.4 | 2 | 0.5 | 3 | 0.6667 | 12 | 0.5 | 3 | 0.6667 | 4 | 0.25 | 2 | 0.5 | 2 | 0.5 | 11 | 0.4545 |
| 12_8399767 | 2 | 1 | 1 | 0 | NA | NA | 1 | 0 | 4 | 0.5 | 1 | 1 | NA | NA | 1 | 0 | 1 | 0 | 3 | 0.3333 |
| 12_8399834 | 4 | 0.5 | 1 | 1 | NA | NA | 2 | 1 | 7 | 0.7143 | 1 | 0 | 1 | 0 | NA | NA | 1 | 1 | 3 | 0.3333 |
| 12_8399844 | 5 | 0.2 | 1 | 0 | NA | NA | 1 | 0 | 7 | 0.1429 | NA | NA | 1 | 0 | 1 | 0 | 1 | 1 | 3 | 0.3333 |
| 12_8399849 | 4 | 0.5 | 1 | 0 | NA | NA | 2 | 0.5 | 7 | 0.4286 | 1 | 0 | 1 | 0 | 1 | 1 | 1 | 1 | 4 | 0.5 |
| 12_8399872 | 4 | 0.5 | 2 | 0 | NA | NA | 2 | 0 | 8 | 0.25 | 1 | 0 | 1 | 0 | 1 | 1 | 1 | 1 | 4 | 0.5 |
| 12_8399877 | 5 | 1 | NA | NA | NA | NA | 2 | 1 | 7 | 1 | 1 | 1 | 1 | 1 | 1 | 1 | 1 | 1 | 4 | 1 |
| 12_8399881 | 4 | 1 | 2 | 1 | NA | NA | 2 | 1 | 8 | 1 | 1 | 1 | 1 | 1 | 1 | 1 | 1 | 1 | 4 | 1 |
| 12_8399886 | 4 | 1 | 2 | 1 | NA | NA | 2 | 1 | 8 | 1 | 1 | 1 | 1 | 1 | 1 | 1 | 1 | 1 | 4 | 1 |
| 12_8399906 | 5 | 0.2 | 2 | 0 | NA | NA | 2 | 0.5 | 9 | 0.2222 | 1 | 0 | 1 | 0 | 1 | 0 | 1 | 1 | 4 | 0.25 |
| 12_8399909 | 4 | 0.25 | 2 | 0 | NA | NA | 2 | 0 | 8 | 0.125 | 2 | 0.5 | 1 | 0 | 1 | 0 | 1 | 0 | 5 | 0.2 |
| 12_8399913 | 3 | 1 | 2 | 1 | NA | NA | 2 | 1 | 7 | 1 | NA | NA | 2 | 0.5 | 1 | 1 | NA | NA | 3 | 0.6667 |
| 12_8399918 | 6 | 0.8333 | 2 | 1 | NA | NA | 2 | 1 | 10 | 0.9 | 2 | 1 | 2 | 1 | 1 | 1 | 1 | 1 | 6 | 1 |
| 12_8399919 | 6 | 0.3333 | 2 | 0 | NA | NA | 2 | 0.5 | 10 | 0.3 | 1 | 0 | NA | NA | 1 | 0 | 1 | 1 | 3 | 0.3333 |
| 12_8399925 | 6 | 0 | 2 | 0 | NA | NA | 2 | 0.5 | 10 | 0.1 | 2 | 0.5 | 1 | 0 | 1 | 0 | 1 | 0 | 5 | 0.2 |
| 12_8399943 | 5 | 0.2 | 3 | 0.3333 | NA | NA | 2 | 0 | 10 | 0.2 | 3 | 0.3333 | 2 | 0 | NA | NA | 1 | 0 | 6 | 0.1667 |
| 12_8399959 | 6 | 1 | 4 | 1 | 1 | 1 | 1 | 1 | 12 | 1 | 4 | 1 | 3 | 1 | 1 | 1 | 1 | 1 | 9 | 1 |
| 12_8403932 | NA | NA | NA | NA | 2 | 0.5 | 2 | 1 | 4 | 0.75 | 1 | 0 | 1 | 1 | 1 | 1 | NA | NA | 3 | 0.6667 |
| 12_8403942 | NA | NA | 1 | 1 | 3 | 0.6667 | 2 | 0.5 | 6 | 0.6667 | 2 | 1 | 1 | 1 | 1 | 1 | NA | NA | 4 | 1 |
| 12_8403981 | NA | NA | 1 | 1 | 4 | 0 | 2 | 1 | 7 | 0.4286 | 1 | 1 | 1 | 1 | 1 | 1 | NA | NA | 3 | 1 |
| 12_8404017 | NA | NA | 1 | 0 | 3 | 0.3333 | 2 | 0.5 | 6 | 0.3333 | 2 | 1 | 2 | 0 | 1 | 0 | 3 | 1 | 8 | 0.625 |
| 12_8404029 | NA | NA | NA | NA | 5 | 0.4 | 3 | 0 | 8 | 0.25 | 2 | 0.5 | 2 | 0 | 1 | 0 | 3 | 0.3333 | 8 | 0.25 |
| 12_8405644 | 2 | 0 | 1 | 0 | 1 | 1 | 2 | 0 | 6 | 0.1667 | 1 | 1 | 1 | 0 | NA | NA | 2 | 0 | 4 | 0.25 |
| 12_8406360 | 6 | 0 | 5 | 0 | 1 | 1 | 2 | 0.5 | 14 | 0.1429 | 1 | 0 | 2 | 0.5 | 1 | 0 | 2 | 0.5 | 6 | 0.3333 |
| 12_8406370 | 7 | 1 | 5 | 0.8 | 1 | 1 | 3 | 1 | 16 | 0.9375 | 2 | 1 | 2 | 1 | 1 | 1 | 3 | 1 | 8 | 1 |
| 12_8406371 | 7 | 0.8571 | 4 | 0.75 | 1 | 1 | 3 | 1 | 15 | 0.8667 | 2 | 0.5 | 2 | 1 | 1 | 0 | 3 | 1 | 8 | 0.75 |
| 12_8406409 | 6 | 0.6667 | 4 | 0.75 | 1 | 1 | 2 | 1 | 13 | 0.7692 | 2 | 1 | 2 | 1 | 1 | 0 | 3 | 0.6667 | 8 | 0.75 |
| 12_8406411 | 4 | 0.25 | 1 | 0 | 1 | 0 | 3 | 0.6667 | 9 | 0.3333 | 2 | 0 | 2 | 0 | 1 | 1 | 3 | 0 | 8 | 0.125 |
| 12_8406423 | 3 | 0.6667 | 5 | 0.4 | 1 | 0 | 3 | 0 | 12 | 0.3333 | 2 | 0 | 2 | 0 | NA | NA | 4 | 0.5 | 8 | 0.25 |
| 12_8406433 | 7 | 0.1429 | 6 | 0 | 1 | 0 | 3 | 0 | 17 | 0.0588 | 3 | 0.3333 | 1 | 0 | NA | NA | 4 | 0.25 | 8 | 0.25 |
| 12_8410606 | NA | NA | 2 | 1 | NA | NA | NA | NA | 2 | 1 | NA | NA | NA | NA | NA | NA | 1 | 1 | 1 | 1 |
| 12_8410613 | NA | NA | 2 | 1 | NA | NA | NA | NA | 2 | 1 | NA | NA | NA | NA | NA | NA | 1 | 1 | 1 | 1 |
| 12_8410621 | NA | NA | 1 | 1 | NA | NA | NA | NA | 1 | 1 | NA | NA | NA | NA | NA | NA | 1 | 1 | 1 | 1 |
| 12_8410657 | NA | NA | 2 | 0.5 | NA | NA | NA | NA | 2 | 0.5 | NA | NA | NA | NA | NA | NA | 1 | 1 | 1 | 1 |
| 12_8410658 | NA | NA | 2 | 0.5 | NA | NA | NA | NA | 2 | 0.5 | NA | NA | NA | NA | NA | NA | 1 | 1 | 1 | 1 |
| 12_8410660 | NA | NA | 2 | 1 | NA | NA | NA | NA | 2 | 1 | NA | NA | NA | NA | NA | NA | 1 | 1 | 1 | 1 |
| 12_8415599 | NA | NA | 1 | 1 | NA | NA | NA | NA | 1 | 1 | NA | NA | 1 | 1 | 1 | 1 | 2 | 1 | 4 | 1 |
| 12_8415711 | NA | NA | 1 | 1 | NA | NA | NA | NA | 1 | 1 | NA | NA | 1 | 1 | 1 | 1 | 3 | 0.3333 | 5 | 0.6 |
| 12_8415726 | NA | NA | 1 | 1 | NA | NA | NA | NA | 1 | 1 | NA | NA | NA | NA | NA | NA | 3 | 0.6667 | 3 | 0.6667 |
| 12_8419016 | 17 | 0.2941 | 11 | 0.3636 | 9 | 0.2222 | 8 | 0.25 | 45 | 0.2889 | 3 | 0 | 11 | 0.4545 | 5 | 0.2 | 10 | 0.1 | 29 | 0.2414 |
| 12_8419617 | 2 | 1 | 1 | 1 | 4 | 0.5 | 4 | 0.5 | 11 | 0.6364 | 4 | 0 | 3 | 0 | 1 | 0 | 3 | 0.3333 | 11 | 0.0909 |
| 12_8419636 | 1 | 0 | 2 | 0 | 3 | 0 | 4 | 0.25 | 10 | 0.1 | 3 | 0.3333 | 1 | 0 | 2 | 0 | 3 | 0 | 9 | 0.1111 |
| 12_8419799 | 1 | 1 | 1 | 0 | NA | NA | 4 | 0.25 | 6 | 0.3333 | 4 | 0 | 2 | 0 | 2 | 0.5 | 3 | 0 | 11 | 0.0909 |
| 12_8419847 | NA | NA | 1 | 0 | NA | NA | 3 | 1 | 4 | 0.75 | 4 | 0.5 | 1 | 1 | NA | NA | 1 | 1 | 6 | 0.6667 |
| 12_8419852 | 1 | 1 | 1 | 1 | NA | NA | 4 | 1 | 6 | 1 | 4 | 1 | 1 | 1 | NA | NA | NA | NA | 5 | 1 |
| 12_8419875 | 1 | 1 | 1 | 1 | NA | NA | 4 | 1 | 6 | 1 | 4 | 1 | 1 | 1 | NA | NA | 1 | 0 | 6 | 0.8333 |
| 12_8419877 | 1 | 1 | 1 | 1 | NA | NA | 4 | 1 | 6 | 1 | 4 | 0.75 | 1 | 1 | 1 | 1 | 1 | 1 | 7 | 0.8571 |
| 12_8419902 | NA | NA | 1 | 1 | NA | NA | 3 | 1 | 4 | 1 | 2 | 1 | NA | NA | 1 | 1 | 1 | 1 | 4 | 1 |
| 12_8419909 | NA | NA | 1 | 1 | NA | NA | 3 | 1 | 4 | 1 | 2 | 1 | NA | NA | 1 | 1 | 1 | 1 | 4 | 1 |
| 12_8430510 | 4 | 0.25 | 4 | 0 | 4 | 0.25 | 5 | 0.2 | 17 | 0.1765 | 4 | 0 | 1 | 0 | 6 | 0.3333 | 5 | 0.4 | 16 | 0.25 |
| 12_8438990 | 2 | 0 | 2 | 0 | 1 | 1 | 4 | 0 | 9 | 0.1111 | 3 | 0.6667 | 3 | 0 | 2 | 0 | 2 | 0.5 | 10 | 0.3 |
| 12_8439009 | 2 | 1 | 1 | 1 | 1 | 1 | 5 | 0.8 | 9 | 0.8889 | 2 | 1 | 3 | 1 | 2 | 1 | 3 | 0.6667 | 10 | 0.9 |
| 12_8439017 | 2 | 0.5 | 2 | 0 | 1 | 0 | 4 | 0 | 9 | 0.1111 | 3 | 0 | 2 | 0 | 2 | 0 | 3 | 0.6667 | 10 | 0.2 |

| | | | | | | | | | | | | | | | | | | | | |
|---|---|---|---|---|---|---|---|---|---|---|---|---|---|---|---|---|---|---|---|---|
| 12_843921 | NA | NA | 3 | 0.3333 | 2 | 0 | 1 | 0 | 6 | 0.1667 | NA | NA | 1 | 1 | 3 | 0 | 1 | 0 | 5 | 0.2 |
| 12_844016 | NA | NA | 1 | 1 | 1 | 0 | NA | NA | 2 | 0.5 | NA | NA | NA | NA | 1 | 1 | 1 | 1 | 2 | 1 |
| 12_844031 | NA | NA | 1 | 0 | 1 | 1 | NA | NA | 2 | 0.5 | NA | NA | 1 | 1 | NA | NA | 1 | 0 | 2 | 0.5 |
| 12_844209 | 5 | 0.2 | 4 | 0.25 | 2 | 0 | 2 | 0 | 13 | 0.1538 | 2 | 0.5 | 1 | 0 | 1 | 0 | 2 | 1 | 6 | 0.5 |
| 12_844237 | 4 | 0.25 | 3 | 0 | 3 | 0.6667 | 1 | 1 | 11 | 0.3636 | NA | NA | 2 | 0 | 2 | 0 | 1 | 1 | 5 | 0.2 |
| 12_844237 | 5 | 0.2 | 3 | 0 | 3 | 0 | 1 | 0 | 12 | 0.0833 | 2 | 0 | 3 | 0.3333 | 1 | 0 | 1 | 0 | 7 | 0.1429 |
| 12_844238 | 5 | 0.8 | 3 | 1 | 2 | 1 | 1 | 1 | 11 | 0.9091 | 2 | 0.5 | 3 | 0 | 2 | 0 | 1 | 1 | 8 | 0.25 |
| 12_844240 | 5 | 0.8 | 2 | 1 | 3 | 0 | 1 | 1 | 11 | 0.6364 | 2 | 1 | 2 | 0.5 | 2 | 0.5 | 1 | 0 | 7 | 0.5714 |
| 12_844240 | 5 | 0.4 | 2 | 0.5 | 3 | 0.6667 | 1 | 0 | 11 | 0.4545 | 2 | 0.5 | 2 | 0.5 | 2 | 0.5 | 1 | 1 | 7 | 0.5714 |
| 12_844644 | 2 | 1 | 2 | 1 | NA | NA | 1 | 1 | 5 | 1 | NA | NA | 2 | 1 | 3 | 0.6667 | 1 | 0 | 6 | 0.6667 |
| 12_844647 | 2 | 1 | 1 | 0 | 1 | 1 | 1 | 0 | 5 | 0.6 | NA | NA | 3 | 0.3333 | 4 | 0.75 | 1 | 1 | 8 | 0.625 |
| 12_844647 | 1 | 1 | 2 | 0 | NA | NA | 1 | 1 | 4 | 0.5 | NA | NA | 3 | 0.6667 | 3 | 0.3333 | 1 | 1 | 7 | 0.5714 |
| 12_844650 | 2 | 1 | 2 | 1 | 1 | 1 | 1 | 1 | 6 | 1 | NA | NA | 3 | 1 | 4 | 1 | 1 | 1 | 8 | 1 |
| 12_844652 | 3 | 0 | 1 | 0 | 1 | 1 | 1 | 0 | 6 | 0.1667 | 1 | 0 | 3 | 0.3333 | 4 | 0.25 | NA | NA | 8 | 0.25 |
| 12_844652 | 2 | 0.5 | 1 | 1 | 1 | 1 | 1 | 0 | 5 | 0.6 | NA | NA | 1 | 1 | 3 | 0 | 1 | 0 | 5 | 0.2 |
| 12_844737 | NA | NA | NA | NA | NA | NA | 1 | 1 | 1 | 1 | NA | NA | 1 | 0 | NA | NA | 1 | 1 | 2 | 0.5 |
| 12_844781 | 14 | 0.2143 | 11 | 0.1818 | 10 | 0.3 | 5 | 0.2 | 40 | 0.225 | 7 | 0.2857 | 5 | 0 | 9 | 0.1111 | 5 | 0.2 | 26 | 0.1538 |
| 12_844790 | 5 | 0.6 | 7 | 0.2857 | 8 | 0.75 | 1 | 0 | 21 | 0.5238 | 5 | 0.6 | 3 | 0.6667 | 4 | 0.5 | 2 | 0.5 | 14 | 0.5714 |
| 12_844792 | 12 | 0.1667 | 9 | 0.2222 | 10 | 0 | 3 | 0 | 34 | 0.1176 | 7 | 0 | 3 | 0.3333 | 8 | 0.125 | 4 | 0 | 22 | 0.0909 |
| 12_844864 | 9 | 0.4444 | 4 | 0 | 8 | 0.25 | 3 | 0 | 24 | 0.25 | 5 | 0.2 | 2 | 0 | 5 | 0.2 | 4 | 0.5 | 16 | 0.25 |
| 12_844878 | NA | NA | 1 | 0 | NA | NA | 3 | 0.6667 | 4 | 0.5 | NA | NA | NA | NA | 1 | 1 | 2 | 0 | 3 | 0.3333 |
| 12_844880 | NA | NA | NA | NA | NA | NA | 2 | 1 | 2 | 1 | NA | NA | NA | NA | 1 | 1 | 2 | 1 | 3 | 1 |
| 12_845764 | 3 | 0 | 3 | 0 | 2 | 0.5 | 3 | 0.3333 | 11 | 0.1818 | 4 | 0 | 7 | 0.1429 | 2 | 0.5 | 2 | 0.5 | 15 | 0.2 |
| 12_845765 | 3 | 0.3333 | 4 | 0.75 | 4 | 1 | 4 | 0.5 | 15 | 0.6667 | 5 | 0.4 | 7 | 0.7143 | 1 | 0 | 2 | 0.5 | 15 | 0.5333 |
| 12_845765 | 5 | 0.4 | 4 | 0.5 | 5 | 0.2 | 5 | 0.2 | 19 | 0.3158 | 5 | 0.6 | 7 | 0.5714 | 2 | 0.5 | 2 | 0 | 16 | 0.5 |
| 12_845766 | 5 | 0.6 | 4 | 0.75 | 5 | 0.6 | 5 | 0.6 | 19 | 0.6316 | 6 | 0.8333 | 7 | 0.7143 | 2 | 1 | 2 | 1 | 17 | 0.8235 |
| 12_845768 | 3 | 1 | 4 | 1 | 4 | 1 | 5 | 0.8 | 16 | 0.9375 | 5 | 0.8 | 6 | 0.6667 | 2 | 1 | 2 | 1 | 15 | 0.8 |
| 12_845769 | 4 | 0 | 4 | 0.25 | 5 | 0.4 | 5 | 0.2 | 18 | 0.2222 | 6 | 0.1667 | 6 | 0 | 1 | 0 | 2 | 0.5 | 15 | 0.1333 |
| 12_845769 | 5 | 0.2 | 4 | 0.5 | 5 | 0.4 | 5 | 1 | 19 | 0.5263 | 5 | 0 | 7 | 0.2857 | 2 | 0.5 | 2 | 0 | 16 | 0.1875 |
| 12_845769 | 3 | 1 | 2 | 1 | 4 | 1 | 2 | 0 | 11 | 0.8182 | 4 | 1 | 6 | 0.8333 | 1 | 0 | 2 | 1 | 13 | 0.8462 |
| 12_845769 | 5 | 0.6 | 3 | 0 | 5 | 0.2 | 4 | 0.5 | 17 | 0.3529 | 6 | 0.3333 | 7 | 0.5714 | 2 | 0.5 | 2 | 0 | 17 | 0.4118 |
| 12_845770 | 3 | 0 | 4 | 0.25 | 5 | 0.2 | 5 | 0 | 17 | 0.1176 | 6 | 0 | 8 | 0.125 | 2 | 0.5 | 2 | 0 | 18 | 0.1111 |
| 12_845770 | 5 | 0.8 | 4 | 0.75 | 5 | 0.6 | 5 | 0.8 | 19 | 0.7368 | 5 | 0.8 | 8 | 0.875 | 2 | 1 | 2 | 1 | 17 | 0.8824 |
| 12_845771 | 4 | 0 | 4 | 0.5 | 5 | 0.2 | 4 | 0 | 17 | 0.1765 | 5 | 0.6 | 8 | 0.25 | 2 | 0 | 2 | 0.5 | 17 | 0.3529 |
| 12_845771 | 5 | 0.8 | 4 | 0.75 | 5 | 1 | 5 | 0.4 | 19 | 0.7368 | 6 | 0.6667 | 8 | 0.875 | 2 | 0.5 | 2 | 1 | 18 | 0.7778 |
| 12_845772 | 5 | 1 | 2 | 1 | 4 | 0 | 6 | 1 | 17 | 0.7647 | 6 | 0.3333 | 8 | 0.875 | 2 | 1 | 1 | 1 | 17 | 0.7059 |
| 12_845773 | 3 | 0 | 3 | 0 | 4 | 0 | 6 | 0.1667 | 16 | 0.0625 | 5 | 0.2 | 8 | 0.375 | 2 | 0.5 | 2 | 0.5 | 17 | 0.3529 |
| 12_846494 | NA | NA | 1 | 0 | 1 | 1 | 1 | 0 | 3 | 0.3333 | NA | NA | NA | NA | NA | NA | 2 | 0.5 | 2 | 0.5 |
| 12_846495 | NA | NA | 1 | 1 | 1 | 1 | 1 | 1 | 3 | 1 | NA | NA | NA | NA | 1 | 1 | 1 | 0 | 2 | 0.5 |
| 12_846495 | NA | NA | 1 | 1 | 1 | 1 | 1 | 1 | 3 | 1 | NA | NA | NA | NA | 1 | 1 | 2 | 0.5 | 3 | 0.6667 |
| 12_846496 | NA | NA | 1 | 1 | 1 | 1 | NA | NA | 2 | 1 | NA | NA | NA | NA | NA | NA | 2 | 0.5 | 2 | 0.5 |
| 12_847206 | NA | NA | NA | NA | 1 | 1 | NA | NA | 1 | 1 | NA | NA | 1 | 1 | NA | NA | NA | NA | 1 | 1 |
| 12_847206 | NA | NA | NA | NA | 1 | 1 | 1 | 0 | 2 | 0.5 | NA | NA | 1 | 1 | NA | NA | NA | NA | 1 | 1 |
| 12_847208 | NA | NA | NA | NA | 1 | 1 | 1 | 1 | 2 | 1 | NA | NA | 1 | 1 | NA | NA | NA | NA | 1 | 1 |
| 12_847239 | NA | NA | NA | NA | 1 | 1 | NA | NA | 1 | 1 | NA | NA | 2 | 0.5 | NA | NA | NA | NA | 2 | 0.5 |
| 12_847239 | 1 | 0 | NA | NA | 1 | 1 | NA | NA | 2 | 0.5 | NA | NA | 2 | 0.5 | 1 | 0 | NA | NA | 3 | 0.3333 |
| 12_847240 | 17 | 0.1176 | 9 | 0 | 11 | 0.0909 | 14 | 0.2143 | 51 | 0.1176 | 7 | 0 | 21 | 0.0952 | 21 | 0.1905 | 10 | 0.2 | 59 | 0.1356 |
| 12_847242 | 16 | 0.375 | 9 | 0.2222 | 15 | 0.2667 | 25 | 0.24 | 65 | 0.2769 | 3 | 0 | 24 | 0.4583 | 22 | 0.3182 | 9 | 0.3333 | 58 | 0.3621 |
| 12_847242 | 19 | 0.2632 | 11 | 0.2727 | 19 | 0.2105 | 26 | 0.3077 | 75 | 0.2667 | 6 | 0.6667 | 27 | 0.3704 | 24 | 0.375 | 10 | 0.4 | 67 | 0.403 |
| 12_847246 | 16 | 0.125 | 11 | 0.0909 | 15 | 0.0667 | 22 | 0 | 64 | 0.0625 | 4 | 0 | 22 | 0.1364 | 19 | 0.0526 | 8 | 0.125 | 53 | 0.0943 |
| 12_847246 | 18 | 0.0556 | 11 | 0 | 17 | 0.0588 | 24 | 0.0833 | 70 | 0.0571 | 7 | 0 | 21 | 0 | 22 | 0.0455 | 10 | 0.2 | 60 | 0.05 |
| 12_847246 | 17 | 0.1176 | 9 | 0.2222 | 14 | 0 | 19 | 0.4211 | 59 | 0.2034 | 6 | 0.5 | 16 | 0.125 | 20 | 0.2 | 9 | 0.1111 | 51 | 0.1961 |
| 12_847247 | 16 | 0.0625 | 8 | 0.125 | 17 | 0.1176 | 22 | 0.2273 | 63 | 0.1429 | 7 | 0.4286 | 24 | 0.375 | 21 | 0.0476 | 9 | 0.2222 | 61 | 0.2459 |
| 12_847249 | 18 | 0.1667 | 10 | 0.1 | 15 | 0.1333 | 22 | 0.1364 | 65 | 0.1385 | 6 | 0.1667 | 20 | 0.25 | 14 | 0.0714 | 8 | 0.375 | 48 | 0.2083 |
| 12_847251 | 14 | 0.2143 | 8 | 0.25 | 13 | 0.0769 | 22 | 0.2727 | 57 | 0.2105 | 6 | 0.1667 | 21 | 0.2381 | 17 | 0.1765 | 10 | 0.2 | 54 | 0.2037 |
| 12_847252 | 18 | 0.3889 | 6 | 0.3333 | 13 | 0.3077 | 22 | 0.1364 | 59 | 0.2712 | 5 | 0 | 19 | 0.3684 | 15 | 0.1333 | 10 | 0.3 | 49 | 0.2449 |
| 12_847254 | 12 | 0.8333 | 11 | 0.9091 | 14 | 0.9286 | 14 | 1 | 51 | 0.9216 | 6 | 1 | 16 | 1 | 13 | 0.7692 | 8 | 1 | 43 | 0.9302 |
| 12_847254 | 16 | 0.75 | 10 | 1 | 10 | 0.8 | 20 | 0.8 | 56 | 0.8214 | 5 | 0.8 | 19 | 0.8421 | 14 | 0.6429 | 9 | 0.7778 | 47 | 0.766 |
| 12_847256 | 13 | 0.1538 | 11 | 0.0909 | 12 | 0.0833 | 14 | 0.2857 | 50 | 0.16 | 5 | 0.2 | 16 | 0.1875 | 15 | 0.0667 | 6 | 0 | 42 | 0.119 |
| 12_847273 | 14 | 0.1429 | 8 | 0 | 11 | 0.4545 | 15 | 0.2667 | 48 | 0.2292 | 4 | 0 | 11 | 0.0909 | 12 | 0.25 | 5 | 0 | 32 | 0.125 |
| 12_848027 | 1 | 1 | NA | NA | 3 | 0.3333 | 1 | 0 | 5 | 0.4 | NA | NA | 1 | 0 | 2 | 0.5 | NA | NA | 3 | 0.3333 |
| 12_848209 | 1 | 0 | 3 | 0.3333 | 1 | 0 | NA | NA | 5 | 0.2 | NA | NA | 2 | 0 | NA | NA | 3 | 0.3333 | 5 | 0.2 |
| 12_848213 | 10 | 0 | 9 | 0.1111 | 6 | 0.1667 | 10 | 0 | 35 | 0.0571 | 5 | 0 | 5 | 0.4 | 11 | 0.2727 | 9 | 0.1111 | 30 | 0.2 |
| 12_848214 | 10 | 0.1 | 9 | 0.1111 | 5 | 0 | 11 | 0.2727 | 35 | 0.1429 | 4 | 0.25 | 6 | 0.3333 | 10 | 0.4 | 9 | 0.2222 | 29 | 0.3103 |
| 12_849624 | NA | NA | NA | NA | NA | NA | 1 | 1 | 1 | 1 | NA | NA | NA | NA | 1 | 1 | NA | NA | 1 | 1 |

| | | | | | | | | | | | | | | | | | | | | |
|---|---|---|---|---|---|---|---|---|---|---|---|---|---|---|---|---|---|---|---|---|
| 12_8496298 | NA | NA | 1 | 1 | 1 | 0 | 1 | 0 | 3 | 0.3333 | NA | NA | NA | NA | 1 | 1 | NA | NA | 1 | 1 |
| 12_8496521 | 27 | 0.2593 | 13 | 0.3077 | 27 | 0.3333 | 24 | 0.375 | 91 | 0.3187 | 12 | 0.25 | 27 | 0.1111 | 27 | 0.4074 | 17 | 0.4118 | 83 | 0.2892 |
| 12_8496531 | 25 | 0.28 | 20 | 0.25 | 24 | 0.125 | 21 | 0.381 | 90 | 0.2556 | 15 | 0.1333 | 29 | 0.2069 | 26 | 0.2692 | 17 | 0.0588 | 87 | 0.1839 |
| 12_8496566 | 17 | 0.5882 | 15 | 0.4667 | 22 | 0.4545 | 23 | 0.4348 | 77 | 0.4805 | 10 | 0.4 | 15 | 0.2667 | 19 | 0.5789 | 17 | 0.4706 | 61 | 0.4426 |
| 12_8496569 | 21 | 0.2857 | 16 | 0.5 | 21 | 0.6667 | 19 | 0.6316 | 77 | 0.5195 | 13 | 0.4615 | 23 | 0.4348 | 23 | 0.6087 | 17 | 0.3529 | 76 | 0.4737 |
| 12_8496570 | 19 | 0.9474 | 12 | 1 | 19 | 0.9474 | 21 | 1 | 71 | 0.9718 | 12 | 1 | 21 | 0.8571 | 24 | 1 | 13 | 1 | 70 | 0.9571 |
| 12_8496575 | 24 | 0.5 | 18 | 0.4444 | 26 | 0.5385 | 23 | 0.4783 | 91 | 0.4945 | 15 | 0.8 | 23 | 0.4783 | 24 | 0.4167 | 20 | 0.65 | 82 | 0.561 |
| 12_8496576 | 21 | 0.5238 | 15 | 0.3333 | 27 | 0.5556 | 21 | 0.5238 | 84 | 0.5 | 15 | 0.6667 | 23 | 0.4783 | 26 | 0.6154 | 21 | 0.6667 | 85 | 0.6 |
| 12_8496584 | 25 | 0.56 | 17 | 0.2353 | 25 | 0.36 | 25 | 0.24 | 92 | 0.3587 | 15 | 0.2667 | 23 | 0.3043 | 27 | 0.3704 | 18 | 0.2778 | 83 | 0.3133 |
| 12_8496592 | 22 | 0.2273 | 16 | 0.0625 | 21 | 0.1905 | 23 | 0.0435 | 82 | 0.1341 | 12 | 0.25 | 22 | 0.2727 | 25 | 0.2 | 17 | 0.0588 | 76 | 0.1974 |
| 12_8496614 | 21 | 0.5714 | 13 | 0.6923 | 23 | 0.5652 | 21 | 0.5714 | 78 | 0.5897 | 13 | 0.4615 | 21 | 0.5238 | 20 | 0.6 | 17 | 0.7059 | 71 | 0.5775 |
| 12_8496619 | 21 | 0.2857 | 12 | 0.3333 | 23 | 0.3043 | 18 | 0.3889 | 74 | 0.3243 | 10 | 0.1 | 17 | 0.3529 | 17 | 0.3529 | 14 | 0.5 | 58 | 0.3448 |
| 12_8496623 | 23 | 0.9565 | 14 | 1 | 24 | 0.9167 | 19 | 0.9474 | 80 | 0.95 | 13 | 0.9231 | 21 | 1 | 21 | 0.9048 | 17 | 0.8824 | 72 | 0.9306 |
| 12_8496628 | 16 | 0.375 | 11 | 0.3636 | 20 | 0.4 | 19 | 0.3684 | 66 | 0.3788 | 12 | 0.4167 | 18 | 0.5 | 20 | 0.3 | 15 | 0.5333 | 65 | 0.4308 |
| 12_8496634 | 18 | 0.5556 | 12 | 0.5 | 19 | 0.4737 | 16 | 0.625 | 65 | 0.5385 | 8 | 0.75 | 15 | 0.4667 | 12 | 0.6667 | 16 | 0.5625 | 51 | 0.5882 |
| 12_8496635 | 18 | 0.6667 | 12 | 0.5833 | 18 | 0.2778 | 15 | 0.4667 | 63 | 0.4921 | 11 | 0.4545 | 18 | 0.5556 | 15 | 0.5333 | 15 | 0.4667 | 59 | 0.5085 |
| 12_8496642 | 16 | 0.9375 | 8 | 1 | 20 | 1 | 15 | 0.8 | 59 | 0.9322 | 11 | 0.7273 | 15 | 1 | 17 | 0.8824 | 15 | 0.9333 | 58 | 0.8966 |
| 12_8496643 | 17 | 0.5294 | 11 | 0.8182 | 22 | 0.5909 | 21 | 0.5714 | 71 | 0.6056 | 12 | 0.4167 | 17 | 0.7647 | 20 | 0.55 | 12 | 0.5833 | 61 | 0.5902 |
| 12_8496647 | 21 | 0.0476 | 12 | 0.0833 | 22 | 0.0909 | 19 | 0.0526 | 74 | 0.0676 | 13 | 0.1538 | 20 | 0.05 | 18 | 0 | 17 | 0.0588 | 68 | 0.0588 |
| 12_8496667 | 18 | 0.7778 | 8 | 0.875 | 19 | 0.9474 | 16 | 0.6875 | 61 | 0.8197 | 11 | 0.8182 | 19 | 0.8947 | 16 | 0.9375 | 14 | 0.9286 | 60 | 0.9 |
| 12_8496668 | 18 | 0.2778 | 10 | 0.2 | 20 | 0.2 | 17 | 0.2353 | 65 | 0.2308 | 10 | 0.3 | 18 | 0.1111 | 16 | 0.1875 | 14 | 0.1429 | 58 | 0.1724 |
| 12_8496671 | 18 | 1 | 9 | 1 | 18 | 0.9444 | 16 | 1 | 61 | 0.9836 | 12 | 1 | 18 | 1 | 17 | 1 | 14 | 1 | 61 | 1 |
| 12_8496684 | 16 | 0.6875 | 11 | 0.2727 | 21 | 0.1429 | 19 | 0.3158 | 67 | 0.3433 | 9 | 0.4444 | 16 | 0.4375 | 16 | 0.3125 | 14 | 0.2143 | 55 | 0.3455 |
| 12_8496693 | 15 | 1 | 9 | 1 | 17 | 0.9412 | 14 | 0.9286 | 55 | 0.9636 | 8 | 1 | 17 | 0.8824 | 16 | 0.9375 | 14 | 0.9286 | 55 | 0.9273 |
| 12_8496694 | 19 | 0.1579 | 10 | 0.1 | 20 | 0.15 | 20 | 0.35 | 69 | 0.2029 | 10 | 0.2 | 17 | 0.2353 | 16 | 0.1875 | 14 | 0.5 | 57 | 0.2807 |
| 12_8496701 | 17 | 0.2353 | 10 | 0.2 | 20 | 0.05 | 20 | 0.15 | 67 | 0.1493 | 11 | 0.0909 | 21 | 0.2857 | 17 | 0.1765 | 16 | 0.1875 | 65 | 0.2 |
| 12_8496720 | 20 | 0.85 | 9 | 1 | 21 | 0.9048 | 19 | 0.7895 | 69 | 0.8696 | 9 | 0.8889 | 19 | 0.8421 | 16 | 0.875 | 15 | 1 | 59 | 0.8983 |
| 12_8499802 | 10 | 0.1 | 4 | 0.25 | 4 | 0.25 | 5 | 0.2 | 23 | 0.1739 | 4 | 0.25 | 5 | 0 | 8 | 0.125 | 5 | 0 | 22 | 0.0909 |
| 12_8506159 | 1 | 0 | NA | NA | 1 | 1 | NA | NA | 2 | 0.5 | 1 | 1 | 2 | 1 | NA | NA | NA | NA | 3 | 1 |
| 12_8506164 | 1 | 1 | NA | NA | 1 | 0 | NA | NA | 2 | 0.5 | 1 | 1 | 2 | 0 | NA | NA | NA | NA | 3 | 0.3333 |
| 12_8506196 | 1 | 1 | 2 | 0 | NA | NA | NA | NA | 3 | 0.3333 | 1 | 1 | 3 | 0 | NA | NA | NA | NA | 4 | 0.25 |
| 12_8506209 | 1 | 1 | 2 | 0.5 | 1 | 1 | NA | NA | 4 | 0.75 | 1 | 0 | 3 | 0.6667 | NA | NA | NA | NA | 4 | 0.5 |
| 12_8506224 | 1 | 1 | 2 | 0.5 | 2 | 1 | NA | NA | 5 | 0.8 | NA | NA | 3 | 1 | NA | NA | NA | NA | 3 | 1 |
| 12_8506229 | NA | NA | 2 | 0.5 | 2 | 0 | NA | NA | 4 | 0.25 | 1 | 0 | 3 | 0.3333 | NA | NA | NA | NA | 4 | 0.25 |
| 12_8506231 | 1 | 1 | 1 | 1 | 2 | 1 | NA | NA | 4 | 1 | 1 | 1 | 2 | 1 | NA | NA | NA | NA | 3 | 1 |
| 12_8507572 | 1 | 0 | 2 | 0.5 | 2 | 0 | NA | NA | 5 | 0.2 | 1 | 0 | 2 | 0.5 | NA | NA | NA | NA | 3 | 0.3333 |
| 12_8507598 | 1 | 1 | 2 | 0.5 | 2 | 0.5 | NA | NA | 5 | 0.6 | 1 | 0 | 2 | 1 | 2 | 0.5 | NA | NA | 5 | 0.6 |
| 12_8507607 | 1 | 1 | 1 | 0 | 2 | 0.5 | NA | NA | 4 | 0.5 | 1 | 0 | 2 | 0 | 2 | 0.5 | NA | NA | 5 | 0.2 |
| 12_8507608 | 1 | 1 | 1 | 1 | 2 | 0 | NA | NA | 4 | 0.5 | 1 | 1 | 2 | 1 | 2 | 1 | NA | NA | 5 | 1 |
| 12_8507636 | NA | NA | 2 | 0.5 | 3 | 0 | NA | NA | 5 | 0.2 | 1 | 1 | 2 | 0.5 | 3 | 1 | 1 | 1 | 7 | 0.8571 |
| 12_8507691 | 1 | 0 | 3 | 0.6667 | 3 | 0.3333 | 1 | 0 | 8 | 0.375 | 1 | 1 | 2 | 0 | 3 | 0.3333 | 2 | 0 | 8 | 0.25 |
| 12_8507692 | 1 | 0 | 4 | 0.25 | 3 | 0 | 1 | 0 | 9 | 0.1111 | 1 | 0 | 2 | 0.5 | 3 | 0 | 2 | 0.5 | 8 | 0.25 |
| 12_8507711 | 1 | 0 | 3 | 0.3333 | 3 | 0 | NA | NA | 7 | 0.1429 | 1 | 1 | 1 | 0 | 2 | 0 | 2 | 0 | 6 | 0.1667 |
| 12_8507718 | 1 | 1 | 4 | 0.25 | 4 | 0.5 | 1 | 1 | 10 | 0.5 | 1 | 0 | 2 | 0 | 3 | 0.6667 | 1 | 0 | 7 | 0.2857 |
| 12_8507733 | 1 | 0 | 4 | 0.75 | 3 | 0.3333 | 1 | 0 | 9 | 0.4444 | 1 | 1 | 3 | 0.6667 | 3 | 0.3333 | 2 | 0 | 9 | 0.4444 |
| 12_8507744 | 1 | 0 | 3 | 1 | 3 | 0.3333 | 1 | 1 | 8 | 0.625 | 1 | 1 | 3 | 0.6667 | 3 | 0.6667 | 2 | 0.5 | 9 | 0.6667 |
| 12_8507749 | 1 | 0 | 4 | 0.75 | 3 | 0.6667 | 2 | 0.5 | 10 | 0.6 | 1 | 1 | 3 | 0.3333 | 2 | 1 | 2 | 0 | 8 | 0.5 |
| 12_8507757 | NA | NA | 3 | 0 | 3 | 0.6667 | 2 | 0 | 8 | 0.25 | 1 | 1 | 2 | 0 | 1 | 1 | 2 | 0 | 6 | 0.3333 |
| 12_8508759 | 2 | 0.5 | 4 | 0.75 | 4 | 0 | 4 | 0 | 14 | 0.2857 | 2 | 0.5 | 5 | 0.2 | 5 | 0 | 3 | 0 | 15 | 0.1333 |
| 12_8510447 | 1 | 0 | 3 | 0.3333 | 4 | 0 | 1 | 0 | 9 | 0.1111 | NA | NA | 2 | 0 | 5 | 0.4 | 1 | 0 | 8 | 0.25 |
| 12_8510455 | 2 | 0.5 | 3 | 0 | 4 | 0.25 | 1 | 0 | 10 | 0.2 | NA | NA | 4 | 0.75 | 4 | 0.25 | 2 | 0.5 | 10 | 0.5 |
| 12_8510480 | 2 | 0.5 | 3 | 0 | 2 | 0.5 | 1 | 0 | 8 | 0.25 | NA | NA | 3 | 0.3333 | 5 | 0 | 2 | 0.5 | 10 | 0.2 |
| 12_8510672 | 1 | 0 | 2 | 0.5 | NA | NA | NA | NA | 3 | 0.3333 | NA | NA | 1 | 0 | 1 | 1 | 1 | 0 | 3 | 0.3333 |
| 12_8510998 | 1 | 1 | 1 | 1 | 1 | 1 | 1 | 1 | 4 | 1 | NA | NA | 1 | 1 | NA | NA | 2 | 0.5 | 3 | 0.6667 |
| 12_8511716 | 4 | 0 | 9 | 0.1111 | 8 | 0.125 | 11 | 0 | 32 | 0.0625 | 3 | 0.3333 | 5 | 0 | 8 | 0.125 | 9 | 0 | 25 | 0.08 |
| 12_8511839 | 1 | 0 | 1 | 1 | NA | NA | 3 | 0.3333 | 5 | 0.4 | 1 | 1 | NA | NA | 1 | 0 | 1 | 1 | 3 | 0.6667 |
| 12_8512020 | 1 | 0 | 1 | 1 | NA | NA | 2 | 0 | 4 | 0.25 | 2 | 0 | NA | NA | 3 | 0.3333 | 1 | 0 | 6 | 0.1667 |
| 12_8512084 | 2 | 1 | 2 | 1 | NA | NA | 2 | 0.5 | 6 | 0.8333 | 2 | 0.5 | NA | NA | 3 | 1 | 1 | 1 | 6 | 0.8333 |
| 12_8512088 | NA | NA | 2 | 0.5 | NA | NA | 2 | 0 | 4 | 0.25 | 1 | 0 | NA | NA | 2 | 0.5 | NA | NA | 3 | 0.3333 |
| 12_8512092 | 2 | 1 | 2 | 0.5 | NA | NA | 2 | 0.5 | 6 | 0.6667 | 2 | 1 | NA | NA | 4 | 0.5 | 1 | 1 | 7 | 0.7143 |
| 12_8512096 | 2 | 0 | 1 | 1 | NA | NA | 2 | 0 | 5 | 0.2 | 2 | 0 | NA | NA | 3 | 0.3333 | 1 | 0 | 6 | 0.1667 |
| 12_8512097 | 2 | 0.5 | 1 | 0 | NA | NA | 2 | 0.5 | 5 | 0.4 | 2 | 1 | NA | NA | 4 | 0.75 | 1 | 1 | 7 | 0.8571 |
| 12_8512099 | 2 | 1 | 2 | 1 | NA | NA | 2 | 1 | 6 | 1 | 2 | 1 | NA | NA | 4 | 0.75 | 1 | 1 | 7 | 0.8571 |
| 12_8512101 | 2 | 1 | 2 | 0.5 | NA | NA | 2 | 0 | 6 | 0.5 | 1 | 0 | NA | NA | 3 | 0.3333 | 1 | 1 | 5 | 0.4 |
| 12_8512103 | 2 | 0 | 2 | 0.5 | NA | NA | 2 | 0.5 | 6 | 0.3333 | 2 | 0.5 | NA | NA | 4 | 0.75 | 1 | 1 | 7 | 0.7143 |

| | | | | | | | | | | | | | | | | | | | | |
|---|---|---|---|---|---|---|---|---|---|---|---|---|---|---|---|---|---|---|---|---|
| 12_8512105 | 2 | 0.5 | 2 | 0.5 | NA | NA | 2 | 1 | 6 | 0.6667 | 2 | 1 | NA | NA | 4 | 0.75 | 1 | 1 | 7 | 0.8571 |
| 12_8512107 | 2 | 1 | 2 | 0 | NA | NA | 2 | 1 | 6 | 0.6667 | 1 | 0 | NA | NA | 4 | 0.25 | 1 | 1 | 6 | 0.3333 |
| 12_8512117 | 2 | 1 | 1 | 1 | NA | NA | 2 | 1 | 5 | 1 | 2 | 1 | NA | NA | 4 | 1 | NA | NA | 6 | 1 |
| 12_8512122 | 2 | 0.5 | 2 | 0.5 | NA | NA | 2 | 0.5 | 6 | 0.5 | 2 | 0.5 | 1 | 0 | 4 | 0.25 | 2 | 0.5 | 9 | 0.3333 |
| 12_8513294 | NA | NA | NA | NA | NA | NA | 2 | 1 | 2 | 1 | NA | NA | 1 | 0 | 2 | 1 | 1 | 0 | 4 | 0.5 |
| 12_8513300 | NA | NA | 1 | 1 | NA | NA | 2 | 1 | 3 | 1 | NA | NA | 1 | 0 | 2 | 1 | 1 | 0 | 4 | 0.5 |
| 12_8513323 | NA | NA | 1 | 1 | NA | NA | 1 | 1 | 2 | 1 | NA | NA | 1 | 0 | 1 | 1 | 1 | 0 | 3 | 0.3333 |
| 12_8513324 | NA | NA | 1 | 1 | NA | NA | 2 | 0.5 | 3 | 0.6667 | NA | NA | 1 | 1 | 1 | 1 | 1 | 0 | 3 | 0.6667 |
| 12_8513386 | NA | NA | NA | NA | NA | NA | 1 | 1 | 1 | 1 | NA | NA | 1 | 1 | 1 | 1 | 1 | 0 | 3 | 0.6667 |
| 12_8513428 | NA | NA | 1 | 0 | NA | NA | 3 | 0.3333 | 4 | 0.25 | NA | NA | 1 | 1 | 2 | 0 | NA | NA | 3 | 0.3333 |
| 12_8513444 | NA | NA | 1 | 0 | NA | NA | 3 | 0.3333 | 4 | 0.25 | NA | NA | 1 | 1 | 2 | 0 | 1 | 0 | 4 | 0.25 |
| 12_8513450 | NA | NA | NA | NA | NA | NA | 2 | 1 | 2 | 1 | NA | NA | 1 | 1 | 2 | 1 | 1 | 1 | 4 | 1 |
| 12_8513466 | NA | NA | 1 | 1 | NA | NA | 3 | 0.6667 | 4 | 0.75 | NA | NA | 1 | 1 | 2 | 1 | 1 | 1 | 4 | 1 |
| 12_8513472 | NA | NA | 1 | 0 | NA | NA | 2 | 0.5 | 3 | 0.3333 | NA | NA | 1 | 1 | 2 | 0.5 | NA | NA | 3 | 0.6667 |
| 12_8513474 | NA | NA | 1 | 0 | NA | NA | 2 | 1 | 3 | 0.6667 | NA | NA | NA | NA | 1 | 0 | 1 | 1 | 2 | 0.5 |
| 12_8513487 | NA | NA | 1 | 1 | NA | NA | 2 | 1 | 3 | 1 | NA | NA | 1 | 1 | 2 | 1 | NA | NA | 3 | 1 |
| 12_8513488 | NA | NA | 1 | 1 | NA | NA | 2 | 0 | 3 | 0.3333 | NA | NA | 1 | 0 | 2 | 0.5 | NA | NA | 3 | 0.3333 |
| 12_8513493 | NA | NA | 1 | 0 | NA | NA | 3 | 1 | 4 | 0.75 | NA | NA | 1 | 1 | 2 | 1 | NA | NA | 3 | 1 |
| 12_8513509 | NA | NA | 1 | 0 | NA | NA | 3 | 1 | 4 | 0.75 | NA | NA | NA | NA | 2 | 1 | NA | NA | 2 | 1 |
| 12_8513634 | NA | NA | NA | NA | 1 | 0 | 2 | 0.5 | 3 | 0.3333 | NA | NA | NA | NA | 1 | 1 | 1 | 1 | 2 | 1 |
| 12_8517946 | 14 | 0.2143 | 14 | 0 | 8 | 0 | 9 | 0 | 45 | 0.0667 | 9 | 0.1111 | 15 | 0.1333 | 16 | 0.0625 | 9 | 0 | 49 | 0.0816 |
| 12_8517956 | 14 | 0.7143 | 13 | 0.5385 | 9 | 0.4444 | 8 | 0.875 | 44 | 0.6364 | 7 | 0.7143 | 14 | 0.6429 | 16 | 0.6875 | 10 | 0.6 | 47 | 0.6596 |
| 12_8517972 | 15 | 0.4667 | 12 | 0.5 | 8 | 0.25 | 11 | 0.4545 | 46 | 0.4348 | 8 | 0 | 13 | 0.3077 | 16 | 0.3125 | 10 | 0.5 | 47 | 0.2979 |
| 12_8517975 | 13 | 0.1538 | 13 | 0.1538 | 8 | 0.375 | 8 | 0.375 | 42 | 0.2381 | 9 | 0.2222 | 15 | 0.2667 | 13 | 0.3077 | 11 | 0.1818 | 48 | 0.25 |
| 12_8517987 | 17 | 0.1176 | 14 | 0.2857 | 12 | 0.0833 | 10 | 0 | 53 | 0.1321 | 9 | 0 | 15 | 0.2 | 16 | 0.0625 | 12 | 0.0833 | 52 | 0.0962 |
| 12_8518027 | 20 | 0.05 | 15 | 0.0667 | 11 | 0 | 11 | 0.1818 | 57 | 0.0702 | 11 | 0.0909 | 13 | 0.2308 | 17 | 0.0588 | 11 | 0.0909 | 52 | 0.1154 |
| 12_8518034 | 19 | 0.3684 | 14 | 0.3571 | 10 | 0.4 | 11 | 0.4545 | 54 | 0.3889 | 10 | 0.1 | 15 | 0.4 | 18 | 0.1111 | 12 | 0.0833 | 55 | 0.1818 |
| 12_8518082 | 18 | 0.1111 | 18 | 0.2778 | 15 | 0 | 13 | 0.0769 | 64 | 0.125 | 10 | 0.4 | 19 | 0.3684 | 22 | 0.2273 | 14 | 0.3571 | 65 | 0.3231 |
| 12_8518239 | 1 | 0 | 1 | 1 | NA | NA | NA | NA | 2 | 0.5 | NA | NA | 1 | 0 | NA | NA | 2 | 0.5 | 3 | 0.3333 |
| 12_8518256 | 1 | 0 | 1 | 0 | 5 | 0 | 7 | 0.1429 | 14 | 0.0714 | 3 | 0 | 4 | 0.25 | 1 | 0 | 4 | 0.25 | 12 | 0.1667 |
| 12_8518273 | 1 | 0 | 1 | 1 | 5 | 0.8 | 6 | 0.5 | 13 | 0.6154 | 3 | 1 | 3 | 0.6667 | 1 | 1 | 3 | 1 | 10 | 0.9 |
| 12_8518274 | 1 | 0 | 1 | 0 | 5 | 0 | 6 | 0.3333 | 13 | 0.1538 | 3 | 0 | 3 | 0 | 1 | 0 | 4 | 0.25 | 11 | 0.0909 |
| 12_8518277 | NA | NA | 1 | 0 | 5 | 0.6 | 5 | 0.6 | 11 | 0.5455 | 3 | 1 | 4 | 1 | 1 | 1 | 3 | 0.6667 | 11 | 0.9091 |
| 12_8518282 | 1 | 0 | 1 | 0 | 4 | 0.25 | 5 | 0.2 | 11 | 0.1818 | 3 | 0.3333 | 3 | 0.3333 | 1 | 0 | 4 | 0 | 11 | 0.1818 |
| 12_8518290 | NA | NA | 1 | 0 | 3 | 0.6667 | 4 | 0.5 | 8 | 0.5 | 3 | 0.6667 | 3 | 0.6667 | 1 | 1 | 4 | 0.25 | 11 | 0.5455 |
| 12_8518291 | NA | NA | 1 | 0 | 4 | 0.5 | 5 | 0.4 | 10 | 0.4 | 3 | 1 | 3 | 0.6667 | NA | NA | 4 | 0.25 | 10 | 0.6 |
| 12_8518300 | NA | NA | 1 | 0 | 4 | 0.25 | 5 | 0.2 | 10 | 0.2 | 3 | 0.3333 | 4 | 0.5 | NA | NA | 3 | 0.3333 | 10 | 0.4 |
| 12_8518306 | NA | NA | NA | NA | 4 | 0.5 | 4 | 1 | 8 | 0.75 | 3 | 0.6667 | 3 | 1 | NA | NA | 4 | 1 | 10 | 0.9 |
| 12_8518310 | NA | NA | 1 | 1 | 3 | 0 | 4 | 0.75 | 8 | 0.5 | 2 | 0 | 4 | 0.25 | NA | NA | 2 | 1 | 8 | 0.375 |
| 12_8518313 | NA | NA | 1 | 1 | 2 | 0.5 | 3 | 1 | 6 | 0.8333 | 2 | 0.5 | 3 | 0.6667 | NA | NA | 3 | 1 | 8 | 0.75 |
| 12_8518626 | NA | NA | 1 | 0 | 3 | 0.3333 | 5 | 0.4 | 9 | 0.3333 | 1 | 1 | 3 | 0.3333 | NA | NA | 1 | 0 | 5 | 0.4 |
| 12_8521346 | NA | NA | NA | NA | 2 | 1 | 1 | 1 | 3 | 1 | 2 | 1 | NA | NA | NA | NA | NA | NA | 2 | 1 |
| 12_8521356 | NA | NA | NA | NA | 2 | 1 | 1 | 1 | 3 | 1 | 1 | 1 | NA | NA | NA | NA | NA | NA | 1 | 1 |
| 12_8521372 | NA | NA | NA | NA | 2 | 1 | NA | NA | 2 | 1 | 1 | 1 | NA | NA | NA | NA | NA | NA | 1 | 1 |
| 12_8521373 | NA | NA | NA | NA | 2 | 0.5 | 1 | 1 | 3 | 0.6667 | 1 | 1 | NA | NA | NA | NA | NA | NA | 1 | 1 |
| 12_8526643 | 5 | 0.2 | 6 | 0 | 2 | 0 | 3 | 0 | 16 | 0.0625 | 6 | 0 | 3 | 0 | 3 | 0 | 6 | 0.1667 | 18 | 0.0556 |
| 12_8527306 | 6 | 0.8333 | 11 | 0.6364 | 4 | 0.5 | 4 | 0.5 | 25 | 0.64 | 9 | 0.6667 | 9 | 0.8889 | 6 | 0.8333 | 8 | 1 | 32 | 0.8438 |
| 12_8527315 | 4 | 0.25 | 8 | 0.25 | 4 | 0.25 | 4 | 0.25 | 20 | 0.25 | 11 | 0.0909 | 9 | 0.1111 | 3 | 0.3333 | 6 | 0.1667 | 29 | 0.1379 |
| 12_8531301 | 1 | 1 | NA | NA | NA | NA | 1 | 1 | 2 | 1 | NA | NA | NA | NA | 4 | 0.5 | 1 | 1 | 5 | 0.6 |
| 12_8531341 | NA | NA | NA | NA | NA | NA | 1 | 1 | 1 | 1 | NA | NA | NA | NA | 1 | 1 | 1 | 0 | 2 | 0.5 |
| 12_8531412 | 1 | 1 | NA | NA | NA | NA | 1 | 0 | 2 | 0.5 | NA | NA | NA | NA | 2 | 0.5 | 1 | 1 | 3 | 0.6667 |
| 12_8531437 | 1 | 1 | NA | NA | NA | NA | NA | NA | 1 | 1 | NA | NA | NA | NA | 2 | 1 | 1 | 1 | 3 | 1 |
| 12_8531487 | NA | NA | NA | NA | NA | NA | 1 | 1 | 1 | 1 | NA | NA | NA | NA | 2 | 0.5 | 1 | 0 | 3 | 0.3333 |
| 12_8531931 | 6 | 0 | 10 | 0.3 | 9 | 0 | 4 | 0 | 29 | 0.1034 | 3 | 0 | 3 | 0.3333 | 4 | 0.25 | 4 | 0.25 | 14 | 0.2143 |
| 12_8531940 | 5 | 0 | 9 | 0.3333 | 9 | 0.2222 | 3 | 0.3333 | 26 | 0.2308 | 3 | 0.3333 | 3 | 0.3333 | 5 | 0.2 | 4 | 0.25 | 15 | 0.2667 |
| 12_8533671 | 1 | 1 | NA | NA | NA | NA | NA | NA | 1 | 1 | 1 | 1 | NA | NA | NA | NA | NA | NA | 1 | 1 |
| 12_8533684 | 1 | 1 | NA | NA | NA | NA | NA | NA | 1 | 1 | 1 | 1 | NA | NA | NA | NA | NA | NA | 1 | 1 |
| 12_8534364 | 1 | 1 | NA | NA | NA | NA | NA | NA | 1 | 1 | 1 | 1 | NA | NA | NA | NA | 1 | 1 | 2 | 1 |
| 12_8534369 | 1 | 1 | NA | NA | NA | NA | NA | NA | 1 | 1 | 1 | 1 | NA | NA | NA | NA | 1 | 1 | 2 | 1 |
| 12_8534397 | 1 | 1 | NA | NA | NA | NA | NA | NA | 1 | 1 | 1 | 1 | NA | NA | NA | NA | 1 | 1 | 2 | 1 |
| 12_8534665 | 5 | 0 | 7 | 0 | 8 | 0.125 | 5 | 0.2 | 25 | 0.08 | 10 | 0 | 5 | 0 | 6 | 0 | 10 | 0.2 | 31 | 0.0645 |
| 12_8534746 | 1 | 0 | 3 | 0 | 6 | 0.5 | 4 | 0.5 | 14 | 0.3571 | 5 | 0 | 1 | 0 | 4 | 0.25 | 5 | 0.2 | 15 | 0.1333 |
| 12_8534791 | 3 | 1 | 5 | 0.6 | 5 | 0.8 | 4 | 1 | 17 | 0.8235 | 3 | 1 | 4 | 1 | 4 | 0.5 | 5 | 1 | 16 | 0.875 |
| 12_8534835 | 2 | 0.5 | 5 | 0 | 5 | 0.2 | 4 | 0.25 | 16 | 0.1875 | 3 | 0 | 5 | 0.2 | 3 | 0 | 3 | 0 | 14 | 0.0714 |
| 12_8534853 | 3 | 1 | 4 | 1 | 4 | 1 | 4 | 1 | 15 | 1 | 2 | 1 | 3 | 0.6667 | NA | NA | 1 | 0 | 6 | 0.6667 |

| | | | | | | | | | | | | | | | | | | | | |
|---|---|---|---|---|---|---|---|---|---|---|---|---|---|---|---|---|---|---|---|---|
| 12_8534854 | 3 | 0.3333 | 5 | 0.4 | 5 | 0.2 | 4 | 0.5 | 17 | 0.3529 | 3 | 0.6667 | 4 | 0.5 | 1 | 1 | 4 | 0.5 | 12 | 0.5833 |
| 12_8534854 | 3 | 0.3333 | 5 | 0.2 | 5 | 0 | 4 | 0 | 17 | 0.1176 | 3 | 0 | 2 | 0 | 1 | 0 | 3 | 0.3333 | 9 | 0.1111 |
| 12_8534860 | 3 | 0.6667 | 5 | 0.4 | 4 | 0.5 | 3 | 0.6667 | 15 | 0.5333 | 3 | 0.3333 | 3 | 0.3333 | 1 | 0 | 3 | 0.3333 | 10 | 0.3 |
| 12_8535231 | 3 | 0 | 3 | 0.3333 | 5 | 0 | 3 | 0 | 14 | 0.0714 | 3 | 0.3333 | 3 | 0 | 1 | 0 | 3 | 0 | 10 | 0.1 |
| 12_8538818 | 16 | 0 | 14 | 0.1429 | 12 | 0.0833 | 11 | 0.0909 | 53 | 0.0755 | 9 | 0.2222 | 16 | 0.0625 | 11 | 0.1818 | 10 | 0 | 46 | 0.1087 |
| 12_8538865 | 6 | 0.1667 | 6 | 0 | 6 | 0.1667 | 5 | 0.2 | 23 | 0.1304 | 6 | 0 | 8 | 0 | 5 | 0.6 | 7 | 0 | 26 | 0.1154 |
| 12_8538973 | 11 | 0.0909 | 12 | 0.1667 | 9 | 0.5556 | 9 | 0.4444 | 41 | 0.2927 | 9 | 0.1111 | 13 | 0.2308 | 6 | 0.1667 | 10 | 0.2 | 38 | 0.1842 |
| 12_8538979 | 17 | 0.0588 | 14 | 0.0714 | 10 | 0.1 | 10 | 0.1 | 51 | 0.0784 | 9 | 0 | 14 | 0.0714 | 7 | 0.1429 | 10 | 0.1 | 40 | 0.075 |
| 12_8539000 | 13 | 1 | 13 | 0.9231 | 9 | 1 | 11 | 1 | 46 | 0.9783 | 8 | 0.875 | 14 | 0.9286 | 7 | 1 | 12 | 1 | 41 | 0.9512 |
| 12_8539002 | 13 | 1 | 12 | 1 | 8 | 1 | 11 | 0.8182 | 44 | 0.9545 | 10 | 1 | 13 | 1 | 6 | 1 | 11 | 1 | 40 | 1 |
| 12_8539008 | 10 | 1 | 10 | 1 | 7 | 0.8571 | 9 | 0.8889 | 36 | 0.9444 | 10 | 0.8 | 13 | 1 | 6 | 1 | 11 | 0.8182 | 40 | 0.9 |
| 12_8539015 | 11 | 0.4545 | 11 | 0.3636 | 5 | 0.6 | 4 | 0.25 | 31 | 0.4194 | 7 | 0.4286 | 10 | 0.5 | 2 | 0.5 | 10 | 0.3 | 29 | 0.4138 |
| 12_8539026 | 10 | 0.9 | 10 | 0.5 | 4 | 1 | 7 | 0.5714 | 31 | 0.7097 | 7 | 0.7143 | 8 | 0.875 | 5 | 0.4 | 6 | 0.3333 | 26 | 0.6154 |
| 12_8552275 | 3 | 0.3333 | 5 | 0.2 | 3 | 0 | NA | NA | 11 | 0.1818 | 3 | 0 | 2 | 0 | 3 | 0.3333 | 2 | 0 | 10 | 0.1 |
| 12_8552290 | 1 | 1 | 6 | 0.1667 | 3 | 0.3333 | NA | NA | 10 | 0.3 | 3 | 0 | 2 | 0 | 3 | 1 | 2 | 0 | 10 | 0.3 |
| 12_8552294 | 2 | 0.5 | 6 | 0.1667 | 2 | 0 | NA | NA | 10 | 0.2 | 4 | 0.25 | 2 | 0 | 3 | 0 | 1 | 0 | 10 | 0.1 |
| 12_8552296 | 2 | 0.5 | 6 | 0.8333 | 3 | 1 | NA | NA | 11 | 0.8182 | 3 | 0.3333 | 1 | 1 | 3 | 1 | 1 | 0 | 8 | 0.625 |
| 12_8552299 | 2 | 1 | 6 | 0.6667 | 3 | 0.3333 | NA | NA | 11 | 0.6364 | 4 | 0.75 | 2 | 0.5 | 3 | 0.6667 | 2 | 0.5 | 11 | 0.6364 |
| 12_8552313 | 1 | 0 | 8 | 0.5 | 3 | 0 | NA | NA | 12 | 0.3333 | 1 | 1 | 3 | 0 | 2 | 0.5 | 1 | 1 | 7 | 0.4286 |
| 12_8552334 | 3 | 0.3333 | 8 | 0.25 | 4 | 0.5 | NA | NA | 15 | 0.3333 | 4 | 0.5 | 3 | 0 | 3 | 1 | 3 | 1 | 13 | 0.6154 |
| 12_8552338 | 3 | 1 | 9 | 1 | 4 | 1 | NA | NA | 16 | 1 | 3 | 1 | 3 | 1 | 3 | 1 | 3 | 1 | 12 | 1 |
| 12_8552372 | 3 | 0.3333 | 10 | 0.2 | 4 | 0.25 | NA | NA | 17 | 0.2353 | 4 | 0 | 3 | 0.3333 | 4 | 0 | 3 | 0 | 14 | 0.0714 |
| 12_8557207 | NA | NA | NA | NA | 1 | 1 | NA | NA | 1 | 1 | NA | NA | 1 | 1 | NA | NA | 1 | 0 | 2 | 0.5 |
| 12_8564451 | NA | NA | 2 | 1 | 3 | 1 | NA | NA | 5 | 1 | NA | NA | NA | NA | NA | NA | 2 | 1 | 2 | 1 |
| 12_8564462 | 1 | 1 | 1 | 1 | 3 | 0.6667 | NA | NA | 5 | 0.8 | NA | NA | NA | NA | NA | NA | 2 | 1 | 2 | 1 |
| 12_8564462 | 1 | 1 | 2 | 0.5 | 3 | 0.6667 | NA | NA | 6 | 0.6667 | NA | NA | NA | NA | NA | NA | 1 | 1 | 1 | 1 |
| 12_8564469 | 1 | 1 | 1 | 1 | 3 | 1 | NA | NA | 5 | 1 | NA | NA | NA | NA | NA | NA | 2 | 1 | 2 | 1 |
| 12_8564474 | 1 | 0 | 1 | 1 | 3 | 0.6667 | NA | NA | 5 | 0.6 | NA | NA | NA | NA | NA | NA | 2 | 1 | 2 | 1 |
| 12_8564505 | 1 | 0 | 1 | 0 | 3 | 0.3333 | NA | NA | 5 | 0.2 | NA | NA | NA | NA | NA | NA | 2 | 0.5 | 2 | 0.5 |
| 12_8564519 | 1 | 1 | 1 | 1 | 3 | 1 | NA | NA | 5 | 1 | NA | NA | NA | NA | NA | NA | 2 | 0.5 | 2 | 0.5 |
| 12_8564527 | NA | NA | 1 | 1 | 1 | 1 | NA | NA | 2 | 1 | NA | NA | NA | NA | NA | NA | 1 | 1 | 1 | 1 |
| 12_8564532 | NA | NA | 1 | 1 | 2 | 0.5 | NA | NA | 3 | 0.6667 | NA | NA | NA | NA | NA | NA | 1 | 1 | 1 | 1 |
| 12_8564584 | NA | NA | NA | NA | 2 | 0.5 | NA | NA | 2 | 0.5 | NA | NA | NA | NA | NA | NA | 2 | 0.5 | 2 | 0.5 |
| 12_8565501 | 5 | 0 | 12 | 0.1667 | 2 | 0 | 2 | 0 | 21 | 0.0952 | 5 | 0.2 | 1 | 0 | 3 | 0.3333 | 6 | 0 | 15 | 0.1333 |
| 12_8568573 | 3 | 0.3333 | 12 | 0.75 | 4 | 0.25 | 1 | 0 | 20 | 0.55 | 7 | 0.4286 | 2 | 1 | 4 | 0 | 4 | 0.25 | 17 | 0.3529 |
| 12_8568583 | 3 | 1 | 7 | 0.8571 | 2 | 1 | 3 | 1 | 15 | 0.9333 | 3 | 0.6667 | 2 | 1 | 2 | 1 | 5 | 0.8 | 12 | 0.8333 |
| 12_8568588 | 4 | 0.25 | 13 | 0.0769 | 5 | 0.2 | 2 | 0 | 24 | 0.125 | 5 | 0 | 1 | 0 | 3 | 0 | 6 | 0.3333 | 15 | 0.1333 |
| 12_8568592 | 4 | 0.75 | 16 | 1 | 5 | 1 | 3 | 1 | 28 | 0.9643 | 7 | 0.5714 | 3 | 1 | 5 | 0.8 | 6 | 0.6667 | 21 | 0.7143 |
| 12_8568593 | 5 | 0.4 | 16 | 0.375 | 6 | 0.1667 | 4 | 0.5 | 31 | 0.3548 | 7 | 0.1429 | 3 | 0.3333 | 5 | 0.2 | 8 | 0.125 | 23 | 0.1739 |
| 12_8568608 | 6 | 0.3333 | 15 | 0.5333 | 5 | 0.2 | 4 | 0.25 | 30 | 0.4 | 7 | 0.2857 | 3 | 0.3333 | 5 | 0.4 | 7 | 0.2857 | 22 | 0.3182 |
| 12_8568614 | 5 | 0.6 | 12 | 0.8333 | 7 | 0.5714 | 3 | 0.6667 | 27 | 0.7037 | 5 | 0.6 | 2 | 0.5 | 3 | 1 | 6 | 0.6667 | 16 | 0.6875 |
| 12_8568615 | 7 | 1 | 14 | 0.8571 | 8 | 1 | 3 | 1 | 32 | 0.9375 | 6 | 1 | 3 | 1 | 4 | 1 | 8 | 1 | 21 | 1 |
| 12_8568629 | 8 | 1 | 14 | 1 | 7 | 1 | 4 | 1 | 33 | 1 | 6 | 1 | 3 | 1 | 3 | 1 | 8 | 0.875 | 20 | 0.95 |
| 12_8568649 | 10 | 0.5 | 18 | 0.7222 | 8 | 1 | 4 | 1 | 40 | 0.75 | 7 | 0.7143 | 4 | 0.25 | 6 | 0.6667 | 9 | 0.8889 | 26 | 0.6923 |
| 12_8587138 | 2 | 0 | 1 | 0 | 1 | 1 | 2 | 0 | 6 | 0.1667 | 1 | 1 | 1 | 0 | 5 | 0.6 | 1 | 1 | 8 | 0.625 |
| 12_8587155 | 3 | 0 | 1 | 1 | 1 | 1 | 1 | 1 | 6 | 0.5 | 2 | 1 | 1 | 0 | 4 | 1 | 1 | 1 | 8 | 0.875 |
| 12_8596586 | NA | NA | 2 | 0.5 | 1 | 0 | NA | NA | 3 | 0.3333 | 2 | 0 | NA | NA | 2 | 0.5 | 1 | 1 | 5 | 0.4 |
| 12_8610364 | 3 | 0.6667 | 2 | 0.5 | 1 | 0 | 5 | 0.4 | 11 | 0.4545 | NA | NA | 7 | 0.4286 | 3 | 0.3333 | 2 | 0.5 | 12 | 0.4167 |
| 12_8610404 | 3 | 1 | 2 | 1 | 1 | 1 | 5 | 0.6 | 11 | 0.8182 | NA | NA | 4 | 0.25 | 3 | 0.3333 | 2 | 0.5 | 9 | 0.3333 |
| 12_8610427 | 3 | 0.6667 | 2 | 0.5 | 1 | 1 | 4 | 0.25 | 10 | 0.5 | NA | NA | 4 | 0.5 | 3 | 0.3333 | 2 | 0.5 | 9 | 0.4444 |
| 12_8610440 | 3 | 0.3333 | 1 | 1 | NA | NA | 5 | 0.6 | 9 | 0.5556 | NA | NA | 2 | 1 | 2 | 1 | 2 | 0.5 | 6 | 0.8333 |
| 12_8610451 | 3 | 0.3333 | 2 | 0 | 1 | 0 | 5 | 0.4 | 11 | 0.2727 | NA | NA | 4 | 0.25 | 4 | 0 | 2 | 0 | 10 | 0.1 |
| 12_8610464 | 3 | 0.3333 | NA | NA | 1 | 0 | 4 | 0 | 8 | 0.125 | NA | NA | 4 | 0 | 4 | 0 | 2 | 0.5 | 10 | 0.1 |
| 12_8610470 | 4 | 0.25 | 1 | 0 | 1 | 0 | 3 | 0 | 9 | 0.1111 | NA | NA | 3 | 0 | 3 | 0.6667 | 2 | 0 | 8 | 0.25 |
| 12_8612436 | 2 | 0 | 2 | 0 | 1 | 1 | 4 | 0.5 | 9 | 0.3333 | NA | NA | 2 | 1 | 2 | 0.5 | 1 | 0 | 5 | 0.6 |
| 12_8612461 | 2 | 0 | NA | NA | 1 | 1 | 3 | 0.3333 | 6 | 0.3333 | NA | NA | 2 | 0.5 | 2 | 1 | 1 | 1 | 5 | 0.8 |
| 12_8612466 | 2 | 1 | 2 | 0 | NA | NA | 2 | 1 | 6 | 0.6667 | NA | NA | 2 | 0 | 2 | 0.5 | 1 | 1 | 5 | 0.4 |
| 12_8612493 | 2 | 0.5 | 1 | 1 | NA | NA | 2 | 0.5 | 5 | 0.6 | NA | NA | 1 | 1 | 2 | 1 | 1 | 1 | 4 | 1 |
| 12_8620915 | 3 | 0 | 1 | 1 | 4 | 0 | 3 | 0.6667 | 11 | 0.2727 | 1 | 0 | 3 | 0 | 4 | 0.25 | 3 | 0.3333 | 11 | 0.1818 |
| 12_8620919 | NA | NA | 1 | 0 | 3 | 0.3333 | 3 | 0 | 7 | 0.1429 | 2 | 0 | 3 | 0 | 4 | 0.25 | 3 | 0 | 12 | 0.0833 |
| 12_8633581 | NA | NA | 1 | 1 | NA | NA | NA | NA | 1 | 1 | 3 | 0.6667 | 2 | 0.5 | 2 | 1 | 2 | 1 | 9 | 0.7778 |
| 12_8633817 | 3 | 0.3333 | 1 | 1 | 1 | 0 | 1 | 0 | 6 | 0.3333 | 5 | 0.4 | 1 | 0 | 2 | 0.5 | 6 | 0.5 | 14 | 0.4286 |
| 12_8674895 | NA | NA | 1 | 1 | NA | NA | NA | NA | 1 | 1 | 1 | 1 | 1 | 0 | NA | NA | 1 | 0 | 3 | 0.3333 |
| 12_8692820 | 6 | 0 | 9 | 0.1111 | 11 | 0.2727 | 8 | 0.125 | 34 | 0.1471 | 8 | 0.125 | 9 | 0.2222 | 8 | 0 | 13 | 0 | 38 | 0.0789 |
| 12_8692835 | 3 | 0.3333 | 8 | 0.25 | 6 | 0 | 5 | 0 | 22 | 0.1364 | 2 | 0 | 10 | 0.3 | 5 | 0.2 | 12 | 0.0833 | 29 | 0.1724 |

| | | | | | | | | | | | | | | | | | | | | |
|---|---|---|---|---|---|---|---|---|---|---|---|---|---|---|---|---|---|---|---|---|
| 12_869283 | 2 | 0 | NA | NA | 1 | 1 | 1 | 0 | 4 | 0.25 | 2 | 1 | 1 | 0 | NA | NA | 2 | 0 | 5 | 0.4 |
| 12_869285 | 5 | 0.4 | 8 | 0.5 | 8 | 0.25 | 4 | 0.25 | 25 | 0.36 | 6 | 0.5 | 8 | 0.625 | 6 | 0.8333 | 12 | 0.3333 | 32 | 0.5312 |
| 12_869286 | 5 | 0 | 5 | 0.2 | 8 | 0.25 | 5 | 0.2 | 23 | 0.1739 | 5 | 0.2 | 9 | 0 | 4 | 0.25 | 7 | 0 | 25 | 0.08 |
| 12_869286 | 3 | 0 | 8 | 0.125 | 5 | 0.2 | 5 | 0 | 21 | 0.0952 | 4 | 0 | 4 | 0.5 | 3 | 0.3333 | 9 | 0.2222 | 20 | 0.25 |
| 12_869286 | 5 | 1 | 6 | 1 | 9 | 1 | 5 | 1 | 25 | 1 | 5 | 1 | 9 | 1 | 4 | 1 | 11 | 1 | 29 | 1 |
| 12_869287 | 3 | 1 | 6 | 1 | 6 | 1 | 2 | 1 | 17 | 1 | 6 | 1 | 9 | 0.7778 | 4 | 1 | 9 | 1 | 28 | 0.9286 |
| 12_869288 | 3 | 0.6667 | 5 | 0.6 | 5 | 0.4 | 2 | 0.5 | 15 | 0.5333 | 3 | 0.6667 | 7 | 0.7143 | 3 | 0 | 9 | 0.6667 | 22 | 0.5909 |
| 12_869289 | 3 | 1 | 4 | 0.25 | 4 | 0.25 | 3 | 0.6667 | 14 | 0.5 | 3 | 0.6667 | 3 | 0 | 3 | 0.3333 | 5 | 0.4 | 14 | 0.3571 |
| 12_869289 | 4 | 1 | 6 | 1 | 7 | 1 | 4 | 0.75 | 21 | 0.9524 | 4 | 1 | 7 | 1 | 4 | 1 | 9 | 0.7778 | 24 | 0.9167 |
| 12_869289 | 4 | 0.75 | 6 | 0.8333 | 5 | 1 | 4 | 0.75 | 19 | 0.8421 | 4 | 1 | 7 | 0.7143 | 4 | 1 | 8 | 0.875 | 23 | 0.8696 |
| 12_869290 | 4 | 0.25 | 6 | 0.1667 | 6 | 0.1667 | 4 | 0.5 | 20 | 0.25 | 4 | 0.25 | 7 | 0.1429 | 4 | 0 | 5 | 0.2 | 20 | 0.15 |
| 12_869291 | 3 | 1 | 3 | 0.3333 | 6 | 0.5 | 3 | 0.6667 | 15 | 0.6 | 4 | 0.5 | 6 | 0.5 | 3 | 1 | 6 | 0.8333 | 19 | 0.6842 |
| 12_869292 | 3 | 0.3333 | 6 | 0.5 | 6 | 0.5 | 4 | 0.25 | 19 | 0.4211 | 4 | 0 | 6 | 0.8333 | 3 | 0.6667 | 5 | 0.4 | 18 | 0.5 |
| 12_869293 | 1 | 1 | 3 | 0.6667 | 5 | 0.8 | 4 | 0.75 | 13 | 0.7692 | 2 | 1 | 6 | 0.6667 | 3 | 1 | 5 | 0.2 | 16 | 0.625 |
| 12_869293 | 3 | 1 | 6 | 1 | 5 | 1 | 4 | 1 | 18 | 1 | 2 | 1 | 6 | 1 | 3 | 1 | 6 | 1 | 17 | 1 |
| 12_869294 | 3 | 1 | 5 | 1 | 5 | 1 | 4 | 1 | 17 | 1 | 3 | 1 | 3 | 1 | 4 | 1 | 5 | 1 | 15 | 1 |
| 12_869295 | 2 | 1 | 4 | 0.5 | 3 | 0.6667 | 2 | 1 | 11 | 0.7273 | 3 | 0.6667 | 2 | 1 | 4 | 0.5 | 4 | 0 | 13 | 0.4615 |
| 12_869299 | 2 | 0 | 1 | 0 | 4 | 0.25 | 4 | 0.5 | 11 | 0.2727 | 3 | 0 | 3 | 0 | 2 | 0 | 5 | 0.2 | 13 | 0.0769 |
| 12_869305 | 1 | 1 | 2 | 0.5 | 1 | 0 | 2 | 1 | 6 | 0.6667 | 3 | 1 | 2 | 1 | 1 | 0 | 5 | 0.4 | 11 | 0.6364 |
| 12_869307 | NA | NA | 1 | 1 | 1 | 0 | 4 | 0.25 | 6 | 0.3333 | 2 | 0 | 2 | 0.5 | 1 | 1 | 5 | 0.4 | 10 | 0.4 |
| 12_869310 | NA | NA | 2 | 0.5 | NA | NA | 3 | 0 | 5 | 0.2 | 1 | 1 | 2 | 1 | 1 | 0 | 2 | 0 | 6 | 0.5 |
| 12_869311 | NA | NA | 1 | 1 | NA | NA | 2 | 0.5 | 3 | 0.6667 | 2 | 1 | 1 | 1 | 1 | 1 | 2 | 1 | 6 | 1 |
| 12_870085 | 5 | 0 | 4 | 0 | 5 | 0.2 | 6 | 0.1667 | 20 | 0.1 | 3 | 0 | 3 | 0.3333 | 5 | 0.6 | 4 | 0.25 | 15 | 0.3333 |
| 12_870086 | 8 | 0.875 | 5 | 0.4 | 6 | 0.8333 | 8 | 0.625 | 27 | 0.7037 | 3 | 0 | 5 | 0.6 | 7 | 0.8571 | 4 | 1 | 19 | 0.6842 |
| 12_870087 | 8 | 0.25 | 6 | 0.8333 | 4 | 0.75 | 8 | 0.75 | 26 | 0.6154 | 3 | 0.6667 | 5 | 0.6 | 6 | 0.8333 | 5 | 0.8 | 19 | 0.7368 |
| 12_870087 | 9 | 0 | 6 | 0.5 | 5 | 0 | 10 | 0.1 | 30 | 0.1333 | 4 | 0 | 3 | 0 | 6 | 0.1667 | 4 | 0 | 17 | 0.0588 |
| 12_870087 | 8 | 0.25 | 6 | 0 | 5 | 0.2 | 8 | 0.25 | 27 | 0.1852 | 4 | 0.5 | 5 | 0.4 | 6 | 0.5 | 5 | 0.4 | 20 | 0.45 |
| 12_870088 | 6 | 0.3333 | 4 | 0.5 | 5 | 0.6 | 8 | 0.75 | 23 | 0.5652 | 4 | 0.5 | 4 | 0.25 | 6 | 0.6667 | 4 | 0.5 | 18 | 0.5 |
| 12_870088 | 8 | 0.25 | 6 | 0.5 | 6 | 0.1667 | 10 | 0.1 | 30 | 0.2333 | 3 | 0.3333 | 4 | 0.5 | 6 | 0.1667 | 5 | 0 | 18 | 0.2222 |
| 12_870088 | 10 | 0.2 | 7 | 0.1429 | 5 | 0 | 9 | 0.2222 | 31 | 0.1613 | 4 | 0 | 5 | 0.2 | 7 | 0.1429 | 4 | 0.25 | 20 | 0.15 |
| 12_870089 | 7 | 0.4286 | 6 | 0.3333 | 5 | 0.4 | 11 | 0.4545 | 29 | 0.4138 | 4 | 0.75 | 5 | 0.6 | 5 | 0.2 | 4 | 0.25 | 18 | 0.4444 |
| 12_870089 | 9 | 0.7778 | 7 | 1 | 5 | 1 | 10 | 0.9 | 31 | 0.9032 | 3 | 1 | 4 | 1 | 6 | 1 | 5 | 1 | 18 | 1 |
| 12_870089 | 7 | 1 | 6 | 0.6667 | 6 | 1 | 8 | 1 | 27 | 0.9259 | 4 | 1 | 5 | 1 | 5 | 1 | 6 | 0.8333 | 20 | 0.95 |
| 12_870090 | 9 | 1 | 7 | 0.8571 | 6 | 0.8333 | 9 | 0.6667 | 31 | 0.8387 | 3 | 0.6667 | 4 | 1 | 7 | 0.8571 | 5 | 0.8 | 19 | 0.8421 |
| 12_870091 | 11 | 0.0909 | 7 | 0.1429 | 5 | 0.2 | 12 | 0 | 35 | 0.0857 | 5 | 0 | 5 | 0.2 | 7 | 0 | 6 | 0.3333 | 23 | 0.1304 |
| 12_870091 | 10 | 0.2 | 6 | 0.6667 | 6 | 0.3333 | 8 | 0.375 | 30 | 0.3667 | 4 | 0 | 4 | 0.5 | 5 | 0.2 | 6 | 0.1667 | 19 | 0.2105 |
| 12_870092 | 11 | 0.3636 | 5 | 0 | 5 | 0.6 | 11 | 0 | 32 | 0.2188 | 4 | 0.25 | 5 | 0.4 | 6 | 0.1667 | 6 | 0.1667 | 21 | 0.2381 |
| 12_870094 | 10 | 0.4 | 8 | 0.375 | 5 | 0.6 | 11 | 0.0909 | 34 | 0.3235 | 7 | 0 | 5 | 0.4 | 8 | 0.125 | 6 | 0.1667 | 26 | 0.1538 |
| 12_870109 | 12 | 0 | 10 | 0.1 | 10 | 0.2 | 13 | 0.0769 | 45 | 0.0889 | 10 | 0.2 | 10 | 0 | 10 | 0 | 8 | 0.25 | 38 | 0.1053 |
| 12_870110 | 10 | 0 | 10 | 0.1 | 11 | 0.2727 | 17 | 0 | 48 | 0.0833 | 6 | 0.5 | 9 | 0.1111 | 9 | 0.3333 | 9 | 0.1111 | 33 | 0.2424 |
| 12_870112 | 12 | 0.25 | 8 | 0.75 | 10 | 0.3 | 16 | 0.625 | 46 | 0.4783 | 6 | 0.6667 | 9 | 0.2222 | 7 | 0.5714 | 9 | 0.4444 | 31 | 0.4516 |
| 12_870112 | 13 | 0.0769 | 9 | 0 | 10 | 0.2 | 16 | 0 | 48 | 0.0625 | 8 | 0.25 | 6 | 0.1667 | 10 | 0.1 | 8 | 0.125 | 32 | 0.1562 |
| 12_870112 | 12 | 0.0833 | 10 | 0.2 | 11 | 0.0909 | 16 | 0.0625 | 49 | 0.102 | 8 | 0 | 10 | 0.1 | 10 | 0.1 | 11 | 0.1818 | 39 | 0.1026 |
| 12_870116 | 14 | 0.0714 | 12 | 0.3333 | 10 | 0.1 | 17 | 0.1765 | 53 | 0.1698 | 7 | 0 | 10 | 0.3 | 12 | 0.1667 | 11 | 0.2727 | 40 | 0.2 |
| 12_870116 | 13 | 0 | 13 | 0.2308 | 10 | 0.1 | 18 | 0.1111 | 54 | 0.1111 | 8 | 0.375 | 11 | 0.1818 | 10 | 0.2 | 13 | 0.1538 | 42 | 0.2143 |
| 12_870117 | 14 | 0.5 | 11 | 0.2727 | 8 | 0.5 | 17 | 0.5882 | 50 | 0.48 | 8 | 0.5 | 8 | 0.5 | 12 | 0.5833 | 12 | 0.5 | 40 | 0.525 |
| 12_870120 | 20 | 0.2 | 15 | 0.3333 | 12 | 0.1667 | 20 | 0.1 | 67 | 0.194 | 13 | 0.3846 | 11 | 0.3636 | 14 | 0.2143 | 12 | 0.4167 | 50 | 0.34 |
| 12_870301 | 1 | 1 | NA | NA | NA | NA | NA | NA | 1 | 1 | NA | NA | 1 | 1 | 2 | 1 | 1 | 1 | 4 | 1 |
| 12_870302 | 1 | 1 | NA | NA | 1 | 1 | NA | NA | 2 | 1 | NA | NA | 1 | 1 | 2 | 1 | 1 | 1 | 4 | 1 |
| 12_870348 | 1 | 1 | NA | NA | NA | NA | NA | NA | 1 | 1 | 1 | 0 | 2 | 1 | NA | NA | NA | NA | 3 | 0.6667 |
| 12_870351 | 2 | 0.5 | NA | NA | 1 | 0 | NA | NA | 3 | 0.3333 | 1 | 0 | 2 | 0.5 | NA | NA | 1 | 0 | 4 | 0.25 |
| 12_871361 | 1 | 0 | 3 | 0.6667 | 2 | 0.5 | NA | NA | 6 | 0.5 | NA | NA | 3 | 0.3333 | 8 | 0.125 | NA | NA | 11 | 0.1818 |
| 12_871362 | 1 | 1 | 2 | 1 | 3 | 1 | NA | NA | 6 | 1 | NA | NA | 3 | 0.6667 | 7 | 0.5714 | 1 | 1 | 11 | 0.6364 |
| 12_871362 | 1 | 1 | 3 | 0.6667 | 3 | 0.6667 | NA | NA | 7 | 0.7143 | NA | NA | 3 | 0.6667 | 9 | 0.7778 | 1 | 1 | 13 | 0.7692 |
| 12_871362 | 1 | 0 | 4 | 0 | 4 | 0.25 | NA | NA | 9 | 0.1111 | 1 | 0 | 4 | 0 | 9 | 0.1111 | 2 | 0 | 16 | 0.0625 |
| 12_871364 | 1 | 0 | 3 | 0.3333 | 4 | 0 | NA | NA | 8 | 0.125 | 1 | 1 | 5 | 0 | 7 | 0 | 2 | 0.5 | 15 | 0.1333 |
| 12_871364 | 1 | 1 | 4 | 1 | 4 | 0.5 | NA | NA | 9 | 0.7778 | 1 | 1 | 5 | 0.4 | 8 | 0.375 | 2 | 1 | 16 | 0.5 |
| 12_871366 | 1 | 0 | 4 | 0.25 | 2 | 0.5 | NA | NA | 7 | 0.2857 | 1 | 0 | 5 | 0.4 | 9 | 0.1111 | 2 | 0 | 17 | 0.1765 |
| 12_871367 | 2 | 0 | 3 | 0.3333 | 3 | 0.3333 | NA | NA | 8 | 0.25 | 1 | 1 | 5 | 0 | 7 | 0.2857 | NA | NA | 13 | 0.2308 |
| 12_871383 | 3 | 0 | 2 | 0 | 1 | 1 | 3 | 0.3333 | 9 | 0.2222 | NA | NA | 4 | 0 | 6 | 0.1667 | 2 | 0.5 | 12 | 0.1667 |
| 12_871385 | 3 | 0 | 1 | 0 | 1 | 1 | 3 | 0 | 8 | 0.125 | NA | NA | 5 | 0 | 6 | 0 | 3 | 0.3333 | 14 | 0.0714 |
| 12_871538 | 5 | 0 | 4 | 0.25 | 1 | 0 | 2 | 0 | 12 | 0.0833 | 3 | 0 | NA | NA | NA | NA | 5 | 0.4 | 8 | 0.25 |
| 12_871552 | 4 | 0 | 3 | 0.3333 | 1 | 1 | NA | NA | 8 | 0.25 | 2 | 0.5 | NA | NA | NA | NA | 3 | 0 | 5 | 0.2 |
| 12_871558 | 4 | 1 | 3 | 1 | 2 | 1 | NA | NA | 9 | 1 | 1 | 1 | NA | NA | NA | NA | 2 | 1 | 3 | 1 |
| 12_871558 | 5 | 0.4 | 3 | 0.3333 | 2 | 0 | NA | NA | 10 | 0.3 | 2 | 0.5 | NA | NA | NA | NA | 2 | 0.5 | 4 | 0.5 |

| | | | | | | | | | | | | | | | | | | | | |
|---|---|---|---|---|---|---|---|---|---|---|---|---|---|---|---|---|---|---|---|---|
| 12_871558 | 3 | 0.6667 | 3 | 0.6667 | 2 | 0 | NA | NA | 8 | 0.5 | 2 | 0.5 | NA | NA | NA | NA | 1 | 1 | 3 | 0.6667 |
| 12_871559 | 3 | 0.3333 | 3 | 0.6667 | 2 | 0 | NA | NA | 8 | 0.375 | 2 | 0.5 | NA | NA | NA | NA | 2 | 0 | 4 | 0.25 |
| 12_871559 | 4 | 1 | 3 | 0.6667 | 2 | 0 | NA | NA | 9 | 0.6667 | 1 | 1 | NA | NA | NA | NA | 2 | 0 | 3 | 0.3333 |
| 12_871559 | 4 | 0.75 | 3 | 1 | 2 | 0 | NA | NA | 9 | 0.6667 | 2 | 1 | NA | NA | NA | NA | 2 | 0.5 | 4 | 0.75 |
| 12_872105 | 2 | 0.5 | 4 | 0.5 | 5 | 0.4 | 4 | 0 | 15 | 0.3333 | 3 | 0 | 3 | 0 | 3 | 0.3333 | 3 | 0.3333 | 12 | 0.1667 |
| 12_872108 | 3 | 0.3333 | 5 | 0.4 | 5 | 0.2 | 4 | 0 | 17 | 0.2353 | 3 | 0.3333 | 3 | 0 | 3 | 0 | 3 | 0.3333 | 12 | 0.1667 |
| 12_872110 | 3 | 0 | 5 | 0.2 | 5 | 0.2 | 4 | 0 | 17 | 0.1176 | 2 | 0 | 3 | 0.6667 | 2 | 1 | 3 | 0 | 10 | 0.4 |
| 12_872114 | 4 | 0.5 | 2 | 1 | 5 | 0.2 | 3 | 0.6667 | 14 | 0.5 | 4 | 0.5 | 2 | 0.5 | 1 | 0 | 2 | 0.5 | 9 | 0.4444 |
| 12_872133 | 3 | 0 | 6 | 0 | 2 | 0 | 6 | 0.1667 | 17 | 0.0588 | 1 | 0 | 2 | 0.5 | 3 | 0 | 11 | 0 | 17 | 0.0588 |
| 12_872159 | 2 | 0.5 | 2 | 0.5 | NA | NA | 5 | 0 | 9 | 0.2222 | NA | NA | 2 | 0 | 3 | 0 | 4 | 0.5 | 9 | 0.2222 |
| 12_872171 | NA | NA | 1 | 1 | NA | NA | 1 | 0 | 2 | 0.5 | NA | NA | 1 | 0 | NA | NA | 2 | 0.5 | 3 | 0.3333 |
| 12_872172 | NA | NA | 1 | 1 | NA | NA | 2 | 0.5 | 3 | 0.6667 | NA | NA | 1 | 1 | NA | NA | 2 | 1 | 3 | 1 |
| 12_872173 | NA | NA | 1 | 1 | NA | NA | 2 | 0 | 3 | 0.3333 | NA | NA | NA | NA | NA | NA | 2 | 1 | 2 | 1 |
| 12_872188 | NA | NA | NA | NA | NA | NA | 1 | 1 | 1 | 1 | NA | NA | 1 | 1 | NA | NA | 1 | 1 | 2 | 1 |
| 12_872486 | NA | NA | NA | NA | NA | NA | 1 | 1 | 1 | 1 | NA | NA | 1 | 1 | NA | NA | NA | NA | 1 | 1 |
| 12_872856 | 6 | 0 | 2 | 0 | 11 | 0.3636 | 6 | 0.1667 | 25 | 0.2 | 7 | 0.1429 | 5 | 0.2 | 11 | 0.1818 | 4 | 0.25 | 27 | 0.1852 |
| 12_872857 | 7 | 0.2857 | 4 | 0.25 | 12 | 0.5833 | 7 | 0.4286 | 30 | 0.4333 | 8 | 0.375 | 6 | 0.3333 | 11 | 0.3636 | 5 | 0.4 | 30 | 0.3667 |
| 12_872860 | 4 | 1 | 3 | 0.6667 | 8 | 0 | 4 | 0.25 | 19 | 0.3684 | 3 | 0 | 4 | 0.5 | 9 | 0.4444 | 4 | 0 | 20 | 0.3 |
| 12_872860 | 6 | 0 | 3 | 0 | 12 | 0.25 | 6 | 0.3333 | 27 | 0.1852 | 8 | 0.125 | 6 | 0 | 12 | 0.0833 | 4 | 0 | 30 | 0.0667 |
| 12_872861 | 6 | 0.8333 | 3 | 0.6667 | 12 | 0.8333 | 4 | 0.75 | 25 | 0.8 | 8 | 0.375 | 5 | 0.4 | 8 | 0.75 | 2 | 1 | 23 | 0.5652 |
| 12_872861 | 6 | 0 | 3 | 0.3333 | 9 | 0.4444 | 4 | 0 | 22 | 0.2273 | 7 | 0 | 5 | 0.2 | 10 | 0.5 | 4 | 0 | 26 | 0.2308 |
| 12_872861 | 5 | 0.6 | 3 | 0 | 10 | 0.3 | 5 | 0.2 | 23 | 0.3043 | 7 | 0.1429 | 6 | 0.1667 | 10 | 0.2 | 4 | 0 | 27 | 0.1481 |
| 12_872862 | 5 | 0.6 | 4 | 0 | 10 | 0.7 | 5 | 0.8 | 24 | 0.5833 | 8 | 0.5 | 6 | 0.1667 | 11 | 0.4545 | 4 | 0.25 | 29 | 0.3793 |
| 12_872862 | 4 | 0.5 | 4 | 0.25 | 11 | 0.6364 | 4 | 0.5 | 23 | 0.5217 | 7 | 0.1429 | 4 | 0 | 10 | 0.4 | 4 | 0.75 | 25 | 0.32 |
| 12_872863 | 4 | 0.75 | 4 | 0.75 | 9 | 0.8889 | 4 | 1 | 21 | 0.8571 | 8 | 1 | 5 | 0.8 | 12 | 0.9167 | 4 | 1 | 29 | 0.931 |
| 12_872864 | 5 | 1 | 4 | 0.75 | 10 | 0.8 | 6 | 0.6667 | 25 | 0.8 | 8 | 0.875 | 5 | 0.8 | 11 | 0.9091 | 4 | 1 | 28 | 0.8929 |
| 12_872864 | 4 | 0.5 | 2 | 1 | 10 | 1 | 5 | 0.6 | 21 | 0.8095 | 7 | 0.7143 | 5 | 0.6 | 8 | 0.25 | 4 | 0.5 | 24 | 0.5 |
| 12_873205 | 21 | 0.0952 | 25 | 0.28 | 17 | 0.2353 | 19 | 0.2632 | 82 | 0.2195 | 16 | 0.125 | 20 | 0 | 20 | 0.1 | 12 | 0.3333 | 68 | 0.1176 |
| 12_873205 | 19 | 0.1579 | 26 | 0.0769 | 18 | 0.1667 | 20 | 0.05 | 83 | 0.1084 | 15 | 0.0667 | 18 | 0.1111 | 21 | 0.0952 | 13 | 0.1538 | 67 | 0.1045 |
| 12_873206 | 22 | 0.2273 | 25 | 0.04 | 14 | 0.1429 | 21 | 0.0952 | 82 | 0.122 | 16 | 0 | 18 | 0.1667 | 20 | 0.2 | 13 | 0.0769 | 67 | 0.1194 |
| 12_873206 | 16 | 0.0625 | 16 | 0.25 | 5 | 0 | 12 | 0.0833 | 49 | 0.1224 | 8 | 0.25 | 15 | 0.0667 | 8 | 0.25 | 6 | 0 | 37 | 0.1351 |
| 12_873535 | 1 | 1 | 1 | 0 | NA | NA | NA | NA | 2 | 0.5 | 2 | 0.5 | 1 | 0 | NA | NA | 1 | 1 | 4 | 0.5 |
| 12_873565 | NA | NA | 3 | 1 | NA | NA | 3 | 0.6667 | 6 | 0.8333 | 1 | 1 | 1 | 1 | 3 | 0.6667 | 3 | 0.6667 | 8 | 0.75 |
| 12_873566 | NA | NA | 3 | 1 | 1 | 1 | 3 | 1 | 7 | 1 | 1 | 1 | 2 | 1 | 3 | 1 | 4 | 0.5 | 10 | 0.8 |
| 12_874756 | NA | NA | 2 | 1 | NA | NA | NA | NA | 2 | 1 | 1 | 1 | 1 | 1 | NA | NA | NA | NA | 2 | 1 |
| 12_874757 | 1 | 0 | 3 | 0.6667 | 1 | 0 | 1 | 0 | 6 | 0.3333 | 1 | 1 | 1 | 1 | NA | NA | 2 | 1 | 4 | 1 |
| 12_874757 | NA | NA | 4 | 0.75 | NA | NA | 1 | 1 | 5 | 0.8 | 1 | 1 | 1 | 0 | NA | NA | 1 | 0 | 3 | 0.3333 |
| 12_874757 | 1 | 0 | 4 | 1 | NA | NA | NA | NA | 5 | 0.8 | 1 | 0 | 1 | 1 | NA | NA | 2 | 1 | 4 | 0.75 |
| 12_874758 | 1 | 1 | 4 | 0.25 | 1 | 0 | 1 | 1 | 7 | 0.4286 | 1 | 1 | 1 | 1 | NA | NA | 2 | 0.5 | 4 | 0.75 |
| 12_874966 | 20 | 0.1 | 14 | 0.0714 | 12 | 0.0833 | 23 | 0.1304 | 69 | 0.1014 | 13 | 0.0769 | 20 | 0.1 | 10 | 0.1 | 17 | 0.1765 | 60 | 0.1167 |
| 12_874970 | 13 | 0.6154 | 8 | 0.75 | 10 | 0.8 | 6 | 0.8333 | 37 | 0.7297 | 11 | 0.7273 | 14 | 0.7857 | 10 | 0.6 | 16 | 0.6875 | 51 | 0.7059 |
| 12_874970 | 12 | 0.25 | 6 | 0.1667 | 8 | 0.125 | 6 | 0.3333 | 32 | 0.2188 | 10 | 0.3 | 13 | 0.1538 | 9 | 0 | 18 | 0.1667 | 50 | 0.16 |
| 12_874971 | 12 | 0.1667 | 10 | 0.3 | 11 | 0.2727 | 7 | 0.1429 | 40 | 0.225 | 10 | 0.3 | 11 | 0.2727 | 8 | 0.5 | 22 | 0.1818 | 51 | 0.2745 |
| 12_874972 | 4 | 0 | 7 | 0.2857 | 11 | 0 | 7 | 0.2857 | 29 | 0.1379 | 7 | 0 | 15 | 0 | 7 | 0.1429 | 17 | 0.1176 | 46 | 0.0652 |
| 12_874972 | 13 | 0.7692 | 10 | 0.7 | 9 | 0.7778 | 8 | 0.75 | 40 | 0.75 | 11 | 0.4545 | 17 | 0.6471 | 8 | 0.5 | 21 | 0.5238 | 57 | 0.5439 |
| 12_874972 | 13 | 0.2308 | 11 | 0.2727 | 11 | 0.2727 | 8 | 0.25 | 43 | 0.2558 | 10 | 0.6 | 17 | 0.1765 | 10 | 0.3 | 22 | 0.3636 | 59 | 0.339 |
| 12_874973 | 12 | 0.0833 | 10 | 0.1 | 11 | 0.0909 | 8 | 0 | 41 | 0.0732 | 9 | 0.3333 | 15 | 0.1333 | 9 | 0.1111 | 20 | 0.05 | 53 | 0.1321 |
| 12_874973 | 11 | 0.0909 | 8 | 0 | 11 | 0.2727 | 8 | 0.25 | 38 | 0.1579 | 8 | 0.375 | 11 | 0.0909 | 10 | 0.1 | 18 | 0.1667 | 47 | 0.1702 |
| 12_883355 | 1 | 1 | NA | NA | NA | NA | NA | NA | 1 | 1 | NA | NA | NA | NA | NA | NA | 1 | 1 | 1 | 1 |
| 12_883356 | 1 | 1 | NA | NA | NA | NA | NA | NA | 1 | 1 | NA | NA | NA | NA | NA | NA | 1 | 1 | 1 | 1 |
| 12_883356 | 1 | 1 | NA | NA | NA | NA | NA | NA | 1 | 1 | NA | NA | NA | NA | NA | NA | 1 | 1 | 1 | 1 |
| 12_883358 | 1 | 1 | NA | NA | NA | NA | NA | NA | 1 | 1 | NA | NA | NA | NA | 1 | 0 | 1 | 1 | 2 | 0.5 |
| 12_883358 | 1 | 1 | NA | NA | NA | NA | NA | NA | 1 | 1 | NA | NA | NA | NA | 1 | 1 | 1 | 0 | 2 | 0.5 |
| 12_884278 | 7 | 0.2857 | 5 | 0 | 4 | 0.25 | 3 | 0.3333 | 19 | 0.2105 | 3 | 0 | 4 | 0.5 | 5 | 0.2 | 6 | 0.1667 | 18 | 0.2222 |
| 12_884278 | 7 | 0.2857 | 5 | 0 | 4 | 0.25 | 4 | 0 | 20 | 0.15 | 5 | 0 | 3 | 0.3333 | 5 | 0.2 | 4 | 0.25 | 17 | 0.1765 |
| 12_884282 | 9 | 0.3333 | 6 | 0.1667 | 5 | 0.2 | 5 | 0.2 | 25 | 0.24 | 4 | 0.25 | 3 | 0 | 4 | 0 | 4 | 0 | 15 | 0.0667 |
| 12_884282 | 6 | 1 | 6 | 0.8333 | 3 | 1 | 5 | 0.8 | 20 | 0.9 | 5 | 0.6 | 4 | 1 | 4 | 0.5 | 5 | 0.8 | 18 | 0.7222 |
| 12_884282 | 8 | 0.125 | 5 | 0 | 1 | 0 | 4 | 0.25 | 18 | 0.1111 | 4 | 0.5 | 4 | 0.5 | 3 | 0 | 4 | 0 | 15 | 0.2667 |
| 12_884283 | 9 | 0.4444 | 7 | 0.1429 | 5 | 0.2 | 6 | 0.5 | 27 | 0.3333 | 5 | 0.4 | 6 | 0.8333 | 5 | 0.2 | 5 | 0.4 | 21 | 0.4762 |
| 12_884283 | 11 | 0.1818 | 8 | 0 | 6 | 0 | 6 | 0 | 31 | 0.0645 | 6 | 0.3333 | 6 | 0 | 5 | 0 | 4 | 0 | 21 | 0.0952 |
| 12_897794 | 2 | 0 | NA | NA | 1 | 0 | 2 | 0.5 | 5 | 0.2 | 2 | 0 | 3 | 0.3333 | NA | NA | 1 | 0 | 6 | 0.1667 |
| 12_897794 | 3 | 1 | NA | NA | 1 | 1 | 2 | 1 | 6 | 1 | 2 | 1 | 3 | 0.3333 | NA | NA | 1 | 1 | 6 | 0.6667 |
| 12_897795 | 1 | 1 | NA | NA | 1 | 0 | 1 | 0 | 3 | 0.3333 | 2 | 0 | 2 | 0 | NA | NA | 1 | 1 | 5 | 0.2 |
| 12_897795 | 3 | 0.6667 | NA | NA | 1 | 0 | 2 | 0.5 | 6 | 0.5 | 2 | 1 | 1 | 0 | NA | NA | 1 | 0 | 4 | 0.5 |
| 12_897795 | 2 | 0 | NA | NA | NA | NA | 1 | 1 | 3 | 0.3333 | 1 | 0 | 3 | 0.3333 | NA | NA | 1 | 1 | 5 | 0.4 |

| | | | | | | | | | | | | | | | | | | | | |
|---|---|---|---|---|---|---|---|---|---|---|---|---|---|---|---|---|---|---|---|---|
| 12_8977976 | 3 | 1 | NA | NA | 1 | 1 | 2 | 0.5 | 6 | 0.8333 | 2 | 1 | 3 | 0.6667 | NA | NA | 2 | 0.5 | 7 | 0.7143 |
| 12_8978006 | 2 | 0.5 | NA | NA | NA | NA | 1 | 0 | 3 | 0.3333 | NA | NA | 1 | 0 | NA | NA | 2 | 1 | 3 | 0.6667 |
| 12_8978010 | 3 | 0.6667 | 1 | 1 | 1 | 1 | 1 | 1 | 6 | 0.8333 | 2 | 0.5 | 3 | 1 | 1 | 1 | 2 | 0.5 | 8 | 0.75 |
| 12_8978013 | 3 | 0.3333 | NA | NA | 1 | 0 | 2 | 0 | 6 | 0.1667 | 1 | 0 | 2 | 0.5 | 1 | 0 | 2 | 0 | 6 | 0.1667 |
| 12_9017140 | 6 | 0 | 2 | 0.5 | 9 | 0.1111 | 8 | 0.125 | 25 | 0.12 | 5 | 0 | 9 | 0 | 8 | 0.125 | 6 | 0.1667 | 28 | 0.0714 |
| 12_9029837 | 3 | 0.3333 | 2 | 0.5 | 3 | 0.3333 | 5 | 0 | 13 | 0.2308 | 2 | 0 | 7 | 0.2857 | 5 | 0 | 5 | 0 | 19 | 0.1053 |
| 12_9029843 | 3 | 0.6667 | 2 | 1 | 5 | 0.6 | 7 | 0.7143 | 17 | 0.7059 | 2 | 1 | 5 | 0.8 | 7 | 0.4286 | 5 | 0.4 | 19 | 0.5789 |
| 12_9029850 | 3 | 0.6667 | 1 | 1 | 4 | 0.75 | 4 | 0.75 | 12 | 0.75 | 2 | 1 | 7 | 1 | 8 | 1 | 5 | 1 | 22 | 1 |
| 12_9029855 | 3 | 0.3333 | 1 | 0 | 5 | 0.2 | 2 | 0 | 11 | 0.1818 | 2 | 0 | 5 | 0 | 7 | 0.1429 | 5 | 0.4 | 19 | 0.1579 |
| 12_9029863 | 5 | 0.8 | 1 | 1 | 11 | 1 | 10 | 1 | 27 | 0.963 | 4 | 1 | 9 | 0.8889 | 9 | 0.8889 | 7 | 0.8571 | 29 | 0.8966 |
| 12_9029863 | 5 | 0.4 | 3 | 0.3333 | 10 | 0 | 11 | 0.1818 | 29 | 0.1724 | 4 | 0 | 9 | 0.1111 | 10 | 0.4 | 7 | 0.1429 | 30 | 0.2 |
| 12_9029865 | 5 | 0 | 1 | 0 | 11 | 0.3636 | 10 | 0.1 | 27 | 0.1852 | 4 | 0.5 | 9 | 0.1111 | 8 | 0.125 | 7 | 0.2857 | 28 | 0.2143 |
| 12_9029866 | 3 | 0.3333 | 1 | 1 | 6 | 0.5 | 9 | 0.7778 | 19 | 0.6316 | 2 | 0.5 | 8 | 0.75 | 8 | 0.75 | 5 | 0.8 | 23 | 0.7391 |
| 12_9029876 | 4 | 0 | 2 | 0 | 9 | 0.2222 | 8 | 0.125 | 23 | 0.1304 | 3 | 0 | 7 | 0.4286 | 7 | 0.2857 | 5 | 0.2 | 22 | 0.2727 |
| 12_9029882 | 4 | 0.5 | 3 | 0.3333 | 10 | 0 | 9 | 0.3333 | 26 | 0.2308 | 3 | 0.3333 | 6 | 0.1667 | 9 | 0.1111 | 6 | 0.1667 | 24 | 0.1667 |
| 12_9029883 | 5 | 0.4 | 5 | 0.8 | 9 | 0.4444 | 9 | 0.7778 | 28 | 0.6071 | 3 | 0.6667 | 7 | 0.7143 | 7 | 0.5714 | 7 | 0.4286 | 24 | 0.5833 |
| 12_9029907 | 8 | 0.25 | 5 | 0.4 | 16 | 0.125 | 12 | 0.3333 | 41 | 0.2439 | 4 | 0 | 12 | 0 | 14 | 0.2143 | 10 | 0 | 40 | 0.075 |
| 12_9029915 | 9 | 0 | 6 | 0.1667 | 14 | 0.1429 | 11 | 0.0909 | 40 | 0.1 | 4 | 0.25 | 14 | 0.1429 | 9 | 0.2222 | 9 | 0.1111 | 36 | 0.1667 |
| 12_9029915 | 9 | 0 | 6 | 0 | 15 | 0.0667 | 12 | 0.1667 | 42 | 0.0714 | 4 | 0 | 13 | 0.0769 | 14 | 0.1429 | 10 | 0.1 | 41 | 0.0976 |
| 12_9029916 | 9 | 1 | 6 | 1 | 16 | 1 | 12 | 1 | 43 | 1 | 5 | 1 | 14 | 0.7143 | 16 | 0.9375 | 11 | 0.9091 | 46 | 0.8696 |
| 12_9029918 | 8 | 0.125 | 6 | 0 | 12 | 0 | 12 | 0.0833 | 38 | 0.0526 | 5 | 0.2 | 13 | 0.1538 | 12 | 0 | 11 | 0.0909 | 41 | 0.0976 |
| 12_9029921 | 8 | 1 | 7 | 1 | 15 | 1 | 12 | 0.9167 | 42 | 0.9762 | 4 | 1 | 13 | 1 | 16 | 1 | 11 | 0.9091 | 44 | 0.9773 |
| 12_9029925 | 7 | 0.2857 | 7 | 0.4286 | 17 | 0.1765 | 12 | 0.5 | 43 | 0.3256 | 5 | 0 | 14 | 0.2857 | 16 | 0.125 | 12 | 0.3333 | 47 | 0.2128 |
| 12_9029926 | 7 | 0.2857 | 3 | 0 | 15 | 0 | 13 | 0.1538 | 38 | 0.1053 | 4 | 0 | 10 | 0.1 | 10 | 0.3 | 11 | 0.2727 | 35 | 0.2 |
| 12_9029933 | 8 | 1 | 7 | 1 | 16 | 1 | 12 | 1 | 43 | 1 | 6 | 1 | 13 | 1 | 16 | 1 | 12 | 1 | 47 | 1 |
| 12_9029947 | 9 | 0 | 8 | 0 | 16 | 0.1875 | 11 | 0.0909 | 44 | 0.0909 | 6 | 0.1667 | 14 | 0 | 15 | 0.0667 | 11 | 0.0909 | 46 | 0.0652 |
| 12_9029950 | 4 | 0.5 | 6 | 0.8333 | 9 | 0.6667 | 9 | 0.5556 | 28 | 0.6429 | 4 | 0.75 | 9 | 0.5556 | 9 | 0.6667 | 10 | 0.2 | 32 | 0.5 |
| 12_9029958 | 9 | 0.1111 | 6 | 0 | 15 | 0.0667 | 12 | 0.25 | 42 | 0.119 | 5 | 0 | 12 | 0.25 | 15 | 0 | 11 | 0 | 43 | 0.0698 |
| 12_9029981 | 13 | 0.3077 | 9 | 0.2222 | 18 | 0.1111 | 13 | 0.3077 | 53 | 0.2264 | 7 | 0.2857 | 14 | 0.4286 | 17 | 0.3529 | 12 | 0.5 | 50 | 0.4 |
| 12_9029986 | 15 | 0.4667 | 9 | 0.2222 | 17 | 0.2941 | 15 | 0.3333 | 56 | 0.3393 | 7 | 0.1429 | 16 | 0.25 | 16 | 0.125 | 11 | 0.1818 | 50 | 0.18 |
| 12_9029988 | 13 | 0.4615 | 6 | 0.3333 | 15 | 0.4 | 6 | 0.6667 | 40 | 0.45 | 6 | 0.3333 | 13 | 0.4615 | 13 | 0.5385 | 8 | 0.25 | 40 | 0.425 |
| 12_9030084 | 13 | 0.0769 | 15 | 0.1333 | 18 | 0.1111 | 14 | 0.2143 | 60 | 0.1333 | 10 | 0 | 17 | 0.1176 | 19 | 0.1053 | 13 | 0.0769 | 59 | 0.0847 |
| 12_9030114 | 16 | 0.25 | 14 | 0.0714 | 25 | 0.04 | 19 | 0.0526 | 74 | 0.0946 | 11 | 0.0909 | 20 | 0.1 | 22 | 0.0909 | 19 | 0 | 72 | 0.0694 |
| 12_9030130 | 19 | 0.5789 | 17 | 0.4118 | 25 | 0.44 | 18 | 0.6111 | 79 | 0.5063 | 11 | 0.6364 | 21 | 0.3333 | 20 | 0.35 | 17 | 0.5294 | 69 | 0.4348 |
| 12_9030142 | 21 | 0.2381 | 21 | 0.2857 | 29 | 0.3103 | 20 | 0.35 | 91 | 0.2967 | 16 | 0.25 | 24 | 0.3333 | 22 | 0.3182 | 19 | 0.2632 | 81 | 0.2963 |
| 12_9030151 | 14 | 0.1429 | 12 | 0.0833 | 17 | 0.2941 | 13 | 0.6154 | 56 | 0.2857 | 10 | 0.3 | 22 | 0.1364 | 18 | 0.3333 | 11 | 0.1818 | 61 | 0.2295 |
| 12_9069152 | 7 | 0.4286 | 13 | 0.5385 | 11 | 0.4545 | 13 | 0.4615 | 44 | 0.4773 | 11 | 0.4545 | 6 | 0.5 | 6 | 0.3333 | 7 | 0.5714 | 30 | 0.4667 |
| 12_9069185 | 5 | 0 | 10 | 0.1 | 6 | 0 | 13 | 0.0769 | 34 | 0.0588 | 8 | 0.125 | 5 | 0 | 8 | 0.25 | 8 | 0 | 29 | 0.1034 |
| 12_9069208 | 5 | 0.2 | 11 | 0.1818 | 7 | 0.5714 | 12 | 0.3333 | 35 | 0.3143 | 8 | 0.5 | 6 | 0.1667 | 3 | 0.3333 | 8 | 0.375 | 25 | 0.36 |
| 12_9069247 | 3 | 0.3333 | 10 | 0.5 | 7 | 0.4286 | 7 | 0.4286 | 27 | 0.4444 | 7 | 0.4286 | 7 | 0.5714 | 4 | 0.25 | 5 | 0 | 23 | 0.3478 |
| 12_9069277 | 2 | 0.5 | 6 | 0 | 5 | 0 | 6 | 0 | 19 | 0.0526 | 7 | 0.1429 | 2 | 0 | 3 | 0 | 1 | 0 | 13 | 0.0769 |
| 12_9069331 | 3 | 0 | 8 | 0 | 4 | 0.25 | 6 | 0.3333 | 21 | 0.1429 | 6 | 0 | 4 | 0 | 3 | 0.3333 | 3 | 0 | 16 | 0.0625 |
| 12_9069380 | 3 | 0 | 8 | 0.125 | 4 | 0.25 | 6 | 0 | 21 | 0.0952 | 8 | 0.125 | 5 | 0 | 4 | 0 | 3 | 0 | 20 | 0.05 |
| 12_9069398 | 3 | 0 | 6 | 0 | 3 | 0.6667 | 5 | 0 | 17 | 0.1176 | 8 | 0.25 | 6 | 0.5 | 2 | 0.5 | 2 | 0.5 | 18 | 0.3889 |
| 12_9069436 | 3 | 0.3333 | 8 | 0.25 | 4 | 0.5 | 6 | 0.5 | 21 | 0.381 | 8 | 0.25 | 3 | 0 | 4 | 0.25 | 3 | 1 | 18 | 0.3333 |
| 12_9069440 | 3 | 1 | 6 | 0.6667 | 4 | 1 | 6 | 1 | 19 | 0.8947 | 7 | 1 | 3 | 1 | 4 | 1 | 1 | 1 | 15 | 1 |
| 12_9069445 | 3 | 0 | 6 | 0.1667 | 4 | 0.25 | 6 | 0.3333 | 19 | 0.2105 | 7 | 0.1429 | 4 | 0 | 3 | 0.3333 | 2 | 0.5 | 16 | 0.1875 |
| 12_9069467 | 2 | 0.5 | 7 | 0 | 4 | 0 | 5 | 0 | 18 | 0.0556 | 6 | 0 | 3 | 0.3333 | 3 | 0 | 2 | 0 | 14 | 0.0714 |
| 12_9069501 | 1 | 0 | 6 | 0.5 | 3 | 0 | 1 | 1 | 11 | 0.3636 | 6 | 0.3333 | 1 | 0 | 1 | 0 | 2 | 0.5 | 10 | 0.3 |
| 12_9069515 | 2 | 0 | 6 | 0.3333 | 3 | 0.6667 | 4 | 0 | 15 | 0.2667 | 6 | 0.5 | 3 | 0.3333 | 1 | 0 | NA | NA | 10 | 0.4 |
| 12_9069537 | 2 | 0.5 | 4 | 0.75 | 3 | 0.6667 | 4 | 0.5 | 13 | 0.6154 | 7 | 1 | 2 | 1 | 2 | 1 | 2 | 0.5 | 13 | 0.9231 |
| 12_9069562 | 1 | 0 | 6 | 0.3333 | 2 | 0 | 4 | 0 | 13 | 0.1538 | 7 | 0.1429 | 3 | 0.3333 | 2 | 0 | 2 | 0 | 14 | 0.1429 |
| 12_9069570 | 2 | 0.5 | 4 | 1 | 2 | 0.5 | 3 | 1 | 11 | 0.8182 | 6 | 0.6667 | 3 | 1 | 2 | 0.5 | 1 | 1 | 12 | 0.75 |
| 12_9069586 | 2 | 1 | 6 | 0.8333 | 1 | 1 | 4 | 0.5 | 13 | 0.7692 | 5 | 0.4 | 3 | 0.6667 | 2 | 1 | 2 | 0 | 12 | 0.5 |
| 12_9069586 | 2 | 0.5 | 5 | 0.6 | 1 | 0 | 3 | 0.6667 | 11 | 0.5455 | 5 | 1 | 3 | 0 | 2 | 1 | NA | NA | 10 | 0.7 |
| 12_9069595 | 2 | 0 | 6 | 0.1667 | 1 | 0 | 4 | 0 | 13 | 0.0769 | 5 | 0.4 | 3 | 0 | 2 | 0.5 | 2 | 0 | 12 | 0.25 |
| 12_9069600 | 2 | 1 | 6 | 0.8333 | 1 | 0 | 3 | 0.3333 | 12 | 0.6667 | 5 | 0.6 | 3 | 1 | 2 | 0.5 | NA | NA | 10 | 0.7 |
| 12_9069604 | 2 | 0.5 | 5 | 0.2 | 1 | 0 | 4 | 0.5 | 12 | 0.3333 | 6 | 0.1667 | 2 | 0 | 2 | 0.5 | 2 | 0.5 | 12 | 0.25 |
| 12_9069605 | 2 | 0 | 5 | 0.2 | 1 | 0 | 4 | 0 | 12 | 0.0833 | 5 | 0 | 3 | 0.3333 | 2 | 0 | 2 | 0 | 12 | 0.0833 |
| 12_9069611 | 1 | 1 | 3 | 1 | 1 | 1 | 3 | 1 | 8 | 1 | 3 | 1 | 1 | 1 | 2 | 1 | NA | NA | 6 | 1 |
| 12_9069612 | 1 | 1 | 3 | 1 | 1 | 1 | 1 | 1 | 6 | 1 | 4 | 1 | 3 | 1 | 1 | 1 | NA | NA | 8 | 1 |
| 12_9069634 | 1 | 1 | 3 | 1 | 1 | 1 | 3 | 1 | 8 | 1 | 4 | 1 | 2 | 1 | 2 | 0.5 | 1 | 1 | 9 | 0.8889 |
| 12_9069642 | 2 | 0.5 | 4 | 0 | 1 | 0 | 3 | 0.3333 | 10 | 0.2 | 5 | 0.4 | 3 | 0 | 2 | 0.5 | 1 | 0 | 11 | 0.2727 |
| 12_9069647 | 2 | 1 | 4 | 0.75 | 1 | 1 | 3 | 1 | 10 | 0.9 | 5 | 1 | 3 | 1 | 1 | 1 | 1 | 1 | 10 | 1 |
| 12_9069653 | 2 | 0.5 | 3 | 0 | 1 | 0 | 3 | 0 | 9 | 0.1111 | 5 | 0.2 | 3 | 0.3333 | 2 | 0.5 | 1 | 0 | 11 | 0.2727 |

| | | | | | | | | | | | | | | | | | | | | |
|---|---|---|---|---|---|---|---|---|---|---|---|---|---|---|---|---|---|---|---|---|
| 12_906966 | 2 | 1 | 3 | 1 | 1 | 0 | 3 | 0.3333 | 9 | 0.6667 | 4 | 0.5 | 2 | 1 | 2 | 0.5 | 1 | 1 | 9 | 0.6667 |
| 12_907050 | NA | NA | 2 | 0.5 | NA | NA | 3 | 0.3333 | 5 | 0.4 | 2 | 1 | 2 | 0 | 1 | 0 | 1 | 0 | 6 | 0.3333 |
| 12_907054 | 1 | 0 | 3 | 0.6667 | NA | NA | 3 | 0.3333 | 7 | 0.4286 | 2 | 0.5 | 1 | 1 | 1 | 1 | 1 | 1 | 5 | 0.8 |
| 12_907149 | 2 | 1 | NA | NA | NA | NA | NA | NA | 2 | 1 | NA | NA | 1 | 0 | 2 | 0.5 | 1 | 1 | 4 | 0.5 |
| 12_907149 | 2 | 0.5 | NA | NA | NA | NA | NA | NA | 2 | 0.5 | NA | NA | 1 | 1 | 2 | 1 | 1 | 1 | 4 | 1 |
| 12_909661 | 1 | 0 | NA | NA | 1 | 1 | 2 | 1 | 4 | 0.75 | 2 | 1 | NA | NA | 1 | 0 | 1 | 1 | 4 | 0.75 |
| 12_915060 | NA | NA | 1 | 1 | 1 | 1 | NA | NA | 2 | 1 | NA | NA | 1 | 1 | NA | NA | NA | NA | 1 | 1 |
| 12_915061 | NA | NA | 1 | 1 | 1 | 0 | NA | NA | 2 | 0.5 | NA | NA | 1 | 1 | NA | NA | NA | NA | 1 | 1 |
| 12_915062 | NA | NA | 1 | 1 | 1 | 1 | NA | NA | 2 | 1 | NA | NA | 1 | 1 | NA | NA | NA | NA | 1 | 1 |
| 12_915064 | NA | NA | 1 | 0 | 1 | 1 | NA | NA | 2 | 0.5 | NA | NA | 1 | 1 | NA | NA | NA | NA | 1 | 1 |
| 12_915066 | NA | NA | 1 | 1 | 1 | 1 | NA | NA | 2 | 1 | NA | NA | 1 | 1 | NA | NA | NA | NA | 1 | 1 |
| 12_915071 | NA | NA | 1 | 1 | 1 | 1 | NA | NA | 2 | 1 | NA | NA | 1 | 1 | NA | NA | NA | NA | 1 | 1 |
| 12_915072 | NA | NA | 1 | 1 | 1 | 1 | NA | NA | 2 | 1 | NA | NA | 1 | 1 | NA | NA | NA | NA | 1 | 1 |
| 12_915072 | NA | NA | 1 | 1 | NA | NA | NA | NA | 1 | 1 | NA | NA | 1 | 1 | NA | NA | NA | NA | 1 | 1 |
| 12_915073 | NA | NA | NA | NA | 1 | 1 | NA | NA | 1 | 1 | NA | NA | 1 | 1 | NA | NA | NA | NA | 1 | 1 |
| 12_915226 | 4 | 0 | 4 | 0.25 | 3 | 0 | 6 | 0.1667 | 17 | 0.1176 | 6 | 0.1667 | 3 | 0 | NA | NA | 6 | 0 | 15 | 0.0667 |
| 12_915229 | 5 | 0 | 4 | 0 | 4 | 0 | 4 | 0.25 | 17 | 0.0588 | 7 | 0.1429 | 2 | 0.5 | 2 | 0 | 8 | 0.125 | 19 | 0.1579 |
| 12_915243 | 3 | 0.3333 | 1 | 0 | 4 | 0 | 1 | 0 | 9 | 0.1111 | 2 | 0 | 1 | 0 | 1 | 0 | 5 | 0.2 | 9 | 0.1111 |
| 12_915261 | 1 | 1 | 1 | 1 | 1 | 1 | 2 | 0.5 | 5 | 0.8 | 2 | 0.5 | NA | NA | NA | NA | NA | NA | 2 | 0.5 |
| 12_915263 | NA | NA | 1 | 1 | 1 | 0 | 1 | 0 | 3 | 0.3333 | 2 | 0.5 | NA | NA | NA | NA | NA | NA | 2 | 0.5 |
| 12_915264 | NA | NA | NA | NA | 1 | 1 | 2 | 0.5 | 3 | 0.6667 | 2 | 0.5 | 2 | 1 | NA | NA | NA | NA | 4 | 0.75 |
| 12_915264 | NA | NA | NA | NA | NA | NA | 2 | 1 | 2 | 1 | 2 | 1 | 2 | 1 | NA | NA | NA | NA | 4 | 1 |
| 12_916001 | 1 | 1 | 2 | 0.5 | 1 | 0 | NA | NA | 4 | 0.5 | 1 | 1 | 1 | 0 | NA | NA | NA | NA | 2 | 0.5 |
| 12_917729 | 14 | 0.0714 | 26 | 0.0385 | 15 | 0.0667 | 12 | 0.0833 | 67 | 0.0597 | 10 | 0.2 | 15 | 0.1333 | 14 | 0.2143 | 14 | 0.0714 | 53 | 0.1509 |
| 12_917730 | 13 | 0.0769 | 23 | 0.1739 | 15 | 0.2667 | 14 | 0.1429 | 65 | 0.1692 | 10 | 0 | 16 | 0.1875 | 14 | 0.5714 | 16 | 0.0625 | 56 | 0.2143 |
| 12_917732 | 11 | 0.4545 | 22 | 0.8182 | 9 | 0.8889 | 11 | 0.9091 | 53 | 0.7736 | 10 | 0.3 | 16 | 0.75 | 12 | 0.5833 | 13 | 0.5385 | 51 | 0.5686 |
| 12_917733 | 13 | 0 | 20 | 0.1 | 11 | 0.0909 | 12 | 0.1667 | 56 | 0.0893 | 9 | 0.2222 | 16 | 0.3125 | 11 | 0.2727 | 14 | 0.2143 | 50 | 0.26 |
| 12_917733 | 2 | 0 | NA | NA | 1 | 1 | 2 | 1 | 5 | 0.6 | NA | NA | 1 | 1 | NA | NA | NA | NA | 1 | 1 |
| 12_917733 | NA | NA | 3 | 0.3333 | 1 | 0 | 1 | 1 | 5 | 0.4 | NA | NA | 2 | 1 | NA | NA | 1 | 1 | 3 | 1 |
| 12_917733 | 1 | 1 | 1 | 1 | 1 | 1 | NA | NA | 3 | 1 | 1 | 1 | NA | NA | NA | NA | NA | NA | 1 | 1 |
| 12_917734 | 9 | 0.3333 | 16 | 0.0625 | 8 | 0 | 11 | 0 | 44 | 0.0909 | 9 | 0 | 15 | 0.1333 | 9 | 0.1111 | 13 | 0.1538 | 46 | 0.1087 |
| 12_917734 | 1 | 0 | 1 | 1 | 1 | 0 | 1 | 0 | 4 | 0.25 | 1 | 1 | NA | NA | NA | NA | NA | NA | 1 | 1 |
| 12_917749 | 4 | 0 | 16 | 0.0625 | 8 | 0.125 | 7 | 0.1429 | 35 | 0.0857 | 8 | 0 | 12 | 0.1667 | 9 | 0.1111 | 11 | 0.0909 | 40 | 0.1 |
| 12_917750 | 4 | 0 | 15 | 0.1333 | 7 | 0.1429 | 7 | 0 | 33 | 0.0909 | 6 | 0 | 9 | 0.2222 | 7 | 0 | 10 | 0.1 | 32 | 0.0938 |
| 12_917754 | 5 | 0 | 19 | 0.1053 | 8 | 0.125 | 8 | 0.125 | 40 | 0.1 | 4 | 0 | 11 | 0.1818 | 10 | 0.1 | 8 | 0 | 33 | 0.0909 |
| 12_917755 | 4 | 1 | 13 | 0.9231 | 9 | 0.6667 | 6 | 1 | 32 | 0.875 | 4 | 1 | 6 | 0.8333 | 8 | 1 | 7 | 0.5714 | 25 | 0.84 |
| 12_917756 | 6 | 1 | 16 | 0.875 | 10 | 0.9 | 10 | 0.9 | 42 | 0.9048 | 6 | 0.8333 | 12 | 0.9167 | 9 | 1 | 11 | 0.9091 | 38 | 0.9211 |
| 12_917756 | 6 | 0.3333 | 16 | 0.1875 | 8 | 0.125 | 9 | 0.2222 | 39 | 0.2051 | 5 | 0 | 13 | 0.1538 | 9 | 0.1111 | 11 | 0.5455 | 38 | 0.2368 |
| 12_917756 | 6 | 0.3333 | 14 | 0.3571 | 5 | 0.4 | 7 | 0.4286 | 32 | 0.375 | 2 | 0.5 | 9 | 0.4444 | 8 | 0.375 | 10 | 0.4 | 29 | 0.4138 |
| 12_917757 | 6 | 0.8333 | 14 | 0.7857 | 8 | 0.875 | 10 | 0.6 | 38 | 0.7632 | 6 | 0.3333 | 12 | 0.75 | 10 | 0.8 | 11 | 0.7273 | 39 | 0.6923 |
| 12_917757 | 7 | 0.2857 | 15 | 0.8 | 10 | 0.6 | 9 | 0.5556 | 41 | 0.6098 | 6 | 0.8333 | 12 | 0.6667 | 9 | 0.7778 | 12 | 0.5833 | 39 | 0.6923 |
| 12_917758 | 4 | 1 | 14 | 0.9286 | 7 | 1 | 6 | 0.8333 | 31 | 0.9355 | 6 | 1 | 8 | 1 | 8 | 1 | 11 | 1 | 33 | 1 |
| 12_917759 | 5 | 0.2 | 13 | 0.0769 | 10 | 0.2 | 8 | 0.125 | 36 | 0.1389 | 6 | 0 | 10 | 0.1 | 10 | 0 | 10 | 0.2 | 36 | 0.0833 |
| 12_917759 | 7 | 1 | 15 | 0.6 | 9 | 0.5556 | 9 | 1 | 40 | 0.75 | 6 | 0.8333 | 10 | 0.5 | 9 | 0.5556 | 11 | 1 | 36 | 0.7222 |
| 12_917759 | 4 | 1 | 12 | 0.75 | 8 | 0.875 | 8 | 1 | 32 | 0.875 | 4 | 0.75 | 10 | 0.9 | 9 | 1 | 11 | 1 | 34 | 0.9412 |
| 12_917760 | 6 | 0.6667 | 14 | 0.7143 | 8 | 1 | 6 | 0.6667 | 34 | 0.7647 | 6 | 0.8333 | 9 | 0.7778 | 9 | 0.6667 | 10 | 0.8 | 34 | 0.7647 |
| 12_917761 | 5 | 0.2 | 13 | 0 | 5 | 0.2 | 8 | 0 | 31 | 0.0645 | 6 | 0.1667 | 10 | 0.1 | 8 | 0.25 | 9 | 0.2222 | 33 | 0.1818 |
| 12_917761 | 6 | 0.8333 | 11 | 0.8182 | 5 | 1 | 8 | 0.875 | 30 | 0.8667 | 6 | 0.8333 | 10 | 0.9 | 9 | 0.8889 | 7 | 1 | 32 | 0.9062 |
| 12_917761 | 5 | 0.2 | 15 | 0.7333 | 6 | 0.6667 | 8 | 0.75 | 34 | 0.6471 | 6 | 0.6667 | 9 | 0.6667 | 9 | 0.7778 | 11 | 0.5455 | 35 | 0.6571 |
| 12_917761 | 2 | 0.5 | 11 | 0.2727 | 3 | 0.6667 | 5 | 0.4 | 21 | 0.381 | 2 | 0 | 8 | 0.25 | 5 | 0 | 8 | 0.25 | 23 | 0.1739 |
| 12_917763 | 6 | 1 | 13 | 0.8462 | 5 | 0.8 | 7 | 1 | 31 | 0.9032 | 5 | 0.6 | 11 | 0.9091 | 8 | 0.625 | 11 | 0.6364 | 35 | 0.7143 |
| 12_917763 | 6 | 0.5 | 13 | 0.5385 | 5 | 0.4 | 7 | 0.1429 | 31 | 0.4194 | 4 | 0.75 | 10 | 0.5 | 9 | 0.5556 | 10 | 0.4 | 33 | 0.5152 |
| 12_917764 | 5 | 0.2 | 14 | 0.1429 | 5 | 0.2 | 6 | 0.1667 | 30 | 0.1667 | 6 | 0.1667 | 10 | 0 | 8 | 0.125 | 9 | 0 | 33 | 0.0606 |
| 12_917764 | 6 | 0.1667 | 14 | 0.1429 | 5 | 0.4 | 5 | 0 | 30 | 0.1667 | 7 | 0.1429 | 8 | 0.375 | 9 | 0.3333 | 8 | 0.125 | 32 | 0.25 |
| 12_917766 | 4 | 0 | 13 | 0.1538 | 4 | 0 | 5 | 0.2 | 26 | 0.1154 | 5 | 0.4 | 7 | 0.4286 | 9 | 0.3333 | 7 | 0 | 28 | 0.2857 |
| 12_917767 | 6 | 0.1667 | 12 | 0.3333 | 2 | 1 | 6 | 0.6667 | 26 | 0.4231 | 6 | 0 | 10 | 0.3 | 9 | 0.5556 | 8 | 0.625 | 33 | 0.3939 |
| 12_917767 | 5 | 0 | 10 | 0 | 4 | 0 | 6 | 0.3333 | 25 | 0.08 | 3 | 0 | 7 | 0.1429 | 8 | 0.125 | 8 | 0.25 | 26 | 0.1538 |
| 12_917998 | 3 | 0 | 7 | 0.1429 | 2 | 0 | 3 | 0 | 15 | 0.0667 | 1 | 0 | 2 | 0.5 | 2 | 0 | 3 | 0 | 8 | 0.125 |
| 12_957453 | NA | NA | 1 | 0 | 3 | 0.3333 | 2 | 1 | 6 | 0.5 | NA | NA | 1 | 1 | 1 | 1 | 1 | 1 | 3 | 1 |
| 12_957454 | NA | NA | 1 | 1 | 4 | 0.25 | 2 | 1 | 7 | 0.5714 | NA | NA | 1 | 1 | 1 | 0 | 1 | 1 | 3 | 0.6667 |
| 12_957455 | NA | NA | 1 | 1 | 4 | 1 | 2 | 0.5 | 7 | 0.8571 | NA | NA | 1 | 1 | 1 | 1 | NA | NA | 2 | 1 |
| 12_957456 | NA | NA | 2 | 0.5 | 3 | 0.6667 | 1 | 1 | 6 | 0.6667 | NA | NA | 1 | 1 | 1 | 1 | NA | NA | 2 | 1 |
| 12_957457 | NA | NA | 2 | 1 | 4 | 1 | 2 | 1 | 8 | 1 | NA | NA | NA | NA | 1 | 1 | 1 | 1 | 2 | 1 |
| 12_957457 | NA | NA | 2 | 1 | 3 | 0.6667 | 1 | 0 | 6 | 0.6667 | NA | NA | 1 | 1 | 1 | 1 | 1 | 0 | 3 | 0.6667 |
| 12_957459 | NA | NA | 2 | 0 | 3 | 0.3333 | 1 | 0 | 6 | 0.1667 | NA | NA | 1 | 1 | 1 | 0 | 1 | 0 | 3 | 0.3333 |

| | | | | | | | | | | | | | | | | | | | | |
|---|---|---|---|---|---|---|---|---|---|---|---|---|---|---|---|---|---|---|---|---|
| 12_957462( | NA | NA | 1 | 0 | 4 | 0.25 | 2 | 0.5 | 7 | 0.2857 | NA | NA | 1 | 1 | 1 | 0 | NA | NA | 2 | 0.5 |
| 12_957464! | NA | NA | 1 | 0 | 3 | 0 | 2 | 0.5 | 6 | 0.1667 | NA | NA | 2 | 0.5 | 1 | 0 | 1 | 0 | 4 | 0.25 |
| 12_957466( | NA | NA | 2 | 1 | 4 | 1 | 2 | 1 | 8 | 1 | NA | NA | 2 | 1 | 1 | 1 | 1 | 1 | 4 | 1 |
| 12_957468 | NA | NA | NA | NA | 4 | 0.25 | 1 | 0 | 5 | 0.2 | NA | NA | 2 | 0.5 | 1 | 1 | 1 | 0 | 4 | 0.5 |
| 12_957469 | NA | NA | 2 | 0 | 4 | 0.75 | 2 | 0.5 | 8 | 0.5 | NA | NA | 2 | 0 | 2 | 0 | 2 | 1 | 6 | 0.3333 |
| 12_957470 | NA | NA | 1 | 1 | 3 | 1 | 2 | 1 | 6 | 1 | NA | NA | 2 | 1 | 1 | 1 | 2 | 1 | 5 | 1 |
| 12_957472 | NA | NA | 2 | 0.5 | 5 | 0.4 | NA | NA | 7 | 0.4286 | NA | NA | 1 | 1 | 1 | 0 | 1 | 0 | 3 | 0.3333 |
| 12_957474! | NA | NA | 2 | 1 | 4 | 1 | 2 | 0.5 | 8 | 0.875 | 1 | 1 | 1 | 1 | 2 | 0.5 | 1 | 1 | 5 | 0.8 |
| 12_957474 | NA | NA | 1 | 0 | 5 | 0.2 | 2 | 0 | 8 | 0.125 | 1 | 0 | 2 | 0.5 | 2 | 0 | 2 | 0 | 7 | 0.1429 |
| 12_957475 | NA | NA | 2 | 1 | 3 | 1 | 2 | 1 | 7 | 1 | 1 | 1 | 1 | 1 | 2 | 1 | 1 | 1 | 5 | 1 |
| 12_957475 | NA | NA | 1 | 1 | 5 | 1 | 2 | 1 | 8 | 1 | 1 | 1 | 2 | 1 | 2 | 1 | 2 | 1 | 7 | 1 |
| 12_957476 | NA | NA | 1 | 1 | 6 | 0.6667 | 2 | 1 | 9 | 0.7778 | 1 | 1 | 2 | 1 | 1 | 1 | 2 | 0 | 6 | 0.6667 |
| 12_957476( | NA | NA | 2 | 1 | 4 | 0.75 | 2 | 1 | 8 | 0.875 | NA | NA | NA | NA | 2 | 0.5 | 1 | 1 | 3 | 0.6667 |
| 12_957477 | NA | NA | 2 | 1 | 5 | 0.6 | 2 | 1 | 9 | 0.7778 | 1 | 1 | 2 | 1 | 2 | 0.5 | 2 | 1 | 7 | 0.8571 |
| 12_957478( | 1 | 0 | 1 | 0 | 7 | 0.1429 | 2 | 0 | 11 | 0.0909 | 2 | 0 | 2 | 0 | 2 | 0 | 2 | 0.5 | 8 | 0.125 |
| 12_957483 | 1 | 0 | 1 | 1 | 7 | 0.1429 | 2 | 0 | 11 | 0.1818 | 1 | 0 | 2 | 0.5 | 2 | 0.5 | 3 | 0 | 8 | 0.25 |
| 12_957487 | 1 | 0 | 3 | 0.6667 | 6 | 0.1667 | 3 | 1 | 13 | 0.4615 | 2 | 0.5 | 3 | 0 | 2 | 0 | 3 | 0.6667 | 10 | 0.3 |
| 12_957500( | 4 | 0 | 2 | 0.5 | 3 | 0.3333 | 3 | 0.3333 | 12 | 0.25 | 1 | 0 | 4 | 0 | 4 | 0.25 | 5 | 0 | 14 | 0.0714 |
| 12_957518! | 8 | 0.5 | 6 | 0.3333 | 5 | 0.4 | 6 | 0.5 | 25 | 0.44 | 4 | 0.25 | 5 | 0.4 | 8 | 0.5 | 7 | 0.4286 | 24 | 0.4167 |
| 12_981683 | NA | NA | NA | NA | 1 | 1 | NA | NA | 1 | 1 | NA | NA | NA | NA | NA | NA | 1 | 1 | 1 | 1 |
| 12_981687 | 4 | 0.5 | 5 | 0.4 | 2 | 0.5 | 2 | 0.5 | 13 | 0.4615 | 4 | 0.75 | 6 | 0.1667 | 5 | 0.2 | 8 | 0.375 | 23 | 0.3478 |
| 12_981691( | 3 | 0 | 6 | 0.1667 | 3 | 0 | 4 | 0.25 | 16 | 0.125 | 4 | 0.25 | 4 | 0 | 4 | 0 | 7 | 0.2857 | 19 | 0.1579 |
| 12_981700 | 3 | 0.6667 | 6 | 0.1667 | 4 | 0.25 | 4 | 0.25 | 17 | 0.2941 | 4 | 0.25 | 6 | 0.3333 | 3 | 0 | 7 | 0.4286 | 20 | 0.3 |
| 12_982415! | NA | NA | NA | NA | NA | NA | 1 | 1 | 1 | 1 | NA | NA | NA | NA | NA | NA | 1 | 1 | 1 | 1 |
| 12_982416 | NA | NA | NA | NA | NA | NA | 1 | 1 | 1 | 1 | NA | NA | NA | NA | NA | NA | 1 | 1 | 1 | 1 |
| 12_982420! | NA | NA | NA | NA | NA | NA | 1 | 1 | 1 | 1 | NA | NA | NA | NA | NA | NA | 1 | 1 | 1 | 1 |
| 12_982422 | NA | NA | NA | NA | NA | NA | 1 | 1 | 1 | 1 | NA | NA | NA | NA | NA | NA | 1 | 1 | 1 | 1 |
| 12_982423! | NA | NA | NA | NA | NA | NA | 1 | 1 | 1 | 1 | NA | NA | NA | NA | NA | NA | 1 | 1 | 1 | 1 |
| 12_982427 | NA | NA | NA | NA | NA | NA | 1 | 1 | 1 | 1 | NA | NA | NA | NA | NA | NA | 1 | 1 | 1 | 1 |
| 12_984010 | 1 | 1 | NA | NA | NA | NA | NA | NA | 1 | 1 | NA | NA | NA | NA | 1 | 1 | NA | NA | 1 | 1 |
| 12_984010! | 1 | 1 | NA | NA | NA | NA | NA | NA | 1 | 1 | NA | NA | NA | NA | 1 | 1 | NA | NA | 1 | 1 |
| 12_984011( | 1 | 1 | NA | NA | NA | NA | NA | NA | 1 | 1 | NA | NA | NA | NA | 1 | 1 | NA | NA | 1 | 1 |
| 12_984011 | 1 | 1 | NA | NA | NA | NA | NA | NA | 1 | 1 | NA | NA | NA | NA | 1 | 1 | NA | NA | 1 | 1 |
| 12_984012! | 1 | 1 | NA | NA | NA | NA | NA | NA | 1 | 1 | NA | NA | NA | NA | 1 | 1 | NA | NA | 1 | 1 |
| 12_986354 | 3 | 0.3333 | 2 | 0 | 5 | 0 | 3 | 0 | 13 | 0.0769 | 2 | 0.5 | 1 | 0 | 1 | 0 | 2 | 0 | 6 | 0.1667 |
| 12_986355( | 2 | 0.5 | 1 | 0 | 4 | 1 | 3 | 0 | 10 | 0.5 | 2 | 1 | NA | NA | 1 | 0 | 2 | 0.5 | 5 | 0.6 |
| 12_986357 | 3 | 0.3333 | 2 | 0 | 5 | 0.4 | 3 | 0 | 13 | 0.2308 | 2 | 0.5 | 1 | 1 | 3 | 0.3333 | 1 | 0 | 7 | 0.4286 |
| 12_986357 | 2 | 1 | 2 | 1 | 5 | 1 | 2 | 1 | 11 | 1 | 2 | 1 | 1 | 1 | 3 | 1 | 2 | 0.5 | 8 | 0.875 |
| 12_986357! | 3 | 0.6667 | 2 | 0.5 | 5 | 0.4 | 3 | 0.3333 | 13 | 0.4615 | 1 | 1 | 1 | 1 | 2 | 0 | 2 | 0.5 | 6 | 0.5 |
| 12_986358 | 2 | 0.5 | 2 | 0.5 | 4 | 1 | 2 | 1 | 10 | 0.8 | 1 | 1 | 1 | 1 | 3 | 0.6667 | 2 | 1 | 7 | 0.8571 |
| 12_986359 | 1 | 1 | 2 | 0.5 | 5 | 0 | 3 | 0.6667 | 11 | 0.3636 | 1 | 0 | 1 | 1 | 3 | 0.3333 | 1 | 0 | 6 | 0.3333 |
| 12_986359! | 1 | 0 | 1 | 0 | 4 | 0.5 | 3 | 0.3333 | 9 | 0.3333 | 1 | 0 | 1 | 1 | 3 | 0 | 2 | 0.5 | 7 | 0.2857 |
| 12_986359 | 2 | 0 | 2 | 1 | 3 | 0 | 1 | 0 | 8 | 0.25 | 1 | 0 | 1 | 0 | 2 | 0 | 2 | 0.5 | 6 | 0.1667 |
| 12_986431 | 1 | 0 | 1 | 1 | 1 | 1 | NA | NA | 3 | 0.6667 | NA | NA | NA | NA | NA | NA | 1 | 1 | 1 | 1 |
| 12_986443 | 1 | 0 | 1 | 0 | 1 | 0 | 1 | 1 | 4 | 0.25 | NA | NA | NA | NA | NA | NA | 1 | 1 | 1 | 1 |
| 12_986445( | 1 | 0 | NA | NA | 1 | 1 | 1 | 0 | 3 | 0.3333 | NA | NA | NA | NA | NA | NA | 1 | 1 | 1 | 1 |
| 12_986447 | 1 | 0 | 1 | 1 | 1 | 1 | 1 | 1 | 4 | 0.75 | NA | NA | NA | NA | NA | NA | 1 | 1 | 1 | 1 |
| 12_986545 | 1 | 1 | NA | NA | 1 | 1 | 1 | 1 | 3 | 1 | NA | NA | NA | NA | NA | NA | 1 | 1 | 1 | 1 |
| 12_986545! | NA | NA | NA | NA | 1 | 1 | 1 | 1 | 2 | 1 | NA | NA | NA | NA | NA | NA | 1 | 1 | 1 | 1 |
| 12_986547 | 1 | 1 | NA | NA | 1 | 0 | 1 | 1 | 3 | 0.6667 | NA | NA | NA | NA | NA | NA | 1 | 1 | 1 | 1 |
| 12_986549( | NA | NA | NA | NA | NA | NA | 1 | 1 | 1 | 1 | NA | NA | NA | NA | NA | NA | 1 | 1 | 1 | 1 |
| 12_986553 | 1 | 1 | NA | NA | NA | NA | 1 | 1 | 2 | 1 | NA | NA | NA | NA | NA | NA | 1 | 1 | 1 | 1 |
| 12_986553 | 1 | 1 | NA | NA | NA | NA | 1 | 0 | 2 | 0.5 | NA | NA | NA | NA | NA | NA | 1 | 1 | 1 | 1 |
| 12_987249 | 7 | 0.1429 | 9 | 0.4444 | 5 | 0 | 4 | 0.25 | 25 | 0.24 | 3 | 0.3333 | 5 | 0.2 | 5 | 0.2 | 4 | 0.25 | 17 | 0.2353 |
| 12_987250 | 7 | 0.4286 | 10 | 0 | 7 | 0.2857 | 4 | 0 | 28 | 0.1786 | 3 | 0.6667 | 6 | 0 | 6 | 0.1667 | 4 | 0 | 19 | 0.1579 |
| 12_987464( | 3 | 1 | 3 | 0.3333 | 1 | 1 | 4 | 0 | 11 | 0.4545 | 1 | 1 | 2 | 0.5 | 3 | 0.6667 | 3 | 1 | 9 | 0.7778 |
| 12_987534( | 8 | 0.625 | 5 | 0 | 5 | 0 | 5 | 0.2 | 23 | 0.2609 | 2 | 0 | 3 | 0.3333 | 2 | 0 | 4 | 0 | 11 | 0.0909 |
| 12_987554 | 3 | 0.3333 | 4 | 0 | 2 | 0 | 6 | 0 | 15 | 0.0667 | 1 | 0 | 4 | 0 | 9 | 0.1111 | 3 | 0 | 17 | 0.0588 |
| 12_987556( | 4 | 0.5 | 4 | 0.25 | 2 | 0.5 | 6 | 0.1667 | 16 | 0.3125 | 1 | 0 | 4 | 0.25 | 7 | 0 | 3 | 0.6667 | 15 | 0.2 |
| 12_987556! | 4 | 0.5 | 3 | 0.3333 | 2 | 0.5 | 6 | 0.5 | 15 | 0.4667 | 1 | 0 | 3 | 0.6667 | 7 | 0.4286 | 3 | 0.3333 | 14 | 0.4286 |
| 12_987556( | 3 | 0.6667 | 4 | 0.25 | 2 | 0.5 | 6 | 0.5 | 15 | 0.4667 | NA | NA | 4 | 0.75 | 6 | 0.3333 | 3 | 0.3333 | 13 | 0.4615 |
| 12_987556 | 3 | 0 | 4 | 0.25 | 2 | 0 | 6 | 0.5 | 15 | 0.2667 | NA | NA | 4 | 0.75 | 6 | 0.1667 | 3 | 1 | 13 | 0.5385 |
| 12_987557 | 2 | 0 | 3 | 0.6667 | 2 | 0 | 6 | 0 | 13 | 0.1538 | 1 | 0 | 3 | 0.3333 | 5 | 0 | 3 | 0 | 12 | 0.0833 |
| 12_987559( | 3 | 0.6667 | 4 | 0.25 | 2 | 0.5 | 6 | 0 | 15 | 0.2667 | 1 | 0 | 5 | 0 | 7 | 0.1429 | 1 | 0 | 14 | 0.0714 |
| 12_987563 | 1 | 0 | 2 | 0.5 | 1 | 0 | 2 | 0 | 6 | 0.1667 | 1 | 0 | 1 | 0 | 3 | 0.6667 | 2 | 0.5 | 7 | 0.4286 |

| | | | | | | | | | | | | | | | | | | | | |
|---|---|---|---|---|---|---|---|---|---|---|---|---|---|---|---|---|---|---|---|---|
| 12_987565( | 3 | 1 | 3 | 1 | 2 | 1 | 4 | 0.25 | 12 | 0.75 | 1 | 0 | 2 | 0.5 | 3 | 0.3333 | 2 | 0.5 | 8 | 0.375 |
| 12_987607 | 2 | 0 | 1 | 0 | 1 | 0 | 2 | 0.5 | 6 | 0.1667 | NA | NA | 3 | 0.3333 | 2 | 0 | 2 | 0 | 7 | 0.1429 |
| 12_989489 | 1 | 0 | 3 | 0.6667 | 5 | 0.2 | 1 | 1 | 10 | 0.4 | NA | NA | 3 | 0.3333 | 2 | 0 | 3 | 0 | 8 | 0.125 |
| 12_989489 | 1 | 0 | 4 | 0.75 | 6 | 0.1667 | 1 | 0 | 12 | 0.3333 | NA | NA | 4 | 0 | 2 | 0 | 3 | 0.3333 | 9 | 0.1111 |
| 12_989490( | 1 | 0 | 4 | 0.5 | 6 | 0 | 1 | 0 | 12 | 0.1667 | NA | NA | 4 | 0.25 | 2 | 1 | 3 | 0.3333 | 9 | 0.4444 |
| 12_991565 | 6 | 0.8333 | 5 | 0.8 | 6 | 1 | 10 | 0.7 | 27 | 0.8148 | 5 | 0.6 | 5 | 1 | 5 | 0.6 | 3 | 1 | 18 | 0.7778 |
| 12_991566 | 5 | 0.6 | 5 | 0.4 | 6 | 0.1667 | 9 | 0 | 25 | 0.24 | 3 | 0 | 5 | 0.4 | 4 | 0 | 3 | 0.3333 | 15 | 0.2 |
| 12_991571 | 4 | 0.5 | 5 | 0.2 | 7 | 0.7143 | 8 | 0.375 | 24 | 0.4583 | 3 | 0.6667 | 4 | 0 | 5 | 0.4 | 2 | 1 | 14 | 0.4286 |
| 12_991572 | 2 | 1 | 4 | 1 | 4 | 1 | 6 | 1 | 16 | 1 | 2 | 1 | 4 | 1 | 5 | 1 | 1 | 1 | 12 | 1 |
| 12_991572 | 3 | 0.6667 | 4 | 0.5 | 6 | 0.6667 | 5 | 0.6 | 18 | 0.6111 | 3 | 0.6667 | 3 | 0.3333 | 4 | 0.5 | 1 | 1 | 11 | 0.5455 |
| 12_991572 | 3 | 0 | 5 | 0.2 | 5 | 0.4 | 8 | 0.25 | 21 | 0.2381 | 2 | 0 | 3 | 0.3333 | 4 | 0 | 2 | 1 | 11 | 0.2727 |
| 12_991574 | 4 | 0.25 | 4 | 0 | 6 | 0.1667 | 7 | 0.2857 | 21 | 0.1905 | 3 | 0.3333 | 3 | 0 | 5 | 0 | 1 | 0 | 12 | 0.0833 |
| 12_991574 | 4 | 0.5 | 4 | 0.75 | 4 | 0.25 | 8 | 0 | 20 | 0.3 | 2 | 0 | 3 | 0.3333 | 5 | 0.4 | 2 | 0.5 | 12 | 0.3333 |
| 12_991575( | 4 | 0.75 | 3 | 0.3333 | 5 | 0.8 | 6 | 1 | 18 | 0.7778 | 2 | 0.5 | 3 | 0.3333 | 5 | 0.8 | 2 | 1 | 12 | 0.6667 |
| 12_991576 | 4 | 0.5 | 4 | 0 | 5 | 0 | 6 | 0 | 19 | 0.1053 | 3 | 0 | 2 | 0.5 | 4 | 0 | 1 | 0 | 10 | 0.1 |
| 12_991578 | 4 | 0 | 3 | 0 | 4 | 0 | 8 | 0.125 | 19 | 0.0526 | 3 | 0 | 3 | 0.3333 | 5 | 0 | 2 | 0 | 13 | 0.0769 |
| 12_991588 | 1 | 0 | 1 | 0 | 3 | 0.3333 | 4 | 0 | 9 | 0.1111 | 2 | 0 | 2 | 0 | 5 | 0.2 | 1 | 0 | 10 | 0.1 |
| 12_991609 | NA | NA | NA | NA | NA | NA | 1 | 1 | 1 | 1 | NA | NA | 1 | 1 | 2 | 0 | 1 | 1 | 4 | 0.5 |
| 12_998578 | 1 | 1 | 1 | 0 | NA | NA | NA | NA | 2 | 0.5 | 1 | 0 | NA | NA | NA | NA | 2 | 1 | 3 | 0.6667 |
| 12_998579 | 1 | 1 | 2 | 1 | NA | NA | NA | NA | 3 | 1 | 1 | 1 | NA | NA | NA | NA | 2 | 0.5 | 3 | 0.6667 |
| 12_100027( | NA | NA | NA | NA | 1 | 0 | 1 | 1 | 2 | 0.5 | NA | NA | NA | NA | 3 | 0.6667 | 1 | 0 | 4 | 0.5 |
| 12_100027( | NA | NA | NA | NA | 1 | 1 | 1 | 1 | 2 | 1 | NA | NA | NA | NA | 3 | 1 | 1 | 1 | 4 | 1 |
| 12_100027( | NA | NA | NA | NA | 1 | 1 | 1 | 1 | 2 | 1 | NA | NA | NA | NA | 2 | 1 | NA | NA | 2 | 1 |
| 12_100027 | NA | NA | NA | NA | 1 | 1 | 1 | 1 | 2 | 1 | NA | NA | NA | NA | 1 | 1 | 1 | 1 | 2 | 1 |
| 12_100027 | NA | NA | NA | NA | 1 | 0 | 1 | 1 | 2 | 0.5 | NA | NA | NA | NA | NA | NA | 1 | 1 | 1 | 1 |
| 12_100027( | NA | NA | NA | NA | 1 | 1 | 1 | 0 | 2 | 0.5 | NA | NA | NA | NA | 3 | 0.3333 | 1 | 0 | 4 | 0.25 |
| 12_100027( | NA | NA | NA | NA | 1 | 0 | 1 | 1 | 2 | 0.5 | NA | NA | NA | NA | 3 | 0.6667 | 1 | 1 | 4 | 0.75 |
| 12_100027 | NA | NA | NA | NA | 1 | 1 | NA | NA | 1 | 1 | NA | NA | NA | NA | 3 | 1 | 1 | 1 | 4 | 1 |
| 12_100027 | NA | NA | NA | NA | 1 | 1 | NA | NA | 1 | 1 | NA | NA | NA | NA | 2 | 1 | NA | NA | 2 | 1 |
| 12_100028( | NA | NA | NA | NA | 1 | 1 | 1 | 0 | 2 | 0.5 | NA | NA | NA | NA | 2 | 0.5 | 1 | 0 | 3 | 0.3333 |
| 12_100028 | NA | NA | 2 | 0.5 | 1 | 1 | 1 | 0 | 4 | 0.5 | NA | NA | NA | NA | 3 | 0.3333 | 1 | 0 | 4 | 0.25 |
| 12_100028 | NA | NA | 1 | 1 | 1 | 1 | 1 | 1 | 3 | 1 | NA | NA | NA | NA | 3 | 1 | 1 | 0 | 4 | 0.75 |
| 12_100028 | NA | NA | 2 | 0.5 | 1 | 0 | 1 | 0 | 4 | 0.25 | NA | NA | NA | NA | 3 | 0.3333 | 1 | 0 | 4 | 0.25 |
| 12_100028 | NA | NA | 2 | 0.5 | 1 | 0 | NA | NA | 3 | 0.3333 | NA | NA | NA | NA | 1 | 0 | 1 | 1 | 2 | 0.5 |
| 12_100028 | NA | NA | 2 | 1 | 1 | 1 | 1 | 1 | 4 | 1 | NA | NA | NA | NA | 3 | 1 | 1 | 1 | 4 | 1 |
| 12_100028( | NA | NA | 2 | 0 | 1 | 1 | 1 | 1 | 4 | 0.5 | NA | NA | NA | NA | 3 | 0.6667 | NA | NA | 3 | 0.6667 |
| 12_100086 | 26 | 0.0769 | 28 | 0.0357 | 25 | 0.04 | 28 | 0.0714 | 107 | 0.0561 | 25 | 0 | 22 | 0.0909 | 23 | 0.0435 | 19 | 0.1053 | 89 | 0.0562 |
| 12_100091 | 2 | 0 | 2 | 0.5 | 2 | 0 | 2 | 1 | 8 | 0.375 | 4 | 0.5 | NA | NA | 3 | 0 | 3 | 0 | 10 | 0.2 |
| 12_100096 | 1 | 1 | 1 | 0 | 2 | 0.5 | 2 | 0 | 6 | 0.3333 | 4 | 0.25 | NA | NA | 3 | 0.3333 | 3 | 0.3333 | 10 | 0.3 |
| 12_100096 | 1 | 0 | 1 | 0 | 2 | 0 | 1 | 1 | 5 | 0.2 | 3 | 0.3333 | NA | NA | 2 | 0.5 | 3 | 0 | 8 | 0.25 |
| 12_100097( | 1 | 1 | 1 | 1 | 2 | 1 | 2 | 1 | 6 | 1 | 4 | 1 | NA | NA | 3 | 1 | 3 | 1 | 10 | 1 |
| 12_100097( | NA | NA | 1 | 1 | 2 | 1 | 2 | 1 | 5 | 1 | 2 | 1 | NA | NA | 3 | 1 | 3 | 1 | 8 | 1 |
| 12_100097 | 1 | 1 | 1 | 1 | 2 | 1 | 2 | 1 | 6 | 1 | 4 | 0.75 | NA | NA | 3 | 0.3333 | 3 | 1 | 10 | 0.7 |
| 12_100097 | 1 | 1 | NA | NA | 2 | 1 | 1 | 1 | 4 | 1 | 4 | 1 | NA | NA | 2 | 1 | 2 | 0.5 | 8 | 0.875 |
| 12_100097 | 1 | 0 | 1 | 1 | 2 | 0.5 | 2 | 0.5 | 6 | 0.5 | 4 | 0.75 | NA | NA | 3 | 0.3333 | 3 | 0.3333 | 10 | 0.5 |
| 12_100097 | NA | NA | 1 | 0 | 1 | 1 | 1 | 0 | 3 | 0.3333 | 3 | 0.3333 | NA | NA | 1 | 0 | 2 | 0 | 6 | 0.1667 |
| 12_100098( | NA | NA | 1 | 1 | 1 | 1 | 1 | 1 | 3 | 1 | 3 | 0.6667 | NA | NA | 1 | 1 | 2 | 1 | 6 | 0.8333 |
| 12_100098 | NA | NA | 1 | 1 | NA | NA | 1 | 1 | 2 | 1 | 4 | 0 | NA | NA | 2 | 0 | 3 | 0.3333 | 9 | 0.1111 |
| 12_100110 | NA | NA | NA | NA | NA | NA | 1 | 1 | 1 | 1 | 1 | 0 | NA | NA | NA | NA | 2 | 0.5 | 3 | 0.3333 |
| 12_100110 | NA | NA | NA | NA | NA | NA | 1 | 1 | 1 | 1 | 1 | 1 | NA | NA | NA | NA | 1 | 1 | 2 | 1 |
| 12_100172 | 265 | 0.1736 | 263 | 0.1825 | 238 | 0.1849 | 267 | 0.1498 | 1033 | 0.1723 | 215 | 0.1814 | 272 | 0.1728 | 262 | 0.1718 | 241 | 0.1992 | 990 | 0.1808 |
| 12_100173 | 306 | 0.0817 | 302 | 0.0596 | 272 | 0.0662 | 287 | 0.0488 | 1167 | 0.0643 | 245 | 0.0367 | 290 | 0.0966 | 302 | 0.0861 | 277 | 0.0722 | 1114 | 0.0745 |
| 12_100173 | 275 | 0.3855 | 262 | 0.4046 | 216 | 0.4259 | 249 | 0.3855 | 1002 | 0.3992 | 214 | 0.4112 | 252 | 0.4087 | 260 | 0.4385 | 224 | 0.4911 | 950 | 0.4368 |
| 12_100173 | 309 | 0.1068 | 306 | 0.0948 | 277 | 0.1191 | 296 | 0.1115 | 1188 | 0.1077 | 254 | 0.1535 | 290 | 0.1138 | 308 | 0.1071 | 290 | 0.1034 | 1142 | 0.1182 |
| 12_100262 | NA | NA | 1 | 1 | NA | NA | NA | NA | 1 | 1 | NA | NA | NA | NA | 2 | 1 | NA | NA | 2 | 1 |
| 12_100263 | NA | NA | 1 | 1 | NA | NA | NA | NA | 1 | 1 | NA | NA | NA | NA | 2 | 1 | NA | NA | 2 | 1 |
| 12_100263 | NA | NA | 1 | 1 | NA | NA | NA | NA | 1 | 1 | NA | NA | NA | NA | 2 | 0.5 | NA | NA | 2 | 0.5 |
| 12_100263 | NA | NA | 1 | 1 | NA | NA | NA | NA | 1 | 1 | NA | NA | NA | NA | 2 | 1 | NA | NA | 2 | 1 |
| 12_100263 | NA | NA | 1 | 1 | NA | NA | NA | NA | 1 | 1 | NA | NA | NA | NA | 2 | 1 | NA | NA | 2 | 1 |
| 12_100263 | NA | NA | 1 | 1 | NA | NA | NA | NA | 1 | 1 | NA | NA | NA | NA | 2 | 1 | NA | NA | 2 | 1 |
| 12_100267 | 22 | 0.8636 | 25 | 0.96 | 16 | 0.875 | 21 | 0.9048 | 84 | 0.9048 | 18 | 0.8889 | 15 | 0.8667 | 20 | 1 | 21 | 0.8571 | 74 | 0.9054 |
| 12_100267 | 14 | 0.3571 | 17 | 0.6471 | 11 | 0.5455 | 15 | 0.6 | 57 | 0.5439 | 10 | 0.2 | 10 | 0.6 | 16 | 0.4375 | 14 | 0.5 | 50 | 0.44 |
| 12_100267 | 22 | 0.0909 | 20 | 0.05 | 11 | 0.1818 | 14 | 0.0714 | 67 | 0.0896 | 17 | 0.0588 | 12 | 0.0833 | 16 | 0.125 | 16 | 0.125 | 61 | 0.0984 |
| 12_100267 | 16 | 0.4375 | 16 | 0.5 | 9 | 0 | 17 | 0.2353 | 58 | 0.3276 | 13 | 0.6154 | 13 | 0.2308 | 16 | 0.1875 | 12 | 0.1667 | 54 | 0.2963 |
| 12_100514 | 4 | 0.25 | NA | NA | 4 | 0.5 | 5 | 0.2 | 13 | 0.3077 | 4 | 0.25 | 6 | 0.5 | 9 | 0.6667 | 1 | 1 | 20 | 0.55 |

| | | | | | | | | | | | | | | | | | | | | |
|---|---|---|---|---|---|---|---|---|---|---|---|---|---|---|---|---|---|---|---|---|
| 12_100515 | 3 | 0.6667 | NA | NA | 3 | 0.3333 | 4 | 0 | 10 | 0.3 | 2 | 0.5 | 4 | 0 | 11 | 0.4545 | 2 | 0 | 19 | 0.3158 |
| 12_100515 | 3 | 0.3333 | NA | NA | 3 | 0.3333 | 4 | 0 | 10 | 0.2 | 1 | 0 | 3 | 0 | 11 | 0.3636 | 2 | 0 | 17 | 0.2353 |
| 12_100515 | 3 | 0.3333 | NA | NA | 3 | 0.6667 | 4 | 0.75 | 10 | 0.6 | 2 | 0 | 2 | 0.5 | 10 | 0.4 | 1 | 0 | 15 | 0.3333 |
| 12_100516 | 6 | 0.3333 | 2 | 0.5 | 1 | 0 | 5 | 0 | 14 | 0.2143 | 2 | 0.5 | 3 | 0.6667 | 7 | 0.1429 | 1 | 0 | 13 | 0.3077 |
| 12_100516 | 6 | 0.6667 | 2 | 0.5 | 1 | 1 | 3 | 0.3333 | 12 | 0.5833 | 2 | 0.5 | 3 | 0.3333 | 7 | 0.2857 | 2 | 1 | 14 | 0.4286 |
| 12_100516 | 7 | 0.4286 | 2 | 0 | 1 | 0 | 5 | 0 | 15 | 0.2 | 2 | 0.5 | 3 | 0 | 8 | 0 | 2 | 0.5 | 15 | 0.1333 |
| 12_100516 | 7 | 0.7143 | 2 | 1 | NA | NA | 3 | 0.6667 | 12 | 0.75 | 2 | 1 | 1 | 1 | 7 | 0.7143 | 1 | 1 | 11 | 0.8182 |
| 12_100517 | 4 | 1 | 1 | 1 | 1 | 1 | 5 | 1 | 11 | 1 | 2 | 1 | 2 | 1 | 6 | 1 | 2 | 0.5 | 12 | 0.9167 |
| 12_100517 | 5 | 0.4 | 1 | 0 | 1 | 0 | 4 | 0.25 | 11 | 0.2727 | 2 | 0.5 | 2 | 0 | 6 | 0 | 2 | 1 | 12 | 0.25 |
| 12_100517 | 4 | 0 | 1 | 1 | 1 | 0 | 4 | 0.25 | 10 | 0.2 | 2 | 0 | 2 | 0 | 6 | 0.1667 | 2 | 0 | 12 | 0.0833 |
| 12_100517 | 5 | 1 | 1 | 1 | 1 | 1 | 5 | 1 | 12 | 1 | 1 | 1 | 2 | 1 | 5 | 1 | 1 | 1 | 9 | 1 |
| 12_100517 | 3 | 0 | 1 | 0 | NA | NA | 5 | 0.2 | 9 | 0.1111 | 2 | 0.5 | 2 | 0.5 | 5 | 0.2 | 2 | 0 | 11 | 0.2727 |
| 12_100517 | 5 | 0.4 | 1 | 0 | 1 | 0 | 4 | 0.5 | 11 | 0.3636 | 2 | 1 | 1 | 1 | 5 | 0.8 | 2 | 0 | 10 | 0.7 |
| 12_100517 | 5 | 1 | 1 | 1 | 1 | 1 | 4 | 1 | 11 | 1 | 2 | 1 | 2 | 1 | 5 | 1 | 1 | 1 | 10 | 1 |
| 12_100517 | 3 | 0 | 1 | 0 | NA | NA | 5 | 0.4 | 9 | 0.2222 | 2 | 0 | 2 | 0 | 6 | 0.1667 | 2 | 0 | 12 | 0.0833 |
| 12_100517 | 3 | 1 | 1 | 1 | NA | NA | 4 | 0.75 | 8 | 0.875 | 2 | 1 | 2 | 1 | 5 | 0.8 | 2 | 1 | 11 | 0.9091 |
| 12_100517 | 3 | 0 | 1 | 0 | 1 | 1 | 4 | 0.5 | 9 | 0.3333 | 2 | 0.5 | 1 | 1 | 6 | 0 | 1 | 0 | 10 | 0.2 |
| 12_100517 | 2 | 0.5 | NA | NA | NA | NA | 3 | 1 | 5 | 0.8 | 1 | 0 | 1 | 1 | 3 | 1 | NA | NA | 5 | 0.8 |
| 12_100517 | 2 | 1 | 1 | 1 | 1 | 1 | 4 | 0.75 | 8 | 0.875 | 2 | 0.5 | 2 | 1 | 6 | 0.8333 | 2 | 1 | 12 | 0.8333 |
| 12_100517 | 2 | 0 | 1 | 0 | 1 | 0 | 3 | 0.3333 | 7 | 0.1429 | 2 | 0 | 2 | 0 | 6 | 0.1667 | 2 | 0 | 12 | 0.0833 |
| 12_100524 | 3 | 0.3333 | NA | NA | 1 | 0 | 3 | 0 | 7 | 0.1429 | 2 | 0 | 1 | 1 | 3 | 0 | 1 | 0 | 7 | 0.1429 |
| 12_100843 | 1 | 0 | NA | NA | NA | NA | 1 | 1 | 2 | 0.5 | 2 | 0.5 | NA | NA | NA | NA | NA | NA | 2 | 0.5 |
| 12_100843 | NA | NA | NA | NA | NA | NA | 1 | 1 | 1 | 1 | 2 | 0.5 | NA | NA | NA | NA | NA | NA | 2 | 0.5 |
| 12_100844 | NA | NA | NA | NA | NA | NA | 1 | 1 | 1 | 1 | 2 | 1 | NA | NA | NA | NA | NA | NA | 2 | 1 |
| 12_100844 | NA | NA | NA | NA | NA | NA | 1 | 1 | 1 | 1 | 2 | 1 | NA | NA | NA | NA | NA | NA | 2 | 1 |
| 12_100844 | NA | NA | NA | NA | NA | NA | 1 | 1 | 1 | 1 | 1 | 1 | NA | NA | NA | NA | NA | NA | 1 | 1 |
| 12_100877 | 2 | 0 | 4 | 0.5 | 3 | 0.6667 | 8 | 0.25 | 17 | 0.3529 | NA | NA | 4 | 0.5 | 5 | 0.6 | 2 | 0 | 11 | 0.4545 |
| 12_100878 | 2 | 0 | 3 | 0 | 3 | 0.3333 | 7 | 0.1429 | 15 | 0.1333 | NA | NA | 6 | 0.1667 | 5 | 0.4 | 2 | 0 | 13 | 0.2308 |
| 12_100878 | 4 | 0 | 4 | 0.25 | 2 | 0 | 7 | 0.1429 | 17 | 0.1176 | NA | NA | 4 | 0 | 3 | 0.3333 | 2 | 0 | 9 | 0.1111 |
| 12_100878 | 4 | 0.5 | 3 | 0.6667 | 3 | 0.3333 | 8 | 0.5 | 18 | 0.5 | NA | NA | 6 | 0.6667 | 3 | 0.6667 | 1 | 0 | 10 | 0.6 |
| 12_100879 | 4 | 0.25 | 3 | 0 | 2 | 0.5 | 8 | 0.625 | 17 | 0.4118 | NA | NA | 5 | 0.4 | 3 | 0 | 1 | 1 | 9 | 0.3333 |
| 12_100879 | 3 | 0.3333 | 3 | 0.3333 | 3 | 0.3333 | 7 | 0.2857 | 16 | 0.3125 | NA | NA | 3 | 0.3333 | 4 | 0.25 | 1 | 0 | 8 | 0.25 |
| 12_100879 | 3 | 0 | 3 | 0 | 2 | 0 | 7 | 0.2857 | 15 | 0.1333 | NA | NA | 5 | 0.2 | 4 | 0 | 1 | 0 | 10 | 0.1 |
| 12_100879 | 4 | 0.75 | 2 | 0.5 | 3 | 0.3333 | 6 | 0.6667 | 15 | 0.6 | NA | NA | 4 | 0.75 | 4 | 0.75 | NA | NA | 8 | 0.75 |
| 12_100879 | 4 | 1 | 2 | 0.5 | 3 | 1 | 5 | 0.8 | 14 | 0.8571 | NA | NA | 3 | 0.6667 | 4 | 1 | 1 | 1 | 8 | 0.875 |
| 12_100879 | 1 | 1 | NA | NA | 2 | 0 | 5 | 0 | 8 | 0.125 | NA | NA | 3 | 0.6667 | 1 | 0 | 1 | 1 | 5 | 0.6 |
| 12_100879 | 4 | 1 | 2 | 0.5 | 3 | 0.6667 | 7 | 0.5714 | 16 | 0.6875 | NA | NA | 3 | 0.6667 | 4 | 0.75 | 1 | 0 | 8 | 0.625 |
| 12_100879 | 3 | 0 | 2 | 0.5 | 3 | 0.3333 | 7 | 0 | 15 | 0.1333 | NA | NA | 3 | 0.3333 | 3 | 0 | NA | NA | 6 | 0.1667 |
| 12_100879 | 3 | 0.3333 | 2 | 0.5 | 3 | 0 | 5 | 0 | 13 | 0.1538 | NA | NA | 3 | 0.3333 | 4 | 0.25 | 1 | 1 | 8 | 0.375 |
| 12_100879 | 4 | 0.75 | 2 | 0.5 | 3 | 0.3333 | 6 | 0.3333 | 15 | 0.4667 | NA | NA | 3 | 0 | 4 | 0.25 | 1 | 0 | 8 | 0.125 |
| 12_101006 | 1 | 0 | 3 | 0.3333 | 5 | 0 | 5 | 0 | 14 | 0.0714 | 5 | 0 | 6 | 0.1667 | 5 | 0.2 | 5 | 0 | 21 | 0.0952 |
| 12_101006 | 2 | 0.5 | 4 | 0.5 | 4 | 0.75 | 3 | 0.3333 | 13 | 0.5385 | 2 | 0 | 6 | 1 | 5 | 0.8 | 6 | 0.5 | 19 | 0.6842 |
| 12_101006 | 2 | 0 | 6 | 0.8333 | 4 | 0.25 | 9 | 0.3333 | 21 | 0.4286 | 5 | 0.4 | 8 | 0.25 | 5 | 0.4 | 6 | 0.5 | 24 | 0.375 |
| 12_101006 | 1 | 1 | 5 | 0.4 | 4 | 0.25 | 7 | 0.1429 | 17 | 0.2941 | 5 | 0.4 | 7 | 0.8571 | 3 | 0.3333 | 4 | 0.5 | 19 | 0.5789 |
| 12_101006 | 2 | 0.5 | 6 | 0.5 | 5 | 0.8 | 7 | 0.8571 | 20 | 0.7 | 5 | 0.6 | 8 | 0.375 | 5 | 0.8 | 5 | 0.4 | 23 | 0.5217 |
| 12_101006 | 2 | 1 | 6 | 0.8333 | 4 | 0.75 | 8 | 0.5 | 20 | 0.7 | 4 | 0.25 | 6 | 0.8333 | 4 | 0.25 | 5 | 0.8 | 19 | 0.5789 |
| 12_101006 | 1 | 1 | 6 | 1 | 4 | 1 | 7 | 0.8571 | 18 | 0.9444 | 3 | 0.3333 | 5 | 1 | 2 | 0.5 | 6 | 0.8333 | 16 | 0.75 |
| 12_101006 | 2 | 0 | 6 | 0.3333 | 3 | 1 | 6 | 0.5 | 17 | 0.4706 | 4 | 0.75 | 5 | 1 | 4 | 1 | 4 | 0.5 | 17 | 0.8235 |
| 12_101006 | 1 | 1 | 6 | 0.8333 | 4 | 0.75 | 9 | 1 | 20 | 0.9 | 2 | 0.5 | 4 | 1 | 3 | 1 | 6 | 1 | 15 | 0.9333 |
| 12_101006 | 1 | 0 | 6 | 0 | 4 | 0 | 9 | 0.1111 | 20 | 0.05 | 3 | 0 | 6 | 0 | 3 | 0 | 6 | 0.3333 | 18 | 0.1111 |
| 12_101006 | 1 | 1 | 4 | 0.5 | 4 | 0.75 | 6 | 0.6667 | 15 | 0.6667 | 2 | 0.5 | 5 | 0.8 | 3 | 0 | 6 | 0.6667 | 16 | 0.5625 |
| 12_101006 | 1 | 0 | 5 | 0.8 | 2 | 0 | 8 | 0.875 | 16 | 0.6875 | 2 | 1 | 4 | 0.75 | 2 | 1 | 3 | 0.3333 | 11 | 0.7273 |
| 12_101006 | 1 | 1 | 5 | 0.2 | 3 | 0.3333 | 8 | 0 | 17 | 0.1765 | 2 | 0.5 | 5 | 0.2 | 2 | 0 | 5 | 0 | 14 | 0.1429 |
| 12_101008 | 1 | 0 | 6 | 0 | 4 | 0.25 | 7 | 0 | 18 | 0.0556 | 3 | 0 | 6 | 0 | 3 | 0 | 5 | 0.2 | 17 | 0.0588 |
| 12_101021 | 1 | 0 | 5 | 0 | 3 | 0.3333 | 4 | 0 | 13 | 0.0769 | 2 | 0 | 2 | 0 | NA | NA | 2 | 0.5 | 6 | 0.1667 |
| 12_101024 | NA | NA | 5 | 1 | 2 | 1 | 3 | 1 | 10 | 1 | 1 | 1 | 2 | 1 | NA | NA | NA | NA | 3 | 1 |
| 12_101024 | NA | NA | 3 | 1 | 2 | 1 | 3 | 1 | 8 | 1 | NA | NA | NA | NA | NA | NA | 1 | 1 | 1 | 1 |
| 12_101025 | NA | NA | 5 | 0.4 | 3 | 0 | 4 | 0.5 | 12 | 0.3333 | 1 | 0 | 2 | 1 | NA | NA | NA | NA | 3 | 0.6667 |
| 12_101049 | NA | NA | 5 | 0.8 | 3 | 1 | 2 | 1 | 10 | 0.9 | 1 | 1 | 2 | 1 | NA | NA | 1 | 0 | 4 | 0.75 |
| 12_101049 | NA | NA | 3 | 0.3333 | 3 | 0 | 3 | 1 | 9 | 0.4444 | NA | NA | 1 | 1 | NA | NA | NA | NA | 1 | 1 |
| 12_101716 | 5 | 0.6 | NA | NA | NA | NA | 1 | 1 | 6 | 0.6667 | NA | NA | 2 | 1 | NA | NA | 2 | 0.5 | 4 | 0.75 |
| 12_101716 | 5 | 0.8 | NA | NA | NA | NA | 1 | 1 | 6 | 0.8333 | NA | NA | 2 | 1 | NA | NA | 2 | 1 | 4 | 1 |
| 12_101716 | 5 | 0.4 | NA | NA | NA | NA | 1 | 1 | 6 | 0.5 | NA | NA | 2 | 1 | NA | NA | 2 | 1 | 4 | 1 |
| 12_101716 | 3 | 0.3333 | NA | NA | NA | NA | 1 | 0 | 4 | 0.25 | NA | NA | 2 | 0.5 | NA | NA | 2 | 0 | 4 | 0.25 |
| 12_101800 | 2 | 1 | 5 | 0.4 | 2 | 1 | 11 | 1 | 20 | 0.85 | NA | NA | 3 | 1 | 5 | 1 | 4 | 1 | 12 | 1 |

| | | | | | | | | | | | | | | | | | | | | |
|---|---|---|---|---|---|---|---|---|---|---|---|---|---|---|---|---|---|---|---|---|
| 12_1018008 | 3 | 0 | 4 | 0.25 | 2 | 0.5 | 8 | 0.25 | 17 | 0.2353 | 1 | 0 | 3 | 0.3333 | 4 | 0 | 4 | 0.5 | 12 | 0.25 |
| 12_1018010 | 3 | 0.3333 | 3 | 0.6667 | 1 | 0 | 11 | 0.1818 | 18 | 0.2778 | NA | NA | 3 | 0.3333 | 4 | 0.25 | 4 | 0.5 | 11 | 0.3636 |
| 12_1018011 | 2 | 1 | 5 | 1 | 2 | 1 | 9 | 0.8889 | 18 | 0.9444 | 1 | 1 | 3 | 1 | 3 | 1 | 3 | 1 | 10 | 1 |
| 12_1018012 | 2 | 1 | 6 | 1 | 1 | 1 | 8 | 1 | 17 | 1 | 1 | 1 | 2 | 1 | 4 | 0.5 | 4 | 1 | 11 | 0.8182 |
| 12_1018012 | 3 | 0.6667 | 6 | 0.1667 | 1 | 0 | 9 | 0.5556 | 19 | 0.4211 | 1 | 1 | 2 | 0.5 | 3 | 0.6667 | 4 | 0.25 | 10 | 0.5 |
| 12_1018013 | 3 | 0.3333 | 5 | 0.4 | 1 | 0 | 10 | 0.1 | 19 | 0.2105 | 1 | 1 | 3 | 0 | 4 | 0 | 4 | 0 | 12 | 0.0833 |
| 12_1018014 | 3 | 1 | 5 | 0.2 | 1 | 1 | 10 | 0.4 | 19 | 0.4737 | NA | NA | 3 | 1 | 4 | 0.5 | 4 | 0 | 11 | 0.4545 |
| 12_1018015 | 3 | 0.3333 | 5 | 0.2 | 1 | 0 | 9 | 0.2222 | 18 | 0.2222 | 1 | 0 | 2 | 0 | 5 | 0 | 4 | 0.25 | 12 | 0.0833 |
| 12_1018015 | 3 | 0.3333 | 6 | 0 | NA | NA | 10 | 0.1 | 19 | 0.1053 | NA | NA | 2 | 0.5 | 4 | 0.25 | 3 | 0.6667 | 9 | 0.4444 |
| 12_1018016 | 3 | 1 | 5 | 0 | NA | NA | 6 | 0.8333 | 14 | 0.5714 | 1 | 1 | 1 | 0 | 3 | 0.6667 | 4 | 0 | 9 | 0.3333 |
| 12_1018018 | 2 | 0 | 6 | 0.1667 | NA | NA | 8 | 0.375 | 16 | 0.25 | 1 | 1 | 1 | 0 | 4 | 0 | 4 | 0.25 | 10 | 0.2 |
| 12_1018037 | NA | NA | 3 | 0.6667 | NA | NA | 4 | 0.25 | 7 | 0.4286 | NA | NA | 1 | 0 | 3 | 0 | 3 | 0.3333 | 7 | 0.1429 |
| 12_1018037 | 2 | 0.5 | 2 | 1 | NA | NA | 4 | 0.25 | 8 | 0.5 | NA | NA | 1 | 0 | 2 | 0.5 | 3 | 0.3333 | 6 | 0.3333 |
| 12_1018037 | 2 | 0 | 3 | 0.3333 | NA | NA | 4 | 0 | 9 | 0.1111 | NA | NA | 1 | 0 | 2 | 0.5 | 3 | 0 | 6 | 0.1667 |
| 12_1018038 | 2 | 0.5 | 2 | 0 | NA | NA | 4 | 0.5 | 8 | 0.375 | NA | NA | 1 | 0 | 2 | 0.5 | 3 | 0 | 6 | 0.1667 |
| 12_1018039 | 2 | 0.5 | 2 | 0.5 | NA | NA | 4 | 0.5 | 8 | 0.5 | NA | NA | NA | NA | 2 | 1 | 2 | 0 | 4 | 0.5 |
| 12_1018497 | 1 | 0 | 1 | 1 | NA | NA | 2 | 1 | 4 | 0.75 | NA | NA | NA | NA | NA | NA | 2 | 0.5 | 2 | 0.5 |
| 12_1018499 | NA | NA | 1 | 1 | NA | NA | 2 | 1 | 3 | 1 | NA | NA | NA | NA | NA | NA | 1 | 1 | 1 | 1 |
| 12_1018502 | 1 | 1 | NA | NA | NA | NA | NA | NA | 1 | 1 | NA | NA | NA | NA | NA | NA | 1 | 1 | 1 | 1 |
| 12_1018522 | 1 | 1 | 1 | 0 | NA | NA | NA | NA | 2 | 0.5 | NA | NA | NA | NA | NA | NA | 1 | 1 | 1 | 1 |
| 12_1018853 | 3 | 0 | 2 | 0 | 7 | 0.2857 | 3 | 0 | 15 | 0.1333 | 2 | 0.5 | 3 | 0.6667 | 5 | 0.6 | 4 | 0.75 | 14 | 0.6429 |
| 12_1018854 | 3 | 1 | 2 | 0 | 4 | 1 | 3 | 1 | 12 | 0.8333 | 1 | 1 | 3 | 0.6667 | 4 | 1 | 4 | 0.75 | 12 | 0.8333 |
| 12_1018854 | 2 | 0 | 1 | 0 | 1 | 0 | 1 | 1 | 5 | 0.2 | 1 | 1 | 1 | 0 | 3 | 1 | 2 | 0 | 7 | 0.5714 |
| 12_1019562 | 2 | 1 | 1 | 1 | 1 | 1 | 2 | 0.5 | 6 | 0.8333 | 2 | 0 | 1 | 1 | 1 | 0 | 4 | 0.25 | 8 | 0.25 |
| 12_1019562 | 2 | 0 | 1 | 0 | 1 | 0 | 2 | 0.5 | 6 | 0.1667 | 2 | 0 | 1 | 0 | 2 | 0.5 | 4 | 0 | 9 | 0.1111 |
| 12_1019669 | 3 | 0 | 1 | 0 | 2 | 0.5 | 3 | 0 | 9 | 0.1111 | 2 | 0 | 1 | 0 | 2 | 0 | 7 | 0.4286 | 12 | 0.25 |
| 12_1019694 | 3 | 0.3333 | 2 | 0.5 | 3 | 0.3333 | 2 | 0 | 10 | 0.3 | 3 | 0 | 1 | 0 | 4 | 0 | 9 | 0.1111 | 17 | 0.0588 |
| 12_1019694 | 2 | 0.5 | 1 | 1 | 2 | 0 | 2 | 0.5 | 7 | 0.4286 | 4 | 0.25 | 1 | 1 | 3 | 0 | 5 | 0.8 | 13 | 0.4615 |
| 12_1019694 | 3 | 0.3333 | 2 | 1 | 3 | 0 | 3 | 0.6667 | 11 | 0.4545 | 4 | 0.75 | 1 | 1 | 4 | 1 | 8 | 0.625 | 17 | 0.7647 |
| 12_1019695 | 3 | 0 | 3 | 0.6667 | 3 | 0 | 3 | 0 | 12 | 0.1667 | 3 | 0.3333 | 1 | 0 | 5 | 0 | 9 | 0.3333 | 18 | 0.2222 |
| 12_1019698 | 3 | 0.3333 | 3 | 0.6667 | 1 | 0 | 3 | 1 | 10 | 0.6 | 2 | 0.5 | 1 | 0 | 3 | 0.6667 | 6 | 0.8333 | 12 | 0.6667 |
| 12_1019699 | 4 | 0.5 | 3 | 0 | 1 | 0 | 3 | 0.6667 | 11 | 0.3636 | 4 | 0.5 | 1 | 0 | 4 | 0.5 | 8 | 0.375 | 17 | 0.4118 |
| 12_1019699 | 3 | 0.6667 | 4 | 0.5 | 2 | 0 | 3 | 0.3333 | 12 | 0.4167 | 3 | 0.6667 | 1 | 1 | 5 | 0.8 | 6 | 0.8333 | 15 | 0.8 |
| 12_1019700 | 1 | 0 | 4 | 0.25 | 1 | 1 | 1 | 1 | 7 | 0.4286 | 3 | 0 | 1 | 0 | 5 | 0.2 | 2 | 0.5 | 11 | 0.1818 |
| 12_1019703 | 2 | 1 | 3 | 0.6667 | NA | NA | 1 | 0 | 6 | 0.6667 | 2 | 0.5 | 1 | 0 | 4 | 1 | 3 | 0.6667 | 10 | 0.7 |
| 12_1019712 | 2 | 0.5 | 3 | 0.3333 | 1 | 0 | 1 | 0 | 7 | 0.2857 | 5 | 0 | NA | NA | 4 | 0.5 | 4 | 0 | 13 | 0.1538 |
| 12_1019713 | 2 | 0 | 3 | 0 | 2 | 0.5 | NA | NA | 7 | 0.1429 | 6 | 0.3333 | NA | NA | 5 | 0 | 4 | 0.25 | 15 | 0.2 |
| 12_1019714 | 2 | 0.5 | 3 | 0 | 2 | 1 | 1 | 0 | 8 | 0.375 | 6 | 0.1667 | NA | NA | 4 | 0 | 2 | 0 | 12 | 0.0833 |
| 12_1019715 | 3 | 0.3333 | 4 | 0.25 | 3 | 0 | NA | NA | 10 | 0.2 | 4 | 1 | NA | NA | 4 | 0 | 2 | 0 | 10 | 0.4 |
| 12_1020941 | 1 | 1 | 1 | 1 | NA | NA | 2 | 0.5 | 4 | 0.75 | 1 | 0 | 1 | 1 | NA | NA | 1 | 0 | 3 | 0.3333 |
| 12_1020944 | 1 | 1 | NA | NA | NA | NA | 1 | 1 | 2 | 1 | 1 | 1 | 1 | 1 | NA | NA | 1 | 1 | 3 | 1 |
| 12_1020945 | 1 | 1 | NA | NA | NA | NA | NA | NA | 1 | 1 | 1 | 0 | 1 | 1 | NA | NA | 1 | 1 | 3 | 0.6667 |
| 12_1020946 | 1 | 1 | 1 | 1 | NA | NA | 1 | 1 | 3 | 1 | 1 | 1 | 1 | 1 | NA | NA | NA | NA | 2 | 1 |
| 12_1020950 | NA | NA | NA | NA | NA | NA | 1 | 1 | 1 | 1 | 1 | 1 | 1 | 0 | NA | NA | 1 | 1 | 3 | 0.6667 |
| 12_1020950 | NA | NA | NA | NA | NA | NA | 1 | 1 | 1 | 1 | 1 | 1 | 1 | 1 | NA | NA | NA | NA | 2 | 1 |
| 12_1020954 | 4 | 0 | NA | NA | 1 | 1 | 2 | 0 | 7 | 0.1429 | NA | NA | 5 | 0.4 | 1 | 0 | 4 | 0.25 | 10 | 0.3 |
| 12_1020956 | 5 | 0.4 | NA | NA | 2 | 0 | 2 | 0.5 | 9 | 0.3333 | NA | NA | 5 | 0.4 | 2 | 0.5 | 3 | 0.6667 | 10 | 0.5 |
| 12_1020956 | NA | NA | NA | NA | NA | NA | 1 | 1 | 1 | 1 | 1 | 1 | 1 | 1 | NA | NA | NA | NA | 2 | 1 |
| 12_1020958 | NA | NA | NA | NA | NA | NA | 1 | 1 | 1 | 1 | 1 | 0 | 1 | 1 | NA | NA | NA | NA | 2 | 0.5 |
| 12_1022327 | 7 | 0 | 3 | 0.3333 | 4 | 0 | 4 | 0.5 | 18 | 0.1667 | 1 | 0 | 8 | 0 | 4 | 0 | 6 | 0.1667 | 19 | 0.0526 |
| 12_1022327 | 8 | 0.125 | 2 | 0.5 | 6 | 0 | 4 | 0 | 20 | 0.1 | 1 | 1 | 6 | 0.3333 | 5 | 0.2 | 8 | 0.25 | 20 | 0.3 |
| 12_1022329 | 10 | 0.8 | 3 | 0.6667 | 7 | 0.8571 | 3 | 1 | 23 | 0.8261 | 1 | 1 | 8 | 1 | 6 | 1 | 9 | 0.8889 | 24 | 0.9583 |
| 12_1022330 | 9 | 0.8889 | 3 | 1 | 8 | 0.625 | 5 | 1 | 25 | 0.84 | 1 | 1 | 8 | 0.875 | 5 | 0.8 | 9 | 0.8889 | 23 | 0.8696 |
| 12_1022331 | 9 | 0.2222 | 2 | 0.5 | 6 | 0 | 4 | 0.25 | 21 | 0.1905 | 1 | 0 | 5 | 0 | 5 | 0.4 | 9 | 0.2222 | 20 | 0.2 |
| 12_1022331 | 12 | 0.8333 | 2 | 1 | 8 | 0.75 | 5 | 1 | 27 | 0.8519 | 1 | 1 | 7 | 1 | 7 | 0.8571 | 8 | 0.75 | 23 | 0.8696 |
| 12_1022331 | 12 | 0.3333 | 3 | 0 | 8 | 0.375 | 5 | 0.2 | 28 | 0.2857 | 1 | 0 | 7 | 0.4286 | 7 | 0.2857 | 9 | 0.3333 | 24 | 0.3333 |
| 12_1022331 | 7 | 0.1429 | 2 | 0 | 8 | 0.125 | 4 | 0.25 | 21 | 0.1429 | 1 | 0 | 8 | 0.25 | 8 | 0.125 | 6 | 0.1667 | 23 | 0.1739 |
| 12_1022332 | 11 | 0.0909 | 2 | 0 | 8 | 0.125 | 4 | 0.25 | 25 | 0.12 | 1 | 0 | 8 | 0.25 | 8 | 0.125 | 10 | 0.1 | 27 | 0.1481 |
| 12_1022333 | 10 | 0.5 | 2 | 0.5 | 6 | 0.1667 | 4 | 0.75 | 22 | 0.4545 | 1 | 1 | 6 | 0.5 | 8 | 0.5 | 8 | 0.375 | 23 | 0.4783 |
| 12_1022333 | 8 | 0.5 | 2 | 1 | 7 | 0.7143 | 4 | 1 | 21 | 0.7143 | NA | NA | 5 | 0.8 | 8 | 0.875 | 9 | 0.5556 | 22 | 0.7273 |
| 12_1022334 | 8 | 0.125 | 2 | 0.5 | 1 | 0 | 3 | 0 | 14 | 0.1429 | 1 | 0 | 5 | 0.2 | 6 | 0.1667 | 5 | 0 | 17 | 0.1176 |
| 12_1022335 | 8 | 0.625 | 2 | 1 | 1 | 1 | 4 | 1 | 15 | 0.8 | 1 | 0 | 4 | 1 | 6 | 0.8333 | 5 | 0.8 | 16 | 0.8125 |
| 12_1022336 | 7 | 0.1429 | 2 | 0 | 1 | 1 | 4 | 0.5 | 14 | 0.2857 | 1 | 0 | 5 | 0.2 | 5 | 0 | 6 | 0.1667 | 17 | 0.1176 |
| 12_1022337 | 5 | 0.2 | 2 | 0.5 | NA | NA | 4 | 1 | 11 | 0.5455 | 1 | 1 | 5 | 0.2 | 4 | 0.75 | 5 | 0.2 | 15 | 0.4 |
| 12_1022338 | 6 | 0.3333 | 2 | 0 | 2 | 0.5 | 4 | 0.25 | 14 | 0.2857 | 1 | 1 | 6 | 0.1667 | 6 | 0.5 | 6 | 0.6667 | 19 | 0.4737 |

| | | | | | | | | | | | | | | | | | | | | |
|---|---|---|---|---|---|---|---|---|---|---|---|---|---|---|---|---|---|---|---|---|
| 12_1022339 | 7 | 0.1429 | 1 | 1 | 2 | 0 | 4 | 0 | 14 | 0.1429 | 1 | 0 | 3 | 0.3333 | 6 | 0.1667 | 5 | 0.2 | 15 | 0.2 |
| 12_1022354 | 9 | 0.2222 | 4 | 0.5 | 4 | 0 | 5 | 0.2 | 22 | 0.2273 | 1 | 0 | 6 | 0 | 8 | 0.125 | 5 | 0 | 20 | 0.05 |
| 12_1022375 | 15 | 0.1333 | 3 | 0.6667 | 9 | 0.2222 | 9 | 0.1111 | 36 | 0.1944 | 9 | 0.1111 | 11 | 0.1818 | 12 | 0 | 6 | 0.3333 | 38 | 0.1316 |
| 12_1022376 | 17 | 0.2353 | 4 | 0.25 | 10 | 0.2 | 10 | 0.3 | 41 | 0.2439 | 7 | 0.1429 | 12 | 0.1667 | 9 | 0.1111 | 6 | 0 | 34 | 0.1176 |
| 12_1022377 | 21 | 0.4286 | 4 | 0.5 | 11 | 0.4545 | 12 | 0.5 | 48 | 0.4583 | 8 | 0.625 | 14 | 0.5714 | 13 | 0.6154 | 4 | 0.75 | 39 | 0.6154 |
| 12_1022377 | 17 | 0.1176 | 5 | 0 | 11 | 0.1818 | 9 | 0.2222 | 42 | 0.1429 | 7 | 0 | 14 | 0.1429 | 14 | 0.2143 | 5 | 0.4 | 40 | 0.175 |
| 12_1022378 | 21 | 0.3333 | 4 | 0.25 | 10 | 0.1 | 13 | 0.4615 | 48 | 0.3125 | 8 | 0.375 | 13 | 0.2308 | 15 | 0.2667 | 8 | 0.25 | 44 | 0.2727 |
| 12_1022379 | 21 | 0.0952 | 5 | 0.4 | 11 | 0.0909 | 11 | 0.0909 | 48 | 0.125 | 5 | 0 | 15 | 0.1333 | 14 | 0.1429 | 6 | 0 | 40 | 0.1 |
| 12_1022380 | 22 | 0.2273 | 4 | 0.25 | 12 | 0.1667 | 12 | 0 | 50 | 0.16 | 5 | 0.2 | 18 | 0.1667 | 15 | 0.2667 | 6 | 0.5 | 44 | 0.25 |
| 12_1022381 | 25 | 0.88 | 4 | 1 | 12 | 1 | 14 | 0.8571 | 55 | 0.9091 | 9 | 1 | 17 | 0.9412 | 15 | 0.9333 | 6 | 1 | 47 | 0.9574 |
| 12_1022382 | 21 | 0.381 | 8 | 0.25 | 12 | 0.3333 | 12 | 0.3333 | 53 | 0.3396 | 6 | 0.3333 | 15 | 0.4 | 14 | 0.3571 | 8 | 0.5 | 43 | 0.3953 |
| 12_1022382 | 26 | 0.1538 | 7 | 0 | 13 | 0.2308 | 15 | 0.0667 | 61 | 0.1311 | 9 | 0.2222 | 17 | 0.2353 | 16 | 0.125 | 7 | 0.1429 | 49 | 0.1837 |
| 12_1022386 | 9 | 0 | 2 | 0.5 | 3 | 0 | 5 | 0 | 19 | 0.0526 | 3 | 0 | 8 | 0.125 | 9 | 0.1111 | 3 | 0 | 23 | 0.087 |
| 12_1022391 | 3 | 0 | 1 | 0 | 1 | 1 | 3 | 0 | 8 | 0.125 | 2 | 0 | 4 | 0 | 5 | 0.2 | 2 | 0.5 | 13 | 0.1538 |
| 12_1022393 | 4 | 1 | 1 | 1 | 2 | 0.5 | 3 | 1 | 10 | 0.9 | 1 | 1 | 6 | 0.8333 | 5 | 0.8 | 3 | 1 | 15 | 0.8667 |
| 12_1022395 | 4 | 0.5 | 2 | 0 | 3 | 0 | 4 | 0.75 | 13 | 0.3846 | 1 | 0 | 6 | 0.3333 | 6 | 0.1667 | 3 | 0 | 16 | 0.1875 |
| 12_1023353 | NA | NA | 2 | 1 | NA | NA | NA | NA | 2 | 1 | 1 | 1 | NA | NA | NA | NA | NA | NA | 1 | 1 |
| 12_1023561 | 1 | 0 | 3 | 0.3333 | NA | NA | 1 | 0 | 5 | 0.2 | 1 | 0 | 1 | 1 | NA | NA | NA | NA | 2 | 0.5 |
| 12_1023563 | 1 | 1 | 3 | 0.3333 | NA | NA | 1 | 0 | 5 | 0.4 | 1 | 1 | 1 | 0 | NA | NA | NA | NA | 2 | 0.5 |
| 12_1023564 | 1 | 1 | 3 | 1 | NA | NA | 1 | 1 | 5 | 1 | 1 | 1 | NA | NA | NA | NA | NA | NA | 1 | 1 |
| 12_1023567 | 1 | 1 | 2 | 1 | NA | NA | 1 | 1 | 4 | 1 | 1 | 1 | 1 | 1 | NA | NA | NA | NA | 2 | 1 |
| 12_1023569 | 1 | 1 | 3 | 1 | NA | NA | 1 | 1 | 5 | 1 | 1 | 1 | 1 | 1 | NA | NA | NA | NA | 2 | 1 |
| 12_1023569 | 1 | 1 | 3 | 1 | NA | NA | 1 | 1 | 5 | 1 | 1 | 1 | 1 | 1 | NA | NA | NA | NA | 2 | 1 |
| 12_1023606 | 24 | 0.2083 | 12 | 0.25 | 22 | 0.2727 | 19 | 0.3684 | 77 | 0.2727 | 14 | 0.2857 | 14 | 0.1429 | 17 | 0.2941 | 19 | 0.3684 | 64 | 0.2812 |
| 12_1023607 | 15 | 0.7333 | 9 | 0.6667 | 17 | 0.6471 | 17 | 0.7059 | 58 | 0.6897 | 13 | 0.8462 | 13 | 0.6154 | 14 | 0.7143 | 17 | 0.5294 | 57 | 0.6667 |
| 12_1024384 | 2 | 0 | 1 | 1 | NA | NA | 1 | 1 | 4 | 0.5 | NA | NA | 1 | 1 | NA | NA | 1 | 1 | 2 | 1 |
| 12_1024384 | 2 | 1 | 2 | 1 | NA | NA | 2 | 1 | 6 | 1 | NA | NA | NA | NA | 1 | 1 | 3 | 0.3333 | 4 | 0.5 |
| 12_1024384 | 1 | 0 | 2 | 1 | 1 | 0 | 1 | 0 | 5 | 0.4 | NA | NA | 1 | 1 | 1 | 1 | 3 | 1 | 5 | 1 |
| 12_1024385 | 2 | 0.5 | 3 | 0.6667 | 1 | 1 | 1 | 1 | 7 | 0.7143 | NA | NA | NA | NA | 2 | 1 | 2 | 1 | 4 | 1 |
| 12_1025280 | 19 | 0.5789 | 15 | 0.6667 | 13 | 0.4615 | 14 | 0.6429 | 61 | 0.5902 | 23 | 0.6087 | 11 | 0.5455 | 8 | 0.5 | 10 | 0.7 | 52 | 0.5962 |
| 12_1025280 | 24 | 0.125 | 21 | 0 | 18 | 0.1111 | 22 | 0.1364 | 85 | 0.0941 | 23 | 0.087 | 20 | 0 | 9 | 0.2222 | 15 | 0.0667 | 67 | 0.0746 |
| 12_1025281 | 22 | 0.0909 | 18 | 0.0556 | 17 | 0.1765 | 23 | 0.087 | 80 | 0.1 | 19 | 0.1579 | 19 | 0 | 9 | 0 | 16 | 0.0625 | 63 | 0.0635 |
| 12_1025288 | 23 | 0.2174 | 23 | 0.0435 | 19 | 0.1053 | 23 | 0.1304 | 88 | 0.125 | 25 | 0.08 | 20 | 0.2 | 11 | 0.0909 | 15 | 0.2667 | 71 | 0.1549 |
| 12_1025289 | 15 | 0 | 13 | 0.2308 | 15 | 0 | 14 | 0 | 57 | 0.0526 | 18 | 0.0556 | 17 | 0.1765 | 9 | 0.1111 | 13 | 0.0769 | 57 | 0.1053 |
| 12_1025290 | 23 | 0.1304 | 24 | 0.0833 | 24 | 0 | 20 | 0.2 | 91 | 0.0989 | 21 | 0.1429 | 22 | 0.1818 | 13 | 0.0769 | 13 | 0 | 69 | 0.1159 |
| 12_1025291 | 29 | 0.1724 | 30 | 0.1667 | 21 | 0.0476 | 25 | 0.16 | 105 | 0.1429 | 27 | 0.037 | 20 | 0.2 | 13 | 0.0769 | 14 | 0 | 74 | 0.0811 |
| 12_1025292 | 27 | 0.1481 | 34 | 0.2353 | 28 | 0.1786 | 27 | 0.1111 | 116 | 0.1724 | 30 | 0.2 | 20 | 0.2 | 16 | 0.125 | 16 | 0.0625 | 82 | 0.1585 |
| 12_1025310 | 25 | 0.2 | 33 | 0.1515 | 23 | 0.1304 | 25 | 0.16 | 106 | 0.1604 | 21 | 0.0476 | 17 | 0.2941 | 13 | 0.1538 | 20 | 0.25 | 71 | 0.1831 |
| 12_1025312 | 26 | 0 | 41 | 0.0976 | 29 | 0.1379 | 30 | 0.1 | 126 | 0.0873 | 32 | 0.0625 | 24 | 0.0833 | 18 | 0.2778 | 20 | 0.05 | 94 | 0.1064 |
| 12_1025349 | 10 | 0.2 | 19 | 0.2105 | 10 | 0.1 | 8 | 0.375 | 47 | 0.2128 | 15 | 0.3333 | 4 | 0 | 7 | 0.2857 | 7 | 0.2857 | 33 | 0.2727 |
| 12_1025349 | 9 | 0.1111 | 14 | 0.2857 | 10 | 0.4 | 7 | 0.4286 | 40 | 0.3 | 13 | 0.3846 | 3 | 0 | 8 | 0 | 7 | 0.1429 | 31 | 0.1935 |
| 12_1025352 | 8 | 0 | 16 | 0.0625 | 11 | 0.0909 | 7 | 0.1429 | 42 | 0.0714 | 12 | 0.0833 | 4 | 0 | 6 | 0.1667 | 6 | 0 | 28 | 0.0714 |
| 12_1025355 | 7 | 0.2857 | 12 | 0.25 | 9 | 0.4444 | 8 | 0.375 | 36 | 0.3333 | 10 | 0.3 | 4 | 0.5 | 5 | 0.4 | 3 | 0.3333 | 22 | 0.3636 |
| 12_1025355 | 8 | 0.25 | 13 | 0 | 9 | 0.1111 | 6 | 0 | 36 | 0.0833 | 10 | 0.1 | 3 | 0 | 4 | 0.25 | 3 | 0 | 20 | 0.1 |
| 12_1025363 | 5 | 1 | 6 | 0.8333 | 7 | 0.7143 | 7 | 0.5714 | 25 | 0.76 | 8 | 0.875 | 2 | 0.5 | 3 | 0.3333 | 2 | 1 | 15 | 0.7333 |
| 12_1025363 | 8 | 0.5 | 9 | 0.3333 | 7 | 0.5714 | 7 | 0.7143 | 31 | 0.5161 | 8 | 1 | 3 | 1 | 4 | 0 | 3 | 0 | 18 | 0.6111 |
| 12_1025364 | 8 | 0.75 | 9 | 1 | 7 | 1 | 6 | 0.8333 | 30 | 0.9 | 7 | 1 | 3 | 1 | 5 | 1 | 2 | 0.5 | 17 | 0.9412 |
| 12_1025365 | 8 | 0.625 | 8 | 0.5 | 4 | 0.5 | 4 | 0.5 | 24 | 0.5417 | 7 | 0.2857 | 2 | 0 | 3 | 0.3333 | 3 | 0.3333 | 15 | 0.2667 |
| 12_1025365 | 8 | 0.5 | 9 | 0.6667 | 7 | 0.4286 | 6 | 0.6667 | 30 | 0.5667 | 7 | 0.2857 | 3 | 0 | 4 | 0.5 | 3 | 0.3333 | 17 | 0.2941 |
| 12_1027083 | 2 | 0 | 7 | 0.1429 | 10 | 0.1 | 4 | 0.25 | 23 | 0.1304 | 3 | 0 | 2 | 0 | 2 | 0.5 | 7 | 0.2857 | 14 | 0.2143 |
| 12_1027084 | 3 | 0.3333 | 9 | 0.1111 | 10 | 0 | 4 | 0 | 26 | 0.0769 | 5 | 0 | 3 | 0.3333 | 3 | 0 | 6 | 0 | 17 | 0.0588 |
| 12_1027085 | 3 | 0.6667 | 8 | 0.375 | 10 | 0.3 | 4 | 0.5 | 25 | 0.4 | 4 | 0.25 | 3 | 0 | 3 | 0.3333 | 6 | 0.3333 | 16 | 0.25 |
| 12_1027086 | 3 | 0 | 8 | 0.25 | 9 | 0 | 4 | 0 | 24 | 0.0833 | 5 | 0.4 | 3 | 0 | 3 | 0 | 6 | 0 | 17 | 0.1176 |
| 12_1027087 | 3 | 0.3333 | 6 | 0.8333 | 8 | 0.75 | 4 | 0.25 | 21 | 0.619 | 4 | 1 | 3 | 0.3333 | 3 | 0.6667 | 3 | 0.3333 | 13 | 0.6154 |
| 12_1027087 | 2 | 1 | 6 | 0.6667 | 8 | 0.875 | 4 | 0.5 | 20 | 0.75 | 4 | 1 | 2 | 1 | 2 | 1 | 6 | 0.6667 | 14 | 0.8571 |
| 12_1027088 | 3 | 0 | 8 | 0.375 | 6 | 0.1667 | 3 | 0 | 20 | 0.2 | 5 | 0 | 2 | 0 | 1 | 0 | 5 | 0.2 | 13 | 0.0769 |
| 12_1027092 | 3 | 1 | 6 | 1 | 4 | 1 | 2 | 0.5 | 15 | 0.9333 | 3 | 1 | 3 | 0.6667 | 2 | 1 | 4 | 1 | 12 | 0.9167 |
| 12_1027093 | 3 | 1 | 7 | 1 | 3 | 1 | 3 | 1 | 16 | 1 | 5 | 1 | 1 | 1 | 3 | 1 | 3 | 1 | 12 | 1 |
| 12_1027093 | 2 | 0.5 | 7 | 0.1429 | 4 | 0.25 | 1 | 0 | 14 | 0.2143 | 4 | 0 | 2 | 0.5 | 2 | 0.5 | 4 | 0.5 | 12 | 0.3333 |
| 12_1027093 | 3 | 0.3333 | 4 | 1 | 3 | 1 | 3 | 1 | 13 | 0.8462 | 4 | 1 | 3 | 1 | 3 | 1 | 4 | 1 | 14 | 1 |
| 12_1027095 | 2 | 0 | 5 | 0.2 | 4 | 0 | 3 | 0.3333 | 14 | 0.1429 | 3 | 0 | 3 | 0.6667 | 3 | 0 | 5 | 0.2 | 14 | 0.2143 |
| 12_1027095 | 3 | 0.3333 | 6 | 0.3333 | 4 | 0.25 | 3 | 0.6667 | 16 | 0.375 | 4 | 0.5 | 3 | 0.3333 | 3 | 0.6667 | 5 | 0.4 | 15 | 0.4667 |
| 12_1027109 | 3 | 1 | NA | NA | 2 | 1 | NA | NA | 5 | 1 | NA | NA | 5 | 0.8 | 4 | 1 | 3 | 0.6667 | 12 | 0.8333 |
| 12_1027109 | 2 | 1 | NA | NA | 2 | 0.5 | NA | NA | 4 | 0.75 | NA | NA | 5 | 0.4 | 3 | 0.3333 | 2 | 0.5 | 10 | 0.4 |
| 12_1027110 | 3 | 1 | NA | NA | 3 | 1 | NA | NA | 6 | 1 | NA | NA | 4 | 1 | 4 | 1 | 4 | 1 | 12 | 1 |

| | | | | | | | | | | | | | | | | | | | | |
|---|---|---|---|---|---|---|---|---|---|---|---|---|---|---|---|---|---|---|---|---|
| 12_102711( | 2 | 1 | NA | NA | 3 | 1 | NA | NA | 5 | 1 | NA | NA | 5 | 1 | 4 | 1 | 4 | 1 | 13 | 1 |
| 12_102711' | 1 | 1 | NA | NA | 2 | 0 | NA | NA | 3 | 0.3333 | NA | NA | 3 | 0.3333 | 2 | 0 | 4 | 0.25 | 9 | 0.2222 |
| 12_102711: | 3 | 0.6667 | NA | NA | 3 | 0 | NA | NA | 6 | 0.3333 | NA | NA | 4 | 0.5 | 3 | 0 | 3 | 0 | 10 | 0.2 |
| 12_102711: | 3 | 1 | NA | NA | 1 | 1 | NA | NA | 4 | 1 | NA | NA | 2 | 0.5 | 2 | 1 | 3 | 0.6667 | 7 | 0.7143 |
| 12_102711: | 3 | 1 | NA | NA | 3 | 0.6667 | NA | NA | 6 | 0.8333 | NA | NA | 2 | 1 | 3 | 1 | 3 | 0.6667 | 8 | 0.875 |
| 12_102711: | 2 | 1 | NA | NA | 3 | 1 | NA | NA | 5 | 1 | NA | NA | 3 | 0.6667 | 3 | 1 | 4 | 1 | 10 | 0.9 |
| 12_102711· | 2 | 1 | NA | NA | 1 | 1 | NA | NA | 3 | 1 | NA | NA | 4 | 0.75 | 4 | 1 | 4 | 1 | 12 | 0.9167 |
| 12_102720( | NA | NA | 3 | 1 | 1 | 0 | 1 | 1 | 5 | 0.8 | 2 | 1 | 3 | 0.6667 | 3 | 1 | NA | NA | 8 | 0.875 |
| 12_102720( | NA | NA | 4 | 0 | 1 | 0 | 1 | 1 | 6 | 0.1667 | 3 | 0.3333 | 2 | 0 | 2 | 0 | 1 | 0 | 8 | 0.125 |
| 12_102720' | NA | NA | 4 | 0.75 | 1 | 1 | NA | NA | 5 | 0.8 | 2 | 1 | 3 | 0.6667 | 2 | 1 | NA | NA | 7 | 0.8571 |
| 12_102720' | NA | NA | 4 | 0.25 | 1 | 1 | 1 | 0 | 6 | 0.3333 | 3 | 0 | 2 | 0.5 | 3 | 0.3333 | 1 | 0 | 9 | 0.2222 |
| 12_102720: | NA | NA | 4 | 0.25 | 1 | 0 | 1 | 0 | 6 | 0.1667 | 2 | 0.5 | 3 | 1 | 2 | 0.5 | 1 | 0 | 8 | 0.625 |
| 12_102720: | NA | NA | 4 | 0.75 | 1 | 1 | 1 | 1 | 6 | 0.8333 | 3 | 1 | 3 | 1 | 3 | 1 | 1 | 1 | 10 | 1 |
| 12_102720: | NA | NA | 3 | 0.6667 | 1 | 1 | 1 | 1 | 5 | 0.8 | 3 | 1 | 3 | 1 | 2 | 1 | 1 | 1 | 9 | 1 |
| 12_102720: | NA | NA | 3 | 0.3333 | 1 | 0 | 1 | 1 | 5 | 0.4 | 2 | 0.5 | 3 | 0.3333 | 3 | 0.3333 | 1 | 0 | 9 | 0.3333 |
| 12_102742' | 7 | 0.7143 | 2 | 0 | 4 | 0.25 | 7 | 0.1429 | 20 | 0.35 | 7 | 0 | 12 | 0.3333 | 5 | 0.4 | 4 | 0.25 | 28 | 0.25 |
| 12_102742: | 5 | 0 | 2 | 0 | 5 | 0.2 | 8 | 0.375 | 20 | 0.2 | 8 | 0.25 | 11 | 0.1818 | 5 | 0.2 | 4 | 0 | 28 | 0.1786 |
| 12_102742! | 6 | 0.5 | 2 | 0 | 4 | 0.75 | 9 | 0.5556 | 21 | 0.5238 | 7 | 0.2857 | 10 | 0.7 | 5 | 0.4 | 3 | 0.6667 | 25 | 0.52 |
| 12_102742{ | 7 | 0.2857 | 2 | 0.5 | 3 | 0 | 7 | 0.4286 | 19 | 0.3158 | 8 | 0.125 | 11 | 0.2727 | 6 | 0.1667 | 6 | 0.1667 | 31 | 0.1935 |
| 12_102742! | 5 | 0.2 | 2 | 0 | 4 | 0 | 8 | 0 | 19 | 0.0526 | 8 | 0 | 7 | 0.4286 | 5 | 0.4 | 4 | 0.25 | 24 | 0.25 |
| 12_102749( | 4 | 0.5 | 3 | 0 | 3 | 0.3333 | 3 | 0 | 13 | 0.2308 | NA | NA | 4 | 0 | 2 | 0.5 | 1 | 0 | 7 | 0.1429 |
| 12_102750( | 1 | 1 | 3 | 0.6667 | 1 | 0 | 3 | 0 | 8 | 0.375 | NA | NA | 1 | 0 | 2 | 0.5 | 1 | 1 | 4 | 0.5 |
| 12_102751( | 1 | 1 | 1 | 0 | NA | NA | 3 | 0.6667 | 5 | 0.6 | NA | NA | NA | NA | 1 | 1 | 1 | 1 | 2 | 1 |
| 12_102751! | 1 | 0 | 2 | 1 | NA | NA | 3 | 0.3333 | 6 | 0.5 | NA | NA | NA | NA | 1 | 1 | 1 | 0 | 2 | 0.5 |
| 12_102751! | 1 | 1 | 2 | 0.5 | NA | NA | 3 | 1 | 6 | 0.8333 | NA | NA | NA | NA | 1 | 1 | 1 | 1 | 2 | 1 |
| 12_102751! | NA | NA | 2 | 0.5 | NA | NA | 2 | 0 | 4 | 0.25 | NA | NA | NA | NA | 1 | 1 | NA | NA | 1 | 1 |
| 12_102751! | NA | NA | 2 | 0.5 | NA | NA | 3 | 1 | 5 | 0.8 | NA | NA | NA | NA | 1 | 1 | 1 | 1 | 2 | 1 |
| 12_102751( | 1 | 1 | 1 | 1 | NA | NA | 3 | 0.6667 | 5 | 0.8 | NA | NA | NA | NA | 1 | 1 | 1 | 1 | 2 | 1 |
| 12_102751( | 1 | 1 | 2 | 0.5 | NA | NA | 2 | 0.5 | 5 | 0.6 | NA | NA | NA | NA | 1 | 1 | NA | NA | 1 | 1 |
| 12_102751{ | NA | NA | 2 | 0.5 | NA | NA | 2 | 1 | 4 | 0.75 | NA | NA | NA | NA | 1 | 1 | 1 | 1 | 2 | 1 |
| 12_102751! | 1 | 1 | 2 | 1 | NA | NA | 3 | 0.6667 | 6 | 0.8333 | NA | NA | NA | NA | 1 | 1 | 1 | 0 | 2 | 0.5 |
| 12_102752( | 1 | 1 | 2 | 1 | NA | NA | 3 | 1 | 6 | 1 | NA | NA | NA | NA | 1 | 1 | 1 | 1 | 2 | 1 |
| 12_102754! | NA | NA | NA | NA | NA | NA | 2 | 1 | 2 | 1 | NA | NA | NA | NA | 1 | 1 | 1 | 1 | 2 | 1 |
| 12_102804' | NA | NA | NA | NA | NA | NA | 2 | 0.5 | 2 | 0.5 | NA | NA | NA | NA | 1 | 1 | NA | NA | 1 | 1 |
| 12_102804: | NA | NA | NA | NA | NA | NA | 1 | 1 | 1 | 1 | NA | NA | NA | NA | 1 | 1 | NA | NA | 1 | 1 |
| 12_102804: | NA | NA | NA | NA | NA | NA | 1 | 1 | 1 | 1 | NA | NA | NA | NA | 1 | 1 | NA | NA | 1 | 1 |
| 12_102804· | NA | NA | NA | NA | NA | NA | 1 | 1 | 1 | 1 | NA | NA | NA | NA | 1 | 1 | NA | NA | 1 | 1 |
| 12_103074{ | 3 | 0 | 1 | 0 | 2 | 0.5 | 2 | 0 | 8 | 0.125 | 4 | 0.25 | 1 | 0 | 4 | 0 | 1 | 0 | 10 | 0.1 |
| 12_103078· | 3 | 0.3333 | 2 | 0 | 1 | 0 | 2 | 0 | 8 | 0.125 | 4 | 0 | 1 | 0 | 4 | 0.25 | 2 | 0 | 11 | 0.0909 |
| 12_103078( | 3 | 0.6667 | 2 | 0.5 | 2 | 0 | 2 | 0 | 9 | 0.3333 | 4 | 0.25 | NA | NA | 5 | 0.6 | 1 | 0 | 10 | 0.4 |
| 12_103078: | 3 | 0 | 2 | 0.5 | 2 | 0 | 2 | 0.5 | 9 | 0.2222 | 4 | 0.25 | 1 | 1 | 5 | 0.2 | 1 | 0 | 11 | 0.2727 |
| 12_103078: | 3 | 0 | 2 | 0.5 | 2 | 0 | 2 | 0 | 9 | 0.1111 | 4 | 0 | 1 | 0 | 5 | 0.4 | 2 | 0.5 | 12 | 0.25 |
| 12_103078{ | 3 | 0 | 1 | 0 | 3 | 0 | 2 | 0.5 | 9 | 0.1111 | 2 | 0.5 | NA | NA | 3 | 0 | 1 | 0 | 6 | 0.1667 |
| 12_103078! | 3 | 0.6667 | 2 | 0.5 | 3 | 1 | 2 | 0 | 10 | 0.6 | 4 | 0.5 | 1 | 1 | 4 | 0.5 | 2 | 0 | 11 | 0.4545 |
| 12_103078! | 1 | 0 | 1 | 0 | 3 | 0.6667 | 2 | 0 | 7 | 0.2857 | 4 | 0.75 | 1 | 1 | 4 | 0.5 | 1 | 0 | 10 | 0.6 |
| 12_103079( | 4 | 1 | 2 | 0.5 | 3 | 0.6667 | 2 | 0.5 | 11 | 0.7273 | 4 | 1 | 1 | 1 | 4 | 0.75 | 2 | 0.5 | 11 | 0.8182 |
| 12_103079' | 4 | 0.75 | 2 | 1 | 2 | 0.5 | 2 | 1 | 10 | 0.8 | 4 | 0.75 | 2 | 1 | 6 | 0.8333 | 2 | 1 | 14 | 0.8571 |
| 12_103079: | 3 | 0 | 1 | 0 | 3 | 0.6667 | 2 | 0 | 9 | 0.2222 | 3 | 0.3333 | 3 | 0 | 6 | 0.3333 | 2 | 0.5 | 14 | 0.2857 |
| 12_103079: | 3 | 0 | 1 | 0 | 4 | 0.25 | 1 | 0 | 9 | 0.1111 | 4 | 0 | 3 | 0.3333 | 6 | 0 | 3 | 0 | 16 | 0.0625 |
| 12_103149{ | 2 | 1 | 4 | 1 | 3 | 1 | 3 | 1 | 12 | 1 | 1 | 1 | 1 | 1 | 6 | 1 | 3 | 1 | 11 | 1 |
| 12_103149! | 4 | 0.25 | 6 | 0.1667 | 4 | 0 | 3 | 0 | 17 | 0.1176 | 4 | 0 | 4 | 0 | 7 | 0.1429 | 6 | 0.1667 | 21 | 0.0952 |
| 12_103149! | 5 | 0 | 5 | 0.2 | 5 | 0 | 6 | 0.1667 | 21 | 0.0952 | 4 | 0 | 3 | 0 | 8 | 0 | 3 | 0.3333 | 18 | 0.0556 |
| 12_103150( | 6 | 0.1667 | 7 | 0.2857 | 1 | 1 | 5 | 0.2 | 19 | 0.2632 | 3 | 0.6667 | 5 | 0 | 7 | 0.2857 | 6 | 0 | 21 | 0.1905 |
| 12_103150' | 4 | 0.25 | 7 | 0.5714 | 3 | 0 | 7 | 0 | 21 | 0.2381 | 4 | 0 | 3 | 0.3333 | 4 | 0.25 | 5 | 0 | 16 | 0.125 |
| 12_103161( | 2 | 1 | NA | NA | NA | NA | 5 | 0.4 | 7 | 0.5714 | 4 | 0.5 | 1 | 0 | 5 | 0.8 | NA | NA | 10 | 0.6 |
| 12_103161' | 1 | 0 | NA | NA | NA | NA | 5 | 0.2 | 6 | 0.1667 | 4 | 0 | 2 | 0 | 5 | 0.2 | 1 | 0 | 12 | 0.0833 |
| 12_103162{ | NA | NA | NA | NA | NA | NA | 5 | 0.4 | 5 | 0.4 | 1 | 1 | 1 | 0 | 4 | 0 | NA | NA | 6 | 0.1667 |
| 12_103163( | NA | NA | NA | NA | NA | NA | 4 | 0.5 | 4 | 0.5 | 2 | 1 | 1 | 0 | 4 | 1 | NA | NA | 7 | 0.8571 |
| 12_103163( | NA | NA | NA | NA | NA | NA | 3 | 0.6667 | 3 | 0.6667 | 2 | 0.5 | 1 | 1 | 4 | 0.75 | NA | NA | 7 | 0.7143 |
| 12_103163: | NA | NA | NA | NA | NA | NA | 3 | 0.6667 | 3 | 0.6667 | 2 | 1 | 1 | 1 | 4 | 0.75 | 2 | 0 | 9 | 0.6667 |
| 12_103163· | NA | NA | NA | NA | NA | NA | 3 | 0.3333 | 3 | 0.3333 | 2 | 0.5 | 1 | 1 | 5 | 0.8 | 2 | 0.5 | 10 | 0.7 |
| 12_103163· | NA | NA | NA | NA | NA | NA | 2 | 0.5 | 2 | 0.5 | 2 | 0 | 1 | 0 | 5 | 0.4 | 1 | 0 | 9 | 0.2222 |
| 12_103163· | NA | NA | NA | NA | NA | NA | 3 | 0.3333 | 3 | 0.3333 | 2 | 0.5 | 1 | 0 | 4 | 0.25 | 2 | 0 | 9 | 0.2222 |
| 12_103370( | 2 | 0 | 1 | 0 | 1 | 0 | 2 | 0.5 | 6 | 0.1667 | 2 | 0 | 1 | 0 | 2 | 0 | 2 | 0.5 | 7 | 0.1429 |
| 12_103370( | 2 | 0 | 1 | 0 | 1 | 0 | 1 | 1 | 5 | 0.2 | 1 | 0 | 1 | 0 | 2 | 0.5 | 1 | 0 | 5 | 0.2 |

| | | | | | | | | | | | | | | | | | | | | |
|---|---|---|---|---|---|---|---|---|---|---|---|---|---|---|---|---|---|---|---|---|
| 12_1033707 | 2 | 1 | 1 | 1 | 1 | 1 | 2 | 1 | 6 | 1 | 2 | 1 | 1 | 1 | 2 | 0.5 | 1 | 1 | 6 | 0.8333 |
| 12_1033707 | 1 | 0 | 1 | 1 | 1 | 1 | 2 | 0 | 5 | 0.4 | 2 | 0 | 1 | 1 | 2 | 0 | 2 | 0 | 7 | 0.1429 |
| 12_1033707 | 2 | 1 | 1 | 0 | 1 | 0 | 1 | 1 | 5 | 0.6 | 2 | 0 | 1 | 1 | 2 | 0.5 | NA | NA | 5 | 0.4 |
| 12_1033708 | 2 | 1 | 1 | 1 | 1 | 1 | 2 | 1 | 6 | 1 | 2 | 0.5 | 1 | 1 | 1 | 1 | 2 | 0.5 | 6 | 0.6667 |
| 12_1033708 | 1 | 1 | 1 | 1 | 1 | 1 | 2 | 1 | 5 | 1 | 3 | 1 | NA | NA | NA | NA | 2 | 1 | 5 | 1 |
| 12_1033708 | 2 | 1 | 1 | 0 | 1 | 1 | 2 | 0.5 | 6 | 0.6667 | 2 | 0.5 | 1 | 1 | 2 | 0 | 1 | 1 | 6 | 0.5 |
| 12_1033709 | 2 | 1 | 1 | 1 | 1 | 1 | 2 | 0.5 | 6 | 0.8333 | 3 | 0 | 1 | 0 | 2 | 0.5 | 2 | 0.5 | 8 | 0.25 |
| 12_1033709 | 1 | 0 | 1 | 1 | 1 | 1 | 2 | 0.5 | 5 | 0.6 | 3 | 0.3333 | NA | NA | 2 | 0.5 | 1 | 0 | 6 | 0.3333 |
| 12_1033710 | 2 | 0.5 | 1 | 0 | NA | NA | 2 | 1 | 5 | 0.6 | 3 | 0.6667 | 1 | 0 | 2 | 1 | 2 | 0.5 | 8 | 0.625 |
| 12_1033712 | 1 | 1 | 1 | 1 | NA | NA | 2 | 0.5 | 4 | 0.75 | 2 | 0 | 1 | 0 | 2 | 0.5 | 2 | 0.5 | 7 | 0.2857 |
| 12_1033721 | 2 | 0 | 1 | 0 | 1 | 1 | 1 | 0 | 5 | 0.2 | 3 | 0.3333 | 2 | 0 | 2 | 0 | 3 | 0 | 10 | 0.1 |
| 12_1033723 | NA | NA | 2 | 0.5 | 1 | 1 | 1 | 0 | 4 | 0.5 | 2 | 0 | 3 | 0.3333 | 1 | 0 | 2 | 0.5 | 8 | 0.25 |
| 12_1033742 | 11 | 0.5455 | 16 | 0.25 | 10 | 0.1 | 24 | 0.4167 | 61 | 0.3443 | 21 | 0.2857 | 9 | 0.3333 | 11 | 0.1818 | 6 | 0.5 | 47 | 0.2979 |
| 12_1033742 | 23 | 0.1304 | 25 | 0.16 | 20 | 0.05 | 31 | 0.2581 | 99 | 0.1616 | 32 | 0.25 | 14 | 0.0714 | 19 | 0.1579 | 13 | 0.0769 | 78 | 0.1667 |
| 12_1033743 | 21 | 0.0476 | 26 | 0.0769 | 19 | 0 | 28 | 0.25 | 94 | 0.1064 | 33 | 0.2121 | 13 | 0.1538 | 18 | 0.1111 | 13 | 0.0769 | 77 | 0.1558 |
| 12_1033744 | 20 | 0.9 | 24 | 0.875 | 16 | 1 | 24 | 0.9167 | 84 | 0.9167 | 23 | 0.8696 | 12 | 0.8333 | 16 | 0.75 | 10 | 0.9 | 61 | 0.8361 |
| 12_1033745 | 22 | 0.0455 | 26 | 0 | 18 | 0.1111 | 28 | 0.0714 | 94 | 0.0532 | 25 | 0.2 | 13 | 0 | 18 | 0.1667 | 11 | 0.0909 | 67 | 0.1343 |
| 12_1033837 | 20 | 0.6 | 18 | 0.6667 | 17 | 0.7059 | 23 | 0.6957 | 78 | 0.6667 | 18 | 0.6667 | 11 | 0.5455 | 14 | 0.6429 | 12 | 0.8333 | 55 | 0.6727 |
| 12_1033839 | 17 | 0.1176 | 17 | 0.0588 | 14 | 0.0714 | 22 | 0.0455 | 70 | 0.0714 | 18 | 0.0556 | 11 | 0.0909 | 14 | 0 | 9 | 0.1111 | 52 | 0.0577 |
| 12_1033839 | 17 | 0.1765 | 17 | 0.1765 | 17 | 0.1176 | 22 | 0.1364 | 73 | 0.1507 | 17 | 0.1765 | 10 | 0.3 | 13 | 0.0769 | 10 | 0.1 | 50 | 0.16 |
| 12_1033839 | 16 | 0.1875 | 17 | 0.0588 | 14 | 0.2143 | 18 | 0.1111 | 65 | 0.1385 | 16 | 0.125 | 9 | 0 | 13 | 0 | 8 | 0.25 | 46 | 0.087 |
| 12_1033852 | 19 | 0.3158 | 15 | 0.2667 | 14 | 0.0714 | 20 | 0.2 | 68 | 0.2206 | 11 | 0 | 8 | 0.375 | 13 | 0.1538 | 8 | 0.25 | 40 | 0.175 |
| 12_1033854 | 18 | 0.6111 | 16 | 0.4375 | 12 | 0.25 | 19 | 0.4211 | 65 | 0.4462 | 12 | 0.3333 | 9 | 0.5556 | 10 | 0.8 | 7 | 0.8571 | 38 | 0.6053 |
| 12_1033855 | 17 | 0.2941 | 14 | 0 | 9 | 0.1111 | 17 | 0.1765 | 57 | 0.1579 | 11 | 0.0909 | 8 | 0.125 | 11 | 0.0909 | 7 | 0.1429 | 37 | 0.1081 |
| 12_1033855 | 15 | 0.2667 | 15 | 0.3333 | 11 | 0.3636 | 19 | 0.1579 | 60 | 0.2667 | 11 | 0.2727 | 8 | 0.25 | 9 | 0.5556 | 7 | 0.2857 | 35 | 0.3429 |
| 12_1033855 | 13 | 0.5385 | 14 | 0.9286 | 8 | 0.75 | 16 | 0.5 | 51 | 0.6667 | 10 | 0.9 | 5 | 0.6 | 10 | 0.6 | 8 | 0.5 | 33 | 0.6667 |
| 12_1033855 | 14 | 0.3571 | 14 | 0.2143 | 9 | 0.1111 | 18 | 0.2778 | 55 | 0.2545 | 12 | 0.3333 | 7 | 0.4286 | 11 | 0.4545 | 6 | 0.3333 | 36 | 0.3889 |
| 12_1033855 | 17 | 0.2941 | 17 | 0.4706 | 10 | 0.3 | 17 | 0.2941 | 61 | 0.3443 | 11 | 0.2727 | 8 | 0.25 | 12 | 0.4167 | 7 | 0.4286 | 38 | 0.3421 |
| 12_1033901 | 13 | 0.0769 | 14 | 0.2143 | 10 | 0.1 | 19 | 0.2105 | 56 | 0.1607 | 11 | 0.0909 | 6 | 0 | 11 | 0 | 8 | 0.125 | 36 | 0.0556 |
| 12_1033902 | 14 | 0.9286 | 17 | 0.8235 | 10 | 0.7 | 17 | 0.9412 | 58 | 0.8621 | 10 | 0.9 | 7 | 0.4286 | 11 | 0.7273 | 7 | 1 | 35 | 0.7714 |
| 12_1033905 | 11 | 0.2727 | 10 | 0.1 | 6 | 0.1667 | 11 | 0.1818 | 38 | 0.1842 | 6 | 0.5 | 5 | 0 | 8 | 0.375 | 6 | 0.6667 | 25 | 0.4 |
| 12_1033907 | 9 | 0.3333 | 10 | 0.7 | 7 | 0.2857 | 10 | 0.6 | 36 | 0.5 | 6 | 0.5 | 3 | 0.6667 | 9 | 0.5556 | 6 | 0.8333 | 24 | 0.625 |
| 12_1035793 | 4 | 0.5 | 4 | 0.75 | 2 | 0 | 7 | 0.5714 | 17 | 0.5294 | NA | NA | 4 | 0.25 | 5 | 0.4 | 3 | 1 | 12 | 0.5 |
| 12_1035794 | 5 | 0 | 4 | 0.25 | 3 | 0 | 8 | 0 | 20 | 0.05 | NA | NA | 4 | 0.5 | 7 | 0.4286 | 3 | 1 | 14 | 0.5714 |
| 12_1035798 | 6 | 0.3333 | 5 | 0.2 | 5 | 0 | 11 | 0.2727 | 27 | 0.2222 | 1 | 0 | 8 | 0 | 7 | 0.1429 | 2 | 0 | 18 | 0.0556 |
| 12_1039612 | 1 | 0 | NA | NA | NA | NA | 1 | 1 | 2 | 0.5 | NA | NA | 1 | 1 | NA | NA | NA | NA | 1 | 1 |
| 12_1039628 | 2 | 1 | 1 | 1 | 3 | 0.6667 | 2 | 1 | 8 | 0.875 | 1 | 1 | 3 | 0.6667 | 4 | 0.75 | 1 | 1 | 9 | 0.7778 |
| 12_1039629 | 8 | 0.5 | 4 | 0.75 | 6 | 0.6667 | 5 | 0.8 | 23 | 0.6522 | 3 | 1 | 8 | 0.625 | 7 | 0.8571 | 4 | 1 | 22 | 0.8182 |
| 12_1039630 | 5 | 0.2 | 4 | 0 | 7 | 0 | 3 | 0 | 19 | 0.0526 | 2 | 0 | 6 | 0 | 5 | 0.2 | 2 | 0.5 | 15 | 0.1333 |
| 12_1039632 | 8 | 1 | 3 | 1 | 7 | 0.8571 | 4 | 1 | 22 | 0.9545 | 1 | 1 | 9 | 1 | 6 | 1 | 2 | 1 | 18 | 1 |
| 12_1039643 | NA | NA | NA | NA | NA | NA | 1 | 1 | 1 | 1 | 1 | 1 | NA | NA | NA | NA | NA | NA | 1 | 1 |
| 12_1039644 | 7 | 0 | 2 | 0 | 6 | 0 | 5 | 0.2 | 20 | 0.05 | 1 | 0 | 8 | 0 | 3 | 0 | 2 | 0.5 | 14 | 0.0714 |
| 12_1039676 | 5 | 0.4 | 1 | 1 | 7 | 0.5714 | 3 | 0.3333 | 16 | 0.5 | 1 | 1 | 9 | 0.3333 | 4 | 0.75 | 1 | 1 | 15 | 0.5333 |
| 12_1039745 | 6 | 0 | 1 | 0 | 5 | 0.2 | 4 | 0.25 | 16 | 0.125 | 2 | 0 | 8 | 0.125 | 4 | 0 | 1 | 0 | 15 | 0.0667 |
| 12_1039746 | 3 | 1 | 1 | 0 | 3 | 0.3333 | 4 | 0.75 | 11 | 0.6364 | 2 | 0.5 | 6 | 0.3333 | 4 | 0 | 1 | 0 | 13 | 0.2308 |
| 12_1039748 | 7 | 0.8571 | 2 | 1 | 3 | 0.6667 | 3 | 1 | 15 | 0.8667 | 2 | 1 | 8 | 0.875 | 5 | 1 | 1 | 1 | 16 | 0.9375 |
| 12_1039748 | 7 | 0.1429 | 2 | 0 | 4 | 0 | 3 | 0 | 16 | 0.0625 | 2 | 0 | 7 | 0.1429 | 5 | 0 | 1 | 0 | 15 | 0.0667 |
| 12_1043750 | NA | NA | 2 | 0.5 | 1 | 1 | 2 | 0.5 | 5 | 0.6 | 3 | 1 | 1 | 1 | 1 | 0 | NA | NA | 5 | 0.8 |
| 12_1043750 | NA | NA | 5 | 0.4 | 1 | 1 | 2 | 0.5 | 8 | 0.5 | 4 | 0.25 | 2 | 0.5 | 1 | 1 | NA | NA | 7 | 0.4286 |
| 12_1043752 | NA | NA | 5 | 0.2 | 2 | 0 | 2 | 0 | 9 | 0.1111 | 4 | 0.25 | 2 | 0 | 1 | 0 | NA | NA | 7 | 0.1429 |
| 12_1043753 | NA | NA | 5 | 0.8 | 2 | 1 | 1 | 0 | 8 | 0.75 | 4 | 0.75 | 2 | 1 | 1 | 1 | NA | NA | 7 | 0.8571 |
| 12_1043754 | NA | NA | 4 | 0 | 1 | 1 | 2 | 0.5 | 7 | 0.2857 | 5 | 0 | 2 | 0.5 | 1 | 1 | NA | NA | 8 | 0.25 |
| 12_1043754 | NA | NA | 6 | 0.3333 | NA | NA | 2 | 1 | 8 | 0.5 | 5 | 0.4 | 1 | 1 | 1 | 0 | NA | NA | 7 | 0.4286 |
| 12_1043755 | NA | NA | 6 | 0.3333 | 2 | 0 | 1 | 0 | 9 | 0.2222 | 3 | 0.3333 | 2 | 0 | NA | NA | NA | NA | 5 | 0.2 |
| 12_1043755 | NA | NA | 7 | 0.4286 | 1 | 1 | 1 | 0 | 9 | 0.4444 | 5 | 0.6 | 2 | 0 | 1 | 0 | NA | NA | 8 | 0.375 |
| 12_1043774 | NA | NA | 6 | 0.1667 | 3 | 0 | 2 | 0 | 11 | 0.0909 | 5 | 0.2 | 2 | 0 | 1 | 0 | 1 | 0 | 9 | 0.1111 |
| 12_1043802 | NA | NA | 10 | 0.1 | 3 | 0 | 3 | 0 | 16 | 0.0625 | 5 | 0 | 2 | 0.5 | 1 | 0 | 1 | 0 | 9 | 0.1111 |
| 12_1046540 | 5 | 0.4 | NA | NA | 1 | 0 | 3 | 0.6667 | 9 | 0.4444 | 1 | 0 | 3 | 0.6667 | 1 | 0 | 5 | 0.6 | 10 | 0.5 |
| 12_1046542 | 2 | 0.5 | 1 | 0 | 3 | 0 | 3 | 0 | 9 | 0.1111 | 1 | 0 | 5 | 0 | 2 | 0.5 | 4 | 0 | 12 | 0.0833 |
| 12_1046543 | 4 | 0 | 1 | 0 | 2 | 0 | 4 | 0.25 | 11 | 0.0909 | 1 | 0 | 3 | 0.3333 | 1 | 0 | 3 | 0.3333 | 8 | 0.25 |
| 12_1046543 | 4 | 0.25 | 1 | 0 | 1 | 0 | 3 | 0.3333 | 9 | 0.2222 | 1 | 0 | 3 | 0.3333 | 2 | 0 | 3 | 0.3333 | 9 | 0.2222 |
| 12_1046544 | 2 | 0.5 | 1 | 0 | 3 | 0 | 3 | 0.3333 | 9 | 0.2222 | 1 | 0 | NA | NA | NA | NA | 2 | 1 | 3 | 0.6667 |
| 12_1046546 | 3 | 0.3333 | 1 | 0 | 1 | 0 | 4 | 0 | 9 | 0.1111 | 1 | 0 | 3 | 0.3333 | 1 | 0 | 3 | 0.6667 | 8 | 0.375 |
| 12_1046547 | 3 | 0.3333 | 1 | 0 | 1 | 0 | 1 | 0 | 6 | 0.1667 | 1 | 1 | 2 | 0 | 1 | 1 | 3 | 0.6667 | 7 | 0.5714 |
| 12_1046551 | 2 | 0.5 | NA | NA | NA | NA | 2 | 0.5 | 4 | 0.5 | 1 | 0 | 4 | 0 | 1 | 1 | 2 | 0.5 | 8 | 0.25 |

| | | | | | | | | | | | | | | | | | | | | |
|---|---|---|---|---|---|---|---|---|---|---|---|---|---|---|---|---|---|---|---|---|
| 12_104655 | 2 | 0.5 | NA | NA | NA | NA | 3 | 0.6667 | 5 | 0.6 | NA | NA | 4 | 0.5 | 1 | 1 | 3 | 0.3333 | 8 | 0.5 |
| 12_104655 | 2 | 0 | NA | NA | 1 | 0 | 3 | 0.3333 | 6 | 0.1667 | 1 | 1 | 4 | 0 | 1 | 0 | 3 | 0 | 9 | 0.1111 |
| 12_104657 | NA | NA | NA | NA | 1 | 1 | 1 | 0 | 2 | 0.5 | NA | NA | 2 | 0 | 1 | 1 | 2 | 0 | 5 | 0.2 |
| 12_104657 | 2 | 1 | 1 | 0 | 3 | 0.6667 | 1 | 1 | 7 | 0.7143 | 1 | 0 | 2 | 0.5 | NA | NA | 3 | 0.3333 | 6 | 0.3333 |
| 12_104660 | 2 | 0.5 | 1 | 0 | 3 | 0.3333 | NA | NA | 6 | 0.3333 | 1 | 1 | 2 | 0.5 | NA | NA | 3 | 0.6667 | 6 | 0.6667 |
| 12_104662 | 1 | 1 | 1 | 1 | 3 | 0.6667 | NA | NA | 5 | 0.8 | 1 | 1 | 2 | 1 | NA | NA | 3 | 0.6667 | 6 | 0.8333 |
| 12_104662 | 1 | 0 | 1 | 1 | 2 | 0.5 | NA | NA | 4 | 0.5 | 1 | 1 | 2 | 0 | NA | NA | 3 | 0.3333 | 6 | 0.3333 |
| 12_104662 | 1 | 1 | 1 | 1 | 3 | 0.6667 | NA | NA | 5 | 0.8 | 1 | 1 | 2 | 1 | NA | NA | 2 | 0.5 | 5 | 0.8 |
| 12_104663 | 1 | 1 | 1 | 1 | 3 | 0.6667 | NA | NA | 5 | 0.8 | 1 | 1 | 2 | 1 | NA | NA | 3 | 0.3333 | 6 | 0.6667 |
| 12_104663 | 1 | 0 | 1 | 0 | 3 | 0.3333 | NA | NA | 5 | 0.2 | 1 | 0 | 2 | 0.5 | NA | NA | 1 | 0 | 4 | 0.25 |
| 12_104668 | NA | NA | NA | NA | 2 | 1 | NA | NA | 2 | 1 | 1 | 1 | 1 | 1 | NA | NA | 2 | 0.5 | 4 | 0.75 |
| 12_104668 | NA | NA | NA | NA | 2 | 1 | NA | NA | 2 | 1 | 1 | 1 | 1 | 1 | NA | NA | 2 | 1 | 4 | 1 |
| 12_104669 | NA | NA | NA | NA | 2 | 1 | NA | NA | 2 | 1 | 1 | 1 | NA | NA | NA | NA | 2 | 0.5 | 3 | 0.6667 |
| 12_104669 | NA | NA | NA | NA | 2 | 1 | NA | NA | 2 | 1 | 1 | 1 | NA | NA | NA | NA | 2 | 0.5 | 3 | 0.6667 |
| 12_104718 | 1 | 1 | NA | NA | NA | NA | NA | NA | 1 | 1 | 2 | 0.5 | NA | NA | NA | NA | NA | NA | 2 | 0.5 |
| 12_104718 | 1 | 1 | 2 | 0.5 | 1 | 1 | NA | NA | 4 | 0.75 | 2 | 1 | NA | NA | NA | NA | NA | NA | 2 | 1 |
| 12_104718 | 1 | 1 | 2 | 0.5 | 1 | 1 | NA | NA | 4 | 0.75 | 1 | 1 | NA | NA | NA | NA | NA | NA | 1 | 1 |
| 12_104729 | 6 | 0.3333 | 5 | 0 | 4 | 0.5 | 7 | 0 | 22 | 0.1818 | 3 | 0.6667 | 2 | 0 | 3 | 0.3333 | 4 | 0.25 | 12 | 0.3333 |
| 12_104729 | 6 | 0.6667 | 3 | 0.3333 | 3 | 0.6667 | 8 | 0.375 | 20 | 0.5 | 3 | 0.6667 | 1 | 1 | 2 | 0 | 4 | 0.75 | 10 | 0.6 |
| 12_104729 | 4 | 0.5 | 2 | 0 | 2 | 0.5 | 6 | 0.3333 | 14 | 0.3571 | 3 | 0 | NA | NA | 2 | 0 | 2 | 0.5 | 7 | 0.1429 |
| 12_104729 | 6 | 0 | 2 | 0 | 3 | 0.3333 | 7 | 0.4286 | 18 | 0.2222 | 3 | 0 | 1 | 1 | 2 | 0 | 3 | 0.3333 | 9 | 0.2222 |
| 12_104729 | 5 | 0.4 | 2 | 0.5 | 3 | 0 | 7 | 0.2857 | 17 | 0.2941 | 3 | 0 | 1 | 1 | 1 | 0 | 3 | 1 | 8 | 0.5 |
| 12_104730 | 6 | 0.3333 | 2 | 0 | 2 | 0.5 | 4 | 0 | 14 | 0.2143 | 4 | 0.25 | NA | NA | 2 | 0.5 | 3 | 0.3333 | 9 | 0.3333 |
| 12_104730 | 4 | 0.5 | 2 | 0 | 3 | 0.3333 | 5 | 0.2 | 14 | 0.2857 | 4 | 0.5 | 1 | 0 | 2 | 0 | 3 | 0.3333 | 10 | 0.3 |
| 12_104732 | 2 | 0.5 | NA | NA | 2 | 0.5 | 4 | 0.5 | 8 | 0.5 | 4 | 0.25 | 1 | 0 | 2 | 0 | 1 | 0 | 8 | 0.125 |
| 12_104732 | 1 | 0 | NA | NA | 2 | 0.5 | 3 | 0.6667 | 6 | 0.5 | 4 | 0.75 | 1 | 1 | 1 | 1 | 1 | 0 | 7 | 0.7143 |
| 12_104732 | 1 | 0 | NA | NA | 2 | 1 | 3 | 0.6667 | 6 | 0.6667 | 3 | 0.6667 | 1 | 1 | 1 | 0 | 1 | 0 | 6 | 0.5 |
| 12_104732 | 1 | 0 | NA | NA | 1 | 0 | 3 | 0.3333 | 5 | 0.2 | 4 | 0.25 | 1 | 0 | 2 | 0 | 1 | 0 | 8 | 0.125 |
| 12_104732 | 1 | 0 | NA | NA | 2 | 0.5 | 2 | 0 | 5 | 0.2 | 4 | 0.5 | NA | NA | 2 | 0.5 | 1 | 0 | 7 | 0.4286 |
| 12_104732 | 1 | 0 | NA | NA | 2 | 0.5 | 3 | 0.3333 | 6 | 0.3333 | 4 | 0.5 | 1 | 0 | 2 | 0.5 | 1 | 0 | 8 | 0.375 |
| 12_104732 | 1 | 1 | NA | NA | 2 | 1 | 3 | 0.3333 | 6 | 0.6667 | 3 | 0.3333 | 1 | 0 | 2 | 0.5 | 1 | 0 | 7 | 0.2857 |
| 12_104735 | NA | NA | 6 | 0 | 2 | 0 | 2 | 0.5 | 10 | 0.1 | NA | NA | 3 | 0 | 4 | 0.25 | 1 | 0 | 8 | 0.125 |
| 12_104737 | NA | NA | 3 | 0.3333 | 1 | 0 | NA | NA | 4 | 0.25 | NA | NA | 1 | 1 | NA | NA | 1 | 1 | 2 | 1 |
| 12_104738 | 1 | 1 | 1 | 1 | 1 | 1 | NA | NA | 3 | 1 | 1 | 0 | 1 | 1 | 1 | 0 | 1 | 1 | 4 | 0.5 |
| 12_104748 | 2 | 0.5 | 2 | 0.5 | 1 | 1 | NA | NA | 5 | 0.6 | NA | NA | NA | NA | 1 | 1 | NA | NA | 1 | 1 |
| 12_104748 | 1 | 1 | 1 | 1 | 1 | 1 | NA | NA | 3 | 1 | NA | NA | NA | NA | 1 | 1 | NA | NA | 1 | 1 |
| 12_105006 | 16 | 0 | 13 | 0.1538 | 14 | 0.2857 | 15 | 0.0667 | 58 | 0.1207 | 6 | 0.1667 | 11 | 0.0909 | 10 | 0.1 | 12 | 0.0833 | 39 | 0.1026 |
| 12_105006 | 13 | 0.6923 | 9 | 0.5556 | 8 | 0.5 | 13 | 0.3846 | 43 | 0.5349 | 7 | 0.5714 | 6 | 0.6667 | 10 | 0.3 | 9 | 0.3333 | 32 | 0.4375 |
| 12_105006 | 15 | 0.3333 | 12 | 0.1667 | 11 | 0.0909 | 15 | 0 | 53 | 0.1509 | 7 | 0.4286 | 10 | 0.3 | 9 | 0.1111 | 10 | 0.3 | 36 | 0.2778 |
| 12_105006 | 8 | 0 | 9 | 0.2222 | 8 | 0.375 | 10 | 0.1 | 35 | 0.1714 | 3 | 0.3333 | 3 | 0.6667 | 7 | 0.1429 | 8 | 0.125 | 21 | 0.2381 |
| 12_105007 | 17 | 0.2353 | 12 | 0.1667 | 13 | 0.0769 | 17 | 0.1765 | 59 | 0.1695 | 9 | 0 | 13 | 0.1538 | 8 | 0 | 9 | 0.3333 | 39 | 0.1282 |
| 12_105303 | NA | NA | 2 | 0.5 | 3 | 0.3333 | 1 | 0 | 6 | 0.3333 | 2 | 0 | 3 | 0.3333 | 2 | 0 | NA | NA | 7 | 0.1429 |
| 12_105347 | 1 | 0 | 2 | 0 | 2 | 0.5 | 2 | 1 | 7 | 0.4286 | 1 | 1 | 5 | 0.8 | 3 | 0 | 1 | 0 | 10 | 0.5 |
| 12_105347 | 1 | 0 | 2 | 0 | 3 | 0.3333 | 1 | 0 | 7 | 0.1429 | 1 | 1 | 5 | 0.4 | 2 | 0 | 1 | 0 | 9 | 0.3333 |
| 12_105347 | NA | NA | 1 | 1 | NA | NA | 1 | 1 | 2 | 1 | 1 | 1 | NA | NA | 1 | 1 | 1 | 1 | 3 | 1 |
| 12_105347 | 1 | 0 | 2 | 0 | 3 | 0 | 2 | 0.5 | 8 | 0.125 | 1 | 0 | 5 | 0.2 | 3 | 0.3333 | 1 | 0 | 10 | 0.2 |
| 12_105347 | 1 | 1 | 2 | 0.5 | 2 | 0.5 | 2 | 1 | 7 | 0.7143 | 1 | 0 | 6 | 0 | 2 | 0.5 | 1 | 1 | 10 | 0.2 |
| 12_105348 | 1 | 1 | 2 | 1 | 2 | 1 | 2 | 1 | 7 | 1 | 1 | 1 | 5 | 1 | 3 | 1 | 1 | 0 | 10 | 0.9 |
| 12_105348 | 1 | 0 | 1 | 0 | 1 | 0 | 1 | 1 | 4 | 0.25 | 1 | 1 | 4 | 0.75 | 3 | 0.3333 | 1 | 1 | 9 | 0.6667 |
| 12_105348 | 1 | 0 | 2 | 0 | 3 | 0.3333 | 2 | 0.5 | 8 | 0.25 | 1 | 1 | 5 | 0.8 | 2 | 0.5 | 1 | 1 | 9 | 0.7778 |
| 12_105348 | 1 | 1 | 2 | 1 | 3 | 1 | 2 | 1 | 8 | 1 | 1 | 1 | 5 | 1 | 3 | 1 | 1 | 1 | 10 | 1 |
| 12_105348 | 1 | 0 | 2 | 0 | 3 | 0.3333 | 3 | 0 | 9 | 0.1111 | 1 | 0 | 5 | 0.2 | 3 | 0 | 1 | 0 | 10 | 0.1 |
| 12_105348 | 1 | 1 | 2 | 0.5 | 2 | 0.5 | 3 | 0 | 8 | 0.375 | NA | NA | 5 | 0.2 | 3 | 0.3333 | 1 | 0 | 9 | 0.2222 |
| 12_105348 | 1 | 0 | 1 | 0 | 4 | 0.25 | 4 | 0 | 10 | 0.1 | NA | NA | 5 | 0.4 | 2 | 0 | NA | NA | 7 | 0.2857 |
| 12_105348 | NA | NA | 2 | 0 | 3 | 0.6667 | 3 | 0.6667 | 8 | 0.5 | NA | NA | 4 | 1 | 5 | 1 | 1 | 1 | 10 | 1 |
| 12_105348 | 1 | 1 | 2 | 1 | 3 | 0 | 4 | 0 | 10 | 0.3 | NA | NA | 4 | 0.25 | 4 | 0.5 | 1 | 1 | 9 | 0.4444 |
| 12_105349 | 1 | 0 | 5 | 0 | 5 | 0.2 | 4 | 0 | 15 | 0.0667 | 2 | 0 | 7 | 0.1429 | 5 | 0 | 1 | 0 | 15 | 0.0667 |
| 12_105349 | 1 | 0 | 5 | 0 | 5 | 0.2 | 4 | 0 | 15 | 0.0667 | 3 | 0 | 5 | 0.4 | 4 | 0 | 2 | 0 | 14 | 0.1429 |
| 12_105349 | 1 | 0 | 5 | 0 | 4 | 0.25 | 3 | 0.3333 | 13 | 0.1538 | 3 | 0 | 5 | 0.4 | 4 | 0.25 | 1 | 1 | 13 | 0.3077 |
| 12_105349 | 2 | 0 | 6 | 0.1667 | 4 | 0.25 | 5 | 0.2 | 17 | 0.1765 | 3 | 0.3333 | 6 | 0 | 6 | 0.3333 | 2 | 0.5 | 17 | 0.2353 |
| 12_105350 | 1 | 0 | 5 | 0.2 | 6 | 0.1667 | 3 | 0 | 15 | 0.1333 | 4 | 0 | 8 | 0.25 | 6 | 0.1667 | 2 | 0.5 | 20 | 0.2 |
| 12_105378 | 1 | 1 | NA | NA | NA | NA | 1 | 1 | 2 | 1 | NA | NA | NA | NA | 1 | 1 | NA | NA | 1 | 1 |
| 12_105378 | 1 | 0 | NA | NA | NA | NA | 1 | 1 | 2 | 0.5 | NA | NA | NA | NA | 1 | 1 | NA | NA | 1 | 1 |
| 12_105453 | 2 | 0.5 | NA | NA | 1 | 0 | 1 | 0 | 4 | 0.25 | 2 | 1 | 1 | 1 | NA | NA | 4 | 0.5 | 7 | 0.7143 |
| 12_105454 | 1 | 1 | NA | NA | NA | NA | 1 | 0 | 2 | 0.5 | 2 | 0 | 1 | 0 | NA | NA | 2 | 0.5 | 5 | 0.2 |

| | | | | | | | | | | | | | | | | | | | | |
|---|---|---|---|---|---|---|---|---|---|---|---|---|---|---|---|---|---|---|---|---|
| 12_105454 | 2 | 1 | NA | NA | 1 | 1 | 1 | 1 | 4 | 1 | 3 | 1 | 1 | 1 | NA | NA | 3 | 0.6667 | 7 | 0.8571 |
| 12_105454 | 2 | 1 | NA | NA | 1 | 1 | 1 | 1 | 4 | 1 | 1 | 1 | 1 | 1 | NA | NA | 3 | 0.6667 | 5 | 0.8 |
| 12_105454 | 3 | 0 | NA | NA | 2 | 0.5 | 1 | 0 | 6 | 0.1667 | 3 | 0.3333 | NA | NA | NA | NA | 2 | 0.5 | 5 | 0.4 |
| 12_105454 | 3 | 0.3333 | NA | NA | 1 | 1 | 2 | 1 | 6 | 0.6667 | 2 | 1 | 1 | 1 | NA | NA | 3 | 0.3333 | 6 | 0.6667 |
| 12_105454 | 3 | 0.3333 | NA | NA | 1 | 0 | 2 | 0 | 6 | 0.1667 | NA | NA | 1 | 0 | NA | NA | 2 | 0.5 | 3 | 0.3333 |
| 12_105454 | 3 | 0.3333 | NA | NA | 2 | 0 | 4 | 0.75 | 9 | 0.4444 | 3 | 0 | 1 | 0 | NA | NA | 3 | 1 | 7 | 0.4286 |
| 12_105454 | 2 | 1 | NA | NA | 2 | 1 | 4 | 1 | 8 | 1 | 2 | 0.5 | NA | NA | NA | NA | 2 | 1 | 4 | 0.75 |
| 12_105454 | 2 | 1 | NA | NA | 1 | 1 | 2 | 0 | 5 | 0.6 | 2 | 0.5 | NA | NA | NA | NA | 3 | 1 | 5 | 0.8 |
| 12_105455 | 3 | 0.3333 | 1 | 0 | 2 | 0 | 4 | 0 | 10 | 0.1 | 2 | 0 | 1 | 0 | 1 | 1 | 2 | 0 | 6 | 0.1667 |
| 12_105457 | 20 | 0.1 | 18 | 0 | 16 | 0.0625 | 29 | 0.1379 | 83 | 0.0843 | 10 | 0.1 | 22 | 0.1364 | 24 | 0.0417 | 18 | 0 | 74 | 0.0676 |
| 12_105458 | 23 | 0.0435 | 21 | 0.2381 | 18 | 0.0556 | 33 | 0.1515 | 95 | 0.1263 | 15 | 0.2 | 24 | 0.125 | 26 | 0.2692 | 23 | 0.0435 | 88 | 0.1591 |
| 12_105458 | 23 | 0.1739 | 22 | 0.1818 | 19 | 0.1053 | 33 | 0.0909 | 97 | 0.134 | 19 | 0.2105 | 21 | 0.1429 | 24 | 0.2083 | 24 | 0.0417 | 88 | 0.1477 |
| 12_105458 | 25 | 0.04 | 19 | 0.1579 | 15 | 0.0667 | 33 | 0.1818 | 92 | 0.1196 | 14 | 0.0714 | 19 | 0.1053 | 23 | 0.3043 | 26 | 0.0769 | 82 | 0.1463 |
| 12_105458 | 25 | 0.92 | 22 | 1 | 19 | 0.9474 | 40 | 0.85 | 106 | 0.9151 | 13 | 0.9231 | 23 | 0.8696 | 28 | 1 | 28 | 0.9286 | 92 | 0.9348 |
| 12_105458 | 30 | 0.0667 | 26 | 0.0769 | 19 | 0.1053 | 41 | 0.0244 | 116 | 0.0603 | 19 | 0.1053 | 27 | 0.0741 | 33 | 0.0909 | 28 | 0.0357 | 107 | 0.0748 |
| 12_105458 | 21 | 0.9048 | 18 | 0.7222 | 13 | 0.7692 | 29 | 0.7931 | 81 | 0.8025 | 17 | 0.8235 | 17 | 0.8235 | 19 | 0.9474 | 17 | 0.8235 | 70 | 0.8571 |
| 12_105458 | 21 | 0.5714 | 26 | 0.7308 | 19 | 0.6316 | 35 | 0.6571 | 101 | 0.6535 | 15 | 0.8 | 22 | 0.7727 | 19 | 0.7895 | 19 | 0.6316 | 75 | 0.7467 |
| 12_105458 | 24 | 0.625 | 24 | 0.625 | 21 | 0.6667 | 38 | 0.7632 | 107 | 0.6822 | 16 | 0.75 | 26 | 0.8462 | 26 | 0.7692 | 25 | 0.84 | 93 | 0.8065 |
| 12_105458 | 27 | 0.3333 | 27 | 0.4444 | 20 | 0.4 | 36 | 0.3333 | 110 | 0.3727 | 18 | 0.5 | 26 | 0.5769 | 28 | 0.4286 | 25 | 0.44 | 97 | 0.4845 |
| 12_105458 | 20 | 0.1 | 23 | 0.1739 | 14 | 0.2857 | 28 | 0.0714 | 85 | 0.1412 | 12 | 0 | 16 | 0.1875 | 21 | 0.1905 | 18 | 0.1111 | 67 | 0.1343 |
| 12_105458 | 30 | 0.2 | 24 | 0.2917 | 21 | 0.1905 | 38 | 0.2105 | 113 | 0.2212 | 17 | 0.1765 | 24 | 0.0833 | 31 | 0.2581 | 26 | 0.1923 | 98 | 0.1837 |
| 12_105458 | 26 | 0.5 | 22 | 0.3182 | 18 | 0.3333 | 33 | 0.3636 | 99 | 0.3838 | 16 | 0.375 | 25 | 0.36 | 28 | 0.3571 | 27 | 0.4074 | 96 | 0.375 |
| 12_105458 | 30 | 0.1333 | 22 | 0.0455 | 22 | 0.0455 | 36 | 0.1111 | 110 | 0.0909 | 16 | 0 | 24 | 0.0833 | 29 | 0.069 | 26 | 0.0769 | 95 | 0.0632 |
| 12_105458 | 25 | 0.72 | 24 | 0.75 | 21 | 0.619 | 36 | 0.8056 | 106 | 0.7358 | 16 | 0.625 | 23 | 0.8261 | 32 | 0.625 | 30 | 0.9 | 101 | 0.7525 |
| 12_105459 | 20 | 0.35 | 20 | 0.2 | 21 | 0.1905 | 34 | 0.1765 | 95 | 0.2211 | 15 | 0.1333 | 26 | 0.4231 | 23 | 0.3913 | 25 | 0.32 | 89 | 0.3371 |
| 12_105459 | 27 | 0.2963 | 20 | 0.2 | 20 | 0.15 | 39 | 0.2821 | 106 | 0.2453 | 13 | 0.3077 | 22 | 0.1364 | 31 | 0.2258 | 27 | 0.1481 | 93 | 0.1935 |
| 12_105459 | 31 | 0.0968 | 24 | 0.0833 | 20 | 0.15 | 46 | 0.0435 | 121 | 0.0826 | 18 | 0.1111 | 24 | 0.125 | 32 | 0.1562 | 32 | 0.125 | 106 | 0.1321 |
| 12_105459 | 11 | 0.3636 | 16 | 0.25 | 12 | 0.5 | 19 | 0.3684 | 58 | 0.3621 | 9 | 0.3333 | 10 | 0 | 14 | 0.0714 | 17 | 0.4706 | 50 | 0.24 |
| 12_105580 | 3 | 0.3333 | 3 | 0 | 2 | 0.5 | 2 | 0.5 | 10 | 0.3 | 3 | 0.6667 | 3 | 0 | 2 | 0.5 | 6 | 0.3333 | 14 | 0.3571 |
| 12_105580 | 2 | 0 | 3 | 0.3333 | 2 | 0 | 2 | 0 | 9 | 0.1111 | 3 | 0 | 3 | 0 | 2 | 0.5 | 5 | 0 | 13 | 0.0769 |
| 12_105581 | 1 | 0 | 3 | 0.3333 | 2 | 0 | 2 | 0 | 8 | 0.125 | 3 | 0.6667 | 2 | 0 | 1 | 0 | 5 | 0.4 | 11 | 0.3636 |
| 12_105582 | 1 | 0 | 3 | 0.3333 | 3 | 0 | 2 | 0 | 9 | 0.1111 | 3 | 0.3333 | 2 | 0 | 1 | 0 | 4 | 0 | 10 | 0.1 |
| 12_105583 | 4 | 0.5 | 3 | 0.3333 | 3 | 0 | 5 | 0.2 | 15 | 0.2667 | 4 | 0.25 | 4 | 0.5 | 2 | 0.5 | 7 | 0 | 17 | 0.2353 |
| 12_105583 | 4 | 0 | 4 | 0 | 2 | 0 | 5 | 0.2 | 15 | 0.0667 | 5 | 0 | 4 | 0 | 1 | 0 | 8 | 0.25 | 18 | 0.1111 |
| 12_105583 | 4 | 0.5 | 3 | 0.3333 | 2 | 0.5 | 3 | 0 | 12 | 0.3333 | 6 | 0.3333 | 4 | 0.5 | 2 | 0.5 | 5 | 0.4 | 17 | 0.4118 |
| 12_105583 | 2 | 0 | 3 | 0.3333 | 3 | 1 | 6 | 0.3333 | 14 | 0.4286 | 4 | 1 | 3 | 0.6667 | 2 | 0 | 4 | 0.25 | 13 | 0.5385 |
| 12_105665 | 1 | 1 | 4 | 0 | 8 | 0 | 5 | 0 | 18 | 0.0556 | 4 | 0 | 4 | 0 | 3 | 0.3333 | 2 | 0 | 13 | 0.0769 |
| 12_105665 | NA | NA | 2 | 0 | 6 | 0.1667 | 6 | 0.3333 | 14 | 0.2143 | 2 | 0 | 3 | 0.3333 | 4 | 0 | 3 | 0.3333 | 12 | 0.1667 |
| 12_105666 | 2 | 0 | 3 | 0.3333 | 7 | 0 | 6 | 0.1667 | 18 | 0.1111 | 4 | 0.25 | 4 | 0 | 3 | 0 | 3 | 0 | 14 | 0.0714 |
| 12_105667 | 2 | 1 | 3 | 0.6667 | 8 | 0.75 | 7 | 0.8571 | 20 | 0.8 | 4 | 0.75 | 4 | 0.75 | 5 | 1 | 4 | 0.5 | 17 | 0.7647 |
| 12_105667 | 1 | 1 | 2 | 1 | 4 | 1 | 6 | 0.6667 | 13 | 0.8462 | 4 | 0.75 | 4 | 0.5 | 5 | 0.6 | 2 | 1 | 15 | 0.6667 |
| 12_105667 | 2 | 0.5 | 4 | 0 | 6 | 0 | 8 | 0 | 20 | 0.05 | 5 | 0.2 | 6 | 0 | 6 | 0.3333 | 5 | 0 | 22 | 0.1364 |
| 12_105667 | 2 | 0.5 | 4 | 0.25 | 7 | 0.1429 | 8 | 0.25 | 21 | 0.2381 | 5 | 0.4 | 7 | 0 | 6 | 0 | 5 | 0.4 | 23 | 0.1739 |
| 12_105667 | 2 | 0.5 | 4 | 0.5 | 6 | 0 | 8 | 0.125 | 20 | 0.2 | 4 | 0.25 | 7 | 0.4286 | 6 | 0.1667 | 4 | 0.5 | 21 | 0.3333 |
| 12_105667 | 2 | 0.5 | 5 | 0 | 6 | 0 | 8 | 0.25 | 21 | 0.1429 | 5 | 0.4 | 7 | 0.1429 | 6 | 0.3333 | 5 | 0.2 | 23 | 0.2609 |
| 12_105667 | 2 | 1 | 3 | 0 | 5 | 0.6 | 6 | 0.6667 | 16 | 0.5625 | 4 | 0.25 | 7 | 0.5714 | 5 | 0.4 | 4 | 0.5 | 20 | 0.45 |
| 12_105668 | 1 | 0 | 5 | 0 | 3 | 0.3333 | 6 | 0.3333 | 15 | 0.2 | 4 | 0 | 6 | 0.3333 | 1 | 0 | 3 | 0 | 14 | 0.1429 |
| 12_105668 | NA | NA | 4 | 0 | 4 | 0.25 | 5 | 0.2 | 13 | 0.1538 | 4 | 0.25 | 6 | 0 | 3 | 0 | 1 | 0 | 14 | 0.0714 |
| 12_105668 | 1 | 0 | NA | NA | NA | NA | 4 | 0.25 | 5 | 0.2 | 1 | 0 | NA | NA | 1 | 1 | NA | NA | 2 | 0.5 |
| 12_105803 | 4 | 0.5 | 10 | 0.4 | 8 | 0 | 10 | 0.5 | 32 | 0.3438 | 2 | 1 | 3 | 0 | 12 | 0.3333 | 7 | 0.2857 | 24 | 0.3333 |
| 12_105803 | 6 | 0.6667 | 13 | 0.9231 | 10 | 0.7 | 11 | 0.6364 | 40 | 0.75 | 3 | 0.3333 | 4 | 0.75 | 13 | 0.6154 | 8 | 0.875 | 28 | 0.6786 |
| 12_105803 | 5 | 0.2 | 12 | 0.0833 | 7 | 0.1429 | 9 | 0.1111 | 33 | 0.1212 | 2 | 0.5 | 4 | 0.25 | 12 | 0.1667 | 7 | 0.5714 | 25 | 0.32 |
| 12_105803 | 5 | 0.4 | 14 | 0.1429 | 9 | 0.2222 | 11 | 0.5455 | 39 | 0.3077 | 3 | 0.6667 | 5 | 0.4 | 12 | 0.1667 | 8 | 0.125 | 28 | 0.25 |
| 12_105803 | 6 | 0.6667 | 14 | 0.2857 | 10 | 0.1 | 9 | 0.3333 | 39 | 0.3077 | 2 | 0.5 | 4 | 0.25 | 11 | 0.3636 | 9 | 0.2222 | 26 | 0.3077 |
| 12_105803 | 4 | 0.25 | 11 | 0.3636 | 9 | 0.5556 | 11 | 0.6364 | 35 | 0.4857 | 3 | 0 | 5 | 0.2 | 11 | 0.4545 | 8 | 0.375 | 27 | 0.3333 |
| 12_105803 | 5 | 0.4 | 10 | 0.1 | 6 | 0 | 7 | 0 | 28 | 0.1071 | 3 | 0 | 4 | 0 | 12 | 0 | 9 | 0.2222 | 28 | 0.0714 |
| 12_105803 | 7 | 0.1429 | 9 | 0.1111 | 5 | 0 | 8 | 0.25 | 29 | 0.1379 | 2 | 0 | 4 | 0 | 8 | 0 | 7 | 0.2857 | 21 | 0.0952 |
| 12_105803 | 7 | 0 | 13 | 0.1538 | 11 | 0.0909 | 9 | 0.1111 | 40 | 0.1 | 2 | 0.5 | 4 | 0 | 14 | 0.0714 | 9 | 0.3333 | 29 | 0.1724 |
| 12_105803 | 8 | 0.25 | 13 | 0.1538 | 10 | 0.2 | 11 | 0.2727 | 42 | 0.2143 | 3 | 0.3333 | 5 | 0.4 | 13 | 0.1538 | 9 | 0.2222 | 30 | 0.2333 |
| 12_105803 | 8 | 0.125 | 14 | 0.1429 | 13 | 0.0769 | 11 | 0.0909 | 46 | 0.1087 | 4 | 0.25 | 4 | 0 | 14 | 0 | 9 | 0.1111 | 31 | 0.0645 |
| 12_105803 | 7 | 0 | 13 | 0.3846 | 10 | 0.2 | 12 | 0.4167 | 42 | 0.2857 | 4 | 0.25 | 4 | 0 | 15 | 0.2667 | 8 | 0.375 | 31 | 0.2581 |
| 12_105804 | 8 | 0.375 | 16 | 0.25 | 12 | 0.25 | 13 | 0.3077 | 49 | 0.2857 | 5 | 0.6 | 6 | 0.1667 | 12 | 0.1667 | 10 | 0.3 | 33 | 0.2727 |
| 12_105805 | 12 | 0.25 | 18 | 0 | 15 | 0.1333 | 16 | 0.25 | 61 | 0.1475 | 5 | 0.2 | 8 | 0.375 | 17 | 0.1765 | 9 | 0.5556 | 39 | 0.3077 |
| 12_105805 | 12 | 0.0833 | 19 | 0.1053 | 16 | 0.0625 | 16 | 0 | 63 | 0.0635 | 6 | 0.1667 | 7 | 0.4286 | 18 | 0.1111 | 11 | 0.2727 | 42 | 0.2143 |
| 12_106042 | 1 | 1 | NA | NA | 1 | 1 | 2 | 0.5 | 4 | 0.75 | NA | NA | 1 | 1 | 2 | 1 | 3 | 1 | 6 | 1 |

| | | | | | | | | | | | | | | | | | | | | |
|---|---|---|---|---|---|---|---|---|---|---|---|---|---|---|---|---|---|---|---|---|
| 12_1060425 | 1 | 1 | NA | NA | 1 | 0 | 1 | 0 | 3 | 0.3333 | NA | NA | 1 | 0 | 2 | 0 | 3 | 0.3333 | 6 | 0.1667 |
| 12_1060425 | 1 | 0 | NA | NA | 1 | 1 | 2 | 0.5 | 4 | 0.5 | NA | NA | 1 | 0 | 1 | 1 | 2 | 1 | 4 | 0.75 |
| 12_1060425 | 1 | 1 | NA | NA | NA | NA | 2 | 1 | 3 | 1 | NA | NA | 1 | 1 | 2 | 1 | 2 | 1 | 5 | 1 |
| 12_1060427 | 1 | 1 | NA | NA | 1 | 1 | 2 | 0 | 4 | 0.5 | NA | NA | 1 | 0 | 2 | 0.5 | 3 | 0.6667 | 6 | 0.5 |
| 12_1078771 | 7 | 0 | 11 | 0.0909 | 7 | 0 | 11 | 0.0909 | 36 | 0.0556 | 7 | 0.1429 | 7 | 0 | 13 | 0.0769 | 7 | 0 | 34 | 0.0588 |
| 12_1078771 | 2 | 0 | 10 | 0 | 6 | 0.1667 | 9 | 0.2222 | 27 | 0.1111 | 7 | 0 | 7 | 0 | 13 | 0.1538 | 7 | 0.1429 | 34 | 0.0882 |
| 12_1078772 | 8 | 0.5 | 9 | 0.3333 | 8 | 0 | 11 | 0.0909 | 36 | 0.2222 | 8 | 0.125 | 8 | 0.25 | 15 | 0.2667 | 7 | 0.2857 | 38 | 0.2368 |
| 12_1078774 | 5 | 0.8 | 7 | 1 | 5 | 1 | 7 | 1 | 24 | 0.9583 | 6 | 1 | 9 | 0.8889 | 10 | 1 | 7 | 0.7143 | 32 | 0.9062 |
| 12_1078774 | 4 | 0.5 | 8 | 0.75 | 4 | 0.25 | 6 | 0.1667 | 22 | 0.4545 | 5 | 0.4 | 9 | 0.4444 | 10 | 0.7 | 6 | 0.1667 | 30 | 0.4667 |
| 12_1078786 | 4 | 0 | 5 | 0.6 | 3 | 0 | 7 | 0.5714 | 19 | 0.3684 | 3 | 0.3333 | 6 | 0 | 4 | 0.75 | 4 | 0.25 | 17 | 0.2941 |
| 12_1078793 | 10 | 0.3 | 13 | 0.3077 | 12 | 0.1667 | 14 | 0.2143 | 49 | 0.2449 | 12 | 0.4167 | 17 | 0.2353 | 16 | 0.3125 | 12 | 0.25 | 57 | 0.2982 |
| 12_1078793 | 12 | 0.1667 | 16 | 0.25 | 14 | 0.1429 | 16 | 0.125 | 58 | 0.1724 | 12 | 0.0833 | 20 | 0.2 | 23 | 0.0435 | 12 | 0.25 | 67 | 0.1343 |
| 12_1078796 | 8 | 0.25 | 11 | 0.2727 | 8 | 0.25 | 12 | 0.25 | 39 | 0.2564 | 8 | 0.375 | 16 | 0.375 | 16 | 0.25 | 11 | 0.2727 | 51 | 0.3137 |
| 12_1078796 | 5 | 0.2 | 13 | 0.2308 | 3 | 0.3333 | 10 | 0.4 | 31 | 0.2903 | 7 | 0 | 18 | 0.1667 | 17 | 0.0588 | 9 | 0.5556 | 51 | 0.1765 |
| 12_1078797 | 8 | 0.125 | 15 | 0 | 8 | 0.125 | 16 | 0.0625 | 47 | 0.0638 | 9 | 0 | 21 | 0.0952 | 21 | 0.1429 | 10 | 0 | 61 | 0.082 |
| 12_1078797 | 7 | 0 | 10 | 0.2 | 6 | 0.1667 | 11 | 0.0909 | 34 | 0.1176 | 9 | 0.3333 | 16 | 0.1875 | 19 | 0.1053 | 9 | 0.3333 | 53 | 0.2075 |
| 12_1078797 | 8 | 0.125 | 15 | 0.0667 | 10 | 0.3 | 16 | 0.0625 | 49 | 0.1224 | 9 | 0.3333 | 20 | 0.35 | 20 | 0.4 | 10 | 0.5 | 59 | 0.3898 |
| 12_1078798 | 7 | 0.5714 | 12 | 0.4167 | 8 | 0.5 | 13 | 0.6923 | 40 | 0.55 | 8 | 0.625 | 19 | 0.5789 | 18 | 0.5556 | 7 | 0.4286 | 52 | 0.5577 |
| 12_1078799 | 7 | 0.7143 | 14 | 0.7857 | 10 | 0.7 | 14 | 0.7857 | 45 | 0.7556 | 9 | 0.6667 | 20 | 0.85 | 18 | 0.8333 | 10 | 0.9 | 57 | 0.8246 |
| 12_1078799 | 2 | 0.5 | 9 | 0.1111 | 6 | 0.1667 | 8 | 0.25 | 25 | 0.2 | 5 | 0 | 10 | 0.2 | 11 | 0.3636 | 7 | 0.1429 | 33 | 0.2121 |
| 12_1078799 | 2 | 0 | NA | NA | 2 | 0.5 | NA | NA | 4 | 0.25 | 1 | 0 | 3 | 0 | 1 | 0 | 1 | 1 | 6 | 0.1667 |
| 12_1078800 | 3 | 0.6667 | 12 | 0.6667 | 8 | 0.625 | 8 | 0.75 | 31 | 0.6774 | 4 | 1 | 17 | 0.7647 | 13 | 0.8462 | 8 | 0.875 | 42 | 0.8333 |
| 12_1078801 | 6 | 0.3333 | 12 | 0.25 | 9 | 0.4444 | 13 | 0.4615 | 40 | 0.375 | 6 | 0.1667 | 18 | 0.3333 | 17 | 0.4118 | 9 | 0.3333 | 50 | 0.34 |
| 12_1078803 | 6 | 0.6667 | 15 | 0.4667 | 8 | 0.5 | 11 | 0.5455 | 40 | 0.525 | 8 | 0.625 | 17 | 0.6471 | 19 | 0.3684 | 9 | 0.7778 | 53 | 0.566 |
| 12_1078803 | 6 | 0.1667 | 14 | 0 | 10 | 0.1 | 10 | 0.1 | 40 | 0.075 | 5 | 0.4 | 18 | 0.1667 | 14 | 0.0714 | 5 | 0.2 | 42 | 0.1667 |
| 12_1078804 | 5 | 0.4 | 9 | 0 | 10 | 0.3 | 10 | 0.1 | 34 | 0.1765 | 6 | 0.1667 | 15 | 0.2667 | 13 | 0.1538 | 6 | 0.3333 | 40 | 0.225 |
| 12_1078805 | 6 | 0.1667 | 14 | 0.0714 | 9 | 0 | 13 | 0.2308 | 42 | 0.119 | 7 | 0.4286 | 16 | 0.125 | 15 | 0.0667 | 9 | 0.3333 | 47 | 0.1915 |
| 12_1078806 | 5 | 0 | 13 | 0 | 9 | 0.2222 | 11 | 0.2727 | 38 | 0.1316 | 7 | 0.1429 | 18 | 0.2778 | 14 | 0.2857 | 7 | 0 | 46 | 0.2174 |
| 12_1078808 | 4 | 0.25 | 12 | 0.25 | 8 | 0.25 | 8 | 0.25 | 32 | 0.25 | 6 | 0.1667 | 13 | 0.1538 | 11 | 0 | 6 | 0.1667 | 36 | 0.1111 |
| 12_1078811 | 5 | 0.4 | 9 | 0.2222 | 7 | 0.2857 | 8 | 0 | 29 | 0.2069 | 6 | 0.3333 | 12 | 0.4167 | 11 | 0.3636 | 6 | 0 | 35 | 0.3143 |
| 12_1078814 | 4 | 0 | 8 | 0 | 7 | 0 | 6 | 0.3333 | 25 | 0.08 | 4 | 0.5 | 10 | 0.1 | 8 | 0.375 | 3 | 0.3333 | 25 | 0.28 |
| 12_1078815 | 5 | 0.2 | 7 | 0.2857 | 6 | 0.1667 | 7 | 0.5714 | 25 | 0.32 | 5 | 0.4 | 9 | 0.1111 | 8 | 0 | 4 | 0 | 26 | 0.1154 |
| 12_1078815 | 5 | 1 | 6 | 1 | 6 | 0.8333 | 5 | 1 | 22 | 0.9545 | 2 | 1 | 8 | 0.875 | 8 | 0.875 | 3 | 1 | 21 | 0.9048 |
| 12_1078815 | 5 | 0.4 | 6 | 0.5 | 6 | 0.5 | 7 | 0.4286 | 24 | 0.4583 | 4 | 0.25 | 9 | 0.5556 | 9 | 0.8889 | 4 | 1 | 26 | 0.6923 |
| 12_1078816 | 4 | 0.5 | 6 | 0.8333 | 4 | 1 | 7 | 0.5714 | 21 | 0.7143 | 5 | 1 | 8 | 0.75 | 9 | 1 | 4 | 1 | 26 | 0.9231 |
| 12_1078816 | 4 | 0.75 | 6 | 0.6667 | 6 | 0.3333 | 5 | 0.4 | 21 | 0.5238 | 5 | 0.6 | 8 | 0.5 | 8 | 0.625 | 2 | 1 | 23 | 0.6087 |
| 12_1078817 | 5 | 1 | 6 | 1 | 6 | 1 | 5 | 1 | 22 | 1 | 3 | 1 | 8 | 0.875 | 7 | 1 | 3 | 1 | 21 | 0.9524 |
| 12_1078819 | 5 | 1 | 5 | 1 | 6 | 1 | 6 | 1 | 22 | 1 | 5 | 1 | 8 | 1 | 7 | 1 | 4 | 1 | 24 | 1 |
| 12_1078820 | 5 | 0 | 4 | 0.25 | 4 | 0.25 | 5 | 0.6 | 18 | 0.2778 | 3 | 0 | 8 | 0.375 | 7 | 0.1429 | 3 | 0 | 21 | 0.1905 |
| 12_1078821 | 5 | 0.4 | 4 | 0.75 | 6 | 0.3333 | 6 | 0.1667 | 21 | 0.381 | 5 | 0 | 5 | 0.2 | 6 | 0.3333 | 4 | 0 | 20 | 0.15 |
| 12_1078821 | 4 | 0.5 | 6 | 0.3333 | 6 | 0.3333 | 6 | 0.6667 | 22 | 0.4545 | 5 | 0.4 | 7 | 0.4286 | 6 | 0.3333 | 4 | 0.5 | 22 | 0.4091 |
| 12_1078822 | 5 | 0 | 5 | 0.2 | 6 | 0.1667 | 7 | 0.4286 | 23 | 0.2174 | 5 | 0.4 | 8 | 0.125 | 8 | 0.375 | 4 | 0.5 | 25 | 0.32 |
| 12_1078826 | 4 | 1 | 5 | 1 | 4 | 0.75 | 7 | 0.8571 | 20 | 0.9 | 4 | 0.5 | 7 | 1 | 5 | 0.8 | 2 | 1 | 18 | 0.8333 |
| 12_1078831 | 3 | 0.3333 | 3 | 0 | 4 | 0.5 | 4 | 0.25 | 14 | 0.2857 | 2 | 0 | 6 | 0.1667 | 3 | 0.3333 | 2 | 0.5 | 13 | 0.2308 |
| 12_1078832 | 3 | 0.3333 | 5 | 0.4 | 4 | 0.75 | 5 | 0.4 | 17 | 0.4706 | 3 | 0.3333 | 7 | 0.2857 | 3 | 0.6667 | 1 | 0 | 14 | 0.3571 |
| 12_1078832 | 3 | 1 | 4 | 1 | 4 | 1 | 4 | 1 | 15 | 1 | 3 | 1 | 6 | 0.8333 | 2 | 1 | 2 | 1 | 13 | 0.9231 |
| 12_1078832 | 2 | 1 | 5 | 0.6 | 5 | 1 | 5 | 0.4 | 17 | 0.7059 | 3 | 0.3333 | 6 | 0.8333 | 2 | 1 | 2 | 0.5 | 13 | 0.6923 |
| 12_1078833 | 2 | 0.5 | 4 | 0.5 | 5 | 0.6 | 4 | 1 | 15 | 0.6667 | 3 | 0.3333 | 6 | 0.1667 | 2 | 0 | 2 | 0 | 13 | 0.1538 |
| 12_1078834 | 1 | 0 | 2 | 0 | 3 | 0 | 4 | 0.25 | 10 | 0.1 | 3 | 0 | 7 | 0.1429 | 2 | 0 | 1 | 0 | 13 | 0.0769 |
| 12_1078834 | 1 | 0 | 5 | 0.2 | 5 | 0.4 | 5 | 0.6 | 16 | 0.375 | 3 | 0.6667 | 6 | 0.3333 | 1 | 1 | 2 | 0 | 12 | 0.4167 |
| 12_1078835 | 2 | 0 | 5 | 0.2 | 5 | 0 | 5 | 0 | 17 | 0.0588 | 4 | 0 | 5 | 0 | 2 | 1 | 2 | 0 | 13 | 0.1538 |
| 12_1079559 | 2 | 0.5 | 5 | 0.2 | 4 | 0.25 | 4 | 0 | 15 | 0.2 | 2 | 0 | 4 | 0.25 | 2 | 0 | 2 | 0 | 10 | 0.1 |
| 12_1079559 | 1 | 1 | 5 | 0.2 | 3 | 0.3333 | 2 | 0.5 | 11 | 0.3636 | 3 | 0.3333 | 2 | 0.5 | NA | NA | 2 | 0 | 7 | 0.2857 |
| 12_1080731 | 19 | 0.7895 | 14 | 0.7857 | 15 | 0.8667 | 18 | 0.7222 | 66 | 0.7879 | 17 | 0.8235 | 19 | 1 | 20 | 0.95 | 18 | 0.8333 | 74 | 0.9054 |
| 12_1080732 | 20 | 0.1 | 13 | 0.4615 | 17 | 0.2941 | 25 | 0.4 | 75 | 0.3067 | 21 | 0.2857 | 20 | 0.35 | 23 | 0.4348 | 22 | 0.4091 | 86 | 0.3721 |
| 12_1080734 | 20 | 0.3 | 10 | 0.6 | 13 | 0.2308 | 17 | 0.4118 | 60 | 0.3667 | 16 | 0.75 | 13 | 0.3846 | 21 | 0.4286 | 20 | 0.4 | 70 | 0.4857 |
| 12_1080735 | 20 | 0.35 | 12 | 0.4167 | 15 | 0.4667 | 21 | 0.5238 | 68 | 0.4412 | 19 | 0.5263 | 15 | 0.4667 | 17 | 0.4118 | 19 | 0.4211 | 70 | 0.4571 |
| 12_1080736 | 19 | 0.7895 | 10 | 0.8 | 16 | 0.6875 | 20 | 0.85 | 65 | 0.7846 | 18 | 0.8333 | 16 | 0.875 | 21 | 0.8571 | 20 | 0.95 | 75 | 0.88 |
| 12_1080737 | 16 | 0.4375 | 12 | 0.1667 | 15 | 0.6 | 17 | 0.4118 | 60 | 0.4167 | 15 | 0.4 | 15 | 0.4 | 18 | 0.5556 | 18 | 0.6111 | 66 | 0.5 |
| 12_1080737 | 18 | 1 | 11 | 0.9091 | 12 | 1 | 17 | 0.8235 | 58 | 0.931 | 17 | 0.8235 | 15 | 0.8667 | 19 | 0.8947 | 16 | 0.875 | 67 | 0.8657 |
| 12_1080845 | 9 | 0 | 8 | 0.125 | 10 | 0 | 10 | 0.1 | 37 | 0.0541 | 10 | 0.2 | 12 | 0.0833 | 13 | 0.0769 | 12 | 0.0833 | 47 | 0.1064 |
| 12_1081699 | 16 | 0.125 | 9 | 0.3333 | 9 | 0.2222 | 21 | 0.4762 | 55 | 0.3091 | 16 | 0.375 | 20 | 0.3 | 16 | 0.5 | 13 | 0.2308 | 65 | 0.3538 |
| 12_1081709 | 13 | 0.1538 | 6 | 0.1667 | 3 | 0 | 15 | 0.0667 | 37 | 0.1081 | 7 | 0 | 13 | 0.1538 | 10 | 0 | 8 | 0 | 38 | 0.0526 |
| 12_1081715 | 4 | 0.25 | 5 | 0 | 1 | 0 | 7 | 0 | 17 | 0.0588 | 5 | 0.2 | 4 | 0 | 1 | 0 | 3 | 0 | 13 | 0.0769 |
| 12_1081716 | 7 | 0.2857 | 4 | 0 | 2 | 0 | 9 | 0.1111 | 22 | 0.1364 | 5 | 0 | 3 | 0 | 2 | 1 | 4 | 0 | 14 | 0.1429 |

| | | | | | | | | | | | | | | | | | | | | |
|---|---|---|---|---|---|---|---|---|---|---|---|---|---|---|---|---|---|---|---|---|
| 12_108172 | 7 | 0.7143 | 4 | 0.25 | 1 | 1 | 9 | 0.5556 | 21 | 0.5714 | 4 | 0.75 | 3 | 0.6667 | 2 | 0.5 | 4 | 0.75 | 13 | 0.6923 |
| 12_108172 | 3 | 0.3333 | 2 | 0.5 | NA | NA | 4 | 0.25 | 9 | 0.3333 | 4 | 0.5 | 2 | 0.5 | 2 | 0.5 | 2 | 0.5 | 10 | 0.5 |
| 12_108172 | 3 | 0.3333 | 4 | 0.25 | NA | NA | 6 | 0.1667 | 13 | 0.2308 | 5 | 0 | 1 | 0 | 1 | 1 | 2 | 0 | 9 | 0.1111 |
| 12_108172 | 3 | 1 | 1 | 1 | 1 | 1 | 7 | 1 | 12 | 1 | 5 | 1 | 3 | 0.6667 | NA | NA | 4 | 1 | 12 | 0.9167 |
| 12_108172 | 7 | 0.4286 | 4 | 0.25 | NA | NA | 8 | 0.5 | 19 | 0.4211 | 5 | 0.2 | 3 | 0.3333 | 2 | 1 | 3 | 0.3333 | 13 | 0.3846 |
| 12_108172 | 6 | 0.8333 | 3 | 0.6667 | NA | NA | 7 | 0.8571 | 16 | 0.8125 | 4 | 1 | 2 | 0.5 | 1 | 1 | 4 | 0.75 | 11 | 0.8182 |
| 12_108172 | 7 | 0.2857 | 4 | 0 | NA | NA | 8 | 0.125 | 19 | 0.1579 | 6 | 0.1667 | 4 | 0 | 2 | 0 | 4 | 0 | 16 | 0.0625 |
| 12_108172 | 5 | 1 | 4 | 1 | NA | NA | 5 | 1 | 14 | 1 | 6 | 0.8333 | 2 | 1 | 1 | 1 | 4 | 1 | 13 | 0.9231 |
| 12_108294 | 2 | 1 | NA | NA | NA | NA | NA | NA | 2 | 1 | NA | NA | NA | NA | 1 | 1 | NA | NA | 1 | 1 |
| 12_108294 | 2 | 0.5 | NA | NA | NA | NA | NA | NA | 2 | 0.5 | NA | NA | NA | NA | 1 | 1 | NA | NA | 1 | 1 |
| 12_108294 | 2 | 1 | NA | NA | NA | NA | NA | NA | 2 | 1 | NA | NA | NA | NA | 2 | 0.5 | NA | NA | 2 | 0.5 |
| 12_108294 | 2 | 0.5 | NA | NA | NA | NA | NA | NA | 2 | 0.5 | NA | NA | NA | NA | 2 | 1 | NA | NA | 2 | 1 |
| 12_108294 | 2 | 1 | NA | NA | NA | NA | NA | NA | 2 | 1 | NA | NA | NA | NA | 2 | 0.5 | NA | NA | 2 | 0.5 |
| 12_108295 | NA | NA | NA | NA | NA | NA | 2 | 0.5 | 2 | 0.5 | 1 | 1 | 1 | 0 | 2 | 0 | NA | NA | 4 | 0.25 |
| 12_108327 | 13 | 0.1538 | 11 | 0.0909 | 14 | 0.1429 | 10 | 0 | 48 | 0.1042 | 13 | 0 | 19 | 0.1053 | 15 | 0.1333 | 8 | 0 | 55 | 0.0727 |
| 12_108328 | 22 | 0.0909 | 19 | 0.2105 | 24 | 0.1667 | 18 | 0.2222 | 83 | 0.1687 | 16 | 0.0625 | 26 | 0.1538 | 23 | 0 | 19 | 0.1053 | 84 | 0.0833 |
| 12_108504 | 5 | 0.6 | 3 | 0.6667 | 2 | 0 | 3 | 0.6667 | 13 | 0.5385 | NA | NA | 4 | 1 | 3 | 0.3333 | 2 | 1 | 9 | 0.7778 |
| 12_108504 | 4 | 0.25 | 3 | 0 | 2 | 0 | 2 | 0 | 11 | 0.0909 | NA | NA | 4 | 0 | 2 | 0 | 5 | 0.2 | 11 | 0.0909 |
| 12_108504 | 5 | 0 | 3 | 0.6667 | 2 | 0 | 3 | 0 | 13 | 0.1538 | NA | NA | 5 | 0 | 3 | 0 | 4 | 0.25 | 12 | 0.0833 |
| 12_108504 | 4 | 0.5 | 3 | 0.3333 | 2 | 0 | 3 | 0 | 12 | 0.25 | NA | NA | 4 | 0.25 | 2 | 0 | 1 | 0 | 7 | 0.1429 |
| 12_108504 | 4 | 0 | 3 | 0.3333 | 2 | 0.5 | 3 | 0.3333 | 12 | 0.25 | NA | NA | 5 | 0.4 | 3 | 0.3333 | 5 | 0.2 | 13 | 0.3077 |
| 12_108505 | 5 | 0.6 | 2 | 0.5 | 2 | 0 | 1 | 1 | 10 | 0.5 | 1 | 0 | 3 | 1 | 4 | 0.25 | 4 | 0.75 | 12 | 0.5833 |
| 12_108505 | 6 | 0.3333 | 3 | 0.3333 | 2 | 0 | 3 | 0.3333 | 14 | 0.2857 | 1 | 0 | 7 | 0.4286 | 3 | 0.3333 | 5 | 0.2 | 16 | 0.3125 |
| 12_108505 | 5 | 0.4 | 3 | 0 | 2 | 0 | 3 | 0 | 13 | 0.1538 | 1 | 0 | 4 | 0.25 | 4 | 0 | 4 | 0.25 | 13 | 0.1538 |
| 12_108505 | 6 | 0.1667 | 3 | 0 | 2 | 0 | 2 | 0 | 13 | 0.0769 | NA | NA | 5 | 0 | 4 | 0.25 | 4 | 0.25 | 13 | 0.1538 |
| 12_108505 | 5 | 0.6 | 2 | 0 | 2 | 0 | 3 | 0.6667 | 12 | 0.4167 | NA | NA | 5 | 0.6 | 4 | 0 | 5 | 0.8 | 14 | 0.5 |
| 12_108505 | 3 | 0.6667 | 3 | 0.3333 | 1 | 0 | 2 | 0 | 9 | 0.3333 | NA | NA | 7 | 0.7143 | 4 | 0.5 | 5 | 0.8 | 16 | 0.6875 |
| 12_108505 | 6 | 0.1667 | 2 | 0 | 2 | 0 | 3 | 0.3333 | 13 | 0.1538 | 1 | 0 | 7 | 0 | 4 | 0.5 | 4 | 0 | 16 | 0.125 |
| 12_108505 | 4 | 0.25 | 3 | 0.6667 | 2 | 0.5 | 3 | 0 | 12 | 0.3333 | 1 | 1 | 7 | 0.4286 | 4 | 0 | 5 | 0.2 | 17 | 0.2941 |
| 12_108505 | 5 | 0.4 | 2 | 0 | 2 | 0 | 3 | 0 | 12 | 0.1667 | 1 | 1 | 7 | 0.2857 | 4 | 0.25 | 4 | 0 | 16 | 0.25 |
| 12_108505 | 6 | 0.1667 | 3 | 0 | 2 | 0 | 4 | 0 | 15 | 0.0667 | NA | NA | 7 | 0 | 4 | 0.25 | 6 | 0.3333 | 17 | 0.1765 |
| 12_108650 | NA | NA | NA | NA | NA | NA | 1 | 1 | 1 | 1 | 1 | 1 | 1 | 0 | NA | NA | NA | NA | 2 | 0.5 |
| 12_108654 | 31 | 0.129 | 21 | 0 | 18 | 0.1667 | 39 | 0.0769 | 109 | 0.0917 | 11 | 0 | 32 | 0.2188 | 22 | 0.2273 | 19 | 0.0526 | 84 | 0.1548 |
| 12_108654 | 25 | 0.12 | 22 | 0.0455 | 18 | 0.1111 | 39 | 0.0769 | 104 | 0.0865 | 13 | 0 | 30 | 0.0667 | 26 | 0.1923 | 17 | 0.2353 | 86 | 0.1279 |
| 12_108656 | 1 | 1 | NA | NA | 1 | 0 | NA | NA | 2 | 0.5 | NA | NA | NA | NA | 2 | 0.5 | 1 | 1 | 3 | 0.6667 |
| 12_108656 | 1 | 1 | NA | NA | 1 | 0 | NA | NA | 2 | 0.5 | NA | NA | NA | NA | 1 | 1 | 1 | 0 | 2 | 0.5 |
| 12_108658 | NA | NA | NA | NA | 1 | 1 | NA | NA | 1 | 1 | NA | NA | NA | NA | 1 | 1 | 1 | 0 | 2 | 0.5 |
| 12_108658 | NA | NA | NA | NA | 1 | 1 | NA | NA | 1 | 1 | NA | NA | NA | NA | NA | NA | 1 | 1 | 1 | 1 |
| 12_108658 | NA | NA | NA | NA | 1 | 1 | NA | NA | 1 | 1 | NA | NA | NA | NA | NA | NA | 1 | 1 | 1 | 1 |
| 12_108757 | NA | NA | 2 | 0 | NA | NA | 1 | 1 | 3 | 0.3333 | NA | NA | 1 | 1 | 1 | 0 | 1 | 1 | 3 | 0.6667 |
| 12_108758 | NA | NA | 2 | 0.5 | 6 | 0.3333 | NA | NA | 8 | 0.375 | 4 | 0.75 | 4 | 0.5 | 3 | 0.3333 | 4 | 0.5 | 15 | 0.5333 |
| 12_108758 | NA | NA | 3 | 1 | 1 | 0 | NA | NA | 4 | 0.75 | 1 | 0 | 2 | 0 | 2 | 0.5 | 1 | 0 | 6 | 0.1667 |
| 12_108760 | NA | NA | 2 | 0.5 | 1 | 0 | 1 | 0 | 4 | 0.25 | 3 | 0.3333 | 1 | 0 | 2 | 0.5 | 2 | 0 | 8 | 0.25 |
| 12_108781 | 5 | 0.6 | 5 | 0.8 | 4 | 0.75 | 3 | 0.6667 | 17 | 0.7059 | 6 | 0.6667 | 2 | 1 | 5 | 0.8 | 2 | 0.5 | 15 | 0.7333 |
| 12_108781 | 8 | 0.75 | 7 | 1 | 7 | 0.8571 | 6 | 1 | 28 | 0.8929 | 6 | 0.6667 | 4 | 1 | 6 | 1 | 3 | 0.6667 | 19 | 0.8421 |
| 12_108781 | 7 | 0.1429 | 8 | 0.25 | 6 | 0 | 7 | 0.4286 | 28 | 0.2143 | 5 | 0 | 4 | 0 | 4 | 0.25 | 3 | 0 | 16 | 0.0625 |
| 12_108781 | 7 | 0.1429 | 8 | 0.125 | 5 | 0 | 5 | 0.2 | 25 | 0.12 | 6 | 0.1667 | 3 | 0 | 7 | 0.5714 | 3 | 0 | 19 | 0.2632 |
| 12_108781 | 7 | 0.2857 | 6 | 0.6667 | 7 | 0.7143 | 6 | 0.6667 | 26 | 0.5769 | 6 | 0.6667 | 4 | 1 | 7 | 0.7143 | 2 | 1 | 19 | 0.7895 |
| 12_108781 | 7 | 0.4286 | 7 | 0.5714 | 6 | 0.5 | 6 | 0.3333 | 26 | 0.4615 | 6 | 0.5 | 4 | 0.75 | 7 | 0.4286 | 3 | 0.3333 | 20 | 0.5 |
| 12_108781 | 8 | 1 | 7 | 1 | 7 | 0.8571 | 5 | 0.6 | 27 | 0.8889 | 5 | 1 | 4 | 1 | 6 | 1 | 3 | 1 | 18 | 1 |
| 12_108783 | 3 | 0.3333 | 2 | 0 | 2 | 0 | 2 | 1 | 9 | 0.3333 | 1 | 0 | 2 | 0 | 4 | 0.25 | 1 | 0 | 8 | 0.125 |
| 12_108783 | 3 | 0 | 1 | 0 | 2 | 0.5 | 1 | 1 | 7 | 0.2857 | 1 | 1 | 3 | 0.6667 | 4 | 0.75 | 1 | 0 | 9 | 0.6667 |
| 12_108783 | 2 | 0 | 1 | 0 | 1 | 0 | 2 | 0.5 | 6 | 0.1667 | 2 | 0 | 3 | 0 | 2 | 0.5 | NA | NA | 7 | 0.1429 |
| 12_108785 | 4 | 0 | 4 | 0 | 5 | 0.2 | 4 | 0 | 17 | 0.0588 | 4 | 0.25 | 4 | 0 | 4 | 0 | 3 | 0 | 15 | 0.0667 |
| 12_108785 | 6 | 0.3333 | 2 | 0.5 | 5 | 0.4 | 4 | 1 | 17 | 0.5294 | 5 | 0.8 | 5 | 0.6 | 5 | 0.6 | 4 | 0.5 | 19 | 0.6316 |
| 12_108791 | 3 | 0.3333 | 5 | 0.2 | 4 | 0 | 6 | 0.1667 | 18 | 0.1667 | 1 | 0 | 2 | 0 | 3 | 0.3333 | 1 | 0 | 7 | 0.1429 |
| 12_108792 | 4 | 0.25 | 6 | 0 | 3 | 0 | 6 | 0.5 | 19 | 0.2105 | 3 | 0.3333 | 3 | 0.6667 | 3 | 0.6667 | 1 | 0 | 10 | 0.5 |
| 12_108792 | NA | NA | 3 | 0.6667 | 2 | 1 | 5 | 1 | 10 | 0.9 | 1 | 1 | 3 | 0.6667 | 1 | 1 | NA | NA | 5 | 0.8 |
| 12_108792 | 4 | 0.75 | 5 | 0.4 | 3 | 0.6667 | 6 | 0.5 | 18 | 0.5556 | 2 | 1 | 3 | 1 | 3 | 1 | 1 | 1 | 9 | 1 |
| 12_108792 | 5 | 0.2 | 4 | 0 | 3 | 0.3333 | 4 | 0 | 16 | 0.125 | 3 | 0.3333 | 3 | 0 | 2 | 0 | 1 | 0 | 9 | 0.1111 |
| 12_108792 | 4 | 0.25 | 4 | 0.5 | 2 | 0.5 | 6 | 1 | 16 | 0.625 | 2 | 1 | 3 | 0.6667 | 2 | 1 | 1 | 1 | 8 | 0.875 |
| 12_108792 | 3 | 0 | 4 | 0.25 | 3 | 0 | 6 | 0.1667 | 16 | 0.125 | 3 | 0 | 3 | 0.3333 | 3 | 0 | 1 | 0 | 10 | 0.1 |
| 12_108792 | 4 | 0.25 | 4 | 0.5 | 2 | 0 | 5 | 0 | 15 | 0.2 | 3 | 0.3333 | 1 | 0 | 3 | 0.3333 | 1 | 0 | 8 | 0.25 |
| 12_108792 | 2 | 0.5 | 2 | 0.5 | 1 | 1 | 3 | 0.6667 | 8 | 0.625 | 1 | 1 | 1 | 1 | 2 | 1 | 1 | 1 | 5 | 1 |
| 12_108792 | 4 | 0.25 | 4 | 0.25 | 3 | 0.6667 | 6 | 0.5 | 17 | 0.4118 | 3 | 0.6667 | 1 | 0 | 3 | 0.3333 | NA | NA | 7 | 0.4286 |

| | | | | | | | | | | | | | | | | | | | | |
|---|---|---|---|---|---|---|---|---|---|---|---|---|---|---|---|---|---|---|---|---|
| 12_108793 | 2 | 0 | 2 | 0 | 1 | 0 | 6 | 0.1667 | 11 | 0.0909 | 3 | 0 | 2 | 0.5 | 3 | 0 | 1 | 0 | 9 | 0.1111 |
| 12_108793 | 3 | 0.6667 | 2 | 1 | 2 | 1 | 6 | 1 | 13 | 0.9231 | 3 | 0.6667 | 2 | 1 | 3 | 1 | 1 | 1 | 9 | 0.8889 |
| 12_108793 | 3 | 0.3333 | 1 | 0 | 1 | 1 | 4 | 0.75 | 9 | 0.5556 | 2 | 1 | 2 | 0.5 | 3 | 0.6667 | 1 | 1 | 8 | 0.75 |
| 12_108793 | 2 | 0 | 1 | 0 | 2 | 0.5 | 6 | 0.5 | 11 | 0.3636 | 1 | 1 | 1 | 0 | 3 | 1 | 1 | 0 | 6 | 0.6667 |
| 12_108793 | 3 | 0.3333 | 1 | 0 | 2 | 0 | 4 | 0.75 | 10 | 0.4 | 2 | 0.5 | 2 | 0.5 | 3 | 0.6667 | NA | NA | 7 | 0.5714 |
| 12_108826 | 14 | 0.0714 | 14 | 0.2857 | 9 | 0 | 17 | 0.1765 | 54 | 0.1481 | 9 | 0.2222 | 7 | 0.1429 | 8 | 0.25 | 16 | 0.25 | 40 | 0.225 |
| 12_108826 | 15 | 0.0667 | 15 | 0.1333 | 10 | 0 | 14 | 0.0714 | 54 | 0.0741 | 8 | 0.375 | 9 | 0 | 12 | 0.3333 | 19 | 0.1053 | 48 | 0.1875 |
| 12_108826 | 15 | 0.0667 | 14 | 0.0714 | 8 | 0.125 | 14 | 0.1429 | 51 | 0.098 | 6 | 0 | 5 | 0.6 | 8 | 0.25 | 17 | 0.1176 | 36 | 0.1944 |
| 12_108827 | 11 | 0.1818 | 12 | 0.6667 | 7 | 0.7143 | 14 | 0.0714 | 44 | 0.3636 | 6 | 0.3333 | 4 | 0.5 | 9 | 0.1111 | 14 | 0.2143 | 33 | 0.2424 |
| 12_108832 | 10 | 0 | 11 | 0.0909 | 4 | 0 | 8 | 0.125 | 33 | 0.0606 | 7 | 0.1429 | 6 | 0 | 8 | 0 | 12 | 0.0833 | 33 | 0.0606 |
| 12_108833 | 8 | 0 | 8 | 0.125 | 2 | 0 | 7 | 0.5714 | 25 | 0.2 | 6 | 0 | 6 | 0.3333 | 6 | 0.3333 | 10 | 0 | 28 | 0.1429 |
| 12_108841 | 11 | 0 | 9 | 0.2222 | 2 | 0.5 | 6 | 0.1667 | 28 | 0.1429 | 6 | 0.3333 | 6 | 0 | 5 | 0.2 | 13 | 0.1538 | 30 | 0.1667 |
| 12_108864 | NA | NA | 2 | 0.5 | NA | NA | 1 | 0 | 3 | 0.3333 | 1 | 0 | NA | NA | NA | NA | 1 | 1 | 2 | 0.5 |
| 12_108864 | 1 | 0 | 3 | 0.3333 | NA | NA | 1 | 1 | 5 | 0.4 | 1 | 0 | NA | NA | NA | NA | 1 | 1 | 2 | 0.5 |
| 12_108864 | 1 | 1 | 3 | 1 | NA | NA | 1 | 1 | 5 | 1 | NA | NA | NA | NA | NA | NA | 1 | 1 | 1 | 1 |
| 12_108864 | 1 | 1 | 1 | 1 | NA | NA | 1 | 1 | 3 | 1 | 1 | 0 | NA | NA | NA | NA | 1 | 1 | 2 | 0.5 |
| 12_108864 | 1 | 1 | 3 | 0.6667 | NA | NA | NA | NA | 4 | 0.75 | 1 | 1 | NA | NA | NA | NA | 1 | 1 | 2 | 1 |
| 12_108864 | 1 | 1 | 3 | 0 | NA | NA | 1 | 0 | 5 | 0.2 | 1 | 1 | NA | NA | NA | NA | 1 | 0 | 2 | 0.5 |
| 12_108993 | 2 | 0 | 4 | 0.25 | NA | NA | 1 | 0 | 7 | 0.1429 | 2 | 0.5 | NA | NA | 1 | 0 | 1 | 0 | 4 | 0.25 |
| 12_108994 | 2 | 0.5 | 3 | 0.6667 | NA | NA | 1 | 0 | 6 | 0.5 | 2 | 0.5 | NA | NA | NA | NA | 1 | 1 | 3 | 0.6667 |
| 12_108998 | 14 | 0.0714 | 17 | 0 | 14 | 0.1429 | 17 | 0.1765 | 62 | 0.0968 | 16 | 0.0625 | 22 | 0.1364 | 24 | 0.1667 | 11 | 0 | 73 | 0.1096 |
| 12_108998 | 17 | 0.2353 | 21 | 0.1905 | 16 | 0.125 | 18 | 0.1667 | 72 | 0.1806 | 14 | 0.0714 | 20 | 0.35 | 29 | 0.1724 | 11 | 0 | 74 | 0.1757 |
| 12_108998 | 12 | 0.0833 | 13 | 0.0769 | 15 | 0 | 20 | 0.05 | 60 | 0.05 | 16 | 0.125 | 16 | 0.125 | 26 | 0.0385 | 10 | 0.1 | 68 | 0.0882 |
| 12_108998 | 15 | 0.7333 | 12 | 0.6667 | 16 | 0.6875 | 16 | 0.4375 | 59 | 0.6271 | 14 | 0.6429 | 19 | 0.7368 | 28 | 0.75 | 9 | 0.7778 | 70 | 0.7286 |
| 12_108998 | 15 | 0.6 | 14 | 0.5714 | 13 | 0.6154 | 18 | 0.3333 | 60 | 0.5167 | 12 | 0.3333 | 18 | 0.6667 | 25 | 0.52 | 11 | 0.5455 | 66 | 0.5303 |
| 12_108999 | 10 | 0.6 | 7 | 0.4286 | 9 | 0.5556 | 14 | 0.5 | 40 | 0.525 | 10 | 0.6 | 14 | 0.5 | 16 | 0.5 | 6 | 0.8333 | 46 | 0.5652 |
| 12_108999 | 10 | 1 | 11 | 0.9091 | 11 | 1 | 16 | 1 | 48 | 0.9792 | 8 | 1 | 18 | 1 | 20 | 0.9 | 8 | 1 | 54 | 0.963 |
| 12_109428 | 7 | 0.1429 | 1 | 0 | 3 | 0 | 6 | 0 | 17 | 0.0588 | NA | NA | 5 | 0 | 7 | 0.1429 | 2 | 0 | 14 | 0.0714 |
| 12_109429 | 4 | 0.5 | 1 | 0 | 2 | 0.5 | 6 | 0.8333 | 13 | 0.6154 | NA | NA | 5 | 0.6 | 6 | 0.3333 | 2 | 0 | 13 | 0.3846 |
| 12_109429 | 7 | 1 | 1 | 1 | 3 | 1 | 7 | 1 | 18 | 1 | NA | NA | 6 | 1 | 8 | 1 | 2 | 0.5 | 16 | 0.9375 |
| 12_109525 | 3 | 0.3333 | 8 | 0.375 | 3 | 0.3333 | 7 | 0.5714 | 21 | 0.4286 | 3 | 0.3333 | 3 | 0.3333 | 10 | 0.4 | 9 | 0.5556 | 25 | 0.44 |
| 12_109525 | 3 | 0.3333 | 9 | 0.2222 | 4 | 0.25 | 7 | 0.2857 | 23 | 0.2609 | 3 | 0 | 3 | 0.3333 | 9 | 0.1111 | 9 | 0.1111 | 24 | 0.125 |
| 12_109525 | 3 | 0 | 7 | 0.2857 | 2 | 0.5 | 6 | 0.3333 | 18 | 0.2778 | 3 | 0 | 2 | 0 | 8 | 0.25 | 7 | 0 | 20 | 0.1 |
| 12_109525 | 3 | 0 | 7 | 0.1429 | 4 | 0 | 6 | 0.1667 | 20 | 0.1 | 3 | 0 | 2 | 0 | 7 | 0.1429 | 6 | 0.3333 | 18 | 0.1667 |
| 12_109525 | 2 | 1 | 8 | 1 | 4 | 0.75 | 4 | 1 | 18 | 0.9444 | 2 | 0.5 | 2 | 0.5 | 8 | 1 | 6 | 0.6667 | 18 | 0.7778 |
| 12_109525 | 2 | 0.5 | 7 | 0.5714 | 2 | 0.5 | 5 | 0.4 | 16 | 0.5 | 1 | 1 | 3 | 0.6667 | 7 | 0.4286 | 5 | 0.4 | 16 | 0.5 |
| 12_109525 | 1 | 1 | 9 | 0.7778 | 3 | 0.3333 | 6 | 0.5 | 19 | 0.6316 | 2 | 0.5 | 1 | 1 | 6 | 1 | 9 | 0.5556 | 18 | 0.7222 |
| 12_109525 | 3 | 1 | 8 | 1 | 4 | 1 | 7 | 0.8571 | 22 | 0.9545 | 3 | 1 | 3 | 0.3333 | 8 | 0.75 | 8 | 1 | 22 | 0.8182 |
| 12_109525 | 3 | 1 | 9 | 1 | 4 | 1 | 5 | 1 | 21 | 1 | 1 | 0 | 4 | 1 | 8 | 1 | 8 | 1 | 21 | 0.9524 |
| 12_109526 | NA | NA | 2 | 0 | NA | NA | 3 | 0.3333 | 5 | 0.2 | 1 | 0 | NA | NA | 1 | 0 | 2 | 1 | 4 | 0.5 |
| 12_109527 | 1 | 0 | 1 | 0 | 2 | 0 | 4 | 0.5 | 8 | 0.25 | 1 | 0 | 2 | 0.5 | 2 | 0 | 4 | 0.25 | 9 | 0.2222 |
| 12_109529 | 5 | 0 | 5 | 0 | 5 | 0.4 | 8 | 0 | 23 | 0.087 | 7 | 0.5714 | 8 | 0.25 | 6 | 0.1667 | 8 | 0 | 29 | 0.2414 |
| 12_109530 | 4 | 0 | 1 | 0 | 1 | 1 | 1 | 0 | 7 | 0.1429 | 1 | 0 | 3 | 0 | NA | NA | 5 | 0.6 | 9 | 0.3333 |
| 12_109530 | 3 | 1 | 2 | 0.5 | 1 | 1 | 1 | 1 | 7 | 0.8571 | 1 | 0 | 3 | 0 | NA | NA | 5 | 0.4 | 9 | 0.2222 |
| 12_109530 | 2 | 0 | 3 | 0.3333 | NA | NA | 2 | 0 | 7 | 0.1429 | 1 | 0 | 3 | 0.3333 | NA | NA | 7 | 0.2857 | 11 | 0.2727 |
| 12_109530 | 4 | 0 | 5 | 0.4 | 2 | 0 | 4 | 0.5 | 15 | 0.2667 | 1 | 1 | 2 | 0 | 2 | 0 | 9 | 0.3333 | 14 | 0.2857 |
| 12_109532 | 13 | 0.1538 | 10 | 0 | 10 | 0.2 | 14 | 0 | 47 | 0.0851 | 3 | 0 | 15 | 0.1333 | 9 | 0 | 12 | 0 | 39 | 0.0513 |
| 12_109533 | 24 | 0.0417 | 14 | 0.0714 | 17 | 0.0588 | 15 | 0.0667 | 70 | 0.0571 | 4 | 0 | 25 | 0.04 | 17 | 0.2353 | 14 | 0.2143 | 60 | 0.1333 |
| 12_109533 | 17 | 0.3529 | 12 | 0.5 | 13 | 0.1538 | 14 | 0.1429 | 56 | 0.2857 | 3 | 1 | 18 | 0.2222 | 16 | 0.375 | 14 | 0.2143 | 51 | 0.3137 |
| 12_109534 | 24 | 0.125 | 18 | 0.0556 | 18 | 0.1111 | 23 | 0 | 83 | 0.0723 | 5 | 0.2 | 17 | 0.1176 | 22 | 0.0455 | 15 | 0 | 59 | 0.0678 |
| 12_109534 | 28 | 0.1071 | 24 | 0.125 | 23 | 0.2609 | 27 | 0.1852 | 102 | 0.1667 | 10 | 0.1 | 26 | 0.3462 | 23 | 0.3478 | 21 | 0.1905 | 80 | 0.275 |
| 12_109535 | 8 | 0 | 10 | 0.2 | 8 | 0.125 | 16 | 0.0625 | 42 | 0.0952 | 2 | 0 | 12 | 0.0833 | 14 | 0 | 9 | 0.1111 | 37 | 0.0541 |
| 12_109536 | 13 | 0 | 8 | 0.25 | 8 | 0 | 17 | 0.0588 | 46 | 0.0652 | 5 | 0 | 11 | 0 | 16 | 0.125 | 16 | 0.1875 | 48 | 0.1042 |
| 12_109537 | 16 | 0.0625 | 14 | 0.2143 | 11 | 0.2727 | 20 | 0.1 | 61 | 0.1475 | 6 | 0 | 15 | 0.1333 | 24 | 0.1667 | 18 | 0.2778 | 63 | 0.1746 |
| 12_109537 | 17 | 0.1176 | 21 | 0 | 18 | 0.1667 | 21 | 0.1905 | 77 | 0.1169 | 8 | 0 | 20 | 0.05 | 38 | 0.1579 | 30 | 0.3 | 96 | 0.1667 |
| 12_109537 | 19 | 0.2105 | 22 | 0.1364 | 19 | 0.1053 | 26 | 0.0385 | 86 | 0.1163 | 8 | 0.25 | 19 | 0.1053 | 40 | 0.025 | 32 | 0.1875 | 99 | 0.1111 |
| 12_109537 | 20 | 0.05 | 24 | 0.2083 | 20 | 0.05 | 27 | 0.1852 | 91 | 0.1319 | 9 | 0.1111 | 23 | 0.2609 | 35 | 0.0857 | 30 | 0.2 | 97 | 0.1649 |
| 12_109537 | 17 | 0.4706 | 18 | 0.5556 | 19 | 0.4737 | 25 | 0.4 | 79 | 0.4684 | 7 | 0.4286 | 18 | 0.6111 | 36 | 0.6944 | 28 | 0.6429 | 89 | 0.6404 |
| 12_109537 | 14 | 0.4286 | 14 | 0.5714 | 20 | 0.75 | 22 | 0.7273 | 70 | 0.6429 | 8 | 0.75 | 16 | 0.6875 | 33 | 0.6061 | 23 | 0.6087 | 80 | 0.6375 |
| 12_109537 | 18 | 0.6667 | 17 | 0.7647 | 19 | 0.5789 | 21 | 0.619 | 75 | 0.6533 | 8 | 0.375 | 17 | 0.6471 | 36 | 0.7222 | 24 | 0.625 | 85 | 0.6471 |
| 12_109537 | 17 | 0.9412 | 22 | 0.7273 | 22 | 0.9091 | 23 | 0.913 | 84 | 0.869 | 8 | 1 | 20 | 0.8 | 32 | 0.7812 | 31 | 0.8387 | 91 | 0.8242 |
| 12_109613 | 4 | 0.25 | 6 | 0.1667 | 3 | 0.3333 | 5 | 0 | 18 | 0.1667 | NA | NA | 3 | 0.3333 | 4 | 0.5 | NA | NA | 7 | 0.4286 |
| 12_109627 | 11 | 0.6364 | 10 | 0.4 | 13 | 0.2308 | 18 | 0.3333 | 52 | 0.3846 | 7 | 0.1429 | 9 | 0.2222 | 12 | 0.0833 | 7 | 0.4286 | 35 | 0.2 |
| 12_109627 | 10 | 0 | 11 | 0.5455 | 13 | 0.1538 | 16 | 0.375 | 50 | 0.28 | 7 | 0.5714 | 10 | 0.3 | 12 | 0.4167 | 7 | 0.4286 | 36 | 0.4167 |
| 12_109627 | 11 | 0.6364 | 11 | 0.8182 | 12 | 0.8333 | 18 | 0.8889 | 52 | 0.8077 | 9 | 0.4444 | 9 | 0.5556 | 12 | 0.5 | 7 | 0.7143 | 37 | 0.5405 |

| | | | | | | | | | | | | | | | | | | | | |
|---|---|---|---|---|---|---|---|---|---|---|---|---|---|---|---|---|---|---|---|---|
| 12_109627 | NA | NA | 1 | 1 | NA | NA | NA | NA | 1 | 1 | NA | NA | NA | NA | NA | NA | 1 | 1 | 1 | 1 |
| 12_109627 | 7 | 1 | 10 | 0.5 | 10 | 0.7 | 14 | 0.6429 | 41 | 0.6829 | 6 | 0.3333 | 7 | 1 | 10 | 0.7 | 7 | 0.5714 | 30 | 0.6667 |
| 12_109627 | 9 | 0.2222 | 8 | 0.25 | 10 | 0.2 | 14 | 0.1429 | 41 | 0.1951 | 7 | 0 | 8 | 0.125 | 13 | 0.0769 | 5 | 0.2 | 33 | 0.0909 |
| 12_109627 | 10 | 0 | 12 | 0.1667 | 13 | 0.0769 | 20 | 0.05 | 55 | 0.0727 | 9 | 0.1111 | 9 | 0 | 12 | 0.1667 | 9 | 0.1111 | 39 | 0.1026 |
| 12_109627 | 11 | 0 | 12 | 0.0833 | 11 | 0.0909 | 19 | 0.1579 | 53 | 0.0943 | 8 | 0 | 10 | 0.3 | 12 | 0 | 8 | 0.125 | 38 | 0.1053 |
| 12_109627 | 10 | 0.1 | 14 | 0.0714 | 15 | 0.1333 | 20 | 0.1 | 59 | 0.1017 | 9 | 0.3333 | 10 | 0.1 | 13 | 0.0769 | 9 | 0 | 41 | 0.122 |
| 12_109627 | 11 | 0.0909 | 14 | 0.3571 | 14 | 0.5 | 19 | 0.3158 | 58 | 0.3276 | 8 | 0.125 | 10 | 0 | 14 | 0.3571 | 9 | 0.3333 | 41 | 0.2195 |
| 12_109627 | 9 | 0.8889 | 12 | 0.9167 | 9 | 1 | 14 | 1 | 44 | 0.9545 | 7 | 1 | 6 | 0.8333 | 10 | 0.8 | 7 | 0.7143 | 30 | 0.8333 |
| 12_109715 | 4 | 0 | 5 | 0 | 6 | 0.1667 | 9 | 0.1111 | 24 | 0.0833 | 3 | 0 | 2 | 0.5 | 9 | 0.1111 | 5 | 0.4 | 19 | 0.2105 |
| 12_110246 | NA | NA | NA | NA | NA | NA | 1 | 1 | 1 | 1 | NA | NA | 1 | 1 | NA | NA | NA | NA | 1 | 1 |
| 12_110537 | 12 | 0.0833 | 7 | 0.1429 | 11 | 0.1818 | 14 | 0.1429 | 44 | 0.1364 | 4 | 0 | 8 | 0 | 7 | 0.1429 | 11 | 0.0909 | 30 | 0.0667 |
| 12_110537 | 11 | 0.2727 | 9 | 0.2222 | 10 | 0.2 | 13 | 0.3077 | 43 | 0.2558 | 7 | 0.2857 | 9 | 0.1111 | 8 | 0 | 15 | 0 | 39 | 0.0769 |
| 12_110547 | 4 | 0.25 | 6 | 0.1667 | 8 | 0 | 7 | 0.1429 | 25 | 0.12 | 3 | 0.3333 | 8 | 0.125 | 6 | 0 | 3 | 0.3333 | 20 | 0.15 |
| 12_110547 | 8 | 0.125 | 5 | 0 | 9 | 0.3333 | 8 | 0.25 | 30 | 0.2 | 5 | 0.2 | 8 | 0 | 4 | 0.5 | 5 | 0 | 22 | 0.1364 |
| 12_110549 | 6 | 0 | 12 | 0.0833 | 12 | 0.0833 | 8 | 0 | 38 | 0.0526 | 7 | 0.1429 | 11 | 0.1818 | 6 | 0 | 5 | 0.2 | 29 | 0.1379 |
| 12_110549 | 7 | 0 | 12 | 0.1667 | 11 | 0.0909 | 8 | 0 | 38 | 0.0789 | 4 | 0 | 7 | 0 | 5 | 0 | 3 | 0.3333 | 19 | 0.0526 |
| 12_110565 | 2 | 1 | 1 | 1 | NA | NA | NA | NA | 3 | 1 | NA | NA | 1 | 1 | 1 | 0 | NA | NA | 2 | 0.5 |
| 12_110565 | 2 | 0.5 | 1 | 0 | NA | NA | NA | NA | 3 | 0.3333 | NA | NA | 1 | 1 | 1 | 0 | NA | NA | 2 | 0.5 |
| 12_110565 | 2 | 0.5 | 1 | 1 | NA | NA | NA | NA | 3 | 0.6667 | NA | NA | 1 | 1 | 1 | 0 | NA | NA | 2 | 0.5 |
| 12_110616 | 6 | 0 | 6 | 0 | 2 | 0 | 3 | 0.3333 | 17 | 0.0588 | 4 | 0 | 6 | 0.1667 | 5 | 0 | 2 | 0 | 17 | 0.0588 |
| 12_110633 | 35 | 0.1714 | 33 | 0.1818 | 38 | 0.2368 | 27 | 0.3333 | 133 | 0.2256 | 28 | 0 | 30 | 0.3667 | 21 | 0.1905 | 21 | 0.2857 | 100 | 0.21 |
| 12_110633 | 40 | 0.5 | 33 | 0.5455 | 35 | 0.5143 | 30 | 0.7333 | 138 | 0.5652 | 32 | 0.4688 | 31 | 0.4839 | 19 | 0.6842 | 21 | 0.381 | 103 | 0.4951 |
| 12_110633 | 37 | 0.9189 | 37 | 0.9459 | 40 | 0.95 | 31 | 0.871 | 145 | 0.9241 | 35 | 0.9429 | 36 | 0.9722 | 27 | 0.963 | 23 | 0.9565 | 121 | 0.9587 |
| 12_110657 | 102 | 0.1275 | 158 | 0.0949 | 127 | 0.1654 | 118 | 0.1186 | 505 | 0.1248 | 67 | 0.1194 | 113 | 0.0885 | 92 | 0.1304 | 136 | 0.1103 | 408 | 0.1103 |
| 12_110658 | 96 | 0.0833 | 146 | 0.0548 | 117 | 0.0427 | 108 | 0.0463 | 467 | 0.0557 | 58 | 0.0517 | 111 | 0.0901 | 90 | 0.0222 | 132 | 0.0455 | 391 | 0.0537 |
| 12_110658 | 62 | 0.0645 | 104 | 0.0673 | 83 | 0.0723 | 88 | 0.0795 | 337 | 0.0712 | 42 | 0.0238 | 83 | 0.0602 | 65 | 0.0769 | 91 | 0.0879 | 281 | 0.0676 |
| 12_110659 | 80 | 0.2 | 125 | 0.192 | 98 | 0.2245 | 96 | 0.1979 | 399 | 0.203 | 46 | 0.1304 | 95 | 0.1789 | 73 | 0.1507 | 114 | 0.1842 | 328 | 0.1677 |
| 12_110661 | 62 | 0.129 | 98 | 0.102 | 73 | 0.1096 | 79 | 0.1772 | 312 | 0.1282 | 39 | 0.1795 | 72 | 0.1111 | 58 | 0.2069 | 98 | 0.1327 | 267 | 0.1498 |
| 12_110759 | 1 | 0 | 5 | 0.2 | NA | NA | 3 | 0.3333 | 9 | 0.2222 | 2 | 0.5 | 1 | 0 | 1 | 0 | 1 | 0 | 5 | 0.2 |
| 12_110759 | 1 | 0 | 5 | 0.2 | NA | NA | 3 | 0.6667 | 9 | 0.3333 | 3 | 0 | 1 | 0 | 1 | 1 | 1 | 0 | 6 | 0.1667 |
| 12_110759 | NA | NA | 4 | 0.25 | NA | NA | 2 | 0.5 | 6 | 0.3333 | 3 | 0 | 1 | 1 | 1 | 0 | 1 | 1 | 6 | 0.3333 |
| 12_110760 | NA | NA | 3 | 0.6667 | NA | NA | 3 | 1 | 6 | 0.8333 | 3 | 0.3333 | 1 | 1 | 1 | 1 | 1 | 1 | 6 | 0.6667 |
| 12_110760 | NA | NA | 5 | 0.4 | NA | NA | 2 | 0.5 | 7 | 0.4286 | 3 | 0.3333 | 3 | 0.6667 | 1 | 1 | 1 | 1 | 8 | 0.625 |
| 12_110760 | NA | NA | 5 | 0.2 | NA | NA | 3 | 0.3333 | 8 | 0.25 | 3 | 0 | 2 | 1 | 1 | 0 | 1 | 1 | 7 | 0.4286 |
| 12_110760 | NA | NA | 5 | 0.4 | NA | NA | 3 | 1 | 8 | 0.625 | 2 | 0.5 | 3 | 0.3333 | 1 | 1 | 1 | 1 | 7 | 0.5714 |
| 12_110760 | 1 | 0 | 5 | 0.6 | NA | NA | 3 | 0.3333 | 9 | 0.4444 | 3 | 1 | 3 | 0.6667 | 1 | 1 | 1 | 0 | 8 | 0.75 |
| 12_110760 | 1 | 0 | 5 | 0 | NA | NA | 2 | 0.5 | 8 | 0.125 | 4 | 0 | 2 | 0.5 | 1 | 0 | 1 | 1 | 8 | 0.25 |
| 12_110760 | NA | NA | 4 | 0.75 | NA | NA | 3 | 1 | 7 | 0.8571 | 3 | 1 | 3 | 0.6667 | 1 | 0 | 1 | 1 | 8 | 0.75 |
| 12_110760 | 1 | 0 | 5 | 0.4 | NA | NA | 3 | 0 | 9 | 0.2222 | 4 | 0.25 | 3 | 0.3333 | 1 | 0 | 1 | 0 | 9 | 0.2222 |
| 12_110760 | 1 | 0 | 5 | 1 | NA | NA | 3 | 1 | 9 | 0.8889 | 4 | 0.75 | 2 | 0.5 | 1 | 1 | 1 | 1 | 8 | 0.75 |
| 12_110760 | NA | NA | 3 | 0.6667 | NA | NA | 2 | 1 | 5 | 0.8 | 3 | 1 | 2 | 1 | 1 | 1 | 1 | 1 | 7 | 1 |
| 12_110760 | 1 | 0 | 5 | 0.4 | NA | NA | 3 | 0.3333 | 9 | 0.3333 | 4 | 0.5 | 3 | 0.3333 | 1 | 0 | 1 | 1 | 9 | 0.4444 |
| 12_110760 | NA | NA | 5 | 1 | NA | NA | 3 | 0.3333 | 8 | 0.75 | 4 | 0.75 | 2 | 1 | 1 | 0 | 1 | 1 | 8 | 0.75 |
| 12_110760 | 1 | 0 | 3 | 0.3333 | NA | NA | 1 | 0 | 5 | 0.2 | 3 | 0.3333 | NA | NA | 1 | 0 | 1 | 0 | 5 | 0.2 |
| 12_110760 | NA | NA | 4 | 0.5 | NA | NA | 2 | 1 | 6 | 0.6667 | 4 | 1 | 3 | 1 | 1 | 1 | 1 | 1 | 9 | 1 |
| 12_110760 | 1 | 0 | 5 | 0.4 | NA | NA | 3 | 1 | 9 | 0.5556 | 4 | 0.75 | 3 | 1 | 2 | 0 | 1 | 1 | 10 | 0.7 |
| 12_110764 | 1 | 0 | NA | NA | NA | NA | 2 | 0.5 | 3 | 0.3333 | 1 | 1 | 1 | 1 | NA | NA | 3 | 0.3333 | 5 | 0.6 |
| 12_110769 | 1 | 0 | 2 | 0.5 | 2 | 0.5 | 1 | 0 | 6 | 0.1667 | NA | NA | 3 | 0.6667 | NA | NA | 1 | 0 | 4 | 0.5 |
| 12_110770 | NA | NA | 2 | 0.5 | 2 | 0.5 | 1 | 0 | 5 | 0.4 | 1 | 0 | 3 | 0 | NA | NA | 1 | 1 | 5 | 0.2 |
| 12_110770 | 2 | 0 | 3 | 0 | 3 | 0.3333 | 2 | 0.5 | 10 | 0.2 | 1 | 0 | 3 | 0.3333 | NA | NA | 1 | 0 | 5 | 0.2 |
| 12_110770 | 2 | 0 | 3 | 0.3333 | 2 | 0.5 | 2 | 0 | 9 | 0.2222 | 1 | 0 | 4 | 0.25 | NA | NA | 1 | 0 | 6 | 0.1667 |
| 12_110770 | 1 | 1 | NA | NA | NA | NA | NA | NA | 1 | 1 | 3 | 0.6667 | 2 | 0 | 1 | 1 | NA | NA | 6 | 0.5 |
| 12_110770 | 1 | 1 | NA | NA | NA | NA | NA | NA | 1 | 1 | 3 | 1 | 1 | 1 | NA | NA | NA | NA | 4 | 1 |
| 12_110770 | 1 | 1 | NA | NA | NA | NA | NA | NA | 1 | 1 | 1 | 1 | 1 | 1 | 1 | 1 | NA | NA | 3 | 1 |
| 12_110770 | 1 | 1 | NA | NA | NA | NA | NA | NA | 1 | 1 | 3 | 1 | NA | NA | 1 | 1 | NA | NA | 4 | 1 |
| 12_110770 | 1 | 1 | NA | NA | NA | NA | NA | NA | 1 | 1 | 1 | 1 | 1 | 1 | 1 | 1 | NA | NA | 3 | 1 |
| 12_110771 | 1 | 1 | NA | NA | NA | NA | NA | NA | 1 | 1 | 3 | 1 | 1 | 1 | 1 | 1 | NA | NA | 5 | 1 |
| 12_110771 | 1 | 1 | NA | NA | 1 | 1 | NA | NA | 2 | 1 | 3 | 1 | 1 | 1 | 1 | 1 | NA | NA | 5 | 1 |
| 12_110771 | 1 | 1 | NA | NA | NA | NA | NA | NA | 1 | 1 | 3 | 0.3333 | 1 | 0 | 1 | 0 | NA | NA | 5 | 0.2 |
| 12_110771 | 1 | 1 | NA | NA | 1 | 0 | NA | NA | 2 | 0.5 | 3 | 0.3333 | 1 | 1 | 1 | 0 | NA | NA | 5 | 0.4 |
| 12_110771 | 1 | 1 | NA | NA | NA | NA | NA | NA | 1 | 1 | 3 | 0.6667 | 1 | 1 | 1 | 1 | NA | NA | 5 | 0.8 |
| 12_110771 | 1 | 1 | NA | NA | 1 | 0 | NA | NA | 2 | 0.5 | 3 | 1 | 1 | 1 | NA | NA | NA | NA | 4 | 1 |
| 12_110772 | 1 | 1 | NA | NA | NA | NA | NA | NA | 1 | 1 | 2 | 1 | 1 | 1 | NA | NA | NA | NA | 3 | 1 |
| 12_110774 | 1 | 1 | 3 | 0.6667 | 2 | 0.5 | 2 | 0.5 | 8 | 0.625 | 1 | 0 | 1 | 1 | 1 | 1 | 2 | 1 | 5 | 0.8 |
| 12_110776 | NA | NA | NA | NA | 1 | 1 | NA | NA | 1 | 1 | NA | NA | NA | NA | 1 | 0 | 1 | 1 | 2 | 0.5 |

| | | | | | | | | | | | | | | | | | | | | |
|---|---|---|---|---|---|---|---|---|---|---|---|---|---|---|---|---|---|---|---|---|
| 12_1108219 | 1 | 1 | NA | NA | NA | NA | NA | NA | 1 | 1 | 1 | 1 | 1 | 1 | NA | NA | 1 | 1 | 3 | 1 |
| 12_1108222 | 1 | 1 | NA | NA | NA | NA | NA | NA | 1 | 1 | NA | NA | 1 | 1 | 1 | 1 | 1 | 1 | 3 | 1 |
| 12_1108347 | 1 | 0 | 1 | 1 | 1 | 0 | NA | NA | 3 | 0.3333 | 2 | 0.5 | 1 | 1 | 2 | 1 | 2 | 0 | 7 | 0.5714 |
| 12_1108349 | 1 | 0 | 2 | 0.5 | 1 | 1 | NA | NA | 4 | 0.5 | 1 | 1 | 2 | 1 | 2 | 0.5 | 2 | 1 | 7 | 0.8571 |
| 12_1108350 | 1 | 0 | 1 | 0 | 1 | 1 | NA | NA | 3 | 0.3333 | 1 | 1 | 2 | 1 | 2 | 0.5 | 2 | 0 | 7 | 0.5714 |
| 12_1108352 | 1 | 1 | 1 | 1 | 2 | 1 | NA | NA | 4 | 1 | 2 | 1 | 2 | 1 | 2 | 1 | 2 | 1 | 8 | 1 |
| 12_1108352 | 1 | 1 | 2 | 1 | 1 | 1 | NA | NA | 4 | 1 | 1 | 1 | 1 | 1 | 1 | 1 | 1 | 1 | 4 | 1 |
| 12_1108353 | 1 | 1 | 2 | 0.5 | 2 | 0.5 | NA | NA | 5 | 0.6 | 1 | 0 | 2 | 0.5 | 2 | 0.5 | 2 | 0 | 7 | 0.2857 |
| 12_1108353 | 1 | 1 | 2 | 1 | 2 | 1 | NA | NA | 5 | 1 | 2 | 1 | 2 | 1 | 2 | 1 | 2 | 1 | 8 | 1 |
| 12_1108398 | 6 | 1 | 3 | 0.6667 | 5 | 0.8 | 7 | 0.8571 | 21 | 0.8571 | 5 | 1 | 9 | 0.8889 | 4 | 0.5 | 6 | 1 | 24 | 0.875 |
| 12_1108399 | 7 | 0.1429 | 2 | 0.5 | 5 | 0 | 7 | 0.2857 | 21 | 0.1905 | 5 | 0.2 | 8 | 0.375 | 2 | 0 | 4 | 0.25 | 19 | 0.2632 |
| 12_1108400 | 5 | 0 | 4 | 0.25 | 5 | 0.4 | 8 | 0.5 | 22 | 0.3182 | 5 | 0 | 10 | 0.2 | 4 | 0.25 | 5 | 0.6 | 24 | 0.25 |
| 12_1108401 | 5 | 0.4 | 4 | 0.5 | 3 | 1 | 6 | 0.5 | 18 | 0.5556 | 4 | 0.75 | 10 | 0.3 | 4 | 0.75 | 4 | 0.75 | 22 | 0.5455 |
| 12_1108401 | 5 | 0 | 4 | 0.5 | 5 | 0 | 6 | 0 | 20 | 0.1 | 5 | 0 | 8 | 0.375 | 4 | 0.5 | 4 | 0.25 | 21 | 0.2857 |
| 12_1108402 | 4 | 0.25 | 2 | 0 | 4 | 0 | 8 | 0 | 18 | 0.0556 | 5 | 0.4 | 7 | 0 | 4 | 0 | 5 | 0 | 21 | 0.0952 |
| 12_1108402 | 5 | 0.4 | 3 | 0.3333 | 4 | 0.5 | 6 | 0.5 | 18 | 0.4444 | 5 | 0.4 | 7 | 0.4286 | 4 | 0 | 5 | 0 | 21 | 0.2381 |
| 12_1108403 | 4 | 0.75 | 3 | 0.6667 | 5 | 1 | 5 | 0.8 | 17 | 0.8235 | 3 | 0.6667 | 5 | 1 | 4 | 1 | 3 | 0.6667 | 15 | 0.8667 |
| 12_1108403 | 6 | 0.3333 | 4 | 0.25 | 5 | 0.2 | 7 | 0.1429 | 22 | 0.2273 | 6 | 0.3333 | 6 | 0.1667 | 4 | 0 | 4 | 0 | 20 | 0.15 |
| 12_1108460 | 1 | 1 | 2 | 1 | NA | NA | NA | NA | 3 | 1 | 1 | 1 | 1 | 1 | NA | NA | NA | NA | 2 | 1 |
| 12_1108504 | 5 | 0.2 | 5 | 0 | 3 | 0 | 7 | 0 | 20 | 0.05 | 8 | 0 | 6 | 0.1667 | 3 | 0 | 4 | 0.25 | 21 | 0.0952 |
| 12_1108557 | 1 | 0 | NA | NA | NA | NA | 1 | 1 | 2 | 0.5 | 1 | 1 | NA | NA | NA | NA | NA | NA | 1 | 1 |
| 12_1108979 | 1 | 1 | 1 | 1 | NA | NA | NA | NA | 2 | 1 | NA | NA | 1 | 1 | NA | NA | NA | NA | 1 | 1 |
| 12_1108980 | 1 | 1 | 1 | 1 | NA | NA | NA | NA | 2 | 1 | NA | NA | 1 | 1 | NA | NA | NA | NA | 1 | 1 |
| 12_1108994 | 1 | 1 | 1 | 0 | NA | NA | NA | NA | 2 | 0.5 | NA | NA | 1 | 1 | NA | NA | NA | NA | 1 | 1 |
| 12_1108995 | 1 | 1 | 1 | 1 | NA | NA | NA | NA | 2 | 1 | NA | NA | 1 | 1 | NA | NA | NA | NA | 1 | 1 |
| 12_1108997 | 1 | 1 | 1 | 0 | NA | NA | NA | NA | 2 | 0.5 | NA | NA | 1 | 1 | NA | NA | NA | NA | 1 | 1 |
| 12_1108999 | 1 | 1 | 1 | 0 | NA | NA | NA | NA | 2 | 0.5 | NA | NA | 2 | 0.5 | NA | NA | NA | NA | 2 | 0.5 |
| 12_1108999 | 1 | 0 | 1 | 1 | NA | NA | NA | NA | 2 | 0.5 | NA | NA | 1 | 1 | NA | NA | NA | NA | 1 | 1 |
| 12_1109355 | 4 | 0.75 | 2 | 0.5 | NA | NA | NA | NA | 6 | 0.6667 | NA | NA | 4 | 1 | NA | NA | NA | NA | 4 | 1 |
| 12_1109356 | 4 | 0.75 | 1 | 0 | NA | NA | NA | NA | 5 | 0.6 | NA | NA | 4 | 1 | NA | NA | NA | NA | 4 | 1 |
| 12_1109356 | 4 | 0.75 | 1 | 1 | NA | NA | NA | NA | 5 | 0.8 | NA | NA | 2 | 1 | NA | NA | NA | NA | 2 | 1 |
| 12_1109357 | 4 | 0.25 | 2 | 0.5 | NA | NA | NA | NA | 6 | 0.3333 | NA | NA | 4 | 0.25 | 1 | 1 | NA | NA | 5 | 0.4 |
| 12_1109367 | 6 | 0 | 2 | 0 | 3 | 0.3333 | 2 | 0 | 13 | 0.0769 | 1 | 0 | 4 | 0.25 | 2 | 0 | 2 | 0 | 9 | 0.1111 |
| 12_1109378 | 7 | 0.2857 | 5 | 0 | 4 | 0 | 2 | 0 | 18 | 0.1111 | 3 | 0 | 6 | 0 | 3 | 0.6667 | 4 | 0 | 16 | 0.125 |
| 12_1109428 | 1 | 0 | 2 | 0.5 | 2 | 0 | NA | NA | 5 | 0.2 | 1 | 0 | 2 | 1 | 2 | 0 | 4 | 0.75 | 9 | 0.5556 |
| 12_1109429 | 1 | 0 | 2 | 0.5 | 2 | 0.5 | NA | NA | 5 | 0.4 | 1 | 0 | 2 | 1 | 2 | 0.5 | 3 | 0.3333 | 8 | 0.5 |
| 12_1109429 | 2 | 0 | 2 | 1 | 2 | 1 | NA | NA | 6 | 0.6667 | 1 | 1 | 2 | 1 | 2 | 0.5 | 4 | 0.75 | 9 | 0.7778 |
| 12_1109429 | 2 | 0.5 | 1 | 1 | 2 | 1 | NA | NA | 5 | 0.8 | NA | NA | 1 | 1 | 1 | 1 | 2 | 1 | 4 | 1 |
| 12_1109429 | 2 | 0.5 | 2 | 0 | 1 | 1 | NA | NA | 5 | 0.4 | 1 | 1 | 2 | 0 | 2 | 0.5 | 3 | 0 | 8 | 0.25 |
| 12_1109430 | 2 | 1 | 2 | 1 | 1 | 1 | NA | NA | 5 | 1 | 1 | 1 | 2 | 0.5 | 2 | 1 | 3 | 1 | 8 | 0.875 |
| 12_1109431 | 2 | 0.5 | 2 | 0.5 | 2 | 0.5 | NA | NA | 6 | 0.5 | 1 | 0 | 1 | 1 | 2 | 1 | 3 | 0.6667 | 7 | 0.7143 |
| 12_1109431 | 1 | 1 | 1 | 1 | 2 | 1 | NA | NA | 4 | 1 | 1 | 1 | 1 | 1 | 2 | 1 | 2 | 1 | 6 | 1 |
| 12_1109432 | 2 | 0 | 1 | 0 | 2 | 0.5 | NA | NA | 5 | 0.2 | 1 | 1 | 2 | 0 | 2 | 0 | 4 | 0.25 | 9 | 0.2222 |
| 12_1109435 | 2 | 1 | 2 | 1 | 2 | 0.5 | NA | NA | 6 | 0.8333 | 1 | 0 | NA | NA | 1 | 1 | 3 | 0.6667 | 5 | 0.6 |
| 12_1109437 | 2 | 1 | 2 | 0.5 | 1 | 1 | NA | NA | 5 | 0.8 | 1 | 0 | NA | NA | 1 | 1 | 4 | 1 | 6 | 0.8333 |
| 12_1110783 | NA | NA | 3 | 0.3333 | 1 | 1 | NA | NA | 4 | 0.5 | NA | NA | 2 | 0.5 | 1 | 0 | 1 | 1 | 4 | 0.5 |
| 12_1110783 | NA | NA | 3 | 0.3333 | 1 | 1 | NA | NA | 4 | 0.5 | NA | NA | 2 | 0 | 1 | 1 | 1 | 0 | 4 | 0.25 |
| 12_1110784 | NA | NA | 4 | 0.25 | 1 | 1 | NA | NA | 5 | 0.4 | NA | NA | 2 | 0.5 | 1 | 0 | 1 | 0 | 4 | 0.25 |
| 12_1110784 | NA | NA | 4 | 0.5 | 1 | 1 | NA | NA | 5 | 0.6 | NA | NA | 2 | 0.5 | 1 | 0 | 1 | 1 | 4 | 0.5 |
| 12_1110786 | NA | NA | 4 | 1 | NA | NA | NA | NA | 4 | 1 | NA | NA | 2 | 1 | NA | NA | 1 | 1 | 3 | 1 |
| 12_1110789 | NA | NA | 4 | 1 | NA | NA | NA | NA | 4 | 1 | NA | NA | 2 | 1 | NA | NA | 1 | 1 | 3 | 1 |
| 12_1110790 | NA | NA | 3 | 0.3333 | 1 | 1 | NA | NA | 4 | 0.5 | NA | NA | 3 | 0.3333 | 1 | 1 | 1 | 1 | 5 | 0.6 |
| 12_1110796 | NA | NA | 2 | 1 | NA | NA | NA | NA | 2 | 1 | 1 | 0 | 1 | 1 | 1 | 0 | NA | NA | 3 | 0.3333 |
| 12_1110807 | NA | NA | 5 | 0.2 | 3 | 0 | 1 | 0 | 9 | 0.1111 | 2 | 0 | 4 | 0 | 1 | 0 | 2 | 0.5 | 9 | 0.1111 |
| 12_1110808 | NA | NA | 5 | 0.6 | 3 | 0.6667 | NA | NA | 8 | 0.625 | 2 | 0 | 2 | 0 | 1 | 1 | 2 | 1 | 7 | 0.4286 |
| 12_1110809 | NA | NA | 7 | 0.2857 | 3 | 0.3333 | 1 | 1 | 11 | 0.3636 | 2 | 0 | 4 | 0 | 1 | 0 | 2 | 0.5 | 9 | 0.1111 |
| 12_1110814 | NA | NA | 5 | 0.2 | 3 | 0 | 1 | 0 | 9 | 0.1111 | 2 | 0 | 3 | 0.6667 | 1 | 0 | 2 | 0 | 8 | 0.25 |
| 12_1112280 | 1 | 0 | 1 | 0 | 1 | 1 | NA | NA | 3 | 0.3333 | NA | NA | 2 | 0.5 | 1 | 0 | 1 | 0 | 4 | 0.25 |
| 12_1112281 | NA | NA | 1 | 1 | 1 | 1 | NA | NA | 2 | 1 | NA | NA | 2 | 1 | 2 | 1 | 1 | 1 | 5 | 1 |
| 12_1112282 | 1 | 1 | 1 | 1 | 1 | 1 | NA | NA | 3 | 1 | NA | NA | 2 | 1 | 2 | 0.5 | 1 | 1 | 5 | 0.8 |
| 12_1112284 | 1 | 0 | 2 | 0.5 | NA | NA | NA | NA | 3 | 0.3333 | NA | NA | 1 | 1 | 2 | 1 | 1 | 0 | 4 | 0.75 |
| 12_1112285 | 1 | 0 | 2 | 1 | 1 | 0 | NA | NA | 4 | 0.5 | NA | NA | 2 | 1 | 2 | 1 | 1 | 1 | 5 | 1 |
| 12_1112286 | NA | NA | 2 | 1 | 1 | 1 | NA | NA | 3 | 1 | NA | NA | 2 | 1 | 1 | 1 | 1 | 1 | 4 | 1 |
| 12_1112291 | 1 | 0 | 3 | 0.3333 | 2 | 0.5 | NA | NA | 6 | 0.3333 | NA | NA | 1 | 1 | 2 | 1 | NA | NA | 3 | 1 |
| 12_1112333 | 1 | 0 | 3 | 0 | 4 | 0.25 | 2 | 0.5 | 10 | 0.2 | 2 | 0 | 3 | 0 | 4 | 0.25 | 2 | 0 | 11 | 0.0909 |

| | | | | | | | | | | | | | | | | | | | | |
|---|---|---|---|---|---|---|---|---|---|---|---|---|---|---|---|---|---|---|---|---|
| 12_1112425 | 1 | 1 | 1 | 1 | NA | NA | 2 | 0.5 | 4 | 0.75 | 1 | 0 | 2 | 0 | 1 | 0 | 1 | 1 | 5 | 0.2 |
| 12_1112596 | 18 | 0.2222 | 10 | 0.2 | 13 | 0.2308 | 14 | 0.0714 | 55 | 0.1818 | 8 | 0.25 | 14 | 0.1429 | 16 | 0.125 | 6 | 0 | 44 | 0.1364 |
| 12_1112597 | 14 | 0.1429 | 9 | 0.1111 | 11 | 0 | 7 | 0 | 41 | 0.0732 | 7 | 0.1429 | 13 | 0 | 15 | 0.2 | 3 | 0 | 38 | 0.1053 |
| 12_1112597 | 16 | 0.625 | 11 | 0.7273 | 9 | 0.8889 | 12 | 0.6667 | 48 | 0.7083 | 8 | 0.75 | 11 | 0.7273 | 14 | 0.7143 | 7 | 1 | 40 | 0.775 |
| 12_1112598 | 20 | 0.2 | 13 | 0 | 14 | 0.1429 | 13 | 0.2308 | 60 | 0.15 | 7 | 0.1429 | 17 | 0.1765 | 21 | 0.0952 | 8 | 0 | 53 | 0.1132 |
| 12_1112601 | 16 | 0.8125 | 13 | 0.6923 | 13 | 0.9231 | 13 | 0.8462 | 55 | 0.8182 | 6 | 0.6667 | 12 | 0.8333 | 20 | 0.8 | 6 | 0.6667 | 44 | 0.7727 |
| 12_1112602 | 13 | 0.1538 | 13 | 0.0769 | 12 | 0 | 14 | 0.2143 | 52 | 0.1154 | 6 | 0 | 14 | 0.1429 | 21 | 0.0952 | 6 | 0.1667 | 47 | 0.1064 |
| 12_1112602 | 15 | 0.3333 | 10 | 0.4 | 11 | 0.1818 | 13 | 0.1538 | 49 | 0.2653 | 6 | 0 | 12 | 0.0833 | 17 | 0.1765 | 4 | 0 | 39 | 0.1026 |
| 12_1112603 | 10 | 0.2 | 9 | 0.5556 | 11 | 0.6364 | 10 | 0.7 | 40 | 0.525 | 6 | 0.3333 | 10 | 0.3 | 17 | 0.4706 | 5 | 0.6 | 38 | 0.4211 |
| 12_1112603 | 11 | 0.1818 | 9 | 0.3333 | 11 | 0 | 12 | 0.0833 | 43 | 0.1395 | 6 | 0.1667 | 9 | 0.1111 | 19 | 0.3158 | 6 | 0.3333 | 40 | 0.25 |
| 12_1112606 | 12 | 0 | 10 | 0.2 | 11 | 0 | 11 | 0.0909 | 44 | 0.0682 | 5 | 0 | 10 | 0.3 | 14 | 0.2143 | 6 | 0 | 35 | 0.1714 |
| 12_1112655 | 11 | 0.2727 | 8 | 0.125 | 8 | 0 | 11 | 0.1818 | 38 | 0.1579 | 5 | 0.2 | 9 | 0.3333 | 10 | 0 | 5 | 0.2 | 29 | 0.1724 |
| 12_1112655 | 9 | 0.7778 | 9 | 0.8889 | 8 | 0.875 | 10 | 0.9 | 36 | 0.8611 | 4 | 1 | 9 | 0.8889 | 9 | 0.8889 | 5 | 0.8 | 27 | 0.8889 |
| 12_1112657 | 10 | 1 | 7 | 0.8571 | 7 | 0.8571 | 12 | 0.6667 | 36 | 0.8333 | 4 | 1 | 8 | 0.625 | 11 | 1 | 4 | 0.5 | 27 | 0.8148 |
| 12_1112658 | 11 | 0.8182 | 8 | 0.375 | 7 | 0.2857 | 12 | 0.4167 | 38 | 0.5 | 4 | 0.25 | 8 | 0.375 | 11 | 0.6364 | 4 | 0.75 | 27 | 0.5185 |
| 12_1112658 | 8 | 0.75 | 7 | 0.7143 | 5 | 0.8 | 11 | 0.8182 | 31 | 0.7742 | 2 | 1 | 6 | 0.6667 | 11 | 0.9091 | 5 | 1 | 24 | 0.875 |
| 12_1112875 | 4 | 0.25 | 2 | 0 | 2 | 0 | 4 | 0 | 12 | 0.0833 | 1 | 0 | 5 | 0 | 4 | 0 | 2 | 0.5 | 12 | 0.0833 |
| 12_1113438 | 3 | 0.3333 | NA | NA | 1 | 0 | 2 | 0 | 6 | 0.1667 | 1 | 0 | 1 | 0 | 3 | 0.3333 | 1 | 0 | 6 | 0.1667 |
| 12_1113439 | 2 | 0 | NA | NA | NA | NA | 1 | 1 | 3 | 0.3333 | NA | NA | NA | NA | 3 | 0.3333 | NA | NA | 3 | 0.3333 |
| 12_1114161 | 1 | 1 | NA | NA | 1 | 1 | NA | NA | 2 | 1 | 2 | 1 | 1 | 1 | NA | NA | 1 | 0 | 4 | 0.75 |
| 12_1114170 | 1 | 0 | 1 | 0 | 1 | 0 | 3 | 0.3333 | 6 | 0.1667 | 2 | 0 | 1 | 1 | 1 | 0 | NA | NA | 4 | 0.25 |
| 12_1114171 | 1 | 1 | 1 | 1 | NA | NA | 2 | 0.5 | 4 | 0.75 | 2 | 0 | 1 | 0 | 1 | 1 | NA | NA | 4 | 0.25 |
| 12_1114513 | 1 | 1 | NA | NA | 1 | 1 | 2 | 1 | 4 | 1 | NA | NA | NA | NA | NA | NA | 1 | 1 | 1 | 1 |
| 12_1114513 | 1 | 1 | NA | NA | 1 | 1 | 2 | 1 | 4 | 1 | NA | NA | NA | NA | NA | NA | 1 | 1 | 1 | 1 |
| 12_1114514 | 1 | 1 | NA | NA | 1 | 1 | 2 | 0.5 | 4 | 0.75 | NA | NA | NA | NA | NA | NA | 1 | 1 | 1 | 1 |
| 12_1114520 | 1 | 0 | NA | NA | 1 | 1 | 3 | 0 | 5 | 0.2 | NA | NA | NA | NA | NA | NA | 1 | 1 | 1 | 1 |
| 12_1114609 | NA | NA | NA | NA | 1 | 1 | NA | NA | 1 | 1 | 1 | 1 | 1 | 1 | 1 | 0 | 1 | 1 | 4 | 0.75 |
| 12_1114610 | NA | NA | NA | NA | 1 | 1 | NA | NA | 1 | 1 | NA | NA | NA | NA | 1 | 1 | NA | NA | 1 | 1 |
| 12_1114611 | NA | NA | NA | NA | 1 | 1 | NA | NA | 1 | 1 | 1 | 1 | 1 | 0 | 1 | 1 | 1 | 0 | 4 | 0.5 |
| 12_1114611 | NA | NA | NA | NA | 1 | 1 | NA | NA | 1 | 1 | 1 | 1 | 1 | 1 | 1 | 0 | 1 | 1 | 4 | 0.75 |
| 12_1114612 | NA | NA | NA | NA | 1 | 1 | NA | NA | 1 | 1 | 1 | 1 | 1 | 1 | 1 | 0 | 1 | 1 | 4 | 0.75 |
| 12_1114612 | NA | NA | NA | NA | 1 | 1 | NA | NA | 1 | 1 | 1 | 0 | 1 | 0 | 1 | 1 | 1 | 1 | 4 | 0.5 |
| 12_1114614 | NA | NA | NA | NA | 1 | 1 | NA | NA | 1 | 1 | 1 | 0 | 1 | 1 | 1 | 0 | 1 | 0 | 4 | 0.25 |
| 12_1115059 | 24 | 0.0417 | 13 | 0 | 18 | 0.1111 | 14 | 0.1429 | 69 | 0.0725 | 8 | 0 | 23 | 0.087 | 17 | 0.0588 | 12 | 0 | 60 | 0.05 |
| 12_1115110 | 25 | 0.12 | 19 | 0.1053 | 23 | 0 | 25 | 0.12 | 92 | 0.087 | 9 | 0.2222 | 25 | 0.12 | 18 | 0.2222 | 19 | 0.1053 | 71 | 0.1549 |
| 12_1115113 | 33 | 0.2424 | 25 | 0.32 | 26 | 0.1538 | 30 | 0.1667 | 114 | 0.2193 | 10 | 0 | 27 | 0.1111 | 27 | 0.1852 | 19 | 0.1579 | 83 | 0.1325 |
| 12_1115113 | 30 | 0.2333 | 19 | 0.5263 | 19 | 0.3158 | 23 | 0.3478 | 91 | 0.3407 | 9 | 0.1111 | 22 | 0.4091 | 20 | 0.25 | 12 | 0.5 | 63 | 0.3333 |
| 12_1115114 | 27 | 0.1111 | 20 | 0.15 | 21 | 0.1429 | 25 | 0.2 | 93 | 0.1505 | 8 | 0 | 26 | 0.0769 | 21 | 0.0952 | 13 | 0.2308 | 68 | 0.1029 |
| 12_1115114 | 26 | 0.2692 | 16 | 0.4375 | 17 | 0.4118 | 21 | 0.4762 | 80 | 0.3875 | 8 | 0.125 | 22 | 0.2727 | 20 | 0.25 | 14 | 0.5 | 64 | 0.2969 |
| 12_1115122 | 16 | 0.5 | 11 | 0.4545 | 18 | 0.3889 | 24 | 0.3333 | 69 | 0.4058 | 11 | 0.3636 | 20 | 0.3 | 15 | 0.6 | 11 | 0.4545 | 57 | 0.4211 |
| 12_1115122 | 17 | 0 | 10 | 0.3 | 17 | 0.1176 | 22 | 0.1818 | 66 | 0.1364 | 10 | 0.2 | 19 | 0.1579 | 17 | 0.1176 | 12 | 0.1667 | 58 | 0.1552 |
| 12_1115390 | 1 | 1 | NA | NA | NA | NA | NA | NA | 1 | 1 | 1 | 1 | NA | NA | NA | NA | 1 | 1 | 2 | 1 |
| 12_1115393 | 1 | 1 | NA | NA | NA | NA | 1 | 0 | 2 | 0.5 | 1 | 0 | NA | NA | NA | NA | 1 | 1 | 2 | 0.5 |
| 12_1115394 | 1 | 1 | NA | NA | NA | NA | NA | NA | 1 | 1 | 1 | 1 | NA | NA | NA | NA | 1 | 0 | 2 | 0.5 |
| 12_1115395 | 1 | 1 | NA | NA | NA | NA | NA | NA | 1 | 1 | NA | NA | NA | NA | NA | NA | 1 | 1 | 1 | 1 |
| 12_1116217 | 10 | 0.2 | 10 | 0.2 | 11 | 0.1818 | 7 | 0 | 38 | 0.1579 | 9 | 0 | 12 | 0.0833 | 5 | 0.2 | 9 | 0.1111 | 35 | 0.0857 |
| 12_1116219 | 10 | 0.1 | 8 | 0 | 12 | 0.3333 | 7 | 0.1429 | 37 | 0.1622 | 9 | 0 | 9 | 0.2222 | 7 | 0.1429 | 7 | 0 | 32 | 0.0938 |
| 12_1116219 | 13 | 0.4615 | 10 | 0.7 | 12 | 0.6667 | 8 | 0.625 | 43 | 0.6047 | 8 | 0.875 | 12 | 0.5833 | 7 | 0.7143 | 8 | 0.5 | 35 | 0.6571 |
| 12_1116219 | 13 | 0.5385 | 11 | 0.6364 | 11 | 0.4545 | 8 | 0.625 | 43 | 0.5581 | 8 | 0.625 | 11 | 0.6364 | 7 | 0.7143 | 9 | 0.5556 | 35 | 0.6286 |
| 12_1116358 | 519 | 0.2197 | 390 | 0.2128 | 410 | 0.2683 | 439 | 0.2073 | 1758 | 0.2264 | 312 | 0.234 | 418 | 0.2297 | 446 | 0.1951 | 356 | 0.2725 | 1532 | 0.2304 |
| 12_1116381 | 2 | 0 | NA | NA | 1 | 0 | 1 | 1 | 4 | 0.25 | 1 | 0 | 4 | 0.25 | 2 | 0 | NA | NA | 7 | 0.1429 |
| 12_1116446 | 1 | 1 | 1 | 1 | NA | NA | 2 | 1 | 4 | 1 | 2 | 0.5 | 5 | 0.4 | 1 | 1 | 2 | 0.5 | 10 | 0.5 |
| 12_1116452 | 1 | 1 | 1 | 1 | 1 | 1 | 1 | 0 | 4 | 0.75 | 2 | 0 | 5 | 0.2 | 2 | 0.5 | 2 | 0 | 11 | 0.1818 |
| 12_1116452 | 1 | 1 | NA | NA | 1 | 1 | 2 | 1 | 4 | 1 | 1 | 1 | 6 | 0.5 | 2 | 1 | 1 | 0 | 10 | 0.6 |
| 12_1116473 | 1 | 0 | 2 | 0.5 | 1 | 0 | 3 | 0 | 7 | 0.1429 | 2 | 0.5 | 7 | 0 | 2 | 0 | 2 | 0 | 13 | 0.0769 |
| 12_1116492 | NA | NA | 3 | 1 | 1 | 0 | 4 | 0.75 | 8 | 0.75 | 2 | 1 | 5 | 1 | 2 | 1 | 2 | 1 | 11 | 1 |
| 12_1116492 | 2 | 0.5 | 3 | 0 | 1 | 1 | 4 | 0.75 | 10 | 0.5 | 2 | 1 | 7 | 0.5714 | 2 | 1 | 2 | 0.5 | 13 | 0.6923 |
| 12_1116493 | 2 | 0 | 1 | 0 | NA | NA | 3 | 0.6667 | 6 | 0.3333 | 2 | 0.5 | 4 | 0 | 1 | 1 | 2 | 0 | 9 | 0.2222 |
| 12_1116498 | 2 | 0 | 1 | 0 | 1 | 0 | 3 | 0.3333 | 7 | 0.1429 | 3 | 0 | 6 | 0 | 1 | 1 | 2 | 0 | 12 | 0.0833 |
| 12_1116501 | 2 | 0 | 3 | 0.6667 | 1 | 1 | 1 | 0 | 7 | 0.4286 | 3 | 0.6667 | 5 | 0.8 | 1 | 1 | 2 | 1 | 11 | 0.8182 |
| 12_1116760 | 7 | 0.7143 | 4 | 0.75 | 6 | 0.1667 | 6 | 0.3333 | 23 | 0.4783 | 3 | 1 | 3 | 0.3333 | 6 | 0.6667 | 7 | 0.7143 | 19 | 0.6842 |
| 12_1116765 | 14 | 0.7857 | 6 | 0.5 | 8 | 0.75 | 13 | 0.8462 | 41 | 0.7561 | 6 | 0.8333 | 8 | 0.625 | 15 | 0.8 | 12 | 0.8333 | 41 | 0.7805 |
| 12_1116765 | 15 | 0.1333 | 5 | 0 | 6 | 0.5 | 12 | 0.0833 | 38 | 0.1579 | 3 | 0 | 8 | 0.25 | 15 | 0.0667 | 7 | 0.1429 | 33 | 0.1212 |
| 12_1116765 | 13 | 0.0769 | 4 | 0.5 | 5 | 0.2 | 14 | 0.2143 | 36 | 0.1944 | 5 | 0.2 | 8 | 0.375 | 14 | 0.2143 | 12 | 0 | 39 | 0.1795 |
| 12_1116760 | 12 | 0.6667 | 5 | 0.4 | 5 | 0.8 | 9 | 0.5556 | 31 | 0.6129 | 3 | 0.6667 | 5 | 0.6 | 10 | 0.5 | 6 | 0.3333 | 24 | 0.5 |

| | | | | | | | | | | | | | | | | | | | | |
|---|---|---|---|---|---|---|---|---|---|---|---|---|---|---|---|---|---|---|---|---|
| 12_111676 | 10 | 0.8 | 3 | 1 | 4 | 1 | 9 | 1 | 26 | 0.9231 | 4 | 1 | 4 | 1 | 10 | 1 | 8 | 0.875 | 26 | 0.9615 |
| 12_111676 | 15 | 0.4 | 6 | 0.1667 | 6 | 0.3333 | 14 | 0.4286 | 41 | 0.3659 | 4 | 0.5 | 6 | 0.1667 | 9 | 0.5556 | 7 | 0.1429 | 26 | 0.3462 |
| 12_111676 | 12 | 0.3333 | 4 | 0.5 | 6 | 0.5 | 14 | 0.6429 | 36 | 0.5 | 4 | 0.5 | 5 | 0.6 | 11 | 0.3636 | 9 | 0.6667 | 29 | 0.5172 |
| 12_111676 | 8 | 0.625 | 5 | 0.6 | 7 | 0.5714 | 11 | 1 | 31 | 0.7419 | 4 | 0.75 | 4 | 1 | 11 | 0.9091 | 9 | 0.6667 | 28 | 0.8214 |
| 12_111677 | 14 | 0.0714 | 6 | 0.3333 | 7 | 0 | 13 | 0.0769 | 40 | 0.1 | 6 | 0.1667 | 6 | 0 | 11 | 0 | 11 | 0.0909 | 34 | 0.0588 |
| 12_111677 | 13 | 1 | 6 | 0.8333 | 6 | 0.5 | 14 | 0.6429 | 39 | 0.7692 | 7 | 0.5714 | 5 | 0.6 | 12 | 0.5833 | 11 | 0.8182 | 35 | 0.6571 |
| 12_111677 | 15 | 0.0667 | 6 | 0.1667 | 7 | 0.1429 | 13 | 0.1538 | 41 | 0.122 | 7 | 0.2857 | 6 | 0 | 12 | 0.1667 | 11 | 0.1818 | 36 | 0.1667 |
| 12_111677 | 14 | 0.2143 | 5 | 0.2 | 7 | 0.1429 | 13 | 0.1538 | 39 | 0.1795 | 6 | 0.1667 | 6 | 0 | 11 | 0.1818 | 8 | 0.375 | 31 | 0.1935 |
| 12_111677 | 12 | 1 | 5 | 1 | 7 | 0.8571 | 13 | 1 | 37 | 0.973 | 5 | 1 | 5 | 1 | 10 | 1 | 9 | 1 | 29 | 1 |
| 12_111677 | 12 | 0.1667 | 4 | 0.25 | 7 | 0.1429 | 13 | 0.2308 | 36 | 0.1944 | 6 | 0 | 4 | 0.25 | 10 | 0.3 | 11 | 0.1818 | 31 | 0.1935 |
| 12_111677 | 11 | 0.8182 | 5 | 1 | 5 | 1 | 11 | 0.7273 | 32 | 0.8438 | 6 | 0.6667 | 5 | 0.8 | 10 | 0.9 | 8 | 0.75 | 29 | 0.7931 |
| 12_111677 | 5 | 0.2 | 4 | 0 | 4 | 0.25 | 7 | 0 | 20 | 0.1 | 3 | 0 | 3 | 0.3333 | 7 | 0 | 6 | 0 | 19 | 0.0526 |
| 12_111677 | 14 | 1 | 4 | 1 | 6 | 1 | 9 | 1 | 33 | 1 | 7 | 1 | 5 | 1 | 12 | 0.8333 | 8 | 1 | 32 | 0.9375 |
| 12_111677 | 14 | 0.2143 | 5 | 0 | 7 | 0 | 13 | 0.2308 | 39 | 0.1538 | 6 | 0 | 6 | 0.1667 | 11 | 0.1818 | 11 | 0.1818 | 34 | 0.1471 |
| 12_111677 | 13 | 0.5385 | 5 | 0.2 | 5 | 0.6 | 11 | 0.2727 | 34 | 0.4118 | 6 | 0.3333 | 6 | 0.3333 | 11 | 0.3636 | 10 | 0.4 | 33 | 0.3636 |
| 12_111677 | 13 | 0.1538 | 5 | 0.2 | 6 | 0.1667 | 11 | 0.1818 | 35 | 0.1714 | 5 | 0.2 | 5 | 0 | 11 | 0.0909 | 10 | 0 | 31 | 0.0645 |
| 12_111677 | 13 | 0.7692 | 5 | 0.8 | 6 | 0.6667 | 9 | 0.8889 | 33 | 0.7879 | 5 | 1 | 5 | 0.8 | 10 | 0.6 | 9 | 0.7778 | 29 | 0.7586 |
| 12_111677 | 13 | 0.1538 | 4 | 0 | 5 | 0 | 11 | 0 | 33 | 0.0606 | 6 | 0 | 5 | 0.2 | 5 | 0 | 10 | 0.1 | 26 | 0.0769 |
| 12_111700 | 5 | 0 | 1 | 1 | 6 | 0 | 7 | 0 | 19 | 0.0526 | 2 | 0 | 6 | 0.1667 | 3 | 0 | 3 | 0 | 14 | 0.0714 |
| 12_111700 | 3 | 0.3333 | 2 | 0.5 | 6 | 0.3333 | 7 | 0.4286 | 18 | 0.3889 | 2 | 1 | 5 | 0.4 | 3 | 0.3333 | 4 | 0.5 | 14 | 0.5 |
| 12_111761 | NA | NA | NA | NA | NA | NA | 1 | 1 | 1 | 1 | NA | NA | 2 | 0 | 2 | 0 | 2 | 0.5 | 6 | 0.1667 |
| 12_111770 | 2 | 0 | 1 | 0 | 3 | 0.3333 | 3 | 0.3333 | 9 | 0.2222 | 1 | 1 | 2 | 0 | 1 | 1 | 1 | 0 | 5 | 0.4 |
| 12_111770 | 1 | 1 | 1 | 1 | 3 | 0.3333 | 2 | 0 | 7 | 0.4286 | NA | NA | 2 | 0 | 1 | 1 | 1 | 1 | 4 | 0.5 |
| 12_111770 | 1 | 0 | 1 | 0 | 2 | 1 | 2 | 0.5 | 6 | 0.5 | 1 | 1 | 1 | 1 | 1 | 0 | 1 | 0 | 4 | 0.5 |
| 12_111770 | 1 | 0 | 1 | 1 | 2 | 1 | 2 | 0 | 6 | 0.5 | NA | NA | 1 | 1 | NA | NA | 1 | 0 | 2 | 0.5 |
| 12_111770 | 1 | 0 | 1 | 0 | 2 | 0 | 3 | 0.6667 | 7 | 0.2857 | 1 | 0 | 2 | 0.5 | 1 | 0 | 1 | 0 | 5 | 0.2 |
| 12_111770 | 1 | 1 | 1 | 1 | 2 | 0 | 2 | 1 | 6 | 0.6667 | 1 | 0 | 2 | 0.5 | NA | NA | 1 | 0 | 4 | 0.25 |
| 12_111770 | 1 | 0 | NA | NA | 2 | 1 | 2 | 1 | 5 | 0.8 | 1 | 0 | 2 | 0 | NA | NA | 1 | 1 | 4 | 0.25 |
| 12_111795 | 10 | 0.4 | 13 | 0.1538 | 8 | 0.25 | 10 | 0.1 | 41 | 0.2195 | 11 | 0.1818 | 10 | 0.2 | 10 | 0.5 | 9 | 0.5556 | 40 | 0.35 |
| 12_111795 | 3 | 0.6667 | 9 | 0.3333 | 7 | 0.2857 | 4 | 0.5 | 23 | 0.3913 | 5 | 0.2 | 9 | 0.3333 | 5 | 0.4 | 8 | 0.375 | 27 | 0.3333 |
| 12_111795 | 6 | 1 | 10 | 0.3 | 7 | 0.4286 | 9 | 0.4444 | 32 | 0.5 | 6 | 0.5 | 8 | 0.125 | 8 | 0.375 | 10 | 0.4 | 32 | 0.3438 |
| 12_111795 | 8 | 0.375 | 10 | 0.3 | 9 | 0.2222 | 10 | 0.4 | 37 | 0.3243 | 8 | 0.125 | 11 | 0.3636 | 9 | 0.1111 | 10 | 0.3 | 38 | 0.2368 |
| 12_111795 | 8 | 0.25 | 9 | 0.2222 | 9 | 0 | 8 | 0.375 | 34 | 0.2059 | 8 | 0.125 | 10 | 0.1 | 9 | 0.3333 | 7 | 0.1429 | 34 | 0.1765 |
| 12_111795 | 8 | 0.375 | 8 | 0.375 | 8 | 0 | 8 | 0.125 | 32 | 0.2188 | 5 | 0 | 9 | 0.4444 | 7 | 0.1429 | 9 | 0.3333 | 30 | 0.2667 |
| 12_111795 | 8 | 0.125 | 5 | 0 | 5 | 0 | 6 | 0.3333 | 24 | 0.125 | 5 | 0.2 | 6 | 0 | 8 | 0.125 | 6 | 0.1667 | 25 | 0.12 |
| 12_111799 | 7 | 0.1429 | 7 | 0.1429 | 6 | 0.5 | 8 | 0.25 | 28 | 0.25 | 2 | 0 | 8 | 0.25 | 5 | 0.4 | 7 | 0.4286 | 22 | 0.3182 |
| 12_111799 | 8 | 0.375 | 7 | 0.2857 | 4 | 0.25 | 8 | 0.375 | 27 | 0.3333 | 1 | 0 | 6 | 0.5 | 4 | 0 | 7 | 0.1429 | 18 | 0.2222 |
| 12_111799 | 6 | 1 | 5 | 1 | 6 | 0.8333 | 9 | 1 | 26 | 0.9615 | 3 | 1 | 7 | 1 | 4 | 1 | 7 | 1 | 21 | 1 |
| 12_111799 | 5 | 0.2 | 5 | 0.4 | 4 | 0 | 8 | 0.125 | 22 | 0.1818 | 2 | 0 | 5 | 0 | 3 | 0.3333 | 6 | 0 | 16 | 0.0625 |
| 12_111799 | 6 | 0.5 | 4 | 1 | 6 | 0.8333 | 9 | 0.8889 | 25 | 0.8 | 3 | 1 | 7 | 1 | 5 | 1 | 7 | 0.8571 | 22 | 0.9545 |
| 12_111800 | 6 | 0.5 | 3 | 0.6667 | 6 | 0.3333 | 7 | 0.2857 | 22 | 0.4091 | 3 | 0 | 6 | 0.1667 | 5 | 0.2 | 6 | 0.3333 | 20 | 0.2 |
| 12_111801 | 6 | 0 | 1 | 0 | 4 | 0.25 | 8 | 0 | 19 | 0.0526 | 2 | 0.5 | 4 | 0 | 3 | 0.6667 | 4 | 0 | 13 | 0.2308 |
| 12_111801 | 6 | 0.5 | 3 | 0.6667 | 5 | 0 | 8 | 0.375 | 22 | 0.3636 | 3 | 0.3333 | 5 | 0.6 | 4 | 0.75 | 4 | 0.25 | 16 | 0.5 |
| 12_111801 | 4 | 0.25 | 2 | 1 | 4 | 0.75 | 4 | 0.25 | 14 | 0.5 | 1 | 0 | 3 | 0.6667 | 3 | 0 | 3 | 0 | 10 | 0.2 |
| 12_111802 | 5 | 0.2 | 1 | 0 | 5 | 0 | 5 | 0 | 16 | 0.0625 | 2 | 0 | 5 | 0 | 4 | 0 | 2 | 0.5 | 13 | 0.0769 |
| 12_111925 | NA | NA | 1 | 1 | NA | NA | NA | NA | 1 | 1 | NA | NA | 1 | 1 | NA | NA | NA | NA | 1 | 1 |
| 12_111925 | NA | NA | 1 | 1 | NA | NA | NA | NA | 1 | 1 | NA | NA | 1 | 1 | NA | NA | NA | NA | 1 | 1 |
| 12_111925 | NA | NA | 1 | 1 | NA | NA | NA | NA | 1 | 1 | NA | NA | 1 | 1 | NA | NA | NA | NA | 1 | 1 |
| 12_112147 | 2 | 0.5 | NA | NA | NA | NA | 1 | 0 | 3 | 0.3333 | NA | NA | 1 | 1 | NA | NA | NA | NA | 1 | 1 |
| 12_112466 | 24 | 0.125 | 27 | 0.1852 | 11 | 0.3636 | 12 | 0.3333 | 74 | 0.2162 | 19 | 0.1579 | 11 | 0.2727 | 19 | 0.2632 | 15 | 0.0667 | 64 | 0.1875 |
| 12_112467 | 23 | 0.0435 | 29 | 0.1034 | 13 | 0.0769 | 17 | 0.0588 | 82 | 0.0732 | 20 | 0.05 | 14 | 0 | 24 | 0.125 | 19 | 0 | 77 | 0.0519 |
| 12_112467 | 18 | 0.1667 | 26 | 0.0385 | 8 | 0.25 | 14 | 0.2857 | 66 | 0.1515 | 15 | 0 | 13 | 0.3846 | 22 | 0 | 12 | 0.1667 | 62 | 0.1129 |
| 12_112468 | 14 | 0.5714 | 17 | 0.6471 | 13 | 0.6923 | 11 | 0.7273 | 55 | 0.6545 | 15 | 0.7333 | 12 | 0.5833 | 16 | 0.875 | 13 | 0.6923 | 56 | 0.7321 |
| 12_112468 | 18 | 0 | 14 | 0.0714 | 12 | 0.0833 | 11 | 0.0909 | 55 | 0.0545 | 14 | 0.0714 | 14 | 0.1429 | 16 | 0.125 | 9 | 0.2222 | 53 | 0.1321 |
| 12_112577 | 34 | 0.0882 | 24 | 0.1667 | 25 | 0.08 | 31 | 0.0323 | 114 | 0.0877 | 13 | 0.1538 | 39 | 0.1282 | 32 | 0.1875 | 17 | 0.1765 | 101 | 0.1584 |
| 12_112578 | 39 | 0.2308 | 24 | 0.2083 | 29 | 0.1379 | 29 | 0.069 | 121 | 0.1653 | 13 | 0.1538 | 41 | 0.2195 | 35 | 0.2286 | 18 | 0.2222 | 107 | 0.215 |
| 12_112578 | 39 | 0.2564 | 23 | 0 | 29 | 0.069 | 30 | 0.2667 | 121 | 0.1653 | 12 | 0.4167 | 40 | 0.175 | 34 | 0.2059 | 19 | 0.2105 | 105 | 0.219 |
| 12_112578 | 39 | 0.7436 | 22 | 0.7727 | 26 | 0.8462 | 25 | 0.76 | 112 | 0.7768 | 13 | 0.4615 | 40 | 0.65 | 31 | 0.7097 | 20 | 0.75 | 104 | 0.6635 |
| 12_112578 | 41 | 0.2439 | 30 | 0.1333 | 27 | 0.2593 | 30 | 0.2333 | 128 | 0.2188 | 13 | 0.2308 | 42 | 0.1905 | 36 | 0.1667 | 18 | 0.1667 | 109 | 0.1835 |
| 12_112578 | 18 | 0.4444 | 19 | 0.5263 | 21 | 0.7143 | 18 | 0.6111 | 76 | 0.5789 | 9 | 0.5556 | 30 | 0.7 | 24 | 0.4583 | 14 | 0.5714 | 77 | 0.5844 |
| 12_112578 | 38 | 0.3158 | 27 | 0.2593 | 30 | 0.2667 | 31 | 0.2258 | 126 | 0.2698 | 16 | 0.125 | 40 | 0.225 | 32 | 0.25 | 19 | 0.1579 | 107 | 0.2056 |
| 12_112578 | 2 | 0 | 1 | 0 | 4 | 0 | 5 | 0.2 | 12 | 0.0833 | NA | NA | 3 | 0.3333 | 3 | 0 | 3 | 0 | 9 | 0.1111 |
| 12_112579 | 2 | 0 | 1 | 1 | 3 | 0 | 5 | 0 | 11 | 0.0909 | NA | NA | 6 | 0.1667 | 5 | 0.2 | 2 | 0 | 13 | 0.1538 |
| 12_112579 | NA | NA | 1 | 0 | 5 | 0.4 | 4 | 0.5 | 10 | 0.4 | NA | NA | 7 | 0.5714 | 2 | 0 | 2 | 0 | 11 | 0.3636 |
| 12_112579 | 3 | 0.6667 | NA | NA | 5 | 0.2 | 5 | 0.4 | 13 | 0.3846 | NA | NA | 6 | 1 | 4 | 1 | 2 | 0.5 | 12 | 0.9167 |

| | | | | | | | | | | | | | | | | | | | | |
|---|---|---|---|---|---|---|---|---|---|---|---|---|---|---|---|---|---|---|---|---|
| 12_112579 | 3 | 0 | 1 | 0 | 5 | 0 | 7 | 0.1429 | 16 | 0.0625 | NA | NA | 7 | 0 | 5 | 0 | 2 | 0.5 | 14 | 0.0714 |
| 12_112579 | 1 | 1 | 1 | 0 | 7 | 0.4286 | 7 | 0.2857 | 16 | 0.375 | NA | NA | 7 | 0.2857 | 5 | 0.8 | 5 | 0.4 | 17 | 0.4706 |
| 12_112579 | 4 | 0.25 | 1 | 0 | 4 | 0 | 4 | 0 | 13 | 0.0769 | NA | NA | 6 | 0 | 6 | 0 | 7 | 0.1429 | 19 | 0.0526 |
| 12_112580 | 4 | 0.25 | NA | NA | 4 | 0.25 | 2 | 0.5 | 10 | 0.3 | NA | NA | 5 | 0.8 | 3 | 1 | 4 | 0.5 | 12 | 0.75 |
| 12_112580 | 4 | 0.25 | 2 | 0 | 5 | 0 | 6 | 0 | 17 | 0.0588 | NA | NA | 7 | 0 | 6 | 0.3333 | 8 | 0.25 | 21 | 0.1905 |
| 12_112580 | 3 | 0.6667 | 2 | 0.5 | 4 | 0.5 | 5 | 0.6 | 14 | 0.5714 | NA | NA | 5 | 0.2 | 4 | 0.5 | 8 | 0.625 | 17 | 0.4706 |
| 12_112580 | 3 | 0.3333 | 1 | 0 | 6 | 0 | 7 | 0 | 17 | 0.0588 | NA | NA | 7 | 0.1429 | 6 | 0.5 | 8 | 0.375 | 21 | 0.3333 |
| 12_112580 | 3 | 0.3333 | 2 | 0.5 | 5 | 0.4 | 3 | 0.3333 | 13 | 0.3846 | NA | NA | 6 | 0.5 | 5 | 0.4 | 5 | 0.6 | 16 | 0.5 |
| 12_112581 | 44 | 0.5455 | 28 | 0.5357 | 29 | 0.4483 | 32 | 0.625 | 133 | 0.5414 | 14 | 0.3571 | 41 | 0.7805 | 41 | 0.5122 | 25 | 0.56 | 121 | 0.595 |
| 12_112581 | 44 | 0.0909 | 24 | 0.0417 | 33 | 0.1515 | 38 | 0.2632 | 139 | 0.1439 | 15 | 0.1333 | 50 | 0.1 | 45 | 0.1556 | 26 | 0.0385 | 136 | 0.1103 |
| 12_112581 | 44 | 0.0909 | 29 | 0.1379 | 34 | 0.1176 | 39 | 0.1282 | 146 | 0.1164 | 18 | 0.1667 | 47 | 0.0638 | 46 | 0.1522 | 27 | 0.2222 | 138 | 0.1377 |
| 12_112581 | 42 | 0.2381 | 28 | 0.4286 | 30 | 0.3 | 38 | 0.2632 | 138 | 0.2971 | 17 | 0.2941 | 49 | 0.1837 | 47 | 0.2766 | 25 | 0.36 | 138 | 0.2609 |
| 12_112620 | 27 | 0.3333 | 18 | 0.1111 | 18 | 0.5556 | 23 | 0.3478 | 86 | 0.3372 | 23 | 0.2609 | 16 | 0.25 | 22 | 0.3636 | 18 | 0.2778 | 79 | 0.2911 |
| 12_112620 | 24 | 0.625 | 13 | 0.6923 | 21 | 0.7143 | 21 | 0.7619 | 79 | 0.6962 | 18 | 0.6667 | 14 | 0.9286 | 20 | 0.85 | 12 | 0.75 | 64 | 0.7969 |
| 12_112620 | 24 | 0 | 16 | 0.125 | 19 | 0 | 18 | 0.1667 | 77 | 0.0649 | 18 | 0.1667 | 14 | 0.1429 | 20 | 0.05 | 14 | 0.2143 | 66 | 0.1364 |
| 12_112620 | 23 | 0.087 | 15 | 0.0667 | 16 | 0.0625 | 19 | 0.0526 | 73 | 0.0685 | 16 | 0 | 14 | 0.0714 | 17 | 0.1765 | 14 | 0.0714 | 61 | 0.082 |
| 12_112621 | 24 | 0.3333 | 17 | 0.4118 | 20 | 0.45 | 20 | 0.3 | 81 | 0.3704 | 21 | 0.3333 | 16 | 0.0625 | 21 | 0.381 | 16 | 0.3125 | 74 | 0.2838 |
| 12_112623 | 14 | 0.1429 | 11 | 0.0909 | 10 | 0.1 | 12 | 0.5833 | 47 | 0.234 | 13 | 0.1538 | 10 | 0.2 | 12 | 0.1667 | 7 | 0.1429 | 42 | 0.1667 |
| 12_112701 | 3 | 0 | 3 | 0 | 4 | 0 | 1 | 1 | 11 | 0.0909 | 4 | 0 | 2 | 0 | 4 | 0.5 | 3 | 0 | 13 | 0.1538 |
| 12_112701 | 3 | 0.3333 | 4 | 0 | 5 | 0 | 3 | 0 | 15 | 0.0667 | 4 | 0 | 3 | 0 | 5 | 0.4 | 6 | 0.1667 | 18 | 0.1667 |
| 12_112702 | 4 | 0.5 | 6 | 0.5 | 6 | 0.5 | 3 | 0.3333 | 19 | 0.4737 | 6 | 0.6667 | 3 | 0 | 3 | 0.3333 | 2 | 0.5 | 14 | 0.4286 |
| 12_112702 | 4 | 0.25 | 5 | 0.2 | 5 | 0.2 | 4 | 0 | 18 | 0.1667 | 6 | 0.1667 | 3 | 0.6667 | 4 | 0.5 | 5 | 0.2 | 18 | 0.3333 |
| 12_112702 | 4 | 1 | 7 | 0.8571 | 6 | 0.5 | 2 | 1 | 19 | 0.7895 | 7 | 0.5714 | 2 | 1 | 4 | 1 | 6 | 1 | 19 | 0.8421 |
| 12_112702 | 4 | 0.5 | 7 | 0.1429 | 7 | 0.4286 | 3 | 0 | 21 | 0.2857 | 6 | 0 | 2 | 0.5 | 4 | 0.25 | 6 | 0.5 | 18 | 0.2778 |
| 12_112702 | 4 | 0.25 | 6 | 0.1667 | 6 | 0.3333 | 4 | 0 | 20 | 0.2 | 7 | 0.4286 | 2 | 0 | 5 | 0.2 | 5 | 0 | 19 | 0.2105 |
| 12_112703 | 4 | 1 | 6 | 0.6667 | 7 | 0.5714 | 4 | 0.75 | 21 | 0.7143 | 7 | 0.7143 | 3 | 0.6667 | 5 | 0.8 | 6 | 0.5 | 21 | 0.6667 |
| 12_112703 | 3 | 0 | 6 | 0.5 | 5 | 0.2 | 2 | 0.5 | 16 | 0.3125 | 6 | 0.3333 | 3 | 0.6667 | 4 | 0.75 | 5 | 0.6 | 18 | 0.5556 |
| 12_112704 | 5 | 0.8 | 9 | 0.3333 | 6 | 0.3333 | 4 | 0.75 | 24 | 0.5 | 8 | 0.5 | 3 | 0.6667 | 5 | 0.6 | 7 | 0.4286 | 23 | 0.5217 |
| 12_112704 | 5 | 1 | 10 | 0.6 | 8 | 0.5 | 6 | 0.5 | 29 | 0.6207 | 7 | 0.7143 | 2 | 0.5 | 7 | 0.8571 | 7 | 0.7143 | 23 | 0.7391 |
| 12_112704 | 4 | 0.25 | 11 | 0.1818 | 7 | 0.1429 | 6 | 0.1667 | 28 | 0.1786 | 8 | 0.5 | 4 | 0.25 | 7 | 0.1429 | 7 | 0 | 26 | 0.2308 |
| 12_112704 | 4 | 0.5 | 9 | 0.3333 | 6 | 0.8333 | 5 | 0.6 | 24 | 0.5417 | 3 | 0.6667 | 3 | 0.3333 | 5 | 0.6 | 4 | 0.5 | 15 | 0.5333 |
| 12_112704 | 5 | 0.2 | 10 | 0.2 | 8 | 0 | 6 | 0 | 29 | 0.1034 | 8 | 0.25 | 3 | 0.6667 | 7 | 0 | 7 | 0.2857 | 25 | 0.24 |
| 12_112705 | 5 | 0.6 | 10 | 0.6 | 8 | 0.625 | 8 | 0.875 | 31 | 0.6774 | 8 | 0.625 | 3 | 0.6667 | 8 | 0.75 | 9 | 0.5556 | 28 | 0.6429 |
| 12_112705 | 5 | 1 | 7 | 0.4286 | 8 | 0.875 | 7 | 0.5714 | 27 | 0.7037 | 8 | 0.625 | 4 | 0.25 | 8 | 0.75 | 9 | 0.4444 | 29 | 0.5517 |
| 12_112705 | 5 | 0 | 10 | 0.3 | 6 | 0 | 8 | 0.25 | 29 | 0.1724 | 5 | 0.2 | 2 | 0 | 5 | 0.2 | 7 | 0.2857 | 19 | 0.2105 |
| 12_112705 | 5 | 0.2 | 10 | 0.4 | 8 | 0.375 | 8 | 0.25 | 31 | 0.3226 | 9 | 0.2222 | 4 | 0.25 | 7 | 0.5714 | 11 | 0.2727 | 31 | 0.3226 |
| 12_112705 | 5 | 0.2 | 10 | 0 | 9 | 0 | 8 | 0.125 | 32 | 0.0625 | 8 | 0 | 4 | 0.25 | 9 | 0.3333 | 11 | 0.0909 | 32 | 0.1562 |
| 12_112705 | 5 | 0.8 | 10 | 0.9 | 8 | 1 | 5 | 1 | 28 | 0.9286 | 10 | 1 | 4 | 1 | 8 | 1 | 10 | 0.9 | 32 | 0.9688 |
| 12_112705 | 6 | 0.5 | 11 | 0.4545 | 5 | 0.4 | 5 | 0.4 | 27 | 0.4444 | 10 | 0.2 | 5 | 0.4 | 10 | 0.4 | 10 | 0.2 | 35 | 0.2857 |
| 12_112705 | 7 | 0.2857 | 11 | 0.6364 | 9 | 0.6667 | 9 | 0.3333 | 36 | 0.5 | 9 | 0.5556 | 5 | 0.4 | 9 | 0.5556 | 11 | 0.5455 | 34 | 0.5294 |
| 12_112705 | 7 | 0.7143 | 11 | 1 | 7 | 0.8571 | 8 | 0.625 | 33 | 0.8182 | 8 | 0.875 | 5 | 0.6 | 9 | 0.5556 | 8 | 0.75 | 30 | 0.7 |
| 12_112705 | 8 | 0.875 | 8 | 0.875 | 6 | 0.8333 | 6 | 0.8333 | 28 | 0.8571 | 7 | 0.8571 | 4 | 0.25 | 6 | 1 | 10 | 0.8 | 27 | 0.7778 |
| 12_112705 | 8 | 0.625 | 12 | 0.9167 | 8 | 0.75 | 9 | 0.8889 | 37 | 0.8108 | 9 | 0.8889 | 4 | 1 | 11 | 0.9091 | 10 | 0.7 | 34 | 0.8529 |
| 12_112711 | 11 | 0.9091 | 15 | 0.9333 | 11 | 0.9091 | 11 | 0.7273 | 48 | 0.875 | 14 | 0.9286 | 8 | 0.625 | 11 | 0.8182 | 19 | 0.8947 | 52 | 0.8462 |
| 12_112711 | 14 | 0 | 19 | 0.1579 | 13 | 0 | 11 | 0.0909 | 57 | 0.0702 | 14 | 0.0714 | 8 | 0.125 | 14 | 0.0714 | 19 | 0.4211 | 55 | 0.2 |
| 12_112711 | 7 | 0 | 11 | 0.0909 | 6 | 0 | 9 | 0.2222 | 33 | 0.0909 | 9 | 0.2222 | 6 | 0.5 | 11 | 0.1818 | 15 | 0.1333 | 41 | 0.2195 |
| 12_112711 | 12 | 0.6667 | 21 | 0.7143 | 12 | 0.5833 | 11 | 0.8182 | 56 | 0.6964 | 18 | 0.5556 | 9 | 0.6667 | 11 | 0.6364 | 15 | 0.2667 | 53 | 0.5094 |
| 12_112711 | 14 | 0.7857 | 22 | 0.5455 | 14 | 0.7143 | 13 | 0.7692 | 63 | 0.6825 | 19 | 0.6842 | 9 | 1 | 12 | 0.9167 | 19 | 0.8421 | 59 | 0.8305 |
| 12_112712 | 13 | 0.6154 | 24 | 0.5833 | 13 | 0.5385 | 11 | 0.5455 | 61 | 0.5738 | 19 | 0.6316 | 8 | 0.5 | 14 | 0.5714 | 17 | 0.5294 | 58 | 0.569 |
| 12_112712 | 15 | 0.2 | 21 | 0.1429 | 13 | 0.1538 | 14 | 0.2857 | 63 | 0.1905 | 23 | 0.087 | 9 | 0.2222 | 18 | 0.2222 | 18 | 0 | 68 | 0.1176 |
| 12_112714 | 1 | 1 | NA | NA | NA | NA | NA | NA | 1 | 1 | NA | NA | 1 | 1 | NA | NA | NA | NA | 1 | 1 |
| 12_112714 | 1 | 1 | NA | NA | NA | NA | NA | NA | 1 | 1 | NA | NA | 1 | 1 | NA | NA | NA | NA | 1 | 1 |
| 12_112735 | NA | NA | 3 | 0 | 2 | 0.5 | 1 | 1 | 6 | 0.3333 | NA | NA | NA | NA | 3 | 1 | NA | NA | 3 | 1 |
| 12_112735 | NA | NA | 3 | 0.3333 | 2 | 0.5 | 1 | 0 | 6 | 0.3333 | 1 | 1 | NA | NA | 3 | 0 | NA | NA | 4 | 0.25 |
| 12_112735 | NA | NA | 3 | 0.6667 | 1 | 1 | 2 | 0.5 | 6 | 0.6667 | NA | NA | 1 | 1 | 3 | 1 | NA | NA | 4 | 1 |
| 12_112772 | 2 | 0.5 | NA | NA | 1 | 1 | 2 | 0 | 5 | 0.4 | 1 | 1 | 1 | 0 | 1 | 1 | NA | NA | 3 | 0.6667 |
| 12_112773 | 3 | 0.6667 | NA | NA | 1 | 0 | 2 | 0.5 | 6 | 0.5 | 1 | 0 | 2 | 0.5 | 1 | 1 | NA | NA | 4 | 0.5 |
| 12_112779 | 4 | 0.25 | NA | NA | 1 | 0 | 2 | 1 | 7 | 0.4286 | 1 | 1 | 2 | 0 | 1 | 0 | NA | NA | 4 | 0.25 |
| 12_112783 | 2 | 0.5 | 2 | 0.5 | 2 | 0.5 | 1 | 0 | 7 | 0.4286 | NA | NA | 1 | 1 | 1 | 0 | NA | NA | 2 | 0.5 |
| 12_112784 | 1 | 1 | 1 | 1 | 2 | 1 | NA | NA | 4 | 1 | NA | NA | 1 | 1 | 1 | 1 | NA | NA | 2 | 1 |
| 12_112784 | 1 | 1 | 1 | 1 | 1 | 1 | NA | NA | 3 | 1 | NA | NA | 1 | 0 | 1 | 1 | NA | NA | 2 | 0.5 |
| 12_112785 | 1 | 0 | 1 | 1 | 1 | 1 | NA | NA | 3 | 0.6667 | NA | NA | 1 | 1 | 1 | 1 | NA | NA | 2 | 1 |
| 12_112785 | 1 | 0 | 1 | 0 | 2 | 1 | NA | NA | 4 | 0.5 | NA | NA | 1 | 1 | 1 | 1 | NA | NA | 2 | 1 |
| 12_112785 | 1 | 0 | 1 | 1 | 2 | 1 | NA | NA | 4 | 0.75 | NA | NA | 1 | 1 | 1 | 1 | NA | NA | 2 | 1 |
| 12_112785 | 1 | 0 | NA | NA | 2 | 1 | NA | NA | 3 | 0.6667 | NA | NA | 1 | 1 | 1 | 1 | NA | NA | 2 | 1 |

| | | | | | | | | | | | | | | | | | | | | |
|---|---|---|---|---|---|---|---|---|---|---|---|---|---|---|---|---|---|---|---|---|
| 12_1127951 | 1 | 1 | NA | NA | NA | NA | NA | NA | 1 | 1 | NA | NA | NA | NA | 1 | 1 | NA | NA | 1 | 1 |
| 12_1128716 | 6 | 0 | 1 | 0 | 4 | 0.5 | 4 | 0 | 15 | 0.1333 | 2 | 0 | 5 | 0.4 | 2 | 0 | 1 | 0 | 10 | 0.2 |
| 12_1128718 | 6 | 0 | 1 | 0 | 4 | 0.25 | 4 | 0.25 | 15 | 0.1333 | 3 | 0.3333 | 4 | 0.25 | 2 | 0 | 2 | 0.5 | 11 | 0.2727 |
| 12_1128719 | 5 | 0.2 | 1 | 1 | 4 | 0.25 | 4 | 0 | 14 | 0.2143 | 3 | 0.3333 | 6 | 0.1667 | 2 | 0.5 | 2 | 0.5 | 13 | 0.3077 |
| 12_1128719 | 6 | 0.6667 | 1 | 1 | 3 | 1 | 4 | 1 | 14 | 0.8571 | 3 | 0.3333 | 6 | 1 | 2 | 1 | 1 | 0 | 12 | 0.75 |
| 12_1128720 | 5 | 0.2 | 1 | 0 | 4 | 0.25 | 4 | 0.25 | 14 | 0.2143 | 3 | 0.3333 | 6 | 0 | 2 | 0 | NA | NA | 11 | 0.0909 |
| 12_1128724 | 6 | 0.8333 | NA | NA | 5 | 0.6 | 3 | 1 | 14 | 0.7857 | 2 | 0.5 | 6 | 0.8333 | 2 | 0.5 | 1 | 1 | 11 | 0.7273 |
| 12_1128724 | 4 | 0.25 | NA | NA | 4 | 0.5 | 3 | 0.6667 | 11 | 0.4545 | 2 | 0 | 5 | 0.6 | 2 | 0.5 | 1 | 1 | 10 | 0.5 |
| 12_1128725 | 4 | 0.75 | NA | NA | 4 | 0.75 | 2 | 0.5 | 10 | 0.7 | 2 | 0.5 | 6 | 0.8333 | 1 | 1 | 1 | 1 | 10 | 0.8 |
| 12_1128726 | 6 | 0.1667 | NA | NA | 4 | 0 | 3 | 0 | 13 | 0.0769 | 2 | 0.5 | 5 | 0 | 1 | 0 | 1 | 0 | 9 | 0.1111 |
| 12_1128726 | 6 | 0.3333 | NA | NA | 4 | 0.25 | 3 | 0.6667 | 13 | 0.3846 | 2 | 0.5 | 5 | 0.4 | 2 | 0 | 1 | 0 | 10 | 0.3 |
| 12_1128727 | NA | NA | NA | NA | 2 | 1 | 1 | 1 | 3 | 1 | 1 | 1 | 3 | 0.6667 | NA | NA | NA | NA | 4 | 0.75 |
| 12_1128727 | 6 | 0.6667 | NA | NA | 3 | 1 | 2 | 1 | 11 | 0.8182 | 2 | 1 | 6 | 0.6667 | 1 | 0 | 1 | 0 | 10 | 0.6 |
| 12_1128728 | 5 | 0.8 | NA | NA | 4 | 0.25 | 2 | 0 | 11 | 0.4545 | 1 | 1 | 5 | 0.8 | 2 | 0 | 1 | 0 | 9 | 0.5556 |
| 12_1128729 | 3 | 0.3333 | NA | NA | 4 | 0.25 | 3 | 0 | 10 | 0.2 | 2 | 0.5 | 6 | 0 | 2 | 0 | 1 | 0 | 11 | 0.0909 |
| 12_1128984 | 3 | 0.3333 | 1 | 0 | 1 | 1 | 1 | 0 | 6 | 0.3333 | 2 | 0.5 | 4 | 0.5 | NA | NA | 1 | 1 | 7 | 0.5714 |
| 12_1129193 | 1 | 1 | NA | NA | 2 | 0 | 1 | 1 | 4 | 0.5 | 4 | 0.5 | NA | NA | 1 | 1 | NA | NA | 5 | 0.6 |
| 12_1129237 | 2 | 0 | 5 | 0.4 | 2 | 1 | 7 | 0.1429 | 16 | 0.3125 | 2 | 1 | 2 | 0.5 | 1 | 0 | 1 | 1 | 6 | 0.6667 |
| 12_1129238 | 2 | 0.5 | 4 | 0.75 | 3 | 0 | 7 | 0.8571 | 16 | 0.625 | 1 | 0 | 2 | 0.5 | 1 | 1 | 1 | 1 | 5 | 0.6 |
| 12_1129239 | 1 | 1 | 4 | 0.5 | 1 | 0 | 7 | 0.5714 | 13 | 0.5385 | 1 | 0 | 2 | 0.5 | 1 | 0 | NA | NA | 4 | 0.25 |
| 12_1129413 | 11 | 0.8182 | 6 | 0.6667 | 6 | 0.5 | 3 | 0.6667 | 26 | 0.6923 | 3 | 0.6667 | 4 | 1 | 4 | 0.75 | 4 | 0.75 | 15 | 0.8 |
| 12_1129413 | 18 | 0.1111 | 6 | 0.1667 | 6 | 0.1667 | 4 | 0 | 34 | 0.1176 | 6 | 0 | 5 | 0 | 8 | 0.25 | 7 | 0 | 26 | 0.0769 |
| 12_1129414 | 15 | 0.7333 | 6 | 0.6667 | 6 | 0.8333 | 5 | 0.8 | 32 | 0.75 | 6 | 0.8333 | 4 | 1 | 7 | 0.7143 | 7 | 0.8571 | 24 | 0.8333 |
| 12_1129414 | 17 | 0.2941 | 6 | 0.5 | 6 | 0.1667 | 5 | 0.4 | 34 | 0.3235 | 6 | 0.5 | 6 | 0 | 8 | 0.125 | 7 | 0.5714 | 27 | 0.2963 |
| 12_1129416 | 15 | 0.2667 | 7 | 0.5714 | 6 | 0.1667 | 6 | 0 | 34 | 0.2647 | 6 | 0.1667 | 6 | 0 | 8 | 0.125 | 6 | 0.1667 | 26 | 0.1154 |
| 12_1129416 | 17 | 0.0588 | 5 | 0.2 | 6 | 0.1667 | 4 | 0.25 | 32 | 0.125 | 5 | 0 | 6 | 0.3333 | 6 | 0.1667 | 4 | 0 | 21 | 0.1429 |
| 12_1129417 | 16 | 0.75 | 5 | 0.8 | 5 | 1 | 5 | 1 | 31 | 0.8387 | 5 | 1 | 5 | 1 | 7 | 0.8571 | 6 | 1 | 23 | 0.9565 |
| 12_1129418 | 8 | 0.75 | 4 | 0.25 | 3 | 0.6667 | 5 | 0.6 | 20 | 0.6 | 4 | 0.75 | 4 | 0.75 | 6 | 0.6667 | 2 | 0 | 16 | 0.625 |
| 12_1129419 | 17 | 0.0588 | 7 | 0 | 6 | 0.3333 | 6 | 0 | 36 | 0.0833 | 6 | 0 | 7 | 0.1429 | 8 | 0.25 | 7 | 0 | 28 | 0.1071 |
| 12_1129419 | 16 | 0.375 | 6 | 0.5 | 7 | 0.4286 | 6 | 0.5 | 35 | 0.4286 | 7 | 0.4286 | 7 | 0.5714 | 9 | 0.3333 | 7 | 0.4286 | 30 | 0.4333 |
| 12_1129420 | 14 | 0.2857 | 6 | 0 | 7 | 0.1429 | 5 | 0.2 | 32 | 0.1875 | 6 | 0.5 | 2 | 0 | 8 | 0.25 | 6 | 0 | 22 | 0.2273 |
| 12_1129421 | 13 | 1 | 6 | 1 | 7 | 0.8571 | 5 | 1 | 31 | 0.9677 | 7 | 1 | 5 | 1 | 11 | 1 | 4 | 1 | 27 | 1 |
| 12_1129421 | 14 | 0.7143 | 6 | 0.5 | 6 | 0.6667 | 6 | 0.3333 | 32 | 0.5938 | 5 | 0.6 | 6 | 0.5 | 6 | 0.1667 | 7 | 0.7143 | 24 | 0.5 |
| 12_1129421 | 16 | 1 | 6 | 1 | 6 | 0.8333 | 6 | 0.8333 | 34 | 0.9412 | 7 | 1 | 7 | 1 | 9 | 0.8889 | 7 | 1 | 30 | 0.9667 |
| 12_1129422 | 17 | 0.1176 | 6 | 0.3333 | 6 | 0.5 | 6 | 0.3333 | 35 | 0.2571 | 8 | 0 | 6 | 0.1667 | 10 | 0.5 | 9 | 0.1111 | 33 | 0.2121 |
| 12_1129422 | 16 | 0.1875 | 6 | 0.3333 | 6 | 0 | 5 | 0.2 | 33 | 0.1818 | 8 | 0.125 | 7 | 0.2857 | 7 | 0 | 7 | 0.5714 | 29 | 0.2414 |
| 12_1129423 | 15 | 0.6 | 6 | 0.6667 | 7 | 0.4286 | 6 | 1 | 34 | 0.6471 | 7 | 0.5714 | 6 | 0.5 | 8 | 0.5 | 8 | 0.375 | 29 | 0.4828 |
| 12_1129423 | 15 | 0.0667 | 9 | 0 | 7 | 0.1429 | 6 | 0.1667 | 37 | 0.0811 | 8 | 0.25 | 9 | 0 | 10 | 0.2 | 7 | 0 | 34 | 0.1176 |
| 12_1129423 | 18 | 0 | 8 | 0.125 | 7 | 0 | 5 | 0.2 | 38 | 0.0526 | 9 | 0.1111 | 8 | 0.125 | 10 | 0 | 7 | 0 | 34 | 0.0588 |
| 12_1129424 | 18 | 0.2222 | 8 | 0.125 | 8 | 0.375 | 6 | 0.5 | 40 | 0.275 | 10 | 0.3 | 10 | 0.4 | 11 | 0.2727 | 8 | 0.375 | 39 | 0.3333 |
| 12_1129425 | 10 | 0.1 | 5 | 0.2 | 4 | 0.25 | 4 | 0.25 | 23 | 0.1739 | 6 | 0.3333 | 5 | 0 | 7 | 0.4286 | 6 | 0.1667 | 24 | 0.25 |
| 12_1129425 | 5 | 0 | NA | NA | 1 | 1 | 2 | 1 | 8 | 0.375 | NA | NA | 1 | 0 | 3 | 0.3333 | 1 | 1 | 5 | 0.4 |
| 12_1129425 | 5 | 1 | NA | NA | 1 | 1 | 2 | 1 | 8 | 1 | NA | NA | 1 | 1 | 1 | 1 | 1 | 1 | 3 | 1 |
| 12_1129427 | 5 | 0.4 | NA | NA | 2 | 0.5 | 3 | 1 | 10 | 0.6 | NA | NA | NA | NA | 3 | 0.6667 | 2 | 0.5 | 5 | 0.6 |
| 12_1129429 | 4 | 0 | NA | NA | 2 | 1 | 3 | 0.3333 | 9 | 0.3333 | NA | NA | 2 | 1 | 1 | 0 | 2 | 0.5 | 5 | 0.6 |
| 12_1129429 | 4 | 0.25 | NA | NA | 1 | 0 | 3 | 0 | 8 | 0.125 | NA | NA | 1 | 0 | 3 | 0 | 1 | 1 | 5 | 0.2 |
| 12_1130352 | 8 | 0.25 | 5 | 0.6 | 9 | 0.3333 | 8 | 0.5 | 30 | 0.4 | 2 | 0.5 | 7 | 0.7143 | 6 | 0.8333 | 9 | 0.5556 | 24 | 0.6667 |
| 12_1130353 | 7 | 0.8571 | 6 | 0.6667 | 5 | 1 | 3 | 0.6667 | 21 | 0.8095 | 1 | 0 | 6 | 1 | 3 | 1 | 7 | 0.7143 | 17 | 0.8235 |
| 12_1130353 | 10 | 0.4 | 10 | 0.1 | 8 | 0.375 | 10 | 0.4 | 38 | 0.3158 | 1 | 0 | 8 | 0.5 | 7 | 0.7143 | 10 | 0.7 | 26 | 0.6154 |
| 12_1130353 | 9 | 0.7778 | 10 | 0.5 | 9 | 0.6667 | 9 | 0.6667 | 37 | 0.6486 | 2 | 1 | 7 | 0.2857 | 7 | 0.4286 | 10 | 0.1 | 26 | 0.3077 |
| 12_1130353 | 9 | 0.4444 | 6 | 0.5 | 7 | 0.2857 | 8 | 0.375 | 30 | 0.4 | 2 | 0 | 7 | 0.4286 | 6 | 0.5 | 10 | 0.4 | 25 | 0.4 |
| 12_1130353 | 7 | 0.8571 | 8 | 0.75 | 6 | 0.6667 | 8 | 0.875 | 29 | 0.7931 | 2 | 0.5 | 7 | 0.7143 | 5 | 0.6 | 10 | 0.8 | 24 | 0.7083 |
| 12_1130355 | 9 | 0.2222 | 11 | 0.3636 | 10 | 0.4 | 7 | 0.4286 | 37 | 0.3514 | 3 | 0.3333 | 7 | 0.4286 | 6 | 0.5 | 7 | 0.5714 | 23 | 0.4783 |
| 12_1130377 | 15 | 0.6667 | 16 | 0.6875 | 11 | 0.6364 | 10 | 0.6 | 52 | 0.6538 | 8 | 0.875 | 11 | 0.8182 | 9 | 0.8889 | 16 | 0.875 | 44 | 0.8636 |
| 12_1130378 | 17 | 0.1765 | 11 | 0.0909 | 10 | 0.1 | 10 | 0.3 | 48 | 0.1667 | 9 | 0.1111 | 10 | 0 | 7 | 0.2857 | 14 | 0.0714 | 40 | 0.1 |
| 12_1131002 | 8 | 0.125 | 10 | 0.1 | 4 | 0 | 12 | 0.0833 | 34 | 0.0882 | 4 | 0.25 | 13 | 0.2308 | 7 | 0.1429 | 8 | 0.125 | 32 | 0.1875 |
| 12_1131003 | 8 | 0.125 | 10 | 0.2 | 4 | 0 | 13 | 0.3077 | 35 | 0.2 | 3 | 0.3333 | 12 | 0.25 | 6 | 0 | 5 | 0.2 | 26 | 0.1923 |
| 12_1131004 | 9 | 0.3333 | 10 | 0.6 | 2 | 1 | 12 | 0.4167 | 33 | 0.4848 | 5 | 0.8 | 13 | 0.3077 | 7 | 0.7143 | 7 | 0.8571 | 32 | 0.5938 |
| 12_1131089 | 41 | 0.122 | 35 | 0.1714 | 25 | 0.12 | 44 | 0.0455 | 145 | 0.1103 | 29 | 0.069 | 32 | 0.125 | 34 | 0.2647 | 26 | 0.2308 | 121 | 0.1736 |
| 12_1131517 | 3 | 0 | 1 | 0 | 3 | 0 | 2 | 0.5 | 9 | 0.1111 | 2 | 0.5 | 1 | 0 | 1 | 0 | 5 | 0.2 | 9 | 0.2222 |
| 12_1131517 | 3 | 0.3333 | 2 | 0.5 | 2 | 0 | 3 | 0.3333 | 10 | 0.3 | 2 | 0 | 1 | 1 | 1 | 0 | 5 | 0.4 | 9 | 0.3333 |
| 12_1131517 | 3 | 1 | 1 | 1 | 3 | 1 | 3 | 1 | 10 | 1 | 2 | 1 | 1 | 1 | NA | NA | 6 | 0.8333 | 9 | 0.8889 |
| 12_1131520 | 2 | 0.5 | 2 | 0.5 | 2 | 0 | 2 | 0 | 8 | 0.25 | 2 | 0 | 1 | 1 | 1 | 0 | 7 | 0.1429 | 11 | 0.1818 |
| 12_1131521 | 2 | 0.5 | 2 | 0.5 | 4 | 0.5 | 2 | 0.5 | 10 | 0.5 | 1 | 0 | 1 | 0 | 1 | 0 | 7 | 0.4286 | 10 | 0.3 |
| 12_1131522 | 3 | 1 | 2 | 1 | 4 | 1 | 4 | 0.75 | 13 | 0.9231 | 3 | 0.6667 | 1 | 1 | NA | NA | 8 | 0.875 | 12 | 0.8333 |

| | | | | | | | | | | | | | | | | | | | | |
|---|---|---|---|---|---|---|---|---|---|---|---|---|---|---|---|---|---|---|---|---|
| 12_113152 | 3 | 1 | 1 | 1 | 4 | 1 | 3 | 1 | 11 | 1 | 2 | 1 | 1 | 1 | 1 | 1 | 6 | 0.8333 | 10 | 0.9 |
| 12_113152 | 4 | 1 | 1 | 1 | 5 | 1 | 4 | 1 | 14 | 1 | 4 | 1 | NA | NA | NA | NA | 4 | 1 | 8 | 1 |
| 12_113152 | 3 | 0.3333 | NA | NA | 4 | 0 | 3 | 0 | 10 | 0.1 | 3 | 0 | 2 | 0 | 1 | 1 | 7 | 0 | 13 | 0.0769 |
| 12_113152 | 2 | 1 | 2 | 1 | 5 | 1 | 4 | 1 | 13 | 1 | 4 | 1 | 2 | 1 | 1 | 1 | 7 | 1 | 14 | 1 |
| 12_113152 | 4 | 0.5 | 2 | 0.5 | 3 | 0.3333 | 3 | 0 | 12 | 0.3333 | 2 | 0 | 1 | 1 | NA | NA | 7 | 0.2857 | 10 | 0.3 |
| 12_113152 | 4 | 1 | 2 | 1 | 5 | 0.8 | 2 | 1 | 13 | 0.9231 | 4 | 1 | 2 | 1 | 2 | 1 | 7 | 1 | 15 | 1 |
| 12_113152 | 5 | 1 | 1 | 1 | 5 | 0.6 | 3 | 0.6667 | 14 | 0.7857 | 4 | 1 | 1 | 1 | 2 | 0.5 | 5 | 0.8 | 12 | 0.8333 |
| 12_113152 | 5 | 1 | 1 | 1 | 6 | 1 | 3 | 1 | 15 | 1 | 4 | 1 | 1 | 1 | 2 | 0.5 | 8 | 1 | 15 | 0.9333 |
| 12_113153 | 5 | 0.6 | NA | NA | 4 | 0.5 | 3 | 0.6667 | 12 | 0.5833 | 3 | 1 | 2 | 0.5 | 2 | 0.5 | 8 | 0.75 | 15 | 0.7333 |
| 12_113153 | 4 | 1 | NA | NA | 5 | 1 | 4 | 1 | 13 | 1 | 4 | 1 | 2 | 1 | 2 | 1 | 7 | 0.7143 | 15 | 0.8667 |
| 12_113222 | 1 | 0 | 32 | 0.0625 | 2 | 0.5 | NA | NA | 35 | 0.0857 | 7 | 0.1429 | NA | NA | NA | NA | 2 | 0 | 9 | 0.1111 |
| 12_113250 | NA | NA | 35 | 0.2571 | 2 | 0.5 | 1 | 0 | 38 | 0.2632 | 20 | 0.2 | NA | NA | NA | NA | 2 | 0.5 | 22 | 0.2273 |
| 12_113251 | NA | NA | 31 | 0.6774 | 1 | 1 | 1 | 0 | 33 | 0.6667 | 14 | 0.5 | NA | NA | NA | NA | 2 | 0.5 | 16 | 0.5 |
| 12_113251 | NA | NA | 29 | 0.4138 | 1 | 0 | 1 | 0 | 31 | 0.3871 | 14 | 0.5714 | NA | NA | NA | NA | 2 | 0 | 16 | 0.5 |
| 12_113269 | 2 | 0.5 | 86 | 0.314 | 14 | 0.2857 | NA | NA | 102 | 0.3137 | 4 | 0 | NA | NA | 5 | 0.6 | 36 | 0.3611 | 45 | 0.3556 |
| 12_113270 | 2 | 0.5 | 73 | 0.4658 | 12 | 0.1667 | NA | NA | 87 | 0.4253 | 4 | 0.5 | NA | NA | 2 | 1 | 29 | 0.4483 | 35 | 0.4857 |
| 12_113271 | 1 | 0 | 70 | 0.0857 | 11 | 0 | NA | NA | 82 | 0.0732 | 4 | 0 | NA | NA | 5 | 0 | 31 | 0.1935 | 40 | 0.15 |
| 12_113323 | 1 | 1 | 3 | 0 | 1 | 0 | NA | NA | 5 | 0.2 | 1 | 1 | NA | NA | NA | NA | NA | NA | 1 | 1 |
| 12_113382 | 1 | 1 | 2 | 0 | 1 | 0 | NA | NA | 4 | 0.25 | 3 | 0.3333 | NA | NA | 2 | 0 | 1 | 1 | 6 | 0.3333 |
| 12_113384 | 3 | 0 | 11 | 0.0909 | 4 | 0 | 2 | 0.5 | 20 | 0.1 | 7 | 0 | NA | NA | 2 | 0 | 10 | 0.2 | 19 | 0.1053 |
| 12_113384 | 3 | 0.3333 | 13 | 0 | 6 | 0.1667 | 2 | 0 | 24 | 0.0833 | 11 | 0.0909 | NA | NA | 4 | 0.25 | 9 | 0.2222 | 24 | 0.1667 |
| 12_113384 | 4 | 0 | 13 | 0 | 6 | 0.3333 | 2 | 0 | 25 | 0.08 | 12 | 0.0833 | NA | NA | 4 | 0 | 10 | 0.1 | 26 | 0.0769 |
| 12_113386 | 2 | 0 | 13 | 0.1538 | 5 | 0.2 | 1 | 0 | 21 | 0.1429 | 7 | 0.2857 | NA | NA | 3 | 0.3333 | 9 | 0 | 19 | 0.1579 |
| 12_113392 | NA | NA | 8 | 0.125 | 4 | 0.5 | NA | NA | 12 | 0.25 | 3 | 0.3333 | NA | NA | NA | NA | 31 | 0.0645 | 34 | 0.0882 |
| 12_113393 | 1 | 0 | 1 | 1 | NA | NA | 1 | 1 | 3 | 0.6667 | 2 | 1 | NA | NA | NA | NA | 1 | 1 | 3 | 1 |
| 12_116022 | 1 | 0 | 1 | 1 | 2 | 0.5 | 3 | 1 | 7 | 0.7143 | 1 | 1 | NA | NA | 1 | 1 | 1 | 1 | 3 | 1 |
| 12_116022 | NA | NA | 1 | 0 | 3 | 0 | 3 | 0.3333 | 7 | 0.1429 | 1 | 0 | NA | NA | 1 | 1 | 1 | 0 | 3 | 0.3333 |
| 12_116022 | 1 | 0 | 1 | 1 | 3 | 0.3333 | 2 | 1 | 7 | 0.5714 | 1 | 0 | NA | NA | 1 | 1 | 1 | 1 | 3 | 0.6667 |
| 12_116023 | 1 | 0 | 1 | 1 | 3 | 0.6667 | 3 | 0.6667 | 8 | 0.625 | 1 | 1 | NA | NA | 1 | 0 | 1 | 1 | 3 | 0.6667 |
| 12_116024 | 2 | 1 | 4 | 1 | 4 | 1 | 3 | 1 | 13 | 1 | 1 | 1 | 2 | 1 | NA | NA | 1 | 1 | 4 | 1 |
| 12_116170 | NA | NA | 1 | 1 | 1 | 0 | 1 | 0 | 3 | 0.3333 | NA | NA | 1 | 0 | 1 | 1 | NA | NA | 2 | 0.5 |
| 12_116333 | 1 | 0 | 7 | 0 | NA | NA | 3 | 0.3333 | 11 | 0.0909 | 2 | 0 | 3 | 0 | 4 | 0.5 | 5 | 0.2 | 14 | 0.2143 |
| 12_116333 | 2 | 0 | 6 | 0 | NA | NA | 3 | 0.3333 | 11 | 0.0909 | 2 | 0 | 2 | 0 | 3 | 0 | 3 | 0.6667 | 10 | 0.2 |
| 12_116333 | 2 | 0 | 5 | 0.2 | 1 | 0 | 3 | 0.3333 | 11 | 0.1818 | 2 | 0 | 2 | 0 | 3 | 0.6667 | 5 | 0.4 | 12 | 0.3333 |
| 12_116334 | 2 | 1 | 5 | 0.6 | 1 | 1 | 4 | 0.5 | 12 | 0.6667 | 2 | 1 | 4 | 1 | 2 | 0.5 | 4 | 1 | 12 | 0.9167 |
| 12_116334 | 2 | 0 | 6 | 0.1667 | 1 | 0 | 3 | 0 | 12 | 0.0833 | 2 | 0 | 4 | 0.25 | 2 | 0 | 5 | 0.8 | 13 | 0.3846 |
| 12_116334 | 2 | 0 | 4 | 0 | 2 | 0.5 | 3 | 0 | 11 | 0.0909 | 2 | 0 | 3 | 0.3333 | 4 | 0 | 4 | 0.5 | 13 | 0.2308 |
| 12_116339 | 1 | 0 | 2 | 0.5 | 2 | 0 | 3 | 0 | 8 | 0.125 | 3 | 0 | 2 | 0 | 3 | 0.3333 | 2 | 0 | 10 | 0.1 |
| 12_116343 | 5 | 0.4 | 3 | 0.3333 | 4 | 0 | 10 | 0 | 22 | 0.1364 | 5 | 0.2 | 3 | 0 | 9 | 0.2222 | 7 | 0.2857 | 24 | 0.2083 |
| 12_116343 | 8 | 0.625 | 3 | 0.3333 | 4 | 0.5 | 10 | 0.4 | 25 | 0.48 | 3 | 0.3333 | 2 | 0 | 8 | 0.5 | 7 | 0.7143 | 20 | 0.5 |
| 12_116344 | 6 | 0.6667 | 1 | 1 | 2 | 1 | 12 | 1 | 21 | 0.9048 | 2 | 1 | 1 | 1 | 8 | 1 | 7 | 0.8571 | 18 | 0.9444 |
| 12_116344 | 8 | 0.25 | 3 | 0 | 4 | 0.25 | 12 | 0.0833 | 27 | 0.1481 | 4 | 0 | 2 | 1 | 9 | 0 | 7 | 0.5714 | 22 | 0.2727 |
| 12_116344 | 7 | 0.2857 | 3 | 0 | 4 | 0.25 | 7 | 0.2857 | 21 | 0.2381 | 3 | 0.3333 | 3 | 0.3333 | 5 | 0 | 7 | 0.1429 | 18 | 0.1667 |
| 12_116344 | 8 | 0.625 | 3 | 0.3333 | 4 | 0.25 | 10 | 0.5 | 25 | 0.48 | 4 | 1 | 2 | 1 | 7 | 0.7143 | 8 | 0.625 | 21 | 0.7619 |
| 12_116344 | 6 | 0.3333 | 5 | 0.4 | 3 | 0.3333 | 11 | 0.7273 | 25 | 0.52 | 4 | 0.5 | 3 | 0.6667 | 8 | 0.625 | 7 | 0.7143 | 22 | 0.6364 |
| 12_116344 | 8 | 0.875 | 7 | 0.8571 | 4 | 1 | 10 | 0.9 | 29 | 0.8966 | 5 | 0.6 | 3 | 1 | 8 | 1 | 8 | 0.875 | 24 | 0.875 |
| 12_116344 | 8 | 0.75 | 5 | 0.4 | 4 | 0.5 | 11 | 0.6364 | 28 | 0.6071 | 4 | 0.5 | 3 | 0.6667 | 7 | 0.7143 | 9 | 0.5556 | 23 | 0.6087 |
| 12_116353 | NA | NA | 3 | 0.6667 | 1 | 0 | NA | NA | 4 | 0.5 | 1 | 1 | NA | NA | NA | NA | NA | NA | 1 | 1 |
| 12_116353 | NA | NA | 3 | 1 | 1 | 1 | NA | NA | 4 | 1 | 1 | 1 | NA | NA | NA | NA | NA | NA | 1 | 1 |
| 12_116353 | NA | NA | 3 | 1 | 1 | 0 | NA | NA | 4 | 0.75 | 1 | 0 | NA | NA | 1 | 1 | NA | NA | 2 | 0.5 |
| 12_116354 | 1 | 1 | 3 | 0.3333 | 1 | 0 | NA | NA | 5 | 0.4 | 1 | 1 | 1 | 0 | 1 | 0 | 1 | 0 | 4 | 0.25 |
| 12_116354 | NA | NA | 2 | 1 | 1 | 0 | NA | NA | 3 | 0.6667 | 1 | 1 | 1 | 0 | 1 | 0 | 1 | 1 | 4 | 0.5 |
| 12_116354 | 1 | 0 | 3 | 0.3333 | 1 | 0 | NA | NA | 5 | 0.2 | 2 | 1 | 1 | 0 | 1 | 0 | 1 | 0 | 5 | 0.4 |
| 12_116362 | 1 | 1 | NA | NA | NA | NA | NA | NA | 1 | 1 | 1 | 0 | 1 | 1 | 1 | 1 | NA | NA | 3 | 0.6667 |
| 12_116363 | 1 | 1 | NA | NA | NA | NA | NA | NA | 1 | 1 | 1 | 0 | 1 | 1 | 1 | 1 | NA | NA | 3 | 0.6667 |
| 12_117652 | 2 | 1 | NA | NA | 1 | 0 | 3 | 0.6667 | 6 | 0.6667 | 2 | 0 | 1 | 0 | 3 | 0 | 2 | 0.5 | 8 | 0.125 |
| 12_117677 | 8 | 0.375 | 4 | 0.25 | 1 | 0 | 4 | 0 | 17 | 0.2353 | 4 | 0.25 | 2 | 1 | 6 | 0.3333 | 3 | 0.3333 | 15 | 0.4 |
| 12_117678 | 1 | 1 | 1 | 1 | NA | NA | NA | NA | 2 | 1 | 1 | 1 | NA | NA | NA | NA | NA | NA | 1 | 1 |
| 12_117719 | 10 | 0.7 | 5 | 1 | 2 | 0 | 4 | 0.5 | 21 | 0.6667 | 6 | 0.3333 | 5 | 0.6 | 6 | 0.6667 | 5 | 0.8 | 22 | 0.5909 |
| 12_117720 | 6 | 0 | 3 | 0 | 3 | 0 | 5 | 0.2 | 17 | 0.0588 | 3 | 0.3333 | 5 | 0 | 3 | 0.3333 | 5 | 0 | 16 | 0.125 |
| 12_117720 | 10 | 0.9 | 5 | 1 | 2 | 1 | 6 | 0.8333 | 23 | 0.913 | 5 | 1 | 5 | 1 | 4 | 0.75 | 6 | 1 | 20 | 0.95 |
| 12_117720 | 12 | 0.9167 | 6 | 0.8333 | 3 | 1 | 6 | 1 | 27 | 0.9259 | 7 | 0.8571 | 5 | 1 | 6 | 1 | 6 | 0.6667 | 24 | 0.875 |
| 12_117720 | 7 | 0.8571 | 4 | 1 | 3 | 0.6667 | 2 | 0 | 16 | 0.75 | 5 | 1 | 5 | 1 | 4 | 1 | 4 | 0.75 | 18 | 0.9444 |
| 12_117720 | 9 | 0.8889 | 6 | 1 | 2 | 0.5 | 6 | 0.8333 | 23 | 0.8696 | 5 | 1 | 5 | 1 | 6 | 1 | 6 | 0.8333 | 22 | 0.9545 |
| 12_117720 | 11 | 0.2727 | 4 | 0 | 3 | 0 | 4 | 0.5 | 22 | 0.2273 | 5 | 0.4 | 5 | 0 | 5 | 0 | 5 | 0.4 | 20 | 0.2 |

| | | | | | | | | | | | | | | | | | | | | |
|---|---|---|---|---|---|---|---|---|---|---|---|---|---|---|---|---|---|---|---|---|
| 12_1177204 | 11 | 0.3636 | 6 | 0.3333 | 3 | 0 | 6 | 0 | 26 | 0.2308 | 6 | 0.1667 | 4 | 0 | 6 | 0 | 6 | 0.1667 | 22 | 0.0909 |
| 12_1177204 | 12 | 0.9167 | 6 | 0.6667 | 3 | 1 | 6 | 1 | 27 | 0.8889 | 5 | 0.6 | 5 | 0.8 | 6 | 1 | 5 | 0.6 | 21 | 0.7619 |
| 12_1177205 | 10 | 0.2 | 6 | 0 | 3 | 0.6667 | 7 | 0.1429 | 26 | 0.1923 | 6 | 0.3333 | 5 | 0 | 7 | 0 | 6 | 0.3333 | 24 | 0.1667 |
| 12_1177209 | 11 | 0.5455 | 5 | 0.6 | 4 | 0.75 | 8 | 0.625 | 28 | 0.6071 | 6 | 1 | 5 | 0.6 | 7 | 0.8571 | 7 | 0.8571 | 25 | 0.84 |
| 12_1177215 | 11 | 0.1818 | 8 | 0.25 | 4 | 0.5 | 10 | 0 | 33 | 0.1818 | 6 | 0.1667 | 5 | 0.2 | 9 | 0 | 6 | 0 | 26 | 0.0769 |
| 12_1177227 | 18 | 0.0556 | 13 | 0.0769 | 7 | 0.1429 | 17 | 0.1176 | 55 | 0.0909 | 10 | 0.2 | 10 | 0.1 | 13 | 0.0769 | 10 | 0 | 43 | 0.093 |
| 12_1177230 | 18 | 0.2222 | 12 | 0.1667 | 5 | 0.2 | 12 | 0.25 | 47 | 0.2128 | 7 | 0.2857 | 10 | 0.1 | 14 | 0.3571 | 6 | 0.5 | 37 | 0.2973 |
| 12_1177231 | 19 | 0.6842 | 12 | 0.3333 | 4 | 0.75 | 15 | 0.4667 | 50 | 0.54 | 10 | 0.7 | 9 | 0.6667 | 14 | 0.8571 | 7 | 0.5714 | 40 | 0.725 |
| 12_1177232 | 21 | 0.0476 | 12 | 0 | 7 | 0.2857 | 16 | 0.125 | 56 | 0.0893 | 13 | 0.2308 | 10 | 0.1 | 13 | 0.0769 | 7 | 0.1429 | 43 | 0.1395 |
| 12_1177232 | 21 | 0.1429 | 13 | 0.0769 | 7 | 0.1429 | 15 | 0.0667 | 56 | 0.1071 | 10 | 0.3 | 10 | 0.1 | 13 | 0.0769 | 6 | 0 | 39 | 0.1282 |
| 12_1181963 | 2 | 0 | 1 | 1 | 2 | 0 | NA | NA | 5 | 0.2 | 2 | 0 | 1 | 1 | 2 | 0 | 1 | 0 | 6 | 0.1667 |
| 12_1181963 | 1 | 1 | 1 | 0 | 2 | 0.5 | NA | NA | 4 | 0.5 | 2 | 0 | 1 | 0 | 2 | 0 | 1 | 1 | 6 | 0.1667 |
| 12_1181964 | 1 | 1 | 1 | 0 | 2 | 0.5 | NA | NA | 4 | 0.5 | NA | NA | 1 | 0 | 1 | 1 | 1 | 1 | 3 | 0.6667 |
| 12_1181964 | 2 | 1 | 1 | 1 | 2 | 1 | NA | NA | 5 | 1 | NA | NA | 1 | 1 | 2 | 1 | 1 | 1 | 4 | 1 |
| 12_1181965 | 2 | 0.5 | 1 | 1 | 2 | 1 | NA | NA | 5 | 0.8 | 1 | 1 | NA | NA | 2 | 0.5 | 1 | 1 | 4 | 0.75 |
| 12_1181966 | 2 | 1 | 1 | 0 | 2 | 0 | NA | NA | 5 | 0.4 | 1 | 1 | 1 | 1 | 2 | 0.5 | NA | NA | 4 | 0.75 |
| 12_1181971 | 1 | 1 | 1 | 1 | 1 | 0 | NA | NA | 3 | 0.6667 | NA | NA | NA | NA | 2 | 0.5 | NA | NA | 2 | 0.5 |
| 12_1181996 | 3 | 0 | 3 | 0 | 4 | 0.25 | 2 | 0.5 | 12 | 0.1667 | 2 | 0 | 1 | 0 | NA | NA | 4 | 0.5 | 7 | 0.2857 |
| 12_1181998 | 2 | 0.5 | 3 | 0.3333 | 5 | 0.4 | 3 | 0 | 13 | 0.3077 | 2 | 0 | 1 | 0 | NA | NA | 3 | 0.3333 | 6 | 0.1667 |
| 12_1182015 | 1 | 1 | 3 | 0.3333 | 3 | 0.6667 | 3 | 1 | 10 | 0.7 | 1 | 1 | 1 | 1 | NA | NA | 3 | 1 | 5 | 1 |
| 12_1182016 | 1 | 0 | 1 | 0 | 4 | 0.25 | 3 | 0 | 9 | 0.1111 | 2 | 0 | 1 | 0 | NA | NA | 3 | 0.3333 | 6 | 0.1667 |
| 12_1182018 | 1 | 1 | 2 | 1 | 3 | 0.6667 | 2 | 1 | 8 | 0.875 | 1 | 0 | NA | NA | NA | NA | 3 | 1 | 4 | 0.75 |
| 12_1182019 | 1 | 1 | 2 | 0.5 | 1 | 1 | 1 | 1 | 5 | 0.8 | 1 | 1 | 1 | 1 | NA | NA | 2 | 1 | 4 | 1 |
| 12_1182019 | 1 | 1 | 2 | 0.5 | 2 | 1 | 1 | 1 | 6 | 0.8333 | 1 | 0 | NA | NA | NA | NA | 1 | 1 | 2 | 0.5 |
| 12_1182020 | NA | NA | 2 | 0.5 | 1 | 1 | 2 | 1 | 5 | 0.8 | 1 | 1 | 1 | 1 | NA | NA | 3 | 1 | 5 | 1 |
| 12_1182624 | NA | NA | 1 | 1 | 1 | 1 | NA | NA | 2 | 1 | 1 | 1 | NA | NA | NA | NA | 1 | 1 | 2 | 1 |
| 12_1182626 | NA | NA | 1 | 0 | NA | NA | 2 | 0.5 | 3 | 0.3333 | NA | NA | NA | NA | NA | NA | 2 | 0.5 | 2 | 0.5 |
| 12_1182631 | NA | NA | NA | NA | NA | NA | 2 | 1 | 2 | 1 | NA | NA | NA | NA | NA | NA | 2 | 0.5 | 2 | 0.5 |
| 12_1182632 | NA | NA | NA | NA | NA | NA | 2 | 0.5 | 2 | 0.5 | NA | NA | NA | NA | NA | NA | 2 | 1 | 2 | 1 |
| 12_1182632 | NA | NA | NA | NA | NA | NA | 2 | 0.5 | 2 | 0.5 | NA | NA | NA | NA | NA | NA | 2 | 0.5 | 2 | 0.5 |
| 12_1182633 | NA | NA | NA | NA | NA | NA | 2 | 1 | 2 | 1 | NA | NA | NA | NA | NA | NA | 1 | 1 | 1 | 1 |
| 12_1188126 | 1 | 1 | 1 | 1 | 1 | 1 | NA | NA | 3 | 1 | NA | NA | NA | NA | NA | NA | 1 | 1 | 1 | 1 |
| 12_1188129 | NA | NA | 1 | 1 | 1 | 1 | NA | NA | 2 | 1 | NA | NA | NA | NA | NA | NA | 1 | 1 | 1 | 1 |
| 12_1188130 | NA | NA | 2 | 1 | 2 | 1 | NA | NA | 4 | 1 | NA | NA | NA | NA | NA | NA | 1 | 1 | 1 | 1 |
| 12_1188137 | 1 | 1 | 2 | 1 | 2 | 1 | NA | NA | 5 | 1 | NA | NA | NA | NA | NA | NA | 1 | 1 | 1 | 1 |
| 12_1188138 | 1 | 1 | 2 | 1 | 2 | 1 | NA | NA | 5 | 1 | NA | NA | NA | NA | NA | NA | 1 | 1 | 1 | 1 |
| 12_1188139 | 1 | 0 | 1 | 1 | 1 | 1 | NA | NA | 3 | 0.6667 | NA | NA | NA | NA | NA | NA | 1 | 1 | 1 | 1 |
| 12_1188139 | 1 | 1 | 2 | 1 | 2 | 1 | NA | NA | 5 | 1 | NA | NA | NA | NA | NA | NA | 1 | 1 | 1 | 1 |
| 12_1188140 | 1 | 1 | 2 | 1 | 2 | 1 | NA | NA | 5 | 1 | NA | NA | NA | NA | NA | NA | 1 | 1 | 1 | 1 |
| 12_1191224 | 3 | 0 | 8 | 0 | 10 | 0.1 | 10 | 0.1 | 31 | 0.0645 | 5 | 0 | 4 | 0 | 8 | 0.125 | 6 | 0.1667 | 23 | 0.087 |
| 12_1191250 | 2 | 0 | 6 | 0 | 7 | 0.2857 | 8 | 0 | 23 | 0.087 | 2 | 0 | 1 | 0 | 4 | 0.25 | 4 | 0.25 | 11 | 0.1818 |
| 12_1191250 | 2 | 0 | 7 | 0.1429 | 6 | 0 | 7 | 0.1429 | 22 | 0.0909 | 2 | 0 | NA | NA | 4 | 0 | 4 | 0.25 | 10 | 0.1 |
| 12_1191255 | 1 | 1 | 4 | 0.5 | 5 | 0.2 | 5 | 0 | 15 | 0.2667 | 1 | 1 | NA | NA | 4 | 0.25 | 3 | 0 | 8 | 0.25 |
| 12_1191261 | 1 | 0 | 6 | 0 | 5 | 0.2 | 5 | 0.2 | 17 | 0.1176 | 2 | 0.5 | 1 | 1 | 4 | 0.25 | 3 | 0.3333 | 10 | 0.4 |
| 12_1191263 | NA | NA | 6 | 0.8333 | 5 | 0.4 | 4 | 0.25 | 15 | 0.5333 | NA | NA | 1 | 0 | 4 | 0.25 | 2 | 1 | 7 | 0.4286 |
| 12_1191264 | NA | NA | 6 | 0.1667 | 5 | 0.2 | 4 | 1 | 15 | 0.4 | 1 | 0 | 1 | 0 | 4 | 0.5 | 2 | 0 | 8 | 0.25 |
| 12_1191266 | NA | NA | 6 | 0.5 | 3 | 1 | 3 | 0.6667 | 12 | 0.6667 | NA | NA | 1 | 1 | 4 | 0.75 | 1 | 0 | 6 | 0.6667 |
| 12_1191267 | NA | NA | 3 | 0 | 4 | 0.5 | 2 | 0.5 | 9 | 0.3333 | NA | NA | 1 | 0 | 4 | 0.25 | 2 | 1 | 7 | 0.4286 |
| 12_1191267 | NA | NA | 6 | 0.1667 | 3 | 0 | 3 | 0 | 12 | 0.0833 | 1 | 0 | 1 | 1 | 4 | 0 | 2 | 0 | 8 | 0.125 |
| 12_1191268 | NA | NA | 3 | 1 | 2 | 1 | 3 | 1 | 8 | 1 | NA | NA | 1 | 1 | 2 | 1 | NA | NA | 3 | 1 |
| 12_1191268 | NA | NA | 5 | 0.2 | 5 | 0.2 | 3 | 0 | 13 | 0.1538 | 1 | 0 | 1 | 1 | 4 | 0.25 | 2 | 0 | 8 | 0.25 |
| 12_1191346 | NA | NA | 4 | 0.5 | 3 | 0.3333 | 3 | 0.3333 | 10 | 0.4 | NA | NA | 1 | 0 | 4 | 0.25 | 1 | 1 | 6 | 0.3333 |
| 12_1191346 | NA | NA | 4 | 0 | 3 | 0.3333 | 2 | 0 | 9 | 0.1111 | 1 | 1 | 1 | 0 | 4 | 0.25 | 1 | 0 | 7 | 0.2857 |
| 12_1192015 | NA | NA | 4 | 0 | 2 | 0 | 1 | 1 | 7 | 0.1429 | NA | NA | 1 | 0 | 3 | 0.3333 | 1 | 0 | 5 | 0.2 |
| 12_1192016 | NA | NA | 2 | 0 | 1 | 0 | 1 | 1 | 4 | 0.25 | NA | NA | 2 | 0.5 | 3 | 1 | 1 | 1 | 6 | 0.8333 |
| 12_1192016 | NA | NA | 3 | 0 | NA | NA | 1 | 1 | 4 | 0.25 | NA | NA | 2 | 0.5 | 2 | 0 | 1 | 0 | 5 | 0.2 |
| 12_1192017 | NA | NA | 1 | 0 | 1 | 0 | 1 | 1 | 3 | 0.3333 | NA | NA | 1 | 1 | NA | NA | 1 | 1 | 2 | 1 |
| 12_1192018 | NA | NA | 1 | 1 | NA | NA | 1 | 1 | 2 | 1 | NA | NA | 1 | 0 | 1 | 1 | 1 | 0 | 3 | 0.3333 |
| 12_1199096 | NA | NA | NA | NA | 2 | 0.5 | 1 | 1 | 3 | 0.6667 | NA | NA | 1 | 1 | NA | NA | NA | NA | 1 | 1 |
| 12_1199098 | NA | NA | 1 | 0 | 3 | 0 | 2 | 0.5 | 6 | 0.1667 | NA | NA | 1 | 0 | NA | NA | 1 | 1 | 2 | 0.5 |
| 12_1199099 | NA | NA | 1 | 1 | 3 | 0.6667 | 1 | 1 | 5 | 0.8 | NA | NA | 2 | 1 | NA | NA | 1 | 0 | 3 | 0.6667 |
| 12_1199101 | NA | NA | 1 | 1 | 3 | 1 | 3 | 1 | 7 | 1 | NA | NA | 1 | 1 | 1 | 1 | 2 | 1 | 4 | 1 |
| 12_1199101 | NA | NA | 1 | 0 | 3 | 1 | 2 | 1 | 6 | 0.8333 | NA | NA | 2 | 1 | 1 | 0 | 2 | 1 | 5 | 0.8 |
| 12_1199103 | NA | NA | 1 | 1 | 2 | 0 | 2 | 0 | 5 | 0.2 | NA | NA | 2 | 0.5 | 1 | 0 | 2 | 0 | 5 | 0.2 |
| 12_1199823 | NA | NA | 1 | 1 | 3 | 1 | 2 | 1 | 6 | 1 | NA | NA | 3 | 0.6667 | 1 | 1 | 2 | 0.5 | 6 | 0.6667 |

| | | | | | | | | | | | | | | | | | | | | |
|---|---|---|---|---|---|---|---|---|---|---|---|---|---|---|---|---|---|---|---|---|
| 12_119982 | NA | NA | 1 | 1 | 2 | 1 | 2 | 0.5 | 5 | 0.8 | NA | NA | 2 | 1 | 1 | 1 | 2 | 1 | 5 | 1 |
| 12_119983 | NA | NA | 2 | 0 | 2 | 0.5 | 3 | 0.3333 | 7 | 0.2857 | 1 | 1 | 3 | 0.3333 | 2 | 0 | 3 | 0.3333 | 9 | 0.3333 |
| 13_354221 | 1 | 1 | NA | NA | NA | NA | NA | NA | 1 | 1 | NA | NA | 1 | 1 | NA | NA | NA | NA | 1 | 1 |
| 13_354237 | 1 | 1 | NA | NA | NA | NA | 1 | 1 | 2 | 1 | NA | NA | 1 | 1 | NA | NA | NA | NA | 1 | 1 |
| 13_354239 | 1 | 1 | NA | NA | NA | NA | 1 | 1 | 2 | 1 | NA | NA | 1 | 1 | NA | NA | NA | NA | 1 | 1 |
| 13_360632 | 25 | 0.12 | 37 | 0.0541 | 34 | 0 | 27 | 0.0741 | 123 | 0.0569 | 11 | 0 | 29 | 0.069 | 22 | 0.0909 | 33 | 0.0303 | 95 | 0.0526 |
| 13_361517 | 38 | 0.1053 | 46 | 0.1087 | 43 | 0.1163 | 36 | 0.0556 | 163 | 0.0982 | 20 | 0.15 | 33 | 0.1515 | 31 | 0.0645 | 33 | 0.1818 | 117 | 0.1368 |
| 13_361548 | 39 | 0.1538 | 53 | 0.2264 | 50 | 0.18 | 40 | 0.15 | 182 | 0.1813 | 20 | 0.15 | 35 | 0.1429 | 29 | 0.1379 | 40 | 0.2 | 124 | 0.1613 |
| 13_361585 | 3 | 0.3333 | 4 | 0.25 | 3 | 0 | 5 | 0 | 15 | 0.1333 | 1 | 1 | 1 | 0 | 2 | 0 | NA | NA | 4 | 0.25 |
| 13_361636 | 2 | 0 | 3 | 0 | 3 | 0.3333 | 6 | 0.1667 | 14 | 0.1429 | 2 | 0 | 1 | 1 | 2 | 0 | 5 | 0 | 10 | 0.1 |
| 13_361730 | 2 | 0 | 1 | 0 | 1 | 0 | 1 | 1 | 5 | 0.2 | 1 | 1 | 2 | 0 | 2 | 0 | 1 | 0 | 6 | 0.1667 |
| 13_362360 | 2 | 0.5 | 3 | 0 | 1 | 0 | 7 | 0.5714 | 13 | 0.3846 | NA | NA | 1 | 0 | 1 | 1 | 3 | 0.6667 | 5 | 0.6 |
| 13_362380 | 1 | 1 | 3 | 1 | NA | NA | 5 | 0.8 | 9 | 0.8889 | NA | NA | 1 | 0 | 1 | 1 | 3 | 1 | 5 | 0.8 |
| 13_362382 | 2 | 1 | 3 | 0.6667 | 1 | 1 | 4 | 1 | 10 | 0.9 | NA | NA | 1 | 0 | 1 | 0 | 2 | 0.5 | 4 | 0.25 |
| 13_362419 | NA | NA | 3 | 0.6667 | 1 | 1 | 3 | 0.6667 | 7 | 0.7143 | NA | NA | 1 | 1 | 1 | 0 | 3 | 0.6667 | 5 | 0.6 |
| 13_362430 | 1 | 0 | 3 | 0.6667 | 1 | 1 | 4 | 1 | 9 | 0.7778 | NA | NA | 1 | 0 | 1 | 0 | 3 | 0.3333 | 5 | 0.2 |
| 13_362447 | 2 | 0 | 3 | 0.3333 | 1 | 0 | 3 | 0.3333 | 9 | 0.2222 | NA | NA | 1 | 0 | 1 | 0 | 3 | 0.3333 | 5 | 0.2 |
| 13_362459 | 2 | 0 | 3 | 0.3333 | 1 | 0 | 3 | 0.3333 | 9 | 0.2222 | NA | NA | NA | NA | 1 | 1 | 2 | 0 | 3 | 0.3333 |
| 13_362469 | 1 | 1 | 2 | 0.5 | NA | NA | 3 | 0.3333 | 6 | 0.5 | NA | NA | 1 | 1 | 1 | 1 | 2 | 0 | 4 | 0.5 |
| 13_362477 | 2 | 1 | 3 | 0.3333 | 1 | 1 | 3 | 0.3333 | 9 | 0.5556 | NA | NA | 1 | 1 | 1 | 1 | 2 | 0 | 4 | 0.5 |
| 13_362485 | 2 | 0 | 3 | 0 | 1 | 1 | 2 | 0.5 | 8 | 0.25 | NA | NA | NA | NA | 1 | 0 | 3 | 0.3333 | 4 | 0.25 |
| 13_362491 | 2 | 0 | 3 | 0.3333 | 1 | 1 | 3 | 0.3333 | 9 | 0.3333 | NA | NA | NA | NA | NA | NA | 2 | 1 | 2 | 1 |
| 13_362529 | 2 | 0 | 3 | 0 | 1 | 1 | 3 | 0 | 9 | 0.1111 | NA | NA | NA | NA | 1 | 1 | 3 | 0 | 4 | 0.25 |
| 13_362640 | 1 | 0 | 3 | 0.6667 | 1 | 0 | NA | NA | 5 | 0.4 | NA | NA | NA | NA | 1 | 1 | 2 | 0 | 3 | 0.3333 |
| 13_363185 | NA | NA | 3 | 0.6667 | 1 | 1 | 1 | 0 | 5 | 0.6 | NA | NA | NA | NA | 1 | 0 | 1 | 1 | 2 | 0.5 |
| 13_363189 | NA | NA | 3 | 0.6667 | 1 | 1 | 1 | 1 | 5 | 0.8 | NA | NA | NA | NA | 1 | 1 | 1 | 0 | 2 | 0.5 |
| 13_363194 | NA | NA | 3 | 0 | NA | NA | 1 | 1 | 4 | 0.25 | NA | NA | NA | NA | 1 | 0 | 1 | 1 | 2 | 0.5 |
| 13_363475 | NA | NA | 2 | 0.5 | 1 | 0 | 1 | 0 | 4 | 0.25 | NA | NA | NA | NA | 1 | 1 | 1 | 0 | 2 | 0.5 |
| 13_363488 | NA | NA | 2 | 0.5 | 1 | 1 | 1 | 1 | 4 | 0.75 | NA | NA | NA | NA | 1 | 1 | NA | NA | 1 | 1 |
| 13_363512 | NA | NA | 3 | 0.3333 | 1 | 1 | 1 | 0 | 5 | 0.4 | NA | NA | NA | NA | NA | NA | 1 | 1 | 1 | 1 |
| 13_370024 | 8 | 0.375 | 6 | 0.5 | 3 | 0 | 7 | 0 | 24 | 0.25 | 5 | 0 | 3 | 0.6667 | 2 | 0.5 | 5 | 0.6 | 15 | 0.4 |
| 13_370040 | 8 | 0.25 | 8 | 0.125 | 4 | 0 | 7 | 0 | 27 | 0.1111 | 5 | 0.2 | 3 | 0 | 4 | 0.5 | 5 | 0.2 | 17 | 0.2353 |
| 13_370041 | 6 | 0.1667 | 3 | 0.3333 | 4 | 0.25 | 5 | 0 | 18 | 0.1667 | 3 | 0.3333 | 2 | 0 | 1 | 0 | 4 | 0 | 10 | 0.1 |
| 13_370058 | 9 | 0.1111 | 7 | 0.2857 | 4 | 0.25 | 7 | 0 | 27 | 0.1481 | 4 | 0 | 3 | 0.3333 | 3 | 0.3333 | 5 | 0 | 15 | 0.1333 |
| 13_370063 | 8 | 0.25 | 6 | 0.3333 | 4 | 0.25 | 5 | 0.6 | 23 | 0.3478 | 6 | 0.8333 | 2 | 1 | 4 | 0.25 | 5 | 0.2 | 17 | 0.5294 |
| 13_370074 | 6 | 0.5 | 7 | 0.7143 | 3 | 1 | 4 | 0.75 | 20 | 0.7 | 5 | 0.6 | 3 | 1 | 5 | 0.8 | 4 | 0.5 | 17 | 0.7059 |
| 13_370074 | 8 | 0.125 | 6 | 0.5 | 2 | 0.5 | 5 | 0.4 | 21 | 0.3333 | 5 | 0.6 | 3 | 0 | 3 | 0.3333 | 5 | 0.6 | 16 | 0.4375 |
| 13_370092 | 9 | 0.1111 | 8 | 0.25 | 5 | 0 | 7 | 0 | 29 | 0.1034 | 6 | 0.1667 | 2 | 0 | 4 | 0 | 5 | 0 | 17 | 0.0588 |
| 13_370361 | 1 | 1 | 1 | 1 | NA | NA | 1 | 1 | 3 | 1 | NA | NA | 1 | 1 | NA | NA | NA | NA | 1 | 1 |
| 13_393288 | 7 | 0.1429 | 6 | 0 | 1 | 0 | 6 | 0 | 20 | 0.05 | 1 | 0 | 4 | 0.5 | 6 | 0.5 | 3 | 0.3333 | 14 | 0.4286 |
| 13_393408 | 7 | 1 | 3 | 1 | NA | NA | 4 | 1 | 14 | 1 | 2 | 1 | 2 | 1 | 2 | 1 | 4 | 1 | 10 | 1 |
| 13_393411 | 6 | 0.1667 | 3 | 0.3333 | NA | NA | 2 | 0.5 | 11 | 0.2727 | NA | NA | 2 | 0.5 | 2 | 0 | 3 | 0.3333 | 7 | 0.2857 |
| 13_393417 | 7 | 0.2857 | 4 | 0 | NA | NA | 4 | 0.25 | 15 | 0.2 | 1 | 0 | 3 | 0.6667 | 2 | 0 | 3 | 0 | 9 | 0.2222 |
| 13_394339 | 4 | 0.25 | 1 | 1 | NA | NA | 2 | 0 | 7 | 0.2857 | 1 | 1 | 2 | 0.5 | 1 | 0 | 3 | 0.3333 | 7 | 0.4286 |
| 13_591490 | 1 | 0 | 4 | 0.25 | NA | NA | NA | NA | 5 | 0.2 | 1 | 0 | 1 | 0 | NA | NA | 2 | 1 | 4 | 0.5 |
| 13_591493 | 2 | 0.5 | 4 | 1 | NA | NA | NA | NA | 6 | 0.8333 | 1 | 1 | 1 | 1 | NA | NA | 2 | 0.5 | 4 | 0.75 |
| 13_591496 | 1 | 0 | 4 | 0.25 | NA | NA | NA | NA | 5 | 0.2 | NA | NA | 1 | 0 | NA | NA | 2 | 0.5 | 3 | 0.3333 |
| 13_591498 | 1 | 1 | 2 | 1 | NA | NA | NA | NA | 3 | 1 | NA | NA | 1 | 1 | NA | NA | 1 | 1 | 2 | 1 |
| 13_591500 | 2 | 0.5 | 3 | 0.6667 | NA | NA | NA | NA | 5 | 0.6 | 1 | 1 | NA | NA | NA | NA | 1 | 1 | 2 | 1 |
| 13_591502 | 2 | 0.5 | 2 | 0.5 | NA | NA | NA | NA | 4 | 0.5 | 1 | 1 | NA | NA | NA | NA | 1 | 0 | 2 | 0.5 |
| 13_591520 | 1 | 0 | 3 | 1 | NA | NA | 1 | 0 | 5 | 0.6 | 1 | 0 | 1 | 0 | NA | NA | 2 | 0.5 | 4 | 0.25 |
| 13_591749 | 2 | 0.5 | 4 | 1 | NA | NA | 1 | 1 | 7 | 0.8571 | 1 | 1 | 1 | 1 | 1 | 1 | 2 | 1 | 5 | 1 |
| 13_591760 | 2 | 0 | 4 | 0.5 | 1 | 1 | 2 | 0 | 9 | 0.3333 | 2 | 0 | 1 | 0 | 1 | 1 | 2 | 0 | 6 | 0.1667 |
| 13_591851 | 3 | 0 | 4 | 0.25 | 3 | 0 | 2 | 0 | 12 | 0.0833 | 3 | 0.3333 | 1 | 0 | 2 | 0 | 2 | 0 | 8 | 0.125 |
| 13_591867 | 3 | 0.3333 | 4 | 0.25 | 3 | 0.3333 | 2 | 0 | 12 | 0.25 | 3 | 0.3333 | 1 | 1 | 2 | 0 | 2 | 0 | 8 | 0.25 |
| 13_591880 | 3 | 0 | 4 | 0.25 | 2 | 0 | 1 | 0 | 10 | 0.1 | 4 | 0 | 1 | 1 | 1 | 1 | 2 | 0 | 8 | 0.25 |
| 13_592008 | 2 | 0 | 6 | 0 | 5 | 0.2 | 2 | 0 | 15 | 0.0667 | 5 | 0 | 2 | 0 | 4 | 0.25 | 1 | 0 | 12 | 0.0833 |
| 13_592012 | 2 | 0 | 6 | 0.1667 | 5 | 0 | 3 | 0 | 16 | 0.0625 | 5 | 0.2 | 2 | 0 | 3 | 0 | 2 | 0 | 12 | 0.0833 |
| 13_592018 | 3 | 0 | 8 | 0.25 | 6 | 0 | 3 | 0.3333 | 20 | 0.15 | 5 | 0.2 | 2 | 0 | 3 | 0 | 2 | 0 | 12 | 0.0833 |
| 13_662993 | 5 | 0.4 | 4 | 0.5 | 10 | 0 | 3 | 0.3333 | 22 | 0.2273 | 3 | 0 | NA | NA | 11 | 0.3636 | 1 | 0 | 15 | 0.2667 |
| 13_663132 | 27 | 0.1481 | 28 | 0.0357 | 19 | 0.0526 | 22 | 0.0909 | 96 | 0.0833 | 24 | 0 | 18 | 0.0556 | 30 | 0.0667 | 21 | 0.1429 | 93 | 0.0645 |
| 13_663144 | 26 | 0.1923 | 30 | 0.3 | 17 | 0.1765 | 18 | 0.2222 | 91 | 0.2308 | 20 | 0.25 | 18 | 0.2222 | 28 | 0.2143 | 20 | 0.2 | 86 | 0.2209 |
| 13_862030 | NA | NA | NA | NA | 1 | 1 | NA | NA | 1 | 1 | NA | NA | NA | NA | 2 | 0.5 | NA | NA | 2 | 0.5 |
| 13_881755 | NA | NA | 1 | 1 | 2 | 0.5 | 2 | 1 | 5 | 0.8 | NA | NA | 1 | 1 | 2 | 1 | NA | NA | 3 | 1 |
| 13_881757 | NA | NA | 1 | 0 | 1 | 1 | 2 | 0.5 | 4 | 0.5 | NA | NA | 1 | 1 | NA | NA | NA | NA | 1 | 1 |

| | | | | | | | | | | | | | | | | | | | | |
|---|---|---|---|---|---|---|---|---|---|---|---|---|---|---|---|---|---|---|---|---|
| 13_881763 | NA | NA | 1 | 0 | 3 | 0.3333 | 2 | 0.5 | 6 | 0.3333 | NA | NA | 1 | 1 | 2 | 1 | NA | NA | 3 | 1 |
| 13_881774 | NA | NA | 1 | 0 | 3 | 0.6667 | 2 | 1 | 6 | 0.6667 | NA | NA | 1 | 1 | 2 | 1 | NA | NA | 3 | 1 |
| 13_881783 | NA | NA | 1 | 0 | 1 | 0 | 1 | 1 | 3 | 0.3333 | NA | NA | 1 | 1 | 1 | 1 | NA | NA | 2 | 1 |
| 13_881814 | NA | NA | 1 | 0 | 3 | 0.3333 | 2 | 0 | 6 | 0.1667 | 1 | 0 | 1 | 0 | 2 | 0.5 | NA | NA | 4 | 0.25 |
| 13_881822 | NA | NA | 1 | 1 | 3 | 0.3333 | 2 | 1 | 6 | 0.6667 | 1 | 1 | 1 | 1 | 1 | 1 | NA | NA | 3 | 1 |
| 13_885330 | 5 | 0.2 | 4 | 0 | 5 | 0 | 3 | 0.3333 | 17 | 0.1176 | 4 | 0.25 | 7 | 0.1429 | 4 | 0 | 7 | 0 | 22 | 0.0909 |
| 13_885541 | 7 | 0.1429 | 5 | 0.4 | 9 | 0.2222 | 9 | 0.3333 | 30 | 0.2667 | 10 | 0.2 | 10 | 0 | 9 | 0.1111 | 9 | 0 | 38 | 0.0789 |
| 13_885553 | 8 | 0 | 4 | 0.25 | 8 | 0.25 | 7 | 0 | 27 | 0.1111 | 10 | 0.1 | 10 | 0 | 7 | 0.1429 | 7 | 0 | 34 | 0.0588 |
| 13_885581 | 7 | 0.8571 | 5 | 1 | 8 | 1 | 10 | 1 | 30 | 0.9667 | 9 | 1 | 9 | 1 | 7 | 1 | 8 | 0.75 | 33 | 0.9394 |
| 13_889958 | 3 | 0.3333 | 2 | 0 | 5 | 0 | 7 | 0.1429 | 17 | 0.1176 | 3 | 0 | 6 | 0 | 1 | 0 | 5 | 0.2 | 15 | 0.0667 |
| 13_889960 | 3 | 0 | 2 | 0.5 | 4 | 0.25 | 9 | 0 | 18 | 0.1111 | 3 | 0 | 6 | 0.1667 | 2 | 0 | 5 | 0 | 16 | 0.0625 |
| 13_890212 | 2 | 0.5 | 3 | 0.6667 | 3 | 0 | 3 | 0 | 11 | 0.2727 | NA | NA | 1 | 1 | 1 | 0 | 2 | 1 | 4 | 0.75 |
| 13_894148 | 5 | 0 | 4 | 0 | 2 | 0.5 | 5 | 0 | 16 | 0.0625 | 8 | 0.125 | 4 | 0.25 | 4 | 0 | 3 | 0 | 19 | 0.1053 |
| 13_902222 | 2 | 1 | NA | NA | 3 | 1 | 1 | 1 | 6 | 1 | 1 | 1 | NA | NA | 1 | 1 | NA | NA | 2 | 1 |
| 13_902285 | 6 | 0 | 3 | 0.3333 | 8 | 0.125 | 5 | 0 | 22 | 0.0909 | 3 | 0 | 7 | 0.2857 | 6 | 0.3333 | 3 | 0 | 19 | 0.2105 |
| 13_902290 | 6 | 0.3333 | 3 | 0.3333 | 7 | 0.4286 | 5 | 0.4 | 21 | 0.381 | 3 | 0.3333 | 8 | 0.25 | 5 | 0 | 2 | 1 | 18 | 0.2778 |
| 13_902304 | 6 | 0 | 3 | 0.6667 | 7 | 0 | 5 | 0.6 | 21 | 0.2381 | 3 | 0.3333 | 8 | 0.375 | 5 | 0.2 | 3 | 0.6667 | 19 | 0.3684 |
| 13_902305 | 6 | 0 | 1 | 0 | 7 | 0 | 5 | 0.4 | 19 | 0.1053 | 3 | 0 | 8 | 0.25 | 1 | 1 | 2 | 0 | 14 | 0.2143 |
| 13_902312 | 5 | 1 | 2 | 0.5 | 7 | 0.8571 | 5 | 0.4 | 19 | 0.7368 | 1 | 1 | 3 | 0.3333 | 5 | 0.6 | 4 | 0.5 | 13 | 0.5385 |
| 13_902314 | 5 | 0.6 | 3 | 0.6667 | 6 | 0 | 4 | 0.25 | 18 | 0.3333 | 2 | 0.5 | 6 | 0 | 3 | 0 | 4 | 0.25 | 15 | 0.1333 |
| 13_902316 | 6 | 0 | 3 | 0 | 7 | 0.1429 | 4 | 0.5 | 20 | 0.15 | 2 | 0 | 7 | 0.2857 | 5 | 0 | 4 | 0.25 | 18 | 0.1667 |
| 13_922077 | 2 | 0.5 | NA | NA | 2 | 0 | 3 | 0.3333 | 7 | 0.2857 | 1 | 0 | 2 | 0 | 5 | 0.4 | 1 | 1 | 9 | 0.3333 |
| 13_922083 | 2 | 0.5 | 2 | 0 | 3 | 0 | 3 | 0.3333 | 10 | 0.2 | 2 | 0 | 5 | 0.2 | 5 | 0.2 | 1 | 0 | 13 | 0.1538 |
| 13_922094 | 2 | 0 | 2 | 0 | 4 | 0 | 3 | 0.6667 | 11 | 0.1818 | 2 | 0 | 4 | 0 | 5 | 0.2 | 2 | 0.5 | 13 | 0.1538 |
| 13_922133 | 2 | 0.5 | 3 | 0.3333 | 4 | 0.25 | 3 | 0 | 12 | 0.25 | 1 | 0 | 6 | 0 | 3 | 0 | 2 | 1 | 12 | 0.1667 |
| 13_922146 | 3 | 0.3333 | 1 | 0 | 5 | 0.8 | 4 | 0.25 | 13 | 0.4615 | 2 | 0.5 | 6 | 0.5 | 4 | 0.25 | 2 | 0.5 | 14 | 0.4286 |
| 13_922147 | 3 | 0 | 3 | 0 | 6 | 0.5 | 4 | 0.25 | 16 | 0.25 | 2 | 0.5 | 7 | 0 | 5 | 0.2 | 2 | 0 | 16 | 0.125 |
| 13_922148 | 3 | 0.6667 | 3 | 0 | 5 | 0.2 | 4 | 0 | 15 | 0.2 | 2 | 0 | 5 | 0.2 | 5 | 0.2 | 1 | 1 | 13 | 0.2308 |
| 13_922227 | 4 | 0 | 5 | 0.4 | 6 | 0.1667 | 5 | 0.2 | 20 | 0.2 | 3 | 0 | 7 | 0.5714 | 5 | 0.4 | 7 | 0.1429 | 22 | 0.3182 |
| 13_922230 | 5 | 0 | 5 | 0.2 | 4 | 0 | 5 | 0.4 | 19 | 0.1579 | 3 | 0 | 6 | 0.5 | 4 | 0.25 | 6 | 0 | 19 | 0.2105 |
| 13_922231 | 5 | 0.2 | 5 | 0 | 5 | 0 | 5 | 0 | 20 | 0.05 | 2 | 0 | 6 | 0.1667 | 4 | 0 | 7 | 0.1429 | 19 | 0.1053 |
| 13_922244 | 5 | 0.2 | 3 | 0.3333 | 5 | 0.2 | 5 | 0.4 | 18 | 0.2778 | 3 | 0 | 5 | 0.4 | 4 | 0.75 | 7 | 0 | 19 | 0.2632 |
| 13_922275 | 4 | 0 | 5 | 0 | 6 | 0 | 5 | 0.2 | 20 | 0.05 | 3 | 0 | 7 | 0.1429 | 4 | 0.25 | 5 | 0 | 19 | 0.1053 |
| 13_922290 | 6 | 0.1667 | 5 | 0 | 6 | 0 | 7 | 0.4286 | 24 | 0.1667 | 3 | 0 | 6 | 0.3333 | 6 | 0.3333 | 8 | 0.125 | 23 | 0.2174 |
| 13_922347 | 5 | 0.4 | 6 | 0.5 | 6 | 0 | 7 | 0.4286 | 24 | 0.3333 | 3 | 0.3333 | 7 | 0.4286 | 5 | 0.2 | 7 | 0.5714 | 22 | 0.4091 |
| 13_922361 | 8 | 0.125 | 6 | 0.1667 | 9 | 0.1111 | 9 | 0.1111 | 32 | 0.125 | 3 | 0.3333 | 5 | 0.2 | 5 | 0 | 9 | 0.1111 | 22 | 0.1364 |
| 13_922380 | 7 | 0.2857 | 6 | 0 | 9 | 0.1111 | 10 | 0.3 | 32 | 0.1875 | 4 | 0.5 | 7 | 0.7143 | 7 | 0.2857 | 9 | 0.2222 | 27 | 0.4074 |
| 13_922386 | 7 | 0.4286 | 6 | 0.5 | 10 | 0.4 | 9 | 0.2222 | 32 | 0.375 | 3 | 0.3333 | 8 | 0.125 | 7 | 0 | 8 | 0.125 | 26 | 0.1154 |
| 13_922409 | 7 | 0 | 4 | 0 | 9 | 0 | 9 | 0.2222 | 29 | 0.069 | 4 | 0.25 | 8 | 0.125 | 7 | 0 | 5 | 0 | 24 | 0.0833 |
| 13_922431 | 7 | 0.4286 | 5 | 0 | 12 | 0.0833 | 9 | 0.1111 | 33 | 0.1515 | 7 | 0.2857 | 6 | 0.1667 | 7 | 0.2857 | 7 | 0.4286 | 27 | 0.2963 |
| 13_932278 | 3 | 0 | 10 | 0.2 | 6 | 0.1667 | 3 | 0.3333 | 22 | 0.1818 | 10 | 0.3 | 3 | 0 | 7 | 0 | 2 | 0.5 | 22 | 0.1818 |
| 13_968723 | 7 | 0.1429 | 4 | 0.25 | 4 | 0 | 2 | 0 | 17 | 0.1176 | 2 | 0 | 4 | 0.25 | 3 | 0 | 2 | 0 | 11 | 0.0909 |
| 13_971585 | 7 | 0.4286 | 3 | 0 | 5 | 0.2 | 3 | 0.6667 | 18 | 0.3333 | 2 | 0 | 5 | 0.2 | 4 | 0 | 1 | 0 | 12 | 0.0833 |
| 13_971588 | 7 | 0.7143 | 3 | 0.6667 | 6 | 0.6667 | 4 | 1 | 20 | 0.75 | 3 | 0.3333 | 5 | 0.8 | 4 | 1 | 3 | 0.3333 | 15 | 0.6667 |
| 13_971595 | 7 | 0.4286 | 3 | 0.3333 | 6 | 0.5 | 2 | 1 | 18 | 0.5 | 2 | 0.5 | 1 | 1 | 3 | 0.3333 | 1 | 0 | 7 | 0.4286 |
| 13_971609 | 8 | 1 | 4 | 1 | 5 | 0.8 | 2 | 1 | 19 | 0.9474 | 3 | 1 | 5 | 1 | 4 | 0.5 | 3 | 1 | 15 | 0.8667 |
| 13_971615 | 6 | 0.3333 | 3 | 0.3333 | 8 | 0.5 | 3 | 0.6667 | 20 | 0.45 | 3 | 0 | 5 | 0.6 | 5 | 0.6 | 3 | 0 | 16 | 0.375 |
| 13_971616 | 7 | 0.7143 | 3 | 0.3333 | 5 | 0.6 | 3 | 0.3333 | 18 | 0.5556 | 2 | 1 | 4 | 0.5 | 4 | 0.5 | 3 | 0.6667 | 13 | 0.6154 |
| 13_971628 | 9 | 0.8889 | 5 | 1 | 10 | 0.9 | 2 | 1 | 26 | 0.9231 | 3 | 1 | 3 | 1 | 4 | 1 | 3 | 1 | 13 | 1 |
| 13_971632 | 8 | 0.875 | 4 | 1 | 9 | 0.8889 | 2 | 1 | 23 | 0.913 | 2 | 0.5 | 6 | 0.6667 | 4 | 1 | 1 | 1 | 13 | 0.7692 |
| 13_971722 | 6 | 0.3333 | 5 | 0 | 6 | 0 | 2 | 0 | 19 | 0.1053 | 3 | 0.6667 | 8 | 0 | 8 | 0.25 | 3 | 0 | 22 | 0.1818 |
| 13_971842 | 9 | 0.4444 | 10 | 0.3 | 8 | 0.625 | 5 | 1 | 32 | 0.5312 | 4 | 0.5 | 9 | 0.4444 | 10 | 0.5 | 2 | 0.5 | 25 | 0.48 |
| 13_971864 | 8 | 0.25 | 10 | 0.4 | 8 | 0.375 | 8 | 0.25 | 34 | 0.3235 | 3 | 0.3333 | 7 | 0.2857 | 8 | 0.25 | 1 | 0 | 19 | 0.2632 |
| 13_971885 | 8 | 0.125 | 7 | 0 | 8 | 0.125 | 8 | 0.125 | 31 | 0.0968 | 4 | 0.25 | 8 | 0 | 11 | 0.0909 | 3 | 0.3333 | 26 | 0.1154 |
| 13_973506 | 19 | 0.2105 | 30 | 0.0333 | 22 | 0.0909 | 25 | 0.16 | 96 | 0.1146 | 16 | 0 | 22 | 0.0909 | 35 | 0.0857 | 15 | 0.1333 | 88 | 0.0795 |
| 13_973536 | 17 | 0.4118 | 29 | 0.3448 | 21 | 0.381 | 24 | 0.2917 | 91 | 0.3516 | 11 | 0.5455 | 21 | 0.2381 | 31 | 0.3226 | 12 | 0.25 | 75 | 0.32 |
| 13_973544 | 16 | 0.3125 | 22 | 0.3636 | 20 | 0.45 | 23 | 0.1304 | 81 | 0.3086 | 13 | 0.0769 | 20 | 0.3 | 30 | 0.1667 | 14 | 0.0714 | 77 | 0.1688 |
| 13_973653 | 13 | 0 | 23 | 0.087 | 12 | 0.3333 | 18 | 0.2222 | 66 | 0.1515 | 9 | 0.2222 | 16 | 0.0625 | 23 | 0.2174 | 8 | 0.125 | 56 | 0.1607 |
| 13_973657 | 14 | 0.3571 | 20 | 0.35 | 15 | 0.4667 | 18 | 0.2778 | 67 | 0.3582 | 8 | 0.125 | 14 | 0.1429 | 23 | 0.2174 | 10 | 0.5 | 55 | 0.2364 |
| 13_973665 | 9 | 0.1111 | 17 | 0.1765 | 10 | 0 | 14 | 0.0714 | 50 | 0.1 | 6 | 0.1667 | 13 | 0.0769 | 16 | 0.125 | 10 | 0.2 | 45 | 0.1333 |
| 13_973921 | 7 | 0.1429 | 20 | 0.2 | 15 | 0.1333 | 11 | 0.0909 | 53 | 0.1509 | 6 | 0.1667 | 11 | 0.2727 | 15 | 0.2667 | 7 | 0 | 39 | 0.2051 |
| 13_973955 | 12 | 0.0833 | 16 | 0.3125 | 13 | 0.3846 | 12 | 0.4167 | 53 | 0.3019 | 5 | 0.4 | 14 | 0.3571 | 18 | 0.1667 | 10 | 0.4 | 47 | 0.2979 |
| 13_974035 | 2 | 0.5 | 3 | 0 | 6 | 0.1667 | 5 | 0.2 | 16 | 0.1875 | 3 | 0 | 2 | 0.5 | 7 | 0 | 3 | 0.3333 | 15 | 0.1333 |
| 13_974555 | NA | NA | NA | NA | NA | NA | 1 | 1 | 1 | 1 | NA | NA | 4 | 0.25 | 1 | 0 | NA | NA | 5 | 0.2 |
| 13_992702 | 1 | 1 | 2 | 1 | NA | NA | NA | NA | 3 | 1 | NA | NA | NA | NA | NA | NA | 1 | 1 | 1 | 1 |

| | | | | | | | | | | | | | | | | | | | | |
|---|---|---|---|---|---|---|---|---|---|---|---|---|---|---|---|---|---|---|---|---|
| 13_9927846 | NA | NA | 1 | 1 | NA | NA | NA | NA | 1 | 1 | NA | NA | NA | NA | NA | NA | 1 | 1 | 1 | 1 |
| 13_9928595 | NA | NA | 1 | 1 | NA | NA | NA | NA | 1 | 1 | NA | NA | NA | NA | NA | NA | 1 | 1 | 1 | 1 |
| 13_1156844 | 2 | 1 | 5 | 0 | 4 | 0.5 | 3 | 0.6667 | 14 | 0.4286 | 5 | 0.6 | 2 | 1 | 3 | 0.3333 | 6 | 0.5 | 16 | 0.5625 |
| 13_1156848 | 3 | 0.3333 | 5 | 0 | 2 | 0 | 4 | 0.25 | 14 | 0.1429 | 5 | 0.4 | 3 | 0 | 4 | 0 | 6 | 0.1667 | 18 | 0.1667 |
| 13_1156854 | 5 | 0.8 | 5 | 0.4 | 4 | 0.75 | 5 | 0.8 | 19 | 0.6842 | 3 | 1 | 2 | 0.5 | 4 | 0.75 | 6 | 0.8333 | 15 | 0.8 |
| 13_1156859 | 5 | 0.8 | 5 | 0.2 | 4 | 0.25 | 5 | 0.8 | 19 | 0.5263 | 5 | 0.6 | 3 | 0.3333 | 4 | 1 | 6 | 0.6667 | 18 | 0.6667 |
| 13_1156865 | 3 | 0.3333 | 4 | 0 | 4 | 0 | 5 | 0.2 | 16 | 0.125 | 5 | 0 | 2 | 0.5 | 4 | 0.5 | 6 | 0 | 17 | 0.1765 |
| 13_1156903 | 5 | 0.6 | 2 | 0 | 3 | 1 | 4 | 1 | 14 | 0.7143 | 3 | 0.6667 | 2 | 1 | 3 | 0.6667 | 6 | 0.8333 | 14 | 0.7857 |
| 13_1156907 | 5 | 0.6 | 4 | 0.75 | 2 | 1 | 3 | 0.3333 | 14 | 0.6429 | 5 | 0.2 | 2 | 0.5 | 3 | 0.6667 | 6 | 0.3333 | 16 | 0.375 |
| 13_1156908 | 5 | 0 | 5 | 0 | 4 | 0.25 | 3 | 0 | 17 | 0.0588 | 4 | 0 | 1 | 1 | 2 | 0.5 | 6 | 0 | 13 | 0.1538 |
| 13_1156916 | 4 | 0 | 3 | 0.3333 | 4 | 0.25 | 4 | 0.5 | 15 | 0.2667 | 5 | 0 | 2 | 0 | 2 | 0.5 | 4 | 0.75 | 13 | 0.3077 |
| 13_1156916 | 4 | 0.25 | 5 | 0.4 | 4 | 0.25 | 4 | 0.25 | 17 | 0.2941 | 5 | 0.2 | 2 | 0 | 3 | 0 | 6 | 0.5 | 16 | 0.25 |
| 13_1156943 | 3 | 0.6667 | 4 | 0 | 3 | 0.6667 | 4 | 0.25 | 14 | 0.3571 | 5 | 0.4 | 2 | 1 | 3 | 0.6667 | 4 | 0.25 | 14 | 0.5 |
| 13_1160036 | 4 | 0 | 2 | 0 | 2 | 0 | 3 | 0.3333 | 11 | 0.0909 | 3 | 0 | 1 | 0 | 3 | 0.3333 | 4 | 0 | 11 | 0.0909 |
| 13_1160972 | 3 | 0 | 2 | 0 | 2 | 0 | 2 | 0.5 | 9 | 0.1111 | 3 | 0.3333 | 1 | 1 | 2 | 0.5 | 3 | 0 | 9 | 0.3333 |
| 13_1160984 | 4 | 0.75 | 2 | 0.5 | 2 | 0.5 | 2 | 0 | 10 | 0.5 | 1 | 0 | 1 | 0 | 3 | 0.6667 | 3 | 0.6667 | 8 | 0.5 |
| 13_1160999 | 4 | 0 | 2 | 0 | 2 | 0 | 2 | 1 | 10 | 0.2 | 3 | 0.6667 | 1 | 1 | 2 | 0 | 1 | 1 | 7 | 0.5714 |
| 13_1161024 | 3 | 0.3333 | 2 | 0.5 | 2 | 0 | 2 | 0 | 9 | 0.2222 | 3 | 0 | 1 | 1 | 2 | 1 | 2 | 0.5 | 8 | 0.5 |
| 13_1161025 | 3 | 0.3333 | 2 | 0.5 | 1 | 0 | 2 | 0 | 8 | 0.25 | 3 | 0.3333 | 1 | 0 | 1 | 0 | 2 | 0 | 7 | 0.1429 |
| 13_1161059 | 1 | 1 | 1 | 1 | 2 | 0 | 2 | 0 | 6 | 0.3333 | 3 | 0.3333 | NA | NA | 2 | 0 | 2 | 0 | 7 | 0.1429 |
| 13_116106 | 2 | 0.5 | 1 | 0 | 2 | 0 | 2 | 0 | 7 | 0.1429 | 3 | 0.6667 | NA | NA | 2 | 0 | 2 | 0 | 7 | 0.2857 |
| 13_1161064 | 2 | 0 | 1 | 1 | 2 | 1 | 2 | 1 | 7 | 0.7143 | 3 | 1 | NA | NA | 2 | 0 | 2 | 1 | 7 | 0.7143 |
| 13_1161068 | 2 | 0.5 | 1 | 0 | 2 | 0 | 2 | 0.5 | 7 | 0.2857 | 2 | 0 | NA | NA | 2 | 0.5 | 2 | 0 | 6 | 0.1667 |
| 13_1161075 | 2 | 1 | 1 | 1 | 1 | 1 | 2 | 1 | 6 | 1 | 3 | 0.3333 | NA | NA | 2 | 0.5 | 2 | 0 | 7 | 0.2857 |
| 13_1228442 | 4 | 0.25 | 7 | 0.4286 | 1 | 1 | NA | NA | 12 | 0.4167 | 5 | 0.4 | 7 | 0.2857 | 8 | 0.25 | 7 | 0 | 27 | 0.2222 |
| 13_124548 | 2 | 0 | 2 | 0 | 3 | 0.3333 | NA | NA | 7 | 0.1429 | NA | NA | 1 | 0 | 2 | 0.5 | 2 | 0 | 5 | 0.2 |
| 13_124551 | NA | NA | 2 | 0 | 3 | 0.3333 | 1 | 0 | 6 | 0.1667 | NA | NA | 2 | 0.5 | 2 | 0 | 1 | 0 | 5 | 0.2 |
| 13_124557 | 1 | 1 | 1 | 0 | 1 | 0 | 1 | 0 | 4 | 0.25 | NA | NA | 2 | 0 | 2 | 0 | 2 | 0.5 | 6 | 0.1667 |
| 13_1245579 | 1 | 1 | 1 | 0 | NA | NA | 1 | 0 | 3 | 0.3333 | NA | NA | 2 | 0.5 | 2 | 1 | 2 | 0 | 6 | 0.5 |
| 13_126218 | 1 | 1 | NA | NA | NA | NA | 2 | 0 | 3 | 0.3333 | 2 | 1 | 1 | 0 | 3 | 0.6667 | NA | NA | 6 | 0.6667 |
| 13_1262183 | 1 | 1 | NA | NA | NA | NA | 2 | 1 | 3 | 1 | 2 | 1 | 1 | 1 | 2 | 0.5 | NA | NA | 5 | 0.8 |
| 13_1262185 | 1 | 1 | NA | NA | NA | NA | 2 | 0.5 | 3 | 0.6667 | 2 | 1 | 1 | 0 | 3 | 0 | NA | NA | 6 | 0.3333 |
| 13_1262188 | 1 | 1 | NA | NA | NA | NA | 2 | 0 | 3 | 0.3333 | 2 | 1 | 1 | 0 | 3 | 0 | NA | NA | 6 | 0.3333 |
| 13_1262369 | 1 | 0 | NA | NA | NA | NA | 3 | 0.3333 | 4 | 0.25 | 2 | 0.5 | 4 | 0.5 | 2 | 0 | 2 | 0 | 10 | 0.3 |
| 13_1262369 | 2 | 0.5 | NA | NA | NA | NA | 3 | 0.3333 | 5 | 0.4 | 2 | 1 | 4 | 0 | 2 | 0 | 1 | 0 | 9 | 0.2222 |
| 13_1263052 | 1 | 1 | 1 | 0 | 1 | 0 | 1 | 0 | 4 | 0.25 | 1 | 1 | NA | NA | NA | NA | NA | NA | 1 | 1 |
| 13_1340642 | 1 | 0 | 3 | 0.3333 | NA | NA | 1 | 0 | 5 | 0.2 | 2 | 0.5 | NA | NA | NA | NA | NA | NA | 2 | 0.5 |
| 13_1353312 | 1 | 0 | 2 | 0 | NA | NA | 2 | 1 | 5 | 0.4 | 1 | 0 | 3 | 0.3333 | NA | NA | 1 | 1 | 5 | 0.4 |
| 13_1353379 | NA | NA | 1 | 0 | NA | NA | 2 | 1 | 3 | 0.6667 | 2 | 0 | 3 | 1 | 2 | 0.5 | 1 | 1 | 8 | 0.625 |
| 13_1356772 | NA | NA | NA | NA | NA | NA | 2 | 0.5 | 2 | 0.5 | 1 | 1 | 3 | 0.6667 | 3 | 0.3333 | 1 | 0 | 8 | 0.5 |
| 13_1395167 | 2 | 0.5 | NA | NA | 3 | 0 | 3 | 0 | 8 | 0.125 | 2 | 0 | 1 | 0 | 5 | 0 | 3 | 0.3333 | 11 | 0.0909 |
| 13_1395183 | 3 | 0.3333 | NA | NA | 4 | 0 | 3 | 0.3333 | 10 | 0.2 | 4 | 0 | 1 | 0 | 8 | 0 | 4 | 0.5 | 17 | 0.1176 |
| 13_1395206 | 3 | 0 | NA | NA | 4 | 0.5 | 4 | 0.25 | 11 | 0.2727 | 4 | 0.25 | 2 | 0 | 6 | 0 | 4 | 0 | 16 | 0.0625 |
| 13_1395216 | 3 | 0.6667 | NA | NA | 3 | 0.6667 | 4 | 1 | 10 | 0.8 | 4 | 1 | 3 | 0.3333 | 8 | 0.625 | 5 | 0.8 | 20 | 0.7 |
| 13_1395232 | 4 | 1 | NA | NA | 3 | 0.6667 | 4 | 1 | 11 | 0.9091 | 4 | 0.75 | 3 | 0.3333 | 8 | 0.875 | 4 | 1 | 19 | 0.7895 |
| 13_1395240 | 3 | 0.3333 | 1 | 0 | 3 | 0.6667 | 3 | 0 | 10 | 0.3 | 4 | 0.75 | 3 | 0.3333 | 7 | 0.1429 | 5 | 0.6 | 19 | 0.4211 |
| 13_1395247 | 4 | 0 | 1 | 0 | 3 | 0.6667 | 4 | 0.25 | 12 | 0.25 | 3 | 0.6667 | 2 | 0.5 | 5 | 0.4 | 5 | 0.6 | 15 | 0.5333 |
| 13_139526 | 3 | 0.3333 | NA | NA | 3 | 0.3333 | 4 | 0.25 | 10 | 0.3 | 3 | 0 | 2 | 0 | 7 | 0.2857 | 2 | 0 | 14 | 0.1429 |
| 13_1395266 | 3 | 1 | 1 | 0 | 3 | 0.6667 | 3 | 0.6667 | 10 | 0.7 | 3 | 1 | 2 | 0.5 | 7 | 1 | 3 | 0.3333 | 15 | 0.8 |
| 13_1395267 | 3 | 1 | 1 | 0 | 4 | 0.75 | 4 | 0.75 | 12 | 0.75 | 4 | 1 | 1 | 0 | 8 | 0.625 | 4 | 0.75 | 17 | 0.7059 |
| 13_139527 | 4 | 0 | 1 | 1 | 4 | 0 | 3 | 0 | 12 | 0.0833 | 4 | 0.25 | 2 | 0 | 7 | 0.1429 | 5 | 0.8 | 18 | 0.3333 |
| 13_1395273 | 2 | 0.5 | NA | NA | 4 | 0 | 4 | 0.25 | 10 | 0.2 | 4 | 0.5 | 3 | 0.3333 | 9 | 0.1111 | 5 | 0.2 | 21 | 0.2381 |
| 13_139530 | 6 | 0 | 1 | 0 | 4 | 0.25 | 8 | 0.125 | 19 | 0.1053 | 6 | 0.1667 | 3 | 0.3333 | 9 | 0.2222 | 6 | 0.1667 | 24 | 0.2083 |
| 13_1400047 | 2 | 0 | 9 | 0.1111 | 4 | 0 | 3 | 0 | 18 | 0.0556 | 3 | 0 | 1 | 0 | 2 | 0.5 | 4 | 0 | 10 | 0.1 |
| 13_1400090 | 3 | 0 | 12 | 0.1667 | 5 | 0 | 2 | 0 | 22 | 0.0909 | 3 | 0.3333 | 6 | 0 | 2 | 0 | 5 | 0 | 16 | 0.0625 |
| 13_1417236 | 1 | 0 | 2 | 0 | 2 | 0.5 | 1 | 1 | 6 | 0.3333 | 3 | 0.3333 | 1 | 0 | 1 | 1 | 2 | 0.5 | 7 | 0.4286 |
| 13_141729 | NA | NA | 2 | 0 | 3 | 0.6667 | 1 | 0 | 6 | 0.3333 | 3 | 0.3333 | 2 | 0.5 | 1 | 0 | 2 | 0.5 | 8 | 0.375 |
| 13_1417295 | 1 | 0 | 2 | 0 | 4 | 0.75 | 1 | 0 | 8 | 0.375 | 2 | 1 | 1 | 0 | 1 | 0 | 2 | 0.5 | 6 | 0.5 |
| 13_1417298 | 1 | 0 | 3 | 0 | 4 | 0 | 1 | 1 | 9 | 0.1111 | 2 | 0.5 | 1 | 0 | 1 | 0 | 2 | 0.5 | 6 | 0.3333 |
| 13_1417302 | 1 | 1 | 2 | 0 | 2 | 1 | NA | NA | 5 | 0.6 | 2 | 0.5 | 1 | 0 | 1 | 1 | 1 | 0 | 5 | 0.4 |
| 13_1417305 | 1 | 0 | 2 | 0 | 4 | 0.25 | 1 | 0 | 8 | 0.125 | 2 | 0.5 | 2 | 0 | 1 | 0 | 1 | 1 | 6 | 0.3333 |
| 13_1417327 | 1 | 0 | 3 | 0.3333 | 4 | 0.5 | 1 | 0 | 9 | 0.3333 | 2 | 0.5 | 2 | 0 | 1 | 0 | 2 | 0 | 7 | 0.1429 |
| 13_141734 | 1 | 0 | 2 | 0 | 4 | 0.25 | 1 | 0 | 8 | 0.125 | 2 | 0.5 | 1 | 0 | 1 | 0 | 2 | 1 | 6 | 0.5 |
| 13_1417810 | NA | NA | NA | NA | NA | NA | 1 | 1 | 1 | 1 | NA | NA | NA | NA | NA | NA | 1 | 1 | 1 | 1 |
| 13_1419762 | 2 | 0.5 | 2 | 1 | NA | NA | NA | NA | 4 | 0.75 | NA | NA | NA | NA | NA | NA | 1 | 1 | 1 | 1 |

| | | | | | | | | | | | | | | | | | | | | |
|---|---|---|---|---|---|---|---|---|---|---|---|---|---|---|---|---|---|---|---|---|
| 13_141976 | 2 | 0.5 | 2 | 0 | NA | NA | NA | NA | 4 | 0.25 | NA | NA | NA | NA | NA | NA | 1 | 1 | 1 | 1 |
| 13_141977 | 2 | 0.5 | 2 | 0 | NA | NA | NA | NA | 4 | 0.25 | NA | NA | NA | NA | NA | NA | 1 | 1 | 1 | 1 |
| 13_141978 | 2 | 0.5 | 2 | 0 | NA | NA | NA | NA | 4 | 0.25 | NA | NA | NA | NA | NA | NA | 1 | 1 | 1 | 1 |
| 13_142264 | 3 | 0 | 2 | 0 | 3 | 0.3333 | 3 | 0 | 11 | 0.0909 | 5 | 0.2 | 3 | 0 | 3 | 0 | 7 | 0 | 18 | 0.0556 |
| 13_142265 | 4 | 0 | 2 | 0.5 | 3 | 0 | 3 | 0 | 12 | 0.0833 | 5 | 0 | 3 | 0.3333 | 3 | 0 | 6 | 0 | 17 | 0.0588 |
| 13_142268 | 5 | 0.2 | 2 | 0 | 3 | 0 | 3 | 0 | 13 | 0.0769 | 5 | 0 | 3 | 0 | 5 | 0 | 7 | 0.1429 | 20 | 0.05 |
| 13_142274 | 4 | 0.25 | NA | NA | 3 | 0.6667 | 3 | 0.3333 | 10 | 0.4 | 3 | 0.6667 | 3 | 0.3333 | 3 | 0 | 5 | 0.2 | 14 | 0.2857 |
| 13_142275 | 3 | 0.3333 | NA | NA | 3 | 0.6667 | 2 | 1 | 8 | 0.625 | 5 | 0.8 | 3 | 0.6667 | 2 | 0.5 | 6 | 0.5 | 16 | 0.625 |
| 13_142283 | 6 | 0.1667 | 2 | 0 | 3 | 0.3333 | 2 | 0 | 13 | 0.1538 | 4 | 0 | 3 | 0 | 5 | 0 | 5 | 0.4 | 17 | 0.1176 |
| 13_142284 | 9 | 0 | NA | NA | 5 | 0.2 | 2 | 0 | 16 | 0.0625 | 5 | 0.2 | 3 | 0 | 3 | 0 | 5 | 0 | 16 | 0.0625 |
| 13_142285 | 5 | 0.4 | NA | NA | 4 | 0.5 | 2 | 0 | 11 | 0.3636 | 4 | 0.5 | 1 | 0 | 5 | 0.6 | 4 | 0.5 | 14 | 0.5 |
| 13_142286 | 9 | 0.1111 | 2 | 0 | 5 | 0 | 1 | 0 | 17 | 0.0588 | 3 | 0.3333 | 2 | 0 | 6 | 0 | 3 | 0 | 14 | 0.0714 |
| 13_142287 | 8 | 0 | 2 | 0.5 | 5 | 0.2 | 2 | 0 | 17 | 0.1176 | 5 | 0.4 | 2 | 0.5 | 6 | 0.1667 | 6 | 0 | 19 | 0.2105 |
| 13_142287 | 9 | 0.7778 | 2 | 1 | 4 | 0.5 | 2 | 1 | 17 | 0.7647 | 5 | 0.8 | 2 | 1 | 6 | 0.8333 | 6 | 0.8333 | 19 | 0.8421 |
| 13_142288 | 8 | 0.375 | 2 | 0.5 | 4 | 0.5 | 2 | 1 | 16 | 0.5 | 3 | 0.6667 | 2 | 0 | 7 | 0.4286 | 4 | 0.5 | 16 | 0.4375 |
| 13_142288 | 7 | 0.2857 | 2 | 0 | 4 | 0.5 | 2 | 0.5 | 15 | 0.3333 | 3 | 0.6667 | 1 | 0 | 6 | 0.5 | 5 | 0.2 | 15 | 0.4 |
| 13_142288 | 7 | 1 | 1 | 1 | 4 | 0.75 | 2 | 1 | 14 | 0.9286 | 4 | 1 | 1 | 1 | 5 | 1 | 6 | 1 | 16 | 1 |
| 13_142290 | 8 | 0.125 | 2 | 0 | 4 | 0 | 1 | 0 | 15 | 0.0667 | 4 | 0 | 2 | 0.5 | 5 | 0.2 | 3 | 0 | 14 | 0.1429 |
| 13_143615 | 1 | 1 | 2 | 1 | 1 | 1 | 6 | 0.8333 | 10 | 0.9 | NA | NA | 1 | 1 | 3 | 0.3333 | 1 | 1 | 5 | 0.6 |
| 13_143615 | 1 | 0 | 2 | 0 | 1 | 0 | 6 | 0.1667 | 10 | 0.1 | NA | NA | 1 | 0 | 2 | 0.5 | 1 | 0 | 4 | 0.25 |
| 13_143617 | 1 | 1 | 3 | 0.6667 | NA | NA | 7 | 0.4286 | 11 | 0.5455 | NA | NA | 1 | 0 | 3 | 0.3333 | 2 | 0 | 6 | 0.1667 |
| 13_143619 | 1 | 0 | 3 | 0.3333 | 1 | 0 | 8 | 0.125 | 13 | 0.1538 | NA | NA | 1 | 0 | 4 | 0 | 3 | 0.3333 | 8 | 0.125 |
| 13_143620 | 1 | 0 | 3 | 0.6667 | 1 | 1 | 8 | 0.5 | 13 | 0.5385 | NA | NA | 1 | 1 | 4 | 0.75 | 2 | 1 | 7 | 0.8571 |
| 13_143620 | 1 | 0 | 4 | 0.75 | NA | NA | 9 | 0.8889 | 14 | 0.7857 | NA | NA | 1 | 0 | 4 | 0.5 | 3 | 0 | 8 | 0.25 |
| 13_143621 | 1 | 0 | 3 | 0.3333 | NA | NA | 9 | 0 | 13 | 0.0769 | NA | NA | 1 | 0 | 4 | 0 | 3 | 0.3333 | 8 | 0.125 |
| 13_143621 | 1 | 1 | 2 | 0.5 | 1 | 0 | 7 | 0.8571 | 11 | 0.7273 | NA | NA | 1 | 1 | 1 | 1 | 3 | 0.6667 | 5 | 0.8 |
| 13_143622 | 1 | 0 | 3 | 0.3333 | 1 | 0 | 8 | 0.25 | 13 | 0.2308 | NA | NA | 1 | 0 | 4 | 0 | 2 | 0.5 | 7 | 0.1429 |
| 13_143624 | 3 | 0 | 3 | 0 | 1 | 0 | 9 | 0.1111 | 16 | 0.0625 | NA | NA | 2 | 0 | 4 | 0 | 3 | 0.6667 | 9 | 0.2222 |
| 13_143624 | 2 | 1 | 4 | 0.75 | 1 | 1 | 7 | 0.8571 | 14 | 0.8571 | NA | NA | 1 | 1 | 4 | 0.5 | 3 | 0.6667 | 8 | 0.625 |
| 13_143723 | 3 | 1 | 5 | 0.8 | 3 | 1 | 12 | 0.75 | 23 | 0.8261 | 1 | 1 | 4 | 0.5 | 5 | 0.2 | 4 | 0.25 | 14 | 0.3571 |
| 13_143723 | 3 | 0.3333 | 5 | 0.4 | 3 | 0.6667 | 12 | 0.0833 | 23 | 0.2609 | 1 | 0 | 4 | 0 | 6 | 0.1667 | 4 | 0.25 | 15 | 0.1333 |
| 13_143724 | 3 | 0 | 5 | 0.2 | 3 | 0 | 11 | 0.1818 | 22 | 0.1364 | 1 | 1 | 4 | 0 | 6 | 0.3333 | 4 | 0.25 | 15 | 0.2667 |
| 13_143724 | 2 | 0 | 5 | 0.6 | 2 | 0 | 12 | 0.4167 | 21 | 0.381 | NA | NA | 3 | 0.3333 | 6 | 0.3333 | 4 | 0.25 | 13 | 0.3077 |
| 13_143725 | 3 | 1 | 7 | 0.8571 | 3 | 1 | 10 | 0.9 | 23 | 0.913 | 1 | 1 | 4 | 0.75 | 6 | 0.6667 | 3 | 0.6667 | 14 | 0.7143 |
| 13_143726 | 3 | 0 | 5 | 0.4 | 3 | 0 | 11 | 0.0909 | 22 | 0.1364 | 1 | 1 | 4 | 0.25 | 5 | 0.2 | 2 | 0.5 | 12 | 0.3333 |
| 13_143729 | 3 | 0 | 8 | 0.125 | 3 | 0.3333 | 14 | 0.2143 | 28 | 0.1786 | 1 | 1 | 4 | 0 | 5 | 0.2 | 4 | 0 | 14 | 0.1429 |
| 13_143729 | 3 | 0.3333 | 7 | 0.2857 | 3 | 0.3333 | 14 | 0.4286 | 27 | 0.3704 | 2 | 1 | 4 | 0.25 | 6 | 0.3333 | 4 | 0 | 16 | 0.3125 |
| 13_143732 | 3 | 0 | 6 | 0.1667 | 6 | 0 | 16 | 0.0625 | 31 | 0.0645 | 3 | 0 | 5 | 0 | 6 | 0 | 4 | 0.25 | 18 | 0.0556 |
| 13_143732 | 1 | 0 | 7 | 0.2857 | 5 | 0.2 | 16 | 0.0625 | 29 | 0.1379 | 3 | 0.3333 | 4 | 0.25 | 5 | 0 | 5 | 0.2 | 17 | 0.1765 |
| 13_143737 | 4 | 0.25 | 8 | 0.25 | 6 | 0.1667 | 16 | 0.1875 | 34 | 0.2059 | 4 | 0.25 | 5 | 0.2 | 7 | 0 | 5 | 0 | 21 | 0.0952 |
| 13_144043 | 2 | 0 | 2 | 0.5 | 2 | 0.5 | NA | NA | 6 | 0.3333 | NA | NA | 4 | 0.25 | 2 | 0 | 2 | 0 | 8 | 0.125 |
| 13_144045 | 1 | 1 | 2 | 0.5 | 3 | 0.6667 | NA | NA | 6 | 0.6667 | NA | NA | 4 | 0.75 | NA | NA | NA | NA | 4 | 0.75 |
| 13_144047 | 1 | 0 | 2 | 0 | 3 | 0.6667 | NA | NA | 6 | 0.3333 | NA | NA | 4 | 0.5 | 1 | 0 | 2 | 0.5 | 7 | 0.4286 |
| 13_144050 | 1 | 1 | NA | NA | 2 | 0.5 | NA | NA | 3 | 0.6667 | NA | NA | 2 | 1 | NA | NA | 2 | 0.5 | 4 | 0.75 |
| 13_144051 | 1 | 1 | 2 | 1 | 1 | 1 | NA | NA | 4 | 1 | NA | NA | 3 | 0.6667 | 1 | 1 | 1 | 1 | 5 | 0.8 |
| 13_144052 | 1 | 0 | 2 | 0.5 | NA | NA | NA | NA | 3 | 0.3333 | NA | NA | 3 | 1 | 1 | 1 | 1 | 0 | 5 | 0.8 |
| 13_144906 | NA | NA | 2 | 1 | 1 | 1 | NA | NA | 3 | 1 | NA | NA | 1 | 1 | NA | NA | NA | NA | 1 | 1 |
| 13_144908 | NA | NA | 1 | 1 | 1 | 0 | NA | NA | 2 | 0.5 | NA | NA | 1 | 1 | NA | NA | NA | NA | 1 | 1 |
| 13_144910 | NA | NA | 1 | 1 | 1 | 1 | NA | NA | 2 | 1 | NA | NA | 1 | 1 | NA | NA | NA | NA | 1 | 1 |
| 13_144911 | NA | NA | 1 | 1 | 1 | 0 | NA | NA | 2 | 0.5 | NA | NA | 1 | 1 | NA | NA | NA | NA | 1 | 1 |
| 13_147850 | 3 | 0 | 4 | 0.25 | 6 | 0.6667 | 6 | 0.3333 | 19 | 0.3684 | 5 | 0.4 | 4 | 0.5 | 4 | 0.25 | 4 | 0.5 | 17 | 0.4118 |
| 13_147851 | 4 | 0.25 | 5 | 0.4 | 3 | 1 | 6 | 0.5 | 18 | 0.5 | 4 | 0.75 | 4 | 0.75 | 4 | 0.5 | 3 | 0.3333 | 15 | 0.6 |
| 13_148110 | 2 | 0 | 1 | 0 | 2 | 0.5 | 2 | 0.5 | 7 | 0.2857 | 2 | 0 | 3 | 0 | 1 | 0 | 1 | 1 | 7 | 0.1429 |
| 13_148110 | 2 | 1 | 1 | 1 | 2 | 1 | 2 | 1 | 7 | 1 | 2 | 1 | 4 | 1 | 2 | 1 | 1 | 1 | 9 | 1 |
| 13_148111 | 2 | 0.5 | 1 | 0 | 3 | 0.6667 | 2 | 1 | 8 | 0.625 | 1 | 1 | 3 | 0.6667 | 1 | 1 | 1 | 1 | 6 | 0.8333 |
| 13_148112 | 2 | 0.5 | 1 | 0 | 3 | 0.6667 | 1 | 1 | 7 | 0.5714 | 2 | 0.5 | 4 | 0.25 | 2 | 1 | 1 | 1 | 9 | 0.5556 |
| 13_148114 | 2 | 0 | 1 | 0 | 3 | 0.6667 | 2 | 0.5 | 8 | 0.375 | 2 | 0 | 4 | 0.25 | 2 | 0 | 1 | 1 | 9 | 0.2222 |
| 13_148117 | 1 | 1 | 1 | 0 | 2 | 0 | 2 | 0 | 6 | 0.1667 | 2 | 0.5 | 4 | 0 | 2 | 0 | 1 | 0 | 9 | 0.1111 |
| 13_148118 | 2 | 0 | 1 | 1 | 3 | 0.3333 | 2 | 0.5 | 8 | 0.375 | 2 | 0 | 4 | 0 | 2 | 0.5 | 1 | 1 | 9 | 0.2222 |
| 13_148118 | 3 | 0.3333 | 1 | 1 | 3 | 0.3333 | 2 | 0.5 | 9 | 0.4444 | 3 | 0 | 4 | 1 | 2 | 0.5 | 1 | 0 | 10 | 0.5 |
| 13_148119 | 3 | 0.6667 | 1 | 1 | 2 | 1 | 2 | 1 | 8 | 0.875 | 3 | 1 | 4 | 1 | 2 | 1 | 1 | 1 | 10 | 1 |
| 13_148120 | 2 | 0 | 3 | 1 | 3 | 0.6667 | 2 | 0.5 | 10 | 0.6 | 3 | 0.3333 | 4 | 0.25 | 2 | 0.5 | 1 | 1 | 10 | 0.4 |
| 13_148120 | 2 | 1 | 3 | 1 | 2 | 1 | 2 | 1 | 9 | 1 | 3 | 1 | 4 | 1 | 2 | 1 | 1 | 1 | 10 | 1 |
| 13_148121 | 3 | 0 | 2 | 0.5 | 1 | 0 | 2 | 1 | 8 | 0.375 | 2 | 0.5 | 3 | 0 | 2 | 1 | 1 | 1 | 8 | 0.5 |
| 13_148121 | 3 | 0.6667 | 3 | 0 | 3 | 0.3333 | 2 | 0 | 11 | 0.2727 | 3 | 0.6667 | 5 | 0 | 2 | 1 | 1 | 0 | 11 | 0.3636 |

| | | | | | | | | | | | | | | | | | | | | |
|---|---|---|---|---|---|---|---|---|---|---|---|---|---|---|---|---|---|---|---|---|
| 13_1481223 | 2 | 1 | 2 | 1 | 3 | 1 | 2 | 1 | 9 | 1 | 2 | 0.5 | 4 | 0.75 | 1 | 1 | 1 | 1 | 8 | 0.75 |
| 13_1481226 | 2 | 1 | 1 | 1 | 1 | 1 | 2 | 1 | 6 | 1 | 3 | 1 | 2 | 1 | 1 | 1 | 1 | 1 | 7 | 1 |
| 13_1481238 | 3 | 0 | 3 | 0.6667 | 3 | 0.6667 | 2 | 0.5 | 11 | 0.4545 | 4 | 0.25 | 4 | 0.25 | 2 | 0 | 1 | 0 | 11 | 0.1818 |
| 13_1481250 | 4 | 0.25 | 3 | 0 | 2 | 0 | 2 | 0 | 11 | 0.0909 | 4 | 0 | 6 | 0 | 1 | 1 | NA | NA | 11 | 0.0909 |
| 13_1481262 | 5 | 0.2 | 2 | 0 | 3 | 0 | 2 | 0 | 12 | 0.0833 | 4 | 0 | 6 | 0.3333 | 2 | 0 | 1 | 0 | 13 | 0.1538 |
| 13_1481265 | 4 | 0.25 | 2 | 0 | 3 | 0 | 2 | 0.5 | 11 | 0.1818 | 4 | 0.25 | 6 | 0.5 | 2 | 0.5 | 1 | 0 | 13 | 0.3846 |
| 13_1563703 | 1 | 1 | NA | NA | NA | NA | NA | NA | 1 | 1 | 1 | 1 | 1 | 1 | NA | NA | 1 | 1 | 3 | 1 |
| 13_1563735 | 1 | 1 | NA | NA | NA | NA | NA | NA | 1 | 1 | 1 | 1 | NA | NA | NA | NA | 1 | 1 | 2 | 1 |
| 13_1563737 | 1 | 1 | 1 | 1 | NA | NA | NA | NA | 2 | 1 | 1 | 1 | NA | NA | NA | NA | 1 | 0 | 2 | 0.5 |
| 13_1563756 | 1 | 1 | NA | NA | NA | NA | NA | NA | 1 | 1 | 1 | 1 | NA | NA | NA | NA | NA | NA | 1 | 1 |
| 13_1563758 | 1 | 1 | NA | NA | NA | NA | NA | NA | 1 | 1 | 1 | 1 | NA | NA | NA | NA | NA | NA | 1 | 1 |
| 13_1590419 | NA | NA | 1 | 0 | 1 | 1 | 1 | 1 | 3 | 0.6667 | 1 | 0 | NA | NA | 1 | 1 | NA | NA | 2 | 0.5 |
| 13_1590450 | NA | NA | 1 | 1 | 1 | 1 | 1 | 1 | 3 | 1 | 1 | 1 | NA | NA | NA | NA | NA | NA | 1 | 1 |
| 13_1594360 | 2 | 1 | 2 | 1 | 1 | 1 | 1 | 1 | 6 | 1 | 3 | 0.6667 | 1 | 1 | 1 | 0 | 3 | 0.3333 | 8 | 0.5 |
| 13_1594371 | 2 | 0.5 | 1 | 1 | 1 | 0 | 1 | 0 | 5 | 0.4 | 3 | 0.3333 | 1 | 0 | NA | NA | 2 | 0 | 6 | 0.1667 |
| 13_1594375 | 2 | 1 | 2 | 1 | NA | NA | 1 | 1 | 5 | 1 | 3 | 1 | 1 | 1 | 1 | 1 | 2 | 0.5 | 7 | 0.8571 |
| 13_1594380 | 2 | 0.5 | 2 | 0.5 | 1 | 1 | 1 | 0 | 6 | 0.5 | 2 | 0.5 | 1 | 0 | 1 | 1 | 3 | 0.3333 | 7 | 0.4286 |
| 13_1594384 | 2 | 1 | 1 | 1 | 1 | 1 | NA | NA | 4 | 1 | 3 | 1 | 1 | 1 | 1 | 1 | 3 | 0.6667 | 8 | 0.875 |
| 13_1594385 | 1 | 0 | 2 | 0.5 | NA | NA | 1 | 0 | 4 | 0.25 | 2 | 1 | 1 | 1 | 1 | 1 | 1 | 0 | 5 | 0.8 |
| 13_1594390 | 2 | 0.5 | 2 | 0 | 1 | 0 | 1 | 0 | 6 | 0.1667 | 3 | 0.3333 | 1 | 0 | 1 | 0 | 2 | 0.5 | 7 | 0.2857 |
| 13_1594397 | NA | NA | 1 | 1 | NA | NA | 1 | 0 | 2 | 0.5 | 3 | 1 | 1 | 1 | NA | NA | 2 | 0.5 | 6 | 0.8333 |
| 13_1594416 | 1 | 1 | 1 | 1 | 1 | 1 | NA | NA | 3 | 1 | NA | NA | 1 | 1 | 1 | 1 | 3 | 0.6667 | 5 | 0.8 |
| 13_1594482 | 1 | 0 | 1 | 1 | 1 | 0 | 3 | 0.3333 | 6 | 0.3333 | 1 | 0 | NA | NA | 1 | 1 | 3 | 0 | 5 | 0.2 |
| 13_1594816 | NA | NA | 1 | 1 | 1 | 0 | NA | NA | 2 | 0.5 | 1 | 0 | NA | NA | NA | NA | 1 | 1 | 2 | 0.5 |
| 13_1594830 | NA | NA | 1 | 1 | 1 | 0 | NA | NA | 2 | 0.5 | 1 | 0 | NA | NA | NA | NA | 1 | 1 | 2 | 0.5 |
| 13_1594894 | NA | NA | 1 | 1 | NA | NA | NA | NA | 1 | 1 | 1 | 0 | NA | NA | NA | NA | 1 | 1 | 2 | 0.5 |
| 13_1594900 | NA | NA | 1 | 1 | NA | NA | NA | NA | 1 | 1 | 1 | 0 | NA | NA | NA | NA | 1 | 1 | 2 | 0.5 |
| 13_1594924 | NA | NA | 1 | 1 | NA | NA | NA | NA | 1 | 1 | 1 | 1 | NA | NA | NA | NA | NA | NA | 1 | 1 |
| 13_1594931 | NA | NA | 1 | 1 | NA | NA | NA | NA | 1 | 1 | 1 | 1 | NA | NA | NA | NA | NA | NA | 1 | 1 |
| 13_1595576 | 2 | 0 | 1 | 1 | NA | NA | NA | NA | 3 | 0.3333 | 1 | 0 | 1 | 1 | 1 | 0 | 1 | 0 | 4 | 0.25 |
| 13_1595604 | 1 | 1 | 1 | 0 | NA | NA | 1 | 1 | 3 | 0.6667 | 1 | 1 | NA | NA | 1 | 1 | 1 | 0 | 3 | 0.6667 |
| 13_1595608 | 1 | 1 | NA | NA | NA | NA | 1 | 1 | 2 | 1 | 1 | 1 | 1 | 1 | NA | NA | 1 | 1 | 3 | 1 |
| 13_1595612 | 1 | 0 | 1 | 1 | NA | NA | 1 | 0 | 3 | 0.3333 | 1 | 1 | 1 | 0 | 1 | 0 | NA | NA | 3 | 0.3333 |
| 13_1595613 | 1 | 1 | 1 | 1 | NA | NA | 1 | 0 | 3 | 0.6667 | 1 | 0 | 1 | 1 | 1 | 1 | NA | NA | 3 | 0.6667 |
| 13_1595631 | 1 | 0 | 2 | 1 | NA | NA | 1 | 1 | 4 | 0.75 | NA | NA | 1 | 1 | 1 | 0 | NA | NA | 2 | 0.5 |
| 13_1598926 | 1 | 1 | NA | NA | NA | NA | 1 | 1 | 2 | 1 | 1 | 1 | 2 | 1 | NA | NA | NA | NA | 3 | 1 |
| 13_1598955 | 2 | 1 | NA | NA | NA | NA | 1 | 1 | 3 | 1 | 1 | 1 | 2 | 1 | NA | NA | 1 | 1 | 4 | 1 |
| 13_1598986 | NA | NA | NA | NA | NA | NA | 1 | 1 | 1 | 1 | NA | NA | 2 | 1 | NA | NA | NA | NA | 2 | 1 |
| 13_1599015 | 1 | 1 | NA | NA | NA | NA | NA | NA | 1 | 1 | NA | NA | 1 | 1 | NA | NA | NA | NA | 1 | 1 |
| 13_1599018 | 1 | 1 | NA | NA | NA | NA | NA | NA | 1 | 1 | NA | NA | 2 | 0.5 | NA | NA | NA | NA | 2 | 0.5 |
| 13_1599046 | 1 | 1 | NA | NA | NA | NA | NA | NA | 1 | 1 | NA | NA | 2 | 0.5 | NA | NA | NA | NA | 2 | 0.5 |
| 13_1611955 | 2 | 0.5 | 1 | 1 | NA | NA | 4 | 1 | 7 | 0.8571 | 1 | 1 | NA | NA | 1 | 1 | NA | NA | 2 | 1 |
| 13_1611970 | 1 | 1 | 1 | 0 | NA | NA | 4 | 0.75 | 6 | 0.6667 | 1 | 1 | NA | NA | 1 | 1 | NA | NA | 2 | 1 |
| 13_1787154 | 1 | 1 | 2 | 0 | 1 | 1 | NA | NA | 4 | 0.5 | 1 | 1 | NA | NA | NA | NA | NA | NA | 1 | 1 |
| 13_1787159 | 1 | 0 | 1 | 0 | 1 | 1 | 1 | 0 | 4 | 0.25 | 2 | 0.5 | NA | NA | NA | NA | NA | NA | 2 | 0.5 |
| 13_1787167 | 1 | 1 | NA | NA | NA | NA | NA | NA | 1 | 1 | 2 | 1 | 1 | 1 | NA | NA | 1 | 1 | 4 | 1 |
| 13_1787171 | NA | NA | 2 | 1 | NA | NA | 1 | 1 | 3 | 1 | 2 | 1 | 1 | 1 | NA | NA | 1 | 1 | 4 | 1 |
| 13_1787174 | 1 | 0 | 2 | 0.5 | 1 | 0 | 1 | 0 | 5 | 0.2 | 3 | 0.6667 | 1 | 0 | NA | NA | 1 | 0 | 5 | 0.4 |
| 13_1787179 | 1 | 1 | 1 | 1 | 1 | 1 | 1 | 0 | 4 | 0.75 | 3 | 1 | 1 | 0 | NA | NA | 1 | 0 | 5 | 0.6 |
| 13_1789103 | 4 | 0.5 | 2 | 0.5 | NA | NA | NA | NA | 6 | 0.5 | 3 | 0.3333 | NA | NA | NA | NA | NA | NA | 3 | 0.3333 |
| 13_1789195 | 2 | 0.5 | NA | NA | NA | NA | NA | NA | 2 | 0.5 | 1 | 1 | NA | NA | NA | NA | NA | NA | 1 | 1 |
| 13_1789228 | 5 | 0.2 | 6 | 0 | 4 | 0.25 | 5 | 0.2 | 20 | 0.15 | 2 | 0 | 6 | 0.1667 | 2 | 0 | 2 | 0 | 12 | 0.0833 |
| 13_1789232 | 6 | 0 | 3 | 0 | 3 | 0.3333 | 4 | 0.25 | 16 | 0.125 | 1 | 0 | 6 | 0.1667 | 2 | 1 | 4 | 0 | 13 | 0.2308 |
| 13_1789256 | 6 | 1 | 6 | 1 | 3 | 1 | 4 | 1 | 19 | 1 | 2 | 1 | 6 | 1 | 1 | 0 | 3 | 1 | 12 | 0.9167 |
| 13_1789260 | 4 | 1 | 6 | 1 | 2 | 1 | 3 | 1 | 15 | 1 | 2 | 1 | 5 | 0.8 | 1 | 0 | 2 | 0.5 | 10 | 0.7 |
| 13_1789269 | 5 | 0.8 | 5 | 1 | 3 | 0.3333 | 3 | 1 | 16 | 0.8125 | 2 | 1 | 6 | 0.5 | 1 | 1 | 3 | 1 | 12 | 0.75 |
| 13_1789272 | 5 | 1 | 6 | 0.6667 | 2 | 0.5 | 2 | 0.5 | 15 | 0.7333 | 2 | 1 | 6 | 0.3333 | 1 | 1 | 3 | 0.6667 | 12 | 0.5833 |
| 13_1789274 | 5 | 0.8 | 6 | 0.8333 | 3 | 1 | 2 | 1 | 16 | 0.875 | 1 | 1 | 5 | 1 | 1 | 0 | 3 | 1 | 10 | 0.9 |
| 13_1789275 | 4 | 1 | 4 | 1 | 3 | 0.6667 | 2 | 1 | 13 | 0.9231 | 1 | 1 | 6 | 0.6667 | NA | NA | 2 | 1 | 9 | 0.7778 |
| 13_1789279 | 3 | 0.3333 | 4 | 0 | 2 | 0 | 1 | 0 | 10 | 0.1 | 1 | 0 | 4 | 0 | 1 | 0 | 2 | 0.5 | 8 | 0.125 |
| 13_1789280 | 3 | 0 | 3 | 0.3333 | 1 | 0 | 3 | 0.6667 | 10 | 0.3 | 2 | 0.5 | 6 | 0 | 1 | 1 | 1 | 0 | 10 | 0.2 |
| 13_1789286 | 4 | 0 | 6 | 0 | 2 | 0 | 2 | 0.5 | 14 | 0.0714 | 1 | 1 | 6 | 0.1667 | 1 | 0 | 2 | 0.5 | 10 | 0.3 |
| 13_1789291 | 4 | 0.75 | 5 | 0.6 | 3 | 1 | 3 | 1 | 15 | 0.8 | 2 | 0 | 6 | 0.5 | 1 | 1 | 2 | 0.5 | 11 | 0.4545 |
| 13_1789295 | 4 | 1 | 4 | 1 | 2 | 1 | 2 | 1 | 12 | 1 | NA | NA | 5 | 1 | 1 | 1 | 2 | 1 | 8 | 1 |
| 13_1789300 | 1 | 0 | 4 | 0.25 | 2 | 0.5 | 1 | 0 | 8 | 0.25 | 2 | 0.5 | 6 | 0.1667 | 1 | 0 | 2 | 0.5 | 11 | 0.2727 |

| | | | | | | | | | | | | | | | | | | | | |
|---|---|---|---|---|---|---|---|---|---|---|---|---|---|---|---|---|---|---|---|---|
| 13_1789307 | 3 | 0.3333 | 5 | 0.2 | 1 | 1 | 2 | 0.5 | 11 | 0.3636 | 2 | 0 | 2 | 0.5 | NA | NA | 2 | 0 | 6 | 0.1667 |
| 13_1789317 | 4 | 0.75 | 5 | 1 | 1 | 1 | 3 | 1 | 13 | 0.9231 | 2 | 1 | 5 | 0.8 | NA | NA | 2 | 1 | 9 | 0.8889 |
| 13_1789318 | 2 | 0 | 5 | 1 | 2 | 1 | 2 | 1 | 11 | 0.8182 | 2 | 0.5 | 4 | 0.75 | 1 | 0 | 2 | 1 | 9 | 0.6667 |
| 13_1789331 | 1 | 1 | 4 | 0.75 | 1 | 1 | NA | NA | 6 | 0.8333 | NA | NA | 4 | 1 | 1 | 1 | NA | NA | 5 | 1 |
| 13_1789335 | 4 | 0.5 | 6 | 0.8333 | 2 | 1 | 2 | 1 | 14 | 0.7857 | 1 | 1 | 5 | 1 | 1 | 1 | 2 | 1 | 9 | 1 |
| 13_1789336 | 3 | 0.6667 | 6 | 0.6667 | 2 | 0 | 3 | 0.6667 | 14 | 0.5714 | NA | NA | 5 | 0.6 | 1 | 1 | 2 | 1 | 8 | 0.75 |
| 13_1789338 | 3 | 0 | 4 | 0 | 2 | 0 | 2 | 0.5 | 11 | 0.0909 | NA | NA | 2 | 0.5 | NA | NA | 2 | 0 | 4 | 0.25 |
| 13_1789406 | 4 | 0.25 | 5 | 0 | 1 | 0 | 3 | 0 | 13 | 0.0769 | 1 | 1 | 6 | 0.1667 | 1 | 0 | 2 | 0 | 10 | 0.2 |
| 13_1789411 | 4 | 0 | 5 | 0.2 | 1 | 0 | 2 | 0 | 12 | 0.0833 | 1 | 0 | 4 | 0.25 | 1 | 0 | 2 | 0 | 8 | 0.125 |
| 13_1789413 | 4 | 0.25 | 5 | 0 | 1 | 0 | 3 | 0 | 13 | 0.0769 | 1 | 0 | 5 | 0.2 | 1 | 0 | 2 | 0 | 9 | 0.1111 |
| 13_1789426 | 3 | 0.6667 | 1 | 1 | 1 | 0 | 2 | 0.5 | 7 | 0.5714 | NA | NA | 5 | 0.4 | 1 | 0 | NA | NA | 6 | 0.3333 |
| 13_1789428 | 2 | 0 | NA | NA | 1 | 1 | 1 | 0 | 4 | 0.25 | NA | NA | 3 | 0.3333 | 1 | 0 | NA | NA | 4 | 0.25 |
| 13_1794720 | 2 | 0.5 | 1 | 0 | NA | NA | 1 | 1 | 4 | 0.5 | 1 | 1 | 1 | 0 | NA | NA | 1 | 1 | 3 | 0.6667 |
| 13_1794725 | 1 | 1 | 1 | 1 | NA | NA | NA | NA | 2 | 1 | 1 | 0 | 4 | 0.5 | NA | NA | 1 | 1 | 6 | 0.5 |
| 13_1797832 | 2 | 0 | NA | NA | NA | NA | 1 | 1 | 3 | 0.3333 | NA | NA | 3 | 0.6667 | NA | NA | NA | NA | 3 | 0.6667 |
| 13_1797839 | 2 | 1 | NA | NA | NA | NA | 1 | 1 | 3 | 1 | 1 | 0 | 2 | 1 | NA | NA | NA | NA | 3 | 0.6667 |
| 13_1797852 | NA | NA | NA | NA | NA | NA | 1 | 1 | 1 | 1 | 1 | 1 | 1 | 0 | NA | NA | NA | NA | 2 | 0.5 |
| 13_1805636 | 6 | 0.6667 | 5 | 0.4 | 2 | 1 | 6 | 0.5 | 19 | 0.5789 | 3 | 0.6667 | 7 | 0.5714 | 4 | 1 | 10 | 0.7 | 24 | 0.7083 |
| 13_1805682 | 4 | 0.5 | 3 | 1 | 1 | 1 | 3 | 0.6667 | 11 | 0.7273 | 2 | 0 | 7 | 0.7143 | 3 | 0.3333 | 3 | 0.6667 | 15 | 0.5333 |
| 13_1805688 | 4 | 0.5 | 3 | 1 | 1 | 1 | 3 | 1 | 11 | 0.8182 | 1 | 1 | 7 | 1 | 3 | 1 | 3 | 1 | 14 | 1 |
| 13_1816353 | 1 | 0 | 5 | 0 | 2 | 0 | 3 | 0.3333 | 11 | 0.0909 | 3 | 0 | 2 | 0 | 4 | 0.25 | 1 | 0 | 10 | 0.1 |
| 13_1816354 | 3 | 0 | 6 | 0.6667 | 2 | 0 | 3 | 0.3333 | 14 | 0.3571 | 3 | 0.6667 | 2 | 0.5 | 3 | 0 | 1 | 1 | 9 | 0.4444 |
| 13_1816387 | 2 | 0 | 5 | 0.4 | 2 | 0 | 3 | 0 | 12 | 0.1667 | 1 | 1 | 2 | 0.5 | 3 | 0 | 2 | 0.5 | 8 | 0.375 |
| 13_1816399 | 3 | 0 | 3 | 0.3333 | 2 | 0 | 2 | 0 | 10 | 0.1 | 3 | 0 | 3 | 0.3333 | 2 | 0 | 2 | 0.5 | 10 | 0.2 |
| 13_1816409 | 2 | 0 | 3 | 0.3333 | 1 | 0 | 1 | 1 | 7 | 0.2857 | 3 | 1 | 3 | 0.3333 | 2 | 1 | 2 | 1 | 10 | 0.8 |
| 13_1816412 | 1 | 1 | 4 | 0 | 2 | 0 | 2 | 0 | 9 | 0.1111 | 3 | 0 | 2 | 0.5 | 2 | 0 | 2 | 0 | 9 | 0.1111 |
| 13_1816415 | 3 | 0 | 4 | 0 | 2 | 0 | 2 | 0.5 | 11 | 0.0909 | 3 | 0.3333 | 3 | 0.3333 | 2 | 0 | 1 | 0 | 9 | 0.2222 |
| 13_1816419 | 3 | 1 | 3 | 1 | 2 | 1 | 2 | 0.5 | 10 | 0.9 | 3 | 1 | 3 | 1 | 2 | 1 | 2 | 1 | 10 | 1 |
| 13_1816429 | 3 | 0 | 2 | 0 | 2 | 0.5 | 2 | 0 | 9 | 0.1111 | 3 | 0.6667 | 3 | 0.3333 | 2 | 0 | 2 | 0 | 10 | 0.3 |
| 13_1905041 | 14 | 0.2857 | 7 | 0.1429 | 16 | 0.1875 | 18 | 0.2222 | 55 | 0.2182 | 9 | 0.1111 | 9 | 0 | 15 | 0.1333 | 15 | 0.0667 | 48 | 0.0833 |
| 13_1905042 | 9 | 0.1111 | 4 | 0.25 | 13 | 0.1538 | 15 | 0.0667 | 41 | 0.122 | 12 | 0.0833 | 8 | 0 | 12 | 0 | 13 | 0.1538 | 45 | 0.0667 |
| 13_1905064 | 15 | 0.8 | 12 | 0.6667 | 20 | 0.75 | 23 | 0.6087 | 70 | 0.7 | 18 | 0.6667 | 12 | 0.9167 | 19 | 0.7895 | 18 | 0.5 | 67 | 0.7015 |
| 13_1905067 | 8 | 0.625 | 10 | 0.8 | 21 | 0.8571 | 17 | 1 | 56 | 0.8571 | 14 | 0.9286 | 11 | 0.9091 | 17 | 0.8824 | 13 | 0.6923 | 55 | 0.8545 |
| 13_1905069 | 12 | 0.4167 | 8 | 0.125 | 15 | 0.2667 | 17 | 0.3529 | 52 | 0.3077 | 16 | 0.3125 | 13 | 0.3077 | 15 | 0.2667 | 15 | 0.0667 | 59 | 0.2373 |
| 13_1905083 | 13 | 0.4615 | 10 | 0.7 | 17 | 0.5882 | 15 | 0.6 | 55 | 0.5818 | 14 | 0.2857 | 12 | 0.25 | 12 | 0.6667 | 12 | 0.5 | 50 | 0.42 |
| 13_1905084 | 15 | 0.3333 | 7 | 0.2857 | 17 | 0.3529 | 17 | 0.2353 | 56 | 0.3036 | 13 | 0.3077 | 12 | 0.4167 | 14 | 0.2143 | 14 | 0.4286 | 53 | 0.3396 |
| 13_1933395 | 16 | 0.0625 | 16 | 0 | 15 | 0 | 13 | 0.1538 | 60 | 0.05 | 20 | 0.1 | 12 | 0.0833 | 6 | 0 | 16 | 0.0625 | 54 | 0.0741 |
| 13_1933399 | 12 | 0.0833 | 15 | 0.1333 | 12 | 0.0833 | 10 | 0 | 49 | 0.0816 | 15 | 0.0667 | 11 | 0 | 6 | 0.1667 | 9 | 0.1111 | 41 | 0.0732 |
| 13_1933404 | 16 | 0.25 | 15 | 0.2 | 14 | 0.2143 | 11 | 0.2727 | 56 | 0.2321 | 18 | 0.3333 | 11 | 0.4545 | 7 | 0.1429 | 16 | 0.25 | 52 | 0.3077 |
| 13_1933411 | 15 | 1 | 17 | 0.9412 | 12 | 0.9167 | 11 | 1 | 55 | 0.9636 | 20 | 0.8 | 9 | 1 | 7 | 0.7143 | 12 | 0.75 | 48 | 0.8125 |
| 13_1933416 | 13 | 0.3077 | 12 | 0.3333 | 10 | 0.4 | 9 | 0.4444 | 44 | 0.3636 | 12 | 0.3333 | 9 | 0.2222 | 3 | 0.3333 | 11 | 0.2727 | 35 | 0.2857 |
| 13_1933426 | 10 | 0.7 | 14 | 0.5714 | 11 | 0.2727 | 11 | 0.4545 | 46 | 0.5 | 15 | 0.4667 | 9 | 0.6667 | 5 | 0.6 | 13 | 0.4615 | 42 | 0.5238 |
| 13_1933431 | 14 | 0.3571 | 18 | 0.3889 | 13 | 0.1538 | 11 | 0.2727 | 56 | 0.3036 | 18 | 0.1111 | 12 | 0.25 | 7 | 0.1429 | 15 | 0.4 | 52 | 0.2308 |
| 13_1933432 | 18 | 0.1667 | 17 | 0.1176 | 14 | 0.2857 | 11 | 0.0909 | 60 | 0.1667 | 18 | 0.0556 | 12 | 0.25 | 7 | 0 | 16 | 0.0625 | 53 | 0.0943 |
| 13_1933446 | 19 | 0.1579 | 22 | 0.2273 | 13 | 0.0769 | 16 | 0.1875 | 70 | 0.1714 | 19 | 0.1053 | 13 | 0 | 7 | 0.1429 | 22 | 0.1364 | 61 | 0.0984 |
| 13_1933446 | 17 | 0 | 21 | 0.0476 | 12 | 0.1667 | 14 | 0.1429 | 64 | 0.0781 | 17 | 0.0588 | 12 | 0 | 6 | 0.1667 | 22 | 0.1364 | 57 | 0.0877 |
| 13_1933452 | 15 | 0.2667 | 19 | 0.2632 | 10 | 0.3 | 15 | 0.5333 | 59 | 0.339 | 18 | 0.3333 | 10 | 0.2 | 8 | 0.375 | 22 | 0.1364 | 58 | 0.2414 |
| 13_1933455 | 20 | 0.15 | 22 | 0.1364 | 13 | 0.0769 | 16 | 0.3125 | 71 | 0.169 | 18 | 0.0556 | 12 | 0.0833 | 7 | 0.1429 | 21 | 0.0952 | 58 | 0.0862 |
| 13_1933485 | 17 | 0.4118 | 21 | 0.2381 | 20 | 0.15 | 16 | 0.3125 | 74 | 0.2703 | 21 | 0.1905 | 13 | 0.0769 | 7 | 0.1429 | 21 | 0.0952 | 62 | 0.129 |
| 13_1933613 | 1 | 0 | 2 | 1 | 2 | 0.5 | 1 | 0 | 6 | 0.5 | 1 | 1 | 2 | 0 | NA | NA | NA | NA | 3 | 0.3333 |
| 13_1933626 | NA | NA | 2 | 0.5 | 2 | 0 | NA | NA | 4 | 0.25 | 1 | 1 | 2 | 0.5 | NA | NA | NA | NA | 3 | 0.6667 |
| 13_1933636 | 1 | 1 | 2 | 1 | 1 | 1 | 1 | 1 | 5 | 1 | 1 | 1 | 2 | 1 | NA | NA | NA | NA | 3 | 1 |
| 13_1933646 | 1 | 1 | 1 | 0 | 2 | 0.5 | 1 | 0 | 5 | 0.4 | 1 | 1 | 2 | 0 | NA | NA | NA | NA | 3 | 0.3333 |
| 13_1933664 | 1 | 0 | 1 | 1 | 2 | 0.5 | 1 | 1 | 5 | 0.6 | 1 | 0 | 2 | 0.5 | NA | NA | NA | NA | 3 | 0.3333 |
| 13_1933677 | 1 | 1 | 1 | 0 | 2 | 0 | 1 | 1 | 5 | 0.4 | 1 | 1 | 2 | 0 | NA | NA | NA | NA | 3 | 0.3333 |
| 13_1933684 | 1 | 0 | 2 | 0.5 | 2 | 0 | 1 | 0 | 6 | 0.1667 | 1 | 0 | 2 | 0.5 | NA | NA | NA | NA | 3 | 0.3333 |
| 13_1933690 | 1 | 1 | 1 | 0 | 2 | 0.5 | 1 | 1 | 5 | 0.6 | 1 | 1 | 1 | 1 | NA | NA | NA | NA | 2 | 1 |
| 13_1933703 | 1 | 1 | 2 | 0.5 | 2 | 1 | 1 | 1 | 6 | 0.8333 | 1 | 1 | 2 | 0.5 | NA | NA | NA | NA | 3 | 0.6667 |
| 13_1933721 | 1 | 0 | 2 | 0 | 2 | 0 | 1 | 1 | 6 | 0.1667 | 1 | 0 | 2 | 0.5 | NA | NA | NA | NA | 3 | 0.3333 |
| 13_1954205 | 6 | 0.3333 | 7 | 0.5714 | 11 | 0.5455 | 13 | 0.6923 | 37 | 0.5676 | 9 | 0.7778 | 13 | 0.7692 | 13 | 0.8462 | 6 | 0.8333 | 41 | 0.8049 |
| 13_1954219 | 4 | 0 | 8 | 0 | 12 | 0.25 | 10 | 0.1 | 34 | 0.1176 | 11 | 0.1818 | 14 | 0.3571 | 10 | 0 | 5 | 0.4 | 40 | 0.225 |
| 13_1956061 | 5 | 0.6 | 5 | 0 | 10 | 0.1 | 12 | 0.4167 | 32 | 0.2812 | 9 | 0.1111 | 10 | 0.2 | 7 | 0.2857 | 6 | 0.1667 | 32 | 0.1875 |
| 13_1977652 | 24 | 0 | 13 | 0.1538 | 29 | 0.1034 | 14 | 0 | 80 | 0.0625 | 14 | 0.0714 | 15 | 0.1333 | 8 | 0 | 10 | 0.1 | 47 | 0.0851 |
| 13_1977675 | 20 | 0.5 | 9 | 0.3333 | 29 | 0.4138 | 14 | 0.5 | 72 | 0.4444 | 10 | 0.2 | 12 | 0.1667 | 5 | 0.2 | 9 | 0.2222 | 36 | 0.1944 |
| 13_1977718 | 21 | 0.2381 | 11 | 0.5455 | 26 | 0.2308 | 14 | 0.2857 | 72 | 0.2917 | 9 | 0.1111 | 15 | 0.4 | 11 | 0.3636 | 6 | 0.1667 | 41 | 0.2927 |

| | | | | | | | | | | | | | | | | | | | | |
|---|---|---|---|---|---|---|---|---|---|---|---|---|---|---|---|---|---|---|---|---|
| 13_1977724 | 24 | 0.0833 | 8 | 0.125 | 25 | 0.24 | 12 | 0.1667 | 69 | 0.1594 | 10 | 0.2 | 15 | 0.2 | 10 | 0.1 | 6 | 0.5 | 41 | 0.2195 |
| 13_1977726 | 25 | 0.56 | 10 | 0.8 | 27 | 0.6296 | 15 | 0.8667 | 77 | 0.6753 | 12 | 0.5 | 16 | 0.625 | 12 | 0.5833 | 7 | 0.7143 | 47 | 0.5957 |
| 13_1977728 | 26 | 0.1154 | 11 | 0.0909 | 26 | 0.0769 | 14 | 0.0714 | 77 | 0.0909 | 12 | 0.0833 | 15 | 0.1333 | 10 | 0.3 | 7 | 0 | 44 | 0.1364 |
| 13_1977731 | 21 | 0.2857 | 7 | 0 | 22 | 0.5 | 10 | 0.3 | 60 | 0.3333 | 9 | 0 | 10 | 0.2 | 5 | 0.8 | 6 | 0.3333 | 30 | 0.2667 |
| 13_1977732 | 24 | 0.1667 | 10 | 0.1 | 26 | 0.1154 | 11 | 0.0909 | 71 | 0.1268 | 11 | 0.2727 | 15 | 0.1333 | 7 | 0.2857 | 7 | 0.1429 | 40 | 0.2 |
| 13_1977739 | 24 | 0.75 | 11 | 0.3636 | 26 | 0.4231 | 14 | 0.5714 | 75 | 0.5467 | 10 | 0.4 | 13 | 0.6154 | 11 | 0.1818 | 7 | 0.5714 | 41 | 0.439 |
| 13_1977745 | 19 | 0.1579 | 10 | 0.1 | 22 | 0.1364 | 10 | 0 | 61 | 0.1148 | 10 | 0.1 | 10 | 0.2 | 8 | 0.125 | 6 | 0 | 34 | 0.1176 |
| 13_1977871 | 19 | 0.6842 | 11 | 0.3636 | 21 | 0.619 | 12 | 0.5833 | 63 | 0.5873 | 10 | 0.6 | 11 | 0.2727 | 6 | 0.5 | 6 | 0 | 33 | 0.3636 |
| 13_1980368 | 15 | 0 | 11 | 0.0909 | 20 | 0.1 | 6 | 0 | 52 | 0.0577 | 11 | 0.0909 | 13 | 0.2308 | 8 | 0.125 | 5 | 0 | 37 | 0.1351 |
| 13_2078944 | 19 | 0.4737 | 20 | 0.2 | 14 | 0.0714 | 21 | 0.1429 | 74 | 0.2297 | 9 | 0.2222 | 24 | 0.125 | 19 | 0.2632 | 19 | 0.1579 | 71 | 0.1831 |
| 13_2078945 | 16 | 0.625 | 17 | 0.7059 | 14 | 0.5714 | 21 | 0.6667 | 68 | 0.6471 | 10 | 0.2 | 26 | 0.6154 | 18 | 0.5556 | 14 | 0.7143 | 68 | 0.5588 |
| 13_2078946 | 16 | 0.1875 | 12 | 0.0833 | 10 | 0.2 | 16 | 0.0625 | 54 | 0.1296 | 5 | 0 | 16 | 0.0625 | 16 | 0.1875 | 8 | 0.25 | 45 | 0.1333 |
| 13_2078947 | 15 | 0.6667 | 15 | 0.7333 | 14 | 0.8571 | 21 | 0.5714 | 65 | 0.6923 | 9 | 0.6667 | 18 | 0.6667 | 15 | 0.6 | 15 | 0.9333 | 57 | 0.7193 |
| 13_2137640 | 5 | 0.8 | 5 | 0.2 | 3 | 0.3333 | 3 | 0.3333 | 16 | 0.4375 | 5 | 0 | 8 | 0.25 | 1 | 1 | 6 | 0 | 20 | 0.15 |
| 13_2137642 | 4 | 0 | 5 | 0 | 4 | 0.5 | 4 | 0 | 17 | 0.1176 | 5 | 0 | 7 | 0.2857 | 1 | 1 | 5 | 0.2 | 18 | 0.2222 |
| 13_2137652 | 5 | 0.2 | 6 | 0 | 4 | 0.25 | 3 | 0.3333 | 18 | 0.1667 | 6 | 0.1667 | 9 | 0.4444 | 1 | 0 | 4 | 0.5 | 20 | 0.35 |
| 13_2137654 | 3 | 0 | 5 | 0.4 | 4 | 0 | 2 | 0 | 14 | 0.1429 | 5 | 0 | 9 | 0 | 1 | 1 | 6 | 0.1667 | 21 | 0.0952 |
| 13_2137657 | 4 | 1 | 6 | 0.6667 | 4 | 0.5 | 3 | 0.6667 | 17 | 0.7059 | 4 | 1 | 7 | 0.8571 | 1 | 1 | 3 | 1 | 15 | 0.9333 |
| 13_2154786 | 1 | 1 | 1 | 1 | 1 | 0 | 1 | 1 | 4 | 0.75 | NA | NA | NA | NA | 1 | 1 | 1 | 1 | 2 | 1 |
| 13_2154805 | 1 | 0 | 1 | 1 | 1 | 0 | 1 | 1 | 4 | 0.5 | NA | NA | NA | NA | 1 | 0 | 1 | 1 | 2 | 0.5 |
| 13_2155293 | NA | NA | 1 | 1 | 1 | 0 | 1 | 1 | 3 | 0.6667 | NA | NA | NA | NA | 1 | 1 | 1 | 1 | 2 | 1 |
| 13_2155310 | 2 | 1 | 1 | 1 | 1 | 1 | 1 | 1 | 5 | 1 | NA | NA | NA | NA | 1 | 1 | 1 | 1 | 2 | 1 |
| 13_2155321 | 2 | 1 | NA | NA | 1 | 1 | 1 | 1 | 4 | 1 | NA | NA | NA | NA | 1 | 1 | 1 | 1 | 2 | 1 |
| 13_2155331 | 2 | 0.5 | 1 | 0 | 1 | 0 | 1 | 0 | 5 | 0.2 | NA | NA | NA | NA | 1 | 1 | 1 | 1 | 2 | 1 |
| 13_2155359 | NA | NA | 1 | 0 | 1 | 1 | 1 | 1 | 3 | 0.6667 | 1 | 1 | NA | NA | 1 | 1 | NA | NA | 2 | 1 |
| 13_2155362 | 2 | 1 | 1 | 1 | 1 | 1 | 1 | 1 | 5 | 1 | 1 | 1 | NA | NA | 1 | 1 | 1 | 1 | 3 | 1 |
| 13_2155366 | 3 | 1 | NA | NA | 1 | 0 | 1 | 1 | 5 | 0.8 | 1 | 1 | NA | NA | 1 | 1 | 1 | 0 | 3 | 0.6667 |
| 13_2155367 | 1 | 1 | 1 | 1 | 1 | 0 | 1 | 0 | 4 | 0.5 | 1 | 1 | NA | NA | NA | NA | 1 | 0 | 2 | 0.5 |
| 13_2155385 | 3 | 0.6667 | 1 | 1 | NA | NA | NA | NA | 4 | 0.75 | 1 | 1 | NA | NA | 1 | 0 | 1 | 1 | 3 | 0.6667 |
| 13_2155395 | 2 | 0 | 1 | 1 | 1 | 0 | 1 | 0 | 5 | 0.2 | 1 | 1 | NA | NA | 1 | 0 | 1 | 0 | 3 | 0.3333 |
| 13_2157160 | 3 | 0 | 4 | 0.25 | 3 | 0 | 6 | 0.1667 | 16 | 0.125 | 5 | 0.4 | 5 | 0 | NA | NA | 4 | 0.5 | 14 | 0.2857 |
| 13_2157165 | 2 | 0 | 4 | 0.5 | 3 | 0.6667 | 5 | 0.4 | 14 | 0.4286 | 4 | 0.75 | 4 | 0.5 | NA | NA | 5 | 0.4 | 13 | 0.5385 |
| 13_2157177 | 1 | 0 | 4 | 0.25 | 3 | 0 | 4 | 0.25 | 12 | 0.1667 | 5 | 0.2 | 5 | 0.8 | NA | NA | 5 | 0.4 | 15 | 0.4667 |
| 13_2157179 | 3 | 0.3333 | 4 | 0.5 | 1 | 1 | 6 | 0.1667 | 14 | 0.3571 | 5 | 0.2 | 4 | 0.25 | NA | NA | 3 | 0.6667 | 12 | 0.3333 |
| 13_2157187 | 3 | 0.3333 | 3 | 0.6667 | 3 | 0.3333 | 6 | 1 | 15 | 0.6667 | 4 | 0.5 | 4 | 0.75 | NA | NA | 3 | 0.3333 | 11 | 0.5455 |
| 13_2157194 | 2 | 0.5 | 3 | 0.3333 | 2 | 1 | 5 | 0.6 | 12 | 0.5833 | 4 | 0 | 4 | 0.5 | NA | NA | 5 | 0.2 | 13 | 0.2308 |
| 13_2157197 | 2 | 1 | 2 | 1 | 1 | 1 | 4 | 1 | 9 | 1 | 4 | 0.75 | 1 | 1 | NA | NA | 5 | 1 | 10 | 0.9 |
| 13_2157203 | 3 | 0 | NA | NA | 3 | 0.3333 | 5 | 0 | 11 | 0.0909 | 2 | 0 | 3 | 0 | NA | NA | 5 | 0.2 | 10 | 0.1 |
| 13_2157214 | 1 | 0 | 1 | 0 | 2 | 1 | 3 | 0.3333 | 7 | 0.4286 | 2 | 0.5 | 3 | 0 | NA | NA | 4 | 0.25 | 9 | 0.2222 |
| 13_2157226 | 2 | 0 | 1 | 0 | 3 | 0.3333 | 5 | 0.2 | 11 | 0.1818 | 3 | 0.6667 | 4 | 0.25 | NA | NA | 5 | 0.2 | 12 | 0.3333 |
| 13_2157229 | 3 | 0.3333 | 1 | 0 | 3 | 0.3333 | 6 | 0.6667 | 13 | 0.4615 | 3 | 0.3333 | 4 | 0.5 | NA | NA | 5 | 0.4 | 12 | 0.4167 |
| 13_2157230 | 3 | 0 | 1 | 0 | 3 | 0.3333 | 4 | 0.25 | 11 | 0.1818 | 3 | 0 | 4 | 0 | NA | NA | 5 | 0.2 | 12 | 0.0833 |
| 13_2157242 | 2 | 0.5 | 1 | 1 | 3 | 0.6667 | 6 | 1 | 12 | 0.8333 | 3 | 1 | 2 | 0.5 | NA | NA | 5 | 0.8 | 10 | 0.8 |
| 13_2157270 | 2 | 0 | 1 | 0 | 3 | 0.3333 | 5 | 0.2 | 11 | 0.1818 | 3 | 0 | 3 | 0 | NA | NA | 4 | 0.25 | 10 | 0.1 |
| 13_2157284 | 2 | 0 | 1 | 0 | 2 | 0 | 6 | 0.1667 | 11 | 0.0909 | 3 | 0.3333 | 3 | 0 | NA | NA | 5 | 0 | 11 | 0.0909 |
| 13_2158063 | 2 | 0 | 1 | 0 | 1 | 0 | 4 | 0.25 | 8 | 0.125 | NA | NA | 1 | 1 | NA | NA | 3 | 0 | 4 | 0.25 |
| 13_2158065 | 2 | 0.5 | 1 | 1 | 1 | 1 | 4 | 0.25 | 8 | 0.5 | NA | NA | NA | NA | NA | NA | 2 | 0.5 | 2 | 0.5 |
| 13_2160690 | 1 | 1 | 1 | 1 | NA | NA | 1 | 1 | 3 | 1 | NA | NA | 1 | 1 | NA | NA | 1 | 0 | 2 | 0.5 |
| 13_2160747 | NA | NA | 1 | 1 | NA | NA | 1 | 1 | 2 | 1 | NA | NA | 1 | 1 | NA | NA | 1 | 1 | 2 | 1 |
| 13_2162690 | 11 | 0.0909 | 3 | 0 | 4 | 0 | 8 | 0.25 | 26 | 0.1154 | 2 | 0 | 15 | 0.2 | 8 | 0.125 | 6 | 0.1667 | 31 | 0.1613 |
| 13_2162698 | 7 | 0.4286 | 4 | 0 | 3 | 0.6667 | 5 | 0.2 | 19 | 0.3158 | 1 | 0 | 12 | 0.1667 | 10 | 0.1 | 4 | 0.25 | 27 | 0.1481 |
| 13_2162701 | 11 | 0.3636 | 3 | 0 | 3 | 0.3333 | 9 | 0.3333 | 26 | 0.3077 | 2 | 0 | 13 | 0.2308 | 8 | 0.25 | 6 | 0.3333 | 29 | 0.2414 |
| 13_2162734 | 8 | 0.125 | 1 | 0 | NA | NA | 6 | 0 | 15 | 0.0667 | NA | NA | 11 | 0.1818 | 5 | 0.2 | 3 | 0.3333 | 19 | 0.2105 |
| 13_2162768 | 6 | 0.3333 | 1 | 1 | 2 | 0 | 6 | 0.3333 | 15 | 0.3333 | 1 | 0 | 10 | 0.3 | 6 | 0.1667 | 3 | 0.6667 | 20 | 0.3 |
| 13_2162783 | 7 | 0.2857 | 1 | 0 | 2 | 1 | 5 | 0.2 | 15 | 0.3333 | 1 | 0 | 11 | 0.1818 | 7 | 0.1429 | 2 | 0.5 | 21 | 0.1905 |
| 13_2162790 | 6 | 0.1667 | 1 | 0 | 1 | 0 | 1 | 0 | 9 | 0.1111 | NA | NA | 9 | 0.1111 | 5 | 0 | 2 | 0.5 | 16 | 0.125 |
| 13_2162792 | 6 | 0.6667 | 1 | 0 | 2 | 0.5 | 3 | 0.6667 | 12 | 0.5833 | 1 | 0 | 11 | 0.1818 | 5 | 0.4 | 2 | 0.5 | 19 | 0.2632 |
| 13_2162852 | 4 | 0.75 | 1 | 1 | 2 | 1 | 2 | 1 | 9 | 0.8889 | 1 | 1 | 8 | 0.875 | 3 | 1 | 2 | 1 | 14 | 0.9286 |
| 13_2162853 | 5 | 0.2 | 1 | 0 | 2 | 0 | 1 | 1 | 9 | 0.2222 | 1 | 0 | 9 | 0.2222 | 3 | 0 | 1 | 0 | 14 | 0.1429 |
| 13_2162866 | 4 | 0.75 | 1 | 1 | 2 | 0.5 | 2 | 1 | 9 | 0.7778 | 1 | 1 | 9 | 0.6667 | 1 | 1 | 2 | 1 | 13 | 0.7692 |
| 13_2162895 | 2 | 1 | 1 | 1 | 1 | 1 | 3 | 1 | 7 | 1 | 1 | 1 | 9 | 0.8889 | 3 | 1 | 1 | 1 | 14 | 0.9286 |
| 13_2162903 | 2 | 1 | NA | NA | NA | NA | 3 | 1 | 5 | 1 | NA | NA | 6 | 1 | 2 | 1 | 1 | 1 | 9 | 1 |
| 13_2162911 | 2 | 1 | 1 | 1 | NA | NA | 3 | 0.6667 | 6 | 0.8333 | 1 | 1 | 4 | 1 | 2 | 0.5 | 1 | 1 | 8 | 0.875 |
| 13_2162912 | 4 | 0 | 1 | 1 | 2 | 0.5 | 3 | 1 | 10 | 0.5 | 1 | 1 | 9 | 0.8889 | 3 | 0.6667 | 1 | 0 | 14 | 0.7857 |
| 13_2162927 | 2 | 0.5 | 1 | 0 | 1 | 0 | 2 | 0.5 | 6 | 0.3333 | 1 | 0 | 7 | 0 | 3 | 0.3333 | NA | NA | 11 | 0.0909 |

| | | | | | | | | | | | | | | | | | | | | |
|---|---|---|---|---|---|---|---|---|---|---|---|---|---|---|---|---|---|---|---|---|
| 13_2162935 | 2 | 0 | 1 | 1 | 1 | 1 | 3 | 1 | 7 | 0.7143 | 1 | 1 | 8 | 0.875 | 3 | 0.6667 | NA | NA | 12 | 0.8333 |
| 13_2162941 | 2 | 0 | 1 | 0 | 1 | 0 | 3 | 0.3333 | 7 | 0.1429 | 1 | 0 | 7 | 0 | 3 | 0 | 1 | 1 | 12 | 0.0833 |
| 13_2162950 | 3 | 0.3333 | 1 | 1 | 1 | 1 | 3 | 1 | 8 | 0.75 | 1 | 0 | 7 | 0.5714 | 3 | 0.6667 | 1 | 1 | 12 | 0.5833 |
| 13_2162970 | 3 | 0 | 1 | 0 | NA | NA | 3 | 0.3333 | 7 | 0.1429 | 1 | 0 | 6 | 0 | 2 | 0.5 | 1 | 1 | 10 | 0.2 |
| 13_2162979 | 3 | 0.6667 | 1 | 0 | 1 | 1 | 3 | 1 | 8 | 0.75 | 1 | 1 | 7 | 0.5714 | 2 | 0.5 | 1 | 1 | 11 | 0.6364 |
| 13_2162980 | 3 | 1 | 1 | 1 | 1 | 1 | 3 | 1 | 8 | 1 | 1 | 1 | 6 | 0.8333 | 2 | 0.5 | 1 | 1 | 10 | 0.8 |
| 13_2162981 | 3 | 1 | 1 | 1 | 1 | 1 | 2 | 1 | 7 | 1 | 1 | 1 | 7 | 1 | 2 | 1 | 1 | 1 | 11 | 1 |
| 13_2162985 | NA | NA | NA | NA | 1 | 1 | NA | NA | 1 | 1 | 1 | 1 | 2 | 0.5 | NA | NA | NA | NA | 3 | 0.6667 |
| 13_2162991 | 3 | 0.3333 | 1 | 0 | 1 | 0 | 3 | 0 | 8 | 0.125 | 1 | 0 | 8 | 0.375 | 2 | 0 | 1 | 0 | 12 | 0.25 |
| 13_2162995 | 3 | 0.3333 | 1 | 0 | 1 | 1 | 2 | 0 | 7 | 0.2857 | 1 | 0 | 8 | 0.125 | 2 | 0 | 1 | 0 | 12 | 0.0833 |
| 13_2162995 | 3 | 0.3333 | 1 | 0 | 1 | 0 | 1 | 0 | 6 | 0.1667 | 1 | 0 | 6 | 0.1667 | 2 | 0 | 1 | 1 | 10 | 0.2 |
| 13_2163240 | 1 | 1 | NA | NA | 1 | 1 | 1 | 0 | 3 | 0.6667 | 1 | 0 | 3 | 0 | NA | NA | 1 | 1 | 5 | 0.2 |
| 13_2166884 | 1 | 1 | NA | NA | NA | NA | NA | NA | 1 | 1 | NA | NA | NA | NA | 2 | 0.5 | NA | NA | 2 | 0.5 |
| 13_2166905 | 1 | 1 | NA | NA | NA | NA | NA | NA | 1 | 1 | NA | NA | NA | NA | 2 | 0.5 | NA | NA | 2 | 0.5 |
| 13_2166906 | 1 | 1 | NA | NA | NA | NA | NA | NA | 1 | 1 | NA | NA | NA | NA | 1 | 1 | NA | NA | 1 | 1 |
| 13_2166914 | 1 | 1 | NA | NA | NA | NA | NA | NA | 1 | 1 | NA | NA | NA | NA | 1 | 1 | NA | NA | 1 | 1 |
| 13_2166932 | 1 | 1 | NA | NA | NA | NA | NA | NA | 1 | 1 | NA | NA | NA | NA | 2 | 1 | NA | NA | 2 | 1 |
| 13_2166938 | 1 | 1 | NA | NA | NA | NA | NA | NA | 1 | 1 | NA | NA | NA | NA | 2 | 0.5 | NA | NA | 2 | 0.5 |
| 13_2168787 | NA | NA | 1 | 1 | NA | NA | NA | NA | 1 | 1 | NA | NA | NA | NA | 1 | 1 | NA | NA | 1 | 1 |
| 13_2168856 | NA | NA | 1 | 1 | NA | NA | NA | NA | 1 | 1 | NA | NA | NA | NA | 1 | 1 | NA | NA | 1 | 1 |
| 13_2168870 | NA | NA | 1 | 1 | NA | NA | NA | NA | 1 | 1 | NA | NA | NA | NA | 1 | 1 | NA | NA | 1 | 1 |
| 13_2169963 | NA | NA | 5 | 0.2 | 2 | 0.5 | 1 | 0 | 8 | 0.25 | 2 | 0.5 | NA | NA | 1 | 0 | NA | NA | 3 | 0.3333 |
| 13_2169970 | NA | NA | 2 | 0.5 | 2 | 0.5 | NA | NA | 4 | 0.5 | 3 | 0.3333 | 1 | 0 | 1 | 1 | 1 | 0 | 6 | 0.3333 |
| 13_2169980 | NA | NA | 4 | 1 | 2 | 0.5 | 1 | 1 | 7 | 0.8571 | 3 | 0.6667 | 1 | 1 | 1 | 1 | 1 | 0 | 6 | 0.6667 |
| 13_2169998 | NA | NA | 4 | 0.25 | 3 | 0 | 1 | 0 | 8 | 0.125 | 3 | 0.6667 | NA | NA | 1 | 0 | 1 | 0 | 5 | 0.4 |
| 13_2170014 | NA | NA | 3 | 0.3333 | 1 | 1 | 1 | 0 | 5 | 0.4 | 2 | 1 | NA | NA | NA | NA | 1 | 0 | 3 | 0.6667 |
| 13_2192435 | NA | NA | 1 | 1 | NA | NA | NA | NA | 1 | 1 | NA | NA | NA | NA | NA | NA | 1 | 1 | 1 | 1 |
| 13_2192438 | NA | NA | 1 | 1 | NA | NA | NA | NA | 1 | 1 | NA | NA | NA | NA | NA | NA | 1 | 1 | 1 | 1 |
| 13_2192440 | NA | NA | 1 | 1 | NA | NA | NA | NA | 1 | 1 | NA | NA | NA | NA | NA | NA | 1 | 1 | 1 | 1 |
| 13_2192446 | NA | NA | 1 | 1 | NA | NA | NA | NA | 1 | 1 | NA | NA | NA | NA | NA | NA | 1 | 1 | 1 | 1 |
| 13_2192450 | NA | NA | 1 | 1 | NA | NA | NA | NA | 1 | 1 | NA | NA | NA | NA | NA | NA | 1 | 1 | 1 | 1 |
| 13_2193154 | 2 | 1 | 1 | 1 | NA | NA | NA | NA | 3 | 1 | NA | NA | NA | NA | 1 | 1 | 1 | 0 | 2 | 0.5 |
| 13_2193158 | 2 | 1 | NA | NA | NA | NA | NA | NA | 2 | 1 | NA | NA | NA | NA | 1 | 1 | 1 | 1 | 2 | 1 |
| 13_2193275 | 2 | 1 | NA | NA | NA | NA | NA | NA | 2 | 1 | NA | NA | NA | NA | NA | NA | 1 | 1 | 1 | 1 |
| 13_2222508 | NA | NA | 1 | 0 | NA | NA | 2 | 0.5 | 3 | 0.3333 | 2 | 0 | 1 | 1 | NA | NA | NA | NA | 3 | 0.3333 |
| 13_2280381 | 1 | 1 | 1 | 1 | NA | NA | NA | NA | 2 | 1 | NA | NA | NA | NA | 1 | 1 | NA | NA | 1 | 1 |
| 13_2280396 | 1 | 1 | NA | NA | NA | NA | NA | NA | 1 | 1 | NA | NA | NA | NA | 1 | 1 | NA | NA | 1 | 1 |
| 13_2280400 | 1 | 1 | NA | NA | NA | NA | NA | NA | 1 | 1 | NA | NA | NA | NA | 1 | 1 | NA | NA | 1 | 1 |
| 13_2353756 | 3 | 0.3333 | 3 | 0 | 2 | 0 | 2 | 0 | 10 | 0.1 | 1 | 0 | 5 | 0 | 2 | 0.5 | 5 | 0 | 13 | 0.0769 |
| 13_2353761 | 4 | 0.25 | 3 | 0 | 1 | 0 | 2 | 0.5 | 10 | 0.2 | NA | NA | 3 | 0.3333 | 1 | 1 | 5 | 0.2 | 9 | 0.3333 |
| 13_2353951 | 2 | 0.5 | 2 | 0 | 1 | 0 | 1 | 0 | 6 | 0.1667 | NA | NA | 3 | 0.3333 | 5 | 0 | 4 | 0.25 | 12 | 0.1667 |
| 13_2353987 | 1 | 0 | 2 | 0.5 | 1 | 1 | NA | NA | 4 | 0.5 | 1 | 0 | NA | NA | 3 | 0.3333 | 5 | 0.4 | 9 | 0.3333 |
| 13_2354005 | 1 | 1 | 2 | 0.5 | 1 | 0 | 1 | 0 | 5 | 0.4 | 1 | 0 | 3 | 0 | 4 | 0 | 4 | 0.5 | 12 | 0.1667 |
| 13_2354006 | 1 | 1 | 2 | 1 | 1 | 0 | NA | NA | 4 | 0.75 | NA | NA | 2 | 1 | 4 | 1 | 4 | 1 | 10 | 1 |
| 13_2354014 | 1 | 1 | 3 | 0 | 2 | 0 | 1 | 1 | 7 | 0.2857 | 1 | 0 | 3 | 0 | 5 | 0.4 | 5 | 0.4 | 14 | 0.2857 |
| 13_2354030 | 1 | 1 | 3 | 0.3333 | 2 | 0.5 | 1 | 0 | 7 | 0.4286 | 1 | 1 | 3 | 0.3333 | 2 | 1 | 3 | 1 | 9 | 0.7778 |
| 13_2354033 | 1 | 1 | 3 | 1 | 2 | 0.5 | 1 | 1 | 7 | 0.8571 | 1 | 0 | 3 | 1 | 5 | 0.4 | 5 | 0.8 | 14 | 0.6429 |
| 13_2354037 | 1 | 1 | 3 | 1 | 2 | 0.5 | 1 | 1 | 7 | 0.8571 | 1 | 1 | 3 | 1 | 5 | 0.8 | 5 | 1 | 14 | 0.9286 |
| 13_2354041 | 1 | 1 | 2 | 0.5 | 2 | 0.5 | NA | NA | 5 | 0.6 | NA | NA | 3 | 0.6667 | 3 | 0.6667 | 3 | 0.3333 | 9 | 0.5556 |
| 13_2354045 | 1 | 1 | 3 | 1 | 2 | 0.5 | NA | NA | 6 | 0.8333 | 1 | 1 | 3 | 1 | 5 | 1 | 4 | 0.75 | 13 | 0.9231 |
| 13_2354060 | 1 | 1 | 2 | 0.5 | NA | NA | NA | NA | 3 | 0.6667 | 1 | 1 | 2 | 0 | 4 | 0.75 | 4 | 1 | 11 | 0.7273 |
| 13_2354063 | 1 | 0 | 3 | 0.6667 | 2 | 0.5 | 1 | 0 | 7 | 0.4286 | 1 | 0 | 3 | 0 | 3 | 0 | 5 | 0.4 | 12 | 0.1667 |
| 13_2354070 | 1 | 1 | 3 | 1 | 2 | 1 | 1 | 1 | 7 | 1 | 1 | 1 | 3 | 1 | 5 | 1 | 4 | 1 | 13 | 1 |
| 13_2354072 | 1 | 0 | 2 | 0.5 | 2 | 0.5 | 1 | 1 | 6 | 0.5 | 1 | 1 | 3 | 1 | 5 | 1 | 4 | 1 | 13 | 1 |
| 13_2354079 | 1 | 1 | 3 | 1 | 2 | 1 | 1 | 1 | 7 | 1 | 1 | 1 | 3 | 1 | 5 | 0.8 | 3 | 1 | 12 | 0.9167 |
| 13_2354095 | 1 | 0 | 2 | 0.5 | 2 | 0.5 | 1 | 1 | 6 | 0.5 | 1 | 0 | 3 | 0.6667 | 5 | 0.4 | 5 | 0.2 | 14 | 0.3571 |
| 13_2354110 | 1 | 1 | 3 | 1 | 2 | 1 | NA | NA | 6 | 1 | 1 | 1 | 3 | 1 | 4 | 1 | 5 | 1 | 13 | 1 |
| 13_2354125 | 1 | 0 | 3 | 0.3333 | 2 | 0 | NA | NA | 6 | 0.1667 | NA | NA | 2 | 0 | 4 | 0.25 | 5 | 0.4 | 11 | 0.2727 |
| 13_2354135 | 1 | 0 | 3 | 0.3333 | 2 | 0.5 | NA | NA | 6 | 0.3333 | NA | NA | 1 | 0 | 4 | 0.75 | 5 | 0.8 | 10 | 0.7 |
| 13_2354136 | 1 | 0 | 3 | 0.6667 | 2 | 1 | NA | NA | 6 | 0.6667 | 1 | 1 | 1 | 1 | 4 | 1 | 4 | 1 | 10 | 1 |
| 13_2354152 | NA | NA | 3 | 0 | 2 | 0.5 | NA | NA | 5 | 0.2 | NA | NA | 1 | 0 | 5 | 0.4 | 4 | 0.25 | 10 | 0.3 |
| 13_2354155 | NA | NA | 2 | 0.5 | 1 | 1 | NA | NA | 3 | 0.6667 | 1 | 0 | 1 | 0 | 1 | 0 | 5 | 0.8 | 8 | 0.5 |
| 13_2354167 | 1 | 0 | 1 | 1 | 2 | 1 | NA | NA | 4 | 0.75 | 1 | 1 | NA | NA | 3 | 0.6667 | 4 | 0.75 | 8 | 0.75 |
| 13_2354176 | 1 | 0 | 3 | 0.3333 | 2 | 0.5 | NA | NA | 6 | 0.3333 | 1 | 0 | 2 | 0.5 | 5 | 0.2 | 4 | 0.5 | 12 | 0.3333 |
| 13_2360303 | NA | NA | NA | NA | 1 | 1 | NA | NA | 1 | 1 | 1 | 1 | NA | NA | NA | NA | NA | NA | 1 | 1 |

| | | | | | | | | | | | | | | | | | | | | |
|---|---|---|---|---|---|---|---|---|---|---|---|---|---|---|---|---|---|---|---|---|
| 13_236032 | NA | NA | NA | NA | 1 | 1 | NA | NA | 1 | 1 | 1 | 1 | NA | NA | NA | NA | NA | NA | 1 | 1 |
| 13_236058 | 4 | 0.75 | 1 | 1 | 1 | 1 | 1 | 1 | 7 | 0.8571 | 4 | 0.75 | 6 | 1 | 2 | 0.5 | 3 | 1 | 15 | 0.8667 |
| 13_236058 | 4 | 1 | 1 | 0 | 3 | 0.3333 | NA | NA | 8 | 0.625 | 4 | 0.5 | 5 | 0.4 | 1 | 0 | 3 | 0 | 13 | 0.3077 |
| 13_236058 | 6 | 1 | 1 | 1 | 3 | 0.6667 | 1 | 1 | 11 | 0.9091 | 4 | 1 | 5 | 0.8 | 3 | 1 | 3 | 1 | 15 | 0.9333 |
| 13_236059 | 5 | 0.2 | 1 | 1 | 2 | 0 | 1 | 0 | 9 | 0.2222 | 4 | 0.5 | 4 | 0.25 | 1 | 0 | 3 | 0.3333 | 12 | 0.3333 |
| 13_236059 | 2 | 0.5 | NA | NA | 2 | 0 | NA | NA | 4 | 0.25 | 1 | 0 | 3 | 0.3333 | 2 | 0.5 | 1 | 0 | 7 | 0.2857 |
| 13_236061 | 6 | 1 | 1 | 1 | 3 | 1 | 1 | 1 | 11 | 1 | 4 | 1 | 3 | 1 | 3 | 1 | 3 | 1 | 13 | 1 |
| 13_236061 | 5 | 0 | NA | NA | 3 | 0.3333 | 1 | 0 | 9 | 0.1111 | 2 | 0 | 2 | 0 | 2 | 0 | 2 | 0.5 | 8 | 0.125 |
| 13_236062 | 5 | 0.2 | 1 | 0 | 3 | 0.3333 | 1 | 0 | 10 | 0.2 | 4 | 0.25 | 3 | 0 | 3 | 0 | 1 | 0 | 11 | 0.0909 |
| 13_236063 | 6 | 1 | 1 | 1 | 3 | 1 | NA | NA | 10 | 1 | 4 | 1 | 3 | 0.6667 | 3 | 1 | 1 | 1 | 11 | 0.9091 |
| 13_236063 | 5 | 0.2 | 1 | 0 | 3 | 0.3333 | 1 | 1 | 10 | 0.3 | 4 | 0.25 | 4 | 0.5 | 2 | 1 | 1 | 1 | 11 | 0.5455 |
| 13_236064 | 5 | 1 | 1 | 1 | 2 | 1 | 1 | 1 | 9 | 1 | NA | NA | 4 | 0.75 | 3 | 0.6667 | 2 | 0.5 | 9 | 0.6667 |
| 13_236064 | NA | NA | NA | NA | 1 | 1 | NA | NA | 1 | 1 | 1 | 1 | NA | NA | NA | NA | NA | NA | 1 | 1 |
| 13_237692 | NA | NA | NA | NA | NA | NA | 2 | 0.5 | 2 | 0.5 | 1 | 1 | 2 | 0.5 | 1 | 0 | 1 | 0 | 5 | 0.4 |
| 13_237693 | NA | NA | NA | NA | NA | NA | 2 | 0.5 | 2 | 0.5 | 1 | 0 | 1 | 1 | NA | NA | NA | NA | 2 | 0.5 |
| 13_237694 | NA | NA | NA | NA | NA | NA | 2 | 1 | 2 | 1 | 1 | 0 | NA | NA | 1 | 1 | NA | NA | 2 | 0.5 |
| 13_237697 | NA | NA | NA | NA | NA | NA | 2 | 1 | 2 | 1 | 1 | 1 | NA | NA | 1 | 1 | 1 | 1 | 3 | 1 |
| 13_238684 | 24 | 0.625 | 14 | 0.5 | 17 | 0.5294 | 15 | 0.2667 | 70 | 0.5 | 17 | 0.3529 | 16 | 0.75 | 18 | 0.7778 | 16 | 0.4375 | 67 | 0.5821 |
| 13_238762 | 24 | 0.2083 | 16 | 0.375 | 17 | 0.1765 | 14 | 0.2143 | 71 | 0.2394 | 15 | 0.1333 | 18 | 0.3333 | 19 | 0.2632 | 16 | 0.0625 | 68 | 0.2059 |
| 13_238763 | 25 | 0.12 | 18 | 0 | 17 | 0.1176 | 16 | 0.1875 | 76 | 0.1053 | 17 | 0.0588 | 19 | 0.0526 | 19 | 0.1053 | 16 | 0 | 71 | 0.0563 |
| 13_238878 | 3 | 0 | 1 | 1 | 1 | 0 | 1 | 0 | 6 | 0.1667 | NA | NA | 1 | 1 | NA | NA | NA | NA | 1 | 1 |
| 13_238879 | 4 | 0.25 | 1 | 0 | 1 | 1 | 1 | 0 | 7 | 0.2857 | 1 | 1 | 1 | 1 | NA | NA | NA | NA | 2 | 1 |
| 13_238881 | 2 | 0.5 | NA | NA | NA | NA | 1 | 0 | 3 | 0.3333 | NA | NA | 1 | 1 | NA | NA | NA | NA | 1 | 1 |
| 13_238882 | 3 | 1 | 1 | 1 | NA | NA | 1 | 1 | 5 | 1 | NA | NA | 1 | 1 | NA | NA | NA | NA | 1 | 1 |
| 13_238882 | 3 | 0 | 1 | 1 | NA | NA | 1 | 0 | 5 | 0.2 | NA | NA | 1 | 1 | NA | NA | NA | NA | 1 | 1 |
| 13_238882 | 4 | 0.25 | 1 | 1 | NA | NA | 1 | 0 | 6 | 0.3333 | NA | NA | 1 | 1 | NA | NA | NA | NA | 1 | 1 |
| 13_238883 | 4 | 0.25 | 1 | 1 | NA | NA | NA | NA | 5 | 0.4 | NA | NA | 1 | 1 | NA | NA | NA | NA | 1 | 1 |
| 13_239235 | 9 | 0.7778 | 2 | 1 | 1 | 0 | 5 | 0.8 | 17 | 0.7647 | 2 | 1 | 1 | 1 | 4 | 0.5 | 2 | 0.5 | 9 | 0.6667 |
| 13_239235 | 8 | 0.625 | 2 | 0.5 | 2 | 1 | 5 | 0.8 | 17 | 0.7059 | 2 | 1 | 1 | 1 | 4 | 1 | 2 | 0.5 | 9 | 0.8889 |
| 13_239236 | 7 | 0.8571 | 1 | 1 | 2 | 1 | 4 | 0.75 | 14 | 0.8571 | 1 | 1 | 1 | 1 | 3 | 0.6667 | 2 | 0.5 | 7 | 0.7143 |
| 13_239237 | 10 | 0.9 | 2 | 1 | 2 | 1 | 5 | 1 | 19 | 0.9474 | 2 | 1 | 1 | 1 | 3 | 1 | 1 | 1 | 7 | 1 |
| 13_239237 | 10 | 1 | 2 | 1 | 2 | 1 | 5 | 1 | 19 | 1 | 2 | 1 | 1 | 1 | 3 | 1 | 2 | 1 | 8 | 1 |
| 13_239239 | 10 | 0.9 | 2 | 0.5 | 2 | 1 | 4 | 1 | 18 | 0.8889 | 2 | 1 | 1 | 1 | 3 | 1 | 1 | 1 | 7 | 1 |
| 13_239239 | 6 | 0.6667 | 3 | 0.3333 | 2 | 0 | 3 | 0.6667 | 14 | 0.5 | 1 | 1 | 1 | 0 | 2 | 1 | 1 | 0 | 5 | 0.6 |
| 13_239240 | 8 | 0.875 | 2 | 1 | 2 | 0.5 | 3 | 1 | 15 | 0.8667 | 1 | 1 | NA | NA | 4 | 1 | 2 | 1 | 7 | 1 |
| 13_239240 | 10 | 1 | 3 | 1 | 3 | 0.6667 | 4 | 1 | 20 | 0.95 | 1 | 1 | 1 | 1 | 4 | 1 | 2 | 1 | 8 | 1 |
| 13_239240 | 10 | 0.5 | 2 | 0 | 3 | 0.3333 | 3 | 0.6667 | 18 | 0.4444 | 2 | 1 | 1 | 0 | 2 | 0.5 | 2 | 0.5 | 7 | 0.5714 |
| 13_248852 | 3 | 0 | 7 | 0.2857 | 3 | 0.3333 | 5 | 0.6 | 18 | 0.3333 | NA | NA | 6 | 0.8333 | 4 | 0.5 | 4 | 0.5 | 14 | 0.6429 |
| 13_248854 | 3 | 0.6667 | 6 | 0.8333 | 3 | 1 | 6 | 0.8333 | 18 | 0.8333 | NA | NA | 5 | 0.6 | 5 | 1 | 3 | 0.6667 | 13 | 0.7692 |
| 13_248854 | 4 | 1 | 7 | 1 | 3 | 0.6667 | 6 | 0.6667 | 20 | 0.85 | NA | NA | 5 | 1 | 4 | 1 | 5 | 0.8 | 14 | 0.9286 |
| 13_248854 | 4 | 0 | 7 | 0.1429 | 3 | 0 | 3 | 0 | 17 | 0.0588 | NA | NA | 5 | 0 | 7 | 0.1429 | 4 | 0.25 | 16 | 0.125 |
| 13_248856 | 4 | 1 | 7 | 0.8571 | 2 | 1 | 6 | 0.8333 | 19 | 0.8947 | NA | NA | 7 | 1 | 7 | 0.8571 | 5 | 0.6 | 19 | 0.8421 |
| 13_248856 | 4 | 0 | 7 | 0.1429 | 3 | 0.3333 | 6 | 0 | 20 | 0.1 | NA | NA | 6 | 0.1667 | 6 | 0 | 4 | 0 | 16 | 0.0625 |
| 13_248887 | 3 | 0 | 1 | 1 | NA | NA | NA | NA | 4 | 0.25 | NA | NA | 5 | 0.6 | 1 | 0 | 1 | 1 | 7 | 0.5714 |
| 13_248888 | 2 | 0.5 | 1 | 1 | NA | NA | NA | NA | 3 | 0.6667 | NA | NA | 5 | 0.8 | 1 | 1 | NA | NA | 6 | 0.8333 |
| 13_248888 | 3 | 1 | 1 | 1 | NA | NA | NA | NA | 4 | 1 | NA | NA | 5 | 1 | 1 | 1 | 1 | 1 | 7 | 1 |
| 13_248888 | 2 | 0.5 | 1 | 1 | NA | NA | NA | NA | 3 | 0.6667 | NA | NA | 5 | 0.6 | 1 | 1 | 1 | 1 | 7 | 0.7143 |
| 13_248891 | 3 | 0.6667 | 1 | 1 | NA | NA | NA | NA | 4 | 0.75 | NA | NA | 5 | 0.4 | 1 | 1 | 1 | 1 | 7 | 0.5714 |
| 13_248891 | 3 | 0 | 1 | 1 | NA | NA | NA | NA | 4 | 0.25 | 1 | 0 | 5 | 0.6 | 1 | 0 | 1 | 0 | 8 | 0.375 |
| 13_248892 | 3 | 0 | 1 | 1 | NA | NA | NA | NA | 4 | 0.25 | 1 | 0 | 4 | 0 | 2 | 0.5 | 1 | 0 | 8 | 0.125 |
| 13_248894 | 1 | 1 | 1 | 1 | NA | NA | NA | NA | 2 | 1 | 1 | 0 | 5 | 0.4 | 2 | 0.5 | 1 | 1 | 9 | 0.4444 |
| 13_248895 | 2 | 0.5 | 1 | 1 | NA | NA | NA | NA | 3 | 0.6667 | 1 | 1 | 5 | 0.6 | 2 | 1 | 1 | 1 | 9 | 0.7778 |
| 13_249156 | 2 | 1 | 8 | 0.375 | 4 | 0 | 7 | 0.5714 | 21 | 0.4286 | NA | NA | 4 | 0.25 | 5 | 0.6 | 6 | 0.1667 | 15 | 0.3333 |
| 13_249158 | 2 | 0.5 | 8 | 0.75 | 5 | 0.8 | 8 | 1 | 23 | 0.8261 | 1 | 0 | 5 | 0.8 | 8 | 1 | 7 | 1 | 21 | 0.9048 |
| 13_249158 | 2 | 0 | 7 | 0 | 4 | 0.25 | 7 | 0.4286 | 20 | 0.2 | 1 | 0 | 4 | 0.25 | 7 | 0.1429 | 7 | 0.2857 | 19 | 0.2105 |
| 13_249158 | 2 | 0 | 9 | 0 | 5 | 0.2 | 7 | 0.1429 | 23 | 0.087 | 1 | 0 | 5 | 0 | 7 | 0.1429 | 7 | 0 | 20 | 0.05 |
| 13_249170 | 5 | 0 | 10 | 0 | 8 | 0.25 | 5 | 0.2 | 28 | 0.1071 | 2 | 0 | 4 | 0.25 | 8 | 0.125 | 4 | 0.25 | 18 | 0.1667 |
| 13_249171 | 4 | 0.25 | 5 | 0 | 3 | 0 | 6 | 0 | 18 | 0.0556 | 2 | 0 | 5 | 0 | 6 | 0 | 7 | 0.1429 | 20 | 0.05 |
| 13_249173 | 7 | 0 | 11 | 0.0909 | 9 | 0.1111 | 8 | 0 | 35 | 0.0571 | 3 | 0 | 12 | 0 | 11 | 0.0909 | 9 | 0.1111 | 35 | 0.0571 |
| 13_249853 | 10 | 0.9 | 8 | 1 | 4 | 1 | 8 | 1 | 30 | 0.9667 | 8 | 0.625 | 4 | 1 | 2 | 1 | 10 | 1 | 24 | 0.875 |
| 13_249853 | 7 | 1 | 7 | 0.8571 | 4 | 1 | 7 | 1 | 25 | 0.96 | 10 | 1 | 4 | 1 | 5 | 1 | 10 | 1 | 29 | 1 |
| 13_249854 | 11 | 0.3636 | 7 | 0.1429 | 4 | 0.5 | 6 | 0.3333 | 28 | 0.3214 | 8 | 0.375 | 2 | 0 | 4 | 0 | 7 | 0.1429 | 21 | 0.1905 |
| 13_249859 | 14 | 0.2857 | 8 | 0.125 | 6 | 0 | 10 | 0.1 | 38 | 0.1579 | 11 | 0 | 6 | 0.1667 | 4 | 0.25 | 12 | 0.0833 | 33 | 0.0909 |
| 13_249872 | 26 | 0.0769 | 17 | 0.1176 | 19 | 0.0526 | 23 | 0.0435 | 85 | 0.0706 | 17 | 0.0588 | 13 | 0 | 14 | 0.0714 | 28 | 0.0714 | 72 | 0.0556 |
| 13_249968 | 32 | 1 | 21 | 1 | 24 | 0.9583 | 28 | 1 | 105 | 0.9905 | 21 | 0.9048 | 13 | 0.9231 | 18 | 1 | 33 | 0.9394 | 85 | 0.9412 |

| | | | | | | | | | | | | | | | | | | | | |
|---|---|---|---|---|---|---|---|---|---|---|---|---|---|---|---|---|---|---|---|---|
| 13_249968 | 28 | 0.2857 | 20 | 0.2 | 24 | 0.375 | 25 | 0.2 | 97 | 0.268 | 20 | 0.45 | 13 | 0.2308 | 12 | 0.5 | 35 | 0.2286 | 80 | 0.325 |
| 13_249968 | 34 | 0.0882 | 21 | 0.3333 | 23 | 0.1304 | 29 | 0.1034 | 107 | 0.1495 | 22 | 0.1818 | 15 | 0 | 17 | 0.2353 | 36 | 0.1389 | 90 | 0.1444 |
| 13_249969 | 24 | 0.8333 | 15 | 0.8667 | 19 | 0.8947 | 25 | 0.84 | 83 | 0.8554 | 14 | 0.7143 | 13 | 0.8462 | 14 | 0.6429 | 30 | 0.8 | 71 | 0.7606 |
| 13_249969 | 32 | 0.8438 | 22 | 0.9545 | 22 | 0.9091 | 27 | 0.8889 | 103 | 0.8932 | 20 | 1 | 14 | 0.8571 | 19 | 0.9474 | 37 | 0.8649 | 90 | 0.9111 |
| 13_249970 | 32 | 0.75 | 22 | 0.8182 | 22 | 0.8636 | 31 | 0.8387 | 107 | 0.8131 | 21 | 0.9048 | 14 | 0.7857 | 16 | 0.625 | 39 | 0.8462 | 90 | 0.8111 |
| 13_249975 | 35 | 0.1714 | 25 | 0.08 | 25 | 0 | 31 | 0.1935 | 116 | 0.1207 | 23 | 0.087 | 17 | 0.1176 | 21 | 0.0476 | 40 | 0.125 | 101 | 0.099 |
| 13_250252 | 5 | 0.2 | 1 | 1 | NA | NA | 4 | 0 | 10 | 0.2 | 3 | 0 | 1 | 0 | 1 | 1 | 2 | 0.5 | 7 | 0.2857 |
| 13_250252 | 6 | 1 | 2 | 1 | NA | NA | 4 | 1 | 12 | 1 | 3 | 0.6667 | 2 | 1 | 1 | 1 | 2 | 1 | 8 | 0.875 |
| 13_250254 | 6 | 0.3333 | 2 | 0.5 | NA | NA | 4 | 0 | 12 | 0.25 | 3 | 0 | 2 | 0.5 | 1 | 0 | 2 | 0 | 8 | 0.125 |
| 13_250255 | 6 | 0 | 1 | 0 | 1 | 0 | 4 | 0.25 | 12 | 0.0833 | 3 | 0 | 2 | 0.5 | 1 | 0 | 2 | 0 | 8 | 0.125 |
| 13_250409 | 1 | 1 | NA | NA | NA | NA | NA | NA | 1 | 1 | NA | NA | NA | NA | 2 | 0.5 | NA | NA | 2 | 0.5 |
| 13_250837 | 2 | 1 | 1 | 1 | 1 | 1 | 2 | 1 | 6 | 1 | NA | NA | NA | NA | 1 | 1 | 1 | 0 | 2 | 0.5 |
| 13_250838 | 3 | 0.6667 | NA | NA | 1 | 1 | 2 | 0 | 6 | 0.5 | NA | NA | NA | NA | 2 | 0.5 | 1 | 1 | 3 | 0.6667 |
| 13_250838 | 2 | 0 | 1 | 1 | 1 | 0 | 2 | 0.5 | 6 | 0.3333 | NA | NA | NA | NA | 2 | 0.5 | 1 | 1 | 3 | 0.6667 |
| 13_250838 | 3 | 0.6667 | 1 | 0 | 1 | 0 | 2 | 0 | 7 | 0.2857 | NA | NA | NA | NA | 2 | 0.5 | 1 | 0 | 3 | 0.3333 |
| 13_250839 | 3 | 1 | 1 | 1 | 1 | 1 | 2 | 1 | 7 | 1 | NA | NA | NA | NA | 2 | 1 | 1 | 1 | 3 | 1 |
| 13_250841 | 3 | 0 | 2 | 0 | 1 | 1 | 2 | 0 | 8 | 0.125 | NA | NA | NA | NA | 2 | 0.5 | 1 | 0 | 3 | 0.3333 |
| 13_250843 | 3 | 0.3333 | 1 | 1 | 1 | 0 | 2 | 0.5 | 7 | 0.4286 | NA | NA | NA | NA | 2 | 0.5 | 1 | 0 | 3 | 0.3333 |
| 13_250844 | 3 | 1 | 2 | 0.5 | NA | NA | 2 | 1 | 7 | 0.8571 | NA | NA | NA | NA | 2 | 1 | 1 | 1 | 3 | 1 |
| 13_250844 | 2 | 1 | 2 | 1 | 3 | 1 | 2 | 1 | 9 | 1 | NA | NA | NA | NA | 1 | 1 | 1 | 1 | 2 | 1 |
| 13_250844 | 2 | 1 | 1 | 0 | 2 | 0.5 | 2 | 0 | 7 | 0.4286 | NA | NA | NA | NA | 1 | 1 | NA | NA | 1 | 1 |
| 13_250844 | 2 | 0 | 1 | 1 | 2 | 1 | 2 | 0 | 7 | 0.4286 | NA | NA | NA | NA | NA | NA | 1 | 1 | 1 | 1 |
| 13_250844 | 3 | 0 | NA | NA | 2 | 0 | 2 | 0.5 | 7 | 0.1429 | NA | NA | NA | NA | 2 | 0.5 | 1 | 0 | 3 | 0.3333 |
| 13_250845 | 3 | 0.3333 | 2 | 0.5 | 2 | 0.5 | 2 | 0 | 9 | 0.3333 | 1 | 0 | NA | NA | 2 | 0.5 | 1 | 0 | 4 | 0.25 |
| 13_250846 | 3 | 1 | 2 | 1 | 1 | 1 | 2 | 1 | 8 | 1 | 1 | 1 | NA | NA | 2 | 0.5 | 1 | 0 | 4 | 0.5 |
| 13_250846 | 3 | 1 | 2 | 1 | 2 | 1 | 2 | 1 | 9 | 1 | 1 | 0 | NA | NA | 2 | 1 | 1 | 1 | 4 | 0.75 |
| 13_250846 | 3 | 1 | 2 | 1 | 1 | 0 | 1 | 1 | 7 | 0.8571 | 1 | 0 | NA | NA | 2 | 1 | NA | NA | 3 | 0.6667 |
| 13_250847 | 3 | 0.3333 | 2 | 1 | 3 | 0.3333 | NA | NA | 8 | 0.5 | NA | NA | NA | NA | 2 | 0.5 | NA | NA | 2 | 0.5 |
| 13_250850 | 2 | 1 | 2 | 1 | 1 | 1 | 3 | 1 | 8 | 1 | NA | NA | NA | NA | 2 | 1 | 1 | 1 | 3 | 1 |
| 13_250850 | 3 | 0.6667 | 2 | 1 | 2 | 0.5 | 2 | 0.5 | 9 | 0.6667 | NA | NA | NA | NA | 2 | 1 | 1 | 1 | 3 | 1 |
| 13_250852 | 3 | 1 | 2 | 1 | 1 | 1 | 3 | 1 | 9 | 1 | 1 | 1 | NA | NA | 2 | 1 | 1 | 1 | 4 | 1 |
| 13_250855 | 2 | 1 | 1 | 1 | 1 | 1 | 2 | 0.5 | 6 | 0.8333 | 2 | 0.5 | NA | NA | 2 | 1 | 2 | 0.5 | 6 | 0.6667 |
| 13_250856 | 1 | 0 | 1 | 1 | NA | NA | 3 | 0 | 5 | 0.2 | 2 | 0.5 | NA | NA | 3 | 0.3333 | 3 | 0.3333 | 8 | 0.375 |
| 13_250859 | 2 | 0 | 1 | 0 | 1 | 0 | 3 | 0.6667 | 7 | 0.2857 | 2 | 0.5 | NA | NA | 3 | 0.3333 | 3 | 0 | 8 | 0.25 |
| 13_250859 | 3 | 1 | 1 | 1 | 1 | 1 | 3 | 0.6667 | 8 | 0.875 | 2 | 1 | NA | NA | 3 | 1 | 2 | 0 | 7 | 0.7143 |
| 13_250860 | 3 | 0 | 2 | 0.5 | NA | NA | 3 | 0 | 8 | 0.125 | 1 | 1 | NA | NA | 2 | 0 | 2 | 0 | 5 | 0.2 |
| 13_250860 | 3 | 0 | 2 | 0 | NA | NA | 3 | 0.6667 | 8 | 0.25 | 2 | 0 | NA | NA | 2 | 0 | 3 | 0.3333 | 7 | 0.1429 |
| 13_250917 | 9 | 0.1111 | 9 | 0.1111 | 11 | 0 | 14 | 0.0714 | 43 | 0.0698 | 4 | 0.25 | 13 | 0 | 9 | 0.1111 | 5 | 0.2 | 31 | 0.0968 |
| 13_250918 | 9 | 0.3333 | 9 | 0.1111 | 8 | 0.125 | 18 | 0.2778 | 44 | 0.2273 | 4 | 0 | 13 | 0.2308 | 8 | 0.125 | 5 | 0.2 | 30 | 0.1667 |
| 13_250921 | 6 | 0 | 9 | 0 | 8 | 0 | 19 | 0.1579 | 42 | 0.0714 | 4 | 0.25 | 10 | 0.1 | 8 | 0 | 5 | 0 | 27 | 0.0741 |
| 13_250922 | 8 | 0.125 | 9 | 0.3333 | 10 | 0.2 | 20 | 0.3 | 47 | 0.2553 | 5 | 0.2 | 10 | 0.1 | 10 | 0.2 | 5 | 0.4 | 30 | 0.2 |
| 13_250930 | 7 | 0.1429 | 5 | 0.4 | 7 | 0.1429 | 10 | 0.2 | 29 | 0.2069 | 4 | 0.5 | 6 | 0 | 2 | 0 | 4 | 0.25 | 16 | 0.1875 |
| 13_250943 | 5 | 0.2 | 2 | 0.5 | 5 | 0.4 | 12 | 0.4167 | 24 | 0.375 | 3 | 0.6667 | 3 | 0.6667 | 5 | 0.2 | 1 | 0 | 12 | 0.4167 |
| 13_251219 | NA | NA | 1 | 1 | NA | NA | NA | NA | 1 | 1 | NA | NA | NA | NA | NA | NA | 1 | 1 | 1 | 1 |
| 13_251717 | 5 | 0.4 | 5 | 0 | 6 | 0 | NA | NA | 16 | 0.125 | 1 | 0 | 3 | 0.3333 | 7 | 0.2857 | 3 | 0.3333 | 14 | 0.2857 |
| 13_251719 | 5 | 0.4 | 5 | 0.4 | 7 | 0.2857 | 1 | 0 | 18 | 0.3333 | 1 | 0 | 3 | 0.6667 | 7 | 0.2857 | 2 | 0 | 13 | 0.3077 |
| 13_251722 | 6 | 0 | 4 | 0.5 | 7 | 0.1429 | 2 | 0 | 19 | 0.1579 | 2 | 0 | 2 | 0.5 | 7 | 0.2857 | 2 | 0.5 | 13 | 0.3077 |
| 13_251725 | 3 | 0 | 6 | 0.1667 | 4 | 0.5 | 2 | 0 | 15 | 0.2 | 2 | 0 | 2 | 0.5 | 5 | 0.6 | 3 | 0 | 12 | 0.3333 |
| 13_251725 | 6 | 0 | 5 | 0.8 | 3 | 0.6667 | 2 | 0 | 16 | 0.375 | NA | NA | 2 | 0.5 | 7 | 0.1429 | 3 | 0.3333 | 12 | 0.25 |
| 13_251726 | 5 | 0.2 | 6 | 0.3333 | 7 | 0.2857 | 2 | 0 | 20 | 0.25 | 1 | 0 | 2 | 0 | 7 | 0.2857 | 3 | 0 | 13 | 0.1538 |
| 13_251728 | 4 | 0 | 3 | 0 | 4 | 0 | 2 | 0.5 | 13 | 0.0769 | NA | NA | 2 | 0.5 | 6 | 0 | 2 | 0 | 10 | 0.1 |
| 13_251739 | 4 | 1 | 5 | 0.8 | 8 | 0.875 | 2 | 0.5 | 19 | 0.8421 | 1 | 1 | 2 | 0.5 | 4 | 1 | 3 | 0.6667 | 10 | 0.8 |
| 13_251739 | 4 | 0.5 | 5 | 0.8 | 6 | 0.6667 | 3 | 0.6667 | 18 | 0.6667 | 1 | 1 | 2 | 0.5 | 4 | 0.75 | 4 | 0.75 | 11 | 0.7273 |
| 13_251739 | 4 | 0.25 | 5 | 0 | 7 | 0.1429 | 3 | 0 | 19 | 0.1053 | 1 | 1 | 2 | 0 | 4 | 0 | 3 | 0 | 10 | 0.1 |
| 13_251740 | 4 | 0.25 | 5 | 0.8 | 7 | 0.4286 | 3 | 0 | 19 | 0.4211 | 1 | 0 | 2 | 0 | 4 | 0.75 | 4 | 0.75 | 11 | 0.5455 |
| 13_251740 | 4 | 0.5 | 4 | 0.75 | 5 | 0.6 | 1 | 0 | 14 | 0.5714 | 1 | 1 | NA | NA | 2 | 0.5 | 3 | 0.6667 | 6 | 0.6667 |
| 13_251741 | 3 | 0 | 5 | 0 | 6 | 0.5 | 2 | 0 | 16 | 0.1875 | 1 | 0 | 2 | 0 | 4 | 0 | 4 | 0.25 | 11 | 0.0909 |
| 13_251741 | 3 | 0.3333 | 4 | 0.5 | 7 | 0.1429 | 1 | 0 | 15 | 0.2667 | 1 | 0 | 2 | 0.5 | 4 | 0.25 | 3 | 0.6667 | 10 | 0.4 |
| 13_251741 | 1 | 0 | 2 | 0.5 | 3 | 0 | 3 | 0 | 9 | 0.1111 | 2 | 0 | 4 | 0.25 | 5 | 0.2 | NA | NA | 11 | 0.1818 |
| 13_251742 | 4 | 0 | 5 | 0 | 6 | 0.5 | 3 | 0 | 18 | 0.1667 | 1 | 0 | 2 | 0.5 | 4 | 0 | 4 | 0.25 | 11 | 0.1818 |
| 13_251743 | 3 | 0.6667 | 4 | 1 | 3 | 0.3333 | 6 | 1 | 16 | 0.8125 | 2 | 0.5 | 4 | 0.75 | 5 | 0.6 | 1 | 1 | 12 | 0.6667 |
| 13_251745 | 3 | 0 | 2 | 0.5 | 3 | 0 | 6 | 0.3333 | 14 | 0.2143 | 1 | 0 | 3 | 0.6667 | 4 | 0 | 1 | 0 | 9 | 0.2222 |
| 13_251748 | 4 | 0 | 4 | 0.25 | 4 | 0 | 2 | 0 | 14 | 0.0714 | NA | NA | 1 | 1 | 4 | 0.25 | 2 | 0.5 | 7 | 0.4286 |
| 13_251752 | 3 | 0 | 3 | 0.3333 | 4 | 0.25 | 2 | 0 | 12 | 0.1667 | NA | NA | 3 | 0 | 3 | 0.6667 | 3 | 0 | 9 | 0.2222 |
| 13_251767 | 2 | 0 | 2 | 0 | 2 | 0 | 2 | 0.5 | 8 | 0.125 | NA | NA | 1 | 1 | 2 | 0.5 | 1 | 0 | 4 | 0.5 |

| | | | | | | | | | | | | | | | | | | | | |
|---|---|---|---|---|---|---|---|---|---|---|---|---|---|---|---|---|---|---|---|---|
| 13_2517673 | 2 | 0 | 2 | 0.5 | 3 | 0 | 2 | 0 | 9 | 0.1111 | NA | NA | 1 | 1 | 2 | 0 | 1 | 0 | 4 | 0.25 |
| 13_2517691 | 2 | 0 | 2 | 0.5 | 3 | 0 | NA | NA | 7 | 0.1429 | NA | NA | 1 | 0 | 2 | 0.5 | 1 | 0 | 4 | 0.25 |
| 13_2517699 | 2 | 0.5 | 2 | 0 | 3 | 0 | 3 | 0 | 10 | 0.1 | NA | NA | 1 | 1 | 3 | 0.3333 | 1 | 0 | 5 | 0.4 |
| 13_2517704 | 2 | 0 | 2 | 0 | 3 | 0.3333 | 3 | 0 | 10 | 0.1 | NA | NA | 1 | 0 | 3 | 0.3333 | 1 | 0 | 5 | 0.2 |
| 13_2517708 | 2 | 0 | 2 | 1 | 3 | 0.6667 | 3 | 0.3333 | 10 | 0.5 | NA | NA | 1 | 0 | 2 | 0.5 | 1 | 1 | 4 | 0.5 |
| 13_2543608 | 44 | 0.0682 | 58 | 0.0517 | 65 | 0.1077 | 56 | 0.0536 | 223 | 0.0717 | 44 | 0.1818 | 65 | 0.0769 | 49 | 0.0204 | 39 | 0.0256 | 197 | 0.0761 |
| 13_2543649 | 44 | 0.0909 | 34 | 0.1176 | 47 | 0.0851 | 41 | 0.122 | 166 | 0.1024 | 28 | 0.0714 | 46 | 0.087 | 38 | 0.0263 | 32 | 0.125 | 144 | 0.0764 |
| 13_2543739 | 36 | 0.0278 | 38 | 0.0526 | 50 | 0.06 | 50 | 0.06 | 174 | 0.0517 | 29 | 0.069 | 55 | 0.0727 | 33 | 0.0303 | 36 | 0.0556 | 153 | 0.0588 |
| 13_2543744 | 37 | 0.2162 | 32 | 0.2188 | 45 | 0.1333 | 47 | 0.1064 | 161 | 0.1615 | 26 | 0.1154 | 52 | 0.1731 | 28 | 0.1786 | 35 | 0.1429 | 141 | 0.156 |
| 13_2543758 | 32 | 0.2812 | 37 | 0.3784 | 43 | 0.2326 | 42 | 0.3333 | 154 | 0.3052 | 24 | 0.25 | 48 | 0.2292 | 31 | 0.2581 | 27 | 0.4074 | 130 | 0.2769 |
| 13_2543761 | 33 | 0.303 | 33 | 0.4545 | 39 | 0.3333 | 46 | 0.3261 | 151 | 0.351 | 25 | 0.2 | 52 | 0.25 | 26 | 0.2308 | 32 | 0.25 | 135 | 0.237 |
| 13_2543769 | 26 | 0.1154 | 20 | 0 | 32 | 0.0625 | 38 | 0.0263 | 116 | 0.0517 | 16 | 0.0625 | 34 | 0.0882 | 24 | 0.0417 | 24 | 0.0833 | 98 | 0.0714 |
| 13_2543769 | 31 | 0.6452 | 31 | 0.6774 | 42 | 0.6667 | 41 | 0.5366 | 145 | 0.6276 | 21 | 0.7143 | 48 | 0.7083 | 26 | 0.7308 | 35 | 0.6286 | 130 | 0.6923 |
| 13_2544626 | 33 | 0.1212 | 31 | 0 | 40 | 0 | 46 | 0.1087 | 150 | 0.06 | 24 | 0 | 49 | 0.1224 | 28 | 0.1071 | 32 | 0.0625 | 133 | 0.0827 |
| 13_2913291 | 4 | 0 | 6 | 0.1667 | 2 | 0 | 3 | 0 | 15 | 0.0667 | 1 | 1 | 7 | 0.1429 | 2 | 0 | 1 | 0 | 11 | 0.1818 |
| 13_2913425 | 1 | 0 | 4 | 0.5 | 2 | 0.5 | 2 | 0.5 | 9 | 0.4444 | NA | NA | 5 | 0.8 | 2 | 1 | NA | NA | 7 | 0.8571 |
| 13_2913433 | 2 | 0 | 4 | 0 | 2 | 0 | 2 | 0.5 | 10 | 0.1 | NA | NA | 4 | 0.25 | 2 | 0 | NA | NA | 6 | 0.1667 |
| 13_2913435 | 1 | 1 | 4 | 0.75 | 2 | 0.5 | 2 | 1 | 9 | 0.7778 | NA | NA | 5 | 0.8 | 2 | 1 | NA | NA | 7 | 0.8571 |
| 13_2913442 | 1 | 0 | 1 | 0 | 2 | 0.5 | 2 | 0 | 6 | 0.1667 | NA | NA | 5 | 0.2 | 3 | 0 | NA | NA | 8 | 0.125 |
| 13_2913450 | 1 | 0 | 3 | 1 | 1 | 1 | 2 | 0 | 7 | 0.5714 | NA | NA | 5 | 0.6 | 3 | 0.6667 | NA | NA | 8 | 0.625 |
| 13_2913465 | NA | NA | 2 | 1 | 1 | 1 | 2 | 1 | 5 | 1 | NA | NA | 4 | 1 | 2 | 1 | NA | NA | 6 | 1 |
| 13_2913466 | NA | NA | 2 | 0.5 | 1 | 0 | 2 | 0 | 5 | 0.2 | NA | NA | 4 | 0.75 | 1 | 1 | NA | NA | 5 | 0.8 |
| 13_2913478 | NA | NA | 2 | 1 | 1 | 0 | 2 | 0.5 | 5 | 0.6 | NA | NA | 4 | 0.75 | 1 | 1 | NA | NA | 5 | 0.8 |
| 13_2913482 | 1 | 0 | 2 | 0.5 | 1 | 0 | 2 | 0.5 | 6 | 0.3333 | NA | NA | 4 | 0.5 | 1 | 1 | NA | NA | 5 | 0.6 |
| 13_2913483 | NA | NA | 2 | 1 | 1 | 1 | 2 | 1 | 5 | 1 | NA | NA | 4 | 1 | NA | NA | NA | NA | 4 | 1 |
| 13_2913494 | NA | NA | 1 | 0 | NA | NA | 2 | 0.5 | 3 | 0.3333 | NA | NA | 3 | 0.3333 | 1 | 0 | NA | NA | 4 | 0.25 |
| 13_2913519 | NA | NA | NA | NA | 1 | 1 | 1 | 1 | 2 | 1 | NA | NA | 3 | 0.6667 | NA | NA | NA | NA | 3 | 0.6667 |
| 13_2913523 | NA | NA | 1 | 1 | 1 | 1 | 2 | 1 | 4 | 1 | NA | NA | 3 | 1 | 1 | 1 | NA | NA | 4 | 1 |
| 13_2913527 | 1 | 1 | 1 | 1 | NA | NA | 1 | 1 | 3 | 1 | NA | NA | 3 | 1 | NA | NA | NA | NA | 3 | 1 |
| 13_2913538 | NA | NA | 1 | 1 | NA | NA | 1 | 1 | 2 | 1 | NA | NA | 1 | 1 | 1 | 1 | NA | NA | 2 | 1 |
| 13_2913576 | NA | NA | NA | NA | 1 | 1 | 2 | 1 | 3 | 1 | NA | NA | NA | NA | 1 | 1 | NA | NA | 1 | 1 |
| 13_2913581 | NA | NA | NA | NA | 1 | 0 | 2 | 0.5 | 3 | 0.3333 | NA | NA | 1 | 1 | 1 | 0 | NA | NA | 2 | 0.5 |
| 13_2913670 | NA | NA | NA | NA | 1 | 0 | 2 | 0.5 | 3 | 0.3333 | NA | NA | NA | NA | 1 | 1 | NA | NA | 1 | 1 |
| 13_2921249 | NA | NA | 1 | 1 | NA | NA | NA | NA | 1 | 1 | NA | NA | 1 | 1 | NA | NA | 2 | 0.5 | 3 | 0.6667 |
| 13_2922511 | 1 | 0 | 1 | 0 | NA | NA | 3 | 0.3333 | 5 | 0.2 | NA | NA | NA | NA | 1 | 1 | NA | NA | 1 | 1 |
| 13_2922526 | 2 | 0 | 1 | 0 | NA | NA | 3 | 0.6667 | 6 | 0.3333 | NA | NA | NA | NA | 1 | 1 | NA | NA | 1 | 1 |
| 13_2950942 | NA | NA | 1 | 0 | 1 | 1 | 3 | 0.6667 | 5 | 0.6 | NA | NA | 2 | 0.5 | 2 | 1 | 3 | 1 | 7 | 0.8571 |
| 13_2950947 | 1 | 1 | 1 | 1 | 2 | 1 | 3 | 1 | 7 | 1 | NA | NA | 1 | 1 | 2 | 1 | 3 | 1 | 6 | 1 |
| 13_2950949 | 1 | 0 | 1 | 0 | 2 | 0.5 | 3 | 0.3333 | 7 | 0.2857 | NA | NA | 2 | 0 | 1 | 0 | 3 | 0.3333 | 6 | 0.1667 |
| 13_2950950 | 1 | 0 | 1 | 0 | 2 | 0.5 | 3 | 0.6667 | 7 | 0.4286 | NA | NA | 2 | 1 | 2 | 0.5 | 2 | 0 | 6 | 0.5 |
| 13_2950960 | NA | NA | 2 | 0 | 2 | 1 | 3 | 0.6667 | 7 | 0.5714 | NA | NA | 2 | 0.5 | 2 | 0.5 | 2 | 1 | 6 | 0.6667 |
| 13_2950983 | NA | NA | 2 | 0.5 | 1 | 0 | 2 | 0.5 | 5 | 0.4 | NA | NA | 2 | 0 | 2 | 0 | 2 | 0.5 | 6 | 0.1667 |
| 13_2950992 | NA | NA | 2 | 0 | 1 | 0 | 3 | 0.3333 | 6 | 0.1667 | NA | NA | 2 | 0 | 1 | 1 | 3 | 0 | 6 | 0.1667 |
| 13_2950993 | NA | NA | 2 | 1 | 1 | 1 | 3 | 1 | 6 | 1 | NA | NA | 1 | 1 | 2 | 0.5 | 3 | 1 | 6 | 0.8333 |
| 13_2950996 | NA | NA | 2 | 0.5 | 1 | 0 | 2 | 0.5 | 5 | 0.4 | NA | NA | 2 | 0.5 | 1 | 0 | 3 | 0.3333 | 6 | 0.3333 |
| 13_2951003 | NA | NA | 1 | 0 | 1 | 0 | 2 | 1 | 4 | 0.5 | NA | NA | 2 | 1 | 1 | 1 | 3 | 1 | 6 | 1 |
| 13_3009312 | NA | NA | 1 | 1 | 1 | 1 | 1 | 0 | 3 | 0.6667 | 4 | 0.75 | 1 | 0 | 1 | 1 | 3 | 0.3333 | 9 | 0.5556 |
| 13_3009323 | NA | NA | 1 | 1 | 1 | 1 | 1 | 1 | 3 | 1 | 5 | 1 | 1 | 1 | 1 | 1 | 1 | 1 | 8 | 1 |
| 13_3009327 | NA | NA | 1 | 0 | 1 | 1 | NA | NA | 2 | 0.5 | 4 | 1 | 1 | 1 | 1 | 1 | 2 | 0.5 | 8 | 0.875 |
| 13_3009354 | NA | NA | 1 | 1 | NA | NA | 1 | 0 | 2 | 0.5 | 4 | 0.5 | 1 | 0 | 1 | 0 | 1 | 1 | 7 | 0.4286 |
| 13_3009357 | NA | NA | 1 | 1 | 1 | 1 | 1 | 1 | 3 | 1 | 5 | 1 | 1 | 1 | 1 | 1 | 1 | 1 | 8 | 1 |
| 13_3009375 | NA | NA | 1 | 1 | 1 | 1 | 1 | 0 | 3 | 0.6667 | 5 | 0.6 | 1 | 1 | NA | NA | 2 | 0.5 | 8 | 0.625 |
| 13_3009387 | NA | NA | 1 | 1 | 1 | 1 | 1 | 1 | 3 | 1 | 4 | 1 | 2 | 0.5 | 1 | 1 | 2 | 1 | 9 | 0.8889 |
| 13_3009390 | NA | NA | 1 | 1 | 1 | 1 | NA | NA | 2 | 1 | 4 | 1 | 1 | 1 | NA | NA | 2 | 1 | 7 | 1 |
| 13_3009398 | NA | NA | 1 | 1 | 1 | 1 | 1 | 1 | 3 | 1 | 1 | 1 | 2 | 0.5 | 1 | 1 | 1 | 1 | 5 | 0.8 |
| 13_3009401 | NA | NA | 1 | 0 | 1 | 1 | 1 | 1 | 3 | 0.6667 | 3 | 1 | NA | NA | 1 | 1 | 2 | 0.5 | 6 | 0.8333 |
| 13_3009402 | NA | NA | 1 | 1 | 1 | 1 | 1 | 1 | 3 | 1 | 3 | 0.6667 | 2 | 1 | 1 | 1 | 2 | 1 | 8 | 0.875 |
| 13_3009405 | NA | NA | 1 | 0 | 1 | 1 | NA | NA | 2 | 0.5 | 4 | 0 | 2 | 1 | 1 | 1 | 1 | 0 | 8 | 0.375 |
| 13_3072394 | NA | NA | NA | NA | 1 | 1 | NA | NA | 1 | 1 | NA | NA | 2 | 0.5 | NA | NA | NA | NA | 2 | 0.5 |
| 13_3072425 | NA | NA | NA | NA | 1 | 1 | NA | NA | 1 | 1 | NA | NA | 2 | 1 | NA | NA | NA | NA | 2 | 1 |
| 13_3072439 | NA | NA | 2 | 0.5 | 1 | 0 | 1 | 0 | 4 | 0.25 | 1 | 0 | 4 | 0 | 3 | 0.3333 | 1 | 0 | 9 | 0.1111 |
| 13_3072470 | 26 | 0.8077 | 33 | 0.7576 | 28 | 0.9286 | 33 | 0.8182 | 120 | 0.825 | 30 | 0.7333 | 33 | 0.8485 | 43 | 0.8605 | 36 | 0.9444 | 142 | 0.8521 |
| 13_3072494 | 29 | 0.1034 | 33 | 0.0606 | 28 | 0.1786 | 36 | 0.0833 | 126 | 0.1032 | 33 | 0.1515 | 34 | 0.0882 | 45 | 0.0889 | 35 | 0.0857 | 147 | 0.102 |
| 13_3072500 | 24 | 0.0417 | 30 | 0.0667 | 26 | 0 | 32 | 0.125 | 112 | 0.0625 | 26 | 0.0385 | 30 | 0.1333 | 36 | 0.0833 | 30 | 0.0667 | 122 | 0.082 |
| 13_3072504 | 23 | 0.1739 | 28 | 0.2857 | 25 | 0.2 | 29 | 0.3103 | 105 | 0.2476 | 23 | 0.1739 | 30 | 0.2 | 38 | 0.2368 | 33 | 0.1818 | 124 | 0.2016 |

| | | | | | | | | | | | | | | | | | | | | |
|---|---|---|---|---|---|---|---|---|---|---|---|---|---|---|---|---|---|---|---|---|
| 13_3072518 | 17 | 0.3529 | 23 | 0.1739 | 21 | 0.3333 | 27 | 0.3333 | 88 | 0.2955 | 23 | 0.1304 | 18 | 0.1667 | 30 | 0.2333 | 26 | 0.4615 | 97 | 0.2577 |
| 13_3089283 | 2 | 1 | 2 | 0.5 | 2 | 0.5 | 1 | 1 | 7 | 0.7143 | NA | NA | 1 | 0 | NA | NA | 2 | 0.5 | 3 | 0.3333 |
| 13_3089293 | 1 | 0 | 2 | 0.5 | 1 | 1 | NA | NA | 4 | 0.5 | 1 | 0 | 1 | 0 | NA | NA | 2 | 0.5 | 4 | 0.25 |
| 13_3089329 | 2 | 0 | 4 | 0.25 | 3 | 0 | 1 | 0 | 10 | 0.1 | 2 | 0.5 | 1 | 0 | NA | NA | 2 | 0 | 5 | 0.2 |
| 13_3089349 | 2 | 0.5 | 2 | 1 | 3 | 0 | 1 | 1 | 8 | 0.5 | 2 | 0 | 1 | 1 | NA | NA | 2 | 0.5 | 5 | 0.4 |
| 13_3089472 | 4 | 0 | 7 | 0.1429 | 6 | 0.3333 | 3 | 0.3333 | 20 | 0.2 | 2 | 0.5 | 2 | 0.5 | 1 | 0 | 4 | 0.5 | 9 | 0.4444 |
| 13_3099873 | 3 | 0 | 3 | 0 | 5 | 0.2 | 6 | 0 | 17 | 0.0588 | 2 | 0 | 4 | 0 | 7 | 0.1429 | 3 | 0 | 16 | 0.0625 |
| 13_3099875 | 4 | 0.25 | 3 | 0.3333 | 4 | 0 | 5 | 0 | 16 | 0.125 | 2 | 0 | 3 | 0 | 9 | 0.1111 | 2 | 0 | 16 | 0.0625 |
| 13_3099924 | 5 | 0 | 1 | 0 | 5 | 0.2 | 3 | 0 | 14 | 0.0714 | 2 | 0 | 5 | 0.2 | 9 | 0 | 3 | 0 | 19 | 0.0526 |
| 13_3174303 | NA | NA | NA | NA | 1 | 1 | NA | NA | 1 | 1 | NA | NA | NA | NA | 1 | 1 | NA | NA | 1 | 1 |
| 13_3174345 | NA | NA | NA | NA | 1 | 1 | NA | NA | 1 | 1 | NA | NA | 1 | 1 | 1 | 1 | 1 | 0 | 3 | 0.6667 |
| 13_3174390 | NA | NA | NA | NA | 1 | 1 | NA | NA | 1 | 1 | NA | NA | 1 | 1 | 1 | 0 | 1 | 0 | 3 | 0.3333 |
| 13_3174408 | NA | NA | NA | NA | 1 | 1 | NA | NA | 1 | 1 | NA | NA | 1 | 1 | 1 | 0 | 1 | 1 | 3 | 0.6667 |
| 13_3199097 | 2 | 0 | 1 | 1 | NA | NA | NA | NA | 3 | 0.3333 | 1 | 0 | NA | NA | 2 | 0.5 | NA | NA | 3 | 0.3333 |
| 13_3199105 | 1 | 1 | 1 | 0 | NA | NA | NA | NA | 2 | 0.5 | 1 | 1 | NA | NA | 1 | 0 | 1 | 0 | 3 | 0.3333 |
| 13_3199110 | 2 | 0.5 | 1 | 1 | NA | NA | NA | NA | 3 | 0.6667 | 1 | 0 | NA | NA | 2 | 1 | 1 | 1 | 4 | 0.75 |
| 13_3199110 | 2 | 0.5 | 1 | 1 | NA | NA | NA | NA | 3 | 0.6667 | 1 | 1 | NA | NA | 1 | 1 | 1 | 1 | 3 | 1 |
| 13_3199141 | 1 | 1 | 1 | 1 | NA | NA | NA | NA | 2 | 1 | 1 | 0 | 1 | 1 | 2 | 0.5 | 1 | 1 | 5 | 0.6 |
| 13_3199154 | 1 | 1 | 1 | 1 | NA | NA | NA | NA | 2 | 1 | 1 | 0 | 1 | 0 | 2 | 0.5 | 1 | 1 | 5 | 0.4 |
| 13_3199166 | 2 | 0 | 1 | 1 | NA | NA | NA | NA | 3 | 0.3333 | 1 | 0 | 1 | 0 | 2 | 0.5 | 2 | 0 | 6 | 0.1667 |
| 13_3199187 | 2 | 0.5 | 1 | 1 | NA | NA | NA | NA | 3 | 0.6667 | 1 | 0 | 2 | 0.5 | 2 | 0 | 2 | 0.5 | 7 | 0.2857 |
| 13_3199269 | 3 | 0.3333 | 2 | 0.5 | NA | NA | NA | NA | 5 | 0.4 | 2 | 0.5 | 2 | 1 | 2 | 0.5 | 2 | 0.5 | 8 | 0.625 |
| 13_3199283 | 3 | 1 | 2 | 1 | NA | NA | 1 | 1 | 6 | 1 | 2 | 1 | 2 | 1 | 2 | 1 | 2 | 1 | 8 | 1 |
| 13_3199290 | 3 | 0.3333 | 2 | 0 | NA | NA | 1 | 0 | 6 | 0.1667 | 2 | 0.5 | 2 | 1 | 2 | 0 | 2 | 0 | 8 | 0.375 |
| 13_3199293 | 4 | 0.25 | 2 | 0.5 | NA | NA | 1 | 0 | 7 | 0.2857 | 2 | 0 | 2 | 0.5 | 2 | 0 | 2 | 0 | 8 | 0.125 |
| 13_3199295 | 4 | 0.5 | 2 | 0.5 | NA | NA | 1 | 0 | 7 | 0.4286 | 2 | 0 | 1 | 1 | 2 | 0 | 1 | 0 | 6 | 0.1667 |
| 13_3199298 | 5 | 0.2 | 3 | 0.6667 | NA | NA | 1 | 0 | 9 | 0.3333 | 1 | 0 | 2 | 0.5 | 2 | 0.5 | 2 | 0 | 7 | 0.2857 |
| 13_3199306 | 4 | 0.5 | 2 | 0.5 | NA | NA | 1 | 1 | 7 | 0.5714 | 2 | 0 | 2 | 0 | 2 | 0.5 | 2 | 1 | 8 | 0.375 |
| 13_3199310 | 3 | 0.3333 | 1 | 0 | NA | NA | 1 | 1 | 5 | 0.4 | 2 | 1 | 1 | 0 | 2 | 1 | 2 | 1 | 7 | 0.8571 |
| 13_3199311 | 5 | 0.4 | 2 | 0.5 | NA | NA | 1 | 0 | 8 | 0.375 | 2 | 1 | 2 | 0.5 | 2 | 0.5 | 2 | 0 | 8 | 0.5 |
| 13_3199314 | 5 | 1 | 3 | 1 | NA | NA | NA | NA | 8 | 1 | 2 | 1 | 2 | 1 | 2 | 1 | 2 | 1 | 8 | 1 |
| 13_3199318 | 4 | 1 | 2 | 1 | NA | NA | 1 | 1 | 7 | 1 | 2 | 1 | 2 | 1 | 2 | 1 | 2 | 1 | 8 | 1 |
| 13_3199331 | 3 | 1 | 3 | 1 | 1 | 1 | 1 | 1 | 8 | 1 | 1 | 1 | 2 | 1 | 2 | 1 | 2 | 1 | 7 | 1 |
| 13_3199337 | 4 | 0.25 | 3 | 0.6667 | 1 | 0 | 1 | 1 | 9 | 0.4444 | 2 | 1 | 1 | 1 | 2 | 1 | 2 | 0.5 | 7 | 0.8571 |
| 13_3200374 | 4 | 0 | 3 | 0 | 1 | 0 | 1 | 1 | 9 | 0.1111 | 3 | 0 | 2 | 0 | 3 | 0.3333 | 1 | 0 | 9 | 0.1111 |
| 13_3200381 | 4 | 0.5 | 1 | 0 | NA | NA | 1 | 1 | 6 | 0.5 | 2 | 1 | 2 | 0.5 | 3 | 0 | 2 | 0.5 | 9 | 0.4444 |
| 13_3298475 | NA | NA | NA | NA | 1 | 1 | NA | NA | 1 | 1 | NA | NA | NA | NA | 1 | 1 | NA | NA | 1 | 1 |
| 13_3298568 | 1 | 0 | NA | NA | 1 | 1 | NA | NA | 2 | 0.5 | NA | NA | NA | NA | 1 | 1 | NA | NA | 1 | 1 |
| 13_3298572 | NA | NA | NA | NA | 1 | 1 | NA | NA | 1 | 1 | NA | NA | NA | NA | 1 | 1 | NA | NA | 1 | 1 |
| 13_3298577 | NA | NA | NA | NA | 1 | 1 | NA | NA | 1 | 1 | NA | NA | NA | NA | 1 | 1 | NA | NA | 1 | 1 |
| 13_3298691 | 4 | 0.25 | 4 | 0 | 1 | 0 | 2 | 0 | 11 | 0.0909 | 2 | 0 | 4 | 0 | 6 | 0.3333 | 2 | 0 | 14 | 0.1429 |
| 13_3300026 | 6 | 0.1667 | 6 | 0 | 3 | 0 | 2 | 0 | 17 | 0.0588 | 3 | 0 | 6 | 0 | 7 | 0 | 4 | 0.25 | 20 | 0.05 |
| 13_3300031 | 5 | 0.6 | 5 | 1 | 4 | 0.5 | 2 | 1 | 16 | 0.75 | 3 | 0.6667 | 6 | 0.5 | 6 | 0.5 | 4 | 0.75 | 19 | 0.5789 |
| 13_3300608 | 9 | 0.1111 | 9 | 0.2222 | 6 | 0 | 4 | 0 | 28 | 0.1071 | 5 | 0 | 7 | 0.2857 | 11 | 0 | 7 | 0.2857 | 30 | 0.1333 |
| 13_3300608 | 10 | 0.3 | 7 | 0.2857 | 6 | 0.1667 | 4 | 0.25 | 27 | 0.2593 | 5 | 0 | 7 | 0.2857 | 10 | 0.4 | 5 | 0.4 | 27 | 0.2963 |
| 13_3300614 | 9 | 0.5556 | 9 | 0.2222 | 6 | 0.6667 | 4 | 0.75 | 28 | 0.5 | 4 | 0 | 7 | 0 | 10 | 0.3 | 7 | 0.1429 | 28 | 0.1429 |
| 13_3300622 | 6 | 0.3333 | 10 | 0.1 | 3 | 0.6667 | 4 | 0.25 | 23 | 0.2609 | 6 | 0.5 | 6 | 0.5 | 10 | 0 | 7 | 0.5714 | 29 | 0.3448 |
| 13_3300623 | 9 | 0 | 9 | 0 | 6 | 0.3333 | 4 | 0 | 28 | 0.0714 | 6 | 0 | 6 | 0 | 12 | 0.0833 | 8 | 0.125 | 32 | 0.0625 |
| 13_3410225 | 6 | 0.1667 | 7 | 0.4286 | 3 | 0 | 6 | 0.3333 | 22 | 0.2727 | 6 | 0.5 | 5 | 0.4 | 2 | 0 | 9 | 0.3333 | 22 | 0.3636 |
| 13_3421661 | NA | NA | 1 | 1 | 1 | 1 | NA | NA | 2 | 1 | NA | NA | 1 | 1 | 1 | 1 | NA | NA | 2 | 1 |
| 13_3421673 | 1 | 0 | 1 | 0 | 1 | 1 | NA | NA | 3 | 0.3333 | NA | NA | 1 | 0 | 1 | 1 | NA | NA | 2 | 0.5 |
| 13_3421679 | 1 | 1 | NA | NA | 1 | 0 | NA | NA | 2 | 0.5 | NA | NA | 1 | 0 | 1 | 1 | NA | NA | 2 | 0.5 |
| 13_3421684 | 1 | 1 | 1 | 0 | 1 | 1 | NA | NA | 3 | 0.6667 | NA | NA | 1 | 1 | 1 | 1 | NA | NA | 2 | 1 |
| 13_3421689 | 1 | 1 | 1 | 1 | 1 | 1 | NA | NA | 3 | 1 | NA | NA | 1 | 1 | 1 | 1 | NA | NA | 2 | 1 |
| 13_3421719 | 2 | 1 | 1 | 1 | 1 | 1 | NA | NA | 4 | 1 | NA | NA | 1 | 1 | 1 | 1 | NA | NA | 2 | 1 |
| 13_3421761 | 2 | 0 | 1 | 1 | 1 | 0 | NA | NA | 4 | 0.25 | NA | NA | 1 | 1 | 1 | 0 | NA | NA | 2 | 0.5 |
| 13_3421857 | 1 | 1 | NA | NA | NA | NA | NA | NA | 1 | 1 | NA | NA | NA | NA | 1 | 1 | NA | NA | 1 | 1 |
| 13_3425789 | 5 | 0.8 | 6 | 0.8333 | 4 | 1 | 5 | 0.8 | 20 | 0.85 | 8 | 0.875 | 4 | 0.75 | 2 | 1 | 5 | 0.8 | 19 | 0.8421 |
| 13_3425796 | 4 | 0.75 | 4 | 0.75 | 3 | 1 | 4 | 1 | 15 | 0.8667 | 6 | 0.6667 | 4 | 0.25 | 2 | 0.5 | 4 | 0.75 | 16 | 0.5625 |
| 13_3425798 | 4 | 0.25 | 3 | 0.6667 | 4 | 0 | 3 | 0 | 14 | 0.2143 | 6 | 0 | 3 | 0.3333 | 2 | 0 | 3 | 0.3333 | 14 | 0.1429 |
| 13_3425841 | 102 | 0.549 | 121 | 0.5702 | 100 | 0.58 | 57 | 0.5614 | 380 | 0.5658 | 75 | 0.5733 | 66 | 0.6061 | 57 | 0.5439 | 77 | 0.6234 | 275 | 0.5891 |
| 13_3425844 | 94 | 0.7021 | 114 | 0.5789 | 98 | 0.5714 | 61 | 0.4754 | 367 | 0.5913 | 74 | 0.527 | 63 | 0.5714 | 55 | 0.6727 | 67 | 0.6269 | 259 | 0.5946 |
| 13_3425889 | 82 | 0.0854 | 97 | 0.0309 | 68 | 0.0441 | 53 | 0.0377 | 300 | 0.05 | 58 | 0.2241 | 58 | 0.1034 | 47 | 0.0638 | 70 | 0.0429 | 233 | 0.1073 |
| 13_3425902 | 75 | 0.2667 | 86 | 0.2093 | 69 | 0.2174 | 49 | 0.1837 | 279 | 0.2222 | 54 | 0.1481 | 54 | 0.3333 | 44 | 0.1364 | 67 | 0.194 | 219 | 0.2055 |
| 13_3425918 | 75 | 0.3333 | 92 | 0.3261 | 69 | 0.4203 | 46 | 0.1957 | 282 | 0.3298 | 56 | 0.4464 | 58 | 0.4138 | 43 | 0.3256 | 66 | 0.3939 | 223 | 0.3991 |

| | | | | | | | | | | | | | | | | | | | | |
|---|---|---|---|---|---|---|---|---|---|---|---|---|---|---|---|---|---|---|---|---|
| 13_3425920 | 82 | 0.1707 | 95 | 0.1474 | 68 | 0.1029 | 52 | 0.1923 | 297 | 0.1515 | 51 | 0.1176 | 60 | 0.1333 | 46 | 0.1522 | 71 | 0.1972 | 228 | 0.1535 |
| 13_3425929 | 59 | 0.1525 | 71 | 0.169 | 58 | 0.0862 | 37 | 0.1892 | 225 | 0.1467 | 41 | 0.0976 | 42 | 0.0952 | 32 | 0.2188 | 46 | 0.1087 | 161 | 0.1242 |
| 13_3425934 | 79 | 0.1899 | 95 | 0.1789 | 69 | 0.1014 | 55 | 0.1091 | 298 | 0.151 | 56 | 0.1786 | 62 | 0.1935 | 45 | 0.1778 | 70 | 0.1571 | 233 | 0.176 |
| 13_3425940 | 51 | 0.7059 | 68 | 0.5882 | 49 | 0.6327 | 37 | 0.5946 | 205 | 0.6293 | 42 | 0.5714 | 38 | 0.5526 | 31 | 0.4194 | 47 | 0.5745 | 158 | 0.538 |
| 13_3425944 | 65 | 0.5231 | 81 | 0.4321 | 58 | 0.5517 | 47 | 0.5745 | 251 | 0.51 | 42 | 0.5238 | 57 | 0.5614 | 38 | 0.5789 | 56 | 0.6786 | 193 | 0.5907 |
| 13_3431178 | 9 | 0 | 26 | 0.0385 | 25 | 0.12 | 17 | 0.0588 | 77 | 0.0649 | 20 | 0.1 | 19 | 0.1579 | 19 | 0.0526 | 16 | 0 | 74 | 0.0811 |
| 13_3431189 | 5 | 0.2 | 13 | 0 | 23 | 0.3043 | 17 | 0.2941 | 58 | 0.2241 | 16 | 0.125 | 16 | 0.25 | 17 | 0.1765 | 14 | 0 | 63 | 0.1429 |
| 13_3431202 | 8 | 0.5 | 21 | 0.4762 | 22 | 0.2727 | 21 | 0.381 | 72 | 0.3889 | 18 | 0.6111 | 16 | 0.25 | 15 | 0.5333 | 14 | 0.5714 | 63 | 0.4921 |
| 13_3431216 | 8 | 0.25 | 23 | 0.0435 | 21 | 0.0476 | 19 | 0 | 71 | 0.0563 | 17 | 0.0588 | 17 | 0.0588 | 16 | 0.1875 | 14 | 0.0714 | 64 | 0.0938 |
| 13_3431226 | 8 | 0 | 22 | 0.4091 | 14 | 0.2143 | 12 | 0.0833 | 56 | 0.2321 | 16 | 0.25 | 17 | 0.2941 | 13 | 0.0769 | 14 | 0.3571 | 60 | 0.25 |
| 13_3431229 | 5 | 0.6 | 16 | 0.3125 | 18 | 0.2222 | 12 | 0.25 | 51 | 0.2941 | 13 | 0.3077 | 13 | 0.1538 | 10 | 0 | 11 | 0.2727 | 47 | 0.1915 |
| 13_3431234 | 8 | 0.5 | 19 | 0.3158 | 20 | 0.25 | 15 | 0.2 | 62 | 0.2903 | 18 | 0.2222 | 17 | 0.2353 | 13 | 0.3846 | 16 | 0.375 | 64 | 0.2969 |
| 13_3431240 | 3 | 0.3333 | 13 | 0.5385 | 15 | 0.6667 | 12 | 0.4167 | 43 | 0.5349 | 16 | 0.625 | 13 | 0.6923 | 7 | 0.7143 | 9 | 0.6667 | 45 | 0.6667 |
| 13_3431245 | 6 | 0.5 | 15 | 0.4667 | 14 | 0.6429 | 12 | 0.5833 | 47 | 0.5532 | 14 | 0.2857 | 13 | 0.4615 | 9 | 0.4444 | 13 | 0.5385 | 49 | 0.4286 |
| 13_3436323 | 5 | 0.6 | NA | NA | 1 | 0 | 5 | 0.6 | 11 | 0.5455 | 1 | 1 | 5 | 0.6 | 2 | 0 | 2 | 0 | 10 | 0.4 |
| 13_3436372 | 5 | 0.6 | 3 | 0 | 3 | 0 | 10 | 0.4 | 21 | 0.3333 | 4 | 0.5 | 11 | 0.1818 | 7 | 0.2857 | 5 | 0.2 | 27 | 0.2593 |
| 13_3436378 | 6 | 0.1667 | 4 | 0.25 | 3 | 0.3333 | 11 | 0.0909 | 24 | 0.1667 | 8 | 0.25 | 12 | 0.0833 | 8 | 0.25 | 7 | 0 | 35 | 0.1429 |
| 13_3436413 | 4 | 0 | 4 | 0.25 | 2 | 0 | 9 | 0.1111 | 19 | 0.1053 | 6 | 0.1667 | 12 | 0.0833 | 8 | 0 | 7 | 0 | 33 | 0.0606 |
| 13_3436461 | 4 | 0.5 | 3 | 0 | 2 | 0 | 10 | 0.5 | 19 | 0.3684 | 6 | 0 | 9 | 0.2222 | 8 | 0 | 5 | 0.4 | 28 | 0.1429 |
| 13_3436493 | 3 | 0.3333 | 3 | 0.3333 | 2 | 0 | 7 | 0.5714 | 15 | 0.4 | 5 | 0.6 | 8 | 0.375 | 7 | 0.2857 | 5 | 0.2 | 25 | 0.36 |
| 13_3436494 | 4 | 0 | 3 | 0.3333 | 2 | 0 | 8 | 0 | 17 | 0.0588 | 5 | 0.4 | 9 | 0 | 8 | 0 | 5 | 0.2 | 27 | 0.1111 |
| 13_3436518 | 4 | 0.25 | 2 | 0 | 2 | 0 | 8 | 0.125 | 16 | 0.125 | 5 | 0 | 10 | 0.1 | 7 | 0 | 5 | 0.2 | 27 | 0.0741 |
| 13_3436531 | 3 | 0 | 2 | 0 | 2 | 0.5 | 8 | 0.375 | 15 | 0.2667 | 6 | 0.1667 | 10 | 0.3 | 7 | 0.2857 | 6 | 0.6667 | 29 | 0.3448 |
| 13_3436542 | 3 | 1 | 2 | 0 | 2 | 0.5 | 7 | 0.7143 | 14 | 0.6429 | 5 | 0.8 | 9 | 0.7778 | 6 | 0.5 | 3 | 0.3333 | 23 | 0.6522 |
| 13_3436543 | 3 | 1 | 2 | 1 | 1 | 0 | 8 | 0.875 | 14 | 0.8571 | 5 | 0.8 | 10 | 1 | 7 | 0.7143 | 4 | 0.75 | 26 | 0.8462 |
| 13_3436557 | 2 | 0.5 | 2 | 0 | 2 | 0.5 | 5 | 0.6 | 11 | 0.4545 | 4 | 0.5 | 7 | 0.5714 | 6 | 0.3333 | 4 | 0.75 | 21 | 0.5238 |
| 13_3436568 | 2 | 0 | 2 | 0 | NA | NA | 6 | 0.3333 | 10 | 0.2 | 2 | 0 | 8 | 0 | 7 | 0.1429 | 3 | 0.3333 | 20 | 0.1 |
| 13_3436599 | 1 | 1 | 1 | 1 | 1 | 0 | 5 | 0.6 | 8 | 0.625 | 2 | 0.5 | 4 | 1 | 2 | 1 | 5 | 0.4 | 13 | 0.6923 |
| 13_3436607 | 1 | 1 | 1 | 0 | 1 | 0 | 6 | 0.5 | 9 | 0.4444 | 2 | 0.5 | 5 | 0.4 | 5 | 0.8 | 4 | 0.75 | 16 | 0.625 |
| 13_3436608 | 1 | 1 | 2 | 0.5 | 1 | 1 | 7 | 0.7143 | 11 | 0.7273 | 2 | 0.5 | 9 | 0.6667 | 2 | 1 | 5 | 0.4 | 18 | 0.6111 |
| 13_3436643 | 2 | 1 | 2 | 1 | 2 | 1 | 8 | 0.875 | 14 | 0.9286 | 3 | 1 | 6 | 1 | 3 | 0.6667 | 4 | 1 | 16 | 0.9375 |
| 13_3436646 | 2 | 1 | 1 | 1 | 2 | 1 | 7 | 1 | 12 | 1 | 3 | 1 | 5 | 1 | 2 | 1 | 4 | 1 | 14 | 1 |
| 13_3436649 | 2 | 0.5 | 2 | 0.5 | 1 | 0 | 8 | 0.375 | 13 | 0.3846 | 3 | 0 | 5 | 0 | 3 | 0.6667 | 4 | 0.75 | 15 | 0.3333 |
| 13_3436651 | 2 | 0 | 2 | 0 | 2 | 0.5 | 8 | 0 | 14 | 0.0714 | 3 | 0.3333 | 4 | 0 | 3 | 0 | 4 | 0 | 14 | 0.0714 |
| 13_3436654 | 2 | 1 | 2 | 1 | 2 | 0.5 | 8 | 1 | 14 | 0.9286 | 1 | 1 | 6 | 1 | 4 | 0.75 | 4 | 1 | 15 | 0.9333 |
| 13_3436658 | 2 | 0.5 | 2 | 0 | 2 | 0 | 7 | 0 | 13 | 0.0769 | 1 | 0 | 5 | 0 | 4 | 0 | 4 | 0.25 | 14 | 0.0714 |
| 13_3436672 | 2 | 1 | 2 | 1 | 2 | 1 | 8 | 1 | 14 | 1 | 2 | 1 | 5 | 1 | 4 | 0.75 | 3 | 1 | 14 | 0.9286 |
| 13_3436676 | 1 | 0 | 1 | 0 | 1 | 0 | 6 | 0.6667 | 9 | 0.4444 | 2 | 0.5 | 4 | 0 | 3 | 0.6667 | 4 | 0.75 | 13 | 0.4615 |
| 13_3436681 | 2 | 1 | 1 | 1 | 2 | 1 | 5 | 1 | 10 | 1 | 2 | 1 | 4 | 1 | 4 | 1 | 4 | 1 | 14 | 1 |
| 13_3436682 | 2 | 0.5 | 2 | 1 | 2 | 0.5 | 7 | 0.2857 | 13 | 0.4615 | 2 | 0 | 5 | 0.2 | 4 | 0.25 | 3 | 0 | 14 | 0.1429 |
| 13_3436685 | 2 | 0 | 2 | 0 | 2 | 0.5 | 6 | 0 | 12 | 0.0833 | 2 | 0 | 5 | 0.4 | 4 | 0 | 2 | 0.5 | 13 | 0.2308 |
| 13_3436692 | 2 | 0.5 | 2 | 0.5 | 2 | 0.5 | 6 | 0.3333 | 12 | 0.4167 | 2 | 0 | 5 | 0.4 | 2 | 1 | 3 | 0.3333 | 12 | 0.4167 |
| 13_3436694 | 2 | 1 | 2 | 0.5 | 2 | 0.5 | 7 | 0.4286 | 13 | 0.5385 | 2 | 1 | 4 | 0.75 | 4 | 0.75 | 3 | 0.6667 | 13 | 0.7692 |
| 13_3436701 | 2 | 1 | 2 | 1 | 2 | 1 | 7 | 1 | 13 | 1 | 2 | 1 | 5 | 1 | 2 | 1 | 1 | 1 | 10 | 1 |
| 13_3436709 | 1 | 0 | 1 | 1 | 1 | 0 | 3 | 0 | 6 | 0.1667 | 2 | 0.5 | 3 | 0.6667 | 2 | 0 | 2 | 0.5 | 9 | 0.4444 |
| 13_3436712 | 2 | 0 | 2 | 0 | 1 | 0 | 6 | 0.1667 | 11 | 0.0909 | 2 | 0.5 | 5 | 0 | 3 | 0 | 2 | 0 | 12 | 0.0833 |
| 13_3452832 | 2 | 0.5 | 1 | 0 | NA | NA | NA | NA | 3 | 0.3333 | 1 | 1 | 1 | 0 | 1 | 1 | 1 | 1 | 4 | 0.75 |
| 13_3452841 | 2 | 0.5 | 1 | 1 | 1 | 0 | 1 | 0 | 5 | 0.4 | 1 | 1 | 2 | 0 | 3 | 0.3333 | 1 | 0 | 7 | 0.2857 |
| 13_3452843 | 2 | 1 | 1 | 0 | 1 | 1 | 1 | 1 | 5 | 0.8 | 1 | 1 | 2 | 0.5 | 3 | 1 | 1 | 1 | 7 | 0.8571 |
| 13_3452854 | 2 | 0.5 | 1 | 1 | NA | NA | NA | NA | 3 | 0.6667 | 1 | 1 | 1 | 0 | 3 | 0.6667 | NA | NA | 5 | 0.6 |
| 13_3452896 | 2 | 0 | 1 | 1 | 1 | 1 | NA | NA | 4 | 0.5 | 1 | 1 | 1 | 1 | 1 | 1 | NA | NA | 3 | 1 |
| 13_3452916 | 1 | 1 | 1 | 1 | NA | NA | NA | NA | 2 | 1 | NA | NA | NA | NA | NA | NA | 1 | 1 | 1 | 1 |
| 13_3452936 | 1 | 1 | NA | NA | NA | NA | NA | NA | 1 | 1 | NA | NA | NA | NA | NA | NA | 1 | 1 | 1 | 1 |
| 13_3452947 | NA | NA | 1 | 1 | NA | NA | NA | NA | 1 | 1 | NA | NA | 1 | 1 | NA | NA | 1 | 1 | 2 | 1 |
| 13_3452951 | 1 | 1 | 1 | 1 | NA | NA | NA | NA | 2 | 1 | NA | NA | 1 | 1 | NA | NA | 1 | 1 | 2 | 1 |
| 13_3452964 | 1 | 1 | 1 | 1 | NA | NA | NA | NA | 2 | 1 | NA | NA | 1 | 1 | NA | NA | 1 | 1 | 2 | 1 |
| 13_3453043 | 1 | 1 | 1 | 1 | NA | NA | NA | NA | 2 | 1 | NA | NA | 1 | 1 | NA | NA | 1 | 1 | 2 | 1 |
| 13_3481234 | 1 | 1 | NA | NA | NA | NA | NA | NA | 1 | 1 | NA | NA | NA | NA | 1 | 1 | NA | NA | 1 | 1 |
| 13_3505773 | NA | NA | NA | NA | 1 | 1 | 2 | 0.5 | 3 | 0.6667 | 1 | 0 | 1 | 1 | NA | NA | NA | NA | 2 | 0.5 |
| 13_3505784 | NA | NA | NA | NA | 1 | 1 | 2 | 1 | 3 | 1 | 1 | 0 | 1 | 1 | NA | NA | NA | NA | 2 | 0.5 |
| 13_3505795 | NA | NA | NA | NA | 1 | 1 | 1 | 1 | 2 | 1 | 1 | 0 | 1 | 1 | NA | NA | NA | NA | 2 | 0.5 |
| 13_3505798 | NA | NA | NA | NA | 1 | 1 | 1 | 1 | 2 | 1 | 1 | 0 | 1 | 1 | NA | NA | NA | NA | 2 | 0.5 |
| 13_3506746 | 1 | 0 | 3 | 0 | 5 | 0.4 | 3 | 0.3333 | 12 | 0.25 | 1 | 0 | 1 | 0 | 6 | 0.1667 | 2 | 0.5 | 10 | 0.2 |
| 13_3506769 | 1 | 0 | 3 | 0.3333 | 5 | 0 | 3 | 0 | 12 | 0.0833 | 1 | 0 | 1 | 0 | 6 | 0.3333 | 2 | 0 | 10 | 0.2 |
| 13_3506776 | 1 | 0 | 2 | 0 | 5 | 0 | 3 | 0.3333 | 11 | 0.0909 | 1 | 0 | 1 | 0 | 6 | 0.3333 | 1 | 1 | 9 | 0.3333 |

| | | | | | | | | | | | | | | | | | | | | |
|---|---|---|---|---|---|---|---|---|---|---|---|---|---|---|---|---|---|---|---|---|
| 13_350683 | NA | NA | 1 | 0 | 3 | 0.3333 | 2 | 0 | 6 | 0.1667 | 1 | 1 | 1 | 0 | 3 | 0.3333 | 2 | 0 | 7 | 0.2857 |
| 13_350854 | 1 | 1 | 3 | 0 | 8 | 0.125 | 5 | 0.2 | 17 | 0.1765 | 3 | 0 | 5 | 0.2 | 8 | 0.125 | 4 | 0 | 20 | 0.1 |
| 13_350854 | 1 | 1 | 6 | 1 | 8 | 0.875 | 5 | 0.8 | 20 | 0.9 | 3 | 1 | 5 | 1 | 8 | 0.625 | 4 | 0.75 | 20 | 0.8 |
| 13_350854 | 1 | 0 | 5 | 0.2 | 7 | 0.2857 | 4 | 0.5 | 17 | 0.2941 | 3 | 0.3333 | 6 | 0.1667 | 9 | 0 | 4 | 0.25 | 22 | 0.1364 |
| 13_350855 | 1 | 1 | 5 | 0.2 | 8 | 0.125 | 5 | 0 | 19 | 0.1579 | 4 | 0.25 | 6 | 0 | 9 | 0.3333 | 4 | 0.25 | 23 | 0.2174 |
| 13_350856 | NA | NA | 5 | 0.8 | 5 | 0.8 | 5 | 0.8 | 15 | 0.8 | 3 | 1 | 6 | 1 | 9 | 0.8889 | 3 | 0.6667 | 21 | 0.9048 |
| 13_350856 | 1 | 1 | 5 | 0.4 | 8 | 0.375 | 4 | 0.5 | 18 | 0.4444 | 4 | 0.5 | 5 | 0.2 | 10 | 0.3 | 4 | 0.25 | 23 | 0.3043 |
| 13_350863 | 2 | 0 | 4 | 0 | 6 | 0.1667 | 2 | 0.5 | 14 | 0.1429 | NA | NA | 5 | 0 | 8 | 0 | 2 | 0.5 | 15 | 0.0667 |
| 13_350864 | 3 | 0 | 6 | 0 | 6 | 0.1667 | 2 | 0.5 | 17 | 0.1176 | 1 | 0 | 5 | 0.2 | 8 | 0.25 | 2 | 0 | 16 | 0.1875 |
| 13_350891 | 1 | 0 | 5 | 0.2 | 2 | 0 | 1 | 1 | 9 | 0.2222 | NA | NA | 4 | 0.5 | 4 | 0 | 1 | 0 | 9 | 0.2222 |
| 13_350895 | 1 | 0 | 4 | 0.25 | 2 | 0 | 1 | 0 | 8 | 0.125 | NA | NA | 4 | 0.25 | 4 | 0 | 1 | 0 | 9 | 0.1111 |
| 13_350898 | 2 | 0.5 | 5 | 0.2 | 4 | 0.25 | 3 | 0 | 14 | 0.2143 | 2 | 0 | 4 | 0.25 | 6 | 0.3333 | 3 | 0 | 15 | 0.2 |
| 13_350940 | NA | NA | NA | NA | NA | NA | 1 | 1 | 1 | 1 | NA | NA | 1 | 0 | NA | NA | 1 | 1 | 2 | 0.5 |
| 13_351618 | 9 | 0.2222 | 6 | 0.5 | 9 | 0.3333 | 9 | 0.4444 | 33 | 0.3636 | 7 | 0.4286 | 11 | 0.2727 | 11 | 0.4545 | 5 | 0.6 | 34 | 0.4118 |
| 13_360566 | NA | NA | NA | NA | 1 | 1 | NA | NA | 1 | 1 | 2 | 1 | NA | NA | 1 | 1 | 2 | 1 | 5 | 1 |
| 13_360577 | NA | NA | 2 | 0 | 1 | 0 | 2 | 0.5 | 5 | 0.2 | 3 | 0 | 1 | 0 | 2 | 1 | 2 | 0 | 8 | 0.25 |
| 13_360807 | NA | NA | 1 | 1 | 1 | 1 | NA | NA | 2 | 1 | 1 | 1 | NA | NA | 1 | 1 | NA | NA | 2 | 1 |
| 13_361525 | 1 | 1 | 1 | 1 | 2 | 0.5 | 1 | 1 | 5 | 0.8 | 1 | 0 | 5 | 0.4 | 1 | 0 | NA | NA | 7 | 0.2857 |
| 13_361525 | 1 | 1 | 1 | 1 | 2 | 0 | 2 | 0 | 6 | 0.3333 | NA | NA | 5 | 0.4 | 1 | 0 | NA | NA | 6 | 0.3333 |
| 13_361525 | 1 | 0 | 1 | 0 | 2 | 0.5 | 2 | 0 | 6 | 0.1667 | 1 | 0 | 5 | 0.2 | 1 | 0 | NA | NA | 7 | 0.1429 |
| 13_361526 | NA | NA | 1 | 0 | 1 | 1 | 2 | 0 | 4 | 0.25 | 1 | 1 | 5 | 0.2 | 1 | 0 | NA | NA | 7 | 0.2857 |
| 13_361529 | 2 | 0 | 1 | 1 | 2 | 0.5 | 4 | 0.25 | 9 | 0.3333 | 1 | 1 | 3 | 0.3333 | 1 | 1 | NA | NA | 5 | 0.6 |
| 13_361529 | 1 | 1 | 1 | 1 | 2 | 1 | 4 | 0.25 | 8 | 0.625 | 1 | 1 | 5 | 0.8 | 1 | 0 | NA | NA | 7 | 0.7143 |
| 13_361532 | 3 | 0.3333 | 1 | 1 | 2 | 0 | 5 | 0 | 11 | 0.1818 | 1 | 1 | 5 | 0.2 | 1 | 0 | NA | NA | 7 | 0.2857 |
| 13_361533 | 3 | 0.3333 | 1 | 0 | 2 | 0 | 5 | 0 | 11 | 0.0909 | 1 | 1 | 6 | 0.5 | 1 | 1 | NA | NA | 8 | 0.625 |
| 13_361533 | 3 | 1 | 1 | 1 | 2 | 1 | 5 | 1 | 11 | 1 | 1 | 1 | 4 | 0.75 | 2 | 1 | NA | NA | 7 | 0.8571 |
| 13_361535 | 3 | 0.3333 | 2 | 0 | 2 | 0 | 5 | 0.2 | 12 | 0.1667 | 1 | 0 | 4 | 0.5 | 1 | 1 | NA | NA | 6 | 0.5 |
| 13_361536 | 3 | 1 | 2 | 0.5 | 3 | 0.3333 | 4 | 0.25 | 12 | 0.5 | 1 | 0 | 4 | 0.5 | 2 | 0.5 | NA | NA | 7 | 0.4286 |
| 13_361536 | 2 | 1 | 2 | 1 | 3 | 0.3333 | 3 | 0.6667 | 10 | 0.7 | 1 | 1 | 5 | 0.8 | 1 | 1 | NA | NA | 7 | 0.8571 |
| 13_361538 | 2 | 0.5 | 2 | 0 | 3 | 0 | 4 | 0.25 | 11 | 0.1818 | NA | NA | 5 | 0.8 | 2 | 0.5 | NA | NA | 7 | 0.7143 |
| 13_361539 | 3 | 1 | 2 | 0.5 | 2 | 1 | 4 | 1 | 11 | 0.9091 | 1 | 1 | 4 | 0.75 | 2 | 0 | NA | NA | 7 | 0.5714 |
| 13_361539 | 2 | 1 | 2 | 1 | 3 | 0.6667 | 4 | 1 | 11 | 0.9091 | 1 | 1 | 4 | 0.5 | 1 | 1 | NA | NA | 6 | 0.6667 |
| 13_361539 | 2 | 1 | 2 | 1 | 3 | 1 | 4 | 1 | 11 | 1 | 1 | 1 | 5 | 1 | 2 | 1 | NA | NA | 8 | 1 |
| 13_361539 | 3 | 0.6667 | 2 | 1 | 3 | 0.3333 | 4 | 0.75 | 12 | 0.6667 | 1 | 1 | 5 | 0.6 | 2 | 0.5 | NA | NA | 8 | 0.625 |
| 13_361635 | 10 | 0 | 9 | 0.1111 | 9 | 0 | 8 | 0.125 | 36 | 0.0556 | 9 | 0.3333 | 9 | 0 | 13 | 0.1538 | 13 | 0.4615 | 44 | 0.25 |
| 13_361636 | 9 | 0.8889 | 7 | 1 | 5 | 1 | 8 | 1 | 29 | 0.9655 | 8 | 1 | 11 | 0.9091 | 11 | 1 | 12 | 0.8333 | 42 | 0.9286 |
| 13_361637 | 4 | 0 | 6 | 0.1667 | 5 | 0.2 | 6 | 0 | 21 | 0.0952 | 7 | 0 | 6 | 0.5 | 10 | 0.5 | 10 | 0.3 | 33 | 0.3333 |
| 13_363889 | NA | NA | 1 | 1 | NA | NA | 1 | 0 | 2 | 0.5 | 1 | 0 | 1 | 0 | 4 | 0 | 4 | 0.5 | 10 | 0.2 |
| 13_363889 | 1 | 0 | 1 | 1 | NA | NA | 1 | 0 | 3 | 0.3333 | 1 | 0 | 2 | 1 | 4 | 0.75 | 4 | 0.5 | 11 | 0.6364 |
| 13_363890 | NA | NA | 1 | 1 | NA | NA | 1 | 1 | 2 | 1 | 1 | 0 | 2 | 0 | 3 | 0 | 4 | 0.5 | 10 | 0.2 |
| 13_367214 | 2 | 0.5 | 4 | 0.25 | 1 | 1 | 1 | 0 | 8 | 0.375 | 2 | 0 | 2 | 0 | 4 | 0.25 | 2 | 0 | 10 | 0.1 |
| 13_369103 | 5 | 0 | 14 | 0.2857 | 7 | 0.4286 | 1 | 0 | 27 | 0.2593 | 9 | 0.2222 | 1 | 0 | NA | NA | 5 | 0 | 15 | 0.1333 |
| 13_369107 | 6 | 0.1667 | 12 | 0.1667 | 6 | 0.1667 | 2 | 0 | 26 | 0.1538 | 7 | 0.2857 | 1 | 1 | 1 | 0 | 4 | 0.25 | 13 | 0.3077 |
| 13_369107 | 7 | 0 | 19 | 0.1053 | 7 | 0.1429 | 2 | 0 | 35 | 0.0857 | 6 | 0 | 2 | 0 | 1 | 0 | 6 | 0.1667 | 15 | 0.0667 |
| 13_381330 | NA | NA | NA | NA | NA | NA | 1 | 1 | 1 | 1 | NA | NA | NA | NA | 1 | 1 | NA | NA | 1 | 1 |
| 13_381331 | NA | NA | NA | NA | NA | NA | 1 | 1 | 1 | 1 | NA | NA | NA | NA | 1 | 1 | NA | NA | 1 | 1 |
| 13_381332 | NA | NA | NA | NA | NA | NA | 2 | 0.5 | 2 | 0.5 | NA | NA | NA | NA | 1 | 1 | NA | NA | 1 | 1 |
| 13_381333 | 1 | 1 | NA | NA | NA | NA | 2 | 1 | 3 | 1 | NA | NA | NA | NA | 1 | 1 | NA | NA | 1 | 1 |
| 13_381334 | 1 | 1 | NA | NA | NA | NA | 2 | 1 | 3 | 1 | NA | NA | NA | NA | 1 | 1 | NA | NA | 1 | 1 |
| 13_381355 | 2 | 1 | NA | NA | NA | NA | 2 | 1 | 4 | 1 | 2 | 0.5 | NA | NA | 2 | 1 | NA | NA | 4 | 0.75 |
| 13_381357 | NA | NA | 1 | 1 | NA | NA | 2 | 1 | 3 | 1 | 2 | 1 | NA | NA | 2 | 0.5 | NA | NA | 4 | 0.75 |
| 13_381559 | 2 | 0 | 7 | 0 | 6 | 0.1667 | 4 | 0 | 19 | 0.0526 | 2 | 0 | 3 | 0 | 1 | 0 | 4 | 0.25 | 10 | 0.1 |
| 13_381560 | 1 | 1 | 6 | 0.5 | 6 | 0.3333 | 5 | 0 | 18 | 0.3333 | NA | NA | 2 | 0 | 1 | 0 | 3 | 0.6667 | 6 | 0.3333 |
| 13_381565 | 1 | 1 | 6 | 0 | 4 | 0 | 4 | 0 | 15 | 0.0667 | NA | NA | 2 | 0 | 1 | 1 | 2 | 0 | 5 | 0.2 |
| 13_381622 | 6 | 0 | 13 | 0.0769 | 7 | 0 | 13 | 0.0769 | 39 | 0.0513 | 9 | 0 | 9 | 0 | 15 | 0.2 | 8 | 0.125 | 41 | 0.0976 |
| 13_381625 | 8 | 0.625 | 9 | 0.4444 | 9 | 0.3333 | 14 | 0.3571 | 40 | 0.425 | 8 | 0.25 | 9 | 0 | 15 | 0.4 | 13 | 0.0769 | 45 | 0.2 |
| 13_381629 | 8 | 0.375 | 6 | 0.1667 | 7 | 0.4286 | 13 | 0.3846 | 34 | 0.3529 | 10 | 0.3 | 7 | 0.2857 | 15 | 0.4667 | 10 | 0.2 | 42 | 0.3333 |
| 13_381634 | 6 | 0 | 10 | 0.2 | 5 | 0 | 9 | 0.2222 | 30 | 0.1333 | 4 | 0 | 3 | 0 | 15 | 0.1333 | 10 | 0.1 | 32 | 0.0938 |
| 13_381635 | 7 | 0.4286 | 9 | 0.5556 | 4 | 1 | 11 | 0.8182 | 31 | 0.6774 | 8 | 0.75 | 5 | 0.4 | 16 | 0.6875 | 10 | 0.7 | 39 | 0.6667 |
| 13_381876 | NA | NA | NA | NA | 1 | 1 | 1 | 1 | 2 | 1 | NA | NA | NA | NA | 1 | 1 | NA | NA | 1 | 1 |
| 13_381877 | NA | NA | NA | NA | 1 | 1 | 1 | 0 | 2 | 0.5 | NA | NA | NA | NA | 1 | 1 | NA | NA | 1 | 1 |
| 13_381881 | NA | NA | NA | NA | 1 | 1 | 1 | 1 | 2 | 1 | NA | NA | NA | NA | 1 | 1 | NA | NA | 1 | 1 |
| 13_381881 | NA | NA | NA | NA | 1 | 0 | 1 | 1 | 2 | 0.5 | NA | NA | NA | NA | 1 | 1 | NA | NA | 1 | 1 |
| 13_381882 | NA | NA | NA | NA | 1 | 1 | NA | NA | 1 | 1 | NA | NA | NA | NA | 1 | 1 | NA | NA | 1 | 1 |
| 13_381883 | NA | NA | NA | NA | 1 | 0 | 1 | 1 | 2 | 0.5 | NA | NA | NA | NA | 1 | 1 | NA | NA | 1 | 1 |

| | | | | | | | | | | | | | | | | | | | | |
|---|---|---|---|---|---|---|---|---|---|---|---|---|---|---|---|---|---|---|---|---|
| 13_3818852 | NA | NA | NA | NA | 1 | 1 | 1 | 0 | 2 | 0.5 | NA | NA | NA | NA | 1 | 1 | NA | NA | 1 | 1 |
| 13_3818873 | NA | NA | NA | NA | 1 | 1 | 1 | 0 | 2 | 0.5 | NA | NA | NA | NA | 1 | 1 | NA | NA | 1 | 1 |
| 13_3818890 | NA | NA | NA | NA | 1 | 1 | NA | NA | 1 | 1 | NA | NA | NA | NA | 1 | 1 | NA | NA | 1 | 1 |
| 13_3818906 | NA | NA | NA | NA | 1 | 1 | NA | NA | 1 | 1 | NA | NA | NA | NA | 1 | 1 | NA | NA | 1 | 1 |
| 13_3818910 | NA | NA | NA | NA | 1 | 0 | 1 | 1 | 2 | 0.5 | NA | NA | NA | NA | 1 | 1 | NA | NA | 1 | 1 |
| 13_3818919 | NA | NA | NA | NA | 1 | 1 | 1 | 0 | 2 | 0.5 | NA | NA | NA | NA | 1 | 1 | NA | NA | 1 | 1 |
| 13_3820714 | NA | NA | NA | NA | NA | NA | 1 | 1 | 1 | 1 | NA | NA | NA | NA | NA | NA | 1 | 1 | 1 | 1 |
| 13_3820719 | NA | NA | NA | NA | NA | NA | 1 | 1 | 1 | 1 | NA | NA | NA | NA | NA | NA | 1 | 1 | 1 | 1 |
| 13_3824499 | 2 | 1 | 1 | 1 | 4 | 0.75 | NA | NA | 7 | 0.8571 | 2 | 0.5 | 2 | 0.5 | 4 | 1 | 3 | 1 | 11 | 0.8182 |
| 13_3824502 | 1 | 1 | 1 | 0 | 4 | 0.5 | NA | NA | 6 | 0.5 | 2 | 0 | 2 | 0 | 4 | 0.25 | 2 | 0.5 | 10 | 0.2 |
| 13_3824511 | 2 | 0.5 | 1 | 0 | 3 | 0.3333 | NA | NA | 6 | 0.3333 | 2 | 0 | 2 | 0.5 | 4 | 0.25 | 1 | 1 | 9 | 0.3333 |
| 13_3824512 | 2 | 0 | 1 | 1 | 2 | 0 | NA | NA | 5 | 0.2 | NA | NA | 2 | 0.5 | 3 | 0 | 2 | 0.5 | 7 | 0.2857 |
| 13_3824515 | 2 | 0 | 1 | 0 | 4 | 0.25 | NA | NA | 7 | 0.1429 | 2 | 0.5 | 2 | 0 | 3 | 0.6667 | 2 | 0 | 9 | 0.3333 |
| 13_3824519 | 2 | 0 | NA | NA | 2 | 0.5 | NA | NA | 4 | 0.25 | 2 | 0 | 2 | 0 | 3 | 0.3333 | 3 | 0 | 10 | 0.1 |
| 13_3824536 | 2 | 0.5 | 1 | 0 | 4 | 0.25 | NA | NA | 7 | 0.2857 | 2 | 0 | 2 | 0.5 | 4 | 0.25 | 2 | 0 | 10 | 0.2 |
| 13_3868322 | 2 | 1 | 1 | 1 | 1 | 1 | 1 | 1 | 5 | 1 | 1 | 1 | NA | NA | NA | NA | 2 | 0.5 | 3 | 0.6667 |
| 13_3868447 | 10 | 0.5 | 7 | 0.4286 | 10 | 0.6 | 9 | 0.4444 | 36 | 0.5 | 7 | 0.5714 | 10 | 0.3 | 7 | 0.5714 | 10 | 0.5 | 34 | 0.4706 |
| 13_3868456 | 8 | 0 | 6 | 0.1667 | 8 | 0.375 | 10 | 0 | 32 | 0.125 | 6 | 0.3333 | 7 | 0 | 6 | 0.1667 | 10 | 0 | 29 | 0.1034 |
| 13_3868471 | 8 | 0.75 | 6 | 0.5 | 10 | 0.7 | 8 | 0.625 | 32 | 0.6562 | 4 | 0.5 | 8 | 0.75 | 4 | 0.75 | 9 | 0.6667 | 25 | 0.68 |
| 13_3868524 | 6 | 0.6667 | 6 | 0.6667 | 7 | 0.1429 | 9 | 0.2222 | 28 | 0.3929 | 4 | 0.5 | 5 | 0.4 | 5 | 0.2 | 10 | 0.4 | 24 | 0.375 |
| 13_3868527 | 3 | 0.6667 | 4 | 0.25 | 4 | 0.75 | 5 | 0.6 | 16 | 0.5625 | 3 | 0.3333 | 2 | 0.5 | 1 | 0 | 5 | 0.8 | 11 | 0.5455 |
| 13_3868541 | 5 | 1 | 6 | 0.8333 | 5 | 0.8 | 10 | 0.8 | 26 | 0.8462 | 4 | 1 | 6 | 1 | 5 | 1 | 7 | 1 | 22 | 1 |
| 13_3868556 | 5 | 0.8 | 3 | 0.3333 | 3 | 0 | 5 | 0 | 16 | 0.3125 | NA | NA | 4 | 0 | 3 | 0.3333 | 8 | 0.5 | 15 | 0.3333 |
| 13_3878678 | 5 | 0 | 5 | 0.2 | 7 | 0 | NA | NA | 17 | 0.0588 | NA | NA | 4 | 0 | 6 | 0 | 2 | 0.5 | 12 | 0.0833 |
| 13_3878719 | 5 | 0.2 | 5 | 0.2 | 6 | 0 | 2 | 0 | 18 | 0.1111 | 1 | 0 | 3 | 0 | 4 | 0.25 | 1 | 0 | 9 | 0.1111 |
| 13_3878728 | 5 | 0.2 | 5 | 0.2 | 7 | 0.1429 | 2 | 0 | 19 | 0.1579 | 1 | 0 | 4 | 0.5 | 5 | 0 | 2 | 0 | 12 | 0.1667 |
| 13_3878735 | 5 | 0.4 | 2 | 0 | 7 | 0.2857 | 2 | 0 | 16 | 0.25 | 1 | 0 | 4 | 0 | 6 | 0.3333 | 2 | 0 | 13 | 0.1538 |
| 13_3878759 | 4 | 0.5 | 3 | 0.3333 | 6 | 0 | 1 | 0 | 14 | 0.2143 | 1 | 1 | 3 | 0 | 4 | 0.5 | 1 | 0 | 9 | 0.3333 |
| 13_3878764 | 3 | 0.3333 | 5 | 0 | 7 | 0 | 2 | 0 | 17 | 0.0588 | 1 | 0 | 4 | 0 | 4 | 0 | 2 | 0.5 | 11 | 0.0909 |
| 13_3878769 | 4 | 0.25 | 5 | 0.2 | 7 | 0.2857 | 2 | 0 | 18 | 0.2222 | 1 | 0 | 4 | 0 | 4 | 0.25 | 2 | 0.5 | 11 | 0.1818 |
| 13_3878780 | 3 | 1 | 4 | 0.25 | 4 | 0.5 | 2 | 1 | 13 | 0.6154 | 1 | 1 | NA | NA | 5 | 0.6 | 1 | 0 | 7 | 0.5714 |
| 13_3878782 | 2 | 1 | 3 | 1 | 4 | 1 | 2 | 0.5 | 11 | 0.9091 | 1 | 1 | 3 | 0.3333 | 6 | 1 | 1 | 1 | 11 | 0.8182 |
| 13_3878785 | 4 | 0.25 | 5 | 0.2 | 6 | 0 | 2 | 0 | 17 | 0.1176 | 1 | 0 | 3 | 0.3333 | 6 | 0.1667 | 2 | 0 | 12 | 0.1667 |
| 13_3883009 | 5 | 0 | 8 | 0 | 5 | 0.4 | 11 | 0.1818 | 29 | 0.1379 | 8 | 0.125 | 5 | 0.4 | 5 | 0.2 | 5 | 0.2 | 23 | 0.2174 |
| 13_3883016 | 5 | 0.2 | 8 | 0.625 | 6 | 0.6667 | 9 | 0.2222 | 28 | 0.4286 | 8 | 0.375 | 3 | 0.3333 | 5 | 0 | 4 | 0.75 | 20 | 0.35 |
| 13_3911635 | 2 | 0.5 | 3 | 1 | 1 | 1 | 2 | 0.5 | 8 | 0.75 | NA | NA | 1 | 1 | 3 | 0 | 2 | 0 | 6 | 0.1667 |
| 13_3911645 | 1 | 1 | 3 | 1 | 1 | 1 | 4 | 0.5 | 9 | 0.7778 | NA | NA | 1 | 1 | 2 | 1 | 3 | 0 | 6 | 0.5 |
| 13_3911646 | 2 | 0.5 | 2 | 0.5 | NA | NA | 3 | 1 | 7 | 0.7143 | NA | NA | 1 | 1 | 2 | 1 | 3 | 1 | 6 | 1 |
| 13_3911648 | 1 | 1 | 2 | 0.5 | 1 | 0 | 1 | 1 | 5 | 0.6 | NA | NA | NA | NA | 2 | 0 | 1 | 1 | 3 | 0.3333 |
| 13_3911673 | 2 | 0.5 | 3 | 1 | 1 | 1 | 3 | 0.6667 | 9 | 0.7778 | NA | NA | 1 | 1 | 1 | 1 | 4 | 0.75 | 6 | 0.8333 |
| 13_3911675 | 2 | 1 | 3 | 1 | 1 | 1 | 3 | 1 | 9 | 1 | NA | NA | 1 | 1 | NA | NA | 4 | 0.75 | 5 | 0.8 |
| 13_4107130 | 1 | 0 | NA | NA | NA | NA | 2 | 0.5 | 3 | 0.3333 | NA | NA | 3 | 0 | 1 | 0 | 1 | 1 | 5 | 0.2 |
| 13_4107156 | 2 | 0.5 | NA | NA | NA | NA | 3 | 0 | 5 | 0.2 | NA | NA | 2 | 0.5 | 1 | 0 | 1 | 1 | 4 | 0.5 |
| 13_4107160 | 2 | 0.5 | NA | NA | NA | NA | 3 | 1 | 5 | 0.8 | NA | NA | 3 | 0.3333 | 1 | 1 | 1 | 1 | 5 | 0.6 |
| 13_4107180 | 2 | 1 | NA | NA | NA | NA | 3 | 1 | 5 | 1 | NA | NA | 3 | 0.3333 | 1 | 1 | 1 | 1 | 5 | 0.6 |
| 13_4107193 | 2 | 0.5 | NA | NA | NA | NA | 3 | 0.6667 | 5 | 0.6 | NA | NA | 2 | 0 | 1 | 1 | 1 | 0 | 4 | 0.25 |
| 13_4107203 | 2 | 1 | NA | NA | NA | NA | 3 | 1 | 5 | 1 | NA | NA | 3 | 1 | 1 | 1 | 2 | 1 | 6 | 1 |
| 13_4107216 | 2 | 1 | NA | NA | NA | NA | 3 | 1 | 5 | 1 | NA | NA | 3 | 1 | 1 | 1 | 2 | 1 | 6 | 1 |
| 13_4107224 | 2 | 0 | NA | NA | NA | NA | 2 | 0.5 | 4 | 0.25 | NA | NA | 3 | 0.3333 | 1 | 1 | 1 | 1 | 5 | 0.6 |
| 13_4111017 | 1 | 1 | NA | NA | NA | NA | NA | NA | 1 | 1 | NA | NA | 1 | 1 | NA | NA | 1 | 1 | 2 | 1 |
| 13_4111022 | 1 | 1 | 1 | 1 | NA | NA | NA | NA | 2 | 1 | NA | NA | 1 | 0 | NA | NA | 1 | 1 | 2 | 0.5 |
| 13_4111025 | 1 | 1 | 1 | 1 | NA | NA | NA | NA | 2 | 1 | NA | NA | 1 | 1 | NA | NA | 1 | 1 | 2 | 1 |
| 13_4111027 | 1 | 1 | 1 | 1 | NA | NA | NA | NA | 2 | 1 | NA | NA | 1 | 0 | NA | NA | 1 | 1 | 2 | 0.5 |
| 13_4111075 | 1 | 1 | NA | NA | NA | NA | NA | NA | 1 | 1 | NA | NA | 1 | 1 | NA | NA | NA | NA | 1 | 1 |
| 13_4111078 | 1 | 1 | NA | NA | NA | NA | NA | NA | 1 | 1 | NA | NA | 1 | 1 | NA | NA | NA | NA | 1 | 1 |
| 13_4111089 | 1 | 1 | 1 | 1 | NA | NA | NA | NA | 2 | 1 | NA | NA | 1 | 1 | NA | NA | NA | NA | 1 | 1 |
| 13_4111160 | 3 | 0 | NA | NA | 2 | 0.5 | 2 | 0 | 7 | 0.1429 | NA | NA | 3 | 0.3333 | 2 | 0 | 2 | 0.5 | 7 | 0.2857 |
| 13_4111171 | 3 | 1 | NA | NA | 2 | 1 | NA | NA | 5 | 1 | NA | NA | 3 | 1 | 2 | 1 | 2 | 1 | 7 | 1 |
| 13_4111175 | 2 | 0.5 | NA | NA | 1 | 0 | 3 | 0.6667 | 6 | 0.5 | NA | NA | 3 | 0.3333 | 2 | 0.5 | 2 | 0.5 | 7 | 0.4286 |
| 13_4111177 | 3 | 0 | NA | NA | 2 | 0.5 | 2 | 0 | 7 | 0.1429 | NA | NA | 2 | 0.5 | 2 | 0.5 | 2 | 0 | 6 | 0.3333 |
| 13_4111188 | 3 | 1 | NA | NA | 2 | 1 | 3 | 1 | 8 | 1 | NA | NA | 3 | 1 | 2 | 1 | 2 | 1 | 7 | 1 |
| 13_4111190 | 3 | 0.3333 | NA | NA | 2 | 1 | 3 | 1 | 8 | 0.75 | NA | NA | 3 | 0.6667 | 2 | 1 | 1 | 1 | 6 | 0.8333 |
| 13_4111214 | 1 | 1 | NA | NA | 1 | 1 | 3 | 1 | 5 | 1 | NA | NA | 2 | 1 | 1 | 1 | 1 | 0 | 4 | 0.75 |
| 13_4111236 | 3 | 1 | NA | NA | 1 | 1 | 3 | 0.6667 | 7 | 0.8571 | NA | NA | 3 | 1 | 1 | 0 | 2 | 0.5 | 6 | 0.6667 |
| 13_4111244 | 3 | 0.6667 | NA | NA | 2 | 1 | 3 | 0.3333 | 8 | 0.625 | NA | NA | 3 | 0.3333 | 2 | 0.5 | 2 | 0.5 | 7 | 0.4286 |

| | | | | | | | | | | | | | | | | | | | | |
|---|---|---|---|---|---|---|---|---|---|---|---|---|---|---|---|---|---|---|---|---|
| 13_411124 | 3 | 0.3333 | NA | NA | 2 | 0 | 3 | 0 | 8 | 0.125 | NA | NA | 3 | 0.3333 | 2 | 0 | 2 | 0 | 7 | 0.1429 |
| 13_411124 | 3 | 0.3333 | NA | NA | 2 | 0 | 2 | 0.5 | 7 | 0.2857 | NA | NA | 3 | 1 | 1 | 1 | NA | NA | 4 | 1 |
| 13_411125 | 3 | 0.6667 | NA | NA | 2 | 1 | 3 | 0.3333 | 8 | 0.625 | NA | NA | 3 | 0.3333 | 2 | 0.5 | 2 | 0 | 7 | 0.2857 |
| 13_411125 | 1 | 0 | NA | NA | 1 | 1 | 3 | 0.6667 | 5 | 0.6 | NA | NA | 2 | 1 | 1 | 1 | 2 | 1 | 5 | 1 |
| 13_411125 | 3 | 0.3333 | NA | NA | 2 | 0.5 | 3 | 0.3333 | 8 | 0.375 | NA | NA | 2 | 0.5 | 2 | 0 | 2 | 0 | 6 | 0.1667 |
| 13_411126 | 3 | 0.3333 | NA | NA | 2 | 0.5 | 3 | 0.3333 | 8 | 0.375 | NA | NA | 2 | 0.5 | 3 | 1 | 2 | 1 | 7 | 0.8571 |
| 13_411127 | 4 | 1 | NA | NA | 2 | 1 | 2 | 1 | 8 | 1 | NA | NA | 3 | 1 | 3 | 1 | 2 | 1 | 8 | 1 |
| 13_411139 | 6 | 0.3333 | 1 | 0 | 2 | 0 | 4 | 0 | 13 | 0.1538 | NA | NA | 4 | 0.25 | 6 | 0 | 2 | 0 | 12 | 0.0833 |
| 13_411141 | 7 | 0 | 1 | 1 | 2 | 0.5 | 4 | 0 | 14 | 0.1429 | 1 | 0 | 5 | 0 | 6 | 0.1667 | 3 | 0 | 15 | 0.0667 |
| 13_411750 | 1 | 0 | NA | NA | NA | NA | 1 | 1 | 2 | 0.5 | NA | NA | NA | NA | NA | NA | 2 | 0.5 | 2 | 0.5 |
| 13_411759 | 12 | 0.1667 | 10 | 0.1 | 16 | 0 | 10 | 0.1 | 48 | 0.0833 | 12 | 0 | 18 | 0.1667 | 13 | 0.1538 | 11 | 0.2727 | 54 | 0.1481 |
| 13_413386 | 1 | 1 | 1 | 1 | 1 | 0 | NA | NA | 3 | 0.6667 | NA | NA | NA | NA | 2 | 1 | 2 | 0.5 | 4 | 0.75 |
| 13_414262 | 6 | 0 | 12 | 0.0833 | 10 | 0 | 9 | 0.1111 | 37 | 0.0541 | 8 | 0 | 4 | 0 | 13 | 0.0769 | 14 | 0.0714 | 39 | 0.0513 |
| 13_414263 | 7 | 0.1429 | 11 | 0 | 10 | 0.2 | 10 | 0.2 | 38 | 0.1316 | 6 | 0.3333 | 5 | 0.4 | 13 | 0.0769 | 13 | 0.0769 | 37 | 0.1622 |
| 13_414263 | 5 | 0.8 | 10 | 0.6 | 6 | 0.8333 | 6 | 0.8333 | 27 | 0.7407 | 3 | 0.3333 | 3 | 0.3333 | 9 | 0.6667 | 5 | 0.4 | 20 | 0.5 |
| 13_414264 | 5 | 1 | 11 | 1 | 10 | 0.8 | 8 | 0.875 | 34 | 0.9118 | 7 | 0.8571 | 4 | 0.75 | 11 | 0.9091 | 14 | 0.8571 | 36 | 0.8611 |
| 13_414265 | 7 | 0 | 12 | 0.0833 | 14 | 0.1429 | 10 | 0 | 43 | 0.0698 | 9 | 0 | 7 | 0.1429 | 14 | 0.0714 | 15 | 0.0667 | 45 | 0.0667 |
| 13_414267 | 7 | 0.2857 | 15 | 0.0667 | 14 | 0.2143 | 10 | 0.1 | 46 | 0.1522 | 9 | 0 | 6 | 0 | 14 | 0.0714 | 15 | 0.1333 | 44 | 0.0682 |
| 13_414268 | 7 | 0.1429 | 11 | 0 | 11 | 0.0909 | 9 | 0 | 38 | 0.0526 | 7 | 0.2857 | 6 | 0.6667 | 8 | 0.125 | 12 | 0.1667 | 33 | 0.2727 |
| 13_414269 | 6 | 0.8333 | 15 | 0.7333 | 13 | 0.7692 | 10 | 0.5 | 44 | 0.7045 | 8 | 0.375 | 6 | 0.8333 | 14 | 0.6429 | 14 | 0.7143 | 42 | 0.6429 |
| 13_414270 | 7 | 0.5714 | 17 | 0.4706 | 9 | 0.7778 | 11 | 0.3636 | 44 | 0.5227 | 8 | 0.375 | 5 | 0.8 | 14 | 0.3571 | 15 | 0.4667 | 42 | 0.4524 |
| 13_414271 | 7 | 0.4286 | 15 | 0.4 | 12 | 0.6667 | 8 | 0.125 | 42 | 0.4286 | 9 | 0.2222 | 7 | 0.5714 | 13 | 0.3077 | 15 | 0.6 | 44 | 0.4318 |
| 13_414278 | 8 | 0.25 | 11 | 0.0909 | 11 | 0.1818 | 9 | 0.3333 | 39 | 0.2051 | 8 | 0.25 | 2 | 0 | 14 | 0.1429 | 13 | 0.1538 | 37 | 0.1622 |
| 13_414279 | 11 | 0.1818 | 17 | 0.0588 | 16 | 0.1875 | 11 | 0 | 55 | 0.1091 | 9 | 0.1111 | 7 | 0 | 20 | 0.15 | 18 | 0.0556 | 54 | 0.0926 |
| 13_414290 | 19 | 0.2632 | 19 | 0.2105 | 20 | 0 | 19 | 0 | 77 | 0.1169 | 19 | 0.0526 | 14 | 0.2143 | 19 | 0.1053 | 22 | 0.0455 | 74 | 0.0946 |
| 13_414297 | 26 | 0.4231 | 24 | 0.2917 | 29 | 0.2759 | 21 | 0.4762 | 100 | 0.36 | 25 | 0.36 | 15 | 0.5333 | 25 | 0.44 | 27 | 0.4074 | 92 | 0.4239 |
| 13_414298 | 29 | 0.4138 | 26 | 0.3846 | 29 | 0.4828 | 24 | 0.4167 | 108 | 0.4259 | 26 | 0.3077 | 18 | 0.4444 | 26 | 0.3077 | 29 | 0.3448 | 99 | 0.3434 |
| 13_414637 | 1 | 0 | NA | NA | 1 | 1 | 1 | 0 | 3 | 0.3333 | 1 | 1 | 1 | 0 | 1 | 1 | 1 | 0 | 4 | 0.5 |
| 13_414638 | 1 | 1 | NA | NA | 1 | 1 | 1 | 1 | 3 | 1 | 1 | 1 | 1 | 1 | 1 | 1 | 1 | 1 | 4 | 1 |
| 13_414638 | 2 | 0.5 | NA | NA | 1 | 0 | 1 | 1 | 4 | 0.5 | 2 | 1 | 1 | 0 | 2 | 1 | NA | NA | 5 | 0.8 |
| 13_414640 | 2 | 0.5 | NA | NA | 1 | 0 | NA | NA | 3 | 0.3333 | 1 | 0 | NA | NA | 2 | 0.5 | NA | NA | 3 | 0.3333 |
| 13_414641 | 2 | 1 | NA | NA | 1 | 1 | 1 | 1 | 4 | 1 | 1 | 1 | 1 | 1 | 2 | 1 | NA | NA | 4 | 1 |
| 13_414641 | NA | NA | NA | NA | 1 | 1 | 1 | 1 | 2 | 1 | 2 | 1 | 1 | 1 | 1 | 1 | NA | NA | 4 | 1 |
| 13_414642 | 2 | 1 | NA | NA | 1 | 1 | 1 | 0 | 4 | 0.75 | 2 | 1 | 1 | 1 | 2 | 1 | NA | NA | 5 | 1 |
| 13_414643 | 1 | 1 | NA | NA | 1 | 1 | NA | NA | 2 | 1 | 2 | 1 | 1 | 1 | 2 | 1 | NA | NA | 5 | 1 |
| 13_414648 | 2 | 1 | NA | NA | 1 | 1 | NA | NA | 3 | 1 | 2 | 0.5 | 1 | 1 | NA | NA | 1 | 0 | 4 | 0.5 |
| 13_414650 | 1 | 0 | NA | NA | 1 | 1 | 1 | 0 | 3 | 0.3333 | 1 | 1 | NA | NA | NA | NA | NA | NA | 1 | 1 |
| 13_414651 | 2 | 1 | NA | NA | 1 | 1 | 1 | 1 | 4 | 1 | 2 | 1 | 1 | 1 | NA | NA | NA | NA | 3 | 1 |
| 13_414653 | 2 | 0.5 | NA | NA | NA | NA | 1 | 0 | 3 | 0.3333 | 2 | 0.5 | 1 | 1 | NA | NA | NA | NA | 3 | 0.6667 |
| 13_414653 | 2 | 1 | NA | NA | 1 | 1 | 1 | 1 | 4 | 1 | 2 | 1 | 1 | 1 | NA | NA | NA | NA | 3 | 1 |
| 13_414653 | 2 | 1 | NA | NA | 1 | 1 | 1 | 1 | 4 | 1 | 1 | 0 | 1 | 1 | NA | NA | NA | NA | 2 | 0.5 |
| 13_414653 | 2 | 1 | NA | NA | 1 | 1 | NA | NA | 3 | 1 | 2 | 1 | 1 | 1 | NA | NA | NA | NA | 3 | 1 |
| 13_414694 | 2 | 1 | NA | NA | 1 | 1 | 1 | 1 | 4 | 1 | 2 | 0 | 1 | 1 | NA | NA | NA | NA | 3 | 0.3333 |
| 13_414699 | 1 | 1 | NA | NA | NA | NA | NA | NA | 1 | 1 | 2 | 1 | 1 | 0 | NA | NA | NA | NA | 3 | 0.6667 |
| 13_414700 | 1 | 1 | NA | NA | NA | NA | NA | NA | 1 | 1 | 1 | 1 | 1 | 1 | NA | NA | NA | NA | 2 | 1 |
| 13_414922 | 1 | 1 | NA | NA | NA | NA | NA | NA | 1 | 1 | NA | NA | 1 | 1 | NA | NA | NA | NA | 1 | 1 |
| 13_417880 | NA | NA | NA | NA | 1 | 1 | 2 | 1 | 3 | 1 | NA | NA | NA | NA | NA | NA | 1 | 1 | 1 | 1 |
| 13_417881 | NA | NA | NA | NA | 1 | 1 | 1 | 1 | 2 | 1 | NA | NA | NA | NA | 1 | 1 | NA | NA | 1 | 1 |
| 13_417882 | NA | NA | NA | NA | 1 | 1 | NA | NA | 1 | 1 | NA | NA | NA | NA | 1 | 1 | 1 | 1 | 2 | 1 |
| 13_417886 | NA | NA | 1 | 1 | 1 | 1 | 2 | 0.5 | 4 | 0.75 | NA | NA | 1 | 1 | 1 | 0 | 1 | 1 | 3 | 0.6667 |
| 13_417899 | 1 | 0 | 1 | 1 | 2 | 0 | 4 | 0 | 8 | 0.125 | 1 | 0 | 2 | 0.5 | 1 | 0 | 1 | 0 | 5 | 0.2 |
| 13_417901 | 1 | 0 | 1 | 0 | 2 | 1 | 4 | 0 | 8 | 0.25 | 1 | 0 | 2 | 0.5 | 1 | 0 | 1 | 0 | 5 | 0.2 |
| 13_417926 | 3 | 0.3333 | 4 | 0 | 6 | 0 | 6 | 0 | 19 | 0.0526 | 2 | 0 | 6 | 0 | 7 | 0.1429 | 1 | 0 | 16 | 0.0625 |
| 13_417928 | 2 | 0 | 3 | 0 | 7 | 0.1429 | 5 | 0.2 | 17 | 0.1176 | 3 | 0 | 5 | 0.6 | 3 | 0.3333 | 1 | 1 | 12 | 0.4167 |
| 13_417946 | 12 | 0.1667 | 8 | 0.25 | 11 | 0.2727 | 7 | 0.4286 | 38 | 0.2632 | 6 | 0 | 9 | 0.5556 | 15 | 0.1333 | 5 | 0.6 | 35 | 0.2857 |
| 13_417946 | 8 | 1 | 6 | 1 | 7 | 1 | 4 | 1 | 25 | 1 | 6 | 1 | 4 | 1 | 9 | 1 | 2 | 1 | 21 | 1 |
| 13_417946 | 12 | 0.5833 | 8 | 1 | 11 | 0.5455 | 8 | 0.625 | 39 | 0.6667 | 9 | 0.3333 | 9 | 1 | 15 | 0.6 | 5 | 0.4 | 38 | 0.6053 |
| 13_420638 | 1 | 1 | NA | NA | NA | NA | NA | NA | 1 | 1 | NA | NA | 1 | 1 | NA | NA | NA | NA | 1 | 1 |
| 13_423094 | 1 | 1 | NA | NA | NA | NA | NA | NA | 1 | 1 | 2 | 1 | NA | NA | NA | NA | NA | NA | 2 | 1 |
| 13_423097 | 1 | 1 | 1 | 0 | NA | NA | NA | NA | 2 | 0.5 | 2 | 1 | NA | NA | NA | NA | NA | NA | 2 | 1 |
| 13_423103 | 1 | 0 | 1 | 0 | 2 | 0.5 | NA | NA | 4 | 0.25 | 2 | 0.5 | NA | NA | NA | NA | NA | NA | 2 | 0.5 |
| 13_423108 | 1 | 1 | 1 | 0 | 2 | 0.5 | NA | NA | 4 | 0.5 | 2 | 0.5 | NA | NA | NA | NA | NA | NA | 2 | 0.5 |
| 13_423110 | 1 | 0 | 1 | 0 | 2 | 0.5 | 1 | 0 | 5 | 0.2 | 2 | 1 | NA | NA | NA | NA | NA | NA | 2 | 1 |
| 13_423112 | 1 | 0 | 1 | 0 | 2 | 0.5 | 1 | 0 | 5 | 0.2 | 2 | 0.5 | NA | NA | NA | NA | NA | NA | 2 | 0.5 |
| 13_423117 | 1 | 1 | 1 | 0 | NA | NA | 1 | 1 | 3 | 0.6667 | 2 | 0.5 | NA | NA | NA | NA | NA | NA | 2 | 0.5 |

| | | | | | | | | | | | | | | | | | | | | |
|---|---|---|---|---|---|---|---|---|---|---|---|---|---|---|---|---|---|---|---|---|
| 13_4231222 | 2 | 0.5 | 1 | 1 | 2 | 0.5 | 1 | 1 | 6 | 0.6667 | 2 | 0.5 | NA | NA | NA | NA | NA | NA | 2 | 0.5 |
| 13_4231265 | 1 | 1 | 1 | 1 | 1 | 1 | 1 | 1 | 4 | 1 | 2 | 0.5 | NA | NA | NA | NA | NA | NA | 2 | 0.5 |
| 13_4231279 | 1 | 1 | 1 | 0 | 2 | 0 | 1 | 0 | 5 | 0.2 | 2 | 0.5 | NA | NA | NA | NA | NA | NA | 2 | 0.5 |
| 13_4231284 | 1 | 1 | NA | NA | 2 | 0.5 | NA | NA | 3 | 0.6667 | 2 | 0.5 | NA | NA | NA | NA | NA | NA | 2 | 0.5 |
| 13_4231291 | 2 | 0.5 | 1 | 0 | 2 | 0 | 1 | 0 | 6 | 0.1667 | 1 | 1 | NA | NA | NA | NA | NA | NA | 1 | 1 |
| 13_4231321 | 1 | 1 | NA | NA | 2 | 0.5 | NA | NA | 3 | 0.6667 | 3 | 0.6667 | NA | NA | NA | NA | NA | NA | 3 | 0.6667 |
| 13_4231327 | 2 | 0.5 | 1 | 1 | 1 | 0 | 1 | 0 | 5 | 0.4 | 3 | 0.3333 | NA | NA | NA | NA | NA | NA | 3 | 0.3333 |
| 13_4231339 | 2 | 0 | 1 | 0 | 1 | 1 | 2 | 1 | 6 | 0.5 | 3 | 0.3333 | NA | NA | NA | NA | NA | NA | 3 | 0.3333 |
| 13_4231353 | 2 | 0.5 | 1 | 0 | 2 | 1 | 2 | 0 | 7 | 0.4286 | 3 | 0.3333 | NA | NA | NA | NA | NA | NA | 3 | 0.3333 |
| 13_4231360 | 1 | 0 | 1 | 0 | 2 | 0 | 2 | 0.5 | 6 | 0.1667 | 3 | 0.3333 | NA | NA | NA | NA | 1 | 0 | 4 | 0.25 |
| 13_4231362 | 2 | 0.5 | NA | NA | 2 | 1 | 2 | 1 | 6 | 0.8333 | 1 | 1 | NA | NA | NA | NA | 1 | 0 | 2 | 0.5 |
| 13_4231370 | 2 | 1 | 1 | 0 | 2 | 0 | 2 | 0 | 7 | 0.2857 | 2 | 0 | NA | NA | NA | NA | 1 | 1 | 3 | 0.3333 |
| 13_4233516 | 2 | 1 | 1 | 1 | 2 | 1 | 3 | 1 | 8 | 1 | 3 | 1 | NA | NA | NA | NA | 1 | 1 | 4 | 1 |
| 13_4233522 | 2 | 1 | 1 | 0 | 2 | 0.5 | 3 | 0.6667 | 8 | 0.625 | 4 | 0.75 | NA | NA | NA | NA | 1 | 1 | 5 | 0.8 |
| 13_4233523 | 2 | 0.5 | NA | NA | 1 | 0 | 3 | 0.3333 | 6 | 0.3333 | 4 | 0.25 | NA | NA | NA | NA | 1 | 0 | 5 | 0.2 |
| 13_4233706 | 2 | 1 | 1 | 1 | 2 | 1 | 2 | 1 | 7 | 1 | 2 | 1 | NA | NA | NA | NA | NA | NA | 2 | 1 |
| 13_4233710 | 2 | 0 | 1 | 1 | 2 | 0.5 | 2 | 0.5 | 7 | 0.4286 | 3 | 0.6667 | NA | NA | NA | NA | NA | NA | 3 | 0.6667 |
| 13_4233719 | 1 | 0 | 1 | 1 | NA | NA | NA | NA | 2 | 0.5 | 2 | 1 | NA | NA | NA | NA | NA | NA | 2 | 1 |
| 13_4233732 | 2 | 1 | 1 | 1 | 2 | 1 | 3 | 1 | 8 | 1 | 3 | 1 | NA | NA | NA | NA | 1 | 1 | 4 | 1 |
| 13_4233737 | 1 | 0 | 1 | 0 | 2 | 0.5 | 2 | 0 | 6 | 0.1667 | 2 | 1 | NA | NA | NA | NA | NA | NA | 2 | 1 |
| 13_4233751 | 2 | 1 | 1 | 0 | 2 | 1 | 3 | 0.6667 | 8 | 0.75 | 5 | 0.8 | NA | NA | NA | NA | NA | NA | 5 | 0.8 |
| 13_4233758 | 2 | 0.5 | 1 | 0 | 1 | 1 | 3 | 1 | 7 | 0.7143 | 4 | 0.75 | NA | NA | NA | NA | 1 | 1 | 5 | 0.8 |
| 13_4233765 | 2 | 0 | NA | NA | 2 | 0.5 | 3 | 1 | 7 | 0.5714 | 3 | 0.6667 | NA | NA | NA | NA | 1 | 1 | 4 | 0.75 |
| 13_4233776 | 1 | 1 | 1 | 1 | 1 | 1 | 3 | 1 | 6 | 1 | 4 | 1 | NA | NA | NA | NA | 1 | 1 | 5 | 1 |
| 13_4233788 | 1 | 1 | NA | NA | 2 | 0 | 4 | 0 | 7 | 0.1429 | 3 | 0.3333 | NA | NA | NA | NA | 1 | 0 | 4 | 0.25 |
| 13_4233796 | 2 | 1 | 1 | 1 | 2 | 0.5 | 2 | 1 | 7 | 0.8571 | 3 | 0.6667 | NA | NA | NA | NA | 1 | 1 | 4 | 0.75 |
| 13_4233807 | 1 | 1 | 1 | 0 | 1 | 1 | 4 | 0.75 | 7 | 0.7143 | 5 | 0.4 | NA | NA | NA | NA | NA | NA | 5 | 0.4 |
| 13_4233810 | 1 | 1 | 1 | 1 | 2 | 1 | 4 | 1 | 8 | 1 | 3 | 1 | NA | NA | NA | NA | 1 | 1 | 4 | 1 |
| 13_4246061 | NA | NA | NA | NA | 1 | 1 | NA | NA | 1 | 1 | 2 | 1 | NA | NA | NA | NA | 1 | 1 | 3 | 1 |
| 13_4246082 | 1 | 1 | 1 | 1 | 1 | 1 | NA | NA | 3 | 1 | 2 | 1 | NA | NA | NA | NA | 1 | 1 | 3 | 1 |
| 13_4246085 | 1 | 1 | 1 | 0 | 1 | 1 | NA | NA | 3 | 0.6667 | 2 | 0 | NA | NA | NA | NA | 1 | 1 | 3 | 0.3333 |
| 13_4284033 | 1 | 1 | NA | NA | NA | NA | NA | NA | 1 | 1 | NA | NA | 2 | 1 | 3 | 0.3333 | 1 | 1 | 6 | 0.6667 |
| 13_4328871 | 35 | 0.3714 | 30 | 0.2333 | 31 | 0.1935 | 28 | 0.2143 | 124 | 0.2581 | 26 | 0.3077 | 38 | 0.2368 | 48 | 0.2292 | 28 | 0.2857 | 140 | 0.2571 |
| 13_4328876 | 24 | 0.9167 | 24 | 0.9167 | 26 | 0.8462 | 22 | 0.9545 | 96 | 0.9062 | 19 | 1 | 25 | 0.92 | 32 | 0.9375 | 23 | 0.913 | 99 | 0.9394 |
| 13_4328898 | 38 | 0.3158 | 36 | 0.3611 | 37 | 0.3784 | 33 | 0.3636 | 144 | 0.3542 | 26 | 0.3846 | 43 | 0.4186 | 57 | 0.4737 | 30 | 0.3333 | 156 | 0.4167 |
| 13_4328908 | 36 | 0.8056 | 27 | 0.9259 | 39 | 0.9231 | 30 | 0.8667 | 132 | 0.8788 | 21 | 0.9524 | 37 | 0.973 | 49 | 0.9184 | 26 | 0.9615 | 133 | 0.9474 |
| 13_4328917 | 37 | 0.027 | 38 | 0.0789 | 37 | 0.027 | 30 | 0.1333 | 142 | 0.0634 | 26 | 0 | 43 | 0.093 | 55 | 0.0727 | 32 | 0 | 156 | 0.0513 |
| 13_4329017 | 55 | 0.0364 | 42 | 0.0476 | 55 | 0.0364 | 41 | 0.0976 | 193 | 0.0518 | 29 | 0.1724 | 63 | 0.0476 | 66 | 0.0606 | 33 | 0.0303 | 191 | 0.0681 |
| 13_4329185 | 85 | 0.2 | 69 | 0.087 | 72 | 0.1389 | 77 | 0.2078 | 303 | 0.1617 | 44 | 0.1136 | 96 | 0.0729 | 101 | 0.1188 | 61 | 0.1475 | 302 | 0.1093 |
| 13_4330528 | 4 | 0 | 3 | 0.3333 | 2 | 0 | 7 | 0 | 16 | 0.0625 | 5 | 0 | 1 | 0 | 4 | 0.25 | 5 | 0 | 15 | 0.0667 |
| 13_4334898 | 12 | 0 | 8 | 0.125 | 6 | 0.3333 | 7 | 0.1429 | 33 | 0.1212 | 8 | 0.375 | 6 | 0 | 13 | 0.3077 | 8 | 0.375 | 35 | 0.2857 |
| 13_4334898 | 11 | 0.8182 | 6 | 0.8333 | 7 | 0.8571 | 7 | 1 | 31 | 0.871 | 7 | 0.5714 | 7 | 0.8571 | 8 | 0.875 | 7 | 0.7143 | 29 | 0.7586 |
| 13_4334961 | 9 | 0.3333 | 8 | 0.125 | 5 | 0.4 | 7 | 0.1429 | 29 | 0.2414 | 4 | 0.25 | 5 | 0.2 | 10 | 0.1 | 6 | 0 | 25 | 0.12 |
| 13_4334965 | 9 | 0.3333 | 6 | 0.3333 | 5 | 0.2 | 8 | 0.125 | 28 | 0.25 | 4 | 0.75 | 4 | 0.5 | 9 | 0.2222 | 4 | 0.25 | 21 | 0.381 |
| 13_4335018 | 6 | 0 | 8 | 0 | 3 | 0.3333 | 7 | 0.2857 | 24 | 0.125 | 4 | 0 | 4 | 0.5 | 10 | 0.2 | 5 | 0 | 23 | 0.1739 |
| 13_4335046 | 6 | 0.1667 | 5 | 0 | 2 | 0 | 8 | 0.125 | 21 | 0.0952 | 2 | 0 | 3 | 0.3333 | 8 | 0.125 | 5 | 0.2 | 18 | 0.1667 |
| 13_4335068 | 6 | 0.1667 | 1 | 0 | 3 | 0 | 5 | 0.6 | 15 | 0.2667 | 1 | 0 | 2 | 0 | 10 | 0.1 | 4 | 0.25 | 17 | 0.1176 |
| 13_4335071 | 6 | 0.1667 | 4 | 0.25 | 3 | 0 | 6 | 0 | 19 | 0.1053 | 2 | 0 | 2 | 0 | 10 | 0.1 | 5 | 0 | 19 | 0.0526 |
| 13_4335378 | NA | NA | 1 | 1 | NA | NA | NA | NA | 1 | 1 | NA | NA | NA | NA | 5 | 1 | NA | NA | 5 | 1 |
| 13_4354827 | 2 | 0.5 | NA | NA | NA | NA | NA | NA | 2 | 0.5 | 3 | 0 | 1 | 1 | 2 | 0 | 1 | 0 | 7 | 0.1429 |
| 13_4354828 | 2 | 1 | NA | NA | NA | NA | NA | NA | 2 | 1 | 3 | 0.3333 | 1 | 1 | 2 | 0 | NA | NA | 6 | 0.3333 |
| 13_4354852 | 1 | 1 | NA | NA | NA | NA | NA | NA | 1 | 1 | 3 | 1 | 1 | 1 | 1 | 1 | 1 | 1 | 6 | 1 |
| 13_4356080 | NA | NA | NA | NA | NA | NA | 1 | 1 | 1 | 1 | NA | NA | NA | NA | 1 | 1 | 1 | 1 | 2 | 1 |
| 13_4356095 | NA | NA | NA | NA | NA | NA | 1 | 1 | 1 | 1 | NA | NA | NA | NA | 1 | 0 | 1 | 1 | 2 | 0.5 |
| 13_4358841 | 3 | 0 | 6 | 0.1667 | 4 | 0 | 3 | 0.3333 | 16 | 0.125 | 4 | 0.25 | 7 | 0 | 4 | 0.25 | 8 | 0 | 23 | 0.087 |
| 13_4363450 | 1 | 0 | 5 | 0.2 | NA | NA | 4 | 0 | 10 | 0.1 | 2 | 0.5 | 3 | 0.3333 | 1 | 0 | 2 | 0.5 | 8 | 0.375 |
| 13_4369466 | 1 | 1 | 1 | 0 | NA | NA | 1 | 0 | 3 | 0.3333 | 1 | 1 | 1 | 0 | NA | NA | 3 | 0.3333 | 5 | 0.4 |
| 13_4369489 | 2 | 0.5 | 1 | 1 | 1 | 0 | NA | NA | 4 | 0.5 | 2 | 0 | 1 | 1 | NA | NA | 2 | 0 | 5 | 0.2 |
| 13_4369500 | 2 | 0 | 1 | 1 | 1 | 0 | NA | NA | 4 | 0.25 | 1 | 1 | 1 | 0 | NA | NA | 3 | 0.6667 | 5 | 0.6 |
| 13_4369513 | 1 | 0 | 1 | 1 | NA | NA | NA | NA | 2 | 0.5 | 1 | 0 | 4 | 0.25 | NA | NA | 3 | 0.6667 | 8 | 0.375 |
| 13_4382160 | NA | NA | 2 | 1 | NA | NA | NA | NA | 2 | 1 | NA | NA | 1 | 1 | NA | NA | NA | NA | 1 | 1 |
| 13_4382161 | NA | NA | 1 | 1 | NA | NA | NA | NA | 1 | 1 | NA | NA | 1 | 1 | NA | NA | 1 | 1 | 2 | 1 |
| 13_4382176 | NA | NA | 1 | 1 | NA | NA | NA | NA | 1 | 1 | NA | NA | 1 | 1 | NA | NA | 1 | 1 | 2 | 1 |
| 13_4382196 | NA | NA | 1 | 1 | NA | NA | NA | NA | 1 | 1 | NA | NA | 1 | 1 | NA | NA | 1 | 1 | 2 | 1 |
| 13_4382217 | NA | NA | 1 | 1 | NA | NA | NA | NA | 1 | 1 | NA | NA | 1 | 1 | NA | NA | 1 | 1 | 2 | 1 |

| | | | | | | | | | | | | | | | | | | | | |
|---|---|---|---|---|---|---|---|---|---|---|---|---|---|---|---|---|---|---|---|---|
| 13_4382412 | NA | NA | 1 | 1 | 1 | 0 | NA | NA | 2 | 0.5 | NA | NA | 1 | 0 | NA | NA | 2 | 1 | 3 | 0.6667 |
| 13_4395550 | 1 | 1 | NA | NA | NA | NA | 1 | 1 | 2 | 1 | NA | NA | 2 | 1 | NA | NA | 1 | 1 | 3 | 1 |
| 13_4395572 | 3 | 0.3333 | NA | NA | NA | NA | NA | NA | 3 | 0.3333 | 1 | 1 | 2 | 1 | NA | NA | 1 | 0 | 4 | 0.75 |
| 13_4395595 | 2 | 0.5 | NA | NA | NA | NA | 1 | 0 | 3 | 0.3333 | 1 | 1 | 2 | 0.5 | NA | NA | 1 | 0 | 4 | 0.5 |
| 13_4395600 | 2 | 1 | NA | NA | 1 | 1 | 1 | 1 | 4 | 1 | 1 | 1 | 1 | 1 | NA | NA | 1 | 1 | 3 | 1 |
| 13_4395627 | 2 | 1 | NA | NA | 1 | 1 | 1 | 1 | 4 | 1 | 1 | 1 | 3 | 1 | NA | NA | 1 | 1 | 5 | 1 |
| 13_4395629 | 2 | 0.5 | NA | NA | 1 | 0 | 1 | 1 | 4 | 0.5 | NA | NA | 2 | 0.5 | NA | NA | 1 | 0 | 3 | 0.3333 |
| 13_4395632 | 3 | 0.6667 | NA | NA | 1 | 1 | 1 | 1 | 5 | 0.8 | 1 | 1 | 2 | 1 | NA | NA | NA | NA | 3 | 1 |
| 13_4395634 | 3 | 1 | NA | NA | 1 | 1 | 1 | 1 | 5 | 1 | 1 | 1 | 3 | 1 | NA | NA | 1 | 1 | 5 | 1 |
| 13_4395637 | 3 | 1 | NA | NA | 1 | 1 | NA | NA | 4 | 1 | 1 | 1 | 3 | 1 | NA | NA | 1 | 1 | 5 | 1 |
| 13_4505583 | NA | NA | 2 | 0.5 | NA | NA | 1 | 1 | 3 | 0.6667 | 1 | 0 | 1 | 1 | 1 | 1 | NA | NA | 3 | 0.6667 |
| 13_4505586 | NA | NA | 1 | 0 | NA | NA | 1 | 1 | 2 | 0.5 | 1 | 1 | NA | NA | 1 | 1 | NA | NA | 2 | 1 |
| 13_4505593 | 1 | 0 | 1 | 0 | NA | NA | 1 | 1 | 3 | 0.3333 | 1 | 0 | 2 | 0.5 | 1 | 0 | NA | NA | 4 | 0.25 |
| 13_4505612 | 1 | 1 | 2 | 0.5 | NA | NA | NA | NA | 3 | 0.6667 | 1 | 0 | 1 | 1 | NA | NA | NA | NA | 2 | 0.5 |
| 13_4505623 | 1 | 0 | 2 | 0 | 1 | 1 | 1 | 0 | 5 | 0.2 | 1 | 1 | 3 | 1 | 1 | 0 | NA | NA | 5 | 0.8 |
| 13_4505977 | 1 | 1 | 2 | 0.5 | 1 | 0 | 2 | 0.5 | 6 | 0.5 | 2 | 1 | 2 | 1 | 1 | 1 | NA | NA | 5 | 1 |
| 13_4507402 | 3 | 0 | 3 | 0 | 4 | 0 | 5 | 0.2 | 15 | 0.0667 | 3 | 0.3333 | 4 | 0 | 3 | 0 | 2 | 0 | 12 | 0.0833 |
| 13_4507410 | 2 | 0 | 3 | 0.3333 | 3 | 0.3333 | 4 | 0.25 | 12 | 0.25 | 2 | 0.5 | 4 | 0.25 | 2 | 0 | 2 | 0.5 | 10 | 0.3 |
| 13_4507487 | 2 | 1 | 1 | 1 | 1 | 0 | 2 | 1 | 6 | 0.8333 | 1 | 1 | 5 | 1 | 3 | 1 | 3 | 1 | 12 | 1 |
| 13_4507494 | 2 | 1 | 1 | 1 | 4 | 1 | 1 | 1 | 8 | 1 | NA | NA | 4 | 1 | 4 | 0.5 | 2 | 0.5 | 10 | 0.7 |
| 13_4507567 | 18 | 0.5 | 20 | 0.15 | 15 | 0.1333 | 17 | 0.1176 | 70 | 0.2286 | 10 | 0.3 | 13 | 0.0769 | 19 | 0.1579 | 10 | 0.2 | 52 | 0.1731 |
| 13_4507569 | 17 | 0.4706 | 19 | 0.5789 | 18 | 0.5 | 17 | 0.4118 | 71 | 0.493 | 12 | 0.5 | 14 | 0.4286 | 14 | 0.3571 | 11 | 0.5455 | 51 | 0.451 |
| 13_4507582 | 12 | 0.25 | 13 | 0.4615 | 8 | 0.25 | 9 | 0.2222 | 42 | 0.3095 | 8 | 0.125 | 6 | 0 | 11 | 0.0909 | 6 | 0.1667 | 31 | 0.0968 |
| 13_4556201 | 1 | 0 | 3 | 0.6667 | 2 | 1 | NA | NA | 6 | 0.6667 | NA | NA | NA | NA | 2 | 0.5 | 2 | 0.5 | 4 | 0.5 |
| 13_4556397 | 2 | 0 | 3 | 0.3333 | 2 | 0 | NA | NA | 7 | 0.1429 | NA | NA | NA | NA | 2 | 0.5 | 4 | 0 | 6 | 0.1667 |
| 13_4556405 | 2 | 0.5 | 3 | 0.3333 | 1 | 0 | NA | NA | 6 | 0.3333 | NA | NA | NA | NA | 2 | 1 | 3 | 1 | 5 | 1 |
| 13_4556410 | 2 | 0.5 | 3 | 0.6667 | 2 | 0.5 | NA | NA | 7 | 0.5714 | NA | NA | NA | NA | 2 | 1 | 2 | 1 | 4 | 1 |
| 13_4556419 | 3 | 0.3333 | 1 | 1 | 2 | 1 | NA | NA | 6 | 0.6667 | NA | NA | NA | NA | 1 | 1 | 3 | 0.6667 | 4 | 0.75 |
| 13_4556420 | 2 | 0.5 | 3 | 0.3333 | 2 | 0 | NA | NA | 7 | 0.2857 | NA | NA | NA | NA | 2 | 1 | 3 | 0.3333 | 5 | 0.6 |
| 13_4556420 | 3 | 1 | 3 | 0.6667 | 2 | 0.5 | NA | NA | 8 | 0.75 | NA | NA | NA | NA | 2 | 1 | 4 | 0.75 | 6 | 0.8333 |
| 13_4556421 | 3 | 0.6667 | NA | NA | 2 | 1 | NA | NA | 5 | 0.8 | NA | NA | NA | NA | 1 | 1 | 2 | 1 | 3 | 1 |
| 13_4569942 | 12 | 0.0833 | 5 | 0.6 | 10 | 0.2 | 15 | 0.2 | 42 | 0.2143 | 4 | 0.25 | 18 | 0.1667 | 12 | 0.0833 | 13 | 0.0769 | 47 | 0.1277 |
| 13_4569964 | 12 | 0.9167 | 6 | 1 | 13 | 1 | 14 | 1 | 45 | 0.9778 | 4 | 1 | 18 | 1 | 10 | 1 | 11 | 0.8182 | 43 | 0.9535 |
| 13_4569967 | 9 | 0.3333 | 5 | 0.6 | 12 | 0.5833 | 11 | 0.2727 | 37 | 0.4324 | 4 | 0.25 | 19 | 0.4737 | 12 | 0.4167 | 12 | 0.6667 | 47 | 0.4894 |
| 13_4569974 | 11 | 0.7273 | 3 | 1 | 10 | 0.8 | 13 | 0.7692 | 37 | 0.7838 | 5 | 1 | 18 | 0.8333 | 10 | 0.8 | 8 | 0.625 | 41 | 0.8049 |
| 13_4569982 | 7 | 0.8571 | 4 | 1 | 11 | 0.9091 | 7 | 1 | 29 | 0.931 | 4 | 1 | 15 | 1 | 8 | 0.875 | 9 | 0.8889 | 36 | 0.9444 |
| 13_4569985 | 1 | 0 | 1 | 1 | 1 | 1 | 2 | 1 | 5 | 0.8 | 1 | 1 | 1 | 0 | 2 | 0.5 | 4 | 0.25 | 8 | 0.375 |
| 13_4570362 | 2 | 0 | 9 | 0 | 7 | 0.2857 | 7 | 0 | 25 | 0.08 | 8 | 0.125 | 13 | 0.0769 | 12 | 0.0833 | 7 | 0 | 40 | 0.075 |
| 13_4570519 | 1 | 0 | 2 | 0 | 1 | 1 | 5 | 0 | 9 | 0.1111 | 2 | 0 | 8 | 0.125 | 7 | 0.1429 | 4 | 0.5 | 21 | 0.1905 |
| 13_4570595 | 1 | 0 | 2 | 0 | 1 | 1 | 2 | 0 | 6 | 0.1667 | 1 | 0 | 2 | 0.5 | 2 | 0 | 2 | 0 | 7 | 0.1429 |
| 13_4570629 | NA | NA | 2 | 1 | 1 | 1 | 2 | 1 | 5 | 1 | NA | NA | 2 | 0.5 | 1 | 0 | 1 | 1 | 4 | 0.5 |
| 13_4570715 | NA | NA | 1 | 1 | NA | NA | 1 | 1 | 2 | 1 | NA | NA | 1 | 1 | 1 | 0 | NA | NA | 2 | 0.5 |
| 13_4570716 | NA | NA | NA | NA | NA | NA | 1 | 1 | 1 | 1 | NA | NA | NA | NA | 1 | 1 | 1 | 1 | 2 | 1 |
| 13_4570720 | NA | NA | 1 | 1 | NA | NA | 1 | 1 | 2 | 1 | NA | NA | 1 | 1 | 1 | 1 | NA | NA | 2 | 1 |
| 13_4570728 | NA | NA | NA | NA | NA | NA | 1 | 1 | 1 | 1 | NA | NA | NA | NA | 1 | 0 | 1 | 1 | 2 | 0.5 |
| 13_4570732 | NA | NA | 1 | 1 | NA | NA | 1 | 1 | 2 | 1 | NA | NA | 1 | 0 | 1 | 1 | NA | NA | 2 | 0.5 |
| 13_4572170 | NA | NA | NA | NA | NA | NA | 1 | 1 | 1 | 1 | NA | NA | NA | NA | 1 | 1 | NA | NA | 1 | 1 |
| 13_4611952 | 1 | 1 | NA | NA | NA | NA | NA | NA | 1 | 1 | NA | NA | 1 | 1 | 2 | 0.5 | 3 | 1 | 6 | 0.8333 |
| 13_4611974 | 1 | 1 | NA | NA | NA | NA | NA | NA | 1 | 1 | NA | NA | 1 | 1 | 2 | 1 | 3 | 1 | 6 | 1 |
| 13_4611980 | 1 | 1 | NA | NA | NA | NA | NA | NA | 1 | 1 | NA | NA | 1 | 1 | 2 | 1 | 3 | 1 | 6 | 1 |
| 13_4658270 | 1 | 0 | 2 | 0 | 3 | 0.3333 | 1 | 1 | 7 | 0.2857 | 2 | 0 | 1 | 0 | 2 | 0.5 | 2 | 0 | 7 | 0.1429 |
| 13_4658307 | 4 | 0.25 | 1 | 0 | 2 | 0 | 3 | 0.3333 | 10 | 0.2 | 4 | 0 | 1 | 0 | 3 | 0 | 3 | 0.3333 | 11 | 0.0909 |
| 13_4658369 | 4 | 0.25 | 3 | 0 | 3 | 0 | 4 | 0.25 | 14 | 0.1429 | 4 | 0 | 1 | 1 | 9 | 0.1111 | 5 | 0 | 19 | 0.1053 |
| 13_4658436 | 4 | 1 | 6 | 0.5 | 3 | 0.6667 | 3 | 0 | 16 | 0.5625 | 4 | 0.75 | 1 | 1 | 9 | 0.8889 | 6 | 0.8333 | 20 | 0.85 |
| 13_4658440 | 5 | 0.6 | 4 | 0.25 | 1 | 1 | 5 | 0.8 | 15 | 0.6 | 4 | 1 | 1 | 1 | 9 | 0.5556 | 3 | 0.6667 | 17 | 0.7059 |
| 13_4680270 | 15 | 0.0667 | 19 | 0 | 26 | 0.0769 | 34 | 0.0882 | 94 | 0.0638 | 12 | 0.1667 | 19 | 0 | 22 | 0.0909 | 20 | 0.2 | 73 | 0.1096 |
| 13_4680284 | 15 | 0.2 | 21 | 0.1429 | 25 | 0.08 | 30 | 0.1 | 91 | 0.1209 | 12 | 0.0833 | 21 | 0.0952 | 24 | 0.0417 | 17 | 0.1765 | 74 | 0.0946 |
| 13_4682907 | 8 | 0.375 | 1 | 0 | 1 | 0 | 5 | 0 | 15 | 0.2 | 2 | 0 | 3 | 0.6667 | 3 | 0 | NA | NA | 8 | 0.25 |
| 13_4682908 | 10 | 0.2 | 2 | 0 | 2 | 0 | 5 | 0.2 | 19 | 0.1579 | 5 | 0 | 6 | 0 | 5 | 0.2 | NA | NA | 16 | 0.0625 |
| 13_4682910 | 8 | 0 | 1 | 0 | 2 | 0.5 | 4 | 0 | 15 | 0.0667 | 4 | 0 | 6 | 0.1667 | 5 | 0 | NA | NA | 15 | 0.0667 |
| 13_4682917 | 9 | 0.2222 | 2 | 0 | 2 | 0.5 | 5 | 0.2 | 18 | 0.2222 | 4 | 0 | 4 | 0.25 | 5 | 0 | NA | NA | 13 | 0.0769 |
| 13_4682927 | 10 | 0.1 | 2 | 0.5 | 3 | 0 | 4 | 0.5 | 19 | 0.2105 | 3 | 0.6667 | 4 | 0.25 | 4 | 0.25 | NA | NA | 11 | 0.3636 |
| 13_4682929 | 11 | 0.1818 | 2 | 0 | 3 | 0 | 5 | 0.2 | 21 | 0.1429 | 5 | 0 | 6 | 0 | 5 | 0.2 | 1 | 0 | 17 | 0.0588 |
| 13_4682949 | 10 | 0.7 | 2 | 0 | 3 | 0.6667 | 5 | 0.2 | 20 | 0.5 | 5 | 0.2 | 5 | 0.2 | 6 | 0.3333 | NA | NA | 16 | 0.25 |
| 13_4683015 | 18 | 0.3333 | 7 | 0.7143 | 13 | 0.6154 | 10 | 0.5 | 48 | 0.5 | 7 | 0.2857 | 11 | 0.6364 | 13 | 0.4615 | 10 | 0.5 | 41 | 0.4878 |

| | | | | | | | | | | | | | | | | | | | | |
|---|---|---|---|---|---|---|---|---|---|---|---|---|---|---|---|---|---|---|---|---|
| 13_4683039 | 20 | 0.2 | 9 | 0.3333 | 15 | 0.2667 | 13 | 0 | 57 | 0.193 | 8 | 0.375 | 11 | 0.4545 | 10 | 0.3 | 10 | 0.1 | 39 | 0.3077 |
| 13_4683061 | 20 | 0.25 | 9 | 0.5556 | 17 | 0.5882 | 14 | 0.5 | 60 | 0.45 | 10 | 0.4 | 10 | 0.6 | 12 | 0.5833 | 13 | 0.6154 | 45 | 0.5556 |
| 13_4683109 | 24 | 0.2083 | 9 | 0.2222 | 23 | 0.1304 | 17 | 0.1765 | 73 | 0.1781 | 8 | 0.125 | 14 | 0.2857 | 17 | 0.1176 | 12 | 0.1667 | 51 | 0.1765 |
| 13_4683113 | 23 | 0.2609 | 9 | 0.3333 | 20 | 0.3 | 16 | 0.125 | 68 | 0.25 | 9 | 0.1111 | 16 | 0.1875 | 17 | 0.4118 | 13 | 0.3077 | 55 | 0.2727 |
| 13_4683121 | 21 | 0.0476 | 11 | 0.0909 | 23 | 0 | 18 | 0.1111 | 73 | 0.0548 | 11 | 0.0909 | 14 | 0 | 18 | 0.1111 | 11 | 0.0909 | 54 | 0.0741 |
| 13_4683137 | 18 | 0.1111 | 8 | 0.25 | 23 | 0.087 | 18 | 0 | 67 | 0.0896 | 11 | 0 | 20 | 0.25 | 21 | 0 | 12 | 0.1667 | 64 | 0.1094 |
| 13_4683431 | 4 | 0 | 6 | 0.3333 | 4 | 0 | 6 | 0 | 20 | 0.1 | 2 | 0 | 12 | 0.0833 | 7 | 0.1429 | 8 | 0 | 29 | 0.069 |
| 13_4683442 | 4 | 0 | 6 | 0.1667 | 3 | 0.3333 | 4 | 0.25 | 17 | 0.1765 | 2 | 0 | 11 | 0.2727 | 6 | 0.6667 | 8 | 0.375 | 27 | 0.3704 |
| 13_4683442 | 4 | 0.75 | 5 | 0 | 3 | 0.3333 | 6 | 0.5 | 18 | 0.3889 | 1 | 1 | 11 | 0.4545 | 5 | 0.4 | 8 | 0.25 | 25 | 0.4 |
| 13_4683443 | 4 | 0 | 5 | 0 | 4 | 0.25 | 5 | 0.4 | 18 | 0.1667 | 2 | 0.5 | 10 | 0.1 | 7 | 0.2857 | 8 | 0.375 | 27 | 0.2593 |
| 13_4683452 | 4 | 0.75 | 6 | 0.8333 | 4 | 1 | 6 | 1 | 20 | 0.9 | 2 | 1 | 9 | 0.8889 | 7 | 1 | 8 | 1 | 26 | 0.9615 |
| 13_4683465 | 4 | 0.25 | 5 | 0.8 | 4 | 0.5 | 6 | 0.6667 | 19 | 0.5789 | 2 | 0.5 | 10 | 0.2 | 8 | 0.625 | 7 | 0.7143 | 27 | 0.4815 |
| 13_4683587 | 1 | 1 | NA | NA | NA | NA | NA | NA | 1 | 1 | 1 | 1 | 1 | 0 | NA | NA | NA | NA | 2 | 0.5 |
| 13_4683655 | 3 | 0.3333 | 4 | 0 | 2 | 0 | 5 | 0 | 14 | 0.0714 | 1 | 0 | 7 | 0.2857 | 5 | 0 | 8 | 0.25 | 21 | 0.1905 |
| 13_4683672 | 2 | 0 | 4 | 0.5 | 2 | 0 | 3 | 0 | 11 | 0.1818 | 1 | 0 | 7 | 0.1429 | 4 | 0.5 | 6 | 0.3333 | 18 | 0.2778 |
| 13_4683757 | 2 | 0 | 4 | 0 | 3 | 0 | 6 | 0.1667 | 15 | 0.0667 | 1 | 0 | 8 | 0.125 | 7 | 0.1429 | 7 | 0 | 23 | 0.087 |
| 13_4683767 | 2 | 0 | 3 | 0.3333 | 2 | 0 | 3 | 0 | 10 | 0.1 | 1 | 0 | 7 | 0 | 5 | 0.2 | 6 | 0 | 19 | 0.0526 |
| 13_4684033 | 1 | 1 | 3 | 0.6667 | 3 | 1 | 6 | 0.6667 | 13 | 0.7692 | NA | NA | 6 | 0.3333 | 5 | 0.8 | 6 | 0.8333 | 17 | 0.6471 |
| 13_4684045 | 1 | 0 | 3 | 1 | 3 | 1 | 6 | 1 | 13 | 0.9231 | 1 | 1 | 7 | 1 | 8 | 0.875 | 4 | 0.75 | 20 | 0.9 |
| 13_4684048 | 2 | 0 | 3 | 0.3333 | 2 | 0 | 5 | 0.4 | 12 | 0.25 | 1 | 0 | 8 | 0.125 | 5 | 0 | 5 | 0 | 19 | 0.0526 |
| 13_4684076 | 2 | 1 | 2 | 0.5 | 3 | 1 | 6 | 0.6667 | 13 | 0.7692 | NA | NA | 6 | 1 | 7 | 1 | 4 | 1 | 17 | 1 |
| 13_4684079 | 1 | 1 | 1 | 1 | 3 | 1 | 5 | 1 | 10 | 1 | 1 | 1 | 7 | 0.7143 | 6 | 0.6667 | 4 | 1 | 18 | 0.7778 |
| 13_4684082 | 2 | 0.5 | 1 | 1 | 2 | 1 | 6 | 0.8333 | 11 | 0.8182 | NA | NA | 7 | 0.7143 | 7 | 0.5714 | 5 | 0.8 | 19 | 0.6842 |
| 13_4684089 | 1 | 0 | 2 | 0 | 3 | 0.3333 | 6 | 0.1667 | 12 | 0.1667 | 1 | 0 | 5 | 0 | 7 | 0 | 5 | 0.2 | 18 | 0.0556 |
| 13_4684713 | 2 | 1 | 1 | 0 | 3 | 0.3333 | 5 | 0.4 | 11 | 0.4545 | 1 | 1 | 6 | 0.6667 | 6 | 0.3333 | 4 | 0 | 17 | 0.4118 |
| 13_4684720 | 2 | 0 | 1 | 0 | 3 | 0 | 5 | 0.2 | 11 | 0.0909 | 1 | 0 | 6 | 0 | 6 | 0 | 5 | 0.2 | 18 | 0.0556 |
| 13_4684727 | 2 | 0 | 1 | 0 | 2 | 0.5 | 5 | 0.2 | 10 | 0.2 | 1 | 0 | 6 | 0.3333 | 6 | 0.6667 | 5 | 1 | 18 | 0.6111 |
| 13_4684730 | 2 | 1 | 1 | 1 | 3 | 0.3333 | 5 | 0.6 | 11 | 0.6364 | 1 | 1 | 6 | 0.5 | 6 | 0.8333 | 5 | 0.6 | 18 | 0.6667 |
| 13_4684734 | 1 | 0 | 1 | 0 | 3 | 0.3333 | 4 | 0.25 | 9 | 0.2222 | 1 | 0 | 7 | 0.4286 | 6 | 0 | 5 | 0.4 | 19 | 0.2632 |
| 13_4684735 | 1 | 0 | 1 | 1 | 3 | 1 | 4 | 0.5 | 9 | 0.6667 | 1 | 0 | 5 | 1 | 4 | 0.75 | 5 | 0.6 | 15 | 0.7333 |
| 13_4684735 | 2 | 0.5 | 1 | 0 | 3 | 0.3333 | 4 | 0.25 | 10 | 0.3 | 1 | 0 | 5 | 0.6 | 3 | 0.3333 | 4 | 0.75 | 13 | 0.5385 |
| 13_4684736 | 2 | 1 | 1 | 1 | 2 | 0 | 3 | 0 | 8 | 0.375 | 1 | 0 | 7 | 0.4286 | 6 | 0.6667 | 4 | 0.75 | 18 | 0.5556 |
| 13_4684741 | 1 | 0 | 1 | 0 | 2 | 0 | 4 | 0.5 | 8 | 0.25 | 1 | 0 | 7 | 0 | 6 | 0.5 | 4 | 0 | 18 | 0.1667 |
| 13_4690284 | 2 | 0 | 2 | 0 | 3 | 0 | 3 | 0.6667 | 10 | 0.2 | NA | NA | 4 | 0.5 | 2 | 0 | 1 | 1 | 7 | 0.4286 |
| 13_4690291 | 2 | 0 | 4 | 0 | 4 | 0.5 | 3 | 0.6667 | 13 | 0.3077 | NA | NA | 5 | 0 | 3 | 0 | 1 | 1 | 9 | 0.1111 |
| 13_4690297 | 2 | 0 | 4 | 0.75 | 4 | 0.25 | 3 | 0 | 13 | 0.3077 | NA | NA | 5 | 0.2 | 3 | 0.3333 | 1 | 0 | 9 | 0.2222 |
| 13_4690303 | 1 | 0 | 3 | 0 | 4 | 0.5 | 3 | 0.6667 | 11 | 0.3636 | NA | NA | 4 | 0.5 | 3 | 0.6667 | 1 | 0 | 8 | 0.5 |
| 13_4690325 | NA | NA | 3 | 0.3333 | 3 | 0.6667 | 2 | 0.5 | 8 | 0.5 | NA | NA | 4 | 1 | 2 | 0.5 | NA | NA | 6 | 0.8333 |
| 13_4690329 | NA | NA | 4 | 0.25 | 3 | 1 | 2 | 0.5 | 9 | 0.5556 | NA | NA | 4 | 0.25 | 2 | 0.5 | NA | NA | 6 | 0.3333 |
| 13_4690347 | NA | NA | 4 | 0.25 | 3 | 0 | 2 | 0 | 9 | 0.1111 | NA | NA | 4 | 0 | 2 | 1 | NA | NA | 6 | 0.3333 |
| 13_4690348 | NA | NA | 4 | 0.25 | 3 | 0.6667 | NA | NA | 7 | 0.4286 | NA | NA | 3 | 0 | 1 | 1 | NA | NA | 4 | 0.25 |
| 13_4690362 | NA | NA | 3 | 1 | 2 | 1 | 2 | 1 | 7 | 1 | NA | NA | 3 | 1 | 1 | 1 | NA | NA | 4 | 1 |
| 13_4690376 | NA | NA | 4 | 0.25 | 3 | 0.3333 | 1 | 0 | 8 | 0.25 | NA | NA | 2 | 0.5 | 1 | 1 | NA | NA | 3 | 0.6667 |
| 13_4690380 | NA | NA | 3 | 1 | 3 | 0.3333 | 2 | 0 | 8 | 0.5 | NA | NA | 3 | 0 | 2 | 0.5 | NA | NA | 5 | 0.2 |
| 13_4690380 | NA | NA | 3 | 1 | 3 | 1 | 2 | 0.5 | 8 | 0.875 | NA | NA | 3 | 0.6667 | 2 | 1 | NA | NA | 5 | 0.8 |
| 13_4690402 | NA | NA | 3 | 0.6667 | 2 | 1 | 2 | 0.5 | 7 | 0.7143 | NA | NA | 3 | 0.6667 | 2 | 1 | NA | NA | 5 | 0.8 |
| 13_4690562 | NA | NA | 3 | 0.3333 | 2 | 0 | 2 | 0 | 7 | 0.1429 | NA | NA | 3 | 0.6667 | 2 | 0.5 | NA | NA | 5 | 0.6 |
| 13_4690569 | NA | NA | 3 | 0.6667 | 2 | 0 | 2 | 1 | 7 | 0.5714 | NA | NA | 1 | 1 | 2 | 0.5 | NA | NA | 3 | 0.6667 |
| 13_4690573 | NA | NA | 3 | 0.3333 | 3 | 0.6667 | 2 | 0.5 | 8 | 0.5 | NA | NA | 2 | 0.5 | 1 | 0 | NA | NA | 3 | 0.3333 |
| 13_4690582 | NA | NA | 3 | 0.6667 | 2 | 1 | 2 | 1 | 7 | 0.8571 | NA | NA | 1 | 0 | 1 | 1 | NA | NA | 2 | 0.5 |
| 13_4690584 | NA | NA | 3 | 1 | 2 | 0.5 | 2 | 1 | 7 | 0.8571 | NA | NA | 2 | 1 | 1 | 1 | NA | NA | 3 | 1 |
| 13_4690594 | NA | NA | 3 | 0 | 3 | 0.3333 | 1 | 0 | 7 | 0.1429 | NA | NA | 2 | 0.5 | 1 | 0 | NA | NA | 3 | 0.3333 |
| 13_4690601 | NA | NA | 2 | 0.5 | 3 | 0 | 2 | 1 | 7 | 0.4286 | NA | NA | 1 | 1 | NA | NA | NA | NA | 1 | 1 |
| 13_4690620 | NA | NA | 2 | 0.5 | 3 | 0.3333 | 2 | 0 | 7 | 0.2857 | NA | NA | 2 | 0.5 | 1 | 0 | NA | NA | 3 | 0.3333 |
| 13_4716619 | 6 | 0.1667 | 3 | 1 | 3 | 0.3333 | 2 | 0.5 | 14 | 0.4286 | 5 | 0.4 | 1 | 1 | 1 | 0 | 3 | 0 | 10 | 0.3 |
| 13_4716667 | 5 | 0.4 | 3 | 0.3333 | 2 | 1 | 2 | 0 | 12 | 0.4167 | 3 | 0.3333 | 1 | 0 | NA | NA | 2 | 0 | 6 | 0.1667 |
| 13_4716674 | 6 | 1 | 3 | 1 | 3 | 1 | 1 | 1 | 13 | 1 | 4 | 1 | 1 | 1 | NA | NA | 2 | 1 | 7 | 1 |
| 13_4716692 | 6 | 1 | 2 | 1 | 3 | 1 | 2 | 1 | 13 | 1 | 3 | 1 | 1 | 1 | NA | NA | 2 | 1 | 6 | 1 |
| 13_4716696 | 6 | 0.1667 | 2 | 0.5 | 3 | 0 | 2 | 0 | 13 | 0.1538 | 3 | 0.3333 | 1 | 1 | NA | NA | 1 | 0 | 5 | 0.4 |
| 13_4716697 | 5 | 0.4 | NA | NA | 3 | 0.6667 | 2 | 0.5 | 10 | 0.5 | 3 | 0.6667 | 1 | 1 | NA | NA | 2 | 0 | 6 | 0.5 |
| 13_4716738 | 3 | 1 | 1 | 1 | 3 | 1 | 2 | 1 | 9 | 1 | 2 | 1 | 1 | 1 | NA | NA | 1 | 1 | 4 | 1 |
| 13_4716753 | 4 | 1 | 1 | 1 | 3 | 0.3333 | 2 | 1 | 10 | 0.8 | 3 | 1 | 1 | 1 | NA | NA | 1 | 1 | 5 | 1 |
| 13_4716755 | 4 | 1 | 1 | 1 | 3 | 1 | 1 | 1 | 9 | 1 | 3 | 1 | 1 | 1 | NA | NA | 1 | 1 | 5 | 1 |
| 13_4716791 | 3 | 0 | 1 | 0 | 1 | 0 | 1 | 1 | 6 | 0.1667 | 1 | 0 | 1 | 1 | NA | NA | 1 | 0 | 3 | 0.3333 |
| 13_4717803 | 1 | 1 | 1 | 1 | 1 | 1 | 1 | 0 | 4 | 0.75 | NA | NA | NA | NA | 2 | 1 | 2 | 0.5 | 4 | 0.75 |

| | | | | | | | | | | | | | | | | | | | | |
|---|---|---|---|---|---|---|---|---|---|---|---|---|---|---|---|---|---|---|---|---|
| 13_4717867 | 1 | 0 | 1 | 1 | 2 | 0 | NA | NA | 4 | 0.25 | NA | NA | NA | NA | 1 | 0 | 3 | 0.3333 | 4 | 0.25 |
| 13_4723644 | 2 | 1 | 1 | 0 | 1 | 1 | 1 | 1 | 5 | 0.8 | 1 | 1 | 1 | 0 | NA | NA | 1 | 1 | 3 | 0.6667 |
| 13_4723645 | 2 | 0.5 | 1 | 1 | 1 | 1 | 3 | 0.3333 | 7 | 0.5714 | 1 | 1 | 1 | 0 | NA | NA | NA | NA | 2 | 0.5 |
| 13_4723652 | 1 | 1 | 1 | 0 | NA | NA | 3 | 0 | 5 | 0.2 | 1 | 0 | 1 | 0 | NA | NA | 1 | 1 | 3 | 0.3333 |
| 13_4723851 | 8 | 0.375 | 16 | 0.125 | 9 | 0.1111 | 11 | 0.0909 | 44 | 0.1591 | 2 | 0 | 6 | 0 | 10 | 0.4 | 9 | 0 | 27 | 0.1481 |
| 13_4723919 | 8 | 0 | 14 | 0 | 9 | 0 | 16 | 0.25 | 47 | 0.0851 | 2 | 0 | 4 | 0 | 10 | 0.2 | 8 | 0.125 | 24 | 0.125 |
| 13_4727037 | NA | NA | 1 | 1 | NA | NA | NA | NA | 1 | 1 | 1 | 1 | NA | NA | NA | NA | NA | NA | 1 | 1 |
| 13_4727049 | NA | NA | 1 | 1 | NA | NA | NA | NA | 1 | 1 | 1 | 1 | NA | NA | NA | NA | NA | NA | 1 | 1 |
| 13_4739949 | 1 | 1 | NA | NA | NA | NA | NA | NA | 1 | 1 | 1 | 1 | NA | NA | 1 | 1 | NA | NA | 2 | 1 |
| 13_4740060 | 1 | 1 | 2 | 0.5 | 2 | 0 | 2 | 0.5 | 7 | 0.4286 | 4 | 0.25 | NA | NA | 2 | 0.5 | 2 | 0.5 | 8 | 0.375 |
| 13_4740096 | 4 | 1 | 2 | 1 | 2 | 0.5 | 2 | 0.5 | 10 | 0.8 | 4 | 0.75 | 1 | 1 | 2 | 1 | 3 | 1 | 10 | 0.9 |
| 13_4740111 | 4 | 0.75 | 1 | 1 | 2 | 0.5 | 2 | 1 | 9 | 0.7778 | 4 | 0.5 | 1 | 1 | 2 | 0 | 3 | 1 | 10 | 0.6 |
| 13_4740114 | 2 | 0.5 | 2 | 1 | 2 | 0.5 | 1 | 0 | 7 | 0.5714 | 4 | 0.75 | 1 | 1 | 1 | 1 | 2 | 1 | 8 | 0.875 |
| 13_4740124 | 4 | 0.25 | 1 | 0 | 1 | 0 | 2 | 0 | 8 | 0.125 | 5 | 0.4 | 1 | 0 | 1 | 0 | 3 | 0 | 10 | 0.2 |
| 13_4841469 | 9 | 0.1111 | 14 | 0.1429 | 11 | 0.1818 | 19 | 0.1053 | 53 | 0.1321 | 13 | 0 | 16 | 0 | 20 | 0.05 | 16 | 0.25 | 65 | 0.0769 |
| 13_4841492 | 10 | 0.1 | 12 | 0.1667 | 14 | 0.0714 | 18 | 0 | 54 | 0.0741 | 12 | 0 | 17 | 0.0588 | 19 | 0.0526 | 18 | 0.1111 | 66 | 0.0606 |
| 13_4841660 | 16 | 0.1875 | 14 | 0.2857 | 17 | 0.2353 | 22 | 0.1364 | 69 | 0.2029 | 13 | 0.2308 | 16 | 0.1875 | 22 | 0.2727 | 23 | 0.1739 | 74 | 0.2162 |
| 13_4841677 | 19 | 0.1579 | 16 | 0.1875 | 22 | 0.0909 | 26 | 0.0769 | 83 | 0.1205 | 13 | 0.2308 | 20 | 0.3 | 32 | 0.2188 | 25 | 0.16 | 90 | 0.2222 |
| 13_4841683 | 15 | 0.2 | 17 | 0.1176 | 21 | 0.2381 | 26 | 0.1923 | 79 | 0.1899 | 11 | 0.1818 | 18 | 0.4444 | 33 | 0.3636 | 21 | 0.2857 | 83 | 0.3373 |
| 13_4841753 | 11 | 0.3636 | 11 | 0.0909 | 12 | 0.1667 | 11 | 0.1818 | 45 | 0.2 | 9 | 0.4444 | 17 | 0.0588 | 18 | 0.1111 | 13 | 0.2308 | 57 | 0.1754 |
| 13_4841755 | 9 | 0.2222 | 10 | 0.1 | 12 | 0.3333 | 8 | 0 | 39 | 0.1795 | 9 | 0.3333 | 15 | 0.2 | 17 | 0.0588 | 15 | 0.0667 | 56 | 0.1429 |
| 13_4841768 | 12 | 0.3333 | 7 | 0.4286 | 11 | 0.1818 | 12 | 0.3333 | 42 | 0.3095 | 8 | 0.375 | 21 | 0.3333 | 15 | 0.3333 | 12 | 0.1667 | 56 | 0.3036 |
| 13_4841769 | 8 | 0.125 | 8 | 0.125 | 11 | 0.0909 | 11 | 0 | 38 | 0.0789 | 8 | 0 | 18 | 0.1111 | 16 | 0.1875 | 13 | 0 | 55 | 0.0909 |
| 13_4841770 | 12 | 0.9167 | 11 | 0.9091 | 12 | 0.75 | 11 | 0.8182 | 46 | 0.8478 | 6 | 0.8333 | 19 | 0.7368 | 16 | 0.8125 | 14 | 0.7857 | 55 | 0.7818 |
| 13_4841864 | 6 | 0.5 | 10 | 0.2 | 9 | 0.3333 | 3 | 0 | 28 | 0.2857 | 5 | 0.4 | 11 | 0.6364 | 9 | 0.2222 | 11 | 0.2727 | 36 | 0.3889 |
| 13_4841873 | 5 | 0.2 | 9 | 0.1111 | 10 | 0.5 | 2 | 0.5 | 26 | 0.3077 | 4 | 0.75 | 11 | 0.2727 | 7 | 0 | 7 | 0.4286 | 29 | 0.3103 |
| 13_4863669 | 3 | 0.6667 | 1 | 0 | NA | NA | 1 | 0 | 5 | 0.4 | 4 | 0.75 | 2 | 0.5 | 1 | 1 | NA | NA | 7 | 0.7143 |
| 13_4863672 | 3 | 0.6667 | 1 | 0 | NA | NA | 1 | 0 | 5 | 0.4 | 4 | 0 | 2 | 0.5 | 1 | 0 | NA | NA | 7 | 0.1429 |
| 13_4864278 | 1 | 1 | 1 | 1 | NA | NA | NA | NA | 2 | 1 | 1 | 0 | 1 | 1 | NA | NA | NA | NA | 2 | 0.5 |
| 13_4864289 | 1 | 1 | 1 | 1 | NA | NA | NA | NA | 2 | 1 | NA | NA | 1 | 1 | NA | NA | NA | NA | 1 | 1 |
| 13_4864294 | 1 | 1 | 1 | 1 | NA | NA | NA | NA | 2 | 1 | NA | NA | 1 | 1 | NA | NA | NA | NA | 1 | 1 |
| 13_4864311 | 1 | 1 | 1 | 1 | NA | NA | NA | NA | 2 | 1 | NA | NA | 1 | 1 | NA | NA | NA | NA | 1 | 1 |
| 13_4873153 | 5 | 0 | 5 | 0.2 | 4 | 0.5 | 4 | 0.25 | 18 | 0.2222 | 2 | 0.5 | 6 | 0.3333 | 1 | 0 | 2 | 0.5 | 11 | 0.3636 |
| 13_4873302 | 2 | 1 | 2 | 1 | 1 | 1 | 2 | 0 | 7 | 0.7143 | NA | NA | NA | NA | NA | NA | 1 | 1 | 1 | 1 |
| 13_4873319 | NA | NA | 3 | 0 | 1 | 0 | 3 | 0.3333 | 7 | 0.1429 | 1 | 1 | 1 | 0 | NA | NA | 2 | 1 | 4 | 0.75 |
| 13_4873329 | 1 | 0 | 3 | 0 | 2 | 0.5 | 3 | 0 | 9 | 0.1111 | 1 | 0 | 2 | 0 | NA | NA | 2 | 0.5 | 5 | 0.2 |
| 13_4873951 | 1 | 1 | 1 | 0 | 2 | 1 | 3 | 1 | 7 | 0.8571 | 1 | 0 | 2 | 0.5 | NA | NA | 1 | 1 | 4 | 0.5 |
| 13_4873957 | 2 | 0.5 | 2 | 1 | 2 | 1 | 2 | 0.5 | 8 | 0.75 | 1 | 0 | 2 | 0.5 | NA | NA | 1 | 1 | 4 | 0.5 |
| 13_4873981 | 2 | 0.5 | 2 | 1 | 2 | 1 | 3 | 1 | 9 | 0.8889 | NA | NA | 2 | 0.5 | NA | NA | 1 | 1 | 3 | 0.6667 |
| 13_4873984 | 2 | 1 | 2 | 0 | 1 | 1 | 2 | 1 | 7 | 0.7143 | NA | NA | 2 | 0.5 | NA | NA | 1 | 1 | 3 | 0.6667 |
| 13_4873987 | 1 | 1 | 1 | 1 | 1 | 1 | 3 | 0.6667 | 6 | 0.8333 | NA | NA | 1 | 1 | NA | NA | 1 | 1 | 2 | 1 |
| 13_4874001 | 2 | 0.5 | 2 | 1 | NA | NA | 2 | 0.5 | 6 | 0.6667 | NA | NA | 1 | 0 | NA | NA | 1 | 1 | 2 | 0.5 |
| 13_4874030 | 2 | 0.5 | 2 | 0 | NA | NA | 1 | 1 | 5 | 0.4 | NA | NA | 1 | 0 | NA | NA | 1 | 1 | 2 | 0.5 |
| 13_4874036 | 2 | 0 | 2 | 0 | NA | NA | 2 | 0.5 | 6 | 0.1667 | NA | NA | 1 | 0 | NA | NA | 1 | 1 | 2 | 0.5 |
| 13_4895550 | 4 | 0.5 | 14 | 0.5 | 4 | 0.5 | 7 | 0.4286 | 29 | 0.4828 | 8 | 0.25 | 3 | 0.3333 | 12 | 0.3333 | 3 | 0 | 26 | 0.2692 |
| 13_4895649 | 5 | 0 | 8 | 0.25 | 3 | 0 | 5 | 0.4 | 21 | 0.1905 | 4 | 0.25 | 3 | 0.3333 | 8 | 0.25 | 5 | 0 | 20 | 0.2 |
| 13_4895665 | 4 | 0.5 | 9 | 1 | 1 | 1 | 5 | 1 | 19 | 0.8947 | 4 | 0.75 | 3 | 1 | 8 | 0.875 | 6 | 1 | 21 | 0.9048 |
| 13_4895679 | 3 | 0 | 8 | 0.375 | 1 | 0 | 3 | 0 | 15 | 0.2 | 4 | 0.5 | 3 | 0 | 8 | 0.25 | 5 | 0.4 | 20 | 0.3 |
| 13_4895684 | 5 | 0 | 9 | 0.2222 | 2 | 0 | 5 | 0 | 21 | 0.0952 | 5 | 0.2 | 3 | 0.3333 | 11 | 0.0909 | 6 | 0 | 25 | 0.12 |
| 13_4895686 | 5 | 0.8 | 11 | 0.7273 | 2 | 0.5 | 5 | 1 | 23 | 0.7826 | 5 | 0.8 | 3 | 0.3333 | 11 | 0.9091 | 5 | 1 | 24 | 0.8333 |
| 13_4903416 | 6 | 0.3333 | 11 | 0.3636 | 12 | 0.1667 | 9 | 0.3333 | 38 | 0.2895 | 12 | 0.5833 | 16 | 0.3125 | 12 | 0.6667 | 3 | 0.3333 | 43 | 0.4884 |
| 13_4903437 | 3 | 1 | 10 | 0.6 | 12 | 0.4167 | 7 | 0.4286 | 32 | 0.5312 | 13 | 0.6923 | 14 | 0.5714 | 12 | 0.8333 | 3 | 1 | 42 | 0.7143 |
| 13_4906376 | 2 | 0 | 4 | 0 | 6 | 0.5 | 6 | 0 | 18 | 0.1667 | 8 | 0 | 5 | 0.2 | 5 | 0.2 | 3 | 0.3333 | 21 | 0.1429 |
| 13_4907535 | 1 | 0 | 2 | 0 | 4 | 0 | 5 | 0.2 | 12 | 0.0833 | 4 | 0 | 2 | 0 | 4 | 0.25 | 2 | 0 | 12 | 0.0833 |
| 13_4912128 | NA | NA | 1 | 0 | 5 | 0.2 | 2 | 0 | 8 | 0.125 | 2 | 0 | 2 | 0 | 2 | 0.5 | 1 | 1 | 7 | 0.2857 |
| 13_4912261 | 2 | 1 | 2 | 0 | 3 | 0.3333 | 4 | 0.25 | 11 | 0.3636 | 2 | 0 | 4 | 0.25 | 4 | 0 | 3 | 1 | 13 | 0.3077 |
| 13_4919130 | 1 | 0 | 4 | 0.5 | 1 | 0 | 6 | 0.3333 | 12 | 0.3333 | 3 | 0 | 2 | 0 | 6 | 0.3333 | 2 | 0 | 13 | 0.1538 |
| 13_4919140 | 1 | 0 | 3 | 0.6667 | 2 | 0.5 | 5 | 0.4 | 11 | 0.4545 | 4 | 0.5 | 2 | 0 | 5 | 0.6 | 3 | 0.3333 | 14 | 0.4286 |
| 13_4921033 | 5 | 0.2 | 6 | 0.1667 | 2 | 0.5 | 7 | 0.2857 | 20 | 0.25 | 3 | 0.3333 | 8 | 0.25 | 10 | 0.3 | 4 | 0.5 | 25 | 0.32 |
| 13_4921036 | 5 | 0 | 7 | 0.4286 | 3 | 0 | 7 | 0 | 22 | 0.1364 | 4 | 0.5 | 8 | 0 | 10 | 0.1 | 5 | 0 | 27 | 0.1111 |
| 13_4921045 | 5 | 0.2 | 5 | 0.4 | 2 | 0 | 6 | 0.1667 | 18 | 0.2222 | 4 | 0.25 | 8 | 0.125 | 10 | 0.3 | 4 | 0.5 | 26 | 0.2692 |
| 13_4921051 | 5 | 0.8 | 4 | 0.5 | 3 | 0 | 7 | 0.8571 | 19 | 0.6316 | 3 | 0.6667 | 6 | 0.6667 | 8 | 0.5 | 4 | 0.5 | 21 | 0.5714 |
| 13_4921059 | 5 | 1 | 5 | 0.6 | 3 | 0.6667 | 7 | 0.7143 | 20 | 0.75 | 4 | 0.75 | 5 | 1 | 10 | 1 | 4 | 0.25 | 23 | 0.8261 |
| 13_4921074 | 4 | 0.5 | 5 | 1 | 2 | 0.5 | 7 | 0.7143 | 18 | 0.7222 | 2 | 0.5 | 7 | 0.7143 | 10 | 0.9 | 4 | 0.75 | 23 | 0.7826 |
| 13_4921078 | 4 | 0.75 | 4 | 0 | 2 | 0.5 | 7 | 0 | 17 | 0.2353 | 1 | 0 | 6 | 0 | 9 | 0.2222 | 4 | 0 | 20 | 0.1 |

| | | | | | | | | | | | | | | | | | | | | |
|---|---|---|---|---|---|---|---|---|---|---|---|---|---|---|---|---|---|---|---|---|
| 13_492108 | 5 | 0.2 | 5 | 0 | 2 | 0 | 5 | 0.4 | 17 | 0.1765 | 2 | 0.5 | 7 | 0 | 10 | 0.2 | 3 | 0 | 22 | 0.1364 |
| 13_492108 | 5 | 0.4 | 4 | 0.25 | 2 | 0.5 | 5 | 0.2 | 16 | 0.3125 | 2 | 0 | 6 | 0 | 8 | 0.25 | 2 | 0.5 | 18 | 0.1667 |
| 13_492143 | 4 | 0.25 | 2 | 0.5 | 2 | 0 | 3 | 0.6667 | 11 | 0.3636 | 1 | 0 | 6 | 0.3333 | 6 | 0.5 | 2 | 1 | 15 | 0.4667 |
| 13_492143 | 4 | 0.25 | 4 | 0.25 | 1 | 0 | 3 | 0 | 12 | 0.1667 | 1 | 0 | 6 | 0.1667 | 6 | 0 | 3 | 0 | 16 | 0.0625 |
| 13_492144 | 2 | 0 | 4 | 0.5 | 2 | 0.5 | 3 | 1 | 11 | 0.5455 | 1 | 1 | 6 | 0.3333 | 6 | 0.5 | 3 | 0 | 16 | 0.375 |
| 13_492145 | 3 | 0.3333 | 2 | 0 | 2 | 0 | 3 | 0 | 10 | 0.1 | NA | NA | 5 | 0 | 6 | 0.3333 | 2 | 0 | 13 | 0.1538 |
| 13_492146 | 3 | 0.6667 | 4 | 1 | 1 | 1 | 1 | 1 | 9 | 0.8889 | 1 | 1 | 5 | 1 | 7 | 0.7143 | 2 | 0.5 | 15 | 0.8 |
| 13_492147 | 3 | 0 | 3 | 0 | 2 | 0 | 2 | 0.5 | 10 | 0.1 | 1 | 0 | 5 | 0 | 7 | 0.1429 | 2 | 0 | 15 | 0.0667 |
| 13_492147 | 2 | 0 | 4 | 0.25 | 2 | 0 | 1 | 0 | 9 | 0.1111 | 1 | 0 | 4 | 0 | 7 | 0 | 1 | 1 | 13 | 0.0769 |
| 13_492147 | 2 | 0 | 4 | 0 | 2 | 0.5 | 2 | 0.5 | 10 | 0.2 | 1 | 1 | 5 | 0.2 | 7 | 0.1429 | 2 | 0 | 15 | 0.2 |
| 13_492147 | 2 | 0.5 | 3 | 0.6667 | 2 | 1 | 2 | 1 | 9 | 0.7778 | 1 | 1 | 5 | 0.6 | 7 | 0.8571 | 2 | 1 | 15 | 0.8 |
| 13_492148 | 1 | 1 | 4 | 0.75 | 2 | 0.5 | 1 | 0 | 8 | 0.625 | 1 | 0 | 4 | 0.5 | 6 | 0.5 | 2 | 0.5 | 13 | 0.4615 |
| 13_492148 | 1 | 0 | 3 | 0 | 2 | 0.5 | 2 | 0 | 8 | 0.125 | 1 | 0 | 5 | 0 | 5 | 0.2 | 2 | 0 | 13 | 0.0769 |
| 13_492299 | 1 | 0 | NA | NA | 1 | 0 | 1 | 1 | 3 | 0.3333 | 1 | 0 | 1 | 1 | 3 | 0 | 1 | 0 | 6 | 0.1667 |
| 13_492300 | 1 | 0 | 1 | 0 | 2 | 0 | 1 | 1 | 5 | 0.2 | 1 | 1 | 1 | 1 | 3 | 0.3333 | 1 | 0 | 6 | 0.5 |
| 13_493019 | NA | NA | 1 | 1 | NA | NA | 1 | 1 | 2 | 1 | NA | NA | 1 | 1 | NA | NA | NA | NA | 1 | 1 |
| 13_493491 | 3 | 0.6667 | 3 | 0 | 7 | 0.1429 | 4 | 0.75 | 17 | 0.3529 | 5 | 0.4 | 3 | 0.3333 | 3 | 0.6667 | 3 | 1 | 14 | 0.5714 |
| 13_493492 | 2 | 1 | 9 | 0.1111 | 8 | 0.125 | 7 | 0.1429 | 26 | 0.1923 | 7 | 0 | 3 | 0.6667 | 6 | 0.1667 | 6 | 0 | 22 | 0.1364 |
| 13_493492 | 3 | 0 | 9 | 0.3333 | 8 | 0.375 | 6 | 0.1667 | 26 | 0.2692 | 6 | 0.1667 | 1 | 0 | 5 | 0.2 | 6 | 0.1667 | 18 | 0.1667 |
| 13_493492 | 4 | 0 | 9 | 0 | 8 | 0.25 | 7 | 0 | 28 | 0.0714 | 7 | 0.1429 | 3 | 0 | 5 | 0.2 | 5 | 0 | 20 | 0.1 |
| 13_493492 | 2 | 0 | 7 | 0.1429 | 6 | 0 | 5 | 0 | 20 | 0.05 | 6 | 0.1667 | 3 | 0.3333 | 5 | 0 | 4 | 0 | 18 | 0.1111 |
| 13_493493 | 4 | 1 | 8 | 0.75 | 8 | 0.75 | 8 | 1 | 28 | 0.8571 | 8 | 0.875 | 3 | 0.6667 | 6 | 0.5 | 6 | 0.6667 | 23 | 0.6957 |
| 13_493493 | 2 | 1 | 6 | 0.8333 | 4 | 1 | 4 | 0.75 | 16 | 0.875 | 3 | 1 | 2 | 1 | 4 | 1 | 4 | 1 | 13 | 1 |
| 13_493493 | 5 | 0.8 | 8 | 0.75 | 8 | 0.75 | 8 | 1 | 29 | 0.8276 | 8 | 0.625 | 3 | 0.6667 | 5 | 0.6 | 6 | 0.6667 | 22 | 0.6364 |
| 13_493494 | 5 | 0.4 | 9 | 0.1111 | 8 | 0.125 | 8 | 0.375 | 30 | 0.2333 | 8 | 0 | 3 | 0.3333 | 6 | 0.3333 | 6 | 0.5 | 23 | 0.2609 |
| 13_493494 | 4 | 0.75 | 8 | 0.25 | 8 | 0.5 | 7 | 0.4286 | 27 | 0.4444 | 7 | 0.2857 | 3 | 0.3333 | 5 | 0 | 6 | 0.3333 | 21 | 0.2381 |
| 13_493495 | 5 | 1 | 9 | 0.4444 | 8 | 0.75 | 8 | 0.875 | 30 | 0.7333 | 7 | 0.8571 | 3 | 1 | 6 | 0.6667 | 5 | 0.6 | 21 | 0.7619 |
| 13_493495 | 4 | 1 | 8 | 0.375 | 6 | 0.6667 | 6 | 0.3333 | 24 | 0.5417 | 7 | 0.5714 | 2 | 1 | 4 | 0.75 | 2 | 0 | 15 | 0.6 |
| 13_493843 | 2 | 0.5 | NA | NA | NA | NA | NA | NA | 2 | 0.5 | NA | NA | NA | NA | 1 | 1 | NA | NA | 1 | 1 |
| 13_495327 | 1 | 1 | 1 | 1 | NA | NA | NA | NA | 2 | 1 | 1 | 0 | 1 | 1 | NA | NA | NA | NA | 2 | 0.5 |
| 13_495332 | NA | NA | 2 | 1 | NA | NA | 1 | 1 | 3 | 1 | 1 | 1 | NA | NA | NA | NA | 1 | 0 | 2 | 0.5 |
| 13_495366 | 2 | 1 | 2 | 0.5 | NA | NA | NA | NA | 4 | 0.75 | 1 | 0 | 2 | 1 | NA | NA | 1 | 0 | 4 | 0.5 |
| 13_495368 | 2 | 0.5 | 1 | 1 | NA | NA | 1 | 0 | 4 | 0.5 | NA | NA | 2 | 0.5 | NA | NA | 2 | 0 | 4 | 0.25 |
| 13_495369 | 2 | 1 | 2 | 1 | NA | NA | NA | NA | 4 | 1 | NA | NA | 2 | 1 | NA | NA | NA | NA | 2 | 1 |
| 13_495377 | 1 | 0 | 4 | 0.5 | NA | NA | 1 | 0 | 6 | 0.3333 | NA | NA | 3 | 0.3333 | NA | NA | 3 | 0.3333 | 6 | 0.3333 |
| 13_495379 | 2 | 0 | 3 | 0.3333 | NA | NA | 1 | 0 | 6 | 0.1667 | NA | NA | 3 | 0.3333 | NA | NA | 2 | 0 | 5 | 0.2 |
| 13_495382 | 2 | 0.5 | 3 | 0.3333 | NA | NA | 1 | 0 | 6 | 0.3333 | NA | NA | 3 | 0 | NA | NA | 3 | 0.3333 | 6 | 0.1667 |
| 13_495389 | 2 | 0 | 5 | 0.2 | NA | NA | 2 | 0 | 9 | 0.1111 | 1 | 0 | 3 | 0 | NA | NA | 4 | 0.25 | 8 | 0.125 |
| 13_495392 | 2 | 0 | 4 | 0 | 1 | 0 | 2 | 0.5 | 9 | 0.1111 | 2 | 0 | 4 | 0 | NA | NA | 4 | 0.25 | 10 | 0.1 |
| 13_495397 | 2 | 0.5 | 4 | 0.5 | 1 | 1 | 2 | 0.5 | 9 | 0.5556 | 2 | 1 | 3 | 0.6667 | NA | NA | 5 | 0.4 | 10 | 0.6 |
| 13_495397 | 2 | 1 | 5 | 1 | 3 | 0.6667 | 3 | 1 | 13 | 0.9231 | 3 | 0.3333 | 4 | 0.75 | NA | NA | 5 | 0.8 | 12 | 0.6667 |
| 13_495404 | 2 | 0 | 8 | 0.125 | 3 | 0 | 3 | 0 | 16 | 0.0625 | 5 | 0 | 5 | 0.2 | 1 | 0 | 4 | 0 | 15 | 0.0667 |
| 13_495738 | 2 | 0.5 | NA | NA | NA | NA | NA | NA | 2 | 0.5 | NA | NA | 1 | 1 | NA | NA | NA | NA | 1 | 1 |
| 13_495742 | 2 | 1 | NA | NA | NA | NA | NA | NA | 2 | 1 | NA | NA | 1 | 1 | NA | NA | NA | NA | 1 | 1 |
| 13_495745 | 1 | 1 | NA | NA | NA | NA | NA | NA | 1 | 1 | NA | NA | 1 | 1 | NA | NA | NA | NA | 1 | 1 |
| 13_496151 | 3 | 1 | 4 | 1 | 2 | 1 | 9 | 1 | 18 | 1 | NA | NA | 1 | 1 | 4 | 1 | 2 | 1 | 7 | 1 |
| 13_496152 | 3 | 0 | 5 | 0.2 | 3 | 0 | 8 | 0.125 | 19 | 0.1053 | NA | NA | 2 | 0 | 5 | 0.2 | 2 | 0 | 9 | 0.1111 |
| 13_496155 | 3 | 0.3333 | 4 | 0 | 3 | 0 | 10 | 0.1 | 20 | 0.1 | NA | NA | 2 | 0 | 6 | 0.1667 | 2 | 0 | 10 | 0.1 |
| 13_496156 | 3 | 0.3333 | 4 | 0.25 | 3 | 0 | 9 | 0 | 19 | 0.1053 | NA | NA | 2 | 0 | 5 | 0.4 | 2 | 0 | 9 | 0.2222 |
| 13_496157 | 3 | 0.6667 | 3 | 0 | 2 | 0 | 7 | 0.2857 | 15 | 0.2667 | NA | NA | 2 | 0 | 6 | 0.3333 | 2 | 0 | 10 | 0.2 |
| 13_496163 | 4 | 0 | 1 | 0 | 5 | 0 | 10 | 0.2 | 20 | 0.1 | 4 | 0 | 2 | 0 | 8 | 0.125 | 2 | 0.5 | 16 | 0.125 |
| 13_497434 | NA | NA | 1 | 1 | NA | NA | NA | NA | 1 | 1 | NA | NA | NA | NA | 1 | 1 | NA | NA | 1 | 1 |
| 13_497435 | NA | NA | 1 | 1 | NA | NA | NA | NA | 1 | 1 | NA | NA | NA | NA | 1 | 1 | NA | NA | 1 | 1 |
| 13_497438 | NA | NA | 1 | 1 | NA | NA | NA | NA | 1 | 1 | NA | NA | NA | NA | 1 | 1 | NA | NA | 1 | 1 |
| 13_498147 | 2 | 0.5 | NA | NA | 1 | 0 | 1 | 0 | 4 | 0.25 | NA | NA | 1 | 1 | 1 | 0 | NA | NA | 2 | 0.5 |
| 13_498148 | 2 | 0 | NA | NA | 1 | 0 | 1 | 1 | 4 | 0.25 | NA | NA | 1 | 0 | 1 | 1 | NA | NA | 2 | 0.5 |
| 13_498148 | 2 | 0 | 1 | 0 | 1 | 0 | 1 | 1 | 5 | 0.2 | NA | NA | 1 | 1 | 1 | 0 | NA | NA | 2 | 0.5 |
| 13_498151 | 2 | 0.5 | 1 | 1 | 1 | 1 | 1 | 1 | 5 | 0.8 | NA | NA | 1 | 1 | 1 | 1 | NA | NA | 2 | 1 |
| 13_498151 | 2 | 1 | 1 | 0 | 1 | 1 | 1 | 1 | 5 | 0.8 | NA | NA | 1 | 0 | 1 | 1 | NA | NA | 2 | 0.5 |
| 13_498158 | 3 | 0.3333 | 1 | 1 | 1 | 0 | 1 | 1 | 6 | 0.5 | NA | NA | 1 | 1 | 1 | 1 | NA | NA | 2 | 1 |
| 13_498162 | 3 | 0 | 1 | 0 | 1 | 1 | 1 | 0 | 6 | 0.1667 | NA | NA | 3 | 0.3333 | 1 | 0 | NA | NA | 4 | 0.25 |
| 13_498320 | 5 | 0 | 1 | 0 | 3 | 0 | 3 | 0.6667 | 12 | 0.1667 | NA | NA | 3 | 0 | 1 | 0 | 1 | 1 | 5 | 0.2 |
| 13_498465 | 3 | 0 | 1 | 0 | 2 | 0.5 | 1 | 0 | 7 | 0.1429 | 3 | 0 | 1 | 0 | 1 | 0 | 2 | 0.5 | 7 | 0.1429 |
| 13_505874 | 1 | 1 | 1 | 0 | 1 | 0 | 1 | 0 | 4 | 0.25 | 1 | 1 | 1 | 0 | 1 | 0 | NA | NA | 3 | 0.3333 |
| 13_505876 | 1 | 1 | 1 | 1 | 1 | 1 | NA | NA | 3 | 1 | 1 | 1 | 1 | 1 | 1 | 1 | NA | NA | 3 | 1 |

| | | | | | | | | | | | | | | | | | | | | |
|---|---|---|---|---|---|---|---|---|---|---|---|---|---|---|---|---|---|---|---|---|
| 13_512533 | 1 | 1 | NA | NA | NA | NA | NA | NA | 1 | 1 | NA | NA | 1 | 1 | 1 | 1 | 1 | 1 | 3 | 1 |
| 13_512540 | NA | NA | NA | NA | NA | NA | 1 | 1 | 1 | 1 | NA | NA | 1 | 1 | 1 | 1 | NA | NA | 2 | 1 |
| 13_512544 | NA | NA | NA | NA | NA | NA | 1 | 1 | 1 | 1 | NA | NA | 1 | 1 | 1 | 1 | NA | NA | 2 | 1 |
| 13_512545 | NA | NA | NA | NA | NA | NA | 1 | 1 | 1 | 1 | NA | NA | 1 | 1 | 1 | 1 | NA | NA | 2 | 1 |
| 13_512935 | 9 | 0.6667 | 20 | 0.55 | 12 | 0.5 | 18 | 0.4444 | 59 | 0.5254 | 7 | 0.5714 | 8 | 0.5 | 10 | 0.6 | 11 | 0.7273 | 36 | 0.6111 |
| 13_512935 | 10 | 0.3 | 20 | 0.1 | 14 | 0.1429 | 21 | 0.2381 | 65 | 0.1846 | 7 | 0.2857 | 9 | 0.1111 | 9 | 0.1111 | 11 | 0.0909 | 36 | 0.1389 |
| 13_512935 | 9 | 0.5556 | 21 | 0.4762 | 13 | 0.6154 | 19 | 0.5789 | 62 | 0.5484 | 7 | 0.8571 | 9 | 0.6667 | 8 | 0.75 | 9 | 0.5556 | 33 | 0.697 |
| 13_512983 | 9 | 0.2222 | 22 | 0.0909 | 14 | 0.1429 | 19 | 0.0526 | 64 | 0.1094 | 7 | 0.1429 | 8 | 0 | 9 | 0.1111 | 12 | 0.1667 | 36 | 0.1111 |
| 13_512984 | 8 | 0.375 | 21 | 0.0476 | 14 | 0.4286 | 19 | 0.2632 | 62 | 0.2419 | 6 | 0.1667 | 9 | 0.2222 | 10 | 0.5 | 10 | 0.3 | 35 | 0.3143 |
| 13_512985 | 11 | 0 | 23 | 0.0435 | 15 | 0 | 21 | 0.1429 | 70 | 0.0571 | 8 | 0.125 | 11 | 0.0909 | 12 | 0.1667 | 12 | 0 | 43 | 0.093 |
| 13_513030 | 9 | 0.4444 | 16 | 0.125 | 11 | 0.3636 | 14 | 0.3571 | 50 | 0.3 | 5 | 0.4 | 9 | 0 | 5 | 0.2 | 3 | 0.3333 | 22 | 0.1818 |
| 13_513030 | 12 | 1 | 21 | 1 | 14 | 0.9286 | 22 | 0.9545 | 69 | 0.971 | 9 | 1 | 11 | 0.8182 | 12 | 0.9167 | 12 | 1 | 44 | 0.9318 |
| 13_513032 | 13 | 0.6923 | 18 | 0.6667 | 14 | 0.7143 | 22 | 0.5455 | 67 | 0.6418 | 7 | 0.2857 | 9 | 0.3333 | 12 | 0.5 | 10 | 0.8 | 38 | 0.5 |
| 13_513032 | 12 | 0.9167 | 20 | 0.8 | 14 | 0.8571 | 24 | 0.7917 | 70 | 0.8286 | 6 | 0.8333 | 8 | 0.75 | 12 | 0.75 | 11 | 0.9091 | 37 | 0.8108 |
| 13_513032 | 14 | 0.1429 | 21 | 0.0476 | 13 | 0.0769 | 25 | 0.16 | 73 | 0.1096 | 9 | 0 | 11 | 0.0909 | 12 | 0.0833 | 10 | 0.2 | 42 | 0.0952 |
| 13_513032 | 15 | 1 | 19 | 0.8947 | 15 | 1 | 24 | 0.875 | 73 | 0.9315 | 9 | 1 | 11 | 0.8182 | 13 | 0.9231 | 12 | 0.9167 | 45 | 0.9111 |
| 13_513033 | 12 | 0.1667 | 19 | 0.1053 | 14 | 0.0714 | 19 | 0.0526 | 64 | 0.0938 | 8 | 0.125 | 10 | 0.1 | 11 | 0 | 10 | 0.1 | 39 | 0.0769 |
| 13_513033 | 14 | 0.4286 | 20 | 0.45 | 11 | 0.6364 | 21 | 0.5714 | 66 | 0.5152 | 9 | 0.7778 | 7 | 0.2857 | 12 | 0.5 | 8 | 0.5 | 36 | 0.5278 |
| 13_513033 | 7 | 0.1429 | 21 | 0.0476 | 10 | 0.2 | 19 | 0.0526 | 57 | 0.0877 | 6 | 0.1667 | 9 | 0.1111 | 11 | 0 | 9 | 0.2222 | 35 | 0.1143 |
| 13_513041 | 11 | 0.0909 | 26 | 0.1154 | 20 | 0.25 | 28 | 0.3214 | 85 | 0.2118 | 11 | 0.0909 | 16 | 0.125 | 14 | 0.1429 | 15 | 0.0667 | 56 | 0.1071 |
| 13_513042 | 14 | 0.0714 | 25 | 0.08 | 19 | 0.1579 | 32 | 0.25 | 90 | 0.1556 | 12 | 0.0833 | 16 | 0.125 | 15 | 0.2667 | 14 | 0.1429 | 57 | 0.1579 |
| 13_513060 | 17 | 0 | 38 | 0.1579 | 32 | 0 | 40 | 0.175 | 127 | 0.1024 | 18 | 0.2222 | 23 | 0.0435 | 26 | 0.0769 | 20 | 0.1 | 87 | 0.1034 |
| 13_513061 | NA | NA | 1 | 1 | NA | NA | NA | NA | 1 | 1 | NA | NA | NA | NA | NA | NA | 1 | 1 | 1 | 1 |
| 13_515733 | 2 | 0.5 | NA | NA | NA | NA | NA | NA | 2 | 0.5 | NA | NA | NA | NA | NA | NA | 1 | 1 | 1 | 1 |
| 13_515740 | 2 | 1 | NA | NA | NA | NA | NA | NA | 2 | 1 | NA | NA | NA | NA | NA | NA | 1 | 1 | 1 | 1 |
| 13_515741 | 2 | 1 | NA | NA | 1 | 1 | NA | NA | 3 | 1 | NA | NA | NA | NA | NA | NA | 1 | 1 | 1 | 1 |
| 13_515741 | 2 | 1 | NA | NA | 1 | 1 | NA | NA | 3 | 1 | NA | NA | NA | NA | NA | NA | 1 | 1 | 1 | 1 |
| 13_515747 | 2 | 0.5 | NA | NA | 1 | 1 | 1 | 0 | 4 | 0.5 | NA | NA | 1 | 1 | NA | NA | 1 | 1 | 2 | 1 |
| 13_515749 | 3 | 1 | NA | NA | 1 | 1 | 1 | 0 | 5 | 0.8 | NA | NA | 1 | 1 | NA | NA | 1 | 1 | 2 | 1 |
| 13_515749 | 3 | 1 | NA | NA | 1 | 1 | 1 | 1 | 5 | 1 | NA | NA | 1 | 1 | NA | NA | 1 | 1 | 2 | 1 |
| 13_515750 | 3 | 1 | 1 | 1 | 1 | 1 | 1 | 1 | 6 | 1 | NA | NA | NA | NA | NA | NA | 1 | 1 | 1 | 1 |
| 13_515752 | 3 | 0.6667 | 1 | 0 | 1 | 0 | 1 | 1 | 6 | 0.5 | NA | NA | 1 | 1 | NA | NA | 1 | 1 | 2 | 1 |
| 13_515753 | 3 | 1 | 1 | 1 | 1 | 1 | 1 | 1 | 6 | 1 | NA | NA | 1 | 1 | NA | NA | 1 | 1 | 2 | 1 |
| 13_515753 | 3 | 0.3333 | 1 | 1 | 1 | 0 | 1 | 0 | 6 | 0.3333 | NA | NA | 1 | 1 | NA | NA | 1 | 0 | 2 | 0.5 |
| 13_515753 | 3 | 0.6667 | 1 | 0 | 1 | 0 | 1 | 0 | 6 | 0.3333 | NA | NA | 1 | 1 | NA | NA | NA | NA | 1 | 1 |
| 13_515754 | 3 | 1 | 1 | 1 | 1 | 1 | 1 | 0 | 6 | 0.8333 | NA | NA | 1 | 1 | NA | NA | NA | NA | 1 | 1 |
| 13_515754 | 3 | 0.6667 | 1 | 0 | 1 | 1 | 1 | 1 | 6 | 0.6667 | NA | NA | 1 | 1 | NA | NA | 1 | 0 | 2 | 0.5 |
| 13_515758 | 3 | 0.6667 | 1 | 1 | 1 | 1 | NA | NA | 5 | 0.8 | NA | NA | 1 | 1 | NA | NA | 1 | 1 | 2 | 1 |
| 13_515779 | 8 | 0 | 8 | 0.125 | 13 | 0 | 10 | 0.2 | 39 | 0.0769 | 7 | 0.1429 | 8 | 0 | 9 | 0.3333 | 6 | 0.5 | 30 | 0.2333 |
| 13_515784 | 9 | 0 | 10 | 0.2 | 13 | 0.0769 | 11 | 0.0909 | 43 | 0.093 | 6 | 0 | 10 | 0 | 6 | 0 | 5 | 0.4 | 27 | 0.0741 |
| 13_515785 | 10 | 0.5 | 9 | 0.1111 | 9 | 0.1111 | 11 | 0.2727 | 39 | 0.2564 | 7 | 0 | 9 | 0.1111 | 7 | 0 | 6 | 0.3333 | 29 | 0.1034 |
| 13_515785 | 8 | 0.5 | 6 | 0.6667 | 11 | 0.3636 | 11 | 0.6364 | 36 | 0.5278 | 7 | 0.5714 | 10 | 0.7 | 4 | 0.25 | 4 | 0.75 | 25 | 0.6 |
| 13_515786 | 10 | 0.4 | 8 | 0.625 | 12 | 0.25 | 10 | 0.3 | 40 | 0.375 | 7 | 0.2857 | 8 | 0.375 | 6 | 0.3333 | 6 | 0.5 | 27 | 0.3704 |
| 13_515786 | 10 | 0.2 | 8 | 0.125 | 13 | 0.1538 | 11 | 0 | 42 | 0.119 | 7 | 0.5714 | 9 | 0.1111 | 5 | 0.2 | 4 | 0 | 25 | 0.24 |
| 13_515786 | 6 | 0.3333 | 5 | 0.6 | 12 | 0.25 | 10 | 0.6 | 33 | 0.4242 | 5 | 0.8 | 5 | 0.6 | 5 | 0.4 | 5 | 1 | 20 | 0.7 |
| 13_515787 | 9 | 0.6667 | 8 | 1 | 10 | 0.6 | 9 | 0.7778 | 36 | 0.75 | 5 | 0.8 | 9 | 0.7778 | 4 | 0.75 | 4 | 1 | 22 | 0.8182 |
| 13_515787 | 10 | 0 | 7 | 0.1429 | 10 | 0 | 11 | 0.0909 | 38 | 0.0526 | 5 | 0 | 8 | 0 | 3 | 0 | 4 | 0.25 | 20 | 0.05 |
| 13_515788 | 7 | 0.1429 | 8 | 0.25 | 12 | 0.0833 | 8 | 0.375 | 35 | 0.2 | 6 | 0.1667 | 6 | 0.3333 | 4 | 0.25 | 5 | 0.8 | 21 | 0.381 |
| 13_515789 | 9 | 0.4444 | 7 | 0.2857 | 8 | 0.375 | 9 | 0.2222 | 33 | 0.3333 | 4 | 0.25 | 9 | 0.4444 | 5 | 0.4 | 5 | 0.8 | 23 | 0.4783 |
| 13_515789 | 8 | 0.375 | 8 | 0.5 | 8 | 0.375 | 7 | 0.7143 | 31 | 0.4839 | 4 | 0.25 | 8 | 0.625 | 3 | 0.3333 | 4 | 1 | 19 | 0.5789 |
| 13_515791 | 8 | 0.125 | 4 | 0 | 8 | 0 | 6 | 0.3333 | 26 | 0.1154 | 5 | 0 | 6 | 0.5 | 3 | 0.6667 | 5 | 0.4 | 19 | 0.3684 |
| 13_515791 | 5 | 0.2 | 5 | 0.4 | 6 | 0.3333 | 6 | 0.6667 | 22 | 0.4091 | 3 | 0.6667 | 7 | 0.4286 | 1 | 1 | 4 | 0.75 | 15 | 0.6 |
| 13_515794 | 5 | 0.2 | 7 | 0.2857 | 10 | 0.1 | 7 | 0 | 29 | 0.1379 | 5 | 0.2 | 5 | 0.2 | 4 | 0.25 | 4 | 0 | 18 | 0.1667 |
| 13_515800 | 6 | 0.3333 | 7 | 0.1429 | 9 | 0 | 5 | 0.2 | 27 | 0.1481 | 5 | 0.2 | 6 | 0 | 2 | 0 | 4 | 0 | 17 | 0.0588 |
| 13_515810 | 4 | 0.5 | 3 | 0.3333 | 5 | 0 | 3 | 0 | 15 | 0.2 | 4 | 0 | 3 | 0 | 3 | 0.3333 | NA | NA | 10 | 0.1 |
| 13_515837 | 1 | 0 | 2 | 0.5 | 4 | 0 | 3 | 0 | 10 | 0.1 | 3 | 0 | 3 | 0 | 3 | 0.3333 | NA | NA | 9 | 0.1111 |
| 13_515838 | 1 | 0 | 2 | 0 | 3 | 0.3333 | 2 | 0.5 | 8 | 0.25 | 2 | 0.5 | 3 | 0.3333 | 3 | 0 | NA | NA | 8 | 0.25 |
| 13_517215 | NA | NA | NA | NA | NA | NA | 1 | 1 | 1 | 1 | 2 | 1 | 1 | 1 | NA | NA | NA | NA | 3 | 1 |
| 13_517884 | NA | NA | 1 | 1 | NA | NA | NA | NA | 1 | 1 | NA | NA | NA | NA | 1 | 1 | NA | NA | 1 | 1 |
| 13_518374 | NA | NA | 2 | 0 | 3 | 0.6667 | 1 | 0 | 6 | 0.3333 | NA | NA | NA | NA | 2 | 1 | 1 | 1 | 3 | 1 |
| 13_518374 | 2 | 1 | 2 | 1 | 3 | 0.6667 | 1 | 0 | 8 | 0.75 | NA | NA | 1 | 1 | 3 | 0.3333 | 1 | 1 | 5 | 0.6 |
| 13_518374 | 2 | 0.5 | 3 | 0.3333 | 3 | 0 | 1 | 0 | 9 | 0.2222 | NA | NA | 1 | 0 | 3 | 0.3333 | 1 | 1 | 5 | 0.4 |
| 13_518375 | 2 | 1 | 3 | 0.3333 | 2 | 0.5 | 1 | 0 | 8 | 0.5 | NA | NA | 1 | 0 | 2 | 0.5 | NA | NA | 3 | 0.3333 |
| 13_518398 | 5 | 0 | 3 | 0 | 3 | 0 | 3 | 0.3333 | 14 | 0.0714 | 7 | 0.1429 | 4 | 0 | 5 | 0 | 2 | 0 | 18 | 0.0556 |
| 13_518399 | 3 | 0.3333 | 3 | 0.6667 | 2 | 0 | 3 | 0 | 11 | 0.2727 | 6 | 0.5 | 5 | 0.2 | 3 | 0.3333 | 2 | 0.5 | 16 | 0.375 |

| | | | | | | | | | | | | | | | | | | | | |
|---|---|---|---|---|---|---|---|---|---|---|---|---|---|---|---|---|---|---|---|---|
| 13_5184008 | 3 | 0 | 2 | 0 | 2 | 0 | 3 | 0.3333 | 10 | 0.1 | 6 | 0 | 4 | 0.25 | 5 | 0 | 2 | 0 | 17 | 0.0588 |
| 13_5184023 | 2 | 0.5 | 3 | 1 | 2 | 0 | 3 | 0 | 10 | 0.4 | 6 | 0.5 | 3 | 0.3333 | 5 | 0.4 | 2 | 0 | 16 | 0.375 |
| 13_5185545 | 1 | 1 | 2 | 0.5 | NA | NA | 3 | 0.6667 | 6 | 0.6667 | 2 | 1 | 1 | 1 | 2 | 1 | 2 | 1 | 7 | 1 |
| 13_5185564 | 1 | 0 | 4 | 0.25 | NA | NA | 3 | 0 | 8 | 0.125 | 1 | 0 | 1 | 0 | 1 | 0 | 2 | 0.5 | 5 | 0.2 |
| 13_5185567 | 1 | 1 | 2 | 0.5 | NA | NA | 3 | 1 | 6 | 0.8333 | 2 | 0.5 | NA | NA | 2 | 1 | 2 | 1 | 6 | 0.8333 |
| 13_5185588 | NA | NA | 4 | 0.25 | NA | NA | 3 | 0.3333 | 7 | 0.2857 | 3 | 0.3333 | NA | NA | 2 | 0 | 1 | 0 | 6 | 0.1667 |
| 13_5185626 | NA | NA | 4 | 1 | NA | NA | 3 | 0.6667 | 7 | 0.8571 | 2 | 1 | 2 | 0.5 | 2 | 0.5 | 1 | 1 | 7 | 0.7143 |
| 13_5185664 | 2 | 1 | 4 | 1 | NA | NA | 2 | 1 | 8 | 1 | 2 | 0.5 | 1 | 0 | 1 | 1 | 1 | 1 | 5 | 0.6 |
| 13_5185668 | 1 | 1 | 4 | 1 | NA | NA | 1 | 1 | 6 | 1 | 2 | 1 | 2 | 1 | 2 | 0.5 | 1 | 1 | 7 | 0.8571 |
| 13_5185670 | 1 | 0 | 4 | 0.25 | NA | NA | 2 | 0.5 | 7 | 0.2857 | 2 | 0.5 | 2 | 1 | 2 | 0.5 | 1 | 0 | 7 | 0.5714 |
| 13_5185672 | NA | NA | 4 | 1 | NA | NA | 1 | 1 | 5 | 1 | 2 | 1 | 1 | 1 | 1 | 1 | 1 | 1 | 5 | 1 |
| 13_5185673 | 1 | 0 | 4 | 0.25 | NA | NA | 2 | 0 | 7 | 0.1429 | 2 | 0 | 2 | 0.5 | 1 | 0 | 1 | 1 | 6 | 0.3333 |
| 13_5185677 | 1 | 0 | 3 | 0.6667 | NA | NA | 2 | 0 | 6 | 0.3333 | 2 | 0 | 2 | 0 | 2 | 0 | 1 | 1 | 7 | 0.1429 |
| 13_5185686 | 1 | 0 | 2 | 1 | NA | NA | 2 | 0.5 | 5 | 0.6 | 1 | 1 | 1 | 1 | 1 | 1 | 1 | 1 | 4 | 1 |
| 13_5185694 | 1 | 1 | 3 | 1 | NA | NA | 2 | 1 | 6 | 1 | 1 | 1 | 2 | 1 | 1 | 1 | 1 | 1 | 5 | 1 |
| 13_5200202 | 4 | 0.5 | 4 | 0.5 | 3 | 1 | 7 | 0.5714 | 18 | 0.6111 | 6 | 0.5 | 2 | 0.5 | 4 | 0.25 | 2 | 0.5 | 14 | 0.4286 |
| 13_5200211 | 4 | 0.5 | 4 | 0.75 | 3 | 0.3333 | 3 | 1 | 14 | 0.6429 | 4 | 0.75 | NA | NA | 1 | 1 | 3 | 0.3333 | 8 | 0.625 |
| 13_5200218 | 4 | 0.5 | 4 | 0 | 2 | 0 | 6 | 0 | 16 | 0.125 | 6 | 0.1667 | 2 | 0 | 4 | 0.25 | 1 | 1 | 13 | 0.2308 |
| 13_5200220 | 4 | 0.5 | 5 | 0.4 | 3 | 0.3333 | 7 | 0.2857 | 19 | 0.3684 | 6 | 0.8333 | 2 | 0 | 3 | 0.6667 | 3 | 0.3333 | 14 | 0.5714 |
| 13_5200230 | 4 | 0 | 5 | 0 | 3 | 0 | 6 | 0.1667 | 18 | 0.0556 | 6 | 0 | 1 | 0 | 4 | 0.25 | 3 | 0.3333 | 14 | 0.1429 |
| 13_5200235 | 4 | 0 | 4 | 0.75 | 4 | 1 | 7 | 0.5714 | 19 | 0.5789 | 7 | 0.5714 | 1 | 0 | 5 | 0.2 | 3 | 0 | 16 | 0.3125 |
| 13_5200240 | 4 | 0.5 | 4 | 0.5 | 4 | 0.5 | 7 | 0.8571 | 19 | 0.6316 | 7 | 0.7143 | 2 | 0 | 5 | 0.6 | 3 | 0.6667 | 17 | 0.5882 |
| 13_5200249 | 3 | 0 | 6 | 0.3333 | 2 | 0 | 7 | 0 | 18 | 0.1111 | 6 | 0.1667 | 1 | 0 | 5 | 0 | 3 | 0.3333 | 15 | 0.1333 |
| 13_5265757 | NA | NA | 1 | 1 | NA | NA | NA | NA | 1 | 1 | NA | NA | NA | NA | NA | NA | 1 | 1 | 1 | 1 |
| 13_5265862 | 39 | 0.5897 | 34 | 0.5882 | 25 | 0.52 | 27 | 0.3333 | 125 | 0.52 | 26 | 0.5 | 29 | 0.5517 | 30 | 0.5 | 28 | 0.6071 | 113 | 0.5398 |
| 13_5265865 | 34 | 0.5882 | 35 | 0.6857 | 25 | 0.8 | 25 | 0.76 | 119 | 0.6975 | 24 | 0.7917 | 23 | 0.6957 | 23 | 0.7826 | 20 | 0.85 | 90 | 0.7778 |
| 13_5265865 | 1 | 1 | NA | NA | 1 | 1 | NA | NA | 2 | 1 | NA | NA | 1 | 1 | NA | NA | NA | NA | 1 | 1 |
| 13_5265866 | 22 | 0.5909 | 31 | 0.5484 | 19 | 0.5263 | 20 | 0.55 | 92 | 0.5543 | 21 | 0.619 | 17 | 0.5294 | 19 | 0.3684 | 22 | 0.4091 | 79 | 0.481 |
| 13_5265911 | 31 | 0.2903 | 31 | 0.4516 | 22 | 0.3182 | 27 | 0.1852 | 111 | 0.3153 | 19 | 0.4737 | 20 | 0.6 | 25 | 0.44 | 21 | 0.3333 | 85 | 0.4588 |
| 13_5265914 | 30 | 0.1333 | 33 | 0.0909 | 23 | 0.2609 | 28 | 0.0714 | 114 | 0.1316 | 20 | 0.15 | 17 | 0.1176 | 26 | 0.1538 | 21 | 0.0952 | 84 | 0.131 |
| 13_5266047 | 18 | 0.2222 | 25 | 0.44 | 13 | 0.7692 | 20 | 0.5 | 76 | 0.4605 | 13 | 0.6154 | 15 | 0.3333 | 14 | 0.5714 | 14 | 0.5 | 56 | 0.5 |
| 13_5266103 | 16 | 0.125 | 17 | 0.1176 | 10 | 0 | 19 | 0.2105 | 62 | 0.129 | 12 | 0.0833 | 11 | 0.0909 | 10 | 0.2 | 11 | 0 | 44 | 0.0909 |
| 13_5266128 | 16 | 0.3125 | 15 | 0.2667 | 10 | 0.4 | 19 | 0.1053 | 60 | 0.25 | 11 | 0.2727 | 10 | 0.3 | 10 | 0.2 | 11 | 0.2727 | 42 | 0.2619 |
| 13_5266141 | 14 | 0.1429 | 15 | 0.1333 | 10 | 0 | 17 | 0 | 56 | 0.0714 | 8 | 0.125 | 9 | 0 | 9 | 0.1111 | 10 | 0.2 | 36 | 0.1111 |
| 13_5266142 | 13 | 0.3846 | 15 | 0.3333 | 9 | 0.5556 | 16 | 0.25 | 53 | 0.3585 | 9 | 0.2222 | 9 | 0.5556 | 9 | 0.2222 | 10 | 0.3 | 37 | 0.3243 |
| 13_5266144 | 17 | 0.5882 | 15 | 0.6667 | 7 | 0.8571 | 16 | 0.625 | 55 | 0.6545 | 10 | 0.7 | 7 | 0.8571 | 9 | 0.8889 | 10 | 0.9 | 36 | 0.8333 |
| 13_5266160 | 12 | 0.1667 | 14 | 0.2143 | 8 | 0.125 | 11 | 0.0909 | 45 | 0.1556 | 7 | 0 | 9 | 0.2222 | 8 | 0 | 8 | 0.375 | 32 | 0.1562 |
| 13_5266164 | 15 | 0.2667 | 12 | 0.0833 | 9 | 0.2222 | 10 | 0.2 | 46 | 0.1957 | 5 | 0 | 9 | 0.1111 | 7 | 0 | 9 | 0.1111 | 30 | 0.0667 |
| 13_5266178 | 15 | 0.2667 | 16 | 0.25 | 10 | 0.1 | 16 | 0.25 | 57 | 0.2281 | 7 | 0.2857 | 7 | 0.2857 | 7 | 0 | 9 | 0.1111 | 30 | 0.1667 |
| 13_5266181 | 15 | 0.2667 | 6 | 0.3333 | 7 | 0.8571 | 14 | 0.6429 | 42 | 0.5 | 6 | 0.6667 | 7 | 0.5714 | 7 | 0.5714 | 3 | 1 | 23 | 0.6522 |
| 13_5266182 | 11 | 1 | 9 | 0.8889 | 5 | 0.8 | 8 | 0.875 | 33 | 0.9091 | 7 | 1 | 4 | 0.75 | 6 | 0.8333 | 5 | 1 | 22 | 0.9091 |
| 13_5266201 | 14 | 0.2857 | 12 | 0.4167 | 7 | 0.2857 | 13 | 0.3846 | 46 | 0.3478 | 7 | 0.4286 | 8 | 0.125 | 4 | 0.25 | 7 | 0.4286 | 26 | 0.3077 |
| 13_5266221 | 10 | 0.6 | 12 | 0.6667 | 6 | 0.5 | 12 | 0.8333 | 40 | 0.675 | 6 | 0.8333 | 4 | 0.5 | 4 | 0.75 | 7 | 1 | 21 | 0.8095 |
| 13_5268462 | 3 | 0.6667 | 7 | 0.1429 | 2 | 0 | 4 | 0.25 | 16 | 0.25 | 1 | 1 | 1 | 0 | 1 | 0 | 1 | 1 | 4 | 0.5 |
| 13_5306121 | NA | NA | NA | NA | 1 | 1 | NA | NA | 1 | 1 | 1 | 1 | 1 | 0 | NA | NA | 1 | 1 | 3 | 0.6667 |
| 13_5306183 | NA | NA | NA | NA | 1 | 1 | NA | NA | 1 | 1 | 1 | 0 | 1 | 1 | NA | NA | 1 | 0 | 3 | 0.3333 |
| 13_5312137 | NA | NA | 1 | 0 | 3 | 0.3333 | 3 | 0.6667 | 7 | 0.4286 | 1 | 1 | NA | NA | 4 | 0.75 | 2 | 0.5 | 7 | 0.7143 |
| 13_5312144 | NA | NA | 1 | 0 | 3 | 1 | 3 | 1 | 7 | 0.8571 | 1 | 1 | NA | NA | 3 | 1 | 3 | 0.6667 | 7 | 0.8571 |
| 13_5312152 | NA | NA | 1 | 1 | 3 | 1 | 3 | 1 | 7 | 1 | 1 | 1 | NA | NA | 4 | 1 | 3 | 0.6667 | 8 | 0.875 |
| 13_5312169 | NA | NA | NA | NA | 3 | 1 | 3 | 0.6667 | 6 | 0.8333 | 1 | 1 | NA | NA | 3 | 1 | 1 | 1 | 5 | 1 |
| 13_5312171 | NA | NA | NA | NA | 3 | 1 | 2 | 0.5 | 5 | 0.8 | 1 | 0 | NA | NA | 4 | 0.25 | 1 | 0 | 6 | 0.1667 |
| 13_5312174 | NA | NA | NA | NA | 3 | 0.6667 | 1 | 0 | 4 | 0.5 | 1 | 0 | NA | NA | 3 | 0.6667 | 1 | 1 | 5 | 0.6 |
| 13_5312182 | NA | NA | NA | NA | 3 | 1 | 3 | 1 | 6 | 1 | 1 | 1 | NA | NA | 2 | 1 | 1 | 0 | 4 | 0.75 |
| 13_5312185 | NA | NA | NA | NA | 3 | 0.3333 | 2 | 0 | 5 | 0.2 | 1 | 0 | NA | NA | 4 | 0.25 | 2 | 0.5 | 7 | 0.2857 |
| 13_5312187 | NA | NA | NA | NA | 3 | 0 | 3 | 0.6667 | 6 | 0.3333 | NA | NA | NA | NA | 3 | 0.3333 | 2 | 1 | 5 | 0.6 |
| 13_5312208 | NA | NA | NA | NA | 3 | 0.3333 | 2 | 0 | 5 | 0.2 | 1 | 0 | NA | NA | 3 | 0.3333 | 2 | 0 | 6 | 0.1667 |
| 13_5312245 | NA | NA | NA | NA | 2 | 0 | 1 | 1 | 3 | 0.3333 | 1 | 0 | NA | NA | 2 | 0.5 | 2 | 0.5 | 5 | 0.4 |
| 13_5312248 | NA | NA | NA | NA | 2 | 0 | 2 | 0.5 | 4 | 0.25 | 1 | 1 | NA | NA | 3 | 0.6667 | 2 | 0.5 | 6 | 0.6667 |
| 13_5312281 | NA | NA | NA | NA | 1 | 1 | 1 | 1 | 2 | 1 | 1 | 0 | NA | NA | 3 | 0.6667 | NA | NA | 4 | 0.5 |
| 13_5348694 | 7 | 0 | 4 | 0 | 6 | 0.1667 | 2 | 0 | 19 | 0.0526 | 5 | 0.2 | NA | NA | 6 | 0 | 7 | 0.1429 | 18 | 0.1111 |
| 13_5348717 | 7 | 0 | 4 | 0.25 | 5 | 0 | 4 | 0 | 20 | 0.05 | 5 | 0.2 | NA | NA | 7 | 0 | 8 | 0 | 20 | 0.05 |
| 13_5348734 | 6 | 0 | 5 | 0 | 7 | 0.2857 | 4 | 0.25 | 22 | 0.1364 | 4 | 0.25 | NA | NA | 7 | 0 | 9 | 0 | 20 | 0.05 |
| 13_5348746 | 8 | 0.375 | 5 | 0.6 | 7 | 0.7143 | 4 | 0.75 | 24 | 0.5833 | 5 | 0.4 | NA | NA | 7 | 0.7143 | 9 | 0.6667 | 21 | 0.619 |
| 13_5348756 | 7 | 0.5714 | 4 | 0.75 | 5 | 0.6 | 4 | 0.75 | 20 | 0.65 | 4 | 0.75 | NA | NA | 7 | 0.7143 | 9 | 0.7778 | 20 | 0.75 |
| 13_5348760 | 7 | 0.7143 | 6 | 0.6667 | 6 | 0.8333 | 4 | 0.75 | 23 | 0.7391 | 5 | 0.8 | 1 | 0 | 7 | 0.5714 | 8 | 0.75 | 21 | 0.6667 |

| | | | | | | | | | | | | | | | | | | | | |
|---|---|---|---|---|---|---|---|---|---|---|---|---|---|---|---|---|---|---|---|---|
| 13_534876 | 7 | 0.7143 | 5 | 0.6 | 4 | 1 | 4 | 1 | 20 | 0.8 | 4 | 1 | 1 | 1 | 6 | 0.6667 | 8 | 0.875 | 19 | 0.8421 |
| 13_534876 | 5 | 0.2 | 6 | 0.1667 | 6 | 0.5 | 3 | 0 | 20 | 0.25 | 3 | 0 | NA | NA | 6 | 0 | 7 | 0.1429 | 16 | 0.0625 |
| 13_542048 | 1 | 0 | 2 | 0 | 1 | 1 | 2 | 0 | 6 | 0.1667 | NA | NA | 5 | 0.4 | 4 | 0 | 1 | 1 | 10 | 0.3 |
| 13_542053 | 18 | 0.1111 | 15 | 0.3333 | 21 | 0.2381 | 30 | 0.2 | 84 | 0.2143 | 12 | 0.1667 | 22 | 0.1818 | 31 | 0.2581 | 20 | 0.25 | 85 | 0.2235 |
| 13_542054 | 21 | 0.4762 | 17 | 0.6471 | 23 | 0.4783 | 33 | 0.7273 | 94 | 0.5957 | 15 | 0.7333 | 26 | 0.5385 | 34 | 0.7059 | 20 | 0.75 | 95 | 0.6737 |
| 13_542058 | 22 | 0.0909 | 14 | 0.1429 | 27 | 0.1111 | 29 | 0.1379 | 92 | 0.1196 | 9 | 0 | 22 | 0.2727 | 29 | 0.2414 | 21 | 0.2857 | 81 | 0.2346 |
| 13_542062 | 16 | 0 | 15 | 0.2 | 28 | 0.25 | 27 | 0.1111 | 86 | 0.1512 | 9 | 0.4444 | 22 | 0.2727 | 32 | 0.2188 | 21 | 0.381 | 84 | 0.2976 |
| 13_542062 | 16 | 0.3125 | 14 | 0.3571 | 28 | 0.2143 | 28 | 0.2143 | 86 | 0.2558 | 11 | 0.2727 | 20 | 0.4 | 31 | 0.3548 | 19 | 0.4211 | 81 | 0.3704 |
| 13_542065 | 11 | 0.0909 | 10 | 0.1 | 22 | 0.0455 | 29 | 0.0345 | 72 | 0.0556 | 9 | 0 | 20 | 0.15 | 24 | 0.0833 | 13 | 0.2308 | 66 | 0.1212 |
| 13_542066 | 14 | 0.2143 | 9 | 0.3333 | 20 | 0.1 | 25 | 0.16 | 68 | 0.1765 | 7 | 0.1429 | 19 | 0.0526 | 24 | 0.125 | 12 | 0.0833 | 62 | 0.0968 |
| 13_542066 | 14 | 1 | 9 | 1 | 20 | 1 | 26 | 1 | 69 | 1 | 11 | 1 | 22 | 1 | 25 | 1 | 15 | 1 | 73 | 1 |
| 13_542067 | 14 | 0.2857 | 9 | 0.4444 | 19 | 0.1053 | 28 | 0.1786 | 70 | 0.2143 | 9 | 0.3333 | 23 | 0.3043 | 23 | 0.2609 | 15 | 0.4667 | 70 | 0.3286 |
| 13_542067 | 12 | 0.4167 | 10 | 0.4 | 19 | 0.3158 | 20 | 0.6 | 61 | 0.4426 | 7 | 0.2857 | 17 | 0.5882 | 20 | 0.3 | 14 | 0.5 | 58 | 0.431 |
| 13_542067 | 13 | 0.4615 | 9 | 0.7778 | 16 | 0.625 | 21 | 0.7143 | 59 | 0.6441 | 7 | 0.8571 | 15 | 0.5333 | 18 | 0.6667 | 14 | 0.5714 | 54 | 0.6296 |
| 13_542067 | 15 | 1 | 9 | 1 | 21 | 1 | 26 | 1 | 71 | 1 | 10 | 1 | 21 | 1 | 26 | 0.9615 | 15 | 1 | 72 | 0.9861 |
| 13_542068 | 11 | 0.4545 | 8 | 0.5 | 15 | 0.5333 | 21 | 0.619 | 55 | 0.5455 | 8 | 0.625 | 17 | 0.7059 | 17 | 0.5882 | 14 | 0.7143 | 56 | 0.6607 |
| 13_542069 | 15 | 0.9333 | 10 | 0.9 | 17 | 1 | 25 | 1 | 67 | 0.9701 | 8 | 1 | 20 | 1 | 23 | 1 | 17 | 1 | 68 | 1 |
| 13_542070 | 15 | 0.3333 | 11 | 0.2727 | 18 | 0.4444 | 27 | 0.4074 | 71 | 0.3803 | 7 | 0.4286 | 20 | 0.5 | 23 | 0.3478 | 13 | 0.6154 | 63 | 0.4603 |
| 13_542070 | 11 | 1 | 10 | 1 | 18 | 1 | 24 | 0.9583 | 63 | 0.9841 | 6 | 1 | 20 | 1 | 25 | 1 | 15 | 0.8667 | 66 | 0.9697 |
| 13_542071 | 12 | 1 | 9 | 1 | 13 | 1 | 21 | 1 | 55 | 1 | 4 | 1 | 20 | 1 | 19 | 1 | 13 | 1 | 56 | 1 |
| 13_542071 | 13 | 1 | 8 | 1 | 20 | 0.95 | 25 | 0.96 | 66 | 0.9697 | 6 | 1 | 20 | 1 | 23 | 0.9565 | 15 | 1 | 64 | 0.9844 |
| 13_542072 | 12 | 0.0833 | 10 | 0.3 | 15 | 0.1333 | 22 | 0.3636 | 59 | 0.2373 | 5 | 0 | 19 | 0.1579 | 21 | 0.1429 | 14 | 0.2857 | 59 | 0.1695 |
| 13_542073 | 5 | 1 | 6 | 1 | 9 | 1 | 12 | 1 | 32 | 1 | 3 | 1 | 13 | 1 | 17 | 0.9412 | 9 | 0.8889 | 42 | 0.9524 |
| 13_542073 | 14 | 0.2857 | 10 | 0.4 | 15 | 0.2667 | 22 | 0.3182 | 61 | 0.3115 | 4 | 0.25 | 20 | 0.15 | 22 | 0.3636 | 14 | 0.2857 | 60 | 0.2667 |
| 13_542074 | 15 | 0.1333 | 10 | 0 | 17 | 0.1176 | 21 | 0.0952 | 63 | 0.0952 | 5 | 0 | 19 | 0.2105 | 24 | 0.0833 | 14 | 0.2143 | 62 | 0.1452 |
| 13_542074 | 6 | 0.5 | 7 | 0.4286 | 12 | 0.5 | 14 | 0.5 | 39 | 0.4872 | 3 | 0 | 12 | 0.4167 | 15 | 0.7333 | 9 | 0.6667 | 39 | 0.5641 |
| 13_542076 | 13 | 0.7692 | 8 | 0.625 | 15 | 0.6667 | 21 | 0.7619 | 57 | 0.7193 | 5 | 0.4 | 17 | 0.6471 | 24 | 0.5417 | 12 | 0.5833 | 58 | 0.569 |
| 13_542076 | 10 | 0 | 9 | 0.1111 | 15 | 0.1333 | 20 | 0.05 | 54 | 0.0741 | 5 | 0.2 | 17 | 0.0588 | 23 | 0 | 12 | 0.0833 | 57 | 0.0526 |
| 13_542077 | 11 | 1 | 9 | 1 | 12 | 1 | 20 | 0.95 | 52 | 0.9808 | 4 | 1 | 13 | 1 | 23 | 1 | 12 | 1 | 52 | 1 |
| 13_542078 | 10 | 0.4 | 7 | 0.7143 | 9 | 0.6667 | 11 | 0.6364 | 37 | 0.5946 | 4 | 1 | 10 | 0.7 | 13 | 0.7692 | 8 | 0.75 | 35 | 0.7714 |
| 13_542079 | 8 | 0.25 | 8 | 0.25 | 7 | 0.4286 | 7 | 0.1429 | 30 | 0.2667 | 4 | 0.25 | 12 | 0.25 | 14 | 0.2857 | 7 | 0.4286 | 37 | 0.2973 |
| 13_542081 | 11 | 0.9091 | 8 | 1 | 10 | 0.8 | 17 | 1 | 46 | 0.9348 | 4 | 1 | 13 | 0.9231 | 16 | 0.9375 | 11 | 1 | 44 | 0.9545 |
| 13_542081 | 9 | 0.6667 | 8 | 0.5 | 7 | 0.8571 | 16 | 0.6875 | 40 | 0.675 | 4 | 1 | 11 | 0.8182 | 16 | 0.8125 | 10 | 0.8 | 41 | 0.8293 |
| 13_542083 | 6 | 0.1667 | 6 | 0.1667 | 7 | 0 | 11 | 0.1818 | 30 | 0.1333 | 2 | 0 | 7 | 0.1429 | 14 | 0.0714 | 7 | 0.2857 | 30 | 0.1333 |
| 13_542083 | 3 | 0 | 4 | 0 | 3 | 0.3333 | 13 | 0.1538 | 23 | 0.1304 | 4 | 0 | 6 | 0.3333 | 13 | 0.1538 | 5 | 0 | 28 | 0.1429 |
| 13_542084 | 9 | 0.2222 | 5 | 0.2 | 7 | 0.1429 | 16 | 0.1875 | 37 | 0.1892 | 4 | 0.25 | 5 | 0.2 | 11 | 0.3636 | 4 | 0.25 | 24 | 0.2917 |
| 13_542085 | 9 | 0.5556 | 5 | 0.2 | 6 | 0.5 | 16 | 0.5625 | 36 | 0.5 | 4 | 0 | 6 | 0 | 13 | 0.1538 | 4 | 0.25 | 27 | 0.1111 |
| 13_542085 | 8 | 1 | 5 | 0.6 | 5 | 0.8 | 16 | 0.75 | 34 | 0.7941 | 4 | 0.75 | 4 | 1 | 13 | 0.5385 | 3 | 1 | 24 | 0.7083 |
| 13_542262 | NA | NA | 1 | 0 | 1 | 1 | NA | NA | 2 | 0.5 | NA | NA | 1 | 1 | NA | NA | 2 | 0.5 | 3 | 0.6667 |
| 13_542605 | 6 | 0.3333 | 3 | 0 | 6 | 0.1667 | 10 | 0.1 | 25 | 0.16 | 5 | 0 | 10 | 0.3 | 9 | 0.3333 | 16 | 0.3125 | 40 | 0.275 |
| 13_542610 | 8 | 0.125 | 4 | 0 | 10 | 0.2 | 11 | 0 | 33 | 0.0909 | 5 | 0 | 11 | 0.1818 | 8 | 0 | 17 | 0.0588 | 41 | 0.0732 |
| 13_542611 | 8 | 0.125 | 4 | 0 | 10 | 0 | 10 | 0.3 | 32 | 0.125 | 6 | 0.1667 | 12 | 0.25 | 10 | 0.1 | 17 | 0.2353 | 45 | 0.2 |
| 13_542612 | 7 | 0.4286 | 4 | 0.5 | 9 | 0 | 11 | 0.1818 | 31 | 0.2258 | 4 | 0 | 10 | 0.3 | 8 | 0.125 | 15 | 0.2667 | 37 | 0.2162 |
| 13_542613 | 7 | 0 | 3 | 0 | 10 | 0.2 | 10 | 0.2 | 30 | 0.1333 | 6 | 0.1667 | 13 | 0.3077 | 9 | 0.2222 | 16 | 0.1875 | 44 | 0.2273 |
| 13_542614 | 8 | 0.25 | 4 | 0 | 8 | 0 | 9 | 0.1111 | 29 | 0.1034 | 6 | 0 | 11 | 0.0909 | 10 | 0 | 17 | 0.2353 | 44 | 0.1136 |
| 13_542615 | 6 | 0.3333 | 4 | 1 | 8 | 0.625 | 10 | 0.4 | 28 | 0.5357 | 8 | 0.625 | 12 | 0.5833 | 8 | 0.625 | 16 | 0.25 | 44 | 0.4773 |
| 13_542617 | 9 | 0.4444 | 5 | 0.8 | 11 | 0.4545 | 11 | 0.2727 | 36 | 0.4444 | 7 | 0.2857 | 12 | 0.5 | 10 | 0.3 | 18 | 0.3333 | 47 | 0.3617 |
| 13_542617 | 7 | 0.5714 | 6 | 0.8333 | 7 | 0.7143 | 8 | 0.875 | 28 | 0.75 | 5 | 1 | 9 | 0.7778 | 7 | 0.7143 | 12 | 0.5833 | 33 | 0.7273 |
| 13_542618 | 8 | 1 | 6 | 1 | 10 | 0.8 | 9 | 0.8889 | 33 | 0.9091 | 6 | 1 | 11 | 1 | 9 | 1 | 17 | 0.9412 | 43 | 0.9767 |
| 13_542619 | 8 | 0.125 | 4 | 0.25 | 10 | 0 | 9 | 0.2222 | 31 | 0.129 | 7 | 0 | 13 | 0.3077 | 10 | 0.3 | 16 | 0.25 | 46 | 0.2391 |
| 13_542619 | 8 | 0 | 5 | 0.4 | 9 | 0 | 12 | 0.1667 | 34 | 0.1176 | 6 | 0.1667 | 14 | 0.3571 | 7 | 0.1429 | 18 | 0.3333 | 45 | 0.2889 |
| 13_542619 | 8 | 0 | 6 | 0.1667 | 11 | 0.2727 | 12 | 0.3333 | 37 | 0.2162 | 7 | 0.2857 | 15 | 0.3333 | 10 | 0.2 | 20 | 0.3 | 52 | 0.2885 |
| 13_543678 | NA | NA | NA | NA | NA | NA | 1 | 1 | 1 | 1 | 1 | 1 | 1 | 1 | NA | NA | NA | NA | 2 | 1 |
| 13_543678 | NA | NA | NA | NA | NA | NA | 1 | 1 | 1 | 1 | 1 | 0 | 1 | 1 | NA | NA | NA | NA | 2 | 0.5 |
| 13_543680 | NA | NA | NA | NA | NA | NA | 1 | 1 | 1 | 1 | 1 | 1 | NA | NA | NA | NA | NA | NA | 1 | 1 |
| 13_543685 | NA | NA | NA | NA | NA | NA | 1 | 1 | 1 | 1 | 1 | 0 | 1 | 1 | NA | NA | NA | NA | 2 | 0.5 |
| 13_543685 | NA | NA | NA | NA | NA | NA | 1 | 1 | 1 | 1 | 1 | 1 | 1 | 1 | NA | NA | NA | NA | 2 | 1 |
| 13_543687 | NA | NA | NA | NA | NA | NA | 1 | 1 | 1 | 1 | 1 | 0 | 1 | 1 | NA | NA | NA | NA | 2 | 0.5 |
| 13_543689 | NA | NA | NA | NA | NA | NA | 1 | 1 | 1 | 1 | NA | NA | 1 | 1 | NA | NA | NA | NA | 1 | 1 |
| 13_543878 | NA | NA | NA | NA | NA | NA | 2 | 1 | 2 | 1 | 1 | 1 | 1 | 1 | NA | NA | NA | NA | 2 | 1 |
| 13_543895 | NA | NA | 2 | 1 | NA | NA | 2 | 0.5 | 4 | 0.75 | 1 | 1 | 1 | 1 | NA | NA | NA | NA | 2 | 1 |
| 13_543896 | NA | NA | 2 | 0.5 | NA | NA | 2 | 1 | 4 | 0.75 | 1 | 1 | 1 | 1 | NA | NA | NA | NA | 2 | 1 |
| 13_543900 | NA | NA | 2 | 0.5 | NA | NA | 2 | 0 | 4 | 0.25 | NA | NA | 1 | 1 | NA | NA | NA | NA | 1 | 1 |
| 13_545274 | 9 | 0.3333 | 15 | 0.1333 | 12 | 0 | 14 | 0.1429 | 50 | 0.14 | 12 | 0.1667 | 9 | 0.2222 | 13 | 0.0769 | 8 | 0.25 | 42 | 0.1667 |
| 13_545275 | 10 | 0.4 | 13 | 0.5385 | 12 | 0.6667 | 15 | 0.4667 | 50 | 0.52 | 15 | 0.3333 | 11 | 0.2727 | 13 | 0.1538 | 9 | 0.2222 | 48 | 0.25 |

| | | | | | | | | | | | | | | | | | | | | |
|---|---|---|---|---|---|---|---|---|---|---|---|---|---|---|---|---|---|---|---|---|
| 13_5452755 | 7 | 0 | 12 | 0.25 | 11 | 0.4545 | 15 | 0 | 45 | 0.1778 | 14 | 0.2143 | 10 | 0.1 | 12 | 0.4167 | 8 | 0.375 | 44 | 0.2727 |
| 13_5452767 | 19 | 0.1579 | 17 | 0.1176 | 16 | 0.1875 | 18 | 0.1111 | 70 | 0.1429 | 18 | 0.1667 | 15 | 0.1333 | 17 | 0.1765 | 13 | 0 | 63 | 0.127 |
| 13_5452804 | 36 | 0.0556 | 36 | 0.0833 | 33 | 0.1515 | 41 | 0.0244 | 146 | 0.0753 | 34 | 0.1176 | 44 | 0.0909 | 41 | 0.0732 | 29 | 0.2414 | 148 | 0.1216 |
| 13_5452810 | 37 | 0.2703 | 38 | 0.2105 | 35 | 0.2571 | 39 | 0.2308 | 149 | 0.2416 | 34 | 0.2059 | 44 | 0.2045 | 39 | 0.1538 | 27 | 0.1111 | 144 | 0.1736 |
| 13_5452815 | 41 | 0.2683 | 40 | 0.25 | 36 | 0.25 | 44 | 0.2045 | 161 | 0.2422 | 36 | 0.2778 | 44 | 0.1136 | 44 | 0.25 | 30 | 0.2 | 154 | 0.2078 |
| 13_5452817 | 41 | 0.2683 | 38 | 0.2105 | 36 | 0.25 | 42 | 0.2143 | 157 | 0.2357 | 31 | 0.1613 | 44 | 0.2045 | 39 | 0.1795 | 28 | 0.3214 | 142 | 0.2113 |
| 13_5452848 | 37 | 0.9459 | 37 | 0.9459 | 37 | 0.973 | 46 | 0.9348 | 157 | 0.949 | 33 | 0.9697 | 42 | 0.9286 | 43 | 0.9767 | 30 | 0.9667 | 148 | 0.9595 |
| 13_5452850 | 44 | 0.0455 | 47 | 0.0213 | 39 | 0.0769 | 49 | 0.0612 | 179 | 0.0503 | 35 | 0.0571 | 49 | 0.1429 | 46 | 0.0217 | 37 | 0.027 | 167 | 0.0659 |
| 13_5452854 | 39 | 0 | 39 | 0.1282 | 36 | 0.0833 | 50 | 0.08 | 164 | 0.0732 | 33 | 0.0303 | 54 | 0.0741 | 39 | 0.0769 | 31 | 0.1613 | 157 | 0.0828 |
| 13_5452856 | 39 | 0.0513 | 42 | 0.1667 | 40 | 0.05 | 50 | 0.12 | 171 | 0.0994 | 34 | 0.0588 | 46 | 0.1304 | 41 | 0.1463 | 35 | 0.0571 | 156 | 0.1026 |
| 13_5452876 | 42 | 0.7857 | 39 | 0.7692 | 37 | 0.7027 | 49 | 0.7347 | 167 | 0.7485 | 34 | 0.7647 | 50 | 0.8 | 44 | 0.7955 | 36 | 0.8333 | 164 | 0.7988 |
| 13_5452882 | 36 | 0.9167 | 41 | 0.9024 | 32 | 0.875 | 49 | 0.9388 | 158 | 0.9114 | 25 | 0.96 | 44 | 0.8636 | 40 | 0.9 | 33 | 0.9091 | 142 | 0.9014 |
| 13_5452887 | 46 | 0.3478 | 47 | 0.234 | 42 | 0.2381 | 55 | 0.3273 | 190 | 0.2895 | 39 | 0.2821 | 54 | 0.2407 | 49 | 0.3265 | 38 | 0.3421 | 180 | 0.2944 |
| 13_5452892 | 44 | 0.2273 | 44 | 0.2273 | 42 | 0.2619 | 47 | 0.2128 | 177 | 0.2316 | 38 | 0.2632 | 57 | 0.2982 | 47 | 0.1915 | 38 | 0.1842 | 180 | 0.2389 |
| 13_5452894 | 38 | 0.3421 | 37 | 0.4054 | 29 | 0.3448 | 44 | 0.2955 | 148 | 0.3446 | 31 | 0.3548 | 44 | 0.2273 | 48 | 0.4375 | 28 | 0.5357 | 151 | 0.3775 |
| 13_5453252 | 8 | 0.125 | 4 | 0 | 5 | 0 | 8 | 0.125 | 25 | 0.08 | 2 | 0 | 6 | 0.1667 | 6 | 0 | 3 | 0 | 17 | 0.0588 |
| 13_5453598 | 2 | 0.5 | 1 | 0 | 2 | 0.5 | 3 | 1 | 8 | 0.625 | 3 | 0.3333 | 2 | 0 | 1 | 0 | 1 | 1 | 7 | 0.2857 |
| 13_5453603 | 2 | 0 | 2 | 0 | 2 | 0 | 3 | 0.3333 | 9 | 0.1111 | 2 | 0.5 | 2 | 0 | 1 | 0 | 1 | 0 | 6 | 0.1667 |
| 13_5460770 | 3 | 0.6667 | 3 | 0.3333 | NA | NA | 5 | 0.8 | 11 | 0.6364 | 4 | 1 | 5 | 0.4 | 7 | 0.4286 | 2 | 1 | 18 | 0.6111 |
| 13_5460777 | 4 | 0.25 | 3 | 0.3333 | NA | NA | 4 | 0.25 | 11 | 0.2727 | 3 | 0 | 5 | 0 | 6 | 0.5 | 3 | 0.3333 | 17 | 0.2353 |
| 13_5460782 | 3 | 0.6667 | 1 | 1 | NA | NA | 4 | 0.75 | 8 | 0.75 | 2 | 0.5 | 4 | 0.75 | 5 | 0.8 | 3 | 0.3333 | 14 | 0.6429 |
| 13_5460784 | 4 | 1 | 3 | 1 | NA | NA | 5 | 0.6 | 12 | 0.8333 | 4 | 0.75 | 5 | 1 | 7 | 0.7143 | 3 | 0.6667 | 19 | 0.7895 |
| 13_5460790 | 3 | 0 | 2 | 1 | NA | NA | 4 | 0.5 | 9 | 0.4444 | 3 | 0 | 4 | 0.25 | 7 | 0.2857 | NA | NA | 14 | 0.2143 |
| 13_5460792 | 4 | 0.25 | 3 | 0 | NA | NA | 3 | 0 | 10 | 0.1 | 4 | 0.75 | 5 | 0.2 | 7 | 0.2857 | 3 | 0.6667 | 19 | 0.4211 |
| 13_5467190 | 1 | 1 | 1 | 0 | NA | NA | 2 | 1 | 4 | 0.75 | NA | NA | 1 | 1 | NA | NA | NA | NA | 1 | 1 |
| 13_5467413 | 3 | 1 | 1 | 1 | NA | NA | 3 | 1 | 7 | 1 | NA | NA | 1 | 1 | NA | NA | NA | NA | 1 | 1 |
| 13_5467436 | 3 | 0.6667 | 1 | 1 | NA | NA | 3 | 0.6667 | 7 | 0.7143 | NA | NA | 2 | 1 | NA | NA | NA | NA | 2 | 1 |
| 13_5467441 | 4 | 1 | 1 | 1 | NA | NA | 1 | 1 | 6 | 1 | NA | NA | 2 | 0.5 | NA | NA | NA | NA | 2 | 0.5 |
| 13_5467620 | 3 | 0.6667 | 1 | 1 | NA | NA | 3 | 0.3333 | 7 | 0.5714 | NA | NA | 2 | 0.5 | NA | NA | NA | NA | 2 | 0.5 |
| 13_5467622 | 4 | 1 | 1 | 1 | NA | NA | 2 | 1 | 7 | 1 | NA | NA | 2 | 1 | NA | NA | NA | NA | 2 | 1 |
| 13_5468777 | NA | NA | 1 | 1 | NA | NA | 1 | 0 | 2 | 0.5 | NA | NA | 1 | 1 | NA | NA | NA | NA | 1 | 1 |
| 13_5468801 | NA | NA | 1 | 1 | NA | NA | 1 | 0 | 2 | 0.5 | NA | NA | 1 | 1 | NA | NA | NA | NA | 1 | 1 |
| 13_5469840 | 4 | 0.75 | 1 | 1 | 1 | 0 | 4 | 0.75 | 10 | 0.7 | 1 | 1 | 1 | 1 | 1 | 1 | NA | NA | 3 | 1 |
| 13_5469845 | 4 | 0.25 | 1 | 0 | 1 | 0 | 5 | 0.4 | 11 | 0.2727 | 1 | 1 | 1 | 0 | 1 | 0 | NA | NA | 3 | 0.3333 |
| 13_5469858 | 4 | 0.25 | 1 | 0 | 1 | 0 | 6 | 0 | 12 | 0.0833 | 1 | 1 | 1 | 0 | 1 | 0 | NA | NA | 3 | 0.3333 |
| 13_5469886 | 3 | 0 | NA | NA | 1 | 1 | 2 | 0 | 6 | 0.1667 | 1 | 1 | 1 | 0 | 1 | 0 | NA | NA | 3 | 0.3333 |
| 13_5469901 | 4 | 0 | 1 | 1 | 1 | 1 | 7 | 0.1429 | 13 | 0.2308 | 1 | 1 | 2 | 0.5 | 1 | 0 | NA | NA | 4 | 0.5 |
| 13_5469933 | 25 | 0.08 | 32 | 0.125 | 25 | 0 | 14 | 0.0714 | 96 | 0.0729 | 23 | 0.087 | 26 | 0 | 12 | 0.0833 | 27 | 0.0741 | 88 | 0.0568 |
| 13_5470006 | 15 | 0.0667 | 18 | 0.0556 | 17 | 0.0588 | 6 | 0.1667 | 56 | 0.0714 | 18 | 0.1111 | 15 | 0.1333 | 8 | 0.125 | 17 | 0.2941 | 58 | 0.1724 |
| 13_5470022 | 17 | 0.0588 | 23 | 0.0435 | 20 | 0.15 | 10 | 0.2 | 70 | 0.1 | 19 | 0.1053 | 18 | 0.0556 | 12 | 0.0833 | 20 | 0.15 | 69 | 0.1014 |
| 13_5472871 | 1 | 0 | 1 | 0 | 2 | 0 | 3 | 0.3333 | 7 | 0.1429 | 3 | 0 | 1 | 0 | 2 | 0 | 2 | 0.5 | 8 | 0.125 |
| 13_5472964 | 2 | 0 | 4 | 0.25 | 4 | 0 | 3 | 0 | 13 | 0.0769 | 3 | 0.3333 | 1 | 0 | 4 | 0 | 2 | 0 | 10 | 0.1 |
| 13_5472970 | 2 | 0.5 | 4 | 0 | 4 | 0 | 3 | 0 | 13 | 0.0769 | 3 | 0.3333 | 1 | 0 | 4 | 0 | 2 | 0 | 10 | 0.1 |
| 13_5472980 | 2 | 0.5 | 4 | 0.5 | 4 | 0 | 4 | 0.5 | 14 | 0.3571 | 4 | 0 | 1 | 1 | 4 | 0.25 | 2 | 0 | 11 | 0.1818 |
| 13_5472981 | 2 | 0 | 3 | 0 | 4 | 0.25 | 4 | 0 | 13 | 0.0769 | 4 | 0.25 | 1 | 0 | 5 | 0 | 2 | 0 | 12 | 0.0833 |
| 13_5472983 | 1 | 0 | 4 | 0.25 | 4 | 0.25 | 1 | 1 | 10 | 0.3 | 4 | 0.25 | 1 | 0 | 3 | 0 | NA | NA | 8 | 0.125 |
| 13_5472987 | 2 | 0 | 2 | 0.5 | 4 | 0.25 | 4 | 0 | 12 | 0.1667 | 4 | 0.5 | 1 | 0 | 5 | 0.2 | 2 | 0.5 | 12 | 0.3333 |
| 13_5472990 | 1 | 1 | 4 | 0.75 | 4 | 0.75 | 4 | 1 | 13 | 0.8462 | 3 | 1 | 1 | 1 | 5 | 0.8 | 2 | 1 | 11 | 0.9091 |
| 13_5472993 | 1 | 0 | 4 | 0.5 | 3 | 0.3333 | 4 | 0.5 | 12 | 0.4167 | 4 | 0.5 | 1 | 0 | 4 | 0.25 | 3 | 0.3333 | 12 | 0.3333 |
| 13_5472996 | 1 | 0 | 3 | 0.6667 | 2 | 0 | 4 | 0.5 | 10 | 0.4 | 4 | 0 | NA | NA | 4 | 0.25 | 2 | 0.5 | 10 | 0.2 |
| 13_5473001 | 2 | 0 | 5 | 0.8 | 3 | 0.3333 | 4 | 0.75 | 14 | 0.5714 | 4 | 0.5 | NA | NA | 4 | 1 | 3 | 1 | 11 | 0.8182 |
| 13_5473002 | 2 | 0 | 5 | 0.2 | 3 | 0 | 4 | 0.25 | 14 | 0.1429 | 4 | 0 | 1 | 0 | 5 | 0 | 3 | 0.3333 | 13 | 0.0769 |
| 13_5473027 | 2 | 0.5 | 5 | 0.6 | 4 | 0.25 | 2 | 0.5 | 13 | 0.4615 | 5 | 0.8 | 1 | 0 | 7 | 0.4286 | 3 | 1 | 16 | 0.625 |
| 13_5473028 | 2 | 0 | 3 | 0 | 4 | 0 | 4 | 0.25 | 13 | 0.0769 | 2 | 0 | 1 | 0 | 7 | 0.1429 | 3 | 0 | 13 | 0.0769 |
| 13_5473030 | 2 | 0 | 4 | 0.25 | 4 | 0 | 3 | 0 | 13 | 0.0769 | 5 | 0.2 | 1 | 0 | 7 | 0.1429 | 3 | 0 | 16 | 0.125 |
| 13_5473039 | 1 | 1 | 4 | 0.25 | 5 | 0.4 | 4 | 0.25 | 14 | 0.3571 | 6 | 0.6667 | NA | NA | 7 | 0.7143 | 3 | 0.6667 | 16 | 0.6875 |
| 13_5473045 | 2 | 0 | 5 | 0 | 5 | 0 | 4 | 0.25 | 16 | 0.0625 | 6 | 0 | 1 | 0 | 7 | 0.1429 | 3 | 0 | 17 | 0.0588 |
| 13_5473277 | 6 | 0.6667 | 3 | 0.6667 | 5 | 0.4 | 3 | 0.6667 | 17 | 0.5882 | 3 | 0.6667 | 4 | 0.5 | 1 | 0 | 6 | 0.5 | 14 | 0.5 |
| 13_5473285 | 4 | 0.25 | 3 | 0 | 4 | 0.25 | 2 | 0 | 13 | 0.1538 | 3 | 0 | 6 | 0.1667 | 1 | 0 | 5 | 0.2 | 15 | 0.1333 |
| 13_5473288 | 5 | 0.8 | 3 | 1 | 5 | 0.2 | 1 | 0 | 14 | 0.5714 | 3 | 0.3333 | 7 | 0.7143 | 1 | 1 | 4 | 0.5 | 15 | 0.6 |
| 13_5473294 | 6 | 0.1667 | 4 | 0.25 | 4 | 0.25 | 2 | 0.5 | 16 | 0.25 | 3 | 0 | 7 | 0.2857 | 1 | 1 | 7 | 0.1429 | 18 | 0.2222 |
| 13_5473309 | 4 | 0.5 | 4 | 0.5 | 4 | 0.5 | 3 | 0.3333 | 15 | 0.4667 | 3 | 0.3333 | 7 | 0.4286 | NA | NA | 6 | 0.5 | 16 | 0.4375 |
| 13_5473875 | 39 | 0.1026 | 25 | 0.12 | 24 | 0.0417 | 25 | 0.08 | 113 | 0.0885 | 25 | 0.16 | 25 | 0.08 | 24 | 0.0417 | 19 | 0.0526 | 93 | 0.086 |
| 13_5473877 | 38 | 0.1316 | 24 | 0.4167 | 22 | 0.4091 | 28 | 0.3214 | 112 | 0.2946 | 23 | 0.1304 | 24 | 0.3333 | 25 | 0.2 | 18 | 0.4444 | 90 | 0.2667 |
| 13_5473879 | 37 | 0.5135 | 25 | 0.4 | 18 | 0.4444 | 26 | 0.5385 | 106 | 0.4811 | 23 | 0.4348 | 27 | 0.4815 | 25 | 0.6 | 15 | 0.5333 | 90 | 0.5111 |

| | | | | | | | | | | | | | | | | | | | | |
|---|---|---|---|---|---|---|---|---|---|---|---|---|---|---|---|---|---|---|---|---|
| 13_547388 | 31 | 0.0645 | 16 | 0.25 | 19 | 0.0526 | 20 | 0.25 | 86 | 0.1395 | 15 | 0.3333 | 12 | 0 | 15 | 0.2 | 11 | 0.0909 | 53 | 0.1698 |
| 13_547408 | 12 | 0.1667 | 9 | 0.2222 | 10 | 0.3 | 8 | 0.125 | 39 | 0.2051 | 7 | 0.2857 | 15 | 0.2667 | 6 | 0 | 3 | 0 | 31 | 0.1935 |
| 13_547414 | 24 | 0.125 | 40 | 0.125 | 30 | 0.0667 | 23 | 0.1304 | 117 | 0.1111 | 24 | 0.125 | 35 | 0.1143 | 29 | 0.1379 | 23 | 0.1739 | 111 | 0.1351 |
| 13_547415 | 26 | 0.1923 | 47 | 0.1277 | 41 | 0.122 | 26 | 0.0769 | 140 | 0.1286 | 28 | 0.1071 | 38 | 0.1842 | 32 | 0.2812 | 28 | 0.1429 | 126 | 0.1825 |
| 13_547416 | 17 | 0.0588 | 38 | 0.0263 | 38 | 0.1579 | 20 | 0.1 | 113 | 0.0885 | 25 | 0.12 | 34 | 0.1765 | 24 | 0.0833 | 21 | 0.1429 | 104 | 0.1346 |
| 13_547422 | NA | NA | 4 | 0.25 | 1 | 0 | 2 | 0 | 7 | 0.1429 | 1 | 0 | NA | NA | 1 | 1 | 1 | 0 | 3 | 0.3333 |
| 13_548089 | 4 | 1 | 5 | 1 | 3 | 0.6667 | 2 | 1 | 14 | 0.9286 | 2 | 1 | 1 | 1 | 2 | 1 | 3 | 1 | 8 | 1 |
| 13_548089 | 5 | 0.8 | 6 | 0.5 | 3 | 0.6667 | 3 | 0 | 17 | 0.5294 | 2 | 1 | 3 | 1 | 2 | 0 | 3 | 0.6667 | 10 | 0.7 |
| 13_548089 | 3 | 0.6667 | 6 | 0.6667 | 3 | 0.6667 | 3 | 0.6667 | 15 | 0.6667 | 2 | 0.5 | 2 | 0.5 | 1 | 0 | 3 | 1 | 8 | 0.625 |
| 13_548089 | 5 | 0.4 | 5 | 0 | 2 | 0 | 2 | 0 | 14 | 0.1429 | 2 | 0 | 2 | 0 | 1 | 1 | 2 | 0 | 7 | 0.1429 |
| 13_548090 | 3 | 0 | 5 | 0.6 | 2 | 0 | 1 | 1 | 11 | 0.3636 | 1 | 1 | 2 | 0 | 1 | 0 | 3 | 0.6667 | 7 | 0.4286 |
| 13_548090 | 3 | 0 | 5 | 0 | 2 | 0.5 | 2 | 0.5 | 12 | 0.1667 | NA | NA | 3 | 0.6667 | 2 | 0 | 2 | 0.5 | 7 | 0.4286 |
| 13_548090 | 3 | 0.3333 | 2 | 0.5 | NA | NA | 1 | 0 | 6 | 0.3333 | 2 | 1 | 1 | 1 | 2 | 0.5 | 2 | 0.5 | 7 | 0.7143 |
| 13_548091 | 7 | 0.1429 | 6 | 0.1667 | 2 | 0.5 | 3 | 0.3333 | 18 | 0.2222 | 1 | 0 | 3 | 0 | 3 | 0.3333 | 3 | 0.6667 | 10 | 0.3 |
| 13_548092 | 6 | 0.5 | 6 | 0.5 | 3 | 0 | 3 | 0.3333 | 18 | 0.3889 | 2 | 1 | 3 | 0.6667 | 2 | 0.5 | 3 | 0.3333 | 10 | 0.6 |
| 13_548092 | 7 | 0.7143 | 5 | 0.4 | 2 | 1 | 3 | 0.6667 | 17 | 0.6471 | 1 | 1 | 2 | 1 | NA | NA | 4 | 0.75 | 7 | 0.8571 |
| 13_548093 | 8 | 1 | 4 | 1 | 3 | 1 | 3 | 1 | 18 | 1 | 1 | 1 | 2 | 1 | 3 | 1 | 4 | 1 | 10 | 1 |
| 13_548093 | 7 | 0.2857 | 4 | 0.5 | 3 | 0.6667 | 1 | 0 | 15 | 0.4 | 2 | 0 | 3 | 0 | 3 | 0 | 3 | 0.6667 | 11 | 0.1818 |
| 13_548094 | 7 | 0 | 6 | 0.1667 | 1 | 0 | 3 | 0 | 17 | 0.0588 | 1 | 1 | 2 | 0 | 2 | 0.5 | 4 | 0 | 9 | 0.2222 |
| 13_548098 | 5 | 0.6 | 5 | 1 | NA | NA | 2 | 0.5 | 12 | 0.75 | 1 | 1 | 1 | 1 | 3 | 0.6667 | 5 | 0.4 | 10 | 0.6 |
| 13_548100 | 6 | 0.8333 | 5 | 0.4 | NA | NA | 2 | 0.5 | 13 | 0.6154 | 1 | 1 | 1 | 1 | 1 | 0 | 5 | 0.6 | 8 | 0.625 |
| 13_548100 | 6 | 0.8333 | 5 | 0.8 | NA | NA | 3 | 0.3333 | 14 | 0.7143 | NA | NA | 1 | 1 | 2 | 1 | 5 | 1 | 8 | 1 |
| 13_548101 | 4 | 0.25 | 5 | 0.2 | NA | NA | 2 | 0 | 11 | 0.1818 | NA | NA | 1 | 0 | 2 | 1 | 4 | 0.75 | 7 | 0.7143 |
| 13_548102 | 4 | 0.5 | 3 | 0.3333 | NA | NA | 3 | 0.6667 | 10 | 0.5 | NA | NA | 1 | 0 | 3 | 0 | 3 | 0.6667 | 7 | 0.2857 |
| 13_548111 | 5 | 0.2 | 6 | 0 | 2 | 0 | 5 | 0.2 | 18 | 0.1111 | 1 | 0 | 3 | 0 | 2 | 0 | 4 | 0.25 | 10 | 0.1 |
| 13_548111 | 5 | 0.4 | 6 | 0.3333 | 2 | 0.5 | 4 | 0 | 17 | 0.2941 | 1 | 1 | 3 | 0.3333 | 4 | 0.75 | 6 | 0.8333 | 14 | 0.7143 |
| 13_548111 | 6 | 0.1667 | 6 | 0 | 2 | 0.5 | 5 | 0 | 19 | 0.1053 | 1 | 0 | 3 | 0.3333 | 4 | 0 | 7 | 0 | 15 | 0.0667 |
| 13_548112 | 6 | 0.6667 | 6 | 0.6667 | 1 | 0 | 5 | 0.6 | 18 | 0.6111 | 1 | 0 | 3 | 0.6667 | 4 | 0.75 | 7 | 0.5714 | 15 | 0.6 |
| 13_548113 | 6 | 0 | 7 | 0.1429 | 2 | 0 | 5 | 0 | 20 | 0.05 | 1 | 0 | 3 | 0.3333 | 3 | 0 | 7 | 0.1429 | 14 | 0.1429 |
| 13_548116 | 5 | 0.2 | 7 | 0.1429 | 3 | 0 | 5 | 0 | 20 | 0.1 | 2 | 0 | 3 | 0 | 5 | 0.2 | 8 | 0.125 | 18 | 0.1111 |
| 13_548116 | 6 | 0.1667 | 8 | 0 | 3 | 0 | 5 | 0.6 | 22 | 0.1818 | 1 | 0 | 2 | 0 | 5 | 0.2 | 7 | 0.1429 | 15 | 0.1333 |
| 13_548274 | 18 | 0.2222 | 21 | 0.2381 | 18 | 0.0556 | 11 | 0.0909 | 68 | 0.1618 | 12 | 0.25 | 22 | 0.2273 | 14 | 0.1429 | 19 | 0.2105 | 67 | 0.209 |
| 13_548278 | 19 | 0.6316 | 20 | 0.85 | 17 | 0.8235 | 14 | 0.5 | 70 | 0.7143 | 11 | 0.5455 | 23 | 0.3913 | 17 | 0.6471 | 28 | 0.6786 | 79 | 0.5696 |
| 13_548285 | 14 | 0 | 14 | 0 | 10 | 0.1 | 10 | 0.2 | 48 | 0.0625 | 7 | 0 | 21 | 0.0952 | 11 | 0.0909 | 16 | 0.125 | 55 | 0.0909 |
| 13_548291 | 13 | 0.1538 | 18 | 0.2778 | 9 | 0.2222 | 10 | 0.1 | 50 | 0.2 | 10 | 0.2 | 18 | 0.0556 | 11 | 0.1818 | 17 | 0.4706 | 56 | 0.2321 |
| 13_548293 | 12 | 0.1667 | 18 | 0 | 10 | 0.1 | 8 | 0.25 | 48 | 0.1042 | 9 | 0.1111 | 17 | 0.0588 | 10 | 0.1 | 15 | 0.2 | 51 | 0.1176 |
| 13_548294 | 6 | 0.1667 | 8 | 0.375 | 9 | 0.2222 | 2 | 1 | 25 | 0.32 | 8 | 0 | 16 | 0.1875 | 9 | 0.4444 | 13 | 0.3846 | 46 | 0.2609 |
| 13_548850 | 6 | 0.8333 | 15 | 0.2667 | 13 | 0.3077 | 7 | 0.2857 | 41 | 0.3659 | 9 | 0.6667 | 12 | 0.4167 | 8 | 0.375 | 16 | 0.1875 | 45 | 0.3778 |
| 13_548851 | 6 | 0 | 17 | 0.2353 | 13 | 0.1538 | 6 | 0 | 42 | 0.1429 | 10 | 0.3 | 12 | 0.0833 | 7 | 0 | 16 | 0 | 45 | 0.0889 |
| 13_548851 | 7 | 0.2857 | 15 | 0.1333 | 13 | 0.4615 | 7 | 0 | 42 | 0.2381 | 10 | 0.3 | 12 | 0.5 | 7 | 0.4286 | 16 | 0.3125 | 45 | 0.3778 |
| 13_548852 | 6 | 0.1667 | 15 | 0.2667 | 13 | 0.1538 | 6 | 0.1667 | 40 | 0.2 | 9 | 0 | 11 | 0.1818 | 8 | 0 | 12 | 0 | 40 | 0.05 |
| 13_548852 | 6 | 1 | 13 | 0.9231 | 10 | 0.9 | 6 | 0.8333 | 35 | 0.9143 | 9 | 1 | 7 | 1 | 6 | 1 | 15 | 0.9333 | 37 | 0.973 |
| 13_548858 | 6 | 0.6667 | 10 | 0.6 | 6 | 0.3333 | 3 | 0.6667 | 25 | 0.56 | 7 | 0.5714 | 7 | 0.5714 | 5 | 0.8 | 10 | 0.5 | 29 | 0.5862 |
| 13_548861 | 5 | 1 | 13 | 1 | 9 | 1 | 4 | 1 | 31 | 1 | 8 | 1 | 7 | 1 | 6 | 1 | 11 | 1 | 32 | 1 |
| 13_548862 | 5 | 0.2 | 13 | 0.2308 | 8 | 0.25 | 4 | 0.5 | 30 | 0.2667 | 8 | 0.125 | 7 | 0.2857 | 6 | 0.6667 | 11 | 0.5455 | 32 | 0.4062 |
| 13_548863 | 4 | 1 | 12 | 0.9167 | 5 | 0.8 | 3 | 1 | 24 | 0.9167 | 6 | 1 | 6 | 1 | 5 | 1 | 8 | 1 | 25 | 1 |
| 13_548863 | 5 | 0.4 | 14 | 0.2857 | 6 | 0 | 4 | 0.25 | 29 | 0.2414 | 6 | 0.1667 | 7 | 0 | 5 | 0 | 10 | 0.3 | 28 | 0.1429 |
| 13_548866 | 4 | 0.25 | 10 | 0.4 | 4 | 0.25 | 3 | 0 | 21 | 0.2857 | 4 | 0.25 | 7 | 0.2857 | 5 | 0.4 | 8 | 0.25 | 24 | 0.2917 |
| 13_548869 | 5 | 0.4 | 10 | 0.4 | 6 | 0.5 | 4 | 1 | 25 | 0.52 | 6 | 0.1667 | 6 | 0.1667 | 4 | 0 | 7 | 0.2857 | 23 | 0.1739 |
| 13_548869 | 5 | 0.4 | 12 | 0.3333 | 6 | 0.3333 | 4 | 0.5 | 27 | 0.3704 | 7 | 0.7143 | 6 | 0.3333 | 5 | 0.2 | 8 | 0.25 | 26 | 0.3846 |
| 13_548871 | 3 | 0.6667 | 10 | 0.7 | 6 | 0.5 | 4 | 0.25 | 23 | 0.5652 | 5 | 0.6 | 5 | 0.4 | 4 | 0.5 | 9 | 0.7778 | 23 | 0.6087 |
| 13_548872 | 4 | 0 | 12 | 0.25 | 6 | 0.6667 | 4 | 0.25 | 26 | 0.3077 | 6 | 0.3333 | 4 | 0.25 | 4 | 0.25 | 9 | 0.5556 | 23 | 0.3913 |
| 13_548872 | 4 | 0 | 11 | 0.1818 | 6 | 0 | 4 | 0 | 25 | 0.08 | 5 | 0 | 5 | 0.2 | 5 | 0.2 | 8 | 0.125 | 23 | 0.1304 |
| 13_548872 | 4 | 0.25 | 12 | 0.1667 | 5 | 0 | 4 | 0 | 25 | 0.12 | 6 | 0.6667 | 4 | 0 | 5 | 0 | 6 | 0.1667 | 21 | 0.2381 |
| 13_548873 | 3 | 0 | 10 | 0.2 | 5 | 0.6 | 4 | 0.75 | 22 | 0.3636 | 6 | 0.5 | 5 | 0 | 5 | 0.4 | 8 | 0.25 | 24 | 0.2917 |
| 13_548873 | 3 | 0 | 6 | 0.3333 | 3 | 1 | 3 | 1 | 15 | 0.5333 | 4 | 0.5 | 5 | 0.4 | 4 | 0 | 8 | 0.625 | 21 | 0.4286 |
| 13_548874 | 4 | 0.5 | 9 | 0.1111 | 3 | 0.3333 | 4 | 0.25 | 20 | 0.25 | 6 | 0.3333 | 4 | 0.5 | 4 | 0.25 | 7 | 0.4286 | 21 | 0.381 |
| 13_548875 | 3 | 0 | 7 | 0.2857 | 3 | 0 | 2 | 0 | 15 | 0.1333 | 5 | 0.6 | 4 | 0 | 3 | 0 | 4 | 0.25 | 16 | 0.25 |
| 13_548875 | 4 | 0.25 | 10 | 0.1 | 4 | 0 | 3 | 0.3333 | 21 | 0.1429 | 6 | 0.1667 | 5 | 0.2 | 5 | 0 | 8 | 0 | 24 | 0.0833 |
| 13_548876 | 3 | 0 | 9 | 0.1111 | 2 | 0 | 3 | 0 | 17 | 0.0588 | 4 | 0 | 3 | 0 | 4 | 0.25 | 5 | 0 | 16 | 0.0625 |
| 13_548876 | 4 | 1 | 10 | 1 | 3 | 1 | 4 | 1 | 21 | 1 | 4 | 1 | 5 | 1 | 5 | 1 | 7 | 1 | 21 | 1 |
| 13_548877 | 4 | 0 | 9 | 0.1111 | 2 | 0 | 4 | 0 | 19 | 0.0526 | 6 | 0 | 4 | 0 | 5 | 0.2 | 7 | 0.1429 | 22 | 0.0909 |
| 13_548878 | 4 | 1 | 8 | 1 | 1 | 0 | 3 | 1 | 16 | 0.9375 | 5 | 1 | 4 | 0.75 | 5 | 1 | 6 | 0.8333 | 20 | 0.9 |
| 13_548879 | 4 | 0.5 | 9 | 0 | 2 | 0 | 4 | 0 | 19 | 0.1053 | 6 | 0.1667 | 5 | 0 | 5 | 0 | 7 | 0.1429 | 23 | 0.087 |
| 13_548880 | 3 | 0 | 8 | 0 | 2 | 0 | 3 | 0.3333 | 16 | 0.0625 | 6 | 0.1667 | 3 | 0.3333 | 5 | 0 | 7 | 0 | 21 | 0.0952 |

| | | | | | | | | | | | | | | | | | | | | |
|---|---|---|---|---|---|---|---|---|---|---|---|---|---|---|---|---|---|---|---|---|
| 13_5488808 | 3 | 0 | 8 | 0 | 2 | 1 | 1 | 0 | 14 | 0.1429 | 6 | 0.5 | 3 | 0.3333 | 4 | 0 | 7 | 0 | 20 | 0.2 |
| 13_5488810 | 3 | 0.3333 | 9 | 0.1111 | 1 | 0 | 1 | 0 | 14 | 0.1429 | 4 | 0 | 1 | 0 | 4 | 0.25 | 6 | 0 | 15 | 0.0667 |
| 13_5488819 | 3 | 0.6667 | 9 | 0.8889 | 2 | 0.5 | 2 | 1 | 16 | 0.8125 | 4 | 1 | 3 | 1 | 5 | 0.8 | 5 | 0.8 | 17 | 0.8824 |
| 13_5488821 | 3 | 0.6667 | 8 | 0.25 | 2 | 0 | 2 | 0 | 15 | 0.2667 | 5 | 0.2 | 2 | 0 | 5 | 0.8 | 5 | 0.2 | 17 | 0.3529 |
| 13_5488826 | 3 | 0.6667 | 8 | 0.875 | 2 | 1 | 1 | 0 | 14 | 0.7857 | 5 | 0.8 | 2 | 1 | 5 | 0.4 | 5 | 1 | 17 | 0.7647 |
| 13_5488833 | 3 | 0.6667 | 9 | 0.4444 | 2 | 0.5 | NA | NA | 14 | 0.5 | 5 | 1 | 1 | 0 | 4 | 1 | 6 | 0.8333 | 16 | 0.875 |
| 13_5490708 | 39 | 0.2051 | 58 | 0.1034 | 49 | 0.0612 | 31 | 0.1613 | 177 | 0.1243 | 36 | 0.1111 | 28 | 0.1786 | 30 | 0.0667 | 32 | 0.0938 | 126 | 0.1111 |
| 13_5494420 | 1 | 1 | 2 | 0.5 | 1 | 1 | NA | NA | 4 | 0.75 | NA | NA | 1 | 1 | 1 | 0 | NA | NA | 2 | 0.5 |
| 13_5515273 | 8 | 0.125 | 4 | 0.25 | 5 | 0 | 4 | 0.25 | 21 | 0.1429 | 7 | 0 | 3 | 0 | 3 | 0.3333 | 3 | 0 | 16 | 0.0625 |
| 13_5515283 | 19 | 0.2105 | 14 | 0.4286 | 10 | 0.2 | 12 | 0.25 | 55 | 0.2727 | 17 | 0.2353 | 12 | 0.4167 | 9 | 0.4444 | 6 | 0 | 44 | 0.2955 |
| 13_5515289 | 9 | 0 | 9 | 0.1111 | 11 | 0 | 10 | 0.1 | 39 | 0.0513 | 14 | 0.0714 | 12 | 0.0833 | 7 | 0 | 5 | 0 | 38 | 0.0526 |
| 13_5515303 | 14 | 0.2143 | 9 | 0.1111 | 11 | 0.0909 | 10 | 0 | 44 | 0.1136 | 18 | 0.2222 | 6 | 0 | 6 | 0.1667 | 6 | 0.1667 | 36 | 0.1667 |
| 13_5517632 | 1 | 0 | 3 | 0.3333 | NA | NA | 1 | 0 | 5 | 0.2 | NA | NA | 2 | 0 | 3 | 0.3333 | 2 | 0 | 7 | 0.1429 |
| 13_5517636 | 1 | 0 | 3 | 0 | NA | NA | 2 | 0.5 | 6 | 0.1667 | NA | NA | 2 | 0 | 2 | 0.5 | 2 | 0 | 6 | 0.1667 |
| 13_5517639 | NA | NA | 3 | 0.3333 | NA | NA | 2 | 0.5 | 5 | 0.4 | NA | NA | 2 | 0.5 | 1 | 0 | 2 | 0 | 5 | 0.2 |
| 13_5517640 | NA | NA | 3 | 0.6667 | NA | NA | 2 | 0.5 | 5 | 0.6 | NA | NA | 1 | 1 | 2 | 0.5 | 2 | 0 | 5 | 0.4 |
| 13_5523410 | 1 | 1 | NA | NA | 2 | 0.5 | 2 | 0.5 | 5 | 0.6 | NA | NA | 1 | 1 | NA | NA | 1 | 1 | 2 | 1 |
| 13_5528018 | 4 | 0.75 | 2 | 0 | 2 | 0 | NA | NA | 8 | 0.375 | 1 | 1 | NA | NA | NA | NA | NA | NA | 1 | 1 |
| 13_5528023 | 4 | 0.25 | 1 | 1 | 1 | 1 | NA | NA | 6 | 0.5 | 2 | 0.5 | NA | NA | NA | NA | NA | NA | 2 | 0.5 |
| 13_5528026 | 5 | 0.6 | 2 | 0 | 2 | 1 | NA | NA | 9 | 0.5556 | 1 | 1 | NA | NA | NA | NA | NA | NA | 1 | 1 |
| 13_5533477 | 4 | 0.25 | 2 | 0 | NA | NA | 1 | 0 | 7 | 0.1429 | NA | NA | NA | NA | NA | NA | 3 | 0.3333 | 3 | 0.3333 |
| 13_5533480 | 4 | 0.25 | 1 | 0 | NA | NA | 1 | 0 | 6 | 0.1667 | NA | NA | NA | NA | NA | NA | 3 | 0.3333 | 3 | 0.3333 |
| 13_5533481 | 3 | 0 | 2 | 0.5 | NA | NA | 1 | 1 | 6 | 0.3333 | NA | NA | NA | NA | NA | NA | 3 | 1 | 3 | 1 |
| 13_5533486 | 4 | 0.5 | 2 | 0 | NA | NA | 1 | 1 | 7 | 0.4286 | NA | NA | NA | NA | NA | NA | 3 | 0.3333 | 3 | 0.3333 |
| 13_5533497 | 3 | 0 | 2 | 0.5 | NA | NA | 1 | 0 | 6 | 0.1667 | NA | NA | NA | NA | NA | NA | 3 | 0.6667 | 3 | 0.6667 |
| 13_5533508 | 4 | 0.5 | 2 | 0.5 | NA | NA | 1 | 0 | 7 | 0.4286 | NA | NA | NA | NA | NA | NA | 3 | 0.3333 | 3 | 0.3333 |
| 13_5533537 | 3 | 0 | 2 | 0 | NA | NA | 1 | 1 | 6 | 0.1667 | NA | NA | NA | NA | NA | NA | 3 | 0.3333 | 3 | 0.3333 |
| 13_5533563 | 4 | 0 | 1 | 1 | NA | NA | 1 | 0 | 6 | 0.1667 | NA | NA | NA | NA | NA | NA | 3 | 0.3333 | 3 | 0.3333 |
| 13_5533571 | 3 | 0.6667 | 2 | 1 | NA | NA | 1 | 1 | 6 | 0.8333 | NA | NA | NA | NA | NA | NA | 2 | 1 | 2 | 1 |
| 13_5539538 | NA | NA | NA | NA | NA | NA | 1 | 1 | 1 | 1 | 1 | 0 | 3 | 0.3333 | 1 | 1 | NA | NA | 5 | 0.4 |
| 13_5539547 | NA | NA | NA | NA | NA | NA | 1 | 1 | 1 | 1 | 1 | 1 | 3 | 0 | 1 | 0 | NA | NA | 5 | 0.2 |
| 13_5539551 | NA | NA | NA | NA | NA | NA | 1 | 1 | 1 | 1 | 1 | 1 | 3 | 0.6667 | 1 | 1 | NA | NA | 5 | 0.8 |
| 13_5539553 | NA | NA | NA | NA | NA | NA | 1 | 1 | 1 | 1 | 1 | 1 | 3 | 1 | 1 | 1 | NA | NA | 5 | 1 |
| 13_5539574 | NA | NA | NA | NA | NA | NA | 1 | 1 | 1 | 1 | 1 | 1 | 3 | 0.6667 | 1 | 0 | NA | NA | 5 | 0.6 |
| 13_5546011 | 3 | 0.6667 | 1 | 0 | 2 | 0.5 | 4 | 0.25 | 10 | 0.4 | 3 | 0.3333 | 7 | 0 | 5 | 0 | 2 | 0 | 17 | 0.0588 |
| 13_5546032 | 2 | 1 | 2 | 1 | 1 | 1 | 4 | 1 | 9 | 1 | 3 | 1 | 8 | 1 | 4 | 1 | 1 | 1 | 16 | 1 |
| 13_5546032 | 2 | 1 | 1 | 1 | 1 | 0 | 4 | 0.75 | 8 | 0.75 | 2 | 1 | 6 | 0.5 | 4 | 1 | 2 | 0 | 14 | 0.6429 |
| 13_5546036 | 1 | 1 | 2 | 0.5 | 2 | 0 | 4 | 1 | 9 | 0.6667 | 3 | 1 | 6 | 0.8333 | 5 | 0.8 | 1 | 1 | 15 | 0.8667 |
| 13_5546037 | 3 | 0 | 2 | 0.5 | 2 | 0.5 | 4 | 0 | 11 | 0.1818 | 3 | 0.3333 | 6 | 0.1667 | 5 | 0.8 | 2 | 0.5 | 16 | 0.4375 |
| 13_5546041 | 2 | 0 | 1 | 1 | 2 | 0 | 3 | 0 | 8 | 0.125 | 1 | 0 | 7 | 0.1429 | 4 | 0.25 | 2 | 0 | 14 | 0.1429 |
| 13_5546044 | 2 | 0 | 1 | 1 | 2 | 0 | 4 | 0 | 9 | 0.1111 | 3 | 0.3333 | 7 | 0 | 6 | 0.1667 | 1 | 0 | 17 | 0.1176 |
| 13_5546052 | 4 | 0 | 2 | 0 | 1 | 0 | 4 | 0.5 | 11 | 0.1818 | 3 | 0 | 8 | 0.125 | 6 | 0.1667 | 2 | 0 | 19 | 0.1053 |
| 13_5546053 | 3 | 0.6667 | 1 | 0 | 2 | 0.5 | 2 | 0.5 | 8 | 0.5 | 2 | 1 | 3 | 1 | 2 | 0.5 | NA | NA | 7 | 0.8571 |
| 13_5546054 | 2 | 0.5 | 2 | 0.5 | 2 | 0 | 4 | 0.25 | 10 | 0.3 | 3 | 0 | 4 | 0 | 6 | 0.3333 | 1 | 0 | 14 | 0.1429 |
| 13_5546062 | 4 | 0 | 2 | 0.5 | 2 | 1 | 4 | 0.25 | 12 | 0.3333 | 2 | 0.5 | 8 | 0.375 | 6 | 0.1667 | 2 | 0 | 18 | 0.2778 |
| 13_5546063 | 4 | 0 | 3 | 0 | 2 | 0 | 3 | 0.3333 | 12 | 0.0833 | 3 | 0 | 8 | 0 | 6 | 0.1667 | 2 | 0.5 | 19 | 0.1053 |
| 13_5546071 | 4 | 1 | 3 | 0.3333 | 2 | 1 | 4 | 1 | 13 | 0.8462 | 3 | 1 | 7 | 1 | 6 | 0.6667 | 1 | 1 | 17 | 0.8824 |
| 13_5546079 | 3 | 0.3333 | 2 | 0 | 2 | 0 | 4 | 0 | 11 | 0.0909 | 3 | 0 | 8 | 0.25 | 6 | 0.3333 | 2 | 0 | 19 | 0.2105 |
| 13_5546080 | 3 | 0.6667 | 3 | 0 | 2 | 0 | 4 | 0 | 12 | 0.1667 | 3 | 0.3333 | 7 | 0 | 5 | 0 | 2 | 0 | 17 | 0.0588 |
| 13_5546084 | 3 | 0 | 3 | 0 | 2 | 0 | 5 | 0.2 | 13 | 0.0769 | 2 | 1 | 7 | 0.2857 | 5 | 0.2 | NA | NA | 14 | 0.3571 |
| 13_5546116 | 4 | 0 | 3 | 0 | 3 | 0 | 5 | 0.2 | 15 | 0.0667 | 3 | 0 | 8 | 0.125 | 7 | 0 | 2 | 0 | 20 | 0.05 |
| 13_5546117 | 3 | 0.3333 | 3 | 0.3333 | 3 | 0.6667 | 3 | 0 | 12 | 0.3333 | 3 | 0.3333 | 7 | 0.1429 | 6 | 0.5 | 1 | 1 | 17 | 0.3529 |
| 13_5546122 | 4 | 1 | 2 | 1 | 2 | 1 | 3 | 1 | 11 | 1 | 3 | 1 | 8 | 1 | 7 | 1 | 1 | 1 | 19 | 1 |
| 13_5546137 | 4 | 1 | 3 | 1 | 4 | 1 | 5 | 1 | 16 | 1 | 3 | 1 | 8 | 1 | 5 | 1 | 2 | 1 | 18 | 1 |
| 13_5546142 | 4 | 1 | 3 | 1 | 3 | 1 | 5 | 0.8 | 15 | 0.9333 | 2 | 0.5 | 5 | 1 | 4 | 1 | 1 | 1 | 12 | 0.9167 |
| 13_5546143 | 3 | 0.6667 | 2 | 0 | 4 | 0 | 5 | 0.2 | 14 | 0.2143 | 4 | 0.75 | 6 | 0.6667 | 4 | 0.5 | 2 | 0 | 16 | 0.5625 |
| 13_5546156 | 2 | 0.5 | 1 | 0 | 3 | 0 | 4 | 0 | 10 | 0.1 | 4 | 0 | 7 | 0 | 7 | 0.1429 | 2 | 0 | 20 | 0.05 |
| 13_5546168 | 4 | 1 | 3 | 1 | 3 | 1 | 3 | 1 | 13 | 1 | NA | NA | 6 | 1 | 1 | 1 | NA | NA | 7 | 1 |
| 13_5546169 | 4 | 0 | 4 | 0 | 4 | 0 | 4 | 0.25 | 16 | 0.0625 | 1 | 1 | 5 | 0.2 | 2 | 1 | 1 | 0 | 9 | 0.4444 |
| 13_5546170 | 5 | 1 | 3 | 0.6667 | 5 | 1 | 3 | 1 | 16 | 0.9375 | 3 | 1 | 5 | 1 | 4 | 1 | NA | NA | 12 | 1 |
| 13_5546172 | 3 | 0.3333 | 5 | 0.4 | 4 | 0 | 4 | 0 | 16 | 0.1875 | 3 | 0.6667 | 6 | 0.1667 | 4 | 0.25 | 2 | 0.5 | 15 | 0.3333 |
| 13_5546176 | 4 | 0.25 | 3 | 0 | 4 | 0.25 | 4 | 0.25 | 15 | 0.2 | 2 | 0 | 7 | 0.1429 | 5 | 0.2 | 2 | 0 | 16 | 0.125 |
| 13_5546178 | 5 | 0.2 | 4 | 0 | 4 | 0.25 | 4 | 0 | 17 | 0.1176 | 3 | 0 | 7 | 0.1429 | 5 | 0 | 3 | 0 | 18 | 0.0556 |
| 13_5546208 | 3 | 1 | 1 | 1 | 1 | 0 | 2 | 1 | 7 | 0.8571 | 1 | 0 | 2 | 0 | 1 | 1 | 2 | 1 | 6 | 0.5 |
| 13_5546481 | 7 | 0.1429 | 2 | 0 | 3 | 0 | 5 | 0 | 17 | 0.0588 | 1 | 0 | 8 | 0.125 | 2 | 0 | 3 | 0 | 14 | 0.0714 |

| | | | | | | | | | | | | | | | | | | | | |
|---|---|---|---|---|---|---|---|---|---|---|---|---|---|---|---|---|---|---|---|---|
| 13_5546772 | 12 | 0.3333 | 4 | 0 | 5 | 0 | 6 | 0 | 27 | 0.1481 | 7 | 0.1429 | 9 | 0.1111 | 5 | 0 | 10 | 0.1 | 31 | 0.0968 |
| 13_5546779 | 13 | 0.7692 | 6 | 1 | 5 | 0.4 | 4 | 0.75 | 28 | 0.75 | 9 | 0.7778 | 9 | 0.8889 | 5 | 0.4 | 13 | 0.5385 | 36 | 0.6667 |
| 13_5546790 | 15 | 0.1333 | 5 | 0 | 5 | 0 | 5 | 0.6 | 30 | 0.1667 | 9 | 0.1111 | 10 | 0.2 | 5 | 0.6 | 13 | 0.0769 | 37 | 0.1892 |
| 13_5546791 | 17 | 0.2941 | 6 | 0 | 5 | 0 | 6 | 0.3333 | 34 | 0.2059 | 9 | 0.3333 | 9 | 0.3333 | 7 | 0.1429 | 13 | 0.1538 | 38 | 0.2368 |
| 13_5546856 | 16 | 0.0625 | 6 | 0.1667 | 2 | 0 | 6 | 0 | 30 | 0.0667 | 8 | 0.125 | 9 | 0.1111 | 6 | 0 | 11 | 0.0909 | 34 | 0.0882 |
| 13_5547018 | NA | NA | 2 | 1 | 1 | 1 | NA | NA | 3 | 1 | NA | NA | 1 | 0 | 1 | 1 | 2 | 1 | 4 | 0.75 |
| 13_5547018 | NA | NA | 1 | 1 | 1 | 1 | 1 | 0 | 3 | 0.6667 | NA | NA | NA | NA | 1 | 1 | 2 | 0 | 3 | 0.3333 |
| 13_5547021 | NA | NA | 2 | 0.5 | 1 | 0 | 1 | 1 | 4 | 0.5 | NA | NA | 1 | 1 | NA | NA | 2 | 1 | 3 | 1 |
| 13_5547082 | 30 | 0 | 35 | 0.1429 | 31 | 0.0968 | 37 | 0.0811 | 133 | 0.0827 | 24 | 0.1667 | 27 | 0.037 | 24 | 0.1667 | 20 | 0.15 | 95 | 0.1263 |
| 13_5549181 | 8 | 0.375 | 6 | 0.1667 | 5 | 0.2 | 8 | 0.25 | 27 | 0.2593 | 9 | 0.4444 | 4 | 0.25 | 4 | 0.25 | 3 | 0.3333 | 20 | 0.35 |
| 13_5549186 | 7 | 0.1429 | 5 | 0 | 5 | 0.2 | 6 | 0.5 | 23 | 0.2174 | 9 | 0.1111 | 4 | 0 | 4 | 0.25 | 4 | 0.5 | 21 | 0.1905 |
| 13_5549189 | 8 | 0.25 | 6 | 0.3333 | 6 | 0.3333 | 8 | 0.25 | 28 | 0.2857 | 9 | 0 | 4 | 0.25 | 4 | 0 | 4 | 0.25 | 21 | 0.0952 |
| 13_5549198 | 7 | 0 | 6 | 0 | 6 | 0.3333 | 8 | 0.125 | 27 | 0.1111 | 8 | 0.375 | 4 | 0 | 4 | 0 | 4 | 0 | 20 | 0.15 |
| 13_5549252 | 4 | 0.5 | 5 | 0 | 5 | 0.2 | 5 | 0.2 | 19 | 0.2105 | 7 | 0 | 2 | 0 | 4 | 0 | 4 | 0.25 | 17 | 0.0588 |
| 13_5549291 | 4 | 0 | 5 | 0.4 | 5 | 0.4 | 7 | 0.1429 | 21 | 0.2381 | 7 | 0.1429 | 2 | 0 | 4 | 0 | 3 | 0 | 16 | 0.0625 |
| 13_5549382 | 4 | 0.75 | 6 | 0.1667 | 5 | 0.2 | 5 | 0.2 | 20 | 0.3 | 5 | 0 | 4 | 0.25 | 5 | 0.2 | 4 | 0.25 | 18 | 0.1667 |
| 13_5549411 | 4 | 0 | 4 | 0 | 3 | 0 | 3 | 0.6667 | 14 | 0.1429 | 4 | 0.5 | 5 | 0 | 4 | 0 | 3 | 0 | 16 | 0.125 |
| 13_5549428 | 4 | 0.25 | 5 | 0 | 3 | 0 | 5 | 0.2 | 17 | 0.1176 | 5 | 0 | 5 | 0 | 3 | 0.3333 | 3 | 0.3333 | 16 | 0.125 |
| 13_5549460 | 7 | 0.1429 | 4 | 0.25 | 5 | 0.2 | 3 | 0 | 19 | 0.1579 | 6 | 0.1667 | 5 | 0.4 | 3 | 0 | 6 | 0.1667 | 20 | 0.2 |
| 13_5549463 | 7 | 0.8571 | 6 | 0.5 | 5 | 0.8 | 3 | 0.6667 | 21 | 0.7143 | 4 | 0.75 | 2 | 1 | 4 | 0.5 | 5 | 0.8 | 15 | 0.7333 |
| 13_5549466 | 6 | 0.1667 | 6 | 0 | 5 | 0 | 3 | 0 | 20 | 0.05 | 6 | 0 | 5 | 0.2 | 3 | 0 | 6 | 0 | 20 | 0.05 |
| 13_5553194 | 4 | 0.75 | 10 | 0.3 | 8 | 0.375 | 7 | 0.2857 | 29 | 0.3793 | 6 | 0.5 | 12 | 0.5 | 14 | 0.6429 | 11 | 0.3636 | 43 | 0.5116 |
| 13_5560777 | 5 | 0.8 | 6 | 0.8333 | 4 | 0.25 | 6 | 0.6667 | 21 | 0.6667 | 7 | 0.7143 | 6 | 1 | 8 | 0.5 | 4 | 0.5 | 25 | 0.68 |
| 13_5560784 | 5 | 0.4 | 9 | 0.2222 | 4 | 0.25 | 5 | 0.2 | 23 | 0.2609 | 8 | 0.125 | 5 | 0.4 | 9 | 0.1111 | 5 | 0.4 | 27 | 0.2222 |
| 13_5560812 | 5 | 0.2 | 9 | 0.2222 | 5 | 0.2 | 6 | 0 | 25 | 0.16 | 9 | 0.2222 | 6 | 0.1667 | 9 | 0.2222 | 6 | 0.1667 | 30 | 0.2 |
| 13_5560814 | 6 | 0.8333 | 8 | 0.625 | 4 | 0.5 | 4 | 0.75 | 22 | 0.6818 | 8 | 0.625 | 6 | 0.5 | 9 | 0.2222 | 7 | 0.7143 | 30 | 0.5 |
| 13_5560823 | 6 | 0.1667 | 10 | 0.3 | 5 | 0 | 6 | 0.3333 | 27 | 0.2222 | 10 | 0.3 | 6 | 0.3333 | 11 | 0.3636 | 6 | 0.3333 | 33 | 0.3333 |
| 13_5560857 | 6 | 0 | 7 | 0 | 8 | 0.25 | 6 | 0 | 27 | 0.0741 | 9 | 0.1111 | 5 | 0 | 12 | 0.0833 | 8 | 0.125 | 34 | 0.0882 |
| 13_5561998 | 4 | 0.5 | 3 | 0.6667 | 6 | 0.1667 | 3 | 0.3333 | 16 | 0.375 | 3 | 0.6667 | 10 | 0.3 | 3 | 0.6667 | 7 | 0.4286 | 23 | 0.4348 |
| 13_5562041 | 3 | 0 | 4 | 0.5 | 5 | 0.2 | 2 | 0 | 14 | 0.2143 | 4 | 0.5 | 5 | 0 | 3 | 0.6667 | 6 | 0.3333 | 18 | 0.3333 |
| 13_5562049 | 4 | 0.75 | 5 | 0.6 | 5 | 0.4 | 2 | 0 | 16 | 0.5 | 4 | 0.75 | 10 | 0.6 | 2 | 0.5 | 6 | 0.8333 | 22 | 0.6818 |
| 13_5562059 | 5 | 0.2 | 5 | 0 | 7 | 0 | 3 | 0 | 20 | 0.05 | 3 | 0.3333 | 8 | 0 | 4 | 0 | 8 | 0.125 | 23 | 0.087 |
| 13_5562070 | 5 | 0.6 | 5 | 0 | 6 | 0.1667 | 3 | 0 | 19 | 0.2105 | 4 | 0.5 | 8 | 0.125 | 4 | 0.25 | 7 | 0.1429 | 23 | 0.2174 |
| 13_5562078 | 4 | 0.75 | 6 | 0.6667 | 6 | 0.8333 | 2 | 0 | 18 | 0.6667 | 2 | 1 | 10 | 0.7 | 5 | 1 | 7 | 0.8571 | 24 | 0.8333 |
| 13_5562083 | 3 | 0.3333 | 5 | 0.4 | 3 | 0.6667 | 2 | 0.5 | 13 | 0.4615 | 3 | 1 | 6 | 0.8333 | 5 | 0.2 | 4 | 0.75 | 18 | 0.6667 |
| 13_5562084 | 5 | 0.2 | 6 | 0.3333 | 5 | 0.2 | 3 | 0 | 19 | 0.2105 | 4 | 0.25 | 10 | 0 | 6 | 0.3333 | 8 | 0.125 | 28 | 0.1429 |
| 13_5562087 | 4 | 1 | 4 | 1 | 2 | 1 | 2 | 1 | 12 | 1 | 3 | 1 | 6 | 1 | 4 | 1 | 8 | 0.875 | 21 | 0.9524 |
| 13_5562169 | 25 | 0.04 | 20 | 0.1 | 12 | 0 | 26 | 0.0769 | 83 | 0.0602 | 23 | 0.087 | 28 | 0.1071 | 29 | 0.0345 | 20 | 0.05 | 100 | 0.07 |
| 13_5562182 | 25 | 0.16 | 25 | 0.24 | 15 | 0.2667 | 26 | 0.1154 | 91 | 0.1868 | 26 | 0.3846 | 27 | 0.2222 | 29 | 0.1724 | 27 | 0.3333 | 109 | 0.2752 |
| 13_5562183 | 30 | 0.1667 | 22 | 0.2727 | 16 | 0.1875 | 30 | 0.3 | 98 | 0.2347 | 25 | 0.36 | 30 | 0.3333 | 27 | 0.1481 | 26 | 0.1538 | 108 | 0.25 |
| 13_5562190 | 25 | 0.12 | 24 | 0.0833 | 17 | 0.1176 | 30 | 0.0667 | 96 | 0.0938 | 25 | 0.28 | 26 | 0.1538 | 28 | 0.1429 | 26 | 0.1538 | 105 | 0.181 |
| 13_5562298 | 24 | 0.0833 | 17 | 0 | 14 | 0 | 25 | 0.16 | 80 | 0.075 | 15 | 0.0667 | 19 | 0.0526 | 19 | 0.0526 | 17 | 0.0588 | 70 | 0.0571 |
| 13_5562320 | 22 | 0.6818 | 16 | 0.6875 | 12 | 0.6667 | 25 | 0.72 | 75 | 0.6933 | 13 | 0.7692 | 22 | 0.6818 | 21 | 0.7619 | 19 | 0.7895 | 75 | 0.7467 |
| 13_5567109 | 2 | 1 | NA | NA | NA | NA | 3 | 0.6667 | 5 | 0.8 | 1 | 1 | NA | NA | 1 | 1 | NA | NA | 2 | 1 |
| 13_5567114 | 3 | 0 | NA | NA | NA | NA | 3 | 0.3333 | 6 | 0.1667 | 1 | 1 | NA | NA | 1 | 1 | NA | NA | 2 | 1 |
| 13_5567120 | 1 | 1 | NA | NA | NA | NA | 3 | 0 | 4 | 0.25 | 1 | 1 | NA | NA | 1 | 0 | NA | NA | 2 | 0.5 |
| 13_5567135 | 1 | 1 | NA | NA | NA | NA | 3 | 0.6667 | 4 | 0.75 | 1 | 1 | NA | NA | 1 | 0 | NA | NA | 2 | 0.5 |
| 13_5567139 | 2 | 1 | NA | NA | NA | NA | 3 | 0.6667 | 5 | 0.8 | 1 | 1 | NA | NA | NA | NA | NA | NA | 1 | 1 |
| 13_5567836 | 5 | 0.2 | 2 | 0 | 3 | 0.3333 | 3 | 1 | 13 | 0.3846 | 1 | 1 | 3 | 0.3333 | 3 | 0.6667 | 2 | 0.5 | 9 | 0.5556 |
| 13_5567841 | 5 | 0.6 | 2 | 0.5 | 4 | 0.75 | 3 | 0.3333 | 14 | 0.5714 | 1 | 0 | 5 | 0.4 | 4 | 0.5 | 3 | 0.3333 | 13 | 0.3846 |
| 13_5567843 | 6 | 0 | 1 | 0 | 3 | 0 | 4 | 0.25 | 14 | 0.0714 | 1 | 0 | 5 | 0.2 | 4 | 0.25 | 3 | 0 | 13 | 0.1538 |
| 13_5568713 | 4 | 0.25 | 11 | 0.0909 | 4 | 0 | 5 | 0 | 24 | 0.0833 | 3 | 0.3333 | 5 | 0.2 | 4 | 0 | 6 | 0 | 18 | 0.1111 |
| 13_5568724 | 4 | 0 | 9 | 0.1111 | 4 | 0 | 2 | 0 | 19 | 0.0526 | 3 | 0.3333 | 4 | 0 | 5 | 0.4 | 7 | 0.1429 | 19 | 0.2105 |
| 13_5568728 | 4 | 0.5 | 11 | 0.6364 | 3 | 0.6667 | 3 | 0.3333 | 21 | 0.5714 | 3 | 0.3333 | 3 | 0.3333 | 5 | 1 | 7 | 0.7143 | 18 | 0.6667 |
| 13_5568729 | 5 | 1 | 12 | 0.8333 | 4 | 0.75 | 5 | 1 | 26 | 0.8846 | 3 | 0.6667 | 4 | 0.5 | 6 | 1 | 6 | 1 | 19 | 0.8421 |
| 13_5569044 | 6 | 0 | 9 | 0 | 4 | 0.25 | 5 | 0.2 | 24 | 0.0833 | 3 | 0 | 5 | 0 | 4 | 0.25 | 7 | 0 | 19 | 0.0526 |
| 13_5569071 | 5 | 0.8 | 9 | 0.8889 | 4 | 0.5 | 5 | 1 | 23 | 0.8261 | 3 | 0.6667 | 4 | 0.75 | 5 | 1 | 7 | 0.8571 | 19 | 0.8421 |
| 13_5569076 | 5 | 0.2 | 7 | 0.1429 | 2 | 0.5 | 5 | 0.2 | 19 | 0.2105 | 2 | 0 | 4 | 0 | 5 | 0 | 6 | 0.1667 | 17 | 0.0588 |
| 13_5569079 | 5 | 0.2 | 9 | 0.1111 | 4 | 0 | 5 | 0 | 23 | 0.087 | 2 | 0.5 | 4 | 0 | 5 | 0.6 | 7 | 0 | 18 | 0.2222 |
| 13_5569082 | 5 | 0 | 9 | 0.2222 | 4 | 0.5 | 5 | 0.2 | 23 | 0.2174 | 2 | 0 | 4 | 0.25 | 4 | 0 | 6 | 0.1667 | 16 | 0.125 |
| 13_5569095 | 4 | 0.25 | 8 | 0.125 | 4 | 0.25 | 5 | 0.4 | 21 | 0.2381 | 2 | 0 | 4 | 0.25 | 4 | 0.25 | 6 | 0 | 16 | 0.125 |
| 13_5569097 | 5 | 0 | 7 | 0.2857 | 4 | 0 | 5 | 0 | 21 | 0.0952 | 1 | 0 | 3 | 0 | 5 | 0.2 | 7 | 0.2857 | 16 | 0.1875 |
| 13_5569112 | 5 | 0.2 | 9 | 0.6667 | 4 | 0.5 | 5 | 0.6 | 23 | 0.5217 | 2 | 0 | 3 | 1 | 3 | 0.3333 | 7 | 0.1429 | 15 | 0.3333 |
| 13_5569118 | 2 | 0.5 | 5 | 0.8 | 3 | 0.6667 | 3 | 1 | 13 | 0.7692 | 1 | 1 | 3 | 0.6667 | 1 | 1 | 6 | 0.6667 | 11 | 0.7273 |
| 13_5569120 | 4 | 0.25 | 9 | 0.3333 | 4 | 0 | 5 | 0 | 22 | 0.1818 | 3 | 0 | 3 | 0 | 4 | 0.25 | 6 | 0 | 16 | 0.0625 |

| | | | | | | | | | | | | | | | | | | | | |
|---|---|---|---|---|---|---|---|---|---|---|---|---|---|---|---|---|---|---|---|---|
| 13_5569127 | 3 | 0.3333 | 9 | 0.3333 | 2 | 1 | 4 | 0.5 | 18 | 0.4444 | 3 | 0.6667 | 3 | 0.6667 | 4 | 0.25 | 7 | 0.4286 | 17 | 0.4706 |
| 13_5569128 | 3 | 0 | 8 | 0 | 3 | 0 | 5 | 0.2 | 19 | 0.0526 | 3 | 0 | 3 | 0 | 4 | 0.25 | 7 | 0 | 17 | 0.0588 |
| 13_5569133 | 4 | 0 | 9 | 0 | 2 | 0.5 | 5 | 0 | 20 | 0.05 | 3 | 0 | 3 | 0 | 4 | 0 | 7 | 0.1429 | 17 | 0.0588 |
| 13_5569136 | 3 | 0.6667 | 7 | 1 | 3 | 1 | 3 | 1 | 16 | 0.9375 | 3 | 0.3333 | 3 | 0.6667 | 4 | 0.75 | 7 | 0.5714 | 17 | 0.5882 |
| 13_5569142 | 4 | 0.75 | 7 | 0.7143 | 3 | 0.3333 | 5 | 0.6 | 19 | 0.6316 | 3 | 0.3333 | 2 | 1 | 2 | 0 | 6 | 0.8333 | 13 | 0.6154 |
| 13_5570394 | 2 | 0 | 2 | 0 | 1 | 0 | 2 | 0.5 | 7 | 0.1429 | 1 | 1 | NA | NA | 1 | 0 | 3 | 0.6667 | 5 | 0.6 |
| 13_5574486 | 14 | 0.8571 | 8 | 0.75 | 10 | 0.7 | 13 | 0.9231 | 45 | 0.8222 | 8 | 0.875 | 10 | 0.9 | 9 | 0.8889 | 11 | 0.8182 | 38 | 0.8684 |
| 13_5574490 | 16 | 0.375 | 9 | 0.5556 | 9 | 0.1111 | 14 | 0.4286 | 48 | 0.375 | 7 | 0.2857 | 15 | 0.4667 | 9 | 0.6667 | 10 | 0.4 | 41 | 0.4634 |
| 13_5574493 | 11 | 0.1818 | 8 | 0 | 5 | 0 | 11 | 0.1818 | 35 | 0.1143 | 5 | 0.4 | 12 | 0.0833 | 12 | 0 | 10 | 0.2 | 39 | 0.1282 |
| 13_5574500 | 16 | 0.6875 | 9 | 0.6667 | 6 | 0.5 | 11 | 0.9091 | 42 | 0.7143 | 6 | 0.8333 | 13 | 0.6923 | 7 | 0.5714 | 10 | 0.6 | 36 | 0.6667 |
| 13_5574538 | 27 | 0.0741 | 20 | 0.15 | 13 | 0.0769 | 17 | 0.1765 | 77 | 0.1169 | 22 | 0.1818 | 22 | 0.1364 | 21 | 0 | 17 | 0.1176 | 82 | 0.1098 |
| 13_5575714 | 2 | 1 | 4 | 1 | NA | NA | 6 | 1 | 12 | 1 | 2 | 1 | 5 | 1 | 4 | 1 | 1 | 1 | 12 | 1 |
| 13_5575719 | 3 | 0.3333 | 3 | 0.3333 | 1 | 0 | 5 | 0 | 12 | 0.1667 | 2 | 0.5 | 5 | 0.2 | 5 | 0.4 | 1 | 0 | 13 | 0.3077 |
| 13_5575727 | 4 | 0 | 5 | 0 | 1 | 0 | 7 | 0.1429 | 17 | 0.0588 | 2 | 0 | 6 | 0.1667 | 4 | 0 | 2 | 0.5 | 14 | 0.1429 |
| 13_5575735 | 4 | 0.25 | 5 | 0 | 2 | 0.5 | 7 | 0.1429 | 18 | 0.1667 | 2 | 0 | 6 | 0.1667 | 5 | 0 | 2 | 0 | 15 | 0.0667 |
| 13_5575738 | 4 | 0.75 | 5 | 0.2 | 2 | 0 | 7 | 0.5714 | 18 | 0.4444 | NA | NA | 4 | 0.5 | 3 | 0.3333 | 1 | 0 | 8 | 0.375 |
| 13_5575739 | 2 | 0 | 3 | 0.3333 | 1 | 0 | 6 | 0.1667 | 12 | 0.1667 | 2 | 0 | 5 | 0.2 | 5 | 0 | 2 | 0 | 14 | 0.0714 |
| 13_5575749 | 3 | 0.6667 | 5 | 1 | 1 | 1 | 7 | 0.8571 | 16 | 0.875 | 2 | 0.5 | 5 | 0.6 | 4 | 1 | 2 | 1 | 13 | 0.7692 |
| 13_5575752 | 4 | 0.5 | 5 | 0.6 | 2 | 1 | 7 | 0.4286 | 18 | 0.5556 | 2 | 1 | 4 | 1 | 4 | 0.5 | 1 | 1 | 11 | 0.8182 |
| 13_5575759 | 4 | 0.5 | 5 | 0.2 | 2 | 0.5 | 6 | 0.5 | 17 | 0.4118 | 2 | 0.5 | 5 | 0.8 | 5 | 0.6 | 2 | 1 | 14 | 0.7143 |
| 13_5575786 | 4 | 1 | 5 | 0.8 | 1 | 1 | 6 | 0.5 | 16 | 0.75 | 2 | 0.5 | 5 | 0.2 | 4 | 0.5 | 1 | 0 | 12 | 0.3333 |
| 13_5575807 | 4 | 1 | 4 | 0.5 | 2 | 0.5 | 6 | 0.5 | 16 | 0.625 | 2 | 0.5 | 4 | 0.5 | 4 | 0.25 | 2 | 0.5 | 12 | 0.4167 |
| 13_5575810 | 4 | 0.75 | 5 | 0.2 | 4 | 0.5 | 5 | 0.6 | 18 | 0.5 | 2 | 0.5 | 6 | 0.5 | 4 | 0.25 | NA | NA | 12 | 0.4167 |
| 13_5577255 | 4 | 0 | 1 | 0 | NA | NA | 6 | 0.5 | 11 | 0.2727 | 5 | 0.4 | 2 | 0.5 | 2 | 0 | 4 | 0.25 | 13 | 0.3077 |
| 13_5577269 | 4 | 0.25 | 2 | 0.5 | 1 | 0 | 6 | 0.3333 | 13 | 0.3077 | 3 | 0.6667 | 2 | 0 | 5 | 0.2 | 4 | 0 | 14 | 0.2143 |
| 13_5577269 | 7 | 1 | 2 | 1 | 1 | 1 | 6 | 1 | 16 | 1 | 4 | 1 | 3 | 0.6667 | 4 | 0.75 | 3 | 0.6667 | 14 | 0.7857 |
| 13_5577978 | 9 | 1 | 2 | 1 | 1 | 1 | 6 | 1 | 18 | 1 | 5 | 1 | 3 | 1 | 6 | 1 | 5 | 1 | 19 | 1 |
| 13_5577982 | 10 | 0.4 | 3 | 0 | 2 | 0.5 | 6 | 0.3333 | 21 | 0.3333 | 6 | 0.3333 | 5 | 0.2 | 5 | 0.4 | 6 | 0.5 | 22 | 0.3636 |
| 13_5578000 | 6 | 0.1667 | 3 | 0 | 3 | 0 | 5 | 0.2 | 17 | 0.1176 | 6 | 0 | 4 | 0 | 5 | 0.4 | 5 | 0 | 20 | 0.1 |
| 13_5578014 | 10 | 0.1 | 4 | 0.25 | 5 | 0.4 | 8 | 0 | 27 | 0.1481 | 7 | 0.1429 | 5 | 0.6 | 7 | 0.2857 | 5 | 0 | 24 | 0.25 |
| 13_5582845 | 1 | 0 | 6 | 0.1667 | 2 | 0 | 5 | 0 | 14 | 0.0714 | 5 | 0.2 | 5 | 0.2 | 1 | 0 | 2 | 0 | 13 | 0.1538 |
| 13_5582879 | 3 | 0.3333 | 11 | 0.0909 | 5 | 0.2 | 4 | 0 | 23 | 0.1304 | 7 | 0.1429 | 5 | 0 | 2 | 0 | 3 | 0 | 17 | 0.0588 |
| 13_5584705 | 2 | 0 | 9 | 0.1111 | 3 | 0.6667 | 9 | 0.5556 | 23 | 0.3478 | 1 | 1 | 9 | 0.2222 | 4 | 1 | 6 | 0.3333 | 20 | 0.45 |
| 13_5584707 | 2 | 0 | 7 | 0.1429 | 2 | 0 | 5 | 0 | 16 | 0.0625 | 1 | 0 | 8 | 0.25 | 3 | 0.3333 | 4 | 0 | 16 | 0.1875 |
| 13_5584711 | 2 | 1 | 5 | 1 | 2 | 1 | 7 | 1 | 16 | 1 | 1 | 1 | 5 | 1 | 3 | 1 | 6 | 1 | 15 | 1 |
| 13_5584713 | 2 | 0 | 7 | 0 | 3 | 0 | 8 | 0.375 | 20 | 0.15 | 1 | 0 | 9 | 0.1111 | 3 | 0 | 2 | 0 | 15 | 0.0667 |
| 13_5584717 | 2 | 0 | 6 | 0.3333 | 4 | 0 | 8 | 0.125 | 20 | 0.15 | 1 | 0 | 8 | 0.25 | 5 | 0 | 6 | 0 | 20 | 0.1 |
| 13_5584719 | 2 | 1 | 8 | 0.125 | 4 | 0 | 10 | 0.2 | 24 | 0.2083 | 2 | 0.5 | 8 | 0.375 | 5 | 0 | 5 | 0.4 | 20 | 0.3 |
| 13_5584722 | 1 | 0 | 6 | 0.3333 | 4 | 0 | 8 | 0.5 | 19 | 0.3158 | 1 | 1 | 8 | 0.25 | 5 | 0.4 | 3 | 0.3333 | 17 | 0.3529 |
| 13_5584726 | 2 | 0 | 9 | 0.1111 | 5 | 0 | 9 | 0.2222 | 25 | 0.12 | 3 | 0.6667 | 9 | 0.2222 | 4 | 0.5 | 6 | 0.5 | 22 | 0.4091 |
| 13_5584730 | 1 | 1 | 9 | 0.2222 | 4 | 0 | 9 | 0.1111 | 23 | 0.1739 | 3 | 0.3333 | 8 | 0 | 4 | 0 | 5 | 0.2 | 20 | 0.1 |
| 13_5584754 | 1 | 1 | 8 | 0.625 | 4 | 0.25 | 10 | 0.8 | 23 | 0.6522 | 3 | 0.6667 | 8 | 0.375 | 5 | 0.8 | 5 | 0.6 | 21 | 0.5714 |
| 13_5584773 | 3 | 0.3333 | 9 | 0.3333 | 3 | 0 | 11 | 0.7273 | 26 | 0.4615 | 3 | 1 | 9 | 0.5556 | 5 | 0.6 | 6 | 0.5 | 23 | 0.6087 |
| 13_5584790 | 2 | 0 | 10 | 0.3 | 2 | 0 | 12 | 0.25 | 26 | 0.2308 | 4 | 0.25 | 9 | 0.1111 | 6 | 0.3333 | 6 | 0.1667 | 25 | 0.2 |
| 13_5584868 | 5 | 0 | 10 | 0.3 | 6 | 0.3333 | 12 | 0.3333 | 33 | 0.2727 | 4 | 0.25 | 9 | 0 | 8 | 0.25 | 10 | 0 | 31 | 0.0968 |
| 13_5584946 | 6 | 0.3333 | 9 | 0.3333 | 7 | 0.2857 | 14 | 0.2857 | 36 | 0.3056 | 5 | 0.4 | 9 | 0.3333 | 7 | 0.2857 | 9 | 0.5556 | 30 | 0.4 |
| 13_5584972 | 5 | 1 | 8 | 1 | 9 | 1 | 14 | 0.9286 | 36 | 0.9722 | 4 | 1 | 9 | 0.7778 | 5 | 1 | 12 | 0.8333 | 30 | 0.8667 |
| 13_5584981 | 6 | 0.1667 | 9 | 0.3333 | 9 | 0.2222 | 15 | 0.3333 | 39 | 0.2821 | 5 | 0 | 10 | 0.1 | 8 | 0.25 | 11 | 0.2727 | 34 | 0.1765 |
| 13_5584988 | 6 | 0 | 7 | 0.1429 | 8 | 0.25 | 14 | 0.2143 | 35 | 0.1714 | 4 | 0 | 7 | 0 | 7 | 0 | 9 | 0.6667 | 27 | 0.2222 |
| 13_5584990 | 7 | 1 | 8 | 1 | 9 | 1 | 16 | 1 | 40 | 1 | 3 | 1 | 9 | 1 | 7 | 1 | 10 | 1 | 29 | 1 |
| 13_5584999 | 7 | 0.8571 | 9 | 1 | 9 | 1 | 17 | 1 | 42 | 0.9762 | 7 | 1 | 9 | 0.8889 | 9 | 1 | 11 | 1 | 36 | 0.9722 |
| 13_5585004 | 8 | 0.625 | 10 | 0.8 | 9 | 0.2222 | 14 | 0.6429 | 41 | 0.5854 | 7 | 0.5714 | 9 | 0.3333 | 8 | 0.625 | 11 | 0.6364 | 35 | 0.5429 |
| 13_5585006 | 6 | 0.1667 | 8 | 0 | 6 | 0.3333 | 15 | 0.1333 | 35 | 0.1429 | 5 | 0.2 | 4 | 0 | 6 | 0.1667 | 10 | 0.1 | 25 | 0.12 |
| 13_5585110 | 11 | 0.1818 | 16 | 0 | 13 | 0.2308 | 21 | 0.0476 | 61 | 0.0984 | 7 | 0 | 12 | 0 | 15 | 0.1333 | 19 | 0.1053 | 53 | 0.0755 |
| 13_5585120 | 11 | 0.0909 | 17 | 0.0588 | 13 | 0 | 19 | 0.1053 | 60 | 0.0667 | 10 | 0 | 13 | 0 | 16 | 0.0625 | 16 | 0.125 | 55 | 0.0545 |
| 13_5586580 | 15 | 0.0667 | 7 | 0 | 6 | 0.3333 | 13 | 0.1538 | 41 | 0.122 | 8 | 0 | 10 | 0.1 | 9 | 0.2222 | 7 | 0 | 34 | 0.0882 |
| 13_5586585 | 14 | 0.0714 | 10 | 0.3 | 6 | 0.3333 | 14 | 0.1429 | 44 | 0.1818 | 8 | 0.125 | 12 | 0.1667 | 9 | 0 | 10 | 0.1 | 39 | 0.1026 |
| 13_5587355 | 18 | 0.5 | 10 | 0.8 | 7 | 0.7143 | 15 | 0.6 | 50 | 0.62 | 8 | 0.75 | 14 | 0.7143 | 9 | 0.8889 | 15 | 0.9333 | 46 | 0.8261 |
| 13_5587367 | 19 | 0 | 11 | 0.1818 | 8 | 0.125 | 13 | 0.1538 | 51 | 0.098 | 12 | 0.1667 | 15 | 0 | 9 | 0 | 14 | 0.2143 | 50 | 0.1 |
| 13_5587368 | 14 | 0.7143 | 10 | 0.7 | 6 | 0.5 | 14 | 0.5 | 44 | 0.6136 | 9 | 0.4444 | 12 | 0.5833 | 7 | 0.5714 | 13 | 0.5385 | 41 | 0.5366 |
| 13_5587368 | 10 | 0.2 | 7 | 0.2857 | 3 | 0.6667 | 15 | 0.2667 | 35 | 0.2857 | 6 | 0.6667 | 7 | 0.2857 | 8 | 0.125 | 8 | 0.125 | 29 | 0.2759 |
| 13_5587372 | 18 | 0.8889 | 14 | 0.7857 | 8 | 0.875 | 16 | 1 | 56 | 0.8929 | 12 | 1 | 15 | 0.8667 | 11 | 0.9091 | 13 | 0.9231 | 51 | 0.9216 |
| 13_5587375 | 19 | 0.8421 | 12 | 0.8333 | 8 | 0.75 | 16 | 0.875 | 55 | 0.8364 | 13 | 0.9231 | 13 | 0.9231 | 11 | 0.8182 | 16 | 0.9375 | 53 | 0.9057 |
| 13_5587379 | 18 | 0.6111 | 14 | 0.7857 | 7 | 0.8571 | 16 | 0.5625 | 55 | 0.6727 | 11 | 0.7273 | 16 | 0.8125 | 11 | 0.9091 | 15 | 0.8 | 53 | 0.8113 |
| 13_5587379 | 1 | 1 | NA | NA | NA | NA | 1 | 1 | 2 | 1 | 1 | 1 | NA | NA | NA | NA | NA | NA | 1 | 1 |

| | | | | | | | | | | | | | | | | | | | | |
|---|---|---|---|---|---|---|---|---|---|---|---|---|---|---|---|---|---|---|---|---|
| 13_558738 | 19 | 0.1053 | 12 | 0.25 | 9 | 0.4444 | 14 | 0.5 | 54 | 0.2963 | 9 | 0.3333 | 12 | 0.3333 | 10 | 0.2 | 14 | 0.2857 | 45 | 0.2889 |
| 13_559264 | 1 | 1 | NA | NA | NA | NA | NA | NA | 1 | 1 | NA | NA | 1 | 1 | NA | NA | 1 | 1 | 2 | 1 |
| 13_562219 | 7 | 0.1429 | 7 | 0.1429 | 12 | 0 | 7 | 0.1429 | 33 | 0.0909 | 5 | 0 | 13 | 0.1538 | 7 | 0.1429 | 7 | 0.1429 | 32 | 0.125 |
| 13_562221 | 9 | 0 | 8 | 0.25 | 12 | 0.0833 | 9 | 0.2222 | 38 | 0.1316 | 6 | 0.1667 | 17 | 0.0588 | 9 | 0 | 7 | 0 | 39 | 0.0513 |
| 13_564375 | 22 | 0.1818 | 41 | 0.122 | 30 | 0.0667 | 40 | 0.25 | 133 | 0.1579 | 30 | 0.4333 | 32 | 0.2188 | 34 | 0.0882 | 25 | 0.2 | 121 | 0.2314 |
| 13_566850 | 1 | 1 | NA | NA | 1 | 1 | NA | NA | 2 | 1 | 1 | 1 | NA | NA | NA | NA | 1 | 1 | 2 | 1 |
| 13_566850 | 1 | 1 | NA | NA | 1 | 1 | NA | NA | 2 | 1 | 1 | 0 | NA | NA | NA | NA | 1 | 1 | 2 | 0.5 |
| 13_566851 | 1 | 1 | NA | NA | 1 | 1 | NA | NA | 2 | 1 | 1 | 1 | NA | NA | NA | NA | 1 | 1 | 2 | 1 |
| 13_566871 | 2 | 0.5 | 1 | 0 | 1 | 0 | 1 | 0 | 5 | 0.2 | 1 | 0 | NA | NA | 2 | 0 | 1 | 1 | 4 | 0.25 |
| 13_566886 | 1 | 0 | 1 | 1 | NA | NA | 1 | 0 | 3 | 0.3333 | 1 | 0 | NA | NA | 3 | 0.6667 | 1 | 0 | 5 | 0.4 |
| 13_566886 | 2 | 0 | 2 | 0 | 1 | 0 | 1 | 1 | 6 | 0.1667 | 1 | 0 | NA | NA | 3 | 0.3333 | 1 | 0 | 5 | 0.2 |
| 13_566887 | 2 | 0.5 | 2 | 1 | 1 | 1 | 1 | 1 | 6 | 0.8333 | 1 | 1 | NA | NA | 3 | 1 | 1 | 1 | 5 | 1 |
| 13_566887 | 2 | 0.5 | 1 | 1 | NA | NA | 1 | 1 | 4 | 0.75 | 1 | 1 | NA | NA | 2 | 1 | 1 | 1 | 4 | 1 |
| 13_566887 | 1 | 0 | 1 | 1 | 1 | 1 | 1 | 0 | 4 | 0.5 | 1 | 1 | NA | NA | 3 | 1 | 1 | 0 | 5 | 0.8 |
| 13_567870 | 2 | 1 | NA | NA | NA | NA | 2 | 1 | 4 | 1 | 1 | 1 | NA | NA | NA | NA | NA | NA | 1 | 1 |
| 13_568713 | 1 | 1 | 2 | 1 | 1 | 1 | NA | NA | 4 | 1 | 1 | 1 | 1 | 1 | 1 | 0 | NA | NA | 3 | 0.6667 |
| 13_568713 | 1 | 0 | 1 | 0 | 1 | 1 | NA | NA | 3 | 0.3333 | 2 | 0.5 | 1 | 1 | NA | NA | NA | NA | 3 | 0.6667 |
| 13_568713 | NA | NA | 2 | 1 | 1 | 0 | NA | NA | 3 | 0.6667 | 2 | 0.5 | 1 | 1 | 1 | 0 | NA | NA | 4 | 0.5 |
| 13_568715 | 1 | 0 | 3 | 0.3333 | 1 | 1 | NA | NA | 5 | 0.4 | 2 | 0 | 1 | 0 | 1 | 1 | NA | NA | 4 | 0.25 |
| 13_568836 | 1 | 0 | 3 | 0 | 1 | 1 | 1 | 0 | 6 | 0.1667 | 2 | 0.5 | 1 | 1 | NA | NA | NA | NA | 3 | 0.6667 |
| 13_568853 | 1 | 1 | 3 | 1 | 2 | 1 | NA | NA | 6 | 1 | 3 | 0.6667 | 1 | 1 | NA | NA | NA | NA | 4 | 0.75 |
| 13_568857 | 3 | 1 | 3 | 0 | 2 | 0.5 | 1 | 0 | 9 | 0.4444 | 3 | 0 | 1 | 1 | 2 | 0.5 | NA | NA | 6 | 0.3333 |
| 13_568860 | 3 | 0.6667 | 2 | 0 | 2 | 0.5 | 2 | 0 | 9 | 0.3333 | 3 | 0 | 1 | 0 | 2 | 1 | 1 | 1 | 7 | 0.4286 |
| 13_568891 | 5 | 0 | 3 | 0.3333 | 2 | 0 | 4 | 0 | 14 | 0.0714 | 3 | 0.3333 | 1 | 0 | 3 | 0.3333 | 2 | 0 | 9 | 0.2222 |
| 13_568978 | 1 | 1 | 3 | 0.6667 | 1 | 1 | NA | NA | 5 | 0.8 | NA | NA | 1 | 1 | 2 | 0 | NA | NA | 3 | 0.3333 |
| 13_568979 | 2 | 1 | 2 | 0.5 | NA | NA | NA | NA | 4 | 0.75 | NA | NA | 1 | 1 | 2 | 0 | NA | NA | 3 | 0.3333 |
| 13_568982 | 2 | 0.5 | 3 | 0.3333 | NA | NA | NA | NA | 5 | 0.4 | NA | NA | 1 | 1 | 1 | 0 | NA | NA | 2 | 0.5 |
| 13_569002 | 2 | 0.5 | NA | NA | NA | NA | 1 | 1 | 3 | 0.6667 | 1 | 1 | NA | NA | NA | NA | NA | NA | 1 | 1 |
| 13_569004 | 2 | 1 | NA | NA | NA | NA | 1 | 1 | 3 | 1 | 1 | 1 | NA | NA | NA | NA | NA | NA | 1 | 1 |
| 13_569005 | 1 | 1 | NA | NA | NA | NA | 1 | 1 | 2 | 1 | 1 | 1 | NA | NA | NA | NA | NA | NA | 1 | 1 |
| 13_569214 | 3 | 1 | 1 | 0 | 1 | 1 | 2 | 0.5 | 7 | 0.7143 | NA | NA | 2 | 1 | 3 | 0.6667 | NA | NA | 5 | 0.8 |
| 13_569215 | 3 | 0.3333 | 1 | 0 | 1 | 0 | 3 | 0.3333 | 8 | 0.25 | NA | NA | 4 | 0.25 | 3 | 0 | 1 | 1 | 8 | 0.25 |
| 13_569215 | 3 | 1 | 1 | 1 | 1 | 0 | 3 | 0.6667 | 8 | 0.75 | NA | NA | 3 | 1 | 3 | 1 | 1 | 1 | 7 | 1 |
| 13_569703 | 1 | 0 | NA | NA | NA | NA | 1 | 1 | 2 | 0.5 | NA | NA | 1 | 1 | NA | NA | 1 | 1 | 2 | 1 |
| 13_570351 | NA | NA | NA | NA | NA | NA | 1 | 1 | 1 | 1 | NA | NA | 1 | 1 | NA | NA | 1 | 1 | 2 | 1 |
| 13_570353 | 2 | 1 | NA | NA | NA | NA | 1 | 1 | 3 | 1 | NA | NA | 1 | 1 | NA | NA | 1 | 1 | 2 | 1 |
| 13_570353 | 2 | 1 | NA | NA | NA | NA | 1 | 0 | 3 | 0.6667 | NA | NA | 1 | 1 | NA | NA | 1 | 1 | 2 | 1 |
| 13_570356 | 2 | 1 | NA | NA | NA | NA | 1 | 1 | 3 | 1 | NA | NA | NA | NA | NA | NA | 1 | 1 | 1 | 1 |
| 13_575742 | 32 | 0.1562 | 27 | 0.1111 | 22 | 0.0909 | 23 | 0.087 | 104 | 0.1154 | 20 | 0.25 | 24 | 0 | 13 | 0.3077 | 13 | 0.1538 | 70 | 0.1571 |
| 13_575767 | 18 | 0.0556 | 17 | 0.1765 | 8 | 0 | 15 | 0 | 58 | 0.069 | 17 | 0.1176 | 19 | 0.0526 | 9 | 0.1111 | 11 | 0.2727 | 56 | 0.125 |
| 13_575769 | 8 | 1 | 15 | 1 | 5 | 1 | 11 | 1 | 39 | 1 | 13 | 0.9231 | 18 | 0.9444 | 3 | 1 | 12 | 0.9167 | 46 | 0.9348 |
| 13_575769 | 11 | 0.0909 | 12 | 0.1667 | 5 | 0.2 | 12 | 0.0833 | 40 | 0.125 | 14 | 0.1429 | 13 | 0.2308 | 5 | 0.2 | 10 | 0.1 | 42 | 0.1667 |
| 13_575770 | 12 | 0.4167 | 15 | 0.7333 | 5 | 0.6 | 12 | 0.4167 | 44 | 0.5455 | 11 | 0.6364 | 17 | 0.5882 | 8 | 0.5 | 11 | 0.6364 | 47 | 0.5957 |
| 13_575770 | 13 | 0.9231 | 16 | 0.6875 | 6 | 0.5 | 13 | 0.9231 | 48 | 0.7917 | 14 | 0.7857 | 18 | 0.7222 | 8 | 0.875 | 12 | 0.6667 | 52 | 0.75 |
| 13_575772 | 12 | 0.1667 | 13 | 0.3846 | 6 | 0.3333 | 11 | 0 | 42 | 0.2143 | 13 | 0.4615 | 16 | 0.125 | 6 | 0.1667 | 12 | 0.1667 | 47 | 0.234 |
| 13_581481 | 2 | 0 | 3 | 0.3333 | NA | NA | NA | NA | 5 | 0.2 | 1 | 1 | 3 | 0 | 3 | 0.3333 | 2 | 0.5 | 9 | 0.3333 |
| 13_581482 | 2 | 0.5 | 3 | 0.3333 | NA | NA | 1 | 0 | 6 | 0.3333 | 1 | 1 | 3 | 0 | 3 | 0.3333 | 2 | 1 | 9 | 0.4444 |
| 13_581482 | 2 | 0.5 | 3 | 0 | NA | NA | NA | NA | 5 | 0.2 | 1 | 0 | 3 | 0.3333 | 3 | 0.3333 | 2 | 0.5 | 9 | 0.3333 |
| 13_581483 | 2 | 0.5 | 4 | 0 | 1 | 0 | 2 | 0 | 9 | 0.1111 | 1 | 0 | 4 | 0.25 | 3 | 0.3333 | 2 | 0.5 | 10 | 0.3 |
| 13_581483 | 3 | 0 | 7 | 0.2857 | 2 | 0 | 5 | 0 | 17 | 0.1176 | 3 | 0 | 8 | 0 | 4 | 0.25 | 7 | 0.1429 | 22 | 0.0909 |
| 13_581485 | 4 | 1 | 4 | 0.5 | 3 | 1 | 4 | 0.75 | 15 | 0.8 | 4 | 0.75 | 7 | 0.5714 | 4 | 1 | 6 | 1 | 21 | 0.8095 |
| 13_581485 | 3 | 1 | 6 | 1 | 3 | 0.6667 | 7 | 0.7143 | 19 | 0.8421 | 4 | 0.75 | 6 | 1 | 4 | 0.75 | 5 | 1 | 19 | 0.8947 |
| 13_581485 | 4 | 0 | 6 | 0 | 3 | 0 | 5 | 0.2 | 18 | 0.0556 | 5 | 0.2 | 8 | 0.375 | 4 | 0.75 | 7 | 0 | 24 | 0.2917 |
| 13_581492 | 3 | 0 | 4 | 0 | 2 | 0.5 | 4 | 0 | 13 | 0.0769 | 3 | 0 | 7 | 0.1429 | 4 | 0 | 5 | 0 | 19 | 0.0526 |
| 13_581499 | 2 | 0 | 3 | 0.3333 | 2 | 0.5 | 3 | 0.3333 | 10 | 0.3 | 3 | 0.3333 | 5 | 0.2 | 1 | 0 | 4 | 0.5 | 13 | 0.3077 |
| 13_581500 | 2 | 0 | 4 | 0.25 | 2 | 0 | 3 | 0 | 11 | 0.0909 | 3 | 0.3333 | 4 | 0 | 2 | 0 | 4 | 0 | 13 | 0.0769 |
| 13_581503 | 3 | 0 | 2 | 0 | 1 | 0 | 3 | 0.3333 | 9 | 0.1111 | 2 | 0 | 3 | 0.3333 | 1 | 0 | 3 | 0 | 9 | 0.1111 |
| 13_581512 | 1 | 0 | 1 | 0 | 2 | 0.5 | 3 | 0 | 7 | 0.1429 | 1 | 0 | 3 | 0.3333 | 2 | 0 | 2 | 0 | 8 | 0.125 |
| 13_581512 | 1 | 0 | 1 | 0 | 2 | 1 | 3 | 0 | 7 | 0.2857 | 1 | 1 | 3 | 0.3333 | 2 | 0.5 | 3 | 0.3333 | 9 | 0.4444 |
| 13_581515 | 1 | 1 | 1 | 1 | 1 | 1 | 1 | 1 | 4 | 1 | 1 | 1 | 3 | 0.6667 | 1 | 1 | 1 | 0 | 6 | 0.6667 |
| 13_581515 | NA | NA | 1 | 0 | 1 | 0 | 2 | 0.5 | 4 | 0.25 | 1 | 1 | 3 | 0 | 1 | 0 | 1 | 0 | 6 | 0.1667 |
| 13_581517 | 1 | 0 | 1 | 0 | 1 | 1 | 2 | 0 | 5 | 0.2 | NA | NA | 2 | 0 | 1 | 0 | 1 | 1 | 4 | 0.25 |
| 13_581517 | 1 | 1 | 1 | 1 | 1 | 1 | 1 | 1 | 4 | 1 | 1 | 1 | 2 | 1 | 1 | 1 | 1 | 0 | 5 | 0.8 |
| 13_581517 | NA | NA | 1 | 0 | 1 | 1 | 2 | 0 | 4 | 0.25 | 1 | 1 | 2 | 0.5 | 1 | 0 | 1 | 0 | 5 | 0.4 |
| 13_581519 | 1 | 1 | 1 | 0 | 1 | 0 | 2 | 0.5 | 5 | 0.4 | 1 | 1 | 1 | 1 | 1 | 1 | 1 | 0 | 4 | 0.75 |

| | | | | | | | | | | | | | | | | | | | | |
|---|---|---|---|---|---|---|---|---|---|---|---|---|---|---|---|---|---|---|---|---|
| 13_581522 | NA | NA | 1 | 0 | 1 | 1 | 1 | 0 | 3 | 0.3333 | 1 | 1 | 1 | 0 | 1 | 0 | NA | NA | 3 | 0.3333 |
| 13_581523 | 1 | 1 | 1 | 1 | 1 | 1 | 1 | 1 | 4 | 1 | 1 | 1 | 1 | 1 | 1 | 1 | NA | NA | 3 | 1 |
| 13_581524 | 1 | 1 | NA | NA | NA | NA | 1 | 1 | 2 | 1 | 1 | 1 | 1 | 1 | NA | NA | NA | NA | 2 | 1 |
| 13_581524 | NA | NA | 1 | 1 | 1 | 1 | NA | NA | 2 | 1 | 1 | 1 | 1 | 0 | NA | NA | 1 | 0 | 3 | 0.3333 |
| 13_581525 | NA | NA | NA | NA | 1 | 1 | 1 | 1 | 2 | 1 | 1 | 1 | NA | NA | 1 | 1 | NA | NA | 2 | 1 |
| 13_581526 | 1 | 1 | 1 | 0 | 1 | 1 | 1 | 0 | 4 | 0.5 | 1 | 1 | NA | NA | NA | NA | 1 | 0 | 2 | 0.5 |
| 13_581527 | 1 | 0 | 1 | 1 | 1 | 0 | 1 | 1 | 4 | 0.5 | 1 | 1 | NA | NA | NA | NA | NA | NA | 1 | 1 |
| 13_581528 | 1 | 1 | NA | NA | 1 | 1 | 1 | 1 | 3 | 1 | NA | NA | NA | NA | NA | NA | 1 | 1 | 1 | 1 |
| 13_582737 | 15 | 0.0667 | 26 | 0.1154 | 22 | 0.2727 | 21 | 0.0476 | 84 | 0.131 | 19 | 0.1579 | 21 | 0.0476 | 17 | 0.0588 | 19 | 0.0526 | 76 | 0.0789 |
| 13_582739 | 18 | 0.0556 | 27 | 0.1481 | 30 | 0.0333 | 26 | 0.0385 | 101 | 0.0693 | 19 | 0.1579 | 17 | 0 | 18 | 0 | 26 | 0.0769 | 80 | 0.0625 |
| 13_582740 | 17 | 0 | 33 | 0.2121 | 34 | 0.0882 | 29 | 0.1379 | 113 | 0.1239 | 20 | 0.15 | 27 | 0.1481 | 21 | 0.0476 | 26 | 0.1154 | 94 | 0.117 |
| 13_582741 | 18 | 0.5556 | 33 | 0.4242 | 33 | 0.4242 | 27 | 0.3333 | 111 | 0.4234 | 19 | 0.3684 | 24 | 0.2917 | 16 | 0.1875 | 25 | 0.36 | 84 | 0.3095 |
| 13_582742 | 11 | 0.1818 | 24 | 0.125 | 26 | 0.1923 | 22 | 0.0455 | 83 | 0.1325 | 14 | 0.0714 | 24 | 0.25 | 15 | 0.0667 | 20 | 0.25 | 73 | 0.1781 |
| 13_582743 | 14 | 0.0714 | 27 | 0.037 | 20 | 0 | 19 | 0.1053 | 80 | 0.05 | 13 | 0 | 21 | 0 | 16 | 0.125 | 20 | 0.15 | 70 | 0.0714 |
| 13_582748 | 16 | 0.625 | 34 | 0.8529 | 24 | 0.8333 | 20 | 0.7 | 94 | 0.7766 | 15 | 0.7333 | 18 | 0.8889 | 17 | 0.8824 | 24 | 0.75 | 74 | 0.8108 |
| 13_582749 | 7 | 0 | 11 | 0.2727 | 8 | 0.125 | 4 | 0 | 30 | 0.1333 | 7 | 0.1429 | 8 | 0.125 | 5 | 0 | 13 | 0.0769 | 33 | 0.0909 |
| 13_582750 | 7 | 1 | 7 | 1 | 7 | 1 | 6 | 1 | 27 | 1 | 5 | 1 | 5 | 1 | 3 | 1 | 11 | 1 | 24 | 1 |
| 13_582750 | 7 | 1 | 9 | 1 | 7 | 1 | 5 | 1 | 28 | 1 | 6 | 1 | 6 | 1 | 3 | 1 | 11 | 1 | 26 | 1 |
| 13_582750 | 7 | 1 | 9 | 1 | 7 | 1 | 6 | 1 | 29 | 1 | 4 | 1 | 6 | 1 | 2 | 1 | 13 | 1 | 25 | 1 |
| 13_582752 | 7 | 0.5714 | 7 | 0.4286 | 4 | 1 | 4 | 0.75 | 22 | 0.6364 | 4 | 0.5 | 5 | 0.6 | 2 | 1 | 6 | 0.6667 | 17 | 0.6471 |
| 13_582760 | 4 | 0.5 | 7 | 0.8571 | 3 | 0.3333 | 3 | 0.3333 | 17 | 0.5882 | 3 | 0.6667 | 3 | 0.3333 | 2 | 1 | 5 | 0.8 | 13 | 0.6923 |
| 13_582761 | 5 | 0 | 6 | 0.1667 | 5 | 0 | 3 | 0.3333 | 19 | 0.1053 | 2 | 0 | 4 | 0.25 | 3 | 0 | 7 | 0.1429 | 16 | 0.125 |
| 13_583112 | 4 | 0.25 | 4 | 0.5 | 3 | 0 | 4 | 0 | 15 | 0.2 | 2 | 0 | 3 | 0.3333 | 4 | 0 | 3 | 0.3333 | 12 | 0.1667 |
| 13_583113 | 4 | 0 | 4 | 0 | 4 | 0.25 | 4 | 0.25 | 16 | 0.125 | 2 | 0.5 | 5 | 0 | 6 | 0 | 3 | 0 | 16 | 0.0625 |
| 13_583113 | 2 | 0 | 3 | 0.3333 | 4 | 0 | 2 | 0 | 11 | 0.0909 | 1 | 0 | 5 | 0.2 | 4 | 0 | 2 | 0 | 12 | 0.0833 |
| 13_583114 | 4 | 1 | 4 | 0.75 | 4 | 1 | 3 | 1 | 15 | 0.9333 | 1 | 1 | 4 | 1 | 6 | 0.8333 | 2 | 1 | 13 | 0.9231 |
| 13_583115 | 4 | 0 | 4 | 0 | 3 | 0.3333 | 3 | 0 | 14 | 0.0714 | 2 | 0.5 | 3 | 0 | 5 | 0.2 | 2 | 0 | 12 | 0.1667 |
| 13_583115 | 1 | 0 | 1 | 1 | 3 | 0.6667 | 3 | 0.3333 | 8 | 0.5 | 1 | 1 | 3 | 0.6667 | 4 | 0.5 | 1 | 1 | 9 | 0.6667 |
| 13_583118 | 1 | 1 | NA | NA | NA | NA | NA | NA | 1 | 1 | NA | NA | NA | NA | NA | NA | 1 | 1 | 1 | 1 |
| 13_583118 | 1 | 1 | NA | NA | NA | NA | 1 | 0 | 2 | 0.5 | NA | NA | NA | NA | NA | NA | 1 | 1 | 1 | 1 |
| 13_583255 | 16 | 0 | 14 | 0.1429 | 10 | 0 | 13 | 0.1538 | 53 | 0.0755 | 17 | 0.0588 | 21 | 0.0476 | 14 | 0.1429 | 14 | 0 | 66 | 0.0606 |
| 13_584354 | NA | NA | NA | NA | 1 | 1 | NA | NA | 1 | 1 | NA | NA | NA | NA | 1 | 1 | NA | NA | 1 | 1 |
| 13_584359 | NA | NA | NA | NA | 1 | 1 | NA | NA | 1 | 1 | NA | NA | 1 | 1 | 2 | 0.5 | NA | NA | 3 | 0.6667 |
| 13_585294 | 1 | 1 | 1 | 1 | NA | NA | 2 | 0 | 4 | 0.5 | 1 | 1 | NA | NA | 3 | 0.3333 | NA | NA | 4 | 0.5 |
| 13_585295 | 5 | 0.2 | 1 | 1 | 1 | 0 | 6 | 0 | 13 | 0.1538 | 2 | 0.5 | NA | NA | 4 | 0 | 1 | 0 | 7 | 0.1429 |
| 13_585296 | 3 | 1 | 2 | 1 | 1 | 1 | 5 | 1 | 11 | 1 | 3 | 1 | NA | NA | 3 | 1 | 2 | 1 | 8 | 1 |
| 13_585296 | 4 | 1 | 1 | 1 | 1 | 1 | 5 | 1 | 11 | 1 | 2 | 1 | NA | NA | 3 | 1 | NA | NA | 5 | 1 |
| 13_585394 | 35 | 0.2 | 31 | 0.3226 | 29 | 0.3793 | 27 | 0.3704 | 122 | 0.3115 | 21 | 0.4286 | 31 | 0.1613 | 22 | 0.2727 | 27 | 0.3333 | 101 | 0.2871 |
| 13_585395 | 37 | 0.1622 | 34 | 0.0588 | 29 | 0.069 | 26 | 0.1923 | 126 | 0.119 | 21 | 0.2857 | 35 | 0.1429 | 27 | 0.2222 | 27 | 0.2222 | 110 | 0.2091 |
| 13_585396 | 31 | 0.0968 | 26 | 0.1154 | 24 | 0.0417 | 22 | 0.2273 | 103 | 0.1165 | 18 | 0.2222 | 32 | 0.0938 | 19 | 0.3684 | 25 | 0.24 | 94 | 0.2128 |
| 13_585399 | 73 | 0.0548 | 68 | 0.0735 | 60 | 0.1 | 67 | 0.0746 | 268 | 0.0746 | 49 | 0.102 | 62 | 0.0323 | 56 | 0.0893 | 51 | 0.0784 | 218 | 0.0734 |
| 13_585559 | 2 | 1 | 1 | 1 | 3 | 1 | 2 | 1 | 8 | 1 | 2 | 0.5 | NA | NA | 1 | 1 | 2 | 1 | 5 | 0.8 |
| 13_585559 | 2 | 0.5 | 1 | 0 | 4 | 0.25 | 2 | 1 | 9 | 0.4444 | 1 | 0 | NA | NA | 1 | 0 | 2 | 0.5 | 4 | 0.25 |
| 13_585559 | 2 | 0 | 1 | 1 | 4 | 1 | 3 | 1 | 10 | 0.8 | 1 | 1 | NA | NA | 1 | 1 | 2 | 1 | 4 | 1 |
| 13_585559 | 2 | 1 | 1 | 1 | 3 | 1 | 3 | 1 | 9 | 1 | 2 | 0.5 | 1 | 1 | 1 | 1 | 2 | 1 | 6 | 0.8333 |
| 13_585564 | 1 | 0 | 1 | 1 | 3 | 0.6667 | 4 | 1 | 9 | 0.7778 | 1 | 1 | 1 | 1 | NA | NA | 2 | 0.5 | 4 | 0.75 |
| 13_585566 | 2 | 1 | 1 | 0 | 4 | 0.5 | 3 | 0.3333 | 10 | 0.5 | 2 | 0.5 | 1 | 0 | 1 | 0 | 2 | 0.5 | 6 | 0.3333 |
| 13_585567 | 2 | 1 | 1 | 1 | 4 | 1 | 4 | 1 | 11 | 1 | 2 | 1 | 1 | 1 | 1 | 1 | 2 | 1 | 6 | 1 |
| 13_585567 | 2 | 0 | 1 | 1 | 2 | 0.5 | 3 | 0.3333 | 8 | 0.375 | 2 | 0.5 | NA | NA | NA | NA | 2 | 0 | 4 | 0.25 |
| 13_585569 | 2 | 0.5 | 1 | 1 | 4 | 0.75 | 3 | 0.6667 | 10 | 0.7 | 2 | 0 | NA | NA | NA | NA | 2 | 1 | 4 | 0.5 |
| 13_585570 | 2 | 0 | 1 | 1 | 5 | 0 | 4 | 0.75 | 12 | 0.3333 | 2 | 0.5 | 1 | 0 | 1 | 0 | 2 | 0.5 | 6 | 0.3333 |
| 13_585570 | 2 | 1 | 1 | 1 | 4 | 1 | 3 | 1 | 10 | 1 | 2 | 1 | 1 | 1 | 1 | 1 | 1 | 1 | 5 | 1 |
| 13_585571 | 2 | 1 | 1 | 1 | 5 | 1 | 3 | 1 | 11 | 1 | 2 | 1 | NA | NA | 1 | 1 | 2 | 1 | 5 | 1 |
| 13_585572 | 2 | 0.5 | 1 | 0 | 5 | 0.2 | 1 | 0 | 9 | 0.2222 | 1 | 1 | 1 | 1 | 1 | 1 | 2 | 0.5 | 5 | 0.8 |
| 13_585572 | 2 | 0.5 | NA | NA | 4 | 0.25 | 1 | 0 | 7 | 0.2857 | 1 | 0 | NA | NA | 1 | 0 | 1 | 1 | 3 | 0.3333 |
| 13_585573 | 1 | 0 | 1 | 1 | 5 | 0.4 | 2 | 0 | 9 | 0.3333 | 1 | 0 | NA | NA | 1 | 0 | 1 | 1 | 3 | 0.3333 |
| 13_585574 | 2 | 0.5 | 1 | 1 | 5 | 1 | 4 | 1 | 12 | 0.9167 | 1 | 1 | 1 | 1 | 2 | 1 | 1 | 1 | 5 | 1 |
| 13_585575 | 2 | 0 | 1 | 1 | 5 | 0 | 3 | 0.6667 | 11 | 0.2727 | 2 | 0 | NA | NA | 2 | 1 | 2 | 0.5 | 6 | 0.5 |
| 13_585575 | 2 | 0 | 1 | 0 | 3 | 0 | 3 | 0.3333 | 9 | 0.1111 | 2 | 0.5 | NA | NA | 1 | 0 | 1 | 1 | 4 | 0.5 |
| 13_585583 | NA | NA | NA | NA | 1 | 1 | 1 | 1 | 2 | 1 | NA | NA | NA | NA | 1 | 1 | NA | NA | 1 | 1 |
| 13_589562 | 15 | 0.3333 | 9 | 0.2222 | 12 | 0.5 | 13 | 0.3077 | 49 | 0.3469 | 8 | 0.25 | 10 | 0.5 | 8 | 0.625 | 7 | 0.2857 | 33 | 0.4242 |
| 13_589565 | 17 | 0.1176 | 10 | 0.4 | 12 | 0.1667 | 13 | 0.0769 | 52 | 0.1731 | 12 | 0.1667 | 14 | 0.1429 | 8 | 0.25 | 7 | 0.2857 | 41 | 0.1951 |
| 13_589565 | 12 | 0.5833 | 8 | 0.375 | 9 | 0.5556 | 9 | 0.6667 | 38 | 0.5526 | 13 | 0.3077 | 9 | 0.5556 | 3 | 0.3333 | 6 | 0.1667 | 31 | 0.3548 |
| 13_589565 | 19 | 0.2632 | 15 | 0.2 | 16 | 0.125 | 18 | 0.1111 | 68 | 0.1765 | 15 | 0.2 | 12 | 0.25 | 12 | 0 | 12 | 0.1667 | 51 | 0.1569 |
| 13_590799 | 21 | 0.1429 | 22 | 0.2273 | 18 | 0.3333 | 21 | 0.0952 | 82 | 0.1951 | 23 | 0.3043 | 18 | 0.3889 | 15 | 0.2667 | 16 | 0.0625 | 72 | 0.2639 |

| | | | | | | | | | | | | | | | | | | | | |
|---|---|---|---|---|---|---|---|---|---|---|---|---|---|---|---|---|---|---|---|---|
| 13_5907994 | 21 | 0.2381 | 22 | 0.2727 | 15 | 0.4 | 24 | 0.3333 | 82 | 0.3049 | 24 | 0.375 | 20 | 0.45 | 16 | 0.4375 | 14 | 0.4286 | 74 | 0.4189 |
| 13_5908004 | 24 | 0.0833 | 24 | 0.0833 | 15 | 0.0667 | 22 | 0.0455 | 85 | 0.0706 | 24 | 0.0417 | 17 | 0.1176 | 18 | 0.1111 | 13 | 0.0769 | 72 | 0.0833 |
| 13_5908453 | 24 | 0.0417 | 25 | 0.08 | 20 | 0.2 | 15 | 0.0667 | 84 | 0.0952 | 13 | 0 | 18 | 0.0556 | 17 | 0.1765 | 11 | 0.0909 | 59 | 0.0847 |
| 13_5914221 | NA | NA | 1 | 1 | 1 | 0 | NA | NA | 2 | 0.5 | 2 | 0 | NA | NA | 2 | 0.5 | 1 | 0 | 5 | 0.2 |
| 13_5927612 | 4 | 0.25 | 1 | 0 | 2 | 0.5 | 4 | 0.5 | 11 | 0.3636 | 4 | 0.25 | 1 | 1 | 2 | 0 | 7 | 0.4286 | 14 | 0.3571 |
| 13_5927621 | 5 | 0.8 | 2 | 1 | 3 | 0.3333 | 4 | 0.25 | 14 | 0.5714 | 5 | 0.6 | 3 | 0.3333 | 1 | 1 | 6 | 0.5 | 15 | 0.5333 |
| 13_5927622 | 3 | 0 | NA | NA | 3 | 0.3333 | 1 | 0 | 7 | 0.1429 | 5 | 0 | 2 | 0 | 2 | 1 | 4 | 0.25 | 13 | 0.2308 |
| 13_5927623 | 2 | 0.5 | 2 | 0.5 | 3 | 1 | 1 | 1 | 8 | 0.75 | 5 | 0.6 | NA | NA | 3 | 0.3333 | 5 | 0.2 | 13 | 0.3846 |
| 13_5927627 | 5 | 0 | 1 | 0 | 1 | 1 | 1 | 0 | 8 | 0.125 | 5 | 0 | 2 | 0 | 2 | 0 | 5 | 0.4 | 14 | 0.1429 |
| 13_5927629 | 5 | 0.2 | 2 | 0 | 2 | 0 | 4 | 0.5 | 13 | 0.2308 | 5 | 0.2 | 3 | 1 | 3 | 0.3333 | 6 | 0.3333 | 17 | 0.4118 |
| 13_5927633 | 4 | 1 | 1 | 1 | 1 | 1 | 4 | 1 | 10 | 1 | 4 | 1 | 3 | 1 | 3 | 1 | 6 | 1 | 16 | 1 |
| 13_5927634 | 2 | 0.5 | 2 | 0.5 | 1 | 0 | 3 | 0.3333 | 8 | 0.375 | 3 | 0.3333 | 2 | 0.5 | 2 | 0 | 4 | 1 | 11 | 0.5455 |
| 13_5927650 | 4 | 0.75 | NA | NA | 2 | 1 | 3 | 0.3333 | 9 | 0.6667 | 3 | 1 | 3 | 1 | 2 | 1 | 5 | 1 | 13 | 1 |
| 13_5927651 | 3 | 0.3333 | 2 | 1 | 3 | 1 | 4 | 0.25 | 12 | 0.5833 | 4 | 0.75 | 4 | 0.75 | 2 | 1 | 5 | 0 | 15 | 0.5333 |
| 13_5927655 | 5 | 0.6 | 2 | 1 | 2 | 0.5 | 2 | 0.5 | 11 | 0.6364 | 6 | 0.1667 | 4 | 0.25 | 3 | 0.3333 | 6 | 0.1667 | 19 | 0.2105 |
| 13_5927664 | 4 | 0.5 | 2 | 1 | 2 | 1 | 4 | 0.5 | 12 | 0.6667 | 6 | 0.5 | 3 | 1 | 3 | 1 | 4 | 0.25 | 16 | 0.625 |
| 13_5927680 | 4 | 0.5 | 2 | 0 | 4 | 0.25 | 4 | 0 | 14 | 0.2143 | 6 | 0.3333 | 3 | 0.3333 | 3 | 0 | 7 | 0.2857 | 19 | 0.2632 |
| 13_5927690 | 4 | 0 | 2 | 0.5 | 5 | 0.6 | 5 | 0.2 | 16 | 0.3125 | 7 | 0.1429 | 2 | 0.5 | 3 | 1 | 7 | 0.4286 | 19 | 0.4211 |
| 13_5927701 | 4 | 0.75 | 2 | 1 | 4 | 1 | 5 | 0.6 | 15 | 0.8 | 5 | 0.8 | 4 | 0.75 | 2 | 1 | 7 | 0.8571 | 18 | 0.8333 |
| 13_5927702 | 3 | 0.3333 | 2 | 0.5 | 3 | 0 | 4 | 0.5 | 12 | 0.3333 | 4 | 0.25 | 2 | 0.5 | 2 | 0.5 | 5 | 0.6 | 13 | 0.4615 |
| 13_5927705 | 4 | 0.75 | 2 | 1 | 5 | 0.2 | 5 | 0 | 16 | 0.375 | 7 | 0.1429 | 2 | 0.5 | 3 | 0.6667 | 6 | 0.3333 | 18 | 0.3333 |
| 13_5927707 | 4 | 0.75 | 2 | 1 | 2 | 0 | 5 | 0.2 | 13 | 0.4615 | 5 | 0.6 | 1 | 1 | 1 | 1 | 4 | 0.75 | 11 | 0.7273 |
| 13_5927709 | 4 | 0.5 | 2 | 0.5 | 5 | 0.2 | 5 | 0 | 16 | 0.25 | 7 | 0.2857 | 4 | 0.75 | 3 | 0 | 6 | 0.1667 | 20 | 0.3 |
| 13_5927710 | 4 | 0.75 | 2 | 1 | 5 | 0.2 | 4 | 0.5 | 15 | 0.5333 | 7 | 0.1429 | 4 | 0.25 | 3 | 0.6667 | 6 | 0.8333 | 20 | 0.45 |
| 13_5927712 | 4 | 1 | 2 | 0.5 | 4 | 0.75 | 4 | 0.75 | 14 | 0.7857 | 5 | 0.8 | 2 | 1 | 3 | 1 | 4 | 0.5 | 14 | 0.7857 |
| 13_5927725 | 5 | 0.8 | NA | NA | 4 | 0.25 | 4 | 0 | 13 | 0.3846 | 6 | 0.6667 | 4 | 0.75 | 2 | 1 | 6 | 0.6667 | 18 | 0.7222 |
| 13_5927730 | 3 | 0.6667 | 1 | 1 | 4 | 1 | 3 | 0.6667 | 11 | 0.8182 | 6 | 1 | 3 | 1 | 3 | 1 | 7 | 1 | 19 | 1 |
| 13_5927735 | 4 | 0 | 2 | 0.5 | 5 | 0.4 | 4 | 0 | 15 | 0.2 | 7 | 0.2857 | 4 | 0.25 | 3 | 0 | 7 | 0.1429 | 21 | 0.1905 |
| 13_5927742 | 5 | 0.6 | 2 | 1 | 4 | 1 | 4 | 0 | 15 | 0.6 | 7 | 0.2857 | 4 | 0.75 | 3 | 0.3333 | 9 | 0.4444 | 23 | 0.4348 |
| 13_5927760 | 6 | 0 | 3 | 0 | 5 | 0.2 | 4 | 0 | 18 | 0.0556 | 7 | 0.2857 | 4 | 0.25 | 3 | 0.3333 | 9 | 0.2222 | 23 | 0.2609 |
| 13_5927924 | 3 | 0.3333 | 2 | 0 | 3 | 0 | 5 | 0 | 13 | 0.0769 | 2 | 0 | 5 | 0.2 | 1 | 0 | 5 | 0 | 13 | 0.0769 |
| 13_5927926 | 3 | 0 | 2 | 0.5 | 4 | 0 | 5 | 0 | 14 | 0.0714 | 2 | 0 | 6 | 0 | 1 | 0 | 5 | 0.2 | 14 | 0.0714 |
| 13_5928114 | 8 | 0.375 | 7 | 0.4286 | 3 | 0.3333 | 6 | 0.6667 | 24 | 0.4583 | 5 | 0.4 | 7 | 0.2857 | 7 | 0.2857 | 10 | 0.6 | 29 | 0.4138 |
| 13_5928160 | 22 | 0.0455 | 17 | 0.0588 | 13 | 0 | 18 | 0.2222 | 70 | 0.0857 | 13 | 0.0769 | 13 | 0.0769 | 16 | 0.125 | 24 | 0.0833 | 66 | 0.0909 |
| 13_5959744 | 15 | 0.1333 | 13 | 0.3846 | 11 | 0.0909 | 18 | 0.2778 | 57 | 0.2281 | 10 | 0.1 | 6 | 0 | 13 | 0.2308 | 6 | 0.1667 | 35 | 0.1429 |
| 13_5959783 | 15 | 0.4667 | 15 | 0.3333 | 12 | 0.5 | 21 | 0.3333 | 63 | 0.3968 | 10 | 0.4 | 5 | 0.6 | 16 | 0.5 | 8 | 0.125 | 39 | 0.4103 |
| 13_5959787 | 17 | 0.5882 | 18 | 0.6111 | 10 | 0.7 | 22 | 0.6364 | 67 | 0.6269 | 10 | 0.2 | 5 | 1 | 15 | 0.3333 | 9 | 0.3333 | 39 | 0.3846 |
| 13_5959796 | 18 | 0.1111 | 16 | 0.125 | 11 | 0 | 19 | 0.1053 | 64 | 0.0938 | 10 | 0 | 4 | 0 | 12 | 0.1667 | 7 | 0.1429 | 33 | 0.0909 |
| 13_5959798 | 17 | 0.4118 | 18 | 0.5 | 12 | 0.5 | 19 | 0.4211 | 66 | 0.4545 | 9 | 0.6667 | 4 | 0.5 | 13 | 0.0769 | 8 | 0.5 | 34 | 0.3824 |
| 13_5959815 | 5 | 0.2 | 6 | 0 | 3 | 0.6667 | 9 | 0.1111 | 23 | 0.1739 | 3 | 0 | NA | NA | 4 | 0 | 3 | 0.3333 | 10 | 0.1 |
| 13_5959829 | 4 | 0 | 7 | 0.1429 | 4 | 0 | 7 | 0.2857 | 22 | 0.1364 | 3 | 0 | 1 | 0 | 3 | 0 | 3 | 0.3333 | 10 | 0.1 |
| 13_5959833 | 3 | 0 | 3 | 0 | 4 | 0 | 8 | 0.25 | 18 | 0.1111 | 2 | 0.5 | NA | NA | 3 | 0 | 2 | 0 | 7 | 0.1429 |
| 13_5959838 | 4 | 0 | 7 | 0.4286 | 4 | 0 | 7 | 0.1429 | 22 | 0.1818 | 3 | 0.3333 | 1 | 0 | 4 | 0 | 3 | 0.3333 | 11 | 0.1818 |
| 13_5959840 | 2 | 0 | 6 | 0.1667 | 4 | 0 | 4 | 0.25 | 16 | 0.125 | 2 | 0 | 1 | 1 | 1 | 0 | 3 | 0.3333 | 7 | 0.2857 |
| 13_5959856 | 3 | 0 | 7 | 0.4286 | 4 | 1 | 5 | 1 | 19 | 0.6316 | 3 | 0.6667 | 1 | 1 | 4 | 0.5 | 2 | 0 | 10 | 0.5 |
| 13_5959866 | 4 | 0 | 6 | 0.1667 | 4 | 0.25 | 6 | 0.3333 | 20 | 0.2 | 3 | 0.3333 | 1 | 1 | 3 | 0.3333 | 3 | 0 | 10 | 0.3 |
| 13_5959872 | 5 | 0.6 | 7 | 0.4286 | 4 | 0.5 | 5 | 0.8 | 21 | 0.5714 | 3 | 0.3333 | 1 | 1 | 4 | 0.75 | 3 | 0 | 11 | 0.4545 |
| 13_5960199 | 1 | 1 | NA | NA | 2 | 0.5 | NA | NA | 3 | 0.6667 | NA | NA | NA | NA | 1 | 1 | 2 | 1 | 3 | 1 |
| 13_5960215 | 2 | 0 | NA | NA | 2 | 0.5 | 2 | 0 | 6 | 0.1667 | 1 | 0 | NA | NA | 1 | 1 | NA | NA | 2 | 0.5 |
| 13_5960220 | 2 | 0.5 | 1 | 0 | 4 | 0 | 2 | 0 | 9 | 0.1111 | 1 | 1 | NA | NA | 2 | 0.5 | 2 | 1 | 5 | 0.8 |
| 13_5960238 | 2 | 1 | 1 | 0 | 4 | 1 | 2 | 0.5 | 9 | 0.7778 | 1 | 1 | NA | NA | 1 | 1 | 2 | 1 | 4 | 1 |
| 13_5960242 | 2 | 0 | 1 | 0 | 4 | 0.25 | 2 | 0 | 9 | 0.1111 | 1 | 1 | NA | NA | 1 | 0 | 2 | 1 | 4 | 0.75 |
| 13_5964798 | 4 | 0.5 | 7 | 0.4286 | 6 | 0.6667 | 6 | 0.6667 | 23 | 0.5652 | 4 | 0.5 | 2 | 1 | 7 | 0.5714 | 4 | 0.75 | 17 | 0.6471 |
| 13_5964803 | 4 | 0.5 | 8 | 0 | 7 | 0 | 7 | 0 | 26 | 0.0769 | 3 | 0 | 2 | 0 | 6 | 0 | 3 | 0.3333 | 14 | 0.0714 |
| 13_5964817 | 4 | 1 | 5 | 1 | 6 | 1 | 7 | 0.8571 | 22 | 0.9545 | 3 | 0.6667 | 2 | 0.5 | 5 | 1 | 4 | 1 | 14 | 0.8571 |
| 13_5964832 | 4 | 0.75 | 4 | 0.75 | 7 | 0.4286 | 6 | 0.5 | 21 | 0.5714 | 2 | 1 | 2 | 1 | 4 | 0.75 | 2 | 0.5 | 10 | 0.8 |
| 13_5964850 | 3 | 0 | 6 | 0.3333 | 7 | 0.4286 | 5 | 0.2 | 21 | 0.2857 | 3 | 0.3333 | 2 | 0 | 2 | 0 | 3 | 0 | 10 | 0.1 |
| 13_5978229 | 6 | 0.5 | 5 | 0.2 | 3 | 0.3333 | 5 | 0.2 | 19 | 0.3158 | 4 | 0 | 9 | 0.1111 | 9 | 0.1111 | 10 | 0.2 | 32 | 0.125 |
| 13_5978239 | 6 | 0 | 5 | 0 | 3 | 0.3333 | 5 | 0 | 19 | 0.0526 | 4 | 0 | 8 | 0.25 | 8 | 0.125 | 10 | 0 | 30 | 0.1 |
| 13_5978243 | 6 | 0.5 | 5 | 1 | 3 | 0.6667 | 4 | 0.5 | 18 | 0.6667 | 4 | 0.5 | 5 | 0.4 | 9 | 0.6667 | 9 | 0.4444 | 27 | 0.5185 |
| 13_5978252 | 8 | 0.125 | 5 | 0 | 3 | 0 | 5 | 0.2 | 21 | 0.0952 | 5 | 0.2 | 9 | 0 | 11 | 0 | 9 | 0.1111 | 34 | 0.0588 |
| 13_5978293 | 2 | 0.5 | 1 | 0 | 1 | 0 | 1 | 0 | 5 | 0.2 | 3 | 0.3333 | 2 | 0.5 | 1 | 0 | 1 | 0 | 7 | 0.2857 |
| 13_5978300 | 3 | 0 | 3 | 0.3333 | 1 | 0 | NA | NA | 7 | 0.1429 | 3 | 0.3333 | 3 | 0 | 1 | 0 | NA | NA | 7 | 0.1429 |
| 13_5978307 | 3 | 1 | 3 | 0.6667 | 1 | 1 | 1 | 1 | 8 | 0.875 | 3 | 0.3333 | 2 | 0.5 | 1 | 1 | NA | NA | 6 | 0.5 |
| 13_5978376 | 1 | 1 | 2 | 0.5 | 1 | 0 | NA | NA | 4 | 0.5 | 3 | 0.6667 | 1 | 1 | 1 | 1 | NA | NA | 5 | 0.8 |

| | | | | | | | | | | | | | | | | | | | | |
|---|---|---|---|---|---|---|---|---|---|---|---|---|---|---|---|---|---|---|---|---|
| 13_597844 | 1 | 0 | 2 | 1 | 1 | 0 | NA | NA | 4 | 0.5 | 3 | 0.3333 | 1 | 0 | NA | NA | NA | NA | 4 | 0.25 |
| 13_597849 | 1 | 0 | 2 | 0.5 | 1 | 1 | NA | NA | 4 | 0.5 | 1 | 1 | 1 | 1 | NA | NA | NA | NA | 2 | 1 |
| 13_597850 | 1 | 1 | 2 | 0.5 | 1 | 1 | NA | NA | 4 | 0.75 | 2 | 1 | 1 | 0 | NA | NA | NA | NA | 3 | 0.6667 |
| 13_599034 | 35 | 0.2286 | 28 | 0.2143 | 30 | 0.1667 | 39 | 0.1795 | 132 | 0.197 | 26 | 0.1923 | 29 | 0.1379 | 35 | 0.2286 | 31 | 0.129 | 121 | 0.1736 |
| 13_599038 | 39 | 0.0769 | 33 | 0.2727 | 34 | 0.3235 | 44 | 0.2273 | 150 | 0.22 | 26 | 0.0385 | 29 | 0.1724 | 41 | 0.3171 | 32 | 0.3125 | 128 | 0.2266 |
| 13_599041 | 35 | 0.0857 | 25 | 0.12 | 30 | 0.0667 | 39 | 0.0256 | 129 | 0.0698 | 23 | 0.0435 | 27 | 0.0741 | 36 | 0.1111 | 26 | 0.0385 | 112 | 0.0714 |
| 13_599044 | 25 | 0.44 | 23 | 0.3913 | 25 | 0.6 | 24 | 0.75 | 97 | 0.5464 | 22 | 0.4545 | 17 | 0.4118 | 28 | 0.5357 | 16 | 0.25 | 83 | 0.4337 |
| 13_599044 | 37 | 0.6216 | 29 | 0.6207 | 28 | 0.5357 | 33 | 0.5758 | 127 | 0.5906 | 22 | 0.7273 | 24 | 0.4583 | 35 | 0.6571 | 27 | 0.6667 | 108 | 0.6296 |
| 13_599044 | 20 | 0.3 | 20 | 0.5 | 23 | 0.6957 | 21 | 0.5714 | 84 | 0.5238 | 18 | 0.2778 | 21 | 0.5714 | 27 | 0.7037 | 20 | 0.75 | 86 | 0.593 |
| 13_599044 | 38 | 0.2895 | 28 | 0.2143 | 29 | 0.3793 | 38 | 0.3947 | 133 | 0.3233 | 24 | 0.25 | 24 | 0.25 | 37 | 0.3514 | 28 | 0.25 | 113 | 0.2832 |
| 13_599046 | 30 | 0.8333 | 25 | 0.92 | 27 | 0.8519 | 34 | 0.9706 | 116 | 0.8966 | 18 | 0.9444 | 19 | 0.9474 | 27 | 0.9259 | 22 | 0.8182 | 86 | 0.907 |
| 13_599047 | 36 | 0.1389 | 24 | 0.0833 | 31 | 0.0323 | 36 | 0.1944 | 127 | 0.1181 | 21 | 0.0476 | 23 | 0.0435 | 34 | 0.2059 | 23 | 0.1304 | 101 | 0.1188 |
| 13_599047 | 30 | 0.9333 | 26 | 0.9231 | 28 | 0.9286 | 30 | 0.9333 | 114 | 0.9298 | 20 | 0.85 | 19 | 0.8947 | 30 | 0.9 | 24 | 0.875 | 93 | 0.8817 |
| 13_599187 | NA | NA | NA | NA | NA | NA | 1 | 1 | 1 | 1 | NA | NA | 1 | 1 | NA | NA | NA | NA | 1 | 1 |
| 13_599197 | 2 | 0 | 6 | 0.3333 | 4 | 0 | 7 | 0 | 19 | 0.1053 | 1 | 0 | 5 | 0 | 4 | 0.25 | 3 | 0 | 13 | 0.0769 |
| 13_599197 | 2 | 0.5 | 7 | 0 | 3 | 0 | 7 | 0.2857 | 19 | 0.1579 | 2 | 0 | 4 | 0.75 | 5 | 0.6 | 4 | 0.5 | 15 | 0.5333 |
| 13_599198 | 3 | 0 | 5 | 0 | 4 | 0 | 7 | 0.2857 | 19 | 0.1053 | 2 | 0 | 5 | 0.2 | 5 | 0 | 3 | 0 | 15 | 0.0667 |
| 13_599199 | 3 | 0 | 7 | 0.5714 | 4 | 0.25 | 4 | 0.25 | 18 | 0.3333 | 2 | 0.5 | 6 | 0.1667 | 5 | 0.8 | 4 | 0.75 | 17 | 0.5294 |
| 13_599200 | 3 | 0.3333 | 4 | 1 | 4 | 1 | 4 | 0.25 | 15 | 0.6667 | NA | NA | 6 | 0.6667 | 4 | 0.75 | 4 | 0.75 | 14 | 0.7143 |
| 13_599200 | 2 | 0 | 7 | 0.1429 | 3 | 0 | 6 | 0.1667 | 18 | 0.1111 | 1 | 0 | 5 | 0.2 | 2 | 0 | 4 | 0 | 12 | 0.0833 |
| 13_599201 | 2 | 0.5 | 5 | 1 | 4 | 1 | 4 | 0.75 | 15 | 0.8667 | 1 | 1 | 5 | 1 | 3 | 1 | 2 | 1 | 11 | 1 |
| 13_599201 | 2 | 0.5 | 5 | 1 | 3 | 0.3333 | 4 | 0.25 | 14 | 0.5714 | 2 | 0.5 | 6 | 0.6667 | 4 | 0.5 | 4 | 0.25 | 16 | 0.5 |
| 13_599203 | 3 | 0 | 5 | 0.2 | 3 | 0.3333 | 5 | 0 | 16 | 0.125 | 2 | 0 | 6 | 0 | 5 | 0.2 | 2 | 0 | 15 | 0.0667 |
| 13_599205 | 2 | 0.5 | 5 | 0.2 | 3 | 0 | 4 | 0 | 14 | 0.1429 | NA | NA | 4 | 0.25 | 5 | 0.2 | 3 | 0.3333 | 12 | 0.25 |
| 13_599205 | 3 | 0 | 5 | 0.2 | 2 | 0 | 4 | 0 | 14 | 0.0714 | NA | NA | 4 | 0.25 | 3 | 0 | 4 | 0.25 | 11 | 0.1818 |
| 13_599298 | 2 | 0.5 | 5 | 0 | 3 | 0 | 4 | 0.25 | 14 | 0.1429 | NA | NA | 3 | 0 | 3 | 0 | 3 | 0.3333 | 9 | 0.1111 |
| 13_599299 | 3 | 0 | 5 | 0.2 | 3 | 0 | 3 | 0 | 14 | 0.0714 | NA | NA | 5 | 0 | 4 | 0.5 | 4 | 0 | 13 | 0.1538 |
| 13_599475 | 1 | 1 | 2 | 0 | 1 | 1 | 3 | 0.3333 | 7 | 0.4286 | NA | NA | 1 | 0 | NA | NA | 4 | 0.25 | 5 | 0.2 |
| 13_599476 | 1 | 0 | 2 | 0.5 | 1 | 0 | 3 | 0 | 7 | 0.1429 | NA | NA | 1 | 0 | NA | NA | 4 | 0.25 | 5 | 0.2 |
| 13_599479 | 2 | 0 | 2 | 0 | 1 | 0 | 3 | 0.6667 | 8 | 0.25 | NA | NA | 1 | 0 | NA | NA | 3 | 0.3333 | 4 | 0.25 |
| 13_599479 | 2 | 0.5 | 1 | 1 | 1 | 1 | 2 | 1 | 6 | 0.8333 | NA | NA | 1 | 1 | NA | NA | 1 | 1 | 2 | 1 |
| 13_599483 | 1 | 1 | 1 | 0 | 1 | 0 | 2 | 0.5 | 5 | 0.4 | NA | NA | 1 | 0 | NA | NA | 3 | 0.6667 | 4 | 0.5 |
| 13_599695 | 1 | 1 | 3 | 1 | NA | NA | 1 | 1 | 5 | 1 | NA | NA | 1 | 1 | NA | NA | 1 | 1 | 2 | 1 |
| 13_599696 | 1 | 0 | 3 | 0.6667 | NA | NA | 2 | 0 | 6 | 0.3333 | NA | NA | NA | NA | NA | NA | 1 | 1 | 1 | 1 |
| 13_599697 | 1 | 1 | 3 | 1 | 1 | 0 | 2 | 1 | 7 | 0.8571 | NA | NA | 1 | 1 | NA | NA | 2 | 1 | 3 | 1 |
| 13_603223 | 3 | 0.3333 | NA | NA | 4 | 0.5 | 2 | 1 | 9 | 0.5556 | 1 | 1 | NA | NA | 1 | 1 | NA | NA | 2 | 1 |
| 13_603227 | 4 | 0.25 | 1 | 1 | 4 | 0 | 2 | 0.5 | 11 | 0.2727 | 2 | 0 | NA | NA | 1 | 1 | NA | NA | 3 | 0.3333 |
| 13_603232 | 3 | 1 | 2 | 1 | NA | NA | 2 | 1 | 7 | 1 | 1 | 1 | NA | NA | NA | NA | NA | NA | 1 | 1 |
| 13_603233 | 3 | 0 | NA | NA | NA | NA | 2 | 0.5 | 5 | 0.2 | 1 | 1 | NA | NA | NA | NA | NA | NA | 1 | 1 |
| 13_603233 | 3 | 1 | 2 | 0.5 | NA | NA | 2 | 0.5 | 7 | 0.7143 | 1 | 1 | NA | NA | NA | NA | NA | NA | 1 | 1 |
| 13_603237 | 2 | 1 | 2 | 1 | NA | NA | 1 | 0 | 5 | 0.8 | 1 | 1 | NA | NA | NA | NA | NA | NA | 1 | 1 |
| 13_603240 | 1 | 1 | 1 | 0 | NA | NA | 1 | 1 | 3 | 0.6667 | 1 | 1 | NA | NA | NA | NA | NA | NA | 1 | 1 |
| 13_603241 | 2 | 1 | NA | NA | NA | NA | 1 | 0 | 3 | 0.6667 | 1 | 1 | NA | NA | NA | NA | NA | NA | 1 | 1 |
| 13_603242 | 1 | 1 | NA | NA | NA | NA | 1 | 1 | 2 | 1 | 1 | 1 | NA | NA | NA | NA | NA | NA | 1 | 1 |
| 13_603245 | 1 | 1 | 1 | 0 | NA | NA | 1 | 0 | 3 | 0.3333 | 1 | 1 | NA | NA | NA | NA | NA | NA | 1 | 1 |
| 13_603247 | 2 | 0.5 | 1 | 1 | NA | NA | 1 | 0 | 4 | 0.5 | 1 | 1 | NA | NA | NA | NA | NA | NA | 1 | 1 |
| 13_603247 | 2 | 1 | 1 | 1 | NA | NA | 1 | 0 | 4 | 0.75 | 1 | 1 | NA | NA | NA | NA | NA | NA | 1 | 1 |
| 13_609086 | 8 | 0.75 | 8 | 0.5 | 3 | 0.6667 | 9 | 0.4444 | 28 | 0.5714 | 5 | 0.6 | 7 | 0.5714 | 6 | 0.6667 | 7 | 0.7143 | 25 | 0.64 |
| 13_609086 | 8 | 1 | 11 | 0.9091 | 3 | 1 | 10 | 1 | 32 | 0.9688 | 4 | 1 | 8 | 1 | 6 | 1 | 7 | 0.8571 | 25 | 0.96 |
| 13_609087 | 4 | 0.25 | 5 | 0.6 | 3 | 0 | 9 | 0.6667 | 21 | 0.4762 | 2 | 0 | 5 | 0.2 | 4 | 0.25 | 7 | 0.1429 | 18 | 0.1667 |
| 13_609088 | 9 | 0.6667 | 10 | 0.4 | 3 | 0.6667 | 12 | 0.4167 | 34 | 0.5 | 5 | 0.6 | 8 | 0.5 | 8 | 0.875 | 7 | 0.5714 | 28 | 0.6429 |
| 13_609090 | 8 | 0.625 | 9 | 1 | 4 | 1 | 10 | 1 | 31 | 0.9032 | 5 | 1 | 6 | 0.6667 | 8 | 1 | 9 | 0.8889 | 28 | 0.8929 |
| 13_609091 | 6 | 0.1667 | 8 | 0 | 2 | 0 | 12 | 0.5 | 28 | 0.25 | 4 | 0.25 | 7 | 0.2857 | 6 | 0.1667 | 6 | 0.1667 | 23 | 0.2174 |
| 13_609091 | 10 | 0.9 | 11 | 1 | 4 | 1 | 13 | 1 | 38 | 0.9737 | 4 | 1 | 7 | 1 | 8 | 1 | 10 | 0.9 | 29 | 0.9655 |
| 13_609092 | 7 | 0 | 10 | 0 | 4 | 0.25 | 15 | 0.1333 | 36 | 0.0833 | 5 | 0.2 | 9 | 0.1111 | 6 | 0 | 10 | 0.2 | 30 | 0.1333 |
| 13_609093 | 10 | 0.4 | 11 | 0.3636 | 3 | 0 | 15 | 0.4667 | 39 | 0.3846 | 3 | 0.3333 | 8 | 0.5 | 11 | 0.2727 | 8 | 0.25 | 30 | 0.3333 |
| 13_609093 | 6 | 1 | 6 | 1 | 2 | 1 | 11 | 1 | 25 | 1 | 2 | 1 | 8 | 1 | 6 | 1 | 8 | 1 | 24 | 1 |
| 13_609093 | 9 | 1 | 5 | 1 | 2 | 1 | 12 | 1 | 28 | 1 | 3 | 1 | 6 | 1 | 8 | 0.875 | 5 | 1 | 22 | 0.9545 |
| 13_609094 | 9 | 0.7778 | 10 | 1 | 4 | 0.5 | 15 | 1 | 38 | 0.8947 | 4 | 1 | 8 | 1 | 9 | 0.8889 | 9 | 0.8889 | 30 | 0.9333 |
| 13_609095 | 7 | 0.4286 | 5 | 0.6 | 3 | 0.6667 | 12 | 0.5 | 27 | 0.5185 | 3 | 0.3333 | 7 | 0.7143 | 10 | 0.8 | 4 | 0.75 | 24 | 0.7083 |
| 13_609095 | 11 | 0.9091 | 12 | 0.9167 | 7 | 1 | 13 | 1 | 43 | 0.9535 | 5 | 1 | 10 | 0.8 | 7 | 1 | 9 | 1 | 31 | 0.9355 |
| 13_609097 | 6 | 0.6667 | 8 | 1 | 6 | 1 | 11 | 1 | 31 | 0.9355 | 3 | 1 | 6 | 0.8333 | 5 | 1 | 5 | 1 | 19 | 0.9474 |
| 13_609099 | 11 | 0.4545 | 11 | 0.6364 | 7 | 0.2857 | 15 | 0.4667 | 44 | 0.4773 | 3 | 1 | 9 | 0.3333 | 9 | 0.4444 | 10 | 0.5 | 31 | 0.4839 |
| 13_609101 | 15 | 0.0667 | 11 | 0 | 8 | 0.125 | 17 | 0.1176 | 51 | 0.0784 | 7 | 0.1429 | 13 | 0 | 11 | 0.0909 | 11 | 0.2727 | 42 | 0.119 |
| 13_609102 | 15 | 0.0667 | 13 | 0.0769 | 6 | 0.1667 | 16 | 0 | 50 | 0.06 | 7 | 0.1429 | 13 | 0.1538 | 10 | 0.1 | 11 | 0.1818 | 41 | 0.1463 |

| | | | | | | | | | | | | | | | | | | | | |
|---|---|---|---|---|---|---|---|---|---|---|---|---|---|---|---|---|---|---|---|---|
| 13_609103 | 16 | 0.0625 | 12 | 0 | 8 | 0 | 19 | 0.1053 | 55 | 0.0545 | 7 | 0.1429 | 13 | 0.1538 | 12 | 0 | 13 | 0 | 45 | 0.0667 |
| 13_609105 | 17 | 0.2941 | 15 | 0.3333 | 11 | 0.3636 | 25 | 0.24 | 68 | 0.2941 | 7 | 0.5714 | 12 | 0.1667 | 13 | 0.4615 | 15 | 0.2 | 47 | 0.3191 |
| 13_621992 | NA | NA | NA | NA | 1 | 1 | NA | NA | 1 | 1 | NA | NA | NA | NA | 1 | 1 | 1 | 1 | 2 | 1 |
| 13_621993 | NA | NA | NA | NA | 1 | 1 | NA | NA | 1 | 1 | NA | NA | NA | NA | 1 | 1 | 1 | 1 | 2 | 1 |
| 13_621996 | NA | NA | NA | NA | 1 | 1 | NA | NA | 1 | 1 | NA | NA | NA | NA | 1 | 1 | 1 | 0 | 2 | 0.5 |
| 13_621996 | NA | NA | NA | NA | 1 | 1 | NA | NA | 1 | 1 | NA | NA | NA | NA | 1 | 1 | 1 | 1 | 2 | 1 |
| 13_621997 | NA | NA | NA | NA | 1 | 1 | NA | NA | 1 | 1 | NA | NA | NA | NA | 1 | 1 | 1 | 1 | 2 | 1 |
| 13_625165 | NA | NA | 1 | 1 | NA | NA | NA | NA | 1 | 1 | NA | NA | NA | NA | 1 | 1 | NA | NA | 1 | 1 |
| 13_626013 | 2 | 0.5 | NA | NA | 2 | 0.5 | NA | NA | 4 | 0.5 | NA | NA | NA | NA | 1 | 1 | NA | NA | 1 | 1 |
| 13_626016 | 2 | 1 | NA | NA | 1 | 1 | 2 | 1 | 5 | 1 | NA | NA | NA | NA | 2 | 1 | NA | NA | 2 | 1 |
| 13_626028 | 1 | 1 | NA | NA | 1 | 1 | 1 | 1 | 3 | 1 | NA | NA | NA | NA | 2 | 1 | NA | NA | 2 | 1 |
| 13_626028 | 2 | 0 | NA | NA | 1 | 1 | 1 | 0 | 4 | 0.25 | NA | NA | NA | NA | 2 | 0.5 | NA | NA | 2 | 0.5 |
| 13_626028 | 1 | 1 | NA | NA | 1 | 1 | 1 | 0 | 3 | 0.6667 | NA | NA | NA | NA | 2 | 0.5 | NA | NA | 2 | 0.5 |
| 13_626647 | 1 | 1 | NA | NA | NA | NA | NA | NA | 1 | 1 | NA | NA | 1 | 1 | 1 | 0 | 2 | 0.5 | 4 | 0.5 |
| 13_626648 | 1 | 1 | NA | NA | NA | NA | NA | NA | 1 | 1 | NA | NA | 1 | 1 | 2 | 1 | 2 | 1 | 5 | 1 |
| 13_626648 | 1 | 1 | NA | NA | NA | NA | NA | NA | 1 | 1 | NA | NA | 1 | 0 | 2 | 0 | 2 | 0.5 | 5 | 0.2 |
| 13_626649 | 1 | 1 | NA | NA | NA | NA | NA | NA | 1 | 1 | NA | NA | 1 | 1 | 2 | 1 | 1 | 1 | 4 | 1 |
| 13_627828 | NA | NA | 1 | 1 | NA | NA | NA | NA | 1 | 1 | NA | NA | NA | NA | NA | NA | 1 | 1 | 1 | 1 |
| 13_628201 | 1 | 0 | NA | NA | NA | NA | 1 | 1 | 2 | 0.5 | 1 | 1 | NA | NA | 1 | 0 | 2 | 0 | 4 | 0.25 |
| 13_628203 | 2 | 1 | NA | NA | NA | NA | NA | NA | 2 | 1 | NA | NA | NA | NA | 1 | 1 | 1 | 1 | 2 | 1 |
| 13_628204 | 2 | 0.5 | NA | NA | NA | NA | NA | NA | 2 | 0.5 | NA | NA | NA | NA | 1 | 1 | 2 | 1 | 3 | 1 |
| 13_628205 | 2 | 0.5 | NA | NA | NA | NA | NA | NA | 2 | 0.5 | NA | NA | NA | NA | 1 | 1 | 2 | 0.5 | 3 | 0.6667 |
| 13_628205 | 2 | 1 | NA | NA | NA | NA | NA | NA | 2 | 1 | NA | NA | NA | NA | 1 | 1 | 2 | 1 | 3 | 1 |
| 13_628212 | 2 | 1 | NA | NA | NA | NA | NA | NA | 2 | 1 | NA | NA | NA | NA | 1 | 1 | 1 | 1 | 2 | 1 |
| 13_628215 | 2 | 0.5 | NA | NA | NA | NA | NA | NA | 2 | 0.5 | NA | NA | NA | NA | 1 | 1 | 1 | 1 | 2 | 1 |
| 13_634470 | NA | NA | 2 | 0 | 2 | 1 | 2 | 0.5 | 6 | 0.5 | 1 | 1 | 1 | 1 | 1 | 1 | 1 | 1 | 4 | 1 |
| 13_634470 | NA | NA | 2 | 0 | 1 | 1 | 2 | 0.5 | 5 | 0.4 | NA | NA | 1 | 1 | 1 | 1 | 1 | 0 | 3 | 0.6667 |
| 13_634471 | NA | NA | 2 | 0 | 2 | 1 | 2 | 0.5 | 6 | 0.5 | 1 | 1 | 1 | 1 | NA | NA | 1 | 0 | 3 | 0.6667 |
| 13_634472 | NA | NA | 1 | 0 | 2 | 1 | 1 | 0 | 4 | 0.5 | 1 | 1 | NA | NA | 1 | 1 | 1 | 0 | 3 | 0.6667 |
| 13_634486 | 1 | 1 | 2 | 0.5 | NA | NA | 2 | 1 | 5 | 0.8 | NA | NA | NA | NA | 1 | 0 | 2 | 1 | 3 | 0.6667 |
| 13_634497 | 1 | 1 | 5 | 1 | NA | NA | 6 | 1 | 12 | 1 | NA | NA | 4 | 1 | 2 | 1 | 2 | 1 | 8 | 1 |
| 13_634497 | 1 | 1 | 4 | 1 | NA | NA | 3 | 0.6667 | 8 | 0.875 | NA | NA | 3 | 0.6667 | 1 | 1 | 3 | 0.6667 | 7 | 0.7143 |
| 13_634498 | 1 | 0 | 5 | 0.6 | NA | NA | 4 | 1 | 10 | 0.7 | NA | NA | 3 | 1 | 2 | 1 | 2 | 1 | 7 | 1 |
| 13_634498 | 1 | 1 | 5 | 0.8 | NA | NA | 4 | 1 | 10 | 0.9 | NA | NA | 4 | 0.5 | 2 | 0 | 3 | 0.3333 | 9 | 0.3333 |
| 13_634498 | 1 | 1 | 5 | 0.6 | NA | NA | 3 | 0.6667 | 9 | 0.6667 | NA | NA | 2 | 1 | 1 | 1 | 3 | 0.3333 | 6 | 0.6667 |
| 13_634498 | NA | NA | NA | NA | NA | NA | 1 | 1 | 1 | 1 | NA | NA | NA | NA | 1 | 1 | NA | NA | 1 | 1 |
| 13_634499 | 1 | 0 | 6 | 0.6667 | NA | NA | 5 | 0.2 | 12 | 0.4167 | NA | NA | 5 | 0.8 | 2 | 1 | 2 | 1 | 9 | 0.8889 |
| 13_634502 | 2 | 1 | 4 | 1 | NA | NA | 3 | 1 | 9 | 1 | NA | NA | 4 | 1 | 2 | 1 | 1 | 1 | 7 | 1 |
| 13_634503 | 1 | 1 | 5 | 0.8 | NA | NA | 3 | 0.6667 | 9 | 0.7778 | 1 | 1 | 4 | 0.25 | 2 | 0.5 | 1 | 1 | 8 | 0.5 |
| 13_634503 | 1 | 1 | 4 | 0.75 | NA | NA | 5 | 0.2 | 10 | 0.5 | 1 | 0 | 2 | 0 | 2 | 0.5 | 1 | 1 | 6 | 0.3333 |
| 13_634504 | 1 | 0 | 5 | 0.8 | NA | NA | 5 | 0.8 | 11 | 0.7273 | 1 | 1 | 4 | 0.25 | 2 | 1 | 1 | 0 | 8 | 0.5 |
| 13_634504 | 2 | 0.5 | 2 | 0.5 | NA | NA | 4 | 0.5 | 8 | 0.5 | 1 | 0 | 3 | 0.6667 | 1 | 1 | 1 | 1 | 6 | 0.6667 |
| 13_634505 | 2 | 0.5 | 5 | 0.6 | NA | NA | 4 | 0.5 | 11 | 0.5455 | 1 | 0 | 3 | 0.6667 | 1 | 1 | 1 | 1 | 6 | 0.6667 |
| 13_634505 | 2 | 0.5 | 4 | 1 | NA | NA | 4 | 1 | 10 | 0.9 | NA | NA | 3 | 1 | 2 | 0.5 | 1 | 1 | 6 | 0.8333 |
| 13_634506 | 1 | 0 | 5 | 0 | NA | NA | 3 | 0.3333 | 9 | 0.1111 | 1 | 0 | 3 | 0.3333 | 3 | 0.3333 | 1 | 1 | 8 | 0.375 |
| 13_634601 | 1 | 0 | NA | NA | 3 | 0.3333 | NA | NA | 4 | 0.25 | 2 | 0.5 | 1 | 0 | NA | NA | NA | NA | 3 | 0.3333 |
| 13_634603 | 1 | 0 | NA | NA | 3 | 0.3333 | NA | NA | 4 | 0.25 | 2 | 0.5 | 1 | 0 | NA | NA | NA | NA | 3 | 0.3333 |
| 13_636563 | 6 | 0.1667 | 12 | 0.1667 | 1 | 0 | 7 | 0 | 26 | 0.1154 | 4 | 0.25 | 6 | 0 | 4 | 0.25 | 9 | 0 | 23 | 0.087 |
| 13_636566 | 5 | 0.4 | 9 | 0.2222 | 2 | 0 | 7 | 0.1429 | 23 | 0.2174 | 4 | 0.5 | 5 | 0.4 | 3 | 0 | 10 | 0.1 | 22 | 0.2273 |
| 13_636588 | 3 | 0 | 6 | 0.1667 | 1 | 0 | 4 | 0.25 | 14 | 0.1429 | 2 | 0.5 | 2 | 0 | 3 | 0 | 2 | 0.5 | 9 | 0.2222 |
| 13_636588 | 3 | 0.3333 | 5 | 0 | 1 | 0 | 5 | 0 | 14 | 0.0714 | 2 | 0.5 | 2 | 0 | 2 | 0 | 2 | 0.5 | 8 | 0.25 |
| 13_636594 | 3 | 0.3333 | 3 | 0.6667 | 1 | 1 | 4 | 0.75 | 11 | 0.6364 | 2 | 0.5 | 1 | 1 | 4 | 0.5 | 3 | 1 | 10 | 0.7 |
| 13_636594 | 3 | 0.6667 | 2 | 0.5 | 1 | 0 | 4 | 0.75 | 10 | 0.6 | 1 | 1 | 1 | 0 | 2 | 1 | 3 | 1 | 7 | 0.8571 |
| 13_636595 | 1 | 1 | 4 | 0.25 | 1 | 1 | 3 | 0.6667 | 9 | 0.5556 | 2 | 0 | 1 | 0 | 2 | 0.5 | 1 | 0 | 6 | 0.1667 |
| 13_636595 | 2 | 0.5 | 4 | 1 | 1 | 0 | 4 | 0.75 | 11 | 0.7273 | 1 | 1 | 1 | 0 | 3 | 1 | 2 | 1 | 7 | 0.8571 |
| 13_636596 | 1 | 1 | 4 | 0.75 | 1 | 1 | 2 | 0.5 | 8 | 0.75 | 1 | 1 | 1 | 1 | 2 | 1 | 3 | 1 | 7 | 1 |
| 13_636597 | 2 | 0 | 5 | 0.2 | 1 | 0 | 4 | 0.25 | 12 | 0.1667 | 1 | 1 | 1 | 0 | 4 | 0.5 | 1 | 0 | 7 | 0.4286 |
| 13_636598 | 2 | 0 | 3 | 1 | 1 | 1 | 3 | 1 | 9 | 0.7778 | 1 | 1 | 1 | 1 | 4 | 0.5 | 1 | 1 | 7 | 0.7143 |
| 13_636803 | 2 | 0 | 1 | 1 | NA | NA | 1 | 0 | 4 | 0.25 | 1 | 0 | NA | NA | 3 | 0.3333 | 1 | 0 | 5 | 0.2 |
| 13_640197 | NA | NA | NA | NA | 2 | 0.5 | 2 | 1 | 4 | 0.75 | 1 | 0 | NA | NA | 1 | 1 | NA | NA | 2 | 0.5 |
| 13_640197 | NA | NA | NA | NA | 1 | 1 | 1 | 0 | 2 | 0.5 | 1 | 0 | NA | NA | 1 | 1 | NA | NA | 2 | 0.5 |
| 13_640199 | NA | NA | NA | NA | 2 | 0.5 | 1 | 1 | 3 | 0.6667 | 1 | 0 | NA | NA | 1 | 1 | NA | NA | 2 | 0.5 |
| 13_640199 | NA | NA | NA | NA | 2 | 1 | 2 | 1 | 4 | 1 | 1 | 1 | NA | NA | NA | NA | NA | NA | 1 | 1 |
| 13_640200 | NA | NA | NA | NA | 2 | 0.5 | 2 | 1 | 4 | 0.75 | 1 | 1 | NA | NA | 1 | 1 | NA | NA | 2 | 1 |
| 13_640200 | NA | NA | 1 | 1 | 2 | 0 | 2 | 1 | 5 | 0.6 | 1 | 0 | NA | NA | 1 | 1 | NA | NA | 2 | 0.5 |

| | | | | | | | | | | | | | | | | | | | | |
|---|---|---|---|---|---|---|---|---|---|---|---|---|---|---|---|---|---|---|---|---|
| 13_640202 | NA | NA | 1 | 1 | 3 | 0 | 1 | 0 | 5 | 0.2 | 1 | 0 | NA | NA | 1 | 1 | NA | NA | 2 | 0.5 |
| 13_640205 | NA | NA | 1 | 1 | 2 | 0.5 | 2 | 1 | 5 | 0.8 | 1 | 1 | NA | NA | 1 | 1 | NA | NA | 2 | 1 |
| 13_640464 | NA | NA | 1 | 0 | 3 | 0.6667 | 3 | 0.3333 | 7 | 0.4286 | 1 | 0 | NA | NA | 1 | 1 | NA | NA | 2 | 0.5 |
| 13_640467 | NA | NA | 1 | 0 | 3 | 0.3333 | 2 | 0 | 6 | 0.1667 | 1 | 1 | NA | NA | 1 | 1 | NA | NA | 2 | 1 |
| 13_640467 | NA | NA | 1 | 1 | 2 | 0.5 | 2 | 0 | 5 | 0.4 | 1 | 1 | NA | NA | 1 | 0 | NA | NA | 2 | 0.5 |
| 13_640469 | NA | NA | 1 | 0 | 2 | 0.5 | 1 | 0 | 4 | 0.25 | 1 | 1 | 1 | 0 | 1 | 0 | NA | NA | 3 | 0.3333 |
| 13_640470 | NA | NA | 1 | 1 | 3 | 0.6667 | 3 | 0 | 7 | 0.4286 | 1 | 1 | 1 | 0 | 1 | 0 | NA | NA | 3 | 0.3333 |
| 13_640471 | NA | NA | 1 | 1 | 3 | 0.6667 | 3 | 1 | 7 | 0.8571 | 1 | 1 | 1 | 0 | 1 | 1 | NA | NA | 3 | 0.6667 |
| 13_640471 | NA | NA | 1 | 1 | 3 | 0.3333 | 2 | 1 | 6 | 0.6667 | 1 | 0 | 1 | 1 | 1 | 0 | NA | NA | 3 | 0.3333 |
| 13_640472 | NA | NA | 1 | 0 | 3 | 0.3333 | NA | NA | 4 | 0.25 | 1 | 0 | 2 | 0.5 | 1 | 0 | NA | NA | 4 | 0.25 |
| 13_640473 | NA | NA | 1 | 1 | 3 | 0.6667 | 2 | 1 | 6 | 0.8333 | NA | NA | 2 | 1 | 1 | 1 | NA | NA | 3 | 1 |
| 13_640473 | NA | NA | 1 | 1 | 2 | 1 | 2 | 1 | 5 | 1 | 1 | 0 | 1 | 0 | 1 | 1 | NA | NA | 3 | 0.3333 |
| 13_640474 | NA | NA | 1 | 0 | 3 | 0 | 2 | 0.5 | 6 | 0.1667 | 1 | 0 | 2 | 0.5 | 1 | 0 | NA | NA | 4 | 0.25 |
| 13_640475 | NA | NA | 1 | 1 | 3 | 0.3333 | 2 | 0.5 | 6 | 0.5 | 1 | 0 | 2 | 0.5 | 1 | 0 | NA | NA | 4 | 0.25 |
| 13_643100 | 11 | 0.0909 | 21 | 0 | 8 | 0.125 | 20 | 0.15 | 60 | 0.0833 | 10 | 0.1 | 11 | 0.0909 | 11 | 0.0909 | 9 | 0 | 41 | 0.0732 |
| 13_643331 | 14 | 0.2857 | 33 | 0.1515 | 15 | 0.1333 | 26 | 0.1538 | 88 | 0.1705 | 20 | 0.35 | 20 | 0.25 | 25 | 0.2 | 20 | 0.15 | 85 | 0.2353 |
| 13_643332 | 15 | 0.0667 | 33 | 0.1212 | 19 | 0.0526 | 27 | 0.1481 | 94 | 0.1064 | 21 | 0.0476 | 24 | 0.0417 | 28 | 0.1071 | 22 | 0.0455 | 95 | 0.0632 |
| 13_643332 | 14 | 0 | 29 | 0.0345 | 17 | 0.1176 | 25 | 0.12 | 85 | 0.0706 | 19 | 0.3158 | 21 | 0.0476 | 23 | 0.0435 | 20 | 0.1 | 83 | 0.1205 |
| 13_643333 | 10 | 0 | 25 | 0.16 | 20 | 0.25 | 22 | 0.1818 | 77 | 0.1688 | 18 | 0.2222 | 22 | 0.0909 | 25 | 0.28 | 21 | 0.381 | 86 | 0.2442 |
| 13_643338 | 11 | 0.4545 | 32 | 0.3438 | 22 | 0.3182 | 25 | 0.36 | 90 | 0.3556 | 15 | 0.5333 | 23 | 0.3478 | 25 | 0.4 | 24 | 0.3333 | 87 | 0.3908 |
| 13_643342 | 14 | 0.1429 | 28 | 0.1786 | 21 | 0.1429 | 31 | 0.1613 | 94 | 0.1596 | 21 | 0 | 21 | 0.0476 | 28 | 0.0714 | 23 | 0.1739 | 93 | 0.0753 |
| 13_645118 | 16 | 0 | 17 | 0.1176 | 17 | 0.0588 | 20 | 0.1 | 70 | 0.0714 | 14 | 0.0714 | 13 | 0 | 12 | 0 | 21 | 0.0952 | 60 | 0.05 |
| 13_645118 | 13 | 0.1538 | 17 | 0.1176 | 15 | 0.2 | 18 | 0.1667 | 63 | 0.1587 | 16 | 0.3125 | 13 | 0.2308 | 11 | 0 | 21 | 0.0476 | 61 | 0.1475 |
| 13_645118 | 16 | 0.4375 | 18 | 0.5 | 16 | 0.4375 | 17 | 0.5882 | 67 | 0.4925 | 15 | 0.4 | 10 | 0.8 | 10 | 0.3 | 21 | 0.6667 | 56 | 0.5536 |
| 13_645119 | 9 | 0 | 17 | 0.1176 | 17 | 0.0588 | 13 | 0.1538 | 56 | 0.0893 | 13 | 0.0769 | 10 | 0 | 10 | 0.1 | 20 | 0.1 | 53 | 0.0755 |
| 13_645148 | 8 | 0 | 12 | 0.0833 | 11 | 0 | 14 | 0.1429 | 45 | 0.0667 | 10 | 0.1 | 8 | 0 | 9 | 0.2222 | 19 | 0.1053 | 46 | 0.1087 |
| 13_645867 | NA | NA | NA | NA | NA | NA | 3 | 0.6667 | 3 | 0.6667 | 3 | 1 | NA | NA | 2 | 1 | 3 | 0.6667 | 8 | 0.875 |
| 13_645869 | NA | NA | NA | NA | 1 | 1 | 2 | 1 | 3 | 1 | 3 | 1 | NA | NA | 3 | 0.6667 | 3 | 1 | 9 | 0.8889 |
| 13_645869 | NA | NA | NA | NA | NA | NA | 1 | 1 | 1 | 1 | 2 | 1 | NA | NA | 3 | 0.6667 | 3 | 0 | 8 | 0.5 |
| 13_645870 | NA | NA | NA | NA | 1 | 1 | 2 | 1 | 3 | 1 | 3 | 1 | NA | NA | 3 | 0.3333 | 3 | 0.6667 | 9 | 0.6667 |
| 13_645871 | NA | NA | NA | NA | 1 | 1 | 2 | 0.5 | 3 | 0.6667 | 3 | 0.6667 | NA | NA | 3 | 0.3333 | 3 | 0 | 9 | 0.3333 |
| 13_645872 | NA | NA | NA | NA | 1 | 1 | 2 | 0.5 | 3 | 0.6667 | 2 | 1 | NA | NA | 3 | 1 | 3 | 0.6667 | 8 | 0.875 |
| 13_645874 | NA | NA | NA | NA | 1 | 1 | 1 | 1 | 2 | 1 | 2 | 1 | NA | NA | 4 | 0.5 | 1 | 0 | 7 | 0.5714 |
| 13_648848 | 4 | 0.25 | 1 | 0 | NA | NA | 4 | 0.25 | 9 | 0.2222 | NA | NA | 2 | 0.5 | 2 | 1 | 2 | 0 | 6 | 0.5 |
| 13_648857 | 3 | 0.3333 | NA | NA | NA | NA | 3 | 1 | 6 | 0.6667 | NA | NA | 2 | 0.5 | 3 | 0.6667 | 2 | 1 | 7 | 0.7143 |
| 13_648860 | 3 | 0.3333 | 1 | 0 | NA | NA | 4 | 0 | 8 | 0.125 | 1 | 1 | 1 | 1 | 3 | 0.3333 | 2 | 0.5 | 7 | 0.5714 |
| 13_648860 | 3 | 0.6667 | 1 | 0 | NA | NA | 3 | 0.3333 | 7 | 0.4286 | 1 | 0 | 2 | 0.5 | 3 | 0 | 2 | 0.5 | 8 | 0.25 |
| 13_648862 | 1 | 1 | 1 | 1 | NA | NA | 1 | 1 | 3 | 1 | 1 | 1 | 1 | 0 | 3 | 0.6667 | 2 | 1 | 7 | 0.7143 |
| 13_648863 | 3 | 1 | 1 | 1 | NA | NA | 4 | 0.5 | 8 | 0.75 | 1 | 1 | 1 | 1 | 3 | 1 | 2 | 0.5 | 7 | 0.8571 |
| 13_648863 | 3 | 0.3333 | 1 | 0 | NA | NA | 4 | 0.75 | 8 | 0.5 | 1 | 0 | 2 | 1 | 3 | 0 | 2 | 1 | 8 | 0.5 |
| 13_648863 | 1 | 1 | 1 | 0 | NA | NA | 4 | 0.5 | 6 | 0.5 | 1 | 1 | 1 | 0 | 3 | 0.3333 | 2 | 0.5 | 7 | 0.4286 |
| 13_652130 | 6 | 0.1667 | 16 | 0.0625 | 8 | 0 | 7 | 0.1429 | 37 | 0.0811 | 7 | 0 | 5 | 0 | 10 | 0.1 | 14 | 0.1429 | 36 | 0.0833 |
| 13_652132 | 7 | 0.5714 | 13 | 0.6154 | 6 | 0.5 | 8 | 0.875 | 34 | 0.6471 | 6 | 0.6667 | 7 | 0.5714 | 7 | 0.5714 | 11 | 0.6364 | 31 | 0.6129 |
| 13_652133 | 7 | 0.1429 | 15 | 0.0667 | 6 | 0.1667 | 7 | 0.2857 | 35 | 0.1429 | 5 | 0 | 7 | 0.4286 | 8 | 0.125 | 11 | 0.2727 | 31 | 0.2258 |
| 13_652143 | 5 | 0 | 10 | 0.1 | 3 | 0.3333 | 3 | 0.6667 | 21 | 0.1905 | 2 | 0 | 7 | 0.4286 | 7 | 0 | 7 | 0 | 23 | 0.1304 |
| 13_652145 | 5 | 1 | 9 | 0.7778 | 2 | 0.5 | 3 | 1 | 19 | 0.8421 | 1 | 1 | 7 | 1 | 6 | 0.8333 | 7 | 1 | 21 | 0.9524 |
| 13_652145 | 5 | 0 | 5 | 0.2 | 2 | 0 | 4 | 0 | 16 | 0.0625 | 2 | 0 | 5 | 0 | 6 | 0 | 6 | 0.1667 | 19 | 0.0526 |
| 13_652157 | 6 | 0.3333 | 9 | 0.1111 | 2 | 0 | 4 | 0.25 | 21 | 0.1905 | 2 | 0.5 | 5 | 0.2 | 7 | 0 | 4 | 0.25 | 18 | 0.1667 |
| 13_654442 | NA | NA | NA | NA | NA | NA | 1 | 1 | 1 | 1 | 1 | 1 | NA | NA | NA | NA | NA | NA | 1 | 1 |
| 13_654511 | 4 | 0 | 7 | 0.2857 | 1 | 0 | 2 | 0.5 | 14 | 0.2143 | 2 | 0 | 2 | 0.5 | 3 | 0.3333 | 2 | 0 | 9 | 0.2222 |
| 13_654515 | 4 | 0.5 | 7 | 0.4286 | 1 | 0 | 3 | 0.3333 | 15 | 0.4 | 4 | 0 | 2 | 0 | 5 | 0.2 | 2 | 0 | 13 | 0.0769 |
| 13_654515 | 5 | 1 | 6 | 0.8333 | 1 | 0 | 3 | 1 | 15 | 0.8667 | 4 | 1 | 2 | 1 | 4 | 1 | 2 | 1 | 12 | 1 |
| 13_654517 | 5 | 0.2 | 7 | 0.1429 | 1 | 0 | 3 | 0.3333 | 16 | 0.1875 | 4 | 0 | 3 | 0 | 5 | 0.4 | 2 | 0 | 14 | 0.1429 |
| 13_670603 | NA | NA | 2 | 0.5 | 2 | 0.5 | 4 | 0 | 8 | 0.25 | NA | NA | 1 | 0 | 2 | 0.5 | 4 | 0.75 | 7 | 0.5714 |
| 13_670780 | NA | NA | NA | NA | 4 | 0.5 | 4 | 0.25 | 8 | 0.375 | NA | NA | 1 | 0 | 3 | 0.6667 | 3 | 1 | 7 | 0.7143 |
| 13_670780 | NA | NA | NA | NA | 4 | 0.25 | 3 | 0.3333 | 7 | 0.2857 | NA | NA | 1 | 0 | 3 | 0.6667 | 3 | 0.6667 | 7 | 0.5714 |
| 13_670806 | NA | NA | NA | NA | NA | NA | 1 | 1 | 1 | 1 | NA | NA | NA | NA | 1 | 1 | NA | NA | 1 | 1 |
| 13_670812 | 1 | 0 | 1 | 1 | NA | NA | NA | NA | 2 | 0.5 | 1 | 1 | 1 | 0 | NA | NA | NA | NA | 2 | 0.5 |
| 13_671452 | 8 | 1 | 6 | 0.5 | 12 | 0.8333 | 12 | 0.5 | 38 | 0.7105 | 8 | 0.75 | 7 | 0.2857 | 3 | 0 | 5 | 0.6 | 23 | 0.4783 |
| 13_671452 | 8 | 0.25 | 6 | 0.1667 | 10 | 0.3 | 12 | 0.1667 | 36 | 0.2222 | 6 | 0.1667 | 5 | 0.4 | 3 | 0.6667 | 5 | 0.4 | 19 | 0.3684 |
| 13_671452 | 8 | 0.25 | 6 | 0.1667 | 12 | 0.0833 | 13 | 0.0769 | 39 | 0.1282 | 7 | 0.2857 | 9 | 0.1111 | 5 | 0.4 | 6 | 0.1667 | 27 | 0.2222 |
| 13_671452 | 9 | 0 | 5 | 0 | 11 | 0.0909 | 14 | 0.2143 | 39 | 0.1026 | 6 | 0 | 9 | 0.3333 | 3 | 0 | 7 | 0 | 25 | 0.12 |
| 13_671453 | 8 | 0.75 | 7 | 0.7143 | 6 | 0.6667 | 12 | 0.4167 | 33 | 0.6061 | 6 | 0.1667 | 6 | 0.5 | 3 | 1 | 7 | 0.7143 | 22 | 0.5455 |
| 13_671454 | 9 | 0.4444 | 6 | 0.1667 | 11 | 0.4545 | 12 | 0.25 | 38 | 0.3421 | 7 | 0.1429 | 7 | 0.2857 | 5 | 0.2 | 7 | 0.1429 | 26 | 0.1923 |
| 13_671455 | 8 | 0.5 | 7 | 0.1429 | 11 | 0.0909 | 12 | 0.5 | 38 | 0.3158 | 8 | 0.5 | 5 | 0.4 | 4 | 0.25 | 5 | 0.4 | 22 | 0.4091 |

| | | | | | | | | | | | | | | | | | | | | |
|---|---|---|---|---|---|---|---|---|---|---|---|---|---|---|---|---|---|---|---|---|
| 13_671457 | 10 | 0.9 | 7 | 0.7143 | 12 | 0.5 | 14 | 0.7143 | 43 | 0.6977 | 6 | 1 | 8 | 1 | 5 | 0.8 | 7 | 0.8571 | 26 | 0.9231 |
| 13_671458 | 10 | 0 | 8 | 0.125 | 12 | 0 | 14 | 0.1429 | 44 | 0.0682 | 8 | 0 | 8 | 0 | 5 | 0.2 | 7 | 0.1429 | 28 | 0.0714 |
| 13_671458 | 9 | 0.5556 | 7 | 0.2857 | 7 | 0.2857 | 9 | 0.1111 | 32 | 0.3125 | 4 | 1 | 6 | 0.1667 | 3 | 0.3333 | 6 | 0.1667 | 19 | 0.3684 |
| 13_671868 | NA | NA | NA | NA | 1 | 1 | 1 | 1 | 2 | 1 | NA | NA | NA | NA | NA | NA | 1 | 1 | 1 | 1 |
| 13_671869 | NA | NA | NA | NA | 1 | 1 | 1 | 0 | 2 | 0.5 | NA | NA | NA | NA | NA | NA | 1 | 1 | 1 | 1 |
| 13_671877 | NA | NA | NA | NA | 1 | 1 | 1 | 0 | 2 | 0.5 | NA | NA | NA | NA | NA | NA | 1 | 1 | 1 | 1 |
| 13_671881 | NA | NA | NA | NA | NA | NA | 1 | 1 | 1 | 1 | NA | NA | NA | NA | NA | NA | 1 | 1 | 1 | 1 |
| 13_671881 | NA | NA | NA | NA | 1 | 1 | 1 | 1 | 2 | 1 | NA | NA | NA | NA | NA | NA | 1 | 1 | 1 | 1 |
| 13_671883 | NA | NA | NA | NA | 1 | 0 | 1 | 1 | 2 | 0.5 | NA | NA | NA | NA | NA | NA | 1 | 1 | 1 | 1 |
| 13_671888 | NA | NA | NA | NA | 1 | 1 | 1 | 1 | 2 | 1 | NA | NA | NA | NA | NA | NA | 1 | 1 | 1 | 1 |
| 13_671895 | NA | NA | NA | NA | NA | NA | 1 | 1 | 1 | 1 | NA | NA | NA | NA | NA | NA | 1 | 1 | 1 | 1 |
| 13_671898 | NA | NA | NA | NA | 1 | 0 | 1 | 1 | 2 | 0.5 | NA | NA | NA | NA | NA | NA | 1 | 1 | 1 | 1 |
| 13_671901 | NA | NA | NA | NA | 1 | 1 | 1 | 1 | 2 | 1 | NA | NA | NA | NA | NA | NA | 1 | 1 | 1 | 1 |
| 13_672176 | NA | NA | NA | NA | 1 | 1 | NA | NA | 1 | 1 | NA | NA | NA | NA | 1 | 1 | NA | NA | 1 | 1 |
| 13_672179 | NA | NA | NA | NA | 1 | 1 | NA | NA | 1 | 1 | NA | NA | NA | NA | 1 | 1 | NA | NA | 1 | 1 |
| 13_672185 | NA | NA | NA | NA | 1 | 1 | NA | NA | 1 | 1 | NA | NA | NA | NA | 1 | 1 | NA | NA | 1 | 1 |
| 13_673299 | NA | NA | NA | NA | 1 | 1 | NA | NA | 1 | 1 | NA | NA | NA | NA | 1 | 1 | NA | NA | 1 | 1 |
| 13_673302 | NA | NA | NA | NA | 1 | 1 | NA | NA | 1 | 1 | NA | NA | NA | NA | 1 | 1 | NA | NA | 1 | 1 |
| 13_673302 | NA | NA | NA | NA | 1 | 1 | NA | NA | 1 | 1 | 1 | 1 | NA | NA | 1 | 0 | NA | NA | 2 | 0.5 |
| 13_673305 | NA | NA | NA | NA | 1 | 1 | 1 | 1 | 2 | 1 | 1 | 1 | NA | NA | NA | NA | NA | NA | 1 | 1 |
| 13_673306 | NA | NA | NA | NA | 1 | 0 | 1 | 1 | 2 | 0.5 | 2 | 1 | NA | NA | NA | NA | NA | NA | 2 | 1 |
| 13_673308 | NA | NA | NA | NA | 1 | 1 | 1 | 0 | 2 | 0.5 | 2 | 0.5 | NA | NA | 1 | 0 | NA | NA | 3 | 0.3333 |
| 13_673750 | 2 | 0.5 | NA | NA | NA | NA | 1 | 0 | 3 | 0.3333 | 1 | 0 | 1 | 1 | NA | NA | NA | NA | 2 | 0.5 |
| 13_673750 | 2 | 1 | NA | NA | NA | NA | 1 | 1 | 3 | 1 | 1 | 1 | 1 | 1 | NA | NA | NA | NA | 2 | 1 |
| 13_673750 | 2 | 1 | NA | NA | NA | NA | 1 | 0 | 3 | 0.6667 | 1 | 1 | NA | NA | NA | NA | NA | NA | 1 | 1 |
| 13_673757 | 1 | 1 | NA | NA | NA | NA | 1 | 1 | 2 | 1 | 1 | 1 | 1 | 0 | NA | NA | NA | NA | 2 | 0.5 |
| 13_673757 | 1 | 0 | NA | NA | NA | NA | 1 | 1 | 2 | 0.5 | 2 | 0 | 1 | 1 | NA | NA | NA | NA | 3 | 0.3333 |
| 13_673758 | 1 | 0 | NA | NA | NA | NA | 1 | 1 | 2 | 0.5 | 1 | 1 | 1 | 1 | NA | NA | NA | NA | 2 | 1 |
| 13_673760 | 1 | 0 | NA | NA | NA | NA | 1 | 1 | 2 | 0.5 | 1 | 1 | 1 | 1 | NA | NA | NA | NA | 2 | 1 |
| 13_673760 | 1 | 1 | NA | NA | NA | NA | 1 | 1 | 2 | 1 | 1 | 1 | 1 | 0 | NA | NA | NA | NA | 2 | 0.5 |
| 13_673761 | 1 | 1 | NA | NA | NA | NA | 1 | 1 | 2 | 1 | 1 | 1 | 1 | 0 | NA | NA | NA | NA | 2 | 0.5 |
| 13_674030 | 1 | 0 | NA | NA | NA | NA | 1 | 1 | 2 | 0.5 | 1 | 1 | 1 | 1 | NA | NA | NA | NA | 2 | 1 |
| 13_674030 | 1 | 1 | NA | NA | NA | NA | 1 | 0 | 2 | 0.5 | 1 | 1 | 1 | 0 | NA | NA | NA | NA | 2 | 0.5 |
| 13_674031 | 1 | 0 | NA | NA | NA | NA | 1 | 1 | 2 | 0.5 | NA | NA | 1 | 1 | NA | NA | NA | NA | 1 | 1 |
| 13_674032 | NA | NA | NA | NA | NA | NA | 1 | 1 | 1 | 1 | 1 | 1 | NA | NA | NA | NA | NA | NA | 1 | 1 |
| 13_674033 | NA | NA | NA | NA | NA | NA | 1 | 1 | 1 | 1 | 1 | 1 | 1 | 1 | NA | NA | NA | NA | 2 | 1 |
| 13_674045 | NA | NA | NA | NA | NA | NA | 1 | 1 | 1 | 1 | 1 | 1 | NA | NA | NA | NA | NA | NA | 1 | 1 |
| 13_674046 | NA | NA | NA | NA | NA | NA | 1 | 1 | 1 | 1 | 1 | 1 | NA | NA | NA | NA | NA | NA | 1 | 1 |
| 13_675379 | NA | NA | NA | NA | 1 | 0 | 1 | 1 | 2 | 0.5 | 1 | 1 | NA | NA | NA | NA | NA | NA | 1 | 1 |
| 13_675384 | NA | NA | NA | NA | NA | NA | 1 | 1 | 1 | 1 | 1 | 1 | NA | NA | NA | NA | NA | NA | 1 | 1 |
| 13_675887 | 1 | 0 | 3 | 0 | NA | NA | 3 | 0.3333 | 7 | 0.1429 | 1 | 0 | 4 | 0.25 | NA | NA | 4 | 0 | 9 | 0.1111 |
| 13_675890 | 1 | 0 | 1 | 0 | NA | NA | 2 | 0.5 | 4 | 0.25 | 1 | 0 | 4 | 0.25 | NA | NA | 4 | 0 | 9 | 0.1111 |
| 13_675895 | NA | NA | 1 | 1 | NA | NA | 2 | 0 | 3 | 0.3333 | 1 | 1 | 4 | 0.5 | NA | NA | 3 | 0 | 8 | 0.375 |
| 13_675899 | NA | NA | 1 | 0 | NA | NA | 1 | 1 | 2 | 0.5 | 1 | 0 | 3 | 0.3333 | NA | NA | 3 | 0.3333 | 7 | 0.2857 |
| 13_675901 | NA | NA | 1 | 1 | NA | NA | 2 | 0.5 | 3 | 0.6667 | 2 | 1 | 3 | 0.6667 | NA | NA | 5 | 0.8 | 10 | 0.8 |
| 13_675902 | NA | NA | 1 | 1 | NA | NA | 2 | 1 | 3 | 1 | 1 | 1 | 4 | 0.25 | NA | NA | 4 | 1 | 9 | 0.6667 |
| 13_675902 | NA | NA | 1 | 1 | NA | NA | 2 | 1 | 3 | 1 | 1 | 1 | 4 | 1 | NA | NA | 4 | 1 | 9 | 1 |
| 13_675903 | 1 | 1 | 1 | 1 | NA | NA | 2 | 1 | 4 | 1 | 2 | 1 | 4 | 1 | NA | NA | 3 | 1 | 9 | 1 |
| 13_675903 | 1 | 1 | 1 | 1 | NA | NA | 2 | 1 | 4 | 1 | 2 | 1 | 4 | 0.75 | NA | NA | 5 | 1 | 11 | 0.9091 |
| 13_675908 | NA | NA | NA | NA | NA | NA | 1 | 1 | 1 | 1 | 1 | 1 | 3 | 0.6667 | NA | NA | 4 | 1 | 8 | 0.875 |
| 13_675912 | NA | NA | NA | NA | NA | NA | 2 | 0.5 | 2 | 0.5 | 1 | 1 | 2 | 1 | NA | NA | 4 | 0.75 | 7 | 0.8571 |
| 13_675912 | NA | NA | 1 | 1 | NA | NA | 2 | 1 | 3 | 1 | 1 | 1 | 3 | 1 | NA | NA | 3 | 1 | 7 | 1 |
| 13_675913 | NA | NA | 1 | 0 | NA | NA | 2 | 0.5 | 3 | 0.3333 | 1 | 0 | 4 | 0.75 | NA | NA | 3 | 1 | 8 | 0.75 |
| 13_676196 | 3 | 0.3333 | 1 | 0 | 1 | 1 | 2 | 0 | 7 | 0.2857 | NA | NA | NA | NA | 1 | 0 | 3 | 0.3333 | 4 | 0.25 |
| 13_676199 | 3 | 0.6667 | 1 | 0 | 1 | 1 | 4 | 0.5 | 9 | 0.5556 | NA | NA | NA | NA | NA | NA | 3 | 0.6667 | 3 | 0.6667 |
| 13_676200 | 3 | 0 | 2 | 0.5 | 3 | 0.6667 | 4 | 0.5 | 12 | 0.4167 | NA | NA | NA | NA | NA | NA | 3 | 0.3333 | 3 | 0.3333 |
| 13_676205 | 4 | 0.25 | 2 | 0 | 2 | 0 | 4 | 0.25 | 12 | 0.1667 | NA | NA | NA | NA | 2 | 0 | 4 | 0.25 | 6 | 0.1667 |
| 13_676206 | 5 | 0.2 | 2 | 0.5 | 2 | 0 | 5 | 0.4 | 14 | 0.2857 | NA | NA | NA | NA | 2 | 0.5 | 5 | 0.4 | 7 | 0.4286 |
| 13_676207 | 2 | 0.5 | 2 | 0.5 | 1 | 1 | 3 | 1 | 8 | 0.75 | NA | NA | NA | NA | 1 | 0 | 3 | 1 | 4 | 0.75 |
| 13_676209 | 3 | 1 | 1 | 1 | 2 | 1 | 4 | 0.75 | 10 | 0.9 | NA | NA | NA | NA | 1 | 0 | 3 | 0.6667 | 4 | 0.5 |
| 13_676210 | 2 | 1 | 1 | 1 | 2 | 1 | 3 | 1 | 8 | 1 | NA | NA | NA | NA | 2 | 1 | 5 | 0.8 | 7 | 0.8571 |
| 13_676213 | 3 | 0 | 1 | 1 | 2 | 0.5 | 3 | 0.3333 | 9 | 0.3333 | NA | NA | NA | NA | 2 | 0.5 | 3 | 0.3333 | 5 | 0.4 |
| 13_676214 | 3 | 0 | NA | NA | 1 | 1 | NA | NA | 4 | 0.25 | NA | NA | NA | NA | 2 | 0.5 | 5 | 0.2 | 7 | 0.2857 |
| 13_676216 | 2 | 1 | NA | NA | 2 | 1 | 3 | 1 | 7 | 1 | NA | NA | NA | NA | NA | NA | 3 | 1 | 3 | 1 |
| 13_676216 | 3 | 0 | NA | NA | 2 | 0.5 | 3 | 0.3333 | 8 | 0.25 | NA | NA | NA | NA | 1 | 0 | 4 | 0.5 | 5 | 0.4 |

| | | | | | | | | | | | | | | | | | | | | |
|---|---|---|---|---|---|---|---|---|---|---|---|---|---|---|---|---|---|---|---|---|
| 13_6762187 | 2 | 1 | NA | NA | 2 | 1 | 2 | 1 | 6 | 1 | NA | NA | NA | NA | 1 | 1 | 4 | 1 | 5 | 1 |
| 13_6762197 | 2 | 0.5 | NA | NA | 2 | 0.5 | 1 | 0 | 5 | 0.4 | NA | NA | NA | NA | 1 | 1 | 3 | 0.6667 | 4 | 0.75 |
| 13_6762199 | 2 | 1 | NA | NA | 2 | 1 | 3 | 1 | 7 | 1 | NA | NA | NA | NA | 1 | 1 | 4 | 0.75 | 5 | 0.8 |
| 13_6762211 | 2 | 1 | NA | NA | 2 | 0.5 | 3 | 0.3333 | 7 | 0.5714 | NA | NA | NA | NA | 2 | 0 | 4 | 0.25 | 6 | 0.1667 |
| 13_6762225 | 2 | 0.5 | NA | NA | 2 | 0 | 2 | 0 | 6 | 0.1667 | NA | NA | NA | NA | 1 | 0 | 4 | 0.5 | 5 | 0.4 |
| 13_6762241 | 2 | 1 | NA | NA | 2 | 0.5 | 3 | 0.3333 | 7 | 0.5714 | NA | NA | NA | NA | 2 | 0.5 | 3 | 0.3333 | 5 | 0.4 |
| 13_6762255 | 2 | 1 | NA | NA | 2 | 0.5 | 2 | 0 | 6 | 0.5 | NA | NA | NA | NA | 1 | 0 | 4 | 0.5 | 5 | 0.4 |
| 13_6762290 | NA | NA | NA | NA | 2 | 0.5 | 5 | 0.4 | 7 | 0.4286 | NA | NA | NA | NA | 1 | 0 | 3 | 0.6667 | 4 | 0.5 |
| 13_6763866 | 1 | 1 | 3 | 0.6667 | 1 | 0 | NA | NA | 5 | 0.6 | NA | NA | 1 | 1 | 1 | 0 | NA | NA | 2 | 0.5 |
| 13_6763868 | 1 | 1 | 3 | 1 | 1 | 1 | NA | NA | 5 | 1 | NA | NA | 1 | 1 | NA | NA | NA | NA | 1 | 1 |
| 13_6763890 | NA | NA | 3 | 0.3333 | 1 | 0 | NA | NA | 4 | 0.25 | NA | NA | NA | NA | 1 | 1 | NA | NA | 1 | 1 |
| 13_6763904 | NA | NA | 2 | 0.5 | 1 | 0 | NA | NA | 3 | 0.3333 | NA | NA | NA | NA | 1 | 1 | NA | NA | 1 | 1 |
| 13_6763915 | NA | NA | 3 | 1 | 1 | 1 | NA | NA | 4 | 1 | NA | NA | NA | NA | 1 | 1 | NA | NA | 1 | 1 |
| 13_6763940 | NA | NA | 1 | 1 | NA | NA | NA | NA | 1 | 1 | NA | NA | NA | NA | 1 | 1 | NA | NA | 1 | 1 |
| 13_6763957 | NA | NA | 3 | 1 | NA | NA | NA | NA | 3 | 1 | NA | NA | NA | NA | 1 | 1 | NA | NA | 1 | 1 |
| 13_6766395 | 2 | 0 | 1 | 1 | NA | NA | 5 | 0.4 | 8 | 0.375 | NA | NA | 1 | 0 | 1 | 1 | NA | NA | 2 | 0.5 |
| 13_6766401 | 2 | 1 | 1 | 1 | NA | NA | 5 | 1 | 8 | 1 | NA | NA | NA | NA | 1 | 1 | NA | NA | 1 | 1 |
| 13_6766414 | 1 | 0 | 1 | 1 | NA | NA | 1 | 1 | 3 | 0.6667 | NA | NA | 1 | 1 | NA | NA | NA | NA | 1 | 1 |
| 13_6766424 | 2 | 1 | 2 | 0.5 | NA | NA | 2 | 1 | 6 | 0.8333 | NA | NA | 1 | 1 | 1 | 1 | NA | NA | 2 | 1 |
| 13_6766426 | 2 | 1 | 1 | 1 | NA | NA | 4 | 0.5 | 7 | 0.7143 | NA | NA | 1 | 1 | 1 | 1 | NA | NA | 2 | 1 |
| 13_6766430 | 2 | 0.5 | 2 | 1 | NA | NA | 5 | 0.4 | 9 | 0.5556 | NA | NA | NA | NA | 1 | 1 | NA | NA | 1 | 1 |
| 13_6766443 | 2 | 1 | 1 | 1 | NA | NA | 5 | 1 | 8 | 1 | NA | NA | 1 | 1 | 1 | 1 | NA | NA | 2 | 1 |
| 13_6766449 | 2 | 0.5 | 2 | 1 | NA | NA | 5 | 0.2 | 9 | 0.4444 | NA | NA | 1 | 1 | 1 | 0 | NA | NA | 2 | 0.5 |
| 13_6766463 | 1 | 1 | 1 | 1 | NA | NA | 5 | 1 | 7 | 1 | NA | NA | 1 | 1 | 1 | 1 | NA | NA | 2 | 1 |
| 13_6766473 | NA | NA | 1 | 1 | NA | NA | 4 | 0.5 | 5 | 0.6 | NA | NA | NA | NA | 1 | 1 | NA | NA | 1 | 1 |
| 13_6766475 | 1 | 1 | 1 | 1 | NA | NA | 4 | 0.75 | 6 | 0.8333 | NA | NA | 1 | 1 | 1 | 1 | NA | NA | 2 | 1 |
| 13_6766477 | 1 | 0 | 1 | 0 | NA | NA | 4 | 0.25 | 6 | 0.1667 | NA | NA | 1 | 0 | 1 | 1 | NA | NA | 2 | 0.5 |
| 13_6766495 | 1 | 1 | 1 | 1 | NA | NA | 3 | 0.6667 | 5 | 0.8 | NA | NA | NA | NA | 1 | 1 | NA | NA | 1 | 1 |
| 13_6766515 | 1 | 0 | 1 | 0 | NA | NA | 5 | 1 | 7 | 0.7143 | NA | NA | 1 | 1 | 1 | 1 | NA | NA | 2 | 1 |
| 13_6766525 | 1 | 1 | 1 | 1 | NA | NA | 3 | 0.3333 | 5 | 0.6 | NA | NA | 1 | 1 | NA | NA | NA | NA | 1 | 1 |
| 13_6766526 | 1 | 0 | 1 | 1 | NA | NA | 4 | 0.5 | 6 | 0.5 | NA | NA | 1 | 0 | 1 | 1 | NA | NA | 2 | 0.5 |
| 13_6766538 | 1 | 0 | 1 | 0 | NA | NA | 4 | 0.25 | 6 | 0.1667 | NA | NA | 1 | 1 | 1 | 0 | NA | NA | 2 | 0.5 |
| 13_6766557 | 1 | 1 | NA | NA | NA | NA | 4 | 0.75 | 5 | 0.8 | NA | NA | 1 | 0 | 1 | 1 | NA | NA | 2 | 0.5 |
| 13_6766590 | 1 | 1 | NA | NA | NA | NA | 3 | 1 | 4 | 1 | NA | NA | 1 | 1 | 1 | 1 | NA | NA | 2 | 1 |
| 13_6768715 | 1 | 0 | 1 | 1 | NA | NA | 1 | 0 | 3 | 0.3333 | NA | NA | 2 | 0.5 | NA | NA | 1 | 0 | 3 | 0.3333 |
| 13_6768740 | 1 | 1 | 1 | 1 | NA | NA | 3 | 0 | 5 | 0.4 | NA | NA | 3 | 0.3333 | 1 | 0 | 1 | 0 | 5 | 0.2 |
| 13_6768741 | 1 | 1 | 1 | 1 | NA | NA | 3 | 0 | 5 | 0.4 | NA | NA | 4 | 0.25 | 1 | 0 | 1 | 0 | 6 | 0.1667 |
| 13_6768754 | 1 | 1 | NA | NA | NA | NA | 1 | 1 | 2 | 1 | NA | NA | 4 | 0.5 | NA | NA | NA | NA | 4 | 0.5 |
| 13_6768759 | 1 | 1 | NA | NA | NA | NA | 3 | 0.6667 | 4 | 0.75 | NA | NA | 3 | 1 | 1 | 1 | NA | NA | 4 | 1 |
| 13_6768761 | 1 | 1 | NA | NA | NA | NA | 3 | 0.6667 | 4 | 0.75 | NA | NA | 4 | 1 | 1 | 1 | 1 | 1 | 6 | 1 |
| 13_6768773 | NA | NA | NA | NA | NA | NA | 3 | 0.3333 | 3 | 0.3333 | NA | NA | 4 | 0.25 | 1 | 0 | NA | NA | 5 | 0.2 |
| 13_6768775 | NA | NA | NA | NA | NA | NA | 3 | 0.6667 | 3 | 0.6667 | NA | NA | 4 | 0.5 | 1 | 0 | 1 | 0 | 6 | 0.3333 |
| 13_6768777 | NA | NA | NA | NA | NA | NA | 2 | 0.5 | 2 | 0.5 | NA | NA | 4 | 0.25 | 1 | 0 | NA | NA | 5 | 0.2 |
| 13_6768821 | 1 | 0 | NA | NA | NA | NA | 3 | 0.3333 | 4 | 0.25 | NA | NA | 3 | 0.3333 | 1 | 1 | NA | NA | 4 | 0.5 |
| 13_6768833 | 1 | 0 | NA | NA | NA | NA | 3 | 0.3333 | 4 | 0.25 | NA | NA | 2 | 0.5 | 1 | 0 | NA | NA | 3 | 0.3333 |
| 13_6768840 | NA | NA | NA | NA | NA | NA | 3 | 1 | 3 | 1 | NA | NA | 2 | 1 | 1 | 1 | NA | NA | 3 | 1 |
| 13_6768843 | NA | NA | NA | NA | NA | NA | 3 | 1 | 3 | 1 | NA | NA | 1 | 1 | 1 | 1 | NA | NA | 2 | 1 |
| 13_6768852 | 1 | 1 | NA | NA | NA | NA | 3 | 0.3333 | 4 | 0.5 | NA | NA | 2 | 0.5 | NA | NA | NA | NA | 2 | 0.5 |
| 13_6768862 | 1 | 1 | NA | NA | NA | NA | 2 | 0.5 | 3 | 0.6667 | NA | NA | 2 | 1 | NA | NA | NA | NA | 2 | 1 |
| 13_6768889 | 1 | 1 | NA | NA | NA | NA | 3 | 1 | 4 | 1 | NA | NA | 2 | 1 | NA | NA | NA | NA | 2 | 1 |
| 13_6768921 | 1 | 1 | NA | NA | NA | NA | 3 | 1 | 4 | 1 | NA | NA | 2 | 1 | NA | NA | NA | NA | 2 | 1 |
| 13_6768948 | 1 | 1 | NA | NA | NA | NA | 3 | 1 | 4 | 1 | NA | NA | 2 | 1 | NA | NA | NA | NA | 2 | 1 |
| 13_6768977 | 1 | 1 | NA | NA | NA | NA | 3 | 1 | 4 | 1 | NA | NA | 1 | 1 | NA | NA | NA | NA | 1 | 1 |
| 13_6768990 | 1 | 1 | NA | NA | NA | NA | 3 | 1 | 4 | 1 | NA | NA | 2 | 1 | NA | NA | NA | NA | 2 | 1 |
| 13_6769053 | 1 | 1 | NA | NA | NA | NA | 2 | 1 | 3 | 1 | NA | NA | 2 | 0.5 | NA | NA | NA | NA | 2 | 0.5 |
| 13_6769055 | NA | NA | NA | NA | NA | NA | 3 | 0.6667 | 3 | 0.6667 | NA | NA | 1 | 1 | NA | NA | NA | NA | 1 | 1 |
| 13_6769069 | NA | NA | NA | NA | NA | NA | 3 | 1 | 3 | 1 | NA | NA | 1 | 1 | NA | NA | NA | NA | 1 | 1 |
| 13_6776607 | 1 | 1 | NA | NA | 2 | 0.5 | NA | NA | 3 | 0.6667 | NA | NA | 1 | 0 | 1 | 1 | NA | NA | 2 | 0.5 |
| 13_6776611 | 1 | 0 | NA | NA | 2 | 0.5 | NA | NA | 3 | 0.3333 | NA | NA | 1 | 1 | 1 | 0 | NA | NA | 2 | 0.5 |
| 13_6776632 | NA | NA | NA | NA | 2 | 1 | NA | NA | 2 | 1 | NA | NA | 1 | 1 | NA | NA | NA | NA | 1 | 1 |
| 13_6776647 | NA | NA | NA | NA | 1 | 1 | NA | NA | 1 | 1 | NA | NA | NA | NA | 1 | 1 | NA | NA | 1 | 1 |
| 13_6776652 | NA | NA | NA | NA | 2 | 0.5 | NA | NA | 2 | 0.5 | NA | NA | 1 | 1 | 1 | 0 | NA | NA | 2 | 0.5 |
| 13_6776655 | NA | NA | NA | NA | 2 | 1 | NA | NA | 2 | 1 | NA | NA | 1 | 1 | 1 | 1 | NA | NA | 2 | 1 |
| 13_6776668 | NA | NA | NA | NA | 2 | 0.5 | NA | NA | 2 | 0.5 | NA | NA | 1 | 1 | 1 | 1 | NA | NA | 2 | 1 |
| 13_6776709 | NA | NA | NA | NA | 2 | 1 | NA | NA | 2 | 1 | NA | NA | 1 | 1 | NA | NA | NA | NA | 1 | 1 |

| | | | | | | | | | | | | | | | | | | | | |
|---|---|---|---|---|---|---|---|---|---|---|---|---|---|---|---|---|---|---|---|---|
| 13_6776751 | NA | NA | NA | NA | 2 | 0.5 | NA | NA | 2 | 0.5 | NA | NA | 1 | 1 | NA | NA | NA | NA | 1 | 1 |
| 13_6776808 | NA | NA | NA | NA | 2 | 1 | NA | NA | 2 | 1 | NA | NA | 1 | 1 | NA | NA | NA | NA | 1 | 1 |
| 13_6776888 | NA | NA | NA | NA | 2 | 0.5 | NA | NA | 2 | 0.5 | NA | NA | 1 | 1 | NA | NA | NA | NA | 1 | 1 |
| 13_6780697 | NA | NA | NA | NA | 1 | 1 | NA | NA | 1 | 1 | NA | NA | NA | NA | 1 | 0 | 1 | 1 | 2 | 0.5 |
| 13_6780788 | NA | NA | NA | NA | 2 | 0.5 | NA | NA | 2 | 0.5 | NA | NA | 1 | 1 | NA | NA | 1 | 0 | 2 | 0.5 |
| 13_6781550 | NA | NA | 3 | 0.6667 | 1 | 0 | NA | NA | 4 | 0.5 | 1 | 0 | NA | NA | 1 | 0 | 1 | 1 | 3 | 0.3333 |
| 13_6781560 | NA | NA | 4 | 0.25 | 3 | 0.3333 | NA | NA | 7 | 0.2857 | 1 | 1 | NA | NA | 1 | 0 | 1 | 0 | 3 | 0.3333 |
| 13_6781582 | NA | NA | 4 | 1 | 2 | 1 | NA | NA | 6 | 1 | 1 | 1 | NA | NA | 1 | 1 | NA | NA | 2 | 1 |
| 13_6781588 | NA | NA | 3 | 0.3333 | 2 | 0 | NA | NA | 5 | 0.2 | 2 | 0 | NA | NA | 1 | 1 | 1 | 1 | 4 | 0.5 |
| 13_6781589 | NA | NA | 4 | 0 | 2 | 0.5 | NA | NA | 6 | 0.1667 | 2 | 0 | NA | NA | 1 | 1 | 1 | 0 | 4 | 0.25 |
| 13_6781597 | NA | NA | 3 | 1 | 1 | 1 | NA | NA | 4 | 1 | 2 | 1 | NA | NA | 1 | 1 | NA | NA | 3 | 1 |
| 13_6781602 | NA | NA | 4 | 1 | 1 | 1 | NA | NA | 5 | 1 | 2 | 0.5 | NA | NA | 1 | 1 | 1 | 1 | 4 | 0.75 |
| 13_6781618 | NA | NA | 4 | 0.5 | 2 | 0.5 | NA | NA | 6 | 0.5 | 2 | 0 | NA | NA | 1 | 0 | 1 | 1 | 4 | 0.25 |
| 13_6781630 | NA | NA | 2 | 1 | 2 | 0.5 | NA | NA | 4 | 0.75 | 2 | 0 | NA | NA | 1 | 1 | NA | NA | 3 | 0.3333 |
| 13_6781630 | NA | NA | 1 | 1 | NA | NA | NA | NA | 1 | 1 | NA | NA | NA | NA | NA | NA | 1 | 1 | 1 | 1 |
| 13_6781639 | NA | NA | 4 | 1 | 2 | 1 | NA | NA | 6 | 1 | 2 | 1 | NA | NA | 1 | 0 | NA | NA | 3 | 0.6667 |
| 13_6781644 | NA | NA | 3 | 1 | 1 | 0 | NA | NA | 4 | 0.75 | 2 | 1 | NA | NA | 1 | 0 | NA | NA | 3 | 0.6667 |
| 13_6781659 | NA | NA | 2 | 1 | 2 | 0.5 | NA | NA | 4 | 0.75 | 1 | 0 | NA | NA | 1 | 1 | NA | NA | 2 | 0.5 |
| 13_6781661 | NA | NA | 2 | 1 | 2 | 0.5 | NA | NA | 4 | 0.75 | 2 | 1 | NA | NA | 1 | 1 | NA | NA | 3 | 1 |
| 13_6781664 | NA | NA | 4 | 0.75 | 2 | 1 | NA | NA | 6 | 0.8333 | 2 | 0.5 | NA | NA | 1 | 1 | NA | NA | 3 | 0.6667 |
| 13_6781667 | NA | NA | 4 | 0.5 | 2 | 0.5 | NA | NA | 6 | 0.5 | 2 | 0.5 | NA | NA | 1 | 0 | NA | NA | 3 | 0.3333 |
| 13_6781684 | NA | NA | 4 | 1 | 2 | 1 | NA | NA | 6 | 1 | 2 | 1 | NA | NA | 1 | 1 | NA | NA | 3 | 1 |
| 13_6781719 | NA | NA | 3 | 0.3333 | 2 | 1 | NA | NA | 5 | 0.6 | 2 | 1 | NA | NA | 1 | 0 | NA | NA | 3 | 0.6667 |
| 13_6781742 | NA | NA | 3 | 1 | 2 | 1 | NA | NA | 5 | 1 | 2 | 0.5 | NA | NA | 1 | 1 | NA | NA | 3 | 0.6667 |
| 13_6781751 | NA | NA | 3 | 1 | 2 | 1 | NA | NA | 5 | 1 | 2 | 1 | NA | NA | 1 | 1 | NA | NA | 3 | 1 |
| 13_6781765 | NA | NA | 1 | 1 | 1 | 1 | NA | NA | 2 | 1 | 2 | 1 | NA | NA | 1 | 1 | NA | NA | 3 | 1 |
| 13_6781799 | NA | NA | 3 | 0.6667 | 1 | 1 | NA | NA | 4 | 0.75 | 2 | 0.5 | NA | NA | 1 | 1 | NA | NA | 3 | 0.6667 |
| 13_6781810 | NA | NA | 3 | 1 | 1 | 1 | NA | NA | 4 | 1 | 1 | 1 | NA | NA | 1 | 0 | NA | NA | 2 | 0.5 |
| 13_6781850 | NA | NA | 3 | 0.6667 | 2 | 0 | NA | NA | 5 | 0.4 | 1 | 1 | NA | NA | 1 | 0 | NA | NA | 2 | 0.5 |
| 13_6781865 | NA | NA | 3 | 1 | 1 | 1 | NA | NA | 4 | 1 | 1 | 1 | NA | NA | 1 | 1 | NA | NA | 2 | 1 |
| 13_6781875 | NA | NA | 1 | 1 | NA | NA | NA | NA | 1 | 1 | 1 | 1 | NA | NA | 1 | 0 | NA | NA | 2 | 0.5 |
| 13_6781880 | NA | NA | 1 | 1 | NA | NA | NA | NA | 1 | 1 | 1 | 1 | NA | NA | 1 | 0 | NA | NA | 2 | 0.5 |
| 13_6781908 | NA | NA | 1 | 1 | NA | NA | NA | NA | 1 | 1 | 1 | 0 | NA | NA | 1 | 1 | NA | NA | 2 | 0.5 |
| 13_6785369 | 1 | 1 | NA | NA | 1 | 1 | 1 | 0 | 3 | 0.6667 | NA | NA | NA | NA | 1 | 1 | 2 | 0 | 3 | 0.3333 |
| 13_6785381 | 1 | 0 | NA | NA | 1 | 1 | NA | NA | 2 | 0.5 | NA | NA | NA | NA | 1 | 0 | 2 | 0.5 | 3 | 0.3333 |
| 13_6785391 | 1 | 1 | NA | NA | 1 | 0 | NA | NA | 2 | 0.5 | NA | NA | NA | NA | 1 | 0 | 2 | 0.5 | 3 | 0.3333 |
| 13_6785419 | 1 | 1 | NA | NA | NA | NA | NA | NA | 1 | 1 | NA | NA | NA | NA | 1 | 1 | 2 | 1 | 3 | 1 |
| 13_6785426 | 1 | 1 | NA | NA | 1 | 1 | NA | NA | 2 | 1 | NA | NA | NA | NA | 1 | 1 | 2 | 1 | 3 | 1 |
| 13_6785471 | 1 | 1 | NA | NA | NA | NA | NA | NA | 1 | 1 | NA | NA | NA | NA | 1 | 0 | 1 | 1 | 2 | 0.5 |
| 13_6785491 | 1 | 1 | NA | NA | NA | NA | NA | NA | 1 | 1 | NA | NA | NA | NA | NA | NA | 1 | 1 | 1 | 1 |
| 13_6785496 | 1 | 1 | NA | NA | NA | NA | NA | NA | 1 | 1 | NA | NA | NA | NA | 1 | 1 | 1 | 1 | 2 | 1 |
| 13_6785535 | 1 | 1 | NA | NA | NA | NA | NA | NA | 1 | 1 | NA | NA | NA | NA | 1 | 1 | NA | NA | 1 | 1 |
| 13_6785538 | 1 | 1 | NA | NA | NA | NA | NA | NA | 1 | 1 | NA | NA | NA | NA | 1 | 1 | NA | NA | 1 | 1 |
| 13_6785541 | 1 | 1 | NA | NA | NA | NA | NA | NA | 1 | 1 | NA | NA | NA | NA | 1 | 1 | NA | NA | 1 | 1 |
| 13_6789602 | 2 | 0 | 1 | 1 | NA | NA | 1 | 1 | 4 | 0.5 | 2 | 0.5 | 1 | 0 | NA | NA | 1 | 1 | 4 | 0.5 |
| 13_6789605 | 2 | 0.5 | 1 | 1 | NA | NA | 1 | 1 | 4 | 0.75 | 2 | 1 | 1 | 1 | NA | NA | 1 | 1 | 4 | 1 |
| 13_6789606 | 2 | 0.5 | 1 | 0 | NA | NA | 1 | 0 | 4 | 0.25 | 2 | 0 | 1 | 1 | NA | NA | 1 | 0 | 4 | 0.25 |
| 13_6789611 | 2 | 0.5 | 1 | 1 | NA | NA | 1 | 1 | 4 | 0.75 | 2 | 1 | 1 | 1 | NA | NA | 1 | 0 | 4 | 0.75 |
| 13_6789615 | 2 | 0.5 | NA | NA | NA | NA | NA | NA | 2 | 0.5 | 1 | 1 | NA | NA | NA | NA | 1 | 0 | 2 | 0.5 |
| 13_6789631 | NA | NA | 1 | 1 | NA | NA | 1 | 1 | 2 | 1 | 1 | 1 | 1 | 1 | NA | NA | NA | NA | 2 | 1 |
| 13_6789634 | 1 | 1 | 1 | 0 | NA | NA | 1 | 0 | 3 | 0.3333 | 2 | 0.5 | 1 | 0 | NA | NA | 1 | 0 | 4 | 0.25 |
| 13_6789635 | 1 | 0 | 1 | 0 | NA | NA | 1 | 1 | 3 | 0.3333 | 1 | 0 | 1 | 1 | NA | NA | 1 | 1 | 3 | 0.6667 |
| 13_6789643 | 1 | 1 | 1 | 1 | NA | NA | NA | NA | 2 | 1 | 2 | 0.5 | NA | NA | NA | NA | 1 | 0 | 3 | 0.3333 |
| 13_6789670 | 1 | 1 | NA | NA | NA | NA | NA | NA | 1 | 1 | 2 | 1 | NA | NA | NA | NA | 1 | 1 | 3 | 1 |
| 13_6789689 | 1 | 1 | 1 | 1 | NA | NA | NA | NA | 2 | 1 | 1 | 1 | NA | NA | NA | NA | 1 | 1 | 2 | 1 |
| 13_6789713 | 1 | 0 | 1 | 1 | NA | NA | NA | NA | 2 | 0.5 | 2 | 0.5 | NA | NA | NA | NA | 1 | 1 | 3 | 0.6667 |
| 13_6789722 | NA | NA | 1 | 1 | NA | NA | NA | NA | 1 | 1 | 1 | 1 | NA | NA | NA | NA | 1 | 1 | 2 | 1 |
| 13_6789753 | NA | NA | 1 | 1 | NA | NA | NA | NA | 1 | 1 | 2 | 1 | NA | NA | NA | NA | 1 | 1 | 3 | 1 |
| 13_6790600 | NA | NA | NA | NA | 1 | 1 | NA | NA | 1 | 1 | 1 | 1 | NA | NA | NA | NA | NA | NA | 1 | 1 |
| 13_6790614 | NA | NA | NA | NA | 1 | 1 | NA | NA | 1 | 1 | 1 | 1 | NA | NA | NA | NA | NA | NA | 1 | 1 |
| 13_6790683 | NA | NA | NA | NA | 1 | 1 | NA | NA | 1 | 1 | 1 | 1 | NA | NA | NA | NA | NA | NA | 1 | 1 |
| 13_6790683 | NA | NA | NA | NA | 1 | 1 | NA | NA | 1 | 1 | 1 | 1 | NA | NA | NA | NA | NA | NA | 1 | 1 |
| 13_6790695 | NA | NA | NA | NA | 1 | 1 | NA | NA | 1 | 1 | 1 | 1 | NA | NA | NA | NA | NA | NA | 1 | 1 |
| 13_6793460 | NA | NA | NA | NA | 1 | 1 | 1 | 1 | 2 | 1 | NA | NA | 1 | 1 | NA | NA | NA | NA | 1 | 1 |
| 13_6793481 | NA | NA | NA | NA | 1 | 1 | 1 | 1 | 2 | 1 | NA | NA | 1 | 1 | NA | NA | NA | NA | 1 | 1 |

| | | | | | | | | | | | | | | | | | | | | |
|---|---|---|---|---|---|---|---|---|---|---|---|---|---|---|---|---|---|---|---|---|
| 13_679348! | NA | NA | NA | NA | 1 | 1 | 2 | 0.5 | 3 | 0.6667 | NA | NA | 1 | 1 | NA | NA | NA | NA | 1 | 1 |
| 13_679350! | NA | NA | NA | NA | 1 | 1 | 2 | 1 | 3 | 1 | NA | NA | 1 | 1 | NA | NA | NA | NA | 1 | 1 |
| 13_679354( | NA | NA | NA | NA | 1 | 1 | 2 | 1 | 3 | 1 | NA | NA | 1 | 1 | NA | NA | NA | NA | 1 | 1 |
| 13_679355( | NA | NA | NA | NA | NA | NA | 2 | 1 | 2 | 1 | NA | NA | 1 | 1 | NA | NA | NA | NA | 1 | 1 |
| 13_679361 | NA | NA | NA | NA | NA | NA | 2 | 1 | 2 | 1 | NA | NA | 1 | 1 | NA | NA | NA | NA | 1 | 1 |
| 13_679365 | NA | NA | NA | NA | 1 | 1 | 2 | 1 | 3 | 1 | NA | NA | 1 | 1 | NA | NA | NA | NA | 1 | 1 |
| 13_679367 | NA | NA | NA | NA | 1 | 0 | 2 | 0.5 | 3 | 0.3333 | NA | NA | 1 | 1 | NA | NA | NA | NA | 1 | 1 |
| 13_679367 | NA | NA | NA | NA | 1 | 0 | 2 | 0.5 | 3 | 0.3333 | NA | NA | 1 | 1 | NA | NA | NA | NA | 1 | 1 |
| 13_679734 | NA | NA | 2 | 1 | NA | NA | 1 | 1 | 3 | 1 | NA | NA | 1 | 1 | NA | NA | NA | NA | 1 | 1 |
| 13_679735! | NA | NA | 2 | 1 | NA | NA | 1 | 1 | 3 | 1 | NA | NA | 1 | 1 | NA | NA | NA | NA | 1 | 1 |
| 13_679737! | NA | NA | 2 | 1 | NA | NA | 1 | 1 | 3 | 1 | NA | NA | 1 | 1 | NA | NA | NA | NA | 1 | 1 |
| 13_679741 | NA | NA | 2 | 1 | NA | NA | 1 | 1 | 3 | 1 | NA | NA | 1 | 1 | NA | NA | NA | NA | 1 | 1 |
| 13_679743( | NA | NA | 2 | 0.5 | NA | NA | 1 | 1 | 3 | 0.6667 | NA | NA | 1 | 1 | NA | NA | 1 | 0 | 2 | 0.5 |
| 13_679743 | NA | NA | 2 | 0.5 | NA | NA | 2 | 0.5 | 4 | 0.5 | NA | NA | 1 | 1 | NA | NA | NA | NA | 1 | 1 |
| 13_679745( | NA | NA | 2 | 0.5 | NA | NA | 1 | 1 | 3 | 0.6667 | NA | NA | NA | NA | NA | NA | 1 | 1 | 1 | 1 |
| 13_679746( | NA | NA | 2 | 1 | NA | NA | 2 | 0.5 | 4 | 0.75 | NA | NA | 1 | 1 | NA | NA | 1 | 1 | 2 | 1 |
| 13_679746 | NA | NA | 2 | 0.5 | NA | NA | 2 | 0.5 | 4 | 0.5 | NA | NA | 1 | 1 | NA | NA | 1 | 0 | 2 | 0.5 |
| 13_679747 | NA | NA | 2 | 1 | NA | NA | 2 | 1 | 4 | 1 | NA | NA | 1 | 1 | NA | NA | NA | NA | 1 | 1 |
| 13_679748 | NA | NA | 2 | 1 | NA | NA | 1 | 1 | 3 | 1 | NA | NA | 1 | 1 | 1 | 1 | NA | NA | 2 | 1 |
| 13_679749 | NA | NA | 2 | 0.5 | NA | NA | 2 | 0.5 | 4 | 0.5 | NA | NA | 1 | 1 | 1 | 1 | 1 | 1 | 3 | 1 |
| 13_679749 | NA | NA | 2 | 1 | NA | NA | 2 | 1 | 4 | 1 | NA | NA | 1 | 1 | 1 | 1 | NA | NA | 2 | 1 |
| 13_679750 | NA | NA | 1 | 1 | NA | NA | 2 | 0.5 | 3 | 0.6667 | NA | NA | 1 | 0 | 2 | 0.5 | NA | NA | 3 | 0.3333 |
| 13_687096! | NA | NA | 3 | 0 | NA | NA | 1 | 1 | 4 | 0.25 | 1 | 1 | 4 | 0 | 1 | 0 | NA | NA | 6 | 0.1667 |
| 13_687097 | NA | NA | 3 | 0.3333 | NA | NA | 1 | 0 | 4 | 0.25 | 1 | 0 | 4 | 0 | 1 | 1 | NA | NA | 6 | 0.1667 |
| 13_687100( | NA | NA | 4 | 0 | 1 | 0 | 1 | 1 | 6 | 0.1667 | 1 | 1 | 3 | 0.6667 | 1 | 1 | NA | NA | 5 | 0.8 |
| 13_687101 | NA | NA | 3 | 0.3333 | 1 | 0 | 1 | 1 | 5 | 0.4 | 1 | 1 | 3 | 0.3333 | 1 | 1 | NA | NA | 5 | 0.6 |
| 13_687102! | NA | NA | 1 | 1 | 1 | 1 | 1 | 1 | 3 | 1 | 1 | 1 | 2 | 1 | 1 | 1 | NA | NA | 4 | 1 |
| 13_687104( | NA | NA | 3 | 0.3333 | 1 | 0 | 1 | 0 | 5 | 0.2 | 1 | 0 | 3 | 0.3333 | 1 | 0 | NA | NA | 5 | 0.2 |
| 13_687104 | NA | NA | 3 | 0.3333 | NA | NA | 1 | 0 | 4 | 0.25 | 1 | 1 | 3 | 0.3333 | NA | NA | NA | NA | 4 | 0.5 |
| 13_687233 | NA | NA | 1 | 1 | NA | NA | NA | NA | 1 | 1 | 1 | 0 | 1 | 1 | NA | NA | NA | NA | 2 | 0.5 |
| 13_687326( | NA | NA | 1 | 1 | 1 | 0 | NA | NA | 2 | 0.5 | NA | NA | 1 | 1 | NA | NA | NA | NA | 1 | 1 |
| 13_687326( | NA | NA | NA | NA | 1 | 1 | NA | NA | 1 | 1 | NA | NA | 1 | 1 | NA | NA | NA | NA | 1 | 1 |
| 13_687328 | NA | NA | 1 | 1 | 1 | 1 | NA | NA | 2 | 1 | NA | NA | 1 | 1 | NA | NA | NA | NA | 1 | 1 |
| 13_687402( | 1 | 1 | 1 | 1 | NA | NA | 1 | 0 | 3 | 0.6667 | NA | NA | 5 | 0.2 | 3 | 0.6667 | 1 | 0 | 9 | 0.3333 |
| 13_687611 | 2 | 0 | 3 | 0 | 1 | 0 | 2 | 0.5 | 8 | 0.125 | 2 | 0.5 | 1 | 0 | NA | NA | NA | NA | 3 | 0.3333 |
| 13_687613 | 2 | 1 | 4 | 0 | 1 | 1 | 2 | 1 | 9 | 0.5556 | 1 | 1 | 1 | 0 | NA | NA | NA | NA | 2 | 0.5 |
| 13_687632 | NA | NA | NA | NA | 1 | 1 | NA | NA | 1 | 1 | 1 | 1 | NA | NA | NA | NA | NA | NA | 1 | 1 |
| 13_687685 | 6 | 0.6667 | 8 | 0.25 | 13 | 0.5385 | 8 | 0.375 | 35 | 0.4571 | 7 | 0.4286 | 7 | 0.2857 | 18 | 0.3333 | 13 | 0.2308 | 45 | 0.3111 |
| 13_687687 | 8 | 0.875 | 10 | 0.7 | 13 | 0.5385 | 7 | 1 | 38 | 0.7368 | 11 | 0.5455 | 6 | 0.6667 | 17 | 0.4706 | 14 | 0.4286 | 48 | 0.5 |
| 13_687687 | 8 | 0.375 | 7 | 0.2857 | 10 | 0.3 | 7 | 0.2857 | 32 | 0.3125 | 9 | 0.2222 | 7 | 0.4286 | 11 | 0.4545 | 12 | 0.4167 | 39 | 0.3846 |
| 13_696461 | 2 | 1 | 7 | 0.4286 | 3 | 0.3333 | 2 | 0 | 14 | 0.4286 | 6 | 0.6667 | 5 | 0.2 | 4 | 1 | 8 | 0.375 | 23 | 0.5217 |
| 13_696461 | 2 | 0 | 7 | 0.1429 | 3 | 0 | 3 | 0.3333 | 15 | 0.1333 | 6 | 0 | 5 | 0.2 | 6 | 0.3333 | 6 | 0.1667 | 23 | 0.1739 |
| 13_696463! | NA | NA | 7 | 1 | 3 | 1 | 1 | 1 | 11 | 1 | 6 | 1 | 3 | 1 | 3 | 1 | 4 | 1 | 16 | 1 |
| 13_696463! | 2 | 1 | 7 | 1 | 3 | 1 | 2 | 1 | 14 | 1 | 5 | 1 | 4 | 1 | 2 | 1 | 5 | 1 | 16 | 1 |
| 13_696465 | 2 | 1 | 7 | 0.8571 | 3 | 0.6667 | 3 | 1 | 15 | 0.8667 | 6 | 1 | 3 | 1 | 2 | 0.5 | 6 | 0.8333 | 17 | 0.8824 |
| 13_696465( | 2 | 1 | 7 | 0.5714 | 2 | 1 | 3 | 0.6667 | 14 | 0.7143 | 6 | 0.8333 | 3 | 0.3333 | 3 | 0.6667 | 6 | 0.6667 | 18 | 0.6667 |
| 13_696465 | 2 | 1 | 7 | 0.7143 | 3 | 1 | 3 | 0.6667 | 15 | 0.8 | 4 | 0.75 | 3 | 0.3333 | 2 | 0 | 5 | 0.6 | 14 | 0.5 |
| 13_696465 | 2 | 0.5 | 7 | 0.2857 | 3 | 0.3333 | 3 | 0.6667 | 15 | 0.4 | 6 | 0.6667 | 2 | 0 | 2 | 0.5 | 5 | 0.2 | 15 | 0.4 |
| 13_696466 | 2 | 0.5 | 6 | 0.3333 | 3 | 0 | 3 | 0 | 14 | 0.2143 | 6 | 0.1667 | 3 | 0 | 2 | 0 | 6 | 0 | 17 | 0.0588 |
| 13_696467( | 2 | 0 | 7 | 0.4286 | 3 | 0.3333 | 3 | 0.6667 | 15 | 0.4 | 6 | 0.5 | 3 | 0.3333 | 2 | 0.5 | 5 | 0.2 | 16 | 0.375 |
| 13_696467 | 2 | 0.5 | 5 | 0.6 | 3 | 1 | 3 | 0.3333 | 13 | 0.6154 | 6 | 0.6667 | 3 | 0.3333 | 5 | 0.6 | 5 | 0.6 | 19 | 0.5789 |
| 13_696469( | 1 | 1 | 7 | 0.2857 | 1 | 1 | 3 | 0.6667 | 12 | 0.5 | 6 | 0.1667 | 3 | 0.3333 | 4 | 0.25 | 5 | 0 | 18 | 0.1667 |
| 13_696470 | 1 | 0 | 6 | 0.3333 | 1 | 0 | 3 | 0 | 11 | 0.1818 | 6 | 0 | 3 | 0.3333 | 3 | 0 | 6 | 0.1667 | 18 | 0.1111 |
| 13_696471 | 1 | 1 | 5 | 0 | 1 | 1 | 3 | 0 | 10 | 0.2 | 6 | 0.3333 | 2 | 0 | NA | NA | 6 | 0.5 | 14 | 0.3571 |
| 13_696472 | 1 | 1 | 4 | 0.25 | 1 | 0 | 3 | 0 | 9 | 0.2222 | 4 | 0 | 3 | 0 | 3 | 0 | 5 | 0.2 | 15 | 0.0667 |
| 13_696476 | NA | NA | 7 | 1 | 1 | 1 | 1 | 1 | 9 | 1 | 4 | 1 | 2 | 1 | 3 | 1 | 4 | 1 | 13 | 1 |
| 13_696477 | NA | NA | 7 | 0.8571 | 1 | 1 | 1 | 0 | 9 | 0.7778 | 3 | 0.6667 | 1 | 1 | 3 | 1 | 4 | 1 | 11 | 0.9091 |
| 13_696480 | NA | NA | 5 | 0.6 | NA | NA | 1 | 1 | 6 | 0.6667 | 2 | 0 | 2 | 0.5 | 2 | 0.5 | 2 | 0.5 | 8 | 0.375 |
| 13_696480 | NA | NA | 6 | 0.1667 | 1 | 1 | 1 | 1 | 8 | 0.375 | 2 | 0 | 2 | 0.5 | 3 | 0 | 4 | 0.5 | 11 | 0.2727 |
| 13_696480( | NA | NA | 6 | 0.3333 | 1 | 1 | 1 | 1 | 8 | 0.5 | 2 | 0.5 | NA | NA | 3 | 0.6667 | 4 | 1 | 9 | 0.7778 |
| 13_696481 | NA | NA | 6 | 0.3333 | 1 | 0 | 2 | 0 | 9 | 0.2222 | 2 | 1 | 1 | 0 | 1 | 0 | 4 | 0.5 | 8 | 0.5 |
| 13_696551 | NA | NA | 5 | 0.2 | NA | NA | NA | NA | 5 | 0.2 | 1 | 0 | 1 | 0 | 1 | 0 | 3 | 0.3333 | 6 | 0.1667 |
| 13_697789! | 1 | 1 | NA | NA | NA | NA | NA | NA | 1 | 1 | NA | NA | NA | NA | 1 | 1 | 1 | 0 | 2 | 0.5 |
| 13_697817! | 5 | 0.8 | 9 | 0.8889 | 3 | 0.6667 | 4 | 1 | 21 | 0.8571 | 2 | 1 | 4 | 0.75 | 2 | 1 | 7 | 0.4286 | 15 | 0.6667 |
| 13_697820 | 6 | 0.6667 | 9 | 0.5556 | 5 | 0.6 | 6 | 0.6667 | 26 | 0.6154 | 2 | 0.5 | 6 | 0.1667 | 2 | 0 | 6 | 0.3333 | 16 | 0.25 |

| | | | | | | | | | | | | | | | | | | | | |
|---|---|---|---|---|---|---|---|---|---|---|---|---|---|---|---|---|---|---|---|---|
| 13_6978224 | 6 | 0.6667 | 6 | 0.6667 | 3 | 0.6667 | 4 | 0.5 | 19 | 0.6316 | 2 | 1 | 3 | 0.6667 | 4 | 0.5 | 5 | 0.6 | 14 | 0.6429 |
| 13_6978229 | 8 | 0.125 | 12 | 0 | 4 | 0 | 7 | 0.1429 | 31 | 0.0645 | 4 | 0.25 | 6 | 0.1667 | 4 | 0 | 7 | 0 | 21 | 0.0952 |
| 13_698084 | 1 | 1 | NA | NA | 1 | 1 | NA | NA | 2 | 1 | NA | NA | NA | NA | NA | NA | 1 | 1 | 1 | 1 |
| 13_698087 | 1 | 1 | NA | NA | NA | NA | NA | NA | 1 | 1 | 1 | 1 | NA | NA | NA | NA | 1 | 1 | 2 | 1 |
| 13_698089 | 1 | 1 | NA | NA | 1 | 1 | NA | NA | 2 | 1 | 1 | 1 | NA | NA | NA | NA | 1 | 0 | 2 | 0.5 |
| 13_698090 | 1 | 1 | NA | NA | 1 | 0 | NA | NA | 2 | 0.5 | 1 | 1 | NA | NA | NA | NA | 1 | 0 | 2 | 0.5 |
| 13_698090 | 1 | 1 | NA | NA | NA | NA | NA | NA | 1 | 1 | 1 | 1 | NA | NA | NA | NA | NA | NA | 1 | 1 |
| 13_698510 | 23 | 0.2174 | 33 | 0.0909 | 27 | 0.2222 | 14 | 0.0714 | 97 | 0.1546 | 29 | 0.1724 | 7 | 0.4286 | 13 | 0.1538 | 24 | 0.2083 | 73 | 0.2055 |
| 13_698512 | 20 | 0.55 | 43 | 0.6744 | 34 | 0.6176 | 17 | 0.7647 | 114 | 0.6491 | 31 | 0.4839 | 8 | 0.375 | 12 | 0.75 | 26 | 0.5385 | 77 | 0.5325 |
| 13_698512 | 24 | 0.375 | 39 | 0.4872 | 33 | 0.4848 | 14 | 0.4286 | 110 | 0.4545 | 30 | 0.2667 | 8 | 0.375 | 10 | 0.3 | 23 | 0.3478 | 71 | 0.3099 |
| 13_698512 | 21 | 0.5238 | 34 | 0.5294 | 24 | 0.4167 | 15 | 0.2667 | 94 | 0.4574 | 27 | 0.3704 | 6 | 0.1667 | 10 | 0.5 | 23 | 0.5652 | 66 | 0.4394 |
| 13_698512 | 21 | 0.7143 | 30 | 0.5667 | 24 | 0.625 | 13 | 0.6154 | 88 | 0.625 | 25 | 0.76 | 7 | 0.7143 | 8 | 0.625 | 21 | 0.5714 | 61 | 0.6721 |
| 13_698513 | 17 | 0.7059 | 38 | 0.6316 | 23 | 0.5652 | 15 | 0.6667 | 93 | 0.6344 | 24 | 0.5 | 7 | 0.7143 | 10 | 0.8 | 23 | 0.5652 | 64 | 0.5938 |
| 13_698514 | 22 | 0.2727 | 35 | 0.2857 | 28 | 0.1071 | 15 | 0.4667 | 100 | 0.26 | 28 | 0.3571 | 4 | 0.5 | 10 | 0.6 | 22 | 0.3636 | 64 | 0.4062 |
| 13_698515 | 19 | 0.2105 | 41 | 0.2439 | 29 | 0.0345 | 15 | 0 | 104 | 0.1442 | 29 | 0.069 | 8 | 0.125 | 10 | 0.1 | 23 | 0.2174 | 70 | 0.1286 |
| 13_698516 | 17 | 0.2353 | 41 | 0.3415 | 24 | 0.2083 | 13 | 0.1538 | 95 | 0.2632 | 25 | 0.28 | 7 | 0.2857 | 11 | 0.1818 | 23 | 0.3043 | 66 | 0.2727 |
| 13_698516 | 17 | 0 | 41 | 0.1707 | 29 | 0.1379 | 15 | 0.0667 | 102 | 0.1176 | 30 | 0.1 | 8 | 0.125 | 11 | 0.2727 | 24 | 0.2083 | 73 | 0.1644 |
| 13_698518 | 17 | 0.2353 | 39 | 0.2564 | 26 | 0.3462 | 13 | 0.2308 | 95 | 0.2737 | 26 | 0.2308 | 7 | 0.4286 | 10 | 0 | 20 | 0.2 | 63 | 0.2063 |
| 13_698519 | 9 | 0.3333 | 31 | 0.129 | 18 | 0.0556 | 12 | 0.0833 | 70 | 0.1286 | 23 | 0.1304 | 7 | 0 | 4 | 0.25 | 16 | 0 | 50 | 0.08 |
| 13_698519 | 15 | 0.4667 | 31 | 0.4194 | 21 | 0.619 | 11 | 0.3636 | 78 | 0.4744 | 25 | 0.32 | 6 | 0.8333 | 8 | 0.75 | 16 | 0.25 | 55 | 0.4182 |
| 13_698520 | 15 | 1 | 27 | 1 | 21 | 1 | 10 | 1 | 73 | 1 | 23 | 1 | 7 | 1 | 8 | 1 | 15 | 1 | 53 | 1 |
| 13_698520 | 15 | 0.8 | 35 | 0.8571 | 24 | 0.8333 | 12 | 0.75 | 86 | 0.8256 | 29 | 0.8621 | 9 | 0.7778 | 10 | 0.8 | 23 | 0.913 | 71 | 0.8592 |
| 13_698520 | 15 | 0.6667 | 30 | 0.4667 | 21 | 0.5714 | 10 | 0.8 | 76 | 0.5789 | 25 | 0.76 | 9 | 0.7778 | 9 | 0.3333 | 20 | 0.65 | 63 | 0.6667 |
| 13_698869 | 12 | 0.3333 | 27 | 0.3333 | 20 | 0.2 | 9 | 0.2222 | 68 | 0.2794 | 21 | 0.2381 | 5 | 0.2 | 8 | 0.125 | 19 | 0.2632 | 53 | 0.2264 |
| 13_698869 | 12 | 0.9167 | 27 | 0.7778 | 21 | 0.7619 | 10 | 0.6 | 70 | 0.7714 | 19 | 0.8421 | 7 | 0.8571 | 10 | 0.6 | 21 | 0.9048 | 57 | 0.8246 |
| 13_698869 | 12 | 0.0833 | 23 | 0.087 | 19 | 0.4211 | 10 | 0.2 | 64 | 0.2031 | 19 | 0.3158 | 6 | 0.3333 | 8 | 0.125 | 16 | 0.125 | 49 | 0.2245 |
| 13_698872 | 5 | 0.2 | 10 | 0 | 11 | 0.0909 | 5 | 0 | 31 | 0.0645 | 13 | 0 | 3 | 0 | 7 | 0.2857 | 9 | 0 | 32 | 0.0625 |
| 13_698877 | 9 | 0.2222 | 15 | 0.1333 | 12 | 0.0833 | 6 | 0 | 42 | 0.119 | 10 | 0.2 | 3 | 0.3333 | 3 | 0 | 12 | 0.0833 | 28 | 0.1429 |
| 13_699638 | NA | NA | 10 | 0.1 | 6 | 0.3333 | 6 | 0 | 22 | 0.1364 | 9 | 0.3333 | 11 | 0.0909 | 7 | 0.1429 | 4 | 0 | 31 | 0.1613 |
| 13_699638 | NA | NA | 7 | 0.5714 | 4 | 0.25 | 3 | 0 | 14 | 0.3571 | 6 | 0.5 | 5 | 0.4 | 6 | 0.5 | 2 | 0 | 19 | 0.4211 |
| 13_699639 | NA | NA | 9 | 0.6667 | 6 | 1 | 4 | 0.75 | 19 | 0.7895 | 7 | 0.8571 | 10 | 0.5 | 6 | 0.6667 | 2 | 0.5 | 25 | 0.64 |
| 13_707375 | 3 | 0.3333 | 7 | 0.2857 | 4 | 0.25 | 4 | 0.25 | 18 | 0.2778 | 5 | 0 | 7 | 0.1429 | 8 | 0.25 | 12 | 0 | 32 | 0.0938 |
| 13_707375 | 3 | 0 | 6 | 0 | 3 | 0 | 2 | 0.5 | 14 | 0.0714 | 4 | 0 | 4 | 0 | 6 | 0.1667 | 8 | 0.125 | 22 | 0.0909 |
| 13_707377 | 3 | 0.6667 | 5 | 0.6 | 4 | 0.25 | 3 | 0 | 15 | 0.4 | 6 | 0.1667 | 8 | 0 | 7 | 0.1429 | 11 | 0.2727 | 32 | 0.1562 |
| 13_707506 | 1 | 0 | 3 | 1 | NA | NA | 3 | 0.6667 | 7 | 0.7143 | 6 | 0.6667 | 5 | 1 | 6 | 0.6667 | 6 | 0.6667 | 23 | 0.7391 |
| 13_707506 | 1 | 0 | 4 | 0 | 1 | 1 | 3 | 0.3333 | 9 | 0.2222 | 6 | 0.1667 | 5 | 0.2 | 6 | 0.1667 | 8 | 0.25 | 25 | 0.2 |
| 13_707507 | 2 | 0.5 | 3 | 0.3333 | 2 | 0.5 | 3 | 0.3333 | 10 | 0.4 | 5 | 0.2 | 4 | 0 | 6 | 0.3333 | 6 | 0.1667 | 21 | 0.1905 |
| 13_707508 | 2 | 1 | 3 | 1 | 2 | 1 | 4 | 1 | 11 | 1 | 5 | 1 | 5 | 1 | 6 | 1 | 8 | 1 | 24 | 1 |
| 13_707508 | 1 | 0 | 3 | 0.6667 | 2 | 0.5 | 3 | 0.3333 | 9 | 0.4444 | 4 | 0.5 | 3 | 0.3333 | 4 | 0.5 | 5 | 0.2 | 16 | 0.375 |
| 13_707512 | 1 | 1 | 1 | 0 | 2 | 0.5 | 3 | 0.3333 | 7 | 0.4286 | 5 | 0.4 | 5 | 0.2 | 2 | 0.5 | 6 | 0.8333 | 18 | 0.5 |
| 13_707514 | 1 | 0 | 1 | 0 | 1 | 0 | 3 | 0.3333 | 6 | 0.1667 | 4 | 0.5 | 4 | 0.25 | 4 | 0.25 | 4 | 0.5 | 16 | 0.375 |
| 13_707519 | 1 | 1 | 2 | 1 | 1 | 1 | 1 | 0 | 5 | 0.8 | 3 | 1 | 4 | 1 | 3 | 1 | 3 | 1 | 13 | 1 |
| 13_707523 | 2 | 1 | 1 | 1 | 2 | 1 | 1 | 1 | 6 | 1 | 3 | 1 | 3 | 0.6667 | 3 | 1 | 1 | 1 | 10 | 0.9 |
| 13_707527 | 1 | 0 | 1 | 0 | 2 | 0 | 1 | 1 | 5 | 0.2 | 3 | 0.3333 | 3 | 0 | 1 | 0 | 2 | 0.5 | 9 | 0.2222 |
| 13_707531 | 1 | 0 | 1 | 0 | 1 | 1 | NA | NA | 3 | 0.3333 | 3 | 0.6667 | 2 | 0 | 1 | 0 | 1 | 1 | 7 | 0.4286 |
| 13_707532 | 2 | 1 | 1 | 0 | NA | NA | NA | NA | 3 | 0.6667 | 2 | 0 | 3 | 0.6667 | 1 | 1 | NA | NA | 6 | 0.5 |
| 13_707533 | 1 | 1 | 1 | 1 | 1 | 0 | NA | NA | 3 | 0.6667 | 2 | 1 | 3 | 0 | 1 | 0 | 1 | 1 | 7 | 0.4286 |
| 13_707536 | 1 | 1 | 1 | 1 | 1 | 1 | NA | NA | 3 | 1 | 3 | 0.6667 | 3 | 0.3333 | NA | NA | 1 | 1 | 7 | 0.5714 |
| 13_707537 | 2 | 1 | NA | NA | 1 | 0 | NA | NA | 3 | 0.6667 | 2 | 0.5 | 3 | 1 | 1 | 1 | NA | NA | 6 | 0.8333 |
| 13_707538 | 2 | 0 | 1 | 1 | 1 | 0 | NA | NA | 4 | 0.25 | 3 | 0 | 2 | 0.5 | NA | NA | NA | NA | 5 | 0.2 |
| 13_707539 | 1 | 1 | 1 | 1 | 1 | 1 | NA | NA | 3 | 1 | 2 | 1 | 3 | 1 | 1 | 1 | NA | NA | 6 | 1 |
| 13_707539 | 1 | 0 | 1 | 0 | 1 | 1 | NA | NA | 3 | 0.3333 | 3 | 0.6667 | 2 | 0 | 1 | 0 | NA | NA | 6 | 0.3333 |
| 13_707540 | 1 | 0 | NA | NA | 1 | 1 | NA | NA | 2 | 0.5 | 2 | 0.5 | 2 | 0 | 1 | 0 | NA | NA | 5 | 0.2 |
| 13_707541 | 2 | 1 | 1 | 1 | 1 | 1 | NA | NA | 4 | 1 | 1 | 0 | 2 | 0.5 | 1 | 1 | NA | NA | 4 | 0.5 |
| 13_707542 | 2 | 0.5 | NA | NA | 1 | 0 | NA | NA | 3 | 0.3333 | 2 | 0.5 | 2 | 0 | 1 | 0 | NA | NA | 5 | 0.2 |
| 13_707544 | NA | NA | 1 | 1 | 1 | 1 | NA | NA | 2 | 1 | 1 | 1 | 2 | 1 | 1 | 0 | NA | NA | 4 | 0.75 |
| 13_707546 | 1 | 1 | NA | NA | 1 | 0 | NA | NA | 2 | 0.5 | 1 | 0 | 2 | 1 | 1 | 0 | NA | NA | 4 | 0.5 |
| 13_707547 | NA | NA | NA | NA | 1 | 1 | NA | NA | 1 | 1 | 1 | 1 | 2 | 0 | 1 | 1 | NA | NA | 4 | 0.5 |
| 13_710290 | NA | NA | NA | NA | NA | NA | 1 | 1 | 1 | 1 | NA | NA | NA | NA | 1 | 1 | NA | NA | 1 | 1 |
| 13_710291 | NA | NA | NA | NA | NA | NA | 1 | 1 | 1 | 1 | NA | NA | NA | NA | 1 | 1 | NA | NA | 1 | 1 |
| 13_710292 | NA | NA | NA | NA | NA | NA | 1 | 1 | 1 | 1 | NA | NA | NA | NA | 1 | 1 | NA | NA | 1 | 1 |
| 13_710293 | NA | NA | NA | NA | NA | NA | 1 | 1 | 1 | 1 | NA | NA | NA | NA | 1 | 1 | NA | NA | 1 | 1 |
| 13_710299 | NA | NA | NA | NA | NA | NA | 1 | 1 | 1 | 1 | NA | NA | NA | NA | 1 | 1 | NA | NA | 1 | 1 |
| 13_734796 | 5 | 0.2 | 9 | 0.1111 | 6 | 0.3333 | 7 | 0.1429 | 27 | 0.1852 | 3 | 0.3333 | 8 | 0.25 | 8 | 0.25 | 7 | 0.2857 | 26 | 0.2692 |
| 13_734797 | 2 | 0.5 | 5 | 1 | 5 | 0.8 | 7 | 0.4286 | 19 | 0.6842 | 3 | 0.6667 | 8 | 0.75 | 6 | 0.6667 | 4 | 1 | 21 | 0.7619 |

| | | | | | | | | | | | | | | | | | | | | |
|---|---|---|---|---|---|---|---|---|---|---|---|---|---|---|---|---|---|---|---|---|
| 13_734798 | 5 | 0.2 | 9 | 0.2222 | 7 | 0.1429 | 10 | 0.2 | 31 | 0.1935 | 4 | 0.5 | 11 | 0.4545 | 8 | 0.375 | 6 | 0.5 | 29 | 0.4483 |
| 13_736626 | 3 | 0.3333 | 2 | 0.5 | 3 | 0.6667 | 4 | 0.75 | 12 | 0.5833 | NA | NA | 2 | 0.5 | 1 | 0 | 1 | 0 | 4 | 0.25 |
| 13_736626 | 4 | 0.75 | 2 | 0.5 | 4 | 1 | 2 | 1 | 12 | 0.8333 | 1 | 1 | 3 | 1 | 1 | 1 | 1 | 1 | 6 | 1 |
| 13_736626 | 4 | 1 | 1 | 1 | 3 | 1 | 2 | 1 | 10 | 1 | NA | NA | 3 | 0.6667 | NA | NA | 1 | 1 | 4 | 0.75 |
| 13_736627 | 5 | 0.6 | 2 | 1 | 3 | 0 | 3 | 0.3333 | 13 | 0.4615 | NA | NA | 3 | 0.3333 | 1 | 0 | 2 | 0 | 6 | 0.1667 |
| 13_736630 | 5 | 0.2 | NA | NA | 3 | 0 | 3 | 0.3333 | 11 | 0.1818 | 1 | 0 | 3 | 0 | 1 | 0 | 2 | 0.5 | 7 | 0.1429 |
| 13_736633 | 5 | 0 | 2 | 0 | 5 | 0.2 | 2 | 0.5 | 14 | 0.1429 | 1 | 0 | 2 | 0 | 1 | 1 | 1 | 0 | 5 | 0.2 |
| 13_736633 | 4 | 0.25 | 1 | 0 | 5 | 0.4 | 3 | 0 | 13 | 0.2308 | 1 | 0 | 3 | 0 | 1 | 1 | 1 | 0 | 6 | 0.1667 |
| 13_736642 | 7 | 0.2857 | 5 | 0.4 | 9 | 0 | 2 | 0 | 23 | 0.1739 | 2 | 0 | 3 | 0 | NA | NA | 4 | 0.5 | 9 | 0.2222 |
| 13_736644 | 8 | 0.25 | 6 | 0.1667 | 9 | 0.4444 | 5 | 0.6 | 28 | 0.3571 | 3 | 0 | 3 | 0.3333 | 1 | 0 | 5 | 0.6 | 12 | 0.3333 |
| 13_737684 | 14 | 0.3571 | 17 | 0.1765 | 8 | 0.75 | 12 | 0.5833 | 51 | 0.4118 | 7 | 0.5714 | 16 | 0.375 | 16 | 0.5625 | 9 | 0.3333 | 48 | 0.4583 |
| 13_737686 | 19 | 0.5789 | 24 | 0.6667 | 13 | 0.6923 | 17 | 0.4706 | 73 | 0.6027 | 10 | 0.4 | 19 | 0.4211 | 26 | 0.5385 | 18 | 0.6667 | 73 | 0.5205 |
| 13_739630 | 1 | 1 | NA | NA | NA | NA | NA | NA | 1 | 1 | NA | NA | NA | NA | NA | NA | 1 | 1 | 1 | 1 |
| 13_739631 | 1 | 1 | NA | NA | NA | NA | NA | NA | 1 | 1 | NA | NA | NA | NA | NA | NA | 1 | 1 | 1 | 1 |
| 13_739638 | 1 | 1 | NA | NA | NA | NA | NA | NA | 1 | 1 | NA | NA | NA | NA | NA | NA | 1 | 1 | 1 | 1 |
| 13_739639 | 1 | 1 | NA | NA | NA | NA | NA | NA | 1 | 1 | NA | NA | NA | NA | NA | NA | 1 | 1 | 1 | 1 |
| 13_739670 | 1 | 1 | 2 | 0.5 | NA | NA | NA | NA | 3 | 0.6667 | NA | NA | 1 | 1 | NA | NA | NA | NA | 1 | 1 |
| 13_741084 | 8 | 0.125 | 7 | 0.4286 | 6 | 0.3333 | 7 | 0.2857 | 28 | 0.2857 | 4 | 0 | 2 | 1 | 4 | 0.5 | 6 | 0 | 16 | 0.25 |
| 13_741084 | 8 | 0.25 | 6 | 0 | 4 | 0.5 | 7 | 0 | 25 | 0.16 | 4 | 0 | 2 | 0 | 4 | 0.25 | 5 | 0 | 15 | 0.0667 |
| 13_741085 | 4 | 0.25 | 7 | 0.1429 | 4 | 0 | 5 | 0.2 | 20 | 0.15 | 2 | 0.5 | 2 | 0 | 3 | 0.3333 | 3 | 0.6667 | 10 | 0.4 |
| 13_741088 | 6 | 0.1667 | 5 | 0 | 6 | 0.1667 | 5 | 0 | 22 | 0.0909 | 2 | 0 | 1 | 0 | 4 | 0.25 | 5 | 0.2 | 12 | 0.1667 |
| 13_741563 | 1 | 1 | NA | NA | NA | NA | 4 | 0.75 | 5 | 0.8 | 2 | 1 | NA | NA | NA | NA | NA | NA | 2 | 1 |
| 13_741564 | 1 | 1 | NA | NA | NA | NA | 4 | 0.75 | 5 | 0.8 | 2 | 1 | NA | NA | NA | NA | NA | NA | 2 | 1 |
| 13_741566 | NA | NA | NA | NA | NA | NA | 4 | 0.75 | 4 | 0.75 | 1 | 1 | NA | NA | NA | NA | NA | NA | 1 | 1 |
| 13_741567 | NA | NA | NA | NA | NA | NA | 4 | 0.75 | 4 | 0.75 | 1 | 1 | NA | NA | NA | NA | NA | NA | 1 | 1 |
| 13_741569 | NA | NA | NA | NA | NA | NA | 3 | 0.6667 | 3 | 0.6667 | 2 | 0.5 | NA | NA | NA | NA | NA | NA | 2 | 0.5 |
| 13_741569 | NA | NA | NA | NA | NA | NA | 4 | 0.5 | 4 | 0.5 | 2 | 0.5 | NA | NA | NA | NA | NA | NA | 2 | 0.5 |
| 13_741694 | 5 | 0 | 4 | 0 | 2 | 0 | 6 | 0.1667 | 17 | 0.0588 | 3 | 0.3333 | 2 | 0 | 3 | 0 | 3 | 0 | 11 | 0.0909 |
| 13_741694 | 4 | 0.5 | 4 | 0.5 | 1 | 0 | 6 | 0.3333 | 15 | 0.4 | 3 | 0 | 2 | 0.5 | 3 | 0.3333 | 3 | 0.3333 | 11 | 0.2727 |
| 13_741695 | 4 | 0.5 | 4 | 0.75 | 2 | 0 | 6 | 0.3333 | 16 | 0.4375 | 3 | 0 | 1 | 0 | 3 | 0.3333 | 4 | 0.25 | 11 | 0.1818 |
| 13_741794 | 4 | 0.25 | 5 | 0.6 | 4 | 0.5 | 9 | 0.6667 | 22 | 0.5455 | 4 | 0.75 | 2 | 0 | 3 | 0.6667 | 7 | 0.8571 | 16 | 0.6875 |
| 13_741795 | 4 | 1 | 4 | 1 | 4 | 1 | 9 | 0.7778 | 21 | 0.9048 | 4 | 0.75 | 2 | 0.5 | 4 | 1 | 6 | 1 | 16 | 0.875 |
| 13_741795 | 5 | 0.8 | 6 | 0.8333 | 3 | 1 | 10 | 0.8 | 24 | 0.8333 | 4 | 0.75 | 2 | 1 | 4 | 0.75 | 7 | 1 | 17 | 0.8824 |
| 13_741796 | 5 | 0.6 | 5 | 0.6 | 4 | 0.5 | 8 | 0.375 | 22 | 0.5 | 4 | 0.75 | 1 | 0 | 4 | 0.5 | 6 | 0.8333 | 15 | 0.6667 |
| 13_741797 | 5 | 1 | 7 | 1 | 4 | 0.5 | 9 | 0.8889 | 25 | 0.88 | 4 | 1 | 2 | 0.5 | 4 | 1 | 7 | 1 | 17 | 0.9412 |
| 13_741798 | 5 | 0.4 | 8 | 0.125 | 4 | 0.25 | 9 | 0 | 26 | 0.1538 | 4 | 0 | 2 | 0 | 5 | 0 | 7 | 0.2857 | 18 | 0.1111 |
| 13_741799 | 5 | 0 | 9 | 0.1111 | 4 | 0.25 | 10 | 0 | 28 | 0.0714 | 5 | 0 | 2 | 0 | 4 | 0 | 7 | 0.1429 | 18 | 0.0556 |
| 13_741801 | 5 | 0.2 | 10 | 0.2 | 5 | 0 | 12 | 0.1667 | 32 | 0.1562 | 4 | 0.5 | 2 | 0 | 5 | 0.8 | 7 | 0.4286 | 18 | 0.5 |
| 13_741801 | 4 | 0.25 | 9 | 0 | 5 | 0.2 | 11 | 0.0909 | 29 | 0.1034 | 5 | 0.2 | 2 | 0 | 6 | 0 | 6 | 0.1667 | 19 | 0.1053 |
| 13_741801 | 2 | 0 | 8 | 0.875 | 4 | 0.75 | 11 | 0.4545 | 25 | 0.6 | 4 | 1 | NA | NA | 4 | 1 | 7 | 0.8571 | 15 | 0.9333 |
| 13_741802 | 5 | 0 | 10 | 0.1 | 4 | 0 | 10 | 0.1 | 29 | 0.069 | 5 | 0 | 3 | 0 | 4 | 0.25 | 7 | 0.1429 | 19 | 0.1053 |
| 13_741802 | 5 | 0.2 | 12 | 0.5 | 6 | 0.3333 | 11 | 0.2727 | 34 | 0.3529 | 4 | 0.5 | 3 | 0.3333 | 6 | 0.5 | 8 | 0.5 | 21 | 0.4762 |
| 13_741802 | 4 | 0.25 | 10 | 0.6 | 6 | 0.5 | 11 | 0.4545 | 31 | 0.4839 | 4 | 0.25 | 3 | 0 | 7 | 0.7143 | 7 | 0.7143 | 21 | 0.5238 |
| 13_741814 | 10 | 0.1 | 21 | 0.2857 | 7 | 0 | 12 | 0 | 50 | 0.14 | 13 | 0 | 6 | 0.1667 | 9 | 0 | 13 | 0.1538 | 41 | 0.0732 |
| 13_741814 | 10 | 0.4 | 27 | 0.2222 | 7 | 0 | 17 | 0.2353 | 61 | 0.2295 | 12 | 0.3333 | 8 | 0.25 | 11 | 0.4545 | 12 | 0.3333 | 43 | 0.3488 |
| 13_741815 | 10 | 0.3 | 26 | 0.1538 | 6 | 0 | 18 | 0.0556 | 60 | 0.1333 | 12 | 0.0833 | 9 | 0.1111 | 9 | 0 | 12 | 0.0833 | 42 | 0.0714 |
| 13_741816 | 10 | 0.6 | 24 | 0.5417 | 6 | 0.5 | 17 | 0.4706 | 57 | 0.5263 | 11 | 0.5455 | 6 | 0.6667 | 9 | 0.5556 | 11 | 0.6364 | 37 | 0.5946 |
| 13_741836 | 27 | 0.0741 | 48 | 0.0417 | 27 | 0.037 | 29 | 0.069 | 131 | 0.0534 | 28 | 0.1071 | 27 | 0.1111 | 22 | 0.2273 | 25 | 0.12 | 102 | 0.1373 |
| 13_743195 | 9 | 0.2222 | 8 | 0.125 | 8 | 0.125 | 10 | 0.1 | 35 | 0.1429 | 4 | 0.25 | 4 | 0 | 3 | 0 | 3 | 0 | 14 | 0.0714 |
| 13_743198 | 7 | 0.1429 | 6 | 0.1667 | 7 | 0.1429 | 11 | 0.0909 | 31 | 0.129 | 5 | 0.4 | 2 | 0.5 | 4 | 0 | 5 | 0.2 | 16 | 0.25 |
| 13_743407 | 3 | 0 | 6 | 0 | 7 | 0 | 12 | 0.1667 | 28 | 0.0714 | 1 | 0 | 1 | 0 | 8 | 0 | 7 | 0.1429 | 17 | 0.0588 |
| 13_743407 | 2 | 0 | 5 | 0.2 | 7 | 0 | 9 | 0.1111 | 23 | 0.087 | 1 | 1 | 1 | 0 | 7 | 0.1429 | 6 | 0 | 15 | 0.1333 |
| 13_743408 | 3 | 0.3333 | 5 | 0 | 7 | 0.1429 | 11 | 0.1818 | 26 | 0.1538 | 1 | 1 | 1 | 1 | 7 | 0 | 6 | 0.1667 | 15 | 0.2 |
| 13_743410 | 3 | 1 | 3 | 0.6667 | 3 | 0.6667 | 10 | 0.7 | 19 | 0.7368 | 1 | 0 | NA | NA | 4 | 1 | 4 | 0.5 | 9 | 0.6667 |
| 13_743410 | 2 | 0.5 | 4 | 0.5 | 7 | 1 | 9 | 1 | 22 | 0.8636 | 1 | 1 | 2 | 1 | 3 | 1 | 6 | 0.1667 | 12 | 0.5833 |
| 13_743410 | 2 | 0 | 5 | 0.6 | 5 | 0.4 | 7 | 0.5714 | 19 | 0.4737 | NA | NA | 2 | 1 | 5 | 0.2 | 6 | 0.6667 | 13 | 0.5385 |
| 13_743410 | NA | NA | 1 | 0 | 1 | 1 | 1 | 1 | 3 | 0.6667 | NA | NA | NA | NA | 1 | 0 | 1 | 1 | 2 | 0.5 |
| 13_743411 | 2 | 0 | 6 | 0.3333 | 6 | 0.1667 | 9 | 0 | 23 | 0.1304 | 1 | 0 | 2 | 0 | 7 | 0.5714 | 7 | 0 | 17 | 0.2353 |
| 13_743412 | 3 | 1 | 3 | 0.3333 | 5 | 1 | 10 | 0.9 | 21 | 0.8571 | NA | NA | 2 | 1 | 5 | 0.8 | 6 | 0.6667 | 13 | 0.7692 |
| 13_743412 | 2 | 0.5 | 2 | 0 | 7 | 0 | 11 | 0.0909 | 22 | 0.0909 | NA | NA | 2 | 0 | 7 | 0.1429 | 3 | 0.3333 | 12 | 0.1667 |
| 13_743412 | 3 | 1 | 4 | 0.75 | 7 | 1 | 9 | 0.8889 | 23 | 0.913 | NA | NA | 2 | 1 | 7 | 0.8571 | 6 | 0.6667 | 15 | 0.8 |
| 13_744518 | 19 | 0.2105 | 17 | 0.1176 | 21 | 0.0952 | 14 | 0.0714 | 71 | 0.1268 | 18 | 0.0556 | 18 | 0.1667 | 21 | 0.2381 | 14 | 0.2857 | 71 | 0.1831 |
| 13_744519 | 16 | 0.0625 | 17 | 0 | 19 | 0.1053 | 13 | 0.0769 | 65 | 0.0615 | 16 | 0.0625 | 15 | 0 | 20 | 0.1 | 16 | 0.1875 | 67 | 0.0896 |
| 13_744520 | 18 | 0.2222 | 16 | 0.375 | 18 | 0.3333 | 14 | 0.2143 | 66 | 0.2879 | 17 | 0.1176 | 17 | 0.1765 | 20 | 0.25 | 15 | 0.2667 | 69 | 0.2029 |
| 13_744704 | 35 | 0.1429 | 36 | 0.0556 | 30 | 0.1 | 41 | 0.122 | 142 | 0.1056 | 31 | 0.129 | 28 | 0.0714 | 42 | 0.119 | 36 | 0.1667 | 137 | 0.1241 |

| | | | | | | | | | | | | | | | | | | | | |
|---|---|---|---|---|---|---|---|---|---|---|---|---|---|---|---|---|---|---|---|---|
| 13_744705 | 34 | 0.5882 | 32 | 0.6562 | 33 | 0.6667 | 33 | 0.6364 | 132 | 0.6364 | 30 | 0.7667 | 27 | 0.4815 | 42 | 0.6667 | 35 | 0.7143 | 134 | 0.6642 |
| 13_745047 | 2 | 1 | 2 | 1 | 2 | 1 | 5 | 1 | 11 | 1 | 5 | 1 | 4 | 0.75 | 2 | 1 | 2 | 1 | 13 | 0.9231 |
| 13_745047 | 3 | 0.3333 | 3 | 0.3333 | 1 | 1 | 2 | 0.5 | 9 | 0.4444 | 2 | 0.5 | 4 | 0.25 | 2 | 0.5 | 3 | 0 | 11 | 0.2727 |
| 13_745048 | 2 | 0.5 | 3 | 0 | 2 | 0 | 5 | 0.2 | 12 | 0.1667 | 6 | 0.1667 | 5 | 0 | 2 | 0 | 4 | 0.25 | 17 | 0.1176 |
| 13_745049 | 3 | 0.3333 | 3 | 0 | 2 | 0 | 5 | 0 | 13 | 0.0769 | 5 | 0.4 | 4 | 0.25 | 2 | 0 | 4 | 0.25 | 15 | 0.2667 |
| 13_745050 | 3 | 0.6667 | 3 | 1 | 2 | 1 | 4 | 0.75 | 12 | 0.8333 | 5 | 0.6 | 3 | 0.3333 | 2 | 1 | 2 | 0.5 | 12 | 0.5833 |
| 13_745051 | 2 | 0 | NA | NA | 1 | 0 | 5 | 0.6 | 8 | 0.375 | 5 | 0 | 2 | 0 | 1 | 0 | 2 | 0.5 | 10 | 0.1 |
| 13_745051 | 3 | 1 | 3 | 0.6667 | 2 | 0.5 | 5 | 0.6 | 13 | 0.6923 | 6 | 0.8333 | 2 | 1 | 2 | 0.5 | 2 | 0.5 | 12 | 0.75 |
| 13_745051 | 3 | 1 | 1 | 1 | 1 | 1 | 3 | 1 | 8 | 1 | 5 | 0.8 | 2 | 1 | 1 | 1 | 2 | 1 | 10 | 0.9 |
| 13_745052 | 2 | 0 | 2 | 0 | 1 | 1 | 5 | 0.2 | 10 | 0.2 | 5 | 0.2 | 2 | 0 | 2 | 0.5 | NA | NA | 9 | 0.2222 |
| 13_745053 | 3 | 0.3333 | 3 | 0 | 2 | 1 | 5 | 0.8 | 13 | 0.5385 | 5 | 0.6 | 2 | 1 | 2 | 0.5 | 2 | 0.5 | 11 | 0.6364 |
| 13_745058 | 1 | 1 | NA | NA | 4 | 0 | 2 | 0 | 7 | 0.1429 | 5 | 0 | 1 | 1 | 1 | 0 | 1 | 0 | 8 | 0.125 |
| 13_745060 | 1 | 1 | NA | NA | 4 | 0 | 3 | 0 | 8 | 0.125 | 4 | 0.5 | 1 | 1 | 1 | 1 | 1 | 0 | 7 | 0.5714 |
| 13_745075 | 2 | 0 | 5 | 0 | 4 | 0.25 | 2 | 0 | 13 | 0.0769 | 4 | 0.25 | 1 | 0 | 1 | 1 | 3 | 0.3333 | 9 | 0.3333 |
| 13_745087 | NA | NA | 1 | 1 | NA | NA | 2 | 0 | 3 | 0.3333 | NA | NA | 2 | 0 | 2 | 0.5 | 3 | 0.3333 | 7 | 0.2857 |
| 13_745088 | NA | NA | 1 | 1 | NA | NA | 2 | 1 | 3 | 1 | 1 | 1 | 2 | 1 | 2 | 1 | 3 | 0.6667 | 8 | 0.875 |
| 13_745088 | NA | NA | 2 | 1 | NA | NA | 1 | 1 | 3 | 1 | 1 | 0 | 2 | 0.5 | 2 | 1 | 3 | 1 | 8 | 0.75 |
| 13_745090 | NA | NA | 2 | 1 | NA | NA | 2 | 1 | 4 | 1 | 1 | 1 | 2 | 1 | 3 | 1 | 2 | 1 | 8 | 1 |
| 13_745090 | NA | NA | 1 | 0 | NA | NA | 1 | 1 | 2 | 0.5 | NA | NA | 2 | 0.5 | 3 | 0.3333 | NA | NA | 5 | 0.4 |
| 13_745091 | NA | NA | 1 | 1 | NA | NA | 1 | 1 | 2 | 1 | 1 | 1 | 1 | 1 | 1 | 1 | NA | NA | 3 | 1 |
| 13_745092 | NA | NA | 2 | 0.5 | NA | NA | 1 | 1 | 3 | 0.6667 | NA | NA | 2 | 0 | 3 | 0.3333 | 3 | 1 | 8 | 0.5 |
| 13_745092 | NA | NA | 1 | 1 | NA | NA | 1 | 1 | 2 | 1 | 1 | 1 | 2 | 1 | 2 | 1 | NA | NA | 5 | 1 |
| 13_745096 | 1 | 1 | 3 | 0.3333 | NA | NA | 2 | 0.5 | 6 | 0.5 | 1 | 0 | 3 | 0 | 4 | 0.25 | 5 | 0 | 13 | 0.0769 |
| 13_745098 | 1 | 1 | 3 | 0.3333 | 1 | 0 | 3 | 0 | 8 | 0.25 | 1 | 1 | 2 | 0 | 3 | 0 | 3 | 0 | 9 | 0.1111 |
| 13_745098 | 1 | 1 | 4 | 0.25 | 1 | 0 | 3 | 0 | 9 | 0.2222 | 1 | 0 | 3 | 0.3333 | 5 | 0 | 5 | 0.2 | 14 | 0.1429 |
| 13_745114 | 1 | 1 | 2 | 0.5 | 1 | 1 | NA | NA | 4 | 0.75 | NA | NA | 1 | 1 | 3 | 1 | NA | NA | 4 | 1 |
| 13_745114 | NA | NA | 1 | 1 | 1 | 0 | NA | NA | 2 | 0.5 | NA | NA | NA | NA | 3 | 0.3333 | NA | NA | 3 | 0.3333 |
| 13_745116 | NA | NA | 2 | 1 | 1 | 1 | NA | NA | 3 | 1 | NA | NA | 1 | 1 | 2 | 0.5 | NA | NA | 3 | 0.6667 |
| 13_745117 | 1 | 1 | 2 | 1 | NA | NA | NA | NA | 3 | 1 | NA | NA | 1 | 1 | 3 | 0.3333 | NA | NA | 4 | 0.5 |
| 13_745121 | 1 | 1 | 2 | 0.5 | 1 | 0 | NA | NA | 4 | 0.5 | NA | NA | 1 | 0 | 2 | 0.5 | NA | NA | 3 | 0.3333 |
| 13_745124 | 1 | 0 | 2 | 0.5 | 2 | 0 | NA | NA | 5 | 0.2 | 1 | 0 | 1 | 1 | 3 | 0.3333 | NA | NA | 5 | 0.4 |
| 13_745125 | 3 | 0.3333 | 2 | 0 | 2 | 0 | NA | NA | 7 | 0.1429 | NA | NA | 1 | 0 | 3 | 0.3333 | NA | NA | 4 | 0.25 |
| 13_745126 | 2 | 1 | 2 | 1 | 2 | 0.5 | NA | NA | 6 | 0.8333 | NA | NA | 1 | 1 | 2 | 1 | NA | NA | 3 | 1 |
| 13_745176 | NA | NA | NA | NA | 1 | 1 | NA | NA | 1 | 1 | NA | NA | 1 | 1 | 1 | 1 | 1 | 1 | 3 | 1 |
| 13_745199 | NA | NA | NA | NA | 1 | 0 | 1 | 1 | 2 | 0.5 | NA | NA | NA | NA | 1 | 1 | NA | NA | 1 | 1 |
| 13_745201 | NA | NA | NA | NA | 1 | 1 | 1 | 0 | 2 | 0.5 | NA | NA | NA | NA | 1 | 1 | NA | NA | 1 | 1 |
| 13_745207 | NA | NA | NA | NA | NA | NA | 1 | 1 | 1 | 1 | NA | NA | NA | NA | 1 | 1 | NA | NA | 1 | 1 |
| 13_745933 | NA | NA | 1 | 1 | NA | NA | NA | NA | 1 | 1 | NA | NA | 1 | 1 | NA | NA | NA | NA | 1 | 1 |
| 13_745935 | 2 | 0 | 1 | 1 | 2 | 0 | NA | NA | 5 | 0.2 | 1 | 1 | 2 | 0 | 1 | 0 | 1 | 0 | 5 | 0.2 |
| 13_745935 | 2 | 0.5 | 1 | 1 | 2 | 1 | NA | NA | 5 | 0.8 | 1 | 1 | 3 | 1 | 1 | 1 | 1 | 1 | 6 | 1 |
| 13_745937 | 2 | 0 | NA | NA | 2 | 0.5 | NA | NA | 4 | 0.25 | NA | NA | 3 | 0 | 1 | 1 | NA | NA | 4 | 0.25 |
| 13_745937 | 2 | 1 | 1 | 1 | 2 | 1 | NA | NA | 5 | 1 | NA | NA | 3 | 1 | 1 | 1 | 1 | 1 | 5 | 1 |
| 13_745939 | 1 | 1 | 1 | 0 | 2 | 0.5 | NA | NA | 4 | 0.5 | 1 | 1 | 2 | 0 | 1 | 1 | NA | NA | 4 | 0.5 |
| 13_745939 | 1 | 1 | 1 | 1 | 2 | 1 | NA | NA | 4 | 1 | NA | NA | 3 | 1 | 1 | 0 | NA | NA | 4 | 0.75 |
| 13_745939 | 1 | 1 | 1 | 1 | 2 | 1 | NA | NA | 4 | 1 | 1 | 1 | 3 | 1 | 1 | 1 | NA | NA | 5 | 1 |
| 13_745939 | 1 | 1 | 1 | 1 | 2 | 1 | NA | NA | 4 | 1 | 1 | 1 | 3 | 1 | 1 | 1 | NA | NA | 5 | 1 |
| 13_745939 | 1 | 0 | 1 | 1 | 2 | 1 | NA | NA | 4 | 0.75 | 1 | 1 | 3 | 0.6667 | 1 | 0 | NA | NA | 5 | 0.6 |
| 13_745940 | 1 | 1 | 1 | 1 | 2 | 1 | NA | NA | 4 | 1 | NA | NA | 3 | 1 | NA | NA | NA | NA | 3 | 1 |
| 13_745946 | 1 | 1 | 1 | 1 | 2 | 0 | NA | NA | 4 | 0.5 | NA | NA | 2 | 0.5 | 1 | 0 | NA | NA | 3 | 0.3333 |
| 13_745947 | NA | NA | 1 | 1 | 2 | 0.5 | NA | NA | 3 | 0.6667 | NA | NA | 1 | 1 | 1 | 1 | NA | NA | 2 | 1 |
| 13_745951 | 1 | 1 | 1 | 1 | 2 | 0 | NA | NA | 4 | 0.5 | NA | NA | 1 | 1 | 1 | 0 | NA | NA | 2 | 0.5 |
| 13_745953 | 1 | 1 | 1 | 1 | 2 | 1 | NA | NA | 4 | 1 | NA | NA | 1 | 1 | 1 | 1 | NA | NA | 2 | 1 |
| 13_745964 | 1 | 0 | 1 | 1 | 2 | 0 | NA | NA | 4 | 0.25 | NA | NA | 1 | 1 | NA | NA | NA | NA | 1 | 1 |
| 13_746282 | 2 | 1 | 1 | 1 | 2 | 0 | 2 | 1 | 7 | 0.7143 | 1 | 1 | NA | NA | 2 | 1 | 1 | 0 | 4 | 0.75 |
| 13_746284 | 2 | 1 | 1 | 0 | 3 | 0.3333 | 2 | 0.5 | 8 | 0.5 | 1 | 1 | NA | NA | 1 | 0 | 1 | 1 | 3 | 0.6667 |
| 13_746285 | 3 | 0.6667 | 1 | 1 | 3 | 0.3333 | 2 | 0.5 | 9 | 0.5556 | 1 | 1 | NA | NA | 2 | 0 | 2 | 0.5 | 5 | 0.4 |
| 13_746290 | 2 | 1 | 1 | 1 | 3 | 1 | 2 | 0.5 | 8 | 0.875 | NA | NA | NA | NA | 2 | 0.5 | 1 | 0 | 3 | 0.3333 |
| 13_746996 | 1 | 1 | NA | NA | 1 | 1 | NA | NA | 2 | 1 | 1 | 1 | NA | NA | NA | NA | NA | NA | 1 | 1 |
| 13_747000 | 1 | 1 | 2 | 0.5 | 2 | 1 | NA | NA | 5 | 0.8 | 1 | 1 | 1 | 0 | 2 | 1 | 1 | 1 | 5 | 0.8 |
| 13_747001 | 1 | 1 | 1 | 0 | 2 | 0.5 | NA | NA | 4 | 0.5 | 1 | 0 | 1 | 0 | 2 | 0.5 | 1 | 0 | 5 | 0.2 |
| 13_747002 | 1 | 0 | 2 | 0 | 1 | 1 | NA | NA | 4 | 0.25 | 1 | 1 | 1 | 0 | 1 | 1 | 1 | 0 | 4 | 0.5 |
| 13_747003 | 1 | 1 | 2 | 1 | 2 | 1 | NA | NA | 5 | 1 | 1 | 1 | 1 | 1 | 2 | 0.5 | 1 | 0 | 5 | 0.6 |
| 13_747004 | 1 | 1 | 2 | 1 | 2 | 1 | NA | NA | 5 | 1 | 1 | 1 | 1 | 1 | 1 | 1 | 1 | 1 | 4 | 1 |
| 13_747004 | 1 | 1 | 2 | 0.5 | 2 | 1 | NA | NA | 5 | 0.8 | 1 | 1 | 1 | 1 | 2 | 1 | 1 | 1 | 5 | 1 |
| 13_747005 | NA | NA | 2 | 0.5 | 2 | 0 | NA | NA | 4 | 0.25 | 1 | 0 | 1 | 1 | 1 | 0 | 1 | 0 | 4 | 0.25 |

| | | | | | | | | | | | | | | | | | | | | |
|---|---|---|---|---|---|---|---|---|---|---|---|---|---|---|---|---|---|---|---|---|
| 13_7470075 | 1 | 1 | 2 | 0.5 | 1 | 0 | NA | NA | 4 | 0.5 | 1 | 1 | 1 | 0 | 1 | 0 | 1 | 1 | 4 | 0.5 |
| 13_7470079 | 1 | 1 | 2 | 0.5 | 1 | 1 | NA | NA | 4 | 0.75 | 1 | 1 | 1 | 1 | NA | NA | 1 | 0 | 3 | 0.6667 |
| 13_7470106 | 1 | 0 | 2 | 0 | 1 | 1 | NA | NA | 4 | 0.25 | 1 | 1 | 1 | 0 | NA | NA | 1 | 0 | 3 | 0.3333 |
| 13_7470112 | NA | NA | 2 | 0.5 | 1 | 1 | NA | NA | 3 | 0.6667 | 1 | 1 | 1 | 1 | NA | NA | 1 | 1 | 3 | 1 |
| 13_7470122 | NA | NA | 1 | 0 | 1 | 1 | NA | NA | 2 | 0.5 | 1 | 1 | 1 | 1 | NA | NA | 1 | 1 | 3 | 1 |
| 13_7470164 | 1 | 1 | 1 | 1 | 1 | 1 | NA | NA | 3 | 1 | 1 | 1 | NA | NA | NA | NA | 1 | 1 | 2 | 1 |
| 13_7470175 | 1 | 1 | NA | NA | 1 | 0 | NA | NA | 2 | 0.5 | 1 | 0 | NA | NA | NA | NA | 1 | 1 | 2 | 0.5 |
| 13_7470177 | 1 | 0 | 1 | 1 | 1 | 0 | NA | NA | 3 | 0.3333 | 1 | 0 | 1 | 1 | NA | NA | 1 | 0 | 3 | 0.3333 |
| 13_7470180 | NA | NA | 1 | 1 | 1 | 1 | NA | NA | 2 | 1 | 1 | 1 | 1 | 1 | NA | NA | 1 | 1 | 3 | 1 |
| 13_7470221 | 1 | 0 | 1 | 0 | 1 | 1 | NA | NA | 3 | 0.3333 | 1 | 0 | 1 | 1 | NA | NA | NA | NA | 2 | 0.5 |
| 13_7470229 | 1 | 1 | 1 | 1 | 1 | 1 | NA | NA | 3 | 1 | 1 | 1 | 1 | 0 | NA | NA | NA | NA | 2 | 0.5 |
| 13_7475316 | NA | NA | 2 | 0 | 1 | 1 | 1 | 0 | 4 | 0.25 | NA | NA | NA | NA | 1 | 1 | 2 | 0 | 3 | 0.3333 |
| 13_7475330 | 1 | 0 | 1 | 0 | 1 | 1 | 1 | 0 | 4 | 0.25 | 1 | 0 | NA | NA | 1 | 0 | 2 | 1 | 4 | 0.5 |
| 13_7475344 | 1 | 1 | 1 | 0 | 1 | 1 | 1 | 1 | 4 | 0.75 | 1 | 1 | NA | NA | 1 | 1 | 2 | 1 | 4 | 1 |
| 13_7475346 | 1 | 0 | 1 | 1 | 1 | 1 | 1 | 0 | 4 | 0.5 | 1 | 0 | NA | NA | 1 | 1 | 1 | 0 | 3 | 0.3333 |
| 13_7475347 | 1 | 1 | NA | NA | 1 | 1 | 1 | 1 | 3 | 1 | 1 | 1 | NA | NA | 1 | 1 | NA | NA | 2 | 1 |
| 13_7475349 | 1 | 1 | 1 | 0 | 1 | 1 | 1 | 1 | 4 | 0.75 | 1 | 1 | NA | NA | 1 | 1 | 1 | 1 | 3 | 1 |
| 13_7475360 | 1 | 0 | 1 | 0 | 1 | 1 | 1 | 0 | 4 | 0.25 | 1 | 0 | NA | NA | 1 | 1 | NA | NA | 2 | 0.5 |
| 13_7475363 | 1 | 1 | 1 | 1 | 1 | 0 | 1 | 1 | 4 | 0.75 | 1 | 0 | NA | NA | 1 | 1 | 1 | 1 | 3 | 0.6667 |
| 13_7475387 | 1 | 0 | 1 | 1 | 1 | 0 | 1 | 1 | 4 | 0.5 | 1 | 1 | NA | NA | 1 | 1 | 1 | 1 | 3 | 1 |
| 13_7475429 | 1 | 0 | 1 | 1 | 1 | 0 | NA | NA | 3 | 0.3333 | 1 | 0 | NA | NA | NA | NA | 1 | 1 | 2 | 0.5 |
| 13_7475487 | 1 | 0 | 1 | 1 | 1 | 1 | 1 | 1 | 4 | 0.75 | 1 | 1 | NA | NA | 1 | 0 | 1 | 1 | 3 | 0.6667 |
| 13_7475567 | NA | NA | 1 | 1 | NA | NA | 1 | 0 | 2 | 0.5 | 1 | 1 | NA | NA | 1 | 0 | 1 | 0 | 3 | 0.3333 |
| 13_7483887 | NA | NA | 2 | 0 | 1 | 1 | 2 | 0.5 | 5 | 0.4 | NA | NA | 2 | 1 | 1 | 1 | 1 | 1 | 4 | 1 |
| 13_7483900 | NA | NA | 1 | 0 | 1 | 1 | 2 | 0 | 4 | 0.25 | 1 | 0 | 1 | 1 | 2 | 0 | 1 | 0 | 5 | 0.2 |
| 13_7483911 | NA | NA | 1 | 0 | NA | NA | 2 | 0.5 | 3 | 0.3333 | 1 | 0 | 1 | 0 | 2 | 0 | 1 | 1 | 5 | 0.2 |
| 13_7523819 | NA | NA | 1 | 0 | 1 | 0 | 2 | 1 | 4 | 0.5 | 1 | 1 | NA | NA | 1 | 0 | 6 | 0.5 | 8 | 0.5 |
| 13_7527819 | 2 | 0.5 | 5 | 0 | 4 | 0 | 2 | 0 | 13 | 0.0769 | 2 | 0 | 1 | 0 | 2 | 0.5 | 9 | 0 | 14 | 0.0714 |
| 13_7528016 | 2 | 0 | 5 | 0.6 | 4 | 0.5 | 2 | 0.5 | 13 | 0.4615 | 2 | 0.5 | 1 | 1 | 1 | 1 | 10 | 0.2 | 14 | 0.3571 |
| 13_7528022 | 2 | 0 | 3 | 0.3333 | 3 | 0 | 2 | 0 | 10 | 0.1 | 2 | 0 | 1 | 0 | 2 | 0 | 8 | 0.125 | 13 | 0.0769 |
| 13_7528032 | 3 | 0 | 6 | 0.1667 | 4 | 0 | 2 | 0.5 | 15 | 0.1333 | 2 | 0 | 1 | 0 | 2 | 0 | 9 | 0.2222 | 14 | 0.1429 |
| 13_7528034 | 3 | 1 | 6 | 0.5 | 3 | 0.6667 | 3 | 1 | 15 | 0.7333 | 2 | 0.5 | 1 | 1 | 1 | 1 | 10 | 0.9 | 14 | 0.8571 |
| 13_7528041 | 3 | 1 | 5 | 1 | 4 | 1 | 3 | 1 | 15 | 1 | 2 | 1 | 1 | 1 | 2 | 1 | 8 | 1 | 13 | 1 |
| 13_7528069 | 1 | 0 | 5 | 0 | 4 | 0.25 | 3 | 0 | 13 | 0.0769 | 2 | 0 | 1 | 0 | 2 | 0 | 7 | 0.1429 | 12 | 0.0833 |
| 13_7528211 | 2 | 0 | 5 | 0.2 | 4 | 0 | 2 | 0 | 13 | 0.0769 | 2 | 0 | NA | NA | 1 | 0 | 7 | 0.2857 | 10 | 0.2 |
| 13_7528251 | 2 | 0 | 7 | 0.2857 | 4 | 0 | 2 | 0 | 15 | 0.1333 | 2 | 0 | 1 | 0 | 4 | 0 | 10 | 0.2 | 17 | 0.1176 |
| 13_7528266 | 2 | 0 | 7 | 0.1429 | 4 | 0 | 3 | 0.6667 | 16 | 0.1875 | 2 | 0 | 1 | 0 | 4 | 0 | 10 | 0.3 | 17 | 0.1765 |
| 13_7591188 | 1 | 0 | 2 | 1 | 1 | 1 | NA | NA | 4 | 0.75 | NA | NA | NA | NA | 2 | 0.5 | 2 | 1 | 4 | 0.75 |
| 13_7591200 | 1 | 0 | 2 | 1 | 1 | 1 | NA | NA | 4 | 0.75 | NA | NA | NA | NA | 2 | 0.5 | 2 | 1 | 4 | 0.75 |
| 13_7599751 | 5 | 1 | 5 | 1 | 6 | 1 | 2 | 1 | 18 | 1 | 4 | 1 | 2 | 1 | 3 | 1 | 4 | 1 | 13 | 1 |
| 13_7599755 | 7 | 0.1429 | 7 | 0.1429 | 7 | 0.2857 | 5 | 0.2 | 26 | 0.1923 | 3 | 0.3333 | 2 | 0 | 4 | 0.25 | 5 | 0 | 14 | 0.1429 |
| 13_7599790 | 4 | 1 | 4 | 0.75 | 5 | 1 | 4 | 0.5 | 17 | 0.8235 | 4 | 0.5 | 2 | 1 | 3 | 1 | 4 | 1 | 13 | 0.8462 |
| 13_7601793 | 6 | 0 | 3 | 0.3333 | 2 | 0 | 2 | 0 | 13 | 0.0769 | 1 | 1 | 1 | 0 | 1 | 1 | 1 | 0 | 4 | 0.5 |
| 13_7601988 | 5 | 0.2 | 3 | 0.3333 | 1 | 0 | 2 | 0 | 11 | 0.1818 | 1 | 1 | 1 | 0 | 1 | 0 | 2 | 0 | 5 | 0.2 |
| 13_7602026 | 5 | 0 | 2 | 0 | 2 | 0.5 | NA | NA | 9 | 0.1111 | 1 | 1 | 1 | 0 | 1 | 0 | 2 | 0 | 5 | 0.2 |
| 13_7602032 | 6 | 0.1667 | 2 | 0.5 | 2 | 0.5 | 2 | 0.5 | 12 | 0.3333 | 1 | 1 | 1 | 1 | 1 | 1 | 2 | 0.5 | 5 | 0.8 |
| 13_7605887 | 2 | 0 | 2 | 0.5 | 1 | 0 | 1 | 0 | 6 | 0.1667 | 1 | 0 | 1 | 0 | 1 | 0 | 2 | 0.5 | 5 | 0.2 |
| 13_7614717 | 2 | 0.5 | 2 | 0.5 | NA | NA | NA | NA | 4 | 0.5 | 1 | 1 | NA | NA | 1 | 0 | 1 | 0 | 3 | 0.3333 |
| 13_7614742 | 1 | 1 | 2 | 1 | NA | NA | 1 | 0 | 4 | 0.75 | 1 | 0 | NA | NA | 1 | 0 | 1 | 1 | 3 | 0.3333 |
| 13_7614746 | 1 | 1 | 2 | 1 | NA | NA | 1 | 0 | 4 | 0.75 | 1 | 0 | NA | NA | 1 | 0 | 1 | 1 | 3 | 0.3333 |
| 13_7615099 | 4 | 0.75 | 2 | 1 | 1 | 0 | 7 | 0.7143 | 14 | 0.7143 | 1 | 0 | 1 | 0 | 5 | 0.6 | 2 | 0 | 9 | 0.3333 |
| 13_7615101 | 2 | 0 | 2 | 0 | 1 | 0 | 7 | 0.1429 | 12 | 0.0833 | 2 | 0.5 | NA | NA | 4 | 0.25 | 2 | 0 | 8 | 0.25 |
| 13_7615102 | 4 | 0.25 | 2 | 0.5 | 1 | 0 | 6 | 0.6667 | 13 | 0.4615 | 1 | 0 | NA | NA | 5 | 0.4 | 3 | 0 | 9 | 0.2222 |
| 13_7615111 | 4 | 0.25 | 2 | 0 | 1 | 0 | 7 | 0.4286 | 14 | 0.2857 | 2 | 0 | NA | NA | 5 | 0 | 3 | 0.3333 | 10 | 0.1 |
| 13_7615119 | 4 | 0.5 | 1 | 1 | 1 | 0 | 5 | 0.2 | 11 | 0.3636 | 2 | 0 | NA | NA | 4 | 0.75 | 2 | 0.5 | 8 | 0.5 |
| 13_7616584 | 1 | 0 | 1 | 1 | NA | NA | 1 | 1 | 3 | 0.6667 | NA | NA | NA | NA | 1 | 1 | NA | NA | 1 | 1 |
| 13_7616635 | 2 | 0.5 | 1 | 1 | NA | NA | 1 | 0 | 4 | 0.5 | 2 | 1 | NA | NA | 2 | 1 | NA | NA | 4 | 1 |
| 13_7616636 | 2 | 0 | 1 | 1 | NA | NA | 4 | 0.5 | 7 | 0.4286 | 2 | 0 | NA | NA | 2 | 0.5 | NA | NA | 4 | 0.25 |
| 13_7616658 | 2 | 1 | 1 | 1 | NA | NA | 3 | 0.3333 | 6 | 0.6667 | 1 | 1 | NA | NA | 2 | 1 | NA | NA | 3 | 1 |
| 13_7616660 | 1 | 1 | 1 | 1 | NA | NA | 2 | 0.5 | 4 | 0.75 | 1 | 1 | NA | NA | 2 | 0 | NA | NA | 3 | 0.3333 |
| 13_7620355 | NA | NA | 1 | 1 | 1 | 1 | NA | NA | 2 | 1 | 2 | 0.5 | 2 | 0 | 1 | 0 | NA | NA | 5 | 0.2 |
| 13_7620356 | NA | NA | 1 | 1 | 1 | 1 | NA | NA | 2 | 1 | 1 | 1 | 2 | 1 | 1 | 1 | NA | NA | 4 | 1 |
| 13_7620364 | NA | NA | 1 | 1 | 1 | 1 | NA | NA | 2 | 1 | 1 | 1 | 2 | 1 | 1 | 1 | NA | NA | 4 | 1 |
| 13_7620365 | NA | NA | 1 | 1 | 1 | 1 | NA | NA | 2 | 1 | 1 | 1 | 1 | 1 | 1 | 1 | NA | NA | 3 | 1 |
| 13_7653173 | NA | NA | 2 | 1 | NA | NA | NA | NA | 2 | 1 | NA | NA | NA | NA | 1 | 1 | NA | NA | 1 | 1 |

| | | | | | | | | | | | | | | | | | | | | |
|---|---|---|---|---|---|---|---|---|---|---|---|---|---|---|---|---|---|---|---|---|
| 13_765317 | NA | NA | 1 | 1 | NA | NA | NA | NA | 1 | 1 | NA | NA | 1 | 1 | 1 | 1 | NA | NA | 2 | 1 |
| 13_765320 | 1 | 1 | 1 | 1 | NA | NA | NA | NA | 2 | 1 | NA | NA | 1 | 0 | 1 | 1 | NA | NA | 2 | 0.5 |
| 13_765320 | 1 | 1 | 2 | 0.5 | NA | NA | NA | NA | 3 | 0.6667 | NA | NA | 1 | 0 | 1 | 1 | NA | NA | 2 | 0.5 |
| 13_765320 | 1 | 1 | 2 | 1 | NA | NA | NA | NA | 3 | 1 | NA | NA | 1 | 0 | 1 | 1 | NA | NA | 2 | 0.5 |
| 13_765322 | 1 | 1 | NA | NA | NA | NA | NA | NA | 1 | 1 | NA | NA | NA | NA | 1 | 1 | NA | NA | 1 | 1 |
| 13_765324 | 1 | 1 | 1 | 0 | NA | NA | NA | NA | 2 | 0.5 | NA | NA | 1 | 0 | 1 | 1 | NA | NA | 2 | 0.5 |
| 13_771706 | NA | NA | NA | NA | 1 | 1 | NA | NA | 1 | 1 | NA | NA | NA | NA | 1 | 1 | NA | NA | 1 | 1 |
| 13_771782 | 18 | 0 | 12 | 0.0833 | 11 | 0 | 18 | 0.1111 | 59 | 0.0508 | 4 | 0.25 | 15 | 0.0667 | 15 | 0.0667 | 17 | 0.1176 | 51 | 0.098 |
| 13_771803 | 7 | 0.2857 | 2 | 0.5 | 3 | 0.3333 | 3 | 0 | 15 | 0.2667 | 1 | 1 | 3 | 0 | 4 | 1 | 2 | 0.5 | 10 | 0.6 |
| 13_771804 | 6 | 0 | 2 | 0 | 4 | 0.25 | 3 | 0.6667 | 15 | 0.2 | 1 | 0 | 3 | 0 | 5 | 0.2 | 2 | 0 | 11 | 0.0909 |
| 13_771805 | 7 | 0.5714 | 2 | 0.5 | 5 | 0.4 | 2 | 0.5 | 16 | 0.5 | 1 | 1 | 2 | 0 | 5 | 0.6 | 3 | 0.6667 | 11 | 0.5455 |
| 13_771806 | 7 | 0 | 2 | 0.5 | 4 | 0.25 | 4 | 0.5 | 17 | 0.2353 | 1 | 1 | 3 | 0 | 3 | 0.3333 | 3 | 0 | 10 | 0.2 |
| 13_771917 | 2 | 0.5 | 1 | 0 | 2 | 0 | 4 | 0 | 9 | 0.1111 | NA | NA | 4 | 0.25 | 5 | 0.2 | NA | NA | 9 | 0.2222 |
| 13_771918 | 2 | 0 | 2 | 0.5 | 2 | 0.5 | 6 | 0.1667 | 12 | 0.25 | NA | NA | 3 | 0 | 5 | 0.6 | 1 | 0 | 9 | 0.3333 |
| 13_771920 | 2 | 0.5 | 1 | 0 | 1 | 0 | 5 | 0 | 9 | 0.1111 | NA | NA | 4 | 0 | 4 | 0.25 | 1 | 1 | 9 | 0.2222 |
| 13_771920 | 1 | 1 | 2 | 0.5 | 2 | 0 | 6 | 0.5 | 11 | 0.4545 | NA | NA | 4 | 0.5 | 5 | 0 | 1 | 0 | 10 | 0.2 |
| 13_771920 | 1 | 0 | 2 | 0 | 2 | 1 | 6 | 0.3333 | 11 | 0.3636 | NA | NA | 5 | 0.2 | 4 | 0 | 1 | 1 | 10 | 0.2 |
| 13_772830 | 1 | 1 | NA | NA | NA | NA | 2 | 0.5 | 3 | 0.6667 | NA | NA | NA | NA | 1 | 1 | 1 | 1 | 2 | 1 |
| 13_772833 | 1 | 1 | NA | NA | NA | NA | 2 | 0.5 | 3 | 0.6667 | NA | NA | NA | NA | 1 | 1 | 1 | 0 | 2 | 0.5 |
| 13_773410 | NA | NA | NA | NA | NA | NA | 1 | 1 | 1 | 1 | NA | NA | 1 | 1 | NA | NA | NA | NA | 1 | 1 |
| 13_783122 | 3 | 1 | 4 | 1 | 2 | 1 | 4 | 1 | 13 | 1 | 2 | 1 | 4 | 0.75 | 4 | 0.75 | 5 | 0.6 | 15 | 0.7333 |
| 13_783123 | 3 | 0 | 4 | 0 | 2 | 0.5 | 4 | 0.25 | 13 | 0.1538 | 2 | 0 | 3 | 0 | 2 | 0 | 4 | 0.75 | 11 | 0.2727 |
| 13_783126 | 3 | 0 | 5 | 0.4 | 4 | 0 | 5 | 0 | 17 | 0.1176 | 2 | 0 | 4 | 0.25 | 4 | 0.25 | 6 | 0.1667 | 16 | 0.1875 |
| 13_783133 | 4 | 0.5 | 4 | 0 | 3 | 0.3333 | 4 | 0 | 15 | 0.2 | 3 | 0 | 4 | 0 | 5 | 0.2 | 4 | 0.25 | 16 | 0.125 |
| 13_783374 | 4 | 0 | 8 | 0.25 | 3 | 0 | 4 | 0.25 | 19 | 0.1579 | 6 | 0.3333 | 2 | 0 | NA | NA | 6 | 0 | 14 | 0.1429 |
| 13_783374 | 5 | 1 | 8 | 1 | 3 | 1 | 4 | 1 | 20 | 1 | 8 | 0.875 | 2 | 1 | 1 | 1 | 6 | 0.8333 | 17 | 0.8824 |
| 13_783375 | 5 | 1 | 6 | 1 | 3 | 1 | 5 | 1 | 19 | 1 | 7 | 1 | 2 | 0.5 | 2 | 1 | 4 | 1 | 15 | 0.9333 |
| 13_783375 | 4 | 0.25 | 9 | 0.6667 | 3 | 1 | 6 | 0.3333 | 22 | 0.5455 | 7 | 0.1429 | 2 | 0.5 | 2 | 0.5 | 4 | 0.25 | 15 | 0.2667 |
| 13_783376 | 5 | 0.8 | 8 | 0.5 | 3 | 0.6667 | 6 | 0.5 | 22 | 0.5909 | 7 | 0.5714 | 2 | 0.5 | 1 | 1 | 5 | 0.6 | 15 | 0.6 |
| 13_783376 | 5 | 0.4 | 9 | 0.5556 | 2 | 0.5 | 6 | 0.5 | 22 | 0.5 | 7 | 0.4286 | 2 | 0 | 2 | 0.5 | 6 | 0.8333 | 17 | 0.5294 |
| 13_783377 | 5 | 0.8 | 5 | 0.4 | 2 | 0.5 | 6 | 0.3333 | 18 | 0.5 | 5 | 0.6 | 2 | 0.5 | 2 | 0 | 3 | 0.3333 | 12 | 0.4167 |
| 13_783377 | 5 | 1 | 6 | 0.8333 | 3 | 0.3333 | 5 | 1 | 19 | 0.8421 | 6 | 1 | 2 | 1 | 2 | 1 | 5 | 1 | 15 | 1 |
| 13_783377 | 5 | 1 | 7 | 1 | 3 | 1 | 6 | 0.8333 | 21 | 0.9524 | 6 | 1 | 2 | 0.5 | 2 | 1 | 6 | 1 | 16 | 0.9375 |
| 13_783378 | 5 | 1 | 9 | 0.8889 | 3 | 1 | 6 | 0.8333 | 23 | 0.913 | 7 | 1 | 1 | 1 | 2 | 1 | 6 | 0.6667 | 16 | 0.875 |
| 13_783378 | 4 | 1 | 7 | 0.8571 | 1 | 1 | 5 | 1 | 17 | 0.9412 | 4 | 1 | NA | NA | 2 | 1 | 4 | 1 | 10 | 1 |
| 13_783379 | 4 | 0 | 7 | 0.1429 | 2 | 0 | 4 | 0 | 17 | 0.0588 | 5 | 0.2 | 1 | 0 | 1 | 0 | 4 | 0.25 | 11 | 0.1818 |
| 13_783379 | 4 | 0.5 | 7 | 0 | 3 | 0 | 5 | 0 | 19 | 0.1053 | 6 | 0.1667 | 1 | 0 | 2 | 0.5 | 5 | 0 | 14 | 0.1429 |
| 13_783433 | 4 | 0.25 | 7 | 0 | 3 | 0 | 6 | 0 | 20 | 0.05 | 7 | 0 | 1 | 0 | 1 | 1 | 5 | 0 | 14 | 0.0714 |
| 13_783433 | 4 | 0.25 | 6 | 0.3333 | 3 | 0 | 6 | 0 | 19 | 0.1579 | 5 | 0.2 | 1 | 0 | 1 | 0 | 5 | 0.2 | 12 | 0.1667 |
| 13_783434 | 4 | 0.25 | 6 | 0.5 | 2 | 0 | 6 | 0.3333 | 18 | 0.3333 | 6 | 0.1667 | 1 | 0 | 1 | 0 | 4 | 0 | 12 | 0.0833 |
| 13_783435 | 3 | 0.6667 | 6 | 0.3333 | 3 | 0.3333 | 5 | 0.2 | 17 | 0.3529 | 4 | 0 | NA | NA | 1 | 1 | 5 | 0.6 | 10 | 0.4 |
| 13_783463 | 3 | 0 | 4 | 0.25 | 2 | 0.5 | 2 | 0.5 | 11 | 0.2727 | 3 | 0.6667 | NA | NA | 1 | 0 | 5 | 0 | 9 | 0.2222 |
| 13_783463 | 3 | 1 | 3 | 0.3333 | 3 | 0.3333 | 2 | 0.5 | 11 | 0.5455 | 4 | 0 | NA | NA | 1 | 0 | 4 | 0.25 | 9 | 0.1111 |
| 13_783464 | 3 | 0.6667 | 4 | 0.25 | 2 | 1 | 1 | 1 | 10 | 0.6 | 3 | 0.3333 | NA | NA | 1 | 0 | 3 | 0.3333 | 7 | 0.2857 |
| 13_810316 | 3 | 0 | 3 | 0 | 2 | 0 | 4 | 0.25 | 12 | 0.0833 | 1 | 0 | 1 | 0 | 2 | 0.5 | 1 | 0 | 5 | 0.2 |
| 13_810415 | 8 | 0.125 | 7 | 0.1429 | 4 | 0 | 4 | 0 | 23 | 0.087 | 4 | 0.25 | 3 | 0 | 6 | 0.1667 | 2 | 0 | 15 | 0.1333 |
| 13_810415 | 6 | 1 | 6 | 0.6667 | 4 | 1 | 3 | 1 | 19 | 0.8947 | 3 | 1 | 3 | 1 | 6 | 1 | 2 | 1 | 14 | 1 |
| 13_810416 | 6 | 0.1667 | 6 | 0.6667 | 4 | 0.25 | 4 | 0.25 | 20 | 0.35 | 4 | 0 | 2 | 0 | 6 | 0.3333 | 3 | 0.6667 | 15 | 0.2667 |
| 13_810417 | 5 | 0.8 | 3 | 1 | 3 | 0.6667 | 3 | 0.6667 | 14 | 0.7857 | 4 | 1 | 4 | 0.75 | 6 | 0.6667 | 2 | 1 | 16 | 0.8125 |
| 13_810418 | 7 | 0.4286 | 6 | 0.1667 | 4 | 0.25 | 3 | 0 | 20 | 0.25 | 3 | 0.3333 | 3 | 0 | 5 | 0.2 | 4 | 0.25 | 15 | 0.2 |
| 13_810419 | 8 | 1 | 5 | 1 | 4 | 1 | 4 | 0.75 | 21 | 0.9524 | 3 | 0.6667 | 4 | 1 | 5 | 1 | 4 | 1 | 16 | 0.9375 |
| 13_810419 | 7 | 0 | 6 | 0.1667 | 4 | 0.5 | 2 | 0 | 19 | 0.1579 | 2 | 0 | 4 | 0 | 5 | 0 | 3 | 0.3333 | 14 | 0.0714 |
| 13_810419 | 7 | 1 | 6 | 1 | 4 | 1 | 4 | 1 | 21 | 1 | 4 | 1 | 2 | 0.5 | 7 | 0.8571 | 3 | 1 | 16 | 0.875 |
| 13_810420 | 6 | 0.6667 | 6 | 0.6667 | 4 | 0.5 | 3 | 0.6667 | 19 | 0.6316 | 3 | 0.3333 | 3 | 0.6667 | 5 | 0.4 | 3 | 1 | 14 | 0.5714 |
| 13_810421 | 8 | 1 | 6 | 1 | 4 | 1 | 4 | 1 | 22 | 1 | 4 | 1 | 4 | 1 | 6 | 1 | 4 | 1 | 18 | 1 |
| 13_810422 | 8 | 0.25 | 6 | 0.3333 | 4 | 0 | 4 | 0 | 22 | 0.1818 | 4 | 0 | 4 | 0.5 | 5 | 0.2 | 3 | 0.3333 | 16 | 0.25 |
| 13_810422 | 8 | 0 | 8 | 0.375 | 3 | 0 | 4 | 0.5 | 23 | 0.2174 | 3 | 0.6667 | 4 | 0.75 | 6 | 0.5 | 4 | 0.25 | 17 | 0.5294 |
| 13_810423 | 8 | 1 | 7 | 1 | 4 | 1 | 5 | 1 | 24 | 1 | 4 | 1 | 4 | 1 | 5 | 0.8 | 4 | 0.75 | 17 | 0.8824 |
| 13_810423 | 7 | 0.5714 | 6 | 0.3333 | 4 | 0.5 | 5 | 0.6 | 22 | 0.5 | 4 | 0.5 | 4 | 0.25 | 6 | 0.3333 | 5 | 0.4 | 19 | 0.3684 |
| 13_811766 | 1 | 1 | NA | NA | NA | NA | NA | NA | 1 | 1 | NA | NA | NA | NA | NA | NA | 1 | 1 | 1 | 1 |
| 13_811788 | 1 | 1 | NA | NA | NA | NA | NA | NA | 1 | 1 | NA | NA | NA | NA | NA | NA | 1 | 1 | 1 | 1 |
| 13_811790 | 1 | 1 | NA | NA | NA | NA | NA | NA | 1 | 1 | NA | NA | NA | NA | NA | NA | 1 | 1 | 1 | 1 |
| 13_818206 | 1 | 0 | 1 | 1 | NA | NA | 4 | 0.25 | 6 | 0.3333 | 3 | 0.3333 | 2 | 0 | 1 | 1 | 3 | 0.6667 | 9 | 0.4444 |
| 13_818234 | NA | NA | 1 | 1 | 3 | 0.3333 | NA | NA | 4 | 0.5 | NA | NA | NA | NA | 1 | 1 | NA | NA | 1 | 1 |
| 13_818236 | 1 | 1 | NA | NA | 1 | 1 | NA | NA | 2 | 1 | NA | NA | NA | NA | 1 | 1 | NA | NA | 1 | 1 |

| | | | | | | | | | | | | | | | | | | | | |
|---|---|---|---|---|---|---|---|---|---|---|---|---|---|---|---|---|---|---|---|---|
| 13_818237 | 1 | 1 | 1 | 1 | 1 | 1 | NA | NA | 3 | 1 | NA | NA | NA | NA | 1 | 1 | NA | NA | 1 | 1 |
| 13_818597 | NA | NA | NA | NA | 1 | 1 | 4 | 0.5 | 5 | 0.6 | 1 | 1 | NA | NA | NA | NA | 1 | 0 | 2 | 0.5 |
| 13_818982 | 1 | 0 | NA | NA | NA | NA | 3 | 0.3333 | 4 | 0.25 | NA | NA | NA | NA | NA | NA | 3 | 0.3333 | 3 | 0.3333 |
| 13_818982 | NA | NA | NA | NA | 1 | 1 | 3 | 0.6667 | 4 | 0.75 | NA | NA | NA | NA | NA | NA | 2 | 0.5 | 2 | 0.5 |
| 13_818982 | 1 | 0 | NA | NA | 1 | 1 | 5 | 0.8 | 7 | 0.7143 | NA | NA | NA | NA | NA | NA | 3 | 0.6667 | 3 | 0.6667 |
| 13_818982 | 1 | 0 | NA | NA | NA | NA | 3 | 0.6667 | 4 | 0.5 | NA | NA | NA | NA | NA | NA | 3 | 0.3333 | 3 | 0.3333 |
| 13_819011 | 4 | 0 | 6 | 0.1667 | 2 | 0.5 | 6 | 0.5 | 18 | 0.2778 | 3 | 0.3333 | 3 | 0.3333 | 3 | 0.3333 | 2 | 0.5 | 11 | 0.3636 |
| 13_819448 | 14 | 0.2857 | 10 | 0.3 | 14 | 0.2857 | 17 | 0.3529 | 55 | 0.3091 | 7 | 0 | 11 | 0.1818 | 18 | 0.2222 | 16 | 0.125 | 52 | 0.1538 |
| 13_838105 | 10 | 0.2 | 13 | 0.1538 | 8 | 0.125 | 9 | 0.2222 | 40 | 0.175 | 3 | 0 | 4 | 0.25 | 4 | 0 | 7 | 0.2857 | 18 | 0.1667 |
| 13_838106 | 6 | 0.8333 | 10 | 0.6 | 8 | 0.625 | 7 | 0.4286 | 31 | 0.6129 | 2 | 0 | 4 | 0.75 | 4 | 0.5 | 4 | 0.75 | 14 | 0.5714 |
| 13_838116 | 10 | 0.1 | 17 | 0.1765 | 8 | 0.375 | 16 | 0.1875 | 51 | 0.1961 | 3 | 0.3333 | 4 | 0.5 | 7 | 0.1429 | 12 | 0.25 | 26 | 0.2692 |
| 13_838140 | 23 | 0.1739 | 31 | 0.1935 | 20 | 0.2 | 19 | 0.1579 | 93 | 0.1828 | 13 | 0.1538 | 12 | 0 | 9 | 0.2222 | 20 | 0.25 | 54 | 0.1667 |
| 13_838150 | 22 | 0.1818 | 35 | 0.2 | 21 | 0.1905 | 19 | 0.4211 | 97 | 0.2371 | 19 | 0.1579 | 13 | 0.2308 | 8 | 0.125 | 12 | 0.1667 | 52 | 0.1731 |
| 13_838150 | 26 | 0.1923 | 39 | 0.2564 | 25 | 0.12 | 15 | 0.2 | 105 | 0.2 | 20 | 0.1 | 15 | 0.2 | 10 | 0.1 | 13 | 0 | 58 | 0.1034 |
| 13_838865 | 1 | 1 | 2 | 0 | 1 | 1 | 1 | 1 | 5 | 0.6 | NA | NA | NA | NA | 2 | 0.5 | 1 | 0 | 3 | 0.3333 |
| 13_838866 | 2 | 1 | 2 | 0.5 | 1 | 1 | 1 | 1 | 6 | 0.8333 | 1 | 1 | NA | NA | 3 | 1 | 1 | 1 | 5 | 1 |
| 13_838867 | 2 | 1 | 1 | 1 | 1 | 1 | 1 | 1 | 5 | 1 | 1 | 1 | NA | NA | 5 | 1 | 1 | 1 | 7 | 1 |
| 13_838867 | 1 | 1 | NA | NA | 1 | 1 | 1 | 0 | 3 | 0.6667 | NA | NA | NA | NA | 2 | 1 | 1 | 1 | 3 | 1 |
| 13_838868 | 2 | 0.5 | 1 | 0 | 2 | 1 | 1 | 1 | 6 | 0.6667 | 1 | 0 | NA | NA | 3 | 0.6667 | 1 | 1 | 5 | 0.6 |
| 13_838870 | 2 | 1 | 6 | 0.3333 | 4 | 0.25 | 3 | 0.6667 | 15 | 0.4667 | 4 | 1 | 1 | 1 | 5 | 0.4 | 4 | 0.5 | 14 | 0.6429 |
| 13_838870 | 2 | 1 | 4 | 0.75 | 4 | 0.5 | 3 | 1 | 13 | 0.7692 | 4 | 0.5 | NA | NA | 5 | 0.4 | 3 | 0.3333 | 12 | 0.4167 |
| 13_838870 | 2 | 0.5 | 6 | 0.1667 | 4 | 0 | 4 | 0 | 16 | 0.125 | 4 | 0 | 1 | 0 | 8 | 0.125 | 4 | 0 | 17 | 0.0588 |
| 13_838875 | 2 | 0 | 7 | 0.4286 | 4 | 0.5 | 3 | 0 | 16 | 0.3125 | 5 | 0.2 | 2 | 0.5 | 9 | 0.2222 | 3 | 0.3333 | 19 | 0.2632 |
| 13_838876 | 2 | 0 | 5 | 0 | 5 | 0.4 | 3 | 0 | 15 | 0.1333 | 4 | 0 | 2 | 0 | 8 | 0.25 | 3 | 0 | 17 | 0.1176 |
| 13_838877 | 2 | 0 | 5 | 0.2 | 4 | 0 | 3 | 0 | 14 | 0.0714 | 5 | 0.2 | 1 | 0 | 8 | 0.25 | 4 | 0.25 | 18 | 0.2222 |
| 13_838878 | 2 | 0.5 | 6 | 0 | 5 | 0 | 3 | 0 | 16 | 0.0625 | 4 | 0 | 1 | 0 | 9 | 0 | 4 | 0.25 | 18 | 0.0556 |
| 13_838889 | 3 | 0.6667 | 9 | 0.3333 | 7 | 0.4286 | 6 | 0.3333 | 25 | 0.4 | 4 | 0.25 | 3 | 0.3333 | 10 | 0.2 | 4 | 0.75 | 21 | 0.3333 |
| 13_844428 | 3 | 0.6667 | 4 | 0.5 | 2 | 0.5 | 2 | 0.5 | 11 | 0.5455 | 3 | 0 | 2 | 0.5 | 2 | 0.5 | 3 | 0.6667 | 10 | 0.4 |
| 13_844428 | 4 | 0 | 4 | 0 | 2 | 0.5 | 1 | 0 | 11 | 0.0909 | 3 | 0.3333 | 1 | 0 | 2 | 0 | 1 | 0 | 7 | 0.1429 |
| 13_844429 | 2 | 1 | 3 | 1 | 2 | 1 | 2 | 1 | 9 | 1 | 3 | 1 | 1 | 1 | 2 | 0.5 | 3 | 1 | 9 | 0.8889 |
| 13_844429 | 3 | 0.3333 | 3 | 0.6667 | 1 | 1 | 1 | 0 | 8 | 0.5 | 3 | 0.6667 | 2 | 0 | 2 | 0.5 | 3 | 0.6667 | 10 | 0.5 |
| 13_844432 | 2 | 0 | 1 | 1 | 2 | 0.5 | 1 | 0 | 6 | 0.3333 | 1 | 0 | 2 | 0.5 | 3 | 0 | 2 | 0 | 8 | 0.125 |
| 13_844433 | 3 | 0.3333 | 4 | 0 | 2 | 0 | 2 | 0 | 11 | 0.0909 | 3 | 0.3333 | 2 | 0.5 | 2 | 0.5 | 3 | 0.3333 | 10 | 0.4 |
| 13_844434 | 3 | 0 | 3 | 0.3333 | 1 | 0 | 2 | 0 | 9 | 0.1111 | 3 | 0 | 2 | 0 | 3 | 0.3333 | 3 | 0 | 11 | 0.0909 |
| 13_853613 | 8 | 0 | 13 | 0.1538 | 4 | 0 | 11 | 0 | 36 | 0.0556 | 8 | 0 | 10 | 0.1 | 10 | 0 | 10 | 0.1 | 38 | 0.0526 |
| 13_853625 | NA | NA | 1 | 0 | 1 | 1 | NA | NA | 2 | 0.5 | 1 | 1 | NA | NA | NA | NA | NA | NA | 1 | 1 |
| 13_853635 | 3 | 0.6667 | 4 | 0.75 | 6 | 0 | 6 | 0.1667 | 19 | 0.3158 | 5 | 0 | 5 | 0.4 | 4 | 0 | 6 | 0.3333 | 20 | 0.2 |
| 13_853636 | 3 | 0 | 4 | 0.25 | 8 | 0.5 | 6 | 0.3333 | 21 | 0.3333 | 5 | 0 | 6 | 0.1667 | 4 | 0.5 | 6 | 0.1667 | 21 | 0.1905 |
| 13_853636 | 3 | 0.6667 | 6 | 0.8333 | 8 | 0.625 | 4 | 0.75 | 21 | 0.7143 | 4 | 1 | 6 | 0.3333 | 3 | 0 | 5 | 0.6 | 18 | 0.5 |
| 13_853636 | 3 | 0 | 5 | 0.4 | 8 | 0.25 | 5 | 0.4 | 21 | 0.2857 | 4 | 0 | 6 | 0.1667 | 3 | 0.3333 | 6 | 0 | 19 | 0.1053 |
| 13_853637 | 3 | 0 | 4 | 0.25 | 7 | 0 | 5 | 0.2 | 19 | 0.1053 | 5 | 0 | 6 | 0 | 3 | 0 | 5 | 0.2 | 19 | 0.0526 |
| 13_853637 | 3 | 0.6667 | 5 | 0 | 6 | 0.6667 | 5 | 0.6 | 19 | 0.4737 | 3 | 0.3333 | 4 | 0.25 | 3 | 0.3333 | 4 | 0.5 | 14 | 0.3571 |
| 13_853637 | 3 | 0.6667 | 3 | 0.3333 | 7 | 0.8571 | 6 | 0.5 | 19 | 0.6316 | 5 | 0.8 | 4 | 0.75 | 2 | 0.5 | 6 | 0.6667 | 17 | 0.7059 |
| 13_853637 | 2 | 0 | 4 | 0 | 8 | 0 | 4 | 0.25 | 18 | 0.0556 | 4 | 0.25 | 3 | 0 | 2 | 0 | 5 | 0.2 | 14 | 0.1429 |
| 13_853637 | 3 | 0 | 4 | 0 | 7 | 0.2857 | 6 | 0.3333 | 20 | 0.2 | 5 | 0 | 6 | 0.1667 | 3 | 0 | 6 | 0.3333 | 20 | 0.15 |
| 13_853638 | 3 | 0 | 3 | 0 | 4 | 0.5 | 6 | 0 | 16 | 0.125 | 4 | 0.75 | 4 | 0 | 2 | 0.5 | 6 | 0.1667 | 16 | 0.3125 |
| 13_853638 | 3 | 0.3333 | 4 | 0.75 | 4 | 0.75 | 6 | 0.8333 | 17 | 0.7059 | 4 | 0.75 | 5 | 0.6 | 2 | 1 | 5 | 0.8 | 16 | 0.75 |
| 13_853639 | 3 | 1 | 2 | 1 | 5 | 1 | 4 | 1 | 14 | 1 | 5 | 1 | 6 | 0.8333 | 3 | 1 | 6 | 1 | 20 | 0.95 |
| 13_853639 | 3 | 0.6667 | 4 | 0.75 | 5 | 1 | 3 | 1 | 15 | 0.8667 | 4 | 0.75 | 6 | 0.8333 | 2 | 1 | 6 | 0.8333 | 18 | 0.8333 |
| 13_853639 | 3 | 0 | 2 | 0.5 | 5 | 0.8 | 5 | 0.6 | 15 | 0.5333 | 5 | 0.2 | 5 | 0.8 | 3 | 0.3333 | 6 | 0.8333 | 19 | 0.5789 |
| 13_853639 | 3 | 0 | 4 | 0.75 | 6 | 0.5 | 3 | 0 | 16 | 0.375 | 4 | 0 | 4 | 0 | 3 | 0.6667 | 6 | 0.5 | 17 | 0.2941 |
| 13_853642 | 1 | 0 | 2 | 0 | 5 | 0 | 5 | 0.2 | 13 | 0.0769 | 2 | 0.5 | 5 | 0 | 2 | 0.5 | 3 | 0 | 12 | 0.1667 |
| 13_853642 | 3 | 0 | 4 | 0.25 | 4 | 0.5 | 6 | 0.6667 | 17 | 0.4118 | 4 | 0.75 | 6 | 0.1667 | 3 | 0.6667 | 5 | 0.2 | 18 | 0.3889 |
| 13_853650 | 2 | 0 | 5 | 0 | 4 | 0 | 4 | 0.25 | 15 | 0.0667 | 3 | 0 | 4 | 0.25 | NA | NA | 5 | 0 | 12 | 0.0833 |
| 13_854366 | 2 | 1 | NA | NA | 1 | 1 | 4 | 0.75 | 7 | 0.8571 | NA | NA | 2 | 1 | NA | NA | 2 | 1 | 4 | 1 |
| 13_889701 | 2 | 0 | 1 | 0 | 5 | 0.2 | 4 | 0 | 12 | 0.0833 | 5 | 0 | 5 | 0 | 1 | 0 | 2 | 0.5 | 13 | 0.0769 |
| 13_894680 | 1 | 1 | 2 | 1 | 4 | 1 | 7 | 0.8571 | 14 | 0.9286 | 5 | 0.6 | 4 | 1 | 2 | 1 | 4 | 1 | 15 | 0.8667 |
| 13_894680 | 3 | 0.3333 | 3 | 0.3333 | 9 | 0 | 10 | 0 | 25 | 0.08 | 9 | 0 | 6 | 0 | 3 | 0.3333 | 8 | 0.25 | 26 | 0.1154 |
| 13_894680 | 3 | 0.3333 | 3 | 0.6667 | 6 | 0.3333 | 10 | 0.5 | 22 | 0.4545 | 7 | 0.4286 | 4 | 0 | 3 | 0.6667 | 4 | 0.75 | 18 | 0.4444 |
| 13_894709 | 1 | 1 | NA | NA | 1 | 0 | NA | NA | 2 | 0.5 | NA | NA | NA | NA | 1 | 0 | 2 | 0.5 | 3 | 0.3333 |
| 13_897497 | 1 | 0 | 2 | 0.5 | 1 | 0 | 1 | 0 | 5 | 0.2 | 1 | 0 | 2 | 0 | 2 | 0.5 | 1 | 0 | 6 | 0.1667 |
| 13_897534 | 1 | 1 | 1 | 1 | NA | NA | 1 | 1 | 3 | 1 | 1 | 1 | 1 | 0 | 5 | 1 | 1 | 1 | 8 | 0.875 |
| 13_897535 | 1 | 0 | 3 | 0 | 1 | 1 | 1 | 1 | 6 | 0.3333 | NA | NA | 2 | 0 | 5 | 0.4 | 1 | 0 | 8 | 0.25 |
| 13_897563 | 1 | 0 | 3 | 0 | 1 | 1 | NA | NA | 5 | 0.2 | 1 | 1 | 1 | 0 | 3 | 0.3333 | 1 | 0 | 6 | 0.3333 |
| 13_897567 | NA | NA | 3 | 0.3333 | 1 | 0 | 3 | 0 | 7 | 0.1429 | NA | NA | 2 | 0 | 5 | 0.2 | 1 | 0 | 8 | 0.125 |

| | | | | | | | | | | | | | | | | | | | | |
|---|---|---|---|---|---|---|---|---|---|---|---|---|---|---|---|---|---|---|---|---|
| 13_898037 | 1 | 0 | 2 | 0 | 2 | 0.5 | NA | NA | 5 | 0.2 | NA | NA | 1 | 0 | 2 | 0.5 | 2 | 0 | 5 | 0.2 |
| 13_898037 | 2 | 0 | 2 | 0.5 | 1 | 0 | NA | NA | 5 | 0.2 | NA | NA | 3 | 0 | 2 | 0.5 | 2 | 0.5 | 7 | 0.2857 |
| 13_898038 | 2 | 0.5 | 2 | 0 | 1 | 0 | NA | NA | 5 | 0.2 | NA | NA | 3 | 0.3333 | 2 | 0.5 | 2 | 0 | 7 | 0.2857 |
| 13_898046 | 1 | 0 | 1 | 1 | 1 | 0 | NA | NA | 3 | 0.3333 | NA | NA | 1 | 0 | 3 | 0.3333 | 1 | 1 | 5 | 0.4 |
| 13_898046 | 2 | 1 | 3 | 1 | 1 | 1 | NA | NA | 6 | 1 | NA | NA | 1 | 1 | 2 | 1 | 2 | 1 | 5 | 1 |
| 13_898047 | NA | NA | 2 | 0.5 | 1 | 1 | NA | NA | 3 | 0.6667 | NA | NA | 1 | 0 | 1 | 1 | 2 | 0 | 4 | 0.25 |
| 13_898048 | 2 | 0.5 | 1 | 0 | 1 | 0 | NA | NA | 4 | 0.25 | NA | NA | 2 | 1 | 2 | 0.5 | 1 | 1 | 5 | 0.8 |
| 13_898050 | NA | NA | 2 | 1 | 1 | 1 | NA | NA | 3 | 1 | NA | NA | 1 | 1 | 1 | 1 | 1 | 1 | 3 | 1 |
| 13_898052 | 1 | 1 | 3 | 1 | 1 | 1 | NA | NA | 5 | 1 | NA | NA | 1 | 1 | 2 | 1 | 1 | 1 | 4 | 1 |
| 13_898054 | 1 | 1 | 3 | 0.6667 | 2 | 0 | NA | NA | 6 | 0.5 | NA | NA | 2 | 0.5 | 2 | 1 | 2 | 0.5 | 6 | 0.6667 |
| 13_898054 | 1 | 1 | 3 | 0.6667 | 2 | 0.5 | NA | NA | 6 | 0.6667 | NA | NA | 2 | 0.5 | 2 | 0.5 | 2 | 1 | 6 | 0.6667 |
| 13_898364 | NA | NA | NA | NA | 1 | 1 | NA | NA | 1 | 1 | NA | NA | 1 | 1 | NA | NA | NA | NA | 1 | 1 |
| 13_898368 | NA | NA | 1 | 1 | 1 | 1 | NA | NA | 2 | 1 | NA | NA | 1 | 1 | NA | NA | NA | NA | 1 | 1 |
| 13_898372 | NA | NA | 1 | 1 | NA | NA | NA | NA | 1 | 1 | NA | NA | 1 | 1 | NA | NA | NA | NA | 1 | 1 |
| 13_898375 | NA | NA | 1 | 1 | NA | NA | NA | NA | 1 | 1 | NA | NA | 1 | 1 | NA | NA | NA | NA | 1 | 1 |
| 13_898378 | NA | NA | 1 | 1 | NA | NA | NA | NA | 1 | 1 | NA | NA | 1 | 1 | NA | NA | NA | NA | 1 | 1 |
| 13_898384 | NA | NA | 1 | 1 | NA | NA | NA | NA | 1 | 1 | NA | NA | 1 | 1 | NA | NA | NA | NA | 1 | 1 |
| 13_898392 | NA | NA | 1 | 1 | NA | NA | NA | NA | 1 | 1 | NA | NA | 1 | 1 | NA | NA | NA | NA | 1 | 1 |
| 13_898515 | NA | NA | 3 | 0.6667 | NA | NA | NA | NA | 3 | 0.6667 | NA | NA | 1 | 1 | 2 | 0.5 | 1 | 1 | 4 | 0.75 |
| 13_898519 | NA | NA | 3 | 0.3333 | NA | NA | NA | NA | 3 | 0.3333 | NA | NA | 1 | 0 | NA | NA | 1 | 1 | 2 | 0.5 |
| 13_898527 | NA | NA | 3 | 0.6667 | NA | NA | NA | NA | 3 | 0.6667 | NA | NA | 1 | 1 | NA | NA | 1 | 0 | 2 | 0.5 |
| 13_898536 | NA | NA | 1 | 1 | NA | NA | NA | NA | 1 | 1 | NA | NA | 1 | 1 | NA | NA | NA | NA | 1 | 1 |
| 13_899974 | 2 | 1 | NA | NA | NA | NA | 3 | 0.3333 | 5 | 0.6 | 1 | 1 | 2 | 1 | NA | NA | 2 | 0.5 | 5 | 0.8 |
| 13_902490 | 10 | 0.2 | 3 | 0 | 4 | 0.25 | 8 | 0 | 25 | 0.12 | 3 | 0.3333 | 3 | 0 | 5 | 0.2 | 4 | 0 | 15 | 0.1333 |
| 13_902525 | 13 | 0.8462 | 5 | 0.8 | 3 | 0.6667 | 9 | 0.8889 | 30 | 0.8333 | 4 | 1 | 5 | 0.6 | 6 | 0.5 | 5 | 0.8 | 20 | 0.7 |
| 13_902526 | 11 | 0.9091 | 4 | 1 | 4 | 1 | 10 | 0.9 | 29 | 0.931 | 4 | 1 | 5 | 0.8 | 6 | 0.8333 | 7 | 0.7143 | 22 | 0.8182 |
| 13_902526 | 11 | 0.1818 | 4 | 0 | 4 | 0 | 11 | 0 | 30 | 0.0667 | 3 | 0 | 5 | 0.4 | 6 | 0 | 8 | 0.25 | 22 | 0.1818 |
| 13_902526 | 12 | 0 | 5 | 0.4 | 4 | 0.5 | 10 | 0.1 | 31 | 0.1613 | 3 | 0 | 5 | 0.2 | 6 | 0.1667 | 8 | 0.125 | 22 | 0.1364 |
| 13_902527 | 11 | 0.6364 | 4 | 1 | 4 | 0.75 | 10 | 0.6 | 29 | 0.6897 | 3 | 0.6667 | 5 | 0.8 | 3 | 0.3333 | 9 | 0.7778 | 20 | 0.7 |
| 13_902528 | 9 | 0.8889 | 3 | 0.6667 | 2 | 1 | 5 | 1 | 19 | 0.8947 | 3 | 1 | 4 | 0.75 | 3 | 1 | 6 | 0.8333 | 16 | 0.875 |
| 13_902529 | 9 | 0.1111 | 2 | 0 | 2 | 0 | 5 | 0 | 18 | 0.0556 | 2 | 0 | 4 | 0 | 2 | 0.5 | 5 | 0 | 13 | 0.0769 |
| 13_902529 | 7 | 0 | 3 | 0.3333 | 2 | 0 | 5 | 0.2 | 17 | 0.1176 | 3 | 0 | 4 | 0 | 3 | 0.3333 | 5 | 0.4 | 15 | 0.2 |
| 13_902531 | 5 | 0 | NA | NA | NA | NA | 6 | 0.1667 | 11 | 0.0909 | 2 | 0.5 | NA | NA | 2 | 0.5 | 2 | 0 | 6 | 0.3333 |
| 13_910316 | 2 | 0.5 | 2 | 0.5 | 5 | 0.4 | 1 | 1 | 10 | 0.5 | 3 | 0.6667 | 1 | 1 | NA | NA | 2 | 0.5 | 6 | 0.6667 |
| 13_910317 | 1 | 0 | 3 | 0.3333 | 7 | 0.2857 | 1 | 1 | 12 | 0.3333 | 3 | 0.3333 | 1 | 1 | NA | NA | 2 | 0.5 | 6 | 0.5 |
| 13_910318 | 1 | 0 | 3 | 0 | 7 | 0.4286 | 1 | 1 | 12 | 0.3333 | 1 | 1 | NA | NA | NA | NA | 2 | 0 | 3 | 0.3333 |
| 13_910469 | 1 | 1 | 3 | 0 | 4 | 0.25 | NA | NA | 8 | 0.25 | 1 | 0 | NA | NA | NA | NA | 1 | 1 | 2 | 0.5 |
| 13_910470 | 1 | 0 | 1 | 0 | 2 | 1 | NA | NA | 4 | 0.5 | NA | NA | NA | NA | NA | NA | 1 | 1 | 1 | 1 |
| 13_910843 | 3 | 0.6667 | 3 | 0.6667 | NA | NA | 1 | 0 | 7 | 0.5714 | 2 | 0.5 | 1 | 0 | 2 | 0.5 | 3 | 0.3333 | 8 | 0.375 |
| 13_918061 | NA | NA | NA | NA | NA | NA | 1 | 1 | 1 | 1 | NA | NA | NA | NA | 1 | 1 | NA | NA | 1 | 1 |
| 13_918064 | NA | NA | NA | NA | NA | NA | 1 | 1 | 1 | 1 | NA | NA | NA | NA | 1 | 1 | NA | NA | 1 | 1 |
| 13_918070 | NA | NA | NA | NA | NA | NA | 1 | 1 | 1 | 1 | NA | NA | NA | NA | 1 | 1 | NA | NA | 1 | 1 |
| 13_918072 | NA | NA | NA | NA | NA | NA | 1 | 1 | 1 | 1 | NA | NA | NA | NA | 1 | 1 | NA | NA | 1 | 1 |
| 13_918490 | 12 | 0.8333 | 11 | 0.8182 | 10 | 1 | 11 | 0.9091 | 44 | 0.8864 | 9 | 1 | 14 | 0.7857 | 7 | 1 | 8 | 1 | 38 | 0.9211 |
| 13_918491 | 16 | 0.8125 | 16 | 0.875 | 12 | 0.9167 | 13 | 0.8462 | 57 | 0.8596 | 10 | 0.6 | 14 | 1 | 12 | 0.6667 | 11 | 0.8182 | 47 | 0.7872 |
| 13_918491 | 17 | 0.8235 | 14 | 0.7143 | 15 | 0.8667 | 13 | 0.7692 | 59 | 0.7966 | 12 | 0.5 | 15 | 0.8667 | 14 | 0.7857 | 11 | 0.6364 | 52 | 0.7115 |
| 13_918503 | 1 | 0 | 2 | 1 | 1 | 1 | 2 | 0 | 6 | 0.5 | NA | NA | 1 | 0 | 3 | 0 | 4 | 0.25 | 8 | 0.125 |
| 13_918507 | 1 | 0 | 2 | 0.5 | 2 | 0.5 | 1 | 0 | 6 | 0.3333 | 1 | 0 | 2 | 0.5 | 5 | 0 | 4 | 0.25 | 12 | 0.1667 |
| 13_918519 | 1 | 0 | 3 | 0.3333 | 2 | 0 | 4 | 0 | 10 | 0.1 | 2 | 1 | 2 | 0 | 5 | 0.6 | 6 | 0.3333 | 15 | 0.4667 |
| 13_918530 | 3 | 0 | 3 | 0 | 3 | 0.3333 | 6 | 0 | 15 | 0.0667 | 3 | 0.3333 | 3 | 0.3333 | 5 | 0.2 | 5 | 0 | 16 | 0.1875 |
| 13_918532 | 3 | 0.3333 | 3 | 0 | 2 | 0.5 | 5 | 0.2 | 13 | 0.2308 | 2 | 0 | 4 | 0 | 5 | 0 | 5 | 0.4 | 16 | 0.125 |
| 13_918538 | 4 | 0 | 2 | 0 | 5 | 0 | 7 | 0.1429 | 18 | 0.0556 | 4 | 0.25 | 5 | 0 | 6 | 0 | 5 | 0 | 20 | 0.05 |
| 13_918563 | NA | NA | 1 | 1 | 1 | 1 | 1 | 1 | 3 | 1 | 1 | 1 | NA | NA | NA | NA | NA | NA | 1 | 1 |
| 13_918563 | NA | NA | 1 | 1 | 1 | 1 | 1 | 1 | 3 | 1 | 1 | 1 | NA | NA | NA | NA | NA | NA | 1 | 1 |
| 13_919448 | 2 | 0.5 | 2 | 0 | 1 | 0 | 6 | 0 | 11 | 0.0909 | 6 | 0.1667 | NA | NA | 3 | 0 | 1 | 0 | 10 | 0.1 |
| 13_919539 | 1 | 0 | 1 | 0 | 2 | 1 | 2 | 0 | 6 | 0.3333 | 2 | 0.5 | NA | NA | NA | NA | 1 | 0 | 3 | 0.3333 |
| 13_919540 | 1 | 0 | 2 | 0 | 2 | 0.5 | 3 | 0.3333 | 8 | 0.25 | 3 | 0 | NA | NA | 3 | 0.3333 | 1 | 0 | 7 | 0.1429 |
| 13_920288 | 10 | 0.1 | 4 | 0 | 6 | 0.5 | 11 | 0.3636 | 31 | 0.2581 | 3 | 0.3333 | 4 | 0 | 8 | 0.125 | 3 | 0 | 18 | 0.1111 |
| 13_920297 | 4 | 0 | 7 | 0 | 8 | 0.125 | 8 | 0.125 | 27 | 0.0741 | 3 | 0 | 5 | 0.2 | 8 | 0.25 | 2 | 0 | 18 | 0.1667 |
| 13_920317 | 2 | 0 | 2 | 0.5 | 4 | 0.25 | 6 | 0 | 14 | 0.1429 | 2 | 0.5 | 3 | 0 | 8 | 0 | NA | NA | 13 | 0.0769 |
| 13_920319 | 3 | 0 | 1 | 0 | 3 | 0 | 4 | 0.25 | 11 | 0.0909 | 2 | 0 | 4 | 0 | 8 | 0.25 | NA | NA | 14 | 0.1429 |
| 13_920322 | 2 | 0.5 | 2 | 0 | 4 | 0 | 4 | 0 | 12 | 0.0833 | 1 | 0 | 2 | 0.5 | 8 | 0.25 | NA | NA | 11 | 0.2727 |
| 13_920323 | 2 | 0 | 2 | 0 | 5 | 0.2 | 4 | 0.25 | 13 | 0.1538 | 1 | 0 | 3 | 0.3333 | 10 | 0.3 | NA | NA | 14 | 0.2857 |
| 13_920323 | 2 | 0 | 2 | 0 | 5 | 0.2 | 4 | 0 | 13 | 0.0769 | NA | NA | 3 | 0 | 8 | 0.125 | NA | NA | 11 | 0.0909 |
| 13_920323 | 3 | 0.3333 | 2 | 0 | 5 | 0.4 | 3 | 0.3333 | 13 | 0.3077 | NA | NA | 4 | 0.5 | 10 | 0.3 | NA | NA | 14 | 0.3571 |

| | | | | | | | | | | | | | | | | | | | | |
|---|---|---|---|---|---|---|---|---|---|---|---|---|---|---|---|---|---|---|---|---|
| 13_920324 | 3 | 1 | NA | NA | 3 | 1 | 4 | 0.5 | 10 | 0.8 | NA | NA | 2 | 1 | 9 | 0.7778 | NA | NA | 11 | 0.8182 |
| 13_920324 | 1 | 1 | 1 | 1 | 2 | 1 | 3 | 0.3333 | 7 | 0.7143 | NA | NA | 3 | 1 | 5 | 0.4 | NA | NA | 8 | 0.625 |
| 13_923485 | 2 | 0.5 | 2 | 1 | 6 | 1 | 2 | 0.5 | 12 | 0.8333 | 1 | 1 | 2 | 1 | 3 | 1 | 2 | 1 | 8 | 1 |
| 13_923486 | 4 | 0 | 1 | 0 | 5 | 0.2 | 1 | 0 | 11 | 0.0909 | 1 | 0 | 3 | 0.3333 | 1 | 1 | 2 | 0.5 | 7 | 0.4286 |
| 13_923486 | 4 | 0 | 2 | 0 | 5 | 0.6 | 2 | 0 | 13 | 0.2308 | 1 | 0 | 3 | 0.3333 | 2 | 0 | 2 | 0.5 | 8 | 0.25 |
| 13_923487 | 4 | 0.25 | 2 | 0 | 4 | 0 | 2 | 0.5 | 12 | 0.1667 | NA | NA | 3 | 1 | 2 | 0 | 2 | 0.5 | 7 | 0.5714 |
| 13_923488 | 4 | 0 | 1 | 0 | 3 | 0 | 1 | 1 | 9 | 0.1111 | NA | NA | 3 | 0.6667 | 2 | 1 | 2 | 0 | 7 | 0.5714 |
| 13_923489 | 4 | 0.75 | 1 | 0 | 3 | 0.3333 | 2 | 0.5 | 10 | 0.5 | NA | NA | 3 | 1 | 3 | 0.6667 | 2 | 0.5 | 8 | 0.75 |
| 13_923489 | 4 | 0.5 | 1 | 0 | 4 | 0.75 | NA | NA | 9 | 0.5556 | NA | NA | 3 | 0 | 3 | 0.6667 | 2 | 0 | 8 | 0.25 |
| 13_924917 | NA | NA | 1 | 1 | NA | NA | NA | NA | 1 | 1 | 1 | 0 | 1 | 0 | 1 | 0 | 2 | 0.5 | 5 | 0.2 |
| 13_925213 | NA | NA | 1 | 1 | 2 | 0 | NA | NA | 3 | 0.3333 | 1 | 0 | 1 | 1 | 1 | 0 | 2 | 0.5 | 5 | 0.4 |
| 13_925213 | NA | NA | 2 | 1 | 2 | 1 | NA | NA | 4 | 1 | 1 | 1 | 1 | 1 | 1 | 0 | 1 | 1 | 4 | 0.75 |
| 13_925748 | NA | NA | 2 | 1 | NA | NA | NA | NA | 2 | 1 | NA | NA | 2 | 1 | 3 | 0.6667 | 1 | 1 | 6 | 0.8333 |
| 13_925750 | NA | NA | 3 | 0.6667 | NA | NA | NA | NA | 3 | 0.6667 | NA | NA | 2 | 0.5 | 3 | 0.6667 | 1 | 0 | 6 | 0.5 |
| 13_925751 | NA | NA | 4 | 0.75 | NA | NA | NA | NA | 4 | 0.75 | NA | NA | 2 | 1 | 3 | 0.6667 | 2 | 0.5 | 7 | 0.7143 |
| 13_925753 | NA | NA | 2 | 1 | NA | NA | NA | NA | 2 | 1 | NA | NA | 2 | 1 | 3 | 0.6667 | 2 | 1 | 7 | 0.8571 |
| 13_925754 | NA | NA | 3 | 0.6667 | NA | NA | NA | NA | 3 | 0.6667 | NA | NA | 2 | 1 | 3 | 0 | 2 | 1 | 7 | 0.5714 |
| 13_925754 | NA | NA | 3 | 0.6667 | NA | NA | NA | NA | 3 | 0.6667 | NA | NA | 2 | 1 | 3 | 0 | 3 | 0.3333 | 8 | 0.375 |
| 13_925756 | NA | NA | 4 | 1 | NA | NA | NA | NA | 4 | 1 | NA | NA | 1 | 1 | 2 | 1 | 2 | 1 | 5 | 1 |
| 13_925757 | NA | NA | 5 | 0.4 | NA | NA | NA | NA | 5 | 0.4 | NA | NA | 2 | 1 | 2 | 1 | 3 | 0.3333 | 7 | 0.7143 |
| 13_925757 | NA | NA | 5 | 1 | NA | NA | NA | NA | 5 | 1 | NA | NA | 1 | 1 | 3 | 1 | 2 | 0.5 | 6 | 0.8333 |
| 13_925758 | NA | NA | 3 | 0.6667 | NA | NA | NA | NA | 3 | 0.6667 | NA | NA | 1 | 1 | 1 | 1 | 2 | 0.5 | 4 | 0.75 |
| 13_925760 | NA | NA | 5 | 0.2 | NA | NA | NA | NA | 5 | 0.2 | NA | NA | 2 | 0 | 3 | 0 | 2 | 0.5 | 7 | 0.1429 |
| 13_925761 | NA | NA | 5 | 1 | NA | NA | NA | NA | 5 | 1 | NA | NA | 1 | 1 | 3 | 1 | 3 | 1 | 7 | 1 |
| 13_925763 | NA | NA | 5 | 0.4 | NA | NA | NA | NA | 5 | 0.4 | NA | NA | 2 | 1 | 2 | 0.5 | 3 | 0 | 7 | 0.4286 |
| 13_925766 | 1 | 1 | 3 | 0 | NA | NA | NA | NA | 4 | 0.25 | NA | NA | NA | NA | 3 | 0.3333 | 3 | 0 | 6 | 0.1667 |
| 13_925766 | 1 | 1 | 6 | 1 | NA | NA | NA | NA | 7 | 1 | NA | NA | 1 | 1 | 3 | 1 | 3 | 1 | 7 | 1 |
| 13_925779 | 5 | 0 | 6 | 0.5 | 1 | 0 | 5 | 0.8 | 17 | 0.4118 | 1 | 0 | 5 | 0.6 | 3 | 0.3333 | 3 | 0.3333 | 12 | 0.4167 |
| 13_926243 | 2 | 0 | NA | NA | 3 | 0.3333 | 1 | 0 | 6 | 0.1667 | 1 | 0 | 3 | 0 | 1 | 0 | 3 | 0.3333 | 8 | 0.125 |
| 13_926482 | NA | NA | 1 | 0 | 2 | 1 | 1 | 0 | 4 | 0.5 | NA | NA | NA | NA | 1 | 1 | 2 | 1 | 3 | 1 |
| 13_926579 | NA | NA | NA | NA | 2 | 0.5 | NA | NA | 2 | 0.5 | NA | NA | NA | NA | NA | NA | 2 | 1 | 2 | 1 |
| 13_926582 | NA | NA | NA | NA | 2 | 1 | NA | NA | 2 | 1 | NA | NA | NA | NA | NA | NA | 2 | 0.5 | 2 | 0.5 |
| 13_926583 | NA | NA | NA | NA | 1 | 1 | NA | NA | 1 | 1 | NA | NA | NA | NA | NA | NA | 2 | 1 | 2 | 1 |
| 13_926583 | NA | NA | NA | NA | 2 | 0.5 | NA | NA | 2 | 0.5 | NA | NA | NA | NA | NA | NA | 2 | 0.5 | 2 | 0.5 |
| 13_928445 | 5 | 0.2 | 3 | 0 | 1 | 0 | 3 | 0 | 12 | 0.0833 | 2 | 0 | 2 | 0 | 2 | 0 | 3 | 0.3333 | 9 | 0.1111 |
| 13_928460 | 7 | 0.1429 | 4 | 0 | 3 | 0 | 2 | 0 | 16 | 0.0625 | 3 | 0 | 4 | 0.25 | 1 | 0 | 5 | 0 | 13 | 0.0769 |
| 13_928476 | 8 | 0.375 | 4 | 0.5 | 3 | 0 | 6 | 0.1667 | 21 | 0.2857 | 7 | 0.1429 | 5 | 0.4 | 2 | 1 | 7 | 0.1429 | 21 | 0.2857 |
| 13_928476 | 9 | 0.2222 | 8 | 0.125 | 3 | 0 | 6 | 0.1667 | 26 | 0.1538 | 6 | 0.1667 | 5 | 0.6 | 3 | 1 | 8 | 0.25 | 22 | 0.4091 |
| 13_928476 | 7 | 0.1429 | 9 | 0 | 4 | 0.25 | 7 | 0 | 27 | 0.0741 | 7 | 0.1429 | 6 | 0.1667 | 3 | 0 | 7 | 0 | 23 | 0.087 |
| 13_928477 | 10 | 0.2 | 9 | 0.4444 | 5 | 0 | 5 | 0.4 | 29 | 0.2759 | 7 | 0.2857 | 6 | 0.5 | 4 | 0.5 | 7 | 0.4286 | 24 | 0.4167 |
| 13_928478 | 6 | 0.1667 | 4 | 0.25 | 5 | 0 | 4 | 0 | 19 | 0.1053 | 5 | 0 | 6 | 0 | 4 | 0.5 | 7 | 0.1429 | 22 | 0.1364 |
| 13_929894 | NA | NA | NA | NA | NA | NA | 2 | 1 | 2 | 1 | NA | NA | NA | NA | NA | NA | 2 | 1 | 2 | 1 |
| 13_929903 | NA | NA | 1 | 0 | 3 | 0 | 3 | 0.3333 | 7 | 0.1429 | 2 | 0 | NA | NA | 2 | 0 | 5 | 0.2 | 9 | 0.1111 |
| 13_929925 | 1 | 0 | 5 | 0 | 5 | 0.2 | 2 | 0 | 13 | 0.0769 | 1 | 1 | 1 | 0 | 5 | 0 | 7 | 0.1429 | 14 | 0.1429 |
| 13_929932 | 1 | 0 | 6 | 0.1667 | 3 | 0 | 3 | 0.3333 | 13 | 0.1538 | 1 | 1 | 2 | 0 | 3 | 0 | 4 | 0 | 10 | 0.1 |
| 13_929941 | 1 | 0 | 6 | 0 | 3 | 0.3333 | 3 | 0 | 13 | 0.0769 | NA | NA | 2 | 0.5 | 4 | 0 | 5 | 0 | 11 | 0.0909 |
| 13_931775 | 2 | 0.5 | 1 | 0 | NA | NA | NA | NA | 3 | 0.3333 | 1 | 0 | 2 | 1 | NA | NA | NA | NA | 3 | 0.6667 |
| 13_932834 | 1 | 1 | 6 | 0.8333 | 2 | 1 | 4 | 0.5 | 13 | 0.7692 | 2 | 0.5 | 1 | 1 | 4 | 0.75 | NA | NA | 7 | 0.7143 |
| 13_932835 | 1 | 0 | 6 | 0.6667 | 2 | 0.5 | 5 | 0 | 14 | 0.3571 | 2 | 0 | 3 | 0.6667 | 3 | 0.3333 | 2 | 0 | 10 | 0.3 |
| 13_932835 | 1 | 0 | 6 | 0.1667 | 2 | 0 | 5 | 0 | 14 | 0.0714 | 1 | 0 | 3 | 0 | 3 | 0.3333 | 2 | 0 | 9 | 0.1111 |
| 13_932836 | NA | NA | 7 | 0.1429 | 1 | 0 | 4 | 0.25 | 12 | 0.1667 | 2 | 0 | 3 | 0 | 3 | 0.3333 | 2 | 0 | 10 | 0.1 |
| 13_934404 | 1 | 0 | 2 | 0.5 | 2 | 0 | 3 | 0 | 8 | 0.125 | 1 | 0 | 2 | 0 | 1 | 1 | 2 | 0 | 6 | 0.1667 |
| 13_934409 | 3 | 1 | 4 | 1 | 2 | 1 | 7 | 0.2857 | 16 | 0.6875 | 1 | 0 | 5 | 0.4 | 2 | 0 | 1 | 0 | 9 | 0.2222 |
| 13_934412 | 6 | 0.8333 | 7 | 0.7143 | 6 | 0.6667 | 8 | 0.5 | 27 | 0.6667 | 2 | 0.5 | 9 | 0.5556 | 4 | 0.5 | 8 | 0.5 | 23 | 0.5217 |
| 13_934412 | 6 | 0.6667 | 7 | 0.7143 | 6 | 0.8333 | 13 | 0.9231 | 32 | 0.8125 | 2 | 1 | 9 | 0.8889 | 5 | 0.6 | 8 | 0.75 | 24 | 0.7917 |
| 13_934413 | 7 | 0.4286 | 7 | 0.4286 | 6 | 0.1667 | 13 | 0.3846 | 33 | 0.3636 | 4 | 0.25 | 11 | 0.3636 | 7 | 0.7143 | 12 | 0.4167 | 34 | 0.4412 |
| 13_934414 | 7 | 0.7143 | 7 | 0.4286 | 7 | 0.4286 | 10 | 0.4 | 31 | 0.4839 | 3 | 0.3333 | 10 | 0.4 | 7 | 0.4286 | 9 | 0.5556 | 29 | 0.4483 |
| 13_934415 | 7 | 0.5714 | 7 | 0.4286 | 7 | 0.2857 | 12 | 0.5833 | 33 | 0.4848 | 4 | 0.25 | 13 | 0.5385 | 7 | 0.4286 | 12 | 0.4167 | 36 | 0.4444 |
| 13_934416 | 4 | 0.75 | 8 | 0.375 | 3 | 1 | 8 | 0.375 | 23 | 0.5217 | 3 | 0.6667 | 5 | 0.4 | 3 | 0.6667 | 7 | 0.4286 | 18 | 0.5 |
| 13_934416 | 8 | 0.5 | 8 | 0.5 | 5 | 0.4 | 8 | 0.625 | 29 | 0.5172 | 3 | 0.6667 | 10 | 0.4 | 7 | 0.5714 | 9 | 0.3333 | 29 | 0.4483 |
| 13_934416 | 6 | 0.3333 | 8 | 0.25 | 7 | 0.2857 | 6 | 0.1667 | 27 | 0.2593 | 3 | 0.3333 | 10 | 0.2 | 5 | 0.2 | 5 | 0.4 | 23 | 0.2609 |
| 13_934927 | 1 | 0 | 2 | 0.5 | 3 | 0.3333 | 2 | 0 | 8 | 0.25 | 4 | 0 | 3 | 0 | 3 | 0.3333 | 2 | 0 | 12 | 0.0833 |
| 13_934931 | 1 | 0 | 3 | 0.3333 | 3 | 0.3333 | 2 | 0.5 | 9 | 0.3333 | 3 | 0 | 3 | 0 | 2 | 0 | 2 | 0.5 | 10 | 0.1 |
| 13_934934 | 2 | 0.5 | 5 | 0 | 4 | 0 | 1 | 0 | 12 | 0.0833 | 5 | 0 | 3 | 0 | 5 | 0.2 | 3 | 0.3333 | 16 | 0.125 |
| 13_934962 | 3 | 0 | 7 | 0.1429 | 8 | 0 | 2 | 0 | 20 | 0.05 | 7 | 0 | 8 | 0.125 | 6 | 0 | 4 | 0.25 | 25 | 0.08 |

| | | | | | | | | | | | | | | | | | | | | |
|---|---|---|---|---|---|---|---|---|---|---|---|---|---|---|---|---|---|---|---|---|
| 13_934964( | 3 | 0 | 7 | 0.1429 | 6 | 0 | 2 | 0 | 18 | 0.0556 | 8 | 0.125 | 6 | 0 | 4 | 0 | 3 | 0.3333 | 21 | 0.0952 |
| 13_935030( | 1 | 1 | 2 | 1 | 1 | 1 | NA | NA | 4 | 1 | NA | NA | NA | NA | NA | NA | 4 | 1 | 4 | 1 |
| 13_935030! | 1 | 1 | 2 | 1 | NA | NA | NA | NA | 3 | 1 | NA | NA | NA | NA | NA | NA | 4 | 1 | 4 | 1 |
| 13_935031' | 1 | 1 | 1 | 1 | NA | NA | NA | NA | 2 | 1 | NA | NA | NA | NA | NA | NA | 4 | 1 | 4 | 1 |
| 13_935032: | 1 | 1 | 2 | 1 | 1 | 1 | NA | NA | 4 | 1 | NA | NA | NA | NA | NA | NA | 5 | 1 | 5 | 1 |
| 13_935386! | 11 | 0.2727 | 11 | 0 | 7 | 0.2857 | 10 | 0.1 | 39 | 0.1538 | 3 | 0 | 12 | 0 | 8 | 0.25 | 9 | 0.1111 | 32 | 0.0938 |
| 13_935386! | 10 | 1 | 10 | 0.7 | 6 | 0.6667 | 9 | 1 | 35 | 0.8571 | 2 | 0 | 11 | 0.9091 | 9 | 0.5556 | 7 | 1 | 29 | 0.7586 |
| 13_935387! | 11 | 1 | 13 | 0.9231 | 8 | 0.875 | 12 | 0.9167 | 44 | 0.9318 | 3 | 0.6667 | 14 | 1 | 7 | 0.7143 | 8 | 1 | 32 | 0.9062 |
| 13_935389: | 11 | 0.0909 | 14 | 0 | 9 | 0 | 12 | 0.1667 | 46 | 0.0652 | 2 | 0 | 14 | 0.1429 | 8 | 0 | 9 | 0.1111 | 33 | 0.0909 |
| 13_935389. | 10 | 0 | 13 | 0.0769 | 7 | 0 | 9 | 0.3333 | 39 | 0.1026 | 3 | 0 | 12 | 0 | 7 | 0.2857 | 10 | 0.2 | 32 | 0.125 |
| 13_935390( | 10 | 0.1 | 14 | 0 | 8 | 0 | 12 | 0.1667 | 44 | 0.0682 | 3 | 0 | 17 | 0.0588 | 8 | 0.125 | 9 | 0 | 37 | 0.0541 |
| 13_935395: | 11 | 0.0909 | 15 | 0.0667 | 9 | 0.1111 | 13 | 0.0769 | 48 | 0.0833 | 5 | 0.2 | 19 | 0 | 10 | 0.2 | 13 | 0.0769 | 47 | 0.0851 |
| 13_935397( | 13 | 0.1538 | 17 | 0.5294 | 10 | 0.1 | 13 | 0.0769 | 53 | 0.2453 | 5 | 0.4 | 19 | 0.2632 | 9 | 0.3333 | 14 | 0.3571 | 47 | 0.3191 |
| 13_935397: | 12 | 0.3333 | 15 | 0.3333 | 11 | 0.3636 | 16 | 0.1875 | 54 | 0.2963 | 7 | 0.4286 | 21 | 0.381 | 11 | 0.2727 | 14 | 0.2857 | 53 | 0.3396 |
| 13_935398. | 9 | 0.5556 | 14 | 0.5 | 11 | 0.3636 | 14 | 0.4286 | 48 | 0.4583 | 5 | 0.2 | 17 | 0.7647 | 8 | 0.375 | 12 | 0.6667 | 42 | 0.5952 |
| 13_935398! | 9 | 0.2222 | 17 | 0.2353 | 10 | 0.2 | 13 | 0.1538 | 49 | 0.2041 | 5 | 0 | 20 | 0.15 | 9 | 0.4444 | 12 | 0.3333 | 46 | 0.2391 |
| 13_935413: | 12 | 0.0833 | 12 | 0.0833 | 5 | 0 | 6 | 0 | 35 | 0.0571 | 4 | 0 | 14 | 0.0714 | 8 | 0 | 7 | 0.1429 | 33 | 0.0606 |
| 13_935413. | 14 | 0.5 | 10 | 0.1 | 3 | 0.3333 | 8 | 0.125 | 35 | 0.2857 | 4 | 0.25 | 15 | 0.2 | 10 | 0.2 | 9 | 0 | 38 | 0.1579 |
| 13_935683! | 2 | 0 | 3 | 0.3333 | 1 | 0 | 1 | 0 | 7 | 0.1429 | 1 | 0 | 3 | 0 | 1 | 0 | 4 | 0.25 | 9 | 0.1111 |
| 13_935687' | 1 | 0 | 2 | 0 | 1 | 0 | 1 | 1 | 5 | 0.2 | 1 | 0 | NA | NA | 1 | 0 | 4 | 0.5 | 6 | 0.3333 |
| 13_935707' | 6 | 0 | 1 | 0 | 2 | 0.5 | 1 | 0 | 10 | 0.1 | 4 | 0 | 4 | 0.25 | 3 | 0 | 4 | 0 | 15 | 0.0667 |
| 13_935711: | 5 | 0 | 1 | 0 | 2 | 0 | 1 | 1 | 9 | 0.1111 | 2 | 0.5 | 3 | 0 | 3 | 0 | 2 | 0.5 | 10 | 0.2 |
| 13_935711! | 5 | 0.2 | NA | NA | 2 | 0 | 1 | 0 | 8 | 0.125 | 3 | 0 | 2 | 0 | 3 | 0 | 3 | 0.3333 | 11 | 0.0909 |
| 13_935713! | 5 | 0 | 1 | 0 | 2 | 0 | 1 | 1 | 9 | 0.1111 | 2 | 0 | 2 | 0 | 3 | 0 | 4 | 0.25 | 11 | 0.0909 |
| 13_935714! | 5 | 0.4 | 1 | 1 | 2 | 0 | 1 | 0 | 9 | 0.3333 | 2 | 1 | 2 | 0.5 | 2 | 0.5 | 4 | 0.25 | 10 | 0.5 |
| 13_935715( | 5 | 0.2 | 1 | 1 | 2 | 0 | 1 | 0 | 9 | 0.2222 | 2 | 0.5 | 2 | 0.5 | 2 | 0 | NA | NA | 6 | 0.3333 |
| 13_935715' | 5 | 0.4 | 1 | 0 | 2 | 1 | 1 | 1 | 9 | 0.5556 | 2 | 0.5 | 2 | 0.5 | 2 | 0.5 | 4 | 0.5 | 10 | 0.5 |
| 13_935715( | 3 | 0.3333 | 1 | 1 | 2 | 1 | 1 | 1 | 7 | 0.7143 | 1 | 1 | 2 | 1 | NA | NA | 2 | 0.5 | 5 | 0.8 |
| 13_935716: | 4 | 1 | NA | NA | 2 | 0.5 | NA | NA | 6 | 0.8333 | 2 | 1 | 1 | 1 | NA | NA | 4 | 1 | 7 | 1 |
| 13_935717: | 1 | 0 | 1 | 0 | 2 | 0.5 | 1 | 0 | 5 | 0.2 | 2 | 0 | 2 | 0 | 1 | 0 | 4 | 0.25 | 9 | 0.1111 |
| 13_935718' | 4 | 0.75 | 1 | 1 | 2 | 0.5 | 1 | 1 | 8 | 0.75 | 1 | 0 | 2 | 0.5 | 1 | 1 | 4 | 0.25 | 8 | 0.375 |
| 13_935718! | 4 | 0.25 | NA | NA | 1 | 0 | 1 | 1 | 6 | 0.3333 | 2 | 1 | 2 | 0 | 1 | 0 | 3 | 0.6667 | 8 | 0.5 |
| 13_935761! | 4 | 0.75 | 1 | 1 | 2 | 0 | 1 | 1 | 8 | 0.625 | 2 | 0.5 | 1 | 1 | 1 | 1 | 4 | 1 | 8 | 0.875 |
| 13_935776: | 4 | 0.25 | 1 | 1 | 2 | 0.5 | 1 | 1 | 8 | 0.5 | 2 | 0 | 1 | 0 | NA | NA | 4 | 0.25 | 7 | 0.1429 |
| 13_935777( | 4 | 0.5 | 1 | 1 | 2 | 1 | 1 | 1 | 8 | 0.75 | 2 | 1 | 1 | 1 | NA | NA | 4 | 0.75 | 7 | 0.8571 |
| 13_935778! | 4 | 1 | 1 | 1 | 2 | 0.5 | NA | NA | 7 | 0.8571 | 2 | 1 | NA | NA | NA | NA | 3 | 1 | 5 | 1 |
| 13_936352' | 4 | 0.75 | NA | NA | 1 | 0 | NA | NA | 5 | 0.6 | 2 | 1 | NA | NA | NA | NA | 2 | 0 | 4 | 0.5 |
| 13_940770. | 7 | 0 | 14 | 0.2857 | 2 | 0 | 3 | 0 | 26 | 0.1538 | 2 | 0.5 | 2 | 0 | 8 | 0.25 | 3 | 0 | 15 | 0.2 |
| 13_940770: | 7 | 0 | 13 | 0.1538 | 2 | 0 | 3 | 0 | 25 | 0.08 | 2 | 0 | 1 | 0 | 7 | 0 | 3 | 0.3333 | 13 | 0.0769 |
| 13_940771! | 8 | 0.625 | 11 | 1 | 2 | 1 | 2 | 1 | 23 | 0.8696 | 2 | 1 | 3 | 0.6667 | 6 | 0.6667 | 2 | 0.5 | 13 | 0.6923 |
| 13_940775( | 8 | 0 | 14 | 0.0714 | 3 | 0.3333 | 4 | 0 | 29 | 0.069 | 3 | 0.3333 | 5 | 0.2 | 9 | 0.1111 | 4 | 0 | 21 | 0.1429 |
| 13_940780( | 8 | 0 | 11 | 0.1818 | 5 | 0 | 5 | 0 | 29 | 0.069 | 5 | 0 | 5 | 0 | 7 | 0 | 3 | 0.3333 | 20 | 0.05 |
| 13_940790. | 9 | 0 | 16 | 0.1875 | 6 | 0.1667 | 8 | 0 | 39 | 0.1026 | 7 | 0.2857 | 6 | 0 | 12 | 0.1667 | 7 | 0 | 32 | 0.125 |
| 13_940790( | 13 | 0 | 14 | 0.2857 | 6 | 0.1667 | 8 | 0.125 | 41 | 0.1463 | 7 | 0.2857 | 5 | 0 | 14 | 0.3571 | 7 | 0.2857 | 33 | 0.2727 |
| 13_943287! | NA | NA | NA | NA | 1 | 1 | NA | NA | 1 | 1 | NA | NA | NA | NA | 1 | 1 | NA | NA | 1 | 1 |
| 13_943288: | NA | NA | NA | NA | 1 | 1 | NA | NA | 1 | 1 | NA | NA | NA | NA | 1 | 1 | NA | NA | 1 | 1 |
| 13_943290( | 1 | 0 | NA | NA | 1 | 1 | NA | NA | 2 | 0.5 | NA | NA | NA | NA | 1 | 1 | NA | NA | 1 | 1 |
| 13_943304: | 1 | 1 | NA | NA | 1 | 0 | NA | NA | 2 | 0.5 | NA | NA | NA | NA | 1 | 1 | NA | NA | 1 | 1 |
| 13_943304( | 1 | 1 | NA | NA | 1 | 1 | NA | NA | 2 | 1 | NA | NA | NA | NA | 1 | 1 | NA | NA | 1 | 1 |
| 13_943305( | 1 | 1 | NA | NA | 2 | 1 | NA | NA | 3 | 1 | NA | NA | NA | NA | 1 | 1 | NA | NA | 1 | 1 |
| 13_943306! | 1 | 1 | NA | NA | 2 | 0.5 | NA | NA | 3 | 0.6667 | NA | NA | NA | NA | 1 | 1 | NA | NA | 1 | 1 |
| 13_943307( | 1 | 1 | NA | NA | 2 | 0 | NA | NA | 3 | 0.3333 | NA | NA | NA | NA | 1 | 1 | NA | NA | 1 | 1 |
| 13_943307! | 1 | 1 | NA | NA | 2 | 1 | NA | NA | 3 | 1 | NA | NA | NA | NA | 1 | 1 | NA | NA | 1 | 1 |
| 13_943308( | 1 | 0 | NA | NA | 2 | 1 | NA | NA | 3 | 0.6667 | NA | NA | NA | NA | 1 | 1 | NA | NA | 1 | 1 |
| 13_943316. | 2 | 0.5 | 1 | 0 | 2 | 0.5 | NA | NA | 5 | 0.4 | NA | NA | 1 | 1 | 1 | 0 | NA | NA | 2 | 0.5 |
| 13_943317( | 1 | 1 | 3 | 0 | 1 | 1 | NA | NA | 5 | 0.4 | NA | NA | 1 | 1 | 1 | 0 | NA | NA | 2 | 0.5 |
| 13_943352( | NA | NA | 1 | 1 | 2 | 1 | 1 | 1 | 4 | 1 | NA | NA | NA | NA | 1 | 1 | NA | NA | 1 | 1 |
| 13_943382( | 31 | 0.0645 | 33 | 0.0909 | 38 | 0.1053 | 48 | 0.1042 | 150 | 0.0933 | 28 | 0.1429 | 34 | 0.0588 | 32 | 0.125 | 29 | 0.069 | 123 | 0.0976 |
| 13_943401: | 14 | 0.1429 | 19 | 0 | 16 | 0 | 22 | 0.0909 | 71 | 0.0563 | 16 | 0 | 16 | 0.0625 | 19 | 0.1053 | 17 | 0.1765 | 68 | 0.0882 |
| 13_943404: | 16 | 1 | 20 | 0.95 | 23 | 0.8696 | 30 | 0.8667 | 89 | 0.9101 | 24 | 0.9583 | 20 | 0.85 | 28 | 0.9286 | 21 | 1 | 93 | 0.9355 |
| 13_943405! | 15 | 0.2 | 18 | 0.3889 | 16 | 0.1875 | 23 | 0.3043 | 72 | 0.2778 | 20 | 0.35 | 15 | 0.4667 | 19 | 0.3684 | 13 | 0.3846 | 67 | 0.3881 |
| 13_943406( | 18 | 0.5556 | 14 | 0.4286 | 18 | 0.5 | 24 | 0.3333 | 74 | 0.4459 | 20 | 0.45 | 17 | 0.4118 | 28 | 0.5714 | 19 | 0.5263 | 84 | 0.5 |
| 13_944934! | 2 | 0 | 3 | 0.3333 | NA | NA | 1 | 1 | 6 | 0.3333 | NA | NA | 1 | 1 | NA | NA | 2 | 0.5 | 3 | 0.6667 |
| 13_944935! | 1 | 1 | 1 | 1 | NA | NA | 1 | 1 | 3 | 1 | NA | NA | 1 | 0 | NA | NA | 2 | 0.5 | 3 | 0.3333 |
| 13_944944( | 1 | 1 | 1 | 1 | NA | NA | 1 | 1 | 3 | 1 | NA | NA | NA | NA | NA | NA | 2 | 1 | 2 | 1 |

| | | | | | | | | | | | | | | | | | | | | |
|---|---|---|---|---|---|---|---|---|---|---|---|---|---|---|---|---|---|---|---|---|
| 13_9449448 | 1 | 1 | 2 | 1 | NA | NA | 1 | 1 | 4 | 1 | NA | NA | NA | NA | NA | NA | 2 | 0.5 | 2 | 0.5 |
| 13_9449452 | 1 | 1 | 1 | 1 | NA | NA | 1 | 1 | 3 | 1 | NA | NA | NA | NA | NA | NA | 2 | 0.5 | 2 | 0.5 |
| 13_9449460 | 1 | 0 | 1 | 1 | NA | NA | 1 | 1 | 3 | 0.6667 | NA | NA | NA | NA | NA | NA | 2 | 1 | 2 | 1 |
| 13_9449469 | 1 | 1 | 2 | 1 | NA | NA | NA | NA | 3 | 1 | NA | NA | NA | NA | NA | NA | 2 | 1 | 2 | 1 |
| 13_9449515 | 1 | 0 | 1 | 1 | NA | NA | NA | NA | 2 | 0.5 | NA | NA | NA | NA | NA | NA | 1 | 1 | 1 | 1 |
| 13_9449528 | 1 | 1 | 1 | 1 | NA | NA | NA | NA | 2 | 1 | NA | NA | NA | NA | NA | NA | 2 | 1 | 2 | 1 |
| 13_9461363 | 2 | 0 | 6 | 0.1667 | NA | NA | 6 | 0 | 14 | 0.0714 | 1 | 0 | 4 | 0 | 4 | 0.25 | 3 | 0 | 12 | 0.0833 |
| 13_9470228 | 3 | 1 | 5 | 0.6 | 2 | 0.5 | 5 | 0.6 | 15 | 0.6667 | 3 | 1 | 4 | 0.75 | 6 | 0.1667 | 6 | 0.8333 | 19 | 0.6316 |
| 13_9470256 | 3 | 0 | 6 | 0.1667 | 3 | 0 | 3 | 0.6667 | 15 | 0.2 | 3 | 0.3333 | 5 | 0.4 | 5 | 0.4 | 6 | 0 | 19 | 0.2632 |
| 13_9513368 | NA | NA | 2 | 0.5 | 1 | 1 | 3 | 0 | 6 | 0.3333 | 4 | 0.25 | 3 | 0 | 1 | 0 | 3 | 0.3333 | 11 | 0.1818 |
| 13_9515820 | NA | NA | 4 | 0.25 | 4 | 0.75 | 3 | 0.3333 | 11 | 0.4545 | 3 | 0.6667 | 4 | 0.25 | 5 | 0.2 | 4 | 0.5 | 16 | 0.375 |
| 13_9515831 | NA | NA | 4 | 0.25 | 4 | 0 | 3 | 0 | 11 | 0.0909 | 5 | 0 | 5 | 0.2 | 6 | 0 | 4 | 0.25 | 20 | 0.1 |
| 13_9515862 | NA | NA | 4 | 0 | 4 | 0.25 | 3 | 0.6667 | 11 | 0.2727 | 5 | 0.2 | 4 | 0.25 | 7 | 0.1429 | 4 | 0.75 | 20 | 0.3 |
| 13_9515887 | NA | NA | 1 | 1 | 3 | 0 | 3 | 0 | 7 | 0.1429 | 5 | 0 | 3 | 0 | 6 | 0 | 4 | 0.25 | 18 | 0.0556 |
| 13_9516129 | 15 | 0.2 | 11 | 0.0909 | 14 | 0.3571 | 13 | 0.2308 | 53 | 0.2264 | 4 | 0.5 | 16 | 0.25 | 10 | 0 | 13 | 0.0769 | 43 | 0.1628 |
| 13_9516134 | 14 | 0.2143 | 10 | 0.1 | 13 | 0.1538 | 12 | 0.1667 | 49 | 0.1633 | 6 | 0.1667 | 16 | 0.3125 | 13 | 0.1538 | 12 | 0.0833 | 47 | 0.1915 |
| 13_9516162 | 16 | 0.1875 | 9 | 0 | 13 | 0.1538 | 11 | 0.1818 | 49 | 0.1429 | 4 | 0 | 17 | 0.2941 | 10 | 0.3 | 15 | 0.1333 | 46 | 0.2174 |
| 13_9516206 | 11 | 0 | 7 | 0.1429 | 7 | 0.1429 | 11 | 0 | 36 | 0.0556 | 2 | 0.5 | 15 | 0.1333 | 11 | 0.1818 | 10 | 0 | 38 | 0.1316 |
| 13_9516212 | 15 | 0 | 5 | 0 | 11 | 0.2727 | 10 | 0 | 41 | 0.0732 | 2 | 0 | 14 | 0.1429 | 8 | 0 | 11 | 0.0909 | 35 | 0.0857 |
| 13_9516233 | 11 | 0.1818 | 6 | 0.1667 | 8 | 0 | 11 | 0.2727 | 36 | 0.1667 | 3 | 0.3333 | 14 | 0.2143 | 10 | 0 | 13 | 0.2308 | 40 | 0.175 |
| 13_9516235 | 10 | 0.3 | 7 | 0.1429 | 8 | 0.125 | 10 | 0.1 | 35 | 0.1714 | 2 | 0 | 15 | 0.3333 | 9 | 0.3333 | 11 | 0 | 37 | 0.2162 |
| 13_9516236 | 12 | 0.4167 | 7 | 0.2857 | 11 | 0.1818 | 11 | 0.2727 | 41 | 0.2927 | 2 | 0 | 14 | 0.3571 | 10 | 0.3 | 11 | 0.6364 | 37 | 0.4054 |
| 13_9516243 | 11 | 0.0909 | 7 | 0.2857 | 11 | 0.3636 | 10 | 0.5 | 39 | 0.3077 | 3 | 0 | 15 | 0.1333 | 8 | 0.375 | 11 | 0.0909 | 37 | 0.1622 |
| 13_9516254 | 11 | 0.9091 | 5 | 0.8 | 9 | 0.7778 | 9 | 0.7778 | 34 | 0.8235 | 3 | 0.6667 | 14 | 0.8571 | 6 | 0.8333 | 9 | 0.6667 | 32 | 0.7812 |
| 13_9516255 | 12 | 0.0833 | 6 | 0 | 12 | 0.0833 | 11 | 0.0909 | 41 | 0.0732 | 4 | 0 | 14 | 0 | 10 | 0.1 | 12 | 0.0833 | 40 | 0.05 |
| 13_9516260 | 13 | 0.2308 | 7 | 0.2857 | 10 | 0.7 | 14 | 0.5714 | 44 | 0.4545 | 4 | 0.25 | 14 | 0.5 | 8 | 0.25 | 13 | 0.3077 | 39 | 0.359 |
| 13_9516264 | 14 | 0.0714 | 6 | 0 | 11 | 0.0909 | 11 | 0.1818 | 42 | 0.0952 | 5 | 0 | 11 | 0.1818 | 11 | 0.1818 | 12 | 0.0833 | 39 | 0.1282 |
| 13_9547271 | 2 | 0.5 | 2 | 0 | 4 | 0.5 | 1 | 0 | 9 | 0.3333 | 4 | 0.25 | 3 | 0 | 1 | 0 | 2 | 0 | 10 | 0.1 |
| 13_9547303 | 3 | 1 | 2 | 1 | 3 | 0.6667 | 1 | 0 | 9 | 0.7778 | 3 | 1 | 2 | 0.5 | 1 | 1 | 2 | 0.5 | 8 | 0.75 |
| 13_9547320 | 3 | 0.3333 | 1 | 0 | 5 | 0.4 | 1 | 0 | 10 | 0.3 | 3 | 1 | 2 | 1 | NA | NA | 2 | 0.5 | 7 | 0.8571 |
| 13_9547323 | 3 | 0.6667 | 1 | 1 | 6 | 0.6667 | 1 | 1 | 11 | 0.7273 | 4 | 0.5 | 3 | 0.3333 | NA | NA | 2 | 0.5 | 9 | 0.4444 |
| 13_9547335 | 3 | 0 | 2 | 0 | 6 | 0.3333 | 1 | 0 | 12 | 0.1667 | 3 | 0 | 3 | 0.3333 | NA | NA | 2 | 0 | 8 | 0.125 |
| 13_9547342 | 3 | 0.6667 | 3 | 1 | 5 | 1 | 1 | 1 | 12 | 0.9167 | 2 | 1 | 3 | 0.6667 | NA | NA | 2 | 1 | 7 | 0.8571 |
| 13_9547351 | 3 | 0.3333 | 3 | 0.3333 | 5 | 0.2 | 1 | 0 | 12 | 0.25 | 4 | 0 | 3 | 0.6667 | NA | NA | 1 | 0 | 8 | 0.25 |
| 13_9547355 | 3 | 0 | 3 | 0.3333 | 6 | 0.1667 | 1 | 0 | 13 | 0.1538 | 2 | 1 | 3 | 0 | NA | NA | 1 | 0 | 6 | 0.3333 |
| 13_9548732 | 10 | 0.3 | 9 | 0.3333 | 11 | 0.2727 | 5 | 0.4 | 35 | 0.3143 | 6 | 0 | 6 | 0.3333 | 12 | 0.25 | 7 | 0.4286 | 31 | 0.2581 |
| 13_9548766 | 11 | 0.0909 | 8 | 0.125 | 10 | 0.1 | 4 | 0 | 33 | 0.0909 | 6 | 0 | 5 | 0 | 13 | 0.1538 | 7 | 0.1429 | 31 | 0.0968 |
| 13_9548772 | 9 | 0 | 9 | 0.1111 | 7 | 0.1429 | 3 | 0 | 28 | 0.0714 | 6 | 0 | 3 | 0 | 9 | 0 | 6 | 0.3333 | 24 | 0.0833 |
| 13_9548787 | 9 | 0.3333 | 9 | 0.1111 | 10 | 0 | 4 | 0.5 | 32 | 0.1875 | 5 | 0.4 | 5 | 0.4 | 12 | 0.25 | 5 | 0 | 27 | 0.2593 |
| 13_9548791 | 5 | 0.4 | 7 | 0.7143 | 6 | 0 | 3 | 0 | 21 | 0.3333 | 5 | 0.8 | 1 | 0 | 8 | 0.625 | 5 | 0.8 | 19 | 0.6842 |
| 13_9548802 | 9 | 0.6667 | 8 | 0.75 | 10 | 0.8 | 3 | 0.6667 | 30 | 0.7333 | 5 | 0.8 | 3 | 1 | 12 | 0.9167 | 5 | 0.8 | 25 | 0.88 |
| 13_9548805 | 6 | 0.3333 | 8 | 0.5 | 9 | 0.1111 | 4 | 0 | 27 | 0.2593 | 6 | 0.1667 | 4 | 0 | 11 | 0.1818 | 4 | 0.5 | 25 | 0.2 |
| 13_9548810 | 9 | 0.1111 | 6 | 0 | 10 | 0.2 | 3 | 0.3333 | 28 | 0.1429 | 6 | 0.1667 | 3 | 0 | 11 | 0 | 4 | 0.5 | 24 | 0.125 |
| 13_9556813 | 3 | 0 | 5 | 0.2 | 2 | 0 | 3 | 0.3333 | 13 | 0.1538 | 4 | 0.25 | 4 | 0 | 2 | 0 | 2 | 0 | 12 | 0.0833 |
| 13_9556824 | 4 | 0.5 | 7 | 0.4286 | 2 | 0.5 | 3 | 1 | 16 | 0.5625 | 4 | 0.25 | 4 | 0.25 | 2 | 0.5 | 3 | 0.3333 | 13 | 0.3077 |
| 13_9556832 | 4 | 0 | 4 | 0.25 | 2 | 0 | 3 | 0 | 13 | 0.0769 | 3 | 0.3333 | 4 | 0 | 2 | 0 | 3 | 0 | 12 | 0.0833 |
| 13_9576449 | 2 | 0.5 | NA | NA | 1 | 0 | NA | NA | 3 | 0.3333 | NA | NA | 4 | 0 | 1 | 0 | 3 | 0.3333 | 8 | 0.125 |
| 13_9609682 | 2 | 0 | 2 | 0 | 3 | 0.3333 | 3 | 0.3333 | 10 | 0.2 | 2 | 0 | 7 | 0 | 5 | 0 | 4 | 0.25 | 18 | 0.0556 |
| 13_9609750 | 3 | 0 | 2 | 0 | 2 | 0.5 | 3 | 0.3333 | 10 | 0.2 | 2 | 0 | 3 | 0.3333 | 6 | 0 | NA | NA | 11 | 0.0909 |
| 13_9609768 | 3 | 0.3333 | 2 | 0 | 2 | 0 | 3 | 0 | 10 | 0.1 | 2 | 0 | 3 | 0 | 5 | 0.2 | 1 | 0 | 11 | 0.0909 |
| 13_9609768 | 3 | 0 | 2 | 0 | 2 | 0.5 | 3 | 0 | 10 | 0.1 | 1 | 0 | 3 | 0 | 4 | 0.25 | 1 | 0 | 9 | 0.1111 |
| 13_9609778 | 3 | 0 | 3 | 0 | 2 | 0 | 3 | 0.3333 | 11 | 0.0909 | 2 | 0 | 4 | 0 | 6 | 0.1667 | 1 | 0 | 13 | 0.0769 |
| 13_9609783 | 3 | 0.6667 | 3 | 0.6667 | 1 | 1 | 3 | 1 | 10 | 0.8 | 2 | 0 | 2 | 0.5 | 5 | 0.8 | 1 | 1 | 10 | 0.6 |
| 13_9622495 | 2 | 1 | 1 | 0 | 1 | 0 | 2 | 0.5 | 6 | 0.5 | 1 | 0 | 1 | 1 | 2 | 1 | 1 | 0 | 5 | 0.6 |
| 13_9622497 | 1 | 0 | 1 | 0 | 1 | 1 | 2 | 0.5 | 5 | 0.4 | 1 | 0 | 1 | 0 | 2 | 0.5 | NA | NA | 4 | 0.25 |
| 13_9622525 | 1 | 1 | 1 | 1 | NA | NA | 1 | 1 | 3 | 1 | 1 | 1 | NA | NA | 2 | 0.5 | NA | NA | 3 | 0.6667 |
| 13_9622534 | 1 | 1 | 1 | 0 | 1 | 0 | 2 | 0 | 5 | 0.2 | 1 | 1 | NA | NA | 2 | 0 | 1 | 0 | 4 | 0.25 |
| 13_9655112 | 3 | 0 | 1 | 0 | 2 | 0.5 | 8 | 0.125 | 14 | 0.1429 | 5 | 0.4 | 3 | 0.3333 | 1 | 1 | 4 | 0.25 | 13 | 0.3846 |
| 13_9655117 | 4 | 0.75 | 1 | 1 | 2 | 1 | 9 | 0.5556 | 16 | 0.6875 | 5 | 1 | 3 | 1 | 1 | 1 | 3 | 1 | 12 | 1 |
| 13_9655120 | 4 | 0.5 | 1 | 0 | 2 | 0 | 9 | 0.3333 | 16 | 0.3125 | 5 | 0 | 4 | 0.25 | 1 | 0 | 2 | 1 | 12 | 0.25 |
| 13_9655123 | 4 | 0.75 | 1 | 1 | 2 | 1 | 8 | 0.75 | 15 | 0.8 | 5 | 1 | 4 | 0.5 | 1 | 0 | 5 | 0.6 | 15 | 0.6667 |
| 13_9655126 | 4 | 0.25 | 2 | 0 | 2 | 1 | 8 | 0.375 | 16 | 0.375 | 5 | 0.4 | 4 | 0.25 | 1 | 1 | 5 | 0.2 | 15 | 0.3333 |
| 13_9655133 | 2 | 0 | 2 | 0.5 | 2 | 0.5 | 7 | 0 | 13 | 0.1538 | 3 | 0 | 4 | 0 | 1 | 0 | 4 | 0.25 | 12 | 0.0833 |
| 13_9655142 | 1 | 1 | 1 | 1 | NA | NA | 4 | 0.25 | 6 | 0.5 | 2 | 1 | 2 | 0.5 | 1 | 1 | 3 | 0 | 8 | 0.5 |
| 13_9655149 | 3 | 0.6667 | 2 | 0 | 1 | 1 | 7 | 0 | 13 | 0.2308 | 3 | 0.3333 | 2 | 0 | 2 | 0 | 2 | 0 | 9 | 0.1111 |

| | | | | | | | | | | | | | | | | | | | | |
|---|---|---|---|---|---|---|---|---|---|---|---|---|---|---|---|---|---|---|---|---|
| 13_9655186 | 4 | 0.5 | 2 | 0 | NA | NA | 5 | 0 | 11 | 0.1818 | 3 | 0.6667 | 3 | 0 | 4 | 0.25 | 3 | 0.3333 | 13 | 0.3077 |
| 13_9655319 | 4 | 0.25 | 1 | 0 | 1 | 0 | 2 | 0 | 8 | 0.125 | NA | NA | 2 | 0.5 | 5 | 0 | NA | NA | 7 | 0.1429 |
| 13_9655339 | 4 | 0 | 1 | 0 | 1 | 1 | 2 | 0 | 8 | 0.125 | NA | NA | 2 | 0.5 | 5 | 0 | 1 | 0 | 8 | 0.125 |
| 13_9661745 | NA | NA | NA | NA | 1 | 0 | 3 | 0.3333 | 4 | 0.25 | NA | NA | NA | NA | 3 | 0.3333 | 1 | 0 | 4 | 0.25 |
| 13_9661766 | NA | NA | NA | NA | 2 | 0.5 | 3 | 0 | 5 | 0.2 | NA | NA | NA | NA | 3 | 0.3333 | 3 | 0.3333 | 6 | 0.3333 |
| 13_9661775 | NA | NA | NA | NA | 2 | 1 | 3 | 1 | 5 | 1 | NA | NA | NA | NA | 2 | 0.5 | 2 | 1 | 4 | 0.75 |
| 13_9661836 | NA | NA | NA | NA | 2 | 1 | 2 | 0.5 | 4 | 0.75 | NA | NA | NA | NA | 2 | 0.5 | 2 | 0.5 | 4 | 0.5 |
| 13_9661853 | NA | NA | NA | NA | 2 | 0.5 | 3 | 0.3333 | 5 | 0.4 | NA | NA | NA | NA | 2 | 0 | 2 | 0.5 | 4 | 0.25 |
| 13_9661867 | NA | NA | NA | NA | 1 | 1 | 2 | 1 | 3 | 1 | NA | NA | NA | NA | 2 | 0.5 | 2 | 0.5 | 4 | 0.5 |
| 13_9662005 | NA | NA | NA | NA | 2 | 0 | 3 | 0.6667 | 5 | 0.4 | NA | NA | NA | NA | 1 | 1 | 2 | 0.5 | 3 | 0.6667 |
| 13_9662012 | NA | NA | NA | NA | 2 | 1 | 2 | 0.5 | 4 | 0.75 | NA | NA | NA | NA | 2 | 0.5 | 2 | 0.5 | 4 | 0.5 |
| 13_9662019 | NA | NA | NA | NA | 2 | 0.5 | 2 | 0.5 | 4 | 0.5 | NA | NA | NA | NA | 2 | 0.5 | 2 | 0.5 | 4 | 0.5 |
| 13_9662030 | NA | NA | NA | NA | 2 | 1 | 2 | 0.5 | 4 | 0.75 | NA | NA | NA | NA | 1 | 0 | 2 | 1 | 3 | 0.6667 |
| 13_9662048 | NA | NA | NA | NA | 2 | 0.5 | 1 | 1 | 3 | 0.6667 | NA | NA | NA | NA | 2 | 0.5 | 1 | 0 | 3 | 0.3333 |
| 13_9662049 | NA | NA | NA | NA | 2 | 1 | 2 | 0.5 | 4 | 0.75 | NA | NA | NA | NA | 2 | 0.5 | 2 | 0.5 | 4 | 0.5 |
| 13_9662052 | NA | NA | NA | NA | 2 | 0 | 2 | 0.5 | 4 | 0.25 | NA | NA | NA | NA | 1 | 0 | 2 | 0.5 | 3 | 0.3333 |
| 13_9662059 | NA | NA | NA | NA | 2 | 1 | NA | NA | 2 | 1 | NA | NA | NA | NA | NA | NA | 1 | 1 | 1 | 1 |
| 13_9662079 | NA | NA | NA | NA | 2 | 0.5 | 2 | 0 | 4 | 0.25 | NA | NA | NA | NA | 2 | 0 | 2 | 0.5 | 4 | 0.25 |
| 13_9663323 | NA | NA | NA | NA | 1 | 0 | 1 | 1 | 2 | 0.5 | NA | NA | NA | NA | 1 | 0 | 1 | 1 | 2 | 0.5 |
| 13_9677373 | 1 | 0 | 3 | 0 | NA | NA | 2 | 0.5 | 6 | 0.1667 | NA | NA | 3 | 0.3333 | 2 | 0 | NA | NA | 5 | 0.2 |
| 13_9677392 | 1 | 0 | 3 | 0.6667 | NA | NA | 2 | 0 | 6 | 0.3333 | 1 | 0 | 3 | 0.3333 | 3 | 0.6667 | 1 | 1 | 8 | 0.5 |
| 13_9677419 | 1 | 0 | 3 | 0.3333 | NA | NA | 2 | 0.5 | 6 | 0.3333 | 1 | 1 | NA | NA | 3 | 0.3333 | 1 | 1 | 5 | 0.6 |
| 13_9677649 | 7 | 0 | 15 | 0.0667 | 5 | 0.2 | 4 | 0 | 31 | 0.0645 | 4 | 0.25 | 5 | 0.2 | 7 | 0 | 6 | 0.3333 | 22 | 0.1818 |
| 13_9678578 | 7 | 0.1429 | 11 | 0.0909 | 13 | 0.0769 | 13 | 0.0769 | 44 | 0.0909 | 8 | 0.125 | 7 | 0.1429 | 7 | 0.1429 | 8 | 0.25 | 30 | 0.1667 |
| 13_9678579 | 7 | 0.4286 | 11 | 0.5455 | 12 | 0.6667 | 12 | 0.5833 | 42 | 0.5714 | 5 | 0.6 | 8 | 0.375 | 8 | 0.125 | 9 | 0.5556 | 30 | 0.4 |
| 13_9678598 | 11 | 0 | 13 | 0.0769 | 12 | 0.1667 | 11 | 0.0909 | 47 | 0.0851 | 8 | 0 | 12 | 0.25 | 8 | 0 | 9 | 0.3333 | 37 | 0.1622 |
| 13_9678673 | 10 | 0.1 | 10 | 0.1 | 12 | 0.1667 | 9 | 0 | 41 | 0.0976 | 6 | 0 | 10 | 0.1 | 8 | 0.125 | 8 | 0 | 32 | 0.0625 |
| 13_9678678 | 8 | 0.25 | 9 | 0.2222 | 11 | 0.3636 | 7 | 0.4286 | 35 | 0.3143 | 6 | 0.1667 | 9 | 0.2222 | 8 | 0.125 | 7 | 0.2857 | 30 | 0.2 |
| 13_9678689 | 7 | 0.7143 | 9 | 0.4444 | 10 | 0.8 | 6 | 0.6667 | 32 | 0.6562 | 5 | 0.4 | 8 | 0.625 | 6 | 0.5 | 7 | 0.7143 | 26 | 0.5769 |
| 13_9678691 | 9 | 0.6667 | 10 | 0.3 | 11 | 0.5455 | 8 | 0.625 | 38 | 0.5263 | 6 | 0.3333 | 9 | 0.5556 | 6 | 0.5 | 8 | 0.25 | 29 | 0.4138 |
| 13_9678698 | 9 | 0.8889 | 10 | 1 | 9 | 0.7778 | 7 | 1 | 35 | 0.9143 | 4 | 0.75 | 8 | 0.875 | 8 | 1 | 8 | 0.875 | 28 | 0.8929 |
| 13_9678704 | 8 | 0.875 | 8 | 0.875 | 10 | 0.8 | 9 | 1 | 35 | 0.8857 | 5 | 1 | 8 | 0.875 | 5 | 1 | 8 | 1 | 26 | 0.9615 |
| 13_9678707 | 5 | 0.2 | 8 | 0 | 9 | 0.1111 | 7 | 0.2857 | 29 | 0.1379 | 5 | 0 | 8 | 0.5 | 7 | 0.1429 | 6 | 0 | 26 | 0.1923 |
| 13_9708636 | 1 | 1 | 2 | 0 | 1 | 0 | NA | NA | 4 | 0.25 | NA | NA | 1 | 1 | NA | NA | NA | NA | 1 | 1 |
| 13_9708655 | 1 | 1 | 2 | 0.5 | 1 | 1 | NA | NA | 4 | 0.75 | NA | NA | 1 | 1 | NA | NA | NA | NA | 1 | 1 |
| 13_9708656 | 1 | 0 | 3 | 0.6667 | 1 | 0 | NA | NA | 5 | 0.4 | NA | NA | 1 | 1 | NA | NA | NA | NA | 1 | 1 |
| 13_9726302 | 5 | 0.4 | 8 | 0.25 | 2 | 0 | 4 | 0 | 19 | 0.2105 | 2 | 0.5 | 4 | 0 | 6 | 0.1667 | 4 | 0 | 16 | 0.125 |
| 13_9726306 | 5 | 0.8 | 6 | 0.8333 | 1 | 1 | 3 | 0.6667 | 15 | 0.8 | 2 | 0.5 | 4 | 0.75 | 6 | 0.8333 | 4 | 0.75 | 16 | 0.75 |
| 13_9726310 | 4 | 0 | 6 | 0.3333 | 1 | 0 | 3 | 0 | 14 | 0.1429 | 1 | 0 | 2 | 0 | 5 | 0 | 4 | 0.25 | 12 | 0.0833 |
| 13_9726312 | 5 | 0.2 | 8 | 0.75 | 2 | 1 | 4 | 0.75 | 19 | 0.6316 | 1 | 1 | 3 | 1 | 6 | 1 | 4 | 0.5 | 14 | 0.8571 |
| 13_9726320 | 4 | 0 | 7 | 0.5714 | 4 | 0 | 4 | 0 | 19 | 0.2105 | 2 | 0.5 | 3 | 0 | 5 | 0.2 | 5 | 0.4 | 15 | 0.2667 |
| 13_9726328 | 5 | 0 | 8 | 0.25 | 3 | 0 | 4 | 0.25 | 20 | 0.15 | 2 | 0 | 4 | 0.25 | 5 | 0.2 | 5 | 0.2 | 16 | 0.1875 |
| 13_9726331 | 5 | 0.4 | 7 | 0.2857 | 4 | 0.25 | 4 | 0.25 | 20 | 0.3 | 2 | 0.5 | 3 | 0 | 5 | 0 | 5 | 0.2 | 15 | 0.1333 |
| 13_9726335 | 5 | 0 | 7 | 0.1429 | 3 | 0 | 4 | 0.75 | 19 | 0.2105 | 2 | 0.5 | 3 | 0 | 5 | 0.4 | 4 | 0.25 | 14 | 0.2857 |
| 13_9726337 | 4 | 0 | 5 | 0.2 | 4 | 0.25 | 4 | 0 | 17 | 0.1176 | 1 | 0 | 4 | 0 | 4 | 0.25 | 4 | 0 | 13 | 0.0769 |
| 13_9726340 | 1 | 1 | 5 | 1 | 3 | 1 | 4 | 0.75 | 13 | 0.9231 | 2 | 1 | 2 | 1 | 3 | 1 | 4 | 0.75 | 11 | 0.9091 |
| 13_9726347 | 5 | 0 | 7 | 0 | 4 | 0 | 4 | 0.25 | 20 | 0.05 | 2 | 0 | 4 | 0.25 | 5 | 0 | 5 | 0 | 16 | 0.0625 |
| 13_9726360 | 5 | 0.4 | 7 | 0.2857 | 4 | 0 | 4 | 0.5 | 20 | 0.3 | 1 | 0 | 3 | 0.6667 | 6 | 0 | 5 | 0.2 | 15 | 0.2 |
| 13_9726361 | 5 | 0.2 | 4 | 0 | 2 | 0 | 4 | 0 | 15 | 0.0667 | 2 | 0 | 1 | 1 | 5 | 0 | 3 | 0.3333 | 11 | 0.1818 |
| 13_9726363 | 5 | 0.8 | 8 | 0.875 | 4 | 1 | 3 | 1 | 20 | 0.9 | 1 | 1 | 2 | 1 | 6 | 0.5 | 5 | 0.8 | 14 | 0.7143 |
| 13_9726366 | 5 | 0.6 | 8 | 0.75 | 3 | 1 | 3 | 1 | 19 | 0.7895 | 2 | 0.5 | 3 | 1 | 6 | 0.5 | 3 | 0.3333 | 14 | 0.5714 |
| 13_9726374 | 4 | 0.25 | 6 | 0.1667 | 3 | 0.3333 | 2 | 0 | 15 | 0.2 | 1 | 0 | 3 | 0 | 4 | 0.25 | 3 | 0 | 11 | 0.0909 |
| 13_9726376 | 5 | 0.2 | 8 | 0 | 2 | 0 | 2 | 0 | 17 | 0.0588 | 2 | 0 | 3 | 0 | 6 | 0.1667 | 4 | 0 | 15 | 0.0667 |
| 13_9726385 | 5 | 0 | 8 | 0.25 | 3 | 0.3333 | 3 | 0 | 19 | 0.1579 | 1 | 0 | 4 | 0.5 | 5 | 0.4 | 3 | 0 | 13 | 0.3077 |
| 13_9726386 | 5 | 0 | 6 | 0.1667 | 2 | 0.5 | 3 | 0 | 16 | 0.125 | NA | NA | 4 | 0.25 | 5 | 0 | 3 | 0.3333 | 12 | 0.1667 |
| 13_9726432 | 4 | 0 | 8 | 0.125 | 2 | 0 | 2 | 0.5 | 16 | 0.125 | 2 | 0 | 4 | 0.25 | 5 | 0.2 | 3 | 0.3333 | 14 | 0.2143 |
| 13_9726572 | 4 | 0 | 9 | 0.3333 | 7 | 0.2857 | 6 | 0.1667 | 26 | 0.2308 | 2 | 0.5 | 3 | 0 | 6 | 0.3333 | 4 | 0.25 | 15 | 0.2667 |
| 13_9726578 | 5 | 0 | 9 | 0.1111 | 6 | 0.1667 | 5 | 0 | 25 | 0.08 | 2 | 0.5 | 4 | 0.25 | 6 | 0.1667 | 4 | 0.5 | 16 | 0.3125 |
| 13_9726743 | 1 | 1 | 2 | 0 | 2 | 0 | 1 | 0 | 6 | 0.1667 | NA | NA | NA | NA | 1 | 1 | 3 | 0.3333 | 4 | 0.5 |
| 13_9726745 | 1 | 0 | 1 | 0 | 3 | 0.6667 | 1 | 0 | 6 | 0.3333 | NA | NA | NA | NA | 1 | 0 | 3 | 0.3333 | 4 | 0.25 |
| 13_9730890 | 2 | 1 | 1 | 1 | NA | NA | NA | NA | 3 | 1 | NA | NA | NA | NA | 1 | 1 | NA | NA | 1 | 1 |
| 13_9731216 | 5 | 0.8 | 2 | 1 | 1 | 1 | 1 | 0 | 9 | 0.7778 | 3 | 0 | 3 | 0.3333 | NA | NA | NA | NA | 6 | 0.1667 |
| 13_9731234 | 5 | 0.8 | 4 | 0.5 | 1 | 1 | 1 | 1 | 11 | 0.7273 | 2 | 0.5 | 3 | 0 | NA | NA | 1 | 1 | 6 | 0.3333 |
| 13_9731254 | 4 | 0.75 | 2 | 1 | NA | NA | 2 | 0.5 | 8 | 0.75 | 3 | 0.6667 | 3 | 0 | NA | NA | 1 | 0 | 7 | 0.2857 |
| 13_9731264 | 4 | 0.25 | NA | NA | 1 | 1 | 2 | 1 | 7 | 0.5714 | 3 | 0.3333 | 3 | 0 | NA | NA | 2 | 0 | 8 | 0.125 |

| | | | | | | | | | | | | | | | | | | | | |
|---|---|---|---|---|---|---|---|---|---|---|---|---|---|---|---|---|---|---|---|---|
| 13_973126 | 4 | 0 | 2 | 1 | 1 | 0 | 1 | 0 | 8 | 0.25 | 3 | 0 | 3 | 0 | NA | NA | 2 | 1 | 8 | 0.25 |
| 13_973157 | 1 | 1 | 1 | 1 | NA | NA | NA | NA | 2 | 1 | NA | NA | NA | NA | NA | NA | 1 | 1 | 1 | 1 |
| 13_973159 | 1 | 1 | 1 | 1 | NA | NA | NA | NA | 2 | 1 | NA | NA | NA | NA | NA | NA | 2 | 1 | 2 | 1 |
| 13_973161 | 1 | 1 | 1 | 1 | NA | NA | NA | NA | 2 | 1 | NA | NA | NA | NA | NA | NA | 1 | 1 | 1 | 1 |
| 13_973161 | 1 | 1 | 1 | 1 | NA | NA | NA | NA | 2 | 1 | NA | NA | NA | NA | NA | NA | 1 | 1 | 1 | 1 |
| 13_973162 | 1 | 1 | 1 | 0 | NA | NA | NA | NA | 2 | 0.5 | NA | NA | 1 | 0 | NA | NA | 2 | 0.5 | 3 | 0.3333 |
| 13_973163 | 1 | 1 | 1 | 1 | NA | NA | NA | NA | 2 | 1 | NA | NA | 1 | 1 | NA | NA | 2 | 1 | 3 | 1 |
| 13_973169 | 1 | 0 | 2 | 0.5 | NA | NA | NA | NA | 3 | 0.3333 | NA | NA | NA | NA | NA | NA | 2 | 0.5 | 2 | 0.5 |
| 13_973815 | 1 | 0 | 8 | 0.125 | 2 | 0 | 5 | 0 | 16 | 0.0625 | 5 | 0.2 | 4 | 0 | 5 | 0 | 3 | 0.6667 | 17 | 0.1765 |
| 13_973816 | 1 | 0 | 11 | 0.4545 | 2 | 0.5 | 5 | 0.4 | 19 | 0.4211 | 6 | 0.3333 | 6 | 0.1667 | 4 | 0 | 3 | 0.3333 | 19 | 0.2105 |
| 13_973817 | 1 | 1 | 10 | 0.3 | 2 | 0 | 6 | 0.3333 | 19 | 0.3158 | 6 | 0.3333 | 5 | 0.6 | 4 | 0.5 | 3 | 0.6667 | 18 | 0.5 |
| 13_973817 | 1 | 0 | 10 | 0.1 | 2 | 0 | 6 | 0.3333 | 19 | 0.1579 | 6 | 0 | 6 | 0 | 5 | 0.2 | 2 | 0 | 19 | 0.0526 |
| 13_973819 | NA | NA | 11 | 0.1818 | 2 | 0 | 5 | 0.4 | 18 | 0.2222 | 7 | 0.8571 | 4 | 0.5 | 3 | 0 | 2 | 0.5 | 16 | 0.5625 |
| 13_973820 | 1 | 0 | 10 | 0.1 | 2 | 0 | 7 | 0.1429 | 20 | 0.1 | 7 | 0.1429 | 5 | 0.4 | 5 | 0 | 5 | 0.4 | 22 | 0.2273 |
| 13_973821 | 1 | 0 | 11 | 0.2727 | 2 | 0.5 | 7 | 0 | 21 | 0.1905 | 8 | 0.125 | 5 | 0.2 | 5 | 0.2 | 2 | 0.5 | 20 | 0.2 |
| 13_973821 | NA | NA | 8 | 0.375 | 2 | 0.5 | 5 | 0.8 | 15 | 0.5333 | 6 | 0.6667 | 6 | 0.8333 | 5 | 0.8 | 4 | 0.75 | 21 | 0.7619 |
| 13_973822 | 1 | 0 | 9 | 0.1111 | 2 | 0 | 6 | 0.1667 | 18 | 0.1111 | 8 | 0 | 7 | 0.1429 | 5 | 0.2 | 6 | 0 | 26 | 0.0769 |
| 13_973823 | 1 | 0 | 10 | 0.1 | 1 | 0 | 4 | 0.25 | 16 | 0.125 | 7 | 0.4286 | 6 | 0 | 4 | 0.25 | 5 | 0.6 | 22 | 0.3182 |
| 13_973824 | 1 | 0 | 11 | 0.5455 | 2 | 0.5 | 7 | 0.7143 | 21 | 0.5714 | 7 | 0.8571 | 6 | 0.5 | 4 | 0.75 | 6 | 0.6667 | 23 | 0.6957 |
| 13_973824 | 1 | 1 | 9 | 0.8889 | 1 | 1 | 6 | 1 | 17 | 0.9412 | 7 | 1 | 4 | 1 | 3 | 1 | 4 | 1 | 18 | 1 |
| 13_973826 | 2 | 1 | 10 | 0.9 | 2 | 1 | 8 | 1 | 22 | 0.9545 | 11 | 1 | 9 | 1 | 6 | 1 | 6 | 1 | 32 | 1 |
| 13_973827 | 3 | 0 | 12 | 0.5 | 3 | 0.3333 | 5 | 0.2 | 23 | 0.3478 | 11 | 0.2727 | 7 | 0.5714 | 5 | 0.4 | 6 | 0.1667 | 29 | 0.3448 |
| 13_973827 | 3 | 1 | 13 | 1 | 3 | 1 | 8 | 1 | 27 | 1 | 11 | 1 | 7 | 1 | 5 | 1 | 5 | 1 | 28 | 1 |
| 13_973828 | 3 | 0 | 12 | 0.1667 | 3 | 0 | 8 | 0 | 26 | 0.0769 | 10 | 0.2 | 8 | 0.375 | 7 | 0 | 6 | 0 | 31 | 0.1613 |
| 13_973830 | 3 | 0.6667 | 13 | 0.6154 | 2 | 1 | 8 | 0.625 | 26 | 0.6538 | 15 | 0.6 | 9 | 0.8889 | 5 | 1 | 9 | 0.4444 | 38 | 0.6842 |
| 13_973830 | 3 | 1 | 10 | 0.8 | 3 | 1 | 6 | 0.6667 | 22 | 0.8182 | 13 | 0.7692 | 8 | 1 | 6 | 0.6667 | 7 | 0.5714 | 34 | 0.7647 |
| 13_973830 | 2 | 0.5 | 11 | 0.0909 | 2 | 0 | 7 | 0 | 22 | 0.0909 | 15 | 0.0667 | 8 | 0 | 6 | 0.1667 | 9 | 0 | 38 | 0.0526 |
| 13_973831 | 3 | 0.3333 | 13 | 0.4615 | 3 | 0 | 8 | 0.375 | 27 | 0.3704 | 13 | 0.3077 | 10 | 0.2 | 7 | 0.5714 | 9 | 0.4444 | 39 | 0.359 |
| 13_973831 | 2 | 1 | 10 | 1 | 4 | 1 | 7 | 1 | 23 | 1 | 14 | 1 | 10 | 1 | 7 | 1 | 10 | 1 | 41 | 1 |
| 13_973871 | 1 | 0 | 13 | 0.1538 | 5 | 0.4 | 7 | 0 | 26 | 0.1538 | 14 | 0.1429 | 12 | 0.25 | 6 | 0.1667 | 10 | 0 | 42 | 0.1429 |
| 13_973871 | 2 | 1 | 11 | 0.8182 | 3 | 1 | 7 | 0.8571 | 23 | 0.8696 | 11 | 0.9091 | 6 | 1 | 2 | 1 | 8 | 0.75 | 27 | 0.8889 |
| 13_973872 | 3 | 1 | 12 | 0.9167 | 4 | 1 | 7 | 1 | 26 | 0.9615 | 13 | 0.7692 | 10 | 1 | 5 | 0.8 | 8 | 1 | 36 | 0.8889 |
| 13_973872 | 3 | 0.3333 | 14 | 0.8571 | 4 | 0.5 | 6 | 0.8333 | 27 | 0.7407 | 14 | 0.7143 | 8 | 0.75 | 9 | 0.6667 | 8 | 0.5 | 39 | 0.6667 |
| 13_973875 | 3 | 0 | 20 | 0.15 | 9 | 0.3333 | 11 | 0.2727 | 43 | 0.2093 | 22 | 0.1818 | 12 | 0.5 | 11 | 0.0909 | 15 | 0.2667 | 60 | 0.25 |
| 13_973875 | 4 | 1 | 22 | 0.8182 | 10 | 1 | 10 | 1 | 46 | 0.913 | 23 | 0.8261 | 14 | 1 | 10 | 0.9 | 13 | 0.9231 | 60 | 0.9 |
| 13_973875 | 4 | 0.25 | 20 | 0.3 | 8 | 0.125 | 11 | 0.1818 | 43 | 0.2326 | 20 | 0.15 | 13 | 0.1538 | 11 | 0.0909 | 14 | 0.1429 | 58 | 0.1379 |
| 13_973875 | 4 | 0.75 | 19 | 0.7895 | 9 | 0.4444 | 11 | 0.7273 | 43 | 0.6977 | 21 | 0.5714 | 14 | 0.6429 | 11 | 0.3636 | 13 | 0.4615 | 59 | 0.5254 |
| 13_973876 | 4 | 0.75 | 21 | 0.619 | 10 | 0.5 | 10 | 0.4 | 45 | 0.5556 | 22 | 0.5 | 12 | 0.25 | 10 | 0.3 | 15 | 0.5333 | 59 | 0.4237 |
| 13_973878 | 5 | 0.4 | 26 | 0.3846 | 9 | 0.5556 | 11 | 0.5455 | 51 | 0.451 | 24 | 0.375 | 16 | 0.5 | 9 | 0.2222 | 19 | 0.3684 | 68 | 0.3824 |
| 13_973879 | 16 | 0.125 | 36 | 0.0278 | 19 | 0.1053 | 16 | 0 | 87 | 0.0575 | 30 | 0.1 | 19 | 0.1053 | 18 | 0.1111 | 36 | 0 | 103 | 0.068 |
| 13_973880 | 18 | 0.1667 | 38 | 0.0789 | 15 | 0 | 15 | 0.1333 | 86 | 0.093 | 27 | 0.037 | 20 | 0.05 | 16 | 0.25 | 31 | 0.0645 | 94 | 0.0851 |
| 13_973880 | 16 | 0.0625 | 36 | 0.1111 | 18 | 0.1111 | 17 | 0.0588 | 87 | 0.092 | 29 | 0.1034 | 17 | 0 | 15 | 0.0667 | 35 | 0.1143 | 96 | 0.0833 |
| 13_973883 | 21 | 0.1429 | 40 | 0.225 | 18 | 0.1111 | 17 | 0.2941 | 96 | 0.1979 | 31 | 0.1935 | 22 | 0.0455 | 20 | 0.2 | 34 | 0.1471 | 107 | 0.1495 |
| 13_973887 | 28 | 0.1429 | 53 | 0.0943 | 25 | 0.16 | 19 | 0.2105 | 125 | 0.136 | 39 | 0.1538 | 28 | 0.2143 | 21 | 0.1429 | 38 | 0.2105 | 126 | 0.1825 |
| 13_976724 | NA | NA | NA | NA | 3 | 1 | 1 | 1 | 4 | 1 | NA | NA | NA | NA | 1 | 1 | NA | NA | 1 | 1 |
| 13_976724 | NA | NA | NA | NA | 3 | 1 | 1 | 1 | 4 | 1 | NA | NA | NA | NA | 1 | 1 | NA | NA | 1 | 1 |
| 13_976724 | NA | NA | NA | NA | 1 | 1 | 1 | 1 | 2 | 1 | NA | NA | NA | NA | 1 | 1 | NA | NA | 1 | 1 |
| 13_976726 | NA | NA | NA | NA | 3 | 0.3333 | 1 | 0 | 4 | 0.25 | NA | NA | NA | NA | 1 | 1 | NA | NA | 1 | 1 |
| 13_976726 | NA | NA | NA | NA | 2 | 0.5 | 1 | 1 | 3 | 0.6667 | NA | NA | NA | NA | 1 | 1 | 1 | 0 | 2 | 0.5 |
| 13_976730 | NA | NA | NA | NA | 2 | 1 | 1 | 1 | 3 | 1 | NA | NA | NA | NA | 1 | 1 | NA | NA | 1 | 1 |
| 13_976730 | NA | NA | NA | NA | 1 | 0 | 1 | 1 | 2 | 0.5 | NA | NA | NA | NA | 1 | 1 | NA | NA | 1 | 1 |
| 13_976733 | NA | NA | NA | NA | 3 | 0.3333 | 1 | 0 | 4 | 0.25 | NA | NA | NA | NA | 1 | 1 | NA | NA | 1 | 1 |
| 13_976741 | NA | NA | NA | NA | 1 | 0 | 1 | 1 | 2 | 0.5 | NA | NA | NA | NA | 1 | 1 | NA | NA | 1 | 1 |
| 13_976741 | NA | NA | NA | NA | 3 | 1 | 1 | 1 | 4 | 1 | NA | NA | NA | NA | 1 | 1 | NA | NA | 1 | 1 |
| 13_976743 | NA | NA | NA | NA | 3 | 0.6667 | 1 | 1 | 4 | 0.75 | NA | NA | NA | NA | 1 | 1 | NA | NA | 1 | 1 |
| 13_976750 | NA | NA | NA | NA | 3 | 0.3333 | 1 | 1 | 4 | 0.5 | NA | NA | NA | NA | 1 | 1 | 1 | 0 | 2 | 0.5 |
| 13_980361 | 1 | 0 | 3 | 0 | 3 | 0.3333 | NA | NA | 7 | 0.1429 | 1 | 1 | 1 | 0 | 1 | 1 | NA | NA | 3 | 0.6667 |
| 13_980659 | 1 | 0 | 5 | 0.4 | 5 | 0.2 | 1 | 0 | 12 | 0.25 | 2 | 1 | NA | NA | 1 | 0 | 1 | 0 | 4 | 0.5 |
| 13_980660 | 1 | 0 | 6 | 0 | 5 | 0.2 | NA | NA | 12 | 0.0833 | 3 | 0.3333 | 1 | 0 | 1 | 0 | 1 | 0 | 6 | 0.1667 |
| 13_980660 | 1 | 1 | 5 | 0.6 | 5 | 0.6 | NA | NA | 11 | 0.6364 | 3 | 0 | 1 | 0 | 1 | 0 | 1 | 1 | 6 | 0.1667 |
| 13_980660 | NA | NA | 6 | 0.6667 | 5 | 0.8 | 1 | 1 | 12 | 0.75 | 4 | 0.75 | 1 | 0 | 1 | 0 | 1 | 1 | 7 | 0.5714 |
| 13_980661 | NA | NA | 4 | 0 | 4 | 0 | 1 | 1 | 9 | 0.1111 | 3 | 0.3333 | 1 | 0 | 1 | 0 | 1 | 0 | 6 | 0.1667 |
| 13_980661 | NA | NA | 6 | 0.8333 | 6 | 1 | 1 | 1 | 13 | 0.9231 | 4 | 0.75 | 1 | 1 | 1 | 1 | 1 | 1 | 7 | 0.8571 |
| 13_980662 | 1 | 0 | 6 | 0.1667 | 6 | 0 | 1 | 0 | 14 | 0.0714 | 4 | 0.25 | 1 | 0 | 1 | 0 | NA | NA | 6 | 0.1667 |
| 13_980748 | NA | NA | 2 | 0.5 | 1 | 0 | NA | NA | 3 | 0.3333 | NA | NA | NA | NA | 1 | 1 | NA | NA | 1 | 1 |

| | | | | | | | | | | | | | | | | | | | | |
|---|---|---|---|---|---|---|---|---|---|---|---|---|---|---|---|---|---|---|---|---|
| 13_980755 | 1 | 0 | 3 | 0.3333 | 1 | 1 | NA | NA | 5 | 0.4 | 1 | 1 | 1 | 1 | 1 | 0 | 1 | 1 | 4 | 0.75 |
| 13_980763 | NA | NA | 6 | 0.3333 | 1 | 0 | NA | NA | 7 | 0.2857 | 3 | 0.6667 | 2 | 0.5 | 1 | 1 | 3 | 0.3333 | 9 | 0.5556 |
| 13_982318 | 1 | 1 | NA | NA | 1 | 0 | NA | NA | 2 | 0.5 | 1 | 1 | 1 | 1 | 1 | 1 | 1 | 1 | 4 | 1 |
| 13_983833 | 1 | 0 | 2 | 0 | 1 | 0 | 1 | 1 | 5 | 0.2 | 3 | 0 | 1 | 0 | 4 | 0.25 | 2 | 0 | 10 | 0.1 |
| 13_983843 | 2 | 0.5 | 3 | 0.3333 | 1 | 0 | 1 | 1 | 7 | 0.4286 | 3 | 0.3333 | NA | NA | 4 | 0 | 2 | 0 | 9 | 0.1111 |
| 13_983850 | 2 | 0 | 5 | 0.4 | 2 | 0 | 3 | 0 | 12 | 0.1667 | 4 | 0 | 2 | 0.5 | 4 | 0.25 | 2 | 0 | 12 | 0.1667 |
| 13_983851 | 2 | 0.5 | 3 | 1 | 2 | 1 | 2 | 1 | 9 | 0.8889 | 3 | 1 | 2 | 0.5 | 4 | 0.75 | 2 | 1 | 11 | 0.8182 |
| 13_983853 | 3 | 0.6667 | 5 | 0.8 | 2 | 0 | 2 | 0.5 | 12 | 0.5833 | 4 | 1 | 2 | 0.5 | 4 | 0.5 | 2 | 0.5 | 12 | 0.6667 |
| 13_983854 | 3 | 0.3333 | 4 | 0 | 2 | 0.5 | 2 | 0 | 11 | 0.1818 | 4 | 0 | 1 | 1 | 3 | 0.3333 | 2 | 1 | 10 | 0.4 |
| 13_983855 | 2 | 1 | 4 | 0.5 | 1 | 1 | 2 | 1 | 9 | 0.7778 | 4 | 1 | 2 | 1 | 3 | 1 | NA | NA | 9 | 1 |
| 13_983855 | 3 | 1 | 3 | 1 | 2 | 1 | 2 | 1 | 10 | 1 | 4 | 1 | 2 | 1 | 3 | 1 | 2 | 0.5 | 11 | 0.9091 |
| 13_983856 | 2 | 0.5 | 5 | 0.6 | 2 | 1 | 1 | 0 | 10 | 0.6 | 4 | 0.25 | 1 | 1 | 2 | 0.5 | 2 | 0.5 | 9 | 0.4444 |
| 13_983856 | 3 | 0.3333 | 5 | 0.6 | 1 | 0 | NA | NA | 9 | 0.4444 | 3 | 0.3333 | 2 | 0.5 | 3 | 0.3333 | 1 | 0 | 9 | 0.3333 |
| 13_983856 | 2 | 0.5 | 4 | 0.75 | 2 | 0.5 | 1 | 1 | 9 | 0.6667 | 3 | 0 | 1 | 1 | 2 | 1 | 2 | 0.5 | 8 | 0.5 |
| 13_983857 | 3 | 0.6667 | 5 | 1 | 2 | 1 | 2 | 0 | 12 | 0.75 | 4 | 1 | 2 | 1 | 3 | 0.3333 | 1 | 1 | 10 | 0.8 |
| 13_983857 | 3 | 0.6667 | 5 | 0.8 | 2 | 1 | 2 | 1 | 12 | 0.8333 | 4 | 0.75 | 2 | 1 | 3 | 0.6667 | 2 | 1 | 11 | 0.8182 |
| 13_983857 | 2 | 0 | 3 | 0.3333 | 1 | 0 | 1 | 0 | 7 | 0.1429 | 3 | 0.6667 | 2 | 0.5 | 3 | 0 | 1 | 1 | 9 | 0.4444 |
| 13_983858 | 3 | 0.3333 | 5 | 0.4 | 1 | 1 | 2 | 0.5 | 11 | 0.4545 | 4 | 0 | 1 | 0 | 2 | 0.5 | NA | NA | 7 | 0.1429 |
| 13_984027 | 5 | 0.4 | 3 | 0.6667 | 5 | 0.8 | 3 | 1 | 16 | 0.6875 | 2 | 1 | 3 | 1 | 3 | 0.3333 | 1 | 0 | 9 | 0.6667 |
| 13_984465 | 36 | 0.0556 | 42 | 0.1429 | 40 | 0.075 | 30 | 0.0667 | 148 | 0.0878 | 37 | 0.0811 | 31 | 0.129 | 39 | 0.1026 | 39 | 0.0513 | 146 | 0.089 |
| 13_984465 | 35 | 0.0286 | 40 | 0.025 | 39 | 0.0256 | 25 | 0.2 | 139 | 0.0576 | 38 | 0.0526 | 26 | 0.0769 | 35 | 0.0571 | 40 | 0.075 | 139 | 0.0647 |
| 13_984583 | 1 | 0 | 2 | 0 | 1 | 1 | 1 | 1 | 5 | 0.4 | NA | NA | NA | NA | 1 | 1 | NA | NA | 1 | 1 |
| 13_984924 | NA | NA | NA | NA | NA | NA | 1 | 1 | 1 | 1 | 1 | 1 | NA | NA | NA | NA | NA | NA | 1 | 1 |
| 13_984927 | NA | NA | NA | NA | NA | NA | 1 | 1 | 1 | 1 | 1 | 1 | NA | NA | NA | NA | NA | NA | 1 | 1 |
| 13_988609 | 6 | 0.1667 | 5 | 0 | 3 | 0 | 3 | 0 | 17 | 0.0588 | 4 | 0.25 | 1 | 0 | 4 | 0.25 | 3 | 0 | 12 | 0.1667 |
| 13_988615 | 2 | 0 | 2 | 0 | 2 | 0 | 3 | 0.3333 | 9 | 0.1111 | 1 | 0 | 1 | 0 | 3 | 0 | 1 | 1 | 6 | 0.1667 |
| 13_988616 | 4 | 0 | 1 | 0 | 1 | 1 | 5 | 0 | 11 | 0.0909 | 2 | 0 | 1 | 1 | 4 | 0 | NA | NA | 7 | 0.1429 |
| 13_992137 | NA | NA | NA | NA | NA | NA | 1 | 1 | 1 | 1 | NA | NA | NA | NA | 1 | 1 | NA | NA | 1 | 1 |
| 13_993855 | 1 | 1 | 1 | 1 | 1 | 0 | NA | NA | 3 | 0.6667 | NA | NA | NA | NA | 2 | 0.5 | 1 | 0 | 3 | 0.3333 |
| 13_993859 | 1 | 1 | 1 | 1 | 1 | 1 | NA | NA | 3 | 1 | NA | NA | NA | NA | 2 | 0.5 | 1 | 1 | 3 | 0.6667 |
| 13_993860 | 1 | 1 | 1 | 1 | 1 | 1 | NA | NA | 3 | 1 | NA | NA | NA | NA | 2 | 1 | NA | NA | 2 | 1 |
| 13_993861 | 1 | 1 | 1 | 1 | 1 | 1 | NA | NA | 3 | 1 | NA | NA | NA | NA | 2 | 0.5 | NA | NA | 2 | 0.5 |
| 13_993863 | 1 | 0 | 1 | 1 | 1 | 1 | NA | NA | 3 | 0.6667 | NA | NA | NA | NA | 2 | 0.5 | 1 | 0 | 3 | 0.3333 |
| 13_993865 | 1 | 1 | 1 | 0 | 1 | 1 | NA | NA | 3 | 0.6667 | NA | NA | NA | NA | 2 | 0.5 | 1 | 0 | 3 | 0.3333 |
| 13_993866 | NA | NA | 1 | 1 | 1 | 1 | NA | NA | 2 | 1 | NA | NA | NA | NA | 1 | 0 | 1 | 1 | 2 | 0.5 |
| 13_993868 | 1 | 1 | 1 | 1 | 1 | 0 | NA | NA | 3 | 0.6667 | NA | NA | NA | NA | 2 | 1 | 1 | 1 | 3 | 1 |
| 13_994562 | 5 | 0 | 9 | 0.1111 | 8 | 0.125 | 7 | 0 | 29 | 0.069 | 10 | 0 | 6 | 0 | 7 | 0.4286 | 6 | 0.1667 | 29 | 0.1379 |
| 13_994563 | 4 | 0.25 | 8 | 0.625 | 7 | 0.4286 | 7 | 0.2857 | 26 | 0.4231 | 10 | 0.2 | 5 | 0.4 | 4 | 0.5 | 4 | 0.5 | 23 | 0.3478 |
| 13_994563 | 4 | 0.25 | 8 | 0.125 | 8 | 0.125 | 5 | 0 | 25 | 0.12 | 8 | 0 | 6 | 0.1667 | 6 | 0.1667 | 4 | 0 | 24 | 0.0833 |
| 13_994564 | 4 | 0.75 | 6 | 1 | 7 | 1 | 6 | 1 | 23 | 0.9565 | 9 | 0.8889 | 5 | 1 | 5 | 1 | 2 | 0.5 | 21 | 0.9048 |
| 13_994566 | 2 | 0 | 2 | 0.5 | 7 | 0 | 7 | 0.8571 | 18 | 0.3889 | 5 | 0.8 | 3 | 0.6667 | 5 | 0.4 | 3 | 0 | 16 | 0.5 |
| 13_994567 | 3 | 0.6667 | 6 | 0.5 | 8 | 0.5 | 7 | 0.7143 | 24 | 0.5833 | 9 | 0.3333 | 5 | 0.6 | 7 | 0.7143 | 3 | 0.6667 | 24 | 0.5417 |
| 13_994567 | 4 | 0 | 6 | 0.3333 | 7 | 0 | 6 | 0 | 23 | 0.087 | 9 | 0.2222 | 5 | 0 | 7 | 0.1429 | 3 | 0 | 24 | 0.125 |
| 13_995513 | 4 | 0.5 | 5 | 0 | 2 | 0 | 4 | 0 | 15 | 0.1333 | 5 | 0.2 | 2 | 0 | 1 | 0 | 9 | 0.2222 | 17 | 0.1765 |
| 13_995515 | 4 | 0.25 | 5 | 0.4 | 2 | 0 | 5 | 0.2 | 16 | 0.25 | 5 | 0.4 | 2 | 0 | 1 | 0 | 8 | 0.375 | 16 | 0.3125 |
| 13_995516 | 4 | 0.5 | 5 | 0 | 2 | 0 | 4 | 0.75 | 15 | 0.3333 | 5 | 0.4 | 2 | 0.5 | 1 | 0 | 8 | 0.25 | 16 | 0.3125 |
| 13_995516 | 4 | 0 | 4 | 0.75 | 2 | 0.5 | 5 | 0.6 | 15 | 0.4667 | 3 | 0.3333 | 2 | 0.5 | 1 | 0 | 9 | 0.7778 | 15 | 0.6 |
| 13_995516 | 4 | 0.25 | 6 | 0.1667 | 2 | 0 | 5 | 0.4 | 17 | 0.2353 | 4 | 0 | 3 | 0 | 1 | 0 | 9 | 0.3333 | 17 | 0.1765 |
| 13_995516 | 4 | 0.75 | 6 | 0.3333 | 2 | 0.5 | 5 | 0.4 | 17 | 0.4706 | 4 | 0 | 3 | 0.3333 | 1 | 0 | 8 | 0.375 | 16 | 0.25 |
| 13_995518 | 4 | 0.75 | 5 | 0.4 | 2 | 0.5 | 4 | 0.25 | 15 | 0.4667 | 5 | 0.6 | 2 | 0.5 | 1 | 0 | 7 | 0.4286 | 15 | 0.4667 |
| 13_995518 | 4 | 1 | 5 | 0.8 | 1 | 1 | 4 | 1 | 14 | 0.9286 | 4 | 1 | 3 | 1 | 1 | 1 | 7 | 1 | 15 | 1 |
| 13_995519 | 4 | 0 | 6 | 0.1667 | 1 | 0 | 5 | 0 | 16 | 0.0625 | 5 | 0 | 3 | 0 | 1 | 0 | 9 | 0.1111 | 18 | 0.0556 |
| 13_995584 | 65 | 0.4308 | 83 | 0.4096 | 69 | 0.4058 | 69 | 0.5507 | 286 | 0.4476 | 68 | 0.4412 | 67 | 0.4179 | 47 | 0.5319 | 55 | 0.6 | 237 | 0.4895 |
| 13_995584 | 1 | 1 | 6 | 0.5 | 3 | 0.3333 | 1 | 1 | 11 | 0.5455 | 4 | 0.5 | 4 | 1 | 5 | 0.6 | 1 | 0 | 14 | 0.6429 |
| 13_995584 | 1 | 1 | 1 | 0 | 1 | 0 | 1 | 0 | 4 | 0.25 | NA | NA | 1 | 0 | NA | NA | 1 | 1 | 2 | 0.5 |
| 13_995584 | 64 | 0.8438 | 91 | 0.9011 | 76 | 0.8421 | 68 | 0.8971 | 299 | 0.8729 | 72 | 0.9167 | 68 | 0.8676 | 58 | 0.8448 | 62 | 0.9032 | 260 | 0.8846 |
| 13_995584 | 63 | 0.2698 | 89 | 0.2921 | 77 | 0.3377 | 67 | 0.3433 | 296 | 0.3108 | 72 | 0.2361 | 70 | 0.4 | 56 | 0.2679 | 53 | 0.1698 | 251 | 0.2749 |
| 13_995584 | 1 | 0 | 4 | 0.5 | 1 | 0 | 2 | 0 | 8 | 0.25 | 2 | 0 | 1 | 1 | 3 | 0.3333 | 4 | 0 | 10 | 0.2 |
| 13_995585 | 62 | 0.5806 | 84 | 0.6667 | 70 | 0.4857 | 63 | 0.6349 | 279 | 0.595 | 62 | 0.5806 | 70 | 0.6286 | 55 | 0.7091 | 57 | 0.5614 | 244 | 0.6189 |
| 13_995586 | 61 | 0.2131 | 74 | 0.2297 | 63 | 0.2222 | 60 | 0.1167 | 258 | 0.1977 | 60 | 0.2 | 68 | 0.2206 | 46 | 0.2174 | 51 | 0.1373 | 225 | 0.1956 |
| 13_995602 | 29 | 0.069 | 44 | 0.0909 | 39 | 0.2821 | 35 | 0.1714 | 147 | 0.1565 | 34 | 0.0882 | 31 | 0.2258 | 27 | 0.1111 | 31 | 0.1935 | 123 | 0.1545 |
| 13_995602 | 26 | 0.4231 | 50 | 0.52 | 44 | 0.4773 | 35 | 0.6571 | 155 | 0.5226 | 32 | 0.4688 | 39 | 0.5128 | 28 | 0.4643 | 36 | 0.5833 | 135 | 0.5111 |
| 13_995603 | 27 | 0.1481 | 37 | 0.2432 | 40 | 0.125 | 33 | 0.1515 | 137 | 0.1679 | 34 | 0.1471 | 38 | 0.2895 | 28 | 0.1429 | 31 | 0.2903 | 131 | 0.2214 |
| 13_995603 | 30 | 0.5333 | 47 | 0.5745 | 44 | 0.7045 | 34 | 0.5588 | 155 | 0.6 | 35 | 0.5429 | 39 | 0.6923 | 24 | 0.5417 | 33 | 0.7273 | 131 | 0.6336 |
| 13_995604 | 30 | 0.0667 | 45 | 0.0667 | 45 | 0.0667 | 34 | 0 | 154 | 0.0519 | 36 | 0.1389 | 35 | 0.0286 | 31 | 0.0323 | 33 | 0.0909 | 135 | 0.0741 |

| | | | | | | | | | | | | | | | | | | | | |
|---|---|---|---|---|---|---|---|---|---|---|---|---|---|---|---|---|---|---|---|---|
| 13_9956068 | 28 | 0.1429 | 41 | 0.2195 | 37 | 0.1892 | 34 | 0.1765 | 140 | 0.1857 | 29 | 0.2759 | 28 | 0.1786 | 24 | 0.125 | 28 | 0.3214 | 109 | 0.2294 |
| 13_9956072 | 1 | 1 | NA | NA | 1 | 0 | NA | NA | 2 | 0.5 | 3 | 0 | 2 | 0.5 | 4 | 0 | 1 | 0 | 10 | 0.1 |
| 13_9956089 | 18 | 0.6667 | 23 | 0.7391 | 22 | 0.7727 | 21 | 0.619 | 84 | 0.7024 | 20 | 0.7 | 22 | 0.8636 | 13 | 1 | 21 | 0.8571 | 76 | 0.8421 |
| 13_9956091 | 28 | 0.6429 | 37 | 0.5135 | 28 | 0.6786 | 31 | 0.5806 | 124 | 0.5968 | 29 | 0.4483 | 23 | 0.5217 | 25 | 0.52 | 22 | 0.6364 | 99 | 0.5253 |
| 13_9956134 | 23 | 0.1739 | 35 | 0.0286 | 24 | 0.0417 | 30 | 0.1333 | 112 | 0.0893 | 24 | 0.0833 | 26 | 0.0385 | 22 | 0.0909 | 20 | 0.1 | 92 | 0.0761 |
| 13_9956135 | 23 | 0.2609 | 35 | 0.2 | 23 | 0.1304 | 29 | 0.2069 | 110 | 0.2 | 23 | 0.087 | 22 | 0.5455 | 19 | 0.1579 | 21 | 0.1429 | 85 | 0.2353 |
| 13_9956152 | 21 | 0.2857 | 32 | 0.1562 | 22 | 0.0909 | 29 | 0.1724 | 104 | 0.1731 | 20 | 0.15 | 23 | 0.3043 | 17 | 0.2941 | 18 | 0.1667 | 78 | 0.2308 |
| 13_9956153 | 20 | 0.15 | 26 | 0.4231 | 18 | 0.0556 | 28 | 0.25 | 92 | 0.2391 | 19 | 0.1053 | 17 | 0.1765 | 16 | 0 | 16 | 0.25 | 68 | 0.1324 |
| 13_9956157 | 19 | 0.2105 | 29 | 0.1034 | 21 | 0.0952 | 27 | 0.1481 | 96 | 0.1354 | 18 | 0.0556 | 23 | 0.087 | 17 | 0.0588 | 18 | 0.0556 | 76 | 0.0658 |
| 13_9956192 | 17 | 0.1176 | 25 | 0 | 18 | 0.1111 | 25 | 0.12 | 85 | 0.0824 | 18 | 0.0556 | 21 | 0.0952 | 17 | 0 | 14 | 0.2143 | 70 | 0.0857 |
| 13_9956202 | 15 | 0.1333 | 24 | 0.125 | 18 | 0.0556 | 26 | 0.0385 | 83 | 0.0843 | 18 | 0.1111 | 17 | 0.1176 | 14 | 0.1429 | 10 | 0 | 59 | 0.1017 |
| 13_9956205 | 16 | 0.1875 | 19 | 0.2105 | 16 | 0.1875 | 19 | 0.1053 | 70 | 0.1714 | 18 | 0.1667 | 12 | 0.1667 | 13 | 0.2308 | 9 | 0.1111 | 52 | 0.1731 |
| 13_9956213 | 8 | 1 | 16 | 0.8125 | 8 | 0.625 | 13 | 0.6923 | 45 | 0.7778 | 10 | 0.8 | 6 | 0.8333 | 6 | 0.6667 | 9 | 0.7778 | 31 | 0.7742 |
| 13_9956220 | 10 | 0.9 | 17 | 0.8235 | 11 | 0.9091 | 17 | 0.8824 | 55 | 0.8727 | 16 | 0.9375 | 9 | 0.7778 | 9 | 0.7778 | 9 | 0.7778 | 43 | 0.8372 |
| 13_9956225 | 12 | 0.25 | 21 | 0.2381 | 12 | 0.1667 | 17 | 0.1176 | 62 | 0.1935 | 16 | 0.1875 | 11 | 0.1818 | 9 | 0.2222 | 9 | 0.2222 | 45 | 0.2 |
| 13_9956228 | 12 | 0.9167 | 17 | 1 | 13 | 1 | 15 | 1 | 57 | 0.9825 | 12 | 1 | 10 | 1 | 10 | 0.9 | 8 | 1 | 40 | 0.975 |
| 13_9956231 | 12 | 0.0833 | 20 | 0.05 | 14 | 0 | 21 | 0.1429 | 67 | 0.0746 | 18 | 0.0556 | 13 | 0.0769 | 13 | 0.0769 | 11 | 0 | 55 | 0.0545 |
| 13_9956232 | 10 | 0.7 | 21 | 0.7619 | 13 | 0.7692 | 22 | 0.9091 | 66 | 0.803 | 16 | 0.875 | 13 | 0.8462 | 11 | 1 | 9 | 0.5556 | 49 | 0.8367 |
| 13_9956390 | 10 | 0 | 19 | 0.1579 | 12 | 0.0833 | 19 | 0.2105 | 60 | 0.1333 | 14 | 0.0714 | 13 | 0 | 12 | 0.0833 | 10 | 0.1 | 49 | 0.0612 |
| 13_9956613 | 9 | 0.3333 | 13 | 0.2308 | 9 | 0.3333 | 17 | 0.1765 | 48 | 0.25 | 11 | 0.2727 | 10 | 0.2 | 12 | 0.3333 | 6 | 0.5 | 39 | 0.3077 |
| 13_9956625 | 7 | 0.8571 | 14 | 0.4286 | 10 | 0.4 | 14 | 0.4286 | 45 | 0.4889 | 12 | 0.4167 | 9 | 0.5556 | 12 | 0.3333 | 7 | 0.4286 | 40 | 0.425 |
| 13_9956680 | 4 | 0.25 | 11 | 0.1818 | 5 | 0.8 | 11 | 0.2727 | 31 | 0.3226 | 9 | 0.3333 | 7 | 0.1429 | 9 | 0.2222 | 6 | 0.1667 | 31 | 0.2258 |
| 13_9956713 | 5 | 0.4 | 11 | 0.2727 | 6 | 0 | 10 | 0.3 | 32 | 0.25 | 10 | 0.4 | 6 | 0.1667 | 8 | 0 | 5 | 0.6 | 29 | 0.2759 |
| 13_9956730 | 3 | 0 | 9 | 0.5556 | 6 | 0.5 | 10 | 0.1 | 28 | 0.3214 | 10 | 0.3 | 6 | 0.5 | 6 | 0.3333 | 4 | 0.75 | 26 | 0.4231 |
| 13_9956764 | 4 | 0.75 | 6 | 1 | 5 | 1 | 9 | 0.8889 | 24 | 0.9167 | 8 | 0.875 | 5 | 1 | 4 | 0.75 | 5 | 1 | 22 | 0.9091 |
| 13_9956766 | 3 | 0.6667 | 5 | 0.4 | 4 | 0.5 | 7 | 0.1429 | 19 | 0.3684 | 6 | 0.8333 | 5 | 0.4 | 4 | 0.5 | 4 | 0.25 | 19 | 0.5263 |
| 13_9956775 | 3 | 1 | 2 | 1 | 4 | 1 | 5 | 1 | 14 | 1 | 7 | 0.8571 | 4 | 1 | 3 | 1 | 2 | 1 | 16 | 0.9375 |
| 13_9956832 | 2 | 0 | 3 | 0 | 5 | 0.4 | 5 | 0 | 15 | 0.1333 | 6 | 0 | 4 | 0 | 5 | 0.2 | 3 | 0.3333 | 18 | 0.1111 |
| 13_9956850 | 2 | 0.5 | 2 | 0 | 4 | 0 | 5 | 0 | 13 | 0.0769 | 4 | 0.25 | 3 | 0.3333 | 4 | 0.25 | 2 | 0 | 13 | 0.2308 |
| 13_9956857 | 2 | 0.5 | 2 | 0.5 | 4 | 0.25 | 5 | 0 | 13 | 0.2308 | 5 | 0 | 4 | 0.25 | 4 | 0.25 | 3 | 0.3333 | 16 | 0.1875 |
| 13_9956881 | 2 | 0.5 | 1 | 0 | 1 | 0 | 3 | 0.3333 | 7 | 0.2857 | 5 | 0.4 | 4 | 0.75 | 4 | 0.25 | 2 | 0.5 | 15 | 0.4667 |
| 13_9956898 | 2 | 0.5 | 1 | 1 | 1 | 1 | 3 | 1 | 7 | 0.8571 | 5 | 1 | 3 | 0.6667 | 1 | 1 | 3 | 0.6667 | 12 | 0.8333 |
| 13_9957021 | 1 | 1 | NA | NA | 1 | 1 | NA | NA | 2 | 1 | 3 | 0.6667 | 1 | 1 | NA | NA | 1 | 1 | 5 | 0.8 |
| 13_9957026 | 2 | 0 | 1 | 1 | 1 | 1 | 1 | 0 | 5 | 0.4 | 2 | 0.5 | 2 | 1 | NA | NA | 2 | 1 | 6 | 0.8333 |
| 13_9957121 | 1 | 1 | NA | NA | 1 | 0 | 2 | 0.5 | 4 | 0.5 | 3 | 0.6667 | 2 | 1 | NA | NA | 2 | 1 | 7 | 0.8571 |
| 13_9957139 | 1 | 1 | NA | NA | 1 | 0 | 2 | 0.5 | 4 | 0.5 | 2 | 0.5 | 2 | 0.5 | NA | NA | 2 | 1 | 6 | 0.6667 |
| 13_9957155 | 1 | 0 | NA | NA | 1 | 0 | 2 | 1 | 4 | 0.5 | 3 | 0.3333 | 2 | 0.5 | NA | NA | 2 | 1 | 7 | 0.5714 |
| 13_9957180 | 1 | 1 | NA | NA | 1 | 0 | 1 | 1 | 3 | 0.6667 | 2 | 0.5 | 2 | 1 | NA | NA | 2 | 1 | 6 | 0.8333 |
| 13_9957204 | 1 | 1 | NA | NA | 1 | 1 | 2 | 1 | 4 | 1 | 1 | 1 | 2 | 1 | NA | NA | 1 | 1 | 4 | 1 |
| 13_1000852 | 4 | 0.25 | 4 | 0 | 6 | 0 | 1 | 0 | 15 | 0.0667 | 3 | 0 | 2 | 1 | 4 | 0.25 | 3 | 0 | 12 | 0.25 |
| 13_1000852 | 2 | 0.5 | 3 | 0.3333 | 5 | 0.2 | 1 | 1 | 11 | 0.3636 | 3 | 0.3333 | 2 | 1 | 5 | 0.4 | 3 | 0.3333 | 13 | 0.4615 |
| 13_1000853 | 5 | 0.4 | 3 | 0.6667 | 6 | 0.3333 | 1 | 0 | 15 | 0.4 | 3 | 0 | 3 | 0.6667 | 5 | 0.6 | 4 | 0.25 | 15 | 0.4 |
| 13_1001547 | 1 | 0 | 6 | 0.1667 | 5 | 0 | 1 | 0 | 13 | 0.0769 | 3 | 0 | 2 | 0.5 | 5 | 0.2 | 3 | 0 | 13 | 0.1538 |
| 13_1001554 | 2 | 0 | 5 | 0 | 4 | 0.25 | 1 | 0 | 12 | 0.0833 | 3 | 0 | 2 | 0 | 3 | 0 | 3 | 0.3333 | 11 | 0.0909 |
| 13_1001555 | 2 | 0 | 5 | 0.2 | 4 | 0 | 1 | 0 | 12 | 0.0833 | 3 | 0.3333 | 2 | 0 | 5 | 0 | 2 | 0 | 12 | 0.0833 |
| 13_1001556 | 2 | 0.5 | 5 | 0 | 4 | 0 | 1 | 0 | 12 | 0.0833 | 2 | 0 | 2 | 0 | 5 | 0.2 | 2 | 0 | 11 | 0.0909 |
| 13_1001565 | 2 | 0 | 3 | 0.3333 | 2 | 0.5 | 1 | 0 | 8 | 0.25 | 2 | 0 | 1 | 1 | 3 | 0.3333 | 1 | 0 | 7 | 0.2857 |
| 13_1001565 | 2 | 0.5 | 2 | 1 | 2 | 1 | NA | NA | 6 | 0.8333 | 2 | 1 | 1 | 0 | 4 | 0.5 | 1 | 1 | 8 | 0.625 |
| 13_1001567 | 2 | 0 | 3 | 0.3333 | 1 | 0 | 1 | 0 | 7 | 0.1429 | 1 | 1 | 1 | 0 | 5 | 0.4 | NA | NA | 7 | 0.4286 |
| 13_1001602 | 1 | 1 | 1 | 0 | 1 | 0 | NA | NA | 3 | 0.3333 | 2 | 0 | 1 | 1 | 2 | 0 | 1 | 0 | 6 | 0.1667 |
| 13_1001863 | 1 | 0 | NA | NA | 1 | 1 | NA | NA | 2 | 0.5 | NA | NA | 1 | 0 | 2 | 0.5 | 1 | 1 | 4 | 0.5 |
| 13_1002739 | 2 | 0 | 2 | 0 | 4 | 0.25 | 2 | 0 | 10 | 0.1 | 4 | 0.5 | 3 | 0 | 4 | 0 | 5 | 0.2 | 16 | 0.1875 |
| 13_1005966 | NA | NA | 2 | 0 | 3 | 0 | 4 | 0.25 | 9 | 0.1111 | 1 | 0 | 1 | 0 | 5 | 0.2 | NA | NA | 7 | 0.1429 |
| 13_1005966 | NA | NA | 2 | 1 | 2 | 1 | 3 | 0.6667 | 7 | 0.8571 | 2 | 1 | 1 | 1 | 5 | 0.8 | NA | NA | 8 | 0.875 |
| 13_1005968 | NA | NA | 2 | 0.5 | 3 | 0.6667 | 4 | 1 | 9 | 0.7778 | NA | NA | 1 | 1 | 3 | 0.3333 | NA | NA | 4 | 0.5 |
| 13_1006054 | NA | NA | 2 | 0 | 4 | 0.25 | 3 | 0 | 9 | 0.1111 | NA | NA | 1 | 1 | 3 | 0.3333 | 2 | 0 | 6 | 0.3333 |
| 13_1006351 | 1 | 1 | 1 | 0 | NA | NA | NA | NA | 2 | 0.5 | 1 | 0 | 1 | 0 | NA | NA | 1 | 1 | 3 | 0.3333 |
| 13_1006352 | 1 | 1 | 2 | 1 | NA | NA | NA | NA | 3 | 1 | 1 | 1 | 1 | 1 | NA | NA | 1 | 1 | 3 | 1 |
| 13_1006352 | 1 | 1 | 2 | 1 | NA | NA | NA | NA | 3 | 1 | 1 | 0 | 1 | 1 | NA | NA | 1 | 1 | 3 | 0.6667 |
| 13_1006761 | 1 | 1 | NA | NA | NA | NA | NA | NA | 1 | 1 | 1 | 0 | 1 | 1 | NA | NA | 1 | 0 | 3 | 0.3333 |
| 13_1006762 | 1 | 1 | NA | NA | NA | NA | NA | NA | 1 | 1 | 1 | 1 | 1 | 0 | NA | NA | 1 | 1 | 3 | 0.6667 |
| 13_1006762 | 1 | 1 | NA | NA | NA | NA | NA | NA | 1 | 1 | 1 | 0 | 1 | 1 | NA | NA | NA | NA | 2 | 0.5 |
| 13_1007302 | NA | NA | 1 | 1 | 2 | 0.5 | NA | NA | 3 | 0.6667 | NA | NA | NA | NA | NA | NA | 1 | 1 | 1 | 1 |
| 13_1007303 | NA | NA | 2 | 1 | 2 | 0 | NA | NA | 4 | 0.5 | NA | NA | NA | NA | NA | NA | 1 | 1 | 1 | 1 |
| 13_1007538 | 1 | 1 | 5 | 0.6 | NA | NA | 2 | 0.5 | 8 | 0.625 | 3 | 1 | NA | NA | 2 | 1 | 3 | 0.6667 | 8 | 0.875 |

| | | | | | | | | | | | | | | | | | | | | |
|---|---|---|---|---|---|---|---|---|---|---|---|---|---|---|---|---|---|---|---|---|
| 13_100754 | 1 | 1 | 4 | 0.5 | NA | NA | 1 | 1 | 6 | 0.6667 | 2 | 1 | NA | NA | 1 | 1 | 3 | 0.3333 | 6 | 0.6667 |
| 13_100792 | 9 | 0.1111 | 9 | 0 | 13 | 0.1538 | 7 | 0.2857 | 38 | 0.1316 | 12 | 0.0833 | 16 | 0.3125 | 13 | 0.1538 | 18 | 0 | 59 | 0.1356 |
| 13_100792 | 6 | 0.1667 | 9 | 0.3333 | 13 | 0.4615 | 11 | 0.1818 | 39 | 0.3077 | 10 | 0.2 | 16 | 0.3125 | 12 | 0.25 | 17 | 0.1176 | 55 | 0.2182 |
| 13_100792 | 8 | 0 | 9 | 0.2222 | 13 | 0.0769 | 10 | 0.1 | 40 | 0.1 | 12 | 0 | 18 | 0.2222 | 10 | 0.4 | 17 | 0 | 57 | 0.1404 |
| 13_100792 | 9 | 0.5556 | 8 | 0.375 | 15 | 0.3333 | 11 | 0.3636 | 43 | 0.3953 | 12 | 0.1667 | 19 | 0.2632 | 12 | 0.4167 | 16 | 0.1875 | 59 | 0.2542 |
| 13_100802 | 4 | 0.5 | 1 | 0 | 5 | 0.2 | 2 | 0.5 | 12 | 0.3333 | 3 | 0 | 2 | 0.5 | 3 | 0 | 2 | 0.5 | 10 | 0.2 |
| 13_100833 | 3 | 0 | 5 | 0 | 4 | 0.25 | 6 | 0.1667 | 18 | 0.1111 | 1 | 0 | 2 | 0.5 | 4 | 0 | 2 | 0 | 9 | 0.1111 |
| 13_100833 | 3 | 0 | 5 | 0.2 | 3 | 0 | 8 | 0 | 19 | 0.0526 | 2 | 0 | 3 | 0 | 4 | 0.25 | 3 | 0 | 12 | 0.0833 |
| 13_100833 | 3 | 0 | 4 | 0 | 2 | 1 | 7 | 0.2857 | 16 | 0.25 | 1 | 1 | 3 | 0 | 4 | 0.25 | 3 | 0.6667 | 11 | 0.3636 |
| 13_100839 | 4 | 0 | NA | NA | 2 | 1 | 2 | 1 | 8 | 0.5 | 2 | 0 | 2 | 0.5 | 2 | 0 | 2 | 0 | 8 | 0.125 |
| 13_100872 | 10 | 0.2 | 10 | 0.2 | 20 | 0.15 | 13 | 0.4615 | 53 | 0.2453 | 12 | 0.3333 | 18 | 0.3889 | 12 | 0.3333 | 11 | 0.1818 | 53 | 0.3208 |
| 13_100939 | 4 | 0.25 | 7 | 0.5714 | 5 | 0.2 | 11 | 0.1818 | 27 | 0.2963 | 8 | 0.25 | 5 | 0.2 | 3 | 0.6667 | 2 | 1 | 18 | 0.3889 |
| 13_100939 | 2 | 1 | 7 | 0.7143 | 3 | 0.3333 | 13 | 0.5385 | 25 | 0.6 | 7 | 0.2857 | 5 | 0.6 | 4 | 0.5 | 2 | 0.5 | 18 | 0.4444 |
| 13_100939 | 3 | 0.3333 | 7 | 0 | 5 | 0.2 | 11 | 0.3636 | 26 | 0.2308 | 7 | 0.2857 | 7 | 0.2857 | 4 | 0.25 | 2 | 0 | 20 | 0.25 |
| 13_100939 | 2 | 0.5 | 4 | 1 | 3 | 0.3333 | 7 | 0.5714 | 16 | 0.625 | 5 | 0.4 | 4 | 0.75 | 3 | 0.6667 | 2 | 0 | 14 | 0.5 |
| 13_100939 | NA | NA | 1 | 1 | NA | NA | NA | NA | 1 | 1 | NA | NA | 1 | 1 | NA | NA | NA | NA | 1 | 1 |
| 13_100939 | 3 | 0.3333 | 5 | 0 | 2 | 0.5 | 6 | 0.3333 | 16 | 0.25 | 4 | 0.25 | 4 | 0.75 | 2 | 0.5 | 1 | 0 | 11 | 0.4545 |
| 13_100949 | 3 | 0.3333 | 2 | 0 | 5 | 0.4 | 11 | 0.0909 | 21 | 0.1905 | 5 | 0.2 | 3 | 0 | 3 | 0 | 1 | 0 | 12 | 0.0833 |
| 13_101820 | 1 | 0 | NA | NA | 2 | 0 | 2 | 0.5 | 5 | 0.2 | 1 | 0 | NA | NA | 1 | 1 | 3 | 0.6667 | 5 | 0.6 |
| 13_101820 | 3 | 0.3333 | NA | NA | 4 | 0 | 6 | 0.3333 | 13 | 0.2308 | 1 | 1 | 5 | 0 | 4 | 0 | 7 | 0.1429 | 17 | 0.1176 |
| 13_101820 | 3 | 0.6667 | NA | NA | 4 | 0.75 | 6 | 0.8333 | 13 | 0.7692 | 1 | 0 | 5 | 0.8 | 4 | 1 | 7 | 0.8571 | 17 | 0.8235 |
| 13_101820 | 3 | 0.6667 | NA | NA | 5 | 0 | 6 | 0.3333 | 14 | 0.2857 | 1 | 1 | 5 | 0.2 | 4 | 0 | 7 | 0.8571 | 17 | 0.4706 |
| 13_101820 | 2 | 0 | NA | NA | 5 | 0.2 | 7 | 0.1429 | 14 | 0.1429 | 1 | 0 | 4 | 0 | 4 | 0 | 7 | 0.1429 | 16 | 0.0625 |
| 13_101820 | 2 | 1 | NA | NA | 4 | 1 | 6 | 0.8333 | 12 | 0.9167 | 1 | 1 | 5 | 1 | 4 | 1 | 8 | 0.875 | 18 | 0.9444 |
| 13_101820 | 3 | 0.3333 | NA | NA | 5 | 1 | 7 | 0.7143 | 15 | 0.7333 | NA | NA | 4 | 0.75 | 4 | 1 | 8 | 1 | 16 | 0.9375 |
| 13_101820 | 2 | 0 | NA | NA | 5 | 0 | 6 | 0.1667 | 13 | 0.0769 | NA | NA | 5 | 0 | 3 | 0 | 8 | 0.125 | 16 | 0.0625 |
| 13_101820 | 1 | 1 | NA | NA | 5 | 1 | 4 | 1 | 10 | 1 | NA | NA | 4 | 1 | 4 | 1 | 6 | 1 | 14 | 1 |
| 13_101821 | 1 | 1 | NA | NA | 5 | 1 | 6 | 0.6667 | 12 | 0.8333 | 1 | 1 | 3 | 0.6667 | 4 | 1 | 8 | 0.5 | 16 | 0.6875 |
| 13_101821 | 1 | 1 | NA | NA | 3 | 0.6667 | 4 | 0.75 | 8 | 0.75 | 1 | 1 | 5 | 0.8 | 3 | 1 | 6 | 0.8333 | 15 | 0.8667 |
| 13_101821 | 1 | 0 | NA | NA | 3 | 0.3333 | 3 | 0.6667 | 7 | 0.4286 | 1 | 1 | 4 | 0.75 | 3 | 0.3333 | 7 | 0.1429 | 15 | 0.4 |
| 13_101821 | 1 | 1 | NA | NA | 3 | 1 | 5 | 0.6 | 9 | 0.7778 | 1 | 1 | 5 | 1 | 3 | 1 | 8 | 0.875 | 17 | 0.9412 |
| 13_101821 | 1 | 0 | NA | NA | 3 | 0.3333 | 3 | 0.3333 | 7 | 0.2857 | NA | NA | 4 | 0.5 | 3 | 1 | 5 | 1 | 12 | 0.8333 |
| 13_101821 | 1 | 1 | NA | NA | 3 | 0.3333 | 3 | 0.3333 | 7 | 0.4286 | 1 | 0 | 5 | 0.6 | 3 | 0.6667 | 6 | 0.6667 | 15 | 0.6 |
| 13_102869 | 2 | 0 | 5 | 0.4 | 3 | 0 | 3 | 0.3333 | 13 | 0.2308 | 5 | 0 | 1 | 0 | 2 | 0.5 | 1 | 1 | 9 | 0.2222 |
| 13_102869 | 2 | 0.5 | 4 | 0 | 2 | 0 | 3 | 0 | 11 | 0.0909 | 3 | 0.3333 | 1 | 0 | 2 | 0 | 1 | 0 | 7 | 0.1429 |
| 13_102869 | 2 | 0.5 | 5 | 0 | 2 | 0 | 3 | 0 | 12 | 0.0833 | 4 | 0.25 | 1 | 0 | 2 | 0 | 1 | 1 | 8 | 0.25 |
| 13_102870 | 2 | 0 | 4 | 0.25 | 2 | 0.5 | 3 | 0 | 11 | 0.1818 | 2 | 0.5 | 1 | 0 | 2 | 0 | NA | NA | 5 | 0.2 |
| 13_102871 | 1 | 0 | 3 | 0.3333 | 2 | 0.5 | 2 | 0.5 | 8 | 0.375 | 2 | 0.5 | 1 | 1 | 1 | 1 | NA | NA | 4 | 0.75 |
| 13_102871 | 2 | 1 | 3 | 1 | 2 | 1 | 3 | 1 | 10 | 1 | 2 | 1 | NA | NA | 1 | 1 | NA | NA | 3 | 1 |
| 13_102871 | 2 | 0.5 | 4 | 0.75 | 2 | 1 | 3 | 1 | 11 | 0.8182 | 2 | 1 | 1 | 1 | 1 | 1 | NA | NA | 4 | 1 |
| 13_102871 | 2 | 0 | 4 | 0.5 | 2 | 0 | 2 | 0.5 | 10 | 0.3 | 1 | 1 | 1 | 1 | 1 | 0 | NA | NA | 3 | 0.6667 |
| 13_102871 | 1 | 0 | 4 | 0.5 | 1 | 1 | 2 | 0 | 8 | 0.375 | 2 | 0.5 | 1 | 1 | 1 | 0 | NA | NA | 4 | 0.5 |
| 13_102871 | 1 | 1 | 4 | 1 | 1 | 1 | 1 | 1 | 7 | 1 | 2 | 1 | 1 | 1 | NA | NA | NA | NA | 3 | 1 |
| 13_102871 | NA | NA | 2 | 0.5 | 1 | 0 | 1 | 0 | 4 | 0.25 | 1 | 1 | 1 | 0 | 1 | 1 | NA | NA | 3 | 0.6667 |
| 13_102871 | 1 | 1 | 2 | 1 | NA | NA | NA | NA | 3 | 1 | NA | NA | 1 | 1 | NA | NA | NA | NA | 1 | 1 |
| 13_102871 | 1 | 0 | 4 | 1 | 1 | 1 | 2 | 1 | 8 | 0.875 | 2 | 1 | 1 | 1 | NA | NA | NA | NA | 3 | 1 |
| 13_102871 | 2 | 1 | 2 | 0.5 | 1 | 1 | 2 | 0.5 | 7 | 0.7143 | 1 | 0 | 1 | 1 | NA | NA | NA | NA | 2 | 0.5 |
| 13_102872 | 2 | 1 | 4 | 1 | 1 | 1 | 1 | 1 | 8 | 1 | 2 | 1 | 1 | 1 | NA | NA | NA | NA | 3 | 1 |
| 13_102872 | 2 | 0 | 4 | 0.5 | NA | NA | 2 | 0.5 | 8 | 0.375 | 1 | 1 | 1 | 0 | NA | NA | NA | NA | 2 | 0.5 |
| 13_102872 | 2 | 1 | 4 | 0.75 | 1 | 0 | 2 | 0.5 | 9 | 0.6667 | 2 | 1 | 1 | 1 | NA | NA | NA | NA | 3 | 1 |
| 13_102872 | 1 | 1 | 3 | 1 | 1 | 1 | 2 | 1 | 7 | 1 | 1 | 1 | NA | NA | NA | NA | NA | NA | 1 | 1 |
| 13_102872 | 2 | 0 | 4 | 0 | 1 | 0 | 2 | 0.5 | 9 | 0.1111 | 2 | 0.5 | 1 | 0 | NA | NA | NA | NA | 3 | 0.3333 |
| 13_102873 | 2 | 0.5 | 4 | 0.5 | NA | NA | 1 | 0 | 7 | 0.4286 | 2 | 0.5 | 1 | 1 | NA | NA | NA | NA | 3 | 0.6667 |
| 13_102873 | 1 | 1 | 2 | 1 | NA | NA | 2 | 0.5 | 5 | 0.8 | 2 | 0.5 | 1 | 0 | NA | NA | NA | NA | 3 | 0.3333 |
| 13_102873 | 1 | 1 | 2 | 0.5 | NA | NA | 2 | 0.5 | 5 | 0.6 | 1 | 1 | 1 | 1 | NA | NA | NA | NA | 2 | 1 |
| 13_102873 | 1 | 1 | 3 | 0.6667 | NA | NA | 2 | 0 | 6 | 0.5 | 2 | 0.5 | 1 | 0 | NA | NA | NA | NA | 3 | 0.3333 |
| 13_102873 | 1 | 1 | 3 | 1 | NA | NA | 2 | 1 | 6 | 1 | 2 | 0.5 | 1 | 0 | NA | NA | NA | NA | 3 | 0.3333 |
| 13_102873 | NA | NA | 3 | 1 | NA | NA | 2 | 1 | 5 | 1 | 1 | 1 | 1 | 1 | NA | NA | NA | NA | 2 | 1 |
| 13_102874 | 1 | 1 | 2 | 0.5 | NA | NA | 2 | 0.5 | 5 | 0.6 | 1 | 1 | 1 | 0 | NA | NA | NA | NA | 2 | 0.5 |
| 13_102874 | 1 | 1 | 3 | 1 | NA | NA | 2 | 1 | 6 | 1 | 1 | 1 | 1 | 1 | NA | NA | NA | NA | 2 | 1 |
| 13_102874 | 1 | 1 | 3 | 1 | NA | NA | 1 | 1 | 5 | 1 | 1 | 1 | 1 | 1 | NA | NA | NA | NA | 2 | 1 |
| 13_102875 | NA | NA | 2 | 1 | NA | NA | 1 | 0 | 3 | 0.6667 | 1 | 1 | NA | NA | NA | NA | NA | NA | 1 | 1 |
| 13_103876 | 1 | 0 | 1 | 0 | 2 | 0.5 | NA | NA | 4 | 0.25 | NA | NA | 3 | 0.3333 | 2 | 0 | 2 | 0 | 7 | 0.1429 |
| 13_103878 | 2 | 0 | 9 | 0 | 9 | 0 | 6 | 0.3333 | 26 | 0.0769 | 1 | 0 | 10 | 0.3 | 7 | 0.1429 | 4 | 0.5 | 22 | 0.2727 |
| 13_103878 | 2 | 0.5 | 8 | 0.25 | 8 | 0.125 | 5 | 0 | 23 | 0.1739 | NA | NA | 7 | 0 | 6 | 0 | 3 | 0.3333 | 16 | 0.0625 |

| | | | | | | | | | | | | | | | | | | | | |
|---|---|---|---|---|---|---|---|---|---|---|---|---|---|---|---|---|---|---|---|---|
| 13_103879 | 3 | 0 | 6 | 0.1667 | 6 | 0.1667 | 6 | 0.3333 | 21 | 0.1905 | NA | NA | 3 | 0 | 3 | 0.3333 | 4 | 0.25 | 10 | 0.2 |
| 13_103879 | 1 | 1 | 7 | 0.8571 | 6 | 0.8333 | 6 | 1 | 20 | 0.9 | 1 | 1 | 6 | 1 | 4 | 1 | 2 | 1 | 13 | 1 |
| 13_103879 | NA | NA | 5 | 0.2 | 7 | 0.2857 | 3 | 0 | 15 | 0.2 | 1 | 0 | 5 | 0.2 | 6 | 0 | 4 | 0.25 | 16 | 0.125 |
| 13_103879 | 2 | 1 | 6 | 0.6667 | 9 | 0.7778 | 5 | 1 | 22 | 0.8182 | 1 | 1 | 6 | 0.8333 | 6 | 0.8333 | 4 | 1 | 17 | 0.8824 |
| 13_103879 | 1 | 1 | 7 | 1 | 8 | 1 | 6 | 1 | 22 | 1 | 1 | 1 | 6 | 1 | 5 | 1 | 4 | 1 | 16 | 1 |
| 13_103879 | 1 | 1 | 7 | 0.4286 | 7 | 0.4286 | 5 | 0.6 | 20 | 0.5 | 1 | 0 | 6 | 0.8333 | 5 | 0.6 | 4 | 0.5 | 16 | 0.625 |
| 13_103879 | 2 | 0 | 7 | 0.1429 | 5 | 0.2 | 5 | 0.4 | 19 | 0.2105 | 1 | 0 | 4 | 0.25 | 4 | 0.25 | 3 | 0 | 12 | 0.1667 |
| 13_103880 | 1 | 1 | 6 | 0.6667 | 7 | 0.8571 | 3 | 1 | 17 | 0.8235 | 1 | 0 | 4 | 1 | 5 | 1 | 4 | 0.75 | 14 | 0.8571 |
| 13_103880 | 2 | 1 | 6 | 0.6667 | 8 | 0.875 | 6 | 0.6667 | 22 | 0.7727 | 1 | 1 | 5 | 0.6 | 5 | 0.8 | 4 | 1 | 15 | 0.8 |
| 13_103955 | 9 | 0.1111 | 7 | 0.1429 | 10 | 0.1 | 17 | 0.1176 | 43 | 0.1163 | 7 | 0 | 7 | 0 | 13 | 0.2308 | 9 | 0 | 36 | 0.0833 |
| 13_103955 | 5 | 0.2 | 8 | 0 | 11 | 0.0909 | 16 | 0.25 | 40 | 0.15 | 8 | 0 | 10 | 0.2 | 16 | 0.375 | 7 | 0.2857 | 41 | 0.2439 |
| 13_103955 | 7 | 0 | 9 | 0.1111 | 10 | 0 | 17 | 0.1765 | 43 | 0.093 | 8 | 0 | 11 | 0.0909 | 12 | 0.0833 | 7 | 0.2857 | 38 | 0.1053 |
| 13_103956 | 4 | 0 | 7 | 0.2857 | 8 | 0.25 | 13 | 0.3077 | 32 | 0.25 | 8 | 0.25 | 7 | 0.2857 | 11 | 0.3636 | 6 | 0.5 | 32 | 0.3438 |
| 13_103956 | 3 | 0 | 6 | 0.1667 | 8 | 0 | 10 | 0.1 | 27 | 0.0741 | 6 | 0.1667 | 5 | 0 | 9 | 0.1111 | 6 | 0 | 26 | 0.0769 |
| 13_103957 | 4 | 1 | 7 | 0.8571 | 9 | 0.8889 | 11 | 0.8182 | 31 | 0.871 | 7 | 0.4286 | 9 | 0.7778 | 12 | 0.9167 | 7 | 0.5714 | 35 | 0.7143 |
| 13_103957 | 5 | 1 | 6 | 0.8333 | 8 | 0.875 | 12 | 0.8333 | 31 | 0.871 | 8 | 0.875 | 8 | 1 | 11 | 1 | 6 | 0.8333 | 33 | 0.9394 |
| 13_103957 | 5 | 0 | 6 | 0.1667 | 9 | 0 | 13 | 0.1538 | 33 | 0.0909 | 9 | 0.1111 | 8 | 0 | 10 | 0.2 | 7 | 0 | 34 | 0.0882 |
| 13_103957 | 5 | 0.6 | 7 | 0.4286 | 9 | 0.4444 | 13 | 0.2308 | 34 | 0.3824 | 9 | 0 | 7 | 0.1429 | 9 | 0.1111 | 7 | 0.1429 | 32 | 0.0938 |
| 13_103957 | 3 | 0.3333 | 5 | 0.4 | 7 | 0.2857 | 11 | 0.4545 | 26 | 0.3846 | 5 | 0 | 7 | 0.5714 | 9 | 0.4444 | 7 | 0.4286 | 28 | 0.3929 |
| 13_103957 | 4 | 0.25 | 7 | 0.5714 | 7 | 0.7143 | 10 | 0.3 | 28 | 0.4643 | 5 | 0.4 | 7 | 0.1429 | 8 | 0.25 | 8 | 0.5 | 28 | 0.3214 |
| 13_103957 | 4 | 0.25 | 4 | 0 | 7 | 0 | 9 | 0.1111 | 24 | 0.0833 | 7 | 0.4286 | 4 | 0 | 8 | 0.125 | 4 | 0.25 | 23 | 0.2174 |
| 13_103957 | 2 | 0 | 4 | 0 | 7 | 0.5714 | 9 | 0.3333 | 22 | 0.3182 | 5 | 0.4 | 6 | 0.5 | 6 | 0.1667 | 5 | 0.6 | 22 | 0.4091 |
| 13_103957 | 3 | 0.3333 | 2 | 0.5 | NA | NA | 2 | 0.5 | 7 | 0.4286 | 2 | 0 | 2 | 0.5 | 1 | 1 | 1 | 0 | 6 | 0.3333 |
| 13_103970 | 3 | 1 | 6 | 0.3333 | 3 | 0 | 10 | 0.7 | 22 | 0.5455 | 4 | 0.25 | 4 | 0.5 | 9 | 0.1111 | 6 | 1 | 23 | 0.4348 |
| 13_103970 | 4 | 0 | 3 | 0.6667 | 6 | 0 | 9 | 0.2222 | 22 | 0.1818 | 5 | 0 | 4 | 0 | 5 | 0.2 | 6 | 0.1667 | 20 | 0.1 |
| 13_103970 | 3 | 0.6667 | 6 | 0.8333 | 5 | 0.6 | 12 | 0.75 | 26 | 0.7308 | 6 | 1 | 3 | 1 | 8 | 0.5 | 6 | 0.8333 | 23 | 0.7826 |
| 13_103970 | 2 | 1 | 5 | 0.8 | 4 | 0.75 | 11 | 0.7273 | 22 | 0.7727 | 6 | 0.6667 | 3 | 1 | 6 | 0.6667 | 6 | 0.5 | 21 | 0.6667 |
| 13_103970 | 3 | 0 | 6 | 0.3333 | 6 | 0.1667 | 12 | 0.1667 | 27 | 0.1852 | 6 | 0.1667 | 4 | 0.25 | 9 | 0.1111 | 7 | 0.2857 | 26 | 0.1923 |
| 13_103970 | 3 | 0.6667 | 5 | 0.2 | 6 | 0 | 12 | 0 | 26 | 0.1154 | 4 | 0.25 | 4 | 0.25 | 9 | 0.2222 | 3 | 0 | 20 | 0.2 |
| 13_103970 | 2 | 0 | 4 | 0 | 6 | 0.1667 | 10 | 0.2 | 22 | 0.1364 | 6 | 0.3333 | 3 | 0.3333 | 9 | 0.1111 | 6 | 0.6667 | 24 | 0.3333 |
| 13_103970 | 2 | 1 | 5 | 0.6 | 4 | 0.5 | 8 | 0.625 | 19 | 0.6316 | 6 | 0.8333 | 2 | 1 | 9 | 0.3333 | 5 | 0.6 | 22 | 0.5909 |
| 13_103970 | 3 | 1 | 3 | 1 | 4 | 1 | 8 | 0.875 | 18 | 0.9444 | 3 | 0.3333 | 3 | 1 | 6 | 0.5 | 6 | 0.6667 | 18 | 0.6111 |
| 13_103970 | 3 | 0 | 5 | 0 | 5 | 0.4 | 9 | 0.1111 | 22 | 0.1364 | 4 | 0 | 4 | 0 | 9 | 0.2222 | 5 | 0 | 22 | 0.0909 |
| 13_103970 | 1 | 0 | 4 | 0 | 4 | 0 | 5 | 0.2 | 14 | 0.0714 | 4 | 0.75 | 4 | 0.25 | 7 | 0.1429 | 6 | 0 | 21 | 0.2381 |
| 13_103970 | 1 | 1 | 4 | 0.75 | 3 | 1 | 8 | 0.125 | 16 | 0.5 | 3 | 0.6667 | 4 | 0.5 | 9 | 0.3333 | 6 | 0.6667 | 22 | 0.5 |
| 13_103971 | 2 | 0 | 5 | 0.8 | 4 | 0.75 | 8 | 0.5 | 19 | 0.5789 | 4 | 1 | 4 | 0.75 | 7 | 0.7143 | 5 | 0.8 | 20 | 0.8 |
| 13_103971 | 2 | 1 | 5 | 0.4 | 2 | 0 | 5 | 0.2 | 14 | 0.3571 | 4 | 0 | 3 | 0 | 9 | 0.2222 | 2 | 0.5 | 18 | 0.1667 |
| 13_103971 | 2 | 0 | 5 | 0 | 5 | 0.2 | 5 | 0.2 | 17 | 0.1176 | 3 | 0 | 3 | 0 | 8 | 0.125 | 6 | 0 | 20 | 0.05 |
| 13_103971 | 2 | 0.5 | 5 | 0.4 | 3 | 0.6667 | 6 | 0.5 | 16 | 0.5 | 4 | 0.75 | 4 | 0.5 | 8 | 0.25 | 5 | 0.2 | 21 | 0.381 |
| 13_103971 | 2 | 0.5 | 5 | 0.6 | 4 | 0.25 | 6 | 0.3333 | 17 | 0.4118 | 4 | 0 | 3 | 0 | 8 | 0.125 | 5 | 0.4 | 20 | 0.15 |
| 13_103971 | 2 | 1 | 4 | 0.5 | 4 | 1 | 7 | 0.4286 | 17 | 0.6471 | 4 | 0.75 | 2 | 0.5 | 8 | 0.25 | 5 | 0.8 | 19 | 0.5263 |
| 13_103971 | 2 | 0.5 | 5 | 0 | 4 | 0.25 | 7 | 0 | 18 | 0.1111 | 3 | 0 | 3 | 0.6667 | 8 | 0.125 | 6 | 0.5 | 20 | 0.3 |
| 13_104057 | 1 | 0 | 1 | 1 | 1 | 0 | 1 | 1 | 4 | 0.5 | NA | NA | NA | NA | 1 | 0 | 3 | 0.6667 | 4 | 0.5 |
| 13_104159 | NA | NA | NA | NA | 2 | 0.5 | 3 | 0 | 5 | 0.2 | 1 | 0 | 3 | 0.3333 | 3 | 0.3333 | 3 | 0 | 10 | 0.2 |
| 13_104159 | 2 | 0.5 | 1 | 0 | 2 | 0.5 | 4 | 0 | 9 | 0.2222 | 1 | 0 | 3 | 0 | 3 | 0.6667 | 3 | 0.3333 | 10 | 0.3 |
| 13_104160 | 1 | 0 | 1 | 0 | 2 | 0.5 | 4 | 0.25 | 8 | 0.25 | 1 | 1 | 3 | 0.3333 | 2 | 0.5 | 4 | 0.75 | 10 | 0.6 |
| 13_104160 | 2 | 0.5 | 1 | 0 | 2 | 0 | 1 | 0 | 6 | 0.1667 | 1 | 0 | 3 | 0 | 1 | 0 | 4 | 0.25 | 9 | 0.1111 |
| 13_104161 | 2 | 0.5 | 1 | 1 | 1 | 0 | 4 | 0.5 | 8 | 0.5 | 1 | 1 | 1 | 1 | 1 | 1 | 3 | 0 | 6 | 0.5 |
| 13_104161 | 1 | 0 | NA | NA | 3 | 0 | 5 | 0.2 | 9 | 0.1111 | 1 | 1 | 1 | 0 | 1 | 0 | 4 | 0 | 7 | 0.1429 |
| 13_104187 | 1 | 1 | 1 | 1 | 2 | 0.5 | 4 | 1 | 8 | 0.875 | 2 | 0.5 | 2 | 1 | NA | NA | 3 | 0.6667 | 7 | 0.7143 |
| 13_104187 | 1 | 0 | 1 | 0 | 1 | 0 | 4 | 0.25 | 7 | 0.1429 | 2 | 0 | 1 | 1 | NA | NA | 2 | 0 | 5 | 0.2 |
| 13_104187 | 1 | 0 | 1 | 1 | NA | NA | 3 | 0.3333 | 5 | 0.4 | 2 | 0.5 | 2 | 0.5 | NA | NA | 2 | 0 | 6 | 0.3333 |
| 13_104278 | 16 | 0 | 17 | 0.0588 | 18 | 0.1667 | 18 | 0.0556 | 69 | 0.0725 | 13 | 0.0769 | 11 | 0.0909 | 13 | 0.1538 | 16 | 0.125 | 53 | 0.1132 |
| 13_104278 | 17 | 0.0588 | 19 | 0.0526 | 20 | 0.15 | 20 | 0 | 76 | 0.0658 | 14 | 0.2143 | 12 | 0.0833 | 16 | 0 | 17 | 0.2941 | 59 | 0.1525 |
| 13_104278 | 16 | 0 | 22 | 0.0909 | 25 | 0.04 | 19 | 0.1053 | 82 | 0.061 | 14 | 0.0714 | 12 | 0.1667 | 17 | 0.0588 | 17 | 0.0588 | 60 | 0.0833 |
| 13_104310 | NA | NA | 1 | 1 | 1 | 0 | NA | NA | 2 | 0.5 | 1 | 1 | NA | NA | NA | NA | NA | NA | 1 | 1 |
| 13_104310 | NA | NA | 1 | 1 | 1 | 1 | NA | NA | 2 | 1 | 1 | 1 | NA | NA | NA | NA | NA | NA | 1 | 1 |
| 13_104310 | NA | NA | 1 | 1 | 1 | 1 | NA | NA | 2 | 1 | 1 | 1 | NA | NA | NA | NA | NA | NA | 1 | 1 |
| 13_104311 | NA | NA | 1 | 1 | 1 | 0 | NA | NA | 2 | 0.5 | 1 | 1 | NA | NA | NA | NA | NA | NA | 1 | 1 |
| 13_104311 | NA | NA | 1 | 1 | 1 | 0 | NA | NA | 2 | 0.5 | 1 | 1 | NA | NA | NA | NA | NA | NA | 1 | 1 |
| 13_104314 | 12 | 0.1667 | 7 | 0 | 8 | 0 | 7 | 0.2857 | 34 | 0.1176 | 5 | 0.4 | 4 | 0 | 3 | 0.3333 | 6 | 0.3333 | 18 | 0.2778 |
| 13_104323 | 9 | 0.2222 | 2 | 0.5 | 4 | 0.25 | 5 | 0.6 | 20 | 0.35 | 5 | 0.2 | 8 | 0.375 | 8 | 0.375 | 2 | 0 | 23 | 0.3043 |
| 13_104324 | 7 | 0.2857 | 4 | 0 | 5 | 0 | 6 | 0.3333 | 22 | 0.1818 | 5 | 0.2 | 8 | 0.375 | 8 | 0 | 2 | 0 | 23 | 0.1739 |
| 13_104337 | 10 | 0.2 | 4 | 0 | 8 | 0 | 8 | 0 | 30 | 0.0667 | 5 | 0.2 | 8 | 0 | 6 | 0 | 5 | 0.2 | 24 | 0.0833 |
| 13_104337 | 10 | 0.9 | 4 | 0.25 | 6 | 0.3333 | 6 | 0.6667 | 26 | 0.6154 | 5 | 0.4 | 7 | 0.4286 | 5 | 0.6 | 5 | 0.4 | 22 | 0.4545 |

| | | | | | | | | | | | | | | | | | | | | |
|---|---|---|---|---|---|---|---|---|---|---|---|---|---|---|---|---|---|---|---|---|
| 13_104338( | 11 | 0.1818 | 8 | 0.125 | 9 | 0.3333 | 6 | 0.1667 | 34 | 0.2059 | 6 | 0.5 | 9 | 0.3333 | 11 | 0.4545 | 5 | 0.2 | 31 | 0.3871 |
| 13_104339( | 13 | 0.0769 | 10 | 0.1 | 10 | 0.2 | 8 | 0 | 41 | 0.0976 | 7 | 0 | 10 | 0.1 | 13 | 0.1538 | 6 | 0 | 36 | 0.0833 |
| 13_104341! | NA | NA | NA | NA | NA | NA | 2 | 0.5 | 2 | 0.5 | 2 | 0.5 | 3 | 0.3333 | 2 | 0.5 | 1 | 0 | 8 | 0.375 |
| 13_104341! | 1 | 1 | 1 | 1 | NA | NA | 3 | 0.6667 | 5 | 0.8 | 2 | 1 | 3 | 1 | 1 | 1 | 1 | 1 | 7 | 1 |
| 13_104356( | 1 | 1 | 1 | 1 | NA | NA | 2 | 1 | 4 | 1 | 2 | 0.5 | 2 | 1 | 1 | 1 | 1 | 1 | 6 | 0.8333 |
| 13_104356 | 1 | 1 | 1 | 0 | NA | NA | 1 | 0 | 3 | 0.3333 | 2 | 0 | 2 | 0.5 | 1 | 0 | 1 | 0 | 6 | 0.1667 |
| 13_104356 | 1 | 0 | 1 | 0 | NA | NA | 2 | 0.5 | 4 | 0.25 | 2 | 0.5 | 2 | 0.5 | 1 | 0 | 1 | 1 | 6 | 0.5 |
| 13_104356 | 1 | 1 | 1 | 1 | NA | NA | 1 | 0 | 3 | 0.6667 | 2 | 0.5 | 1 | 1 | NA | NA | 1 | 0 | 4 | 0.5 |
| 13_104356 | 1 | 1 | 1 | 0 | NA | NA | 1 | 0 | 3 | 0.3333 | 1 | 0 | 2 | 0 | 1 | 1 | NA | NA | 4 | 0.25 |
| 13_104356 | 1 | 0 | 1 | 1 | NA | NA | 1 | 1 | 3 | 0.6667 | 2 | 1 | 2 | 1 | 1 | 0 | 1 | 1 | 6 | 0.8333 |
| 13_104356 | 1 | 1 | 1 | 1 | NA | NA | 2 | 0.5 | 4 | 0.75 | NA | NA | 2 | 1 | 1 | 1 | NA | NA | 3 | 1 |
| 13_104365 | 1 | 1 | NA | NA | NA | NA | 1 | 0 | 2 | 0.5 | 1 | 0 | 2 | 1 | 1 | 0 | 2 | 0 | 6 | 0.3333 |
| 13_104768 | 2 | 0 | 3 | 0.3333 | 2 | 0.5 | 1 | 0 | 8 | 0.25 | NA | NA | 1 | 0 | 3 | 0 | 2 | 0.5 | 6 | 0.1667 |
| 13_104974( | 2 | 0.5 | 1 | 0 | NA | NA | 1 | 1 | 4 | 0.5 | 1 | 1 | 3 | 0.6667 | 2 | 1 | 3 | 1 | 9 | 0.8889 |
| 13_104974 | 4 | 0.25 | 1 | 1 | 2 | 0.5 | 1 | 0 | 8 | 0.375 | 1 | 0 | 4 | 0.25 | 2 | 1 | 1 | 1 | 8 | 0.5 |
| 13_104974 | 4 | 1 | 1 | 1 | 3 | 1 | 3 | 1 | 11 | 1 | 1 | 1 | 4 | 1 | 2 | 1 | 3 | 1 | 10 | 1 |
| 13_104974 | 4 | 0.25 | 2 | 0 | 3 | 0.3333 | 3 | 0 | 12 | 0.1667 | 1 | 0 | 5 | 0.2 | 1 | 1 | 4 | 0.25 | 11 | 0.2727 |
| 13_105083 | 8 | 0.125 | 14 | 0.1429 | 10 | 0 | 18 | 0.0556 | 50 | 0.08 | 4 | 0.25 | 11 | 0 | 12 | 0 | 17 | 0.1176 | 44 | 0.0682 |
| 13_105084 | 9 | 0.3333 | 20 | 0.1 | 21 | 0.1429 | 19 | 0.1053 | 69 | 0.1449 | 4 | 0 | 17 | 0.1176 | 18 | 0.0556 | 17 | 0.2353 | 56 | 0.125 |
| 13_105085 | 5 | 0 | 16 | 0.125 | 15 | 0.2667 | 16 | 0.1875 | 52 | 0.1731 | 3 | 0 | 14 | 0.2143 | 12 | 0.0833 | 13 | 0.2308 | 42 | 0.1667 |
| 13_105085 | 4 | 0 | 13 | 0.6154 | 12 | 0.3333 | 15 | 0.4 | 44 | 0.4091 | 4 | 0.5 | 12 | 0.3333 | 13 | 0.2308 | 12 | 0.3333 | 41 | 0.3171 |
| 13_105086 | 3 | 0.3333 | 13 | 0.2308 | 14 | 0.1429 | 14 | 0.2857 | 44 | 0.2273 | 3 | 0.3333 | 12 | 0.3333 | 14 | 0.2857 | 14 | 0.1429 | 43 | 0.2558 |
| 13_105086 | 4 | 0 | 12 | 0 | 15 | 0.1333 | 15 | 0.0667 | 46 | 0.0652 | 3 | 0 | 11 | 0 | 13 | 0 | 13 | 0.3077 | 40 | 0.1 |
| 13_105086 | 3 | 0.3333 | 11 | 0.8182 | 7 | 0.2857 | 10 | 0.3 | 31 | 0.4839 | 2 | 0 | 6 | 0.6667 | 9 | 0.5556 | 11 | 0.5455 | 28 | 0.5357 |
| 13_105087 | 2 | 0 | 13 | 0.1538 | 10 | 0.1 | 16 | 0.0625 | 41 | 0.0976 | 2 | 0.5 | 8 | 0 | 10 | 0.1 | 7 | 0 | 27 | 0.0741 |
| 13_105087 | 1 | 1 | 13 | 0.2308 | 9 | 0.5556 | 15 | 0.3333 | 38 | 0.3684 | 1 | 0 | 8 | 0.25 | 10 | 0.4 | 6 | 0.3333 | 25 | 0.32 |
| 13_105087 | 1 | 0 | 9 | 0.2222 | 8 | 0 | 12 | 0.0833 | 30 | 0.1 | 2 | 0 | 7 | 0.4286 | 7 | 0 | 8 | 0.125 | 24 | 0.1667 |
| 13_105087 | 3 | 0 | 13 | 0.0769 | 9 | 0.1111 | 13 | 0.1538 | 38 | 0.1053 | 2 | 0 | 10 | 0 | 10 | 0.2 | 9 | 0 | 31 | 0.0645 |
| 13_105087 | 2 | 0 | 13 | 0.0769 | 9 | 0.1111 | 11 | 0.0909 | 35 | 0.0857 | 2 | 0 | 9 | 0.1111 | 10 | 0.1 | 9 | 0.1111 | 30 | 0.1 |
| 13_105119! | NA | NA | NA | NA | 2 | 1 | NA | NA | 2 | 1 | NA | NA | NA | NA | 3 | 0.3333 | NA | NA | 3 | 0.3333 |
| 13_105190 | 1 | 0 | 7 | 0.2857 | 6 | 0.1667 | 9 | 0 | 23 | 0.1304 | NA | NA | 4 | 0.25 | 2 | 0 | 4 | 0.5 | 10 | 0.3 |
| 13_105267 | NA | NA | 2 | 0 | 3 | 0.6667 | 5 | 0.6 | 10 | 0.5 | NA | NA | 2 | 0.5 | NA | NA | 1 | 0 | 3 | 0.3333 |
| 13_105386 | NA | NA | 2 | 0 | 7 | 0.2857 | 4 | 0 | 13 | 0.1538 | 1 | 0 | 1 | 1 | NA | NA | 2 | 0 | 4 | 0.25 |
| 13_105386 | NA | NA | 1 | 0 | 4 | 0.75 | 2 | 0.5 | 7 | 0.5714 | NA | NA | 2 | 0 | NA | NA | 2 | 1 | 4 | 0.5 |
| 13_105386 | NA | NA | 1 | 0 | 4 | 0.75 | 1 | 0 | 6 | 0.5 | 1 | 1 | 1 | 1 | NA | NA | 2 | 0.5 | 4 | 0.75 |
| 13_105386 | NA | NA | 1 | 1 | 4 | 0 | 1 | 0 | 6 | 0.1667 | 1 | 0 | 1 | 0 | NA | NA | 1 | 1 | 3 | 0.3333 |
| 13_105386 | NA | NA | 1 | 0 | 4 | 1 | NA | NA | 5 | 0.8 | 1 | 1 | NA | NA | NA | NA | 2 | 1 | 3 | 1 |
| 13_105386 | NA | NA | 1 | 1 | 4 | 0.25 | NA | NA | 5 | 0.4 | 1 | 0 | 1 | 0 | NA | NA | 2 | 0.5 | 4 | 0.25 |
| 13_105386 | NA | NA | 1 | 1 | 3 | 1 | NA | NA | 4 | 1 | NA | NA | 1 | 1 | NA | NA | 2 | 1 | 3 | 1 |
| 13_105387( | NA | NA | 1 | 0 | 5 | 0.2 | NA | NA | 6 | 0.1667 | NA | NA | 1 | 0 | NA | NA | 2 | 0.5 | 3 | 0.3333 |
| 13_105581 | 1 | 0 | NA | NA | 2 | 0 | 1 | 1 | 4 | 0.25 | NA | NA | 1 | 1 | 1 | 0 | NA | NA | 2 | 0.5 |
| 13_105581 | 1 | 1 | NA | NA | 2 | 1 | 1 | 0 | 4 | 0.75 | NA | NA | 1 | 1 | 1 | 0 | NA | NA | 2 | 0.5 |
| 13_105581 | 1 | 1 | NA | NA | 1 | 1 | 1 | 0 | 3 | 0.6667 | NA | NA | 1 | 1 | 1 | 1 | NA | NA | 2 | 1 |
| 13_105582 | 1 | 0 | NA | NA | 2 | 0.5 | 1 | 0 | 4 | 0.25 | NA | NA | 1 | 0 | 1 | 1 | NA | NA | 2 | 0.5 |
| 13_105582 | 1 | 1 | NA | NA | 2 | 1 | 1 | 1 | 4 | 1 | NA | NA | 1 | 1 | 1 | 1 | NA | NA | 2 | 1 |
| 13_105582 | NA | NA | NA | NA | 1 | 1 | 1 | 1 | 2 | 1 | NA | NA | 1 | 1 | 1 | 1 | NA | NA | 2 | 1 |
| 13_105582 | 1 | 1 | NA | NA | 2 | 1 | 1 | 1 | 4 | 1 | NA | NA | 1 | 1 | 1 | 1 | NA | NA | 2 | 1 |
| 13_105582 | 1 | 1 | NA | NA | 2 | 0.5 | 1 | 1 | 4 | 0.75 | NA | NA | 1 | 1 | 1 | 1 | NA | NA | 2 | 1 |
| 13_105582 | 1 | 1 | NA | NA | NA | NA | 1 | 1 | 2 | 1 | NA | NA | 1 | 1 | NA | NA | NA | NA | 1 | 1 |
| 13_105582 | 1 | 1 | NA | NA | 2 | 0.5 | 1 | 1 | 4 | 0.75 | NA | NA | 1 | 1 | 1 | 0 | NA | NA | 2 | 0.5 |
| 13_105582( | 1 | 1 | NA | NA | 1 | 1 | 1 | 1 | 3 | 1 | NA | NA | 1 | 1 | 1 | 1 | NA | NA | 2 | 1 |
| 13_105582 | 1 | 1 | NA | NA | 2 | 1 | 1 | 1 | 4 | 1 | NA | NA | 1 | 1 | NA | NA | NA | NA | 1 | 1 |
| 13_105582 | 1 | 1 | NA | NA | 2 | 0.5 | 1 | 0 | 4 | 0.5 | NA | NA | 1 | 1 | 1 | 0 | NA | NA | 2 | 0.5 |
| 13_105582 | 2 | 1 | NA | NA | 3 | 1 | 1 | 1 | 6 | 1 | NA | NA | 1 | 1 | 1 | 1 | NA | NA | 2 | 1 |
| 13_105584( | 3 | 0.6667 | NA | NA | 3 | 0.6667 | 1 | 1 | 7 | 0.7143 | NA | NA | NA | NA | 1 | 1 | 2 | 0 | 3 | 0.3333 |
| 13_105584 | 3 | 1 | NA | NA | 3 | 0.6667 | NA | NA | 6 | 0.8333 | NA | NA | 1 | 1 | 1 | 0 | 3 | 0.6667 | 5 | 0.6 |
| 13_105584 | 2 | 0 | NA | NA | 2 | 0.5 | NA | NA | 4 | 0.25 | NA | NA | 1 | 0 | 1 | 0 | 2 | 0.5 | 4 | 0.25 |
| 13_105584 | 3 | 0.3333 | NA | NA | 2 | 0 | NA | NA | 5 | 0.2 | NA | NA | 1 | 0 | 1 | 0 | 2 | 0.5 | 4 | 0.25 |
| 13_106931 | 5 | 0 | 12 | 0.1667 | 8 | 0.25 | 11 | 0 | 36 | 0.1111 | 1 | 0 | 5 | 0.2 | 3 | 0 | 7 | 0.1429 | 16 | 0.125 |
| 13_106932 | 3 | 0.3333 | 12 | 0 | 8 | 0.125 | 12 | 0.0833 | 35 | 0.0857 | 2 | 0 | 4 | 0 | NA | NA | 8 | 0.125 | 14 | 0.0714 |
| 13_107083 | 2 | 0.5 | NA | NA | NA | NA | 1 | 1 | 3 | 0.6667 | NA | NA | NA | NA | NA | NA | 1 | 1 | 1 | 1 |
| 13_107083 | 2 | 1 | NA | NA | NA | NA | 1 | 1 | 3 | 1 | NA | NA | NA | NA | NA | NA | 1 | 1 | 1 | 1 |
| 13_107084( | 2 | 0.5 | 2 | 0.5 | 3 | 0 | 3 | 0.6667 | 10 | 0.4 | 1 | 0 | 2 | 0.5 | 3 | 0 | 3 | 0.6667 | 9 | 0.3333 |
| 13_107095 | 3 | 0 | 2 | 0 | 2 | 0 | 5 | 0.4 | 12 | 0.1667 | NA | NA | 3 | 0 | 4 | 0.5 | 4 | 0 | 11 | 0.1818 |
| 13_107097 | 13 | 0.0769 | 11 | 0.0909 | 10 | 0 | 9 | 0.1111 | 43 | 0.0698 | 4 | 0 | 7 | 0.1429 | 12 | 0.25 | 3 | 0 | 26 | 0.1538 |

| | | | | | | | | | | | | | | | | | | | | |
|---|---|---|---|---|---|---|---|---|---|---|---|---|---|---|---|---|---|---|---|---|
| 13_107098 | 10 | 0.1 | 10 | 0.4 | 9 | 0.4444 | 7 | 0.1429 | 36 | 0.2778 | 5 | 0 | 4 | 0.25 | 10 | 0.1 | 3 | 0.3333 | 22 | 0.1364 |
| 13_107098 | 9 | 0.1111 | 11 | 0.0909 | 10 | 0.3 | 8 | 0.25 | 38 | 0.1842 | 4 | 0 | 7 | 0.1429 | 13 | 0.3846 | 3 | 0.3333 | 27 | 0.2593 |
| 13_107098 | 9 | 0.4444 | 14 | 0.2857 | 8 | 0.5 | 10 | 0.2 | 41 | 0.3415 | 7 | 0.7143 | 8 | 0.5 | 12 | 0.6667 | 4 | 0.5 | 31 | 0.6129 |
| 13_107098 | 10 | 0 | 13 | 0.1538 | 9 | 0.2222 | 9 | 0 | 41 | 0.0976 | 9 | 0.1111 | 7 | 0.2857 | 12 | 0.0833 | 5 | 0 | 33 | 0.1212 |
| 13_107159 | NA | NA | 1 | 1 | NA | NA | 1 | 0 | 2 | 0.5 | 1 | 1 | NA | NA | NA | NA | NA | NA | 1 | 1 |
| 13_107160 | NA | NA | 1 | 1 | NA | NA | 1 | 0 | 2 | 0.5 | 1 | 1 | NA | NA | NA | NA | NA | NA | 1 | 1 |
| 13_107160 | NA | NA | 1 | 1 | NA | NA | 1 | 0 | 2 | 0.5 | 1 | 1 | NA | NA | NA | NA | NA | NA | 1 | 1 |
| 13_107550 | 3 | 0 | 6 | 0.1667 | NA | NA | 5 | 0.6 | 14 | 0.2857 | 2 | 0.5 | 3 | 0 | 1 | 0 | 3 | 0 | 9 | 0.1111 |
| 13_107862 | 4 | 0.25 | 2 | 0 | 1 | 0 | 7 | 0 | 14 | 0.0714 | 3 | 0.3333 | 9 | 0.1111 | 6 | 0 | NA | NA | 18 | 0.1111 |
| 13_107862 | 3 | 0 | 2 | 0 | 1 | 1 | 5 | 0.6 | 11 | 0.3636 | 4 | 0.75 | 10 | 0.4 | 3 | 0.3333 | 1 | 0 | 18 | 0.4444 |
| 13_107863 | 2 | 1 | 2 | 1 | 1 | 1 | 5 | 1 | 10 | 1 | 3 | 1 | 11 | 0.9091 | 4 | 1 | 1 | 1 | 19 | 0.9474 |
| 13_107863 | 3 | 0.6667 | 2 | 1 | 1 | 1 | 5 | 0.4 | 11 | 0.6364 | 4 | 0.5 | 12 | 0.5833 | 5 | 0.8 | 1 | 1 | 22 | 0.6364 |
| 13_107863 | 1 | 1 | 1 | 1 | 1 | 1 | 1 | 1 | 4 | 1 | 3 | 0.6667 | 6 | 1 | 2 | 0.5 | 2 | 1 | 13 | 0.8462 |
| 13_107863 | 3 | 0.6667 | 2 | 1 | 1 | 1 | 4 | 1 | 10 | 0.9 | 4 | 1 | 11 | 0.9091 | 4 | 1 | NA | NA | 19 | 0.9474 |
| 13_107863 | 3 | 1 | 1 | 0 | 1 | 1 | 4 | 1 | 9 | 0.8889 | 4 | 0.75 | 11 | 1 | 4 | 1 | 2 | 1 | 21 | 0.9524 |
| 13_107863 | 3 | 1 | 2 | 0 | 1 | 1 | 3 | 1 | 9 | 0.7778 | 2 | 1 | 9 | 0.8889 | 3 | 1 | 1 | 1 | 15 | 0.9333 |
| 13_107863 | 2 | 1 | 2 | 1 | 1 | 1 | 4 | 1 | 9 | 1 | 3 | 1 | 9 | 0.8889 | 3 | 1 | 1 | 1 | 16 | 0.9375 |
| 13_107863 | 2 | 1 | 1 | 0 | 1 | 1 | 4 | 1 | 8 | 0.875 | 4 | 1 | 11 | 1 | 4 | 1 | 1 | 1 | 20 | 1 |
| 13_107863 | 2 | 0 | 1 | 0 | 1 | 0 | 4 | 0.25 | 8 | 0.125 | 4 | 0.5 | 10 | 0 | 3 | 0 | 1 | 0 | 18 | 0.1111 |
| 13_107863 | 3 | 0.6667 | 1 | 1 | 1 | 1 | 4 | 0.5 | 9 | 0.6667 | 4 | 1 | 7 | 0.7143 | 3 | 0.3333 | 1 | 1 | 15 | 0.7333 |
| 13_107863 | 2 | 0 | 2 | 0.5 | 1 | 0 | 4 | 1 | 9 | 0.5556 | 4 | 0.5 | 10 | 0.4 | 2 | 1 | 1 | 0 | 17 | 0.4706 |
| 13_107863 | 3 | 1 | 1 | 0 | 1 | 1 | 3 | 1 | 8 | 0.875 | 4 | 0.75 | 7 | 1 | 4 | 0.75 | 1 | 0 | 16 | 0.8125 |
| 13_107924 | 1 | 0 | 1 | 1 | NA | NA | 1 | 1 | 3 | 0.6667 | 2 | 0.5 | 5 | 1 | 2 | 1 | NA | NA | 9 | 0.8889 |
| 13_107924 | 2 | 1 | 1 | 1 | NA | NA | 1 | 1 | 4 | 1 | 2 | 0.5 | 5 | 0.8 | 1 | 1 | NA | NA | 8 | 0.75 |
| 13_108184 | 2 | 0 | 6 | 0.5 | 2 | 0 | 8 | 0.75 | 18 | 0.5 | 1 | 0 | 4 | 0.75 | 4 | 0.75 | 7 | 0.5714 | 16 | 0.625 |
| 13_108184 | 1 | 1 | 5 | 1 | 1 | 1 | 5 | 0.8 | 12 | 0.9167 | 2 | 1 | 5 | 0.8 | 2 | 1 | 5 | 0.8 | 14 | 0.8571 |
| 13_108184 | 1 | 1 | 6 | 1 | 3 | 1 | 7 | 1 | 17 | 1 | 2 | 0.5 | 5 | 1 | 4 | 1 | 8 | 1 | 19 | 0.9474 |
| 13_108184 | 2 | 0.5 | 4 | 0.25 | 3 | 0 | 6 | 0.1667 | 15 | 0.2 | 2 | 0.5 | 5 | 0.4 | 4 | 0 | 7 | 0.2857 | 18 | 0.2778 |
| 13_108184 | 1 | 0 | 5 | 0.2 | 2 | 0 | 8 | 0 | 16 | 0.0625 | 1 | 0 | 2 | 0 | 4 | 0 | 7 | 0.1429 | 14 | 0.0714 |
| 13_108184 | 2 | 0.5 | 6 | 0 | 2 | 0.5 | 7 | 0.1429 | 17 | 0.1765 | NA | NA | 4 | 0.5 | 5 | 0.2 | 7 | 0.4286 | 16 | 0.375 |
| 13_108184 | 2 | 1 | 6 | 0.6667 | 3 | 0.3333 | 8 | 0.5 | 19 | 0.5789 | 1 | 1 | 4 | 0.75 | 5 | 0.6 | 7 | 0.2857 | 17 | 0.5294 |
| 13_108184 | NA | NA | 6 | 0.1667 | 3 | 0 | 6 | 0.1667 | 15 | 0.1333 | 1 | 0 | 4 | 0 | 5 | 0 | 7 | 0.1429 | 17 | 0.0588 |
| 13_108184 | 1 | 0 | 4 | 0 | 3 | 0.3333 | 6 | 0 | 14 | 0.0714 | 1 | 0 | 4 | 0.25 | 5 | 0 | 7 | 0.1429 | 17 | 0.1176 |
| 13_108189 | 9 | 0.1111 | 6 | 0.1667 | 9 | 0 | 9 | 0 | 33 | 0.0606 | 9 | 0.1111 | 8 | 0 | 8 | 0 | 3 | 0.3333 | 28 | 0.0714 |
| 13_108189 | 10 | 0.1 | 6 | 0.3333 | 9 | 0.2222 | 9 | 0 | 34 | 0.1471 | 7 | 0 | 7 | 0.2857 | 7 | 0.4286 | 4 | 0 | 25 | 0.2 |
| 13_108189 | 10 | 0 | 5 | 0.6 | 8 | 0 | 9 | 0.1111 | 32 | 0.125 | 9 | 0.2222 | 8 | 0.125 | 8 | 0.125 | 1 | 0 | 26 | 0.1538 |
| 13_108330 | 2 | 0.5 | 1 | 1 | 5 | 0.6 | 2 | 0.5 | 10 | 0.6 | NA | NA | 1 | 0 | 2 | 0.5 | 2 | 0.5 | 5 | 0.4 |
| 13_108330 | 3 | 0 | 1 | 1 | 6 | 0.1667 | 2 | 0 | 12 | 0.1667 | NA | NA | 1 | 0 | 2 | 0.5 | 2 | 0 | 5 | 0.2 |
| 13_108419 | NA | NA | 3 | 0.6667 | 3 | 1 | 1 | 0 | 7 | 0.7143 | NA | NA | NA | NA | 1 | 1 | NA | NA | 1 | 1 |
| 13_108419 | NA | NA | 1 | 1 | NA | NA | 1 | 1 | 2 | 1 | NA | NA | NA | NA | 1 | 1 | NA | NA | 1 | 1 |
| 13_108423 | NA | NA | NA | NA | 2 | 1 | NA | NA | 2 | 1 | NA | NA | 1 | 1 | NA | NA | NA | NA | 1 | 1 |
| 13_110087 | 3 | 0.3333 | 2 | 0 | 2 | 0 | 4 | 0.25 | 11 | 0.1818 | 1 | 0 | 3 | 0.3333 | 1 | 0 | 4 | 0.25 | 9 | 0.2222 |
| 13_110087 | 3 | 0 | 4 | 0.75 | NA | NA | 5 | 0.6 | 12 | 0.5 | 1 | 1 | 4 | 0 | NA | NA | 4 | 0.25 | 9 | 0.2222 |
| 13_110087 | 3 | 1 | 4 | 0.5 | 1 | 1 | 5 | 0.8 | 13 | 0.7692 | NA | NA | 4 | 1 | NA | NA | 4 | 0.5 | 8 | 0.75 |
| 13_110087 | 3 | 0 | 3 | 0 | 1 | 0 | 5 | 0.2 | 12 | 0.0833 | 1 | 0 | 4 | 0.25 | 1 | 0 | 4 | 0.25 | 10 | 0.2 |
| 13_110087 | 3 | 1 | 2 | 0 | 2 | 1 | 4 | 0.75 | 11 | 0.7273 | 1 | 1 | 3 | 1 | 2 | 0.5 | 4 | 0.5 | 10 | 0.7 |
| 13_110087 | 2 | 0.5 | 2 | 1 | 2 | 0.5 | 3 | 0.6667 | 9 | 0.6667 | 1 | 1 | 4 | 1 | 1 | 1 | 3 | 0.3333 | 9 | 0.7778 |
| 13_110087 | 3 | 0.3333 | 3 | 0.3333 | 2 | 1 | 5 | 0.6 | 13 | 0.5385 | 1 | 1 | 3 | 0.6667 | 2 | 0.5 | 4 | 0.5 | 10 | 0.6 |
| 13_110087 | 3 | 0 | 3 | 0.3333 | 1 | 0 | 5 | 0 | 12 | 0.0833 | 1 | 0 | 4 | 0 | 2 | 0 | 4 | 0.25 | 11 | 0.0909 |
| 13_110087 | 3 | 1 | 2 | 0.5 | 1 | 0 | 5 | 0.2 | 11 | 0.4545 | 1 | 0 | 2 | 0 | 1 | 0 | 3 | 0.3333 | 7 | 0.1429 |
| 13_110088 | 1 | 1 | 2 | 0.5 | 2 | 0.5 | 5 | 0.8 | 10 | 0.7 | 1 | 1 | 4 | 0.25 | 1 | 1 | 2 | 0 | 8 | 0.375 |
| 13_110088 | 2 | 1 | 3 | 0 | 1 | 0 | 2 | 0 | 8 | 0.25 | NA | NA | 1 | 0 | 1 | 0 | 2 | 0.5 | 4 | 0.25 |
| 13_110088 | 3 | 0 | 1 | 0 | 1 | 0 | 2 | 0.5 | 7 | 0.1429 | 1 | 0 | 1 | 1 | 1 | 1 | 3 | 0 | 6 | 0.3333 |
| 13_110088 | 2 | 0.5 | 3 | 1 | 2 | 1 | 6 | 0.8333 | 13 | 0.8462 | 1 | 1 | 4 | 0.75 | 2 | 0.5 | 2 | 1 | 9 | 0.7778 |
| 13_110088 | 1 | 0 | 3 | 0.6667 | 2 | 1 | 4 | 0.75 | 10 | 0.7 | 1 | 1 | 4 | 0.5 | 4 | 1 | 4 | 0.75 | 13 | 0.7692 |
| 13_110088 | 2 | 0.5 | 3 | 0.6667 | 2 | 0.5 | 5 | 0.6 | 12 | 0.5833 | 1 | 0 | 4 | 0.25 | 4 | 0.5 | 4 | 0.25 | 13 | 0.3077 |
| 13_110088 | 2 | 1 | NA | NA | 2 | 1 | NA | NA | 4 | 1 | 1 | 1 | 3 | 1 | NA | NA | 3 | 1 | 7 | 1 |
| 13_110088 | 2 | 0 | 4 | 0.25 | 4 | 0 | 5 | 0.2 | 15 | 0.1333 | 1 | 0 | 6 | 0.3333 | 3 | 0.3333 | 3 | 0.3333 | 13 | 0.3077 |
| 13_110088 | 2 | 0.5 | 3 | 0.6667 | 3 | 0.6667 | 3 | 0.6667 | 11 | 0.6364 | 1 | 1 | 6 | 0.8333 | 2 | 0.5 | 3 | 0.6667 | 12 | 0.75 |
| 13_110089 | 3 | 0.3333 | 3 | 0.3333 | 4 | 0.5 | 3 | 0.3333 | 13 | 0.3846 | 3 | 0.3333 | 6 | 0.6667 | 1 | 0 | 4 | 0.75 | 14 | 0.5714 |
| 13_110089 | 2 | 0 | 3 | 0.3333 | 4 | 0.5 | 5 | 0.2 | 14 | 0.2857 | 3 | 0.6667 | 8 | 0.75 | 2 | 0 | 4 | 0.5 | 17 | 0.5882 |
| 13_110089 | 3 | 0.3333 | 2 | 0 | 3 | 0 | 3 | 0 | 11 | 0.0909 | 2 | 0 | 6 | 0.1667 | 1 | 0 | 3 | 0.3333 | 12 | 0.1667 |
| 13_110089 | 3 | 0.3333 | 4 | 0 | 4 | 0.5 | 4 | 0.5 | 15 | 0.3333 | 3 | 0 | 6 | 0.3333 | 2 | 0 | 4 | 0 | 15 | 0.1333 |
| 13_110090 | 3 | 0.3333 | 3 | 0 | 4 | 0 | 5 | 0.2 | 15 | 0.1333 | 4 | 0.5 | 6 | 0 | 3 | 0 | 4 | 0.25 | 17 | 0.1765 |
| 13_110091 | 4 | 0.25 | 6 | 0.3333 | 6 | 0 | 6 | 0.1667 | 22 | 0.1818 | 6 | 0 | 6 | 0.8333 | 4 | 0.25 | 5 | 0 | 21 | 0.2857 |

| | | | | | | | | | | | | | | | | | | | | |
|---|---|---|---|---|---|---|---|---|---|---|---|---|---|---|---|---|---|---|---|---|
| 13_1100918 | 4 | 0.25 | 8 | 0.125 | 6 | 0 | 7 | 0.2857 | 25 | 0.16 | 6 | 0 | 6 | 0.1667 | 4 | 0 | 4 | 0 | 20 | 0.05 |
| 13_1100922 | 5 | 0.2 | 8 | 0 | 7 | 0.1429 | 7 | 0.1429 | 27 | 0.1111 | 5 | 0 | 7 | 0 | 5 | 0.4 | 5 | 0 | 22 | 0.0909 |
| 13_1101910 | 38 | 0.0789 | 38 | 0.0526 | 26 | 0.1154 | 24 | 0.0417 | 126 | 0.0714 | 14 | 0.1429 | 23 | 0.087 | 20 | 0.05 | 27 | 0.037 | 84 | 0.0714 |
| 13_1101911 | 33 | 0.1818 | 39 | 0.0769 | 22 | 0.2273 | 22 | 0.1364 | 116 | 0.1466 | 13 | 0.3077 | 19 | 0.2105 | 23 | 0.1739 | 25 | 0.16 | 80 | 0.2 |
| 13_1101911 | 33 | 0.0909 | 36 | 0.1389 | 25 | 0.04 | 20 | 0.25 | 114 | 0.1228 | 14 | 0.0714 | 18 | 0.2222 | 18 | 0.2222 | 28 | 0.1429 | 78 | 0.1667 |
| 13_1101915 | 32 | 0.0625 | 43 | 0.093 | 24 | 0 | 21 | 0.0476 | 120 | 0.0583 | 14 | 0.1429 | 22 | 0.0455 | 21 | 0.0476 | 28 | 0.1071 | 85 | 0.0824 |
| 13_1101916 | 27 | 0.2963 | 40 | 0.35 | 21 | 0.4762 | 20 | 0.5 | 108 | 0.3889 | 13 | 0.3077 | 20 | 0.4 | 18 | 0.2222 | 23 | 0.3478 | 74 | 0.3243 |
| 13_1101944 | 18 | 0.1667 | 15 | 0.0667 | 14 | 0.1429 | 9 | 0 | 56 | 0.1071 | 7 | 0.1429 | 12 | 0.25 | 5 | 0.4 | 14 | 0.0714 | 38 | 0.1842 |
| 13_1101945 | 16 | 0.0625 | 13 | 0.3077 | 12 | 0 | 7 | 0.1429 | 48 | 0.125 | 6 | 0 | 13 | 0 | 5 | 0.2 | 12 | 0.0833 | 36 | 0.0556 |
| 13_1101946 | 15 | 0.2 | 12 | 0.1667 | 9 | 0 | 8 | 0 | 44 | 0.1136 | 6 | 0 | 9 | 0 | 3 | 0 | 11 | 0.1818 | 29 | 0.069 |
| 13_1101951 | 13 | 0.4615 | 8 | 0.5 | 9 | 0.4444 | 5 | 0.4 | 35 | 0.4571 | 5 | 0.4 | 5 | 0.4 | 4 | 0.5 | 10 | 0.5 | 24 | 0.4583 |
| 13_1101952 | 11 | 0.1818 | 9 | 0.1111 | 8 | 0.125 | 7 | 0.1429 | 35 | 0.1429 | 5 | 0 | 10 | 0.4 | 5 | 0.2 | 7 | 0.1429 | 27 | 0.2222 |
| 13_1101965 | 4 | 0 | 3 | 0.3333 | 4 | 0 | 4 | 0.25 | 15 | 0.1333 | 3 | 0 | 6 | 0 | 3 | 0.3333 | 4 | 0 | 16 | 0.0625 |
| 13_1101965 | 7 | 0.2857 | 4 | 0 | 3 | 0 | 4 | 0 | 18 | 0.1111 | 4 | 0.25 | 7 | 0.2857 | 4 | 0.5 | 4 | 0.25 | 19 | 0.3158 |
| 13_1101975 | 3 | 0 | 6 | 0.1667 | 3 | 0 | 2 | 0 | 14 | 0.0714 | 3 | 0 | 3 | 0.3333 | 2 | 0 | 1 | 0 | 9 | 0.1111 |
| 13_1101978 | 3 | 0.3333 | 6 | 0.3333 | 3 | 0 | 1 | 0 | 13 | 0.2308 | 3 | 0 | 3 | 0.3333 | 2 | 0.5 | 1 | 0 | 9 | 0.2222 |
| 13_1101980 | 2 | 0 | 4 | 0.75 | 5 | 0.2 | 2 | 0.5 | 13 | 0.3846 | 2 | 0.5 | 3 | 0.3333 | 2 | 1 | 2 | 0.5 | 9 | 0.5556 |
| 13_1105366 | 34 | 0.1176 | 21 | 0.1905 | 19 | 0.0526 | 27 | 0 | 101 | 0.0891 | 17 | 0.1176 | 32 | 0.2188 | 31 | 0.129 | 21 | 0.1429 | 101 | 0.1584 |
| 13_1105367 | 31 | 0.3871 | 21 | 0.5714 | 20 | 0.35 | 29 | 0.3103 | 101 | 0.396 | 16 | 0.5 | 29 | 0.4828 | 33 | 0.3636 | 22 | 0.3182 | 100 | 0.41 |
| 13_1105367 | 30 | 0.2333 | 20 | 0.15 | 18 | 0.1667 | 27 | 0.1852 | 95 | 0.1895 | 19 | 0 | 28 | 0.1786 | 28 | 0.1429 | 21 | 0.2381 | 96 | 0.1458 |
| 13_1105367 | 24 | 0.375 | 21 | 0.1905 | 15 | 0.1333 | 24 | 0.1667 | 84 | 0.2262 | 16 | 0.3125 | 27 | 0.2593 | 27 | 0.4074 | 20 | 0.55 | 90 | 0.3778 |
| 13_1105368 | 34 | 0.9412 | 26 | 1 | 20 | 0.9 | 31 | 0.9355 | 111 | 0.9459 | 18 | 0.9444 | 32 | 0.9688 | 36 | 0.9722 | 25 | 1 | 111 | 0.973 |
| 13_1105368 | 34 | 0.4412 | 22 | 0.5455 | 21 | 0.381 | 27 | 0.4074 | 104 | 0.4423 | 16 | 0.4375 | 35 | 0.4571 | 37 | 0.4324 | 26 | 0.2692 | 114 | 0.4035 |
| 13_1105369 | 32 | 0.5938 | 22 | 0.5455 | 17 | 0.7059 | 24 | 0.4583 | 95 | 0.5684 | 17 | 0.4706 | 32 | 0.5938 | 39 | 0.641 | 25 | 0.44 | 113 | 0.5575 |
| 13_1105369 | 32 | 0.3125 | 26 | 0.3462 | 21 | 0.1429 | 32 | 0.2188 | 111 | 0.2613 | 18 | 0.0556 | 32 | 0.4375 | 39 | 0.2821 | 27 | 0.2222 | 116 | 0.2759 |
| 13_1105369 | 30 | 0.8333 | 23 | 0.7391 | 19 | 0.8421 | 25 | 0.84 | 97 | 0.8144 | 19 | 0.8947 | 29 | 0.7931 | 33 | 0.9394 | 24 | 0.875 | 105 | 0.8762 |
| 13_1105370 | 32 | 0.9688 | 25 | 0.96 | 22 | 0.9545 | 33 | 0.9091 | 112 | 0.9464 | 20 | 1 | 37 | 0.8919 | 37 | 1 | 25 | 0.96 | 119 | 0.958 |
| 13_1105370 | 28 | 0.6071 | 22 | 0.7273 | 21 | 0.8095 | 26 | 0.7308 | 97 | 0.7113 | 17 | 0.6471 | 28 | 0.6071 | 29 | 0.4138 | 25 | 0.48 | 99 | 0.5253 |
| 13_1105370 | 27 | 0.963 | 18 | 0.9444 | 22 | 0.9545 | 31 | 0.9677 | 98 | 0.9592 | 14 | 0.9286 | 30 | 0.9333 | 32 | 0.9688 | 21 | 0.9048 | 97 | 0.9381 |
| 13_1105371 | 29 | 0.3448 | 24 | 0.375 | 23 | 0.3913 | 27 | 0.4444 | 103 | 0.3883 | 21 | 0.3333 | 32 | 0.5 | 36 | 0.4167 | 26 | 0.5 | 115 | 0.4435 |
| 13_1115624 | 4 | 0.75 | 3 | 0 | 6 | 0 | 4 | 0 | 17 | 0.1765 | 3 | 0.3333 | 9 | 0.6667 | 7 | 0.2857 | 4 | 0.75 | 23 | 0.5217 |
| 13_1115625 | 4 | 0.25 | 4 | 0.25 | 6 | 0.3333 | 4 | 0 | 18 | 0.2222 | 3 | 0.3333 | 8 | 0 | 8 | 0.25 | 6 | 0.1667 | 25 | 0.16 |
| 13_1115626 | 3 | 0.3333 | 4 | 0 | 6 | 0 | 4 | 0 | 17 | 0.0588 | 1 | 0 | 9 | 0.2222 | 8 | 0 | 6 | 0 | 24 | 0.0833 |
| 13_1115627 | 9 | 0.6667 | 11 | 0.4545 | 14 | 0.8571 | 10 | 0.7 | 44 | 0.6818 | 5 | 0.6 | 10 | 0.3 | 10 | 0.4 | 10 | 0.4 | 35 | 0.4 |
| 13_1115628 | 12 | 1 | 13 | 1 | 14 | 0.9286 | 11 | 0.9091 | 50 | 0.96 | 6 | 1 | 12 | 1 | 13 | 0.9231 | 14 | 0.8571 | 45 | 0.9333 |
| 13_1115628 | 9 | 0.3333 | 14 | 0.3571 | 14 | 0.4286 | 11 | 0.2727 | 48 | 0.3542 | 6 | 0.5 | 11 | 0.5455 | 11 | 0.1818 | 11 | 0.4545 | 39 | 0.4103 |
| 13_1115628 | 11 | 0.5455 | 13 | 0.6923 | 12 | 0.6667 | 12 | 0.5833 | 48 | 0.625 | 7 | 0.5714 | 14 | 0.4286 | 11 | 0.3636 | 11 | 0.5455 | 43 | 0.4651 |
| 13_1115629 | 9 | 0.2222 | 9 | 0 | 8 | 0 | 7 | 0 | 33 | 0.0606 | 6 | 0 | 15 | 0.0667 | 10 | 0.1 | 8 | 0.125 | 39 | 0.0769 |
| 13_1115629 | 8 | 0.875 | 10 | 0.9 | 11 | 1 | 10 | 1 | 39 | 0.9487 | 7 | 0.8571 | 12 | 0.75 | 10 | 0.8 | 11 | 0.8182 | 40 | 0.8 |
| 13_1115630 | 10 | 0.1 | 12 | 0 | 11 | 0.1818 | 11 | 0 | 44 | 0.0682 | 6 | 0.1667 | 15 | 0.0667 | 11 | 0.0909 | 11 | 0.2727 | 43 | 0.1395 |
| 13_1115631 | 9 | 0.1111 | 11 | 0.1818 | 11 | 0 | 11 | 0.0909 | 42 | 0.0952 | 7 | 0.2857 | 15 | 0 | 13 | 0.1538 | 11 | 0 | 46 | 0.087 |
| 13_1116240 | 5 | 0.8 | 7 | 0.1429 | 7 | 0.4286 | 7 | 0.7143 | 26 | 0.5 | 4 | 0.75 | 9 | 0.4444 | 6 | 0.5 | 2 | 1 | 21 | 0.5714 |
| 13_1116240 | 6 | 0.6667 | 7 | 0.5714 | 7 | 0.2857 | 7 | 1 | 27 | 0.6296 | 4 | 0.75 | 13 | 0.8462 | 8 | 0.75 | 2 | 1 | 27 | 0.8148 |
| 13_1116240 | 5 | 0.6 | 9 | 0.5556 | 7 | 0.5714 | 6 | 0.5 | 27 | 0.5556 | 3 | 0 | 13 | 0.2308 | 6 | 0.8333 | 2 | 1 | 24 | 0.4167 |
| 13_1116241 | 5 | 0.2 | 9 | 0.1111 | 6 | 0 | 7 | 0 | 27 | 0.0741 | 4 | 0.25 | 9 | 0.2222 | 9 | 0.3333 | 2 | 0 | 24 | 0.25 |
| 13_1116241 | 3 | 0.6667 | 6 | 0 | 7 | 0.1429 | 5 | 0.2 | 21 | 0.1905 | 4 | 0 | 9 | 0.2222 | 8 | 0.25 | 2 | 0 | 23 | 0.1739 |
| 13_1116241 | 3 | 0 | 8 | 0.25 | 3 | 0.3333 | 4 | 0.25 | 18 | 0.2222 | 3 | 0.3333 | 9 | 0.1111 | 5 | 0.2 | 2 | 0 | 19 | 0.1579 |
| 13_1116241 | 6 | 0.1667 | 7 | 0.1429 | 7 | 0 | 6 | 0 | 26 | 0.0769 | 4 | 0.25 | 12 | 0 | 9 | 0.1111 | 2 | 0 | 27 | 0.0741 |
| 13_1116241 | 6 | 0.5 | 5 | 0.2 | 3 | 0.6667 | 4 | 0.5 | 18 | 0.4444 | 3 | 0.6667 | 8 | 0.125 | 5 | 0.2 | 1 | 1 | 17 | 0.2941 |
| 13_1116244 | 6 | 0.5 | 6 | 0.6667 | 5 | 0.4 | 4 | 0.5 | 21 | 0.5238 | 4 | 0.25 | 12 | 0.5 | 6 | 0.6667 | 1 | 0 | 23 | 0.4783 |
| 13_1116245 | 5 | 0.6 | 5 | 0 | 5 | 0.6 | 6 | 0.1667 | 21 | 0.3333 | 3 | 0.3333 | 12 | 0.5833 | 7 | 0.4286 | 2 | 0 | 24 | 0.4583 |
| 13_1116247 | 6 | 0 | 5 | 0 | 3 | 0 | 5 | 0.2 | 19 | 0.0526 | 2 | 0 | 8 | 0.125 | 5 | 0 | 1 | 0 | 16 | 0.0625 |
| 13_1117298 | 1 | 1 | 1 | 0 | 1 | 0 | NA | NA | 3 | 0.3333 | 1 | 1 | NA | NA | 1 | 1 | NA | NA | 2 | 1 |
| 13_1118515 | 1 | 1 | NA | NA | NA | NA | NA | NA | 1 | 1 | 1 | 1 | NA | NA | 1 | 1 | NA | NA | 2 | 1 |
| 13_1118516 | 1 | 1 | NA | NA | NA | NA | NA | NA | 1 | 1 | 1 | 1 | NA | NA | 1 | 0 | 1 | 1 | 3 | 0.6667 |
| 13_1118518 | 1 | 1 | NA | NA | NA | NA | NA | NA | 1 | 1 | NA | NA | NA | NA | 1 | 1 | 1 | 1 | 2 | 1 |
| 13_1118518 | 1 | 1 | NA | NA | NA | NA | NA | NA | 1 | 1 | 1 | 1 | NA | NA | NA | NA | 1 | 1 | 2 | 1 |
| 13_1118520 | 1 | 1 | NA | NA | NA | NA | NA | NA | 1 | 1 | 1 | 1 | NA | NA | 1 | 1 | 1 | 1 | 3 | 1 |
| 13_1118522 | 1 | 1 | 1 | 0 | NA | NA | NA | NA | 2 | 0.5 | 1 | 1 | NA | NA | 1 | 0 | 1 | 1 | 3 | 0.6667 |
| 13_1118524 | 1 | 1 | 1 | 0 | NA | NA | NA | NA | 2 | 0.5 | 1 | 1 | 1 | 0 | 1 | 1 | 1 | 1 | 4 | 0.75 |
| 13_1118526 | 1 | 1 | 1 | 0 | NA | NA | NA | NA | 2 | 0.5 | 1 | 0 | 1 | 0 | 1 | 0 | 1 | 1 | 4 | 0.25 |
| 13_1118537 | 1 | 0 | 3 | 0.3333 | NA | NA | NA | NA | 4 | 0.25 | 1 | 1 | 2 | 0 | 5 | 0.2 | 1 | 1 | 9 | 0.3333 |
| 13_1118540 | 2 | 0 | 4 | 0 | NA | NA | 1 | 1 | 7 | 0.1429 | 2 | 0 | 1 | 0 | 5 | 0.2 | 1 | 0 | 9 | 0.1111 |
| 13_1118851 | 1 | 1 | NA | NA | NA | NA | NA | NA | 1 | 1 | NA | NA | 1 | 1 | NA | NA | NA | NA | 1 | 1 |
| 13_1118853 | 1 | 1 | NA | NA | NA | NA | NA | NA | 1 | 1 | NA | NA | 1 | 1 | NA | NA | NA | NA | 1 | 1 |

| | | | | | | | | | | | | | | | | | | | | |
|---|---|---|---|---|---|---|---|---|---|---|---|---|---|---|---|---|---|---|---|---|
| 13_112504 | NA | NA | NA | NA | NA | NA | 1 | 1 | 1 | 1 | NA | NA | NA | NA | 1 | 1 | 1 | 1 | 2 | 1 |
| 13_112504 | NA | NA | NA | NA | NA | NA | 1 | 1 | 1 | 1 | NA | NA | NA | NA | NA | NA | 1 | 1 | 1 | 1 |
| 13_112504 | NA | NA | NA | NA | NA | NA | 1 | 1 | 1 | 1 | NA | NA | NA | NA | 1 | 1 | 1 | 1 | 2 | 1 |
| 13_112504 | NA | NA | NA | NA | NA | NA | 1 | 1 | 1 | 1 | NA | NA | NA | NA | 1 | 1 | 1 | 0 | 2 | 0.5 |
| 13_112640 | 1 | 0 | 2 | 0.5 | 1 | 0 | 2 | 0.5 | 6 | 0.3333 | 2 | 0 | 7 | 0.2857 | 6 | 0 | 4 | 0 | 19 | 0.1053 |
| 13_112640 | 3 | 0 | 2 | 0.5 | NA | NA | 2 | 0 | 7 | 0.1429 | NA | NA | 5 | 0.6 | 7 | 0.2857 | 3 | 0.6667 | 15 | 0.4667 |
| 13_112640 | 4 | 0.5 | 2 | 0 | 1 | 0 | 3 | 0.3333 | 10 | 0.3 | 2 | 0.5 | 8 | 0.25 | 6 | 0.3333 | 4 | 1 | 20 | 0.45 |
| 13_112640 | 3 | 1 | 2 | 0.5 | 1 | 0 | 3 | 1 | 9 | 0.7778 | 2 | 1 | 8 | 0.75 | 7 | 1 | 3 | 1 | 20 | 0.9 |
| 13_112640 | 5 | 1 | 2 | 1 | 1 | 1 | 3 | 0.6667 | 11 | 0.9091 | 2 | 1 | 8 | 0.875 | 7 | 1 | 4 | 1 | 21 | 0.9524 |
| 13_112640 | 5 | 0.8 | 2 | 1 | NA | NA | 3 | 1 | 10 | 0.9 | 3 | 1 | 9 | 0.6667 | 6 | 1 | 3 | 1 | 21 | 0.8571 |
| 13_112641 | 5 | 0.4 | 2 | 0.5 | 1 | 0 | 2 | 0 | 10 | 0.3 | 3 | 0.3333 | 8 | 0.375 | 6 | 0.1667 | 4 | 0 | 21 | 0.2381 |
| 13_112641 | 5 | 0.2 | 2 | 0 | 1 | 0 | 3 | 0.6667 | 11 | 0.2727 | 2 | 0.5 | 9 | 0.3333 | 7 | 0.1429 | 4 | 0 | 22 | 0.2273 |
| 13_112641 | NA | NA | 1 | 1 | 1 | 1 | 2 | 1 | 4 | 1 | 2 | 1 | 7 | 0.8571 | 4 | 0.75 | 1 | 1 | 14 | 0.8571 |
| 13_112641 | 5 | 0.8 | NA | NA | 1 | 1 | 3 | 0.6667 | 9 | 0.7778 | 3 | 0.3333 | 5 | 0.8 | 5 | 0.6 | 2 | 0 | 15 | 0.5333 |
| 13_112641 | 5 | 1 | 2 | 0.5 | 1 | 1 | 4 | 0.75 | 12 | 0.8333 | 3 | 0.6667 | 11 | 0.8182 | 9 | 1 | 4 | 0.75 | 27 | 0.8519 |
| 13_112641 | 5 | 1 | 2 | 1 | 1 | 0 | 4 | 1 | 12 | 0.9167 | 2 | 0.5 | 11 | 0.8182 | 7 | 0.8571 | 5 | 1 | 25 | 0.84 |
| 13_112641 | 5 | 0.4 | 2 | 0 | NA | NA | 4 | 0.25 | 11 | 0.2727 | 2 | 0.5 | 10 | 0.4 | 7 | 0.4286 | 3 | 0 | 22 | 0.3636 |
| 13_112641 | 4 | 1 | 1 | 1 | NA | NA | 3 | 0.6667 | 8 | 0.875 | 3 | 0.6667 | 10 | 0.9 | 8 | 1 | 4 | 0.75 | 25 | 0.88 |
| 13_112641 | 5 | 0.4 | 2 | 0 | 2 | 0 | 3 | 0.3333 | 12 | 0.25 | 2 | 0 | 12 | 0.0833 | 9 | 0.2222 | 5 | 0.2 | 28 | 0.1429 |
| 13_112641 | 5 | 0 | 2 | 0 | 2 | 0.5 | 5 | 0 | 14 | 0.0714 | 4 | 0.25 | 11 | 0.0909 | 8 | 0.25 | 5 | 0.2 | 28 | 0.1786 |
| 13_112641 | 5 | 1 | 1 | 1 | 1 | 1 | 5 | 1 | 12 | 1 | 4 | 1 | 12 | 0.9167 | 8 | 1 | 4 | 1 | 28 | 0.9643 |
| 13_112641 | 5 | 0.2 | 2 | 1 | 2 | 0 | 5 | 0.6 | 14 | 0.4286 | 4 | 0.5 | 9 | 0.3333 | 8 | 0.5 | 5 | 0.4 | 26 | 0.4231 |
| 13_112641 | 5 | 0.4 | 2 | 0 | 1 | 1 | 2 | 0 | 10 | 0.3 | 2 | 0 | 9 | 0.2222 | 6 | 0 | 4 | 0 | 21 | 0.0952 |
| 13_112641 | 2 | 0.5 | 1 | 1 | 1 | 1 | 5 | 0.6 | 9 | 0.6667 | 4 | 1 | 9 | 0.6667 | 8 | 0.875 | 2 | 0.5 | 23 | 0.7826 |
| 13_112641 | 4 | 1 | 1 | 1 | 2 | 1 | 4 | 1 | 11 | 1 | 3 | 1 | 11 | 1 | 9 | 1 | 4 | 1 | 27 | 1 |
| 13_112641 | 4 | 1 | 1 | 0 | 2 | 0.5 | 4 | 0.75 | 11 | 0.7273 | 3 | 0.6667 | 11 | 0.9091 | 7 | 0.5714 | 5 | 0.6 | 26 | 0.7308 |
| 13_112641 | 3 | 0.3333 | 1 | 0 | 2 | 0 | 6 | 0.3333 | 12 | 0.25 | 4 | 0 | 10 | 0 | 8 | 0.25 | 5 | 0 | 27 | 0.0741 |
| 13_112642 | 5 | 0.4 | 1 | 0 | 2 | 0 | 5 | 0.4 | 13 | 0.3077 | 4 | 0.75 | 8 | 0.5 | 9 | 0.5556 | 5 | 0.6 | 26 | 0.5769 |
| 13_112642 | 4 | 0.25 | 1 | 0 | 2 | 0 | 3 | 0 | 10 | 0.1 | 4 | 0.25 | 11 | 0.2727 | 6 | 0.3333 | 5 | 0.4 | 26 | 0.3077 |
| 13_112642 | 4 | 0.75 | 1 | 1 | 1 | 1 | 6 | 0.6667 | 12 | 0.75 | 3 | 1 | 8 | 1 | 9 | 0.6667 | 5 | 1 | 25 | 0.88 |
| 13_112642 | 5 | 0.2 | 1 | 1 | 2 | 0 | 6 | 0.8333 | 14 | 0.5 | 5 | 0 | 10 | 0.4 | 9 | 0.2222 | 3 | 0.6667 | 27 | 0.2963 |
| 13_112642 | 7 | 0.1429 | 1 | 1 | 2 | 0 | 7 | 0.1429 | 17 | 0.1765 | 5 | 0 | 12 | 0.0833 | 14 | 0.0714 | 8 | 0.25 | 39 | 0.1026 |
| 13_112642 | 7 | 0.2857 | 2 | 0.5 | 2 | 0.5 | 8 | 0.25 | 19 | 0.3158 | 5 | 0 | 12 | 0.0833 | 14 | 0.0714 | 8 | 0.25 | 39 | 0.1026 |
| 13_112642 | 8 | 0.375 | 2 | 0.5 | 2 | 0 | 6 | 0.3333 | 18 | 0.3333 | 5 | 0.4 | 10 | 0.6 | 13 | 0.6154 | 7 | 0.8571 | 35 | 0.6286 |
| 13_112643 | 10 | 0.2 | 2 | 0.5 | 2 | 0 | 8 | 0.125 | 22 | 0.1818 | 5 | 0 | 13 | 0.1538 | 14 | 0.1429 | 9 | 0 | 41 | 0.0976 |
| 13_112643 | NA | NA | NA | NA | NA | NA | 1 | 1 | 1 | 1 | 2 | 0 | 2 | 0.5 | 3 | 0 | 2 | 0 | 9 | 0.1111 |
| 13_112644 | NA | NA | NA | NA | NA | NA | 3 | 0.3333 | 3 | 0.3333 | 2 | 0 | 2 | 0.5 | 4 | 0 | 5 | 0.2 | 13 | 0.1538 |
| 13_112644 | 2 | 0.5 | NA | NA | NA | NA | 2 | 0 | 4 | 0.25 | 2 | 0 | 3 | 0.3333 | 4 | 0.5 | 4 | 0.75 | 13 | 0.4615 |
| 13_112868 | NA | NA | 1 | 1 | 3 | 0 | 1 | 0 | 5 | 0.2 | 1 | 0 | 3 | 0.3333 | 1 | 0 | 1 | 0 | 6 | 0.1667 |
| 13_112868 | NA | NA | 1 | 0 | 3 | 0.3333 | 1 | 1 | 5 | 0.4 | NA | NA | 3 | 0.3333 | 1 | 1 | 1 | 0 | 5 | 0.4 |
| 13_112868 | NA | NA | 1 | 0 | 2 | 0.5 | 1 | 0 | 4 | 0.25 | 1 | 0 | 3 | 0.3333 | 1 | 0 | 1 | 0 | 6 | 0.1667 |
| 13_112868 | NA | NA | 1 | 1 | 3 | 1 | 1 | 1 | 5 | 1 | 1 | 1 | 3 | 0.6667 | 1 | 1 | 1 | 0 | 6 | 0.6667 |
| 13_112874 | 1 | 1 | 1 | 1 | 1 | 0 | 1 | 0 | 4 | 0.5 | NA | NA | 2 | 1 | 1 | 0 | 1 | 0 | 4 | 0.5 |
| 13_112874 | 1 | 1 | 2 | 0 | 1 | 0 | 1 | 0 | 5 | 0.2 | NA | NA | 1 | 0 | 1 | 1 | 1 | 0 | 3 | 0.3333 |
| 13_112874 | 1 | 1 | 1 | 1 | 1 | 1 | 1 | 1 | 4 | 1 | NA | NA | 2 | 1 | 1 | 1 | NA | NA | 3 | 1 |
| 13_112874 | 1 | 1 | 2 | 0.5 | 2 | 0 | 1 | 0 | 6 | 0.3333 | NA | NA | 2 | 0.5 | 1 | 1 | 1 | 0 | 4 | 0.5 |
| 13_112874 | 1 | 0 | 2 | 0.5 | 2 | 0.5 | 2 | 0 | 7 | 0.2857 | NA | NA | 2 | 0.5 | 1 | 0 | 1 | 0 | 4 | 0.25 |
| 13_112874 | NA | NA | 2 | 1 | 2 | 1 | 2 | 1 | 6 | 1 | NA | NA | 2 | 1 | NA | NA | NA | NA | 2 | 1 |
| 13_112875 | 1 | 1 | 2 | 0.5 | 3 | 0 | 2 | 0.5 | 8 | 0.375 | NA | NA | 2 | 1 | 1 | 0 | 1 | 0 | 4 | 0.5 |
| 13_113004 | 4 | 0.25 | 3 | 0 | 3 | 0 | 2 | 0.5 | 12 | 0.1667 | 2 | 0 | 2 | 0 | 13 | 0 | 2 | 0.5 | 19 | 0.0526 |
| 13_113004 | 5 | 0.8 | 2 | 1 | 1 | 1 | 1 | 1 | 9 | 0.8889 | 2 | 1 | 2 | 0.5 | 10 | 0.7 | 2 | 1 | 16 | 0.75 |
| 13_113004 | 9 | 0 | 6 | 0.3333 | 3 | 0 | 7 | 0 | 25 | 0.08 | 2 | 0 | 3 | 0 | 10 | 0 | 3 | 0.3333 | 18 | 0.0556 |
| 13_113083 | NA | NA | 3 | 0.6667 | NA | NA | 3 | 1 | 6 | 0.8333 | NA | NA | 1 | 1 | 2 | 1 | NA | NA | 3 | 1 |
| 13_113096 | 1 | 0 | 5 | 0.2 | NA | NA | 3 | 0 | 9 | 0.1111 | 1 | 1 | 2 | 0 | 2 | 0 | NA | NA | 5 | 0.2 |
| 13_113101 | 2 | 0 | 4 | 0.5 | NA | NA | 4 | 0.25 | 10 | 0.3 | NA | NA | 2 | 1 | 3 | 0.3333 | NA | NA | 5 | 0.6 |
| 13_113105 | 1 | 0 | 9 | 0.3333 | NA | NA | 1 | 0 | 11 | 0.2727 | NA | NA | 2 | 0.5 | 4 | 0.25 | NA | NA | 6 | 0.3333 |
| 13_113728 | 1 | 0 | 3 | 0 | 1 | 0 | 2 | 0.5 | 7 | 0.1429 | 3 | 0.3333 | 2 | 0.5 | 3 | 0.3333 | 3 | 0 | 11 | 0.2727 |
| 13_113729 | NA | NA | 3 | 0.3333 | 1 | 0 | 2 | 0.5 | 6 | 0.3333 | 3 | 0 | 2 | 0 | 1 | 0 | 2 | 0.5 | 8 | 0.125 |
| 13_113729 | NA | NA | 3 | 0.3333 | 1 | 0 | 1 | 0 | 5 | 0.2 | 2 | 0 | 1 | 0 | 2 | 0.5 | 3 | 0 | 8 | 0.125 |
| 13_113729 | NA | NA | 1 | 1 | 1 | 0 | 2 | 0.5 | 4 | 0.5 | 2 | 0 | 2 | 0 | 2 | 0 | 3 | 0.3333 | 9 | 0.1111 |
| 13_113730 | NA | NA | 1 | 1 | 1 | 0 | 2 | 0 | 4 | 0.25 | 2 | 0.5 | 2 | 0 | 1 | 0 | 2 | 0 | 7 | 0.1429 |
| 13_113730 | NA | NA | NA | NA | 1 | 0 | 2 | 0.5 | 3 | 0.3333 | 1 | 0 | 1 | 1 | 1 | 0 | NA | NA | 3 | 0.3333 |
| 13_113730 | NA | NA | NA | NA | 1 | 0 | 2 | 0.5 | 3 | 0.3333 | 2 | 0.5 | 2 | 0.5 | 1 | 0 | NA | NA | 5 | 0.4 |
| 13_113730 | NA | NA | NA | NA | 1 | 0 | 1 | 1 | 2 | 0.5 | 2 | 0 | 2 | 0.5 | 1 | 0 | NA | NA | 5 | 0.2 |
| 13_113731 | NA | NA | NA | NA | 2 | 1 | 3 | 0.6667 | 5 | 0.8 | 2 | 0.5 | 2 | 1 | 1 | 1 | NA | NA | 5 | 0.8 |

| | | | | | | | | | | | | | | | | | | | | |
|---|---|---|---|---|---|---|---|---|---|---|---|---|---|---|---|---|---|---|---|---|
| 13_113731 | NA | NA | NA | NA | 2 | 0.5 | NA | NA | 2 | 0.5 | 2 | 1 | 2 | 1 | 1 | 1 | NA | NA | 5 | 1 |
| 13_113731 | NA | NA | 1 | 1 | 2 | 1 | 3 | 1 | 6 | 1 | 2 | 1 | 2 | 1 | NA | NA | NA | NA | 4 | 1 |
| 13_113753 | NA | NA | NA | NA | 2 | 1 | 4 | 1 | 6 | 1 | 2 | 0.5 | 3 | 0.6667 | NA | NA | NA | NA | 5 | 0.6 |
| 13_113753 | NA | NA | NA | NA | NA | NA | 1 | 1 | 1 | 1 | 2 | 0 | 3 | 0.3333 | NA | NA | NA | NA | 5 | 0.2 |
| 13_113753 | NA | NA | NA | NA | 2 | 0 | 3 | 0.3333 | 5 | 0.2 | 1 | 0 | 2 | 0.5 | NA | NA | NA | NA | 3 | 0.3333 |
| 13_113753 | NA | NA | NA | NA | 2 | 0.5 | 2 | 0 | 4 | 0.25 | NA | NA | 2 | 1 | NA | NA | NA | NA | 2 | 1 |
| 13_113799 | NA | NA | 1 | 1 | NA | NA | NA | NA | 1 | 1 | 2 | 0.5 | NA | NA | NA | NA | NA | NA | 2 | 0.5 |
| 13_113799 | NA | NA | 1 | 1 | NA | NA | NA | NA | 1 | 1 | 1 | 1 | NA | NA | NA | NA | NA | NA | 1 | 1 |
| 13_113799 | NA | NA | 1 | 1 | NA | NA | NA | NA | 1 | 1 | 2 | 0.5 | NA | NA | NA | NA | NA | NA | 2 | 0.5 |
| 13_113799 | NA | NA | 1 | 1 | NA | NA | NA | NA | 1 | 1 | 1 | 1 | NA | NA | NA | NA | NA | NA | 1 | 1 |
| 13_113800 | NA | NA | 1 | 1 | NA | NA | NA | NA | 1 | 1 | 1 | 1 | NA | NA | NA | NA | NA | NA | 1 | 1 |
| 13_114104 | 1 | 0 | NA | NA | NA | NA | 1 | 1 | 2 | 0.5 | NA | NA | 3 | 0.3333 | 1 | 0 | 1 | 0 | 5 | 0.2 |
| 13_114291 | 1 | 1 | NA | NA | NA | NA | 1 | 0 | 2 | 0.5 | NA | NA | 4 | 0.75 | 1 | 0 | 1 | 0 | 6 | 0.5 |
| 13_114291 | 1 | 1 | NA | NA | 1 | 1 | 1 | 1 | 3 | 1 | NA | NA | 4 | 1 | 1 | 1 | 2 | 1 | 7 | 1 |
| 13_114291 | 1 | 1 | NA | NA | 1 | 1 | 1 | 1 | 3 | 1 | NA | NA | 3 | 1 | 1 | 1 | 2 | 1 | 6 | 1 |
| 13_114291 | 1 | 1 | NA | NA | 1 | 1 | 1 | 1 | 3 | 1 | NA | NA | 3 | 1 | 1 | 1 | NA | NA | 4 | 1 |
| 13_114291 | 1 | 1 | NA | NA | 1 | 0 | 1 | 1 | 3 | 0.6667 | NA | NA | 4 | 1 | 1 | 1 | 1 | 1 | 6 | 1 |
| 13_114291 | 1 | 1 | NA | NA | 1 | 1 | 1 | 1 | 3 | 1 | NA | NA | 4 | 1 | 1 | 1 | 1 | 1 | 6 | 1 |
| 13_114291 | NA | NA | NA | NA | 1 | 1 | 1 | 0 | 2 | 0.5 | NA | NA | 5 | 0.4 | 1 | 1 | 2 | 0.5 | 8 | 0.5 |
| 13_114291 | NA | NA | NA | NA | 1 | 1 | 1 | 0 | 2 | 0.5 | NA | NA | 3 | 0.6667 | 2 | 0.5 | 1 | 1 | 6 | 0.6667 |
| 13_114292 | 1 | 1 | NA | NA | 1 | 0 | 1 | 0 | 3 | 0.3333 | NA | NA | 5 | 0.4 | 3 | 0.3333 | 2 | 0 | 10 | 0.3 |
| 13_114293 | 2 | 0.5 | 1 | 1 | 2 | 0.5 | 3 | 0.6667 | 8 | 0.625 | 1 | 0 | 3 | 1 | 6 | 0.5 | 3 | 1 | 13 | 0.6923 |
| 13_114293 | 4 | 0.5 | 1 | 1 | 2 | 0.5 | 3 | 0.6667 | 10 | 0.6 | 1 | 1 | 5 | 0.6 | 6 | 0.5 | 5 | 0.6 | 17 | 0.5882 |
| 13_114293 | 5 | 0 | 4 | 0 | 7 | 0.4286 | 3 | 0 | 19 | 0.1579 | 4 | 0 | 10 | 0.1 | 8 | 0.375 | 11 | 0.1818 | 33 | 0.1818 |
| 13_114294 | NA | NA | NA | NA | NA | NA | 1 | 1 | 1 | 1 | NA | NA | 1 | 1 | NA | NA | NA | NA | 1 | 1 |
| 13_114298 | NA | NA | NA | NA | NA | NA | 1 | 1 | 1 | 1 | 1 | 1 | 2 | 0.5 | 1 | 1 | NA | NA | 4 | 0.75 |
| 13_114298 | NA | NA | NA | NA | NA | NA | 1 | 1 | 1 | 1 | 1 | 1 | 2 | 1 | 1 | 0 | NA | NA | 4 | 0.75 |
| 13_114298 | NA | NA | NA | NA | NA | NA | 1 | 1 | 1 | 1 | 1 | 1 | 1 | 1 | 1 | 1 | NA | NA | 3 | 1 |
| 13_114424 | 2 | 0 | 2 | 0 | 1 | 0 | 2 | 0.5 | 7 | 0.1429 | NA | NA | 2 | 0.5 | 1 | 1 | 4 | 0 | 7 | 0.2857 |
| 13_114424 | 2 | 0.5 | 2 | 0 | 1 | 0 | 2 | 0.5 | 7 | 0.2857 | NA | NA | 2 | 0.5 | 1 | 0 | 4 | 0.25 | 7 | 0.2857 |
| 13_114425 | 26 | 0.0385 | 25 | 0 | 29 | 0.1724 | 29 | 0 | 109 | 0.055 | 19 | 0 | 28 | 0.0357 | 20 | 0.15 | 16 | 0.0625 | 83 | 0.0602 |
| 13_114964 | 8 | 0.25 | 14 | 0.2857 | 6 | 0.1667 | 12 | 0.3333 | 40 | 0.275 | 6 | 0.1667 | 10 | 0.3 | 10 | 0 | 6 | 0.1667 | 32 | 0.1562 |
| 13_114965 | 7 | 0 | 14 | 0.1429 | 7 | 0.1429 | 12 | 0 | 40 | 0.075 | 6 | 0 | 12 | 0.1667 | 11 | 0.2727 | 7 | 0 | 36 | 0.1389 |
| 13_115224 | 5 | 0 | 6 | 0 | 5 | 0.2 | 4 | 0.75 | 20 | 0.2 | 4 | 0.25 | 2 | 0 | NA | NA | 6 | 0 | 12 | 0.0833 |
| 13_115225 | 5 | 1 | 6 | 0.8333 | 4 | 1 | 5 | 1 | 20 | 0.95 | 4 | 0.5 | 2 | 0.5 | NA | NA | 6 | 0.8333 | 12 | 0.6667 |
| 13_115225 | 4 | 0.25 | 3 | 1 | 4 | 0.75 | 3 | 0.6667 | 14 | 0.6429 | 3 | 1 | 1 | 0 | NA | NA | 2 | 1 | 6 | 0.8333 |
| 13_115225 | 4 | 1 | 3 | 1 | 3 | 1 | 3 | 1 | 13 | 1 | 2 | 1 | 2 | 0.5 | NA | NA | 4 | 1 | 8 | 0.875 |
| 13_115225 | 4 | 1 | 6 | 1 | 5 | 1 | 4 | 1 | 19 | 1 | 2 | 1 | 2 | 0 | NA | NA | 6 | 1 | 10 | 0.8 |
| 13_115225 | 4 | 1 | 5 | 1 | 4 | 1 | 4 | 1 | 17 | 1 | 2 | 1 | 2 | 1 | NA | NA | 6 | 1 | 10 | 1 |
| 13_115225 | 5 | 0.2 | 6 | 0 | 4 | 0 | 4 | 0.25 | 19 | 0.1053 | 2 | 0 | 2 | 0 | NA | NA | 5 | 0.4 | 9 | 0.2222 |
| 13_115225 | 4 | 0 | 4 | 0.25 | 4 | 0 | 3 | 0.6667 | 15 | 0.2 | NA | NA | 1 | 0 | NA | NA | 6 | 0.1667 | 7 | 0.1429 |
| 13_115226 | 4 | 1 | 4 | 0.5 | 3 | 1 | 3 | 0.6667 | 14 | 0.7857 | NA | NA | NA | NA | NA | NA | 4 | 0.75 | 4 | 0.75 |
| 13_115226 | 4 | 0.25 | 3 | 0 | 3 | 0.3333 | 3 | 0 | 13 | 0.1538 | NA | NA | 1 | 1 | NA | NA | 5 | 0.2 | 6 | 0.3333 |
| 13_115226 | 4 | 0 | 5 | 0.6 | 4 | 0 | 2 | 0 | 15 | 0.2 | NA | NA | 1 | 0 | NA | NA | 5 | 0.4 | 6 | 0.3333 |
| 13_115226 | 4 | 0.25 | 5 | 0.2 | 4 | 0 | 3 | 0.6667 | 16 | 0.25 | NA | NA | 2 | 0.5 | NA | NA | 4 | 0 | 6 | 0.1667 |
| 13_115226 | 3 | 0 | 5 | 0.4 | 4 | 0 | 3 | 0.3333 | 15 | 0.2 | NA | NA | 1 | 1 | NA | NA | 4 | 0.5 | 5 | 0.6 |
| 13_115226 | 1 | 0 | 4 | 0.25 | 1 | 0 | 3 | 0.6667 | 9 | 0.3333 | NA | NA | 2 | 0.5 | NA | NA | 4 | 0 | 6 | 0.1667 |
| 13_116435 | 3 | 0.3333 | 1 | 0 | 1 | 1 | 5 | 0.4 | 10 | 0.4 | 2 | 0 | 5 | 0 | 4 | 0.5 | 1 | 0 | 12 | 0.1667 |
| 13_116435 | 3 | 0.3333 | NA | NA | 1 | 1 | 4 | 0.5 | 8 | 0.5 | 2 | 0.5 | 4 | 0.25 | 4 | 0.5 | 1 | 1 | 11 | 0.4545 |
| 13_116435 | 3 | 0.3333 | 1 | 0 | 1 | 1 | 4 | 0.75 | 9 | 0.5556 | 2 | 1 | 5 | 0.2 | 5 | 0.4 | 1 | 1 | 13 | 0.4615 |
| 13_116435 | 3 | 0.6667 | 1 | 1 | 1 | 1 | 4 | 1 | 9 | 0.8889 | 2 | 1 | 4 | 0.75 | 4 | 0.25 | 1 | 1 | 11 | 0.6364 |
| 13_116435 | 3 | 0.3333 | 1 | 1 | 1 | 0 | 5 | 0.4 | 10 | 0.4 | 2 | 0 | 5 | 0.4 | 3 | 0 | 1 | 1 | 11 | 0.2727 |
| 13_116435 | 2 | 0 | 1 | 0 | 1 | 0 | 4 | 0.25 | 8 | 0.125 | 1 | 0 | 4 | 0 | 4 | 0.25 | NA | NA | 9 | 0.1111 |
| 13_116435 | 3 | 0 | 1 | 0 | 1 | 0 | 5 | 0.4 | 10 | 0.2 | 1 | 0 | 5 | 0.2 | 3 | 0.3333 | 1 | 0 | 10 | 0.2 |
| 13_116435 | 3 | 0 | NA | NA | 1 | 0 | 4 | 0.25 | 8 | 0.125 | 2 | 0 | 5 | 0.4 | 4 | 0.5 | 1 | 1 | 12 | 0.4167 |
| 13_116435 | 3 | 0.3333 | 1 | 0 | 1 | 0 | 5 | 0.8 | 10 | 0.5 | 2 | 0.5 | 5 | 0.2 | 4 | 0.25 | 1 | 1 | 12 | 0.3333 |
| 13_116435 | 3 | 1 | 1 | 1 | NA | NA | 5 | 0.8 | 9 | 0.8889 | 2 | 1 | 5 | 0.6 | 2 | 1 | NA | NA | 9 | 0.7778 |
| 13_116436 | 2 | 0 | 1 | 0 | NA | NA | 4 | 0.75 | 7 | 0.4286 | 2 | 0.5 | 4 | 0.5 | 4 | 0.25 | 1 | 1 | 11 | 0.4545 |
| 13_117031 | 2 | 0.5 | NA | NA | 2 | 0 | 3 | 0.3333 | 7 | 0.2857 | NA | NA | 1 | 1 | NA | NA | 3 | 0 | 4 | 0.25 |
| 13_117031 | 2 | 0.5 | NA | NA | 2 | 0 | 2 | 0.5 | 6 | 0.3333 | NA | NA | 1 | 1 | NA | NA | 3 | 0.3333 | 4 | 0.5 |
| 13_117031 | 2 | 0 | 1 | 0 | 2 | 0 | 3 | 0.6667 | 8 | 0.25 | NA | NA | 1 | 0 | NA | NA | 3 | 0.3333 | 4 | 0.25 |
| 13_117031 | 2 | 0 | 1 | 1 | 2 | 0.5 | 3 | 0.6667 | 8 | 0.5 | NA | NA | 1 | 1 | NA | NA | 3 | 1 | 4 | 1 |
| 13_117031 | 2 | 0.5 | NA | NA | NA | NA | 2 | 0.5 | 4 | 0.5 | NA | NA | NA | NA | NA | NA | 2 | 0.5 | 2 | 0.5 |
| 13_117031 | 2 | 0.5 | 1 | 1 | 2 | 0.5 | 3 | 0.6667 | 8 | 0.625 | NA | NA | 1 | 1 | NA | NA | 3 | 1 | 4 | 1 |
| 13_117031 | 2 | 0.5 | 1 | 0 | 2 | 0.5 | 3 | 0.6667 | 8 | 0.5 | NA | NA | 1 | 0 | NA | NA | 3 | 0.6667 | 4 | 0.5 |

| | | | | | | | | | | | | | | | | | | | | |
|---|---|---|---|---|---|---|---|---|---|---|---|---|---|---|---|---|---|---|---|---|
| 13_117409 | 1 | 0 | NA | NA | 3 | 0.6667 | NA | NA | 4 | 0.5 | 1 | 0 | 3 | 0.6667 | 2 | 0 | 2 | 0.5 | 8 | 0.375 |
| 13_117409 | 1 | 0 | NA | NA | 5 | 0.4 | 1 | 0 | 7 | 0.2857 | 3 | 0 | 4 | 0 | 3 | 0 | 3 | 0.3333 | 13 | 0.0769 |
| 13_117410 | 3 | 0 | 1 | 0 | 5 | 0.4 | 2 | 0 | 11 | 0.1818 | 3 | 0.6667 | 3 | 0 | 2 | 1 | 3 | 0.3333 | 11 | 0.4545 |
| 13_117414 | 1 | 1 | NA | NA | 2 | 0 | 1 | 0 | 4 | 0.25 | 1 | 0 | NA | NA | 1 | 1 | NA | NA | 2 | 0.5 |
| 13_117736 | 1 | 0 | NA | NA | 1 | 1 | NA | NA | 2 | 0.5 | NA | NA | 2 | 0.5 | 1 | 1 | NA | NA | 3 | 0.6667 |
| 13_117737 | 1 | 1 | NA | NA | 1 | 0 | NA | NA | 2 | 0.5 | NA | NA | 2 | 0.5 | 1 | 0 | NA | NA | 3 | 0.3333 |
| 13_117737 | 1 | 1 | NA | NA | 1 | 0 | NA | NA | 2 | 0.5 | NA | NA | 2 | 0.5 | 1 | 0 | NA | NA | 3 | 0.3333 |
| 13_117737 | 1 | 1 | NA | NA | NA | NA | NA | NA | 1 | 1 | NA | NA | 2 | 0.5 | 1 | 0 | NA | NA | 3 | 0.3333 |
| 13_117737 | 1 | 1 | NA | NA | 1 | 1 | NA | NA | 2 | 1 | NA | NA | 2 | 0.5 | 1 | 1 | NA | NA | 3 | 0.6667 |
| 13_117737 | 1 | 1 | NA | NA | 1 | 1 | NA | NA | 2 | 1 | NA | NA | 2 | 0.5 | 1 | 0 | NA | NA | 3 | 0.3333 |
| 13_117793 | 2 | 0.5 | 1 | 1 | 1 | 1 | 2 | 0.5 | 6 | 0.6667 | 1 | 0 | NA | NA | 2 | 1 | NA | NA | 3 | 0.6667 |
| 13_117793 | 2 | 1 | 2 | 1 | 2 | 1 | 1 | 1 | 7 | 1 | 1 | 1 | NA | NA | 2 | 1 | NA | NA | 3 | 1 |
| 13_117793 | 2 | 1 | 1 | 0 | 2 | 1 | 2 | 1 | 7 | 0.8571 | 1 | 1 | NA | NA | 2 | 1 | 1 | 1 | 4 | 1 |
| 13_117793 | 2 | 0.5 | 2 | 0.5 | 2 | 0 | 2 | 0.5 | 8 | 0.375 | 1 | 0 | NA | NA | 2 | 0.5 | 1 | 1 | 4 | 0.5 |
| 13_117793 | 2 | 1 | 2 | 1 | 2 | 0 | 1 | 0 | 7 | 0.5714 | 1 | 1 | NA | NA | 2 | 1 | 1 | 1 | 4 | 1 |
| 13_117793 | 2 | 0.5 | 2 | 1 | 1 | 1 | 2 | 0 | 7 | 0.5714 | 1 | 0 | NA | NA | 1 | 1 | 1 | 0 | 3 | 0.3333 |
| 13_118111 | 4 | 0.75 | 4 | 0.25 | 3 | 0 | 2 | 0 | 13 | 0.3077 | 2 | 0 | NA | NA | 4 | 0 | 3 | 0.3333 | 9 | 0.1111 |
| 13_118112 | 3 | 1 | 4 | 1 | 3 | 1 | 2 | 1 | 12 | 1 | 3 | 1 | NA | NA | 4 | 1 | 2 | 1 | 9 | 1 |
| 13_118112 | 4 | 0.75 | 5 | 0.4 | 3 | 0 | 2 | 0 | 14 | 0.3571 | 3 | 0.3333 | 1 | 0 | 3 | 0.3333 | 2 | 0.5 | 9 | 0.3333 |
| 13_118112 | 4 | 0.25 | 4 | 0 | 3 | 0.6667 | 2 | 0.5 | 13 | 0.3077 | 2 | 1 | 1 | 0 | 3 | 0.6667 | 3 | 0.6667 | 9 | 0.6667 |
| 13_118112 | 3 | 0.3333 | 5 | 0.4 | 4 | 0.75 | 2 | 0 | 14 | 0.4286 | 3 | 0 | 1 | 0 | 4 | 0.25 | 3 | 0.6667 | 11 | 0.2727 |
| 13_118112 | 4 | 0.25 | 4 | 0.25 | 4 | 0 | 2 | 0.5 | 14 | 0.2143 | 3 | 0.6667 | 1 | 0 | 4 | 0.25 | 3 | 0.3333 | 11 | 0.3636 |
| 13_118112 | 4 | 0.75 | 4 | 0.75 | 3 | 1 | 2 | 1 | 13 | 0.8462 | 3 | 0.6667 | 2 | 1 | 3 | 0.6667 | 3 | 1 | 11 | 0.8182 |
| 13_118112 | 4 | 0.25 | 5 | 0 | 3 | 0 | 2 | 0 | 14 | 0.0714 | 3 | 0.3333 | 2 | 0 | 4 | 0 | 2 | 0 | 11 | 0.0909 |
| 13_118112 | 3 | 0.3333 | 4 | 1 | 2 | 0 | 1 | 0 | 10 | 0.5 | 2 | 0 | 2 | 1 | 4 | 0.5 | 1 | 1 | 9 | 0.5556 |
| 13_118112 | 3 | 0.3333 | 3 | 0.3333 | 4 | 0 | 2 | 0 | 12 | 0.1667 | 3 | 0 | 1 | 0 | 4 | 0 | 2 | 0.5 | 10 | 0.1 |
| 13_118112 | 3 | 1 | 5 | 0.8 | 4 | 1 | 2 | 1 | 14 | 0.9286 | 3 | 1 | 2 | 1 | 5 | 0.8 | 3 | 1 | 13 | 0.9231 |
| 13_118112 | 4 | 0 | 2 | 0 | 3 | 0.3333 | 1 | 0 | 10 | 0.1 | 2 | 0 | 1 | 1 | 5 | 0 | 2 | 0 | 10 | 0.1 |
| 13_118113 | 3 | 0.6667 | 4 | 0.25 | 4 | 0 | 1 | 0 | 12 | 0.25 | 2 | 0 | 1 | 0 | 4 | 0.25 | 3 | 0.3333 | 10 | 0.2 |
| 13_118113 | 2 | 1 | 4 | 0.75 | 6 | 1 | 1 | 1 | 13 | 0.9231 | 2 | 0.5 | 2 | 1 | 3 | 1 | 1 | 1 | 8 | 0.875 |
| 13_118113 | 3 | 0.3333 | 4 | 0 | 6 | 0 | 2 | 0 | 15 | 0.0667 | 2 | 0 | 2 | 0.5 | 3 | 0 | 3 | 0 | 10 | 0.1 |
| 13_118113 | 2 | 0 | 5 | 0 | 6 | 0 | 2 | 0.5 | 15 | 0.0667 | 3 | 0 | 2 | 0 | 4 | 0 | 4 | 0.25 | 13 | 0.0769 |
| 13_118114 | 2 | 1 | 3 | 1 | 5 | 0.6 | NA | NA | 10 | 0.8 | 2 | 1 | 1 | 1 | 4 | 0.75 | 3 | 1 | 10 | 0.9 |
| 13_118114 | 2 | 0.5 | 4 | 0 | 6 | 0 | NA | NA | 12 | 0.0833 | 2 | 0 | 2 | 0 | 4 | 0.25 | 4 | 0 | 12 | 0.0833 |
| 13_118114 | 2 | 0 | 4 | 0.25 | 6 | 0 | 1 | 0 | 13 | 0.0769 | 2 | 0 | 3 | 0 | 4 | 0 | 4 | 0.25 | 13 | 0.0769 |
| 13_118115 | 1 | 1 | 1 | 0 | 7 | 0 | 1 | 0 | 10 | 0.1 | 2 | 0 | 3 | 0.3333 | 4 | 0.5 | 4 | 0 | 13 | 0.2308 |
| 13_118115 | 1 | 0 | 1 | 0 | 7 | 0.1429 | 1 | 0 | 10 | 0.1 | 1 | 0 | 3 | 0 | 2 | 0.5 | 4 | 0 | 10 | 0.1 |
| 13_118116 | 1 | 1 | 1 | 0 | 2 | 0 | NA | NA | 4 | 0.25 | 2 | 0 | 3 | 0.3333 | 2 | 0 | 3 | 0 | 10 | 0.1 |
| 13_118116 | 1 | 0 | 2 | 0 | 9 | 0.2222 | 1 | 0 | 13 | 0.1538 | 2 | 0.5 | 4 | 0 | 2 | 0 | 4 | 0 | 12 | 0.0833 |
| 13_118117 | 1 | 1 | 1 | 1 | 9 | 0.5556 | 2 | 0.5 | 13 | 0.6154 | 3 | 0.3333 | 5 | 0.4 | 3 | 0 | 4 | 0.5 | 15 | 0.3333 |
| 13_118117 | NA | NA | 2 | 1 | 8 | 0.375 | 1 | 0 | 11 | 0.4545 | 2 | 1 | 4 | 1 | 3 | 0.3333 | 4 | 0.75 | 13 | 0.7692 |
| 13_118117 | 1 | 0 | 2 | 0.5 | 10 | 0.7 | 3 | 0.3333 | 16 | 0.5625 | 2 | 1 | 3 | 0.6667 | 2 | 0.5 | 5 | 0.4 | 12 | 0.5833 |
| 13_118117 | 1 | 1 | 2 | 1 | 12 | 0.9167 | 3 | 0.6667 | 18 | 0.8889 | 2 | 1 | 2 | 1 | 2 | 0.5 | 5 | 1 | 11 | 0.9091 |
| 13_118117 | 1 | 0 | 3 | 0.6667 | 12 | 0.5833 | 3 | 1 | 19 | 0.6316 | 3 | 0.6667 | 5 | 1 | 2 | 0 | 5 | 0.8 | 15 | 0.7333 |
| 13_118516 | 6 | 0.1667 | 4 | 0 | 1 | 0 | 2 | 0 | 13 | 0.0769 | 3 | 0.3333 | 6 | 0.1667 | 7 | 0 | 2 | 0 | 18 | 0.1111 |
| 13_118516 | 8 | 0.75 | 6 | 0.5 | 1 | 0 | 4 | 0.5 | 19 | 0.5789 | 7 | 0.7143 | 6 | 0.6667 | 11 | 0.7273 | 2 | 1 | 26 | 0.7308 |
| 13_118516 | 7 | 0.2857 | 6 | 0.1667 | 1 | 1 | 4 | 0.75 | 18 | 0.3889 | 8 | 0.375 | 6 | 0.3333 | 11 | 0.2727 | 2 | 0 | 27 | 0.2963 |
| 13_118517 | 7 | 0.4286 | 5 | 0 | NA | NA | 4 | 0 | 16 | 0.1875 | 6 | 0.1667 | 6 | 0 | 10 | 0.2 | 2 | 0 | 24 | 0.125 |
| 13_118517 | 4 | 1 | 5 | 1 | 1 | 1 | 2 | 1 | 12 | 1 | 5 | 0.8 | 6 | 1 | 9 | 0.8889 | 2 | 1 | 22 | 0.9091 |
| 13_118523 | NA | NA | NA | NA | NA | NA | 1 | 1 | 1 | 1 | NA | NA | NA | NA | 2 | 1 | NA | NA | 2 | 1 |
| 13_118523 | NA | NA | NA | NA | NA | NA | 1 | 1 | 1 | 1 | NA | NA | NA | NA | 2 | 1 | NA | NA | 2 | 1 |
| 13_118851 | NA | NA | 1 | 1 | 1 | 1 | NA | NA | 2 | 1 | NA | NA | 1 | 1 | NA | NA | NA | NA | 1 | 1 |
| 13_118851 | NA | NA | 1 | 1 | NA | NA | NA | NA | 1 | 1 | NA | NA | 1 | 1 | NA | NA | NA | NA | 1 | 1 |
| 13_118851 | NA | NA | 1 | 1 | NA | NA | NA | NA | 1 | 1 | NA | NA | 1 | 1 | NA | NA | NA | NA | 1 | 1 |
| 13_118925 | NA | NA | 2 | 1 | NA | NA | NA | NA | 2 | 1 | NA | NA | NA | NA | 1 | 1 | 1 | 1 | 2 | 1 |
| 13_118925 | NA | NA | 3 | 0.6667 | NA | NA | NA | NA | 3 | 0.6667 | NA | NA | 1 | 1 | 1 | 1 | NA | NA | 2 | 1 |
| 13_118925 | NA | NA | 3 | 0.6667 | NA | NA | NA | NA | 3 | 0.6667 | NA | NA | NA | NA | 1 | 1 | NA | NA | 1 | 1 |
| 13_118925 | NA | NA | 3 | 0.3333 | NA | NA | NA | NA | 3 | 0.3333 | NA | NA | 1 | 1 | 1 | 0 | 1 | 0 | 3 | 0.3333 |
| 13_118926 | NA | NA | 3 | 1 | NA | NA | NA | NA | 3 | 1 | NA | NA | 1 | 1 | 1 | 1 | 1 | 1 | 3 | 1 |
| 13_119472 | 4 | 1 | 3 | 0.3333 | 3 | 0.6667 | 4 | 0.75 | 14 | 0.7143 | 1 | 1 | NA | NA | 3 | 0.3333 | 3 | 1 | 7 | 0.7143 |
| 13_119472 | 6 | 0.5 | 3 | 0.6667 | 2 | 0.5 | 4 | 0.25 | 15 | 0.4667 | 1 | 1 | NA | NA | 2 | 1 | 5 | 0 | 8 | 0.375 |
| 13_119565 | NA | NA | NA | NA | 4 | 0.25 | 3 | 0.3333 | 7 | 0.2857 | 3 | 0.3333 | 1 | 0 | 6 | 0 | 3 | 0 | 13 | 0.0769 |
| 13_119599 | 1 | 0 | 4 | 0.25 | 1 | 0 | NA | NA | 6 | 0.1667 | 1 | 0 | 1 | 1 | 1 | 0 | 5 | 0.6 | 8 | 0.5 |
| 13_119602 | 1 | 0 | 2 | 0.5 | NA | NA | NA | NA | 3 | 0.3333 | 1 | 0 | NA | NA | 1 | 1 | 4 | 0 | 6 | 0.1667 |
| 13_119606 | 1 | 0 | 4 | 0.5 | 1 | 0 | NA | NA | 6 | 0.3333 | 2 | 0 | 3 | 0.6667 | 2 | 0.5 | 5 | 0.2 | 12 | 0.3333 |

| | | | | | | | | | | | | | | | | | | | | |
|---|---|---|---|---|---|---|---|---|---|---|---|---|---|---|---|---|---|---|---|---|
| 13_119606 | 1 | 1 | 4 | 1 | 1 | 1 | NA | NA | 6 | 1 | 2 | 1 | 4 | 0.25 | 3 | 0.6667 | 4 | 0.5 | 13 | 0.5385 |
| 13_119606 | 1 | 0 | 4 | 0.5 | NA | NA | NA | NA | 5 | 0.4 | 2 | 0 | 2 | 0.5 | 2 | 0 | 3 | 0 | 9 | 0.1111 |
| 13_119606 | 1 | 1 | 3 | 0 | 1 | 0 | NA | NA | 5 | 0.2 | 2 | 0 | 4 | 0.25 | 3 | 0 | 3 | 0 | 12 | 0.0833 |
| 13_119606 | 2 | 1 | 4 | 0.75 | 1 | 1 | NA | NA | 7 | 0.8571 | 2 | 0.5 | 4 | 0.75 | 3 | 0.3333 | 3 | 0 | 12 | 0.4167 |
| 13_119606 | 1 | 0 | 3 | 0.6667 | 1 | 0 | NA | NA | 5 | 0.4 | 1 | 0 | 3 | 1 | 4 | 0 | 3 | 0.3333 | 11 | 0.3636 |
| 13_119606 | 2 | 1 | 4 | 0.75 | 1 | 0 | NA | NA | 7 | 0.7143 | 1 | 1 | 3 | 1 | 2 | 0.5 | 3 | 0.3333 | 9 | 0.6667 |
| 13_119606 | 2 | 0 | 4 | 0.25 | 1 | 0 | NA | NA | 7 | 0.1429 | 2 | 0.5 | 4 | 0 | 4 | 0.25 | 5 | 0 | 15 | 0.1333 |
| 13_119606 | 2 | 0 | 4 | 0.25 | 1 | 1 | NA | NA | 7 | 0.2857 | 2 | 0 | 3 | 0 | 3 | 0.3333 | 3 | 0 | 11 | 0.0909 |
| 13_119622 | 3 | 0.3333 | 6 | 0.6667 | NA | NA | 2 | 0.5 | 11 | 0.5455 | 1 | 1 | 5 | 0 | 10 | 0.2 | 7 | 0.2857 | 23 | 0.2174 |
| 13_119622 | 4 | 0 | 9 | 0.2222 | 1 | 1 | 2 | 0 | 16 | 0.1875 | 2 | 1 | 6 | 0.1667 | 9 | 0.2222 | 8 | 0.5 | 25 | 0.36 |
| 13_119623 | NA | NA | NA | NA | NA | NA | 1 | 1 | 1 | 1 | NA | NA | NA | NA | 1 | 1 | NA | NA | 1 | 1 |
| 13_119623 | NA | NA | NA | NA | NA | NA | 1 | 1 | 1 | 1 | NA | NA | NA | NA | 1 | 1 | NA | NA | 1 | 1 |
| 13_119623 | NA | NA | NA | NA | NA | NA | 1 | 1 | 1 | 1 | NA | NA | NA | NA | 1 | 1 | NA | NA | 1 | 1 |
| 13_119624 | NA | NA | NA | NA | NA | NA | 1 | 1 | 1 | 1 | NA | NA | NA | NA | 1 | 1 | NA | NA | 1 | 1 |
| 13_119624 | NA | NA | NA | NA | NA | NA | 1 | 1 | 1 | 1 | NA | NA | NA | NA | 1 | 1 | NA | NA | 1 | 1 |
| 13_120068 | 2 | 1 | 3 | 1 | 7 | 1 | 2 | 1 | 14 | 1 | 2 | 1 | 4 | 1 | 7 | 1 | 4 | 1 | 17 | 1 |
| 13_120068 | 3 | 0.3333 | 3 | 0.6667 | 8 | 0.5 | 1 | 1 | 15 | 0.5333 | 2 | 0.5 | 4 | 0.5 | 6 | 0.3333 | 4 | 0 | 16 | 0.3125 |
| 13_120068 | NA | NA | 1 | 0 | 5 | 0.6 | 2 | 1 | 8 | 0.625 | NA | NA | 3 | 0.6667 | 7 | 0.4286 | 2 | 0 | 12 | 0.4167 |
| 13_120068 | 1 | 0 | 3 | 0.3333 | 8 | 0.75 | 1 | 0 | 13 | 0.5385 | 2 | 1 | 4 | 1 | 6 | 0.8333 | 3 | 1 | 15 | 0.9333 |
| 13_120070 | 3 | 1 | 3 | 0.6667 | 8 | 0.375 | 2 | 1 | 16 | 0.625 | 2 | 1 | 3 | 1 | 5 | 0.8 | 3 | 0.6667 | 13 | 0.8462 |
| 13_120070 | 1 | 0 | 2 | 0.5 | 6 | 0.1667 | 1 | 1 | 10 | 0.3 | 2 | 0 | 2 | 0.5 | 4 | 0.25 | 2 | 0.5 | 10 | 0.3 |
| 13_120070 | 3 | 0.3333 | 2 | 0 | 7 | 0 | 2 | 0.5 | 14 | 0.1429 | 2 | 0 | 4 | 0 | 6 | 0.1667 | 2 | 0 | 14 | 0.0714 |
| 13_120071 | 1 | 0 | 3 | 0.3333 | 8 | 0.5 | 2 | 0.5 | 14 | 0.4286 | 2 | 0.5 | 4 | 0.25 | 7 | 0.5714 | 1 | 1 | 14 | 0.5 |
| 13_120071 | 2 | 0 | 3 | 0 | 9 | 0 | 2 | 0.5 | 16 | 0.0625 | 3 | 0 | 4 | 0 | 8 | 0.125 | 4 | 0 | 19 | 0.0526 |
| 13_120071 | 2 | 0 | 3 | 0.3333 | 13 | 0.0769 | 2 | 0 | 20 | 0.1 | 2 | 0 | 4 | 0 | 8 | 0.125 | 4 | 0.25 | 18 | 0.1111 |
| 13_120071 | 3 | 0.3333 | 3 | 0.6667 | 12 | 0.6667 | 2 | 0.5 | 20 | 0.6 | 2 | 0.5 | 3 | 0.6667 | 6 | 0.1667 | 4 | 0.25 | 15 | 0.3333 |
| 13_120071 | 2 | 0.5 | 3 | 0 | 11 | 0 | 1 | 0 | 17 | 0.0588 | 2 | 0 | 2 | 0 | 6 | 0.3333 | 3 | 0 | 13 | 0.1538 |
| 13_120071 | 1 | 0 | 3 | 0 | 12 | 0.0833 | 1 | 0 | 17 | 0.0588 | 2 | 0.5 | 4 | 0.25 | 8 | 0.125 | 3 | 0 | 17 | 0.1765 |
| 13_120071 | 3 | 0 | 3 | 0.3333 | 13 | 0.1538 | 2 | 0 | 21 | 0.1429 | 3 | 0.3333 | 4 | 0.25 | 8 | 0.375 | 4 | 0.25 | 19 | 0.3158 |
| 13_120073 | NA | NA | NA | NA | 1 | 1 | NA | NA | 1 | 1 | NA | NA | NA | NA | 1 | 1 | NA | NA | 1 | 1 |
| 13_120073 | NA | NA | NA | NA | 1 | 1 | NA | NA | 1 | 1 | NA | NA | NA | NA | 1 | 1 | NA | NA | 1 | 1 |
| 13_120074 | NA | NA | NA | NA | 1 | 1 | NA | NA | 1 | 1 | NA | NA | NA | NA | 1 | 1 | NA | NA | 1 | 1 |
| 13_120161 | 10 | 0.6 | 12 | 0.4167 | 10 | 0.4 | 10 | 0.4 | 42 | 0.4524 | 8 | 0.375 | 9 | 0.2222 | 7 | 0.4286 | 12 | 0.5 | 36 | 0.3889 |
| 13_120161 | 8 | 0.25 | 13 | 0.4615 | 8 | 0.25 | 10 | 0.1 | 39 | 0.2821 | 7 | 0.2857 | 10 | 0.3 | 8 | 0.375 | 12 | 0.5833 | 37 | 0.4054 |
| 13_120161 | 10 | 0.1 | 12 | 0.25 | 9 | 0.1111 | 10 | 0 | 41 | 0.122 | 7 | 0.1429 | 10 | 0 | 7 | 0.4286 | 10 | 0.4 | 34 | 0.2353 |
| 13_120161 | 10 | 0.4 | 14 | 0.4286 | 8 | 0.125 | 9 | 0.2222 | 41 | 0.3171 | 8 | 0.25 | 10 | 0.2 | 8 | 0.625 | 12 | 0.4167 | 38 | 0.3684 |
| 13_120168 | 17 | 0.2353 | 19 | 0.6316 | 22 | 0.2273 | 15 | 0.4 | 73 | 0.3699 | 9 | 0.4444 | 14 | 0.2143 | 11 | 0.4545 | 16 | 0.25 | 50 | 0.32 |
| 13_120168 | 19 | 0.1579 | 19 | 0.1579 | 17 | 0.2353 | 15 | 0.2 | 70 | 0.1857 | 12 | 0.25 | 14 | 0.1429 | 10 | 0.2 | 17 | 0.2353 | 53 | 0.2075 |
| 13_120168 | 19 | 0 | 21 | 0 | 22 | 0.0455 | 14 | 0.2143 | 76 | 0.0526 | 11 | 0.2727 | 15 | 0.1333 | 11 | 0 | 17 | 0 | 54 | 0.0926 |
| 13_120168 | 17 | 0.1176 | 17 | 0.1176 | 20 | 0.15 | 17 | 0.1176 | 71 | 0.1268 | 11 | 0.1818 | 11 | 0 | 10 | 0.1 | 13 | 0.0769 | 45 | 0.0889 |
| 13_120168 | 26 | 0.0385 | 28 | 0.0714 | 24 | 0.0417 | 19 | 0.1053 | 97 | 0.0619 | 12 | 0.1667 | 16 | 0 | 14 | 0.0714 | 19 | 0.0526 | 61 | 0.0656 |
| 13_120173 | 4 | 0.5 | 5 | 0.6 | 5 | 0.6 | 9 | 0.2222 | 23 | 0.4348 | 3 | 0 | 2 | 1 | 3 | 0.3333 | 1 | 1 | 9 | 0.4444 |
| 13_120173 | 4 | 0 | 4 | 0 | 5 | 0.2 | 7 | 0 | 20 | 0.05 | 2 | 0.5 | 1 | 0 | 4 | 0 | 1 | 0 | 8 | 0.125 |
| 13_120173 | 5 | 0.4 | 5 | 0 | 5 | 0.4 | 9 | 0.1111 | 24 | 0.2083 | 3 | 0.6667 | 1 | 1 | 5 | 0.2 | 1 | 0 | 10 | 0.4 |
| 13_120173 | 4 | 1 | 3 | 0.6667 | 4 | 1 | 6 | 0.8333 | 17 | 0.8824 | 3 | 1 | 1 | 1 | 4 | 1 | 1 | 1 | 9 | 1 |
| 13_120173 | 4 | 0.25 | 4 | 0.25 | 3 | 0.6667 | 6 | 0 | 17 | 0.2353 | 3 | 0.6667 | 2 | 0.5 | 5 | 0.2 | 1 | 0 | 11 | 0.3636 |
| 13_120173 | 4 | 0.5 | 4 | 0.25 | 4 | 0.5 | 7 | 0.2857 | 19 | 0.3684 | 2 | 0 | 2 | 1 | 4 | 0.5 | NA | NA | 8 | 0.5 |
| 13_120173 | 3 | 1 | 2 | 0.5 | 2 | 1 | 5 | 0.8 | 12 | 0.8333 | 2 | 1 | 2 | 1 | 4 | 1 | NA | NA | 8 | 1 |
| 13_120173 | 2 | 0 | 2 | 0.5 | 5 | 0 | 7 | 0.5714 | 16 | 0.3125 | 3 | 0.3333 | 1 | 1 | 3 | 0 | NA | NA | 7 | 0.2857 |
| 13_120174 | 5 | 0.2 | 5 | 0.2 | 5 | 0.2 | 7 | 0 | 22 | 0.1364 | 3 | 0 | 2 | 0.5 | 4 | 0 | 1 | 0 | 10 | 0.1 |
| 13_120174 | 4 | 0.5 | 4 | 1 | 4 | 0.5 | 5 | 0.4 | 17 | 0.5882 | 3 | 1 | 2 | 0.5 | 3 | 0.6667 | 1 | 1 | 9 | 0.7778 |
| 13_120174 | 3 | 1 | 2 | 1 | 4 | 1 | 6 | 1 | 15 | 1 | 2 | 0.5 | 1 | 1 | 3 | 1 | NA | NA | 6 | 0.8333 |
| 13_120174 | 5 | 1 | 4 | 1 | 4 | 1 | 4 | 0.75 | 17 | 0.9412 | 2 | 0.5 | 1 | 0 | 4 | 1 | 1 | 1 | 8 | 0.75 |
| 13_120175 | 5 | 0 | 3 | 0.3333 | 3 | 0 | 7 | 0 | 18 | 0.0556 | 1 | 0 | 2 | 0.5 | 4 | 0 | 1 | 0 | 8 | 0.125 |
| 13_120189 | 3 | 0.3333 | 5 | 0.4 | 4 | 0 | 6 | 0.6667 | 18 | 0.3889 | 2 | 0.5 | 1 | 0 | 4 | 0 | 1 | 0 | 8 | 0.125 |
| 13_120189 | 2 | 1 | 4 | 0.5 | 1 | 0 | 6 | 0.1667 | 13 | 0.3846 | 2 | 0.5 | 2 | 0.5 | 4 | 0.25 | 1 | 0 | 9 | 0.3333 |
| 13_120189 | 2 | 0 | 3 | 0.3333 | 3 | 0.3333 | 4 | 0 | 12 | 0.1667 | 2 | 0.5 | 1 | 0 | 3 | 0 | NA | NA | 6 | 0.1667 |
| 13_120189 | 3 | 0 | 4 | 0.5 | 4 | 0 | 5 | 0.2 | 16 | 0.1875 | 1 | 0 | 1 | 0 | 4 | 0.25 | 1 | 1 | 7 | 0.2857 |
| 13_120189 | 3 | 0.3333 | 5 | 0.4 | 4 | 0 | 5 | 0.2 | 17 | 0.2353 | 1 | 1 | 2 | 0 | 4 | 0 | 1 | 0 | 8 | 0.125 |
| 13_120189 | 3 | 0 | 2 | 0 | 4 | 0.25 | 6 | 0 | 15 | 0.0667 | 2 | 0.5 | 1 | 0 | 3 | 0 | 1 | 0 | 7 | 0.1429 |
| 13_120189 | 1 | 0 | 3 | 0 | 4 | 0.5 | 5 | 0.2 | 13 | 0.2308 | 2 | 0 | 1 | 1 | 2 | 0 | 1 | 0 | 6 | 0.1667 |
| 13_120189 | 3 | 0 | 3 | 0 | 4 | 0.25 | 6 | 0 | 16 | 0.0625 | 2 | 0 | NA | NA | 3 | 0.3333 | 1 | 0 | 6 | 0.1667 |
| 13_120189 | 3 | 0.3333 | 4 | 0.5 | 2 | 1 | 4 | 0.5 | 13 | 0.5385 | 2 | 0.5 | 1 | 0 | 2 | 0 | 1 | 0 | 6 | 0.1667 |
| 13_120189 | 1 | 0 | 3 | 0 | 4 | 0.25 | 5 | 0.2 | 13 | 0.1538 | 2 | 0.5 | 1 | 0 | 2 | 1 | 1 | 0 | 6 | 0.5 |
| 13_120189 | 2 | 1 | 2 | 0.5 | 4 | 1 | 3 | 1 | 11 | 0.9091 | 2 | 1 | 1 | 1 | 2 | 0.5 | 1 | 1 | 6 | 0.8333 |

| | | | | | | | | | | | | | | | | | | | | |
|---|---|---|---|---|---|---|---|---|---|---|---|---|---|---|---|---|---|---|---|---|
| 13_1201898 | 2 | 0.5 | 4 | 1 | 4 | 1 | 3 | 1 | 13 | 0.9231 | 2 | 1 | 1 | 0 | 2 | 1 | 1 | 1 | 6 | 0.8333 |
| 13_1201898 | 2 | 0.5 | 4 | 0.25 | 3 | 0 | 5 | 0.2 | 14 | 0.2143 | 2 | 0 | 1 | 1 | 1 | 0 | 1 | 0 | 5 | 0.2 |
| 13_1201899 | 2 | 0.5 | 2 | 1 | 4 | 0.75 | 4 | 0.75 | 12 | 0.75 | 2 | 1 | 1 | 1 | 2 | 0.5 | NA | NA | 5 | 0.8 |
| 13_1202268 | 5 | 0.2 | 3 | 0 | 7 | 0 | 2 | 0 | 17 | 0.0588 | 3 | 0 | 10 | 0.1 | 5 | 0 | 3 | 0.3333 | 21 | 0.0952 |
| 13_1202277 | 2 | 0 | 3 | 1 | 3 | 0.3333 | NA | NA | 8 | 0.5 | 2 | 0.5 | 4 | 0.5 | 4 | 0.25 | 1 | 0 | 11 | 0.3636 |
| 13_1202278 | 1 | 1 | 4 | 0.5 | 7 | 0.7143 | NA | NA | 12 | 0.6667 | 2 | 1 | 6 | 0.8333 | 4 | 0.5 | 2 | 0.5 | 14 | 0.7143 |
| 13_1202278 | 2 | 0.5 | 3 | 0 | 5 | 0.4 | NA | NA | 10 | 0.3 | 1 | 0 | 5 | 0.6 | 2 | 0.5 | 1 | 0 | 9 | 0.4444 |
| 13_1202279 | 3 | 1 | 4 | 0.75 | 8 | 0.25 | NA | NA | 15 | 0.5333 | 2 | 0.5 | 7 | 0.1429 | 3 | 0.6667 | 2 | 1 | 14 | 0.4286 |
| 13_1202279 | 3 | 0 | 4 | 0.5 | 7 | 0.4286 | NA | NA | 14 | 0.3571 | 1 | 0 | 7 | 0.4286 | 3 | 0.3333 | 1 | 1 | 12 | 0.4167 |
| 13_1202280 | 2 | 0.5 | 4 | 1 | 4 | 1 | NA | NA | 10 | 0.9 | NA | NA | 3 | 0.6667 | 2 | 1 | NA | NA | 5 | 0.8 |
| 13_1202281 | 3 | 0.6667 | 2 | 0 | 6 | 0.5 | NA | NA | 11 | 0.4545 | 1 | 0 | 7 | 0.1429 | 3 | 0.3333 | 1 | 0 | 12 | 0.1667 |
| 13_1202282 | 3 | 0 | NA | NA | 7 | 0.2857 | NA | NA | 10 | 0.2 | 2 | 0 | 7 | 0.2857 | 1 | 0 | 1 | 0 | 11 | 0.1818 |
| 13_1202282 | 2 | 0 | 2 | 0.5 | 3 | 0.3333 | NA | NA | 7 | 0.2857 | 2 | 0 | 6 | 0.3333 | 3 | 0.6667 | 2 | 0.5 | 13 | 0.3846 |
| 13_1202283 | 2 | 1 | 1 | 1 | 4 | 1 | NA | NA | 7 | 1 | 2 | 1 | 5 | 0.8 | 3 | 1 | 2 | 0.5 | 12 | 0.8333 |
| 13_1202283 | 3 | 0.3333 | 3 | 0 | 6 | 0.3333 | NA | NA | 12 | 0.25 | 2 | 0 | 5 | 0.6 | 3 | 0 | 2 | 0 | 12 | 0.25 |
| 13_1202283 | 3 | 0 | 3 | 0 | 6 | 0.1667 | NA | NA | 12 | 0.0833 | 2 | 0 | 5 | 0 | 3 | 0 | 2 | 0.5 | 12 | 0.0833 |
| 13_1202283 | 3 | 0 | 1 | 0 | 7 | 0.2857 | NA | NA | 11 | 0.1818 | 2 | 0 | 4 | 0.5 | 3 | 0 | 1 | 1 | 10 | 0.3 |
| 13_1202284 | 3 | 0.6667 | 2 | 0.5 | 4 | 0.5 | NA | NA | 9 | 0.5556 | 2 | 1 | 3 | 0.6667 | 2 | 0.5 | 1 | 1 | 8 | 0.75 |
| 13_1202373 | 2 | 0.5 | 3 | 0.6667 | 6 | 0.5 | NA | NA | 11 | 0.5455 | 3 | 0.6667 | 4 | 0.25 | 2 | 0 | 2 | 1 | 11 | 0.4545 |
| 13_1202373 | 2 | 0.5 | 2 | 0 | 5 | 0 | NA | NA | 9 | 0.1111 | NA | NA | 5 | 0 | 3 | 0.3333 | 1 | 0 | 9 | 0.1111 |
| 13_1202373 | 3 | 0.6667 | 5 | 0.8 | 6 | 0.8333 | NA | NA | 14 | 0.7857 | 2 | 1 | 4 | 1 | 3 | 1 | 3 | 1 | 12 | 1 |
| 13_1202375 | 3 | 0.3333 | 5 | 0.2 | 4 | 0 | NA | NA | 12 | 0.1667 | 1 | 0 | 4 | 0.5 | 2 | 0.5 | 2 | 0 | 9 | 0.3333 |
| 13_1202375 | 1 | 0 | 3 | 0 | 4 | 0.25 | NA | NA | 8 | 0.125 | 1 | 0 | 2 | 0.5 | 1 | 0 | NA | NA | 4 | 0.25 |
| 13_1202375 | 1 | 1 | 3 | 1 | 3 | 1 | NA | NA | 7 | 1 | 1 | 1 | 2 | 0.5 | 1 | 1 | NA | NA | 4 | 0.75 |
| 13_1202376 | 1 | 0 | 3 | 0.6667 | 4 | 0.25 | NA | NA | 8 | 0.375 | 1 | 1 | 2 | 0 | 2 | 0 | NA | NA | 5 | 0.2 |
| 13_1202376 | 2 | 0 | 5 | 0.4 | 6 | 0.5 | NA | NA | 13 | 0.3846 | 2 | 0 | 5 | 0.2 | 3 | 0.3333 | 1 | 0 | 11 | 0.1818 |
| 13_1202376 | 2 | 0 | 4 | 1 | 5 | 0.4 | NA | NA | 11 | 0.5455 | 2 | 0.5 | 5 | 0.2 | 3 | 0.3333 | 1 | 0 | 11 | 0.2727 |
| 13_1202379 | 3 | 0 | 5 | 0 | 7 | 0.1429 | NA | NA | 15 | 0.0667 | 1 | 0 | 5 | 0.2 | 3 | 0 | 2 | 0 | 11 | 0.0909 |
| 14_3936907 | 1 | 0 | 2 | 0 | 3 | 0.6667 | NA | NA | 6 | 0.3333 | 2 | 0.5 | NA | NA | 1 | 1 | NA | NA | 3 | 0.6667 |
| 14_3936958 | 1 | 1 | 1 | 0 | 3 | 0 | NA | NA | 5 | 0.2 | 2 | 0.5 | NA | NA | 1 | 0 | NA | NA | 3 | 0.3333 |
| 14_3936968 | 1 | 1 | 1 | 0 | 2 | 0.5 | NA | NA | 4 | 0.5 | 1 | 1 | NA | NA | 1 | 1 | NA | NA | 2 | 1 |
| 14_3936980 | 1 | 1 | 1 | 0 | 3 | 0.3333 | NA | NA | 5 | 0.4 | 2 | 0.5 | NA | NA | 1 | 1 | NA | NA | 3 | 0.6667 |
| 14_4189349 | 8 | 0.375 | 18 | 0.2222 | 8 | 0.5 | 15 | 0.4 | 49 | 0.3469 | 8 | 0.375 | 9 | 0.4444 | 7 | 0.2857 | 10 | 0.4 | 34 | 0.3824 |
| 14_4189511 | 9 | 0.2222 | 16 | 0.375 | 13 | 0.2308 | 14 | 0.2143 | 52 | 0.2692 | 9 | 0.3333 | 5 | 0.2 | 5 | 0.4 | 13 | 0.3846 | 32 | 0.3438 |
| 14_4189526 | 14 | 0.0714 | 17 | 0.0588 | 13 | 0.0769 | 16 | 0.3125 | 60 | 0.1333 | 9 | 0.3333 | 11 | 0.1818 | 8 | 0.125 | 13 | 0.3846 | 41 | 0.2683 |
| 14_4189620 | 18 | 0.3333 | 20 | 0.3 | 15 | 0.3333 | 18 | 0.6667 | 71 | 0.4085 | 10 | 0.3 | 10 | 0.3 | 9 | 0.5556 | 16 | 0.375 | 45 | 0.3778 |
| 14_4189702 | 14 | 0.0714 | 23 | 0.087 | 16 | 0.0625 | 17 | 0.1176 | 70 | 0.0857 | 9 | 0.1111 | 10 | 0 | 10 | 0.2 | 16 | 0.0625 | 45 | 0.0889 |
| 14_4189718 | 13 | 0.2308 | 17 | 0.3529 | 11 | 0.0909 | 15 | 0.2667 | 56 | 0.25 | 8 | 0.125 | 7 | 0.4286 | 10 | 0.3 | 15 | 0.4 | 40 | 0.325 |
| 14_4191673 | 2 | 0.5 | NA | NA | 1 | 0 | 1 | 0 | 4 | 0.25 | 1 | 1 | 3 | 0 | NA | NA | 1 | 0 | 5 | 0.2 |
| 14_4200730 | 12 | 0.3333 | 18 | 0.3333 | 16 | 0.375 | 9 | 0.3333 | 55 | 0.3455 | 10 | 0.2 | 18 | 0.3333 | 10 | 0.2 | 7 | 0 | 45 | 0.2222 |
| 14_4254284 | 5 | 0 | 3 | 0.3333 | 4 | 0 | 6 | 0 | 18 | 0.0556 | 1 | 0 | 6 | 0 | 8 | 0 | 3 | 0.3333 | 18 | 0.0556 |
| 14_4254327 | 6 | 0.1667 | 3 | 0.3333 | 8 | 0.125 | 9 | 0 | 26 | 0.1154 | 4 | 0 | 6 | 0 | 8 | 0.25 | 8 | 0.125 | 26 | 0.1154 |
| 14_4254404 | 5 | 0.2 | 3 | 0.3333 | 8 | 0.5 | 7 | 0.4286 | 23 | 0.3913 | 3 | 0 | 4 | 0.25 | 9 | 0.3333 | 7 | 0.4286 | 23 | 0.3043 |
| 14_4254486 | 9 | 0.2222 | 4 | 0.5 | 7 | 0.1429 | 8 | 0.25 | 28 | 0.25 | 3 | 0 | 7 | 0 | 7 | 0.1429 | 4 | 0.25 | 21 | 0.0952 |
| 14_4254545 | 11 | 0.2727 | 3 | 0.3333 | 8 | 0.125 | 7 | 0.4286 | 29 | 0.2759 | 5 | 0.2 | 6 | 0.3333 | 9 | 0 | 7 | 0 | 27 | 0.1111 |
| 14_4254636 | 10 | 1 | 4 | 1 | 4 | 1 | 9 | 0.6667 | 27 | 0.8889 | 3 | 1 | 6 | 0.8333 | 8 | 0.875 | 5 | 0.8 | 22 | 0.8636 |
| 14_4254653 | 7 | 0.2857 | 4 | 0.25 | 7 | 0.1429 | 8 | 0 | 26 | 0.1538 | 4 | 0 | 7 | 0.1429 | 8 | 0.125 | 8 | 0.375 | 27 | 0.1852 |
| 14_4254699 | 8 | 1 | 5 | 1 | 8 | 1 | 6 | 0.8333 | 27 | 0.963 | 5 | 1 | 6 | 1 | 10 | 0.8 | 8 | 1 | 29 | 0.931 |
| 14_4254710 | 6 | 0.8333 | 4 | 0.5 | 7 | 1 | 6 | 1 | 23 | 0.8696 | 4 | 1 | 3 | 1 | 6 | 1 | 7 | 1 | 20 | 1 |
| 14_4254772 | 10 | 0.3 | 5 | 0 | 9 | 0.2222 | 8 | 0.25 | 32 | 0.2188 | 5 | 0 | 6 | 0.1667 | 9 | 0.2222 | 8 | 0.125 | 28 | 0.1429 |
| 14_4257909 | 10 | 0 | 8 | 0 | 8 | 0.25 | 9 | 0.2222 | 35 | 0.1143 | 5 | 0 | 5 | 0.4 | 12 | 0.1667 | 9 | 0 | 31 | 0.129 |
| 14_4261612 | 9 | 0.6667 | 9 | 0.7778 | 6 | 0.5 | 11 | 0.5455 | 35 | 0.6286 | 5 | 0.2 | 6 | 0.5 | 10 | 0.5 | 7 | 0.4286 | 28 | 0.4286 |
| 14_4262883 | 13 | 0.7692 | 10 | 0.7 | 9 | 0.7778 | 11 | 0.7273 | 43 | 0.7442 | 6 | 0.5 | 6 | 0.8333 | 14 | 0.7857 | 10 | 0.6 | 36 | 0.6944 |
| 14_4262984 | 9 | 0.3333 | 8 | 0.75 | 8 | 0.75 | 7 | 0.5714 | 32 | 0.5938 | 5 | 0.6 | 6 | 1 | 10 | 0.5 | 10 | 0.5 | 31 | 0.6129 |
| 14_4263021 | 11 | 0.1818 | 11 | 0.0909 | 9 | 0 | 8 | 0 | 39 | 0.0769 | 4 | 0 | 6 | 0.1667 | 15 | 0 | 12 | 0.0833 | 37 | 0.0541 |
| 14_4263027 | 12 | 0.9167 | 8 | 1 | 8 | 1 | 9 | 0.8889 | 37 | 0.9459 | 5 | 1 | 5 | 1 | 15 | 0.8667 | 9 | 0.8889 | 34 | 0.9118 |
| 14_4263100 | 12 | 0.0833 | 10 | 0.1 | 9 | 0 | 10 | 0.1 | 41 | 0.0732 | 5 | 0.2 | 8 | 0 | 13 | 0 | 13 | 0.0769 | 39 | 0.0513 |
| 14_4263160 | 12 | 0.1667 | 10 | 0.2 | 9 | 0.1111 | 11 | 0 | 42 | 0.119 | 5 | 0.4 | 6 | 0.3333 | 14 | 0 | 10 | 0 | 35 | 0.1143 |
| 14_4263346 | 13 | 0.1538 | 11 | 0.6364 | 10 | 0.1 | 10 | 0.5 | 44 | 0.3409 | 5 | 0.2 | 8 | 0.125 | 17 | 0.2353 | 13 | 0.1538 | 43 | 0.186 |
| 14_4263392 | 13 | 0.1538 | 11 | 0.0909 | 11 | 0 | 11 | 0.0909 | 46 | 0.087 | 7 | 0.2857 | 6 | 0 | 12 | 0.0833 | 13 | 0.2308 | 38 | 0.1579 |
| 14_4263743 | 12 | 0 | 13 | 0.0769 | 10 | 0.2 | 13 | 0.0769 | 48 | 0.0833 | 8 | 0.25 | 8 | 0.25 | 18 | 0 | 14 | 0 | 48 | 0.0833 |
| 14_4265309 | 12 | 0.1667 | 17 | 0.0588 | 14 | 0.2143 | 19 | 0.0526 | 62 | 0.1129 | 10 | 0.1 | 11 | 0 | 21 | 0.0476 | 14 | 0.2143 | 56 | 0.0893 |
| 14_4265399 | NA | NA | NA | NA | 1 | 0 | 1 | 1 | 2 | 0.5 | NA | NA | NA | NA | NA | NA | 2 | 0.5 | 2 | 0.5 |
| 14_4432119 | 4 | 0.25 | 4 | 0 | 3 | 0 | 3 | 0 | 14 | 0.0714 | 3 | 0 | 4 | 0 | 2 | 0 | 4 | 0.25 | 13 | 0.0769 |
| 14_4432153 | 4 | 0.25 | 2 | 0 | 4 | 0.5 | 2 | 0 | 12 | 0.25 | 3 | 0 | 4 | 0.25 | 2 | 0.5 | 4 | 0.25 | 13 | 0.2308 |

| | | | | | | | | | | | | | | | | | | | | |
|---|---|---|---|---|---|---|---|---|---|---|---|---|---|---|---|---|---|---|---|---|
| 14_443215 | 4 | 0 | 4 | 0.5 | 4 | 0.75 | 4 | 0.5 | 16 | 0.4375 | 3 | 0 | 3 | 0 | 1 | 0 | 4 | 0.75 | 11 | 0.2727 |
| 14_443218 | 5 | 0 | 3 | 0.3333 | 3 | 0 | 4 | 0 | 15 | 0.0667 | 3 | 0 | 4 | 0 | 2 | 0.5 | 4 | 0.25 | 13 | 0.1538 |
| 14_443241 | 5 | 0 | 4 | 0.25 | 4 | 0.25 | 4 | 0.25 | 17 | 0.1765 | 3 | 0.3333 | 4 | 0.25 | 1 | 1 | 4 | 0 | 12 | 0.25 |
| 14_443247 | 3 | 0 | 4 | 0.25 | 4 | 0.75 | 4 | 0 | 15 | 0.2667 | 3 | 0.3333 | 4 | 0.5 | 1 | 0 | 4 | 0.25 | 12 | 0.3333 |
| 14_443254 | 3 | 0.3333 | 4 | 0.25 | 4 | 0.5 | 3 | 0 | 14 | 0.2857 | 3 | 0.6667 | 4 | 0.25 | 2 | 0.5 | 4 | 0.5 | 13 | 0.4615 |
| 14_443266 | 3 | 0.3333 | 2 | 0 | 3 | 0.3333 | 3 | 0.6667 | 11 | 0.3636 | 3 | 1 | 3 | 0.3333 | 2 | 0.5 | 3 | 0.3333 | 11 | 0.5455 |
| 14_443267 | 3 | 0 | 3 | 0.3333 | 4 | 0.25 | 3 | 0 | 13 | 0.1538 | 3 | 0.3333 | 4 | 0 | 2 | 0 | 3 | 0 | 12 | 0.0833 |
| 14_443272 | 2 | 0.5 | 3 | 1 | 4 | 1 | 4 | 0.75 | 13 | 0.8462 | 2 | 1 | 4 | 0.75 | 2 | 1 | 2 | 1 | 10 | 0.9 |
| 14_443281 | 3 | 0 | 3 | 0 | 3 | 0.6667 | 4 | 0 | 13 | 0.1538 | 2 | 0 | 4 | 0.5 | 2 | 0.5 | 3 | 0.3333 | 11 | 0.3636 |
| 14_443283 | 3 | 0.6667 | 2 | 1 | 4 | 0.75 | 3 | 1 | 12 | 0.8333 | 2 | 1 | 2 | 0.5 | 2 | 0.5 | 3 | 0.6667 | 9 | 0.6667 |
| 14_443285 | 3 | 0.3333 | 3 | 0.6667 | 4 | 0.5 | 4 | 0 | 14 | 0.3571 | 2 | 0 | 4 | 0.25 | 1 | 0 | 3 | 0 | 10 | 0.1 |
| 14_443295 | 3 | 0 | 2 | 0.5 | 4 | 0.5 | 3 | 0.3333 | 12 | 0.3333 | 1 | 1 | 2 | 1 | 1 | 1 | 3 | 0.6667 | 7 | 0.8571 |
| 14_443312 | NA | NA | NA | NA | 1 | 1 | NA | NA | 1 | 1 | 1 | 1 | 1 | 1 | NA | NA | 3 | 0.6667 | 5 | 0.8 |
| 14_443314 | 3 | 0 | 3 | 0.6667 | 4 | 0.75 | 3 | 0.6667 | 13 | 0.5385 | 2 | 1 | 3 | 0.3333 | 2 | 1 | 3 | 0.6667 | 10 | 0.7 |
| 14_443315 | 2 | 0 | 3 | 0.3333 | 4 | 0.75 | 2 | 0.5 | 11 | 0.4545 | 2 | 1 | 1 | 0 | 1 | 0 | 2 | 0 | 6 | 0.3333 |
| 14_443325 | 2 | 1 | 3 | 1 | 3 | 1 | 3 | 0.6667 | 11 | 0.9091 | 2 | 1 | 1 | 1 | 2 | 1 | 3 | 1 | 8 | 1 |
| 14_443327 | 2 | 0.5 | 3 | 0.3333 | 4 | 0 | 3 | 0.3333 | 12 | 0.25 | 2 | 0 | 2 | 0.5 | 2 | 0.5 | 3 | 0.3333 | 9 | 0.3333 |
| 14_443335 | 2 | 0 | 1 | 0 | 3 | 0 | 4 | 0.25 | 10 | 0.1 | 1 | 0 | 3 | 0.3333 | 2 | 0 | 3 | 0.3333 | 9 | 0.2222 |
| 14_443336 | 1 | 0 | 2 | 1 | 2 | 0.5 | 3 | 0.3333 | 8 | 0.5 | 2 | 0.5 | 2 | 0.5 | 2 | 0 | 3 | 0.6667 | 9 | 0.4444 |
| 14_443338 | 2 | 0 | 3 | 0.6667 | 4 | 0.25 | 3 | 0 | 12 | 0.25 | 2 | 0 | 3 | 0.3333 | 2 | 0.5 | 3 | 0.3333 | 10 | 0.3 |
| 14_443348 | 3 | 0.3333 | 2 | 0.5 | 1 | 1 | 3 | 0 | 9 | 0.3333 | 2 | 0 | 3 | 0.3333 | 2 | 0.5 | 3 | 0.3333 | 10 | 0.3 |
| 14_443349 | 3 | 0 | 3 | 0.3333 | 2 | 0.5 | 3 | 0.6667 | 11 | 0.3636 | 2 | 1 | 2 | 0 | 2 | 0.5 | 3 | 0.3333 | 9 | 0.4444 |
| 14_443359 | 2 | 0 | 3 | 0 | 3 | 0.6667 | 2 | 0 | 10 | 0.2 | 1 | 0 | 3 | 0 | 2 | 0 | 2 | 1 | 8 | 0.25 |
| 14_443380 | 3 | 0 | 3 | 0 | 3 | 0.3333 | 3 | 0 | 12 | 0.0833 | 2 | 0 | 3 | 0.3333 | 2 | 0 | 2 | 0 | 9 | 0.1111 |
| 14_443387 | 1 | 0 | 2 | 0 | 2 | 0.5 | 3 | 0.3333 | 8 | 0.25 | 2 | 0.5 | 1 | 0 | 2 | 1 | NA | NA | 5 | 0.6 |
| 14_443394 | 2 | 0 | 2 | 1 | 1 | 0 | 2 | 0 | 7 | 0.2857 | 2 | 0 | 2 | 0 | 2 | 0.5 | 2 | 1 | 8 | 0.375 |
| 14_443410 | 2 | 0.5 | 2 | 0 | 2 | 0 | 2 | 1 | 8 | 0.375 | 2 | 0 | 2 | 0.5 | 2 | 0 | 2 | 1 | 8 | 0.375 |
| 14_443431 | 2 | 0.5 | 2 | 0 | 2 | 0 | 1 | 1 | 7 | 0.2857 | 2 | 0.5 | 1 | 0 | 2 | 0 | 2 | 1 | 7 | 0.4286 |
| 14_443435 | 2 | 0.5 | 2 | 0.5 | 2 | 0.5 | 2 | 1 | 8 | 0.625 | 2 | 0.5 | 1 | 0 | 1 | 0 | 2 | 1 | 6 | 0.5 |
| 14_443459 | 2 | 0 | 2 | 0 | 2 | 0 | 1 | 1 | 7 | 0.1429 | 2 | 0.5 | 1 | 0 | 2 | 0 | 1 | 0 | 6 | 0.1667 |
| 14_443460 | 2 | 0 | 2 | 0 | 2 | 0 | 2 | 0.5 | 8 | 0.125 | 2 | 0.5 | 1 | 0 | 2 | 0 | 1 | 0 | 6 | 0.1667 |
| 14_443463 | 1 | 1 | 2 | 0.5 | 2 | 1 | 2 | 1 | 7 | 0.8571 | 2 | 0.5 | 1 | 1 | 2 | 0.5 | 1 | 1 | 6 | 0.6667 |
| 14_443511 | 2 | 0 | 1 | 1 | 2 | 0.5 | 1 | 1 | 6 | 0.5 | 2 | 0.5 | 1 | 0 | 1 | 0 | 1 | 0 | 5 | 0.2 |
| 14_443540 | 1 | 0 | 1 | 1 | 3 | 0.6667 | 1 | 1 | 6 | 0.6667 | 2 | 1 | NA | NA | 1 | 1 | 1 | 1 | 4 | 1 |
| 14_443589 | 1 | 1 | NA | NA | 1 | 1 | 2 | 0.5 | 4 | 0.75 | 2 | 1 | 1 | 0 | 1 | 1 | 1 | 1 | 5 | 0.8 |
| 14_443602 | 1 | 1 | NA | NA | 1 | 1 | 2 | 1 | 4 | 1 | 2 | 1 | 1 | 0 | 1 | 1 | 1 | 1 | 5 | 0.8 |
| 14_443603 | 1 | 1 | NA | NA | 1 | 1 | 1 | 1 | 3 | 1 | 2 | 1 | 1 | 1 | 1 | 1 | 1 | 1 | 5 | 1 |
| 14_443620 | 1 | 1 | NA | NA | 1 | 0 | NA | NA | 2 | 0.5 | 2 | 1 | 1 | 0 | 1 | 1 | NA | NA | 4 | 0.75 |
| 14_565071 | 1 | 1 | NA | NA | 1 | 1 | NA | NA | 2 | 1 | NA | NA | NA | NA | 1 | 1 | 1 | 1 | 2 | 1 |
| 14_589468 | 4 | 0.75 | 4 | 0.75 | 3 | 1 | 4 | 1 | 15 | 0.8667 | 2 | 1 | 4 | 0.5 | 6 | 0.8333 | 1 | 1 | 13 | 0.7692 |
| 14_589475 | 5 | 0.4 | 3 | 0 | 4 | 0.25 | 5 | 0.2 | 17 | 0.2353 | 3 | 1 | 5 | 0.6 | 7 | 0.4286 | 2 | 0.5 | 17 | 0.5882 |
| 14_603758 | 11 | 0.5455 | 5 | 0.4 | 8 | 0.75 | 8 | 0.75 | 32 | 0.625 | 5 | 0.4 | 17 | 0.8824 | 15 | 0.4667 | 9 | 0.5556 | 46 | 0.6304 |
| 14_603758 | 11 | 0 | 5 | 0.2 | 7 | 0.2857 | 10 | 0.1 | 33 | 0.1212 | 6 | 0.3333 | 17 | 0.1765 | 14 | 0.0714 | 7 | 0 | 44 | 0.1364 |
| 14_603765 | 10 | 0.2 | 5 | 0 | 6 | 0.1667 | 8 | 0.125 | 29 | 0.1379 | 6 | 0.1667 | 11 | 0 | 14 | 0.0714 | 6 | 0 | 37 | 0.0541 |
| 14_603769 | 11 | 0.2727 | 5 | 0 | 8 | 0 | 10 | 0.3 | 34 | 0.1765 | 6 | 0.3333 | 17 | 0.1176 | 15 | 0.2 | 10 | 0.2 | 48 | 0.1875 |
| 14_603772 | 10 | 0.3 | 5 | 0.6 | 7 | 0.1429 | 10 | 0.1 | 32 | 0.25 | 6 | 0.1667 | 19 | 0.3158 | 14 | 0.2143 | 10 | 0.3 | 49 | 0.2653 |
| 14_603778 | 11 | 0.8182 | 4 | 1 | 5 | 1 | 9 | 1 | 29 | 0.931 | 5 | 1 | 19 | 0.8421 | 13 | 0.8462 | 9 | 0.8889 | 46 | 0.8696 |
| 14_603781 | 11 | 0.6364 | 6 | 0.6667 | 7 | 0.4286 | 9 | 0.6667 | 33 | 0.6061 | 6 | 0.3333 | 18 | 0.6111 | 15 | 0.5333 | 10 | 0.6 | 49 | 0.551 |
| 14_603787 | 11 | 0.3636 | 5 | 0.2 | 7 | 0.5714 | 9 | 0.4444 | 32 | 0.4062 | 5 | 0.6 | 17 | 0.1765 | 13 | 0.6154 | 10 | 0.5 | 45 | 0.4222 |
| 14_603800 | 11 | 0.1818 | 6 | 0.3333 | 8 | 0.125 | 9 | 0.3333 | 34 | 0.2353 | 6 | 0.3333 | 18 | 0.1667 | 13 | 0.0769 | 9 | 0.2222 | 46 | 0.1739 |
| 14_603857 | 9 | 0.2222 | 3 | 0.3333 | 6 | 0.1667 | 10 | 0.2 | 28 | 0.2143 | 6 | 0.1667 | 18 | 0.1667 | 13 | 0.1538 | 10 | 0.1 | 47 | 0.1489 |
| 14_603869 | 10 | 0 | 5 | 0.2 | 6 | 0 | 9 | 0.2222 | 30 | 0.1 | 7 | 0 | 21 | 0.0952 | 16 | 0.0625 | 11 | 0.1818 | 55 | 0.0909 |
| 14_603872 | 10 | 0 | 5 | 0.2 | 9 | 0 | 10 | 0.1 | 34 | 0.0588 | 6 | 0.3333 | 21 | 0.0952 | 13 | 0 | 12 | 0.1667 | 52 | 0.1154 |
| 14_603927 | 7 | 0.4286 | 20 | 0.2 | 13 | 0.1538 | 6 | 0.1667 | 46 | 0.2174 | 5 | 0 | 13 | 0.0769 | 8 | 0 | 9 | 0.1111 | 35 | 0.0571 |
| 14_603937 | 9 | 0 | 16 | 0.4375 | 13 | 0.2308 | 4 | 0 | 42 | 0.2381 | 3 | 0.3333 | 12 | 0.4167 | 9 | 0.3333 | 11 | 0.1818 | 35 | 0.3143 |
| 14_605983 | 7 | 0 | 6 | 0.1667 | 7 | 0.1429 | 2 | 0 | 22 | 0.0909 | 2 | 0 | 5 | 0.2 | 3 | 0 | 6 | 0 | 16 | 0.0625 |
| 14_610443 | 1 | 0 | 2 | 0 | 4 | 0.25 | NA | NA | 7 | 0.1429 | 2 | 0 | 2 | 0.5 | 1 | 0 | 5 | 0 | 10 | 0.1 |
| 14_610451 | NA | NA | 2 | 0.5 | 2 | 0.5 | NA | NA | 4 | 0.5 | 2 | 0 | 3 | 0.3333 | 1 | 0 | 4 | 0.25 | 10 | 0.2 |
| 14_615848 | 4 | 0.25 | 3 | 0.3333 | 2 | 0 | 6 | 0 | 15 | 0.1333 | NA | NA | 4 | 0 | 3 | 0.3333 | 4 | 0 | 11 | 0.0909 |
| 14_615855 | 4 | 0 | 2 | 0.5 | 3 | 0 | 6 | 0 | 15 | 0.0667 | NA | NA | 4 | 0 | 3 | 0 | 2 | 0.5 | 9 | 0.1111 |
| 14_615877 | 3 | 0.3333 | 3 | 0 | 4 | 0 | 5 | 0.2 | 15 | 0.1333 | NA | NA | 3 | 0.3333 | 3 | 0 | 3 | 0 | 9 | 0.1111 |
| 14_618874 | 1 | 1 | NA | NA | 1 | 1 | NA | NA | 2 | 1 | NA | NA | 1 | 1 | NA | NA | NA | NA | 1 | 1 |
| 14_618878 | 1 | 1 | NA | NA | 1 | 0 | NA | NA | 2 | 0.5 | NA | NA | 1 | 1 | NA | NA | NA | NA | 1 | 1 |
| 14_619181 | 3 | 0 | 2 | 0 | 4 | 0 | 5 | 0.2 | 14 | 0.0714 | NA | NA | 4 | 0.75 | 2 | 0 | 1 | 0 | 7 | 0.4286 |
| 14_622709 | NA | NA | NA | NA | 1 | 1 | NA | NA | 1 | 1 | 1 | 1 | NA | NA | NA | NA | NA | NA | 1 | 1 |

| | | | | | | | | | | | | | | | | | | | | |
|---|---|---|---|---|---|---|---|---|---|---|---|---|---|---|---|---|---|---|---|---|
| 14_6227390 | NA | NA | NA | NA | 1 | 1 | 1 | 1 | 2 | 1 | 1 | 1 | NA | NA | NA | NA | NA | NA | 1 | 1 |
| 14_6227648 | NA | NA | NA | NA | 1 | 1 | 1 | 1 | 2 | 1 | 1 | 0 | NA | NA | NA | NA | 1 | 1 | 2 | 0.5 |
| 14_6227913 | NA | NA | NA | NA | 1 | 1 | 1 | 1 | 2 | 1 | 1 | 1 | NA | NA | NA | NA | 1 | 1 | 2 | 1 |
| 14_6228821 | NA | NA | NA | NA | 1 | 1 | 1 | 1 | 2 | 1 | 1 | 1 | NA | NA | NA | NA | 1 | 1 | 2 | 1 |
| 14_6228933 | NA | NA | NA | NA | 1 | 1 | 1 | 1 | 2 | 1 | NA | NA | NA | NA | NA | NA | 1 | 1 | 1 | 1 |
| 14_6229035 | NA | NA | NA | NA | 1 | 1 | 1 | 1 | 2 | 1 | NA | NA | NA | NA | NA | NA | 1 | 1 | 1 | 1 |
| 14_7111133 | 1 | 1 | 1 | 1 | 2 | 1 | 1 | 1 | 5 | 1 | 2 | 1 | 3 | 1 | 2 | 1 | NA | NA | 7 | 1 |
| 14_7111373 | 1 | 1 | 3 | 1 | 2 | 1 | 1 | 1 | 7 | 1 | 2 | 1 | 3 | 1 | 2 | 1 | NA | NA | 7 | 1 |
| 14_7111415 | 1 | 0 | 3 | 1 | 2 | 0.5 | 1 | 1 | 7 | 0.7143 | 2 | 1 | 3 | 0.6667 | 2 | 0.5 | NA | NA | 7 | 0.7143 |
| 14_7111514 | 1 | 1 | 3 | 1 | 2 | 1 | 1 | 1 | 7 | 1 | 2 | 1 | 3 | 1 | 2 | 1 | NA | NA | 7 | 1 |
| 14_7111607 | 1 | 1 | 3 | 1 | 2 | 1 | 1 | 1 | 7 | 1 | 2 | 1 | 2 | 1 | 2 | 1 | NA | NA | 6 | 1 |
| 14_7111654 | 1 | 1 | 2 | 0 | 2 | 0 | 1 | 0 | 6 | 0.1667 | 2 | 0.5 | 3 | 0 | 2 | 0 | NA | NA | 7 | 0.1429 |
| 14_7111736 | 1 | 0 | 3 | 0.3333 | 1 | 0 | 1 | 0 | 6 | 0.1667 | 2 | 0 | 3 | 0 | 2 | 0.5 | NA | NA | 7 | 0.1429 |
| 14_7111759 | NA | NA | 2 | 1 | 2 | 0.5 | 1 | 1 | 5 | 0.8 | 2 | 0.5 | 2 | 1 | 1 | 1 | NA | NA | 5 | 0.8 |
| 14_7111766 | 1 | 1 | 3 | 0.6667 | 2 | 0.5 | 1 | 0 | 7 | 0.5714 | 2 | 0.5 | 2 | 0.5 | 2 | 1 | NA | NA | 6 | 0.6667 |
| 14_7111919 | NA | NA | 3 | 0.6667 | 1 | 1 | 1 | 0 | 5 | 0.6 | 1 | 0 | 3 | 0.3333 | 2 | 0.5 | NA | NA | 6 | 0.3333 |
| 14_7112065 | NA | NA | 3 | 0.6667 | 2 | 0.5 | 1 | 0 | 6 | 0.5 | 1 | 1 | 2 | 1 | 2 | 0.5 | NA | NA | 5 | 0.8 |
| 14_7670452 | NA | NA | 6 | 0 | 3 | 0.6667 | 6 | 0.3333 | 15 | 0.2667 | 2 | 0.5 | 3 | 0.3333 | NA | NA | 3 | 0 | 8 | 0.25 |
| 14_7671131 | NA | NA | 6 | 0.1667 | 4 | 0 | 5 | 0 | 15 | 0.0667 | 2 | 0 | 5 | 0 | NA | NA | 2 | 0.5 | 9 | 0.1111 |
| 14_8362673 | 4 | 0 | 1 | 0 | 2 | 0.5 | 3 | 0 | 10 | 0.1 | 1 | 1 | 3 | 0.6667 | 3 | 0.6667 | NA | NA | 7 | 0.7143 |
| 14_8362950 | 4 | 1 | 2 | 0.5 | 3 | 1 | 2 | 0 | 11 | 0.7273 | 1 | 1 | 3 | 1 | 2 | 1 | NA | NA | 6 | 1 |
| 14_8363156 | 3 | 0.3333 | 2 | 0 | 3 | 0 | 1 | 0 | 9 | 0.1111 | 1 | 0 | 3 | 0 | 1 | 1 | NA | NA | 5 | 0.2 |
| 14_8363315 | 3 | 0 | 2 | 0 | 1 | 0 | 3 | 0.3333 | 9 | 0.1111 | 1 | 1 | 2 | 0 | 2 | 0 | NA | NA | 5 | 0.2 |
| 14_8364256 | 2 | 0.5 | 2 | 0.5 | NA | NA | 4 | 0.25 | 8 | 0.375 | 1 | 1 | 2 | 0 | 2 | 0.5 | NA | NA | 5 | 0.4 |
| 14_8364378 | 2 | 0 | 2 | 0.5 | NA | NA | 4 | 0.25 | 8 | 0.25 | NA | NA | 1 | 0 | 1 | 1 | NA | NA | 2 | 0.5 |
| 14_8365814 | 2 | 0.5 | 1 | 1 | NA | NA | 2 | 1 | 5 | 0.8 | 1 | 1 | 2 | 0 | 1 | 0 | NA | NA | 4 | 0.25 |
| 14_8366010 | 2 | 0 | 1 | 1 | NA | NA | 3 | 0 | 6 | 0.1667 | 1 | 1 | 1 | 0 | 1 | 0 | NA | NA | 3 | 0.3333 |
| 14_8366197 | 1 | 1 | 1 | 1 | NA | NA | 2 | 1 | 4 | 1 | 1 | 1 | NA | NA | 1 | 1 | NA | NA | 2 | 1 |
| 14_8366248 | 2 | 0.5 | 1 | 1 | NA | NA | 2 | 1 | 5 | 0.8 | 1 | 0 | NA | NA | 1 | 1 | NA | NA | 2 | 0.5 |
| 14_8366364 | 2 | 0 | NA | NA | NA | NA | 3 | 0.6667 | 5 | 0.4 | 1 | 1 | 1 | 0 | 1 | 1 | NA | NA | 3 | 0.6667 |
| 14_8369069 | 2 | 0 | 1 | 1 | NA | NA | 2 | 0 | 5 | 0.2 | 1 | 1 | NA | NA | NA | NA | NA | NA | 1 | 1 |
| 14_8369278 | 2 | 1 | NA | NA | NA | NA | 1 | 1 | 3 | 1 | 1 | 1 | NA | NA | NA | NA | NA | NA | 1 | 1 |
| 14_8369298 | 1 | 1 | 1 | 0 | NA | NA | 2 | 0.5 | 4 | 0.5 | 1 | 1 | NA | NA | NA | NA | NA | NA | 1 | 1 |
| 14_8369313 | 1 | 1 | NA | NA | NA | NA | 2 | 0.5 | 3 | 0.6667 | 1 | 1 | NA | NA | NA | NA | NA | NA | 1 | 1 |
| 14_8369604 | 2 | 0.5 | 1 | 0 | NA | NA | 2 | 1 | 5 | 0.6 | 1 | 1 | NA | NA | NA | NA | NA | NA | 1 | 1 |
| 14_8520508 | 12 | 0.1667 | 15 | 0.2 | 14 | 0.1429 | 7 | 0.1429 | 48 | 0.1667 | 8 | 0 | 13 | 0.2308 | 10 | 0.2 | 8 | 0 | 39 | 0.1282 |
| 14_8855959 | 9 | 0.1111 | 7 | 0 | 10 | 0.7 | 13 | 0.3846 | 39 | 0.3333 | 5 | 0.2 | 17 | 0.1765 | 11 | 0.5455 | 8 | 0.375 | 41 | 0.3171 |
| 14_8855979 | 18 | 0.2222 | 13 | 0 | 8 | 0 | 20 | 0.1 | 59 | 0.1017 | 7 | 0 | 21 | 0.0476 | 17 | 0.0588 | 14 | 0.0714 | 59 | 0.0508 |
| 14_8861128 | 16 | 0.125 | 12 | 0.25 | 6 | 0.1667 | 19 | 0.2632 | 53 | 0.2075 | 7 | 0.5714 | 20 | 0.25 | 16 | 0.375 | 14 | 0.2143 | 57 | 0.3158 |
| 14_9015648 | 8 | 0.125 | 7 | 0.1429 | 1 | 0 | 9 | 0 | 25 | 0.08 | 4 | 0 | 9 | 0.1111 | 9 | 0.1111 | 3 | 0 | 25 | 0.08 |
| 14_9649337 | NA | NA | 1 | 0 | NA | NA | 3 | 0.3333 | 4 | 0.25 | 3 | 0.3333 | NA | NA | 1 | 0 | 5 | 0 | 9 | 0.1111 |
| 14_9763980 | 2 | 0 | 6 | 0 | 4 | 0.25 | 3 | 0.6667 | 15 | 0.2 | 2 | 0 | 2 | 0 | 4 | 0 | 7 | 0.1429 | 15 | 0.0667 |
| 14_1009611 | 16 | 0.375 | 22 | 0.2727 | 18 | 0.1667 | 13 | 0.5385 | 69 | 0.3188 | 20 | 0.35 | 14 | 0.3571 | 14 | 0.4286 | 11 | 0.3636 | 59 | 0.3729 |
| 14_1012717 | NA | NA | 2 | 0.5 | 1 | 0 | NA | NA | 3 | 0.3333 | NA | NA | 1 | 1 | NA | NA | NA | NA | 1 | 1 |
| 14_1012724 | 1 | 1 | 2 | 1 | 1 | 0 | NA | NA | 4 | 0.75 | NA | NA | 1 | 1 | NA | NA | NA | NA | 1 | 1 |
| 14_1012732 | 1 | 1 | 1 | 1 | 1 | 0 | NA | NA | 3 | 0.6667 | NA | NA | 1 | 1 | NA | NA | NA | NA | 1 | 1 |
| 14_1016705 | 2 | 0.5 | 5 | 0.2 | 2 | 0 | 3 | 0.3333 | 12 | 0.25 | 3 | 0.3333 | 1 | 0 | 2 | 0 | 1 | 0 | 7 | 0.1429 |
| 14_1016797 | 1 | 0 | 7 | 0.4286 | 3 | 0.3333 | 4 | 0.5 | 15 | 0.4 | 2 | 0.5 | 1 | 0 | 2 | 0.5 | 3 | 0 | 8 | 0.25 |
| 14_1016800 | 1 | 0 | 5 | 0.2 | 5 | 0 | 4 | 0 | 15 | 0.0667 | 3 | 0 | 1 | 0 | 2 | 0 | 3 | 0.3333 | 9 | 0.1111 |
| 14_1022790 | 9 | 0.1111 | 6 | 0 | 3 | 0.6667 | 6 | 0.1667 | 24 | 0.1667 | 9 | 0.2222 | 2 | 0 | 6 | 0 | 4 | 0.25 | 21 | 0.1429 |
| 14_1022969 | 4 | 0 | 1 | 1 | 1 | 0 | 2 | 0 | 8 | 0.125 | 5 | 0.4 | 1 | 0 | 2 | 0.5 | NA | NA | 8 | 0.375 |
| 14_1023180 | 5 | 0 | 8 | 0.125 | 5 | 0 | 2 | 0 | 20 | 0.05 | 7 | 0.1429 | 5 | 0 | 2 | 0 | 6 | 0.1667 | 20 | 0.1 |
| 14_1023184 | 4 | 0 | 9 | 0.2222 | 5 | 0 | 2 | 0.5 | 20 | 0.15 | 7 | 0.1429 | 4 | 0 | 1 | 0 | 6 | 0 | 18 | 0.0556 |
| 14_1031548 | 2 | 0.5 | 3 | 0 | 1 | 0 | NA | NA | 6 | 0.1667 | 2 | 0.5 | 4 | 0.5 | NA | NA | 1 | 1 | 7 | 0.5714 |
| 14_1031549 | 2 | 0.5 | 3 | 0 | 2 | 0 | NA | NA | 7 | 0.1429 | 2 | 0.5 | 3 | 0.3333 | NA | NA | 1 | 0 | 6 | 0.3333 |
| 14_1031554 | 2 | 0.5 | 3 | 0.3333 | 2 | 0 | NA | NA | 7 | 0.2857 | 2 | 0 | 4 | 0.25 | NA | NA | 1 | 0 | 7 | 0.1429 |
| 14_1031556 | 1 | 1 | 2 | 0.5 | 2 | 0 | NA | NA | 5 | 0.4 | 2 | 0 | 3 | 1 | NA | NA | 1 | 1 | 6 | 0.6667 |
| 14_1031557 | 2 | 0 | 3 | 0.3333 | 2 | 0 | NA | NA | 7 | 0.1429 | 2 | 0 | 3 | 0.6667 | NA | NA | 1 | 0 | 6 | 0.3333 |
| 14_1031558 | 2 | 1 | 2 | 1 | 2 | 0 | NA | NA | 6 | 0.6667 | 1 | 0 | 3 | 0.3333 | NA | NA | 1 | 1 | 5 | 0.4 |
| 14_1031562 | 2 | 1 | 3 | 1 | 2 | 1 | NA | NA | 7 | 1 | 2 | 1 | 2 | 1 | NA | NA | 1 | 1 | 5 | 1 |
| 14_1031568 | 2 | 0.5 | 3 | 0 | 1 | 1 | NA | NA | 6 | 0.3333 | 2 | 1 | 3 | 0.3333 | NA | NA | 1 | 0 | 6 | 0.5 |
| 14_1031576 | 1 | 1 | 3 | 0.3333 | NA | NA | NA | NA | 4 | 0.5 | 2 | 0 | 4 | 0.25 | NA | NA | 1 | 1 | 7 | 0.2857 |
| 14_1031583 | 2 | 1 | 3 | 1 | NA | NA | NA | NA | 5 | 1 | 1 | 0 | 3 | 0.6667 | NA | NA | 1 | 1 | 5 | 0.6 |
| 14_1031604 | 2 | 0 | 3 | 0.3333 | NA | NA | NA | NA | 5 | 0.2 | 1 | 0 | 4 | 0.25 | NA | NA | 1 | 0 | 6 | 0.1667 |
| 14_1367014 | 5 | 0.2 | 2 | 1 | 6 | 0.1667 | 4 | 0 | 17 | 0.2353 | 5 | 0.2 | 2 | 0 | 3 | 0 | NA | NA | 10 | 0.1 |

| | | | | | | | | | | | | | | | | | | | | |
|---|---|---|---|---|---|---|---|---|---|---|---|---|---|---|---|---|---|---|---|---|
| 14_1395874 | NA | NA | 2 | 1 | NA | NA | 1 | 0 | 3 | 0.6667 | 1 | 1 | NA | NA | 2 | 1 | 1 | 0 | 4 | 0.75 |
| 14_1403814 | NA | NA | 2 | 0.5 | NA | NA | 1 | 0 | 3 | 0.3333 | 1 | 1 | 1 | 0 | 4 | 0.25 | 1 | 0 | 7 | 0.2857 |
| 14_1417115 | 55 | 0.2727 | 45 | 0.4444 | 35 | 0.3429 | 30 | 0.3 | 165 | 0.3394 | 34 | 0.3824 | 37 | 0.4054 | 45 | 0.2444 | 47 | 0.2766 | 163 | 0.319 |
| 14_1417128 | 53 | 0.4151 | 44 | 0.5455 | 31 | 0.4839 | 27 | 0.5185 | 155 | 0.4839 | 36 | 0.5 | 35 | 0.5429 | 44 | 0.5682 | 46 | 0.413 | 161 | 0.5031 |
| 14_1417136 | 60 | 0.0833 | 43 | 0.0465 | 37 | 0.1892 | 36 | 0.0833 | 176 | 0.0966 | 37 | 0.0811 | 35 | 0.0286 | 44 | 0.1136 | 47 | 0.0851 | 163 | 0.0798 |
| 14_1417151 | 60 | 0.1167 | 46 | 0.2174 | 40 | 0.125 | 37 | 0.1892 | 183 | 0.1585 | 38 | 0.2105 | 39 | 0.1282 | 53 | 0.1698 | 49 | 0.1633 | 179 | 0.1676 |
| 14_1417179 | 24 | 0.0417 | 13 | 0.1538 | 11 | 0 | 11 | 0 | 59 | 0.0508 | 17 | 0.0588 | 8 | 0 | 15 | 0 | 18 | 0.1111 | 58 | 0.0517 |
| 14_1417215 | 52 | 0.1346 | 52 | 0.2115 | 39 | 0.2308 | 32 | 0.1562 | 175 | 0.1829 | 44 | 0.1818 | 33 | 0.3636 | 43 | 0.2558 | 47 | 0.2766 | 167 | 0.2635 |
| 14_1417220 | 62 | 0.3387 | 62 | 0.2419 | 43 | 0.2093 | 38 | 0.2895 | 205 | 0.2732 | 49 | 0.3265 | 39 | 0.4103 | 55 | 0.2 | 50 | 0.38 | 193 | 0.3212 |
| 14_1417316 | 87 | 0.1034 | 64 | 0.0312 | 51 | 0.0392 | 54 | 0.0185 | 256 | 0.0547 | 64 | 0.0625 | 51 | 0.0196 | 67 | 0.1194 | 61 | 0.0656 | 243 | 0.07 |
| 14_1417336 | 87 | 0.092 | 67 | 0.1642 | 54 | 0.1296 | 48 | 0.2292 | 256 | 0.1445 | 63 | 0.0952 | 49 | 0.2245 | 66 | 0.0758 | 65 | 0.1692 | 243 | 0.1358 |
| 14_1424642 | 8 | 1 | 2 | 1 | 9 | 0.5556 | 1 | 1 | 20 | 0.8 | 4 | 0.5 | 5 | 1 | 3 | 1 | 5 | 1 | 17 | 0.8824 |
| 14_1424675 | 8 | 0.125 | 3 | 0.3333 | 9 | 0.1111 | 2 | 0 | 22 | 0.1364 | 3 | 0 | 7 | 0.1429 | 5 | 0 | 5 | 0.4 | 20 | 0.15 |
| 14_1424684 | 6 | 1 | 3 | 1 | 6 | 1 | 2 | 1 | 17 | 1 | 3 | 1 | 4 | 1 | 4 | 1 | 4 | 1 | 15 | 1 |
| 14_1424685 | 5 | 0.4 | 3 | 0 | 5 | 0.2 | 2 | 0 | 15 | 0.2 | 3 | 0.6667 | 4 | 0.25 | 5 | 0.2 | 4 | 0.25 | 16 | 0.3125 |
| 14_1424687 | 7 | 0 | 3 | 0.3333 | 6 | 0 | 2 | 0.5 | 18 | 0.1111 | 2 | 0.5 | 6 | 0 | 6 | 0.3333 | 5 | 0 | 19 | 0.1579 |
| 14_1424690 | 6 | 0.3333 | 2 | 0 | 6 | 0.3333 | 1 | 0 | 15 | 0.2667 | 2 | 0 | 6 | 0.1667 | 6 | 0.1667 | 5 | 0 | 19 | 0.1053 |
| 14_1424692 | 6 | 0.1667 | 3 | 0 | 4 | 0.5 | 1 | 1 | 14 | 0.2857 | 1 | 0 | 5 | 0 | 4 | 0.25 | 4 | 0 | 14 | 0.0714 |
| 14_1424693 | 7 | 1 | 3 | 1 | 5 | 1 | 2 | 1 | 17 | 1 | 3 | 1 | 6 | 0.8333 | 5 | 1 | 3 | 1 | 17 | 0.9412 |
| 14_1424695 | 6 | 0.3333 | 3 | 0.3333 | 6 | 0 | 1 | 0 | 16 | 0.1875 | 2 | 1 | 3 | 0.6667 | 3 | 0.6667 | 5 | 0.2 | 13 | 0.5385 |
| 14_1424707 | 6 | 0.3333 | 3 | 0 | 6 | 0.6667 | 2 | 0.5 | 17 | 0.4118 | 3 | 0.3333 | 5 | 0.4 | 4 | 0.25 | 4 | 0.25 | 16 | 0.3125 |
| 14_1424709 | 6 | 0.8333 | 3 | 1 | 6 | 0.6667 | 2 | 0.5 | 17 | 0.7647 | 2 | 1 | 6 | 0.5 | 5 | 1 | 4 | 0.5 | 17 | 0.7059 |
| 14_1424710 | 5 | 1 | 3 | 1 | 4 | 1 | 2 | 1 | 14 | 1 | 3 | 1 | 5 | 1 | 5 | 1 | 2 | 1 | 15 | 1 |
| 14_1424729 | 5 | 0 | 3 | 0 | 5 | 0.2 | 2 | 0 | 15 | 0.0667 | 2 | 0 | 4 | 0.25 | 3 | 0.3333 | 3 | 0 | 12 | 0.1667 |
| 14_1424731 | 4 | 0.75 | 2 | 0.5 | 4 | 0 | 1 | 0 | 11 | 0.3636 | 2 | 0 | 4 | 0.75 | 5 | 0.4 | 3 | 0.6667 | 14 | 0.5 |
| 14_1424736 | 4 | 0.5 | 3 | 0.6667 | 5 | 1 | 1 | 1 | 13 | 0.7692 | 1 | 1 | 5 | 0.8 | 5 | 0.8 | 3 | 1 | 14 | 0.8571 |
| 14_1424738 | 4 | 0 | 2 | 0.5 | 5 | 0 | NA | NA | 11 | 0.0909 | NA | NA | 3 | 0.3333 | 4 | 0 | 4 | 0 | 11 | 0.0909 |
| 14_1424738 | 5 | 0.6 | 2 | 1 | 4 | 0.75 | 1 | 1 | 12 | 0.75 | 1 | 1 | 4 | 0.75 | 5 | 0.4 | 3 | 0.6667 | 13 | 0.6154 |
| 14_1424739 | 3 | 0.6667 | 3 | 1 | 6 | 1 | 1 | 1 | 13 | 0.9231 | 2 | 1 | 4 | 1 | 4 | 1 | 4 | 0.75 | 14 | 0.9286 |
| 14_1424742 | 5 | 0.2 | 3 | 0 | 6 | 0.3333 | 2 | 0 | 16 | 0.1875 | 1 | 0 | 4 | 0.25 | 4 | 0.5 | 4 | 0 | 13 | 0.2308 |
| 14_1424746 | 4 | 0.25 | 3 | 0 | 5 | 0.2 | 2 | 0.5 | 14 | 0.2143 | 2 | 0.5 | 5 | 0 | 5 | 0 | 3 | 0 | 15 | 0.0667 |
| 14_1424755 | 4 | 0 | 3 | 0 | 6 | 0.3333 | 2 | 0 | 15 | 0.1333 | 2 | 0 | 4 | 0 | 5 | 0.4 | 4 | 0.25 | 15 | 0.2 |
| 14_1430344 | 15 | 0.4667 | 26 | 0.6538 | 34 | 0.7647 | 24 | 0.625 | 99 | 0.6566 | 17 | 0.7059 | 23 | 0.7826 | 24 | 0.6667 | 32 | 0.8125 | 96 | 0.75 |
| 14_1430348 | 16 | 0.4375 | 24 | 0.4167 | 29 | 0.4828 | 21 | 0.3333 | 90 | 0.4222 | 16 | 0.375 | 18 | 0.4444 | 23 | 0.3478 | 29 | 0.3448 | 86 | 0.3721 |
| 14_1430351 | 16 | 0.0625 | 29 | 0.069 | 35 | 0.1143 | 22 | 0 | 102 | 0.0686 | 17 | 0.1176 | 22 | 0.0455 | 26 | 0.0769 | 33 | 0.0303 | 98 | 0.0612 |
| 14_1430355 | 15 | 0.1333 | 27 | 0.5556 | 28 | 0.1786 | 17 | 0.3529 | 87 | 0.3218 | 15 | 0.6 | 21 | 0.4286 | 24 | 0.3333 | 31 | 0.2903 | 91 | 0.3846 |
| 14_1430501 | 3 | 0.3333 | 5 | 0 | 2 | 0 | 1 | 0 | 11 | 0.0909 | 2 | 0 | 1 | 1 | 6 | 0.6667 | 3 | 0.3333 | 12 | 0.5 |
| 14_1430507 | NA | NA | 2 | 0 | 2 | 0.5 | 1 | 0 | 5 | 0.2 | 2 | 0 | 2 | 0 | 5 | 0 | 5 | 0.4 | 14 | 0.1429 |
| 14_1430508 | 2 | 1 | 4 | 0.75 | 2 | 0 | 1 | 1 | 9 | 0.6667 | 2 | 0.5 | 2 | 0.5 | 6 | 0.8333 | 5 | 0.6 | 15 | 0.6667 |
| 14_1430566 | 2 | 1 | 3 | 0 | 2 | 0.5 | 1 | 0 | 8 | 0.375 | 1 | 0 | 2 | 0.5 | 3 | 0 | 3 | 1 | 9 | 0.4444 |
| 14_1437906 | 7 | 0 | 7 | 0 | 1 | 0 | 4 | 0.25 | 19 | 0.0526 | 4 | 0.25 | 8 | 0 | 6 | 0 | 4 | 0.25 | 22 | 0.0909 |
| 14_1437916 | 5 | 0 | 4 | 0.5 | NA | NA | 4 | 0.25 | 13 | 0.2308 | 4 | 0.75 | 6 | 0.3333 | 4 | 0.25 | 2 | 0.5 | 16 | 0.4375 |
| 14_1437927 | 5 | 0.4 | 6 | 0 | 2 | 1 | 4 | 0 | 17 | 0.2353 | 2 | 1 | 3 | 0 | 8 | 0 | 2 | 0 | 15 | 0.1333 |
| 14_1437945 | 5 | 1 | 7 | 1 | 2 | 1 | 4 | 1 | 18 | 1 | 2 | 1 | 6 | 1 | 9 | 1 | 2 | 1 | 19 | 1 |
| 14_1437948 | 7 | 0.8571 | 9 | 0.8889 | 2 | 1 | 5 | 0.8 | 23 | 0.8696 | 1 | 1 | 6 | 0.8333 | 7 | 0.7143 | 2 | 1 | 16 | 0.8125 |
| 14_1437967 | 8 | 0.625 | 9 | 0.7778 | 1 | 1 | 5 | 0.4 | 23 | 0.6522 | 2 | 1 | 6 | 0.6667 | 8 | 0.75 | 2 | 0.5 | 18 | 0.7222 |
| 14_1437971 | 7 | 0.4286 | 9 | 0.2222 | 2 | 0 | 5 | 0.2 | 23 | 0.2609 | 2 | 1 | 5 | 0 | 5 | 0.2 | 2 | 0.5 | 14 | 0.2857 |
| 14_1437981 | 7 | 0.7143 | 8 | 0.875 | 2 | 0.5 | 4 | 1 | 21 | 0.8095 | 1 | 1 | 6 | 1 | 7 | 0.8571 | 1 | 1 | 15 | 0.9333 |
| 14_1441308 | 5 | 0.2 | 7 | 0 | 2 | 0 | 3 | 0 | 17 | 0.0588 | 7 | 0 | 6 | 0 | 3 | 0.3333 | 9 | 0.1111 | 25 | 0.08 |
| 14_1441319 | 7 | 0.8571 | 8 | 0.625 | 4 | 1 | 3 | 0.3333 | 22 | 0.7273 | 9 | 0.8889 | 6 | 0.5 | 5 | 0.6 | 12 | 0.6667 | 32 | 0.6875 |
| 14_1441321 | 7 | 0.2857 | 8 | 0.25 | 4 | 0 | 3 | 0 | 22 | 0.1818 | 10 | 0.1 | 7 | 0.4286 | 4 | 0.25 | 10 | 0.1 | 31 | 0.1935 |
| 14_1441323 | 6 | 1 | 8 | 0.875 | 5 | 0.8 | 4 | 1 | 23 | 0.913 | 10 | 0.9 | 7 | 0.8571 | 5 | 1 | 11 | 0.8182 | 33 | 0.8788 |
| 14_1441334 | 7 | 1 | 7 | 1 | 2 | 1 | 1 | 1 | 17 | 1 | 10 | 0.8 | 6 | 0.8333 | 4 | 1 | 8 | 0.875 | 28 | 0.8571 |
| 14_1441336 | 8 | 0.125 | 9 | 0 | 5 | 0.2 | 2 | 0 | 24 | 0.0833 | 11 | 0.2727 | 7 | 0.1429 | 4 | 0.25 | 9 | 0.2222 | 31 | 0.2258 |
| 14_1441346 | 8 | 0.625 | 7 | 0.5714 | 5 | 0.6 | 2 | 1 | 22 | 0.6364 | 8 | 0.5 | 6 | 0.5 | 5 | 0.8 | 8 | 0.625 | 27 | 0.5926 |
| 14_1441357 | 5 | 0.8 | 7 | 0.5714 | 3 | 0.6667 | 2 | 1 | 17 | 0.7059 | 10 | 0.6 | 4 | 0.75 | 4 | 0.75 | 11 | 0.8182 | 29 | 0.7241 |
| 14_1441358 | 6 | 0.1667 | 7 | 0 | 4 | 0.5 | 2 | 0 | 19 | 0.1579 | 10 | 0.4 | 7 | 0.1429 | 5 | 0.2 | 10 | 0.1 | 32 | 0.2188 |
| 14_1441366 | 6 | 0.8333 | 5 | 0.8 | 3 | 0.3333 | 2 | 0.5 | 16 | 0.6875 | 10 | 0.5 | 6 | 0.8333 | 4 | 1 | 11 | 0.8182 | 31 | 0.7419 |
| 14_1441377 | 8 | 0.875 | 8 | 0.625 | 3 | 1 | 3 | 0.6667 | 22 | 0.7727 | 9 | 0.7778 | 6 | 0.8333 | 4 | 1 | 9 | 0.4444 | 28 | 0.7143 |
| 14_1441379 | 9 | 1 | 9 | 0.8889 | 2 | 1 | 5 | 0.8 | 25 | 0.92 | 10 | 1 | 7 | 1 | 4 | 1 | 11 | 1 | 32 | 1 |
| 14_1441389 | 5 | 0.2 | 5 | 0.2 | 1 | 0 | 2 | 0 | 13 | 0.1538 | 5 | 0.2 | 3 | 0 | 2 | 0 | 4 | 0.5 | 14 | 0.2143 |
| 14_1441390 | 11 | 0.1818 | 10 | 0.1 | 3 | 0 | 6 | 0 | 30 | 0.1 | 13 | 0 | 7 | 0.2857 | 4 | 0.25 | 10 | 0.3 | 34 | 0.1765 |
| 14_1441391 | 9 | 0.2222 | 9 | 0.1111 | 3 | 0 | 5 | 0.4 | 26 | 0.1923 | 13 | 0.3846 | 5 | 0.2 | 5 | 0 | 9 | 0 | 32 | 0.1875 |
| 14_1441395 | 9 | 0.2222 | 7 | 0.5714 | 3 | 0 | 5 | 0.2 | 24 | 0.2917 | 12 | 0.5833 | 7 | 0.1429 | 6 | 0.3333 | 7 | 0.5714 | 32 | 0.4375 |
| 14_1441413 | 9 | 0.8889 | 5 | 0.8 | 3 | 0.6667 | 5 | 0.6 | 22 | 0.7727 | 11 | 0.8182 | 7 | 0.8571 | 4 | 1 | 7 | 0.8571 | 29 | 0.8621 |

| | | | | | | | | | | | | | | | | | | | | |
|---|---|---|---|---|---|---|---|---|---|---|---|---|---|---|---|---|---|---|---|---|
| 14_145183( | 3 | 0 | 4 | 0.25 | 2 | 0 | 1 | 0 | 10 | 0.1 | 2 | 0 | NA | NA | 1 | 0 | 6 | 0.3333 | 9 | 0.2222 |
| 14_145186( | 3 | 0.3333 | 2 | 0 | 2 | 0 | 1 | 0 | 8 | 0.125 | 2 | 0 | NA | NA | 1 | 1 | 4 | 0.5 | 7 | 0.4286 |
| 14_145187( | 3 | 0.6667 | 2 | 0 | 1 | 1 | NA | NA | 6 | 0.5 | 2 | 0.5 | NA | NA | 1 | 0 | 5 | 0.4 | 8 | 0.375 |
| 14_145189( | 3 | 0.3333 | 2 | 0.5 | 2 | 1 | 1 | 1 | 8 | 0.625 | 2 | 0.5 | NA | NA | 1 | 0 | 3 | 0.6667 | 6 | 0.5 |
| 14_145189! | 2 | 0 | 1 | 0 | 2 | 0.5 | 1 | 0 | 6 | 0.1667 | 2 | 0.5 | NA | NA | 1 | 0 | 5 | 0.2 | 8 | 0.25 |
| 14_145190( | 2 | 0.5 | 1 | 0 | 2 | 0.5 | 1 | 1 | 6 | 0.5 | 2 | 1 | NA | NA | 1 | 1 | 5 | 0.6 | 8 | 0.75 |
| 14_145190( | 1 | 1 | NA | NA | 2 | 0.5 | 1 | 1 | 4 | 0.75 | 1 | 0 | NA | NA | NA | NA | 4 | 0.25 | 5 | 0.2 |
| 14_145190! | 2 | 0.5 | 1 | 0 | 1 | 1 | 1 | 1 | 5 | 0.6 | 2 | 0.5 | NA | NA | 1 | 1 | 4 | 0.5 | 7 | 0.5714 |
| 14_145191( | 3 | 1 | 2 | 0 | 2 | 1 | 1 | 1 | 8 | 0.75 | 1 | 1 | NA | NA | 1 | 0 | 5 | 0.6 | 7 | 0.5714 |
| 14_145191: | 1 | 1 | 2 | 0 | 1 | 0 | 1 | 0 | 5 | 0.2 | NA | NA | NA | NA | 1 | 0 | 4 | 0.25 | 5 | 0.2 |
| 14_145640: | 1 | 1 | NA | NA | NA | NA | NA | NA | 1 | 1 | NA | NA | NA | NA | 1 | 1 | NA | NA | 1 | 1 |
| 14_152463( | NA | NA | NA | NA | NA | NA | 1 | 1 | 1 | 1 | NA | NA | NA | NA | NA | NA | 1 | 1 | 1 | 1 |
| 14_152469! | NA | NA | NA | NA | NA | NA | 1 | 1 | 1 | 1 | NA | NA | NA | NA | NA | NA | 1 | 1 | 1 | 1 |
| 14_153858: | NA | NA | 1 | 1 | 1 | 1 | 1 | 1 | 3 | 1 | NA | NA | 1 | 1 | NA | NA | NA | NA | 1 | 1 |
| 14_153862! | NA | NA | 1 | 1 | 1 | 1 | 1 | 1 | 3 | 1 | NA | NA | 1 | 1 | NA | NA | NA | NA | 1 | 1 |
| 14_154633! | NA | NA | 1 | 1 | NA | NA | NA | NA | 1 | 1 | NA | NA | NA | NA | NA | NA | 1 | 1 | 1 | 1 |
| 14_154634( | NA | NA | 1 | 1 | NA | NA | NA | NA | 1 | 1 | NA | NA | NA | NA | NA | NA | 1 | 1 | 1 | 1 |
| 14_157331! | 2 | 1 | 1 | 1 | 1 | 1 | NA | NA | 4 | 1 | 1 | 1 | NA | NA | 2 | 1 | NA | NA | 3 | 1 |
| 14_157333' | 1 | 1 | 2 | 0.5 | 1 | 0 | NA | NA | 4 | 0.5 | 1 | 0 | NA | NA | 2 | 0.5 | NA | NA | 3 | 0.3333 |
| 14_157334: | 2 | 1 | 2 | 1 | 1 | 1 | NA | NA | 5 | 1 | 1 | 1 | NA | NA | 2 | 0 | NA | NA | 3 | 0.3333 |
| 14_157334! | 1 | 0 | 2 | 0.5 | 1 | 0 | NA | NA | 4 | 0.25 | 1 | 1 | NA | NA | 2 | 0.5 | NA | NA | 3 | 0.6667 |
| 14_157338: | 2 | 1 | 2 | 1 | 1 | 1 | NA | NA | 5 | 1 | 1 | 1 | NA | NA | 3 | 0.6667 | NA | NA | 4 | 0.75 |
| 14_157338! | 2 | 1 | 2 | 0.5 | 1 | 1 | NA | NA | 5 | 0.8 | 1 | 1 | NA | NA | 3 | 0 | NA | NA | 4 | 0.25 |
| 14_165303 | NA | NA | 1 | 1 | NA | NA | NA | NA | 1 | 1 | NA | NA | NA | NA | 1 | 1 | NA | NA | 1 | 1 |
| 14_170309( | NA | NA | 1 | 1 | NA | NA | NA | NA | 1 | 1 | 1 | 1 | NA | NA | NA | NA | NA | NA | 1 | 1 |
| 14_170310 | NA | NA | 1 | 1 | NA | NA | NA | NA | 1 | 1 | 1 | 1 | NA | NA | NA | NA | NA | NA | 1 | 1 |
| 14_170312! | NA | NA | 1 | 1 | NA | NA | NA | NA | 1 | 1 | 1 | 1 | NA | NA | NA | NA | NA | NA | 1 | 1 |
| 14_170314< | NA | NA | 1 | 1 | NA | NA | NA | NA | 1 | 1 | 1 | 1 | NA | NA | NA | NA | NA | NA | 1 | 1 |
| 14_182650! | NA | NA | NA | NA | 1 | 1 | NA | NA | 1 | 1 | NA | NA | 1 | 1 | NA | NA | NA | NA | 1 | 1 |
| 14_182651' | NA | NA | NA | NA | 1 | 1 | NA | NA | 1 | 1 | NA | NA | 1 | 1 | NA | NA | NA | NA | 1 | 1 |
| 14_182651< | NA | NA | NA | NA | 1 | 1 | NA | NA | 1 | 1 | NA | NA | 1 | 1 | NA | NA | NA | NA | 1 | 1 |
| 14_182657! | NA | NA | NA | NA | 1 | 1 | NA | NA | 1 | 1 | NA | NA | 1 | 1 | NA | NA | NA | NA | 1 | 1 |
| 14_186393! | NA | NA | NA | NA | 1 | 1 | NA | NA | 1 | 1 | NA | NA | NA | NA | NA | NA | 1 | 1 | 1 | 1 |
| 14_186399! | NA | NA | NA | NA | 1 | 1 | NA | NA | 1 | 1 | NA | NA | NA | NA | NA | NA | 1 | 1 | 1 | 1 |
| 14_192360: | 1 | 1 | 1 | 1 | 2 | 1 | 2 | 1 | 6 | 1 | NA | NA | NA | NA | NA | NA | 1 | 1 | 1 | 1 |
| 14_193931( | NA | NA | NA | NA | 1 | 1 | 2 | 1 | 3 | 1 | NA | NA | 1 | 1 | NA | NA | NA | NA | 1 | 1 |
| 14_193933! | NA | NA | NA | NA | 1 | 0 | 1 | 1 | 2 | 0.5 | NA | NA | 1 | 1 | NA | NA | NA | NA | 1 | 1 |
| 14_193935! | NA | NA | NA | NA | 1 | 1 | 1 | 0 | 2 | 0.5 | NA | NA | 1 | 1 | NA | NA | NA | NA | 1 | 1 |
| 14_193938: | NA | NA | NA | NA | 1 | 1 | 2 | 1 | 3 | 1 | NA | NA | 1 | 1 | NA | NA | NA | NA | 1 | 1 |
| 14_196145: | NA | NA | NA | NA | 1 | 1 | NA | NA | 1 | 1 | NA | NA | NA | NA | NA | NA | 2 | 1 | 2 | 1 |
| 14_196147 | NA | NA | NA | NA | 1 | 1 | NA | NA | 1 | 1 | NA | NA | NA | NA | NA | NA | 2 | 1 | 2 | 1 |
| 14_198617< | NA | NA | NA | NA | 1 | 1 | NA | NA | 1 | 1 | NA | NA | NA | NA | 1 | 1 | 1 | 1 | 2 | 1 |
| 14_198620< | 9 | 0 | 17 | 0.2941 | 18 | 0.0556 | 10 | 0 | 54 | 0.1111 | 13 | 0.0769 | 13 | 0.1538 | 14 | 0.1429 | 12 | 0 | 52 | 0.0962 |
| 14_198621 | 10 | 0.2 | 17 | 0.1176 | 17 | 0.1176 | 13 | 0.1538 | 57 | 0.1404 | 13 | 0 | 16 | 0.25 | 12 | 0.1667 | 11 | 0 | 52 | 0.1154 |
| 14_199237! | 6 | 0 | 5 | 0.4 | 1 | 0 | 7 | 0 | 19 | 0.1053 | 4 | 0 | 7 | 0 | 7 | 0.2857 | 4 | 0 | 22 | 0.0909 |
| 14_199255! | 4 | 0 | 4 | 0 | 4 | 0 | 8 | 0.5 | 20 | 0.2 | 6 | 0 | 5 | 0.2 | 6 | 0.1667 | 6 | 0.3333 | 23 | 0.1739 |
| 14_199257! | 6 | 0.6667 | 6 | 0.5 | 2 | 0.5 | 8 | 0.75 | 22 | 0.6364 | 6 | 0.3333 | 5 | 0.8 | 6 | 0.5 | 6 | 0.8333 | 23 | 0.6087 |
| 14_199258: | 5 | 0.8 | 6 | 0.8333 | 2 | 1 | 8 | 0.75 | 21 | 0.8095 | 7 | 0.4286 | 4 | 1 | 4 | 0.5 | 5 | 0.6 | 20 | 0.6 |
| 14_201272: | 2 | 1 | 2 | 1 | 1 | 1 | 2 | 1 | 7 | 1 | 2 | 1 | 4 | 1 | 2 | 1 | 1 | 1 | 9 | 1 |
| 14_201272 | 2 | 0.5 | 2 | 0.5 | 1 | 0 | 1 | 0 | 6 | 0.3333 | 1 | 1 | 4 | 0.25 | 2 | 0.5 | 1 | 1 | 8 | 0.5 |
| 14_201273: | 2 | 1 | NA | NA | NA | NA | 2 | 1 | 4 | 1 | NA | NA | 4 | 0.75 | 1 | 1 | NA | NA | 5 | 0.8 |
| 14_201273: | 2 | 0 | 2 | 0.5 | 1 | 0 | 2 | 0 | 7 | 0.1429 | 2 | 0.5 | 4 | 0.25 | 2 | 0.5 | NA | NA | 8 | 0.375 |
| 14_201273< | 2 | 1 | 2 | 1 | 1 | 1 | 2 | 1 | 7 | 1 | 2 | 1 | 2 | 1 | NA | NA | NA | NA | 4 | 1 |
| 14_201274: | 2 | 0.5 | 2 | 0 | 1 | 0 | 1 | 0 | 6 | 0.1667 | 1 | 0 | 4 | 0 | 2 | 0.5 | 1 | 0 | 8 | 0.125 |
| 14_201275: | 2 | 1 | 1 | 1 | 1 | 0 | 2 | 0 | 6 | 0.5 | 2 | 0.5 | 4 | 0.5 | 2 | 0.5 | 1 | 0 | 9 | 0.4444 |
| 14_201275! | 1 | 1 | 2 | 1 | 1 | 1 | 1 | 1 | 5 | 1 | 2 | 1 | 3 | 1 | 2 | 1 | 1 | 1 | 8 | 1 |
| 14_201275! | 2 | 1 | 2 | 1 | 1 | 1 | 1 | 1 | 6 | 1 | 2 | 1 | 3 | 1 | 2 | 1 | 1 | 1 | 8 | 1 |
| 14_203664< | 3 | 1 | 1 | 0 | 1 | 0 | 1 | 1 | 6 | 0.6667 | 1 | 0 | 1 | 1 | NA | NA | NA | NA | 2 | 0.5 |
| 14_203665! | 2 | 1 | 1 | 1 | NA | NA | 2 | 1 | 5 | 1 | 1 | 1 | 2 | 0.5 | NA | NA | NA | NA | 3 | 0.6667 |
| 14_203665! | 2 | 0.5 | 1 | 0 | 1 | 0 | 3 | 0.6667 | 7 | 0.4286 | 1 | 1 | 3 | 0.3333 | 1 | 1 | NA | NA | 5 | 0.6 |
| 14_203666! | 3 | 0.6667 | NA | NA | NA | NA | 3 | 0.3333 | 6 | 0.5 | 1 | 1 | 2 | 0 | 1 | 0 | NA | NA | 4 | 0.25 |
| 14_203667' | 3 | 0.6667 | 1 | 1 | 1 | 1 | 3 | 0.3333 | 8 | 0.625 | 1 | 1 | 3 | 0.6667 | NA | NA | NA | NA | 4 | 0.75 |
| 14_203705: | 3 | 0.3333 | 4 | 0.25 | 3 | 0 | 2 | 0 | 12 | 0.1667 | 6 | 0.1667 | 5 | 0.4 | 10 | 0.1 | 3 | 0.6667 | 24 | 0.25 |
| 14_203707! | 5 | 1 | 5 | 0.8 | 2 | 0.5 | 2 | 1 | 14 | 0.8571 | 5 | 0.8 | 5 | 1 | 12 | 0.9167 | 3 | 0.6667 | 25 | 0.88 |
| 14_203709: | 5 | 0 | 4 | 0.25 | 2 | 0 | 2 | 0 | 13 | 0.0769 | 5 | 0 | 4 | 0 | 12 | 0.1667 | 5 | 0 | 26 | 0.0769 |

| | | | | | | | | | | | | | | | | | | | | |
|---|---|---|---|---|---|---|---|---|---|---|---|---|---|---|---|---|---|---|---|---|
| 14_2041801 | NA | NA | 1 | 0 | 2 | 0 | 2 | 0.5 | 5 | 0.2 | 1 | 0 | 3 | 0.3333 | 2 | 0.5 | 1 | 0 | 7 | 0.2857 |
| 14_2041801 | NA | NA | 1 | 0 | 1 | 0 | 3 | 0.6667 | 5 | 0.4 | NA | NA | 3 | 0 | 2 | 0 | 1 | 1 | 6 | 0.1667 |
| 14_2042582 | NA | NA | NA | NA | 1 | 1 | NA | NA | 1 | 1 | 1 | 0 | 1 | 0 | NA | NA | 4 | 0.5 | 6 | 0.3333 |
| 14_2042623 | NA | NA | 1 | 0 | 1 | 0 | 1 | 1 | 3 | 0.3333 | 1 | 0 | 1 | 0 | 2 | 0 | 4 | 0.5 | 8 | 0.25 |
| 14_2043322 | 1 | 1 | 6 | 0.1667 | 3 | 0.6667 | 2 | 1 | 12 | 0.5 | 3 | 1 | 1 | 1 | 1 | 0 | 2 | 0.5 | 7 | 0.7143 |
| 14_2043324 | 1 | 1 | 5 | 0.2 | 3 | 0 | 3 | 0 | 12 | 0.1667 | 4 | 0.25 | NA | NA | 1 | 0 | 2 | 0.5 | 7 | 0.2857 |
| 14_2043332 | 1 | 0 | 6 | 0.5 | 3 | 0.3333 | 3 | 1 | 13 | 0.5385 | 4 | 0.5 | 1 | 1 | 1 | 0 | 2 | 1 | 8 | 0.625 |
| 14_2043347 | 2 | 0 | 6 | 0.1667 | 3 | 0.3333 | 2 | 0 | 13 | 0.1538 | 5 | 0.4 | 1 | 0 | 1 | 0 | 2 | 0.5 | 9 | 0.3333 |
| 14_2043551 | 3 | 0.3333 | 1 | 0 | 3 | 0.6667 | 1 | 0 | 8 | 0.375 | 1 | 1 | NA | NA | 2 | 0 | 1 | 1 | 4 | 0.5 |
| 14_2043813 | 3 | 0.6667 | 2 | 0 | 3 | 0 | 1 | 0 | 9 | 0.2222 | 1 | 1 | 1 | 0 | 1 | 0 | 1 | 0 | 4 | 0.25 |
| 14_2043817 | 4 | 0.25 | 2 | 0 | 4 | 0 | NA | NA | 10 | 0.1 | 3 | 0.3333 | 1 | 0 | 1 | 1 | NA | NA | 5 | 0.4 |
| 14_2043819 | 4 | 0.25 | 2 | 0 | 4 | 0.25 | 1 | 0 | 11 | 0.1818 | 2 | 0.5 | 1 | 1 | 1 | 1 | 1 | 1 | 5 | 0.8 |
| 14_2050554 | 11 | 0 | 8 | 0.125 | 6 | 0.1667 | 2 | 0 | 27 | 0.0741 | 3 | 0 | NA | NA | 4 | 0.25 | 5 | 0 | 12 | 0.0833 |
| 14_2050643 | 11 | 0.3636 | 8 | 0.25 | 8 | 0.375 | 8 | 0.25 | 35 | 0.3143 | 5 | 0.2 | 1 | 1 | 6 | 0.1667 | 7 | 0.2857 | 19 | 0.2632 |
| 14_2055015 | 47 | 0.0851 | 60 | 0.1167 | 55 | 0.0727 | 43 | 0.0233 | 205 | 0.078 | 47 | 0.1064 | 55 | 0.0909 | 44 | 0.0909 | 49 | 0.0408 | 195 | 0.0821 |
| 14_2055034 | 46 | 0.087 | 58 | 0.1034 | 52 | 0.1346 | 38 | 0.0526 | 194 | 0.0979 | 45 | 0.2 | 49 | 0.1224 | 43 | 0.1163 | 45 | 0.1111 | 182 | 0.1374 |
| 14_2055038 | 41 | 0.0488 | 58 | 0.1207 | 46 | 0.1739 | 38 | 0.0789 | 183 | 0.1093 | 43 | 0.2093 | 50 | 0.18 | 38 | 0.2105 | 44 | 0.1364 | 175 | 0.1829 |
| 14_2055119 | 29 | 0.1034 | 41 | 0.0732 | 34 | 0.0882 | 35 | 0.0857 | 139 | 0.0863 | 31 | 0.129 | 44 | 0.1591 | 35 | 0.1143 | 28 | 0.0714 | 138 | 0.1232 |
| 14_2055121 | 31 | 0.0645 | 40 | 0.05 | 31 | 0.0968 | 33 | 0.0303 | 135 | 0.0593 | 30 | 0.1333 | 43 | 0.0233 | 33 | 0.0303 | 31 | 0.0968 | 137 | 0.0657 |
| 14_2055138 | 31 | 0.9677 | 41 | 0.9756 | 33 | 1 | 34 | 0.9412 | 139 | 0.9712 | 29 | 0.9655 | 43 | 0.9302 | 33 | 1 | 29 | 0.9655 | 134 | 0.9627 |
| 14_2055150 | 23 | 0.9565 | 32 | 0.9062 | 20 | 0.95 | 25 | 0.88 | 100 | 0.92 | 23 | 0.9565 | 29 | 0.8621 | 25 | 0.88 | 19 | 0.9474 | 96 | 0.9062 |
| 14_2055153 | 27 | 0.8519 | 37 | 0.8919 | 34 | 0.9118 | 34 | 0.8824 | 132 | 0.8864 | 26 | 1 | 42 | 0.9286 | 32 | 0.875 | 28 | 0.9286 | 128 | 0.9297 |
| 14_2055157 | 27 | 0.8148 | 38 | 0.4737 | 28 | 0.6429 | 32 | 0.5312 | 125 | 0.6 | 25 | 0.6 | 39 | 0.641 | 32 | 0.5625 | 23 | 0.4783 | 119 | 0.5798 |
| 14_2055158 | 24 | 0.0833 | 30 | 0.0667 | 25 | 0 | 25 | 0.12 | 104 | 0.0673 | 18 | 0.1667 | 35 | 0.0571 | 28 | 0.0357 | 21 | 0.0952 | 102 | 0.0784 |
| 14_2059976 | 3 | 0.3333 | 3 | 0.6667 | 2 | 0 | 2 | 0.5 | 10 | 0.4 | 3 | 0.6667 | 3 | 0 | 3 | 0.3333 | 2 | 0.5 | 11 | 0.3636 |
| 14_2059977 | 7 | 0 | 5 | 0.4 | 4 | 0.75 | 5 | 0.2 | 21 | 0.2857 | 3 | 0 | 3 | 0.3333 | 5 | 0.2 | 3 | 0 | 14 | 0.1429 |
| 14_2059992 | 2 | 0 | 4 | 0 | 4 | 0.25 | 5 | 0 | 15 | 0.0667 | 3 | 0 | 3 | 0.3333 | 3 | 0 | 1 | 0 | 10 | 0.1 |
| 14_2059992 | 1 | 1 | 1 | 0 | NA | NA | NA | NA | 2 | 0.5 | NA | NA | NA | NA | 2 | 0 | 2 | 0.5 | 4 | 0.25 |
| 14_2060041 | 4 | 0.5 | 1 | 0 | 1 | 0 | 6 | 0.3333 | 12 | 0.3333 | 3 | 0 | 3 | 0 | 4 | 0.25 | NA | NA | 10 | 0.1 |
| 14_2060044 | 4 | 0.5 | 2 | 0 | 2 | 0 | 6 | 0 | 14 | 0.1429 | 2 | 0.5 | 3 | 0.3333 | 4 | 0.5 | 1 | 0 | 10 | 0.4 |
| 14_2061101 | 4 | 0.5 | 1 | 0 | 1 | 1 | 5 | 0 | 11 | 0.2727 | 3 | 0 | 2 | 0.5 | 3 | 0.6667 | 1 | 0 | 9 | 0.3333 |
| 14_2061104 | 4 | 0.25 | 1 | 0 | 2 | 0 | 4 | 0 | 11 | 0.0909 | 3 | 0 | 1 | 1 | 2 | 0.5 | 1 | 0 | 7 | 0.2857 |
| 14_2061120 | 4 | 1 | 1 | 1 | 1 | 1 | 5 | 1 | 11 | 1 | 2 | 1 | 2 | 1 | 1 | 1 | 1 | 1 | 6 | 1 |
| 14_2061124 | 4 | 1 | 1 | 1 | 1 | 1 | 6 | 0.3333 | 12 | 0.6667 | 2 | 0 | 1 | 1 | 3 | 0.3333 | 1 | 1 | 7 | 0.4286 |
| 14_2061130 | 4 | 0.25 | 1 | 1 | 1 | 0 | 6 | 0.5 | 12 | 0.4167 | 2 | 0.5 | 2 | 0.5 | 3 | 0.3333 | 1 | 0 | 8 | 0.375 |
| 14_2061146 | 4 | 0.5 | 1 | 0 | 1 | 0 | 3 | 0.3333 | 9 | 0.3333 | 1 | 1 | 2 | 0 | 3 | 0 | 1 | 0 | 7 | 0.1429 |
| 14_2061158 | 4 | 1 | 1 | 1 | 1 | 0 | 5 | 0.8 | 11 | 0.8182 | 1 | 1 | 1 | 1 | 2 | 1 | 1 | 1 | 5 | 1 |
| 14_2061164 | 4 | 0 | 1 | 1 | NA | NA | 5 | 0.6 | 10 | 0.4 | 1 | 0 | 2 | 0.5 | 2 | 0.5 | 1 | 1 | 6 | 0.5 |
| 14_2061192 | 2 | 0.5 | 1 | 0 | NA | NA | 3 | 0 | 6 | 0.1667 | 1 | 0 | 1 | 0 | 2 | 0.5 | 1 | 0 | 5 | 0.2 |
| 14_2061202 | 2 | 1 | 1 | 1 | NA | NA | 4 | 1 | 7 | 1 | 1 | 1 | 1 | 1 | 2 | 0.5 | NA | NA | 4 | 0.75 |
| 14_2061203 | 1 | 1 | 1 | 0 | NA | NA | 4 | 0.5 | 6 | 0.5 | 1 | 0 | NA | NA | 2 | 1 | NA | NA | 3 | 0.6667 |
| 14_2074360 | 1 | 1 | NA | NA | NA | NA | NA | NA | 1 | 1 | NA | NA | 1 | 0 | 2 | 0.5 | NA | NA | 3 | 0.3333 |
| 14_2074381 | 4 | 0.75 | 5 | 0.8 | 7 | 0.5714 | 5 | 1 | 21 | 0.7619 | 4 | 1 | 5 | 0.6 | 7 | 1 | 4 | 1 | 20 | 0.9 |
| 14_2074393 | 5 | 0.6 | 5 | 0.8 | 8 | 0.75 | 5 | 0.2 | 23 | 0.6087 | 5 | 0.8 | 6 | 0.6667 | 8 | 0.625 | 4 | 1 | 23 | 0.7391 |
| 14_2074393 | 1 | 1 | NA | NA | NA | NA | NA | NA | 1 | 1 | NA | NA | 1 | 1 | NA | NA | NA | NA | 1 | 1 |
| 14_2074395 | 1 | 1 | NA | NA | NA | NA | NA | NA | 1 | 1 | NA | NA | 1 | 0 | 1 | 1 | NA | NA | 2 | 0.5 |
| 14_2074404 | 1 | 1 | NA | NA | NA | NA | NA | NA | 1 | 1 | NA | NA | NA | NA | 1 | 1 | NA | NA | 1 | 1 |
| 14_2074413 | 1 | 1 | NA | NA | NA | NA | NA | NA | 1 | 1 | NA | NA | 1 | 1 | NA | NA | NA | NA | 1 | 1 |
| 14_2074440 | 1 | 1 | NA | NA | NA | NA | NA | NA | 1 | 1 | NA | NA | 1 | 1 | 1 | 1 | NA | NA | 2 | 1 |
| 14_2074473 | 10 | 0.7 | 9 | 0.7778 | 9 | 0.5556 | 8 | 0.625 | 36 | 0.6667 | 6 | 0.6667 | 8 | 0.875 | 9 | 0.6667 | 5 | 1 | 28 | 0.7857 |
| 14_2074484 | 10 | 0.5 | 9 | 0.6667 | 6 | 0.6667 | 7 | 1 | 32 | 0.6875 | 8 | 1 | 7 | 0.8571 | 8 | 0.875 | 3 | 1 | 26 | 0.9231 |
| 14_2074543 | NA | NA | 2 | 0 | 1 | 1 | NA | NA | 3 | 0.3333 | NA | NA | 2 | 1 | NA | NA | 2 | 0 | 4 | 0.5 |
| 14_2074594 | NA | NA | 3 | 0.6667 | 2 | 0 | 2 | 0 | 7 | 0.2857 | NA | NA | 1 | 1 | 3 | 0 | 5 | 0.4 | 9 | 0.3333 |
| 14_2074602 | NA | NA | 3 | 1 | 1 | 1 | 2 | 1 | 6 | 1 | NA | NA | 1 | 1 | 3 | 1 | 4 | 1 | 8 | 1 |
| 14_2074608 | NA | NA | 3 | 0.6667 | 1 | 0 | 2 | 1 | 6 | 0.6667 | NA | NA | 2 | 1 | 3 | 1 | 5 | 0.4 | 10 | 0.7 |
| 14_2074612 | NA | NA | 3 | 0.6667 | 2 | 0.5 | 1 | 1 | 6 | 0.6667 | NA | NA | 2 | 0 | 2 | 0.5 | 3 | 0.6667 | 7 | 0.4286 |
| 14_2074616 | NA | NA | 3 | 1 | 2 | 1 | 1 | 1 | 6 | 1 | NA | NA | 1 | 1 | 3 | 1 | 4 | 1 | 8 | 1 |
| 14_2074616 | NA | NA | 3 | 1 | 2 | 1 | 2 | 0.5 | 7 | 0.8571 | NA | NA | 2 | 1 | 3 | 1 | 5 | 0.6 | 10 | 0.8 |
| 14_2074632 | 1 | 1 | 2 | 0 | 1 | 0 | 1 | 0 | 5 | 0.2 | NA | NA | 2 | 0 | 3 | 0.3333 | 4 | 0 | 9 | 0.1111 |
| 14_2074633 | NA | NA | NA | NA | 1 | 1 | NA | NA | 1 | 1 | NA | NA | NA | NA | NA | NA | 1 | 1 | 1 | 1 |
| 14_2074637 | 1 | 0 | 2 | 0.5 | 1 | 1 | 2 | 0 | 6 | 0.3333 | NA | NA | NA | NA | 2 | 0 | 5 | 0.4 | 7 | 0.2857 |
| 14_2074642 | 1 | 1 | 3 | 1 | 1 | 1 | 1 | 0 | 6 | 0.8333 | NA | NA | 2 | 0 | 2 | 0.5 | 4 | 0.75 | 8 | 0.5 |
| 14_2074842 | 1 | 1 | 4 | 0 | 2 | 1 | 1 | 0 | 8 | 0.375 | NA | NA | 1 | 0 | 3 | 0.3333 | 4 | 0.25 | 8 | 0.25 |
| 14_2074845 | 1 | 0 | 2 | 0 | 2 | 1 | 1 | 1 | 6 | 0.5 | NA | NA | 1 | 1 | 3 | 0.6667 | 3 | 1 | 7 | 0.8571 |
| 14_2074855 | 1 | 1 | 2 | 0.5 | 2 | 0.5 | 2 | 1 | 7 | 0.7143 | NA | NA | 1 | 0 | 3 | 0.6667 | 4 | 0.75 | 8 | 0.625 |

| | | | | | | | | | | | | | | | | | | | | |
|---|---|---|---|---|---|---|---|---|---|---|---|---|---|---|---|---|---|---|---|---|
| 14_2074873 | 1 | 0 | 2 | 0.5 | 2 | 0.5 | 2 | 0.5 | 7 | 0.4286 | NA | NA | NA | NA | 3 | 0 | 3 | 0.6667 | 6 | 0.3333 |
| 14_2075278 | NA | NA | NA | NA | NA | NA | 2 | 0.5 | 2 | 0.5 | NA | NA | 1 | 0 | NA | NA | 1 | 1 | 2 | 0.5 |
| 14_2075432 | 7 | 0.8571 | 8 | 0.875 | 15 | 0.8667 | 10 | 0.8 | 40 | 0.85 | 7 | 0.7143 | 8 | 0.75 | 11 | 0.8182 | 8 | 0.875 | 34 | 0.7941 |
| 14_2076425 | 2 | 0.5 | 4 | 0 | NA | NA | 1 | 0 | 7 | 0.1429 | 1 | 0 | 3 | 0.3333 | 1 | 0 | 1 | 1 | 6 | 0.3333 |
| 14_2076458 | 2 | 1 | 4 | 0.25 | NA | NA | 1 | 1 | 7 | 0.5714 | 1 | 0 | 4 | 0.5 | NA | NA | NA | NA | 5 | 0.4 |
| 14_2076465 | 2 | 0.5 | 4 | 0.5 | NA | NA | 1 | 0 | 7 | 0.4286 | 1 | 1 | 4 | 1 | NA | NA | NA | NA | 5 | 1 |
| 14_2076487 | 2 | 0 | 3 | 0.3333 | NA | NA | 1 | 0 | 6 | 0.1667 | 1 | 0 | 2 | 1 | 1 | 0 | NA | NA | 4 | 0.5 |
| 14_2077156 | 5 | 0 | 9 | 0 | 1 | 0 | 4 | 0.5 | 19 | 0.1053 | 2 | 0 | 7 | 0 | 4 | 0 | 6 | 0.1667 | 19 | 0.0526 |
| 14_2077344 | 8 | 0.5 | 11 | 0.3636 | 4 | 0.25 | 8 | 0.25 | 31 | 0.3548 | 4 | 0.25 | 11 | 0.3636 | 7 | 0.4286 | 9 | 0.1111 | 31 | 0.2903 |
| 14_2077349 | 9 | 0.8889 | 15 | 0.7333 | 5 | 0.8 | 8 | 0.625 | 37 | 0.7568 | 4 | 0.5 | 11 | 0.8182 | 6 | 0.8333 | 9 | 0.6667 | 30 | 0.7333 |
| 14_2078506 | 17 | 0.1765 | 17 | 0.1176 | 21 | 0.1429 | 20 | 0.05 | 75 | 0.12 | 17 | 0.1765 | 11 | 0.2727 | 16 | 0.0625 | 19 | 0.1579 | 63 | 0.1587 |
| 14_2078512 | 16 | 0.25 | 14 | 0.5 | 20 | 0.45 | 16 | 0.4375 | 66 | 0.4091 | 15 | 0.4667 | 12 | 0.5833 | 14 | 0.5 | 18 | 0.4444 | 59 | 0.4915 |
| 14_2078513 | 17 | 0.3529 | 17 | 0.4706 | 19 | 0.5263 | 16 | 0.25 | 69 | 0.4058 | 17 | 0.4706 | 12 | 0.5833 | 16 | 0.25 | 17 | 0.4706 | 62 | 0.4355 |
| 14_2078532 | 15 | 0.0667 | 17 | 0.1176 | 18 | 0.1667 | 17 | 0.0588 | 67 | 0.1045 | 17 | 0.1176 | 12 | 0.0833 | 16 | 0.1875 | 16 | 0.125 | 61 | 0.1311 |
| 14_2078542 | 17 | 0.1765 | 18 | 0 | 18 | 0 | 17 | 0.0588 | 70 | 0.0571 | 18 | 0.1667 | 11 | 0.2727 | 15 | 0.1333 | 15 | 0 | 59 | 0.1356 |
| 14_2078570 | 12 | 0.25 | 10 | 0.1 | 18 | 0.1667 | 12 | 0.1667 | 52 | 0.1731 | 11 | 0.0909 | 11 | 0 | 13 | 0.2308 | 10 | 0.2 | 45 | 0.1333 |
| 14_2078571 | 15 | 0.2667 | 17 | 0.1765 | 16 | 0.4375 | 15 | 0.2667 | 63 | 0.2857 | 16 | 0.3125 | 12 | 0.3333 | 13 | 0.3077 | 16 | 0.1875 | 57 | 0.2807 |
| 14_2078577 | 13 | 0.1538 | 13 | 0.0769 | 13 | 0.1538 | 15 | 0.2 | 54 | 0.1481 | 17 | 0.1176 | 9 | 0.2222 | 12 | 0 | 11 | 0 | 49 | 0.0816 |
| 14_2078578 | 11 | 0.2727 | 11 | 0.0909 | 12 | 0.25 | 13 | 0.2308 | 47 | 0.2128 | 15 | 0.2667 | 10 | 0.5 | 9 | 0.2222 | 12 | 0.1667 | 46 | 0.2826 |
| 14_2078599 | 10 | 0.2 | 10 | 0.1 | 17 | 0.2941 | 14 | 0.1429 | 51 | 0.1961 | 12 | 0.5 | 11 | 0.2727 | 14 | 0.2143 | 13 | 0.0769 | 50 | 0.26 |
| 14_2078628 | 9 | 1 | 14 | 0.9286 | 14 | 1 | 15 | 0.8 | 52 | 0.9231 | 17 | 1 | 11 | 0.8182 | 13 | 0.9231 | 15 | 1 | 56 | 0.9464 |
| 14_2078629 | 9 | 1 | 13 | 0.7692 | 13 | 0.8462 | 14 | 0.9286 | 49 | 0.8776 | 14 | 1 | 12 | 0.8333 | 12 | 0.8333 | 14 | 0.6429 | 52 | 0.8269 |
| 14_2078636 | 9 | 0.2222 | 12 | 0.0833 | 14 | 0.2143 | 15 | 0.2 | 50 | 0.18 | 14 | 0.2143 | 11 | 0.2727 | 10 | 0.1 | 14 | 0 | 49 | 0.1429 |
| 14_2078640 | 10 | 0.1 | 11 | 0.0909 | 14 | 0 | 16 | 0.125 | 51 | 0.0784 | 15 | 0 | 11 | 0.0909 | 12 | 0.25 | 12 | 0 | 50 | 0.08 |
| 14_2078642 | 7 | 0.7143 | 13 | 0.6923 | 11 | 1 | 15 | 0.8667 | 46 | 0.8261 | 11 | 1 | 11 | 0.7273 | 12 | 0.75 | 11 | 1 | 45 | 0.8667 |
| 14_2078648 | 8 | 0.75 | 8 | 0.875 | 11 | 1 | 8 | 0.875 | 35 | 0.8857 | 12 | 0.75 | 10 | 0.6 | 5 | 0.2 | 9 | 0.8889 | 36 | 0.6667 |
| 14_2078651 | 9 | 0.8889 | 13 | 1 | 12 | 1 | 12 | 0.8333 | 46 | 0.9348 | 13 | 1 | 11 | 1 | 12 | 0.9167 | 12 | 0.9167 | 48 | 0.9583 |
| 14_2078658 | 4 | 1 | 6 | 0.1667 | 5 | 0.8 | 4 | 0.75 | 19 | 0.6316 | 6 | 0.8333 | 4 | 0.75 | 6 | 0.8333 | 7 | 0.7143 | 23 | 0.7826 |
| 14_2089247 | NA | NA | NA | NA | 1 | 1 | NA | NA | 1 | 1 | NA | NA | 1 | 1 | NA | NA | NA | NA | 1 | 1 |
| 14_2089278 | NA | NA | NA | NA | 1 | 1 | NA | NA | 1 | 1 | NA | NA | 1 | 1 | NA | NA | NA | NA | 1 | 1 |
| 14_2107061 | 1 | 0 | 1 | 1 | NA | NA | NA | NA | 2 | 0.5 | 4 | 0 | 5 | 0 | 3 | 0.3333 | 2 | 0 | 14 | 0.0714 |
| 14_2107078 | 1 | 1 | 1 | 0 | 1 | 0 | NA | NA | 3 | 0.3333 | 4 | 0 | 4 | 0.5 | 3 | 0.3333 | 2 | 0.5 | 13 | 0.3077 |
| 14_2108178 | 4 | 0.25 | 5 | 0.4 | 3 | 0 | 3 | 0.3333 | 15 | 0.2667 | 5 | 0.2 | 5 | 0 | 6 | 0.6667 | 2 | 1 | 18 | 0.3889 |
| 14_2108185 | 3 | 0 | 6 | 0.1667 | 2 | 0 | 3 | 0 | 14 | 0.0714 | 4 | 0 | 5 | 0 | 4 | 0.25 | 2 | 0 | 15 | 0.0667 |
| 14_2108193 | 2 | 0.5 | 3 | 0.6667 | 2 | 0.5 | 3 | 0.6667 | 10 | 0.6 | 2 | 0.5 | 4 | 0.5 | 5 | 0.4 | 1 | 0 | 12 | 0.4167 |
| 14_2108197 | 3 | 0.3333 | 6 | 0.1667 | 2 | 0 | 3 | 0.3333 | 14 | 0.2143 | 4 | 0 | 6 | 0.1667 | 6 | 0.5 | 2 | 0.5 | 18 | 0.2778 |
| 14_2108204 | 1 | 0 | 4 | 0 | 3 | 0.3333 | 2 | 0 | 10 | 0.1 | 2 | 0 | 5 | 0 | 5 | 0.2 | 1 | 0 | 13 | 0.0769 |
| 14_2108215 | 1 | 0 | 5 | 0 | 2 | 0 | 4 | 0.5 | 12 | 0.1667 | 3 | 0.3333 | 6 | 0 | 6 | 0.3333 | 2 | 1 | 17 | 0.2941 |
| 14_2108293 | 3 | 0 | 7 | 0.1429 | 3 | 0.3333 | 6 | 0.1667 | 19 | 0.1579 | 5 | 0 | 5 | 0.2 | 8 | 0 | 2 | 0 | 20 | 0.05 |
| 14_2108545 | 2 | 0 | 4 | 0.25 | 8 | 0.125 | 3 | 0 | 17 | 0.1176 | 3 | 0.3333 | NA | NA | 2 | 0 | 7 | 0 | 12 | 0.0833 |
| 14_2108551 | 3 | 0.3333 | 4 | 0 | 6 | 0.1667 | 3 | 0 | 16 | 0.125 | 3 | 0.3333 | NA | NA | 2 | 0 | 9 | 0.1111 | 14 | 0.1429 |
| 14_2108602 | 4 | 0.25 | 6 | 0.1667 | 7 | 0.1429 | 4 | 0 | 21 | 0.1429 | 3 | 0 | 2 | 0 | 3 | 0.3333 | 8 | 0 | 16 | 0.0625 |
| 14_2108632 | 3 | 0.3333 | 6 | 0.5 | 5 | 0.2 | 4 | 0.25 | 18 | 0.3333 | 2 | 0 | 1 | 0 | 3 | 1 | 7 | 0.4286 | 13 | 0.4615 |
| 14_2108648 | 4 | 0 | 6 | 0.1667 | 6 | 0 | 4 | 0 | 20 | 0.05 | 2 | 0 | 1 | 0 | 3 | 0.3333 | 9 | 0.1111 | 15 | 0.1333 |
| 14_2108726 | 2 | 0 | 5 | 0.2 | 7 | 0.1429 | 4 | 0 | 18 | 0.1111 | 1 | 0 | 1 | 0 | 3 | 0 | 9 | 0.1111 | 14 | 0.0714 |
| 14_2149834 | 9 | 0.1111 | 17 | 0 | 15 | 0.1333 | 13 | 0 | 54 | 0.0556 | 8 | 0.25 | 4 | 0 | 16 | 0 | 11 | 0.0909 | 39 | 0.0769 |
| 14_2149852 | 9 | 0.1111 | 15 | 0.4 | 15 | 0.0667 | 12 | 0 | 51 | 0.1569 | 9 | 0.3333 | 2 | 0 | 15 | 0.3333 | 8 | 0.25 | 34 | 0.2941 |
| 14_2154570 | NA | NA | 1 | 0 | 1 | 1 | NA | NA | 2 | 0.5 | 1 | 0 | NA | NA | 1 | 1 | NA | NA | 2 | 0.5 |
| 14_2171866 | 2 | 0 | 3 | 0 | 2 | 0 | 1 | 1 | 8 | 0.125 | NA | NA | 3 | 0.3333 | 2 | 0 | NA | NA | 5 | 0.2 |
| 14_2171871 | 2 | 0.5 | 2 | 0 | 2 | 0 | 1 | 0 | 7 | 0.1429 | NA | NA | 3 | 0.3333 | 3 | 0.3333 | NA | NA | 6 | 0.3333 |
| 14_2171899 | 1 | 1 | 5 | 0.4 | 2 | 1 | 1 | 0 | 9 | 0.5556 | NA | NA | 5 | 0.2 | 4 | 0 | 1 | 0 | 10 | 0.1 |
| 14_2171910 | 2 | 0 | 5 | 0.2 | 2 | 0 | 1 | 0 | 10 | 0.1 | NA | NA | 5 | 0.2 | 4 | 0 | 1 | 0 | 10 | 0.1 |
| 14_2171914 | 2 | 1 | 3 | 0.3333 | 1 | 0 | 1 | 1 | 7 | 0.5714 | NA | NA | 4 | 0.75 | 4 | 0.25 | 1 | 1 | 9 | 0.5556 |
| 14_2171921 | 2 | 1 | 3 | 0.3333 | 1 | 0 | 1 | 0 | 7 | 0.4286 | NA | NA | 5 | 0.6 | 4 | 0.75 | 1 | 1 | 10 | 0.7 |
| 14_2171941 | NA | NA | 5 | 0.2 | 2 | 1 | 1 | 0 | 8 | 0.375 | NA | NA | 4 | 0 | 4 | 0.5 | 1 | 0 | 9 | 0.2222 |
| 14_2171944 | 2 | 0 | 5 | 0.2 | 2 | 0 | 1 | 0 | 10 | 0.1 | NA | NA | 4 | 0.25 | 3 | 0.3333 | 1 | 0 | 8 | 0.25 |
| 14_2171961 | 3 | 1 | 5 | 0.4 | 2 | 0.5 | 1 | 0 | 11 | 0.5455 | NA | NA | 5 | 1 | 4 | 0.5 | 2 | 0.5 | 11 | 0.7273 |
| 14_2171972 | 3 | 0 | 5 | 0 | 2 | 0.5 | 1 | 0 | 11 | 0.0909 | NA | NA | 6 | 0.1667 | 4 | 0 | 1 | 0 | 11 | 0.0909 |
| 14_2171975 | 3 | 1 | 5 | 0.4 | 2 | 1 | NA | NA | 10 | 0.7 | NA | NA | 5 | 0.4 | 4 | 0.5 | 2 | 0.5 | 11 | 0.4545 |
| 14_2171976 | 3 | 0 | 4 | 0 | 1 | 1 | 1 | 0 | 9 | 0.1111 | NA | NA | 6 | 0 | 5 | 0.4 | 1 | 0 | 12 | 0.1667 |
| 14_2171977 | 2 | 0 | 4 | 0.25 | 1 | 0 | 1 | 0 | 8 | 0.125 | NA | NA | 5 | 0.2 | 4 | 0.5 | 1 | 0 | 10 | 0.3 |
| 14_2171986 | 2 | 0.5 | 2 | 0 | 1 | 0 | 1 | 1 | 6 | 0.3333 | NA | NA | 5 | 0.2 | 3 | 0 | 3 | 0.3333 | 11 | 0.1818 |
| 14_2172017 | 2 | 0 | 5 | 0.2 | 2 | 0.5 | NA | NA | 9 | 0.2222 | NA | NA | 5 | 0 | 4 | 0.25 | 3 | 0.3333 | 12 | 0.1667 |
| 14_2172024 | 3 | 0 | 5 | 0 | 2 | 0.5 | 1 | 0 | 11 | 0.0909 | NA | NA | 6 | 0 | 5 | 0.2 | 3 | 0 | 14 | 0.0714 |
| 14_2172025 | 1 | 1 | 3 | 0.3333 | 1 | 0 | 1 | 0 | 6 | 0.3333 | NA | NA | 2 | 0.5 | 4 | 0.5 | 1 | 0 | 7 | 0.4286 |

| | | | | | | | | | | | | | | | | | | | | |
|---|---|---|---|---|---|---|---|---|---|---|---|---|---|---|---|---|---|---|---|---|
| 14_217203 | 3 | 0.6667 | 3 | 0.3333 | 2 | 0.5 | 1 | 1 | 9 | 0.5556 | NA | NA | 5 | 0.4 | 3 | 0.6667 | 3 | 0.3333 | 11 | 0.4545 |
| 14_217204 | 3 | 1 | 5 | 0.8 | 2 | 1 | 1 | 1 | 11 | 0.9091 | NA | NA | 4 | 1 | 5 | 1 | 2 | 1 | 11 | 1 |
| 14_217204 | 3 | 0.3333 | 6 | 0 | 2 | 0 | 1 | 0 | 12 | 0.0833 | NA | NA | 7 | 0 | 5 | 0.4 | 3 | 0.6667 | 15 | 0.2667 |
| 14_217205 | 2 | 1 | 4 | 1 | 2 | 1 | 1 | 1 | 9 | 1 | NA | NA | 7 | 0.8571 | 5 | 0.8 | 4 | 0.75 | 16 | 0.8125 |
| 14_217205 | 3 | 1 | 6 | 1 | 2 | 0.5 | 1 | 1 | 12 | 0.9167 | NA | NA | 5 | 1 | 5 | 1 | 4 | 1 | 14 | 1 |
| 14_217206 | 3 | 1 | 6 | 0.1667 | 2 | 0 | 1 | 0 | 12 | 0.3333 | NA | NA | 6 | 0.3333 | 5 | 0.6 | 4 | 0 | 15 | 0.3333 |
| 14_217206 | 2 | 1 | 6 | 0.8333 | 2 | 1 | 1 | 0 | 11 | 0.8182 | NA | NA | 6 | 0.3333 | 5 | 0 | 4 | 0.25 | 15 | 0.2 |
| 14_217206 | 3 | 0 | 4 | 0.25 | 2 | 0 | 1 | 0 | 10 | 0.1 | NA | NA | 7 | 0.1429 | 4 | 0.25 | 3 | 0 | 14 | 0.1429 |
| 14_217206 | 3 | 0.6667 | 3 | 0.6667 | 2 | 0.5 | 1 | 1 | 9 | 0.6667 | NA | NA | 5 | 0.6 | 4 | 0.25 | 4 | 0.75 | 13 | 0.5385 |
| 14_217206 | 2 | 1 | 4 | 1 | 2 | 1 | NA | NA | 8 | 1 | NA | NA | 4 | 0.75 | 4 | 1 | 4 | 1 | 12 | 0.9167 |
| 14_217207 | 2 | 1 | 5 | 0.8 | 1 | 0 | 1 | 1 | 9 | 0.7778 | NA | NA | 7 | 0.8571 | 3 | 0.6667 | 3 | 1 | 13 | 0.8462 |
| 14_217208 | 3 | 0 | 5 | 0.2 | 2 | 0.5 | 1 | 0 | 11 | 0.1818 | NA | NA | 7 | 0.1429 | 7 | 0 | 4 | 0 | 18 | 0.0556 |
| 14_217208 | 3 | 1 | 6 | 0.6667 | 2 | 0 | 1 | 0 | 12 | 0.5833 | NA | NA | 8 | 0.375 | 5 | 0.6 | 4 | 0.25 | 17 | 0.4118 |
| 14_217208 | 3 | 0.3333 | 6 | 0.5 | NA | NA | 1 | 1 | 10 | 0.5 | NA | NA | 8 | 0.5 | 5 | 0.2 | 4 | 0.5 | 17 | 0.4118 |
| 14_217210 | 4 | 0.75 | 5 | 0.4 | 1 | 1 | 1 | 0 | 11 | 0.5455 | NA | NA | 8 | 0.75 | 7 | 0.7143 | 4 | 0.75 | 19 | 0.7368 |
| 14_217212 | 4 | 1 | 6 | 1 | 2 | 1 | 1 | 1 | 13 | 1 | NA | NA | 9 | 1 | 10 | 0.9 | 4 | 1 | 23 | 0.9565 |
| 14_217215 | 4 | 0 | 6 | 0.1667 | 2 | 0 | 1 | 0 | 13 | 0.0769 | 1 | 0 | 7 | 0.1429 | 9 | 0.1111 | 2 | 0 | 19 | 0.1053 |
| 14_217220 | 5 | 0.6 | 10 | 0.5 | 2 | 0.5 | 1 | 0 | 18 | 0.5 | 1 | 1 | 9 | 0.4444 | 12 | 0.6667 | 8 | 0.25 | 30 | 0.5 |
| 14_218254 | NA | NA | 1 | 1 | NA | NA | NA | NA | 1 | 1 | 2 | 0.5 | NA | NA | NA | NA | NA | NA | 2 | 0.5 |
| 14_218430 | NA | NA | 1 | 1 | NA | NA | NA | NA | 1 | 1 | 1 | 1 | NA | NA | NA | NA | NA | NA | 1 | 1 |
| 14_218482 | 3 | 0.3333 | 13 | 0.1538 | 5 | 0 | 10 | 0.2 | 31 | 0.1613 | 4 | 0 | 7 | 0.1429 | 11 | 0.3636 | 8 | 0.375 | 30 | 0.2667 |
| 14_218972 | 5 | 0 | 6 | 0.1667 | 2 | 0 | 5 | 0 | 18 | 0.0556 | 1 | 0 | 3 | 0 | 4 | 0 | 4 | 0.25 | 12 | 0.0833 |
| 14_218976 | 1 | 1 | 1 | 0 | NA | NA | NA | NA | 2 | 0.5 | NA | NA | 1 | 1 | NA | NA | 2 | 1 | 3 | 1 |
| 14_218977 | 6 | 0.1667 | 7 | 0.4286 | 2 | 0 | 3 | 0 | 18 | 0.2222 | 1 | 1 | 3 | 0.3333 | 1 | 0 | 3 | 0 | 8 | 0.25 |
| 14_220345 | 4 | 0 | 5 | 0.4 | 2 | 0.5 | 3 | 0 | 14 | 0.2143 | 3 | 0 | 11 | 0.2727 | 6 | 0.1667 | 2 | 0 | 22 | 0.1818 |
| 14_220346 | 4 | 1 | 4 | 1 | 2 | 1 | 4 | 0.5 | 14 | 0.8571 | 3 | 0.6667 | 10 | 0.8 | 7 | 0.5714 | 3 | 1 | 23 | 0.7391 |
| 14_220346 | 2 | 0 | 4 | 0.25 | 2 | 0 | 4 | 0.25 | 12 | 0.1667 | 2 | 0.5 | 12 | 0.25 | 7 | 0.4286 | 4 | 0 | 25 | 0.28 |
| 14_220346 | 4 | 0.75 | 4 | 1 | 2 | 1 | 5 | 0.6 | 15 | 0.8 | 2 | 1 | 10 | 0.6 | 5 | 0.8 | 4 | 0.75 | 21 | 0.7143 |
| 14_220347 | 4 | 1 | 4 | 1 | 3 | 1 | 4 | 1 | 15 | 1 | 2 | 1 | 8 | 0.875 | 6 | 1 | 3 | 0.6667 | 19 | 0.8947 |
| 14_220347 | 4 | 1 | 2 | 1 | 2 | 1 | 4 | 0.25 | 12 | 0.75 | 2 | 0.5 | 10 | 0.3 | 6 | 0.5 | 3 | 1 | 21 | 0.4762 |
| 14_220347 | 2 | 0 | 3 | 0 | 1 | 0 | 5 | 0.4 | 11 | 0.1818 | 2 | 0.5 | 8 | 0.25 | 6 | 0 | 4 | 0.25 | 20 | 0.2 |
| 14_220349 | 3 | 1 | 3 | 1 | 2 | 1 | 4 | 1 | 12 | 1 | 2 | 1 | 9 | 1 | 5 | 1 | 3 | 0.6667 | 19 | 0.9474 |
| 14_220349 | 3 | 1 | 3 | 1 | 2 | 1 | 4 | 1 | 12 | 1 | 2 | 1 | 8 | 1 | 7 | 0.8571 | 4 | 1 | 21 | 0.9524 |
| 14_220350 | 2 | 0.5 | 3 | 1 | 3 | 0 | 4 | 0.75 | 12 | 0.5833 | 2 | 0.5 | 7 | 0.5714 | 8 | 0.75 | 3 | 1 | 20 | 0.7 |
| 14_220351 | 3 | 1 | 3 | 1 | 2 | 1 | 5 | 0.8 | 13 | 0.9231 | 2 | 1 | 10 | 0.9 | 8 | 1 | 3 | 1 | 23 | 0.9565 |
| 14_220351 | 3 | 1 | 3 | 1 | 3 | 1 | 5 | 1 | 14 | 1 | 2 | 1 | 10 | 1 | 7 | 1 | 2 | 1 | 21 | 1 |
| 14_220352 | 3 | 0.3333 | 1 | 0 | 1 | 0 | 3 | 0.3333 | 8 | 0.25 | 1 | 0 | 9 | 0.2222 | 8 | 0.25 | 3 | 1 | 21 | 0.3333 |
| 14_220352 | 2 | 1 | 3 | 0.6667 | 3 | 1 | 3 | 1 | 11 | 0.9091 | 1 | 1 | 10 | 0.9 | 8 | 0.875 | 2 | 1 | 21 | 0.9048 |
| 14_220352 | 3 | 0.6667 | 2 | 0.5 | 1 | 0 | 4 | 0 | 10 | 0.3 | 1 | 0 | 12 | 0.3333 | 7 | 0.5714 | 3 | 0.6667 | 23 | 0.4348 |
| 14_220356 | 3 | 0.3333 | 3 | 0.6667 | 3 | 0.3333 | 5 | 1 | 14 | 0.6429 | 1 | 1 | 11 | 0.6364 | 7 | 0.2857 | 3 | 0.3333 | 22 | 0.5 |
| 14_220356 | 3 | 0 | 3 | 0.3333 | 3 | 0.3333 | 5 | 0 | 14 | 0.1429 | 1 | 0 | 9 | 0.1111 | 6 | 0 | 3 | 0 | 19 | 0.0526 |
| 14_220358 | 3 | 0.3333 | 3 | 0.3333 | 1 | 1 | 3 | 0.3333 | 10 | 0.4 | 1 | 0 | 10 | 0.1 | 6 | 0.5 | 3 | 0 | 20 | 0.2 |
| 14_220359 | 3 | 1 | 3 | 1 | 3 | 1 | 3 | 1 | 12 | 1 | 1 | 1 | 8 | 1 | 5 | 1 | 2 | 1 | 16 | 1 |
| 14_220362 | 2 | 1 | 2 | 1 | 2 | 0 | 1 | 1 | 7 | 0.7143 | 1 | 1 | 6 | 0.3333 | 5 | 0.8 | 3 | 0.3333 | 15 | 0.5333 |
| 14_220365 | 2 | 0.5 | 2 | 0.5 | 2 | 0 | 2 | 0 | 8 | 0.25 | NA | NA | 6 | 0 | 4 | 0 | 3 | 0.3333 | 13 | 0.0769 |
| 14_220373 | 2 | 0 | 2 | 1 | 2 | 0.5 | 2 | 0 | 8 | 0.375 | 1 | 0 | 4 | 0.5 | 2 | 0.5 | 3 | 0.3333 | 10 | 0.4 |
| 14_220388 | 1 | 1 | NA | NA | NA | NA | NA | NA | 1 | 1 | NA | NA | NA | NA | 1 | 1 | NA | NA | 1 | 1 |
| 14_220390 | 1 | 1 | NA | NA | NA | NA | NA | NA | 1 | 1 | NA | NA | NA | NA | 1 | 1 | NA | NA | 1 | 1 |
| 14_220395 | 1 | 1 | NA | NA | NA | NA | NA | NA | 1 | 1 | NA | NA | NA | NA | 1 | 1 | NA | NA | 1 | 1 |
| 14_220395 | 1 | 1 | NA | NA | NA | NA | NA | NA | 1 | 1 | NA | NA | NA | NA | 1 | 1 | NA | NA | 1 | 1 |
| 14_233427 | 9 | 0.2222 | 16 | 0.125 | 7 | 0.1429 | 15 | 0.0667 | 47 | 0.1277 | 12 | 0.1667 | 10 | 0 | 6 | 0 | 8 | 0 | 36 | 0.0556 |
| 14_233481 | NA | NA | 2 | 0 | 2 | 0 | 3 | 0.3333 | 7 | 0.1429 | 4 | 0.25 | 4 | 0 | NA | NA | 2 | 0 | 10 | 0.1 |
| 14_233487 | NA | NA | 2 | 0.5 | 2 | 1 | 2 | 0.5 | 6 | 0.6667 | 1 | 0 | 4 | 0.25 | 1 | 0 | 1 | 0 | 7 | 0.1429 |
| 14_233490 | NA | NA | 2 | 0 | 1 | 1 | 2 | 0 | 5 | 0.2 | 2 | 1 | 3 | 0 | NA | NA | 2 | 0.5 | 7 | 0.4286 |
| 14_233498 | NA | NA | 1 | 1 | 1 | 1 | 1 | 0 | 3 | 0.6667 | 2 | 1 | 2 | 0 | 2 | 0 | 1 | 0 | 7 | 0.2857 |
| 14_233499 | NA | NA | 1 | 0 | 2 | 0 | 1 | 1 | 4 | 0.25 | 2 | 1 | 2 | 0 | 2 | 0 | 1 | 0 | 7 | 0.2857 |
| 14_233584 | NA | NA | NA | NA | 1 | 0 | 1 | 1 | 2 | 0.5 | 1 | 0 | NA | NA | NA | NA | 2 | 0.5 | 3 | 0.3333 |
| 14_241840 | 4 | 0.5 | 5 | 0 | 1 | 0 | 2 | 0 | 12 | 0.1667 | 3 | 0.3333 | 4 | 0.25 | 4 | 0.25 | 3 | 0.3333 | 14 | 0.2857 |
| 14_241847 | 2 | 0 | 5 | 0.2 | 3 | 0 | 2 | 0 | 12 | 0.0833 | 1 | 0 | 3 | 0 | 3 | 0 | 2 | 0.5 | 9 | 0.1111 |
| 14_241847 | 2 | 0.5 | 4 | 0.25 | 3 | 0.6667 | 2 | 1 | 11 | 0.5455 | NA | NA | 3 | 0.3333 | 3 | 0.3333 | 2 | 0 | 8 | 0.25 |
| 14_241847 | 2 | 0.5 | 5 | 0.2 | 3 | 0 | 2 | 0 | 12 | 0.1667 | 1 | 0 | 4 | 0.25 | 3 | 0 | 2 | 0 | 10 | 0.1 |
| 14_241851 | 1 | 0 | 3 | 0 | 1 | 0 | 1 | 1 | 6 | 0.1667 | 1 | 0 | 3 | 0 | 3 | 0.3333 | 1 | 0 | 8 | 0.125 |
| 14_241851 | 2 | 0.5 | 5 | 0.4 | 3 | 0.3333 | 2 | 0.5 | 12 | 0.4167 | NA | NA | 4 | 0.5 | 3 | 0 | 1 | 0 | 8 | 0.25 |
| 14_241852 | 2 | 0.5 | 4 | 0.5 | 3 | 0.6667 | 2 | 0.5 | 11 | 0.5455 | 1 | 1 | 4 | 1 | 3 | 0.6667 | 1 | 0 | 9 | 0.7778 |
| 14_241853 | 2 | 1 | 5 | 0.8 | 3 | 0.3333 | 3 | 0.6667 | 13 | 0.6923 | 1 | 0 | 3 | 1 | 2 | 1 | 1 | 1 | 7 | 0.8571 |

| | | | | | | | | | | | | | | | | | | | | |
|---|---|---|---|---|---|---|---|---|---|---|---|---|---|---|---|---|---|---|---|---|
| 14_2418540 | 1 | 1 | 3 | 1 | 2 | 1 | 2 | 0 | 8 | 0.75 | 1 | 1 | 4 | 1 | NA | NA | 1 | 1 | 6 | 1 |
| 14_2418541 | 2 | 0.5 | 5 | 0 | 3 | 0 | 2 | 0 | 12 | 0.0833 | 1 | 0 | 4 | 0.25 | 1 | 0 | 1 | 0 | 7 | 0.1429 |
| 14_2418545 | 2 | 0.5 | 3 | 0.6667 | 3 | 1 | 3 | 1 | 11 | 0.8182 | 1 | 1 | 3 | 0 | 1 | 1 | NA | NA | 5 | 0.4 |
| 14_2418551 | 2 | 1 | 5 | 0.6 | 3 | 0.3333 | 3 | 0.6667 | 13 | 0.6154 | 1 | 1 | 4 | 0 | 1 | 1 | NA | NA | 6 | 0.3333 |
| 14_2418554 | 1 | 0 | 3 | 0.6667 | 3 | 0.3333 | 1 | 1 | 8 | 0.5 | 1 | 1 | 3 | 0.6667 | 1 | 1 | 1 | 1 | 6 | 0.8333 |
| 14_2418559 | 2 | 0.5 | 4 | 0.25 | 2 | 0.5 | 2 | 0 | 10 | 0.3 | 1 | 1 | 3 | 0.3333 | 1 | 0 | 1 | 0 | 6 | 0.3333 |
| 14_2418567 | 2 | 0 | 5 | 0.2 | 1 | 0 | 2 | 0 | 10 | 0.1 | 1 | 0 | 4 | 0.25 | NA | NA | 1 | 0 | 6 | 0.1667 |
| 14_2420814 | NA | NA | 1 | 1 | 1 | 1 | 1 | 1 | 3 | 1 | 1 | 1 | 2 | 0 | NA | NA | NA | NA | 3 | 0.3333 |
| 14_2420823 | 1 | 1 | 1 | 0 | 1 | 0 | NA | NA | 3 | 0.3333 | 1 | 0 | 2 | 0.5 | NA | NA | NA | NA | 3 | 0.3333 |
| 14_2420832 | 1 | 1 | 1 | 1 | 1 | 1 | NA | NA | 3 | 1 | 1 | 1 | 1 | 1 | NA | NA | NA | NA | 2 | 1 |
| 14_2420836 | 1 | 1 | 1 | 1 | 1 | 0 | 1 | 0 | 4 | 0.5 | 1 | 1 | 1 | 0 | NA | NA | NA | NA | 2 | 0.5 |
| 14_2420844 | NA | NA | 1 | 1 | 1 | 1 | 1 | 1 | 3 | 1 | 1 | 1 | 2 | 0.5 | NA | NA | NA | NA | 3 | 0.6667 |
| 14_2420852 | NA | NA | 1 | 0 | 1 | 1 | 1 | 1 | 3 | 0.6667 | 1 | 1 | 2 | 1 | NA | NA | NA | NA | 3 | 1 |
| 14_2420862 | NA | NA | 1 | 1 | 1 | 1 | NA | NA | 2 | 1 | 1 | 0 | 2 | 0.5 | NA | NA | NA | NA | 3 | 0.3333 |
| 14_2420876 | NA | NA | 1 | 0 | 1 | 1 | 1 | 1 | 3 | 0.6667 | 1 | 0 | 1 | 1 | NA | NA | NA | NA | 2 | 0.5 |
| 14_2448888 | 1 | 1 | 1 | 0 | NA | NA | NA | NA | 2 | 0.5 | 1 | 0 | NA | NA | NA | NA | 1 | 1 | 2 | 0.5 |
| 14_2449886 | 4 | 0.25 | 2 | 0.5 | 1 | 0 | 1 | 0 | 8 | 0.25 | 1 | 0 | 1 | 0 | NA | NA | 3 | 0.3333 | 5 | 0.2 |
| 14_2449896 | 3 | 0.3333 | 3 | 1 | 1 | 1 | 1 | 0 | 8 | 0.625 | 1 | 0 | NA | NA | NA | NA | 2 | 1 | 3 | 0.6667 |
| 14_2449899 | 4 | 0.75 | 3 | 1 | NA | NA | 1 | 1 | 8 | 0.875 | 1 | 1 | NA | NA | NA | NA | 3 | 0 | 4 | 0.25 |
| 14_2449907 | 4 | 1 | 2 | 1 | 1 | 1 | 1 | 1 | 8 | 1 | 1 | 1 | 1 | 1 | NA | NA | 2 | 1 | 4 | 1 |
| 14_2449911 | 2 | 0 | 3 | 0.3333 | 1 | 0 | 1 | 0 | 7 | 0.1429 | 1 | 0 | 1 | 1 | NA | NA | 3 | 0.3333 | 5 | 0.4 |
| 14_2454120 | NA | NA | 1 | 0 | NA | NA | 1 | 1 | 2 | 0.5 | NA | NA | NA | NA | 1 | 1 | NA | NA | 1 | 1 |
| 14_2545298 | NA | NA | 1 | 1 | NA | NA | NA | NA | 1 | 1 | 1 | 1 | NA | NA | NA | NA | NA | NA | 1 | 1 |
| 14_2545307 | NA | NA | 1 | 1 | NA | NA | NA | NA | 1 | 1 | 1 | 1 | NA | NA | NA | NA | NA | NA | 1 | 1 |
| 14_2566344 | NA | NA | 4 | 0 | 4 | 0.25 | 2 | 0 | 10 | 0.1 | 3 | 0 | 4 | 0 | 1 | 0 | 3 | 0.3333 | 11 | 0.0909 |
| 14_2566346 | NA | NA | 3 | 0.3333 | 3 | 0 | 2 | 0.5 | 8 | 0.25 | 2 | 0.5 | 3 | 0 | 1 | 0 | 2 | 0 | 8 | 0.125 |
| 14_2566357 | 1 | 0 | 4 | 0.5 | 2 | 0.5 | 2 | 1 | 9 | 0.5556 | 2 | 1 | 3 | 0.6667 | NA | NA | 2 | 0.5 | 7 | 0.7143 |
| 14_2566373 | 2 | 1 | 3 | 0.6667 | 2 | 0.5 | 2 | 0.5 | 9 | 0.6667 | 4 | 0.5 | 4 | 0.25 | 1 | 0 | 4 | 0.5 | 13 | 0.3846 |
| 14_2566442 | 1 | 0 | 4 | 0.25 | 6 | 0 | 2 | 0.5 | 13 | 0.1538 | 3 | 0 | 4 | 0.25 | 1 | 0 | 2 | 0 | 10 | 0.1 |
| 14_2566448 | 2 | 0.5 | 3 | 0 | 6 | 0 | 3 | 0 | 14 | 0.0714 | 4 | 0.25 | 3 | 0 | 1 | 0 | 4 | 0 | 12 | 0.0833 |
| 14_2566511 | 2 | 0 | 8 | 0.125 | 5 | 0 | 4 | 0 | 19 | 0.0526 | 4 | 0.25 | 5 | 0 | 4 | 0 | 5 | 0.2 | 18 | 0.1111 |
| 14_2566552 | 4 | 0.5 | 11 | 0.2727 | 4 | 0.25 | 3 | 0 | 22 | 0.2727 | 6 | 0.1667 | 7 | 0.4286 | 2 | 0.5 | 7 | 0.1429 | 22 | 0.2727 |
| 14_2566560 | 4 | 1 | 11 | 1 | 5 | 0.8 | 3 | 0.6667 | 23 | 0.913 | 6 | 0.8333 | 6 | 1 | 4 | 1 | 5 | 0.8 | 21 | 0.9048 |
| 14_2566562 | 4 | 1 | 10 | 0.6 | 4 | 0.25 | 2 | 0 | 20 | 0.55 | 5 | 0 | 6 | 0.6667 | 3 | 0.6667 | 7 | 0.2857 | 21 | 0.381 |
| 14_2566575 | 3 | 0 | 12 | 0.1667 | 4 | 0 | 6 | 0.1667 | 25 | 0.12 | 6 | 0 | 7 | 0.1429 | 4 | 0.25 | 7 | 0.1429 | 24 | 0.125 |
| 14_2566595 | 3 | 0.6667 | 12 | 0.3333 | 6 | 0.5 | 5 | 0.2 | 26 | 0.3846 | 5 | 0.2 | 7 | 0.2857 | 4 | 0.5 | 8 | 0.125 | 24 | 0.25 |
| 14_2566644 | 6 | 0.1667 | 16 | 0.25 | 6 | 0.3333 | 8 | 0.375 | 36 | 0.2778 | 8 | 0.375 | 7 | 0.2857 | 6 | 0.1667 | 10 | 0.1 | 31 | 0.2258 |
| 14_2566652 | 5 | 0 | 16 | 0.1875 | 6 | 0.3333 | 8 | 0.25 | 35 | 0.2 | 9 | 0.3333 | 8 | 0.125 | 6 | 0.1667 | 10 | 0.1 | 33 | 0.1818 |
| 14_2570000 | 5 | 0.6 | 8 | 0.375 | 10 | 0.5 | 4 | 0.25 | 27 | 0.4444 | 3 | 0.3333 | 2 | 1 | 4 | 0.25 | 4 | 0.25 | 13 | 0.3846 |
| 14_2570004 | 5 | 0.2 | 8 | 0 | 9 | 0 | 5 | 0.2 | 27 | 0.0741 | 4 | 0.5 | 2 | 0 | 4 | 0 | 4 | 0.25 | 14 | 0.2143 |
| 14_2570009 | 5 | 0.6 | 5 | 0.8 | 8 | 0.25 | 5 | 0.6 | 23 | 0.5217 | 4 | 1 | 2 | 0.5 | 5 | 0.2 | 4 | 0.25 | 15 | 0.4667 |
| 14_2570027 | 3 | 1 | 5 | 1 | 8 | 0.875 | 5 | 0.4 | 21 | 0.8095 | 1 | 0 | 3 | 0.6667 | 6 | 1 | 4 | 0.5 | 14 | 0.7143 |
| 14_2570031 | 5 | 0 | 5 | 0.2 | 9 | 0.1111 | 5 | 0 | 24 | 0.0833 | 4 | 0.25 | 3 | 0 | 5 | 0 | 6 | 0.1667 | 18 | 0.1111 |
| 14_2570041 | 4 | 0 | 8 | 0 | 9 | 0.2222 | 5 | 0 | 26 | 0.0769 | 5 | 0.2 | 3 | 0 | 4 | 0 | 5 | 0 | 17 | 0.0588 |
| 14_2570072 | 5 | 0 | 6 | 0 | 11 | 0.1818 | 5 | 0 | 27 | 0.0741 | 5 | 0 | 4 | 0 | 6 | 0.5 | 5 | 0.2 | 20 | 0.2 |
| 14_2571207 | 7 | 0.1429 | 13 | 0 | 11 | 0.0909 | 5 | 0.2 | 36 | 0.0833 | 10 | 0.1 | 4 | 0 | 7 | 0.2857 | 7 | 0 | 28 | 0.1071 |
| 14_2571213 | 8 | 0 | 12 | 0.0833 | 9 | 0.1111 | 5 | 0 | 34 | 0.0588 | 8 | 0.125 | 4 | 0 | 7 | 0 | 7 | 0.2857 | 26 | 0.1154 |
| 14_2571245 | 8 | 0.75 | 12 | 1 | 9 | 1 | 5 | 0.6 | 34 | 0.8824 | 8 | 0.875 | 6 | 0.8333 | 6 | 0.8333 | 7 | 0.7143 | 27 | 0.8148 |
| 14_2571257 | 8 | 0.125 | 11 | 0 | 10 | 0.1 | 5 | 0 | 34 | 0.0588 | 8 | 0.125 | 6 | 0 | 7 | 0.1429 | 6 | 0 | 27 | 0.0741 |
| 14_2571262 | 8 | 0.125 | 11 | 0.1818 | 8 | 0.125 | 4 | 0.5 | 31 | 0.1935 | 9 | 0.2222 | 6 | 0.3333 | 6 | 0 | 4 | 0.25 | 25 | 0.2 |
| 14_2571391 | 3 | 0 | 7 | 0.1429 | 4 | 0 | 1 | 1 | 15 | 0.1333 | 5 | 0.2 | 1 | 0 | 4 | 0 | 3 | 0.3333 | 13 | 0.1538 |
| 14_2571400 | 5 | 0 | 8 | 0 | 3 | 0 | 3 | 0.3333 | 19 | 0.0526 | 5 | 0.2 | 2 | 0 | 4 | 0 | 4 | 0 | 15 | 0.0667 |
| 14_2571434 | 5 | 0.2 | 7 | 0 | 2 | 0 | 3 | 0 | 17 | 0.0588 | 7 | 0 | 3 | 0.3333 | 3 | 0 | 3 | 0 | 16 | 0.0625 |
| 14_2571435 | 5 | 0.2 | 8 | 0.5 | 2 | 0 | 2 | 0 | 17 | 0.2941 | 7 | 0.7143 | 3 | 0.3333 | 3 | 0.3333 | 3 | 0.3333 | 16 | 0.5 |
| 14_2571486 | 4 | 0.25 | 5 | 0 | 2 | 0 | 2 | 0.5 | 13 | 0.1538 | 5 | 0 | 1 | 0 | 4 | 0 | 2 | 0.5 | 12 | 0.0833 |
| 14_2571492 | 2 | 0.5 | 4 | 0.25 | 2 | 0.5 | 3 | 0.6667 | 11 | 0.4545 | 5 | 0.4 | 1 | 0 | 3 | 0.6667 | 3 | 0.6667 | 12 | 0.5 |
| 14_2571525 | NA | NA | 3 | 1 | 1 | 1 | 2 | 1 | 6 | 1 | 3 | 0.6667 | 1 | 0 | 2 | 0 | 2 | 1 | 8 | 0.5 |
| 14_2571539 | NA | NA | NA | NA | 1 | 1 | 1 | 0 | 2 | 0.5 | 1 | 1 | 1 | 1 | 3 | 1 | NA | NA | 5 | 1 |
| 14_2571553 | 2 | 0.5 | 2 | 0.5 | 2 | 0 | 2 | 0 | 8 | 0.25 | 1 | 1 | 1 | 0 | 2 | 0.5 | NA | NA | 4 | 0.5 |
| 14_2588694 | 4 | 0 | 2 | 0 | 3 | 0.3333 | 2 | 0 | 11 | 0.0909 | 2 | 0 | 1 | 0 | 2 | 0 | 8 | 0.375 | 13 | 0.2308 |
| 14_2588857 | NA | NA | 2 | 0 | 1 | 1 | NA | NA | 3 | 0.3333 | 1 | 1 | 1 | 0 | NA | NA | NA | NA | 2 | 0.5 |
| 14_2588868 | NA | NA | 2 | 0.5 | 1 | 0 | NA | NA | 3 | 0.3333 | 2 | 0 | 1 | 1 | NA | NA | NA | NA | 3 | 0.3333 |
| 14_2592464 | 10 | 0.8 | 13 | 0.6154 | 8 | 0.625 | 4 | 0.5 | 35 | 0.6571 | 4 | 0.75 | 6 | 0.8333 | 4 | 0.75 | 3 | 0.6667 | 17 | 0.7647 |
| 14_2592467 | 8 | 0.125 | 8 | 0.125 | 8 | 0.25 | 2 | 0 | 26 | 0.1538 | 4 | 0.25 | 7 | 0 | 4 | 0.25 | 3 | 0 | 18 | 0.1111 |
| 14_2592471 | 11 | 0.0909 | 13 | 0.1538 | 7 | 0.1429 | 4 | 0 | 35 | 0.1143 | 3 | 0 | 8 | 0.125 | 3 | 0 | 3 | 0.6667 | 17 | 0.1765 |

| | | | | | | | | | | | | | | | | | | | | |
|---|---|---|---|---|---|---|---|---|---|---|---|---|---|---|---|---|---|---|---|---|
| 14_2592528 | 9 | 0.1111 | 9 | 0.1111 | 7 | 0 | 4 | 0 | 29 | 0.069 | 4 | 0 | 8 | 0.125 | 1 | 0 | 3 | 0 | 16 | 0.0625 |
| 14_2613474 | 14 | 0.0714 | 13 | 0.2308 | 17 | 0 | 34 | 0.0882 | 78 | 0.0897 | 8 | 0.125 | 23 | 0.1739 | 33 | 0.0909 | 21 | 0.0952 | 85 | 0.1176 |
| 14_2613541 | 14 | 0.2143 | 13 | 0.0769 | 16 | 0.1875 | 32 | 0.1562 | 75 | 0.16 | 6 | 0.1667 | 21 | 0.1905 | 29 | 0.2069 | 17 | 0 | 73 | 0.1507 |
| 14_2614855 | 7 | 0.4286 | 9 | 0.3333 | 14 | 0 | 25 | 0.12 | 55 | 0.1636 | 4 | 0.25 | 13 | 0.3077 | 26 | 0.2308 | 13 | 0.3077 | 56 | 0.2679 |
| 14_2614856 | 7 | 0.4286 | 7 | 0.2857 | 12 | 0.25 | 24 | 0.4583 | 50 | 0.38 | 6 | 0.3333 | 12 | 0.4167 | 25 | 0.56 | 12 | 0.25 | 55 | 0.4364 |
| 14_2625464 | 4 | 0.5 | 3 | 0 | 5 | 1 | 9 | 0.1111 | 21 | 0.381 | 2 | 1 | 2 | 0 | 7 | 0.1429 | 4 | 0.5 | 15 | 0.3333 |
| 14_2625480 | 4 | 0.5 | 7 | 0.2857 | 6 | 0.3333 | 12 | 0.25 | 29 | 0.3103 | 2 | 0 | 6 | 0.1667 | 13 | 0.2308 | 5 | 0.6 | 26 | 0.2692 |
| 14_2625491 | 4 | 1 | 5 | 1 | 6 | 1 | 12 | 1 | 27 | 1 | 1 | 1 | 5 | 1 | 14 | 0.9286 | 4 | 1 | 24 | 0.9583 |
| 14_2625493 | 3 | 1 | 2 | 1 | 5 | 0.8 | 13 | 0.8462 | 23 | 0.8696 | 1 | 1 | 7 | 1 | 13 | 0.8462 | 5 | 0.8 | 26 | 0.8846 |
| 14_2625500 | 1 | 0 | 3 | 0 | 2 | 0.5 | 10 | 0.2 | 16 | 0.1875 | 2 | 0 | 6 | 0.1667 | 10 | 0.3 | 4 | 0.75 | 22 | 0.3182 |
| 14_2625503 | 3 | 1 | 7 | 0.2857 | 6 | 0.5 | 12 | 0.5833 | 28 | 0.5357 | 2 | 0 | 7 | 0.4286 | 13 | 0.3846 | 5 | 0 | 27 | 0.2963 |
| 14_2625504 | 3 | 1 | 4 | 1 | 4 | 1 | 11 | 0.8182 | 22 | 0.9091 | 2 | 0.5 | 6 | 1 | 11 | 1 | 4 | 1 | 23 | 0.9565 |
| 14_2625508 | 2 | 0 | 6 | 0.1667 | 4 | 0 | 13 | 0.0769 | 25 | 0.08 | 2 | 0.5 | 3 | 0.6667 | 10 | 0.2 | 4 | 0 | 19 | 0.2632 |
| 14_2625514 | 3 | 0 | 5 | 0 | 5 | 0.4 | 14 | 0.4286 | 27 | 0.2963 | 2 | 0.5 | 6 | 0.6667 | 11 | 0.2727 | 2 | 0 | 21 | 0.381 |
| 14_2625539 | 3 | 0.3333 | 3 | 0.3333 | 5 | 0.2 | 7 | 0.1429 | 18 | 0.2222 | 1 | 1 | 5 | 0.2 | 8 | 0.375 | 2 | 0 | 16 | 0.3125 |
| 14_2634924 | 2 | 0.5 | 2 | 0 | 2 | 0.5 | 2 | 0.5 | 8 | 0.375 | 3 | 0 | 1 | 0 | 1 | 1 | NA | NA | 5 | 0.2 |
| 14_2634928 | 1 | 0 | 2 | 0.5 | 2 | 0.5 | 2 | 0 | 7 | 0.2857 | 3 | 0.3333 | 1 | 0 | 1 | 0 | NA | NA | 5 | 0.2 |
| 14_2634929 | 2 | 0 | 2 | 0.5 | 1 | 0 | 2 | 0 | 7 | 0.1429 | 3 | 0 | 1 | 0 | 1 | 1 | NA | NA | 5 | 0.2 |
| 14_2635064 | 3 | 0.6667 | 1 | 1 | NA | NA | 4 | 0.5 | 8 | 0.625 | NA | NA | NA | NA | 3 | 0.3333 | 3 | 0 | 6 | 0.1667 |
| 14_2635074 | 2 | 0 | 2 | 0.5 | NA | NA | 4 | 0.75 | 8 | 0.5 | NA | NA | 1 | 1 | 2 | 1 | 3 | 0.6667 | 6 | 0.8333 |
| 14_2635085 | 3 | 0.3333 | 2 | 0.5 | NA | NA | 4 | 0 | 9 | 0.2222 | NA | NA | 1 | 0 | 3 | 0.6667 | 3 | 1 | 7 | 0.7143 |
| 14_2635086 | 3 | 0.6667 | 2 | 0 | NA | NA | 4 | 0 | 9 | 0.2222 | NA | NA | 1 | 1 | 3 | 1 | 2 | 0.5 | 6 | 0.8333 |
| 14_2635091 | 3 | 0.6667 | 2 | 1 | NA | NA | 4 | 0 | 9 | 0.4444 | NA | NA | 1 | 0 | 3 | 1 | 3 | 1 | 7 | 0.8571 |
| 14_2635095 | 3 | 0.6667 | 2 | 0.5 | NA | NA | 4 | 0 | 9 | 0.3333 | NA | NA | 1 | 0 | 4 | 0.5 | 3 | 0.3333 | 8 | 0.375 |
| 14_2635105 | 2 | 0.5 | 2 | 0.5 | NA | NA | 4 | 0 | 8 | 0.25 | NA | NA | NA | NA | 3 | 0.6667 | 3 | 0.6667 | 6 | 0.6667 |
| 14_2635111 | 2 | 1 | 2 | 0.5 | NA | NA | 2 | 0.5 | 6 | 0.6667 | NA | NA | NA | NA | 4 | 0 | 3 | 0.3333 | 7 | 0.1429 |
| 14_2637710 | 28 | 0.0714 | 21 | 0.0476 | 20 | 0.1 | 32 | 0.0625 | 101 | 0.0693 | 26 | 0.0769 | 17 | 0.0588 | 12 | 0.0833 | 17 | 0.0588 | 72 | 0.0694 |
| 14_2637712 | 25 | 0.16 | 22 | 0.3636 | 17 | 0.2941 | 29 | 0.1724 | 93 | 0.2366 | 23 | 0.2174 | 18 | 0.2778 | 11 | 0.0909 | 15 | 0.4667 | 67 | 0.2687 |
| 14_2637716 | 25 | 0.08 | 19 | 0.2105 | 20 | 0.15 | 31 | 0.129 | 95 | 0.1368 | 25 | 0.16 | 17 | 0.2353 | 12 | 0.1667 | 15 | 0.4667 | 69 | 0.2464 |
| 14_2637929 | 5 | 0.2 | NA | NA | 1 | 0 | 3 | 0 | 9 | 0.1111 | 2 | 0.5 | 3 | 0 | 1 | 0 | 2 | 0 | 8 | 0.125 |
| 14_2638627 | 3 | 0 | 3 | 0.3333 | 6 | 0 | 3 | 0 | 15 | 0.0667 | 5 | 0 | 1 | 0 | 2 | 0.5 | 3 | 0 | 11 | 0.0909 |
| 14_2638653 | 4 | 0.5 | 3 | 0.6667 | 5 | 0 | 1 | 0 | 13 | 0.3077 | 4 | 0.25 | NA | NA | 1 | 1 | 3 | 0 | 8 | 0.25 |
| 14_2638658 | 4 | 0.5 | 2 | 0 | 5 | 0 | 2 | 0 | 13 | 0.1538 | 5 | 0.2 | 1 | 0 | 2 | 0 | 3 | 0 | 11 | 0.0909 |
| 14_2638664 | 3 | 0.6667 | 2 | 0 | 4 | 0.5 | 2 | 0 | 11 | 0.3636 | 4 | 0.5 | NA | NA | 1 | 1 | 3 | 0.3333 | 8 | 0.5 |
| 14_2638669 | 4 | 0.25 | 1 | 0 | 4 | 0 | 2 | 0 | 11 | 0.0909 | 5 | 0 | NA | NA | 2 | 0.5 | 3 | 0 | 10 | 0.1 |
| 14_2638679 | 3 | 0.6667 | NA | NA | 2 | 0 | 1 | 0 | 6 | 0.3333 | 5 | 0.4 | NA | NA | 1 | 0 | 3 | 0 | 9 | 0.2222 |
| 14_2638681 | 4 | 1 | NA | NA | 4 | 1 | 2 | 0.5 | 10 | 0.9 | 5 | 0.8 | NA | NA | 1 | 1 | 3 | 0.3333 | 9 | 0.6667 |
| 14_2638689 | 3 | 0 | NA | NA | 4 | 0.25 | 2 | 0.5 | 9 | 0.2222 | 3 | 0 | NA | NA | 2 | 0 | 3 | 0.3333 | 8 | 0.125 |
| 14_2638695 | 3 | 0 | 1 | 0 | 3 | 1 | 2 | 0 | 9 | 0.3333 | 4 | 0.25 | NA | NA | 2 | 0 | 3 | 0.3333 | 9 | 0.2222 |
| 14_2638702 | 3 | 0.6667 | 1 | 1 | 4 | 0.5 | 2 | 0.5 | 10 | 0.6 | 5 | 1 | NA | NA | 2 | 0.5 | 3 | 1 | 10 | 0.9 |
| 14_2638707 | 3 | 0 | 1 | 0 | 4 | 0.25 | 2 | 0 | 10 | 0.1 | 5 | 0.2 | NA | NA | 2 | 0 | 3 | 0.6667 | 10 | 0.3 |
| 14_2638948 | 2 | 0 | 1 | 0 | 3 | 0.3333 | 2 | 0 | 8 | 0.125 | 4 | 0.25 | NA | NA | 2 | 0.5 | 2 | 0.5 | 8 | 0.375 |
| 14_2638978 | 2 | 0 | 1 | 0 | 3 | 0.3333 | 1 | 0 | 7 | 0.1429 | 3 | 0.3333 | NA | NA | 1 | 1 | 3 | 0.3333 | 7 | 0.4286 |
| 14_2638979 | 2 | 0 | 1 | 0 | 3 | 0.3333 | 1 | 0 | 7 | 0.1429 | 3 | 0 | NA | NA | 2 | 0 | 3 | 0.3333 | 8 | 0.125 |
| 14_2638983 | 2 | 1 | 1 | 1 | 3 | 1 | 1 | 1 | 7 | 1 | 3 | 0.6667 | NA | NA | 1 | 1 | 3 | 1 | 7 | 0.8571 |
| 14_2639062 | 1 | 1 | NA | NA | 2 | 0 | 1 | 0 | 4 | 0.25 | NA | NA | NA | NA | 1 | 1 | 2 | 0.5 | 3 | 0.6667 |
| 14_2639080 | NA | NA | NA | NA | NA | NA | 1 | 1 | 1 | 1 | NA | NA | NA | NA | NA | NA | 1 | 1 | 1 | 1 |
| 14_2639107 | NA | NA | NA | NA | 1 | 1 | 1 | 1 | 2 | 1 | NA | NA | NA | NA | NA | NA | 1 | 1 | 1 | 1 |
| 14_2639119 | NA | NA | NA | NA | 1 | 0 | 1 | 1 | 2 | 0.5 | NA | NA | NA | NA | NA | NA | 1 | 1 | 1 | 1 |
| 14_2664104 | 2 | 0.5 | 9 | 0 | 6 | 0 | 7 | 0.1429 | 24 | 0.0833 | 1 | 0 | 4 | 0.25 | 4 | 0 | 2 | 0 | 11 | 0.0909 |
| 14_2664128 | 1 | 0 | 9 | 0.3333 | 7 | 0.1429 | 10 | 0 | 27 | 0.1481 | 1 | 0 | 3 | 0 | 4 | 0.25 | 2 | 0 | 10 | 0.1 |
| 14_2664312 | 2 | 0 | 6 | 0.1667 | 3 | 0 | 3 | 0 | 14 | 0.0714 | 1 | 0 | 1 | 0 | 2 | 0 | 2 | 0.5 | 6 | 0.1667 |
| 14_2664503 | 4 | 0.5 | 9 | 0.4444 | 3 | 0 | 4 | 0.25 | 20 | 0.35 | 2 | 1 | 5 | 0 | 5 | 0.2 | 2 | 0 | 14 | 0.2143 |
| 14_2664540 | 3 | 0 | 7 | 0.1429 | 2 | 0.5 | 3 | 0 | 15 | 0.1333 | 2 | 0.5 | 4 | 0 | 3 | 0.3333 | 2 | 0.5 | 11 | 0.2727 |
| 14_2664554 | 4 | 0.25 | 7 | 0.5714 | 2 | 1 | 4 | 0.25 | 17 | 0.4706 | 1 | 1 | 6 | 0.6667 | 2 | 0.5 | 1 | 1 | 10 | 0.7 |
| 14_2664556 | 4 | 0.25 | 6 | 0.3333 | 3 | 0.3333 | 4 | 0.75 | 17 | 0.4118 | 2 | 0.5 | 7 | 0.1429 | 3 | 0.3333 | 2 | 1 | 14 | 0.3571 |
| 14_2664556 | 3 | 1 | 7 | 0.8571 | 3 | 0.6667 | 4 | 0.75 | 17 | 0.8235 | 1 | 1 | 6 | 0.6667 | 2 | 1 | 2 | 1 | 11 | 0.8182 |
| 14_2664568 | 4 | 0.75 | 5 | 0.4 | 3 | 0.3333 | 4 | 0.5 | 16 | 0.5 | 2 | 1 | 6 | 0.3333 | 3 | 0 | 1 | 0 | 12 | 0.3333 |
| 14_2664582 | 3 | 0 | 7 | 0.2857 | 3 | 0 | 3 | 0.3333 | 16 | 0.1875 | 2 | 0 | 7 | 0.2857 | 3 | 0 | 2 | 0.5 | 14 | 0.2143 |
| 14_2664794 | 2 | 0.5 | 7 | 0.1429 | 3 | 0 | 2 | 0 | 14 | 0.1429 | 2 | 0.5 | 8 | 0 | 3 | 0 | 2 | 0 | 15 | 0.0667 |
| 14_2712370 | 6 | 1 | 11 | 0.9091 | 5 | 1 | 7 | 0.7143 | 29 | 0.8966 | 5 | 1 | 2 | 1 | 6 | 1 | 7 | 1 | 20 | 1 |
| 14_2712374 | 6 | 0.6667 | 11 | 0.3636 | 6 | 0.5 | 6 | 0.5 | 29 | 0.4828 | 5 | 0.8 | 2 | 1 | 4 | 0.5 | 8 | 0.375 | 19 | 0.5789 |
| 14_2712375 | 7 | 0.8571 | 9 | 1 | 7 | 0.8571 | 8 | 0.75 | 31 | 0.871 | 4 | 1 | 2 | 1 | 5 | 1 | 10 | 0.9 | 21 | 0.9524 |
| 14_2712381 | 6 | 0.3333 | 12 | 0.0833 | 8 | 0.5 | 7 | 0.2857 | 33 | 0.2727 | 4 | 0.25 | 2 | 0 | 7 | 0.1429 | 12 | 0.1667 | 25 | 0.16 |
| 14_2712382 | 7 | 0.2857 | 8 | 0 | 6 | 0 | 7 | 0 | 28 | 0.0714 | 4 | 0 | 2 | 0 | 7 | 0.1429 | 10 | 0.1 | 23 | 0.087 |

| | | | | | | | | | | | | | | | | | | | | |
|---|---|---|---|---|---|---|---|---|---|---|---|---|---|---|---|---|---|---|---|---|
| 14_271238 | 7 | 0.5714 | 9 | 0.6667 | 7 | 0.2857 | 8 | 0.375 | 31 | 0.4839 | 3 | 0.6667 | 2 | 1 | 7 | 0.8571 | 10 | 0.7 | 22 | 0.7727 |
| 14_271238 | 7 | 0.2857 | 11 | 0.4545 | 7 | 0.7143 | 8 | 0.625 | 33 | 0.5152 | 5 | 0.8 | 2 | 0.5 | 7 | 0.7143 | 9 | 0.8889 | 23 | 0.7826 |
| 14_271239 | 6 | 0.6667 | 10 | 0.9 | 7 | 1 | 4 | 0.75 | 27 | 0.8519 | 4 | 0.75 | 2 | 1 | 6 | 0.6667 | 6 | 1 | 18 | 0.8333 |
| 14_271240 | 5 | 1 | 8 | 1 | 7 | 1 | 6 | 1 | 26 | 1 | 4 | 0.75 | 1 | 1 | 4 | 1 | 8 | 1 | 17 | 0.9412 |
| 14_271241 | 7 | 0.2857 | 12 | 0.3333 | 7 | 0.5714 | 9 | 0.1111 | 35 | 0.3143 | 6 | 0.3333 | 2 | 0 | 7 | 0.4286 | 10 | 0.4 | 25 | 0.36 |
| 14_271241 | 7 | 0 | 10 | 0 | 7 | 0.1429 | 9 | 0.1111 | 33 | 0.0606 | 6 | 0 | 2 | 0.5 | 6 | 0 | 11 | 0.1818 | 25 | 0.12 |
| 14_271241 | 7 | 0 | 12 | 0.1667 | 6 | 0.1667 | 8 | 0 | 33 | 0.0909 | 3 | 0 | 2 | 0 | 6 | 0 | 12 | 0.1667 | 23 | 0.087 |
| 14_271242 | 9 | 0.6667 | 11 | 0.6364 | 7 | 0.2857 | 9 | 0.5556 | 36 | 0.5556 | 5 | 0 | 2 | 1 | 6 | 0.6667 | 13 | 0.6154 | 26 | 0.5385 |
| 14_271246 | 9 | 0.4444 | 12 | 0.25 | 8 | 0.375 | 10 | 0.2 | 39 | 0.3077 | 4 | 0.25 | 2 | 0.5 | 6 | 0 | 13 | 0 | 25 | 0.08 |
| 14_271247 | 9 | 0.2222 | 12 | 0.25 | 7 | 0 | 7 | 0.7143 | 35 | 0.2857 | 5 | 0.2 | 1 | 1 | 5 | 0.4 | 12 | 0.1667 | 23 | 0.2609 |
| 14_271248 | 11 | 0.0909 | 15 | 0 | 9 | 0.3333 | 11 | 0 | 46 | 0.087 | 6 | 0 | 2 | 0.5 | 7 | 0 | 14 | 0.0714 | 29 | 0.069 |
| 14_271249 | 10 | 0.3 | 14 | 0.0714 | 8 | 0.25 | 9 | 0.1111 | 41 | 0.1707 | 7 | 0.2857 | 2 | 0 | 7 | 0.2857 | 14 | 0.1429 | 30 | 0.2 |
| 14_271249 | 11 | 0 | 14 | 0.0714 | 6 | 0.1667 | 9 | 0.1111 | 40 | 0.075 | 7 | 0.2857 | 2 | 0 | 7 | 0 | 15 | 0.2667 | 31 | 0.1935 |
| 14_271249 | 11 | 0.8182 | 14 | 0.7857 | 9 | 0.8889 | 10 | 0.8 | 44 | 0.8182 | 6 | 0.6667 | 2 | 1 | 6 | 1 | 15 | 1 | 29 | 0.931 |
| 14_271249 | 11 | 0.9091 | 11 | 0.9091 | 8 | 1 | 11 | 0.9091 | 41 | 0.9268 | 4 | 1 | 2 | 0.5 | 7 | 1 | 15 | 0.9333 | 28 | 0.9286 |
| 14_271250 | 11 | 0.6364 | 16 | 0.4375 | 6 | 0.8333 | 11 | 0.6364 | 44 | 0.5909 | 6 | 0.5 | 2 | 0.5 | 7 | 0.4286 | 13 | 0.6154 | 28 | 0.5357 |
| 14_271256 | 7 | 0.5714 | 15 | 0.2667 | 9 | 0.3333 | 10 | 0.5 | 41 | 0.3902 | 7 | 0.4286 | 2 | 0 | 8 | 0.25 | 10 | 0.2 | 27 | 0.2593 |
| 14_271468 | NA | NA | NA | NA | NA | NA | 1 | 1 | 1 | 1 | NA | NA | 1 | 1 | NA | NA | 1 | 1 | 2 | 1 |
| 14_271834 | 2 | 0.5 | NA | NA | 1 | 0 | NA | NA | 3 | 0.3333 | NA | NA | NA | NA | 1 | 0 | 3 | 0.3333 | 4 | 0.25 |
| 14_271862 | 2 | 1 | NA | NA | 2 | 1 | NA | NA | 4 | 1 | NA | NA | 2 | 0.5 | 1 | 0 | 2 | 0.5 | 5 | 0.4 |
| 14_271908 | 1 | 1 | NA | NA | NA | NA | 1 | 0 | 2 | 0.5 | 2 | 0 | 2 | 0 | NA | NA | 1 | 1 | 5 | 0.2 |
| 14_271937 | 2 | 0.5 | NA | NA | 3 | 0.3333 | 1 | 1 | 6 | 0.5 | 1 | 1 | 1 | 1 | 1 | 0 | 2 | 0.5 | 5 | 0.6 |
| 14_271938 | NA | NA | 1 | 1 | NA | NA | NA | NA | 1 | 1 | NA | NA | NA | NA | NA | NA | 2 | 1 | 2 | 1 |
| 14_271940 | 2 | 0.5 | NA | NA | 3 | 0.3333 | 1 | 0 | 6 | 0.3333 | 1 | 0 | 2 | 1 | 1 | 1 | 4 | 0.25 | 8 | 0.5 |
| 14_271941 | 2 | 0.5 | NA | NA | 3 | 0.6667 | 1 | 0 | 6 | 0.5 | 1 | 0 | 2 | 0 | 1 | 1 | 3 | 0.3333 | 7 | 0.2857 |
| 14_271941 | 2 | 1 | NA | NA | 3 | 1 | 1 | 0 | 6 | 0.8333 | 1 | 1 | 2 | 1 | 1 | 1 | 4 | 1 | 8 | 1 |
| 14_271941 | 2 | 0.5 | NA | NA | 3 | 0.6667 | 1 | 0 | 6 | 0.5 | 1 | 1 | 2 | 1 | 1 | 1 | 4 | 0.5 | 8 | 0.75 |
| 14_271942 | 2 | 1 | NA | NA | 2 | 1 | 1 | 0 | 5 | 0.8 | 1 | 1 | 2 | 1 | 1 | 0 | 4 | 0.25 | 8 | 0.5 |
| 14_271942 | 1 | 1 | NA | NA | 3 | 0 | 1 | 0 | 5 | 0.2 | 1 | 0 | 2 | 0.5 | 1 | 0 | 4 | 0.5 | 8 | 0.375 |
| 14_272027 | 3 | 0.6667 | 1 | 0 | 3 | 0 | 1 | 1 | 8 | 0.375 | NA | NA | 1 | 1 | 3 | 0 | NA | NA | 4 | 0.25 |
| 14_272027 | 3 | 0.6667 | 1 | 0 | 3 | 0.3333 | 1 | 0 | 8 | 0.375 | 1 | 1 | 1 | 1 | 3 | 0.3333 | NA | NA | 5 | 0.6 |
| 14_272027 | 3 | 0.6667 | 1 | 1 | 3 | 0.6667 | 1 | 1 | 8 | 0.75 | 1 | 1 | 1 | 1 | 3 | 0.6667 | NA | NA | 5 | 0.8 |
| 14_272028 | 3 | 0.6667 | 1 | 1 | 3 | 0 | 1 | 0 | 8 | 0.375 | 1 | 0 | 1 | 1 | 3 | 0.3333 | NA | NA | 5 | 0.4 |
| 14_272030 | 2 | 0 | NA | NA | 3 | 0.3333 | 1 | 0 | 6 | 0.1667 | 2 | 0 | 1 | 1 | 2 | 0 | 4 | 0.25 | 9 | 0.2222 |
| 14_272030 | 3 | 0.6667 | 1 | 1 | 4 | 0.75 | 1 | 1 | 9 | 0.7778 | 2 | 0 | 2 | 1 | 3 | 0.6667 | 4 | 1 | 11 | 0.7273 |
| 14_272031 | 2 | 0 | 1 | 0 | 2 | 0 | 1 | 1 | 6 | 0.1667 | 1 | 0 | 2 | 0 | 3 | 0 | 4 | 0.25 | 10 | 0.1 |
| 14_272031 | 3 | 0.3333 | 1 | 0 | 4 | 0 | 1 | 0 | 9 | 0.1111 | 2 | 0 | 2 | 0 | 4 | 0.25 | 4 | 0 | 12 | 0.0833 |
| 14_272032 | 3 | 0.3333 | 1 | 0 | 4 | 0.5 | 1 | 1 | 9 | 0.4444 | 1 | 1 | 1 | 1 | 2 | 1 | 3 | 1 | 7 | 1 |
| 14_272033 | 3 | 0.3333 | 1 | 1 | 4 | 0.5 | 1 | 1 | 9 | 0.5556 | 2 | 1 | 2 | 1 | 2 | 0.5 | 1 | 1 | 7 | 0.8571 |
| 14_272034 | 3 | 0.6667 | 1 | 1 | 4 | 1 | 1 | 1 | 9 | 0.8889 | 2 | 1 | 2 | 1 | 4 | 0.75 | 4 | 0.75 | 12 | 0.8333 |
| 14_272042 | 3 | 0 | 2 | 0.5 | 4 | 0 | 1 | 0 | 10 | 0.1 | 2 | 1 | 3 | 0 | 3 | 0.3333 | 4 | 0.25 | 12 | 0.3333 |
| 14_272052 | 5 | 0.6 | 6 | 0 | 5 | 0.2 | 1 | 0 | 17 | 0.2353 | 2 | 0 | 4 | 0.25 | 3 | 0 | 6 | 0 | 15 | 0.0667 |
| 14_272054 | 3 | 0.3333 | 6 | 0 | 4 | 0 | 1 | 0 | 14 | 0.0714 | 2 | 0 | 6 | 0.1667 | 4 | 0 | 5 | 0.2 | 17 | 0.1176 |
| 14_272087 | 1 | 1 | 1 | 1 | 1 | 0 | NA | NA | 3 | 0.6667 | 1 | 1 | 1 | 0 | 2 | 1 | 1 | 1 | 5 | 0.8 |
| 14_272087 | 1 | 1 | 2 | 0 | 2 | 0.5 | NA | NA | 5 | 0.4 | 1 | 0 | NA | NA | 2 | 1 | 2 | 1 | 5 | 0.8 |
| 14_272087 | 1 | 0 | 2 | 0.5 | 1 | 1 | NA | NA | 4 | 0.5 | 1 | 1 | 1 | 1 | 2 | 0 | 2 | 0.5 | 6 | 0.5 |
| 14_272224 | 1 | 0 | 2 | 0.5 | 2 | 0 | NA | NA | 5 | 0.2 | 1 | 0 | NA | NA | 2 | 1 | 1 | 0 | 4 | 0.5 |
| 14_272224 | 1 | 1 | 2 | 0 | 2 | 0 | NA | NA | 5 | 0.2 | 1 | 0 | NA | NA | 2 | 0.5 | 1 | 0 | 4 | 0.25 |
| 14_272225 | 1 | 0 | 1 | 0 | 2 | 0.5 | NA | NA | 4 | 0.25 | 1 | 1 | NA | NA | 2 | 0.5 | 1 | 0 | 4 | 0.5 |
| 14_272225 | 1 | 1 | 2 | 0.5 | 1 | 0 | NA | NA | 4 | 0.5 | 1 | 1 | NA | NA | 2 | 0 | NA | NA | 3 | 0.3333 |
| 14_273749 | 1 | 0 | NA | NA | 1 | 1 | 1 | 1 | 3 | 0.6667 | NA | NA | NA | NA | 1 | 0 | 1 | 1 | 2 | 0.5 |
| 14_273749 | 1 | 0 | NA | NA | 1 | 0 | 1 | 1 | 3 | 0.3333 | NA | NA | NA | NA | 1 | 0 | 1 | 1 | 2 | 0.5 |
| 14_273751 | 1 | 1 | NA | NA | NA | NA | 1 | 1 | 2 | 1 | NA | NA | NA | NA | 2 | 0.5 | 1 | 0 | 3 | 0.3333 |
| 14_274990 | NA | NA | 3 | 1 | NA | NA | 1 | 1 | 4 | 1 | NA | NA | 1 | 0 | 2 | 1 | 3 | 0.6667 | 6 | 0.6667 |
| 14_276015 | NA | NA | NA | NA | 1 | 1 | 4 | 1 | 5 | 1 | NA | NA | 2 | 1 | 2 | 1 | NA | NA | 4 | 1 |
| 14_276015 | 1 | 1 | NA | NA | 1 | 1 | 4 | 1 | 6 | 1 | NA | NA | 2 | 1 | 1 | 0 | NA | NA | 3 | 0.6667 |
| 14_276016 | 1 | 1 | NA | NA | 1 | 0 | 4 | 0.75 | 6 | 0.6667 | NA | NA | 2 | 0 | 2 | 0.5 | NA | NA | 4 | 0.25 |
| 14_276018 | NA | NA | NA | NA | 1 | 1 | 4 | 0.75 | 5 | 0.8 | 1 | 0 | 2 | 0.5 | 2 | 0 | NA | NA | 5 | 0.2 |
| 14_276019 | 1 | 1 | NA | NA | 1 | 1 | 5 | 0.8 | 7 | 0.8571 | 1 | 0 | 2 | 0.5 | 2 | 0.5 | NA | NA | 5 | 0.4 |
| 14_281978 | 3 | 1 | 5 | 1 | 7 | 1 | 8 | 1 | 23 | 1 | 3 | 1 | 9 | 1 | 12 | 1 | 11 | 1 | 35 | 1 |
| 14_281979 | 4 | 1 | 8 | 1 | 9 | 1 | 6 | 0.8333 | 27 | 0.963 | 4 | 1 | 10 | 1 | 12 | 0.8333 | 14 | 1 | 40 | 0.95 |
| 14_281979 | 3 | 1 | 7 | 0.7143 | 8 | 0.875 | 8 | 0.625 | 26 | 0.7692 | 4 | 1 | 10 | 1 | 11 | 0.8182 | 15 | 0.9333 | 40 | 0.925 |
| 14_281980 | 4 | 0.25 | 4 | 0 | 6 | 0.3333 | 6 | 0.1667 | 20 | 0.2 | 2 | 0 | 5 | 0.4 | 8 | 0.25 | 8 | 0.125 | 23 | 0.2174 |
| 14_281980 | 3 | 0.6667 | 8 | 1 | 8 | 0.625 | 10 | 0.2 | 29 | 0.5862 | 4 | 0.5 | 11 | 0.4545 | 11 | 0.5455 | 14 | 0.7143 | 40 | 0.575 |
| 14_281981 | 4 | 0 | 7 | 0.1429 | 6 | 0.1667 | 10 | 0.1 | 27 | 0.1111 | 1 | 0 | 8 | 0 | 9 | 0.2222 | 14 | 0 | 32 | 0.0625 |

| | | | | | | | | | | | | | | | | | | | | |
|---|---|---|---|---|---|---|---|---|---|---|---|---|---|---|---|---|---|---|---|---|
| 14_2819814 | 3 | 0.3333 | 6 | 0.1667 | 6 | 0.1667 | 8 | 0.25 | 23 | 0.2174 | 3 | 0.6667 | 7 | 0.4286 | 11 | 0 | 10 | 0.4 | 31 | 0.2903 |
| 14_2819821 | 4 | 0.25 | 5 | 0 | 9 | 0.2222 | 10 | 0.1 | 28 | 0.1429 | 4 | 0.75 | 7 | 0.1429 | 10 | 0 | 11 | 0.1818 | 32 | 0.1875 |
| 14_2819824 | 1 | 0 | 6 | 0.1667 | 8 | 0 | 10 | 0.1 | 25 | 0.08 | 5 | 0.2 | 11 | 0.1818 | 14 | 0.0714 | 13 | 0.1538 | 43 | 0.1395 |
| 14_2819826 | 4 | 0 | 7 | 0.1429 | 7 | 0.1429 | 9 | 0 | 27 | 0.0741 | 3 | 0 | 11 | 0.1818 | 12 | 0.0833 | 13 | 0 | 39 | 0.0769 |
| 14_2819839 | 4 | 0 | 8 | 0.125 | 10 | 0.1 | 9 | 0 | 31 | 0.0645 | 5 | 0.4 | 13 | 0.0769 | 14 | 0.1429 | 15 | 0.0667 | 47 | 0.1277 |
| 14_2819902 | 9 | 0 | 12 | 0.0833 | 9 | 0.1111 | 16 | 0.0625 | 46 | 0.0652 | 7 | 0 | 13 | 0.0769 | 18 | 0.1111 | 16 | 0.125 | 54 | 0.0926 |
| 14_2819969 | 11 | 0.0909 | 17 | 0.0588 | 10 | 0 | 20 | 0.05 | 58 | 0.0517 | 7 | 0.1429 | 12 | 0.0833 | 21 | 0.0476 | 21 | 0.0476 | 61 | 0.0656 |
| 14_2820027 | 11 | 0.8182 | 21 | 0.8571 | 15 | 0.4667 | 20 | 0.6 | 67 | 0.6866 | 13 | 0.6923 | 15 | 0.6 | 23 | 0.6087 | 25 | 0.64 | 76 | 0.6316 |
| 14_2824037 | NA | NA | 1 | 0 | NA | NA | 1 | 1 | 2 | 0.5 | 2 | 0.5 | NA | NA | NA | NA | NA | NA | 2 | 0.5 |
| 14_2824452 | NA | NA | 1 | 1 | NA | NA | 2 | 1 | 3 | 1 | 2 | 1 | NA | NA | 1 | 1 | NA | NA | 3 | 1 |
| 14_2824467 | NA | NA | 1 | 1 | NA | NA | 2 | 1 | 3 | 1 | 3 | 1 | NA | NA | 1 | 1 | NA | NA | 4 | 1 |
| 14_2824479 | NA | NA | 1 | 1 | NA | NA | 2 | 0.5 | 3 | 0.6667 | 2 | 0 | NA | NA | 1 | 1 | NA | NA | 3 | 0.3333 |
| 14_2824493 | NA | NA | 1 | 0 | NA | NA | 2 | 0.5 | 3 | 0.3333 | 3 | 0.3333 | NA | NA | 1 | 0 | NA | NA | 4 | 0.25 |
| 14_2824505 | NA | NA | 1 | 1 | NA | NA | 2 | 0 | 3 | 0.3333 | 3 | 0.6667 | NA | NA | NA | NA | NA | NA | 3 | 0.6667 |
| 14_2824601 | NA | NA | 1 | 1 | NA | NA | 1 | 1 | 2 | 1 | 2 | 0.5 | NA | NA | 1 | 0 | NA | NA | 3 | 0.3333 |
| 14_2824605 | NA | NA | 1 | 0 | NA | NA | 1 | 1 | 2 | 0.5 | 3 | 0.3333 | NA | NA | 1 | 0 | NA | NA | 4 | 0.25 |
| 14_2824680 | 2 | 0 | 2 | 0.5 | NA | NA | 3 | 0 | 7 | 0.1429 | 3 | 0.3333 | NA | NA | 1 | 0 | NA | NA | 4 | 0.25 |
| 14_2862708 | 17 | 0.3529 | 9 | 0 | 11 | 0.2727 | 22 | 0.1818 | 59 | 0.2203 | 8 | 0.125 | 26 | 0.0385 | 28 | 0.1071 | 17 | 0.1176 | 79 | 0.0886 |
| 14_2862710 | 15 | 0.4667 | 9 | 0 | 9 | 0.2222 | 19 | 0.0526 | 52 | 0.1923 | 7 | 0.1429 | 23 | 0.0435 | 28 | 0.2857 | 16 | 0.1875 | 74 | 0.1757 |
| 14_2862717 | 17 | 0.4706 | 7 | 0.1429 | 10 | 0.3 | 15 | 0.4667 | 49 | 0.3878 | 7 | 0.4286 | 19 | 0.1053 | 25 | 0.32 | 17 | 0.4118 | 68 | 0.2941 |
| 14_2940951 | NA | NA | 2 | 1 | 1 | 1 | NA | NA | 3 | 1 | NA | NA | NA | NA | 1 | 1 | NA | NA | 1 | 1 |
| 14_2940975 | NA | NA | 2 | 1 | 1 | 1 | NA | NA | 3 | 1 | NA | NA | NA | NA | 1 | 1 | NA | NA | 1 | 1 |
| 14_2971275 | 4 | 0 | 3 | 0 | 6 | 0.5 | 7 | 0.1429 | 20 | 0.2 | 9 | 0 | 7 | 0.2857 | 6 | 0 | 6 | 0 | 28 | 0.0714 |
| 14_2971277 | 6 | 0 | 4 | 0.25 | 6 | 0 | 7 | 0.1429 | 23 | 0.087 | 11 | 0.0909 | 8 | 0 | 4 | 0 | 5 | 0.2 | 28 | 0.0714 |
| 14_2971279 | 4 | 0.25 | 4 | 0.75 | 5 | 0.2 | 7 | 0.2857 | 20 | 0.35 | 11 | 0.1818 | 9 | 0.3333 | 7 | 0.2857 | 4 | 0.5 | 31 | 0.2903 |
| 14_2971291 | 6 | 1 | 4 | 0.75 | 4 | 0.5 | 7 | 0.4286 | 21 | 0.6667 | 10 | 0.7 | 9 | 0.5556 | 5 | 0.4 | 7 | 0.7143 | 31 | 0.6129 |
| 14_2971350 | NA | NA | 2 | 0.5 | NA | NA | NA | NA | 2 | 0.5 | NA | NA | 1 | 0 | NA | NA | 2 | 0.5 | 3 | 0.3333 |
| 14_2971498 | 7 | 0 | 7 | 0.1429 | 5 | 0.2 | 6 | 0.6667 | 25 | 0.24 | 16 | 0.1875 | 10 | 0 | 5 | 0.2 | 6 | 0.5 | 37 | 0.1892 |
| 14_2971509 | 6 | 0.5 | 7 | 0.2857 | 6 | 0.1667 | 8 | 0.125 | 27 | 0.2593 | 17 | 0.5294 | 10 | 0.2 | 7 | 0.7143 | 9 | 0.3333 | 43 | 0.4419 |
| 14_2976210 | 1 | 0 | NA | NA | 2 | 1 | 1 | 1 | 4 | 0.75 | 2 | 1 | 2 | 1 | 3 | 1 | 2 | 1 | 9 | 1 |
| 14_2976227 | 2 | 0.5 | NA | NA | 1 | 0 | 1 | 0 | 4 | 0.25 | 1 | 1 | 2 | 0.5 | 2 | 0 | 2 | 0.5 | 7 | 0.4286 |
| 14_2976233 | 2 | 0.5 | NA | NA | 2 | 0.5 | 1 | 0 | 5 | 0.4 | 3 | 0.6667 | 2 | 1 | 3 | 0.6667 | 2 | 0.5 | 10 | 0.7 |
| 14_2976235 | 2 | 0 | NA | NA | 2 | 0.5 | NA | NA | 4 | 0.25 | 3 | 0.3333 | 2 | 0 | 3 | 0 | 2 | 0 | 10 | 0.1 |
| 14_2976238 | 2 | 0.5 | NA | NA | 2 | 0 | 1 | 0 | 5 | 0.2 | 3 | 0.6667 | 2 | 0 | 3 | 0.3333 | 2 | 0.5 | 10 | 0.4 |
| 14_2976240 | 2 | 0.5 | NA | NA | 2 | 0 | 1 | 0 | 5 | 0.2 | 3 | 0.3333 | 2 | 0.5 | 2 | 0 | 2 | 0.5 | 9 | 0.3333 |
| 14_2976256 | 3 | 0.6667 | NA | NA | 2 | 0.5 | 1 | 0 | 6 | 0.5 | 3 | 0.6667 | 1 | 1 | 2 | 1 | 2 | 0.5 | 8 | 0.75 |
| 14_2976259 | 2 | 0.5 | NA | NA | 1 | 1 | 1 | 1 | 4 | 0.75 | 3 | 1 | 2 | 1 | 1 | 1 | 2 | 0.5 | 8 | 0.875 |
| 14_2976297 | 2 | 0.5 | NA | NA | 2 | 0.5 | 1 | 0 | 5 | 0.4 | 2 | 0.5 | 2 | 1 | 2 | 0.5 | 2 | 0.5 | 8 | 0.625 |
| 14_2976310 | 3 | 0.3333 | 1 | 1 | 2 | 0 | 3 | 0.6667 | 9 | 0.4444 | 1 | 0 | 2 | 0 | 4 | 0.25 | 2 | 0 | 9 | 0.1111 |
| 14_2976351 | 3 | 0 | 1 | 0 | 2 | 0 | 3 | 0.3333 | 9 | 0.1111 | 1 | 0 | 1 | 1 | 5 | 0 | 1 | 0 | 8 | 0.125 |
| 14_2976391 | 3 | 1 | 1 | 1 | 2 | 1 | 3 | 0 | 9 | 0.6667 | 1 | 1 | 2 | 1 | 3 | 1 | 2 | 0.5 | 8 | 0.875 |
| 14_2976406 | 1 | 1 | 1 | 0 | 2 | 1 | 2 | 1 | 6 | 0.8333 | 1 | 0 | 1 | 1 | 3 | 1 | 2 | 0.5 | 7 | 0.7143 |
| 14_2976408 | 1 | 0 | 1 | 1 | 2 | 0.5 | 2 | 0 | 6 | 0.3333 | 1 | 0 | 1 | 0 | 3 | 0 | 2 | 1 | 7 | 0.2857 |
| 14_2976412 | 2 | 1 | 1 | 1 | 2 | 0.5 | 2 | 0.5 | 7 | 0.7143 | 1 | 1 | 1 | 1 | 2 | 0.5 | 2 | 1 | 6 | 0.8333 |
| 14_2976413 | 2 | 1 | 1 | 1 | 1 | 1 | 2 | 1 | 6 | 1 | NA | NA | 1 | 1 | 1 | 1 | 2 | 1 | 4 | 1 |
| 14_2976428 | 1 | 0 | 1 | 0 | 1 | 0 | 2 | 0.5 | 5 | 0.2 | 1 | 1 | NA | NA | 3 | 0 | 1 | 1 | 5 | 0.4 |
| 14_2976481 | 2 | 0.5 | 1 | 1 | 2 | 0.5 | 1 | 1 | 6 | 0.6667 | 1 | 0 | NA | NA | 3 | 0.6667 | 2 | 1 | 6 | 0.6667 |
| 14_2976488 | 2 | 0.5 | 1 | 1 | 1 | 0 | 2 | 0.5 | 6 | 0.5 | 1 | 1 | NA | NA | 2 | 0.5 | 1 | 0 | 4 | 0.5 |
| 14_3024524 | 1 | 0 | 1 | 0 | 1 | 0 | 2 | 0.5 | 5 | 0.2 | NA | NA | NA | NA | 1 | 1 | NA | NA | 1 | 1 |
| 14_3024546 | 1 | 1 | 1 | 1 | 1 | 1 | 1 | 1 | 4 | 1 | NA | NA | 1 | 1 | 1 | 1 | 1 | 1 | 3 | 1 |
| 14_3024553 | 1 | 1 | 1 | 0 | 1 | 0 | 2 | 1 | 5 | 0.6 | NA | NA | 1 | 0 | 1 | 1 | 1 | 0 | 3 | 0.3333 |
| 14_3024562 | 1 | 1 | 1 | 1 | 1 | 1 | 2 | 1 | 5 | 1 | NA | NA | 1 | 0 | 1 | 1 | NA | NA | 2 | 0.5 |
| 14_3024567 | 1 | 1 | 1 | 1 | 2 | 1 | 2 | 1 | 6 | 1 | NA | NA | 1 | 1 | 1 | 1 | 1 | 1 | 3 | 1 |
| 14_3024574 | 1 | 1 | 1 | 1 | 2 | 1 | 2 | 1 | 6 | 1 | NA | NA | NA | NA | NA | NA | 1 | 1 | 1 | 1 |
| 14_3024592 | 1 | 1 | 1 | 1 | 1 | 1 | 2 | 1 | 5 | 1 | NA | NA | 1 | 1 | 1 | 1 | 1 | 1 | 3 | 1 |
| 14_3024603 | 1 | 1 | 1 | 0 | 2 | 0.5 | 2 | 0.5 | 6 | 0.5 | NA | NA | 2 | 0.5 | 1 | 0 | 1 | 0 | 4 | 0.25 |
| 14_3024604 | 1 | 0 | 1 | 1 | 2 | 0 | 2 | 1 | 6 | 0.5 | NA | NA | 2 | 1 | 1 | 1 | 1 | 0 | 4 | 0.75 |
| 14_3024624 | 1 | 1 | 1 | 1 | 2 | 1 | 1 | 1 | 5 | 1 | NA | NA | 2 | 1 | 1 | 1 | 1 | 0 | 4 | 0.75 |
| 14_3024827 | 1 | 0 | 3 | 0.3333 | 1 | 0 | 3 | 0 | 8 | 0.125 | NA | NA | 2 | 1 | 2 | 0 | 2 | 0 | 6 | 0.3333 |
| 14_3024834 | 1 | 1 | 3 | 0.3333 | 2 | 0.5 | 4 | 0.5 | 10 | 0.5 | NA | NA | 1 | 0 | 2 | 0 | 2 | 0.5 | 5 | 0.2 |
| 14_3024837 | 2 | 1 | 2 | 0 | 2 | 1 | 4 | 0.25 | 10 | 0.5 | NA | NA | 2 | 0 | 2 | 0.5 | 1 | 0 | 5 | 0.2 |
| 14_3024849 | 2 | 1 | 2 | 0.5 | 2 | 0.5 | 5 | 0.6 | 11 | 0.6364 | NA | NA | 2 | 0 | 2 | 0.5 | 2 | 0 | 6 | 0.1667 |
| 14_3024860 | 3 | 0 | 3 | 0.3333 | 1 | 1 | 3 | 0 | 10 | 0.2 | NA | NA | 2 | 0 | 2 | 0.5 | 2 | 0.5 | 6 | 0.3333 |
| 14_3024869 | 2 | 0.5 | 3 | 0.3333 | 2 | 0.5 | 5 | 0.2 | 12 | 0.3333 | NA | NA | 1 | 0 | 2 | 0.5 | 2 | 0 | 5 | 0.2 |
| 14_3031765 | NA | NA | 1 | 1 | 1 | 1 | 2 | 1 | 4 | 1 | NA | NA | 1 | 0 | 3 | 0.6667 | 1 | 1 | 5 | 0.6 |

| | | | | | | | | | | | | | | | | | | | | |
|---|---|---|---|---|---|---|---|---|---|---|---|---|---|---|---|---|---|---|---|---|
| 14_303178 | 3 | 1 | 2 | 0.5 | 1 | 1 | 1 | 1 | 7 | 0.8571 | NA | NA | 1 | 1 | 3 | 0.6667 | 2 | 1 | 6 | 0.8333 |
| 14_304125 | NA | NA | NA | NA | 1 | 1 | 1 | 0 | 2 | 0.5 | NA | NA | 1 | 1 | NA | NA | NA | NA | 1 | 1 |
| 14_304126 | NA | NA | NA | NA | 1 | 1 | 1 | 0 | 2 | 0.5 | NA | NA | 1 | 1 | NA | NA | NA | NA | 1 | 1 |
| 14_305397 | 1 | 1 | 4 | 0.75 | 3 | 1 | NA | NA | 8 | 0.875 | NA | NA | 1 | 1 | 3 | 1 | 2 | 1 | 6 | 1 |
| 14_305398 | 1 | 0 | 5 | 0.2 | 3 | 0 | 1 | 0 | 10 | 0.1 | NA | NA | 1 | 1 | 4 | 0 | 2 | 0 | 7 | 0.1429 |
| 14_305399 | 1 | 0 | 5 | 0.4 | 3 | 0.3333 | 1 | 0 | 10 | 0.3 | NA | NA | 1 | 1 | 4 | 0.25 | 2 | 1 | 7 | 0.5714 |
| 14_305400 | 1 | 1 | 4 | 1 | 3 | 1 | 1 | 1 | 9 | 1 | NA | NA | NA | NA | 4 | 1 | 2 | 1 | 6 | 1 |
| 14_305400 | 1 | 1 | 5 | 1 | 2 | 1 | 1 | 1 | 9 | 1 | NA | NA | NA | NA | 4 | 0.75 | 2 | 1 | 6 | 0.8333 |
| 14_305401 | 1 | 0 | 5 | 0.4 | 4 | 0.5 | NA | NA | 10 | 0.4 | NA | NA | 1 | 1 | 5 | 0.4 | 2 | 1 | 8 | 0.625 |
| 14_305401 | 1 | 1 | 5 | 1 | 5 | 1 | 1 | 1 | 12 | 1 | NA | NA | 1 | 1 | 4 | 1 | 1 | 1 | 6 | 1 |
| 14_305401 | 1 | 0 | 5 | 0.4 | 5 | 0.2 | 1 | 1 | 12 | 0.3333 | NA | NA | NA | NA | 4 | 0.25 | 2 | 0 | 6 | 0.1667 |
| 14_305402 | 1 | 1 | 4 | 1 | 5 | 0.8 | 1 | 1 | 11 | 0.9091 | NA | NA | 1 | 1 | 5 | 0.6 | 2 | 1 | 8 | 0.75 |
| 14_305414 | 4 | 0 | 4 | 0.25 | 1 | 0 | NA | NA | 9 | 0.1111 | 1 | 1 | NA | NA | 5 | 0 | 5 | 0 | 11 | 0.0909 |
| 14_305415 | 1 | 0 | 4 | 0.25 | NA | NA | NA | NA | 5 | 0.2 | 1 | 1 | NA | NA | NA | NA | 1 | 0 | 2 | 0.5 |
| 14_305432 | 2 | 0 | 2 | 1 | NA | NA | NA | NA | 4 | 0.5 | 2 | 0.5 | 1 | 1 | NA | NA | 1 | 0 | 4 | 0.5 |
| 14_305437 | 1 | 0 | 3 | 0.3333 | 1 | 0 | NA | NA | 5 | 0.2 | 2 | 1 | 1 | 0 | NA | NA | 1 | 0 | 4 | 0.5 |
| 14_305965 | 2 | 0.5 | 3 | 0.6667 | 2 | 0 | 1 | 1 | 8 | 0.5 | 2 | 0.5 | NA | NA | NA | NA | 3 | 0.6667 | 5 | 0.6 |
| 14_305966 | 1 | 1 | 2 | 1 | 2 | 0.5 | NA | NA | 5 | 0.8 | 2 | 0 | NA | NA | NA | NA | 3 | 0.3333 | 5 | 0.2 |
| 14_305967 | 2 | 1 | 3 | 1 | 1 | 1 | 1 | 0 | 7 | 0.8571 | 1 | 1 | NA | NA | NA | NA | 1 | 1 | 2 | 1 |
| 14_305968 | 1 | 0 | 3 | 0.3333 | 1 | 0 | 1 | 0 | 6 | 0.1667 | 1 | 0 | NA | NA | NA | NA | 4 | 0.25 | 5 | 0.2 |
| 14_305970 | 2 | 0.5 | 3 | 0.3333 | 1 | 1 | 1 | 1 | 7 | 0.5714 | 2 | 0.5 | 1 | 0 | NA | NA | 3 | 0.6667 | 6 | 0.5 |
| 14_305977 | 2 | 0.5 | 2 | 0 | 1 | 0 | 1 | 0 | 6 | 0.1667 | 2 | 0.5 | 2 | 0 | 2 | 0.5 | 2 | 0 | 8 | 0.25 |
| 14_305988 | 4 | 0.25 | 2 | 0 | 1 | 0 | 1 | 0 | 8 | 0.125 | 2 | 0 | 2 | 0 | 3 | 0.3333 | 2 | 0 | 9 | 0.1111 |
| 14_306030 | 2 | 0.5 | 2 | 0.5 | NA | NA | NA | NA | 4 | 0.5 | 1 | 0 | NA | NA | 1 | 1 | NA | NA | 2 | 0.5 |
| 14_306305 | 2 | 0 | 3 | 0.3333 | NA | NA | 2 | 0 | 7 | 0.1429 | 2 | 0 | 3 | 0.3333 | 2 | 0 | 1 | 0 | 8 | 0.125 |
| 14_306306 | 2 | 0.5 | 2 | 0 | NA | NA | 1 | 0 | 5 | 0.2 | 3 | 0 | 4 | 0.25 | 1 | 0 | NA | NA | 8 | 0.125 |
| 14_307042 | 1 | 0 | 4 | 0.25 | 1 | 0 | 1 | 0 | 7 | 0.1429 | NA | NA | 3 | 0 | 1 | 0 | 1 | 1 | 5 | 0.2 |
| 14_307093 | 1 | 1 | 4 | 1 | 1 | 1 | 1 | 1 | 7 | 1 | NA | NA | 3 | 0.6667 | 1 | 1 | NA | NA | 4 | 0.75 |
| 14_307093 | 1 | 0 | 4 | 0.5 | 1 | 0 | 1 | 0 | 7 | 0.2857 | NA | NA | 3 | 0.3333 | 1 | 0 | 2 | 0.5 | 6 | 0.3333 |
| 14_307093 | 1 | 1 | 4 | 0.5 | 1 | 0 | 1 | 1 | 7 | 0.5714 | NA | NA | 1 | 0 | 1 | 1 | 1 | 1 | 3 | 0.6667 |
| 14_307097 | 1 | 1 | 4 | 0.5 | 2 | 1 | 1 | 1 | 8 | 0.75 | NA | NA | 2 | 0 | 1 | 1 | 2 | 0.5 | 5 | 0.4 |
| 14_307106 | NA | NA | 1 | 1 | NA | NA | NA | NA | 1 | 1 | NA | NA | NA | NA | NA | NA | 1 | 1 | 1 | 1 |
| 14_307220 | 43 | 0.6279 | 22 | 0.7727 | 29 | 0.5172 | 25 | 0.68 | 119 | 0.6387 | 16 | 0.625 | 27 | 0.7037 | 24 | 0.625 | 25 | 0.68 | 92 | 0.663 |
| 14_307236 | NA | NA | 1 | 0 | 1 | 1 | NA | NA | 2 | 0.5 | NA | NA | NA | NA | 1 | 0 | 3 | 0.3333 | 4 | 0.25 |
| 14_307321 | NA | NA | 1 | 0 | 1 | 0 | 5 | 0.2 | 7 | 0.1429 | 2 | 0 | NA | NA | 4 | 0.25 | 2 | 0.5 | 8 | 0.25 |
| 14_307323 | NA | NA | 6 | 0.1667 | 3 | 0 | 6 | 0 | 15 | 0.0667 | 3 | 0 | 3 | 0 | 6 | 0.1667 | 7 | 0 | 19 | 0.0526 |
| 14_307325 | NA | NA | 2 | 0.5 | 1 | 1 | 4 | 0.75 | 7 | 0.7143 | 1 | 1 | NA | NA | 4 | 1 | 4 | 1 | 9 | 1 |
| 14_307325 | NA | NA | 1 | 0 | 2 | 0 | 4 | 0.25 | 7 | 0.1429 | NA | NA | NA | NA | 5 | 0.2 | 4 | 0.5 | 9 | 0.3333 |
| 14_307327 | NA | NA | 1 | 1 | 2 | 0 | 5 | 0.6 | 8 | 0.5 | 1 | 0 | NA | NA | 2 | 0.5 | 3 | 0.3333 | 6 | 0.3333 |
| 14_307327 | NA | NA | 2 | 0 | 1 | 0 | 5 | 0.4 | 8 | 0.25 | NA | NA | NA | NA | 4 | 0.25 | 2 | 1 | 6 | 0.5 |
| 14_307327 | NA | NA | 2 | 0 | 2 | 0 | 5 | 0.4 | 9 | 0.2222 | 1 | 0 | NA | NA | 4 | 0.25 | 3 | 0.6667 | 8 | 0.375 |
| 14_307328 | 1 | 0 | 7 | 0.1429 | 3 | 0 | 5 | 0 | 16 | 0.0625 | 2 | 0 | 2 | 0 | 9 | 0.1111 | 7 | 0 | 20 | 0.05 |
| 14_307335 | 1 | 1 | 6 | 0.5 | 4 | 0.5 | 6 | 0.5 | 17 | 0.5294 | 3 | 0 | 2 | 0.5 | 6 | 0.5 | 6 | 0.3333 | 17 | 0.3529 |
| 14_307336 | 1 | 1 | 7 | 0.8571 | 4 | 1 | 6 | 0.6667 | 18 | 0.8333 | 2 | 1 | 2 | 0.5 | 5 | 0.6 | 5 | 0.8 | 14 | 0.7143 |
| 14_307336 | 1 | 0 | 6 | 0.1667 | 3 | 0 | 4 | 0.25 | 14 | 0.1429 | 3 | 0 | 1 | 0 | 6 | 0 | 2 | 0.5 | 12 | 0.0833 |
| 14_307337 | 1 | 0 | 7 | 0.1429 | 4 | 0 | 4 | 0.5 | 16 | 0.1875 | 3 | 0 | 3 | 0 | 9 | 0 | 6 | 0.5 | 21 | 0.1429 |
| 14_307338 | 1 | 0 | 8 | 0.25 | 3 | 0 | 6 | 0.1667 | 18 | 0.1667 | 4 | 0 | 2 | 0 | 8 | 0.125 | 6 | 0.6667 | 20 | 0.25 |
| 14_307338 | 1 | 1 | 8 | 0.75 | 3 | 0.6667 | 6 | 0.6667 | 18 | 0.7222 | 3 | 1 | 3 | 0.6667 | 9 | 0.5556 | 7 | 0.7143 | 22 | 0.6818 |
| 14_307338 | 1 | 0 | 7 | 0.1429 | 4 | 0.5 | 5 | 0.4 | 17 | 0.2941 | 4 | 0.5 | 3 | 0.3333 | 9 | 0 | 7 | 0.1429 | 23 | 0.1739 |
| 14_307338 | NA | NA | 8 | 0.125 | 3 | 0 | 5 | 0.2 | 16 | 0.125 | 2 | 0 | 3 | 0.3333 | 5 | 0 | 5 | 0 | 15 | 0.0667 |
| 14_307338 | 1 | 1 | 9 | 0.1111 | 4 | 0 | 6 | 0.5 | 20 | 0.25 | 4 | 0 | 3 | 0 | 8 | 0.125 | 8 | 0.125 | 23 | 0.087 |
| 14_307835 | 2 | 1 | 3 | 0.3333 | NA | NA | 2 | 0 | 7 | 0.4286 | NA | NA | 1 | 0 | 1 | 1 | 2 | 0 | 4 | 0.25 |
| 14_307835 | 2 | 0 | 3 | 0.6667 | NA | NA | 2 | 0 | 7 | 0.2857 | NA | NA | 1 | 1 | 1 | 0 | 2 | 0 | 4 | 0.25 |
| 14_307893 | 2 | 1 | 4 | 0.75 | NA | NA | 2 | 0 | 8 | 0.625 | NA | NA | 1 | 0 | 1 | 1 | 1 | 0 | 3 | 0.3333 |
| 14_307894 | 2 | 1 | 4 | 0.75 | NA | NA | 3 | 0.6667 | 9 | 0.7778 | NA | NA | NA | NA | 1 | 1 | 2 | 0.5 | 3 | 0.6667 |
| 14_307895 | 2 | 1 | 4 | 0.75 | NA | NA | 3 | 0.6667 | 9 | 0.7778 | NA | NA | 1 | 1 | 1 | 1 | 2 | 1 | 4 | 1 |
| 14_307896 | 2 | 0.5 | 4 | 0 | NA | NA | 3 | 0.3333 | 9 | 0.2222 | NA | NA | 1 | 1 | 1 | 0 | 2 | 0.5 | 4 | 0.5 |
| 14_307896 | 2 | 0 | 4 | 0.25 | NA | NA | 2 | 0 | 8 | 0.125 | NA | NA | 1 | 0 | NA | NA | 1 | 1 | 2 | 0.5 |
| 14_308393 | 5 | 0.4 | 3 | 0 | 2 | 0.5 | 2 | 1 | 12 | 0.4167 | NA | NA | 2 | 0.5 | 5 | 0.8 | 3 | 0.3333 | 10 | 0.6 |
| 14_308408 | 6 | 0.3333 | 4 | 0.25 | 3 | 0 | 1 | 0 | 14 | 0.2143 | NA | NA | 2 | 0.5 | 4 | 0 | 3 | 0.3333 | 9 | 0.2222 |
| 14_308409 | 5 | 1 | 5 | 0.4 | 3 | 0.3333 | 2 | 0.5 | 15 | 0.6 | 1 | 1 | 2 | 0.5 | 6 | 0.6667 | 5 | 0.6 | 14 | 0.6429 |
| 14_308411 | 7 | 0.4286 | 6 | 0.1667 | 4 | 0.25 | 3 | 0 | 20 | 0.25 | NA | NA | 2 | 0.5 | 6 | 0.5 | 5 | 0.4 | 13 | 0.4615 |
| 14_308420 | 7 | 0.2857 | 4 | 0.5 | 4 | 0.25 | 5 | 0 | 20 | 0.25 | 1 | 0 | 2 | 0.5 | 5 | 0.4 | 5 | 0.4 | 13 | 0.3846 |
| 14_308426 | 7 | 0.1429 | 6 | 0.5 | 5 | 0 | 5 | 0 | 23 | 0.1739 | 1 | 0 | 3 | 0.3333 | 4 | 0.25 | 5 | 0 | 13 | 0.1538 |
| 14_308433 | 8 | 0.375 | 7 | 0.2857 | 7 | 0.2857 | 10 | 0.2 | 32 | 0.2812 | 3 | 0 | 3 | 0.3333 | 6 | 0.6667 | 7 | 0.2857 | 19 | 0.3684 |

| | | | | | | | | | | | | | | | | | | | | |
|---|---|---|---|---|---|---|---|---|---|---|---|---|---|---|---|---|---|---|---|---|
| 14_308931 | 25 | 0.24 | 43 | 0.186 | 32 | 0.2188 | 30 | 0.3333 | 130 | 0.2385 | 41 | 0.2439 | 49 | 0.3469 | 32 | 0.1875 | 41 | 0.2439 | 163 | 0.2638 |
| 14_308933 | 25 | 0.84 | 41 | 0.8537 | 27 | 0.9259 | 27 | 0.7407 | 120 | 0.8417 | 41 | 0.9268 | 47 | 0.8723 | 35 | 0.8286 | 40 | 0.9 | 163 | 0.8834 |
| 14_308936 | 15 | 0.2 | 22 | 0.3182 | 20 | 0.3 | 23 | 0.1304 | 80 | 0.2375 | 19 | 0.2632 | 22 | 0.2727 | 14 | 0.2857 | 23 | 0.2609 | 78 | 0.2692 |
| 14_308937 | 16 | 0.125 | 32 | 0.4062 | 21 | 0.381 | 20 | 0.25 | 89 | 0.3146 | 22 | 0.2727 | 31 | 0.3226 | 21 | 0.1429 | 31 | 0.3548 | 105 | 0.2857 |
| 14_308938 | 16 | 0.5 | 30 | 0.7667 | 17 | 0.7059 | 25 | 0.68 | 88 | 0.6818 | 24 | 0.5833 | 27 | 0.5926 | 16 | 0.8125 | 27 | 0.8889 | 94 | 0.7128 |
| 14_308938 | 19 | 0.3684 | 34 | 0.5 | 19 | 0.4737 | 24 | 0.5417 | 96 | 0.4792 | 24 | 0.5833 | 35 | 0.4857 | 21 | 0.4286 | 28 | 0.3929 | 108 | 0.4722 |
| 14_308940 | 13 | 0.1538 | 27 | 0.0741 | 15 | 0 | 24 | 0.0417 | 79 | 0.0633 | 22 | 0.0909 | 30 | 0.1667 | 17 | 0.1176 | 28 | 0.1429 | 97 | 0.134 |
| 14_308987 | 5 | 1 | 11 | 0.9091 | 4 | 1 | 11 | 0.5455 | 31 | 0.8065 | 7 | 0.5714 | 9 | 0.7778 | 4 | 0.5 | 10 | 0.8 | 30 | 0.7 |
| 14_308987 | 7 | 0 | 12 | 0 | 5 | 0.2 | 11 | 0.0909 | 35 | 0.0571 | 7 | 0 | 8 | 0 | 7 | 0.1429 | 10 | 0.2 | 32 | 0.0938 |
| 14_308991 | 5 | 0.4 | 10 | 0.2 | 5 | 0 | 10 | 0.1 | 30 | 0.1667 | 4 | 0.25 | 1 | 0 | 5 | 0 | 10 | 0.1 | 20 | 0.1 |
| 14_308992 | 3 | 0.6667 | 9 | 0.6667 | 1 | 0 | 10 | 0.7 | 23 | 0.6522 | 4 | 1 | 5 | 0.6 | 5 | 0.8 | 9 | 0.7778 | 23 | 0.7826 |
| 14_308995 | 4 | 0.25 | 9 | 0.7778 | 5 | 0.6 | 8 | 0.375 | 26 | 0.5385 | 3 | 0.6667 | 4 | 0.5 | 6 | 0.6667 | 7 | 0.4286 | 20 | 0.55 |
| 14_309059 | 1 | 0 | NA | NA | 3 | 0.6667 | 3 | 0 | 7 | 0.2857 | 1 | 0 | 5 | 0.2 | 3 | 0 | 3 | 0 | 12 | 0.0833 |
| 14_309062 | 2 | 0 | 1 | 0 | 3 | 0.3333 | 3 | 0 | 9 | 0.1111 | 1 | 0 | 5 | 0 | 2 | 0.5 | 2 | 0.5 | 10 | 0.2 |
| 14_309521 | NA | NA | 1 | 1 | NA | NA | NA | NA | 1 | 1 | NA | NA | NA | NA | 1 | 1 | NA | NA | 1 | 1 |
| 14_309598 | NA | NA | 1 | 1 | 1 | 1 | NA | NA | 2 | 1 | NA | NA | NA | NA | 3 | 1 | 1 | 1 | 4 | 1 |
| 14_309598 | NA | NA | 1 | 1 | 1 | 0 | NA | NA | 2 | 0.5 | NA | NA | NA | NA | 3 | 0.3333 | 1 | 0 | 4 | 0.25 |
| 14_309726 | NA | NA | 1 | 1 | NA | NA | NA | NA | 1 | 1 | NA | NA | NA | NA | 2 | 1 | 1 | 1 | 3 | 1 |
| 14_309727 | NA | NA | 1 | 1 | NA | NA | NA | NA | 1 | 1 | NA | NA | NA | NA | 2 | 1 | 1 | 1 | 3 | 1 |
| 14_309729 | NA | NA | 1 | 1 | NA | NA | NA | NA | 1 | 1 | NA | NA | NA | NA | 2 | 1 | 1 | 1 | 3 | 1 |
| 14_309793 | 7 | 0.1429 | 9 | 0 | 11 | 0.0909 | 5 | 0 | 32 | 0.0625 | 6 | 0 | 10 | 0 | 10 | 0.2 | 7 | 0 | 33 | 0.0606 |
| 14_309794 | 8 | 0.125 | 9 | 0.2222 | 8 | 0.375 | 4 | 0.25 | 29 | 0.2414 | 8 | 0.25 | 9 | 0.2222 | 13 | 0.4615 | 10 | 0.2 | 40 | 0.3 |
| 14_309808 | 7 | 0.1429 | 16 | 0.0625 | 12 | 0.0833 | 9 | 0.1111 | 44 | 0.0909 | 13 | 0.1538 | 14 | 0.2857 | 13 | 0.1538 | 14 | 0.2857 | 54 | 0.2222 |
| 14_309809 | 9 | 0.2222 | 15 | 0.1333 | 12 | 0.1667 | 9 | 0 | 45 | 0.1333 | 15 | 0 | 16 | 0.125 | 10 | 0 | 14 | 0.1429 | 55 | 0.0727 |
| 14_309809 | 8 | 0.375 | 15 | 0.2 | 11 | 0.2727 | 7 | 0 | 41 | 0.2195 | 14 | 0.2857 | 16 | 0.25 | 14 | 0.2143 | 16 | 0.125 | 60 | 0.2167 |
| 14_309809 | 8 | 0.375 | 12 | 0.5833 | 8 | 0.375 | 7 | 0.2857 | 35 | 0.4286 | 13 | 0.3077 | 14 | 0.6429 | 13 | 0.5385 | 11 | 0.2727 | 51 | 0.451 |
| 14_309810 | 8 | 0.375 | 15 | 0.2 | 12 | 0.25 | 9 | 0.4444 | 44 | 0.2955 | 13 | 0.3077 | 15 | 0.4 | 14 | 0.3571 | 16 | 0.3125 | 58 | 0.3448 |
| 14_309811 | 6 | 0.1667 | 15 | 0.3333 | 10 | 0.2 | 10 | 0.5 | 41 | 0.3171 | 17 | 0.1765 | 14 | 0.0714 | 15 | 0.2 | 12 | 0.25 | 58 | 0.1724 |
| 14_309813 | 10 | 0.1 | 15 | 0.2 | 13 | 0 | 10 | 0 | 48 | 0.0833 | 17 | 0.1176 | 15 | 0.0667 | 17 | 0.0588 | 14 | 0.1429 | 63 | 0.0952 |
| 14_309813 | 8 | 0.375 | 16 | 0.375 | 13 | 0.2308 | 10 | 0.4 | 47 | 0.3404 | 17 | 0.2353 | 16 | 0.25 | 16 | 0.4375 | 17 | 0.1176 | 66 | 0.2576 |
| 14_309814 | 9 | 0.4444 | 12 | 0.4167 | 11 | 0.2727 | 8 | 0.375 | 40 | 0.375 | 10 | 0.2 | 15 | 0.2 | 12 | 0.3333 | 12 | 0.3333 | 49 | 0.2653 |
| 14_309814 | 9 | 0.2222 | 15 | 0.0667 | 13 | 0.2308 | 11 | 0.1818 | 48 | 0.1667 | 15 | 0.1333 | 15 | 0.0667 | 17 | 0.1765 | 16 | 0.3125 | 63 | 0.1746 |
| 14_309831 | 3 | 0 | 1 | 0 | 3 | 0.6667 | NA | NA | 7 | 0.2857 | 1 | 1 | NA | NA | 1 | 0 | 1 | 1 | 3 | 0.6667 |
| 14_309869 | 3 | 0.3333 | 1 | 0 | 1 | 0 | NA | NA | 5 | 0.2 | 1 | 0 | 1 | 0 | NA | NA | 1 | 1 | 3 | 0.3333 |
| 14_310928 | 6 | 0.3333 | 5 | 0 | 2 | 0 | 10 | 0 | 23 | 0.087 | 2 | 0 | 6 | 0.1667 | 2 | 0 | 7 | 0.1429 | 17 | 0.1176 |
| 14_310929 | 6 | 0.5 | 4 | 0.25 | 3 | 0.3333 | 10 | 0.5 | 23 | 0.4348 | 1 | 0 | 7 | 0.2857 | 3 | 1 | 6 | 0.6667 | 17 | 0.5294 |
| 14_310929 | 6 | 1 | 4 | 1 | 3 | 1 | 8 | 0.875 | 21 | 0.9524 | 2 | 1 | 7 | 0.8571 | 3 | 1 | 7 | 1 | 19 | 0.9474 |
| 14_310929 | 6 | 0.5 | 5 | 0.8 | 3 | 1 | 11 | 0.5455 | 25 | 0.64 | 2 | 1 | 7 | 0.7143 | 3 | 0.6667 | 6 | 1 | 18 | 0.8333 |
| 14_310932 | 4 | 0.25 | 5 | 0.2 | 3 | 0.3333 | 6 | 0.1667 | 18 | 0.2222 | 4 | 0 | 3 | 0.3333 | 4 | 0.5 | 7 | 0.1429 | 18 | 0.2222 |
| 14_310937 | 4 | 0 | 4 | 0 | 2 | 0.5 | 7 | 0 | 17 | 0.0588 | 4 | 0 | 4 | 0 | 4 | 0.25 | 7 | 0 | 19 | 0.0526 |
| 14_310958 | 11 | 0.0909 | 4 | 0.25 | 10 | 0.2 | 3 | 0 | 28 | 0.1429 | 4 | 0 | 4 | 0.5 | 4 | 0 | 4 | 0.25 | 16 | 0.1875 |
| 14_310963 | 3 | 0.3333 | 1 | 0 | 3 | 0 | NA | NA | 7 | 0.1429 | 3 | 0 | 3 | 0 | 3 | 0 | 3 | 0.3333 | 12 | 0.0833 |
| 14_310966 | 3 | 0.3333 | 1 | 0 | 3 | 0.3333 | 1 | 0 | 8 | 0.25 | 2 | 1 | 1 | 0 | 1 | 0 | 2 | 0 | 6 | 0.3333 |
| 14_310968 | 2 | 1 | 1 | 1 | 3 | 0.3333 | 1 | 1 | 7 | 0.7143 | 2 | 1 | 2 | 0.5 | 1 | 1 | 2 | 1 | 7 | 0.8571 |
| 14_310974 | 1 | 1 | NA | NA | 1 | 1 | NA | NA | 2 | 1 | NA | NA | NA | NA | 1 | 1 | NA | NA | 1 | 1 |
| 14_310974 | 1 | 1 | NA | NA | 1 | 1 | NA | NA | 2 | 1 | NA | NA | NA | NA | 1 | 1 | NA | NA | 1 | 1 |
| 14_310975 | 1 | 1 | NA | NA | NA | NA | NA | NA | 1 | 1 | NA | NA | NA | NA | 1 | 1 | NA | NA | 1 | 1 |
| 14_310981 | 2 | 0.5 | NA | NA | 1 | 1 | NA | NA | 3 | 0.6667 | NA | NA | NA | NA | 1 | 1 | NA | NA | 1 | 1 |
| 14_310982 | 2 | 0.5 | NA | NA | NA | NA | NA | NA | 2 | 0.5 | NA | NA | NA | NA | 1 | 1 | NA | NA | 1 | 1 |
| 14_310983 | 2 | 0.5 | NA | NA | 1 | 1 | NA | NA | 3 | 0.6667 | NA | NA | NA | NA | 1 | 1 | NA | NA | 1 | 1 |
| 14_311000 | 1 | 1 | NA | NA | NA | NA | NA | NA | 1 | 1 | NA | NA | 1 | 0 | 1 | 1 | NA | NA | 2 | 0.5 |
| 14_311007 | 1 | 1 | NA | NA | NA | NA | NA | NA | 1 | 1 | NA | NA | 1 | 0 | 1 | 1 | NA | NA | 2 | 0.5 |
| 14_311016 | 1 | 1 | NA | NA | NA | NA | NA | NA | 1 | 1 | NA | NA | 1 | 0 | 2 | 0.5 | NA | NA | 3 | 0.3333 |
| 14_312012 | 2 | 0 | 2 | 0 | 2 | 0 | 5 | 0.2 | 11 | 0.0909 | 3 | 0.3333 | 2 | 0 | 4 | 0.25 | 3 | 0 | 12 | 0.1667 |
| 14_312015 | 2 | 0 | 2 | 0.5 | 2 | 0 | 5 | 0.6 | 11 | 0.3636 | 2 | 0.5 | 3 | 0 | 4 | 0.5 | 3 | 0 | 12 | 0.25 |
| 14_312015 | 1 | 0 | 1 | 1 | 2 | 0 | 4 | 0 | 8 | 0.125 | 2 | 0.5 | 3 | 0.3333 | 3 | 0 | 2 | 0 | 10 | 0.2 |
| 14_312016 | 2 | 0.5 | 1 | 1 | 2 | 1 | 5 | 0.8 | 10 | 0.8 | 3 | 1 | 3 | 0 | 3 | 0.6667 | 2 | 0.5 | 11 | 0.5455 |
| 14_312016 | 2 | 1 | 1 | 1 | 2 | 0.5 | 5 | 0.6 | 10 | 0.7 | 2 | 1 | 3 | 0.6667 | 3 | 0.6667 | NA | NA | 8 | 0.75 |
| 14_312155 | 1 | 0 | NA | NA | 1 | 0 | 4 | 0.25 | 6 | 0.1667 | 2 | 0.5 | 3 | 0 | 4 | 0 | 1 | 0 | 10 | 0.1 |
| 14_312156 | 1 | 0 | NA | NA | 2 | 0.5 | 3 | 0.3333 | 6 | 0.3333 | 1 | 1 | 2 | 0.5 | 4 | 0.5 | 1 | 0 | 8 | 0.5 |
| 14_312156 | 1 | 1 | NA | NA | 2 | 1 | 2 | 1 | 5 | 1 | 2 | 1 | 3 | 0 | 4 | 0.5 | 1 | 0 | 10 | 0.4 |
| 14_312156 | 1 | 0 | NA | NA | 2 | 0.5 | 2 | 0.5 | 5 | 0.4 | 2 | 0 | 2 | 1 | 3 | 0 | 1 | 0 | 8 | 0.25 |
| 14_312156 | 1 | 1 | NA | NA | 2 | 0.5 | 2 | 0.5 | 5 | 0.6 | 1 | 0 | 3 | 0.6667 | 2 | 0 | 1 | 0 | 7 | 0.2857 |
| 14_312157 | 1 | 1 | NA | NA | 2 | 0.5 | 2 | 1 | 5 | 0.8 | 1 | 0 | 2 | 0.5 | 4 | 0.75 | NA | NA | 7 | 0.5714 |
| 14_312286 | NA | NA | NA | NA | 1 | 1 | 1 | 0 | 2 | 0.5 | 1 | 1 | 2 | 1 | 1 | 0 | NA | NA | 4 | 0.75 |

| | | | | | | | | | | | | | | | | | | | | |
|---|---|---|---|---|---|---|---|---|---|---|---|---|---|---|---|---|---|---|---|---|
| 14_3123069 | NA | NA | 1 | 0 | 1 | 0 | 3 | 0.6667 | 5 | 0.4 | 1 | 0 | 2 | 0 | 3 | 0 | 3 | 0.3333 | 9 | 0.1111 |
| 14_3123144 | 1 | 0 | 1 | 0 | 2 | 0 | 3 | 0.3333 | 7 | 0.1429 | 1 | 0 | 4 | 0.25 | 4 | 0 | 3 | 0 | 12 | 0.0833 |
| 14_3138885 | 2 | 0.5 | 1 | 1 | 2 | 0.5 | NA | NA | 5 | 0.6 | 1 | 1 | 1 | 1 | NA | NA | NA | NA | 2 | 1 |
| 14_3138892 | 3 | 0.6667 | 1 | 1 | 2 | 1 | 1 | 1 | 7 | 0.8571 | 1 | 1 | 2 | 1 | NA | NA | NA | NA | 3 | 1 |
| 14_3138901 | 4 | 1 | 1 | 1 | 2 | 1 | 1 | 1 | 8 | 1 | 1 | 1 | 2 | 1 | NA | NA | NA | NA | 3 | 1 |
| 14_3138927 | 4 | 0.25 | 1 | 0 | 3 | 0 | NA | NA | 8 | 0.125 | 1 | 0 | 2 | 0.5 | 1 | 0 | NA | NA | 4 | 0.25 |
| 14_3138930 | 4 | 1 | 1 | 1 | 3 | 1 | NA | NA | 8 | 1 | 1 | 0 | 2 | 1 | 1 | 1 | NA | NA | 4 | 0.75 |
| 14_3138944 | 5 | 0.4 | 1 | 1 | 3 | 0.3333 | 1 | 1 | 10 | 0.5 | NA | NA | 2 | 0.5 | 1 | 1 | NA | NA | 3 | 0.6667 |
| 14_3138957 | 5 | 0.8 | 1 | 1 | 1 | 1 | 1 | 1 | 8 | 0.875 | 1 | 0 | 3 | 1 | 1 | 1 | NA | NA | 5 | 0.8 |
| 14_3138968 | 4 | 0.5 | 1 | 1 | 3 | 0.6667 | 1 | 1 | 9 | 0.6667 | 1 | 0 | 3 | 1 | 1 | 1 | NA | NA | 5 | 0.8 |
| 14_3138970 | 5 | 0.4 | 2 | 0.5 | 2 | 0 | 1 | 0 | 10 | 0.3 | 1 | 0 | 3 | 0.6667 | NA | NA | NA | NA | 4 | 0.5 |
| 14_3138975 | 2 | 0.5 | 2 | 0.5 | 2 | 0.5 | NA | NA | 6 | 0.5 | 2 | 0 | 3 | 0 | 1 | 1 | NA | NA | 6 | 0.1667 |
| 14_3138975 | 4 | 0.5 | 1 | 1 | 1 | 1 | NA | NA | 6 | 0.6667 | 2 | 0.5 | 2 | 0 | 1 | 1 | NA | NA | 5 | 0.4 |
| 14_3138978 | 5 | 0 | 2 | 0 | 3 | 0.3333 | 1 | 0 | 11 | 0.0909 | 2 | 0.5 | 3 | 0 | 1 | 0 | NA | NA | 6 | 0.1667 |
| 14_3138981 | 3 | 1 | 2 | 0.5 | 2 | 1 | 1 | 1 | 8 | 0.875 | 1 | 1 | 3 | 1 | 1 | 1 | NA | NA | 5 | 1 |
| 14_3138987 | 2 | 1 | 2 | 1 | 1 | 1 | NA | NA | 5 | 1 | 2 | 0.5 | 3 | 1 | NA | NA | NA | NA | 5 | 0.8 |
| 14_3139002 | 2 | 1 | 2 | 1 | 1 | 1 | 1 | 0 | 6 | 0.8333 | 2 | 0.5 | 2 | 0.5 | 1 | 1 | NA | NA | 5 | 0.6 |
| 14_3139003 | 3 | 0.3333 | 2 | 0 | 2 | 0.5 | 1 | 0 | 8 | 0.25 | 1 | 0 | 3 | 0.6667 | 1 | 0 | NA | NA | 5 | 0.4 |
| 14_3142129 | 4 | 0 | 5 | 0.4 | 3 | 0 | 8 | 0 | 20 | 0.1 | 3 | 0 | 6 | 0.1667 | 5 | 0 | 5 | 0 | 19 | 0.0526 |
| 14_3142135 | 5 | 0.8 | 5 | 0.8 | 2 | 0.5 | 6 | 0.5 | 18 | 0.6667 | 3 | 0.3333 | 6 | 0.8333 | 6 | 0.8333 | 6 | 0.8333 | 21 | 0.7619 |
| 14_3142154 | 4 | 0.25 | 2 | 0.5 | 4 | 0 | 6 | 0.1667 | 16 | 0.1875 | 1 | 0 | 5 | 0.8 | 5 | 0 | 5 | 0.4 | 16 | 0.375 |
| 14_3142155 | 4 | 0 | 4 | 0.25 | 2 | 0 | 4 | 0.25 | 14 | 0.1429 | 2 | 1 | 4 | 0.25 | 2 | 0 | 4 | 0.75 | 12 | 0.5 |
| 14_3142207 | 2 | 0 | 5 | 0 | 3 | 0 | 6 | 0.1667 | 16 | 0.0625 | 1 | 0 | 5 | 0 | 5 | 0.2 | 3 | 0.3333 | 14 | 0.1429 |
| 14_3142220 | 2 | 0.5 | 5 | 0 | 2 | 0 | 6 | 0.1667 | 15 | 0.1333 | 1 | 0 | 3 | 0 | 5 | 0.2 | 3 | 0.6667 | 12 | 0.25 |
| 14_3142241 | 2 | 0 | 2 | 0 | 2 | 0 | 5 | 0.6 | 11 | 0.2727 | 1 | 0 | 2 | 0.5 | 5 | 0 | 3 | 0.3333 | 11 | 0.1818 |
| 14_3142296 | 2 | 0.5 | 4 | 0.25 | 2 | 0 | 5 | 0.2 | 13 | 0.2308 | 1 | 0 | 4 | 0 | 4 | 0.5 | 3 | 0 | 12 | 0.1667 |
| 14_3142300 | 2 | 1 | 3 | 0 | 1 | 0 | 5 | 0.4 | 11 | 0.3636 | 1 | 0 | 4 | 0.25 | 3 | 0.3333 | 3 | 0 | 11 | 0.1818 |
| 14_3142306 | 2 | 0 | 3 | 0.3333 | 2 | 0 | 4 | 0.5 | 11 | 0.2727 | NA | NA | 4 | 0 | 2 | 0.5 | 3 | 0 | 9 | 0.1111 |
| 14_3142326 | 2 | 0.5 | 4 | 0.25 | 2 | 0 | 4 | 0 | 12 | 0.1667 | 1 | 0 | 3 | 0.3333 | 3 | 0 | 2 | 0 | 9 | 0.1111 |
| 14_3142331 | 2 | 0.5 | 2 | 0 | 2 | 0 | 6 | 0 | 12 | 0.0833 | 1 | 0 | 4 | 0 | 3 | 0.3333 | 3 | 0 | 11 | 0.0909 |
| 14_3142354 | 2 | 0 | 4 | 0.25 | 2 | 0 | 5 | 0.2 | 13 | 0.1538 | 1 | 0 | 4 | 0.25 | 3 | 0.3333 | 3 | 0.3333 | 11 | 0.2727 |
| 14_3142369 | 2 | 0.5 | 4 | 0.5 | 2 | 0 | 6 | 0.5 | 14 | 0.4286 | 1 | 1 | 4 | 0.75 | 3 | 1 | 3 | 0.3333 | 11 | 0.7273 |
| 14_3142376 | 2 | 0.5 | 4 | 0 | 2 | 0 | 5 | 0 | 13 | 0.0769 | 1 | 0 | 3 | 0 | 3 | 0 | 2 | 0.5 | 9 | 0.1111 |
| 14_3142377 | 2 | 1 | 4 | 1 | 2 | 1 | 4 | 1 | 12 | 1 | NA | NA | 3 | 0.6667 | 3 | 1 | 2 | 0.5 | 8 | 0.75 |
| 14_3142379 | 2 | 1 | 4 | 1 | 2 | 1 | 5 | 0.8 | 13 | 0.9231 | NA | NA | 2 | 0.5 | 3 | 0.6667 | 2 | 0 | 7 | 0.4286 |
| 14_3142383 | 3 | 0 | 5 | 0.2 | 2 | 0 | 5 | 0 | 15 | 0.0667 | 2 | 0 | 4 | 0 | 5 | 0 | 5 | 0.4 | 16 | 0.125 |
| 14_3142393 | 2 | 0 | 3 | 0 | 2 | 0 | 3 | 0.3333 | 10 | 0.1 | 2 | 0.5 | 3 | 0 | 4 | 0 | 4 | 0 | 13 | 0.0769 |
| 14_3142394 | 2 | 0 | 5 | 0 | 2 | 0.5 | 5 | 0.4 | 14 | 0.2143 | 2 | 0 | 4 | 0.5 | 5 | 0 | 5 | 0.4 | 16 | 0.25 |
| 14_3142399 | 2 | 0 | 4 | 0.25 | 1 | 0 | 6 | 0 | 13 | 0.0769 | 2 | 0 | 4 | 0.25 | 5 | 0.2 | 5 | 0.2 | 16 | 0.1875 |
| 14_3142411 | 1 | 0 | 5 | 0.6 | 1 | 1 | 6 | 0.5 | 13 | 0.5385 | 2 | 1 | 2 | 0.5 | 4 | 0.75 | 4 | 0.75 | 12 | 0.75 |
| 14_3142415 | 2 | 0 | 5 | 0.2 | 2 | 0 | 5 | 0.8 | 14 | 0.3571 | 2 | 0.5 | 4 | 0.5 | 4 | 0.5 | 4 | 0.5 | 14 | 0.5 |
| 14_3142416 | 2 | 0 | 5 | 0.2 | 2 | 0 | 6 | 0 | 15 | 0.0667 | 1 | 0 | 4 | 0.25 | 3 | 0 | 4 | 0 | 12 | 0.0833 |
| 14_3178380 | NA | NA | NA | NA | 1 | 1 | NA | NA | 1 | 1 | 1 | 0 | NA | NA | NA | NA | 1 | 1 | 2 | 0.5 |
| 14_3178399 | NA | NA | NA | NA | 1 | 1 | NA | NA | 1 | 1 | 1 | 1 | NA | NA | NA | NA | 1 | 0 | 2 | 0.5 |
| 14_3180266 | 3 | 0.3333 | 2 | 1 | 3 | 1 | 3 | 0.6667 | 11 | 0.7273 | 3 | 0.6667 | 1 | 1 | 2 | 0.5 | 3 | 0.3333 | 9 | 0.5556 |
| 14_3180268 | 3 | 1 | 2 | 1 | 3 | 1 | 4 | 0.5 | 12 | 0.8333 | 2 | 1 | NA | NA | 2 | 0.5 | 2 | 0 | 6 | 0.5 |
| 14_3180297 | 3 | 0 | 2 | 1 | 1 | 0 | 2 | 0.5 | 8 | 0.375 | 2 | 0.5 | 1 | 0 | 2 | 0.5 | 1 | 0 | 6 | 0.3333 |
| 14_3180361 | 3 | 0.3333 | 1 | 0 | 2 | 0.5 | 2 | 1 | 8 | 0.5 | 2 | 0 | 1 | 0 | 4 | 0.25 | 1 | 0 | 8 | 0.125 |
| 14_3180389 | 3 | 0 | 1 | 0 | 2 | 0.5 | 1 | 0 | 7 | 0.1429 | 3 | 0 | 1 | 0 | 4 | 0.25 | 1 | 0 | 9 | 0.1111 |
| 14_3180487 | 2 | 0.5 | 5 | 0 | 5 | 0.4 | 3 | 0 | 15 | 0.2 | 2 | 0 | 2 | 0 | 7 | 0.1429 | 2 | 1 | 13 | 0.2308 |
| 14_3180490 | 3 | 1 | 5 | 1 | 6 | 1 | 3 | 1 | 17 | 1 | 2 | 1 | 2 | 1 | 9 | 1 | 2 | 1 | 15 | 1 |
| 14_3180496 | 3 | 1 | 4 | 1 | 6 | 0.8333 | 4 | 1 | 17 | 0.9412 | 3 | 0.6667 | 4 | 1 | 10 | 1 | 2 | 1 | 19 | 0.9474 |
| 14_3180498 | 3 | 0.3333 | 4 | 0 | 5 | 0 | 4 | 0 | 16 | 0.0625 | 2 | 0 | 3 | 0 | 11 | 0.0909 | 2 | 0 | 18 | 0.0556 |
| 14_3180501 | 3 | 0.6667 | 2 | 0 | 3 | 0.3333 | 2 | 1 | 10 | 0.5 | 2 | 0.5 | 3 | 0 | 9 | 0.2222 | NA | NA | 14 | 0.2143 |
| 14_3182426 | 5 | 0.6 | 3 | 0.3333 | 4 | 0.75 | 1 | 1 | 13 | 0.6154 | 1 | 0 | NA | NA | 1 | 0 | 2 | 1 | 4 | 0.5 |
| 14_3182428 | 2 | 0 | 3 | 0 | 4 | 0.25 | 1 | 0 | 10 | 0.1 | 1 | 1 | NA | NA | 2 | 0 | 1 | 1 | 4 | 0.5 |
| 14_3182429 | 5 | 0.6 | 3 | 0.3333 | 2 | 0.5 | 1 | 0 | 11 | 0.4545 | NA | NA | NA | NA | 1 | 0 | 2 | 1 | 3 | 0.6667 |
| 14_3182440 | 5 | 0.6 | 3 | 0.3333 | 4 | 0.25 | 1 | 1 | 13 | 0.4615 | 1 | 0 | NA | NA | 2 | 0 | 2 | 1 | 5 | 0.4 |
| 14_3182445 | 4 | 0.25 | 3 | 0 | 4 | 0 | 1 | 0 | 12 | 0.0833 | 1 | 0 | 1 | 0 | 2 | 0.5 | 2 | 0 | 6 | 0.1667 |
| 14_3182448 | 5 | 0.2 | 3 | 0.3333 | 1 | 0 | 1 | 1 | 10 | 0.3 | 1 | 0 | 1 | 0 | 2 | 0.5 | 2 | 0 | 6 | 0.1667 |
| 14_3182454 | 6 | 0.1667 | 3 | 0 | 4 | 0.25 | 1 | 1 | 14 | 0.2143 | 1 | 1 | 1 | 0 | 1 | 0 | 1 | 0 | 4 | 0.25 |
| 14_3182455 | 5 | 0.6 | 3 | 0.6667 | 4 | 0.5 | 1 | 1 | 13 | 0.6154 | 1 | 0 | 1 | 1 | 2 | 1 | 2 | 0.5 | 6 | 0.6667 |
| 14_3182483 | 6 | 0.8333 | 3 | 0.6667 | 2 | 1 | 2 | 1 | 13 | 0.8462 | 1 | 1 | NA | NA | 2 | 1 | 2 | 1 | 5 | 1 |
| 14_3182498 | 5 | 0.8 | 4 | 1 | 3 | 1 | 2 | 1 | 14 | 0.9286 | 3 | 0.6667 | 1 | 1 | 3 | 0.6667 | 2 | 1 | 9 | 0.7778 |
| 14_3182506 | 3 | 0 | 3 | 0 | 5 | 0.2 | 1 | 1 | 12 | 0.1667 | 3 | 0 | 1 | 0 | 3 | 0.3333 | 2 | 0.5 | 9 | 0.2222 |

| | | | | | | | | | | | | | | | | | | | | |
|---|---|---|---|---|---|---|---|---|---|---|---|---|---|---|---|---|---|---|---|---|
| 14_318406 | 1 | 0 | 2 | 0.5 | NA | NA | NA | NA | 3 | 0.3333 | 1 | 1 | NA | NA | NA | NA | NA | NA | 1 | 1 |
| 14_318411 | 3 | 0.6667 | 2 | 0.5 | NA | NA | NA | NA | 5 | 0.6 | 1 | 1 | NA | NA | NA | NA | NA | NA | 1 | 1 |
| 14_318980 | 6 | 0 | 9 | 0.2222 | 6 | 0.1667 | 10 | 0.3 | 31 | 0.1935 | 6 | 0.3333 | 8 | 0 | 9 | 0.4444 | 5 | 0 | 28 | 0.2143 |
| 14_318981 | 5 | 0.4 | 8 | 0.125 | 6 | 0.1667 | 9 | 0.4444 | 28 | 0.2857 | 6 | 0.1667 | 7 | 0.1429 | 10 | 0 | 4 | 0 | 27 | 0.0741 |
| 14_318982 | 7 | 0.1429 | 9 | 0 | 6 | 0.3333 | 10 | 0 | 32 | 0.0938 | 6 | 0.3333 | 8 | 0.125 | 10 | 0.2 | 5 | 0 | 29 | 0.1724 |
| 14_318982 | 7 | 0 | 9 | 0.1111 | 5 | 0 | 10 | 0.4 | 31 | 0.1613 | 6 | 0.1667 | 8 | 0.5 | 10 | 0 | 5 | 0.2 | 29 | 0.2069 |
| 14_318983 | 7 | 0.2857 | 10 | 0.1 | 6 | 0.1667 | 10 | 0 | 33 | 0.1212 | 6 | 0 | 7 | 0 | 9 | 0 | 5 | 0.4 | 27 | 0.0741 |
| 14_318984 | 7 | 0.8571 | 8 | 0.5 | 6 | 0 | 10 | 0.3 | 31 | 0.4194 | 6 | 0.3333 | 7 | 0.5714 | 10 | 0.4 | 5 | 0.4 | 28 | 0.4286 |
| 14_318985 | 7 | 0.1429 | 11 | 0.4545 | 6 | 0.1667 | 9 | 0.3333 | 33 | 0.303 | 5 | 0.2 | 8 | 0.25 | 10 | 0.3 | 4 | 0 | 27 | 0.2222 |
| 14_318985 | 7 | 0 | 11 | 0.2727 | 6 | 0.1667 | 11 | 0.3636 | 35 | 0.2286 | 5 | 0.4 | 8 | 0.25 | 9 | 0.2222 | 4 | 0.5 | 26 | 0.3077 |
| 14_318985 | 7 | 0.1429 | 10 | 0 | 5 | 0 | 11 | 0.1818 | 33 | 0.0909 | 6 | 0.1667 | 7 | 0.1429 | 10 | 0.1 | 6 | 0 | 29 | 0.1034 |
| 14_318986 | 6 | 0.1667 | 9 | 0 | 6 | 0.1667 | 11 | 0.0909 | 32 | 0.0938 | 5 | 0.2 | 7 | 0.1429 | 9 | 0.2222 | 5 | 0 | 26 | 0.1538 |
| 14_318998 | 7 | 0.5714 | 12 | 0.1667 | 8 | 0 | 11 | 0.1818 | 38 | 0.2105 | 7 | 0.2857 | 9 | 0 | 11 | 0.3636 | 8 | 0.25 | 35 | 0.2286 |
| 14_318999 | 6 | 0.1667 | 11 | 0.2727 | 6 | 0.3333 | 6 | 0 | 29 | 0.2069 | 6 | 0.1667 | 8 | 0.125 | 7 | 0 | 4 | 0.25 | 25 | 0.12 |
| 14_319004 | 6 | 0 | 15 | 0.0667 | 11 | 0 | 13 | 0.1538 | 45 | 0.0667 | 8 | 0.125 | 8 | 0.125 | 13 | 0 | 7 | 0 | 36 | 0.0556 |
| 14_319020 | NA | NA | 1 | 1 | NA | NA | NA | NA | 1 | 1 | NA | NA | 1 | 1 | 1 | 0 | NA | NA | 2 | 0.5 |
| 14_319023 | 15 | 1 | 14 | 1 | 13 | 0.9231 | 14 | 0.9286 | 56 | 0.9643 | 12 | 1 | 13 | 0.8462 | 14 | 0.9286 | 17 | 0.9412 | 56 | 0.9286 |
| 14_319023 | 14 | 0.6429 | 17 | 0.7059 | 12 | 0.6667 | 16 | 0.5 | 59 | 0.6271 | 12 | 0.5833 | 12 | 0.5 | 14 | 0.3571 | 18 | 0.9444 | 56 | 0.625 |
| 14_319024 | 12 | 0.25 | 15 | 0.4 | 11 | 0.4545 | 11 | 0.1818 | 49 | 0.3265 | 8 | 0.5 | 10 | 0.2 | 15 | 0.1333 | 13 | 0.0769 | 46 | 0.1957 |
| 14_319024 | 12 | 0.4167 | 16 | 0.625 | 13 | 0.7692 | 16 | 0.375 | 57 | 0.5439 | 11 | 0.6364 | 11 | 0.5455 | 12 | 0.5 | 17 | 0.4118 | 51 | 0.5098 |
| 14_319025 | 14 | 0.0714 | 18 | 0.0556 | 14 | 0.0714 | 16 | 0.1875 | 62 | 0.0968 | 12 | 0.0833 | 15 | 0 | 15 | 0 | 19 | 0.1579 | 61 | 0.0656 |
| 14_319309 | NA | NA | 2 | 1 | 3 | 0.3333 | 1 | 1 | 6 | 0.6667 | 1 | 1 | 1 | 1 | NA | NA | 4 | 0.75 | 6 | 0.8333 |
| 14_319309 | 1 | 0 | 2 | 0 | 3 | 0.3333 | 1 | 0 | 7 | 0.1429 | 1 | 1 | 1 | 1 | NA | NA | 4 | 0.5 | 6 | 0.6667 |
| 14_319311 | 1 | 0 | 2 | 0.5 | 2 | 0 | 1 | 1 | 6 | 0.3333 | 1 | 0 | 1 | 0 | NA | NA | 4 | 0.25 | 6 | 0.1667 |
| 14_319311 | 1 | 0 | 2 | 0.5 | 2 | 0 | 1 | 1 | 6 | 0.3333 | 1 | 0 | 1 | 0 | NA | NA | 5 | 0.2 | 7 | 0.1429 |
| 14_319311 | NA | NA | 2 | 0.5 | 1 | 0 | 1 | 0 | 4 | 0.25 | 1 | 0 | 1 | 0 | NA | NA | 5 | 0.4 | 7 | 0.2857 |
| 14_319311 | 1 | 0 | 2 | 0 | 1 | 1 | 1 | 0 | 5 | 0.2 | 1 | 1 | NA | NA | NA | NA | 5 | 0.4 | 6 | 0.5 |
| 14_319312 | 1 | 0 | 2 | 0.5 | 2 | 0.5 | 1 | 0 | 6 | 0.3333 | 1 | 0 | 1 | 0 | NA | NA | 3 | 0.3333 | 5 | 0.2 |
| 14_319313 | 1 | 0 | 1 | 0 | 1 | 1 | 1 | 0 | 4 | 0.25 | 1 | 1 | 1 | 0 | NA | NA | 4 | 0.25 | 6 | 0.3333 |
| 14_319314 | 1 | 1 | 2 | 0 | 3 | 1 | 1 | 0 | 7 | 0.5714 | 1 | 0 | 2 | 0.5 | NA | NA | 4 | 0 | 7 | 0.1429 |
| 14_319315 | 1 | 0 | 2 | 0.5 | 3 | 0 | 1 | 0 | 7 | 0.1429 | 1 | 1 | 1 | 0 | NA | NA | 5 | 0 | 7 | 0.1429 |
| 14_319315 | 1 | 0 | 1 | 0 | 2 | 0 | 1 | 1 | 5 | 0.2 | 1 | 0 | 2 | 0 | NA | NA | 5 | 0.2 | 8 | 0.125 |
| 14_320182 | 4 | 0 | 3 | 0 | 6 | 0 | 1 | 1 | 14 | 0.0714 | 3 | 0.3333 | 1 | 0 | 1 | 0 | 7 | 0.1429 | 12 | 0.1667 |
| 14_320182 | 3 | 0.3333 | 3 | 0.3333 | 7 | 0 | 1 | 0 | 14 | 0.1429 | 3 | 0 | 3 | 0.3333 | 1 | 0 | 6 | 0 | 13 | 0.0769 |
| 14_320183 | 4 | 0 | 3 | 0 | 9 | 0.2222 | 1 | 0 | 17 | 0.1176 | 3 | 0.6667 | 2 | 0 | 1 | 0 | 6 | 0.1667 | 12 | 0.25 |
| 14_320183 | 3 | 0.3333 | 2 | 0.5 | 8 | 0.25 | 1 | 0 | 14 | 0.2857 | 3 | 0 | 2 | 0 | 1 | 1 | 6 | 0.1667 | 12 | 0.1667 |
| 14_320183 | 3 | 0.6667 | 3 | 0.6667 | 8 | 0.625 | 1 | 1 | 15 | 0.6667 | 2 | 1 | 3 | 0.6667 | 1 | 1 | 7 | 0.5714 | 13 | 0.6923 |
| 14_320184 | 2 | 0 | 2 | 1 | 7 | 0.4286 | 1 | 0 | 12 | 0.4167 | 3 | 0.3333 | 3 | 0 | 1 | 0 | 6 | 0.1667 | 13 | 0.1538 |
| 14_320295 | 1 | 1 | NA | NA | NA | NA | NA | NA | 1 | 1 | NA | NA | NA | NA | 1 | 1 | NA | NA | 1 | 1 |
| 14_320696 | 9 | 0.3333 | 9 | 0.2222 | 11 | 0 | 15 | 0.0667 | 44 | 0.1364 | 13 | 0.2308 | 16 | 0 | 7 | 0.1429 | 15 | 0 | 51 | 0.0784 |
| 14_320696 | 10 | 0.1 | 7 | 0.1429 | 12 | 0.1667 | 14 | 0.2143 | 43 | 0.1628 | 13 | 0.0769 | 16 | 0.125 | 7 | 0.1429 | 14 | 0.3571 | 50 | 0.18 |
| 14_321308 | NA | NA | 1 | 1 | NA | NA | 2 | 0.5 | 3 | 0.6667 | NA | NA | 3 | 0.6667 | NA | NA | 2 | 1 | 5 | 0.8 |
| 14_321849 | NA | NA | 1 | 1 | 1 | 0 | NA | NA | 2 | 0.5 | NA | NA | NA | NA | 2 | 0.5 | 2 | 0 | 4 | 0.25 |
| 14_321851 | NA | NA | NA | NA | 1 | 1 | NA | NA | 1 | 1 | 1 | 1 | NA | NA | 4 | 1 | 2 | 0 | 7 | 0.7143 |
| 14_322919 | NA | NA | 1 | 1 | NA | NA | 1 | 1 | 2 | 1 | NA | NA | 1 | 0 | NA | NA | 1 | 1 | 2 | 0.5 |
| 14_322987 | NA | NA | 2 | 0.5 | NA | NA | 3 | 0.3333 | 5 | 0.4 | 1 | 1 | 2 | 0.5 | NA | NA | 1 | 0 | 4 | 0.5 |
| 14_322988 | NA | NA | 2 | 1 | NA | NA | 4 | 0.75 | 6 | 0.8333 | 2 | 1 | 1 | 1 | NA | NA | 1 | 0 | 4 | 0.75 |
| 14_322988 | NA | NA | 2 | 0.5 | NA | NA | 4 | 0.75 | 6 | 0.6667 | 2 | 0.5 | 2 | 1 | NA | NA | 1 | 0 | 5 | 0.6 |
| 14_322989 | NA | NA | 3 | 1 | NA | NA | 4 | 0.5 | 7 | 0.7143 | 2 | 1 | 2 | 1 | NA | NA | 1 | 1 | 5 | 1 |
| 14_322990 | NA | NA | 2 | 1 | NA | NA | 2 | 0.5 | 4 | 0.75 | 2 | 0.5 | 2 | 1 | NA | NA | NA | NA | 4 | 0.75 |
| 14_322993 | NA | NA | 3 | 0.3333 | NA | NA | 4 | 0.25 | 7 | 0.2857 | 2 | 0 | 2 | 0.5 | NA | NA | 1 | 0 | 5 | 0.2 |
| 14_323001 | 1 | 0 | 4 | 0.25 | 1 | 0 | 2 | 0.5 | 8 | 0.25 | 3 | 0 | 2 | 0.5 | 1 | 0 | NA | NA | 6 | 0.1667 |
| 14_323001 | NA | NA | 4 | 0.25 | 1 | 0 | 3 | 0 | 8 | 0.125 | 2 | 0 | 3 | 0.3333 | 1 | 0 | 1 | 0 | 7 | 0.1429 |
| 14_323018 | 3 | 0 | 3 | 0.3333 | NA | NA | 4 | 0 | 10 | 0.1 | 2 | 0 | 4 | 0.25 | 1 | 0 | NA | NA | 7 | 0.1429 |
| 14_323018 | 1 | 0 | 3 | 0.6667 | NA | NA | 4 | 0 | 8 | 0.25 | 1 | 0 | 4 | 0 | 1 | 1 | NA | NA | 6 | 0.1667 |
| 14_323019 | 3 | 0.3333 | 5 | 0.4 | NA | NA | 2 | 1 | 10 | 0.5 | 2 | 0.5 | 2 | 0 | 1 | 1 | NA | NA | 5 | 0.4 |
| 14_325389 | 1 | 1 | NA | NA | NA | NA | NA | NA | 1 | 1 | NA | NA | NA | NA | NA | NA | 1 | 1 | 1 | 1 |
| 14_325392 | 1 | 1 | NA | NA | NA | NA | NA | NA | 1 | 1 | NA | NA | NA | NA | NA | NA | 1 | 1 | 1 | 1 |
| 14_325393 | 1 | 1 | NA | NA | NA | NA | NA | NA | 1 | 1 | NA | NA | NA | NA | NA | NA | 1 | 1 | 1 | 1 |
| 14_325396 | 1 | 1 | NA | NA | NA | NA | NA | NA | 1 | 1 | NA | NA | NA | NA | NA | NA | 1 | 1 | 1 | 1 |
| 14_326599 | 2 | 0.5 | 1 | 0 | NA | NA | NA | NA | 3 | 0.3333 | 1 | 1 | NA | NA | 1 | 0 | NA | NA | 2 | 0.5 |
| 14_326600 | 2 | 1 | 1 | 1 | NA | NA | NA | NA | 3 | 1 | 1 | 1 | 1 | 1 | 1 | 1 | NA | NA | 3 | 1 |
| 14_326602 | 2 | 0.5 | 1 | 1 | NA | NA | NA | NA | 3 | 0.6667 | 1 | 1 | 1 | 1 | 1 | 0 | NA | NA | 3 | 0.6667 |
| 14_330887 | 1 | 1 | 1 | 1 | NA | NA | NA | NA | 2 | 1 | NA | NA | NA | NA | 1 | 1 | NA | NA | 1 | 1 |
| 14_330890 | 1 | 1 | 1 | 0 | NA | NA | NA | NA | 2 | 0.5 | NA | NA | NA | NA | 1 | 1 | NA | NA | 1 | 1 |

| | | | | | | | | | | | | | | | | | | | | |
|---|---|---|---|---|---|---|---|---|---|---|---|---|---|---|---|---|---|---|---|---|
| 14_330917 | NA | NA | 1 | 1 | NA | NA | NA | NA | 1 | 1 | NA | NA | NA | NA | 1 | 1 | NA | NA | 1 | 1 |
| 14_331001 | 3 | 0 | 9 | 0.1111 | 9 | 0.1111 | 6 | 0 | 27 | 0.0741 | 3 | 0.6667 | 6 | 0.1667 | 3 | 0 | 7 | 0.1429 | 19 | 0.2105 |
| 14_331029 | 3 | 0 | 10 | 0.1 | 8 | 0.125 | 3 | 0 | 24 | 0.0833 | 6 | 0.3333 | 6 | 0 | 6 | 0.1667 | 7 | 0.1429 | 25 | 0.16 |
| 14_331031 | 3 | 0.3333 | 8 | 0.125 | 5 | 0 | 5 | 0 | 21 | 0.0952 | 2 | 0.5 | 6 | 0 | 5 | 0 | 5 | 0.2 | 18 | 0.1111 |
| 14_331032 | 3 | 0.3333 | 9 | 0 | 4 | 0 | 4 | 0 | 20 | 0.05 | 2 | 0.5 | 4 | 0.25 | 5 | 0.2 | 6 | 0.1667 | 17 | 0.2353 |
| 14_331033 | 3 | 0 | 13 | 0.0769 | 5 | 0.2 | 5 | 0 | 26 | 0.0769 | 4 | 0 | 6 | 0 | 6 | 0.1667 | 8 | 0.125 | 24 | 0.0833 |
| 14_331040 | 3 | 0.3333 | 11 | 0.1818 | 5 | 0.4 | 4 | 0.25 | 23 | 0.2609 | 3 | 0 | 4 | 0 | 5 | 0.2 | 6 | 0 | 18 | 0.0556 |
| 14_331108 | NA | NA | 6 | 0.3333 | 5 | 0 | 1 | 0 | 12 | 0.1667 | 1 | 0 | 2 | 0 | 3 | 0.3333 | 2 | 0 | 8 | 0.125 |
| 14_336565 | NA | NA | NA | NA | NA | NA | 1 | 1 | 1 | 1 | 2 | 1 | NA | NA | 1 | 1 | 1 | 1 | 4 | 1 |
| 14_336565 | NA | NA | NA | NA | NA | NA | 1 | 1 | 1 | 1 | 2 | 0.5 | NA | NA | 2 | 0.5 | 1 | 0 | 5 | 0.4 |
| 14_339542 | NA | NA | NA | NA | 1 | 1 | NA | NA | 1 | 1 | 1 | 1 | NA | NA | NA | NA | 1 | 1 | 2 | 1 |
| 14_339547 | NA | NA | NA | NA | 1 | 1 | NA | NA | 1 | 1 | 1 | 1 | NA | NA | NA | NA | 1 | 1 | 2 | 1 |
| 14_340657 | 34 | 0.2353 | 52 | 0.0962 | 29 | 0.069 | 40 | 0.15 | 155 | 0.1355 | 23 | 0 | 39 | 0.0513 | 30 | 0.0333 | 30 | 0.2333 | 122 | 0.082 |
| 14_340657 | 32 | 0.6562 | 52 | 0.5962 | 32 | 0.6562 | 41 | 0.5854 | 157 | 0.6178 | 25 | 0.48 | 42 | 0.7143 | 33 | 0.6667 | 33 | 0.5455 | 133 | 0.6165 |
| 14_340664 | 33 | 0.0606 | 40 | 0.175 | 23 | 0.2609 | 36 | 0.1111 | 132 | 0.1439 | 19 | 0.1579 | 38 | 0.2368 | 33 | 0.2424 | 30 | 0.2667 | 120 | 0.2333 |
| 14_340664 | 33 | 0.0303 | 40 | 0.05 | 27 | 0.0741 | 38 | 0.0789 | 138 | 0.058 | 22 | 0.0455 | 40 | 0.075 | 33 | 0.0606 | 30 | 0.0333 | 125 | 0.056 |
| 14_340665 | 39 | 0.1795 | 43 | 0.2558 | 26 | 0.1923 | 34 | 0.1471 | 142 | 0.1972 | 21 | 0.1429 | 29 | 0.1724 | 36 | 0.0556 | 26 | 0.1538 | 112 | 0.125 |
| 14_340666 | 39 | 0.0256 | 44 | 0.0682 | 27 | 0.037 | 38 | 0.0789 | 148 | 0.0541 | 23 | 0.087 | 37 | 0.0811 | 32 | 0.125 | 28 | 0.1071 | 120 | 0.1 |
| 14_340674 | 4 | 0.25 | 1 | 0 | 1 | 0 | 1 | 0 | 7 | 0.1429 | NA | NA | 1 | 0 | 1 | 0 | 3 | 0.3333 | 5 | 0.2 |
| 14_341261 | NA | NA | NA | NA | NA | NA | 1 | 1 | 1 | 1 | NA | NA | NA | NA | 2 | 0.5 | NA | NA | 2 | 0.5 |
| 14_341357 | 3 | 0.6667 | 7 | 0.7143 | 7 | 0.4286 | 5 | 0.8 | 22 | 0.6364 | 6 | 0.6667 | 2 | 0.5 | 3 | 0.3333 | 5 | 0.8 | 16 | 0.625 |
| 14_341357 | 5 | 0 | 7 | 0.1429 | 7 | 0.1429 | 8 | 0.125 | 27 | 0.1111 | 6 | 0 | 4 | 0 | 4 | 0 | 7 | 0.2857 | 21 | 0.0952 |
| 14_341358 | 5 | 0 | 9 | 0 | 6 | 0.5 | 8 | 0.375 | 28 | 0.2143 | 6 | 0.5 | 4 | 0 | 5 | 0.2 | 7 | 0.2857 | 22 | 0.2727 |
| 14_341362 | 5 | 0 | 8 | 0.25 | 4 | 0.25 | 6 | 0.3333 | 23 | 0.2174 | 5 | 0 | 4 | 0 | 5 | 0.4 | 7 | 0.2857 | 21 | 0.1905 |
| 14_341362 | 5 | 0.8 | 8 | 0.875 | 4 | 0.75 | 5 | 1 | 22 | 0.8636 | 6 | 0.8333 | 3 | 1 | 4 | 0.75 | 7 | 1 | 20 | 0.9 |
| 14_341362 | 5 | 1 | 7 | 1 | 3 | 1 | 6 | 1 | 21 | 1 | 6 | 1 | 3 | 1 | 2 | 1 | 4 | 1 | 15 | 1 |
| 14_341363 | 5 | 0.6 | 7 | 0.7143 | 5 | 0.8 | 7 | 0.1429 | 24 | 0.5417 | 5 | 0.2 | 4 | 0.25 | 4 | 0.25 | 5 | 0.6 | 18 | 0.3333 |
| 14_341363 | 5 | 0.4 | 8 | 0.25 | 5 | 0.4 | 4 | 0.25 | 22 | 0.3182 | 6 | 0.5 | 1 | 1 | 3 | 0.3333 | 5 | 0.2 | 15 | 0.4 |
| 14_341364 | 4 | 0 | 7 | 0 | 4 | 0.75 | 6 | 0.1667 | 21 | 0.1905 | 5 | 0.2 | 3 | 0.6667 | 3 | 0 | 3 | 0.3333 | 14 | 0.2857 |
| 14_341364 | 4 | 0.5 | 7 | 0.7143 | 4 | 0.5 | 5 | 0.4 | 20 | 0.55 | 5 | 1 | 3 | 0.6667 | 3 | 0.6667 | 4 | 0.75 | 15 | 0.8 |
| 14_341364 | 3 | 1 | 4 | 1 | 3 | 1 | 6 | 1 | 16 | 1 | 4 | 1 | 2 | 1 | 2 | 0.5 | 3 | 0.6667 | 11 | 0.8182 |
| 14_341364 | 4 | 1 | 5 | 1 | 4 | 0.75 | 5 | 1 | 18 | 0.9444 | 3 | 0.6667 | 1 | 0 | 3 | 0.6667 | 4 | 1 | 11 | 0.7273 |
| 14_341364 | 3 | 0 | 8 | 0 | 3 | 0.3333 | 4 | 0.25 | 18 | 0.1111 | 3 | 0.3333 | 3 | 0.3333 | 2 | 0 | 3 | 0 | 11 | 0.1818 |
| 14_341365 | 3 | 0.3333 | 8 | 0.5 | 4 | 0.5 | 5 | 0.4 | 20 | 0.45 | 4 | 0.25 | 2 | 1 | 1 | 0 | 5 | 0.4 | 12 | 0.4167 |
| 14_341365 | 3 | 0.3333 | 8 | 0.125 | 4 | 0.25 | 5 | 0.4 | 20 | 0.25 | 5 | 0 | 3 | 0 | 3 | 0.3333 | 5 | 0 | 16 | 0.0625 |
| 14_341365 | 3 | 0 | 8 | 0.375 | 4 | 0.5 | 7 | 0.1429 | 22 | 0.2727 | 5 | 0 | 3 | 0 | 4 | 0 | 5 | 0.2 | 17 | 0.0588 |
| 14_341447 | 1 | 1 | 7 | 0.4286 | 3 | 0.3333 | 6 | 0.3333 | 17 | 0.4118 | 4 | 0 | 3 | 0.3333 | 1 | 0 | 3 | 0.3333 | 11 | 0.1818 |
| 14_342501 | 1 | 0 | 1 | 1 | NA | NA | NA | NA | 2 | 0.5 | NA | NA | 3 | 0.3333 | NA | NA | NA | NA | 3 | 0.3333 |
| 14_342501 | 1 | 1 | 1 | 1 | NA | NA | NA | NA | 2 | 1 | NA | NA | 3 | 1 | NA | NA | NA | NA | 3 | 1 |
| 14_342501 | 1 | 1 | 1 | 0 | NA | NA | NA | NA | 2 | 0.5 | NA | NA | 2 | 0.5 | NA | NA | NA | NA | 2 | 0.5 |
| 14_342505 | 1 | 1 | 1 | 1 | NA | NA | NA | NA | 2 | 1 | NA | NA | 2 | 0.5 | NA | NA | NA | NA | 2 | 0.5 |
| 14_342507 | 1 | 0 | 1 | 1 | NA | NA | NA | NA | 2 | 0.5 | NA | NA | 2 | 0.5 | NA | NA | NA | NA | 2 | 0.5 |
| 14_342508 | 1 | 1 | 1 | 1 | NA | NA | NA | NA | 2 | 1 | NA | NA | 1 | 1 | NA | NA | NA | NA | 1 | 1 |
| 14_342508 | 1 | 0 | 1 | 1 | NA | NA | NA | NA | 2 | 0.5 | NA | NA | 2 | 0.5 | NA | NA | NA | NA | 2 | 0.5 |
| 14_342509 | 1 | 1 | 1 | 0 | NA | NA | NA | NA | 2 | 0.5 | NA | NA | 3 | 0.3333 | NA | NA | NA | NA | 3 | 0.3333 |
| 14_342510 | 1 | 1 | NA | NA | NA | NA | NA | NA | 1 | 1 | NA | NA | 2 | 0.5 | NA | NA | NA | NA | 2 | 0.5 |
| 14_342511 | 1 | 0 | 1 | 1 | NA | NA | NA | NA | 2 | 0.5 | NA | NA | 3 | 0.6667 | NA | NA | NA | NA | 3 | 0.6667 |
| 14_342511 | 1 | 1 | 1 | 1 | NA | NA | NA | NA | 2 | 1 | NA | NA | 2 | 1 | NA | NA | NA | NA | 2 | 1 |
| 14_343955 | 1 | 1 | NA | NA | NA | NA | NA | NA | 1 | 1 | NA | NA | 2 | 0.5 | 1 | 1 | NA | NA | 3 | 0.6667 |
| 14_343956 | 1 | 1 | NA | NA | NA | NA | NA | NA | 1 | 1 | NA | NA | 2 | 0.5 | NA | NA | NA | NA | 2 | 0.5 |
| 14_343990 | 1 | 1 | 1 | 1 | 1 | 1 | 4 | 0.25 | 7 | 0.5714 | NA | NA | 1 | 1 | NA | NA | 1 | 1 | 2 | 1 |
| 14_343990 | NA | NA | NA | NA | 1 | 1 | NA | NA | 1 | 1 | 1 | 1 | NA | NA | NA | NA | 1 | 1 | 2 | 1 |
| 14_344137 | 3 | 0.3333 | 3 | 0 | 2 | 0 | 6 | 0 | 14 | 0.0714 | NA | NA | 1 | 1 | NA | NA | 2 | 0 | 3 | 0.3333 |
| 14_344138 | 2 | 0.5 | 3 | 1 | 2 | 1 | 7 | 1 | 14 | 0.9286 | 1 | 1 | 1 | 1 | NA | NA | 2 | 1 | 4 | 1 |
| 14_344139 | 3 | 0.3333 | 3 | 0.6667 | 2 | 1 | 7 | 0.7143 | 15 | 0.6667 | 1 | 1 | 1 | 0 | NA | NA | 2 | 0.5 | 4 | 0.5 |
| 14_344141 | 3 | 0 | 3 | 0 | 2 | 0.5 | 7 | 0 | 15 | 0.0667 | 1 | 1 | NA | NA | NA | NA | 2 | 0 | 3 | 0.3333 |
| 14_344142 | 2 | 0.5 | 2 | 1 | 2 | 1 | 5 | 1 | 11 | 0.9091 | 2 | 1 | 1 | 0 | NA | NA | 2 | 1 | 5 | 0.8 |
| 14_344144 | 1 | 0 | 3 | 0.3333 | 2 | 0 | 5 | 0.2 | 11 | 0.1818 | 2 | 0 | 1 | 1 | NA | NA | 2 | 0.5 | 5 | 0.4 |
| 14_344145 | 3 | 0 | 3 | 0 | 3 | 0 | 7 | 0.1429 | 16 | 0.0625 | 2 | 0.5 | NA | NA | NA | NA | 2 | 0 | 4 | 0.25 |
| 14_344676 | 3 | 0 | 3 | 0 | 5 | 0.8 | 5 | 0.2 | 16 | 0.3125 | 2 | 0.5 | 3 | 0.3333 | 2 | 1 | 4 | 0.5 | 11 | 0.5455 |
| 14_344677 | 4 | 0 | 4 | 0.5 | 5 | 0.2 | 7 | 0 | 20 | 0.15 | 2 | 0.5 | 3 | 0 | 6 | 0.1667 | 4 | 0 | 15 | 0.1333 |
| 14_344677 | 4 | 0 | 4 | 0 | 5 | 0.2 | 6 | 0.6667 | 19 | 0.2632 | 2 | 0.5 | 2 | 0 | 4 | 0.25 | 3 | 0.3333 | 11 | 0.2727 |
| 14_344678 | 4 | 0.5 | 4 | 0.25 | 4 | 0.25 | 7 | 0.2857 | 19 | 0.3158 | 2 | 0.5 | 2 | 0 | 6 | 0.6667 | 3 | 0.6667 | 13 | 0.5385 |
| 14_344680 | 3 | 0 | 4 | 0.75 | 5 | 0.4 | 6 | 0 | 18 | 0.2778 | 2 | 0 | 1 | 0 | 6 | 0.3333 | 4 | 0 | 13 | 0.1538 |
| 14_344692 | 2 | 0 | 4 | 0.25 | 5 | 0 | 6 | 0 | 17 | 0.0588 | 3 | 0 | 2 | 0 | 6 | 0.1667 | 2 | 0 | 13 | 0.0769 |

| | | | | | | | | | | | | | | | | | | | | |
|---|---|---|---|---|---|---|---|---|---|---|---|---|---|---|---|---|---|---|---|---|
| 14_3446939 | 1 | 0 | 5 | 0 | 5 | 0.2 | 5 | 0 | 16 | 0.0625 | 3 | 0.3333 | 1 | 0 | 5 | 0 | 2 | 0 | 11 | 0.0909 |
| 14_3517210 | 6 | 0.3333 | 1 | 0 | 4 | 0 | 3 | 0.3333 | 14 | 0.2143 | 4 | 0 | 6 | 0.1667 | 2 | 0.5 | 3 | 0.3333 | 15 | 0.2 |
| 14_3517218 | 4 | 0.25 | 2 | 0.5 | 3 | 0 | 5 | 0.4 | 14 | 0.2857 | 3 | 0.3333 | 8 | 0.25 | 3 | 0 | 2 | 0.5 | 16 | 0.25 |
| 14_3529155 | 6 | 0 | 5 | 0.2 | 4 | 0 | 6 | 0.3333 | 21 | 0.1429 | 9 | 0.2222 | 7 | 0 | 10 | 0 | 5 | 0 | 31 | 0.0645 |
| 14_3530256 | 6 | 0.8333 | 5 | 1 | 3 | 1 | 7 | 1 | 21 | 0.9524 | 9 | 0.8889 | 10 | 0.9 | 10 | 1 | 3 | 1 | 32 | 0.9375 |
| 14_3530265 | 7 | 1 | 4 | 0.75 | 4 | 1 | 8 | 0.875 | 23 | 0.913 | 11 | 0.8182 | 9 | 1 | 11 | 1 | 4 | 1 | 35 | 0.9429 |
| 14_3530274 | 8 | 0.875 | 4 | 0.75 | 5 | 0.6 | 8 | 0.625 | 25 | 0.72 | 12 | 0.5 | 10 | 0.5 | 6 | 0.6667 | 5 | 0.4 | 33 | 0.5152 |
| 14_3530283 | 6 | 0.1667 | 5 | 0 | 4 | 0 | 9 | 0.1111 | 24 | 0.0833 | 12 | 0 | 11 | 0 | 11 | 0.1818 | 4 | 0 | 38 | 0.0526 |
| 14_3530289 | 6 | 0.8333 | 2 | 1 | 6 | 0.6667 | 7 | 0.7143 | 21 | 0.7619 | 10 | 1 | 8 | 0.75 | 9 | 0.8889 | 3 | 0.3333 | 30 | 0.8333 |
| 14_3530293 | 6 | 0.3333 | 6 | 0.1667 | 4 | 0 | 7 | 0 | 23 | 0.1304 | 7 | 0.1429 | 9 | 0.2222 | 12 | 0.25 | 3 | 0.3333 | 31 | 0.2258 |
| 14_3530301 | 4 | 0.75 | 5 | 1 | 4 | 0.75 | 7 | 0.8571 | 20 | 0.85 | 10 | 0.8 | 6 | 0.6667 | 5 | 1 | 4 | 1 | 25 | 0.84 |
| 14_3530302 | 6 | 0.6667 | 5 | 0.8 | 4 | 0.25 | 9 | 1 | 24 | 0.75 | 10 | 0.9 | 7 | 0.7143 | 8 | 0.875 | 4 | 0.75 | 29 | 0.8276 |
| 14_3530358 | 2 | 0.5 | 3 | 0.3333 | NA | NA | 3 | 0.3333 | 8 | 0.375 | 3 | 0.3333 | 3 | 0 | 1 | 1 | NA | NA | 7 | 0.2857 |
| 14_3530419 | 2 | 0 | 2 | 0 | 1 | 0 | 5 | 0.2 | 10 | 0.1 | 3 | 0 | 3 | 0.3333 | 1 | 0 | 1 | 0 | 8 | 0.125 |
| 14_3659769 | 15 | 0.1333 | 9 | 0.2222 | 4 | 0 | 10 | 0.4 | 38 | 0.2105 | 5 | 0 | 9 | 0.1111 | 12 | 0.1667 | 5 | 0.2 | 31 | 0.129 |
| 14_3659802 | 14 | 0.1429 | 9 | 0.1111 | 3 | 0 | 9 | 0.2222 | 35 | 0.1429 | 3 | 0 | 8 | 0 | 14 | 0.2143 | 5 | 0 | 30 | 0.1 |
| 14_3660002 | 8 | 0.25 | 5 | 0.6 | 2 | 0 | 7 | 0.2857 | 22 | 0.3182 | 1 | 0 | 5 | 0.4 | 9 | 0.2222 | 1 | 0 | 16 | 0.25 |
| 14_3660104 | 7 | 0.1429 | 2 | 0.5 | 2 | 0 | 8 | 0.375 | 19 | 0.2632 | 3 | 0.6667 | 3 | 0.3333 | 8 | 0.25 | 1 | 0 | 15 | 0.3333 |
| 14_3660106 | 8 | 0.875 | 4 | 0.75 | 2 | 1 | 7 | 0.8571 | 21 | 0.8571 | NA | NA | 3 | 0.6667 | 7 | 1 | 1 | 1 | 11 | 0.9091 |
| 14_3660109 | 3 | 0.3333 | 4 | 0.5 | 2 | 0.5 | 7 | 0.1429 | 16 | 0.3125 | 1 | 0 | 3 | 0.3333 | 7 | 0.2857 | 1 | 1 | 12 | 0.3333 |
| 14_3660116 | 6 | 0.1667 | 4 | 0 | 2 | 0 | 8 | 0.25 | 20 | 0.15 | 3 | 0.6667 | 2 | 0 | 8 | 0.375 | 1 | 0 | 14 | 0.3571 |
| 14_3660123 | 4 | 0.75 | 3 | 1 | 1 | 1 | 7 | 0.7143 | 15 | 0.8 | 2 | 0.5 | NA | NA | 6 | 0.6667 | 2 | 0 | 10 | 0.5 |
| 14_3660129 | 5 | 0.6 | 4 | 0.25 | 2 | 0 | 7 | 0.1429 | 18 | 0.2778 | 3 | 0.6667 | 2 | 0.5 | 8 | 0.375 | 1 | 1 | 14 | 0.5 |
| 14_3660134 | 5 | 0 | 2 | 0.5 | 1 | 1 | 7 | 0 | 15 | 0.1333 | NA | NA | 1 | 0 | 7 | 0.1429 | 2 | 0 | 10 | 0.1 |
| 14_3660135 | 6 | 0.6667 | 3 | 0.6667 | 1 | 1 | 6 | 0.8333 | 16 | 0.75 | 2 | 1 | 2 | 1 | 5 | 1 | NA | NA | 9 | 1 |
| 14_3660139 | 5 | 0 | 3 | 0 | 2 | 0.5 | 7 | 0.2857 | 17 | 0.1765 | 3 | 0 | 2 | 0 | 7 | 0.4286 | 2 | 0.5 | 14 | 0.2857 |
| 14_3660140 | 7 | 0.7143 | 4 | 1 | 2 | 1 | 7 | 0.8571 | 20 | 0.85 | 2 | 1 | 2 | 1 | 8 | 0.875 | 1 | 1 | 13 | 0.9231 |
| 14_3660141 | 6 | 0.1667 | 1 | 0 | 2 | 0.5 | 6 | 0.1667 | 15 | 0.2 | 2 | 0 | 2 | 0 | 7 | 0.2857 | NA | NA | 11 | 0.1818 |
| 14_3660145 | 7 | 0.5714 | 4 | 0.25 | 2 | 0 | 8 | 0.25 | 21 | 0.3333 | 3 | 0 | 2 | 0 | 7 | 0 | 2 | 0.5 | 14 | 0.0714 |
| 14_3660146 | 6 | 1 | 3 | 0.6667 | 2 | 1 | 6 | 0.8333 | 17 | 0.8824 | 3 | 1 | 2 | 1 | 7 | 1 | 1 | 1 | 13 | 1 |
| 14_3660150 | 6 | 0.1667 | 4 | 0 | 2 | 0 | 8 | 0 | 20 | 0.05 | 3 | 0.3333 | 2 | 0.5 | 8 | 0 | 1 | 0 | 14 | 0.1429 |
| 14_3660155 | 7 | 0 | 4 | 0 | 2 | 0 | 8 | 0.375 | 21 | 0.1429 | 3 | 0 | 2 | 0 | 8 | 0.125 | 2 | 0.5 | 15 | 0.1333 |
| 14_3660162 | 7 | 0 | 4 | 0.25 | 2 | 0 | 7 | 0.2857 | 20 | 0.15 | 3 | 0 | 2 | 0 | 8 | 0.25 | 2 | 0 | 15 | 0.1333 |
| 14_3666191 | 3 | 0 | 1 | 0 | 1 | 1 | 4 | 0.5 | 9 | 0.3333 | 1 | 0 | 2 | 0.5 | 4 | 0.5 | 2 | 0.5 | 9 | 0.4444 |
| 14_3666192 | 3 | 0 | 1 | 1 | 1 | 0 | 4 | 0 | 9 | 0.1111 | 1 | 1 | 2 | 0.5 | 4 | 0.25 | 1 | 0 | 8 | 0.375 |
| 14_3666205 | 4 | 0.25 | 1 | 0 | 1 | 1 | 4 | 0 | 10 | 0.2 | 1 | 0 | 2 | 0 | 4 | 0.25 | 1 | 0 | 8 | 0.125 |
| 14_3666208 | 2 | 0.5 | 1 | 0 | 1 | 0 | 4 | 0 | 8 | 0.125 | 1 | 0 | 2 | 1 | 4 | 0.25 | 2 | 0 | 9 | 0.3333 |
| 14_3666209 | 3 | 0.3333 | 1 | 0 | 1 | 0 | 4 | 0 | 9 | 0.1111 | 1 | 0 | 2 | 0 | 4 | 0.25 | 2 | 0 | 9 | 0.1111 |
| 14_3666211 | 2 | 0 | 1 | 0 | 1 | 0 | 4 | 0.25 | 8 | 0.125 | 1 | 0 | 1 | 0 | 4 | 0.25 | 2 | 0 | 8 | 0.125 |
| 14_3666233 | 2 | 1 | 1 | 1 | 1 | 0 | 2 | 1 | 6 | 0.8333 | 1 | 1 | 2 | 1 | 4 | 0.75 | 2 | 1 | 9 | 0.8889 |
| 14_3666261 | 1 | 0 | NA | NA | NA | NA | 4 | 0.25 | 5 | 0.2 | NA | NA | 2 | 0 | 4 | 0.25 | 1 | 0 | 7 | 0.1429 |
| 14_3666270 | 1 | 1 | 1 | 1 | NA | NA | 3 | 0.3333 | 5 | 0.6 | 1 | 1 | 2 | 0.5 | 2 | 1 | 1 | 1 | 6 | 0.8333 |
| 14_3666274 | 1 | 0 | 1 | 0 | NA | NA | 3 | 0.3333 | 5 | 0.2 | 1 | 0 | 2 | 0.5 | 2 | 0.5 | 1 | 0 | 6 | 0.3333 |
| 14_3666291 | 1 | 0 | 1 | 0 | NA | NA | 4 | 0.5 | 6 | 0.3333 | 1 | 0 | 2 | 0.5 | 3 | 0.3333 | 1 | 0 | 7 | 0.2857 |
| 14_3666294 | 1 | 0 | NA | NA | NA | NA | 4 | 0.25 | 5 | 0.2 | 1 | 1 | 2 | 1 | 4 | 1 | 1 | 0 | 8 | 0.875 |
| 14_3666306 | NA | NA | NA | NA | NA | NA | 4 | 0.25 | 4 | 0.25 | 1 | 0 | 2 | 0.5 | 4 | 0.5 | 1 | 0 | 8 | 0.375 |
| 14_3666307 | 1 | 1 | NA | NA | NA | NA | 4 | 0.5 | 5 | 0.6 | 1 | 1 | 2 | 0.5 | 3 | 1 | 1 | 0 | 7 | 0.7143 |
| 14_3680001 | 2 | 0.5 | 4 | 0 | 2 | 0.5 | NA | NA | 8 | 0.25 | 4 | 0 | NA | NA | 1 | 1 | 2 | 0 | 7 | 0.1429 |
| 14_3680049 | 1 | 0 | 2 | 0.5 | 2 | 0.5 | NA | NA | 5 | 0.4 | 2 | 0.5 | NA | NA | 1 | 0 | 2 | 1 | 5 | 0.6 |
| 14_3680087 | 1 | 1 | 3 | 0 | 2 | 0.5 | NA | NA | 6 | 0.3333 | 4 | 1 | NA | NA | 1 | 0 | 2 | 0.5 | 7 | 0.7143 |
| 14_3680123 | 1 | 1 | 4 | 0.25 | 2 | 0.5 | NA | NA | 7 | 0.4286 | 2 | 1 | NA | NA | 1 | 1 | 1 | 1 | 4 | 1 |
| 14_3680127 | 1 | 1 | 4 | 1 | 1 | 1 | NA | NA | 6 | 1 | 4 | 1 | NA | NA | 1 | 1 | 2 | 0.5 | 7 | 0.8571 |
| 14_3680160 | 1 | 1 | 3 | 1 | NA | NA | NA | NA | 4 | 1 | 2 | 1 | NA | NA | 1 | 1 | 1 | 1 | 4 | 1 |
| 14_3680185 | 1 | 1 | 3 | 0.6667 | 1 | 1 | NA | NA | 5 | 0.8 | 3 | 0.6667 | NA | NA | 1 | 1 | NA | NA | 4 | 0.75 |
| 14_3680248 | 1 | 0 | 2 | 0.5 | 1 | 0 | NA | NA | 4 | 0.25 | 3 | 0.3333 | NA | NA | 1 | 0 | 1 | 0 | 5 | 0.2 |
| 14_3684262 | 6 | 0.1667 | 2 | 0.5 | 3 | 0.3333 | 7 | 0.4286 | 18 | 0.3333 | 8 | 0.125 | 3 | 0.3333 | 3 | 0.6667 | 2 | 1 | 16 | 0.375 |
| 14_3684334 | 63 | 0.0635 | 60 | 0.0833 | 57 | 0.1579 | 69 | 0.1159 | 249 | 0.1044 | 48 | 0.2292 | 49 | 0.0816 | 51 | 0.1765 | 51 | 0.1176 | 199 | 0.1508 |
| 14_3684353 | 52 | 0.0385 | 50 | 0.16 | 43 | 0.0465 | 53 | 0.1887 | 198 | 0.1111 | 36 | 0.0278 | 38 | 0.1579 | 44 | 0.0909 | 43 | 0.1163 | 161 | 0.0994 |
| 14_3809249 | 1 | 1 | 3 | 0.6667 | NA | NA | NA | NA | 4 | 0.75 | 1 | 1 | 3 | 1 | 1 | 1 | 3 | 1 | 8 | 1 |
| 14_3809250 | 1 | 1 | 5 | 1 | NA | NA | NA | NA | 6 | 1 | 1 | 1 | 3 | 1 | 1 | 1 | 3 | 1 | 8 | 1 |
| 14_3809260 | 1 | 1 | 3 | 0.6667 | NA | NA | NA | NA | 4 | 0.75 | 1 | 1 | 3 | 1 | 1 | 1 | 2 | 1 | 7 | 1 |
| 14_3809262 | 1 | 0 | 5 | 0.6 | NA | NA | NA | NA | 6 | 0.5 | 1 | 1 | 4 | 0.25 | 1 | 0 | 3 | 0.3333 | 9 | 0.3333 |
| 14_3809268 | 1 | 1 | 3 | 1 | NA | NA | NA | NA | 4 | 1 | 1 | 1 | 4 | 1 | 1 | 1 | 1 | 1 | 7 | 1 |
| 14_3809277 | NA | NA | 3 | 1 | NA | NA | NA | NA | 3 | 1 | NA | NA | 2 | 1 | NA | NA | 2 | 1 | 4 | 1 |
| 14_3809289 | 1 | 1 | 5 | 1 | NA | NA | NA | NA | 6 | 1 | 1 | 1 | 4 | 1 | 1 | 1 | 3 | 1 | 9 | 1 |

| | | | | | | | | | | | | | | | | | | | | |
|---|---|---|---|---|---|---|---|---|---|---|---|---|---|---|---|---|---|---|---|---|
| 14_3809293 | 1 | 1 | 5 | 0.2 | NA | NA | NA | NA | 6 | 0.3333 | 1 | 1 | 2 | 0.5 | 1 | 1 | 3 | 0.3333 | 7 | 0.5714 |
| 14_3919583 | NA | NA | NA | NA | 1 | 1 | 1 | 1 | 2 | 1 | 1 | 0 | NA | NA | 1 | 0 | 1 | 1 | 3 | 0.3333 |
| 14_3919592 | NA | NA | NA | NA | 1 | 0 | 1 | 1 | 2 | 0.5 | 1 | 1 | NA | NA | 2 | 0.5 | 1 | 0 | 4 | 0.5 |
| 14_3919601 | NA | NA | NA | NA | 1 | 0 | 1 | 1 | 2 | 0.5 | 1 | 0 | NA | NA | 1 | 0 | 1 | 1 | 3 | 0.3333 |
| 14_3919609 | NA | NA | NA | NA | 1 | 1 | 1 | 1 | 2 | 1 | 1 | 1 | NA | NA | 1 | 1 | 1 | 1 | 3 | 1 |
| 14_4062886 | 3 | 0 | 2 | 0.5 | 3 | 0 | 2 | 0 | 10 | 0.1 | 4 | 0.25 | 2 | 0 | 1 | 0 | 3 | 0 | 10 | 0.1 |
| 14_4062907 | 4 | 0.25 | NA | NA | 3 | 0 | 1 | 0 | 8 | 0.125 | 4 | 0 | 2 | 0.5 | 1 | 0 | 2 | 0 | 9 | 0.1111 |
| 14_4062908 | 5 | 0.2 | 1 | 0 | 1 | 0 | 2 | 0 | 9 | 0.1111 | 4 | 0.25 | 2 | 1 | 1 | 0 | 2 | 0 | 9 | 0.3333 |
| 14_4062938 | 2 | 0.5 | 1 | 0 | 3 | 0 | 1 | 0 | 7 | 0.1429 | 4 | 0 | 1 | 1 | 1 | 1 | 1 | 0 | 7 | 0.2857 |
| 14_4062960 | 2 | 0.5 | 1 | 1 | 3 | 1 | 1 | 1 | 7 | 0.8571 | 4 | 0.25 | 2 | 0.5 | 1 | 1 | NA | NA | 7 | 0.4286 |
| 14_4062975 | 2 | 1 | 1 | 1 | 1 | 1 | 1 | 1 | 5 | 1 | 4 | 0.75 | 1 | 1 | 1 | 1 | NA | NA | 6 | 0.8333 |
| 14_4062976 | NA | NA | NA | NA | 2 | 0.5 | 1 | 0 | 3 | 0.3333 | 4 | 0.25 | 2 | 0.5 | 1 | 0 | NA | NA | 7 | 0.2857 |
| 14_4062979 | 2 | 1 | 1 | 1 | 2 | 0.5 | 1 | 1 | 6 | 0.8333 | 3 | 1 | 2 | 1 | 1 | 1 | NA | NA | 6 | 1 |
| 14_4313252 | NA | NA | 1 | 0 | NA | NA | 3 | 0.3333 | 4 | 0.25 | NA | NA | 1 | 0 | 2 | 0.5 | NA | NA | 3 | 0.3333 |
| 14_4313257 | NA | NA | 1 | 0 | NA | NA | 2 | 0.5 | 3 | 0.3333 | NA | NA | 1 | 0 | 2 | 1 | NA | NA | 3 | 0.6667 |
| 14_4314646 | NA | NA | NA | NA | NA | NA | 2 | 1 | 2 | 1 | NA | NA | NA | NA | 1 | 1 | NA | NA | 1 | 1 |
| 14_4314661 | 2 | 0.5 | NA | NA | NA | NA | 2 | 0.5 | 4 | 0.5 | 1 | 1 | 1 | 0 | 2 | 1 | NA | NA | 4 | 0.75 |
| 14_4316269 | NA | NA | 1 | 1 | 1 | 1 | 2 | 1 | 4 | 1 | NA | NA | NA | NA | 1 | 1 | NA | NA | 1 | 1 |
| 14_4395253 | NA | NA | NA | NA | 1 | 1 | 2 | 1 | 3 | 1 | 1 | 1 | NA | NA | NA | NA | 2 | 1 | 3 | 1 |
| 14_4395260 | NA | NA | NA | NA | 1 | 1 | 2 | 1 | 3 | 1 | 2 | 1 | NA | NA | NA | NA | 2 | 1 | 4 | 1 |
| 14_4537120 | 1 | 0 | 2 | 1 | NA | NA | 3 | 0.6667 | 6 | 0.6667 | 1 | 0 | 1 | 0 | 3 | 0.3333 | 3 | 1 | 8 | 0.5 |
| 14_4537169 | 1 | 0 | 1 | 0 | NA | NA | 3 | 0.3333 | 5 | 0.2 | 1 | 0 | 1 | 1 | 3 | 0 | 3 | 0 | 8 | 0.125 |
| 14_4537209 | 1 | 0 | 1 | 1 | NA | NA | 3 | 0 | 5 | 0.2 | 1 | 0 | 1 | 1 | 3 | 0 | 2 | 0 | 7 | 0.1429 |
| 14_4537224 | 1 | 0 | 1 | 0 | 2 | 1 | 3 | 0 | 7 | 0.2857 | 1 | 0 | 1 | 1 | 5 | 0.4 | 3 | 0.3333 | 10 | 0.4 |
| 14_4537241 | 2 | 0.5 | 4 | 0 | 6 | 0.3333 | 12 | 0 | 24 | 0.125 | 4 | 0.5 | 7 | 0.2857 | 14 | 0.2143 | 4 | 0.25 | 29 | 0.2759 |
| 14_4537264 | 6 | 0.3333 | 7 | 0.4286 | 8 | 0.375 | 12 | 0.1667 | 33 | 0.303 | 4 | 0 | 8 | 0.125 | 12 | 0.1667 | 5 | 0.4 | 29 | 0.1724 |
| 14_4537273 | 6 | 0 | 5 | 0.2 | 8 | 0.125 | 12 | 0.1667 | 31 | 0.129 | 3 | 0 | 8 | 0.25 | 11 | 0.3636 | 5 | 0 | 27 | 0.2222 |
| 14_4537309 | 5 | 1 | 5 | 0.6 | 8 | 0.75 | 10 | 0.9 | 28 | 0.8214 | 4 | 1 | 7 | 0.8571 | 11 | 0.8182 | 5 | 0.8 | 27 | 0.8519 |
| 14_4537310 | 4 | 0.25 | 4 | 0 | 6 | 0.5 | 11 | 0 | 25 | 0.16 | 3 | 0.3333 | 8 | 0.25 | 10 | 0.4 | 5 | 0 | 26 | 0.2692 |
| 14_4537320 | 6 | 0.1667 | 6 | 0.3333 | 8 | 0.125 | 12 | 0.3333 | 32 | 0.25 | 4 | 0.75 | 7 | 0.2857 | 12 | 0.25 | 5 | 0.4 | 28 | 0.3571 |
| 14_4537322 | 2 | 0 | 6 | 0.3333 | 6 | 0.1667 | 9 | 0.4444 | 23 | 0.3043 | 3 | 0.3333 | 6 | 0 | 11 | 0.2727 | 5 | 0.4 | 25 | 0.24 |
| 14_4544855 | 1 | 1 | 1 | 1 | NA | NA | NA | NA | 2 | 1 | NA | NA | NA | NA | 1 | 1 | NA | NA | 1 | 1 |
| 14_4544857 | 1 | 1 | 1 | 0 | NA | NA | NA | NA | 2 | 0.5 | NA | NA | NA | NA | 1 | 1 | NA | NA | 1 | 1 |
| 14_4544863 | 4 | 0.25 | 4 | 0.25 | 4 | 0 | 4 | 0 | 16 | 0.125 | 1 | 0 | 2 | 0 | 5 | 0.2 | 2 | 0 | 10 | 0.1 |
| 14_4545081 | 4 | 0.25 | 4 | 0 | 3 | 0 | 4 | 0 | 15 | 0.0667 | 1 | 0 | 1 | 1 | 4 | 0 | 2 | 0 | 8 | 0.125 |
| 14_4545845 | 1 | 0 | NA | NA | NA | NA | 2 | 0.5 | 3 | 0.3333 | NA | NA | NA | NA | 2 | 0.5 | NA | NA | 2 | 0.5 |
| 14_4548385 | 6 | 0.6667 | 3 | 0.6667 | 1 | 0 | 1 | 1 | 11 | 0.6364 | 2 | 1 | 2 | 0 | 4 | 1 | NA | NA | 8 | 0.75 |
| 14_4551102 | NA | NA | NA | NA | 1 | 1 | NA | NA | 1 | 1 | NA | NA | 1 | 1 | NA | NA | NA | NA | 1 | 1 |
| 14_4551106 | NA | NA | NA | NA | 1 | 1 | NA | NA | 1 | 1 | NA | NA | 1 | 1 | NA | NA | NA | NA | 1 | 1 |
| 14_4551737 | 5 | 0.2 | 3 | 0 | 1 | 0 | NA | NA | 9 | 0.1111 | 1 | 0 | 3 | 0.3333 | 4 | 0 | NA | NA | 8 | 0.125 |
| 14_4551744 | 6 | 1 | 4 | 0.75 | NA | NA | 1 | 1 | 11 | 0.9091 | 2 | 1 | 3 | 1 | 4 | 1 | NA | NA | 9 | 1 |
| 14_4551745 | 4 | 0.25 | 3 | 0.3333 | 1 | 0 | 2 | 1 | 10 | 0.4 | 1 | 0 | 3 | 0 | 5 | 0.4 | NA | NA | 9 | 0.2222 |
| 14_4551752 | 5 | 0 | 3 | 0.3333 | 1 | 0 | 2 | 0.5 | 11 | 0.1818 | 2 | 0 | 2 | 0 | 3 | 0.3333 | NA | NA | 7 | 0.1429 |
| 14_4551753 | 6 | 1 | 4 | 1 | 1 | 1 | 2 | 1 | 13 | 1 | 1 | 1 | 3 | 1 | 3 | 1 | NA | NA | 7 | 1 |
| 14_4551756 | 5 | 0.8 | 3 | 1 | 1 | 1 | 2 | 1 | 11 | 0.9091 | 1 | 1 | 2 | 1 | 5 | 1 | NA | NA | 8 | 1 |
| 14_4551769 | 4 | 0.25 | 4 | 0.25 | 1 | 0 | 2 | 0 | 11 | 0.1818 | 2 | 0.5 | 3 | 0 | 4 | 0 | NA | NA | 9 | 0.1111 |
| 14_4551775 | 5 | 0.2 | 3 | 0.3333 | 1 | 1 | 1 | 0 | 10 | 0.3 | 2 | 0.5 | 3 | 0 | 4 | 0.25 | NA | NA | 9 | 0.2222 |
| 14_4551777 | 5 | 0.6 | 4 | 0.25 | 1 | 0 | 2 | 0.5 | 12 | 0.4167 | 2 | 0.5 | 3 | 0 | 5 | 0.4 | NA | NA | 10 | 0.3 |
| 14_4551794 | 6 | 0.1667 | 3 | 0.3333 | 1 | 0 | 2 | 0.5 | 12 | 0.25 | NA | NA | 2 | 0 | 5 | 0.2 | NA | NA | 7 | 0.1429 |
| 14_4551803 | 6 | 0.1667 | 4 | 0.25 | 1 | 0 | 1 | 0 | 12 | 0.1667 | NA | NA | 3 | 0 | 6 | 0.3333 | 1 | 1 | 10 | 0.3 |
| 14_4551819 | 6 | 0 | 3 | 0.3333 | 1 | 1 | 2 | 0.5 | 12 | 0.25 | 1 | 0 | 3 | 0.3333 | 5 | 1 | NA | NA | 9 | 0.6667 |
| 14_4551821 | 7 | 0.5714 | 4 | 1 | 1 | 1 | 2 | 1 | 14 | 0.7857 | 1 | 1 | 3 | 0.3333 | 4 | 0.75 | NA | NA | 8 | 0.625 |
| 14_4551823 | 4 | 0.25 | 3 | 0.3333 | 1 | 0 | 2 | 0.5 | 10 | 0.3 | 1 | 0 | 3 | 0 | 6 | 0.1667 | 1 | 0 | 11 | 0.0909 |
| 14_4551825 | 6 | 0 | 5 | 0.2 | 1 | 0 | 1 | 0 | 13 | 0.0769 | 2 | 0 | 3 | 0.3333 | 5 | 0 | 1 | 0 | 11 | 0.0909 |
| 14_4551831 | 6 | 0.5 | 5 | 0.2 | 1 | 1 | 2 | 0 | 14 | 0.3571 | 2 | 0 | 3 | 0.3333 | 5 | 0.2 | 1 | 0 | 11 | 0.1818 |
| 14_4551844 | 8 | 0.25 | 5 | 0.2 | 1 | 0 | 2 | 0 | 16 | 0.1875 | 3 | 0.3333 | 3 | 0 | 6 | 0 | 1 | 0 | 13 | 0.0769 |
| 14_4551892 | 7 | 0.1429 | 5 | 0.2 | 1 | 0 | 3 | 0 | 16 | 0.125 | 2 | 0 | 3 | 0 | 4 | 0.5 | 1 | 1 | 10 | 0.3 |
| 14_4551897 | 7 | 0.2857 | 4 | 0.25 | 1 | 1 | 2 | 0 | 14 | 0.2857 | 2 | 0.5 | 2 | 0 | 5 | 0.2 | 1 | 0 | 10 | 0.2 |
| 14_4552133 | 4 | 0.25 | 1 | 1 | 5 | 0.2 | 4 | 0.25 | 14 | 0.2857 | 5 | 0 | 5 | 0 | 5 | 0.4 | 7 | 0.2857 | 22 | 0.1818 |
| 14_4552340 | 4 | 0 | 2 | 0 | 4 | 0 | 5 | 0.2 | 15 | 0.0667 | 5 | 0.2 | 4 | 0.75 | 3 | 0 | 8 | 0.25 | 20 | 0.3 |
| 14_4558614 | 7 | 0 | 14 | 0.2143 | 13 | 0.0769 | 12 | 0.5 | 46 | 0.2174 | 10 | 0 | 10 | 0 | 15 | 0.2 | 19 | 0.0526 | 54 | 0.0741 |
| 14_4558687 | 5 | 0 | 4 | 0.25 | 9 | 0.1111 | NA | NA | 18 | 0.1111 | 3 | 0 | 4 | 0 | 5 | 0 | 8 | 0.125 | 20 | 0.05 |
| 14_4561119 | 1 | 1 | 1 | 0 | 2 | 1 | NA | NA | 4 | 0.75 | NA | NA | 1 | 1 | NA | NA | 1 | 1 | 2 | 1 |
| 14_4561129 | 1 | 1 | 1 | 0 | 2 | 1 | NA | NA | 4 | 0.75 | NA | NA | 1 | 1 | NA | NA | 1 | 0 | 2 | 0.5 |
| 14_4561407 | 2 | 0.5 | NA | NA | 2 | 0 | 3 | 0 | 7 | 0.1429 | 1 | 1 | 2 | 0 | 1 | 0 | 1 | 0 | 5 | 0.2 |

| | | | | | | | | | | | | | | | | | | | | |
|---|---|---|---|---|---|---|---|---|---|---|---|---|---|---|---|---|---|---|---|---|
| 14_4561472 | 2 | 1 | 1 | 0 | 3 | 0.6667 | 2 | 0 | 8 | 0.5 | NA | NA | 4 | 0.5 | 1 | 0 | 1 | 1 | 6 | 0.5 |
| 14_4561531 | 2 | 0.5 | 1 | 0 | 3 | 0 | 2 | 0 | 8 | 0.125 | 2 | 0 | 4 | 0 | 4 | 0 | 4 | 0.25 | 14 | 0.0714 |
| 14_4561539 | 2 | 0 | NA | NA | 3 | 0.3333 | 2 | 0 | 7 | 0.1429 | 1 | 0 | 4 | 0.5 | 3 | 0.3333 | 4 | 0 | 12 | 0.25 |
| 14_4561575 | 1 | 0 | 1 | 0 | 2 | 0.5 | 3 | 0.6667 | 7 | 0.4286 | 1 | 0 | 3 | 0 | 2 | 0 | 3 | 0.3333 | 9 | 0.1111 |
| 14_4561590 | 2 | 0 | 1 | 1 | 3 | 0 | 3 | 0 | 9 | 0.1111 | 1 | 0 | 2 | 0 | 3 | 0.3333 | 3 | 0.3333 | 9 | 0.2222 |
| 14_4569710 | 9 | 0.1111 | 9 | 0 | 5 | 0.4 | 7 | 0.1429 | 30 | 0.1333 | 3 | 0 | 4 | 0.25 | 5 | 0.4 | 5 | 0 | 17 | 0.1765 |
| 14_4569721 | 11 | 0.0909 | 9 | 0 | 4 | 0.25 | 7 | 0.1429 | 31 | 0.0968 | 4 | 0 | 5 | 0.2 | 7 | 0.1429 | 5 | 0.2 | 21 | 0.1429 |
| 14_4571354 | 9 | 0.2222 | 7 | 0 | 3 | 0 | 5 | 0.2 | 24 | 0.125 | 2 | 0 | 4 | 0 | 3 | 0.3333 | 2 | 0.5 | 11 | 0.1818 |
| 14_4582621 | 2 | 0.5 | 1 | 1 | 2 | 0.5 | 1 | 1 | 6 | 0.6667 | 2 | 0.5 | NA | NA | 2 | 0.5 | 4 | 0 | 8 | 0.25 |
| 14_4582759 | 2 | 1 | 1 | 0 | NA | NA | 1 | 0 | 4 | 0.5 | 2 | 0.5 | NA | NA | 1 | 0 | 2 | 0 | 5 | 0.2 |
| 14_4583088 | 2 | 0 | NA | NA | 3 | 0.3333 | 2 | 0 | 7 | 0.1429 | NA | NA | 1 | 1 | 2 | 0 | NA | NA | 3 | 0.3333 |
| 14_4583244 | 6 | 0 | 4 | 0.25 | 4 | 0 | 5 | 0.4 | 19 | 0.1579 | 3 | 0 | 2 | 0 | 3 | 0 | 5 | 0.2 | 13 | 0.0769 |
| 14_4583246 | 6 | 0 | 3 | 0 | 6 | 0 | 5 | 0.2 | 20 | 0.05 | 2 | 0 | 5 | 0.4 | 2 | 1 | 4 | 0 | 13 | 0.3077 |
| 14_4583258 | 7 | 0.1429 | 4 | 0.25 | 8 | 0.25 | 6 | 0.1667 | 25 | 0.2 | 4 | 0.25 | 7 | 0.1429 | 5 | 0.2 | 4 | 0.25 | 20 | 0.2 |
| 14_4583269 | 8 | 0.375 | 3 | 0.3333 | 7 | 0 | 6 | 0.3333 | 24 | 0.25 | 4 | 0.5 | 8 | 0.5 | 6 | 0.1667 | 5 | 0 | 23 | 0.3043 |
| 14_4583284 | 9 | 0.4444 | 4 | 0.5 | 8 | 0.5 | 5 | 0.8 | 26 | 0.5385 | 4 | 0.5 | 7 | 0.5714 | 3 | 0.6667 | 5 | 0.6 | 19 | 0.5789 |
| 14_4583288 | 11 | 0.5455 | 3 | 0.6667 | 9 | 0.3333 | 6 | 0.6667 | 29 | 0.5172 | 4 | 1 | 7 | 0.7143 | 4 | 1 | 4 | 1 | 19 | 0.8947 |
| 14_4583293 | 6 | 1 | 2 | 0.5 | 7 | 0.7143 | 4 | 0.75 | 19 | 0.7895 | 1 | 1 | 6 | 1 | 3 | 0.6667 | 5 | 1 | 15 | 0.9333 |
| 14_4583298 | 11 | 0 | 2 | 0 | 9 | 0 | 5 | 0.4 | 27 | 0.0741 | 4 | 0.25 | 9 | 0 | 2 | 0 | 2 | 0.5 | 17 | 0.1176 |
| 14_4583912 | 10 | 0 | 4 | 0.25 | 6 | 0 | 4 | 0.25 | 24 | 0.0833 | 4 | 0.25 | 8 | 0.125 | 3 | 0 | 2 | 0 | 17 | 0.1176 |
| 14_4584805 | 5 | 0 | 1 | 0 | 6 | 0.1667 | 3 | 0 | 15 | 0.0667 | 3 | 0 | 8 | 0.375 | 3 | 0 | 1 | 1 | 15 | 0.2667 |
| 14_4586711 | 5 | 0.2 | 1 | 0 | 4 | 0 | 2 | 0 | 12 | 0.0833 | 3 | 0.3333 | 7 | 0 | 1 | 0 | 1 | 0 | 12 | 0.0833 |
| 14_4586718 | 5 | 0.2 | 1 | 1 | 4 | 0.25 | 2 | 0 | 12 | 0.25 | 3 | 0.3333 | 7 | 0 | 1 | 0 | 1 | 0 | 12 | 0.0833 |
| 14_4589512 | 2 | 1 | 1 | 0 | 2 | 0.5 | 1 | 1 | 6 | 0.6667 | 2 | 0.5 | 3 | 1 | NA | NA | NA | NA | 5 | 0.8 |
| 14_4662118 | NA | NA | 1 | 1 | NA | NA | 1 | 1 | 2 | 1 | 1 | 1 | NA | NA | NA | NA | 1 | 1 | 2 | 1 |
| 14_4662141 | NA | NA | 1 | 1 | NA | NA | 1 | 1 | 2 | 1 | NA | NA | NA | NA | NA | NA | 1 | 1 | 1 | 1 |
| 14_4662152 | NA | NA | 1 | 1 | NA | NA | 1 | 1 | 2 | 1 | 1 | 1 | NA | NA | NA | NA | 1 | 1 | 2 | 1 |
| 14_4662162 | NA | NA | 1 | 0 | NA | NA | 1 | 1 | 2 | 0.5 | NA | NA | NA | NA | NA | NA | 1 | 1 | 1 | 1 |
| 14_4701348 | 22 | 0.1818 | 15 | 0.2667 | 21 | 0.2381 | 19 | 0.1579 | 77 | 0.2078 | 17 | 0.1765 | 20 | 0.2 | 25 | 0.08 | 20 | 0.1 | 82 | 0.1341 |
| 14_4701355 | 21 | 0 | 16 | 0.125 | 16 | 0.0625 | 18 | 0.1111 | 71 | 0.0704 | 17 | 0.0588 | 21 | 0.0476 | 22 | 0.1364 | 22 | 0.0455 | 82 | 0.0732 |
| 14_4701362 | 23 | 0.1304 | 16 | 0.125 | 13 | 0.3077 | 18 | 0.2778 | 70 | 0.2 | 18 | 0.1667 | 22 | 0.0909 | 26 | 0.1154 | 17 | 0.2353 | 83 | 0.1446 |
| 14_4701381 | 22 | 0.2727 | 16 | 0.3125 | 15 | 0.4 | 18 | 0.1111 | 71 | 0.2676 | 18 | 0.3333 | 20 | 0.3 | 22 | 0.1818 | 20 | 0.35 | 80 | 0.2875 |
| 14_4701385 | 19 | 0.0526 | 14 | 0.1429 | 11 | 0.1818 | 14 | 0 | 58 | 0.0862 | 13 | 0 | 19 | 0.0526 | 21 | 0.0952 | 16 | 0.125 | 69 | 0.0725 |
| 14_4704878 | 10 | 0 | 12 | 0.1667 | 5 | 0 | 16 | 0.0625 | 43 | 0.0698 | 10 | 0 | 10 | 0 | 13 | 0.3077 | 6 | 0.1667 | 39 | 0.1282 |
| 14_4704899 | 7 | 0.1429 | 12 | 0.0833 | 5 | 0.2 | 16 | 0 | 40 | 0.075 | 10 | 0 | 9 | 0 | 9 | 0.1111 | 5 | 0.2 | 33 | 0.0606 |
| 14_4704920 | 7 | 0.2857 | 11 | 0 | 6 | 0.1667 | 13 | 0.0769 | 37 | 0.1081 | 9 | 0 | 8 | 0 | 11 | 0.1818 | 6 | 0 | 34 | 0.0588 |
| 14_4708370 | 3 | 0 | 2 | 0.5 | 4 | 0 | 4 | 0 | 13 | 0.0769 | 4 | 0.25 | 5 | 0 | 5 | 0 | 4 | 0.25 | 18 | 0.1111 |
| 14_4709125 | 2 | 0.5 | 2 | 0.5 | 3 | 0.6667 | 3 | 0.6667 | 10 | 0.6 | 4 | 0.25 | 6 | 0.5 | 4 | 0.75 | 2 | 1 | 16 | 0.5625 |
| 14_4709129 | 4 | 0.75 | 2 | 0.5 | 2 | 0 | 6 | 0.6667 | 14 | 0.5714 | 4 | 0.5 | 7 | 0.8571 | 5 | 0.4 | 4 | 0.75 | 20 | 0.65 |
| 14_4709132 | 4 | 0.5 | 2 | 0.5 | 2 | 0 | 5 | 0.6 | 13 | 0.4615 | 5 | 0.8 | 6 | 0.6667 | 4 | 0.5 | 3 | 0.3333 | 18 | 0.6111 |
| 14_4709145 | 4 | 1 | 2 | 1 | 4 | 0.75 | 6 | 0.6667 | 16 | 0.8125 | 3 | 1 | 7 | 0.7143 | 5 | 0.8 | 3 | 0.6667 | 18 | 0.7778 |
| 14_4709146 | 4 | 0 | 2 | 0 | 5 | 0 | 5 | 0.2 | 16 | 0.0625 | 5 | 0 | 7 | 0.1429 | 5 | 0 | 4 | 0.75 | 21 | 0.1905 |
| 14_4709156 | 2 | 0 | 2 | 0 | 4 | 0.25 | 6 | 0.3333 | 14 | 0.2143 | 4 | 0 | 5 | 0.2 | 3 | 0 | 3 | 0.6667 | 15 | 0.2 |
| 14_4709186 | NA | NA | NA | NA | NA | NA | 1 | 1 | 1 | 1 | NA | NA | 1 | 0 | NA | NA | 2 | 1 | 3 | 0.6667 |
| 14_4733949 | 1 | 0 | NA | NA | 1 | 1 | NA | NA | 2 | 0.5 | NA | NA | 1 | 0 | 1 | 0 | 1 | 1 | 3 | 0.3333 |
| 14_4752728 | NA | NA | NA | NA | 1 | 1 | 2 | 1 | 3 | 1 | 2 | 1 | NA | NA | 1 | 0 | 1 | 1 | 4 | 0.75 |
| 14_4752733 | NA | NA | NA | NA | 1 | 1 | 2 | 1 | 3 | 1 | 1 | 0 | NA | NA | 1 | 1 | 1 | 1 | 3 | 0.6667 |
| 14_4752743 | NA | NA | NA | NA | 1 | 1 | 2 | 1 | 3 | 1 | 2 | 0.5 | NA | NA | 1 | 1 | 1 | 1 | 4 | 0.75 |
| 14_4752765 | NA | NA | NA | NA | 1 | 1 | 1 | 0 | 2 | 0.5 | 1 | 0 | NA | NA | 1 | 0 | 1 | 1 | 3 | 0.3333 |
| 14_4752768 | NA | NA | NA | NA | 1 | 1 | 1 | 1 | 2 | 1 | 3 | 0.6667 | NA | NA | 1 | 1 | 1 | 1 | 5 | 0.8 |
| 14_4752770 | NA | NA | NA | NA | 1 | 1 | 2 | 1 | 3 | 1 | 3 | 1 | NA | NA | 1 | 1 | NA | NA | 4 | 1 |
| 14_4752793 | NA | NA | NA | NA | 1 | 1 | 2 | 1 | 3 | 1 | 2 | 1 | NA | NA | 1 | 1 | 1 | 1 | 4 | 1 |
| 14_4752814 | NA | NA | NA | NA | 1 | 1 | 2 | 1 | 3 | 1 | 3 | 0.6667 | NA | NA | 1 | 1 | 1 | 1 | 5 | 0.8 |
| 14_4752815 | NA | NA | NA | NA | 1 | 0 | 2 | 0.5 | 3 | 0.3333 | 3 | 0.6667 | NA | NA | NA | NA | 1 | 1 | 4 | 0.75 |
| 14_4752848 | NA | NA | NA | NA | 1 | 0 | 2 | 0.5 | 3 | 0.3333 | 3 | 0 | NA | NA | 1 | 1 | 1 | 0 | 5 | 0.2 |
| 14_4753197 | NA | NA | NA | NA | 2 | 0 | 4 | 0.25 | 6 | 0.1667 | 3 | 0 | 1 | 1 | 3 | 0 | 3 | 0 | 10 | 0.1 |
| 14_4754800 | 1 | 1 | 3 | 0.3333 | NA | NA | 3 | 0.6667 | 7 | 0.5714 | 2 | 0.5 | 1 | 1 | 4 | 0.75 | NA | NA | 7 | 0.7143 |
| 14_4754805 | 1 | 1 | 3 | 0.3333 | 1 | 0 | 3 | 0.3333 | 8 | 0.375 | 2 | 0 | 1 | 1 | 4 | 0.25 | 1 | 0 | 8 | 0.25 |
| 14_4754809 | NA | NA | 3 | 0.6667 | NA | NA | 3 | 0.6667 | 6 | 0.6667 | NA | NA | 1 | 0 | 3 | 0.3333 | NA | NA | 4 | 0.25 |
| 14_4754811 | 1 | 1 | 3 | 0.3333 | 1 | 1 | 3 | 0.6667 | 8 | 0.625 | 1 | 0 | 1 | 1 | 4 | 0.75 | 1 | 0 | 7 | 0.5714 |
| 14_4754813 | NA | NA | 3 | 1 | 1 | 1 | 3 | 0.6667 | 7 | 0.8571 | 1 | 0 | 1 | 1 | 4 | 1 | 1 | 1 | 7 | 0.8571 |
| 14_4754829 | 1 | 1 | 3 | 0.3333 | 1 | 0 | 3 | 0.3333 | 8 | 0.375 | 1 | 0 | NA | NA | 3 | 0.3333 | 1 | 0 | 5 | 0.2 |
| 14_4754830 | 1 | 1 | 3 | 1 | 1 | 1 | 3 | 1 | 8 | 1 | 1 | 1 | 1 | 1 | 3 | 1 | 1 | 1 | 6 | 1 |
| 14_4754844 | 1 | 0 | 3 | 0.3333 | 1 | 0 | 3 | 0.3333 | 8 | 0.25 | 2 | 0 | 2 | 0 | 3 | 0.3333 | 1 | 0 | 8 | 0.125 |
| 14_4755112 | NA | NA | NA | NA | NA | NA | 1 | 1 | 1 | 1 | NA | NA | 1 | 1 | NA | NA | NA | NA | 1 | 1 |

| | | | | | | | | | | | | | | | | | | | | |
|---|---|---|---|---|---|---|---|---|---|---|---|---|---|---|---|---|---|---|---|---|
| 14_4755699 | 2 | 1 | 3 | 1 | 1 | 1 | 3 | 1 | 9 | 1 | 2 | 1 | NA | NA | 2 | 1 | 1 | 1 | 5 | 1 |
| 14_4755748 | 2 | 0 | 3 | 0.3333 | 1 | 0 | 3 | 0.3333 | 9 | 0.2222 | 2 | 0 | 1 | 0 | 3 | 0 | 2 | 0.5 | 8 | 0.125 |
| 14_4755764 | 2 | 0.5 | 3 | 0.3333 | 2 | 0.5 | 3 | 0.6667 | 10 | 0.5 | 3 | 0.3333 | 1 | 0 | 3 | 0 | 1 | 0 | 8 | 0.125 |
| 14_4755842 | 1 | 0 | 1 | 1 | 2 | 0.5 | 4 | 0.75 | 8 | 0.625 | 2 | 0 | NA | NA | 3 | 1 | 1 | 1 | 6 | 0.6667 |
| 14_4755869 | 2 | 0 | 3 | 0 | 3 | 0.3333 | 5 | 0.2 | 13 | 0.1538 | 3 | 0 | 2 | 0 | 4 | 0.25 | 2 | 0 | 11 | 0.0909 |
| 14_4755926 | 4 | 0.25 | 2 | 0 | 5 | 0.2 | 3 | 0 | 14 | 0.1429 | 2 | 0 | 5 | 0.2 | 4 | 0 | 2 | 0.5 | 13 | 0.1538 |
| 14_4755937 | 3 | 1 | 3 | 0 | 3 | 0.3333 | 5 | 0 | 14 | 0.2857 | 1 | 0 | 4 | 0.25 | 2 | 0 | 2 | 0.5 | 9 | 0.2222 |
| 14_4755948 | 2 | 1 | 3 | 0 | 4 | 0.5 | 5 | 1 | 14 | 0.6429 | 3 | 0.3333 | 4 | 0.25 | 4 | 1 | 2 | 1 | 13 | 0.6154 |
| 14_4755953 | 3 | 1 | 2 | 1 | 4 | 1 | 5 | 1 | 14 | 1 | 3 | 1 | 4 | 1 | 4 | 1 | 1 | 1 | 12 | 1 |
| 14_4755957 | 4 | 0.75 | 3 | 1 | 3 | 1 | 4 | 1 | 14 | 0.9286 | 2 | 1 | 3 | 1 | 5 | 0.8 | 2 | 1 | 12 | 0.9167 |
| 14_4756001 | 4 | 0.5 | 3 | 0 | 4 | 0 | 8 | 0 | 19 | 0.1053 | 3 | 0 | 6 | 0.1667 | 6 | 0 | 2 | 0 | 17 | 0.0588 |
| 14_4756020 | 5 | 0.4 | 3 | 0.3333 | 4 | 0.5 | 9 | 0.3333 | 21 | 0.381 | 3 | 0.6667 | 6 | 0.6667 | 7 | 0.5714 | 2 | 0.5 | 18 | 0.6111 |
| 14_4776670 | 4 | 0 | 7 | 0.1429 | 5 | 0.2 | 2 | 0 | 18 | 0.1111 | NA | NA | 4 | 0.25 | 3 | 0.3333 | 4 | 0 | 11 | 0.1818 |
| 14_4776688 | 3 | 0.6667 | 6 | 0.1667 | 3 | 0 | 2 | 0.5 | 14 | 0.2857 | NA | NA | 4 | 0.25 | 1 | 0 | 4 | 0.5 | 9 | 0.3333 |
| 14_4776695 | 4 | 1 | 7 | 1 | 5 | 1 | 2 | 0.5 | 18 | 0.9444 | 2 | 1 | 5 | 0.8 | 2 | 1 | 4 | 0.75 | 13 | 0.8462 |
| 14_4776698 | 2 | 0.5 | 6 | 0.3333 | 1 | 0 | 2 | 0.5 | 11 | 0.3636 | 2 | 0 | 3 | 0.3333 | 2 | 0 | 3 | 0 | 10 | 0.1 |
| 14_4776721 | 3 | 0 | 7 | 0.5714 | 5 | 0.2 | 1 | 1 | 16 | 0.375 | 2 | 0 | 5 | 0.2 | 3 | 0.6667 | 4 | 0.25 | 14 | 0.2857 |
| 14_4776732 | 4 | 0.75 | 7 | 0.8571 | 4 | 0.5 | 1 | 0 | 16 | 0.6875 | 2 | 1 | 4 | 0.5 | 2 | 0 | 2 | 1 | 10 | 0.6 |
| 14_4776744 | 4 | 0.25 | 6 | 0.3333 | 4 | 0.25 | 2 | 0 | 16 | 0.25 | 1 | 0 | 5 | 0.2 | 2 | 0.5 | 5 | 0.2 | 13 | 0.2308 |
| 14_4776748 | 3 | 0.3333 | 5 | 0.2 | 4 | 0.25 | 1 | 0 | 13 | 0.2308 | 2 | 0 | 6 | 0 | 3 | 0.3333 | 3 | 0 | 14 | 0.0714 |
| 14_4776763 | 5 | 0 | 7 | 0.1429 | 5 | 0.2 | 2 | 0 | 19 | 0.1053 | 2 | 0 | 6 | 0.1667 | 4 | 0 | 5 | 0 | 17 | 0.0588 |
| 14_4776769 | 3 | 0 | 8 | 0.125 | 4 | 0.5 | 2 | 0 | 17 | 0.1765 | 1 | 0 | 3 | 0.3333 | 3 | 0 | 6 | 0 | 13 | 0.0769 |
| 14_4776778 | 5 | 0.6 | 6 | 0.8333 | 7 | 0.8571 | 2 | 1 | 20 | 0.8 | 1 | 1 | 8 | 0.875 | 3 | 1 | 1 | 1 | 13 | 0.9231 |
| 14_4776794 | 5 | 0 | 8 | 0.125 | 8 | 0.25 | 2 | 0 | 23 | 0.1304 | 2 | 0 | 7 | 0.1429 | 5 | 0.2 | 6 | 0 | 20 | 0.1 |
| 14_4776800 | 5 | 0.2 | 4 | 0.25 | 7 | 0.2857 | 2 | 0 | 18 | 0.2222 | 2 | 0 | 6 | 0.1667 | 2 | 0 | 5 | 0 | 15 | 0.0667 |
| 14_4776805 | 4 | 0.5 | 6 | 0 | 7 | 0.2857 | 2 | 0.5 | 19 | 0.2632 | 2 | 0 | 7 | 0.4286 | 5 | 0.2 | 5 | 0 | 19 | 0.2105 |
| 14_4776806 | 5 | 0 | 9 | 0 | 8 | 0.25 | 2 | 0 | 24 | 0.0833 | 2 | 0.5 | 8 | 0.125 | 5 | 0 | 5 | 0.2 | 20 | 0.15 |
| 14_4776826 | 4 | 1 | 8 | 1 | 8 | 1 | 3 | 1 | 23 | 1 | 1 | 1 | 8 | 1 | 5 | 1 | 5 | 1 | 19 | 1 |
| 14_4776848 | 5 | 0.2 | 9 | 0.1111 | 7 | 0.2857 | 4 | 0 | 25 | 0.16 | 2 | 0 | 6 | 0 | 5 | 0.2 | 6 | 0.6667 | 19 | 0.2632 |
| 14_4776864 | 5 | 0.4 | 8 | 0.375 | 7 | 0.1429 | 4 | 0.25 | 24 | 0.2917 | 2 | 1 | 7 | 0.2857 | 5 | 0.4 | 7 | 0.2857 | 21 | 0.381 |
| 14_4776874 | 5 | 0.2 | 6 | 0.5 | 10 | 0.4 | 1 | 0 | 22 | 0.3636 | 2 | 0.5 | 8 | 0.375 | 4 | 0.75 | 4 | 1 | 18 | 0.6111 |
| 14_4776875 | 6 | 0.6667 | 11 | 0.6364 | 11 | 0.8182 | 3 | 0.6667 | 31 | 0.7097 | 2 | 0 | 8 | 0.625 | 5 | 0.6 | 7 | 0.8571 | 22 | 0.6364 |
| 14_4776876 | 6 | 0.3333 | 10 | 0 | 10 | 0.1 | 4 | 0 | 30 | 0.1 | 2 | 0 | 8 | 0.375 | 5 | 0 | 7 | 0.2857 | 22 | 0.2273 |
| 14_4776882 | 3 | 0.3333 | 9 | 0.4444 | 10 | 0.8 | 4 | 0.5 | 26 | 0.5769 | 2 | 0.5 | 7 | 0.8571 | 3 | 0.6667 | 6 | 0.6667 | 18 | 0.7222 |
| 14_4776885 | 5 | 1 | 9 | 1 | 10 | 1 | 5 | 1 | 29 | 1 | 2 | 1 | 7 | 1 | 4 | 1 | 5 | 1 | 18 | 1 |
| 14_4776893 | 4 | 1 | 11 | 0.9091 | 8 | 1 | 4 | 1 | 27 | 0.963 | 3 | 1 | 7 | 1 | 5 | 1 | 5 | 1 | 20 | 1 |
| 14_4776896 | 5 | 1 | 10 | 1 | 9 | 1 | 3 | 1 | 27 | 1 | 3 | 1 | 7 | 1 | 3 | 1 | 4 | 1 | 17 | 1 |
| 14_4776912 | 5 | 1 | 8 | 1 | 7 | 0.8571 | 4 | 0.25 | 24 | 0.8333 | 2 | 0.5 | 6 | 0.6667 | 4 | 1 | 4 | 1 | 16 | 0.8125 |
| 14_4776929 | 5 | 0 | 6 | 0.1667 | 11 | 0.3636 | 4 | 0 | 26 | 0.1923 | 3 | 0.3333 | 5 | 0.2 | 4 | 0.25 | 6 | 0.6667 | 18 | 0.3889 |
| 14_4776932 | 5 | 0 | 10 | 0 | 12 | 0.1667 | 6 | 0.3333 | 33 | 0.1212 | 2 | 0 | 8 | 0.125 | 4 | 0.25 | 7 | 0.1429 | 21 | 0.1429 |
| 14_4777857 | 3 | 0.3333 | 5 | 0.4 | 3 | 0 | 3 | 0 | 14 | 0.2143 | 3 | 0 | 7 | 0.2857 | 2 | 0 | 2 | 0.5 | 14 | 0.2143 |
| 14_4777885 | 3 | 0 | 7 | 0.7143 | 2 | 0.5 | 5 | 0 | 17 | 0.3529 | 3 | 0.3333 | 4 | 0.5 | 2 | 1 | NA | NA | 9 | 0.5556 |
| 14_4777901 | 1 | 0 | 5 | 0 | 2 | 0 | 5 | 0.2 | 13 | 0.0769 | 4 | 0.5 | 4 | 0.5 | 2 | 0 | 1 | 0 | 11 | 0.3636 |
| 14_4777902 | 1 | 0 | 5 | 0.2 | 2 | 1 | 2 | 1 | 10 | 0.5 | 3 | 0 | 4 | 0.5 | 1 | 0 | NA | NA | 8 | 0.25 |
| 14_4777915 | 2 | 0 | 5 | 0.2 | 2 | 0 | 4 | 0.25 | 13 | 0.1538 | 4 | 0.25 | 4 | 0 | 2 | 0 | 1 | 0 | 11 | 0.0909 |
| 14_4777929 | 2 | 0 | 6 | 0.1667 | 2 | 0 | 4 | 0 | 14 | 0.0714 | 2 | 0 | 4 | 0.25 | 2 | 0 | NA | NA | 8 | 0.125 |
| 14_4778028 | 1 | 1 | 4 | 0.75 | 1 | 1 | 3 | 0.3333 | 9 | 0.6667 | NA | NA | 3 | 1 | 1 | 0 | NA | NA | 4 | 0.75 |
| 14_4778030 | 1 | 1 | 4 | 0.5 | NA | NA | 3 | 0.3333 | 8 | 0.5 | NA | NA | 3 | 0.6667 | 1 | 0 | NA | NA | 4 | 0.5 |
| 14_4778047 | 1 | 0 | 4 | 0 | 1 | 0 | 3 | 0.6667 | 9 | 0.2222 | NA | NA | 3 | 0.6667 | 1 | 0 | NA | NA | 4 | 0.5 |
| 14_4778048 | 1 | 0 | 4 | 0 | 1 | 1 | 3 | 0.3333 | 9 | 0.2222 | NA | NA | 3 | 0.6667 | 1 | 0 | NA | NA | 4 | 0.5 |
| 14_4790133 | NA | NA | 2 | 0 | 4 | 0.25 | 7 | 0.2857 | 13 | 0.2308 | 1 | 0 | 1 | 0 | NA | NA | 7 | 0.1429 | 9 | 0.1111 |
| 14_4790145 | NA | NA | 3 | 0 | 4 | 0.5 | 7 | 0.1429 | 14 | 0.2143 | 1 | 0 | 2 | 0 | NA | NA | 9 | 0.2222 | 12 | 0.1667 |
| 14_4790149 | NA | NA | 3 | 0.6667 | 4 | 0.75 | 7 | 0.8571 | 14 | 0.7857 | 1 | 1 | 2 | 0.5 | NA | NA | 8 | 0.75 | 11 | 0.7273 |
| 14_4790152 | NA | NA | 2 | 0.5 | 3 | 0.6667 | 7 | 0.2857 | 12 | 0.4167 | 1 | 0 | 2 | 1 | NA | NA | 6 | 0.1667 | 9 | 0.3333 |
| 14_4790156 | NA | NA | 3 | 0 | 5 | 0.2 | 7 | 0.2857 | 15 | 0.2 | 1 | 0 | 3 | 0 | NA | NA | 8 | 0.25 | 12 | 0.1667 |
| 14_4797173 | NA | NA | NA | NA | NA | NA | 1 | 1 | 1 | 1 | 1 | 0 | NA | NA | 1 | 0 | 2 | 0.5 | 4 | 0.25 |
| 14_4797348 | NA | NA | 2 | 0.5 | NA | NA | 2 | 0 | 4 | 0.25 | 1 | 0 | NA | NA | 1 | 0 | 4 | 0.25 | 6 | 0.1667 |
| 14_4797366 | 1 | 0 | 8 | 0 | 6 | 0 | 12 | 0.1667 | 27 | 0.0741 | 3 | 0 | 7 | 0 | 5 | 0 | 11 | 0.2727 | 26 | 0.1154 |
| 14_4797400 | 6 | 0.1667 | 7 | 0.1429 | 1 | 0 | 13 | 0.2308 | 27 | 0.1852 | 6 | 0.5 | 7 | 0.1429 | 9 | 0.4444 | 14 | 0.3571 | 36 | 0.3611 |
| 14_4849379 | 1 | 1 | NA | NA | 1 | 1 | 2 | 1 | 4 | 1 | 1 | 1 | 2 | 1 | 1 | 1 | NA | NA | 4 | 1 |
| 14_4849385 | 2 | 0 | NA | NA | 1 | 1 | 2 | 0.5 | 5 | 0.4 | 1 | 1 | 1 | 0 | 1 | 0 | NA | NA | 3 | 0.3333 |
| 14_4849399 | 2 | 0.5 | NA | NA | 1 | 0 | 1 | 0 | 4 | 0.25 | 1 | 1 | 2 | 1 | NA | NA | NA | NA | 3 | 1 |
| 14_4849400 | 2 | 1 | NA | NA | 1 | 1 | 1 | 1 | 4 | 1 | 1 | 1 | 3 | 1 | 1 | 1 | NA | NA | 5 | 1 |
| 14_4849403 | 2 | 1 | NA | NA | 1 | 1 | 2 | 1 | 5 | 1 | NA | NA | 3 | 1 | 1 | 1 | NA | NA | 4 | 1 |
| 14_4849414 | 2 | 0.5 | NA | NA | 1 | 0 | 2 | 1 | 5 | 0.6 | 1 | 1 | 3 | 0.6667 | 1 | 0 | NA | NA | 5 | 0.6 |

| | | | | | | | | | | | | | | | | | | | | |
|---|---|---|---|---|---|---|---|---|---|---|---|---|---|---|---|---|---|---|---|---|
| 14_4849768 | 3 | 0 | 3 | 0 | NA | NA | 4 | 0.25 | 10 | 0.1 | 1 | 0 | 5 | 0.4 | 4 | 0.75 | NA | NA | 10 | 0.5 |
| 14_4874941 | 1 | 1 | NA | NA | NA | NA | 2 | 0 | 3 | 0.3333 | 1 | 1 | 1 | 1 | 2 | 0 | NA | NA | 4 | 0.5 |
| 14_4874946 | 1 | 1 | NA | NA | NA | NA | 2 | 0 | 3 | 0.3333 | 1 | 1 | 1 | 0 | 2 | 0.5 | NA | NA | 4 | 0.5 |
| 14_4874958 | 1 | 1 | NA | NA | NA | NA | 2 | 1 | 3 | 1 | 1 | 1 | 1 | 1 | 2 | 1 | NA | NA | 4 | 1 |
| 14_4874987 | 1 | 1 | NA | NA | NA | NA | 2 | 1 | 3 | 1 | NA | NA | 1 | 1 | 2 | 0 | NA | NA | 3 | 0.3333 |
| 14_4874989 | 1 | 1 | NA | NA | NA | NA | 2 | 1 | 3 | 1 | 1 | 1 | 1 | 1 | 2 | 1 | NA | NA | 4 | 1 |
| 14_4875134 | 1 | 1 | 1 | 1 | NA | NA | 4 | 0.5 | 6 | 0.6667 | 1 | 1 | 3 | 0.6667 | 1 | 1 | 1 | 1 | 6 | 0.8333 |
| 14_4875136 | 1 | 1 | 1 | 1 | NA | NA | 4 | 0.25 | 6 | 0.5 | 1 | 0 | 3 | 1 | 1 | 1 | NA | NA | 5 | 0.8 |
| 14_4875137 | 1 | 0 | 1 | 1 | NA | NA | 3 | 0 | 5 | 0.2 | 1 | 0 | 3 | 0 | NA | NA | 1 | 1 | 5 | 0.2 |
| 14_4875165 | 1 | 1 | 1 | 0 | NA | NA | 3 | 0 | 5 | 0.2 | 1 | 0 | 3 | 0.3333 | 1 | 0 | 1 | 0 | 6 | 0.1667 |
| 14_4875183 | NA | NA | 1 | 1 | 1 | 0 | 3 | 0 | 5 | 0.2 | 1 | 0 | 3 | 0.3333 | 1 | 0 | 1 | 0 | 6 | 0.1667 |
| 14_4875237 | 1 | 0 | NA | NA | 1 | 1 | 3 | 0 | 5 | 0.2 | 1 | 1 | 4 | 0 | 1 | 0 | 1 | 0 | 7 | 0.1429 |
| 14_4924991 | 3 | 0 | 5 | 0.2 | 7 | 0.1429 | 3 | 0.3333 | 18 | 0.1667 | 3 | 0.6667 | 7 | 0 | 11 | 0.1818 | 2 | 0 | 23 | 0.1739 |
| 14_4925028 | 4 | 0.25 | 6 | 0.1667 | 6 | 0 | 3 | 0 | 19 | 0.1053 | 3 | 0 | 7 | 0.5714 | 12 | 0.1667 | 3 | 0 | 25 | 0.24 |
| 14_4925115 | 3 | 0.3333 | 4 | 0 | 5 | 0 | 1 | 0 | 13 | 0.0769 | 2 | 0 | 7 | 0.1429 | 8 | 0 | 2 | 0 | 19 | 0.0526 |
| 14_4925137 | 7 | 0.2857 | 5 | 0.6 | 6 | 0.3333 | NA | NA | 18 | 0.3889 | 4 | 0.25 | 9 | 0.2222 | 7 | 0.1429 | 5 | 0.6 | 25 | 0.28 |
| 14_4925146 | 7 | 0.8571 | 5 | 0.8 | 6 | 0.6667 | 1 | 1 | 19 | 0.7895 | 3 | 1 | 8 | 0.875 | 8 | 0.625 | 5 | 1 | 24 | 0.8333 |
| 14_4925151 | 6 | 0.5 | 4 | 1 | 5 | 0.8 | 1 | 1 | 16 | 0.75 | 3 | 1 | 7 | 1 | 8 | 0.875 | 3 | 1 | 21 | 0.9524 |
| 14_4925165 | 7 | 0.4286 | 3 | 1 | 6 | 0 | 1 | 0 | 17 | 0.3529 | 4 | 0.5 | 7 | 0.4286 | 7 | 0.1429 | 4 | 0.5 | 22 | 0.3636 |
| 14_4925168 | 6 | 0.1667 | 2 | 0 | 6 | 0 | 1 | 0 | 15 | 0.0667 | 3 | 0.3333 | 8 | 0.125 | 7 | 0.2857 | 5 | 0.2 | 23 | 0.2174 |
| 14_4925169 | 6 | 0.1667 | 4 | 0.5 | 6 | 0.1667 | 1 | 0 | 17 | 0.2353 | 4 | 0.5 | 6 | 0.6667 | 6 | 0.5 | 3 | 0.6667 | 19 | 0.5789 |
| 14_4925288 | 7 | 0 | 4 | 0 | 4 | 0.25 | 1 | 0 | 16 | 0.0625 | 3 | 0.3333 | 7 | 0.4286 | 6 | 0.1667 | 4 | 0 | 20 | 0.25 |
| 14_4925301 | 6 | 0.1667 | 3 | 0 | 4 | 0.25 | NA | NA | 13 | 0.1538 | 2 | 0 | 6 | 0 | 7 | 0.2857 | 5 | 0.2 | 20 | 0.15 |
| 14_4931113 | NA | NA | 3 | 0.3333 | 2 | 0.5 | 1 | 1 | 6 | 0.5 | NA | NA | 2 | 1 | 1 | 0 | 3 | 0.6667 | 6 | 0.6667 |
| 14_4931133 | NA | NA | 2 | 1 | 2 | 1 | NA | NA | 4 | 1 | NA | NA | 2 | 1 | 1 | 1 | 1 | 1 | 4 | 1 |
| 14_4931139 | NA | NA | 2 | 0.5 | 1 | 0 | NA | NA | 3 | 0.3333 | NA | NA | 1 | 0 | 1 | 1 | 1 | 1 | 3 | 0.6667 |
| 14_4931149 | NA | NA | 3 | 0.3333 | 1 | 0 | 1 | 0 | 5 | 0.2 | NA | NA | 2 | 0.5 | 1 | 1 | 2 | 0 | 5 | 0.4 |
| 14_4941034 | NA | NA | 1 | 1 | 2 | 0 | 1 | 0 | 4 | 0.25 | 3 | 0.3333 | 2 | 1 | 3 | 0.3333 | NA | NA | 8 | 0.5 |
| 14_4941043 | NA | NA | 1 | 1 | 2 | 1 | 1 | 0 | 4 | 0.75 | 4 | 1 | 2 | 0.5 | 3 | 1 | NA | NA | 9 | 0.8889 |
| 14_4941045 | 1 | 1 | 1 | 0 | 2 | 0.5 | NA | NA | 4 | 0.5 | NA | NA | 1 | 0 | 1 | 1 | NA | NA | 2 | 0.5 |
| 14_4941050 | 1 | 1 | 1 | 1 | 2 | 0.5 | NA | NA | 4 | 0.75 | 3 | 0.3333 | 2 | 1 | 3 | 1 | NA | NA | 8 | 0.75 |
| 14_4941058 | NA | NA | 1 | 1 | 2 | 1 | 1 | 1 | 4 | 1 | 4 | 1 | 2 | 1 | 2 | 1 | NA | NA | 8 | 1 |
| 14_4941076 | 1 | 1 | 1 | 1 | 2 | 1 | 1 | 1 | 5 | 1 | 5 | 1 | 2 | 1 | 3 | 1 | NA | NA | 10 | 1 |
| 14_4942512 | 1 | 1 | 1 | 0 | 2 | 0.5 | 1 | 0 | 5 | 0.4 | 6 | 0.3333 | 2 | 0.5 | 5 | 0.2 | 2 | 0 | 15 | 0.2667 |
| 14_4942514 | 2 | 0 | 2 | 0.5 | 1 | 0 | 1 | 0 | 6 | 0.1667 | 6 | 0.3333 | 3 | 0 | 4 | 0 | 2 | 0.5 | 15 | 0.2 |
| 14_4942585 | 2 | 0.5 | 2 | 0 | 1 | 0 | 1 | 0 | 6 | 0.1667 | 4 | 0.25 | 4 | 0.25 | 4 | 0.25 | 1 | 0 | 13 | 0.2308 |
| 14_4942815 | 8 | 0 | 9 | 0 | 6 | 0 | 5 | 0.4 | 28 | 0.0714 | 8 | 0.25 | 8 | 0.125 | 8 | 0 | 5 | 0 | 29 | 0.1034 |
| 14_4942818 | 8 | 0.375 | 5 | 0.6 | 5 | 0.2 | 5 | 0.2 | 23 | 0.3478 | 4 | 0.25 | 7 | 0.8571 | 6 | 0.5 | 2 | 1 | 19 | 0.6316 |
| 14_4953624 | 1 | 1 | 1 | 0 | 5 | 0.2 | 3 | 0.3333 | 10 | 0.3 | NA | NA | 1 | 0 | 4 | 0.25 | 1 | 0 | 6 | 0.1667 |
| 14_4955107 | NA | NA | 2 | 0.5 | 5 | 0.2 | 3 | 0 | 10 | 0.2 | NA | NA | 1 | 0 | 3 | 0.6667 | 2 | 0 | 6 | 0.3333 |
| 14_4955119 | NA | NA | 3 | 1 | 5 | 1 | 3 | 1 | 11 | 1 | NA | NA | 1 | 1 | 3 | 1 | 2 | 1 | 6 | 1 |
| 14_4955121 | NA | NA | 3 | 1 | 4 | 0.25 | 2 | 0.5 | 9 | 0.5556 | NA | NA | 1 | 1 | 1 | 1 | 2 | 0.5 | 4 | 0.75 |
| 14_4955624 | NA | NA | 3 | 0.3333 | 3 | 0.6667 | 2 | 0 | 8 | 0.375 | NA | NA | 1 | 1 | 2 | 1 | 1 | 0 | 4 | 0.75 |
| 14_4955624 | NA | NA | 3 | 0.6667 | 4 | 0 | 3 | 0.6667 | 10 | 0.4 | NA | NA | 1 | 0 | 3 | 0.3333 | 1 | 0 | 5 | 0.2 |
| 14_4955635 | NA | NA | 3 | 1 | 5 | 0.6 | 3 | 0.6667 | 11 | 0.7273 | NA | NA | 1 | 1 | 3 | 1 | 1 | 0 | 5 | 0.8 |
| 14_4955641 | NA | NA | 3 | 0.3333 | 5 | 0.2 | 3 | 1 | 11 | 0.4545 | NA | NA | 1 | 1 | 3 | 0 | NA | NA | 4 | 0.25 |
| 14_4991944 | NA | NA | 5 | 0.8 | 2 | 1 | 1 | 1 | 8 | 0.875 | NA | NA | 2 | 1 | 3 | 0.6667 | 1 | 1 | 6 | 0.8333 |
| 14_4991970 | 1 | 1 | 5 | 0.6 | 1 | 0 | 2 | 0.5 | 9 | 0.5556 | 1 | 1 | 2 | 0.5 | 3 | 0.6667 | 1 | 1 | 7 | 0.7143 |
| 14_4991979 | 1 | 1 | 5 | 0.8 | 4 | 0.75 | 2 | 0.5 | 12 | 0.75 | NA | NA | 1 | 1 | 2 | 1 | 1 | 0 | 4 | 0.75 |
| 14_4998894 | NA | NA | 4 | 0.5 | 4 | 0.5 | 2 | 0.5 | 10 | 0.5 | NA | NA | 2 | 0.5 | 2 | 0.5 | 1 | 1 | 5 | 0.6 |
| 14_4998902 | NA | NA | 3 | 0.6667 | 4 | 0.75 | 2 | 0.5 | 9 | 0.6667 | NA | NA | 1 | 0 | 3 | 0.3333 | 1 | 1 | 5 | 0.4 |
| 14_5001038 | NA | NA | 3 | 0.6667 | 3 | 0.6667 | 2 | 0 | 8 | 0.5 | NA | NA | NA | NA | 3 | 0.3333 | 1 | 1 | 4 | 0.5 |
| 14_5001063 | NA | NA | 3 | 0.3333 | 3 | 0.3333 | 2 | 1 | 8 | 0.5 | NA | NA | 1 | 1 | 3 | 0.3333 | 1 | 0 | 5 | 0.4 |
| 14_5001065 | NA | NA | 2 | 0.5 | 2 | 1 | 1 | 1 | 5 | 0.8 | NA | NA | 1 | 1 | 1 | 1 | 1 | 1 | 3 | 1 |
| 14_5001068 | NA | NA | 2 | 1 | 3 | 0.6667 | 2 | 0.5 | 7 | 0.7143 | NA | NA | 1 | 1 | 3 | 1 | 1 | 1 | 5 | 1 |
| 14_5001084 | NA | NA | NA | NA | 2 | 0.5 | NA | NA | 2 | 0.5 | NA | NA | 1 | 0 | 2 | 0.5 | 1 | 1 | 4 | 0.5 |
| 14_5001092 | NA | NA | 1 | 0 | 3 | 0.3333 | 2 | 0 | 6 | 0.1667 | NA | NA | 1 | 0 | 2 | 0.5 | 1 | 1 | 4 | 0.5 |
| 14_5001116 | NA | NA | 1 | 0 | 2 | 1 | 2 | 0.5 | 5 | 0.6 | NA | NA | 1 | 1 | 1 | 1 | 1 | 1 | 3 | 1 |
| 14_5001140 | NA | NA | 1 | 0 | 2 | 0.5 | 2 | 0.5 | 5 | 0.4 | NA | NA | NA | NA | 1 | 1 | 1 | 1 | 2 | 1 |
| 14_5100312 | 12 | 0.1667 | 4 | 0 | 4 | 0 | 3 | 0 | 23 | 0.087 | 5 | 0.2 | 3 | 0 | 6 | 0 | 5 | 0 | 19 | 0.0526 |
| 14_5100314 | 8 | 0.375 | 4 | 0 | 5 | 0.4 | 3 | 0.3333 | 20 | 0.3 | 3 | 0 | 3 | 0 | 3 | 0.3333 | 5 | 0 | 14 | 0.0714 |
| 14_5100315 | 7 | 0.8571 | 2 | 1 | 3 | 1 | 3 | 1 | 15 | 0.9333 | 2 | 0 | 2 | 1 | 3 | 1 | 2 | 1 | 9 | 0.7778 |
| 14_5100316 | 11 | 0.2727 | 4 | 0.5 | 5 | 0.6 | 3 | 0.6667 | 23 | 0.4348 | 5 | 0.4 | 3 | 0.3333 | 5 | 0.8 | 5 | 0.6 | 18 | 0.5556 |
| 14_5100322 | 12 | 0.25 | 3 | 0 | 5 | 0.2 | 3 | 0.6667 | 23 | 0.2609 | 5 | 0.2 | 3 | 0.6667 | 5 | 0.4 | 5 | 0.6 | 18 | 0.4444 |
| 14_5100328 | 12 | 0.1667 | 3 | 0 | 4 | 0 | 1 | 0 | 20 | 0.1 | 3 | 0.3333 | 3 | 0 | 3 | 0 | 5 | 0 | 14 | 0.0714 |

| | | | | | | | | | | | | | | | | | | | | |
|---|---|---|---|---|---|---|---|---|---|---|---|---|---|---|---|---|---|---|---|---|
| 14_510033 | 11 | 0.1818 | 4 | 0 | 5 | 0 | 3 | 0.6667 | 23 | 0.1739 | 5 | 0.6 | 3 | 0 | 4 | 0.5 | 5 | 0.2 | 17 | 0.3529 |
| 14_511180 | NA | NA | NA | NA | 1 | 1 | 2 | 0.5 | 3 | 0.6667 | NA | NA | NA | NA | NA | NA | 1 | 1 | 1 | 1 |
| 14_511181 | NA | NA | NA | NA | NA | NA | 2 | 0.5 | 2 | 0.5 | NA | NA | NA | NA | NA | NA | 1 | 1 | 1 | 1 |
| 14_511530 | NA | NA | 1 | 1 | NA | NA | NA | NA | 1 | 1 | NA | NA | NA | NA | NA | NA | 1 | 1 | 1 | 1 |
| 14_511637 | 6 | 0.1667 | 2 | 0 | 9 | 0.2222 | 6 | 0 | 23 | 0.1304 | 5 | 0 | 9 | 0.1111 | 3 | 0 | 6 | 0.1667 | 23 | 0.087 |
| 14_511640 | 8 | 0.375 | 4 | 0.25 | 9 | 0.1111 | 7 | 0.5714 | 28 | 0.3214 | 5 | 0.6 | 10 | 0.1 | 4 | 0.5 | 8 | 0.25 | 27 | 0.2963 |
| 14_511640 | 8 | 1 | 4 | 1 | 9 | 1 | 7 | 1 | 28 | 1 | 5 | 1 | 9 | 0.7778 | 5 | 1 | 8 | 1 | 27 | 0.9259 |
| 14_511641 | 8 | 0.75 | 5 | 0.4 | 10 | 0.4 | 7 | 0.1429 | 30 | 0.4333 | 5 | 0.4 | 9 | 0.4444 | 6 | 0.3333 | 4 | 0.75 | 24 | 0.4583 |
| 14_511642 | 8 | 0.875 | 5 | 1 | 11 | 1 | 7 | 1 | 31 | 0.9677 | 5 | 0.8 | 8 | 1 | 5 | 1 | 8 | 1 | 26 | 0.9615 |
| 14_511648 | 8 | 0.125 | 12 | 0.3333 | 9 | 0.1111 | 14 | 0.1429 | 43 | 0.186 | 11 | 0 | 18 | 0.1111 | 19 | 0.1579 | 10 | 0.2 | 58 | 0.1207 |
| 14_511649 | 8 | 0.125 | 12 | 0.1667 | 7 | 0.2857 | 11 | 0.0909 | 38 | 0.1579 | 11 | 0 | 16 | 0.0625 | 19 | 0.2105 | 11 | 0.1818 | 57 | 0.1228 |
| 14_511652 | 6 | 0.1667 | 8 | 0.5 | 8 | 0.375 | 12 | 0.4167 | 34 | 0.3824 | 9 | 0.5556 | 10 | 0.9 | 15 | 0.4 | 8 | 0.25 | 42 | 0.5238 |
| 14_511653 | 8 | 0.25 | 11 | 0.1818 | 6 | 0 | 14 | 0 | 39 | 0.1026 | 10 | 0 | 16 | 0.1875 | 20 | 0 | 10 | 0.4 | 56 | 0.125 |
| 14_511889 | 1 | 0 | 4 | 0.25 | 3 | 0.3333 | NA | NA | 8 | 0.25 | 4 | 0 | 2 | 0 | 5 | 0.4 | 3 | 0.6667 | 14 | 0.2857 |
| 14_511890 | 1 | 0 | 4 | 0.5 | 2 | 0 | NA | NA | 7 | 0.2857 | 4 | 1 | 2 | 0 | 4 | 0.25 | 1 | 1 | 11 | 0.5455 |
| 14_511892 | 1 | 0 | 2 | 0.5 | 3 | 0.3333 | NA | NA | 6 | 0.3333 | 1 | 1 | 2 | 0 | 5 | 0.2 | 2 | 0 | 10 | 0.2 |
| 14_511893 | 1 | 1 | 4 | 0.5 | 2 | 0 | NA | NA | 7 | 0.4286 | 4 | 0 | 2 | 0 | 3 | 0.3333 | 2 | 0 | 11 | 0.0909 |
| 14_511893 | 1 | 1 | 4 | 0.75 | 3 | 0.3333 | NA | NA | 8 | 0.625 | 4 | 0.75 | 2 | 0.5 | 5 | 0.8 | 4 | 0 | 15 | 0.5333 |
| 14_512019 | 1 | 1 | NA | NA | NA | NA | NA | NA | 1 | 1 | 1 | 0 | NA | NA | 1 | 0 | 1 | 1 | 3 | 0.3333 |
| 14_513424 | 3 | 0 | 4 | 0.25 | 6 | 0 | 6 | 0.1667 | 19 | 0.1053 | 3 | 0 | 2 | 0 | 4 | 0.25 | 8 | 0 | 17 | 0.0588 |
| 14_513425 | 4 | 0.75 | 6 | 0.8333 | 6 | 1 | 6 | 1 | 22 | 0.9091 | 2 | 0.5 | 2 | 1 | 2 | 0.5 | 5 | 0.8 | 11 | 0.7273 |
| 14_513426 | 3 | 0 | 5 | 0.6 | 6 | 1 | 6 | 1 | 20 | 0.75 | 2 | 1 | 2 | 1 | 4 | 0.75 | 8 | 1 | 16 | 0.9375 |
| 14_513429 | 4 | 0.25 | 7 | 0.2857 | 5 | 0.4 | 9 | 0.1111 | 25 | 0.24 | 5 | 0.2 | 2 | 0.5 | 4 | 0.25 | 7 | 0.1429 | 18 | 0.2222 |
| 14_521320 | 1 | 0 | NA | NA | NA | NA | 1 | 1 | 2 | 0.5 | 2 | 0 | NA | NA | 4 | 0.25 | 4 | 0 | 10 | 0.1 |
| 14_521324 | 1 | 0 | NA | NA | NA | NA | 2 | 0.5 | 3 | 0.3333 | 1 | 0 | NA | NA | 4 | 0 | 4 | 0.5 | 9 | 0.2222 |
| 14_521326 | 2 | 0.5 | NA | NA | 1 | 1 | 2 | 1 | 5 | 0.8 | 3 | 0.6667 | NA | NA | 5 | 0.4 | 2 | 1 | 10 | 0.6 |
| 14_521328 | 2 | 0 | NA | NA | 1 | 0 | 3 | 0.3333 | 6 | 0.1667 | 3 | 0 | NA | NA | 4 | 0 | 5 | 0.2 | 12 | 0.0833 |
| 14_521330 | 2 | 0.5 | NA | NA | 1 | 0 | 3 | 0 | 6 | 0.1667 | 3 | 0 | NA | NA | 5 | 0.2 | 4 | 0 | 12 | 0.0833 |
| 14_521330 | 2 | 0.5 | NA | NA | 1 | 1 | 3 | 0.6667 | 6 | 0.6667 | 3 | 0.3333 | NA | NA | 5 | 0.4 | 4 | 0.75 | 12 | 0.5 |
| 14_521331 | 2 | 1 | NA | NA | 1 | 1 | 3 | 1 | 6 | 1 | 2 | 0.5 | NA | NA | 4 | 0.75 | 5 | 0.6 | 11 | 0.6364 |
| 14_521331 | 1 | 0 | NA | NA | 1 | 0 | 3 | 0.3333 | 5 | 0.2 | 3 | 0 | NA | NA | 5 | 0.2 | 5 | 0 | 13 | 0.0769 |
| 14_521428 | 124 | 0.121 | 114 | 0.1316 | 87 | 0.1609 | 107 | 0.1402 | 432 | 0.1366 | 105 | 0.1429 | 81 | 0.1481 | 82 | 0.1585 | 107 | 0.1402 | 375 | 0.1467 |
| 14_521436 | 102 | 0.2255 | 96 | 0.3125 | 90 | 0.2 | 106 | 0.2547 | 394 | 0.2487 | 101 | 0.2772 | 82 | 0.2683 | 77 | 0.2857 | 92 | 0.2935 | 352 | 0.2812 |
| 14_522265 | NA | NA | NA | NA | 1 | 1 | NA | NA | 1 | 1 | 1 | 1 | NA | NA | NA | NA | NA | NA | 1 | 1 |
| 14_522266 | NA | NA | NA | NA | 1 | 1 | NA | NA | 1 | 1 | 2 | 0.5 | 1 | 1 | NA | NA | NA | NA | 3 | 0.6667 |
| 14_522267 | NA | NA | NA | NA | 1 | 1 | NA | NA | 1 | 1 | 2 | 0.5 | 1 | 0 | NA | NA | NA | NA | 3 | 0.3333 |
| 14_522270 | NA | NA | NA | NA | 1 | 1 | NA | NA | 1 | 1 | 2 | 0 | 2 | 0.5 | NA | NA | NA | NA | 4 | 0.25 |
| 14_522348 | 1 | 0 | NA | NA | 1 | 1 | NA | NA | 2 | 0.5 | 2 | 0 | 2 | 0.5 | NA | NA | NA | NA | 4 | 0.25 |
| 14_522428 | 2 | 0 | 1 | 0 | 2 | 0 | 3 | 0.3333 | 8 | 0.125 | 2 | 1 | 4 | 0.25 | NA | NA | 1 | 0 | 7 | 0.4286 |
| 14_522428 | 2 | 0 | 1 | 0 | 2 | 0 | 3 | 0.3333 | 8 | 0.125 | 2 | 0.5 | 4 | 0.25 | NA | NA | 1 | 0 | 7 | 0.2857 |
| 14_522438 | 3 | 0 | 5 | 0 | 2 | 0.5 | 6 | 0 | 16 | 0.0625 | 4 | 0 | 5 | 0.2 | 5 | 0 | 1 | 0 | 15 | 0.0667 |
| 14_522442 | 4 | 0 | 3 | 0 | 3 | 0.3333 | 7 | 0 | 17 | 0.0588 | 2 | 0 | 6 | 0.1667 | 3 | 0 | 1 | 0 | 12 | 0.0833 |
| 14_522461 | 2 | 0 | 3 | 0 | 2 | 0.5 | 5 | 0.4 | 12 | 0.25 | 1 | 0 | 3 | 0.6667 | 3 | 0 | NA | NA | 7 | 0.2857 |
| 14_522463 | 2 | 0.5 | 2 | 0.5 | 3 | 0.6667 | 7 | 0.1429 | 14 | 0.3571 | 3 | 0.3333 | 4 | 0.5 | 3 | 0.6667 | NA | NA | 10 | 0.5 |
| 14_522464 | 2 | 1 | 2 | 1 | 2 | 0 | 5 | 1 | 11 | 0.8182 | 3 | 0.6667 | 4 | 0.25 | 4 | 0.75 | NA | NA | 11 | 0.5455 |
| 14_522464 | 2 | 0 | 3 | 0 | 2 | 0 | 6 | 0.1667 | 13 | 0.0769 | 3 | 0.3333 | 4 | 0 | 4 | 0 | NA | NA | 11 | 0.0909 |
| 14_522469 | 3 | 0 | 2 | 1 | 2 | 0 | 5 | 0.8 | 12 | 0.5 | 3 | 0.3333 | 2 | 0.5 | 3 | 1 | NA | NA | 8 | 0.625 |
| 14_522470 | 2 | 0 | 3 | 0.3333 | 1 | 1 | 4 | 0.75 | 10 | 0.5 | 2 | 0.5 | 2 | 0.5 | 3 | 0.3333 | NA | NA | 7 | 0.4286 |
| 14_522470 | 2 | 0.5 | 3 | 0 | 2 | 0 | 5 | 0.2 | 12 | 0.1667 | 3 | 0 | 2 | 0.5 | 3 | 0 | NA | NA | 8 | 0.125 |
| 14_523980 | 9 | 0.2222 | 5 | 0 | 5 | 0 | 6 | 0.1667 | 25 | 0.12 | 7 | 0.1429 | 4 | 0 | 6 | 0.1667 | 9 | 0.2222 | 26 | 0.1538 |
| 14_523985 | 8 | 0.25 | 4 | 0.25 | 3 | 0.3333 | 4 | 0 | 19 | 0.2105 | 5 | 0.2 | 2 | 0 | 8 | 0.25 | 10 | 0.3 | 25 | 0.24 |
| 14_523988 | 7 | 0.5714 | 5 | 0.4 | 4 | 0 | 3 | 0.3333 | 19 | 0.3684 | 6 | 0.6667 | 3 | 1 | 7 | 0.8571 | 8 | 0.625 | 24 | 0.75 |
| 14_523989 | 7 | 0.4286 | 5 | 0.4 | 4 | 0.5 | 3 | 0.6667 | 19 | 0.4737 | 5 | 0.8 | 3 | 1 | 7 | 0.4286 | 7 | 0.2857 | 22 | 0.5455 |
| 14_523990 | 5 | 0.4 | 4 | 0.75 | 3 | 0.3333 | 2 | 0 | 14 | 0.4286 | 5 | 0.8 | 2 | 1 | 3 | 1 | 8 | 0.75 | 18 | 0.8333 |
| 14_523991 | 7 | 1 | 6 | 0.8333 | 4 | 1 | 2 | 1 | 19 | 0.9474 | 4 | 1 | 3 | 1 | 5 | 1 | 9 | 0.7778 | 21 | 0.9048 |
| 14_524356 | 4 | 0.25 | 10 | 0.4 | 9 | 0.2222 | 6 | 0.3333 | 29 | 0.3103 | 6 | 0.1667 | 4 | 0 | 3 | 0 | 3 | 0 | 16 | 0.0625 |
| 14_524357 | 3 | 0.3333 | 9 | 0.4444 | 9 | 0.4444 | 6 | 0.6667 | 27 | 0.4815 | 5 | 0.6 | 4 | 0.75 | 5 | 0.8 | 3 | 0.3333 | 17 | 0.6471 |
| 14_524358 | 3 | 0 | 9 | 0.1111 | 9 | 0.1111 | 5 | 0 | 26 | 0.0769 | 5 | 0.2 | 3 | 0.6667 | 4 | 0.25 | 2 | 0 | 14 | 0.2857 |
| 14_524358 | 2 | 0.5 | 5 | 0.2 | 4 | 0.5 | 3 | 0 | 14 | 0.2857 | 4 | 1 | 2 | 0 | 4 | 0.25 | 1 | 0 | 11 | 0.4545 |
| 14_524358 | 1 | 0 | NA | NA | NA | NA | 1 | 1 | 2 | 0.5 | NA | NA | NA | NA | NA | NA | 1 | 1 | 1 | 1 |
| 14_524359 | 2 | 0 | 7 | 0.1429 | 4 | 0 | 3 | 0 | 16 | 0.0625 | 4 | 0 | 2 | 0 | 3 | 0.3333 | 2 | 0 | 11 | 0.0909 |
| 14_524359 | 3 | 0 | 9 | 0.2222 | 9 | 0.2222 | 6 | 0.3333 | 27 | 0.2222 | 5 | 0.8 | 2 | 0.5 | 3 | 0.3333 | 3 | 0.3333 | 13 | 0.5385 |
| 14_524359 | 3 | 1 | 7 | 1 | 8 | 1 | 6 | 1 | 24 | 1 | 5 | 1 | 3 | 1 | 2 | 1 | 3 | 1 | 13 | 1 |
| 14_524360 | 4 | 0.25 | 9 | 0.3333 | 8 | 0.375 | 6 | 0.5 | 27 | 0.3704 | 6 | 0 | 3 | 0 | 3 | 0 | 3 | 0.3333 | 15 | 0.0667 |
| 14_524360 | 3 | 0 | 8 | 0 | 9 | 0.2222 | 7 | 0.2857 | 27 | 0.1481 | 6 | 0.5 | 3 | 0.3333 | 4 | 0 | 3 | 0 | 16 | 0.25 |

| | | | | | | | | | | | | | | | | | | | | |
|---|---|---|---|---|---|---|---|---|---|---|---|---|---|---|---|---|---|---|---|---|
| 14_524361 | 3 | 0.6667 | 10 | 1 | 6 | 1 | 6 | 1 | 25 | 0.96 | 5 | 1 | 3 | 1 | 4 | 1 | 3 | 1 | 15 | 1 |
| 14_524363 | 3 | 0.3333 | 7 | 0.4286 | 8 | 0.375 | 5 | 0.8 | 23 | 0.4783 | 3 | 0 | 2 | 0 | 2 | 0 | 3 | 0.3333 | 10 | 0.1 |
| 14_524423 | 1 | 1 | 4 | 0.75 | 4 | 0.25 | 3 | 0.6667 | 12 | 0.5833 | 2 | 1 | 1 | 1 | NA | NA | 1 | 0 | 4 | 0.75 |
| 14_524423 | 1 | 0 | 3 | 0 | 4 | 0 | 2 | 0.5 | 10 | 0.1 | 2 | 0 | 1 | 0 | NA | NA | 1 | 1 | 4 | 0.25 |
| 14_524424 | 1 | 1 | 4 | 0 | 3 | 0.6667 | 3 | 0 | 11 | 0.2727 | 1 | 1 | 1 | 1 | NA | NA | NA | NA | 2 | 1 |
| 14_524424 | 1 | 0 | 4 | 0.25 | 3 | 0.6667 | 3 | 0.6667 | 11 | 0.4545 | 2 | 0 | 1 | 1 | NA | NA | 1 | 0 | 4 | 0.25 |
| 14_524427 | NA | NA | 2 | 0.5 | 3 | 0 | 2 | 0 | 7 | 0.1429 | 2 | 1 | 1 | 0 | NA | NA | 1 | 0 | 4 | 0.5 |
| 14_524427 | 1 | 1 | 3 | 1 | 3 | 1 | 2 | 0.5 | 9 | 0.8889 | NA | NA | NA | NA | NA | NA | 1 | 1 | 1 | 1 |
| 14_524993 | 7 | 0 | 2 | 0 | 5 | 0 | 4 | 0.25 | 18 | 0.0556 | 6 | 0.3333 | 9 | 0 | 4 | 0.25 | 8 | 0 | 27 | 0.1111 |
| 14_525231 | 2 | 0 | 3 | 0.3333 | 1 | 0 | 2 | 0 | 8 | 0.125 | 1 | 1 | 2 | 0.5 | 4 | 0.25 | 1 | 0 | 8 | 0.375 |
| 14_525232 | 2 | 0.5 | 4 | 0 | 2 | 0 | 3 | 0 | 11 | 0.0909 | 2 | 0 | 3 | 0.6667 | 4 | 0 | NA | NA | 9 | 0.2222 |
| 14_525242 | 4 | 0.25 | 4 | 0 | 2 | 0 | 3 | 0 | 13 | 0.0769 | 2 | 0 | 2 | 0.5 | 3 | 0 | 2 | 0 | 9 | 0.1111 |
| 14_525243 | 3 | 0.3333 | 4 | 0.25 | 2 | 0 | 3 | 0.3333 | 12 | 0.25 | 2 | 0.5 | 3 | 0.3333 | 4 | 0.25 | 2 | 0.5 | 11 | 0.3636 |
| 14_525248 | 4 | 0 | 3 | 0 | 2 | 0.5 | 3 | 0 | 12 | 0.0833 | 3 | 0 | 4 | 0 | 3 | 0 | 5 | 0.2 | 15 | 0.0667 |
| 14_525249 | 3 | 0.3333 | 4 | 0 | 2 | 0 | 2 | 0 | 11 | 0.0909 | 1 | 0 | 3 | 0 | 4 | 0.25 | 4 | 0.5 | 12 | 0.25 |
| 14_525295 | 6 | 0 | 7 | 0.1429 | 3 | 0 | 2 | 0 | 18 | 0.0556 | 5 | 0.2 | 4 | 0.75 | 7 | 0.4286 | 5 | 0.2 | 21 | 0.381 |
| 14_525308 | 5 | 0 | 7 | 0.1429 | 5 | 0.2 | 2 | 0 | 19 | 0.1053 | 3 | 0 | 5 | 0 | 6 | 0.1667 | 5 | 0 | 19 | 0.0526 |
| 14_525323 | 1 | 1 | 2 | 0 | 4 | 0.5 | NA | NA | 7 | 0.4286 | 2 | 0.5 | 4 | 0.5 | 4 | 0.5 | 2 | 0.5 | 12 | 0.5 |
| 14_525487 | 5 | 0 | 7 | 0.1429 | 7 | 0.1429 | 12 | 0 | 31 | 0.0645 | 5 | 0 | 9 | 0.1111 | 8 | 0.125 | 6 | 0 | 28 | 0.0714 |
| 14_525491 | 5 | 0.4 | 7 | 0.5714 | 5 | 0.4 | 8 | 0.625 | 25 | 0.52 | 5 | 0.6 | 6 | 0.8333 | 7 | 0.5714 | 5 | 0.2 | 23 | 0.5652 |
| 14_525491 | 3 | 1 | 6 | 0.6667 | 5 | 1 | 8 | 0.75 | 22 | 0.8182 | 4 | 0.5 | 6 | 0.5 | 5 | 1 | 4 | 0.75 | 19 | 0.6842 |
| 14_525493 | 4 | 0.25 | 7 | 0.7143 | 4 | 0.75 | 8 | 0.625 | 23 | 0.6087 | 3 | 0.6667 | 5 | 0.4 | 7 | 0.2857 | 5 | 0.8 | 20 | 0.5 |
| 14_525493 | 2 | 0 | 6 | 0 | 4 | 0 | 8 | 0.25 | 20 | 0.1 | 3 | 0 | 2 | 0 | 6 | 0.1667 | 3 | 0 | 14 | 0.0714 |
| 14_525494 | 4 | 1 | 6 | 1 | 3 | 0.6667 | 6 | 1 | 19 | 0.9474 | 2 | 1 | 6 | 1 | 6 | 1 | 3 | 1 | 17 | 1 |
| 14_525494 | 3 | 1 | 8 | 0.25 | 4 | 0.25 | 7 | 0.1429 | 22 | 0.3182 | 3 | 0 | 6 | 0.1667 | 6 | 0.3333 | 4 | 0.25 | 19 | 0.2105 |
| 14_525494 | 2 | 0.5 | 3 | 0.3333 | 4 | 0.5 | 6 | 0.8333 | 15 | 0.6 | 2 | 1 | 5 | 0.6 | 5 | 0.6 | 3 | 0.3333 | 15 | 0.6 |
| 14_525495 | 3 | 0 | 5 | 0 | 2 | 0.5 | 7 | 0 | 17 | 0.0588 | 3 | 0 | 3 | 0.3333 | 7 | 0 | 5 | 0 | 18 | 0.0556 |
| 14_525495 | 4 | 0.5 | 7 | 0.2857 | 3 | 0.6667 | 7 | 0.2857 | 21 | 0.381 | 3 | 0 | 5 | 0 | 6 | 0.3333 | 4 | 0 | 18 | 0.1111 |
| 14_525496 | 2 | 1 | 7 | 0.1429 | 1 | 0 | 6 | 0.3333 | 16 | 0.3125 | 3 | 0.3333 | 5 | 0 | 4 | 0 | 5 | 0.4 | 17 | 0.1765 |
| 14_525496 | 4 | 1 | 7 | 0.8571 | 3 | 1 | 5 | 1 | 19 | 0.9474 | 3 | 1 | 4 | 1 | 6 | 0.8333 | 4 | 1 | 17 | 0.9412 |
| 14_525496 | 4 | 0.75 | 8 | 0.875 | 3 | 0.6667 | 7 | 0.5714 | 22 | 0.7273 | 3 | 0.6667 | 6 | 0.8333 | 7 | 1 | 5 | 0.8 | 21 | 0.8571 |
| 14_525497 | 4 | 1 | 8 | 0.875 | 4 | 1 | 7 | 0.8571 | 23 | 0.913 | 3 | 1 | 6 | 0.8333 | 5 | 1 | 5 | 1 | 19 | 0.9474 |
| 14_525497 | 4 | 1 | 7 | 0.7143 | 3 | 1 | 6 | 0.6667 | 20 | 0.8 | 2 | 0.5 | 4 | 1 | 6 | 0.8333 | 5 | 0.4 | 17 | 0.7059 |
| 14_525500 | 3 | 0 | 7 | 0 | 2 | 0.5 | 5 | 0 | 17 | 0.0588 | 2 | 0.5 | 4 | 0 | 5 | 0.2 | 3 | 0 | 14 | 0.1429 |
| 14_525501 | 3 | 0 | 8 | 0.25 | 2 | 0 | 5 | 0.6 | 18 | 0.2778 | 2 | 0.5 | 3 | 0 | 5 | 0 | 4 | 0 | 14 | 0.0714 |
| 14_525504 | 4 | 0.25 | 7 | 0 | 2 | 0 | 6 | 0 | 19 | 0.0526 | 1 | 0 | 3 | 0 | 6 | 0 | 5 | 0.2 | 15 | 0.0667 |
| 14_525505 | 4 | 0.75 | 7 | 0.2857 | 1 | 1 | 5 | 0.4 | 17 | 0.4706 | 1 | 1 | 3 | 0 | 2 | 0 | 3 | 0 | 9 | 0.1111 |
| 14_525505 | 4 | 0 | 7 | 0.1429 | 2 | 0 | 5 | 0 | 18 | 0.0556 | 1 | 0 | 3 | 0 | 5 | 0.2 | 5 | 0 | 14 | 0.0714 |
| 14_525507 | 3 | 1 | 7 | 0.8571 | 2 | 0.5 | 4 | 1 | 16 | 0.875 | 1 | 1 | 3 | 1 | 6 | 1 | 5 | 1 | 15 | 1 |
| 14_525511 | 3 | 0 | 5 | 0.4 | 1 | 0 | 4 | 0.5 | 13 | 0.3077 | 1 | 0 | NA | NA | 4 | 0.25 | 4 | 0.25 | 9 | 0.2222 |
| 14_525513 | 2 | 1 | 5 | 0.2 | 1 | 0 | 2 | 0.5 | 10 | 0.4 | 1 | 0 | 1 | 1 | 5 | 0.6 | 4 | 0.5 | 11 | 0.5455 |
| 14_525514 | 2 | 0.5 | 6 | 0.5 | 1 | 0 | 2 | 0.5 | 11 | 0.4545 | 1 | 1 | 1 | 1 | 4 | 0.25 | 3 | 0.6667 | 9 | 0.5556 |
| 14_525521 | 2 | 1 | 3 | 1 | NA | NA | 1 | 1 | 6 | 1 | 1 | 1 | NA | NA | 3 | 1 | 2 | 1 | 6 | 1 |
| 14_525522 | 2 | 1 | 3 | 1 | NA | NA | 2 | 1 | 7 | 1 | 1 | 1 | 1 | 1 | 4 | 1 | 2 | 1 | 8 | 1 |
| 14_525524 | 1 | 0 | 3 | 0.3333 | NA | NA | 1 | 1 | 5 | 0.4 | 1 | 0 | 1 | 0 | 3 | 0.6667 | 1 | 0 | 6 | 0.3333 |
| 14_525525 | 1 | 1 | 2 | 0 | NA | NA | 2 | 0 | 5 | 0.2 | 1 | 0 | 1 | 1 | 3 | 0.3333 | 1 | 1 | 6 | 0.5 |
| 14_545003 | 11 | 0.1818 | 2 | 0 | 4 | 0 | 5 | 0.2 | 22 | 0.1364 | 7 | 0.1429 | 4 | 0 | 3 | 0 | 4 | 0 | 18 | 0.0556 |
| 14_545053 | 14 | 0.2143 | 4 | 0 | 5 | 0.2 | 4 | 0.25 | 27 | 0.1852 | 6 | 0.1667 | 5 | 0.4 | 4 | 0 | 9 | 0.3333 | 24 | 0.25 |
| 14_545053 | 17 | 0.1176 | 4 | 0.25 | 7 | 0 | 5 | 0 | 33 | 0.0909 | 6 | 0.1667 | 4 | 0 | 7 | 0 | 9 | 0.1111 | 26 | 0.0769 |
| 14_545054 | 14 | 0.0714 | 6 | 0 | 6 | 0.1667 | 5 | 0.2 | 31 | 0.0968 | 7 | 0.2857 | 5 | 0 | 6 | 0 | 8 | 0.5 | 26 | 0.2308 |
| 14_545055 | 9 | 0.4444 | 4 | 0 | 2 | 1 | 1 | 1 | 16 | 0.4375 | 4 | 0 | 3 | 0.3333 | 4 | 1 | 5 | 0.6 | 16 | 0.5 |
| 14_545055 | 15 | 0.2 | 6 | 0.1667 | 7 | 0 | 5 | 0.2 | 33 | 0.1515 | 7 | 0.1429 | 5 | 0.4 | 8 | 0.125 | 10 | 0.2 | 30 | 0.2 |
| 14_545058 | 12 | 0.0833 | 4 | 0.25 | 6 | 0.5 | 4 | 0 | 26 | 0.1923 | 9 | 0 | 4 | 0.25 | 7 | 0.1429 | 7 | 0.2857 | 27 | 0.1481 |
| 14_545305 | NA | NA | 2 | 0.5 | 1 | 0 | 2 | 0.5 | 5 | 0.4 | 1 | 0 | 1 | 0 | NA | NA | 1 | 1 | 3 | 0.3333 |
| 14_545310 | NA | NA | 2 | 0.5 | NA | NA | NA | NA | 2 | 0.5 | 1 | 1 | 1 | 1 | NA | NA | NA | NA | 2 | 1 |
| 14_545314 | NA | NA | 2 | 0.5 | NA | NA | NA | NA | 2 | 0.5 | 1 | 1 | 1 | 1 | NA | NA | 1 | 0 | 3 | 0.6667 |
| 14_545314 | NA | NA | 2 | 0.5 | NA | NA | NA | NA | 2 | 0.5 | 1 | 1 | 1 | 1 | NA | NA | 1 | 1 | 3 | 1 |
| 14_545314 | NA | NA | 2 | 0.5 | NA | NA | NA | NA | 2 | 0.5 | 1 | 1 | 1 | 1 | NA | NA | 1 | 1 | 3 | 1 |
| 14_545318 | NA | NA | 2 | 0.5 | NA | NA | NA | NA | 2 | 0.5 | 1 | 1 | 1 | 0 | NA | NA | 1 | 1 | 3 | 0.6667 |
| 14_546059 | 5 | 0.6 | 6 | 0.1667 | NA | NA | 4 | 0 | 15 | 0.2667 | 5 | 0.8 | 7 | 0.5714 | 4 | 0 | 7 | 0.2857 | 23 | 0.4348 |
| 14_546060 | 7 | 0.7143 | 9 | 0.8889 | 2 | 1 | 8 | 0.875 | 26 | 0.8462 | 8 | 1 | 6 | 0.6667 | 11 | 0.8182 | 9 | 0.7778 | 34 | 0.8235 |
| 14_546060 | 2 | 0.5 | 4 | 0.5 | 2 | 0.5 | 4 | 0 | 12 | 0.3333 | 5 | 0.2 | 8 | 0.25 | 9 | 0.3333 | 7 | 0 | 29 | 0.2069 |
| 14_546061 | 1 | 0 | 3 | 0.3333 | 1 | 0 | NA | NA | 5 | 0.2 | NA | NA | 3 | 0 | 1 | 1 | NA | NA | 4 | 0.25 |
| 14_546671 | 31 | 0.1613 | 15 | 0.2 | 18 | 0.2222 | 21 | 0.2381 | 85 | 0.2 | 17 | 0.2353 | 23 | 0.2609 | 16 | 0.3125 | 18 | 0.1111 | 74 | 0.2297 |
| 14_546778 | 2 | 1 | NA | NA | 1 | 1 | 2 | 0.5 | 5 | 0.8 | NA | NA | 1 | 0 | 1 | 1 | NA | NA | 2 | 0.5 |

| | | | | | | | | | | | | | | | | | | | | |
|---|---|---|---|---|---|---|---|---|---|---|---|---|---|---|---|---|---|---|---|---|
| 14_5467793 | 1 | 1 | NA | NA | 2 | 1 | 2 | 0.5 | 5 | 0.8 | NA | NA | 1 | 1 | 1 | 1 | NA | NA | 2 | 1 |
| 14_5467816 | 2 | 0.5 | NA | NA | 2 | 1 | 2 | 0.5 | 6 | 0.6667 | NA | NA | 1 | 1 | 1 | 1 | NA | NA | 2 | 1 |
| 14_5467887 | 1 | 1 | NA | NA | 2 | 1 | 2 | 0 | 5 | 0.6 | NA | NA | 1 | 0 | 1 | 1 | NA | NA | 2 | 0.5 |
| 14_5467984 | 3 | 0 | NA | NA | 1 | 0 | 1 | 1 | 5 | 0.2 | 1 | 1 | 3 | 0.3333 | NA | NA | 2 | 0 | 6 | 0.3333 |
| 14_5470540 | 6 | 1 | 5 | 1 | 2 | 1 | 1 | 1 | 14 | 1 | 4 | 1 | 1 | 1 | 2 | 1 | 4 | 0.75 | 11 | 0.9091 |
| 14_5470548 | 6 | 0.1667 | 6 | 0.3333 | 2 | 0 | 2 | 0 | 16 | 0.1875 | 3 | 0 | 1 | 0 | 1 | 0 | 5 | 0.4 | 10 | 0.2 |
| 14_5470553 | 4 | 1 | 4 | 1 | 2 | 1 | 1 | 1 | 11 | 1 | 4 | 1 | 1 | 1 | 1 | 1 | 3 | 1 | 9 | 1 |
| 14_5470557 | 6 | 0 | 5 | 0 | 2 | 0 | 2 | 0.5 | 15 | 0.0667 | 5 | 0.2 | 1 | 1 | 2 | 0.5 | 5 | 0.4 | 13 | 0.3846 |
| 14_5470564 | 6 | 0.1667 | 6 | 0 | 2 | 0.5 | 2 | 0.5 | 16 | 0.1875 | 5 | 0.6 | 1 | 0 | 2 | 0 | 5 | 0.2 | 13 | 0.3077 |
| 14_5470574 | 2 | 0.5 | 4 | 0.75 | 1 | 1 | 2 | 0.5 | 9 | 0.6667 | 4 | 0.75 | 1 | 1 | NA | NA | 4 | 0.75 | 9 | 0.7778 |
| 14_5470575 | 5 | 0 | 4 | 0 | 3 | 0.3333 | 2 | 0 | 14 | 0.0714 | 5 | 0.2 | 1 | 0 | 2 | 0 | 4 | 0 | 12 | 0.0833 |
| 14_5470579 | 6 | 0.1667 | 3 | 0 | 2 | 1 | 2 | 1 | 13 | 0.3846 | 6 | 0.1667 | 1 | 0 | 2 | 0.5 | 5 | 0 | 14 | 0.1429 |
| 14_5470584 | 5 | 1 | 5 | 0.8 | 2 | 1 | 2 | 1 | 14 | 0.9286 | 5 | 1 | 1 | 1 | 1 | 1 | 5 | 0.8 | 12 | 0.9167 |
| 14_5470605 | 6 | 0.1667 | 5 | 0 | 4 | 0 | 2 | 0 | 17 | 0.0588 | 4 | 0.25 | 1 | 1 | 2 | 0 | 5 | 0 | 12 | 0.1667 |
| 14_5470608 | 5 | 0.4 | 5 | 0.2 | 3 | 0.6667 | 2 | 0 | 15 | 0.3333 | 5 | 0.4 | 1 | 0 | 3 | 1 | 5 | 0.2 | 14 | 0.4286 |
| 14_5470610 | 6 | 0.6667 | 4 | 0.75 | 4 | 1 | 2 | 1 | 16 | 0.8125 | 5 | 1 | 1 | 1 | 3 | 1 | 6 | 0.6667 | 15 | 0.8667 |
| 14_5470687 | 6 | 0.1667 | 5 | 0 | 5 | 0 | 3 | 0 | 19 | 0.0526 | 5 | 0 | NA | NA | 3 | 0 | 7 | 0.2857 | 15 | 0.1333 |
| 14_5470718 | 7 | 0.1429 | 6 | 0 | 7 | 0.4286 | 3 | 0 | 23 | 0.1739 | 5 | 0 | 1 | 0 | 3 | 0 | 8 | 0.125 | 17 | 0.0588 |
| 14_5470719 | 5 | 0.2 | 4 | 0.5 | 3 | 0.6667 | 3 | 0.6667 | 15 | 0.4667 | 4 | 0.25 | 1 | 0 | 2 | 0.5 | 4 | 0.25 | 11 | 0.2727 |
| 14_5470731 | 7 | 0.1429 | 7 | 0.4286 | 7 | 0.2857 | 3 | 0.3333 | 24 | 0.2917 | 6 | 0 | 1 | 0 | 3 | 0 | 6 | 0.1667 | 16 | 0.0625 |
| 14_5470743 | 7 | 0.2857 | 6 | 0 | 7 | 0.2857 | 3 | 0 | 23 | 0.1739 | 4 | 0.25 | NA | NA | 3 | 0 | 9 | 0.3333 | 16 | 0.25 |
| 14_5470751 | 6 | 0.1667 | 6 | 0.1667 | 5 | 0.4 | 3 | 0.3333 | 20 | 0.25 | 5 | 0 | 1 | 0 | 3 | 0 | 10 | 0.1 | 19 | 0.0526 |
| 14_5470766 | 6 | 0.1667 | 8 | 0 | 6 | 0.1667 | 3 | 0 | 23 | 0.087 | 7 | 0.2857 | 1 | 0 | 3 | 0 | 10 | 0.1 | 21 | 0.1429 |
| 14_5471748 | 5 | 0.4 | 3 | 0 | 2 | 0.5 | 13 | 0.2308 | 23 | 0.2609 | NA | NA | 10 | 0.5 | 11 | 0.4545 | 5 | 0.4 | 26 | 0.4615 |
| 14_5471749 | 2 | 0.5 | 3 | 0 | 2 | 0.5 | 12 | 0.1667 | 19 | 0.2105 | NA | NA | 10 | 0.2 | 10 | 0 | 4 | 0.25 | 24 | 0.125 |
| 14_5471750 | 4 | 0 | 2 | 0 | 1 | 1 | 11 | 0.1818 | 18 | 0.1667 | NA | NA | 10 | 0.1 | 12 | 0.0833 | 5 | 0 | 27 | 0.0741 |
| 14_5471755 | 5 | 0.4 | 2 | 0 | 1 | 0 | 9 | 0.2222 | 17 | 0.2353 | NA | NA | 10 | 0.2 | 13 | 0.0769 | 3 | 0.3333 | 26 | 0.1538 |
| 14_5471760 | 4 | 0.75 | 3 | 0.3333 | 2 | 1 | 12 | 0.25 | 21 | 0.4286 | NA | NA | 10 | 0.6 | 14 | 0.5 | 5 | 0.6 | 29 | 0.5517 |
| 14_5471762 | 4 | 0.5 | 3 | 0 | 1 | 0 | 11 | 0.0909 | 19 | 0.1579 | NA | NA | 10 | 0.1 | 12 | 0.0833 | 4 | 0 | 26 | 0.0769 |
| 14_5471770 | 2 | 1 | 3 | 1 | 2 | 1 | 15 | 0.9333 | 22 | 0.9545 | NA | NA | 10 | 1 | 11 | 1 | 5 | 1 | 26 | 1 |
| 14_5471776 | 3 | 0.3333 | 3 | 0 | 2 | 0 | 13 | 0.1538 | 21 | 0.1429 | NA | NA | 13 | 0.0769 | 12 | 0.0833 | 5 | 0 | 30 | 0.0667 |
| 14_5474474 | 12 | 0.0833 | 8 | 0.25 | 3 | 0.6667 | 4 | 0.25 | 27 | 0.2222 | 5 | 0.2 | 7 | 0 | 4 | 0 | 3 | 0.3333 | 19 | 0.1053 |
| 14_5474484 | 9 | 0 | 11 | 0.5455 | 3 | 0.3333 | 5 | 0.2 | 28 | 0.2857 | 6 | 0 | 9 | 0.3333 | 6 | 0.3333 | 5 | 0.2 | 26 | 0.2308 |
| 14_5474489 | 10 | 0.4 | 10 | 0.4 | 4 | 0 | 5 | 0.4 | 29 | 0.3448 | 5 | 0.2 | 9 | 0.3333 | 6 | 0.1667 | 5 | 0.2 | 25 | 0.24 |
| 14_5474490 | 9 | 0.3333 | 6 | 0.1667 | 2 | 0.5 | 5 | 0.4 | 22 | 0.3182 | 3 | 0.6667 | 7 | 0.2857 | 5 | 0.4 | 4 | 0.5 | 19 | 0.4211 |
| 14_5474507 | 10 | 0.5 | 7 | 1 | 4 | 1 | 5 | 0.6 | 26 | 0.7308 | 4 | 0.75 | 8 | 0.75 | 5 | 0.4 | 4 | 0.75 | 21 | 0.6667 |
| 14_5482162 | NA | NA | NA | NA | 1 | 1 | NA | NA | 1 | 1 | NA | NA | 1 | 1 | NA | NA | NA | NA | 1 | 1 |
| 14_5482178 | NA | NA | NA | NA | 1 | 1 | NA | NA | 1 | 1 | NA | NA | NA | NA | 1 | 1 | NA | NA | 1 | 1 |
| 14_5485168 | 39 | 0.0769 | 40 | 0.05 | 24 | 0.0833 | 30 | 0.0333 | 133 | 0.0602 | 24 | 0 | 32 | 0.0938 | 20 | 0.05 | 25 | 0.08 | 101 | 0.0594 |
| 14_5485169 | 42 | 0.0952 | 46 | 0 | 25 | 0.04 | 33 | 0.1515 | 146 | 0.0685 | 27 | 0.1111 | 38 | 0.1053 | 27 | 0.0741 | 27 | 0.1481 | 119 | 0.1092 |
| 14_5486921 | NA | NA | 5 | 0.2 | 3 | 0.3333 | NA | NA | 8 | 0.25 | 1 | 0 | 2 | 0.5 | 1 | 0 | 2 | 0 | 6 | 0.1667 |
| 14_5486933 | NA | NA | 5 | 0.4 | 3 | 0.3333 | NA | NA | 8 | 0.375 | 1 | 0 | 2 | 0.5 | NA | NA | 2 | 0 | 5 | 0.2 |
| 14_5486974 | NA | NA | 5 | 0.6 | 1 | 0 | NA | NA | 6 | 0.5 | 1 | 0 | 1 | 1 | NA | NA | 2 | 0 | 4 | 0.25 |
| 14_5486985 | NA | NA | 5 | 0.6 | 1 | 0 | NA | NA | 6 | 0.5 | 1 | 0 | 2 | 0.5 | NA | NA | 2 | 0 | 5 | 0.2 |
| 14_5487981 | 13 | 0 | 15 | 0.1333 | 9 | 0 | 14 | 0.1429 | 51 | 0.0784 | 7 | 0.1429 | 5 | 0 | 12 | 0 | 13 | 0.1538 | 37 | 0.0811 |
| 14_5487983 | 7 | 0.1429 | 13 | 0.0769 | 8 | 0 | 12 | 0.0833 | 40 | 0.075 | 6 | 0 | 5 | 0 | 9 | 0.2222 | 8 | 0 | 28 | 0.0714 |
| 14_5488005 | 12 | 0.1667 | 13 | 0.1538 | 8 | 0.125 | 14 | 0.0714 | 47 | 0.1277 | 6 | 0.1667 | 5 | 0 | 10 | 0.1 | 12 | 0.25 | 33 | 0.1515 |
| 14_5488715 | NA | NA | 1 | 1 | NA | NA | NA | NA | 1 | 1 | NA | NA | NA | NA | 1 | 1 | NA | NA | 1 | 1 |
| 14_5488723 | NA | NA | 1 | 1 | NA | NA | NA | NA | 1 | 1 | NA | NA | NA | NA | 1 | 1 | NA | NA | 1 | 1 |
| 14_5488729 | NA | NA | 1 | 1 | NA | NA | NA | NA | 1 | 1 | NA | NA | NA | NA | 1 | 1 | NA | NA | 1 | 1 |
| 14_5488746 | NA | NA | 1 | 1 | NA | NA | NA | NA | 1 | 1 | NA | NA | NA | NA | 1 | 1 | NA | NA | 1 | 1 |
| 14_5488776 | NA | NA | 1 | 1 | NA | NA | NA | NA | 1 | 1 | NA | NA | NA | NA | 1 | 1 | NA | NA | 1 | 1 |
| 14_5492439 | NA | NA | 2 | 0.5 | NA | NA | 1 | 1 | 3 | 0.6667 | NA | NA | 1 | 1 | 2 | 1 | 1 | 1 | 4 | 1 |
| 14_5492733 | 1 | 0 | 9 | 0.1111 | 1 | 0 | 2 | 0 | 13 | 0.0769 | 5 | 0.2 | 5 | 0 | 5 | 0 | 4 | 0 | 19 | 0.0526 |
| 14_5492744 | 1 | 0 | 10 | 0.3 | 1 | 1 | 3 | 0.3333 | 15 | 0.3333 | 4 | 0 | 5 | 0 | 6 | 0.3333 | 4 | 0.5 | 19 | 0.2105 |
| 14_5492768 | 1 | 1 | 9 | 1 | 1 | 1 | 5 | 1 | 16 | 1 | 5 | 1 | 5 | 1 | 5 | 1 | 3 | 1 | 18 | 1 |
| 14_5492773 | 1 | 1 | 9 | 0.5556 | 1 | 0 | 4 | 0.5 | 15 | 0.5333 | 5 | 0.4 | 5 | 0 | 6 | 0.6667 | 3 | 0.6667 | 19 | 0.4211 |
| 14_5492774 | 1 | 0 | 9 | 0.3333 | 1 | 1 | 4 | 0.25 | 15 | 0.3333 | 5 | 0.4 | 4 | 0.25 | 6 | 0.5 | 4 | 0.5 | 19 | 0.4211 |
| 14_5492778 | NA | NA | 8 | 0.75 | 1 | 0 | 1 | 1 | 10 | 0.7 | 2 | 1 | 4 | 0.75 | 3 | 0.3333 | NA | NA | 9 | 0.6667 |
| 14_5492787 | 1 | 0 | 10 | 0.1 | 1 | 0 | 5 | 0.2 | 17 | 0.1176 | 5 | 0 | 5 | 0.2 | 5 | 0 | 4 | 0 | 19 | 0.0526 |
| 14_5492790 | 1 | 0 | 9 | 0.7778 | 1 | 0 | 5 | 0.8 | 16 | 0.6875 | 3 | 0.3333 | 3 | 0.3333 | 5 | 0.8 | 3 | 0.6667 | 14 | 0.5714 |
| 14_5492796 | 1 | 1 | 10 | 1 | 1 | 1 | 6 | 1 | 18 | 1 | 5 | 1 | 5 | 1 | 6 | 1 | 4 | 1 | 20 | 1 |
| 14_5492802 | NA | NA | 9 | 0.6667 | 1 | 1 | 3 | 0.6667 | 13 | 0.6923 | 6 | 1 | 4 | 0.75 | 5 | 0.6 | 3 | 1 | 18 | 0.8333 |
| 14_5492805 | 1 | 0 | 10 | 0.7 | 1 | 0 | 6 | 0.3333 | 18 | 0.5 | 6 | 0.3333 | 4 | 0.25 | 6 | 0.5 | 4 | 0.75 | 20 | 0.45 |
| 14_5496027 | 6 | 0.3333 | 5 | 0 | 5 | 0.4 | 4 | 0 | 20 | 0.2 | 6 | 0.1667 | 7 | 0.2857 | 8 | 0.25 | 3 | 0.6667 | 24 | 0.2917 |

| | | | | | | | | | | | | | | | | | | | | |
|---|---|---|---|---|---|---|---|---|---|---|---|---|---|---|---|---|---|---|---|---|
| 14_5496029 | 6 | 0 | 5 | 0.4 | 4 | 0 | 3 | 0 | 18 | 0.1111 | 6 | 0 | 6 | 0.1667 | 8 | 0.125 | 3 | 0 | 23 | 0.087 |
| 14_5496049 | 3 | 0.3333 | 4 | 0 | 3 | 0.3333 | 4 | 0.25 | 14 | 0.2143 | 6 | 0.5 | 6 | 0.1667 | 8 | 0.375 | 5 | 0 | 25 | 0.28 |
| 14_5496090 | 3 | 0.6667 | 4 | 0 | 3 | 0.3333 | 3 | 0.3333 | 13 | 0.3077 | 5 | 0.2 | 6 | 0 | 5 | 0.4 | 4 | 0.25 | 20 | 0.2 |
| 14_5496107 | 4 | 0 | 4 | 0 | 3 | 0 | 3 | 0.3333 | 14 | 0.0714 | 6 | 0.5 | 7 | 0.1429 | 7 | 0.2857 | 4 | 0 | 24 | 0.25 |
| 14_5501882 | 2 | 1 | 1 | 1 | 1 | 1 | 1 | 1 | 5 | 1 | NA | NA | 3 | 0.6667 | 2 | 1 | 1 | 0 | 6 | 0.6667 |
| 14_5501895 | 2 | 0 | 2 | 0.5 | 1 | 1 | NA | NA | 5 | 0.4 | 1 | 0 | 3 | 0.3333 | 2 | 1 | 2 | 0 | 8 | 0.375 |
| 14_5501896 | 2 | 0 | 2 | 0.5 | 1 | 0 | NA | NA | 5 | 0.2 | NA | NA | 2 | 0 | 2 | 0.5 | 2 | 0 | 6 | 0.1667 |
| 14_5501919 | 2 | 0 | 2 | 0 | 1 | 1 | NA | NA | 5 | 0.2 | NA | NA | 3 | 0.6667 | 2 | 0.5 | NA | NA | 5 | 0.6 |
| 14_5509311 | 2 | 0 | 4 | 0.25 | 4 | 0 | 3 | 0.3333 | 13 | 0.1538 | 1 | 0 | 1 | 1 | 3 | 0 | 2 | 0.5 | 7 | 0.2857 |
| 14_5509378 | 2 | 1 | 3 | 0.3333 | 2 | 1 | 2 | 0.5 | 9 | 0.6667 | 1 | 0 | 3 | 1 | 2 | 0 | 2 | 0 | 8 | 0.375 |
| 14_5509388 | 1 | 1 | 2 | 1 | 2 | 1 | 2 | 1 | 7 | 1 | 1 | 1 | 2 | 1 | 3 | 0.6667 | 1 | 1 | 7 | 0.8571 |
| 14_5509400 | 1 | 0 | 2 | 0 | 2 | 0.5 | 2 | 0.5 | 7 | 0.2857 | 1 | 0 | 3 | 0 | 2 | 0.5 | 2 | 0 | 8 | 0.125 |
| 14_5509409 | NA | NA | 3 | 0.6667 | 2 | 1 | 2 | 1 | 7 | 0.8571 | 1 | 1 | 2 | 1 | 2 | 1 | 1 | 1 | 6 | 1 |
| 14_5509415 | NA | NA | 2 | 1 | 2 | 1 | 1 | 1 | 5 | 1 | 1 | 1 | 1 | 1 | 1 | 1 | 1 | 1 | 4 | 1 |
| 14_5509423 | 1 | 1 | 2 | 0.5 | 1 | 1 | NA | NA | 4 | 0.75 | 1 | 0 | 2 | 1 | 2 | 0.5 | 1 | 0 | 6 | 0.5 |
| 14_5509430 | NA | NA | 3 | 0 | 2 | 1 | 1 | 0 | 6 | 0.3333 | 1 | 0 | 2 | 1 | 3 | 0 | 1 | 0 | 7 | 0.2857 |
| 14_5509431 | 1 | 1 | 2 | 1 | 2 | 1 | 1 | 1 | 6 | 1 | NA | NA | 2 | 1 | 3 | 1 | 1 | 1 | 6 | 1 |
| 14_5509435 | 1 | 0 | 1 | 1 | 2 | 0 | 1 | 0 | 5 | 0.2 | 1 | 0 | 2 | 0.5 | 2 | 0 | 1 | 0 | 6 | 0.1667 |
| 14_5509453 | 1 | 1 | 2 | 1 | 2 | 0.5 | 1 | 1 | 6 | 0.8333 | NA | NA | 1 | 1 | 1 | 1 | 1 | 1 | 3 | 1 |
| 14_5509455 | 1 | 1 | 1 | 1 | 2 | 1 | 1 | 1 | 5 | 1 | 1 | 1 | 1 | 1 | 2 | 1 | 1 | 1 | 5 | 1 |
| 14_5509471 | 1 | 0 | 2 | 0.5 | 1 | 0 | 1 | 0 | 5 | 0.2 | 1 | 1 | 1 | 0 | 2 | 0 | 1 | 1 | 5 | 0.4 |
| 14_5509505 | 1 | 1 | 2 | 1 | 1 | 1 | 1 | 1 | 5 | 1 | 1 | 1 | NA | NA | 2 | 0.5 | 1 | 1 | 4 | 0.75 |
| 14_5509513 | 1 | 0 | 2 | 1 | NA | NA | 1 | 1 | 4 | 0.75 | 1 | 0 | NA | NA | 2 | 0 | 1 | 1 | 4 | 0.25 |
| 14_5509515 | 1 | 1 | 2 | 0.5 | 1 | 1 | 1 | 0 | 5 | 0.6 | 1 | 1 | 1 | 1 | 2 | 0.5 | 1 | 1 | 5 | 0.8 |
| 14_5509528 | 1 | 1 | 2 | 1 | 1 | 1 | 1 | 1 | 5 | 1 | 1 | 1 | 1 | 1 | 2 | 1 | 1 | 1 | 5 | 1 |
| 14_5509547 | 1 | 0 | 2 | 0.5 | 1 | 0 | NA | NA | 4 | 0.25 | 1 | 0 | NA | NA | 2 | 0.5 | NA | NA | 3 | 0.3333 |
| 14_5511113 | NA | NA | 2 | 0.5 | 1 | 1 | 1 | 1 | 4 | 0.75 | 2 | 1 | 2 | 0.5 | 2 | 1 | 1 | 1 | 7 | 0.8571 |
| 14_5511116 | NA | NA | 2 | 0 | 1 | 0 | 1 | 1 | 4 | 0.25 | 2 | 0 | 2 | 0.5 | 2 | 0 | 1 | 1 | 7 | 0.2857 |
| 14_5511137 | NA | NA | 2 | 0.5 | 1 | 0 | NA | NA | 3 | 0.3333 | 2 | 0 | 2 | 0.5 | 2 | 0.5 | 1 | 0 | 7 | 0.2857 |
| 14_5512186 | NA | NA | NA | NA | NA | NA | 2 | 0.5 | 2 | 0.5 | NA | NA | 3 | 0 | 1 | 1 | NA | NA | 4 | 0.25 |
| 14_5512209 | NA | NA | NA | NA | 1 | 1 | NA | NA | 1 | 1 | NA | NA | 2 | 1 | 2 | 1 | NA | NA | 4 | 1 |
| 14_5512519 | 11 | 0.1818 | 13 | 0.0769 | 5 | 0.2 | 9 | 0 | 38 | 0.1053 | 6 | 0.1667 | 15 | 0.0667 | 4 | 0.5 | 10 | 0 | 35 | 0.1143 |
| 14_5513490 | 4 | 0 | 6 | 0.1667 | NA | NA | 2 | 0.5 | 12 | 0.1667 | 2 | 0 | NA | NA | 4 | 0 | 2 | 0.5 | 8 | 0.125 |
| 14_5513567 | 5 | 0.2 | 8 | 0.25 | 4 | 0.25 | 5 | 0.6 | 22 | 0.3182 | 7 | 0.1429 | 4 | 0.75 | 3 | 0 | 5 | 0.4 | 19 | 0.3158 |
| 14_5513568 | 4 | 0.25 | 12 | 0.3333 | 4 | 0.25 | 6 | 0.5 | 26 | 0.3462 | 10 | 0.1 | 6 | 0.3333 | 10 | 0.2 | 6 | 0.1667 | 32 | 0.1875 |
| 14_5513576 | 5 | 0.4 | 16 | 0.9375 | 2 | 0 | 5 | 1 | 28 | 0.7857 | 10 | 0.5 | 5 | 0.6 | 7 | 0.8571 | 7 | 0.5714 | 29 | 0.6207 |
| 14_5514418 | 1 | 0 | 2 | 0 | NA | NA | 2 | 0.5 | 5 | 0.2 | 2 | 0 | 2 | 0.5 | 2 | 0 | NA | NA | 6 | 0.1667 |
| 14_5514461 | 80 | 0.0875 | 95 | 0.0737 | 73 | 0.0411 | 88 | 0.0909 | 336 | 0.0744 | 74 | 0.0811 | 91 | 0.1648 | 88 | 0.0455 | 65 | 0.1077 | 318 | 0.1006 |
| 14_5514668 | NA | NA | 2 | 0.5 | 1 | 0 | NA | NA | 3 | 0.3333 | 1 | 0 | 4 | 0.25 | NA | NA | NA | NA | 5 | 0.2 |
| 14_5514973 | 43 | 0.0233 | 35 | 0.0571 | 26 | 0.0769 | 46 | 0.1304 | 150 | 0.0733 | 36 | 0.0556 | 61 | 0.0492 | 37 | 0.1081 | 34 | 0.0588 | 168 | 0.0655 |
| 14_5514974 | 46 | 0.1087 | 35 | 0.1429 | 29 | 0.0345 | 47 | 0.0851 | 157 | 0.0955 | 34 | 0.0588 | 45 | 0.0667 | 36 | 0.0833 | 37 | 0.1081 | 152 | 0.0789 |
| 14_5514974 | 51 | 0.0196 | 41 | 0.0488 | 32 | 0.0938 | 56 | 0.1429 | 180 | 0.0778 | 44 | 0.0455 | 58 | 0.0862 | 42 | 0.0476 | 40 | 0.05 | 184 | 0.0598 |
| 14_5515429 | NA | NA | 2 | 0.5 | 1 | 1 | 1 | 1 | 4 | 0.75 | NA | NA | NA | NA | 3 | 0.3333 | 1 | 0 | 4 | 0.25 |
| 14_5515731 | NA | NA | 3 | 0 | 1 | 1 | NA | NA | 4 | 0.25 | 1 | 0 | NA | NA | 3 | 0.3333 | 1 | 0 | 5 | 0.2 |
| 14_5515748 | NA | NA | 4 | 1 | 1 | 1 | NA | NA | 5 | 1 | 2 | 1 | NA | NA | 3 | 0.6667 | 2 | 1 | 7 | 0.8571 |
| 14_5515853 | NA | NA | 3 | 0.3333 | 1 | 0 | NA | NA | 4 | 0.25 | 1 | 1 | NA | NA | 1 | 0 | NA | NA | 2 | 0.5 |
| 14_5516318 | NA | NA | 2 | 0 | NA | NA | 1 | 1 | 3 | 0.3333 | NA | NA | NA | NA | 1 | 1 | 2 | 0.5 | 3 | 0.6667 |
| 14_5516319 | NA | NA | 2 | 1 | 1 | 0 | 1 | 1 | 4 | 0.75 | NA | NA | NA | NA | 1 | 0 | 2 | 0.5 | 3 | 0.3333 |
| 14_5517665 | 6 | 0 | 17 | 0 | 18 | 0.2222 | 9 | 0.2222 | 50 | 0.12 | 3 | 0 | 16 | 0.0625 | 8 | 0.125 | 10 | 0.1 | 37 | 0.0811 |
| 14_5522698 | NA | NA | 1 | 1 | NA | NA | 2 | 0.5 | 3 | 0.6667 | NA | NA | NA | NA | 1 | 1 | NA | NA | 1 | 1 |
| 14_5529776 | 1 | 1 | 1 | 0 | 3 | 0 | 4 | 0 | 9 | 0.1111 | NA | NA | 3 | 0.3333 | 3 | 0.3333 | NA | NA | 6 | 0.3333 |
| 14_5529916 | 3 | 0 | 3 | 0 | 4 | 0.25 | 6 | 0.5 | 16 | 0.25 | NA | NA | 6 | 0.5 | 8 | 0.375 | 2 | 0.5 | 16 | 0.4375 |
| 14_5529941 | 4 | 0.25 | 2 | 0 | 5 | 0 | 6 | 0 | 17 | 0.0588 | 1 | 0 | 8 | 0.375 | 7 | 0.1429 | 1 | 1 | 17 | 0.2941 |
| 14_5529942 | 4 | 0 | 4 | 0.25 | 7 | 0.1429 | 7 | 0 | 22 | 0.0909 | 1 | 0 | 7 | 0.1429 | 8 | 0 | NA | NA | 16 | 0.0625 |
| 14_5529948 | 3 | 1 | 3 | 1 | 4 | 1 | 7 | 0.8571 | 17 | 0.9412 | NA | NA | 6 | 0.8333 | 8 | 0.625 | 1 | 0 | 15 | 0.6667 |
| 14_5529976 | 2 | 0.5 | 2 | 0 | 5 | 0.6 | 4 | 0.25 | 13 | 0.3846 | 1 | 0 | 6 | 0.5 | 6 | 0.5 | 1 | 0 | 14 | 0.4286 |
| 14_5530010 | 2 | 1 | 4 | 0.5 | 3 | 0 | 6 | 0.5 | 15 | 0.4667 | NA | NA | 4 | 0.5 | 3 | 0 | 1 | 1 | 8 | 0.375 |
| 14_5530043 | 2 | 0 | 4 | 0.25 | 3 | 0.3333 | 6 | 0.1667 | 15 | 0.2 | NA | NA | 5 | 0.4 | 3 | 0.3333 | 1 | 0 | 9 | 0.3333 |
| 14_5530062 | 2 | 0.5 | 2 | 0 | 2 | 0 | 6 | 0.5 | 12 | 0.3333 | 1 | 0 | 4 | 0.75 | 2 | 0 | 1 | 0 | 8 | 0.375 |
| 14_5530079 | 1 | 1 | 3 | 0 | 2 | 0.5 | 5 | 0.4 | 11 | 0.3636 | 1 | 0 | 5 | 0.2 | 1 | 0 | NA | NA | 7 | 0.1429 |
| 14_5530089 | 1 | 0 | 2 | 0 | 2 | 0 | 6 | 0.1667 | 11 | 0.0909 | 1 | 0 | 5 | 0 | 2 | 0.5 | 1 | 0 | 9 | 0.1111 |
| 14_5530092 | 1 | 1 | 3 | 1 | 2 | 0 | 6 | 0.5 | 12 | 0.5833 | NA | NA | 5 | 1 | 3 | 0.3333 | 1 | 0 | 9 | 0.6667 |
| 14_5530143 | 1 | 0 | 2 | 0.5 | 2 | 0 | 6 | 0.1667 | 11 | 0.1818 | NA | NA | 4 | 0 | 2 | 0.5 | 1 | 0 | 7 | 0.1429 |
| 14_5530153 | 1 | 0 | 3 | 0 | 2 | 0 | 6 | 0.1667 | 12 | 0.0833 | 1 | 0 | 4 | 0.25 | 2 | 0.5 | 1 | 0 | 8 | 0.25 |
| 14_5530169 | 1 | 1 | 2 | 1 | 2 | 0.5 | 4 | 1 | 9 | 0.8889 | 1 | 1 | 3 | 1 | 1 | 1 | 1 | 1 | 6 | 1 |

| | | | | | | | | | | | | | | | | | | | | |
|---|---|---|---|---|---|---|---|---|---|---|---|---|---|---|---|---|---|---|---|---|
| 14_553017! | 1 | 1 | 3 | 0.6667 | 2 | 0 | 5 | 0.2 | 11 | 0.3636 | 1 | 0 | 3 | 0.3333 | 2 | 0 | NA | NA | 6 | 0.1667 |
| 14_553023' | 1 | 0 | 3 | 1 | 1 | 1 | 6 | 0.8333 | 11 | 0.8182 | 1 | 1 | 3 | 1 | 3 | 0.6667 | NA | NA | 7 | 0.8571 |
| 14_553023( | 1 | 0 | 3 | 0.3333 | 1 | 0 | 6 | 0.1667 | 11 | 0.1818 | 1 | 0 | 2 | 0.5 | 2 | 0.5 | NA | NA | 5 | 0.4 |
| 14_553024: | 1 | 0 | 3 | 0 | 1 | 0 | 6 | 0.1667 | 11 | 0.0909 | NA | NA | 3 | 0.3333 | 3 | 0 | NA | NA | 6 | 0.1667 |
| 14_553026( | 1 | 0 | 2 | 0 | 1 | 0 | 4 | 0.25 | 8 | 0.125 | NA | NA | 3 | 0.3333 | 1 | 0 | NA | NA | 4 | 0.25 |
| 14_553032! | 1 | 0 | 1 | 0 | 1 | 0 | 4 | 0.5 | 7 | 0.2857 | 1 | 0 | 3 | 0.3333 | 2 | 0.5 | NA | NA | 6 | 0.3333 |
| 14_553035( | 1 | 0 | NA | NA | NA | NA | 5 | 0.4 | 6 | 0.3333 | 1 | 1 | 2 | 0 | 1 | 1 | NA | NA | 4 | 0.5 |
| 14_553035' | 1 | 1 | 1 | 0 | 1 | 0 | 5 | 0.2 | 8 | 0.25 | NA | NA | 2 | 0.5 | 1 | 0 | NA | NA | 3 | 0.3333 |
| 14_553035‹ | NA | NA | 1 | 0 | 1 | 0 | 5 | 0.6 | 7 | 0.4286 | 1 | 0 | 2 | 0.5 | 1 | 0 | NA | NA | 4 | 0.25 |
| 14_553037‹ | 1 | 1 | 1 | 0 | 2 | 1 | 5 | 0.6 | 9 | 0.6667 | 1 | 0 | 1 | 1 | 1 | 1 | NA | NA | 3 | 0.6667 |
| 14_553040‹ | 1 | 1 | 1 | 1 | 2 | 1 | 4 | 1 | 8 | 1 | 1 | 1 | 1 | 1 | 2 | 1 | NA | NA | 4 | 1 |
| 14_553041( | NA | NA | NA | NA | NA | NA | 4 | 0.5 | 4 | 0.5 | NA | NA | 1 | 1 | 1 | 0 | NA | NA | 2 | 0.5 |
| 14_553044! | 1 | 1 | 1 | 1 | NA | NA | 5 | 0.4 | 7 | 0.5714 | 1 | 1 | NA | NA | 1 | 1 | NA | NA | 2 | 1 |
| 14_553124: | 1 | 1 | NA | NA | NA | NA | NA | NA | 1 | 1 | NA | NA | NA | NA | NA | NA | 1 | 1 | 1 | 1 |
| 14_553216! | NA | NA | 1 | 1 | NA | NA | NA | NA | 1 | 1 | NA | NA | NA | NA | 1 | 1 | NA | NA | 1 | 1 |
| 14_553327: | 1 | 0 | NA | NA | 1 | 0 | 3 | 0.3333 | 5 | 0.2 | 3 | 0.3333 | NA | NA | 3 | 0 | 2 | 0 | 8 | 0.125 |
| 14_553328' | 1 | 1 | 1 | 1 | 1 | 0 | 2 | 0.5 | 5 | 0.6 | 3 | 0.6667 | 2 | 0 | 3 | 0.6667 | 3 | 1 | 11 | 0.6364 |
| 14_553328‹ | 1 | 1 | 1 | 0 | 1 | 0 | 3 | 0.3333 | 6 | 0.3333 | 3 | 0.3333 | 2 | 0 | 3 | 0 | 3 | 0 | 11 | 0.0909 |
| 14_553331! | 1 | 1 | NA | NA | 1 | 0 | 3 | 0.6667 | 5 | 0.6 | 4 | 0.5 | 2 | 0.5 | 2 | 0.5 | 2 | 0.5 | 10 | 0.5 |
| 14_553333: | NA | NA | NA | NA | 1 | 1 | 2 | 0 | 3 | 0.3333 | 3 | 0.6667 | 1 | 0 | 2 | 0.5 | 2 | 1 | 8 | 0.625 |
| 14_553333! | 1 | 1 | 1 | 0 | 1 | 0 | 3 | 0.6667 | 6 | 0.5 | 4 | 0.5 | 1 | 1 | 4 | 0.25 | 3 | 0.6667 | 12 | 0.5 |
| 14_553333( | 1 | 1 | 1 | 0 | 1 | 1 | 3 | 0.6667 | 6 | 0.6667 | 4 | 0.5 | 1 | 1 | 4 | 0.25 | 2 | 0.5 | 11 | 0.4545 |
| 14_553349( | 1 | 1 | 2 | 0.5 | 1 | 1 | 3 | 1 | 7 | 0.8571 | 2 | 1 | 2 | 1 | 3 | 1 | 3 | 0.6667 | 10 | 0.9 |
| 14_553349! | 1 | 1 | 3 | 1 | 1 | 1 | 4 | 0.75 | 9 | 0.8889 | 3 | 0.6667 | 3 | 0.6667 | 4 | 0.75 | 3 | 1 | 13 | 0.7692 |
| 14_553349! | NA | NA | 3 | 0.3333 | 1 | 0 | 4 | 0.25 | 8 | 0.25 | 4 | 0 | 4 | 0.25 | 4 | 0.25 | 4 | 0 | 16 | 0.125 |
| 14_553350: | 1 | 0 | 1 | 0 | 1 | 1 | 5 | 0.4 | 8 | 0.375 | 4 | 0.25 | 4 | 0.25 | 3 | 0.6667 | 2 | 0.5 | 13 | 0.3846 |
| 14_553356( | 2 | 0.5 | 2 | 0 | 2 | 0.5 | 6 | 0 | 12 | 0.1667 | 4 | 0.25 | 4 | 0.5 | 4 | 0.25 | 3 | 0 | 15 | 0.2667 |
| 14_553360: | 1 | 0 | 3 | 0.6667 | 2 | 0 | 6 | 0.1667 | 12 | 0.25 | 4 | 0.25 | 3 | 0.3333 | 5 | 0.6 | 3 | 0 | 15 | 0.3333 |
| 14_553361! | 1 | 1 | 3 | 0.3333 | 2 | 0 | 4 | 0.25 | 10 | 0.3 | 4 | 0.5 | 5 | 0.4 | 4 | 0.5 | 2 | 0 | 15 | 0.4 |
| 14_553363: | 2 | 1 | 4 | 0.25 | 2 | 0 | 4 | 0.5 | 12 | 0.4167 | 2 | 0.5 | 5 | 0.4 | 5 | 0.4 | 4 | 0.25 | 16 | 0.375 |
| 14_553363( | 3 | 0.6667 | 3 | 1 | 2 | 0 | 6 | 0.6667 | 14 | 0.6429 | 5 | 0.6 | 5 | 0.4 | 5 | 0.8 | 4 | 0.5 | 19 | 0.5789 |
| 14_553446( | 42 | 0.0714 | 40 | 0.075 | 37 | 0.1892 | 50 | 0.16 | 169 | 0.1243 | 36 | 0.0278 | 44 | 0.0909 | 49 | 0.102 | 46 | 0.1739 | 175 | 0.1029 |
| 14_553446' | 38 | 0.2105 | 35 | 0.2 | 36 | 0.0556 | 46 | 0.1304 | 155 | 0.1484 | 32 | 0.0625 | 42 | 0.119 | 46 | 0.1522 | 45 | 0.1111 | 165 | 0.1152 |
| 14_553446! | 37 | 0.2432 | 31 | 0.1613 | 31 | 0.2258 | 36 | 0.1389 | 135 | 0.1926 | 26 | 0.2308 | 31 | 0.129 | 39 | 0.2821 | 37 | 0.2432 | 133 | 0.2256 |
| 14_553449: | 37 | 0.3784 | 36 | 0.5278 | 31 | 0.3871 | 49 | 0.3878 | 153 | 0.4183 | 34 | 0.1765 | 39 | 0.4872 | 46 | 0.2826 | 39 | 0.4103 | 158 | 0.3418 |
| 14_553449' | 38 | 0.8421 | 34 | 0.7941 | 34 | 0.9118 | 46 | 0.8696 | 152 | 0.8553 | 38 | 0.9211 | 40 | 0.85 | 47 | 0.7872 | 42 | 0.9286 | 167 | 0.8683 |
| 14_553470: | 27 | 0.037 | 28 | 0.1071 | 23 | 0.1739 | 34 | 0.1176 | 112 | 0.1071 | 19 | 0.2105 | 30 | 0.0667 | 32 | 0.125 | 24 | 0.125 | 105 | 0.1238 |
| 14_553470! | 24 | 0.125 | 21 | 0.0476 | 23 | 0.1739 | 37 | 0.0541 | 105 | 0.0952 | 18 | 0.1111 | 32 | 0.0312 | 27 | 0.1481 | 27 | 0.1111 | 104 | 0.0962 |
| 14_553471! | 23 | 0.5217 | 24 | 0.4583 | 24 | 0.625 | 36 | 0.3611 | 107 | 0.4766 | 21 | 0.4286 | 32 | 0.5 | 26 | 0.4231 | 33 | 0.4848 | 112 | 0.4643 |
| 14_553509: | 29 | 0.1379 | 23 | 0.1304 | 25 | 0.24 | 32 | 0.0938 | 109 | 0.1468 | 23 | 0.2174 | 27 | 0.037 | 30 | 0.1333 | 29 | 0.069 | 109 | 0.1101 |
| 14_553511‹ | 25 | 0.08 | 21 | 0.0952 | 24 | 0.0417 | 33 | 0.1515 | 103 | 0.0971 | 22 | 0.0909 | 21 | 0.0476 | 26 | 0.1538 | 22 | 0.3636 | 91 | 0.1648 |
| 14_553512‹ | 25 | 0.28 | 23 | 0.2174 | 22 | 0.1818 | 35 | 0.2 | 105 | 0.219 | 20 | 0.35 | 29 | 0.2414 | 24 | 0.125 | 25 | 0.12 | 98 | 0.2041 |
| 14_553515: | 27 | 0.2963 | 22 | 0.1818 | 21 | 0.0952 | 29 | 0.2069 | 99 | 0.202 | 21 | 0.0952 | 25 | 0.32 | 24 | 0.25 | 20 | 0.3 | 90 | 0.2444 |
| 14_553524( | 23 | 0.1739 | 18 | 0.1667 | 19 | 0.1579 | 26 | 0.2308 | 86 | 0.186 | 18 | 0.0556 | 22 | 0.1818 | 18 | 0.0556 | 11 | 0 | 69 | 0.087 |
| 14_553525! | 27 | 0.1852 | 21 | 0.1905 | 18 | 0.3333 | 28 | 0.2143 | 94 | 0.2234 | 20 | 0.15 | 23 | 0.3043 | 23 | 0.3478 | 15 | 0.0667 | 81 | 0.2346 |
| 14_553535! | 21 | 0.0952 | 18 | 0.0556 | 17 | 0.1176 | 23 | 0.087 | 79 | 0.0886 | 14 | 0.0714 | 18 | 0.0556 | 16 | 0 | 15 | 0.1333 | 63 | 0.0635 |
| 14_557276‹ | NA | NA | 3 | 1 | 2 | 0 | 1 | 0 | 6 | 0.5 | 2 | 0.5 | NA | NA | NA | NA | NA | NA | 2 | 0.5 |
| 14_557392: | NA | NA | 3 | 0.3333 | 1 | 1 | 1 | 0 | 5 | 0.4 | 1 | 1 | NA | NA | 2 | 0 | NA | NA | 3 | 0.3333 |
| 14_557392( | 1 | 0 | 3 | 0.3333 | 1 | 0 | NA | NA | 5 | 0.2 | 1 | 1 | NA | NA | 2 | 0.5 | NA | NA | 3 | 0.6667 |
| 14_557454! | 3 | 0 | 6 | 0.1667 | 3 | 0.3333 | 7 | 0.2857 | 19 | 0.2105 | 3 | 0.3333 | 1 | 0 | 3 | 0.6667 | 4 | 0.25 | 11 | 0.3636 |
| 14_557455‹ | 1 | 0 | 3 | 0 | 2 | 0.5 | 6 | 0.6667 | 12 | 0.4167 | 3 | 0.6667 | NA | NA | 2 | 0.5 | 3 | 0.3333 | 8 | 0.5 |
| 14_557548! | 13 | 0.0769 | 22 | 0.0455 | 10 | 0.4 | 18 | 0.1111 | 63 | 0.127 | 24 | 0.0833 | 18 | 0.0556 | 18 | 0.1111 | 19 | 0.0526 | 79 | 0.0759 |
| 14_557858: | 1 | 0 | 3 | 0 | 2 | 0 | 1 | 1 | 7 | 0.1429 | 3 | 0 | 3 | 0.3333 | 2 | 0.5 | 2 | 0 | 10 | 0.2 |
| 14_557858! | 1 | 1 | 3 | 0 | 4 | 0 | 3 | 0 | 11 | 0.0909 | 5 | 0 | 2 | 0.5 | 2 | 0 | 3 | 0 | 12 | 0.0833 |
| 14_557860( | 1 | 1 | NA | NA | 4 | 0.25 | 3 | 0.6667 | 8 | 0.5 | 5 | 0.4 | 2 | 0 | 1 | 1 | 3 | 0.3333 | 11 | 0.3636 |
| 14_557860‹ | 1 | 0 | 3 | 0 | 2 | 0 | 3 | 0.6667 | 9 | 0.2222 | 5 | 0 | 2 | 0 | 1 | 0 | 2 | 0.5 | 10 | 0.1 |
| 14_557860( | 1 | 0 | 3 | 0.6667 | 3 | 0 | 3 | 0 | 10 | 0.2 | 5 | 0.4 | 3 | 0.3333 | 2 | 0 | 3 | 0.3333 | 13 | 0.3077 |
| 14_557860! | 1 | 1 | 3 | 0.6667 | 4 | 0.75 | 3 | 1 | 11 | 0.8182 | 6 | 0.6667 | 2 | 0 | 2 | 0.5 | 2 | 0.5 | 12 | 0.5 |
| 14_557861! | 1 | 1 | 3 | 0.3333 | 3 | 0.3333 | 3 | 0.3333 | 10 | 0.4 | 6 | 0.1667 | 3 | 0.6667 | 2 | 0 | 2 | 0 | 13 | 0.2308 |
| 14_557862! | NA | NA | 2 | 0.5 | 2 | 1 | 4 | 0.75 | 8 | 0.75 | 5 | 0.8 | 3 | 0.6667 | 1 | 1 | 2 | 1 | 11 | 0.8182 |
| 14_557862! | NA | NA | 2 | 0.5 | 3 | 0 | 3 | 0 | 8 | 0.125 | 7 | 0.2857 | 3 | 0.3333 | 2 | 0 | 2 | 0 | 14 | 0.2143 |
| 14_557863: | 1 | 1 | NA | NA | 2 | 0.5 | 2 | 1 | 5 | 0.8 | 5 | 0.6 | 1 | 0 | 1 | 1 | 2 | 0.5 | 9 | 0.5556 |
| 14_557864( | 1 | 1 | NA | NA | 1 | 0 | NA | NA | 2 | 0.5 | NA | NA | 1 | 1 | NA | NA | 1 | 1 | 2 | 1 |
| 14_557880‹ | NA | NA | NA | NA | NA | NA | 1 | 1 | 1 | 1 | NA | NA | NA | NA | NA | NA | 2 | 0.5 | 2 | 0.5 |
| 14_558069' | 9 | 0.3333 | 11 | 0.2727 | 10 | 0.2 | 14 | 0.3571 | 44 | 0.2955 | 3 | 0.3333 | 8 | 0.375 | 7 | 0.4286 | 6 | 0.1667 | 24 | 0.3333 |

| | | | | | | | | | | | | | | | | | | | | |
|---|---|---|---|---|---|---|---|---|---|---|---|---|---|---|---|---|---|---|---|---|
| 14_558072 | 7 | 0.1429 | 12 | 0.1667 | 10 | 0 | 14 | 0.0714 | 43 | 0.093 | 5 | 0 | 10 | 0.1 | 9 | 0 | 7 | 0.1429 | 31 | 0.0645 |
| 14_558073 | 10 | 0.3 | 13 | 0.1538 | 11 | 0.0909 | 15 | 0.0667 | 49 | 0.1429 | 8 | 0 | 10 | 0.2 | 9 | 0.1111 | 7 | 0.2857 | 34 | 0.1471 |
| 14_558073 | 5 | 0.4 | 13 | 0.6923 | 11 | 0.4545 | 14 | 0.3571 | 43 | 0.4884 | 6 | 1 | 10 | 0.8 | 7 | 0.7143 | 7 | 0.7143 | 30 | 0.8 |
| 14_558073 | 10 | 0.7 | 11 | 0.5455 | 10 | 0.7 | 15 | 0.0667 | 46 | 0.4565 | 8 | 0.625 | 11 | 0.3636 | 10 | 0.4 | 8 | 0.375 | 37 | 0.4324 |
| 14_558074 | 9 | 0.4444 | 12 | 0.4167 | 10 | 0.3 | 15 | 0.2 | 46 | 0.3261 | 10 | 0.3 | 13 | 0.1538 | 10 | 0.4 | 7 | 0.7143 | 40 | 0.35 |
| 14_558188 | 9 | 0 | 9 | 0.1111 | 7 | 0.1429 | 7 | 0 | 32 | 0.0625 | 9 | 0.2222 | 9 | 0.5556 | 8 | 0.25 | 4 | 0 | 30 | 0.3 |
| 14_558189 | 10 | 0 | 13 | 0.1538 | 8 | 0 | 9 | 0 | 40 | 0.05 | 12 | 0.1667 | 10 | 0.2 | 8 | 0.25 | 7 | 0 | 37 | 0.1622 |
| 14_558190 | 2 | 0 | 8 | 0 | 2 | 0 | 5 | 0.2 | 17 | 0.0588 | 1 | 0 | 5 | 0 | 6 | 0 | 3 | 0.3333 | 15 | 0.0667 |
| 14_558190 | 3 | 0 | 8 | 0.125 | 5 | 0.2 | 7 | 0.1429 | 23 | 0.1304 | 1 | 0 | 7 | 0.1429 | 8 | 0 | 2 | 0 | 18 | 0.0556 |
| 14_558191 | 3 | 0.3333 | 7 | 0.4286 | 4 | 0.5 | 8 | 0.625 | 22 | 0.5 | NA | NA | 5 | 0.4 | 8 | 0.375 | 2 | 0.5 | 15 | 0.4 |
| 14_558410 | 2 | 0 | 2 | 0 | 4 | 0.25 | NA | NA | 8 | 0.125 | 3 | 0.3333 | 2 | 0 | 2 | 0 | 2 | 0 | 9 | 0.1111 |
| 14_558465 | 1 | 1 | 5 | 1 | 1 | 0 | NA | NA | 7 | 0.8571 | 2 | 1 | NA | NA | NA | NA | 2 | 0.5 | 4 | 0.75 |
| 14_558467 | 1 | 1 | 5 | 1 | 1 | 1 | NA | NA | 7 | 1 | 2 | 1 | NA | NA | NA | NA | 2 | 0.5 | 4 | 0.75 |
| 14_558467 | 1 | 0 | 5 | 0 | 1 | 1 | NA | NA | 7 | 0.1429 | 2 | 0.5 | NA | NA | NA | NA | 2 | 0 | 4 | 0.25 |
| 14_558468 | 1 | 0 | 5 | 0.2 | 2 | 0 | NA | NA | 8 | 0.125 | 2 | 0.5 | NA | NA | NA | NA | 2 | 1 | 4 | 0.75 |
| 14_558469 | 1 | 0 | 5 | 0.4 | 2 | 0.5 | NA | NA | 8 | 0.375 | 2 | 0.5 | NA | NA | NA | NA | 2 | 0 | 4 | 0.25 |
| 14_558632 | 15 | 0.8 | 12 | 0.9167 | 11 | 0.9091 | 9 | 0.6667 | 47 | 0.8298 | 5 | 0.4 | 8 | 0.5 | 6 | 0.8333 | 5 | 0.8 | 24 | 0.625 |
| 14_558633 | 12 | 0.4167 | 7 | 0.2857 | 11 | 0.1818 | 7 | 0.2857 | 37 | 0.2973 | 4 | 0.25 | 7 | 0.4286 | 6 | 0.5 | 6 | 0.5 | 23 | 0.4348 |
| 14_558633 | 15 | 0.1333 | 9 | 0.1111 | 10 | 0.1 | 9 | 0 | 43 | 0.093 | 4 | 0 | 6 | 0.1667 | 5 | 0.2 | 7 | 0.1429 | 22 | 0.1364 |
| 14_558735 | 5 | 0.2 | 6 | 0.3333 | 5 | 0.4 | 5 | 0.2 | 21 | 0.2857 | 5 | 0.2 | 2 | 0 | 2 | 0.5 | 6 | 0 | 15 | 0.1333 |
| 14_558736 | 6 | 0.8333 | 4 | 0.75 | 5 | 0.2 | 4 | 0.75 | 19 | 0.6316 | 5 | 0.6 | 4 | 0.75 | 4 | 0.75 | 7 | 0.5714 | 20 | 0.65 |
| 14_558737 | 5 | 0.2 | 5 | 0.6 | 5 | 0.2 | 3 | 0.3333 | 18 | 0.3333 | 4 | 0 | 3 | 0.3333 | 3 | 0.6667 | 8 | 0.25 | 18 | 0.2778 |
| 14_558738 | 4 | 0.25 | 6 | 0.1667 | 5 | 0.2 | 4 | 0.25 | 19 | 0.2105 | 4 | 0 | 3 | 0.6667 | 3 | 0.6667 | 8 | 0 | 18 | 0.2222 |
| 14_558787 | 4 | 0 | 3 | 0 | 3 | 0.3333 | 1 | 0 | 11 | 0.0909 | 2 | 0 | 1 | 0 | 3 | 0 | 6 | 0.3333 | 12 | 0.1667 |
| 14_558787 | 5 | 0 | 6 | 0 | 3 | 0.3333 | 1 | 1 | 15 | 0.1333 | 1 | 0 | 3 | 0.3333 | 3 | 0 | 6 | 0 | 13 | 0.0769 |
| 14_558789 | 5 | 1 | 4 | 1 | 3 | 0.3333 | 2 | 0.5 | 14 | 0.7857 | 2 | 1 | 2 | 0.5 | 3 | 1 | 6 | 1 | 13 | 0.9231 |
| 14_558789 | 5 | 0.4 | 5 | 1 | 3 | 0.3333 | 1 | 1 | 14 | 0.6429 | 2 | 0.5 | 2 | 0.5 | 2 | 1 | 6 | 0.5 | 12 | 0.5833 |
| 14_558802 | 5 | 0 | 5 | 0.2 | 3 | 0 | 2 | 0 | 15 | 0.0667 | 2 | 0.5 | 3 | 0 | 2 | 0 | 5 | 0.4 | 12 | 0.25 |
| 14_558955 | 4 | 0.25 | 9 | 0.1111 | 5 | 0 | 4 | 0 | 22 | 0.0909 | 4 | 0 | 3 | 0 | 4 | 0 | 6 | 0.1667 | 17 | 0.0588 |
| 14_558956 | 3 | 0 | 8 | 0 | 4 | 0 | 5 | 0.2 | 20 | 0.05 | 4 | 0.25 | 3 | 0 | 3 | 0 | 4 | 0 | 14 | 0.0714 |
| 14_558957 | 5 | 0.2 | 8 | 0.25 | 3 | 0.3333 | 5 | 0 | 21 | 0.1905 | 3 | 0.3333 | 4 | 0.5 | 3 | 0 | 7 | 0 | 17 | 0.1765 |
| 14_558957 | 5 | 0.4 | 8 | 0.25 | 5 | 0.4 | 6 | 0.3333 | 24 | 0.3333 | 5 | 0.6 | 3 | 0 | 5 | 0.2 | 7 | 0.2857 | 20 | 0.3 |
| 14_558964 | 3 | 0.3333 | 7 | 0.4286 | 4 | 0.25 | 4 | 0.5 | 18 | 0.3889 | 2 | 0.5 | 2 | 0 | 4 | 0.25 | 7 | 0.7143 | 15 | 0.4667 |
| 14_558964 | 5 | 0.2 | 9 | 0.3333 | 6 | 0.1667 | 5 | 0.8 | 25 | 0.36 | 2 | 0 | 4 | 0.25 | 4 | 0.5 | 5 | 0.6 | 15 | 0.4 |
| 14_559158 | 3 | 0.3333 | 4 | 0 | 2 | 0 | 2 | 0 | 11 | 0.0909 | 3 | 0 | 3 | 0 | 5 | 0.2 | 5 | 0.4 | 16 | 0.1875 |
| 14_559160 | 2 | 0 | 4 | 0.25 | 2 | 0.5 | 2 | 0 | 10 | 0.2 | 3 | 0 | 2 | 0.5 | 7 | 0.1429 | 4 | 0.5 | 16 | 0.25 |
| 14_559176 | NA | NA | 3 | 0 | 1 | 0 | 5 | 0.2 | 9 | 0.1111 | 1 | 0 | 1 | 0 | 2 | 0.5 | 1 | 0 | 5 | 0.2 |
| 14_559177 | NA | NA | 2 | 0 | NA | NA | 3 | 0.6667 | 5 | 0.4 | 1 | 1 | 2 | 0.5 | 2 | 0.5 | 1 | 0 | 6 | 0.5 |
| 14_559177 | NA | NA | 3 | 0 | 1 | 1 | 3 | 0 | 7 | 0.1429 | NA | NA | 1 | 0 | 2 | 0.5 | 1 | 1 | 4 | 0.5 |
| 14_559535 | 2 | 0.5 | 4 | 0.5 | 4 | 0.25 | 6 | 0.3333 | 16 | 0.375 | 1 | 1 | 2 | 1 | 4 | 0.5 | 3 | 0.6667 | 10 | 0.7 |
| 14_559537 | 4 | 0.25 | 4 | 0.5 | 7 | 0.1429 | 9 | 0.6667 | 24 | 0.4167 | 4 | 1 | 6 | 0.1667 | 8 | 0.25 | 5 | 0.6 | 23 | 0.4348 |
| 14_559538 | 6 | 0.6667 | 4 | 0.75 | 7 | 0.5714 | 7 | 0.7143 | 24 | 0.6667 | 3 | 1 | 5 | 0.4 | 7 | 0.7143 | 5 | 0.8 | 20 | 0.7 |
| 14_559579 | 2 | 0 | 2 | 0 | 5 | 0.2 | 6 | 0 | 15 | 0.0667 | 2 | 0.5 | 3 | 0 | 3 | 0 | 2 | 0 | 10 | 0.1 |
| 14_559836 | NA | NA | 2 | 0.5 | NA | NA | 2 | 0 | 4 | 0.25 | 2 | 0 | NA | NA | 2 | 1 | NA | NA | 4 | 0.5 |
| 14_559912 | 3 | 0.3333 | 4 | 0 | 1 | 0 | 4 | 0.25 | 12 | 0.1667 | 2 | 0 | 1 | 1 | 2 | 0.5 | NA | NA | 5 | 0.4 |
| 14_559912 | 3 | 0.6667 | 4 | 0.5 | 1 | 1 | 4 | 0.75 | 12 | 0.6667 | NA | NA | 1 | 1 | 2 | 0.5 | NA | NA | 3 | 0.6667 |
| 14_559913 | 3 | 0.6667 | 5 | 0.2 | NA | NA | 3 | 0.6667 | 11 | 0.4545 | 1 | 1 | 1 | 0 | 3 | 1 | NA | NA | 5 | 0.8 |
| 14_559918 | 1 | 1 | 1 | 1 | 2 | 0.5 | 1 | 1 | 5 | 0.8 | 2 | 0.5 | NA | NA | 1 | 0 | 3 | 0.6667 | 6 | 0.5 |
| 14_561101 | 1 | 0 | NA | NA | 1 | 1 | NA | NA | 2 | 0.5 | NA | NA | NA | NA | 1 | 1 | NA | NA | 1 | 1 |
| 14_561111 | 1 | 1 | NA | NA | 1 | 1 | NA | NA | 2 | 1 | NA | NA | NA | NA | NA | NA | 1 | 1 | 1 | 1 |
| 14_561324 | NA | NA | NA | NA | 1 | 1 | NA | NA | 1 | 1 | 1 | 1 | 1 | 1 | 2 | 1 | 1 | 0 | 5 | 0.8 |
| 14_561337 | NA | NA | NA | NA | 3 | 1 | NA | NA | 3 | 1 | 1 | 1 | 2 | 1 | 2 | 1 | 1 | 1 | 6 | 1 |
| 14_561338 | NA | NA | NA | NA | 3 | 0.3333 | NA | NA | 3 | 0.3333 | NA | NA | 1 | 1 | 1 | 0 | 1 | 0 | 3 | 0.3333 |
| 14_561346 | NA | NA | 1 | 1 | 2 | 0 | NA | NA | 3 | 0.3333 | NA | NA | NA | NA | 1 | 1 | NA | NA | 1 | 1 |
| 14_561349 | 14 | 0.0714 | 10 | 0.4 | 7 | 0.1429 | 18 | 0.2222 | 49 | 0.2041 | 14 | 0.1429 | 14 | 0.1429 | 10 | 0.2 | 9 | 0 | 47 | 0.1277 |
| 14_561351 | 15 | 0.3333 | 7 | 0.5714 | 7 | 0.4286 | 17 | 0.3529 | 46 | 0.3913 | 14 | 0.6429 | 15 | 0.1333 | 10 | 0.3 | 9 | 0 | 48 | 0.2917 |
| 14_561351 | 1 | 1 | NA | NA | 3 | 0 | NA | NA | 4 | 0.25 | 1 | 1 | NA | NA | 1 | 0 | 1 | 0 | 3 | 0.3333 |
| 14_561352 | 1 | 1 | NA | NA | 3 | 0.3333 | NA | NA | 4 | 0.5 | NA | NA | NA | NA | 1 | 1 | NA | NA | 1 | 1 |
| 14_561353 | 1 | 1 | NA | NA | 3 | 0.6667 | NA | NA | 4 | 0.75 | 1 | 1 | NA | NA | 1 | 1 | 1 | 0 | 3 | 0.6667 |
| 14_561353 | 12 | 0.25 | 9 | 0.7778 | 5 | 0.6 | 17 | 0.2941 | 43 | 0.4186 | 15 | 0.4 | 10 | 0.4 | 10 | 0.3 | 9 | 0.5556 | 44 | 0.4091 |
| 14_561354 | 1 | 1 | NA | NA | 2 | 0 | NA | NA | 3 | 0.3333 | 1 | 1 | NA | NA | NA | NA | NA | NA | 1 | 1 |
| 14_561356 | NA | NA | 3 | 0.6667 | 1 | 1 | 2 | 0.5 | 6 | 0.6667 | 4 | 1 | 2 | 1 | 4 | 1 | NA | NA | 10 | 1 |
| 14_561356 | NA | NA | 2 | 0.5 | 1 | 1 | 1 | 0 | 4 | 0.5 | 4 | 0 | 3 | 0.6667 | 3 | 0 | NA | NA | 10 | 0.2 |
| 14_561357 | 9 | 0 | 7 | 0.1429 | 3 | 0.3333 | 13 | 0.0769 | 32 | 0.0938 | 14 | 0.2143 | 11 | 0.1818 | 10 | 0.1 | 10 | 0.1 | 45 | 0.1556 |
| 14_561358 | 2 | 0 | 1 | 1 | 2 | 0 | 5 | 0.2 | 10 | 0.2 | 4 | 0.75 | 5 | 0.2 | 5 | 0.4 | 2 | 1 | 16 | 0.5 |

| | | | | | | | | | | | | | | | | | | | | |
|---|---|---|---|---|---|---|---|---|---|---|---|---|---|---|---|---|---|---|---|---|
| 14_561359 | NA | NA | 2 | 1 | 1 | 1 | 2 | 0.5 | 5 | 0.8 | 4 | 0.75 | 3 | 1 | 4 | 0.75 | NA | NA | 11 | 0.8182 |
| 14_561360 | NA | NA | 3 | 0 | 1 | 0 | 2 | 0.5 | 6 | 0.1667 | 4 | 0.5 | 3 | 0 | 4 | 0.5 | 1 | 0 | 12 | 0.3333 |
| 14_567643 | 1 | 1 | NA | NA | NA | NA | 1 | 1 | 2 | 1 | NA | NA | NA | NA | 1 | 1 | 1 | 1 | 2 | 1 |
| 14_567644 | 1 | 0 | NA | NA | NA | NA | 1 | 1 | 2 | 0.5 | NA | NA | NA | NA | NA | NA | 1 | 1 | 1 | 1 |
| 14_567898 | 1 | 1 | NA | NA | NA | NA | NA | NA | 1 | 1 | NA | NA | NA | NA | NA | NA | 1 | 1 | 1 | 1 |
| 14_567906 | 1 | 1 | NA | NA | NA | NA | NA | NA | 1 | 1 | NA | NA | NA | NA | NA | NA | 1 | 1 | 1 | 1 |
| 14_567907 | 1 | 1 | NA | NA | NA | NA | NA | NA | 1 | 1 | NA | NA | NA | NA | NA | NA | 1 | 1 | 1 | 1 |
| 14_569114 | 4 | 0 | 4 | 0 | 5 | 0 | 4 | 0.25 | 17 | 0.0588 | 2 | 0 | 6 | 0.1667 | 3 | 0 | 1 | 0 | 12 | 0.0833 |
| 14_569116 | 4 | 0.75 | 3 | 1 | 5 | 0.6 | 4 | 0.5 | 16 | 0.6875 | 2 | 1 | 6 | 0.6667 | 3 | 0.6667 | 1 | 1 | 12 | 0.75 |
| 14_569118 | 4 | 0 | 4 | 0.25 | 7 | 0.4286 | 3 | 0.6667 | 18 | 0.3333 | 2 | 0 | 7 | 0.7143 | 3 | 0.3333 | 1 | 1 | 13 | 0.5385 |
| 14_569118 | 3 | 0.3333 | 2 | 0 | 6 | 0 | 3 | 0 | 14 | 0.0714 | 2 | 0.5 | 5 | 0 | 3 | 0 | 1 | 0 | 11 | 0.0909 |
| 14_569352 | 2 | 1 | 1 | 1 | 1 | 1 | 2 | 1 | 6 | 1 | NA | NA | 3 | 0.6667 | 1 | 1 | 1 | 1 | 5 | 0.8 |
| 14_569606 | 4 | 0.25 | 5 | 0 | 3 | 0 | NA | NA | 12 | 0.0833 | 3 | 0 | 4 | 0.25 | 2 | 0 | 7 | 0 | 16 | 0.0625 |
| 14_569607 | 3 | 0 | 6 | 0.1667 | 3 | 0 | NA | NA | 12 | 0.0833 | 2 | 0.5 | 4 | 0 | 2 | 0 | 4 | 0.5 | 12 | 0.25 |
| 14_570153 | 2 | 0 | 5 | 0.4 | 1 | 0 | NA | NA | 8 | 0.25 | 2 | 1 | 1 | 0 | NA | NA | 3 | 0.3333 | 6 | 0.5 |
| 14_570307 | 8 | 0.125 | 2 | 0 | 2 | 0 | NA | NA | 12 | 0.0833 | 1 | 0 | 5 | 0 | 1 | 0 | 5 | 0.4 | 12 | 0.1667 |
| 14_570309 | 7 | 0.4286 | 2 | 0.5 | 1 | 0 | NA | NA | 10 | 0.4 | 1 | 0 | 4 | 0.25 | 1 | 1 | 5 | 0.6 | 11 | 0.4545 |
| 14_570310 | 7 | 0.2857 | 2 | 0 | 1 | 0 | NA | NA | 10 | 0.2 | 1 | 0 | 3 | 0 | 1 | 0 | 4 | 0.25 | 9 | 0.1111 |
| 14_570311 | 7 | 0.7143 | 2 | 0.5 | 2 | 1 | NA | NA | 11 | 0.7273 | 1 | 0 | 3 | 1 | NA | NA | 4 | 0.75 | 8 | 0.75 |
| 14_570311 | 8 | 0.25 | 2 | 0 | 2 | 0 | NA | NA | 12 | 0.1667 | 1 | 0 | 4 | 0.25 | 1 | 0 | 4 | 0 | 10 | 0.1 |
| 14_570312 | 7 | 0.7143 | 2 | 0 | 2 | 0.5 | NA | NA | 11 | 0.5455 | 1 | 0 | 3 | 0.3333 | 1 | 1 | 3 | 0.3333 | 8 | 0.375 |
| 14_570314 | 8 | 1 | 2 | 1 | 2 | 1 | NA | NA | 12 | 1 | 1 | 1 | 4 | 0.75 | 1 | 1 | 3 | 1 | 9 | 0.8889 |
| 14_570314 | 8 | 1 | 2 | 0.5 | 1 | 1 | NA | NA | 11 | 0.9091 | 1 | 1 | 4 | 1 | 1 | 1 | 3 | 0.6667 | 9 | 0.8889 |
| 14_570315 | 7 | 0.2857 | 1 | 0 | 2 | 0.5 | NA | NA | 10 | 0.3 | 1 | 0 | NA | NA | NA | NA | 2 | 0.5 | 3 | 0.3333 |
| 14_570317 | 7 | 0.1429 | 2 | 0 | 2 | 0.5 | NA | NA | 11 | 0.1818 | 1 | 0 | 2 | 0.5 | 1 | 0 | 4 | 0 | 8 | 0.125 |
| 14_570317 | 6 | 0.8333 | 2 | 1 | 2 | 1 | NA | NA | 10 | 0.9 | 1 | 0 | 2 | 1 | 1 | 1 | 2 | 1 | 6 | 0.8333 |
| 14_570341 | 6 | 0 | NA | NA | 2 | 0.5 | NA | NA | 8 | 0.125 | 1 | 0 | 2 | 1 | 1 | 0 | 2 | 0 | 6 | 0.3333 |
| 14_570419 | 6 | 0.1667 | 1 | 0 | 2 | 0.5 | NA | NA | 9 | 0.2222 | 1 | 0 | 1 | 0 | NA | NA | 4 | 0.25 | 6 | 0.1667 |
| 14_570419 | 6 | 0.8333 | 1 | 0 | 2 | 0 | NA | NA | 9 | 0.5556 | 1 | 0 | 1 | 0 | NA | NA | 4 | 0.5 | 6 | 0.3333 |
| 14_570420 | 6 | 0.1667 | 1 | 0 | 2 | 0 | NA | NA | 9 | 0.1111 | 1 | 0 | NA | NA | NA | NA | 3 | 0.3333 | 4 | 0.25 |
| 14_570420 | 5 | 0.8 | NA | NA | 1 | 1 | NA | NA | 6 | 0.8333 | 1 | 1 | NA | NA | NA | NA | 4 | 0.75 | 5 | 0.8 |
| 14_570421 | 4 | 0.5 | 1 | 1 | 1 | 0 | NA | NA | 6 | 0.5 | 1 | 1 | NA | NA | NA | NA | 4 | 0.25 | 5 | 0.4 |
| 14_570497 | NA | NA | 1 | 1 | NA | NA | 1 | 1 | 2 | 1 | NA | NA | 2 | 1 | NA | NA | NA | NA | 2 | 1 |
| 14_570498 | NA | NA | 1 | 1 | NA | NA | 1 | 0 | 2 | 0.5 | NA | NA | 3 | 0.3333 | NA | NA | NA | NA | 3 | 0.3333 |
| 14_570499 | NA | NA | 1 | 0 | NA | NA | 1 | 1 | 2 | 0.5 | NA | NA | 2 | 1 | NA | NA | NA | NA | 2 | 1 |
| 14_570499 | NA | NA | 1 | 0 | NA | NA | 1 | 1 | 2 | 0.5 | NA | NA | 3 | 0.6667 | NA | NA | NA | NA | 3 | 0.6667 |
| 14_570500 | NA | NA | 1 | 1 | NA | NA | 1 | 0 | 2 | 0.5 | NA | NA | 3 | 0.3333 | NA | NA | NA | NA | 3 | 0.3333 |
| 14_570503 | NA | NA | 1 | 0 | NA | NA | 1 | 1 | 2 | 0.5 | NA | NA | 3 | 0 | NA | NA | 1 | 1 | 4 | 0.25 |
| 14_570503 | NA | NA | 1 | 1 | NA | NA | 1 | 1 | 2 | 1 | NA | NA | 3 | 1 | NA | NA | 1 | 1 | 4 | 1 |
| 14_570504 | NA | NA | 1 | 1 | NA | NA | 1 | 0 | 2 | 0.5 | NA | NA | 2 | 0 | NA | NA | 1 | 1 | 3 | 0.3333 |
| 14_571401 | 3 | 0 | 2 | 0 | 2 | 0.5 | 4 | 0.25 | 11 | 0.1818 | 2 | 0.5 | 3 | 0 | 2 | 0.5 | 5 | 0 | 12 | 0.1667 |
| 14_571403 | 3 | 0 | 4 | 0.25 | 2 | 0 | 3 | 0.3333 | 12 | 0.1667 | 2 | 0 | 3 | 0 | 2 | 0.5 | 4 | 0.75 | 11 | 0.3636 |
| 14_571404 | 3 | 0 | 4 | 0 | 2 | 0.5 | 3 | 0 | 12 | 0.0833 | 3 | 0 | 4 | 0 | 2 | 0.5 | 5 | 0.6 | 14 | 0.2857 |
| 14_571404 | 3 | 0.3333 | 4 | 0 | 2 | 0 | 4 | 0.5 | 13 | 0.2308 | 2 | 0.5 | 4 | 0 | 2 | 0 | 3 | 0 | 11 | 0.0909 |
| 14_571405 | 3 | 0 | 4 | 0.25 | 2 | 0 | 4 | 0.25 | 13 | 0.1538 | 3 | 0 | 4 | 0 | 2 | 0 | 5 | 0.4 | 14 | 0.1429 |
| 14_571405 | 3 | 0 | 4 | 0.25 | 2 | 0 | 4 | 0.25 | 13 | 0.1538 | 3 | 1 | 4 | 0.25 | 1 | 0 | 5 | 0.2 | 13 | 0.3846 |
| 14_571405 | 3 | 0.3333 | 3 | 0 | 2 | 0 | 4 | 0 | 12 | 0.0833 | 2 | 0 | 4 | 0.25 | 3 | 0 | 5 | 0.2 | 14 | 0.1429 |
| 14_571406 | 2 | 0.5 | 3 | 1 | 2 | 0 | 1 | 1 | 8 | 0.625 | 1 | 0 | 2 | 0.5 | 2 | 0.5 | 3 | 0.3333 | 8 | 0.375 |
| 14_571406 | 3 | 0.3333 | 2 | 0 | 2 | 0 | 4 | 0.5 | 11 | 0.2727 | 3 | 0 | 4 | 0.5 | 3 | 0.6667 | 4 | 0.75 | 14 | 0.5 |
| 14_571407 | 3 | 1 | 3 | 1 | 2 | 1 | 4 | 1 | 12 | 1 | 3 | 0.6667 | 3 | 1 | 2 | 1 | 5 | 0.6 | 13 | 0.7692 |
| 14_571408 | 3 | 0.6667 | 3 | 0 | 2 | 1 | 3 | 0.6667 | 11 | 0.5455 | 3 | 0.6667 | 5 | 1 | 3 | 0.3333 | 4 | 1 | 15 | 0.8 |
| 14_571408 | 3 | 0.3333 | 4 | 0.25 | 2 | 0 | 4 | 0.25 | 13 | 0.2308 | 3 | 0.3333 | 4 | 0.5 | 3 | 0.3333 | 5 | 0.2 | 15 | 0.3333 |
| 14_571409 | 3 | 0.6667 | 2 | 1 | 2 | 0.5 | 3 | 0.6667 | 10 | 0.7 | 2 | 0 | 5 | 0.6 | 1 | 0 | 3 | 1 | 11 | 0.5455 |
| 14_571944 | 5 | 0.2 | 7 | 0.4286 | 4 | 0 | 5 | 0 | 21 | 0.1905 | 6 | 0.1667 | 9 | 0.2222 | 7 | 0.2857 | 7 | 0.2857 | 29 | 0.2414 |
| 14_571973 | 4 | 0.5 | 10 | 0.7 | 9 | 0.4444 | 7 | 0.4286 | 30 | 0.5333 | 7 | 0.7143 | 9 | 0.5556 | 6 | 0.3333 | 10 | 0.6 | 32 | 0.5625 |
| 14_571973 | 5 | 0.6 | 10 | 0.5 | 8 | 0.75 | 5 | 1 | 28 | 0.6786 | 6 | 1 | 9 | 0.6667 | 6 | 0.8333 | 6 | 0.8333 | 27 | 0.8148 |
| 14_571973 | 5 | 0.2 | 9 | 0.2222 | 9 | 0.3333 | 7 | 0.1429 | 30 | 0.2333 | 6 | 0.1667 | 9 | 0.2222 | 7 | 0 | 8 | 0.5 | 30 | 0.2333 |
| 14_571973 | 5 | 0.6 | 10 | 0.4 | 8 | 0.5 | 9 | 0.4444 | 32 | 0.4688 | 7 | 0.2857 | 8 | 0.25 | 7 | 0.5714 | 10 | 0.3 | 32 | 0.3438 |
| 14_571984 | 4 | 0.25 | 5 | 0 | 4 | 0 | 3 | 0.3333 | 16 | 0.125 | 2 | 0 | 3 | 0.3333 | NA | NA | 2 | 0 | 7 | 0.1429 |
| 14_571991 | 3 | 0.3333 | 5 | 0.4 | 4 | 0 | 4 | 0 | 16 | 0.1875 | 2 | 0 | 3 | 0.3333 | 2 | 0.5 | 2 | 0 | 9 | 0.2222 |
| 14_571995 | 5 | 0 | 4 | 0.25 | 4 | 0.25 | 4 | 0 | 17 | 0.1176 | NA | NA | 4 | 0.75 | 1 | 0 | 2 | 0 | 7 | 0.4286 |
| 14_573362 | 9 | 0.3333 | 3 | 0.3333 | NA | NA | 3 | 1 | 15 | 0.4667 | 1 | 1 | NA | NA | NA | NA | 1 | 1 | 2 | 1 |
| 14_573376 | 7 | 1 | 2 | 1 | NA | NA | 1 | 1 | 10 | 1 | 1 | 1 | NA | NA | NA | NA | NA | NA | 1 | 1 |
| 14_573376 | 7 | 1 | 1 | 1 | NA | NA | 1 | 1 | 9 | 1 | 1 | 1 | NA | NA | NA | NA | NA | NA | 1 | 1 |
| 14_573608 | 28 | 0.1429 | 23 | 0.2174 | 21 | 0.1429 | 17 | 0.1176 | 89 | 0.1573 | 12 | 0.3333 | 10 | 0.1 | 12 | 0.1667 | 15 | 0.2667 | 49 | 0.2245 |

| | | | | | | | | | | | | | | | | | | | | |
|---|---|---|---|---|---|---|---|---|---|---|---|---|---|---|---|---|---|---|---|---|
| 14_5736089 | 19 | 0.4211 | 18 | 0.5556 | 21 | 0.4286 | 12 | 0.4167 | 70 | 0.4571 | 11 | 0.5455 | 10 | 0.4 | 11 | 0.3636 | 14 | 0.3571 | 46 | 0.413 |
| 14_5736099 | 22 | 0.1364 | 21 | 0.0952 | 19 | 0 | 12 | 0.0833 | 74 | 0.0811 | 11 | 0.1818 | 7 | 0.1429 | 10 | 0 | 13 | 0.0769 | 41 | 0.0976 |
| 14_5736107 | 24 | 0 | 19 | 0.1053 | 21 | 0.0952 | 12 | 0 | 76 | 0.0526 | 13 | 0.2308 | 10 | 0.3 | 10 | 0.1 | 13 | 0 | 46 | 0.1522 |
| 14_5736117 | 21 | 0.0952 | 21 | 0.0952 | 16 | 0.0625 | 15 | 0.0667 | 73 | 0.0822 | 12 | 0.3333 | 7 | 0.2857 | 10 | 0.2 | 14 | 0.0714 | 43 | 0.2093 |
| 14_5736122 | 21 | 0.2381 | 22 | 0.5 | 17 | 0.1765 | 17 | 0.3529 | 77 | 0.3247 | 12 | 0.25 | 9 | 0.5556 | 10 | 0.3 | 13 | 0.4615 | 44 | 0.3864 |
| 14_5736123 | 18 | 0 | 18 | 0.2778 | 17 | 0.1176 | 12 | 0.3333 | 65 | 0.1692 | 11 | 0.4545 | 8 | 0.375 | 7 | 0.2857 | 10 | 0.1 | 36 | 0.3056 |
| 14_5736136 | 20 | 0.25 | 21 | 0.2381 | 18 | 0.3333 | 14 | 0.7143 | 73 | 0.3562 | 10 | 0.4 | 10 | 0.7 | 9 | 0.5556 | 12 | 0.25 | 41 | 0.4634 |
| 14_5736158 | 18 | 0.1111 | 16 | 0.125 | 15 | 0 | 10 | 0.2 | 59 | 0.1017 | 6 | 0.1667 | 8 | 0 | 7 | 0.1429 | 13 | 0.1538 | 34 | 0.1176 |
| 14_5736172 | 21 | 0.9048 | 20 | 0.9 | 13 | 0.9231 | 13 | 0.8462 | 67 | 0.8955 | 8 | 1 | 9 | 1 | 7 | 0.8571 | 13 | 1 | 37 | 0.973 |
| 14_5736190 | 20 | 0.25 | 18 | 0.3889 | 19 | 0.3684 | 12 | 0.25 | 69 | 0.3188 | 12 | 0.3333 | 8 | 0.375 | 8 | 0.5 | 12 | 0.1667 | 40 | 0.325 |
| 14_5736191 | 23 | 0.087 | 20 | 0.15 | 20 | 0.1 | 15 | 0.2667 | 78 | 0.141 | 12 | 0.0833 | 8 | 0.375 | 8 | 0.125 | 13 | 0.0769 | 41 | 0.1463 |
| 14_5736195 | 17 | 0.3529 | 14 | 0.7143 | 16 | 0.625 | 10 | 0.4 | 57 | 0.5263 | 9 | 0.4444 | 5 | 0.4 | 3 | 0.6667 | 9 | 0.5556 | 26 | 0.5 |
| 14_5736196 | 22 | 0.5455 | 19 | 0.5789 | 18 | 0.5 | 15 | 0.6 | 74 | 0.5541 | 10 | 0.5 | 8 | 0.625 | 7 | 0.4286 | 13 | 0.7692 | 38 | 0.6053 |
| 14_5736211 | 20 | 0.85 | 20 | 0.7 | 17 | 0.7647 | 12 | 0.75 | 69 | 0.7681 | 12 | 0.75 | 7 | 1 | 6 | 0.8333 | 14 | 0.5 | 39 | 0.7179 |
| 14_5736225 | 18 | 0.2222 | 16 | 0.0625 | 17 | 0.0588 | 10 | 0.3 | 61 | 0.1475 | 11 | 0.1818 | 4 | 0 | 6 | 0 | 12 | 0.25 | 33 | 0.1515 |
| 14_5775492 | NA | NA | 1 | 1 | 2 | 0.5 | 1 | 0 | 4 | 0.5 | 2 | 0 | 3 | 0.3333 | 2 | 1 | 1 | 0 | 8 | 0.375 |
| 14_5775725 | NA | NA | NA | NA | 1 | 1 | NA | NA | 1 | 1 | NA | NA | NA | NA | NA | NA | 1 | 1 | 1 | 1 |
| 14_5776172 | 4 | 1 | 6 | 0.5 | 3 | 1 | 6 | 1 | 19 | 0.8421 | 7 | 0.8571 | 2 | 1 | 4 | 1 | 6 | 1 | 19 | 0.9474 |
| 14_5776181 | 10 | 0 | 10 | 0.3 | 3 | 0 | 9 | 0.1111 | 32 | 0.125 | 8 | 0 | 5 | 0.2 | 6 | 0.1667 | 8 | 0.125 | 27 | 0.1111 |
| 14_5776191 | 7 | 0.1429 | 8 | 0.5 | 4 | 0.5 | 11 | 0.6364 | 30 | 0.4667 | 6 | 0.3333 | 3 | 0.3333 | 3 | 0.6667 | 6 | 0.3333 | 18 | 0.3889 |
| 14_5776197 | 6 | 0.1667 | 7 | 0.1429 | 1 | 1 | 5 | 0.4 | 19 | 0.2632 | 6 | 0.3333 | 3 | 0.6667 | 3 | 0.3333 | 4 | 0 | 16 | 0.3125 |
| 14_5776204 | 11 | 0.9091 | 10 | 1 | 4 | 1 | 10 | 0.9 | 35 | 0.9429 | 6 | 1 | 5 | 1 | 6 | 1 | 9 | 1 | 26 | 1 |
| 14_5776206 | 4 | 1 | 3 | 0.6667 | 2 | 1 | 6 | 0.8333 | 15 | 0.8667 | 5 | 0.8 | 2 | 1 | 4 | 1 | 7 | 1 | 18 | 0.9444 |
| 14_5776208 | 6 | 0.5 | 4 | 0 | 2 | 0.5 | 5 | 0 | 17 | 0.2353 | 3 | 1 | 4 | 0.5 | 5 | 0.4 | 7 | 0.4286 | 19 | 0.5263 |
| 14_5776210 | 6 | 0.5 | 6 | 0.5 | 4 | 0.75 | 7 | 0.2857 | 23 | 0.4783 | 5 | 0.4 | 4 | 0.75 | 3 | 0 | 6 | 0.1667 | 18 | 0.3333 |
| 14_5776217 | 7 | 0.5714 | 9 | 0.1111 | 2 | 0.5 | 9 | 0.4444 | 27 | 0.3704 | 8 | 0.5 | 4 | 0.75 | 7 | 0.4286 | 7 | 0.5714 | 26 | 0.5385 |
| 14_5776219 | 9 | 0.2222 | 7 | 0 | 1 | 0 | 9 | 0.4444 | 26 | 0.2308 | 6 | 0.1667 | 5 | 0.2 | 7 | 0 | 6 | 0 | 24 | 0.0833 |
| 14_5776223 | 3 | 1 | 9 | 0.7778 | 3 | 1 | 7 | 1 | 22 | 0.9091 | 7 | 1 | 3 | 1 | 3 | 1 | 6 | 1 | 19 | 1 |
| 14_5776234 | 6 | 0.6667 | 8 | 0.5 | 3 | 0.6667 | 9 | 0.4444 | 26 | 0.5385 | 5 | 0.6 | 2 | 0 | 6 | 0.6667 | 5 | 0.6 | 18 | 0.5556 |
| 14_5778003 | 3 | 0.6667 | 4 | 0.75 | 4 | 0 | NA | NA | 11 | 0.4545 | 5 | 0.8 | 1 | 0 | 1 | 1 | 1 | 1 | 8 | 0.75 |
| 14_5778005 | 4 | 0.25 | 5 | 0.6 | 4 | 0.75 | NA | NA | 13 | 0.5385 | 5 | 0.6 | 1 | 1 | 2 | 0.5 | 1 | 0 | 9 | 0.5556 |
| 14_5778010 | 5 | 0.2 | 5 | 1 | 3 | 0.3333 | NA | NA | 13 | 0.5385 | 4 | 0.25 | 2 | 0.5 | 3 | 0.3333 | 1 | 1 | 10 | 0.4 |
| 14_5778020 | 4 | 0.5 | 4 | 1 | 3 | 0.3333 | NA | NA | 11 | 0.6364 | 3 | 0.6667 | 1 | 1 | 3 | 1 | 1 | 1 | 8 | 0.875 |
| 14_5778028 | 5 | 0 | 5 | 0.2 | 4 | 0 | NA | NA | 14 | 0.0714 | 4 | 0.25 | 2 | 0.5 | 3 | 0 | 1 | 0 | 10 | 0.2 |
| 14_5778040 | 4 | 0 | 5 | 0 | 3 | 0.6667 | NA | NA | 12 | 0.1667 | 4 | 0.25 | NA | NA | 3 | 0.3333 | 1 | 0 | 8 | 0.25 |
| 14_5778717 | 12 | 0.3333 | 9 | 0.4444 | 11 | 0.4545 | 8 | 0.125 | 40 | 0.35 | 2 | 0 | 6 | 0.5 | 9 | 0.4444 | 5 | 0.4 | 22 | 0.4091 |
| 14_5778721 | 14 | 0.3571 | 10 | 0.1 | 11 | 0.5455 | 7 | 0.4286 | 42 | 0.3571 | 1 | 0 | 7 | 0.5714 | 8 | 0.375 | 4 | 0.75 | 20 | 0.5 |
| 14_5778722 | 10 | 0.2 | 9 | 0.1111 | 7 | 0.4286 | 6 | 0 | 32 | 0.1875 | 1 | 0 | 4 | 0 | 8 | 0.25 | 1 | 0 | 14 | 0.1429 |
| 14_5778723 | 12 | 0.3333 | 10 | 0.5 | 9 | 0.4444 | 7 | 0.4286 | 38 | 0.4211 | 1 | 0 | 4 | 0 | 5 | 0.2 | 4 | 0.5 | 14 | 0.2143 |
| 14_5781579 | 4 | 0 | 2 | 0 | 1 | 1 | 4 | 0.25 | 11 | 0.1818 | NA | NA | 2 | 0.5 | 2 | 0 | 2 | 0 | 6 | 0.1667 |
| 14_5781926 | 2 | 1 | 1 | 1 | NA | NA | 2 | 0.5 | 5 | 0.8 | NA | NA | NA | NA | 1 | 1 | NA | NA | 1 | 1 |
| 14_5781960 | 2 | 1 | 1 | 0 | NA | NA | 2 | 0.5 | 5 | 0.6 | NA | NA | NA | NA | 1 | 1 | NA | NA | 1 | 1 |
| 14_5782836 | 1 | 1 | 1 | 1 | NA | NA | 1 | 1 | 3 | 1 | NA | NA | NA | NA | 1 | 1 | NA | NA | 1 | 1 |
| 14_5792834 | 7 | 0.5714 | 4 | 0.25 | 3 | 0.6667 | 2 | 0 | 16 | 0.4375 | 2 | 1 | 1 | 0 | 5 | 0.2 | 2 | 0.5 | 10 | 0.4 |
| 14_5792849 | 5 | 0.8 | 2 | 0.5 | 3 | 0.6667 | 1 | 1 | 11 | 0.7273 | NA | NA | 1 | 0 | 3 | 1 | 1 | 1 | 5 | 0.8 |
| 14_5792850 | 3 | 1 | 3 | 1 | 3 | 1 | 1 | 1 | 10 | 1 | 2 | 1 | 1 | 1 | 4 | 1 | 2 | 1 | 9 | 1 |
| 14_5792858 | 5 | 0.6 | 4 | 0.5 | 2 | 0.5 | 1 | 0 | 12 | 0.5 | 2 | 0 | 1 | 1 | 3 | 0 | 1 | 1 | 7 | 0.2857 |
| 14_5792860 | 5 | 1 | 4 | 1 | 3 | 1 | 2 | 1 | 14 | 1 | 1 | 1 | 1 | 1 | 5 | 1 | 2 | 1 | 9 | 1 |
| 14_5792868 | 4 | 0.5 | 3 | 0.6667 | 3 | 0.6667 | 1 | 0 | 11 | 0.5455 | 1 | 1 | 1 | 0 | 3 | 0 | NA | NA | 5 | 0.2 |
| 14_5792869 | 6 | 0.1667 | 4 | 0.25 | 4 | 0 | 1 | 1 | 15 | 0.2 | 2 | 0 | 1 | 1 | 5 | 0 | 1 | 0 | 9 | 0.1111 |
| 14_5792878 | 6 | 1 | 4 | 1 | 3 | 1 | 1 | 1 | 14 | 1 | 2 | 1 | 1 | 1 | 4 | 1 | NA | NA | 7 | 1 |
| 14_5792880 | 5 | 0.6 | 4 | 0.25 | 4 | 0 | 1 | 0 | 14 | 0.2857 | 2 | 0 | 1 | 0 | 5 | 0.4 | 1 | 0 | 9 | 0.2222 |
| 14_5792893 | 4 | 0.5 | 4 | 0.5 | 3 | 0.3333 | 1 | 1 | 12 | 0.5 | 2 | 0.5 | 1 | 0 | 6 | 0.8333 | 1 | 1 | 10 | 0.7 |
| 14_5792899 | 5 | 0.2 | 4 | 0 | 4 | 0 | 1 | 0 | 14 | 0.0714 | 2 | 0 | 1 | 0 | 5 | 0 | 1 | 1 | 9 | 0.1111 |
| 14_5792900 | 6 | 1 | 4 | 1 | 4 | 0.75 | 1 | 1 | 15 | 0.9333 | 1 | 1 | 1 | 1 | 5 | 0.8 | 1 | 1 | 8 | 0.875 |
| 14_5792910 | 6 | 0 | 4 | 0 | 4 | 0 | 1 | 1 | 15 | 0.0667 | 2 | 0 | 1 | 0 | 6 | 0.1667 | 1 | 0 | 10 | 0.1 |
| 14_5793440 | 4 | 1 | 4 | 0.25 | 4 | 0.5 | 1 | 0 | 13 | 0.5385 | 2 | 0.5 | 1 | 0 | 4 | 0.25 | 1 | 1 | 8 | 0.375 |
| 14_5793461 | 4 | 1 | 3 | 0.6667 | 2 | 1 | NA | NA | 9 | 0.8889 | 1 | 1 | NA | NA | 5 | 1 | 1 | 1 | 7 | 1 |
| 14_5793463 | 3 | 1 | 3 | 0 | 2 | 0 | NA | NA | 8 | 0.375 | 1 | 0 | NA | NA | 4 | 0.5 | 1 | 1 | 6 | 0.5 |
| 14_5793464 | 4 | 0 | 3 | 0.3333 | 2 | 0 | NA | NA | 9 | 0.1111 | 1 | 0 | NA | NA | 3 | 0.3333 | 1 | 0 | 5 | 0.2 |
| 14_5793470 | 4 | 0.25 | 3 | 0 | 3 | 0.6667 | NA | NA | 10 | 0.3 | 2 | 0.5 | NA | NA | 5 | 0.2 | 1 | 0 | 8 | 0.25 |
| 14_5793478 | 4 | 0.75 | 3 | 1 | 1 | 1 | NA | NA | 8 | 0.875 | 2 | 0.5 | NA | NA | 3 | 0.6667 | 1 | 0 | 6 | 0.5 |
| 14_5793483 | 4 | 1 | 3 | 0 | 1 | 0 | NA | NA | 8 | 0.5 | 1 | 1 | NA | NA | 3 | 0.6667 | 1 | 0 | 5 | 0.6 |
| 14_5793486 | 4 | 0.75 | 3 | 0 | 2 | 0 | NA | NA | 9 | 0.3333 | 1 | 1 | NA | NA | 4 | 0.25 | 1 | 0 | 6 | 0.3333 |
| 14_5793701 | 3 | 0.3333 | 2 | 0.5 | 2 | 1 | NA | NA | 7 | 0.5714 | NA | NA | NA | NA | 3 | 0.3333 | 1 | 1 | 4 | 0.5 |

| | | | | | | | | | | | | | | | | | | | | |
|---|---|---|---|---|---|---|---|---|---|---|---|---|---|---|---|---|---|---|---|---|
| 14_579370 | 4 | 0.75 | 3 | 0.6667 | 3 | 0.6667 | NA | NA | 10 | 0.7 | 1 | 1 | NA | NA | 2 | 0 | 1 | 0 | 4 | 0.25 |
| 14_579371 | 4 | 1 | 2 | 1 | 2 | 1 | NA | NA | 8 | 1 | NA | NA | NA | NA | 3 | 1 | NA | NA | 3 | 1 |
| 14_579373 | 3 | 0 | 3 | 0 | 2 | 0.5 | NA | NA | 8 | 0.125 | NA | NA | NA | NA | 3 | 0.6667 | NA | NA | 3 | 0.6667 |
| 14_579378 | 2 | 1 | 1 | 1 | NA | NA | NA | NA | 3 | 1 | NA | NA | NA | NA | 1 | 1 | NA | NA | 1 | 1 |
| 14_579714 | 2 | 1 | 2 | 1 | 1 | 1 | NA | NA | 5 | 1 | NA | NA | NA | NA | 3 | 1 | NA | NA | 3 | 1 |
| 14_579715 | 1 | 1 | 2 | 0 | 1 | 0 | NA | NA | 4 | 0.25 | NA | NA | NA | NA | 3 | 0.3333 | NA | NA | 3 | 0.3333 |
| 14_579716 | 3 | 1 | 2 | 0.5 | 1 | 1 | NA | NA | 6 | 0.8333 | NA | NA | NA | NA | 2 | 1 | NA | NA | 2 | 1 |
| 14_579716 | 3 | 0.3333 | 2 | 0.5 | 1 | 1 | NA | NA | 6 | 0.5 | NA | NA | NA | NA | 2 | 1 | NA | NA | 2 | 1 |
| 14_579717 | 3 | 1 | 1 | 1 | NA | NA | NA | NA | 4 | 1 | NA | NA | NA | NA | 2 | 0.5 | NA | NA | 2 | 0.5 |
| 14_579718 | 2 | 0 | 2 | 0 | 1 | 1 | NA | NA | 5 | 0.2 | NA | NA | NA | NA | 2 | 1 | NA | NA | 2 | 1 |
| 14_580394 | 10 | 0.3 | 12 | 0.25 | 12 | 0.0833 | 14 | 0.0714 | 48 | 0.1667 | 11 | 0.0909 | 9 | 0.3333 | 13 | 0.3846 | 9 | 0.1111 | 42 | 0.2381 |
| 14_580395 | 16 | 0.1875 | 12 | 0.1667 | 16 | 0.25 | 19 | 0.1053 | 63 | 0.1746 | 16 | 0.1875 | 13 | 0.3077 | 15 | 0.1333 | 15 | 0.2 | 59 | 0.2034 |
| 14_580395 | 13 | 0 | 12 | 0.0833 | 15 | 0.0667 | 17 | 0.0588 | 57 | 0.0526 | 14 | 0.0714 | 11 | 0.0909 | 18 | 0.0556 | 16 | 0 | 59 | 0.0508 |
| 14_580396 | 12 | 0.5 | 13 | 0.4615 | 12 | 0.6667 | 13 | 0.6923 | 50 | 0.58 | 13 | 0.4615 | 10 | 0.4 | 12 | 0.6667 | 15 | 0.6667 | 50 | 0.56 |
| 14_580422 | 2 | 0 | 1 | 1 | 1 | 1 | NA | NA | 4 | 0.5 | 1 | 1 | NA | NA | 1 | 0 | 1 | 1 | 3 | 0.6667 |
| 14_580643 | 7 | 1 | 10 | 1 | 10 | 1 | 8 | 1 | 35 | 1 | 6 | 1 | 7 | 1 | 7 | 1 | 9 | 1 | 29 | 1 |
| 14_580643 | 6 | 0.1667 | 9 | 0.1111 | 9 | 0 | 7 | 0 | 31 | 0.0645 | 6 | 0.3333 | 7 | 0 | 7 | 0 | 9 | 0 | 29 | 0.069 |
| 14_580644 | 2 | 1 | 2 | 1 | 1 | 1 | NA | NA | 5 | 1 | 1 | 1 | NA | NA | 1 | 1 | NA | NA | 2 | 1 |
| 14_580645 | 2 | 1 | 1 | 1 | 1 | 1 | NA | NA | 4 | 1 | 1 | 1 | NA | NA | 2 | 1 | NA | NA | 3 | 1 |
| 14_580650 | 2 | 0.5 | 2 | 0 | 1 | 0 | NA | NA | 5 | 0.2 | 1 | 0 | 1 | 0 | 3 | 0.3333 | NA | NA | 5 | 0.2 |
| 14_580653 | 2 | 0.5 | 2 | 0 | 1 | 0 | 1 | 0 | 6 | 0.1667 | 1 | 0 | 1 | 0 | 3 | 0.3333 | NA | NA | 5 | 0.2 |
| 14_580656 | 2 | 0 | 2 | 1 | 1 | 0 | 1 | 1 | 6 | 0.5 | 1 | 0 | 1 | 0 | 3 | 0.3333 | NA | NA | 5 | 0.2 |
| 14_580703 | 2 | 0 | 2 | 0 | 1 | 0 | 5 | 0.2 | 10 | 0.1 | 2 | 0 | 3 | 0 | 9 | 0.1111 | 4 | 0.25 | 18 | 0.1111 |
| 14_580712 | 1 | 0 | 5 | 0.4 | 2 | 0 | 4 | 0 | 12 | 0.1667 | 2 | 0 | 1 | 0 | 8 | 0.125 | 4 | 0 | 15 | 0.0667 |
| 14_580719 | NA | NA | 3 | 0.3333 | 2 | 0 | 6 | 0.1667 | 11 | 0.1818 | NA | NA | 1 | 0 | 10 | 0.1 | 4 | 0.5 | 15 | 0.2 |
| 14_580720 | 1 | 0 | 4 | 0 | 2 | 0 | 4 | 0.25 | 11 | 0.0909 | NA | NA | 1 | 0 | 9 | 0 | 3 | 0.3333 | 13 | 0.0769 |
| 14_580721 | 1 | 0 | 6 | 0 | 2 | 0 | 6 | 0.3333 | 15 | 0.1333 | NA | NA | 1 | 1 | 11 | 0.2727 | 4 | 0.25 | 16 | 0.3125 |
| 14_580723 | 1 | 0 | 5 | 0.2 | 2 | 0.5 | 6 | 0.1667 | 14 | 0.2143 | NA | NA | 1 | 0 | 10 | 0.2 | 5 | 0.4 | 16 | 0.25 |
| 14_580723 | NA | NA | 3 | 0 | 2 | 0 | 6 | 0.6667 | 11 | 0.3636 | NA | NA | 1 | 0 | 6 | 0.5 | 3 | 0.3333 | 10 | 0.4 |
| 14_580723 | 1 | 0 | 4 | 0.5 | 2 | 0.5 | 7 | 0.4286 | 14 | 0.4286 | NA | NA | 1 | 1 | 10 | 0.6 | 4 | 0.5 | 15 | 0.6 |
| 14_580727 | 1 | 0 | 4 | 0.25 | 2 | 0 | 5 | 0.4 | 12 | 0.25 | NA | NA | NA | NA | 7 | 0.1429 | 4 | 0.25 | 11 | 0.1818 |
| 14_580738 | NA | NA | 2 | 0.5 | 1 | 1 | 2 | 1 | 5 | 0.8 | NA | NA | 1 | 1 | 7 | 0.8571 | 6 | 0.5 | 14 | 0.7143 |
| 14_580740 | NA | NA | 3 | 1 | 1 | 0 | 2 | 1 | 6 | 0.8333 | NA | NA | 1 | 0 | 5 | 0.6 | 6 | 0.5 | 12 | 0.5 |
| 14_581546 | 3 | 0 | 4 | 0 | 1 | 1 | 2 | 0 | 10 | 0.1 | 3 | 0.3333 | 2 | 0 | 3 | 0 | 2 | 0 | 10 | 0.1 |
| 14_581546 | 2 | 0 | 3 | 0.6667 | NA | NA | 1 | 0 | 6 | 0.3333 | 2 | 0.5 | 2 | 1 | 2 | 1 | 2 | 1 | 8 | 0.875 |
| 14_583099 | NA | NA | 1 | 0 | 1 | 1 | 1 | 0 | 3 | 0.3333 | NA | NA | NA | NA | 1 | 1 | NA | NA | 1 | 1 |
| 14_583105 | NA | NA | 1 | 0 | 1 | 1 | 1 | 1 | 3 | 0.6667 | 1 | 1 | 1 | 1 | NA | NA | NA | NA | 2 | 1 |
| 14_583106 | NA | NA | NA | NA | 1 | 1 | 1 | 0 | 2 | 0.5 | 2 | 1 | 1 | 1 | 1 | 1 | NA | NA | 4 | 1 |
| 14_583107 | NA | NA | 1 | 0 | 1 | 1 | NA | NA | 2 | 0.5 | 2 | 0 | 1 | 1 | 1 | 0 | NA | NA | 4 | 0.25 |
| 14_583469 | 2 | 1 | 2 | 0.5 | 3 | 1 | 1 | 0 | 8 | 0.75 | 2 | 0.5 | 2 | 1 | 1 | 1 | NA | NA | 5 | 0.8 |
| 14_583469 | 2 | 0 | 2 | 0 | 3 | 0.3333 | 1 | 0 | 8 | 0.125 | 2 | 0 | 2 | 0 | 1 | 1 | 1 | 0 | 6 | 0.1667 |
| 14_583469 | 2 | 0.5 | 2 | 0.5 | 3 | 0.6667 | 1 | 0 | 8 | 0.5 | 2 | 0.5 | 2 | 0 | 1 | 1 | 1 | 0 | 6 | 0.3333 |
| 14_583469 | 2 | 1 | 2 | 0.5 | 2 | 1 | 1 | 1 | 7 | 0.8571 | 2 | 0.5 | 3 | 0.6667 | 1 | 1 | 1 | 1 | 7 | 0.7143 |
| 14_583470 | 2 | 1 | 2 | 0.5 | 3 | 1 | NA | NA | 7 | 0.8571 | 2 | 1 | 2 | 0.5 | 1 | 1 | 1 | 1 | 6 | 0.8333 |
| 14_583470 | 2 | 0.5 | 2 | 0.5 | 2 | 1 | 1 | 0 | 7 | 0.5714 | 2 | 1 | 1 | 1 | 1 | 0 | 1 | 1 | 5 | 0.8 |
| 14_583471 | 2 | 1 | 2 | 1 | 3 | 0.3333 | 1 | 0 | 8 | 0.625 | 2 | 0.5 | 3 | 0.6667 | 2 | 1 | 1 | 1 | 8 | 0.75 |
| 14_583473 | 2 | 0 | 2 | 0 | 2 | 0.5 | NA | NA | 6 | 0.1667 | 1 | 0 | 3 | 0 | 2 | 0.5 | 1 | 0 | 7 | 0.1429 |
| 14_583473 | 2 | 0 | 3 | 0.3333 | 4 | 0.5 | 1 | 0 | 10 | 0.3 | 2 | 0 | 2 | 0 | 2 | 0.5 | 1 | 0 | 7 | 0.1429 |
| 14_583475 | 2 | 0 | 3 | 0.6667 | 3 | 0.6667 | 1 | 0 | 9 | 0.4444 | 2 | 0.5 | 2 | 0.5 | 1 | 1 | 1 | 1 | 6 | 0.6667 |
| 14_583492 | 1 | 0 | 1 | 0 | 2 | 0.5 | 2 | 0 | 6 | 0.1667 | 1 | 0 | 2 | 0 | 1 | 0 | 2 | 0.5 | 6 | 0.1667 |
| 14_594594 | 1 | 1 | NA | NA | 1 | 1 | NA | NA | 2 | 1 | NA | NA | NA | NA | 1 | 0 | 1 | 1 | 2 | 0.5 |
| 14_594728 | 5 | 0.4 | 2 | 1 | 4 | 1 | 4 | 0.5 | 15 | 0.6667 | 2 | 0 | 3 | 1 | 2 | 0 | 5 | 0.6 | 12 | 0.5 |
| 14_594728 | 4 | 0.25 | 2 | 1 | 6 | 0.3333 | 5 | 0 | 17 | 0.2941 | 1 | 0 | 3 | 0.3333 | 1 | 0 | 4 | 0 | 9 | 0.1111 |
| 14_594741 | 7 | 0 | 4 | 0.25 | 6 | 0.1667 | 6 | 0 | 23 | 0.087 | 3 | 0.3333 | 5 | 0.2 | 3 | 0 | 4 | 0.25 | 15 | 0.2 |
| 14_594741 | 8 | 0.125 | 6 | 0.1667 | 7 | 0.2857 | 6 | 0.1667 | 27 | 0.1852 | 3 | 0 | 5 | 0.4 | 3 | 0 | 6 | 0 | 17 | 0.1176 |
| 14_597354 | 1 | 1 | NA | NA | NA | NA | NA | NA | 1 | 1 | NA | NA | NA | NA | 1 | 1 | NA | NA | 1 | 1 |
| 14_597845 | 2 | 0.5 | 4 | 0 | 4 | 0.5 | 4 | 0.5 | 14 | 0.3571 | 4 | 0.5 | NA | NA | 2 | 0.5 | 3 | 0.6667 | 9 | 0.5556 |
| 14_597847 | 5 | 0.6 | 4 | 0.5 | 4 | 0.25 | 5 | 0.2 | 18 | 0.3889 | 3 | 0.3333 | NA | NA | 5 | 0.2 | 2 | 0 | 10 | 0.2 |
| 14_597848 | 3 | 0 | 2 | 1 | 3 | 0 | 4 | 0 | 12 | 0.1667 | 3 | 0 | NA | NA | 3 | 0.3333 | 2 | 0.5 | 8 | 0.25 |
| 14_597848 | 4 | 0.25 | 1 | 0 | 3 | 0 | 2 | 0 | 10 | 0.1 | 3 | 0 | NA | NA | 1 | 1 | 3 | 0 | 7 | 0.1429 |
| 14_597848 | 4 | 0 | 4 | 0 | 4 | 0.25 | 4 | 0 | 16 | 0.0625 | 3 | 0.3333 | NA | NA | 3 | 0 | 3 | 0 | 9 | 0.1111 |
| 14_597849 | 5 | 0.2 | 4 | 0.25 | 4 | 0 | 4 | 0.25 | 17 | 0.1765 | 3 | 0.3333 | NA | NA | 3 | 0 | 3 | 0.3333 | 9 | 0.2222 |
| 14_597852 | 3 | 0 | 3 | 0 | 4 | 0.25 | 2 | 0 | 12 | 0.0833 | 1 | 0 | NA | NA | 3 | 0.3333 | 3 | 0.3333 | 7 | 0.2857 |
| 14_597856 | 5 | 0.4 | 4 | 0.5 | 4 | 0.75 | 3 | 0 | 16 | 0.4375 | 3 | 0 | 1 | 1 | 3 | 1 | 5 | 0.4 | 12 | 0.5 |
| 14_597857 | 4 | 0.25 | 4 | 0.25 | 3 | 0 | 4 | 0.25 | 15 | 0.2 | 3 | 0 | 1 | 0 | 3 | 0 | 5 | 0.2 | 12 | 0.0833 |

| | | | | | | | | | | | | | | | | | | | | |
|---|---|---|---|---|---|---|---|---|---|---|---|---|---|---|---|---|---|---|---|---|
| 14_5979704 | 3 | 0.6667 | 5 | 0.6 | NA | NA | 2 | 0.5 | 10 | 0.6 | 4 | 0.5 | 2 | 0.5 | 4 | 0.75 | 3 | 0.6667 | 13 | 0.6154 |
| 14_5979711 | 3 | 0 | 4 | 0.25 | 1 | 0 | NA | NA | 8 | 0.125 | 4 | 0.25 | 2 | 0 | 3 | 0 | 2 | 0 | 11 | 0.0909 |
| 14_5979727 | 2 | 0 | 5 | 0.2 | 1 | 1 | 3 | 0 | 11 | 0.1818 | 3 | 0 | 2 | 0 | 3 | 0 | 4 | 0.25 | 12 | 0.0833 |
| 14_5979728 | 1 | 0 | 5 | 0.8 | NA | NA | NA | NA | 6 | 0.6667 | 1 | 0 | 1 | 0 | 3 | 0.6667 | 1 | 1 | 6 | 0.5 |
| 14_5979728 | NA | NA | NA | NA | NA | NA | 2 | 1 | 2 | 1 | NA | NA | NA | NA | NA | NA | 1 | 1 | 1 | 1 |
| 14_5979745 | 2 | 0.5 | 5 | 0.8 | 1 | 1 | 2 | 1 | 10 | 0.8 | 3 | 0.3333 | 1 | 1 | 3 | 0 | 4 | 0.75 | 11 | 0.4545 |
| 14_5979750 | 2 | 1 | 4 | 0.25 | NA | NA | 2 | 1 | 8 | 0.625 | 3 | 0.3333 | 1 | 0 | 3 | 0.3333 | 3 | 0.6667 | 10 | 0.4 |
| 14_5979767 | 2 | 0.5 | 2 | 1 | 1 | 1 | 1 | 0 | 6 | 0.6667 | 1 | 1 | 1 | 1 | 1 | 0 | 2 | 1 | 5 | 0.8 |
| 14_5979767 | 2 | 0 | 4 | 0.25 | 1 | 0 | 2 | 0.5 | 9 | 0.2222 | 1 | 1 | NA | NA | 3 | 0.3333 | 2 | 0 | 6 | 0.3333 |
| 14_5979771 | 1 | 0 | 4 | 1 | 1 | 0 | 2 | 1 | 8 | 0.75 | 3 | 0.3333 | 1 | 1 | 3 | 0.3333 | 4 | 0.75 | 11 | 0.5455 |
| 14_5979783 | 1 | 0 | 5 | 0.2 | NA | NA | 2 | 0 | 8 | 0.125 | 4 | 0 | 1 | 1 | 3 | 0.3333 | 4 | 0 | 12 | 0.1667 |
| 14_5979786 | 1 | 0 | 4 | 0.75 | NA | NA | 1 | 1 | 6 | 0.6667 | 3 | 0.6667 | 1 | 1 | 3 | 0.3333 | 4 | 0.75 | 11 | 0.6364 |
| 14_5979790 | 2 | 0.5 | 3 | 0.6667 | 1 | 0 | 2 | 1 | 8 | 0.625 | 1 | 1 | 1 | 0 | 2 | 0 | 3 | 1 | 7 | 0.5714 |
| 14_5979801 | 2 | 1 | 5 | 0.6 | 1 | 1 | 2 | 0.5 | 10 | 0.7 | 3 | 0.6667 | 1 | 1 | 3 | 1 | 4 | 0.75 | 11 | 0.8182 |
| 14_5979804 | 2 | 0.5 | 4 | 0.5 | 1 | 0 | 2 | 0 | 9 | 0.3333 | 4 | 0 | 1 | 0 | 2 | 0 | 4 | 0.25 | 11 | 0.0909 |
| 14_5979806 | 2 | 0.5 | 4 | 1 | 1 | 1 | 1 | 0 | 8 | 0.75 | 1 | 1 | 1 | 1 | 2 | 1 | 3 | 1 | 7 | 1 |
| 14_5979808 | 1 | 1 | 5 | 0.4 | 1 | 1 | 2 | 1 | 9 | 0.6667 | 2 | 0 | 1 | 1 | 2 | 0.5 | 3 | 0.3333 | 8 | 0.375 |
| 14_5979828 | 2 | 0 | 3 | 0.3333 | 1 | 0 | 1 | 0 | 7 | 0.1429 | 2 | 0 | NA | NA | 2 | 0 | 3 | 0.3333 | 7 | 0.1429 |
| 14_5979874 | 1 | 0 | 1 | 1 | NA | NA | 1 | 0 | 3 | 0.3333 | 1 | 0 | 1 | 0 | 2 | 0.5 | 1 | 0 | 5 | 0.2 |
| 14_5980154 | 5 | 0.4 | 8 | 0.375 | 4 | 0.5 | 3 | 0.6667 | 20 | 0.45 | 1 | 0 | 2 | 0 | 2 | 0.5 | 6 | 0.3333 | 11 | 0.2727 |
| 14_5980156 | 4 | 0.75 | 8 | 0.125 | 4 | 0.75 | 3 | 0.3333 | 19 | 0.4211 | 1 | 1 | 3 | 0.3333 | 2 | 0 | 5 | 0.8 | 11 | 0.5455 |
| 14_5980162 | 4 | 0.25 | 8 | 0.375 | 3 | 0.6667 | 2 | 0 | 17 | 0.3529 | 1 | 0 | 3 | 0.3333 | 2 | 0 | 6 | 0.1667 | 12 | 0.1667 |
| 14_5980165 | 4 | 0.25 | 6 | 0.1667 | 3 | 0.6667 | 2 | 0 | 15 | 0.2667 | 1 | 0 | 2 | 0 | 1 | 0 | 7 | 0.1429 | 11 | 0.0909 |
| 14_5980190 | 3 | 1 | 6 | 1 | 1 | 1 | 3 | 0.6667 | 13 | 0.9231 | 1 | 0 | 3 | 0.6667 | 2 | 0.5 | 5 | 1 | 11 | 0.7273 |
| 14_5980688 | NA | NA | 1 | 1 | NA | NA | 2 | 0.5 | 3 | 0.6667 | NA | NA | 1 | 0 | 3 | 0.3333 | 1 | 1 | 5 | 0.4 |
| 14_5980689 | NA | NA | 1 | 1 | NA | NA | 2 | 0.5 | 3 | 0.6667 | NA | NA | 1 | 0 | 3 | 1 | 1 | 0 | 5 | 0.6 |
| 14_5980695 | NA | NA | 1 | 1 | NA | NA | 2 | 0.5 | 3 | 0.6667 | NA | NA | 1 | 0 | 3 | 0.6667 | 1 | 1 | 5 | 0.6 |
| 14_5980697 | NA | NA | 1 | 1 | NA | NA | 1 | 1 | 2 | 1 | NA | NA | 1 | 1 | 2 | 1 | NA | NA | 3 | 1 |
| 14_5980721 | NA | NA | 1 | 1 | NA | NA | 1 | 1 | 2 | 1 | NA | NA | 1 | 1 | 3 | 1 | 1 | 1 | 5 | 1 |
| 14_5980744 | NA | NA | 1 | 1 | NA | NA | 2 | 0 | 3 | 0.3333 | NA | NA | 1 | 0 | 3 | 0.3333 | 1 | 0 | 5 | 0.2 |
| 14_5980748 | NA | NA | NA | NA | NA | NA | 2 | 1 | 2 | 1 | NA | NA | 1 | 1 | 2 | 0.5 | 1 | 1 | 4 | 0.75 |
| 14_5980777 | NA | NA | 1 | 1 | NA | NA | 2 | 1 | 3 | 1 | NA | NA | 1 | 0 | 3 | 0.6667 | 1 | 1 | 5 | 0.6 |
| 14_5980788 | NA | NA | 1 | 1 | NA | NA | 2 | 1 | 3 | 1 | NA | NA | NA | NA | 1 | 1 | NA | NA | 1 | 1 |
| 14_5980791 | NA | NA | 1 | 0 | NA | NA | 2 | 0.5 | 3 | 0.3333 | NA | NA | NA | NA | 3 | 0.6667 | NA | NA | 3 | 0.6667 |
| 14_5980807 | NA | NA | 1 | 1 | NA | NA | 2 | 0 | 3 | 0.3333 | NA | NA | NA | NA | 3 | 0 | 1 | 1 | 4 | 0.25 |
| 14_5980817 | NA | NA | 1 | 1 | NA | NA | 2 | 0 | 3 | 0.3333 | NA | NA | NA | NA | 3 | 0.3333 | 1 | 1 | 4 | 0.5 |
| 14_5980831 | NA | NA | 1 | 0 | NA | NA | 2 | 1 | 3 | 0.6667 | NA | NA | NA | NA | 2 | 0.5 | 1 | 1 | 3 | 0.6667 |
| 14_5980854 | NA | NA | NA | NA | NA | NA | 2 | 0.5 | 2 | 0.5 | NA | NA | NA | NA | 3 | 0.3333 | NA | NA | 3 | 0.3333 |
| 14_5980859 | NA | NA | 1 | 0 | NA | NA | 2 | 0.5 | 3 | 0.3333 | NA | NA | NA | NA | 1 | 1 | NA | NA | 1 | 1 |
| 14_5980867 | NA | NA | 1 | 1 | NA | NA | 2 | 0 | 3 | 0.3333 | NA | NA | NA | NA | 3 | 0.6667 | 1 | 0 | 4 | 0.5 |
| 14_5980879 | NA | NA | 1 | 1 | NA | NA | 1 | 1 | 2 | 1 | NA | NA | NA | NA | 2 | 1 | NA | NA | 2 | 1 |
| 14_5980895 | NA | NA | 1 | 1 | NA | NA | 1 | 1 | 2 | 1 | NA | NA | NA | NA | 2 | 0.5 | NA | NA | 2 | 0.5 |
| 14_5980923 | NA | NA | 1 | 1 | NA | NA | 1 | 1 | 2 | 1 | NA | NA | NA | NA | 3 | 0.6667 | NA | NA | 3 | 0.6667 |
| 14_5980925 | NA | NA | 1 | 1 | NA | NA | 1 | 1 | 2 | 1 | NA | NA | NA | NA | 3 | 1 | 1 | 0 | 4 | 0.75 |
| 14_5980926 | NA | NA | 1 | 1 | NA | NA | 1 | 1 | 2 | 1 | NA | NA | NA | NA | 2 | 0 | 1 | 1 | 3 | 0.3333 |
| 14_5982227 | NA | NA | NA | NA | 1 | 1 | NA | NA | 1 | 1 | NA | NA | NA | NA | NA | NA | 2 | 0.5 | 2 | 0.5 |
| 14_5982349 | 5 | 0 | 8 | 0 | 2 | 0.5 | 3 | 0 | 18 | 0.0556 | 3 | 0 | 3 | 0.3333 | 5 | 0.2 | 6 | 0 | 17 | 0.1176 |
| 14_5982365 | 6 | 0.3333 | 9 | 0.4444 | 3 | 0 | 5 | 0 | 23 | 0.2609 | 3 | 0.3333 | 3 | 0.3333 | 5 | 0.2 | 8 | 0.125 | 19 | 0.2105 |
| 14_5982371 | 4 | 0 | 7 | 0.1429 | 2 | 0 | 5 | 0.2 | 18 | 0.1111 | 3 | 0.3333 | 3 | 0.3333 | 3 | 0 | 5 | 0.2 | 14 | 0.2143 |
| 14_5982381 | 4 | 0 | 9 | 0.2222 | 1 | 0 | 3 | 0 | 17 | 0.1176 | 3 | 0.6667 | 3 | 0.3333 | 5 | 0 | 7 | 0 | 18 | 0.1667 |
| 14_5982384 | 5 | 0.2 | 6 | 0.3333 | 2 | 0 | 2 | 0 | 15 | 0.2 | 2 | 0.5 | 3 | 0 | 6 | 0.3333 | 6 | 0.1667 | 17 | 0.2353 |
| 14_5982385 | 5 | 0.6 | 8 | 0.875 | 2 | 1 | 4 | 0.75 | 19 | 0.7895 | 2 | 0.5 | 3 | 0.6667 | 6 | 1 | 6 | 0.3333 | 17 | 0.6471 |
| 14_5986739 | 1 | 1 | NA | NA | 2 | 1 | 1 | 1 | 4 | 1 | 1 | 1 | NA | NA | 1 | 1 | 1 | 0 | 3 | 0.6667 |
| 14_5986751 | 1 | 1 | NA | NA | 2 | 1 | NA | NA | 3 | 1 | NA | NA | NA | NA | 1 | 1 | 1 | 1 | 2 | 1 |
| 14_5986762 | 1 | 1 | NA | NA | 2 | 1 | NA | NA | 3 | 1 | 1 | 1 | NA | NA | 2 | 1 | 1 | 1 | 4 | 1 |
| 14_5986765 | 1 | 1 | NA | NA | 1 | 0 | 1 | 1 | 3 | 0.6667 | 1 | 1 | NA | NA | 2 | 1 | 1 | 1 | 4 | 1 |
| 14_5986767 | 1 | 1 | NA | NA | 1 | 1 | NA | NA | 2 | 1 | 1 | 1 | NA | NA | 1 | 0 | NA | NA | 2 | 0.5 |
| 14_5986767 | 1 | 0 | NA | NA | 2 | 0 | 1 | 1 | 4 | 0.25 | 1 | 0 | NA | NA | 2 | 0 | 1 | 1 | 4 | 0.25 |
| 14_5986777 | 1 | 1 | NA | NA | 2 | 1 | 1 | 1 | 4 | 1 | 1 | 1 | NA | NA | 1 | 1 | 2 | 1 | 4 | 1 |
| 14_5986807 | 1 | 0 | NA | NA | 1 | 1 | NA | NA | 2 | 0.5 | 1 | 1 | NA | NA | 2 | 1 | 2 | 0.5 | 5 | 0.8 |
| 14_5987666 | 1 | 1 | 1 | 0 | 2 | 0.5 | 1 | 0 | 5 | 0.4 | NA | NA | NA | NA | 1 | 0 | 2 | 1 | 3 | 0.6667 |
| 14_5988107 | 1 | 0 | NA | NA | 1 | 1 | 1 | 0 | 3 | 0.3333 | 1 | 0 | 1 | 0 | NA | NA | 1 | 1 | 3 | 0.3333 |
| 14_5988314 | 1 | 1 | NA | NA | 1 | 0 | 1 | 0 | 3 | 0.3333 | NA | NA | 1 | 0 | NA | NA | 1 | 1 | 2 | 0.5 |
| 14_5988336 | 1 | 1 | NA | NA | 1 | 0 | 1 | 0 | 3 | 0.3333 | NA | NA | 1 | 0 | 1 | 0 | 1 | 1 | 3 | 0.3333 |
| 14_5988338 | 1 | 1 | NA | NA | 1 | 0 | 1 | 1 | 3 | 0.6667 | NA | NA | 1 | 1 | 1 | 0 | 1 | 0 | 3 | 0.3333 |

| | | | | | | | | | | | | | | | | | | | | |
|---|---|---|---|---|---|---|---|---|---|---|---|---|---|---|---|---|---|---|---|---|
| 14_598838( | 1 | 0 | NA | NA | 1 | 1 | 1 | 0 | 3 | 0.3333 | NA | NA | 1 | 0 | NA | NA | 1 | 1 | 2 | 0.5 |
| 14_598849 | 1 | 1 | NA | NA | 1 | 0 | NA | NA | 2 | 0.5 | NA | NA | 1 | 1 | NA | NA | 1 | 0 | 2 | 0.5 |
| 14_604558 | NA | NA | 4 | 0.5 | 1 | 0 | 5 | 0.2 | 10 | 0.3 | NA | NA | 2 | 0 | 4 | 0.25 | 6 | 0 | 12 | 0.0833 |
| 14_604565 | NA | NA | 4 | 0.25 | 1 | 0 | 3 | 0.3333 | 8 | 0.25 | NA | NA | 2 | 0.5 | 4 | 0 | 5 | 0.4 | 11 | 0.2727 |
| 14_604570 | 4 | 0 | 4 | 0.25 | 4 | 0 | 6 | 0.1667 | 18 | 0.1111 | 3 | 0.3333 | 6 | 0 | 4 | 0.5 | 3 | 0.3333 | 16 | 0.25 |
| 14_604573( | 4 | 0.75 | 2 | 0.5 | 3 | 1 | 7 | 0.8571 | 16 | 0.8125 | 1 | 0 | 3 | 0.3333 | 5 | 0.6 | 2 | 0.5 | 11 | 0.4545 |
| 14_604573( | 4 | 0 | 2 | 1 | 3 | 0.6667 | 6 | 0.3333 | 15 | 0.4 | 2 | 1 | 5 | 0.4 | 4 | 0.25 | 3 | 0 | 14 | 0.3571 |
| 14_604573 | 4 | 0 | 4 | 0 | 3 | 0 | 6 | 0.1667 | 17 | 0.0588 | 2 | 0 | 5 | 0.2 | 4 | 0 | 2 | 0 | 13 | 0.0769 |
| 14_604573 | 3 | 0 | 4 | 0.75 | 3 | 0.3333 | 8 | 0.5 | 18 | 0.4444 | 2 | 0 | 5 | 0.6 | 4 | 0.75 | 3 | 0.3333 | 14 | 0.5 |
| 14_604574 | 4 | 0.25 | 4 | 0.25 | 3 | 0 | 8 | 0 | 19 | 0.1053 | 2 | 0 | 6 | 0.1667 | 3 | 0 | 3 | 0 | 14 | 0.0714 |
| 14_604575 | 4 | 0.5 | 3 | 0.6667 | 3 | 0.3333 | 4 | 0.25 | 14 | 0.4286 | 2 | 0 | 3 | 0.3333 | 3 | 0 | 2 | 0 | 10 | 0.1 |
| 14_605378 | 7 | 0.1429 | 14 | 0.0714 | 8 | 0.125 | 15 | 0.2667 | 44 | 0.1591 | 9 | 0.2222 | 9 | 0.1111 | 9 | 0.2222 | 15 | 0.1333 | 42 | 0.1667 |
| 14_605378 | 7 | 0.2857 | 14 | 0.0714 | 8 | 0.125 | 13 | 0.1538 | 42 | 0.1429 | 11 | 0.2727 | 9 | 0.2222 | 8 | 0 | 14 | 0.2857 | 42 | 0.2143 |
| 14_605379 | 7 | 0.1429 | 15 | 0.1333 | 8 | 0 | 16 | 0.1875 | 46 | 0.1304 | 10 | 0 | 9 | 0.2222 | 8 | 0.375 | 15 | 0 | 42 | 0.119 |
| 14_605380 | 7 | 0.1429 | 15 | 0.2667 | 8 | 0.375 | 16 | 0.3125 | 46 | 0.2826 | 12 | 0.0833 | 8 | 0.125 | 10 | 0.2 | 16 | 0.0625 | 46 | 0.1087 |
| 14_605383 | 8 | 0.125 | 15 | 0 | 8 | 0.375 | 15 | 0.3333 | 46 | 0.1957 | 11 | 0.1818 | 13 | 0.1538 | 9 | 0 | 16 | 0.0625 | 49 | 0.102 |
| 14_606141 | 1 | 0 | 1 | 1 | NA | NA | NA | NA | 2 | 0.5 | NA | NA | NA | NA | NA | NA | 1 | 1 | 1 | 1 |
| 14_606141 | 1 | 1 | 1 | 1 | NA | NA | NA | NA | 2 | 1 | NA | NA | NA | NA | NA | NA | 1 | 1 | 1 | 1 |
| 14_606142 | 1 | 1 | 1 | 1 | NA | NA | NA | NA | 2 | 1 | NA | NA | NA | NA | NA | NA | 1 | 1 | 1 | 1 |
| 14_606143 | NA | NA | 1 | 1 | NA | NA | NA | NA | 1 | 1 | NA | NA | NA | NA | NA | NA | 1 | 1 | 1 | 1 |
| 14_606149 | NA | NA | 1 | 1 | NA | NA | NA | NA | 1 | 1 | NA | NA | NA | NA | NA | NA | 1 | 1 | 1 | 1 |
| 14_606152 | 1 | 1 | 1 | 1 | NA | NA | NA | NA | 2 | 1 | NA | NA | NA | NA | NA | NA | 1 | 1 | 1 | 1 |
| 14_606155 | NA | NA | 1 | 1 | NA | NA | NA | NA | 1 | 1 | NA | NA | NA | NA | NA | NA | 1 | 1 | 1 | 1 |
| 14_606168 | NA | NA | NA | NA | 2 | 0.5 | NA | NA | 2 | 0.5 | NA | NA | 2 | 0 | NA | NA | 1 | 1 | 3 | 0.3333 |
| 14_606168 | NA | NA | NA | NA | 2 | 1 | NA | NA | 2 | 1 | NA | NA | 2 | 0.5 | NA | NA | 1 | 0 | 3 | 0.3333 |
| 14_606170 | NA | NA | NA | NA | 2 | 0 | 1 | 1 | 3 | 0.3333 | 2 | 0 | 3 | 0 | NA | NA | 2 | 0.5 | 7 | 0.1429 |
| 14_606170 | NA | NA | NA | NA | 2 | 0.5 | 1 | 1 | 3 | 0.6667 | 2 | 0 | 3 | 0.3333 | NA | NA | 2 | 0.5 | 7 | 0.2857 |
| 14_606171 | NA | NA | NA | NA | 2 | 0 | 1 | 1 | 3 | 0.3333 | 3 | 0.3333 | 3 | 0.3333 | 1 | 0 | 2 | 0.5 | 9 | 0.3333 |
| 14_606172 | NA | NA | NA | NA | 1 | 1 | 1 | 1 | 2 | 1 | 2 | 1 | 2 | 0 | 1 | 0 | 2 | 0 | 7 | 0.2857 |
| 14_606175 | NA | NA | NA | NA | 2 | 1 | 2 | 1 | 4 | 1 | 2 | 1 | 3 | 1 | 2 | 1 | 2 | 1 | 9 | 1 |
| 14_606176 | NA | NA | NA | NA | 3 | 0.6667 | 1 | 1 | 4 | 0.75 | 2 | 0.5 | 3 | 0.3333 | 1 | 0 | 2 | 0.5 | 8 | 0.375 |
| 14_606176 | NA | NA | NA | NA | 3 | 0 | 2 | 0.5 | 5 | 0.2 | 3 | 0 | 3 | 0.3333 | 1 | 1 | 2 | 0 | 9 | 0.2222 |
| 14_606178 | NA | NA | NA | NA | 2 | 0.5 | 2 | 1 | 4 | 0.75 | 3 | 1 | 2 | 0.5 | 2 | 1 | 2 | 1 | 9 | 0.8889 |
| 14_606178 | NA | NA | NA | NA | 2 | 1 | 1 | 1 | 3 | 1 | 1 | 1 | 3 | 0.6667 | 1 | 1 | 2 | 1 | 7 | 0.8571 |
| 14_606181 | NA | NA | NA | NA | 3 | 0 | 2 | 0.5 | 5 | 0.2 | 2 | 0 | 3 | 0 | 2 | 0 | 2 | 0.5 | 9 | 0.1111 |
| 14_606182 | NA | NA | NA | NA | 3 | 1 | 2 | 1 | 5 | 1 | 2 | 0.5 | 2 | 1 | 1 | 1 | 2 | 1 | 7 | 0.8571 |
| 14_606182 | NA | NA | NA | NA | 2 | 1 | 2 | 0.5 | 4 | 0.75 | 3 | 0 | 2 | 0.5 | 1 | 1 | 2 | 0.5 | 8 | 0.375 |
| 14_606216 | 2 | 0 | 2 | 1 | 2 | 0 | 7 | 0.2857 | 13 | 0.3077 | 1 | 0 | NA | NA | 3 | 0 | 1 | 1 | 5 | 0.2 |
| 14_606225 | 3 | 0 | 2 | 0.5 | 1 | 0 | 5 | 0.4 | 11 | 0.2727 | 3 | 0.3333 | 1 | 0 | 1 | 0 | 1 | 0 | 6 | 0.1667 |
| 14_606242 | 7 | 0 | 3 | 0.3333 | 4 | 0.5 | 8 | 0 | 22 | 0.1364 | 3 | 0 | 1 | 0 | 3 | 0.3333 | 4 | 0 | 11 | 0.0909 |
| 14_606252 | 8 | 0.5 | 6 | 0.1667 | 5 | 0.4 | 9 | 0.3333 | 28 | 0.3571 | 3 | 0 | 2 | 0.5 | 4 | 0.5 | 3 | 0.6667 | 12 | 0.4167 |
| 14_606255 | 6 | 0.3333 | 6 | 0.3333 | 4 | 0.75 | 8 | 0.125 | 24 | 0.3333 | 4 | 0 | 2 | 1 | 3 | 0.3333 | 4 | 0.25 | 13 | 0.3077 |
| 14_609293 | NA | NA | 1 | 1 | NA | NA | NA | NA | 1 | 1 | NA | NA | NA | NA | 1 | 1 | NA | NA | 1 | 1 |
| 14_609296 | NA | NA | 2 | 1 | NA | NA | NA | NA | 2 | 1 | NA | NA | NA | NA | 1 | 1 | NA | NA | 1 | 1 |
| 14_609296 | NA | NA | 2 | 1 | NA | NA | NA | NA | 2 | 1 | NA | NA | NA | NA | 1 | 1 | NA | NA | 1 | 1 |
| 14_609938 | NA | NA | 1 | 0 | 1 | 1 | NA | NA | 2 | 0.5 | 1 | 0 | 1 | 1 | 1 | 0 | NA | NA | 3 | 0.3333 |
| 14_609986 | NA | NA | 1 | 1 | 1 | 1 | NA | NA | 2 | 1 | 1 | 1 | 2 | 1 | 4 | 1 | 1 | 1 | 8 | 1 |
| 14_609987 | NA | NA | 1 | 1 | NA | NA | NA | NA | 1 | 1 | 1 | 1 | 2 | 1 | 5 | 0.8 | 1 | 1 | 9 | 0.8889 |
| 14_609988 | NA | NA | 1 | 1 | 1 | 0 | NA | NA | 2 | 0.5 | 1 | 0 | 1 | 1 | 4 | 0 | 1 | 0 | 7 | 0.1429 |
| 14_609988 | NA | NA | 1 | 1 | 2 | 0.5 | NA | NA | 3 | 0.6667 | 1 | 1 | 2 | 1 | 5 | 0.6 | NA | NA | 8 | 0.75 |
| 14_609991 | 1 | 1 | 1 | 0 | 2 | 0.5 | 1 | 1 | 5 | 0.6 | 1 | 1 | 2 | 0 | 7 | 0.2857 | 1 | 1 | 11 | 0.3636 |
| 14_609992 | 1 | 1 | 1 | 0 | 2 | 0 | 2 | 0.5 | 6 | 0.3333 | 1 | 1 | 2 | 0 | 7 | 0.1429 | 1 | 1 | 11 | 0.2727 |
| 14_609992 | 1 | 0 | 1 | 1 | 2 | 0.5 | NA | NA | 4 | 0.5 | 1 | 0 | 2 | 0 | 7 | 0.2857 | 1 | 0 | 11 | 0.1818 |
| 14_609992 | 1 | 0 | NA | NA | 1 | 0 | 2 | 0.5 | 4 | 0.25 | 1 | 0 | 2 | 0.5 | 8 | 0.375 | 1 | 1 | 12 | 0.4167 |
| 14_609993 | 1 | 1 | 1 | 1 | 1 | 1 | 2 | 1 | 5 | 1 | 1 | 1 | 2 | 1 | 9 | 1 | 1 | 1 | 13 | 1 |
| 14_609993 | 1 | 1 | 1 | 0 | 2 | 0 | 2 | 0 | 6 | 0.1667 | 1 | 1 | 2 | 0 | 10 | 0.1 | 2 | 0 | 15 | 0.1333 |
| 14_609994 | 1 | 0 | 1 | 1 | 1 | 0 | 2 | 1 | 5 | 0.6 | 1 | 1 | 1 | 1 | 9 | 0.6667 | 1 | 1 | 12 | 0.75 |
| 14_609995 | 1 | 0 | 1 | 0 | 2 | 0 | 2 | 0.5 | 6 | 0.1667 | NA | NA | 2 | 0.5 | 8 | 0.625 | 2 | 0.5 | 12 | 0.5833 |
| 14_609995 | 1 | 1 | NA | NA | 2 | 0 | 2 | 0.5 | 5 | 0.4 | NA | NA | 2 | 0.5 | 3 | 0 | 2 | 0 | 7 | 0.1429 |
| 14_609995 | 1 | 1 | 1 | 0 | 1 | 1 | 2 | 1 | 5 | 0.8 | 1 | 1 | 2 | 1 | 9 | 0.8889 | 2 | 1 | 14 | 0.9286 |
| 14_609998 | NA | NA | 1 | 1 | 3 | 0.3333 | 1 | 1 | 5 | 0.6 | 2 | 1 | 3 | 1 | 7 | 0.8571 | 2 | 0.5 | 14 | 0.8571 |
| 14_609998 | 1 | 1 | 2 | 1 | 3 | 1 | 1 | 1 | 7 | 1 | 2 | 1 | 3 | 1 | 9 | 1 | 3 | 1 | 17 | 1 |
| 14_610005 | 2 | 0 | 2 | 0 | 3 | 0.3333 | 3 | 0.3333 | 10 | 0.2 | NA | NA | 3 | 0.3333 | 7 | 0 | 2 | 0 | 12 | 0.0833 |
| 14_610009 | 2 | 0.5 | 1 | 1 | 4 | 0.25 | 1 | 0 | 8 | 0.375 | 1 | 1 | 3 | 0.3333 | 3 | 0.3333 | 1 | 0 | 8 | 0.375 |
| 14_610011 | 3 | 1 | 2 | 1 | 4 | 0.5 | 2 | 0.5 | 11 | 0.7273 | 1 | 0 | 3 | 0 | 4 | 1 | 1 | 0 | 9 | 0.4444 |

| | | | | | | | | | | | | | | | | | | | | |
|---|---|---|---|---|---|---|---|---|---|---|---|---|---|---|---|---|---|---|---|---|
| 14_6100148 | 4 | 0.5 | 2 | 0.5 | 3 | 0 | 3 | 0.6667 | 12 | 0.4167 | 1 | 1 | 4 | 0.25 | 5 | 0.6 | 1 | 0 | 11 | 0.4545 |
| 14_6114088 | 4 | 0.5 | 5 | 0.4 | 5 | 0.4 | 2 | 0 | 16 | 0.375 | 3 | 0.6667 | 2 | 0.5 | 7 | 0.5714 | 3 | 0.3333 | 15 | 0.5333 |
| 14_6114100 | 2 | 0 | 6 | 0.1667 | 5 | 0 | 2 | 0 | 15 | 0.0667 | 3 | 0.3333 | 1 | 0 | 8 | 0 | 2 | 0 | 14 | 0.0714 |
| 14_6114103 | 3 | 1 | 6 | 0.6667 | 4 | 1 | 2 | 1 | 15 | 0.8667 | 3 | 0.6667 | NA | NA | 7 | 0.7143 | 2 | 0.5 | 12 | 0.6667 |
| 14_6114426 | NA | NA | 1 | 1 | NA | NA | NA | NA | 1 | 1 | NA | NA | NA | NA | 1 | 1 | NA | NA | 1 | 1 |
| 14_6114428 | NA | NA | 1 | 1 | NA | NA | NA | NA | 1 | 1 | NA | NA | NA | NA | 1 | 1 | NA | NA | 1 | 1 |
| 14_6120145 | 5 | 0.2 | 7 | 0 | 4 | 0 | 4 | 0.5 | 20 | 0.15 | 7 | 0.1429 | 4 | 0.75 | 10 | 0.1 | 5 | 0 | 26 | 0.1923 |
| 14_6120159 | 5 | 1 | 7 | 0.7143 | 3 | 1 | 4 | 0.75 | 19 | 0.8421 | 8 | 0.75 | 6 | 1 | 11 | 0.7273 | 5 | 1 | 30 | 0.8333 |
| 14_6120167 | 4 | 0.5 | 7 | 0 | 2 | 1 | 4 | 0.25 | 17 | 0.2941 | 8 | 0.25 | 4 | 0 | 11 | 0.1818 | 5 | 0.2 | 28 | 0.1786 |
| 14_6120172 | 3 | 0 | 5 | 0.2 | 4 | 0 | 4 | 0 | 16 | 0.0625 | 6 | 0 | 2 | 0 | 8 | 0 | 4 | 0.25 | 20 | 0.05 |
| 14_6120181 | 4 | 0.75 | 6 | 0 | 3 | 0 | 3 | 0 | 16 | 0.1875 | 6 | 0 | 3 | 0 | 10 | 0.4 | 4 | 1 | 23 | 0.3478 |
| 14_6120182 | 4 | 0 | 7 | 0 | 4 | 0 | 4 | 0.25 | 19 | 0.0526 | 8 | 0.125 | 3 | 0 | 10 | 0.1 | 5 | 0 | 26 | 0.0769 |
| 14_6120194 | 4 | 1 | 7 | 0.2857 | 3 | 0.3333 | 4 | 0.25 | 18 | 0.4444 | 6 | 0.3333 | 3 | 0.6667 | 11 | 0.1818 | 5 | 0 | 25 | 0.24 |
| 14_6120198 | 4 | 0.25 | 6 | 0 | 3 | 0 | 3 | 0 | 16 | 0.0625 | 6 | 0 | 3 | 0.3333 | 10 | 0.1 | 4 | 0.25 | 23 | 0.1304 |
| 14_6120199 | 4 | 1 | 5 | 0.6 | 4 | 1 | 4 | 0.25 | 17 | 0.7059 | 7 | 0.8571 | 4 | 0.75 | 9 | 0.4444 | 5 | 0.6 | 25 | 0.64 |
| 14_6120221 | 3 | 0 | 7 | 0.2857 | 4 | 0.5 | 3 | 0.3333 | 17 | 0.2941 | 6 | 0.1667 | 2 | 0.5 | 8 | 0.25 | 3 | 0 | 19 | 0.2105 |
| 14_6120250 | 3 | 0.3333 | 7 | 0.1429 | 4 | 0 | 3 | 0 | 17 | 0.1176 | 7 | 0.1429 | 3 | 0 | 9 | 0.1111 | 4 | 0.25 | 23 | 0.1304 |
| 14_6120254 | 4 | 0.25 | 6 | 0 | 4 | 0 | 2 | 0 | 16 | 0.0625 | 6 | 0.1667 | 2 | 0 | 9 | 0.1111 | 2 | 0 | 19 | 0.1053 |
| 14_6120263 | 4 | 0.5 | 6 | 0.3333 | 4 | 0 | 2 | 1 | 16 | 0.375 | 5 | 0 | 2 | 0.5 | 8 | 0.375 | 3 | 0.3333 | 18 | 0.2778 |
| 14_6120269 | 4 | 0.5 | 7 | 0.4286 | 3 | 0.3333 | 3 | 0 | 17 | 0.3529 | 7 | 0 | 3 | 0 | 10 | 0.2 | 4 | 0.25 | 24 | 0.125 |
| 14_6120279 | 4 | 0 | 4 | 0.25 | 3 | 0.3333 | 3 | 0 | 14 | 0.1429 | 4 | 0 | 3 | 0 | 8 | 0.125 | 3 | 0 | 18 | 0.0556 |
| 14_6120293 | 4 | 0.25 | 5 | 0.2 | 3 | 0 | 3 | 0 | 15 | 0.1333 | 6 | 0.3333 | 2 | 0.5 | 9 | 0.2222 | 3 | 0 | 20 | 0.25 |
| 14_6120312 | 4 | 0.25 | 6 | 0.3333 | 3 | 0 | 3 | 0 | 16 | 0.1875 | 5 | 0 | 1 | 0 | 9 | 0.1111 | 3 | 0.6667 | 18 | 0.1667 |
| 14_6120313 | 4 | 0.25 | 7 | 0.1429 | 2 | 0 | 3 | 0.3333 | 16 | 0.1875 | 6 | 0.1667 | 2 | 0 | 8 | 0 | 3 | 0.3333 | 19 | 0.1053 |
| 14_6120352 | 5 | 0.8 | 6 | 0.6667 | 2 | 1 | 3 | 0.6667 | 16 | 0.75 | 5 | 0.8 | 3 | 0.3333 | 7 | 0.5714 | 2 | 0.5 | 17 | 0.5882 |
| 14_6120353 | 2 | 0 | 4 | 0.5 | 2 | 0.5 | 3 | 0.6667 | 11 | 0.4545 | 4 | 0.5 | 3 | 0.3333 | 7 | 0.4286 | 2 | 1 | 16 | 0.5 |
| 14_6120360 | 3 | 0 | 7 | 0.2857 | 1 | 0 | 3 | 0 | 14 | 0.1429 | 5 | 0.2 | 3 | 0 | 7 | 0 | 2 | 0 | 17 | 0.0588 |
| 14_6120364 | 4 | 0.75 | 5 | 1 | 2 | 1 | 3 | 0.6667 | 14 | 0.8571 | 2 | 1 | 3 | 1 | 7 | 0.7143 | 2 | 1 | 14 | 0.8571 |
| 14_6120367 | 2 | 0 | 7 | 0.1429 | 1 | 0 | 3 | 0.3333 | 13 | 0.1538 | 5 | 0.4 | 3 | 0.6667 | 7 | 0.1429 | 2 | 0 | 17 | 0.2941 |
| 14_6120369 | 4 | 0.75 | 5 | 0.6 | 2 | 0.5 | 3 | 0.6667 | 14 | 0.6429 | 3 | 0 | 3 | 0.6667 | 8 | 0.5 | 2 | 0.5 | 16 | 0.4375 |
| 14_6120371 | 3 | 0.3333 | 6 | 0.1667 | 2 | 1 | 3 | 0 | 14 | 0.2857 | 5 | 0.4 | 2 | 0.5 | 7 | 0.4286 | 2 | 0 | 16 | 0.375 |
| 14_6120371 | 3 | 1 | 4 | 1 | 1 | 0 | 3 | 0.6667 | 11 | 0.8182 | 3 | 0.6667 | 1 | 1 | 7 | 0.8571 | 2 | 1 | 13 | 0.8462 |
| 14_6120380 | 2 | 0 | 5 | 0.2 | 2 | 0 | 3 | 0.6667 | 12 | 0.25 | 4 | 0 | 2 | 0.5 | 7 | 0 | 1 | 1 | 14 | 0.1429 |
| 14_6120380 | 3 | 0 | 6 | 0.3333 | 2 | 0 | 2 | 0 | 13 | 0.1538 | 4 | 0.25 | 1 | 0 | 8 | 0 | 1 | 0 | 14 | 0.0714 |
| 14_6120384 | 4 | 0.25 | 6 | 0.8333 | 1 | 0 | 3 | 1 | 14 | 0.6429 | 5 | 0.6 | 3 | 0.6667 | 6 | 0.1667 | 1 | 0 | 15 | 0.4 |
| 14_6120834 | 3 | 0.3333 | 5 | 0.6 | 2 | 0.5 | 3 | 0.3333 | 13 | 0.4615 | 5 | 0.6 | 3 | 1 | 7 | 0.4286 | 2 | 0.5 | 17 | 0.5882 |
| 14_6120840 | 4 | 0 | 4 | 0.25 | 2 | 0 | 3 | 0 | 13 | 0.0769 | 5 | 0.2 | 3 | 0 | 8 | 0.125 | 2 | 0.5 | 18 | 0.1667 |
| 14_6120845 | 5 | 0.4 | 4 | 0.25 | 2 | 0 | 3 | 0.3333 | 14 | 0.2857 | 4 | 0 | 3 | 0 | 8 | 0.5 | 2 | 0.5 | 17 | 0.2941 |
| 14_6120855 | 5 | 0.6 | 5 | 0.8 | 2 | 0.5 | 2 | 0.5 | 14 | 0.6429 | 3 | 0 | 3 | 0.6667 | 8 | 0.875 | 2 | 0.5 | 16 | 0.625 |
| 14_6144546 | NA | NA | NA | NA | NA | NA | 1 | 1 | 1 | 1 | NA | NA | NA | NA | 1 | 1 | 1 | 0 | 2 | 0.5 |
| 14_6144554 | NA | NA | NA | NA | NA | NA | 1 | 1 | 1 | 1 | NA | NA | NA | NA | 1 | 1 | 1 | 0 | 2 | 0.5 |
| 14_6144582 | NA | NA | NA | NA | NA | NA | 1 | 1 | 1 | 1 | NA | NA | NA | NA | 1 | 1 | 1 | 1 | 2 | 1 |
| 14_6144586 | NA | NA | NA | NA | NA | NA | 1 | 1 | 1 | 1 | NA | NA | NA | NA | 1 | 1 | 1 | 1 | 2 | 1 |
| 14_6144597 | NA | NA | NA | NA | NA | NA | 1 | 1 | 1 | 1 | NA | NA | NA | NA | 1 | 1 | 1 | 0 | 2 | 0.5 |
| 14_6144603 | NA | NA | NA | NA | NA | NA | 1 | 1 | 1 | 1 | NA | NA | NA | NA | 1 | 1 | 1 | 1 | 2 | 1 |
| 14_6144616 | NA | NA | NA | NA | NA | NA | 1 | 1 | 1 | 1 | NA | NA | NA | NA | 1 | 1 | NA | NA | 1 | 1 |
| 14_6144634 | NA | NA | NA | NA | NA | NA | 2 | 1 | 2 | 1 | NA | NA | NA | NA | 1 | 1 | 1 | 0 | 2 | 0.5 |
| 14_6144668 | NA | NA | NA | NA | NA | NA | 2 | 1 | 2 | 1 | NA | NA | NA | NA | 1 | 1 | 2 | 1 | 3 | 1 |
| 14_6144682 | NA | NA | NA | NA | NA | NA | 2 | 1 | 2 | 1 | NA | NA | NA | NA | 1 | 0 | 2 | 0.5 | 3 | 0.3333 |
| 14_6144687 | NA | NA | NA | NA | NA | NA | 2 | 1 | 2 | 1 | NA | NA | NA | NA | 1 | 1 | 2 | 0.5 | 3 | 0.6667 |
| 14_6144727 | NA | NA | NA | NA | NA | NA | 1 | 1 | 1 | 1 | NA | NA | NA | NA | 1 | 1 | 2 | 0.5 | 3 | 0.6667 |
| 14_6144736 | NA | NA | NA | NA | NA | NA | 2 | 0.5 | 2 | 0.5 | NA | NA | NA | NA | 1 | 1 | 1 | 0 | 2 | 0.5 |
| 14_6144741 | NA | NA | NA | NA | NA | NA | 2 | 1 | 2 | 1 | NA | NA | NA | NA | 1 | 1 | 2 | 1 | 3 | 1 |
| 14_6144744 | NA | NA | NA | NA | NA | NA | 2 | 1 | 2 | 1 | NA | NA | NA | NA | 1 | 1 | 2 | 1 | 3 | 1 |
| 14_6144751 | NA | NA | NA | NA | NA | NA | 2 | 0.5 | 2 | 0.5 | NA | NA | NA | NA | 1 | 1 | 2 | 1 | 3 | 1 |
| 14_6144756 | NA | NA | NA | NA | NA | NA | 2 | 1 | 2 | 1 | NA | NA | NA | NA | 1 | 1 | 2 | 1 | 3 | 1 |
| 14_6144765 | NA | NA | 1 | 0 | NA | NA | 2 | 1 | 3 | 0.6667 | NA | NA | NA | NA | 1 | 1 | 2 | 1 | 3 | 1 |
| 14_6144766 | NA | NA | 1 | 1 | NA | NA | 2 | 0.5 | 3 | 0.6667 | NA | NA | NA | NA | 1 | 1 | 2 | 0.5 | 3 | 0.6667 |
| 14_6144779 | NA | NA | 1 | 1 | NA | NA | 1 | 1 | 2 | 1 | NA | NA | NA | NA | 1 | 1 | 2 | 0.5 | 3 | 0.6667 |
| 14_6144798 | NA | NA | 1 | 0 | NA | NA | 2 | 0.5 | 3 | 0.3333 | NA | NA | NA | NA | 1 | 0 | 2 | 0.5 | 3 | 0.3333 |
| 14_6144808 | NA | NA | 1 | 1 | NA | NA | 2 | 0.5 | 3 | 0.6667 | NA | NA | NA | NA | NA | NA | 1 | 1 | 1 | 1 |
| 14_6144810 | NA | NA | 1 | 1 | NA | NA | 2 | 0.5 | 3 | 0.6667 | NA | NA | NA | NA | 1 | 1 | 1 | 0 | 2 | 0.5 |
| 14_6144829 | NA | NA | 1 | 1 | NA | NA | 1 | 1 | 2 | 1 | NA | NA | NA | NA | 2 | 1 | 2 | 0 | 4 | 0.5 |
| 14_6144834 | NA | NA | 1 | 1 | NA | NA | 1 | 0 | 2 | 0.5 | NA | NA | NA | NA | 2 | 1 | 2 | 1 | 4 | 1 |
| 14_6144859 | NA | NA | 1 | 0 | NA | NA | 2 | 1 | 3 | 0.6667 | NA | NA | NA | NA | 2 | 1 | 1 | 1 | 3 | 1 |

| | | | | | | | | | | | | | | | | | | | | |
|---|---|---|---|---|---|---|---|---|---|---|---|---|---|---|---|---|---|---|---|---|
| 14_614487 | NA | NA | 1 | 1 | NA | NA | 2 | 1 | 3 | 1 | NA | NA | NA | NA | 3 | 1 | 2 | 1 | 5 | 1 |
| 14_614489 | NA | NA | 1 | 1 | NA | NA | 2 | 1 | 3 | 1 | NA | NA | NA | NA | 2 | 1 | 2 | 1 | 4 | 1 |
| 14_614489 | NA | NA | 1 | 0 | NA | NA | 2 | 0.5 | 3 | 0.3333 | NA | NA | NA | NA | 3 | 0.3333 | 2 | 0 | 5 | 0.2 |
| 14_614498 | 1 | 0 | 2 | 0 | NA | NA | 6 | 0.3333 | 9 | 0.2222 | NA | NA | 1 | 0 | 4 | 0.5 | 3 | 0.3333 | 8 | 0.375 |
| 14_614500 | 1 | 1 | 3 | 0.6667 | NA | NA | 5 | 0.8 | 9 | 0.7778 | NA | NA | 1 | 1 | 4 | 0.75 | 3 | 0.6667 | 8 | 0.75 |
| 14_614500 | 1 | 1 | 2 | 1 | 1 | 1 | 6 | 1 | 10 | 1 | NA | NA | 1 | 1 | 2 | 0.5 | 3 | 1 | 6 | 0.8333 |
| 14_614501 | 1 | 0 | 2 | 0 | 1 | 1 | 5 | 0 | 9 | 0.1111 | NA | NA | 1 | 1 | 5 | 0.2 | 3 | 0.6667 | 9 | 0.4444 |
| 14_614502 | NA | NA | 2 | 0 | 1 | 1 | 4 | 0.25 | 7 | 0.2857 | NA | NA | 1 | 0 | 6 | 0.8333 | 3 | 1 | 10 | 0.8 |
| 14_614502 | 1 | 0 | 2 | 0 | 1 | 0 | 7 | 0.1429 | 11 | 0.0909 | NA | NA | 1 | 0 | 5 | 0.2 | 3 | 0 | 9 | 0.1111 |
| 14_614503 | NA | NA | 3 | 0.3333 | NA | NA | 6 | 0 | 9 | 0.1111 | NA | NA | 1 | 0 | 6 | 0.6667 | 3 | 0.3333 | 10 | 0.5 |
| 14_614504 | 1 | 0 | 3 | 0 | 1 | 0 | 7 | 0.2857 | 12 | 0.1667 | NA | NA | 1 | 0 | 5 | 0.2 | 3 | 0 | 9 | 0.1111 |
| 14_614504 | 1 | 1 | 2 | 0 | 1 | 0 | 3 | 0.3333 | 7 | 0.2857 | NA | NA | 1 | 1 | 6 | 0.1667 | 3 | 0 | 10 | 0.2 |
| 14_614507 | 1 | 1 | 3 | 0.6667 | 1 | 1 | 7 | 0.7143 | 12 | 0.75 | NA | NA | 1 | 1 | 5 | 0.6 | 3 | 0.6667 | 9 | 0.6667 |
| 14_614508 | 1 | 0 | 4 | 0.25 | 1 | 0 | 4 | 0 | 10 | 0.1 | NA | NA | 2 | 0 | 6 | 0.1667 | 3 | 0 | 11 | 0.0909 |
| 14_614512 | 1 | 0 | 2 | 0 | NA | NA | 9 | 0.6667 | 12 | 0.5 | NA | NA | 3 | 0.3333 | 5 | 0.4 | 4 | 0.75 | 12 | 0.5 |
| 14_614512 | 1 | 0 | 5 | 0 | 1 | 0 | 10 | 0.2 | 17 | 0.1176 | NA | NA | 3 | 0 | 6 | 0.1667 | 4 | 0 | 13 | 0.0769 |
| 14_614513 | 2 | 0.5 | 5 | 0 | 2 | 0 | 10 | 0 | 19 | 0.0526 | NA | NA | 3 | 0 | 6 | 0.1667 | 3 | 0 | 12 | 0.0833 |
| 14_614515 | 2 | 1 | 5 | 1 | 2 | 1 | 8 | 1 | 17 | 1 | NA | NA | 3 | 1 | 6 | 0.8333 | 4 | 1 | 13 | 0.9231 |
| 14_614515 | 2 | 0 | 5 | 0.2 | 2 | 0 | 8 | 0.375 | 17 | 0.2353 | NA | NA | 2 | 0.5 | 7 | 0.1429 | 3 | 0.3333 | 12 | 0.25 |
| 14_614516 | 2 | 0 | 5 | 0.2 | 3 | 0 | 10 | 0.3 | 20 | 0.2 | NA | NA | 3 | 0.6667 | 8 | 0.25 | 4 | 0.5 | 15 | 0.4 |
| 14_614516 | 2 | 1 | 5 | 0.4 | 3 | 0 | 11 | 0.2727 | 21 | 0.3333 | NA | NA | 3 | 0.6667 | 7 | 0.1429 | 4 | 0.5 | 14 | 0.3571 |
| 14_614518 | 2 | 0.5 | 6 | 0.5 | 3 | 1 | 10 | 0.8 | 21 | 0.7143 | NA | NA | 2 | 1 | 8 | 0.875 | 4 | 1 | 14 | 0.9286 |
| 14_614518 | 2 | 0.5 | 6 | 0.1667 | 4 | 0 | 9 | 0 | 21 | 0.0952 | NA | NA | 3 | 0 | 8 | 0 | 4 | 0.25 | 15 | 0.0667 |
| 14_614518 | 1 | 0 | 4 | 0 | 4 | 0.25 | 8 | 0 | 17 | 0.0588 | NA | NA | 2 | 0 | 6 | 0.1667 | 3 | 0 | 11 | 0.0909 |
| 14_614519 | 1 | 0 | 6 | 0.1667 | 4 | 0 | 12 | 0.0833 | 23 | 0.087 | NA | NA | 3 | 0 | 8 | 0.125 | 3 | 0 | 14 | 0.0714 |
| 14_614519 | 1 | 1 | 5 | 0.8 | 3 | 1 | 5 | 0.6 | 14 | 0.7857 | NA | NA | 3 | 1 | 7 | 0.8571 | 3 | 1 | 13 | 0.9231 |
| 14_614520 | 2 | 0 | 7 | 0.1429 | 4 | 0 | 9 | 0.1111 | 22 | 0.0909 | NA | NA | 2 | 0.5 | 7 | 0.5714 | 5 | 0.2 | 14 | 0.4286 |
| 14_614520 | 2 | 0 | 7 | 0.2857 | 3 | 0.6667 | 10 | 0.4 | 22 | 0.3636 | NA | NA | 3 | 0.6667 | 6 | 0.6667 | 3 | 0.6667 | 12 | 0.6667 |
| 14_614520 | 2 | 1 | 6 | 0.8333 | 4 | 0.75 | 8 | 0.875 | 20 | 0.85 | NA | NA | 3 | 0.3333 | 6 | 0.6667 | 5 | 1 | 14 | 0.7143 |
| 14_614520 | 2 | 1 | 7 | 0.8571 | 4 | 0.75 | 12 | 0.9167 | 25 | 0.88 | NA | NA | 3 | 1 | 5 | 1 | 5 | 0.8 | 13 | 0.9231 |
| 14_614521 | 2 | 1 | 5 | 0.8 | 4 | 1 | 10 | 1 | 21 | 0.9524 | NA | NA | 3 | 1 | 8 | 0.875 | 4 | 1 | 15 | 0.9333 |
| 14_614521 | 1 | 1 | 7 | 0.7143 | 4 | 0.75 | 11 | 0.9091 | 23 | 0.8261 | NA | NA | 3 | 1 | 6 | 0.6667 | 4 | 1 | 13 | 0.8462 |
| 14_614522 | 2 | 0 | 7 | 0.1429 | 3 | 0 | 12 | 0.0833 | 24 | 0.0833 | NA | NA | 3 | 0 | 6 | 0.3333 | 5 | 0.4 | 14 | 0.2857 |
| 14_614522 | 2 | 0 | 6 | 0.3333 | 4 | 0.25 | 12 | 0.1667 | 24 | 0.2083 | NA | NA | 3 | 0.3333 | 7 | 0.2857 | 5 | 0 | 15 | 0.2 |
| 14_614523 | 2 | 0 | 6 | 0.1667 | 4 | 0 | 11 | 0.0909 | 23 | 0.087 | NA | NA | 4 | 0 | 7 | 0 | 5 | 0.2 | 16 | 0.0625 |
| 14_614525 | 2 | 0.5 | 6 | 0.1667 | 3 | 0 | 9 | 0.2222 | 20 | 0.2 | NA | NA | 2 | 0.5 | 7 | 0 | 3 | 0 | 12 | 0.0833 |
| 14_614526 | 3 | 0 | 7 | 0.1429 | 6 | 0 | 12 | 0.0833 | 28 | 0.0714 | NA | NA | 4 | 0.25 | 7 | 0 | 5 | 0 | 16 | 0.0625 |
| 14_614526 | 3 | 0.6667 | 6 | 0.3333 | 6 | 0.5 | 13 | 0.6154 | 28 | 0.5357 | NA | NA | 4 | 0.75 | 7 | 0.2857 | 6 | 0.5 | 17 | 0.4706 |
| 14_615726 | NA | NA | 1 | 1 | NA | NA | NA | NA | 1 | 1 | NA | NA | 1 | 1 | NA | NA | NA | NA | 1 | 1 |
| 14_615728 | NA | NA | 1 | 1 | NA | NA | NA | NA | 1 | 1 | NA | NA | 1 | 1 | NA | NA | NA | NA | 1 | 1 |
| 14_615729 | NA | NA | 1 | 1 | NA | NA | NA | NA | 1 | 1 | NA | NA | 1 | 1 | NA | NA | NA | NA | 1 | 1 |
| 14_615785 | 2 | 0.5 | 6 | 0.3333 | 3 | 0 | 2 | 0.5 | 13 | 0.3077 | 3 | 0 | 4 | 0 | 4 | 0.5 | 6 | 0.1667 | 17 | 0.1765 |
| 14_615786 | 4 | 0.25 | 3 | 0.3333 | 5 | 0.2 | 1 | 0 | 13 | 0.2308 | 2 | 0 | 5 | 0.4 | 3 | 0 | 6 | 0.3333 | 16 | 0.25 |
| 14_616031 | 10 | 0 | 8 | 0.125 | 14 | 0 | 5 | 0.2 | 37 | 0.0541 | 6 | 0 | 7 | 0 | 13 | 0.0769 | 11 | 0.0909 | 37 | 0.0541 |
| 14_616045 | 8 | 0.25 | 6 | 0.3333 | 12 | 0.3333 | 6 | 0.6667 | 32 | 0.375 | 4 | 0.25 | 5 | 0.6 | 10 | 0.4 | 9 | 0.3333 | 28 | 0.3929 |
| 14_616045 | 10 | 0.2 | 6 | 0.1667 | 11 | 0.0909 | 6 | 0.3333 | 33 | 0.1818 | 5 | 0.2 | 4 | 0 | 11 | 0.2727 | 11 | 0.4545 | 31 | 0.2903 |
| 14_616045 | 10 | 0.2 | 5 | 0.2 | 11 | 0 | 6 | 0 | 32 | 0.0938 | 4 | 0 | 4 | 0.25 | 10 | 0.1 | 10 | 0.3 | 28 | 0.1786 |
| 14_616045 | 10 | 0.5 | 6 | 0.3333 | 10 | 0.2 | 6 | 0.5 | 32 | 0.375 | 4 | 0.75 | 3 | 0.6667 | 8 | 0.125 | 11 | 0.5455 | 26 | 0.4615 |
| 14_616046 | 7 | 0.1429 | 6 | 0.1667 | 11 | 0 | 5 | 0 | 29 | 0.069 | 3 | 0 | 4 | 0.25 | 10 | 0 | 6 | 0.5 | 23 | 0.1739 |
| 14_616047 | 9 | 0.1111 | 6 | 0.3333 | 11 | 0.2727 | 6 | 0.1667 | 32 | 0.2188 | 4 | 0 | 3 | 0.3333 | 9 | 0.2222 | 11 | 0.4545 | 27 | 0.2963 |
| 14_616048 | 8 | 0.625 | 5 | 0.6 | 8 | 0.875 | 3 | 0.6667 | 24 | 0.7083 | 3 | 1 | 3 | 1 | 11 | 0.6364 | 7 | 0.8571 | 24 | 0.7917 |
| 14_616049 | 10 | 1 | 6 | 1 | 11 | 1 | 6 | 1 | 33 | 1 | 3 | 1 | 4 | 1 | 10 | 1 | 10 | 0.9 | 27 | 0.963 |
| 14_617721 | 7 | 0.1429 | 11 | 0.2727 | 10 | 0.2 | 18 | 0.2778 | 46 | 0.2391 | 3 | 0 | 3 | 0 | 5 | 0.4 | 7 | 0.4286 | 18 | 0.2778 |
| 14_617724 | 10 | 0 | 10 | 0.1 | 11 | 0.1818 | 15 | 0.2 | 46 | 0.1304 | 5 | 0.2 | 6 | 0 | 5 | 0 | 4 | 0 | 20 | 0.05 |
| 14_617725 | 11 | 0.2727 | 8 | 0.125 | 10 | 0.1 | 18 | 0.3889 | 47 | 0.2553 | 6 | 0.3333 | 7 | 0.4286 | 6 | 0.1667 | 6 | 0.5 | 25 | 0.36 |
| 14_617725 | 11 | 0.4545 | 9 | 0.4444 | 11 | 0.3636 | 16 | 0.5625 | 47 | 0.4681 | 6 | 0.3333 | 8 | 0.125 | 6 | 0.3333 | 5 | 0.4 | 25 | 0.28 |
| 14_618408 | NA | NA | 4 | 0.5 | NA | NA | 1 | 1 | 5 | 0.6 | 1 | 0 | 1 | 0 | 3 | 1 | NA | NA | 5 | 0.6 |
| 14_618409 | NA | NA | 4 | 0.5 | NA | NA | 1 | 0 | 5 | 0.4 | 1 | 0 | 1 | 0 | 3 | 0.3333 | NA | NA | 5 | 0.2 |
| 14_618410 | NA | NA | 3 | 1 | NA | NA | 1 | 1 | 4 | 1 | 1 | 1 | 1 | 1 | 3 | 1 | NA | NA | 5 | 1 |
| 14_618411 | NA | NA | 3 | 0 | NA | NA | 1 | 1 | 4 | 0.25 | 1 | 1 | 1 | 1 | 3 | 0.6667 | NA | NA | 5 | 0.8 |
| 14_618422 | NA | NA | 1 | 1 | 4 | 1 | 1 | 1 | 6 | 1 | 4 | 1 | 1 | 1 | 4 | 1 | 1 | 0 | 10 | 0.9 |
| 14_618423 | NA | NA | 1 | 1 | 5 | 0.2 | 1 | 0 | 7 | 0.2857 | 3 | 0.3333 | NA | NA | 5 | 0.2 | 1 | 0 | 9 | 0.2222 |
| 14_618424 | NA | NA | 1 | 0 | 4 | 0.25 | 1 | 0 | 6 | 0.1667 | 3 | 0.6667 | NA | NA | 5 | 0.4 | 1 | 0 | 9 | 0.4444 |
| 14_618425 | NA | NA | 1 | 0 | 4 | 0.75 | NA | NA | 5 | 0.6 | 3 | 0.3333 | 1 | 1 | 4 | 0.5 | 1 | 1 | 9 | 0.5556 |
| 14_618427 | NA | NA | 1 | 0 | 6 | 0.1667 | 1 | 0 | 8 | 0.125 | 4 | 0.25 | 1 | 0 | 5 | 0 | 1 | 0 | 11 | 0.0909 |

| | | | | | | | | | | | | | | | | | | | | |
|---|---|---|---|---|---|---|---|---|---|---|---|---|---|---|---|---|---|---|---|---|
| 14_618427 | NA | NA | 1 | 0 | 7 | 0.1429 | 1 | 1 | 9 | 0.2222 | 4 | 0.75 | 1 | 0 | 4 | 0.25 | NA | NA | 9 | 0.4444 |
| 14_618428 | NA | NA | NA | NA | 7 | 1 | 1 | 1 | 8 | 1 | 3 | 1 | 1 | 1 | 2 | 1 | 1 | 1 | 7 | 1 |
| 14_618428 | NA | NA | 1 | 1 | 7 | 0.4286 | 1 | 1 | 9 | 0.5556 | 5 | 0 | NA | NA | 5 | 0.2 | 1 | 0 | 11 | 0.0909 |
| 14_618428 | NA | NA | NA | NA | 5 | 1 | 1 | 1 | 6 | 1 | 5 | 0.8 | 1 | 1 | 4 | 1 | 1 | 1 | 11 | 0.9091 |
| 14_618430 | NA | NA | 1 | 1 | 7 | 1 | 1 | 1 | 9 | 1 | 5 | 1 | 1 | 1 | 5 | 1 | 1 | 1 | 12 | 1 |
| 14_618430 | NA | NA | 1 | 0 | 8 | 0.25 | 1 | 1 | 10 | 0.3 | 5 | 0.2 | 1 | 0 | 5 | 0.4 | 1 | 0 | 12 | 0.25 |
| 14_618430 | NA | NA | 1 | 1 | 7 | 0.7143 | 1 | 0 | 9 | 0.6667 | 3 | 0.6667 | 1 | 0 | 4 | 0.5 | 1 | 0 | 9 | 0.4444 |
| 14_618431 | NA | NA | 1 | 0 | 5 | 0.6 | 1 | 0 | 7 | 0.4286 | 5 | 0.2 | 1 | 0 | 5 | 0.2 | 1 | 0 | 12 | 0.1667 |
| 14_618431 | NA | NA | 1 | 0 | 4 | 0.5 | 1 | 0 | 6 | 0.3333 | 2 | 0.5 | NA | NA | 4 | 0 | NA | NA | 6 | 0.1667 |
| 14_618431 | NA | NA | 1 | 0 | 6 | 0.6667 | 1 | 1 | 8 | 0.625 | 5 | 0.4 | 1 | 1 | 4 | 1 | 1 | 1 | 11 | 0.7273 |
| 14_618433 | NA | NA | 1 | 1 | 4 | 0.5 | 1 | 0 | 6 | 0.5 | 4 | 1 | 1 | 1 | 3 | 0.3333 | 1 | 0 | 9 | 0.6667 |
| 14_625140 | NA | NA | 3 | 0.3333 | 1 | 0 | 2 | 0.5 | 6 | 0.3333 | NA | NA | 1 | 0 | 1 | 1 | NA | NA | 2 | 0.5 |
| 14_625141 | NA | NA | 3 | 0 | 1 | 0 | 2 | 0.5 | 6 | 0.1667 | NA | NA | NA | NA | 1 | 1 | 1 | 0 | 2 | 0.5 |
| 14_625142 | NA | NA | 4 | 0.75 | 1 | 1 | 2 | 0.5 | 7 | 0.7143 | NA | NA | 1 | 1 | 1 | 1 | 1 | 1 | 3 | 1 |
| 14_625143 | NA | NA | 2 | 0.5 | 1 | 1 | 2 | 0 | 5 | 0.4 | NA | NA | 1 | 1 | 1 | 0 | 1 | 1 | 3 | 0.6667 |
| 14_625144 | NA | NA | 3 | 0.6667 | 1 | 0 | 2 | 1 | 6 | 0.6667 | NA | NA | 1 | 0 | 1 | 1 | 1 | 1 | 3 | 0.6667 |
| 14_626101 | NA | NA | 1 | 0 | 1 | 1 | NA | NA | 2 | 0.5 | NA | NA | NA | NA | NA | NA | 1 | 1 | 1 | 1 |
| 14_626101 | NA | NA | 2 | 0.5 | 1 | 0 | NA | NA | 3 | 0.3333 | NA | NA | NA | NA | NA | NA | 1 | 1 | 1 | 1 |
| 14_628249 | NA | NA | 1 | 1 | NA | NA | NA | NA | 1 | 1 | NA | NA | NA | NA | 1 | 1 | NA | NA | 1 | 1 |
| 14_628251 | NA | NA | 1 | 1 | NA | NA | NA | NA | 1 | 1 | NA | NA | NA | NA | 1 | 1 | NA | NA | 1 | 1 |
| 14_628252 | NA | NA | 1 | 1 | NA | NA | NA | NA | 1 | 1 | NA | NA | NA | NA | 1 | 1 | NA | NA | 1 | 1 |
| 14_628436 | NA | NA | 2 | 1 | 1 | 0 | NA | NA | 3 | 0.6667 | 1 | 1 | NA | NA | 2 | 0.5 | NA | NA | 3 | 0.6667 |
| 14_628436 | NA | NA | 2 | 0.5 | 1 | 1 | NA | NA | 3 | 0.6667 | 1 | 0 | NA | NA | 1 | 1 | NA | NA | 2 | 0.5 |
| 14_628436 | NA | NA | 2 | 1 | 1 | 0 | NA | NA | 3 | 0.6667 | 1 | 0 | NA | NA | 2 | 1 | NA | NA | 3 | 0.6667 |
| 14_628438 | NA | NA | 2 | 1 | 1 | 1 | NA | NA | 3 | 1 | 1 | 1 | NA | NA | 1 | 1 | NA | NA | 2 | 1 |
| 14_628438 | NA | NA | 2 | 0.5 | 1 | 0 | NA | NA | 3 | 0.3333 | 1 | 0 | NA | NA | 2 | 0.5 | NA | NA | 3 | 0.3333 |
| 14_628439 | NA | NA | 2 | 0 | 1 | 1 | NA | NA | 3 | 0.3333 | 1 | 1 | NA | NA | 2 | 0 | NA | NA | 3 | 0.3333 |
| 14_628440 | NA | NA | NA | NA | 2 | 1 | NA | NA | 2 | 1 | 1 | 1 | NA | NA | 2 | 1 | NA | NA | 3 | 1 |
| 14_628441 | NA | NA | 2 | 0.5 | 2 | 0 | NA | NA | 4 | 0.25 | 1 | 1 | NA | NA | 2 | 0.5 | NA | NA | 3 | 0.6667 |
| 14_628442 | NA | NA | 1 | 1 | 1 | 0 | NA | NA | 2 | 0.5 | 1 | 1 | NA | NA | 1 | 1 | NA | NA | 2 | 1 |
| 14_628442 | NA | NA | 2 | 0.5 | 2 | 0.5 | NA | NA | 4 | 0.5 | 1 | 1 | NA | NA | NA | NA | NA | NA | 1 | 1 |
| 14_628446 | NA | NA | 3 | 1 | 3 | 0.6667 | NA | NA | 6 | 0.8333 | 1 | 1 | 1 | 1 | 2 | 0.5 | 1 | 1 | 5 | 0.8 |
| 14_628446 | NA | NA | 4 | 0 | 4 | 0.25 | NA | NA | 8 | 0.125 | 1 | 1 | 1 | 0 | 3 | 0 | NA | NA | 5 | 0.2 |
| 14_628447 | NA | NA | 3 | 0.3333 | 4 | 0.25 | NA | NA | 7 | 0.2857 | 1 | 1 | 1 | 0 | 3 | 0.3333 | 1 | 0 | 6 | 0.3333 |
| 14_628447 | NA | NA | 4 | 0.25 | 4 | 0 | NA | NA | 8 | 0.125 | 1 | 0 | 1 | 0 | 3 | 0.6667 | 1 | 1 | 6 | 0.5 |
| 14_628449 | NA | NA | 4 | 0.5 | 4 | 0.75 | NA | NA | 8 | 0.625 | 1 | 1 | 1 | 1 | 3 | 1 | 1 | 1 | 6 | 1 |
| 14_628450 | NA | NA | 2 | 0.5 | NA | NA | NA | NA | 2 | 0.5 | 1 | 1 | 1 | 0 | 2 | 1 | 1 | 1 | 5 | 0.8 |
| 14_628451 | NA | NA | 5 | 0.4 | 2 | 0 | NA | NA | 7 | 0.2857 | 1 | 0 | 1 | 0 | 3 | 0.6667 | NA | NA | 5 | 0.4 |
| 14_628452 | NA | NA | 5 | 0.6 | 1 | 0 | NA | NA | 6 | 0.5 | 1 | 0 | 1 | 0 | 3 | 0.6667 | 1 | 0 | 6 | 0.3333 |
| 14_628453 | NA | NA | 5 | 0.4 | 3 | 0.6667 | NA | NA | 8 | 0.5 | 1 | 0 | 1 | 0 | 3 | 1 | 2 | 0.5 | 7 | 0.5714 |
| 14_628457 | NA | NA | 3 | 0.6667 | 3 | 1 | NA | NA | 6 | 0.8333 | 2 | 1 | 1 | 1 | 2 | 1 | 2 | 1 | 7 | 1 |
| 14_628457 | NA | NA | 5 | 1 | 4 | 1 | NA | NA | 9 | 1 | 3 | 1 | 2 | 1 | 3 | 1 | 2 | 1 | 10 | 1 |
| 14_628457 | NA | NA | 5 | 1 | 4 | 0.75 | NA | NA | 9 | 0.8889 | 3 | 1 | 2 | 1 | 3 | 1 | 2 | 1 | 10 | 1 |
| 14_629309 | 2 | 0 | 6 | 0.8333 | 1 | 0 | 2 | 0.5 | 11 | 0.5455 | 2 | 0 | 2 | 0.5 | 1 | 0 | NA | NA | 5 | 0.2 |
| 14_629342 | 1 | 1 | 4 | 0.75 | NA | NA | 2 | 1 | 7 | 0.8571 | 3 | 1 | 1 | 1 | 2 | 0.5 | NA | NA | 6 | 0.8333 |
| 14_631939 | NA | NA | 1 | 1 | NA | NA | NA | NA | 1 | 1 | NA | NA | NA | NA | NA | NA | 1 | 1 | 1 | 1 |
| 14_631945 | NA | NA | 1 | 1 | NA | NA | NA | NA | 1 | 1 | NA | NA | 1 | 1 | 1 | 1 | 1 | 1 | 3 | 1 |
| 14_631945 | NA | NA | 1 | 1 | NA | NA | NA | NA | 1 | 1 | NA | NA | 1 | 1 | 1 | 1 | 2 | 1 | 4 | 1 |
| 14_631946 | NA | NA | 1 | 1 | NA | NA | NA | NA | 1 | 1 | 1 | 1 | 1 | 1 | 1 | 1 | 2 | 0 | 5 | 0.6 |
| 14_631951 | 1 | 0 | 1 | 1 | NA | NA | NA | NA | 2 | 0.5 | 1 | 0 | 1 | 1 | 1 | 0 | 2 | 0.5 | 5 | 0.4 |
| 14_631953 | 1 | 0 | 1 | 1 | NA | NA | NA | NA | 2 | 0.5 | 1 | 1 | NA | NA | NA | NA | 2 | 0 | 3 | 0.3333 |
| 14_631961 | 1 | 0 | 1 | 0 | 1 | 0 | 1 | 1 | 4 | 0.25 | NA | NA | 1 | 0 | 1 | 0 | 1 | 1 | 3 | 0.3333 |
| 14_631982 | 3 | 0 | 2 | 0.5 | 1 | 0 | 3 | 0.3333 | 9 | 0.2222 | 2 | 0.5 | 4 | 0 | 1 | 1 | 2 | 0.5 | 9 | 0.3333 |
| 14_631983 | 3 | 0 | 3 | 0.3333 | 1 | 0 | 2 | 0 | 9 | 0.1111 | 2 | 0.5 | 4 | 0.25 | 1 | 1 | 2 | 0 | 9 | 0.3333 |
| 14_631984 | 2 | 0.5 | 2 | 0 | 1 | 0 | NA | NA | 5 | 0.2 | 1 | 0 | 1 | 1 | 1 | 0 | 2 | 0.5 | 5 | 0.4 |
| 14_631985 | 4 | 0.25 | 3 | 0.3333 | 1 | 0 | 3 | 0.6667 | 11 | 0.3636 | 2 | 0.5 | 4 | 0.5 | 1 | 1 | 2 | 0 | 9 | 0.4444 |
| 14_631985 | 3 | 0.3333 | 4 | 0.75 | 1 | 1 | 3 | 0.6667 | 11 | 0.6364 | 2 | 1 | 4 | 0.75 | 1 | 1 | 3 | 0.6667 | 10 | 0.8 |
| 14_631987 | 1 | 1 | 3 | 1 | 1 | 1 | 1 | 1 | 6 | 1 | 2 | 1 | 4 | 1 | NA | NA | 3 | 0.3333 | 9 | 0.7778 |
| 14_631987 | 3 | 0 | 4 | 0.5 | 1 | 0 | 3 | 0 | 11 | 0.1818 | 2 | 0.5 | 4 | 0.25 | 1 | 0 | 4 | 0.25 | 11 | 0.2727 |
| 14_631987 | 2 | 0 | 4 | 0.75 | NA | NA | 3 | 0.3333 | 9 | 0.4444 | 2 | 0.5 | 3 | 0.6667 | 1 | 0 | 3 | 0.6667 | 9 | 0.5556 |
| 14_633797 | 67 | 0.3134 | 59 | 0.0847 | 67 | 0.209 | 56 | 0.3036 | 249 | 0.2289 | 66 | 0.2424 | 50 | 0.22 | 59 | 0.1695 | 52 | 0.3462 | 227 | 0.2423 |
| 14_633799 | 71 | 0.0563 | 71 | 0.0986 | 79 | 0.0886 | 64 | 0.0312 | 285 | 0.0702 | 67 | 0.1194 | 57 | 0.1404 | 59 | 0.0847 | 55 | 0.1818 | 238 | 0.1303 |
| 14_633800 | 71 | 0.2817 | 77 | 0.1688 | 76 | 0.2105 | 59 | 0.2373 | 283 | 0.2226 | 71 | 0.2254 | 51 | 0.1373 | 60 | 0.2167 | 51 | 0.1373 | 233 | 0.1845 |
| 14_633801 | 78 | 0.2949 | 76 | 0.1974 | 81 | 0.2099 | 65 | 0.2 | 300 | 0.2267 | 78 | 0.3333 | 64 | 0.3125 | 61 | 0.2623 | 55 | 0.3091 | 258 | 0.3062 |
| 14_633801 | 64 | 0.1406 | 69 | 0.058 | 75 | 0.0933 | 55 | 0.0545 | 263 | 0.0875 | 70 | 0.0857 | 55 | 0.0909 | 59 | 0.1186 | 54 | 0.1481 | 238 | 0.1092 |

| | | | | | | | | | | | | | | | | | | | | |
|---|---|---|---|---|---|---|---|---|---|---|---|---|---|---|---|---|---|---|---|---|
| 14_6338013 | 71 | 0.6761 | 70 | 0.6571 | 76 | 0.5921 | 58 | 0.6379 | 275 | 0.64 | 68 | 0.6029 | 55 | 0.7455 | 54 | 0.6481 | 50 | 0.8 | 227 | 0.6916 |
| 14_6338015 | 33 | 0.4848 | 40 | 0.55 | 53 | 0.4717 | 32 | 0.5312 | 158 | 0.5063 | 38 | 0.5263 | 32 | 0.5312 | 36 | 0.3611 | 30 | 0.5 | 136 | 0.4779 |
| 14_6338016 | 76 | 0.1711 | 76 | 0.1184 | 80 | 0.1625 | 61 | 0.1311 | 293 | 0.1468 | 73 | 0.137 | 57 | 0.1228 | 57 | 0.1579 | 58 | 0.0862 | 245 | 0.1265 |
| 14_6338050 | 73 | 0.137 | 70 | 0.1143 | 80 | 0.1 | 66 | 0.1212 | 289 | 0.1176 | 79 | 0.1266 | 58 | 0.069 | 60 | 0.1167 | 58 | 0.0862 | 255 | 0.102 |
| 14_6338068 | 6 | 0 | 8 | 0 | 4 | 0 | 2 | 0.5 | 20 | 0.05 | 1 | 0 | 4 | 0.25 | 1 | 0 | 4 | 0 | 10 | 0.1 |
| 14_6338078 | 4 | 0 | 8 | 0.25 | 4 | 0.25 | 2 | 0.5 | 18 | 0.2222 | 2 | 0.5 | 2 | 0.5 | 1 | 0 | 4 | 0.25 | 9 | 0.3333 |
| 14_6338392 | 11 | 0.1818 | 17 | 0.0588 | 16 | 0.0625 | 18 | 0 | 62 | 0.0645 | 9 | 0.1111 | 10 | 0.1 | 19 | 0.0526 | 9 | 0 | 47 | 0.0638 |
| 14_6338401 | 11 | 0.2727 | 17 | 0.4118 | 20 | 0.2 | 18 | 0.4444 | 66 | 0.3333 | 9 | 0.2222 | 11 | 0.1818 | 17 | 0.4706 | 10 | 0.4 | 47 | 0.3404 |
| 14_6338406 | 11 | 0.3636 | 14 | 0.4286 | 16 | 0.375 | 16 | 0.3125 | 57 | 0.3684 | 9 | 0.3333 | 8 | 0.25 | 16 | 0.3125 | 10 | 0.4 | 43 | 0.3256 |
| 14_6338407 | 11 | 0.5455 | 16 | 0.75 | 17 | 0.7059 | 15 | 0.4667 | 59 | 0.6271 | 6 | 0.8333 | 7 | 0.2857 | 16 | 0.5 | 8 | 0.75 | 37 | 0.5676 |
| 14_6338409 | 12 | 0.75 | 13 | 0.6154 | 17 | 0.7059 | 17 | 0.7647 | 59 | 0.7119 | 8 | 0.875 | 10 | 0.8 | 14 | 0.9286 | 9 | 0.7778 | 41 | 0.8537 |
| 14_6338420 | 10 | 0.1 | 14 | 0.0714 | 15 | 0.0667 | 15 | 0.2 | 54 | 0.1111 | 8 | 0.25 | 8 | 0.25 | 17 | 0.3529 | 9 | 0.3333 | 42 | 0.3095 |
| 14_6338422 | 11 | 0.2727 | 16 | 0.125 | 19 | 0.1579 | 19 | 0.3158 | 65 | 0.2154 | 10 | 0.1 | 11 | 0.0909 | 19 | 0.1579 | 8 | 0.25 | 48 | 0.1458 |
| 14_6368962 | NA | NA | NA | NA | 1 | 1 | NA | NA | 1 | 1 | NA | NA | NA | NA | NA | NA | 1 | 1 | 1 | 1 |
| 14_6375773 | 3 | 0.3333 | 3 | 0 | 6 | 0.3333 | 2 | 0 | 14 | 0.2143 | 5 | 0 | 5 | 0.2 | 6 | 0 | NA | NA | 16 | 0.0625 |
| 14_6375794 | 3 | 0.3333 | 2 | 0 | 6 | 0 | 1 | 0 | 12 | 0.0833 | 5 | 0.2 | 3 | 0 | 5 | 0 | NA | NA | 13 | 0.0769 |
| 14_6375796 | 3 | 0 | 3 | 0 | 6 | 0.1667 | 2 | 0 | 14 | 0.0714 | 5 | 0 | 5 | 0.2 | 7 | 0 | 1 | 0 | 18 | 0.0556 |
| 14_6375814 | 17 | 0.2941 | 18 | 0.2222 | 12 | 0.0833 | 10 | 0.3 | 57 | 0.2281 | 13 | 0.1538 | 9 | 0 | 13 | 0.0769 | 13 | 0.3846 | 48 | 0.1667 |
| 14_6375840 | 14 | 0.5 | 13 | 0.4615 | 13 | 0.3077 | 10 | 0.6 | 50 | 0.46 | 11 | 0.2727 | 7 | 0.4286 | 14 | 0.2857 | 10 | 0.5 | 42 | 0.3571 |
| 14_6375962 | 13 | 0.1538 | 15 | 0 | 11 | 0.0909 | 5 | 0.2 | 44 | 0.0909 | 9 | 0.1111 | 5 | 0 | 14 | 0.0714 | 5 | 0 | 33 | 0.0606 |
| 14_6376069 | 10 | 0 | 7 | 0.4286 | 11 | 0 | 6 | 0 | 34 | 0.0882 | 5 | 0 | 4 | 0.5 | 11 | 0 | 4 | 0 | 24 | 0.0833 |
| 14_6376075 | 9 | 0.1111 | 7 | 0.1429 | 10 | 0.1 | 4 | 0 | 30 | 0.1 | 3 | 0.3333 | 3 | 0 | 12 | 0.0833 | 2 | 0 | 20 | 0.1 |
| 14_6376076 | 6 | 0.3333 | 5 | 0.6 | 7 | 0.1429 | 3 | 0.3333 | 21 | 0.3333 | 5 | 0.6 | 3 | 0.3333 | 6 | 0 | 4 | 0.5 | 18 | 0.3333 |
| 14_6376105 | 8 | 0.75 | 3 | 0.3333 | 7 | 0.7143 | 4 | 1 | 22 | 0.7273 | 4 | 0.75 | 2 | 1 | 9 | 0.6667 | 4 | 0.75 | 19 | 0.7368 |
| 14_6376107 | 10 | 0.1 | 8 | 0 | 9 | 0.1111 | 2 | 0 | 29 | 0.069 | 5 | 0 | 4 | 0 | 11 | 0.1818 | 4 | 0.25 | 24 | 0.125 |
| 14_6376141 | 8 | 0.375 | 8 | 0.625 | 9 | 0.6667 | 4 | 0.5 | 29 | 0.5517 | 4 | 0.75 | 5 | 0.4 | 8 | 0.375 | 2 | 0.5 | 19 | 0.4737 |
| 14_6376147 | 10 | 0.6 | 8 | 0.5 | 10 | 0.5 | 5 | 0.4 | 33 | 0.5152 | 4 | 0.75 | 5 | 0.2 | 10 | 0.6 | 3 | 0.3333 | 22 | 0.5 |
| 14_6378112 | 5 | 0 | 2 | 0.5 | 7 | 0 | 5 | 0.2 | 19 | 0.1053 | 2 | 0 | 1 | 0 | 4 | 0.5 | 2 | 0.5 | 9 | 0.3333 |
| 14_6384435 | NA | NA | 1 | 1 | NA | NA | NA | NA | 1 | 1 | 1 | 0 | NA | NA | 2 | 0.5 | NA | NA | 3 | 0.3333 |
| 14_6384444 | NA | NA | 1 | 1 | NA | NA | NA | NA | 1 | 1 | 1 | 1 | NA | NA | 2 | 1 | NA | NA | 3 | 1 |
| 14_6390109 | NA | NA | 1 | 1 | NA | NA | NA | NA | 1 | 1 | 1 | 0 | 1 | 1 | NA | NA | NA | NA | 2 | 0.5 |
| 14_6390232 | NA | NA | 1 | 1 | NA | NA | NA | NA | 1 | 1 | 1 | 0 | 1 | 1 | NA | NA | NA | NA | 2 | 0.5 |
| 14_6390271 | NA | NA | 1 | 1 | NA | NA | NA | NA | 1 | 1 | 2 | 0.5 | 1 | 1 | NA | NA | NA | NA | 3 | 0.6667 |
| 14_6390288 | NA | NA | 1 | 1 | NA | NA | NA | NA | 1 | 1 | 2 | 0.5 | NA | NA | NA | NA | NA | NA | 2 | 0.5 |
| 14_6390307 | NA | NA | 1 | 1 | NA | NA | NA | NA | 1 | 1 | 1 | 1 | 1 | 1 | NA | NA | NA | NA | 2 | 1 |
| 14_6390330 | NA | NA | 1 | 1 | NA | NA | NA | NA | 1 | 1 | 2 | 0.5 | NA | NA | NA | NA | NA | NA | 2 | 0.5 |
| 14_6390344 | NA | NA | 1 | 1 | NA | NA | NA | NA | 1 | 1 | 2 | 0.5 | NA | NA | NA | NA | NA | NA | 2 | 0.5 |
| 14_6408651 | 1 | 1 | NA | NA | 1 | 1 | NA | NA | 2 | 1 | NA | NA | NA | NA | 2 | 1 | 1 | 1 | 3 | 1 |
| 14_6408652 | 1 | 1 | NA | NA | 1 | 0 | NA | NA | 2 | 0.5 | NA | NA | NA | NA | 2 | 1 | 1 | 0 | 3 | 0.6667 |
| 14_6408657 | NA | NA | NA | NA | 1 | 1 | NA | NA | 1 | 1 | NA | NA | NA | NA | 2 | 1 | 1 | 1 | 3 | 1 |
| 14_6408676 | NA | NA | NA | NA | 1 | 1 | NA | NA | 1 | 1 | NA | NA | NA | NA | 2 | 1 | 1 | 1 | 3 | 1 |
| 14_6409263 | NA | NA | NA | NA | 2 | 0.5 | 5 | 0.4 | 7 | 0.4286 | NA | NA | 4 | 0.25 | 5 | 0 | 2 | 0 | 11 | 0.0909 |
| 14_6410362 | NA | NA | 1 | 1 | 2 | 0 | NA | NA | 3 | 0.3333 | NA | NA | 1 | 0 | NA | NA | 1 | 1 | 2 | 0.5 |
| 14_6436038 | 2 | 0 | 4 | 0.25 | NA | NA | 5 | 1 | 11 | 0.0909 | 2 | 0 | 2 | 0 | 3 | 0 | 6 | 0.5 | 13 | 0.2308 |
| 14_6436052 | 3 | 0.6667 | 3 | 0.6667 | NA | NA | 5 | 1 | 11 | 0.8182 | 2 | 1 | 2 | 1 | 2 | 1 | 6 | 0.8333 | 12 | 0.9167 |
| 14_6482853 | 4 | 0 | 15 | 0 | 11 | 0.0909 | 9 | 0.1111 | 39 | 0.0513 | 6 | 0 | 9 | 0.1111 | 10 | 0.1 | 6 | 0 | 31 | 0.0645 |
| 14_6482880 | 6 | 0.3333 | 14 | 0.2857 | 9 | 0.2222 | 11 | 0.2727 | 40 | 0.275 | 5 | 0.2 | 10 | 0.4 | 9 | 0.3333 | 4 | 0.25 | 28 | 0.3214 |
| 14_6482882 | 5 | 0.2 | 12 | 0 | 11 | 0.0909 | 8 | 0.125 | 36 | 0.0833 | 6 | 0 | 8 | 0.25 | 6 | 0 | 4 | 0 | 24 | 0.0833 |
| 14_6483048 | 9 | 0.1111 | 19 | 0.3158 | 10 | 0.1 | 8 | 0.125 | 46 | 0.1957 | 5 | 0.2 | 10 | 0.1 | 8 | 0.375 | 8 | 0.125 | 31 | 0.1935 |
| 14_6483073 | 9 | 0 | 21 | 0.0952 | 14 | 0.2143 | 8 | 0.25 | 52 | 0.1346 | 7 | 0 | 11 | 0 | 9 | 0 | 9 | 0.2222 | 36 | 0.0556 |
| 14_6483090 | 7 | 0.1429 | 25 | 0.32 | 14 | 0.2143 | 7 | 0.7143 | 53 | 0.3208 | 6 | 0.1667 | 11 | 0.1818 | 9 | 0 | 10 | 0.2 | 36 | 0.1389 |
| 14_6504056 | 1 | 0 | NA | NA | 1 | 0 | 1 | 1 | 3 | 0.3333 | NA | NA | NA | NA | 1 | 1 | NA | NA | 1 | 1 |
| 14_6504090 | 1 | 0 | NA | NA | 1 | 1 | 1 | 1 | 3 | 0.6667 | NA | NA | NA | NA | 1 | 1 | NA | NA | 1 | 1 |
| 14_6504100 | 1 | 1 | NA | NA | 1 | 1 | NA | NA | 2 | 1 | NA | NA | NA | NA | 1 | 1 | NA | NA | 1 | 1 |
| 14_6504376 | 1 | 1 | NA | NA | 3 | 1 | 1 | 1 | 5 | 1 | NA | NA | NA | NA | 1 | 1 | 1 | 0 | 2 | 0.5 |
| 14_6504377 | 1 | 1 | NA | NA | 1 | 1 | NA | NA | 2 | 1 | NA | NA | NA | NA | 1 | 1 | 2 | 0.5 | 3 | 0.6667 |
| 14_6504384 | 1 | 1 | NA | NA | 3 | 1 | 1 | 1 | 5 | 1 | NA | NA | NA | NA | 1 | 1 | 3 | 0.6667 | 4 | 0.75 |
| 14_6504396 | 1 | 1 | NA | NA | 3 | 1 | 1 | 1 | 5 | 1 | NA | NA | NA | NA | 1 | 1 | 2 | 1 | 3 | 1 |
| 14_6504406 | 1 | 1 | NA | NA | 1 | 1 | 1 | 0 | 3 | 0.6667 | NA | NA | NA | NA | 1 | 1 | 3 | 0 | 4 | 0.25 |
| 14_6504412 | 1 | 0 | NA | NA | 2 | 1 | 1 | 1 | 4 | 0.75 | NA | NA | NA | NA | 1 | 0 | 3 | 1 | 4 | 0.75 |
| 14_6504413 | 1 | 1 | NA | NA | 3 | 0.6667 | NA | NA | 4 | 0.75 | NA | NA | NA | NA | 1 | 1 | 3 | 0.3333 | 4 | 0.5 |
| 14_6504432 | 1 | 0 | NA | NA | 3 | 0.3333 | 1 | 0 | 5 | 0.2 | NA | NA | NA | NA | 1 | 1 | 3 | 1 | 4 | 1 |
| 14_6505005 | 3 | 0 | 12 | 0.0833 | 8 | 0.25 | 4 | 0 | 27 | 0.1111 | 10 | 0.1 | 3 | 0 | 9 | 0 | 4 | 0.25 | 26 | 0.0769 |
| 14_6505072 | 2 | 0 | 11 | 0.1818 | 6 | 0.1667 | NA | NA | 19 | 0.1579 | 5 | 0 | 2 | 0 | 8 | 0.25 | 3 | 0.3333 | 18 | 0.1667 |
| 14_6505073 | 2 | 0 | 13 | 0.2308 | 7 | 0.5714 | NA | NA | 22 | 0.3182 | 7 | 0.2857 | 4 | 0.25 | 8 | 0.125 | 3 | 0.3333 | 22 | 0.2273 |

| | | | | | | | | | | | | | | | | | | | | |
|---|---|---|---|---|---|---|---|---|---|---|---|---|---|---|---|---|---|---|---|---|
| 14_6505112 | 2 | 0.5 | 9 | 0 | 5 | 0 | NA | NA | 16 | 0.0625 | 5 | 0.2 | 4 | 0 | 7 | 0.1429 | 2 | 0 | 18 | 0.1111 |
| 14_6505120 | 2 | 0 | 9 | 0.1111 | 5 | 0 | NA | NA | 16 | 0.0625 | 6 | 0.3333 | 4 | 0.25 | 7 | 0 | 2 | 0 | 19 | 0.1579 |
| 14_6527684 | 2 | 0 | 2 | 0 | NA | NA | 5 | 0.2 | 9 | 0.1111 | 2 | 0 | NA | NA | 3 | 0 | 4 | 0.25 | 9 | 0.1111 |
| 14_6527693 | 2 | 0.5 | 1 | 1 | NA | NA | 4 | 0 | 7 | 0.2857 | 1 | 1 | NA | NA | 2 | 0.5 | 4 | 0.25 | 7 | 0.4286 |
| 14_6527700 | 1 | 0 | 2 | 0.5 | NA | NA | 4 | 0 | 7 | 0.1429 | 2 | 0 | NA | NA | 3 | 0.3333 | 5 | 0.2 | 10 | 0.2 |
| 14_6527709 | 2 | 0 | 2 | 0 | NA | NA | 4 | 0.25 | 8 | 0.125 | 1 | 0 | NA | NA | 3 | 0.3333 | 4 | 0.5 | 8 | 0.375 |
| 14_6527716 | 1 | 0 | NA | NA | NA | NA | 4 | 0.75 | 5 | 0.6 | 2 | 1 | NA | NA | 1 | 1 | 4 | 0.25 | 7 | 0.5714 |
| 14_6528999 | 17 | 0.1176 | 11 | 0.0909 | 10 | 0 | 13 | 0 | 51 | 0.0588 | 11 | 0.1818 | 5 | 0 | 6 | 0.1667 | 12 | 0.1667 | 34 | 0.1471 |
| 14_6529000 | 19 | 0.2105 | 8 | 0.375 | 10 | 0.4 | 10 | 0.3 | 47 | 0.2979 | 13 | 0.4615 | 5 | 0 | 7 | 0.1429 | 12 | 0.4167 | 37 | 0.3243 |
| 14_6529041 | 11 | 0 | 12 | 0.1667 | 10 | 0.1 | 9 | 0.1111 | 42 | 0.0952 | 8 | 0.25 | 5 | 0 | 6 | 0.1667 | 9 | 0.1111 | 28 | 0.1429 |
| 14_6529149 | 6 | 0.8333 | 6 | 0.5 | 4 | 0.5 | 4 | 0.75 | 20 | 0.65 | 4 | 0.5 | 1 | 1 | 3 | 0.3333 | 3 | 1 | 11 | 0.6364 |
| 14_6529179 | 8 | 0 | 9 | 0.1111 | 5 | 0 | 3 | 0.3333 | 25 | 0.08 | 5 | 0.2 | 2 | 0 | 5 | 0 | 5 | 0 | 17 | 0.0588 |
| 14_6529181 | 8 | 0.625 | 7 | 0.5714 | 4 | 0.75 | 4 | 0.75 | 23 | 0.6522 | 5 | 0.8 | 2 | 0.5 | 3 | 1 | 3 | 0.6667 | 13 | 0.7692 |
| 14_6529183 | 9 | 0.1111 | 8 | 0.125 | 4 | 0.25 | 4 | 0.25 | 25 | 0.16 | 4 | 0.75 | 2 | 0.5 | 4 | 0 | 4 | 0.5 | 14 | 0.4286 |
| 14_6529212 | 8 | 0.875 | 7 | 0.5714 | 4 | 0.25 | 3 | 1 | 22 | 0.6818 | 5 | 0.4 | 1 | 0 | 3 | 0.3333 | 5 | 0.8 | 14 | 0.5 |
| 14_6529221 | 8 | 0.25 | 7 | 0 | 5 | 0.2 | 2 | 0.5 | 22 | 0.1818 | 4 | 0.25 | 1 | 0 | 3 | 0 | 4 | 0.75 | 12 | 0.3333 |
| 14_6529225 | 8 | 0.125 | 7 | 0.1429 | 3 | 0 | 4 | 0.25 | 22 | 0.1364 | 5 | 0.6 | 1 | 0 | 2 | 0 | 4 | 0 | 12 | 0.25 |
| 14_6531305 | 8 | 0 | 6 | 0.1667 | 3 | 0 | 1 | 0 | 18 | 0.0556 | 4 | 0 | NA | NA | 3 | 0 | 4 | 0.25 | 11 | 0.0909 |
| 14_6531306 | 8 | 0.625 | 5 | 0.6 | 3 | 0 | 1 | 1 | 17 | 0.5294 | 4 | 0.5 | NA | NA | 3 | 0.6667 | 4 | 0.25 | 11 | 0.4545 |
| 14_6531309 | 6 | 0.5 | 6 | 0 | 3 | 0.3333 | 1 | 0 | 16 | 0.25 | 2 | 0.5 | NA | NA | 3 | 0 | 4 | 0.5 | 9 | 0.3333 |
| 14_6531313 | 7 | 1 | 6 | 0.8333 | 3 | 0.6667 | NA | NA | 16 | 0.875 | 3 | 0.3333 | NA | NA | 3 | 0.6667 | 4 | 0.25 | 10 | 0.4 |
| 14_6531314 | 6 | 0.5 | 6 | 0.5 | 4 | 0.5 | NA | NA | 16 | 0.5 | 3 | 0 | NA | NA | 3 | 0 | 2 | 1 | 8 | 0.25 |
| 14_6531328 | 5 | 1 | 4 | 1 | 4 | 1 | NA | NA | 13 | 1 | 4 | 1 | NA | NA | 3 | 0.6667 | 4 | 1 | 11 | 0.9091 |
| 14_6531335 | 8 | 0.25 | 5 | 0 | 4 | 0.5 | NA | NA | 17 | 0.2353 | 2 | 0 | NA | NA | 3 | 0.3333 | 4 | 0.25 | 9 | 0.2222 |
| 14_6531336 | 6 | 0.6667 | 4 | 0.5 | 4 | 0.5 | NA | NA | 14 | 0.5714 | 2 | 1 | NA | NA | 3 | 0.3333 | 2 | 0.5 | 7 | 0.5714 |
| 14_6531338 | 3 | 1 | 1 | 1 | 4 | 0.75 | NA | NA | 8 | 0.875 | 2 | 1 | NA | NA | 2 | 1 | 2 | 1 | 6 | 1 |
| 14_6531339 | 4 | 0.75 | 3 | 1 | 4 | 1 | NA | NA | 11 | 0.9091 | 2 | 0.5 | NA | NA | 2 | 0.5 | 4 | 0.25 | 8 | 0.375 |
| 14_6531344 | 8 | 0.625 | 4 | 0.5 | 3 | 0 | NA | NA | 15 | 0.4667 | 2 | 0.5 | NA | NA | 3 | 0.3333 | 3 | 0.6667 | 8 | 0.5 |
| 14_6531346 | 6 | 1 | 3 | 0.6667 | 4 | 1 | NA | NA | 13 | 0.9231 | 3 | 0.6667 | NA | NA | 2 | 0.5 | 4 | 0.5 | 9 | 0.5556 |
| 14_6531346 | 8 | 0.5 | 3 | 0.3333 | 3 | 0.6667 | NA | NA | 14 | 0.5 | 2 | 1 | NA | NA | 3 | 0.3333 | 3 | 0 | 8 | 0.375 |
| 14_6531347 | 7 | 0.1429 | 4 | 0 | 4 | 0.5 | NA | NA | 15 | 0.2 | 3 | 0 | NA | NA | 3 | 0.6667 | 3 | 0 | 9 | 0.2222 |
| 14_6531349 | 5 | 1 | 3 | 0.6667 | 3 | 0.3333 | NA | NA | 11 | 0.7273 | 3 | 0.3333 | NA | NA | 1 | 0 | 3 | 0.6667 | 7 | 0.4286 |
| 14_6531357 | 6 | 1 | 3 | 0.6667 | 4 | 1 | NA | NA | 13 | 0.9231 | 2 | 1 | NA | NA | 3 | 1 | 4 | 1 | 9 | 1 |
| 14_6531361 | 7 | 1 | 1 | 1 | 3 | 0.6667 | NA | NA | 11 | 0.9091 | 2 | 1 | NA | NA | 3 | 0 | 2 | 0 | 7 | 0.2857 |
| 14_6531367 | 7 | 0.1429 | 2 | 0 | 4 | 0.25 | NA | NA | 13 | 0.1538 | 2 | 0 | NA | NA | 2 | 0.5 | 1 | 0 | 5 | 0.2 |
| 14_6531401 | 7 | 0.4286 | 3 | 0 | 4 | 0 | NA | NA | 14 | 0.2143 | 3 | 0.3333 | NA | NA | 2 | 0 | 3 | 0.3333 | 8 | 0.25 |
| 14_6531433 | 7 | 0 | 3 | 0 | 4 | 0.25 | NA | NA | 14 | 0.0714 | 2 | 0.5 | NA | NA | 2 | 0 | 2 | 0 | 6 | 0.1667 |
| 14_6531437 | 7 | 0.2857 | 2 | 0 | 4 | 0.5 | NA | NA | 13 | 0.3077 | 2 | 0.5 | NA | NA | 1 | 1 | 3 | 1 | 6 | 0.8333 |
| 14_6531454 | 7 | 0.5714 | 2 | 0.5 | 4 | 0.5 | NA | NA | 13 | 0.5385 | 2 | 0.5 | NA | NA | 2 | 0 | 3 | 0.3333 | 7 | 0.2857 |
| 14_6531518 | 5 | 0.8 | 1 | 1 | 4 | 0.5 | NA | NA | 10 | 0.7 | 3 | 0.6667 | NA | NA | 1 | 1 | 3 | 0.6667 | 7 | 0.7143 |
| 14_6531524 | 2 | 1 | 1 | 1 | 1 | 1 | NA | NA | 4 | 1 | 1 | 1 | NA | NA | 1 | 1 | 2 | 0.5 | 4 | 0.75 |
| 14_6531527 | 5 | 0 | 2 | 0 | 3 | 0.3333 | NA | NA | 10 | 0.1 | 3 | 1 | NA | NA | 1 | 0 | 3 | 0.6667 | 7 | 0.7143 |
| 14_6531536 | 6 | 0.1667 | 2 | 0.5 | 3 | 0 | NA | NA | 11 | 0.1818 | 3 | 0.3333 | NA | NA | NA | NA | 3 | 0.3333 | 6 | 0.3333 |
| 14_6531551 | 5 | 1 | 2 | 1 | 3 | 0.6667 | NA | NA | 10 | 0.9 | 2 | 1 | NA | NA | NA | NA | 3 | 1 | 5 | 1 |
| 14_6534675 | NA | NA | NA | NA | 2 | 1 | NA | NA | 2 | 1 | 1 | 1 | NA | NA | NA | NA | NA | NA | 1 | 1 |
| 14_6543883 | NA | NA | NA | NA | NA | NA | 1 | 1 | 1 | 1 | NA | NA | 1 | 1 | 1 | 0 | 1 | 0 | 3 | 0.3333 |
| 14_6544020 | 2 | 0.5 | 2 | 0.5 | 1 | 0 | 1 | 0 | 6 | 0.3333 | NA | NA | 1 | 1 | 2 | 0.5 | 1 | 0 | 4 | 0.5 |
| 14_6547270 | 1 | 0 | 1 | 0 | 1 | 1 | NA | NA | 3 | 0.3333 | NA | NA | 1 | 1 | 1 | 0 | 3 | 0 | 5 | 0.2 |
| 14_6547274 | 1 | 1 | NA | NA | NA | NA | NA | NA | 1 | 1 | NA | NA | 1 | 1 | 1 | 1 | 1 | 1 | 3 | 1 |
| 14_6547298 | 2 | 0.5 | 1 | 0 | NA | NA | NA | NA | 3 | 0.3333 | NA | NA | 1 | 0 | 1 | 0 | 3 | 0.3333 | 5 | 0.2 |
| 14_6547317 | 2 | 1 | 1 | 1 | NA | NA | NA | NA | 3 | 1 | NA | NA | 1 | 1 | 1 | 1 | 3 | 1 | 5 | 1 |
| 14_6549041 | 2 | 0.5 | 1 | 0 | NA | NA | NA | NA | 3 | 0.3333 | NA | NA | NA | NA | 1 | 0 | 3 | 0.3333 | 4 | 0.25 |
| 14_6549054 | 1 | 1 | 1 | 1 | NA | NA | NA | NA | 2 | 1 | NA | NA | NA | NA | 1 | 1 | 2 | 1 | 3 | 1 |
| 14_6549069 | 2 | 0.5 | 1 | 0 | NA | NA | NA | NA | 3 | 0.3333 | NA | NA | NA | NA | 1 | 0 | 2 | 0.5 | 3 | 0.3333 |
| 14_6555886 | 2 | 1 | 2 | 1 | NA | NA | 1 | 1 | 5 | 1 | NA | NA | 2 | 0.5 | 2 | 1 | 2 | 1 | 6 | 0.8333 |
| 14_6555893 | 2 | 0.5 | 1 | 0 | NA | NA | NA | NA | 3 | 0.3333 | NA | NA | 2 | 0.5 | 2 | 0 | 2 | 0 | 6 | 0.1667 |
| 14_6576805 | 2 | 1 | 5 | 0 | 7 | 0.1429 | 7 | 0.1429 | 21 | 0.1905 | 7 | 0 | 7 | 0.2857 | 4 | 0 | 6 | 0.1667 | 24 | 0.125 |
| 14_6576822 | 6 | 0 | 10 | 0.1 | 9 | 0.1111 | 6 | 0.1667 | 31 | 0.0968 | 8 | 0 | 11 | 0.1818 | 7 | 0 | 10 | 0 | 36 | 0.0556 |
| 14_6576828 | 4 | 0 | 9 | 0.1111 | 8 | 0.25 | 6 | 0 | 27 | 0.1111 | 6 | 0.1667 | 10 | 0.1 | 5 | 0.2 | 8 | 0 | 29 | 0.1034 |
| 14_6576829 | 6 | 0.1667 | 9 | 0.4444 | 9 | 0.5556 | 8 | 0.625 | 32 | 0.4688 | 8 | 0.375 | 11 | 0.0909 | 7 | 0.7143 | 10 | 0.4 | 36 | 0.3611 |
| 14_6576856 | 6 | 0.1667 | 6 | 0.1667 | 7 | 0.4286 | 6 | 0.1667 | 25 | 0.24 | 7 | 0.1429 | 6 | 0.1667 | 6 | 0 | 9 | 0.5556 | 28 | 0.25 |
| 14_6576856 | 6 | 0 | 7 | 0 | 8 | 0.125 | 7 | 0.1429 | 28 | 0.0714 | 7 | 0.4286 | 6 | 0.1667 | 7 | 0 | 10 | 0 | 30 | 0.1333 |
| 14_6576857 | 6 | 0.3333 | 9 | 0.1111 | 8 | 0.125 | 7 | 0 | 30 | 0.1333 | 7 | 0.1429 | 7 | 0.2857 | 6 | 0 | 10 | 0.1 | 30 | 0.1333 |
| 14_6576859 | 6 | 0.3333 | 8 | 0.375 | 7 | 0 | 5 | 0.4 | 26 | 0.2692 | 7 | 0.2857 | 6 | 0.5 | 5 | 0.4 | 9 | 0.1111 | 27 | 0.2963 |
| 14_6576871 | 6 | 0 | 9 | 0 | 7 | 0 | 8 | 0.25 | 30 | 0.0667 | 7 | 0 | 7 | 0.1429 | 6 | 0.1667 | 10 | 0 | 30 | 0.0667 |

| | | | | | | | | | | | | | | | | | | | | |
|---|---|---|---|---|---|---|---|---|---|---|---|---|---|---|---|---|---|---|---|---|
| 14_657688 | 4 | 0 | 6 | 0.1667 | 4 | 0 | 5 | 0 | 19 | 0.0526 | 6 | 0 | 7 | 0.4286 | 6 | 0 | 9 | 0.1111 | 28 | 0.1429 |
| 14_661027 | 4 | 0.25 | 7 | 0.5714 | 3 | 0.3333 | 6 | 0.6667 | 20 | 0.5 | 1 | 1 | 3 | 0 | 4 | 0.5 | 3 | 1 | 11 | 0.5455 |
| 14_661027 | 5 | 0.6 | 6 | 0.5 | 3 | 0.6667 | 8 | 0.625 | 22 | 0.5909 | 1 | 1 | 1 | 1 | 4 | 0.5 | 3 | 0.6667 | 9 | 0.6667 |
| 14_661028 | 3 | 0.6667 | 5 | 0.8 | 3 | 1 | 5 | 1 | 16 | 0.875 | NA | NA | 2 | 1 | 4 | 0.5 | 2 | 1 | 8 | 0.75 |
| 14_661028 | 5 | 0 | 6 | 0.3333 | 4 | 0 | 8 | 0.25 | 23 | 0.1739 | 1 | 0 | 3 | 0 | 4 | 0.5 | 3 | 0.3333 | 11 | 0.2727 |
| 14_661028 | 5 | 0.2 | 4 | 0.25 | 4 | 0 | 8 | 0.25 | 21 | 0.1905 | NA | NA | 3 | 0 | 3 | 0.3333 | 2 | 0 | 8 | 0.125 |
| 14_661029 | 5 | 1 | 7 | 0.7143 | 3 | 1 | 8 | 0.5 | 23 | 0.7391 | 1 | 1 | 3 | 1 | 3 | 1 | 3 | 0.6667 | 10 | 0.9 |
| 14_661029 | 6 | 1 | 7 | 1 | 3 | 1 | 8 | 1 | 24 | 1 | 1 | 1 | 3 | 1 | 4 | 1 | 3 | 1 | 11 | 1 |
| 14_661030 | 6 | 0.1667 | 7 | 0 | 4 | 0 | 9 | 0.5556 | 26 | 0.2308 | 1 | 1 | 3 | 0 | 4 | 0 | 3 | 0.3333 | 11 | 0.1818 |
| 14_661036 | 6 | 0.1667 | 6 | 0 | 5 | 0 | 10 | 0.2 | 27 | 0.1111 | 3 | 0 | 4 | 0 | 4 | 0.25 | 2 | 0 | 13 | 0.0769 |
| 14_661038 | 6 | 0 | 7 | 0.1429 | 7 | 0.1429 | 9 | 0.2222 | 29 | 0.1379 | 3 | 0 | 3 | 0 | 4 | 0.5 | 3 | 0.3333 | 13 | 0.2308 |
| 14_661570 | 7 | 0.7143 | 5 | 0.4 | 5 | 1 | 3 | 0.6667 | 20 | 0.7 | 1 | 0 | 3 | 0.6667 | 5 | 0.4 | 3 | 0.3333 | 12 | 0.4167 |
| 14_661570 | 5 | 0.4 | 5 | 0 | 3 | 0 | 2 | 0 | 15 | 0.1333 | NA | NA | 1 | 0 | 4 | 0 | 2 | 0.5 | 7 | 0.1429 |
| 14_661570 | 3 | 0.3333 | 4 | 1 | 4 | 1 | 2 | 1 | 13 | 0.8462 | 1 | 0 | 3 | 1 | 4 | 1 | 3 | 1 | 11 | 0.9091 |
| 14_661571 | 6 | 0.6667 | 5 | 0.8 | 4 | 0.25 | 3 | 0.6667 | 18 | 0.6111 | 1 | 0 | 3 | 0 | 5 | 0.2 | 3 | 0.6667 | 12 | 0.25 |
| 14_661571 | 6 | 1 | 5 | 0.8 | 3 | 1 | 2 | 1 | 16 | 0.9375 | 1 | 1 | 2 | 1 | 2 | 1 | 3 | 0.6667 | 8 | 0.875 |
| 14_661571 | 7 | 0.1429 | 5 | 0 | 5 | 0 | 2 | 0 | 19 | 0.0526 | 1 | 0 | 3 | 0 | 4 | 0.25 | 3 | 0 | 11 | 0.0909 |
| 14_661572 | 7 | 0.4286 | 5 | 0 | 4 | 0 | 3 | 0 | 19 | 0.1579 | 1 | 1 | 3 | 0.3333 | 3 | 0 | 2 | 0 | 9 | 0.2222 |
| 14_661572 | 3 | 0 | 4 | 0.25 | 3 | 0 | NA | NA | 10 | 0.1 | 1 | 1 | 3 | 0 | 1 | 0 | 2 | 0.5 | 7 | 0.2857 |
| 14_661573 | 5 | 0.2 | 4 | 0.25 | 5 | 0 | 3 | 0.3333 | 17 | 0.1765 | NA | NA | 3 | 0.6667 | 3 | 0 | 3 | 0.6667 | 9 | 0.4444 |
| 14_661574 | 5 | 1 | 4 | 1 | 5 | 0.8 | 2 | 1 | 16 | 0.9375 | 1 | 1 | 3 | 1 | 3 | 1 | 3 | 1 | 10 | 1 |
| 14_661580 | 7 | 0 | 1 | 0 | 4 | 0 | 3 | 0.6667 | 15 | 0.1333 | 1 | 1 | 2 | 0 | 3 | 0.3333 | 3 | 0.3333 | 9 | 0.3333 |
| 14_661584 | 4 | 0.25 | 2 | 0.5 | 4 | 0 | 2 | 0.5 | 12 | 0.25 | 1 | 0 | NA | NA | 1 | 0 | 1 | 1 | 3 | 0.3333 |
| 14_661584 | 5 | 0.4 | 2 | 1 | 5 | 0.2 | 3 | 0.3333 | 15 | 0.4 | 1 | 1 | 1 | 0 | 2 | 0.5 | 2 | 0.5 | 6 | 0.5 |
| 14_661585 | 4 | 0.25 | 1 | 1 | 5 | 0.2 | 3 | 0.3333 | 13 | 0.3077 | 1 | 1 | 1 | 0 | 2 | 0 | 2 | 0.5 | 6 | 0.3333 |
| 14_661585 | 3 | 0 | 1 | 1 | 5 | 0.4 | 3 | 0.6667 | 12 | 0.4167 | 1 | 1 | 1 | 1 | 2 | 0.5 | 2 | 0.5 | 6 | 0.6667 |
| 14_661586 | 2 | 0 | 1 | 0 | 5 | 0.2 | 3 | 0 | 11 | 0.0909 | 1 | 0 | 1 | 0 | 2 | 0.5 | 2 | 0 | 6 | 0.1667 |
| 14_661682 | 3 | 0 | 1 | 0 | 4 | 0.25 | 3 | 0 | 11 | 0.0909 | 1 | 0 | 1 | 0 | 2 | 0 | 2 | 0.5 | 6 | 0.1667 |
| 14_661683 | NA | NA | 1 | 0 | 4 | 0.75 | 3 | 0.3333 | 8 | 0.5 | 2 | 0.5 | 1 | 0 | 2 | 0.5 | 2 | 0.5 | 7 | 0.4286 |
| 14_661686 | 2 | 0.5 | 1 | 0 | 4 | 0.75 | 2 | 0.5 | 9 | 0.5556 | 1 | 0 | 1 | 0 | 2 | 0 | 2 | 0.5 | 6 | 0.1667 |
| 14_661688 | 2 | 0 | 1 | 1 | 3 | 0.3333 | 2 | 0 | 8 | 0.25 | 1 | 0 | 1 | 0 | 2 | 1 | 2 | 0 | 6 | 0.3333 |
| 14_661688 | 1 | 0 | 1 | 0 | 2 | 0.5 | 3 | 0.3333 | 7 | 0.2857 | 1 | 1 | 1 | 0 | 2 | 1 | 1 | 1 | 5 | 0.8 |
| 14_661688 | 2 | 0 | 1 | 0 | 4 | 0.75 | 3 | 0.6667 | 10 | 0.5 | 1 | 0 | 1 | 0 | 2 | 0.5 | 2 | 0 | 6 | 0.1667 |
| 14_661689 | 2 | 0 | NA | NA | 3 | 0 | 3 | 0.3333 | 8 | 0.125 | 1 | 0 | 1 | 1 | 2 | 0.5 | 1 | 1 | 5 | 0.6 |
| 14_661689 | 2 | 1 | 1 | 0 | 4 | 0.25 | 3 | 0.3333 | 10 | 0.4 | 1 | 0 | 1 | 0 | 2 | 1 | 1 | 0 | 5 | 0.4 |
| 14_661690 | 2 | 0.5 | 1 | 1 | 3 | 1 | 3 | 1 | 9 | 0.8889 | 1 | 0 | 1 | 1 | 2 | 1 | 1 | 1 | 5 | 0.8 |
| 14_661690 | 1 | 1 | 1 | 1 | 4 | 0.75 | 2 | 1 | 8 | 0.875 | 1 | 1 | 1 | 1 | 2 | 1 | 1 | 1 | 5 | 1 |
| 14_661691 | 2 | 1 | 1 | 0 | 3 | 0 | 2 | 0 | 8 | 0.25 | 1 | 0 | 1 | 1 | 2 | 0 | 1 | 0 | 5 | 0.2 |
| 14_662131 | 138 | 0.2681 | 159 | 0.2579 | 131 | 0.1985 | 126 | 0.2302 | 554 | 0.2401 | 132 | 0.2652 | 119 | 0.1513 | 124 | 0.2419 | 113 | 0.2124 | 488 | 0.2193 |
| 14_662131 | 138 | 0.1014 | 159 | 0.0629 | 136 | 0.0588 | 132 | 0.0606 | 565 | 0.0708 | 140 | 0.0929 | 128 | 0.1094 | 135 | 0.0815 | 121 | 0.0909 | 524 | 0.0935 |
| 14_663221 | 6 | 0.3333 | 3 | 0 | 3 | 0 | 4 | 0 | 16 | 0.125 | 5 | 0.2 | 3 | 0.3333 | 5 | 0.2 | 7 | 0.1429 | 20 | 0.2 |
| 14_663870 | NA | NA | NA | NA | 1 | 1 | NA | NA | 1 | 1 | NA | NA | 2 | 1 | NA | NA | NA | NA | 2 | 1 |
| 14_663886 | NA | NA | NA | NA | 1 | 1 | NA | NA | 1 | 1 | NA | NA | 2 | 0.5 | 1 | 1 | NA | NA | 3 | 0.6667 |
| 14_663889 | NA | NA | NA | NA | 1 | 1 | NA | NA | 1 | 1 | NA | NA | 1 | 1 | 1 | 0 | NA | NA | 2 | 0.5 |
| 14_663908 | 62 | 0.0645 | 62 | 0.1613 | 59 | 0.1186 | 45 | 0.1333 | 228 | 0.1184 | 48 | 0.1042 | 54 | 0.2407 | 64 | 0.0938 | 57 | 0.1053 | 223 | 0.1345 |
| 14_663909 | 49 | 0.4082 | 47 | 0.3191 | 58 | 0.4828 | 34 | 0.5 | 188 | 0.4255 | 39 | 0.5385 | 45 | 0.5333 | 46 | 0.4783 | 49 | 0.4898 | 179 | 0.5084 |
| 14_663909 | 62 | 0.4516 | 57 | 0.6316 | 60 | 0.6667 | 48 | 0.4792 | 227 | 0.5595 | 45 | 0.6444 | 49 | 0.4286 | 64 | 0.6094 | 61 | 0.6885 | 219 | 0.5982 |
| 14_663909 | 56 | 0.0893 | 48 | 0.1458 | 55 | 0.0727 | 43 | 0.1395 | 202 | 0.1089 | 40 | 0.1 | 41 | 0.1707 | 60 | 0.0833 | 55 | 0.2 | 196 | 0.1378 |
| 14_663911 | 53 | 0.0755 | 43 | 0.1395 | 50 | 0.08 | 41 | 0.0488 | 187 | 0.0856 | 37 | 0.0811 | 43 | 0.0698 | 48 | 0.0625 | 47 | 0.0851 | 175 | 0.0743 |
| 14_663911 | 54 | 0.8519 | 57 | 0.7544 | 54 | 0.7963 | 40 | 0.725 | 205 | 0.7854 | 43 | 0.8372 | 50 | 0.78 | 58 | 0.8276 | 54 | 0.8333 | 205 | 0.8195 |
| 14_663912 | 53 | 0.1509 | 46 | 0.087 | 51 | 0.0588 | 37 | 0.1622 | 187 | 0.1123 | 37 | 0.0811 | 42 | 0.0714 | 53 | 0.2453 | 53 | 0.0943 | 185 | 0.1297 |
| 14_663913 | 53 | 0.8679 | 52 | 0.9038 | 52 | 0.7692 | 34 | 0.6471 | 191 | 0.8115 | 40 | 0.95 | 48 | 0.8542 | 54 | 0.7407 | 54 | 0.8889 | 196 | 0.852 |
| 14_663913 | 49 | 0.3061 | 46 | 0.3913 | 47 | 0.383 | 36 | 0.2778 | 178 | 0.3427 | 38 | 0.2895 | 40 | 0.425 | 50 | 0.26 | 48 | 0.3542 | 176 | 0.3295 |
| 14_663913 | 55 | 0.3818 | 53 | 0.2642 | 53 | 0.1698 | 42 | 0.3095 | 203 | 0.2808 | 43 | 0.3256 | 50 | 0.44 | 60 | 0.3833 | 55 | 0.2909 | 208 | 0.3606 |
| 14_663915 | 51 | 0.098 | 42 | 0.0952 | 45 | 0.0667 | 32 | 0.0312 | 170 | 0.0765 | 36 | 0.0278 | 42 | 0.1429 | 48 | 0.1667 | 50 | 0.04 | 176 | 0.0966 |
| 14_665465 | 11 | 0.2727 | 12 | 0.1667 | 14 | 0.1429 | 16 | 0.3125 | 53 | 0.2264 | 10 | 0.3 | 16 | 0.1875 | 15 | 0.1333 | 12 | 0.25 | 53 | 0.2075 |
| 14_665466 | 14 | 0.7143 | 9 | 0.4444 | 14 | 0.8571 | 13 | 0.7692 | 50 | 0.72 | 8 | 0.625 | 16 | 0.6875 | 13 | 0.4615 | 13 | 0.6154 | 50 | 0.6 |
| 14_665467 | 13 | 0.6154 | 12 | 0.6667 | 12 | 0.5833 | 14 | 0.7143 | 51 | 0.6471 | 9 | 0.4444 | 14 | 0.7143 | 12 | 0.25 | 11 | 0.8182 | 46 | 0.5652 |
| 14_665473 | 16 | 0.4375 | 16 | 0.1875 | 17 | 0.3529 | 21 | 0.3333 | 70 | 0.3286 | 15 | 0.2667 | 17 | 0.1765 | 17 | 0.2353 | 15 | 0.2667 | 64 | 0.2344 |
| 14_665475 | 15 | 0.2 | 13 | 0.0769 | 19 | 0.2632 | 18 | 0.2222 | 65 | 0.2 | 14 | 0.2857 | 18 | 0.1667 | 18 | 0.1111 | 9 | 0.4444 | 59 | 0.2203 |
| 14_665476 | 14 | 0.2857 | 17 | 0.1765 | 18 | 0.2778 | 22 | 0.4545 | 71 | 0.3099 | 12 | 0.1667 | 19 | 0.2632 | 15 | 0.3333 | 12 | 0.3333 | 58 | 0.2759 |
| 14_665479 | 19 | 0.2105 | 18 | 0.0556 | 18 | 0.1667 | 22 | 0.2273 | 77 | 0.1688 | 15 | 0.2667 | 21 | 0.1429 | 15 | 0.2667 | 17 | 0.1765 | 68 | 0.2059 |
| 14_665480 | 11 | 0.0909 | 16 | 0.25 | 16 | 0.1875 | 21 | 0.1905 | 64 | 0.1875 | 14 | 0.1429 | 19 | 0 | 15 | 0.0667 | 16 | 0.1875 | 64 | 0.0938 |
| 14_665484 | 24 | 0 | 15 | 0.2 | 22 | 0.0455 | 26 | 0.0769 | 87 | 0.069 | 12 | 0.25 | 25 | 0 | 17 | 0.0588 | 21 | 0.0476 | 75 | 0.0667 |
| 14_665487 | 24 | 0.125 | 18 | 0.1667 | 22 | 0.2727 | 25 | 0.16 | 89 | 0.1798 | 18 | 0.1111 | 25 | 0.2 | 20 | 0.2 | 19 | 0.2632 | 82 | 0.1951 |

| | | | | | | | | | | | | | | | | | | | | |
|---|---|---|---|---|---|---|---|---|---|---|---|---|---|---|---|---|---|---|---|---|
| 14_665988 | 28 | 0.3929 | 29 | 0.4138 | 29 | 0.5172 | 16 | 0.5 | 102 | 0.451 | 28 | 0.5 | 25 | 0.56 | 27 | 0.3704 | 22 | 0.3636 | 102 | 0.451 |
| 14_665988 | 27 | 0.0741 | 23 | 0.1739 | 22 | 0.2727 | 11 | 0.0909 | 83 | 0.1566 | 26 | 0.1923 | 21 | 0.1429 | 24 | 0.25 | 15 | 0.2 | 86 | 0.1977 |
| 14_665988 | 26 | 0.1923 | 23 | 0.1304 | 24 | 0.125 | 17 | 0.0588 | 90 | 0.1333 | 25 | 0.12 | 20 | 0.1 | 21 | 0.2381 | 19 | 0.2105 | 85 | 0.1647 |
| 14_665989 | 24 | 0.0417 | 26 | 0.0769 | 26 | 0.1154 | 15 | 0 | 91 | 0.0659 | 29 | 0.1034 | 22 | 0.1818 | 27 | 0.0741 | 23 | 0.0435 | 101 | 0.099 |
| 14_669707 | NA | NA | NA | NA | 1 | 1 | NA | NA | 1 | 1 | NA | NA | 1 | 0 | NA | NA | 1 | 1 | 2 | 0.5 |
| 14_669765 | NA | NA | 1 | 1 | 3 | 0 | 2 | 0 | 6 | 0.1667 | 1 | 0 | 2 | 0.5 | 1 | 0 | 1 | 0 | 5 | 0.2 |
| 14_669766 | 1 | 0 | 1 | 1 | 2 | 0.5 | 2 | 0 | 6 | 0.3333 | 2 | 1 | 2 | 1 | 1 | 1 | 1 | 0 | 6 | 0.8333 |
| 14_669766 | 2 | 0 | 1 | 1 | 1 | 0 | 2 | 0.5 | 6 | 0.3333 | 1 | 1 | 1 | 1 | 1 | 1 | NA | NA | 3 | 1 |
| 14_669767 | NA | NA | NA | NA | 3 | 0.3333 | 2 | 0 | 5 | 0.2 | 2 | 0 | 2 | 0 | 1 | 1 | 1 | 0 | 6 | 0.1667 |
| 14_669771 | 2 | 0.5 | 1 | 1 | 4 | 0.75 | 3 | 0.6667 | 10 | 0.7 | 2 | 1 | 2 | 0.5 | 3 | 0.3333 | 1 | 1 | 8 | 0.625 |
| 14_670427 | 6 | 0.3333 | 10 | 0.2 | 8 | 0.125 | 6 | 0.1667 | 30 | 0.2 | 8 | 0.125 | 9 | 0.2222 | 7 | 0.2857 | 14 | 0.4286 | 38 | 0.2895 |
| 14_670992 | NA | NA | NA | NA | 1 | 1 | NA | NA | 1 | 1 | NA | NA | NA | NA | NA | NA | 1 | 1 | 1 | 1 |
| 14_671519 | 1 | 1 | NA | NA | 1 | 1 | NA | NA | 2 | 1 | NA | NA | NA | NA | 1 | 1 | NA | NA | 1 | 1 |
| 14_671534 | 2 | 1 | NA | NA | 2 | 1 | 2 | 0.5 | 6 | 0.8333 | NA | NA | NA | NA | 2 | 1 | 1 | 1 | 3 | 1 |
| 14_671535 | 3 | 1 | NA | NA | 2 | 0.5 | 3 | 0.6667 | 8 | 0.75 | 1 | 0 | 1 | 0 | 1 | 1 | 1 | 1 | 4 | 0.5 |
| 14_671535 | 3 | 0.6667 | NA | NA | NA | NA | 3 | 0.6667 | 6 | 0.6667 | 1 | 1 | 1 | 0 | 2 | 1 | NA | NA | 4 | 0.75 |
| 14_671536 | 3 | 0.6667 | NA | NA | 2 | 1 | 3 | 0.6667 | 8 | 0.75 | 1 | 1 | 2 | 1 | 2 | 1 | 1 | 1 | 6 | 1 |
| 14_671537 | 2 | 1 | NA | NA | 1 | 1 | 2 | 0.5 | 5 | 0.8 | 1 | 0 | 2 | 1 | 2 | 1 | 1 | 0 | 6 | 0.6667 |
| 14_671538 | 3 | 0.6667 | NA | NA | 2 | 1 | 3 | 0.6667 | 8 | 0.75 | 1 | 1 | 2 | 0.5 | 2 | 1 | 1 | 1 | 6 | 0.8333 |
| 14_671538 | 2 | 1 | NA | NA | 2 | 1 | 1 | 1 | 5 | 1 | 1 | 1 | NA | NA | 2 | 1 | 1 | 1 | 4 | 1 |
| 14_671542 | 3 | 0.3333 | NA | NA | 2 | 1 | 3 | 0.3333 | 8 | 0.5 | 1 | 1 | 3 | 0.6667 | 2 | 1 | 2 | 0 | 8 | 0.625 |
| 14_671543 | 4 | 0.5 | NA | NA | 2 | 0 | 3 | 0 | 9 | 0.2222 | 1 | 0 | 3 | 0.3333 | 2 | 0.5 | 3 | 0.3333 | 9 | 0.3333 |
| 14_671543 | 3 | 0 | NA | NA | 2 | 0.5 | 3 | 0.3333 | 8 | 0.25 | NA | NA | 3 | 0.3333 | 4 | 0 | 3 | 0 | 10 | 0.1 |
| 14_671544 | 2 | 0.5 | NA | NA | 2 | 0.5 | 1 | 1 | 5 | 0.6 | 1 | 1 | 3 | 0.3333 | 2 | 0.5 | 2 | 1 | 8 | 0.625 |
| 14_671546 | 3 | 1 | NA | NA | 3 | 1 | 5 | 0.4 | 11 | 0.7273 | 4 | 0.25 | 3 | 0.3333 | 3 | 0.3333 | NA | NA | 10 | 0.3 |
| 14_671547 | 5 | 1 | 1 | 1 | 3 | 1 | 5 | 1 | 14 | 1 | 4 | 1 | 3 | 1 | 4 | 1 | 3 | 1 | 14 | 1 |
| 14_671547 | 5 | 1 | 1 | 0 | 3 | 1 | 5 | 1 | 14 | 0.9286 | 3 | 0.6667 | 4 | 1 | 4 | 0.75 | 3 | 1 | 14 | 0.8571 |
| 14_671548 | 5 | 1 | 1 | 1 | 3 | 1 | 5 | 1 | 14 | 1 | 2 | 1 | 1 | 1 | 3 | 0.6667 | 2 | 1 | 8 | 0.875 |
| 14_671548 | 5 | 0 | 1 | 1 | 3 | 0 | 4 | 0 | 13 | 0.0769 | 3 | 0.6667 | 4 | 0.5 | 4 | 0 | 4 | 0.25 | 15 | 0.3333 |
| 14_671549 | 5 | 0 | NA | NA | 3 | 0 | 5 | 0.2 | 13 | 0.0769 | 3 | 0 | 3 | 0.6667 | 3 | 0 | 4 | 0.5 | 13 | 0.3077 |
| 14_671550 | 5 | 0 | NA | NA | 2 | 0.5 | 5 | 0.4 | 12 | 0.25 | 3 | 0 | 2 | 0.5 | 4 | 0 | 3 | 0.6667 | 12 | 0.25 |
| 14_671556 | 5 | 0.2 | 2 | 0.5 | 4 | 0.25 | 6 | 0.1667 | 17 | 0.2353 | 3 | 0 | 4 | 0.25 | 5 | 0.4 | 5 | 0 | 17 | 0.1765 |
| 14_671562 | 5 | 0.6 | 3 | 1 | 4 | 0.5 | 6 | 0.3333 | 18 | 0.5556 | 3 | 0.6667 | 4 | 0.5 | 6 | 0.6667 | 7 | 0.7143 | 20 | 0.65 |
| 14_671565 | 5 | 0.2 | 4 | 0.25 | 5 | 0 | 6 | 0.3333 | 20 | 0.2 | 3 | 0.3333 | 4 | 0.5 | 6 | 0.1667 | 7 | 0.4286 | 20 | 0.35 |
| 14_671566 | 6 | 0.3333 | 3 | 0.3333 | 2 | 1 | 6 | 0.1667 | 17 | 0.3529 | 2 | 0.5 | 4 | 0.25 | 5 | 0.2 | 7 | 0.7143 | 18 | 0.4444 |
| 14_671567 | 6 | 0.1667 | 5 | 0.2 | 4 | 0.75 | 6 | 0.3333 | 21 | 0.3333 | 3 | 0.6667 | 4 | 0.5 | 6 | 0 | 7 | 0.1429 | 20 | 0.25 |
| 14_671570 | 4 | 0.5 | 4 | 0.25 | 4 | 0 | 3 | 0.3333 | 15 | 0.2667 | 4 | 0 | 6 | 0.1667 | 7 | 0 | 7 | 0.1429 | 24 | 0.0833 |
| 14_672228 | 9 | 0.2222 | 16 | 0.125 | 14 | 0.2143 | 7 | 0.1429 | 46 | 0.1739 | 8 | 0.375 | 11 | 0.1818 | 14 | 0.0714 | 6 | 0.3333 | 39 | 0.2051 |
| 14_672229 | 10 | 0.1 | 11 | 0.1818 | 11 | 0 | 9 | 0.1111 | 41 | 0.0976 | 6 | 0.1667 | 9 | 0 | 9 | 0 | 9 | 0.2222 | 33 | 0.0909 |
| 14_672232 | 8 | 0 | 15 | 0.0667 | 8 | 0 | 7 | 0.1429 | 38 | 0.0526 | 8 | 0.125 | 8 | 0.125 | 10 | 0 | 6 | 0 | 32 | 0.0625 |
| 14_672232 | 7 | 0.4286 | 13 | 0.2308 | 11 | 0.1818 | 6 | 0.5 | 37 | 0.2973 | 6 | 0.1667 | 12 | 0.1667 | 8 | 0.25 | 8 | 0.375 | 34 | 0.2353 |
| 14_672241 | 7 | 0.1429 | 14 | 0 | 10 | 0.2 | 7 | 0 | 38 | 0.0789 | 6 | 0 | 11 | 0.1818 | 8 | 0.125 | 7 | 0 | 32 | 0.0938 |
| 14_672522 | 3 | 0.6667 | 3 | 1 | 1 | 1 | 1 | 1 | 8 | 0.875 | 1 | 0 | 2 | 1 | NA | NA | NA | NA | 3 | 0.6667 |
| 14_672523 | 2 | 1 | 3 | 1 | 1 | 1 | 1 | 1 | 7 | 1 | 1 | 1 | 2 | 1 | NA | NA | NA | NA | 3 | 1 |
| 14_672524 | 1 | 1 | 3 | 1 | 1 | 0 | 1 | 1 | 6 | 0.8333 | 1 | 1 | 1 | 1 | NA | NA | NA | NA | 2 | 1 |
| 14_672525 | 2 | 1 | 3 | 1 | 1 | 1 | 1 | 1 | 7 | 1 | 1 | 1 | 2 | 1 | NA | NA | NA | NA | 3 | 1 |
| 14_672525 | 2 | 0.5 | 2 | 0 | 1 | 0 | NA | NA | 5 | 0.2 | 1 | 0 | 2 | 0.5 | NA | NA | NA | NA | 3 | 0.3333 |
| 14_672526 | 2 | 1 | 2 | 0 | 1 | 1 | 1 | 1 | 6 | 0.6667 | 1 | 0 | 2 | 1 | NA | NA | NA | NA | 3 | 0.6667 |
| 14_672527 | 1 | 0 | 2 | 0.5 | 1 | 1 | 1 | 0 | 5 | 0.4 | NA | NA | 2 | 0.5 | NA | NA | NA | NA | 2 | 0.5 |
| 14_672529 | 2 | 0.5 | 1 | 0 | 2 | 0 | 1 | 0 | 6 | 0.1667 | 1 | 0 | 2 | 1 | NA | NA | NA | NA | 3 | 0.6667 |
| 14_672533 | 13 | 0.7692 | 18 | 0.8333 | 27 | 1 | 15 | 1 | 73 | 0.9178 | 15 | 0.9333 | 28 | 0.9643 | 27 | 1 | 19 | 1 | 89 | 0.9775 |
| 14_672535 | 13 | 1 | 17 | 0.9412 | 21 | 1 | 13 | 0.9231 | 64 | 0.9688 | 11 | 1 | 25 | 0.92 | 26 | 1 | 16 | 1 | 78 | 0.9744 |
| 14_672535 | 4 | 1 | 8 | 0.875 | 13 | 0.9231 | 11 | 1 | 36 | 0.9444 | 5 | 0.8 | 17 | 0.9412 | 15 | 0.8667 | 6 | 0.8333 | 43 | 0.8837 |
| 14_672535 | 12 | 0.8333 | 13 | 1 | 20 | 0.9 | 13 | 1 | 58 | 0.931 | 8 | 0.875 | 24 | 0.7917 | 22 | 0.8182 | 12 | 0.9167 | 66 | 0.8333 |
| 14_672535 | 12 | 0.3333 | 15 | 0.5333 | 21 | 0.5714 | 15 | 0.5333 | 63 | 0.5079 | 13 | 0.6154 | 26 | 0.3077 | 21 | 0.7143 | 18 | 0.4444 | 78 | 0.5 |
| 14_672537 | 13 | 0.2308 | 18 | 0.0556 | 24 | 0.25 | 14 | 0.3571 | 69 | 0.2174 | 14 | 0.2857 | 24 | 0.375 | 24 | 0.5417 | 15 | 0.3333 | 77 | 0.4026 |
| 14_672538 | 14 | 0.0714 | 17 | 0.2353 | 26 | 0.1923 | 14 | 0.0714 | 71 | 0.1549 | 14 | 0.2143 | 24 | 0.1667 | 26 | 0.1154 | 15 | 0.2 | 79 | 0.1646 |
| 14_672538 | 14 | 0.5714 | 16 | 0.25 | 22 | 0.3182 | 15 | 0.2667 | 67 | 0.3433 | 10 | 0.4 | 27 | 0.4815 | 24 | 0.5 | 15 | 0.5333 | 76 | 0.4868 |
| 14_672539 | 14 | 0 | 16 | 0.125 | 24 | 0.0417 | 15 | 0.0667 | 69 | 0.058 | 12 | 0.0833 | 26 | 0.1538 | 22 | 0.0909 | 16 | 0.125 | 76 | 0.1184 |
| 14_679780 | NA | NA | 1 | 1 | 2 | 1 | NA | NA | 3 | 1 | NA | NA | NA | NA | 2 | 1 | 1 | 0 | 3 | 0.6667 |
| 14_679780 | 1 | 1 | 1 | 0 | 2 | 0.5 | NA | NA | 4 | 0.5 | NA | NA | NA | NA | 2 | 0 | 2 | 1 | 4 | 0.5 |
| 14_679794 | NA | NA | 4 | 0.25 | 3 | 0.3333 | 1 | 0 | 8 | 0.25 | NA | NA | 3 | 0 | 1 | 0 | 2 | 0.5 | 6 | 0.1667 |
| 14_679900 | NA | NA | 4 | 0.25 | NA | NA | 1 | 1 | 5 | 0.4 | NA | NA | NA | NA | 1 | 1 | NA | NA | 1 | 1 |
| 14_679935 | NA | NA | NA | NA | NA | NA | 1 | 1 | 1 | 1 | NA | NA | NA | NA | 2 | 0.5 | NA | NA | 2 | 0.5 |
| 14_679935 | NA | NA | 1 | 1 | NA | NA | 1 | 1 | 2 | 1 | NA | NA | NA | NA | 1 | 1 | NA | NA | 1 | 1 |

| | | | | | | | | | | | | | | | | | | | | |
|---|---|---|---|---|---|---|---|---|---|---|---|---|---|---|---|---|---|---|---|---|
| 14_6799364 | NA | NA | 1 | 1 | NA | NA | NA | NA | 1 | 1 | NA | NA | NA | NA | 1 | 1 | NA | NA | 1 | 1 |
| 14_6799385 | NA | NA | 1 | 1 | NA | NA | NA | NA | 1 | 1 | NA | NA | NA | NA | 2 | 1 | NA | NA | 2 | 1 |
| 14_6831480 | NA | NA | NA | NA | 1 | 1 | NA | NA | 1 | 1 | NA | NA | NA | NA | NA | NA | 1 | 1 | 1 | 1 |
| 14_6832219 | 34 | 0.1471 | 40 | 0.075 | 30 | 0.0333 | 29 | 0.1034 | 133 | 0.0902 | 22 | 0.0455 | 21 | 0.1905 | 27 | 0 | 19 | 0.1579 | 89 | 0.0899 |
| 14_6832503 | 43 | 0.9302 | 64 | 0.875 | 48 | 0.7292 | 42 | 0.9286 | 197 | 0.8629 | 38 | 0.8684 | 39 | 0.8462 | 41 | 0.9024 | 30 | 0.9 | 148 | 0.8784 |
| 14_6832546 | 23 | 0 | 35 | 0.1714 | 30 | 0.0333 | 28 | 0.0357 | 116 | 0.069 | 16 | 0 | 17 | 0 | 24 | 0.0833 | 20 | 0.1 | 77 | 0.0519 |
| 14_6832563 | 25 | 0.36 | 37 | 0.3243 | 36 | 0.25 | 39 | 0.2308 | 137 | 0.2847 | 23 | 0.3913 | 21 | 0.1905 | 27 | 0.1111 | 22 | 0.3636 | 93 | 0.2581 |
| 14_6832566 | 26 | 0.2308 | 43 | 0.2326 | 39 | 0.1795 | 38 | 0.1579 | 146 | 0.1986 | 25 | 0.24 | 22 | 0.1818 | 29 | 0.1724 | 24 | 0.0833 | 100 | 0.17 |
| 14_6832605 | 28 | 0.1429 | 53 | 0.2075 | 46 | 0.1957 | 35 | 0.3714 | 162 | 0.2284 | 26 | 0.3846 | 24 | 0.1667 | 30 | 0.2667 | 24 | 0.125 | 104 | 0.2404 |
| 14_6832619 | 31 | 0.2581 | 54 | 0.1111 | 50 | 0.18 | 36 | 0.1944 | 171 | 0.1754 | 26 | 0.1538 | 27 | 0.1111 | 32 | 0.25 | 24 | 0.125 | 109 | 0.1651 |
| 14_6835710 | 14 | 0.7857 | 21 | 0.9048 | 17 | 0.7059 | 15 | 0.8667 | 67 | 0.8209 | 9 | 1 | 13 | 0.6923 | 19 | 0.7368 | 5 | 0.4 | 46 | 0.7391 |
| 14_6835717 | 8 | 0.5 | 18 | 0.3889 | 16 | 0.5625 | 13 | 0.4615 | 55 | 0.4727 | 7 | 0.5714 | 12 | 0.6667 | 16 | 0.625 | 5 | 0.2 | 40 | 0.575 |
| 14_6836129 | NA | NA | NA | NA | NA | NA | 2 | 1 | 2 | 1 | NA | NA | NA | NA | 1 | 1 | 2 | 1 | 3 | 1 |
| 14_6836198 | NA | NA | 1 | 1 | 1 | 1 | 2 | 1 | 4 | 1 | NA | NA | 2 | 1 | 2 | 1 | 2 | 1 | 6 | 1 |
| 14_6836202 | NA | NA | 1 | 1 | 1 | 1 | 2 | 1 | 4 | 1 | NA | NA | 2 | 1 | 1 | 1 | 3 | 1 | 6 | 1 |
| 14_6836211 | NA | NA | 1 | 1 | 1 | 1 | 1 | 1 | 3 | 1 | NA | NA | 1 | 0 | 2 | 1 | 3 | 0.6667 | 6 | 0.6667 |
| 14_6836214 | NA | NA | 1 | 1 | NA | NA | 2 | 0.5 | 3 | 0.6667 | NA | NA | 2 | 0.5 | 2 | 1 | 3 | 1 | 7 | 0.8571 |
| 14_6836238 | NA | NA | NA | NA | NA | NA | 1 | 1 | 1 | 1 | NA | NA | 1 | 1 | 2 | 1 | 1 | 1 | 4 | 1 |
| 14_6836311 | 1 | 0 | NA | NA | NA | NA | 2 | 0.5 | 3 | 0.3333 | NA | NA | 1 | 1 | 2 | 0.5 | 2 | 0 | 5 | 0.4 |
| 14_6836337 | NA | NA | NA | NA | NA | NA | 1 | 1 | 1 | 1 | NA | NA | 2 | 1 | NA | NA | NA | NA | 2 | 1 |
| 14_6836368 | NA | NA | NA | NA | NA | NA | 2 | 0.5 | 2 | 0.5 | NA | NA | 1 | 1 | NA | NA | NA | NA | 1 | 1 |
| 14_6836393 | NA | NA | NA | NA | NA | NA | 2 | 0.5 | 2 | 0.5 | NA | NA | 1 | 1 | 1 | 1 | NA | NA | 2 | 1 |
| 14_6836474 | NA | NA | NA | NA | NA | NA | 1 | 1 | 1 | 1 | NA | NA | NA | NA | 1 | 1 | 1 | 0 | 2 | 0.5 |
| 14_6836615 | NA | NA | 3 | 0.3333 | 1 | 0 | 1 | 0 | 5 | 0.2 | 1 | 0 | 4 | 0.25 | 2 | 0 | 1 | 0 | 8 | 0.125 |
| 14_6836701 | NA | NA | 2 | 0 | 2 | 0.5 | 1 | 0 | 5 | 0.2 | NA | NA | 4 | 0.25 | 2 | 0 | 1 | 0 | 7 | 0.1429 |
| 14_6836750 | NA | NA | 3 | 0 | 2 | 0.5 | 1 | 0 | 6 | 0.1667 | NA | NA | 5 | 0.2 | 1 | 0 | 3 | 0 | 9 | 0.1111 |
| 14_6836766 | NA | NA | 4 | 0.25 | 2 | 0.5 | 1 | 0 | 7 | 0.2857 | 1 | 0 | 5 | 0.2 | 2 | 0 | 3 | 0.6667 | 11 | 0.2727 |
| 14_6836771 | NA | NA | 4 | 0.25 | 1 | 0 | 1 | 0 | 6 | 0.1667 | 1 | 1 | 6 | 0 | 2 | 0.5 | 3 | 0.6667 | 12 | 0.3333 |
| 14_6836795 | NA | NA | 4 | 0 | 1 | 1 | 1 | 0 | 6 | 0.1667 | 1 | 0 | 5 | 0.2 | 2 | 0 | 3 | 0 | 11 | 0.0909 |
| 14_6836800 | NA | NA | 4 | 0.25 | 2 | 0 | 1 | 0 | 7 | 0.1429 | 1 | 1 | 4 | 0.25 | 2 | 0 | 3 | 0 | 10 | 0.2 |
| 14_6836802 | NA | NA | 4 | 0.25 | 2 | 0.5 | 1 | 1 | 7 | 0.4286 | 1 | 0 | 5 | 0.2 | 1 | 0 | 3 | 0.6667 | 10 | 0.3 |
| 14_6836814 | NA | NA | 3 | 0 | 2 | 0 | 2 | 0.5 | 7 | 0.1429 | 1 | 0 | 4 | 0.25 | 2 | 0 | 3 | 0 | 10 | 0.1 |
| 14_6836819 | NA | NA | 4 | 0.5 | 2 | 1 | 1 | 1 | 7 | 0.7143 | 1 | 1 | 6 | 0.6667 | 2 | 0.5 | 2 | 1 | 11 | 0.7273 |
| 14_6836828 | 1 | 0 | 4 | 0 | 2 | 1 | 4 | 0.5 | 11 | 0.3636 | 1 | 0 | 6 | 0.3333 | 3 | 0 | 6 | 0 | 16 | 0.125 |
| 14_6836831 | 1 | 0 | 3 | 0.6667 | 3 | 0.3333 | 3 | 0 | 10 | 0.3 | 1 | 0 | 4 | 0.5 | 2 | 0 | 4 | 0 | 11 | 0.1818 |
| 14_6836834 | 1 | 0 | 3 | 1 | 1 | 1 | 3 | 0.3333 | 8 | 0.625 | 1 | 1 | 6 | 0.6667 | 3 | 0.6667 | 4 | 0.75 | 14 | 0.7143 |
| 14_6836836 | 1 | 0 | 3 | 0 | 3 | 0 | 4 | 0.25 | 11 | 0.0909 | 1 | 0 | 5 | 0.2 | 3 | 0 | 6 | 0.1667 | 15 | 0.1333 |
| 14_6836857 | 1 | 1 | 5 | 1 | 3 | 0.6667 | 4 | 0.25 | 13 | 0.6923 | 1 | 1 | 7 | 0.7143 | 3 | 0.6667 | 6 | 0.5 | 17 | 0.6471 |
| 14_6836869 | 1 | 0 | 4 | 0.25 | 3 | 0 | 4 | 0 | 12 | 0.0833 | 1 | 0 | 6 | 0.1667 | 2 | 0 | 5 | 0 | 14 | 0.0714 |
| 14_6836875 | 1 | 1 | 4 | 0.75 | 2 | 0 | 5 | 0.6 | 12 | 0.5833 | 1 | 0 | 7 | 0.4286 | 3 | 0 | 5 | 0 | 16 | 0.1875 |
| 14_6948060 | 4 | 0 | 2 | 0 | NA | NA | 4 | 0.25 | 10 | 0.1 | NA | NA | 5 | 0.2 | 4 | 0 | 2 | 0.5 | 11 | 0.1818 |
| 14_6948068 | 4 | 0.75 | 2 | 0 | NA | NA | 4 | 0.5 | 10 | 0.5 | NA | NA | 5 | 0.8 | 2 | 0.5 | 1 | 1 | 8 | 0.75 |
| 14_6948075 | 4 | 0.25 | 1 | 0 | NA | NA | 3 | 0 | 8 | 0.125 | NA | NA | 4 | 0 | 4 | 0.5 | 1 | 1 | 9 | 0.3333 |
| 14_6948105 | 3 | 0.6667 | 2 | 0.5 | NA | NA | 4 | 0.25 | 9 | 0.4444 | NA | NA | 5 | 0.4 | 4 | 0.5 | 1 | 0 | 10 | 0.4 |
| 14_6948181 | 1 | 1 | 2 | 0.5 | NA | NA | 3 | 0 | 6 | 0.3333 | NA | NA | 3 | 0 | 4 | 0.25 | 1 | 0 | 8 | 0.125 |
| 14_6948208 | 1 | 1 | 1 | 1 | NA | NA | 3 | 0.6667 | 5 | 0.8 | NA | NA | 1 | 0 | 4 | 0.25 | 1 | 0 | 6 | 0.1667 |
| 14_6948211 | 1 | 1 | 2 | 0 | NA | NA | 1 | 0 | 4 | 0.25 | NA | NA | 1 | 1 | 3 | 0 | 1 | 0 | 5 | 0.2 |
| 14_6948229 | 1 | 1 | NA | NA | NA | NA | 2 | 1 | 3 | 1 | NA | NA | 1 | 1 | 3 | 1 | NA | NA | 4 | 1 |
| 14_6948260 | 1 | 1 | 1 | 0 | NA | NA | 3 | 0.3333 | 5 | 0.4 | NA | NA | 1 | 1 | 3 | 0.6667 | NA | NA | 4 | 0.75 |
| 14_6948272 | 1 | 1 | 1 | 1 | NA | NA | 2 | 1 | 4 | 1 | NA | NA | 1 | 1 | 3 | 1 | 1 | 1 | 5 | 1 |
| 14_6948274 | NA | NA | NA | NA | NA | NA | 2 | 1 | 2 | 1 | NA | NA | NA | NA | 2 | 1 | 1 | 1 | 3 | 1 |
| 14_6948636 | 1 | 1 | NA | NA | NA | NA | NA | NA | 1 | 1 | NA | NA | 1 | 1 | 1 | 0 | 1 | 0 | 3 | 0.3333 |
| 14_6948641 | 1 | 1 | NA | NA | NA | NA | 1 | 0 | 2 | 0.5 | NA | NA | 2 | 1 | NA | NA | NA | NA | 2 | 1 |
| 14_6948649 | 1 | 1 | NA | NA | NA | NA | 1 | 1 | 2 | 1 | NA | NA | 2 | 0 | NA | NA | 1 | 1 | 3 | 0.3333 |
| 14_6948651 | 1 | 1 | NA | NA | NA | NA | 1 | 1 | 2 | 1 | NA | NA | 2 | 1 | NA | NA | 1 | 0 | 3 | 0.6667 |
| 14_6948674 | 1 | 1 | NA | NA | NA | NA | NA | NA | 1 | 1 | NA | NA | 2 | 0.5 | NA | NA | NA | NA | 2 | 0.5 |
| 14_6959915 | 3 | 0 | 9 | 0.3333 | 2 | 0 | 2 | 0 | 16 | 0.1875 | 1 | 1 | 5 | 0.2 | 10 | 0.1 | NA | NA | 16 | 0.1875 |
| 14_6959920 | 3 | 0 | 10 | 0.1 | NA | NA | 5 | 0 | 18 | 0.0556 | 1 | 0 | 7 | 0.1429 | 10 | 0 | NA | NA | 18 | 0.0556 |
| 14_6959921 | 2 | 0 | 10 | 0.2 | 3 | 0 | 5 | 0 | 20 | 0.1 | 1 | 0 | 7 | 0.5714 | 11 | 0.1818 | NA | NA | 19 | 0.3158 |
| 14_6959924 | 3 | 0 | 9 | 0 | 3 | 0.3333 | 3 | 0 | 18 | 0.0556 | 1 | 0 | 6 | 0.1667 | 11 | 0 | NA | NA | 18 | 0.0556 |
| 14_6960319 | 1 | 1 | 11 | 0.2727 | 2 | 0.5 | 6 | 0.5 | 20 | 0.4 | 4 | 0.5 | 10 | 0.4 | 9 | 0.4444 | 2 | 0 | 25 | 0.4 |
| 14_6960321 | 3 | 0 | 13 | 0.1538 | 3 | 0 | 6 | 0.5 | 25 | 0.2 | 4 | 0.25 | 10 | 0 | 12 | 0.0833 | 3 | 0 | 29 | 0.069 |
| 14_6960333 | 2 | 0.5 | 11 | 0.1818 | 4 | 0.5 | 5 | 0.4 | 22 | 0.3182 | 3 | 0 | 12 | 0.4167 | 11 | 0.3636 | 3 | 0 | 29 | 0.3103 |
| 14_6960343 | 2 | 0 | 9 | 0 | 4 | 0.25 | 5 | 0.2 | 20 | 0.1 | 2 | 0 | 9 | 0.3333 | 9 | 0.1111 | 3 | 0 | 23 | 0.1739 |
| 14_6960348 | 4 | 0.25 | 12 | 0.1667 | 3 | 0 | 5 | 0.2 | 24 | 0.1667 | 4 | 0 | 12 | 0.1667 | 12 | 0.25 | 3 | 0.6667 | 31 | 0.2258 |

| | | | | | | | | | | | | | | | | | | | | |
|---|---|---|---|---|---|---|---|---|---|---|---|---|---|---|---|---|---|---|---|---|
| 14_696035 | NA | NA | 3 | 1 | NA | NA | 2 | 1 | 5 | 1 | NA | NA | 1 | 1 | 1 | 1 | NA | NA | 2 | 1 |
| 14_696036 | 5 | 0.2 | 13 | 0 | 4 | 0.5 | 6 | 0 | 28 | 0.1071 | 6 | 0.1667 | 13 | 0.0769 | 13 | 0.0769 | 3 | 0 | 35 | 0.0857 |
| 14_696038 | NA | NA | 1 | 1 | NA | NA | 3 | 1 | 4 | 1 | NA | NA | 1 | 1 | NA | NA | NA | NA | 1 | 1 |
| 14_696038 | NA | NA | 3 | 1 | NA | NA | 3 | 1 | 6 | 1 | NA | NA | 1 | 1 | 1 | 1 | NA | NA | 2 | 1 |
| 14_696040 | NA | NA | 3 | 0.6667 | NA | NA | 3 | 0.6667 | 6 | 0.6667 | NA | NA | 1 | 1 | NA | NA | NA | NA | 1 | 1 |
| 14_698173 | 5 | 0.2 | 3 | 0 | 1 | 0 | 5 | 0 | 14 | 0.0714 | 3 | 0 | 8 | 0 | 6 | 0.1667 | 2 | 0 | 19 | 0.0526 |
| 14_698174 | 5 | 0.2 | 1 | 0 | 1 | 0 | 5 | 0.4 | 12 | 0.25 | 2 | 0 | 6 | 0.3333 | 5 | 0.4 | 2 | 0.5 | 15 | 0.3333 |
| 14_698174 | 5 | 0.4 | 3 | 0.3333 | 1 | 0 | 5 | 0.4 | 14 | 0.3571 | 3 | 0 | 6 | 0.1667 | 5 | 0.4 | 2 | 0 | 16 | 0.1875 |
| 14_698174 | 6 | 0.1667 | 3 | 0.3333 | 1 | 0 | 5 | 0.2 | 15 | 0.2 | 4 | 0 | 8 | 0.125 | 6 | 0.1667 | 2 | 0 | 20 | 0.1 |
| 14_698174 | 4 | 0 | 2 | 0.5 | NA | NA | 5 | 0 | 11 | 0.0909 | 3 | 0.3333 | 6 | 0 | 4 | 0 | 2 | 0 | 15 | 0.0667 |
| 14_698194 | NA | NA | 1 | 0 | NA | NA | 1 | 1 | 2 | 0.5 | NA | NA | NA | NA | NA | NA | 1 | 1 | 1 | 1 |
| 14_698196 | NA | NA | 1 | 0 | NA | NA | 1 | 1 | 2 | 0.5 | NA | NA | NA | NA | 1 | 1 | NA | NA | 1 | 1 |
| 14_698196 | NA | NA | 1 | 1 | NA | NA | 1 | 1 | 2 | 1 | NA | NA | NA | NA | 1 | 1 | NA | NA | 1 | 1 |
| 14_698204 | NA | NA | 1 | 1 | NA | NA | NA | NA | 1 | 1 | NA | NA | NA | NA | 1 | 1 | 1 | 0 | 2 | 0.5 |
| 14_698204 | NA | NA | 1 | 1 | NA | NA | NA | NA | 1 | 1 | NA | NA | NA | NA | NA | NA | 1 | 1 | 1 | 1 |
| 14_698207 | NA | NA | 1 | 1 | NA | NA | NA | NA | 1 | 1 | NA | NA | NA | NA | 1 | 1 | NA | NA | 1 | 1 |
| 14_698207 | NA | NA | 1 | 1 | NA | NA | NA | NA | 1 | 1 | NA | NA | NA | NA | 1 | 1 | NA | NA | 1 | 1 |
| 14_698214 | 7 | 0 | 3 | 0 | 1 | 0 | 6 | 0.1667 | 17 | 0.0588 | 6 | 0.1667 | 9 | 0 | 8 | 0.125 | 3 | 0 | 26 | 0.0769 |
| 14_698215 | 8 | 0.375 | 4 | 0.75 | 1 | 0 | 7 | 0.2857 | 20 | 0.4 | 4 | 0.75 | 10 | 0.3 | 7 | 0.2857 | 3 | 0 | 24 | 0.3333 |
| 14_698216 | 6 | 0.1667 | 2 | 0 | 1 | 0 | 4 | 0.5 | 13 | 0.2308 | 2 | 1 | 9 | 0.2222 | 7 | 0.1429 | 1 | 0 | 19 | 0.2632 |
| 14_698217 | 5 | 0 | 4 | 0.5 | 2 | 0 | 7 | 0.1429 | 18 | 0.1667 | 4 | 0.25 | 8 | 0 | 8 | 0.375 | 2 | 0.5 | 22 | 0.2273 |
| 14_698300 | NA | NA | 2 | 1 | NA | NA | NA | NA | 2 | 1 | NA | NA | NA | NA | NA | NA | 1 | 1 | 1 | 1 |
| 14_698300 | NA | NA | 2 | 1 | NA | NA | NA | NA | 2 | 1 | NA | NA | NA | NA | NA | NA | 1 | 1 | 1 | 1 |
| 14_699349 | 4 | 0.25 | 6 | 0.3333 | 3 | 0 | 4 | 0.5 | 17 | 0.2941 | 3 | 0.3333 | 2 | 0.5 | 2 | 0 | 2 | 0 | 9 | 0.2222 |
| 14_699545 | 7 | 0.8571 | 11 | 0.8182 | 12 | 0.9167 | 12 | 0.8333 | 42 | 0.8571 | 10 | 0.9 | 8 | 0.75 | 8 | 0.875 | 8 | 0.75 | 34 | 0.8235 |
| 14_699546 | 5 | 0 | 14 | 0.0714 | 12 | 0.0833 | 8 | 0 | 39 | 0.0513 | 11 | 0.0909 | 10 | 0 | 6 | 0 | 8 | 0.125 | 35 | 0.0571 |
| 14_699546 | 6 | 0.3333 | 15 | 0.3333 | 14 | 0.5 | 10 | 0.7 | 45 | 0.4667 | 8 | 0.375 | 9 | 0.8889 | 9 | 0.5556 | 8 | 0.5 | 34 | 0.5882 |
| 14_699546 | 7 | 0.4286 | 13 | 0.2308 | 15 | 0.3333 | 10 | 0.3 | 45 | 0.3111 | 11 | 0.4545 | 10 | 0.5 | 9 | 0.5556 | 8 | 0.25 | 38 | 0.4474 |
| 14_699547 | 8 | 0.375 | 12 | 0.6667 | 14 | 0.2857 | 12 | 0.3333 | 46 | 0.413 | 11 | 0.2727 | 8 | 0.25 | 9 | 0.5556 | 11 | 0 | 39 | 0.2564 |
| 14_699547 | 5 | 0.4 | 9 | 0.8889 | 14 | 0.5 | 8 | 0.5 | 36 | 0.5833 | 7 | 0.7143 | 7 | 0.7143 | 7 | 0.8571 | 9 | 0.8889 | 30 | 0.8 |
| 14_699547 | 7 | 0.2857 | 13 | 0.1538 | 12 | 0.1667 | 11 | 0.0909 | 43 | 0.1628 | 10 | 0.2 | 10 | 0.4 | 10 | 0.2 | 11 | 0.2727 | 41 | 0.2683 |
| 14_699552 | 7 | 0.1429 | 12 | 0.0833 | 12 | 0 | 9 | 0 | 40 | 0.05 | 11 | 0.0909 | 10 | 0.1 | 6 | 0 | 9 | 0.2222 | 36 | 0.1111 |
| 14_699552 | 7 | 0 | 11 | 0.2727 | 15 | 0.2667 | 10 | 0.1 | 43 | 0.186 | 9 | 0.2222 | 9 | 0.4444 | 6 | 0.6667 | 9 | 0.3333 | 33 | 0.3939 |
| 14_699552 | 4 | 0.25 | 10 | 0.4 | 8 | 0.5 | 7 | 0.2857 | 29 | 0.3793 | 8 | 0.375 | 7 | 0.4286 | 4 | 1 | 6 | 0.1667 | 25 | 0.44 |
| 14_699554 | 6 | 1 | 11 | 1 | 12 | 0.9167 | 8 | 1 | 37 | 0.973 | 7 | 0.8571 | 9 | 1 | 4 | 1 | 5 | 1 | 25 | 0.96 |
| 14_699556 | 5 | 0 | 9 | 0.2222 | 10 | 0 | 5 | 0.2 | 29 | 0.1034 | 8 | 0.25 | 8 | 0 | 4 | 0.25 | 6 | 0 | 26 | 0.1154 |
| 14_699556 | 6 | 0.6667 | 11 | 0.3636 | 14 | 0.4286 | 9 | 0.5556 | 40 | 0.475 | 8 | 0.25 | 8 | 0.125 | 5 | 0.8 | 8 | 0.875 | 29 | 0.4828 |
| 14_699556 | 4 | 0.5 | 11 | 0.0909 | 6 | 0.1667 | 6 | 0.3333 | 27 | 0.2222 | 6 | 0.3333 | 8 | 0.375 | 3 | 0.3333 | 4 | 0.5 | 21 | 0.381 |
| 14_699924 | 1 | 1 | NA | NA | NA | NA | NA | NA | 1 | 1 | NA | NA | 1 | 0 | 1 | 1 | NA | NA | 2 | 0.5 |
| 14_699924 | 1 | 1 | NA | NA | NA | NA | NA | NA | 1 | 1 | NA | NA | NA | NA | 1 | 1 | NA | NA | 1 | 1 |
| 14_699925 | 1 | 1 | NA | NA | NA | NA | NA | NA | 1 | 1 | NA | NA | 1 | 1 | 1 | 1 | NA | NA | 2 | 1 |
| 14_699928 | 1 | 1 | NA | NA | NA | NA | NA | NA | 1 | 1 | NA | NA | 1 | 1 | 1 | 0 | NA | NA | 2 | 0.5 |
| 14_699928 | 1 | 1 | NA | NA | NA | NA | NA | NA | 1 | 1 | NA | NA | 1 | 1 | 1 | 0 | NA | NA | 2 | 0.5 |
| 14_700225 | 1 | 0 | 5 | 0.6 | 7 | 0.5714 | 2 | 0.5 | 15 | 0.5333 | 4 | 0.5 | 6 | 0.1667 | 8 | 0.375 | 7 | 0.2857 | 25 | 0.32 |
| 14_700243 | 2 | 0.5 | 1 | 1 | 2 | 0 | 1 | 0 | 6 | 0.3333 | 3 | 0.6667 | 4 | 0.25 | 3 | 0.3333 | 4 | 0 | 14 | 0.2857 |
| 14_700245 | 1 | 1 | 1 | 0 | 2 | 0 | NA | NA | 4 | 0.25 | 2 | 1 | 4 | 0.5 | 1 | 1 | 2 | 0.5 | 9 | 0.6667 |
| 14_700246 | 4 | 0.75 | 4 | 1 | 12 | 0.8333 | 3 | 0.6667 | 23 | 0.8261 | 8 | 1 | 3 | 1 | 12 | 0.8333 | 9 | 0.8889 | 32 | 0.9062 |
| 14_700250 | 3 | 1 | 7 | 0.5714 | 10 | 0.6 | 5 | 0.6 | 25 | 0.64 | 9 | 0.6667 | 6 | 0.6667 | 10 | 0.6 | 8 | 0.625 | 33 | 0.6364 |
| 14_700337 | 4 | 0 | 5 | 0 | 6 | 0.1667 | 4 | 0.25 | 19 | 0.1053 | 7 | 0.1429 | 4 | 0 | 11 | 0.0909 | 7 | 0.1429 | 29 | 0.1034 |
| 14_700338 | 4 | 0.25 | 7 | 0.4286 | 6 | 0.5 | 3 | 1 | 20 | 0.5 | 8 | 1 | 4 | 1 | 9 | 0.8889 | 7 | 0.7143 | 28 | 0.8929 |
| 14_700340 | 3 | 0.3333 | 4 | 0.25 | 5 | 0 | 4 | 0 | 16 | 0.125 | 7 | 0 | 3 | 0 | 8 | 0.125 | 7 | 0.1429 | 25 | 0.08 |
| 14_700342 | 3 | 1 | 6 | 0.5 | 4 | 0.5 | 4 | 1 | 17 | 0.7059 | 5 | 1 | 3 | 1 | 7 | 1 | 6 | 0.5 | 21 | 0.8571 |
| 14_700342 | 3 | 0 | 6 | 0.3333 | 4 | 0 | 4 | 0.25 | 17 | 0.1765 | 6 | 0.3333 | 4 | 0 | 8 | 0.125 | 6 | 0 | 24 | 0.125 |
| 14_700343 | 3 | 1 | 5 | 0.6 | 3 | 0.6667 | 3 | 1 | 14 | 0.7857 | 3 | 1 | 4 | 0.5 | 6 | 0.8333 | 6 | 0.8333 | 19 | 0.7895 |
| 14_700343 | 3 | 1 | 4 | 0.75 | 4 | 0.75 | 3 | 1 | 14 | 0.8571 | 5 | 0.8 | 3 | 1 | 6 | 0.8333 | 6 | 0.6667 | 20 | 0.8 |
| 14_703163 | 2 | 0.5 | 2 | 1 | 2 | 0.5 | 5 | 0.2 | 11 | 0.4545 | 2 | 0 | NA | NA | 4 | 0.25 | 3 | 1 | 9 | 0.4444 |
| 14_703164 | 6 | 0 | 2 | 0.5 | 4 | 0.5 | 6 | 0.3333 | 18 | 0.2778 | 2 | 0.5 | 1 | 0 | 4 | 0.25 | 4 | 0.75 | 11 | 0.4545 |
| 14_703167 | 6 | 1 | 3 | 1 | 5 | 1 | 7 | 1 | 21 | 1 | 2 | 1 | 1 | 1 | 5 | 1 | 4 | 1 | 12 | 1 |
| 14_703167 | 3 | 0 | 2 | 1 | 5 | 0.2 | 7 | 0.4286 | 17 | 0.3529 | 1 | 1 | 1 | 0 | 5 | 0.2 | 3 | 0.3333 | 10 | 0.3 |
| 14_703168 | 5 | 0.8 | 3 | 0.6667 | 4 | 1 | 4 | 1 | 16 | 0.875 | 1 | 1 | 1 | 1 | 4 | 1 | 4 | 1 | 10 | 1 |
| 14_703168 | 7 | 0.5714 | 1 | 1 | 4 | 1 | 6 | 0.5 | 18 | 0.6667 | 2 | 1 | NA | NA | 4 | 1 | 3 | 0.3333 | 9 | 0.7778 |
| 14_703168 | 8 | 0 | 2 | 0.5 | 5 | 0.4 | 7 | 0.2857 | 22 | 0.2273 | 2 | 0 | 1 | 1 | 4 | 0 | 3 | 0 | 10 | 0.1 |
| 14_703176 | 7 | 1 | 3 | 1 | 4 | 1 | 1 | 1 | 15 | 1 | 2 | 1 | 1 | 1 | 3 | 1 | 3 | 1 | 9 | 1 |
| 14_703176 | 9 | 1 | 3 | 1 | 5 | 0.8 | 5 | 0.8 | 22 | 0.9091 | 2 | 1 | 1 | 1 | 3 | 1 | 3 | 1 | 9 | 1 |
| 14_703179 | 8 | 0.125 | 3 | 0 | 7 | 0.1429 | 8 | 0.125 | 26 | 0.1154 | 2 | 0 | NA | NA | 6 | 0 | 4 | 0.25 | 12 | 0.0833 |

| | | | | | | | | | | | | | | | | | | | | |
|---|---|---|---|---|---|---|---|---|---|---|---|---|---|---|---|---|---|---|---|---|
| 14_7031804 | 8 | 0 | 5 | 0.2 | 8 | 0.125 | 9 | 0.3333 | 30 | 0.1667 | 3 | 0.3333 | 1 | 0 | 6 | 0.1667 | 7 | 0 | 17 | 0.1176 |
| 14_7031894 | 10 | 0 | 5 | 0.2 | 12 | 0.0833 | 11 | 0 | 38 | 0.0526 | 3 | 0 | 2 | 0 | 6 | 0.1667 | 9 | 0 | 20 | 0.05 |
| 14_7031899 | 11 | 0.0909 | 5 | 0 | 11 | 0.0909 | 12 | 0.0833 | 39 | 0.0769 | 4 | 0 | 3 | 0 | 6 | 0.5 | 9 | 0.1111 | 22 | 0.1818 |
| 14_7031917 | 11 | 0.6364 | 6 | 0.8333 | 13 | 0.6923 | 12 | 0.6667 | 42 | 0.6905 | 4 | 0.25 | 3 | 0.6667 | 7 | 0.7143 | 8 | 0.375 | 22 | 0.5 |
| 14_7031964 | 13 | 0.2308 | 6 | 0.3333 | 13 | 0.1538 | 18 | 0.3889 | 50 | 0.28 | 8 | 0 | 8 | 0.375 | 11 | 0.3636 | 10 | 0.1 | 37 | 0.2162 |
| 14_7031970 | 13 | 0.4615 | 12 | 0.5 | 17 | 0.3529 | 18 | 0.4444 | 60 | 0.4333 | 9 | 0.3333 | 9 | 0.4444 | 13 | 0.4615 | 12 | 0.5 | 43 | 0.4419 |
| 14_7037482 | 1 | 1 | 1 | 1 | 1 | 1 | 4 | 0.25 | 7 | 0.5714 | 1 | 1 | NA | NA | 2 | 1 | 1 | 0 | 4 | 0.75 |
| 14_7037506 | 2 | 0 | 1 | 0 | 1 | 0 | 5 | 0.2 | 9 | 0.1111 | NA | NA | 1 | 0 | 1 | 0 | 1 | 1 | 3 | 0.3333 |
| 14_7037509 | 1 | 0 | 1 | 0 | 1 | 0 | 5 | 0.2 | 8 | 0.125 | 1 | 0 | 1 | 0 | 2 | 0.5 | 1 | 1 | 5 | 0.4 |
| 14_7037512 | 2 | 0 | NA | NA | NA | NA | 5 | 0.6 | 7 | 0.4286 | 1 | 0 | 1 | 0 | 2 | 0.5 | 1 | 0 | 5 | 0.2 |
| 14_7037617 | 8 | 0.125 | 11 | 0.0909 | 8 | 0 | 6 | 0 | 33 | 0.0606 | 10 | 0.1 | 9 | 0 | 9 | 0.2222 | 11 | 0.0909 | 39 | 0.1026 |
| 14_7037621 | 7 | 0 | 12 | 0 | 7 | 0.1429 | 7 | 0.1429 | 33 | 0.0606 | 8 | 0.25 | 9 | 0 | 8 | 0.25 | 10 | 0.1 | 35 | 0.1429 |
| 14_7037951 | 4 | 0 | 6 | 0.1667 | 6 | 0 | 4 | 0 | 20 | 0.05 | 5 | 0 | 5 | 0.4 | 8 | 0.25 | 4 | 0 | 22 | 0.1818 |
| 14_7039217 | NA | NA | NA | NA | 1 | 1 | 1 | 1 | 2 | 1 | 1 | 1 | NA | NA | NA | NA | NA | NA | 1 | 1 |
| 14_7039219 | NA | NA | NA | NA | 1 | 1 | NA | NA | 1 | 1 | 2 | 1 | NA | NA | NA | NA | NA | NA | 2 | 1 |
| 14_7039228 | NA | NA | NA | NA | 1 | 1 | 1 | 1 | 2 | 1 | 2 | 1 | NA | NA | NA | NA | NA | NA | 2 | 1 |
| 14_7039237 | NA | NA | NA | NA | 1 | 1 | 1 | 1 | 2 | 1 | 2 | 1 | NA | NA | NA | NA | NA | NA | 2 | 1 |
| 14_7039488 | NA | NA | NA | NA | 1 | 1 | 1 | 1 | 2 | 1 | 2 | 1 | NA | NA | NA | NA | NA | NA | 2 | 1 |
| 14_7039493 | NA | NA | NA | NA | 1 | 0 | 1 | 1 | 2 | 0.5 | 2 | 1 | NA | NA | NA | NA | NA | NA | 2 | 1 |
| 14_7039499 | NA | NA | NA | NA | 1 | 1 | 1 | 0 | 2 | 0.5 | 2 | 0.5 | NA | NA | NA | NA | NA | NA | 2 | 0.5 |
| 14_7040188 | 4 | 0.5 | 3 | 0.6667 | 3 | 0 | 6 | 0.1667 | 16 | 0.3125 | 3 | 0 | 2 | 0 | 3 | 0.3333 | 1 | 0 | 9 | 0.1111 |
| 14_7040190 | 5 | 0 | 3 | 0.3333 | 3 | 0.3333 | 6 | 0.1667 | 17 | 0.1765 | 3 | 0 | 2 | 0 | 3 | 0.3333 | NA | NA | 8 | 0.125 |
| 14_7040195 | 6 | 0.5 | 2 | 0.5 | 1 | 0 | 4 | 0.5 | 13 | 0.4615 | 2 | 0.5 | 1 | 0 | 3 | 1 | 1 | 0 | 7 | 0.5714 |
| 14_7040202 | 7 | 0.1429 | 3 | 0 | 3 | 0 | 6 | 0 | 19 | 0.0526 | 3 | 0 | 2 | 0 | 4 | 0.25 | 1 | 0 | 10 | 0.1 |
| 14_7040208 | 5 | 0.4 | 3 | 0.3333 | 3 | 0.3333 | 5 | 0.2 | 16 | 0.3125 | 2 | 0.5 | 2 | 0.5 | 3 | 0 | NA | NA | 7 | 0.2857 |
| 14_7041794 | 4 | 0 | 3 | 0.3333 | 5 | 0 | 2 | 0 | 14 | 0.0714 | 5 | 0.2 | 5 | 0.2 | 3 | 0.3333 | 2 | 0.5 | 15 | 0.2667 |
| 14_7041826 | 5 | 0.2 | 3 | 0.6667 | 5 | 0 | 2 | 0 | 15 | 0.2 | 4 | 0 | 4 | 0 | 3 | 0.3333 | 1 | 0 | 12 | 0.0833 |
| 14_7041837 | 5 | 0.4 | 3 | 0.3333 | 3 | 0 | 2 | 0 | 13 | 0.2308 | 4 | 0 | 4 | 0.25 | 3 | 0 | 2 | 0 | 13 | 0.0769 |
| 14_7045487 | 1 | 1 | NA | NA | NA | NA | NA | NA | 1 | 1 | NA | NA | NA | NA | NA | NA | 2 | 0.5 | 2 | 0.5 |
| 14_7045553 | 3 | 0.3333 | 3 | 0 | NA | NA | 3 | 0 | 9 | 0.1111 | 1 | 0 | 3 | 0.3333 | 5 | 0.4 | 4 | 0.25 | 13 | 0.3077 |
| 14_7045555 | 3 | 0 | 4 | 0 | NA | NA | 3 | 0.3333 | 10 | 0.1 | 1 | 0 | 3 | 0.3333 | 5 | 0 | 4 | 0 | 13 | 0.0769 |
| 14_7054282 | 4 | 0.25 | 2 | 0.5 | 1 | 0 | 1 | 0 | 8 | 0.25 | 2 | 0.5 | 1 | 0 | 3 | 0 | 4 | 0 | 10 | 0.1 |
| 14_7054303 | 3 | 0.3333 | 5 | 0.2 | 1 | 0 | 1 | 0 | 10 | 0.2 | 2 | 0 | 1 | 0 | 3 | 0.3333 | 4 | 0 | 10 | 0.1 |
| 14_7054307 | 4 | 0.5 | 4 | 0.25 | 1 | 1 | 1 | 0 | 10 | 0.4 | 2 | 0 | 1 | 0 | 3 | 0 | 4 | 0.25 | 10 | 0.1 |
| 14_7054384 | 3 | 1 | 4 | 0.5 | NA | NA | 1 | 1 | 8 | 0.75 | NA | NA | NA | NA | 3 | 0.6667 | 4 | 0.5 | 7 | 0.5714 |
| 14_7054390 | 3 | 0 | 4 | 0.25 | NA | NA | 1 | 0 | 8 | 0.125 | 1 | 0 | NA | NA | 2 | 0.5 | 4 | 0.25 | 7 | 0.2857 |
| 14_7054398 | 2 | 1 | 2 | 1 | 1 | 1 | 1 | 1 | 6 | 1 | 1 | 1 | 1 | 1 | 2 | 1 | 3 | 1 | 7 | 1 |
| 14_7054399 | 3 | 1 | 2 | 1 | NA | NA | NA | NA | 5 | 1 | 1 | 1 | NA | NA | 1 | 1 | 2 | 1 | 4 | 1 |
| 14_7054404 | 3 | 1 | 4 | 0.75 | 1 | 1 | 1 | 1 | 9 | 0.8889 | 1 | 1 | NA | NA | 3 | 1 | 3 | 1 | 7 | 1 |
| 14_7054408 | 3 | 1 | 3 | 1 | 1 | 1 | NA | NA | 7 | 1 | 1 | 1 | 1 | 1 | 2 | 1 | 3 | 0.6667 | 7 | 0.8571 |
| 14_7054411 | 2 | 0.5 | 2 | 1 | NA | NA | 1 | 0 | 5 | 0.6 | 1 | 0 | NA | NA | 2 | 1 | 3 | 0.6667 | 6 | 0.6667 |
| 14_7067635 | 5 | 0 | 1 | 0 | 3 | 0 | 5 | 0.2 | 14 | 0.0714 | 1 | 1 | 4 | 0 | 2 | 0 | 2 | 0 | 9 | 0.1111 |
| 14_7067663 | 5 | 0.4 | 1 | 0 | 1 | 0 | 4 | 0 | 11 | 0.1818 | 1 | 0 | 4 | 0.25 | NA | NA | 2 | 1 | 7 | 0.4286 |
| 14_7067673 | 5 | 0.2 | 1 | 0 | 2 | 0 | 5 | 0.8 | 13 | 0.3846 | 1 | 1 | 2 | 1 | 2 | 0 | 2 | 0 | 7 | 0.4286 |
| 14_7067689 | 3 | 0.6667 | 1 | 1 | 1 | 0 | 3 | 1 | 8 | 0.75 | 1 | 1 | 2 | 1 | 2 | 0.5 | 1 | 1 | 6 | 0.8333 |
| 14_7069477 | 2 | 0 | 2 | 1 | 1 | 1 | 4 | 0.5 | 9 | 0.5556 | NA | NA | 2 | 0 | 1 | 0 | 3 | 0.6667 | 6 | 0.3333 |
| 14_7070427 | 45 | 0.0444 | 33 | 0.0606 | 36 | 0.0278 | 40 | 0.125 | 154 | 0.0649 | 27 | 0.0741 | 50 | 0.08 | 40 | 0.05 | 39 | 0.0256 | 156 | 0.0577 |
| 14_7070437 | 42 | 0.7143 | 33 | 0.697 | 35 | 0.8286 | 40 | 0.7 | 150 | 0.7333 | 26 | 0.6923 | 44 | 0.7955 | 39 | 0.8205 | 33 | 0.9394 | 142 | 0.8169 |
| 14_7070445 | 45 | 1 | 38 | 0.9474 | 45 | 1 | 45 | 1 | 173 | 0.9884 | 27 | 1 | 52 | 0.9808 | 45 | 0.9333 | 43 | 0.9535 | 167 | 0.9641 |
| 14_7070452 | 44 | 0.2045 | 42 | 0.119 | 44 | 0.2045 | 41 | 0.122 | 171 | 0.1637 | 27 | 0.0741 | 52 | 0.1346 | 49 | 0.0204 | 43 | 0.186 | 171 | 0.1053 |
| 14_7070455 | 46 | 0.9783 | 41 | 0.9512 | 44 | 0.9545 | 49 | 0.9592 | 180 | 0.9611 | 37 | 0.9459 | 57 | 0.9825 | 53 | 1 | 43 | 0.9535 | 190 | 0.9737 |
| 14_7070462 | 51 | 0.0392 | 46 | 0.1957 | 44 | 0.1364 | 50 | 0.14 | 191 | 0.1257 | 32 | 0.125 | 53 | 0.1321 | 51 | 0.0196 | 44 | 0.1364 | 180 | 0.1 |
| 14_7070477 | 43 | 1 | 47 | 0.9149 | 41 | 1 | 42 | 1 | 173 | 0.9769 | 35 | 1 | 51 | 1 | 45 | 0.9778 | 42 | 1 | 173 | 0.9942 |
| 14_7070483 | 55 | 0.0727 | 58 | 0.069 | 52 | 0.1538 | 60 | 0.1167 | 225 | 0.1022 | 42 | 0.0952 | 67 | 0.0896 | 53 | 0.1509 | 49 | 0.1429 | 211 | 0.1185 |
| 14_7070494 | 51 | 0.1569 | 46 | 0.2391 | 41 | 0.1707 | 54 | 0.1296 | 192 | 0.1719 | 39 | 0.2051 | 53 | 0.1321 | 44 | 0.2727 | 40 | 0.175 | 176 | 0.1932 |
| 14_7070559 | 73 | 0.5342 | 61 | 0.5246 | 60 | 0.5833 | 60 | 0.6 | 254 | 0.5591 | 46 | 0.6304 | 82 | 0.622 | 66 | 0.6061 | 57 | 0.614 | 251 | 0.6175 |
| 14_7070568 | 71 | 0.2535 | 64 | 0.2031 | 61 | 0.2623 | 59 | 0.3898 | 255 | 0.2745 | 47 | 0.3191 | 78 | 0.3333 | 67 | 0.3433 | 54 | 0.4444 | 246 | 0.3577 |
| 14_7070595 | 69 | 0.2029 | 56 | 0.125 | 63 | 0.1587 | 65 | 0.2 | 253 | 0.1739 | 51 | 0.1765 | 70 | 0.1286 | 65 | 0.1846 | 57 | 0.2105 | 243 | 0.1728 |
| 14_7071227 | 2 | 0.5 | 4 | 0 | 2 | 0 | 2 | 0 | 10 | 0.1 | 1 | 1 | 2 | 0 | 1 | 1 | 5 | 0.2 | 9 | 0.3333 |
| 14_7071239 | 2 | 0.5 | 4 | 0.5 | 1 | 0 | 1 | 0 | 8 | 0.375 | 1 | 0 | 3 | 0.6667 | 1 | 1 | 4 | 0.25 | 9 | 0.4444 |
| 14_7071240 | 2 | 0.5 | 4 | 0.25 | 2 | 0.5 | 2 | 1 | 10 | 0.5 | NA | NA | 3 | 0.6667 | 1 | 1 | 5 | 0.4 | 9 | 0.5556 |
| 14_7071246 | 2 | 0 | 4 | 0.25 | 2 | 0 | 2 | 0 | 10 | 0.1 | NA | NA | 3 | 0.3333 | 1 | 0 | 4 | 0 | 8 | 0.125 |
| 14_7071249 | 1 | 0 | 3 | 0.3333 | NA | NA | 1 | 0 | 5 | 0.2 | NA | NA | 2 | 0.5 | 1 | 1 | 3 | 0.3333 | 6 | 0.5 |
| 14_7071249 | 1 | 0 | 4 | 0.25 | 2 | 0 | 2 | 0 | 9 | 0.1111 | NA | NA | 3 | 0.3333 | 1 | 0 | 4 | 0.25 | 8 | 0.25 |
| 14_7076834 | 2 | 0.5 | 9 | 0.2222 | 4 | 0 | 6 | 0 | 21 | 0.1429 | 6 | 0 | 2 | 0 | 4 | 0.5 | 4 | 0.25 | 16 | 0.1875 |

| | | | | | | | | | | | | | | | | | | | | |
|---|---|---|---|---|---|---|---|---|---|---|---|---|---|---|---|---|---|---|---|---|
| 14_707686 | 3 | 0 | 10 | 0.1 | 6 | 0.5 | 8 | 0.25 | 27 | 0.2222 | 9 | 0.1111 | 4 | 0 | 4 | 0 | 5 | 0.4 | 22 | 0.1364 |
| 14_707686 | 2 | 0 | 7 | 0.2857 | 7 | 0 | 6 | 0.1667 | 22 | 0.1364 | 9 | 0.3333 | 3 | 0 | 4 | 0.25 | 4 | 0.5 | 20 | 0.3 |
| 14_707709 | 2 | 0 | 6 | 0.1667 | 2 | 0 | 1 | 0 | 11 | 0.0909 | 2 | 0 | 2 | 0.5 | 2 | 0 | 2 | 0.5 | 8 | 0.25 |
| 14_707737 | 4 | 0 | 10 | 0.1 | 13 | 0.2308 | 11 | 0.0909 | 38 | 0.1316 | 8 | 0.125 | 8 | 0.25 | 6 | 0.1667 | 5 | 0.2 | 27 | 0.1852 |
| 14_707738 | 6 | 0.5 | 12 | 0.4167 | 13 | 0.4615 | 12 | 0.4167 | 43 | 0.4419 | 10 | 0.8 | 9 | 0.4444 | 6 | 0.5 | 7 | 0.5714 | 32 | 0.5938 |
| 14_707740 | 5 | 0.8 | 8 | 1 | 10 | 0.7 | 9 | 0.6667 | 32 | 0.7812 | 6 | 1 | 4 | 1 | 3 | 0.6667 | 7 | 0.8571 | 20 | 0.9 |
| 14_707740 | 8 | 0.875 | 13 | 0.6154 | 15 | 0.7333 | 11 | 0.6364 | 47 | 0.7021 | 10 | 0.8 | 9 | 0.6667 | 4 | 0.75 | 8 | 0.75 | 31 | 0.7419 |
| 14_707741 | 5 | 0.4 | 11 | 0.1818 | 12 | 0.3333 | 9 | 0.6667 | 37 | 0.3784 | 11 | 0.2727 | 7 | 0.7143 | 6 | 0.5 | 6 | 0.5 | 30 | 0.4667 |
| 14_707741 | 7 | 0.2857 | 10 | 0.2 | 14 | 0 | 12 | 0.4167 | 43 | 0.2093 | 10 | 0.1 | 9 | 0.3333 | 4 | 0 | 7 | 0.2857 | 30 | 0.2 |
| 14_707742 | 7 | 0.4286 | 12 | 0.75 | 12 | 0.6667 | 12 | 0.6667 | 43 | 0.6512 | 8 | 0.75 | 9 | 0.6667 | 6 | 0.8333 | 6 | 0.8333 | 29 | 0.7586 |
| 14_707744 | 9 | 0.6667 | 12 | 0.75 | 14 | 0.5714 | 13 | 0.6923 | 48 | 0.6667 | 12 | 0.25 | 11 | 0.8182 | 8 | 0.5 | 8 | 0.375 | 39 | 0.4872 |
| 14_707744 | 8 | 0.125 | 9 | 0.1111 | 12 | 0.0833 | 10 | 0 | 39 | 0.0769 | 11 | 0.0909 | 11 | 0 | 7 | 0.1429 | 8 | 0 | 37 | 0.0541 |
| 14_707744 | 7 | 0.7143 | 14 | 0.7143 | 14 | 0.6429 | 11 | 0.6364 | 46 | 0.6739 | 10 | 0.6 | 9 | 0.4444 | 8 | 0.375 | 5 | 0.8 | 32 | 0.5312 |
| 14_707746 | 6 | 0 | 9 | 0.4444 | 10 | 0.3 | 12 | 0.25 | 37 | 0.2703 | 11 | 0.2727 | 8 | 0.25 | 10 | 0.2 | 6 | 0.1667 | 35 | 0.2286 |
| 14_707749 | 5 | 0.6 | 11 | 1 | 11 | 0.9091 | 10 | 0.8 | 37 | 0.8649 | 5 | 0.8 | 9 | 0.8889 | 3 | 0.3333 | 7 | 0.7143 | 24 | 0.75 |
| 14_707749 | 6 | 0.6667 | 10 | 1 | 11 | 0.7273 | 10 | 0.7 | 37 | 0.7838 | 7 | 0.8571 | 7 | 0.7143 | 3 | 0.6667 | 5 | 1 | 22 | 0.8182 |
| 14_707750 | 5 | 0 | 8 | 0.625 | 10 | 0.8 | 6 | 1 | 29 | 0.6552 | 8 | 0.5 | 7 | 0.8571 | 4 | 0.75 | 6 | 0.6667 | 25 | 0.68 |
| 14_707752 | 5 | 0 | 12 | 0 | 12 | 0.4167 | 10 | 0 | 39 | 0.1282 | 8 | 0 | 9 | 0.1111 | 5 | 0 | 5 | 0.2 | 27 | 0.0741 |
| 14_707753 | 4 | 0.25 | 11 | 0.2727 | 11 | 0.4545 | 9 | 0.4444 | 35 | 0.3714 | 10 | 0.2 | 8 | 0.25 | 4 | 0 | 8 | 0.25 | 30 | 0.2 |
| 14_707754 | 6 | 0.3333 | 13 | 0.3077 | 13 | 0.1538 | 13 | 0.1538 | 45 | 0.2222 | 10 | 0.3 | 10 | 0.4 | 6 | 0.1667 | 7 | 0.4286 | 33 | 0.3333 |
| 14_707756 | 4 | 0.25 | 14 | 0.0714 | 11 | 0.1818 | 9 | 0.1111 | 38 | 0.1316 | 11 | 0 | 9 | 0.2222 | 7 | 0 | 4 | 0.5 | 31 | 0.129 |
| 14_707941 | 1 | 1 | 1 | 0 | NA | NA | NA | NA | 2 | 0.5 | NA | NA | NA | NA | 1 | 1 | NA | NA | 1 | 1 |
| 14_708104 | 1 | 0 | 5 | 0.4 | 1 | 0 | 1 | 0 | 8 | 0.25 | NA | NA | 1 | 1 | 2 | 0.5 | NA | NA | 3 | 0.6667 |
| 14_708170 | 6 | 0 | 10 | 0.1 | 6 | 0 | 2 | 0.5 | 24 | 0.0833 | 3 | 0 | 9 | 0 | 4 | 0.25 | 3 | 0 | 19 | 0.0526 |
| 14_708170 | 5 | 0.2 | 10 | 0.2 | 5 | 0 | 2 | 0.5 | 22 | 0.1818 | 3 | 0.3333 | 8 | 0.25 | 2 | 0 | 3 | 0 | 16 | 0.1875 |
| 14_708171 | 6 | 0.6667 | 10 | 0.6 | 8 | 0.625 | 1 | 0 | 25 | 0.6 | 4 | 0.5 | 10 | 0.6 | 6 | 0.3333 | 4 | 0.25 | 24 | 0.4583 |
| 14_708171 | 6 | 0.1667 | 10 | 0.5 | 7 | 0.4286 | 3 | 0.3333 | 26 | 0.3846 | 3 | 0.3333 | 9 | 0 | 7 | 0.1429 | 3 | 0 | 22 | 0.0909 |
| 14_708172 | 6 | 0 | 14 | 0.1429 | 8 | 0.25 | 3 | 0 | 31 | 0.129 | 3 | 0 | 10 | 0.2 | 6 | 0.1667 | 4 | 0 | 23 | 0.1304 |
| 14_708173 | 7 | 0.4286 | 14 | 0.5714 | 8 | 0.5 | 3 | 0.3333 | 32 | 0.5 | 3 | 0.3333 | 10 | 0.6 | 8 | 0.125 | 4 | 0.25 | 25 | 0.36 |
| 14_708173 | 8 | 0.375 | 11 | 0.3636 | 9 | 0.2222 | 2 | 1 | 30 | 0.3667 | 4 | 0.5 | 9 | 0.1111 | 9 | 0 | 4 | 0.25 | 26 | 0.1538 |
| 14_708173 | 6 | 0.3333 | 14 | 0.3571 | 9 | 0.2222 | 4 | 0.25 | 33 | 0.303 | 5 | 0 | 12 | 0.25 | 6 | 0.1667 | 4 | 0.25 | 27 | 0.1852 |
| 14_708175 | 9 | 0 | 13 | 0.2308 | 9 | 0.4444 | 3 | 0.3333 | 34 | 0.2353 | 5 | 0.4 | 12 | 0.3333 | 9 | 0.1111 | 3 | 0.3333 | 29 | 0.2759 |
| 14_708212 | 5 | 0.2 | 4 | 0.75 | 1 | 0 | 6 | 0.3333 | 16 | 0.375 | 1 | 0 | 6 | 0.3333 | 4 | 0.5 | 3 | 0.3333 | 14 | 0.3571 |
| 14_708213 | 6 | 0 | 4 | 0 | 2 | 0.5 | 5 | 0 | 17 | 0.0588 | 1 | 0 | 4 | 0.25 | 3 | 0.3333 | 4 | 0 | 12 | 0.1667 |
| 14_708215 | 5 | 0.2 | 4 | 0.25 | 2 | 0 | 6 | 0 | 17 | 0.1176 | 1 | 0 | 6 | 0.1667 | 3 | 0 | 4 | 0 | 14 | 0.0714 |
| 14_708216 | 5 | 0 | 3 | 0 | 2 | 0 | 5 | 0.2 | 15 | 0.0667 | 1 | 1 | 6 | 0.3333 | 3 | 0.3333 | 3 | 0 | 13 | 0.3077 |
| 14_708216 | 5 | 0.8 | 3 | 0.6667 | 2 | 0 | 5 | 0.6 | 15 | 0.6 | 1 | 1 | 4 | 0.75 | 3 | 1 | 3 | 0.3333 | 11 | 0.7273 |
| 14_708216 | 6 | 0.3333 | 4 | 0.5 | 2 | 1 | 5 | 0.2 | 17 | 0.4118 | 1 | 1 | 6 | 0.6667 | 3 | 0.3333 | 4 | 0.5 | 14 | 0.5714 |
| 14_708220 | 3 | 0.3333 | 1 | 1 | 2 | 1 | 3 | 0.3333 | 9 | 0.5556 | 1 | 0 | 4 | 0.25 | 2 | 0.5 | 3 | 0.6667 | 10 | 0.4 |
| 14_708220 | 6 | 0.5 | 3 | 0.6667 | 2 | 0 | 4 | 0 | 15 | 0.3333 | 1 | 1 | 6 | 0.3333 | 1 | 0 | 4 | 0.5 | 12 | 0.4167 |
| 14_708221 | 5 | 0.8 | NA | NA | 1 | 1 | 2 | 1 | 8 | 0.875 | 1 | 1 | 4 | 0.75 | 3 | 1 | 1 | 1 | 9 | 0.8889 |
| 14_708221 | 6 | 0.3333 | 2 | 0 | 2 | 0 | 5 | 0 | 15 | 0.1333 | NA | NA | 5 | 0 | 2 | 0.5 | 4 | 0 | 11 | 0.0909 |
| 14_708222 | 5 | 0 | 3 | 0.3333 | 2 | 0 | 5 | 0 | 15 | 0.0667 | 1 | 0 | 5 | 0.2 | 3 | 0 | 3 | 0 | 12 | 0.0833 |
| 14_708222 | 5 | 0.8 | 3 | 1 | 2 | 1 | 3 | 0.6667 | 13 | 0.8462 | 1 | 1 | 6 | 0.6667 | 3 | 0.6667 | 4 | 0.5 | 14 | 0.6429 |
| 14_708223 | 6 | 0.6667 | 2 | 1 | 1 | 0 | 5 | 0.8 | 14 | 0.7143 | 1 | 1 | 3 | 0.3333 | 2 | 0 | 4 | 0.25 | 10 | 0.3 |
| 14_708223 | 5 | 0.2 | 2 | 1 | 2 | 0.5 | 4 | 0.5 | 13 | 0.4615 | 1 | 1 | 5 | 0 | 2 | 1 | 3 | 0 | 11 | 0.2727 |
| 14_708225 | 4 | 1 | 1 | 1 | 2 | 0 | 4 | 0.25 | 11 | 0.5455 | 1 | 1 | 2 | 0.5 | 2 | 0.5 | NA | NA | 5 | 0.6 |
| 14_708235 | 2 | 0 | 2 | 0.5 | 2 | 0 | 3 | 0.3333 | 9 | 0.2222 | NA | NA | 3 | 0 | 2 | 0 | 2 | 0.5 | 7 | 0.1429 |
| 14_708263 | 1 | 0 | 2 | 0.5 | 1 | 0 | 3 | 0 | 7 | 0.1429 | 1 | 0 | 3 | 0.3333 | 2 | 0 | 2 | 0 | 8 | 0.125 |
| 14_708264 | 1 | 0 | 2 | 0.5 | NA | NA | 3 | 0 | 6 | 0.1667 | 1 | 0 | 3 | 0 | 2 | 0 | 2 | 1 | 8 | 0.25 |
| 14_708264 | NA | NA | 1 | 0 | 1 | 1 | 2 | 0 | 4 | 0.25 | 1 | 0 | 4 | 0.25 | NA | NA | 1 | 0 | 6 | 0.1667 |
| 14_708393 | NA | NA | 1 | 1 | 1 | 1 | NA | NA | 2 | 1 | 2 | 1 | NA | NA | 1 | 1 | 2 | 1 | 5 | 1 |
| 14_708396 | NA | NA | 1 | 1 | 2 | 1 | 1 | 1 | 4 | 1 | 2 | 1 | NA | NA | NA | NA | 1 | 1 | 3 | 1 |
| 14_708396 | NA | NA | 2 | 0.5 | 3 | 0.6667 | 1 | 0 | 6 | 0.5 | 2 | 1 | NA | NA | NA | NA | 1 | 1 | 3 | 1 |
| 14_708397 | NA | NA | 2 | 1 | 3 | 1 | 1 | 0 | 6 | 0.8333 | 2 | 0.5 | NA | NA | NA | NA | 1 | 1 | 3 | 0.6667 |
| 14_708398 | NA | NA | NA | NA | 3 | 0.3333 | 1 | 1 | 4 | 0.5 | 2 | 0 | NA | NA | NA | NA | 1 | 1 | 3 | 0.3333 |
| 14_708398 | NA | NA | 2 | 0 | 3 | 0.3333 | 1 | 0 | 6 | 0.1667 | 2 | 1 | NA | NA | NA | NA | 1 | 1 | 3 | 1 |
| 14_708400 | NA | NA | 1 | 1 | 1 | 1 | 1 | 0 | 3 | 0.6667 | 2 | 1 | NA | NA | NA | NA | 1 | 0 | 3 | 0.6667 |
| 14_708402 | NA | NA | 2 | 1 | 3 | 0.6667 | 1 | 1 | 6 | 0.8333 | 2 | 0.5 | NA | NA | NA | NA | NA | NA | 2 | 0.5 |
| 14_708403 | NA | NA | 2 | 0.5 | 3 | 0 | 1 | 0 | 6 | 0.1667 | 2 | 0.5 | NA | NA | NA | NA | 1 | 1 | 3 | 0.6667 |
| 14_708414 | 34 | 0.0588 | 39 | 0.0256 | 25 | 0 | 44 | 0.1136 | 142 | 0.0563 | 35 | 0.1429 | 30 | 0.0333 | 37 | 0.1351 | 31 | 0.0968 | 133 | 0.1053 |
| 14_708415 | 35 | 0.3714 | 35 | 0.2571 | 21 | 0.4762 | 43 | 0.2326 | 134 | 0.3134 | 37 | 0.2432 | 32 | 0.1875 | 39 | 0.2564 | 33 | 0.2727 | 141 | 0.2411 |
| 14_708416 | 20 | 0.35 | 23 | 0.1304 | 12 | 0.5 | 37 | 0.2973 | 92 | 0.2935 | 21 | 0.5714 | 24 | 0.375 | 28 | 0.3214 | 30 | 0.3 | 103 | 0.3786 |
| 14_708417 | 37 | 0.1081 | 36 | 0.1667 | 22 | 0.1364 | 45 | 0.2667 | 140 | 0.1786 | 34 | 0.1765 | 31 | 0.1935 | 39 | 0.1282 | 34 | 0.2353 | 138 | 0.1812 |
| 14_708418 | 34 | 0.1471 | 36 | 0.1111 | 23 | 0.087 | 43 | 0.0465 | 136 | 0.0956 | 37 | 0.0811 | 29 | 0.069 | 36 | 0.0556 | 35 | 0.1429 | 137 | 0.0876 |

| | | | | | | | | | | | | | | | | | | | | |
|---|---|---|---|---|---|---|---|---|---|---|---|---|---|---|---|---|---|---|---|---|
| 14_708418 | 33 | 0.6667 | 35 | 0.6857 | 23 | 0.8261 | 41 | 0.8537 | 132 | 0.7576 | 34 | 0.7941 | 27 | 0.8148 | 37 | 0.8108 | 34 | 0.6176 | 132 | 0.7576 |
| 14_708419 | 31 | 0.6774 | 33 | 0.7576 | 22 | 0.8182 | 36 | 0.9167 | 122 | 0.7951 | 29 | 0.8621 | 22 | 0.9091 | 34 | 0.8529 | 32 | 0.7812 | 117 | 0.8462 |
| 14_708646 | 1 | 1 | NA | NA | NA | NA | NA | NA | 1 | 1 | 1 | 1 | NA | NA | NA | NA | NA | NA | 1 | 1 |
| 14_709021 | 5 | 0.2 | 8 | 0.25 | 5 | 0 | 12 | 0.0833 | 30 | 0.1333 | 5 | 0 | 4 | 0 | 9 | 0.1111 | 4 | 0.25 | 22 | 0.0909 |
| 14_709032 | 4 | 0.25 | 2 | 0 | 2 | 0 | 7 | 0.1429 | 15 | 0.1333 | 4 | 0.25 | 4 | 0 | 4 | 0 | 3 | 0 | 15 | 0.0667 |
| 14_709034 | 2 | 1 | 2 | 0 | 1 | 1 | 6 | 0.1667 | 11 | 0.3636 | 2 | 0 | 3 | 0 | 3 | 1 | 4 | 0.75 | 12 | 0.5 |
| 14_711279 | NA | NA | 2 | 0.5 | 2 | 1 | NA | NA | 4 | 0.75 | NA | NA | 2 | 0 | 1 | 1 | 1 | 1 | 4 | 0.5 |
| 14_711279 | NA | NA | 2 | 1 | 2 | 1 | NA | NA | 4 | 1 | NA | NA | 2 | 0 | 2 | 0.5 | 1 | 0 | 5 | 0.2 |
| 14_711280 | NA | NA | 2 | 0.5 | 2 | 0 | NA | NA | 4 | 0.25 | NA | NA | 2 | 0.5 | 2 | 0 | 1 | 0 | 5 | 0.2 |
| 14_711281 | NA | NA | 2 | 0 | 2 | 1 | 1 | 0 | 5 | 0.4 | NA | NA | 1 | 0 | 2 | 0 | 1 | 1 | 4 | 0.25 |
| 14_711282 | NA | NA | 2 | 1 | 2 | 0 | 1 | 1 | 5 | 0.6 | NA | NA | 2 | 0 | 2 | 0 | 1 | 1 | 5 | 0.2 |
| 14_711282 | NA | NA | 2 | 0.5 | 2 | 0.5 | 1 | 1 | 5 | 0.6 | NA | NA | 1 | 1 | 2 | 0.5 | 1 | 1 | 4 | 0.75 |
| 14_712162 | 4 | 0 | 4 | 0.25 | 3 | 0 | 1 | 0 | 12 | 0.0833 | 1 | 0 | 2 | 1 | 1 | 0 | 3 | 0 | 7 | 0.2857 |
| 14_712163 | 4 | 1 | 5 | 0.8 | 3 | 1 | 1 | 1 | 13 | 0.9231 | 1 | 1 | 3 | 1 | 2 | 1 | 2 | 0.5 | 8 | 0.875 |
| 14_712163 | 4 | 0.25 | 5 | 0.2 | 3 | 0 | 1 | 0 | 13 | 0.1538 | NA | NA | 3 | 0.3333 | 2 | 0 | 3 | 0 | 8 | 0.125 |
| 14_712164 | 3 | 0.6667 | 5 | 1 | 3 | 1 | 1 | 1 | 12 | 0.9167 | 1 | 1 | 3 | 0.6667 | 1 | 0 | 3 | 0.6667 | 8 | 0.625 |
| 14_712164 | 4 | 1 | 5 | 1 | 3 | 0.6667 | 1 | 1 | 13 | 0.9231 | 1 | 1 | 4 | 1 | 1 | 1 | 2 | 0.5 | 8 | 0.875 |
| 14_712164 | 3 | 0 | 5 | 0.2 | 2 | 0 | 1 | 0 | 11 | 0.0909 | NA | NA | 4 | 0 | 2 | 0.5 | 3 | 0 | 9 | 0.1111 |
| 14_712165 | 2 | 1 | 5 | 0.8 | 3 | 0.6667 | 1 | 0 | 11 | 0.7273 | 1 | 1 | 4 | 1 | 2 | 1 | 3 | 1 | 10 | 1 |
| 14_727766 | 5 | 0.4 | 6 | 0.1667 | 3 | 0 | 6 | 0 | 20 | 0.15 | 2 | 0 | 7 | 0 | 2 | 0 | 10 | 0.2 | 21 | 0.0952 |
| 14_727771 | 5 | 0.2 | 4 | 0 | 3 | 0 | 5 | 0.2 | 17 | 0.1176 | 3 | 0 | 6 | 0 | 1 | 0 | 6 | 0.1667 | 16 | 0.0625 |
| 14_727772 | 5 | 0.2 | 5 | 0.6 | 3 | 0.3333 | 6 | 0.5 | 19 | 0.4211 | 2 | 1 | 10 | 0.1 | 1 | 0 | 11 | 0.2727 | 24 | 0.25 |
| 14_727773 | 5 | 0 | 5 | 0 | 3 | 0 | 6 | 0.1667 | 19 | 0.0526 | 3 | 0 | 10 | 0.1 | 2 | 0 | 11 | 0.0909 | 26 | 0.0769 |
| 14_727775 | 5 | 0.2 | 5 | 0.4 | 3 | 0.3333 | 6 | 0 | 19 | 0.2105 | 3 | 0 | 8 | 0.125 | 2 | 0 | 11 | 0.1818 | 24 | 0.125 |
| 14_727777 | 3 | 0.3333 | 5 | 0.6 | 2 | 0.5 | 2 | 0 | 12 | 0.4167 | 2 | 1 | 10 | 0.4 | NA | NA | 6 | 0.3333 | 18 | 0.4444 |
| 14_728221 | 1 | 0 | 3 | 1 | 3 | 1 | 2 | 0.5 | 9 | 0.7778 | 1 | 0 | 3 | 0.3333 | 1 | 1 | 3 | 0 | 8 | 0.25 |
| 14_733810 | NA | NA | 1 | 1 | NA | NA | NA | NA | 1 | 1 | NA | NA | 1 | 1 | NA | NA | NA | NA | 1 | 1 |
| 14_734199 | 3 | 0.3333 | 3 | 0 | 3 | 0 | 2 | 0 | 11 | 0.0909 | 3 | 0.3333 | 4 | 0 | 6 | 0 | 3 | 0 | 16 | 0.0625 |
| 14_734199 | 3 | 0.3333 | 3 | 0 | 2 | 0.5 | 4 | 0 | 12 | 0.1667 | 1 | 0 | 4 | 0 | 5 | 0.2 | 3 | 0.3333 | 13 | 0.1538 |
| 14_734199 | 1 | 0 | 3 | 0.3333 | 2 | 0 | 4 | 0 | 10 | 0.1 | 3 | 0.3333 | 4 | 0.25 | 5 | 0 | 3 | 1 | 15 | 0.3333 |
| 14_734200 | 2 | 0.5 | 3 | 0 | 3 | 0 | 4 | 0.5 | 12 | 0.25 | 2 | 0.5 | 3 | 0.3333 | 4 | 0.5 | 3 | 1 | 12 | 0.5833 |
| 14_734201 | 1 | 1 | NA | NA | 1 | 0 | 1 | 0 | 3 | 0.3333 | 2 | 0.5 | 2 | 0 | 3 | 0 | 2 | 0 | 9 | 0.1111 |
| 14_734201 | 1 | 1 | NA | NA | 1 | 0 | 1 | 0 | 3 | 0.3333 | 1 | 1 | 2 | 0 | 3 | 0 | 1 | 0 | 7 | 0.1429 |
| 14_734341 | 1 | 0 | 1 | 1 | 2 | 0 | 1 | 0 | 5 | 0.2 | 2 | 0 | 1 | 0 | 4 | 0.25 | NA | NA | 7 | 0.1429 |
| 14_734342 | 1 | 1 | 1 | 0 | 2 | 0 | 1 | 0 | 5 | 0.2 | 2 | 0 | 1 | 0 | 4 | 0.25 | NA | NA | 7 | 0.1429 |
| 14_734757 | NA | NA | NA | NA | 2 | 0.5 | NA | NA | 2 | 0.5 | 1 | 0 | NA | NA | 1 | 1 | NA | NA | 2 | 0.5 |
| 14_734758 | NA | NA | NA | NA | 2 | 0.5 | NA | NA | 2 | 0.5 | 1 | 1 | NA | NA | 1 | 1 | NA | NA | 2 | 1 |
| 14_734758 | NA | NA | NA | NA | 2 | 0.5 | NA | NA | 2 | 0.5 | 1 | 0 | NA | NA | 1 | 1 | NA | NA | 2 | 0.5 |
| 14_734764 | NA | NA | NA | NA | 2 | 0.5 | NA | NA | 2 | 0.5 | 1 | 1 | NA | NA | 1 | 1 | NA | NA | 2 | 1 |
| 14_734765 | NA | NA | NA | NA | 2 | 0.5 | NA | NA | 2 | 0.5 | 2 | 0.5 | NA | NA | 1 | 0 | NA | NA | 3 | 0.3333 |
| 14_734766 | NA | NA | NA | NA | 2 | 1 | NA | NA | 2 | 1 | 2 | 1 | NA | NA | 1 | 1 | NA | NA | 3 | 1 |
| 14_734768 | NA | NA | NA | NA | 2 | 0.5 | NA | NA | 2 | 0.5 | 2 | 0.5 | NA | NA | 1 | 1 | NA | NA | 3 | 0.6667 |
| 14_734768 | NA | NA | NA | NA | 2 | 0.5 | NA | NA | 2 | 0.5 | 2 | 1 | NA | NA | 1 | 1 | NA | NA | 3 | 1 |
| 14_734768 | NA | NA | NA | NA | 2 | 0.5 | NA | NA | 2 | 0.5 | 2 | 0.5 | NA | NA | 1 | 0 | NA | NA | 3 | 0.3333 |
| 14_734770 | NA | NA | NA | NA | 2 | 1 | NA | NA | 2 | 1 | 2 | 1 | NA | NA | 1 | 1 | NA | NA | 3 | 1 |
| 14_734771 | NA | NA | NA | NA | 1 | 1 | NA | NA | 1 | 1 | 1 | 1 | NA | NA | NA | NA | NA | NA | 1 | 1 |
| 14_734771 | NA | NA | NA | NA | 2 | 0.5 | NA | NA | 2 | 0.5 | 2 | 0.5 | NA | NA | 1 | 1 | NA | NA | 3 | 0.6667 |
| 14_736005 | 157 | 0.1911 | 141 | 0.1418 | 111 | 0.2072 | 117 | 0.1709 | 526 | 0.1768 | 107 | 0.1121 | 111 | 0.1712 | 119 | 0.1849 | 123 | 0.1382 | 460 | 0.1522 |
| 14_737554 | 7 | 0.4286 | 7 | 0.4286 | 2 | 0.5 | 4 | 0.5 | 20 | 0.45 | 3 | 0.3333 | 1 | 0 | 4 | 0.25 | 3 | 0.3333 | 11 | 0.2727 |
| 14_737554 | 7 | 0.1429 | 8 | 0 | 2 | 0 | 6 | 0.1667 | 23 | 0.087 | 3 | 0 | 2 | 0 | 4 | 0.25 | 2 | 0 | 11 | 0.0909 |
| 14_737555 | 6 | 0 | 6 | 0.5 | 2 | 0 | 6 | 0.3333 | 20 | 0.25 | 3 | 0 | 2 | 0.5 | 5 | 0 | 3 | 0 | 13 | 0.0769 |
| 14_737557 | 7 | 0.5714 | 7 | 0.4286 | 2 | 0.5 | 6 | 0.3333 | 22 | 0.4545 | 3 | 0 | 2 | 0.5 | 3 | 0.3333 | 4 | 0.25 | 12 | 0.25 |
| 14_737557 | 6 | 0.1667 | 8 | 0.25 | 2 | 0 | 6 | 0 | 22 | 0.1364 | 3 | 0.3333 | 2 | 0 | 5 | 0 | 4 | 0 | 14 | 0.0714 |
| 14_737558 | 7 | 0 | 6 | 0 | NA | NA | 4 | 0.25 | 17 | 0.0588 | 2 | 0 | 1 | 1 | 6 | 0 | 3 | 0 | 12 | 0.0833 |
| 14_737580 | 1 | 0 | 1 | 0 | 1 | 1 | NA | NA | 3 | 0.3333 | 2 | 0.5 | NA | NA | NA | NA | 4 | 0.25 | 6 | 0.3333 |
| 14_737581 | NA | NA | 1 | 0 | 1 | 1 | NA | NA | 2 | 0.5 | 2 | 0.5 | NA | NA | NA | NA | 4 | 0.5 | 6 | 0.5 |
| 14_737582 | 1 | 0 | 1 | 0 | 1 | 1 | NA | NA | 3 | 0.3333 | 2 | 0.5 | NA | NA | NA | NA | 4 | 0.25 | 6 | 0.3333 |
| 14_737583 | 1 | 1 | 1 | 0 | 1 | 0 | NA | NA | 3 | 0.3333 | 2 | 0.5 | NA | NA | NA | NA | 3 | 0 | 5 | 0.2 |
| 14_737585 | 1 | 1 | 1 | 0 | 1 | 1 | NA | NA | 3 | 0.6667 | 2 | 0.5 | NA | NA | NA | NA | 3 | 0 | 5 | 0.2 |
| 14_737585 | 1 | 1 | NA | NA | 1 | 1 | NA | NA | 2 | 1 | 1 | 1 | NA | NA | NA | NA | 3 | 1 | 4 | 1 |
| 14_737586 | 1 | 0 | 1 | 1 | 1 | 1 | NA | NA | 3 | 0.6667 | 1 | 0 | NA | NA | NA | NA | 2 | 1 | 3 | 0.6667 |
| 14_737588 | NA | NA | 1 | 1 | 1 | 1 | NA | NA | 2 | 1 | 1 | 1 | NA | NA | NA | NA | 2 | 1 | 3 | 1 |
| 14_737588 | 1 | 0 | 1 | 0 | 1 | 1 | NA | NA | 3 | 0.3333 | 2 | 0.5 | NA | NA | NA | NA | 2 | 1 | 4 | 0.75 |
| 14_737588 | 1 | 1 | 1 | 0 | 1 | 1 | NA | NA | 3 | 0.6667 | 2 | 0 | NA | NA | NA | NA | 1 | 1 | 3 | 0.3333 |
| 14_737589 | 1 | 0 | 1 | 0 | 1 | 1 | NA | NA | 3 | 0.3333 | 2 | 0.5 | NA | NA | NA | NA | 2 | 1 | 4 | 0.75 |

| | | | | | | | | | | | | | | | | | | | | |
|---|---|---|---|---|---|---|---|---|---|---|---|---|---|---|---|---|---|---|---|---|
| 14_737590 | 1 | 0 | 1 | 1 | 1 | 0 | NA | NA | 3 | 0.3333 | 2 | 0.5 | NA | NA | NA | NA | 2 | 0 | 4 | 0.25 |
| 14_738330 | 24 | 0.5417 | 30 | 0.6333 | 30 | 0.5 | 26 | 0.5769 | 110 | 0.5636 | 21 | 0.4286 | 21 | 0.4762 | 29 | 0.5517 | 26 | 0.6923 | 97 | 0.5464 |
| 14_738332 | 20 | 0 | 18 | 0.0556 | 12 | 0.0833 | 17 | 0.1176 | 67 | 0.0597 | 11 | 0.0909 | 13 | 0 | 20 | 0.15 | 15 | 0 | 59 | 0.0678 |
| 14_738334 | 21 | 0.1429 | 17 | 0.1176 | 14 | 0.0714 | 19 | 0.1053 | 71 | 0.1127 | 11 | 0.2727 | 13 | 0.0769 | 20 | 0.15 | 14 | 0.0714 | 58 | 0.1379 |
| 14_748786 | NA | NA | 2 | 0 | 1 | 1 | 6 | 0.5 | 9 | 0.4444 | NA | NA | 1 | 1 | 1 | 0 | 2 | 0 | 4 | 0.25 |
| 14_748788 | NA | NA | 2 | 0.5 | 3 | 1 | 4 | 1 | 9 | 0.8889 | NA | NA | 1 | 1 | 1 | 0 | 2 | 1 | 4 | 0.75 |
| 14_748789 | NA | NA | 2 | 0.5 | 3 | 0.6667 | 4 | 0.75 | 9 | 0.6667 | NA | NA | 1 | 1 | 1 | 1 | 2 | 0.5 | 4 | 0.75 |
| 14_748790 | NA | NA | 1 | 1 | 3 | 0.6667 | 4 | 0.75 | 8 | 0.75 | NA | NA | 1 | 1 | 1 | 0 | 2 | 0.5 | 4 | 0.5 |
| 14_748791 | 1 | 1 | 1 | 1 | 3 | 0.6667 | 5 | 0.4 | 10 | 0.6 | 1 | 0 | NA | NA | NA | NA | 3 | 0.3333 | 4 | 0.25 |
| 14_748791 | NA | NA | NA | NA | 2 | 1 | 6 | 1 | 8 | 1 | 1 | 0 | 1 | 1 | 1 | 1 | 3 | 1 | 6 | 0.8333 |
| 14_748794 | 1 | 1 | 2 | 0 | 2 | 1 | 5 | 0.4 | 10 | 0.5 | 2 | 0 | 1 | 0 | NA | NA | 3 | 0.6667 | 6 | 0.3333 |
| 14_748795 | 1 | 1 | 2 | 0.5 | 3 | 1 | 7 | 1 | 13 | 0.9231 | 1 | 1 | 1 | 1 | 1 | 0 | 3 | 1 | 6 | 0.8333 |
| 14_748795 | NA | NA | 2 | 0.5 | 3 | 0.6667 | 5 | 1 | 10 | 0.8 | 2 | 1 | 1 | 1 | 1 | 0 | 1 | 1 | 5 | 0.8 |
| 14_749431 | 2 | 0.5 | 2 | 0 | 4 | 0.25 | 5 | 0.4 | 13 | 0.3077 | 3 | 0 | 1 | 0 | 1 | 0 | 3 | 0.6667 | 8 | 0.25 |
| 14_749432 | 2 | 0.5 | 3 | 0 | 4 | 0.25 | 7 | 0.1429 | 16 | 0.1875 | 4 | 0 | 1 | 0 | 1 | 1 | 3 | 0 | 9 | 0.1111 |
| 14_749435 | 1 | 1 | 3 | 1 | 3 | 0.3333 | 3 | 1 | 10 | 0.8 | 4 | 0.5 | NA | NA | 2 | 1 | 2 | 1 | 8 | 0.75 |
| 14_749436 | 3 | 0.6667 | 3 | 0.3333 | 5 | 0.4 | 6 | 0.5 | 17 | 0.4706 | 4 | 0 | 1 | 0 | NA | NA | 3 | 0.3333 | 8 | 0.125 |
| 14_749436 | 3 | 0.6667 | 4 | 1 | 5 | 0.8 | 6 | 0.8333 | 18 | 0.8333 | 4 | 1 | 1 | 1 | 2 | 1 | 3 | 0.6667 | 10 | 0.9 |
| 14_749437 | 3 | 0.3333 | 4 | 1 | 5 | 1 | 7 | 0.7143 | 19 | 0.7895 | 6 | 0.6667 | 1 | 1 | 2 | 1 | 4 | 0.75 | 13 | 0.7692 |
| 14_749437 | 3 | 0.3333 | 3 | 0.6667 | 4 | 0.75 | 5 | 0.8 | 15 | 0.6667 | 3 | 1 | 1 | 1 | 2 | 1 | 1 | 1 | 7 | 1 |
| 14_749437 | 3 | 1 | 4 | 0.75 | 4 | 1 | 5 | 1 | 16 | 0.9375 | 5 | 0.6 | 1 | 1 | 1 | 1 | 2 | 1 | 9 | 0.7778 |
| 14_749437 | 3 | 0.6667 | 5 | 0.8 | 4 | 0.5 | 6 | 0.8333 | 18 | 0.7222 | 6 | 0.8333 | 1 | 1 | 1 | 1 | 4 | 0.75 | 12 | 0.8333 |
| 14_749438 | 3 | 1 | 4 | 1 | 4 | 1 | 5 | 1 | 16 | 1 | 6 | 1 | 1 | 1 | 2 | 1 | 4 | 1 | 13 | 1 |
| 14_749438 | 2 | 0.5 | 4 | 0.25 | 3 | 0 | 4 | 0.5 | 13 | 0.3077 | 4 | 0 | NA | NA | 1 | 0 | 2 | 0.5 | 7 | 0.1429 |
| 14_749439 | 2 | 0.5 | 4 | 0.25 | 5 | 0.4 | 5 | 0.2 | 16 | 0.3125 | 5 | 0.2 | NA | NA | 1 | 0 | 4 | 0.25 | 10 | 0.2 |
| 14_749439 | 2 | 0.5 | 4 | 0.5 | 5 | 0.8 | 6 | 0 | 17 | 0.4118 | 5 | 0.4 | 1 | 1 | 2 | 0 | 3 | 0.6667 | 11 | 0.4545 |
| 14_749439 | 3 | 0.6667 | 5 | 0.6 | 5 | 0.8 | 7 | 0.2857 | 20 | 0.55 | 5 | 0.6 | 1 | 1 | 1 | 0 | 4 | 1 | 11 | 0.7273 |
| 14_749439 | 3 | 0 | 4 | 0.25 | 5 | 0.4 | 7 | 0.2857 | 19 | 0.2632 | 3 | 0 | NA | NA | 2 | 0 | 4 | 0.25 | 9 | 0.1111 |
| 14_749446 | 4 | 0.25 | 5 | 0.4 | 6 | 0.3333 | 8 | 0.125 | 23 | 0.2609 | 6 | 0 | NA | NA | 3 | 0.3333 | 3 | 0.3333 | 12 | 0.1667 |
| 14_749449 | 3 | 0 | 4 | 0 | 4 | 0.25 | 5 | 0.2 | 16 | 0.125 | 3 | 0 | 2 | 0 | 4 | 0.25 | 4 | 0 | 13 | 0.0769 |
| 14_749459 | 6 | 0.6667 | 7 | 0.4286 | 7 | 0.4286 | 8 | 0.25 | 28 | 0.4286 | 6 | 0.1667 | 3 | 0.3333 | 5 | 0.6 | 5 | 0 | 19 | 0.2632 |
| 14_749460 | 7 | 0.1429 | 7 | 0 | 9 | 0 | 9 | 0.1111 | 32 | 0.0625 | 4 | 0.25 | 2 | 0 | 6 | 0 | 4 | 0 | 16 | 0.0625 |
| 14_749463 | 10 | 0.1 | 8 | 0.125 | 11 | 0 | 12 | 0.1667 | 41 | 0.0976 | 6 | 0.3333 | 3 | 0 | 7 | 0.1429 | 6 | 0 | 22 | 0.1364 |
| 14_749466 | 9 | 0 | 12 | 0 | 12 | 0.0833 | 13 | 0.2308 | 46 | 0.087 | 6 | 0.1667 | 4 | 0 | 7 | 0 | 7 | 0.1429 | 24 | 0.0833 |
| 14_749873 | NA | NA | 1 | 1 | 1 | 1 | 2 | 0.5 | 4 | 0.75 | NA | NA | 1 | 1 | NA | NA | NA | NA | 1 | 1 |
| 14_749942 | NA | NA | 1 | 1 | 1 | 0 | 1 | 1 | 3 | 0.6667 | 2 | 0.5 | NA | NA | 1 | 1 | 1 | 1 | 4 | 0.75 |
| 14_749943 | NA | NA | 1 | 1 | 1 | 0 | 1 | 0 | 3 | 0.3333 | 2 | 0 | NA | NA | NA | NA | 1 | 1 | 3 | 0.3333 |
| 14_749943 | NA | NA | 1 | 0 | 1 | 1 | 2 | 0.5 | 4 | 0.5 | 2 | 0.5 | NA | NA | 1 | 0 | 1 | 0 | 4 | 0.25 |
| 14_749943 | NA | NA | 1 | 0 | 1 | 1 | 2 | 0 | 4 | 0.25 | 2 | 0.5 | NA | NA | 1 | 0 | 1 | 0 | 4 | 0.25 |
| 14_749944 | NA | NA | 2 | 1 | 1 | 0 | 2 | 1 | 5 | 0.8 | 2 | 1 | NA | NA | 1 | 1 | 1 | 1 | 4 | 1 |
| 14_749944 | NA | NA | 1 | 0 | 1 | 1 | 1 | 0 | 3 | 0.3333 | 2 | 0 | NA | NA | NA | NA | 1 | 1 | 3 | 0.3333 |
| 14_754668 | NA | NA | 1 | 0 | 1 | 1 | 2 | 0.5 | 4 | 0.5 | NA | NA | 1 | 0 | 2 | 1 | NA | NA | 3 | 0.6667 |
| 14_754668 | NA | NA | 1 | 1 | NA | NA | 2 | 0.5 | 3 | 0.6667 | NA | NA | 1 | 0 | 2 | 0.5 | NA | NA | 3 | 0.3333 |
| 14_755681 | 3 | 0.3333 | 3 | 0.3333 | NA | NA | 6 | 0.6667 | 12 | 0.5 | NA | NA | 1 | 1 | 1 | 0 | NA | NA | 2 | 0.5 |
| 14_755694 | 6 | 0.5 | 5 | 0.4 | NA | NA | 8 | 0.25 | 19 | 0.3684 | NA | NA | 1 | 1 | 1 | 0 | 2 | 0 | 4 | 0.25 |
| 14_755694 | 4 | 0.25 | 3 | 0.6667 | NA | NA | 6 | 0.3333 | 13 | 0.3846 | NA | NA | NA | NA | NA | NA | 1 | 1 | 1 | 1 |
| 14_755694 | 6 | 1 | 6 | 0.6667 | NA | NA | 9 | 1 | 21 | 0.9048 | NA | NA | 1 | 1 | 1 | 1 | 2 | 1 | 4 | 1 |
| 14_755694 | 5 | 0.6 | 5 | 0.2 | NA | NA | 7 | 0.4286 | 17 | 0.4118 | NA | NA | 1 | 0 | 1 | 1 | 1 | 0 | 3 | 0.3333 |
| 14_755695 | 6 | 0 | 5 | 0.2 | NA | NA | 8 | 0.25 | 19 | 0.1579 | NA | NA | 1 | 0 | 1 | 1 | 2 | 0 | 4 | 0.25 |
| 14_755811 | 5 | 0.6 | 8 | 0 | 1 | 0 | 7 | 0 | 21 | 0.1429 | NA | NA | 1 | 0 | 2 | 0 | 3 | 0.3333 | 6 | 0.1667 |
| 14_755811 | 6 | 0.5 | 9 | 0.6667 | 1 | 0 | 9 | 0.7778 | 25 | 0.64 | NA | NA | 1 | 0 | 2 | 0.5 | 3 | 1 | 6 | 0.6667 |
| 14_755812 | 6 | 0.3333 | 8 | 0 | 1 | 0 | 9 | 0.5556 | 24 | 0.2917 | NA | NA | 1 | 0 | 2 | 0 | 3 | 0.6667 | 6 | 0.3333 |
| 14_755813 | 5 | 0.6 | 7 | 0.5714 | 3 | 0.6667 | 10 | 0.9 | 25 | 0.72 | NA | NA | 1 | 1 | 2 | 1 | 3 | 0.6667 | 6 | 0.8333 |
| 14_755813 | 6 | 0.6667 | 8 | 0.75 | 3 | 0.6667 | 9 | 0.5556 | 26 | 0.6538 | NA | NA | 1 | 0 | 2 | 0.5 | 3 | 0.6667 | 6 | 0.5 |
| 14_755815 | 6 | 0.6667 | 7 | 0.7143 | 2 | 0.5 | 10 | 0.3 | 25 | 0.52 | NA | NA | 1 | 0 | 2 | 1 | 3 | 1 | 6 | 0.8333 |
| 14_755817 | 6 | 0.3333 | 6 | 0.1667 | 2 | 1 | 5 | 0.2 | 19 | 0.3158 | NA | NA | 1 | 1 | 2 | 0 | 3 | 0.3333 | 6 | 0.3333 |
| 14_755817 | 6 | 0.5 | 8 | 0.5 | 3 | 0.6667 | 9 | 0.3333 | 26 | 0.4615 | 1 | 1 | 1 | 1 | 3 | 0.6667 | 3 | 1 | 8 | 0.875 |
| 14_755818 | 6 | 0.3333 | 9 | 0.3333 | 2 | 0 | 9 | 0.2222 | 26 | 0.2692 | 1 | 1 | NA | NA | 3 | 0 | 4 | 0.25 | 8 | 0.25 |
| 14_757631 | 1 | 1 | NA | NA | NA | NA | NA | NA | 1 | 1 | 1 | 1 | NA | NA | NA | NA | NA | NA | 1 | 1 |
| 14_757633 | 1 | 1 | NA | NA | NA | NA | NA | NA | 1 | 1 | 1 | 1 | NA | NA | NA | NA | NA | NA | 1 | 1 |
| 14_757634 | 1 | 1 | NA | NA | NA | NA | NA | NA | 1 | 1 | 1 | 1 | NA | NA | NA | NA | NA | NA | 1 | 1 |
| 14_757634 | 1 | 1 | NA | NA | NA | NA | NA | NA | 1 | 1 | 1 | 1 | NA | NA | NA | NA | NA | NA | 1 | 1 |
| 14_757635 | 1 | 1 | NA | NA | NA | NA | NA | NA | 1 | 1 | 1 | 1 | NA | NA | NA | NA | NA | NA | 1 | 1 |
| 14_757636 | 1 | 1 | NA | NA | NA | NA | NA | NA | 1 | 1 | 1 | 1 | NA | NA | NA | NA | NA | NA | 1 | 1 |
| 14_757666 | 1 | 1 | NA | NA | NA | NA | 1 | 0 | 2 | 0.5 | 1 | 0 | NA | NA | NA | NA | 1 | 1 | 2 | 0.5 |

| | | | | | | | | | | | | | | | | | | | | |
|---|---|---|---|---|---|---|---|---|---|---|---|---|---|---|---|---|---|---|---|---|
| 14_757667 | 1 | 0 | NA | NA | NA | NA | 1 | 1 | 2 | 0.5 | 1 | 0 | NA | NA | NA | NA | 1 | 1 | 2 | 0.5 |
| 14_757669 | 2 | 1 | NA | NA | NA | NA | 1 | 1 | 3 | 1 | 1 | 1 | NA | NA | NA | NA | 1 | 0 | 2 | 0.5 |
| 14_759400 | 5 | 0.6 | 6 | 0.1667 | 2 | 0.5 | 1 | 0 | 14 | 0.3571 | 2 | 0 | 3 | 0 | 3 | 1 | 6 | 0.3333 | 14 | 0.3571 |
| 14_759400 | 5 | 0.2 | 6 | 0 | 3 | 0.3333 | 1 | 0 | 15 | 0.1333 | 2 | 0.5 | 3 | 0.3333 | 3 | 0.3333 | 7 | 0 | 15 | 0.2 |
| 14_759400 | 4 | 0 | 6 | 0 | 3 | 0.3333 | 1 | 0 | 14 | 0.0714 | 1 | 0 | 1 | 0 | 2 | 0 | 7 | 0.1429 | 11 | 0.0909 |
| 14_759405 | 2 | 0 | 3 | 0.3333 | 2 | 0 | 1 | 1 | 8 | 0.25 | 2 | 0 | 1 | 1 | 2 | 0.5 | 6 | 0.1667 | 11 | 0.2727 |
| 14_759409 | 1 | 1 | 2 | 1 | 2 | 0.5 | NA | NA | 5 | 0.8 | 1 | 1 | 1 | 1 | 2 | 1 | 4 | 1 | 8 | 1 |
| 14_759410 | 2 | 1 | 3 | 0 | NA | NA | NA | NA | 5 | 0.4 | 2 | 0.5 | NA | NA | 2 | 0 | 4 | 0 | 8 | 0.125 |
| 14_759411 | 2 | 0.5 | 3 | 0.3333 | NA | NA | NA | NA | 5 | 0.4 | 2 | 0 | NA | NA | 1 | 1 | 6 | 0.3333 | 9 | 0.3333 |
| 14_759411 | 1 | 1 | 2 | 0.5 | NA | NA | NA | NA | 3 | 0.6667 | 2 | 0.5 | 1 | 1 | 2 | 0.5 | 6 | 0.1667 | 11 | 0.3636 |
| 14_759412 | 1 | 1 | NA | NA | NA | NA | NA | NA | 1 | 1 | 2 | 0.5 | 1 | 1 | NA | NA | 2 | 1 | 5 | 0.8 |
| 14_759413 | NA | NA | 2 | 0.5 | NA | NA | NA | NA | 2 | 0.5 | 2 | 1 | 1 | 1 | 2 | 1 | 4 | 1 | 9 | 1 |
| 14_759414 | 1 | 0 | 2 | 0.5 | NA | NA | NA | NA | 3 | 0.3333 | 1 | 0 | NA | NA | 2 | 0.5 | 5 | 0 | 8 | 0.125 |
| 14_759976 | 1 | 1 | NA | NA | 3 | 0.3333 | 2 | 0 | 6 | 0.3333 | 3 | 0.3333 | 1 | 0 | 4 | 0.25 | 3 | 0 | 11 | 0.1818 |
| 14_759996 | NA | NA | NA | NA | NA | NA | 2 | 1 | 2 | 1 | 2 | 1 | 1 | 1 | 4 | 1 | 1 | 1 | 8 | 1 |
| 14_760000 | NA | NA | NA | NA | NA | NA | 2 | 0.5 | 2 | 0.5 | 2 | 0 | NA | NA | 3 | 0.3333 | 1 | 0 | 6 | 0.1667 |
| 14_760820 | 1 | 0 | NA | NA | NA | NA | 2 | 1 | 3 | 0.6667 | NA | NA | NA | NA | NA | NA | 2 | 0.5 | 2 | 0.5 |
| 14_760821 | 1 | 0 | NA | NA | NA | NA | 2 | 1 | 3 | 0.6667 | NA | NA | NA | NA | NA | NA | 2 | 0.5 | 2 | 0.5 |
| 14_760822 | 1 | 1 | NA | NA | NA | NA | 2 | 1 | 3 | 1 | NA | NA | NA | NA | NA | NA | 2 | 1 | 2 | 1 |
| 14_760823 | NA | NA | NA | NA | NA | NA | 1 | 1 | 1 | 1 | NA | NA | NA | NA | NA | NA | 2 | 0.5 | 2 | 0.5 |
| 14_760823 | 1 | 0 | NA | NA | NA | NA | 1 | 1 | 2 | 0.5 | NA | NA | NA | NA | NA | NA | 1 | 1 | 1 | 1 |
| 14_760825 | 1 | 0 | NA | NA | NA | NA | 1 | 1 | 2 | 0.5 | NA | NA | NA | NA | NA | NA | 2 | 1 | 2 | 1 |
| 14_760826 | NA | NA | NA | NA | NA | NA | 1 | 1 | 1 | 1 | NA | NA | NA | NA | NA | NA | 1 | 1 | 1 | 1 |
| 14_760826 | NA | NA | NA | NA | NA | NA | 1 | 1 | 1 | 1 | NA | NA | NA | NA | NA | NA | 1 | 1 | 1 | 1 |
| 14_760856 | 161 | 0.0559 | 184 | 0.0761 | 179 | 0.0894 | 176 | 0.0455 | 700 | 0.0671 | 150 | 0.0467 | 139 | 0.0432 | 152 | 0.0789 | 162 | 0.0494 | 603 | 0.0547 |
| 14_760856 | 171 | 0.4561 | 186 | 0.4624 | 194 | 0.4794 | 177 | 0.4181 | 728 | 0.4547 | 157 | 0.4522 | 142 | 0.3944 | 155 | 0.5032 | 173 | 0.4682 | 627 | 0.4561 |
| 14_761345 | 1 | 1 | 3 | 0 | 2 | 0 | 1 | 0 | 7 | 0.1429 | 3 | 0.3333 | 1 | 0 | 1 | 0 | 2 | 0.5 | 7 | 0.2857 |
| 14_761347 | 2 | 0.5 | 3 | 0.6667 | 2 | 1 | 1 | 1 | 8 | 0.75 | 2 | 0.5 | 1 | 0 | 1 | 1 | 2 | 0.5 | 6 | 0.5 |
| 14_761347 | 2 | 1 | 3 | 0.6667 | 2 | 1 | NA | NA | 7 | 0.8571 | 2 | 1 | 1 | 0 | 1 | 0 | 1 | 1 | 5 | 0.6 |
| 14_761348 | 2 | 1 | 1 | 0 | 3 | 0.3333 | 1 | 0 | 7 | 0.4286 | 2 | 0.5 | 5 | 0 | NA | NA | 2 | 0.5 | 9 | 0.2222 |
| 14_761348 | 1 | 1 | 2 | 0 | 3 | 0.3333 | 2 | 0.5 | 8 | 0.375 | 2 | 1 | 5 | 0.2 | 1 | 0 | 2 | 1 | 10 | 0.5 |
| 14_761349 | 2 | 0 | 3 | 0.6667 | 3 | 0.3333 | 2 | 0.5 | 10 | 0.4 | 2 | 0.5 | 5 | 0 | 1 | 1 | 2 | 0.5 | 10 | 0.3 |
| 14_762001 | 1 | 0 | 1 | 0 | NA | NA | 2 | 0.5 | 4 | 0.25 | 2 | 1 | 2 | 1 | 2 | 0.5 | 1 | 1 | 7 | 0.8571 |
| 14_762001 | 1 | 0 | 1 | 1 | 1 | 0 | 2 | 0.5 | 5 | 0.4 | 3 | 0 | 2 | 0.5 | 2 | 0 | 1 | 0 | 8 | 0.125 |
| 14_762004 | 2 | 1 | 2 | 1 | 1 | 1 | 2 | 0.5 | 7 | 0.8571 | 3 | 0.6667 | 2 | 1 | 2 | 1 | 1 | 1 | 8 | 0.875 |
| 14_762004 | 1 | 1 | 2 | 1 | 1 | 1 | 2 | 1 | 6 | 1 | 3 | 0.6667 | 2 | 1 | 2 | 1 | NA | NA | 7 | 0.8571 |
| 14_762006 | 2 | 0.5 | 2 | 1 | 1 | 0 | 1 | 0 | 6 | 0.5 | 2 | 0 | NA | NA | NA | NA | 1 | 1 | 3 | 0.3333 |
| 14_762007 | 1 | 0 | 2 | 1 | 1 | 1 | 2 | 1 | 6 | 0.8333 | 3 | 0.6667 | 1 | 1 | 2 | 1 | 1 | 1 | 7 | 0.8571 |
| 14_762007 | 2 | 1 | 2 | 1 | 1 | 1 | 1 | 1 | 6 | 1 | 3 | 1 | 2 | 1 | 1 | 1 | 1 | 1 | 7 | 1 |
| 14_762010 | 2 | 1 | 2 | 1 | 1 | 1 | 1 | 1 | 6 | 1 | 2 | 0.5 | 2 | 1 | 2 | 1 | 2 | 1 | 8 | 0.875 |
| 14_762012 | 3 | 0.6667 | 3 | 1 | 1 | 1 | 1 | 1 | 8 | 0.875 | 2 | 1 | 2 | 1 | 2 | 1 | 3 | 1 | 9 | 1 |
| 14_762012 | 2 | 1 | 3 | 1 | 2 | 1 | 2 | 1 | 9 | 1 | NA | NA | 2 | 0.5 | 2 | 1 | 2 | 0.5 | 6 | 0.6667 |
| 14_762013 | 3 | 0.3333 | 1 | 1 | 1 | 0 | 2 | 0.5 | 7 | 0.4286 | 1 | 0 | 2 | 1 | 1 | 0 | 2 | 0.5 | 6 | 0.5 |
| 14_763239 | 3 | 0 | 5 | 0 | 2 | 0.5 | 7 | 0 | 17 | 0.0588 | 3 | 0 | 2 | 0 | 3 | 0.3333 | 5 | 0 | 13 | 0.0769 |
| 14_763240 | 3 | 0 | 3 | 0 | 2 | 0 | 8 | 0.125 | 16 | 0.0625 | 3 | 0 | 4 | 0 | 5 | 0.2 | 6 | 0 | 18 | 0.0556 |
| 14_763244 | 2 | 0 | 4 | 0.25 | NA | NA | 6 | 0 | 12 | 0.0833 | 4 | 0.25 | 4 | 0 | 4 | 0 | 5 | 0 | 17 | 0.0588 |
| 14_763258 | NA | NA | 2 | 0 | 2 | 0 | 4 | 0.25 | 8 | 0.125 | 5 | 0.2 | 3 | 0.3333 | 3 | 0 | 7 | 0 | 18 | 0.1111 |
| 14_763281 | 1 | 1 | NA | NA | NA | NA | 2 | 0.5 | 3 | 0.6667 | 2 | 0.5 | NA | NA | 2 | 1 | 3 | 0.6667 | 7 | 0.7143 |
| 14_763705 | 3 | 0 | 3 | 0.3333 | 1 | 0 | 2 | 0 | 9 | 0.1111 | 3 | 1 | 1 | 0 | 1 | 0 | 1 | 0 | 6 | 0.5 |
| 14_763722 | 4 | 0.5 | 3 | 1 | 1 | 1 | 2 | 0 | 10 | 0.6 | 3 | 1 | 1 | 1 | 1 | 1 | 1 | 0 | 6 | 0.8333 |
| 14_763723 | 3 | 1 | 3 | 0.6667 | 1 | 1 | 1 | 1 | 8 | 0.875 | 3 | 1 | 1 | 1 | 1 | 1 | NA | NA | 5 | 1 |
| 14_766532 | NA | NA | 3 | 1 | 2 | 1 | 1 | 0 | 6 | 0.8333 | 1 | 1 | NA | NA | 1 | 1 | NA | NA | 2 | 1 |
| 14_766532 | NA | NA | 3 | 1 | 2 | 1 | 2 | 1 | 7 | 1 | 1 | 1 | NA | NA | 1 | 1 | NA | NA | 2 | 1 |
| 14_766535 | NA | NA | 4 | 0.25 | 1 | 0 | 2 | 0 | 7 | 0.1429 | 1 | 1 | NA | NA | 2 | 1 | NA | NA | 3 | 1 |
| 14_766537 | NA | NA | 5 | 0.2 | 3 | 0.3333 | 1 | 0 | 9 | 0.2222 | 2 | 0 | 1 | 0 | 3 | 0.3333 | NA | NA | 6 | 0.1667 |
| 14_766538 | 1 | 1 | 5 | 0.2 | 2 | 1 | 2 | 1 | 10 | 0.6 | 2 | 0.5 | 1 | 1 | 3 | 0.3333 | 1 | 1 | 7 | 0.5714 |
| 14_766538 | 1 | 1 | 5 | 0.2 | 2 | 0.5 | 2 | 0 | 10 | 0.3 | 2 | 0.5 | 1 | 0 | 3 | 0.6667 | NA | NA | 6 | 0.5 |
| 14_766540 | NA | NA | 7 | 0.1429 | 2 | 0.5 | 3 | 0.3333 | 12 | 0.25 | 2 | 0.5 | 1 | 1 | 2 | 0 | 1 | 1 | 6 | 0.5 |
| 14_766540 | NA | NA | 7 | 0.4286 | 3 | 0.3333 | 3 | 0 | 13 | 0.3077 | 3 | 0.6667 | 1 | 1 | 3 | 0.6667 | 1 | 1 | 8 | 0.75 |
| 14_766543 | 1 | 0 | 7 | 0.4286 | 2 | 0.5 | 3 | 0 | 13 | 0.3077 | 3 | 0 | 1 | 0 | 3 | 0.3333 | 1 | 1 | 8 | 0.25 |
| 14_766545 | NA | NA | 5 | 0.6 | 2 | 1 | 1 | 1 | 8 | 0.75 | 2 | 0.5 | NA | NA | 3 | 1 | NA | NA | 5 | 0.8 |
| 14_766547 | 1 | 0 | 4 | 0.5 | 4 | 0 | 3 | 1 | 12 | 0.4167 | 3 | 0 | 2 | 0 | 5 | 0.4 | NA | NA | 10 | 0.2 |
| 14_766548 | 1 | 1 | 7 | 1 | 4 | 1 | 4 | 1 | 16 | 1 | 4 | 1 | 2 | 1 | 4 | 1 | 1 | 0 | 11 | 0.9091 |
| 14_766552 | 1 | 1 | 7 | 0.1429 | 4 | 0.25 | 5 | 0.6 | 17 | 0.3529 | 6 | 0.3333 | 3 | 1 | 5 | 0.2 | 1 | 0 | 15 | 0.4 |
| 14_766553 | 1 | 0 | 7 | 0.1429 | 4 | 0.25 | 4 | 0.25 | 16 | 0.1875 | 8 | 0.25 | 2 | 0.5 | 5 | 0.2 | 2 | 0.5 | 17 | 0.2941 |

| | | | | | | | | | | | | | | | | | | | | |
|---|---|---|---|---|---|---|---|---|---|---|---|---|---|---|---|---|---|---|---|---|
| 14_766553 | 1 | 0 | 6 | 0.5 | 4 | 0.25 | 4 | 0.25 | 15 | 0.3333 | 6 | 0 | 2 | 0 | 4 | 0 | 2 | 0.5 | 14 | 0.0714 |
| 14_766554 | 2 | 1 | 6 | 1 | 3 | 1 | 5 | 1 | 16 | 1 | 7 | 1 | 2 | 1 | 5 | 1 | 2 | 1 | 16 | 1 |
| 14_766556 | 3 | 0.3333 | 6 | 0 | 4 | 0 | 4 | 0.25 | 17 | 0.1176 | 6 | 0.1667 | 2 | 0 | 4 | 0.25 | 3 | 0.3333 | 15 | 0.2 |
| 14_766559 | 5 | 0.6 | 4 | 0.5 | 4 | 0.25 | 5 | 1 | 18 | 0.6111 | 5 | 0.6 | 2 | 1 | 2 | 0 | 2 | 0.5 | 11 | 0.5455 |
| 14_766560 | 5 | 0 | 7 | 0.1429 | 4 | 0 | 6 | 0.1667 | 22 | 0.0909 | 8 | 0 | 3 | 0 | 6 | 0.3333 | 3 | 0 | 20 | 0.1 |
| 14_766563 | 5 | 0.8 | 7 | 0.8571 | 4 | 0.75 | 6 | 1 | 22 | 0.8636 | 8 | 0.5 | 3 | 1 | 6 | 0.5 | 4 | 0.25 | 21 | 0.5238 |
| 14_766563 | 4 | 0.5 | 7 | 0.7143 | 3 | 0.3333 | 6 | 0.3333 | 20 | 0.5 | 5 | 0.2 | 3 | 0.6667 | 4 | 0.75 | 4 | 0.75 | 16 | 0.5625 |
| 14_766775 | 1 | 1 | 1 | 0 | NA | NA | 2 | 0 | 4 | 0.25 | 1 | 1 | 2 | 0.5 | 2 | 0 | 1 | 0 | 6 | 0.3333 |
| 14_766776 | 1 | 1 | 1 | 0 | NA | NA | 2 | 0.5 | 4 | 0.5 | 2 | 1 | NA | NA | 2 | 0.5 | NA | NA | 4 | 0.75 |
| 14_766777 | NA | NA | 1 | 1 | NA | NA | 1 | 1 | 2 | 1 | 1 | 1 | 1 | 1 | 2 | 1 | NA | NA | 4 | 1 |
| 14_766784 | 2 | 0.5 | 1 | 0 | NA | NA | 2 | 0 | 5 | 0.2 | 2 | 0.5 | 2 | 0 | 1 | 0 | 1 | 0 | 6 | 0.1667 |
| 14_767436 | 1 | 1 | 1 | 1 | NA | NA | 2 | 0.5 | 4 | 0.75 | NA | NA | 1 | 1 | NA | NA | NA | NA | 1 | 1 |
| 14_767438 | 1 | 0 | 1 | 0 | NA | NA | 2 | 0.5 | 4 | 0.25 | 1 | 0 | 1 | 1 | NA | NA | 1 | 0 | 3 | 0.3333 |
| 14_767439 | 66 | 0.1364 | 67 | 0.194 | 53 | 0.2075 | 93 | 0.2151 | 279 | 0.19 | 66 | 0.2576 | 77 | 0.3117 | 88 | 0.1477 | 58 | 0.1724 | 289 | 0.2215 |
| 14_767440 | 68 | 0.0294 | 64 | 0.125 | 51 | 0.098 | 90 | 0.0889 | 273 | 0.0842 | 57 | 0.0351 | 74 | 0.0541 | 91 | 0.1099 | 55 | 0.0727 | 277 | 0.0722 |
| 14_767440 | 64 | 0.0781 | 61 | 0.0984 | 45 | 0.0222 | 83 | 0.012 | 253 | 0.0514 | 62 | 0.0968 | 68 | 0.0735 | 91 | 0.022 | 58 | 0.0517 | 279 | 0.0573 |
| 14_767441 | 74 | 0.6081 | 64 | 0.5312 | 53 | 0.7736 | 90 | 0.5333 | 281 | 0.5979 | 64 | 0.5156 | 83 | 0.5542 | 98 | 0.6122 | 59 | 0.4407 | 304 | 0.5428 |
| 14_767441 | 71 | 0.5352 | 65 | 0.5846 | 49 | 0.6122 | 85 | 0.5059 | 270 | 0.5519 | 59 | 0.5763 | 72 | 0.5556 | 91 | 0.6593 | 63 | 0.6032 | 285 | 0.6035 |
| 14_767442 | 79 | 0.8101 | 65 | 0.7538 | 52 | 0.7692 | 89 | 0.7303 | 285 | 0.7649 | 62 | 0.7419 | 81 | 0.7284 | 95 | 0.7579 | 65 | 0.7538 | 303 | 0.7459 |
| 14_767445 | 68 | 0.6618 | 55 | 0.7273 | 49 | 0.7143 | 85 | 0.6941 | 257 | 0.6965 | 58 | 0.6379 | 71 | 0.7887 | 82 | 0.7073 | 53 | 0.6604 | 264 | 0.7045 |
| 14_767445 | 57 | 0.6667 | 53 | 0.6981 | 49 | 0.6327 | 72 | 0.7361 | 231 | 0.6883 | 52 | 0.6538 | 63 | 0.6825 | 75 | 0.7067 | 54 | 0.7407 | 244 | 0.6967 |
| 14_767446 | 98 | 0.4388 | 77 | 0.5455 | 59 | 0.3559 | 110 | 0.4182 | 344 | 0.4419 | 71 | 0.4225 | 102 | 0.4902 | 108 | 0.5463 | 71 | 0.493 | 352 | 0.4943 |
| 14_767447 | 111 | 0.1351 | 84 | 0.131 | 72 | 0.0417 | 120 | 0.0583 | 387 | 0.093 | 85 | 0.1294 | 113 | 0.0973 | 119 | 0.1008 | 84 | 0.0595 | 401 | 0.0973 |
| 14_767448 | 106 | 0.3962 | 83 | 0.5422 | 71 | 0.3521 | 114 | 0.3772 | 374 | 0.4144 | 80 | 0.4 | 103 | 0.4175 | 107 | 0.4393 | 81 | 0.4444 | 371 | 0.4259 |
| 14_767449 | 107 | 0.486 | 78 | 0.5 | 68 | 0.5 | 106 | 0.4057 | 359 | 0.468 | 81 | 0.4938 | 106 | 0.3302 | 109 | 0.3945 | 77 | 0.4156 | 373 | 0.4021 |
| 14_767704 | 27 | 0.8889 | 22 | 0.9091 | 37 | 0.8649 | 38 | 0.8684 | 124 | 0.879 | 20 | 0.85 | 34 | 0.8529 | 22 | 0.9091 | 33 | 0.8788 | 109 | 0.8716 |
| 14_773490 | NA | NA | NA | NA | 2 | 1 | NA | NA | 2 | 1 | NA | NA | NA | NA | 3 | 1 | 1 | 1 | 4 | 1 |
| 14_779589 | 1 | 0 | 3 | 0.3333 | 4 | 0.5 | 2 | 0 | 10 | 0.3 | 3 | 0 | 3 | 0 | 4 | 0.25 | 5 | 0.4 | 15 | 0.2 |
| 14_779589 | 1 | 0 | 3 | 0.3333 | 5 | 0.4 | 2 | 0 | 11 | 0.2727 | 3 | 0.3333 | 3 | 0 | 3 | 0.6667 | 6 | 0.1667 | 15 | 0.2667 |
| 14_779590 | 2 | 1 | 3 | 0.6667 | 5 | 0.4 | NA | NA | 10 | 0.6 | 3 | 0.6667 | 2 | 0.5 | 4 | 0.75 | 5 | 0.2 | 14 | 0.5 |
| 14_780525 | 1 | 1 | NA | NA | NA | NA | NA | NA | 1 | 1 | 1 | 1 | NA | NA | NA | NA | 1 | 0 | 2 | 0.5 |
| 14_780582 | NA | NA | 3 | 0.6667 | NA | NA | NA | NA | 3 | 0.6667 | NA | NA | 3 | 0.3333 | 1 | 0 | 2 | 0 | 6 | 0.1667 |
| 14_780620 | 1 | 1 | 1 | 1 | 1 | 0 | NA | NA | 3 | 0.6667 | NA | NA | 1 | 1 | 1 | 1 | 4 | 0.25 | 6 | 0.5 |
| 14_780634 | 1 | 1 | 1 | 1 | NA | NA | NA | NA | 2 | 1 | NA | NA | NA | NA | 1 | 1 | 1 | 1 | 2 | 1 |
| 14_780636 | 2 | 0.5 | 6 | 0.1667 | 2 | 0.5 | 5 | 0.2 | 15 | 0.2667 | 1 | 0 | NA | NA | 3 | 0 | 3 | 0.3333 | 7 | 0.1429 |
| 14_780638 | 12 | 0.5833 | 17 | 0.5882 | 7 | 0.4286 | 10 | 0.6 | 46 | 0.5652 | 9 | 0.8889 | 6 | 0.8333 | 13 | 0.6154 | 9 | 0.6667 | 37 | 0.7297 |
| 14_787303 | 19 | 0.2105 | 20 | 0.1 | 14 | 0.1429 | 24 | 0.25 | 77 | 0.1818 | 11 | 0.1818 | 23 | 0.087 | 15 | 0.2 | 15 | 0.1333 | 64 | 0.1406 |
| 14_787306 | 18 | 0.1667 | 24 | 0.2083 | 14 | 0 | 25 | 0.12 | 81 | 0.1358 | 10 | 0.1 | 25 | 0.16 | 15 | 0.2667 | 17 | 0.1176 | 67 | 0.1642 |
| 14_787319 | 11 | 0.3636 | 12 | 0 | 7 | 0 | 19 | 0.1053 | 49 | 0.1224 | 6 | 0.1667 | 18 | 0.3333 | 9 | 0.1111 | 11 | 0.2727 | 44 | 0.25 |
| 14_787332 | 9 | 0.2222 | 9 | 0.1111 | 4 | 0 | 16 | 0.0625 | 38 | 0.1053 | 7 | 0 | 21 | 0.1429 | 11 | 0.0909 | 12 | 0.3333 | 51 | 0.1569 |
| 14_787474 | 4 | 0.25 | 9 | 0 | 4 | 0.25 | 13 | 0.1538 | 30 | 0.1333 | 6 | 0.1667 | 16 | 0.25 | 5 | 0.2 | 8 | 0 | 35 | 0.1714 |
| 14_787475 | 5 | 0 | 8 | 0 | 3 | 0 | 14 | 0.2143 | 30 | 0.1 | 4 | 0 | 16 | 0.25 | 4 | 0 | 9 | 0.2222 | 33 | 0.1818 |
| 14_787477 | 6 | 0.1667 | 8 | 0.125 | 4 | 0.75 | 10 | 0.2 | 28 | 0.25 | 5 | 0.2 | 16 | 0.1875 | 5 | 0.2 | 7 | 0 | 33 | 0.1515 |
| 14_787478 | 4 | 0.25 | 5 | 0 | 2 | 1 | 10 | 0.3 | 21 | 0.2857 | 4 | 0 | 12 | 0.25 | 6 | 0.1667 | 6 | 0.1667 | 28 | 0.1786 |
| 14_787479 | 5 | 0.6 | 6 | 0.8333 | 4 | 0.25 | 10 | 0.4 | 25 | 0.52 | 4 | 0 | 16 | 0.375 | 6 | 0.5 | 5 | 0.6 | 31 | 0.3871 |
| 14_787479 | 6 | 0.1667 | 8 | 0 | 4 | 0.25 | 11 | 0.1818 | 29 | 0.1379 | 4 | 0.25 | 14 | 0.0714 | 6 | 0 | 7 | 0.1429 | 31 | 0.0968 |
| 14_787480 | 4 | 1 | 5 | 0.8 | 3 | 1 | 10 | 1 | 22 | 0.9545 | 5 | 1 | 10 | 1 | 4 | 1 | 4 | 1 | 23 | 1 |
| 14_787480 | 3 | 1 | 9 | 1 | 3 | 1 | 11 | 1 | 26 | 1 | 4 | 0.75 | 12 | 1 | 5 | 0.8 | 7 | 1 | 28 | 0.9286 |
| 14_787481 | 5 | 0.2 | 7 | 0.1429 | 4 | 0 | 11 | 0.1818 | 27 | 0.1481 | 5 | 0.2 | 14 | 0.1429 | 6 | 0.1667 | 7 | 0.1429 | 32 | 0.1562 |
| 14_787482 | 5 | 0.8 | 5 | 0.8 | 4 | 1 | 8 | 0.625 | 22 | 0.7727 | 3 | 0.6667 | 14 | 0.4286 | 5 | 0.8 | 7 | 0.7143 | 29 | 0.5862 |
| 14_787484 | 6 | 0.5 | 7 | 0.7143 | 3 | 1 | 9 | 0.3333 | 25 | 0.56 | 2 | 0.5 | 14 | 0.7143 | 5 | 0.6 | 7 | 0.5714 | 28 | 0.6429 |
| 14_787485 | 3 | 0.6667 | 6 | 0.1667 | 2 | 0 | 8 | 0.25 | 19 | 0.2632 | 2 | 0.5 | 11 | 0 | 5 | 0.4 | 7 | 0.2857 | 25 | 0.2 |
| 14_787487 | 5 | 0 | 6 | 0 | 4 | 0.25 | 8 | 0.125 | 23 | 0.087 | 2 | 0 | 11 | 0.1818 | 5 | 0.2 | 7 | 0.1429 | 25 | 0.16 |
| 14_787490 | 4 | 0.5 | 7 | 0.2857 | 3 | 0.6667 | 9 | 0.2222 | 23 | 0.3478 | 2 | 0.5 | 11 | 0.3636 | 5 | 0.8 | 6 | 0.3333 | 24 | 0.4583 |
| 14_787491 | 4 | 0.25 | 5 | 0.8 | 1 | 0 | 6 | 0.8333 | 16 | 0.625 | 2 | 1 | 7 | 0.8571 | 2 | 0 | 6 | 0.8333 | 17 | 0.7647 |
| 14_787492 | 2 | 0 | 5 | 0 | 1 | 0 | 6 | 0.6667 | 14 | 0.2857 | 2 | 0.5 | 10 | 0.4 | 3 | 0 | 4 | 0.25 | 19 | 0.3158 |
| 14_787492 | 3 | 0 | 6 | 0.5 | 2 | 0.5 | 6 | 0 | 17 | 0.2353 | 2 | 0 | 9 | 0.1111 | 4 | 0.25 | 5 | 0 | 20 | 0.1 |
| 14_787494 | 3 | 0.3333 | 6 | 0 | 3 | 0.6667 | 7 | 0.2857 | 19 | 0.2632 | 2 | 0.5 | 10 | 0.2 | 4 | 0.75 | 5 | 0.6 | 21 | 0.4286 |
| 14_787494 | 3 | 0 | 6 | 0 | 3 | 0.3333 | 7 | 0 | 19 | 0.0526 | 1 | 1 | 10 | 0 | 4 | 0.25 | 6 | 0 | 21 | 0.0952 |
| 14_787496 | 3 | 0.3333 | 6 | 0.6667 | 2 | 1 | 7 | 0.8571 | 18 | 0.7222 | 2 | 1 | 11 | 0.6364 | 3 | 0.6667 | 6 | 0.8333 | 22 | 0.7273 |
| 14_787497 | 3 | 0 | 5 | 0 | 3 | 0.3333 | 4 | 0.25 | 15 | 0.1333 | 2 | 0 | 10 | 0.2 | 4 | 0.5 | 6 | 0 | 22 | 0.1818 |
| 14_787498 | 3 | 0 | 5 | 0 | 3 | 0 | 5 | 0.2 | 16 | 0.0625 | 2 | 0 | 10 | 0.3 | 3 | 0.3333 | 6 | 0 | 21 | 0.1905 |
| 14_787498 | 2 | 0 | 5 | 0.6 | 2 | 0.5 | 4 | 0.25 | 13 | 0.3846 | 2 | 0.5 | 8 | 0.25 | 3 | 0.3333 | 5 | 0 | 18 | 0.2222 |
| 14_787502 | 1 | 1 | 4 | 0.5 | 3 | 0.3333 | 2 | 0 | 10 | 0.4 | 1 | 1 | 7 | 0.1429 | 2 | 0.5 | 4 | 0.25 | 14 | 0.2857 |
| 14_787504 | 1 | 0 | 3 | 0 | 3 | 0 | 5 | 0.2 | 12 | 0.0833 | 2 | 0 | 6 | 0.3333 | 1 | 0 | 3 | 0 | 12 | 0.1667 |

| | | | | | | | | | | | | | | | | | | | | |
|---|---|---|---|---|---|---|---|---|---|---|---|---|---|---|---|---|---|---|---|---|
| 14_787504 | 1 | 0 | 4 | 0.25 | 3 | 0 | 4 | 0.5 | 12 | 0.25 | 2 | 0 | 6 | 0.1667 | 2 | 0 | 5 | 0 | 15 | 0.0667 |
| 14_787505 | 1 | 1 | 4 | 0.75 | 3 | 1 | 4 | 1 | 12 | 0.9167 | 1 | 1 | 8 | 0.625 | 2 | 1 | 4 | 1 | 15 | 0.8 |
| 14_787506 | 1 | 1 | 2 | 1 | 2 | 1 | 2 | 1 | 7 | 1 | 1 | 1 | 7 | 0.7143 | 1 | 1 | 3 | 1 | 12 | 0.8333 |
| 14_787506 | 1 | 1 | 4 | 0.75 | 3 | 0.6667 | 4 | 0.5 | 12 | 0.6667 | 2 | 0 | 8 | 0.625 | 2 | 1 | 4 | 0.5 | 16 | 0.5625 |
| 14_787508 | 1 | 0 | 4 | 0.5 | 3 | 1 | 4 | 0.75 | 12 | 0.6667 | 2 | 0.5 | 8 | 0.875 | 2 | 0.5 | 4 | 1 | 16 | 0.8125 |
| 14_787512 | 1 | 0 | 4 | 0.5 | 3 | 0.3333 | 5 | 0 | 13 | 0.2308 | 1 | 0 | 7 | 0.5714 | 2 | 0.5 | 4 | 0.25 | 14 | 0.4286 |
| 14_787512 | 1 | 0 | 3 | 0 | 3 | 0 | 5 | 0.2 | 12 | 0.0833 | 1 | 0 | 7 | 0.4286 | 1 | 0 | 4 | 0 | 13 | 0.2308 |
| 14_787512 | 1 | 0 | 4 | 0.75 | 1 | 1 | 1 | 0 | 7 | 0.5714 | 1 | 1 | 6 | 0.5 | 1 | 1 | 4 | 0.75 | 12 | 0.6667 |
| 14_787514 | 1 | 0 | 4 | 0 | 2 | 0.5 | 5 | 0 | 12 | 0.0833 | 1 | 1 | 6 | 0 | 2 | 0 | 3 | 0 | 12 | 0.0833 |
| 14_787515( | NA | NA | 4 | 1 | 3 | 0 | 4 | 0.75 | 11 | 0.6364 | 1 | 1 | 5 | 0.8 | 1 | 0 | 3 | 0.3333 | 10 | 0.6 |
| 14_787515( | NA | NA | 4 | 0 | 3 | 0.3333 | 5 | 0 | 12 | 0.0833 | 1 | 0 | 6 | 0 | 2 | 0 | 3 | 0.3333 | 12 | 0.0833 |
| 14_787524 | 1 | 0 | 3 | 0.6667 | 3 | 0.3333 | 3 | 0.3333 | 10 | 0.4 | 1 | 0 | 7 | 0.4286 | 1 | 0 | 3 | 0.6667 | 12 | 0.4167 |
| 14_787968 | 11 | 0 | 10 | 0.2 | 12 | 0 | 10 | 0.2 | 43 | 0.093 | 3 | 0.3333 | 9 | 0 | 11 | 0 | 9 | 0.2222 | 32 | 0.0938 |
| 14_789705 | NA | NA | 1 | 1 | NA | NA | NA | NA | 1 | 1 | NA | NA | NA | NA | NA | NA | 1 | 1 | 1 | 1 |
| 14_789707 | NA | NA | 1 | 1 | NA | NA | NA | NA | 1 | 1 | NA | NA | NA | NA | NA | NA | 1 | 1 | 1 | 1 |
| 14_789709 | NA | NA | 1 | 1 | NA | NA | NA | NA | 1 | 1 | NA | NA | NA | NA | NA | NA | 1 | 1 | 1 | 1 |
| 14_789709 | NA | NA | 1 | 1 | NA | NA | NA | NA | 1 | 1 | NA | NA | NA | NA | NA | NA | 1 | 1 | 1 | 1 |
| 14_789712 | NA | NA | 1 | 1 | NA | NA | NA | NA | 1 | 1 | NA | NA | NA | NA | NA | NA | 1 | 1 | 1 | 1 |
| 14_789712 | NA | NA | 1 | 1 | NA | NA | NA | NA | 1 | 1 | NA | NA | NA | NA | NA | NA | 1 | 1 | 1 | 1 |
| 14_789714 | NA | NA | 1 | 1 | NA | NA | NA | NA | 1 | 1 | NA | NA | NA | NA | NA | NA | 1 | 1 | 1 | 1 |
| 14_789725( | NA | NA | 1 | 1 | NA | NA | NA | NA | 1 | 1 | NA | NA | NA | NA | NA | NA | 1 | 1 | 1 | 1 |
| 14_794388 | 4 | 0.5 | 1 | 1 | 1 | 1 | 3 | 0.3333 | 9 | 0.5556 | 1 | 1 | NA | NA | 3 | 0.3333 | 3 | 0.3333 | 7 | 0.4286 |
| 14_794389 | 4 | 0.25 | 1 | 0 | 1 | 1 | 3 | 0 | 9 | 0.2222 | 2 | 0 | NA | NA | 3 | 0.3333 | 4 | 0.25 | 9 | 0.2222 |
| 14_794390 | 4 | 0.5 | 1 | 1 | NA | NA | 3 | 0 | 8 | 0.375 | 2 | 0.5 | NA | NA | 3 | 0 | 1 | 0 | 6 | 0.1667 |
| 14_794391 | 4 | 0.5 | NA | NA | NA | NA | 2 | 0 | 6 | 0.3333 | 2 | 0.5 | NA | NA | 3 | 0 | 1 | 1 | 6 | 0.3333 |
| 14_794394 | 4 | 0.25 | 1 | 0 | NA | NA | 4 | 0 | 9 | 0.1111 | 2 | 0.5 | NA | NA | 4 | 0 | 1 | 0 | 7 | 0.1429 |
| 14_795731( | NA | NA | 2 | 0 | NA | NA | 1 | 1 | 3 | 0.3333 | NA | NA | 1 | 1 | NA | NA | 2 | 0 | 3 | 0.3333 |
| 14_795731 | NA | NA | 2 | 0 | NA | NA | 1 | 1 | 3 | 0.3333 | NA | NA | 1 | 1 | NA | NA | 2 | 0.5 | 3 | 0.6667 |
| 14_796389 | NA | NA | 1 | 1 | 1 | 0 | NA | NA | 2 | 0.5 | 1 | 1 | 1 | 0 | NA | NA | 2 | 0 | 4 | 0.25 |
| 14_796390( | NA | NA | 1 | 1 | 2 | 0 | NA | NA | 3 | 0.3333 | 1 | 0 | 1 | 1 | NA | NA | 3 | 0.6667 | 5 | 0.6 |
| 14_796609 | NA | NA | 2 | 0 | 2 | 0 | 1 | 1 | 5 | 0.2 | 1 | 0 | 1 | 0 | 1 | 0 | 1 | 1 | 4 | 0.25 |
| 14_796639 | 2 | 0 | 1 | 0 | 4 | 0.25 | 4 | 0 | 11 | 0.0909 | 1 | 0 | 4 | 0 | 2 | 0.5 | 1 | 1 | 8 | 0.25 |
| 14_796645 | 1 | 0 | 3 | 0.3333 | 5 | 0.6 | 7 | 0.7143 | 16 | 0.5625 | 1 | 1 | 4 | 0.75 | 3 | 0.6667 | 2 | 0.5 | 10 | 0.7 |
| 14_796648 | 2 | 0 | 4 | 0.75 | 5 | 0.6 | 4 | 0.5 | 15 | 0.5333 | 1 | 1 | 5 | 0.4 | 3 | 0.6667 | 2 | 0.5 | 11 | 0.5455 |
| 14_796648 | 1 | 0 | 4 | 0.75 | 6 | 0.5 | 8 | 0.5 | 19 | 0.5263 | 2 | 0.5 | 6 | 0.6667 | 3 | 0.6667 | 2 | 1 | 13 | 0.6923 |
| 14_796649 | 2 | 0 | 4 | 0.75 | 5 | 0.4 | 4 | 0.5 | 15 | 0.4667 | 1 | 0 | 7 | 0.2857 | 2 | 0.5 | 2 | 0.5 | 12 | 0.3333 |
| 14_796650 | NA | NA | 2 | 1 | 5 | 1 | 5 | 1 | 12 | 1 | 2 | 1 | 8 | 1 | 3 | 1 | 1 | 1 | 14 | 1 |
| 14_796653 | 2 | 0 | 4 | 0.5 | 6 | 0.1667 | 7 | 0.2857 | 19 | 0.2632 | 3 | 0 | 9 | 0.2222 | 4 | 0.75 | 2 | 0 | 18 | 0.2778 |
| 14_796653 | 2 | 0 | 5 | 0.4 | 3 | 0.3333 | 6 | 0 | 16 | 0.1875 | 3 | 0.3333 | 8 | 0.375 | 2 | 0.5 | 2 | 0.5 | 15 | 0.4 |
| 14_796653 | 3 | 0 | 6 | 0.1667 | 5 | 0 | 5 | 0 | 19 | 0.0526 | 3 | 0 | 9 | 0 | 3 | 0.3333 | 1 | 0 | 16 | 0.0625 |
| 14_796654 | 2 | 0.5 | 6 | 1 | 4 | 0.75 | 7 | 0.8571 | 19 | 0.8421 | 3 | 0.6667 | 7 | 0.8571 | 3 | 1 | 2 | 1 | 15 | 0.8667 |
| 14_796654 | 2 | 0 | 3 | 0 | 5 | 0 | 7 | 0.1429 | 17 | 0.0588 | 3 | 0 | 9 | 0.1111 | 3 | 0.3333 | 2 | 0 | 17 | 0.1176 |
| 14_796655 | 3 | 0.3333 | 6 | 0.1667 | 6 | 0.1667 | 7 | 0.1429 | 22 | 0.1818 | 4 | 0.5 | 9 | 0 | 4 | 0.5 | 2 | 0.5 | 19 | 0.2632 |
| 14_796655 | 3 | 0 | 6 | 0.3333 | 4 | 0.25 | 6 | 0 | 19 | 0.1579 | 4 | 0 | 9 | 0.2222 | 4 | 0 | 3 | 0 | 20 | 0.1 |
| 14_796656 | 2 | 1 | 3 | 0.6667 | 4 | 1 | 6 | 1 | 15 | 0.9333 | 3 | 0.6667 | 8 | 0.875 | 4 | 1 | 2 | 1 | 17 | 0.8824 |
| 14_796657 | 4 | 1 | 4 | 0.75 | 6 | 0.6667 | 6 | 0.8333 | 20 | 0.8 | 2 | 0 | 6 | 1 | 3 | 1 | 3 | 1 | 14 | 0.8571 |
| 14_796658 | 4 | 0.75 | 5 | 1 | 6 | 1 | 7 | 1 | 22 | 0.9545 | 4 | 1 | 7 | 1 | 4 | 1 | 2 | 1 | 17 | 1 |
| 14_796659 | 4 | 0.5 | 5 | 0.6 | 8 | 0.5 | 6 | 0.6667 | 23 | 0.5652 | 4 | 0 | 9 | 0.3333 | 4 | 0.25 | 3 | 0.3333 | 20 | 0.25 |
| 14_796660 | 2 | 1 | 6 | 1 | 7 | 1 | 7 | 1 | 22 | 1 | 2 | 1 | 9 | 1 | 3 | 0.6667 | 2 | 1 | 16 | 0.9375 |
| 14_796660 | 4 | 0.25 | 6 | 0.1667 | 7 | 0.1429 | 7 | 0 | 24 | 0.125 | 4 | 0.25 | 9 | 0.2222 | 4 | 0.25 | 3 | 0.3333 | 20 | 0.25 |
| 14_796661 | 4 | 1 | 6 | 1 | 7 | 1 | 6 | 1 | 23 | 1 | 3 | 1 | 9 | 1 | 4 | 1 | 3 | 1 | 19 | 1 |
| 14_796661 | 3 | 1 | 4 | 1 | 4 | 1 | 5 | 0.4 | 16 | 0.8125 | 2 | 0.5 | 3 | 1 | 2 | 1 | 1 | 1 | 8 | 0.875 |
| 14_796661 | 4 | 1 | 6 | 1 | 7 | 1 | 7 | 1 | 24 | 1 | 3 | 1 | 8 | 1 | 4 | 1 | 2 | 0.5 | 17 | 0.9412 |
| 14_796662 | 4 | 0 | 5 | 0.2 | 8 | 0.5 | 5 | 0.2 | 22 | 0.2727 | 3 | 0 | 9 | 0.3333 | 4 | 0 | 3 | 0 | 19 | 0.1579 |
| 14_796663 | 4 | 0 | 6 | 0 | 8 | 0.25 | 8 | 0 | 26 | 0.0769 | 4 | 0 | 9 | 0 | 4 | 0.5 | 3 | 0 | 20 | 0.1 |
| 14_796663 | 4 | 0 | 6 | 0.6667 | 8 | 0.5 | 7 | 0.4286 | 25 | 0.44 | 4 | 0.75 | 9 | 0.4444 | 4 | 0.75 | 3 | 1 | 20 | 0.65 |
| 14_796666 | 2 | 0.5 | 4 | 0.25 | 8 | 0.25 | 6 | 0.3333 | 20 | 0.3 | 2 | 0 | 6 | 0.1667 | 3 | 0.3333 | 3 | 0.3333 | 14 | 0.2143 |
| 14_796668 | 5 | 0 | 7 | 0.1429 | 9 | 0.1111 | 8 | 0.25 | 29 | 0.1379 | 4 | 0 | 8 | 0.125 | 4 | 0.25 | 4 | 0.25 | 20 | 0.15 |
| 14_796670 | 5 | 0.4 | 5 | 0.4 | 7 | 0.4286 | 6 | 0.3333 | 23 | 0.3913 | 2 | 0.5 | 8 | 0.125 | 3 | 0 | 3 | 0.3333 | 16 | 0.1875 |
| 14_796681 | 5 | 0 | 10 | 0.3 | 12 | 0 | 7 | 0 | 34 | 0.0882 | 4 | 0.25 | 15 | 0.0667 | 6 | 0.1667 | 4 | 0 | 29 | 0.1034 |
| 14_796981 | NA | NA | 1 | 0 | 1 | 1 | 3 | 0 | 5 | 0.2 | NA | NA | NA | NA | 1 | 1 | NA | NA | 1 | 1 |
| 14_796985 | 3 | 0.3333 | 9 | 0.3333 | 8 | 0.5 | 18 | 0.2778 | 38 | 0.3421 | 6 | 0.3333 | 8 | 0.375 | 16 | 0.3125 | 10 | 0.7 | 40 | 0.425 |
| 14_796988 | 3 | 0.6667 | 8 | 0.375 | 6 | 0.5 | 11 | 0.1818 | 28 | 0.3571 | 5 | 0.6 | 6 | 0.3333 | 14 | 0.3571 | 8 | 0.625 | 33 | 0.4545 |
| 14_796988 | 3 | 0.3333 | 10 | 0 | 7 | 0 | 13 | 0.2308 | 33 | 0.1212 | 8 | 0 | 8 | 0 | 14 | 0.2857 | 6 | 0 | 36 | 0.1111 |
| 14_796993 | 3 | 1 | 13 | 1 | 5 | 0.8 | 16 | 0.75 | 37 | 0.8649 | 7 | 0.5714 | 7 | 1 | 12 | 0.75 | 10 | 0.8 | 36 | 0.7778 |

| | | | | | | | | | | | | | | | | | | | | |
|---|---|---|---|---|---|---|---|---|---|---|---|---|---|---|---|---|---|---|---|---|
| 14_7969938 | 3 | 0.3333 | 14 | 0 | 6 | 0.1667 | 14 | 0.0714 | 37 | 0.0811 | 6 | 0 | 7 | 0 | 14 | 0 | 11 | 0.1818 | 38 | 0.0526 |
| 14_7969939 | 3 | 0 | 14 | 0 | 6 | 0.1667 | 15 | 0.0667 | 38 | 0.0526 | 6 | 0.1667 | 6 | 0.3333 | 14 | 0 | 10 | 0.4 | 36 | 0.1944 |
| 14_7969947 | 2 | 0 | 7 | 0 | 3 | 0 | 12 | 0.1667 | 24 | 0.0833 | 5 | 0.2 | 4 | 0 | 10 | 0.1 | 7 | 0 | 26 | 0.0769 |
| 14_7971868 | 1 | 0 | 3 | 0 | 2 | 0.5 | 3 | 0 | 9 | 0.1111 | 1 | 0 | NA | NA | 3 | 0.3333 | NA | NA | 4 | 0.25 |
| 14_7981095 | 1 | 1 | NA | NA | NA | NA | 1 | 0 | 2 | 0.5 | NA | NA | NA | NA | 1 | 1 | NA | NA | 1 | 1 |
| 14_7981796 | 1 | 1 | NA | NA | NA | NA | 1 | 1 | 2 | 1 | 1 | 1 | NA | NA | 1 | 1 | NA | NA | 2 | 1 |
| 14_7981804 | 1 | 1 | NA | NA | NA | NA | 1 | 1 | 2 | 1 | 1 | 1 | NA | NA | 1 | 0 | NA | NA | 2 | 0.5 |
| 14_7981818 | 1 | 1 | NA | NA | NA | NA | 1 | 1 | 2 | 1 | NA | NA | NA | NA | 1 | 1 | NA | NA | 1 | 1 |
| 14_7986754 | 9 | 0.1111 | 4 | 0.25 | 7 | 0 | 3 | 0 | 23 | 0.087 | 5 | 0 | NA | NA | 6 | 0 | 4 | 0.25 | 15 | 0.0667 |
| 14_7987233 | 9 | 0.1111 | 5 | 0.2 | 11 | 0 | 1 | 0 | 26 | 0.0769 | 1 | 1 | 3 | 0 | 2 | 0.5 | 6 | 0.1667 | 12 | 0.25 |
| 14_8000494 | 3 | 0.3333 | 1 | 1 | 4 | 0 | 6 | 0.6667 | 14 | 0.4286 | 2 | 1 | NA | NA | 7 | 0.4286 | 6 | 0.5 | 15 | 0.5333 |
| 14_8000495 | 4 | 1 | 3 | 0.6667 | 4 | 0.25 | 9 | 0.7778 | 20 | 0.7 | 2 | 1 | NA | NA | 10 | 0.4 | 8 | 0.75 | 20 | 0.6 |
| 14_8000502 | 4 | 0.5 | 2 | 0 | 4 | 0 | 8 | 0.625 | 18 | 0.3889 | 2 | 0 | NA | NA | 9 | 0.2222 | 9 | 0.4444 | 20 | 0.3 |
| 14_8000518 | 3 | 0.3333 | 3 | 0 | 5 | 0 | 8 | 0 | 19 | 0.0526 | 2 | 0 | NA | NA | 9 | 0 | 8 | 0.125 | 19 | 0.0526 |
| 14_8000632 | 3 | 0 | 3 | 0.6667 | 5 | 0 | 4 | 0 | 15 | 0.1333 | NA | NA | NA | NA | 3 | 0 | 5 | 0.2 | 8 | 0.125 |
| 14_8000661 | 3 | 0.3333 | 2 | 0.5 | 3 | 0.3333 | 6 | 0 | 14 | 0.2143 | 1 | 1 | NA | NA | 7 | 0.4286 | 4 | 0.5 | 12 | 0.5 |
| 14_8000667 | 3 | 0.6667 | 2 | 1 | 4 | 0.5 | 6 | 0.8333 | 15 | 0.7333 | 1 | 0 | NA | NA | 8 | 0.5 | 4 | 0 | 13 | 0.3077 |
| 14_8000676 | 3 | 0.3333 | 2 | 0 | 5 | 0.4 | 5 | 0.4 | 15 | 0.3333 | 1 | 1 | NA | NA | 6 | 0.3333 | 6 | 0.1667 | 13 | 0.3077 |
| 14_8000685 | 2 | 1 | 2 | 0.5 | 4 | 0.5 | 3 | 0.3333 | 11 | 0.5455 | 1 | 1 | NA | NA | 8 | 0.625 | 5 | 0.6 | 14 | 0.6429 |
| 14_8000685 | 3 | 0 | 3 | 0 | 2 | 0 | 3 | 0.3333 | 11 | 0.0909 | 1 | 0 | NA | NA | 4 | 0.25 | 4 | 0.25 | 9 | 0.2222 |
| 14_8000719 | 3 | 0 | 3 | 0 | 4 | 0.25 | 4 | 0 | 14 | 0.0714 | 1 | 0 | NA | NA | 5 | 0.2 | 4 | 0 | 10 | 0.1 |
| 14_8000730 | 3 | 1 | 3 | 1 | 5 | 0.8 | 5 | 0.6 | 16 | 0.8125 | 1 | 1 | NA | NA | 6 | 1 | 3 | 0.6667 | 10 | 0.9 |
| 14_8000730 | 3 | 0 | 1 | 0 | 4 | 0.5 | 3 | 0.3333 | 11 | 0.2727 | 1 | 0 | NA | NA | 3 | 0.6667 | 3 | 0 | 7 | 0.2857 |
| 14_8000738 | 3 | 0 | 3 | 0.3333 | 5 | 0 | 3 | 0 | 14 | 0.0714 | 1 | 0 | NA | NA | 5 | 0.2 | 4 | 0 | 10 | 0.1 |
| 14_8000755 | 2 | 0.5 | 2 | 0 | 3 | 0 | 3 | 0 | 10 | 0.1 | 1 | 1 | NA | NA | 5 | 0 | 2 | 0 | 8 | 0.125 |
| 14_8000760 | 3 | 0 | 3 | 0.6667 | 4 | 0 | 4 | 0.5 | 14 | 0.2857 | 1 | 1 | NA | NA | 5 | 0.2 | 4 | 0.5 | 10 | 0.4 |
| 14_8000762 | 2 | 0.5 | 3 | 0.6667 | 5 | 0.2 | 4 | 0 | 14 | 0.2857 | NA | NA | NA | NA | 5 | 0.4 | 4 | 0.5 | 9 | 0.4444 |
| 14_8000795 | 3 | 0.3333 | 2 | 0.5 | NA | NA | 3 | 0 | 8 | 0.25 | 1 | 0 | NA | NA | 1 | 1 | 3 | 0 | 5 | 0.2 |
| 14_8000797 | 3 | 0 | 3 | 0.3333 | 4 | 0 | 4 | 0.25 | 14 | 0.1429 | 1 | 1 | NA | NA | 4 | 0.25 | 3 | 0.3333 | 8 | 0.375 |
| 14_8000822 | 2 | 0 | NA | NA | 2 | 0.5 | 1 | 0 | 5 | 0.2 | 1 | 0 | NA | NA | 2 | 0 | 3 | 0.3333 | 6 | 0.1667 |
| 14_8468269 | NA | NA | NA | NA | 1 | 1 | 1 | 1 | 2 | 1 | NA | NA | 2 | 0.5 | NA | NA | NA | NA | 2 | 0.5 |
| 14_8468278 | NA | NA | NA | NA | 1 | 1 | 1 | 1 | 2 | 1 | NA | NA | 2 | 1 | NA | NA | NA | NA | 2 | 1 |
| 14_8468337 | NA | NA | NA | NA | NA | NA | 2 | 0.5 | 2 | 0.5 | 1 | 1 | 3 | 1 | 2 | 0.5 | NA | NA | 6 | 0.8333 |
| 14_8468341 | NA | NA | NA | NA | 1 | 1 | 1 | 0 | 2 | 0.5 | 1 | 1 | 3 | 0.6667 | 2 | 0.5 | NA | NA | 6 | 0.6667 |
| 14_8468342 | NA | NA | NA | NA | 1 | 1 | 2 | 0 | 3 | 0.3333 | 1 | 0 | 2 | 0.5 | 2 | 0 | NA | NA | 5 | 0.2 |
| 14_8468395 | 1 | 0 | NA | NA | NA | NA | 3 | 0.6667 | 4 | 0.5 | 1 | 0 | 3 | 0.6667 | 1 | 0 | NA | NA | 5 | 0.4 |
| 14_8468446 | 1 | 1 | NA | NA | 2 | 1 | 4 | 0.25 | 7 | 0.5714 | 1 | 1 | 4 | 0.75 | 2 | 0.5 | NA | NA | 7 | 0.7143 |
| 14_8468451 | NA | NA | NA | NA | 3 | 0.6667 | 5 | 0.2 | 8 | 0.375 | 1 | 0 | 4 | 0.5 | 2 | 0.5 | NA | NA | 7 | 0.4286 |
| 14_8468467 | 1 | 1 | 1 | 0 | 2 | 1 | 5 | 0.8 | 9 | 0.7778 | 1 | 0 | 4 | 1 | 2 | 0.5 | NA | NA | 7 | 0.7143 |
| 14_8468472 | 1 | 0 | 1 | 0 | 2 | 0.5 | 5 | 0.8 | 9 | 0.5556 | 1 | 0 | 3 | 0.3333 | 1 | 1 | NA | NA | 5 | 0.4 |
| 14_8468477 | 1 | 0 | 1 | 0 | 3 | 0 | 6 | 0.1667 | 11 | 0.0909 | 1 | 0 | 4 | 0.5 | 2 | 0.5 | NA | NA | 7 | 0.4286 |
| 14_8468480 | 1 | 0 | 1 | 1 | 2 | 0.5 | 6 | 0.1667 | 10 | 0.3 | 1 | 0 | 3 | 0.3333 | 1 | 0 | 1 | 0 | 6 | 0.1667 |
| 14_8468482 | 1 | 1 | 1 | 0 | 2 | 0.5 | 1 | 0 | 5 | 0.4 | 1 | 1 | 2 | 0 | 2 | 0 | 1 | 0 | 6 | 0.1667 |
| 14_8468495 | 1 | 1 | 1 | 1 | 2 | 1 | 6 | 0.8333 | 10 | 0.9 | 1 | 1 | 4 | 0.75 | 2 | 1 | NA | NA | 7 | 0.8571 |
| 14_8468504 | 1 | 1 | 1 | 1 | 1 | 1 | 5 | 0.8 | 8 | 0.875 | 1 | 1 | 3 | 1 | 2 | 1 | NA | NA | 6 | 1 |
| 14_8468515 | 2 | 0.5 | 1 | 0 | 2 | 0 | 4 | 0.25 | 9 | 0.2222 | 1 | 1 | 3 | 0.3333 | 2 | 0 | NA | NA | 6 | 0.3333 |
| 14_8468519 | 1 | 1 | NA | NA | 4 | 1 | 6 | 1 | 11 | 1 | 1 | 0 | 4 | 1 | 2 | 1 | 1 | 0 | 8 | 0.75 |
| 14_8468536 | 2 | 0 | 1 | 0 | 6 | 0.5 | 4 | 0.75 | 13 | 0.4615 | 1 | 0 | 4 | 0.25 | 2 | 0 | NA | NA | 7 | 0.1429 |
| 14_8468549 | 3 | 0.3333 | NA | NA | 6 | 0 | 6 | 0.1667 | 15 | 0.1333 | 2 | 0 | 4 | 0.25 | 3 | 0 | 1 | 0 | 10 | 0.1 |
| 14_8468556 | 2 | 0 | 1 | 1 | 4 | 0 | 4 | 0 | 11 | 0.0909 | 2 | 0 | 3 | 0 | 2 | 0 | 2 | 0.5 | 9 | 0.1111 |
| 14_8468563 | 3 | 0.6667 | 1 | 0 | 6 | 1 | 4 | 0.5 | 14 | 0.7143 | 1 | 0 | 3 | 0.6667 | 4 | 0.25 | 1 | 1 | 9 | 0.4444 |
| 14_8468565 | 1 | 1 | 1 | 1 | 7 | 0.4286 | 5 | 0.6 | 14 | 0.5714 | 2 | 1 | 3 | 0.3333 | 4 | 0.75 | 1 | 1 | 10 | 0.7 |
| 14_8468599 | 1 | 1 | 2 | 0.5 | 4 | 0.5 | 5 | 0.6 | 12 | 0.5833 | 2 | 0.5 | 4 | 0.5 | 6 | 0.5 | 5 | 0.4 | 17 | 0.4706 |
| 14_8468604 | 3 | 0 | 2 | 0 | 6 | 0.1667 | 9 | 0.4444 | 20 | 0.25 | 2 | 0 | 4 | 0.25 | 5 | 0.6 | 3 | 0.3333 | 14 | 0.3571 |
| 14_8477046 | 1 | 1 | 2 | 1 | 6 | 0.6667 | 8 | 1 | 17 | 0.8824 | 3 | 1 | 6 | 1 | 5 | 1 | 3 | 1 | 17 | 1 |
| 14_8477047 | 2 | 1 | 2 | 1 | 7 | 0.7143 | 5 | 0.8 | 16 | 0.8125 | 3 | 1 | 5 | 0.8 | 3 | 1 | 2 | 0.5 | 13 | 0.8462 |
| 14_8477048 | 4 | 1 | 2 | 1 | 7 | 0.7143 | 8 | 0.875 | 21 | 0.8571 | 3 | 1 | 4 | 1 | 6 | 0.8333 | 5 | 1 | 18 | 0.9444 |
| 14_8477050 | 3 | 1 | 2 | 1 | 6 | 0.8333 | 8 | 1 | 19 | 0.9474 | 3 | 1 | 6 | 1 | 6 | 1 | 6 | 0.8333 | 21 | 0.9524 |
| 14_8477052 | 4 | 0.5 | 2 | 0.5 | 5 | 0.6 | 8 | 0.625 | 19 | 0.5789 | 3 | 0.3333 | 3 | 0.6667 | 6 | 0.5 | 5 | 0.4 | 17 | 0.4706 |
| 14_8477056 | 4 | 0.25 | 2 | 0.5 | 6 | 0 | 7 | 0.2857 | 19 | 0.2105 | 2 | 0.5 | 6 | 0.1667 | 5 | 0.2 | 4 | 0.25 | 17 | 0.2353 |
| 14_8477074 | 3 | 0.6667 | NA | NA | 4 | 0.25 | 4 | 0.75 | 11 | 0.5455 | 3 | 0.6667 | 6 | 0.6667 | 4 | 0 | 4 | 1 | 17 | 0.5882 |
| 14_8477077 | 5 | 0.4 | 2 | 0.5 | 9 | 0.6667 | 7 | 0.4286 | 23 | 0.5217 | 3 | 0.3333 | 6 | 0.6667 | 6 | 1 | 7 | 0.2857 | 22 | 0.5909 |
| 14_8477096 | 5 | 1 | 3 | 1 | 7 | 1 | 9 | 1 | 24 | 1 | 3 | 1 | 6 | 1 | 5 | 1 | 3 | 1 | 17 | 1 |
| 14_8477108 | 5 | 0.4 | 3 | 0.3333 | 7 | 0.4286 | 12 | 0.5833 | 27 | 0.4815 | 2 | 0.5 | 8 | 0.625 | 6 | 0.6667 | 6 | 0.3333 | 22 | 0.5455 |
| 14_8477111 | 5 | 0 | 2 | 0 | 7 | 0.2857 | 7 | 0.1429 | 21 | 0.1429 | 2 | 0.5 | 7 | 0.1429 | 5 | 0 | 3 | 0 | 17 | 0.1176 |

| | | | | | | | | | | | | | | | | | | | | |
|---|---|---|---|---|---|---|---|---|---|---|---|---|---|---|---|---|---|---|---|---|
| 14_847712( | 5 | 0.6 | 2 | 0 | 7 | 0.5714 | 12 | 0.25 | 26 | 0.3846 | 3 | 0.6667 | 8 | 0.5 | 6 | 0.5 | 7 | 0.1429 | 24 | 0.4167 |
| 14_847712! | 4 | 0.25 | 2 | 0 | 5 | 0.2 | 10 | 0 | 21 | 0.0952 | 2 | 0 | 7 | 0.1429 | 5 | 0 | 6 | 0 | 20 | 0.05 |
| 14_847713! | 5 | 0.4 | 4 | 0.25 | 7 | 0.7143 | 12 | 0.25 | 28 | 0.3929 | 3 | 0.6667 | 7 | 0.7143 | 6 | 0.6667 | 7 | 0.2857 | 23 | 0.5652 |
| 14_847714: | 4 | 0.25 | 3 | 0 | 7 | 0.4286 | 9 | 0.1111 | 23 | 0.2174 | 3 | 0.3333 | 7 | 0.5714 | 6 | 0 | 7 | 0.2857 | 23 | 0.3043 |
| 14_847714{ | 5 | 0.2 | 3 | 0.3333 | 6 | 0.3333 | 11 | 0 | 25 | 0.16 | 3 | 0 | 7 | 0 | 4 | 0.25 | 8 | 0.125 | 22 | 0.0909 |
| 14_847715; | 5 | 0 | 4 | 0 | 9 | 0.3333 | 10 | 0 | 28 | 0.1071 | 4 | 0 | 7 | 0.1429 | 7 | 0 | 7 | 0.1429 | 25 | 0.08 |
| 14_847746: | 8 | 0 | 8 | 0.125 | 9 | 0.2222 | 8 | 0 | 33 | 0.0909 | 6 | 0 | 8 | 0.125 | 11 | 0.2727 | 11 | 0.0909 | 36 | 0.1389 |
| 14_847746: | 8 | 0.375 | 7 | 0.2857 | 10 | 0.5 | 11 | 0.4545 | 36 | 0.4167 | 10 | 0.3 | 8 | 0.375 | 11 | 0.4545 | 10 | 0.4 | 39 | 0.3846 |
| 14_847749{ | 8 | 0.25 | 9 | 0.4444 | 16 | 0.3125 | 14 | 0.4286 | 47 | 0.3617 | 13 | 0.1538 | 10 | 0.3 | 14 | 0.2143 | 13 | 0.5385 | 50 | 0.3 |
| 14_877434! | NA | NA | 2 | 0 | 1 | 1 | 2 | 1 | 5 | 0.6 | 1 | 1 | NA | NA | 2 | 0 | NA | NA | 3 | 0.3333 |
| 14_877435! | NA | NA | 4 | 1 | 2 | 1 | 1 | 1 | 7 | 1 | 1 | 1 | NA | NA | 2 | 0.5 | NA | NA | 3 | 0.6667 |
| 14_877437 | NA | NA | 4 | 1 | 2 | 0.5 | 2 | 1 | 8 | 0.875 | 1 | 1 | 1 | 0 | 2 | 0.5 | NA | NA | 4 | 0.5 |
| 14_877437! | NA | NA | 4 | 0.5 | 2 | 0.5 | 2 | 0 | 8 | 0.375 | 1 | 0 | NA | NA | 2 | 0.5 | NA | NA | 3 | 0.3333 |
| 14_877438! | NA | NA | 5 | 0 | 1 | 1 | 2 | 0 | 8 | 0.125 | 1 | 1 | 1 | 0 | 2 | 0 | NA | NA | 4 | 0.25 |
| 14_877827; | 2 | 0.5 | 6 | 0.1667 | 3 | 0.3333 | 2 | 0 | 13 | 0.2308 | 1 | 0 | 5 | 0.2 | 3 | 0.3333 | 2 | 0 | 11 | 0.1818 |
| 14_887043 | 2 | 0.5 | 3 | 1 | 2 | 1 | NA | NA | 7 | 0.8571 | NA | NA | 1 | 1 | 2 | 0.5 | 1 | 1 | 4 | 0.75 |
| 14_887044 | 2 | 0.5 | 4 | 1 | 2 | 1 | NA | NA | 8 | 0.875 | NA | NA | 1 | 1 | 2 | 0.5 | 1 | 1 | 4 | 0.75 |
| 14_887044{ | 1 | 1 | 2 | 1 | 2 | 1 | NA | NA | 5 | 1 | NA | NA | 1 | 1 | 2 | 0.5 | 1 | 1 | 4 | 0.75 |
| 14_887046{ | 2 | 0.5 | 2 | 0 | 1 | 0 | NA | NA | 5 | 0.2 | NA | NA | 1 | 0 | 2 | 0.5 | 1 | 0 | 4 | 0.25 |
| 14_887047( | 1 | 1 | 3 | 0.6667 | 1 | 0 | NA | NA | 5 | 0.6 | NA | NA | 1 | 0 | 2 | 0.5 | 1 | 1 | 4 | 0.5 |
| 14_887047; | 1 | 1 | 3 | 0.6667 | 1 | 1 | NA | NA | 5 | 0.8 | NA | NA | 1 | 1 | 2 | 1 | 1 | 1 | 4 | 1 |
| 14_887048! | 1 | 1 | 2 | 1 | 1 | 0 | NA | NA | 4 | 0.75 | NA | NA | 1 | 1 | 2 | 0.5 | 1 | 0 | 4 | 0.5 |
| 14_887049; | 1 | 1 | 3 | 0.3333 | 1 | 1 | NA | NA | 5 | 0.6 | NA | NA | 1 | 1 | 2 | 1 | 1 | 1 | 4 | 1 |
| 14_887050! | NA | NA | 2 | 0.5 | 1 | 0 | NA | NA | 3 | 0.3333 | NA | NA | 1 | 1 | 2 | 1 | 1 | 1 | 4 | 1 |
| 14_887052: | 1 | 1 | 2 | 1 | 1 | 0 | NA | NA | 4 | 0.75 | NA | NA | 1 | 1 | 2 | 0.5 | 1 | 1 | 4 | 0.75 |
| 14_887052 | NA | NA | 3 | 1 | 1 | 1 | NA | NA | 4 | 1 | NA | NA | 1 | 1 | 2 | 1 | 1 | 1 | 4 | 1 |
| 14_887052! | 1 | 1 | 3 | 0 | NA | NA | NA | NA | 4 | 0.25 | NA | NA | NA | NA | 2 | 0.5 | 1 | 0 | 3 | 0.3333 |
| 14_887053 | 1 | 0 | 2 | 0.5 | 1 | 0 | NA | NA | 4 | 0.25 | NA | NA | 1 | 0 | 2 | 0.5 | 1 | 0 | 4 | 0.25 |
| 14_887053( | 1 | 1 | 2 | 1 | 1 | 1 | NA | NA | 4 | 1 | NA | NA | 1 | 1 | 1 | 1 | 1 | 1 | 3 | 1 |
| 14_887054! | 1 | 1 | 1 | 1 | NA | NA | NA | NA | 2 | 1 | NA | NA | 1 | 1 | 2 | 1 | 1 | 1 | 4 | 1 |
| 14_887055( | 1 | 0 | 2 | 0.5 | 1 | 0 | NA | NA | 4 | 0.25 | NA | NA | 1 | 0 | 2 | 0.5 | 1 | 0 | 4 | 0.25 |
| 14_887055; | 1 | 0 | 2 | 0 | 1 | 1 | NA | NA | 4 | 0.25 | NA | NA | 1 | 0 | 2 | 0.5 | 1 | 1 | 4 | 0.5 |
| 14_887057( | 1 | 1 | 2 | 0 | 1 | 0 | NA | NA | 4 | 0.25 | NA | NA | 1 | 0 | 2 | 0.5 | 1 | 0 | 4 | 0.25 |
| 14_887057( | 1 | 1 | 1 | 0 | 1 | 0 | NA | NA | 3 | 0.3333 | NA | NA | 1 | 0 | 2 | 1 | 1 | 0 | 4 | 0.5 |
| 14_887058( | 1 | 1 | 1 | 1 | 1 | 1 | NA | NA | 3 | 1 | NA | NA | 1 | 1 | 2 | 1 | 1 | 1 | 4 | 1 |
| 14_887058! | 1 | 0 | 2 | 0.5 | 1 | 0 | NA | NA | 4 | 0.25 | NA | NA | 1 | 0 | 1 | 1 | 1 | 0 | 3 | 0.3333 |
| 14_887059( | 1 | 0 | 2 | 0.5 | 1 | 0 | NA | NA | 4 | 0.25 | NA | NA | 1 | 1 | 1 | 0 | 1 | 0 | 3 | 0.3333 |
| 14_887060! | 1 | 1 | 2 | 1 | 1 | 1 | NA | NA | 4 | 1 | NA | NA | 1 | 1 | 1 | 1 | 1 | 1 | 3 | 1 |
| 14_887062 | 1 | 1 | 2 | 0 | NA | NA | NA | NA | 3 | 0.3333 | NA | NA | 1 | 1 | 1 | 0 | 1 | 0 | 3 | 0.3333 |
| 14_887062! | 1 | 1 | 2 | 1 | 1 | 1 | NA | NA | 4 | 1 | NA | NA | 1 | 1 | NA | NA | 1 | 1 | 2 | 1 |
| 14_887063: | NA | NA | 1 | 1 | 1 | 1 | NA | NA | 2 | 1 | NA | NA | 1 | 1 | 1 | 1 | 1 | 1 | 3 | 1 |
| 14_887063( | 1 | 1 | 2 | 1 | 1 | 1 | NA | NA | 4 | 1 | NA | NA | 1 | 0 | 1 | 1 | 1 | 1 | 3 | 0.6667 |
| 14_887064( | NA | NA | 2 | 1 | 1 | 1 | NA | NA | 3 | 1 | NA | NA | 1 | 1 | 1 | 1 | 1 | 1 | 3 | 1 |
| 14_887065 | 1 | 1 | 1 | 1 | 1 | 0 | NA | NA | 3 | 0.6667 | NA | NA | 1 | 1 | 1 | 0 | 1 | 1 | 3 | 0.6667 |
| 14_887065{ | 1 | 0 | 1 | 1 | 1 | 1 | NA | NA | 3 | 0.6667 | NA | NA | 1 | 1 | 1 | 1 | 1 | 1 | 3 | 1 |
| 14_887066{ | NA | NA | 2 | 0 | 1 | 1 | NA | NA | 3 | 0.3333 | NA | NA | 1 | 0 | NA | NA | 1 | 1 | 2 | 0.5 |
| 14_887067; | NA | NA | 1 | 0 | 1 | 1 | NA | NA | 2 | 0.5 | NA | NA | 1 | 1 | NA | NA | 1 | 0 | 2 | 0.5 |
| 14_887069: | NA | NA | 2 | 1 | 1 | 1 | NA | NA | 3 | 1 | NA | NA | 1 | 1 | NA | NA | 1 | 1 | 2 | 1 |
| 14_887070; | NA | NA | 2 | 1 | NA | NA | NA | NA | 2 | 1 | NA | NA | 1 | 1 | NA | NA | NA | NA | 1 | 1 |
| 14_932521( | 6 | 0 | 2 | 0 | 6 | 0.1667 | 2 | 0 | 16 | 0.0625 | NA | NA | 8 | 0.125 | 3 | 0 | 2 | 0 | 13 | 0.0769 |
| 14_932524( | 5 | 0.2 | 2 | 0 | 3 | 0 | 2 | 0 | 12 | 0.0833 | NA | NA | 8 | 0 | 3 | 0 | 3 | 0.3333 | 14 | 0.0714 |
| 14_932525 | 5 | 0.8 | 2 | 0 | 4 | 0.25 | 2 | 0 | 13 | 0.3846 | NA | NA | 8 | 0.25 | 3 | 0.6667 | 3 | 0.3333 | 14 | 0.3571 |
| 14_932526( | 4 | 0.25 | 2 | 0 | 5 | 0.2 | 2 | 0.5 | 13 | 0.2308 | NA | NA | 7 | 0.2857 | 3 | 1 | 2 | 1 | 12 | 0.5833 |
| 14_932526! | 5 | 0.6 | 1 | 1 | 5 | 0.6 | 1 | 0 | 12 | 0.5833 | NA | NA | 6 | 0.3333 | 3 | 0 | 2 | 1 | 11 | 0.3636 |
| 14_932526( | 5 | 1 | 2 | 1 | 4 | 0.75 | 2 | 1 | 13 | 0.9231 | NA | NA | 5 | 1 | 2 | 1 | 2 | 0.5 | 9 | 0.8889 |
| 14_932528: | 3 | 0.3333 | 2 | 0 | 5 | 0 | 2 | 0 | 12 | 0.0833 | NA | NA | 7 | 0 | 1 | 0 | 3 | 0.3333 | 11 | 0.0909 |
| 14_932529( | 5 | 0.8 | 2 | 0.5 | 5 | 0.2 | 2 | 0 | 14 | 0.4286 | NA | NA | 7 | 0.4286 | 2 | 0.5 | NA | NA | 9 | 0.4444 |
| 14_932529: | 5 | 1 | 2 | 1 | 4 | 1 | 2 | 1 | 13 | 1 | NA | NA | 8 | 0.875 | 3 | 1 | 2 | 1 | 13 | 0.9231 |
| 14_932530: | 4 | 1 | 1 | 1 | 4 | 1 | 2 | 1 | 11 | 1 | NA | NA | 7 | 1 | 3 | 1 | 2 | 1 | 12 | 1 |
| 14_932530{ | 5 | 0 | 2 | 0 | 4 | 0.25 | 2 | 0 | 13 | 0.0769 | NA | NA | 7 | 0 | 3 | 0.3333 | 2 | 0.5 | 12 | 0.1667 |
| 14_932531! | 5 | 0.2 | 2 | 0 | 4 | 0 | 2 | 0 | 13 | 0.0769 | NA | NA | 8 | 0.125 | 3 | 0 | 2 | 0 | 13 | 0.0769 |
| 14_940993( | 2 | 1 | 7 | 0.1429 | 6 | 0 | 11 | 0.0909 | 26 | 0.1538 | 1 | 0 | 4 | 0.5 | 7 | 0 | 6 | 0.1667 | 18 | 0.1667 |
| 14_940998( | 1 | 0 | 4 | 0 | 5 | 0.2 | 8 | 0 | 18 | 0.0556 | 1 | 0 | 4 | 0 | 5 | 0.2 | 4 | 0 | 14 | 0.0714 |
| 14_941012 | 2 | 0.5 | 3 | 0 | 2 | 1 | 4 | 0.25 | 11 | 0.3636 | 1 | 0 | 2 | 0 | 5 | 0.4 | 4 | 0.5 | 12 | 0.3333 |
| 14_941232( | 2 | 1 | 1 | 1 | 3 | 0.3333 | NA | NA | 6 | 0.6667 | 1 | 0 | 4 | 0.25 | NA | NA | 3 | 0 | 8 | 0.125 |

| | | | | | | | | | | | | | | | | | | | | |
|---|---|---|---|---|---|---|---|---|---|---|---|---|---|---|---|---|---|---|---|---|
| 14_941234 | 2 | 0 | 1 | 1 | 3 | 0.6667 | 3 | 0.3333 | 9 | 0.4444 | 1 | 1 | 3 | 0.3333 | NA | NA | 3 | 0 | 7 | 0.2857 |
| 14_941235 | 2 | 0.5 | 2 | 1 | 3 | 0.6667 | 2 | 0.5 | 9 | 0.6667 | NA | NA | 3 | 1 | NA | NA | 3 | 1 | 6 | 1 |
| 14_941237 | 2 | 1 | 1 | 1 | 3 | 0.6667 | 3 | 0.6667 | 9 | 0.7778 | NA | NA | 4 | 1 | NA | NA | 3 | 1 | 7 | 1 |
| 14_941237 | 2 | 0 | 1 | 1 | 3 | 0 | 1 | 0 | 7 | 0.1429 | 1 | 1 | 4 | 0.25 | NA | NA | 3 | 0 | 8 | 0.25 |
| 14_941237 | 1 | 1 | 2 | 1 | 3 | 0.6667 | 3 | 0.6667 | 9 | 0.7778 | 1 | 1 | 3 | 0.6667 | NA | NA | 3 | 0.6667 | 7 | 0.7143 |
| 14_941238 | 2 | 0.5 | 2 | 0 | 2 | 0.5 | 3 | 0 | 9 | 0.2222 | 1 | 0 | 4 | 0.25 | NA | NA | 3 | 0 | 8 | 0.125 |
| 14_941238 | 2 | 0.5 | 1 | 1 | 3 | 0 | 3 | 0.3333 | 9 | 0.3333 | 1 | 1 | 4 | 0.5 | NA | NA | 3 | 0.3333 | 8 | 0.5 |
| 14_941240 | 2 | 1 | 2 | 0.5 | 2 | 0.5 | 2 | 0 | 8 | 0.5 | 1 | 1 | 4 | 0.25 | NA | NA | 3 | 0.3333 | 8 | 0.375 |
| 14_941243 | 2 | 0 | 2 | 0.5 | 1 | 0 | 2 | 0 | 7 | 0.1429 | 1 | 0 | 4 | 0.75 | NA | NA | 3 | 0.3333 | 8 | 0.5 |
| 14_941243 | 1 | 1 | 2 | 1 | 2 | 1 | 2 | 0.5 | 7 | 0.8571 | 1 | 1 | 4 | 0.5 | NA | NA | 3 | 0.6667 | 8 | 0.625 |
| 14_941245 | 2 | 0.5 | 2 | 0 | 2 | 0 | 2 | 0 | 8 | 0.125 | 1 | 0 | 4 | 0.5 | NA | NA | 2 | 0 | 7 | 0.2857 |
| 14_941245 | 1 | 1 | 1 | 1 | 2 | 1 | 1 | 1 | 5 | 1 | NA | NA | 3 | 0.6667 | NA | NA | 2 | 0.5 | 5 | 0.6 |
| 14_941248 | 2 | 0.5 | 1 | 0 | 2 | 0 | 2 | 0 | 7 | 0.1429 | 1 | 0 | 4 | 0.25 | NA | NA | 1 | 0 | 6 | 0.1667 |
| 14_941248 | 2 | 1 | 1 | 1 | 2 | 0.5 | 2 | 0.5 | 7 | 0.7143 | 1 | 1 | 3 | 0.6667 | NA | NA | 3 | 0.3333 | 7 | 0.5714 |
| 14_941252 | 1 | 1 | 2 | 1 | 2 | 0.5 | 2 | 0.5 | 7 | 0.7143 | 1 | 1 | 4 | 0.75 | NA | NA | 2 | 1 | 7 | 0.8571 |
| 14_941256 | 1 | 1 | 2 | 1 | NA | NA | 2 | 0.5 | 5 | 0.8 | 1 | 1 | 2 | 1 | NA | NA | 2 | 0.5 | 5 | 0.8 |
| 14_941256 | 1 | 0 | NA | NA | 1 | 1 | 2 | 1 | 4 | 0.75 | 1 | 0 | 2 | 1 | NA | NA | 2 | 1 | 5 | 0.8 |
| 14_941258 | 1 | 1 | 2 | 1 | 1 | 1 | 1 | 1 | 5 | 1 | 1 | 1 | 1 | 0 | NA | NA | 2 | 0.5 | 4 | 0.5 |
| 14_941258 | 1 | 1 | 2 | 0.5 | 1 | 1 | 2 | 0.5 | 6 | 0.6667 | 1 | 0 | NA | NA | NA | NA | 1 | 1 | 2 | 0.5 |
| 14_941258 | NA | NA | NA | NA | 1 | 0 | 2 | 0.5 | 3 | 0.3333 | 1 | 1 | NA | NA | NA | NA | 2 | 0 | 3 | 0.3333 |
| 14_941259 | 1 | 1 | 1 | 1 | 1 | 1 | 2 | 1 | 5 | 1 | 1 | 0 | NA | NA | NA | NA | 2 | 1 | 3 | 0.6667 |
| 14_941260 | 1 | 1 | 1 | 1 | 1 | 1 | 2 | 1 | 5 | 1 | 1 | 1 | NA | NA | NA | NA | 1 | 0 | 2 | 0.5 |
| 14_941261 | 1 | 1 | NA | NA | 1 | 0 | 2 | 1 | 4 | 0.75 | 1 | 1 | NA | NA | NA | NA | 1 | 1 | 2 | 1 |
| 14_941261 | 1 | 1 | NA | NA | 1 | 0 | 1 | 1 | 3 | 0.6667 | 1 | 1 | NA | NA | NA | NA | NA | NA | 1 | 1 |
| 14_980247 | 5 | 0.2 | NA | NA | 6 | 0 | 4 | 0.25 | 15 | 0.1333 | NA | NA | 2 | 0.5 | 4 | 0 | 2 | 0.5 | 8 | 0.25 |
| 14_980266 | 2 | 0.5 | NA | NA | 3 | 0.6667 | 1 | 0 | 6 | 0.5 | NA | NA | 1 | 0 | 2 | 0 | 2 | 0.5 | 5 | 0.2 |
| 14_980266 | 1 | 1 | NA | NA | 4 | 0.75 | NA | NA | 5 | 0.8 | NA | NA | 1 | 1 | 2 | 0.5 | 1 | 0 | 4 | 0.5 |
| 14_980267 | 2 | 1 | NA | NA | 3 | 0.3333 | 2 | 1 | 7 | 0.7143 | NA | NA | 1 | 1 | 1 | 0 | 1 | 1 | 3 | 0.6667 |
| 14_980268 | 2 | 1 | NA | NA | 4 | 0.75 | 2 | 1 | 8 | 0.875 | NA | NA | 1 | 1 | 2 | 1 | 2 | 1 | 5 | 1 |
| 14_980269 | 1 | 0 | NA | NA | 4 | 0.25 | 2 | 0 | 7 | 0.1429 | NA | NA | 1 | 0 | 2 | 0.5 | 1 | 1 | 4 | 0.5 |
| 14_992723 | 20 | 0.05 | 14 | 0.1429 | 11 | 0 | 16 | 0.0625 | 61 | 0.0656 | 8 | 0.125 | 25 | 0.08 | 16 | 0.125 | 19 | 0.2105 | 68 | 0.1324 |
| 14_992738 | 11 | 0 | 13 | 0 | 14 | 0.2143 | 15 | 0.2 | 53 | 0.1132 | 8 | 0 | 19 | 0 | 15 | 0.0667 | 18 | 0.1111 | 60 | 0.05 |
| 14_993127 | 3 | 0.3333 | 1 | 1 | 1 | 1 | NA | NA | 5 | 0.6 | NA | NA | 2 | 0.5 | 1 | 0 | NA | NA | 3 | 0.3333 |
| 14_993128 | 2 | 0.5 | 1 | 0 | 1 | 0 | NA | NA | 4 | 0.25 | NA | NA | 2 | 0.5 | 1 | 1 | NA | NA | 3 | 0.6667 |
| 14_993146 | 2 | 1 | NA | NA | 1 | 0 | NA | NA | 3 | 0.6667 | NA | NA | NA | NA | 1 | 1 | NA | NA | 1 | 1 |
| 14_995393 | 1 | 1 | NA | NA | NA | NA | NA | NA | 1 | 1 | NA | NA | 1 | 1 | NA | NA | 1 | 1 | 2 | 1 |
| 14_995507 | NA | NA | 1 | 1 | NA | NA | 1 | 0 | 2 | 0.5 | 1 | 0 | 1 | 1 | NA | NA | NA | NA | 2 | 0.5 |
| 14_995508 | 1 | 1 | 2 | 1 | NA | NA | 1 | 1 | 4 | 1 | NA | NA | 1 | 1 | NA | NA | 1 | 1 | 2 | 1 |
| 14_995511 | 1 | 0 | 2 | 0.5 | NA | NA | NA | NA | 3 | 0.3333 | 1 | 0 | 1 | 1 | NA | NA | 1 | 0 | 3 | 0.3333 |
| 14_995517 | 2 | 0 | 3 | 0.3333 | NA | NA | 1 | 0 | 6 | 0.1667 | 1 | 0 | 2 | 0 | 1 | 1 | 2 | 0 | 6 | 0.1667 |
| 14_100104 | 6 | 0.6667 | 5 | 0.2 | 4 | 0.25 | 4 | 0.5 | 19 | 0.4211 | 3 | 1 | 3 | 0 | 2 | 0.5 | 2 | 0.5 | 10 | 0.5 |
| 14_100109 | 2 | 0.5 | 2 | 0 | NA | NA | 1 | 0 | 5 | 0.2 | 2 | 0.5 | NA | NA | 2 | 0 | NA | NA | 4 | 0.25 |
| 14_100112 | 2 | 1 | 1 | 0 | NA | NA | NA | NA | 3 | 0.6667 | 2 | 0.5 | NA | NA | 1 | 0 | NA | NA | 3 | 0.3333 |
| 14_100112 | 2 | 0.5 | 1 | 1 | NA | NA | NA | NA | 3 | 0.6667 | 2 | 1 | NA | NA | 1 | 1 | NA | NA | 3 | 1 |
| 14_100112 | 2 | 1 | 1 | 1 | NA | NA | NA | NA | 3 | 1 | 2 | 0.5 | NA | NA | 1 | 1 | NA | NA | 3 | 0.6667 |
| 14_100260 | NA | NA | 1 | 1 | NA | NA | NA | NA | 1 | 1 | 2 | 1 | NA | NA | 1 | 0 | NA | NA | 3 | 0.6667 |
| 14_100260 | 1 | 0 | 1 | 1 | NA | NA | NA | NA | 2 | 0.5 | 2 | 0.5 | NA | NA | NA | NA | NA | NA | 2 | 0.5 |
| 14_100260 | 2 | 0.5 | 1 | 1 | NA | NA | NA | NA | 3 | 0.6667 | 1 | 1 | NA | NA | 1 | 0 | NA | NA | 2 | 0.5 |
| 14_100260 | 2 | 0 | 1 | 1 | NA | NA | NA | NA | 3 | 0.3333 | 2 | 1 | NA | NA | 1 | 1 | NA | NA | 3 | 1 |
| 14_101680 | NA | NA | NA | NA | 2 | 0 | 2 | 0.5 | 4 | 0.25 | NA | NA | 1 | 0 | 1 | 1 | 2 | 0 | 4 | 0.25 |
| 14_101681 | NA | NA | NA | NA | 1 | 0 | 2 | 1 | 3 | 0.6667 | NA | NA | 1 | 0 | 1 | 0 | 1 | 1 | 3 | 0.3333 |
| 14_101681 | NA | NA | NA | NA | 2 | 0 | 2 | 0.5 | 4 | 0.25 | NA | NA | 1 | 1 | 1 | 1 | 1 | 0 | 3 | 0.6667 |
| 14_101681 | NA | NA | NA | NA | 1 | 0 | 2 | 1 | 3 | 0.6667 | NA | NA | NA | NA | 1 | 1 | 1 | 1 | 2 | 1 |
| 14_101682 | NA | NA | NA | NA | 2 | 0.5 | 2 | 1 | 4 | 0.75 | NA | NA | 1 | 1 | 1 | 1 | 1 | 1 | 3 | 1 |
| 14_101846 | NA | NA | NA | NA | NA | NA | 1 | 1 | 1 | 1 | NA | NA | 1 | 1 | NA | NA | NA | NA | 1 | 1 |
| 14_101870 | 1 | 0 | 4 | 0.25 | 5 | 0 | 3 | 0 | 13 | 0.0769 | 2 | 0 | 8 | 0.125 | 5 | 0 | 1 | 0 | 16 | 0.0625 |
| 14_101871 | 9 | 0.4444 | 9 | 0.4444 | 11 | 0.4545 | 8 | 0.25 | 37 | 0.4054 | 6 | 0.3333 | 14 | 0.4286 | 4 | 0.5 | 6 | 0.1667 | 30 | 0.3667 |
| 14_101871 | 9 | 0.1111 | 9 | 0 | 10 | 0.2 | 10 | 0.2 | 38 | 0.1316 | 6 | 0.1667 | 14 | 0.1429 | 4 | 0.25 | 6 | 0.3333 | 30 | 0.2 |
| 14_101871 | 8 | 0.25 | 6 | 0.3333 | 10 | 0.2 | 7 | 0 | 31 | 0.1935 | 6 | 0 | 12 | 0.25 | 2 | 0.5 | 5 | 0.2 | 25 | 0.2 |
| 14_101871 | 8 | 0.25 | 10 | 0.1 | 10 | 0.2 | 9 | 0.6667 | 37 | 0.2973 | 6 | 0.1667 | 12 | 0.25 | 4 | 0.5 | 4 | 0 | 26 | 0.2308 |
| 14_101871 | 7 | 0.5714 | 9 | 0.6667 | 9 | 0.4444 | 8 | 0.25 | 33 | 0.4848 | 6 | 0.5 | 14 | 0.6429 | 3 | 0.3333 | 6 | 0.3333 | 29 | 0.5172 |
| 14_101871 | 7 | 0 | 9 | 0.3333 | 7 | 0.1429 | 7 | 0 | 30 | 0.1333 | 4 | 0.25 | 13 | 0.0769 | 4 | 0 | 7 | 0.1429 | 28 | 0.1071 |
| 14_101871 | 9 | 0.3333 | 9 | 0.4444 | 11 | 0.3636 | 8 | 0.5 | 37 | 0.4054 | 6 | 0.3333 | 15 | 0.3333 | 4 | 0.25 | 6 | 0.8333 | 31 | 0.4194 |
| 14_102122 | 3 | 0.3333 | NA | NA | NA | NA | 2 | 0 | 5 | 0.2 | 1 | 1 | 1 | 0 | 2 | 0.5 | 1 | 1 | 5 | 0.6 |
| 14_102122 | 3 | 0.3333 | 1 | 1 | 1 | 0 | 2 | 0 | 7 | 0.2857 | 1 | 1 | 1 | 0 | 3 | 0 | 1 | 1 | 6 | 0.3333 |

| | | | | | | | | | | | | | | | | | | | | |
|---|---|---|---|---|---|---|---|---|---|---|---|---|---|---|---|---|---|---|---|---|
| 14_1021229 | 3 | 0 | 1 | 0 | 1 | 1 | 2 | 0.5 | 7 | 0.2857 | 1 | 1 | 1 | 0 | 3 | 0.3333 | 1 | 0 | 6 | 0.3333 |
| 14_102124( | 4 | 0.5 | 1 | 1 | NA | NA | 2 | 0 | 7 | 0.4286 | 1 | 0 | 2 | 0.5 | 2 | 0 | 3 | 1 | 8 | 0.5 |
| 14_102124( | 3 | 0.6667 | 1 | 1 | 1 | 1 | 2 | 1 | 7 | 0.8571 | 1 | 1 | 2 | 1 | 3 | 0.3333 | 3 | 0.3333 | 9 | 0.5556 |
| 14_1021248 | 4 | 0.25 | 1 | 0 | 1 | 1 | 2 | 0 | 8 | 0.25 | 1 | 0 | 2 | 0.5 | 4 | 0 | 3 | 0.3333 | 10 | 0.2 |
| 14_1021248 | 4 | 0.75 | 1 | 1 | 1 | 0 | 1 | 1 | 7 | 0.7143 | NA | NA | 2 | 0.5 | 4 | 0.5 | 3 | 0.6667 | 9 | 0.5556 |
| 14_1021248 | 4 | 1 | 1 | 0 | 1 | 1 | 1 | 0 | 7 | 0.7143 | NA | NA | 2 | 0.5 | 1 | 1 | 3 | 1 | 6 | 0.8333 |
| 14_1021249 | 3 | 0.3333 | 1 | 1 | 1 | 1 | 2 | 1 | 7 | 0.7143 | 1 | 1 | 2 | 1 | 3 | 1 | 3 | 1 | 9 | 1 |
| 14_1021375 | 3 | 0 | 1 | 0 | 3 | 0.3333 | 2 | 0.5 | 9 | 0.2222 | 1 | 0 | 3 | 0.3333 | 4 | 0.5 | 6 | 0.3333 | 14 | 0.3571 |
| 14_102137( | 4 | 0.25 | 1 | 0 | 3 | 0 | 2 | 0.5 | 10 | 0.2 | 1 | 0 | 4 | 0 | 5 | 0 | 6 | 0.1667 | 16 | 0.0625 |
| 14_102137( | 4 | 0.25 | 1 | 0 | 3 | 0 | 2 | 0 | 10 | 0.1 | 1 | 0 | 4 | 0 | 5 | 0.4 | 5 | 0.2 | 15 | 0.2 |
| 14_1021377 | 3 | 1 | 1 | 0 | 3 | 1 | 1 | 1 | 8 | 0.875 | 1 | 1 | 3 | 0.6667 | 5 | 0.8 | 4 | 1 | 13 | 0.8462 |
| 14_1021377 | 4 | 0 | 1 | 0 | 4 | 0.5 | 2 | 0 | 11 | 0.1818 | 1 | 0 | 4 | 0 | 5 | 0.2 | 6 | 0.1667 | 16 | 0.125 |
| 14_1021399 | 3 | 0 | 1 | 0 | 3 | 0.3333 | 3 | 0 | 10 | 0.1 | 1 | 0 | 5 | 0.2 | 5 | 0 | 5 | 0 | 16 | 0.0625 |
| 14_102170: | 1 | 0 | NA | NA | 1 | 1 | 2 | 1 | 4 | 0.75 | 1 | 1 | 1 | 0 | NA | NA | 1 | 1 | 3 | 0.6667 |
| 14_102170: | 1 | 0 | NA | NA | NA | NA | 2 | 1 | 3 | 0.6667 | 1 | 0 | 1 | 1 | NA | NA | 1 | 0 | 3 | 0.3333 |
| 14_102170( | 1 | 0 | NA | NA | 2 | 0.5 | 2 | 1 | 5 | 0.6 | 1 | 1 | 1 | 1 | NA | NA | 1 | 1 | 3 | 1 |
| 14_1021716 | 14 | 0.2143 | 10 | 0.1 | 13 | 0.3077 | 19 | 0.2105 | 56 | 0.2143 | 10 | 0.2 | 20 | 0.15 | 20 | 0.15 | 14 | 0.2143 | 64 | 0.1719 |
| 14_1021719 | 11 | 0.6364 | 11 | 0.2727 | 21 | 0.4762 | 18 | 0.3333 | 61 | 0.4262 | 11 | 0.4545 | 20 | 0.4 | 23 | 0.3478 | 13 | 0.3846 | 67 | 0.3881 |
| 14_1032142 | 6 | 0 | 13 | 0.1538 | 5 | 0 | 9 | 0 | 33 | 0.0606 | 8 | 0.25 | 9 | 0.1111 | 8 | 0 | 12 | 0.0833 | 37 | 0.1081 |
| 14_103214: | 6 | 0.1667 | 11 | 0.0909 | 5 | 0.2 | 8 | 0.125 | 30 | 0.1333 | 8 | 0.25 | 8 | 0.5 | 8 | 0.25 | 11 | 0.1818 | 35 | 0.2857 |
| 14_103215: | 8 | 0.125 | 14 | 0.0714 | 10 | 0.1 | 13 | 0.3846 | 45 | 0.1778 | 3 | 0 | 8 | 0.125 | 10 | 0.1 | 11 | 0.0909 | 32 | 0.0938 |
| 14_1032158 | 3 | 0 | 14 | 0.2857 | 8 | 0 | 8 | 0 | 33 | 0.1212 | 1 | 0 | 5 | 0.2 | 6 | 0.1667 | 8 | 0.25 | 20 | 0.2 |
| 14_1032159 | 4 | 0 | 13 | 0.1538 | 9 | 0.1111 | 12 | 0.3333 | 38 | 0.1842 | 2 | 0.5 | 7 | 0.2857 | 6 | 0.3333 | 7 | 0.4286 | 22 | 0.3636 |
| 14_103216' | 2 | 0 | 13 | 0.1538 | 9 | 0.3333 | 12 | 0.1667 | 36 | 0.1944 | 1 | 0 | 5 | 0.2 | 4 | 0.5 | 6 | 0.5 | 16 | 0.375 |
| 14_103216' | 4 | 0.25 | 14 | 0.5714 | 7 | 0.5714 | 9 | 0.1111 | 34 | 0.4118 | 2 | 1 | 5 | 0.4 | 6 | 0.8333 | 5 | 0.4 | 18 | 0.6111 |
| 14_103216' | 2 | 0 | 13 | 0.2308 | 7 | 0.2857 | 11 | 0.2727 | 33 | 0.2424 | 2 | 0.5 | 5 | 0.2 | 6 | 0.3333 | 7 | 0.5714 | 20 | 0.4 |
| 14_1032162 | 4 | 0.25 | 11 | 0.1818 | 8 | 0.375 | 10 | 0.2 | 33 | 0.2424 | 2 | 0.5 | 5 | 0.2 | 8 | 0 | 5 | 0.4 | 20 | 0.2 |
| 14_103216: | 4 | 0 | 14 | 0.2143 | 9 | 0.1111 | 11 | 0.1818 | 38 | 0.1579 | 2 | 1 | 5 | 0 | 7 | 0.2857 | 7 | 0.2857 | 21 | 0.2857 |
| 14_1032164 | 3 | 0 | 12 | 0.0833 | 8 | 0.125 | 8 | 0 | 31 | 0.0645 | 2 | 0 | 4 | 0 | 6 | 0 | 5 | 0.4 | 17 | 0.1176 |
| 14_1032164 | 3 | 0 | 10 | 0.5 | 8 | 0.5 | 11 | 0.5455 | 32 | 0.4688 | 2 | 0.5 | 5 | 0.6 | 6 | 0.5 | 6 | 1 | 19 | 0.6842 |
| 14_1032167 | 2 | 0.5 | 10 | 0 | 4 | 0.25 | 8 | 0.375 | 24 | 0.2083 | 1 | 0 | 5 | 0.2 | 4 | 0 | 5 | 0.2 | 15 | 0.1333 |
| 14_1032169 | 2 | 0 | 14 | 0.0714 | 7 | 0 | 11 | 0.0909 | 34 | 0.0588 | 1 | 0 | 5 | 0.2 | 7 | 0 | 6 | 0.1667 | 19 | 0.1053 |
| 14_1032169 | 2 | 1 | 12 | 0.5 | 6 | 0.6667 | 10 | 0.8 | 30 | 0.6667 | NA | NA | 5 | 0.8 | 6 | 0.6667 | 5 | 0.6 | 16 | 0.6875 |
| 14_103217: | 1 | 1 | 10 | 0.4 | 6 | 0.1667 | 6 | 0.3333 | 23 | 0.3478 | NA | NA | 4 | 0.5 | 5 | 0.2 | 6 | 0.5 | 15 | 0.4 |
| 14_1032180 | 1 | 0 | 7 | 0.1429 | 2 | 0.5 | 3 | 0 | 13 | 0.1538 | NA | NA | 4 | 0 | 3 | 0 | 4 | 0.25 | 11 | 0.0909 |
| 14_103218: | 1 | 1 | 6 | 0.8333 | 2 | 1 | 6 | 0.6667 | 15 | 0.8 | NA | NA | 3 | 1 | 3 | 1 | 3 | 1 | 9 | 1 |
| 14_103218: | 1 | 1 | 4 | 0.25 | 2 | 0 | 5 | 0.2 | 12 | 0.25 | NA | NA | 2 | 0.5 | 3 | 0 | 4 | 0 | 9 | 0.1111 |
| 14_1032184 | 1 | 1 | 4 | 1 | 2 | 1 | 5 | 1 | 12 | 1 | NA | NA | 2 | 1 | 2 | 1 | 4 | 1 | 8 | 1 |
| 14_1032184 | 1 | 0 | 5 | 0.4 | NA | NA | 5 | 0.4 | 11 | 0.3636 | NA | NA | 2 | 0 | 2 | 1 | 4 | 0.75 | 8 | 0.625 |
| 14_1032185 | NA | NA | 3 | 1 | 2 | 1 | 3 | 1 | 8 | 1 | NA | NA | 2 | 1 | 2 | 1 | 3 | 1 | 7 | 1 |
| 14_1032185 | NA | NA | 5 | 1 | 2 | 1 | 2 | 1 | 9 | 1 | NA | NA | 3 | 1 | 1 | 1 | 4 | 1 | 8 | 1 |
| 14_1032186 | 1 | 1 | 5 | 1 | NA | NA | 3 | 1 | 9 | 1 | NA | NA | 1 | 1 | 1 | 1 | 3 | 0.6667 | 5 | 0.8 |
| 14_1032187 | 1 | 0 | 5 | 0 | 2 | 0.5 | 4 | 0 | 12 | 0.0833 | NA | NA | 1 | 1 | 2 | 0 | 3 | 0.3333 | 6 | 0.3333 |
| 14_1032188 | 1 | 0 | 5 | 0.4 | 2 | 1 | 2 | 1 | 10 | 0.6 | NA | NA | 3 | 0.6667 | 3 | 1 | 3 | 0.3333 | 9 | 0.6667 |
| 14_1032189 | NA | NA | 5 | 0.4 | 1 | 0 | 4 | 0.5 | 10 | 0.4 | NA | NA | 2 | 0.5 | 1 | 0 | 2 | 0 | 5 | 0.2 |
| 14_1032189 | 1 | 1 | 4 | 0.75 | 1 | 0 | 2 | 0.5 | 8 | 0.625 | NA | NA | 3 | 0.3333 | 1 | 0 | 2 | 0 | 6 | 0.1667 |
| 14_103219' | NA | NA | 5 | 0.6 | 1 | 1 | 3 | 0.6667 | 9 | 0.6667 | NA | NA | 2 | 1 | 1 | 0 | 3 | 0.6667 | 6 | 0.6667 |
| 14_1032199 | NA | NA | 3 | 1 | 1 | 1 | 2 | 1 | 6 | 1 | NA | NA | 1 | 1 | 1 | 1 | 2 | 0.5 | 4 | 0.75 |
| 14_103219( | NA | NA | 3 | 0 | 1 | 0 | 2 | 0.5 | 6 | 0.1667 | NA | NA | 2 | 0 | 2 | 0.5 | 3 | 0 | 7 | 0.1429 |
| 14_103219( | NA | NA | 3 | 1 | 1 | 0 | 2 | 0.5 | 6 | 0.6667 | NA | NA | 2 | 0.5 | NA | NA | 2 | 0.5 | 4 | 0.5 |
| 14_1032197 | NA | NA | 3 | 0.3333 | 1 | 1 | 2 | 0 | 6 | 0.3333 | NA | NA | 2 | 0 | 1 | 0 | 3 | 0.3333 | 6 | 0.1667 |
| 14_1032204 | NA | NA | 1 | 1 | NA | NA | NA | NA | 1 | 1 | NA | NA | 2 | 1 | NA | NA | 2 | 1 | 4 | 1 |
| 14_1032212 | 1 | 1 | NA | NA | NA | NA | NA | NA | 1 | 1 | NA | NA | 2 | 1 | NA | NA | 2 | 1 | 4 | 1 |
| 14_1032214 | 1 | 0 | 1 | 1 | NA | NA | NA | NA | 2 | 0.5 | NA | NA | 2 | 0.5 | NA | NA | 1 | 1 | 3 | 0.6667 |
| 14_1033091 | NA | NA | 2 | 0.5 | 2 | 0.5 | NA | NA | 4 | 0.5 | 1 | 0 | 2 | 0 | 4 | 0.25 | 4 | 0.25 | 11 | 0.1818 |
| 14_1033092 | NA | NA | 2 | 0.5 | 2 | 0 | NA | NA | 4 | 0.25 | 1 | 0 | 2 | 0 | 4 | 0.25 | 4 | 0.25 | 11 | 0.1818 |
| 14_1033100 | NA | NA | 2 | 1 | 2 | 0.5 | NA | NA | 4 | 0.75 | 1 | 0 | 1 | 0 | 4 | 0.5 | 3 | 0.6667 | 9 | 0.4444 |
| 14_1033108 | NA | NA | 2 | 0.5 | 2 | 0.5 | NA | NA | 4 | 0.5 | 1 | 0 | 2 | 0.5 | 3 | 0.3333 | 3 | 0 | 9 | 0.2222 |
| 14_1033134 | NA | NA | 4 | 1 | 2 | 1 | NA | NA | 6 | 1 | 2 | 1 | 2 | 1 | 5 | 1 | 3 | 1 | 12 | 1 |
| 14_1033134 | NA | NA | 4 | 0 | 2 | 1 | NA | NA | 6 | 0.3333 | 2 | 0.5 | 2 | 0.5 | 5 | 0.4 | 3 | 1 | 12 | 0.5833 |
| 14_1033135 | NA | NA | 4 | 1 | 1 | 1 | NA | NA | 5 | 1 | 2 | 0.5 | 2 | 1 | 5 | 0.8 | 3 | 1 | 12 | 0.8333 |
| 14_1033135 | NA | NA | 4 | 0.5 | 2 | 0.5 | NA | NA | 6 | 0.5 | 2 | 0 | 2 | 0 | 3 | 0.3333 | 3 | 0.6667 | 10 | 0.3 |
| 14_1033136 | NA | NA | 4 | 1 | 2 | 1 | NA | NA | 6 | 1 | 2 | 1 | 2 | 1 | 4 | 1 | 3 | 0.6667 | 11 | 0.9091 |
| 14_1033136 | NA | NA | 2 | 0.5 | 1 | 0 | NA | NA | 3 | 0.3333 | 1 | 1 | 1 | 0 | 2 | 0 | 3 | 0 | 7 | 0.1429 |
| 14_1033138 | NA | NA | 4 | 0.75 | 2 | 1 | 1 | 1 | 7 | 0.8571 | 2 | 1 | 3 | 1 | 4 | 1 | 3 | 1 | 12 | 1 |

| | | | | | | | | | | | | | | | | | | | | |
|---|---|---|---|---|---|---|---|---|---|---|---|---|---|---|---|---|---|---|---|---|
| 14_103314 | 1 | 0 | 3 | 0.3333 | 2 | 0.5 | 1 | 1 | 7 | 0.4286 | 2 | 0.5 | 4 | 0.25 | 5 | 0.6 | 3 | 0.6667 | 14 | 0.5 |
| 14_103314 | 1 | 1 | 5 | 0.2 | 2 | 0.5 | 1 | 0 | 9 | 0.3333 | 2 | 0 | 4 | 0.25 | 4 | 0.25 | 3 | 0.6667 | 13 | 0.3077 |
| 14_103314 | 1 | 0 | 5 | 0.4 | 1 | 0 | 1 | 0 | 8 | 0.25 | 2 | 0 | 4 | 0 | 5 | 0.4 | 3 | 0 | 14 | 0.1429 |
| 14_103320 | 7 | 0.1429 | 15 | 0.3333 | 6 | 0.1667 | 10 | 0.3 | 38 | 0.2632 | 9 | 0.2222 | 7 | 0 | 9 | 0.3333 | 18 | 0.1111 | 43 | 0.1628 |
| 14_103320 | 7 | 0.1429 | 15 | 0.5333 | 7 | 0.4286 | 11 | 0.2727 | 40 | 0.375 | 9 | 0.6667 | 6 | 0.5 | 9 | 0.2222 | 18 | 0.6111 | 42 | 0.5238 |
| 14_103320 | 4 | 1 | 12 | 0.9167 | 6 | 1 | 10 | 1 | 32 | 0.9688 | 8 | 0.875 | 6 | 1 | 7 | 0.7143 | 16 | 1 | 37 | 0.9189 |
| 14_103320 | 6 | 0.8333 | 15 | 0.6 | 5 | 1 | 9 | 0.7778 | 35 | 0.7429 | 8 | 0.75 | 6 | 0.5 | 7 | 0.5714 | 13 | 0.8462 | 34 | 0.7059 |
| 14_103320 | 6 | 0.1667 | 11 | 0.0909 | 6 | 0 | 8 | 0 | 31 | 0.0645 | 8 | 0.25 | 7 | 0 | 5 | 0 | 14 | 0.0714 | 34 | 0.0882 |
| 14_103320 | 5 | 0 | 11 | 0.0909 | 6 | 0.1667 | 9 | 0 | 31 | 0.0645 | 8 | 0.125 | 6 | 0 | 6 | 0.1667 | 17 | 0.0588 | 37 | 0.0811 |
| 14_103320 | 5 | 0.8 | 12 | 0.8333 | 4 | 0.75 | 9 | 0.8889 | 30 | 0.8333 | 7 | 0.8571 | 5 | 0.8 | 7 | 0.8571 | 15 | 0.8 | 34 | 0.8235 |
| 14_103320 | 6 | 0.5 | 9 | 0.5556 | 7 | 0.4286 | 8 | 0.5 | 30 | 0.5 | 6 | 1 | 4 | 0.75 | 3 | 0.3333 | 12 | 0.5 | 25 | 0.64 |
| 14_103320 | 6 | 0.1667 | 10 | 0.4 | 6 | 0.1667 | 8 | 0.375 | 30 | 0.3 | 7 | 0.4286 | 6 | 0.1667 | 4 | 0.25 | 13 | 0.1538 | 30 | 0.2333 |
| 14_103320 | 7 | 0.7143 | 10 | 0.6 | 5 | 0.6 | 8 | 0.75 | 30 | 0.6667 | 8 | 0.875 | 6 | 0.5 | 4 | 0.75 | 14 | 0.3571 | 32 | 0.5625 |
| 14_103332 | 4 | 0.5 | 6 | 1 | 2 | 1 | 6 | 0.5 | 18 | 0.7222 | 4 | 1 | 3 | 0.6667 | 4 | 1 | 8 | 0.75 | 19 | 0.8421 |
| 14_103332 | 6 | 0.1667 | 6 | 0 | 2 | 0 | 8 | 0.125 | 22 | 0.0909 | 7 | 0 | 2 | 0 | 5 | 0.2 | 7 | 0.1429 | 21 | 0.0952 |
| 14_103332 | 6 | 0 | 7 | 0 | 4 | 0 | 9 | 0.3333 | 26 | 0.1154 | 6 | 0.1667 | 3 | 0 | 6 | 0 | 7 | 0.1429 | 22 | 0.0909 |
| 14_103351 | 16 | 0.8125 | 24 | 0.7917 | 13 | 0.8462 | 21 | 0.8095 | 74 | 0.8108 | 9 | 0.7778 | 13 | 0.7692 | 16 | 0.8125 | 22 | 0.7273 | 60 | 0.7667 |
| 14_103351 | 13 | 0 | 16 | 0.0625 | 14 | 0.0714 | 19 | 0.1053 | 62 | 0.0645 | 7 | 0.1429 | 14 | 0.0714 | 13 | 0 | 19 | 0.1053 | 53 | 0.0755 |
| 14_103351 | 14 | 0.9286 | 23 | 0.7826 | 12 | 0.6667 | 22 | 0.7273 | 71 | 0.7746 | 10 | 1 | 16 | 0.9375 | 16 | 0.75 | 22 | 0.9091 | 64 | 0.8906 |
| 14_103351 | 15 | 0.0667 | 23 | 0.1304 | 12 | 0.1667 | 22 | 0.0455 | 72 | 0.0972 | 9 | 0.1111 | 16 | 0.25 | 14 | 0.1429 | 19 | 0.0526 | 58 | 0.1379 |
| 14_103351 | NA | NA | 1 | 1 | NA | NA | NA | NA | 1 | 1 | NA | NA | NA | NA | NA | NA | 1 | 1 | 1 | 1 |
| 14_103351 | 10 | 0.1 | 15 | 0.2 | 9 | 0.2222 | 12 | 0.0833 | 46 | 0.1522 | 7 | 0.1429 | 13 | 0.3077 | 9 | 0.3333 | 19 | 0.1579 | 48 | 0.2292 |
| 14_103351 | 14 | 0.6429 | 22 | 0.4091 | 14 | 0.2143 | 22 | 0.3636 | 72 | 0.4028 | 9 | 0.3333 | 17 | 0.4706 | 15 | 0.7333 | 24 | 0.4583 | 65 | 0.5077 |
| 14_103351 | 15 | 0.6 | 22 | 0.5455 | 13 | 0.5385 | 18 | 0.3889 | 68 | 0.5147 | 8 | 1 | 16 | 0.4375 | 14 | 0.4286 | 25 | 0.48 | 63 | 0.5238 |
| 14_103351 | 13 | 0.6923 | 20 | 0.45 | 11 | 0.6364 | 15 | 0.6 | 59 | 0.5763 | 6 | 0.8333 | 14 | 0.7143 | 12 | 0.5 | 19 | 0.6842 | 51 | 0.6667 |
| 14_103351 | 12 | 0.1667 | 22 | 0.0455 | 13 | 0 | 18 | 0.0556 | 65 | 0.0615 | 7 | 0.1429 | 16 | 0.0625 | 13 | 0.0769 | 23 | 0 | 59 | 0.0508 |
| 14_103371 | 9 | 0.1111 | 13 | 0.1538 | 5 | 0 | 7 | 0.2857 | 34 | 0.1471 | 2 | 0 | 10 | 0.1 | 6 | 0.3333 | 6 | 0.1667 | 24 | 0.1667 |
| 14_103393 | 5 | 0.8 | 4 | 0.75 | 5 | 0.8 | 3 | 0.6667 | 17 | 0.7647 | NA | NA | 4 | 0.75 | 2 | 0.5 | 4 | 0.75 | 10 | 0.7 |
| 14_103393 | 5 | 0 | 4 | 0 | 4 | 0.25 | 3 | 0 | 16 | 0.0625 | NA | NA | 5 | 0.2 | 2 | 0 | 2 | 0.5 | 9 | 0.2222 |
| 14_103393 | 4 | 0.25 | 3 | 0 | 5 | 0 | 3 | 0.3333 | 15 | 0.1333 | NA | NA | 4 | 0 | 3 | 0.3333 | 3 | 0 | 10 | 0.1 |
| 14_103393 | 5 | 0.4 | 4 | 0.5 | 4 | 0.75 | 3 | 0.3333 | 16 | 0.5 | NA | NA | 3 | 0.3333 | 3 | 0.3333 | 1 | 0 | 7 | 0.2857 |
| 14_103393 | 4 | 0.5 | 1 | 1 | 2 | 1 | 2 | 0 | 9 | 0.5556 | NA | NA | 3 | 0.3333 | 2 | 0 | 2 | 0.5 | 7 | 0.2857 |
| 14_103393 | 4 | 0 | 3 | 0.3333 | 4 | 0 | 1 | 0 | 12 | 0.0833 | NA | NA | 4 | 0 | 3 | 0 | 2 | 0.5 | 9 | 0.1111 |
| 14_103393 | 5 | 0.2 | 3 | 0.3333 | 4 | 0.25 | 2 | 0.5 | 14 | 0.2857 | NA | NA | 4 | 0.25 | 2 | 0 | 2 | 0 | 8 | 0.125 |
| 14_103393 | 4 | 0 | 3 | 0 | 4 | 0.25 | 2 | 0 | 13 | 0.0769 | NA | NA | 4 | 0 | 2 | 0 | 2 | 0.5 | 8 | 0.125 |
| 14_103393 | 5 | 0.2 | 3 | 0 | 2 | 0.5 | 2 | 0 | 12 | 0.1667 | NA | NA | 4 | 0.25 | 1 | 0 | 2 | 0 | 7 | 0.1429 |
| 14_103393 | 5 | 0.2 | 3 | 0 | 4 | 0.5 | 2 | 0 | 14 | 0.2143 | NA | NA | 4 | 0.25 | 2 | 0 | 2 | 0.5 | 8 | 0.25 |
| 14_103394 | 5 | 0.8 | 2 | 0 | 3 | 0.3333 | 2 | 0.5 | 12 | 0.5 | NA | NA | 4 | 0.25 | 2 | 1 | 2 | 0 | 8 | 0.375 |
| 14_103678 | 1 | 1 | 1 | 1 | 3 | 1 | 1 | 1 | 6 | 1 | NA | NA | 2 | 0.5 | NA | NA | NA | NA | 2 | 0.5 |
| 14_103679 | 1 | 1 | 1 | 1 | 3 | 1 | 1 | 1 | 6 | 1 | NA | NA | 2 | 1 | NA | NA | NA | NA | 2 | 1 |
| 14_103679 | 1 | 1 | 2 | 0.5 | 2 | 0 | 1 | 1 | 6 | 0.5 | NA | NA | 2 | 0.5 | NA | NA | NA | NA | 2 | 0.5 |
| 14_103679 | 1 | 1 | 2 | 1 | 3 | 0.6667 | NA | NA | 6 | 0.8333 | NA | NA | 1 | 1 | NA | NA | NA | NA | 1 | 1 |
| 14_103679 | 1 | 0 | 2 | 1 | 2 | 0 | 1 | 0 | 6 | 0.3333 | NA | NA | 2 | 0.5 | NA | NA | NA | NA | 2 | 0.5 |
| 14_103679 | 1 | 1 | 2 | 1 | 3 | 0 | 1 | 1 | 7 | 0.5714 | NA | NA | 2 | 0.5 | NA | NA | NA | NA | 2 | 0.5 |
| 14_103680 | 1 | 0 | 1 | 0 | 2 | 0.5 | 1 | 1 | 5 | 0.4 | NA | NA | 2 | 1 | NA | NA | NA | NA | 2 | 1 |
| 14_103680 | 1 | 1 | 2 | 1 | 4 | 1 | 1 | 1 | 8 | 1 | NA | NA | 2 | 1 | NA | NA | NA | NA | 2 | 1 |
| 14_103680 | 1 | 1 | 2 | 0 | 4 | 0.5 | NA | NA | 7 | 0.4286 | NA | NA | 2 | 0.5 | NA | NA | NA | NA | 2 | 0.5 |
| 14_103680 | 1 | 0 | 3 | 0.6667 | 3 | 0.6667 | 1 | 1 | 8 | 0.625 | NA | NA | 2 | 0.5 | NA | NA | NA | NA | 2 | 0.5 |
| 14_103680 | 1 | 1 | 3 | 0 | 4 | 0 | 1 | 1 | 9 | 0.2222 | NA | NA | 2 | 0.5 | NA | NA | NA | NA | 2 | 0.5 |
| 14_103680 | 1 | 1 | 2 | 1 | 3 | 1 | 1 | 1 | 7 | 1 | NA | NA | 2 | 1 | NA | NA | NA | NA | 2 | 1 |
| 14_103680 | 1 | 1 | 3 | 0 | 3 | 0.3333 | NA | NA | 7 | 0.2857 | NA | NA | 2 | 1 | NA | NA | NA | NA | 2 | 1 |
| 14_103686 | 1 | 1 | 2 | 1 | 4 | 1 | 1 | 1 | 8 | 1 | NA | NA | 2 | 0.5 | 1 | 0 | 1 | 1 | 4 | 0.5 |
| 14_103686 | 1 | 0 | 3 | 0.6667 | 4 | 0.25 | 1 | 1 | 9 | 0.4444 | NA | NA | 1 | 1 | 1 | 1 | 1 | 1 | 3 | 1 |
| 14_103687 | 1 | 0 | 1 | 0 | 3 | 0.3333 | 1 | 0 | 6 | 0.1667 | NA | NA | 2 | 0 | NA | NA | 1 | 1 | 3 | 0.3333 |
| 14_103925 | 6 | 0.3333 | 6 | 0 | 5 | 0 | 5 | 0 | 22 | 0.0909 | 2 | 0.5 | 4 | 0.25 | 2 | 0.5 | 8 | 0.375 | 16 | 0.375 |
| 14_103925 | 6 | 0.3333 | 7 | 0.7143 | 4 | 0.25 | 3 | 0.3333 | 20 | 0.45 | 3 | 0.3333 | 4 | 0.25 | 3 | 0.3333 | 9 | 0.3333 | 19 | 0.3158 |
| 14_103941 | 6 | 0.8333 | 9 | 0.6667 | 7 | 1 | 6 | 0.5 | 28 | 0.75 | 4 | 0.75 | 5 | 0.4 | 5 | 1 | 11 | 0.7273 | 25 | 0.72 |
| 14_103941 | 8 | 0.375 | 9 | 0.6667 | 6 | 0.5 | 6 | 0.5 | 29 | 0.5172 | 3 | 0 | 4 | 0.25 | 4 | 0.25 | 9 | 0.6667 | 20 | 0.4 |
| 14_103941 | 9 | 1 | 9 | 1 | 7 | 1 | 6 | 1 | 31 | 1 | 4 | 0.5 | 7 | 0.8571 | 5 | 0.8 | 11 | 0.7273 | 27 | 0.7407 |
| 14_103941 | 9 | 0.8889 | 9 | 0.4444 | 5 | 0.8 | 6 | 0.5 | 29 | 0.6552 | 6 | 0.3333 | 5 | 0.8 | 5 | 0.4 | 10 | 0.5 | 26 | 0.5 |
| 14_103942 | 9 | 0.1111 | 7 | 0 | 9 | 0.3333 | 5 | 0 | 30 | 0.1333 | 6 | 0.1667 | 7 | 0 | 4 | 0.25 | 11 | 0.0909 | 28 | 0.1071 |
| 14_104052 | 3 | 0.3333 | 6 | 0 | 6 | 0.1667 | 4 | 0 | 19 | 0.1053 | 3 | 0 | 10 | 0.1 | 3 | 0 | 7 | 0.1429 | 23 | 0.087 |
| 14_104053 | 2 | 0.5 | 5 | 0.4 | 5 | 0.2 | 5 | 0.2 | 17 | 0.2941 | 4 | 0.5 | 8 | 0.375 | 3 | 0.3333 | 6 | 0.6667 | 21 | 0.4762 |
| 14_104053 | 2 | 0.5 | 6 | 0 | 4 | 0 | 5 | 0 | 17 | 0.0588 | 3 | 0 | 6 | 0.3333 | 3 | 0 | 5 | 0 | 17 | 0.1176 |
| 14_104053 | 2 | 0 | 5 | 0 | 4 | 0 | 5 | 0.2 | 16 | 0.0625 | 2 | 0 | 5 | 0.2 | 3 | 0 | 3 | 0 | 13 | 0.0769 |

| | | | | | | | | | | | | | | | | | | | | |
|---|---|---|---|---|---|---|---|---|---|---|---|---|---|---|---|---|---|---|---|---|
| 14_104054( | 1 | 0 | NA | NA | 3 | 0 | 1 | 1 | 5 | 0.2 | 2 | 0 | 2 | 0.5 | NA | NA | NA | NA | 4 | 0.25 |
| 14_104054' | 2 | 0 | 6 | 0 | 5 | 0 | 3 | 0.3333 | 16 | 0.0625 | 3 | 0.3333 | 7 | 0 | 3 | 0 | 4 | 0 | 17 | 0.0588 |
| 14_104054; | NA | NA | 5 | 0 | 4 | 0.25 | 3 | 0.3333 | 12 | 0.1667 | 1 | 0 | 3 | 0.3333 | 2 | 0 | 2 | 0.5 | 8 | 0.25 |
| 14_104054 | 2 | 0 | 6 | 0.1667 | 5 | 0 | 4 | 0 | 17 | 0.0588 | 1 | 0 | 7 | 0.1429 | 3 | 0 | 3 | 0 | 14 | 0.0714 |
| 14_104054 | 2 | 0 | 5 | 0 | 5 | 0 | 4 | 0.25 | 16 | 0.0625 | 2 | 0 | 6 | 0 | 2 | 0.5 | 3 | 0 | 13 | 0.0769 |
| 14_104054 | 2 | 0 | 5 | 0.4 | 4 | 0.75 | 5 | 0.4 | 16 | 0.4375 | 3 | 0.6667 | 6 | 0.6667 | 3 | 0.3333 | 3 | 0.3333 | 15 | 0.5333 |
| 14_104054 | 1 | 0 | 4 | 0.25 | 3 | 0 | 4 | 0.5 | 12 | 0.25 | 2 | 0 | 5 | 0 | 2 | 0 | 2 | 0.5 | 11 | 0.0909 |
| 14_104080 | 1 | 0 | 1 | 0 | 5 | 0 | 3 | 0.3333 | 10 | 0.1 | 1 | 0 | 1 | 0 | 2 | 0 | 3 | 0.3333 | 7 | 0.1429 |
| 14_104080( | 1 | 1 | 5 | 0.8 | 5 | 0.6 | 4 | 0.25 | 15 | 0.6 | 1 | 0 | 4 | 0.75 | 2 | 1 | 4 | 1 | 11 | 0.8182 |
| 14_104080 | 1 | 1 | 5 | 0.2 | 5 | 0 | 3 | 0.3333 | 14 | 0.2143 | 1 | 0 | 3 | 0.6667 | 1 | 0 | 2 | 0 | 7 | 0.2857 |
| 14_104080 | 1 | 1 | 4 | 0.75 | 5 | 0.8 | 4 | 0.75 | 14 | 0.7857 | 1 | 0 | 3 | 0.3333 | NA | NA | 1 | 1 | 5 | 0.4 |
| 14_104080 | 1 | 0 | 4 | 0.25 | 5 | 0 | 3 | 0 | 13 | 0.0769 | 1 | 1 | 3 | 0 | NA | NA | 2 | 0 | 6 | 0.1667 |
| 14_104715 | 1 | 0 | 1 | 0 | NA | NA | 1 | 1 | 3 | 0.3333 | 1 | 0 | 1 | 0 | 2 | 0.5 | 1 | 0 | 5 | 0.2 |
| 14_104716( | 1 | 0 | NA | NA | NA | NA | 1 | 1 | 2 | 0.5 | 1 | 0 | 1 | 0 | 2 | 0.5 | 2 | 0 | 6 | 0.1667 |
| 14_104716' | 1 | 1 | 1 | 0 | NA | NA | 1 | 1 | 3 | 0.6667 | NA | NA | 1 | 1 | 2 | 0.5 | 2 | 0.5 | 5 | 0.6 |
| 14_104716 | 1 | 1 | NA | NA | NA | NA | 1 | 0 | 2 | 0.5 | NA | NA | 1 | 1 | 1 | 1 | 2 | 0.5 | 4 | 0.75 |
| 14_104716 | NA | NA | 1 | 1 | NA | NA | 1 | 0 | 2 | 0.5 | NA | NA | NA | NA | 2 | 1 | 1 | 0 | 3 | 0.6667 |
| 14_104716 | 1 | 1 | 1 | 1 | NA | NA | 1 | 1 | 3 | 1 | NA | NA | 1 | 0 | 2 | 1 | 1 | 0 | 4 | 0.5 |
| 14_104716 | 1 | 1 | 1 | 1 | NA | NA | NA | NA | 2 | 1 | NA | NA | 1 | 1 | 2 | 1 | NA | NA | 3 | 1 |
| 14_104716( | NA | NA | NA | NA | NA | NA | 1 | 1 | 1 | 1 | NA | NA | 1 | 1 | 1 | 1 | 2 | 0.5 | 4 | 0.75 |
| 14_104716 | 1 | 1 | NA | NA | NA | NA | NA | NA | 1 | 1 | NA | NA | 1 | 1 | 2 | 1 | 2 | 1 | 5 | 1 |
| 14_104716 | 1 | 1 | 1 | 0 | NA | NA | 1 | 0 | 3 | 0.3333 | NA | NA | 1 | 0 | 2 | 0.5 | 2 | 0 | 5 | 0.2 |
| 14_104717( | 1 | 1 | NA | NA | NA | NA | 1 | 0 | 2 | 0.5 | NA | NA | 1 | 1 | 2 | 0.5 | 1 | 0 | 4 | 0.5 |
| 14_104717 | NA | NA | 1 | 1 | NA | NA | 1 | 0 | 2 | 0.5 | NA | NA | NA | NA | 2 | 1 | 1 | 1 | 3 | 1 |
| 14_104717 | NA | NA | 1 | 1 | NA | NA | 1 | 1 | 2 | 1 | NA | NA | NA | NA | 2 | 0.5 | 2 | 0 | 4 | 0.25 |
| 14_104717 | NA | NA | 1 | 0 | NA | NA | 1 | 1 | 2 | 0.5 | NA | NA | NA | NA | 1 | 1 | 2 | 1 | 3 | 1 |
| 14_105344 | 1 | 0 | 6 | 0.1667 | 6 | 0 | 6 | 0.1667 | 19 | 0.1053 | NA | NA | 6 | 0.1667 | 4 | 0.25 | 1 | 0 | 11 | 0.1818 |
| 14_105344 | 1 | 0 | 4 | 0 | 2 | 0 | 5 | 0.2 | 12 | 0.0833 | NA | NA | 5 | 0.2 | 4 | 0 | NA | NA | 9 | 0.1111 |
| 14_105344 | 1 | 1 | 6 | 1 | 6 | 0.6667 | 6 | 0.5 | 19 | 0.7368 | NA | NA | 6 | 0.5 | 5 | 1 | NA | NA | 11 | 0.7273 |
| 14_105345 | 1 | 0 | 6 | 0 | 5 | 0.2 | 4 | 0 | 16 | 0.0625 | NA | NA | 6 | 0.3333 | 5 | 0 | NA | NA | 11 | 0.1818 |
| 14_105347' | NA | NA | 8 | 0.625 | 6 | 0.5 | 4 | 0.25 | 18 | 0.5 | 1 | 0 | 3 | 0.3333 | 4 | 0.5 | 3 | 0.3333 | 11 | 0.3636 |
| 14_105352( | 1 | 1 | 4 | 0 | 1 | 0 | 4 | 0 | 10 | 0.1 | NA | NA | 5 | 0.2 | 3 | 0 | 3 | 0 | 11 | 0.0909 |
| 14_105352 | 1 | 0 | 3 | 0 | 3 | 0 | 3 | 0.3333 | 10 | 0.1 | NA | NA | 4 | 0.25 | 3 | 0.3333 | 3 | 0.3333 | 10 | 0.3 |
| 14_105352 | 1 | 1 | 3 | 1 | 2 | 1 | 2 | 1 | 8 | 1 | NA | NA | 4 | 1 | 2 | 1 | 3 | 0.3333 | 9 | 0.7778 |
| 14_105352( | 1 | 0 | 2 | 0 | 1 | 0 | 3 | 0.3333 | 7 | 0.1429 | NA | NA | 4 | 0.25 | 3 | 0.3333 | 3 | 0 | 10 | 0.2 |
| 14_105415 | 1 | 1 | NA | NA | NA | NA | NA | NA | 1 | 1 | 1 | 1 | NA | NA | 1 | 1 | NA | NA | 2 | 1 |
| 14_105415 | 1 | 1 | NA | NA | NA | NA | NA | NA | 1 | 1 | NA | NA | NA | NA | 1 | 1 | NA | NA | 1 | 1 |
| 14_105415 | 1 | 1 | NA | NA | NA | NA | NA | NA | 1 | 1 | 1 | 0 | NA | NA | 1 | 1 | NA | NA | 2 | 0.5 |
| 14_105415 | 1 | 1 | NA | NA | NA | NA | NA | NA | 1 | 1 | 1 | 1 | NA | NA | 1 | 0 | NA | NA | 2 | 0.5 |
| 14_105537 | NA | NA | NA | NA | 1 | 1 | NA | NA | 1 | 1 | NA | NA | 1 | 1 | NA | NA | NA | NA | 1 | 1 |
| 14_105538' | NA | NA | NA | NA | 1 | 1 | NA | NA | 1 | 1 | NA | NA | 1 | 1 | NA | NA | NA | NA | 1 | 1 |
| 14_105538' | NA | NA | NA | NA | 1 | 1 | NA | NA | 1 | 1 | NA | NA | 1 | 1 | NA | NA | NA | NA | 1 | 1 |
| 14_105858( | 2 | 1 | 2 | 1 | 1 | 1 | 1 | 0 | 6 | 0.8333 | 1 | 1 | 2 | 1 | 2 | 0.5 | 2 | 1 | 7 | 0.8571 |
| 14_105858 | 2 | 0.5 | 1 | 0 | 3 | 0 | 1 | 0 | 7 | 0.1429 | 1 | 1 | 4 | 0 | 2 | 0 | 2 | 0.5 | 9 | 0.2222 |
| 14_105858 | 2 | 1 | 2 | 0.5 | 2 | 1 | 1 | 1 | 7 | 0.8571 | NA | NA | 5 | 0.8 | 2 | 0 | 2 | 1 | 9 | 0.6667 |
| 14_105858 | 2 | 1 | 2 | 0.5 | 3 | 0.6667 | 1 | 1 | 8 | 0.75 | NA | NA | 1 | 1 | 2 | 0 | 2 | 0.5 | 5 | 0.4 |
| 14_105858 | 2 | 1 | 2 | 0 | 3 | 0.3333 | NA | NA | 7 | 0.4286 | NA | NA | 3 | 0.6667 | 1 | 0 | 2 | 0.5 | 6 | 0.5 |
| 14_105859( | 1 | 1 | 2 | 1 | 2 | 1 | 1 | 0 | 6 | 0.8333 | NA | NA | 4 | 0.75 | 1 | 1 | 2 | 1 | 7 | 0.8571 |
| 14_105859' | 2 | 1 | 1 | 0 | 3 | 0.3333 | 1 | 1 | 7 | 0.5714 | NA | NA | 4 | 0.75 | 2 | 0.5 | 2 | 0 | 8 | 0.5 |
| 14_105859 | 2 | 1 | 2 | 1 | 2 | 0 | 1 | 0 | 7 | 0.5714 | NA | NA | 5 | 0.2 | 2 | 0 | 2 | 0.5 | 9 | 0.2222 |
| 14_105859 | 2 | 0 | 2 | 0.5 | 2 | 0.5 | NA | NA | 6 | 0.3333 | NA | NA | 3 | 0.3333 | 2 | 0 | 2 | 0.5 | 7 | 0.2857 |
| 14_105860( | 1 | 1 | 1 | 1 | 2 | 1 | NA | NA | 4 | 1 | NA | NA | 4 | 1 | 2 | 1 | 1 | 1 | 7 | 1 |
| 14_105919 | 89 | 0.3146 | 118 | 0.2627 | 108 | 0.2315 | 92 | 0.3804 | 407 | 0.2924 | 100 | 0.32 | 128 | 0.3047 | 110 | 0.3545 | 104 | 0.3365 | 442 | 0.3281 |
| 14_105919 | 80 | 0.325 | 111 | 0.2973 | 89 | 0.2247 | 85 | 0.2 | 365 | 0.263 | 91 | 0.3407 | 112 | 0.2768 | 94 | 0.234 | 91 | 0.3956 | 388 | 0.3093 |
| 14_106129( | 16 | 0.125 | 11 | 0.1818 | 18 | 0.4444 | 9 | 0.4444 | 54 | 0.2963 | 8 | 0.25 | 21 | 0.3333 | 8 | 0.375 | 9 | 0.3333 | 46 | 0.3261 |
| 14_106129 | 16 | 0.0625 | 10 | 0.2 | 17 | 0.1765 | 9 | 0.1111 | 52 | 0.1346 | 6 | 0.1667 | 18 | 0.1111 | 6 | 0.1667 | 10 | 0.1 | 40 | 0.125 |
| 14_106130 | 10 | 0.1 | 8 | 0.125 | 11 | 0.2727 | 8 | 0.5 | 37 | 0.2432 | 4 | 0.25 | 14 | 0.2143 | 7 | 0.4286 | 10 | 0.4 | 35 | 0.3143 |
| 14_106130 | 12 | 0.3333 | 8 | 0.125 | 10 | 0.2 | 7 | 0.1429 | 37 | 0.2162 | 3 | 0.3333 | 9 | 0.3333 | 6 | 0 | 5 | 0.4 | 23 | 0.2609 |
| 14_106130 | 8 | 1 | 9 | 0.8889 | 11 | 1 | 7 | 1 | 35 | 0.9714 | 3 | 1 | 10 | 1 | 7 | 1 | 8 | 1 | 28 | 1 |
| 14_106130 | 9 | 1 | 7 | 1 | 13 | 1 | 8 | 0.875 | 37 | 0.973 | 4 | 1 | 11 | 1 | 5 | 1 | 10 | 1 | 30 | 1 |
| 14_106130 | 8 | 0.875 | 8 | 1 | 12 | 0.8333 | 7 | 1 | 35 | 0.9143 | 2 | 1 | 10 | 0.6 | 5 | 1 | 10 | 1 | 27 | 0.8519 |
| 14_106130 | 9 | 0.8889 | 6 | 0.6667 | 12 | 0.6667 | 7 | 0.8571 | 34 | 0.7647 | 4 | 0.75 | 12 | 0.5833 | 5 | 1 | 10 | 0.9 | 31 | 0.7742 |
| 14_106130( | 8 | 1 | 7 | 1 | 12 | 1 | 8 | 1 | 35 | 1 | 3 | 1 | 12 | 0.9167 | 6 | 1 | 10 | 1 | 31 | 0.9677 |
| 14_106130 | 8 | 0.375 | 6 | 0.5 | 9 | 0.2222 | 7 | 0.5714 | 30 | 0.4 | 2 | 0 | 9 | 0.3333 | 6 | 0.6667 | 10 | 0.5 | 27 | 0.4444 |
| 14_106130 | 8 | 1 | 8 | 1 | 8 | 1 | 7 | 1 | 31 | 1 | 3 | 1 | 11 | 1 | 6 | 1 | 9 | 1 | 29 | 1 |

| | | | | | | | | | | | | | | | | | | | | |
|---|---|---|---|---|---|---|---|---|---|---|---|---|---|---|---|---|---|---|---|---|
| 14_1061307 | 5 | 0.2 | 5 | 0.2 | 8 | 0.375 | 6 | 0.3333 | 24 | 0.2917 | 3 | 0 | 8 | 0.375 | 3 | 0 | 7 | 0.1429 | 21 | 0.1905 |
| 14_1061308 | 7 | 1 | 9 | 1 | 11 | 0.9091 | 7 | 0.8571 | 34 | 0.9412 | 3 | 0.6667 | 9 | 0.7778 | 5 | 1 | 10 | 0.8 | 27 | 0.8148 |
| 14_1061309 | 8 | 1 | 7 | 0.5714 | 9 | 0.8889 | 7 | 0.7143 | 31 | 0.8065 | 4 | 1 | 10 | 0.5 | 4 | 0.75 | 9 | 0.6667 | 27 | 0.6667 |
| 14_1061340 | 7 | 0.5714 | 6 | 0.5 | 5 | 0.4 | 6 | 0.3333 | 24 | 0.4583 | NA | NA | 6 | 0.3333 | 5 | 1 | 7 | 0.5714 | 18 | 0.6111 |
| 14_1061341 | 5 | 1 | 5 | 1 | 5 | 0.8 | 7 | 0.5714 | 22 | 0.8182 | NA | NA | 4 | 0.75 | 5 | 0.8 | 6 | 0.8333 | 15 | 0.8 |
| 14_1061341 | 6 | 1 | 5 | 1 | 5 | 1 | 6 | 1 | 22 | 1 | NA | NA | 6 | 1 | 5 | 1 | 6 | 1 | 17 | 1 |
| 14_1061343 | 6 | 0.3333 | 3 | 1 | 6 | 0.8333 | 6 | 0.3333 | 21 | 0.5714 | NA | NA | 4 | 0.25 | 4 | 1 | 7 | 0.7143 | 15 | 0.6667 |
| 14_1091469 | NA | NA | NA | NA | 1 | 1 | NA | NA | 1 | 1 | 2 | 0.5 | NA | NA | 1 | 0 | NA | NA | 3 | 0.3333 |
| 14_1091478 | 8 | 0.375 | 9 | 0.2222 | 11 | 0.1818 | 6 | 0.1667 | 34 | 0.2353 | 7 | 0.1429 | 2 | 0 | 5 | 0 | 6 | 0.1667 | 20 | 0.1 |
| 14_1091479 | 8 | 0.25 | 8 | 0 | 10 | 0.1 | 4 | 0.25 | 30 | 0.1333 | 6 | 0.3333 | 3 | 0 | 5 | 0.2 | 6 | 0.1667 | 20 | 0.2 |
| 14_1091479 | 7 | 0.7143 | 6 | 1 | 12 | 1 | 6 | 0.8333 | 31 | 0.9032 | 7 | 0.7143 | 3 | 0.6667 | 5 | 0.6 | 7 | 0.4286 | 22 | 0.5909 |
| 14_1091480 | 7 | 1 | 9 | 1 | 11 | 1 | 6 | 1 | 33 | 1 | 7 | 0.8571 | 2 | 1 | 7 | 0.8571 | 6 | 1 | 22 | 0.9091 |
| 14_1091481 | 6 | 0.1667 | 9 | 0.2222 | 10 | 0.2 | 3 | 0.6667 | 28 | 0.25 | 6 | 0.1667 | 2 | 0 | 5 | 0.2 | 4 | 0 | 17 | 0.1176 |
| 14_1091481 | 8 | 0.25 | 8 | 0.375 | 11 | 0.1818 | 5 | 0.2 | 32 | 0.25 | 6 | 0 | 3 | 0.3333 | 3 | 0.3333 | 5 | 0.4 | 17 | 0.2353 |
| 14_1091481 | 9 | 0.2222 | 9 | 0.6667 | 12 | 0.6667 | 5 | 1 | 35 | 0.6 | 6 | 1 | 2 | 0.5 | 4 | 1 | 4 | 0.75 | 16 | 0.875 |
| 14_1091483 | 7 | 0.4286 | 8 | 0.5 | 10 | 0.5 | 6 | 0.3333 | 31 | 0.4516 | 6 | 0.6667 | 1 | 0 | 3 | 1 | 5 | 0.4 | 15 | 0.6 |
| 14_1091484 | 8 | 0.625 | 6 | 0.8333 | 10 | 0.9 | 6 | 0.6667 | 30 | 0.7667 | 7 | 0.8571 | 1 | 1 | 5 | 1 | 3 | 0.6667 | 16 | 0.875 |
| 14_1091484 | 5 | 0.6 | 7 | 1 | 11 | 0.8182 | 6 | 0.5 | 29 | 0.7586 | 5 | 0.6 | 2 | 0.5 | 5 | 0.8 | 4 | 0.75 | 16 | 0.6875 |
| 14_1091484 | 4 | 1 | 8 | 1 | 9 | 0.7778 | 5 | 1 | 26 | 0.9231 | 6 | 0.8333 | 2 | 1 | 5 | 1 | 6 | 1 | 19 | 0.9474 |
| 14_1091485 | 7 | 1 | 9 | 0.8889 | 11 | 0.8182 | 6 | 1 | 33 | 0.9091 | 7 | 0.8571 | 2 | 1 | 5 | 1 | 6 | 1 | 20 | 0.95 |
| 14_1091485 | 6 | 0 | 9 | 0.2222 | 9 | 0.3333 | 6 | 0.6667 | 30 | 0.3 | 6 | 0.1667 | 2 | 0.5 | 5 | 0 | 5 | 0.4 | 18 | 0.2222 |
| 14_1091486 | 6 | 0.6667 | 9 | 0.3333 | 8 | 0.25 | 6 | 0.3333 | 29 | 0.3793 | 5 | 0.8 | 2 | 0 | 4 | 0.25 | 4 | 0.25 | 15 | 0.4 |
| 14_1091486 | 4 | 0.5 | 8 | 0.875 | 6 | 1 | 4 | 1 | 22 | 0.8636 | 5 | 0.8 | 2 | 0.5 | 5 | 0.8 | 3 | 1 | 15 | 0.8 |
| 14_1091487 | 6 | 0.5 | 8 | 0.375 | 9 | 0 | 5 | 1 | 28 | 0.3929 | 5 | 0.6 | 1 | 1 | 5 | 0.4 | 5 | 0.6 | 16 | 0.5625 |
| 14_1091487 | 5 | 0.2 | 7 | 0.4286 | 6 | 0.5 | 4 | 0.25 | 22 | 0.3636 | 3 | 0.6667 | 2 | 0 | 2 | 0.5 | 4 | 0.5 | 11 | 0.4545 |
| 14_1091488 | 5 | 0.6 | 8 | 0.5 | 6 | 0.6667 | 6 | 0.6667 | 25 | 0.6 | 3 | 1 | 2 | 1 | 5 | 0.6 | 4 | 0.5 | 14 | 0.7143 |
| 14_1091489 | 7 | 0.2857 | 7 | 0.1429 | 9 | 0.1111 | 6 | 0.5 | 29 | 0.2414 | 4 | 0.25 | 2 | 0 | 5 | 0.4 | 5 | 0.6 | 16 | 0.375 |
| 14_1091490 | 4 | 1 | 3 | 0.3333 | 8 | 0.25 | 5 | 0.2 | 20 | 0.4 | 2 | 1 | 2 | 0 | 2 | 0.5 | 5 | 0.6 | 11 | 0.5455 |
| 14_1091491 | 6 | 0.1667 | 6 | 0.1667 | 7 | 0.1429 | 6 | 0.1667 | 25 | 0.16 | 4 | 0 | 2 | 0 | 5 | 0.2 | 5 | 0.4 | 16 | 0.1875 |
| 14_1091491 | 5 | 0.8 | 7 | 0.4286 | 7 | 0.1429 | 6 | 0.6667 | 25 | 0.48 | 4 | 0.5 | 2 | 0 | 4 | 0.5 | 5 | 0.4 | 15 | 0.4 |
| 14_1091492 | 4 | 0 | 5 | 0.2 | 6 | 0.1667 | 6 | 0 | 21 | 0.0952 | 4 | 0 | 2 | 0 | 5 | 0.2 | 4 | 0.25 | 15 | 0.1333 |
| 14_1091494 | 4 | 0.5 | 6 | 0.8333 | 4 | 0.75 | 5 | 0.8 | 19 | 0.7368 | 3 | 0.6667 | 1 | 0 | 6 | 0.8333 | 5 | 0.8 | 15 | 0.7333 |
| 14_1091494 | 4 | 0 | 7 | 0.4286 | 4 | 0.25 | 5 | 0.6 | 20 | 0.35 | 4 | 0.25 | 1 | 1 | 6 | 0.8333 | 4 | 0.5 | 15 | 0.6 |
| 14_1091494 | 5 | 0 | 6 | 0.1667 | 4 | 0.25 | 4 | 0 | 19 | 0.1053 | 4 | 0.25 | 1 | 0 | 2 | 0 | 3 | 0 | 10 | 0.1 |
| 14_1091495 | 5 | 0.2 | 6 | 0 | 5 | 0 | 4 | 0 | 20 | 0.05 | 4 | 0.25 | 1 | 0 | 4 | 0 | 5 | 0 | 14 | 0.0714 |
| 14_1091496 | 4 | 1 | 5 | 1 | 3 | 1 | 4 | 1 | 16 | 1 | 4 | 1 | 1 | 1 | 4 | 1 | 5 | 1 | 14 | 1 |
| 14_1091499 | 4 | 0 | 6 | 0 | 4 | 0.25 | 4 | 0.25 | 18 | 0.1111 | 2 | 0.5 | 1 | 0 | 4 | 0 | 5 | 0 | 12 | 0.0833 |
| 14_1091499 | 4 | 0.25 | 6 | 0.1667 | 4 | 0 | 4 | 0 | 18 | 0.1111 | 2 | 0 | 1 | 0 | 3 | 0.6667 | 4 | 0.25 | 10 | 0.3 |
| 14_1091500 | 3 | 0.3333 | 5 | 0.4 | 3 | 0 | 3 | 0.3333 | 14 | 0.2857 | 1 | 0 | NA | NA | 4 | 0.25 | 5 | 0.4 | 10 | 0.3 |
| 14_1091500 | 2 | 1 | 4 | 0.75 | 1 | 1 | 3 | 1 | 10 | 0.9 | 1 | 1 | 1 | 1 | 5 | 1 | 5 | 1 | 12 | 1 |
| 14_1091501 | 4 | 0 | 6 | 0.1667 | 3 | 0 | 3 | 0 | 16 | 0.0625 | 1 | 0 | 1 | 0 | 5 | 0.2 | 5 | 0 | 12 | 0.0833 |
| 14_1091502 | 3 | 0.3333 | 6 | 1 | 3 | 1 | 3 | 1 | 15 | 0.8667 | 1 | 1 | 1 | 1 | 5 | 1 | 4 | 1 | 11 | 1 |
| 14_1091502 | 3 | 0 | 6 | 0.1667 | 3 | 0 | 3 | 0.3333 | 15 | 0.1333 | 1 | 0 | 1 | 1 | 5 | 0.2 | 4 | 0.25 | 11 | 0.2727 |
| 14_1091503 | 1 | 1 | 4 | 0.25 | 3 | 0.6667 | 2 | 0 | 10 | 0.4 | 1 | 0 | NA | NA | 3 | 0.6667 | 4 | 0.75 | 8 | 0.625 |
| 14_1091504 | 2 | 0 | 5 | 0.4 | 3 | 0 | 3 | 0.3333 | 13 | 0.2308 | 1 | 0 | 1 | 0 | 4 | 0.5 | 2 | 0 | 8 | 0.25 |
| 14_1091504 | 3 | 0 | 2 | 0.5 | 3 | 0.6667 | 2 | 0 | 10 | 0.3 | 1 | 1 | 1 | 1 | 3 | 0.3333 | 2 | 0 | 7 | 0.4286 |
| 14_1091505 | 2 | 1 | 3 | 1 | 2 | 1 | 1 | 1 | 8 | 1 | NA | NA | NA | NA | 3 | 1 | 2 | 1 | 5 | 1 |
| 14_1091506 | 3 | 1 | 5 | 1 | 2 | 1 | 3 | 1 | 13 | 1 | 1 | 1 | 1 | 1 | 3 | 1 | 3 | 1 | 8 | 1 |
| 14_1091507 | 2 | 1 | 3 | 1 | 2 | 1 | 1 | 1 | 8 | 1 | 1 | 1 | 1 | 1 | 4 | 1 | 2 | 1 | 8 | 1 |
| 14_1091508 | 2 | 1 | 4 | 1 | 3 | 0.6667 | 3 | 1 | 12 | 0.9167 | NA | NA | 1 | 0 | 4 | 1 | 3 | 1 | 8 | 0.875 |
| 14_1091509 | 3 | 0 | 3 | 0.6667 | 2 | 0.5 | 2 | 0 | 10 | 0.3 | 1 | 0 | 1 | 0 | 4 | 0.5 | 3 | 0.6667 | 9 | 0.4444 |
| 14_1091512 | 2 | 1 | 4 | 0.25 | 1 | 0 | 2 | 0 | 9 | 0.3333 | 1 | 0 | 1 | 0 | 3 | 0.3333 | 3 | 0 | 8 | 0.125 |
| 14_1091514 | 2 | 0.5 | 3 | 0.6667 | NA | NA | 1 | 0 | 6 | 0.5 | 1 | 1 | 1 | 1 | 2 | 0.5 | 2 | 1 | 6 | 0.8333 |
| 14_1117480 | NA | NA | NA | NA | 1 | 1 | NA | NA | 1 | 1 | NA | NA | 2 | 0 | 1 | 1 | NA | NA | 3 | 0.3333 |
| 14_1119164 | 5 | 0.2 | 2 | 0 | NA | NA | 1 | 0 | 8 | 0.125 | 1 | 0 | NA | NA | 3 | 0.3333 | 1 | 1 | 5 | 0.4 |
| 14_1119165 | 5 | 0.2 | NA | NA | 1 | 0 | 1 | 1 | 7 | 0.2857 | 2 | 0 | NA | NA | 1 | 1 | 1 | 0 | 4 | 0.25 |
| 14_1119165 | 4 | 0 | 1 | 0 | 1 | 0 | 1 | 1 | 7 | 0.1429 | 2 | 0 | NA | NA | 2 | 0.5 | 1 | 0 | 5 | 0.2 |
| 14_1119167 | 5 | 1 | 1 | 1 | 1 | 1 | NA | NA | 7 | 1 | 2 | 1 | NA | NA | 2 | 0.5 | 1 | 1 | 5 | 0.8 |
| 14_1119168 | 4 | 0.75 | 2 | 1 | 1 | 1 | 1 | 1 | 8 | 0.875 | 2 | 1 | NA | NA | 3 | 0.6667 | 1 | 1 | 6 | 0.8333 |
| 14_1119168 | 3 | 0.3333 | NA | NA | NA | NA | 1 | 1 | 4 | 0.5 | 2 | 0.5 | NA | NA | 2 | 1 | 1 | 0 | 5 | 0.6 |
| 14_1119168 | 5 | 0 | 2 | 0 | 1 | 0 | 1 | 1 | 9 | 0.1111 | 2 | 0 | NA | NA | 3 | 0.3333 | 1 | 0 | 6 | 0.1667 |
| 14_1119169 | 5 | 0.4 | 3 | 0 | 1 | 1 | 1 | 1 | 10 | 0.4 | 2 | 1 | NA | NA | 3 | 0.3333 | 1 | 1 | 6 | 0.6667 |
| 14_1119169 | 6 | 1 | 1 | 1 | 1 | 1 | 1 | 1 | 9 | 1 | 3 | 1 | NA | NA | 3 | 1 | 1 | 1 | 7 | 1 |
| 14_1119170 | 5 | 0.2 | 2 | 0 | 1 | 0 | 1 | 0 | 9 | 0.1111 | 3 | 0.3333 | NA | NA | 2 | 0.5 | NA | NA | 5 | 0.4 |
| 14_1119171 | 5 | 1 | 3 | 1 | 1 | 1 | 1 | 1 | 10 | 1 | 3 | 0.6667 | NA | NA | 3 | 1 | 1 | 1 | 7 | 0.8571 |

| | | | | | | | | | | | | | | | | | | | | |
|---|---|---|---|---|---|---|---|---|---|---|---|---|---|---|---|---|---|---|---|---|
| 14_1119172 | 3 | 0.6667 | 4 | 0.75 | 1 | 1 | 1 | 0 | 9 | 0.6667 | 3 | 0.3333 | NA | NA | 3 | 0.6667 | 1 | 1 | 7 | 0.5714 |
| 14_1119172 | 6 | 0.3333 | 4 | 0 | 1 | 1 | 1 | 0 | 12 | 0.25 | 3 | 0.3333 | 1 | 0 | 3 | 0.3333 | 1 | 0 | 8 | 0.25 |
| 14_1119172 | 3 | 0.3333 | 4 | 1 | 1 | 0 | 1 | 0 | 9 | 0.5556 | 3 | 0.3333 | 1 | 1 | 3 | 0.6667 | 1 | 0 | 8 | 0.5 |
| 14_1119173 | 6 | 0.1667 | 4 | 0.5 | 1 | 0 | 1 | 0 | 12 | 0.25 | 3 | 0.3333 | 1 | 0 | 3 | 0.3333 | 1 | 0 | 8 | 0.25 |
| 14_1119173 | 5 | 0 | 4 | 0.25 | 1 | 0 | 1 | 0 | 11 | 0.0909 | 3 | 0 | NA | NA | 3 | 0.3333 | 1 | 0 | 7 | 0.1429 |
| 14_1119174 | 5 | 0.8 | 4 | 1 | 1 | 1 | 1 | 1 | 11 | 0.9091 | 3 | 0.6667 | 1 | 1 | 3 | 1 | 1 | 1 | 8 | 0.875 |
| 14_1119176 | 6 | 0.5 | 4 | 0.25 | 1 | 0 | NA | NA | 11 | 0.3636 | 3 | 0 | 1 | 0 | 2 | 0.5 | 1 | 1 | 7 | 0.2857 |
| 14_1119176 | 5 | 0.2 | 4 | 0 | 1 | 0 | 1 | 0 | 11 | 0.0909 | 3 | 0.3333 | 1 | 1 | 2 | 0 | 1 | 0 | 7 | 0.2857 |
| 14_1119177 | 5 | 0 | 3 | 0 | 1 | 1 | 1 | 0 | 10 | 0.1 | 2 | 0 | 1 | 0 | 3 | 0.3333 | 1 | 0 | 7 | 0.1429 |
| 14_1119178 | 5 | 0.4 | 5 | 1 | 3 | 1 | 1 | 1 | 14 | 0.7857 | 3 | 1 | 1 | 1 | 3 | 1 | 1 | 0 | 8 | 0.875 |
| 14_1119179 | 5 | 0 | 3 | 0.3333 | 3 | 0 | 1 | 0 | 12 | 0.0833 | 3 | 0.6667 | 1 | 0 | 3 | 0.3333 | 1 | 0 | 8 | 0.375 |
| 14_1119180 | 7 | 0 | 4 | 0.25 | 3 | 0 | 1 | 1 | 15 | 0.1333 | 3 | 0.3333 | NA | NA | 3 | 0 | 1 | 0 | 7 | 0.1429 |
| 14_1119181 | 7 | 0.1429 | 5 | 0 | 3 | 0.3333 | 1 | 0 | 16 | 0.125 | 3 | 0.3333 | 2 | 0 | 3 | 0.3333 | 1 | 0 | 9 | 0.2222 |
| 14_1119182 | 6 | 1 | 5 | 0.6 | 2 | 1 | 1 | 1 | 14 | 0.8571 | 3 | 1 | 1 | 1 | 1 | 1 | 1 | 1 | 6 | 1 |
| 14_1119182 | 5 | 0 | 5 | 0 | 3 | 0 | 1 | 1 | 14 | 0.0714 | 3 | 0.3333 | 2 | 0 | 2 | 0 | 1 | 1 | 8 | 0.25 |
| 14_1119183 | 6 | 0.5 | 3 | 0.3333 | 3 | 0.3333 | 1 | 0 | 13 | 0.3846 | 3 | 0.3333 | 1 | 0 | 2 | 1 | 1 | 0 | 7 | 0.4286 |
| 14_1119184 | 7 | 0 | 5 | 0.4 | 3 | 0 | 1 | 0 | 16 | 0.125 | 3 | 0 | 2 | 0 | 3 | 0.3333 | 1 | 0 | 9 | 0.1111 |
| 14_1119186 | 8 | 0.25 | 5 | 0.6 | 3 | 0.6667 | 1 | 0 | 17 | 0.4118 | 3 | 0.3333 | 1 | 1 | 3 | 0.6667 | 1 | 0 | 8 | 0.5 |
| 14_1119187 | 6 | 0.3333 | 4 | 0.25 | 2 | 0 | 1 | 1 | 13 | 0.3077 | 4 | 0.25 | 3 | 0 | 3 | 0 | 1 | 0 | 11 | 0.0909 |
| 14_1119189 | 6 | 0.6667 | 6 | 0.8333 | 3 | 0.3333 | 1 | 1 | 16 | 0.6875 | 4 | 0.75 | 3 | 1 | 3 | 1 | 1 | 1 | 11 | 0.9091 |
| 14_1119189 | 7 | 0.1429 | 6 | 0.3333 | 4 | 0.25 | NA | NA | 17 | 0.2353 | 3 | 0.3333 | 2 | 0 | 5 | 0 | 1 | 0 | 11 | 0.0909 |
| 14_1119189 | 6 | 0.8333 | 6 | 0.8333 | 4 | 0.5 | 1 | 1 | 17 | 0.7647 | 4 | 0.75 | 1 | 1 | 4 | 0.75 | 1 | 1 | 10 | 0.8 |
| 14_1119190 | 6 | 0.1667 | 5 | 0.4 | 2 | 0.5 | 1 | 1 | 14 | 0.3571 | 3 | 0.3333 | 2 | 0.5 | 2 | 0 | 1 | 1 | 8 | 0.375 |
| 14_1119190 | 7 | 0.2857 | 5 | 0.2 | 2 | 0 | 1 | 0 | 15 | 0.2 | 3 | 0 | 4 | 0.5 | 3 | 0.6667 | 1 | 0 | 11 | 0.3636 |
| 14_1119191 | 8 | 0.75 | 4 | 0.25 | 6 | 0.1667 | 1 | 0 | 19 | 0.4211 | 2 | 0 | 3 | 0 | 4 | 0.25 | 1 | 0 | 10 | 0.1 |
| 14_1119191 | 7 | 0.2857 | 5 | 0.4 | 6 | 0.5 | 1 | 0 | 19 | 0.3684 | 1 | 1 | 4 | 0.75 | 4 | 1 | 1 | 1 | 10 | 0.9 |
| 14_1119191 | 8 | 0.25 | 6 | 0.1667 | 4 | 0.25 | 1 | 0 | 19 | 0.2105 | 3 | 0 | 3 | 0.6667 | 3 | 0.3333 | NA | NA | 9 | 0.3333 |
| 14_1119191 | 6 | 0.5 | 5 | 0.4 | 4 | 0.5 | 1 | 1 | 16 | 0.5 | 3 | 1 | 4 | 0.25 | 3 | 0 | 1 | 1 | 11 | 0.4545 |
| 14_1119192 | 10 | 1 | 6 | 1 | 6 | 0.8333 | 1 | 1 | 23 | 0.9565 | 3 | 1 | 4 | 1 | 4 | 1 | 1 | 1 | 12 | 1 |
| 14_1119192 | 6 | 0.1667 | 5 | 0 | 5 | 0 | 1 | 0 | 17 | 0.0588 | 4 | 0.25 | 2 | 0 | 5 | 0.2 | 1 | 1 | 12 | 0.25 |
| 14_1119192 | 6 | 0.3333 | 5 | 0.8 | 4 | 0.25 | 1 | 0 | 16 | 0.4375 | 4 | 0.75 | 2 | 0 | 4 | 1 | 1 | 0 | 11 | 0.6364 |
| 14_1119192 | 6 | 0.1667 | 6 | 0.5 | 4 | 0 | 1 | 0 | 17 | 0.2353 | 3 | 0 | 4 | 0.25 | 4 | 0 | 2 | 0.5 | 13 | 0.1538 |
| 14_1119192 | 7 | 1 | 6 | 0.8333 | 5 | 1 | NA | NA | 18 | 0.9444 | 2 | 1 | 3 | 1 | 3 | 1 | 2 | 1 | 10 | 1 |
| 14_1119192 | 5 | 0.6 | 6 | 0.3333 | 6 | 0.6667 | 1 | 0 | 18 | 0.5 | 4 | 0.25 | 3 | 0.3333 | 3 | 0 | 2 | 0 | 12 | 0.1667 |
| 14_1119193 | 8 | 0.125 | 6 | 0 | 5 | 0.2 | 1 | 0 | 20 | 0.1 | 4 | 0 | 4 | 0 | 4 | 0.25 | 2 | 0.5 | 14 | 0.1429 |
| 14_1119193 | 9 | 0.5556 | 5 | 0.8 | 6 | 0.8333 | 1 | 1 | 21 | 0.7143 | 4 | 1 | 4 | 0.75 | 5 | 1 | 2 | 1 | 15 | 0.9333 |
| 14_1119194 | 9 | 0.1111 | 2 | 0.5 | 5 | 0.6 | 1 | 0 | 17 | 0.2941 | 4 | 0.25 | 3 | 0.3333 | 3 | 0 | 2 | 0.5 | 12 | 0.25 |
| 14_1119194 | 8 | 0.25 | 5 | 0 | 6 | 0.1667 | NA | NA | 19 | 0.1579 | 4 | 0.25 | 4 | 0.5 | 4 | 0.5 | 1 | 1 | 13 | 0.4615 |
| 14_1119194 | 6 | 0.1667 | 4 | 0.25 | 6 | 0.1667 | 1 | 1 | 17 | 0.2353 | 4 | 0.25 | 3 | 0 | 5 | 0 | 1 | 1 | 13 | 0.1538 |
| 14_1119195 | 7 | 0.7143 | 5 | 0.8 | 6 | 1 | 1 | 1 | 19 | 0.8421 | 2 | 1 | 4 | 0.75 | 4 | 1 | 2 | 1 | 12 | 0.9167 |
| 14_1119198 | 7 | 0.5714 | 2 | 0.5 | 6 | 0.6667 | 2 | 1 | 17 | 0.6471 | 3 | 0.6667 | 5 | 0.6 | 5 | 0.4 | 2 | 1 | 15 | 0.6 |
| 14_1119198 | 8 | 0.125 | 4 | 0 | 5 | 0 | 2 | 0 | 19 | 0.0526 | 3 | 0 | 4 | 0 | 5 | 0 | 2 | 0.5 | 14 | 0.0714 |
| 14_1119201 | 10 | 0.5 | 4 | 0.5 | 9 | 0.7778 | 2 | 1 | 25 | 0.64 | 3 | 0.6667 | 6 | 0.6667 | 5 | 0.2 | 2 | 0.5 | 16 | 0.5 |
| 14_1119201 | 10 | 0.1 | 3 | 0.3333 | 8 | 0.125 | 1 | 0 | 22 | 0.1364 | 3 | 0.3333 | 4 | 0 | 5 | 0.2 | 2 | 0 | 14 | 0.1429 |
| 14_1119204 | 11 | 0.0909 | 6 | 0.6667 | 10 | 0.3 | 2 | 0 | 29 | 0.2759 | 6 | 0.8333 | 6 | 0.3333 | 6 | 0.3333 | 3 | 0.3333 | 21 | 0.4762 |
| 14_1119204 | 7 | 0.2857 | 6 | 0.1667 | 10 | 0 | 2 | 0.5 | 25 | 0.16 | 6 | 0.1667 | 6 | 0.5 | 7 | 0 | 4 | 0.5 | 23 | 0.2609 |
| 14_1119204 | 10 | 0.3 | 6 | 0 | 9 | 0.1111 | 2 | 0 | 27 | 0.1481 | 4 | 0 | 5 | 0.2 | 7 | 0 | 3 | 0.3333 | 19 | 0.1053 |
| 14_1156071 | 1 | 1 | 3 | 0 | 5 | 0.4 | 3 | 0.3333 | 12 | 0.3333 | 3 | 0 | 2 | 0 | 3 | 0.6667 | 2 | 0 | 10 | 0.2 |
| 14_1156074 | 2 | 0.5 | 2 | 0 | 5 | 0 | 3 | 0 | 12 | 0.0833 | 3 | 0 | 2 | 1 | 2 | 0.5 | 2 | 0.5 | 9 | 0.4444 |
| 14_1160256 | 1 | 1 | 2 | 0.5 | NA | NA | 1 | 0 | 4 | 0.5 | NA | NA | 1 | 1 | NA | NA | NA | NA | 1 | 1 |
| 14_1160258 | 1 | 0 | 2 | 0.5 | NA | NA | 1 | 0 | 4 | 0.25 | NA | NA | 1 | 1 | NA | NA | NA | NA | 1 | 1 |
| 14_1160258 | 1 | 0 | 1 | 1 | NA | NA | 1 | 0 | 3 | 0.3333 | NA | NA | 1 | 1 | NA | NA | NA | NA | 1 | 1 |
| 14_1160259 | 1 | 1 | 1 | 0 | NA | NA | 1 | 0 | 3 | 0.3333 | NA | NA | 1 | 1 | NA | NA | NA | NA | 1 | 1 |
| 14_1167617 | NA | NA | 1 | 1 | NA | NA | 1 | 0 | 2 | 0.5 | 1 | 0 | 1 | 1 | 1 | 0 | NA | NA | 3 | 0.3333 |
| 14_1167622 | 1 | 1 | 1 | 0 | NA | NA | 2 | 0.5 | 4 | 0.5 | 1 | 0 | 2 | 0.5 | 1 | 0 | NA | NA | 4 | 0.25 |
| 14_1183502 | NA | NA | 1 | 1 | NA | NA | 1 | 1 | 2 | 1 | NA | NA | NA | NA | 1 | 1 | 1 | 1 | 2 | 1 |
| 14_1184000 | 2 | 0 | 2 | 1 | 3 | 0.6667 | NA | NA | 7 | 0.5714 | 4 | 0.5 | 3 | 0.3333 | 2 | 0 | 4 | 0.75 | 13 | 0.4615 |
| 14_1184002 | 2 | 0.5 | 3 | 0.6667 | 4 | 0.75 | NA | NA | 9 | 0.6667 | 4 | 0.75 | 1 | 1 | 1 | 1 | 4 | 0.75 | 10 | 0.8 |
| 14_1184003 | 2 | 0.5 | 4 | 0.75 | 4 | 0 | NA | NA | 10 | 0.4 | 5 | 0.4 | 2 | 0 | 2 | 0 | 4 | 0.25 | 13 | 0.2308 |
| 14_1184003 | 1 | 0 | 3 | 0.3333 | 3 | 0.6667 | NA | NA | 7 | 0.4286 | 3 | 0.6667 | 1 | 1 | 1 | 1 | 5 | 0.4 | 10 | 0.6 |
| 14_1184003 | 2 | 0.5 | 4 | 0.5 | 3 | 0.6667 | NA | NA | 9 | 0.5556 | 4 | 0.5 | 3 | 0.3333 | 3 | 0 | 2 | 1 | 12 | 0.4167 |
| 14_1184004 | 2 | 1 | 4 | 1 | 3 | 0.6667 | NA | NA | 9 | 0.8889 | 5 | 0.6 | 3 | 0.3333 | 2 | 1 | 3 | 0.3333 | 13 | 0.5385 |
| 14_1184004 | 1 | 0 | 3 | 0 | 2 | 0.5 | NA | NA | 6 | 0.1667 | 3 | 0.3333 | 3 | 0 | 3 | 0.3333 | 5 | 0 | 14 | 0.1429 |
| 14_1184006 | 2 | 1 | 4 | 0.75 | 5 | 0.6 | NA | NA | 11 | 0.7273 | 5 | 1 | 4 | 0.5 | 4 | 1 | 5 | 0.8 | 18 | 0.8333 |
| 14_1184714 | NA | NA | NA | NA | 2 | 1 | 3 | 1 | 5 | 1 | 2 | 1 | 1 | 0 | 2 | 1 | 1 | 1 | 6 | 0.8333 |

| | | | | | | | | | | | | | | | | | | | | |
|---|---|---|---|---|---|---|---|---|---|---|---|---|---|---|---|---|---|---|---|---|
| 14_1184719 | NA | NA | NA | NA | 1 | 0 | 3 | 0.3333 | 4 | 0.25 | 1 | 1 | 1 | 0 | 2 | 1 | 1 | 1 | 5 | 0.8 |
| 14_1184721 | NA | NA | NA | NA | 2 | 1 | 3 | 0.6667 | 5 | 0.8 | NA | NA | 1 | 1 | 1 | 1 | 1 | 1 | 3 | 1 |
| 14_1184722 | NA | NA | NA | NA | 2 | 1 | 3 | 0.6667 | 5 | 0.8 | 2 | 1 | 1 | 1 | 2 | 1 | 1 | 1 | 6 | 1 |
| 14_1184722 | NA | NA | NA | NA | 2 | 0.5 | 2 | 0.5 | 4 | 0.5 | 1 | 1 | 1 | 1 | 2 | 1 | 1 | 0 | 5 | 0.8 |
| 14_1184727 | NA | NA | NA | NA | 2 | 1 | 3 | 1 | 5 | 1 | 1 | 1 | NA | NA | NA | NA | 1 | 1 | 2 | 1 |
| 14_1184733 | NA | NA | NA | NA | 2 | 0.5 | 2 | 0 | 4 | 0.25 | 1 | 1 | NA | NA | NA | NA | 1 | 1 | 2 | 1 |
| 14_1184733 | NA | NA | NA | NA | 2 | 0 | 2 | 0.5 | 4 | 0.25 | 1 | 1 | NA | NA | NA | NA | 1 | 0 | 2 | 0.5 |
| 14_1187216 | 2 | 1 | 3 | 1 | 1 | 1 | 2 | 1 | 8 | 1 | NA | NA | 1 | 1 | NA | NA | 1 | 1 | 2 | 1 |
| 14_1187216 | 2 | 1 | 3 | 0.6667 | NA | NA | 2 | 1 | 7 | 0.8571 | NA | NA | 1 | 1 | NA | NA | 1 | 0 | 2 | 0.5 |
| 14_1189438 | NA | NA | 2 | 1 | NA | NA | NA | NA | 2 | 1 | NA | NA | 2 | 1 | 1 | 0 | 1 | 0 | 4 | 0.5 |
| 14_1189440 | 1 | 1 | 2 | 1 | NA | NA | NA | NA | 3 | 1 | NA | NA | 3 | 1 | 1 | 1 | 1 | 0 | 5 | 0.8 |
| 14_1189440 | 1 | 0 | 2 | 1 | NA | NA | NA | NA | 3 | 0.6667 | NA | NA | 3 | 1 | 1 | 0 | 1 | 0 | 5 | 0.6 |
| 14_1189441 | 1 | 1 | 1 | 1 | NA | NA | NA | NA | 2 | 1 | NA | NA | 3 | 1 | NA | NA | 1 | 1 | 4 | 1 |
| 14_1189442 | 1 | 1 | 2 | 1 | NA | NA | NA | NA | 3 | 1 | NA | NA | 3 | 1 | NA | NA | 1 | 1 | 4 | 1 |
| 14_1189443 | 1 | 0 | 2 | 0.5 | NA | NA | NA | NA | 3 | 0.3333 | NA | NA | 3 | 1 | NA | NA | 1 | 1 | 4 | 1 |
| 14_1189444 | NA | NA | 2 | 1 | NA | NA | NA | NA | 2 | 1 | NA | NA | 1 | 1 | 1 | 1 | 1 | 1 | 3 | 1 |
| 14_1189447 | NA | NA | 1 | 1 | NA | NA | NA | NA | 1 | 1 | NA | NA | 1 | 1 | NA | NA | 1 | 0 | 2 | 0.5 |
| 14_1191117 | 15 | 0.2 | 11 | 0.2727 | 7 | 0 | 10 | 0.2 | 43 | 0.186 | 16 | 0.0625 | 13 | 0.1538 | 14 | 0.3571 | 7 | 0 | 50 | 0.16 |
| 14_1191143 | 5 | 0 | 4 | 0.25 | 4 | 0.25 | 3 | 0.3333 | 16 | 0.1875 | NA | NA | 5 | 0 | 3 | 0.3333 | 4 | 0.25 | 12 | 0.1667 |
| 14_1191144 | 6 | 0 | 4 | 0.5 | 4 | 0.25 | 3 | 0 | 17 | 0.1765 | 1 | 1 | 5 | 0.4 | 4 | 0 | 4 | 0 | 14 | 0.2143 |
| 14_1191144 | 6 | 0 | 4 | 0.25 | 3 | 0 | 3 | 0 | 16 | 0.0625 | 1 | 0 | 4 | 0.25 | 3 | 0 | 2 | 0 | 10 | 0.1 |
| 14_1191145 | 6 | 0.5 | 5 | 0.6 | 4 | 0.5 | 4 | 0.75 | 19 | 0.5789 | 1 | 1 | 5 | 0.8 | 3 | 0.6667 | 4 | 0.75 | 13 | 0.7692 |
| 14_1191146 | 6 | 0 | 4 | 0.25 | 5 | 0 | 4 | 0 | 19 | 0.0526 | 1 | 0 | 4 | 0 | 1 | 0 | 4 | 0.5 | 10 | 0.2 |
| 14_1191599 | 1 | 1 | 3 | 0 | NA | NA | 1 | 0 | 5 | 0.2 | NA | NA | NA | NA | 1 | 1 | 1 | 0 | 2 | 0.5 |
| 14_1191600 | 1 | 1 | 3 | 0.6667 | NA | NA | NA | NA | 4 | 0.75 | NA | NA | NA | NA | 1 | 1 | NA | NA | 1 | 1 |
| 14_1192155 | 2 | 0.5 | 4 | 0.25 | 5 | 0 | 1 | 0 | 12 | 0.1667 | 5 | 0.4 | 5 | 0 | 5 | 0.2 | 5 | 0.2 | 20 | 0.2 |
| 14_1192155 | 3 | 0 | 4 | 0 | 4 | 0.25 | 2 | 0 | 13 | 0.0769 | 4 | 0.5 | 6 | 0.3333 | 5 | 0.2 | 5 | 0 | 20 | 0.25 |
| 14_1192156 | 2 | 0.5 | 3 | 0 | 1 | 0 | 2 | 0 | 8 | 0.125 | 2 | 0 | 3 | 0 | 3 | 0 | 5 | 0.2 | 13 | 0.0769 |
| 14_1192156 | 2 | 0.5 | 3 | 0 | 4 | 0 | 2 | 0 | 11 | 0.0909 | 4 | 0 | 4 | 0 | 2 | 0 | 6 | 0.1667 | 16 | 0.0625 |
| 14_1192158 | 2 | 0 | 3 | 0 | 4 | 0.25 | 2 | 0 | 11 | 0.0909 | 4 | 0 | 4 | 0.25 | 3 | 0.3333 | 5 | 0.2 | 16 | 0.1875 |
| 14_1192158 | 2 | 0 | 4 | 0 | 4 | 0.25 | 2 | 0 | 12 | 0.0833 | 4 | 0 | 5 | 0 | 4 | 0.25 | 6 | 0.1667 | 19 | 0.1053 |
| 14_1192164 | 3 | 0.3333 | 2 | 1 | 4 | 0.25 | 2 | 0.5 | 11 | 0.4545 | 5 | 0 | 5 | 0.2 | 5 | 0.4 | 5 | 0.4 | 20 | 0.25 |
| 14_1192186 | 4 | 0.25 | 4 | 0 | 3 | 0 | 2 | 0 | 13 | 0.0769 | 2 | 0 | 3 | 0.6667 | 4 | 0.25 | 6 | 0.5 | 15 | 0.4 |
| 14_1192203 | NA | NA | 1 | 0 | 1 | 0 | 2 | 0.5 | 4 | 0.25 | 1 | 1 | NA | NA | 1 | 0 | 1 | 0 | 3 | 0.3333 |
| 14_1192203 | NA | NA | 2 | 1 | 1 | 1 | 2 | 0.5 | 5 | 0.8 | 1 | 1 | NA | NA | 1 | 1 | 1 | 1 | 3 | 1 |
| 14_1192205 | NA | NA | 2 | 1 | 1 | 0 | 2 | 1 | 5 | 0.8 | 1 | 1 | NA | NA | 1 | 1 | 1 | 0 | 3 | 0.6667 |
| 14_1192205 | NA | NA | 2 | 1 | 1 | 1 | 2 | 1 | 5 | 1 | 1 | 1 | NA | NA | 1 | 1 | 1 | 1 | 3 | 1 |
| 14_1192206 | NA | NA | 2 | 1 | 1 | 0 | 2 | 1 | 5 | 0.8 | NA | NA | NA | NA | 1 | 0 | 1 | 1 | 2 | 0.5 |
| 14_1192206 | NA | NA | 1 | 0 | 1 | 0 | 1 | 1 | 3 | 0.3333 | 1 | 0 | NA | NA | 1 | 1 | 1 | 1 | 3 | 0.6667 |
| 14_1192239 | 1 | 0 | 2 | 0.5 | 2 | 0.5 | 5 | 0 | 10 | 0.2 | 2 | 0.5 | 1 | 0 | 3 | 0 | NA | NA | 6 | 0.1667 |
| 14_1192241 | 2 | 1 | 2 | 0.5 | NA | NA | 5 | 0.8 | 9 | 0.7778 | 2 | 1 | 1 | 1 | 3 | 1 | NA | NA | 6 | 1 |
| 14_1192243 | 1 | 0 | 2 | 0 | 1 | 0 | 5 | 0.4 | 9 | 0.2222 | 2 | 0.5 | NA | NA | 2 | 0 | NA | NA | 4 | 0.25 |
| 14_1201065 | NA | NA | NA | NA | 2 | 0 | 1 | 1 | 3 | 0.3333 | NA | NA | 2 | 1 | NA | NA | NA | NA | 2 | 1 |
| 14_1201066 | NA | NA | 1 | 1 | 2 | 0.5 | 1 | 0 | 4 | 0.5 | NA | NA | 2 | 0.5 | NA | NA | NA | NA | 2 | 0.5 |
| 14_1201066 | NA | NA | 1 | 0 | 2 | 0.5 | 1 | 1 | 4 | 0.5 | NA | NA | 2 | 1 | NA | NA | NA | NA | 2 | 1 |
| 14_1201067 | NA | NA | 1 | 1 | 1 | 0 | 1 | 1 | 3 | 0.6667 | NA | NA | 1 | 1 | NA | NA | NA | NA | 1 | 1 |
| 14_1201068 | NA | NA | 1 | 0 | 2 | 0.5 | NA | NA | 3 | 0.3333 | NA | NA | 2 | 1 | NA | NA | NA | NA | 2 | 1 |
| 14_1201069 | NA | NA | 2 | 1 | 2 | 1 | NA | NA | 4 | 1 | NA | NA | 2 | 1 | NA | NA | NA | NA | 2 | 1 |
| 14_1201069 | NA | NA | 1 | 1 | 2 | 1 | NA | NA | 3 | 1 | NA | NA | 2 | 1 | NA | NA | NA | NA | 2 | 1 |
| 14_1201070 | NA | NA | 2 | 1 | 1 | 1 | NA | NA | 3 | 1 | NA | NA | 2 | 1 | NA | NA | NA | NA | 2 | 1 |
| 14_1201071 | NA | NA | 2 | 1 | 2 | 0.5 | NA | NA | 4 | 0.75 | NA | NA | 3 | 1 | NA | NA | NA | NA | 3 | 1 |
| 14_1201071 | NA | NA | 2 | 0.5 | 2 | 1 | NA | NA | 4 | 0.75 | NA | NA | 3 | 1 | NA | NA | NA | NA | 3 | 1 |
| 14_1201072 | NA | NA | 2 | 0.5 | NA | NA | NA | NA | 2 | 0.5 | NA | NA | 3 | 0.3333 | NA | NA | NA | NA | 3 | 0.3333 |
| 14_1201074 | NA | NA | 2 | 1 | 2 | 1 | NA | NA | 4 | 1 | NA | NA | 3 | 0.6667 | NA | NA | NA | NA | 3 | 0.6667 |
| 14_1201074 | NA | NA | 2 | 0.5 | 1 | 0 | 1 | 0 | 4 | 0.25 | NA | NA | 2 | 0.5 | NA | NA | NA | NA | 2 | 0.5 |
| 14_1201077 | NA | NA | 2 | 0.5 | 2 | 0 | 1 | 0 | 5 | 0.2 | NA | NA | 3 | 0.6667 | NA | NA | NA | NA | 3 | 0.6667 |
| 14_1201094 | NA | NA | 3 | 0.3333 | 3 | 0 | 2 | 0 | 8 | 0.125 | NA | NA | 5 | 0.4 | 1 | 1 | NA | NA | 6 | 0.5 |
| 14_1201103 | NA | NA | 3 | 0 | 5 | 0.2 | 5 | 0.2 | 13 | 0.1538 | NA | NA | 4 | 0.75 | 2 | 0.5 | 1 | 0 | 7 | 0.5714 |
| 14_1201114 | NA | NA | 6 | 0 | 5 | 0.2 | 6 | 0 | 17 | 0.0588 | NA | NA | 5 | 0.2 | 2 | 0.5 | 2 | 0 | 9 | 0.2222 |
| 14_1201120 | 2 | 0 | 7 | 0 | 7 | 0.2857 | 6 | 0 | 22 | 0.0909 | 1 | 0 | 5 | 0.2 | 2 | 0 | 3 | 0 | 11 | 0.0909 |
| 14_1201121 | 1 | 0 | 6 | 0 | 6 | 0.1667 | 7 | 0.1429 | 20 | 0.1 | 1 | 0 | 5 | 0 | 2 | 0 | 3 | 0.3333 | 11 | 0.0909 |
| 14_1201123 | 2 | 0.5 | 3 | 0.3333 | 5 | 0.2 | 6 | 0.5 | 16 | 0.375 | NA | NA | 2 | 0 | 1 | 1 | 3 | 0.6667 | 6 | 0.5 |
| 14_1201123 | 2 | 0 | 6 | 0.1667 | 7 | 0.1429 | 5 | 0.2 | 20 | 0.15 | 1 | 0 | 4 | 0.25 | 2 | 0.5 | 2 | 0.5 | 9 | 0.3333 |
| 14_1201124 | 2 | 0.5 | 6 | 0.1667 | 4 | 0.5 | 5 | 0.6 | 17 | 0.4118 | 1 | 0 | 2 | 0 | 2 | 0.5 | 3 | 0.6667 | 8 | 0.375 |
| 14_1201125 | 1 | 1 | 6 | 1 | 7 | 1 | 8 | 0.875 | 22 | 0.9545 | NA | NA | 5 | 1 | 2 | 1 | 3 | 1 | 10 | 1 |

| | | | | | | | | | | | | | | | | | | | | |
|---|---|---|---|---|---|---|---|---|---|---|---|---|---|---|---|---|---|---|---|---|
| 14_1201126 | 2 | 0.5 | 5 | 0.4 | 7 | 0.8571 | 5 | 1 | 19 | 0.7368 | 1 | 1 | 4 | 0.5 | 1 | 1 | 3 | 1 | 9 | 0.7778 |
| 14_1201126 | 2 | 0.5 | 6 | 0.6667 | 7 | 1 | 8 | 0.875 | 23 | 0.8261 | 1 | 1 | 4 | 0.75 | 2 | 1 | 1 | 1 | 8 | 0.875 |
| 14_1201126 | 2 | 0 | 6 | 0.1667 | 8 | 0.125 | 7 | 0.2857 | 23 | 0.1739 | NA | NA | 5 | 0.2 | 2 | 0 | 3 | 0 | 10 | 0.1 |
| 14_1201126 | 2 | 1 | 7 | 0.8571 | 9 | 0.8889 | 8 | 0.875 | 26 | 0.8846 | 1 | 1 | 4 | 0.75 | 2 | 1 | 3 | 1 | 10 | 0.9 |
| 14_1201127 | 1 | 0 | 3 | 0 | 4 | 1 | 4 | 1 | 12 | 0.6667 | 1 | 1 | 1 | 0 | 1 | 1 | 1 | 0 | 4 | 0.5 |
| 14_1201127 | 2 | 1 | 6 | 1 | 8 | 0.875 | 7 | 0.8571 | 23 | 0.913 | 1 | 0 | 5 | 1 | 2 | 1 | 3 | 1 | 11 | 0.9091 |
| 14_1201128 | 1 | 1 | 6 | 1 | 8 | 1 | 7 | 0.8571 | 22 | 0.9545 | 2 | 1 | 4 | 1 | 1 | 1 | 3 | 1 | 10 | 1 |
| 14_1201129 | 2 | 1 | 7 | 0.8571 | 9 | 0.8889 | 7 | 0.8571 | 25 | 0.88 | 2 | 1 | 5 | 0.8 | 2 | 1 | 2 | 1 | 11 | 0.9091 |
| 14_1201129 | NA | NA | 8 | 0.625 | 4 | 0.25 | 5 | 0.6 | 17 | 0.5294 | 1 | 1 | 3 | 1 | 2 | 1 | 2 | 1 | 8 | 1 |
| 14_1201129 | NA | NA | 7 | 0.5714 | 10 | 0.7 | 8 | 0.625 | 25 | 0.64 | 2 | 0.5 | 4 | 0.25 | 3 | 1 | 3 | 1 | 12 | 0.6667 |
| 14_1201130 | 2 | 0 | 8 | 0.5 | 10 | 0.4 | 8 | 0 | 28 | 0.2857 | 2 | 0.5 | 5 | 0.4 | 3 | 0.3333 | 3 | 0.6667 | 13 | 0.4615 |
| 14_1205623 | 14 | 0.2857 | 16 | 0.5 | 14 | 0.5714 | 13 | 0.4615 | 57 | 0.4561 | 3 | 0.3333 | 16 | 0.625 | 18 | 0.6111 | 7 | 0.4286 | 44 | 0.5682 |
| 14_1205623 | 12 | 0.4167 | 14 | 0.8571 | 13 | 0.5385 | 12 | 0.5 | 51 | 0.5882 | 5 | 0.6 | 15 | 0.6 | 17 | 0.8235 | 6 | 0.5 | 43 | 0.6744 |
| 14_1205623 | 15 | 0.2 | 18 | 0.3333 | 16 | 0.0625 | 13 | 0.3077 | 62 | 0.2258 | 5 | 0.6 | 15 | 0.3333 | 20 | 0.25 | 7 | 0.2857 | 47 | 0.3191 |
| 14_1205624 | 13 | 0.5385 | 18 | 0.5556 | 15 | 0.6667 | 17 | 0.4706 | 63 | 0.5556 | 5 | 0.2 | 18 | 0.5556 | 19 | 0.4737 | 7 | 0.2857 | 49 | 0.449 |
| 14_1205624 | 15 | 0 | 17 | 0.2353 | 17 | 0.1176 | 15 | 0.1333 | 64 | 0.125 | 5 | 0 | 17 | 0.2353 | 14 | 0.2143 | 7 | 0.1429 | 43 | 0.186 |
| 14_1205625 | 13 | 1 | 18 | 0.6111 | 15 | 0.8667 | 15 | 0.6667 | 61 | 0.7705 | 5 | 0.8 | 16 | 0.75 | 19 | 0.7895 | 6 | 0.8333 | 46 | 0.7826 |
| 14_1205625 | 15 | 0.2667 | 18 | 0.2778 | 16 | 0.4375 | 17 | 0.2941 | 66 | 0.3182 | 4 | 0.75 | 19 | 0.4737 | 18 | 0.3889 | 7 | 0.4286 | 48 | 0.4583 |
| 14_1205626 | 14 | 0.2143 | 15 | 0.1333 | 12 | 0.25 | 16 | 0.25 | 57 | 0.2105 | 5 | 0 | 17 | 0 | 19 | 0.1579 | 7 | 0.1429 | 48 | 0.0833 |
| 14_1205626 | 16 | 0.25 | 19 | 0.2105 | 18 | 0.4444 | 15 | 0.1333 | 68 | 0.2647 | 5 | 0.2 | 18 | 0.3889 | 20 | 0.25 | 7 | 0.2857 | 50 | 0.3 |
| 14_1206664 | 23 | 0.087 | 27 | 0.0741 | 25 | 0.12 | 22 | 0.1364 | 97 | 0.1031 | 7 | 0.1429 | 23 | 0.0435 | 27 | 0.1111 | 12 | 0.4167 | 69 | 0.1449 |
| 14_1206664 | 18 | 0.1667 | 23 | 0.2609 | 23 | 0.1739 | 17 | 0 | 81 | 0.1605 | 5 | 0 | 16 | 0.0625 | 19 | 0.0526 | 13 | 0.1538 | 53 | 0.0755 |
| 14_1206668 | 22 | 0.2273 | 24 | 0.3333 | 22 | 0.3182 | 24 | 0.4167 | 92 | 0.3261 | 7 | 0.8571 | 22 | 0.5909 | 28 | 0.4643 | 9 | 0.7778 | 66 | 0.5909 |
| 14_1206669 | 18 | 0.0556 | 24 | 0.2083 | 23 | 0.1304 | 22 | 0 | 87 | 0.1034 | 5 | 0.2 | 20 | 0.15 | 20 | 0.1 | 11 | 0.2727 | 56 | 0.1607 |
| 14_1206671 | 24 | 0.4167 | 28 | 0.4286 | 25 | 0.4 | 28 | 0.2857 | 105 | 0.381 | 7 | 0.2857 | 29 | 0.2414 | 28 | 0.3571 | 14 | 0.4286 | 78 | 0.3205 |
| 14_1206688 | 48 | 0.2083 | 50 | 0.08 | 35 | 0.2 | 44 | 0.1136 | 177 | 0.1469 | 17 | 0.1765 | 45 | 0.1556 | 53 | 0.1698 | 28 | 0.1071 | 143 | 0.1538 |
| 14_1206690 | 47 | 0.0638 | 51 | 0.0588 | 36 | 0.0556 | 51 | 0.0588 | 185 | 0.0595 | 23 | 0 | 47 | 0.0213 | 57 | 0.0526 | 29 | 0.2069 | 156 | 0.0641 |
| 14_1207162 | 1 | 1 | 4 | 0.5 | 1 | 0 | 2 | 0.5 | 8 | 0.5 | 3 | 0 | 1 | 0 | 2 | 0.5 | 4 | 0 | 10 | 0.1 |
| 14_1207162 | 1 | 0 | 3 | 0 | 1 | 1 | 2 | 0 | 7 | 0.1429 | 2 | 0 | 1 | 1 | 1 | 0 | 4 | 0 | 8 | 0.125 |
| 14_1207163 | 1 | 0 | 3 | 0 | 1 | 0 | 2 | 0.5 | 7 | 0.1429 | 2 | 0 | 3 | 0.3333 | 3 | 0.3333 | 1 | 0 | 9 | 0.2222 |
| 14_1207163 | 1 | 0 | 4 | 0.25 | 1 | 0 | 3 | 0 | 9 | 0.1111 | 3 | 0.3333 | 3 | 0.3333 | 3 | 0 | 4 | 0.25 | 13 | 0.2308 |
| 14_1207164 | 2 | 0 | 4 | 0 | 2 | 0 | 2 | 0.5 | 10 | 0.1 | 2 | 0 | 3 | 0.3333 | 3 | 0 | 4 | 0.25 | 12 | 0.1667 |
| 14_1207411 | 1 | 1 | 4 | 0.25 | 2 | 0.5 | 4 | 1 | 11 | 0.6364 | 2 | 0.5 | 2 | 1 | 2 | 1 | 4 | 1 | 10 | 0.9 |
| 14_1207411 | NA | NA | 3 | 1 | 1 | 1 | 4 | 1 | 8 | 1 | 4 | 1 | 1 | 1 | 2 | 1 | 4 | 1 | 11 | 1 |
| 14_1207412 | NA | NA | 3 | 0.3333 | 1 | 0 | 3 | 0 | 7 | 0.1429 | 4 | 0.25 | 1 | 0 | 1 | 0 | 4 | 0.25 | 10 | 0.2 |
| 14_1207412 | 1 | 1 | 5 | 0.8 | 2 | 0.5 | 5 | 0.8 | 13 | 0.7692 | 4 | 0.5 | 3 | 0.3333 | 2 | 1 | 4 | 0.75 | 13 | 0.6154 |
| 14_1207413 | 2 | 0 | 4 | 0.25 | 2 | 0 | 5 | 0 | 13 | 0.0769 | 4 | 0 | 3 | 0 | 3 | 0 | 3 | 0.3333 | 13 | 0.0769 |
| 14_1207415 | 1 | 0 | 7 | 0.2857 | 1 | 0 | 2 | 0.5 | 11 | 0.2727 | 3 | 0.3333 | 4 | 0 | 2 | 0 | 3 | 0 | 12 | 0.0833 |
| 14_1207417 | 2 | 0.5 | 6 | 0 | 2 | 0.5 | 4 | 0.25 | 14 | 0.2143 | 3 | 0.3333 | 4 | 0.25 | 3 | 0 | 5 | 0.4 | 15 | 0.2667 |
| 14_1207419 | 4 | 0 | 6 | 0 | 3 | 0 | 4 | 0.25 | 17 | 0.0588 | 3 | 0 | 4 | 0 | 3 | 0 | 5 | 0.2 | 15 | 0.0667 |
| 14_1207421 | 4 | 0.25 | 8 | 0.125 | 4 | 0.25 | 3 | 0 | 19 | 0.1579 | 3 | 0.6667 | 2 | 0 | 3 | 0.3333 | 6 | 0.3333 | 14 | 0.3571 |
| 14_1207422 | 3 | 0 | 6 | 0.1667 | 3 | 0 | 4 | 0 | 16 | 0.0625 | 3 | 0 | 2 | 0 | 3 | 0 | 6 | 0.1667 | 14 | 0.0714 |
| 14_1207443 | 6 | 0.1667 | 12 | 0 | 6 | 0.1667 | 5 | 0.2 | 29 | 0.1034 | 6 | 0.1667 | 8 | 0.125 | 4 | 0.25 | 9 | 0.1111 | 27 | 0.1481 |
| 14_1207444 | 5 | 1 | 12 | 0.6667 | 5 | 0.8 | 4 | 0.75 | 26 | 0.7692 | 7 | 1 | 8 | 0.5 | 5 | 0.8 | 9 | 0.8889 | 29 | 0.7931 |
| 14_1211841 | 11 | 0.1818 | 10 | 0.4 | 11 | 0 | 7 | 0 | 39 | 0.1538 | 7 | 0 | 13 | 0 | 9 | 0.2222 | 11 | 0.2727 | 40 | 0.125 |
| 14_1211843 | 11 | 0.9091 | 15 | 1 | 12 | 0.9167 | 7 | 0.7143 | 45 | 0.9111 | 12 | 0.9167 | 14 | 0.7857 | 9 | 0.8889 | 9 | 1 | 44 | 0.8864 |
| 14_1211845 | 12 | 0 | 16 | 0.1875 | 12 | 0.1667 | 7 | 0.1429 | 47 | 0.1277 | 12 | 0.25 | 14 | 0 | 12 | 0.0833 | 12 | 0.0833 | 50 | 0.1 |
| 14_1211845 | 12 | 0.4167 | 15 | 0.4 | 12 | 0.5833 | 11 | 0.4545 | 50 | 0.46 | 11 | 0.5455 | 16 | 0.625 | 13 | 0.7692 | 10 | 1 | 50 | 0.72 |
| 14_1211845 | 12 | 0.0833 | 14 | 0.2143 | 10 | 0 | 10 | 0 | 46 | 0.087 | 12 | 0.0833 | 15 | 0.1333 | 12 | 0.1667 | 11 | 0.2727 | 50 | 0.16 |
| 14_1211847 | 10 | 0.3 | 16 | 0.375 | 10 | 0.2 | 9 | 0.2222 | 45 | 0.2889 | 13 | 0.1538 | 16 | 0.125 | 12 | 0.4167 | 9 | 0.2222 | 50 | 0.22 |
| 14_1214814 | NA | NA | NA | NA | NA | NA | 2 | 0.5 | 2 | 0.5 | NA | NA | NA | NA | NA | NA | 1 | 1 | 1 | 1 |
| 14_1214847 | NA | NA | 1 | 1 | 1 | 1 | 1 | 0 | 3 | 0.6667 | NA | NA | 1 | 1 | NA | NA | 3 | 1 | 4 | 1 |
| 14_1214848 | NA | NA | 1 | 1 | 1 | 0 | 1 | 0 | 3 | 0.3333 | NA | NA | 1 | 1 | NA | NA | 3 | 0 | 4 | 0.25 |
| 14_1214858 | 2 | 0 | 2 | 0.5 | 3 | 0 | 1 | 0 | 8 | 0.125 | 1 | 0 | NA | NA | NA | NA | 3 | 0.3333 | 4 | 0.25 |
| 14_1214873 | 2 | 0.5 | 4 | 0.25 | 6 | 0 | 1 | 0 | 13 | 0.1538 | 1 | 1 | 3 | 0.3333 | 1 | 0 | 3 | 0 | 8 | 0.25 |
| 14_1214880 | 2 | 0 | 3 | 0.3333 | 4 | 0 | 1 | 0 | 10 | 0.1 | 1 | 0 | 3 | 0 | NA | NA | 2 | 0.5 | 6 | 0.1667 |
| 14_1214911 | 6 | 0 | 14 | 0.0714 | 11 | 0.0909 | 3 | 0 | 34 | 0.0588 | 7 | 0.1429 | 4 | 0 | 3 | 0 | 3 | 0 | 17 | 0.0588 |
| 14_1215198 | 2 | 0 | 2 | 1 | 5 | 0 | 2 | 0 | 11 | 0.1818 | 4 | 0 | 3 | 0 | 2 | 0 | 1 | 1 | 10 | 0.1 |
| 14_1215198 | 2 | 0 | 5 | 0 | 7 | 0.1429 | 4 | 0.25 | 18 | 0.1111 | 7 | 0 | 7 | 0.1429 | 1 | 0 | 1 | 0 | 16 | 0.0625 |
| 14_1215199 | 1 | 0 | 2 | 0.5 | 8 | 0.125 | 3 | 0 | 14 | 0.1429 | 6 | 0.1667 | 6 | 0 | 1 | 1 | NA | NA | 13 | 0.1538 |
| 14_1215199 | 3 | 0.6667 | 6 | 0.5 | 6 | 1 | 1 | 0 | 16 | 0.6875 | 4 | 0.75 | 3 | 0.6667 | 2 | 1 | 1 | 1 | 10 | 0.8 |
| 14_1215200 | 4 | 0 | 6 | 0.1667 | 6 | 0 | 4 | 0 | 20 | 0.05 | 7 | 0 | 6 | 0.3333 | 1 | 0 | 1 | 0 | 15 | 0.1333 |
| 14_1215201 | 5 | 0.2 | 6 | 0.5 | 8 | 0.5 | 4 | 0.75 | 23 | 0.4783 | 8 | 0.5 | 7 | 0.2857 | 2 | 1 | 1 | 1 | 18 | 0.5 |
| 14_1215202 | 5 | 0.4 | 6 | 0.8333 | 5 | 0.6 | 3 | 0.6667 | 19 | 0.6316 | 7 | 0.7143 | 7 | 0.4286 | 2 | 1 | 2 | 0.5 | 18 | 0.6111 |
| 14_1215202 | 4 | 0.5 | 6 | 0.6667 | 8 | 0.75 | 4 | 0.5 | 22 | 0.6364 | 8 | 0.75 | 7 | 0.5714 | 2 | 1 | 2 | 0.5 | 19 | 0.6842 |

| | | | | | | | | | | | | | | | | | | | | |
|---|---|---|---|---|---|---|---|---|---|---|---|---|---|---|---|---|---|---|---|---|
| 14_1215207 | 3 | 0.3333 | 3 | 0.6667 | 6 | 0 | 3 | 0 | 15 | 0.2 | 5 | 0.2 | 4 | 0 | 1 | 1 | 1 | 0 | 11 | 0.1818 |
| 14_1215208 | 8 | 0.25 | 5 | 0 | 8 | 0 | 2 | 0.5 | 23 | 0.1304 | 10 | 0.1 | 6 | 0 | 2 | 0.5 | 2 | 0.5 | 20 | 0.15 |
| 14_1215242 | 6 | 0 | 10 | 0.1 | 9 | 0.1111 | 7 | 0 | 32 | 0.0625 | 8 | 0 | 6 | 0 | 5 | 0.2 | 12 | 0.25 | 31 | 0.129 |
| 14_1215749 | 2 | 0 | 5 | 0.2 | 4 | 0.75 | 3 | 0.3333 | 14 | 0.3571 | 2 | 0.5 | 2 | 0.5 | 1 | 0 | 5 | 0 | 10 | 0.2 |
| 14_1217796 | 12 | 0.1667 | 13 | 0 | 11 | 0 | 3 | 0 | 39 | 0.0513 | 5 | 0 | 7 | 0.1429 | 4 | 0 | 4 | 0 | 20 | 0.05 |
| 14_1217798 | 11 | 0.1818 | 12 | 0.0833 | 11 | 0 | 4 | 0.25 | 38 | 0.1053 | 6 | 0 | 7 | 0.1429 | 4 | 0 | 4 | 0.5 | 21 | 0.1429 |
| 14_1217798 | 10 | 0.1 | 9 | 0.3333 | 10 | 0 | 4 | 0 | 33 | 0.1212 | 5 | 0 | 6 | 0 | 3 | 0.3333 | 3 | 0.3333 | 17 | 0.1176 |
| 14_1217803 | 8 | 0.125 | 10 | 0 | 7 | 0.1429 | 4 | 0 | 29 | 0.069 | 2 | 0 | 5 | 0.2 | 3 | 0 | 3 | 0 | 13 | 0.0769 |
| 14_1217804 | 8 | 0 | 10 | 0.2 | 7 | 0 | 2 | 0 | 27 | 0.0741 | 2 | 0.5 | 4 | 0 | 3 | 0 | 3 | 0 | 12 | 0.0833 |
| 14_1217807 | 7 | 1 | 10 | 0.9 | 6 | 1 | 4 | 1 | 27 | 0.963 | 1 | 1 | 3 | 0.6667 | 2 | 1 | 4 | 0.5 | 10 | 0.7 |
| 14_1217808 | 8 | 0.75 | 7 | 0.5714 | 6 | 1 | 3 | 0.3333 | 24 | 0.7083 | 2 | 0 | 2 | 0.5 | NA | NA | 4 | 0 | 8 | 0.125 |
| 14_1217809 | 4 | 1 | 6 | 0.8333 | 5 | 0.4 | 3 | 0.3333 | 18 | 0.6667 | 2 | 1 | 3 | 0.6667 | 1 | 0 | 2 | 0 | 8 | 0.5 |
| 14_1222578 | 6 | 0.5 | 7 | 0.5714 | 4 | 0.5 | 3 | 0.6667 | 20 | 0.55 | 8 | 0.625 | 3 | 0.6667 | NA | NA | 6 | 0.5 | 17 | 0.5882 |
| 14_1222581 | 8 | 0.625 | 8 | 0.25 | 4 | 0.25 | 4 | 0 | 24 | 0.3333 | 8 | 0.125 | 3 | 0 | 2 | 0 | 4 | 0.5 | 17 | 0.1765 |
| 14_1223961 | 27 | 0.1852 | 15 | 0.2667 | 15 | 0.1333 | 19 | 0.1053 | 76 | 0.1711 | 14 | 0.2143 | 23 | 0.2174 | 12 | 0.5833 | 15 | 0.1333 | 64 | 0.2656 |
| 14_1223961 | 24 | 0.0833 | 19 | 0.1053 | 15 | 0.2 | 20 | 0.1 | 78 | 0.1154 | 13 | 0.0769 | 21 | 0.0476 | 14 | 0.1429 | 14 | 0 | 62 | 0.0645 |
| 14_1223961 | 29 | 0.1034 | 18 | 0.1667 | 17 | 0.2353 | 20 | 0.05 | 84 | 0.131 | 13 | 0.0769 | 24 | 0.2083 | 13 | 0.1538 | 16 | 0.5 | 66 | 0.2424 |
| 14_1223961 | 27 | 0.037 | 17 | 0.1176 | 16 | 0.0625 | 20 | 0.1 | 80 | 0.075 | 10 | 0.1 | 23 | 0.0435 | 15 | 0.0667 | 16 | 0.1875 | 64 | 0.0938 |
| 14_1223962 | 25 | 0.2 | 17 | 0.1176 | 15 | 0.2667 | 20 | 0.45 | 77 | 0.2597 | 13 | 0.2308 | 26 | 0.3846 | 14 | 0.4286 | 17 | 0.2941 | 70 | 0.3429 |
| 14_1223964 | 28 | 0.1786 | 16 | 0.125 | 18 | 0.1111 | 20 | 0.2 | 82 | 0.1585 | 12 | 0.4167 | 26 | 0.0385 | 15 | 0.2 | 16 | 0.0625 | 69 | 0.1449 |
| 14_1223964 | 31 | 0.2581 | 18 | 0.4444 | 20 | 0.4 | 18 | 0.1111 | 87 | 0.2989 | 15 | 0.4 | 26 | 0.2692 | 14 | 0.2143 | 18 | 0.2778 | 73 | 0.2877 |
| 14_1223964 | 25 | 0.12 | 16 | 0.25 | 15 | 0.2667 | 16 | 0.3125 | 72 | 0.2222 | 15 | 0.3333 | 24 | 0.2083 | 12 | 0.0833 | 15 | 0.2667 | 66 | 0.2273 |
| 14_1226033 | NA | NA | NA | NA | 1 | 0 | 2 | 0.5 | 3 | 0.3333 | NA | NA | 1 | 1 | 1 | 0 | 1 | 0 | 3 | 0.3333 |
| 14_1226034 | NA | NA | NA | NA | 1 | 1 | 2 | 0.5 | 3 | 0.6667 | NA | NA | 1 | 1 | 1 | 0 | NA | NA | 2 | 0.5 |
| 14_1227123 | 1 | 0 | 2 | 0.5 | NA | NA | NA | NA | 3 | 0.3333 | NA | NA | 1 | 0 | NA | NA | 1 | 1 | 2 | 0.5 |
| 14_1227128 | NA | NA | 2 | 0.5 | NA | NA | NA | NA | 2 | 0.5 | NA | NA | 1 | 0 | NA | NA | 1 | 1 | 2 | 0.5 |
| 14_1227129 | 2 | 0 | 2 | 1 | NA | NA | NA | NA | 4 | 0.5 | NA | NA | 1 | 0 | NA | NA | 1 | 1 | 2 | 0.5 |
| 14_1227131 | 2 | 0.5 | 2 | 0.5 | NA | NA | NA | NA | 4 | 0.5 | NA | NA | 1 | 1 | NA | NA | 1 | 0 | 2 | 0.5 |
| 14_1227132 | 1 | 1 | 2 | 1 | 1 | 1 | NA | NA | 4 | 1 | NA | NA | 1 | 1 | NA | NA | 1 | 1 | 2 | 1 |
| 14_1227132 | 1 | 1 | 2 | 1 | 1 | 1 | NA | NA | 4 | 1 | NA | NA | NA | NA | NA | NA | 1 | 1 | 1 | 1 |
| 14_1227134 | 2 | 1 | 2 | 1 | NA | NA | NA | NA | 4 | 1 | NA | NA | 1 | 1 | NA | NA | 1 | 1 | 2 | 1 |
| 14_1227134 | 2 | 1 | 2 | 1 | NA | NA | NA | NA | 4 | 1 | NA | NA | 1 | 1 | NA | NA | 1 | 1 | 2 | 1 |
| 14_1227136 | 2 | 1 | 2 | 1 | 1 | 0 | NA | NA | 5 | 0.8 | NA | NA | 1 | 1 | NA | NA | 1 | 1 | 2 | 1 |
| 14_1227137 | 3 | 1 | 2 | 1 | NA | NA | NA | NA | 5 | 1 | NA | NA | 1 | 0 | NA | NA | 1 | 1 | 2 | 0.5 |
| 14_1227139 | 2 | 0 | 2 | 1 | 1 | 1 | NA | NA | 5 | 0.6 | NA | NA | 1 | 0 | NA | NA | 1 | 1 | 2 | 0.5 |
| 14_1227140 | 3 | 0.3333 | 2 | 0.5 | 1 | 1 | NA | NA | 6 | 0.5 | NA | NA | 1 | 1 | NA | NA | 1 | 1 | 2 | 1 |
| 14_1231284 | 5 | 0 | 9 | 0.2222 | 2 | 0 | 2 | 0 | 18 | 0.1111 | 5 | 0.2 | 7 | 0.1429 | NA | NA | 4 | 0.25 | 16 | 0.1875 |
| 14_1231286 | 4 | 0 | 7 | 0.5714 | 2 | 0.5 | 1 | 1 | 14 | 0.4286 | 4 | 0.5 | 6 | 0.5 | NA | NA | 4 | 0.5 | 14 | 0.5 |
| 14_1231286 | 5 | 0 | 7 | 0.1429 | 2 | 0 | 1 | 0 | 15 | 0.0667 | 4 | 0 | 7 | 0.2857 | NA | NA | 4 | 0 | 15 | 0.1333 |
| 14_1231288 | 1 | 1 | 4 | 0.25 | 2 | 1 | NA | NA | 7 | 0.5714 | 2 | 1 | 5 | 0.8 | NA | NA | 3 | 0.6667 | 10 | 0.8 |
| 14_1231289 | 3 | 0.6667 | 6 | 0.3333 | 2 | 0 | 1 | 0 | 12 | 0.3333 | 4 | 0 | 7 | 0.2857 | NA | NA | 3 | 0.3333 | 14 | 0.2143 |
| 14_1231289 | 2 | 0 | 7 | 0.2857 | 2 | 0 | 1 | 0 | 12 | 0.1667 | 4 | 0.5 | 4 | 0 | NA | NA | 3 | 0 | 11 | 0.1818 |
| 14_1231289 | 3 | 0 | 7 | 0.1429 | 2 | 0 | 1 | 0 | 13 | 0.0769 | 4 | 0 | 6 | 0.1667 | NA | NA | 3 | 0 | 13 | 0.0769 |
| 14_1231291 | 2 | 0.5 | 6 | 0 | 1 | 0 | 1 | 0 | 10 | 0.1 | 4 | 0.5 | 4 | 0.25 | NA | NA | 1 | 1 | 9 | 0.4444 |
| 14_1235143 | 9 | 0.2222 | 10 | 0.1 | 8 | 0.125 | 10 | 0 | 37 | 0.1081 | 3 | 0 | 10 | 0 | 11 | 0 | 6 | 0.3333 | 30 | 0.0667 |
| 14_1235146 | 8 | 0 | 7 | 0.1429 | 7 | 0 | 8 | 0.25 | 30 | 0.1 | 2 | 0 | 11 | 0.0909 | 11 | 0 | 8 | 0.125 | 32 | 0.0625 |
| 14_1235544 | 2 | 0 | NA | NA | 1 | 1 | 1 | 0 | 4 | 0.25 | 1 | 0 | 3 | 0.3333 | 5 | 0 | 4 | 0 | 13 | 0.0769 |
| 14_1242058 | 1 | 1 | 3 | 0.3333 | 4 | 0 | 3 | 0 | 11 | 0.1818 | 1 | 1 | 4 | 0 | 4 | 0 | 2 | 0.5 | 11 | 0.1818 |
| 14_1242068 | 1 | 1 | 2 | 0 | 2 | 0 | 1 | 0 | 6 | 0.1667 | 1 | 0 | 4 | 0.25 | 3 | 0 | 4 | 0 | 12 | 0.0833 |
| 14_1242069 | 1 | 1 | 1 | 1 | 3 | 0.3333 | 1 | 1 | 6 | 0.6667 | 1 | 1 | 2 | 1 | 2 | 1 | 2 | 1 | 7 | 1 |
| 14_1242070 | 1 | 1 | NA | NA | 3 | 0 | 1 | 0 | 5 | 0.2 | 1 | 0 | 3 | 0.6667 | 2 | 0 | 3 | 0.3333 | 9 | 0.3333 |
| 14_1242075 | 1 | 1 | NA | NA | 2 | 0.5 | NA | NA | 3 | 0.6667 | 1 | 1 | 2 | 0.5 | 2 | 0.5 | 3 | 0.3333 | 8 | 0.5 |
| 14_1242116 | 1 | 0 | 2 | 0 | 2 | 0.5 | 1 | 0 | 6 | 0.1667 | 2 | 0 | 1 | 0 | 2 | 0 | 1 | 1 | 6 | 0.1667 |
| 14_1242152 | 1 | 0 | 2 | 0.5 | 4 | 0 | 2 | 0.5 | 9 | 0.2222 | 1 | 0 | 3 | 0.3333 | 1 | 0 | 3 | 0 | 8 | 0.125 |
| 14_1242174 | 3 | 0 | 4 | 0.25 | 4 | 0.5 | 3 | 0 | 14 | 0.2143 | 3 | 0 | 3 | 0 | 5 | 0 | 3 | 0.3333 | 14 | 0.0714 |
| 14_1242174 | 2 | 0 | 4 | 0.75 | 4 | 0.5 | 3 | 0 | 13 | 0.3846 | 3 | 0 | 3 | 0 | 5 | 0 | 4 | 0.5 | 15 | 0.1333 |
| 14_1242174 | 1 | 0 | 4 | 0 | 4 | 0.5 | 2 | 1 | 11 | 0.3636 | 3 | 0 | 3 | 0.3333 | 4 | 0.25 | 3 | 0.3333 | 13 | 0.2308 |
| 14_1242174 | 2 | 0.5 | 3 | 1 | 4 | 0.75 | NA | NA | 9 | 0.7778 | 3 | 1 | 2 | 1 | 3 | 0.6667 | 3 | 1 | 11 | 0.9091 |
| 14_1242175 | 2 | 0 | 4 | 0 | 4 | 0.5 | 3 | 0 | 13 | 0.1538 | 3 | 0.3333 | 3 | 0 | 4 | 0.25 | 4 | 0.25 | 14 | 0.2143 |
| 14_1242175 | 2 | 0 | 4 | 0 | 4 | 0.25 | 2 | 0 | 12 | 0.0833 | 3 | 0.3333 | 2 | 0 | 5 | 0 | 3 | 0 | 13 | 0.0769 |
| 14_1242175 | 3 | 0 | 4 | 0.5 | 4 | 0.75 | 2 | 1 | 13 | 0.5385 | 3 | 0 | 3 | 0.6667 | 3 | 0.6667 | 4 | 1 | 13 | 0.6154 |
| 14_1242176 | 3 | 0.3333 | 3 | 1 | 4 | 0.5 | 3 | 0.6667 | 13 | 0.6154 | 3 | 0.3333 | 3 | 0 | 5 | 0.6 | 3 | 0.3333 | 14 | 0.3571 |
| 14_1242176 | 3 | 1 | 3 | 1 | 4 | 1 | 3 | 1 | 13 | 1 | 3 | 1 | 3 | 1 | 5 | 1 | 3 | 1 | 14 | 1 |
| 14_1242177 | 3 | 0.6667 | 4 | 1 | 4 | 0.75 | 2 | 1 | 13 | 0.8462 | 3 | 1 | 3 | 1 | 4 | 1 | 2 | 1 | 12 | 1 |
| 14_1242177 | 2 | 0 | 3 | 0.3333 | 4 | 0 | 3 | 0 | 12 | 0.0833 | 2 | 0 | 2 | 0 | 3 | 0.3333 | 2 | 0 | 9 | 0.1111 |

| | | | | | | | | | | | | | | | | | | | | |
|---|---|---|---|---|---|---|---|---|---|---|---|---|---|---|---|---|---|---|---|---|
| 15_3309779 | 57 | 0.0526 | 80 | 0.1125 | 60 | 0.1333 | 53 | 0.1321 | 250 | 0.108 | 59 | 0.1186 | 37 | 0 | 44 | 0.1136 | 47 | 0.1915 | 187 | 0.1123 |
| 15_3347791 | NA | NA | 1 | 0 | 2 | 0.5 | 1 | 0 | 4 | 0.25 | NA | NA | NA | NA | 4 | 0.25 | 2 | 0 | 6 | 0.1667 |
| 15_3349007 | NA | NA | 1 | 0 | 2 | 0.5 | 1 | 0 | 4 | 0.25 | NA | NA | NA | NA | 3 | 0.3333 | 1 | 0 | 4 | 0.25 |
| 15_3349169 | NA | NA | 1 | 1 | 2 | 1 | NA | NA | 3 | 1 | NA | NA | NA | NA | 2 | 0.5 | 1 | 0 | 3 | 0.3333 |
| 15_3349403 | NA | NA | 1 | 0 | 2 | 0 | 1 | 1 | 4 | 0.25 | NA | NA | NA | NA | 3 | 0.3333 | 1 | 0 | 4 | 0.25 |
| 15_3349453 | NA | NA | 1 | 1 | 2 | 1 | 1 | 1 | 4 | 1 | NA | NA | NA | NA | 3 | 0.6667 | 1 | 0 | 4 | 0.5 |
| 15_3349552 | NA | NA | 1 | 1 | 2 | 1 | NA | NA | 3 | 1 | NA | NA | NA | NA | 2 | 1 | 1 | 1 | 3 | 1 |
| 15_3349930 | NA | NA | 1 | 1 | 2 | 0.5 | 1 | 0 | 4 | 0.5 | NA | NA | NA | NA | 3 | 1 | 1 | 0 | 4 | 0.75 |
| 15_3350052 | NA | NA | NA | NA | 1 | 1 | NA | NA | 1 | 1 | NA | NA | NA | NA | 2 | 0.5 | 1 | 0 | 3 | 0.3333 |
| 15_3995011 | NA | NA | NA | NA | 1 | 1 | NA | NA | 1 | 1 | NA | NA | 1 | 1 | 1 | 0 | NA | NA | 2 | 0.5 |
| 15_4011881 | 5 | 0 | 10 | 0.2 | 2 | 1 | 10 | 0.1 | 27 | 0.1852 | 4 | 0 | 4 | 0 | 6 | 0.3333 | 9 | 0.1111 | 23 | 0.1304 |
| 15_4013118 | 5 | 0.4 | 12 | 0.0833 | 3 | 0.3333 | 6 | 0.3333 | 26 | 0.2308 | 2 | 0 | 2 | 0.5 | 4 | 0.25 | 9 | 0 | 17 | 0.1176 |
| 15_4013140 | 3 | 0 | 11 | 0.0909 | 3 | 0 | 6 | 0.1667 | 23 | 0.087 | 2 | 0.5 | 2 | 0 | 5 | 0.2 | 8 | 0.25 | 17 | 0.2353 |
| 15_4015006 | 4 | 0 | 2 | 0.5 | 1 | 0 | 4 | 0 | 11 | 0.0909 | 1 | 0 | 1 | 0 | 2 | 0.5 | 6 | 0 | 10 | 0.1 |
| 15_4015405 | 2 | 1 | 3 | 1 | 1 | 1 | 3 | 0.3333 | 9 | 0.7778 | NA | NA | 1 | 1 | 2 | 1 | 6 | 1 | 9 | 1 |
| 15_4015452 | 2 | 0.5 | 2 | 0 | 1 | 0 | 2 | 0.5 | 7 | 0.2857 | NA | NA | NA | NA | 1 | 0 | 4 | 0.5 | 5 | 0.4 |
| 15_4015458 | 3 | 0.6667 | 2 | 1 | 1 | 1 | 2 | 0.5 | 8 | 0.75 | 1 | 0 | 1 | 1 | NA | NA | 4 | 1 | 6 | 0.8333 |
| 15_4015463 | 2 | 0.5 | 2 | 0.5 | 1 | 1 | 3 | 0 | 8 | 0.375 | NA | NA | NA | NA | 1 | 0 | 4 | 0.25 | 5 | 0.2 |
| 15_4015753 | 3 | 0.3333 | 2 | 1 | 1 | 1 | 1 | 0 | 7 | 0.5714 | 1 | 1 | 2 | 0.5 | 1 | 1 | 6 | 0.8333 | 10 | 0.8 |
| 15_4015815 | 3 | 0 | 2 | 0 | 1 | 0 | 3 | 0.3333 | 9 | 0.1111 | 1 | 0 | 2 | 0.5 | NA | NA | 5 | 0 | 8 | 0.125 |
| 15_4183408 | 23 | 0.4783 | 29 | 0.5862 | 13 | 0.3846 | 20 | 0.6 | 85 | 0.5294 | 18 | 0.2778 | 20 | 0.45 | 27 | 0.2963 | 27 | 0.4815 | 92 | 0.3804 |
| 15_4183786 | 28 | 0 | 33 | 0.0909 | 16 | 0.125 | 17 | 0.1765 | 94 | 0.0851 | 21 | 0.0476 | 20 | 0.05 | 30 | 0.0333 | 28 | 0.0714 | 99 | 0.0505 |
| 15_4183886 | 33 | 0.1212 | 35 | 0.1143 | 19 | 0.0526 | 19 | 0 | 106 | 0.0849 | 23 | 0.2609 | 19 | 0 | 33 | 0.1515 | 31 | 0.1935 | 106 | 0.1604 |
| 15_4546196 | 1 | 0 | 1 | 1 | NA | NA | 1 | 0 | 3 | 0.3333 | 1 | 1 | 2 | 0 | 1 | 0 | NA | NA | 4 | 0.25 |
| 15_4546324 | 1 | 1 | 2 | 1 | 1 | 1 | 1 | 1 | 5 | 1 | 1 | 1 | 2 | 1 | 1 | 1 | NA | NA | 4 | 1 |
| 15_4546425 | 1 | 1 | 2 | 0.5 | 1 | 1 | NA | NA | 4 | 0.75 | 1 | 1 | 2 | 1 | 1 | 1 | NA | NA | 4 | 1 |
| 15_4546468 | 1 | 1 | 2 | 0 | 1 | 0 | NA | NA | 4 | 0.25 | 2 | 1 | 2 | 0 | NA | NA | NA | NA | 4 | 0.5 |
| 15_4546560 | 1 | 1 | 2 | 1 | 1 | 1 | 1 | 1 | 5 | 1 | 2 | 1 | 2 | 1 | 1 | 1 | NA | NA | 5 | 1 |
| 15_4546624 | 1 | 1 | 2 | 0.5 | NA | NA | 1 | 0 | 4 | 0.5 | 2 | 0.5 | 1 | 1 | 1 | 0 | NA | NA | 4 | 0.5 |
| 15_4604290 | 2 | 0 | 2 | 0 | 1 | 1 | NA | NA | 5 | 0.2 | 2 | 1 | 2 | 0.5 | 1 | 0 | NA | NA | 5 | 0.6 |
| 15_4604298 | 2 | 0.5 | 2 | 0.5 | 1 | 1 | NA | NA | 5 | 0.6 | 2 | 1 | 2 | 0 | NA | NA | NA | NA | 4 | 0.5 |
| 15_4606868 | 2 | 0 | 3 | 0.3333 | 1 | 0 | 1 | 1 | 7 | 0.2857 | 1 | 0 | 4 | 0.75 | 1 | 0 | NA | NA | 6 | 0.5 |
| 15_4610300 | 5 | 0.2 | 15 | 0.1333 | 3 | 0 | 6 | 0 | 29 | 0.1034 | 3 | 0 | 8 | 0 | 4 | 0.25 | 1 | 0 | 16 | 0.0625 |
| 15_5018072 | 1 | 1 | NA | NA | 1 | 1 | NA | NA | 2 | 1 | NA | NA | 1 | 1 | NA | NA | NA | NA | 1 | 1 |
| 15_5150889 | 2 | 1 | NA | NA | 1 | 1 | NA | NA | 3 | 1 | NA | NA | NA | NA | 1 | 1 | NA | NA | 1 | 1 |
| 15_5150935 | 2 | 0.5 | NA | NA | NA | NA | NA | NA | 2 | 0.5 | NA | NA | NA | NA | 1 | 1 | NA | NA | 1 | 1 |
| 15_5150981 | 2 | 1 | NA | NA | 1 | 1 | NA | NA | 3 | 1 | NA | NA | NA | NA | 1 | 1 | NA | NA | 1 | 1 |
| 15_5150996 | 2 | 1 | NA | NA | 1 | 1 | NA | NA | 3 | 1 | NA | NA | NA | NA | 1 | 1 | NA | NA | 1 | 1 |
| 15_5151174 | 2 | 1 | NA | NA | 1 | 1 | NA | NA | 3 | 1 | NA | NA | NA | NA | 1 | 1 | 1 | 1 | 2 | 1 |
| 15_5151378 | 2 | 0.5 | NA | NA | 1 | 0 | NA | NA | 3 | 0.3333 | NA | NA | NA | NA | 1 | 1 | 1 | 0 | 2 | 0.5 |
| 15_5152190 | 2 | 1 | NA | NA | NA | NA | NA | NA | 2 | 1 | NA | NA | NA | NA | 1 | 1 | 1 | 1 | 2 | 1 |
| 15_5208195 | NA | NA | 3 | 0.3333 | 2 | 0 | 3 | 0 | 8 | 0.125 | NA | NA | 2 | 0 | 2 | 0.5 | 1 | 1 | 5 | 0.4 |
| 15_5208549 | NA | NA | 4 | 0 | 2 | 1 | 5 | 0.4 | 11 | 0.3636 | 3 | 0.3333 | 1 | 0 | 2 | 0.5 | 1 | 0 | 7 | 0.2857 |
| 15_5208655 | NA | NA | 4 | 0.25 | 3 | 0.3333 | 5 | 0.8 | 12 | 0.5 | 3 | 0.6667 | 1 | 1 | 1 | 1 | 1 | 0 | 6 | 0.6667 |
| 15_5208678 | NA | NA | 4 | 0.25 | 2 | 0 | 5 | 0 | 11 | 0.0909 | 3 | 0.3333 | 1 | 0 | 2 | 0 | 1 | 0 | 7 | 0.1429 |
| 15_5209264 | 1 | 0 | 4 | 0 | 3 | 0.3333 | 5 | 0.2 | 13 | 0.1538 | 2 | 0 | 1 | 0 | 3 | 0.3333 | 1 | 0 | 7 | 0.1429 |
| 15_5210011 | 3 | 0.3333 | 5 | 0.2 | 3 | 0 | 5 | 0.2 | 16 | 0.1875 | 3 | 0.6667 | 2 | 0 | 4 | 0 | 1 | 0 | 10 | 0.2 |
| 15_5210096 | 3 | 0 | 5 | 0 | 2 | 0.5 | 5 | 0 | 15 | 0.0667 | 3 | 0 | 2 | 0 | 4 | 0.25 | 1 | 0 | 10 | 0.1 |
| 15_5210401 | 2 | 0 | 5 | 0.6 | NA | NA | 5 | 0.4 | 12 | 0.4167 | 3 | 0 | 1 | 1 | 3 | 0 | 1 | 0 | 8 | 0.125 |
| 15_5210923 | 3 | 0 | 6 | 0.1667 | 2 | 0.5 | 5 | 0 | 16 | 0.125 | 2 | 0.5 | 2 | 0 | 5 | 0 | 1 | 1 | 10 | 0.2 |
| 15_5246681 | 3 | 0 | 9 | 0 | 5 | 0 | 14 | 0.1429 | 31 | 0.0645 | 7 | 0 | 6 | 0 | 5 | 0 | 7 | 0.2857 | 25 | 0.08 |
| 15_5246779 | 4 | 0 | 11 | 0.2727 | 4 | 0 | 13 | 0.2308 | 32 | 0.1875 | 6 | 0 | 4 | 0 | 3 | 0.3333 | 7 | 0.2857 | 20 | 0.15 |
| 15_5246803 | 3 | 1 | 9 | 0.8889 | 4 | 0.75 | 11 | 1 | 27 | 0.9259 | 6 | 1 | 5 | 1 | 5 | 1 | 5 | 1 | 21 | 1 |
| 15_5246809 | 3 | 0.3333 | 10 | 0.7 | 6 | 0.3333 | 9 | 0.5556 | 28 | 0.5357 | 6 | 0.5 | 4 | 0.75 | 5 | 0.8 | 7 | 0.5714 | 22 | 0.6364 |
| 15_5246883 | 4 | 0 | 11 | 0 | 5 | 0.4 | 13 | 0.1538 | 33 | 0.1212 | 7 | 0.1429 | 5 | 0.2 | 4 | 0.25 | 7 | 0 | 23 | 0.1304 |
| 15_5246910 | 4 | 1 | 10 | 1 | 5 | 1 | 14 | 1 | 33 | 1 | 7 | 1 | 6 | 1 | 5 | 1 | 6 | 1 | 24 | 1 |
| 15_5247173 | 4 | 0.75 | 11 | 0.8182 | 5 | 1 | 12 | 0.8333 | 32 | 0.8438 | 6 | 0.5 | 7 | 0.7143 | 6 | 0.8333 | 6 | 0.8333 | 25 | 0.72 |
| 15_5247256 | 5 | 0 | 9 | 0 | 7 | 0.5714 | 11 | 0.4545 | 32 | 0.2812 | 7 | 0 | 6 | 0.1667 | 4 | 0.5 | 6 | 0.3333 | 23 | 0.2174 |
| 15_5247351 | 6 | 0 | 12 | 0.25 | 9 | 0.3333 | 12 | 0 | 39 | 0.1538 | 6 | 0 | 6 | 0 | 6 | 0.1667 | 7 | 0.5714 | 25 | 0.2 |
| 15_5247669 | 6 | 0.1667 | 14 | 0.2857 | 11 | 0 | 15 | 0.2 | 46 | 0.1739 | 7 | 0 | 7 | 0 | 6 | 0.6667 | 7 | 0 | 27 | 0.1481 |
| 15_6465036 | 1 | 1 | 1 | 1 | 1 | 1 | 2 | 1 | 5 | 1 | 1 | 1 | NA | NA | NA | NA | NA | NA | 1 | 1 |
| 15_6465114 | 1 | 0 | 1 | 0 | 1 | 0 | 2 | 0.5 | 5 | 0.2 | 1 | 1 | NA | NA | NA | NA | NA | NA | 1 | 1 |
| 15_6610291 | 1 | 1 | 1 | 0 | NA | NA | NA | NA | 2 | 0.5 | NA | NA | NA | NA | 1 | 1 | NA | NA | 1 | 1 |
| 15_6815960 | NA | NA | 2 | 0.5 | NA | NA | NA | NA | 2 | 0.5 | NA | NA | NA | NA | NA | NA | 1 | 1 | 1 | 1 |
| 15_6816394 | NA | NA | 2 | 1 | NA | NA | NA | NA | 2 | 1 | NA | NA | NA | NA | NA | NA | 1 | 1 | 1 | 1 |

| | | | | | | | | | | | | | | | | | | | | |
|---|---|---|---|---|---|---|---|---|---|---|---|---|---|---|---|---|---|---|---|---|
| 15_6816444 | NA | NA | 2 | 1 | NA | NA | 1 | 0 | 3 | 0.6667 | NA | NA | NA | NA | NA | NA | 1 | 1 | 1 | 1 |
| 15_6825798 | 1 | 1 | 2 | 0.5 | 3 | 0.3333 | 2 | 1 | 8 | 0.625 | NA | NA | 1 | 0 | NA | NA | 1 | 1 | 2 | 0.5 |
| 15_6825866 | 1 | 1 | 3 | 1 | 3 | 1 | 2 | 1 | 9 | 1 | NA | NA | 1 | 1 | NA | NA | 1 | 1 | 2 | 1 |
| 15_6826162 | 2 | 1 | 3 | 0.6667 | 3 | 0.6667 | 2 | 0.5 | 10 | 0.7 | NA | NA | 1 | 0 | 1 | 1 | 1 | 1 | 3 | 0.6667 |
| 15_6826208 | 2 | 0 | 3 | 0.3333 | 3 | 0 | 2 | 0.5 | 10 | 0.2 | NA | NA | 1 | 1 | NA | NA | NA | NA | 1 | 1 |
| 15_6826281 | 2 | 0 | 3 | 0 | 2 | 0 | 2 | 0.5 | 9 | 0.1111 | 1 | 0 | 1 | 0 | 1 | 1 | 1 | 0 | 4 | 0.25 |
| 15_6826502 | 1 | 0 | 1 | 1 | 1 | 1 | 3 | 1 | 6 | 0.8333 | 1 | 1 | 2 | 1 | 1 | 0 | 1 | 0 | 5 | 0.6 |
| 15_6826573 | 2 | 0 | 3 | 0.3333 | 3 | 0 | 3 | 0.3333 | 11 | 0.1818 | 1 | 0 | 2 | 0.5 | 1 | 0 | 1 | 0 | 5 | 0.2 |
| 15_6826646 | 2 | 0 | 3 | 0.3333 | 3 | 0.3333 | 3 | 0.6667 | 11 | 0.3636 | 1 | 0 | 2 | 1 | 1 | 0 | 1 | 0 | 5 | 0.4 |
| 15_6827448 | 1 | 1 | 1 | 0 | 1 | 0 | 1 | 0 | 4 | 0.25 | 1 | 1 | 1 | 0 | 3 | 0.3333 | 1 | 0 | 6 | 0.3333 |
| 15_6843190 | 1 | 0 | 1 | 0 | NA | NA | 1 | 1 | 3 | 0.3333 | NA | NA | 1 | 1 | 1 | 1 | NA | NA | 2 | 1 |
| 15_6844088 | 1 | 1 | NA | NA | 1 | 1 | 1 | 1 | 3 | 1 | 1 | 1 | 1 | 0 | 2 | 1 | NA | NA | 4 | 0.75 |
| 15_6844281 | 1 | 1 | NA | NA | 1 | 0 | 1 | 1 | 3 | 0.6667 | 1 | 1 | 1 | 1 | 2 | 1 | NA | NA | 4 | 1 |
| 15_6844309 | 1 | 1 | NA | NA | 1 | 0 | 1 | 0 | 3 | 0.3333 | 1 | 1 | 1 | 0 | 2 | 0 | NA | NA | 4 | 0.25 |
| 15_6844677 | 1 | 1 | NA | NA | 1 | 0 | 1 | 1 | 3 | 0.6667 | 1 | 1 | 1 | 0 | 1 | 1 | NA | NA | 3 | 0.6667 |
| 15_6844997 | 1 | 1 | NA | NA | 1 | 1 | 1 | 1 | 3 | 1 | 1 | 1 | 1 | 0 | 2 | 1 | NA | NA | 4 | 0.75 |
| 15_6845087 | NA | NA | NA | NA | 1 | 1 | 1 | 1 | 2 | 1 | 1 | 0 | 1 | 1 | 2 | 1 | NA | NA | 4 | 0.75 |
| 15_6845131 | 1 | 1 | NA | NA | 1 | 1 | 1 | 1 | 3 | 1 | 1 | 1 | 1 | 1 | 2 | 1 | NA | NA | 4 | 1 |
| 15_6845150 | 1 | 0 | NA | NA | 1 | 0 | 1 | 1 | 3 | 0.3333 | 1 | 1 | 1 | 0 | 2 | 0.5 | NA | NA | 4 | 0.5 |
| 15_6845189 | 1 | 0 | NA | NA | 1 | 1 | 1 | 1 | 3 | 0.6667 | 1 | 1 | 1 | 1 | 2 | 0.5 | NA | NA | 4 | 0.75 |
| 15_7220502 | NA | NA | 1 | 0 | 2 | 0.5 | 2 | 0 | 5 | 0.2 | NA | NA | NA | NA | 1 | 1 | NA | NA | 1 | 1 |
| 15_7220646 | NA | NA | 1 | 1 | 1 | 1 | 2 | 0.5 | 4 | 0.75 | NA | NA | NA | NA | 1 | 1 | NA | NA | 1 | 1 |
| 15_7220738 | NA | NA | 1 | 1 | 3 | 1 | 2 | 0.5 | 6 | 0.8333 | NA | NA | NA | NA | 1 | 1 | NA | NA | 1 | 1 |
| 15_7220763 | NA | NA | 1 | 0 | 3 | 1 | 2 | 1 | 6 | 0.8333 | NA | NA | NA | NA | 1 | 1 | NA | NA | 1 | 1 |
| 15_7220783 | NA | NA | 1 | 1 | 3 | 0.6667 | 2 | 0.5 | 6 | 0.6667 | NA | NA | NA | NA | 1 | 1 | NA | NA | 1 | 1 |
| 15_7222792 | 2 | 0 | 2 | 0 | 3 | 0.3333 | 3 | 0.3333 | 10 | 0.2 | 1 | 1 | NA | NA | 1 | 0 | NA | NA | 2 | 0.5 |
| 15_7223141 | 2 | 0.5 | 1 | 0 | 6 | 0.3333 | 3 | 0.3333 | 12 | 0.3333 | 1 | 1 | NA | NA | 1 | 0 | NA | NA | 2 | 0.5 |
| 15_7224496 | 5 | 0 | 2 | 0 | 5 | 0 | 3 | 0.3333 | 15 | 0.0667 | 2 | 0.5 | 2 | 0.5 | 5 | 0 | NA | NA | 9 | 0.2222 |
| 15_7224868 | 5 | 0 | 2 | 0 | 8 | 0 | 5 | 0.2 | 20 | 0.05 | 2 | 0 | 4 | 0.25 | 4 | 0 | 1 | 0 | 11 | 0.0909 |
| 15_7225048 | 5 | 0.2 | 2 | 0 | 8 | 0.125 | 5 | 0.4 | 20 | 0.2 | 2 | 0.5 | 4 | 0.25 | 4 | 0 | 1 | 0 | 11 | 0.1818 |
| 15_7225983 | 5 | 0 | 2 | 0 | 8 | 0.125 | 4 | 0 | 19 | 0.0526 | 4 | 0 | 5 | 0 | 5 | 0.2 | 2 | 0 | 16 | 0.0625 |
| 15_7226489 | 6 | 0.1667 | 1 | 0 | 8 | 0.25 | 5 | 0 | 20 | 0.15 | 4 | 0.5 | 5 | 0.2 | 6 | 0.1667 | 1 | 0 | 16 | 0.25 |
| 15_7226580 | 6 | 0.3333 | 2 | 1 | 7 | 0.1429 | 5 | 0.4 | 20 | 0.35 | 4 | 0.25 | 5 | 0.2 | 5 | 0.2 | 1 | 0 | 15 | 0.2 |
| 15_7226782 | 6 | 0 | 3 | 0.3333 | 8 | 0.125 | 5 | 0.2 | 22 | 0.1364 | 4 | 0.75 | 5 | 0.2 | 6 | 0 | 3 | 0.3333 | 18 | 0.2778 |
| 15_7864063 | NA | NA | NA | NA | NA | NA | 1 | 1 | 1 | 1 | NA | NA | NA | NA | 1 | 1 | 1 | 1 | 2 | 1 |
| 15_7864094 | NA | NA | NA | NA | NA | NA | 1 | 1 | 1 | 1 | NA | NA | NA | NA | NA | NA | 1 | 1 | 1 | 1 |
| 15_7864328 | NA | NA | NA | NA | NA | NA | 1 | 1 | 1 | 1 | NA | NA | 1 | 1 | 1 | 1 | 1 | 1 | 3 | 1 |
| 15_7864940 | NA | NA | NA | NA | NA | NA | 1 | 1 | 1 | 1 | NA | NA | 1 | 1 | 1 | 1 | 1 | 1 | 3 | 1 |
| 15_8121799 | 5 | 0 | 3 | 0 | 3 | 0.3333 | 6 | 0.1667 | 17 | 0.1176 | 4 | 0 | 2 | 0.5 | 3 | 0 | 3 | 0 | 12 | 0.0833 |
| 15_8121909 | 6 | 0 | 2 | 0 | 4 | 0.25 | 6 | 0 | 18 | 0.0556 | 3 | 0.3333 | 2 | 0.5 | 4 | 0.25 | 2 | 0.5 | 11 | 0.3636 |
| 15_8122046 | 5 | 0 | 2 | 0 | 4 | 0.25 | 6 | 0.1667 | 17 | 0.1176 | 4 | 0.25 | 2 | 0 | 4 | 0 | 2 | 0 | 12 | 0.0833 |
| 15_8122387 | 4 | 0.25 | 2 | 0 | 2 | 0 | 6 | 0 | 14 | 0.0714 | 4 | 0.25 | 2 | 0.5 | 4 | 0 | 1 | 0 | 11 | 0.1818 |
| 15_8258315 | NA | NA | 1 | 0 | 3 | 0.3333 | 1 | 0 | 5 | 0.2 | NA | NA | NA | NA | 2 | 0.5 | NA | NA | 2 | 0.5 |
| 15_8258405 | 1 | 1 | 1 | 1 | 3 | 0.6667 | NA | NA | 5 | 0.8 | 1 | 0 | NA | NA | 3 | 0.6667 | NA | NA | 4 | 0.5 |
| 15_8258434 | 1 | 1 | 1 | 1 | 3 | 0.3333 | 1 | 0 | 6 | 0.5 | 1 | 1 | 1 | 0 | 3 | 0.3333 | NA | NA | 5 | 0.4 |
| 15_8258664 | 2 | 0.5 | NA | NA | 3 | 0.6667 | 1 | 0 | 6 | 0.5 | 1 | 0 | NA | NA | 1 | 1 | NA | NA | 2 | 0.5 |
| 15_8276438 | 2 | 0 | NA | NA | 2 | 0.5 | 1 | 0 | 5 | 0.2 | 1 | 1 | 2 | 0 | 2 | 0 | NA | NA | 5 | 0.2 |
| 15_8300221 | 3 | 0 | 1 | 1 | 4 | 0.25 | 3 | 0 | 11 | 0.1818 | 5 | 0 | 3 | 0 | 4 | 0.25 | 5 | 0 | 17 | 0.0588 |
| 15_8300341 | 3 | 0.3333 | 2 | 0 | 6 | 0 | 3 | 0.3333 | 14 | 0.1429 | 6 | 0.1667 | 4 | 0 | 6 | 0.1667 | 5 | 0.2 | 21 | 0.1429 |
| 15_8300416 | 3 | 0.3333 | 2 | 0 | 5 | 0.4 | 3 | 0.6667 | 13 | 0.3846 | 6 | 0.6667 | 3 | 0.3333 | 5 | 0.4 | 6 | 0.3333 | 20 | 0.45 |
| 15_8320599 | 3 | 0.3333 | 2 | 0.5 | 2 | 0 | 7 | 0.1429 | 14 | 0.2143 | 2 | 0 | 2 | 0 | 7 | 0.1429 | 4 | 0 | 15 | 0.0667 |
| 15_8320651 | NA | NA | 1 | 1 | NA | NA | NA | NA | 1 | 1 | 1 | 1 | NA | NA | 1 | 1 | 2 | 0.5 | 4 | 0.75 |
| 15_8320720 | 2 | 1 | 2 | 1 | NA | NA | 7 | 1 | 11 | 1 | 2 | 1 | 2 | 0.5 | 7 | 0.8571 | 5 | 1 | 16 | 0.875 |
| 15_8320753 | 2 | 0 | 2 | 1 | NA | NA | 6 | 1 | 10 | 0.8 | 2 | 0 | 2 | 0.5 | 6 | 0.5 | 5 | 0.6 | 15 | 0.4667 |
| 15_8320771 | 3 | 0.3333 | 2 | 0.5 | NA | NA | 5 | 0.6 | 10 | 0.5 | 2 | 0 | 2 | 0 | 7 | 0.2857 | 4 | 0.5 | 15 | 0.2667 |
| 15_8320815 | 3 | 0.6667 | 2 | 1 | NA | NA | 6 | 0.8333 | 11 | 0.8182 | 2 | 1 | 2 | 0 | 6 | 0.8333 | 5 | 1 | 15 | 0.8 |
| 15_8320833 | 3 | 0.6667 | 2 | 0.5 | NA | NA | 6 | 0.3333 | 11 | 0.4545 | 1 | 0 | 2 | 0.5 | 7 | 0.2857 | 4 | 0 | 14 | 0.2143 |
| 15_8320850 | 3 | 0.3333 | 2 | 0 | NA | NA | 6 | 0 | 11 | 0.0909 | 2 | 0 | 2 | 0 | 7 | 0.2857 | 5 | 0 | 16 | 0.125 |
| 15_8372001 | 2 | 1 | NA | NA | NA | NA | NA | NA | 2 | 1 | NA | NA | 1 | 1 | NA | NA | 1 | 1 | 2 | 1 |
| 15_8372103 | 2 | 1 | NA | NA | NA | NA | NA | NA | 2 | 1 | NA | NA | 1 | 0 | NA | NA | 1 | 1 | 2 | 0.5 |
| 15_8379879 | 2 | 1 | NA | NA | NA | NA | 2 | 0.5 | 4 | 0.75 | NA | NA | NA | NA | 1 | 1 | 1 | 1 | 2 | 1 |
| 15_8380048 | 2 | 1 | NA | NA | NA | NA | 3 | 0.6667 | 5 | 0.8 | NA | NA | 1 | 1 | 1 | 1 | 1 | 1 | 3 | 1 |
| 15_8380170 | 2 | 0 | NA | NA | NA | NA | 2 | 0.5 | 4 | 0.25 | NA | NA | 1 | 1 | 1 | 0 | 1 | 1 | 3 | 0.6667 |
| 15_8380379 | 2 | 1 | NA | NA | NA | NA | 3 | 1 | 5 | 1 | NA | NA | 1 | 1 | 1 | 1 | 1 | 1 | 3 | 1 |
| 15_8380841 | 2 | 0.5 | NA | NA | NA | NA | 3 | 0 | 5 | 0.2 | NA | NA | 1 | 1 | 1 | 0 | 1 | 0 | 3 | 0.3333 |

| | | | | | | | | | | | | | | | | | | | | |
|---|---|---|---|---|---|---|---|---|---|---|---|---|---|---|---|---|---|---|---|---|
| 15_838113( | 1 | 1 | NA | NA | NA | NA | 3 | 1 | 4 | 1 | NA | NA | 1 | 1 | 1 | 0 | 1 | 1 | 3 | 0.6667 |
| 15_838827 | 1 | 1 | NA | NA | NA | NA | 2 | 0 | 3 | 0.3333 | NA | NA | 1 | 1 | 1 | 1 | 1 | 0 | 3 | 0.6667 |
| 15_838830( | NA | NA | NA | NA | NA | NA | 3 | 0.3333 | 3 | 0.3333 | NA | NA | 1 | 1 | NA | NA | NA | NA | 1 | 1 |
| 15_838850 | NA | NA | NA | NA | NA | NA | 3 | 0.3333 | 3 | 0.3333 | NA | NA | 1 | 1 | NA | NA | 1 | 1 | 2 | 1 |
| 15_838853( | NA | NA | NA | NA | NA | NA | 3 | 0.6667 | 3 | 0.6667 | NA | NA | 1 | 1 | 1 | 1 | 1 | 0 | 3 | 0.6667 |
| 15_838855 | NA | NA | NA | NA | NA | NA | 3 | 0.3333 | 3 | 0.3333 | NA | NA | 1 | 1 | 1 | 0 | 1 | 1 | 3 | 0.6667 |
| 15_847510 | 1 | 0 | 1 | 1 | NA | NA | 1 | 1 | 3 | 0.6667 | 1 | 1 | NA | NA | 1 | 1 | 1 | 1 | 3 | 1 |
| 15_852947 | NA | NA | NA | NA | NA | NA | 1 | 1 | 1 | 1 | NA | NA | NA | NA | 1 | 1 | 1 | 1 | 2 | 1 |
| 15_852960 | NA | NA | NA | NA | NA | NA | 1 | 1 | 1 | 1 | NA | NA | NA | NA | 1 | 1 | 1 | 1 | 2 | 1 |
| 15_852993( | NA | NA | NA | NA | NA | NA | 1 | 1 | 1 | 1 | NA | NA | NA | NA | 1 | 0 | 1 | 1 | 2 | 0.5 |
| 15_852995( | NA | NA | NA | NA | NA | NA | 1 | 1 | 1 | 1 | NA | NA | NA | NA | 1 | 1 | 1 | 1 | 2 | 1 |
| 15_853059( | NA | NA | NA | NA | NA | NA | 1 | 1 | 1 | 1 | NA | NA | NA | NA | 1 | 0 | 1 | 1 | 2 | 0.5 |
| 15_853214 | NA | NA | 1 | 1 | 1 | 1 | 1 | 1 | 3 | 1 | NA | NA | NA | NA | 1 | 1 | NA | NA | 1 | 1 |
| 15_866366 | 64 | 0.0156 | 58 | 0.1034 | 67 | 0.0597 | 49 | 0.1224 | 238 | 0.0714 | 45 | 0.0889 | 39 | 0.0513 | 53 | 0.0566 | 40 | 0.075 | 177 | 0.0678 |
| 15_866399 | 74 | 0.0946 | 58 | 0.1034 | 69 | 0.1884 | 47 | 0.0638 | 248 | 0.1169 | 44 | 0.1818 | 40 | 0.05 | 53 | 0.1132 | 43 | 0.1163 | 180 | 0.1167 |
| 15_866401 | 62 | 0.2419 | 56 | 0.4464 | 70 | 0.2571 | 50 | 0.24 | 238 | 0.2941 | 50 | 0.4 | 44 | 0.4545 | 53 | 0.3396 | 42 | 0.2857 | 189 | 0.3704 |
| 15_866490 | 45 | 0.5111 | 36 | 0.3889 | 53 | 0.3962 | 33 | 0.3333 | 167 | 0.4132 | 34 | 0.2647 | 23 | 0.3043 | 36 | 0.4167 | 34 | 0.2353 | 127 | 0.3071 |
| 15_866491 | 58 | 0.0517 | 41 | 0.0976 | 58 | 0.0345 | 41 | 0.0244 | 198 | 0.0505 | 32 | 0.125 | 30 | 0 | 43 | 0.0698 | 37 | 0.027 | 142 | 0.0563 |
| 15_866500 | 59 | 0.0508 | 46 | 0.0652 | 59 | 0.1356 | 41 | 0.0244 | 205 | 0.0732 | 36 | 0.0556 | 25 | 0.04 | 44 | 0.0227 | 40 | 0.1 | 145 | 0.0552 |
| 15_866557 | 55 | 0.7091 | 43 | 0.5349 | 60 | 0.6833 | 35 | 0.6571 | 193 | 0.6528 | 34 | 0.6765 | 23 | 0.4783 | 38 | 0.5526 | 35 | 0.6286 | 130 | 0.5923 |
| 15_866561 | 38 | 0.7105 | 36 | 0.75 | 45 | 0.6444 | 31 | 0.6774 | 150 | 0.6933 | 30 | 0.6333 | 23 | 0.6522 | 27 | 0.5185 | 31 | 0.7097 | 111 | 0.6306 |
| 15_866566 | 50 | 0.26 | 43 | 0.2326 | 54 | 0.3148 | 35 | 0.1429 | 182 | 0.2473 | 27 | 0.3704 | 26 | 0.2308 | 35 | 0.3714 | 37 | 0.2162 | 125 | 0.296 |
| 15_866571 | 52 | 0.0385 | 37 | 0.0811 | 49 | 0.0408 | 34 | 0.1471 | 172 | 0.0698 | 34 | 0.0882 | 25 | 0.16 | 34 | 0.0882 | 35 | 0 | 128 | 0.0781 |
| 15_866574 | 48 | 0.1875 | 36 | 0.1944 | 48 | 0.2708 | 29 | 0.069 | 161 | 0.1925 | 30 | 0.2667 | 21 | 0.1905 | 32 | 0.3438 | 32 | 0.2188 | 115 | 0.2609 |
| 15_899752 | 2 | 0.5 | NA | NA | 1 | 0 | NA | NA | 3 | 0.3333 | NA | NA | 1 | 1 | NA | NA | 1 | 0 | 2 | 0.5 |
| 15_899786 | 2 | 0.5 | 1 | 0 | 1 | 0 | NA | NA | 4 | 0.25 | NA | NA | 1 | 0 | NA | NA | 1 | 1 | 2 | 0.5 |
| 15_910842 | 1 | 1 | 1 | 1 | 2 | 0 | 9 | 1 | 13 | 0.8462 | 6 | 0.8333 | 6 | 1 | 7 | 0.2857 | 3 | 0.6667 | 22 | 0.6818 |
| 15_910854 | 1 | 0 | 3 | 0.3333 | 3 | 0 | 10 | 0.1 | 17 | 0.1176 | 8 | 0.25 | 7 | 0 | 7 | 0 | 5 | 0 | 27 | 0.0741 |
| 15_910863( | 1 | 1 | 3 | 0.3333 | 3 | 0 | 10 | 0.4 | 17 | 0.3529 | 8 | 0.5 | 6 | 0.8333 | 7 | 0.5714 | 5 | 0.8 | 26 | 0.6538 |
| 15_910863 | 1 | 1 | 1 | 1 | 3 | 0.3333 | 8 | 0.75 | 13 | 0.6923 | 8 | 0.625 | 7 | 1 | 7 | 0.7143 | 4 | 0.75 | 26 | 0.7692 |
| 15_910924 | 1 | 0 | 2 | 0 | 4 | 0.25 | 8 | 0.125 | 15 | 0.1333 | 7 | 0 | 7 | 0.1429 | 8 | 0.125 | 5 | 0 | 27 | 0.0741 |
| 15_911381 | 1 | 1 | NA | NA | NA | NA | 1 | 0 | 2 | 0.5 | NA | NA | 1 | 1 | NA | NA | 1 | 1 | 2 | 1 |
| 15_919686 | 2 | 0.5 | 3 | 0 | 5 | 0.2 | 1 | 0 | 11 | 0.1818 | 2 | 1 | 3 | 0 | 1 | 0 | 3 | 0 | 9 | 0.2222 |
| 15_919722 | 2 | 0.5 | 1 | 0 | 4 | 0.75 | 1 | 0 | 8 | 0.5 | 2 | 0.5 | 2 | 0 | 1 | 0 | 3 | 0.3333 | 8 | 0.25 |
| 15_919723 | 2 | 0 | 3 | 0 | 6 | 0.3333 | 1 | 0 | 12 | 0.1667 | 2 | 0.5 | 3 | 0.3333 | 1 | 1 | 3 | 0 | 9 | 0.3333 |
| 15_919777 | 2 | 0 | 2 | 0 | 5 | 0.2 | 1 | 0 | 10 | 0.1 | 2 | 0 | 3 | 0.3333 | 1 | 0 | 3 | 0 | 9 | 0.1111 |
| 15_919900 | 2 | 0 | 3 | 0 | 7 | 0.1429 | 3 | 0 | 15 | 0.0667 | 2 | 0.5 | 2 | 0 | 2 | 0 | 6 | 0 | 12 | 0.0833 |
| 15_104470 | 1 | 1 | 3 | 0.3333 | 1 | 0 | 2 | 0.5 | 7 | 0.4286 | NA | NA | NA | NA | 1 | 0 | 3 | 0.3333 | 4 | 0.25 |
| 15_104613 | 1 | 1 | 2 | 0 | NA | NA | 2 | 0 | 5 | 0.2 | 1 | 1 | NA | NA | 1 | 0 | 3 | 0.6667 | 5 | 0.6 |
| 15_104763 | NA | NA | 4 | 0 | 2 | 0.5 | 1 | 0 | 7 | 0.1429 | 2 | 0.5 | 5 | 0 | 3 | 0 | 3 | 0 | 13 | 0.0769 |
| 15_104765 | 2 | 0 | 5 | 0.2 | 5 | 0.4 | 3 | 0.3333 | 15 | 0.2667 | 3 | 0.3333 | 4 | 0.25 | 3 | 0.6667 | 2 | 0.5 | 12 | 0.4167 |
| 15_104766 | 2 | 0 | 5 | 0 | 5 | 0.2 | 3 | 0 | 15 | 0.0667 | 3 | 0 | 5 | 0 | 4 | 0.25 | 3 | 0 | 15 | 0.0667 |
| 15_104934 | 2 | 0.5 | NA | NA | 1 | 0 | 1 | 0 | 4 | 0.25 | 3 | 0.3333 | 1 | 0 | 3 | 0 | 4 | 0.5 | 11 | 0.2727 |
| 15_105751 | NA | NA | 1 | 1 | 1 | 1 | NA | NA | 2 | 1 | NA | NA | 1 | 1 | NA | NA | 1 | 1 | 2 | 1 |
| 15_109950 | 2 | 1 | 2 | 0.5 | 1 | 0 | 2 | 1 | 7 | 0.7143 | 1 | 0 | 3 | 0.3333 | NA | NA | 1 | 0 | 5 | 0.2 |
| 15_109951 | 1 | 0 | 1 | 1 | 1 | 0 | 2 | 0 | 5 | 0.2 | 1 | 1 | 3 | 0 | NA | NA | 1 | 0 | 5 | 0.2 |
| 15_109952 | 2 | 0.5 | 2 | 0 | 2 | 0.5 | 2 | 1 | 8 | 0.5 | 1 | 0 | 3 | 1 | NA | NA | 1 | 0 | 5 | 0.6 |
| 15_109954 | 2 | 0 | 2 | 0 | 2 | 0.5 | 1 | 0 | 7 | 0.1429 | 1 | 0 | 3 | 0.3333 | NA | NA | NA | NA | 4 | 0.25 |
| 15_119831 | 5 | 0 | 14 | 0 | 5 | 0 | 15 | 0.2 | 39 | 0.0769 | 3 | 0 | 7 | 0.1429 | 6 | 0 | 10 | 0.2 | 26 | 0.1154 |
| 15_121851 | 17 | 0 | 36 | 0 | 36 | 0.1944 | 30 | 0.2333 | 119 | 0.1176 | 22 | 0.0909 | 28 | 0.0714 | 51 | 0.2549 | 33 | 0.1212 | 134 | 0.1567 |
| 15_122697( | 2 | 1 | 4 | 0 | 4 | 0.25 | 1 | 1 | 11 | 0.3636 | 1 | 0 | 2 | 0 | 3 | 0 | 3 | 0.3333 | 9 | 0.1111 |
| 15_122697 | NA | NA | 2 | 0 | 3 | 0.6667 | 1 | 1 | 6 | 0.5 | NA | NA | 1 | 1 | 1 | 1 | 2 | 0.5 | 4 | 0.75 |
| 15_122697 | 2 | 0.5 | 4 | 0.25 | 5 | 0.8 | 1 | 0 | 12 | 0.5 | 1 | 1 | 2 | 0.5 | 3 | 0.3333 | 3 | 0.3333 | 9 | 0.4444 |
| 15_122697 | 1 | 1 | 4 | 0.75 | 6 | 1 | 1 | 0 | 12 | 0.8333 | 1 | 1 | 1 | 1 | 3 | 0.3333 | 3 | 0.3333 | 8 | 0.5 |
| 15_122699( | 2 | 0.5 | 2 | 0.5 | 4 | 0 | 1 | 1 | 9 | 0.3333 | 1 | 1 | 2 | 1 | 2 | 0.5 | 3 | 0.3333 | 8 | 0.625 |
| 15_122699 | 2 | 0 | 2 | 0 | 4 | 0 | 1 | 1 | 9 | 0.1111 | 1 | 1 | 2 | 0 | 3 | 0.3333 | 3 | 0.3333 | 9 | 0.3333 |
| 15_122700( | 2 | 1 | 3 | 1 | 6 | 1 | 1 | 0 | 12 | 0.9167 | 1 | 1 | 2 | 1 | 4 | 1 | 3 | 1 | 10 | 1 |
| 15_122701 | 2 | 1 | 4 | 1 | 5 | 1 | 2 | 1 | 13 | 1 | 1 | 0 | 1 | 0 | 4 | 1 | 4 | 1 | 10 | 0.8 |
| 15_122701 | 2 | 1 | 5 | 1 | 6 | 1 | 2 | 1 | 15 | 1 | 1 | 1 | 3 | 1 | 4 | 0.75 | 5 | 1 | 13 | 0.9231 |
| 15_122702( | 2 | 1 | 5 | 0.8 | 6 | 1 | 2 | 0 | 15 | 0.8 | 1 | 1 | 1 | 1 | 3 | 1 | 5 | 0.8 | 10 | 0.9 |
| 15_122702 | 2 | 1 | 3 | 1 | 6 | 1 | 2 | 1 | 13 | 1 | 1 | 1 | 2 | 1 | 4 | 1 | 3 | 0.6667 | 10 | 0.9 |
| 15_122703 | 2 | 0.5 | 5 | 0.6 | 4 | 0.75 | 1 | 0 | 12 | 0.5833 | 2 | 0 | 2 | 0 | 3 | 1 | 5 | 0.2 | 12 | 0.3333 |
| 15_122706 | NA | NA | 4 | 0.5 | 7 | 0.7143 | NA | NA | 11 | 0.6364 | 2 | 1 | 2 | 0.5 | 2 | 0.5 | 1 | 1 | 7 | 0.7143 |
| 15_122709 | 2 | 0.5 | 3 | 1 | 6 | 1 | 2 | 1 | 13 | 0.9231 | 2 | 1 | 2 | 0.5 | 2 | 1 | 4 | 1 | 10 | 0.9 |
| 15_122712 | 2 | 0 | 5 | 0.4 | 7 | 0.4286 | 5 | 0.2 | 19 | 0.3158 | 1 | 0 | 2 | 0 | 4 | 0.25 | 4 | 0 | 11 | 0.0909 |

| | | | | | | | | | | | | | | | | | | | | |
|---|---|---|---|---|---|---|---|---|---|---|---|---|---|---|---|---|---|---|---|---|
| 15_1232170 | 6 | 0.1667 | 11 | 0.3636 | 7 | 0.4286 | 8 | 0 | 32 | 0.25 | 5 | 0.2 | 8 | 0.25 | 6 | 0.3333 | 8 | 0.25 | 27 | 0.2593 |
| 15_1232203 | 7 | 0.2857 | 13 | 0.1538 | 9 | 0.2222 | 10 | 0.3 | 39 | 0.2308 | 7 | 0.2857 | 8 | 0.5 | 9 | 0.3333 | 11 | 0.1818 | 35 | 0.3143 |
| 15_1234955 | 8 | 0.5 | 16 | 0.375 | 8 | 0.5 | 8 | 0.25 | 40 | 0.4 | 8 | 0.625 | 7 | 0.4286 | 7 | 0.4286 | 12 | 0.3333 | 34 | 0.4412 |
| 15_1234964 | 7 | 0 | 14 | 0.0714 | 7 | 0.1429 | 11 | 0 | 39 | 0.0513 | 6 | 0.1667 | 9 | 0.1111 | 6 | 0 | 8 | 0.125 | 29 | 0.1034 |
| 15_1234966 | 10 | 0 | 18 | 0.1667 | 9 | 0.3333 | 12 | 0.0833 | 49 | 0.1429 | 8 | 0 | 8 | 0.125 | 11 | 0 | 11 | 0.0909 | 38 | 0.0526 |
| 15_1234967 | 10 | 1 | 18 | 1 | 9 | 0.8889 | 12 | 1 | 49 | 0.9796 | 7 | 1 | 11 | 1 | 9 | 1 | 12 | 1 | 39 | 1 |
| 15_1234979 | 7 | 0.2857 | 17 | 0.0588 | 9 | 0.3333 | 10 | 0.5 | 43 | 0.2558 | 7 | 0.2857 | 9 | 0.3333 | 10 | 0.2 | 10 | 0.3 | 36 | 0.2778 |
| 15_1234980 | 9 | 0 | 16 | 0.0625 | 10 | 0.1 | 12 | 0.0833 | 47 | 0.0638 | 8 | 0 | 10 | 0.1 | 10 | 0.2 | 13 | 0.0769 | 41 | 0.0976 |
| 15_1234987 | 4 | 0.5 | 11 | 0.6364 | 6 | 1 | 8 | 0.875 | 29 | 0.7586 | 6 | 0.3333 | 7 | 0.5714 | 10 | 0.6 | 11 | 0.5455 | 34 | 0.5294 |
| 15_1234993 | 10 | 0.7 | 15 | 0.6667 | 9 | 0.7778 | 12 | 0.5833 | 46 | 0.6739 | 5 | 1 | 10 | 0.6 | 9 | 0.4444 | 10 | 0.5 | 34 | 0.5882 |
| 15_1235074 | 16 | 0 | 19 | 0.0526 | 11 | 0.0909 | 15 | 0.2 | 61 | 0.082 | 7 | 0 | 10 | 0.2 | 10 | 0.1 | 14 | 0.1429 | 41 | 0.122 |
| 15_1235128 | 13 | 0.0769 | 21 | 0 | 16 | 0.125 | 17 | 0.1765 | 67 | 0.0896 | 12 | 0.0833 | 14 | 0.1429 | 12 | 0.0833 | 14 | 0.1429 | 52 | 0.1154 |
| 15_1236076 | 11 | 0 | 7 | 0.2857 | 7 | 0.1429 | 10 | 0.1 | 35 | 0.1143 | 4 | 0 | 9 | 0 | 12 | 0.0833 | 5 | 0.2 | 30 | 0.0667 |
| 15_1236094 | 12 | 0.8333 | 7 | 0.8571 | 5 | 0.6 | 9 | 0.8889 | 33 | 0.8182 | 5 | 0.8 | 9 | 0.8889 | 11 | 0.7273 | 5 | 0.4 | 30 | 0.7333 |
| 15_1236112 | 14 | 0.0714 | 9 | 0 | 4 | 0 | 12 | 0.0833 | 39 | 0.0513 | 5 | 0 | 9 | 0.2222 | 13 | 0.2308 | 6 | 0 | 33 | 0.1515 |
| 15_1236133 | 10 | 0.5 | 6 | 0 | 3 | 0 | 7 | 0.5714 | 26 | 0.3462 | 3 | 0 | 6 | 0.3333 | 9 | 0.2222 | 5 | 0 | 23 | 0.1739 |
| 15_1236202 | 8 | 0.75 | 4 | 0.75 | 3 | 0.6667 | 6 | 1 | 21 | 0.8095 | 4 | 0.75 | 6 | 0.6667 | 8 | 0.75 | 4 | 1 | 22 | 0.7727 |
| 15_1236204 | 7 | 0 | 6 | 0.1667 | 3 | 0 | 7 | 0.4286 | 23 | 0.1739 | 4 | 0.25 | 6 | 0.1667 | 7 | 0.2857 | 5 | 0 | 22 | 0.1818 |
| 15_1236812 | 5 | 0 | 4 | 0.25 | 3 | 0.3333 | 4 | 0 | 16 | 0.125 | 4 | 0.25 | 5 | 0.2 | 7 | 0 | 2 | 0.5 | 18 | 0.1667 |
| 15_1237280 | 5 | 0.8 | 4 | 0.75 | 2 | 1 | 4 | 1 | 15 | 0.8667 | 3 | 0.6667 | 4 | 1 | 7 | 1 | 3 | 1 | 17 | 0.9412 |
| 15_1237282 | 2 | 1 | 3 | 0.3333 | 2 | 0 | 6 | 0 | 13 | 0.2308 | 3 | 0.6667 | 4 | 0.25 | 2 | 1 | 1 | 0 | 10 | 0.5 |
| 15_1237300 | 5 | 0.8 | 4 | 1 | 2 | 1 | 5 | 0.6 | 16 | 0.8125 | 3 | 0.3333 | 2 | 0.5 | 5 | 0.8 | 3 | 0.6667 | 13 | 0.6154 |
| 15_1237353 | 3 | 0.3333 | 1 | 0 | 1 | 0 | 6 | 0.1667 | 11 | 0.1818 | 2 | 0.5 | 2 | 0 | 5 | 0 | 3 | 0 | 12 | 0.0833 |
| 15_1237361 | 4 | 1 | 1 | 1 | 1 | 0 | 5 | 0.6 | 11 | 0.7273 | 2 | 1 | 2 | 1 | 4 | 0.75 | 3 | 0.6667 | 11 | 0.8182 |
| 15_1237402 | 5 | 0.2 | 2 | 0 | 1 | 0 | 6 | 0.5 | 14 | 0.2857 | 2 | 0 | 1 | 0 | 3 | 0 | 1 | 1 | 7 | 0.1429 |
| 15_1237410 | 5 | 0.2 | 2 | 0 | 1 | 0 | 5 | 0.2 | 13 | 0.1538 | 2 | 0.5 | 1 | 0 | 2 | 0 | 1 | 0 | 6 | 0.1667 |
| 15_1237419 | 4 | 0 | 2 | 0.5 | 1 | 0 | 4 | 0 | 11 | 0.0909 | 2 | 0 | 1 | 0 | 1 | 1 | NA | NA | 4 | 0.25 |
| 15_1237422 | 2 | 0.5 | 1 | 1 | NA | NA | 3 | 0.3333 | 6 | 0.5 | 1 | 0 | 1 | 1 | 1 | 1 | 1 | 0 | 4 | 0.5 |
| 15_1237437 | 5 | 0.2 | 2 | 0 | NA | NA | 5 | 0 | 12 | 0.0833 | 2 | 0 | 1 | 1 | 1 | 1 | 1 | 0 | 5 | 0.4 |
| 15_1237445 | 3 | 0 | 1 | 0 | 1 | 1 | 1 | 0 | 6 | 0.1667 | 2 | 0 | 1 | 0 | 1 | 1 | 1 | 0 | 5 | 0.2 |
| 15_1237455 | 5 | 0 | 2 | 0.5 | 1 | 0 | 3 | 0 | 11 | 0.0909 | 2 | 0.5 | 1 | 0 | 1 | 0 | 1 | 0 | 5 | 0.2 |
| 15_1237461 | 3 | 0.3333 | 2 | 0 | 1 | 0 | 3 | 0 | 9 | 0.1111 | 2 | 0.5 | 1 | 0 | 1 | 0 | NA | NA | 4 | 0.25 |
| 15_1237466 | 3 | 1 | 1 | 0 | 1 | 1 | 1 | 0 | 6 | 0.6667 | 1 | 1 | 1 | 1 | 1 | 1 | 1 | 0 | 4 | 0.75 |
| 15_1237469 | 3 | 0.6667 | 2 | 1 | 1 | 1 | 3 | 1 | 9 | 0.8889 | 2 | 1 | 1 | 0 | 1 | 0 | 1 | 1 | 5 | 0.6 |
| 15_1237487 | 5 | 0.4 | 1 | 1 | NA | NA | 3 | 0 | 9 | 0.3333 | 1 | 1 | NA | NA | 1 | 0 | 1 | 0 | 3 | 0.3333 |
| 15_1237507 | 5 | 0.8 | 1 | 0 | NA | NA | 3 | 0.3333 | 9 | 0.5556 | 1 | 1 | NA | NA | 1 | 0 | 1 | 0 | 3 | 0.3333 |
| 15_1237523 | 5 | 0.2 | 1 | 1 | NA | NA | 3 | 0.3333 | 9 | 0.3333 | NA | NA | NA | NA | 1 | 0 | 1 | 1 | 2 | 0.5 |
| 15_1237524 | 5 | 0.8 | 1 | 1 | NA | NA | 3 | 0.3333 | 9 | 0.6667 | 1 | 0 | NA | NA | NA | NA | 1 | 1 | 2 | 0.5 |
| 15_1237537 | 3 | 0.6667 | 1 | 0 | NA | NA | 2 | 0 | 6 | 0.3333 | 1 | 1 | NA | NA | NA | NA | 1 | 1 | 2 | 1 |
| 15_1237584 | 3 | 1 | NA | NA | NA | NA | 2 | 1 | 5 | 1 | NA | NA | NA | NA | NA | NA | 1 | 1 | 1 | 1 |
| 15_1237585 | 3 | 0.6667 | NA | NA | NA | NA | 3 | 1 | 6 | 0.8333 | 1 | 1 | NA | NA | NA | NA | 1 | 1 | 2 | 1 |
| 15_1237592 | 3 | 0.3333 | NA | NA | NA | NA | 3 | 0.3333 | 6 | 0.3333 | 1 | 1 | NA | NA | NA | NA | 1 | 1 | 2 | 1 |
| 15_1237623 | 3 | 1 | NA | NA | NA | NA | 3 | 0.6667 | 6 | 0.8333 | 1 | 1 | NA | NA | NA | NA | 1 | 1 | 2 | 1 |
| 15_1259239 | 1 | 1 | NA | NA | NA | NA | NA | NA | 1 | 1 | NA | NA | NA | NA | NA | NA | 1 | 1 | 1 | 1 |
| 15_1259271 | 2 | 1 | NA | NA | NA | NA | NA | NA | 2 | 1 | NA | NA | NA | NA | NA | NA | 1 | 1 | 1 | 1 |
| 15_1259289 | 1 | 1 | NA | NA | NA | NA | NA | NA | 1 | 1 | NA | NA | NA | NA | NA | NA | 1 | 1 | 1 | 1 |
| 15_1280864 | 1 | 1 | 3 | 0 | 7 | 0.1429 | 4 | 0.5 | 15 | 0.2667 | 5 | 0.6 | 3 | 0.3333 | 2 | 0 | 5 | 0.2 | 15 | 0.3333 |
| 15_1280881 | 1 | 0 | 3 | 0.3333 | 6 | 0.3333 | 4 | 0.75 | 14 | 0.4286 | 4 | 0.25 | NA | NA | 2 | 0.5 | 6 | 0.1667 | 12 | 0.25 |
| 15_1280995 | 1 | 0 | 1 | 0 | 5 | 0.2 | 5 | 0.2 | 12 | 0.1667 | 3 | 0.3333 | 2 | 0 | 3 | 0 | 3 | 0.3333 | 11 | 0.1818 |
| 15_1281017 | NA | NA | 1 | 0 | 4 | 0 | 6 | 0.1667 | 11 | 0.0909 | 3 | 0 | 2 | 0.5 | 2 | 0 | 3 | 0 | 10 | 0.1 |
| 15_1281032 | NA | NA | 1 | 1 | 4 | 0.25 | 5 | 0.4 | 10 | 0.4 | 3 | 0.6667 | 2 | 1 | 2 | 0.5 | 2 | 0 | 9 | 0.5556 |
| 15_1281042 | NA | NA | 1 | 0 | 3 | 0 | 6 | 0.3333 | 10 | 0.2 | 3 | 0.6667 | 2 | 0 | 2 | 0.5 | 3 | 0 | 10 | 0.3 |
| 15_1281043 | NA | NA | 1 | 1 | 3 | 0.6667 | 5 | 0 | 9 | 0.3333 | 2 | 1 | 2 | 0 | 2 | 0 | 3 | 0 | 9 | 0.2222 |
| 15_1281058 | NA | NA | 1 | 1 | 3 | 1 | 6 | 0.8333 | 10 | 0.9 | 3 | 1 | 2 | 1 | 1 | 1 | 2 | 0.5 | 8 | 0.875 |
| 15_1281067 | NA | NA | 1 | 0 | 2 | 0 | 4 | 0.25 | 7 | 0.1429 | 2 | 0 | 2 | 0.5 | 2 | 0 | 3 | 0 | 9 | 0.1111 |
| 15_1281071 | NA | NA | NA | NA | 2 | 0 | 6 | 0.1667 | 8 | 0.125 | 1 | 0 | 2 | 0 | 2 | 0.5 | 3 | 0 | 8 | 0.125 |
| 15_1281075 | NA | NA | 1 | 0 | 3 | 0.3333 | 6 | 0.5 | 10 | 0.4 | 3 | 0.3333 | 2 | 0 | 2 | 1 | 2 | 0 | 9 | 0.3333 |
| 15_1281076 | NA | NA | 1 | 0 | 3 | 0.3333 | 6 | 0 | 10 | 0.1 | 3 | 0 | 2 | 0.5 | 2 | 0 | 3 | 0 | 10 | 0.1 |
| 15_1281083 | NA | NA | NA | NA | 2 | 0.5 | 6 | 0.6667 | 8 | 0.625 | 1 | 1 | 2 | 0.5 | 2 | 1 | 3 | 0 | 8 | 0.5 |
| 15_1281084 | NA | NA | 1 | 0 | 3 | 0.3333 | 5 | 0.2 | 9 | 0.2222 | 3 | 0.3333 | 2 | 0 | 2 | 0 | 3 | 0 | 10 | 0.1 |
| 15_1281799 | 1 | 0 | 1 | 1 | 2 | 0 | 4 | 0 | 8 | 0.125 | 3 | 0.3333 | 2 | 0 | 2 | 0.5 | NA | NA | 7 | 0.2857 |
| 15_1281803 | 1 | 1 | NA | NA | 1 | 0 | 4 | 0 | 6 | 0.1667 | 2 | 0 | 2 | 0.5 | NA | NA | 1 | 0 | 5 | 0.2 |
| 15_1281823 | NA | NA | 1 | 0 | 2 | 1 | 5 | 0.6 | 8 | 0.625 | 3 | 0.6667 | 1 | 1 | 2 | 1 | 1 | 1 | 7 | 0.8571 |
| 15_1283408 | 2 | 0 | 4 | 0.5 | 2 | 0.5 | 2 | 0.5 | 10 | 0.4 | 2 | 1 | 3 | 1 | 2 | 1 | 1 | 1 | 8 | 1 |
| 15_1293524 | 14 | 0.1429 | 13 | 0.2308 | 12 | 0.25 | 11 | 0.0909 | 50 | 0.18 | 3 | 0.6667 | 9 | 0.2222 | 14 | 0.2857 | 12 | 0.1667 | 38 | 0.2632 |

| | | | | | | | | | | | | | | | | | | | | |
|---|---|---|---|---|---|---|---|---|---|---|---|---|---|---|---|---|---|---|---|---|
| 15_130291 | NA | NA | 1 | 1 | 1 | 0 | NA | NA | 2 | 0.5 | NA | NA | 1 | 1 | 2 | 0.5 | NA | NA | 3 | 0.6667 |
| 15_130296 | NA | NA | 1 | 1 | 1 | 0 | 1 | 1 | 3 | 0.6667 | 1 | 1 | 1 | 0 | 1 | 1 | 1 | 1 | 4 | 0.75 |
| 15_130341 | NA | NA | 1 | 1 | NA | NA | NA | NA | 1 | 1 | NA | NA | 2 | 0.5 | NA | NA | 1 | 1 | 3 | 0.6667 |
| 15_130342 | NA | NA | 1 | 1 | NA | NA | NA | NA | 1 | 1 | NA | NA | 1 | 1 | NA | NA | 1 | 1 | 2 | 1 |
| 15_130342 | NA | NA | 1 | 1 | NA | NA | NA | NA | 1 | 1 | NA | NA | 2 | 1 | NA | NA | NA | NA | 2 | 1 |
| 15_130343 | NA | NA | 1 | 1 | NA | NA | NA | NA | 1 | 1 | NA | NA | 2 | 1 | NA | NA | NA | NA | 2 | 1 |
| 15_130343 | NA | NA | 1 | 1 | NA | NA | NA | NA | 1 | 1 | NA | NA | 1 | 1 | NA | NA | 1 | 1 | 2 | 1 |
| 15_130344 | NA | NA | 1 | 1 | NA | NA | NA | NA | 1 | 1 | NA | NA | 1 | 1 | NA | NA | 1 | 1 | 2 | 1 |
| 15_130344 | NA | NA | 1 | 1 | NA | NA | NA | NA | 1 | 1 | NA | NA | 1 | 1 | NA | NA | 1 | 0 | 2 | 0.5 |
| 15_130345 | NA | NA | 1 | 1 | NA | NA | NA | NA | 1 | 1 | NA | NA | 1 | 1 | NA | NA | 1 | 1 | 2 | 1 |
| 15_130345 | NA | NA | 1 | 1 | NA | NA | NA | NA | 1 | 1 | NA | NA | 2 | 1 | NA | NA | 1 | 1 | 3 | 1 |
| 15_130346 | NA | NA | 1 | 1 | NA | NA | NA | NA | 1 | 1 | NA | NA | NA | NA | NA | NA | 1 | 1 | 1 | 1 |
| 15_167782 | 1 | 1 | NA | NA | 3 | 0.3333 | 1 | 0 | 5 | 0.4 | NA | NA | 2 | 0.5 | NA | NA | 1 | 1 | 3 | 0.6667 |
| 15_168322 | 1 | 1 | NA | NA | 4 | 0 | 2 | 0 | 7 | 0.1429 | 1 | 0 | 3 | 0 | 1 | 0 | 2 | 0.5 | 7 | 0.1429 |
| 15_168559 | 1 | 0 | 1 | 0 | 4 | 0.5 | 3 | 0.6667 | 9 | 0.4444 | NA | NA | 2 | 0.5 | 2 | 1 | 2 | 0.5 | 6 | 0.6667 |
| 15_168560 | 1 | 0 | 2 | 0 | 4 | 0 | 3 | 0.3333 | 10 | 0.1 | 2 | 0 | 1 | 0 | 2 | 0.5 | 2 | 0 | 7 | 0.1429 |
| 15_168560 | 1 | 0 | 2 | 0.5 | 4 | 0.5 | 3 | 0.3333 | 10 | 0.4 | 2 | 0.5 | 2 | 0 | 2 | 0.5 | 2 | 0.5 | 8 | 0.375 |
| 15_168560 | NA | NA | 1 | 1 | 3 | 0.3333 | 2 | 1 | 6 | 0.6667 | 2 | 1 | 2 | 0.5 | 2 | 1 | 1 | 1 | 7 | 0.8571 |
| 15_168561 | 1 | 0 | 2 | 0 | 3 | 0.3333 | 3 | 0.6667 | 9 | 0.3333 | 1 | 0 | 2 | 1 | 2 | 1 | 3 | 0.3333 | 8 | 0.625 |
| 15_168562 | 1 | 1 | 3 | 1 | 4 | 0.75 | 2 | 0.5 | 10 | 0.8 | 2 | 0.5 | 2 | 1 | 2 | 0.5 | 2 | 0.5 | 8 | 0.625 |
| 15_168562 | NA | NA | 1 | 1 | 3 | 1 | 3 | 0.6667 | 7 | 0.8571 | 2 | 1 | 2 | 0.5 | NA | NA | 2 | 1 | 6 | 0.8333 |
| 15_168563 | 1 | 1 | 3 | 1 | 4 | 1 | 2 | 1 | 10 | 1 | 2 | 1 | 1 | 1 | 2 | 1 | 3 | 0.6667 | 8 | 0.875 |
| 15_168563 | 1 | 0 | 2 | 0 | 4 | 0.25 | 2 | 0 | 9 | 0.1111 | 1 | 0 | 2 | 0 | 2 | 0.5 | 2 | 0.5 | 7 | 0.2857 |
| 15_168567 | 1 | 0 | 4 | 0.25 | 3 | 0.3333 | 3 | 0 | 11 | 0.1818 | 1 | 0 | 3 | 0 | 2 | 0 | 4 | 0.25 | 10 | 0.1 |
| 15_168571 | NA | NA | 1 | 1 | 5 | 0.2 | 2 | 0.5 | 8 | 0.375 | NA | NA | 2 | 0 | 1 | 1 | 3 | 0 | 6 | 0.1667 |
| 15_188996 | 5 | 0.6 | 2 | 0 | 2 | 0 | NA | NA | 9 | 0.3333 | 1 | 1 | 3 | 0 | 4 | 0.25 | 4 | 0.5 | 12 | 0.3333 |
| 15_188996 | 5 | 0.2 | 3 | 0 | 2 | 0 | 1 | 0 | 11 | 0.0909 | 1 | 1 | 4 | 0 | 5 | 0 | 4 | 0 | 14 | 0.0714 |
| 15_188996 | 5 | 1 | 3 | 1 | 1 | 1 | 1 | 0 | 10 | 0.9 | 1 | 1 | 4 | 0.75 | 5 | 0.6 | 4 | 1 | 14 | 0.7857 |
| 15_188997 | 4 | 0.75 | 2 | 1 | 2 | 0 | NA | NA | 8 | 0.625 | 1 | 1 | 3 | 1 | 3 | 1 | 4 | 1 | 11 | 1 |
| 15_190133 | 4 | 0.25 | 8 | 0.125 | 3 | 0 | 5 | 0 | 20 | 0.1 | 3 | 0 | 5 | 0.6 | 5 | 0 | 6 | 0.1667 | 19 | 0.2105 |
| 15_190133 | 4 | 0.25 | 9 | 0 | 3 | 0.3333 | 5 | 0.4 | 21 | 0.1905 | 4 | 0 | 6 | 0.5 | 6 | 0.1667 | 6 | 0.1667 | 22 | 0.2273 |
| 15_190134 | 4 | 0.25 | 9 | 0.2222 | 3 | 0 | 5 | 0 | 21 | 0.1429 | 4 | 0.25 | 6 | 0 | 6 | 0.1667 | 6 | 0.1667 | 22 | 0.1364 |
| 15_190134 | 5 | 0.2 | 8 | 0 | 3 | 0 | 5 | 0.2 | 21 | 0.0952 | 4 | 0.25 | 6 | 0.1667 | 6 | 0.3333 | 4 | 0.25 | 20 | 0.25 |
| 15_190135 | 4 | 1 | 8 | 1 | 2 | 1 | 5 | 1 | 19 | 1 | 5 | 1 | 6 | 1 | 6 | 1 | 3 | 1 | 20 | 1 |
| 15_190135 | 5 | 0.6 | 8 | 0.125 | 2 | 0 | 5 | 0.2 | 20 | 0.25 | 5 | 0.2 | 7 | 0.4286 | 6 | 0.1667 | 5 | 0.2 | 23 | 0.2609 |
| 15_190137 | 4 | 1 | 9 | 0.8889 | 3 | 0.6667 | 7 | 0.7143 | 23 | 0.8261 | 5 | 1 | 6 | 0.8333 | 6 | 1 | 5 | 0.6 | 22 | 0.8636 |
| 15_190137 | 5 | 0.6 | 7 | 0.7143 | 3 | 0.6667 | 6 | 0.6667 | 21 | 0.6667 | 3 | 0.6667 | 7 | 1 | 6 | 1 | 5 | 1 | 21 | 0.9524 |
| 15_190138 | 5 | 0.2 | 8 | 0.25 | 2 | 0.5 | 6 | 0.1667 | 21 | 0.2381 | 4 | 0.5 | 5 | 0.4 | 5 | 0.4 | 5 | 0.6 | 19 | 0.4737 |
| 15_190138 | 4 | 1 | 8 | 1 | 3 | 1 | 7 | 1 | 22 | 1 | 5 | 1 | 7 | 1 | 6 | 1 | 6 | 1 | 24 | 1 |
| 15_190140 | 5 | 0.8 | 10 | 0.7 | 2 | 1 | 4 | 1 | 21 | 0.8095 | 5 | 0.8 | 5 | 0.8 | 5 | 1 | 5 | 0.8 | 20 | 0.85 |
| 15_190140 | 2 | 0.5 | 5 | 0.6 | 4 | 0.5 | 1 | 0 | 12 | 0.5 | 3 | 0.3333 | 2 | 0 | 4 | 0.25 | 3 | 0 | 12 | 0.1667 |
| 15_212377 | NA | NA | 2 | 0 | 1 | 1 | 1 | 1 | 4 | 0.5 | 1 | 1 | NA | NA | 1 | 1 | NA | NA | 2 | 1 |
| 15_213590 | NA | NA | 2 | 0 | 1 | 1 | 1 | 1 | 4 | 0.5 | 1 | 1 | NA | NA | 1 | 0 | NA | NA | 2 | 0.5 |
| 15_215861 | NA | NA | 1 | 1 | 1 | 1 | 1 | 1 | 3 | 1 | 1 | 0 | NA | NA | 1 | 1 | NA | NA | 2 | 0.5 |
| 15_215862 | NA | NA | 2 | 0.5 | 1 | 1 | 1 | 1 | 4 | 0.75 | 1 | 0 | NA | NA | 1 | 1 | NA | NA | 2 | 0.5 |
| 15_215864 | NA | NA | 1 | 1 | NA | NA | 1 | 1 | 2 | 1 | 1 | 1 | NA | NA | 1 | 1 | NA | NA | 2 | 1 |
| 15_215864 | NA | NA | 2 | 1 | 1 | 1 | 1 | 0 | 4 | 0.75 | 1 | 1 | NA | NA | 1 | 1 | NA | NA | 2 | 1 |
| 15_215864 | NA | NA | 2 | 0.5 | 1 | 0 | NA | NA | 3 | 0.3333 | 1 | 1 | NA | NA | 1 | 0 | NA | NA | 2 | 0.5 |
| 15_215866 | NA | NA | 2 | 1 | 1 | 0 | 1 | 1 | 4 | 0.75 | NA | NA | NA | NA | 1 | 1 | NA | NA | 1 | 1 |
| 15_215892 | NA | NA | 2 | 0 | 1 | 1 | 1 | 0 | 4 | 0.25 | 3 | 0.3333 | NA | NA | NA | NA | 1 | 0 | 4 | 0.25 |
| 15_231736 | 2 | 0.5 | 1 | 1 | 1 | 1 | 2 | 0.5 | 6 | 0.6667 | NA | NA | 1 | 1 | 1 | 1 | 2 | 0 | 4 | 0.5 |
| 15_231736 | 2 | 1 | 2 | 0.5 | 1 | 1 | 2 | 1 | 7 | 0.8571 | NA | NA | NA | NA | 1 | 1 | 1 | 1 | 2 | 1 |
| 15_231737 | 2 | 0 | 2 | 0.5 | 1 | 1 | 2 | 0.5 | 7 | 0.4286 | NA | NA | 1 | 1 | 1 | 1 | 2 | 1 | 4 | 1 |
| 15_231738 | 2 | 0.5 | 2 | 1 | 1 | 1 | 2 | 1 | 7 | 0.8571 | NA | NA | 1 | 0 | NA | NA | 2 | 0.5 | 3 | 0.3333 |
| 15_232268 | 3 | 0.3333 | 2 | 0 | 1 | 0 | 2 | 0.5 | 8 | 0.25 | NA | NA | 2 | 0 | 1 | 1 | 1 | 0 | 4 | 0.25 |
| 15_234742 | 5 | 0.8 | 2 | 0.5 | 2 | 1 | 3 | 1 | 12 | 0.8333 | NA | NA | 3 | 0.3333 | 3 | 1 | 2 | 0.5 | 8 | 0.625 |
| 15_234742 | 4 | 1 | 2 | 1 | 2 | 0.5 | 3 | 0.6667 | 11 | 0.8182 | NA | NA | 3 | 0.6667 | 3 | 1 | 1 | 0 | 7 | 0.7143 |
| 15_234743 | 4 | 0.25 | 2 | 1 | 3 | 0.6667 | 3 | 0.6667 | 12 | 0.5833 | NA | NA | 2 | 1 | 3 | 0.6667 | 2 | 0.5 | 7 | 0.7143 |
| 15_234743 | 3 | 0.6667 | 2 | 0.5 | 3 | 1 | 3 | 1 | 11 | 0.8182 | NA | NA | 3 | 0.3333 | 3 | 0.3333 | 2 | 0.5 | 8 | 0.375 |
| 15_234743 | 4 | 1 | 2 | 1 | 3 | 0.6667 | 3 | 1 | 12 | 0.9167 | NA | NA | 3 | 1 | 2 | 1 | 3 | 1 | 8 | 1 |
| 15_234744 | 4 | 0.5 | 2 | 1 | 3 | 0.6667 | 3 | 1 | 12 | 0.75 | NA | NA | 3 | 1 | 3 | 1 | 3 | 1 | 9 | 1 |
| 15_234744 | 4 | 1 | 2 | 0.5 | 3 | 1 | 2 | 0.5 | 11 | 0.8182 | NA | NA | 3 | 1 | 3 | 0.6667 | 3 | 1 | 9 | 0.8889 |
| 15_234744 | 4 | 0.5 | 2 | 0 | 2 | 1 | 3 | 0.3333 | 11 | 0.4545 | NA | NA | 1 | 1 | NA | NA | 2 | 0 | 3 | 0.3333 |
| 15_234745 | 5 | 0 | 2 | 0.5 | 3 | 0.3333 | 2 | 0.5 | 12 | 0.25 | NA | NA | 3 | 0.3333 | 3 | 0.3333 | 2 | 0.5 | 8 | 0.375 |
| 15_234746 | 3 | 1 | 2 | 1 | 1 | 1 | 1 | 1 | 7 | 1 | NA | NA | 3 | 1 | 2 | 1 | 2 | 1 | 7 | 1 |

| | | | | | | | | | | | | | | | | | | | | |
|---|---|---|---|---|---|---|---|---|---|---|---|---|---|---|---|---|---|---|---|---|
| 15_2499012 | 1 | 1 | NA | NA | NA | NA | NA | NA | 1 | 1 | NA | NA | NA | NA | NA | NA | 1 | 1 | 1 | 1 |
| 15_2499013 | 1 | 1 | NA | NA | NA | NA | NA | NA | 1 | 1 | NA | NA | NA | NA | NA | NA | 1 | 1 | 1 | 1 |
| 15_2536388 | 64 | 0.0938 | 116 | 0.0862 | 71 | 0.1408 | 49 | 0.1429 | 300 | 0.11 | 76 | 0.0658 | 44 | 0.1818 | 52 | 0.0769 | 66 | 0.1818 | 238 | 0.1218 |
| 15_2536430 | 49 | 0.6531 | 72 | 0.6944 | 45 | 0.6222 | 29 | 0.5172 | 195 | 0.641 | 45 | 0.5778 | 24 | 0.5 | 37 | 0.5946 | 35 | 0.4857 | 141 | 0.5461 |
| 15_2536449 | 52 | 0.9615 | 71 | 1 | 47 | 1 | 32 | 0.9062 | 202 | 0.9752 | 51 | 0.9608 | 24 | 1 | 40 | 0.975 | 44 | 1 | 159 | 0.9811 |
| 15_2536455 | 44 | 1 | 67 | 0.9701 | 41 | 0.9756 | 30 | 0.9667 | 182 | 0.978 | 48 | 0.9792 | 28 | 0.9643 | 40 | 1 | 45 | 0.9556 | 161 | 0.9752 |
| 15_2541494 | NA | NA | NA | NA | NA | NA | 1 | 1 | 1 | 1 | 1 | 0 | 1 | 1 | NA | NA | NA | NA | 2 | 0.5 |
| 15_2578112 | NA | NA | 3 | 0 | NA | NA | 1 | 1 | 4 | 0.25 | 1 | 0 | NA | NA | 1 | 1 | 1 | 0 | 3 | 0.3333 |
| 15_2578153 | 1 | 1 | 3 | 0.6667 | NA | NA | 2 | 1 | 6 | 0.8333 | 1 | 0 | NA | NA | 1 | 0 | 1 | 1 | 3 | 0.3333 |
| 15_2578161 | 1 | 0 | 4 | 0.5 | NA | NA | 2 | 0.5 | 7 | 0.4286 | 1 | 0 | NA | NA | 1 | 0 | 1 | 1 | 3 | 0.3333 |
| 15_2578162 | 1 | 0 | 2 | 0.5 | NA | NA | 2 | 1 | 5 | 0.6 | NA | NA | NA | NA | 1 | 1 | NA | NA | 1 | 1 |
| 15_2578169 | 1 | 1 | 4 | 0.75 | NA | NA | 3 | 0.6667 | 8 | 0.75 | 1 | 0 | NA | NA | 1 | 0 | 1 | 1 | 3 | 0.3333 |
| 15_2580453 | 1 | 0 | 6 | 0.1667 | 1 | 0 | 3 | 0.3333 | 11 | 0.1818 | 3 | 0.3333 | 1 | 0 | 1 | 0 | 1 | 0 | 6 | 0.1667 |
| 15_2580789 | 2 | 0 | 6 | 0.3333 | 1 | 1 | 3 | 0 | 12 | 0.25 | 4 | 0 | 2 | 0.5 | 1 | 0 | 1 | 0 | 8 | 0.125 |
| 15_2580796 | 2 | 0.5 | 7 | 0.5714 | 1 | 0 | 3 | 0.6667 | 13 | 0.5385 | 5 | 0.8 | 2 | 1 | 1 | 1 | 1 | 1 | 9 | 0.8889 |
| 15_2581254 | 3 | 1 | 7 | 0.7143 | 3 | 0.6667 | 3 | 0.6667 | 16 | 0.75 | 5 | 1 | 2 | 0.5 | 1 | 0 | 2 | 1 | 10 | 0.8 |
| 15_2581261 | 2 | 0 | 7 | 0.1429 | 3 | 0 | 3 | 0 | 15 | 0.0667 | 5 | 0 | 2 | 0.5 | 1 | 1 | 2 | 0 | 10 | 0.2 |
| 15_2581262 | 4 | 0.5 | 7 | 0.1429 | 4 | 0.5 | 3 | 0.3333 | 18 | 0.3333 | 5 | 0.8 | 2 | 0 | 1 | 1 | 2 | 1 | 10 | 0.7 |
| 15_2581264 | 4 | 0.25 | 5 | 0 | 3 | 0 | 3 | 0 | 15 | 0.0667 | 3 | 0.3333 | 1 | 0 | 1 | 0 | 2 | 0 | 7 | 0.1429 |
| 15_2581284 | 3 | 0 | 5 | 0.2 | 4 | 0.25 | 2 | 0 | 14 | 0.1429 | 4 | 0.75 | 1 | 0 | 1 | 0 | 2 | 0.5 | 8 | 0.5 |
| 15_2581314 | 6 | 0 | 4 | 0.5 | 4 | 0 | 3 | 0.3333 | 17 | 0.1765 | 4 | 0.25 | 3 | 0 | 1 | 0 | 4 | 0.25 | 12 | 0.1667 |
| 15_2581347 | 5 | 0.2 | 5 | 0 | 4 | 0.75 | 4 | 0 | 18 | 0.2222 | 6 | 0.1667 | 4 | 0.25 | 2 | 0 | 5 | 0.2 | 17 | 0.1765 |
| 15_2581359 | 6 | 0.5 | 7 | 0.7143 | 4 | 0.5 | 5 | 0.4 | 22 | 0.5455 | 5 | 1 | 4 | 0.75 | 3 | 0.6667 | 6 | 0.3333 | 18 | 0.6667 |
| 15_2581360 | 4 | 1 | 4 | 0.75 | 4 | 1 | 4 | 1 | 16 | 0.9375 | 6 | 1 | 3 | 0.6667 | 2 | 0.5 | 7 | 0.8571 | 18 | 0.8333 |
| 15_2581369 | 4 | 0.25 | 5 | 0.2 | 4 | 0.25 | 3 | 0.3333 | 16 | 0.25 | 5 | 0.2 | 4 | 0.5 | 2 | 0 | 7 | 0.2857 | 18 | 0.2778 |
| 15_2597191 | 3 | 0 | 5 | 0 | 6 | 0.1667 | 5 | 0.4 | 19 | 0.1579 | 4 | 0.25 | 7 | 0.1429 | 4 | 0 | 2 | 0 | 17 | 0.1176 |
| 15_2597197 | 3 | 0.6667 | 5 | 0.8 | 6 | 0.8333 | 4 | 1 | 18 | 0.8333 | 5 | 0.8 | 5 | 0.8 | 3 | 0.6667 | 3 | 0.6667 | 16 | 0.75 |
| 15_2597198 | 3 | 0.6667 | 4 | 0.25 | 5 | 0.4 | 4 | 0.25 | 16 | 0.375 | 3 | 0.6667 | 5 | 0.4 | 5 | 0.4 | 2 | 0 | 15 | 0.4 |
| 15_2597199 | 3 | 0.3333 | 4 | 0.25 | 4 | 0.5 | 5 | 0.6 | 16 | 0.4375 | 5 | 0.6 | 5 | 0.6 | 5 | 0.2 | 3 | 0.3333 | 18 | 0.4444 |
| 15_2597200 | 2 | 0.5 | 3 | 0.3333 | 5 | 0.8 | 4 | 0.5 | 14 | 0.5714 | 3 | 0 | 5 | 0.6 | 4 | 0.75 | 1 | 0 | 13 | 0.4615 |
| 15_2597205 | 2 | 1 | 4 | 1 | 6 | 1 | 6 | 1 | 18 | 1 | 6 | 0.8333 | 6 | 1 | 4 | 1 | 1 | 1 | 17 | 0.9412 |
| 15_2597206 | 3 | 0 | 4 | 0 | 4 | 0.25 | 5 | 0.2 | 16 | 0.125 | 5 | 0.2 | 6 | 0.5 | 2 | 0.5 | 3 | 0 | 16 | 0.3125 |
| 15_2597231 | 4 | 0.25 | 5 | 0.2 | 7 | 0.4286 | 8 | 0 | 24 | 0.2083 | 5 | 0 | 7 | 0.5714 | 4 | 0.25 | 4 | 0 | 20 | 0.25 |
| 15_2597236 | 3 | 0 | 5 | 0.2 | 7 | 0.4286 | 7 | 0.2857 | 22 | 0.2727 | 5 | 0 | 7 | 0.2857 | 3 | 0 | 4 | 0.25 | 19 | 0.1579 |
| 15_2597267 | 4 | 0.5 | 6 | 0.3333 | 6 | 0 | 7 | 0.4286 | 23 | 0.3043 | 5 | 0 | 8 | 0.375 | 4 | 0.25 | 4 | 0.25 | 21 | 0.2381 |
| 15_2597351 | 8 | 0.25 | 9 | 0 | 10 | 0.3 | 10 | 0 | 37 | 0.1351 | 6 | 0.1667 | 10 | 0 | 5 | 0 | 5 | 0.2 | 26 | 0.0769 |
| 15_2598471 | 2 | 0.5 | 6 | 0.3333 | 1 | 0 | 3 | 0.6667 | 12 | 0.4167 | 3 | 0.3333 | 2 | 0 | 3 | 0.3333 | 3 | 0 | 11 | 0.1818 |
| 15_2598700 | 6 | 0.5 | 8 | 0.25 | 3 | 0.3333 | 7 | 1 | 24 | 0.5417 | 5 | 0.4 | 5 | 0.2 | 4 | 0.25 | 6 | 0.8333 | 20 | 0.45 |
| 15_2640932 | 2 | 1 | 3 | 1 | 1 | 1 | 1 | 1 | 7 | 1 | 1 | 0 | NA | NA | NA | NA | 1 | 1 | 2 | 0.5 |
| 15_2640936 | 2 | 0.5 | 3 | 0.6667 | 1 | 1 | 1 | 0 | 7 | 0.5714 | 1 | 1 | NA | NA | NA | NA | 1 | 1 | 2 | 1 |
| 15_2640960 | 2 | 1 | 3 | 1 | 1 | 0 | NA | NA | 6 | 0.8333 | NA | NA | NA | NA | 1 | 1 | 1 | 1 | 2 | 1 |
| 15_2746690 | 10 | 0.1 | 17 | 0.3529 | 9 | 0.1111 | 9 | 0.2222 | 45 | 0.2222 | 3 | 0 | 7 | 0.2857 | 8 | 0.125 | 4 | 0.25 | 22 | 0.1818 |
| 15_2746708 | 15 | 0.1333 | 21 | 0.1905 | 8 | 0.5 | 15 | 0 | 59 | 0.1695 | 4 | 0 | 13 | 0.2308 | 11 | 0 | 9 | 0 | 37 | 0.0811 |
| 15_2759333 | 15 | 0.4 | 17 | 0.7059 | 9 | 0.5556 | 19 | 0.5789 | 60 | 0.5667 | 1 | 1 | 9 | 0.4444 | 13 | 0.3077 | 10 | 0.6 | 33 | 0.4545 |
| 15_2759334 | 24 | 0.125 | 32 | 0.0938 | 15 | 0.1333 | 27 | 0.037 | 98 | 0.0918 | 5 | 0.2 | 24 | 0.0833 | 19 | 0.2105 | 19 | 0 | 67 | 0.1045 |
| 15_2759348 | 24 | 0.1667 | 29 | 0.3103 | 16 | 0.1875 | 26 | 0.2692 | 95 | 0.2421 | 6 | 0.3333 | 28 | 0.2857 | 19 | 0.3684 | 21 | 0.1905 | 74 | 0.2838 |
| 15_2759351 | 22 | 0.0455 | 29 | 0.1379 | 16 | 0.0625 | 23 | 0.1739 | 90 | 0.1111 | 5 | 0 | 26 | 0.0385 | 21 | 0.0952 | 19 | 0.0526 | 71 | 0.0563 |
| 15_2759353 | 21 | 0 | 24 | 0.125 | 12 | 0.1667 | 21 | 0.0476 | 78 | 0.0769 | 5 | 0.2 | 24 | 0.1667 | 13 | 0.0769 | 18 | 0.0556 | 60 | 0.1167 |
| 15_2759360 | 20 | 0.85 | 27 | 0.9259 | 12 | 0.9167 | 23 | 0.7826 | 82 | 0.8659 | 4 | 0.75 | 23 | 0.7391 | 17 | 1 | 15 | 0.7333 | 59 | 0.8136 |
| 15_2759362 | 23 | 0.913 | 34 | 1 | 16 | 0.9375 | 27 | 0.963 | 100 | 0.96 | 7 | 1 | 28 | 0.9643 | 20 | 0.95 | 21 | 0.9524 | 76 | 0.9605 |
| 15_2759363 | 24 | 0.4583 | 32 | 0.4375 | 16 | 0.4375 | 27 | 0.3333 | 99 | 0.4141 | 7 | 0.5714 | 27 | 0.5185 | 19 | 0.2632 | 19 | 0.5789 | 72 | 0.4722 |
| 15_2759370 | 25 | 0.32 | 32 | 0.3125 | 14 | 0.3571 | 26 | 0.2692 | 97 | 0.3093 | 8 | 0 | 27 | 0.1111 | 17 | 0.2941 | 19 | 0.5263 | 71 | 0.2535 |
| 15_2759373 | 25 | 0.88 | 25 | 0.96 | 15 | 0.9333 | 27 | 0.8889 | 92 | 0.913 | 8 | 0.875 | 25 | 0.92 | 17 | 1 | 17 | 0.9412 | 67 | 0.9403 |
| 15_2759394 | 21 | 0.1429 | 25 | 0.12 | 16 | 0.3125 | 22 | 0.0909 | 84 | 0.1548 | 8 | 0 | 24 | 0.0417 | 17 | 0.1176 | 18 | 0.1111 | 67 | 0.0746 |
| 15_2759437 | 29 | 0.2069 | 33 | 0.2121 | 19 | 0.1053 | 33 | 0.2121 | 114 | 0.193 | 8 | 0.25 | 25 | 0.16 | 24 | 0.25 | 24 | 0.2083 | 81 | 0.2099 |
| 15_2759479 | 12 | 0.25 | 10 | 0 | 11 | 0.0909 | 20 | 0.1 | 53 | 0.1132 | 8 | 0.125 | 14 | 0.2143 | 11 | 0.1818 | 11 | 0.0909 | 44 | 0.1591 |
| 15_2760607 | 2 | 0.5 | 5 | 0.2 | 1 | 0 | 3 | 0.6667 | 11 | 0.3636 | 3 | 0 | 1 | 0 | 6 | 0 | 5 | 0.2 | 15 | 0.0667 |
| 15_2760621 | 4 | 0.25 | 6 | 0.3333 | 2 | 0 | 3 | 0 | 15 | 0.2 | 3 | 0.3333 | 2 | 0 | 6 | 0 | 4 | 0.25 | 15 | 0.1333 |
| 15_2760623 | 3 | 1 | 4 | 1 | 2 | 0.5 | 3 | 1 | 12 | 0.9167 | 3 | 1 | 1 | 1 | 6 | 0.8333 | 4 | 0.75 | 14 | 0.8571 |
| 15_2760626 | 4 | 0.75 | 5 | 0.2 | 1 | 0 | 3 | 0.6667 | 13 | 0.4615 | 3 | 0.6667 | 1 | 1 | 6 | 0.5 | 3 | 0 | 13 | 0.4615 |
| 15_2760627 | 4 | 0.75 | 5 | 1 | 2 | 1 | 3 | 1 | 14 | 0.9286 | 3 | 1 | 2 | 1 | 5 | 0.8 | 4 | 1 | 14 | 0.9286 |
| 15_2760633 | 3 | 0.6667 | 4 | 0.75 | 2 | 0 | 2 | 0.5 | 11 | 0.5455 | 2 | 1 | 2 | 0 | 4 | 0.75 | 3 | 0.6667 | 11 | 0.6364 |
| 15_2760634 | 4 | 0.25 | 6 | 0.5 | 2 | 0.5 | 2 | 0 | 14 | 0.3571 | 3 | 0.3333 | 1 | 0 | 5 | 0.4 | 3 | 0 | 12 | 0.25 |
| 15_2760643 | 3 | 0.6667 | 6 | 1 | 2 | 1 | 3 | 0.6667 | 14 | 0.8571 | 2 | 1 | 2 | 0.5 | 5 | 1 | 3 | 1 | 12 | 0.9167 |
| 15_2760881 | 3 | 0.3333 | 4 | 0 | 2 | 0 | 3 | 0 | 12 | 0.0833 | 1 | 1 | 1 | 1 | 5 | 0.4 | 3 | 0.6667 | 10 | 0.6 |

| | | | | | | | | | | | | | | | | | | | | |
|---|---|---|---|---|---|---|---|---|---|---|---|---|---|---|---|---|---|---|---|---|
| 15_2765665 | 4 | 0.25 | NA | NA | 3 | 0 | 7 | 0 | 14 | 0.0714 | 4 | 0 | 4 | 0 | 4 | 0 | 3 | 0.3333 | 15 | 0.0667 |
| 15_2765672 | 2 | 0 | NA | NA | 4 | 0.75 | 8 | 0.25 | 14 | 0.3571 | 4 | 0.5 | 4 | 0.25 | 7 | 0.1429 | 1 | 0 | 16 | 0.25 |
| 15_2765674 | 1 | 1 | NA | NA | 3 | 1 | 6 | 0.6667 | 10 | 0.8 | 4 | 0.75 | 4 | 1 | 4 | 0.75 | 2 | 0.5 | 14 | 0.7857 |
| 15_2765675 | 3 | 0 | NA | NA | 2 | 0.5 | 7 | 0.1429 | 12 | 0.1667 | 4 | 0.25 | 5 | 0.2 | 7 | 0 | 3 | 0 | 19 | 0.1053 |
| 15_2765680 | 2 | 0.5 | NA | NA | 4 | 0.75 | 7 | 0.8571 | 13 | 0.7692 | 4 | 0.5 | 5 | 0.6 | 5 | 0.4 | 2 | 1 | 16 | 0.5625 |
| 15_2765691 | 3 | 0.3333 | NA | NA | 3 | 0 | 7 | 0 | 13 | 0.0769 | 3 | 0 | 4 | 0 | 6 | 0.3333 | 3 | 0 | 16 | 0.125 |
| 15_2765694 | 2 | 1 | NA | NA | 3 | 1 | 7 | 1 | 12 | 1 | 2 | 1 | 5 | 0.8 | 5 | 0.4 | 3 | 0.6667 | 15 | 0.6667 |
| 15_2765696 | 3 | 0.3333 | NA | NA | 3 | 0.3333 | 7 | 0.1429 | 13 | 0.2308 | 2 | 0 | 4 | 0.75 | 5 | 0.2 | 3 | 0 | 14 | 0.2857 |
| 15_2765697 | 2 | 0.5 | NA | NA | 3 | 0.3333 | 6 | 0.8333 | 11 | 0.6364 | 4 | 0.25 | 2 | 0.5 | 6 | 0.5 | 2 | 0 | 14 | 0.3571 |
| 15_2765702 | 2 | 0.5 | NA | NA | 3 | 0 | 7 | 0.1429 | 12 | 0.1667 | 4 | 0.25 | 5 | 0.2 | 6 | 0.6667 | 3 | 0.6667 | 18 | 0.4444 |
| 15_2765708 | 2 | 1 | NA | NA | 2 | 0.5 | 7 | 0.7143 | 11 | 0.7273 | 4 | 0.75 | 5 | 0.4 | 4 | 0.5 | 3 | 0.6667 | 16 | 0.5625 |
| 15_2773107 | 17 | 0.1765 | 12 | 0 | 9 | 0 | 10 | 0 | 48 | 0.0625 | 10 | 0.1 | 12 | 0 | 17 | 0.0588 | 10 | 0.1 | 49 | 0.0612 |
| 15_2773146 | 13 | 0.5385 | 12 | 0.1667 | 6 | 0.1667 | 7 | 0.4286 | 38 | 0.3421 | 7 | 0.2857 | 10 | 0.4 | 10 | 0.2 | 9 | 0.3333 | 36 | 0.3056 |
| 15_2773150 | 15 | 0.3333 | 11 | 0.2727 | 5 | 0 | 8 | 0.375 | 39 | 0.2821 | 8 | 0.125 | 12 | 0.1667 | 14 | 0.0714 | 8 | 0 | 42 | 0.0952 |
| 15_2773183 | 11 | 0.2727 | 11 | 0.2727 | 6 | 0.1667 | 5 | 0 | 33 | 0.2121 | 7 | 0.4286 | 10 | 0 | 11 | 0.3636 | 7 | 0.5714 | 35 | 0.3143 |
| 15_2773193 | 11 | 0.4545 | 9 | 0.3333 | 9 | 0.8889 | 5 | 0.6 | 34 | 0.5588 | 5 | 0.6 | 8 | 0.75 | 8 | 0.625 | 7 | 0.7143 | 28 | 0.6786 |
| 15_2773203 | 15 | 0.1333 | 11 | 0.0909 | 9 | 0.2222 | 6 | 0 | 41 | 0.122 | 7 | 0.1429 | 8 | 0 | 13 | 0.0769 | 7 | 0 | 35 | 0.0571 |
| 15_2773214 | 12 | 0.75 | 9 | 0.8889 | 9 | 0.6667 | 5 | 0.8 | 35 | 0.7714 | 6 | 1 | 8 | 0.625 | 10 | 0.8 | 6 | 0.8333 | 30 | 0.8 |
| 15_2773216 | 7 | 1 | 8 | 0.875 | 5 | 1 | 3 | 1 | 23 | 0.9565 | 5 | 1 | 6 | 0.8333 | 9 | 0.8889 | 3 | 1 | 23 | 0.913 |
| 15_2773217 | 8 | 0.875 | 5 | 0.8 | 3 | 1 | 3 | 1 | 19 | 0.8947 | 5 | 1 | 5 | 1 | 6 | 0.8333 | 2 | 1 | 18 | 0.9444 |
| 15_2773224 | 13 | 0.3077 | 6 | 0 | 6 | 0 | 4 | 0 | 29 | 0.1379 | 7 | 0.2857 | 6 | 0.1667 | 8 | 0.125 | 6 | 0.3333 | 27 | 0.2222 |
| 15_2773226 | 5 | 1 | 4 | 1 | NA | NA | 5 | 0.8 | 14 | 0.9286 | 3 | 1 | 2 | 1 | 7 | 1 | 5 | 0.8 | 17 | 0.9412 |
| 15_2773235 | 11 | 0.0909 | 8 | 0 | 8 | 0.25 | 6 | 0 | 33 | 0.0909 | 7 | 0 | 7 | 0.1429 | 11 | 0.1818 | 5 | 0 | 30 | 0.1 |
| 15_2773238 | 7 | 1 | 8 | 1 | 6 | 1 | 6 | 1 | 27 | 1 | 6 | 1 | 7 | 0.7143 | 9 | 1 | 5 | 1 | 27 | 0.9259 |
| 15_2773248 | 4 | 0.25 | 5 | 0.6 | 5 | 0.6 | 4 | 0.25 | 18 | 0.4444 | 4 | 0.75 | 3 | 0.6667 | 2 | 0.5 | 3 | 0.6667 | 12 | 0.6667 |
| 15_2773256 | 10 | 0.2 | 9 | 0.1111 | 8 | 0.25 | 5 | 0.4 | 32 | 0.2188 | 6 | 0.8333 | 7 | 0.2857 | 11 | 0.1818 | 6 | 0 | 30 | 0.3 |
| 15_2773260 | 9 | 0.8889 | 5 | 1 | 5 | 1 | 6 | 1 | 25 | 0.96 | 6 | 1 | 3 | 1 | 7 | 1 | 4 | 0.75 | 20 | 0.95 |
| 15_2773266 | 11 | 0.2727 | 8 | 0.625 | 7 | 1 | 6 | 0.8333 | 32 | 0.625 | 7 | 0.5714 | 5 | 0.6 | 10 | 0.6 | 5 | 0.8 | 27 | 0.6296 |
| 15_2773272 | 8 | 0 | 7 | 0.1429 | 6 | 0.1667 | 5 | 0.4 | 26 | 0.1538 | 6 | 0.1667 | 6 | 0.5 | 9 | 0.3333 | 4 | 0.25 | 25 | 0.32 |
| 15_2773282 | 10 | 0.8 | 7 | 1 | 7 | 0.8571 | 5 | 0.8 | 29 | 0.8621 | 5 | 1 | 4 | 1 | 10 | 0.7 | 5 | 1 | 24 | 0.875 |
| 15_2773284 | 12 | 1 | 7 | 1 | 8 | 1 | 6 | 1 | 33 | 1 | 5 | 1 | 5 | 1 | 10 | 1 | 5 | 1 | 25 | 1 |
| 15_2773294 | 8 | 0.625 | 7 | 0.4286 | 6 | 0.6667 | 4 | 0.75 | 25 | 0.6 | 4 | 0.25 | 3 | 0.3333 | 10 | 0.5 | 4 | 0.5 | 21 | 0.4286 |
| 15_2773306 | 9 | 0.1111 | 7 | 0.1429 | 8 | 0 | 4 | 0.25 | 28 | 0.1071 | 7 | 0.1429 | 4 | 0 | 7 | 0.1429 | 6 | 0 | 24 | 0.0833 |
| 15_2774445 | 1 | 0 | 2 | 1 | NA | NA | NA | NA | 3 | 0.6667 | NA | NA | NA | NA | 1 | 1 | NA | NA | 1 | 1 |
| 15_2774533 | NA | NA | 2 | 0.5 | NA | NA | NA | NA | 2 | 0.5 | NA | NA | NA | NA | 1 | 1 | NA | NA | 1 | 1 |
| 15_2774704 | 3 | 1 | 7 | 0.8571 | 9 | 0.6667 | 4 | 0.75 | 23 | 0.7826 | 10 | 0.7 | 7 | 1 | 8 | 0.875 | 4 | 0.75 | 29 | 0.8276 |
| 15_2774730 | 6 | 0.3333 | 6 | 0.8333 | 8 | 0.75 | 7 | 0.4286 | 27 | 0.5926 | 11 | 0.4545 | 9 | 0.6667 | 8 | 0.75 | 7 | 0.2857 | 35 | 0.5429 |
| 15_2774735 | 4 | 0 | 6 | 0.3333 | 9 | 0 | 9 | 0.2222 | 28 | 0.1429 | 10 | 0.1 | 7 | 0.1429 | 8 | 0 | 7 | 0.1429 | 32 | 0.0938 |
| 15_2774752 | 5 | 0 | 6 | 0.1667 | 7 | 0 | 6 | 0.5 | 24 | 0.1667 | 9 | 0.1111 | 6 | 0 | 6 | 0.1667 | 7 | 0.1429 | 28 | 0.1071 |
| 15_2774762 | 6 | 1 | 7 | 0.8571 | 8 | 1 | 5 | 1 | 26 | 0.9615 | 10 | 1 | 5 | 1 | 7 | 1 | 5 | 1 | 27 | 1 |
| 15_2774775 | 7 | 0.5714 | 8 | 0.75 | 8 | 0.875 | 7 | 0.8571 | 30 | 0.7667 | 9 | 0.8889 | 8 | 0.875 | 5 | 1 | 6 | 0.6667 | 28 | 0.8571 |
| 15_2774776 | 7 | 0.2857 | 8 | 0 | 7 | 0.1429 | 6 | 0.1667 | 28 | 0.1429 | 10 | 0 | 5 | 0.2 | 5 | 0 | 5 | 0.2 | 25 | 0.08 |
| 15_2774807 | 5 | 0.6 | 7 | 0.5714 | 5 | 1 | 8 | 1 | 25 | 0.8 | 7 | 0.8571 | 8 | 0.75 | 7 | 0.8571 | 6 | 0.8333 | 28 | 0.8214 |
| 15_2774826 | 4 | 0.5 | 5 | 1 | 4 | 0.75 | 6 | 1 | 19 | 0.8421 | 6 | 1 | 6 | 0.8333 | 3 | 0.6667 | 3 | 1 | 18 | 0.8889 |
| 15_2774829 | 7 | 0 | 9 | 0 | 5 | 0 | 6 | 0.3333 | 27 | 0.0741 | 9 | 0.1111 | 6 | 0 | 6 | 0.1667 | 4 | 0 | 25 | 0.08 |
| 15_2774829 | 6 | 0 | 7 | 0.2857 | 4 | 0.25 | 5 | 0.2 | 22 | 0.1818 | 7 | 0.2857 | 5 | 0.2 | 4 | 0 | 4 | 0.5 | 20 | 0.25 |
| 15_2774834 | 2 | 0 | 4 | 0.25 | 1 | 0 | 4 | 0 | 11 | 0.0909 | 4 | 0 | 4 | 0 | 4 | 0.25 | 3 | 0 | 15 | 0.0667 |
| 15_2790275 | 1 | 1 | 1 | 1 | 1 | 1 | NA | NA | 3 | 1 | 1 | 0 | 2 | 0.5 | NA | NA | NA | NA | 3 | 0.3333 |
| 15_2790281 | 1 | 1 | 1 | 1 | 2 | 0.5 | 1 | 1 | 5 | 0.8 | 4 | 0.25 | 2 | 1 | 1 | 0 | NA | NA | 7 | 0.4286 |
| 15_2790281 | NA | NA | NA | NA | 2 | 1 | 1 | 1 | 3 | 1 | 3 | 0.3333 | 2 | 1 | 1 | 0 | NA | NA | 6 | 0.5 |
| 15_2790289 | 1 | 0 | 1 | 0 | 1 | 1 | 1 | 1 | 4 | 0.5 | 4 | 0.25 | 2 | 0.5 | 1 | 0 | NA | NA | 7 | 0.2857 |
| 15_3102776 | 11 | 0.8182 | 9 | 0.6667 | 13 | 0.7692 | 5 | 0.8 | 38 | 0.7632 | 15 | 0.8667 | 13 | 0.6154 | 8 | 0.75 | 7 | 0.7143 | 43 | 0.7442 |
| 15_3102777 | 9 | 0.2222 | 11 | 0.3636 | 16 | 0.25 | 9 | 0.2222 | 45 | 0.2667 | 16 | 0.25 | 13 | 0.3846 | 9 | 0.2222 | 12 | 0.5 | 50 | 0.34 |
| 15_3102782 | 14 | 0.0714 | 16 | 0 | 20 | 0.2 | 12 | 0 | 62 | 0.0806 | 22 | 0.0909 | 20 | 0.05 | 12 | 0.1667 | 16 | 0.0625 | 70 | 0.0857 |
| 15_3102784 | 10 | 0.1 | 10 | 0.3 | 15 | 0.0667 | 9 | 0.1111 | 44 | 0.1364 | 15 | 0.4 | 15 | 0.2667 | 9 | 0.2222 | 10 | 0.2 | 49 | 0.2857 |
| 15_3102787 | 11 | 0.6364 | 12 | 0.3333 | 17 | 0.5294 | 11 | 0.4545 | 51 | 0.4902 | 18 | 0.3333 | 19 | 0.4211 | 11 | 0.4545 | 13 | 0.5385 | 61 | 0.4262 |
| 15_3127405 | 2 | 1 | NA | NA | NA | NA | 1 | 0 | 3 | 0.6667 | 1 | 1 | 1 | 1 | 1 | 1 | NA | NA | 3 | 1 |
| 15_3129244 | 9 | 0.2222 | 15 | 0.2 | 11 | 0.1818 | 9 | 0.2222 | 44 | 0.2045 | 3 | 0.3333 | 5 | 0.2 | 8 | 0.25 | 4 | 0 | 20 | 0.2 |
| 15_3129305 | 8 | 0.25 | 13 | 0.0769 | 9 | 0 | 8 | 0 | 38 | 0.0789 | 3 | 0 | 7 | 0 | 7 | 0.1429 | 3 | 0 | 20 | 0.05 |
| 15_3129319 | 8 | 0.125 | 13 | 0.0769 | 9 | 0.2222 | 9 | 0.1111 | 39 | 0.1282 | 3 | 0 | 6 | 0 | 5 | 0.2 | 4 | 0 | 18 | 0.0556 |
| 15_3129340 | 7 | 0.2857 | 13 | 0.0769 | 9 | 0 | 8 | 0.375 | 37 | 0.1622 | 3 | 0 | 7 | 0.2857 | 7 | 0.1429 | 2 | 0 | 19 | 0.1579 |
| 15_3129381 | 7 | 0.5714 | 9 | 0 | 6 | 0.3333 | 7 | 0.1429 | 29 | 0.2414 | 3 | 0.3333 | 4 | 0 | 6 | 0.5 | 2 | 0 | 15 | 0.2667 |
| 15_3129394 | 7 | 0.5714 | 9 | 0.2222 | 6 | 0.5 | 7 | 0.1429 | 29 | 0.3448 | 2 | 0.5 | 3 | 0 | 7 | 0.2857 | 1 | 1 | 13 | 0.3077 |
| 15_3129492 | 5 | 0.4 | 9 | 0.1111 | 3 | 0.3333 | 7 | 0.1429 | 24 | 0.2083 | 2 | 0 | 2 | 0 | 3 | 0.3333 | NA | NA | 7 | 0.1429 |
| 15_3129533 | 4 | 0.75 | 6 | 0.3333 | 3 | 0.3333 | 6 | 0.1667 | 19 | 0.3684 | 2 | 0 | 2 | 0.5 | 1 | 0 | NA | NA | 5 | 0.2 |

| | | | | | | | | | | | | | | | | | | | | |
|---|---|---|---|---|---|---|---|---|---|---|---|---|---|---|---|---|---|---|---|---|
| 15_3129728 | 2 | 0.5 | 2 | 0 | 1 | 1 | 3 | 0.6667 | 8 | 0.5 | 2 | 0.5 | 2 | 0 | NA | NA | 1 | 0 | 5 | 0.2 |
| 15_3129743 | 3 | 0.6667 | 2 | 1 | 1 | 1 | 2 | 0.5 | 8 | 0.75 | NA | NA | 2 | 1 | NA | NA | 1 | 0 | 3 | 0.6667 |
| 15_3129764 | 4 | 0.5 | 2 | 0 | 1 | 1 | 1 | 1 | 8 | 0.5 | 1 | 1 | 2 | 0.5 | NA | NA | 1 | 1 | 4 | 0.75 |
| 15_3129769 | 2 | 1 | 2 | 1 | 1 | 1 | 3 | 1 | 8 | 1 | NA | NA | 2 | 1 | NA | NA | 1 | 0 | 3 | 0.6667 |
| 15_3129778 | 4 | 0.75 | 1 | 1 | 1 | 1 | 2 | 0.5 | 8 | 0.75 | 1 | 1 | 2 | 0.5 | NA | NA | NA | NA | 3 | 0.6667 |
| 15_3129793 | 4 | 1 | 2 | 1 | 1 | 1 | 1 | 1 | 8 | 1 | NA | NA | 1 | 1 | NA | NA | NA | NA | 1 | 1 |
| 15_3129793 | NA | NA | NA | NA | NA | NA | 1 | 1 | 1 | 1 | NA | NA | 1 | 1 | NA | NA | NA | NA | 1 | 1 |
| 15_3129799 | 4 | 0.25 | 1 | 0 | 1 | 0 | 2 | 0.5 | 8 | 0.25 | 1 | 0 | 2 | 0.5 | NA | NA | NA | NA | 3 | 0.3333 |
| 15_3145478 | 1 | 1 | NA | NA | NA | NA | 1 | 0 | 2 | 0.5 | NA | NA | 1 | 0 | NA | NA | 1 | 1 | 2 | 0.5 |
| 15_3145678 | 14 | 0.1429 | 20 | 0.3 | 17 | 0.2941 | 26 | 0.3462 | 77 | 0.2857 | 16 | 0.0625 | 19 | 0.1053 | 19 | 0.2632 | 16 | 0.25 | 70 | 0.1714 |
| 15_3145767 | 15 | 0.2 | 17 | 0.1176 | 15 | 0.1333 | 28 | 0.25 | 75 | 0.1867 | 11 | 0.0909 | 20 | 0.25 | 26 | 0.1923 | 21 | 0.2857 | 78 | 0.2179 |
| 15_3145880 | 16 | 0.3125 | 21 | 0.2381 | 13 | 0.5385 | 24 | 0.2917 | 74 | 0.3243 | 12 | 0.1667 | 15 | 0.3333 | 18 | 0.1111 | 21 | 0.2381 | 66 | 0.2121 |
| 15_3145882 | 15 | 0.3333 | 18 | 0.2222 | 12 | 0.5 | 19 | 0.2632 | 64 | 0.3125 | 10 | 0.6 | 13 | 0.5385 | 17 | 0.4706 | 20 | 0.55 | 60 | 0.5333 |
| 15_3145890 | 12 | 0.25 | 17 | 0 | 10 | 0.1 | 21 | 0.0476 | 60 | 0.0833 | 11 | 0 | 13 | 0.0769 | 11 | 0 | 18 | 0.1111 | 53 | 0.0566 |
| 15_3145895 | 16 | 0.125 | 20 | 0.15 | 13 | 0.1538 | 25 | 0.12 | 74 | 0.1351 | 11 | 0.1818 | 15 | 0.0667 | 21 | 0.1905 | 21 | 0.2381 | 68 | 0.1765 |
| 15_3145896 | 15 | 0.4 | 18 | 0.3333 | 10 | 0.2 | 22 | 0.3636 | 65 | 0.3385 | 11 | 0.4545 | 13 | 0.4615 | 17 | 0.2941 | 20 | 0.55 | 61 | 0.4426 |
| 15_3145901 | 13 | 0.3846 | 17 | 0.5294 | 11 | 0.6364 | 20 | 0.3 | 61 | 0.4426 | 11 | 0.6364 | 15 | 0.4 | 20 | 0.4 | 21 | 0.4762 | 67 | 0.4627 |
| 15_3145922 | 12 | 0.4167 | 17 | 0.2941 | 9 | 0.5556 | 24 | 0.5417 | 62 | 0.4516 | 13 | 0.3077 | 15 | 0.4 | 18 | 0.2778 | 19 | 0.5789 | 65 | 0.4 |
| 15_3145927 | 13 | 1 | 14 | 1 | 8 | 1 | 19 | 0.9474 | 54 | 0.9815 | 11 | 0.9091 | 9 | 0.8889 | 16 | 0.9375 | 16 | 1 | 52 | 0.9423 |
| 15_3159008 | 3 | 0 | 3 | 0.3333 | 2 | 0 | NA | NA | 8 | 0.125 | NA | NA | 1 | 1 | 1 | 0 | 1 | 0 | 3 | 0.3333 |
| 15_3161722 | 1 | 0 | 2 | 0 | NA | NA | 3 | 0.3333 | 6 | 0.1667 | NA | NA | 2 | 0 | NA | NA | 3 | 0.3333 | 5 | 0.2 |
| 15_3161732 | 1 | 1 | 1 | 1 | NA | NA | 1 | 1 | 3 | 1 | NA | NA | 1 | 1 | NA | NA | 1 | 1 | 2 | 1 |
| 15_3161733 | NA | NA | 1 | 1 | NA | NA | 4 | 0.75 | 5 | 0.8 | NA | NA | 3 | 1 | NA | NA | NA | NA | 3 | 1 |
| 15_3161742 | 2 | 1 | 1 | 1 | NA | NA | 3 | 0.6667 | 6 | 0.8333 | NA | NA | 3 | 1 | NA | NA | 3 | 1 | 6 | 1 |
| 15_3161747 | 2 | 1 | 1 | 1 | NA | NA | 4 | 0.75 | 7 | 0.8571 | NA | NA | 3 | 0.6667 | NA | NA | 2 | 0.5 | 5 | 0.6 |
| 15_3161751 | 2 | 0 | 2 | 0.5 | NA | NA | 4 | 1 | 8 | 0.625 | NA | NA | 3 | 0 | NA | NA | 2 | 0.5 | 5 | 0.2 |
| 15_3162059 | NA | NA | 1 | 1 | NA | NA | NA | NA | 1 | 1 | NA | NA | 1 | 1 | 1 | 1 | NA | NA | 2 | 1 |
| 15_3162413 | NA | NA | 1 | 1 | NA | NA | 1 | 1 | 2 | 1 | 1 | 1 | 2 | 1 | 1 | 1 | 1 | 1 | 5 | 1 |
| 15_3162414 | NA | NA | 1 | 1 | NA | NA | 1 | 0 | 2 | 0.5 | 1 | 1 | 2 | 0.5 | 2 | 1 | 1 | 1 | 6 | 0.8333 |
| 15_3162423 | NA | NA | 1 | 1 | NA | NA | 1 | 0 | 2 | 0.5 | NA | NA | 1 | 1 | 3 | 0 | NA | NA | 4 | 0.25 |
| 15_3162427 | NA | NA | 1 | 1 | NA | NA | 1 | 0 | 2 | 0.5 | NA | NA | 1 | 1 | 3 | 1 | NA | NA | 4 | 1 |
| 15_3162463 | NA | NA | 1 | 1 | NA | NA | 1 | 0 | 2 | 0.5 | NA | NA | 1 | 1 | 3 | 0.3333 | NA | NA | 4 | 0.5 |
| 15_3162553 | NA | NA | NA | NA | NA | NA | 1 | 1 | 1 | 1 | NA | NA | NA | NA | 1 | 1 | NA | NA | 1 | 1 |
| 15_3162638 | NA | NA | NA | NA | NA | NA | 1 | 1 | 1 | 1 | NA | NA | NA | NA | 1 | 1 | NA | NA | 1 | 1 |
| 15_3268968 | NA | NA | 1 | 1 | NA | NA | NA | NA | 1 | 1 | NA | NA | 1 | 1 | NA | NA | NA | NA | 1 | 1 |
| 15_3269008 | NA | NA | 1 | 0 | NA | NA | 1 | 1 | 2 | 0.5 | NA | NA | 1 | 1 | NA | NA | NA | NA | 1 | 1 |
| 15_3269029 | NA | NA | 1 | 1 | NA | NA | NA | NA | 1 | 1 | NA | NA | 1 | 1 | NA | NA | NA | NA | 1 | 1 |
| 15_3269238 | 1 | 0 | 1 | 1 | NA | NA | NA | NA | 2 | 0.5 | 1 | 0 | 1 | 0 | 2 | 0.5 | NA | NA | 4 | 0.25 |
| 15_3269505 | 1 | 0 | 2 | 0 | 1 | 1 | 2 | 0.5 | 6 | 0.3333 | 1 | 0 | 2 | 0.5 | 2 | 0 | 2 | 0.5 | 7 | 0.2857 |
| 15_3269506 | 1 | 0 | 1 | 1 | 1 | 1 | 2 | 0.5 | 5 | 0.6 | 1 | 0 | 2 | 0 | 3 | 0.3333 | 2 | 1 | 8 | 0.375 |
| 15_3269514 | 1 | 0 | 2 | 0.5 | 1 | 0 | 2 | 0 | 6 | 0.1667 | 1 | 0 | 2 | 0 | 2 | 0.5 | 1 | 1 | 6 | 0.3333 |
| 15_3269525 | 1 | 1 | 3 | 0.3333 | 1 | 1 | 1 | 0 | 6 | 0.5 | 1 | 0 | 2 | 0.5 | 2 | 1 | 1 | 0 | 6 | 0.5 |
| 15_3269536 | 1 | 1 | 2 | 1 | NA | NA | 2 | 0.5 | 5 | 0.8 | 1 | 1 | 2 | 1 | 2 | 1 | 2 | 1 | 7 | 1 |
| 15_3269551 | 1 | 0 | 3 | 0.3333 | NA | NA | 2 | 0 | 6 | 0.1667 | 1 | 0 | 2 | 0.5 | 3 | 0 | 2 | 0 | 8 | 0.125 |
| 15_3269575 | 1 | 1 | 2 | 0 | 1 | 0 | 2 | 0 | 6 | 0.1667 | 1 | 0 | 2 | 0 | 3 | 0.6667 | 2 | 0 | 8 | 0.25 |
| 15_3279541 | 2 | 0.5 | NA | NA | 1 | 1 | NA | NA | 3 | 0.6667 | 1 | 1 | NA | NA | 1 | 0 | NA | NA | 2 | 0.5 |
| 15_3292114 | 9 | 0.2222 | 4 | 0 | 5 | 0.2 | 10 | 0 | 28 | 0.1071 | 2 | 0 | 4 | 0.25 | 7 | 0 | 7 | 0 | 20 | 0.05 |
| 15_3303272 | 9 | 0.7778 | 6 | 1 | 8 | 0.875 | 13 | 1 | 36 | 0.9167 | 2 | 1 | 6 | 0.5 | 8 | 0.875 | 9 | 0.8889 | 25 | 0.8 |
| 15_3303362 | 11 | 0.2727 | 5 | 0 | 9 | 0.2222 | 14 | 0.2143 | 39 | 0.2051 | 2 | 0 | 7 | 0.1429 | 11 | 0.0909 | 8 | 0.5 | 28 | 0.2143 |
| 15_3321318 | 2 | 0 | 1 | 0 | 1 | 0 | 3 | 0.3333 | 7 | 0.1429 | 1 | 0 | 3 | 0.3333 | 1 | 1 | NA | NA | 5 | 0.4 |
| 15_3321320 | 2 | 0.5 | 2 | 0 | 2 | 0.5 | 3 | 0 | 9 | 0.2222 | 1 | 0 | 3 | 0.3333 | 1 | 1 | NA | NA | 5 | 0.4 |
| 15_3321322 | 2 | 0.5 | 2 | 0 | 2 | 0 | 4 | 0.25 | 10 | 0.2 | 2 | 0.5 | 3 | 0 | 1 | 0 | NA | NA | 6 | 0.1667 |
| 15_3321339 | 2 | 0.5 | 1 | 0 | 1 | 0 | 3 | 0 | 7 | 0.1429 | 1 | 0 | 2 | 0.5 | NA | NA | NA | NA | 3 | 0.3333 |
| 15_3359430 | 4 | 0 | 3 | 0.3333 | 3 | 0 | 3 | 0 | 13 | 0.0769 | 2 | 0 | 4 | 0.25 | 1 | 0 | 2 | 0 | 9 | 0.1111 |
| 15_3359442 | 4 | 0.25 | 3 | 0.3333 | 3 | 0 | 4 | 0 | 14 | 0.1429 | 2 | 0 | 4 | 0.5 | NA | NA | 2 | 0 | 8 | 0.25 |
| 15_3368420 | 8 | 0.75 | 7 | 0.8571 | 6 | 1 | 8 | 0.75 | 29 | 0.8276 | 5 | 0.6 | 6 | 0.5 | 2 | 0 | 8 | 0.375 | 21 | 0.4286 |
| 15_3368433 | 3 | 0.3333 | 8 | 0.875 | 4 | 0.75 | 6 | 0.3333 | 21 | 0.619 | 3 | 0.3333 | 8 | 0.375 | 2 | 0 | 8 | 0.25 | 21 | 0.2857 |
| 15_3368436 | 6 | 0.1667 | 6 | 0.3333 | 4 | 0.5 | 9 | 0.1111 | 25 | 0.24 | 4 | 0.25 | 8 | 0 | 2 | 0 | 7 | 0.1429 | 21 | 0.0952 |
| 15_3368439 | 7 | 0.8571 | 6 | 1 | 5 | 1 | 8 | 0.75 | 26 | 0.8846 | 3 | 0.6667 | 6 | 0.3333 | 2 | 1 | 8 | 0.75 | 19 | 0.6316 |
| 15_3368441 | 5 | 0 | 6 | 0 | 4 | 0 | 9 | 0.3333 | 24 | 0.125 | 4 | 0 | 6 | 0 | 2 | 0 | 7 | 0.1429 | 19 | 0.0526 |
| 15_3368446 | 7 | 0.2857 | 7 | 0.4286 | 5 | 0.8 | 9 | 0 | 28 | 0.3214 | 4 | 0 | 8 | 0.125 | 2 | 0 | 6 | 0 | 20 | 0.05 |
| 15_3368458 | 6 | 0 | 7 | 0.2857 | 5 | 0.2 | 9 | 0 | 27 | 0.1111 | 4 | 0.25 | 7 | 0 | 2 | 0 | 7 | 0.1429 | 20 | 0.1 |
| 15_3368512 | 4 | 0.5 | 4 | 0.25 | 1 | 0 | 6 | 0 | 15 | 0.2 | 2 | 0 | 5 | 0 | 1 | 0 | 6 | 0.3333 | 14 | 0.1429 |
| 15_3368569 | 3 | 0 | 3 | 0 | NA | NA | 6 | 0.1667 | 12 | 0.0833 | 3 | 0 | 6 | 0.1667 | 1 | 0 | 6 | 0.3333 | 16 | 0.1875 |
| 15_3368586 | 3 | 0 | 4 | 0.25 | 2 | 0 | 4 | 0 | 13 | 0.0769 | 3 | 0 | 5 | 0 | 1 | 0 | 6 | 0.1667 | 15 | 0.0667 |

| | | | | | | | | | | | | | | | | | | | | |
|---|---|---|---|---|---|---|---|---|---|---|---|---|---|---|---|---|---|---|---|---|
| 15_3368596 | 3 | 0 | 3 | 0.3333 | 2 | 0 | 4 | 0 | 12 | 0.0833 | 3 | 0.3333 | 4 | 0.5 | 1 | 1 | 6 | 0.3333 | 14 | 0.4286 |
| 15_3368601 | 2 | 0 | 4 | 1 | 1 | 0 | 5 | 0 | 12 | 0.3333 | 3 | 0.6667 | 5 | 0.4 | NA | NA | 6 | 0.6667 | 14 | 0.5714 |
| 15_3368602 | 2 | 0.5 | 3 | 1 | 2 | 0.5 | 5 | 0.8 | 12 | 0.75 | 1 | 1 | 4 | 0.25 | 1 | 1 | 5 | 1 | 11 | 0.7273 |
| 15_3368608 | 2 | 0 | 4 | 0.75 | 2 | 0 | 1 | 1 | 9 | 0.4444 | 3 | 0.3333 | 3 | 0.3333 | 1 | 1 | 4 | 0.75 | 11 | 0.5455 |
| 15_3368612 | 2 | 0 | 4 | 0.5 | 2 | 0 | 4 | 0.25 | 12 | 0.25 | 3 | 0.3333 | 5 | 0 | 1 | 0 | 5 | 0.2 | 14 | 0.1429 |
| 15_3368614 | 2 | 0 | 3 | 1 | 1 | 0 | 4 | 0.25 | 10 | 0.4 | 2 | 0 | 5 | 0.4 | 1 | 0 | 6 | 0.5 | 14 | 0.3571 |
| 15_3368619 | 2 | 0 | 3 | 0.3333 | 1 | 0 | 3 | 0 | 9 | 0.1111 | 2 | 0 | 5 | 0.2 | 1 | 0 | 6 | 0 | 14 | 0.0714 |
| 15_3368621 | 2 | 0 | 2 | 0.5 | 1 | 0 | 4 | 0.25 | 9 | 0.2222 | 2 | 0 | 5 | 0.4 | 1 | 0 | 6 | 0.3333 | 14 | 0.2857 |
| 15_3368627 | 2 | 0.5 | 3 | 0.3333 | 2 | 0 | 4 | 0 | 11 | 0.1818 | 2 | 0 | 5 | 0.2 | 1 | 0 | 3 | 0 | 11 | 0.0909 |
| 15_3368643 | 1 | 0 | 2 | 1 | 1 | 1 | 3 | 0.3333 | 7 | 0.5714 | 1 | 0 | 5 | 1 | 1 | 1 | 3 | 0.6667 | 10 | 0.8 |
| 15_3368649 | 2 | 0 | 1 | 1 | 2 | 0.5 | 2 | 0.5 | 7 | 0.4286 | 1 | 0 | 3 | 0.6667 | 1 | 1 | 4 | 0.25 | 9 | 0.4444 |
| 15_3368653 | 2 | 0 | 2 | 0.5 | 2 | 0.5 | 3 | 0.3333 | 9 | 0.3333 | 1 | 1 | 4 | 0 | 1 | 1 | 3 | 0 | 9 | 0.2222 |
| 15_3368660 | 2 | 0.5 | 2 | 0 | 2 | 0.5 | 2 | 0.5 | 8 | 0.375 | 1 | 1 | 4 | 0.75 | 1 | 1 | 3 | 0.3333 | 9 | 0.6667 |
| 15_3368661 | 2 | 1 | 2 | 1 | 2 | 1 | 3 | 1 | 9 | 1 | 1 | 1 | 4 | 1 | 1 | 1 | 4 | 1 | 10 | 1 |
| 15_3368668 | 2 | 1 | 2 | 1 | 2 | 1 | 3 | 0.6667 | 9 | 0.8889 | 1 | 1 | 1 | 1 | 1 | 0 | 5 | 1 | 8 | 0.875 |
| 15_3368675 | 2 | 1 | 2 | 0 | 2 | 0 | 3 | 1 | 9 | 0.5556 | NA | NA | 2 | 1 | 1 | 0 | 3 | 0 | 6 | 0.3333 |
| 15_3368684 | 2 | 1 | 2 | 0.5 | 2 | 1 | 3 | 0.6667 | 9 | 0.7778 | 1 | 1 | 3 | 1 | 1 | 1 | 3 | 0.3333 | 8 | 0.75 |
| 15_3368692 | 2 | 0 | 1 | 1 | 2 | 1 | 2 | 0 | 7 | 0.4286 | 1 | 0 | 3 | 0.3333 | 1 | 1 | 3 | 0 | 8 | 0.25 |
| 15_3368711 | 2 | 0.5 | 2 | 0 | 2 | 0 | 2 | 0.5 | 8 | 0.25 | NA | NA | 1 | 0 | 1 | 0 | 3 | 0.3333 | 5 | 0.2 |
| 15_3368712 | 1 | 1 | NA | NA | NA | NA | 1 | 1 | 2 | 1 | 1 | 1 | NA | NA | 1 | 1 | 2 | 1 | 4 | 1 |
| 15_3368732 | 2 | 0.5 | 1 | 1 | 1 | 1 | 2 | 0.5 | 6 | 0.6667 | 1 | 0 | 2 | 0.5 | 1 | 0 | 3 | 0.3333 | 7 | 0.2857 |
| 15_3368734 | NA | NA | NA | NA | 1 | 1 | NA | NA | 1 | 1 | 1 | 1 | NA | NA | NA | NA | NA | NA | 1 | 1 |
| 15_3368748 | 1 | 0 | 1 | 1 | NA | NA | NA | NA | 2 | 0.5 | 1 | 0 | NA | NA | 1 | 1 | 2 | 0 | 4 | 0.25 |
| 15_3368771 | 1 | 1 | 1 | 0 | 1 | 1 | NA | NA | 3 | 0.6667 | NA | NA | NA | NA | 1 | 0 | 3 | 0.3333 | 4 | 0.25 |
| 15_3383649 | NA | NA | 1 | 1 | NA | NA | 1 | 1 | 2 | 1 | NA | NA | NA | NA | 1 | 1 | NA | NA | 1 | 1 |
| 15_3383653 | NA | NA | 1 | 1 | NA | NA | 1 | 1 | 2 | 1 | NA | NA | NA | NA | 1 | 1 | NA | NA | 1 | 1 |
| 15_3408226 | 1 | 1 | NA | NA | NA | NA | NA | NA | 1 | 1 | 1 | 1 | NA | NA | NA | NA | NA | NA | 1 | 1 |
| 15_3408242 | 1 | 1 | NA | NA | NA | NA | NA | NA | 1 | 1 | 1 | 1 | NA | NA | NA | NA | NA | NA | 1 | 1 |
| 15_3408247 | 1 | 1 | NA | NA | NA | NA | NA | NA | 1 | 1 | 1 | 1 | NA | NA | NA | NA | NA | NA | 1 | 1 |
| 15_3408253 | 1 | 1 | NA | NA | NA | NA | NA | NA | 1 | 1 | 1 | 1 | NA | NA | NA | NA | NA | NA | 1 | 1 |
| 15_3408276 | 1 | 1 | NA | NA | NA | NA | NA | NA | 1 | 1 | 1 | 1 | NA | NA | NA | NA | NA | NA | 1 | 1 |
| 15_3408294 | 2 | 0 | 1 | 0 | 2 | 0.5 | NA | NA | 5 | 0.2 | NA | NA | 5 | 0.2 | 2 | 0 | 2 | 0 | 9 | 0.1111 |
| 15_3408300 | 3 | 0 | 2 | 0.5 | 2 | 0 | 1 | 0 | 8 | 0.125 | NA | NA | 5 | 0.4 | 2 | 0 | 4 | 0 | 11 | 0.1818 |
| 15_3411817 | 3 | 0 | 5 | 0.2 | 4 | 0.25 | 3 | 0 | 15 | 0.1333 | 3 | 0 | 4 | 0 | 6 | 0.1667 | 4 | 0 | 17 | 0.0588 |
| 15_3414092 | 1 | 1 | 6 | 0.6667 | 7 | 0.7143 | 5 | 0.8 | 19 | 0.7368 | 5 | 0 | 3 | 0 | 7 | 0.4286 | 5 | 0.6 | 20 | 0.3 |
| 15_3414098 | 2 | 0 | 8 | 0.5 | 7 | 0.8571 | 3 | 0.6667 | 20 | 0.6 | 4 | 0.5 | 9 | 0.7778 | 9 | 0.5556 | 5 | 0.6 | 27 | 0.6296 |
| 15_3414102 | 2 | 0 | 7 | 0.1429 | 4 | 0 | 4 | 0 | 17 | 0.0588 | 3 | 0 | 6 | 0.1667 | 9 | 0.2222 | 4 | 0.25 | 22 | 0.1818 |
| 15_3414103 | 2 | 0.5 | 6 | 0.6667 | 3 | 1 | 5 | 0.8 | 16 | 0.75 | 4 | 0.5 | 2 | 0.5 | 4 | 0.75 | 3 | 0.6667 | 13 | 0.6154 |
| 15_3414103 | 3 | 0 | 5 | 0.2 | 5 | 0 | 3 | 0.3333 | 16 | 0.125 | 4 | 0.75 | 6 | 0.6667 | 5 | 0.2 | 3 | 0.3333 | 18 | 0.5 |
| 15_3414111 | 3 | 0 | 8 | 0 | 7 | 0.1429 | 4 | 0.5 | 22 | 0.1364 | 5 | 0.4 | 7 | 0.1429 | 8 | 0.25 | 7 | 0.5714 | 27 | 0.3333 |
| 15_3414116 | 3 | 0.6667 | 7 | 0.8571 | 6 | 0.8333 | 6 | 1 | 22 | 0.8636 | 4 | 0.75 | 8 | 0.75 | 10 | 0.6 | 6 | 0.8333 | 28 | 0.7143 |
| 15_3414120 | 2 | 1 | 7 | 0.7143 | 5 | 1 | 6 | 1 | 20 | 0.9 | 3 | 0.6667 | 6 | 1 | 9 | 0.6667 | 7 | 0.8571 | 25 | 0.8 |
| 15_3423849 | 3 | 0 | 10 | 0.1 | 9 | 0.2222 | 6 | 0.1667 | 28 | 0.1429 | 6 | 0 | 6 | 0.3333 | 11 | 0.2727 | 7 | 0.1429 | 30 | 0.2 |
| 15_3428360 | 10 | 0.5 | 13 | 0.4615 | 11 | 0.2727 | 12 | 0.3333 | 46 | 0.3913 | 9 | 0.3333 | 8 | 0.375 | 16 | 0.5 | 7 | 0.4286 | 40 | 0.425 |
| 15_3428413 | 13 | 0 | 19 | 0.1053 | 16 | 0.1875 | 17 | 0.2941 | 65 | 0.1538 | 12 | 0 | 10 | 0.2 | 16 | 0.25 | 8 | 0.125 | 46 | 0.1522 |
| 15_3434525 | 1 | 1 | 1 | 0 | 2 | 0 | 3 | 0 | 7 | 0.1429 | 1 | 0 | 1 | 0 | 1 | 1 | NA | NA | 3 | 0.3333 |
| 15_3434549 | 2 | 0.5 | 1 | 1 | 2 | 1 | 2 | 1 | 7 | 0.8571 | NA | NA | 1 | 1 | NA | NA | NA | NA | 1 | 1 |
| 15_3434551 | 2 | 0.5 | NA | NA | 2 | 0.5 | 3 | 0.6667 | 7 | 0.5714 | 1 | 1 | 1 | 1 | 1 | 0 | NA | NA | 3 | 0.6667 |
| 15_3434559 | 2 | 0.5 | 1 | 0 | 2 | 1 | 3 | 1 | 8 | 0.75 | 1 | 1 | 1 | 1 | 1 | 0 | NA | NA | 3 | 0.6667 |
| 15_3434569 | 2 | 1 | 1 | 0 | 2 | 0 | 2 | 0.5 | 7 | 0.4286 | 1 | 0 | 2 | 0 | 1 | 1 | NA | NA | 4 | 0.25 |
| 15_3442864 | 4 | 0 | 5 | 0.4 | 2 | 0 | 5 | 0.6 | 16 | 0.3125 | 3 | 0.3333 | 5 | 0.2 | 4 | 0.25 | 1 | 1 | 13 | 0.3077 |
| 15_3442865 | 5 | 0.4 | 5 | 0.4 | 2 | 0.5 | 4 | 0.25 | 16 | 0.375 | 3 | 1 | 5 | 0.4 | 4 | 0.5 | 1 | 0 | 13 | 0.5385 |
| 15_3442875 | 3 | 0.3333 | 2 | 0 | 2 | 0 | 4 | 0 | 11 | 0.0909 | 2 | 0 | 5 | 0 | 3 | 0.3333 | 1 | 0 | 11 | 0.0909 |
| 15_3443601 | 7 | 0 | 6 | 0.1667 | 2 | 0 | 5 | 0 | 20 | 0.05 | 3 | 0.3333 | 6 | 0 | 5 | 0 | 3 | 0 | 17 | 0.0588 |
| 15_3443607 | 8 | 0.375 | 6 | 0.3333 | 3 | 0.3333 | 5 | 0 | 22 | 0.2727 | 4 | 0.25 | 5 | 0.4 | 5 | 0 | 3 | 0.6667 | 17 | 0.2941 |
| 15_3443608 | 6 | 0.6667 | 5 | 0.8 | 3 | 0.6667 | 5 | 0.8 | 19 | 0.7368 | 2 | 0.5 | 4 | 0.5 | 4 | 0.75 | 2 | 0.5 | 12 | 0.5833 |
| 15_3443609 | 8 | 0.125 | 6 | 0 | 2 | 0 | 4 | 0 | 20 | 0.05 | 4 | 0 | 6 | 0.1667 | 5 | 0 | 3 | 0 | 18 | 0.0556 |
| 15_3443630 | 7 | 0 | 4 | 0.25 | 3 | 0.3333 | 5 | 0.2 | 19 | 0.1579 | 5 | 0.2 | 5 | 0.4 | 4 | 0.25 | 4 | 0.75 | 18 | 0.3889 |
| 15_3448445 | 11 | 0.1818 | 10 | 0.3 | 11 | 0.1818 | 12 | 0.25 | 44 | 0.2273 | 14 | 0.2143 | 8 | 0.25 | 6 | 0.1667 | 10 | 0.2 | 38 | 0.2105 |
| 15_3453009 | NA | NA | NA | NA | 1 | 1 | NA | NA | 1 | 1 | NA | NA | NA | NA | NA | NA | 1 | 1 | 1 | 1 |
| 15_3453043 | NA | NA | NA | NA | 1 | 1 | NA | NA | 1 | 1 | NA | NA | NA | NA | NA | NA | 2 | 0.5 | 2 | 0.5 |
| 15_3457515 | NA | NA | 1 | 1 | 1 | 0 | NA | NA | 2 | 0.5 | 1 | 1 | NA | NA | 1 | 0 | 1 | 0 | 3 | 0.3333 |
| 15_3457547 | 1 | 1 | 1 | 1 | 1 | 1 | 1 | 1 | 4 | 1 | 1 | 1 | NA | NA | 1 | 0 | 2 | 0.5 | 4 | 0.5 |
| 15_3483736 | NA | NA | 1 | 0 | NA | NA | 1 | 1 | 2 | 0.5 | NA | NA | NA | NA | 1 | 1 | NA | NA | 1 | 1 |
| 15_3483740 | NA | NA | 1 | 1 | NA | NA | 1 | 1 | 2 | 1 | NA | NA | NA | NA | 1 | 1 | NA | NA | 1 | 1 |

| | | | | | | | | | | | | | | | | | | | | |
|---|---|---|---|---|---|---|---|---|---|---|---|---|---|---|---|---|---|---|---|---|
| 15_348374! | NA | NA | 1 | 1 | NA | NA | 1 | 1 | 2 | 1 | NA | NA | NA | NA | 1 | 1 | NA | NA | 1 | 1 |
| 15_348390: | NA | NA | 1 | 0 | 1 | 1 | 1 | 0 | 3 | 0.3333 | NA | NA | 2 | 0.5 | 1 | 0 | NA | NA | 3 | 0.3333 |
| 15_348391 | NA | NA | 1 | 1 | 1 | 1 | 1 | 1 | 3 | 1 | 1 | 0 | 2 | 1 | 1 | 1 | NA | NA | 4 | 0.75 |
| 15_348392 | NA | NA | 1 | 1 | 1 | 1 | NA | NA | 2 | 1 | 1 | 1 | 2 | 1 | 1 | 1 | 1 | 1 | 5 | 1 |
| 15_348393! | NA | NA | 1 | 1 | 1 | 0 | 1 | 1 | 3 | 0.6667 | 1 | 1 | 2 | 0.5 | 2 | 1 | 1 | 1 | 6 | 0.8333 |
| 15_348394! | NA | NA | NA | NA | 1 | 1 | NA | NA | 1 | 1 | NA | NA | 1 | 1 | 1 | 1 | NA | NA | 2 | 1 |
| 15_348394 | NA | NA | 1 | 0 | NA | NA | 1 | 1 | 2 | 0.5 | 1 | 1 | 2 | 0.5 | 1 | 1 | 1 | 0 | 5 | 0.6 |
| 15_348395( | NA | NA | 1 | 1 | 1 | 0 | 2 | 0 | 4 | 0.25 | 1 | 0 | 3 | 0 | 2 | 0 | 2 | 0.5 | 8 | 0.125 |
| 15_348397! | NA | NA | 1 | 1 | 2 | 1 | 2 | 1 | 5 | 1 | NA | NA | 2 | 1 | 2 | 1 | 2 | 1 | 6 | 1 |
| 15_348398( | NA | NA | 1 | 1 | NA | NA | NA | NA | 1 | 1 | NA | NA | 2 | 0.5 | 2 | 1 | 2 | 1 | 6 | 0.8333 |
| 15_348399: | 1 | 1 | 1 | 1 | 1 | 1 | NA | NA | 3 | 1 | 1 | 1 | 4 | 1 | 1 | 1 | 2 | 1 | 8 | 1 |
| 15_348399 | 1 | 1 | 1 | 1 | 1 | 1 | 2 | 1 | 5 | 1 | 1 | 1 | 4 | 1 | 2 | 1 | 2 | 1 | 9 | 1 |
| 15_348400 | 1 | 0 | 1 | 1 | 1 | 0 | 2 | 0 | 5 | 0.2 | 1 | 0 | 4 | 0 | 2 | 0 | 2 | 0.5 | 9 | 0.1111 |
| 15_348400 | NA | NA | 1 | 1 | 1 | 1 | 2 | 1 | 4 | 1 | NA | NA | 4 | 0.75 | 2 | 1 | 1 | 1 | 7 | 0.8571 |
| 15_348400 | 1 | 1 | 1 | 0 | 1 | 1 | 2 | 1 | 5 | 0.8 | 1 | 1 | 3 | 0.6667 | 2 | 1 | 2 | 0 | 8 | 0.625 |
| 15_348401 | 1 | 1 | 1 | 1 | 1 | 1 | 2 | 1 | 5 | 1 | 1 | 1 | 3 | 1 | 1 | 1 | 2 | 1 | 7 | 1 |
| 15_348402 | 1 | 0 | 1 | 0 | 1 | 0 | 2 | 0.5 | 5 | 0.2 | NA | NA | 4 | 0 | 2 | 0.5 | 2 | 0 | 8 | 0.125 |
| 15_348418 | 2 | 0 | 1 | 0 | 1 | 0 | 2 | 0.5 | 6 | 0.1667 | 2 | 0 | 2 | 0 | 1 | 0 | 2 | 0.5 | 7 | 0.1429 |
| 15_348419( | 3 | 0.6667 | NA | NA | 1 | 1 | 1 | 0 | 5 | 0.6 | 2 | 0.5 | 2 | 0 | 2 | 0.5 | 2 | 0.5 | 8 | 0.375 |
| 15_348420! | 3 | 1 | 1 | 1 | 1 | 1 | 2 | 1 | 7 | 1 | 2 | 1 | 2 | 1 | 2 | 1 | 2 | 1 | 8 | 1 |
| 15_348421 | 1 | 1 | NA | NA | 1 | 1 | 2 | 0.5 | 4 | 0.75 | 2 | 0.5 | 4 | 1 | 2 | 1 | 2 | 1 | 10 | 0.9 |
| 15_348421 | 4 | 0.25 | NA | NA | 2 | 0.5 | 3 | 0.3333 | 9 | 0.3333 | 2 | 0.5 | 4 | 0.25 | 2 | 0 | 2 | 0.5 | 10 | 0.3 |
| 15_348421! | 4 | 1 | 1 | 1 | 1 | 1 | 3 | 1 | 9 | 1 | 2 | 1 | 3 | 1 | 2 | 1 | 2 | 1 | 9 | 1 |
| 15_348421! | 4 | 0.25 | 1 | 0 | 1 | 0 | 3 | 1 | 9 | 0.4444 | 2 | 0.5 | 4 | 0 | 2 | 0.5 | 1 | 0 | 9 | 0.2222 |
| 15_348422! | 3 | 1 | NA | NA | NA | NA | 2 | 1 | 5 | 1 | 2 | 0.5 | 2 | 1 | NA | NA | 2 | 1 | 6 | 0.8333 |
| 15_348423! | 4 | 0.5 | 1 | 0 | 2 | 0.5 | 3 | 0.3333 | 10 | 0.4 | 2 | 0 | 3 | 0.6667 | 2 | 0.5 | 2 | 0 | 9 | 0.3333 |
| 15_348423 | 4 | 0 | 1 | 0 | 2 | 0.5 | 3 | 0 | 10 | 0.1 | 2 | 0 | 3 | 0.3333 | 2 | 0 | 2 | 0.5 | 9 | 0.2222 |
| 15_348424: | 4 | 1 | NA | NA | 3 | 1 | 3 | 1 | 10 | 1 | 2 | 0.5 | 4 | 1 | 3 | 1 | 2 | 1 | 11 | 0.9091 |
| 15_348424! | 3 | 0.3333 | 1 | 0 | 2 | 1 | 2 | 0 | 8 | 0.375 | 1 | 0 | 3 | 0 | 3 | 0.3333 | 2 | 0 | 9 | 0.1111 |
| 15_348425! | 3 | 0 | NA | NA | 3 | 0.3333 | 2 | 0 | 8 | 0.125 | 2 | 0 | 3 | 0 | 3 | 0.3333 | NA | NA | 8 | 0.125 |
| 15_348428 | 3 | 0 | 1 | 1 | 3 | 1 | 2 | 1 | 9 | 0.6667 | 2 | 0.5 | 3 | 1 | 2 | 1 | 2 | 0 | 9 | 0.6667 |
| 15_348759 | 2 | 0.5 | 3 | 0.3333 | 1 | 0 | NA | NA | 6 | 0.3333 | 2 | 0.5 | NA | NA | 2 | 0.5 | 1 | 0 | 5 | 0.4 |
| 15_348760 | 2 | 0.5 | 3 | 0 | 1 | 0 | NA | NA | 6 | 0.1667 | 2 | 0 | NA | NA | 2 | 0.5 | 1 | 0 | 5 | 0.2 |
| 15_348760 | 2 | 1 | 3 | 0.6667 | 1 | 0 | NA | NA | 6 | 0.6667 | 2 | 0.5 | NA | NA | 1 | 0 | NA | NA | 3 | 0.3333 |
| 15_348765( | 1 | 0 | 3 | 0.6667 | NA | NA | NA | NA | 4 | 0.5 | 2 | 0 | NA | NA | 2 | 1 | NA | NA | 4 | 0.5 |
| 15_348766! | 1 | 0 | 2 | 1 | NA | NA | NA | NA | 3 | 0.6667 | 2 | 0.5 | NA | NA | 2 | 0 | NA | NA | 4 | 0.25 |
| 15_348767( | NA | NA | 2 | 0.5 | 1 | 0 | NA | NA | 3 | 0.3333 | NA | NA | NA | NA | 1 | 1 | NA | NA | 1 | 1 |
| 15_348768 | 1 | 1 | 3 | 0 | 1 | 0 | NA | NA | 5 | 0.2 | 2 | 0.5 | NA | NA | NA | NA | NA | NA | 2 | 0.5 |
| 15_349452( | 1 | 1 | 3 | 0 | 1 | 0 | NA | NA | 5 | 0.2 | 2 | 0 | NA | NA | 2 | 0.5 | NA | NA | 4 | 0.25 |
| 15_349452 | 1 | 1 | 3 | 0 | 1 | 0 | NA | NA | 5 | 0.2 | 2 | 0 | NA | NA | 1 | 1 | NA | NA | 3 | 0.3333 |
| 15_359104( | 3 | 0.3333 | 1 | 1 | NA | NA | 1 | 0 | 5 | 0.4 | 3 | 1 | NA | NA | 2 | 1 | NA | NA | 5 | 1 |
| 15_359105 | 5 | 0.2 | 2 | 0 | NA | NA | 1 | 0 | 8 | 0.125 | 3 | 0.3333 | NA | NA | 2 | 0 | NA | NA | 5 | 0.2 |
| 15_359106( | 4 | 0.5 | 2 | 0.5 | NA | NA | 1 | 0 | 7 | 0.4286 | 4 | 0.75 | NA | NA | 2 | 0 | NA | NA | 6 | 0.5 |
| 15_359106! | 5 | 0.4 | 2 | 0.5 | NA | NA | 1 | 1 | 8 | 0.5 | 4 | 0.5 | NA | NA | 2 | 0.5 | NA | NA | 6 | 0.5 |
| 15_359107: | 4 | 0.75 | 2 | 1 | NA | NA | 1 | 1 | 7 | 0.8571 | 2 | 1 | NA | NA | 2 | 0.5 | NA | NA | 4 | 0.75 |
| 15_359107! | 4 | 0.75 | 2 | 1 | NA | NA | 1 | 1 | 7 | 0.8571 | 4 | 1 | NA | NA | 2 | 1 | NA | NA | 6 | 1 |
| 15_359107! | 4 | 0.25 | 2 | 0.5 | NA | NA | 1 | 0 | 7 | 0.2857 | 3 | 1 | NA | NA | 1 | 1 | NA | NA | 4 | 1 |
| 15_359110 | 5 | 0.2 | 1 | 0 | NA | NA | 1 | 0 | 7 | 0.1429 | 3 | 0.3333 | NA | NA | 2 | 0.5 | NA | NA | 5 | 0.4 |
| 15_359300! | 5 | 0.2 | 1 | 1 | 1 | 0 | 2 | 0.5 | 9 | 0.3333 | 4 | 0.25 | NA | NA | 3 | 0.3333 | 1 | 0 | 8 | 0.25 |
| 15_359301 | 5 | 0.2 | 2 | 0.5 | 1 | 0 | 2 | 0 | 10 | 0.2 | 6 | 0.1667 | NA | NA | 3 | 0.3333 | 1 | 0 | 10 | 0.2 |
| 15_359303 | 8 | 0.125 | 2 | 0 | 1 | 1 | 2 | 0.5 | 13 | 0.2308 | 4 | 0.25 | NA | NA | 3 | 0 | 2 | 0 | 9 | 0.1111 |
| 15_359309! | 11 | 0 | 2 | 0.5 | 2 | 0 | 2 | 0 | 17 | 0.0588 | 5 | 0.2 | NA | NA | 5 | 0 | 2 | 0 | 12 | 0.0833 |
| 15_359310! | 9 | 0.3333 | 3 | 0.6667 | 2 | 0 | 2 | 0.5 | 16 | 0.375 | 6 | 0.8333 | NA | NA | 3 | 0.3333 | 1 | 1 | 10 | 0.7 |
| 15_362407! | 4 | 0 | 8 | 0.5 | 6 | 0.1667 | 5 | 0 | 23 | 0.2174 | 2 | 0.5 | 1 | 1 | 5 | 0.2 | NA | NA | 8 | 0.375 |
| 15_362408! | 4 | 0 | 6 | 0.1667 | 6 | 0.5 | 4 | 0.25 | 20 | 0.25 | 1 | 0 | NA | NA | 3 | 0.3333 | NA | NA | 4 | 0.25 |
| 15_362411: | 4 | 0 | 8 | 0.125 | 6 | 0.1667 | 4 | 0.25 | 22 | 0.1364 | 2 | 0.5 | 1 | 0 | 4 | 0 | NA | NA | 7 | 0.1429 |
| 15_362412 | 4 | 0.25 | 8 | 0.125 | 6 | 0 | 3 | 0 | 21 | 0.0952 | 2 | 0 | 1 | 1 | 4 | 0 | NA | NA | 7 | 0.1429 |
| 15_362414! | 3 | 0.3333 | 8 | 0.125 | 6 | 0.1667 | 4 | 0 | 21 | 0.1429 | 1 | 0 | 1 | 0 | 5 | 0.4 | NA | NA | 7 | 0.2857 |
| 15_362415: | 4 | 1 | 8 | 0.875 | 5 | 1 | 2 | 1 | 19 | 0.9474 | 2 | 1 | 1 | 1 | 3 | 1 | NA | NA | 6 | 1 |
| 15_362415 | 4 | 1 | 8 | 1 | 6 | 0.6667 | 4 | 1 | 22 | 0.9091 | 2 | 1 | 1 | 1 | 4 | 1 | NA | NA | 7 | 1 |
| 15_362417: | 2 | 1 | 7 | 0.5714 | 3 | 1 | 3 | 0.3333 | 15 | 0.6667 | 1 | 0 | 1 | 1 | 1 | 0 | NA | NA | 3 | 0.3333 |
| 15_362418: | 3 | 1 | 7 | 0.8571 | 6 | 1 | 4 | 0.5 | 20 | 0.85 | 1 | 1 | 1 | 1 | 4 | 1 | NA | NA | 6 | 1 |
| 15_362418! | 4 | 0.75 | 7 | 0.5714 | 5 | 0.2 | 1 | 1 | 17 | 0.5294 | 2 | 0 | 1 | 0 | 3 | 0.3333 | NA | NA | 6 | 0.1667 |
| 15_362419: | 4 | 1 | 5 | 1 | 5 | 1 | 4 | 1 | 18 | 1 | 2 | 1 | 1 | 1 | 3 | 1 | NA | NA | 6 | 1 |
| 15_362419( | 3 | 0.6667 | 6 | 0.8333 | 3 | 1 | 4 | 0.5 | 16 | 0.75 | 1 | 1 | 1 | 1 | 3 | 0.6667 | NA | NA | 5 | 0.8 |

| | | | | | | | | | | | | | | | | | | | | |
|---|---|---|---|---|---|---|---|---|---|---|---|---|---|---|---|---|---|---|---|---|
| 15_362420 | 4 | 0.75 | 5 | 0.8 | 6 | 0 | 3 | 0.6667 | 18 | 0.5 | 1 | 1 | 1 | 1 | 2 | 0 | NA | NA | 4 | 0.5 |
| 15_362655 | 2 | 0.5 | 3 | 0 | 3 | 0 | 3 | 0 | 11 | 0.0909 | NA | NA | NA | NA | 1 | 1 | NA | NA | 1 | 1 |
| 15_362656 | 3 | 0.6667 | 2 | 1 | 4 | 0.75 | 3 | 0.3333 | 12 | 0.6667 | 1 | 1 | NA | NA | 2 | 0.5 | NA | NA | 3 | 0.6667 |
| 15_362657 | 1 | 1 | 2 | 1 | 3 | 1 | 3 | 0.6667 | 9 | 0.8889 | NA | NA | NA | NA | 2 | 1 | NA | NA | 2 | 1 |
| 15_362657 | 3 | 0.6667 | 2 | 0.5 | 2 | 0 | 3 | 0.3333 | 10 | 0.4 | 1 | 0 | NA | NA | 1 | 1 | NA | NA | 2 | 0.5 |
| 15_362660 | 2 | 1 | NA | NA | 3 | 0.6667 | 3 | 1 | 8 | 0.875 | 1 | 0 | NA | NA | 2 | 1 | NA | NA | 3 | 0.6667 |
| 15_362660 | 1 | 1 | NA | NA | 4 | 1 | 2 | 1 | 7 | 1 | 1 | 1 | NA | NA | 1 | 0 | NA | NA | 2 | 0.5 |
| 15_364781 | 24 | 0.125 | 25 | 0.24 | 25 | 0.28 | 28 | 0.2143 | 102 | 0.2157 | 15 | 0.3333 | 21 | 0.3333 | 18 | 0.2222 | 21 | 0.1905 | 75 | 0.2667 |
| 15_364785 | 25 | 0.08 | 29 | 0.1034 | 22 | 0.0455 | 28 | 0.1786 | 104 | 0.1058 | 15 | 0.1333 | 23 | 0.0435 | 17 | 0.1765 | 20 | 0.15 | 75 | 0.12 |
| 15_364791 | 19 | 0.2632 | 25 | 0.36 | 18 | 0.4444 | 22 | 0.3182 | 84 | 0.3452 | 14 | 0.2857 | 14 | 0.2857 | 15 | 0.3333 | 19 | 0.3684 | 62 | 0.3226 |
| 15_364792 | 21 | 0.2857 | 22 | 0.3182 | 21 | 0.1905 | 23 | 0.2609 | 87 | 0.2644 | 14 | 0.2143 | 19 | 0.1579 | 12 | 0.0833 | 17 | 0.1765 | 62 | 0.1613 |
| 15_364792 | 21 | 0.0476 | 27 | 0.0741 | 20 | 0.15 | 23 | 0.087 | 91 | 0.0879 | 15 | 0.2667 | 20 | 0.25 | 13 | 0.1538 | 14 | 0.1429 | 62 | 0.2097 |
| 15_364793 | 20 | 0.15 | 27 | 0.2222 | 18 | 0.0556 | 25 | 0.16 | 90 | 0.1556 | 14 | 0.1429 | 21 | 0.2381 | 13 | 0.0769 | 20 | 0.15 | 68 | 0.1618 |
| 15_364859 | 15 | 0.1333 | 20 | 0.05 | 19 | 0.2632 | 21 | 0.0952 | 75 | 0.1333 | 11 | 0.1818 | 20 | 0.15 | 13 | 0.1538 | 15 | 0.0667 | 59 | 0.1356 |
| 15_364860 | 10 | 0.2 | 18 | 0.1667 | 15 | 0 | 19 | 0.3158 | 62 | 0.1774 | 7 | 0.1429 | 15 | 0.2667 | 13 | 0.0769 | 12 | 0.1667 | 47 | 0.1702 |
| 15_364861 | 10 | 0.5 | 17 | 0.0588 | 16 | 0.4375 | 21 | 0.381 | 64 | 0.3281 | 9 | 0.2222 | 20 | 0.25 | 11 | 0.4545 | 14 | 0.2143 | 54 | 0.2778 |
| 15_364862 | 11 | 0 | 18 | 0.1667 | 16 | 0.125 | 20 | 0.15 | 65 | 0.1231 | 9 | 0 | 19 | 0.1579 | 11 | 0.0909 | 14 | 0.0714 | 53 | 0.0943 |
| 15_367707 | 14 | 0.2857 | 18 | 0.1667 | 15 | 0.2 | 16 | 0.1875 | 63 | 0.2063 | 15 | 0.2 | 22 | 0.2273 | 13 | 0.2308 | 17 | 0.2941 | 67 | 0.2388 |
| 15_367709 | 10 | 0.3 | 14 | 0 | 11 | 0.0909 | 11 | 0 | 46 | 0.087 | 10 | 0 | 15 | 0.0667 | 9 | 0.3333 | 14 | 0.2143 | 48 | 0.1458 |
| 15_367717 | 105 | 0.181 | 94 | 0.2021 | 72 | 0.1944 | 90 | 0.1667 | 361 | 0.1856 | 56 | 0.3214 | 100 | 0.2 | 92 | 0.2391 | 115 | 0.2522 | 363 | 0.2452 |
| 15_367719 | 158 | 0.1709 | 158 | 0.2405 | 122 | 0.1721 | 168 | 0.1369 | 606 | 0.1799 | 110 | 0.2545 | 172 | 0.1512 | 163 | 0.1534 | 178 | 0.2303 | 623 | 0.1926 |
| 15_367939 | 3 | 0.3333 | 4 | 0 | 4 | 0 | 3 | 0.3333 | 14 | 0.1429 | NA | NA | 5 | 0 | 3 | 0.3333 | 5 | 0.2 | 13 | 0.1538 |
| 15_369967 | 1 | 0 | NA | NA | 4 | 0.5 | 2 | 0 | 7 | 0.2857 | 1 | 0 | 1 | 1 | 1 | 1 | 1 | 1 | 4 | 0.75 |
| 15_369968 | 1 | 0 | NA | NA | 4 | 0 | 1 | 1 | 6 | 0.1667 | 2 | 0 | 1 | 1 | 1 | 0 | 1 | 0 | 5 | 0.2 |
| 15_369969 | 1 | 1 | NA | NA | 4 | 1 | 2 | 0.5 | 7 | 0.8571 | 2 | 1 | 1 | 1 | 1 | 1 | 1 | 1 | 5 | 1 |
| 15_369969 | 1 | 1 | NA | NA | 3 | 1 | 2 | 1 | 6 | 1 | 1 | 1 | 1 | 1 | 1 | 0 | 1 | 0 | 4 | 0.5 |
| 15_369970 | 1 | 1 | NA | NA | 2 | 1 | 1 | 1 | 4 | 1 | 1 | 1 | 1 | 1 | 1 | 1 | 1 | 1 | 4 | 1 |
| 15_369970 | 1 | 0 | NA | NA | 4 | 0.25 | 2 | 0.5 | 7 | 0.2857 | 1 | 0 | 1 | 1 | 1 | 1 | NA | NA | 3 | 0.6667 |
| 15_369971 | NA | NA | NA | NA | 2 | 1 | 2 | 1 | 4 | 1 | 2 | 1 | 1 | 1 | 1 | 1 | 1 | 1 | 5 | 1 |
| 15_369971 | 1 | 0 | NA | NA | 3 | 0.6667 | 1 | 1 | 5 | 0.6 | 1 | 1 | 1 | 0 | 1 | 1 | 1 | 1 | 4 | 0.75 |
| 15_369979 | 1 | 1 | 1 | 1 | 3 | 0 | 2 | 0.5 | 7 | 0.4286 | 2 | 0.5 | 1 | 1 | NA | NA | 2 | 0.5 | 5 | 0.6 |
| 15_369994 | 24 | 0.0833 | 20 | 0.15 | 25 | 0.04 | 28 | 0.2143 | 97 | 0.1237 | 19 | 0.1053 | 32 | 0.0938 | 28 | 0.1071 | 28 | 0.0357 | 107 | 0.0841 |
| 15_370010 | 29 | 0.2414 | 26 | 0.3462 | 19 | 0.4737 | 35 | 0.3714 | 109 | 0.3486 | 15 | 0.5333 | 31 | 0.4839 | 38 | 0.4211 | 33 | 0.5455 | 117 | 0.4872 |
| 15_370610 | NA | NA | NA | NA | 1 | 1 | NA | NA | 1 | 1 | NA | NA | NA | NA | NA | NA | 1 | 1 | 1 | 1 |
| 15_373666 | 31 | 0.5484 | 63 | 0.5873 | 40 | 0.55 | 20 | 0.75 | 154 | 0.5909 | 38 | 0.5789 | 17 | 0.7059 | 18 | 0.5556 | 25 | 0.6 | 98 | 0.602 |
| 15_373666 | 33 | 0.9091 | 57 | 0.9474 | 35 | 0.8571 | 22 | 0.9545 | 147 | 0.9184 | 39 | 0.9744 | 17 | 0.9412 | 17 | 0.9412 | 24 | 0.9167 | 97 | 0.9485 |
| 15_373667 | 32 | 0.6875 | 56 | 0.5893 | 37 | 0.5946 | 20 | 0.8 | 145 | 0.6414 | 32 | 0.5938 | 15 | 0.6667 | 16 | 0.5625 | 20 | 0.6 | 83 | 0.6024 |
| 15_373668 | 33 | 0.5152 | 60 | 0.5167 | 36 | 0.5833 | 22 | 0.6364 | 151 | 0.5497 | 34 | 0.6176 | 17 | 0.7059 | 17 | 0.3529 | 24 | 0.625 | 92 | 0.587 |
| 15_373668 | 28 | 0.1786 | 46 | 0.1739 | 32 | 0.0625 | 13 | 0.2308 | 119 | 0.1513 | 29 | 0.1379 | 14 | 0.2143 | 18 | 0.0556 | 16 | 0.25 | 77 | 0.1558 |
| 15_373669 | 35 | 0.1429 | 59 | 0.2881 | 36 | 0.2778 | 23 | 0.2609 | 153 | 0.2484 | 36 | 0.1667 | 15 | 0.2667 | 20 | 0.45 | 25 | 0.24 | 96 | 0.2604 |
| 15_373669 | 27 | 0.3333 | 56 | 0.1607 | 30 | 0.2 | 22 | 0.2727 | 135 | 0.2222 | 29 | 0.1379 | 14 | 0.3571 | 18 | 0.2222 | 22 | 0.1364 | 83 | 0.1928 |
| 15_373671 | 27 | 0.037 | 47 | 0.0426 | 30 | 0.0667 | 17 | 0.1176 | 121 | 0.0579 | 25 | 0 | 12 | 0 | 14 | 0.0714 | 20 | 0.15 | 71 | 0.0563 |
| 15_373674 | 26 | 0.0769 | 52 | 0.0962 | 31 | 0.1613 | 21 | 0.1905 | 130 | 0.1231 | 23 | 0.1304 | 11 | 0.2727 | 14 | 0.0714 | 17 | 0.0588 | 65 | 0.1231 |
| 15_373677 | 21 | 0.0476 | 48 | 0.0625 | 25 | 0.04 | 17 | 0.1176 | 111 | 0.0631 | 22 | 0.1364 | 9 | 0.1111 | 14 | 0 | 18 | 0.1111 | 63 | 0.0952 |
| 15_373678 | 17 | 0.4118 | 51 | 0.3137 | 28 | 0.4643 | 19 | 0.4211 | 115 | 0.3826 | 19 | 0.3684 | 8 | 0.625 | 13 | 0.3077 | 19 | 0.3684 | 59 | 0.3898 |
| 15_373682 | 15 | 0.2667 | 42 | 0.0952 | 23 | 0.2174 | 16 | 0.25 | 96 | 0.1771 | 17 | 0.1765 | 8 | 0.25 | 11 | 0.1818 | 17 | 0.1765 | 53 | 0.1887 |
| 15_373683 | 23 | 0.4348 | 51 | 0.5686 | 30 | 0.6 | 16 | 0.625 | 120 | 0.5583 | 26 | 0.5 | 11 | 0.5455 | 14 | 0.5 | 20 | 0.55 | 71 | 0.5211 |
| 15_373685 | 19 | 0.2632 | 44 | 0.2955 | 16 | 0.5 | 14 | 0.2857 | 93 | 0.3226 | 20 | 0.25 | 8 | 0.25 | 11 | 0.3636 | 20 | 0.25 | 59 | 0.2712 |
| 15_373685 | 18 | 0.0556 | 40 | 0.275 | 21 | 0.1905 | 14 | 0.1429 | 93 | 0.1935 | 22 | 0.0909 | 7 | 0.2857 | 9 | 0 | 18 | 0.0556 | 56 | 0.0893 |
| 15_373686 | 19 | 0 | 31 | 0.129 | 19 | 0.0526 | 10 | 0 | 79 | 0.0633 | 19 | 0.1053 | 6 | 0.1667 | 9 | 0 | 13 | 0 | 47 | 0.0638 |
| 15_373688 | 16 | 0.375 | 43 | 0.6279 | 22 | 0.5909 | 16 | 0.625 | 97 | 0.5773 | 23 | 0.5652 | 7 | 0.7143 | 10 | 0.7 | 16 | 0.625 | 56 | 0.625 |
| 15_373689 | 19 | 0.0526 | 41 | 0.2683 | 19 | 0.2632 | 14 | 0.2143 | 93 | 0.2151 | 24 | 0.2083 | 7 | 0.1429 | 10 | 0.2 | 15 | 0.2667 | 56 | 0.2143 |
| 15_373689 | 20 | 0.25 | 39 | 0.2051 | 20 | 0.2 | 16 | 0.25 | 95 | 0.2211 | 21 | 0 | 9 | 0.1111 | 10 | 0.1 | 18 | 0.1111 | 58 | 0.069 |
| 15_373690 | 20 | 0 | 38 | 0.1316 | 17 | 0.1176 | 15 | 0.0667 | 90 | 0.0889 | 22 | 0.1818 | 9 | 0 | 10 | 0 | 13 | 0.0769 | 54 | 0.0926 |
| 15_373974 | 15 | 0.8667 | 29 | 0.7931 | 14 | 0.7857 | 13 | 0.6923 | 71 | 0.7887 | 19 | 1 | 7 | 0.8571 | 9 | 0.7778 | 15 | 0.6667 | 50 | 0.84 |
| 15_373974 | 18 | 0.7222 | 37 | 0.6486 | 15 | 1 | 14 | 0.5714 | 84 | 0.7143 | 20 | 0.75 | 7 | 0.7143 | 9 | 0.8889 | 13 | 0.6154 | 49 | 0.7347 |
| 15_373975 | 8 | 0.875 | 21 | 1 | 8 | 0.875 | 6 | 0.8333 | 43 | 0.9302 | 16 | 0.875 | 5 | 0.6 | 6 | 1 | 11 | 0.9091 | 38 | 0.8684 |
| 15_373975 | 16 | 0.6875 | 37 | 0.7297 | 14 | 0.7143 | 13 | 0.6923 | 80 | 0.7125 | 17 | 0.6471 | 9 | 0.5556 | 8 | 1 | 15 | 0.6667 | 49 | 0.6939 |
| 15_373975 | 16 | 0.6875 | 36 | 0.7222 | 13 | 0.4615 | 13 | 0.6154 | 78 | 0.6538 | 19 | 0.7368 | 9 | 0.6667 | 7 | 0.5714 | 16 | 0.6875 | 51 | 0.6863 |
| 15_373976 | 16 | 0.1875 | 36 | 0.2222 | 16 | 0.0625 | 13 | 0.2308 | 81 | 0.1852 | 20 | 0 | 8 | 0.125 | 9 | 0.1111 | 15 | 0.2667 | 52 | 0.1154 |
| 15_379244 | 2 | 0 | 6 | 0 | 3 | 0 | 8 | 0.125 | 19 | 0.0526 | 6 | 0 | 6 | 0 | 4 | 0.25 | 3 | 0 | 19 | 0.0526 |
| 15_379245 | 2 | 0 | 5 | 0 | 4 | 0 | 7 | 0.1429 | 18 | 0.0556 | 6 | 0 | 4 | 0 | 4 | 0.25 | 4 | 0 | 18 | 0.0556 |
| 15_379261 | 1 | 0 | 2 | 0 | 3 | 0.3333 | 5 | 0 | 11 | 0.0909 | 2 | 0 | 4 | 0 | 1 | 0 | 2 | 0.5 | 9 | 0.1111 |
| 15_379276 | 2 | 0 | 1 | 0 | 1 | 1 | 2 | 0.5 | 6 | 0.3333 | 2 | 0.5 | 2 | 0.5 | NA | NA | 1 | 0 | 5 | 0.4 |
| 15_379447 | 2 | 0.5 | 1 | 0 | 1 | 1 | 2 | 0 | 6 | 0.3333 | 2 | 0 | 1 | 0 | 1 | 0 | 1 | 1 | 5 | 0.2 |

| | | | | | | | | | | | | | | | | | | | | |
|---|---|---|---|---|---|---|---|---|---|---|---|---|---|---|---|---|---|---|---|---|
| 15_379972 | 7 | 0.1429 | 4 | 0 | 9 | 0.1111 | 6 | 0 | 26 | 0.0769 | 3 | 0.3333 | 8 | 0.125 | 4 | 0.25 | 4 | 0 | 19 | 0.1579 |
| 15_382946 | 4 | 0 | 4 | 0.25 | 2 | 0 | 2 | 0 | 12 | 0.0833 | 2 | 0 | NA | NA | 1 | 0 | 3 | 0.3333 | 6 | 0.1667 |
| 15_382947 | 6 | 0.1667 | 7 | 0 | 2 | 0 | 4 | 0 | 19 | 0.0526 | 6 | 0 | 1 | 1 | 2 | 0 | 4 | 0 | 13 | 0.0769 |
| 15_382948 | 6 | 0.1667 | 4 | 0.25 | 3 | 0.6667 | 2 | 0.5 | 15 | 0.3333 | 5 | 0.2 | 1 | 1 | 2 | 0 | 4 | 0.5 | 12 | 0.3333 |
| 15_382949 | 4 | 0 | 5 | 0.2 | 3 | 0.3333 | 3 | 0.3333 | 15 | 0.2 | 5 | 0 | 1 | 1 | 2 | 0 | 4 | 0.25 | 12 | 0.1667 |
| 15_382950 | 6 | 0.5 | 5 | 0 | 2 | 0 | 3 | 0 | 16 | 0.1875 | 6 | 0.1667 | 1 | 0 | 2 | 0.5 | 4 | 0.5 | 13 | 0.3077 |
| 15_382951 | 5 | 0.4 | 6 | 0.1667 | 3 | 0.3333 | 3 | 0 | 17 | 0.2353 | 5 | 0.4 | 1 | 1 | 2 | 0 | 4 | 0.25 | 12 | 0.3333 |
| 15_382952 | 5 | 0.4 | 6 | 0.3333 | 3 | 0.3333 | 3 | 0.6667 | 17 | 0.4118 | 6 | 0.1667 | 1 | 0 | 2 | 0.5 | 4 | 0.5 | 13 | 0.3077 |
| 15_382958 | 4 | 1 | 5 | 0.6 | 1 | 1 | 3 | 1 | 13 | 0.8462 | 4 | 0.75 | NA | NA | 1 | 1 | 3 | 1 | 8 | 0.875 |
| 15_382961 | 2 | 1 | 5 | 0.8 | 1 | 1 | 2 | 1 | 10 | 0.9 | 3 | 1 | 1 | 1 | 1 | 0 | 3 | 1 | 8 | 0.875 |
| 15_382961 | 2 | 0 | 5 | 0.8 | 1 | 1 | 3 | 0.6667 | 11 | 0.6364 | 3 | 0.6667 | NA | NA | 1 | 1 | 3 | 0.6667 | 7 | 0.7143 |
| 15_382963 | 2 | 0.5 | 5 | 1 | 2 | 1 | 3 | 0.3333 | 12 | 0.75 | 2 | 1 | 1 | 1 | 1 | 1 | 1 | 1 | 5 | 1 |
| 15_382969 | 4 | 0 | 6 | 0.1667 | 2 | 0 | 3 | 0 | 15 | 0.0667 | 5 | 0.2 | 1 | 1 | 1 | 0 | 4 | 0 | 11 | 0.1818 |
| 15_382970 | 3 | 1 | 5 | 1 | 2 | 1 | 3 | 1 | 13 | 1 | 5 | 1 | 1 | 1 | NA | NA | 4 | 1 | 10 | 1 |
| 15_382973 | 4 | 1 | 6 | 1 | 2 | 1 | 3 | 1 | 15 | 1 | 5 | 1 | 1 | 1 | 1 | 1 | 3 | 1 | 10 | 1 |
| 15_382973 | 3 | 1 | 5 | 0.4 | 1 | 0 | 2 | 0 | 11 | 0.4545 | 5 | 0.6 | 1 | 1 | NA | NA | 4 | 0.75 | 10 | 0.7 |
| 15_382974 | 3 | 0 | 5 | 0.2 | 1 | 0 | 2 | 0 | 11 | 0.0909 | 2 | 0.5 | 1 | 0 | NA | NA | 4 | 0 | 7 | 0.1429 |
| 15_382974 | 4 | 0.75 | 6 | 0.8333 | 2 | 1 | 3 | 1 | 15 | 0.8667 | 3 | 0.6667 | 1 | 0 | 1 | 1 | 4 | 0.75 | 9 | 0.6667 |
| 15_384900 | 13 | 0.2308 | 17 | 0.1765 | 5 | 0.2 | 7 | 0.1429 | 42 | 0.1905 | 7 | 0.1429 | 10 | 0 | 5 | 0 | 19 | 0.1053 | 41 | 0.0732 |
| 15_386920 | 22 | 0.0909 | 38 | 0.1316 | 25 | 0.16 | 20 | 0.05 | 105 | 0.1143 | 18 | 0.0556 | 12 | 0 | 23 | 0.1739 | 31 | 0.1613 | 84 | 0.119 |
| 15_386926 | 18 | 0.1111 | 31 | 0.0645 | 29 | 0.1379 | 21 | 0.1429 | 99 | 0.1111 | 10 | 0.2 | 24 | 0.125 | 24 | 0.2083 | 27 | 0.1852 | 85 | 0.1765 |
| 15_388125 | 10 | 0.6 | 13 | 0.6923 | 17 | 0.5882 | 9 | 0.6667 | 49 | 0.6327 | 16 | 0.5625 | 11 | 0.5455 | 14 | 0.4286 | 18 | 0.5 | 59 | 0.5085 |
| 15_388126 | 9 | 0.3333 | 18 | 0.5 | 13 | 0.4615 | 12 | 0.5 | 52 | 0.4615 | 16 | 0.4375 | 13 | 0.6154 | 14 | 0.4286 | 18 | 0.6111 | 61 | 0.5246 |
| 15_388126 | 13 | 0.6154 | 18 | 0.7222 | 15 | 0.8 | 11 | 0.7273 | 57 | 0.7193 | 14 | 0.6429 | 13 | 0.8462 | 13 | 0.6154 | 14 | 0.9286 | 54 | 0.7593 |
| 15_388127 | 12 | 0.4167 | 17 | 0.5294 | 19 | 0.4211 | 9 | 0.6667 | 57 | 0.4912 | 16 | 0.25 | 14 | 0.1429 | 15 | 0.6 | 17 | 0.2353 | 62 | 0.3065 |
| 15_388128 | 11 | 0.2727 | 19 | 0.5263 | 17 | 0.3529 | 11 | 0.2727 | 58 | 0.3793 | 16 | 0.0625 | 15 | 0.3333 | 16 | 0.125 | 16 | 0.375 | 63 | 0.2222 |
| 15_388129 | 12 | 0.0833 | 19 | 0.2105 | 18 | 0.2778 | 12 | 0.0833 | 61 | 0.1803 | 16 | 0.0625 | 13 | 0.2308 | 17 | 0.2353 | 16 | 0.25 | 62 | 0.1935 |
| 15_388130 | 14 | 0.5714 | 21 | 0.5238 | 16 | 0.8125 | 14 | 0.8571 | 65 | 0.6769 | 16 | 0.5 | 15 | 0.6 | 17 | 0.7059 | 18 | 0.6667 | 66 | 0.6212 |
| 15_388132 | 14 | 0.2857 | 19 | 0.1579 | 17 | 0.1765 | 14 | 0.4286 | 64 | 0.25 | 15 | 0.2667 | 12 | 0.4167 | 16 | 0.3125 | 17 | 0.6471 | 60 | 0.4167 |
| 15_388136 | 15 | 0.4 | 21 | 0.6667 | 19 | 0.6842 | 13 | 0.5385 | 68 | 0.5882 | 16 | 0.25 | 16 | 0.75 | 16 | 0.8125 | 19 | 0.5789 | 67 | 0.597 |
| 15_388137 | 15 | 0 | 21 | 0.0952 | 19 | 0.1053 | 13 | 0.3846 | 68 | 0.1324 | 19 | 0.1053 | 15 | 0.1333 | 17 | 0 | 19 | 0.1053 | 70 | 0.0857 |
| 15_388138 | 17 | 0.0588 | 22 | 0.1364 | 17 | 0.2941 | 14 | 0.0714 | 70 | 0.1429 | 19 | 0.1579 | 16 | 0.125 | 18 | 0.2222 | 19 | 0.2632 | 72 | 0.1944 |
| 15_388138 | 14 | 0 | 21 | 0.3333 | 16 | 0.25 | 13 | 0.0769 | 64 | 0.1875 | 21 | 0.1905 | 16 | 0.3125 | 14 | 0.2143 | 19 | 0.2105 | 70 | 0.2286 |
| 15_388140 | 17 | 0.2353 | 25 | 0.2 | 19 | 0.2105 | 13 | 0.1538 | 74 | 0.2027 | 21 | 0.1429 | 17 | 0.1176 | 20 | 0.2 | 20 | 0.15 | 78 | 0.1538 |
| 15_388140 | 17 | 0.3529 | 22 | 0.2273 | 18 | 0.2778 | 12 | 0.3333 | 69 | 0.2899 | 22 | 0.1364 | 18 | 0.5 | 20 | 0.2 | 21 | 0.2857 | 81 | 0.2716 |
| 15_388145 | 20 | 0.05 | 27 | 0.037 | 24 | 0 | 12 | 0.3333 | 83 | 0.0723 | 21 | 0.0476 | 18 | 0.1667 | 15 | 0.1333 | 21 | 0.0476 | 75 | 0.0933 |
| 15_388145 | 21 | 0 | 21 | 0.0952 | 20 | 0.2 | 9 | 0.1111 | 71 | 0.0986 | 18 | 0.1111 | 18 | 0.0556 | 17 | 0.0588 | 21 | 0.1905 | 74 | 0.1081 |
| 15_388157 | 1 | 1 | 1 | 0 | NA | NA | NA | NA | 2 | 0.5 | 1 | 1 | 1 | 1 | 1 | 1 | 1 | 1 | 4 | 1 |
| 15_388158 | 1 | 1 | 2 | 0 | NA | NA | NA | NA | 3 | 0.3333 | 1 | 1 | 1 | 1 | 1 | 1 | NA | NA | 3 | 1 |
| 15_388159 | 1 | 0 | 2 | 0.5 | NA | NA | NA | NA | 3 | 0.3333 | 1 | 1 | NA | NA | 1 | 1 | 1 | 0 | 3 | 0.6667 |
| 15_388943 | 1 | 1 | NA | NA | 1 | 1 | NA | NA | 2 | 1 | NA | NA | 1 | 1 | NA | NA | NA | NA | 1 | 1 |
| 15_388943 | 1 | 1 | NA | NA | 1 | 1 | NA | NA | 2 | 1 | NA | NA | 1 | 1 | NA | NA | NA | NA | 1 | 1 |
| 15_388943 | 1 | 0 | NA | NA | 1 | 1 | NA | NA | 2 | 0.5 | NA | NA | 1 | 1 | NA | NA | NA | NA | 1 | 1 |
| 15_388985 | 1 | 1 | NA | NA | NA | NA | NA | NA | 1 | 1 | NA | NA | 1 | 1 | NA | NA | NA | NA | 1 | 1 |
| 15_389000 | NA | NA | NA | NA | 1 | 1 | NA | NA | 1 | 1 | NA | NA | 1 | 0 | NA | NA | 1 | 1 | 2 | 0.5 |
| 15_389502 | 6 | 0.5 | 2 | 0.5 | 1 | 1 | 2 | 1 | 11 | 0.6364 | 2 | 0.5 | 1 | 0 | NA | NA | 2 | 0 | 5 | 0.2 |
| 15_389589 | 1 | 0 | 1 | 1 | 1 | 1 | 1 | 0 | 4 | 0.5 | NA | NA | 1 | 0 | 1 | 0 | 1 | 1 | 3 | 0.3333 |
| 15_389590 | 1 | 0 | 1 | 0 | 1 | 1 | 1 | 1 | 4 | 0.5 | NA | NA | NA | NA | 1 | 1 | 1 | 1 | 2 | 1 |
| 15_389590 | 1 | 1 | 1 | 1 | 1 | 1 | NA | NA | 3 | 1 | NA | NA | NA | NA | NA | NA | 1 | 1 | 1 | 1 |
| 15_389592 | 1 | 1 | 1 | 1 | 1 | 1 | 1 | 1 | 4 | 1 | NA | NA | NA | NA | 1 | 1 | 1 | 1 | 2 | 1 |
| 15_393005 | 2 | 0 | 3 | 0 | 3 | 0.6667 | 2 | 0 | 10 | 0.2 | 1 | 0 | 3 | 0 | 2 | 0 | 1 | 1 | 7 | 0.1429 |
| 15_393006 | 1 | 1 | 4 | 0.75 | 3 | 1 | 2 | 0.5 | 10 | 0.8 | NA | NA | 2 | 0.5 | 2 | 0.5 | 1 | 1 | 5 | 0.6 |
| 15_393009 | 1 | 0 | 4 | 0.5 | 4 | 0.75 | 1 | 1 | 10 | 0.6 | 1 | 1 | 3 | 0.3333 | 3 | 0.6667 | 1 | 0 | 8 | 0.5 |
| 15_393009 | 2 | 0.5 | 5 | 0.6 | 5 | 0.2 | 2 | 0.5 | 14 | 0.4286 | 1 | 0 | 3 | 1 | 3 | 0.6667 | 1 | 0 | 8 | 0.625 |
| 15_393009 | 2 | 0.5 | 5 | 1 | 5 | 0.6 | 2 | 0.5 | 14 | 0.7143 | 1 | 0 | 3 | 1 | 3 | 0.3333 | 1 | 1 | 8 | 0.625 |
| 15_393010 | 2 | 0 | 4 | 0 | 5 | 0.4 | 1 | 0 | 12 | 0.1667 | 1 | 0 | 2 | 0 | 2 | 0.5 | NA | NA | 5 | 0.2 |
| 15_395450 | 5 | 0.8 | 7 | 0.8571 | 9 | 0.8889 | 5 | 1 | 26 | 0.8846 | 4 | 1 | 7 | 1 | 6 | 0.8333 | 3 | 1 | 20 | 0.95 |
| 15_395450 | 7 | 0 | 8 | 0 | 9 | 0 | 6 | 0.5 | 30 | 0.1 | 5 | 0.2 | 7 | 0 | 9 | 0.3333 | 3 | 0 | 24 | 0.1667 |
| 15_395451 | 7 | 0.7143 | 7 | 0.4286 | 9 | 0.2222 | 7 | 0.4286 | 30 | 0.4333 | 5 | 0.2 | 7 | 0.5714 | 9 | 0.5556 | 4 | 0.25 | 25 | 0.44 |
| 15_395451 | 7 | 1 | 5 | 0.8 | 8 | 0.875 | 6 | 1 | 26 | 0.9231 | 4 | 1 | 7 | 1 | 7 | 1 | 4 | 1 | 22 | 1 |
| 15_395453 | 5 | 1 | 8 | 0.75 | 9 | 0.6667 | 8 | 0.875 | 30 | 0.8 | 6 | 0.8333 | 8 | 0.75 | 9 | 0.6667 | 4 | 1 | 27 | 0.7778 |
| 15_395453 | 3 | 1 | 7 | 0.7143 | 10 | 0.5 | 7 | 0.5714 | 27 | 0.6296 | 6 | 0.3333 | 7 | 0.4286 | 9 | 1 | 5 | 0.6 | 27 | 0.6296 |
| 15_395455 | 6 | 0.3333 | 9 | 0.4444 | 11 | 0.4545 | 8 | 0.25 | 34 | 0.3824 | 7 | 0.5714 | 9 | 0.3333 | 7 | 0.5714 | 3 | 0 | 26 | 0.4231 |
| 15_395456 | 7 | 0.1429 | 9 | 0.1111 | 11 | 0.0909 | 7 | 0.1429 | 34 | 0.1176 | 6 | 0 | 8 | 0.125 | 9 | 0.2222 | 4 | 0 | 27 | 0.1111 |
| 15_395457 | 8 | 1 | 10 | 0.9 | 11 | 0.9091 | 7 | 1 | 36 | 0.9444 | 6 | 1 | 8 | 0.875 | 7 | 1 | 5 | 1 | 26 | 0.9615 |

| | | | | | | | | | | | | | | | | | | | | |
|---|---|---|---|---|---|---|---|---|---|---|---|---|---|---|---|---|---|---|---|---|
| 15_3954575 | 8 | 0.375 | 9 | 0.2222 | 9 | 0.3333 | 7 | 0.4286 | 33 | 0.3333 | 5 | 0.4 | 8 | 0.25 | 9 | 0.3333 | 6 | 0.5 | 28 | 0.3571 |
| 15_3954578 | 7 | 0.5714 | 8 | 0.25 | 9 | 0.3333 | 5 | 0.2 | 29 | 0.3448 | 5 | 0.2 | 8 | 0.375 | 9 | 0.1111 | 6 | 0.1667 | 28 | 0.2143 |
| 15_3954580 | 6 | 1 | 5 | 1 | 9 | 0.7778 | 7 | 1 | 27 | 0.9259 | 5 | 0.6 | 4 | 1 | 7 | 0.7143 | 3 | 1 | 19 | 0.7895 |
| 15_3954637 | 2 | 0.5 | 5 | 0.2 | 5 | 0 | 3 | 0.6667 | 15 | 0.2667 | NA | NA | 4 | 0.25 | 4 | 0 | 4 | 0 | 12 | 0.0833 |
| 15_3954723 | 3 | 0.3333 | 4 | 0 | 11 | 0.0909 | 4 | 0.5 | 22 | 0.1818 | 3 | 0 | 8 | 0 | 3 | 0 | 4 | 0.25 | 18 | 0.0556 |
| 15_3954728 | 4 | 0.25 | 4 | 0.25 | 8 | 0 | 4 | 0.25 | 20 | 0.15 | 5 | 0.2 | 8 | 0.25 | 4 | 0.5 | 6 | 0.1667 | 23 | 0.2609 |
| 15_3973457 | 5 | 0.2 | 3 | 0.6667 | 6 | 0.5 | 6 | 0.5 | 20 | 0.45 | 5 | 0.4 | 4 | 0.25 | 6 | 0.8333 | 4 | 0.25 | 19 | 0.4737 |
| 15_3973465 | 3 | 0.6667 | NA | NA | 4 | 0.5 | 4 | 0.75 | 11 | 0.6364 | 3 | 1 | 3 | 0.3333 | 3 | 1 | 3 | 0.6667 | 12 | 0.75 |
| 15_3973470 | 4 | 0.5 | 3 | 0.3333 | 6 | 0 | 7 | 0.2857 | 20 | 0.25 | 5 | 0 | 2 | 0 | 4 | 0.75 | 2 | 0 | 13 | 0.2308 |
| 15_3973477 | 3 | 0.6667 | 2 | 0.5 | 5 | 0.4 | 5 | 0.6 | 15 | 0.5333 | 4 | 0.5 | 5 | 0.2 | 7 | 0.1429 | 4 | 0.25 | 20 | 0.25 |
| 15_3973480 | 3 | 0.3333 | 2 | 0 | 5 | 0.2 | 4 | 0.25 | 14 | 0.2143 | 3 | 0 | 4 | 0 | 4 | 0.5 | 4 | 0 | 15 | 0.1333 |
| 15_3973489 | 2 | 1 | 2 | 1 | 6 | 1 | 4 | 0.75 | 14 | 0.9286 | 4 | 1 | 4 | 0.75 | 7 | 0.8571 | 3 | 0.6667 | 18 | 0.8333 |
| 15_3973503 | 2 | 0 | 2 | 0.5 | 5 | 0.2 | 4 | 0.5 | 13 | 0.3077 | 5 | 0 | 5 | 0.2 | 5 | 0.4 | 4 | 0 | 19 | 0.1579 |
| 15_3973535 | 1 | 0 | 1 | 0 | 6 | 0.1667 | 4 | 0 | 12 | 0.0833 | 3 | 0.3333 | 4 | 0 | 4 | 0 | 3 | 0.3333 | 14 | 0.1429 |
| 15_3973539 | 1 | 0 | 2 | 1 | 5 | 1 | 4 | 0.75 | 12 | 0.8333 | 5 | 0.8 | 4 | 1 | 4 | 0.75 | 4 | 1 | 17 | 0.8824 |
| 15_3973540 | 2 | 0.5 | 2 | 0.5 | 6 | 0.6667 | 2 | 1 | 12 | 0.6667 | 5 | 0.8 | 4 | 0.5 | 4 | 0.5 | 3 | 0.6667 | 16 | 0.625 |
| 15_3973547 | 1 | 1 | 2 | 1 | 6 | 1 | 6 | 1 | 15 | 1 | 3 | 1 | 5 | 0.8 | 4 | 1 | 2 | 0.5 | 14 | 0.8571 |
| 15_3973557 | 2 | 0 | 2 | 0 | 6 | 0 | 4 | 0.25 | 14 | 0.0714 | 3 | 0 | 2 | 0 | 3 | 0 | 1 | 1 | 9 | 0.1111 |
| 15_3973558 | 1 | 0 | 2 | 0 | 5 | 0.2 | 5 | 0.2 | 13 | 0.1538 | 3 | 0.3333 | 3 | 0 | 4 | 0.25 | 2 | 0 | 12 | 0.1667 |
| 15_3973562 | 2 | 1 | 2 | 1 | 3 | 1 | 2 | 1 | 9 | 1 | 1 | 1 | 2 | 1 | 3 | 0.6667 | 1 | 1 | 7 | 0.8571 |
| 15_3973563 | 2 | 1 | 2 | 1 | 6 | 1 | 4 | 1 | 14 | 1 | 4 | 1 | 2 | 1 | 4 | 1 | 2 | 1 | 12 | 1 |
| 15_3973570 | 2 | 0.5 | 1 | 1 | 6 | 0.6667 | 5 | 0.8 | 14 | 0.7143 | 4 | 0.25 | 3 | 1 | 3 | 0 | 2 | 0.5 | 12 | 0.4167 |
| 15_3973571 | 2 | 0 | 2 | 0.5 | 6 | 0.1667 | 5 | 0 | 15 | 0.1333 | 4 | 0.25 | 3 | 0 | 4 | 0.25 | 2 | 0 | 13 | 0.1538 |
| 15_3973579 | NA | NA | 1 | 0 | 3 | 0.6667 | 2 | 1 | 6 | 0.6667 | NA | NA | 2 | 0.5 | 4 | 0.5 | 2 | 1 | 8 | 0.625 |
| 15_3973602 | 2 | 0 | 1 | 0 | 6 | 0 | 4 | 0.5 | 13 | 0.1538 | 3 | 0.3333 | 4 | 0.25 | 4 | 0 | 2 | 0 | 13 | 0.1538 |
| 15_3973602 | 1 | 0 | 1 | 1 | 6 | 0.1667 | 5 | 0.4 | 13 | 0.3077 | 3 | 0.3333 | 4 | 0.25 | 4 | 0.25 | 2 | 0.5 | 13 | 0.3077 |
| 15_3973607 | 2 | 0.5 | 1 | 1 | 4 | 0.5 | 5 | 0.6 | 12 | 0.5833 | 3 | 0.6667 | 4 | 0.75 | 3 | 1 | 1 | 0 | 11 | 0.7273 |
| 15_3973954 | 2 | 1 | 1 | 0 | 3 | 0 | 4 | 0 | 10 | 0.2 | 2 | 0 | 3 | 0.3333 | 3 | 0 | 1 | 0 | 9 | 0.1111 |
| 15_3974132 | 1 | 0 | 1 | 1 | 4 | 0 | 3 | 0 | 9 | 0.1111 | 2 | 0 | 3 | 0 | 3 | 0.3333 | 1 | 0 | 9 | 0.1111 |
| 15_3974134 | 1 | 0 | 1 | 1 | 3 | 0.3333 | 3 | 0 | 8 | 0.25 | 2 | 0.5 | 2 | 0.5 | 3 | 0 | 1 | 1 | 8 | 0.375 |
| 15_3974143 | NA | NA | 1 | 0 | 3 | 0.3333 | 3 | 0.6667 | 7 | 0.4286 | 1 | 0 | 2 | 1 | 1 | 0 | 1 | 1 | 5 | 0.6 |
| 15_3974172 | 1 | 1 | 1 | 1 | 4 | 0.25 | 1 | 1 | 7 | 0.5714 | 2 | 0.5 | 3 | 0.3333 | 1 | 1 | 1 | 0 | 7 | 0.4286 |
| 15_3974177 | NA | NA | 1 | 0 | 3 | 0.3333 | 3 | 0.3333 | 7 | 0.2857 | 2 | 0.5 | 4 | 0 | 2 | 0.5 | 1 | 0 | 9 | 0.2222 |
| 15_3974181 | 1 | 1 | 1 | 1 | 3 | 1 | 1 | 1 | 6 | 1 | 2 | 1 | 4 | 1 | 2 | 1 | 1 | 1 | 9 | 1 |
| 15_3974186 | 1 | 0 | 1 | 1 | 4 | 0 | 2 | 1 | 8 | 0.375 | 1 | 1 | 2 | 0.5 | 2 | 0.5 | 1 | 1 | 6 | 0.6667 |
| 15_3974190 | 1 | 0 | 1 | 0 | 4 | 0.5 | 2 | 0 | 8 | 0.25 | 1 | 1 | 4 | 0.25 | 2 | 0 | 1 | 0 | 8 | 0.25 |
| 15_3974199 | 1 | 0 | 1 | 1 | 4 | 1 | 3 | 0.6667 | 9 | 0.7778 | 1 | 1 | 4 | 0.25 | 2 | 0 | 1 | 0 | 8 | 0.25 |
| 15_3974201 | 1 | 1 | 1 | 1 | 2 | 1 | 3 | 1 | 7 | 1 | 1 | 1 | 3 | 1 | 2 | 1 | 1 | 0 | 7 | 0.8571 |
| 15_3974204 | NA | NA | 1 | 1 | 3 | 1 | 2 | 0.5 | 6 | 0.8333 | 1 | 1 | 4 | 0.75 | 1 | 0 | NA | NA | 6 | 0.6667 |
| 15_3974204 | 1 | 1 | NA | NA | 3 | 0 | 3 | 0.3333 | 7 | 0.2857 | 1 | 0 | 2 | 0 | 1 | 0 | 1 | 1 | 5 | 0.2 |
| 15_4168340 | 4 | 0.25 | 8 | 0.625 | 7 | 0.4286 | 5 | 0.4 | 24 | 0.4583 | 3 | 0.3333 | 2 | 1 | 10 | 0.4 | 7 | 0.4286 | 22 | 0.4545 |
| 15_4168348 | 4 | 0.75 | 9 | 0.4444 | 8 | 0.875 | 4 | 0.25 | 25 | 0.6 | 4 | 0.25 | 3 | 0.6667 | 8 | 0.625 | 8 | 0.75 | 23 | 0.6087 |
| 15_4168358 | 4 | 0.25 | 4 | 0.5 | 6 | 0.5 | 3 | 0.6667 | 17 | 0.4706 | 2 | 0.5 | 2 | 0.5 | 5 | 0.4 | 5 | 0.8 | 14 | 0.5714 |
| 15_4168377 | 3 | 0 | 10 | 0.4 | 9 | 0.4444 | 6 | 0.6667 | 28 | 0.4286 | 2 | 0.5 | 4 | 0.75 | 8 | 0.625 | 11 | 0.5455 | 25 | 0.6 |
| 15_4168383 | 5 | 0.6 | 10 | 0.9 | 5 | 0.4 | 4 | 0.5 | 24 | 0.6667 | 4 | 0.5 | 3 | 0.6667 | 11 | 0.8182 | 13 | 0.8462 | 31 | 0.7742 |
| 15_4168383 | 4 | 0 | 12 | 0.25 | 9 | 0.2222 | 6 | 0.1667 | 31 | 0.1935 | 4 | 0.25 | 3 | 0 | 9 | 0.4444 | 12 | 0.25 | 28 | 0.2857 |
| 15_4168390 | 6 | 0 | 12 | 0.1667 | 9 | 0.2222 | 6 | 0 | 33 | 0.1212 | 4 | 0.25 | 4 | 0 | 12 | 0.0833 | 13 | 0.1538 | 33 | 0.1212 |
| 15_4172266 | 15 | 0.3333 | 23 | 0.2174 | 12 | 0.25 | 11 | 0.3636 | 61 | 0.2787 | 7 | 0 | 10 | 0.2 | 21 | 0.3333 | 14 | 0.2143 | 52 | 0.2308 |
| 15_4172269 | 13 | 0 | 22 | 0.0909 | 14 | 0.0714 | 8 | 0 | 57 | 0.0526 | 11 | 0 | 9 | 0.1111 | 18 | 0.1111 | 16 | 0.0625 | 54 | 0.0741 |
| 15_4172271 | 16 | 0.1875 | 21 | 0.1429 | 18 | 0.2778 | 11 | 0.2727 | 66 | 0.2121 | 11 | 0.0909 | 10 | 0.4 | 23 | 0.2174 | 19 | 0.2632 | 63 | 0.2381 |
| 15_4172273 | 17 | 0.8235 | 25 | 0.76 | 17 | 0.8824 | 11 | 0.8182 | 70 | 0.8143 | 10 | 0.7 | 11 | 0.9091 | 22 | 0.8182 | 21 | 0.9048 | 64 | 0.8438 |
| 15_4172428 | 28 | 0.0357 | 34 | 0.0294 | 21 | 0.1429 | 12 | 0.1667 | 95 | 0.0737 | 19 | 0.0526 | 15 | 0.0667 | 26 | 0 | 27 | 0.1111 | 87 | 0.0575 |
| 15_4228891 | NA | NA | 2 | 0 | 2 | 0.5 | 2 | 0.5 | 6 | 0.3333 | 1 | 1 | 1 | 0 | 1 | 0 | NA | NA | 3 | 0.3333 |
| 15_4228975 | NA | NA | 2 | 1 | 1 | 0 | NA | NA | 3 | 0.6667 | NA | NA | 1 | 1 | 1 | 1 | NA | NA | 2 | 1 |
| 15_4229010 | NA | NA | 2 | 1 | NA | NA | NA | NA | 2 | 1 | 1 | 1 | 1 | 1 | 1 | 1 | NA | NA | 3 | 1 |
| 15_4229015 | NA | NA | 2 | 0.5 | 1 | 1 | NA | NA | 3 | 0.6667 | 1 | 1 | 1 | 1 | 1 | 0 | NA | NA | 3 | 0.6667 |
| 15_4229037 | NA | NA | 2 | 1 | 1 | 1 | NA | NA | 3 | 1 | 1 | 1 | 1 | 1 | 1 | 1 | NA | NA | 3 | 1 |
| 15_4253985 | NA | NA | 1 | 1 | NA | NA | NA | NA | 1 | 1 | 1 | 1 | NA | NA | 1 | 1 | NA | NA | 2 | 1 |
| 15_4339092 | 8 | 0.25 | 6 | 0.3333 | 8 | 0.125 | 17 | 0.3529 | 39 | 0.2821 | 11 | 0.1818 | 10 | 0.1 | 15 | 0.3333 | 10 | 0.1 | 46 | 0.1957 |
| 15_4372714 | NA | NA | 1 | 1 | NA | NA | 1 | 0 | 2 | 0.5 | 1 | 1 | 1 | 0 | 1 | 0 | 1 | 0 | 4 | 0.25 |
| 15_4372932 | NA | NA | 1 | 1 | NA | NA | 1 | 1 | 2 | 1 | 1 | 1 | 1 | 1 | NA | NA | NA | NA | 2 | 1 |
| 15_4372992 | 1 | 1 | 1 | 1 | NA | NA | NA | NA | 2 | 1 | 1 | 0 | 2 | 0.5 | NA | NA | NA | NA | 3 | 0.3333 |
| 15_4373010 | NA | NA | 1 | 1 | NA | NA | 1 | 1 | 2 | 1 | 1 | 1 | 2 | 1 | NA | NA | NA | NA | 3 | 1 |
| 15_4373016 | NA | NA | 1 | 1 | NA | NA | 1 | 1 | 2 | 1 | 1 | 1 | 1 | 1 | NA | NA | NA | NA | 2 | 1 |
| 15_4373022 | NA | NA | 1 | 1 | NA | NA | 1 | 0 | 2 | 0.5 | 1 | 0 | 2 | 1 | NA | NA | NA | NA | 3 | 0.6667 |

| | | | | | | | | | | | | | | | | | | | | |
|---|---|---|---|---|---|---|---|---|---|---|---|---|---|---|---|---|---|---|---|---|
| 15_4373035 | NA | NA | NA | NA | NA | NA | 1 | 1 | 1 | 1 | 1 | 1 | 1 | 1 | NA | NA | NA | NA | 2 | 1 |
| 15_4373043 | NA | NA | 1 | 1 | NA | NA | NA | NA | 1 | 1 | 1 | 0 | 2 | 0.5 | NA | NA | NA | NA | 3 | 0.3333 |
| 15_4373051 | NA | NA | NA | NA | NA | NA | 1 | 1 | 1 | 1 | 1 | 1 | 2 | 1 | NA | NA | NA | NA | 3 | 1 |
| 15_4373059 | NA | NA | 1 | 0 | NA | NA | 1 | 1 | 2 | 0.5 | 1 | 0 | 2 | 0.5 | NA | NA | NA | NA | 3 | 0.3333 |
| 15_4373072 | NA | NA | 1 | 0 | NA | NA | 1 | 1 | 2 | 0.5 | 1 | 1 | 1 | 1 | NA | NA | NA | NA | 2 | 1 |
| 15_4373079 | NA | NA | 1 | 1 | NA | NA | 1 | 1 | 2 | 1 | 1 | 0 | 2 | 0.5 | NA | NA | NA | NA | 3 | 0.3333 |
| 15_4373096 | NA | NA | 1 | 1 | NA | NA | 1 | 1 | 2 | 1 | 1 | 0 | 2 | 1 | NA | NA | NA | NA | 3 | 0.6667 |
| 15_4373100 | NA | NA | 1 | 1 | NA | NA | 1 | 1 | 2 | 1 | 1 | 1 | 2 | 0.5 | NA | NA | NA | NA | 3 | 0.6667 |
| 15_4373101 | NA | NA | NA | NA | NA | NA | 1 | 1 | 1 | 1 | NA | NA | 2 | 0.5 | NA | NA | NA | NA | 2 | 0.5 |
| 15_4373120 | NA | NA | 1 | 1 | NA | NA | NA | NA | 1 | 1 | 1 | 0 | 1 | 1 | NA | NA | NA | NA | 2 | 0.5 |
| 15_4406059 | NA | NA | NA | NA | 1 | 1 | NA | NA | 1 | 1 | NA | NA | 1 | 1 | 2 | 1 | NA | NA | 3 | 1 |
| 15_4406066 | NA | NA | NA | NA | 1 | 1 | NA | NA | 1 | 1 | NA | NA | 2 | 1 | 2 | 0.5 | NA | NA | 4 | 0.75 |
| 15_4406100 | NA | NA | NA | NA | 1 | 1 | NA | NA | 1 | 1 | NA | NA | 3 | 1 | 2 | 1 | NA | NA | 5 | 1 |
| 15_4409261 | NA | NA | NA | NA | NA | NA | 1 | 1 | 1 | 1 | NA | NA | 3 | 0.6667 | 3 | 1 | NA | NA | 6 | 0.8333 |
| 15_4409700 | NA | NA | NA | NA | 1 | 1 | 1 | 1 | 2 | 1 | NA | NA | 3 | 1 | 4 | 1 | 1 | 1 | 8 | 1 |
| 15_4409705 | NA | NA | NA | NA | NA | NA | 1 | 1 | 1 | 1 | NA | NA | 2 | 0 | 4 | 0.5 | 1 | 1 | 7 | 0.4286 |
| 15_4409732 | NA | NA | NA | NA | 1 | 0 | 1 | 1 | 2 | 0.5 | NA | NA | 4 | 0 | 4 | 0.25 | 1 | 0 | 9 | 0.1111 |
| 15_4409747 | NA | NA | NA | NA | 1 | 1 | 1 | 1 | 2 | 1 | NA | NA | 4 | 0.75 | 3 | 1 | 1 | 0 | 8 | 0.75 |
| 15_4429929 | 5 | 0 | 4 | 0.25 | 7 | 0.2857 | 6 | 0.1667 | 22 | 0.1818 | 3 | 0 | 5 | 0 | 3 | 0.3333 | 3 | 0.3333 | 14 | 0.1429 |
| 15_4429952 | 6 | 0.3333 | 5 | 0 | 7 | 0.1429 | 5 | 0.2 | 23 | 0.1739 | 2 | 0 | 5 | 0.4 | 2 | 0.5 | 3 | 0 | 12 | 0.25 |
| 15_4429960 | 8 | 0.25 | 4 | 0 | 5 | 0 | 5 | 0 | 22 | 0.0909 | 3 | 0 | 5 | 0 | 3 | 0.3333 | 3 | 0 | 14 | 0.0714 |
| 15_4429963 | 8 | 0.625 | 5 | 0.4 | 7 | 0.7143 | 4 | 0.25 | 24 | 0.5417 | NA | NA | 5 | 0.4 | 3 | 0.6667 | 2 | 0.5 | 10 | 0.5 |
| 15_4429968 | 7 | 0.5714 | 4 | 0.75 | 7 | 1 | 6 | 0.8333 | 24 | 0.7917 | 3 | 0.6667 | 4 | 0.75 | 1 | 1 | 2 | 1 | 10 | 0.8 |
| 15_4429975 | 8 | 1 | 5 | 1 | 6 | 1 | 6 | 1 | 25 | 1 | 2 | 1 | 5 | 1 | 3 | 1 | 3 | 1 | 13 | 1 |
| 15_4429982 | 8 | 0.625 | 5 | 0.4 | 5 | 0.8 | 5 | 0.8 | 23 | 0.6522 | 3 | 0.6667 | 5 | 0.8 | 2 | 0.5 | 3 | 0.6667 | 13 | 0.6923 |
| 15_4429992 | 5 | 0.2 | 2 | 0.5 | 4 | 0.25 | 3 | 0.3333 | 14 | 0.2857 | 3 | 0.6667 | 4 | 0.5 | 2 | 0.5 | 3 | 0.3333 | 12 | 0.5 |
| 15_4450389 | 6 | 1 | 6 | 0.8333 | 5 | 1 | 5 | 0.8 | 22 | 0.9091 | 4 | 1 | 3 | 1 | 5 | 1 | 3 | 1 | 15 | 1 |
| 15_4450395 | 8 | 0.5 | 6 | 0 | 6 | 0.5 | 4 | 0.5 | 24 | 0.375 | 4 | 0 | 3 | 0.3333 | 6 | 0.6667 | 3 | 0.3333 | 16 | 0.375 |
| 15_4450398 | 8 | 0.875 | 4 | 1 | 6 | 1 | 5 | 0.6 | 23 | 0.8696 | 3 | 1 | 3 | 1 | 4 | 1 | 4 | 1 | 14 | 1 |
| 15_4450400 | 7 | 0.7143 | 4 | 0.75 | 6 | 0.8333 | 5 | 0.8 | 22 | 0.7727 | 3 | 1 | 3 | 1 | 5 | 0.6 | 4 | 0.75 | 15 | 0.8 |
| 15_4450402 | 7 | 0.5714 | 5 | 0.2 | 6 | 0.8333 | 5 | 0.2 | 23 | 0.4783 | 3 | 0.3333 | 3 | 0.6667 | 5 | 0.6 | 5 | 0.4 | 16 | 0.5 |
| 15_4453549 | 10 | 0.1 | 18 | 0.1111 | 10 | 0.1 | 14 | 0.2143 | 52 | 0.1346 | 11 | 0.3636 | 10 | 0.1 | 8 | 0.125 | 5 | 0 | 34 | 0.1765 |
| 15_4453553 | 10 | 0.2 | 15 | 0.2 | 9 | 0.3333 | 13 | 0.2308 | 47 | 0.234 | 12 | 0.5 | 7 | 0.5714 | 6 | 0.3333 | 5 | 0 | 30 | 0.4 |
| 15_4453617 | 10 | 1 | 8 | 1 | 7 | 1 | 9 | 1 | 34 | 1 | 9 | 1 | 4 | 1 | 5 | 1 | 3 | 1 | 21 | 1 |
| 15_4453635 | 8 | 0.125 | 10 | 0.4 | 8 | 0.625 | 10 | 0.3 | 36 | 0.3611 | 9 | 0.4444 | 5 | 0.8 | 6 | 0.3333 | 5 | 0.4 | 25 | 0.48 |
| 15_4453640 | 8 | 0.75 | 10 | 1 | 9 | 1 | 11 | 0.6364 | 38 | 0.8421 | 10 | 0.9 | 6 | 0.8333 | 6 | 1 | 4 | 1 | 26 | 0.9231 |
| 15_4453659 | 7 | 0.5714 | 8 | 0.75 | 5 | 0.6 | 10 | 0.4 | 30 | 0.5667 | 7 | 0.4286 | 6 | 0.6667 | 4 | 1 | 3 | 0.6667 | 20 | 0.65 |
| 15_4453667 | 6 | 0.6667 | 8 | 0.5 | 6 | 0.1667 | 9 | 0.3333 | 29 | 0.4138 | 6 | 0.6667 | 3 | 0.3333 | 4 | 0.25 | 2 | 0.5 | 15 | 0.4667 |
| 15_4453675 | 7 | 1 | 10 | 1 | 7 | 1 | 11 | 0.9091 | 35 | 0.9714 | 7 | 0.8571 | 5 | 1 | 5 | 1 | 3 | 1 | 20 | 0.95 |
| 15_4453676 | 5 | 1 | 7 | 1 | 7 | 1 | 8 | 1 | 27 | 1 | 6 | 1 | 5 | 1 | 5 | 1 | 3 | 1 | 19 | 1 |
| 15_4453682 | 7 | 0.4286 | 7 | 0.5714 | 5 | 0.6 | 9 | 0.4444 | 28 | 0.5 | 5 | 0 | 6 | 0.5 | 4 | 0.75 | 3 | 0.6667 | 18 | 0.4444 |
| 15_4453684 | 9 | 0.7778 | 11 | 0.7273 | 7 | 1 | 9 | 0.8889 | 36 | 0.8333 | 7 | 0.5714 | 6 | 0.6667 | 5 | 1 | 2 | 0.5 | 20 | 0.7 |
| 15_4453690 | 8 | 0.25 | 11 | 0 | 6 | 0 | 10 | 0 | 35 | 0.0571 | 6 | 0.1667 | 5 | 0 | 4 | 0.25 | 2 | 0 | 17 | 0.1176 |
| 15_4453699 | 6 | 0.8333 | 7 | 0.5714 | 5 | 0.8 | 10 | 0.6 | 28 | 0.6786 | 7 | 0.7143 | 6 | 0.5 | 6 | 0.6667 | 2 | 0.5 | 21 | 0.619 |
| 15_4466894 | 1 | 0 | 4 | 0.25 | NA | NA | NA | NA | 5 | 0.2 | 2 | 0 | 1 | 1 | 1 | 0 | 1 | 1 | 5 | 0.4 |
| 15_4466909 | 2 | 1 | 4 | 0.75 | 1 | 0 | NA | NA | 7 | 0.7143 | 2 | 1 | 1 | 1 | 1 | 1 | 3 | 0.6667 | 7 | 0.8571 |
| 15_4466912 | 1 | 1 | 3 | 0.6667 | 1 | 0 | NA | NA | 5 | 0.6 | 2 | 0.5 | NA | NA | NA | NA | 3 | 0.3333 | 5 | 0.4 |
| 15_4474484 | NA | NA | 1 | 1 | NA | NA | 1 | 0 | 2 | 0.5 | 1 | 1 | NA | NA | NA | NA | 1 | 1 | 2 | 1 |
| 15_4474509 | NA | NA | 1 | 1 | NA | NA | NA | NA | 1 | 1 | 1 | 1 | NA | NA | NA | NA | 1 | 1 | 2 | 1 |
| 15_4474522 | NA | NA | 1 | 1 | NA | NA | 2 | 0.5 | 3 | 0.6667 | 1 | 1 | NA | NA | NA | NA | 1 | 0 | 2 | 0.5 |
| 15_4474533 | NA | NA | 1 | 0 | NA | NA | 2 | 0.5 | 3 | 0.3333 | 1 | 1 | NA | NA | NA | NA | 1 | 0 | 2 | 0.5 |
| 15_4474557 | NA | NA | 1 | 1 | NA | NA | 3 | 1 | 4 | 1 | 1 | 1 | NA | NA | NA | NA | 1 | 0 | 2 | 0.5 |
| 15_4474559 | NA | NA | 1 | 0 | NA | NA | 2 | 0.5 | 3 | 0.3333 | 1 | 1 | NA | NA | NA | NA | 1 | 1 | 2 | 1 |
| 15_4474580 | NA | NA | 1 | 0 | NA | NA | 3 | 0.6667 | 4 | 0.5 | 1 | 1 | NA | NA | NA | NA | 1 | 1 | 2 | 1 |
| 15_4474613 | NA | NA | 1 | 1 | NA | NA | 3 | 0.3333 | 4 | 0.5 | 1 | 1 | NA | NA | NA | NA | 1 | 0 | 2 | 0.5 |
| 15_4474661 | NA | NA | 1 | 1 | NA | NA | 3 | 1 | 4 | 1 | 1 | 1 | NA | NA | NA | NA | NA | NA | 1 | 1 |
| 15_4474663 | NA | NA | NA | NA | NA | NA | 2 | 0.5 | 2 | 0.5 | 1 | 1 | NA | NA | NA | NA | 1 | 1 | 2 | 1 |
| 15_4474672 | NA | NA | 1 | 0 | NA | NA | 3 | 1 | 4 | 0.75 | 1 | 1 | NA | NA | NA | NA | 1 | 1 | 2 | 1 |
| 15_4474681 | NA | NA | 1 | 1 | NA | NA | 3 | 1 | 4 | 1 | 1 | 1 | NA | NA | NA | NA | 1 | 0 | 2 | 0.5 |
| 15_4474686 | NA | NA | 1 | 1 | NA | NA | 2 | 1 | 3 | 1 | 1 | 1 | NA | NA | NA | NA | NA | NA | 1 | 1 |
| 15_4474701 | NA | NA | 1 | 1 | NA | NA | 3 | 1 | 4 | 1 | 1 | 1 | NA | NA | NA | NA | 1 | 1 | 2 | 1 |
| 15_4474703 | NA | NA | 1 | 1 | NA | NA | 3 | 0.6667 | 4 | 0.75 | NA | NA | NA | NA | NA | NA | 1 | 1 | 1 | 1 |
| 15_4474713 | NA | NA | 1 | 1 | NA | NA | 2 | 1 | 3 | 1 | 1 | 1 | NA | NA | NA | NA | 1 | 0 | 2 | 0.5 |
| 15_4474718 | NA | NA | 1 | 1 | NA | NA | 3 | 1 | 4 | 1 | 1 | 1 | NA | NA | NA | NA | 1 | 1 | 2 | 1 |
| 15_4474720 | NA | NA | 1 | 1 | NA | NA | 2 | 1 | 3 | 1 | NA | NA | NA | NA | NA | NA | 1 | 1 | 1 | 1 |

| | | | | | | | | | | | | | | | | | | | | |
|---|---|---|---|---|---|---|---|---|---|---|---|---|---|---|---|---|---|---|---|---|
| 15_447472 | NA | NA | 1 | 0 | NA | NA | 3 | 0.3333 | 4 | 0.25 | NA | NA | NA | NA | NA | NA | 1 | 1 | 1 | 1 |
| 15_447474 | NA | NA | 1 | 0 | 1 | 0 | 3 | 0.3333 | 5 | 0.2 | 1 | 1 | NA | NA | NA | NA | 1 | 0 | 2 | 0.5 |
| 15_447474 | NA | NA | 1 | 1 | 1 | 1 | 3 | 0.3333 | 5 | 0.6 | NA | NA | NA | NA | NA | NA | 1 | 1 | 1 | 1 |
| 15_447475 | NA | NA | 1 | 1 | 1 | 1 | 2 | 1 | 4 | 1 | 1 | 1 | NA | NA | NA | NA | 1 | 1 | 2 | 1 |
| 15_447475 | NA | NA | 1 | 1 | 1 | 0 | 1 | 1 | 3 | 0.6667 | 1 | 1 | NA | NA | NA | NA | 1 | 1 | 2 | 1 |
| 15_447476 | NA | NA | 1 | 1 | 1 | 1 | 2 | 1 | 4 | 1 | 1 | 1 | NA | NA | NA | NA | 1 | 1 | 2 | 1 |
| 15_449597 | NA | NA | 3 | 0.6667 | NA | NA | NA | NA | 3 | 0.6667 | NA | NA | 1 | 1 | NA | NA | 1 | 0 | 2 | 0.5 |
| 15_449692 | 2 | 0 | 7 | 0.1429 | 4 | 0 | 2 | 0 | 15 | 0.0667 | 3 | 0 | 5 | 0.4 | 1 | 0 | 2 | 0 | 11 | 0.1818 |
| 15_449693 | 2 | 0 | 8 | 0.125 | 3 | 0 | 2 | 0 | 15 | 0.0667 | 2 | 0 | 5 | 0 | 3 | 0.3333 | 2 | 0 | 12 | 0.0833 |
| 15_449694 | 2 | 0 | 7 | 0.2857 | 4 | 0.25 | 1 | 0 | 14 | 0.2143 | 2 | 0.5 | 3 | 0.6667 | 3 | 0.3333 | 2 | 0.5 | 10 | 0.5 |
| 15_449697 | 5 | 0.4 | 8 | 0.375 | 4 | 0 | 3 | 0.3333 | 20 | 0.3 | 2 | 0 | 4 | 0.25 | 2 | 0 | 4 | 0.25 | 12 | 0.1667 |
| 15_449698 | 5 | 0.8 | 9 | 0.7778 | 5 | 1 | 5 | 0.6 | 24 | 0.7917 | 2 | 1 | 4 | 1 | 2 | 0.5 | 4 | 0.25 | 12 | 0.6667 |
| 15_449699 | 4 | 0 | 9 | 0.1111 | 5 | 0 | 5 | 0.4 | 23 | 0.1304 | 2 | 0 | 4 | 0.5 | 3 | 0.3333 | 4 | 0 | 13 | 0.2308 |
| 15_449700 | 3 | 0 | 7 | 0.2857 | 5 | 0 | 5 | 0 | 20 | 0.1 | 1 | 0 | 4 | 0 | 3 | 0 | 4 | 0.25 | 12 | 0.0833 |
| 15_449709 | 2 | 0 | 6 | 0.1667 | 3 | 0 | 2 | 0 | 13 | 0.0769 | 1 | 0 | 3 | 0 | 1 | 1 | 3 | 0 | 8 | 0.125 |
| 15_449714 | 3 | 0.3333 | 8 | 0.125 | 6 | 0 | 4 | 0 | 21 | 0.0952 | 3 | 0.3333 | 5 | 0 | 6 | 0.1667 | 4 | 0 | 18 | 0.1111 |
| 15_449722 | 4 | 0 | 9 | 0.2222 | 8 | 0.25 | 7 | 0.1429 | 28 | 0.1786 | 2 | 0.5 | 7 | 0.1429 | 3 | 0 | 5 | 0.4 | 17 | 0.2353 |
| 15_449724 | 4 | 1 | 7 | 0.8571 | 4 | 1 | 6 | 0.5 | 21 | 0.8095 | 2 | 1 | 6 | 0.8333 | 3 | 1 | 5 | 0.8 | 16 | 0.875 |
| 15_449725 | 5 | 0.4 | 9 | 0.2222 | 8 | 0.25 | 7 | 0 | 29 | 0.2069 | 2 | 0.5 | 7 | 0.2857 | 3 | 0 | 6 | 0.1667 | 18 | 0.2222 |
| 15_449725 | 5 | 0 | 9 | 0 | 8 | 0.25 | 6 | 0.1667 | 28 | 0.1071 | 2 | 0 | 5 | 0 | 3 | 0.3333 | 5 | 0.2 | 15 | 0.1333 |
| 15_449725 | 4 | 0.25 | 6 | 0 | 7 | 0.1429 | 7 | 0 | 24 | 0.0833 | 2 | 0 | 7 | 0.1429 | 2 | 0 | 4 | 0.25 | 15 | 0.1333 |
| 15_449726 | 1 | 0 | 6 | 0.3333 | 4 | 0.25 | 6 | 0.3333 | 17 | 0.2941 | 2 | 0.5 | 4 | 0.25 | 2 | 0 | 4 | 0.5 | 12 | 0.3333 |
| 15_449726 | 2 | 0.5 | 8 | 0.25 | 6 | 0.1667 | 6 | 0.1667 | 22 | 0.2273 | 2 | 0 | 6 | 0 | 3 | 0 | 6 | 0.1667 | 17 | 0.0588 |
| 15_449726 | 2 | 1 | 7 | 0.8571 | 6 | 1 | 5 | 1 | 20 | 0.95 | 2 | 1 | 7 | 0.8571 | 3 | 1 | 6 | 1 | 18 | 0.9444 |
| 15_449727 | 3 | 0.6667 | 8 | 0.5 | 5 | 0.6 | 5 | 0.2 | 21 | 0.4762 | 2 | 0.5 | 6 | 0.3333 | 2 | 1 | 6 | 0.8333 | 16 | 0.625 |
| 15_449763 | 2 | 0 | 8 | 0 | 3 | 0 | 7 | 0.1429 | 20 | 0.05 | 1 | 0 | 6 | 0.1667 | 2 | 0 | 6 | 0 | 15 | 0.0667 |
| 15_449764 | NA | NA | 5 | 0.2 | 1 | 0 | 3 | 0.6667 | 9 | 0.3333 | 1 | 0 | 4 | 0.25 | 2 | 0 | 2 | 0 | 9 | 0.1111 |
| 15_449764 | 2 | 0.5 | 8 | 0.5 | 4 | 0.5 | 5 | 0.4 | 19 | 0.4737 | 1 | 0 | 7 | 0.8571 | 2 | 0.5 | 5 | 0.6 | 15 | 0.6667 |
| 15_449764 | 3 | 0.6667 | 7 | 0.8571 | 3 | 0.6667 | 7 | 0.2857 | 20 | 0.6 | 1 | 1 | 6 | 0.6667 | 2 | 0.5 | 5 | 0.4 | 14 | 0.5714 |
| 15_449767 | 2 | 1 | 8 | 0.875 | 4 | 1 | 6 | 1 | 20 | 0.95 | 1 | 1 | 5 | 0.8 | 2 | 1 | 5 | 0.8 | 13 | 0.8462 |
| 15_449767 | 1 | 1 | 7 | 1 | 4 | 1 | 4 | 1 | 16 | 1 | 1 | 1 | 5 | 0.8 | 2 | 1 | 5 | 1 | 13 | 0.9231 |
| 15_449767 | 2 | 0 | 6 | 0.1667 | 2 | 0.5 | 5 | 0.2 | 15 | 0.2 | 1 | 0 | 6 | 0.6667 | 2 | 0 | 5 | 0.2 | 14 | 0.3571 |
| 15_449777 | 3 | 0.3333 | 8 | 0.125 | 2 | 0 | 6 | 0.1667 | 19 | 0.1579 | 1 | 1 | 6 | 0.5 | 1 | 0 | 5 | 0 | 13 | 0.3077 |
| 15_449779 | 2 | 0 | 3 | 1 | 2 | 1 | 3 | 0.6667 | 10 | 0.7 | 1 | 1 | 5 | 0.6 | 1 | 1 | 5 | 0.4 | 12 | 0.5833 |
| 15_449779 | 2 | 0.5 | 4 | 0 | 2 | 0.5 | 6 | 0.6667 | 14 | 0.4286 | 1 | 0 | 3 | 0.3333 | NA | NA | 3 | 0 | 7 | 0.1429 |
| 15_449780 | 1 | 0 | NA | NA | 1 | 0 | 4 | 0.25 | 6 | 0.1667 | NA | NA | 3 | 0 | NA | NA | 4 | 0.25 | 7 | 0.1429 |
| 15_474455 | 3 | 0 | 1 | 0 | NA | NA | 3 | 0.6667 | 7 | 0.2857 | NA | NA | 4 | 0.5 | 2 | 0 | 2 | 0.5 | 8 | 0.375 |
| 15_474456 | 1 | 0 | NA | NA | 1 | 1 | 1 | 1 | 3 | 0.6667 | NA | NA | 3 | 0.6667 | 1 | 0 | 1 | 0 | 5 | 0.4 |
| 15_474457 | 2 | 1 | 1 | 1 | 1 | 1 | 3 | 1 | 7 | 1 | NA | NA | 4 | 1 | 2 | 1 | 2 | 1 | 8 | 1 |
| 15_474457 | 2 | 0 | 1 | 1 | NA | NA | 3 | 0.3333 | 6 | 0.3333 | NA | NA | 4 | 0.25 | NA | NA | 2 | 1 | 6 | 0.5 |
| 15_474460 | 1 | 0 | 1 | 1 | 1 | 0 | 2 | 0.5 | 5 | 0.4 | NA | NA | 2 | 1 | 1 | 0 | 1 | 1 | 4 | 0.75 |
| 15_505187 | 2 | 0 | 2 | 0 | 2 | 0.5 | 1 | 0 | 7 | 0.1429 | 2 | 0 | 6 | 0 | 3 | 0 | 4 | 0.25 | 15 | 0.0667 |
| 15_505209 | NA | NA | NA | NA | 1 | 1 | NA | NA | 1 | 1 | NA | NA | 3 | 0.3333 | 3 | 0 | NA | NA | 6 | 0.1667 |
| 15_505211 | NA | NA | NA | NA | 1 | 1 | 1 | 1 | 2 | 1 | NA | NA | 3 | 0.6667 | 3 | 1 | 1 | 0 | 7 | 0.7143 |
| 15_505214 | NA | NA | NA | NA | 1 | 1 | 1 | 1 | 2 | 1 | NA | NA | NA | NA | 3 | 0.6667 | NA | NA | 3 | 0.6667 |
| 15_505217 | 1 | 1 | NA | NA | 1 | 1 | 1 | 1 | 3 | 1 | NA | NA | 1 | 0 | 3 | 0.3333 | 3 | 0.6667 | 7 | 0.4286 |
| 15_505219 | 1 | 0 | NA | NA | 1 | 1 | 1 | 1 | 3 | 0.6667 | 1 | 0 | 2 | 0.5 | 3 | 0.3333 | 3 | 0.3333 | 9 | 0.3333 |
| 15_505227 | 1 | 1 | NA | NA | 1 | 1 | 1 | 1 | 3 | 1 | 1 | 1 | 1 | 0 | 1 | 1 | 2 | 0 | 5 | 0.4 |
| 15_505241 | NA | NA | NA | NA | 1 | 1 | NA | NA | 1 | 1 | 2 | 1 | 1 | 1 | NA | NA | 1 | 1 | 4 | 1 |
| 15_505241 | NA | NA | NA | NA | 1 | 1 | NA | NA | 1 | 1 | 2 | 1 | 1 | 1 | NA | NA | NA | NA | 3 | 1 |
| 15_505244 | NA | NA | NA | NA | 1 | 1 | NA | NA | 1 | 1 | 2 | 0.5 | 1 | 1 | NA | NA | 1 | 1 | 4 | 0.75 |
| 15_505246 | NA | NA | NA | NA | 1 | 1 | NA | NA | 1 | 1 | 2 | 0.5 | 1 | 1 | NA | NA | 1 | 1 | 4 | 0.75 |
| 15_516497 | NA | NA | NA | NA | NA | NA | 1 | 1 | 1 | 1 | NA | NA | 1 | 1 | NA | NA | NA | NA | 1 | 1 |
| 15_516500 | 30 | 0.4667 | 33 | 0.3636 | 43 | 0.5581 | 30 | 0.4333 | 136 | 0.4632 | 36 | 0.4167 | 36 | 0.5833 | 33 | 0.4848 | 22 | 0.6364 | 127 | 0.5197 |
| 15_517456 | 1 | 0 | 2 | 0 | 2 | 0.5 | NA | NA | 5 | 0.2 | 1 | 1 | NA | NA | 1 | 0 | NA | NA | 2 | 0.5 |
| 15_517459 | 1 | 0 | 1 | 0 | 2 | 0.5 | NA | NA | 4 | 0.25 | 1 | 0 | NA | NA | 2 | 0.5 | 1 | 1 | 4 | 0.5 |
| 15_517460 | 2 | 1 | 2 | 1 | 2 | 1 | NA | NA | 6 | 1 | 1 | 1 | NA | NA | 1 | 1 | 1 | 1 | 3 | 1 |
| 15_518266 | 15 | 0 | 20 | 0.15 | 14 | 0.0714 | 12 | 0.0833 | 61 | 0.082 | 14 | 0.0714 | 8 | 0.5 | 12 | 0 | 13 | 0.1538 | 47 | 0.1489 |
| 15_518266 | 15 | 0.3333 | 17 | 0.4706 | 13 | 0.3846 | 14 | 0.2857 | 59 | 0.3729 | 16 | 0.25 | 9 | 0.5556 | 15 | 0 | 14 | 0.3571 | 54 | 0.2593 |
| 15_518268 | 15 | 0.0667 | 21 | 0.2381 | 15 | 0 | 14 | 0.0714 | 65 | 0.1077 | 16 | 0.0625 | 7 | 0 | 15 | 0.0667 | 14 | 0.2143 | 52 | 0.0962 |
| 15_518268 | 15 | 0.1333 | 21 | 0.3333 | 13 | 0.2308 | 12 | 0.0833 | 61 | 0.2131 | 15 | 0.0667 | 6 | 0.1667 | 15 | 0.0667 | 13 | 0.2308 | 49 | 0.1224 |
| 15_518270 | 15 | 0.7333 | 20 | 0.65 | 9 | 0.5556 | 9 | 0.3333 | 53 | 0.6038 | 15 | 0.6 | 6 | 0.6667 | 12 | 0.5 | 12 | 0.8333 | 45 | 0.6444 |
| 15_518272 | 14 | 0 | 19 | 0.1053 | 13 | 0.1538 | 13 | 0.3846 | 59 | 0.1525 | 16 | 0.125 | 8 | 0.375 | 15 | 0.2 | 13 | 0.0769 | 52 | 0.1731 |
| 15_519084 | NA | NA | 3 | 0.6667 | NA | NA | 1 | 1 | 4 | 0.75 | 1 | 1 | NA | NA | NA | NA | 1 | 1 | 2 | 1 |
| 15_525935 | 8 | 0.125 | 6 | 0 | 2 | 0.5 | 7 | 0.1429 | 23 | 0.1304 | 6 | 0.1667 | 3 | 0 | 3 | 0.3333 | 6 | 0 | 18 | 0.1111 |

| | | | | | | | | | | | | | | | | | | | | |
|---|---|---|---|---|---|---|---|---|---|---|---|---|---|---|---|---|---|---|---|---|
| 15_5259355 | 7 | 0 | 6 | 0 | 2 | 0.5 | 5 | 0 | 20 | 0.05 | 7 | 0 | 5 | 0.2 | 2 | 0 | 6 | 0 | 20 | 0.05 |
| 15_5292890 | 1 | 0 | NA | NA | 1 | 1 | NA | NA | 2 | 0.5 | NA | NA | 1 | 0 | 1 | 1 | 1 | 1 | 3 | 0.6667 |
| 15_5293190 | 8 | 0.625 | 6 | 0.5 | 7 | 0.4286 | 3 | 0.3333 | 24 | 0.5 | 4 | 0.5 | 4 | 0.25 | 2 | 1 | 1 | 0 | 11 | 0.4545 |
| 15_5293192 | 4 | 0.25 | 5 | 0.2 | 6 | 0.1667 | 1 | 0 | 16 | 0.1875 | 3 | 0 | 1 | 1 | 3 | 0 | 1 | 0 | 8 | 0.125 |
| 15_5293196 | 5 | 0.6 | 7 | 0.7143 | 8 | 0.5 | 3 | 1 | 23 | 0.6522 | 4 | 0.75 | 4 | 0.75 | 3 | 0.6667 | 1 | 0 | 12 | 0.6667 |
| 15_5293207 | 8 | 0.875 | 6 | 1 | 8 | 0.75 | 2 | 0.5 | 24 | 0.8333 | 4 | 0.5 | 3 | 0.3333 | 2 | 0.5 | 1 | 1 | 10 | 0.5 |
| 15_5293212 | 8 | 0.125 | 6 | 0.5 | 8 | 0.25 | 3 | 0.3333 | 25 | 0.28 | 4 | 0.25 | 3 | 0 | 1 | 0 | NA | NA | 8 | 0.125 |
| 15_5293678 | 7 | 0 | 6 | 0.1667 | 5 | 0.2 | 3 | 0 | 21 | 0.0952 | 3 | 0 | 2 | 0.5 | 2 | 0 | 1 | 0 | 8 | 0.125 |
| 15_5320803 | NA | NA | 2 | 1 | 1 | 1 | 2 | 1 | 5 | 1 | 2 | 0.5 | NA | NA | NA | NA | 1 | 0 | 3 | 0.3333 |
| 15_5320807 | 1 | 1 | 2 | 0.5 | 1 | 0 | 2 | 0.5 | 6 | 0.5 | 2 | 0.5 | NA | NA | NA | NA | 1 | 1 | 3 | 0.6667 |
| 15_5320808 | NA | NA | 2 | 0.5 | 1 | 0 | 2 | 0.5 | 5 | 0.4 | 2 | 1 | NA | NA | NA | NA | 1 | 1 | 3 | 1 |
| 15_5320820 | 1 | 1 | 1 | 1 | 1 | 1 | 2 | 0.5 | 5 | 0.8 | 2 | 1 | NA | NA | NA | NA | 1 | 1 | 3 | 1 |
| 15_5320826 | NA | NA | 2 | 1 | 1 | 0 | 2 | 0 | 5 | 0.4 | 2 | 1 | NA | NA | NA | NA | NA | NA | 2 | 1 |
| 15_5320827 | 1 | 0 | 2 | 0.5 | 1 | 0 | 2 | 0 | 6 | 0.1667 | 2 | 0 | NA | NA | NA | NA | 1 | 1 | 3 | 0.3333 |
| 15_5320876 | 1 | 1 | 1 | 0 | 1 | 1 | 2 | 1 | 5 | 0.8 | 2 | 1 | NA | NA | NA | NA | 1 | 1 | 3 | 1 |
| 15_5320883 | 1 | 1 | 2 | 0.5 | 1 | 1 | 2 | 1 | 6 | 0.8333 | 2 | 1 | NA | NA | NA | NA | 1 | 1 | 3 | 1 |
| 15_5331693 | NA | NA | NA | NA | NA | NA | 1 | 1 | 1 | 1 | NA | NA | NA | NA | 1 | 1 | NA | NA | 1 | 1 |
| 15_5331697 | NA | NA | NA | NA | NA | NA | 1 | 1 | 1 | 1 | NA | NA | NA | NA | 1 | 1 | NA | NA | 1 | 1 |
| 15_5331752 | 6 | 0.1667 | 4 | 0.25 | 6 | 0 | 8 | 0 | 24 | 0.0833 | NA | NA | 10 | 0.1 | 7 | 0.1429 | 3 | 0 | 20 | 0.1 |
| 15_5332093 | 1 | 1 | 3 | 0.6667 | 2 | 0 | 5 | 0 | 11 | 0.2727 | NA | NA | 2 | 0 | 3 | 0.3333 | 2 | 1 | 7 | 0.4286 |
| 15_5332098 | 1 | 0 | 3 | 0.6667 | 2 | 0 | 4 | 0.25 | 10 | 0.3 | NA | NA | 2 | 0 | 3 | 0.3333 | 2 | 0.5 | 7 | 0.2857 |
| 15_5332533 | NA | NA | 2 | 1 | NA | NA | NA | NA | 2 | 1 | NA | NA | 3 | 0.6667 | 2 | 1 | 2 | 1 | 7 | 0.8571 |
| 15_5332543 | NA | NA | NA | NA | 1 | 1 | NA | NA | 1 | 1 | NA | NA | 2 | 0.5 | 2 | 1 | 2 | 0.5 | 6 | 0.6667 |
| 15_5332552 | NA | NA | 1 | 0 | 1 | 1 | NA | NA | 2 | 0.5 | NA | NA | 2 | 0 | 2 | 0.5 | 1 | 1 | 5 | 0.4 |
| 15_5446415 | 6 | 0.1667 | 13 | 0.0769 | 9 | 0 | 7 | 0 | 35 | 0.0571 | 7 | 0.2857 | 6 | 0 | 6 | 0 | 10 | 0 | 29 | 0.069 |
| 15_5446531 | 9 | 0.5556 | 23 | 0.0435 | 15 | 0 | 10 | 0 | 57 | 0.1053 | 8 | 0.125 | 10 | 0 | 10 | 0.3 | 16 | 0 | 44 | 0.0909 |
| 15_5446555 | 10 | 0.1 | 21 | 0.0476 | 12 | 0.0833 | 13 | 0 | 56 | 0.0536 | 11 | 0 | 10 | 0.1 | 12 | 0.3333 | 17 | 0.1176 | 50 | 0.14 |
| 15_5461122 | 11 | 0.0909 | 16 | 0.125 | 10 | 0 | 6 | 0.1667 | 43 | 0.093 | 9 | 0.2222 | 3 | 0.3333 | 2 | 0 | 16 | 0.0625 | 30 | 0.1333 |
| 15_5461581 | 12 | 0.0833 | 21 | 0.1429 | 12 | 0.1667 | 7 | 0.1429 | 52 | 0.1346 | 10 | 0.3 | 3 | 0.6667 | 2 | 0 | 13 | 0.1538 | 28 | 0.25 |
| 15_5461584 | 9 | 0 | 19 | 0.1579 | 11 | 0.1818 | 7 | 0.1429 | 46 | 0.1304 | 11 | 0.0909 | 3 | 0 | 2 | 0 | 14 | 0.2857 | 30 | 0.1667 |
| 15_5461647 | 13 | 0.0769 | 28 | 0.1071 | 16 | 0.1875 | 9 | 0 | 66 | 0.1061 | 13 | 0 | 5 | 0.2 | 3 | 0 | 18 | 0.1111 | 39 | 0.0769 |
| 15_5461655 | 14 | 0.2143 | 26 | 0.5385 | 16 | 0.4375 | 7 | 0.2857 | 63 | 0.4127 | 15 | 0.6 | 4 | 0.5 | 3 | 0 | 18 | 0.2778 | 40 | 0.4 |
| 15_5461670 | 12 | 0.4167 | 27 | 0.5926 | 16 | 0.5625 | 8 | 0.375 | 63 | 0.5238 | 14 | 0.7143 | 5 | 0.8 | 3 | 0.6667 | 19 | 0.4737 | 41 | 0.6098 |
| 15_5470240 | 25 | 0.04 | 21 | 0.1429 | 23 | 0.087 | 11 | 0.1818 | 80 | 0.1 | 28 | 0.1429 | 10 | 0.2 | 9 | 0.2222 | 31 | 0.0968 | 78 | 0.141 |
| 15_5487907 | 3 | 0.3333 | 1 | 1 | 5 | 0 | 5 | 0.6 | 14 | 0.3571 | 2 | 0 | 2 | 0 | 5 | 0.4 | 2 | 0.5 | 11 | 0.2727 |
| 15_5487932 | 4 | 0.25 | 2 | 0 | 6 | 0 | 5 | 0 | 17 | 0.0588 | NA | NA | 2 | 0 | 4 | 0 | 2 | 0.5 | 8 | 0.125 |
| 15_5487933 | 5 | 0.2 | 3 | 0 | 6 | 0 | 5 | 0.6 | 19 | 0.2105 | 1 | 1 | 2 | 1 | 4 | 0.5 | 3 | 0 | 10 | 0.5 |
| 15_5487936 | 5 | 0.8 | 2 | 1 | 5 | 0.6 | 4 | 1 | 16 | 0.8125 | 1 | 1 | 2 | 1 | 5 | 1 | 4 | 1 | 12 | 1 |
| 15_5487946 | 4 | 0.75 | 3 | 0.6667 | 6 | 0.6667 | 6 | 1 | 19 | 0.7895 | 1 | 1 | 2 | 1 | 5 | 1 | 4 | 1 | 12 | 1 |
| 15_5487969 | 4 | 0.5 | 3 | 1 | 5 | 0.6 | 4 | 0.75 | 16 | 0.6875 | 1 | 0 | 2 | 1 | 4 | 1 | 1 | 1 | 8 | 0.875 |
| 15_5487974 | 4 | 0.5 | 3 | 0.6667 | 3 | 0 | 4 | 0.75 | 14 | 0.5 | 1 | 1 | 2 | 0.5 | 4 | 1 | 2 | 1 | 9 | 0.8889 |
| 15_5487980 | 3 | 0 | 3 | 0.3333 | 6 | 0 | 5 | 0 | 17 | 0.0588 | NA | NA | 2 | 0.5 | 4 | 0 | 4 | 0.25 | 10 | 0.2 |
| 15_5512229 | 3 | 0 | NA | NA | 1 | 0 | 3 | 0.3333 | 7 | 0.1429 | NA | NA | 1 | 0 | 1 | 1 | NA | NA | 2 | 0.5 |
| 15_5539767 | 11 | 0.2727 | 12 | 0.0833 | 7 | 0 | 8 | 0.25 | 38 | 0.1579 | 5 | 0 | 9 | 0.1111 | 10 | 0 | 4 | 0.5 | 28 | 0.1071 |
| 15_5550207 | NA | NA | NA | NA | NA | NA | 1 | 1 | 1 | 1 | NA | NA | NA | NA | NA | NA | 2 | 1 | 2 | 1 |
| 15_5768492 | NA | NA | 2 | 0.5 | NA | NA | 1 | 0 | 3 | 0.3333 | NA | NA | NA | NA | NA | NA | 2 | 0.5 | 2 | 0.5 |
| 15_5768621 | NA | NA | 2 | 0.5 | 1 | 0 | 3 | 0.3333 | 6 | 0.3333 | NA | NA | NA | NA | NA | NA | 3 | 0.3333 | 3 | 0.3333 |
| 15_5768624 | NA | NA | 2 | 0.5 | 1 | 1 | 3 | 1 | 6 | 0.8333 | NA | NA | NA | NA | NA | NA | 3 | 0.6667 | 3 | 0.6667 |
| 15_5768664 | NA | NA | 2 | 1 | 1 | 1 | 3 | 0.6667 | 6 | 0.8333 | NA | NA | NA | NA | 1 | 1 | 3 | 0.6667 | 4 | 0.75 |
| 15_5768668 | NA | NA | 2 | 0.5 | NA | NA | 3 | 0.3333 | 5 | 0.4 | NA | NA | NA | NA | 1 | 1 | 3 | 1 | 4 | 1 |
| 15_5768690 | NA | NA | 2 | 1 | 1 | 1 | 3 | 0.6667 | 6 | 0.8333 | NA | NA | NA | NA | 1 | 1 | 3 | 1 | 4 | 1 |
| 15_5768700 | NA | NA | 2 | 1 | 2 | 1 | 3 | 0.3333 | 7 | 0.7143 | NA | NA | NA | NA | 1 | 0 | 3 | 0.6667 | 4 | 0.5 |
| 15_5770212 | NA | NA | 1 | 0 | 1 | 0 | 3 | 0.3333 | 5 | 0.2 | NA | NA | NA | NA | 1 | 0 | 3 | 0.3333 | 4 | 0.25 |
| 15_5770226 | NA | NA | 2 | 1 | 3 | 1 | 3 | 1 | 8 | 1 | NA | NA | NA | NA | NA | NA | 3 | 1 | 3 | 1 |
| 15_5773338 | 6 | 0.1667 | 2 | 0 | 3 | 0 | NA | NA | 11 | 0.0909 | 5 | 0 | 8 | 0.25 | 2 | 0.5 | 6 | 0.1667 | 21 | 0.1905 |
| 15_5773346 | 7 | 0.2857 | 3 | 0.3333 | 5 | 0.6 | 1 | 0 | 16 | 0.375 | 5 | 0.6 | 10 | 0.5 | 4 | 1 | 4 | 0.5 | 23 | 0.6087 |
| 15_5773352 | 6 | 1 | 2 | 1 | 4 | 1 | 2 | 1 | 14 | 1 | 6 | 1 | 11 | 0.9091 | 5 | 1 | 5 | 1 | 27 | 0.963 |
| 15_5773430 | 4 | 0.25 | 5 | 0 | 8 | 0.25 | 8 | 0.125 | 25 | 0.16 | 8 | 0 | 12 | 0.0833 | 5 | 0.2 | 8 | 0.125 | 33 | 0.0909 |
| 15_5773439 | 2 | 0.5 | 4 | 0 | 10 | 0.3 | 10 | 0.3 | 26 | 0.2692 | 9 | 0.3333 | 10 | 0.6 | 4 | 0.5 | 7 | 0.2857 | 30 | 0.4333 |
| 15_5773440 | 3 | 0 | 4 | 0.25 | 10 | 0.3 | 10 | 0.2 | 27 | 0.2222 | 11 | 0.0909 | 11 | 0.1818 | 2 | 0 | 6 | 0 | 30 | 0.1 |
| 15_5773451 | 4 | 1 | 5 | 0.4 | 8 | 0.375 | 11 | 0.2727 | 28 | 0.4286 | 10 | 0.1 | 14 | 0.5 | 4 | 0.25 | 6 | 0.3333 | 34 | 0.3235 |
| 15_5773454 | 5 | 0.2 | 5 | 0 | 8 | 0 | 11 | 0.1818 | 29 | 0.1034 | 10 | 0.1 | 12 | 0.0833 | 3 | 0 | 6 | 0.3333 | 31 | 0.129 |
| 15_5773468 | 5 | 1 | 5 | 0.8 | 7 | 0.8571 | 8 | 0.625 | 25 | 0.8 | 8 | 0.75 | 7 | 0.7143 | 4 | 1 | 6 | 0.6667 | 25 | 0.76 |
| 15_5773474 | 4 | 0.75 | 4 | 0.75 | 7 | 0.4286 | 7 | 0.5714 | 22 | 0.5909 | 9 | 0.6667 | 11 | 0.4545 | 4 | 0.25 | 6 | 0.1667 | 30 | 0.4333 |
| 15_5773475 | 5 | 0 | 5 | 0.2 | 9 | 0.2222 | 9 | 0.2222 | 28 | 0.1786 | 8 | 0 | 11 | 0.1818 | 4 | 0.25 | 6 | 0.1667 | 29 | 0.1379 |

| | | | | | | | | | | | | | | | | | | | | |
|---|---|---|---|---|---|---|---|---|---|---|---|---|---|---|---|---|---|---|---|---|
| 15_5773482 | 4 | 0 | 5 | 0 | 9 | 0.1111 | 10 | 0.3 | 28 | 0.1429 | 9 | 0.1111 | 11 | 0 | 5 | 0 | 7 | 0.1429 | 32 | 0.0625 |
| 15_5773491 | 3 | 1 | 3 | 0.6667 | 8 | 0.375 | 9 | 0.7778 | 23 | 0.6522 | 10 | 0.5 | 9 | 0.4444 | 5 | 0.6 | 5 | 0.6 | 29 | 0.5172 |
| 15_5773492 | 3 | 0.3333 | 3 | 0.3333 | 8 | 0.5 | 6 | 0.3333 | 20 | 0.4 | 7 | 0.7143 | 11 | 0.5455 | 4 | 0.75 | 5 | 0.4 | 27 | 0.5926 |
| 15_5773502 | 4 | 0.25 | 6 | 0 | 8 | 0.125 | 12 | 0.1667 | 30 | 0.1333 | 12 | 0.1667 | 13 | 0.4615 | 7 | 0.5714 | 7 | 0.1429 | 39 | 0.3333 |
| 15_5773506 | 4 | 0.75 | 4 | 0.75 | 6 | 0.8333 | 12 | 0.5833 | 26 | 0.6923 | 8 | 0.75 | 11 | 0.8182 | 5 | 0.6 | 6 | 1 | 30 | 0.8 |
| 15_5773512 | 4 | 0 | 6 | 0.1667 | 6 | 0.1667 | 10 | 0.2 | 26 | 0.1538 | 10 | 0.3 | 9 | 0.3333 | 6 | 0.1667 | 7 | 0.1429 | 32 | 0.25 |
| 15_5789755 | 4 | 0 | 4 | 0 | 7 | 0 | 4 | 0.5 | 19 | 0.1053 | 3 | 0.3333 | 6 | 0.1667 | 5 | 0 | 4 | 0 | 18 | 0.1111 |
| 15_5791080 | 7 | 0.1429 | 10 | 0 | 12 | 0 | 7 | 0.1429 | 36 | 0.0556 | 2 | 0 | 13 | 0.2308 | 12 | 0.0833 | 4 | 0.5 | 31 | 0.1935 |
| 15_5791110 | 7 | 0.5714 | 7 | 0.1429 | 7 | 0.4286 | 5 | 0.2 | 26 | 0.3462 | 1 | 0 | 13 | 0.3846 | 9 | 0.4444 | 3 | 0.3333 | 26 | 0.3846 |
| 15_5791147 | 7 | 0.7143 | 6 | 0 | 9 | 0.3333 | 6 | 0.5 | 28 | 0.3929 | NA | NA | 12 | 0.4167 | 9 | 0.4444 | 5 | 0.4 | 26 | 0.4231 |
| 15_5791149 | 7 | 0 | 6 | 0.3333 | 7 | 0.1429 | 5 | 0 | 25 | 0.12 | NA | NA | 12 | 0 | 10 | 0.2 | 5 | 0 | 27 | 0.0741 |
| 15_5791201 | 4 | 0 | 3 | 0 | 6 | 0.3333 | 5 | 0 | 18 | 0.1111 | NA | NA | 10 | 0.2 | 7 | 0.1429 | 5 | 0.2 | 22 | 0.1818 |
| 15_5791203 | 5 | 0.6 | 3 | 0.6667 | 7 | 0.5714 | 6 | 0.8333 | 21 | 0.6667 | NA | NA | 11 | 0.6364 | 9 | 0.4444 | 4 | 0.75 | 24 | 0.5833 |
| 15_5791210 | 3 | 0 | 3 | 0 | 3 | 0 | 5 | 0.2 | 14 | 0.0714 | NA | NA | 12 | 0.1667 | 8 | 0 | 5 | 0 | 25 | 0.08 |
| 15_5791567 | 4 | 0 | 2 | 0 | 6 | 0.1667 | 5 | 0 | 17 | 0.0588 | NA | NA | 11 | 0.0909 | 8 | 0.375 | 4 | 0.25 | 23 | 0.2174 |
| 15_5791578 | 4 | 0.25 | 3 | 0.3333 | 5 | 0 | 5 | 0 | 17 | 0.1176 | NA | NA | 11 | 0 | 9 | 0.2222 | 3 | 0 | 23 | 0.087 |
| 15_5791583 | 4 | 1 | 2 | 1 | 4 | 1 | 4 | 1 | 14 | 1 | NA | NA | 11 | 0.8182 | 9 | 0.8889 | 3 | 1 | 23 | 0.8696 |
| 15_5791590 | 3 | 0 | 3 | 0 | 4 | 0 | 4 | 0.25 | 14 | 0.0714 | NA | NA | 6 | 0.1667 | 5 | 0 | 1 | 0 | 12 | 0.0833 |
| 15_5791590 | 2 | 0.5 | 3 | 0.6667 | 5 | 0.6 | 4 | 0.25 | 14 | 0.5 | NA | NA | 8 | 0.25 | 8 | 0.25 | 3 | 0.6667 | 19 | 0.3158 |
| 15_5791593 | 3 | 0 | 3 | 0.6667 | 4 | 0.75 | 4 | 0.25 | 14 | 0.4286 | NA | NA | 10 | 0.3 | 8 | 0.375 | 4 | 0 | 22 | 0.2727 |
| 15_5791594 | 2 | 1 | 3 | 0.6667 | 3 | 0.6667 | 1 | 0 | 9 | 0.6667 | NA | NA | 9 | 0.6667 | 5 | 0.8 | 3 | 1 | 17 | 0.7647 |
| 15_5791614 | 3 | 0 | 3 | 0.3333 | 5 | 0.4 | 5 | 0 | 16 | 0.1875 | NA | NA | 10 | 0.1 | 7 | 0.1429 | 2 | 0 | 19 | 0.1053 |
| 15_5791626 | 3 | 0 | 3 | 0 | 3 | 0 | 5 | 0.2 | 14 | 0.0714 | NA | NA | 10 | 0.3 | 8 | 0.375 | 1 | 0 | 19 | 0.3158 |
| 15_5791654 | 2 | 0 | 3 | 0.3333 | 5 | 0.2 | 5 | 0.2 | 15 | 0.2 | NA | NA | 9 | 0.2222 | 8 | 0.125 | 1 | 0 | 18 | 0.1667 |
| 15_5791662 | 2 | 0 | 3 | 1 | 4 | 0.25 | 5 | 0.6 | 14 | 0.5 | NA | NA | 8 | 0.25 | 8 | 0.375 | 1 | 0 | 17 | 0.2941 |
| 15_5791677 | 2 | 0 | 3 | 0.3333 | 4 | 0 | 4 | 0 | 13 | 0.0769 | NA | NA | 10 | 0.1 | 8 | 0.125 | 1 | 0 | 19 | 0.1053 |
| 15_5791681 | 2 | 0.5 | 3 | 0.6667 | 4 | 0.75 | 4 | 0.25 | 13 | 0.5385 | NA | NA | 9 | 0.6667 | 7 | 0.8571 | 1 | 0 | 17 | 0.7059 |
| 15_5791683 | 2 | 0.5 | 3 | 1 | 2 | 0 | 3 | 0.6667 | 10 | 0.6 | NA | NA | 7 | 0.5714 | 7 | 0.4286 | 1 | 1 | 15 | 0.5333 |
| 15_5791692 | 2 | 0.5 | 3 | 0 | 4 | 0.25 | 4 | 0.25 | 13 | 0.2308 | NA | NA | 10 | 0.1 | 7 | 0 | 1 | 0 | 18 | 0.0556 |
| 15_5791703 | 2 | 0 | 3 | 0.3333 | 3 | 0 | 3 | 0.3333 | 11 | 0.1818 | NA | NA | 8 | 0.125 | 7 | 0.1429 | 1 | 0 | 16 | 0.125 |
| 15_5791708 | 2 | 1 | 3 | 0 | 3 | 0.6667 | 4 | 0.5 | 12 | 0.5 | NA | NA | 7 | 0.4286 | 7 | 0.4286 | 1 | 0 | 15 | 0.4 |
| 15_5791714 | 1 | 0 | 3 | 0.3333 | 1 | 0 | 3 | 0 | 8 | 0.125 | NA | NA | 4 | 0.25 | 7 | 0 | 1 | 0 | 12 | 0.0833 |
| 15_5791721 | 1 | 0 | 3 | 0.3333 | 3 | 0.6667 | 4 | 0.75 | 11 | 0.5455 | NA | NA | 7 | 0.4286 | 7 | 0.8571 | 1 | 1 | 15 | 0.6667 |
| 15_5791743 | 1 | 1 | 3 | 0.3333 | 3 | 0.6667 | 3 | 0.3333 | 10 | 0.5 | NA | NA | 6 | 0.3333 | 6 | 0.1667 | 1 | 0 | 13 | 0.2308 |
| 15_5791749 | 1 | 0 | 2 | 0 | 3 | 0 | 4 | 0.25 | 10 | 0.1 | NA | NA | 8 | 0 | 7 | 0.4286 | 1 | 0 | 16 | 0.1875 |
| 15_5791770 | 1 | 0 | 3 | 0.3333 | 2 | 0 | 3 | 0 | 9 | 0.1111 | NA | NA | 4 | 0 | 5 | 0.2 | 1 | 0 | 10 | 0.1 |
| 15_5791779 | 1 | 0 | 3 | 0 | 3 | 0 | 4 | 0.25 | 11 | 0.0909 | NA | NA | 7 | 0.1429 | 6 | 0.1667 | 1 | 0 | 14 | 0.1429 |
| 15_5791780 | 1 | 1 | 3 | 0.6667 | 2 | 0.5 | 4 | 0.75 | 10 | 0.7 | NA | NA | 7 | 0.5714 | 7 | 0.4286 | 1 | 0 | 15 | 0.4667 |
| 15_5791782 | 1 | 0 | 2 | 0 | 3 | 0 | 3 | 0.3333 | 9 | 0.1111 | NA | NA | 7 | 0.2857 | 7 | 0.2857 | 1 | 0 | 15 | 0.2667 |
| 15_5791801 | NA | NA | 2 | 0 | 2 | 0.5 | 4 | 0 | 8 | 0.125 | NA | NA | 5 | 0 | 7 | 0.2857 | 1 | 0 | 13 | 0.1538 |
| 15_5791835 | NA | NA | 2 | 0.5 | 3 | 0.6667 | 4 | 0.75 | 9 | 0.6667 | NA | NA | 5 | 0.8 | 7 | 1 | 1 | 1 | 13 | 0.9231 |
| 15_5791837 | NA | NA | 1 | 1 | 3 | 1 | 4 | 1 | 8 | 1 | NA | NA | 7 | 0.8571 | 6 | 0.6667 | 1 | 0 | 14 | 0.7143 |
| 15_5804132 | 1 | 1 | 2 | 0.5 | 1 | 1 | NA | NA | 4 | 0.75 | NA | NA | NA | NA | 1 | 1 | 2 | 0.5 | 3 | 0.6667 |
| 15_5804372 | NA | NA | 2 | 0.5 | 1 | 1 | 1 | 0 | 4 | 0.5 | NA | NA | 1 | 0 | NA | NA | 3 | 0.3333 | 4 | 0.25 |
| 15_5804378 | NA | NA | 1 | 0 | 1 | 1 | 1 | 0 | 3 | 0.3333 | NA | NA | 1 | 1 | NA | NA | 3 | 0.3333 | 4 | 0.5 |
| 15_5804394 | NA | NA | 1 | 0 | 1 | 1 | 1 | 0 | 3 | 0.3333 | NA | NA | 1 | 0 | NA | NA | 3 | 0.3333 | 4 | 0.25 |
| 15_5804443 | NA | NA | 1 | 1 | 1 | 1 | 1 | 1 | 3 | 1 | NA | NA | 1 | 1 | NA | NA | 2 | 0.5 | 3 | 0.6667 |
| 15_5804447 | NA | NA | 1 | 0 | 1 | 0 | 1 | 1 | 3 | 0.3333 | NA | NA | 1 | 1 | NA | NA | 2 | 0 | 3 | 0.3333 |
| 15_5804451 | NA | NA | 1 | 1 | 1 | 1 | NA | NA | 2 | 1 | NA | NA | 1 | 1 | NA | NA | 3 | 0 | 4 | 0.25 |
| 15_5804458 | NA | NA | NA | NA | NA | NA | 1 | 1 | 1 | 1 | NA | NA | 1 | 1 | NA | NA | 2 | 0.5 | 3 | 0.6667 |
| 15_5804463 | NA | NA | 1 | 1 | 1 | 1 | 1 | 1 | 3 | 1 | NA | NA | 1 | 1 | NA | NA | 2 | 1 | 3 | 1 |
| 15_5804467 | NA | NA | 1 | 1 | 1 | 1 | NA | NA | 2 | 1 | NA | NA | NA | NA | NA | NA | 1 | 1 | 1 | 1 |
| 15_5804475 | NA | NA | 1 | 0 | 1 | 1 | 1 | 1 | 3 | 0.6667 | NA | NA | 1 | 1 | NA | NA | 3 | 0.3333 | 4 | 0.5 |
| 15_5804480 | NA | NA | 1 | 1 | 1 | 1 | 1 | 1 | 3 | 1 | NA | NA | 1 | 1 | NA | NA | 3 | 1 | 4 | 1 |
| 15_5804488 | NA | NA | 1 | 0 | 1 | 1 | 1 | 0 | 3 | 0.3333 | NA | NA | 1 | 1 | NA | NA | 3 | 0.3333 | 4 | 0.5 |
| 15_5832669 | 4 | 0.5 | 3 | 0 | 5 | 0.4 | 2 | 0 | 14 | 0.2857 | 1 | 0 | 7 | 0.5714 | 7 | 0.2857 | 4 | 0.75 | 19 | 0.4737 |
| 15_5832700 | 5 | 0.2 | 4 | 0 | 6 | 0.3333 | 2 | 0.5 | 17 | 0.2353 | 3 | 0.3333 | 8 | 0.25 | 8 | 0.125 | 4 | 0 | 23 | 0.1739 |
| 15_5832702 | 5 | 0.6 | 3 | 0.3333 | 5 | 0.2 | 2 | 0 | 15 | 0.3333 | 1 | 0 | 5 | 0.6 | 5 | 0.4 | 4 | 0.5 | 15 | 0.4667 |
| 15_5832703 | 5 | 1 | 4 | 0.5 | 7 | 0.5714 | 2 | 1 | 18 | 0.7222 | 2 | 1 | 6 | 1 | 6 | 1 | 5 | 0.8 | 19 | 0.9474 |
| 15_5832705 | 4 | 0.25 | 3 | 0 | 7 | 0.2857 | 2 | 0 | 16 | 0.1875 | 3 | 0.6667 | 7 | 0.4286 | 8 | 0.5 | 5 | 0.6 | 23 | 0.5217 |
| 15_5832718 | 5 | 0.8 | 3 | 1 | 7 | 1 | 2 | 1 | 17 | 0.9412 | 3 | 0.3333 | 8 | 0.875 | 8 | 0.875 | 5 | 1 | 24 | 0.8333 |
| 15_5832735 | 5 | 0 | 4 | 0 | 5 | 0.2 | 3 | 0 | 17 | 0.0588 | 3 | 0 | 5 | 0 | 7 | 0.1429 | 5 | 0 | 20 | 0.05 |
| 15_5832742 | 4 | 0.5 | 4 | 0.5 | 5 | 0.6 | 2 | 0 | 15 | 0.4667 | 2 | 0 | 6 | 0.5 | 6 | 0.6667 | 4 | 0.25 | 18 | 0.4444 |
| 15_5832745 | 5 | 0.8 | 3 | 0.6667 | 6 | 0.6667 | 3 | 0.6667 | 17 | 0.7059 | 3 | 0.3333 | 5 | 0.8 | 7 | 0.8571 | 4 | 0.25 | 19 | 0.6316 |
| 15_5832757 | 3 | 1 | 3 | 0 | 5 | 0.4 | 3 | 0.6667 | 14 | 0.5 | 3 | 1 | 7 | 0.4286 | 8 | 0.375 | 5 | 0.6 | 23 | 0.5217 |

| | | | | | | | | | | | | | | | | | | | | |
|---|---|---|---|---|---|---|---|---|---|---|---|---|---|---|---|---|---|---|---|---|
| 15_5832766 | 4 | 0.5 | 4 | 0.5 | 6 | 0.8333 | 2 | 0.5 | 16 | 0.625 | 3 | 0.6667 | 7 | 0.7143 | 9 | 0.7778 | 4 | 0.75 | 23 | 0.7391 |
| 15_5832774 | 5 | 0.2 | 4 | 0 | 6 | 0 | 2 | 0.5 | 17 | 0.1176 | 3 | 0 | 9 | 0.1111 | 7 | 0.2857 | 4 | 0 | 23 | 0.1304 |
| 15_5832792 | 4 | 0 | 5 | 0.2 | 7 | 0.2857 | 3 | 0.3333 | 19 | 0.2105 | 3 | 0.3333 | 7 | 0.1429 | 9 | 0.1111 | 5 | 0 | 24 | 0.125 |
| 15_5832799 | 5 | 0.4 | 5 | 0 | 5 | 0 | 3 | 0.3333 | 18 | 0.1667 | 3 | 0.6667 | 8 | 0.125 | 8 | 0.375 | 6 | 0.1667 | 25 | 0.28 |
| 15_5832808 | 4 | 0.25 | 5 | 0 | 7 | 0 | 1 | 0 | 17 | 0.0588 | 2 | 0 | 7 | 0.1429 | 6 | 0 | 4 | 0 | 19 | 0.0526 |
| 15_5832858 | 4 | 0.25 | 7 | 0.1429 | 8 | 0.25 | 3 | 0 | 22 | 0.1818 | 3 | 0 | 4 | 0.25 | 8 | 0 | 8 | 0.125 | 23 | 0.087 |
| 15_5832925 | 8 | 0.125 | 7 | 0 | 7 | 0.5714 | 3 | 0.6667 | 25 | 0.28 | 1 | 1 | 6 | 0.1667 | 8 | 0 | 9 | 0.4444 | 24 | 0.25 |
| 15_5865445 | 6 | 0.5 | 7 | 0.4286 | 10 | 0.6 | 5 | 0.2 | 28 | 0.4643 | 5 | 0 | 7 | 0.5714 | 8 | 0.75 | 9 | 0.4444 | 29 | 0.4828 |
| 15_5865447 | 8 | 0.5 | 7 | 0.7143 | 11 | 0.5455 | 7 | 0.1429 | 33 | 0.4848 | 3 | 0 | 9 | 0.5556 | 8 | 0.375 | 9 | 0.4444 | 29 | 0.4138 |
| 15_5865451 | 7 | 0.2857 | 9 | 0.3333 | 11 | 0.2727 | 7 | 0 | 34 | 0.2353 | 4 | 0.25 | 6 | 0.1667 | 5 | 0.2 | 9 | 0.3333 | 24 | 0.25 |
| 15_5869500 | 3 | 0 | 4 | 0 | 5 | 0.2 | 7 | 0.4286 | 19 | 0.2105 | 3 | 0.3333 | 7 | 0.1429 | 8 | 0 | 5 | 0.2 | 23 | 0.1304 |
| 15_5874505 | 1 | 0 | NA | NA | 2 | 0.5 | 1 | 1 | 4 | 0.5 | 2 | 0.5 | 1 | 1 | 1 | 0 | 1 | 1 | 5 | 0.6 |
| 15_5874506 | 1 | 0 | 1 | 0 | 2 | 0.5 | 1 | 1 | 5 | 0.4 | 2 | 0 | 1 | 0 | 1 | 1 | 1 | 0 | 5 | 0.2 |
| 15_5874512 | 1 | 1 | 1 | 0 | 2 | 1 | 1 | 1 | 5 | 0.8 | 2 | 1 | 1 | 1 | NA | NA | NA | NA | 3 | 1 |
| 15_5874514 | NA | NA | 1 | 1 | 2 | 0.5 | 1 | 1 | 4 | 0.75 | 2 | 0.5 | 1 | 1 | NA | NA | 1 | 0 | 4 | 0.5 |
| 15_5874545 | 1 | 0 | 1 | 0 | 2 | 0.5 | 1 | 1 | 5 | 0.4 | 2 | 1 | 1 | 0 | NA | NA | 1 | 0 | 4 | 0.5 |
| 15_5874546 | 1 | 0 | 1 | 0 | 2 | 0.5 | NA | NA | 4 | 0.25 | 2 | 0 | 1 | 1 | NA | NA | 1 | 0 | 4 | 0.25 |
| 15_5874547 | NA | NA | NA | NA | 2 | 1 | NA | NA | 2 | 1 | 2 | 1 | 1 | 0 | NA | NA | 1 | 1 | 4 | 0.75 |
| 15_5874549 | 1 | 0 | 1 | 1 | 1 | 0 | 1 | 0 | 4 | 0.25 | 3 | 0.3333 | 1 | 1 | 1 | 0 | 1 | 0 | 6 | 0.3333 |
| 15_5874569 | 1 | 1 | 1 | 0 | NA | NA | 1 | 1 | 3 | 0.6667 | 2 | 0.5 | 1 | 1 | NA | NA | 1 | 1 | 4 | 0.75 |
| 15_5874577 | 1 | 0 | 2 | 0.5 | 2 | 0 | 1 | 0 | 6 | 0.1667 | 4 | 0.25 | 1 | 1 | 2 | 0 | 1 | 0 | 8 | 0.25 |
| 15_5874581 | 1 | 1 | 2 | 0.5 | 2 | 0 | 1 | 1 | 6 | 0.5 | 3 | 0.6667 | 1 | 1 | 2 | 0.5 | NA | NA | 6 | 0.6667 |
| 15_5874593 | 1 | 1 | 2 | 0 | 3 | 0.3333 | 1 | 0 | 7 | 0.2857 | 4 | 0 | 1 | 1 | 2 | 0 | 1 | 0 | 8 | 0.125 |
| 15_5874600 | 1 | 0 | 2 | 0 | 3 | 0.3333 | 1 | 0 | 7 | 0.1429 | 3 | 0.3333 | NA | NA | 2 | 0 | 2 | 0 | 7 | 0.1429 |
| 15_5874603 | NA | NA | 1 | 1 | 2 | 1 | 1 | 1 | 4 | 1 | 3 | 1 | 1 | 1 | 2 | 1 | 1 | 1 | 7 | 1 |
| 15_5874610 | 1 | 1 | 1 | 1 | 1 | 1 | NA | NA | 3 | 1 | 4 | 0.75 | NA | NA | 1 | 0 | 2 | 1 | 7 | 0.7143 |
| 15_5877028 | 3 | 0.3333 | 7 | 0.8571 | 2 | 0.5 | 3 | 0.3333 | 15 | 0.6 | 2 | 1 | 6 | 0.5 | 3 | 1 | 3 | 1 | 14 | 0.7857 |
| 15_5877050 | 2 | 1 | 5 | 1 | 3 | 1 | 3 | 1 | 13 | 1 | 4 | 1 | 4 | 1 | 3 | 1 | 3 | 1 | 14 | 1 |
| 15_5877056 | 3 | 0.3333 | 8 | 0.25 | 3 | 0 | 5 | 0.2 | 19 | 0.2105 | 2 | 0 | 6 | 0.6667 | 4 | 0.75 | 3 | 0 | 15 | 0.4667 |
| 15_5877065 | 3 | 0.6667 | 5 | 0.8 | 2 | 0.5 | 4 | 0.75 | 14 | 0.7143 | 3 | 1 | 6 | 0.5 | 3 | 0 | 3 | 0.6667 | 15 | 0.5333 |
| 15_5877068 | 2 | 0 | 7 | 0.2857 | 2 | 0 | 5 | 0 | 16 | 0.125 | 3 | 0 | 6 | 0.1667 | 4 | 0 | 2 | 0 | 15 | 0.0667 |
| 15_5877076 | 3 | 0.6667 | 6 | 0.3333 | 3 | 1 | 5 | 0.8 | 17 | 0.6471 | 2 | 0.5 | 3 | 0.6667 | 4 | 1 | 3 | 1 | 12 | 0.8333 |
| 15_5877084 | 2 | 1 | 4 | 1 | 3 | 1 | 3 | 1 | 12 | 1 | 2 | 0.5 | 3 | 1 | 3 | 1 | 1 | 0 | 9 | 0.7778 |
| 15_5877098 | 4 | 0.75 | 5 | 1 | 2 | 0.5 | 5 | 1 | 16 | 0.875 | 4 | 0.5 | 5 | 1 | 4 | 1 | 3 | 0.6667 | 16 | 0.8125 |
| 15_5877102 | 3 | 0.6667 | 6 | 0.8333 | 4 | 0.75 | 4 | 1 | 17 | 0.8235 | 4 | 1 | 5 | 1 | 4 | 1 | 3 | 0.3333 | 16 | 0.875 |
| 15_5877105 | 4 | 0.5 | 6 | 0.5 | 3 | 0 | 4 | 0 | 17 | 0.2941 | 3 | 0.3333 | 4 | 0.75 | 4 | 0.25 | 3 | 0 | 14 | 0.3571 |
| 15_5877109 | 4 | 0 | 8 | 0.25 | 5 | 0.2 | 5 | 0.2 | 22 | 0.1818 | 4 | 0.25 | 6 | 0.1667 | 5 | 0 | 3 | 0 | 18 | 0.1111 |
| 15_5877114 | 3 | 0.6667 | 7 | 0.5714 | 5 | 0.2 | 5 | 0.8 | 20 | 0.55 | 4 | 0.5 | 6 | 0.6667 | 4 | 1 | 1 | 0 | 15 | 0.6667 |
| 15_5877126 | 4 | 0.5 | 5 | 0.8 | 5 | 0.2 | 5 | 1 | 19 | 0.6316 | 3 | 0.3333 | 6 | 0.5 | 4 | 0.75 | 3 | 0.6667 | 16 | 0.5625 |
| 15_5877135 | 4 | 0.25 | 5 | 0.2 | 5 | 0.2 | 3 | 0 | 17 | 0.1765 | 4 | 0.25 | 6 | 0.3333 | 4 | 0.25 | 2 | 0 | 16 | 0.25 |
| 15_5877149 | 4 | 0 | 5 | 0.2 | 6 | 0 | 4 | 0 | 19 | 0.0526 | 3 | 0.3333 | 6 | 0 | 6 | 0 | 3 | 0.3333 | 18 | 0.1111 |
| 15_5877152 | 4 | 0.5 | 9 | 0.4444 | 6 | 0.1667 | 5 | 0 | 24 | 0.2917 | 4 | 0.25 | 6 | 0 | 7 | 0.1429 | 3 | 0 | 20 | 0.1 |
| 15_5877154 | 4 | 1 | 7 | 1 | 4 | 1 | 4 | 1 | 19 | 1 | 4 | 1 | 6 | 0.8333 | 5 | 1 | 3 | 1 | 18 | 0.9444 |
| 15_5877186 | 4 | 1 | 8 | 0.875 | 4 | 0.5 | 5 | 1 | 21 | 0.8571 | 4 | 1 | 5 | 0.8 | 6 | 1 | 3 | 1 | 18 | 0.9444 |
| 15_5877188 | 4 | 1 | 10 | 1 | 4 | 1 | 5 | 1 | 23 | 1 | 5 | 1 | 5 | 1 | 5 | 1 | 3 | 1 | 18 | 1 |
| 15_5877197 | 5 | 0.6 | 9 | 0.4444 | 4 | 0.75 | 4 | 0.75 | 22 | 0.5909 | 5 | 0.4 | 5 | 0.4 | 6 | 0.5 | 3 | 0.3333 | 19 | 0.4211 |
| 15_5877209 | 4 | 0.5 | 10 | 0.3 | 7 | 0.4286 | 5 | 0.4 | 26 | 0.3846 | 5 | 0.6 | 6 | 0.6667 | 6 | 0.8333 | 3 | 0.6667 | 20 | 0.7 |
| 15_5877233 | NA | NA | NA | NA | NA | NA | 1 | 1 | 1 | 1 | NA | NA | NA | NA | 2 | 1 | NA | NA | 2 | 1 |
| 15_5880043 | 1 | 1 | NA | NA | NA | NA | NA | NA | 1 | 1 | 1 | 1 | NA | NA | NA | NA | NA | NA | 1 | 1 |
| 15_5881368 | 4 | 0.25 | 4 | 0.25 | 5 | 0.2 | 3 | 0 | 16 | 0.1875 | 5 | 0 | 2 | 0 | 1 | 1 | 4 | 0.25 | 12 | 0.1667 |
| 15_5881395 | 4 | 0.25 | 3 | 0 | 4 | 0 | 1 | 0 | 12 | 0.0833 | 3 | 0.3333 | 2 | 0 | NA | NA | 3 | 0 | 8 | 0.125 |
| 15_5881469 | 1 | 0 | 2 | 0 | 2 | 0.5 | 1 | 0 | 6 | 0.1667 | 1 | 1 | 1 | 0 | NA | NA | 3 | 0.6667 | 5 | 0.6 |
| 15_5881523 | 2 | 1 | 4 | 1 | 1 | 1 | NA | NA | 7 | 1 | 1 | 1 | NA | NA | NA | NA | 3 | 1 | 4 | 1 |
| 15_5881548 | 2 | 0 | 3 | 0.3333 | 1 | 0 | NA | NA | 6 | 0.1667 | 1 | 0 | NA | NA | NA | NA | 3 | 0.3333 | 4 | 0.25 |
| 15_5881549 | 3 | 0 | 4 | 0.25 | 1 | 1 | NA | NA | 8 | 0.25 | 1 | 0 | NA | NA | NA | NA | 3 | 0.3333 | 4 | 0.25 |
| 15_5881581 | 3 | 0.3333 | 3 | 0.3333 | 1 | 1 | NA | NA | 7 | 0.4286 | 1 | 0 | NA | NA | NA | NA | 3 | 0.3333 | 4 | 0.25 |
| 15_5918729 | 3 | 0 | 6 | 0.1667 | 1 | 0 | 4 | 0 | 14 | 0.0714 | 4 | 0 | 3 | 0.3333 | 2 | 0 | 5 | 0.2 | 14 | 0.1429 |
| 15_5918731 | 2 | 0 | 6 | 0 | 1 | 0 | 4 | 0.25 | 13 | 0.0769 | 2 | 0 | 3 | 0 | 2 | 0 | 5 | 0.2 | 12 | 0.0833 |
| 15_5918745 | 3 | 1 | 6 | 0.8333 | NA | NA | 5 | 1 | 14 | 0.9286 | 4 | 0.75 | 3 | 0.3333 | 1 | 1 | 5 | 0.8 | 13 | 0.6923 |
| 15_5918752 | 3 | 0 | 6 | 0.1667 | NA | NA | 5 | 0 | 14 | 0.0714 | 4 | 0.25 | 4 | 0 | 2 | 0.5 | 5 | 0.6 | 15 | 0.3333 |
| 15_5918757 | 2 | 0 | 4 | 0.25 | NA | NA | 3 | 0.6667 | 9 | 0.3333 | 2 | 1 | 3 | 0.3333 | 2 | 0 | 3 | 0 | 10 | 0.3 |
| 15_5918758 | 3 | 1 | 6 | 1 | 1 | 1 | 5 | 1 | 15 | 1 | 3 | 1 | 2 | 0.5 | 1 | 1 | 5 | 0.6 | 11 | 0.7273 |
| 15_5918763 | 2 | 0.5 | 7 | 0 | 1 | 0 | 5 | 0 | 15 | 0.0667 | 3 | 0 | 3 | 0 | 2 | 0 | 5 | 0.2 | 13 | 0.0769 |
| 15_5918770 | 3 | 1 | 8 | 0.75 | 1 | 1 | 6 | 1 | 18 | 0.8889 | 4 | 1 | 3 | 0.6667 | 2 | 1 | 5 | 1 | 14 | 0.9286 |
| 15_5918796 | 4 | 0 | 8 | 0.125 | 1 | 0 | 6 | 0 | 19 | 0.0526 | 2 | 1 | 4 | 0 | 2 | 0 | 5 | 0.2 | 13 | 0.2308 |

| | | | | | | | | | | | | | | | | | | | | |
|---|---|---|---|---|---|---|---|---|---|---|---|---|---|---|---|---|---|---|---|---|
| 15_592011 | NA | NA | 1 | 0 | NA | NA | 2 | 0.5 | 3 | 0.3333 | NA | NA | 1 | 0 | NA | NA | 1 | 1 | 2 | 0.5 |
| 15_592027 | NA | NA | 1 | 1 | NA | NA | NA | NA | 1 | 1 | NA | NA | 2 | 0.5 | NA | NA | 1 | 1 | 3 | 0.6667 |
| 15_592028 | 5 | 0.4 | 9 | 0.3333 | 1 | 1 | 6 | 0.5 | 21 | 0.4286 | 7 | 0.2857 | 5 | 0.4 | 3 | 0.6667 | 8 | 0.5 | 23 | 0.4348 |
| 15_592028 | 5 | 0.4 | 8 | 0.375 | NA | NA | 6 | 0.5 | 19 | 0.4211 | 8 | 0.5 | 3 | 0 | 1 | 0 | 8 | 0.25 | 20 | 0.3 |
| 15_592039 | 4 | 0.5 | 6 | 0.6667 | 7 | 0.5714 | 5 | 0.4 | 22 | 0.5455 | 4 | 1 | 5 | 0.6 | 5 | 0.6 | 3 | 1 | 17 | 0.7647 |
| 15_592041 | 5 | 0 | 6 | 0 | 6 | 0.3333 | 4 | 0.25 | 21 | 0.1429 | 4 | 0.5 | 5 | 0.4 | 6 | 0.3333 | 3 | 0.6667 | 18 | 0.4444 |
| 15_592138 | 3 | 0.3333 | 4 | 0 | 2 | 0 | 3 | 0 | 12 | 0.0833 | 2 | 0.5 | 5 | 0 | 6 | 0 | 1 | 0 | 14 | 0.0714 |
| 15_592180 | 3 | 0 | 4 | 0.25 | 3 | 0 | 2 | 0 | 12 | 0.0833 | 3 | 0.3333 | 5 | 0.2 | 3 | 0.3333 | 2 | 0 | 13 | 0.2308 |
| 15_592202 | 1 | 0 | 4 | 0.25 | 2 | 0 | 2 | 0 | 9 | 0.1111 | 3 | 0.3333 | 5 | 0 | 3 | 0 | 1 | 0 | 12 | 0.0833 |
| 15_592419 | NA | NA | 1 | 1 | 3 | 0 | 1 | 0 | 5 | 0.2 | NA | NA | 4 | 0.25 | NA | NA | NA | NA | 4 | 0.25 |
| 15_595263 | 1 | 1 | 1 | 1 | NA | NA | NA | NA | 2 | 1 | NA | NA | NA | NA | 1 | 1 | NA | NA | 1 | 1 |
| 15_595265 | 1 | 1 | 1 | 1 | NA | NA | NA | NA | 2 | 1 | NA | NA | NA | NA | 2 | 1 | NA | NA | 2 | 1 |
| 15_595267 | 1 | 1 | 1 | 0 | NA | NA | NA | NA | 2 | 0.5 | NA | NA | 1 | 1 | 1 | 1 | NA | NA | 2 | 1 |
| 15_595268 | 1 | 0 | 1 | 1 | NA | NA | NA | NA | 2 | 0.5 | NA | NA | 1 | 0 | 1 | 1 | NA | NA | 2 | 0.5 |
| 15_595271 | 1 | 1 | 1 | 0 | NA | NA | NA | NA | 2 | 0.5 | NA | NA | 1 | 1 | 1 | 0 | NA | NA | 2 | 0.5 |
| 15_595272 | 1 | 0 | 1 | 1 | NA | NA | NA | NA | 2 | 0.5 | NA | NA | 1 | 1 | 1 | 1 | NA | NA | 2 | 1 |
| 15_595273 | 1 | 0 | 1 | 1 | NA | NA | NA | NA | 2 | 0.5 | NA | NA | 1 | 1 | 1 | 0 | NA | NA | 2 | 0.5 |
| 15_595276 | 1 | 1 | 1 | 1 | NA | NA | NA | NA | 2 | 1 | NA | NA | 1 | 1 | 2 | 1 | NA | NA | 3 | 1 |
| 15_595278 | 1 | 0 | 1 | 1 | NA | NA | NA | NA | 2 | 0.5 | NA | NA | 1 | 0 | 2 | 1 | NA | NA | 3 | 0.6667 |
| 15_606915 | NA | NA | 1 | 1 | 1 | 0 | 1 | 0 | 3 | 0.3333 | 1 | 0 | 2 | 0 | 2 | 0.5 | NA | NA | 5 | 0.2 |
| 15_606916 | NA | NA | 1 | 1 | NA | NA | 1 | 0 | 2 | 0.5 | 1 | 1 | 2 | 1 | 2 | 0.5 | NA | NA | 5 | 0.8 |
| 15_606918 | NA | NA | 1 | 1 | 1 | 0 | 1 | 0 | 3 | 0.3333 | 1 | 1 | 2 | 0 | 1 | 1 | NA | NA | 4 | 0.5 |
| 15_606918 | NA | NA | NA | NA | 1 | 1 | 1 | 0 | 2 | 0.5 | 1 | 1 | 2 | 0 | 2 | 0.5 | NA | NA | 5 | 0.4 |
| 15_606918 | NA | NA | NA | NA | 1 | 0 | 1 | 1 | 2 | 0.5 | 1 | 1 | 2 | 0 | 2 | 0.5 | NA | NA | 5 | 0.4 |
| 15_606919 | NA | NA | NA | NA | 1 | 0 | 1 | 1 | 2 | 0.5 | 1 | 1 | 2 | 0 | 2 | 0.5 | NA | NA | 5 | 0.4 |
| 15_606920 | NA | NA | NA | NA | 1 | 1 | 1 | 1 | 2 | 1 | 1 | 1 | 2 | 0.5 | 2 | 1 | NA | NA | 5 | 0.8 |
| 15_606939 | NA | NA | NA | NA | 1 | 1 | NA | NA | 1 | 1 | 1 | 1 | 1 | 1 | 2 | 1 | NA | NA | 4 | 1 |
| 15_606946 | NA | NA | NA | NA | 1 | 1 | NA | NA | 1 | 1 | NA | NA | 1 | 1 | 1 | 0 | NA | NA | 2 | 0.5 |
| 15_606948 | NA | NA | NA | NA | 1 | 1 | NA | NA | 1 | 1 | NA | NA | NA | NA | 1 | 1 | NA | NA | 1 | 1 |
| 15_606948 | NA | NA | NA | NA | 1 | 1 | NA | NA | 1 | 1 | NA | NA | 1 | 0 | 1 | 1 | NA | NA | 2 | 0.5 |
| 15_606949 | NA | NA | NA | NA | 1 | 1 | NA | NA | 1 | 1 | NA | NA | 1 | 1 | 1 | 1 | NA | NA | 2 | 1 |
| 15_606954 | NA | NA | NA | NA | 1 | 1 | NA | NA | 1 | 1 | NA | NA | NA | NA | 1 | 1 | NA | NA | 1 | 1 |
| 15_621306 | NA | NA | 1 | 1 | 1 | 0 | NA | NA | 2 | 0.5 | 1 | 0 | NA | NA | 1 | 1 | 1 | 0 | 3 | 0.3333 |
| 15_621306 | NA | NA | 1 | 1 | 1 | 1 | NA | NA | 2 | 1 | 1 | 1 | NA | NA | 1 | 0 | 1 | 0 | 3 | 0.3333 |
| 15_621311 | NA | NA | 1 | 1 | 1 | 1 | 1 | 1 | 3 | 1 | 1 | 0 | NA | NA | 1 | 1 | 1 | 1 | 3 | 0.6667 |
| 15_621313 | NA | NA | 1 | 1 | 1 | 1 | 1 | 1 | 3 | 1 | 1 | 1 | NA | NA | 1 | 1 | 1 | 1 | 3 | 1 |
| 15_621313 | NA | NA | 1 | 1 | 1 | 1 | 1 | 1 | 3 | 1 | 1 | 1 | NA | NA | NA | NA | NA | NA | 1 | 1 |
| 15_621315 | NA | NA | 1 | 1 | 1 | 1 | 1 | 1 | 3 | 1 | 1 | 1 | NA | NA | NA | NA | 1 | 1 | 2 | 1 |
| 15_621317 | NA | NA | 1 | 1 | 1 | 1 | 1 | 1 | 3 | 1 | 1 | 1 | NA | NA | NA | NA | 1 | 1 | 2 | 1 |
| 15_638010 | 14 | 0.6429 | 6 | 0.3333 | 13 | 0.6154 | 12 | 0.5 | 45 | 0.5556 | 2 | 1 | 12 | 0.3333 | 8 | 0.375 | 10 | 0.5 | 32 | 0.4375 |
| 15_638011 | 9 | 0.2222 | 2 | 0 | 8 | 0 | 6 | 0 | 25 | 0.08 | 2 | 0 | 6 | 0.3333 | 5 | 0 | 7 | 0.1429 | 20 | 0.15 |
| 15_638012 | 17 | 1 | 7 | 1 | 13 | 1 | 12 | 1 | 49 | 1 | 2 | 1 | 11 | 0.8182 | 8 | 1 | 10 | 1 | 31 | 0.9355 |
| 15_638013 | 16 | 1 | 5 | 1 | 12 | 1 | 11 | 0.9091 | 44 | 0.9773 | 1 | 1 | 12 | 1 | 5 | 1 | 9 | 1 | 27 | 1 |
| 15_638015 | 15 | 0.9333 | 5 | 1 | 10 | 1 | 10 | 1 | 40 | 0.975 | 1 | 1 | 10 | 0.8 | 7 | 1 | 10 | 0.8 | 28 | 0.8571 |
| 15_638015 | 10 | 0.4 | 5 | 0 | 11 | 0.3636 | 9 | 0 | 35 | 0.2286 | 1 | 0 | 9 | 0.2222 | 7 | 0.5714 | 9 | 0.3333 | 26 | 0.3462 |
| 15_638017 | 15 | 0.9333 | 6 | 1 | 12 | 0.8333 | 9 | 0.7778 | 42 | 0.881 | 1 | 1 | 10 | 0.8 | 8 | 0.75 | 10 | 0.7 | 29 | 0.7586 |
| 15_638018 | 13 | 0.4615 | 5 | 0.2 | 9 | 0.2222 | 10 | 0.5 | 37 | 0.3784 | 1 | 1 | 11 | 0.4545 | 8 | 0.125 | 7 | 0.2857 | 27 | 0.3333 |
| 15_638019 | 12 | 0.0833 | 3 | 0 | 10 | 0.3 | 9 | 0.2222 | 34 | 0.1765 | 1 | 1 | 11 | 0.0909 | 7 | 0.1429 | 9 | 0.2222 | 28 | 0.1786 |
| 15_638019 | 2 | 0.5 | NA | NA | 1 | 0 | 1 | 0 | 4 | 0.25 | NA | NA | 1 | 1 | NA | NA | 1 | 0 | 2 | 0.5 |
| 15_638020 | 13 | 0.1538 | 4 | 0.25 | 10 | 0 | 10 | 0 | 37 | 0.0811 | 1 | 0 | 7 | 0.2857 | 7 | 0.1429 | 9 | 0 | 24 | 0.125 |
| 15_638021 | 12 | 0.75 | 4 | 0.75 | 10 | 0.8 | 7 | 1 | 33 | 0.8182 | 1 | 1 | 10 | 0.7 | 6 | 0.6667 | 9 | 0.6667 | 26 | 0.6923 |
| 15_638022 | 13 | 0.8462 | 3 | 0.3333 | 9 | 0.3333 | 11 | 0.8182 | 36 | 0.6667 | 1 | 1 | 10 | 0.8 | 7 | 0.5714 | 10 | 0.9 | 28 | 0.7857 |
| 15_638022 | 14 | 0.7857 | 4 | 0.75 | 11 | 0.8182 | 10 | 0.6 | 39 | 0.7436 | 1 | 0 | 8 | 0.375 | 7 | 1 | 9 | 1 | 25 | 0.76 |
| 15_638023 | 13 | 0.1538 | 3 | 0.3333 | 10 | 0.4 | 10 | 0.2 | 36 | 0.25 | 1 | 0 | 9 | 0.2222 | 5 | 0 | 7 | 0.2857 | 22 | 0.1818 |
| 15_638033 | 29 | 0.1379 | 14 | 0.0714 | 23 | 0.0435 | 27 | 0.1111 | 93 | 0.0968 | 13 | 0.0769 | 20 | 0.05 | 22 | 0.0455 | 25 | 0.04 | 80 | 0.05 |
| 15_639589 | 1 | 0 | 5 | 0.4 | 2 | 0.5 | 5 | 0 | 13 | 0.2308 | 2 | 0 | 4 | 0 | 1 | 1 | 2 | 0.5 | 9 | 0.2222 |
| 15_639596 | 3 | 0 | 3 | 0 | NA | NA | 5 | 0.2 | 11 | 0.0909 | 1 | 0 | 1 | 0 | 1 | 1 | 2 | 0 | 5 | 0.2 |
| 15_639604 | 5 | 1 | 9 | 0.7778 | 8 | 0.25 | 5 | 0.6 | 27 | 0.6296 | 5 | 0.4 | 3 | 0.6667 | 4 | 0 | 8 | 0.625 | 20 | 0.45 |
| 15_639605 | 6 | 0.3333 | 9 | 0.3333 | 8 | 0.25 | 6 | 0.3333 | 29 | 0.3103 | 5 | 0.2 | 2 | 0 | 6 | 0 | 9 | 0.4444 | 22 | 0.2273 |
| 15_639605 | 5 | 0.6 | 9 | 0.8889 | 9 | 0.7778 | 5 | 1 | 28 | 0.8214 | 4 | 0.5 | 1 | 1 | 6 | 0.5 | 8 | 0.75 | 19 | 0.6316 |
| 15_639605 | 7 | 0.2857 | 9 | 0.3333 | 8 | 0.625 | 4 | 0.25 | 28 | 0.3929 | 5 | 0 | 1 | 0 | 7 | 0.2857 | 8 | 0.125 | 21 | 0.1429 |
| 15_639609 | 7 | 0.2857 | 9 | 0.3333 | 7 | 0.4286 | 4 | 0.25 | 27 | 0.3333 | 4 | 0.5 | 2 | 1 | 5 | 0 | 7 | 0.4286 | 18 | 0.3889 |
| 15_639610 | 7 | 0.7143 | 9 | 0.4444 | 4 | 0.5 | 4 | 0.25 | 24 | 0.5 | 3 | 0.3333 | 2 | 0.5 | 4 | 0 | 6 | 0.3333 | 15 | 0.2667 |
| 15_639611 | 2 | 1 | 4 | 0.75 | 4 | 0.75 | 3 | 1 | 13 | 0.8462 | 4 | 1 | 1 | 0 | 5 | 0.6 | 5 | 0.8 | 15 | 0.7333 |
| 15_639611 | 5 | 0.2 | 8 | 0.125 | 6 | 0 | 3 | 0 | 22 | 0.0909 | 4 | 0 | 2 | 1 | 6 | 0 | 5 | 0 | 17 | 0.1176 |

| | | | | | | | | | | | | | | | | | | | | |
|---|---|---|---|---|---|---|---|---|---|---|---|---|---|---|---|---|---|---|---|---|
| 15_639660 | 2 | 0.5 | 4 | 0 | 3 | 0.6667 | 2 | 0 | 11 | 0.2727 | 4 | 0.25 | 2 | 0 | 5 | 0.4 | 2 | 0.5 | 13 | 0.3077 |
| 15_639661 | 3 | 0.6667 | 7 | 0 | 4 | 0.5 | 3 | 0.6667 | 17 | 0.3529 | 4 | 0.25 | 2 | 0 | 3 | 0.6667 | 2 | 0.5 | 11 | 0.3636 |
| 15_639662 | 3 | 0.6667 | 7 | 0.4286 | 2 | 0.5 | 1 | 0 | 13 | 0.4615 | 2 | 0.5 | 2 | 0 | 5 | 0.6 | 3 | 0.3333 | 12 | 0.4167 |
| 15_639662 | 3 | 0.3333 | 6 | 0.1667 | 3 | 0 | 3 | 0 | 15 | 0.1333 | 4 | 0 | 1 | 0 | 5 | 0.2 | 3 | 0.3333 | 13 | 0.1538 |
| 15_645710 | NA | NA | 3 | 1 | NA | NA | 2 | 1 | 5 | 1 | 1 | 1 | NA | NA | 1 | 1 | 1 | 1 | 3 | 1 |
| 15_645712 | NA | NA | 5 | 0.2 | NA | NA | 2 | 0 | 7 | 0.1429 | 1 | 1 | NA | NA | 1 | 1 | NA | NA | 2 | 1 |
| 15_645713 | NA | NA | 4 | 0.5 | NA | NA | 2 | 0.5 | 6 | 0.5 | 1 | 0 | NA | NA | NA | NA | 1 | 1 | 2 | 0.5 |
| 15_646557 | NA | NA | 4 | 0.25 | NA | NA | 2 | 0.5 | 6 | 0.3333 | NA | NA | NA | NA | 1 | 1 | NA | NA | 1 | 1 |
| 15_647923 | NA | NA | 3 | 1 | NA | NA | NA | NA | 3 | 1 | NA | NA | NA | NA | 1 | 1 | NA | NA | 1 | 1 |
| 15_647932 | NA | NA | 1 | 1 | NA | NA | NA | NA | 1 | 1 | NA | NA | NA | NA | 1 | 1 | NA | NA | 1 | 1 |
| 15_657431 | 3 | 0.6667 | 10 | 0.8 | 5 | 0.6 | 1 | 1 | 19 | 0.7368 | 4 | 0.5 | 3 | 0.3333 | 3 | 0.6667 | 5 | 1 | 15 | 0.6667 |
| 15_657432 | 3 | 0 | 11 | 0.0909 | 6 | 0.1667 | 2 | 0 | 22 | 0.0909 | 4 | 0.25 | 3 | 0 | 4 | 0.25 | 6 | 0.1667 | 17 | 0.1765 |
| 15_657435 | 3 | 0 | 9 | 0.1111 | 7 | 0 | NA | NA | 19 | 0.0526 | 5 | 0 | 3 | 0.6667 | 4 | 0.25 | 4 | 0.25 | 16 | 0.25 |
| 15_658593 | 1 | 0 | 18 | 0.0556 | 3 | 0.3333 | 4 | 0 | 26 | 0.0769 | 8 | 0 | 6 | 0 | 11 | 0 | 4 | 0.5 | 29 | 0.069 |
| 15_658595 | NA | NA | 17 | 0.1176 | 3 | 0.3333 | 5 | 0 | 25 | 0.12 | 8 | 0.25 | 6 | 0 | 11 | 0.0909 | 1 | 0 | 26 | 0.1154 |
| 15_658607 | 1 | 0 | 11 | 0.0909 | 1 | 0 | 3 | 0 | 16 | 0.0625 | 5 | 0 | 2 | 0 | 7 | 0.2857 | 2 | 0 | 16 | 0.125 |
| 15_658616 | NA | NA | 11 | 0.0909 | 1 | 0 | 2 | 0 | 14 | 0.0714 | 4 | 0 | 3 | 0.3333 | 6 | 0 | 2 | 0 | 15 | 0.0667 |
| 15_658616 | NA | NA | 11 | 0.0909 | 1 | 0 | 4 | 0 | 16 | 0.0625 | 5 | 0.2 | 4 | 0 | 6 | 0 | 2 | 0 | 17 | 0.0588 |
| 15_658618 | 1 | 0 | 11 | 0.0909 | NA | NA | 5 | 0 | 17 | 0.0588 | 3 | 0 | 2 | 0 | 6 | 0 | 3 | 0.3333 | 14 | 0.0714 |
| 15_658620 | 2 | 0.5 | 11 | 0.1818 | 1 | 0 | 3 | 0 | 17 | 0.1765 | 4 | 0.25 | 4 | 0.25 | 5 | 0.2 | 2 | 0.5 | 15 | 0.2667 |
| 15_658621 | 2 | 0.5 | 8 | 0.625 | NA | NA | 5 | 0.8 | 15 | 0.6667 | 5 | 0.6 | 5 | 0.8 | 6 | 0.6667 | 2 | 0 | 18 | 0.6111 |
| 15_658622 | 1 | 1 | 8 | 0.625 | NA | NA | 3 | 1 | 12 | 0.75 | 3 | 1 | 4 | 1 | 3 | 0.6667 | 2 | 1 | 12 | 0.9167 |
| 15_658622 | NA | NA | 3 | 0.3333 | NA | NA | 4 | 0 | 7 | 0.1429 | 1 | 0 | 2 | 0.5 | 3 | 0.6667 | 2 | 0 | 8 | 0.375 |
| 15_658623 | NA | NA | 8 | 1 | NA | NA | 4 | 1 | 12 | 1 | 3 | 1 | 3 | 1 | 3 | 1 | 2 | 1 | 11 | 1 |
| 15_658624 | NA | NA | 7 | 0.1429 | NA | NA | 4 | 0.5 | 11 | 0.2727 | NA | NA | 2 | 0.5 | 4 | 0.25 | 2 | 0 | 8 | 0.25 |
| 15_658624 | NA | NA | 7 | 0.5714 | NA | NA | 5 | 0.4 | 12 | 0.5 | 3 | 1 | 3 | 1 | 4 | 0.5 | 2 | 1 | 12 | 0.8333 |
| 15_658624 | 1 | 0 | 9 | 0.1111 | NA | NA | 4 | 0 | 14 | 0.0714 | 3 | 0.3333 | 3 | 0.6667 | 4 | 0 | 2 | 0.5 | 12 | 0.3333 |
| 15_658625 | NA | NA | 7 | 0 | NA | NA | 3 | 0.3333 | 10 | 0.1 | 3 | 0 | 3 | 0.3333 | 4 | 0 | 1 | 0 | 11 | 0.0909 |
| 15_658633 | 1 | 0 | 6 | 0.1667 | NA | NA | 4 | 0.5 | 11 | 0.2727 | 4 | 0.25 | 4 | 0 | 3 | 0 | 2 | 0.5 | 13 | 0.1538 |
| 15_658651 | 2 | 0.5 | 4 | 0.25 | NA | NA | 3 | 0.3333 | 9 | 0.3333 | 1 | 0 | 4 | 0.25 | 1 | 0 | 1 | 0 | 7 | 0.1429 |
| 15_658660 | 1 | 1 | 4 | 0.75 | NA | NA | 2 | 0.5 | 7 | 0.7143 | NA | NA | 2 | 1 | 1 | 0 | NA | NA | 3 | 0.6667 |
| 15_658660 | 2 | 0 | 4 | 0.75 | NA | NA | 3 | 1 | 9 | 0.6667 | 1 | 1 | 2 | 0.5 | 1 | 0 | NA | NA | 4 | 0.5 |
| 15_658663 | 2 | 1 | 4 | 0.75 | NA | NA | 3 | 0 | 9 | 0.5556 | 1 | 0 | 2 | 0.5 | 1 | 1 | NA | NA | 4 | 0.5 |
| 15_658668 | 3 | 0.3333 | 5 | 0 | 1 | 0 | 2 | 0 | 11 | 0.0909 | 1 | 0 | NA | NA | 3 | 0.3333 | 1 | 0 | 5 | 0.2 |
| 15_658669 | 3 | 1 | 3 | 1 | 1 | 1 | 2 | 1 | 9 | 1 | 1 | 1 | NA | NA | 3 | 1 | 1 | 1 | 5 | 1 |
| 15_658669 | 1 | 1 | 4 | 1 | 1 | 1 | 1 | 1 | 7 | 1 | NA | NA | NA | NA | 3 | 1 | 1 | 1 | 4 | 1 |
| 15_658670 | 3 | 0 | 4 | 0.25 | NA | NA | 2 | 0.5 | 9 | 0.2222 | 1 | 1 | 1 | 1 | 3 | 0 | NA | NA | 5 | 0.4 |
| 15_658670 | 3 | 1 | 3 | 1 | NA | NA | 2 | 1 | 8 | 1 | NA | NA | NA | NA | 3 | 1 | 1 | 1 | 4 | 1 |
| 15_658671 | NA | NA | 3 | 1 | NA | NA | 1 | 1 | 4 | 1 | NA | NA | NA | NA | 2 | 1 | 1 | 1 | 3 | 1 |
| 15_658671 | NA | NA | 2 | 1 | NA | NA | 1 | 1 | 3 | 1 | NA | NA | NA | NA | 3 | 1 | 1 | 1 | 4 | 1 |
| 15_658672 | 1 | 0 | 4 | 0.25 | NA | NA | 2 | 0 | 7 | 0.1429 | 1 | 0 | NA | NA | 2 | 0 | 1 | 1 | 4 | 0.25 |
| 15_658672 | NA | NA | 4 | 1 | NA | NA | 2 | 0.5 | 6 | 0.8333 | NA | NA | NA | NA | 3 | 1 | 1 | 1 | 4 | 1 |
| 15_658672 | 1 | 1 | 4 | 1 | NA | NA | 2 | 1 | 7 | 1 | NA | NA | 1 | 1 | 3 | 1 | 1 | 1 | 5 | 1 |
| 15_658673 | NA | NA | 3 | 0.3333 | NA | NA | 2 | 0 | 5 | 0.2 | NA | NA | NA | NA | 2 | 1 | 1 | 1 | 3 | 1 |
| 15_658673 | 1 | 1 | 5 | 0.8 | NA | NA | 2 | 1 | 8 | 0.875 | NA | NA | NA | NA | 3 | 1 | 1 | 1 | 4 | 1 |
| 15_658673 | 1 | 1 | 4 | 0.5 | NA | NA | 2 | 1 | 7 | 0.7143 | NA | NA | NA | NA | 1 | 0 | 1 | 1 | 2 | 0.5 |
| 15_658674 | 1 | 0 | 4 | 0.5 | NA | NA | 2 | 0.5 | 7 | 0.4286 | 1 | 1 | 1 | 0 | 3 | 0.3333 | 1 | 0 | 6 | 0.3333 |
| 15_658674 | 1 | 0 | 4 | 0.25 | NA | NA | 2 | 0 | 7 | 0.1429 | 1 | 0 | 1 | 0 | 2 | 0.5 | 1 | 1 | 5 | 0.4 |
| 15_658675 | 2 | 0.5 | 5 | 0.4 | NA | NA | 2 | 0.5 | 9 | 0.4444 | 1 | 1 | NA | NA | 2 | 1 | 1 | 0 | 4 | 0.75 |
| 15_661579 | NA | NA | 2 | 0.5 | NA | NA | NA | NA | 2 | 0.5 | NA | NA | NA | NA | 2 | 0.5 | 1 | 0 | 3 | 0.3333 |
| 15_664815 | 2 | 0.5 | 2 | 0 | 2 | 0 | 6 | 0.1667 | 12 | 0.1667 | 1 | 0 | 3 | 0.6667 | 4 | 0.75 | 3 | 0.6667 | 11 | 0.6364 |
| 15_664815 | 1 | 0 | 2 | 0.5 | 2 | 1 | 6 | 0.6667 | 11 | 0.6364 | NA | NA | 3 | 0.3333 | 4 | 0.25 | 3 | 0.3333 | 10 | 0.3 |
| 15_664816 | 2 | 1 | 2 | 0.5 | 2 | 1 | 7 | 1 | 13 | 0.9231 | 2 | 0.5 | 4 | 1 | 5 | 1 | 3 | 1 | 14 | 0.9286 |
| 15_664816 | 1 | 1 | 2 | 0.5 | 2 | 1 | 7 | 1 | 12 | 0.9167 | 1 | 1 | 4 | 0.75 | 3 | 1 | 3 | 1 | 11 | 0.9091 |
| 15_664817 | 2 | 0 | 1 | 1 | 2 | 0.5 | 3 | 0.3333 | 8 | 0.375 | 1 | 1 | 3 | 0.6667 | 4 | 0 | 3 | 0.3333 | 11 | 0.3636 |
| 15_664817 | 2 | 0 | 2 | 0 | 1 | 0 | 6 | 0.1667 | 11 | 0.0909 | 2 | 0.5 | 3 | 0.3333 | 5 | 0 | 3 | 0.3333 | 13 | 0.2308 |
| 15_665087 | 1 | 0 | NA | NA | 2 | 1 | 4 | 1 | 7 | 0.8571 | 1 | 1 | 1 | 1 | 4 | 0.5 | 1 | 0 | 7 | 0.5714 |
| 15_665137 | 1 | 1 | 1 | 0 | 1 | 1 | 5 | 0.6 | 8 | 0.625 | 2 | 0.5 | 2 | 0 | 4 | 0.25 | 1 | 0 | 9 | 0.2222 |
| 15_665138 | 1 | 1 | 1 | 1 | 1 | 1 | 5 | 1 | 8 | 1 | 2 | 1 | 2 | 0.5 | 4 | 0.75 | 2 | 0.5 | 10 | 0.7 |
| 15_665138 | 1 | 1 | 1 | 0 | 2 | 0.5 | 6 | 0 | 10 | 0.2 | 3 | 0 | 2 | 0.5 | 5 | 0.2 | 1 | 0 | 11 | 0.1818 |
| 15_665138 | 1 | 0 | 2 | 0 | 2 | 0 | 6 | 0.3333 | 11 | 0.1818 | 2 | 0 | 2 | 0 | 5 | 0.4 | 1 | 0 | 10 | 0.2 |
| 15_665140 | 1 | 0 | 2 | 0 | 2 | 0.5 | 6 | 0.1667 | 11 | 0.1818 | 3 | 0.3333 | 2 | 0 | 5 | 0 | 3 | 0 | 13 | 0.0769 |
| 15_665140 | 1 | 1 | 2 | 0.5 | 2 | 0 | 5 | 1 | 10 | 0.7 | 2 | 1 | 2 | 0.5 | 6 | 0.6667 | 2 | 1 | 12 | 0.75 |
| 15_665141 | NA | NA | 2 | 0 | 2 | 0 | 5 | 0.6 | 9 | 0.3333 | 3 | 0 | 1 | 1 | 4 | 0 | 2 | 0 | 10 | 0.1 |
| 15_665142 | 1 | 1 | 1 | 1 | 2 | 0.5 | 6 | 0.8333 | 10 | 0.8 | 3 | 0.6667 | 2 | 1 | 5 | 0.8 | 3 | 0.3333 | 13 | 0.6923 |

| | | | | | | | | | | | | | | | | | | | | |
|---|---|---|---|---|---|---|---|---|---|---|---|---|---|---|---|---|---|---|---|---|
| 15_6651427 | 1 | 0 | 2 | 0.5 | 1 | 1 | 5 | 0.8 | 9 | 0.6667 | 3 | 0 | 2 | 0.5 | 6 | 0.1667 | 1 | 0 | 12 | 0.1667 |
| 15_6651433 | NA | NA | 2 | 1 | 2 | 0 | 3 | 0.6667 | 7 | 0.5714 | 2 | 0.5 | 1 | 0 | 6 | 0.5 | 3 | 0.3333 | 12 | 0.4167 |
| 15_6651433 | 1 | 0 | 3 | 0 | 2 | 0 | 4 | 0.75 | 10 | 0.3 | 2 | 0.5 | 2 | 0 | 6 | 0.3333 | 3 | 0 | 13 | 0.2308 |
| 15_6651456 | 1 | 0 | 3 | 0 | 2 | 0 | 6 | 0.3333 | 12 | 0.1667 | 3 | 0 | 2 | 0 | 5 | 0.4 | 2 | 0 | 12 | 0.1667 |
| 15_6651468 | 1 | 0 | 2 | 0 | 2 | 0 | 5 | 0.2 | 10 | 0.1 | 3 | 0 | 1 | 0 | 5 | 0 | 1 | 1 | 10 | 0.1 |
| 15_6651485 | 1 | 1 | 2 | 1 | 2 | 0.5 | 4 | 0.5 | 9 | 0.6667 | 3 | 0.3333 | 2 | 0.5 | 6 | 0.5 | 2 | 0.5 | 13 | 0.4615 |
| 15_6651511 | 1 | 0 | 3 | 0 | 2 | 0.5 | 6 | 0.6667 | 12 | 0.4167 | 3 | 0.3333 | 2 | 0.5 | 6 | 0.5 | 3 | 0 | 14 | 0.3571 |
| 15_6651518 | 1 | 0 | 3 | 0 | 2 | 0.5 | 5 | 0 | 11 | 0.0909 | 3 | 0 | 2 | 0 | 7 | 0.1429 | 3 | 0 | 15 | 0.0667 |
| 15_6651601 | 2 | 0.5 | 4 | 0 | NA | NA | 4 | 0 | 10 | 0.1 | NA | NA | 1 | 0 | 3 | 0.6667 | 1 | 0 | 5 | 0.4 |
| 15_6651620 | 2 | 0 | 4 | 0.25 | NA | NA | 3 | 0 | 9 | 0.1111 | NA | NA | 2 | 0 | 4 | 0.25 | 1 | 0 | 7 | 0.1429 |
| 15_6651661 | 2 | 0 | 5 | 0.4 | NA | NA | 2 | 0 | 9 | 0.2222 | NA | NA | 2 | 0 | 4 | 0.5 | 1 | 0 | 7 | 0.2857 |
| 15_6665393 | NA | NA | 5 | 0.4 | NA | NA | NA | NA | 5 | 0.4 | 1 | 1 | NA | NA | NA | NA | NA | NA | 1 | 1 |
| 15_6665397 | NA | NA | 4 | 1 | NA | NA | NA | NA | 4 | 1 | 1 | 1 | NA | NA | NA | NA | NA | NA | 1 | 1 |
| 15_6665411 | NA | NA | 5 | 0.6 | NA | NA | NA | NA | 5 | 0.6 | 1 | 1 | NA | NA | NA | NA | NA | NA | 1 | 1 |
| 15_6665413 | NA | NA | 2 | 0.5 | NA | NA | NA | NA | 2 | 0.5 | 1 | 1 | NA | NA | NA | NA | NA | NA | 1 | 1 |
| 15_6665416 | NA | NA | 3 | 1 | NA | NA | NA | NA | 3 | 1 | 1 | 1 | NA | NA | NA | NA | NA | NA | 1 | 1 |
| 15_6665424 | NA | NA | 4 | 1 | NA | NA | NA | NA | 4 | 1 | 1 | 1 | NA | NA | NA | NA | NA | NA | 1 | 1 |
| 15_6665426 | NA | NA | 4 | 1 | NA | NA | NA | NA | 4 | 1 | 1 | 1 | NA | NA | NA | NA | NA | NA | 1 | 1 |
| 15_6665434 | NA | NA | 2 | 1 | NA | NA | NA | NA | 2 | 1 | 1 | 1 | NA | NA | NA | NA | NA | NA | 1 | 1 |
| 15_6665438 | NA | NA | 5 | 0.4 | NA | NA | NA | NA | 5 | 0.4 | 1 | 1 | NA | NA | NA | NA | NA | NA | 1 | 1 |
| 15_6680164 | 7 | 0 | 24 | 0.125 | 9 | 0 | 8 | 0.125 | 48 | 0.0833 | 16 | 0 | 14 | 0.0714 | 8 | 0.125 | 13 | 0.1538 | 51 | 0.0784 |
| 15_6680183 | 11 | 0.0909 | 25 | 0 | 11 | 0.0909 | 9 | 0.1111 | 56 | 0.0536 | 17 | 0 | 15 | 0.1333 | 8 | 0.125 | 11 | 0.1818 | 51 | 0.098 |
| 15_6680215 | 6 | 0.1667 | 12 | 0.1667 | 9 | 0 | 2 | 0 | 29 | 0.1034 | 9 | 0 | 11 | 0.0909 | 4 | 0 | 6 | 0.1667 | 30 | 0.0667 |
| 15_6680255 | 5 | 0 | 20 | 0.1 | 8 | 0 | 5 | 0.2 | 38 | 0.0789 | 9 | 0.1111 | 8 | 0.125 | 2 | 0 | 8 | 0.375 | 27 | 0.1852 |
| 15_6680260 | 4 | 1 | 13 | 0.7692 | 8 | 0.375 | 4 | 0.5 | 29 | 0.6552 | 10 | 0.6 | 6 | 0.5 | 5 | 0.4 | 6 | 0.6667 | 27 | 0.5556 |
| 15_6680269 | 6 | 0.3333 | 13 | 0.2308 | 8 | 0.25 | 4 | 0.5 | 31 | 0.2903 | 7 | 0.4286 | 5 | 0.4 | 5 | 0.6 | 9 | 0.2222 | 26 | 0.3846 |
| 15_6697505 | 1 | 0 | 2 | 1 | 1 | 0 | 4 | 0.25 | 8 | 0.375 | 3 | 0.6667 | 2 | 0.5 | 5 | 0.6 | 2 | 0.5 | 12 | 0.5833 |
| 15_6697724 | 3 | 0.6667 | 1 | 0 | 1 | 0 | 3 | 0.3333 | 8 | 0.375 | 1 | 1 | 1 | 0 | 2 | 0 | 1 | 1 | 5 | 0.4 |
| 15_6697749 | 1 | 0 | 1 | 1 | 1 | 1 | 1 | 0 | 4 | 0.5 | NA | NA | 1 | 1 | 3 | 0.6667 | 1 | 1 | 5 | 0.8 |
| 15_6697758 | 2 | 0.5 | 1 | 1 | 1 | 1 | 1 | 0 | 5 | 0.6 | 1 | 0 | 1 | 1 | 4 | 0.25 | NA | NA | 6 | 0.3333 |
| 15_6697806 | 2 | 1 | 1 | 0 | 1 | 1 | 1 | 1 | 5 | 0.8 | NA | NA | 1 | 1 | 2 | 0.5 | NA | NA | 3 | 0.6667 |
| 15_6697809 | 1 | 0 | 1 | 0 | 1 | 1 | 1 | 1 | 4 | 0.5 | 1 | 1 | 1 | 0 | 3 | 0 | NA | NA | 5 | 0.2 |
| 15_6697812 | 2 | 0.5 | NA | NA | 1 | 1 | 1 | 0 | 4 | 0.5 | 1 | 1 | 1 | 1 | 1 | 1 | NA | NA | 3 | 1 |
| 15_6697840 | 2 | 1 | NA | NA | NA | NA | 1 | 0 | 3 | 0.6667 | 1 | 1 | 1 | 1 | 1 | 0 | NA | NA | 3 | 0.6667 |
| 15_6697848 | 1 | 1 | 1 | 1 | 2 | 0.5 | 2 | 1 | 6 | 0.8333 | 1 | 0 | 1 | 1 | 1 | 1 | NA | NA | 3 | 0.6667 |
| 15_6698545 | 1 | 1 | NA | NA | 2 | 0 | 2 | 0 | 5 | 0.2 | 1 | 1 | 1 | 0 | 1 | 0 | NA | NA | 3 | 0.3333 |
| 15_6698552 | NA | NA | NA | NA | 1 | 0 | 1 | 1 | 2 | 0.5 | 1 | 1 | 1 | 0 | 1 | 1 | NA | NA | 3 | 0.6667 |
| 15_6698567 | 1 | 0 | NA | NA | 1 | 0 | 2 | 1 | 4 | 0.5 | 1 | 1 | 1 | 0 | NA | NA | NA | NA | 2 | 0.5 |
| 15_6698577 | 1 | 1 | NA | NA | 1 | 1 | 2 | 1 | 4 | 1 | 1 | 1 | 1 | 1 | 1 | 1 | NA | NA | 3 | 1 |
| 15_6698588 | 1 | 1 | NA | NA | 1 | 0 | 2 | 0.5 | 4 | 0.5 | 1 | 1 | 1 | 0 | 1 | 0 | NA | NA | 3 | 0.3333 |
| 15_6698589 | 1 | 0 | NA | NA | NA | NA | 2 | 0.5 | 3 | 0.3333 | 1 | 1 | 1 | 1 | 1 | 0 | NA | NA | 3 | 0.6667 |
| 15_6698594 | 1 | 1 | NA | NA | 1 | 0 | 2 | 0.5 | 4 | 0.5 | 1 | 0 | 1 | 0 | 1 | 1 | NA | NA | 3 | 0.3333 |
| 15_6698599 | 1 | 1 | NA | NA | 1 | 0 | 2 | 0 | 4 | 0.25 | 1 | 1 | 1 | 1 | 1 | 0 | NA | NA | 3 | 0.6667 |
| 15_6795569 | 1 | 0 | 1 | 0 | NA | NA | 3 | 1 | 5 | 0.6 | 1 | 0 | NA | NA | 2 | 0.5 | 2 | 0 | 5 | 0.2 |
| 15_6795598 | 1 | 0 | 1 | 0 | NA | NA | 3 | 0.3333 | 5 | 0.2 | 1 | 0 | NA | NA | 2 | 0.5 | 2 | 0 | 5 | 0.2 |
| 15_6795628 | NA | NA | 1 | 0 | NA | NA | 3 | 0.6667 | 4 | 0.5 | 1 | 1 | NA | NA | 2 | 1 | 2 | 0.5 | 5 | 0.8 |
| 15_6805174 | NA | NA | NA | NA | NA | NA | 2 | 0.5 | 2 | 0.5 | NA | NA | NA | NA | 1 | 1 | NA | NA | 1 | 1 |
| 15_7131151 | 2 | 0 | 4 | 0 | 5 | 0 | 4 | 0.25 | 15 | 0.0667 | 3 | 0.3333 | 6 | 0 | 1 | 0 | 4 | 0 | 14 | 0.0714 |
| 15_7131776 | NA | NA | 1 | 1 | NA | NA | 2 | 0.5 | 3 | 0.6667 | 1 | 1 | 1 | 1 | NA | NA | 3 | 1 | 5 | 1 |
| 15_7131780 | NA | NA | NA | NA | NA | NA | 1 | 1 | 1 | 1 | 1 | 1 | 1 | 0 | NA | NA | 3 | 0.6667 | 5 | 0.6 |
| 15_7131789 | NA | NA | 1 | 1 | NA | NA | 2 | 0.5 | 3 | 0.6667 | 1 | 0 | 1 | 1 | NA | NA | 2 | 1 | 4 | 0.75 |
| 15_7133403 | NA | NA | 1 | 1 | NA | NA | 1 | 1 | 2 | 1 | NA | NA | NA | NA | NA | NA | 2 | 1 | 2 | 1 |
| 15_7133415 | NA | NA | 1 | 1 | NA | NA | 1 | 1 | 2 | 1 | NA | NA | NA | NA | NA | NA | 1 | 1 | 1 | 1 |
| 15_7133502 | NA | NA | 1 | 1 | NA | NA | NA | NA | 1 | 1 | NA | NA | NA | NA | NA | NA | 2 | 0.5 | 2 | 0.5 |
| 15_7133523 | NA | NA | 1 | 1 | NA | NA | NA | NA | 1 | 1 | NA | NA | NA | NA | NA | NA | 2 | 0.5 | 2 | 0.5 |
| 15_7166727 | 2 | 1 | 3 | 1 | 1 | 0 | 2 | 0 | 8 | 0.625 | 2 | 0 | 1 | 0 | 1 | 0 | 2 | 0.5 | 6 | 0.1667 |
| 15_7237397 | 1 | 1 | 1 | 1 | NA | NA | NA | NA | 2 | 1 | 2 | 0 | NA | NA | 2 | 1 | 2 | 0.5 | 6 | 0.5 |
| 15_7237402 | 1 | 0 | 2 | 0.5 | 1 | 0 | NA | NA | 4 | 0.25 | 2 | 0 | NA | NA | 2 | 0.5 | 3 | 0 | 7 | 0.1429 |
| 15_7237430 | 1 | 1 | 2 | 0.5 | 1 | 1 | NA | NA | 4 | 0.75 | 2 | 0 | NA | NA | 2 | 0.5 | 2 | 0.5 | 6 | 0.3333 |
| 15_7237895 | 1 | 0 | 5 | 0.2 | 1 | 0 | 1 | 0 | 8 | 0.125 | 2 | 0 | 1 | 1 | 2 | 0 | 1 | 0 | 6 | 0.1667 |
| 15_7237915 | 1 | 1 | 4 | 0 | 1 | 0 | 1 | 0 | 7 | 0.1429 | 2 | 0 | 1 | 1 | 2 | 0.5 | 1 | 0 | 6 | 0.3333 |
| 15_7238037 | 1 | 0 | 2 | 0.5 | 1 | 1 | 1 | 0 | 5 | 0.4 | NA | NA | NA | NA | 1 | 1 | 1 | 0 | 2 | 0.5 |
| 15_7238223 | 1 | 0 | NA | NA | NA | NA | 1 | 1 | 2 | 0.5 | NA | NA | NA | NA | NA | NA | 1 | 1 | 1 | 1 |
| 15_7238229 | 1 | 1 | NA | NA | NA | NA | 1 | 1 | 2 | 1 | NA | NA | NA | NA | NA | NA | 1 | 1 | 1 | 1 |
| 15_7238236 | 2 | 1 | NA | NA | NA | NA | 1 | 1 | 3 | 1 | NA | NA | NA | NA | NA | NA | 1 | 1 | 1 | 1 |

| | | | | | | | | | | | | | | | | | | | | |
|---|---|---|---|---|---|---|---|---|---|---|---|---|---|---|---|---|---|---|---|---|
| 15_723825 | 1 | 0 | NA | NA | NA | NA | 1 | 1 | 2 | 0.5 | NA | NA | NA | NA | NA | NA | 1 | 1 | 1 | 1 |
| 15_726781 | 19 | 0.1053 | 33 | 0.1212 | 17 | 0.2941 | 22 | 0.0455 | 91 | 0.1319 | 12 | 0.0833 | 18 | 0.0556 | 29 | 0.1034 | 20 | 0.05 | 79 | 0.0759 |
| 15_726784 | 20 | 0.1 | 31 | 0.1935 | 15 | 0.2667 | 23 | 0.1739 | 89 | 0.1798 | 10 | 0.3 | 22 | 0.1818 | 25 | 0.16 | 19 | 0.2105 | 76 | 0.1974 |
| 15_726784 | 17 | 1 | 33 | 0.9394 | 14 | 0.9286 | 23 | 0.9565 | 87 | 0.954 | 12 | 1 | 19 | 0.8947 | 25 | 0.92 | 19 | 0.8421 | 75 | 0.9067 |
| 15_726785 | 8 | 0.75 | 15 | 0.8 | 5 | 0.6 | 11 | 0.9091 | 39 | 0.7949 | 5 | 1 | 14 | 0.8571 | 14 | 0.8571 | 12 | 0.9167 | 45 | 0.8889 |
| 15_726786 | 20 | 1 | 31 | 1 | 12 | 1 | 22 | 1 | 85 | 1 | 12 | 1 | 20 | 1 | 27 | 1 | 18 | 1 | 77 | 1 |
| 15_726787 | 16 | 0.8125 | 33 | 0.7273 | 8 | 0.875 | 22 | 0.8636 | 79 | 0.7975 | 13 | 0.9231 | 19 | 0.7368 | 24 | 0.625 | 20 | 0.9 | 76 | 0.7763 |
| 15_726787 | 17 | 1 | 31 | 0.9032 | 10 | 0.9 | 22 | 0.9091 | 80 | 0.925 | 13 | 1 | 19 | 0.7368 | 26 | 0.8846 | 17 | 0.8824 | 75 | 0.8667 |
| 15_726788 | 18 | 1 | 29 | 1 | 11 | 1 | 21 | 1 | 79 | 1 | 13 | 0.9231 | 20 | 1 | 26 | 0.9615 | 18 | 1 | 77 | 0.974 |
| 15_726788 | 17 | 0.3529 | 25 | 0.24 | 9 | 0.3333 | 18 | 0.3333 | 69 | 0.3043 | 11 | 0.4545 | 16 | 0.25 | 23 | 0.3478 | 18 | 0.2222 | 68 | 0.3088 |
| 15_726789 | 15 | 0.0667 | 28 | 0.0714 | 11 | 0.0909 | 19 | 0.1579 | 73 | 0.0959 | 12 | 0.1667 | 16 | 0.0625 | 18 | 0 | 19 | 0.1579 | 65 | 0.0923 |
| 15_726790 | 13 | 0.3077 | 31 | 0.0323 | 9 | 0.1111 | 17 | 0 | 70 | 0.0857 | 10 | 0 | 17 | 0 | 22 | 0.1364 | 16 | 0.125 | 65 | 0.0769 |
| 15_726790 | 12 | 1 | 25 | 1 | 8 | 1 | 14 | 1 | 59 | 1 | 11 | 1 | 16 | 1 | 20 | 1 | 16 | 1 | 63 | 1 |
| 15_726790 | 16 | 0.375 | 29 | 0.2414 | 8 | 0.375 | 16 | 0.5 | 69 | 0.3478 | 10 | 0.2 | 18 | 0.2778 | 22 | 0.3182 | 17 | 0.3529 | 67 | 0.2985 |
| 15_726790 | 15 | 0.5333 | 31 | 0.6129 | 10 | 0.5 | 18 | 0.6111 | 74 | 0.5811 | 9 | 0.6667 | 17 | 0.5882 | 23 | 0.5652 | 18 | 0.6667 | 67 | 0.6119 |
| 15_726791 | 15 | 0.3333 | 31 | 0.1613 | 10 | 0 | 16 | 0.25 | 72 | 0.1944 | 10 | 0.1 | 17 | 0.2353 | 24 | 0.0417 | 18 | 0.1111 | 69 | 0.1159 |
| 15_726791 | 13 | 0.6154 | 30 | 0.6 | 9 | 0.8889 | 15 | 0.9333 | 67 | 0.7164 | 9 | 0.6667 | 18 | 0.7778 | 21 | 0.6667 | 16 | 0.625 | 64 | 0.6875 |
| 15_726793 | 13 | 1 | 29 | 0.8966 | 9 | 0.8889 | 16 | 0.9375 | 67 | 0.9254 | 9 | 1 | 16 | 0.9375 | 21 | 0.9524 | 15 | 1 | 61 | 0.9672 |
| 15_726795 | 11 | 0.7273 | 20 | 0.9 | 4 | 0.75 | 10 | 0.8 | 45 | 0.8222 | 8 | 0.75 | 12 | 0.9167 | 20 | 0.95 | 13 | 1 | 53 | 0.9245 |
| 15_726795 | 10 | 0.3 | 17 | 0.1176 | 4 | 0.25 | 8 | 0.375 | 39 | 0.2308 | 7 | 0.2857 | 10 | 0.5 | 13 | 0.3077 | 8 | 0.5 | 38 | 0.3947 |
| 15_726795 | 11 | 0.4545 | 26 | 0.1538 | 7 | 0.2857 | 16 | 0.1875 | 60 | 0.2333 | 10 | 0.1 | 14 | 0.3571 | 19 | 0.1579 | 14 | 0.1429 | 57 | 0.193 |
| 15_726796 | 10 | 0.9 | 22 | 0.6818 | 6 | 0.8333 | 10 | 0.7 | 48 | 0.75 | 6 | 0.8333 | 14 | 0.7143 | 14 | 0.7143 | 11 | 0.9091 | 45 | 0.7778 |
| 15_726796 | 8 | 0.5 | 17 | 0.2941 | 6 | 0.5 | 7 | 0.4286 | 38 | 0.3947 | 8 | 0.25 | 9 | 0.5556 | 14 | 0.5 | 12 | 0.3333 | 43 | 0.4186 |
| 15_726797 | 12 | 0.5 | 20 | 0.35 | 7 | 0.5714 | 14 | 0.7143 | 53 | 0.5094 | 9 | 0.6667 | 12 | 0.4167 | 12 | 0.5 | 10 | 0.4 | 43 | 0.4884 |
| 15_726798 | 13 | 0.2308 | 20 | 0.1 | 7 | 0.1429 | 12 | 0.4167 | 52 | 0.2115 | 10 | 0.5 | 9 | 0.2222 | 15 | 0.2 | 12 | 0.3333 | 46 | 0.3043 |
| 15_726798 | 12 | 1 | 25 | 1 | 7 | 1 | 14 | 1 | 58 | 1 | 9 | 1 | 10 | 1 | 19 | 1 | 13 | 1 | 51 | 1 |
| 15_726800 | 12 | 1 | 21 | 1 | 6 | 1 | 12 | 0.9167 | 51 | 0.9804 | 5 | 1 | 8 | 1 | 13 | 1 | 11 | 1 | 37 | 1 |
| 15_726805 | 12 | 0.3333 | 19 | 0.3684 | 6 | 0.3333 | 14 | 0.5714 | 51 | 0.4118 | 9 | 0.4444 | 9 | 0.3333 | 14 | 0.5 | 11 | 0.0909 | 43 | 0.3488 |
| 15_726807 | 8 | 0.875 | 16 | 1 | 5 | 1 | 11 | 1 | 40 | 0.975 | 8 | 0.875 | 9 | 1 | 12 | 1 | 9 | 1 | 38 | 0.9737 |
| 15_726807 | 8 | 0.375 | 9 | 0.4444 | 2 | 0 | 8 | 0.25 | 27 | 0.3333 | 6 | 0.3333 | 7 | 0.1429 | 8 | 0.5 | 8 | 0.25 | 29 | 0.3103 |
| 15_726808 | 4 | 0 | 15 | 0.1333 | 6 | 0.1667 | 11 | 0.2727 | 36 | 0.1667 | 8 | 0.125 | 8 | 0.125 | 12 | 0.0833 | 9 | 0 | 37 | 0.0811 |
| 15_726808 | 8 | 0 | 18 | 0.1667 | 6 | 0 | 12 | 0 | 44 | 0.0682 | 9 | 0.3333 | 7 | 0 | 11 | 0.1818 | 10 | 0 | 37 | 0.1351 |
| 15_726808 | 8 | 0 | 17 | 0.0588 | 5 | 0 | 8 | 0.125 | 38 | 0.0526 | 9 | 0 | 8 | 0.25 | 11 | 0.1818 | 10 | 0 | 38 | 0.1053 |
| 15_726809 | 8 | 0.875 | 14 | 1 | 5 | 1 | 11 | 1 | 38 | 0.9737 | 9 | 1 | 6 | 1 | 8 | 0.875 | 8 | 1 | 31 | 0.9677 |
| 15_726810 | 7 | 0.1429 | 16 | 0.0625 | 5 | 0 | 11 | 0 | 39 | 0.0513 | 8 | 0.25 | 7 | 0.1429 | 8 | 0.125 | 9 | 0 | 32 | 0.125 |
| 15_726811 | 7 | 1 | 10 | 1 | 3 | 1 | 9 | 1 | 29 | 1 | 5 | 1 | 7 | 1 | 9 | 1 | 7 | 1 | 28 | 1 |
| 15_726812 | 7 | 1 | 10 | 0.9 | 4 | 0.75 | 9 | 0.8889 | 30 | 0.9 | 9 | 1 | 8 | 0.875 | 8 | 0.75 | 8 | 1 | 33 | 0.9091 |
| 15_726817 | 6 | 0.1667 | 10 | 0.2 | 3 | 0 | 8 | 0.125 | 27 | 0.1481 | 9 | 0.1111 | 5 | 0.2 | 5 | 0 | 8 | 0 | 27 | 0.0741 |
| 15_729300 | 3 | 0.6667 | NA | NA | 2 | 0.5 | NA | NA | 5 | 0.6 | NA | NA | 4 | 0.5 | NA | NA | 1 | 0 | 5 | 0.4 |
| 15_729655 | 4 | 0 | 4 | 0 | 5 | 0.2 | 6 | 0 | 19 | 0.0526 | 4 | 0 | 5 | 0.2 | 5 | 0.4 | 5 | 0 | 19 | 0.1579 |
| 15_729655 | 6 | 0.1667 | 6 | 0 | 5 | 0.2 | 7 | 0 | 24 | 0.0833 | 4 | 0 | 6 | 0.3333 | 6 | 0 | 6 | 0 | 22 | 0.0909 |
| 15_729655 | 7 | 0.5714 | 8 | 0.25 | 3 | 0.3333 | 7 | 0.4286 | 25 | 0.4 | 3 | 0.3333 | 6 | 0.5 | 6 | 0.8333 | 7 | 0.5714 | 22 | 0.5909 |
| 15_729656 | 5 | 1 | 8 | 1 | 5 | 0.8 | 7 | 1 | 25 | 0.96 | 4 | 1 | 5 | 1 | 3 | 1 | 7 | 1 | 19 | 1 |
| 15_729656 | 7 | 0.2857 | 7 | 0.2857 | 3 | 0 | 5 | 0.4 | 22 | 0.2727 | 4 | 0.25 | 5 | 0.2 | 5 | 0.2 | 6 | 0.3333 | 20 | 0.25 |
| 15_729657 | 4 | 0.25 | 6 | 0.5 | 3 | 0.6667 | 4 | 0.75 | 17 | 0.5294 | 3 | 0.6667 | 4 | 0.25 | 4 | 0.25 | 3 | 0.3333 | 14 | 0.3571 |
| 15_729657 | 6 | 0.3333 | 6 | 0.1667 | 4 | 0.5 | 7 | 0.5714 | 23 | 0.3913 | 4 | 0.5 | 6 | 0.3333 | 5 | 0 | 7 | 0.5714 | 22 | 0.3636 |
| 15_729658 | 5 | 0.2 | 8 | 0 | 5 | 0.2 | 5 | 0 | 23 | 0.087 | 4 | 0.5 | 6 | 0.1667 | 5 | 0 | 7 | 0.1429 | 22 | 0.1818 |
| 15_729658 | 5 | 0.8 | 7 | 0.4286 | 4 | 1 | 4 | 0.75 | 20 | 0.7 | 3 | 1 | 4 | 1 | 3 | 1 | 6 | 0.8333 | 16 | 0.9375 |
| 15_729680 | 2 | 0 | 4 | 0.25 | 4 | 0 | 6 | 0.3333 | 16 | 0.1875 | 2 | 0.5 | 6 | 0.1667 | 5 | 0.2 | 7 | 0.1429 | 20 | 0.2 |
| 15_729680 | 3 | 0 | 5 | 0 | 4 | 0.25 | 6 | 0 | 18 | 0.0556 | 1 | 1 | 5 | 0 | 5 | 0 | 4 | 0.25 | 15 | 0.1333 |
| 15_729681 | 2 | 0 | 5 | 0.4 | 4 | 0.5 | 6 | 0.3333 | 17 | 0.3529 | 1 | 1 | 5 | 0.4 | 4 | 0.25 | 6 | 0.3333 | 16 | 0.375 |
| 15_729693 | 1 | 0 | 4 | 0.25 | 5 | 0 | 4 | 0 | 14 | 0.0714 | 2 | 0 | 4 | 0 | 4 | 0.5 | 5 | 0.4 | 15 | 0.2667 |
| 15_729706 | 5 | 0.2 | 3 | 0 | 4 | 0.25 | 7 | 0 | 19 | 0.1053 | 4 | 0.25 | 7 | 0.1429 | 12 | 0.3333 | 7 | 0.2857 | 30 | 0.2667 |
| 15_729721 | 3 | 0 | 1 | 0 | 2 | 0.5 | 6 | 0.1667 | 12 | 0.1667 | 2 | 0.5 | 6 | 0 | 6 | 0.1667 | 6 | 0.3333 | 20 | 0.2 |
| 15_729730 | 1 | 0 | 2 | 0 | 2 | 0.5 | 4 | 0 | 9 | 0.1111 | 1 | 1 | 4 | 0 | 6 | 0 | 2 | 0.5 | 13 | 0.1538 |
| 15_729733 | 2 | 0.5 | 1 | 0 | 2 | 0.5 | 3 | 0 | 8 | 0.25 | 2 | 0 | 3 | 0.3333 | 6 | 0.1667 | 1 | 0 | 12 | 0.1667 |
| 15_729734 | 4 | 0 | 2 | 0 | 2 | 0.5 | 3 | 0 | 11 | 0.0909 | 2 | 0 | 2 | 0 | 6 | 0.3333 | 3 | 0 | 13 | 0.1538 |
| 15_729738 | 2 | 0 | 1 | 0 | 3 | 0 | 4 | 0.25 | 10 | 0.1 | NA | NA | 3 | 0 | 5 | 0.2 | 4 | 0 | 12 | 0.0833 |
| 15_729773 | NA | NA | 1 | 1 | NA | NA | 1 | 1 | 2 | 1 | NA | NA | 1 | 1 | 3 | 0.6667 | 2 | 0.5 | 6 | 0.6667 |
| 15_729773 | NA | NA | 1 | 0 | NA | NA | 3 | 0.3333 | 4 | 0.25 | NA | NA | 1 | 0 | 3 | 0 | 4 | 0.25 | 8 | 0.125 |
| 15_729773 | NA | NA | 1 | 1 | NA | NA | 3 | 0.3333 | 4 | 0.5 | NA | NA | 1 | 0 | 4 | 0.5 | 3 | 0 | 8 | 0.25 |
| 15_729774 | NA | NA | 1 | 0 | NA | NA | 3 | 0.6667 | 4 | 0.5 | NA | NA | 1 | 0 | 3 | 0.3333 | 3 | 0 | 7 | 0.1429 |
| 15_729774 | NA | NA | 1 | 1 | NA | NA | 1 | 1 | 2 | 1 | NA | NA | 1 | 1 | 4 | 1 | 3 | 1 | 8 | 1 |
| 15_729774 | NA | NA | 1 | 1 | NA | NA | 1 | 0 | 2 | 0.5 | NA | NA | 1 | 1 | 3 | 0.3333 | 3 | 0.3333 | 7 | 0.4286 |
| 15_729776 | NA | NA | 1 | 1 | NA | NA | 2 | 0.5 | 3 | 0.6667 | NA | NA | 1 | 1 | 4 | 0.75 | 4 | 0.5 | 9 | 0.6667 |

| | | | | | | | | | | | | | | | | | | | | |
|---|---|---|---|---|---|---|---|---|---|---|---|---|---|---|---|---|---|---|---|---|
| 15_729777( | NA | NA | 1 | 1 | NA | NA | 2 | 1 | 3 | 1 | NA | NA | 1 | 1 | 2 | 1 | 3 | 0.6667 | 6 | 0.8333 |
| 15_729780( | NA | NA | NA | NA | NA | NA | 2 | 0.5 | 2 | 0.5 | NA | NA | NA | NA | 3 | 0 | 3 | 0.3333 | 6 | 0.1667 |
| 15_729802 | NA | NA | 1 | 1 | 1 | 0 | 2 | 0.5 | 4 | 0.5 | 1 | 0 | 1 | 1 | 3 | 0.3333 | 3 | 1 | 8 | 0.625 |
| 15_729806 | NA | NA | 1 | 1 | 1 | 0 | 2 | 0.5 | 4 | 0.5 | 1 | 0 | 1 | 1 | 3 | 1 | 4 | 0.5 | 9 | 0.6667 |
| 15_729843 | NA | NA | 1 | 1 | 1 | 0 | 2 | 0 | 4 | 0.25 | 1 | 0 | 1 | 1 | 2 | 1 | 3 | 0 | 7 | 0.4286 |
| 15_729845 | NA | NA | 1 | 1 | NA | NA | 1 | 1 | 2 | 1 | 1 | 1 | 1 | 1 | 3 | 1 | 3 | 1 | 8 | 1 |
| 15_729858 | NA | NA | 1 | 1 | 1 | 0 | 2 | 0 | 4 | 0.25 | 1 | 0 | 1 | 1 | 2 | 1 | 3 | 1 | 7 | 0.8571 |
| 15_729878 | NA | NA | 1 | 1 | NA | NA | 1 | 0 | 2 | 0.5 | 1 | 0 | 1 | 1 | 3 | 0.3333 | 4 | 0.75 | 9 | 0.5556 |
| 15_730771 | 4 | 0.25 | 10 | 0.4 | 4 | 0.75 | 5 | 0.2 | 23 | 0.3913 | 10 | 0.3 | 4 | 0.5 | 10 | 0.5 | 9 | 0.2222 | 33 | 0.3636 |
| 15_730771 | 4 | 0.25 | 8 | 0.125 | 4 | 0.25 | 4 | 0 | 20 | 0.15 | 8 | 0.25 | 5 | 0.2 | 8 | 0.125 | 7 | 0.1429 | 28 | 0.1786 |
| 15_730771 | 3 | 0 | 5 | 0 | 2 | 0 | 3 | 0.3333 | 13 | 0.0769 | 5 | 0.2 | 4 | 0 | 6 | 0 | 4 | 0 | 19 | 0.0526 |
| 15_730771 | 11 | 0.6364 | 14 | 0.7143 | 7 | 0.5714 | 7 | 0.4286 | 39 | 0.6154 | 11 | 0.9091 | 10 | 0.8 | 14 | 0.8571 | 14 | 0.7143 | 49 | 0.8163 |
| 15_730772 | 15 | 0.2667 | 16 | 0.125 | 10 | 0.1 | 9 | 0 | 50 | 0.14 | 11 | 0.0909 | 12 | 0.0833 | 16 | 0.1875 | 13 | 0.3077 | 52 | 0.1731 |
| 15_730773 | 15 | 0.6 | 11 | 0.5455 | 4 | 0.75 | 7 | 0.4286 | 37 | 0.5676 | 10 | 0.5 | 11 | 0.5455 | 16 | 0.625 | 12 | 0.6667 | 49 | 0.5918 |
| 15_730773 | 11 | 0.9091 | 13 | 0.7692 | 7 | 1 | 8 | 0.875 | 39 | 0.8718 | 10 | 1 | 9 | 0.8889 | 13 | 0.9231 | 12 | 0.9167 | 44 | 0.9318 |
| 15_730773 | 12 | 0.5833 | 11 | 0.7273 | 10 | 0.7 | 6 | 0.5 | 39 | 0.641 | 11 | 0.6364 | 10 | 0.7 | 15 | 0.9333 | 10 | 0.7 | 46 | 0.7609 |
| 15_730774 | 1 | 0 | 1 | 1 | NA | NA | NA | NA | 2 | 0.5 | 1 | 0 | 1 | 1 | 1 | 0 | 1 | 0 | 4 | 0.25 |
| 15_730774 | 13 | 0.7692 | 15 | 0.8667 | 11 | 0.9091 | 9 | 1 | 48 | 0.875 | 12 | 0.75 | 10 | 0.8 | 16 | 1 | 13 | 0.9231 | 51 | 0.8824 |
| 15_730775 | 14 | 0.5714 | 14 | 0.6429 | 11 | 0.7273 | 9 | 0.5556 | 48 | 0.625 | 12 | 0.6667 | 11 | 0.8182 | 16 | 0.8125 | 13 | 0.8462 | 52 | 0.7885 |
| 15_730775 | 13 | 0.0769 | 14 | 0.1429 | 11 | 0.1818 | 9 | 0 | 47 | 0.1064 | 11 | 0.1818 | 10 | 0.1 | 16 | 0.125 | 13 | 0.1538 | 50 | 0.14 |
| 15_730775 | 12 | 0 | 12 | 0.0833 | 6 | 0 | 8 | 0.125 | 38 | 0.0526 | 11 | 0.0909 | 9 | 0.1111 | 15 | 0.2667 | 10 | 0.1 | 45 | 0.1556 |
| 15_730776 | 12 | 0.5833 | 13 | 0.2308 | 10 | 0.5 | 9 | 0.4444 | 44 | 0.4318 | 10 | 0.5 | 10 | 0.6 | 16 | 0.5625 | 13 | 0.7692 | 49 | 0.6122 |
| 15_730777 | 10 | 0.3 | 10 | 0.3 | 8 | 0.375 | 7 | 0 | 35 | 0.2571 | 7 | 0.4286 | 9 | 0.2222 | 15 | 0.0667 | 11 | 0.2727 | 42 | 0.2143 |
| 15_730779 | 14 | 0.7857 | 9 | 0.7778 | 8 | 0.75 | 6 | 0.5 | 37 | 0.7297 | 5 | 1 | 7 | 0.2857 | 11 | 0.7273 | 9 | 1 | 32 | 0.75 |
| 15_734432 | 1 | 1 | NA | NA | NA | NA | NA | NA | 1 | 1 | NA | NA | 1 | 1 | NA | NA | NA | NA | 1 | 1 |
| 15_734433 | 1 | 1 | NA | NA | NA | NA | NA | NA | 1 | 1 | NA | NA | 1 | 1 | NA | NA | NA | NA | 1 | 1 |
| 15_734434 | 1 | 1 | NA | NA | NA | NA | NA | NA | 1 | 1 | NA | NA | 1 | 1 | NA | NA | NA | NA | 1 | 1 |
| 15_734496 | 3 | 0.3333 | 1 | 0 | 6 | 0.3333 | NA | NA | 10 | 0.3 | 4 | 0 | 3 | 0 | 1 | 0 | 2 | 0.5 | 10 | 0.1 |
| 15_734496 | 3 | 0 | 1 | 0 | 6 | 0.1667 | NA | NA | 10 | 0.1 | 2 | 0 | 2 | 0 | 1 | 0 | 1 | 1 | 6 | 0.1667 |
| 15_734499 | 2 | 1 | NA | NA | 4 | 0 | NA | NA | 6 | 0.3333 | 2 | 0 | 2 | 0 | 1 | 1 | 1 | 1 | 6 | 0.3333 |
| 15_734499 | 2 | 0 | 1 | 0 | 4 | 0.25 | NA | NA | 7 | 0.1429 | 1 | 0 | 3 | 0 | 1 | 0 | 1 | 1 | 6 | 0.1667 |
| 15_734513 | 1 | 1 | 1 | 0 | 5 | 0.4 | NA | NA | 7 | 0.4286 | 1 | 0 | 2 | 0.5 | 1 | 0 | 1 | 0 | 5 | 0.2 |
| 15_734518 | NA | NA | 1 | 0 | 4 | 0.25 | NA | NA | 5 | 0.2 | 1 | 0 | 1 | 1 | NA | NA | 1 | 0 | 3 | 0.3333 |
| 15_734519 | NA | NA | 1 | 1 | 4 | 0 | NA | NA | 5 | 0.2 | 1 | 0 | 2 | 0.5 | 1 | 1 | 1 | 0 | 5 | 0.4 |
| 15_734523 | 3 | 0.6667 | 5 | 1 | 5 | 1 | 3 | 1 | 16 | 0.9375 | 2 | 1 | 3 | 1 | 2 | 1 | 8 | 1 | 15 | 1 |
| 15_734524 | 2 | 0 | 2 | 1 | 4 | 0.25 | 2 | 1 | 10 | 0.5 | NA | NA | 4 | 0.75 | 1 | 0 | 6 | 1 | 11 | 0.8182 |
| 15_734525 | 1 | 0 | 3 | 0.3333 | 5 | 0.4 | 3 | 0 | 12 | 0.25 | 1 | 0 | 3 | 0.3333 | 1 | 0 | 5 | 0.2 | 10 | 0.2 |
| 15_734526 | 3 | 0.3333 | 3 | 0.6667 | 4 | 0.25 | 3 | 0 | 13 | 0.3077 | 2 | 0 | 4 | 0.25 | 2 | 0.5 | 8 | 0.25 | 16 | 0.25 |
| 15_734527 | 2 | 0 | 5 | 0.4 | 4 | 0.5 | 2 | 0 | 13 | 0.3077 | 2 | 0 | 4 | 0.5 | 2 | 0.5 | 8 | 0.25 | 16 | 0.3125 |
| 15_734528 | 3 | 0 | 5 | 0.6 | 4 | 0.25 | 3 | 0.6667 | 15 | 0.4 | 2 | 0.5 | 2 | 1 | 1 | 1 | 8 | 0.25 | 13 | 0.4615 |
| 15_734529 | 2 | 1 | 4 | 0.75 | 5 | 0.8 | 2 | 0.5 | 13 | 0.7692 | 2 | 1 | 3 | 0.6667 | 1 | 1 | 5 | 1 | 11 | 0.9091 |
| 15_734529 | 1 | 0 | 5 | 0.4 | 4 | 0.25 | 2 | 0 | 12 | 0.25 | 2 | 0.5 | 2 | 0 | 1 | 1 | 7 | 0.4286 | 12 | 0.4167 |
| 15_734530 | 2 | 0.5 | 1 | 0 | 5 | 0 | 2 | 0 | 10 | 0.1 | 2 | 0 | 2 | 0 | NA | NA | 4 | 0.25 | 8 | 0.125 |
| 15_734537 | 2 | 0 | 2 | 0.5 | 5 | 0 | 2 | 0 | 11 | 0.0909 | 1 | 1 | 2 | 0 | NA | NA | 3 | 0.6667 | 6 | 0.5 |
| 15_734537 | 2 | 0 | 3 | 0 | 5 | 0.2 | 2 | 0 | 12 | 0.0833 | 2 | 0 | 2 | 0.5 | NA | NA | 4 | 0.25 | 8 | 0.25 |
| 15_734539 | 1 | 0 | 3 | 0 | 5 | 0.2 | 2 | 0 | 11 | 0.0909 | 2 | 0 | 3 | 0.3333 | NA | NA | 5 | 0 | 10 | 0.1 |
| 15_734563 | 2 | 0 | 3 | 0.3333 | 1 | 0 | 1 | 1 | 7 | 0.2857 | 1 | 0 | 3 | 0.3333 | NA | NA | 5 | 0.4 | 9 | 0.3333 |
| 15_734582 | 2 | 0.5 | 4 | 0.5 | 2 | 0.5 | 1 | 0 | 9 | 0.4444 | 1 | 1 | 2 | 0.5 | NA | NA | 4 | 0.25 | 7 | 0.4286 |
| 15_734586 | 2 | 0.5 | 4 | 0.75 | 1 | 0 | 1 | 1 | 8 | 0.625 | 1 | 1 | 2 | 0.5 | NA | NA | 4 | 0.5 | 7 | 0.5714 |
| 15_734773 | 2 | 0.5 | 3 | 0.6667 | 1 | 1 | 1 | 1 | 7 | 0.7143 | NA | NA | NA | NA | NA | NA | 2 | 1 | 2 | 1 |
| 15_734774 | 2 | 1 | 4 | 0.5 | 1 | 0 | 1 | 1 | 8 | 0.625 | NA | NA | 1 | 0 | NA | NA | 2 | 1 | 3 | 0.6667 |
| 15_735973 | 1 | 1 | 2 | 1 | NA | NA | NA | NA | 3 | 1 | NA | NA | NA | NA | NA | NA | 1 | 1 | 1 | 1 |
| 15_736231 | 3 | 0.3333 | 8 | 1 | 5 | 0.8 | 5 | 1 | 21 | 0.8571 | NA | NA | 10 | 0.8 | 2 | 0.5 | 9 | 0.7778 | 21 | 0.7619 |
| 15_736232 | 2 | 0.5 | 9 | 0.4444 | 8 | 0.25 | 5 | 0.2 | 24 | 0.3333 | NA | NA | 11 | 0.2727 | 3 | 0.3333 | 9 | 0.3333 | 23 | 0.3043 |
| 15_736233 | 5 | 0.6 | 10 | 0.5 | 8 | 0.25 | 3 | 0 | 26 | 0.3846 | NA | NA | 9 | 0.5556 | 2 | 0 | 8 | 0.5 | 19 | 0.4737 |
| 15_736233 | 4 | 0.5 | 8 | 0.75 | 8 | 0.75 | 4 | 1 | 24 | 0.75 | NA | NA | 10 | 0.7 | 2 | 0 | 8 | 1 | 20 | 0.75 |
| 15_736234 | 3 | 0 | 8 | 0.125 | 8 | 0.5 | 2 | 1 | 21 | 0.3333 | NA | NA | 9 | 0.2222 | 2 | 0 | 6 | 0.3333 | 17 | 0.2353 |
| 15_736234 | 5 | 1 | 9 | 0.8889 | 8 | 0.875 | 4 | 1 | 26 | 0.9231 | NA | NA | 9 | 0.8889 | 2 | 1 | 7 | 1 | 18 | 0.9444 |
| 15_736236 | 6 | 0 | 9 | 0.2222 | 7 | 0 | 4 | 0 | 26 | 0.0769 | NA | NA | 10 | 0.5 | 3 | 0 | 11 | 0.3636 | 24 | 0.375 |
| 15_736236 | 5 | 1 | 6 | 0.8333 | 7 | 0.8571 | 3 | 0.6667 | 21 | 0.8571 | 1 | 1 | 8 | 0.875 | 3 | 1 | 7 | 1 | 19 | 0.9474 |
| 15_736290 | 7 | 0.5714 | 13 | 0.6154 | 15 | 0.6667 | 13 | 0.6154 | 48 | 0.625 | 6 | 0.5 | 9 | 0.8889 | 6 | 0.6667 | 9 | 0.4444 | 30 | 0.6333 |
| 15_743884 | 1 | 0 | 4 | 0.5 | 3 | 0 | 4 | 0.25 | 12 | 0.25 | 2 | 0 | 4 | 0 | 3 | 0.3333 | 1 | 0 | 10 | 0.1 |
| 15_744008 | 1 | 0 | 5 | 0.6 | 4 | 0.5 | 3 | 0.3333 | 13 | 0.4615 | 3 | 0.6667 | 5 | 0.2 | 2 | 0 | 3 | 0.3333 | 13 | 0.3077 |
| 15_744009 | 1 | 0 | 5 | 0.8 | 4 | 0.75 | 3 | 1 | 13 | 0.7692 | 3 | 1 | 7 | 0.8571 | 3 | 0.6667 | 2 | 1 | 15 | 0.8667 |
| 15_744011( | NA | NA | 6 | 0 | 3 | 0.6667 | 3 | 0.3333 | 12 | 0.25 | 3 | 0 | 7 | 0.1429 | 2 | 0 | 2 | 0.5 | 14 | 0.1429 |

| | | | | | | | | | | | | | | | | | | | | |
|---|---|---|---|---|---|---|---|---|---|---|---|---|---|---|---|---|---|---|---|---|
| 15_7440112 | 1 | 1 | 7 | 0.8571 | 4 | 1 | 3 | 1 | 15 | 0.9333 | 3 | 1 | 8 | 0.875 | 3 | 1 | 4 | 1 | 18 | 0.9444 |
| 15_7440121 | 1 | 1 | 7 | 0.7143 | 3 | 1 | 3 | 0.3333 | 14 | 0.7143 | 2 | 1 | 8 | 0.75 | 3 | 1 | 2 | 1 | 15 | 0.8667 |
| 15_7440123 | NA | NA | 6 | 0.6667 | 4 | 0 | 5 | 0.2 | 15 | 0.3333 | 3 | 0 | 7 | 0.4286 | 3 | 0.3333 | 3 | 0.6667 | 16 | 0.375 |
| 15_7440142 | 1 | 0 | 7 | 0 | 4 | 0.25 | 2 | 0 | 14 | 0.0714 | 2 | 0 | 7 | 0 | 3 | 0.3333 | 3 | 0 | 15 | 0.0667 |
| 15_7440147 | 1 | 0 | 6 | 0.3333 | 4 | 0.5 | 5 | 0.4 | 16 | 0.375 | 3 | 0.6667 | 8 | 0.5 | 3 | 0.3333 | 3 | 0.3333 | 17 | 0.4706 |
| 15_7440163 | 1 | 1 | 6 | 0.6667 | 4 | 0.25 | 4 | 0.5 | 15 | 0.5333 | 3 | 1 | 7 | 0.5714 | 3 | 0.3333 | 3 | 0.3333 | 16 | 0.5625 |
| 15_7445484 | 3 | 0 | 2 | 0.5 | 9 | 0.1111 | 6 | 0 | 20 | 0.1 | 3 | 0.3333 | 6 | 0 | 6 | 0 | 2 | 0 | 17 | 0.0588 |
| 15_7445507 | 5 | 0 | 2 | 0 | 7 | 0.1429 | 5 | 0 | 19 | 0.0526 | 2 | 0 | 7 | 0.1429 | 8 | 0.25 | 3 | 0 | 20 | 0.15 |
| 15_7445510 | 3 | 0 | 2 | 0.5 | 8 | 0 | 6 | 0 | 19 | 0.0526 | 3 | 0 | 6 | 0 | 7 | 0.1429 | 3 | 0 | 19 | 0.0526 |
| 15_7445534 | 4 | 0 | 1 | 0 | 9 | 0.1111 | 4 | 0 | 18 | 0.0556 | 4 | 0 | 5 | 0 | 5 | 0 | 4 | 0.25 | 18 | 0.0556 |
| 15_7445879 | 25 | 0.04 | 25 | 0.2 | 24 | 0.2083 | 20 | 0.15 | 94 | 0.1489 | 18 | 0.1111 | 32 | 0.0625 | 19 | 0.0526 | 22 | 0.0455 | 91 | 0.0659 |
| 15_7445882 | 27 | 0.7037 | 27 | 0.6296 | 27 | 0.5185 | 19 | 0.6316 | 100 | 0.62 | 16 | 0.625 | 30 | 0.5667 | 24 | 0.7917 | 24 | 0.625 | 94 | 0.6489 |
| 15_7445886 | 27 | 0.3333 | 26 | 0.3077 | 32 | 0.2188 | 21 | 0.1905 | 106 | 0.2642 | 16 | 0.5625 | 29 | 0.3103 | 26 | 0.3462 | 21 | 0.3333 | 92 | 0.3696 |
| 15_7445887 | 21 | 0.5238 | 27 | 0.5185 | 29 | 0.5517 | 16 | 0.5625 | 93 | 0.5376 | 15 | 0.7333 | 27 | 0.4815 | 25 | 0.56 | 20 | 0.6 | 87 | 0.5747 |
| 15_7445888 | 21 | 0.8095 | 25 | 0.8 | 31 | 0.9032 | 17 | 0.8824 | 94 | 0.8511 | 17 | 1 | 27 | 0.963 | 19 | 0.9474 | 18 | 0.8333 | 81 | 0.9383 |
| 15_7445891 | 25 | 0.24 | 30 | 0.2 | 24 | 0.2083 | 17 | 0.5882 | 96 | 0.2812 | 15 | 0.2 | 26 | 0.5 | 23 | 0.3478 | 17 | 0.3529 | 81 | 0.3704 |
| 15_7446013 | 2 | 0 | 4 | 0 | 6 | 0.1667 | 1 | 0 | 13 | 0.0769 | 1 | 0 | 2 | 0 | 4 | 0.25 | 3 | 0 | 10 | 0.1 |
| 15_7446070 | 33 | 0.5152 | 40 | 0.675 | 40 | 0.6 | 29 | 0.6897 | 142 | 0.6197 | 22 | 0.7273 | 37 | 0.5676 | 33 | 0.7273 | 27 | 0.6296 | 119 | 0.6555 |
| 15_7446089 | 40 | 0.1 | 46 | 0.1957 | 47 | 0.234 | 29 | 0.2414 | 162 | 0.1914 | 27 | 0.2593 | 44 | 0.2045 | 37 | 0.2703 | 29 | 0.1034 | 137 | 0.2117 |
| 15_7446091 | 34 | 0.5294 | 38 | 0.6579 | 44 | 0.5227 | 27 | 0.6296 | 143 | 0.5804 | 24 | 0.7083 | 44 | 0.6818 | 36 | 0.7222 | 26 | 0.5 | 130 | 0.6615 |
| 15_7446096 | 30 | 0.6 | 40 | 0.6 | 35 | 0.5429 | 21 | 0.4286 | 126 | 0.5556 | 25 | 0.52 | 38 | 0.4474 | 35 | 0.5429 | 25 | 0.48 | 123 | 0.4959 |
| 15_7446101 | 31 | 0.0968 | 36 | 0.1111 | 35 | 0.1714 | 21 | 0.0476 | 123 | 0.1138 | 21 | 0.1905 | 35 | 0.1143 | 33 | 0.1515 | 22 | 0.0455 | 111 | 0.1261 |
| 15_7446102 | 26 | 0.4615 | 34 | 0.4412 | 34 | 0.4706 | 29 | 0.3793 | 123 | 0.439 | 17 | 0.3529 | 36 | 0.5278 | 33 | 0.3939 | 21 | 0.4286 | 107 | 0.4393 |
| 15_7446108 | 34 | 0.6765 | 40 | 0.75 | 42 | 0.6905 | 27 | 0.7037 | 143 | 0.7063 | 24 | 0.875 | 41 | 0.7317 | 36 | 0.8056 | 27 | 0.8519 | 128 | 0.8047 |
| 15_7446116 | 43 | 0.093 | 44 | 0.25 | 47 | 0.234 | 32 | 0.3438 | 166 | 0.2229 | 28 | 0.2857 | 50 | 0.28 | 42 | 0.3095 | 33 | 0.2727 | 153 | 0.2876 |
| 15_7446122 | 43 | 0.1163 | 48 | 0.1042 | 46 | 0.0652 | 33 | 0.0909 | 170 | 0.0941 | 28 | 0.1429 | 48 | 0.0208 | 45 | 0.2 | 35 | 0.1143 | 156 | 0.1154 |
| 15_7446122 | 35 | 0.5429 | 38 | 0.4474 | 44 | 0.4091 | 32 | 0.5312 | 149 | 0.4765 | 30 | 0.5333 | 44 | 0.7727 | 38 | 0.6842 | 31 | 0.6129 | 143 | 0.6643 |
| 15_7454136 | 8 | 0 | 12 | 0.0833 | 6 | 0 | 2 | 0.5 | 28 | 0.0714 | 10 | 0.1 | 6 | 0 | 4 | 0.25 | 9 | 0.1111 | 29 | 0.1034 |
| 15_7454173 | 8 | 0 | 17 | 0.0588 | 6 | 0.1667 | 3 | 0 | 34 | 0.0588 | 12 | 0.0833 | 6 | 0 | 5 | 0 | 12 | 0.25 | 35 | 0.1143 |
| 15_7454263 | 7 | 0.1429 | 15 | 0.2 | 5 | 0 | 3 | 0.3333 | 30 | 0.1667 | 9 | 0.2222 | 5 | 0.4 | 4 | 0.5 | 10 | 0.2 | 28 | 0.2857 |
| 15_7454274 | 6 | 0.3333 | 13 | 0.0769 | 2 | 0 | 3 | 0 | 24 | 0.125 | 8 | 0.125 | 4 | 0 | 3 | 0 | 9 | 0.1111 | 24 | 0.0833 |
| 15_7454286 | 5 | 1 | 10 | 0.9 | 5 | 0.8 | 3 | 1 | 23 | 0.913 | 6 | 0.8333 | 3 | 1 | 2 | 1 | 7 | 1 | 18 | 0.9444 |
| 15_7454296 | 4 | 0.75 | 11 | 1 | 4 | 0.75 | 2 | 1 | 21 | 0.9048 | 7 | 1 | 5 | 1 | 2 | 1 | 7 | 1 | 21 | 1 |
| 15_7454304 | 3 | 0 | 11 | 0.2727 | 5 | 0.2 | 3 | 0.3333 | 22 | 0.2273 | 7 | 0.1429 | 5 | 0 | 3 | 0 | 9 | 0.2222 | 24 | 0.125 |
| 15_7454322 | 3 | 0 | 10 | 0 | 4 | 0.5 | 2 | 0.5 | 19 | 0.1579 | 7 | 0 | 4 | 0 | 2 | 0.5 | 8 | 0.25 | 21 | 0.1429 |
| 15_7454330 | 3 | 0.3333 | 10 | 0.3 | 4 | 0.75 | 3 | 0.6667 | 20 | 0.45 | 8 | 0.25 | 5 | 0.2 | 3 | 0 | 8 | 0.25 | 24 | 0.2083 |
| 15_7454343 | 3 | 0 | 9 | 0 | 3 | 0 | 3 | 0.3333 | 18 | 0.0556 | 8 | 0.125 | 5 | 0 | 3 | 0.3333 | 7 | 0.1429 | 23 | 0.1304 |
| 15_7454357 | 3 | 0 | 8 | 0.25 | 4 | 0 | 3 | 0.3333 | 18 | 0.1667 | 8 | 0.25 | 4 | 0.25 | 3 | 0 | 6 | 0.1667 | 21 | 0.1905 |
| 15_7454367 | 2 | 1 | 5 | 0.4 | 3 | 1 | 2 | 0.5 | 12 | 0.6667 | 7 | 0.4286 | 3 | 0 | 2 | 0.5 | 5 | 0.6 | 17 | 0.4118 |
| 15_7454372 | 2 | 0.5 | 7 | 0.1429 | 3 | 0 | 3 | 0.3333 | 15 | 0.2 | 7 | 0.1429 | 2 | 0.5 | 3 | 0 | 6 | 0.6667 | 18 | 0.3333 |
| 15_7457657 | NA | NA | 1 | 1 | 1 | 1 | 2 | 0.5 | 4 | 0.75 | 3 | 0.6667 | NA | NA | 1 | 0 | NA | NA | 4 | 0.5 |
| 15_7457678 | NA | NA | 1 | 1 | 1 | 1 | 2 | 0.5 | 4 | 0.75 | 3 | 0.3333 | NA | NA | NA | NA | NA | NA | 3 | 0.3333 |
| 15_7457754 | NA | NA | 1 | 1 | NA | NA | 1 | 0 | 2 | 0.5 | 3 | 0.3333 | NA | NA | NA | NA | NA | NA | 3 | 0.3333 |
| 15_7457787 | 1 | 1 | 1 | 1 | 1 | 1 | 1 | 1 | 4 | 1 | 1 | 1 | NA | NA | NA | NA | NA | NA | 1 | 1 |
| 15_7457798 | 1 | 1 | 1 | 0 | 1 | 1 | 1 | 1 | 4 | 0.75 | 2 | 1 | NA | NA | NA | NA | NA | NA | 2 | 1 |
| 15_7457813 | 1 | 1 | 1 | 0 | 1 | 0 | 1 | 0 | 4 | 0.25 | 2 | 1 | NA | NA | NA | NA | NA | NA | 2 | 1 |
| 15_7457847 | 1 | 1 | NA | NA | NA | NA | NA | NA | 1 | 1 | 1 | 1 | NA | NA | NA | NA | NA | NA | 1 | 1 |
| 15_7457863 | 1 | 1 | NA | NA | NA | NA | NA | NA | 1 | 1 | 1 | 1 | NA | NA | NA | NA | NA | NA | 1 | 1 |
| 15_7457864 | 1 | 1 | NA | NA | NA | NA | NA | NA | 1 | 1 | 1 | 1 | NA | NA | NA | NA | NA | NA | 1 | 1 |
| 15_7457871 | 1 | 1 | NA | NA | NA | NA | NA | NA | 1 | 1 | 1 | 1 | NA | NA | NA | NA | NA | NA | 1 | 1 |
| 15_7457893 | 1 | 1 | NA | NA | NA | NA | NA | NA | 1 | 1 | 1 | 1 | NA | NA | NA | NA | NA | NA | 1 | 1 |
| 15_7459541 | 17 | 0.3529 | 12 | 0.0833 | 15 | 0.1333 | 11 | 0.2727 | 55 | 0.2182 | 8 | 0.25 | 17 | 0.1176 | 14 | 0.2143 | 9 | 0.3333 | 48 | 0.2083 |
| 15_7459563 | 16 | 0.3125 | 11 | 0.6364 | 20 | 0.5 | 11 | 0.5455 | 58 | 0.4828 | 6 | 0.1667 | 12 | 0.5 | 13 | 0.6154 | 11 | 0.6364 | 42 | 0.5238 |
| 15_7459576 | 15 | 0.2667 | 12 | 0.25 | 20 | 0.4 | 12 | 0.0833 | 59 | 0.2712 | 11 | 0.2727 | 20 | 0.15 | 15 | 0.2 | 11 | 0.4545 | 57 | 0.2456 |
| 15_7461988 | 25 | 0.68 | 23 | 0.4783 | 25 | 0.68 | 14 | 0.6429 | 87 | 0.6207 | 22 | 0.5455 | 23 | 0.6087 | 15 | 0.3333 | 12 | 0.5 | 72 | 0.5139 |
| 15_7461997 | 25 | 0.16 | 23 | 0.3043 | 32 | 0.2188 | 17 | 0.2941 | 97 | 0.2371 | 26 | 0.2308 | 26 | 0.2692 | 18 | 0.1111 | 21 | 0.2381 | 91 | 0.2198 |
| 15_7462000 | 25 | 0.08 | 23 | 0.1739 | 32 | 0.125 | 14 | 0.0714 | 94 | 0.117 | 29 | 0.1034 | 26 | 0.1538 | 22 | 0.0455 | 21 | 0 | 98 | 0.0816 |
| 15_7462009 | 22 | 0.2273 | 21 | 0.1905 | 26 | 0.1538 | 12 | 0.3333 | 81 | 0.2099 | 26 | 0.3077 | 20 | 0.2 | 19 | 0.0526 | 17 | 0.2353 | 82 | 0.2073 |
| 15_7462023 | 24 | 0.2917 | 24 | 0.0833 | 31 | 0.1935 | 17 | 0.1765 | 96 | 0.1875 | 25 | 0.16 | 26 | 0.0769 | 18 | 0.1667 | 21 | 0.3333 | 90 | 0.1778 |
| 15_7462033 | 20 | 0.05 | 17 | 0.0588 | 20 | 0.2 | 9 | 0.1111 | 66 | 0.1061 | 16 | 0.25 | 17 | 0.1176 | 17 | 0.1176 | 19 | 0.0526 | 69 | 0.1304 |
| 15_7462137 | 14 | 0 | 13 | 0.1538 | 19 | 0.2105 | 8 | 0.25 | 54 | 0.1481 | 8 | 0.125 | 17 | 0.2353 | 11 | 0.0909 | 15 | 0.1333 | 51 | 0.1569 |
| 15_7462179 | 8 | 0.25 | 7 | 0.1429 | 8 | 0.375 | 2 | 0 | 25 | 0.24 | 4 | 0.5 | 4 | 0.25 | 3 | 0 | 7 | 0.2857 | 18 | 0.2778 |
| 15_7462184 | 8 | 0.25 | 8 | 0.375 | 7 | 0.1429 | 2 | 1 | 25 | 0.32 | 3 | 0.3333 | 4 | 0.5 | 2 | 1 | 6 | 0.3333 | 15 | 0.4667 |
| 15_7462191 | 7 | 0 | 8 | 0.25 | 5 | 0 | 1 | 0 | 21 | 0.0952 | 3 | 0.3333 | 3 | 0.3333 | 3 | 0.3333 | 5 | 0.6 | 14 | 0.4286 |
| 15_7462210 | 7 | 0.1429 | 8 | 0.125 | 7 | 0.1429 | 2 | 0 | 24 | 0.125 | 3 | 0 | 2 | 0 | 2 | 0.5 | 4 | 0.25 | 11 | 0.1818 |

| | | | | | | | | | | | | | | | | | | | | |
|---|---|---|---|---|---|---|---|---|---|---|---|---|---|---|---|---|---|---|---|---|
| 15_746225 | 5 | 0.8 | 6 | 1 | 4 | 1 | 2 | 1 | 17 | 0.9412 | 1 | 1 | 2 | 1 | 1 | 1 | 2 | 1 | 6 | 1 |
| 15_746226 | 2 | 0.5 | 4 | 0.5 | 5 | 0.8 | 1 | 1 | 12 | 0.6667 | 1 | 1 | 2 | 1 | 1 | 1 | 3 | 1 | 7 | 1 |
| 15_746228 | 5 | 1 | 3 | 1 | 6 | 1 | 2 | 1 | 16 | 1 | 2 | 1 | 1 | 1 | 1 | 1 | 2 | 1 | 6 | 1 |
| 15_746229 | 6 | 1 | 5 | 1 | 7 | 1 | 1 | 0 | 19 | 0.9474 | 1 | 1 | 1 | 1 | NA | NA | 2 | 1 | 4 | 1 |
| 15_746229 | 5 | 0 | 3 | 0.3333 | 7 | 0 | NA | NA | 15 | 0.0667 | 2 | 0.5 | 1 | 0 | 1 | 0 | 2 | 0.5 | 6 | 0.3333 |
| 15_748304 | 19 | 0.2105 | 27 | 0.1111 | 13 | 0.0769 | 19 | 0.0526 | 78 | 0.1154 | 20 | 0.05 | 14 | 0.1429 | 12 | 0 | 20 | 0.15 | 66 | 0.0909 |
| 15_748305 | 19 | 0.5263 | 29 | 0.4483 | 14 | 0.5 | 18 | 0.6667 | 80 | 0.525 | 21 | 0.4286 | 15 | 0.6667 | 13 | 0.3077 | 21 | 0.4762 | 70 | 0.4714 |
| 15_748308 | 16 | 0.5625 | 27 | 0.2593 | 13 | 0.2308 | 15 | 0.2 | 71 | 0.3099 | 17 | 0.5294 | 12 | 0.3333 | 11 | 0.2727 | 23 | 0.3043 | 63 | 0.3651 |
| 15_748310 | 18 | 0.2778 | 29 | 0.1034 | 15 | 0.2 | 18 | 0.1111 | 80 | 0.1625 | 21 | 0.0476 | 11 | 0 | 16 | 0.125 | 29 | 0.1034 | 77 | 0.0779 |
| 15_748671 | 6 | 0.1667 | 8 | 0 | 6 | 0.3333 | 4 | 0 | 24 | 0.125 | 7 | 0 | 4 | 0 | 5 | 0 | 8 | 0.25 | 24 | 0.0833 |
| 15_749173 | 11 | 0.2727 | 15 | 0.4667 | 10 | 0.1 | 9 | 0.1111 | 45 | 0.2667 | 8 | 0.25 | 12 | 0.3333 | 9 | 0.2222 | 11 | 0 | 40 | 0.2 |
| 15_749173 | 18 | 0.1667 | 20 | 0.1 | 14 | 0 | 18 | 0 | 70 | 0.0714 | 12 | 0.0833 | 19 | 0.1053 | 15 | 0 | 13 | 0.2308 | 59 | 0.1017 |
| 15_754367 | 43 | 0.0698 | 71 | 0.0845 | 57 | 0.0526 | 53 | 0.1132 | 224 | 0.0804 | 44 | 0.0455 | 57 | 0.0702 | 44 | 0.0682 | 50 | 0.04 | 195 | 0.0564 |
| 15_754370 | 46 | 0.4783 | 60 | 0.4167 | 52 | 0.4808 | 47 | 0.4468 | 205 | 0.4537 | 41 | 0.4878 | 48 | 0.5 | 43 | 0.3488 | 45 | 0.5111 | 177 | 0.4633 |
| 15_754370 | 44 | 0.4091 | 67 | 0.5075 | 53 | 0.4717 | 49 | 0.6122 | 213 | 0.5023 | 36 | 0.5 | 53 | 0.4151 | 46 | 0.6087 | 49 | 0.4898 | 184 | 0.5 |
| 15_754371 | 48 | 0.2083 | 74 | 0.2838 | 53 | 0.2453 | 53 | 0.3019 | 228 | 0.2632 | 34 | 0.3824 | 56 | 0.3393 | 45 | 0.3556 | 50 | 0.44 | 185 | 0.3784 |
| 15_754389 | 1 | 0 | 15 | 0.2 | 7 | 0 | 8 | 0 | 31 | 0.0968 | 7 | 0 | 9 | 0 | 8 | 0.125 | 6 | 0.1667 | 30 | 0.0667 |
| 15_754970 | 1 | 1 | NA | NA | NA | NA | NA | NA | 1 | 1 | 1 | 1 | NA | NA | NA | NA | NA | NA | 1 | 1 |
| 15_755289 | 1 | 0 | NA | NA | 3 | 0.3333 | 5 | 0.2 | 9 | 0.2222 | 1 | 0 | 3 | 0 | NA | NA | 2 | 0.5 | 6 | 0.1667 |
| 15_755860 | 2 | 1 | 1 | 0 | 1 | 0 | 4 | 0.75 | 8 | 0.625 | NA | NA | NA | NA | 4 | 0.5 | 2 | 1 | 6 | 0.6667 |
| 15_755866 | 1 | 1 | NA | NA | NA | NA | NA | NA | 1 | 1 | NA | NA | NA | NA | NA | NA | 1 | 1 | 1 | 1 |
| 15_756176 | NA | NA | NA | NA | 1 | 1 | 2 | 0.5 | 3 | 0.6667 | NA | NA | NA | NA | 1 | 0 | 2 | 1 | 3 | 0.6667 |
| 15_756178 | 3 | 0 | NA | NA | 3 | 0.3333 | 4 | 0.75 | 10 | 0.4 | 2 | 0.5 | 1 | 0 | 3 | 0.3333 | 3 | 0.6667 | 9 | 0.4444 |
| 15_756179 | 2 | 0 | NA | NA | 3 | 0.3333 | 4 | 0 | 9 | 0.1111 | 3 | 0 | 1 | 0 | 2 | 0 | 3 | 0.3333 | 9 | 0.1111 |
| 15_756180 | 1 | 0 | NA | NA | 3 | 0 | 4 | 0.25 | 8 | 0.125 | 1 | 1 | 1 | 0 | 3 | 0.3333 | 3 | 0.3333 | 8 | 0.375 |
| 15_756184 | 1 | 1 | NA | NA | 2 | 1 | 3 | 1 | 6 | 1 | 2 | 0.5 | 1 | 0 | 2 | 0.5 | 2 | 0.5 | 7 | 0.4286 |
| 15_756186 | 1 | 1 | NA | NA | 2 | 1 | 3 | 0.6667 | 6 | 0.8333 | 1 | 1 | NA | NA | 2 | 1 | 1 | 1 | 4 | 1 |
| 15_756186 | NA | NA | NA | NA | 3 | 0.6667 | 3 | 0.3333 | 6 | 0.5 | 2 | 0.5 | 1 | 1 | 2 | 1 | 1 | 1 | 6 | 0.8333 |
| 15_756186 | 1 | 1 | NA | NA | 3 | 0.6667 | 4 | 0.5 | 8 | 0.625 | 2 | 0.5 | 1 | 0 | 1 | 0 | 2 | 1 | 6 | 0.5 |
| 15_756188 | NA | NA | NA | NA | 3 | 1 | 3 | 1 | 6 | 1 | 2 | 1 | 1 | 1 | 1 | 0 | 3 | 0.6667 | 7 | 0.7143 |
| 15_756189 | 1 | 1 | NA | NA | 3 | 0 | 3 | 0 | 7 | 0.1429 | 1 | 0 | 1 | 0 | 1 | 1 | 2 | 0.5 | 5 | 0.4 |
| 15_756263 | 4 | 0 | 3 | 0.3333 | 2 | 0 | NA | NA | 9 | 0.1111 | 4 | 0 | 5 | 0.6 | 1 | 0 | 3 | 0.3333 | 13 | 0.3077 |
| 15_756263 | 4 | 0.25 | 5 | 0.2 | 2 | 0 | 3 | 1 | 14 | 0.3571 | 3 | 0.3333 | 6 | 0.1667 | 3 | 0 | 3 | 0.6667 | 15 | 0.2667 |
| 15_756264 | 2 | 0.5 | 3 | 0.6667 | 1 | 0 | 2 | 0.5 | 8 | 0.5 | 5 | 0.4 | 4 | 1 | 2 | 0.5 | 3 | 1 | 14 | 0.7143 |
| 15_757112 | 3 | 0.3333 | 2 | 0 | NA | NA | 3 | 0.6667 | 8 | 0.375 | 3 | 0.3333 | 1 | 1 | 1 | 0 | 2 | 0.5 | 7 | 0.4286 |
| 15_757114 | 2 | 0 | 3 | 0 | NA | NA | 3 | 0.3333 | 8 | 0.125 | 3 | 0 | 1 | 1 | 1 | 1 | 2 | 0.5 | 7 | 0.4286 |
| 15_757116 | NA | NA | 2 | 0.5 | NA | NA | 3 | 0.6667 | 5 | 0.6 | NA | NA | 1 | 0 | 1 | 1 | NA | NA | 2 | 0.5 |
| 15_757118 | 1 | 1 | 2 | 0.5 | NA | NA | 3 | 0.6667 | 6 | 0.6667 | 2 | 0.5 | 1 | 1 | 1 | 1 | 2 | 1 | 6 | 0.8333 |
| 15_757119 | 1 | 1 | 3 | 0 | NA | NA | 3 | 0.3333 | 7 | 0.2857 | 2 | 0 | 1 | 0 | 1 | 0 | 1 | 1 | 5 | 0.2 |
| 15_757123 | 1 | 0 | 2 | 1 | NA | NA | 2 | 0.5 | 5 | 0.6 | 2 | 0 | 1 | 1 | 1 | 1 | 1 | 1 | 5 | 0.6 |
| 15_757659 | 1 | 1 | NA | NA | NA | NA | NA | NA | 1 | 1 | 2 | 0 | 1 | 1 | NA | NA | NA | NA | 3 | 0.3333 |
| 15_757668 | 1 | 0 | NA | NA | 1 | 1 | NA | NA | 2 | 0.5 | NA | NA | 1 | 1 | NA | NA | NA | NA | 1 | 1 |
| 15_757671 | 1 | 1 | NA | NA | 1 | 0 | NA | NA | 2 | 0.5 | NA | NA | 1 | 1 | NA | NA | NA | NA | 1 | 1 |
| 15_757673 | 13 | 0.0769 | 20 | 0.15 | 16 | 0.3125 | 10 | 0.1 | 59 | 0.1695 | 14 | 0.0714 | 17 | 0.1176 | 16 | 0.125 | 11 | 0 | 58 | 0.0862 |
| 15_757677 | 11 | 0.2727 | 20 | 0.3 | 16 | 0.375 | 10 | 0.1 | 57 | 0.2807 | 12 | 0.1667 | 17 | 0.1176 | 15 | 0 | 11 | 0 | 55 | 0.0727 |
| 15_757745 | 4 | 0 | 2 | 0 | 3 | 0 | 4 | 0.25 | 13 | 0.0769 | 3 | 0.3333 | 8 | 0.125 | 3 | 0 | 4 | 0.25 | 18 | 0.1667 |
| 15_757747 | 4 | 0 | 3 | 0.6667 | 4 | 0.25 | 4 | 0.25 | 15 | 0.2667 | 3 | 0 | 5 | 0.4 | 2 | 1 | 2 | 0.5 | 12 | 0.4167 |
| 15_757750 | 4 | 1 | 2 | 1 | 2 | 1 | NA | NA | 8 | 1 | 3 | 1 | 5 | 1 | 2 | 1 | 2 | 1 | 12 | 1 |
| 15_757751 | 3 | 0.6667 | 1 | 1 | 2 | 1 | 1 | 1 | 7 | 0.8571 | 1 | 1 | 5 | 0.8 | 2 | 0.5 | 1 | 1 | 9 | 0.7778 |
| 15_757754 | 3 | 0.6667 | 2 | 0.5 | 2 | 1 | 1 | 1 | 8 | 0.75 | 3 | 1 | 4 | 0.75 | 2 | 0.5 | 1 | 1 | 10 | 0.8 |
| 15_757754 | 3 | 1 | 2 | 1 | 2 | 0.5 | 1 | 0 | 8 | 0.75 | 3 | 0.3333 | 5 | 0.8 | 1 | 1 | 1 | 1 | 10 | 0.7 |
| 15_757756 | 4 | 0.25 | NA | NA | 2 | 0 | NA | NA | 6 | 0.1667 | 3 | 0 | 5 | 0 | 2 | 0.5 | NA | NA | 10 | 0.1 |
| 15_757816 | 1 | 1 | NA | NA | NA | NA | NA | NA | 1 | 1 | NA | NA | NA | NA | 1 | 1 | NA | NA | 1 | 1 |
| 15_757822 | 1 | 1 | NA | NA | NA | NA | NA | NA | 1 | 1 | NA | NA | NA | NA | 1 | 1 | NA | NA | 1 | 1 |
| 15_757824 | 2 | 0 | 2 | 0.5 | NA | NA | 1 | 1 | 5 | 0.4 | 1 | 1 | NA | NA | 1 | 0 | NA | NA | 2 | 0.5 |
| 15_757824 | NA | NA | 2 | 0.5 | NA | NA | NA | NA | 2 | 0.5 | 1 | 1 | NA | NA | NA | NA | NA | NA | 1 | 1 |
| 15_757828 | 2 | 0.5 | 1 | 1 | NA | NA | 1 | 1 | 4 | 0.75 | 1 | 0 | 1 | 1 | 1 | 1 | NA | NA | 3 | 0.6667 |
| 15_757830 | 2 | 1 | 2 | 1 | NA | NA | 1 | 1 | 5 | 1 | NA | NA | 1 | 1 | 1 | 1 | NA | NA | 2 | 1 |
| 15_757830 | 2 | 1 | 2 | 1 | NA | NA | 2 | 0.5 | 6 | 0.8333 | NA | NA | 1 | 1 | 1 | 1 | NA | NA | 2 | 1 |
| 15_757830 | NA | NA | 2 | 1 | NA | NA | 2 | 0.5 | 4 | 0.75 | 1 | 1 | 1 | 1 | 1 | 1 | NA | NA | 3 | 1 |
| 15_757831 | 2 | 0.5 | 2 | 1 | NA | NA | 1 | 1 | 5 | 0.8 | 1 | 1 | 1 | 1 | 1 | 1 | NA | NA | 3 | 1 |
| 15_757831 | 2 | 0 | 2 | 0 | NA | NA | 2 | 0.5 | 6 | 0.1667 | 1 | 1 | 1 | 1 | 1 | 0 | NA | NA | 3 | 0.6667 |
| 15_757834 | 2 | 1 | 1 | 1 | NA | NA | 1 | 1 | 4 | 1 | 1 | 1 | NA | NA | 1 | 0 | NA | NA | 2 | 0.5 |
| 15_757834 | 2 | 0.5 | 2 | 0.5 | NA | NA | 2 | 0.5 | 6 | 0.5 | 1 | 1 | 1 | 0 | NA | NA | NA | NA | 2 | 0.5 |
| 15_757835 | 2 | 0.5 | 2 | 0.5 | NA | NA | 2 | 0.5 | 6 | 0.5 | 1 | 0 | NA | NA | 1 | 1 | NA | NA | 2 | 0.5 |

| | | | | | | | | | | | | | | | | | | | | |
|---|---|---|---|---|---|---|---|---|---|---|---|---|---|---|---|---|---|---|---|---|
| 15_757838 | 1 | 0 | NA | NA | NA | NA | 2 | 0.5 | 3 | 0.3333 | 1 | 1 | NA | NA | 1 | 0 | NA | NA | 2 | 0.5 |
| 15_757838 | NA | NA | 1 | 1 | NA | NA | 2 | 0.5 | 3 | 0.6667 | 1 | 1 | NA | NA | 1 | 1 | NA | NA | 2 | 1 |
| 15_757883 | NA | NA | 5 | 0.2 | 1 | 0 | 6 | 0 | 12 | 0.0833 | 4 | 0 | 2 | 0 | 6 | 0.1667 | 3 | 0.3333 | 15 | 0.1333 |
| 15_757885 | NA | NA | 4 | 0.25 | 2 | 0 | 7 | 0.2857 | 13 | 0.2308 | 3 | 0 | 2 | 0 | 6 | 0 | 3 | 0.3333 | 14 | 0.0714 |
| 15_757886 | NA | NA | 4 | 0.25 | 2 | 0 | 6 | 0.1667 | 12 | 0.1667 | 3 | 0.3333 | 3 | 0 | 5 | 0.2 | 2 | 0 | 13 | 0.1538 |
| 15_757887 | NA | NA | 4 | 0.25 | 1 | 0 | 5 | 0.4 | 10 | 0.3 | 2 | 1 | 3 | 0 | 5 | 0.4 | 1 | 0 | 11 | 0.3636 |
| 15_757965 | 6 | 0.5 | 4 | 0.5 | 4 | 0.5 | 5 | 0.8 | 19 | 0.5789 | 1 | 1 | 7 | 0.7143 | 5 | 0.4 | 3 | 0.3333 | 16 | 0.5625 |
| 15_758190 | 3 | 0.3333 | NA | NA | NA | NA | NA | NA | 3 | 0.3333 | 1 | 1 | NA | NA | 1 | 0 | 1 | 0 | 3 | 0.3333 |
| 15_758191 | 4 | 0.25 | 2 | 0 | 1 | 0 | 1 | 0 | 8 | 0.125 | 1 | 0 | NA | NA | 1 | 0 | 2 | 0.5 | 4 | 0.25 |
| 15_758192 | 3 | 1 | 1 | 1 | 1 | 0 | 1 | 1 | 6 | 0.8333 | 1 | 1 | NA | NA | NA | NA | 2 | 1 | 3 | 1 |
| 15_758195 | 4 | 0.25 | 2 | 1 | 1 | 1 | 1 | 1 | 8 | 0.625 | 1 | 1 | NA | NA | 1 | 1 | 2 | 0.5 | 4 | 0.75 |
| 15_758198 | 4 | 0.75 | 2 | 0.5 | 1 | 1 | 1 | 1 | 8 | 0.75 | 1 | 1 | NA | NA | 1 | 1 | 2 | 0.5 | 4 | 0.75 |
| 15_758200 | 3 | 0.6667 | 2 | 0.5 | NA | NA | 2 | 0.5 | 7 | 0.5714 | 1 | 1 | NA | NA | NA | NA | 2 | 1 | 3 | 1 |
| 15_758202 | 4 | 1 | 2 | 1 | 1 | 1 | 1 | 1 | 8 | 1 | 1 | 1 | NA | NA | 1 | 1 | 2 | 1 | 4 | 1 |
| 15_758206 | 4 | 1 | 1 | 1 | 1 | 1 | 1 | 1 | 7 | 1 | 1 | 1 | NA | NA | 1 | 1 | NA | NA | 2 | 1 |
| 15_758206 | 4 | 0 | 2 | 0 | NA | NA | 2 | 0.5 | 8 | 0.125 | 1 | 0 | NA | NA | 1 | 0 | 1 | 1 | 3 | 0.3333 |
| 15_758207 | 4 | 0.75 | 2 | 1 | NA | NA | 2 | 1 | 8 | 0.875 | 1 | 1 | NA | NA | 1 | 1 | 1 | 1 | 3 | 1 |
| 15_758208 | 2 | 0 | 2 | 0 | 1 | 1 | 1 | 1 | 6 | 0.3333 | 1 | 0 | NA | NA | 1 | 1 | 1 | 0 | 3 | 0.3333 |
| 15_758208 | 4 | 0.75 | 2 | 1 | 1 | 1 | 2 | 1 | 9 | 0.8889 | 1 | 1 | NA | NA | 1 | 1 | 2 | 1 | 4 | 1 |
| 15_758211 | 3 | 0 | 1 | 1 | 1 | 0 | 1 | 0 | 6 | 0.1667 | 1 | 1 | NA | NA | NA | NA | 1 | 0 | 2 | 0.5 |
| 15_758464 | 3 | 1 | 1 | 1 | NA | NA | 2 | 0.5 | 6 | 0.8333 | NA | NA | 6 | 0.8333 | 1 | 1 | 3 | 1 | 10 | 0.9 |
| 15_758465 | 2 | 0.5 | 1 | 1 | NA | NA | 2 | 0.5 | 5 | 0.6 | NA | NA | 6 | 0.6667 | 1 | 1 | 2 | 1 | 9 | 0.7778 |
| 15_758465 | 1 | 1 | 1 | 1 | NA | NA | 1 | 0 | 3 | 0.6667 | NA | NA | 5 | 0.4 | 1 | 0 | NA | NA | 6 | 0.3333 |
| 15_758469 | 3 | 1 | 1 | 1 | NA | NA | 2 | 0.5 | 6 | 0.8333 | NA | NA | 8 | 0.875 | 1 | 1 | 3 | 0.6667 | 12 | 0.8333 |
| 15_758471 | 1 | 1 | 2 | 0.5 | NA | NA | 2 | 1 | 5 | 0.8 | NA | NA | 7 | 0.8571 | NA | NA | 3 | 1 | 10 | 0.9 |
| 15_758471 | 3 | 0.6667 | 2 | 1 | NA | NA | 2 | 0 | 7 | 0.5714 | NA | NA | 7 | 0.5714 | 2 | 0 | 3 | 0.6667 | 12 | 0.5 |
| 15_758472 | 1 | 1 | 2 | 1 | NA | NA | NA | NA | 3 | 1 | NA | NA | 5 | 1 | 1 | 1 | 1 | 1 | 7 | 1 |
| 15_758472 | 1 | 1 | 2 | 0.5 | NA | NA | 1 | 1 | 4 | 0.75 | NA | NA | 6 | 0.5 | 2 | 1 | 2 | 1 | 10 | 0.7 |
| 15_758473 | 1 | 0 | 2 | 0.5 | NA | NA | 1 | 0 | 4 | 0.25 | NA | NA | 5 | 0.6 | 1 | 0 | 2 | 1 | 8 | 0.625 |
| 15_758475 | 1 | 0 | 1 | 1 | NA | NA | 2 | 0 | 4 | 0.25 | NA | NA | 8 | 0.125 | 2 | 0.5 | 2 | 0 | 12 | 0.1667 |
| 15_758520 | 3 | 0.6667 | NA | NA | NA | NA | 1 | 0 | 4 | 0.5 | 2 | 0.5 | NA | NA | 1 | 1 | 1 | 0 | 4 | 0.5 |
| 15_758525 | 3 | 0.6667 | NA | NA | NA | NA | NA | NA | 3 | 0.6667 | 2 | 1 | 1 | 0 | 2 | 0.5 | 2 | 0.5 | 7 | 0.5714 |
| 15_758526 | 4 | 0.5 | NA | NA | NA | NA | NA | NA | 4 | 0.5 | 2 | 0.5 | 1 | 1 | 2 | 0 | 2 | 0.5 | 7 | 0.4286 |
| 15_758527 | 2 | 0.5 | NA | NA | NA | NA | NA | NA | 2 | 0.5 | 2 | 1 | 1 | 0 | 1 | 1 | 2 | 0 | 6 | 0.5 |
| 15_758528 | 3 | 1 | NA | NA | NA | NA | NA | NA | 3 | 1 | 2 | 1 | 1 | 1 | 1 | 1 | 2 | 1 | 6 | 1 |
| 15_758530 | 4 | 0.5 | NA | NA | NA | NA | NA | NA | 4 | 0.5 | 2 | 0.5 | 1 | 0 | 1 | 0 | 2 | 0 | 6 | 0.1667 |
| 15_758531 | 3 | 1 | NA | NA | NA | NA | NA | NA | 3 | 1 | 1 | 1 | 1 | 1 | 1 | 1 | 2 | 0.5 | 5 | 0.8 |
| 15_758532 | 4 | 0.25 | NA | NA | NA | NA | NA | NA | 4 | 0.25 | 2 | 0.5 | 1 | 0 | 1 | 0 | 1 | 0 | 5 | 0.2 |
| 15_758733 | NA | NA | NA | NA | NA | NA | 1 | 1 | 1 | 1 | 1 | 1 | NA | NA | NA | NA | NA | NA | 1 | 1 |
| 15_758735 | NA | NA | NA | NA | NA | NA | 1 | 1 | 1 | 1 | 1 | 1 | NA | NA | NA | NA | NA | NA | 1 | 1 |
| 15_758736 | NA | NA | NA | NA | NA | NA | 1 | 1 | 1 | 1 | 1 | 1 | NA | NA | NA | NA | NA | NA | 1 | 1 |
| 15_758737 | NA | NA | NA | NA | NA | NA | 1 | 1 | 1 | 1 | 1 | 1 | NA | NA | NA | NA | NA | NA | 1 | 1 |
| 15_758739 | NA | NA | NA | NA | NA | NA | 1 | 1 | 1 | 1 | 1 | 1 | NA | NA | NA | NA | NA | NA | 1 | 1 |
| 15_759190 | 5 | 0 | 1 | 0 | 5 | 0 | 6 | 0.1667 | 17 | 0.0588 | 1 | 0 | 3 | 0.3333 | 3 | 0 | 5 | 0.2 | 12 | 0.1667 |
| 15_759192 | 5 | 0.2 | 2 | 0 | 5 | 0 | 8 | 0 | 20 | 0.05 | 1 | 0 | 5 | 0 | 1 | 0 | 5 | 0.2 | 12 | 0.0833 |
| 15_759420 | 23 | 0.0435 | 26 | 0.1154 | 21 | 0.0476 | 28 | 0.0714 | 98 | 0.0714 | 16 | 0.0625 | 11 | 0.0909 | 23 | 0 | 23 | 0.087 | 73 | 0.0548 |
| 15_759421 | 24 | 0.9167 | 27 | 0.963 | 19 | 1 | 29 | 1 | 99 | 0.9697 | 17 | 0.9412 | 10 | 1 | 25 | 0.84 | 24 | 0.9583 | 76 | 0.9211 |
| 15_759421 | 23 | 0.3913 | 25 | 0.4 | 20 | 0.35 | 25 | 0.2 | 93 | 0.3333 | 17 | 0.3529 | 7 | 0.1429 | 24 | 0.2917 | 22 | 0.4545 | 70 | 0.3429 |
| 15_759422 | 22 | 0.7727 | 27 | 0.8519 | 20 | 0.8 | 25 | 0.92 | 94 | 0.8404 | 14 | 0.8571 | 10 | 0.9 | 25 | 0.8 | 23 | 0.913 | 72 | 0.8611 |
| 15_759575 | 1 | 0 | 6 | 0.1667 | 7 | 0.1429 | 1 | 0 | 15 | 0.1333 | 2 | 0 | 7 | 0.1429 | 3 | 0 | 3 | 0.3333 | 15 | 0.1333 |
| 15_759578 | 1 | 0 | 9 | 0.1111 | 5 | 0 | 2 | 0.5 | 17 | 0.1176 | 3 | 0 | 4 | 0.25 | 4 | 0.25 | 3 | 0 | 14 | 0.1429 |
| 15_759579 | 1 | 0 | 10 | 0.3 | 8 | 0.25 | 2 | 0.5 | 21 | 0.2857 | 4 | 0.5 | 5 | 0 | 4 | 0 | 4 | 0.5 | 17 | 0.2353 |
| 15_759582 | 2 | 0 | 7 | 0.4286 | 6 | 0.3333 | 3 | 0 | 18 | 0.2778 | 4 | 0.5 | 4 | 0.5 | 3 | 0.3333 | 3 | 0.3333 | 14 | 0.4286 |
| 15_759584 | 1 | 0 | 8 | 0.125 | 7 | 0.1429 | 4 | 0 | 20 | 0.1 | 3 | 0 | 5 | 0.2 | 3 | 0 | 3 | 0.3333 | 14 | 0.1429 |
| 15_759585 | 2 | 0 | 8 | 0.5 | 6 | 0.5 | 2 | 0.5 | 18 | 0.4444 | 3 | 0.3333 | 4 | 0 | 4 | 0.5 | 2 | 1 | 13 | 0.3846 |
| 15_759586 | 2 | 0 | 7 | 0.2857 | 7 | 0.1429 | 2 | 0.5 | 18 | 0.2222 | 3 | 0 | 4 | 0.25 | 4 | 0.25 | 3 | 0.3333 | 14 | 0.2143 |
| 15_759587 | 1 | 0 | 8 | 0.125 | 7 | 0 | 2 | 0 | 18 | 0.0556 | 2 | 0 | 5 | 0.4 | 2 | 0 | 3 | 0.6667 | 12 | 0.3333 |
| 15_759588 | 2 | 0 | 7 | 0.2857 | 7 | 0.1429 | 2 | 0 | 18 | 0.1667 | 3 | 0 | 4 | 0 | 2 | 0.5 | 3 | 0.3333 | 12 | 0.1667 |
| 15_759591 | NA | NA | 7 | 0 | 5 | 0.2 | 1 | 1 | 13 | 0.1538 | 3 | 0 | 3 | 0.3333 | 1 | 0 | 2 | 0 | 9 | 0.1111 |
| 15_760553 | 17 | 0.1176 | 15 | 0.0667 | 8 | 0.25 | 12 | 0.25 | 52 | 0.1538 | 6 | 0.1667 | 11 | 0 | 12 | 0.0833 | 4 | 0.25 | 33 | 0.0909 |
| 15_760554 | 12 | 0.1667 | 16 | 0.3125 | 7 | 0.2857 | 12 | 0.5833 | 47 | 0.3404 | 7 | 0.1429 | 9 | 0.2222 | 8 | 0 | 4 | 0.25 | 28 | 0.1429 |
| 15_760554 | 14 | 0.6429 | 16 | 0.9375 | 7 | 0.8571 | 12 | 0.5 | 49 | 0.7347 | 7 | 0.4286 | 9 | 0.7778 | 10 | 0.6 | 3 | 1 | 29 | 0.6552 |
| 15_760555 | 12 | 0 | 14 | 0.4286 | 7 | 0.4286 | 11 | 0.3636 | 44 | 0.2955 | 6 | 0 | 9 | 0.3333 | 10 | 0.2 | 4 | 0 | 29 | 0.1724 |
| 15_760555 | 10 | 0.7 | 10 | 1 | 5 | 1 | 11 | 1 | 36 | 0.9167 | 7 | 0.7143 | 7 | 1 | 10 | 0.9 | 4 | 1 | 28 | 0.8929 |
| 15_760555 | 12 | 0 | 14 | 0.0714 | 7 | 0.1429 | 12 | 0.0833 | 45 | 0.0667 | 6 | 0 | 9 | 0.2222 | 11 | 0.0909 | 4 | 0.25 | 30 | 0.1333 |

| | | | | | | | | | | | | | | | | | | | | |
|---|---|---|---|---|---|---|---|---|---|---|---|---|---|---|---|---|---|---|---|---|
| 15_7605593 | 10 | 0.6 | 7 | 0.8571 | 2 | 1 | 5 | 1 | 24 | 0.7917 | 3 | 0.3333 | 5 | 0.8 | 5 | 1 | 2 | 0.5 | 15 | 0.7333 |
| 15_7605593 | 13 | 0.0769 | 11 | 0.1818 | 5 | 0.4 | 10 | 0.1 | 39 | 0.1538 | 5 | 0 | 7 | 0.1429 | 8 | 0.375 | 1 | 0 | 21 | 0.1905 |
| 15_7605613 | 11 | 0.1818 | 12 | 0.0833 | 6 | 0 | 10 | 0.1 | 39 | 0.1026 | 4 | 0.25 | 7 | 0.2857 | 6 | 0 | 2 | 1 | 19 | 0.2632 |
| 15_7605621 | 13 | 0.3077 | 6 | 0.1667 | 6 | 0.3333 | 9 | 0.1111 | 34 | 0.2353 | 7 | 0.1429 | 7 | 0.7143 | 6 | 0.3333 | 1 | 0 | 21 | 0.381 |
| 15_7605621 | 12 | 0.1667 | 5 | 0.2 | 6 | 0.1667 | 8 | 0.125 | 31 | 0.1613 | 7 | 0.4286 | 8 | 0.375 | 7 | 0.2857 | 2 | 0.5 | 24 | 0.375 |
| 15_7605660 | 11 | 0.6364 | 4 | 0.75 | 3 | 1 | 9 | 0.7778 | 27 | 0.7407 | 2 | 0.5 | 7 | 0.8571 | 4 | 1 | 1 | 1 | 14 | 0.8571 |
| 15_7605697 | 9 | 0.2222 | 6 | 0 | 3 | 0 | 8 | 0 | 26 | 0.0769 | 3 | 0 | 8 | 0.5 | 4 | 0 | 2 | 0 | 17 | 0.2353 |
| 15_7605698 | 9 | 0.4444 | 5 | 0.2 | 3 | 1 | 8 | 0.625 | 25 | 0.52 | 3 | 1 | 8 | 0.5 | 4 | 0.5 | 2 | 0.5 | 17 | 0.5882 |
| 15_7605715 | 7 | 0.7143 | 5 | 1 | 4 | 0.75 | 8 | 1 | 24 | 0.875 | 3 | 0.6667 | 6 | 1 | 4 | 1 | 1 | 1 | 14 | 0.9286 |
| 15_7605716 | 9 | 0.6667 | 4 | 0.5 | 4 | 0.25 | 8 | 0.25 | 25 | 0.44 | 3 | 0 | 8 | 0.75 | 4 | 0.25 | 2 | 0.5 | 17 | 0.4706 |
| 15_7605722 | 8 | 0.25 | 5 | 0.2 | 3 | 0.3333 | 8 | 0.125 | 24 | 0.2083 | 3 | 0.3333 | 7 | 0.2857 | 5 | 0.2 | 1 | 1 | 16 | 0.3125 |
| 15_7605758 | 8 | 0.125 | 5 | 0.4 | 3 | 0.6667 | 8 | 0.125 | 24 | 0.25 | 3 | 0.3333 | 7 | 0.2857 | 4 | 0 | 1 | 0 | 15 | 0.2 |
| 15_7605772 | 4 | 0 | 2 | 0 | 2 | 0 | 7 | 0.1429 | 15 | 0.0667 | NA | NA | 3 | 0.3333 | 2 | 0 | 1 | 0 | 6 | 0.1667 |
| 15_7605777 | 9 | 0.2222 | 4 | 0.25 | 2 | 0.5 | 7 | 0.1429 | 22 | 0.2273 | 3 | 0.3333 | 4 | 0.25 | 3 | 0 | 1 | 0 | 11 | 0.1818 |
| 15_7605790 | 9 | 0.4444 | 4 | 1 | 1 | 1 | 7 | 0.7143 | 21 | 0.6667 | 1 | 1 | 5 | 0.6 | 3 | 1 | 2 | 0.5 | 11 | 0.7273 |
| 15_7605796 | 8 | 0.25 | 3 | 0.6667 | 1 | 0 | 7 | 0.1429 | 19 | 0.2632 | 2 | 0 | 6 | 0.3333 | 3 | 0.3333 | 2 | 0 | 13 | 0.2308 |
| 15_7605800 | 9 | 0.2222 | 4 | 0 | 1 | 1 | 7 | 0 | 21 | 0.1429 | 1 | 0 | 6 | 0 | 2 | 0.5 | 2 | 0.5 | 11 | 0.1818 |
| 15_7605806 | 6 | 0.6667 | 2 | 1 | 1 | 1 | 4 | 1 | 13 | 0.8462 | 1 | 1 | 3 | 0 | 2 | 1 | 1 | 1 | 7 | 0.5714 |
| 15_7605813 | 7 | 0.1429 | 4 | 0.5 | NA | NA | 5 | 0.2 | 16 | 0.25 | 2 | 0 | 5 | 0.2 | 3 | 0 | 1 | 0 | 11 | 0.0909 |
| 15_7605847 | 8 | 0.125 | 4 | 0.25 | 1 | 0 | 4 | 0 | 17 | 0.1176 | 2 | 0 | 5 | 0 | 1 | 0 | 2 | 0.5 | 10 | 0.1 |
| 15_7605851 | 6 | 0.8333 | 3 | 1 | 1 | 1 | 3 | 0.6667 | 13 | 0.8462 | 2 | 1 | 3 | 1 | 1 | 1 | NA | NA | 6 | 1 |
| 15_7605858 | 8 | 0.625 | 4 | 0.5 | 1 | 0 | 4 | 0.5 | 17 | 0.5294 | 1 | 0 | 3 | 0.3333 | 1 | 1 | 2 | 0.5 | 7 | 0.4286 |
| 15_7605881 | 5 | 0.6 | 4 | 0.5 | 1 | 1 | 3 | 0.3333 | 13 | 0.5385 | 1 | 0 | 5 | 0.8 | 2 | 0.5 | 1 | 0 | 9 | 0.5556 |
| 15_7605919 | 5 | 0.2 | 3 | 0.3333 | 1 | 1 | 2 | 0.5 | 11 | 0.3636 | NA | NA | 4 | 0 | 2 | 0.5 | 1 | 1 | 7 | 0.2857 |
| 15_7605932 | 5 | 0.2 | 3 | 0.3333 | NA | NA | 2 | 0.5 | 10 | 0.3 | 2 | 0 | 3 | 0.3333 | 1 | 1 | 1 | 0 | 7 | 0.2857 |
| 15_7605951 | 5 | 0.4 | 3 | 0 | NA | NA | 2 | 0.5 | 10 | 0.3 | 1 | 0 | 5 | 0.2 | 2 | 0 | 2 | 0 | 10 | 0.1 |
| 15_7605963 | 5 | 0.2 | 3 | 0.6667 | NA | NA | 2 | 0.5 | 10 | 0.4 | 1 | 0 | 3 | 0.6667 | 1 | 1 | 2 | 0.5 | 7 | 0.5714 |
| 15_7605975 | 4 | 0.25 | 2 | 0.5 | NA | NA | 1 | 0 | 7 | 0.2857 | 2 | 0.5 | 2 | 0.5 | 1 | 0 | 1 | 1 | 6 | 0.5 |
| 15_7605981 | 4 | 0.5 | 3 | 0.3333 | NA | NA | 2 | 0 | 9 | 0.3333 | 2 | 0.5 | 3 | 0.3333 | 1 | 0 | 2 | 0 | 8 | 0.25 |
| 15_7605992 | 4 | 0.75 | 2 | 1 | NA | NA | 2 | 1 | 8 | 0.875 | 2 | 1 | 3 | 0.6667 | NA | NA | 2 | 1 | 7 | 0.8571 |
| 15_7605996 | 3 | 0.6667 | 3 | 1 | NA | NA | 2 | 1 | 8 | 0.875 | 2 | 1 | 3 | 0.6667 | NA | NA | 2 | 1 | 7 | 0.8571 |
| 15_7606010 | 2 | 1 | 1 | 1 | NA | NA | 1 | 1 | 4 | 1 | 2 | 1 | 3 | 1 | 1 | 1 | 2 | 1 | 8 | 1 |
| 15_7606026 | 4 | 1 | 1 | 1 | NA | NA | 1 | 1 | 6 | 1 | 2 | 1 | 3 | 0.3333 | NA | NA | 1 | 0 | 6 | 0.5 |
| 15_7606051 | 4 | 0.75 | 2 | 1 | NA | NA | 2 | 1 | 8 | 0.875 | NA | NA | 3 | 0.6667 | NA | NA | 2 | 1 | 5 | 0.8 |
| 15_7606052 | 4 | 0.5 | 2 | 0.5 | NA | NA | 2 | 0 | 8 | 0.375 | 2 | 1 | 2 | 0 | 1 | 0 | 2 | 0 | 7 | 0.2857 |
| 15_7606068 | 4 | 0.25 | 2 | 1 | NA | NA | 2 | 0.5 | 8 | 0.5 | 2 | 0 | 3 | 0.3333 | 1 | 1 | 2 | 0.5 | 8 | 0.375 |
| 15_7606087 | 2 | 0.5 | NA | NA | NA | NA | 2 | 0 | 4 | 0.25 | 1 | 1 | 1 | 1 | NA | NA | NA | NA | 2 | 1 |
| 15_7606102 | 4 | 0.5 | NA | NA | NA | NA | 2 | 0.5 | 6 | 0.5 | 2 | 0 | 3 | 0.6667 | 1 | 1 | 2 | 0.5 | 8 | 0.5 |
| 15_7606105 | 3 | 0.6667 | NA | NA | NA | NA | 2 | 1 | 5 | 0.8 | 2 | 1 | 2 | 1 | 1 | 1 | 1 | 0 | 6 | 0.8333 |
| 15_7606111 | 4 | 0.25 | NA | NA | NA | NA | 2 | 0 | 6 | 0.1667 | 2 | 0.5 | 1 | 0 | 1 | 0 | 2 | 0 | 6 | 0.1667 |
| 15_7606163 | 2 | 0.5 | NA | NA | NA | NA | NA | NA | 2 | 0.5 | 2 | 0.5 | 2 | 1 | 1 | 1 | 1 | 1 | 6 | 0.8333 |
| 15_7606188 | 3 | 0.3333 | NA | NA | NA | NA | NA | NA | 3 | 0.3333 | 1 | 0 | 2 | 1 | 1 | 0 | 2 | 0.5 | 6 | 0.5 |
| 15_7606191 | 3 | 1 | NA | NA | NA | NA | NA | NA | 3 | 1 | 2 | 1 | 2 | 1 | 1 | 1 | 2 | 0.5 | 7 | 0.8571 |
| 15_7606201 | 3 | 1 | NA | NA | NA | NA | NA | NA | 3 | 1 | 2 | 0 | 1 | 1 | 1 | 0 | 2 | 0.5 | 6 | 0.3333 |
| 15_7606207 | 3 | 0.6667 | NA | NA | NA | NA | NA | NA | 3 | 0.6667 | 2 | 0 | 2 | 0 | NA | NA | 2 | 1 | 6 | 0.3333 |
| 15_7606251 | 2 | 0.5 | NA | NA | NA | NA | NA | NA | 2 | 0.5 | 1 | 0 | 2 | 0 | 1 | 0 | 2 | 0.5 | 6 | 0.1667 |
| 15_7606309 | 1 | 1 | NA | NA | NA | NA | NA | NA | 1 | 1 | 2 | 0.5 | 2 | 0.5 | 1 | 0 | 1 | 0 | 6 | 0.3333 |
| 15_7606336 | 1 | 1 | NA | NA | NA | NA | NA | NA | 1 | 1 | NA | NA | 1 | 1 | NA | NA | 1 | 0 | 2 | 0.5 |
| 15_7606487 | 1 | 1 | NA | NA | NA | NA | NA | NA | 1 | 1 | 1 | 0 | 1 | 1 | NA | NA | NA | NA | 2 | 0.5 |
| 15_7611735 | NA | NA | 3 | 1 | NA | NA | NA | NA | 3 | 1 | NA | NA | NA | NA | 2 | 1 | NA | NA | 2 | 1 |
| 15_7611736 | NA | NA | 2 | 1 | NA | NA | NA | NA | 2 | 1 | NA | NA | NA | NA | 2 | 1 | NA | NA | 2 | 1 |
| 15_7613351 | 32 | 0.0625 | 83 | 0.1566 | 65 | 0.1077 | 33 | 0.2424 | 213 | 0.1408 | 49 | 0.102 | 59 | 0.1017 | 68 | 0.1618 | 47 | 0.1702 | 223 | 0.1345 |
| 15_7613372 | 28 | 0.2143 | 58 | 0.2414 | 53 | 0.3019 | 27 | 0.2222 | 166 | 0.253 | 42 | 0.2857 | 38 | 0.1842 | 52 | 0.2692 | 37 | 0.3514 | 169 | 0.2722 |
| 15_7618263 | 3 | 0 | 3 | 0 | 1 | 0 | 2 | 0.5 | 9 | 0.1111 | 4 | 0.25 | 3 | 0 | 2 | 0 | NA | NA | 9 | 0.1111 |
| 15_7621357 | 6 | 0.5 | 7 | 0 | 7 | 0 | 8 | 0 | 28 | 0.1071 | 6 | 0.3333 | 7 | 0 | 9 | 0 | 7 | 0 | 29 | 0.069 |
| 15_7621364 | 6 | 0.5 | 6 | 0.6667 | 7 | 0.5714 | 8 | 0.625 | 27 | 0.5926 | 6 | 0.6667 | 8 | 0.875 | 10 | 0.7 | 7 | 0.7143 | 31 | 0.7419 |
| 15_7621378 | 4 | 0.75 | 8 | 0.875 | 5 | 0.8 | 7 | 0.5714 | 24 | 0.75 | 6 | 0.6667 | 8 | 0.875 | 9 | 0.7778 | 6 | 0.6667 | 29 | 0.7586 |
| 15_7621386 | 5 | 0 | 8 | 0.375 | 6 | 0 | 7 | 0.1429 | 26 | 0.1538 | 8 | 0 | 8 | 0.25 | 9 | 0.2222 | 7 | 0 | 32 | 0.125 |
| 15_7621391 | 4 | 0.75 | 7 | 0.7143 | 6 | 0.1667 | 7 | 0.8571 | 24 | 0.625 | 8 | 0.75 | 6 | 0.3333 | 9 | 0.6667 | 6 | 0.5 | 29 | 0.5862 |
| 15_7621455 | 1 | 0 | 2 | 0 | 1 | 0 | 2 | 1 | 6 | 0.3333 | 1 | 1 | 3 | 1 | 5 | 0.6 | 2 | 1 | 11 | 0.8182 |
| 15_7621776 | 4 | 0 | 1 | 0 | 4 | 0.25 | 1 | 0 | 10 | 0.1 | 3 | 0.3333 | 2 | 0 | 4 | 0 | 5 | 0.2 | 14 | 0.1429 |
| 15_7621878 | 10 | 0 | 6 | 0.1667 | 8 | 0 | 3 | 0.3333 | 27 | 0.0741 | 9 | 0.2222 | 4 | 0 | 8 | 0 | 7 | 0 | 28 | 0.0714 |
| 15_7623007 | 43 | 0.1395 | 65 | 0.0308 | 36 | 0.0833 | 32 | 0.0625 | 176 | 0.0739 | 45 | 0.0889 | 47 | 0.0638 | 50 | 0 | 47 | 0.0851 | 189 | 0.0582 |
| 15_7623015 | NA | NA | 3 | 0 | 1 | 1 | NA | NA | 4 | 0.25 | 2 | 0.5 | NA | NA | 1 | 0 | NA | NA | 3 | 0.3333 |
| 15_7623132 | 4 | 0 | 4 | 0 | 8 | 0.25 | 8 | 0.125 | 24 | 0.125 | 2 | 0 | 3 | 0 | 5 | 0.2 | 7 | 0.1429 | 17 | 0.1176 |

| | | | | | | | | | | | | | | | | | | | | |
|---|---|---|---|---|---|---|---|---|---|---|---|---|---|---|---|---|---|---|---|---|
| 15_762347 | 4 | 0.75 | 2 | 0 | 2 | 1 | 8 | 0.75 | 16 | 0.6875 | 1 | 0 | 4 | 0.75 | 3 | 0.3333 | 6 | 0.3333 | 14 | 0.4286 |
| 15_762347 | 4 | 0 | 2 | 0 | 2 | 0 | 8 | 0.125 | 16 | 0.0625 | 1 | 0 | 2 | 0.5 | 3 | 0.3333 | 3 | 0.3333 | 9 | 0.3333 |
| 15_762347 | 4 | 0 | 1 | 0 | 2 | 0.5 | 5 | 0 | 12 | 0.0833 | 1 | 0 | 1 | 0 | 2 | 0 | 5 | 0.2 | 9 | 0.1111 |
| 15_762348 | 2 | 0 | NA | NA | 1 | 0 | 3 | 0.3333 | 6 | 0.1667 | NA | NA | 1 | 0 | 1 | 1 | 5 | 0.2 | 7 | 0.2857 |
| 15_762350 | 3 | 0.6667 | NA | NA | 1 | 1 | 2 | 0.5 | 6 | 0.6667 | NA | NA | 2 | 1 | NA | NA | 5 | 0.6 | 7 | 0.7143 |
| 15_762350 | 3 | 0.3333 | NA | NA | 1 | 0 | 2 | 0 | 6 | 0.1667 | NA | NA | 2 | 0 | NA | NA | 5 | 0.2 | 7 | 0.1429 |
| 15_762383 | 15 | 0.9333 | 20 | 0.95 | 13 | 0.9231 | 17 | 0.8824 | 65 | 0.9231 | 6 | 0.8333 | 12 | 0.8333 | 10 | 0.9 | 12 | 0.8333 | 40 | 0.85 |
| 15_762383 | 14 | 0.1429 | 20 | 0.3 | 11 | 0.0909 | 16 | 0.1875 | 61 | 0.1967 | 6 | 0.3333 | 10 | 0 | 9 | 0 | 8 | 0.125 | 33 | 0.0909 |
| 15_762551 | 1 | 1 | 3 | 1 | 1 | 1 | 2 | 1 | 7 | 1 | NA | NA | NA | NA | 1 | 1 | NA | NA | 1 | 1 |
| 15_762551 | 1 | 1 | 2 | 1 | 2 | 1 | 2 | 1 | 7 | 1 | NA | NA | NA | NA | 1 | 1 | NA | NA | 1 | 1 |
| 15_762553 | 1 | 0 | 3 | 0.6667 | 3 | 0.3333 | 2 | 0.5 | 9 | 0.4444 | NA | NA | NA | NA | 1 | 1 | NA | NA | 1 | 1 |
| 15_762554 | 1 | 1 | 4 | 1 | 3 | 0.6667 | 2 | 1 | 10 | 0.9 | NA | NA | NA | NA | 1 | 1 | NA | NA | 1 | 1 |
| 15_763371 | 3 | 0.3333 | 7 | 0.1429 | 2 | 0 | NA | NA | 12 | 0.1667 | 7 | 0 | 3 | 0 | 2 | 0.5 | 3 | 0 | 15 | 0.0667 |
| 15_763382 | 1 | 0 | 4 | 0 | 5 | 0 | 7 | 0.2857 | 17 | 0.1176 | 3 | 0.3333 | 6 | 0 | 4 | 0 | NA | NA | 13 | 0.0769 |
| 15_763435 | NA | NA | NA | NA | 1 | 1 | NA | NA | 1 | 1 | NA | NA | NA | NA | NA | NA | 1 | 1 | 1 | 1 |
| 15_763828 | 4 | 0 | 5 | 0 | 7 | 0.1429 | 2 | 0.5 | 18 | 0.1111 | 2 | 0 | 4 | 0 | 5 | 0 | 2 | 0.5 | 13 | 0.0769 |
| 15_763848 | 5 | 0.2 | 8 | 0 | 9 | 0.4444 | 3 | 0.3333 | 25 | 0.24 | 5 | 0 | 5 | 0.6 | 6 | 0.1667 | 1 | 0 | 17 | 0.2353 |
| 15_763848 | 5 | 0 | 9 | 0 | 9 | 0.2222 | 3 | 0 | 26 | 0.0769 | 4 | 0.5 | 4 | 0.25 | 7 | 0 | 2 | 0 | 17 | 0.1765 |
| 15_763849 | 5 | 0 | 8 | 0.375 | 6 | 0 | 2 | 0 | 21 | 0.1429 | 1 | 0 | 5 | 0.2 | 7 | 0.1429 | 1 | 0 | 14 | 0.1429 |
| 15_763849 | 5 | 0 | 8 | 0.125 | 9 | 0.1111 | 3 | 0 | 25 | 0.08 | 5 | 0 | 5 | 0.2 | 7 | 0 | 2 | 0 | 19 | 0.0526 |
| 15_763849 | 4 | 0.75 | 7 | 0.1429 | 8 | 0.625 | 2 | 0 | 21 | 0.4286 | 2 | 0 | 5 | 0.4 | 7 | 0 | 1 | 1 | 15 | 0.2 |
| 15_763850 | 4 | 1 | 6 | 0.8333 | 8 | 0.875 | 3 | 0.6667 | 21 | 0.8571 | 4 | 1 | 2 | 1 | 5 | 0.8 | 3 | 0.6667 | 14 | 0.8571 |
| 15_763865 | 2 | 0 | 2 | 0 | 6 | 0.1667 | 1 | 0 | 11 | 0.0909 | 3 | 0.3333 | 2 | 0.5 | 5 | 0.2 | 3 | 0 | 13 | 0.2308 |
| 15_764004 | 16 | 0.125 | 22 | 0.3636 | 26 | 0.2308 | 17 | 0.2353 | 81 | 0.2469 | 12 | 0.25 | 8 | 0.375 | 18 | 0.0556 | 16 | 0.125 | 54 | 0.1667 |
| 15_764005 | 13 | 0 | 20 | 0.25 | 21 | 0.0952 | 15 | 0.0667 | 69 | 0.1159 | 12 | 0.25 | 9 | 0 | 11 | 0.0909 | 12 | 0 | 44 | 0.0909 |
| 15_764019 | 9 | 0 | 16 | 0 | 16 | 0.125 | 8 | 0.25 | 49 | 0.0816 | 7 | 0 | 6 | 0 | 12 | 0.0833 | 9 | 0.1111 | 34 | 0.0588 |
| 15_764022 | 8 | 1 | 13 | 0.9231 | 14 | 0.8571 | 12 | 1 | 47 | 0.9362 | 5 | 1 | 7 | 0.7143 | 15 | 0.8667 | 8 | 0.75 | 35 | 0.8286 |
| 15_764025 | 8 | 0.625 | 15 | 0.7333 | 13 | 0.6923 | 12 | 0.4167 | 48 | 0.625 | 9 | 1 | 10 | 0.6 | 18 | 0.7222 | 9 | 0.6667 | 46 | 0.7391 |
| 15_764026 | 5 | 0.2 | 10 | 0.2 | 6 | 0.1667 | 9 | 0.3333 | 30 | 0.2333 | 5 | 0 | 9 | 0.3333 | 16 | 0.125 | 6 | 0.3333 | 36 | 0.1944 |
| 15_764026 | 6 | 0.1667 | 13 | 0.1538 | 9 | 0 | 9 | 0.1111 | 37 | 0.1081 | 4 | 0.25 | 11 | 0.1818 | 16 | 0 | 5 | 0 | 36 | 0.0833 |
| 15_764031 | 7 | 0.1429 | 14 | 0 | 11 | 0.1818 | 12 | 0.0833 | 44 | 0.0909 | 8 | 0.125 | 8 | 0.375 | 13 | 0 | 6 | 0.1667 | 35 | 0.1429 |
| 15_764031 | 7 | 0 | 14 | 0.2143 | 12 | 0.0833 | 12 | 0.1667 | 45 | 0.1333 | 7 | 0.1429 | 8 | 0.125 | 13 | 0 | 6 | 0.1667 | 34 | 0.0882 |
| 15_764033 | 5 | 1 | 11 | 1 | 7 | 1 | 7 | 1 | 30 | 1 | 2 | 1 | 5 | 1 | 7 | 1 | 4 | 1 | 18 | 1 |
| 15_764033 | 5 | 0.4 | 14 | 0.5 | 9 | 0.3333 | 11 | 0.0909 | 39 | 0.3333 | 5 | 0.2 | 8 | 0.875 | 13 | 0.2308 | 5 | 0.4 | 31 | 0.4194 |
| 15_764033 | 5 | 0.2 | 11 | 0 | 10 | 0.4 | 12 | 0.1667 | 38 | 0.1842 | 5 | 0 | 7 | 0.1429 | 9 | 0.1111 | 6 | 0.1667 | 27 | 0.1111 |
| 15_764038 | 3 | 0 | 11 | 0.1818 | 11 | 0 | 11 | 0.2727 | 36 | 0.1389 | 4 | 0 | 6 | 0 | 11 | 0.0909 | 5 | 0.2 | 26 | 0.0769 |
| 15_764060 | 4 | 0.25 | 8 | 0 | 8 | 0.125 | 8 | 0.125 | 28 | 0.1071 | 4 | 0.75 | 3 | 0 | 8 | 0 | 3 | 0.3333 | 18 | 0.2222 |
| 15_764062 | 3 | 0.3333 | 6 | 0 | 3 | 0 | 3 | 0 | 15 | 0.0667 | 4 | 0.25 | 2 | 0 | 1 | 0 | 1 | 0 | 8 | 0.125 |
| 15_764063 | 3 | 0 | 1 | 0 | 2 | 0 | 1 | 1 | 7 | 0.1429 | 3 | 0 | 1 | 0 | 2 | 0 | 2 | 0.5 | 8 | 0.125 |
| 15_764226 | NA | NA | 2 | 0.5 | NA | NA | 3 | 0.3333 | 5 | 0.4 | 3 | 1 | 2 | 1 | 2 | 0.5 | 3 | 0 | 10 | 0.6 |
| 15_764228 | NA | NA | 1 | 1 | NA | NA | 3 | 0 | 4 | 0.25 | 4 | 0 | 3 | 0 | 2 | 0 | 3 | 0.3333 | 12 | 0.0833 |
| 15_764243 | 1 | 0 | 4 | 0.75 | 1 | 1 | NA | NA | 6 | 0.6667 | 1 | 0 | 1 | 1 | 2 | 0 | 2 | 0.5 | 6 | 0.3333 |
| 15_764253 | 3 | 0.6667 | 7 | 0.7143 | 2 | 1 | 1 | 1 | 13 | 0.7692 | 1 | 1 | 1 | 1 | 3 | 1 | NA | NA | 5 | 1 |
| 15_764254 | 3 | 0.3333 | 7 | 0.2857 | 2 | 0 | 1 | 0 | 13 | 0.2308 | 1 | 0 | 1 | 0 | 3 | 1 | NA | NA | 5 | 0.6 |
| 15_764254 | 3 | 0.3333 | 7 | 0.8571 | 2 | 0.5 | NA | NA | 12 | 0.6667 | NA | NA | 1 | 1 | 3 | 0.3333 | NA | NA | 4 | 0.5 |
| 15_764255 | 3 | 0.6667 | 6 | 0.6667 | 2 | 0.5 | 1 | 1 | 12 | 0.6667 | NA | NA | NA | NA | 3 | 0.6667 | NA | NA | 3 | 0.6667 |
| 15_764257 | 1 | 1 | 2 | 1 | NA | NA | NA | NA | 3 | 1 | NA | NA | NA | NA | 1 | 1 | NA | NA | 1 | 1 |
| 15_765083 | 18 | 0.1111 | 21 | 0.0476 | 16 | 0.0625 | 23 | 0.0435 | 78 | 0.0641 | 22 | 0 | 17 | 0.1176 | 20 | 0.05 | 17 | 0.0588 | 76 | 0.0526 |
| 15_765105 | 1 | 1 | NA | NA | NA | NA | NA | NA | 1 | 1 | NA | NA | NA | NA | NA | NA | 1 | 1 | 1 | 1 |
| 15_765281 | 10 | 0.4 | 9 | 0.4444 | 4 | 0.75 | 8 | 0.5 | 31 | 0.4839 | 3 | 1 | 6 | 0.3333 | 3 | 0.6667 | 5 | 0.6 | 17 | 0.5882 |
| 15_765282 | 11 | 0.2727 | 8 | 0.625 | 5 | 0.6 | 7 | 0.5714 | 31 | 0.4839 | 2 | 0.5 | 9 | 0.8889 | 5 | 0.6 | 7 | 0.4286 | 23 | 0.6522 |
| 15_765283 | 11 | 0.0909 | 7 | 0.1429 | 5 | 0.6 | 7 | 0.1429 | 30 | 0.2 | 2 | 0.5 | 10 | 0.2 | 5 | 0.2 | 6 | 0.3333 | 23 | 0.2609 |
| 15_765285 | 9 | 0.1111 | 6 | 0.5 | 3 | 0.6667 | 7 | 0.1429 | 25 | 0.28 | 2 | 0.5 | 8 | 0.375 | 4 | 0 | 5 | 0 | 19 | 0.2105 |
| 15_765290 | 9 | 0.2222 | 6 | 0.1667 | 4 | 0.25 | 6 | 0 | 25 | 0.16 | NA | NA | 8 | 0 | 2 | 0 | 2 | 0.5 | 12 | 0.0833 |
| 15_765290 | 10 | 0 | 4 | 0.5 | 5 | 0.6 | 4 | 0 | 23 | 0.2174 | 1 | 0 | 7 | 0.2857 | 4 | 0.25 | 2 | 0 | 14 | 0.2143 |
| 15_765292 | 10 | 0.4 | 7 | 0 | 4 | 0.75 | 5 | 0.4 | 26 | 0.3462 | NA | NA | 8 | 0.125 | 4 | 0.5 | 5 | 0.4 | 17 | 0.2941 |
| 15_765293 | 10 | 0 | 7 | 0.2857 | 3 | 0 | 4 | 0 | 24 | 0.0833 | 1 | 0 | 8 | 0 | 6 | 0.3333 | 5 | 0.2 | 20 | 0.15 |
| 15_765297 | 9 | 0.7778 | 6 | 0.6667 | 3 | 1 | 2 | 0.5 | 20 | 0.75 | 1 | 0 | 5 | 1 | 3 | 0.6667 | 5 | 0.8 | 14 | 0.7857 |
| 15_765297 | 8 | 0.875 | 4 | 0.75 | 3 | 1 | 4 | 0.75 | 19 | 0.8421 | 1 | 0 | 6 | 0.6667 | 2 | 1 | 5 | 1 | 14 | 0.7857 |
| 15_765298 | 8 | 1 | 6 | 1 | 3 | 1 | 4 | 0.75 | 21 | 0.9524 | 1 | 0 | 4 | 0.75 | 3 | 1 | 5 | 1 | 13 | 0.8462 |
| 15_765298 | 9 | 1 | 4 | 1 | 2 | 1 | 4 | 0.75 | 19 | 0.9474 | 1 | 1 | 5 | 1 | 3 | 1 | 5 | 1 | 14 | 1 |
| 15_765299 | 8 | 1 | 5 | 0.8 | 1 | 1 | 4 | 1 | 18 | 0.9444 | 1 | 1 | 5 | 1 | 2 | 1 | 5 | 0.8 | 13 | 0.9231 |
| 15_765300 | 14 | 0.3571 | 10 | 0.2 | 8 | 0.125 | 11 | 0.4545 | 43 | 0.3023 | 5 | 0.4 | 11 | 0.3636 | 11 | 0.2727 | 13 | 0.3077 | 40 | 0.325 |
| 15_765302 | 11 | 0.2727 | 10 | 0.1 | 12 | 0.1667 | 12 | 0.1667 | 45 | 0.1778 | 4 | 0.5 | 10 | 0.1 | 15 | 0.0667 | 12 | 0.0833 | 41 | 0.122 |
| 15_765302 | 10 | 0.1 | 9 | 0.1111 | 11 | 0 | 8 | 0 | 38 | 0.0526 | 4 | 0 | 8 | 0.125 | 12 | 0.0833 | 9 | 0.1111 | 33 | 0.0909 |

| | | | | | | | | | | | | | | | | | | | | |
|---|---|---|---|---|---|---|---|---|---|---|---|---|---|---|---|---|---|---|---|---|
| 15_765302( | 14 | 0.2857 | 9 | 0.2222 | 12 | 0.1667 | 10 | 0.3 | 45 | 0.2444 | 6 | 0.1667 | 9 | 0.2222 | 14 | 0.1429 | 11 | 0.1818 | 40 | 0.175 |
| 15_765303 | 11 | 0.6364 | 10 | 0.4 | 12 | 0.4167 | 9 | 0.4444 | 42 | 0.4762 | 6 | 0.3333 | 10 | 0.3 | 13 | 0.4615 | 11 | 0.3636 | 40 | 0.375 |
| 15_765304 | 14 | 1 | 10 | 0.6 | 10 | 0.5 | 9 | 0.4444 | 43 | 0.6744 | 5 | 0.8 | 8 | 0.75 | 14 | 0.7857 | 9 | 0.7778 | 36 | 0.7778 |
| 15_765304( | 12 | 0.75 | 8 | 0.5 | 8 | 0.5 | 11 | 0.8182 | 39 | 0.6667 | 6 | 0.8333 | 7 | 0.5714 | 12 | 0.6667 | 9 | 0.6667 | 34 | 0.6765 |
| 15_765304( | 13 | 0.1538 | 8 | 0.375 | 10 | 0.3 | 9 | 0 | 40 | 0.2 | 7 | 0 | 9 | 0.3333 | 11 | 0.2727 | 10 | 0.1 | 37 | 0.1892 |
| 15_765306( | 6 | 0.5 | 12 | 0.4167 | 10 | 0.2 | 11 | 0.5455 | 39 | 0.4103 | 7 | 0.1429 | 8 | 0.5 | 11 | 0.2727 | 11 | 0.3636 | 37 | 0.3243 |
| 15_765312 | 8 | 0.375 | 8 | 0.75 | 4 | 1 | 7 | 1 | 27 | 0.7407 | 1 | 0 | 7 | 0.8571 | 12 | 0.6667 | 4 | 0.5 | 24 | 0.6667 |
| 15_765313 | 6 | 0.5 | 6 | 0.8333 | 5 | 0.8 | 6 | 1 | 23 | 0.7826 | 1 | 1 | 7 | 0.8571 | 12 | 0.75 | 4 | 0.75 | 24 | 0.7917 |
| 15_765313( | 8 | 0.625 | 8 | 0.5 | 6 | 0.8333 | 5 | 1 | 27 | 0.7037 | 1 | 1 | 8 | 0.875 | 11 | 0.7273 | 4 | 1 | 24 | 0.8333 |
| 15_765314( | 7 | 0.1429 | 8 | 0 | 5 | 0 | 5 | 0.2 | 25 | 0.08 | 1 | 0 | 8 | 0.125 | 11 | 0.1818 | 4 | 0.25 | 24 | 0.1667 |
| 15_765315 | 8 | 0 | 5 | 0.4 | 5 | 0 | 6 | 0.1667 | 24 | 0.125 | 1 | 0 | 7 | 0.4286 | 8 | 0.125 | 3 | 0.6667 | 19 | 0.3158 |
| 15_765315( | 5 | 0.8 | 7 | 0.7143 | 5 | 1 | 5 | 1 | 22 | 0.8636 | 1 | 1 | 8 | 0.5 | 10 | 0.9 | 5 | 1 | 24 | 0.7917 |
| 15_765317( | 5 | 0.2 | 8 | 0.125 | 5 | 0.4 | 6 | 0.3333 | 24 | 0.25 | NA | NA | 6 | 0 | 9 | 0.2222 | 4 | 0 | 19 | 0.1053 |
| 15_765317 | 6 | 0.3333 | 5 | 0.4 | 5 | 0.4 | 3 | 0.6667 | 19 | 0.4211 | 1 | 0 | 4 | 0 | 6 | 0.6667 | 4 | 0.75 | 15 | 0.4667 |
| 15_765318( | 5 | 0 | 8 | 0 | 6 | 0.5 | 5 | 0.4 | 24 | 0.2083 | 1 | 0 | 7 | 0.4286 | 11 | 0.1818 | 2 | 0 | 21 | 0.2381 |
| 15_765361! | NA | NA | 1 | 1 | 1 | 0 | 1 | 0 | 3 | 0.3333 | NA | NA | 1 | 1 | NA | NA | 1 | 0 | 2 | 0.5 |
| 15_765424( | 4 | 0.25 | 10 | 0.4 | 6 | 0.5 | 7 | 0.4286 | 27 | 0.4074 | 5 | 0.6 | 8 | 0.375 | 3 | 0 | 6 | 0.3333 | 22 | 0.3636 |
| 15_765429( | 2 | 0 | 10 | 0.5 | 5 | 0.2 | 5 | 0.2 | 22 | 0.3182 | 5 | 0.4 | 6 | 0.6667 | 2 | 0.5 | 2 | 0.5 | 15 | 0.5333 |
| 15_765429 | 2 | 0 | 8 | 0.125 | 5 | 0.2 | 5 | 0.4 | 20 | 0.2 | 4 | 0.5 | 5 | 0.4 | 1 | 1 | 3 | 0.6667 | 13 | 0.5385 |
| 15_765434 | 2 | 0.5 | 6 | 0.1667 | 6 | 0.3333 | 6 | 0.1667 | 20 | 0.25 | 3 | 0.3333 | 6 | 0.5 | 3 | 0.6667 | 4 | 0.25 | 16 | 0.4375 |
| 15_765436( | 2 | 0.5 | 7 | 0.2857 | 6 | 0.3333 | 4 | 0.25 | 19 | 0.3158 | 4 | 0.5 | 5 | 0.2 | 2 | 0.5 | 3 | 0.3333 | 14 | 0.3571 |
| 15_765436( | 1 | 0 | 6 | 0.5 | 3 | 0 | 4 | 0.5 | 14 | 0.3571 | 3 | 1 | 3 | 0 | 3 | 0.3333 | 3 | 0.6667 | 12 | 0.5 |
| 15_765514 | 22 | 0.1364 | 34 | 0 | 27 | 0.1111 | 27 | 0.1111 | 110 | 0.0818 | 13 | 0 | 18 | 0 | 25 | 0.12 | 18 | 0.0556 | 74 | 0.0541 |
| 15_765515 | 30 | 0.2 | 32 | 0.3438 | 34 | 0.2647 | 26 | 0.1923 | 122 | 0.2541 | 18 | 0.1667 | 22 | 0.1364 | 28 | 0.1786 | 24 | 0.1667 | 92 | 0.163 |
| 15_765516 | 30 | 0.5333 | 29 | 0.4138 | 32 | 0.4688 | 24 | 0.3333 | 115 | 0.4435 | 20 | 0.45 | 24 | 0.3333 | 28 | 0.4286 | 24 | 0.2917 | 96 | 0.375 |
| 15_765516 | 30 | 0.1 | 33 | 0 | 38 | 0.0526 | 32 | 0.0938 | 133 | 0.0602 | 22 | 0.0909 | 21 | 0.0952 | 29 | 0.069 | 22 | 0.0909 | 94 | 0.0851 |
| 15_765517( | 13 | 0.1538 | 17 | 0.2353 | 8 | 0.375 | 6 | 0 | 44 | 0.2045 | 11 | 0.2727 | 11 | 0.1818 | 12 | 0.3333 | 13 | 0.0769 | 47 | 0.2128 |
| 15_765521 | 39 | 0.2051 | 35 | 0.0571 | 37 | 0.1892 | 31 | 0.2258 | 142 | 0.169 | 24 | 0.1667 | 27 | 0.1481 | 38 | 0.1316 | 26 | 0.1923 | 115 | 0.1565 |
| 15_765522 | 36 | 0.0278 | 30 | 0.1333 | 35 | 0 | 29 | 0.1379 | 130 | 0.0692 | 24 | 0.0833 | 24 | 0.0417 | 33 | 0.1212 | 24 | 0.0417 | 105 | 0.0762 |
| 15_765744 | 9 | 0.6667 | 5 | 0.6 | 3 | 0.3333 | 2 | 0.5 | 19 | 0.5789 | 5 | 0.6 | 4 | 0.5 | 4 | 0.5 | 8 | 0.5 | 21 | 0.5238 |
| 15_765745 | 8 | 0.125 | 5 | 0 | 4 | 0.25 | NA | NA | 17 | 0.1176 | 5 | 0.6 | 5 | 0.2 | 4 | 0 | 5 | 0.6 | 19 | 0.3684 |
| 15_765746 | 7 | 1 | 2 | 1 | 4 | 1 | 2 | 1 | 15 | 1 | 4 | 1 | 5 | 0.6 | 5 | 1 | 7 | 1 | 21 | 0.9048 |
| 15_765746 | 7 | 0.8571 | 4 | 0.75 | 4 | 0.25 | 1 | 0 | 16 | 0.625 | 4 | 1 | 6 | 0.1667 | 3 | 0.3333 | 6 | 0.5 | 19 | 0.4737 |
| 15_765747 | 9 | 0.1111 | 5 | 0.2 | 3 | 0.3333 | 2 | 0.5 | 19 | 0.2105 | 5 | 0.2 | 3 | 0 | 4 | 0 | 4 | 0 | 16 | 0.0625 |
| 15_765786 | 8 | 0 | 3 | 0 | 5 | 0.2 | 3 | 0 | 19 | 0.0526 | 4 | 0.25 | 4 | 0 | 5 | 0 | 5 | 0.2 | 18 | 0.1111 |
| 15_765786( | 11 | 0.4545 | 7 | 0.5714 | 8 | 0.625 | 4 | 0.25 | 30 | 0.5 | 10 | 0.7 | 5 | 0.6 | 5 | 0.4 | 8 | 0.5 | 28 | 0.5714 |
| 15_765787 | 14 | 0.9286 | 9 | 0.6667 | 9 | 0.7778 | 7 | 0.4286 | 39 | 0.7436 | 11 | 0.8182 | 7 | 0.5714 | 8 | 0.5 | 11 | 0.7273 | 37 | 0.6757 |
| 15_765787 | 12 | 0.1667 | 10 | 0 | 10 | 0 | 6 | 0 | 38 | 0.0526 | 11 | 0.0909 | 6 | 0.3333 | 7 | 0 | 12 | 0.3333 | 36 | 0.1944 |
| 15_765787 | 12 | 0.8333 | 10 | 0.9 | 10 | 0.6 | 5 | 0.6 | 37 | 0.7568 | 8 | 0.625 | 6 | 0.8333 | 6 | 0.6667 | 8 | 0.5 | 28 | 0.6429 |
| 15_765787 | 11 | 0.5455 | 9 | 0.3333 | 9 | 0.2222 | 4 | 0 | 33 | 0.3333 | 8 | 0.375 | 8 | 0.125 | 6 | 0.1667 | 11 | 0.3636 | 33 | 0.2727 |
| 15_765790 | 13 | 0.8462 | 10 | 0.6 | 8 | 0.625 | 6 | 0.3333 | 37 | 0.6486 | 9 | 1 | 7 | 0.8571 | 8 | 0.5 | 11 | 0.4545 | 35 | 0.6857 |
| 15_765790 | 13 | 1 | 10 | 0.9 | 8 | 0.875 | 7 | 1 | 38 | 0.9474 | 11 | 1 | 6 | 1 | 8 | 0.875 | 12 | 0.9167 | 37 | 0.9459 |
| 15_765790 | 11 | 0.7273 | 10 | 0.8 | 7 | 0.7143 | 9 | 0.7778 | 37 | 0.7568 | 10 | 0.6 | 6 | 0.8333 | 6 | 0.6667 | 11 | 0.8182 | 33 | 0.7273 |
| 15_765834 | 1 | 0 | 1 | 0 | 2 | 0 | 4 | 0.25 | 8 | 0.125 | 3 | 0 | 4 | 0 | NA | NA | 4 | 0.25 | 11 | 0.0909 |
| 15_765859 | NA | NA | 1 | 1 | NA | NA | 2 | 1 | 3 | 1 | 1 | 1 | 1 | 1 | 1 | 1 | 1 | 0 | 4 | 0.75 |
| 15_765861 | NA | NA | 1 | 1 | NA | NA | 2 | 0.5 | 3 | 0.6667 | 1 | 1 | 1 | 1 | NA | NA | 2 | 0 | 4 | 0.5 |
| 15_765862( | NA | NA | 1 | 0 | NA | NA | 2 | 1 | 3 | 0.6667 | 1 | 0 | 1 | 0 | 1 | 1 | 2 | 0 | 5 | 0.2 |
| 15_765874 | 1 | 1 | 2 | 0.5 | NA | NA | 4 | 0 | 7 | 0.2857 | 2 | 0 | 1 | 0 | 3 | 0.3333 | 2 | 0.5 | 8 | 0.25 |
| 15_765972 | 3 | 0.6667 | 1 | 0 | 4 | 0.5 | 2 | 0 | 10 | 0.4 | 2 | 0 | 1 | 1 | 2 | 0.5 | 1 | 0 | 6 | 0.3333 |
| 15_765976 | NA | NA | NA | NA | 2 | 0.5 | 2 | 1 | 4 | 0.75 | 2 | 1 | 1 | 1 | 1 | 0 | 1 | 1 | 5 | 0.8 |
| 15_765977 | 1 | 0 | 1 | 0 | 2 | 1 | 2 | 0 | 6 | 0.3333 | 2 | 0.5 | 1 | 0 | 1 | 0 | 1 | 1 | 5 | 0.4 |
| 15_765978 | 1 | 0 | 1 | 0 | 1 | 1 | 2 | 0.5 | 5 | 0.4 | 2 | 0.5 | NA | NA | 1 | 0 | 1 | 0 | 4 | 0.25 |
| 15_765980 | 2 | 1 | 2 | 0.5 | 2 | 1 | 3 | 0.6667 | 9 | 0.7778 | 3 | 0 | NA | NA | 2 | 1 | NA | NA | 5 | 0.4 |
| 15_765983 | 2 | 0 | 2 | 0 | 4 | 0 | 3 | 0.3333 | 11 | 0.0909 | 4 | 0.25 | 1 | 0 | 2 | 0 | NA | NA | 7 | 0.1429 |
| 15_765984 | 3 | 0.3333 | 1 | 1 | 3 | 0.3333 | 3 | 0.6667 | 10 | 0.5 | 3 | 1 | 2 | 1 | 3 | 1 | 1 | 1 | 9 | 1 |
| 15_765985 | 2 | 1 | 1 | 1 | 3 | 1 | 3 | 1 | 9 | 1 | 3 | 0.6667 | 1 | 1 | 1 | 1 | 1 | 1 | 6 | 0.8333 |
| 15_765986( | 3 | 0.6667 | 2 | 1 | 3 | 1 | 3 | 1 | 11 | 0.9091 | 4 | 1 | 2 | 1 | 2 | 1 | 1 | 1 | 9 | 1 |
| 15_765986 | 3 | 1 | 2 | 1 | 4 | 1 | 3 | 1 | 12 | 1 | 4 | 0.75 | 2 | 1 | 2 | 1 | 1 | 1 | 9 | 0.8889 |
| 15_765986 | 3 | 1 | 2 | 1 | 3 | 1 | 2 | 1 | 10 | 1 | 3 | 1 | 1 | 1 | 2 | 1 | 1 | 1 | 7 | 1 |
| 15_765988( | 4 | 1 | 2 | 1 | 3 | 0.3333 | 3 | 1 | 12 | 0.8333 | 4 | 0.75 | 2 | 1 | 3 | 0.6667 | 1 | 1 | 10 | 0.8 |
| 15_765988( | 3 | 0 | 2 | 0.5 | 3 | 0.3333 | 2 | 0 | 10 | 0.2 | 3 | 0 | 2 | 0.5 | 2 | 1 | 1 | 0 | 8 | 0.375 |
| 15_765988 | 4 | 1 | 2 | 1 | 3 | 1 | 4 | 1 | 13 | 1 | 4 | 1 | 1 | 1 | 2 | 1 | NA | NA | 7 | 1 |
| 15_765990 | 3 | 1 | 1 | 0 | 2 | 0 | 3 | 0.3333 | 9 | 0.4444 | 3 | 1 | 1 | 1 | 2 | 0.5 | 1 | 0 | 7 | 0.7143 |
| 15_765993 | NA | NA | NA | NA | NA | NA | 1 | 1 | 1 | 1 | 1 | 1 | NA | NA | NA | NA | NA | NA | 1 | 1 |
| 15_765993 | NA | NA | NA | NA | NA | NA | 1 | 1 | 1 | 1 | 1 | 1 | NA | NA | NA | NA | NA | NA | 1 | 1 |

| | | | | | | | | | | | | | | | | | | | | |
|---|---|---|---|---|---|---|---|---|---|---|---|---|---|---|---|---|---|---|---|---|
| 15_7659964 | 5 | 0.4 | 9 | 0.4444 | 4 | 0 | 4 | 0.25 | 22 | 0.3182 | 5 | 0.2 | 6 | 0.5 | 4 | 0.75 | 1 | 0 | 16 | 0.4375 |
| 15_7659969 | 8 | 0.375 | 8 | 0 | 4 | 0 | 5 | 0.2 | 25 | 0.16 | 6 | 0 | 8 | 0.125 | 5 | 0.2 | 1 | 0 | 20 | 0.1 |
| 15_7659971 | 7 | 1 | 9 | 1 | 4 | 1 | 5 | 1 | 25 | 1 | 6 | 0.8333 | 9 | 1 | 4 | 0.75 | 1 | 0 | 20 | 0.85 |
| 15_7659990 | 7 | 0.4286 | 8 | 0.5 | 4 | 0.75 | 4 | 1 | 23 | 0.6087 | 4 | 0.5 | 7 | 0.4286 | 4 | 0 | NA | NA | 15 | 0.3333 |
| 15_7659995 | 5 | 0.4 | 7 | 0.4286 | 3 | 0 | 5 | 0.8 | 20 | 0.45 | 7 | 0.5714 | 8 | 0.375 | 4 | 0.25 | NA | NA | 19 | 0.4211 |
| 15_7659996 | 6 | 0.8333 | 8 | 1 | 3 | 1 | 5 | 1 | 22 | 0.9545 | 5 | 0.8 | 8 | 0.875 | 2 | 1 | NA | NA | 15 | 0.8667 |
| 15_7660028 | 4 | 0.25 | 7 | 0.5714 | 3 | 1 | 3 | 0.6667 | 17 | 0.5882 | 4 | 1 | 6 | 1 | 2 | 0.5 | NA | NA | 12 | 0.9167 |
| 15_7660048 | 7 | 0.1429 | 7 | 0.1429 | 3 | 0.3333 | 2 | 0 | 19 | 0.1579 | 5 | 0.2 | 6 | 0 | 3 | 0 | NA | NA | 14 | 0.0714 |
| 15_7660581 | 1 | 0 | 3 | 0.3333 | 1 | 0 | 1 | 0 | 6 | 0.1667 | 1 | 0 | 3 | 0 | 3 | 0.3333 | 4 | 0 | 11 | 0.0909 |
| 15_7660606 | NA | NA | 1 | 0 | 2 | 0.5 | 1 | 0 | 4 | 0.25 | NA | NA | 2 | 0.5 | 3 | 0 | 5 | 0.6 | 10 | 0.4 |
| 15_7660617 | NA | NA | 3 | 0.6667 | 1 | 0 | NA | NA | 4 | 0.5 | NA | NA | 2 | 0 | 2 | 0 | 3 | 0.3333 | 7 | 0.1429 |
| 15_7660661 | NA | NA | 2 | 0.5 | 2 | 0.5 | NA | NA | 4 | 0.5 | NA | NA | 4 | 0.5 | 2 | 0.5 | 2 | 0 | 8 | 0.375 |
| 15_7660670 | NA | NA | 3 | 0.3333 | 2 | 0 | NA | NA | 5 | 0.2 | NA | NA | 4 | 0.25 | 2 | 0.5 | 2 | 0.5 | 8 | 0.375 |
| 15_7660673 | NA | NA | 3 | 0.3333 | 2 | 0.5 | NA | NA | 5 | 0.4 | NA | NA | 4 | 0.25 | 2 | 0.5 | 2 | 0.5 | 8 | 0.375 |
| 15_7660841 | 3 | 0.6667 | 7 | 0.5714 | 6 | 0.8333 | 5 | 1 | 21 | 0.7619 | 6 | 0.8333 | 3 | 0.6667 | 5 | 0.8 | 9 | 1 | 23 | 0.8696 |
| 15_7660851 | 3 | 0.3333 | 5 | 0.2 | 5 | 0.2 | 4 | 0 | 17 | 0.1765 | 5 | 0.4 | 1 | 1 | 2 | 0 | 8 | 0.125 | 16 | 0.25 |
| 15_7660855 | 6 | 0.3333 | 6 | 0.1667 | 6 | 0.1667 | 3 | 0.3333 | 21 | 0.2381 | 8 | 0.125 | 2 | 0.5 | 4 | 0.25 | 10 | 0.3 | 24 | 0.25 |
| 15_7660876 | 3 | 0.6667 | 6 | 0.6667 | 5 | 0.4 | 3 | 0.3333 | 17 | 0.5294 | 7 | 0.4286 | 2 | 0.5 | 3 | 0.6667 | 10 | 0.8 | 22 | 0.6364 |
| 15_7660890 | 3 | 0 | 7 | 0.4286 | 5 | 0 | 4 | 0 | 19 | 0.1579 | 8 | 0.75 | 2 | 0.5 | 4 | 1 | 10 | 0.6 | 24 | 0.7083 |
| 15_7661909 | 2 | 0 | 3 | 0.3333 | 2 | 0 | NA | NA | 7 | 0.1429 | 6 | 0.3333 | NA | NA | 1 | 0 | 7 | 0.4286 | 14 | 0.3571 |
| 15_7661921 | 1 | 1 | NA | NA | 2 | 1 | NA | NA | 3 | 1 | 3 | 0.6667 | NA | NA | 1 | 1 | 4 | 1 | 8 | 0.875 |
| 15_7661933 | 2 | 0.5 | 2 | 0.5 | 2 | 0.5 | NA | NA | 6 | 0.5 | 6 | 0.5 | NA | NA | 1 | 1 | 4 | 0.5 | 11 | 0.5455 |
| 15_7661952 | 2 | 0 | 2 | 0.5 | 2 | 0 | NA | NA | 6 | 0.1667 | 4 | 0.25 | NA | NA | NA | NA | 5 | 0.6 | 9 | 0.4444 |
| 15_7662111 | 1 | 1 | 2 | 0 | 2 | 0 | NA | NA | 5 | 0.2 | 2 | 0.5 | NA | NA | NA | NA | 3 | 0.3333 | 5 | 0.4 |
| 15_7662129 | NA | NA | 2 | 0 | 2 | 0.5 | NA | NA | 4 | 0.25 | 4 | 0.5 | NA | NA | NA | NA | 2 | 0.5 | 6 | 0.5 |
| 15_7662171 | 1 | 1 | 1 | 1 | 1 | 1 | NA | NA | 3 | 1 | 2 | 1 | NA | NA | NA | NA | 2 | 1 | 4 | 1 |
| 15_7662175 | 1 | 1 | 2 | 0.5 | 1 | 1 | NA | NA | 4 | 0.75 | 2 | 1 | NA | NA | NA | NA | 1 | 1 | 3 | 1 |
| 15_7662184 | NA | NA | 2 | 0.5 | NA | NA | NA | NA | 2 | 0.5 | 3 | 0.3333 | NA | NA | NA | NA | 3 | 0.3333 | 6 | 0.3333 |
| 15_7673582 | 2 | 0 | 8 | 0.375 | NA | NA | 2 | 0.5 | 12 | 0.3333 | 3 | 0.3333 | 5 | 0 | 4 | 0.5 | 2 | 0 | 14 | 0.2143 |
| 15_7673584 | 4 | 0.75 | 13 | 0.4615 | 3 | 0.3333 | 5 | 0.6 | 25 | 0.52 | 4 | 0.5 | 7 | 1 | 8 | 0.625 | 6 | 0.8333 | 25 | 0.76 |
| 15_7673596 | 5 | 0.2 | 10 | 0.7 | 4 | 0.5 | 5 | 0.8 | 24 | 0.5833 | 3 | 0.3333 | 9 | 0.7778 | 9 | 0.4444 | 6 | 0.6667 | 27 | 0.5926 |
| 15_7673604 | 5 | 1 | 12 | 1 | 4 | 1 | 5 | 1 | 26 | 1 | 1 | 1 | 8 | 1 | 8 | 1 | 6 | 1 | 23 | 1 |
| 15_7673648 | 4 | 0.25 | 10 | 0 | 2 | 0.5 | 3 | 0 | 19 | 0.1053 | 2 | 0 | 5 | 0.2 | 5 | 0 | 4 | 0.25 | 16 | 0.125 |
| 15_7673663 | 5 | 0.6 | 10 | 0.5 | 2 | 0.5 | 4 | 0.75 | 21 | 0.5714 | 2 | 1 | 6 | 0.6667 | 6 | 0.8333 | 5 | 0.6 | 19 | 0.7368 |
| 15_7673668 | 4 | 0 | 10 | 0.1 | 2 | 0 | 5 | 0.4 | 21 | 0.1429 | 2 | 0 | 6 | 0 | 7 | 0.2857 | 4 | 0.25 | 19 | 0.1579 |
| 15_7673677 | 3 | 0.6667 | 7 | 0.4286 | 1 | 0 | 3 | 1 | 14 | 0.5714 | 1 | 0 | 5 | 0.6 | 5 | 0.4 | 4 | 0.25 | 15 | 0.4 |
| 15_7673688 | 4 | 0.75 | 7 | 0.7143 | 1 | 1 | 3 | 1 | 15 | 0.8 | 2 | 0 | 6 | 0.3333 | 4 | 0.5 | 2 | 0 | 14 | 0.2857 |
| 15_7673710 | 2 | 0.5 | 4 | 1 | 2 | 1 | 2 | 0.5 | 10 | 0.8 | 1 | 1 | 6 | 0.3333 | 3 | 1 | 2 | 1 | 12 | 0.6667 |
| 15_7673711 | 4 | 1 | 5 | 1 | 2 | 1 | 3 | 1 | 14 | 1 | 2 | 1 | 5 | 1 | 4 | 1 | 4 | 1 | 15 | 1 |
| 15_7673718 | 4 | 0.5 | 7 | 0.2857 | 1 | 0 | 3 | 0 | 15 | 0.2667 | 2 | 0.5 | 6 | 0.3333 | 3 | 0 | 4 | 0 | 15 | 0.2 |
| 15_7673735 | 4 | 0 | 6 | 0.1667 | 2 | 0 | 3 | 0 | 15 | 0.0667 | 2 | 0 | 5 | 0.2 | 3 | 0 | 4 | 0 | 14 | 0.0714 |
| 15_7673738 | 3 | 1 | 7 | 1 | 2 | 1 | 2 | 1 | 14 | 1 | 2 | 1 | 4 | 1 | 2 | 1 | 3 | 1 | 11 | 1 |
| 15_7673747 | 3 | 0.3333 | 6 | 0.1667 | 1 | 0 | 3 | 0.3333 | 13 | 0.2308 | 2 | 0.5 | 5 | 0.6 | 2 | 0.5 | 4 | 0 | 13 | 0.3846 |
| 15_7673750 | 4 | 1 | 6 | 1 | 2 | 1 | 3 | 1 | 15 | 1 | 2 | 1 | 4 | 0.75 | 2 | 1 | 4 | 1 | 12 | 0.9167 |
| 15_7673753 | 4 | 0.25 | 7 | 0.2857 | 2 | 0 | 3 | 0.3333 | 16 | 0.25 | 1 | 0 | 4 | 0 | 3 | 0.6667 | 4 | 0.25 | 12 | 0.25 |
| 15_7673829 | 4 | 0.5 | 6 | 0.5 | NA | NA | NA | NA | 10 | 0.5 | 1 | 0 | 4 | 0.5 | 2 | 0.5 | 2 | 0.5 | 9 | 0.4444 |
| 15_7673844 | 2 | 0.5 | 4 | 0.5 | 1 | 0 | NA | NA | 7 | 0.4286 | 2 | 0 | 4 | 0.25 | 2 | 0.5 | 1 | 1 | 9 | 0.3333 |
| 15_7673847 | 1 | 0 | 5 | 0.8 | 1 | 0 | NA | NA | 7 | 0.5714 | 2 | 0 | 4 | 0.5 | 2 | 1 | 4 | 0.25 | 12 | 0.4167 |
| 15_7673849 | 1 | 1 | 5 | 0.6 | 1 | 1 | NA | NA | 7 | 0.7143 | 2 | 1 | 4 | 0.75 | 2 | 1 | 4 | 1 | 12 | 0.9167 |
| 15_7673856 | 2 | 0.5 | 5 | 0.4 | NA | NA | NA | NA | 7 | 0.4286 | 1 | 1 | 4 | 0.5 | 2 | 0.5 | 3 | 0.6667 | 10 | 0.6 |
| 15_7675450 | 2 | 0.5 | 2 | 0.5 | NA | NA | NA | NA | 4 | 0.5 | 2 | 1 | 2 | 0 | 2 | 0 | 3 | 0 | 9 | 0.2222 |
| 15_7675506 | 1 | 1 | 1 | 0 | NA | NA | NA | NA | 2 | 0.5 | 2 | 0 | 4 | 0 | 2 | 0.5 | 2 | 0.5 | 10 | 0.2 |
| 15_7675509 | 1 | 1 | 1 | 0 | NA | NA | NA | NA | 2 | 0.5 | 2 | 0 | 4 | 0.25 | 2 | 0.5 | 1 | 1 | 9 | 0.3333 |
| 15_7678472 | 3 | 1 | 1 | 1 | 3 | 0.6667 | 7 | 1 | 14 | 0.9286 | 4 | 1 | 4 | 1 | 6 | 1 | 3 | 1 | 17 | 1 |
| 15_7678476 | 3 | 0.6667 | 1 | 0 | 4 | 0 | 6 | 0.1667 | 14 | 0.2143 | 3 | 0.6667 | 4 | 0.5 | 6 | 0 | 4 | 0 | 17 | 0.2353 |
| 15_7678479 | 1 | 1 | 1 | 1 | 4 | 0.75 | 5 | 0.8 | 11 | 0.8182 | 3 | 0.3333 | 4 | 1 | 5 | 1 | 2 | 0.5 | 14 | 0.7857 |
| 15_7678494 | 2 | 0.5 | 1 | 0 | 3 | 0.3333 | 6 | 0.3333 | 12 | 0.3333 | 2 | 0.5 | 2 | 1 | 6 | 0 | 2 | 0 | 12 | 0.25 |
| 15_7678494 | 4 | 0.5 | 1 | 0 | 4 | 0.5 | 6 | 0.8333 | 15 | 0.6 | 4 | 0.25 | 4 | 0.5 | 3 | 1 | 5 | 0.6 | 16 | 0.5625 |
| 15_7678527 | 3 | 1 | 1 | 1 | 3 | 1 | 5 | 1 | 12 | 1 | 3 | 1 | 2 | 1 | 5 | 1 | 5 | 1 | 15 | 1 |
| 15_7678528 | 5 | 0.4 | 1 | 1 | 4 | 0 | 5 | 0 | 15 | 0.2 | 3 | 0.6667 | 2 | 0.5 | 6 | 0.1667 | 3 | 0.3333 | 14 | 0.3571 |
| 15_7678534 | 5 | 0.6 | 1 | 0 | 5 | 0.4 | 7 | 0.4286 | 18 | 0.4444 | 3 | 0.6667 | 3 | 0.6667 | 5 | 0.6 | 6 | 0.5 | 17 | 0.5882 |
| 15_7678536 | 5 | 1 | NA | NA | 5 | 0.6 | 5 | 1 | 15 | 0.8667 | 2 | 1 | 4 | 1 | 6 | 1 | 6 | 0.8333 | 18 | 0.9444 |
| 15_7678537 | 3 | 1 | NA | NA | 4 | 1 | 5 | 1 | 12 | 1 | 2 | 1 | 3 | 1 | 4 | 1 | 4 | 1 | 13 | 1 |
| 15_7678563 | 2 | 1 | NA | NA | NA | NA | 2 | 0.5 | 4 | 0.75 | NA | NA | 2 | 0.5 | 1 | 1 | 4 | 0.75 | 7 | 0.7143 |
| 15_7683305 | 3 | 0 | 4 | 0 | 6 | 0.1667 | NA | NA | 13 | 0.0769 | 4 | 0.25 | NA | NA | 2 | 0 | 6 | 0 | 12 | 0.0833 |

| | | | | | | | | | | | | | | | | | | | | |
|---|---|---|---|---|---|---|---|---|---|---|---|---|---|---|---|---|---|---|---|---|
| 15_7683455 | 4 | 0.25 | 5 | 0.2 | 6 | 0.8333 | 1 | 0 | 16 | 0.4375 | 4 | 1 | NA | NA | 2 | 1 | 6 | 0.3333 | 12 | 0.6667 |
| 15_7683460 | 1 | 0 | 2 | 0.5 | 4 | 0 | 1 | 1 | 8 | 0.25 | 3 | 0.3333 | NA | NA | 2 | 1 | 4 | 0.5 | 9 | 0.5556 |
| 15_7683471 | 6 | 0.5 | 6 | 0.6667 | 6 | 0.5 | 1 | 1 | 19 | 0.5789 | 4 | 0.5 | NA | NA | 2 | 0 | 6 | 0.6667 | 12 | 0.5 |
| 15_7683482 | 6 | 0.6667 | 7 | 0.8571 | 7 | 0.8571 | 1 | 1 | 21 | 0.8095 | 4 | 0.75 | NA | NA | 2 | 1 | 5 | 0.8 | 11 | 0.8182 |
| 15_7683484 | 5 | 0.2 | 6 | 0.5 | 7 | 0.4286 | 1 | 1 | 19 | 0.4211 | 3 | 0.6667 | NA | NA | 2 | 1 | 5 | 0.8 | 10 | 0.8 |
| 15_7696488 | 13 | 0.3846 | 10 | 0.3 | 7 | 0.2857 | 10 | 0.2 | 40 | 0.3 | 6 | 0.1667 | 7 | 0.8571 | 14 | 0.3571 | 12 | 0.4167 | 39 | 0.4359 |
| 15_7696521 | 15 | 0.1333 | 11 | 0 | 8 | 0.125 | 9 | 0.3333 | 43 | 0.1395 | 6 | 0.1667 | 7 | 0.1429 | 12 | 0.0833 | 11 | 0.1818 | 36 | 0.1389 |
| 15_7696522 | 10 | 0.3 | 7 | 0.1429 | 10 | 0.4 | 11 | 0.5455 | 38 | 0.3684 | 6 | 0 | 6 | 0.3333 | 13 | 0.5385 | 8 | 0.125 | 33 | 0.303 |
| 15_7696531 | 15 | 0.0667 | 11 | 0 | 8 | 0.125 | 13 | 0.1538 | 47 | 0.0851 | 6 | 0 | 8 | 0.25 | 14 | 0.1429 | 12 | 0.1667 | 40 | 0.15 |
| 15_7696568 | 13 | 0.1538 | 8 | 0.125 | 9 | 0 | 7 | 0.1429 | 37 | 0.1081 | 4 | 0.25 | 7 | 0.2857 | 10 | 0.4 | 7 | 0.1429 | 28 | 0.2857 |
| 15_7696586 | 12 | 0.1667 | 8 | 0.125 | 8 | 0.25 | 10 | 0.1 | 38 | 0.1579 | 5 | 0 | 8 | 0.125 | 13 | 0.2308 | 8 | 0.125 | 34 | 0.1471 |
| 15_7696685 | 12 | 0.5 | 6 | 0.6667 | 6 | 0.6667 | 9 | 0.3333 | 33 | 0.5152 | 3 | 0.6667 | 4 | 0 | 9 | 0.5556 | 4 | 0 | 20 | 0.35 |
| 15_7696695 | 11 | 0.5455 | 9 | 0.2222 | 7 | 0.4286 | 8 | 0.25 | 35 | 0.3714 | 3 | 0.3333 | 4 | 0.75 | 9 | 0.4444 | 3 | 0.3333 | 19 | 0.4737 |
| 15_7696828 | 7 | 0.8571 | 5 | 0.4 | 4 | 0.5 | 3 | 0.6667 | 19 | 0.6316 | 4 | 0.75 | 2 | 1 | 6 | 1 | 2 | 1 | 14 | 0.9286 |
| 15_7764537 | 4 | 0.5 | 6 | 0.1667 | 10 | 0 | 5 | 0 | 25 | 0.12 | 6 | 0 | 3 | 0 | 3 | 0.3333 | 3 | 0 | 15 | 0.0667 |
| 15_7764602 | 4 | 0.5 | 5 | 0.4 | 9 | 0.4444 | 6 | 0.5 | 24 | 0.4583 | 5 | 0.8 | 4 | 0.75 | 3 | 0.6667 | 3 | 1 | 15 | 0.8 |
| 15_7779942 | 34 | 0.2353 | 32 | 0.2188 | 29 | 0.2414 | 28 | 0.3214 | 123 | 0.252 | 23 | 0.4783 | 28 | 0.1429 | 25 | 0.16 | 29 | 0.2414 | 105 | 0.2476 |
| 15_7782567 | NA | NA | NA | NA | 1 | 1 | NA | NA | 1 | 1 | NA | NA | NA | NA | NA | NA | 1 | 1 | 1 | 1 |
| 15_7782602 | NA | NA | NA | NA | 1 | 1 | NA | NA | 1 | 1 | NA | NA | NA | NA | 1 | 1 | 1 | 1 | 2 | 1 |
| 15_7782614 | NA | NA | NA | NA | 1 | 0 | 1 | 1 | 2 | 0.5 | NA | NA | NA | NA | 1 | 1 | 1 | 1 | 2 | 1 |
| 15_7782630 | NA | NA | NA | NA | 1 | 0 | 1 | 1 | 2 | 0.5 | NA | NA | NA | NA | 1 | 0 | 1 | 1 | 2 | 0.5 |
| 15_7782688 | NA | NA | NA | NA | 1 | 1 | NA | NA | 1 | 1 | NA | NA | NA | NA | 1 | 1 | 1 | 0 | 2 | 0.5 |
| 15_7782706 | NA | NA | NA | NA | 1 | 1 | NA | NA | 1 | 1 | NA | NA | NA | NA | 1 | 1 | 1 | 1 | 2 | 1 |
| 15_7782726 | NA | NA | NA | NA | 1 | 1 | NA | NA | 1 | 1 | NA | NA | NA | NA | 1 | 1 | 1 | 1 | 2 | 1 |
| 15_7782727 | NA | NA | NA | NA | 1 | 1 | NA | NA | 1 | 1 | NA | NA | NA | NA | 1 | 1 | 1 | 1 | 2 | 1 |
| 15_7782746 | NA | NA | NA | NA | 1 | 1 | NA | NA | 1 | 1 | NA | NA | NA | NA | 1 | 1 | 1 | 1 | 2 | 1 |
| 15_7784738 | 14 | 0 | 10 | 0.2 | 12 | 0.0833 | 9 | 0.1111 | 45 | 0.0889 | 13 | 0.0769 | 6 | 0 | 7 | 0 | 8 | 0.125 | 34 | 0.0588 |
| 15_7787617 | 6 | 0.1667 | 4 | 0.25 | 7 | 0.1429 | 4 | 0.75 | 21 | 0.2857 | 5 | 0.2 | 2 | 0 | 1 | 0 | 1 | 0 | 9 | 0.1111 |
| 15_7787626 | 6 | 0.5 | 3 | 0 | 9 | 0.4444 | 4 | 0.5 | 22 | 0.4091 | 5 | 0.2 | 1 | 0 | 1 | 0 | 1 | 0 | 8 | 0.125 |
| 15_7787631 | 7 | 0.8571 | 2 | 0.5 | 6 | 0.6667 | 2 | 1 | 17 | 0.7647 | 5 | 0.6 | 3 | 0.6667 | 1 | 1 | 1 | 0 | 10 | 0.6 |
| 15_7787639 | 6 | 0.1667 | 2 | 0 | 7 | 0 | 4 | 0 | 19 | 0.0526 | 5 | 0 | 3 | 0.6667 | 1 | 0 | 1 | 0 | 10 | 0.2 |
| 15_7787933 | 1 | 1 | 3 | 0 | 1 | 1 | NA | NA | 5 | 0.4 | 1 | 1 | 3 | 1 | 4 | 0.5 | 2 | 0.5 | 10 | 0.7 |
| 15_7787934 | 1 | 0 | 2 | 0.5 | 1 | 1 | 1 | 0 | 5 | 0.4 | 1 | 0 | 4 | 0.5 | 3 | 0.3333 | 2 | 0 | 10 | 0.3 |
| 15_7787939 | 2 | 1 | 1 | 1 | 1 | 1 | 1 | 1 | 5 | 1 | 1 | 1 | 3 | 1 | 4 | 1 | 2 | 1 | 10 | 1 |
| 15_7787940 | 2 | 0 | 2 | 0.5 | 1 | 0 | 1 | 1 | 6 | 0.3333 | 1 | 0 | 3 | 0.3333 | 4 | 0.25 | 1 | 1 | 9 | 0.3333 |
| 15_7787944 | 2 | 0.5 | 2 | 0 | 1 | 1 | 1 | 0 | 6 | 0.3333 | 1 | 0 | 3 | 0.6667 | 4 | 0.25 | 2 | 0 | 10 | 0.3 |
| 15_7787949 | 2 | 0.5 | 2 | 0 | 1 | 1 | NA | NA | 5 | 0.4 | 1 | 1 | 3 | 0.6667 | 3 | 0.3333 | 2 | 0.5 | 9 | 0.5556 |
| 15_7787980 | 2 | 0 | 2 | 1 | 1 | 0 | 1 | 0 | 6 | 0.3333 | 1 | 0 | 2 | 0 | 4 | 0.25 | 2 | 0 | 9 | 0.1111 |
| 15_7787981 | 2 | 0.5 | 2 | 0 | 1 | 0 | 1 | 0 | 6 | 0.1667 | 1 | 0 | 2 | 0.5 | 4 | 0 | 2 | 0 | 9 | 0.1111 |
| 15_7800330 | 2 | 0.5 | 1 | 1 | 1 | 0 | NA | NA | 4 | 0.5 | 1 | 0 | 2 | 0.5 | 3 | 0 | NA | NA | 6 | 0.1667 |
| 15_7803211 | NA | NA | 1 | 0 | NA | NA | 1 | 1 | 2 | 0.5 | NA | NA | 2 | 0.5 | NA | NA | NA | NA | 2 | 0.5 |
| 15_7804379 | 9 | 0.2222 | 11 | 0.1818 | 11 | 0.3636 | 8 | 0.125 | 39 | 0.2308 | 9 | 0.4444 | 5 | 0.2 | 2 | 0 | 5 | 0.2 | 21 | 0.2857 |
| 15_7804383 | 7 | 0.1429 | 12 | 0.1667 | 9 | 0 | 2 | 0.5 | 30 | 0.1333 | 11 | 0.3636 | 5 | 0 | 5 | 0.4 | 6 | 0 | 27 | 0.2222 |
| 15_7807539 | NA | NA | NA | NA | 1 | 0 | 1 | 1 | 2 | 0.5 | 2 | 0.5 | NA | NA | 1 | 0 | 2 | 0.5 | 5 | 0.4 |
| 15_7828396 | 5 | 0 | 10 | 0.1 | 4 | 0 | 7 | 0.1429 | 26 | 0.0769 | 7 | 0.2857 | 4 | 0 | 3 | 0 | 7 | 0 | 21 | 0.0952 |
| 15_7828963 | 3 | 0.3333 | 7 | 0.1429 | 2 | 0.5 | 5 | 0.4 | 17 | 0.2941 | 7 | 0.4286 | 2 | 0.5 | 2 | 0 | 5 | 0.8 | 16 | 0.5 |
| 15_7828978 | 4 | 0.5 | 7 | 0.5714 | 2 | 0 | 3 | 0.3333 | 16 | 0.4375 | 7 | 0.7143 | 3 | 0.6667 | 2 | 0 | 5 | 0.6 | 17 | 0.5882 |
| 15_7828983 | 4 | 0 | 7 | 0 | 3 | 0 | 6 | 0.3333 | 20 | 0.1 | 7 | 0 | 4 | 0 | 1 | 0 | 6 | 0.1667 | 18 | 0.0556 |
| 15_7829004 | 3 | 0 | 3 | 0 | 3 | 0.6667 | 6 | 0.1667 | 15 | 0.2 | 3 | 0.3333 | 2 | 0 | NA | NA | 5 | 0 | 10 | 0.1 |
| 15_7829802 | 1 | 1 | 2 | 0 | 2 | 0 | NA | NA | 5 | 0.2 | NA | NA | NA | NA | 3 | 0.3333 | 1 | 1 | 4 | 0.5 |
| 15_7832079 | 2 | 0 | 3 | 0 | 1 | 0 | 3 | 0.3333 | 9 | 0.1111 | 1 | 0 | 1 | 0 | 5 | 0 | 1 | 1 | 8 | 0.125 |
| 15_7832193 | 7 | 0 | 9 | 0 | 9 | 0.1111 | 9 | 0.3333 | 34 | 0.1176 | 5 | 0 | 6 | 0.1667 | 10 | 0.1 | 10 | 0.2 | 31 | 0.129 |
| 15_7832273 | 30 | 0.1 | 30 | 0.1667 | 35 | 0.2 | 22 | 0.2273 | 117 | 0.1709 | 22 | 0.1364 | 15 | 0.1333 | 30 | 0.1333 | 20 | 0.25 | 87 | 0.1609 |
| 15_7832285 | 28 | 0.3214 | 29 | 0.1379 | 29 | 0.2069 | 19 | 0.2632 | 105 | 0.2286 | 24 | 0.2917 | 11 | 0.1818 | 25 | 0.28 | 17 | 0.2353 | 77 | 0.2597 |
| 15_7832289 | 26 | 0.2308 | 26 | 0.1538 | 29 | 0.1724 | 19 | 0.0526 | 100 | 0.16 | 24 | 0.375 | 12 | 0.1667 | 22 | 0.1818 | 18 | 0.1667 | 76 | 0.2368 |
| 15_7832322 | 29 | 0.2069 | 31 | 0.3226 | 32 | 0.25 | 22 | 0.3636 | 114 | 0.2807 | 23 | 0.3478 | 17 | 0.1176 | 30 | 0.3667 | 21 | 0.4762 | 91 | 0.3407 |
| 15_7832326 | 32 | 0.2812 | 36 | 0.1944 | 37 | 0.2162 | 26 | 0.1538 | 131 | 0.2137 | 25 | 0.32 | 18 | 0.1667 | 36 | 0.3056 | 24 | 0.4167 | 103 | 0.3107 |
| 15_7832336 | 33 | 0.0909 | 32 | 0.1562 | 28 | 0.1786 | 24 | 0.1667 | 117 | 0.1453 | 22 | 0.1818 | 17 | 0 | 21 | 0.1429 | 19 | 0.3158 | 79 | 0.1646 |
| 15_7842214 | 1 | 1 | 4 | 0.25 | 1 | 0 | NA | NA | 6 | 0.3333 | 2 | 0.5 | 4 | 0 | 1 | 1 | NA | NA | 7 | 0.2857 |
| 15_7842219 | 1 | 0 | 3 | 0.3333 | 1 | 0 | NA | NA | 5 | 0.2 | 2 | 0 | 4 | 0.25 | 1 | 0 | NA | NA | 7 | 0.1429 |
| 15_7842245 | 1 | 0 | 3 | 0.3333 | 1 | 0 | NA | NA | 5 | 0.2 | 2 | 0.5 | 4 | 0 | 1 | 0 | NA | NA | 7 | 0.1429 |
| 15_7842278 | 1 | 0 | 3 | 1 | 1 | 1 | NA | NA | 5 | 0.8 | 1 | 0 | 2 | 1 | 1 | 1 | NA | NA | 4 | 0.75 |
| 15_7842301 | 1 | 1 | 3 | 1 | 1 | 1 | NA | NA | 5 | 1 | 1 | 1 | 3 | 1 | 1 | 1 | NA | NA | 5 | 1 |
| 15_7842318 | 1 | 0 | 3 | 0.3333 | NA | NA | NA | NA | 4 | 0.25 | 2 | 0.5 | 3 | 0.3333 | 1 | 0 | NA | NA | 6 | 0.3333 |
| 15_7842319 | 1 | 1 | 4 | 1 | 1 | 1 | NA | NA | 6 | 1 | 2 | 1 | 1 | 1 | 1 | 1 | NA | NA | 4 | 1 |

| | | | | | | | | | | | | | | | | | | | | |
|---|---|---|---|---|---|---|---|---|---|---|---|---|---|---|---|---|---|---|---|---|
| 15_784232 | 1 | 1 | 3 | 1 | 1 | 1 | NA | NA | 5 | 1 | 3 | 1 | 3 | 1 | 1 | 1 | NA | NA | 7 | 1 |
| 15_784232 | 1 | 1 | 2 | 1 | 1 | 1 | NA | NA | 4 | 1 | 2 | 1 | 2 | 1 | NA | NA | NA | NA | 4 | 1 |
| 15_784233 | 1 | 0 | 3 | 0.3333 | 1 | 0 | NA | NA | 5 | 0.2 | 3 | 0.3333 | 3 | 0.3333 | 1 | 0 | NA | NA | 7 | 0.2857 |
| 15_784233 | NA | NA | 3 | 0.6667 | NA | NA | NA | NA | 3 | 0.6667 | 3 | 0.3333 | 3 | 0.3333 | 1 | 1 | NA | NA | 7 | 0.4286 |
| 15_784233 | NA | NA | 3 | 0.3333 | 1 | 0 | NA | NA | 4 | 0.25 | 2 | 0.5 | 2 | 0 | 1 | 1 | NA | NA | 5 | 0.4 |
| 15_784234 | NA | NA | 3 | 0.6667 | 1 | 0 | NA | NA | 4 | 0.5 | 3 | 0 | 3 | 0.6667 | 1 | 0 | NA | NA | 7 | 0.2857 |
| 15_784234 | NA | NA | 2 | 1 | 1 | 1 | NA | NA | 3 | 1 | 2 | 1 | 2 | 1 | 1 | 1 | NA | NA | 5 | 1 |
| 15_784234 | NA | NA | 3 | 0.6667 | 1 | 0 | NA | NA | 4 | 0.5 | 3 | 0.3333 | 2 | 0.5 | 1 | 0 | NA | NA | 6 | 0.3333 |
| 15_784235 | NA | NA | 3 | 1 | 1 | 1 | NA | NA | 4 | 1 | 3 | 1 | 3 | 1 | 1 | 1 | NA | NA | 7 | 1 |
| 15_784235 | NA | NA | 2 | 0.5 | 1 | 1 | NA | NA | 3 | 0.6667 | 3 | 0.6667 | 3 | 0.6667 | 1 | 1 | NA | NA | 7 | 0.7143 |
| 15_784243 | NA | NA | 2 | 1 | NA | NA | NA | NA | 2 | 1 | 3 | 1 | 1 | 1 | 1 | 1 | NA | NA | 5 | 1 |
| 15_784365 | NA | NA | 1 | 1 | NA | NA | NA | NA | 1 | 1 | NA | NA | NA | NA | NA | NA | 1 | 1 | 1 | 1 |
| 15_784366 | NA | NA | 1 | 1 | NA | NA | NA | NA | 1 | 1 | NA | NA | NA | NA | NA | NA | 1 | 1 | 1 | 1 |
| 15_784366 | NA | NA | 1 | 1 | NA | NA | NA | NA | 1 | 1 | NA | NA | NA | NA | NA | NA | 1 | 1 | 1 | 1 |
| 15_784376 | NA | NA | NA | NA | 2 | 1 | 1 | 1 | 3 | 1 | 1 | 1 | NA | NA | NA | NA | NA | NA | 1 | 1 |
| 15_784376 | NA | NA | NA | NA | 2 | 1 | 1 | 1 | 3 | 1 | 1 | 1 | NA | NA | NA | NA | NA | NA | 1 | 1 |
| 15_784377 | NA | NA | NA | NA | 2 | 1 | NA | NA | 2 | 1 | 1 | 1 | NA | NA | NA | NA | NA | NA | 1 | 1 |
| 15_784378 | NA | NA | NA | NA | 1 | 1 | 1 | 1 | 2 | 1 | 1 | 1 | NA | NA | NA | NA | NA | NA | 1 | 1 |
| 15_784378 | NA | NA | NA | NA | NA | NA | 1 | 1 | 1 | 1 | 1 | 1 | NA | NA | NA | NA | NA | NA | 1 | 1 |
| 15_784391 | NA | NA | 1 | 1 | NA | NA | NA | NA | 1 | 1 | NA | NA | NA | NA | NA | NA | 1 | 1 | 1 | 1 |
| 15_784392 | 1 | 1 | 1 | 0 | NA | NA | NA | NA | 2 | 0.5 | NA | NA | NA | NA | NA | NA | 1 | 1 | 1 | 1 |
| 15_784392 | 1 | 1 | 1 | 1 | NA | NA | NA | NA | 2 | 1 | NA | NA | NA | NA | NA | NA | 1 | 1 | 1 | 1 |
| 15_784393 | 1 | 1 | 1 | 1 | NA | NA | NA | NA | 2 | 1 | NA | NA | NA | NA | NA | NA | 1 | 1 | 1 | 1 |
| 15_784395 | 1 | 1 | NA | NA | NA | NA | NA | NA | 1 | 1 | NA | NA | NA | NA | NA | NA | 1 | 1 | 1 | 1 |
| 15_784451 | 1 | 0 | 1 | 1 | 1 | 1 | NA | NA | 3 | 0.6667 | 1 | 1 | 2 | 0.5 | 1 | 1 | NA | NA | 4 | 0.75 |
| 15_785048 | NA | NA | 1 | 1 | NA | NA | 1 | 0 | 2 | 0.5 | 1 | 1 | NA | NA | NA | NA | NA | NA | 1 | 1 |
| 15_785050 | 1 | 1 | 1 | 1 | 1 | 1 | 1 | 1 | 4 | 1 | 2 | 1 | NA | NA | NA | NA | NA | NA | 2 | 1 |
| 15_785052 | NA | NA | 2 | 0.5 | 1 | 1 | 1 | 1 | 4 | 0.75 | 3 | 0.6667 | NA | NA | NA | NA | NA | NA | 3 | 0.6667 |
| 15_785054 | NA | NA | 2 | 0 | 1 | 1 | 1 | 0 | 4 | 0.25 | 2 | 0.5 | NA | NA | NA | NA | NA | NA | 2 | 0.5 |
| 15_785055 | 1 | 1 | 2 | 0 | NA | NA | 1 | 1 | 4 | 0.5 | 3 | 0.6667 | NA | NA | NA | NA | NA | NA | 3 | 0.6667 |
| 15_785059 | NA | NA | 1 | 1 | 1 | 1 | NA | NA | 2 | 1 | 3 | 1 | NA | NA | NA | NA | NA | NA | 3 | 1 |
| 15_785060 | 1 | 0 | 2 | 1 | 1 | 1 | 2 | 0.5 | 6 | 0.6667 | 4 | 0.25 | NA | NA | NA | NA | NA | NA | 4 | 0.25 |
| 15_785060 | 1 | 1 | 2 | 1 | 1 | 0 | 2 | 1 | 6 | 0.8333 | 4 | 0.75 | NA | NA | NA | NA | NA | NA | 4 | 0.75 |
| 15_785060 | 1 | 0 | 1 | 0 | 1 | 0 | 2 | 0.5 | 5 | 0.2 | 4 | 0.5 | NA | NA | NA | NA | NA | NA | 4 | 0.5 |
| 15_785061 | 1 | 1 | 2 | 0 | 1 | 0 | 2 | 0 | 6 | 0.1667 | 3 | 0.3333 | NA | NA | NA | NA | NA | NA | 3 | 0.3333 |
| 15_785517 | 8 | 0.375 | 14 | 0.5714 | 8 | 0.75 | 5 | 0.2 | 35 | 0.5143 | 10 | 0.9 | 12 | 0.4167 | 7 | 0.7143 | 12 | 0.5 | 41 | 0.6098 |
| 15_785527 | 2 | 0 | 8 | 0.125 | 1 | 0 | 3 | 0.3333 | 14 | 0.1429 | 5 | 0 | 6 | 0 | 2 | 0 | 3 | 0.3333 | 16 | 0.0625 |
| 15_785535 | 4 | 0.25 | 8 | 0.125 | 5 | 0.2 | 2 | 0 | 19 | 0.1579 | 7 | 0.1429 | 8 | 0.25 | 2 | 0 | 6 | 0.1667 | 23 | 0.1739 |
| 15_785541 | 1 | 0 | 6 | 0.6667 | 3 | 0.6667 | 1 | 1 | 11 | 0.6364 | 4 | 0.75 | 3 | 0.6667 | NA | NA | 3 | 0.3333 | 10 | 0.6 |
| 15_785556 | 3 | 1 | 4 | 1 | 2 | 1 | 1 | 0 | 10 | 0.9 | 3 | 1 | 5 | 1 | 1 | 0 | 2 | 0.5 | 11 | 0.8182 |
| 15_785557 | 3 | 0.3333 | 5 | 0.8 | 2 | 0 | 1 | 1 | 11 | 0.5455 | 3 | 0 | 5 | 0.2 | 1 | 0 | 2 | 0 | 11 | 0.0909 |
| 15_785557 | 3 | 1 | 3 | 1 | 1 | 1 | 1 | 0 | 8 | 0.875 | 3 | 0.6667 | 5 | 0.8 | 1 | 1 | 3 | 1 | 12 | 0.8333 |
| 15_785558 | 3 | 0 | 4 | 0 | 1 | 1 | 1 | 0 | 9 | 0.1111 | 4 | 0.25 | 5 | 0 | 1 | 0 | 2 | 0 | 12 | 0.0833 |
| 15_785558 | 2 | 0.5 | 4 | 0.25 | 2 | 0 | 1 | 0 | 9 | 0.2222 | 5 | 0.2 | 5 | 0.4 | 1 | 0 | 3 | 0 | 14 | 0.2143 |
| 15_785558 | 2 | 0.5 | 4 | 0.25 | 2 | 1 | NA | NA | 8 | 0.5 | 5 | 0.4 | 6 | 0.8333 | 1 | 0 | 3 | 0 | 15 | 0.4667 |
| 15_785560 | 1 | 1 | 3 | 1 | NA | NA | 1 | 1 | 5 | 1 | 1 | 1 | 3 | 1 | NA | NA | NA | NA | 4 | 1 |
| 15_785561 | 1 | 1 | 4 | 0.25 | 2 | 0 | 1 | 0 | 8 | 0.25 | 5 | 0.6 | 5 | 0.6 | 1 | 0 | 3 | 0.3333 | 14 | 0.5 |
| 15_785562 | NA | NA | 3 | 0.6667 | 1 | 1 | 1 | 1 | 5 | 0.8 | 4 | 1 | 4 | 1 | NA | NA | 1 | 0 | 9 | 0.8889 |
| 15_785563 | NA | NA | 2 | 0.5 | 2 | 0.5 | 1 | 1 | 5 | 0.6 | 4 | 0.5 | 4 | 0.5 | NA | NA | 3 | 0.6667 | 11 | 0.5455 |
| 15_785563 | 1 | 0 | 3 | 0.3333 | 2 | 0.5 | 1 | 0 | 7 | 0.2857 | 3 | 0 | 4 | 0 | NA | NA | 2 | 0.5 | 9 | 0.1111 |
| 15_785565 | 1 | 1 | 3 | 1 | 2 | 1 | 1 | 1 | 7 | 1 | 5 | 1 | 3 | 1 | NA | NA | 2 | 1 | 10 | 1 |
| 15_785565 | 1 | 1 | 3 | 1 | 2 | 1 | 1 | 1 | 7 | 1 | 4 | 1 | 4 | 1 | NA | NA | 2 | 1 | 10 | 1 |
| 15_785568 | 1 | 1 | 3 | 1 | 2 | 0.5 | 1 | 1 | 7 | 0.8571 | 4 | 0.75 | 4 | 0.5 | NA | NA | 2 | 0 | 10 | 0.5 |
| 15_785569 | 1 | 1 | 3 | 0 | 2 | 0.5 | 1 | 0 | 7 | 0.2857 | 4 | 0.25 | 3 | 0.6667 | NA | NA | 2 | 0 | 9 | 0.3333 |
| 15_785571 | NA | NA | 2 | 0 | 1 | 0 | 1 | 1 | 4 | 0.25 | 2 | 0.5 | 3 | 0.3333 | NA | NA | 2 | 0 | 7 | 0.2857 |
| 15_785575 | 1 | 0 | 1 | 1 | 2 | 0 | 1 | 0 | 5 | 0.2 | 4 | 0 | 1 | 0 | NA | NA | 2 | 0.5 | 7 | 0.1429 |
| 15_785579 | NA | NA | 2 | 0.5 | NA | NA | NA | NA | 2 | 0.5 | 3 | 0.3333 | 1 | 0 | NA | NA | 1 | 0 | 5 | 0.2 |
| 15_785584 | 1 | 0 | 3 | 0.3333 | NA | NA | NA | NA | 4 | 0.25 | 4 | 0.25 | 1 | 0 | NA | NA | 1 | 0 | 6 | 0.1667 |
| 15_785586 | 1 | 1 | 2 | 1 | NA | NA | NA | NA | 3 | 1 | 4 | 1 | 1 | 1 | NA | NA | 2 | 1 | 7 | 1 |
| 15_786594 | 1 | 1 | NA | NA | NA | NA | 1 | 0 | 2 | 0.5 | NA | NA | 1 | 0 | 1 | 1 | NA | NA | 2 | 0.5 |
| 15_786596 | 1 | 0 | NA | NA | NA | NA | 2 | 0.5 | 3 | 0.3333 | NA | NA | 1 | 1 | 1 | 0 | NA | NA | 2 | 0.5 |
| 15_786599 | NA | NA | NA | NA | NA | NA | 1 | 1 | 1 | 1 | NA | NA | NA | NA | 1 | 1 | NA | NA | 1 | 1 |
| 15_786599 | 1 | 1 | NA | NA | NA | NA | 2 | 1 | 3 | 1 | NA | NA | NA | NA | 1 | 1 | NA | NA | 1 | 1 |
| 15_786600 | 1 | 1 | NA | NA | NA | NA | 2 | 1 | 3 | 1 | NA | NA | NA | NA | 1 | 1 | NA | NA | 1 | 1 |
| 15_787315 | 1 | 1 | 2 | 1 | 2 | 1 | 1 | 1 | 6 | 1 | NA | NA | 2 | 1 | 2 | 1 | 2 | 1 | 6 | 1 |

| | | | | | | | | | | | | | | | | | | | | |
|---|---|---|---|---|---|---|---|---|---|---|---|---|---|---|---|---|---|---|---|---|
| 15_787316( | 1 | 0 | 2 | 0 | 2 | 0.5 | NA | NA | 5 | 0.2 | NA | NA | 2 | 0.5 | 2 | 0.5 | 2 | 0 | 6 | 0.3333 |
| 15_787316! | NA | NA | 1 | 1 | NA | NA | 1 | 0 | 2 | 0.5 | NA | NA | 2 | 0 | 1 | 1 | 2 | 1 | 5 | 0.6 |
| 15_787335' | 1 | 1 | 1 | 0 | 2 | 0.5 | 1 | 1 | 5 | 0.6 | NA | NA | 2 | 0.5 | 1 | 1 | 2 | 0.5 | 5 | 0.6 |
| 15_787335· | 1 | 1 | 1 | 0 | 1 | 0 | 1 | 1 | 4 | 0.5 | NA | NA | 2 | 0.5 | 2 | 0.5 | 2 | 0 | 6 | 0.3333 |
| 15_787337! | 1 | 1 | 1 | 1 | 1 | 1 | 2 | 1 | 5 | 1 | NA | NA | 1 | 1 | 2 | 1 | NA | NA | 3 | 1 |
| 15_787341: | 1 | 1 | 1 | 1 | 2 | 0.5 | 2 | 1 | 6 | 0.8333 | NA | NA | 1 | 1 | 2 | 1 | 3 | 1 | 6 | 1 |
| 15_787344' | 1 | 1 | 2 | 0 | 2 | 0 | 2 | 0 | 7 | 0.1429 | NA | NA | 2 | 0.5 | 2 | 0 | 2 | 0 | 6 | 0.1667 |
| 15_787780· | 4 | 0.25 | 8 | 0.125 | 5 | 0 | NA | NA | 17 | 0.1176 | 3 | 0 | 4 | 0.25 | 8 | 0 | 5 | 0.2 | 20 | 0.1 |
| 15_787781: | 8 | 0.625 | 9 | 0.7778 | 9 | 0.5556 | 4 | 0.25 | 30 | 0.6 | 6 | 0.6667 | 7 | 0.2857 | 7 | 0.4286 | 6 | 0.6667 | 26 | 0.5 |
| 15_787782( | 8 | 0.125 | 10 | 0.1 | 10 | 0.1 | 3 | 0 | 31 | 0.0968 | 5 | 0 | 8 | 0 | 9 | 0.1111 | 6 | 0.1667 | 28 | 0.0714 |
| 15_787782! | 8 | 1 | 8 | 0.875 | 7 | 0.8571 | 5 | 1 | 28 | 0.9286 | 4 | 0.75 | 7 | 0.8571 | 8 | 0.875 | 5 | 0.8 | 24 | 0.8333 |
| 15_787784' | 8 | 0.125 | 10 | 0 | 10 | 0.1 | 5 | 0.2 | 33 | 0.0909 | 5 | 0 | 7 | 0 | 8 | 0.25 | 5 | 0 | 25 | 0.08 |
| 15_787784: | 6 | 0 | 6 | 0 | 10 | 0.1 | 4 | 0.25 | 26 | 0.0769 | 5 | 0.2 | 8 | 0.125 | 5 | 0 | 6 | 0 | 24 | 0.0833 |
| 15_787961! | 1 | 0 | 2 | 0 | 1 | 1 | 1 | 0 | 5 | 0.2 | 3 | 0 | 2 | 0 | 3 | 0 | 2 | 0.5 | 10 | 0.1 |
| 15_788026: | 7 | 0.8571 | 32 | 1 | 27 | 1 | 8 | 1 | 74 | 0.9865 | 25 | 0.92 | 28 | 1 | 21 | 1 | 15 | 1 | 89 | 0.9775 |
| 15_788026· | 6 | 1 | 34 | 0.8529 | 26 | 0.8846 | 8 | 0.875 | 74 | 0.8784 | 23 | 0.8696 | 27 | 0.8148 | 23 | 0.913 | 17 | 0.9412 | 90 | 0.8778 |
| 15_788027' | 6 | 0.3333 | 32 | 0.2812 | 23 | 0.3913 | 10 | 0.3 | 71 | 0.3239 | 21 | 0.2381 | 28 | 0.3929 | 23 | 0.3043 | 15 | 0.2667 | 87 | 0.3103 |
| 15_788028' | 8 | 0.125 | 32 | 0.3438 | 28 | 0.2857 | 8 | 0.125 | 76 | 0.2763 | 22 | 0.2727 | 25 | 0.2 | 21 | 0.4762 | 15 | 0.2667 | 83 | 0.3012 |
| 15_788032! | 10 | 0 | 42 | 0.0952 | 33 | 0.2121 | 11 | 0.0909 | 96 | 0.125 | 28 | 0.1071 | 29 | 0.1724 | 27 | 0.1481 | 18 | 0.2778 | 102 | 0.1667 |
| 15_788142· | 13 | 0 | 9 | 0.1111 | 6 | 0 | 4 | 0.25 | 32 | 0.0625 | 7 | 0.1429 | 1 | 0 | 3 | 0.3333 | 4 | 0 | 15 | 0.1333 |
| 15_788247: | 11 | 0.0909 | 6 | 0 | 7 | 0 | 3 | 0.3333 | 27 | 0.0741 | 3 | 0.6667 | 5 | 0.6 | 3 | 0 | 11 | 0.0909 | 22 | 0.2727 |
| 15_788249: | 7 | 1 | 6 | 1 | 5 | 1 | 2 | 1 | 20 | 1 | 4 | 1 | 5 | 0.8 | 2 | 1 | 11 | 0.7273 | 22 | 0.8182 |
| 15_788249( | 9 | 1 | 5 | 1 | 6 | 0.8333 | 3 | 1 | 23 | 0.9565 | 3 | 1 | 4 | 1 | 3 | 1 | 9 | 0.8889 | 19 | 0.9474 |
| 15_788250( | 9 | 0.1111 | 5 | 0.2 | 7 | 0.1429 | 3 | 0.3333 | 24 | 0.1667 | 4 | 0 | 5 | 0.2 | 2 | 0 | 9 | 0 | 20 | 0.05 |
| 15_788250· | 4 | 0.5 | 5 | 1 | 6 | 1 | 3 | 1 | 18 | 0.8889 | 3 | 1 | 5 | 1 | 3 | 1 | 8 | 1 | 19 | 1 |
| 15_788277! | NA | NA | NA | NA | NA | NA | 1 | 1 | 1 | 1 | NA | NA | NA | NA | NA | NA | 1 | 1 | 1 | 1 |
| 15_788772· | 1 | 0 | 3 | 1 | NA | NA | NA | NA | 4 | 0.75 | NA | NA | NA | NA | NA | NA | 1 | 1 | 1 | 1 |
| 15_789128! | 6 | 0.1667 | 1 | 0 | 2 | 0 | 5 | 0 | 14 | 0.0714 | 4 | 0 | 5 | 0.2 | NA | NA | 6 | 0 | 15 | 0.0667 |
| 15_789129: | 5 | 0 | 1 | 1 | 2 | 0 | 2 | 0.5 | 10 | 0.2 | 6 | 0.1667 | 3 | 0 | NA | NA | 4 | 0.25 | 13 | 0.1538 |
| 15_789135( | 9 | 0.1111 | 3 | 0 | 8 | 0.25 | 5 | 0.4 | 25 | 0.2 | 8 | 0.5 | 9 | 0.5556 | 2 | 0.5 | 7 | 0.2857 | 26 | 0.4615 |
| 15_789135: | 8 | 1 | 3 | 1 | 9 | 1 | 7 | 0.8571 | 27 | 0.963 | 9 | 1 | 9 | 0.8889 | 2 | 1 | 8 | 1 | 28 | 0.9643 |
| 15_789136( | 6 | 0.8333 | 4 | 1 | 8 | 1 | 7 | 0.8571 | 25 | 0.92 | 9 | 0.7778 | 8 | 1 | 2 | 1 | 7 | 1 | 26 | 0.9231 |
| 15_789136' | 6 | 1 | 4 | 0.75 | 8 | 1 | 3 | 0.6667 | 21 | 0.9048 | 8 | 1 | 7 | 1 | 1 | 0 | 7 | 1 | 23 | 0.9565 |
| 15_789136· | 8 | 0.75 | 4 | 0.75 | 8 | 1 | 6 | 1 | 26 | 0.8846 | 9 | 0.6667 | 8 | 0.875 | 2 | 1 | 9 | 1 | 28 | 0.8571 |
| 15_789136! | 8 | 0.75 | 4 | 0.25 | 9 | 0.7778 | 5 | 0.2 | 26 | 0.5769 | 8 | 0.625 | 9 | 0.5556 | 2 | 0.5 | 10 | 0.4 | 29 | 0.5172 |
| 15_789136{ | 7 | 1 | 4 | 1 | 10 | 1 | 5 | 1 | 26 | 1 | 9 | 0.8889 | 7 | 1 | 2 | 1 | 9 | 1 | 27 | 0.963 |
| 15_789136! | 8 | 0.125 | 4 | 0 | 10 | 0.2 | 7 | 0.4286 | 29 | 0.2069 | 9 | 0.1111 | 8 | 0.25 | 2 | 0 | 8 | 0.125 | 27 | 0.1481 |
| 15_789138' | 10 | 0.6 | 3 | 0 | 8 | 0.25 | 6 | 0.3333 | 27 | 0.3704 | 8 | 0 | 9 | 0.3333 | 2 | 0 | 7 | 0.1429 | 26 | 0.1538 |
| 15_789138: | 9 | 0.1111 | 3 | 0 | 7 | 0.2857 | 7 | 0.2857 | 26 | 0.1923 | 9 | 0.1111 | 8 | 0 | 1 | 0 | 7 | 0.2857 | 25 | 0.12 |
| 15_789138{ | 10 | 0.5 | 4 | 0 | 8 | 0.125 | 7 | 0 | 29 | 0.2069 | 10 | 0 | 8 | 0.25 | 2 | 0.5 | 10 | 0.1 | 30 | 0.1333 |
| 15_789141: | 9 | 0 | 5 | 0.2 | 7 | 0.1429 | 5 | 0.2 | 26 | 0.1154 | 7 | 0.1429 | 7 | 0 | 1 | 0 | 12 | 0.3333 | 27 | 0.1852 |
| 15_789142( | 12 | 0.0833 | 7 | 0 | 10 | 0.1 | 10 | 0.2 | 39 | 0.1026 | 10 | 0.1 | 11 | 0 | 2 | 0.5 | 14 | 0.1429 | 37 | 0.1081 |
| 15_789144: | 12 | 0 | 6 | 0 | 11 | 0.2727 | 10 | 0 | 39 | 0.0769 | 10 | 0 | 10 | 0.1 | 2 | 0 | 16 | 0.125 | 38 | 0.0789 |
| 15_789215! | 1 | 0 | 1 | 1 | 4 | 0.25 | 1 | 0 | 7 | 0.2857 | 1 | 1 | 2 | 0 | NA | NA | 1 | 0 | 4 | 0.25 |
| 15_789377( | 4 | 0 | 2 | 0.5 | 3 | 0 | 2 | 0 | 11 | 0.0909 | 4 | 0 | 1 | 0 | 2 | 0 | 4 | 0.25 | 11 | 0.0909 |
| 15_789380{ | 4 | 0 | 2 | 0 | 4 | 0 | 3 | 0.3333 | 13 | 0.0769 | 4 | 0 | NA | NA | 2 | 0 | 3 | 0.3333 | 9 | 0.1111 |
| 15_789380! | 4 | 0 | 1 | 0 | 4 | 0 | 3 | 0.3333 | 12 | 0.0833 | 4 | 0.25 | 1 | 0 | 3 | 0 | 4 | 0.25 | 12 | 0.1667 |
| 15_789381: | 2 | 0 | 2 | 0 | 4 | 0.25 | 3 | 0 | 11 | 0.0909 | 3 | 0 | 1 | 0 | 3 | 0.3333 | 4 | 0 | 11 | 0.0909 |
| 15_789738' | 13 | 0.6154 | 13 | 0.7692 | 6 | 0.5 | 8 | 0.75 | 40 | 0.675 | 8 | 0.5 | 11 | 0.5455 | 11 | 0.7273 | 9 | 0.8889 | 39 | 0.6667 |
| 15_789740: | 12 | 0 | 13 | 0.2308 | 5 | 0 | 9 | 0.3333 | 39 | 0.1538 | 7 | 0.1429 | 11 | 0.2727 | 10 | 0.1 | 7 | 0 | 35 | 0.1429 |
| 15_789740! | 16 | 0.125 | 14 | 0.2143 | 7 | 0 | 9 | 0 | 46 | 0.1087 | 8 | 0 | 11 | 0.0909 | 10 | 0 | 9 | 0.1111 | 38 | 0.0526 |
| 15_789785! | 15 | 0.8 | 13 | 0.5385 | 6 | 0.8333 | 10 | 0.7 | 44 | 0.7045 | 5 | 0.8 | 11 | 0.3636 | 9 | 0.4444 | 6 | 1 | 31 | 0.5806 |
| 15_790191{ | 2 | 0.5 | NA | NA | NA | NA | 4 | 0 | 6 | 0.1667 | NA | NA | NA | NA | 1 | 0 | 2 | 0.5 | 3 | 0.3333 |
| 15_790202{ | 3 | 0 | 1 | 1 | 1 | 0 | 2 | 0 | 7 | 0.1429 | NA | NA | 1 | 1 | 1 | 0 | 3 | 0 | 5 | 0.2 |
| 15_790358· | 9 | 0 | 16 | 0.0625 | 10 | 0.3 | 14 | 0.2857 | 49 | 0.1633 | 8 | 0.125 | 12 | 0 | 14 | 0.1429 | 8 | 0.125 | 42 | 0.0952 |
| 15_790396· | 4 | 0 | 2 | 0 | 4 | 0.25 | 3 | 0 | 13 | 0.0769 | 4 | 0.25 | 7 | 0.1429 | 6 | 0 | 1 | 0 | 18 | 0.1111 |
| 15_790396: | 3 | 0 | 3 | 0 | 3 | 0 | 3 | 0.3333 | 12 | 0.0833 | 4 | 0.25 | 8 | 0.125 | 7 | 0 | 2 | 0 | 21 | 0.0952 |
| 15_790403' | 4 | 0 | 1 | 1 | 3 | 0 | 3 | 0 | 11 | 0.0909 | 2 | 0 | 4 | 0 | 3 | 0 | 2 | 0.5 | 11 | 0.0909 |
| 15_790403: | 3 | 0 | 1 | 0 | 3 | 0.3333 | 3 | 0 | 10 | 0.1 | 2 | 0 | 6 | 0.1667 | 4 | 0.25 | 2 | 0 | 14 | 0.1429 |
| 15_790405! | 4 | 0 | 1 | 1 | 2 | 0.5 | 3 | 0.6667 | 10 | 0.4 | 2 | 0.5 | 5 | 0.6 | 2 | 0.5 | 2 | 1 | 11 | 0.6364 |
| 15_790406· | 3 | 0.6667 | NA | NA | 3 | 0.6667 | 3 | 0.3333 | 9 | 0.5556 | 3 | 0.6667 | 5 | 0.6 | 3 | 0.6667 | 1 | 0 | 12 | 0.5833 |
| 15_790406! | 4 | 0.75 | 1 | 1 | 2 | 0 | 3 | 0.3333 | 10 | 0.5 | 3 | 1 | 4 | 0.5 | 2 | 0.5 | 1 | 0 | 10 | 0.6 |
| 15_791329{ | 6 | 0.6667 | 6 | 0 | 3 | 0.3333 | 4 | 0 | 19 | 0.2632 | 4 | 0 | 5 | 0.2 | 5 | 0.4 | 6 | 0.3333 | 20 | 0.25 |
| 15_791330: | 6 | 0.1667 | 5 | 0 | 4 | 0 | 2 | 0 | 17 | 0.0588 | 3 | 0 | 5 | 0.2 | 6 | 0 | 6 | 0.1667 | 20 | 0.1 |
| 15_791331( | 6 | 0 | 4 | 0.25 | 4 | 0 | 4 | 0.25 | 18 | 0.1111 | 3 | 0.3333 | 6 | 0.3333 | 8 | 0.125 | 6 | 0.5 | 23 | 0.3043 |

| | | | | | | | | | | | | | | | | | | | | |
|---|---|---|---|---|---|---|---|---|---|---|---|---|---|---|---|---|---|---|---|---|
| 15_791332( | 6 | 0.6667 | 4 | 0.5 | 5 | 0.8 | 3 | 1 | 18 | 0.7222 | 5 | 0.8 | 5 | 0.2 | 6 | 0.8333 | 4 | 0.5 | 20 | 0.6 |
| 15_791333 | 6 | 0.6667 | 7 | 0.7143 | 7 | 1 | 6 | 0.8333 | 26 | 0.8077 | 4 | 1 | 9 | 0.6667 | 7 | 0.7143 | 4 | 1 | 24 | 0.7917 |
| 15_791334 | 6 | 0.5 | 5 | 0.4 | 6 | 0.5 | 5 | 0.4 | 22 | 0.4545 | 5 | 0.8 | 7 | 0.7143 | 5 | 0.8 | 4 | 0.5 | 21 | 0.7143 |
| 15_791401! | NA | NA | NA | NA | NA | NA | 1 | 1 | 1 | 1 | 1 | 1 | NA | NA | NA | NA | NA | NA | 1 | 1 |
| 15_791403( | NA | NA | NA | NA | NA | NA | 1 | 1 | 1 | 1 | 1 | 1 | NA | NA | NA | NA | NA | NA | 1 | 1 |
| 15_791426 | 2 | 0.5 | 1 | 1 | NA | NA | 1 | 0 | 4 | 0.5 | 1 | 1 | NA | NA | NA | NA | NA | NA | 1 | 1 |
| 15_791706! | NA | NA | 3 | 0.6667 | 3 | 0 | 1 | 1 | 7 | 0.4286 | 4 | 1 | 4 | 0.25 | NA | NA | 1 | 1 | 9 | 0.6667 |
| 15_791707( | NA | NA | 5 | 0 | 4 | 0.25 | 1 | 0 | 10 | 0.1 | 4 | 0 | 4 | 0.25 | 1 | 0 | 1 | 0 | 10 | 0.1 |
| 15_791707( | NA | NA | 4 | 0.25 | 4 | 0.25 | 1 | 1 | 9 | 0.3333 | 4 | 0.25 | 3 | 0.3333 | 1 | 1 | 1 | 1 | 9 | 0.4444 |
| 15_791708 | NA | NA | 4 | 0.75 | 4 | 0.5 | NA | NA | 8 | 0.625 | 4 | 0.25 | 4 | 0.75 | 1 | 1 | 1 | 1 | 10 | 0.6 |
| 15_791718 | NA | NA | 4 | 0 | 2 | 0 | 1 | 1 | 7 | 0.1429 | 4 | 0.25 | 2 | 0 | NA | NA | 1 | 0 | 7 | 0.1429 |
| 15_792423 | NA | NA | NA | NA | NA | NA | 1 | 1 | 1 | 1 | 3 | 0 | NA | NA | 1 | 0 | 1 | 1 | 5 | 0.2 |
| 15_792426 | NA | NA | NA | NA | NA | NA | 1 | 1 | 1 | 1 | 3 | 0.3333 | NA | NA | 1 | 0 | 1 | 1 | 5 | 0.4 |
| 15_792439( | NA | NA | 1 | 0 | NA | NA | 1 | 1 | 2 | 0.5 | NA | NA | 1 | 1 | 1 | 0 | NA | NA | 2 | 0.5 |
| 15_792453 | 12 | 0.0833 | 16 | 0.125 | 9 | 0.1111 | 12 | 0 | 49 | 0.0816 | 5 | 0.4 | 11 | 0.0909 | 16 | 0 | 11 | 0 | 43 | 0.0698 |
| 15_792464( | 9 | 0.1111 | 14 | 0.1429 | 11 | 0 | 11 | 0.0909 | 45 | 0.0889 | 6 | 0.1667 | 9 | 0.3333 | 15 | 0.2 | 10 | 0 | 40 | 0.175 |
| 15_792467 | 5 | 0.4 | 10 | 0.3 | 5 | 0.4 | 8 | 0.5 | 28 | 0.3929 | 5 | 0.6 | 9 | 0.2222 | 8 | 0.75 | 6 | 0.5 | 28 | 0.5 |
| 15_792467 | 8 | 0.75 | 11 | 0.9091 | 6 | 0.8333 | 7 | 1 | 32 | 0.875 | 5 | 0.8 | 9 | 0.6667 | 10 | 0.6 | 9 | 1 | 33 | 0.7576 |
| 15_792468 | 7 | 0.1429 | 12 | 0.4167 | 3 | 0.3333 | 6 | 0.3333 | 28 | 0.3214 | 5 | 0.6 | 10 | 0.2 | 10 | 0.1 | 9 | 0.3333 | 34 | 0.2647 |
| 15_793021 | 4 | 0.25 | 13 | 0.0769 | 10 | 0 | 10 | 0.2 | 37 | 0.1081 | 7 | 0.1429 | 6 | 0.1667 | 7 | 0 | 9 | 0 | 29 | 0.069 |
| 15_793021 | 4 | 0.5 | 14 | 0.7143 | 14 | 0.5 | 11 | 0.7273 | 43 | 0.6279 | 9 | 0.6667 | 7 | 0.4286 | 13 | 0.5385 | 11 | 0.5455 | 40 | 0.55 |
| 15_793023 | 3 | 0.3333 | 13 | 0.0769 | 11 | 0.0909 | 11 | 0.0909 | 38 | 0.1053 | 7 | 0.1429 | 11 | 0.0909 | 11 | 0.0909 | 9 | 0 | 38 | 0.0789 |
| 15_793026( | 3 | 0 | 13 | 0 | 11 | 0 | 11 | 0.1818 | 38 | 0.0526 | 9 | 0.2222 | 10 | 0.1 | 10 | 0 | 10 | 0 | 39 | 0.0769 |
| 15_793030 | 3 | 0 | 13 | 0.3077 | 9 | 0.2222 | 13 | 0.1538 | 38 | 0.2105 | 8 | 0.375 | 8 | 0 | 8 | 0.125 | 11 | 0.2727 | 35 | 0.2 |
| 15_793031 | 3 | 0.6667 | 11 | 0.8182 | 8 | 0.875 | 8 | 0.875 | 30 | 0.8333 | 8 | 0.875 | 9 | 0.8889 | 9 | 0.8889 | 8 | 0.625 | 34 | 0.8235 |
| 15_793032( | 3 | 0 | 12 | 0.25 | 7 | 0 | 8 | 0 | 30 | 0.1 | 9 | 0.1111 | 8 | 0.25 | 7 | 0.1429 | 8 | 0 | 32 | 0.125 |
| 15_793033( | 3 | 0 | 7 | 0.1429 | 6 | 0.1667 | 6 | 0 | 22 | 0.0909 | 6 | 0.5 | 6 | 0 | 6 | 0 | 7 | 0.1429 | 25 | 0.16 |
| 15_793033 | 1 | 0 | 8 | 0.5 | 4 | 0.25 | 7 | 0.4286 | 20 | 0.4 | 6 | 0.1667 | 4 | 0.5 | 5 | 0.8 | 6 | 0.5 | 21 | 0.4762 |
| 15_793033 | 3 | 0.6667 | 12 | 0.6667 | 4 | 0.75 | 8 | 0.5 | 27 | 0.6296 | 6 | 0.5 | 8 | 0.625 | 8 | 0.625 | 8 | 0.375 | 30 | 0.5333 |
| 15_793068 | NA | NA | 1 | 1 | 1 | 1 | 3 | 0.3333 | 5 | 0.6 | NA | NA | NA | NA | 2 | 0.5 | NA | NA | 2 | 0.5 |
| 15_793069 | NA | NA | 2 | 0 | 1 | 1 | 3 | 0.6667 | 6 | 0.5 | NA | NA | NA | NA | 2 | 0.5 | NA | NA | 2 | 0.5 |
| 15_793070 | NA | NA | 2 | 0.5 | NA | NA | 3 | 0.3333 | 5 | 0.4 | NA | NA | NA | NA | 2 | 1 | NA | NA | 2 | 1 |
| 15_793072 | NA | NA | 2 | 0.5 | NA | NA | 1 | 0 | 3 | 0.3333 | NA | NA | NA | NA | 2 | 1 | NA | NA | 2 | 1 |
| 15_793073 | NA | NA | 2 | 0.5 | NA | NA | 2 | 0.5 | 4 | 0.5 | NA | NA | NA | NA | 1 | 1 | NA | NA | 1 | 1 |
| 15_793081! | NA | NA | 2 | 0.5 | NA | NA | 3 | 0 | 5 | 0.2 | NA | NA | NA | NA | 2 | 1 | NA | NA | 2 | 1 |
| 15_793647( | 5 | 0 | 4 | 0 | 5 | 0.4 | 5 | 0 | 19 | 0.1053 | 2 | 0 | 6 | 0 | 1 | 0 | 6 | 0.1667 | 15 | 0.0667 |
| 15_793648 | 4 | 0.25 | 4 | 0 | 7 | 0.1429 | 5 | 0.2 | 20 | 0.15 | 2 | 0.5 | 4 | 0 | 1 | 0 | 5 | 0.2 | 12 | 0.1667 |
| 15_793650 | 4 | 0 | 4 | 0 | 5 | 0.2 | 4 | 0 | 17 | 0.0588 | 2 | 0 | 3 | 0 | 1 | 0 | 4 | 0.25 | 10 | 0.1 |
| 15_793653 | 2 | 0 | 4 | 0.25 | 6 | 0 | 5 | 0.2 | 17 | 0.1176 | 2 | 0 | 4 | 0 | 1 | 0 | 4 | 0.25 | 11 | 0.0909 |
| 15_793654! | 2 | 0.5 | 3 | 0.3333 | 6 | 0.5 | 4 | 0.25 | 15 | 0.4 | 2 | 0.5 | 4 | 0.25 | 1 | 1 | 4 | 0.5 | 11 | 0.4545 |
| 15_793656( | 2 | 0 | 2 | 0 | 6 | 0 | 4 | 0.25 | 14 | 0.0714 | 2 | 0.5 | 3 | 0 | 1 | 1 | 4 | 0 | 10 | 0.2 |
| 15_793659 | NA | NA | 1 | 0 | 4 | 0 | 2 | 0.5 | 7 | 0.1429 | 1 | 0 | 3 | 0 | 1 | 1 | 2 | 0 | 7 | 0.1429 |
| 15_793664( | NA | NA | 1 | 1 | 4 | 0.25 | 4 | 0 | 9 | 0.2222 | NA | NA | 2 | 0 | 1 | 0 | 1 | 1 | 4 | 0.25 |
| 15_793664 | NA | NA | 1 | 0 | 4 | 0.25 | 5 | 0 | 10 | 0.1 | 1 | 0 | 3 | 0 | 1 | 0 | 3 | 0.3333 | 8 | 0.125 |
| 15_793664 | NA | NA | NA | NA | 4 | 0.5 | 5 | 0 | 9 | 0.2222 | NA | NA | 3 | 0.3333 | 1 | 1 | 2 | 0 | 6 | 0.3333 |
| 15_793680 | 15 | 0.0667 | 19 | 0.1053 | 25 | 0.08 | 19 | 0.1053 | 78 | 0.0897 | 15 | 0.0667 | 13 | 0.2308 | 21 | 0.0476 | 13 | 0.2308 | 62 | 0.129 |
| 15_793683 | 23 | 1 | 17 | 1 | 26 | 1 | 19 | 1 | 85 | 1 | 13 | 1 | 18 | 1 | 17 | 1 | 17 | 1 | 65 | 1 |
| 15_793683 | 23 | 0.8696 | 20 | 0.95 | 30 | 0.9 | 18 | 1 | 91 | 0.9231 | 17 | 0.9412 | 21 | 0.9048 | 25 | 0.88 | 18 | 0.8889 | 81 | 0.9012 |
| 15_793684( | 26 | 0.8077 | 17 | 0.8235 | 31 | 0.9032 | 17 | 1 | 91 | 0.8791 | 18 | 0.7778 | 22 | 0.9091 | 25 | 0.88 | 19 | 0.8947 | 84 | 0.869 |
| 15_793684 | 25 | 1 | 19 | 1 | 23 | 1 | 16 | 1 | 83 | 1 | 17 | 1 | 19 | 1 | 22 | 1 | 16 | 1 | 74 | 1 |
| 15_793684 | 17 | 0.0588 | 16 | 0.1875 | 28 | 0.1786 | 17 | 0.2353 | 78 | 0.1667 | 16 | 0.125 | 20 | 0.15 | 23 | 0.2609 | 18 | 0.2222 | 77 | 0.1948 |
| 15_793685( | 25 | 1 | 16 | 1 | 29 | 1 | 16 | 0.9375 | 86 | 0.9884 | 16 | 1 | 19 | 1 | 23 | 1 | 20 | 1 | 78 | 1 |
| 15_793685 | 23 | 1 | 16 | 0.875 | 26 | 0.8846 | 17 | 0.9412 | 82 | 0.9268 | 17 | 1 | 20 | 0.9 | 21 | 0.9524 | 16 | 1 | 74 | 0.9595 |
| 15_793685( | 21 | 0.8571 | 17 | 0.7647 | 26 | 0.5385 | 16 | 0.5625 | 80 | 0.675 | 12 | 0.5833 | 18 | 0.6111 | 20 | 0.75 | 16 | 0.75 | 66 | 0.6818 |
| 15_793685 | 24 | 0.625 | 20 | 0.45 | 25 | 0.6 | 18 | 0.7222 | 87 | 0.5977 | 16 | 0.75 | 21 | 0.5714 | 20 | 0.65 | 18 | 0.5 | 75 | 0.6133 |
| 15_793686 | 23 | 0.9565 | 15 | 1 | 26 | 1 | 15 | 1 | 79 | 0.9873 | 16 | 1 | 18 | 1 | 21 | 0.9524 | 14 | 1 | 69 | 0.9855 |
| 15_793689( | 23 | 0.1739 | 16 | 0.1875 | 24 | 0.1667 | 16 | 0.25 | 79 | 0.1899 | 13 | 0.3077 | 11 | 0.1818 | 18 | 0.1667 | 12 | 0.25 | 54 | 0.2222 |
| 15_793689 | 20 | 1 | 15 | 0.9333 | 22 | 0.9091 | 17 | 1 | 74 | 0.9595 | 12 | 1 | 12 | 0.9167 | 16 | 1 | 14 | 1 | 54 | 0.9815 |
| 15_793691 | 16 | 1 | 14 | 1 | 23 | 1 | 15 | 1 | 68 | 1 | 13 | 1 | 12 | 1 | 13 | 1 | 13 | 1 | 51 | 1 |
| 15_793692 | 16 | 0.1875 | 15 | 0.2 | 21 | 0.2381 | 14 | 0 | 66 | 0.1667 | 11 | 0.0909 | 10 | 0.2 | 15 | 0.2667 | 12 | 0.3333 | 48 | 0.2292 |
| 15_793692( | 14 | 0.0714 | 16 | 0.1875 | 20 | 0.05 | 10 | 0.2 | 60 | 0.1167 | 10 | 0.4 | 8 | 0.25 | 10 | 0.2 | 14 | 0.2857 | 42 | 0.2857 |
| 15_793694 | 11 | 0.1818 | 12 | 0.0833 | 21 | 0.1429 | 14 | 0.4286 | 58 | 0.2069 | 13 | 0.1538 | 9 | 0.3333 | 11 | 0.3636 | 12 | 0.1667 | 45 | 0.2444 |
| 15_793694! | 9 | 0.8889 | 10 | 0.9 | 14 | 0.9286 | 10 | 1 | 43 | 0.9302 | 10 | 0.9 | 6 | 1 | 10 | 1 | 11 | 1 | 37 | 0.973 |
| 15_793695! | 11 | 0 | 11 | 0 | 17 | 0.1765 | 12 | 0.0833 | 51 | 0.0784 | 11 | 0 | 6 | 0 | 8 | 0 | 12 | 0.1667 | 37 | 0.0541 |
| 15_793697 | 11 | 0.2727 | 11 | 0.1818 | 16 | 0.1875 | 12 | 0 | 50 | 0.16 | 12 | 0.1667 | 10 | 0.3 | 8 | 0 | 12 | 0.0833 | 42 | 0.1429 |

| | | | | | | | | | | | | | | | | | | | | |
|---|---|---|---|---|---|---|---|---|---|---|---|---|---|---|---|---|---|---|---|---|
| 15_7936982 | 10 | 0.2 | 10 | 0.1 | 12 | 0.25 | 11 | 0.0909 | 43 | 0.1628 | 11 | 0.0909 | 9 | 0.1111 | 8 | 0.25 | 11 | 0.2727 | 39 | 0.1795 |
| 15_7936997 | 9 | 0.3333 | 10 | 0.2 | 14 | 0.2857 | 7 | 0.2857 | 40 | 0.275 | 10 | 0.3 | 7 | 0.1429 | 9 | 0.4444 | 10 | 0 | 36 | 0.2222 |
| 15_7938943 | 9 | 0.1111 | 8 | 0.125 | 13 | 0 | 9 | 0 | 39 | 0.0513 | 9 | 0.2222 | 5 | 0.2 | 2 | 0 | 9 | 0 | 25 | 0.12 |
| 15_7938945 | 8 | 0.25 | 5 | 0 | 4 | 0 | 9 | 0.1111 | 26 | 0.1154 | 5 | 0 | 4 | 0 | 3 | 0.3333 | 1 | 0 | 13 | 0.0769 |
| 15_7938960 | 7 | 0.1429 | 5 | 0 | 8 | 0.125 | 7 | 0 | 27 | 0.0741 | 5 | 0.4 | 3 | 0 | 2 | 0 | 6 | 0 | 16 | 0.125 |
| 15_7938973 | 5 | 1 | 2 | 1 | 4 | 0.75 | 6 | 0.6667 | 17 | 0.8235 | 3 | 0.6667 | 3 | 0.6667 | NA | NA | 4 | 1 | 10 | 0.8 |
| 15_7938977 | 7 | 0 | 2 | 0.5 | 6 | 0.3333 | 6 | 0.1667 | 21 | 0.1905 | 6 | 0.1667 | 4 | 0.25 | 1 | 0 | 7 | 0 | 18 | 0.1111 |
| 15_7938989 | 2 | 0.5 | 3 | 0 | 6 | 0.1667 | 5 | 0.2 | 16 | 0.1875 | 6 | 0.1667 | 3 | 0 | 1 | 0 | 5 | 0 | 15 | 0.0667 |
| 15_7939540 | NA | NA | NA | NA | 1 | 1 | NA | NA | 1 | 1 | 1 | 0 | NA | NA | 2 | 0 | 2 | 0.5 | 5 | 0.2 |
| 15_7939558 | NA | NA | NA | NA | 1 | 1 | NA | NA | 1 | 1 | 1 | 0 | NA | NA | 2 | 0.5 | 1 | 1 | 4 | 0.5 |
| 15_7941206 | 20 | 0.05 | 23 | 0.3043 | 29 | 0.069 | 15 | 0 | 87 | 0.1149 | 19 | 0 | 30 | 0.0667 | 24 | 0.1667 | 23 | 0.1304 | 96 | 0.0938 |
| 15_7941215 | 18 | 0.0556 | 23 | 0.1304 | 25 | 0.08 | 17 | 0 | 83 | 0.0723 | 19 | 0.0526 | 28 | 0.1071 | 22 | 0.0909 | 22 | 0 | 91 | 0.0659 |
| 15_7941324 | 11 | 0.3636 | 15 | 0.3333 | 15 | 0.1333 | 13 | 0.3077 | 54 | 0.2778 | 12 | 0 | 19 | 0.1579 | 17 | 0.3529 | 13 | 0.0769 | 61 | 0.1639 |
| 15_7941334 | 8 | 0.375 | 15 | 0.2667 | 14 | 0.4286 | 11 | 0.3636 | 48 | 0.3542 | 9 | 0.2222 | 15 | 0.1333 | 16 | 0.25 | 10 | 0.4 | 50 | 0.24 |
| 15_7941371 | 8 | 0.375 | 12 | 0.1667 | 14 | 0.0714 | 8 | 0.125 | 42 | 0.1667 | 7 | 0.2857 | 15 | 0.0667 | 18 | 0.2222 | 12 | 0.0833 | 52 | 0.1538 |
| 15_7941385 | 9 | 0 | 7 | 0.2857 | 11 | 0.2727 | 4 | 0 | 31 | 0.1613 | 8 | 0 | 14 | 0.1429 | 11 | 0 | 3 | 0 | 36 | 0.0556 |
| 15_7941416 | 5 | 0.2 | 9 | 0.6667 | 10 | 0.3 | 3 | 0.3333 | 27 | 0.4074 | 9 | 0.3333 | 12 | 0.1667 | 11 | 0.0909 | 8 | 0.125 | 40 | 0.175 |
| 15_7941419 | 6 | 0 | 11 | 0.3636 | 10 | 0.3 | 2 | 0 | 29 | 0.2414 | 10 | 0 | 12 | 0.0833 | 11 | 0.1818 | 7 | 0.2857 | 40 | 0.125 |
| 15_7941421 | 4 | 0.25 | 8 | 0 | 9 | 0.2222 | 2 | 0 | 23 | 0.1304 | 8 | 0 | 10 | 0 | 9 | 0.2222 | 5 | 0 | 32 | 0.0625 |
| 15_7941426 | 6 | 0 | 9 | 0.2222 | 9 | 0.1111 | 3 | 0 | 27 | 0.1111 | 10 | 0.2 | 11 | 0.0909 | 8 | 0 | 8 | 0.125 | 37 | 0.1081 |
| 15_7941458 | 4 | 0 | 8 | 0.25 | 5 | 0 | 3 | 0.3333 | 20 | 0.15 | 8 | 0.125 | 9 | 0 | 8 | 0.25 | 7 | 0 | 32 | 0.0938 |
| 15_7941469 | 4 | 0.5 | 8 | 0.25 | 5 | 0.6 | 3 | 0.6667 | 20 | 0.45 | 6 | 0.5 | 9 | 0.4444 | 8 | 0.5 | 8 | 0.5 | 31 | 0.4839 |
| 15_7949543 | 1 | 1 | 1 | 0 | NA | NA | 2 | 0.5 | 4 | 0.5 | NA | NA | NA | NA | 1 | 1 | NA | NA | 1 | 1 |
| 15_7949654 | NA | NA | 1 | 0 | NA | NA | 1 | 1 | 2 | 0.5 | 1 | 0 | NA | NA | 1 | 1 | NA | NA | 2 | 0.5 |
| 15_7955129 | 8 | 0.125 | 8 | 0.25 | 3 | 0 | 6 | 0 | 25 | 0.12 | 7 | 0.1429 | 5 | 0.2 | 5 | 0.2 | 8 | 0 | 25 | 0.12 |
| 15_7955198 | 1 | 0 | 1 | 0 | 2 | 0 | 2 | 0.5 | 6 | 0.1667 | 1 | 0 | NA | NA | NA | NA | 2 | 0.5 | 3 | 0.3333 |
| 15_7955210 | 1 | 1 | NA | NA | 2 | 0 | 1 | 0 | 4 | 0.25 | 1 | 0 | 1 | 0 | NA | NA | 2 | 0.5 | 4 | 0.25 |
| 15_7955652 | 19 | 0.0526 | 18 | 0 | 21 | 0.1905 | 17 | 0.2353 | 75 | 0.12 | 13 | 0.2308 | 15 | 0.0667 | 9 | 0 | 16 | 0.1875 | 53 | 0.1321 |
| 15_7955654 | 18 | 0.1667 | 19 | 0.3684 | 20 | 0.3 | 18 | 0.1667 | 75 | 0.2533 | 15 | 0.2667 | 14 | 0.4286 | 9 | 0.2222 | 17 | 0.3529 | 55 | 0.3273 |
| 15_7955656 | 16 | 0.1875 | 16 | 0.375 | 19 | 0.3684 | 18 | 0.6111 | 69 | 0.3913 | 14 | 0.5 | 17 | 0.2353 | 9 | 0.6667 | 14 | 0.4286 | 54 | 0.4259 |
| 15_7955660 | 18 | 0.0556 | 20 | 0.1 | 18 | 0.1111 | 16 | 0.125 | 72 | 0.0972 | 14 | 0.2143 | 17 | 0.1765 | 10 | 0.2 | 12 | 0.0833 | 53 | 0.1698 |
| 15_7955662 | 17 | 0.1176 | 16 | 0.1875 | 21 | 0.1429 | 18 | 0 | 72 | 0.1111 | 13 | 0.2308 | 17 | 0.2941 | 9 | 0.1111 | 14 | 0.2857 | 53 | 0.2453 |
| 15_7955675 | 17 | 0.0588 | 16 | 0 | 18 | 0.1667 | 10 | 0.1 | 61 | 0.082 | 13 | 0.0769 | 16 | 0.125 | 9 | 0 | 13 | 0.0769 | 51 | 0.0784 |
| 15_7955681 | 17 | 0.3529 | 20 | 0.5 | 22 | 0.3636 | 20 | 0.45 | 79 | 0.4177 | 16 | 0.25 | 18 | 0.4444 | 12 | 0.25 | 16 | 0.375 | 62 | 0.3387 |
| 15_7955890 | 7 | 0 | 11 | 0 | 17 | 0.0588 | 14 | 0.1429 | 49 | 0.0612 | 8 | 0 | 9 | 0.2222 | 10 | 0.2 | 10 | 0 | 37 | 0.1081 |
| 15_7956014 | 4 | 0.75 | 5 | 0.6 | 7 | 0.8571 | 5 | 0.6 | 21 | 0.7143 | 3 | 1 | 4 | 1 | 5 | 1 | 7 | 0.8571 | 19 | 0.9474 |
| 15_7956016 | 4 | 0.5 | 5 | 0.2 | 5 | 0.4 | 6 | 0.5 | 20 | 0.4 | 3 | 0.3333 | 5 | 0.4 | 4 | 0.25 | 6 | 0 | 18 | 0.2222 |
| 15_7956034 | 3 | 0.3333 | 4 | 0.25 | 5 | 0.4 | 8 | 0.125 | 20 | 0.25 | 2 | 0 | 4 | 0.5 | 3 | 0 | 4 | 0.25 | 13 | 0.2308 |
| 15_7956034 | 4 | 0.5 | 2 | 0 | 6 | 0.5 | 7 | 0.1429 | 19 | 0.3158 | 2 | 0.5 | 2 | 0 | 3 | 0.6667 | 5 | 0.2 | 12 | 0.3333 |
| 15_7957106 | 3 | 0.3333 | 2 | 0 | 4 | 0.25 | 4 | 0.5 | 13 | 0.3077 | 1 | 0 | 2 | 0 | 2 | 0.5 | 3 | 0 | 8 | 0.125 |
| 15_7957113 | 3 | 0 | 3 | 0.3333 | 4 | 0.25 | 5 | 0.4 | 15 | 0.2667 | 1 | 0 | 2 | 0 | 2 | 0 | 3 | 0.6667 | 8 | 0.25 |
| 15_7957118 | 2 | 1 | 1 | 1 | 5 | 0.6 | 3 | 0.3333 | 11 | 0.6364 | 1 | 0 | NA | NA | 1 | 0 | 2 | 0.5 | 4 | 0.25 |
| 15_7957123 | 2 | 0.5 | 3 | 0.6667 | 4 | 0.5 | 4 | 0.25 | 13 | 0.4615 | 1 | 0 | NA | NA | NA | NA | 2 | 0.5 | 3 | 0.3333 |
| 15_7959196 | 2 | 0.5 | 1 | 0 | 2 | 0 | 3 | 0.3333 | 8 | 0.25 | 2 | 0 | 3 | 0.3333 | 2 | 0 | 1 | 1 | 8 | 0.25 |
| 15_7959199 | 1 | 0 | 2 | 0 | 2 | 0 | 4 | 0.25 | 9 | 0.1111 | 2 | 0 | 3 | 0.3333 | 1 | 0 | 1 | 1 | 7 | 0.2857 |
| 15_7959212 | 3 | 0.3333 | 2 | 0.5 | 3 | 0 | 4 | 0.25 | 12 | 0.25 | 3 | 0 | 3 | 0 | 2 | 0.5 | 1 | 0 | 9 | 0.1111 |
| 15_7960447 | 6 | 0.1667 | 4 | 0 | 5 | 0 | 7 | 0.4286 | 22 | 0.1818 | 8 | 0.25 | 6 | 0.3333 | 4 | 0 | 2 | 0 | 20 | 0.2 |
| 15_7960449 | 7 | 1 | 4 | 0.5 | 5 | 0.4 | 7 | 0.4286 | 23 | 0.6087 | 8 | 0.625 | 6 | 0.5 | 5 | 0.6 | 2 | 0 | 21 | 0.5238 |
| 15_7960563 | 8 | 0.125 | 8 | 0.5 | 6 | 0.1667 | 4 | 0.25 | 26 | 0.2692 | 6 | 0.5 | 4 | 0.25 | 5 | 0.4 | 3 | 0 | 18 | 0.3333 |
| 15_7960891 | 1 | 0 | 3 | 0.3333 | 1 | 0 | 1 | 0 | 6 | 0.1667 | NA | NA | 2 | 0.5 | 1 | 0 | 3 | 0.3333 | 6 | 0.3333 |
| 15_7960924 | 1 | 1 | 2 | 0 | 1 | 0 | 1 | 0 | 5 | 0.2 | NA | NA | 1 | 0 | 1 | 1 | 3 | 0.6667 | 5 | 0.6 |
| 15_7960938 | 1 | 0 | NA | NA | 1 | 1 | 1 | 1 | 3 | 0.6667 | NA | NA | 1 | 1 | 1 | 0 | 3 | 1 | 5 | 0.8 |
| 15_7964581 | 8 | 0.25 | 12 | 0.0833 | 6 | 0 | 12 | 0.1667 | 38 | 0.1316 | 1 | 0 | 6 | 0.3333 | 11 | 0 | 13 | 0.0769 | 31 | 0.0968 |
| 15_7964582 | 11 | 0.3636 | 10 | 0.3 | 7 | 0.4286 | 12 | 0.25 | 40 | 0.325 | 1 | 1 | 7 | 0.1429 | 11 | 0.1818 | 11 | 0.1818 | 30 | 0.2 |
| 15_7964595 | 8 | 0 | 9 | 0.1111 | 5 | 0.2 | 13 | 0.1538 | 35 | 0.1143 | NA | NA | 7 | 0 | 7 | 0.1429 | 10 | 0.2 | 24 | 0.125 |
| 15_7964598 | 9 | 0.2222 | 8 | 0 | 4 | 0.25 | 10 | 0.3 | 31 | 0.1935 | 1 | 0 | 3 | 0 | 10 | 0.1 | 10 | 0.1 | 24 | 0.0833 |
| 15_7964606 | 9 | 0.2222 | 10 | 0.1 | 5 | 0 | 9 | 0.1111 | 33 | 0.1212 | 1 | 1 | 4 | 0.25 | 11 | 0.1818 | 11 | 0.0909 | 27 | 0.1852 |
| 15_7964608 | 9 | 0.5556 | 9 | 0.6667 | 6 | 0.5 | 12 | 0.4167 | 36 | 0.5278 | 1 | 0 | 5 | 0.8 | 11 | 0.2727 | 12 | 0.5833 | 29 | 0.4828 |
| 15_7964622 | 9 | 0.1111 | 10 | 0.2 | 5 | 0.4 | 11 | 0.5455 | 35 | 0.3143 | NA | NA | 4 | 0.25 | 10 | 0.3 | 12 | 0.1667 | 26 | 0.2308 |
| 15_7964627 | 9 | 0.7778 | 6 | 0.5 | 4 | 0.25 | 9 | 0.7778 | 28 | 0.6429 | NA | NA | 4 | 0.5 | 10 | 0.6 | 11 | 0.4545 | 25 | 0.52 |
| 15_7964635 | 7 | 0 | 10 | 0.1 | 6 | 0.1667 | 9 | 0.1111 | 32 | 0.0938 | NA | NA | 4 | 0.25 | 9 | 0.2222 | 10 | 0 | 23 | 0.1304 |
| 15_7967073 | 37 | 0.2432 | 68 | 0.3382 | 44 | 0.4091 | 28 | 0.3929 | 177 | 0.3446 | 58 | 0.4828 | 21 | 0.4286 | 14 | 0.3571 | 35 | 0.4286 | 128 | 0.4453 |
| 15_7967078 | 47 | 0.1064 | 79 | 0.0506 | 53 | 0.0377 | 31 | 0.0645 | 210 | 0.0619 | 68 | 0.0588 | 24 | 0.0417 | 16 | 0.125 | 41 | 0.0488 | 149 | 0.0604 |
| 15_7967087 | 40 | 0.025 | 71 | 0.0282 | 47 | 0.1277 | 28 | 0.1429 | 186 | 0.0699 | 63 | 0.1111 | 21 | 0.0476 | 16 | 0.0625 | 37 | 0.0541 | 137 | 0.0803 |
| 15_7967155 | 36 | 0.0278 | 60 | 0.1667 | 42 | 0.0952 | 21 | 0.1905 | 159 | 0.1195 | 55 | 0.1273 | 19 | 0.1053 | 12 | 0.1667 | 32 | 0.1875 | 118 | 0.1441 |

| | | | | | | | | | | | | | | | | | | | | |
|---|---|---|---|---|---|---|---|---|---|---|---|---|---|---|---|---|---|---|---|---|
| 15_7967175 | 25 | 0.08 | 41 | 0.122 | 32 | 0.0938 | 15 | 0 | 113 | 0.0885 | 44 | 0.0909 | 15 | 0.0667 | 10 | 0.1 | 25 | 0.04 | 94 | 0.0745 |
| 15_7967211 | 30 | 0.0333 | 51 | 0.0196 | 28 | 0.0714 | 17 | 0.2353 | 126 | 0.0635 | 47 | 0.1064 | 14 | 0.0714 | 10 | 0.1 | 26 | 0.0385 | 97 | 0.0825 |
| 15_7967357 | 12 | 0.1667 | 20 | 0.25 | 14 | 0.1429 | 6 | 0.5 | 52 | 0.2308 | 19 | 0.2632 | 9 | 0.2222 | 3 | 0.6667 | 11 | 0.0909 | 42 | 0.2381 |
| 15_7967359 | 18 | 0.1667 | 30 | 0.1 | 19 | 0 | 9 | 0 | 76 | 0.0789 | 28 | 0.0357 | 9 | 0.1111 | 3 | 0 | 12 | 0.0833 | 52 | 0.0577 |
| 15_7967384 | 18 | 0.8889 | 26 | 0.9615 | 18 | 1 | 9 | 0.8889 | 71 | 0.9437 | 28 | 1 | 9 | 0.8889 | 2 | 1 | 11 | 0.9091 | 50 | 0.96 |
| 15_7967404 | 15 | 0.0667 | 23 | 0.3043 | 13 | 0.0769 | 8 | 0.25 | 59 | 0.1864 | 23 | 0.1739 | 6 | 0.1667 | 2 | 0 | 8 | 0.125 | 39 | 0.1538 |
| 15_7967407 | 15 | 0.0667 | 24 | 0.0417 | 18 | 0.0556 | 8 | 0.125 | 65 | 0.0615 | 24 | 0.0833 | 9 | 0.1111 | 3 | 0.3333 | 11 | 0.0909 | 47 | 0.1064 |
| 15_7968835 | 14 | 0.1429 | 16 | 0 | 7 | 0 | 6 | 0.3333 | 43 | 0.093 | 15 | 0.2667 | 7 | 0.1429 | 2 | 0 | 6 | 0 | 30 | 0.1667 |
| 15_7970828 | 38 | 0.2632 | 42 | 0.381 | 32 | 0.2188 | 31 | 0.3548 | 143 | 0.3077 | 13 | 0.3077 | 36 | 0.3611 | 37 | 0.4324 | 26 | 0.4615 | 112 | 0.4018 |
| 15_7970832 | 19 | 0.1053 | 20 | 0.3 | 18 | 0 | 18 | 0 | 75 | 0.1067 | 9 | 0.1111 | 20 | 0 | 17 | 0.0588 | 12 | 0.0833 | 58 | 0.0517 |
| 15_7970852 | 33 | 0.2727 | 41 | 0.2683 | 28 | 0.2857 | 29 | 0.2069 | 131 | 0.2595 | 12 | 0.0833 | 31 | 0.129 | 30 | 0.3667 | 20 | 0.3 | 93 | 0.2366 |
| 15_7970869 | 30 | 0.2 | 33 | 0.0909 | 25 | 0.12 | 23 | 0.1304 | 111 | 0.1351 | 10 | 0.2 | 24 | 0.0833 | 30 | 0.2333 | 16 | 0.1875 | 80 | 0.175 |
| 15_7970882 | 33 | 0.1515 | 33 | 0.2121 | 25 | 0.04 | 25 | 0.12 | 116 | 0.1379 | 11 | 0.0909 | 22 | 0.0909 | 30 | 0.2667 | 17 | 0.1176 | 80 | 0.1625 |
| 15_7970911 | 26 | 0.2692 | 28 | 0.2857 | 22 | 0.4545 | 21 | 0.2857 | 97 | 0.3196 | 10 | 0.3 | 25 | 0.36 | 18 | 0.2778 | 15 | 0.2667 | 68 | 0.3088 |
| 15_7970915 | 20 | 0.35 | 27 | 0.3333 | 15 | 0.3333 | 20 | 0.4 | 82 | 0.3537 | 7 | 0.4286 | 17 | 0.6471 | 18 | 0.5556 | 12 | 0.8333 | 54 | 0.6296 |
| 15_7970964 | 22 | 0.0909 | 27 | 0.1852 | 16 | 0.125 | 22 | 0.0455 | 87 | 0.1149 | 10 | 0 | 22 | 0.1818 | 21 | 0.2381 | 12 | 0.5 | 65 | 0.2308 |
| 15_7971119 | 10 | 0.3 | 17 | 0.1765 | 8 | 0 | 14 | 0.0714 | 49 | 0.1429 | 6 | 0.1667 | 8 | 0.5 | 10 | 0.2 | 5 | 0 | 29 | 0.2414 |
| 15_7971202 | 7 | 0.8571 | 9 | 1 | 5 | 1 | 10 | 0.8 | 31 | 0.9032 | 5 | 0.8 | 5 | 0.8 | 6 | 0.8333 | 3 | 0.6667 | 19 | 0.7895 |
| 15_7971235 | 6 | 0.8333 | 8 | 1 | 2 | 1 | 10 | 1 | 26 | 0.9615 | 4 | 1 | 7 | 1 | 8 | 0.875 | 5 | 1 | 24 | 0.9583 |
| 15_7971241 | 8 | 0.625 | 8 | 0.625 | 2 | 1 | 9 | 0.6667 | 27 | 0.6667 | 2 | 0.5 | 5 | 0.8 | 7 | 0.4286 | 5 | 0.6 | 19 | 0.5789 |
| 15_7971252 | 5 | 0.2 | 9 | 0.1111 | 2 | 0.5 | 9 | 0 | 25 | 0.12 | 3 | 0.3333 | 7 | 0.1429 | 8 | 0.25 | 5 | 0 | 23 | 0.1739 |
| 15_7971280 | 3 | 0 | 5 | 0 | 2 | 0 | 8 | 0.125 | 18 | 0.0556 | 3 | 0 | 5 | 0 | 5 | 0 | 5 | 0.2 | 18 | 0.0556 |
| 15_7971288 | 6 | 0 | 5 | 0.2 | 2 | 0 | 6 | 0.1667 | 19 | 0.1053 | 3 | 0.3333 | 7 | 0 | 4 | 0 | 3 | 0 | 17 | 0.0588 |
| 15_7971289 | 7 | 0.1429 | 8 | 0.25 | 2 | 0 | 9 | 0.1111 | 26 | 0.1538 | 4 | 0 | 7 | 0.1429 | 4 | 0.25 | 4 | 0 | 19 | 0.1053 |
| 15_7971301 | 5 | 0 | 5 | 0 | 1 | 0 | 8 | 0.125 | 19 | 0.0526 | 2 | 0 | 4 | 0 | 3 | 0 | 4 | 0.25 | 13 | 0.0769 |
| 15_7971306 | 5 | 0.4 | 7 | 0.1429 | 2 | 0 | 9 | 0 | 23 | 0.1304 | 2 | 0 | 7 | 0.1429 | 4 | 0 | 4 | 0 | 17 | 0.0588 |
| 15_7971666 | NA | NA | 1 | 0 | 2 | 0 | 2 | 0.5 | 5 | 0.2 | 2 | 1 | 1 | 0 | 2 | 1 | 1 | 1 | 6 | 0.8333 |
| 15_7971686 | NA | NA | 1 | 0 | 1 | 0 | 5 | 0.4 | 7 | 0.2857 | 1 | 0 | NA | NA | 3 | 0.3333 | 2 | 0.5 | 6 | 0.3333 |
| 15_7979932 | NA | NA | NA | NA | NA | NA | 1 | 1 | 1 | 1 | NA | NA | NA | NA | NA | NA | 1 | 1 | 1 | 1 |
| 15_7979938 | NA | NA | NA | NA | NA | NA | 1 | 1 | 1 | 1 | NA | NA | NA | NA | NA | NA | 1 | 1 | 1 | 1 |
| 15_7979955 | NA | NA | NA | NA | NA | NA | 1 | 1 | 1 | 1 | NA | NA | NA | NA | NA | NA | 1 | 1 | 1 | 1 |
| 15_7979972 | NA | NA | NA | NA | NA | NA | 1 | 1 | 1 | 1 | NA | NA | NA | NA | NA | NA | 1 | 1 | 1 | 1 |
| 15_7979990 | NA | NA | NA | NA | NA | NA | 1 | 1 | 1 | 1 | NA | NA | NA | NA | NA | NA | 1 | 1 | 1 | 1 |
| 15_7980009 | 9 | 0 | 10 | 0.1 | 5 | 0 | 10 | 0.3 | 34 | 0.1176 | 4 | 0 | 8 | 0.125 | 4 | 0.25 | 8 | 0.375 | 24 | 0.2083 |
| 15_7980043 | 8 | 0.25 | 8 | 0.25 | 3 | 0.3333 | 5 | 0.2 | 24 | 0.25 | 4 | 0.5 | 7 | 0.1429 | 4 | 0 | 6 | 0.1667 | 21 | 0.1905 |
| 15_7980061 | 9 | 0 | 6 | 0.3333 | 3 | 0 | 6 | 0.1667 | 24 | 0.125 | 3 | 0 | 5 | 0.2 | 3 | 0 | 4 | 0.25 | 15 | 0.1333 |
| 15_7980070 | 9 | 0 | 5 | 0 | 4 | 0 | 8 | 0.25 | 26 | 0.0769 | 2 | 0 | 8 | 0.125 | 3 | 0 | 6 | 0 | 19 | 0.0526 |
| 15_7980171 | 6 | 0.1667 | 7 | 0.1429 | 2 | 0 | 7 | 0.1429 | 22 | 0.1364 | 2 | 0.5 | 6 | 0.3333 | 3 | 0 | 5 | 0.2 | 16 | 0.25 |
| 15_7980172 | 5 | 0 | 6 | 0.1667 | 3 | 0 | 6 | 0.3333 | 20 | 0.15 | 2 | 0.5 | 7 | 0.1429 | 2 | 0.5 | 5 | 0.4 | 16 | 0.3125 |
| 15_7980181 | 6 | 0.3333 | 5 | 0.2 | NA | NA | 5 | 0.4 | 16 | 0.3125 | 2 | 0.5 | 5 | 0.2 | 2 | 0.5 | 4 | 1 | 13 | 0.5385 |
| 15_7980185 | 5 | 0 | 7 | 0.1429 | 3 | 0 | 5 | 0.2 | 20 | 0.1 | 2 | 0 | 5 | 0 | 3 | 0 | 6 | 0.1667 | 16 | 0.0625 |
| 15_7980189 | 2 | 1 | 6 | 0.6667 | 1 | 1 | 3 | 0.3333 | 12 | 0.6667 | 2 | 1 | 5 | 0.6 | 3 | 1 | 3 | 1 | 13 | 0.8462 |
| 15_7980190 | 2 | 0 | 7 | 0 | 3 | 0 | 3 | 0.3333 | 15 | 0.0667 | 2 | 0 | 6 | 0 | 1 | 0 | 1 | 1 | 10 | 0.1 |
| 15_7980192 | 6 | 0.1667 | 6 | 0.1667 | 2 | 0.5 | 6 | 0.5 | 20 | 0.3 | 2 | 0 | 4 | 0 | 3 | 0.3333 | 5 | 0.2 | 14 | 0.1429 |
| 15_7980193 | 5 | 0 | 7 | 0.2857 | 2 | 0.5 | 6 | 0.3333 | 20 | 0.25 | 2 | 0 | 7 | 0.1429 | 2 | 1 | 3 | 0.3333 | 14 | 0.2857 |
| 15_7980196 | 6 | 0.5 | 7 | 0.4286 | 3 | 0.3333 | 5 | 0.6 | 21 | 0.4762 | 2 | 0 | 6 | 0.3333 | 3 | 0 | 5 | 0.4 | 16 | 0.25 |
| 15_7980197 | 5 | 0.2 | 6 | 0.1667 | 1 | 0 | 3 | 0.3333 | 15 | 0.2 | 2 | 0 | 5 | 0.2 | 1 | 1 | 3 | 0 | 11 | 0.1818 |
| 15_7980205 | 5 | 0.2 | 6 | 0.1667 | NA | NA | 3 | 0 | 14 | 0.1429 | 2 | 0.5 | 3 | 0 | 2 | 0 | 4 | 0.25 | 11 | 0.1818 |
| 15_7980208 | 6 | 0.6667 | 6 | 0.3333 | 2 | 0 | 5 | 0.2 | 19 | 0.3684 | 2 | 0.5 | 6 | 0.3333 | 3 | 0.3333 | 5 | 0.6 | 16 | 0.4375 |
| 15_7980211 | 6 | 0.1667 | 7 | 0 | 2 | 0.5 | 6 | 0 | 21 | 0.0952 | 1 | 1 | 5 | 0.2 | 3 | 0.3333 | 4 | 0 | 13 | 0.2308 |
| 15_7980213 | 5 | 1 | 5 | 1 | 2 | 0.5 | 5 | 1 | 17 | 0.9412 | 2 | 1 | 5 | 0.8 | 3 | 1 | 2 | 1 | 12 | 0.9167 |
| 15_7980229 | 4 | 0.25 | 1 | 0 | 3 | 0 | 3 | 0.3333 | 11 | 0.1818 | 2 | 0.5 | 4 | 0 | NA | NA | 1 | 0 | 7 | 0.1429 |
| 15_7980246 | 4 | 0.25 | 2 | 0 | 1 | 0 | 1 | 1 | 8 | 0.25 | 2 | 0 | 2 | 0.5 | NA | NA | 3 | 0.3333 | 7 | 0.2857 |
| 15_7980249 | 4 | 0.25 | 2 | 0 | 2 | 0 | 3 | 0 | 11 | 0.0909 | 2 | 1 | 5 | 0.2 | 2 | 0 | 2 | 0 | 11 | 0.2727 |
| 15_7988064 | 2 | 0.5 | 5 | 0.2 | NA | NA | 2 | 0 | 9 | 0.2222 | 2 | 0 | 2 | 0 | 1 | 0 | 3 | 0.6667 | 8 | 0.25 |
| 15_7988096 | 2 | 1 | 2 | 0.5 | NA | NA | 2 | 1 | 6 | 0.8333 | 1 | 0 | 1 | 1 | 2 | 1 | 2 | 1 | 6 | 0.8333 |
| 15_7988096 | 2 | 1 | 5 | 0 | NA | NA | 2 | 0 | 9 | 0.2222 | 2 | 0.5 | NA | NA | 2 | 0 | 2 | 0 | 6 | 0.1667 |
| 15_7988104 | 2 | 1 | 3 | 0.3333 | NA | NA | 2 | 0.5 | 7 | 0.5714 | 2 | 0.5 | 1 | 1 | 2 | 1 | 2 | 0.5 | 7 | 0.7143 |
| 15_7988105 | 2 | 0 | 5 | 0.2 | NA | NA | 2 | 0 | 9 | 0.1111 | 2 | 1 | 1 | 1 | 2 | 0.5 | 3 | 0.3333 | 8 | 0.625 |
| 15_7988115 | 2 | 0 | 5 | 0.4 | NA | NA | 2 | 0.5 | 9 | 0.3333 | 2 | 0 | 1 | 1 | 1 | 0 | 3 | 0.6667 | 7 | 0.4286 |
| 15_7988118 | 1 | 1 | 4 | 1 | NA | NA | NA | NA | 5 | 1 | 2 | 1 | 1 | 1 | NA | NA | 2 | 1 | 5 | 1 |
| 15_7988119 | 2 | 1 | 3 | 1 | NA | NA | 2 | 1 | 7 | 1 | 2 | 1 | 1 | 1 | 2 | 1 | 3 | 1 | 8 | 1 |
| 15_7988127 | 2 | 0.5 | 3 | 0.6667 | NA | NA | 2 | 0.5 | 7 | 0.5714 | 2 | 0.5 | 1 | 0 | 2 | 0.5 | 2 | 0.5 | 7 | 0.4286 |
| 15_7989848 | NA | NA | 1 | 0 | NA | NA | 1 | 1 | 2 | 0.5 | NA | NA | NA | NA | NA | NA | 1 | 1 | 1 | 1 |
| 15_8000013 | 7 | 0 | 5 | 0.2 | 10 | 0.1 | 9 | 0.1111 | 31 | 0.0968 | 9 | 0.1111 | 9 | 0 | 5 | 0.2 | 8 | 0 | 31 | 0.0645 |

| | | | | | | | | | | | | | | | | | | | | |
|---|---|---|---|---|---|---|---|---|---|---|---|---|---|---|---|---|---|---|---|---|
| 15_8000014 | 8 | 0.125 | 6 | 0.3333 | 9 | 0 | 9 | 0 | 32 | 0.0938 | 9 | 0.1111 | 9 | 0 | 6 | 0.1667 | 9 | 0.2222 | 33 | 0.1212 |
| 15_8000019 | 8 | 0.375 | 6 | 0 | 9 | 0 | 8 | 0.25 | 31 | 0.1613 | 10 | 0.1 | 9 | 0.1111 | 5 | 0 | 8 | 0.375 | 32 | 0.1562 |
| 15_8000037 | 11 | 0.1818 | 6 | 0.3333 | 8 | 0 | 9 | 0.1111 | 34 | 0.1471 | 10 | 0.4 | 9 | 0.4444 | 4 | 0.5 | 7 | 0.1429 | 30 | 0.3667 |
| 15_8000040 | 12 | 0.1667 | 5 | 0 | 10 | 0.1 | 9 | 0.1111 | 36 | 0.1111 | 9 | 0.3333 | 10 | 0.2 | 6 | 0.5 | 9 | 0 | 34 | 0.2353 |
| 15_8000166 | 26 | 0.0385 | 12 | 0.25 | 17 | 0.0588 | 19 | 0.0526 | 74 | 0.0811 | 20 | 0.05 | 18 | 0 | 19 | 0.1053 | 16 | 0.125 | 73 | 0.0685 |
| 15_8000247 | 35 | 0.1143 | 15 | 0 | 26 | 0.0385 | 23 | 0.0435 | 99 | 0.0606 | 21 | 0.0952 | 17 | 0.0588 | 16 | 0.125 | 20 | 0.1 | 74 | 0.0946 |
| 15_8000318 | 50 | 0.26 | 33 | 0.0606 | 36 | 0.1667 | 30 | 0.0333 | 149 | 0.1477 | 33 | 0.1212 | 27 | 0.1852 | 30 | 0.1 | 29 | 0.0345 | 119 | 0.1092 |
| 15_8000359 | 55 | 0.1455 | 48 | 0.0417 | 45 | 0.0889 | 37 | 0.1351 | 185 | 0.1027 | 44 | 0.1136 | 31 | 0.2258 | 24 | 0.0833 | 40 | 0 | 139 | 0.1007 |
| 15_8004441 | 1 | 1 | NA | NA | NA | NA | 1 | 0 | 2 | 0.5 | NA | NA | 2 | 1 | NA | NA | NA | NA | 2 | 1 |
| 15_8004464 | NA | NA | NA | NA | NA | NA | 1 | 1 | 1 | 1 | 1 | 1 | 2 | 0.5 | NA | NA | NA | NA | 3 | 0.6667 |
| 15_8004477 | 1 | 1 | NA | NA | 1 | 1 | NA | NA | 2 | 1 | 1 | 0 | 3 | 0.6667 | 1 | 0 | NA | NA | 5 | 0.4 |
| 15_8004539 | 2 | 0.5 | 1 | 1 | 2 | 1 | 3 | 0.6667 | 8 | 0.75 | 1 | 1 | 3 | 0.3333 | 2 | 0.5 | 2 | 1 | 8 | 0.625 |
| 15_8004543 | 2 | 1 | 1 | 1 | 2 | 1 | 2 | 1 | 7 | 1 | 1 | 1 | 3 | 1 | 2 | 0.5 | 2 | 1 | 8 | 0.875 |
| 15_8004546 | 2 | 0.5 | 1 | 0 | 1 | 1 | 1 | 0 | 5 | 0.4 | 1 | 0 | 4 | 0.25 | 2 | 0.5 | NA | NA | 7 | 0.2857 |
| 15_8004567 | 2 | 1 | NA | NA | 2 | 0.5 | 3 | 0.3333 | 7 | 0.5714 | 1 | 1 | 6 | 1 | 2 | 1 | 2 | 1 | 11 | 1 |
| 15_8004582 | 2 | 1 | 1 | 1 | 2 | 1 | 2 | 1 | 7 | 1 | 1 | 1 | 6 | 1 | 1 | 1 | 2 | 1 | 10 | 1 |
| 15_8004583 | 2 | 0.5 | 1 | 1 | 2 | 1 | 3 | 0.3333 | 8 | 0.625 | 1 | 0 | 5 | 0.6 | 3 | 0 | 1 | 1 | 10 | 0.4 |
| 15_8009525 | 1 | 0 | 5 | 0.4 | 3 | 0.6667 | 1 | 1 | 10 | 0.5 | 1 | 1 | 3 | 0.3333 | 2 | 0 | 1 | 0 | 7 | 0.2857 |
| 15_8009528 | 1 | 1 | 5 | 0.8 | 2 | 0.5 | NA | NA | 8 | 0.75 | NA | NA | 3 | 0.6667 | 2 | 1 | 1 | 1 | 6 | 0.8333 |
| 15_8009532 | NA | NA | 5 | 0.4 | 3 | 0.6667 | NA | NA | 8 | 0.5 | 1 | 1 | 2 | 0.5 | 2 | 0.5 | NA | NA | 5 | 0.6 |
| 15_8009538 | 1 | 1 | 5 | 0.8 | 3 | 0.6667 | 1 | 1 | 10 | 0.8 | 1 | 1 | 3 | 0.3333 | 2 | 1 | 1 | 0 | 7 | 0.5714 |
| 15_8009545 | 1 | 1 | 4 | 0.75 | 4 | 1 | 1 | 1 | 10 | 0.9 | NA | NA | 2 | 1 | 2 | 1 | 1 | 1 | 5 | 1 |
| 15_8009546 | 1 | 0 | 5 | 0.4 | 2 | 1 | 1 | 1 | 9 | 0.5556 | 1 | 0 | 3 | 0.3333 | 2 | 1 | 1 | 0 | 7 | 0.4286 |
| 15_8009557 | 1 | 0 | 4 | 0.25 | 1 | 0 | 1 | 0 | 7 | 0.1429 | NA | NA | 2 | 1 | 2 | 0.5 | NA | NA | 4 | 0.75 |
| 15_8009561 | NA | NA | 4 | 0 | 2 | 1 | 1 | 1 | 7 | 0.4286 | NA | NA | 3 | 1 | 2 | 0.5 | 1 | 0 | 6 | 0.6667 |
| 15_8009601 | 2 | 0.5 | 5 | 0.6 | 3 | 1 | 3 | 1 | 13 | 0.7692 | 2 | 1 | 4 | 0.75 | 2 | 0 | 2 | 0.5 | 10 | 0.6 |
| 15_8009617 | 1 | 0 | 5 | 0.2 | 2 | 0 | 3 | 0 | 11 | 0.0909 | 1 | 1 | 5 | 0.2 | 2 | 0 | 1 | 0 | 9 | 0.2222 |
| 15_8009619 | 2 | 1 | 6 | 0.5 | 2 | 0.5 | 4 | 0.75 | 14 | 0.6429 | 2 | 1 | 4 | 0.25 | 3 | 0.3333 | 1 | 1 | 10 | 0.5 |
| 15_8009628 | 2 | 1 | 6 | 1 | 4 | 1 | 3 | 1 | 15 | 1 | 2 | 1 | 5 | 1 | 4 | 1 | 2 | 1 | 13 | 1 |
| 15_8009637 | 2 | 0 | 7 | 0.4286 | 4 | 0.5 | 3 | 0.3333 | 16 | 0.375 | 3 | 0.3333 | 5 | 0.4 | 3 | 0.3333 | 4 | 0.25 | 15 | 0.3333 |
| 15_8009647 | 3 | 0.3333 | 6 | 0.5 | 4 | 0 | 3 | 0.6667 | 16 | 0.375 | 3 | 1 | 4 | 0.5 | 5 | 0.4 | 4 | 0.5 | 16 | 0.5625 |
| 15_8009652 | 2 | 0.5 | 5 | 0.4 | 2 | 0.5 | 4 | 0 | 13 | 0.3077 | 2 | 0.5 | 5 | 0.8 | 4 | 0.75 | 5 | 0.2 | 16 | 0.5625 |
| 15_8009652 | 3 | 1 | 7 | 1 | 4 | 1 | 3 | 1 | 17 | 1 | 3 | 1 | 5 | 1 | 6 | 1 | 5 | 1 | 19 | 1 |
| 15_8009653 | 3 | 1 | 7 | 1 | 5 | 1 | 4 | 1 | 19 | 1 | 3 | 0.6667 | 7 | 1 | 5 | 0.8 | 3 | 1 | 18 | 0.8889 |
| 15_8009661 | 3 | 1 | 5 | 0.8 | 4 | 1 | 5 | 0.8 | 17 | 0.8824 | 1 | 0 | 6 | 0.6667 | 4 | 0.5 | 5 | 0.8 | 16 | 0.625 |
| 15_8009665 | 3 | 1 | 7 | 1 | 4 | 1 | 3 | 1 | 17 | 1 | 2 | 1 | 4 | 1 | 5 | 1 | 3 | 1 | 14 | 1 |
| 15_8009699 | 4 | 0.5 | 6 | 0.3333 | 7 | 0.4286 | 7 | 0.5714 | 24 | 0.4583 | 3 | 0 | 7 | 0.1429 | 6 | 0.1667 | 6 | 0.1667 | 22 | 0.1364 |
| 15_8009707 | 5 | 0.2 | 8 | 0.25 | 8 | 0.25 | 7 | 0.2857 | 28 | 0.25 | 3 | 0 | 7 | 0.2857 | 9 | 0.3333 | 5 | 0.6 | 24 | 0.3333 |
| 15_8009713 | 6 | 0.8333 | 8 | 0.5 | 9 | 0.4444 | 5 | 0.6 | 28 | 0.5714 | 3 | 1 | 6 | 0.6667 | 5 | 1 | 6 | 0.6667 | 20 | 0.8 |
| 15_8009724 | 5 | 0.2 | 10 | 0.1 | 8 | 0 | 6 | 0.1667 | 29 | 0.1034 | 4 | 0 | 6 | 0.1667 | 7 | 0.1429 | 5 | 0.6 | 22 | 0.2273 |
| 15_8009751 | 6 | 0.1667 | 11 | 0 | 11 | 0 | 10 | 0.2 | 38 | 0.0789 | 4 | 0 | 4 | 0 | 9 | 0.2222 | 7 | 0 | 24 | 0.0833 |
| 15_8009913 | 8 | 0 | 20 | 0.1 | 15 | 0.2 | 12 | 0.0833 | 55 | 0.1091 | 8 | 0.125 | 10 | 0 | 13 | 0.2308 | 11 | 0 | 42 | 0.0952 |
| 15_8009955 | 11 | 0.1818 | 23 | 0.087 | 15 | 0.1333 | 16 | 0.0625 | 65 | 0.1077 | 10 | 0.1 | 13 | 0.2308 | 14 | 0.0714 | 14 | 0.0714 | 51 | 0.1176 |
| 15_8013369 | 1 | 1 | NA | NA | 2 | 0.5 | 1 | 1 | 4 | 0.75 | NA | NA | 1 | 1 | 2 | 1 | 2 | 1 | 5 | 1 |
| 15_8013372 | 1 | 1 | NA | NA | 1 | 1 | 1 | 1 | 3 | 1 | NA | NA | 1 | 1 | 1 | 1 | 2 | 1 | 4 | 1 |
| 15_8013379 | 1 | 1 | NA | NA | 1 | 1 | 1 | 1 | 3 | 1 | NA | NA | 1 | 1 | 2 | 0.5 | 2 | 0.5 | 5 | 0.6 |
| 15_8013385 | 1 | 1 | 1 | 0 | 2 | 0.5 | 1 | 1 | 5 | 0.6 | NA | NA | NA | NA | 2 | 1 | 2 | 1 | 4 | 1 |
| 15_8013387 | 1 | 0 | 1 | 1 | 2 | 0.5 | 1 | 0 | 5 | 0.4 | NA | NA | 1 | 1 | 2 | 0.5 | 2 | 0.5 | 5 | 0.6 |
| 15_8013390 | 1 | 1 | 1 | 0 | 2 | 0 | NA | NA | 4 | 0.25 | NA | NA | NA | NA | 2 | 0.5 | 2 | 0 | 4 | 0.25 |
| 15_8013401 | 1 | 1 | 1 | 1 | 2 | 1 | 1 | 0 | 5 | 0.8 | NA | NA | 1 | 1 | 2 | 0.5 | 3 | 0.3333 | 6 | 0.5 |
| 15_8013413 | 1 | 0 | 1 | 0 | 2 | 0.5 | 1 | 0 | 5 | 0.2 | NA | NA | 1 | 1 | 2 | 0 | 3 | 0.3333 | 6 | 0.3333 |
| 15_8013418 | 1 | 1 | 1 | 1 | 2 | 1 | 1 | 1 | 5 | 1 | NA | NA | 1 | 1 | 2 | 0.5 | 3 | 1 | 6 | 0.8333 |
| 15_8013423 | 1 | 1 | 1 | 0 | 2 | 0 | 1 | 0 | 5 | 0.2 | NA | NA | 1 | 0 | 2 | 0 | 2 | 0.5 | 5 | 0.2 |
| 15_8013424 | 1 | 0 | 1 | 0 | 2 | 0.5 | 1 | 1 | 5 | 0.4 | NA | NA | 1 | 1 | 2 | 0 | 2 | 0 | 5 | 0.2 |
| 15_8013436 | 1 | 0 | NA | NA | 2 | 0.5 | 1 | 0 | 4 | 0.25 | NA | NA | 1 | 0 | 2 | 0 | 1 | 1 | 4 | 0.25 |
| 15_8013484 | 1 | 0 | NA | NA | 2 | 0 | 1 | 1 | 4 | 0.25 | NA | NA | 2 | 0.5 | 2 | 0 | 3 | 0 | 7 | 0.1429 |
| 15_8013641 | 1 | 1 | NA | NA | 2 | 0.5 | 1 | 1 | 4 | 0.75 | 1 | 1 | 2 | 0.5 | 3 | 0.6667 | 1 | 1 | 7 | 0.7143 |
| 15_8013649 | 2 | 0 | 1 | 0 | 2 | 0.5 | 2 | 0.5 | 7 | 0.2857 | 1 | 1 | 3 | 0 | 2 | 0.5 | 2 | 0.5 | 8 | 0.375 |
| 15_8013650 | 2 | 0 | 1 | 0 | 2 | 0.5 | 2 | 0 | 7 | 0.1429 | 1 | 0 | 3 | 0.3333 | 3 | 0 | 3 | 0 | 10 | 0.1 |
| 15_8013751 | 4 | 0.25 | 2 | 0 | 2 | 0.5 | 3 | 0.3333 | 11 | 0.2727 | 3 | 0 | 4 | 0.75 | 6 | 0 | 1 | 0 | 14 | 0.2143 |
| 15_8014099 | 20 | 0.2 | 30 | 0.4 | 13 | 0.4615 | 10 | 0.4 | 73 | 0.3562 | 17 | 0.5294 | 17 | 0.2941 | 17 | 0.3529 | 17 | 0.4118 | 68 | 0.3971 |
| 15_8014497 | 7 | 0.5714 | 7 | 1 | 5 | 0.8 | 7 | 0.8571 | 26 | 0.8077 | 5 | 0.8 | 6 | 1 | 10 | 1 | 5 | 1 | 26 | 0.9615 |
| 15_8014502 | 3 | 0.3333 | 6 | 0.1667 | 5 | 0 | 6 | 0 | 20 | 0.1 | 5 | 0 | 7 | 0 | 8 | 0.125 | 4 | 0.25 | 24 | 0.0833 |
| 15_8014503 | 6 | 1 | 9 | 0.7778 | 6 | 0.8333 | 7 | 1 | 28 | 0.8929 | 5 | 1 | 7 | 1 | 10 | 0.9 | 4 | 1 | 26 | 0.9615 |
| 15_8014507 | 5 | 0.2 | 6 | 0.1667 | 3 | 0 | 8 | 0.25 | 22 | 0.1818 | 3 | 0 | 5 | 0 | 6 | 0.3333 | 4 | 0.25 | 18 | 0.1667 |

| | | | | | | | | | | | | | | | | | | | | |
|---|---|---|---|---|---|---|---|---|---|---|---|---|---|---|---|---|---|---|---|---|
| 15_801451 | 5 | 0.6 | 8 | 0.875 | 7 | 0.8571 | 9 | 0.7778 | 29 | 0.7931 | 4 | 1 | 4 | 0.75 | 8 | 0.75 | 4 | 1 | 20 | 0.85 |
| 15_801451 | 7 | 0.4286 | 8 | 0.125 | 6 | 0.6667 | 9 | 0.8889 | 30 | 0.5333 | 4 | 0.5 | 7 | 0.5714 | 9 | 0.4444 | 4 | 0.5 | 24 | 0.5 |
| 15_802792 | NA | NA | 2 | 0.5 | NA | NA | 1 | 1 | 3 | 0.6667 | 1 | 1 | 2 | 1 | 1 | 1 | 2 | 1 | 6 | 1 |
| 15_802795 | NA | NA | 2 | 0 | NA | NA | 2 | 0.5 | 4 | 0.25 | 1 | 0 | 3 | 0.6667 | 1 | 1 | 1 | 1 | 6 | 0.6667 |
| 15_802797 | NA | NA | 3 | 0.6667 | NA | NA | 3 | 1 | 6 | 0.8333 | 1 | 1 | 2 | 1 | 1 | 1 | 2 | 1 | 6 | 1 |
| 15_802803 | 1 | 0 | 5 | 0.4 | 1 | 0 | 4 | 0 | 11 | 0.1818 | 1 | 0 | 2 | 0 | 1 | 0 | 4 | 0.25 | 8 | 0.125 |
| 15_802807 | 3 | 0 | 6 | 0.3333 | 3 | 0.3333 | 6 | 0.3333 | 18 | 0.2778 | 4 | 0.25 | 6 | 0.1667 | 5 | 0 | 5 | 0 | 20 | 0.1 |
| 15_802809 | 4 | 0.25 | 6 | 0.3333 | 4 | 0.5 | 7 | 0.5714 | 21 | 0.4286 | 3 | 1 | 6 | 0.3333 | 6 | 0.5 | 5 | 0.4 | 20 | 0.5 |
| 15_802819 | 9 | 0.1111 | 11 | 0.3636 | 6 | 0.1667 | 8 | 0.25 | 34 | 0.2353 | 3 | 0 | 8 | 0.125 | 8 | 0.125 | 8 | 0.125 | 27 | 0.1111 |
| 15_802822 | 6 | 0 | 13 | 0.1538 | 8 | 0.125 | 8 | 0 | 35 | 0.0857 | 2 | 0 | 6 | 0.1667 | 5 | 0 | 7 | 0.2857 | 20 | 0.15 |
| 15_802823 | 13 | 0.0769 | 14 | 0.3571 | 10 | 0.2 | 11 | 0.1818 | 48 | 0.2083 | 4 | 0 | 11 | 0.0909 | 6 | 0.1667 | 8 | 0.5 | 29 | 0.2069 |
| 15_802823 | 10 | 0.2 | 13 | 0.4615 | 10 | 0.4 | 10 | 0.6 | 43 | 0.4186 | 4 | 0 | 9 | 0.3333 | 8 | 0.125 | 8 | 0.25 | 29 | 0.2069 |
| 15_808240 | 3 | 0.3333 | 3 | 0 | 3 | 0.3333 | 4 | 0 | 13 | 0.1538 | 2 | 0 | 6 | 0.3333 | 1 | 1 | 2 | 0 | 11 | 0.2727 |
| 15_808244 | 2 | 0.5 | 6 | 0.3333 | 4 | 0.5 | 5 | 0.4 | 17 | 0.4118 | 2 | 0.5 | 8 | 0.375 | 3 | 0 | 2 | 0 | 15 | 0.2667 |
| 15_808247 | 4 | 0 | 9 | 0.1111 | 7 | 0 | 6 | 0.3333 | 26 | 0.1154 | 5 | 0.2 | 9 | 0 | 3 | 0 | 5 | 0.2 | 22 | 0.0909 |
| 15_808531 | 2 | 0 | 3 | 0.6667 | 6 | 0 | 2 | 0.5 | 13 | 0.2308 | 4 | 0.25 | 7 | 0.4286 | 4 | 0 | 3 | 0.3333 | 18 | 0.2778 |
| 15_808922 | 7 | 0.1429 | 4 | 0 | 3 | 0 | 4 | 0 | 18 | 0.0556 | 3 | 0 | 6 | 0 | 4 | 0 | 3 | 0.3333 | 16 | 0.0625 |
| 15_808939 | 8 | 0.125 | 6 | 0 | 4 | 0 | 4 | 0.5 | 22 | 0.1364 | 4 | 0.25 | 7 | 0.1429 | 4 | 0.25 | 5 | 0 | 20 | 0.15 |
| 15_808959 | 8 | 0.125 | 7 | 0.1429 | 4 | 0.75 | 7 | 0.4286 | 26 | 0.3077 | 4 | 0.5 | 9 | 0.2222 | 7 | 0 | 4 | 0 | 24 | 0.1667 |
| 15_808960 | 10 | 0.2 | 8 | 0.125 | 6 | 0.1667 | 11 | 0.1818 | 35 | 0.1714 | 6 | 0.1667 | 9 | 0.1111 | 10 | 0.2 | 5 | 0.6 | 30 | 0.2333 |
| 15_808962 | 6 | 0.1667 | 8 | 0 | 5 | 0.4 | 7 | 0.4286 | 26 | 0.2308 | 4 | 0.25 | 5 | 0.2 | 8 | 0.25 | 4 | 0.25 | 21 | 0.2381 |
| 15_808962 | 7 | 0 | 6 | 0 | 6 | 0.1667 | 11 | 0.0909 | 30 | 0.0667 | 6 | 0.1667 | 9 | 0.3333 | 8 | 0 | 5 | 0 | 28 | 0.1429 |
| 15_808965 | 11 | 0.1818 | 15 | 0.0667 | 11 | 0.1818 | 16 | 0 | 53 | 0.0943 | 8 | 0.125 | 13 | 0.1538 | 14 | 0.0714 | 14 | 0.2143 | 49 | 0.1429 |
| 15_808974 | 11 | 0.2727 | 7 | 0.2857 | 6 | 0.3333 | 11 | 0.3636 | 35 | 0.3143 | 4 | 0.25 | 10 | 0.6 | 9 | 0.3333 | 8 | 0.625 | 31 | 0.4839 |
| 15_808974 | 10 | 0.1 | 6 | 0.3333 | 4 | 0 | 11 | 0.1818 | 31 | 0.1613 | 4 | 0.25 | 9 | 0 | 11 | 0.0909 | 8 | 0.125 | 32 | 0.0938 |
| 15_808974 | 10 | 0.1 | 7 | 0.1429 | 5 | 0 | 11 | 0.0909 | 33 | 0.0909 | 5 | 0 | 10 | 0.1 | 12 | 0.25 | 7 | 0 | 34 | 0.1176 |
| 15_808975 | 10 | 0.5 | 8 | 0.5 | 4 | 0.25 | 11 | 0.4545 | 33 | 0.4545 | 5 | 0.6 | 9 | 0.6667 | 11 | 0.5455 | 7 | 0.8571 | 32 | 0.6562 |
| 15_808975 | 9 | 0.8889 | 7 | 0.7143 | 5 | 1 | 9 | 0.8889 | 30 | 0.8667 | 5 | 1 | 9 | 0.7778 | 9 | 0.8889 | 6 | 0.8333 | 29 | 0.8621 |
| 15_808976 | 8 | 0.875 | 5 | 1 | 5 | 1 | 12 | 1 | 30 | 0.9667 | 5 | 1 | 9 | 0.8889 | 10 | 0.8 | 6 | 1 | 30 | 0.9 |
| 15_808976 | 9 | 0.7778 | 2 | 1 | 3 | 0.6667 | 9 | 0.4444 | 23 | 0.6522 | 4 | 0.75 | 7 | 0.8571 | 9 | 0.8889 | 4 | 1 | 24 | 0.875 |
| 15_808976 | 10 | 0.6 | 7 | 0.4286 | 4 | 0.75 | 10 | 0.4 | 31 | 0.5161 | 4 | 0.5 | 7 | 0.4286 | 9 | 0.5556 | 2 | 0.5 | 22 | 0.5 |
| 15_808976 | 9 | 1 | 6 | 1 | 4 | 1 | 12 | 1 | 31 | 1 | 5 | 0.8 | 8 | 0.875 | 10 | 1 | 5 | 1 | 28 | 0.9286 |
| 15_808977 | 9 | 0.2222 | 3 | 0 | 3 | 0 | 6 | 0.1667 | 21 | 0.1429 | 3 | 0.3333 | 8 | 0.25 | 8 | 0.125 | 5 | 0 | 24 | 0.1667 |
| 15_808977 | 8 | 0.25 | 6 | 0.3333 | 5 | 0.4 | 12 | 0.1667 | 31 | 0.2581 | 5 | 0.4 | 7 | 0.2857 | 11 | 0.1818 | 5 | 0.2 | 28 | 0.25 |
| 15_808978 | 10 | 0.1 | 6 | 0 | 3 | 0.3333 | 10 | 0 | 29 | 0.069 | 5 | 0.2 | 8 | 0.125 | 8 | 0.125 | 6 | 0.1667 | 27 | 0.1481 |
| 15_808978 | 9 | 0 | 7 | 0.4286 | 4 | 0 | 11 | 0.2727 | 31 | 0.1935 | 5 | 0.2 | 7 | 0.1429 | 8 | 0.375 | 5 | 0.4 | 25 | 0.28 |
| 15_809005 | 8 | 0.875 | 6 | 0.8333 | 4 | 0.75 | 10 | 1 | 28 | 0.8929 | 3 | 1 | 7 | 1 | 6 | 0.6667 | 5 | 0.6 | 21 | 0.8095 |
| 15_809005 | 6 | 0.3333 | 6 | 0 | 2 | 0 | 8 | 0.125 | 22 | 0.1364 | 3 | 0 | 6 | 0.1667 | 6 | 0.1667 | 6 | 0.3333 | 21 | 0.1905 |
| 15_809006 | 6 | 0.5 | 6 | 0 | 5 | 0.6 | 7 | 0.7143 | 24 | 0.4583 | 4 | 0 | 7 | 0.4286 | 6 | 0.3333 | 5 | 0.2 | 22 | 0.2727 |
| 15_809008 | 5 | 0.8 | 3 | 0.6667 | 3 | 0.6667 | 9 | 0.2222 | 20 | 0.5 | 4 | 0.25 | 6 | 0.5 | 6 | 0.5 | 3 | 0.3333 | 19 | 0.4211 |
| 15_809009 | 7 | 0.2857 | 4 | 0.25 | 5 | 0.2 | 10 | 0.3 | 26 | 0.2692 | 4 | 0.25 | 7 | 0.1429 | 6 | 0.5 | 5 | 0 | 22 | 0.2273 |
| 15_809009 | 2 | 1 | 1 | 0 | 2 | 0 | 7 | 0.1429 | 12 | 0.25 | 2 | 0.5 | 4 | 0 | 2 | 0 | 3 | 0.6667 | 11 | 0.2727 |
| 15_809009 | 6 | 1 | 4 | 1 | 5 | 1 | 9 | 0.5556 | 24 | 0.8333 | 4 | 1 | 8 | 0.75 | 5 | 0.6 | 5 | 0.6 | 22 | 0.7273 |
| 15_809025 | 5 | 0.2 | 3 | 0 | 3 | 0 | 9 | 0 | 20 | 0.05 | 2 | 0 | 4 | 0 | 3 | 0.3333 | 2 | 0 | 11 | 0.0909 |
| 15_809891 | 2 | 0.5 | NA | NA | 3 | 0.3333 | 5 | 0.4 | 10 | 0.4 | NA | NA | 1 | 1 | 1 | 0 | 1 | 1 | 3 | 0.6667 |
| 15_811214 | 5 | 0 | 3 | 0 | 4 | 0.25 | 3 | 0 | 15 | 0.0667 | 4 | 0 | 3 | 0 | 6 | 0.1667 | 3 | 0.3333 | 16 | 0.125 |
| 15_811216 | 7 | 0.5714 | 3 | 0.3333 | 3 | 1 | 4 | 1 | 17 | 0.7059 | 4 | 0.75 | 3 | 0.6667 | 6 | 0.5 | 2 | 0 | 15 | 0.5333 |
| 15_811220 | 7 | 0.5714 | 5 | 0.8 | 4 | 0.75 | 8 | 1 | 24 | 0.7917 | 2 | 1 | 4 | 1 | 8 | 0.875 | 3 | 1 | 17 | 0.9412 |
| 15_811222 | 8 | 0.875 | 5 | 0.6 | 6 | 0.8333 | 12 | 0.75 | 31 | 0.7742 | 6 | 0.8333 | 7 | 0.8571 | 8 | 0.875 | 3 | 0.6667 | 24 | 0.8333 |
| 15_811222 | 7 | 1 | 4 | 1 | 6 | 1 | 11 | 0.9091 | 28 | 0.9643 | 4 | 0.75 | 4 | 1 | 5 | 1 | 4 | 1 | 17 | 0.9412 |
| 15_811223 | 9 | 0.4444 | 5 | 0.4 | 5 | 0.2 | 10 | 0.1 | 29 | 0.2759 | 6 | 0.1667 | 5 | 0.2 | 8 | 0.25 | 4 | 0 | 23 | 0.1739 |
| 15_811226 | 10 | 0.1 | 6 | 0.3333 | 5 | 0.2 | 13 | 0.2308 | 34 | 0.2059 | 5 | 0.4 | 8 | 0.125 | 9 | 0.2222 | 5 | 0.2 | 27 | 0.2222 |
| 15_811228 | 8 | 0.25 | 3 | 0.6667 | 6 | 0.3333 | 11 | 0.4545 | 28 | 0.3929 | 4 | 0.25 | 5 | 0.6 | 5 | 0.4 | 4 | 0.75 | 18 | 0.5 |
| 15_812035 | 12 | 0 | 6 | 0.1667 | 2 | 0 | 7 | 0.2857 | 27 | 0.1111 | 7 | 0 | 8 | 0.125 | 7 | 0.1429 | 8 | 0 | 30 | 0.0667 |
| 15_812037 | 10 | 0.1 | 6 | 0.1667 | 3 | 0.3333 | 5 | 0.4 | 24 | 0.2083 | 3 | 0.3333 | 3 | 0.6667 | 7 | 0.1429 | 6 | 0.3333 | 19 | 0.3158 |
| 15_812038 | 11 | 0.3636 | 5 | 0 | 3 | 0.3333 | 5 | 0.2 | 24 | 0.25 | 4 | 0.75 | 5 | 0.2 | 8 | 0.25 | 8 | 0.25 | 25 | 0.32 |
| 15_812038 | 12 | 0.1667 | 6 | 0 | 2 | 0 | 6 | 0.3333 | 26 | 0.1538 | 6 | 0 | 5 | 0.4 | 8 | 0 | 7 | 0.4286 | 26 | 0.1923 |
| 15_812479 | 25 | 0.04 | 23 | 0.1739 | 27 | 0.037 | 23 | 0.1739 | 98 | 0.102 | 16 | 0.0625 | 21 | 0 | 28 | 0.1429 | 24 | 0.2083 | 89 | 0.1124 |
| 15_812482 | 30 | 0.1333 | 21 | 0.0476 | 26 | 0.0769 | 24 | 0.125 | 101 | 0.099 | 18 | 0 | 19 | 0.2105 | 26 | 0.1154 | 19 | 0.0526 | 82 | 0.0976 |
| 15_812492 | 16 | 0.125 | 16 | 0.1875 | 16 | 0.3125 | 18 | 0.2222 | 66 | 0.2121 | 8 | 0 | 15 | 0.0667 | 17 | 0.0588 | 13 | 0.0769 | 53 | 0.0566 |
| 15_812494 | 18 | 0.1667 | 14 | 0.0714 | 20 | 0.05 | 19 | 0 | 71 | 0.0704 | 9 | 0 | 18 | 0.0556 | 25 | 0.12 | 16 | 0.125 | 68 | 0.0882 |
| 15_812494 | 18 | 0.2222 | 18 | 0.0556 | 26 | 0.0769 | 28 | 0.2143 | 90 | 0.1444 | 13 | 0.1538 | 20 | 0.25 | 26 | 0.0769 | 19 | 0.2105 | 78 | 0.1667 |
| 15_812496 | 15 | 0.2667 | 14 | 0 | 24 | 0.1667 | 29 | 0.0345 | 82 | 0.1098 | 10 | 0.1 | 21 | 0.0476 | 25 | 0.2 | 19 | 0.2105 | 75 | 0.1467 |
| 15_813372 | NA | NA | NA | NA | NA | NA | 2 | 0.5 | 2 | 0.5 | NA | NA | 1 | 1 | NA | NA | NA | NA | 1 | 1 |
| 15_813373 | NA | NA | NA | NA | NA | NA | 2 | 0.5 | 2 | 0.5 | NA | NA | 1 | 1 | NA | NA | NA | NA | 1 | 1 |

| | | | | | | | | | | | | | | | | | | | | |
|---|---|---|---|---|---|---|---|---|---|---|---|---|---|---|---|---|---|---|---|---|
| 15_8135976 | 5 | 0 | 12 | 0.0833 | 14 | 0.1429 | 9 | 0.3333 | 40 | 0.15 | 10 | 0.4 | 9 | 0.4444 | 14 | 0.1429 | 13 | 0.4615 | 46 | 0.3478 |
| 15_8135982 | 5 | 0.2 | 14 | 0.2857 | 13 | 0.1538 | 10 | 0.7 | 42 | 0.3333 | 8 | 0.5 | 10 | 0.3 | 16 | 0.25 | 13 | 0.5385 | 47 | 0.383 |
| 15_8135993 | 4 | 0.25 | 13 | 0.2308 | 11 | 0 | 5 | 0.6 | 33 | 0.2121 | 6 | 0.1667 | 7 | 0.2857 | 12 | 0.5833 | 7 | 0 | 32 | 0.3125 |
| 15_8146999 | NA | NA | NA | NA | 1 | 1 | 1 | 1 | 2 | 1 | NA | NA | 1 | 1 | NA | NA | NA | NA | 1 | 1 |
| 15_8148556 | 1 | 0 | NA | NA | 1 | 1 | 1 | 1 | 3 | 0.6667 | NA | NA | 1 | 1 | NA | NA | NA | NA | 1 | 1 |
| 15_8153266 | 2 | 0 | 1 | 0 | 3 | 0 | 2 | 1 | 8 | 0.25 | 3 | 0.3333 | 1 | 0 | 6 | 0 | 2 | 0.5 | 12 | 0.1667 |
| 15_8153333 | NA | NA | 1 | 0 | 3 | 0.3333 | 2 | 0 | 6 | 0.1667 | 2 | 0 | 1 | 0 | 2 | 0 | 3 | 0.3333 | 8 | 0.125 |
| 15_8153345 | 2 | 1 | 1 | 1 | 5 | 1 | 1 | 1 | 9 | 1 | 1 | 1 | 2 | 1 | 8 | 0.75 | 4 | 1 | 15 | 0.8667 |
| 15_8153354 | 2 | 0.5 | 1 | 1 | 2 | 0.5 | 2 | 0 | 7 | 0.4286 | 3 | 0.3333 | 2 | 0.5 | 7 | 0.1429 | 3 | 0.3333 | 15 | 0.2667 |
| 15_8153361 | 3 | 1 | NA | NA | 5 | 1 | 2 | 1 | 10 | 1 | 3 | 1 | 1 | 1 | 8 | 0.875 | 3 | 0.6667 | 15 | 0.8667 |
| 15_8153362 | 1 | 0 | 1 | 0 | 6 | 0.1667 | 2 | 0.5 | 10 | 0.2 | 2 | 0 | 2 | 0.5 | 7 | 0.1429 | 2 | 0.5 | 13 | 0.2308 |
| 15_8153368 | 3 | 1 | 1 | 0 | 4 | 0.75 | 1 | 1 | 9 | 0.7778 | 1 | 1 | 2 | 1 | 8 | 0.5 | 4 | 1 | 15 | 0.7333 |
| 15_8153375 | 3 | 0.6667 | 2 | 0.5 | 6 | 0.6667 | 1 | 0 | 12 | 0.5833 | 3 | 0.3333 | 2 | 0 | 9 | 0.3333 | 4 | 0.5 | 18 | 0.3333 |
| 15_8153385 | 2 | 0 | 2 | 0 | 5 | 0 | 2 | 1 | 11 | 0.1818 | 2 | 0 | 2 | 0.5 | 6 | 0 | 4 | 0 | 14 | 0.0714 |
| 15_8153410 | 2 | 0 | 2 | 0.5 | 7 | 0 | 4 | 0 | 15 | 0.0667 | 2 | 0 | 2 | 0 | 5 | 0 | 4 | 0.25 | 13 | 0.0769 |
| 15_8153411 | 3 | 1 | 2 | 1 | 6 | 0.8333 | 3 | 1 | 14 | 0.9286 | 3 | 1 | 2 | 1 | 9 | 0.8889 | 4 | 1 | 18 | 0.9444 |
| 15_8153425 | 2 | 0 | 1 | 0 | 8 | 0.5 | 4 | 0 | 15 | 0.2667 | 3 | 0 | 2 | 0 | 8 | 0.5 | 4 | 0.25 | 17 | 0.2941 |
| 15_8153427 | 2 | 0.5 | NA | NA | 8 | 0.25 | 3 | 0.3333 | 13 | 0.3077 | NA | NA | 1 | 0 | 9 | 0.3333 | 2 | 1 | 12 | 0.4167 |
| 15_8153434 | 2 | 1 | 2 | 0 | 7 | 0.1429 | 4 | 0.25 | 15 | 0.2667 | 3 | 0 | 2 | 0 | 7 | 0.2857 | 4 | 0 | 16 | 0.125 |
| 15_8153445 | 3 | 0.3333 | 2 | 1 | 5 | 0.4 | 4 | 0.5 | 14 | 0.5 | 3 | 0.6667 | 2 | 0.5 | 6 | 0.3333 | 3 | 0.6667 | 14 | 0.5 |
| 15_8153448 | 3 | 1 | 3 | 0 | 8 | 1 | 4 | 0.25 | 18 | 0.6667 | 3 | 0.6667 | 2 | 0.5 | 9 | 0.6667 | 4 | 0.75 | 18 | 0.6667 |
| 15_8153465 | 2 | 1 | 3 | 1 | 6 | 1 | 6 | 1 | 17 | 1 | 2 | 1 | 1 | 1 | 8 | 1 | 4 | 1 | 15 | 1 |
| 15_8153496 | 3 | 0 | 3 | 0.3333 | 9 | 0.2222 | 5 | 0.2 | 20 | 0.2 | 3 | 0 | 2 | 0 | 8 | 0.375 | 5 | 0.2 | 18 | 0.2222 |
| 15_8153498 | 3 | 0.6667 | 3 | 1 | 3 | 1 | 3 | 1 | 12 | 0.9167 | 3 | 1 | 1 | 1 | 7 | 0.8571 | 4 | 1 | 15 | 0.9333 |
| 15_8153501 | 2 | 0.5 | 2 | 0 | 5 | 0 | 4 | 0 | 13 | 0.0769 | 2 | 0 | 1 | 0 | 5 | 0.2 | 3 | 0.3333 | 11 | 0.1818 |
| 15_8153511 | 4 | 0.5 | 4 | 0.75 | 10 | 0.9 | 6 | 0.6667 | 24 | 0.75 | 4 | 0 | 2 | 0.5 | 10 | 0.5 | 5 | 0.6 | 21 | 0.4286 |
| 15_8153512 | 3 | 1 | 4 | 1 | 6 | 1 | 6 | 1 | 19 | 1 | 4 | 1 | 2 | 1 | 10 | 1 | 5 | 1 | 21 | 1 |
| 15_8153514 | 4 | 0.75 | 3 | 0.6667 | 8 | 0.625 | 5 | 0.6 | 20 | 0.65 | 4 | 0.75 | 2 | 0.5 | 11 | 0.6364 | 5 | 0.6 | 22 | 0.6364 |
| 15_8153517 | 4 | 0 | 4 | 0.25 | 10 | 0.1 | 6 | 0 | 24 | 0.0833 | 3 | 0 | 2 | 0 | 10 | 0.1 | 5 | 0.2 | 20 | 0.1 |
| 15_8153518 | 3 | 0 | 4 | 0.5 | 10 | 0.4 | 6 | 0.5 | 23 | 0.3913 | 4 | 0.75 | 2 | 1 | 11 | 0.6364 | 4 | 0.75 | 21 | 0.7143 |
| 15_8153523 | 3 | 0.6667 | 4 | 0.5 | 9 | 0.6667 | 5 | 0.2 | 21 | 0.5238 | 3 | 0.6667 | 2 | 0.5 | 11 | 0.6364 | 4 | 1 | 20 | 0.7 |
| 15_8153528 | 5 | 0.4 | 4 | 0 | 10 | 0 | 7 | 0 | 26 | 0.0769 | 4 | 0 | 2 | 0 | 11 | 0.0909 | 5 | 0.4 | 22 | 0.1364 |
| 15_8153530 | 4 | 1 | 3 | 1 | 7 | 1 | 4 | 1 | 18 | 1 | 4 | 1 | 2 | 1 | 8 | 1 | 3 | 1 | 17 | 1 |
| 15_8153535 | 4 | 0 | 4 | 0 | 11 | 0.3636 | 7 | 0 | 26 | 0.1538 | 4 | 0 | 2 | 0 | 11 | 0.2727 | 4 | 0 | 21 | 0.1429 |
| 15_8153562 | 3 | 0 | 2 | 0 | 4 | 0.25 | 7 | 0 | 16 | 0.0625 | 5 | 0 | NA | NA | 10 | 0.1 | 3 | 0 | 18 | 0.0556 |
| 15_8153589 | 8 | 0.25 | 4 | 0.25 | 10 | 0.1 | 8 | 0 | 30 | 0.1333 | 9 | 0.1111 | 3 | 0.3333 | 10 | 0.1 | 6 | 0.1667 | 28 | 0.1429 |
| 15_8153603 | 6 | 0 | 5 | 0.2 | 10 | 0 | 9 | 0.1111 | 30 | 0.0667 | 8 | 0.25 | 3 | 0 | 11 | 0.3636 | 6 | 0.1667 | 28 | 0.25 |
| 15_8156904 | 6 | 0 | 2 | 0 | 2 | 0 | 6 | 0.1667 | 16 | 0.0625 | 1 | 0 | 4 | 0.25 | 2 | 0 | 4 | 0 | 11 | 0.0909 |
| 15_8156907 | 6 | 0.1667 | 2 | 0 | 2 | 0 | 7 | 0.2857 | 17 | 0.1765 | 1 | 0 | 4 | 0.25 | 3 | 0 | 3 | 0 | 11 | 0.0909 |
| 15_8157125 | 6 | 0 | 2 | 0 | 3 | 0 | 8 | 0.25 | 19 | 0.1053 | 2 | 0 | 5 | 0 | 4 | 0.25 | 4 | 0 | 15 | 0.0667 |
| 15_8157127 | 5 | 0.2 | 3 | 0 | 3 | 0 | 9 | 0.1111 | 20 | 0.1 | 2 | 0 | 6 | 0.1667 | 4 | 0.25 | 4 | 0 | 16 | 0.125 |
| 15_8157128 | 4 | 0.25 | 1 | 0 | 3 | 0 | 9 | 0 | 17 | 0.0588 | NA | NA | 5 | 0 | 4 | 0.25 | 3 | 0 | 12 | 0.0833 |
| 15_8157134 | 5 | 0.4 | 3 | 0.3333 | 2 | 0.5 | 6 | 0.5 | 16 | 0.4375 | NA | NA | 6 | 0.6667 | 3 | 1 | 3 | 1 | 12 | 0.8333 |
| 15_8157140 | 6 | 0 | 3 | 0 | 3 | 0 | 8 | 0.125 | 20 | 0.05 | 1 | 0 | 5 | 0 | 4 | 0 | 4 | 0.25 | 14 | 0.0714 |
| 15_8157141 | 4 | 0.75 | 3 | 0.6667 | 2 | 1 | 8 | 0.75 | 17 | 0.7647 | 1 | 1 | 6 | 0.8333 | 4 | 1 | 3 | 0.6667 | 14 | 0.8571 |
| 15_8157164 | 5 | 0.4 | 2 | 0.5 | 2 | 0.5 | 8 | 0.125 | 17 | 0.2941 | 1 | 0 | 5 | 0.2 | 4 | 0 | 5 | 0.2 | 15 | 0.1333 |
| 15_8157177 | 4 | 0 | 3 | 0.3333 | 3 | 0 | 8 | 0.125 | 18 | 0.1111 | 2 | 0 | 3 | 0 | 4 | 0 | 4 | 0.25 | 13 | 0.0769 |
| 15_8157190 | 6 | 0.3333 | 3 | 0.3333 | 4 | 0.25 | 6 | 0.3333 | 19 | 0.3158 | NA | NA | 5 | 0 | 3 | 0.6667 | 3 | 0.3333 | 11 | 0.2727 |
| 15_8157212 | 7 | 0.8571 | 4 | 1 | 3 | 1 | 8 | 0.75 | 22 | 0.8636 | 1 | 1 | 7 | 0.7143 | 4 | 0.75 | 4 | 1 | 16 | 0.8125 |
| 15_8157224 | 8 | 0.375 | 3 | 0 | 2 | 0 | 9 | 0.4444 | 22 | 0.3182 | 2 | 0.5 | 6 | 0.5 | 4 | 0.5 | 4 | 0.75 | 16 | 0.5625 |
| 15_8157527 | NA | NA | 1 | 1 | NA | NA | NA | NA | 1 | 1 | 1 | 1 | 1 | 0 | NA | NA | 1 | 0 | 3 | 0.3333 |
| 15_8157803 | 3 | 0 | 2 | 0.5 | 1 | 0 | 2 | 0 | 8 | 0.125 | 4 | 0 | 1 | 0 | NA | NA | 1 | 1 | 6 | 0.1667 |
| 15_8157804 | 4 | 0.25 | 3 | 0.3333 | 2 | 0 | 2 | 0 | 11 | 0.1818 | 7 | 0 | 2 | 0 | 1 | 0 | 4 | 0.25 | 14 | 0.0714 |
| 15_8157821 | 4 | 0.5 | 3 | 1 | 2 | 1 | 2 | 0 | 11 | 0.6364 | 3 | 0.3333 | 2 | 0 | 1 | 1 | 4 | 0.25 | 10 | 0.3 |
| 15_8157833 | 3 | 0.6667 | 3 | 0.6667 | 2 | 1 | 1 | 1 | 9 | 0.7778 | 4 | 0.5 | 1 | 0 | 1 | 0 | 3 | 0.6667 | 9 | 0.4444 |
| 15_8157884 | 1 | 1 | 2 | 1 | 1 | 0 | NA | NA | 4 | 0.75 | 1 | 0 | NA | NA | 1 | 1 | 1 | 1 | 3 | 0.6667 |
| 15_8157915 | 1 | 0 | 3 | 0.3333 | NA | NA | NA | NA | 4 | 0.25 | 1 | 0 | NA | NA | 1 | 0 | 1 | 1 | 3 | 0.3333 |
| 15_8157934 | 2 | 0.5 | 3 | 1 | 2 | 0.5 | NA | NA | 7 | 0.7143 | 1 | 1 | NA | NA | 1 | 0 | 1 | 1 | 3 | 0.6667 |
| 15_8158932 | 26 | 0.5385 | 30 | 0.4667 | 19 | 0.4737 | 19 | 0.3158 | 94 | 0.4574 | 28 | 0.6071 | 18 | 0.5 | 28 | 0.5357 | 13 | 0.4615 | 87 | 0.5402 |
| 15_8158934 | 28 | 0.0357 | 28 | 0.0714 | 21 | 0 | 17 | 0.1176 | 94 | 0.0532 | 27 | 0.1481 | 19 | 0.2105 | 28 | 0.1429 | 12 | 0 | 86 | 0.1395 |
| 15_8167782 | 5 | 0 | 23 | 0.3043 | 9 | 0.3333 | 7 | 0.2857 | 44 | 0.2727 | 5 | 0 | 6 | 0.1667 | 9 | 0.3333 | 7 | 0.5714 | 27 | 0.2963 |
| 15_8167785 | 5 | 0 | 23 | 0.1304 | 9 | 0 | 7 | 0 | 44 | 0.0682 | 6 | 0 | 6 | 0 | 9 | 0.2222 | 7 | 0.1429 | 28 | 0.1071 |
| 15_8167941 | 10 | 0.1 | 25 | 0.04 | 14 | 0.0714 | 15 | 0.0667 | 64 | 0.0625 | 11 | 0 | 14 | 0.0714 | 17 | 0.0588 | 5 | 0.2 | 47 | 0.0638 |
| 15_8167986 | 14 | 0 | 23 | 0.1304 | 13 | 0.1538 | 14 | 0.2857 | 64 | 0.1406 | 13 | 0 | 12 | 0.1667 | 15 | 0.2 | 9 | 0.1111 | 49 | 0.1224 |
| 15_8169928 | 21 | 0.1905 | 18 | 0.1111 | 11 | 0.1818 | 15 | 0.2667 | 65 | 0.1846 | 19 | 0.1053 | 16 | 0.1875 | 20 | 0.2 | 14 | 0.2857 | 69 | 0.1884 |

| | | | | | | | | | | | | | | | | | | | | |
|---|---|---|---|---|---|---|---|---|---|---|---|---|---|---|---|---|---|---|---|---|
| 15_817000 | 1 | 0 | 2 | 0.5 | NA | NA | NA | NA | 3 | 0.3333 | 2 | 0.5 | 1 | 1 | 1 | 0 | 1 | 0 | 5 | 0.4 |
| 15_817078 | 32 | 0.125 | 19 | 0.1053 | 12 | 0.0833 | 19 | 0.2105 | 82 | 0.1341 | 16 | 0.125 | 17 | 0.0588 | 25 | 0.12 | 21 | 0.1429 | 79 | 0.1139 |
| 15_817079 | 27 | 0.4444 | 17 | 0.6471 | 12 | 0.5833 | 19 | 0.4211 | 75 | 0.5067 | 17 | 0.6471 | 16 | 0.5 | 26 | 0.6154 | 19 | 0.7368 | 78 | 0.6282 |
| 15_817079 | 31 | 0.6129 | 23 | 0.8261 | 16 | 0.6875 | 21 | 0.7619 | 91 | 0.7143 | 19 | 0.6842 | 25 | 0.64 | 33 | 0.7273 | 23 | 0.5652 | 100 | 0.66 |
| 15_817079 | 35 | 0.6571 | 20 | 0.8 | 15 | 0.7333 | 22 | 0.7727 | 92 | 0.7283 | 20 | 0.65 | 24 | 0.8333 | 27 | 0.7778 | 24 | 0.8333 | 95 | 0.7789 |
| 15_817079 | 33 | 1 | 21 | 1 | 14 | 1 | 22 | 0.9545 | 90 | 0.9889 | 23 | 1 | 25 | 0.92 | 33 | 0.9394 | 24 | 0.9583 | 105 | 0.9524 |
| 15_817080 | 23 | 0.0435 | 22 | 0.0455 | 9 | 0.1111 | 20 | 0.1 | 74 | 0.0676 | 16 | 0 | 19 | 0.1053 | 31 | 0.0645 | 18 | 0.0556 | 84 | 0.0595 |
| 15_817080 | 22 | 0.3182 | 19 | 0.3158 | 10 | 0.4 | 20 | 0.35 | 71 | 0.338 | 14 | 0.2857 | 20 | 0.2 | 28 | 0.3571 | 13 | 0.3846 | 75 | 0.3067 |
| 15_817080 | 32 | 0.2812 | 23 | 0.3478 | 13 | 0.8462 | 21 | 0.5238 | 89 | 0.4382 | 23 | 0.3478 | 18 | 0.3889 | 29 | 0.3103 | 23 | 0.3913 | 93 | 0.3548 |
| 15_817082 | 28 | 0.9286 | 24 | 1 | 16 | 1 | 21 | 0.9524 | 89 | 0.9663 | 25 | 1 | 25 | 1 | 29 | 0.9655 | 23 | 1 | 102 | 0.9902 |
| 15_817083 | 35 | 0 | 26 | 0.1154 | 16 | 0.125 | 23 | 0.1739 | 100 | 0.09 | 23 | 0.1739 | 27 | 0.1111 | 32 | 0.0938 | 21 | 0.0952 | 103 | 0.1165 |
| 15_817083 | 35 | 0.1429 | 24 | 0.125 | 18 | 0.1111 | 22 | 0.0909 | 99 | 0.1212 | 24 | 0.1667 | 29 | 0.069 | 29 | 0.069 | 24 | 0.125 | 106 | 0.1038 |
| 15_817084 | 25 | 0.04 | 18 | 0.0556 | 13 | 0.0769 | 15 | 0.0667 | 71 | 0.0563 | 20 | 0.05 | 16 | 0.125 | 25 | 0.08 | 14 | 0.0714 | 75 | 0.08 |
| 15_817085 | 35 | 0.3143 | 30 | 0.4 | 17 | 0.1765 | 28 | 0.1429 | 110 | 0.2727 | 26 | 0.3077 | 28 | 0.2143 | 31 | 0.1935 | 26 | 0.2692 | 111 | 0.2432 |
| 15_817090 | 40 | 0.075 | 37 | 0.0541 | 30 | 0.1333 | 34 | 0.1176 | 141 | 0.0922 | 30 | 0.1667 | 37 | 0.1351 | 45 | 0.1778 | 30 | 0.0667 | 142 | 0.1408 |
| 15_817091 | 41 | 0.2927 | 31 | 0.3226 | 27 | 0.4074 | 30 | 0.2667 | 129 | 0.3178 | 30 | 0.3 | 33 | 0.3333 | 41 | 0.2683 | 29 | 0.3103 | 133 | 0.3008 |
| 15_817093 | 47 | 0.1489 | 39 | 0.1282 | 36 | 0.1389 | 36 | 0.1944 | 158 | 0.1519 | 35 | 0.1143 | 41 | 0.1463 | 49 | 0.1429 | 33 | 0.1515 | 158 | 0.1392 |
| 15_817094 | 40 | 0.35 | 34 | 0.2647 | 27 | 0.4815 | 30 | 0.2333 | 131 | 0.3282 | 29 | 0.3793 | 41 | 0.3659 | 44 | 0.25 | 30 | 0.3333 | 144 | 0.3264 |
| 15_817095 | 32 | 0.0625 | 30 | 0.0333 | 28 | 0.2143 | 30 | 0.1667 | 120 | 0.1167 | 24 | 0.0417 | 28 | 0.1786 | 47 | 0.1277 | 25 | 0.12 | 124 | 0.121 |
| 15_817100 | 53 | 0.1132 | 53 | 0.1132 | 42 | 0.1667 | 43 | 0.1163 | 191 | 0.1257 | 41 | 0.0976 | 55 | 0.0182 | 63 | 0.2381 | 38 | 0.1053 | 197 | 0.1218 |
| 15_817103 | 60 | 0.0167 | 54 | 0.0741 | 43 | 0.093 | 49 | 0.102 | 206 | 0.068 | 41 | 0.0732 | 56 | 0.0714 | 67 | 0.0299 | 40 | 0.175 | 204 | 0.0784 |
| 15_817103 | 55 | 0.1091 | 48 | 0.1042 | 42 | 0.1667 | 43 | 0.093 | 188 | 0.117 | 39 | 0.1795 | 52 | 0.0577 | 61 | 0.1148 | 39 | 0.0769 | 191 | 0.1047 |
| 15_817103 | 62 | 0.1452 | 55 | 0.1273 | 44 | 0.1591 | 46 | 0.1522 | 207 | 0.1449 | 43 | 0.1628 | 59 | 0.1356 | 63 | 0.1587 | 41 | 0.1463 | 206 | 0.1505 |
| 15_817326 | 2 | 1 | 3 | 0 | 5 | 0 | 4 | 0 | 14 | 0.1429 | NA | NA | 4 | 0 | 5 | 0 | 3 | 0.3333 | 12 | 0.0833 |
| 15_817347 | 2 | 1 | 1 | 1 | 3 | 0.6667 | 3 | 1 | 9 | 0.8889 | NA | NA | 3 | 1 | 3 | 1 | 2 | 1 | 8 | 1 |
| 15_817348 | 3 | 1 | 1 | 1 | 2 | 1 | 2 | 1 | 8 | 1 | NA | NA | 1 | 1 | 3 | 1 | 1 | 1 | 5 | 1 |
| 15_817349 | 2 | 1 | NA | NA | 3 | 0.3333 | 3 | 0.3333 | 8 | 0.5 | NA | NA | 2 | 0.5 | 3 | 0.3333 | 2 | 0.5 | 7 | 0.4286 |
| 15_817352 | 2 | 0 | NA | NA | 1 | 1 | 3 | 1 | 6 | 0.6667 | NA | NA | 3 | 0.3333 | 3 | 0.6667 | 2 | 0.5 | 8 | 0.5 |
| 15_817352 | 3 | 0 | 1 | 1 | 3 | 0.6667 | 2 | 0.5 | 9 | 0.4444 | NA | NA | 2 | 0 | 3 | 0 | 1 | 1 | 6 | 0.1667 |
| 15_817355 | 3 | 1 | 1 | 1 | 3 | 1 | 3 | 1 | 10 | 1 | NA | NA | 3 | 1 | 3 | 0.6667 | 2 | 1 | 8 | 0.875 |
| 15_817355 | 3 | 0.3333 | NA | NA | 3 | 0.3333 | NA | NA | 6 | 0.3333 | NA | NA | 3 | 1 | 2 | 0.5 | 1 | 0 | 6 | 0.6667 |
| 15_817356 | 3 | 0.3333 | 1 | 1 | 3 | 0 | 2 | 0 | 9 | 0.2222 | NA | NA | 3 | 0.3333 | 2 | 0 | 1 | 1 | 6 | 0.3333 |
| 15_817358 | 3 | 0 | 1 | 0 | 3 | 0 | 2 | 0.5 | 9 | 0.1111 | NA | NA | 3 | 0.3333 | 3 | 0 | 1 | 0 | 7 | 0.1429 |
| 15_817496 | 6 | 0 | 11 | 0 | 8 | 0.125 | 8 | 0.125 | 33 | 0.0606 | 11 | 0 | 6 | 0 | 5 | 0 | 9 | 0.2222 | 31 | 0.0645 |
| 15_817496 | 5 | 0 | 9 | 0.2222 | 8 | 0 | 8 | 0.25 | 30 | 0.1333 | 7 | 0 | 3 | 0.3333 | 3 | 0 | 5 | 0.2 | 18 | 0.1111 |
| 15_817497 | 6 | 0.3333 | 12 | 0.4167 | 8 | 0.25 | 7 | 0.1429 | 33 | 0.303 | 10 | 0.6 | 6 | 0.5 | 5 | 0.4 | 9 | 0.3333 | 30 | 0.4667 |
| 15_817498 | 3 | 0 | 12 | 0.3333 | 8 | 0.125 | 7 | 0.1429 | 30 | 0.2 | 12 | 0.0833 | 4 | 0.5 | 5 | 0 | 8 | 0.375 | 29 | 0.2069 |
| 15_817499 | 6 | 0 | 11 | 0.1818 | 6 | 0 | 8 | 0.125 | 31 | 0.0968 | 8 | 0 | 3 | 0.3333 | 5 | 0 | 8 | 0.25 | 24 | 0.125 |
| 15_817989 | 65 | 0.0615 | 56 | 0.125 | 66 | 0.1515 | 69 | 0.1739 | 256 | 0.1289 | 58 | 0.1724 | 57 | 0.1404 | 48 | 0.1458 | 55 | 0.2 | 218 | 0.1651 |
| 15_817992 | 62 | 0.0645 | 64 | 0.1406 | 68 | 0.1176 | 67 | 0.1343 | 261 | 0.1149 | 66 | 0.1818 | 71 | 0.1268 | 53 | 0.1698 | 65 | 0.0923 | 255 | 0.1412 |
| 15_817996 | 2 | 0 | 2 | 0.5 | NA | NA | NA | NA | 4 | 0.25 | 1 | 0 | 1 | 0 | 2 | 0.5 | NA | NA | 4 | 0.25 |
| 15_818003 | 2 | 0 | 7 | 0 | 5 | 0 | 3 | 0.3333 | 17 | 0.0588 | 13 | 0.3077 | 5 | 0 | 6 | 0.1667 | 4 | 0 | 28 | 0.1786 |
| 15_818004 | NA | NA | 9 | 0.5556 | 5 | 0.8 | 3 | 1 | 17 | 0.7059 | 11 | 1 | 5 | 0.8 | 5 | 1 | 6 | 0.8333 | 27 | 0.9259 |
| 15_818005 | 2 | 0.5 | 10 | 0.3 | 6 | 0.1667 | 3 | 0.3333 | 21 | 0.2857 | 14 | 0.2857 | 6 | 0.1667 | 7 | 0.5714 | 7 | 0.2857 | 34 | 0.3235 |
| 15_818007 | 2 | 0 | 8 | 0.25 | 5 | 0.2 | 3 | 0 | 18 | 0.1667 | 13 | 0.1538 | 5 | 0 | 7 | 0.2857 | 3 | 0.3333 | 28 | 0.1786 |
| 15_818015 | 1 | 0 | 8 | 0 | 6 | 0.1667 | 2 | 0.5 | 17 | 0.1176 | 8 | 0.25 | 5 | 0 | 6 | 0.1667 | 3 | 0.3333 | 22 | 0.1818 |
| 15_818095 | 1 | 1 | 1 | 0 | NA | NA | NA | NA | 2 | 0.5 | 3 | 0.6667 | 1 | 0 | 2 | 0.5 | NA | NA | 6 | 0.5 |
| 15_818342 | 11 | 0.5455 | 31 | 0.4194 | 22 | 0.2727 | 20 | 0.55 | 84 | 0.4286 | 16 | 0.3125 | 8 | 0.625 | 19 | 0.4737 | 12 | 0.5833 | 55 | 0.4727 |
| 15_818348 | 19 | 0.2105 | 28 | 0.1429 | 23 | 0.2174 | 20 | 0.1 | 90 | 0.1667 | 15 | 0.2667 | 11 | 0.0909 | 17 | 0.1765 | 12 | 0.0833 | 55 | 0.1636 |
| 15_818353 | 39 | 0.4872 | 30 | 0.5 | 32 | 0.75 | 28 | 0.75 | 129 | 0.6124 | 30 | 0.6333 | 27 | 0.7407 | 36 | 0.4444 | 28 | 0.75 | 121 | 0.6281 |
| 15_818353 | 39 | 0.5128 | 29 | 0.2069 | 29 | 0.4483 | 21 | 0.381 | 118 | 0.3983 | 25 | 0.36 | 24 | 0.125 | 29 | 0.4138 | 26 | 0.3462 | 104 | 0.3173 |
| 15_818353 | 40 | 0.375 | 27 | 0.3704 | 27 | 0.4444 | 25 | 0.52 | 119 | 0.4202 | 26 | 0.4231 | 22 | 0.4091 | 31 | 0.5484 | 25 | 0.44 | 104 | 0.4615 |
| 15_818354 | 33 | 0.0909 | 25 | 0.04 | 26 | 0.1538 | 18 | 0.1667 | 102 | 0.1078 | 17 | 0.1765 | 17 | 0.1176 | 27 | 0.1481 | 24 | 0.1667 | 85 | 0.1529 |
| 15_818355 | 36 | 0.3333 | 27 | 0.2222 | 27 | 0.0741 | 23 | 0.2174 | 113 | 0.2212 | 23 | 0.3043 | 19 | 0.1053 | 23 | 0 | 28 | 0.1429 | 93 | 0.1398 |
| 15_818356 | 42 | 0.1429 | 28 | 0.1071 | 32 | 0.125 | 28 | 0.1429 | 130 | 0.1308 | 24 | 0.0833 | 22 | 0.2273 | 31 | 0.1613 | 28 | 0.2143 | 105 | 0.1714 |
| 15_818357 | 38 | 0.0789 | 28 | 0 | 29 | 0.1034 | 24 | 0.125 | 119 | 0.0756 | 23 | 0.087 | 22 | 0.0455 | 28 | 0.1071 | 27 | 0.1111 | 100 | 0.09 |
| 15_818771 | 7 | 0.1429 | 5 | 0 | 4 | 0 | 2 | 0 | 18 | 0.0556 | 4 | 0 | 1 | 0 | 4 | 0 | 3 | 0.3333 | 12 | 0.0833 |
| 15_818780 | 15 | 0.0667 | 11 | 0.0909 | 12 | 0 | 15 | 0.0667 | 53 | 0.0566 | 11 | 0 | 9 | 0.3333 | 10 | 0 | 8 | 0 | 38 | 0.0789 |
| 15_818783 | 14 | 0.1429 | 11 | 0.1818 | 16 | 0.1875 | 10 | 0.2 | 51 | 0.1765 | 13 | 0.1538 | 8 | 0.125 | 7 | 0.2857 | 8 | 0 | 36 | 0.1389 |
| 15_818785 | 13 | 0.8462 | 11 | 0.8182 | 11 | 0.8182 | 10 | 0.8 | 45 | 0.8222 | 11 | 1 | 8 | 0.875 | 5 | 0.8 | 10 | 0.8 | 34 | 0.8824 |
| 15_818786 | 14 | 0.1429 | 14 | 0.2857 | 15 | 0.2667 | 11 | 0.2727 | 54 | 0.2407 | 12 | 0.1667 | 9 | 0 | 8 | 0.125 | 10 | 0.2 | 39 | 0.1282 |
| 15_818788 | 15 | 0.5333 | 13 | 0.0769 | 17 | 0.4706 | 10 | 0.2 | 55 | 0.3455 | 13 | 0.3077 | 7 | 0.5714 | 9 | 0.3333 | 10 | 0.2 | 39 | 0.3333 |
| 15_818789 | 7 | 0 | 8 | 0.25 | 8 | 0 | 6 | 0 | 29 | 0.069 | 12 | 0.1667 | 4 | 0 | 6 | 0.1667 | 4 | 0 | 26 | 0.1154 |
| 15_818789 | 11 | 0.5455 | 13 | 0.5385 | 15 | 0.6 | 11 | 0.5455 | 50 | 0.56 | 15 | 0.6667 | 3 | 0.6667 | 8 | 0.25 | 9 | 0.6667 | 35 | 0.5714 |
| 15_818789 | 13 | 0 | 12 | 0.0833 | 16 | 0.0625 | 7 | 0.2857 | 48 | 0.0833 | 12 | 0.25 | 4 | 0 | 7 | 0.1429 | 9 | 0.2222 | 32 | 0.1875 |

| | | | | | | | | | | | | | | | | | | | | |
|---|---|---|---|---|---|---|---|---|---|---|---|---|---|---|---|---|---|---|---|---|
| 15_8187901 | 11 | 0.0909 | 14 | 0.1429 | 15 | 0 | 8 | 0.125 | 48 | 0.0833 | 12 | 0 | 5 | 0.2 | 6 | 0 | 9 | 0.2222 | 32 | 0.0938 |
| 15_8192403 | 1 | 0 | 2 | 0 | 1 | 0 | 3 | 0.3333 | 7 | 0.1429 | 3 | 0.3333 | 3 | 0.3333 | 2 | 0 | 3 | 0 | 11 | 0.1818 |
| 15_8192408 | 1 | 1 | 3 | 0.3333 | 1 | 0 | 3 | 0 | 8 | 0.25 | 4 | 0.25 | 3 | 0 | 3 | 0 | 3 | 0 | 13 | 0.0769 |
| 15_8192411 | 1 | 0 | 3 | 0.3333 | 1 | 0 | 3 | 0.6667 | 8 | 0.375 | 4 | 0.25 | 3 | 0.3333 | 3 | 0.3333 | 3 | 0 | 13 | 0.2308 |
| 15_8192573 | 30 | 0.1333 | 22 | 0.0909 | 24 | 0.2083 | 23 | 0.087 | 99 | 0.1313 | 11 | 0.0909 | 20 | 0.25 | 20 | 0.25 | 19 | 0.3158 | 70 | 0.2429 |
| 15_8192593 | 25 | 0.16 | 20 | 0.05 | 25 | 0.08 | 22 | 0.0909 | 92 | 0.0978 | 9 | 0.1111 | 18 | 0 | 15 | 0.1333 | 16 | 0 | 58 | 0.0517 |
| 15_8192604 | 29 | 0.0345 | 23 | 0.1739 | 22 | 0.0455 | 24 | 0 | 98 | 0.0612 | 9 | 0.2222 | 19 | 0.1579 | 15 | 0.0667 | 17 | 0.1765 | 60 | 0.15 |
| 15_8202531 | 1 | 1 | 2 | 1 | 1 | 1 | NA | NA | 4 | 1 | 1 | 1 | 1 | 1 | 1 | 1 | NA | NA | 3 | 1 |
| 15_8202536 | 1 | 0 | 1 | 1 | 1 | 1 | NA | NA | 3 | 0.6667 | 1 | 1 | 1 | 1 | 1 | 1 | NA | NA | 3 | 1 |
| 15_8202539 | 1 | 1 | 1 | 1 | NA | NA | NA | NA | 2 | 1 | NA | NA | 1 | 1 | 1 | 1 | NA | NA | 2 | 1 |
| 15_8202545 | 1 | 1 | 1 | 0 | 1 | 1 | NA | NA | 3 | 0.6667 | NA | NA | 1 | 1 | 2 | 1 | NA | NA | 3 | 1 |
| 15_8202562 | 1 | 0 | 1 | 0 | 1 | 1 | NA | NA | 3 | 0.3333 | 1 | 0 | 1 | 0 | 2 | 0.5 | NA | NA | 4 | 0.25 |
| 15_8202592 | 1 | 1 | 2 | 0.5 | 1 | 0 | 1 | 0 | 5 | 0.4 | 1 | 0 | NA | NA | 3 | 0.3333 | NA | NA | 4 | 0.25 |
| 15_8208500 | NA | NA | 2 | 0.5 | NA | NA | NA | NA | 2 | 0.5 | 1 | 1 | NA | NA | NA | NA | NA | NA | 1 | 1 |
| 15_8208819 | 17 | 0.4706 | 8 | 0.125 | 14 | 0.3571 | 12 | 0.3333 | 51 | 0.3529 | 11 | 0.4545 | 15 | 0.4 | 12 | 0.5 | 14 | 0.5714 | 52 | 0.4808 |
| 15_8208824 | 15 | 0.5333 | 9 | 0.2222 | 12 | 0.25 | 12 | 0.4167 | 48 | 0.375 | 10 | 0.3 | 12 | 0.4167 | 13 | 0.4615 | 13 | 0.7692 | 48 | 0.5 |
| 15_8208830 | 20 | 0.4 | 8 | 0.5 | 15 | 0.4 | 12 | 0.5 | 55 | 0.4364 | 11 | 0.5455 | 15 | 0.4 | 15 | 0.6667 | 15 | 0.7333 | 56 | 0.5893 |
| 15_8208845 | 21 | 0.4762 | 9 | 0.7778 | 14 | 0.6429 | 14 | 0.5 | 58 | 0.569 | 11 | 1 | 14 | 0.6429 | 17 | 0.6471 | 15 | 0.6667 | 57 | 0.7193 |
| 15_8208857 | 20 | 0.5 | 9 | 0.5556 | 15 | 0.3333 | 12 | 0.3333 | 56 | 0.4286 | 12 | 0.5 | 15 | 0.5333 | 14 | 0.4286 | 14 | 0.5 | 55 | 0.4909 |
| 15_8208997 | NA | NA | 1 | 1 | NA | NA | NA | NA | 1 | 1 | 1 | 1 | NA | NA | NA | NA | NA | NA | 1 | 1 |
| 15_8209079 | NA | NA | 1 | 1 | NA | NA | NA | NA | 1 | 1 | 1 | 1 | NA | NA | NA | NA | NA | NA | 1 | 1 |
| 15_8209126 | 3 | 1 | 7 | 1 | 2 | 1 | NA | NA | 12 | 1 | 1 | 0 | 3 | 1 | 3 | 1 | 2 | 0.5 | 9 | 0.7778 |
| 15_8209134 | 3 | 0.6667 | 6 | 0.1667 | 1 | 0 | NA | NA | 10 | 0.3 | 2 | 0.5 | 3 | 0 | 2 | 0 | 2 | 0 | 9 | 0.1111 |
| 15_8209156 | 2 | 0 | 6 | 0.1667 | 2 | 0.5 | NA | NA | 10 | 0.2 | 1 | 0 | 1 | 0 | 1 | 1 | 1 | 1 | 4 | 0.5 |
| 15_8209165 | 3 | 0.6667 | 5 | 0.8 | 2 | 0.5 | NA | NA | 10 | 0.7 | 2 | 1 | 2 | 1 | 2 | 0.5 | 2 | 0.5 | 8 | 0.75 |
| 15_8209173 | 3 | 0 | 6 | 0.1667 | 2 | 0.5 | NA | NA | 11 | 0.1818 | 2 | 0.5 | 2 | 0.5 | 1 | 1 | 1 | 0 | 6 | 0.5 |
| 15_8209659 | 1 | 0 | 3 | 0 | 2 | 1 | NA | NA | 6 | 0.3333 | NA | NA | 1 | 0 | 1 | 1 | NA | NA | 2 | 0.5 |
| 15_8214379 | 5 | 0.4 | 3 | 0.6667 | 1 | 0 | 2 | 0.5 | 11 | 0.4545 | NA | NA | 3 | 0.3333 | NA | NA | 1 | 0 | 4 | 0.25 |
| 15_8214392 | 5 | 0.4 | 3 | 0 | 2 | 0.5 | 3 | 0.6667 | 13 | 0.3846 | 1 | 0 | 3 | 0.3333 | NA | NA | 2 | 0.5 | 6 | 0.3333 |
| 15_8214398 | 5 | 0 | 3 | 0.3333 | 2 | 0.5 | 3 | 0 | 13 | 0.1538 | NA | NA | 3 | 0.3333 | NA | NA | 3 | 0 | 6 | 0.1667 |
| 15_8214419 | 3 | 0 | 3 | 0.3333 | 2 | 0 | 3 | 0.3333 | 11 | 0.1818 | 2 | 1 | 5 | 0.2 | NA | NA | 5 | 0.2 | 12 | 0.3333 |
| 15_8214427 | 6 | 0 | 3 | 0.6667 | 5 | 0.2 | 3 | 0 | 17 | 0.1765 | 3 | 0.3333 | 4 | 0 | 1 | 0 | 7 | 0.1429 | 15 | 0.1333 |
| 15_8214447 | 6 | 0.1667 | 3 | 0.3333 | 6 | 0.1667 | 3 | 0 | 18 | 0.1667 | 2 | 0 | 6 | 0 | 2 | 0.5 | 8 | 0.125 | 18 | 0.1111 |
| 15_8214485 | 12 | 0.1667 | 9 | 0.1111 | 19 | 0.2632 | 10 | 0.4 | 50 | 0.24 | 12 | 0.1667 | 20 | 0.1 | 15 | 0.1333 | 16 | 0.25 | 63 | 0.1587 |
| 15_8217498 | 8 | 0.375 | 15 | 0.2 | 15 | 0.2667 | 12 | 0.3333 | 50 | 0.28 | 13 | 0.1538 | 19 | 0.4737 | 13 | 0.2308 | 14 | 0.2143 | 59 | 0.2881 |
| 15_8217504 | NA | NA | 10 | 0.4 | 5 | 0.2 | 9 | 0.2222 | 24 | 0.2917 | 2 | 0.5 | 8 | 0 | 7 | 0.1429 | 7 | 0.1429 | 24 | 0.125 |
| 15_8217558 | NA | NA | 7 | 0.5714 | 5 | 1 | 6 | 0.5 | 18 | 0.6667 | 4 | 0.75 | 7 | 0.4286 | 4 | 0.5 | 6 | 0.6667 | 21 | 0.5714 |
| 15_8217567 | 3 | 0.3333 | 12 | 0.0833 | 9 | 0.2222 | 9 | 0.1111 | 33 | 0.1515 | 5 | 0.6 | 13 | 0.1538 | 6 | 0.1667 | 8 | 0.125 | 32 | 0.2188 |
| 15_8217760 | 18 | 0.6667 | 31 | 0.4516 | 39 | 0.5128 | 22 | 0.2727 | 110 | 0.4727 | 20 | 0.6 | 27 | 0.5185 | 24 | 0.5 | 20 | 0.5 | 91 | 0.5275 |
| 15_8217778 | 19 | 0.3684 | 30 | 0.3 | 41 | 0.3171 | 23 | 0.2609 | 113 | 0.3097 | 21 | 0.1905 | 28 | 0.2857 | 26 | 0.3077 | 21 | 0.4762 | 96 | 0.3125 |
| 15_8217783 | 17 | 0.0588 | 25 | 0.08 | 33 | 0.1212 | 20 | 0.1 | 95 | 0.0947 | 17 | 0.1176 | 24 | 0.0417 | 24 | 0.2083 | 17 | 0.1176 | 82 | 0.122 |
| 15_8217822 | 19 | 0.3158 | 33 | 0.2121 | 30 | 0.4 | 16 | 0.25 | 98 | 0.2959 | 20 | 0.4 | 26 | 0.3462 | 24 | 0.25 | 17 | 0.4118 | 87 | 0.3448 |
| 15_8217853 | 22 | 0.4545 | 33 | 0.1212 | 41 | 0.1463 | 24 | 0.0833 | 120 | 0.1833 | 19 | 0.1579 | 27 | 0.3704 | 30 | 0.1667 | 23 | 0.2174 | 99 | 0.2323 |
| 15_8217855 | 29 | 0.0345 | 38 | 0.1316 | 49 | 0.0612 | 30 | 0.0333 | 146 | 0.0685 | 24 | 0.2083 | 33 | 0.0606 | 31 | 0.0323 | 27 | 0 | 115 | 0.0696 |
| 15_8217870 | 26 | 0.1154 | 32 | 0.0938 | 44 | 0.1136 | 23 | 0.087 | 125 | 0.104 | 24 | 0.0833 | 27 | 0.037 | 25 | 0.04 | 21 | 0.1429 | 97 | 0.0722 |
| 15_8221389 | 6 | 0.1667 | 1 | 0 | 4 | 0.25 | 3 | 0 | 14 | 0.1429 | 4 | 0.25 | 1 | 0 | 3 | 0.3333 | 5 | 0 | 13 | 0.1538 |
| 15_8221393 | 5 | 0.2 | 1 | 0 | 5 | 0.2 | 3 | 0.3333 | 14 | 0.2143 | 5 | 0.2 | 1 | 0 | 3 | 0.3333 | 5 | 0.2 | 14 | 0.2143 |
| 15_8221394 | 7 | 0.4286 | 2 | 0 | 5 | 0.4 | 3 | 0 | 17 | 0.2941 | 4 | 0.25 | 1 | 0 | 3 | 0 | 6 | 0.5 | 14 | 0.2857 |
| 15_8221395 | 7 | 0.4286 | 2 | 0.5 | 5 | 0.8 | 3 | 0.3333 | 17 | 0.5294 | 3 | 0.3333 | 1 | 1 | 3 | 1 | 6 | 0.5 | 13 | 0.6154 |
| 15_8221400 | 5 | 0.6 | 2 | 1 | 5 | 0.8 | 3 | 0.6667 | 15 | 0.7333 | 4 | 0.25 | 1 | 1 | 2 | 1 | 5 | 0.8 | 12 | 0.6667 |
| 15_8221406 | 6 | 1 | 2 | 1 | 4 | 1 | 2 | 1 | 14 | 1 | 4 | 0.5 | NA | NA | 2 | 1 | 6 | 1 | 12 | 0.8333 |
| 15_8221419 | 4 | 0.5 | 2 | 1 | 5 | 0.6 | 2 | 0.5 | 13 | 0.6154 | 4 | 0.5 | 1 | 0 | 3 | 0.3333 | 6 | 0.1667 | 14 | 0.2857 |
| 15_8221425 | 7 | 0.4286 | 2 | 0.5 | 5 | 0 | 2 | 0 | 16 | 0.25 | 4 | 0 | 1 | 1 | 3 | 0.3333 | 6 | 0 | 14 | 0.1429 |
| 15_8221429 | 7 | 0 | 2 | 0.5 | 4 | 0.5 | 2 | 0.5 | 15 | 0.2667 | 4 | 0 | 1 | 1 | 3 | 0 | 5 | 0.4 | 13 | 0.2308 |
| 15_8221660 | 5 | 0.2 | 2 | 0 | 4 | 0.25 | 2 | 0 | 13 | 0.1538 | 4 | 0 | NA | NA | 3 | 0 | 5 | 0.6 | 12 | 0.25 |
| 15_8224971 | 2 | 1 | 2 | 0.5 | 1 | 0 | NA | NA | 5 | 0.6 | 1 | 1 | 1 | 1 | NA | NA | 2 | 1 | 4 | 1 |
| 15_8224991 | 2 | 0.5 | 1 | 0 | 1 | 0 | 1 | 0 | 5 | 0.2 | NA | NA | 2 | 0.5 | NA | NA | 3 | 0.6667 | 5 | 0.6 |
| 15_8224992 | 2 | 0.5 | 1 | 0 | 1 | 0 | 1 | 0 | 5 | 0.2 | 1 | 0 | 2 | 0 | NA | NA | 1 | 1 | 4 | 0.25 |
| 15_8225026 | 3 | 0.3333 | 2 | 0 | NA | NA | 1 | 0 | 6 | 0.1667 | 1 | 0 | 1 | 0 | NA | NA | 2 | 0.5 | 4 | 0.25 |
| 15_8225031 | 3 | 0.3333 | 2 | 0.5 | 1 | 0 | 1 | 1 | 7 | 0.4286 | 1 | 0 | 1 | 0 | NA | NA | 3 | 0.3333 | 5 | 0.2 |
| 15_8225043 | 3 | 0.3333 | 2 | 0 | 1 | 0 | 1 | 0 | 7 | 0.1429 | NA | NA | 2 | 0.5 | NA | NA | 2 | 0 | 4 | 0.25 |
| 15_8225067 | 3 | 0.3333 | 2 | 0 | 1 | 1 | 1 | 0 | 7 | 0.2857 | 1 | 0 | 2 | 0 | NA | NA | 2 | 1 | 5 | 0.4 |
| 15_8225076 | 3 | 0.3333 | 3 | 0.6667 | NA | NA | 1 | 1 | 7 | 0.5714 | 1 | 1 | 2 | 0.5 | NA | NA | 2 | 0 | 5 | 0.4 |
| 15_8225085 | 3 | 0 | 3 | 0.3333 | 1 | 0 | 1 | 0 | 8 | 0.125 | 1 | 1 | 2 | 0 | NA | NA | 2 | 0.5 | 5 | 0.4 |
| 15_8225108 | 3 | 0 | 3 | 0.3333 | 1 | 0 | 1 | 0 | 8 | 0.125 | 1 | 0 | 2 | 0 | NA | NA | 3 | 0.6667 | 6 | 0.3333 |

| | | | | | | | | | | | | | | | | | | | | |
|---|---|---|---|---|---|---|---|---|---|---|---|---|---|---|---|---|---|---|---|---|
| 15_8225151 | 3 | 0.3333 | 3 | 0 | 1 | 0 | 1 | 0 | 8 | 0.125 | 1 | 0 | 1 | 0 | 1 | 1 | 2 | 0 | 5 | 0.2 |
| 15_8225159 | 3 | 0 | 2 | 0.5 | 1 | 0 | 1 | 0 | 7 | 0.1429 | 1 | 1 | 1 | 0 | NA | NA | 2 | 0 | 4 | 0.25 |
| 15_8225169 | 3 | 0 | 3 | 0.3333 | 1 | 0 | 1 | 0 | 8 | 0.125 | 1 | 1 | 1 | 0 | 2 | 0 | 2 | 0 | 6 | 0.1667 |
| 15_8225556 | 4 | 0.25 | 1 | 1 | 5 | 0.2 | 2 | 0.5 | 12 | 0.3333 | 2 | 0 | 2 | 1 | 1 | 0 | 1 | 0 | 6 | 0.3333 |
| 15_8225582 | 3 | 0 | 1 | 0 | 4 | 0.25 | 2 | 0 | 10 | 0.1 | 3 | 0.3333 | 2 | 0 | 1 | 0 | 3 | 0.3333 | 9 | 0.2222 |
| 15_8225691 | 1 | 1 | 1 | 0 | NA | NA | NA | NA | 2 | 0.5 | 1 | 1 | NA | NA | NA | NA | NA | NA | 1 | 1 |
| 15_8225716 | 1 | 1 | NA | NA | 1 | 1 | NA | NA | 2 | 1 | 1 | 1 | NA | NA | NA | NA | NA | NA | 1 | 1 |
| 15_8225718 | 1 | 1 | 1 | 1 | 1 | 1 | NA | NA | 3 | 1 | 1 | 1 | NA | NA | NA | NA | NA | NA | 1 | 1 |
| 15_8225721 | 1 | 1 | 1 | 0 | NA | NA | NA | NA | 2 | 0.5 | 1 | 1 | NA | NA | NA | NA | NA | NA | 1 | 1 |
| 15_8225734 | 2 | 0 | 2 | 0 | 5 | 0.4 | 2 | 0 | 11 | 0.1818 | 2 | 0.5 | 3 | 0 | 1 | 0 | 1 | 0 | 7 | 0.1429 |
| 15_8225759 | 1 | 1 | 1 | 1 | 1 | 1 | NA | NA | 3 | 1 | 1 | 1 | NA | NA | NA | NA | NA | NA | 1 | 1 |
| 15_8269346 | 6 | 0.3333 | 2 | 0 | 3 | 0 | 4 | 1 | 15 | 0.4 | 4 | 0 | 2 | 0 | 3 | 0.3333 | 6 | 0.3333 | 15 | 0.2 |
| 15_8269354 | 6 | 0 | 3 | 0.3333 | 4 | 0.25 | 4 | 0 | 17 | 0.1176 | 4 | 0.25 | 1 | 0 | 3 | 0 | 4 | 0.25 | 12 | 0.1667 |
| 15_8269377 | 1 | 1 | 2 | 0.5 | 2 | 0.5 | 1 | 0 | 6 | 0.5 | 2 | 0.5 | 1 | 1 | 3 | 0.3333 | 1 | 1 | 7 | 0.5714 |
| 15_8273591 | 2 | 1 | 1 | 1 | 1 | 1 | NA | NA | 4 | 1 | NA | NA | NA | NA | 2 | 1 | NA | NA | 2 | 1 |
| 15_8273598 | 2 | 1 | 1 | 1 | 1 | 0 | NA | NA | 4 | 0.75 | NA | NA | 1 | 1 | 2 | 0.5 | NA | NA | 3 | 0.6667 |
| 15_8273620 | 1 | 1 | NA | NA | 1 | 1 | NA | NA | 2 | 1 | NA | NA | 1 | 1 | 2 | 1 | NA | NA | 3 | 1 |
| 15_8273807 | 1 | 1 | NA | NA | NA | NA | NA | NA | 1 | 1 | NA | NA | NA | NA | 1 | 1 | NA | NA | 1 | 1 |
| 15_8288213 | 1 | 1 | 1 | 0 | NA | NA | NA | NA | 2 | 0.5 | NA | NA | NA | NA | 1 | 1 | 1 | 1 | 2 | 1 |
| 15_8288218 | 1 | 1 | 1 | 1 | NA | NA | NA | NA | 2 | 1 | NA | NA | NA | NA | 1 | 1 | 1 | 1 | 2 | 1 |
| 15_8288224 | NA | NA | 1 | 1 | NA | NA | NA | NA | 1 | 1 | NA | NA | NA | NA | 1 | 0 | 1 | 1 | 2 | 0.5 |
| 15_8288375 | NA | NA | 1 | 1 | NA | NA | NA | NA | 1 | 1 | 1 | 1 | NA | NA | NA | NA | NA | NA | 1 | 1 |
| 15_8288387 | NA | NA | 1 | 1 | NA | NA | NA | NA | 1 | 1 | 1 | 1 | NA | NA | NA | NA | NA | NA | 1 | 1 |
| 15_8288391 | NA | NA | 1 | 1 | NA | NA | NA | NA | 1 | 1 | 1 | 1 | NA | NA | NA | NA | NA | NA | 1 | 1 |
| 15_8299814 | 1 | 1 | NA | NA | NA | NA | NA | NA | 1 | 1 | NA | NA | 1 | 1 | NA | NA | NA | NA | 1 | 1 |
| 15_8299824 | 1 | 1 | NA | NA | NA | NA | NA | NA | 1 | 1 | NA | NA | 1 | 1 | NA | NA | NA | NA | 1 | 1 |
| 15_8299859 | 1 | 1 | NA | NA | NA | NA | NA | NA | 1 | 1 | NA | NA | 1 | 1 | NA | NA | NA | NA | 1 | 1 |
| 15_8299864 | 1 | 1 | NA | NA | NA | NA | NA | NA | 1 | 1 | NA | NA | 1 | 1 | NA | NA | NA | NA | 1 | 1 |
| 15_8299890 | 1 | 1 | NA | NA | NA | NA | NA | NA | 1 | 1 | NA | NA | 1 | 1 | NA | NA | NA | NA | 1 | 1 |
| 15_8299895 | 1 | 1 | NA | NA | NA | NA | NA | NA | 1 | 1 | NA | NA | 1 | 1 | NA | NA | NA | NA | 1 | 1 |
| 15_8299909 | 1 | 1 | NA | NA | NA | NA | NA | NA | 1 | 1 | NA | NA | 1 | 1 | NA | NA | NA | NA | 1 | 1 |
| 15_8299919 | 1 | 1 | NA | NA | NA | NA | NA | NA | 1 | 1 | NA | NA | 1 | 1 | NA | NA | NA | NA | 1 | 1 |
| 15_8300036 | 19 | 0.4737 | 22 | 0.6364 | 17 | 0.4118 | 15 | 0.6 | 73 | 0.5342 | 16 | 0.5625 | 19 | 0.6842 | 16 | 0.375 | 9 | 0.2222 | 60 | 0.5 |
| 15_8300041 | 20 | 0.05 | 19 | 0.2105 | 17 | 0.2353 | 14 | 0.2143 | 70 | 0.1714 | 17 | 0 | 20 | 0.1 | 17 | 0.1765 | 8 | 0 | 62 | 0.0806 |
| 15_8300046 | 15 | 0.3333 | 17 | 0.4706 | 11 | 0.2727 | 12 | 0.4167 | 55 | 0.3818 | 11 | 0.2727 | 19 | 0.1579 | 15 | 0.3333 | 7 | 0.4286 | 52 | 0.2692 |
| 15_8300047 | 18 | 0.8889 | 21 | 0.8571 | 17 | 1 | 17 | 0.9412 | 73 | 0.9178 | 16 | 0.9375 | 18 | 0.9444 | 15 | 0.9333 | 10 | 0.9 | 59 | 0.9322 |
| 15_8300053 | 20 | 0.5 | 22 | 0.3182 | 15 | 0.2667 | 16 | 0.375 | 73 | 0.3699 | 17 | 0.2941 | 18 | 0.2222 | 15 | 0.2667 | 9 | 0.3333 | 59 | 0.2712 |
| 15_8300066 | 18 | 0.1111 | 17 | 0 | 11 | 0.0909 | 11 | 0.1818 | 57 | 0.0877 | 14 | 0 | 19 | 0.0526 | 15 | 0.0667 | 10 | 0.2 | 58 | 0.069 |
| 15_8300087 | 13 | 0.6154 | 16 | 0.625 | 16 | 0.75 | 11 | 0.8182 | 56 | 0.6964 | 15 | 0.8667 | 15 | 0.8667 | 8 | 0.75 | 9 | 0.8889 | 47 | 0.8511 |
| 15_8300095 | 14 | 0.9286 | 11 | 1 | 15 | 0.9333 | 12 | 0.9167 | 52 | 0.9423 | 15 | 0.9333 | 18 | 0.8333 | 11 | 0.8182 | 11 | 0.8182 | 55 | 0.8545 |
| 15_8300112 | 14 | 0 | 16 | 0.1875 | 16 | 0.1875 | 15 | 0.1333 | 61 | 0.1311 | 13 | 0.0769 | 19 | 0.1579 | 10 | 0.1 | 9 | 0.1111 | 51 | 0.1176 |
| 15_8301033 | 14 | 0.0714 | 12 | 0.0833 | 6 | 0.3333 | 19 | 0 | 51 | 0.0784 | 12 | 0.0833 | 20 | 0.1 | 12 | 0 | 11 | 0.1818 | 55 | 0.0909 |
| 15_8301094 | 10 | 0 | 11 | 0.0909 | 6 | 0.1667 | 10 | 0.2 | 37 | 0.1081 | 6 | 0 | 14 | 0.1429 | 8 | 0.125 | 7 | 0 | 35 | 0.0857 |
| 15_8301171 | 7 | 0.4286 | 6 | 0.1667 | 2 | 0 | 13 | 0 | 28 | 0.1429 | 7 | 0.1429 | 11 | 0 | 4 | 0 | 4 | 0.5 | 26 | 0.1154 |
| 15_8301199 | 9 | 0.7778 | 6 | 0.6667 | 2 | 1 | 10 | 0.8 | 27 | 0.7778 | 6 | 0.8333 | 10 | 0.7 | 3 | 0.6667 | 5 | 1 | 24 | 0.7917 |
| 15_8301208 | 7 | 0.1429 | 5 | 0.4 | 1 | 0 | 7 | 0.1429 | 20 | 0.2 | 5 | 0.2 | 10 | 0.1 | 2 | 0 | 4 | 0.5 | 21 | 0.1905 |
| 15_8301216 | 10 | 0 | 7 | 0 | 2 | 0 | 9 | 0.2222 | 28 | 0.0714 | 8 | 0 | 11 | 0.1818 | 4 | 0.5 | 5 | 0 | 28 | 0.1429 |
| 15_8302237 | 8 | 0 | 5 | 0.2 | 1 | 0 | 6 | 0 | 20 | 0.05 | 5 | 0.2 | 8 | 0.125 | 3 | 0 | 3 | 0.6667 | 19 | 0.2105 |
| 15_8303820 | 9 | 0.1111 | 13 | 0.1538 | 12 | 0.1667 | 13 | 0.1538 | 47 | 0.1489 | 12 | 0.1667 | 9 | 0.1111 | 16 | 0.0625 | 9 | 0 | 46 | 0.087 |
| 15_8318422 | 3 | 0.3333 | 2 | 0 | 6 | 0.3333 | 5 | 0.6 | 16 | 0.375 | 3 | 1 | 2 | 0 | 1 | 0 | 5 | 0 | 11 | 0.2727 |
| 15_8318463 | 2 | 0.5 | NA | NA | 5 | 0.2 | 4 | 0.25 | 11 | 0.2727 | 2 | 0 | 2 | 1 | 1 | 1 | 5 | 0.6 | 10 | 0.6 |
| 15_8318483 | 2 | 0 | 1 | 0 | 6 | 0 | 5 | 0.2 | 14 | 0.0714 | 3 | 0.3333 | 3 | 0 | NA | NA | 5 | 0 | 11 | 0.0909 |
| 15_8326057 | 6 | 0.1667 | 9 | 0 | 10 | 0.1 | 9 | 0.2222 | 34 | 0.1176 | 6 | 0.3333 | 10 | 0.1 | 6 | 0.5 | 7 | 0.2857 | 29 | 0.2759 |
| 15_8326060 | 5 | 0 | 10 | 0.1 | 9 | 0.1111 | 10 | 0.4 | 34 | 0.1765 | 5 | 0.2 | 10 | 0.2 | 8 | 0.25 | 5 | 0.2 | 28 | 0.2143 |
| 15_8326075 | 6 | 0 | 12 | 0.0833 | 11 | 0 | 10 | 0.2 | 39 | 0.0769 | 6 | 0.1667 | 10 | 0.2 | 8 | 0.125 | 8 | 0 | 32 | 0.125 |
| 15_8326158 | 5 | 0 | 8 | 0.125 | 7 | 0.1429 | 8 | 0 | 28 | 0.0714 | 6 | 0.1667 | 6 | 0.3333 | 5 | 0 | 8 | 0.25 | 25 | 0.2 |
| 15_8327679 | 1 | 0 | 4 | 0 | 4 | 0 | 6 | 0.1667 | 15 | 0.0667 | 2 | 0 | 4 | 0.25 | 4 | 0 | 5 | 0 | 15 | 0.0667 |
| 15_8336806 | 8 | 0 | 7 | 0.1429 | 7 | 0.1429 | 9 | 0 | 31 | 0.0645 | 5 | 0.2 | 4 | 0 | 9 | 0.1111 | 7 | 0.1429 | 25 | 0.12 |
| 15_8336825 | 6 | 0.6667 | 3 | 0.3333 | 6 | 0.6667 | 5 | 1 | 20 | 0.7 | 5 | 0.6 | 4 | 0.5 | 5 | 0.6 | 5 | 0.8 | 19 | 0.6316 |
| 15_8336835 | 9 | 0.3333 | 4 | 0 | 7 | 0.1429 | 8 | 0 | 28 | 0.1429 | 6 | 0.1667 | 5 | 0 | 6 | 0.1667 | 6 | 0 | 23 | 0.087 |
| 15_8336841 | 8 | 0.125 | 7 | 0 | 7 | 0.1429 | 8 | 0.125 | 30 | 0.1 | 6 | 0 | 5 | 0.2 | 7 | 0.1429 | 6 | 0.1667 | 24 | 0.125 |
| 15_8339499 | 3 | 0 | 1 | 1 | 1 | 0 | 1 | 0 | 6 | 0.1667 | 2 | 0 | 3 | 0 | 4 | 0 | 2 | 0.5 | 11 | 0.0909 |
| 15_8343111 | 3 | 0 | 5 | 0.6 | 3 | 0 | 3 | 0.3333 | 14 | 0.2857 | 5 | 0.2 | 3 | 0 | 3 | 0 | 5 | 0.2 | 16 | 0.125 |
| 15_8343113 | 2 | 0.5 | 5 | 0 | 1 | 0 | 3 | 0 | 11 | 0.0909 | 5 | 0.2 | 3 | 0 | 2 | 0 | 5 | 0.2 | 15 | 0.1333 |
| 15_8343127 | 1 | 1 | 3 | 0.3333 | 3 | 0 | 2 | 0 | 9 | 0.2222 | 5 | 0.2 | 3 | 0.6667 | 3 | 0.3333 | 5 | 0.4 | 16 | 0.375 |

| | | | | | | | | | | | | | | | | | | | | |
|---|---|---|---|---|---|---|---|---|---|---|---|---|---|---|---|---|---|---|---|---|
| 15_834314 | 2 | 1 | 3 | 1 | 3 | 1 | 2 | 1 | 10 | 1 | 6 | 1 | 3 | 1 | 3 | 1 | 5 | 1 | 17 | 1 |
| 15_834314 | 2 | 1 | 2 | 1 | 4 | 1 | 3 | 1 | 11 | 1 | 5 | 1 | 3 | 1 | 3 | 1 | 4 | 1 | 15 | 1 |
| 15_834314 | 2 | 0.5 | 3 | 0.3333 | 3 | 0 | 3 | 0 | 11 | 0.1818 | 5 | 0 | 3 | 0.3333 | 3 | 0 | 3 | 0 | 14 | 0.0714 |
| 15_834315 | 1 | 1 | 2 | 1 | 3 | 1 | 2 | 0.5 | 8 | 0.875 | 4 | 1 | 2 | 0.5 | 2 | 1 | 3 | 1 | 11 | 0.9091 |
| 15_834316 | 1 | 1 | 3 | 1 | 4 | 1 | 3 | 1 | 11 | 1 | 5 | 1 | 3 | 0.6667 | 3 | 1 | 3 | 0.3333 | 14 | 0.7857 |
| 15_834316 | 2 | 0.5 | 3 | 0 | 4 | 0.5 | 2 | 0.5 | 11 | 0.3636 | 5 | 0.4 | 3 | 0.6667 | 3 | 0.3333 | 5 | 0.2 | 16 | 0.375 |
| 15_834317 | 1 | 0 | 2 | 0 | 4 | 0.25 | 3 | 0.3333 | 10 | 0.2 | 6 | 0 | 3 | 0 | 3 | 0.3333 | 3 | 0.6667 | 15 | 0.2 |
| 15_834318 | 2 | 0.5 | 3 | 0.6667 | 4 | 1 | 2 | 0 | 11 | 0.6364 | 6 | 0.5 | 3 | 0 | 3 | 0.6667 | 4 | 0.75 | 16 | 0.5 |
| 15_834395 | 1 | 0 | 2 | 0 | 1 | 1 | 2 | 0 | 6 | 0.1667 | 4 | 0 | 2 | 0.5 | 1 | 0 | 4 | 0 | 11 | 0.0909 |
| 15_834397 | NA | NA | 2 | 1 | NA | NA | 2 | 0 | 4 | 0.5 | 2 | 0.5 | 2 | 0 | 1 | 0 | 2 | 0.5 | 7 | 0.2857 |
| 15_834600 | 2 | 0.5 | 3 | 0.3333 | NA | NA | NA | NA | 5 | 0.4 | 1 | 1 | 2 | 0.5 | 1 | 1 | 1 | 1 | 5 | 0.8 |
| 15_834602 | 2 | 0 | 4 | 0.5 | NA | NA | NA | NA | 6 | 0.3333 | 1 | 0 | 4 | 0.5 | 2 | 0 | 2 | 0.5 | 9 | 0.3333 |
| 15_834605 | 3 | 0 | 4 | 1 | NA | NA | NA | NA | 7 | 0.5714 | 1 | 0 | 3 | 0.3333 | 2 | 0.5 | 2 | 0.5 | 8 | 0.375 |
| 15_834608 | 2 | 1 | 4 | 1 | NA | NA | NA | NA | 6 | 1 | 1 | 1 | 2 | 1 | 2 | 1 | 2 | 0.5 | 7 | 0.8571 |
| 15_834608 | 2 | 0.5 | 3 | 1 | NA | NA | NA | NA | 5 | 0.8 | 1 | 1 | 3 | 1 | 2 | 1 | 2 | 1 | 8 | 1 |
| 15_834610 | 1 | 1 | 3 | 0.6667 | NA | NA | NA | NA | 4 | 0.75 | 1 | 1 | 3 | 0.3333 | 2 | 0.5 | 1 | 1 | 7 | 0.5714 |
| 15_834611 | 1 | 0 | 4 | 0.5 | NA | NA | NA | NA | 5 | 0.4 | 1 | 1 | 3 | 0 | 2 | 0.5 | 1 | 0 | 7 | 0.2857 |
| 15_834877 | 1 | 1 | 2 | 1 | 1 | 1 | 1 | 1 | 5 | 1 | 2 | 1 | NA | NA | 1 | 1 | 1 | 1 | 4 | 1 |
| 15_834880 | 1 | 1 | 3 | 1 | 1 | 1 | NA | NA | 5 | 1 | 1 | 0 | NA | NA | 1 | 1 | 1 | 0 | 3 | 0.3333 |
| 15_834881 | 1 | 1 | 3 | 0.6667 | NA | NA | NA | NA | 4 | 0.75 | 2 | 1 | NA | NA | 1 | 1 | 1 | 0 | 4 | 0.75 |
| 15_834883 | NA | NA | 3 | 0.3333 | 1 | 1 | NA | NA | 4 | 0.5 | NA | NA | NA | NA | 1 | 1 | 1 | 1 | 2 | 1 |
| 15_834886 | NA | NA | 3 | 1 | 1 | 1 | NA | NA | 4 | 1 | 1 | 1 | NA | NA | 1 | 1 | 1 | 1 | 3 | 1 |
| 15_834895 | NA | NA | 2 | 1 | NA | NA | NA | NA | 2 | 1 | NA | NA | NA | NA | 1 | 1 | NA | NA | 1 | 1 |
| 15_834900 | NA | NA | 2 | 0.5 | NA | NA | NA | NA | 2 | 0.5 | NA | NA | NA | NA | 1 | 1 | NA | NA | 1 | 1 |
| 15_834902 | NA | NA | 2 | 0.5 | NA | NA | NA | NA | 2 | 0.5 | NA | NA | NA | NA | 1 | 1 | NA | NA | 1 | 1 |
| 15_834909 | 2 | 0.5 | 1 | 0 | 2 | 1 | NA | NA | 5 | 0.6 | 1 | 0 | NA | NA | NA | NA | 2 | 0.5 | 3 | 0.3333 |
| 15_834909 | 2 | 1 | 3 | 1 | 2 | 1 | NA | NA | 7 | 1 | 1 | 0 | NA | NA | 1 | 1 | 2 | 0.5 | 4 | 0.5 |
| 15_834909 | 1 | 0 | 2 | 1 | 2 | 1 | NA | NA | 5 | 0.8 | NA | NA | NA | NA | 1 | 1 | 2 | 1 | 3 | 1 |
| 15_834910 | 2 | 0.5 | 3 | 1 | 2 | 1 | NA | NA | 7 | 0.8571 | 1 | 0 | NA | NA | 1 | 1 | 1 | 1 | 3 | 0.6667 |
| 15_834911 | 2 | 0.5 | 3 | 1 | 2 | 1 | NA | NA | 7 | 0.8571 | 1 | 1 | NA | NA | NA | NA | 2 | 0.5 | 3 | 0.6667 |
| 15_834912 | 2 | 0.5 | 3 | 1 | 2 | 1 | NA | NA | 7 | 0.8571 | 1 | 1 | NA | NA | 1 | 1 | NA | NA | 2 | 1 |
| 15_834912 | 1 | 0 | 3 | 0 | 2 | 1 | NA | NA | 6 | 0.3333 | 1 | 0 | NA | NA | 1 | 1 | 2 | 0.5 | 4 | 0.5 |
| 15_834915 | 1 | 0 | 1 | 1 | 2 | 0.5 | NA | NA | 4 | 0.5 | 1 | 1 | NA | NA | NA | NA | 1 | 0 | 2 | 0.5 |
| 15_837205 | 5 | 0.4 | 18 | 0.0556 | 6 | 0.1667 | 6 | 0 | 35 | 0.1143 | 17 | 0.1176 | 4 | 0 | 3 | 0 | 5 | 0.4 | 29 | 0.1379 |
| 15_837405 | 8 | 0.125 | 20 | 0.2 | 8 | 0 | 7 | 0 | 43 | 0.1163 | 21 | 0.0952 | 4 | 0.5 | 3 | 0 | 7 | 0 | 35 | 0.1143 |
| 15_837405 | 7 | 0 | 19 | 0.4211 | 8 | 0 | 7 | 0.1429 | 41 | 0.2195 | 19 | 0.1053 | 4 | 0.25 | 3 | 0.3333 | 7 | 0.2857 | 33 | 0.1818 |
| 15_837406 | 9 | 0.1111 | 18 | 0.1111 | 8 | 0.125 | 6 | 0.1667 | 41 | 0.122 | 20 | 0.05 | 2 | 0 | 2 | 0.5 | 8 | 0.125 | 32 | 0.0938 |
| 15_837406 | 7 | 0.8571 | 12 | 0.8333 | 4 | 1 | 3 | 1 | 26 | 0.8846 | 16 | 0.875 | NA | NA | 2 | 0.5 | 3 | 1 | 21 | 0.8571 |
| 15_837406 | 9 | 0 | 22 | 0.1818 | 9 | 0.1111 | 7 | 0.2857 | 47 | 0.1489 | 25 | 0.16 | 5 | 0 | 3 | 0.3333 | 8 | 0.25 | 41 | 0.1707 |
| 15_837407 | 9 | 0.3333 | 20 | 0.4 | 10 | 0.4 | 7 | 0.2857 | 46 | 0.3696 | 24 | 0.5 | 4 | 0 | 3 | 0.6667 | 8 | 0.5 | 39 | 0.4615 |
| 15_837407 | 4 | 0 | 12 | 0.0833 | 7 | 0 | 3 | 0.3333 | 26 | 0.0769 | 14 | 0.1429 | 4 | 0.25 | 3 | 0.3333 | 5 | 0.2 | 26 | 0.1923 |
| 15_837408 | 8 | 0 | 21 | 0.2381 | 8 | 0.25 | 5 | 0 | 42 | 0.1667 | 22 | 0.1818 | 4 | 0.25 | 2 | 0.5 | 8 | 0.125 | 36 | 0.1944 |
| 15_837408 | 8 | 0.375 | 19 | 0.4737 | 7 | 0.2857 | 5 | 0.6 | 39 | 0.4359 | 18 | 0.3333 | 3 | 0.3333 | 3 | 0.6667 | 3 | 0.6667 | 27 | 0.4074 |
| 15_837410 | 6 | 1 | 22 | 0.9091 | 11 | 0.9091 | 6 | 1 | 45 | 0.9333 | 21 | 0.8571 | 3 | 1 | 3 | 1 | 7 | 1 | 34 | 0.9118 |
| 15_837410 | 9 | 1 | 23 | 0.6522 | 11 | 0.4545 | 7 | 0.7143 | 50 | 0.68 | 25 | 0.68 | 5 | 0.6 | 4 | 0.5 | 9 | 0.4444 | 43 | 0.6047 |
| 15_837411 | 8 | 0.25 | 23 | 0.3478 | 12 | 0.3333 | 7 | 0.2857 | 50 | 0.32 | 25 | 0.4 | 5 | 0.2 | 4 | 0.5 | 7 | 0.4286 | 41 | 0.3902 |
| 15_837411 | 7 | 1 | 21 | 0.8571 | 12 | 1 | 7 | 0.7143 | 47 | 0.8936 | 23 | 0.8696 | 4 | 0.75 | 4 | 0.75 | 7 | 0.5714 | 38 | 0.7895 |
| 15_837413 | 10 | 0.6 | 18 | 0.4444 | 12 | 0.4167 | 7 | 0.4286 | 47 | 0.4681 | 24 | 0.5417 | 4 | 0.75 | 6 | 0.5 | 7 | 0.1429 | 41 | 0.4878 |
| 15_837416 | 12 | 0.3333 | 26 | 0.5 | 13 | 0.3077 | 10 | 0.1 | 61 | 0.3607 | 28 | 0.4643 | 7 | 0.1429 | 5 | 0 | 8 | 0.25 | 48 | 0.3333 |
| 15_837421 | 13 | 0.1538 | 30 | 0.1333 | 12 | 0.0833 | 8 | 0.25 | 63 | 0.1429 | 26 | 0.1923 | 7 | 0.1429 | 7 | 0.2857 | 13 | 0.2308 | 53 | 0.2075 |
| 15_837423 | 12 | 0.4167 | 34 | 0.5882 | 17 | 0.5294 | 9 | 0.4444 | 72 | 0.5278 | 36 | 0.6944 | 6 | 0.6667 | 6 | 0.6667 | 12 | 0.5833 | 60 | 0.6667 |
| 15_837423 | 12 | 0.9167 | 31 | 0.8387 | 14 | 0.9286 | 10 | 0.9 | 67 | 0.8806 | 33 | 0.9394 | 7 | 1 | 7 | 1 | 13 | 0.9231 | 60 | 0.95 |
| 15_837424 | 12 | 0.8333 | 31 | 0.7097 | 19 | 0.7368 | 11 | 0.9091 | 73 | 0.7671 | 33 | 0.9394 | 6 | 1 | 9 | 0.7778 | 11 | 0.6364 | 59 | 0.8644 |
| 15_837424 | 12 | 0.6667 | 27 | 0.5556 | 19 | 0.7368 | 10 | 0.9 | 68 | 0.6765 | 27 | 0.7407 | 5 | 1 | 8 | 0.75 | 12 | 0.75 | 52 | 0.7692 |
| 15_837424 | 14 | 0.3571 | 31 | 0.4194 | 19 | 0.2632 | 8 | 0.25 | 72 | 0.3472 | 29 | 0.4483 | 5 | 0.2 | 7 | 0.5714 | 12 | 0.5 | 53 | 0.4528 |
| 15_839606 | 61 | 0.2459 | 61 | 0.3279 | 52 | 0.3269 | 34 | 0.3824 | 208 | 0.3125 | 53 | 0.2264 | 42 | 0.3333 | 42 | 0.3333 | 53 | 0.3019 | 190 | 0.2947 |
| 15_839606 | 55 | 0.1818 | 67 | 0.2388 | 47 | 0.2128 | 30 | 0.1667 | 199 | 0.206 | 48 | 0.25 | 42 | 0.2143 | 42 | 0.0952 | 47 | 0.1489 | 179 | 0.1788 |
| 15_839607 | 57 | 0.0877 | 65 | 0.1846 | 49 | 0.102 | 30 | 0.1667 | 201 | 0.1343 | 54 | 0.1667 | 39 | 0.1282 | 42 | 0.0476 | 49 | 0.102 | 184 | 0.1141 |
| 15_839608 | 58 | 0.6552 | 67 | 0.7015 | 48 | 0.7083 | 32 | 0.7812 | 205 | 0.7024 | 51 | 0.7647 | 44 | 0.8409 | 41 | 0.7073 | 54 | 0.7037 | 190 | 0.7526 |
| 15_839609 | 51 | 0.2941 | 67 | 0.3582 | 43 | 0.3256 | 26 | 0.3462 | 187 | 0.3316 | 45 | 0.3556 | 37 | 0.2703 | 41 | 0.2683 | 53 | 0.2264 | 176 | 0.2784 |
| 15_839611 | 50 | 0.12 | 61 | 0.082 | 40 | 0.1 | 22 | 0.0909 | 173 | 0.0983 | 48 | 0.1042 | 39 | 0.1795 | 37 | 0.1351 | 42 | 0.1429 | 166 | 0.1386 |
| 15_839615 | 32 | 0.375 | 32 | 0.375 | 30 | 0.2 | 22 | 0.3182 | 116 | 0.319 | 25 | 0.24 | 26 | 0.3077 | 29 | 0.4138 | 31 | 0.3871 | 111 | 0.3423 |
| 15_839616 | 34 | 0.5294 | 43 | 0.5581 | 30 | 0.2667 | 24 | 0.4167 | 131 | 0.458 | 33 | 0.3939 | 26 | 0.6538 | 31 | 0.6129 | 40 | 0.6 | 130 | 0.5615 |
| 15_839616 | 30 | 0.5 | 41 | 0.5854 | 25 | 0.84 | 21 | 0.6667 | 117 | 0.6325 | 30 | 0.7333 | 25 | 0.88 | 25 | 0.64 | 39 | 0.7179 | 119 | 0.7395 |
| 15_839617 | 29 | 0.0345 | 39 | 0.0769 | 28 | 0.0357 | 21 | 0.0476 | 117 | 0.0513 | 32 | 0.0625 | 29 | 0.069 | 29 | 0.1379 | 34 | 0.1176 | 124 | 0.0968 |

| | | | | | | | | | | | | | | | | | | | | |
|---|---|---|---|---|---|---|---|---|---|---|---|---|---|---|---|---|---|---|---|---|
| 15_840985 | 3 | 1 | 3 | 1 | 1 | 1 | 3 | 1 | 10 | 1 | 1 | 1 | 1 | 1 | 7 | 1 | 1 | 1 | 10 | 1 |
| 15_840985 | 4 | 1 | 1 | 1 | 2 | 1 | 3 | 1 | 10 | 1 | 1 | 1 | 1 | 0 | 5 | 1 | NA | NA | 7 | 0.8571 |
| 15_840986 | 4 | 0.25 | 3 | 0 | 2 | 1 | 4 | 0.5 | 13 | 0.3846 | 1 | 0 | 1 | 0 | 5 | 0.2 | 1 | 1 | 8 | 0.25 |
| 15_840986 | 5 | 0.6 | 3 | 0.6667 | 2 | 1 | 3 | 0.6667 | 13 | 0.6923 | 1 | 0 | 1 | 0 | 6 | 0.5 | 1 | 1 | 9 | 0.4444 |
| 15_840987 | 5 | 1 | 3 | 1 | 2 | 0.5 | 4 | 1 | 14 | 0.9286 | 2 | 1 | 1 | 0 | 5 | 1 | 1 | 0 | 9 | 0.7778 |
| 15_840987 | 4 | 0.75 | 3 | 0.3333 | 2 | 0.5 | 4 | 0.75 | 13 | 0.6154 | 2 | 0 | 1 | 0 | 6 | 0.1667 | 1 | 0 | 10 | 0.1 |
| 15_840988 | 4 | 0.25 | 2 | 0 | 2 | 0.5 | 3 | 0 | 11 | 0.1818 | 1 | 0 | 1 | 0 | 7 | 0.5714 | 1 | 0 | 10 | 0.4 |
| 15_840989 | 4 | 0.75 | 3 | 0 | 2 | 0 | 4 | 0.75 | 13 | 0.4615 | 2 | 0.5 | 1 | 0 | 5 | 0.2 | 1 | 0 | 9 | 0.2222 |
| 15_840989 | 5 | 0.6 | 2 | 0.5 | 2 | 0.5 | 4 | 0.5 | 13 | 0.5385 | 2 | 0.5 | 1 | 1 | 5 | 0 | 1 | 0 | 9 | 0.2222 |
| 15_840989 | 5 | 0 | 3 | 0.3333 | 2 | 0 | 4 | 0.75 | 14 | 0.2857 | 2 | 0.5 | 1 | 0 | 6 | 0.5 | 1 | 0 | 10 | 0.4 |
| 15_840989 | 5 | 0.2 | 3 | 0 | 2 | 0 | 4 | 0 | 14 | 0.0714 | 2 | 0 | 1 | 0 | 6 | 0.3333 | 1 | 0 | 10 | 0.2 |
| 15_840990 | 5 | 1 | 3 | 0.6667 | 2 | 1 | 1 | 1 | 11 | 0.9091 | 1 | 1 | 1 | 1 | 6 | 0.8333 | 1 | 1 | 9 | 0.8889 |
| 15_840990 | 4 | 0.25 | 3 | 0 | 2 | 1 | 4 | 0.25 | 13 | 0.3077 | 2 | 0 | 1 | 0 | 8 | 0.375 | 1 | 0 | 12 | 0.25 |
| 15_840992 | 4 | 0 | 3 | 0.3333 | 1 | 0 | 4 | 0.25 | 12 | 0.1667 | 1 | 0 | NA | NA | 8 | 0.25 | 1 | 0 | 10 | 0.2 |
| 15_841001 | 6 | 0 | 5 | 0.2 | 2 | 0 | 3 | 0.3333 | 16 | 0.125 | 3 | 0 | 2 | 0 | 7 | 0.1429 | 3 | 0 | 15 | 0.0667 |
| 15_841026 | 1 | 1 | NA | NA | NA | NA | NA | NA | 1 | 1 | NA | NA | NA | NA | 1 | 1 | NA | NA | 1 | 1 |
| 15_841029 | 1 | 1 | NA | NA | NA | NA | NA | NA | 1 | 1 | NA | NA | NA | NA | 1 | 1 | NA | NA | 1 | 1 |
| 15_841032 | 1 | 1 | NA | NA | NA | NA | NA | NA | 1 | 1 | NA | NA | NA | NA | 1 | 1 | NA | NA | 1 | 1 |
| 15_841036 | 1 | 1 | NA | NA | NA | NA | NA | NA | 1 | 1 | NA | NA | NA | NA | 1 | 1 | NA | NA | 1 | 1 |
| 15_841978 | NA | NA | NA | NA | 4 | 0.25 | 3 | 0.3333 | 7 | 0.2857 | 1 | 0 | 5 | 0.4 | 2 | 0 | 2 | 0.5 | 10 | 0.3 |
| 15_841978 | NA | NA | NA | NA | 3 | 0.3333 | 3 | 0.6667 | 6 | 0.5 | 1 | 1 | 3 | 0 | 2 | 0 | 1 | 0 | 7 | 0.1429 |
| 15_841979 | NA | NA | NA | NA | 3 | 0.3333 | 2 | 0.5 | 5 | 0.4 | NA | NA | 5 | 0.2 | 1 | 0 | 2 | 0 | 8 | 0.125 |
| 15_842635 | 6 | 0.1667 | 13 | 0.0769 | 7 | 0 | 2 | 0 | 28 | 0.0714 | 6 | 0 | 3 | 0 | 4 | 0 | 5 | 0.4 | 18 | 0.1111 |
| 15_842636 | 6 | 0.3333 | 12 | 0.25 | 5 | 0 | 2 | 0 | 25 | 0.2 | 5 | 0.6 | 4 | 0.25 | 4 | 0 | 6 | 0.1667 | 19 | 0.2632 |
| 15_842646 | 2 | 0 | 10 | 0.1 | 4 | 0 | 1 | 0 | 17 | 0.0588 | 3 | 0 | 4 | 0 | 3 | 0 | 5 | 0.2 | 15 | 0.0667 |
| 15_842656 | 2 | 0.5 | 9 | 0 | 1 | 0 | 1 | 0 | 13 | 0.0769 | 4 | 0 | 2 | 0.5 | 3 | 0 | 5 | 0 | 14 | 0.0714 |
| 15_842681 | 2 | 0 | 1 | 0 | NA | NA | 1 | 1 | 4 | 0.25 | 1 | 0 | 1 | 0 | 2 | 0.5 | 3 | 0 | 7 | 0.1429 |
| 15_842683 | 2 | 0.5 | 1 | 1 | NA | NA | 2 | 0.5 | 5 | 0.6 | 2 | 0 | NA | NA | 3 | 0.3333 | 4 | 0.5 | 9 | 0.3333 |
| 15_842684 | 1 | 1 | 1 | 1 | NA | NA | 2 | 0.5 | 4 | 0.75 | 2 | 0 | NA | NA | 1 | 0 | 4 | 0.25 | 7 | 0.1429 |
| 15_842684 | 2 | 1 | 1 | 0 | NA | NA | 2 | 0.5 | 5 | 0.6 | 2 | 0.5 | 1 | 0 | NA | NA | 3 | 1 | 6 | 0.6667 |
| 15_842684 | 1 | 1 | 1 | 0 | NA | NA | 2 | 1 | 4 | 0.75 | 1 | 1 | 1 | 1 | NA | NA | 1 | 1 | 3 | 1 |
| 15_844376 | 5 | 0.4 | 5 | 0 | 5 | 0.4 | 3 | 0 | 18 | 0.2222 | 2 | 0 | 2 | 0 | 4 | 0.25 | 7 | 0.1429 | 15 | 0.1333 |
| 15_844377 | 7 | 0.5714 | 7 | 0.1429 | 8 | 0.375 | 5 | 0.2 | 27 | 0.3333 | 3 | 0.3333 | 1 | 0 | 2 | 0.5 | 8 | 0.375 | 14 | 0.3571 |
| 15_844378 | 7 | 0.1429 | 9 | 0.1111 | 8 | 0.125 | 5 | 0 | 29 | 0.1034 | 6 | 0.1667 | 4 | 0 | 6 | 0.1667 | 9 | 0.3333 | 25 | 0.2 |
| 15_844380 | 6 | 0.5 | 9 | 0.4444 | 7 | 0.8571 | 5 | 0.6 | 27 | 0.5926 | 4 | 0.25 | 2 | 0.5 | 5 | 0.4 | 7 | 0.5714 | 18 | 0.4444 |
| 15_844383 | 4 | 0.75 | 5 | 0.4 | 5 | 0.4 | 2 | 0 | 16 | 0.4375 | 4 | 0.5 | 3 | 0.3333 | 3 | 0.3333 | 7 | 0.2857 | 17 | 0.3529 |
| 15_844388 | 5 | 0.2 | 8 | 0 | 6 | 0.3333 | 3 | 0.6667 | 22 | 0.2273 | 3 | 0.3333 | 2 | 0 | 3 | 0.3333 | 6 | 0.3333 | 14 | 0.2857 |
| 15_844388 | 4 | 0.75 | 6 | 0.3333 | 6 | 0.6667 | 3 | 0.6667 | 19 | 0.5789 | 1 | 0 | 2 | 0.5 | 2 | 1 | 5 | 0.4 | 10 | 0.5 |
| 15_844389 | 5 | 0.2 | 6 | 0.3333 | 4 | 0.25 | 3 | 0.6667 | 18 | 0.3333 | NA | NA | 2 | 0 | 2 | 0 | 6 | 0.6667 | 10 | 0.4 |
| 15_844391 | 5 | 0.4 | 5 | 0.6 | 6 | 0.6667 | 2 | 1 | 18 | 0.6111 | 1 | 0 | 1 | 1 | 2 | 0.5 | 5 | 0.6 | 9 | 0.5556 |
| 15_844392 | 3 | 0.3333 | 4 | 0.75 | 5 | 1 | 3 | 1 | 15 | 0.8 | NA | NA | 2 | 0.5 | 1 | 1 | 3 | 0.6667 | 6 | 0.6667 |
| 15_844392 | 5 | 0.2 | 4 | 0.25 | 6 | 0.5 | 2 | 1 | 17 | 0.4118 | NA | NA | 2 | 0 | 1 | 0 | 5 | 0.6 | 8 | 0.375 |
| 15_844393 | 3 | 0.6667 | 4 | 0.5 | 4 | 0.5 | 2 | 0.5 | 13 | 0.5385 | NA | NA | 2 | 0 | 1 | 1 | 3 | 0.3333 | 6 | 0.3333 |
| 15_844395 | 4 | 0.5 | 4 | 0.5 | 5 | 0 | 1 | 0 | 14 | 0.2857 | NA | NA | 1 | 1 | 1 | 0 | 3 | 0.6667 | 5 | 0.6 |
| 15_844397 | 3 | 0.6667 | 4 | 1 | 5 | 1 | 2 | 1 | 14 | 0.9286 | NA | NA | NA | NA | 1 | 1 | 1 | 1 | 2 | 1 |
| 15_844398 | 3 | 0.6667 | 2 | 0.5 | 2 | 1 | 1 | 1 | 8 | 0.75 | NA | NA | 1 | 1 | 1 | 1 | 1 | 1 | 3 | 1 |
| 15_844399 | 3 | 1 | 2 | 1 | 3 | 1 | NA | NA | 8 | 1 | NA | NA | NA | NA | 1 | 1 | NA | NA | 1 | 1 |
| 15_844399 | 4 | 1 | 4 | 1 | 5 | 1 | NA | NA | 13 | 1 | NA | NA | 1 | 1 | 1 | 1 | 1 | 1 | 3 | 1 |
| 15_844399 | 4 | 0.75 | 3 | 0.3333 | 3 | 0.3333 | NA | NA | 10 | 0.5 | NA | NA | 1 | 1 | 1 | 0 | NA | NA | 2 | 0.5 |
| 15_844401 | 3 | 1 | 4 | 1 | 2 | 1 | NA | NA | 9 | 1 | NA | NA | 1 | 1 | 1 | 1 | NA | NA | 2 | 1 |
| 15_844402 | 2 | 0.5 | 3 | 1 | 2 | 0 | NA | NA | 7 | 0.5714 | NA | NA | 1 | 0 | 1 | 1 | NA | NA | 2 | 0.5 |
| 15_844403 | 4 | 0 | 2 | 0.5 | 3 | 0 | NA | NA | 9 | 0.1111 | NA | NA | 1 | 1 | 1 | 0 | NA | NA | 2 | 0.5 |
| 15_844406 | 3 | 1 | 2 | 1 | 2 | 1 | NA | NA | 7 | 1 | NA | NA | 1 | 1 | NA | NA | NA | NA | 1 | 1 |
| 15_844407 | 3 | 0 | 2 | 0 | 2 | 0.5 | NA | NA | 7 | 0.1429 | NA | NA | 1 | 0 | 1 | 1 | NA | NA | 2 | 0.5 |
| 15_844410 | 3 | 0.6667 | 2 | 1 | 2 | 1 | NA | NA | 7 | 0.8571 | NA | NA | 1 | 1 | NA | NA | NA | NA | 1 | 1 |
| 15_844412 | 3 | 0 | 2 | 0.5 | 2 | 1 | NA | NA | 7 | 0.4286 | NA | NA | 1 | 1 | NA | NA | NA | NA | 1 | 1 |
| 15_844413 | 2 | 0 | 1 | 1 | 1 | 0 | NA | NA | 4 | 0.25 | NA | NA | 1 | 1 | NA | NA | NA | NA | 1 | 1 |
| 15_845871 | NA | NA | 2 | 0.5 | 2 | 0 | NA | NA | 4 | 0.25 | 1 | 0 | 1 | 0 | 1 | 1 | 1 | 0 | 4 | 0.25 |
| 15_845871 | NA | NA | 2 | 1 | 2 | 0.5 | 1 | 0 | 5 | 0.6 | 1 | 0 | 1 | 1 | 4 | 0.75 | 1 | 1 | 7 | 0.7143 |
| 15_845872 | NA | NA | 2 | 1 | 2 | 1 | 1 | 1 | 5 | 1 | 1 | 1 | 1 | 0 | 4 | 0.75 | 1 | 1 | 7 | 0.7143 |
| 15_845872 | NA | NA | 2 | 0.5 | 1 | 1 | 2 | 1 | 5 | 0.8 | 1 | 1 | 1 | 1 | 4 | 0.25 | 1 | 1 | 7 | 0.5714 |
| 15_845872 | NA | NA | 2 | 0 | 2 | 0.5 | 2 | 0 | 6 | 0.1667 | 1 | 0 | 1 | 1 | 3 | 0.3333 | 1 | 0 | 6 | 0.3333 |
| 15_845873 | NA | NA | 2 | 1 | 2 | 1 | NA | NA | 4 | 1 | 1 | 1 | 1 | 1 | 4 | 0.75 | 1 | 1 | 7 | 0.8571 |
| 15_845876 | NA | NA | 2 | 1 | 2 | 1 | 2 | 0.5 | 6 | 0.8333 | 2 | 0.5 | 1 | 1 | 5 | 0.4 | 1 | 1 | 9 | 0.5556 |
| 15_845877 | NA | NA | 2 | 1 | 2 | 1 | 2 | 1 | 6 | 1 | 2 | 1 | 2 | 1 | 5 | 0.8 | 1 | 1 | 10 | 0.9 |

| | | | | | | | | | | | | | | | | | | | | |
|---|---|---|---|---|---|---|---|---|---|---|---|---|---|---|---|---|---|---|---|---|
| 15_8467059 | NA | NA | NA | NA | NA | NA | 2 | 1 | 2 | 1 | NA | NA | NA | NA | 2 | 0.5 | 2 | 0 | 4 | 0.25 |
| 15_8467134 | NA | NA | NA | NA | NA | NA | 2 | 0.5 | 2 | 0.5 | NA | NA | NA | NA | 1 | 1 | 1 | 1 | 2 | 1 |
| 15_8467138 | NA | NA | NA | NA | NA | NA | 1 | 1 | 1 | 1 | NA | NA | NA | NA | 1 | 0 | 1 | 1 | 2 | 0.5 |
| 15_8467142 | NA | NA | NA | NA | NA | NA | 3 | 0.6667 | 3 | 0.6667 | NA | NA | NA | NA | 1 | 1 | 1 | 1 | 2 | 1 |
| 15_8481784 | 1 | 1 | 7 | 0.2857 | 3 | 0.3333 | 3 | 0 | 14 | 0.2857 | 2 | 0.5 | 2 | 0 | 3 | 0.6667 | 3 | 0 | 10 | 0.3 |
| 15_8481921 | 1 | 1 | 6 | 0.5 | 3 | 0.6667 | 3 | 0.6667 | 13 | 0.6154 | 2 | 0 | 2 | 0 | 2 | 0.5 | 3 | 0 | 9 | 0.1111 |
| 15_8481927 | 1 | 0 | 7 | 0.2857 | 2 | 0 | 3 | 0 | 13 | 0.1538 | 2 | 0 | 2 | 0.5 | 3 | 0.3333 | 3 | 0 | 10 | 0.2 |
| 15_8481929 | 1 | 1 | 5 | 0.6 | 3 | 1 | 3 | 1 | 12 | 0.8333 | 2 | 1 | 1 | 1 | 3 | 1 | 3 | 1 | 9 | 1 |
| 15_8481946 | NA | NA | 6 | 0.3333 | 2 | 0.5 | 3 | 0 | 11 | 0.2727 | 2 | 1 | 2 | 0.5 | 3 | 0.6667 | 3 | 0.6667 | 10 | 0.7 |
| 15_8482413 | 3 | 0.6667 | 7 | 0.5714 | 4 | 0.25 | 4 | 0.5 | 18 | 0.5 | 4 | 0.5 | 3 | 0.3333 | 6 | 0.3333 | 2 | 0.5 | 15 | 0.4 |
| 15_8482414 | 3 | 0.6667 | 5 | 0.6 | 5 | 0.8 | 3 | 0.6667 | 16 | 0.6875 | 2 | 1 | 3 | 0.6667 | 6 | 0.3333 | 4 | 0.75 | 15 | 0.6 |
| 15_8482415 | 2 | 0 | 7 | 0 | 4 | 0 | 4 | 0.25 | 17 | 0.0588 | 4 | 0 | 3 | 0 | 6 | 0 | 4 | 0.25 | 17 | 0.0588 |
| 15_8482416 | 2 | 1 | 6 | 0.6667 | 5 | 0.8 | 3 | 0.6667 | 16 | 0.75 | 3 | 1 | 3 | 1 | 7 | 1 | 4 | 0.75 | 17 | 0.9412 |
| 15_8482423 | 2 | 1 | NA | NA | 3 | 0.6667 | 4 | 0.75 | 9 | 0.7778 | 3 | 0.6667 | 2 | 0.5 | 3 | 0.6667 | 2 | 1 | 10 | 0.7 |
| 15_8482432 | 1 | 1 | 4 | 1 | 3 | 1 | 3 | 1 | 11 | 1 | 3 | 1 | 2 | 1 | 4 | 1 | 3 | 1 | 12 | 1 |
| 15_8482445 | 1 | 1 | 4 | 0.5 | 3 | 0 | 3 | 0.3333 | 11 | 0.3636 | 3 | 0.3333 | 2 | 1 | 4 | 0.25 | 3 | 0.6667 | 12 | 0.5 |
| 15_8482452 | 1 | 0 | 3 | 0.6667 | 2 | 0 | 3 | 0.6667 | 9 | 0.4444 | 3 | 0.3333 | 2 | 0 | 3 | 0.3333 | 3 | 0.3333 | 11 | 0.2727 |
| 15_8482459 | NA | NA | 3 | 0.3333 | 3 | 0.3333 | 3 | 0.3333 | 9 | 0.3333 | 2 | 1 | 2 | 0 | 2 | 0.5 | 2 | 0 | 8 | 0.375 |
| 15_8482465 | 1 | 1 | 3 | 0.6667 | 3 | 0.6667 | 3 | 0.6667 | 10 | 0.7 | 3 | 0.3333 | 2 | 1 | 4 | 0.5 | 2 | 1 | 11 | 0.6364 |
| 15_8482466 | 1 | 0 | 1 | 0 | 2 | 1 | 2 | 0.5 | 6 | 0.5 | 3 | 0.3333 | 2 | 1 | 2 | 0 | 3 | 0.3333 | 10 | 0.4 |
| 15_8482597 | NA | NA | 3 | 0.6667 | 2 | 0.5 | 4 | 0.5 | 9 | 0.5556 | 4 | 0.5 | 1 | 0 | 3 | 0.6667 | 4 | 0.5 | 12 | 0.5 |
| 15_8482607 | 2 | 0.5 | 4 | 0.25 | 3 | 0 | 5 | 0.2 | 14 | 0.2143 | 4 | 0 | 1 | 0 | 5 | 0 | 5 | 0.2 | 15 | 0.0667 |
| 15_8482671 | 1 | 0 | 4 | 0.5 | 3 | 0 | 3 | 0.3333 | 11 | 0.2727 | 4 | 0.5 | 2 | 0 | 4 | 0 | 5 | 0.6 | 15 | 0.3333 |
| 15_8482681 | 2 | 0 | 4 | 0 | 4 | 0.25 | 6 | 0 | 16 | 0.0625 | 3 | 0 | 3 | 0 | 6 | 0.1667 | 5 | 0 | 17 | 0.0588 |
| 15_8482695 | 2 | 1 | 4 | 1 | 4 | 1 | 3 | 0.3333 | 13 | 0.8462 | 4 | 1 | 3 | 0.6667 | 4 | 0.5 | 3 | 1 | 14 | 0.7857 |
| 15_8482703 | 2 | 0 | 4 | 0.75 | 4 | 0 | 6 | 0.1667 | 16 | 0.25 | 4 | 0.25 | 3 | 0 | 5 | 0 | 5 | 0.2 | 17 | 0.1176 |
| 15_8482825 | 8 | 0 | 8 | 0.5 | 6 | 0 | 4 | 0.25 | 26 | 0.1923 | 6 | 0.5 | 8 | 0.375 | 6 | 0.5 | 4 | 0.5 | 24 | 0.4583 |
| 15_8482843 | 10 | 0.3 | 9 | 0.1111 | 6 | 0 | 7 | 0 | 32 | 0.125 | 7 | 0 | 7 | 0.2857 | 6 | 0.3333 | 6 | 0.1667 | 26 | 0.1923 |
| 15_8482921 | 7 | 0.4286 | 4 | 0.25 | 4 | 0 | 3 | 0 | 18 | 0.2222 | 4 | 0 | 4 | 0 | 1 | 0 | 2 | 0.5 | 11 | 0.0909 |
| 15_8483235 | 4 | 0 | 5 | 0.6 | 7 | 0.4286 | 9 | 0.2222 | 25 | 0.32 | 5 | 0.2 | 7 | 0.2857 | 3 | 0.3333 | 4 | 0.5 | 19 | 0.3158 |
| 15_8483241 | 6 | 0.3333 | 3 | 1 | 5 | 0.8 | 9 | 0.4444 | 23 | 0.5652 | 3 | 0.3333 | 5 | 0 | 3 | 0 | 3 | 0.6667 | 14 | 0.2143 |
| 15_8483284 | 5 | 0.2 | 3 | 0 | 3 | 0 | 7 | 0.1429 | 18 | 0.1111 | 4 | 0.5 | 4 | 0 | 2 | 0 | 2 | 0 | 12 | 0.1667 |
| 15_8483292 | 4 | 0.5 | 4 | 0.75 | 2 | 0.5 | 8 | 0.5 | 18 | 0.5556 | 3 | 0.3333 | 5 | 0.4 | 1 | 0 | 2 | 0.5 | 11 | 0.3636 |
| 15_8483300 | 3 | 1 | 3 | 1 | 2 | 1 | 8 | 0.75 | 16 | 0.875 | 2 | 1 | 4 | 0.75 | 2 | 1 | 1 | 1 | 9 | 0.8889 |
| 15_8483310 | 3 | 0.3333 | 2 | 0 | 4 | 0 | 6 | 0.3333 | 15 | 0.2 | NA | NA | 3 | 0.3333 | 2 | 0 | NA | NA | 5 | 0.2 |
| 15_8483316 | 3 | 0.6667 | 3 | 1 | 3 | 0.6667 | 7 | 1 | 16 | 0.875 | 2 | 1 | 6 | 0.5 | 2 | 1 | 3 | 1 | 13 | 0.7692 |
| 15_8483327 | 2 | 1 | 3 | 0.3333 | 2 | 0.5 | 8 | 0.125 | 15 | 0.3333 | 2 | 0 | 3 | 0 | 1 | 0 | 2 | 0.5 | 8 | 0.125 |
| 15_8493788 | 3 | 0.6667 | 1 | 1 | 2 | 1 | 2 | 0.5 | 8 | 0.75 | NA | NA | NA | NA | 1 | 1 | 3 | 0.6667 | 4 | 0.75 |
| 15_8493798 | 4 | 0.75 | 1 | 0 | 2 | 0.5 | 3 | 0.6667 | 10 | 0.6 | 2 | 0.5 | 2 | 1 | 1 | 1 | 3 | 0.6667 | 8 | 0.75 |
| 15_8494585 | 3 | 1 | 3 | 1 | 2 | 0.5 | 3 | 0.6667 | 11 | 0.8182 | 2 | 0.5 | 1 | 0 | NA | NA | 1 | 0 | 4 | 0.25 |
| 15_8494595 | 4 | 0.75 | 3 | 0.3333 | 2 | 1 | 3 | 0 | 12 | 0.5 | 3 | 0.6667 | 2 | 0.5 | 1 | 1 | 2 | 1 | 8 | 0.75 |
| 15_8494598 | 3 | 1 | 2 | 0.5 | 2 | 0.5 | 3 | 0.6667 | 10 | 0.7 | 3 | 1 | 3 | 1 | 1 | 1 | 2 | 1 | 9 | 1 |
| 15_8494598 | 3 | 0.3333 | 1 | 1 | 2 | 1 | 2 | 0 | 8 | 0.5 | 3 | 0.6667 | 3 | 0.3333 | NA | NA | 1 | 1 | 7 | 0.5714 |
| 15_8494615 | 4 | 0.25 | 5 | 0 | 2 | 0 | 4 | 0 | 15 | 0.0667 | 3 | 0 | 3 | 0 | 1 | 0 | 3 | 0.3333 | 10 | 0.1 |
| 15_8494677 | 6 | 0 | 5 | 0 | 3 | 0 | 5 | 0.2 | 19 | 0.0526 | 4 | 0 | 3 | 0.3333 | 1 | 0 | 3 | 0 | 11 | 0.0909 |
| 15_8513590 | 1 | 1 | 2 | 0.5 | NA | NA | 1 | 1 | 4 | 0.75 | NA | NA | 1 | 0 | 1 | 0 | 2 | 1 | 4 | 0.5 |
| 15_8534720 | 26 | 0.1538 | 43 | 0.3256 | 28 | 0.25 | 34 | 0.0882 | 131 | 0.2137 | 32 | 0.1875 | 30 | 0.2 | 40 | 0.15 | 39 | 0.1026 | 141 | 0.156 |
| 15_8534741 | NA | NA | 7 | 0.4286 | 4 | 0 | 4 | 0.5 | 15 | 0.3333 | 1 | 0 | 2 | 0 | 4 | 0 | 6 | 0.5 | 13 | 0.2308 |
| 15_8534742 | NA | NA | 6 | 0.1667 | 4 | 0 | 4 | 0.5 | 14 | 0.2143 | 1 | 0 | 2 | 0 | 4 | 0 | 6 | 0.1667 | 13 | 0.0769 |
| 15_8534750 | 1 | 0 | 7 | 0.5714 | 4 | 0.5 | 4 | 0.5 | 16 | 0.5 | 1 | 0 | 2 | 0.5 | 3 | 0 | 5 | 0.2 | 11 | 0.1818 |
| 15_8534773 | 2 | 0 | 7 | 0.1429 | 3 | 0 | 4 | 0.5 | 16 | 0.1875 | 1 | 0 | 2 | 0 | 4 | 0 | 7 | 0.1429 | 14 | 0.0714 |
| 15_8534799 | 2 | 0 | 7 | 0 | 4 | 0.25 | 5 | 0.2 | 18 | 0.1111 | 1 | 0 | 3 | 0 | 4 | 0.25 | 7 | 0.1429 | 15 | 0.1333 |
| 15_8542023 | 7 | 0.4286 | 6 | 0.3333 | 4 | 1 | 1 | 1 | 18 | 0.5556 | 10 | 0.4 | 10 | 0.5 | 2 | 0.5 | 8 | 0.25 | 30 | 0.4 |
| 15_8542041 | 8 | 0 | 6 | 0.1667 | 7 | 0 | 2 | 0.5 | 23 | 0.087 | 12 | 0 | 13 | 0.1538 | 2 | 0 | 10 | 0.1 | 37 | 0.0811 |
| 15_8542048 | 8 | 0.125 | 9 | 0.1111 | 6 | 0 | 2 | 0 | 25 | 0.08 | 11 | 0.1818 | 14 | 0 | 3 | 0 | 12 | 0 | 40 | 0.05 |
| 15_8542056 | 7 | 0.2857 | 9 | 0.3333 | 5 | 0.2 | 2 | 0 | 23 | 0.2609 | 13 | 0.1538 | 14 | 0.0714 | 2 | 0 | 11 | 0.3636 | 40 | 0.175 |
| 15_8542073 | 7 | 0.8571 | 9 | 0.6667 | 5 | 0.6 | 2 | 1 | 23 | 0.7391 | 11 | 0.5455 | 11 | 0.4545 | 2 | 0 | 10 | 0.6 | 34 | 0.5 |
| 15_8542076 | 7 | 0.4286 | 9 | 0.6667 | 7 | 0.4286 | 2 | 0.5 | 25 | 0.52 | 10 | 0.3 | 11 | 0.1818 | 3 | 0 | 10 | 0.5 | 34 | 0.2941 |
| 15_8542079 | 8 | 0.875 | 7 | 0.7143 | 6 | 0.5 | 1 | 1 | 22 | 0.7273 | 10 | 0.2 | 10 | 0.3 | 2 | 0.5 | 9 | 0.6667 | 31 | 0.3871 |
| 15_8542093 | 6 | 1 | 8 | 0.875 | 7 | 0.5714 | 2 | 1 | 23 | 0.8261 | 11 | 0.6364 | 14 | 0.8571 | 2 | 1 | 10 | 0.9 | 37 | 0.8108 |
| 15_8542096 | 5 | 1 | 9 | 0.7778 | 4 | 0.75 | 2 | 1 | 20 | 0.85 | 9 | 0.7778 | 11 | 0.6364 | 1 | 1 | 8 | 0.625 | 29 | 0.6897 |
| 15_8542102 | 5 | 0.4 | 9 | 0.4444 | 5 | 0 | 2 | 0 | 21 | 0.2857 | 8 | 0.25 | 12 | 0.3333 | 2 | 0 | 8 | 0.5 | 30 | 0.3333 |
| 15_8542107 | 6 | 0.1667 | 7 | 0.1429 | 6 | 0 | 2 | 0.5 | 21 | 0.1429 | 9 | 0.3333 | 11 | 0.1818 | 2 | 0 | 8 | 0.375 | 30 | 0.2667 |
| 15_8542112 | 5 | 0.8 | 8 | 0.75 | 5 | 0.6 | 2 | 1 | 20 | 0.75 | 8 | 0.5 | 11 | 0.7273 | 2 | 0 | 9 | 0.4444 | 30 | 0.5333 |
| 15_8542117 | 6 | 1 | 9 | 1 | 5 | 1 | 2 | 1 | 22 | 1 | 10 | 1 | 8 | 1 | 2 | 1 | 9 | 1 | 29 | 1 |

| | | | | | | | | | | | | | | | | | | | | |
|---|---|---|---|---|---|---|---|---|---|---|---|---|---|---|---|---|---|---|---|---|
| 15_854212 | 5 | 0.8 | 9 | 0.8889 | 2 | 0.5 | 1 | 0 | 17 | 0.7647 | 9 | 0.6667 | 9 | 0.4444 | 2 | 1 | 8 | 0.75 | 28 | 0.6429 |
| 15_854213 | 4 | 0.5 | 8 | 0.75 | 2 | 0.5 | NA | NA | 14 | 0.6429 | 8 | 0.375 | 9 | 0.5556 | 1 | 0 | 6 | 0.6667 | 24 | 0.5 |
| 15_854213 | 3 | 1 | 9 | 1 | 3 | 1 | 2 | 1 | 17 | 1 | 9 | 1 | 10 | 1 | 2 | 1 | 7 | 1 | 28 | 1 |
| 15_854213 | 4 | 1 | 9 | 1 | 4 | 1 | 2 | 1 | 19 | 1 | 6 | 1 | 10 | 1 | 2 | 1 | 7 | 1 | 25 | 1 |
| 15_854213 | 5 | 0.6 | 7 | 0.1429 | 4 | 0 | 2 | 0 | 18 | 0.2222 | 7 | 0.1429 | 9 | 0.1111 | 1 | 1 | 6 | 0 | 23 | 0.1304 |
| 15_854213 | 5 | 0.2 | 9 | 0.2222 | 4 | 0.5 | 1 | 0 | 19 | 0.2632 | 10 | 0.3 | 10 | 0.4 | 2 | 0 | 6 | 0.5 | 28 | 0.3571 |
| 15_854216 | 4 | 0.75 | 5 | 0.2 | 4 | 0.25 | 1 | 1 | 14 | 0.4286 | 5 | 0.4 | 9 | 0.5556 | 2 | 0 | 1 | 0 | 17 | 0.4118 |
| 15_854218 | 3 | 0.3333 | 7 | 0.4286 | 4 | 0.75 | NA | NA | 14 | 0.5 | 6 | 0.5 | 9 | 0.4444 | 2 | 1 | 3 | 0.6667 | 20 | 0.55 |
| 15_854219 | 1 | 0 | 4 | 0.25 | 3 | 0.3333 | NA | NA | 8 | 0.25 | 6 | 0 | 8 | 0.25 | 1 | 1 | 2 | 0 | 17 | 0.1765 |
| 15_856896 | NA | NA | 1 | 1 | 1 | 1 | NA | NA | 2 | 1 | NA | NA | 1 | 1 | NA | NA | 2 | 1 | 3 | 1 |
| 15_856913 | NA | NA | 2 | 0 | 3 | 0 | 2 | 0.5 | 7 | 0.1429 | 1 | 1 | NA | NA | 3 | 0 | 5 | 0.4 | 9 | 0.3333 |
| 15_856914 | NA | NA | 2 | 0 | 3 | 0.3333 | 2 | 0 | 7 | 0.1429 | 1 | 0 | NA | NA | 2 | 0 | 6 | 0.5 | 9 | 0.3333 |
| 15_856914 | NA | NA | 1 | 0 | 2 | 1 | 1 | 1 | 4 | 0.75 | NA | NA | NA | NA | 2 | 0.5 | 5 | 0.6 | 7 | 0.5714 |
| 15_856914 | NA | NA | 2 | 0 | 3 | 0.3333 | 2 | 1 | 7 | 0.4286 | 1 | 1 | NA | NA | 3 | 0.6667 | 5 | 0.4 | 9 | 0.5556 |
| 15_856915 | NA | NA | 2 | 0 | 2 | 0.5 | 1 | 0 | 5 | 0.2 | 1 | 1 | NA | NA | 3 | 0.3333 | 4 | 0.75 | 8 | 0.625 |
| 15_856917 | NA | NA | 2 | 0 | 3 | 0.3333 | 2 | 0 | 7 | 0.1429 | 1 | 1 | NA | NA | 3 | 0 | 4 | 0 | 8 | 0.125 |
| 15_857408 | NA | NA | NA | NA | 1 | 1 | NA | NA | 1 | 1 | NA | NA | NA | NA | 1 | 1 | NA | NA | 1 | 1 |
| 15_857408 | NA | NA | NA | NA | 1 | 1 | NA | NA | 1 | 1 | NA | NA | NA | NA | 1 | 1 | NA | NA | 1 | 1 |
| 15_857412 | NA | NA | NA | NA | 1 | 1 | NA | NA | 1 | 1 | NA | NA | NA | NA | 1 | 1 | NA | NA | 1 | 1 |
| 15_857415 | NA | NA | NA | NA | 1 | 1 | NA | NA | 1 | 1 | NA | NA | NA | NA | 1 | 1 | NA | NA | 1 | 1 |
| 15_857813 | NA | NA | NA | NA | 2 | 1 | 1 | 0 | 3 | 0.6667 | 2 | 0 | 3 | 0 | 2 | 0.5 | NA | NA | 7 | 0.1429 |
| 15_857815 | NA | NA | NA | NA | 1 | 0 | 2 | 0.5 | 3 | 0.3333 | 2 | 0 | 5 | 0.2 | 4 | 1 | NA | NA | 11 | 0.4545 |
| 15_857832 | NA | NA | 1 | 1 | 1 | 0 | 2 | 1 | 4 | 0.75 | 1 | 1 | NA | NA | NA | NA | NA | NA | 1 | 1 |
| 15_857835 | NA | NA | 1 | 1 | 1 | 1 | 1 | 0 | 3 | 0.6667 | 1 | 1 | NA | NA | NA | NA | NA | NA | 1 | 1 |
| 15_857836 | NA | NA | 1 | 1 | 1 | 1 | 1 | 1 | 3 | 1 | 1 | 1 | NA | NA | NA | NA | NA | NA | 1 | 1 |
| 15_857837 | NA | NA | 1 | 1 | 1 | 1 | NA | NA | 2 | 1 | 1 | 1 | NA | NA | NA | NA | NA | NA | 1 | 1 |
| 15_857843 | NA | NA | 1 | 1 | 1 | 1 | 1 | 0 | 3 | 0.6667 | 1 | 1 | NA | NA | NA | NA | NA | NA | 1 | 1 |
| 15_857844 | NA | NA | 1 | 1 | 1 | 0 | 1 | 0 | 3 | 0.3333 | 1 | 1 | NA | NA | NA | NA | NA | NA | 1 | 1 |
| 15_857845 | NA | NA | NA | NA | 1 | 1 | 1 | 1 | 2 | 1 | 1 | 1 | NA | NA | NA | NA | NA | NA | 1 | 1 |
| 15_857846 | NA | NA | 1 | 1 | 1 | 1 | 1 | 1 | 3 | 1 | 1 | 1 | NA | NA | NA | NA | NA | NA | 1 | 1 |
| 15_857847 | NA | NA | 1 | 1 | 1 | 1 | 1 | 0 | 3 | 0.6667 | 1 | 1 | NA | NA | NA | NA | NA | NA | 1 | 1 |
| 15_857848 | NA | NA | 1 | 0 | 1 | 1 | 1 | 1 | 3 | 0.6667 | 1 | 1 | NA | NA | NA | NA | NA | NA | 1 | 1 |
| 15_860196 | 2 | 1 | 3 | 1 | 3 | 0.6667 | 2 | 1 | 10 | 0.9 | 3 | 0.6667 | 2 | 1 | 2 | 1 | 2 | 1 | 9 | 0.8889 |
| 15_860197 | 2 | 0 | 5 | 0 | 3 | 0.6667 | 2 | 0 | 12 | 0.1667 | 5 | 0 | 3 | 0 | 2 | 0 | 3 | 0.3333 | 13 | 0.0769 |
| 15_860197 | 1 | 1 | 5 | 0.8 | 3 | 1 | 1 | 1 | 10 | 0.9 | 4 | 0.75 | 3 | 0.6667 | 1 | 1 | 3 | 1 | 11 | 0.8182 |
| 15_860197 | 2 | 0.5 | 2 | 0.5 | 1 | 1 | 2 | 1 | 7 | 0.7143 | 5 | 0.8 | 2 | 1 | 2 | 1 | 3 | 1 | 12 | 0.9167 |
| 15_860198 | 2 | 0 | 2 | 0.5 | 3 | 0 | 2 | 0.5 | 9 | 0.2222 | 4 | 0.5 | 3 | 0 | 2 | 0 | 2 | 0 | 11 | 0.1818 |
| 15_860199 | 2 | 0 | 6 | 0.5 | 1 | 0 | 2 | 1 | 11 | 0.4545 | 4 | 0 | 4 | 0.5 | 2 | 1 | 2 | 0.5 | 12 | 0.4167 |
| 15_860201 | 3 | 0.3333 | 6 | 0.5 | 3 | 0.6667 | 2 | 0 | 14 | 0.4286 | 4 | 1 | 4 | 1 | 2 | 1 | 3 | 0.3333 | 13 | 0.8462 |
| 15_860203 | 2 | 0 | 5 | 0 | 2 | 0 | 4 | 0.25 | 13 | 0.0769 | 5 | 0 | 5 | 0.2 | 2 | 0 | 3 | 0.3333 | 15 | 0.1333 |
| 15_860204 | 2 | 0.5 | 6 | 0.3333 | 3 | 0 | 4 | 0.25 | 15 | 0.2667 | 5 | 0.4 | 5 | 0.4 | 2 | 0 | 3 | 0.3333 | 15 | 0.3333 |
| 15_860205 | 3 | 0 | 6 | 0 | 3 | 0 | 4 | 0.25 | 16 | 0.0625 | 4 | 0 | 5 | 0.2 | 1 | 0 | 1 | 0 | 11 | 0.0909 |
| 15_860209 | 3 | 0 | 7 | 0.2857 | 3 | 0 | 7 | 0 | 20 | 0.1 | 3 | 0.3333 | 5 | 0 | 2 | 0 | 2 | 0 | 12 | 0.0833 |
| 15_860215 | 4 | 0.5 | 6 | 0.1667 | 5 | 0.4 | 6 | 0 | 21 | 0.2381 | 5 | 0.2 | 5 | 0.4 | 2 | 0 | 3 | 0 | 15 | 0.2 |
| 15_860217 | 5 | 0.6 | 5 | 0.2 | 7 | 0 | 7 | 0 | 24 | 0.1667 | 5 | 0 | 6 | 0.1667 | 4 | 0 | 3 | 0 | 18 | 0.0556 |
| 15_860234 | 9 | 0 | 11 | 0.1818 | 10 | 0.1 | 8 | 0.375 | 38 | 0.1579 | 6 | 0.1667 | 6 | 0 | 9 | 0.1111 | 1 | 1 | 22 | 0.1364 |
| 15_860239 | 8 | 0.125 | 8 | 0 | 7 | 0.1429 | 4 | 0 | 27 | 0.0741 | 2 | 0 | 5 | 0.2 | 9 | 0.1111 | 3 | 0.3333 | 19 | 0.1579 |
| 15_860242 | 7 | 0.4286 | 9 | 0 | 9 | 0.1111 | 4 | 0.25 | 29 | 0.1724 | 4 | 0 | 4 | 0.25 | 9 | 0.1111 | 3 | 0.6667 | 20 | 0.2 |
| 15_860243 | 7 | 0.1429 | 8 | 0.125 | 8 | 0 | 4 | 0.25 | 27 | 0.1111 | 3 | 0 | 4 | 0 | 8 | 0.125 | 2 | 0 | 17 | 0.0588 |
| 15_860244 | 8 | 0.375 | 8 | 1 | 6 | 0.8333 | 5 | 0.6 | 27 | 0.7037 | 2 | 0.5 | 4 | 0 | 9 | 0.7778 | 3 | 0.6667 | 18 | 0.5556 |
| 15_860245 | 8 | 0.25 | 10 | 0.4 | 9 | 0.1111 | 6 | 0.1667 | 33 | 0.2424 | 2 | 0 | 4 | 0.25 | 9 | 0.1111 | 3 | 0.3333 | 18 | 0.1667 |
| 15_860248 | 4 | 1 | 7 | 0.8571 | 5 | 1 | 4 | 1 | 20 | 0.95 | 1 | 1 | 3 | 1 | 5 | 1 | 2 | 1 | 11 | 1 |
| 15_860249 | 7 | 0.2857 | 8 | 0.25 | 5 | 0.2 | 4 | 0.25 | 24 | 0.25 | 1 | 0 | 3 | 0.3333 | 5 | 0.2 | 2 | 0 | 11 | 0.1818 |
| 15_860250 | 5 | 0.8 | 10 | 0.3 | 7 | 0.1429 | 5 | 0.2 | 27 | 0.3333 | NA | NA | 4 | 0.25 | 4 | 0.75 | 3 | 0.6667 | 11 | 0.5455 |
| 15_860252 | 6 | 0.6667 | 9 | 0.6667 | 6 | 0.8333 | 5 | 0.4 | 26 | 0.6538 | NA | NA | 2 | 1 | 7 | 0.8571 | 2 | 0 | 11 | 0.7273 |
| 15_860253 | 6 | 1 | 9 | 0.8889 | 5 | 1 | 5 | 1 | 25 | 0.96 | NA | NA | 1 | 1 | 5 | 1 | 2 | 1 | 8 | 1 |
| 15_860254 | 6 | 0.5 | 9 | 0 | 6 | 0.1667 | 4 | 0.5 | 25 | 0.24 | NA | NA | 2 | 0.5 | 5 | 0.2 | 3 | 0.3333 | 10 | 0.3 |
| 15_860254 | 7 | 1 | 8 | 0.875 | 6 | 1 | 5 | 1 | 26 | 0.9615 | NA | NA | 2 | 1 | 5 | 1 | 3 | 1 | 10 | 1 |
| 15_860256 | 7 | 0.2857 | 9 | 0.3333 | 5 | 0.4 | 4 | 0.5 | 25 | 0.36 | NA | NA | 2 | 0 | 4 | 1 | 2 | 1 | 8 | 0.75 |
| 15_860256 | 7 | 0.2857 | 9 | 0.2222 | 6 | 0.1667 | 4 | 0.25 | 26 | 0.2308 | NA | NA | 2 | 0 | 5 | 1 | 3 | 0.6667 | 10 | 0.7 |
| 15_860983 | 2 | 1 | NA | NA | 2 | 1 | 1 | 1 | 5 | 1 | 4 | 1 | NA | NA | NA | NA | 1 | 1 | 5 | 1 |
| 15_860984 | 2 | 0.5 | NA | NA | 2 | 1 | 1 | 0 | 5 | 0.6 | 4 | 0.75 | NA | NA | NA | NA | 1 | 1 | 5 | 0.8 |
| 15_863812 | 3 | 0.6667 | 1 | 1 | 2 | 0 | NA | NA | 6 | 0.5 | 2 | 0 | 4 | 0 | 2 | 0.5 | 1 | 0 | 9 | 0.1111 |
| 15_863822 | 4 | 0 | 2 | 0 | 7 | 0.1429 | 3 | 0 | 16 | 0.0625 | 3 | 0 | 4 | 0 | 3 | 0.6667 | 2 | 0 | 12 | 0.1667 |
| 15_874291 | 1 | 1 | NA | NA | NA | NA | NA | NA | 1 | 1 | NA | NA | 1 | 1 | 2 | 0 | NA | NA | 3 | 0.3333 |

| | | | | | | | | | | | | | | | | | | | | |
|---|---|---|---|---|---|---|---|---|---|---|---|---|---|---|---|---|---|---|---|---|
| 15_8750888 | 1 | 1 | 1 | 1 | 1 | 1 | 2 | 0.5 | 5 | 0.8 | 2 | 0.5 | 4 | 1 | 4 | 0.25 | 1 | 0 | 11 | 0.5455 |
| 15_8750934 | 5 | 0.4 | 3 | 0.6667 | 3 | 0.6667 | 1 | 0 | 12 | 0.5 | 4 | 0.5 | 7 | 0.1429 | 6 | 0.3333 | 3 | 0.3333 | 20 | 0.3 |
| 15_8750954 | 7 | 0 | 2 | 0 | 4 | 0.25 | 3 | 0 | 16 | 0.0625 | 5 | 0.2 | 5 | 0 | 7 | 0.2857 | 4 | 0 | 21 | 0.1429 |
| 15_8764159 | 14 | 0.2857 | 15 | 0.4 | 15 | 0.2667 | 13 | 0.1538 | 57 | 0.2807 | 12 | 0.1667 | 19 | 0.3158 | 14 | 0.1429 | 6 | 0.1667 | 51 | 0.2157 |
| 15_8764164 | 14 | 0.3571 | 12 | 0.4167 | 16 | 0.625 | 15 | 0.6667 | 57 | 0.5263 | 7 | 0.4286 | 18 | 0.4444 | 10 | 0.7 | 8 | 0.25 | 43 | 0.4651 |
| 15_8764170 | 15 | 0.9333 | 14 | 0.5714 | 17 | 0.7059 | 15 | 0.6667 | 61 | 0.7213 | 16 | 0.8125 | 24 | 0.75 | 16 | 0.6875 | 10 | 0.9 | 66 | 0.7727 |
| 15_8764171 | 13 | 0.1538 | 14 | 0 | 15 | 0.2 | 10 | 0.2 | 52 | 0.1346 | 11 | 0 | 20 | 0.05 | 16 | 0.1875 | 8 | 0 | 55 | 0.0727 |
| 15_8764181 | 18 | 0.3333 | 15 | 0.0667 | 18 | 0.6111 | 15 | 0.6 | 66 | 0.4091 | 17 | 0.2941 | 21 | 0.3333 | 15 | 0.4 | 10 | 0.4 | 63 | 0.3492 |
| 15_8764186 | 14 | 0.9286 | 13 | 1 | 11 | 1 | 12 | 1 | 50 | 0.98 | 8 | 0.875 | 21 | 1 | 15 | 1 | 10 | 1 | 54 | 0.9815 |
| 15_8764190 | 15 | 1 | 17 | 0.9412 | 17 | 1 | 16 | 1 | 65 | 0.9846 | 14 | 1 | 22 | 1 | 14 | 1 | 10 | 0.8 | 60 | 0.9667 |
| 15_8764195 | 19 | 0.6842 | 15 | 0.4 | 16 | 0.3125 | 15 | 0.4667 | 65 | 0.4769 | 13 | 0.3846 | 23 | 0.4783 | 13 | 0.6154 | 11 | 0.5455 | 60 | 0.5 |
| 15_8764198 | 17 | 1 | 15 | 0.8667 | 18 | 0.8889 | 15 | 1 | 65 | 0.9385 | 12 | 0.9167 | 23 | 0.9565 | 15 | 0.8667 | 12 | 1 | 62 | 0.9355 |
| 15_8764210 | 14 | 0.3571 | 15 | 0.2667 | 13 | 0.3846 | 16 | 0.1875 | 58 | 0.2931 | 11 | 0.0909 | 23 | 0.2609 | 11 | 0.2727 | 12 | 0.3333 | 57 | 0.2456 |
| 15_8764214 | 13 | 0.6154 | 14 | 0.2143 | 15 | 0.4667 | 14 | 0.3571 | 56 | 0.4107 | 14 | 0.5 | 23 | 0.5652 | 13 | 0.4615 | 13 | 0.3846 | 63 | 0.4921 |
| 15_8764217 | 18 | 0.1667 | 14 | 0.2143 | 20 | 0.15 | 14 | 0.1429 | 66 | 0.1667 | 12 | 0.3333 | 24 | 0.2083 | 16 | 0.375 | 13 | 0.0769 | 65 | 0.2462 |
| 15_8764220 | 12 | 0.6667 | 16 | 0.6875 | 17 | 0.8235 | 16 | 0.75 | 61 | 0.7377 | 11 | 0.9091 | 23 | 0.7391 | 12 | 0.5 | 11 | 1 | 57 | 0.7719 |
| 15_8764222 | 16 | 0.5 | 15 | 0.6 | 19 | 0.4211 | 18 | 0.6667 | 68 | 0.5441 | 13 | 0.2308 | 25 | 0.64 | 15 | 0.4667 | 13 | 0.4615 | 66 | 0.4848 |
| 15_8764222 | 16 | 0.0625 | 16 | 0.1875 | 19 | 0.3158 | 17 | 0 | 68 | 0.1471 | 16 | 0.1875 | 26 | 0.1154 | 18 | 0.1667 | 12 | 0.0833 | 72 | 0.1389 |
| 15_8764237 | 15 | 0.5333 | 18 | 0.4444 | 21 | 0.5238 | 18 | 0.6111 | 72 | 0.5278 | 15 | 0.7333 | 27 | 0.6296 | 17 | 0.7059 | 15 | 0.4 | 74 | 0.6216 |
| 15_8764242 | 18 | 0.0556 | 16 | 0 | 19 | 0.3158 | 19 | 0.1053 | 72 | 0.125 | 16 | 0.25 | 22 | 0.1818 | 18 | 0.3889 | 16 | 0.125 | 72 | 0.2361 |
| 15_8764259 | 11 | 0.0909 | 6 | 0.1667 | 8 | 0.25 | 3 | 0.6667 | 28 | 0.2143 | 5 | 0.2 | 10 | 0.3 | 6 | 0.1667 | 5 | 0.6 | 26 | 0.3077 |
| 15_8857150 | 2 | 0 | 10 | 0.1 | 4 | 0 | 2 | 0 | 18 | 0.0556 | NA | NA | 3 | 0.3333 | 3 | 0 | 3 | 0 | 9 | 0.1111 |
| 15_8857240 | 2 | 0.5 | 8 | 0 | 5 | 0 | 2 | 0 | 17 | 0.0588 | 4 | 0.25 | 3 | 0 | 2 | 0 | 6 | 0 | 15 | 0.0667 |
| 15_8857250 | 1 | 0 | 6 | 0 | 3 | 0.6667 | 2 | 0 | 12 | 0.1667 | 3 | 0 | 3 | 0 | 2 | 0 | 5 | 0.2 | 13 | 0.0769 |
| 15_8857251 | 1 | 0 | 6 | 0.1667 | 3 | 0 | 2 | 0 | 12 | 0.0833 | 4 | 0 | 3 | 0.3333 | 2 | 0 | 6 | 0 | 15 | 0.0667 |
| 15_8858513 | 1 | 0 | 5 | 0.4 | NA | NA | NA | NA | 6 | 0.3333 | 1 | 0 | 1 | 0 | 1 | 0 | 3 | 0.3333 | 6 | 0.1667 |
| 15_8861369 | NA | NA | 4 | 0.75 | NA | NA | 1 | 1 | 5 | 0.8 | 1 | 1 | 1 | 1 | NA | NA | 4 | 0.75 | 6 | 0.8333 |
| 15_8861375 | NA | NA | 2 | 0.5 | 1 | 1 | 1 | 1 | 4 | 0.75 | NA | NA | 1 | 0 | 1 | 0 | 3 | 1 | 5 | 0.6 |
| 15_8861392 | NA | NA | 4 | 0 | 1 | 0 | 1 | 1 | 6 | 0.1667 | 1 | 0 | 1 | 0 | 1 | 1 | 2 | 0 | 5 | 0.2 |
| 15_8861404 | NA | NA | 3 | 1 | NA | NA | NA | NA | 3 | 1 | 1 | 1 | 1 | 1 | 1 | 1 | 2 | 0.5 | 5 | 0.8 |
| 15_8861432 | NA | NA | 4 | 0.5 | NA | NA | 1 | 1 | 5 | 0.6 | 1 | 0 | 1 | 1 | 1 | 0 | 2 | 0 | 5 | 0.2 |
| 15_8861436 | NA | NA | 4 | 0.25 | NA | NA | 1 | 0 | 5 | 0.2 | 1 | 0 | 1 | 0 | 1 | 0 | 2 | 0.5 | 5 | 0.2 |
| 15_8861462 | NA | NA | 4 | 1 | NA | NA | 1 | 0 | 5 | 0.8 | 1 | 1 | 1 | 0 | NA | NA | NA | NA | 2 | 0.5 |
| 15_8866585 | NA | NA | NA | NA | 1 | 0 | 1 | 1 | 2 | 0.5 | NA | NA | NA | NA | 1 | 1 | NA | NA | 1 | 1 |
| 15_8866587 | NA | NA | NA | NA | 1 | 0 | 1 | 1 | 2 | 0.5 | 1 | 1 | NA | NA | 1 | 0 | NA | NA | 2 | 0.5 |
| 15_8866701 | NA | NA | 1 | 1 | 1 | 1 | 2 | 1 | 4 | 1 | 2 | 0 | NA | NA | 1 | 1 | NA | NA | 3 | 0.3333 |
| 15_8866705 | NA | NA | NA | NA | 1 | 0 | 3 | 0.6667 | 4 | 0.5 | 2 | 0.5 | NA | NA | 1 | 1 | NA | NA | 3 | 0.6667 |
| 15_8866707 | NA | NA | NA | NA | 1 | 0 | 3 | 0.3333 | 4 | 0.25 | 2 | 0.5 | NA | NA | 1 | 0 | NA | NA | 3 | 0.3333 |
| 15_8866710 | NA | NA | 1 | 1 | NA | NA | 3 | 0.6667 | 4 | 0.75 | 1 | 1 | NA | NA | 1 | 1 | NA | NA | 2 | 1 |
| 15_8866723 | NA | NA | 1 | 0 | 1 | 0 | 3 | 0.3333 | 5 | 0.2 | 1 | 0 | NA | NA | 1 | 1 | NA | NA | 2 | 0.5 |
| 15_8866744 | NA | NA | NA | NA | 1 | 1 | 2 | 0.5 | 3 | 0.6667 | NA | NA | NA | NA | 2 | 1 | NA | NA | 2 | 1 |
| 15_8866746 | NA | NA | 2 | 1 | 1 | 1 | 3 | 0.3333 | 6 | 0.6667 | 2 | 0.5 | NA | NA | 2 | 1 | NA | NA | 4 | 0.75 |
| 15_8866750 | NA | NA | 2 | 1 | 1 | 0 | 3 | 0.3333 | 6 | 0.5 | 2 | 0.5 | NA | NA | 2 | 0 | NA | NA | 4 | 0.25 |
| 15_8866752 | NA | NA | 2 | 0 | 1 | 0 | 3 | 0.3333 | 6 | 0.1667 | 2 | 0.5 | NA | NA | 2 | 0.5 | NA | NA | 4 | 0.5 |
| 15_8866847 | NA | NA | 2 | 0 | 1 | 0 | 3 | 0.3333 | 6 | 0.1667 | 2 | 1 | NA | NA | 2 | 0 | 1 | 0 | 5 | 0.4 |
| 15_8866864 | NA | NA | 2 | 0.5 | NA | NA | 3 | 0.3333 | 5 | 0.4 | 2 | 0 | NA | NA | 1 | 0 | 1 | 1 | 4 | 0.25 |
| 15_8868994 | 2 | 1 | 2 | 0.5 | 2 | 1 | 1 | 1 | 7 | 0.8571 | NA | NA | 1 | 1 | 1 | 1 | NA | NA | 2 | 1 |
| 15_8869002 | 3 | 0.6667 | 2 | 0 | 1 | 0 | 1 | 0 | 7 | 0.2857 | NA | NA | 1 | 1 | 1 | 0 | NA | NA | 2 | 0.5 |
| 15_8869016 | 3 | 0.3333 | 2 | 0 | 2 | 0 | NA | NA | 7 | 0.1429 | NA | NA | 1 | 1 | 1 | 0 | NA | NA | 2 | 0.5 |
| 15_8871071 | 3 | 0.6667 | 7 | 0.2857 | 2 | 0.5 | 6 | 0.5 | 18 | 0.4444 | 5 | 0.4 | 2 | 0.5 | 4 | 0.75 | 4 | 0.75 | 15 | 0.6 |
| 15_8871072 | 4 | 0 | 8 | 0.25 | 2 | 0 | 7 | 0 | 21 | 0.0952 | 6 | 0.1667 | 3 | 0 | 3 | 0 | 5 | 0 | 17 | 0.0588 |
| 15_8871076 | 4 | 0 | 8 | 0.125 | 2 | 0 | 8 | 0.125 | 22 | 0.0909 | 6 | 0.1667 | 2 | 0 | 4 | 0 | 5 | 0 | 17 | 0.0588 |
| 15_8874889 | 6 | 0.8333 | 9 | 0.7778 | 7 | 0.7143 | 6 | 0.6667 | 28 | 0.75 | 4 | 0.5 | 1 | 1 | 4 | 1 | 8 | 0.75 | 17 | 0.7647 |
| 15_8874901 | 4 | 1 | 4 | 1 | 6 | 0.6667 | 6 | 0.8333 | 20 | 0.85 | 3 | 1 | NA | NA | 2 | 1 | 5 | 1 | 10 | 1 |
| 15_8874905 | 8 | 0.125 | 12 | 0.0833 | 6 | 0.1667 | 9 | 0.1111 | 35 | 0.1143 | 4 | 0 | 1 | 0 | 4 | 0.25 | 7 | 0.5714 | 16 | 0.3125 |
| 15_8874907 | 6 | 0.1667 | 9 | 0.2222 | 5 | 0.2 | 7 | 0.2857 | 27 | 0.2222 | 5 | 0.4 | NA | NA | 4 | 0.25 | 7 | 0 | 16 | 0.1875 |
| 15_8874921 | 7 | 0 | 12 | 0.4167 | 8 | 0.125 | 9 | 0.2222 | 36 | 0.2222 | 6 | 0.3333 | 1 | 0 | 4 | 0.25 | 7 | 0.5714 | 18 | 0.3889 |
| 15_8874927 | 7 | 0 | 10 | 0.1 | 8 | 0 | 8 | 0.125 | 33 | 0.0606 | 6 | 0.3333 | 1 | 0 | 3 | 0 | 7 | 0 | 17 | 0.1176 |
| 15_8884116 | 17 | 0.3529 | 15 | 0.7333 | 13 | 0.3077 | 20 | 0.5 | 65 | 0.4769 | 19 | 0.4737 | 16 | 0.5 | 15 | 0.4 | 19 | 0.5263 | 69 | 0.4783 |
| 15_8884120 | 18 | 0.1111 | 15 | 0.4 | 14 | 0.0714 | 21 | 0.0476 | 68 | 0.1471 | 16 | 0.0625 | 19 | 0.1579 | 15 | 0.0667 | 22 | 0.0455 | 72 | 0.0833 |
| 15_8894441 | 1 | 0 | 2 | 0 | 3 | 0.6667 | NA | NA | 6 | 0.3333 | 1 | 0 | 2 | 1 | 2 | 0 | 2 | 0.5 | 7 | 0.4286 |
| 15_8894449 | 4 | 0 | 7 | 0 | 4 | 0.25 | 1 | 0 | 16 | 0.0625 | 1 | 0 | 3 | 0.3333 | 1 | 0 | 2 | 0 | 7 | 0.1429 |
| 15_8895070 | NA | NA | NA | NA | 1 | 1 | 2 | 0 | 3 | 0.3333 | NA | NA | NA | NA | 2 | 0.5 | NA | NA | 2 | 0.5 |
| 15_8895160 | NA | NA | NA | NA | 1 | 1 | 2 | 0.5 | 3 | 0.6667 | NA | NA | NA | NA | 2 | 0.5 | NA | NA | 2 | 0.5 |
| 15_8895183 | 3 | 0.3333 | 6 | 0.5 | 4 | 1 | 2 | 0 | 15 | 0.5333 | NA | NA | 1 | 1 | 2 | 1 | 2 | 0 | 5 | 0.6 |

| | | | | | | | | | | | | | | | | | | | | |
|---|---|---|---|---|---|---|---|---|---|---|---|---|---|---|---|---|---|---|---|---|
| 15_8895241 | 6 | 0.6667 | 9 | 0.4444 | 5 | 0.2 | 3 | 0.6667 | 23 | 0.4783 | 1 | 0 | 5 | 0.2 | 4 | 0.75 | NA | NA | 10 | 0.4 |
| 15_8895263 | 4 | 0.75 | 8 | 0.625 | 4 | 0.75 | 2 | 0 | 18 | 0.6111 | 2 | 0.5 | 4 | 0.5 | 5 | 0.8 | 2 | 1 | 13 | 0.6923 |
| 15_8895273 | 7 | 1 | 7 | 1 | 5 | 0.8 | 3 | 0.6667 | 22 | 0.9091 | 2 | 1 | 4 | 1 | 5 | 1 | 2 | 1 | 13 | 1 |
| 15_8895290 | 7 | 0 | 12 | 0.0833 | 7 | 0.2857 | 3 | 0.6667 | 29 | 0.1724 | 2 | 0.5 | 4 | 0 | 6 | 0 | 2 | 0.5 | 14 | 0.1429 |
| 15_8895294 | 6 | 0.3333 | 9 | 0.1111 | 4 | 0.25 | 3 | 0.3333 | 22 | 0.2273 | 1 | 0 | 2 | 0 | 4 | 0.25 | 2 | 0 | 9 | 0.1111 |
| 15_8895311 | 8 | 0.375 | 12 | 0.3333 | 5 | 0 | 3 | 1 | 28 | 0.3571 | 4 | 0.25 | 5 | 0.6 | 7 | 0.1429 | 4 | 0 | 20 | 0.25 |
| 15_8895398 | 8 | 0 | 14 | 0.1429 | 5 | 0 | 6 | 0 | 33 | 0.0606 | 4 | 0 | 4 | 0.5 | 8 | 0.125 | 3 | 0 | 19 | 0.1579 |
| 15_8895405 | 7 | 0.4286 | 13 | 0.3846 | 7 | 0.2857 | 5 | 0 | 32 | 0.3125 | 4 | 0 | 4 | 0.5 | 9 | 0.5556 | 2 | 0 | 19 | 0.3684 |
| 15_8895410 | 3 | 1 | 7 | 1 | 3 | 1 | 2 | 0.5 | 15 | 0.9333 | 2 | 0.5 | 5 | 0.6 | 1 | 1 | 2 | 1 | 10 | 0.7 |
| 15_8895416 | 5 | 0.8 | 10 | 1 | 6 | 1 | 3 | 0.3333 | 24 | 0.875 | 2 | 0.5 | 5 | 1 | 6 | 0.5 | 2 | 0.5 | 15 | 0.6667 |
| 15_8895449 | 4 | 0.25 | 13 | 0.2308 | 6 | 0.5 | 3 | 0 | 26 | 0.2692 | 3 | 0.6667 | 5 | 0.4 | 7 | 0.2857 | 2 | 0 | 17 | 0.3529 |
| 15_8895678 | NA | NA | 1 | 1 | 1 | 0 | NA | NA | 2 | 0.5 | NA | NA | NA | NA | 1 | 0 | 1 | 1 | 2 | 0.5 |
| 15_8895774 | 4 | 0 | 4 | 0.25 | 2 | 0.5 | 3 | 0 | 13 | 0.1538 | 1 | 1 | NA | NA | 4 | 0 | 2 | 0 | 7 | 0.1429 |
| 15_8897015 | 4 | 0.5 | 8 | 0.125 | 1 | 1 | 1 | 0 | 14 | 0.2857 | 1 | 0 | 2 | 0.5 | 1 | 0 | NA | NA | 4 | 0.25 |
| 15_8897023 | 5 | 0.2 | 5 | 0 | 1 | 0 | 1 | 0 | 12 | 0.0833 | 2 | 0 | 3 | 0.6667 | 1 | 1 | NA | NA | 6 | 0.5 |
| 15_8897032 | 4 | 0 | 6 | 0.1667 | 1 | 0 | 1 | 0 | 12 | 0.0833 | 2 | 0.5 | 3 | 0 | 1 | 0 | NA | NA | 6 | 0.1667 |
| 15_8897043 | 6 | 0.3333 | 8 | 0.125 | 1 | 1 | NA | NA | 15 | 0.2667 | 3 | 0.3333 | 2 | 0.5 | 1 | 0 | NA | NA | 6 | 0.3333 |
| 15_8897044 | 5 | 0.4 | 7 | 0 | NA | NA | NA | NA | 12 | 0.1667 | 2 | 0.5 | 3 | 0.6667 | 1 | 1 | NA | NA | 6 | 0.6667 |
| 15_8897352 | 3 | 0.6667 | 3 | 0 | 4 | 0 | 3 | 0.3333 | 13 | 0.2308 | 3 | 0.3333 | NA | NA | 1 | 0 | 2 | 1 | 6 | 0.5 |
| 15_8897374 | 3 | 0.3333 | 2 | 0 | 3 | 0 | 2 | 0 | 10 | 0.1 | 3 | 0.3333 | 1 | 1 | 1 | 0 | 2 | 0.5 | 7 | 0.4286 |
| 15_8898389 | 6 | 0 | 3 | 0 | 5 | 0 | 2 | 0.5 | 16 | 0.0625 | 3 | 0 | 1 | 0 | 2 | 0 | 2 | 0.5 | 8 | 0.125 |
| 15_8898416 | 6 | 0.3333 | 4 | 0.75 | 6 | 0.3333 | 4 | 0.5 | 20 | 0.45 | 3 | 0.3333 | 2 | 0 | 2 | 0.5 | 2 | 1 | 9 | 0.4444 |
| 15_8898427 | 4 | 0.25 | 2 | 0 | 3 | 0 | 2 | 0 | 11 | 0.0909 | 1 | 0 | NA | NA | 2 | 0 | 2 | 1 | 5 | 0.4 |
| 15_8898427 | 4 | 0.75 | 2 | 0.5 | 3 | 0.6667 | 3 | 0.6667 | 12 | 0.6667 | 1 | 0 | 1 | 1 | 2 | 0.5 | 2 | 1 | 6 | 0.6667 |
| 15_8898440 | 4 | 0.75 | 2 | 0.5 | 3 | 0.6667 | 2 | 0 | 11 | 0.5455 | 1 | 0 | 1 | 1 | 2 | 1 | 2 | 1 | 6 | 0.8333 |
| 15_8898461 | 5 | 0.6 | 1 | 0 | 3 | 0.3333 | 2 | 0 | 11 | 0.3636 | NA | NA | 1 | 1 | 2 | 0.5 | 3 | 0.3333 | 6 | 0.5 |
| 15_8898462 | 3 | 0.3333 | 1 | 0 | 1 | 0 | 2 | 0.5 | 7 | 0.2857 | NA | NA | 1 | 1 | 1 | 1 | 3 | 0.6667 | 5 | 0.8 |
| 15_8898739 | 1 | 0 | 1 | 1 | 2 | 0 | 1 | 0 | 5 | 0.2 | NA | NA | NA | NA | 2 | 0.5 | 2 | 1 | 4 | 0.75 |
| 15_8898761 | NA | NA | NA | NA | 2 | 0 | 1 | 1 | 3 | 0.3333 | NA | NA | NA | NA | 2 | 0.5 | 1 | 1 | 3 | 0.6667 |
| 15_8898762 | NA | NA | NA | NA | 2 | 0.5 | 1 | 1 | 3 | 0.6667 | NA | NA | NA | NA | 2 | 1 | 2 | 0.5 | 4 | 0.75 |
| 15_8898768 | NA | NA | NA | NA | 2 | 1 | 1 | 1 | 3 | 1 | NA | NA | NA | NA | 2 | 1 | 2 | 1 | 4 | 1 |
| 15_8898800 | NA | NA | NA | NA | 2 | 0.5 | 1 | 1 | 3 | 0.6667 | NA | NA | NA | NA | 2 | 0.5 | 2 | 0.5 | 4 | 0.5 |
| 15_8898829 | NA | NA | NA | NA | 1 | 1 | NA | NA | 1 | 1 | NA | NA | NA | NA | 2 | 1 | NA | NA | 2 | 1 |
| 15_8898831 | NA | NA | NA | NA | 1 | 0 | 1 | 1 | 2 | 0.5 | NA | NA | NA | NA | 2 | 0.5 | 1 | 1 | 3 | 0.6667 |
| 15_8901490 | 3 | 0 | 3 | 0.3333 | 1 | 1 | 1 | 0 | 8 | 0.25 | 3 | 0 | 1 | 0 | 1 | 0 | 1 | 1 | 6 | 0.1667 |
| 15_8901491 | 4 | 0.25 | 3 | 0 | 1 | 0 | 1 | 0 | 9 | 0.1111 | 3 | 0.6667 | NA | NA | 1 | 0 | 1 | 0 | 5 | 0.4 |
| 15_8901491 | 4 | 0.25 | 3 | 0.6667 | 1 | 1 | 1 | 0 | 9 | 0.4444 | 3 | 0.6667 | 1 | 1 | 1 | 1 | NA | NA | 5 | 0.8 |
| 15_8901493 | 4 | 0 | 3 | 0.3333 | 1 | 1 | NA | NA | 8 | 0.25 | 3 | 0.3333 | 1 | 0 | 1 | 0 | NA | NA | 5 | 0.2 |
| 15_8901496 | 3 | 0.3333 | 3 | 0 | 1 | 0 | 1 | 0 | 8 | 0.125 | 2 | 0 | 1 | 0 | 1 | 1 | NA | NA | 4 | 0.25 |
| 15_8901507 | 3 | 0.6667 | 2 | 1 | NA | NA | 1 | 0 | 6 | 0.6667 | 2 | 1 | NA | NA | 1 | 0 | NA | NA | 3 | 0.6667 |
| 15_8901511 | 3 | 0.3333 | 2 | 0 | 1 | 0 | 1 | 0 | 7 | 0.1429 | 3 | 0 | 1 | 1 | 1 | 0 | NA | NA | 5 | 0.2 |
| 15_8901517 | 4 | 0 | 3 | 0.3333 | 1 | 0 | NA | NA | 8 | 0.125 | 3 | 0.6667 | 1 | 0 | 1 | 1 | NA | NA | 5 | 0.6 |
| 15_8901520 | 3 | 0.3333 | 3 | 0.3333 | 1 | 0 | 1 | 1 | 8 | 0.375 | 3 | 0.3333 | 1 | 0 | NA | NA | 1 | 1 | 5 | 0.4 |
| 15_8901554 | NA | NA | 1 | 1 | NA | NA | NA | NA | 1 | 1 | NA | NA | NA | NA | 1 | 1 | NA | NA | 1 | 1 |
| 15_8902700 | 1 | 0 | 3 | 0.6667 | 3 | 0.3333 | NA | NA | 7 | 0.4286 | 2 | 1 | NA | NA | NA | NA | NA | NA | 2 | 1 |
| 15_8902756 | 1 | 0 | 4 | 0.25 | 3 | 0.3333 | NA | NA | 8 | 0.25 | 1 | 1 | NA | NA | NA | NA | NA | NA | 1 | 1 |
| 15_8902760 | 1 | 0 | 3 | 0.3333 | 3 | 0.6667 | NA | NA | 7 | 0.4286 | 1 | 1 | NA | NA | NA | NA | NA | NA | 1 | 1 |
| 15_8902781 | 1 | 0 | 3 | 0.3333 | 3 | 0.6667 | NA | NA | 7 | 0.4286 | 1 | 1 | NA | NA | NA | NA | NA | NA | 1 | 1 |
| 15_8902784 | 1 | 0 | 4 | 0.5 | 2 | 0 | NA | NA | 7 | 0.2857 | 1 | 1 | NA | NA | NA | NA | NA | NA | 1 | 1 |
| 15_8903981 | NA | NA | 2 | 0 | 2 | 0.5 | 3 | 0 | 7 | 0.1429 | 2 | 0 | 2 | 0 | 1 | 0 | 4 | 0.25 | 9 | 0.1111 |
| 15_8903990 | NA | NA | 5 | 0 | 6 | 0.1667 | 2 | 0 | 13 | 0.0769 | 4 | 0 | 2 | 0 | 2 | 0.5 | 5 | 0.2 | 13 | 0.1538 |
| 15_8904007 | NA | NA | 3 | 0.3333 | 5 | 0.2 | 4 | 0 | 12 | 0.1667 | 3 | 0 | 3 | 0.3333 | 3 | 0.3333 | 5 | 0.2 | 14 | 0.2143 |
| 15_8904008 | NA | NA | 2 | 1 | 7 | 0.8571 | 5 | 1 | 14 | 0.9286 | 3 | 1 | 3 | 1 | 3 | 0.6667 | 5 | 0.8 | 14 | 0.8571 |
| 15_8904032 | NA | NA | 3 | 1 | 4 | 0.75 | 5 | 0.8 | 12 | 0.8333 | 2 | 0.5 | 3 | 1 | 3 | 0.3333 | 5 | 0.6 | 13 | 0.6154 |
| 15_8905090 | NA | NA | 1 | 0 | NA | NA | 1 | 1 | 2 | 0.5 | NA | NA | NA | NA | NA | NA | 1 | 1 | 1 | 1 |
| 15_8905139 | NA | NA | 1 | 1 | NA | NA | 1 | 1 | 2 | 1 | NA | NA | NA | NA | NA | NA | 1 | 1 | 1 | 1 |
| 15_8905143 | NA | NA | 1 | 1 | NA | NA | 1 | 1 | 2 | 1 | NA | NA | NA | NA | NA | NA | 1 | 1 | 1 | 1 |
| 15_8906772 | NA | NA | NA | NA | 2 | 0.5 | NA | NA | 2 | 0.5 | NA | NA | NA | NA | 2 | 0 | 1 | 1 | 3 | 0.3333 |
| 15_8906902 | 5 | 0.2 | 5 | 0.2 | 6 | 0 | 4 | 0 | 20 | 0.1 | 6 | 0.1667 | 5 | 0 | 8 | 0.25 | 4 | 0 | 23 | 0.1304 |
| 15_8906921 | 4 | 0 | 3 | 0.3333 | 4 | 0.25 | 5 | 0.4 | 16 | 0.25 | 6 | 0 | 5 | 0 | 8 | 0.25 | 5 | 0.4 | 24 | 0.1667 |
| 15_8917061 | 4 | 0.75 | 5 | 0.6 | 7 | 0.7143 | 4 | 0.75 | 20 | 0.7 | 4 | 0.75 | 10 | 0.5 | 6 | 0.5 | 5 | 0.4 | 25 | 0.52 |
| 15_8917097 | 6 | 0 | 9 | 0 | 10 | 0.3 | 4 | 0.25 | 29 | 0.1379 | 9 | 0.2222 | 12 | 0.25 | 6 | 0 | 11 | 0.2727 | 38 | 0.2105 |
| 15_8917101 | 6 | 0 | 9 | 0.1111 | 8 | 0.125 | 3 | 0.3333 | 26 | 0.1154 | 9 | 0.1111 | 13 | 0.0769 | 6 | 0.1667 | 10 | 0.1 | 38 | 0.1053 |
| 15_8917249 | 15 | 0.0667 | 14 | 0.0714 | 11 | 0.0909 | 16 | 0 | 56 | 0.0536 | 11 | 0.0909 | 19 | 0.2632 | 12 | 0 | 13 | 0.1538 | 55 | 0.1455 |
| 15_8917278 | 14 | 0.0714 | 9 | 0.1111 | 15 | 0.0667 | 11 | 0.0909 | 49 | 0.0816 | 11 | 0 | 14 | 0.0714 | 7 | 0.1429 | 10 | 0.1 | 42 | 0.0714 |

| | | | | | | | | | | | | | | | | | | | | |
|---|---|---|---|---|---|---|---|---|---|---|---|---|---|---|---|---|---|---|---|---|
| 15_8917297 | 1 | 1 | 1 | 0 | 1 | 0 | NA | NA | 3 | 0.3333 | 1 | 1 | 2 | 1 | NA | NA | NA | NA | 3 | 1 |
| 15_8917304 | 16 | 0.3125 | 14 | 0.3571 | 15 | 0.8 | 16 | 0.4375 | 61 | 0.4754 | 10 | 0.7 | 17 | 0.3529 | 11 | 0.5455 | 12 | 0.5 | 50 | 0.5 |
| 15_8917312 | 11 | 0.1818 | 11 | 0.4545 | 12 | 0.4167 | 12 | 0.1667 | 46 | 0.3043 | 10 | 0.1 | 16 | 0.125 | 15 | 0.3333 | 10 | 0.2 | 51 | 0.1961 |
| 15_8923537 | 1 | 1 | 2 | 0.5 | 2 | 0 | NA | NA | 5 | 0.4 | 2 | 0.5 | 3 | 0.6667 | 2 | 0 | NA | NA | 7 | 0.4286 |
| 15_8923539 | 2 | 0 | 2 | 0.5 | 2 | 0 | NA | NA | 6 | 0.1667 | 2 | 0 | 4 | 0.5 | 2 | 0 | NA | NA | 8 | 0.25 |
| 15_8923542 | 2 | 1 | 2 | 0 | 1 | 0 | NA | NA | 5 | 0.4 | 2 | 0.5 | 3 | 0 | 2 | 0 | NA | NA | 7 | 0.1429 |
| 15_8923553 | 2 | 1 | 1 | 1 | 1 | 1 | NA | NA | 4 | 1 | 1 | 1 | 3 | 1 | NA | NA | 1 | 1 | 5 | 1 |
| 15_8923582 | 2 | 0 | 2 | 0 | 2 | 0.5 | NA | NA | 6 | 0.1667 | NA | NA | 2 | 1 | 1 | 0 | 1 | 1 | 4 | 0.75 |
| 15_8923590 | 1 | 0 | 2 | 0.5 | 2 | 0 | NA | NA | 5 | 0.2 | NA | NA | 2 | 0 | 1 | 1 | 1 | 0 | 4 | 0.25 |
| 15_8925565 | 5 | 0 | 14 | 0.4286 | 4 | 0.25 | 9 | 0.2222 | 32 | 0.2812 | 9 | 0.2222 | 11 | 0.2727 | 10 | 0.3 | 10 | 0.3 | 40 | 0.275 |
| 15_8925579 | 3 | 0.6667 | 4 | 0.25 | NA | NA | 4 | 0 | 11 | 0.2727 | 2 | 0 | 1 | 0 | 2 | 0.5 | 5 | 0.2 | 10 | 0.2 |
| 15_8925580 | 2 | 0 | 3 | 0 | 1 | 0 | 4 | 0.5 | 10 | 0.2 | 2 | 1 | 1 | 1 | 1 | 1 | 4 | 0.5 | 8 | 0.75 |
| 15_8925588 | 6 | 0.3333 | 1 | 0 | 3 | 0 | 5 | 0.6 | 15 | 0.3333 | NA | NA | 8 | 0.5 | 2 | 0 | 3 | 0.6667 | 13 | 0.4615 |
| 15_8925601 | 6 | 0.5 | 3 | 0.3333 | 3 | 0.3333 | 5 | 0 | 17 | 0.2941 | NA | NA | 6 | 0.3333 | NA | NA | 5 | 0.2 | 11 | 0.2727 |
| 15_8925603 | 6 | 0.5 | 4 | 0.5 | 3 | 1 | 6 | 1 | 19 | 0.7368 | NA | NA | 6 | 0.6667 | 2 | 1 | 8 | 0.625 | 16 | 0.6875 |
| 15_8925603 | 7 | 0.4286 | 4 | 0.25 | 3 | 0.3333 | 8 | 0.125 | 22 | 0.2727 | NA | NA | 6 | 0 | 2 | 0 | 8 | 0.375 | 16 | 0.1875 |
| 15_8925612 | 8 | 0.75 | 4 | 1 | 3 | 0.6667 | 6 | 0.8333 | 21 | 0.8095 | NA | NA | 8 | 1 | 2 | 1 | 7 | 1 | 17 | 1 |
| 15_8925618 | 8 | 0.375 | 4 | 0.25 | 3 | 0 | 6 | 0.6667 | 21 | 0.381 | NA | NA | 7 | 0.1429 | 2 | 0.5 | 6 | 0.3333 | 15 | 0.2667 |
| 15_8925637 | 8 | 0.125 | 4 | 0 | 2 | 0 | 7 | 0.1429 | 21 | 0.0952 | NA | NA | 6 | 0.1667 | 2 | 0.5 | 6 | 0 | 14 | 0.1429 |
| 15_8925643 | 5 | 0.4 | 2 | 1 | 1 | 0 | 3 | 1 | 11 | 0.6364 | NA | NA | 5 | 0.6 | 2 | 1 | 5 | 0.8 | 12 | 0.75 |
| 15_8925650 | 4 | 1 | 3 | 0.6667 | 1 | 1 | 3 | 1 | 11 | 0.9091 | NA | NA | 6 | 1 | 1 | 1 | 5 | 0.6 | 12 | 0.8333 |
| 15_8925651 | 6 | 0.6667 | 2 | 1 | 2 | 1 | 5 | 1 | 15 | 0.8667 | NA | NA | 7 | 1 | NA | NA | 6 | 1 | 13 | 1 |
| 15_8925657 | 7 | 0 | 3 | 0 | 2 | 0 | 7 | 0.2857 | 19 | 0.1053 | NA | NA | 6 | 0 | 2 | 0 | 6 | 0.1667 | 14 | 0.0714 |
| 15_8925660 | 1 | 1 | 4 | 1 | NA | NA | 5 | 1 | 10 | 1 | 4 | 1 | NA | NA | 3 | 0.6667 | 6 | 1 | 13 | 0.9231 |
| 15_8925665 | 3 | 0.3333 | 4 | 0.25 | 1 | 0 | 5 | 0 | 13 | 0.1538 | 2 | 0.5 | 2 | 0.5 | 3 | 0.3333 | 6 | 0.3333 | 13 | 0.3846 |
| 15_8925667 | 3 | 0 | 3 | 0 | 1 | 1 | 5 | 0 | 12 | 0.0833 | 3 | 0.6667 | 2 | 0 | 3 | 0.3333 | 6 | 0 | 14 | 0.2143 |
| 15_8925670 | 2 | 1 | 5 | 0.8 | 1 | 1 | 5 | 1 | 13 | 0.9231 | 3 | 0.6667 | 2 | 1 | 3 | 1 | 7 | 0.5714 | 15 | 0.7333 |
| 15_8925673 | 3 | 0.3333 | 3 | 0 | 1 | 0 | 6 | 0.3333 | 13 | 0.2308 | 3 | 0 | 2 | 0 | 3 | 0.6667 | 6 | 0.1667 | 14 | 0.2143 |
| 15_8925680 | 2 | 0.5 | 3 | 1 | 1 | 1 | 6 | 0.6667 | 12 | 0.75 | 3 | 1 | 2 | 1 | 3 | 0.6667 | 6 | 1 | 14 | 0.9286 |
| 15_8931068 | 4 | 0.25 | 6 | 0.3333 | 4 | 0.5 | 7 | 0.2857 | 21 | 0.3333 | 4 | 0.25 | 4 | 0.25 | 1 | 0 | 2 | 0 | 11 | 0.1818 |
| 15_8931074 | 4 | 0.25 | 5 | 0.2 | 3 | 0.3333 | 4 | 0 | 16 | 0.1875 | 3 | 0.3333 | 5 | 0 | 1 | 0 | 1 | 1 | 10 | 0.2 |
| 15_8933704 | NA | NA | 1 | 1 | NA | NA | NA | NA | 1 | 1 | NA | NA | 1 | 1 | NA | NA | NA | NA | 1 | 1 |
| 15_8933710 | NA | NA | 1 | 1 | NA | NA | NA | NA | 1 | 1 | NA | NA | 1 | 1 | NA | NA | NA | NA | 1 | 1 |
| 15_8933720 | NA | NA | 1 | 1 | NA | NA | NA | NA | 1 | 1 | NA | NA | 1 | 1 | NA | NA | NA | NA | 1 | 1 |
| 15_8933750 | NA | NA | 1 | 1 | NA | NA | NA | NA | 1 | 1 | NA | NA | 1 | 1 | NA | NA | NA | NA | 1 | 1 |
| 15_8934112 | 9 | 0.1111 | 8 | 0 | 4 | 0.25 | 9 | 0 | 30 | 0.0667 | 4 | 0 | 11 | 0 | 6 | 0.3333 | 7 | 0.1429 | 28 | 0.1071 |
| 15_8934141 | 7 | 0.1429 | 7 | 0.1429 | 3 | 0.3333 | 9 | 0.2222 | 26 | 0.1923 | 4 | 0.25 | 10 | 0.1 | 7 | 0.2857 | 7 | 0.4286 | 28 | 0.25 |
| 15_8934156 | 6 | 0.8333 | 9 | 0.4444 | 3 | 1 | 5 | 0.8 | 23 | 0.6957 | 4 | 1 | 10 | 0.8 | 3 | 0.6667 | 7 | 0.8571 | 24 | 0.8333 |
| 15_8934174 | 11 | 0 | 9 | 0 | 3 | 0 | 7 | 0.2857 | 30 | 0.0667 | 1 | 0 | 13 | 0.0769 | 5 | 0 | 6 | 0.3333 | 25 | 0.12 |
| 15_8934184 | 10 | 0.5 | 7 | 0.2857 | 4 | 0.25 | 11 | 0.3636 | 32 | 0.375 | 5 | 0 | 14 | 0.1429 | 7 | 0.1429 | 7 | 0.4286 | 33 | 0.1818 |
| 15_8934198 | 9 | 0.6667 | 16 | 0.875 | 5 | 0.8 | 11 | 0.7273 | 41 | 0.7805 | 6 | 0.8333 | 15 | 0.7333 | 6 | 0.8333 | 8 | 0.625 | 35 | 0.7429 |
| 15_8934202 | 8 | 0.5 | 11 | 0.8182 | 5 | 0.8 | 10 | 0.8 | 34 | 0.7353 | 9 | 0.8889 | 17 | 0.8824 | 8 | 0.875 | 8 | 0.625 | 42 | 0.8333 |
| 15_8934203 | 10 | 0.9 | 13 | 0.8462 | 2 | 1 | 12 | 1 | 37 | 0.9189 | 9 | 1 | 18 | 0.8889 | 7 | 0.7143 | 6 | 1 | 40 | 0.9 |
| 15_8934228 | 10 | 0.2 | 16 | 0.0625 | 6 | 0 | 14 | 0.2143 | 46 | 0.1304 | 11 | 0.0909 | 14 | 0.2857 | 10 | 0.1 | 11 | 0.2727 | 46 | 0.1957 |
| 15_8934598 | 16 | 0.75 | 29 | 0.7931 | 18 | 0.7222 | 20 | 0.75 | 83 | 0.759 | 15 | 0.9333 | 27 | 0.7778 | 20 | 0.85 | 24 | 0.7917 | 86 | 0.8256 |
| 15_8934599 | 16 | 0.3125 | 25 | 0.44 | 13 | 0.3077 | 20 | 0.45 | 74 | 0.3919 | 15 | 0.3333 | 23 | 0.3043 | 19 | 0.4737 | 22 | 0.5 | 79 | 0.4051 |
| 15_8934603 | 19 | 0.2105 | 29 | 0.1379 | 19 | 0.1053 | 22 | 0.0455 | 89 | 0.1236 | 17 | 0.1176 | 26 | 0.1154 | 19 | 0.2105 | 24 | 0.2083 | 86 | 0.1628 |
| 15_8934623 | 8 | 0.25 | 20 | 0 | 15 | 0.2667 | 10 | 0 | 53 | 0.1132 | 14 | 0.0714 | 16 | 0.125 | 16 | 0.125 | 16 | 0.125 | 62 | 0.1129 |
| 15_8935682 | 1 | 0 | 3 | 0 | 2 | 0.5 | 5 | 0 | 11 | 0.0909 | 3 | 0 | 4 | 0 | 7 | 0.1429 | 2 | 0 | 16 | 0.0625 |
| 15_8935687 | 1 | 0 | 4 | 0.25 | 3 | 0 | 5 | 0.4 | 13 | 0.2308 | 2 | 0.5 | 5 | 0.8 | 7 | 0.4286 | 2 | 0 | 16 | 0.5 |
| 15_8935692 | 1 | 0 | 5 | 0.2 | 2 | 0 | 5 | 0.2 | 13 | 0.1538 | 2 | 0.5 | 6 | 0.1667 | 7 | 0.1429 | 3 | 0 | 18 | 0.1667 |
| 15_8935720 | 1 | 0 | 3 | 0.3333 | 2 | 0.5 | 5 | 0 | 11 | 0.1818 | 2 | 0 | 5 | 0.2 | 6 | 0.5 | 1 | 0 | 14 | 0.2857 |
| 15_8935722 | 1 | 1 | 4 | 0.75 | 2 | 0.5 | 3 | 0.6667 | 10 | 0.7 | 2 | 1 | 4 | 0.5 | 4 | 0.75 | 2 | 0.5 | 12 | 0.6667 |
| 15_8935730 | 1 | 0 | 5 | 0.2 | 3 | 0 | 5 | 0.2 | 14 | 0.1429 | 2 | 1 | 7 | 0.4286 | 7 | 0.2857 | 2 | 0 | 18 | 0.3889 |
| 15_8935734 | 1 | 0 | 5 | 0.2 | 3 | 0 | 4 | 0.25 | 13 | 0.1538 | 2 | 0.5 | 7 | 0.1429 | 7 | 0.1429 | 3 | 0 | 19 | 0.1579 |
| 15_8935743 | 1 | 0 | 4 | 0.5 | 3 | 0 | 5 | 0.2 | 13 | 0.2308 | 2 | 0.5 | 6 | 0.3333 | 7 | 0.4286 | 1 | 0 | 16 | 0.375 |
| 15_8935752 | 1 | 1 | 4 | 0 | 3 | 0.3333 | 5 | 0.2 | 13 | 0.2308 | 2 | 0.5 | 7 | 0.2857 | 7 | 0.1429 | 2 | 0.5 | 18 | 0.2778 |
| 15_8935755 | 1 | 0 | 3 | 0.3333 | 3 | 0 | 5 | 0 | 12 | 0.0833 | 1 | 0 | 6 | 0 | 7 | 0.1429 | 2 | 0 | 16 | 0.0625 |
| 15_8935760 | 1 | 0 | 4 | 0.25 | 3 | 0.3333 | 5 | 0.4 | 13 | 0.3077 | 1 | 0 | 7 | 0.2857 | 7 | 0.4286 | 2 | 0 | 17 | 0.2941 |
| 15_8935991 | 1 | 1 | 2 | 1 | 2 | 0 | 3 | 0 | 8 | 0.375 | 1 | 1 | 3 | 0.3333 | 2 | 0.5 | 2 | 1 | 8 | 0.625 |
| 15_8935995 | 1 | 0 | 1 | 1 | 2 | 0 | 3 | 0.3333 | 7 | 0.2857 | 1 | 1 | 2 | 0 | 2 | 0 | 2 | 0 | 7 | 0.1429 |
| 15_8935995 | 1 | 1 | 2 | 0.5 | 2 | 0 | 2 | 1 | 7 | 0.5714 | 1 | 1 | 3 | 0.6667 | 1 | 1 | 2 | 0 | 7 | 0.5714 |
| 15_8936003 | NA | NA | 1 | 1 | 1 | 0 | 3 | 0.6667 | 5 | 0.6 | NA | NA | 1 | 1 | 2 | 1 | 2 | 1 | 5 | 1 |
| 15_8936009 | 1 | 0 | 1 | 0 | 1 | 1 | 3 | 0 | 6 | 0.1667 | 1 | 0 | 3 | 0.3333 | 2 | 0 | 2 | 0.5 | 8 | 0.25 |
| 15_8936020 | NA | NA | 1 | 0 | 2 | 0.5 | 3 | 0 | 6 | 0.1667 | NA | NA | 3 | 0.6667 | 2 | 0 | 2 | 0 | 7 | 0.2857 |

| | | | | | | | | | | | | | | | | | | | | |
|---|---|---|---|---|---|---|---|---|---|---|---|---|---|---|---|---|---|---|---|---|
| 15_893796 | 1 | 1 | NA | NA | NA | NA | NA | NA | 1 | 1 | 1 | 1 | NA | NA | NA | NA | NA | NA | 1 | 1 |
| 15_894318 | 9 | 0.1111 | 2 | 0 | 2 | 0 | 4 | 0 | 17 | 0.0588 | 3 | 0.3333 | 5 | 0.2 | 4 | 0.25 | 2 | 0 | 14 | 0.2143 |
| 15_894320 | 8 | 0 | 3 | 0.3333 | 3 | 0 | 5 | 0.2 | 19 | 0.1053 | 3 | 0.3333 | 4 | 0 | 6 | 0 | 3 | 0 | 16 | 0.0625 |
| 15_894321 | 9 | 1 | 3 | 0.6667 | 2 | 1 | 5 | 1 | 19 | 0.9474 | 3 | 0.6667 | 6 | 0.8333 | 5 | 1 | 3 | 0.6667 | 17 | 0.8235 |
| 15_894324 | 9 | 0.1111 | 3 | 0.3333 | 2 | 0 | 3 | 0 | 17 | 0.1176 | 3 | 0.3333 | 7 | 0.2857 | 7 | 0.1429 | 3 | 0 | 20 | 0.2 |
| 15_894324 | 9 | 0.2222 | 5 | 0 | 4 | 0 | 5 | 0 | 23 | 0.087 | 3 | 0 | 7 | 0.1429 | 4 | 0.25 | 3 | 0 | 17 | 0.1176 |
| 15_894326 | 10 | 0.4 | 5 | 0.2 | 3 | 0 | 3 | 0.6667 | 21 | 0.3333 | 3 | 0 | 6 | 0.5 | 4 | 0.25 | 2 | 1 | 15 | 0.4 |
| 15_894329 | 10 | 0.1 | 6 | 0 | 3 | 0.6667 | 4 | 0.25 | 23 | 0.1739 | 2 | 0 | 7 | 0.4286 | 7 | 0.2857 | 3 | 0.3333 | 19 | 0.3158 |
| 15_894329 | 7 | 0.4286 | 6 | 0.6667 | 4 | 0.25 | 4 | 0.75 | 21 | 0.5238 | 3 | 0.6667 | 2 | 0 | 6 | 0.8333 | 3 | 0.6667 | 14 | 0.6429 |
| 15_894329 | 11 | 0.2727 | 6 | 0.1667 | 4 | 0.25 | 5 | 0.2 | 26 | 0.2308 | 2 | 0.5 | 6 | 0.3333 | 7 | 0.4286 | 3 | 0.3333 | 18 | 0.3889 |
| 15_894330 | 11 | 0.2727 | 6 | 0.1667 | 2 | 1 | 2 | 0 | 21 | 0.2857 | 4 | 0.25 | 6 | 0.3333 | 7 | 0.2857 | 1 | 1 | 18 | 0.3333 |
| 15_894335 | 11 | 0.1818 | 6 | 0 | 4 | 0 | 5 | 0 | 26 | 0.0769 | 4 | 0 | 8 | 0 | 5 | 0.4 | 3 | 0 | 20 | 0.1 |
| 15_894336 | 13 | 0.1538 | 5 | 0 | 5 | 0 | 5 | 0 | 28 | 0.0714 | 5 | 0.6 | 8 | 0 | 9 | 0.2222 | 3 | 0 | 25 | 0.2 |
| 15_894336 | 12 | 0.3333 | 3 | 0.3333 | 5 | 0.2 | 4 | 0.5 | 24 | 0.3333 | 4 | 0.25 | 8 | 0.25 | 8 | 0.25 | 3 | 0 | 23 | 0.2174 |
| 15_894336 | 13 | 0.9231 | 5 | 1 | 4 | 0.75 | 5 | 1 | 27 | 0.9259 | 5 | 1 | 9 | 0.8889 | 9 | 0.8889 | 3 | 1 | 26 | 0.9231 |
| 15_894337 | 13 | 0.4615 | 7 | 0.2857 | 3 | 0.3333 | 5 | 0.6 | 28 | 0.4286 | 5 | 0.2 | 9 | 0.2222 | 10 | 0.5 | 4 | 0.5 | 28 | 0.3571 |
| 15_894337 | 11 | 0.8182 | 5 | 1 | 3 | 0.6667 | 5 | 0.8 | 24 | 0.8333 | 4 | 1 | 4 | 1 | 5 | 0.6 | 4 | 0.75 | 17 | 0.8235 |
| 15_894340 | 12 | 0.5 | 5 | 0.4 | 3 | 0 | 3 | 0 | 23 | 0.3478 | 6 | 0.3333 | 6 | 0.5 | 10 | 0.1 | 2 | 0 | 24 | 0.25 |
| 15_894421 | 15 | 0 | 8 | 0.25 | 4 | 0 | 4 | 0.5 | 31 | 0.129 | 5 | 0 | 9 | 0 | 13 | 0.1538 | 5 | 0 | 32 | 0.0625 |
| 15_894424 | 19 | 0.5263 | 8 | 0.25 | 5 | 0.8 | 4 | 0.5 | 36 | 0.5 | 6 | 0.6667 | 8 | 0.5 | 16 | 0.25 | 6 | 0.6667 | 36 | 0.4444 |
| 15_894424 | 14 | 0.2857 | 10 | 0 | 6 | 0.3333 | 6 | 0.1667 | 36 | 0.1944 | 7 | 0.1429 | 7 | 0 | 16 | 0.1875 | 4 | 0 | 34 | 0.1176 |
| 15_894424 | 10 | 0.4 | 4 | 0.5 | NA | NA | 6 | 1 | 20 | 0.6 | 2 | 1 | 6 | 0.6667 | 10 | 0.8 | 3 | 0.6667 | 21 | 0.7619 |
| 15_894426 | 19 | 0.1579 | 9 | 0.1111 | 6 | 0.1667 | 6 | 0 | 40 | 0.125 | 8 | 0.25 | 10 | 0.2 | 17 | 0.0588 | 7 | 0.2857 | 42 | 0.1667 |
| 15_894427 | 16 | 0.125 | 6 | 0.3333 | 5 | 0.2 | 4 | 0.25 | 31 | 0.1935 | 7 | 0.2857 | 6 | 0.1667 | 14 | 0.0714 | 7 | 0 | 34 | 0.1176 |
| 15_894428 | 18 | 0.0556 | 9 | 0.1111 | 6 | 0 | 7 | 0 | 40 | 0.05 | 7 | 0.1429 | 9 | 0 | 17 | 0.0588 | 5 | 0 | 38 | 0.0526 |
| 15_894438 | 17 | 0.1176 | 12 | 0 | 10 | 0.3 | 11 | 0.0909 | 50 | 0.12 | 8 | 0 | 17 | 0 | 17 | 0.1176 | 9 | 0.1111 | 51 | 0.0588 |
| 15_894439 | 21 | 0.1429 | 11 | 0.1818 | 10 | 0.2 | 11 | 0.0909 | 53 | 0.1509 | 10 | 0 | 17 | 0.1176 | 20 | 0.1 | 10 | 0.2 | 57 | 0.1053 |
| 15_894669 | 1 | 0 | 6 | 0.1667 | 1 | 1 | 4 | 0 | 12 | 0.1667 | 4 | 0.25 | 1 | 0 | 2 | 0.5 | 6 | 0.3333 | 13 | 0.3077 |
| 15_894670 | 1 | 0 | 7 | 0.5714 | 1 | 1 | 3 | 0 | 12 | 0.4167 | 3 | 0.3333 | 2 | 0.5 | 3 | 1 | 6 | 0.3333 | 14 | 0.5 |
| 15_894672 | 1 | 0 | 6 | 0.6667 | 1 | 0 | 5 | 0.6 | 13 | 0.5385 | 2 | 0.5 | 2 | 0.5 | 3 | 0.6667 | 6 | 0.5 | 13 | 0.5385 |
| 15_894674 | 1 | 0 | 7 | 0.1429 | 1 | 0 | 3 | 0 | 12 | 0.0833 | 2 | 0.5 | 1 | 0 | 2 | 0 | 5 | 0 | 10 | 0.1 |
| 15_896615 | NA | NA | 3 | 0 | 2 | 0.5 | 4 | 0 | 9 | 0.1111 | 2 | 0 | 2 | 0.5 | 7 | 0 | 1 | 0 | 12 | 0.0833 |
| 15_896617 | NA | NA | 2 | 0 | 2 | 0.5 | 5 | 0 | 9 | 0.1111 | 1 | 1 | 2 | 0 | 7 | 0 | 1 | 0 | 11 | 0.0909 |
| 15_896623 | NA | NA | 3 | 0 | 2 | 0.5 | 4 | 0.25 | 9 | 0.2222 | 1 | 0 | 2 | 0 | 7 | 0.4286 | 1 | 0 | 11 | 0.2727 |
| 15_896984 | NA | NA | NA | NA | NA | NA | 2 | 1 | 2 | 1 | NA | NA | 2 | 1 | NA | NA | NA | NA | 2 | 1 |
| 15_896985 | NA | NA | NA | NA | NA | NA | 2 | 1 | 2 | 1 | NA | NA | 2 | 1 | NA | NA | NA | NA | 2 | 1 |
| 15_896986 | NA | NA | NA | NA | NA | NA | 1 | 1 | 1 | 1 | NA | NA | 2 | 1 | NA | NA | NA | NA | 2 | 1 |
| 15_901116 | 2 | 0.5 | 2 | 0.5 | NA | NA | 2 | 1 | 6 | 0.6667 | 2 | 1 | 3 | 1 | NA | NA | 2 | 0.5 | 7 | 0.8571 |
| 15_901117 | 2 | 0.5 | 1 | 1 | NA | NA | 2 | 1 | 5 | 0.8 | 2 | 1 | 3 | 0.6667 | 1 | 1 | 1 | 1 | 7 | 0.8571 |
| 15_901118 | 2 | 0 | 2 | 0.5 | 1 | 1 | 3 | 0.6667 | 8 | 0.5 | 2 | 0.5 | 1 | 0 | 1 | 1 | 2 | 0 | 6 | 0.3333 |
| 15_901118 | 2 | 0 | 2 | 0 | NA | NA | 3 | 0.6667 | 7 | 0.2857 | 2 | 0 | 3 | 0 | NA | NA | 2 | 0.5 | 7 | 0.1429 |
| 15_901126 | 2 | 0 | 2 | 0.5 | 1 | 0 | 3 | 0.3333 | 8 | 0.25 | 3 | 0 | 4 | 0 | 2 | 0 | 2 | 0.5 | 11 | 0.0909 |
| 15_901144 | 3 | 0.3333 | 5 | 0 | 4 | 0.25 | 5 | 0.2 | 17 | 0.1765 | 4 | 0.25 | 4 | 0.25 | 4 | 0.25 | 2 | 0 | 14 | 0.2143 |
| 15_901145 | 3 | 0 | 4 | 0.25 | 4 | 0.25 | 5 | 0.4 | 16 | 0.25 | 4 | 0.25 | 4 | 0 | 5 | 0.2 | 3 | 0.3333 | 16 | 0.1875 |
| 15_901166 | 2 | 0.5 | 4 | 0.25 | 5 | 0.4 | 3 | 0.3333 | 14 | 0.3571 | 3 | 1 | 6 | 0.1667 | 5 | 0.8 | 4 | 0.75 | 18 | 0.6111 |
| 15_901168 | 3 | 0 | 6 | 0 | 4 | 0.25 | 2 | 0 | 15 | 0.0667 | 2 | 0.5 | 7 | 0 | 6 | 0.1667 | 4 | 0.5 | 19 | 0.2105 |
| 15_901171 | 6 | 0.1667 | 7 | 0 | 6 | 0.1667 | 5 | 0 | 24 | 0.0833 | 5 | 0.2 | 8 | 0.375 | 6 | 0.3333 | 6 | 0 | 25 | 0.24 |
| 15_901174 | 6 | 0.5 | 8 | 0.125 | 6 | 0.6667 | 5 | 0.2 | 25 | 0.36 | 5 | 0.4 | 9 | 0.3333 | 7 | 0.1429 | 5 | 0.2 | 26 | 0.2692 |
| 15_903730 | 1 | 1 | NA | NA | NA | NA | 3 | 0.3333 | 4 | 0.5 | 1 | 1 | NA | NA | 1 | 0 | NA | NA | 2 | 0.5 |
| 15_903730 | 2 | 0.5 | 1 | 1 | 1 | 1 | 3 | 0.6667 | 7 | 0.7143 | 3 | 0.3333 | NA | NA | 1 | 1 | NA | NA | 4 | 0.5 |
| 15_903731 | 2 | 0.5 | 1 | 0 | 1 | 0 | 2 | 0.5 | 6 | 0.3333 | 3 | 0 | 1 | 0 | 1 | 1 | NA | NA | 5 | 0.2 |
| 15_908183 | 16 | 0.0625 | 17 | 0.1176 | 6 | 0.1667 | 16 | 0.125 | 55 | 0.1091 | 8 | 0 | 8 | 0.125 | 11 | 0.1818 | 18 | 0.1667 | 45 | 0.1333 |
| 15_908184 | 17 | 0.1765 | 16 | 0.0625 | 8 | 0.125 | 18 | 0.1111 | 59 | 0.1186 | 7 | 0.2857 | 10 | 0.2 | 9 | 0.5556 | 17 | 0.2941 | 43 | 0.3256 |
| 15_908199 | 15 | 0 | 16 | 0.125 | 7 | 0 | 17 | 0.0588 | 55 | 0.0545 | 6 | 0.1667 | 7 | 0.2857 | 9 | 0 | 18 | 0.1111 | 40 | 0.125 |
| 15_910327 | 2 | 0 | 1 | 0 | 4 | 0 | 5 | 0.2 | 12 | 0.0833 | 2 | 0 | 2 | 0.5 | 4 | 0.25 | NA | NA | 8 | 0.25 |
| 15_910328 | 2 | 0 | 2 | 0 | 6 | 0.3333 | 3 | 0.6667 | 13 | 0.3077 | 1 | 0 | 1 | 1 | 5 | 0.4 | NA | NA | 7 | 0.4286 |
| 15_910329 | 3 | 0 | 2 | 0.5 | 6 | 0.3333 | 5 | 0 | 16 | 0.1875 | 2 | 0 | NA | NA | 5 | 0.2 | NA | NA | 7 | 0.1429 |
| 15_910330 | 2 | 0 | 2 | 0 | 6 | 0 | 5 | 0.6 | 15 | 0.2 | 2 | 0 | NA | NA | 4 | 0.5 | NA | NA | 6 | 0.3333 |
| 15_910331 | 3 | 1 | 2 | 1 | 6 | 1 | 4 | 1 | 15 | 1 | 2 | 0.5 | 1 | 1 | 5 | 1 | NA | NA | 8 | 0.875 |
| 15_910331 | 2 | 0.5 | 2 | 0.5 | 6 | 0.5 | 4 | 0.5 | 14 | 0.5 | 2 | 0 | 1 | 1 | 4 | 0.75 | NA | NA | 7 | 0.5714 |
| 15_910332 | 2 | 0 | 1 | 1 | 5 | 0 | 5 | 0.4 | 13 | 0.2308 | 1 | 0 | 1 | 1 | 4 | 0 | NA | NA | 6 | 0.1667 |
| 15_910332 | 2 | 0 | 2 | 1 | 6 | 0.3333 | 5 | 0.2 | 15 | 0.3333 | 2 | 1 | NA | NA | 3 | 0.3333 | NA | NA | 5 | 0.6 |
| 15_910730 | 2 | 0.5 | 4 | 0.25 | 2 | 0.5 | 3 | 0 | 11 | 0.2727 | 2 | 0.5 | NA | NA | NA | NA | NA | NA | 2 | 0.5 |
| 15_910750 | 2 | 1 | 4 | 0.75 | 4 | 0.75 | 3 | 1 | 13 | 0.8462 | 2 | 0.5 | NA | NA | 1 | 0 | NA | NA | 3 | 0.3333 |
| 15_910752 | 1 | 1 | 2 | 0.5 | 4 | 0.25 | 2 | 1 | 9 | 0.5556 | 1 | 0 | NA | NA | 1 | 1 | NA | NA | 2 | 0.5 |

| | | | | | | | | | | | | | | | | | | | | |
|---|---|---|---|---|---|---|---|---|---|---|---|---|---|---|---|---|---|---|---|---|
| 15_9107528 | 2 | 0.5 | 4 | 1 | 4 | 0.5 | 2 | 1 | 12 | 0.75 | 1 | 0 | NA | NA | 1 | 1 | NA | NA | 2 | 0.5 |
| 15_9107538 | 1 | 0 | 3 | 0.3333 | 4 | 0.5 | 2 | 1 | 10 | 0.5 | 2 | 0.5 | NA | NA | 1 | 1 | NA | NA | 3 | 0.6667 |
| 15_9107546 | NA | NA | 2 | 1 | 4 | 0.75 | 2 | 0.5 | 8 | 0.75 | 1 | 0 | NA | NA | 1 | 1 | NA | NA | 2 | 0.5 |
| 15_9107557 | 2 | 0 | 3 | 0.3333 | 4 | 0 | 1 | 1 | 10 | 0.2 | 1 | 1 | NA | NA | 1 | 0 | NA | NA | 2 | 0.5 |
| 15_9107576 | 1 | 0 | 3 | 0.6667 | 4 | 0.5 | 2 | 1 | 10 | 0.6 | 1 | 0 | 1 | 1 | NA | NA | NA | NA | 2 | 0.5 |
| 15_9115212 | 21 | 0.6667 | 27 | 0.6296 | 18 | 0.7222 | 20 | 0.75 | 86 | 0.686 | 14 | 0.5 | 31 | 0.7742 | 15 | 0.7333 | 14 | 0.6429 | 74 | 0.6892 |
| 15_9115221 | 19 | 0.1053 | 26 | 0.0769 | 17 | 0.1765 | 19 | 0.2632 | 81 | 0.1481 | 14 | 0.2857 | 28 | 0.0714 | 14 | 0.4286 | 15 | 0.1333 | 71 | 0.1972 |
| 15_9115251 | 17 | 0.0588 | 23 | 0.2174 | 13 | 0.0769 | 17 | 0.1176 | 70 | 0.1286 | 12 | 0.25 | 26 | 0.0385 | 15 | 0.0667 | 14 | 0.0714 | 67 | 0.0896 |
| 15_9115298 | 14 | 0.0714 | 20 | 0.15 | 11 | 0.2727 | 14 | 0.0714 | 59 | 0.1356 | 10 | 0 | 26 | 0.3077 | 12 | 0.1667 | 12 | 0.1667 | 60 | 0.2 |
| 15_9115365 | 12 | 0 | 20 | 0.3 | 13 | 0.1538 | 17 | 0.0588 | 62 | 0.1452 | 6 | 0.1667 | 22 | 0.2727 | 5 | 0.2 | 9 | 0.1111 | 42 | 0.2143 |
| 15_9115391 | 12 | 0.1667 | 16 | 0.125 | 12 | 0.25 | 13 | 0.1538 | 53 | 0.1698 | 5 | 0.2 | 16 | 0.0625 | 6 | 0.1667 | 7 | 0 | 34 | 0.0882 |
| 15_9115536 | 4 | 0.25 | 11 | 0.2727 | 9 | 0.3333 | 10 | 0.4 | 34 | 0.3235 | 6 | 0.5 | 9 | 0.4444 | 4 | 0 | 4 | 0.5 | 23 | 0.3913 |
| 15_9115538 | 7 | 0 | 12 | 0.0833 | 9 | 0.3333 | 10 | 0.1 | 38 | 0.1316 | 6 | 0.3333 | 10 | 0.2 | 4 | 0 | 5 | 0.2 | 25 | 0.2 |
| 15_9115552 | 7 | 0.1429 | 12 | 0.1667 | 8 | 0.625 | 8 | 0.375 | 35 | 0.3143 | 4 | 0.5 | 12 | 0.3333 | 4 | 0 | 5 | 0.2 | 25 | 0.28 |
| 15_9115567 | 4 | 0 | 14 | 0.0714 | 8 | 0.125 | 10 | 0 | 36 | 0.0556 | 4 | 0 | 10 | 0.2 | 3 | 0 | 5 | 0 | 22 | 0.0909 |
| 15_9115575 | 5 | 0 | 9 | 0.2222 | 9 | 0.1111 | 9 | 0.1111 | 32 | 0.125 | 3 | 0.3333 | 8 | 0.25 | 3 | 0.3333 | 4 | 0.25 | 18 | 0.2778 |
| 15_9115609 | 2 | 0 | 14 | 0.1429 | 6 | 0.1667 | 9 | 0.1111 | 31 | 0.129 | 3 | 0 | 6 | 0.1667 | 3 | 0.3333 | 5 | 0.2 | 17 | 0.1765 |
| 15_9115612 | 5 | 0.2 | 12 | 0.3333 | 6 | 0.1667 | 8 | 0.125 | 31 | 0.2258 | 2 | 0 | 8 | 0.25 | 3 | 0 | 4 | 0 | 17 | 0.1176 |
| 15_9115623 | 4 | 0 | 13 | 0.3846 | 7 | 0.7143 | 9 | 0.3333 | 33 | 0.3939 | 3 | 0.6667 | 10 | 0.4 | 3 | 0.3333 | 5 | 0.6 | 21 | 0.4762 |
| 15_9115634 | 3 | 1 | 6 | 0.8333 | 5 | 1 | 6 | 1 | 20 | 0.95 | 3 | 1 | 5 | 1 | 2 | 1 | 3 | 1 | 13 | 1 |
| 15_9115635 | 3 | 0 | 11 | 0.2727 | 6 | 0.1667 | 8 | 0.125 | 28 | 0.1786 | 3 | 0 | 9 | 0.1111 | 3 | 0 | 5 | 0.6 | 20 | 0.2 |
| 15_9115654 | 4 | 1 | 8 | 1 | 5 | 1 | 5 | 1 | 22 | 1 | 3 | 1 | 9 | 1 | 2 | 1 | 3 | 1 | 17 | 1 |
| 15_9116039 | 3 | 0 | 5 | 0.2 | 5 | 0 | 3 | 0 | 16 | 0.0625 | 3 | 0 | 8 | 0.125 | 2 | 0 | 3 | 0 | 16 | 0.0625 |
| 15_9169974 | 6 | 0 | 5 | 0 | 6 | 0.1667 | 11 | 0.0909 | 28 | 0.0714 | 9 | 0.1111 | 6 | 0.1667 | 11 | 0 | 5 | 0 | 31 | 0.0645 |
| 15_9169980 | 6 | 0.1667 | 6 | 0 | 5 | 0.4 | 11 | 0.2727 | 28 | 0.2143 | 10 | 0.2 | 5 | 0.4 | 10 | 0.3 | 5 | 1 | 30 | 0.4 |
| 15_9170019 | 6 | 0 | 6 | 0.1667 | 5 | 0 | 11 | 0.1818 | 28 | 0.1071 | 7 | 0.1429 | 7 | 0.1429 | 11 | 0 | 4 | 0.5 | 29 | 0.1379 |
| 15_9170023 | 1 | 0 | 2 | 0 | 2 | 0.5 | 2 | 0.5 | 7 | 0.2857 | 2 | 0 | 2 | 0 | 2 | 0.5 | 2 | 0 | 8 | 0.125 |
| 15_9223752 | 18 | 0.4444 | 32 | 0.5 | 23 | 0.6522 | 25 | 0.6 | 98 | 0.551 | 12 | 0.75 | 13 | 0.5385 | 22 | 0.7273 | 20 | 0.5 | 67 | 0.6269 |
| 15_9223758 | 16 | 1 | 27 | 1 | 21 | 0.9048 | 26 | 1 | 90 | 0.9778 | 11 | 0.8182 | 15 | 1 | 22 | 1 | 20 | 0.85 | 68 | 0.9265 |
| 15_9223765 | 12 | 0.0833 | 18 | 0.2222 | 17 | 0.1176 | 20 | 0.15 | 67 | 0.1493 | 12 | 0.1667 | 12 | 0.1667 | 17 | 0.1176 | 18 | 0.2778 | 59 | 0.1864 |
| 15_9223775 | 19 | 0.5263 | 28 | 0.6429 | 23 | 0.6087 | 24 | 0.375 | 94 | 0.5426 | 10 | 0.7 | 18 | 0.4444 | 23 | 0.7826 | 20 | 0.7 | 71 | 0.662 |
| 15_9223830 | 23 | 0.087 | 30 | 0.0667 | 23 | 0.087 | 28 | 0.1429 | 104 | 0.0962 | 9 | 0 | 21 | 0 | 25 | 0.12 | 22 | 0.0909 | 77 | 0.0649 |
| 15_9223847 | 21 | 0.0476 | 30 | 0.0333 | 24 | 0.1667 | 26 | 0.1154 | 101 | 0.0891 | 10 | 0.1 | 20 | 0.05 | 19 | 0.1579 | 21 | 0.1429 | 70 | 0.1143 |
| 15_9223859 | 21 | 0.2857 | 33 | 0.2121 | 27 | 0.0741 | 29 | 0.0345 | 110 | 0.1455 | 13 | 0.1538 | 21 | 0.1905 | 23 | 0.1304 | 24 | 0.0417 | 81 | 0.1235 |
| 15_9223923 | 26 | 0.1538 | 39 | 0.1026 | 33 | 0.0303 | 33 | 0.0606 | 131 | 0.084 | 18 | 0.1111 | 22 | 0.0455 | 30 | 0.0667 | 26 | 0.1154 | 96 | 0.0833 |
| 15_9223944 | 29 | 0 | 44 | 0.0909 | 33 | 0.0606 | 37 | 0.1351 | 143 | 0.0769 | 21 | 0.1429 | 25 | 0.08 | 37 | 0.0541 | 26 | 0.0769 | 109 | 0.0826 |
| 15_9223948 | 29 | 0 | 46 | 0.0652 | 34 | 0.0588 | 38 | 0.1053 | 147 | 0.0612 | 22 | 0.0909 | 25 | 0.16 | 34 | 0.0588 | 30 | 0.1667 | 111 | 0.1171 |
| 15_9266786 | NA | NA | 2 | 0.5 | NA | NA | NA | NA | 2 | 0.5 | NA | NA | NA | NA | NA | NA | 1 | 1 | 1 | 1 |
| 15_9266798 | NA | NA | 3 | 0.6667 | NA | NA | NA | NA | 3 | 0.6667 | NA | NA | NA | NA | NA | NA | 1 | 1 | 1 | 1 |
| 15_9266803 | NA | NA | 3 | 0.6667 | NA | NA | NA | NA | 3 | 0.6667 | NA | NA | NA | NA | NA | NA | 2 | 0.5 | 2 | 0.5 |
| 15_9266847 | NA | NA | 3 | 0.6667 | NA | NA | NA | NA | 3 | 0.6667 | NA | NA | NA | NA | 1 | 1 | 2 | 1 | 3 | 1 |
| 15_9266850 | NA | NA | 3 | 0.3333 | NA | NA | NA | NA | 3 | 0.3333 | NA | NA | NA | NA | 1 | 1 | 2 | 0.5 | 3 | 0.6667 |
| 15_9266875 | NA | NA | 2 | 1 | NA | NA | NA | NA | 2 | 1 | NA | NA | NA | NA | 1 | 1 | 2 | 1 | 3 | 1 |
| 15_9266881 | NA | NA | 1 | 1 | NA | NA | NA | NA | 1 | 1 | NA | NA | NA | NA | 1 | 1 | 2 | 1 | 3 | 1 |
| 15_9266882 | NA | NA | 2 | 1 | NA | NA | NA | NA | 2 | 1 | 1 | 1 | NA | NA | 1 | 1 | 2 | 1 | 4 | 1 |
| 15_9266889 | NA | NA | 3 | 0.3333 | NA | NA | NA | NA | 3 | 0.3333 | 1 | 0 | NA | NA | 1 | 1 | 1 | 0 | 3 | 0.3333 |
| 15_9266890 | NA | NA | 3 | 1 | NA | NA | NA | NA | 3 | 1 | 1 | 1 | NA | NA | NA | NA | 2 | 1 | 3 | 1 |
| 15_9266899 | NA | NA | 3 | 0.6667 | NA | NA | NA | NA | 3 | 0.6667 | 1 | 1 | NA | NA | 1 | 1 | 1 | 1 | 3 | 1 |
| 15_9266917 | NA | NA | 3 | 0.6667 | NA | NA | 1 | 0 | 4 | 0.5 | 1 | 0 | NA | NA | 1 | 0 | 2 | 0.5 | 4 | 0.25 |
| 15_9266918 | NA | NA | 3 | 0.6667 | NA | NA | 1 | 1 | 4 | 0.75 | 1 | 1 | NA | NA | 1 | 0 | 2 | 0.5 | 4 | 0.5 |
| 15_9266945 | NA | NA | 2 | 0.5 | NA | NA | NA | NA | 2 | 0.5 | NA | NA | NA | NA | 1 | 1 | 1 | 0 | 2 | 0.5 |
| 15_9266956 | NA | NA | 3 | 1 | NA | NA | NA | NA | 3 | 1 | 1 | 0 | NA | NA | 1 | 1 | 2 | 1 | 4 | 0.75 |
| 15_9314260 | 1 | 1 | NA | NA | NA | NA | 1 | 1 | 2 | 1 | 1 | 1 | NA | NA | NA | NA | 2 | 1 | 3 | 1 |
| 15_9314269 | 1 | 1 | NA | NA | NA | NA | 1 | 1 | 2 | 1 | 1 | 1 | NA | NA | NA | NA | 1 | 0 | 2 | 0.5 |
| 15_9314278 | NA | NA | NA | NA | NA | NA | 1 | 1 | 1 | 1 | 1 | 1 | NA | NA | NA | NA | 2 | 1 | 3 | 1 |
| 15_9318342 | 9 | 0.3333 | 17 | 0.7647 | 14 | 0.5714 | 5 | 0.8 | 45 | 0.6222 | 11 | 0.6364 | 14 | 0.5714 | 12 | 0.75 | 11 | 0.9091 | 48 | 0.7083 |
| 15_9338703 | 2 | 0.5 | 2 | 0.5 | 2 | 0 | 7 | 0.4286 | 13 | 0.3846 | 4 | 0.75 | 5 | 0.8 | 3 | 1 | 6 | 0.3333 | 18 | 0.6667 |
| 15_9339711 | 6 | 0.5 | 14 | 0.5 | 6 | 0.1667 | 6 | 0.3333 | 32 | 0.4062 | 11 | 0.3636 | 9 | 0.2222 | 9 | 0.3333 | 13 | 0.6154 | 42 | 0.4048 |
| 15_9339717 | 7 | 0.1429 | 16 | 0.1875 | 6 | 0 | 6 | 0 | 35 | 0.1143 | 8 | 0.25 | 9 | 0.1111 | 6 | 0.5 | 12 | 0.3333 | 35 | 0.2857 |
| 15_9339730 | 6 | 0 | 15 | 0.1333 | 6 | 0.1667 | 5 | 0 | 32 | 0.0938 | 11 | 0.3636 | 9 | 0.2222 | 5 | 0.2 | 14 | 0.2857 | 39 | 0.2821 |
| 15_9339744 | 5 | 0.2 | 14 | 0.2857 | 6 | 0.1667 | 5 | 0.2 | 30 | 0.2333 | 11 | 0.0909 | 9 | 0.1111 | 6 | 0 | 13 | 0.0769 | 39 | 0.0769 |
| 15_9339764 | 3 | 0 | 13 | 0.1538 | 5 | 0.2 | 4 | 0 | 25 | 0.12 | 7 | 0.1429 | 7 | 0.1429 | 6 | 0.3333 | 11 | 0 | 31 | 0.129 |
| 15_9339783 | 3 | 0.6667 | 14 | 0.8571 | 5 | 0.8 | 4 | 1 | 26 | 0.8462 | 8 | 1 | 8 | 0.625 | 4 | 1 | 11 | 0.9091 | 31 | 0.871 |
| 15_9339797 | 3 | 0 | 10 | 0.1 | 4 | 0.5 | 5 | 0 | 22 | 0.1364 | 8 | 0.125 | 8 | 0 | 4 | 0.25 | 9 | 0.1111 | 29 | 0.1034 |
| 15_9339806 | 2 | 1 | 12 | 0.9167 | 5 | 1 | 4 | 1 | 23 | 0.9565 | 7 | 1 | 8 | 0.875 | 3 | 1 | 9 | 1 | 27 | 0.963 |

| | | | | | | | | | | | | | | | | | | | | |
|---|---|---|---|---|---|---|---|---|---|---|---|---|---|---|---|---|---|---|---|---|
| 15_9339807 | 2 | 0.5 | 9 | 0.3333 | 3 | 0.6667 | 2 | 1 | 16 | 0.5 | 5 | 0.6 | 6 | 0.5 | 4 | 0.75 | 7 | 0.4286 | 22 | 0.5455 |
| 15_9339809 | 2 | 0 | 7 | 0.1429 | 3 | 0.6667 | 3 | 1 | 15 | 0.4 | 6 | 0.8333 | 7 | 0.1429 | 3 | 0.3333 | 5 | 0.8 | 21 | 0.5238 |
| 15_9339827 | 2 | 1 | 10 | 1 | 5 | 1 | 4 | 1 | 21 | 1 | 5 | 1 | 8 | 1 | 2 | 1 | 10 | 1 | 25 | 1 |
| 15_9339829 | 2 | 0 | 6 | 0.3333 | 5 | 0.8 | 4 | 0.25 | 17 | 0.4118 | 6 | 0.3333 | 7 | 0.4286 | 2 | 0 | 10 | 0.2 | 25 | 0.28 |
| 15_9339831 | 2 | 0 | 9 | 0.4444 | 5 | 0 | 4 | 0 | 20 | 0.2 | 6 | 0.1667 | 8 | 0.25 | 2 | 0 | 10 | 0.1 | 26 | 0.1538 |
| 15_9339834 | 2 | 1 | 9 | 1 | 1 | 1 | 2 | 1 | 14 | 1 | 4 | 1 | 7 | 1 | 2 | 1 | 10 | 0.9 | 23 | 0.9565 |
| 15_9339835 | 2 | 1 | 10 | 1 | 5 | 1 | 4 | 1 | 21 | 1 | 6 | 1 | 8 | 1 | 2 | 1 | 10 | 1 | 26 | 1 |
| 15_9339838 | 1 | 1 | 10 | 0.3 | 5 | 0.6 | 4 | 0.5 | 20 | 0.45 | 5 | 0.4 | 8 | 0.375 | 3 | 0.3333 | 9 | 0.7778 | 25 | 0.52 |
| 15_9339843 | 1 | 0 | 8 | 0.25 | 5 | 0.4 | 2 | 0 | 16 | 0.25 | 6 | 0.1667 | 7 | 0.2857 | 3 | 0 | 9 | 0.2222 | 25 | 0.2 |
| 15_9339857 | 1 | 0 | 8 | 0.375 | 4 | 0.25 | 3 | 0.3333 | 16 | 0.3125 | 3 | 0 | 3 | 0.6667 | 2 | 1 | 8 | 0.25 | 16 | 0.375 |
| 15_9339870 | NA | NA | 8 | 0 | 4 | 0.25 | 3 | 0 | 15 | 0.0667 | 4 | 0 | 7 | 0.1429 | 1 | 0 | 8 | 0 | 20 | 0.05 |
| 15_9339872 | 1 | 0 | 8 | 0 | 4 | 0.25 | 2 | 0.5 | 15 | 0.1333 | 5 | 0 | 8 | 0 | 3 | 0 | 9 | 0.3333 | 25 | 0.12 |
| 15_9339872 | 1 | 0 | 9 | 0.6667 | 3 | 0.6667 | 2 | 1 | 15 | 0.6667 | 6 | 0.6667 | 6 | 0.3333 | 3 | 0.6667 | 8 | 0.625 | 23 | 0.5652 |
| 15_9339885 | 1 | 1 | 6 | 1 | 4 | 0.75 | 3 | 0.6667 | 14 | 0.8571 | 5 | 1 | 8 | 0.75 | 3 | 0.6667 | 9 | 0.8889 | 25 | 0.84 |
| 15_9339902 | NA | NA | 5 | 1 | 3 | 0.6667 | 2 | 1 | 10 | 0.9 | 1 | 1 | 2 | 0.5 | 1 | 1 | 6 | 1 | 10 | 0.9 |
| 15_9339907 | NA | NA | 3 | 1 | 1 | 1 | 1 | 1 | 5 | 1 | 2 | 0.5 | 2 | 1 | 1 | 1 | 7 | 0.8571 | 12 | 0.8333 |
| 15_9339908 | 1 | 1 | 3 | 1 | 3 | 1 | 1 | 1 | 8 | 1 | 1 | 1 | 4 | 1 | 2 | 1 | 8 | 0.875 | 15 | 0.9333 |
| 15_9340092 | 1 | 0 | 6 | 0 | 5 | 0.2 | 2 | 0 | 14 | 0.0714 | 4 | 0 | 5 | 0 | 1 | 0 | 8 | 0.125 | 18 | 0.0556 |
| 15_9340097 | 1 | 0 | 7 | 0.5714 | 3 | 0.3333 | 2 | 1 | 13 | 0.5385 | 2 | 1 | 2 | 1 | 2 | 1 | 6 | 0.6667 | 12 | 0.8333 |
| 15_9340103 | 1 | 0 | 6 | 0.3333 | 4 | 0.25 | NA | NA | 11 | 0.2727 | 5 | 0.2 | 5 | 0.2 | 2 | 0 | 7 | 0 | 19 | 0.1053 |
| 15_9340149 | NA | NA | 3 | 0.3333 | 4 | 1 | 2 | 1 | 9 | 0.7778 | 3 | 1 | 2 | 1 | 2 | 0.5 | 5 | 0.6 | 12 | 0.75 |
| 15_9340152 | NA | NA | 5 | 0.4 | 3 | 0.3333 | 2 | 0.5 | 10 | 0.4 | 3 | 0 | 3 | 0.3333 | 2 | 0 | 5 | 0.2 | 13 | 0.1538 |
| 15_9442144 | 1 | 1 | NA | NA | NA | NA | NA | NA | 1 | 1 | NA | NA | NA | NA | NA | NA | 2 | 0.5 | 2 | 0.5 |
| 15_9443871 | 1 | 1 | NA | NA | NA | NA | NA | NA | 1 | 1 | NA | NA | NA | NA | NA | NA | 1 | 1 | 1 | 1 |
| 15_9447612 | 3 | 1 | 12 | 0.5 | 10 | 0.8 | 10 | 0.9 | 35 | 0.7429 | 8 | 0.875 | 3 | 0.6667 | 4 | 0.25 | 4 | 0.5 | 19 | 0.6316 |
| 15_9447630 | 3 | 0.3333 | 11 | 0.1818 | 8 | 0.25 | 5 | 0.2 | 27 | 0.2222 | 8 | 0.375 | 3 | 0 | 6 | 0.1667 | 1 | 0 | 18 | 0.2222 |
| 15_9447632 | 3 | 0 | 14 | 0.0714 | 8 | 0 | 9 | 0.1111 | 34 | 0.0588 | 9 | 0.1111 | 3 | 0.3333 | 5 | 0 | 3 | 0 | 20 | 0.1 |
| 15_9447641 | 3 | 0 | 16 | 0.125 | 8 | 0.125 | 10 | 0 | 37 | 0.0811 | 11 | 0.1818 | 3 | 0.3333 | 5 | 0 | 3 | 0.3333 | 22 | 0.1818 |
| 15_9447653 | 3 | 0.3333 | 16 | 0.0625 | 9 | 0.1111 | 10 | 0 | 38 | 0.0789 | 8 | 0.125 | 3 | 0 | 5 | 0 | 2 | 0 | 18 | 0.0556 |
| 15_9447704 | 2 | 0 | 15 | 0.0667 | 7 | 0 | 8 | 0.125 | 32 | 0.0625 | 8 | 0.125 | 1 | 0 | 4 | 0.25 | 3 | 0 | 16 | 0.125 |
| 15_9447763 | 1 | 1 | 13 | 0.3077 | 8 | 0.5 | 8 | 0.375 | 30 | 0.4 | 4 | 0.75 | 2 | 0 | 2 | 0 | 2 | 0 | 10 | 0.3 |
| 15_9447769 | 1 | 1 | 8 | 0.5 | 6 | 0.5 | 9 | 0.8889 | 24 | 0.6667 | 2 | 1 | 1 | 1 | 2 | 0 | 2 | 1 | 7 | 0.7143 |
| 15_9499233 | 2 | 1 | 1 | 1 | NA | NA | 1 | 1 | 4 | 1 | NA | NA | 1 | 1 | NA | NA | NA | NA | 1 | 1 |
| 15_9499237 | 2 | 1 | 1 | 1 | NA | NA | 1 | 1 | 4 | 1 | NA | NA | 1 | 1 | NA | NA | NA | NA | 1 | 1 |
| 15_9511763 | 43 | 0.8605 | 49 | 0.6327 | 47 | 0.6596 | 24 | 0.7917 | 163 | 0.7239 | 32 | 0.6562 | 28 | 0.7143 | 34 | 0.7059 | 32 | 0.8438 | 126 | 0.7302 |
| 15_9511765 | 53 | 0.6415 | 58 | 0.7414 | 52 | 0.75 | 28 | 0.6786 | 191 | 0.7068 | 40 | 0.575 | 38 | 0.5526 | 40 | 0.55 | 43 | 0.6047 | 161 | 0.5714 |
| 15_9511782 | 37 | 0.0811 | 48 | 0.0417 | 45 | 0.0444 | 23 | 0.087 | 153 | 0.0588 | 27 | 0.1481 | 31 | 0 | 33 | 0.0606 | 37 | 0.0541 | 128 | 0.0625 |
| 15_9511785 | 37 | 1 | 43 | 1 | 35 | 1 | 24 | 0.9583 | 139 | 0.9928 | 20 | 0.9 | 23 | 0.913 | 27 | 0.963 | 29 | 0.931 | 99 | 0.9293 |
| 15_9511792 | 42 | 0.8571 | 48 | 0.8542 | 40 | 0.675 | 25 | 0.92 | 155 | 0.8194 | 29 | 0.8621 | 36 | 0.8889 | 36 | 0.8611 | 43 | 0.8372 | 144 | 0.8611 |
| 15_9552235 | 4 | 0.25 | 1 | 0 | 3 | 1 | 6 | 0.5 | 14 | 0.5 | 6 | 0.5 | 2 | 1 | 6 | 0.5 | 4 | 0.25 | 18 | 0.5 |
| 15_9552249 | 4 | 0.5 | 1 | 0 | 3 | 0.6667 | 4 | 0.5 | 12 | 0.5 | 2 | 0 | 1 | 0 | 2 | 0 | 4 | 0.5 | 9 | 0.2222 |
| 15_9552252 | 3 | 0.3333 | 1 | 0 | 2 | 0 | 6 | 0.1667 | 12 | 0.1667 | 6 | 0.1667 | 2 | 0 | 4 | 0 | 4 | 0 | 16 | 0.0625 |
| 15_9552254 | 4 | 0.5 | 1 | 0 | 3 | 0 | 7 | 0 | 15 | 0.1333 | 6 | 0.1667 | 2 | 0 | 6 | 0.1667 | 4 | 0 | 18 | 0.1111 |
| 15_9552259 | 4 | 1 | NA | NA | 3 | 1 | 6 | 1 | 13 | 1 | 6 | 1 | 1 | 1 | 6 | 0.6667 | 2 | 0.5 | 15 | 0.8 |
| 15_9552260 | 4 | 0 | 1 | 0 | 3 | 0.3333 | 7 | 0.1429 | 15 | 0.1333 | 6 | 0.5 | 2 | 0.5 | 6 | 0.1667 | 3 | 0 | 17 | 0.2941 |
| 15_9552260 | 4 | 0.25 | 1 | 0 | 2 | 0 | 7 | 0 | 14 | 0.0714 | 5 | 0 | 2 | 0.5 | 6 | 0.1667 | 4 | 0.25 | 17 | 0.1765 |
| 15_9552268 | 3 | 0.3333 | 1 | 1 | NA | NA | 6 | 0.5 | 10 | 0.5 | 3 | 1 | 2 | 1 | 4 | 0.75 | 3 | 0.6667 | 12 | 0.8333 |
| 15_9552270 | 3 | 0 | 1 | 0 | 3 | 0.3333 | 7 | 0 | 14 | 0.0714 | 5 | 0 | 2 | 0 | 6 | 0.1667 | 4 | 0 | 17 | 0.0588 |
| 15_9555233 | 1 | 1 | NA | NA | 2 | 0.5 | 3 | 0 | 6 | 0.3333 | 2 | 0.5 | 1 | 0 | 1 | 0 | 2 | 0 | 6 | 0.1667 |
| 15_9555254 | 1 | 0 | NA | NA | 2 | 0 | 2 | 0.5 | 5 | 0.2 | 2 | 1 | NA | NA | NA | NA | 2 | 0.5 | 4 | 0.75 |
| 15_9587066 | 2 | 1 | NA | NA | 2 | 1 | 1 | 1 | 5 | 1 | NA | NA | NA | NA | 1 | 1 | 1 | 1 | 2 | 1 |
| 15_9587075 | 2 | 1 | 1 | 1 | 2 | 0 | 1 | 1 | 6 | 0.6667 | NA | NA | NA | NA | 2 | 0.5 | 2 | 1 | 4 | 0.75 |
| 15_9587087 | 4 | 1 | 1 | 1 | 1 | 1 | NA | NA | 6 | 1 | NA | NA | NA | NA | 2 | 1 | 2 | 1 | 4 | 1 |
| 15_9587095 | 4 | 0 | 3 | 0.3333 | 2 | 0 | 1 | 0 | 10 | 0.1 | NA | NA | NA | NA | 3 | 0.6667 | 2 | 0.5 | 5 | 0.6 |
| 15_9587096 | 4 | 0.25 | 3 | 1 | 2 | 0.5 | 1 | 0 | 10 | 0.5 | NA | NA | NA | NA | 3 | 0.6667 | 2 | 0.5 | 5 | 0.6 |
| 15_9587112 | 4 | 0 | 3 | 0.3333 | 2 | 0.5 | 1 | 0 | 10 | 0.2 | NA | NA | NA | NA | 3 | 0 | 2 | 0.5 | 5 | 0.2 |
| 15_9587126 | 4 | 0.75 | 3 | 1 | 2 | 0 | 1 | 1 | 10 | 0.7 | NA | NA | NA | NA | 3 | 0.6667 | 2 | 0.5 | 5 | 0.6 |
| 15_9587131 | 2 | 1 | 2 | 1 | NA | NA | 1 | 1 | 5 | 1 | NA | NA | NA | NA | 3 | 1 | 2 | 1 | 5 | 1 |
| 15_9587141 | 3 | 1 | 1 | 1 | 2 | 0 | 1 | 1 | 7 | 0.7143 | NA | NA | NA | NA | 3 | 0.6667 | 1 | 1 | 4 | 0.75 |
| 15_9587147 | 3 | 1 | 1 | 1 | 1 | 1 | NA | NA | 5 | 1 | NA | NA | NA | NA | 2 | 1 | NA | NA | 2 | 1 |
| 15_9587164 | 3 | 0.3333 | 3 | 0 | NA | NA | NA | NA | 6 | 0.1667 | NA | NA | NA | NA | 3 | 0.3333 | 2 | 0.5 | 5 | 0.4 |
| 15_9587165 | 3 | 0.3333 | NA | NA | 2 | 0.5 | 1 | 0 | 6 | 0.3333 | NA | NA | NA | NA | 2 | 1 | 1 | 1 | 3 | 1 |
| 15_9587170 | 4 | 0.75 | 2 | 0.5 | 1 | 0 | NA | NA | 7 | 0.5714 | NA | NA | NA | NA | 2 | 0.5 | 2 | 0.5 | 4 | 0.5 |
| 15_9587177 | 4 | 0.5 | 2 | 0.5 | 1 | 1 | 1 | 0 | 8 | 0.5 | NA | NA | NA | NA | 4 | 0.25 | 2 | 0 | 6 | 0.1667 |
| 15_9587198 | 3 | 0.3333 | 3 | 0.3333 | 2 | 0.5 | 1 | 1 | 9 | 0.4444 | NA | NA | NA | NA | 3 | 0.3333 | 2 | 0.5 | 5 | 0.4 |

| | | | | | | | | | | | | | | | | | | | | |
|---|---|---|---|---|---|---|---|---|---|---|---|---|---|---|---|---|---|---|---|---|
| 15_9587202 | 4 | 0.5 | 4 | 1 | 2 | 0 | 1 | 0 | 11 | 0.5455 | NA | NA | NA | NA | 4 | 0.75 | 2 | 0.5 | 6 | 0.6667 |
| 15_9587206 | 4 | 0.25 | 3 | 0.6667 | 2 | 0.5 | 1 | 1 | 10 | 0.5 | NA | NA | NA | NA | 3 | 0.3333 | 1 | 0 | 4 | 0.25 |
| 15_9587212 | 4 | 1 | 3 | 1 | 3 | 0.6667 | 1 | 1 | 11 | 0.9091 | NA | NA | NA | NA | 4 | 1 | 2 | 1 | 6 | 1 |
| 15_9587232 | 5 | 0.6 | 3 | 0.3333 | 3 | 0.3333 | 1 | 1 | 12 | 0.5 | NA | NA | NA | NA | 5 | 0.4 | 1 | 0 | 6 | 0.3333 |
| 15_9587246 | 5 | 0.2 | 4 | 0.25 | 2 | 0 | 1 | 0 | 12 | 0.1667 | NA | NA | NA | NA | 5 | 0.4 | 2 | 1 | 7 | 0.5714 |
| 15_9587261 | 5 | 0.2 | 3 | 0 | 2 | 0 | 1 | 1 | 11 | 0.1818 | NA | NA | NA | NA | 2 | 0.5 | 2 | 0 | 4 | 0.25 |
| 15_9587285 | 5 | 0 | 3 | 0.3333 | 3 | 0 | 1 | 0 | 12 | 0.0833 | NA | NA | NA | NA | 4 | 0.25 | 2 | 0 | 6 | 0.1667 |
| 15_9587311 | 5 | 0 | 4 | 0 | 3 | 0.3333 | 1 | 1 | 13 | 0.1538 | NA | NA | 1 | 0 | 4 | 0 | 1 | 1 | 6 | 0.1667 |
| 15_9587323 | 5 | 0.8 | 4 | 0.75 | 4 | 0.25 | 1 | 1 | 14 | 0.6429 | NA | NA | 1 | 1 | 4 | 0.25 | 2 | 0.5 | 7 | 0.4286 |
| 15_9626792 | 1 | 1 | NA | NA | 1 | 1 | 1 | 1 | 3 | 1 | NA | NA | 4 | 0.5 | 1 | 1 | NA | NA | 5 | 0.6 |
| 15_9626798 | 1 | 1 | 1 | 0 | 1 | 0 | 1 | 0 | 4 | 0.25 | NA | NA | 4 | 0.25 | 1 | 0 | 1 | 0 | 6 | 0.1667 |
| 15_9626805 | 1 | 1 | NA | NA | NA | NA | 1 | 1 | 2 | 1 | NA | NA | 2 | 0 | 1 | 1 | 1 | 1 | 4 | 0.5 |
| 15_9626816 | 1 | 0 | 1 | 0 | 1 | 1 | NA | NA | 3 | 0.3333 | NA | NA | 4 | 0 | 1 | 0 | 1 | 1 | 6 | 0.1667 |
| 15_9626932 | NA | NA | NA | NA | 1 | 1 | NA | NA | 1 | 1 | 1 | 1 | 3 | 1 | 1 | 1 | 2 | 1 | 7 | 1 |
| 15_9626938 | 3 | 0.6667 | 2 | 1 | 2 | 1 | 1 | 1 | 8 | 0.875 | 1 | 1 | 4 | 0.5 | 1 | 1 | 2 | 1 | 8 | 0.75 |
| 15_9626952 | 3 | 0.6667 | 1 | 0 | 1 | 1 | NA | NA | 5 | 0.6 | 1 | 1 | 4 | 0.75 | 1 | 1 | 2 | 0.5 | 8 | 0.75 |
| 15_9626954 | 3 | 0.3333 | 1 | 1 | 2 | 0.5 | 1 | 1 | 7 | 0.5714 | 1 | 0 | 4 | 0.5 | 1 | 1 | 2 | 0.5 | 8 | 0.5 |
| 15_9626973 | 3 | 0 | 2 | 0 | 2 | 0.5 | 1 | 0 | 8 | 0.125 | 1 | 0 | 4 | 0.5 | 2 | 0 | 1 | 1 | 8 | 0.375 |
| 15_9626977 | 3 | 0.6667 | 2 | 0.5 | 1 | 0 | 1 | 0 | 7 | 0.4286 | 1 | 1 | 5 | 0.6 | 2 | 0.5 | 2 | 0 | 10 | 0.5 |
| 15_9626988 | 3 | 0.3333 | 1 | 0 | 2 | 1 | 1 | 0 | 7 | 0.4286 | 1 | 1 | 3 | 0.3333 | 2 | 0 | 1 | 0 | 7 | 0.2857 |
| 15_9626999 | 3 | 0.6667 | 1 | 1 | 2 | 1 | NA | NA | 6 | 0.8333 | 1 | 0 | 3 | 1 | 2 | 0.5 | 3 | 0 | 9 | 0.4444 |
| 15_9627006 | 2 | 1 | 2 | 1 | 2 | 0.5 | 1 | 0 | 7 | 0.7143 | 1 | 1 | 5 | 0.6 | 2 | 1 | 2 | 1 | 10 | 0.8 |
| 15_9627012 | 3 | 0.6667 | 2 | 0.5 | 2 | 1 | 1 | 1 | 8 | 0.75 | 1 | 0 | 4 | 0.75 | NA | NA | 3 | 1 | 8 | 0.75 |
| 15_9627013 | 1 | 0 | NA | NA | 1 | 0 | 1 | 1 | 3 | 0.3333 | 1 | 1 | 1 | 0 | 1 | 1 | 2 | 1 | 5 | 0.8 |
| 15_9627022 | 3 | 1 | 1 | 0 | 2 | 1 | 1 | 1 | 7 | 0.8571 | 1 | 0 | 5 | 0.2 | 2 | 0.5 | 3 | 0.6667 | 11 | 0.3636 |
| 15_9627034 | 3 | 0.3333 | 1 | 1 | 3 | 0.3333 | 1 | 1 | 8 | 0.5 | 1 | 0 | 6 | 0.5 | 2 | 1 | 3 | 1 | 12 | 0.6667 |
| 15_9627041 | 2 | 1 | 1 | 1 | 2 | 0.5 | 1 | 1 | 6 | 0.8333 | 1 | 0 | 5 | 0.8 | 1 | 1 | 3 | 1 | 10 | 0.8 |
| 15_9627044 | 2 | 0.5 | 3 | 1 | 2 | 1 | 1 | 0 | 8 | 0.75 | 1 | 0 | 6 | 0.6667 | 2 | 1 | 2 | 1 | 11 | 0.7273 |
| 15_9627055 | 3 | 0.6667 | 3 | 0.3333 | 3 | 0.3333 | 1 | 0 | 10 | 0.4 | 1 | 1 | 6 | 0 | 2 | 0 | 2 | 0.5 | 11 | 0.1818 |
| 15_9629992 | 3 | 1 | 1 | 1 | 1 | 0 | 1 | 1 | 6 | 0.8333 | 1 | 1 | 5 | 1 | 2 | 1 | 4 | 0.75 | 12 | 0.9167 |
| 15_9629994 | 4 | 0.25 | 2 | 0 | 1 | 0 | NA | NA | 7 | 0.1429 | 1 | 0 | 6 | 0 | 2 | 0 | 4 | 0.25 | 13 | 0.0769 |
| 15_9629995 | 3 | 0.6667 | 2 | 0.5 | 1 | 1 | 1 | 1 | 7 | 0.7143 | NA | NA | 5 | 0.6 | 2 | 1 | 4 | 1 | 11 | 0.8182 |
| 15_9630006 | 3 | 0.3333 | 3 | 0.6667 | 1 | 0 | 1 | 0 | 8 | 0.375 | 1 | 0 | 5 | 0.4 | 3 | 0.3333 | 5 | 0.6 | 14 | 0.4286 |
| 15_9630020 | 3 | 1 | 2 | 1 | 3 | 1 | 1 | 1 | 9 | 1 | NA | NA | 5 | 0.8 | 1 | 1 | 3 | 1 | 9 | 0.8889 |
| 15_9630026 | 4 | 0.5 | 3 | 0.3333 | 2 | 0.5 | 2 | 0.5 | 11 | 0.4545 | 1 | 0 | 6 | 0.1667 | 3 | 0.3333 | 5 | 0 | 15 | 0.1333 |
| 15_9630061 | 4 | 0.25 | 3 | 0 | 3 | 0 | 2 | 0 | 12 | 0.0833 | 1 | 0 | 4 | 0.25 | 3 | 0 | 5 | 0 | 13 | 0.0769 |
| 15_9630081 | 5 | 0.2 | 4 | 0.5 | 3 | 0 | 3 | 0 | 15 | 0.2 | 1 | 0 | 7 | 0.2857 | 2 | 0 | 4 | 0 | 14 | 0.1429 |
| 15_9631238 | NA | NA | 3 | 0.3333 | NA | NA | 1 | 1 | 4 | 0.5 | NA | NA | 3 | 1 | 3 | 0.3333 | NA | NA | 6 | 0.6667 |
| 15_9631262 | 2 | 0 | 5 | 0.2 | NA | NA | 3 | 0.3333 | 10 | 0.2 | 3 | 0.3333 | 4 | 0.25 | 4 | 0.25 | 3 | 0.6667 | 14 | 0.3571 |
| 15_9631264 | 3 | 0 | 6 | 0.5 | 1 | 1 | 3 | 0.6667 | 13 | 0.4615 | 3 | 0.3333 | 6 | 0.5 | 4 | 0 | 2 | 0 | 15 | 0.2667 |
| 15_9631605 | 3 | 0.3333 | 3 | 0.3333 | 1 | 0 | 2 | 0 | 9 | 0.2222 | 2 | 0 | 3 | 0 | 4 | 0.25 | 3 | 0.6667 | 12 | 0.25 |
| 15_9631630 | 3 | 0 | 1 | 0 | 1 | 0 | 1 | 1 | 6 | 0.1667 | 1 | 0 | 1 | 1 | 2 | 0 | 2 | 0.5 | 6 | 0.3333 |
| 15_9631631 | 3 | 1 | 3 | 1 | NA | NA | NA | NA | 6 | 1 | 2 | 0.5 | 2 | 1 | 3 | 1 | 3 | 1 | 10 | 0.9 |
| 15_9631640 | 3 | 0 | 3 | 0 | NA | NA | 2 | 0.5 | 8 | 0.125 | 2 | 1 | 3 | 0 | 2 | 0 | 3 | 0 | 10 | 0.2 |
| 15_9631643 | 2 | 1 | 3 | 1 | 1 | 0 | 2 | 0 | 8 | 0.625 | 2 | 0.5 | 3 | 0 | 2 | 0.5 | 3 | 0.6667 | 10 | 0.4 |
| 15_9631657 | 2 | 0 | 2 | 0.5 | 1 | 1 | 2 | 0.5 | 7 | 0.4286 | 1 | 1 | 3 | 0.3333 | 2 | 0 | 3 | 0.3333 | 9 | 0.3333 |
| 15_9631769 | 2 | 0 | 2 | 0 | NA | NA | 2 | 1 | 6 | 0.3333 | 1 | 0 | 2 | 0.5 | NA | NA | 2 | 1 | 5 | 0.6 |
| 15_9631833 | 1 | 1 | NA | NA | NA | NA | 2 | 0.5 | 3 | 0.6667 | 1 | 0 | 2 | 1 | NA | NA | NA | NA | 3 | 0.6667 |
| 15_9647001 | 12 | 0.5833 | 13 | 0.1538 | 7 | 0.1429 | 11 | 0.1818 | 43 | 0.2791 | 10 | 0.2 | 6 | 0.6667 | 8 | 0.375 | 11 | 0.1818 | 35 | 0.3143 |
| 15_9647031 | 13 | 0.3077 | 13 | 0.4615 | 7 | 0.2857 | 9 | 0.3333 | 42 | 0.3571 | 11 | 0.2727 | 4 | 0.25 | 7 | 0.2857 | 11 | 0.0909 | 33 | 0.2121 |
| 15_9647042 | 10 | 0.2 | 8 | 0.5 | 5 | 0.2 | 7 | 0.1429 | 30 | 0.2667 | 8 | 0.25 | 4 | 0.25 | 6 | 0 | 8 | 0.25 | 26 | 0.1923 |
| 15_9647179 | 4 | 0 | 5 | 0.2 | 3 | 0 | 3 | 0 | 15 | 0.0667 | 6 | 0.3333 | 2 | 0 | 2 | 0 | 8 | 0.125 | 18 | 0.1667 |
| 15_9647229 | 2 | 0.5 | 6 | 0.5 | 3 | 0.6667 | 5 | 0 | 16 | 0.375 | 6 | 0.1667 | 3 | 0.3333 | 1 | 1 | 8 | 0.375 | 18 | 0.3333 |
| 15_9647251 | 5 | 0.4 | 5 | 0 | 3 | 0 | 6 | 0 | 19 | 0.1053 | 5 | 0 | 2 | 0 | 1 | 0 | 8 | 0.125 | 16 | 0.0625 |
| 15_9647255 | 3 | 0 | 4 | 0 | 2 | 0 | 4 | 0.25 | 13 | 0.0769 | 3 | 0 | 2 | 0 | 1 | 0 | 8 | 0.375 | 14 | 0.2143 |
| 15_9647276 | 3 | 0 | 6 | 0 | 1 | 0 | 6 | 0.1667 | 16 | 0.0625 | 4 | 0 | 1 | 0 | 1 | 1 | 6 | 0 | 12 | 0.0833 |
| 15_9647323 | 5 | 0 | 4 | 0.25 | 3 | 0 | 4 | 0 | 16 | 0.0625 | 4 | 0.25 | 1 | 0 | 1 | 0 | 5 | 0 | 11 | 0.0909 |
| 15_9647355 | 5 | 0.2 | 5 | 0.4 | 3 | 0 | 2 | 0 | 15 | 0.2 | 5 | 0 | 1 | 0 | 1 | 0 | 6 | 0.1667 | 13 | 0.0769 |
| 15_9647377 | 2 | 1 | 4 | 0.5 | 1 | 0 | 2 | 0.5 | 9 | 0.5556 | 4 | 0.25 | 1 | 0 | 1 | 1 | 4 | 0.5 | 10 | 0.4 |
| 15_9647386 | 1 | 1 | 4 | 0.5 | 1 | 1 | 2 | 0 | 8 | 0.5 | 5 | 0.2 | 1 | 0 | 1 | 0 | 4 | 0 | 11 | 0.0909 |
| 15_9647399 | 2 | 1 | 5 | 0.8 | 2 | 1 | 2 | 1 | 11 | 0.9091 | 5 | 1 | 1 | 1 | 2 | 1 | 6 | 1 | 14 | 1 |
| 15_9647406 | 2 | 1 | 5 | 0.2 | 2 | 0.5 | 2 | 0 | 11 | 0.3636 | 5 | 0.2 | 1 | 0 | 2 | 0 | 6 | 0.1667 | 14 | 0.1429 |
| 15_9647430 | 4 | 1 | 4 | 0.75 | NA | NA | 2 | 1 | 10 | 0.9 | 4 | 0.75 | 1 | 1 | NA | NA | 3 | 1 | 8 | 0.875 |
| 15_9647441 | 4 | 1 | 4 | 0.75 | 1 | 1 | 3 | 1 | 12 | 0.9167 | 6 | 1 | 1 | 0 | 2 | 1 | 5 | 0.8 | 14 | 0.8571 |
| 15_9647443 | 4 | 0.75 | 5 | 0.8 | 1 | 0 | 3 | 0.3333 | 13 | 0.6154 | 6 | 0.3333 | 1 | 0 | 2 | 1 | 4 | 0.5 | 13 | 0.4615 |

| | | | | | | | | | | | | | | | | | | | | |
|---|---|---|---|---|---|---|---|---|---|---|---|---|---|---|---|---|---|---|---|---|
| 15_964744 | 3 | 0.6667 | 3 | 0.6667 | 1 | 1 | 2 | 1 | 9 | 0.7778 | 4 | 0.5 | 1 | 1 | 2 | 0.5 | 3 | 1 | 10 | 0.7 |
| 15_964744 | 3 | 1 | 4 | 1 | 1 | 1 | 2 | 1 | 10 | 1 | 5 | 0.8 | 1 | 1 | 2 | 1 | 4 | 1 | 12 | 0.9167 |
| 15_964745 | 4 | 0 | 5 | 0.6 | 1 | 0 | 2 | 0 | 12 | 0.25 | 4 | 0 | 1 | 0 | 1 | 0 | 4 | 0.25 | 10 | 0.1 |
| 15_965409 | NA | NA | NA | NA | NA | NA | 1 | 1 | 1 | 1 | 1 | 1 | NA | NA | NA | NA | 1 | 0 | 2 | 0.5 |
| 15_965410 | NA | NA | NA | NA | NA | NA | 1 | 1 | 1 | 1 | 1 | 1 | NA | NA | NA | NA | 1 | 1 | 2 | 1 |
| 15_965416 | 1 | 0 | NA | NA | NA | NA | 1 | 1 | 2 | 0.5 | 1 | 1 | NA | NA | NA | NA | NA | NA | 1 | 1 |
| 15_965421 | NA | NA | NA | NA | NA | NA | 1 | 1 | 1 | 1 | 1 | 1 | NA | NA | NA | NA | NA | NA | 1 | 1 |
| 15_965422 | NA | NA | NA | NA | NA | NA | 1 | 1 | 1 | 1 | 1 | 1 | NA | NA | NA | NA | NA | NA | 1 | 1 |
| 15_965854 | 25 | 0.12 | 31 | 0.0323 | 32 | 0.0625 | 12 | 0 | 100 | 0.06 | 24 | 0.0417 | 16 | 0.0625 | 15 | 0.1333 | 18 | 0.2222 | 73 | 0.1096 |
| 15_965858 | 18 | 0.2222 | 19 | 0.3158 | 22 | 0.1364 | 10 | 0 | 69 | 0.1884 | 17 | 0.1176 | 13 | 0.2308 | 14 | 0.1429 | 18 | 0.3333 | 62 | 0.2097 |
| 15_965859 | 17 | 0.8824 | 22 | 0.9091 | 24 | 0.875 | 10 | 1 | 73 | 0.9041 | 16 | 0.875 | 15 | 1 | 13 | 0.8462 | 18 | 0.9444 | 62 | 0.9194 |
| 15_965860 | 13 | 0.9231 | 24 | 0.9583 | 20 | 0.8 | 7 | 0.7143 | 64 | 0.875 | 15 | 0.8 | 11 | 0.8182 | 15 | 0.7333 | 18 | 0.7222 | 59 | 0.7627 |
| 15_965861 | 15 | 0.6 | 22 | 0.4545 | 23 | 0.6522 | 10 | 0.4 | 70 | 0.5429 | 16 | 0.5625 | 14 | 0.5 | 15 | 0.4 | 16 | 0.4375 | 61 | 0.4754 |
| 15_965862 | 15 | 0.7333 | 21 | 0.7619 | 24 | 0.875 | 8 | 0.75 | 68 | 0.7941 | 14 | 0.8571 | 13 | 0.8462 | 11 | 0.7273 | 17 | 0.7647 | 55 | 0.8 |
| 15_965863 | 12 | 0.75 | 18 | 0.7222 | 23 | 0.7826 | 8 | 0.875 | 61 | 0.7705 | 10 | 0.6 | 10 | 0.7 | 9 | 0.6667 | 16 | 0.875 | 45 | 0.7333 |
| 15_965863 | 14 | 0.1429 | 16 | 0.0625 | 18 | 0.1111 | 9 | 0.1111 | 57 | 0.1053 | 13 | 0.3077 | 8 | 0.125 | 11 | 0 | 18 | 0.1111 | 50 | 0.14 |
| 15_965867 | 11 | 0.1818 | 17 | 0.2941 | 13 | 0.1538 | 4 | 0.25 | 45 | 0.2222 | 4 | 0.25 | 5 | 0.6 | 3 | 0 | 6 | 0 | 18 | 0.2222 |
| 15_965869 | 12 | 0 | 18 | 0.0556 | 20 | 0.1 | 8 | 0.125 | 58 | 0.069 | 9 | 0.1111 | 9 | 0 | 10 | 0.1 | 14 | 0.2143 | 42 | 0.119 |
| 15_965874 | 8 | 0 | 14 | 0.0714 | 16 | 0.0625 | 8 | 0.125 | 46 | 0.0652 | 8 | 0.125 | 9 | 0.2222 | 10 | 0.1 | 11 | 0.0909 | 38 | 0.1316 |
| 15_965874 | 14 | 0.1429 | 15 | 0.6 | 14 | 0.2857 | 7 | 0.5714 | 50 | 0.38 | 7 | 0.5714 | 11 | 0.3636 | 11 | 0.7273 | 14 | 0.3571 | 43 | 0.4884 |
| 15_965875 | 14 | 0.0714 | 19 | 0.0526 | 18 | 0.1111 | 9 | 0 | 60 | 0.0667 | 7 | 0 | 11 | 0.0909 | 12 | 0.0833 | 15 | 0.1333 | 45 | 0.0889 |
| 15_965876 | 14 | 0.1429 | 16 | 0 | 16 | 0.125 | 9 | 0 | 55 | 0.0727 | 6 | 0 | 11 | 0 | 8 | 0 | 14 | 0.2143 | 39 | 0.0769 |
| 15_965877 | 12 | 0.5833 | 13 | 0.4615 | 13 | 0.3846 | 5 | 0.8 | 43 | 0.5116 | 6 | 0.5 | 6 | 0.6667 | 7 | 0.5714 | 13 | 0.9231 | 32 | 0.7188 |
| 15_965878 | 11 | 0.6364 | 16 | 0.75 | 15 | 0.5333 | 7 | 0.5714 | 49 | 0.6327 | 4 | 0.75 | 10 | 0.7 | 8 | 0.625 | 12 | 0.75 | 34 | 0.7059 |
| 15_965973 | 3 | 0 | 12 | 0 | 10 | 0.1 | 6 | 0.1667 | 31 | 0.0645 | 2 | 0 | 7 | 0.4286 | 4 | 0 | 14 | 0.1429 | 27 | 0.1852 |
| 15_971422 | NA | NA | 2 | 0.5 | 3 | 0.3333 | 4 | 0.25 | 9 | 0.3333 | 1 | 1 | NA | NA | 1 | 0 | 3 | 0.6667 | 5 | 0.6 |
| 15_971425 | NA | NA | 1 | 1 | 3 | 0.3333 | 4 | 0.25 | 8 | 0.375 | NA | NA | NA | NA | NA | NA | 3 | 0.6667 | 3 | 0.6667 |
| 15_971426 | NA | NA | 1 | 1 | 3 | 1 | 4 | 0.75 | 8 | 0.875 | NA | NA | NA | NA | NA | NA | 3 | 1 | 3 | 1 |
| 15_971429 | NA | NA | 2 | 0.5 | 3 | 1 | 4 | 0.5 | 9 | 0.6667 | NA | NA | NA | NA | NA | NA | 1 | 1 | 1 | 1 |
| 15_971429 | NA | NA | 2 | 0 | 2 | 0 | 4 | 0.25 | 8 | 0.125 | NA | NA | NA | NA | NA | NA | 1 | 1 | 1 | 1 |
| 15_971429 | NA | NA | 1 | 0 | 2 | 0 | 3 | 0.3333 | 6 | 0.1667 | NA | NA | NA | NA | NA | NA | 3 | 0.3333 | 3 | 0.3333 |
| 15_971429 | NA | NA | 2 | 0.5 | 2 | 1 | 2 | 0 | 6 | 0.5 | NA | NA | NA | NA | NA | NA | 1 | 1 | 1 | 1 |
| 15_971430 | NA | NA | 2 | 0.5 | 3 | 0.3333 | 2 | 0.5 | 7 | 0.4286 | NA | NA | NA | NA | NA | NA | 3 | 0.3333 | 3 | 0.3333 |
| 15_971431 | NA | NA | 2 | 0.5 | 3 | 0.3333 | 3 | 1 | 8 | 0.625 | NA | NA | NA | NA | NA | NA | 2 | 1 | 2 | 1 |
| 15_971433 | NA | NA | 2 | 1 | 1 | 1 | 2 | 0 | 5 | 0.6 | NA | NA | NA | NA | NA | NA | 2 | 0.5 | 2 | 0.5 |
| 15_971434 | NA | NA | 1 | 1 | 3 | 0.3333 | 3 | 1 | 7 | 0.7143 | NA | NA | NA | NA | NA | NA | 2 | 0.5 | 2 | 0.5 |
| 15_971434 | NA | NA | 2 | 0.5 | 2 | 0.5 | 3 | 1 | 7 | 0.7143 | NA | NA | NA | NA | NA | NA | 2 | 1 | 2 | 1 |
| 15_971435 | NA | NA | 1 | 1 | 1 | 1 | 3 | 1 | 5 | 1 | NA | NA | NA | NA | NA | NA | 2 | 1 | 2 | 1 |
| 15_971436 | NA | NA | 1 | 0 | 3 | 0.3333 | 3 | 0.3333 | 7 | 0.2857 | NA | NA | NA | NA | NA | NA | 1 | 1 | 1 | 1 |
| 15_971437 | NA | NA | 2 | 1 | 2 | 1 | 1 | 1 | 5 | 1 | NA | NA | NA | NA | NA | NA | 2 | 1 | 2 | 1 |
| 15_971437 | NA | NA | 2 | 0.5 | 2 | 0 | 3 | 0 | 7 | 0.1429 | NA | NA | NA | NA | NA | NA | 1 | 1 | 1 | 1 |
| 15_971439 | NA | NA | 1 | 1 | 3 | 0.3333 | 3 | 0 | 7 | 0.2857 | NA | NA | NA | NA | NA | NA | 2 | 0.5 | 2 | 0.5 |
| 15_971439 | NA | NA | 1 | 1 | 2 | 1 | 2 | 0 | 5 | 0.6 | NA | NA | NA | NA | NA | NA | 2 | 1 | 2 | 1 |
| 15_971440 | NA | NA | 2 | 1 | 1 | 1 | 2 | 1 | 5 | 1 | NA | NA | NA | NA | NA | NA | 1 | 1 | 1 | 1 |
| 15_971441 | NA | NA | 2 | 1 | 2 | 1 | 2 | 1 | 6 | 1 | NA | NA | NA | NA | NA | NA | 1 | 1 | 1 | 1 |
| 15_971442 | NA | NA | 2 | 0.5 | 2 | 1 | 2 | 1 | 6 | 0.8333 | NA | NA | NA | NA | NA | NA | 1 | 1 | 1 | 1 |
| 15_971444 | NA | NA | 2 | 0 | 2 | 0.5 | 2 | 0 | 6 | 0.1667 | NA | NA | NA | NA | NA | NA | 1 | 1 | 1 | 1 |
| 15_975731 | 2 | 0 | 4 | 0 | 2 | 0 | 2 | 0.5 | 10 | 0.1 | 1 | 0 | 6 | 0 | 6 | 0.3333 | 8 | 0.25 | 21 | 0.1905 |
| 15_976415 | 1 | 0 | NA | NA | NA | NA | 1 | 1 | 2 | 0.5 | 1 | 1 | NA | NA | NA | NA | NA | NA | 1 | 1 |
| 15_976431 | 1 | 1 | 5 | 0.8 | 3 | 1 | 1 | 1 | 10 | 0.9 | 4 | 1 | 3 | 0.6667 | 3 | 0.6667 | 2 | 1 | 12 | 0.8333 |
| 15_976432 | 1 | 1 | 5 | 0.6 | 3 | 0.6667 | 2 | 1 | 11 | 0.7273 | 3 | 0.3333 | 3 | 0.3333 | 3 | 0.3333 | 3 | 1 | 12 | 0.5 |
| 15_976432 | 1 | 1 | 5 | 0.4 | 2 | 0.5 | 2 | 0.5 | 10 | 0.5 | 3 | 0.6667 | 4 | 0 | 2 | 0.5 | 3 | 0.3333 | 12 | 0.3333 |
| 15_976433 | NA | NA | 3 | 0.6667 | 3 | 1 | 2 | 1 | 8 | 0.875 | 4 | 1 | 3 | 1 | 2 | 1 | 3 | 0.6667 | 12 | 0.9167 |
| 15_976434 | 1 | 1 | 4 | 0.25 | 2 | 0.5 | 2 | 0.5 | 9 | 0.4444 | 4 | 0.25 | 3 | 0 | 3 | 0.6667 | 3 | 0.6667 | 13 | 0.3846 |
| 15_976885 | 11 | 0.3636 | 29 | 0.1034 | 14 | 0.2143 | 16 | 0.0625 | 70 | 0.1571 | 17 | 0.1176 | 25 | 0.16 | 20 | 0.1 | 19 | 0.2105 | 81 | 0.1481 |
| 15_976886 | 13 | 0.2308 | 28 | 0.3929 | 14 | 0.4286 | 17 | 0.1176 | 72 | 0.3056 | 15 | 0.4667 | 22 | 0.1818 | 18 | 0.1667 | 19 | 0.3158 | 74 | 0.2703 |
| 15_976891 | 9 | 0.2222 | 27 | 0.1111 | 15 | 0.0667 | 9 | 0.1111 | 60 | 0.1167 | 15 | 0.2 | 24 | 0.0417 | 15 | 0.0667 | 16 | 0 | 70 | 0.0714 |
| 15_976894 | 13 | 0 | 31 | 0.1935 | 11 | 0.1818 | 16 | 0.125 | 71 | 0.1408 | 17 | 0.1176 | 17 | 0.1765 | 19 | 0.1579 | 18 | 0 | 71 | 0.1127 |
| 15_976895 | 10 | 0.2 | 27 | 0.1852 | 11 | 0.0909 | 14 | 0 | 62 | 0.129 | 20 | 0.2 | 16 | 0.25 | 12 | 0.0833 | 17 | 0 | 65 | 0.1385 |
| 15_976905 | 2 | 0.5 | 6 | 0 | 2 | 0 | 4 | 0 | 14 | 0.0714 | 6 | 0 | 2 | 0 | 2 | 0 | 2 | 1 | 12 | 0.1667 |
| 15_976912 | 3 | 0.6667 | 2 | 0.5 | 2 | 0.5 | NA | NA | 7 | 0.5714 | 3 | 0.3333 | 3 | 0.6667 | 1 | 1 | 1 | 1 | 8 | 0.625 |
| 15_978680 | 3 | 0.6667 | 5 | 0.8 | 2 | 0.5 | 4 | 0.75 | 14 | 0.7143 | 3 | 0.6667 | 4 | 0.25 | 3 | 0.3333 | 3 | 0 | 13 | 0.3077 |
| 15_979280 | NA | NA | NA | NA | 1 | 1 | NA | NA | 1 | 1 | NA | NA | NA | NA | 1 | 1 | NA | NA | 1 | 1 |
| 15_979283 | NA | NA | NA | NA | 1 | 1 | NA | NA | 1 | 1 | NA | NA | NA | NA | 1 | 0 | 1 | 1 | 2 | 0.5 |
| 15_979367 | NA | NA | NA | NA | NA | NA | 3 | 0.6667 | 3 | 0.6667 | NA | NA | 1 | 1 | 1 | 0 | NA | NA | 2 | 0.5 |

| | | | | | | | | | | | | | | | | | | | | |
|---|---|---|---|---|---|---|---|---|---|---|---|---|---|---|---|---|---|---|---|---|
| 15_979403 | 1 | 1 | 3 | 0 | 1 | 1 | 1 | 1 | 6 | 0.5 | 1 | 0 | 2 | 0 | 1 | 1 | NA | NA | 4 | 0.25 |
| 15_979405 | 1 | 1 | 3 | 1 | 1 | 1 | 1 | 1 | 6 | 1 | 1 | 1 | 2 | 1 | 1 | 1 | 1 | 1 | 5 | 1 |
| 15_979405 | 1 | 1 | 2 | 1 | 1 | 1 | NA | NA | 4 | 1 | 1 | 1 | 2 | 1 | 1 | 1 | NA | NA | 4 | 1 |
| 15_979406 | 1 | 0 | 3 | 0.3333 | 1 | 0 | 1 | 0 | 6 | 0.1667 | 1 | 0 | 1 | 0 | 1 | 1 | 1 | 0 | 4 | 0.25 |
| 15_979643 | 1 | 1 | 3 | 0.3333 | 1 | 1 | 1 | 0 | 6 | 0.5 | NA | NA | 2 | 0 | 1 | 0 | 1 | 1 | 4 | 0.25 |
| 15_979910 | NA | NA | 1 | 1 | 1 | 1 | NA | NA | 2 | 1 | NA | NA | NA | NA | NA | NA | 1 | 1 | 1 | 1 |
| 15_980301 | 47 | 0.1277 | 47 | 0.0851 | 42 | 0.1429 | 35 | 0.1714 | 171 | 0.1287 | 25 | 0.12 | 39 | 0.1026 | 39 | 0.0769 | 36 | 0.2222 | 139 | 0.1295 |
| 15_980327 | 7 | 0 | 10 | 0.1 | 10 | 0 | 6 | 0.1667 | 33 | 0.0606 | 2 | 0 | 7 | 0.2857 | 5 | 0 | 8 | 0.25 | 22 | 0.1818 |
| 15_980331 | 8 | 0.125 | 9 | 0.1111 | 14 | 0.1429 | 7 | 0.2857 | 38 | 0.1579 | 3 | 0 | 6 | 0 | 5 | 0 | 6 | 0.1667 | 20 | 0.05 |
| 15_980335 | 6 | 0.1667 | 4 | 0 | 11 | 0 | 5 | 0.2 | 26 | 0.0769 | 3 | 0 | 6 | 0 | 3 | 0 | 6 | 0.1667 | 18 | 0.0556 |
| 15_980335 | 6 | 0.8333 | 7 | 0.8571 | 12 | 0.75 | 5 | 0.8 | 30 | 0.8 | 3 | 0.6667 | 7 | 0.7143 | 4 | 0.75 | 6 | 0.6667 | 20 | 0.7 |
| 15_980337 | 6 | 1 | 5 | 0.8 | 13 | 1 | 5 | 1 | 29 | 0.9655 | 3 | 1 | 7 | 1 | 3 | 1 | 3 | 1 | 16 | 1 |
| 15_980338 | 7 | 1 | 8 | 1 | 11 | 1 | 5 | 0.8 | 31 | 0.9677 | 3 | 1 | 6 | 1 | 4 | 1 | 4 | 1 | 17 | 1 |
| 15_980338 | 5 | 0.4 | 4 | 0.25 | 9 | 0.2222 | 4 | 0.5 | 22 | 0.3182 | 2 | 0.5 | 2 | 0.5 | 2 | 0 | 1 | 0 | 7 | 0.2857 |
| 15_980338 | NA | NA | NA | NA | NA | NA | 2 | 1 | 2 | 1 | NA | NA | NA | NA | NA | NA | 1 | 1 | 1 | 1 |
| 15_980338 | 7 | 0.1429 | 8 | 0 | 11 | 0.1818 | 7 | 0.1429 | 33 | 0.1212 | 3 | 0 | 6 | 0 | 4 | 0.25 | 6 | 0.1667 | 19 | 0.1053 |
| 15_980339 | 6 | 1 | 8 | 1 | 10 | 1 | 7 | 1 | 31 | 1 | 3 | 1 | 7 | 1 | 4 | 1 | 5 | 1 | 19 | 1 |
| 15_980339 | 5 | 0 | 7 | 0.1429 | 11 | 0.0909 | 7 | 0.2857 | 30 | 0.1333 | 2 | 0 | 7 | 0.2857 | 4 | 0 | 5 | 0.2 | 18 | 0.1667 |
| 15_980339 | 4 | 0.25 | 6 | 0 | 6 | 0 | 4 | 0 | 20 | 0.05 | 3 | 0 | 5 | 0 | 4 | 0.25 | 5 | 0 | 17 | 0.0588 |
| 15_980339 | 7 | 0.7143 | 8 | 0.875 | 11 | 1 | 6 | 1 | 32 | 0.9062 | 3 | 1 | 7 | 1 | 4 | 1 | 5 | 1 | 19 | 1 |
| 15_980339 | 6 | 0.1667 | 8 | 0 | 11 | 0 | 7 | 0.1429 | 32 | 0.0625 | 3 | 0 | 6 | 0 | 4 | 0 | 5 | 0.2 | 18 | 0.0556 |
| 15_980341 | 4 | 0.75 | 6 | 0.6667 | 8 | 0.875 | 7 | 0.5714 | 25 | 0.72 | 1 | 1 | 5 | 1 | 4 | 1 | 5 | 0.6 | 15 | 0.8667 |
| 15_980341 | 5 | 0.2 | 7 | 0.2857 | 10 | 0.3 | 7 | 0 | 29 | 0.2069 | 2 | 0 | 7 | 0.2857 | 4 | 0.25 | 4 | 0.5 | 17 | 0.2941 |
| 15_980659 | 1 | 0 | 3 | 0.6667 | 2 | 1 | 1 | 0 | 7 | 0.5714 | 1 | 1 | 3 | 0.6667 | 1 | 1 | NA | NA | 5 | 0.8 |
| 15_980662 | 3 | 0.6667 | 4 | 0.75 | 2 | 0.5 | NA | NA | 9 | 0.6667 | 1 | 0 | 2 | 1 | 1 | 1 | NA | NA | 4 | 0.75 |
| 15_980663 | 3 | 1 | 3 | 0.3333 | 2 | 1 | 1 | 1 | 9 | 0.7778 | 1 | 1 | 3 | 0.6667 | 2 | 0.5 | 2 | 0.5 | 8 | 0.625 |
| 15_980666 | 3 | 0.3333 | 4 | 0.5 | 3 | 0.3333 | 1 | 0 | 11 | 0.3636 | 1 | 0 | 3 | 0 | 2 | 0 | 2 | 0.5 | 8 | 0.125 |
| 15_980668 | 3 | 1 | 3 | 1 | 4 | 1 | 1 | 1 | 11 | 1 | NA | NA | 2 | 1 | 3 | 1 | 2 | 0.5 | 7 | 0.8571 |
| 15_980669 | 4 | 0.75 | 4 | 1 | 4 | 0.25 | NA | NA | 12 | 0.6667 | 1 | 1 | 2 | 0 | 4 | 0.75 | 2 | 1 | 9 | 0.6667 |
| 15_980669 | 4 | 1 | 4 | 1 | 4 | 1 | 1 | 1 | 13 | 1 | 1 | 1 | 3 | 1 | 4 | 0.75 | 2 | 1 | 10 | 0.9 |
| 15_980670 | 3 | 1 | 4 | 1 | 4 | 1 | 1 | 1 | 12 | 1 | 1 | 1 | 2 | 1 | 5 | 1 | 2 | 1 | 10 | 1 |
| 15_980670 | 4 | 0.5 | 4 | 0.25 | 4 | 0 | 1 | 1 | 13 | 0.3077 | 1 | 1 | 2 | 0 | 5 | 0.4 | 2 | 0 | 10 | 0.3 |
| 15_980671 | 3 | 1 | 3 | 1 | 3 | 1 | NA | NA | 9 | 1 | NA | NA | NA | NA | 1 | 0 | 2 | 1 | 3 | 0.6667 |
| 15_980675 | 2 | 0.5 | 2 | 0.5 | 3 | 0.3333 | 1 | 1 | 8 | 0.5 | NA | NA | 1 | 0 | 2 | 0.5 | 2 | 0.5 | 5 | 0.4 |
| 15_980675 | 3 | 1 | 3 | 1 | 4 | 0.75 | NA | NA | 10 | 0.9 | NA | NA | 1 | 0 | 2 | 1 | 2 | 1 | 5 | 0.8 |
| 15_980675 | 3 | 0 | 3 | 0.3333 | 4 | 0 | 1 | 0 | 11 | 0.0909 | NA | NA | 1 | 0 | 2 | 0 | 1 | 1 | 4 | 0.25 |
| 15_980675 | 4 | 1 | 3 | 1 | 4 | 0.5 | NA | NA | 11 | 0.8182 | NA | NA | NA | NA | 1 | 0 | 1 | 1 | 2 | 0.5 |
| 15_981008 | NA | NA | 1 | 1 | NA | NA | NA | NA | 1 | 1 | 1 | 1 | NA | NA | NA | NA | 1 | 1 | 2 | 1 |
| 15_981049 | 1 | 1 | 4 | 0.75 | 2 | 0.5 | NA | NA | 7 | 0.7143 | 1 | 1 | 2 | 1 | 2 | 0.5 | 1 | 1 | 6 | 0.8333 |
| 15_981051 | 1 | 1 | 3 | 0.6667 | 2 | 1 | NA | NA | 6 | 0.8333 | 1 | 1 | 1 | 1 | 2 | 1 | 1 | 1 | 5 | 1 |
| 15_981054 | NA | NA | 4 | 0.25 | 2 | 0 | NA | NA | 6 | 0.1667 | 1 | 0 | 2 | 0.5 | 2 | 0.5 | 1 | 1 | 6 | 0.5 |
| 15_981055 | NA | NA | 3 | 0.3333 | 2 | 0.5 | NA | NA | 5 | 0.4 | 1 | 1 | 2 | 0 | 2 | 0.5 | 1 | 1 | 6 | 0.5 |
| 15_981057 | NA | NA | 2 | 0 | 1 | 1 | NA | NA | 3 | 0.3333 | NA | NA | 2 | 0.5 | 2 | 1 | NA | NA | 4 | 0.75 |
| 15_981058 | NA | NA | 4 | 0.75 | 1 | 0 | NA | NA | 5 | 0.6 | 1 | 0 | 1 | 1 | NA | NA | NA | NA | 2 | 0.5 |
| 15_981059 | NA | NA | 3 | 0.3333 | 2 | 0 | NA | NA | 5 | 0.2 | 1 | 0 | 2 | 0.5 | 1 | 0 | 1 | 0 | 5 | 0.2 |
| 15_981060 | NA | NA | 4 | 0.75 | 2 | 1 | NA | NA | 6 | 0.8333 | NA | NA | 1 | 1 | 1 | 1 | 1 | 1 | 3 | 1 |
| 15_981068 | NA | NA | 1 | 1 | NA | NA | NA | NA | 1 | 1 | 1 | 1 | 1 | 0 | 1 | 0 | NA | NA | 3 | 0.3333 |
| 15_981838 | 1 | 0 | NA | NA | NA | NA | 2 | 0.5 | 3 | 0.3333 | NA | NA | 2 | 0.5 | 1 | 0 | NA | NA | 3 | 0.3333 |
| 15_981841 | 1 | 1 | NA | NA | NA | NA | 1 | 1 | 2 | 1 | NA | NA | 3 | 0.6667 | NA | NA | NA | NA | 3 | 0.6667 |
| 15_981863 | NA | NA | NA | NA | 1 | 0 | 1 | 1 | 2 | 0.5 | NA | NA | 2 | 0.5 | NA | NA | NA | NA | 2 | 0.5 |
| 15_981863 | NA | NA | NA | NA | 1 | 1 | 1 | 1 | 2 | 1 | NA | NA | 1 | 1 | NA | NA | NA | NA | 1 | 1 |
| 15_981863 | NA | NA | NA | NA | 1 | 1 | 1 | 1 | 2 | 1 | NA | NA | 2 | 1 | NA | NA | NA | NA | 2 | 1 |
| 15_981865 | NA | NA | NA | NA | 1 | 0 | 1 | 1 | 2 | 0.5 | NA | NA | 2 | 1 | NA | NA | NA | NA | 2 | 1 |
| 15_981866 | NA | NA | NA | NA | 1 | 1 | 1 | 1 | 2 | 1 | NA | NA | 2 | 1 | NA | NA | NA | NA | 2 | 1 |
| 15_981868 | NA | NA | NA | NA | 1 | 1 | 1 | 1 | 2 | 1 | NA | NA | 1 | 1 | NA | NA | NA | NA | 1 | 1 |
| 15_981869 | NA | NA | NA | NA | 1 | 0 | 1 | 1 | 2 | 0.5 | NA | NA | 2 | 0.5 | NA | NA | NA | NA | 2 | 0.5 |
| 15_981919 | NA | NA | NA | NA | 1 | 0 | 1 | 1 | 2 | 0.5 | NA | NA | 1 | 1 | NA | NA | NA | NA | 1 | 1 |
| 15_984168 | 3 | 0.6667 | 10 | 0.6 | 6 | 0.3333 | 7 | 0.5714 | 26 | 0.5385 | 7 | 0.5714 | 5 | 0.6 | 5 | 0.6 | 6 | 0.3333 | 23 | 0.5217 |
| 15_984170 | 6 | 0.6667 | 11 | 0.1818 | 7 | 0.1429 | 13 | 0.3846 | 37 | 0.3243 | 5 | 0.4 | 7 | 0.2857 | 5 | 0.2 | 8 | 0.125 | 25 | 0.24 |
| 15_984170 | 6 | 0.8333 | 11 | 0.7273 | 8 | 0.75 | 13 | 0.7692 | 38 | 0.7632 | 5 | 0.8 | 5 | 0.6 | 5 | 0.6 | 7 | 0.8571 | 22 | 0.7273 |
| 15_984170 | 4 | 0.75 | 10 | 0.5 | 8 | 0.75 | 10 | 0.4 | 32 | 0.5625 | 5 | 0.8 | 5 | 0.6 | 6 | 0.6667 | 6 | 0.6667 | 22 | 0.6818 |
| 15_984170 | 1 | 0 | NA | NA | 1 | 1 | 1 | 1 | 3 | 0.6667 | NA | NA | 2 | 0.5 | NA | NA | 1 | 0 | 3 | 0.3333 |
| 15_984170 | 4 | 0.5 | 10 | 0.2 | 8 | 0.25 | 12 | 0.3333 | 34 | 0.2941 | 6 | 0.1667 | 6 | 0.3333 | 5 | 0.2 | 6 | 0.1667 | 23 | 0.2174 |
| 15_984171 | 5 | 0.2 | 11 | 0.1818 | 8 | 0.125 | 12 | 0.1667 | 36 | 0.1667 | 6 | 0.1667 | 7 | 0.2857 | 6 | 0 | 7 | 0.2857 | 26 | 0.1923 |
| 15_984172 | 5 | 0 | 10 | 0.1 | 7 | 0.1429 | 11 | 0.4545 | 33 | 0.2121 | 5 | 0.2 | 7 | 0.1429 | 6 | 0 | 7 | 0.1429 | 25 | 0.12 |

| | | | | | | | | | | | | | | | | | | | | |
|---|---|---|---|---|---|---|---|---|---|---|---|---|---|---|---|---|---|---|---|---|
| 15_9841726 | 5 | 0.6 | 10 | 0.7 | 7 | 0.7143 | 12 | 0.6667 | 34 | 0.6765 | 4 | 0.75 | 6 | 0.8333 | 4 | 0.75 | 7 | 0.4286 | 21 | 0.6667 |
| 15_9842688 | 5 | 0.2 | 8 | 0 | 4 | 0 | 9 | 0.2222 | 26 | 0.1154 | 4 | 0 | 5 | 0.2 | 6 | 0 | 5 | 0 | 20 | 0.05 |
| 15_9843995 | 1 | 0 | 2 | 0.5 | 1 | 0 | 1 | 0 | 5 | 0.2 | 1 | 0 | 1 | 1 | 3 | 0 | 1 | 0 | 6 | 0.1667 |
| 15_9843998 | NA | NA | 2 | 0 | 1 | 1 | 1 | 0 | 4 | 0.25 | 1 | 0 | 1 | 1 | 3 | 0 | 1 | 0 | 6 | 0.1667 |
| 15_9844066 | NA | NA | NA | NA | 1 | 1 | NA | NA | 1 | 1 | 1 | 1 | 1 | 1 | NA | NA | NA | NA | 2 | 1 |
| 15_9844075 | 1 | 1 | NA | NA | 1 | 1 | NA | NA | 2 | 1 | 1 | 0 | 1 | 1 | NA | NA | NA | NA | 2 | 0.5 |
| 15_9844094 | 1 | 1 | NA | NA | 1 | 1 | NA | NA | 2 | 1 | 1 | 0 | 1 | 1 | NA | NA | NA | NA | 2 | 0.5 |
| 15_9844104 | 1 | 1 | NA | NA | NA | NA | NA | NA | 1 | 1 | 1 | 0 | 1 | 1 | NA | NA | NA | NA | 2 | 0.5 |
| 15_9844106 | 1 | 1 | NA | NA | 1 | 1 | NA | NA | 2 | 1 | 1 | 1 | 1 | 1 | NA | NA | NA | NA | 2 | 1 |
| 15_9844129 | 1 | 0 | NA | NA | 2 | 1 | NA | NA | 3 | 0.6667 | 1 | 1 | 1 | 1 | 1 | 1 | NA | NA | 3 | 1 |
| 15_9844135 | NA | NA | NA | NA | 1 | 1 | NA | NA | 1 | 1 | 1 | 1 | 1 | 1 | 1 | 1 | NA | NA | 3 | 1 |
| 15_9844137 | 1 | 1 | NA | NA | 1 | 0 | NA | NA | 2 | 0.5 | 1 | 1 | 1 | 1 | 1 | 0 | NA | NA | 3 | 0.6667 |
| 15_9844153 | 1 | 1 | NA | NA | 2 | 0.5 | NA | NA | 3 | 0.6667 | 1 | 1 | 1 | 1 | NA | NA | NA | NA | 2 | 1 |
| 15_9844173 | 1 | 1 | NA | NA | 2 | 1 | NA | NA | 3 | 1 | 1 | 1 | 1 | 0 | NA | NA | NA | NA | 2 | 0.5 |
| 15_9844188 | 1 | 1 | NA | NA | 1 | 1 | NA | NA | 2 | 1 | 1 | 1 | 1 | 1 | NA | NA | NA | NA | 2 | 1 |
| 15_9844221 | 1 | 1 | NA | NA | 1 | 1 | NA | NA | 2 | 1 | NA | NA | 1 | 1 | NA | NA | NA | NA | 1 | 1 |
| 15_9848868 | NA | NA | 3 | 0.3333 | 1 | 0 | 1 | 0 | 5 | 0.2 | 1 | 0 | NA | NA | 1 | 0 | 3 | 0.3333 | 5 | 0.2 |
| 15_9848883 | NA | NA | 3 | 0 | 1 | 0 | 1 | 1 | 5 | 0.2 | 1 | 0 | 1 | 1 | NA | NA | 3 | 0.3333 | 5 | 0.4 |
| 15_9848909 | NA | NA | 3 | 0.3333 | 1 | 0 | 1 | 0 | 5 | 0.2 | NA | NA | NA | NA | 1 | 0 | 2 | 0.5 | 3 | 0.3333 |
| 15_9848948 | NA | NA | NA | NA | NA | NA | 1 | 1 | 1 | 1 | NA | NA | NA | NA | 1 | 0 | 1 | 1 | 2 | 0.5 |
| 15_9848969 | NA | NA | NA | NA | NA | NA | 1 | 1 | 1 | 1 | 1 | 0 | NA | NA | 1 | 1 | 1 | 1 | 3 | 0.6667 |
| 15_9848970 | NA | NA | NA | NA | NA | NA | 1 | 1 | 1 | 1 | 1 | 0 | NA | NA | 1 | 0 | 1 | 1 | 3 | 0.3333 |
| 15_9848983 | NA | NA | NA | NA | NA | NA | 1 | 1 | 1 | 1 | 1 | 1 | NA | NA | 1 | 1 | 1 | 1 | 3 | 1 |
| 15_9849002 | NA | NA | NA | NA | NA | NA | 1 | 1 | 1 | 1 | 1 | 1 | NA | NA | 1 | 0 | NA | NA | 2 | 0.5 |
| 15_9849008 | NA | NA | NA | NA | NA | NA | 1 | 1 | 1 | 1 | NA | NA | NA | NA | 1 | 1 | NA | NA | 1 | 1 |
| 15_9854128 | 28 | 0.1429 | 16 | 0.1875 | 18 | 0.2778 | 17 | 0.1765 | 79 | 0.1899 | 15 | 0 | 15 | 0.2667 | 19 | 0.0526 | 27 | 0.2963 | 76 | 0.1711 |
| 15_9854138 | 27 | 0.3704 | 17 | 0.2941 | 19 | 0.1579 | 19 | 0.3158 | 82 | 0.2927 | 18 | 0.1667 | 19 | 0.3684 | 19 | 0.2632 | 29 | 0.4138 | 85 | 0.3176 |
| 15_9854141 | 25 | 0.28 | 16 | 0.5 | 16 | 0.25 | 17 | 0.4706 | 74 | 0.3649 | 19 | 0.5263 | 18 | 0.2778 | 20 | 0.4 | 25 | 0.32 | 82 | 0.378 |
| 15_9854144 | 30 | 0.6333 | 20 | 0.55 | 20 | 0.55 | 15 | 0.3333 | 85 | 0.5412 | 19 | 0.5263 | 18 | 0.5556 | 21 | 0.4286 | 29 | 0.4138 | 87 | 0.4713 |
| 15_9854187 | 32 | 0.25 | 27 | 0.1852 | 24 | 0.125 | 20 | 0.15 | 103 | 0.1845 | 22 | 0.0909 | 26 | 0.2308 | 22 | 0.2727 | 32 | 0.2188 | 102 | 0.2059 |
| 15_9854211 | 16 | 0.75 | 7 | 0.7143 | 8 | 0.625 | 7 | 0.5714 | 38 | 0.6842 | 8 | 1 | 7 | 0.8571 | 6 | 0.5 | 11 | 0.6364 | 32 | 0.75 |
| 15_9854322 | 7 | 0.2857 | 4 | 0.25 | 3 | 0 | 5 | 0 | 19 | 0.1579 | 7 | 0.1429 | 1 | 0 | 4 | 0 | 2 | 0 | 14 | 0.0714 |
| 15_9854328 | 11 | 0.9091 | 8 | 0.75 | 4 | 0.75 | 7 | 0.8571 | 30 | 0.8333 | 7 | 1 | 7 | 1 | 8 | 0.75 | 4 | 0.75 | 26 | 0.8846 |
| 15_9854329 | 12 | 0.1667 | 8 | 0.125 | 5 | 0.2 | 7 | 0 | 32 | 0.125 | 6 | 0.1667 | 5 | 0 | 6 | 0.1667 | 5 | 0.2 | 22 | 0.1364 |
| 15_9854335 | 13 | 0.3846 | 8 | 0.625 | 4 | 0.5 | 6 | 0.3333 | 31 | 0.4516 | 5 | 0.6 | 6 | 0.3333 | 8 | 0.25 | 5 | 0 | 24 | 0.2917 |
| 15_9854337 | 12 | 0.6667 | 7 | 0.4286 | 6 | 0.3333 | 5 | 0.6 | 30 | 0.5333 | 6 | 0.5 | 6 | 0.3333 | 5 | 0.4 | 4 | 0.5 | 21 | 0.4286 |
| 15_9854358 | 12 | 0.4167 | 4 | 0.75 | 6 | 0.8333 | 4 | 0.75 | 26 | 0.6154 | 4 | 0.25 | 6 | 0.6667 | 6 | 0.6667 | 3 | 1 | 19 | 0.6316 |
| 15_9854380 | 8 | 0.125 | 5 | 0 | 5 | 0.6 | 4 | 0.25 | 22 | 0.2273 | 1 | 0 | 6 | 0.5 | 7 | 0.1429 | 3 | 0 | 17 | 0.2353 |
| 15_9856776 | 1 | 0 | NA | NA | 2 | 0.5 | NA | NA | 3 | 0.3333 | NA | NA | 2 | 0.5 | 2 | 0.5 | 2 | 0.5 | 6 | 0.5 |
| 15_9856783 | 1 | 0 | NA | NA | 2 | 1 | NA | NA | 3 | 0.6667 | NA | NA | 2 | 1 | 2 | 1 | 2 | 0.5 | 6 | 0.8333 |
| 15_9856807 | 2 | 1 | NA | NA | 2 | 1 | NA | NA | 4 | 1 | NA | NA | 2 | 1 | 1 | 1 | 1 | 1 | 4 | 1 |
| 15_9856818 | 1 | 0 | NA | NA | 2 | 0.5 | NA | NA | 3 | 0.3333 | NA | NA | 2 | 1 | 1 | 1 | 1 | 0 | 4 | 0.75 |
| 15_9856832 | 2 | 0.5 | NA | NA | NA | NA | NA | NA | 2 | 0.5 | NA | NA | 2 | 1 | 1 | 1 | 1 | 1 | 4 | 1 |
| 15_9856835 | 2 | 1 | NA | NA | 2 | 1 | NA | NA | 4 | 1 | NA | NA | 1 | 1 | 1 | 1 | 1 | 1 | 3 | 1 |
| 15_9862108 | NA | NA | 1 | 1 | NA | NA | 1 | 1 | 2 | 1 | 1 | 1 | 1 | 1 | 1 | 0 | NA | NA | 3 | 0.6667 |
| 15_9863444 | 4 | 0 | 14 | 0.0714 | 8 | 0.125 | 1 | 0 | 27 | 0.0741 | 3 | 0 | 6 | 0 | 4 | 0 | 5 | 0.2 | 18 | 0.0556 |
| 15_9863455 | 2 | 1 | 11 | 0.8182 | 7 | 0.4286 | 1 | 0 | 21 | 0.6667 | 3 | 0.6667 | 6 | 0.6667 | 4 | 0.5 | 4 | 0.75 | 17 | 0.6471 |
| 15_9863522 | 3 | 0.3333 | 11 | 0.0909 | 4 | 0 | 1 | 0 | 19 | 0.1053 | 2 | 0.5 | 4 | 0 | 4 | 0 | 3 | 0 | 13 | 0.0769 |
| 15_9863528 | 2 | 0 | 10 | 0.1 | 3 | 0 | 1 | 0 | 16 | 0.0625 | 3 | 0.3333 | 5 | 0 | 4 | 0 | 3 | 0 | 15 | 0.0667 |
| 15_9863536 | 2 | 0 | 8 | 0.25 | 3 | 0 | 1 | 0 | 14 | 0.1429 | 3 | 0 | 5 | 0.4 | 3 | 0 | 3 | 0 | 14 | 0.1429 |
| 15_9863777 | 46 | 0.0652 | 52 | 0.0577 | 36 | 0.0833 | 60 | 0.1 | 194 | 0.0773 | 34 | 0.0882 | 56 | 0.0893 | 80 | 0.0625 | 36 | 0.0278 | 206 | 0.068 |
| 15_9863824 | 39 | 0.5641 | 46 | 0.4783 | 34 | 0.5294 | 44 | 0.5455 | 163 | 0.5276 | 31 | 0.4839 | 51 | 0.549 | 67 | 0.5672 | 46 | 0.5 | 195 | 0.5333 |
| 15_9870193 | 98 | 0.1122 | 87 | 0.1839 | 74 | 0.1081 | 84 | 0.1786 | 343 | 0.1458 | 96 | 0.1354 | 96 | 0.1458 | 94 | 0.1702 | 71 | 0.1127 | 357 | 0.1429 |
| 15_9870194 | 105 | 0.6952 | 89 | 0.6854 | 85 | 0.6235 | 99 | 0.7172 | 378 | 0.6825 | 111 | 0.7117 | 111 | 0.7387 | 100 | 0.67 | 79 | 0.6962 | 401 | 0.7057 |
| 15_9870199 | 87 | 0.1149 | 74 | 0.2162 | 68 | 0.1912 | 79 | 0.2405 | 308 | 0.1883 | 86 | 0.2791 | 91 | 0.2198 | 83 | 0.2048 | 67 | 0.0896 | 327 | 0.2049 |
| 15_9870225 | 90 | 0.0889 | 85 | 0.0706 | 73 | 0.0411 | 87 | 0.0575 | 335 | 0.0657 | 94 | 0.0851 | 99 | 0.0404 | 92 | 0.0978 | 67 | 0.0149 | 352 | 0.0625 |
| 15_9870225 | 106 | 0.4245 | 93 | 0.4731 | 80 | 0.5125 | 95 | 0.4211 | 374 | 0.4545 | 107 | 0.4953 | 107 | 0.5047 | 98 | 0.3776 | 75 | 0.4 | 387 | 0.4496 |
| 15_9870236 | 102 | 0.6569 | 81 | 0.5926 | 68 | 0.6618 | 92 | 0.6304 | 343 | 0.6356 | 100 | 0.69 | 96 | 0.6562 | 88 | 0.6136 | 68 | 0.6176 | 352 | 0.6477 |
| 15_9870249 | 97 | 0.433 | 81 | 0.5309 | 76 | 0.5526 | 93 | 0.5591 | 347 | 0.5159 | 97 | 0.4742 | 95 | 0.4737 | 87 | 0.5172 | 75 | 0.4267 | 354 | 0.4746 |
| 15_9870252 | 103 | 0.0388 | 86 | 0.0814 | 77 | 0.0649 | 95 | 0.0632 | 361 | 0.0609 | 104 | 0.1058 | 97 | 0.1134 | 94 | 0.0532 | 76 | 0.0395 | 371 | 0.0809 |
| 15_9870264 | 97 | 0.3711 | 83 | 0.4337 | 72 | 0.4583 | 81 | 0.4321 | 333 | 0.4204 | 97 | 0.4639 | 91 | 0.3736 | 86 | 0.3605 | 66 | 0.3485 | 340 | 0.3912 |
| 15_9870267 | 80 | 0.0625 | 63 | 0.0317 | 60 | 0.05 | 71 | 0.0704 | 274 | 0.0547 | 81 | 0.0617 | 77 | 0.039 | 72 | 0.0556 | 60 | 0.0833 | 290 | 0.0586 |
| 15_9870268 | 100 | 0.24 | 78 | 0.1282 | 71 | 0.0845 | 89 | 0.2022 | 338 | 0.1716 | 98 | 0.1633 | 86 | 0.1628 | 85 | 0.1176 | 69 | 0.1739 | 338 | 0.1538 |
| 15_9870279 | 90 | 0.7778 | 76 | 0.6842 | 64 | 0.7812 | 82 | 0.7805 | 312 | 0.7564 | 85 | 0.6824 | 85 | 0.7882 | 80 | 0.7625 | 65 | 0.6923 | 315 | 0.7333 |
| 15_9870282 | 80 | 0.4375 | 66 | 0.5152 | 63 | 0.4603 | 66 | 0.4545 | 275 | 0.4655 | 66 | 0.3485 | 74 | 0.5405 | 72 | 0.3472 | 57 | 0.4561 | 269 | 0.4238 |

| | | | | | | | | | | | | | | | | | | | | |
|---|---|---|---|---|---|---|---|---|---|---|---|---|---|---|---|---|---|---|---|---|
| 15_9870292 | 83 | 0.2651 | 67 | 0.3134 | 61 | 0.3443 | 78 | 0.3077 | 289 | 0.3045 | 73 | 0.2466 | 78 | 0.3333 | 71 | 0.2817 | 59 | 0.322 | 281 | 0.2954 |
| 15_9870319 | 73 | 0.3288 | 63 | 0.3175 | 49 | 0.2245 | 66 | 0.303 | 251 | 0.2988 | 62 | 0.3387 | 67 | 0.2239 | 65 | 0.2308 | 49 | 0.3265 | 243 | 0.2757 |
| 15_9870331 | 72 | 0.3611 | 61 | 0.3115 | 53 | 0.3962 | 66 | 0.3485 | 252 | 0.3532 | 73 | 0.3973 | 66 | 0.2576 | 63 | 0.2857 | 52 | 0.4038 | 254 | 0.3346 |
| 15_9870341 | 61 | 0.918 | 54 | 0.8704 | 38 | 0.8947 | 52 | 0.9231 | 205 | 0.9024 | 61 | 0.9344 | 53 | 0.8868 | 55 | 0.9273 | 42 | 0.9286 | 211 | 0.9194 |
| 15_9870348 | 70 | 0.7 | 60 | 0.7667 | 42 | 0.7143 | 60 | 0.7167 | 232 | 0.7241 | 58 | 0.5517 | 57 | 0.8246 | 56 | 0.7679 | 45 | 0.8 | 216 | 0.7315 |
| 15_9870368 | 46 | 0.6522 | 42 | 0.6429 | 23 | 0.6087 | 46 | 0.6522 | 157 | 0.6433 | 43 | 0.6279 | 43 | 0.6977 | 46 | 0.587 | 31 | 0.6774 | 163 | 0.6442 |
| 15_9870441 | 34 | 0.2941 | 26 | 0.3077 | 20 | 0.2 | 28 | 0.2143 | 108 | 0.2593 | 38 | 0.3421 | 27 | 0.3704 | 34 | 0.2353 | 18 | 0.2778 | 117 | 0.3077 |
| 15_9870450 | 41 | 0.439 | 31 | 0.3226 | 23 | 0.3913 | 30 | 0.2 | 125 | 0.344 | 37 | 0.3784 | 33 | 0.303 | 33 | 0.2424 | 28 | 0.5 | 131 | 0.3511 |
| 15_9870456 | 31 | 0.4839 | 25 | 0.52 | 21 | 0.381 | 27 | 0.4444 | 104 | 0.4615 | 30 | 0.5 | 28 | 0.3214 | 27 | 0.2963 | 21 | 0.4286 | 106 | 0.3868 |
| 15_9870457 | 45 | 0.3333 | 30 | 0.3333 | 26 | 0.5 | 34 | 0.4412 | 135 | 0.3926 | 42 | 0.4048 | 36 | 0.3056 | 31 | 0.2903 | 29 | 0.4483 | 138 | 0.3623 |
| 15_9870462 | 37 | 0.0541 | 26 | 0.1154 | 22 | 0.0909 | 31 | 0.0968 | 116 | 0.0862 | 37 | 0.2162 | 33 | 0.0303 | 27 | 0.1852 | 27 | 0.2222 | 124 | 0.1613 |
| 15_9870474 | 37 | 0.0811 | 21 | 0.1429 | 20 | 0.05 | 30 | 0 | 108 | 0.0648 | 32 | 0.0312 | 23 | 0.0435 | 25 | 0.12 | 23 | 0.0435 | 103 | 0.0583 |
| 15_9870552 | 31 | 0.1613 | 17 | 0.2353 | 16 | 0.0625 | 24 | 0.0417 | 88 | 0.125 | 34 | 0.1176 | 19 | 0.1053 | 17 | 0.0588 | 18 | 0.2222 | 88 | 0.125 |
| 15_9871109 | 9 | 0.4444 | 11 | 0.4545 | 6 | 0.3333 | 11 | 0.2727 | 37 | 0.3784 | 7 | 0.1429 | 6 | 0.1667 | 6 | 0.5 | 7 | 0.4286 | 26 | 0.3077 |
| 15_9871116 | 10 | 0.1 | 10 | 0 | 7 | 0 | 13 | 0.1538 | 40 | 0.075 | 7 | 0.1429 | 7 | 0.2857 | 6 | 0 | 7 | 0.4286 | 27 | 0.2222 |
| 15_9871215 | NA | NA | 1 | 1 | 1 | 1 | NA | NA | 2 | 1 | NA | NA | NA | NA | 2 | 0.5 | 2 | 1 | 4 | 0.75 |
| 15_9871219 | NA | NA | 1 | 1 | 1 | 1 | NA | NA | 2 | 1 | NA | NA | NA | NA | 2 | 1 | 2 | 1 | 4 | 1 |
| 15_9871240 | NA | NA | 1 | 1 | NA | NA | NA | NA | 1 | 1 | NA | NA | NA | NA | 2 | 1 | 2 | 0.5 | 4 | 0.75 |
| 15_9871244 | NA | NA | 1 | 1 | 1 | 1 | NA | NA | 2 | 1 | NA | NA | NA | NA | 2 | 1 | 2 | 0.5 | 4 | 0.75 |
| 15_9871263 | NA | NA | NA | NA | 1 | 1 | NA | NA | 1 | 1 | NA | NA | NA | NA | 2 | 0.5 | NA | NA | 2 | 0.5 |
| 15_9871273 | NA | NA | 1 | 1 | 1 | 1 | NA | NA | 2 | 1 | NA | NA | 1 | 0 | 2 | 0.5 | 1 | 1 | 4 | 0.5 |
| 15_9871333 | 1 | 1 | 1 | 1 | 1 | 1 | NA | NA | 3 | 1 | 1 | 1 | 2 | 0.5 | 2 | 0 | 2 | 1 | 7 | 0.5714 |
| 15_9871336 | 1 | 0 | 1 | 0 | 1 | 1 | NA | NA | 3 | 0.3333 | 1 | 0 | 2 | 0.5 | 2 | 0.5 | 2 | 0.5 | 7 | 0.4286 |
| 15_9871352 | 1 | 1 | 1 | 1 | 1 | 0 | NA | NA | 3 | 0.6667 | 1 | 0 | 2 | 0 | 1 | 0 | 2 | 1 | 6 | 0.3333 |
| 15_9871354 | 1 | 1 | NA | NA | NA | NA | NA | NA | 1 | 1 | 1 | 1 | 1 | 0 | 1 | 1 | 2 | 1 | 5 | 0.8 |
| 15_9871358 | 1 | 1 | NA | NA | 1 | 0 | NA | NA | 2 | 0.5 | 1 | 1 | 2 | 0 | 2 | 0.5 | 1 | 1 | 6 | 0.5 |
| 15_9871372 | 1 | 0 | 1 | 0 | NA | NA | 1 | 1 | 3 | 0.3333 | 1 | 0 | 1 | 0 | 2 | 1 | 2 | 0.5 | 6 | 0.5 |
| 15_9871379 | NA | NA | NA | NA | 1 | 1 | 1 | 1 | 2 | 1 | 1 | 1 | 2 | 1 | 2 | 1 | 2 | 1 | 7 | 1 |
| 15_9871382 | 1 | 1 | 1 | 0 | 1 | 0 | 1 | 1 | 4 | 0.5 | 1 | 0 | 1 | 1 | 2 | 0 | 2 | 1 | 6 | 0.5 |
| 15_9872965 | 8 | 0.125 | 10 | 0.1 | 12 | 0.0833 | 6 | 0 | 36 | 0.0833 | 6 | 0.1667 | 9 | 0.2222 | 4 | 0 | 7 | 0.1429 | 26 | 0.1538 |
| 15_9872980 | 5 | 0 | 12 | 0.3333 | 11 | 0.0909 | 6 | 0.1667 | 34 | 0.1765 | 7 | 0.1429 | 6 | 0.1667 | 3 | 0.3333 | 6 | 0.6667 | 22 | 0.3182 |
| 15_9873083 | 4 | 0.25 | 7 | 0 | 4 | 0 | 3 | 0 | 18 | 0.0556 | 1 | 0 | 4 | 0.5 | 1 | 1 | 2 | 0.5 | 8 | 0.5 |
| 15_9873159 | 5 | 0.2 | 7 | 0.2857 | 4 | 0 | 2 | 0 | 18 | 0.1667 | 3 | 0 | 2 | 0.5 | 3 | 0 | 2 | 1 | 10 | 0.3 |
| 15_9873165 | 6 | 0.1667 | 6 | 0 | 3 | 0 | 1 | 0 | 16 | 0.0625 | 3 | 0 | 3 | 0 | 3 | 0 | 2 | 0.5 | 11 | 0.0909 |
| 15_9873198 | 6 | 0.3333 | 6 | 0.5 | 4 | 0.25 | 2 | 0 | 18 | 0.3333 | 3 | 0.6667 | 3 | 0.6667 | 1 | 0 | 2 | 1 | 9 | 0.6667 |
| 15_9873208 | 4 | 0 | 5 | 0.2 | 4 | 0.25 | 2 | 0 | 15 | 0.1333 | 1 | 0 | 3 | 0.3333 | 2 | 0 | 2 | 0 | 8 | 0.125 |
| 15_9873219 | 6 | 0.1667 | 6 | 0 | 4 | 0.5 | 2 | 0 | 18 | 0.1667 | 3 | 0.3333 | 2 | 0 | 2 | 0 | 2 | 0.5 | 9 | 0.2222 |
| 15_9873222 | 6 | 0 | 5 | 0.2 | 3 | 0 | 2 | 0 | 16 | 0.0625 | 3 | 0.3333 | 3 | 0.6667 | 1 | 0 | 2 | 0.5 | 9 | 0.4444 |
| 15_9873228 | 5 | 0.4 | 5 | 0.2 | 3 | 0.6667 | 2 | 0 | 15 | 0.3333 | 3 | 0 | 2 | 0.5 | 2 | 0.5 | 2 | 0.5 | 9 | 0.3333 |
| 15_9873238 | 3 | 0.3333 | 4 | 0 | 2 | 0 | 1 | 0 | 10 | 0.1 | 3 | 0 | 1 | 1 | 2 | 0 | 1 | 1 | 7 | 0.2857 |
| 15_9873240 | 6 | 0.1667 | 4 | 0.25 | 2 | 0.5 | 2 | 0 | 14 | 0.2143 | 3 | 0.3333 | 1 | 1 | 1 | 0 | 1 | 1 | 6 | 0.5 |
| 15_9873734 | 3 | 1 | 2 | 0.5 | 2 | 0.5 | 1 | 1 | 8 | 0.75 | 2 | 1 | 2 | 0.5 | 1 | 0 | 1 | 1 | 6 | 0.6667 |
| 15_9873737 | 3 | 0.3333 | 3 | 0.6667 | 1 | 0 | 1 | 1 | 8 | 0.5 | 1 | 1 | 2 | 0 | 1 | 1 | 1 | 1 | 5 | 0.6 |
| 15_9873755 | 4 | 0.25 | 3 | 0 | 1 | 0 | 1 | 1 | 9 | 0.2222 | 2 | 0.5 | 2 | 0.5 | 1 | 1 | 1 | 1 | 6 | 0.6667 |
| 15_9873765 | 3 | 0.6667 | 3 | 0.6667 | 1 | 1 | 1 | 1 | 8 | 0.75 | 2 | 1 | 2 | 1 | 1 | 1 | 1 | 1 | 6 | 1 |
| 15_9873777 | 3 | 0.6667 | 3 | 0.6667 | 2 | 1 | 1 | 0 | 9 | 0.6667 | 2 | 0 | 2 | 1 | 1 | 1 | 1 | 0 | 6 | 0.5 |
| 15_9873813 | 2 | 1 | 2 | 1 | NA | NA | 1 | 1 | 5 | 1 | NA | NA | 2 | 1 | 1 | 1 | NA | NA | 3 | 1 |
| 15_9874179 | 2 | 0.5 | 2 | 0.5 | NA | NA | NA | NA | 4 | 0.5 | 1 | 1 | 2 | 1 | 1 | 1 | NA | NA | 4 | 1 |
| 15_9874188 | 4 | 0.5 | 2 | 1 | NA | NA | NA | NA | 6 | 0.6667 | 1 | 1 | 1 | 1 | 1 | 1 | NA | NA | 3 | 1 |
| 15_9874197 | 3 | 1 | 2 | 1 | NA | NA | NA | NA | 5 | 1 | 1 | 1 | 2 | 1 | 1 | 1 | NA | NA | 4 | 1 |
| 15_9874203 | 3 | 0.3333 | 2 | 0.5 | NA | NA | NA | NA | 5 | 0.4 | 1 | 0 | 2 | 0.5 | 1 | 0 | NA | NA | 4 | 0.25 |
| 15_9874253 | 2 | 0 | 1 | 1 | NA | NA | NA | NA | 3 | 0.3333 | 1 | 0 | 2 | 0.5 | 1 | 0 | NA | NA | 4 | 0.25 |
| 15_9874255 | 2 | 0.5 | 1 | 1 | NA | NA | NA | NA | 3 | 0.6667 | 1 | 1 | 2 | 1 | 1 | 0 | NA | NA | 4 | 0.75 |
| 15_9874267 | 1 | 1 | 1 | 1 | NA | NA | NA | NA | 2 | 1 | 1 | 1 | 2 | 1 | 1 | 0 | NA | NA | 4 | 0.75 |
| 15_9874277 | 2 | 0 | 1 | 1 | NA | NA | NA | NA | 3 | 0.3333 | 1 | 1 | 1 | 1 | 1 | 1 | NA | NA | 3 | 1 |
| 15_9874283 | 2 | 1 | NA | NA | NA | NA | NA | NA | 2 | 1 | 1 | 1 | 1 | 1 | 1 | 1 | NA | NA | 3 | 1 |
| 15_9874287 | 1 | 1 | 1 | 1 | NA | NA | NA | NA | 2 | 1 | 1 | 1 | 1 | 1 | 1 | 1 | NA | NA | 3 | 1 |
| 15_9874360 | 1 | 1 | 1 | 0 | NA | NA | NA | NA | 2 | 0.5 | NA | NA | 1 | 1 | 1 | 0 | NA | NA | 2 | 0.5 |
| 15_9874787 | 1 | 1 | 1 | 1 | NA | NA | NA | NA | 2 | 1 | NA | NA | 1 | 1 | 1 | 0 | NA | NA | 2 | 0.5 |
| 15_9876998 | NA | NA | 2 | 1 | 1 | 1 | NA | NA | 3 | 1 | NA | NA | NA | NA | 2 | 1 | NA | NA | 2 | 1 |
| 15_9877012 | NA | NA | 1 | 1 | 1 | 1 | NA | NA | 2 | 1 | NA | NA | NA | NA | 2 | 1 | 1 | 1 | 3 | 1 |
| 15_9877032 | NA | NA | 1 | 1 | 1 | 1 | NA | NA | 2 | 1 | NA | NA | NA | NA | 2 | 1 | 1 | 1 | 3 | 1 |
| 15_9877033 | NA | NA | 1 | 1 | 1 | 1 | NA | NA | 2 | 1 | NA | NA | NA | NA | 1 | 1 | NA | NA | 1 | 1 |
| 15_9877035 | NA | NA | 2 | 1 | NA | NA | NA | NA | 2 | 1 | NA | NA | NA | NA | 2 | 1 | 1 | 1 | 3 | 1 |
| 15_9877042 | NA | NA | 2 | 1 | 1 | 1 | NA | NA | 3 | 1 | NA | NA | NA | NA | 2 | 1 | NA | NA | 2 | 1 |

| | | | | | | | | | | | | | | | | | | | | |
|---|---|---|---|---|---|---|---|---|---|---|---|---|---|---|---|---|---|---|---|---|
| 15_988018 | 2 | 0.5 | NA | NA | NA | NA | NA | NA | 2 | 0.5 | 1 | 1 | NA | NA | NA | NA | NA | NA | 1 | 1 |
| 15_988020 | 3 | 0.3333 | NA | NA | NA | NA | NA | NA | 3 | 0.3333 | 1 | 1 | NA | NA | NA | NA | NA | NA | 1 | 1 |
| 15_988021 | 3 | 0.3333 | NA | NA | NA | NA | NA | NA | 3 | 0.3333 | 1 | 1 | NA | NA | NA | NA | NA | NA | 1 | 1 |
| 15_988297 | 121 | 0.0909 | 166 | 0.0964 | 137 | 0.0803 | 105 | 0.0952 | 529 | 0.0907 | 118 | 0.0847 | 100 | 0.08 | 107 | 0.1589 | 84 | 0.131 | 409 | 0.1125 |
| 15_988302 | 104 | 0.0385 | 123 | 0.0894 | 108 | 0.0463 | 88 | 0.0909 | 423 | 0.0662 | 106 | 0.0566 | 89 | 0.0787 | 76 | 0.0395 | 69 | 0.1014 | 340 | 0.0676 |
| 15_988482 | 163 | 0.3252 | 219 | 0.3288 | 155 | 0.4129 | 137 | 0.4161 | 674 | 0.365 | 175 | 0.3771 | 128 | 0.3438 | 138 | 0.3696 | 146 | 0.2877 | 587 | 0.3458 |
| 15_988486 | 164 | 0.0854 | 229 | 0.0568 | 156 | 0.1154 | 150 | 0.0667 | 699 | 0.0787 | 180 | 0.0444 | 141 | 0.1064 | 133 | 0.1278 | 150 | 0.1 | 604 | 0.0911 |
| 15_990476 | 2 | 0.5 | NA | NA | NA | NA | NA | NA | 2 | 0.5 | 1 | 1 | 1 | 1 | 2 | 0.5 | NA | NA | 4 | 0.75 |
| 15_991101 | 2 | 0 | 4 | 0.25 | NA | NA | 3 | 0 | 9 | 0.1111 | 2 | 0 | 6 | 0.3333 | 2 | 0.5 | 1 | 0 | 11 | 0.2727 |
| 15_991101 | 2 | 1 | 3 | 0.6667 | NA | NA | 3 | 0 | 8 | 0.5 | 2 | 1 | 6 | 0.5 | 3 | 0.6667 | 1 | 0 | 12 | 0.5833 |
| 15_991102 | 2 | 0 | 2 | 0.5 | NA | NA | 2 | 1 | 6 | 0.5 | 1 | 0 | 6 | 0.3333 | 3 | 0 | 1 | 1 | 11 | 0.2727 |
| 15_991102 | 3 | 1 | 4 | 1 | NA | NA | 3 | 1 | 10 | 1 | 2 | 1 | 6 | 1 | 3 | 1 | 1 | 1 | 12 | 1 |
| 15_991104 | 2 | 1 | 5 | 0.8 | NA | NA | 2 | 0.5 | 9 | 0.7778 | 1 | 1 | 5 | 0.8 | 2 | 0.5 | NA | NA | 8 | 0.75 |
| 15_991229 | NA | NA | 1 | 0 | 2 | 0 | 7 | 0.1429 | 10 | 0.1 | 2 | 0 | 5 | 0.2 | 6 | 0 | NA | NA | 13 | 0.0769 |
| 15_991375 | 9 | 0.1111 | 11 | 0.0909 | 12 | 0.0833 | 12 | 0.1667 | 44 | 0.1136 | 7 | 0.2857 | 12 | 0.25 | 10 | 0.3 | 8 | 0 | 37 | 0.2162 |
| 15_991376 | 6 | 0.1667 | 4 | 0.25 | 8 | 0 | 13 | 0.2308 | 31 | 0.1613 | 6 | 0 | 6 | 0.3333 | 6 | 0.3333 | 8 | 0.375 | 26 | 0.2692 |
| 15_991399 | 10 | 0.8 | 10 | 0.6 | 11 | 0.5455 | 10 | 0.6 | 41 | 0.6341 | 5 | 0.8 | 10 | 0.4 | 8 | 0.625 | 8 | 0.375 | 31 | 0.5161 |
| 15_991400 | 10 | 0.2 | 13 | 0.2308 | 13 | 0.1538 | 17 | 0.1176 | 53 | 0.1698 | 9 | 0.2222 | 16 | 0.25 | 14 | 0 | 8 | 0.125 | 47 | 0.1489 |
| 15_991401 | 10 | 1 | 14 | 0.9286 | 12 | 1 | 12 | 0.8333 | 48 | 0.9375 | 9 | 0.7778 | 16 | 1 | 13 | 0.7692 | 7 | 0.8571 | 45 | 0.8667 |
| 15_991401 | 11 | 0 | 12 | 0.4167 | 13 | 0.2308 | 16 | 0.4375 | 52 | 0.2885 | 8 | 0 | 15 | 0.3333 | 15 | 0.2 | 7 | 0.2857 | 45 | 0.2222 |
| 15_991402 | 11 | 0.4545 | 10 | 0.3 | 12 | 0.4167 | 15 | 0.6 | 48 | 0.4583 | 8 | 0.75 | 13 | 0.5385 | 14 | 0.5 | 6 | 0.3333 | 41 | 0.5366 |
| 15_991404 | 10 | 0.2 | 13 | 0 | 14 | 0.1429 | 16 | 0.1875 | 53 | 0.1321 | 10 | 0 | 17 | 0.1176 | 13 | 0.1538 | 9 | 0.3333 | 49 | 0.1429 |
| 15_991449 | 7 | 0 | 13 | 0.0769 | 13 | 0 | 6 | 0.1667 | 39 | 0.0513 | 8 | 0.125 | 8 | 0 | 10 | 0.1 | 8 | 0 | 34 | 0.0588 |
| 15_992048 | 5 | 0.2 | 1 | 0 | 4 | 0 | NA | NA | 10 | 0.1 | 1 | 1 | 1 | 0 | 4 | 0 | NA | NA | 6 | 0.1667 |
| 15_992144 | 5 | 0.2 | 4 | 0.25 | 11 | 0.1818 | 3 | 0.3333 | 23 | 0.2174 | 2 | 0 | 1 | 0 | 3 | 0.6667 | 1 | 0 | 7 | 0.2857 |
| 15_992152 | NA | NA | 1 | 0 | 2 | 0 | 3 | 0.6667 | 6 | 0.3333 | NA | NA | NA | NA | 3 | 0.6667 | NA | NA | 3 | 0.6667 |
| 15_992152 | NA | NA | 1 | 0 | 1 | 1 | 3 | 1 | 5 | 0.8 | NA | NA | NA | NA | 3 | 1 | NA | NA | 3 | 1 |
| 15_992156 | NA | NA | 1 | 1 | 1 | 0 | 1 | 1 | 3 | 0.6667 | NA | NA | NA | NA | 3 | 0.6667 | NA | NA | 3 | 0.6667 |
| 15_992157 | NA | NA | 1 | 1 | NA | NA | 3 | 1 | 4 | 1 | NA | NA | NA | NA | 3 | 1 | NA | NA | 3 | 1 |
| 15_993065 | 61 | 0.1148 | 60 | 0.0667 | 24 | 0.2083 | 52 | 0.0962 | 197 | 0.1066 | 42 | 0.0714 | 42 | 0.1905 | 41 | 0.2683 | 38 | 0.2105 | 163 | 0.184 |
| 15_993068 | 62 | 0.7258 | 64 | 0.8125 | 27 | 0.8148 | 58 | 0.7931 | 211 | 0.782 | 42 | 0.6905 | 41 | 0.6585 | 41 | 0.6098 | 43 | 0.7442 | 167 | 0.6766 |
| 15_993068 | 74 | 0.0541 | 65 | 0.2 | 27 | 0.0741 | 58 | 0.0517 | 224 | 0.0982 | 46 | 0.0652 | 43 | 0.0698 | 43 | 0.1628 | 46 | 0.1304 | 178 | 0.1067 |
| 15_993071 | 73 | 0.274 | 62 | 0.371 | 30 | 0.4 | 56 | 0.3929 | 221 | 0.3484 | 47 | 0.4043 | 42 | 0.3571 | 38 | 0.3947 | 51 | 0.3922 | 178 | 0.3876 |
| 15_993071 | 74 | 0.2838 | 71 | 0.4366 | 28 | 0.3929 | 57 | 0.2632 | 230 | 0.3391 | 53 | 0.2642 | 46 | 0.3261 | 47 | 0.3617 | 51 | 0.3529 | 197 | 0.3249 |
| 15_993074 | 67 | 0.6418 | 60 | 0.6 | 29 | 0.7241 | 56 | 0.6786 | 212 | 0.6509 | 47 | 0.5957 | 46 | 0.7174 | 47 | 0.6383 | 43 | 0.7209 | 183 | 0.6667 |
| 15_993075 | 1 | 1 | 3 | 0.6667 | NA | NA | 2 | 0 | 6 | 0.5 | 2 | 0.5 | 1 | 1 | 1 | 0 | NA | NA | 4 | 0.5 |
| 15_993076 | 77 | 0.1039 | 76 | 0.1974 | 30 | 0.0667 | 68 | 0.1324 | 251 | 0.1355 | 56 | 0.125 | 45 | 0.2 | 48 | 0.0625 | 51 | 0.098 | 200 | 0.12 |
| 15_993077 | 75 | 0.0933 | 68 | 0.0294 | 26 | 0.0769 | 60 | 0.0833 | 229 | 0.0699 | 48 | 0.0833 | 48 | 0.125 | 45 | 0.0444 | 51 | 0.1569 | 192 | 0.1042 |
| 15_993077 | NA | NA | NA | NA | NA | NA | 1 | 1 | 1 | 1 | NA | NA | NA | NA | NA | NA | 2 | 0.5 | 2 | 0.5 |
| 15_993203 | 3 | 0.6667 | 3 | 0.3333 | 5 | 0.2 | 1 | 0 | 12 | 0.3333 | NA | NA | NA | NA | 2 | 0.5 | NA | NA | 2 | 0.5 |
| 15_993204 | NA | NA | 3 | 1 | 6 | 0.1667 | 1 | 1 | 10 | 0.5 | NA | NA | 1 | 0 | 2 | 1 | 1 | 0 | 4 | 0.5 |
| 15_993205 | 2 | 0 | 3 | 0 | 5 | 0.4 | 1 | 1 | 11 | 0.2727 | NA | NA | 1 | 1 | 2 | 0.5 | 1 | 0 | 4 | 0.5 |
| 15_993206 | 2 | 0 | 1 | 0 | 5 | 0.2 | 1 | 0 | 9 | 0.1111 | NA | NA | 1 | 0 | 1 | 0 | 1 | 1 | 3 | 0.3333 |
| 15_993238 | 2 | 1 | 2 | 0.5 | 2 | 0.5 | 1 | 1 | 7 | 0.7143 | NA | NA | 1 | 0 | 1 | 1 | 1 | 1 | 3 | 0.6667 |
| 15_993242 | 2 | 0 | 2 | 0 | 3 | 0.3333 | 1 | 0 | 8 | 0.125 | NA | NA | 1 | 1 | 1 | 0 | 1 | 0 | 3 | 0.3333 |
| 15_993242 | 2 | 0.5 | 2 | 1 | 3 | 1 | 1 | 1 | 8 | 0.875 | NA | NA | 1 | 1 | 1 | 1 | 1 | 0 | 3 | 0.6667 |
| 15_993246 | 2 | 0 | 2 | 0.5 | 2 | 0.5 | 1 | 1 | 7 | 0.4286 | NA | NA | 1 | 0 | 1 | 0 | 1 | 1 | 3 | 0.3333 |
| 15_993246 | 2 | 0.5 | 1 | 1 | 3 | 0.6667 | 1 | 1 | 7 | 0.7143 | NA | NA | 1 | 0 | 1 | 1 | 1 | 0 | 3 | 0.3333 |
| 15_993247 | NA | NA | 4 | 1 | 2 | 1 | 1 | 1 | 7 | 1 | NA | NA | 1 | 0 | 1 | 0 | 1 | 1 | 3 | 0.3333 |
| 15_993255 | 1 | 0 | 2 | 1 | 2 | 1 | 1 | 1 | 6 | 0.8333 | NA | NA | 1 | 1 | 1 | 1 | NA | NA | 2 | 1 |
| 15_993255 | 1 | 0 | 2 | 0.5 | 2 | 0.5 | 1 | 0 | 6 | 0.3333 | NA | NA | 1 | 1 | 1 | 0 | NA | NA | 2 | 0.5 |
| 15_993258 | 1 | 0 | 2 | 0.5 | 2 | 0 | 1 | 0 | 6 | 0.1667 | NA | NA | 1 | 1 | 1 | 0 | NA | NA | 2 | 0.5 |
| 15_993259 | 1 | 1 | 2 | 1 | 2 | 0.5 | 1 | 1 | 6 | 0.8333 | NA | NA | 1 | 1 | 1 | 0 | NA | NA | 2 | 0.5 |
| 15_993680 | NA | NA | 1 | 1 | NA | NA | 1 | 0 | 2 | 0.5 | 1 | 0 | 1 | 0 | 3 | 0 | 1 | 1 | 6 | 0.1667 |
| 15_993681 | NA | NA | 2 | 1 | NA | NA | 1 | 1 | 3 | 1 | 1 | 1 | 1 | 1 | 5 | 0.8 | 1 | 1 | 8 | 0.875 |
| 15_993682 | NA | NA | 2 | 1 | NA | NA | 1 | 1 | 3 | 1 | 1 | 1 | 1 | 1 | 5 | 0.2 | 1 | 1 | 8 | 0.5 |
| 15_993683 | NA | NA | 2 | 0.5 | NA | NA | 1 | 0 | 3 | 0.3333 | 1 | 1 | 1 | 0 | 5 | 0.4 | 1 | 0 | 8 | 0.375 |
| 15_993683 | NA | NA | 2 | 0.5 | NA | NA | 1 | 1 | 3 | 0.6667 | 1 | 1 | 1 | 1 | 4 | 0.75 | 1 | 1 | 7 | 0.8571 |
| 15_993683 | NA | NA | 2 | 1 | NA | NA | 1 | 0 | 3 | 0.6667 | 1 | 0 | 1 | 0 | 5 | 0.6 | 1 | 1 | 8 | 0.5 |
| 15_993683 | NA | NA | 1 | 1 | NA | NA | 1 | 0 | 2 | 0.5 | 1 | 1 | 1 | 0 | 4 | 0 | 1 | 1 | 7 | 0.2857 |
| 15_993685 | NA | NA | NA | NA | NA | NA | 1 | 1 | 1 | 1 | 1 | 1 | 1 | 1 | 4 | 0.75 | 1 | 1 | 7 | 0.8571 |
| 15_993688 | NA | NA | 2 | 1 | NA | NA | 1 | 1 | 3 | 1 | 1 | 1 | 1 | 1 | 5 | 1 | 1 | 1 | 8 | 1 |
| 15_993689 | NA | NA | 2 | 0.5 | NA | NA | 1 | 0 | 3 | 0.3333 | 1 | 1 | 1 | 0 | 4 | 0 | 1 | 0 | 7 | 0.1429 |
| 15_993950 | 10 | 0.1 | 11 | 0.2727 | 6 | 0.5 | 11 | 0.0909 | 38 | 0.2105 | 16 | 0.375 | 17 | 0.2353 | 11 | 0.0909 | 11 | 0.1818 | 55 | 0.2364 |
| 15_993951 | 10 | 0.3 | 9 | 0.5556 | 5 | 0.4 | 11 | 0.2727 | 35 | 0.3714 | 13 | 0.1538 | 15 | 0.1333 | 11 | 0.1818 | 10 | 0.2 | 49 | 0.1633 |

| | | | | | | | | | | | | | | | | | | | | |
|---|---|---|---|---|---|---|---|---|---|---|---|---|---|---|---|---|---|---|---|---|
| 15_993955 | 9 | 0.3333 | 5 | 0.4 | 2 | 0 | 8 | 0.25 | 24 | 0.2917 | 11 | 0.1818 | 13 | 0.1538 | 8 | 0 | 8 | 0 | 40 | 0.1 |
| 15_993961 | 6 | 0 | 6 | 0.1667 | 2 | 0 | 5 | 0 | 19 | 0.0526 | 7 | 0.2857 | 8 | 0.125 | 7 | 0.2857 | 6 | 0.1667 | 28 | 0.2143 |
| 15_993961 | 7 | 0.2857 | 8 | 0.375 | 2 | 0 | 7 | 0.2857 | 24 | 0.2917 | 9 | 0.2222 | 10 | 0.2 | 7 | 0.1429 | 6 | 0.3333 | 32 | 0.2188 |
| 15_993963 | 5 | 0 | 7 | 0.1429 | 2 | 0.5 | 5 | 0.2 | 19 | 0.1579 | 9 | 0.4444 | 9 | 0.2222 | 5 | 0.2 | 8 | 0.25 | 31 | 0.2903 |
| 15_993963 | 9 | 0.2222 | 8 | 0.375 | 4 | 0 | 6 | 0 | 27 | 0.1852 | 9 | 0.1111 | 10 | 0 | 7 | 0 | 8 | 0.125 | 34 | 0.0588 |
| 15_993965 | 5 | 0.2 | 4 | 0.5 | 3 | 0 | 4 | 0.5 | 16 | 0.3125 | 8 | 0.125 | 7 | 0 | 1 | 1 | 4 | 0 | 20 | 0.1 |
| 15_993976 | 32 | 0.125 | 33 | 0.2727 | 31 | 0.2581 | 35 | 0.1429 | 131 | 0.1985 | 23 | 0.2609 | 36 | 0.1944 | 27 | 0.1481 | 38 | 0.3947 | 124 | 0.2581 |
| 15_993979 | 29 | 0.1379 | 37 | 0.1622 | 30 | 0.0667 | 36 | 0.1389 | 132 | 0.1288 | 29 | 0.2759 | 36 | 0.1389 | 29 | 0.1379 | 42 | 0.1429 | 136 | 0.1691 |
| 15_993980 | 25 | 0.04 | 34 | 0.1176 | 28 | 0.0714 | 41 | 0.122 | 128 | 0.0938 | 26 | 0.1154 | 34 | 0.1765 | 23 | 0.0435 | 42 | 0.0952 | 125 | 0.112 |
| 15_995183 | NA | NA | 1 | 0 | NA | NA | 2 | 0.5 | 3 | 0.3333 | NA | NA | NA | NA | 1 | 1 | 1 | 1 | 2 | 1 |
| 15_995186 | NA | NA | 1 | 1 | NA | NA | 1 | 0 | 2 | 0.5 | NA | NA | 1 | 1 | 1 | 1 | 1 | 1 | 3 | 1 |
| 15_995186 | NA | NA | 1 | 1 | NA | NA | 2 | 0 | 3 | 0.3333 | NA | NA | 1 | 1 | 1 | 0 | 1 | 1 | 3 | 0.6667 |
| 15_995186 | NA | NA | 1 | 0 | NA | NA | 2 | 0.5 | 3 | 0.3333 | NA | NA | 1 | 0 | 1 | 1 | 1 | 0 | 3 | 0.3333 |
| 15_995188 | NA | NA | NA | NA | 2 | 0 | 3 | 0.3333 | 5 | 0.2 | NA | NA | 1 | 1 | 2 | 0 | NA | NA | 3 | 0.3333 |
| 15_995189 | 1 | 1 | 1 | 0 | 3 | 0 | 2 | 0.5 | 7 | 0.2857 | NA | NA | 1 | 1 | 2 | 0 | NA | NA | 3 | 0.3333 |
| 15_995241 | NA | NA | 1 | 1 | 2 | 0 | 2 | 0 | 5 | 0.2 | 1 | 0 | NA | NA | 2 | 0.5 | 1 | 0 | 4 | 0.25 |
| 15_995707 | 2 | 0 | 2 | 0 | 2 | 0 | 6 | 0.1667 | 12 | 0.0833 | 1 | 0 | 1 | 0 | 4 | 0.25 | 3 | 0 | 9 | 0.1111 |
| 15_995970 | 4 | 0.25 | 9 | 0.1111 | 11 | 0.2727 | 8 | 0.125 | 32 | 0.1875 | 5 | 0 | 6 | 0.3333 | 7 | 0.4286 | 4 | 0.75 | 22 | 0.3636 |
| 15_995971 | 5 | 0.4 | 10 | 0.2 | 11 | 0.2727 | 9 | 0.1111 | 35 | 0.2286 | 5 | 0 | 6 | 0 | 9 | 0 | 4 | 0.75 | 24 | 0.125 |
| 15_995975 | 6 | 0.3333 | 11 | 0.1818 | 13 | 0.3077 | 10 | 0.4 | 40 | 0.3 | 6 | 0.3333 | 9 | 0.2222 | 10 | 0.2 | 6 | 0.1667 | 31 | 0.2258 |
| 15_995976 | 5 | 0 | 7 | 0.4286 | 8 | 0.5 | 8 | 0.75 | 28 | 0.4643 | 5 | 0.2 | 7 | 0.4286 | 9 | 0.5556 | 4 | 0.5 | 25 | 0.44 |
| 15_995978 | 6 | 0 | 6 | 0 | 9 | 0.2222 | 8 | 0.25 | 29 | 0.1379 | 4 | 0.25 | 9 | 0.1111 | 6 | 0 | 5 | 0 | 24 | 0.0833 |
| 15_995979 | 7 | 0.7143 | 9 | 0.6667 | 14 | 0.7143 | 11 | 0.5455 | 41 | 0.6585 | 7 | 1 | 10 | 0.6 | 9 | 0.5556 | 7 | 0.5714 | 33 | 0.6667 |
| 15_995983 | 7 | 0.2857 | 11 | 0.0909 | 10 | 0.4 | 7 | 0.2857 | 35 | 0.2571 | 4 | 0.25 | 4 | 0 | 7 | 0.1429 | 3 | 0.3333 | 18 | 0.1667 |
| 15_995984 | 7 | 0.1429 | 14 | 0.1429 | 9 | 0.4444 | 9 | 0.1111 | 39 | 0.2051 | 7 | 0.2857 | 3 | 0.3333 | 8 | 0 | 3 | 0 | 21 | 0.1429 |
| 15_995987 | 7 | 0.8571 | 12 | 0.75 | 9 | 0.7778 | 7 | 1 | 35 | 0.8286 | 8 | 0.625 | 5 | 0.4 | 6 | 0.8333 | 4 | 0.5 | 23 | 0.6087 |
| 15_996104 | 24 | 0.125 | 22 | 0.1364 | 18 | 0.2222 | 19 | 0.1579 | 83 | 0.1566 | 14 | 0.0714 | 20 | 0.2 | 12 | 0.1667 | 16 | 0.125 | 62 | 0.1452 |
| 15_996104 | 29 | 0 | 23 | 0.087 | 24 | 0.1667 | 26 | 0 | 102 | 0.0588 | 19 | 0 | 27 | 0.0741 | 15 | 0.1333 | 20 | 0.05 | 81 | 0.0617 |
| 15_996211 | 7 | 0.1429 | 1 | 0 | 7 | 0 | 4 | 0.25 | 19 | 0.1053 | 4 | 0 | 3 | 0 | 4 | 0 | 3 | 0.6667 | 14 | 0.1429 |
| 15_996212 | 7 | 0.5714 | 2 | 0.5 | 8 | 0.375 | 5 | 0.6 | 22 | 0.5 | 6 | 0.6667 | 8 | 0.625 | 4 | 0.25 | 4 | 0.75 | 22 | 0.5909 |
| 15_996212 | 7 | 0.1429 | 1 | 1 | 5 | 0.6 | 4 | 0 | 17 | 0.2941 | 5 | 0 | 8 | 0.125 | 2 | 0.5 | 3 | 0.3333 | 18 | 0.1667 |
| 15_996213 | 8 | 0.375 | 2 | 1 | 6 | 0.5 | 4 | 0.25 | 20 | 0.45 | 4 | 0.5 | 8 | 0.25 | 3 | 0.3333 | 4 | 0.25 | 19 | 0.3158 |
| 15_996213 | 6 | 0.6667 | 1 | 0 | 5 | 0.6 | 4 | 0.25 | 16 | 0.5 | 3 | 0.6667 | 5 | 0.2 | 3 | 0 | 3 | 0.3333 | 14 | 0.2857 |
| 15_996218 | 10 | 0.5 | 4 | 0.25 | 8 | 0.25 | 4 | 0.5 | 26 | 0.3846 | 9 | 0.2222 | 8 | 0.375 | 5 | 0.2 | 4 | 0.25 | 26 | 0.2692 |
| 15_996223 | 12 | 0.0833 | 6 | 0.1667 | 12 | 0.0833 | 8 | 0.125 | 38 | 0.1053 | 8 | 0 | 12 | 0.3333 | 7 | 0.4286 | 6 | 0.1667 | 33 | 0.2424 |
| 15_996225 | 13 | 0 | 5 | 0 | 9 | 0.1111 | 5 | 0.2 | 32 | 0.0625 | 7 | 0 | 14 | 0.0714 | 5 | 0 | 7 | 0.1429 | 33 | 0.0606 |
| 15_996253 | NA | NA | 3 | 0.6667 | 1 | 0 | NA | NA | 4 | 0.5 | NA | NA | 1 | 1 | NA | NA | 1 | 0 | 2 | 0.5 |
| 15_996253 | NA | NA | 2 | 0.5 | 1 | 1 | NA | NA | 3 | 0.6667 | NA | NA | 1 | 1 | NA | NA | NA | NA | 1 | 1 |
| 15_996257 | 16 | 0.375 | 21 | 0.3333 | 15 | 0.2 | 19 | 0.5263 | 71 | 0.3662 | 14 | 0.5714 | 19 | 0.4737 | 24 | 0.5417 | 20 | 0.3 | 77 | 0.4675 |
| 15_996336 | 3 | 0 | 4 | 0.25 | 5 | 0 | 8 | 0.125 | 20 | 0.1 | 8 | 0.125 | 7 | 0.2857 | 6 | 0.1667 | 2 | 0.5 | 23 | 0.2174 |
| 15_996337 | 3 | 1 | 6 | 0.6667 | 5 | 0 | 9 | 0.3333 | 23 | 0.4348 | 9 | 0.3333 | 7 | 0.5714 | 5 | 0.8 | 3 | 0.6667 | 24 | 0.5417 |
| 15_996337 | 4 | 1 | 5 | 1 | 4 | 0.25 | 7 | 0.4286 | 20 | 0.65 | 8 | 0.625 | 8 | 0.75 | 6 | 0.8333 | 4 | 1 | 26 | 0.7692 |
| 15_996339 | 2 | 0 | 5 | 0.2 | 5 | 0.2 | 9 | 0 | 21 | 0.0952 | 6 | 0.1667 | 7 | 0.1429 | 6 | 0 | 3 | 0 | 22 | 0.0909 |
| 15_996340 | 3 | 0.6667 | 5 | 0 | 5 | 0 | 8 | 0.5 | 21 | 0.2857 | 9 | 0.2222 | 7 | 0 | 6 | 0 | 3 | 0 | 25 | 0.08 |
| 15_996340 | 5 | 0.8 | 5 | 1 | 4 | 0.75 | 8 | 0.625 | 22 | 0.7727 | 7 | 0.5714 | 6 | 0.6667 | 7 | 1 | 2 | 0.5 | 22 | 0.7273 |
| 15_996341 | 4 | 1 | 4 | 0.75 | 5 | 0.4 | 8 | 0.5 | 21 | 0.619 | 8 | 0.625 | 7 | 0.5714 | 6 | 0.6667 | 3 | 0 | 24 | 0.5417 |
| 15_996771 | NA | NA | 2 | 0 | 2 | 0.5 | 4 | 0 | 8 | 0.125 | NA | NA | 1 | 1 | 3 | 0.3333 | NA | NA | 4 | 0.5 |
| 15_996772 | NA | NA | 1 | 1 | 2 | 1 | 2 | 1 | 5 | 1 | NA | NA | 1 | 1 | 4 | 1 | NA | NA | 5 | 1 |
| 15_996773 | NA | NA | 2 | 0 | 2 | 0.5 | 4 | 0 | 8 | 0.125 | NA | NA | 1 | 0 | 3 | 0.3333 | NA | NA | 4 | 0.25 |
| 15_996773 | NA | NA | 2 | 0 | 2 | 0.5 | 4 | 0.5 | 8 | 0.375 | NA | NA | 1 | 1 | 1 | 0 | NA | NA | 2 | 0.5 |
| 15_996780 | NA | NA | 2 | 1 | 1 | 1 | 3 | 0 | 6 | 0.5 | NA | NA | NA | NA | 2 | 0.5 | NA | NA | 2 | 0.5 |
| 15_996780 | NA | NA | 2 | 0.5 | 1 | 1 | 3 | 0.3333 | 6 | 0.5 | NA | NA | NA | NA | 3 | 0.3333 | NA | NA | 3 | 0.3333 |
| 15_996784 | NA | NA | 2 | 1 | 1 | 1 | 3 | 0.3333 | 6 | 0.6667 | NA | NA | NA | NA | 2 | 0.5 | NA | NA | 2 | 0.5 |
| 15_996790 | NA | NA | 2 | 1 | NA | NA | 2 | 1 | 4 | 1 | NA | NA | NA | NA | 1 | 1 | NA | NA | 1 | 1 |
| 15_999100 | 3 | 1 | 2 | 1 | 1 | 0 | 1 | 0 | 7 | 0.7143 | NA | NA | NA | NA | 3 | 0.3333 | NA | NA | 3 | 0.3333 |
| 15_999101 | 2 | 1 | 2 | 0.5 | 1 | 1 | 1 | 1 | 6 | 0.8333 | NA | NA | NA | NA | 3 | 1 | NA | NA | 3 | 1 |
| 15_999137 | 3 | 0.3333 | 5 | 0 | 5 | 0 | 3 | 0.3333 | 16 | 0.125 | NA | NA | 4 | 0 | 2 | 0.5 | 3 | 0 | 9 | 0.1111 |
| 15_100113 | 3 | 0.3333 | 1 | 0 | NA | NA | 1 | 0 | 5 | 0.2 | 3 | 0.3333 | 1 | 1 | 3 | 0.3333 | 5 | 0.2 | 12 | 0.3333 |
| 15_100113 | 4 | 1 | 1 | 0 | NA | NA | 2 | 0.5 | 7 | 0.7143 | 3 | 0.6667 | 1 | 0 | 3 | 0.6667 | 5 | 1 | 12 | 0.75 |
| 15_100113 | 3 | 0 | 3 | 0.3333 | NA | NA | 2 | 0.5 | 8 | 0.25 | 3 | 0 | NA | NA | 3 | 0 | 5 | 1 | 11 | 0.4545 |
| 15_100113 | 4 | 0.75 | 2 | 0 | NA | NA | 1 | 0 | 7 | 0.4286 | 3 | 1 | 1 | 0 | 3 | 1 | 5 | 0.6 | 12 | 0.75 |
| 15_100113 | 3 | 1 | 3 | 1 | NA | NA | NA | NA | 6 | 1 | 2 | 1 | 1 | 1 | 3 | 1 | 4 | 1 | 10 | 1 |
| 15_100113 | 1 | 1 | 3 | 0.6667 | NA | NA | 2 | 0.5 | 6 | 0.6667 | 3 | 1 | 1 | 1 | 3 | 0.3333 | 5 | 0.6 | 12 | 0.6667 |
| 15_100113 | 4 | 0.75 | 3 | 0 | NA | NA | 1 | 0 | 8 | 0.375 | 3 | 0.3333 | 1 | 0 | 4 | 0.25 | 6 | 0.3333 | 14 | 0.2857 |
| 15_100113 | 4 | 0.5 | 3 | 1 | 1 | 1 | 1 | 1 | 9 | 0.7778 | 2 | 1 | 2 | 1 | 4 | 0.75 | 5 | 0.4 | 13 | 0.6923 |

| | | | | | | | | | | | | | | | | | | | | |
|---|---|---|---|---|---|---|---|---|---|---|---|---|---|---|---|---|---|---|---|---|
| 15_100114 | 4 | 0.25 | 3 | 0.3333 | 1 | 0 | 1 | 0 | 9 | 0.2222 | 3 | 0.3333 | 2 | 1 | 3 | 0 | 2 | 0.5 | 10 | 0.4 |
| 15_100114 | 6 | 0.1667 | 4 | 0 | 1 | 0 | 1 | 0 | 12 | 0.0833 | 4 | 0.25 | 1 | 1 | 3 | 0.3333 | 3 | 0.3333 | 11 | 0.3636 |
| 15_100114 | 5 | 0 | 4 | 0.5 | 1 | 0 | 1 | 0 | 11 | 0.1818 | 4 | 0 | 2 | 0.5 | 4 | 0.25 | 6 | 0 | 16 | 0.125 |
| 15_100114 | 7 | 0 | 3 | 0.3333 | 3 | 0.6667 | 3 | 0.3333 | 16 | 0.25 | 6 | 0.3333 | 5 | 0.2 | 4 | 0.25 | 6 | 0.3333 | 21 | 0.2857 |
| 15_100116 | 6 | 0.1667 | 4 | 0.5 | 6 | 0.1667 | 6 | 0 | 22 | 0.1818 | 9 | 0.1111 | 8 | 0.125 | 9 | 0.1111 | 10 | 0 | 36 | 0.0833 |
| 15_100117 | 11 | 0.1818 | 4 | 0.75 | 7 | 0.4286 | 8 | 0.375 | 30 | 0.3667 | 10 | 0.2 | 10 | 0.2 | 8 | 0.625 | 10 | 0.2 | 38 | 0.2895 |
| 15_100117 | 10 | 0.1 | 6 | 0.1667 | 6 | 0 | 6 | 0 | 28 | 0.0714 | 10 | 0 | 12 | 0.0833 | 8 | 0.125 | 8 | 0 | 38 | 0.0526 |
| 15_100157 | 1 | 1 | 4 | 0.5 | 2 | 1 | NA | NA | 7 | 0.7143 | 1 | 1 | 2 | 1 | 2 | 1 | 1 | 1 | 6 | 1 |
| 15_100157 | 1 | 1 | 3 | 0.3333 | 2 | 0.5 | NA | NA | 6 | 0.5 | 1 | 1 | 2 | 1 | 1 | 0 | 1 | 1 | 5 | 0.8 |
| 15_100157 | 1 | 1 | 4 | 1 | 2 | 1 | NA | NA | 7 | 1 | 1 | 1 | 2 | 0.5 | 1 | 1 | 1 | 1 | 5 | 0.8 |
| 15_100158 | 1 | 0 | 4 | 0.75 | 2 | 1 | NA | NA | 7 | 0.7143 | NA | NA | 2 | 0.5 | 1 | 1 | 1 | 1 | 4 | 0.75 |
| 15_100158 | 2 | 0 | 4 | 0 | 3 | 0.3333 | NA | NA | 9 | 0.1111 | 1 | 1 | 2 | 0 | 2 | 0 | 1 | 0 | 6 | 0.1667 |
| 15_100211 | 4 | 1 | 5 | 0.8 | 2 | 1 | 1 | 1 | 12 | 0.9167 | 2 | 1 | 6 | 0.6667 | 2 | 1 | 2 | 1 | 12 | 0.8333 |
| 15_100211 | 4 | 0.25 | 5 | 0.4 | 1 | 0 | 2 | 1 | 12 | 0.4167 | 2 | 0 | 6 | 0.5 | 1 | 0 | 1 | 1 | 10 | 0.4 |
| 15_100211 | 4 | 0.25 | 6 | 0 | 2 | 0 | 2 | 0 | 14 | 0.0714 | 1 | 0 | 2 | 0 | 2 | 0 | 3 | 0.6667 | 8 | 0.25 |
| 15_100211 | 4 | 0.25 | 5 | 0.2 | 3 | 0 | 3 | 0.3333 | 15 | 0.2 | 3 | 0 | 6 | 0.1667 | 2 | 0 | 3 | 0.3333 | 14 | 0.1429 |
| 15_100211 | 2 | 0 | 5 | 0.2 | 2 | 0 | 3 | 0 | 12 | 0.0833 | 3 | 0 | 6 | 0.1667 | 3 | 0 | 3 | 0 | 15 | 0.0667 |
| 15_100211 | 3 | 0.3333 | 5 | 0.4 | 3 | 0 | 3 | 0 | 14 | 0.2143 | 2 | 0.5 | 7 | 0 | 3 | 0 | 3 | 0.3333 | 15 | 0.1333 |
| 15_100286 | 4 | 0.5 | 4 | 0.25 | 3 | 0.3333 | NA | NA | 11 | 0.3636 | NA | NA | 2 | 0.5 | 1 | 1 | NA | NA | 3 | 0.6667 |
| 15_100286 | 4 | 0.25 | 4 | 0 | 1 | 1 | 1 | 0 | 10 | 0.2 | NA | NA | 2 | 1 | NA | NA | 1 | 0 | 3 | 0.6667 |
| 15_100290 | 3 | 0 | 4 | 0.25 | NA | NA | 5 | 0 | 12 | 0.0833 | 6 | 0 | 5 | 0.2 | 1 | 0 | 2 | 0 | 14 | 0.0714 |
| 15_100290 | 3 | 0.3333 | 5 | 0.2 | NA | NA | 6 | 0 | 14 | 0.1429 | 6 | 0.3333 | 8 | 0 | 2 | 0.5 | 2 | 0 | 18 | 0.1667 |
| 15_100291 | 2 | 0 | 5 | 0.2 | NA | NA | 4 | 0 | 11 | 0.0909 | 6 | 0 | 7 | 0.2857 | 2 | 0 | 1 | 0 | 16 | 0.125 |
| 15_100291 | 2 | 0.5 | 5 | 0.4 | NA | NA | 2 | 1 | 9 | 0.5556 | 5 | 0.2 | 7 | 0.1429 | 1 | 0 | 1 | 0 | 14 | 0.1429 |
| 15_100291 | 2 | 0 | 3 | 0.3333 | NA | NA | 4 | 0.25 | 9 | 0.2222 | 6 | 0 | 7 | 0.1429 | 1 | 0 | NA | NA | 14 | 0.0714 |
| 15_100291 | 2 | 0.5 | 3 | 0.3333 | NA | NA | 3 | 0.3333 | 8 | 0.375 | 4 | 0.5 | 4 | 1 | NA | NA | NA | NA | 8 | 0.75 |
| 15_100291 | 2 | 0.5 | 3 | 0 | NA | NA | 4 | 0 | 9 | 0.1111 | 6 | 0 | 8 | 0.125 | 1 | 0 | NA | NA | 15 | 0.0667 |
| 15_100291 | 1 | 1 | 3 | 0.6667 | NA | NA | 4 | 1 | 8 | 0.875 | 6 | 0.8333 | 8 | 0.5 | 1 | 1 | NA | NA | 15 | 0.6667 |
| 15_100292 | 1 | 0 | 3 | 0 | NA | NA | 3 | 0.3333 | 7 | 0.1429 | 6 | 0.1667 | 8 | 0.625 | NA | NA | NA | NA | 14 | 0.4286 |
| 15_100292 | NA | NA | 2 | 0.5 | NA | NA | 1 | 0 | 3 | 0.3333 | NA | NA | 5 | 0.6 | NA | NA | NA | NA | 5 | 0.6 |
| 15_100292 | 1 | 1 | 3 | 1 | NA | NA | 2 | 1 | 6 | 1 | 5 | 1 | 6 | 0.8333 | NA | NA | NA | NA | 11 | 0.9091 |
| 15_100292 | 1 | 1 | 3 | 0.3333 | NA | NA | 4 | 0 | 8 | 0.25 | 6 | 0.3333 | 8 | 0.625 | 1 | 0 | NA | NA | 15 | 0.4667 |
| 15_100292 | 1 | 0 | 2 | 0 | NA | NA | 3 | 0.3333 | 6 | 0.1667 | 5 | 0.2 | 6 | 0.5 | 1 | 1 | NA | NA | 12 | 0.4167 |
| 15_100292 | 1 | 1 | 2 | 0.5 | NA | NA | 4 | 0.25 | 7 | 0.4286 | 5 | 0.2 | 6 | 0.3333 | 1 | 0 | NA | NA | 12 | 0.25 |
| 15_100292 | 1 | 0 | 2 | 0.5 | NA | NA | 1 | 1 | 4 | 0.5 | 4 | 0.25 | 6 | 0.1667 | 1 | 0 | NA | NA | 11 | 0.1818 |
| 15_100293 | 6 | 0 | 4 | 0.25 | 3 | 0 | 2 | 0 | 15 | 0.0667 | NA | NA | 8 | 0.375 | 1 | 0 | 1 | 0 | 10 | 0.3 |
| 15_100375 | 6 | 0.1667 | 7 | 0.4286 | 3 | 0 | 5 | 0 | 21 | 0.1905 | 4 | 0.75 | 2 | 0.5 | 6 | 0.5 | 5 | 0.2 | 17 | 0.4706 |
| 15_100375 | 6 | 0.3333 | 5 | 0.4 | 3 | 0.6667 | 5 | 0.2 | 19 | 0.3684 | 4 | 0.5 | 2 | 0.5 | 4 | 0 | 4 | 0.25 | 14 | 0.2857 |
| 15_100376 | 6 | 0 | 7 | 0.2857 | 1 | 0 | 8 | 0.125 | 22 | 0.1364 | 6 | 0.3333 | 2 | 0 | 5 | 0.2 | 6 | 0.3333 | 19 | 0.2632 |
| 15_100377 | 6 | 0.8333 | 5 | 1 | 7 | 1 | 5 | 1 | 23 | 0.9565 | 7 | 1 | 4 | 1 | 4 | 1 | 1 | 1 | 16 | 1 |
| 15_100377 | 4 | 0.25 | 3 | 0.3333 | 6 | 0 | 4 | 0 | 17 | 0.1176 | 6 | 0 | 4 | 0.25 | 4 | 0.25 | 2 | 0 | 16 | 0.125 |
| 15_100377 | 4 | 0.75 | 4 | 0.5 | 3 | 0.6667 | 4 | 0.5 | 15 | 0.6 | 6 | 0.3333 | 2 | 0.5 | 2 | 1 | 1 | 0 | 11 | 0.4545 |
| 15_100377 | 5 | 0.2 | 5 | 0.4 | 8 | 0.25 | 4 | 0.25 | 22 | 0.2727 | 6 | 0.1667 | 3 | 0.3333 | 4 | 0 | 2 | 0 | 15 | 0.1333 |
| 15_100377 | 4 | 0 | 5 | 0 | 6 | 0.1667 | 3 | 0.3333 | 18 | 0.1111 | 7 | 0.1429 | 4 | 0 | 3 | 0 | 2 | 0 | 16 | 0.0625 |
| 15_100378 | 4 | 0 | 5 | 0.4 | 6 | 0.1667 | 3 | 0.3333 | 18 | 0.2222 | 6 | 0.5 | 2 | 0 | 2 | 0.5 | 2 | 0.5 | 12 | 0.4167 |
| 15_100378 | 3 | 0 | 4 | 0.25 | 5 | 0 | 3 | 0 | 15 | 0.0667 | 5 | 0.2 | 1 | 0 | 1 | 1 | 2 | 0 | 9 | 0.2222 |
| 15_100378 | 3 | 0.3333 | 3 | 0 | 4 | 0.25 | 2 | 0 | 12 | 0.1667 | 5 | 0.4 | 1 | 1 | 1 | 0 | 1 | 0 | 8 | 0.375 |
| 15_100378 | 3 | 0.3333 | 3 | 0.3333 | 4 | 1 | 2 | 1 | 12 | 0.6667 | 5 | 1 | 1 | 1 | 1 | 1 | 1 | 1 | 8 | 1 |
| 15_100378 | 3 | 0.6667 | 3 | 0 | 3 | 0.6667 | 1 | 0 | 10 | 0.4 | 4 | 0.5 | NA | NA | NA | NA | 2 | 1 | 6 | 0.6667 |
| 15_100378 | 2 | 0.5 | 3 | 0 | 3 | 0 | 2 | 0 | 10 | 0.1 | 3 | 0.3333 | 1 | 0 | 1 | 0 | 1 | 1 | 6 | 0.3333 |
| 15_100378 | 2 | 1 | 1 | 1 | 3 | 1 | 1 | 1 | 7 | 1 | 4 | 0.75 | 2 | 1 | 1 | 1 | 2 | 1 | 9 | 0.8889 |
| 15_100379 | 1 | 0 | 3 | 0 | 1 | 0 | 1 | 1 | 6 | 0.1667 | 2 | 0.5 | 1 | 0 | 1 | 0 | NA | NA | 4 | 0.25 |
| 15_100379 | 2 | 0 | 3 | 0.3333 | 3 | 0.6667 | 1 | 1 | 9 | 0.4444 | 4 | 0.75 | 2 | 1 | 1 | 0 | 1 | 1 | 8 | 0.75 |
| 15_100379 | 2 | 0 | 3 | 0 | 2 | 0.5 | 2 | 0 | 9 | 0.1111 | 4 | 0.25 | 1 | 0 | 1 | 1 | 2 | 0 | 8 | 0.25 |
| 15_100379 | 2 | 1 | 1 | 1 | 1 | 1 | 1 | 1 | 5 | 1 | 3 | 1 | 2 | 1 | NA | NA | 2 | 1 | 7 | 1 |
| 15_100379 | 2 | 0.5 | 2 | 0 | 2 | 0.5 | 1 | 0 | 7 | 0.2857 | 4 | 0 | 1 | 1 | 1 | 0 | 2 | 0.5 | 8 | 0.25 |
| 15_100379 | 2 | 0.5 | 2 | 0.5 | 2 | 0.5 | NA | NA | 6 | 0.5 | 4 | 0.25 | 1 | 0 | 1 | 0 | 2 | 0.5 | 8 | 0.25 |
| 15_100379 | 2 | 0 | 1 | 0 | 2 | 0.5 | 1 | 0 | 6 | 0.1667 | 4 | 0.25 | 1 | 0 | 1 | 0 | 2 | 0 | 8 | 0.125 |
| 15_100379 | 2 | 1 | 1 | 1 | 2 | 1 | NA | NA | 5 | 1 | 3 | 1 | 1 | 1 | 1 | 1 | 2 | 1 | 7 | 1 |
| 15_100379 | 2 | 1 | 2 | 1 | 2 | 0.5 | 1 | 1 | 7 | 0.8571 | 4 | 0.75 | 2 | 0.5 | 1 | 1 | 2 | 1 | 9 | 0.7778 |
| 15_100379 | 2 | 0.5 | 2 | 1 | 1 | 1 | NA | NA | 5 | 0.8 | 3 | 1 | 1 | 1 | NA | NA | 1 | 1 | 5 | 1 |
| 15_100380 | 2 | 1 | 1 | 1 | 2 | 0.5 | 1 | 1 | 6 | 0.8333 | 3 | 1 | 1 | 1 | 1 | 1 | 1 | 1 | 6 | 1 |
| 15_100380 | 1 | 0 | 1 | 1 | 1 | 1 | NA | NA | 3 | 0.6667 | NA | NA | 1 | 1 | 1 | 1 | 1 | 1 | 3 | 1 |
| 15_100380 | 2 | 1 | 1 | 1 | 2 | 1 | 1 | 1 | 6 | 1 | 1 | 1 | 1 | 1 | 1 | 1 | 1 | 1 | 4 | 1 |
| 15_100380 | 2 | 1 | 1 | 1 | 1 | 1 | NA | NA | 4 | 1 | 1 | 1 | 2 | 1 | 1 | 1 | 1 | 1 | 5 | 1 |

| | | | | | | | | | | | | | | | | | | | | |
|---|---|---|---|---|---|---|---|---|---|---|---|---|---|---|---|---|---|---|---|---|
| 15_1003802 | 2 | 1 | 2 | 1 | 1 | 1 | NA | NA | 5 | 1 | 1 | 1 | 1 | 1 | 1 | 1 | 1 | 1 | 4 | 1 |
| 15_1003803 | 2 | 0 | 2 | 0.5 | 1 | 1 | NA | NA | 5 | 0.4 | 2 | 0.5 | 2 | 0.5 | NA | NA | 1 | 1 | 5 | 0.6 |
| 15_1003803 | 2 | 0.5 | NA | NA | 1 | 0 | NA | NA | 3 | 0.3333 | 2 | 0.5 | 2 | 0.5 | NA | NA | 1 | 0 | 5 | 0.4 |
| 15_1003804 | 2 | 0.5 | 2 | 0.5 | 1 | 0 | NA | NA | 5 | 0.4 | 2 | 0.5 | 1 | 1 | NA | NA | 1 | 1 | 4 | 0.75 |
| 15_1003860 | 2 | 0.5 | 2 | 0 | 1 | 0 | NA | NA | 5 | 0.2 | 2 | 0 | 1 | 0 | NA | NA | 1 | 1 | 4 | 0.25 |
| 15_1004013 | 3 | 0.3333 | 3 | 0 | 1 | 0 | NA | NA | 7 | 0.1429 | 3 | 0 | 2 | 0 | 1 | 0 | 2 | 0.5 | 8 | 0.125 |
| 15_1004734 | 6 | 0 | 10 | 0.2 | 6 | 0.3333 | 3 | 0 | 25 | 0.16 | 11 | 0.0909 | 7 | 0.2857 | 6 | 0 | 11 | 0 | 35 | 0.0857 |
| 15_1004734 | 6 | 0.1667 | 13 | 0.0769 | 6 | 0 | 5 | 0 | 30 | 0.0667 | 10 | 0 | 5 | 0.2 | 5 | 0.2 | 11 | 0 | 31 | 0.0645 |
| 15_1004734 | 6 | 0 | 9 | 0 | 7 | 0.2857 | 5 | 0.4 | 27 | 0.1481 | 10 | 0.1 | 7 | 0 | 6 | 0.1667 | 10 | 0.1 | 33 | 0.0909 |
| 15_1004735 | 4 | 0 | 12 | 0.1667 | 8 | 0 | 4 | 0.25 | 28 | 0.1071 | 9 | 0.2222 | 5 | 0.4 | 6 | 0.1667 | 11 | 0.3636 | 31 | 0.2903 |
| 15_1004736 | 3 | 0 | 9 | 0.5556 | 9 | 0.4444 | 5 | 0.6 | 26 | 0.4615 | 8 | 0.375 | 8 | 0.75 | 4 | 1 | 12 | 0.5833 | 32 | 0.625 |
| 15_1004737 | 5 | 1 | 9 | 1 | 7 | 1 | 5 | 1 | 26 | 1 | 7 | 0.7143 | 8 | 1 | 6 | 0.8333 | 11 | 1 | 32 | 0.9062 |
| 15_1004737 | 4 | 1 | 10 | 0.7 | 7 | 0.8571 | 2 | 1 | 23 | 0.8261 | 9 | 0.4444 | 7 | 0.7143 | 5 | 0.4 | 11 | 0.5455 | 32 | 0.5312 |
| 15_1004737 | 4 | 0.5 | 5 | 0.2 | 5 | 0.2 | 3 | 0 | 17 | 0.2353 | 4 | 0.25 | 5 | 0.2 | 5 | 0.2 | 9 | 0 | 23 | 0.1304 |
| 15_1004739 | 3 | 0.6667 | 5 | 0.6 | 7 | 0.1429 | 3 | 0.6667 | 18 | 0.4444 | 5 | 0.4 | 6 | 0.6667 | 5 | 0.6 | 7 | 0.4286 | 23 | 0.5217 |
| 15_1004744 | 1 | 0 | 6 | 0.5 | 4 | 0.75 | 2 | 0.5 | 13 | 0.5385 | 7 | 0.5714 | 8 | 1 | 7 | 0.7143 | 10 | 0.7 | 32 | 0.75 |
| 15_1004744 | 5 | 0.4 | 9 | 0.2222 | 8 | 0.125 | 4 | 0 | 26 | 0.1923 | 9 | 0.2222 | 8 | 0.25 | 6 | 0.1667 | 10 | 0.2 | 33 | 0.2121 |
| 15_1004748 | 3 | 0.3333 | 7 | 0.4286 | 6 | 0.3333 | 2 | 0 | 18 | 0.3333 | 6 | 0.1667 | 6 | 0.5 | 6 | 0.3333 | 8 | 0.5 | 26 | 0.3846 |
| 15_1004749 | 1 | 0 | 7 | 0.2857 | 6 | 0.1667 | 4 | 0.25 | 18 | 0.2222 | 4 | 0.25 | 6 | 0 | 3 | 0 | 6 | 0.8333 | 19 | 0.3158 |
| 15_1004751 | 2 | 0.5 | 7 | 0.1429 | 6 | 0.1667 | 2 | 0 | 17 | 0.1765 | 4 | 0 | 2 | 0 | 6 | 0 | 7 | 0.1429 | 19 | 0.0526 |
| 15_1004752 | 3 | 0.3333 | 8 | 0.125 | 7 | 0 | 4 | 0 | 22 | 0.0909 | 4 | 0 | 4 | 0 | 6 | 0.1667 | 7 | 0.1429 | 21 | 0.0952 |
| 15_1004752 | 2 | 0 | 5 | 0.6 | 6 | 0.5 | 2 | 0 | 15 | 0.4 | 2 | 0 | 3 | 0 | 5 | 0.2 | 5 | 0.4 | 15 | 0.2 |
| 15_1004754 | 2 | 0.5 | 5 | 0.2 | 4 | 0 | 4 | 0.25 | 15 | 0.2 | 5 | 0.2 | 5 | 0.2 | 4 | 0 | 7 | 0.1429 | 21 | 0.1429 |
| 15_1004756 | 2 | 0 | 3 | 0.6667 | 4 | 0.25 | 3 | 0.3333 | 12 | 0.3333 | 4 | 0 | 2 | 0 | 6 | 0.3333 | 6 | 0.5 | 18 | 0.2778 |
| 15_1004758 | 2 | 0 | 4 | 0.25 | 4 | 0 | 2 | 0 | 12 | 0.0833 | 3 | 0.3333 | 3 | 0 | 5 | 0.2 | 5 | 0 | 16 | 0.125 |
| 15_1004760 | 2 | 0 | 5 | 0 | 2 | 0 | 3 | 0.3333 | 12 | 0.0833 | 2 | 0 | 3 | 0 | 6 | 0.1667 | 5 | 0.2 | 16 | 0.125 |
| 15_1004761 | 1 | 0 | 3 | 0 | 2 | 0 | 2 | 0.5 | 8 | 0.125 | 2 | 0 | 3 | 0 | 5 | 0 | 4 | 0.5 | 14 | 0.1429 |
| 15_1004842 | NA | NA | 1 | 1 | NA | NA | NA | NA | 1 | 1 | 1 | 1 | NA | NA | NA | NA | NA | NA | 1 | 1 |
| 15_1004844 | NA | NA | 1 | 1 | NA | NA | NA | NA | 1 | 1 | 2 | 1 | NA | NA | NA | NA | NA | NA | 2 | 1 |
| 15_1004845 | NA | NA | 1 | 1 | NA | NA | NA | NA | 1 | 1 | 2 | 0.5 | NA | NA | NA | NA | NA | NA | 2 | 0.5 |
| 15_1004845 | NA | NA | 2 | 1 | NA | NA | 3 | 0 | 5 | 0.4 | 2 | 0.5 | 2 | 0 | 5 | 0 | 2 | 0 | 11 | 0.0909 |
| 15_1004846 | NA | NA | 2 | 0.5 | NA | NA | 3 | 0 | 5 | 0.2 | 2 | 1 | 2 | 0 | 3 | 0.3333 | 2 | 0 | 9 | 0.3333 |
| 15_1004988 | 1 | 1 | NA | NA | NA | NA | NA | NA | 1 | 1 | 1 | 1 | NA | NA | NA | NA | NA | NA | 1 | 1 |
| 15_1005195 | NA | NA | 1 | 1 | 1 | 1 | NA | NA | 2 | 1 | NA | NA | 1 | 0 | 1 | 1 | NA | NA | 2 | 0.5 |
| 15_1005898 | NA | NA | 4 | 0.75 | 1 | 1 | 2 | 1 | 7 | 0.8571 | 1 | 1 | NA | NA | 2 | 0.5 | 1 | 0 | 4 | 0.5 |
| 15_1005898 | NA | NA | 4 | 1 | 2 | 1 | 2 | 0.5 | 8 | 0.875 | 1 | 1 | NA | NA | 2 | 0 | 1 | 1 | 4 | 0.5 |
| 15_1005903 | NA | NA | 3 | 0.6667 | 2 | 0 | 1 | 0 | 6 | 0.3333 | 1 | 1 | NA | NA | 1 | 0 | 1 | 1 | 3 | 0.6667 |
| 15_1005909 | NA | NA | 1 | 0 | 1 | 1 | 1 | 1 | 3 | 0.6667 | 1 | 1 | NA | NA | 1 | 1 | 1 | 1 | 3 | 1 |
| 15_1007043 | NA | NA | NA | NA | NA | NA | 1 | 1 | 1 | 1 | NA | NA | NA | NA | 1 | 1 | NA | NA | 1 | 1 |
| 15_1007055 | NA | NA | NA | NA | NA | NA | 1 | 1 | 1 | 1 | NA | NA | NA | NA | 1 | 1 | NA | NA | 1 | 1 |
| 15_1007057 | NA | NA | NA | NA | NA | NA | 1 | 1 | 1 | 1 | NA | NA | NA | NA | 1 | 1 | NA | NA | 1 | 1 |
| 15_1007058 | NA | NA | NA | NA | NA | NA | 1 | 1 | 1 | 1 | NA | NA | NA | NA | 1 | 1 | NA | NA | 1 | 1 |
| 15_1007060 | NA | NA | NA | NA | NA | NA | 1 | 1 | 1 | 1 | NA | NA | NA | NA | 1 | 1 | NA | NA | 1 | 1 |
| 15_1007061 | NA | NA | NA | NA | NA | NA | 1 | 1 | 1 | 1 | NA | NA | NA | NA | 1 | 1 | NA | NA | 1 | 1 |
| 15_1007061 | NA | NA | NA | NA | NA | NA | 1 | 1 | 1 | 1 | NA | NA | NA | NA | 1 | 1 | NA | NA | 1 | 1 |
| 15_1007138 | 1 | 1 | 1 | 1 | 1 | 1 | NA | NA | 3 | 1 | 1 | 1 | 1 | 0 | NA | NA | NA | NA | 2 | 0.5 |
| 15_1007139 | 1 | 0 | 1 | 1 | 1 | 0 | NA | NA | 3 | 0.3333 | 1 | 1 | NA | NA | NA | NA | NA | NA | 1 | 1 |
| 15_1007141 | 2 | 1 | 1 | 1 | 2 | 0.5 | 1 | 1 | 6 | 0.8333 | 2 | 0.5 | NA | NA | 1 | 1 | 1 | 1 | 4 | 0.75 |
| 15_1007143 | 2 | 1 | 1 | 1 | 3 | 1 | 1 | 1 | 7 | 1 | 2 | 1 | NA | NA | 1 | 1 | 1 | 1 | 4 | 1 |
| 15_1007143 | 2 | 0.5 | 1 | 1 | 3 | 0 | 1 | 1 | 7 | 0.4286 | 2 | 0 | NA | NA | 1 | 0 | 1 | 1 | 4 | 0.25 |
| 15_1007144 | 1 | 1 | NA | NA | 3 | 0.3333 | 1 | 1 | 5 | 0.6 | 2 | 1 | NA | NA | 1 | 1 | 1 | 1 | 4 | 1 |
| 15_1007145 | 2 | 0.5 | 1 | 1 | 3 | 0 | 1 | 1 | 7 | 0.4286 | 2 | 0.5 | NA | NA | 1 | 0 | 1 | 0 | 4 | 0.25 |
| 15_1007145 | 2 | 1 | 1 | 0 | 3 | 0.6667 | 1 | 1 | 7 | 0.7143 | 2 | 0.5 | NA | NA | 1 | 1 | 1 | 1 | 4 | 0.75 |
| 15_1007154 | NA | NA | 1 | 1 | NA | NA | NA | NA | 1 | 1 | 1 | 1 | NA | NA | 1 | 0 | 2 | 0.5 | 4 | 0.5 |
| 15_1007159 | NA | NA | 1 | 1 | 1 | 0 | NA | NA | 2 | 0.5 | 2 | 0.5 | NA | NA | 1 | 0 | 2 | 0.5 | 5 | 0.4 |
| 15_1007162 | NA | NA | 1 | 1 | 1 | 0 | NA | NA | 2 | 0.5 | 2 | 0.5 | NA | NA | 1 | 0 | 2 | 0 | 5 | 0.2 |
| 15_1007206 | 1 | 0 | 5 | 0 | 3 | 0 | 5 | 0.2 | 14 | 0.0714 | 4 | 0 | NA | NA | 5 | 0 | 4 | 0.25 | 13 | 0.0769 |
| 15_1007207 | 2 | 0 | 5 | 0 | 3 | 0.3333 | 5 | 0 | 15 | 0.0667 | 4 | 0.25 | NA | NA | 5 | 0 | 4 | 0 | 13 | 0.0769 |
| 15_1007208 | 2 | 0 | 5 | 0 | 4 | 0.25 | 4 | 0 | 15 | 0.0667 | 4 | 0 | NA | NA | 5 | 0.4 | 4 | 0 | 13 | 0.1538 |
| 15_1007213 | 2 | 0 | 7 | 0.1429 | 1 | 0 | 5 | 0.2 | 15 | 0.1333 | 4 | 0 | NA | NA | 4 | 0.25 | 2 | 0 | 10 | 0.1 |
| 15_1007215 | 4 | 0.25 | 7 | 0 | 4 | 0 | 5 | 0.2 | 20 | 0.1 | 4 | 0.25 | NA | NA | 2 | 0 | 5 | 0 | 11 | 0.0909 |
| 15_1009163 | NA | NA | NA | NA | 1 | 0 | 2 | 0.5 | 3 | 0.3333 | 2 | 0.5 | 3 | 0.3333 | 1 | 0 | 1 | 0 | 7 | 0.2857 |
| 15_1009163 | NA | NA | NA | NA | 1 | 0 | 2 | 0.5 | 3 | 0.3333 | 2 | 0.5 | 2 | 0 | NA | NA | 1 | 0 | 5 | 0.2 |
| 15_1009377 | 4 | 0 | 2 | 0.5 | 1 | 0 | 3 | 0 | 10 | 0.1 | 3 | 0 | 3 | 0 | 4 | 0 | 5 | 0.2 | 15 | 0.0667 |
| 15_1009377 | 4 | 0.25 | 2 | 0 | 2 | 1 | 4 | 0.25 | 12 | 0.3333 | 3 | 0 | 4 | 0 | 4 | 0 | 6 | 0.1667 | 17 | 0.0588 |

| | | | | | | | | | | | | | | | | | | | | |
|---|---|---|---|---|---|---|---|---|---|---|---|---|---|---|---|---|---|---|---|---|
| 15_1009377 | 4 | 0.5 | 2 | 0.5 | 2 | 0.5 | 4 | 0.5 | 12 | 0.5 | 3 | 0.3333 | 4 | 0.75 | 4 | 0.25 | 3 | 0 | 14 | 0.3571 |
| 15_1009378 | 4 | 1 | 3 | 1 | 2 | 1 | 4 | 1 | 13 | 1 | 2 | 1 | 4 | 0.75 | 4 | 1 | 5 | 1 | 15 | 0.9333 |
| 15_1009379 | 4 | 0.25 | 4 | 0 | 2 | 0 | 3 | 0.6667 | 13 | 0.2308 | 4 | 0.25 | 4 | 0.25 | 5 | 0.2 | 5 | 0.2 | 18 | 0.2222 |
| 15_1009379 | 6 | 0.8333 | 2 | 0.5 | 2 | 1 | 5 | 0.6 | 15 | 0.7333 | 4 | 0.75 | 3 | 1 | 4 | 0.75 | 4 | 0.5 | 15 | 0.7333 |
| 15_1009381 | 3 | 0 | 2 | 0 | 3 | 0.3333 | 5 | 0 | 13 | 0.0769 | 4 | 0.25 | 2 | 0.5 | 4 | 0.25 | 5 | 0.2 | 15 | 0.2667 |
| 15_1009381 | 3 | 0.3333 | 3 | 0.6667 | 3 | 0.3333 | 4 | 0 | 13 | 0.3077 | 4 | 0.5 | 2 | 0.5 | 1 | 0 | 6 | 0.5 | 13 | 0.4615 |
| 15_1009381 | 6 | 0.3333 | 3 | 0.6667 | 2 | 0 | 5 | 0 | 16 | 0.25 | 2 | 0 | 2 | 0 | 1 | 0 | 6 | 0.1667 | 11 | 0.0909 |
| 15_1009382 | 4 | 0.25 | 3 | 0.6667 | 3 | 0.3333 | 4 | 0.5 | 14 | 0.4286 | 2 | 0 | 3 | 0 | 3 | 0.3333 | 6 | 0.3333 | 14 | 0.2143 |
| 15_1009383 | 7 | 0.8571 | 4 | 1 | 2 | 1 | 5 | 0.8 | 18 | 0.8889 | 4 | 0.75 | 4 | 1 | 2 | 0.5 | 5 | 0.6 | 15 | 0.7333 |
| 15_1009386 | 6 | 0.6667 | 4 | 0.5 | 2 | 1 | 3 | 0.3333 | 15 | 0.6 | 4 | 1 | 4 | 0.25 | 5 | 0.2 | 5 | 0.8 | 18 | 0.5556 |
| 15_1009386 | 3 | 1 | 6 | 1 | 3 | 0.6667 | 4 | 1 | 16 | 0.9375 | 3 | 1 | 4 | 1 | 4 | 1 | 5 | 1 | 16 | 1 |
| 15_1009386 | 7 | 0.1429 | 7 | 0 | 3 | 0.3333 | 5 | 0 | 22 | 0.0909 | 4 | 0.25 | 5 | 0 | 6 | 0.1667 | 6 | 0.5 | 21 | 0.2381 |
| 15_1009386 | 6 | 0 | 6 | 0.3333 | 3 | 0.3333 | 5 | 0 | 20 | 0.15 | 4 | 0.25 | 5 | 0 | 5 | 0 | 4 | 0 | 18 | 0.0556 |
| 15_1009387 | 7 | 0.1429 | 4 | 0 | 3 | 0 | 5 | 0 | 19 | 0.0526 | 4 | 0 | 5 | 0 | 5 | 0.2 | 6 | 0.1667 | 20 | 0.1 |
| 15_1009387 | 6 | 0.1667 | 6 | 0.1667 | 3 | 0.3333 | 5 | 0.4 | 20 | 0.25 | 4 | 0.25 | 3 | 0 | 4 | 0.5 | 3 | 0.6667 | 14 | 0.3571 |
| 15_1009387 | 7 | 0.8571 | 7 | 0.8571 | 2 | 0.5 | 5 | 0.8 | 21 | 0.8095 | 4 | 1 | 5 | 0.8 | 6 | 0.6667 | 6 | 0.8333 | 21 | 0.8095 |
| 15_1009387 | 7 | 1 | 5 | 1 | 3 | 1 | 4 | 1 | 19 | 1 | 3 | 1 | 4 | 1 | 5 | 1 | 5 | 1 | 17 | 1 |
| 15_1009388 | 5 | 0.2 | 6 | 0.1667 | 2 | 0.5 | 4 | 0.25 | 17 | 0.2353 | 4 | 0.25 | 5 | 0.2 | 5 | 0.4 | 5 | 0 | 19 | 0.2105 |
| 15_1009388 | 5 | 0 | 4 | 0 | 3 | 0.3333 | 3 | 0 | 15 | 0.0667 | 3 | 0.3333 | 4 | 0 | 5 | 0 | 6 | 0 | 18 | 0.0556 |
| 15_1009388 | 7 | 0.5714 | 7 | 0.8571 | 4 | 0.5 | 5 | 0.8 | 23 | 0.6957 | 4 | 0.25 | 5 | 0.6 | 6 | 0.5 | 7 | 0.8571 | 22 | 0.5909 |
| 15_1009391 | 8 | 0.75 | 7 | 1 | 4 | 0.5 | 5 | 1 | 24 | 0.8333 | 5 | 0.8 | 4 | 1 | 8 | 0.875 | 7 | 0.8571 | 24 | 0.875 |
| 15_1009393 | 7 | 0.5714 | 4 | 0 | 3 | 0.3333 | 4 | 0.25 | 18 | 0.3333 | 3 | 0.3333 | 4 | 0 | 5 | 0.4 | 5 | 0.4 | 17 | 0.2941 |
| 15_1009395 | 7 | 0.1429 | 9 | 0.2222 | 4 | 0 | 4 | 0.25 | 24 | 0.1667 | 5 | 0.4 | 7 | 0.2857 | 9 | 0.3333 | 7 | 0.1429 | 28 | 0.2857 |
| 15_1009395 | 8 | 0.25 | 8 | 0.125 | 3 | 0.3333 | 4 | 0 | 23 | 0.1739 | 5 | 0.2 | 5 | 0.2 | 9 | 0.3333 | 6 | 0.1667 | 25 | 0.24 |
| 15_1009397 | 6 | 0.1667 | 3 | 0 | 3 | 0 | 3 | 0.3333 | 15 | 0.1333 | 4 | 0 | 3 | 0.6667 | 7 | 0.1429 | 3 | 0 | 17 | 0.1765 |
| 15_1009401 | 4 | 0.25 | 8 | 0 | 4 | 0 | 1 | 0 | 17 | 0.0588 | 4 | 0 | 8 | 0.125 | 3 | 0.3333 | 2 | 0 | 17 | 0.1176 |
| 15_1009402 | 7 | 0.1429 | 11 | 0.0909 | 7 | 0.2857 | 6 | 0.3333 | 31 | 0.1935 | 5 | 0 | 9 | 0.1111 | 9 | 0.3333 | 6 | 0.1667 | 29 | 0.1724 |
| 15_1009406 | 10 | 0.1 | 10 | 0.1 | 7 | 0 | 7 | 0 | 34 | 0.0588 | 6 | 0.1667 | 12 | 0 | 12 | 0.3333 | 7 | 0.1429 | 37 | 0.1622 |
| 15_1009434 | 11 | 0 | 15 | 0.2667 | 12 | 0.0833 | 12 | 0.0833 | 50 | 0.12 | 6 | 0.1667 | 9 | 0 | 12 | 0.0833 | 10 | 0.1 | 37 | 0.0811 |
| 15_1009434 | 12 | 0.75 | 11 | 0.8182 | 12 | 0.8333 | 12 | 0.9167 | 47 | 0.8298 | 5 | 0.8 | 9 | 0.8889 | 9 | 0.7778 | 11 | 0.9091 | 34 | 0.8529 |
| 15_1009434 | 14 | 0.1429 | 16 | 0.125 | 14 | 0.1429 | 14 | 0.2857 | 58 | 0.1724 | 7 | 0 | 8 | 0.125 | 11 | 0.0909 | 12 | 0.0833 | 38 | 0.0789 |
| 15_1009435 | 9 | 0 | 17 | 0.1765 | 13 | 0.0769 | 12 | 0.0833 | 51 | 0.098 | 7 | 0 | 7 | 0.1429 | 9 | 0.2222 | 10 | 0.1 | 33 | 0.1212 |
| 15_1009435 | 14 | 0.4286 | 19 | 0.2105 | 14 | 0.1429 | 13 | 0.3077 | 60 | 0.2667 | 8 | 0.125 | 6 | 0.6667 | 12 | 0.3333 | 13 | 0.4615 | 39 | 0.3846 |
| 15_1010417 | 2 | 1 | 1 | 1 | NA | NA | NA | NA | 3 | 1 | NA | NA | NA | NA | 1 | 1 | 1 | 0 | 2 | 0.5 |
| 15_1010425 | 2 | 1 | 1 | 1 | 1 | 1 | 2 | 1 | 6 | 1 | NA | NA | NA | NA | NA | NA | 1 | 1 | 1 | 1 |
| 15_1010429 | 2 | 1 | 1 | 1 | 1 | 1 | 2 | 0.5 | 6 | 0.8333 | NA | NA | NA | NA | 1 | 1 | NA | NA | 1 | 1 |
| 15_1010430 | 3 | 0.3333 | 1 | 0 | 1 | 0 | 2 | 0.5 | 7 | 0.2857 | NA | NA | NA | NA | 1 | 0 | 1 | 1 | 2 | 0.5 |
| 15_1011087 | 3 | 0 | 2 | 0 | 4 | 0.25 | 1 | 0 | 10 | 0.1 | 3 | 0.3333 | 1 | 0 | 4 | 0 | 2 | 0 | 10 | 0.1 |
| 15_1011088 | 3 | 0.3333 | 1 | 1 | 2 | 1 | 1 | 1 | 7 | 0.7143 | 2 | 0.5 | 1 | 1 | 2 | 1 | 2 | 1 | 7 | 0.8571 |
| 15_1011089 | 3 | 0.6667 | 2 | 0.5 | 4 | 0.5 | 2 | 0 | 11 | 0.4545 | 2 | 1 | 1 | 0 | 5 | 0.8 | 2 | 0 | 10 | 0.6 |
| 15_1011089 | 2 | 1 | 2 | 1 | 3 | 0.6667 | 2 | 1 | 9 | 0.8889 | 2 | 1 | 1 | 1 | 4 | 0.75 | 3 | 0.6667 | 10 | 0.8 |
| 15_1011089 | 4 | 1 | 2 | 1 | 4 | 0.75 | 2 | 1 | 12 | 0.9167 | 3 | 1 | 1 | 1 | 5 | 0.8 | 3 | 1 | 12 | 0.9167 |
| 15_1011089 | 3 | 0 | 2 | 0 | 4 | 0.75 | 2 | 0 | 11 | 0.2727 | 3 | 0.6667 | 1 | 0 | 4 | 0.25 | 2 | 0 | 10 | 0.3 |
| 15_1011089 | 4 | 0.25 | 1 | 0 | 4 | 0.25 | 1 | 1 | 10 | 0.3 | 3 | 0 | NA | NA | 5 | 0.8 | 1 | 1 | 9 | 0.5556 |
| 15_1011091 | 4 | 0.25 | 2 | 0.5 | 4 | 0.5 | NA | NA | 10 | 0.4 | 3 | 0.6667 | 1 | 0 | 5 | 0.6 | 3 | 0.6667 | 12 | 0.5833 |
| 15_1011092 | 4 | 0.5 | 2 | 0.5 | 3 | 1 | 3 | 1 | 12 | 0.75 | 3 | 0.6667 | 1 | 1 | 5 | 1 | 2 | 1 | 11 | 0.9091 |
| 15_1011092 | 4 | 0.25 | 2 | 0.5 | 1 | 1 | 3 | 0 | 10 | 0.3 | 3 | 0.3333 | 1 | 0 | 6 | 0.5 | 2 | 0.5 | 12 | 0.4167 |
| 15_1011092 | 4 | 0.25 | 2 | 0.5 | 4 | 0 | 2 | 0.5 | 12 | 0.25 | 3 | 0.6667 | 1 | 0 | 6 | 0 | 3 | 0.6667 | 13 | 0.3077 |
| 15_1011096 | 4 | 0 | 2 | 0 | 4 | 0.5 | 2 | 0 | 12 | 0.1667 | 3 | 0.3333 | 1 | 1 | 5 | 0.2 | 5 | 0 | 14 | 0.2143 |
| 15_1011097 | 5 | 0.4 | 4 | 0.25 | 3 | 0.6667 | 3 | 0.3333 | 15 | 0.4 | 4 | 0.25 | 1 | 0 | 7 | 0.4286 | 5 | 0.6 | 17 | 0.4118 |
| 15_1011103 | 9 | 0.2222 | 7 | 0.1429 | 8 | 0.125 | 6 | 0 | 30 | 0.1333 | 4 | 0.25 | 3 | 0 | 14 | 0.0714 | 7 | 0.1429 | 28 | 0.1071 |
| 15_1011104 | 8 | 0.375 | 8 | 0.625 | 8 | 0.625 | 6 | 0.6667 | 30 | 0.5667 | 5 | 0.4 | 3 | 0.6667 | 10 | 0.7 | 5 | 0.8 | 23 | 0.6522 |
| 15_1011107 | 11 | 0.2727 | 7 | 0 | 7 | 0 | 5 | 0 | 30 | 0.1 | 6 | 0.1667 | 4 | 0.25 | 13 | 0.3077 | 9 | 0.1111 | 32 | 0.2188 |
| 15_1011107 | 5 | 0 | 5 | 0.4 | 5 | 0.2 | 5 | 0.2 | 20 | 0.2 | 3 | 0 | 3 | 0 | 11 | 0.2727 | 5 | 0 | 22 | 0.1364 |
| 15_1011107 | 10 | 0.6 | 7 | 0.2857 | 8 | 0.375 | 5 | 0.2 | 30 | 0.4 | 7 | 0.4286 | 5 | 0 | 14 | 0.5714 | 6 | 0 | 32 | 0.3438 |
| 15_1011109 | 8 | 0.375 | 9 | 0.3333 | 4 | 0.25 | 6 | 0.1667 | 27 | 0.2963 | 5 | 0.4 | 4 | 0.5 | 12 | 0.4167 | 10 | 0.4 | 31 | 0.4194 |
| 15_1011110 | 12 | 0.5 | 9 | 0.6667 | 11 | 0.7273 | 8 | 0.625 | 40 | 0.625 | 8 | 0.875 | 4 | 1 | 9 | 0.6667 | 10 | 0.8 | 31 | 0.8065 |
| 15_1011111 | 6 | 0 | 6 | 0.1667 | 4 | 0.25 | 5 | 0 | 21 | 0.0952 | 7 | 0.2857 | 5 | 0.4 | 10 | 0.2 | 10 | 0.1 | 32 | 0.2188 |
| 15_1011113 | 13 | 0.1538 | 14 | 0.0714 | 11 | 0 | 8 | 0.125 | 46 | 0.087 | 13 | 0.0769 | 9 | 0.2222 | 16 | 0.125 | 14 | 0.1429 | 52 | 0.1346 |
| 15_1011114 | 12 | 0.8333 | 13 | 0.9231 | 10 | 0.6 | 9 | 0.8889 | 44 | 0.8182 | 11 | 1 | 9 | 1 | 14 | 0.9286 | 13 | 0.9231 | 47 | 0.9574 |
| 15_1011196 | 1 | 0 | 2 | 0 | 1 | 0 | 1 | 1 | 5 | 0.2 | NA | NA | 1 | 0 | 2 | 0.5 | 2 | 0.5 | 5 | 0.4 |
| 15_1011197 | 1 | 1 | 3 | 0 | 2 | 0.5 | NA | NA | 6 | 0.3333 | NA | NA | 1 | 1 | 2 | 0 | 3 | 0 | 6 | 0.1667 |
| 15_1011197 | NA | NA | 2 | 0 | 2 | 0.5 | 1 | 0 | 5 | 0.2 | NA | NA | 2 | 0.5 | 2 | 0.5 | 3 | 0.3333 | 7 | 0.4286 |
| 15_1011199 | 1 | 1 | 2 | 1 | 1 | 1 | 1 | 1 | 5 | 1 | NA | NA | 2 | 0.5 | 2 | 0.5 | 2 | 1 | 6 | 0.6667 |
| 15_1011199 | 1 | 1 | 1 | 1 | 1 | 1 | NA | NA | 3 | 1 | NA | NA | 1 | 1 | 2 | 1 | 2 | 0 | 5 | 0.6 |

| | | | | | | | | | | | | | | | | | | | | |
|---|---|---|---|---|---|---|---|---|---|---|---|---|---|---|---|---|---|---|---|---|
| 15_101119 | 1 | 1 | 2 | 1 | 1 | 1 | NA | NA | 4 | 1 | NA | NA | 2 | 1 | 2 | 1 | 2 | 1 | 6 | 1 |
| 15_101119 | 1 | 1 | 1 | 1 | NA | NA | 1 | 0 | 3 | 0.6667 | NA | NA | 2 | 0.5 | 1 | 1 | 1 | 0 | 4 | 0.5 |
| 15_101129 | 4 | 0.75 | 6 | 0.3333 | 3 | 1 | 6 | 0.3333 | 19 | 0.5263 | 6 | 0.1667 | 1 | 1 | 3 | 0.6667 | 3 | 1 | 13 | 0.5385 |
| 15_101129 | 6 | 0.8333 | 6 | 1 | 2 | 1 | 5 | 0.6 | 19 | 0.8421 | 5 | 1 | 1 | 1 | 3 | 0.6667 | 1 | 1 | 10 | 0.9 |
| 15_101129 | 4 | 0.5 | 7 | 0.4286 | 3 | 0 | 7 | 0.5714 | 21 | 0.4286 | 5 | 0.4 | 1 | 0 | 4 | 0.5 | 3 | 0.3333 | 13 | 0.3846 |
| 15_101129 | 7 | 0.1429 | 8 | 0 | 5 | 0.2 | 8 | 0 | 28 | 0.0714 | 8 | 0.25 | 1 | 0 | 4 | 0 | 4 | 0 | 17 | 0.1176 |
| 15_101130 | 8 | 0.5 | 5 | 0.2 | 6 | 0 | 9 | 0.2222 | 28 | 0.25 | 6 | 0.3333 | 1 | 0 | 8 | 0 | 2 | 0.5 | 17 | 0.1765 |
| 15_101168 | 2 | 0 | 6 | 0.3333 | 5 | 0.2 | 3 | 0 | 16 | 0.1875 | 5 | 0 | 2 | 0.5 | 3 | 0 | 2 | 0 | 12 | 0.0833 |
| 15_101168 | 2 | 1 | 6 | 0.6667 | 4 | 0.5 | 2 | 1 | 14 | 0.7143 | 4 | 0.75 | 2 | 1 | 3 | 0.3333 | 2 | 0.5 | 11 | 0.6364 |
| 15_101168 | 2 | 0 | 7 | 0.2857 | 5 | 0 | 3 | 1 | 17 | 0.2941 | 5 | 0 | 3 | 0.3333 | 3 | 0 | 2 | 0 | 13 | 0.0769 |
| 15_101168 | 2 | 0.5 | 6 | 0.6667 | 4 | 0.5 | 2 | 0.5 | 14 | 0.5714 | 4 | 0.75 | 3 | 0.3333 | 3 | 0.6667 | 3 | 0.3333 | 13 | 0.5385 |
| 15_101168 | 2 | 1 | 7 | 1 | 6 | 0.8333 | 4 | 1 | 19 | 0.9474 | 4 | 1 | 3 | 1 | 4 | 1 | 3 | 0.3333 | 14 | 0.8571 |
| 15_101168 | 2 | 0 | 7 | 0.1429 | 4 | 0 | 3 | 0 | 16 | 0.0625 | 5 | 0.2 | 4 | 0 | 3 | 0 | 3 | 0 | 15 | 0.0667 |
| 15_101172 | 5 | 0 | 11 | 0.1818 | 8 | 0 | 7 | 0 | 31 | 0.0645 | 7 | 0.1429 | 5 | 0 | 8 | 0.125 | 4 | 0 | 24 | 0.0833 |
| 15_101172 | 5 | 0.6 | 9 | 0.6667 | 5 | 0.4 | 7 | 0.5714 | 26 | 0.5769 | 8 | 0.5 | 4 | 0.5 | 9 | 0.3333 | 4 | 0.5 | 25 | 0.44 |
| 15_101172 | 5 | 0.2 | 8 | 0.375 | 7 | 0.2857 | 3 | 0 | 23 | 0.2609 | 4 | 0.5 | 5 | 0.2 | 9 | 0.1111 | 4 | 0.5 | 22 | 0.2727 |
| 15_101172 | 4 | 1 | 10 | 0.9 | 9 | 1 | 7 | 1 | 30 | 0.9667 | 7 | 0.8571 | 5 | 1 | 9 | 1 | 2 | 0.5 | 23 | 0.913 |
| 15_101172 | 6 | 0.3333 | 10 | 0.2 | 9 | 0.3333 | 7 | 0.4286 | 32 | 0.3125 | 8 | 0.5 | 5 | 0.6 | 7 | 0.1429 | 3 | 0.3333 | 23 | 0.3913 |
| 15_101172 | 6 | 0.1667 | 9 | 0.4444 | 10 | 0.2 | 6 | 0.3333 | 31 | 0.2903 | 6 | 0.5 | 5 | 0.2 | 8 | 0.25 | 4 | 0.5 | 23 | 0.3478 |
| 15_101172 | 4 | 0 | 9 | 0.1111 | 10 | 0.3 | 5 | 0.2 | 28 | 0.1786 | 9 | 0.1111 | 3 | 0 | 7 | 0.4286 | 2 | 0.5 | 21 | 0.2381 |
| 15_101172 | 6 | 0.3333 | 11 | 0.1818 | 9 | 0.2222 | 6 | 0.1667 | 32 | 0.2188 | 9 | 0.1111 | 4 | 0 | 7 | 0 | 4 | 0.25 | 24 | 0.0833 |
| 15_101184 | 4 | 0.5 | 4 | 0 | 10 | 0.2 | 4 | 0.25 | 22 | 0.2273 | 5 | 0 | 4 | 0.5 | 2 | 0.5 | 3 | 0 | 14 | 0.2143 |
| 15_101185 | 4 | 0.25 | 4 | 0 | 9 | 0.1111 | 6 | 0.1667 | 23 | 0.1304 | 4 | 0 | 3 | 0.3333 | 3 | 0.3333 | 3 | 0 | 13 | 0.1538 |
| 15_101188 | 6 | 1 | 5 | 1 | 9 | 0.7778 | 6 | 1 | 26 | 0.9231 | 5 | 1 | 4 | 0.5 | 4 | 0.75 | 4 | 0.75 | 17 | 0.7647 |
| 15_101188 | 6 | 0.6667 | 6 | 0.8333 | 10 | 0.2 | 6 | 0.3333 | 28 | 0.4643 | 4 | 0.75 | 5 | 0.2 | 3 | 0 | 5 | 0.4 | 17 | 0.3529 |
| 15_101188 | 6 | 1 | 5 | 0.4 | 10 | 0.7 | 6 | 0.5 | 27 | 0.6667 | 5 | 0.6 | 4 | 0.5 | 3 | 1 | 5 | 0.4 | 17 | 0.5882 |
| 15_101188 | 5 | 0.6 | 5 | 0 | 11 | 0.5455 | 6 | 0.5 | 27 | 0.4444 | 5 | 0.8 | 5 | 0.8 | 4 | 0.5 | 6 | 0.5 | 20 | 0.65 |
| 15_101194 | NA | NA | 1 | 1 | NA | NA | NA | NA | 1 | 1 | NA | NA | 1 | 1 | NA | NA | NA | NA | 1 | 1 |
| 15_101754 | 1 | 1 | NA | NA | 1 | 0 | NA | NA | 2 | 0.5 | NA | NA | NA | NA | 1 | 1 | 1 | 0 | 2 | 0.5 |
| 15_101754 | 1 | 0 | 2 | 0.5 | 2 | 0 | 1 | 0 | 6 | 0.1667 | 1 | 0 | 1 | 0 | 1 | 1 | 1 | 0 | 4 | 0.25 |
| 15_101754 | 1 | 1 | 2 | 1 | 2 | 0.5 | NA | NA | 5 | 0.8 | 1 | 1 | 1 | 1 | 1 | 1 | 1 | 1 | 4 | 1 |
| 15_101767 | NA | NA | NA | NA | NA | NA | 3 | 0.6667 | 3 | 0.6667 | NA | NA | 4 | 0.25 | 1 | 1 | NA | NA | 5 | 0.4 |
| 15_101767 | NA | NA | NA | NA | 1 | 1 | 2 | 1 | 3 | 1 | NA | NA | 4 | 0.75 | 1 | 1 | NA | NA | 5 | 0.8 |
| 15_101768 | NA | NA | NA | NA | NA | NA | 2 | 0.5 | 2 | 0.5 | NA | NA | 2 | 0.5 | NA | NA | NA | NA | 2 | 0.5 |
| 15_101768 | NA | NA | NA | NA | NA | NA | 3 | 0.3333 | 3 | 0.3333 | NA | NA | 3 | 0.3333 | NA | NA | NA | NA | 3 | 0.3333 |
| 15_102003 | 14 | 0 | 13 | 0.0769 | 13 | 0 | 16 | 0.125 | 56 | 0.0536 | 7 | 0.1429 | 19 | 0.1053 | 12 | 0 | 14 | 0 | 52 | 0.0577 |
| 15_102003 | 13 | 0.0769 | 10 | 0 | 12 | 0.0833 | 18 | 0.1111 | 53 | 0.0755 | 6 | 0.1667 | 18 | 0.1111 | 9 | 0 | 11 | 0 | 44 | 0.0682 |
| 15_102003 | 14 | 0.1429 | 10 | 0.1 | 12 | 0.1667 | 15 | 0.2 | 51 | 0.1569 | 7 | 0.1429 | 20 | 0.2 | 9 | 0.2222 | 12 | 0.1667 | 48 | 0.1875 |
| 15_102003 | 16 | 0.0625 | 12 | 0.0833 | 14 | 0.0714 | 17 | 0.0588 | 59 | 0.0678 | 6 | 0 | 19 | 0.1579 | 11 | 0.0909 | 15 | 0 | 51 | 0.0784 |
| 15_102003 | 15 | 0.2 | 11 | 0.1818 | 12 | 0.0833 | 16 | 0.25 | 54 | 0.1852 | 6 | 0.1667 | 18 | 0.1667 | 11 | 0 | 12 | 0.1667 | 47 | 0.1277 |
| 15_102003 | 14 | 0.0714 | 11 | 0.0909 | 8 | 0.25 | 14 | 0.0714 | 47 | 0.1064 | 6 | 0 | 15 | 0.1333 | 11 | 0.1818 | 13 | 0 | 45 | 0.0889 |
| 15_102019 | 1 | 0 | 2 | 1 | 2 | 0.5 | 3 | 1 | 8 | 0.75 | NA | NA | NA | NA | 1 | 1 | 1 | 1 | 2 | 1 |
| 15_102019 | NA | NA | 2 | 0 | 2 | 0 | 3 | 0.6667 | 7 | 0.2857 | NA | NA | NA | NA | 1 | 1 | 1 | 1 | 2 | 1 |
| 15_102020 | 1 | 1 | 2 | 1 | 2 | 1 | 3 | 1 | 8 | 1 | NA | NA | NA | NA | 1 | 1 | 1 | 1 | 2 | 1 |
| 15_102020 | 1 | 0 | 2 | 1 | 2 | 1 | 2 | 0.5 | 7 | 0.7143 | NA | NA | NA | NA | 1 | 1 | 1 | 1 | 2 | 1 |
| 15_102020 | 1 | 0 | 2 | 1 | 2 | 1 | 3 | 0.6667 | 8 | 0.75 | NA | NA | NA | NA | 1 | 1 | 1 | 1 | 2 | 1 |
| 15_102021 | 1 | 0 | 2 | 0.5 | 2 | 0.5 | 3 | 1 | 8 | 0.625 | NA | NA | 2 | 0 | 1 | 1 | 1 | 1 | 4 | 0.5 |
| 15_102021 | NA | NA | 2 | 1 | 1 | 1 | 2 | 1 | 5 | 1 | NA | NA | 2 | 0.5 | 1 | 0 | 1 | 1 | 4 | 0.5 |
| 15_102024 | 3 | 0.6667 | 4 | 0.25 | 2 | 0 | 3 | 0.3333 | 12 | 0.3333 | NA | NA | 1 | 0 | 4 | 0.25 | 1 | 0 | 6 | 0.1667 |
| 15_102025 | 15 | 0.0667 | 12 | 0 | 8 | 0.125 | 10 | 0.1 | 45 | 0.0667 | 14 | 0 | 16 | 0.0625 | 17 | 0.0588 | 9 | 0.1111 | 56 | 0.0536 |
| 15_102052 | NA | NA | NA | NA | 2 | 0.5 | NA | NA | 2 | 0.5 | NA | NA | 1 | 1 | NA | NA | 1 | 0 | 2 | 0.5 |
| 15_102085 | 2 | 0.5 | 3 | 0.6667 | 2 | 0 | 7 | 0.1429 | 14 | 0.2857 | 2 | 0.5 | 2 | 0.5 | 1 | 1 | 2 | 0 | 7 | 0.4286 |
| 15_102085 | 2 | 0 | 2 | 0 | 3 | 0.3333 | 6 | 0.3333 | 13 | 0.2308 | 5 | 0 | 3 | 0.3333 | 3 | 0 | 2 | 0.5 | 13 | 0.1538 |
| 15_102085 | 1 | 1 | 3 | 0 | 2 | 0 | 5 | 0 | 11 | 0.0909 | 6 | 0.1667 | 4 | 0 | 3 | 0 | 2 | 0 | 15 | 0.0667 |
| 15_102086 | 2 | 0 | 1 | 0 | 2 | 0.5 | 6 | 0.1667 | 11 | 0.1818 | 5 | 0 | 4 | 0 | 3 | 0.3333 | 2 | 0 | 14 | 0.0714 |
| 15_102113 | 2 | 1 | 1 | 0 | NA | NA | 3 | 0.6667 | 6 | 0.6667 | NA | NA | 1 | 1 | NA | NA | NA | NA | 1 | 1 |
| 15_102113 | 2 | 0 | 1 | 0 | NA | NA | 3 | 0.6667 | 6 | 0.3333 | NA | NA | 1 | 1 | NA | NA | NA | NA | 1 | 1 |
| 15_102113 | 2 | 1 | 1 | 0 | NA | NA | 3 | 0.3333 | 6 | 0.5 | NA | NA | 1 | 1 | NA | NA | NA | NA | 1 | 1 |
| 15_102121 | 4 | 0.5 | 1 | 1 | NA | NA | 6 | 0 | 11 | 0.2727 | 2 | 0 | 3 | 0.3333 | 1 | 0 | 3 | 0 | 9 | 0.1111 |
| 15_102121 | 4 | 0.25 | 1 | 0 | NA | NA | 6 | 0.1667 | 11 | 0.1818 | 1 | 0 | 3 | 0.3333 | 1 | 0 | 3 | 0.6667 | 8 | 0.375 |
| 15_102121 | 3 | 0.6667 | 1 | 1 | NA | NA | 7 | 0.4286 | 11 | 0.5455 | 2 | 0.5 | 3 | 0.6667 | 2 | 0.5 | 3 | 1 | 10 | 0.7 |
| 15_102121 | 3 | 0.6667 | 1 | 1 | NA | NA | 6 | 1 | 10 | 0.9 | NA | NA | 3 | 0.6667 | 2 | 1 | 3 | 1 | 8 | 0.875 |
| 15_102121 | 2 | 0 | 1 | 0 | NA | NA | 6 | 0.3333 | 9 | 0.2222 | 2 | 0 | 3 | 0 | 2 | 0.5 | 2 | 0 | 9 | 0.1111 |
| 15_102121 | 4 | 0.25 | 1 | 1 | NA | NA | 7 | 0.5714 | 12 | 0.5 | 2 | 0.5 | 3 | 0.6667 | 2 | 0.5 | 3 | 1 | 10 | 0.7 |
| 15_102121 | 3 | 1 | NA | NA | NA | NA | 6 | 1 | 9 | 1 | 1 | 1 | 3 | 1 | 2 | 1 | 1 | 1 | 7 | 1 |

| | | | | | | | | | | | | | | | | | | | | |
|---|---|---|---|---|---|---|---|---|---|---|---|---|---|---|---|---|---|---|---|---|
| 15_1021214 | 4 | 0.5 | NA | NA | NA | NA | 5 | 0.6 | 9 | 0.5556 | 1 | 1 | 3 | 0.3333 | 1 | 0 | 3 | 0 | 8 | 0.25 |
| 15_1021214 | 3 | 1 | 1 | 1 | NA | NA | 7 | 1 | 11 | 1 | 1 | 1 | 3 | 1 | 2 | 1 | 3 | 0.6667 | 9 | 0.8889 |
| 15_1021215 | 3 | 0 | NA | NA | NA | NA | 5 | 0.2 | 8 | 0.125 | 1 | 0 | 3 | 0.6667 | 2 | 0 | 2 | 1 | 8 | 0.5 |
| 15_1021215 | 4 | 0.25 | 1 | 0 | NA | NA | 4 | 0.75 | 9 | 0.4444 | 1 | 1 | 2 | 1 | 2 | 0 | 3 | 0.3333 | 8 | 0.5 |
| 15_1021216 | 3 | 1 | 2 | 1 | NA | NA | 7 | 0.8571 | 12 | 0.9167 | 1 | 1 | 3 | 1 | 2 | 1 | 4 | 0.75 | 10 | 0.9 |
| 15_1021217 | 3 | 0 | 2 | 0 | NA | NA | 6 | 0.3333 | 11 | 0.1818 | 1 | 1 | 4 | 0.25 | 2 | 0.5 | 3 | 0.6667 | 10 | 0.5 |
| 15_1021218 | 4 | 0 | 2 | 0 | NA | NA | 7 | 0.4286 | 13 | 0.2308 | 1 | 1 | 3 | 0.3333 | 3 | 0.3333 | 5 | 0.2 | 12 | 0.3333 |
| 15_1021245 | 1 | 1 | NA | NA | NA | NA | 1 | 0 | 2 | 0.5 | NA | NA | NA | NA | 1 | 1 | NA | NA | 1 | 1 |
| 15_1021885 | NA | NA | 1 | 1 | NA | NA | 2 | 0.5 | 3 | 0.6667 | NA | NA | 1 | 1 | NA | NA | NA | NA | 1 | 1 |
| 15_1021886 | 9 | 0.2222 | 4 | 0 | 5 | 0 | 5 | 0.4 | 23 | 0.1739 | 4 | 0 | 4 | 0.25 | 3 | 0.3333 | 7 | 0.1429 | 18 | 0.1667 |
| 15_1021887 | 10 | 0.9 | 4 | 0.75 | 5 | 0.8 | 4 | 1 | 23 | 0.8696 | 3 | 1 | 5 | 1 | 2 | 0.5 | 9 | 0.8889 | 19 | 0.8947 |
| 15_1021888 | 5 | 1 | 3 | 1 | 4 | 1 | 5 | 0.8 | 17 | 0.9412 | 3 | 1 | 4 | 0.75 | 3 | 1 | 8 | 1 | 18 | 0.9444 |
| 15_1021889 | 9 | 1 | 5 | 1 | 5 | 1 | 6 | 0.8333 | 25 | 0.96 | 4 | 1 | 3 | 1 | 3 | 1 | 10 | 1 | 20 | 1 |
| 15_1021890 | 9 | 1 | 4 | 1 | 3 | 0.6667 | 7 | 0.8571 | 23 | 0.913 | 3 | 1 | 3 | 1 | 3 | 1 | 10 | 0.9 | 19 | 0.9474 |
| 15_1021891 | 9 | 0 | 4 | 0.5 | 4 | 0.25 | 5 | 0 | 22 | 0.1364 | 4 | 0 | 5 | 0.4 | 3 | 0.3333 | 9 | 0.1111 | 21 | 0.1905 |
| 15_1021891 | 11 | 0.2727 | 4 | 0.25 | 5 | 0 | 6 | 0 | 26 | 0.1538 | 4 | 0.25 | 4 | 0 | 2 | 0 | 7 | 0 | 17 | 0.0588 |
| 15_1021892 | 9 | 0.1111 | 4 | 0.75 | 5 | 0.2 | 5 | 0 | 23 | 0.2174 | 4 | 0.25 | 4 | 0 | 2 | 0 | 9 | 0.4444 | 19 | 0.2632 |
| 15_1021892 | 13 | 0.3846 | 5 | 0 | 6 | 0.1667 | 6 | 0.1667 | 30 | 0.2333 | 2 | 0 | 5 | 0.4 | 3 | 0.3333 | 9 | 0.1111 | 19 | 0.2105 |
| 15_1021893 | 13 | 0.1538 | 5 | 0.2 | 4 | 0 | 6 | 0 | 28 | 0.1071 | 4 | 0 | 6 | 0.1667 | 3 | 0.3333 | 10 | 0 | 23 | 0.087 |
| 15_1021894 | 12 | 0.75 | 5 | 0.8 | 5 | 0.8 | 6 | 1 | 28 | 0.8214 | 4 | 0.75 | 4 | 0.75 | 3 | 1 | 9 | 0.8889 | 20 | 0.85 |
| 15_1021896 | 8 | 1 | 5 | 0.8 | 5 | 1 | 4 | 0.75 | 22 | 0.9091 | 4 | 0.75 | 6 | 1 | 4 | 1 | 6 | 0.8333 | 20 | 0.9 |
| 15_1021897 | 12 | 0.1667 | 4 | 0.25 | 5 | 0.2 | 6 | 0.1667 | 27 | 0.1852 | 3 | 0.6667 | 5 | 0 | 1 | 0 | 7 | 0.1429 | 16 | 0.1875 |
| 15_1021899 | 11 | 1 | 9 | 1 | 8 | 1 | 7 | 1 | 35 | 1 | 5 | 1 | 6 | 1 | 5 | 1 | 10 | 1 | 26 | 1 |
| 15_1021899 | 9 | 0.8889 | 6 | 1 | 8 | 0.875 | 5 | 1 | 28 | 0.9286 | 6 | 1 | 3 | 1 | 4 | 1 | 7 | 0.8571 | 20 | 0.95 |
| 15_1021899 | 9 | 0.7778 | 7 | 0.7143 | 5 | 0.6 | 7 | 0.4286 | 28 | 0.6429 | 6 | 0.3333 | 5 | 0.8 | 4 | 0.5 | 9 | 0.7778 | 24 | 0.625 |
| 15_1021900 | 11 | 0.3636 | 8 | 0.25 | 4 | 0.25 | 7 | 0.1429 | 30 | 0.2667 | 6 | 0.1667 | 7 | 0.1429 | 5 | 0 | 9 | 0.1111 | 27 | 0.1111 |
| 15_1021900 | 12 | 0.3333 | 9 | 0.3333 | 6 | 0.5 | 8 | 0.25 | 35 | 0.3429 | 5 | 0.4 | 6 | 0.5 | 5 | 0.6 | 11 | 0.5455 | 27 | 0.5185 |
| 15_1021900 | 15 | 0.9333 | 8 | 0.75 | 7 | 0.8571 | 8 | 0.75 | 38 | 0.8421 | 6 | 1 | 6 | 1 | 5 | 0.8 | 11 | 0.7273 | 28 | 0.8571 |
| 15_1021900 | 12 | 0.9167 | 5 | 1 | 4 | 1 | 8 | 0.625 | 29 | 0.8621 | 4 | 0.75 | 7 | 1 | 3 | 0.6667 | 9 | 0.7778 | 23 | 0.8261 |
| 15_1021901 | 12 | 0.5833 | 7 | 0.8571 | 4 | 0.75 | 6 | 0.5 | 29 | 0.6552 | 2 | 0 | 5 | 0.6 | 4 | 0.75 | 11 | 0.6364 | 22 | 0.5909 |
| 15_1022398 | 52 | 0.3269 | 35 | 0.2857 | 50 | 0.3 | 38 | 0.2105 | 175 | 0.2857 | 47 | 0.4894 | 60 | 0.3 | 38 | 0.3947 | 42 | 0.4762 | 187 | 0.4064 |
| 15_1022405 | 6 | 0.1667 | 9 | 0.4444 | 4 | 0.25 | 5 | 0 | 24 | 0.25 | 3 | 0 | 5 | 0.2 | 2 | 0 | 6 | 0.1667 | 16 | 0.125 |
| 15_1022427 | 6 | 0.3333 | 6 | 0.1667 | 5 | 0.2 | 5 | 0 | 22 | 0.1818 | 3 | 0.3333 | 4 | 0 | 2 | 0 | 6 | 0 | 15 | 0.0667 |
| 15_1022467 | 4 | 0.25 | 6 | 0.1667 | 4 | 0 | 1 | 0 | 15 | 0.1333 | 2 | 0 | 3 | 0 | 1 | 0 | 1 | 1 | 7 | 0.1429 |
| 15_1023174 | NA | NA | NA | NA | 1 | 1 | NA | NA | 1 | 1 | NA | NA | 1 | 1 | NA | NA | NA | NA | 1 | 1 |
| 15_1023394 | NA | NA | 1 | 0 | 1 | 1 | NA | NA | 2 | 0.5 | NA | NA | 2 | 0.5 | NA | NA | NA | NA | 2 | 0.5 |
| 15_1023709 | 6 | 0 | 4 | 0.25 | 3 | 0 | 7 | 0.1429 | 20 | 0.1 | 1 | 0 | 2 | 0 | 2 | 0 | 2 | 0.5 | 7 | 0.1429 |
| 15_1023710 | 5 | 0 | 5 | 0.2 | 2 | 0 | 4 | 0.25 | 16 | 0.125 | NA | NA | 3 | 0.3333 | 4 | 0.25 | 2 | 0 | 9 | 0.2222 |
| 15_1024073 | 19 | 0 | 23 | 0 | 17 | 0.0588 | 18 | 0.2222 | 77 | 0.0649 | 28 | 0.0714 | 21 | 0.0952 | 22 | 0.0909 | 22 | 0.1364 | 93 | 0.0968 |
| 15_1024074 | 2 | 0.5 | 2 | 0.5 | NA | NA | 1 | 1 | 5 | 0.6 | 1 | 1 | 1 | 0 | 3 | 0.6667 | 2 | 0.5 | 7 | 0.5714 |
| 15_1024083 | 8 | 0.5 | 17 | 0.1765 | 7 | 0.2857 | 10 | 0.3 | 42 | 0.2857 | 7 | 0.1429 | 7 | 0.2857 | 8 | 0 | 6 | 0.5 | 28 | 0.2143 |
| 15_1024085 | 13 | 0.0769 | 16 | 0.25 | 10 | 0.3 | 10 | 0.3 | 49 | 0.2245 | 9 | 0.4444 | 8 | 0.25 | 8 | 0.125 | 8 | 0.25 | 33 | 0.2727 |
| 15_1024096 | 7 | 0.5714 | 7 | 0.1429 | 1 | 0 | 9 | 0.5556 | 24 | 0.4167 | 4 | 0.5 | 4 | 0.5 | 4 | 1 | 3 | 0.3333 | 15 | 0.6 |
| 15_1024096 | 9 | 0.1111 | 11 | 0.2727 | 3 | 0.3333 | 9 | 0.1111 | 32 | 0.1875 | 6 | 0 | 6 | 0 | 5 | 0.2 | 5 | 0.2 | 22 | 0.0909 |
| 15_1024098 | 13 | 0.4615 | 11 | 0.5455 | 2 | 1 | 8 | 0.75 | 34 | 0.5882 | 6 | 0.5 | 8 | 0.625 | 5 | 0.6 | 6 | 0.6667 | 25 | 0.6 |
| 15_1024099 | 11 | 0.5455 | 11 | 0.2727 | 3 | 0 | 8 | 0.5 | 33 | 0.3939 | 7 | 0.7143 | 7 | 0.1429 | 5 | 0.8 | 5 | 0.4 | 24 | 0.5 |
| 15_1024135 | 10 | 0 | 2 | 0 | 1 | 0 | 2 | 0.5 | 15 | 0.0667 | 2 | 0 | 3 | 0.3333 | 2 | 0 | 3 | 0 | 10 | 0.1 |
| 15_1024138 | 10 | 0.3 | 3 | 0 | NA | NA | 3 | 0.3333 | 16 | 0.25 | 1 | 0 | 2 | 0 | 2 | 0 | 3 | 0.3333 | 8 | 0.125 |
| 15_1024318 | 7 | 0 | 6 | 0.1667 | 1 | 0 | 2 | 0 | 16 | 0.0625 | 1 | 0 | 4 | 0 | 3 | 0 | 2 | 0.5 | 10 | 0.1 |
| 15_1024319 | 7 | 1 | 7 | 0.5714 | NA | NA | 2 | 1 | 16 | 0.8125 | 1 | 1 | 3 | 1 | 2 | 1 | 2 | 0.5 | 8 | 0.875 |
| 15_1024319 | 8 | 0.25 | 6 | 0.1667 | NA | NA | 1 | 0 | 15 | 0.2 | NA | NA | 4 | 0.25 | 3 | 0.3333 | 2 | 0 | 9 | 0.2222 |
| 15_1024320 | 7 | 0.1429 | 4 | 0 | 1 | 0 | 2 | 0 | 14 | 0.0714 | 1 | 0 | 3 | 0.3333 | 3 | 0 | 2 | 0 | 9 | 0.1111 |
| 15_1024425 | 1 | 1 | NA | NA | NA | NA | 1 | 1 | 2 | 1 | NA | NA | NA | NA | NA | NA | 3 | 1 | 3 | 1 |
| 15_1024427 | 1 | 1 | NA | NA | NA | NA | 1 | 1 | 2 | 1 | NA | NA | NA | NA | NA | NA | 3 | 0.3333 | 3 | 0.3333 |
| 15_1024427 | 1 | 1 | NA | NA | NA | NA | 1 | 1 | 2 | 1 | NA | NA | NA | NA | NA | NA | 3 | 0.6667 | 3 | 0.6667 |
| 15_1025735 | 10 | 0.1 | 8 | 0.125 | 11 | 0.1818 | 10 | 0 | 39 | 0.1026 | 8 | 0 | 7 | 0.1429 | 11 | 0.0909 | 9 | 0 | 35 | 0.0571 |
| 15_1026152 | 10 | 0.4 | 12 | 0 | 5 | 0.2 | 6 | 0.1667 | 33 | 0.1818 | 3 | 0.3333 | 10 | 0.1 | 9 | 0.3333 | 11 | 0.2727 | 33 | 0.2424 |
| 15_1030619 | 3 | 0 | 2 | 1 | 2 | 0 | 2 | 0 | 9 | 0.2222 | 1 | 0 | 2 | 0.5 | 1 | 0 | 2 | 0 | 6 | 0.1667 |
| 15_1030640 | 1 | 1 | NA | NA | NA | NA | NA | NA | 1 | 1 | 1 | 1 | 1 | 1 | 2 | 1 | 1 | 1 | 5 | 1 |
| 15_1030640 | 1 | 0 | 3 | 0 | NA | NA | 1 | 1 | 5 | 0.2 | 1 | 0 | 1 | 0 | 2 | 0 | 1 | 1 | 5 | 0.2 |
| 15_1030641 | NA | NA | 3 | 1 | NA | NA | 1 | 1 | 4 | 1 | 1 | 0 | 1 | 1 | 2 | 0.5 | 1 | 0 | 5 | 0.4 |
| 15_1030644 | NA | NA | 1 | 0 | NA | NA | 1 | 1 | 2 | 0.5 | 1 | 1 | 1 | 1 | 3 | 0.3333 | 1 | 0 | 6 | 0.5 |
| 15_1030645 | NA | NA | 2 | 0 | NA | NA | 1 | 1 | 3 | 0.3333 | 1 | 0 | 1 | 0 | 2 | 0 | 1 | 1 | 5 | 0.2 |
| 15_1030645 | NA | NA | 1 | 1 | NA | NA | 1 | 1 | 2 | 1 | 1 | 1 | 1 | 1 | NA | NA | 1 | 1 | 3 | 1 |
| 15_1031081 | 5 | 0.4 | 6 | 0.3333 | 1 | 1 | 3 | 0 | 15 | 0.3333 | 4 | 0 | 8 | 0.375 | 4 | 0 | 4 | 0.25 | 20 | 0.2 |

| | | | | | | | | | | | | | | | | | | | | |
|---|---|---|---|---|---|---|---|---|---|---|---|---|---|---|---|---|---|---|---|---|
| 15_1031527 | 28 | 0.0714 | 39 | 0.0513 | 23 | 0.1739 | 37 | 0.1622 | 127 | 0.1102 | 10 | 0.1 | 30 | 0.1 | 29 | 0.1379 | 25 | 0.04 | 94 | 0.0957 |
| 15_1031528 | 30 | 0.6 | 44 | 0.6591 | 25 | 0.64 | 42 | 0.5476 | 141 | 0.6099 | 14 | 0.7857 | 34 | 0.6765 | 34 | 0.7353 | 30 | 0.6333 | 112 | 0.6964 |
| 15_1032075 | 15 | 0.0667 | 14 | 0.1429 | 7 | 0.1429 | 8 | 0.125 | 44 | 0.1136 | 10 | 0.3 | 12 | 0.0833 | 12 | 0 | 19 | 0.2105 | 53 | 0.1509 |
| 15_1032079 | 14 | 0.0714 | 14 | 0.2143 | 7 | 0 | 8 | 0 | 43 | 0.093 | 11 | 0.0909 | 11 | 0.1818 | 12 | 0 | 15 | 0.4667 | 49 | 0.2041 |
| 15_1032081 | 18 | 0.1111 | 17 | 0.1765 | 7 | 0 | 9 | 0.2222 | 51 | 0.1373 | 14 | 0.0714 | 14 | 0.2143 | 12 | 0.1667 | 22 | 0.1818 | 62 | 0.1613 |
| 15_1032224 | 18 | 0.2778 | 14 | 0.2143 | 19 | 0.4211 | 9 | 0.2222 | 60 | 0.3 | 11 | 0.1818 | 16 | 0.25 | 8 | 0.125 | 8 | 0.125 | 43 | 0.186 |
| 15_1032230 | 12 | 0 | 13 | 0.0769 | 16 | 0.125 | 6 | 0 | 47 | 0.0638 | 7 | 0.2857 | 9 | 0 | 5 | 0 | 6 | 0 | 27 | 0.0741 |
| 15_1032230 | 11 | 0.5455 | 9 | 0.4444 | 15 | 0.3333 | 10 | 0.7 | 45 | 0.4889 | 7 | 0.2857 | 12 | 0.6667 | 5 | 0.2 | 7 | 0.4286 | 31 | 0.4516 |
| 15_1032529 | 2 | 0.5 | 1 | 1 | 3 | 0.6667 | 1 | 1 | 7 | 0.7143 | NA | NA | NA | NA | 1 | 1 | 2 | 0.5 | 3 | 0.6667 |
| 15_1032530 | 2 | 0.5 | 2 | 1 | 2 | 1 | 1 | 1 | 7 | 0.8571 | NA | NA | NA | NA | 1 | 1 | 1 | 1 | 2 | 1 |
| 15_1032532 | 2 | 0 | 2 | 0.5 | 1 | 0 | 1 | 0 | 6 | 0.1667 | NA | NA | NA | NA | 1 | 1 | 2 | 0 | 3 | 0.3333 |
| 15_1032533 | 1 | 1 | 2 | 0.5 | 1 | 1 | NA | NA | 4 | 0.75 | NA | NA | NA | NA | 1 | 1 | 1 | 1 | 2 | 1 |
| 15_1032534 | 2 | 1 | 2 | 1 | 2 | 0.5 | 1 | 1 | 7 | 0.8571 | NA | NA | NA | NA | 1 | 0 | 1 | 1 | 2 | 0.5 |
| 15_1032534 | 2 | 1 | 1 | 1 | 1 | 1 | 1 | 1 | 5 | 1 | NA | NA | NA | NA | 1 | 0 | 2 | 0.5 | 3 | 0.3333 |
| 15_1032535 | 2 | 1 | 1 | 1 | 2 | 0.5 | NA | NA | 5 | 0.8 | NA | NA | NA | NA | NA | NA | 2 | 0.5 | 2 | 0.5 |
| 15_1032535 | 2 | 1 | 2 | 1 | 2 | 0.5 | 1 | 0 | 7 | 0.7143 | NA | NA | NA | NA | 1 | 0 | 2 | 1 | 3 | 0.6667 |
| 15_1032536 | 2 | 0.5 | 2 | 0.5 | 2 | 0.5 | 1 | 1 | 7 | 0.5714 | NA | NA | NA | NA | NA | NA | 2 | 0.5 | 2 | 0.5 |
| 15_1034061 | 1 | 1 | 1 | 0 | NA | NA | 2 | 0.5 | 4 | 0.5 | 3 | 0 | 4 | 0.75 | NA | NA | 2 | 0.5 | 9 | 0.4444 |
| 15_1034374 | 50 | 0.1 | 24 | 0.0833 | 37 | 0.1351 | 47 | 0.1277 | 158 | 0.1139 | 14 | 0.0714 | 62 | 0.0645 | 46 | 0.1739 | 42 | 0.1429 | 164 | 0.1159 |
| 15_1034388 | 3 | 0.3333 | 3 | 0 | 7 | 0.4286 | 9 | 0.1111 | 22 | 0.2273 | 2 | 0.5 | 4 | 0 | 4 | 0 | 2 | 0 | 12 | 0.0833 |
| 15_1034393 | 27 | 0.037 | 22 | 0.0909 | 27 | 0.1852 | 21 | 0.2381 | 97 | 0.134 | 14 | 0.0714 | 34 | 0.0294 | 35 | 0.1143 | 28 | 0.1071 | 111 | 0.0811 |
| 16_3562306 | NA | NA | 3 | 1 | NA | NA | NA | NA | 3 | 1 | 2 | 0.5 | 1 | 1 | NA | NA | NA | NA | 3 | 0.6667 |
| 16_3562610 | NA | NA | 3 | 0.3333 | NA | NA | 1 | 0 | 4 | 0.25 | 1 | 1 | NA | NA | NA | NA | NA | NA | 1 | 1 |
| 16_3566718 | NA | NA | 4 | 0.75 | 1 | 0 | 1 | 1 | 6 | 0.6667 | 2 | 1 | NA | NA | NA | NA | NA | NA | 2 | 1 |
| 16_3566741 | NA | NA | 3 | 0 | 1 | 1 | 1 | 0 | 5 | 0.2 | 3 | 0.3333 | 1 | 0 | NA | NA | NA | NA | 4 | 0.25 |
| 16_3566784 | NA | NA | 3 | 0.6667 | 1 | 1 | 1 | 1 | 5 | 0.8 | 3 | 1 | NA | NA | NA | NA | NA | NA | 3 | 1 |
| 16_3566843 | NA | NA | 4 | 1 | 1 | 1 | 1 | 1 | 6 | 1 | 3 | 1 | NA | NA | NA | NA | NA | NA | 3 | 1 |
| 16_3566937 | NA | NA | 4 | 0.25 | 1 | 1 | NA | NA | 5 | 0.4 | 2 | 0.5 | NA | NA | NA | NA | NA | NA | 2 | 0.5 |
| 16_3567133 | NA | NA | 4 | 0.5 | NA | NA | 2 | 1 | 6 | 0.6667 | 3 | 0.6667 | NA | NA | NA | NA | NA | NA | 3 | 0.6667 |
| 16_3567301 | NA | NA | 4 | 0.75 | 1 | 1 | 3 | 1 | 8 | 0.875 | 3 | 1 | NA | NA | NA | NA | NA | NA | 3 | 1 |
| 16_3567318 | NA | NA | 4 | 1 | 1 | 1 | 3 | 1 | 8 | 1 | 3 | 1 | NA | NA | NA | NA | NA | NA | 3 | 1 |
| 16_3567356 | NA | NA | 4 | 0.25 | NA | NA | 3 | 0.3333 | 7 | 0.2857 | 2 | 0.5 | NA | NA | NA | NA | NA | NA | 2 | 0.5 |
| 16_3567396 | NA | NA | 5 | 0.8 | 1 | 1 | 3 | 0.3333 | 9 | 0.6667 | 3 | 1 | NA | NA | NA | NA | NA | NA | 3 | 1 |
| 16_3567648 | NA | NA | 3 | 1 | 1 | 1 | 3 | 0.3333 | 7 | 0.7143 | 3 | 0.3333 | NA | NA | NA | NA | NA | NA | 3 | 0.3333 |
| 16_3567715 | NA | NA | 4 | 1 | 1 | 1 | 3 | 1 | 8 | 1 | 2 | 1 | NA | NA | NA | NA | NA | NA | 2 | 1 |
| 16_3567845 | NA | NA | 5 | 0.2 | NA | NA | 2 | 1 | 7 | 0.4286 | 2 | 1 | NA | NA | NA | NA | NA | NA | 2 | 1 |
| 16_3568374 | 1 | 0 | 5 | 0.2 | 1 | 0 | 2 | 0 | 9 | 0.1111 | 3 | 0.3333 | 1 | 0 | 1 | 0 | 2 | 0 | 7 | 0.1429 |
| 16_3568474 | 1 | 1 | 4 | 0.25 | NA | NA | 2 | 0.5 | 7 | 0.4286 | 2 | 0.5 | 1 | 0 | 1 | 1 | 1 | 0 | 5 | 0.4 |
| 16_3665521 | 2 | 0.5 | NA | NA | NA | NA | 1 | 0 | 3 | 0.3333 | 1 | 0 | NA | NA | 1 | 1 | NA | NA | 2 | 0.5 |
| 16_3665556 | 2 | 0 | NA | NA | NA | NA | 1 | 1 | 3 | 0.3333 | 1 | 0 | NA | NA | 1 | 1 | NA | NA | 2 | 0.5 |
| 16_3666091 | 2 | 0.5 | NA | NA | NA | NA | 1 | 0 | 3 | 0.3333 | 1 | 0 | NA | NA | 1 | 1 | NA | NA | 2 | 0.5 |
| 16_3667318 | 1 | 1 | 1 | 0 | NA | NA | 2 | 0.5 | 4 | 0.5 | 1 | 1 | NA | NA | 1 | 0 | NA | NA | 2 | 0.5 |
| 16_3672203 | 1 | 0 | 1 | 1 | NA | NA | 1 | 0 | 3 | 0.3333 | 1 | 1 | NA | NA | 1 | 1 | NA | NA | 2 | 1 |
| 16_3672672 | NA | NA | 1 | 1 | NA | NA | 2 | 0.5 | 3 | 0.6667 | 1 | 1 | NA | NA | 1 | 1 | NA | NA | 2 | 1 |
| 16_3672766 | 3 | 0 | NA | NA | NA | NA | 2 | 1 | 5 | 0.4 | 1 | 1 | NA | NA | 1 | 1 | NA | NA | 2 | 1 |
| 16_3672914 | 2 | 0.5 | 1 | 1 | NA | NA | 1 | 1 | 4 | 0.75 | 1 | 1 | NA | NA | 1 | 1 | NA | NA | 2 | 1 |
| 16_3672952 | 2 | 1 | 1 | 1 | NA | NA | 2 | 1 | 5 | 1 | 1 | 1 | NA | NA | 1 | 1 | NA | NA | 2 | 1 |
| 16_3674322 | 1 | 0 | NA | NA | NA | NA | 2 | 0.5 | 3 | 0.3333 | NA | NA | NA | NA | 1 | 1 | NA | NA | 1 | 1 |
| 16_3680713 | 1 | 0 | NA | NA | NA | NA | 3 | 0.3333 | 4 | 0.25 | NA | NA | 2 | 0 | NA | NA | 2 | 0.5 | 4 | 0.25 |
| 16_3680789 | 1 | 1 | NA | NA | 1 | 0 | 4 | 0 | 6 | 0.1667 | NA | NA | 1 | 0 | NA | NA | 2 | 0.5 | 3 | 0.3333 |
| 16_3680801 | 2 | 1 | 1 | 0 | 1 | 0 | 5 | 0.8 | 9 | 0.6667 | NA | NA | 3 | 1 | NA | NA | 2 | 1 | 5 | 1 |
| 16_3683114 | 1 | 1 | 1 | 1 | 1 | 0 | 5 | 0.8 | 8 | 0.75 | NA | NA | 3 | 0.6667 | NA | NA | 1 | 0 | 4 | 0.5 |
| 16_3683260 | 1 | 0 | 1 | 0 | 1 | 0 | 4 | 0.5 | 7 | 0.2857 | NA | NA | 3 | 0.3333 | NA | NA | 1 | 1 | 4 | 0.5 |
| 16_3683641 | NA | NA | NA | NA | NA | NA | 5 | 1 | 5 | 1 | NA | NA | 2 | 1 | NA | NA | 1 | 1 | 3 | 1 |
| 16_3683725 | 1 | 1 | 1 | 1 | 1 | 1 | 4 | 0.75 | 7 | 0.8571 | NA | NA | 3 | 1 | NA | NA | 1 | 1 | 4 | 1 |
| 16_3683730 | NA | NA | 1 | 0 | 1 | 1 | 4 | 0.75 | 6 | 0.6667 | NA | NA | 2 | 1 | NA | NA | NA | NA | 2 | 1 |
| 16_3683778 | NA | NA | 1 | 1 | 1 | 1 | 5 | 0.8 | 7 | 0.8571 | NA | NA | 3 | 1 | NA | NA | 1 | 1 | 4 | 1 |
| 16_3683799 | 1 | 0 | 1 | 1 | NA | NA | 2 | 1 | 4 | 0.75 | NA | NA | 1 | 1 | NA | NA | 1 | 1 | 2 | 1 |
| 16_3683808 | 1 | 1 | 1 | 1 | 1 | 1 | 5 | 0.8 | 8 | 0.875 | NA | NA | 2 | 0.5 | NA | NA | 1 | 1 | 3 | 0.6667 |
| 16_3683892 | NA | NA | 1 | 0 | 1 | 1 | 5 | 0.2 | 7 | 0.2857 | NA | NA | 2 | 0 | NA | NA | 1 | 1 | 3 | 0.3333 |
| 16_3683987 | 1 | 0 | 1 | 0 | 1 | 1 | 5 | 0 | 8 | 0.125 | NA | NA | 3 | 0.3333 | NA | NA | 1 | 0 | 4 | 0.25 |
| 16_3684044 | 1 | 1 | NA | NA | 1 | 1 | 4 | 1 | 6 | 1 | NA | NA | 3 | 0.6667 | NA | NA | 1 | 1 | 4 | 0.75 |
| 16_3684157 | NA | NA | 1 | 1 | 1 | 1 | 4 | 0.5 | 6 | 0.6667 | NA | NA | 2 | 0.5 | NA | NA | 1 | 1 | 3 | 0.6667 |
| 16_3684279 | 1 | 1 | 1 | 1 | 1 | 1 | 4 | 0.5 | 7 | 0.7143 | NA | NA | 1 | 1 | NA | NA | 1 | 1 | 2 | 1 |
| 16_3684387 | 1 | 1 | NA | NA | 1 | 0 | 5 | 0.6 | 7 | 0.5714 | NA | NA | 2 | 0.5 | NA | NA | 1 | 0 | 3 | 0.3333 |

| | | | | | | | | | | | | | | | | | | | | |
|---|---|---|---|---|---|---|---|---|---|---|---|---|---|---|---|---|---|---|---|---|
| 16_3684427 | 1 | 0 | 1 | 0 | 1 | 1 | 5 | 0.2 | 8 | 0.25 | NA | NA | 3 | 1 | NA | NA | 1 | 0 | 4 | 0.75 |
| 16_3684516 | 1 | 1 | 1 | 1 | 1 | 1 | 4 | 0.5 | 7 | 0.7143 | NA | NA | 3 | 1 | NA | NA | 1 | 1 | 4 | 1 |
| 16_3718630 | 13 | 0.3846 | 13 | 0.1538 | 11 | 0.4545 | 7 | 0.4286 | 44 | 0.3409 | 8 | 0.25 | 10 | 0.5 | 15 | 0.4667 | 11 | 0.1818 | 44 | 0.3636 |
| 16_3719853 | 15 | 0.1333 | 17 | 0 | 9 | 0.1111 | 9 | 0 | 50 | 0.06 | 8 | 0.125 | 12 | 0.0833 | 17 | 0.0588 | 11 | 0 | 48 | 0.0625 |
| 16_3719874 | 14 | 0 | 17 | 0.1176 | 9 | 0.3333 | 7 | 0.1429 | 47 | 0.1277 | 7 | 0.2857 | 13 | 0.1538 | 16 | 0.375 | 9 | 0.1111 | 45 | 0.2444 |
| 16_3721472 | 16 | 0.8125 | 19 | 0.7368 | 13 | 0.6154 | 9 | 1 | 57 | 0.7719 | 10 | 0.9 | 11 | 0.8182 | 16 | 0.8125 | 13 | 0.9231 | 50 | 0.86 |
| 16_3721495 | 14 | 0.7143 | 20 | 0.8 | 14 | 0.7143 | 9 | 0.7778 | 57 | 0.7544 | 9 | 0.6667 | 11 | 0.8182 | 18 | 0.8889 | 11 | 0.7273 | 49 | 0.7959 |
| 16_3721694 | 9 | 0.4444 | 19 | 0.5789 | 10 | 0.4 | 7 | 0.1429 | 45 | 0.4444 | 7 | 0.2857 | 8 | 0.375 | 14 | 0.3571 | 10 | 0.3 | 39 | 0.3333 |
| 16_3721706 | 16 | 0.9375 | 22 | 0.7727 | 14 | 0.7857 | 11 | 0.9091 | 63 | 0.8413 | 11 | 0.7273 | 12 | 0.75 | 18 | 0.9444 | 13 | 0.6154 | 54 | 0.7778 |
| 16_3721753 | 18 | 0.1111 | 22 | 0.0909 | 14 | 0.2143 | 8 | 0.125 | 62 | 0.129 | 10 | 0.2 | 12 | 0.0833 | 15 | 0.2 | 11 | 0.1818 | 48 | 0.1667 |
| 16_3721817 | 16 | 0.5 | 18 | 0.4444 | 14 | 0.6429 | 12 | 0.5833 | 60 | 0.5333 | 11 | 0.3636 | 12 | 0.3333 | 17 | 0.2941 | 12 | 0.5 | 52 | 0.3654 |
| 16_3721939 | 14 | 0.7143 | 20 | 0.65 | 13 | 0.3077 | 9 | 0.4444 | 56 | 0.5536 | 7 | 0.1429 | 8 | 0.375 | 16 | 0.875 | 11 | 0.1818 | 42 | 0.4762 |
| 16_3721948 | 17 | 0.1176 | 20 | 0.25 | 13 | 0.4615 | 11 | 0.1818 | 61 | 0.2459 | 11 | 0.1818 | 11 | 0.2727 | 18 | 0.3889 | 12 | 0.1667 | 52 | 0.2692 |
| 16_3726975 | 1 | 1 | 4 | 0.75 | 3 | 0.6667 | 4 | 0.25 | 12 | 0.5833 | 3 | 0 | 2 | 0 | 2 | 0.5 | 1 | 0 | 8 | 0.125 |
| 16_3758768 | 3 | 0 | 8 | 0.375 | 5 | 0.2 | 7 | 0.1429 | 23 | 0.2174 | 5 | 0.2 | 3 | 0 | 2 | 0 | 3 | 0 | 13 | 0.0769 |
| 16_3758774 | 3 | 0.3333 | 8 | 0.375 | 4 | 0 | 5 | 0.4 | 20 | 0.3 | 5 | 0 | 1 | 1 | 2 | 0.5 | 3 | 0 | 11 | 0.1818 |
| 16_3758836 | 4 | 1 | 10 | 0.7 | 5 | 0.6 | 7 | 0.4286 | 26 | 0.6538 | 5 | 0.4 | 4 | 1 | 2 | 0.5 | 3 | 0.3333 | 14 | 0.5714 |
| 16_3758997 | 5 | 0.2 | 5 | 0 | 4 | 0.5 | 6 | 0 | 20 | 0.15 | 4 | 0.25 | 3 | 0.3333 | 2 | 0 | 3 | 0.3333 | 12 | 0.25 |
| 16_3797021 | NA | NA | 1 | 0 | 3 | 0.3333 | 1 | 1 | 5 | 0.4 | 1 | 0 | 3 | 0 | 4 | 0.5 | 7 | 0.1429 | 15 | 0.2 |
| 16_3797133 | NA | NA | 2 | 1 | 3 | 0.6667 | 1 | 1 | 6 | 0.8333 | 1 | 1 | 4 | 0.5 | 5 | 0.8 | 7 | 1 | 17 | 0.8235 |
| 16_3797349 | NA | NA | 1 | 1 | 3 | 0.3333 | NA | NA | 4 | 0.5 | 1 | 0 | 4 | 1 | 4 | 0.25 | 7 | 0.5714 | 16 | 0.5625 |
| 16_3797533 | NA | NA | 2 | 1 | 3 | 0.6667 | 1 | 1 | 6 | 0.8333 | 1 | 1 | 5 | 0.8 | 5 | 1 | 7 | 1 | 18 | 0.9444 |
| 16_3797578 | NA | NA | 2 | 1 | 2 | 1 | NA | NA | 4 | 1 | 1 | 1 | 5 | 0.6 | 4 | 1 | 8 | 0.75 | 18 | 0.7778 |
| 16_3797584 | NA | NA | 2 | 1 | 2 | 0.5 | 1 | 1 | 5 | 0.8 | 1 | 1 | 5 | 1 | 4 | 0.75 | 8 | 1 | 18 | 0.9444 |
| 16_3797654 | NA | NA | 2 | 0.5 | 3 | 0 | 1 | 0 | 6 | 0.1667 | 1 | 0 | 5 | 0.2 | 5 | 0.4 | 7 | 0.4286 | 18 | 0.3333 |
| 16_3797739 | NA | NA | 2 | 0 | 3 | 0.3333 | 1 | 0 | 6 | 0.1667 | 1 | 0 | 4 | 0 | 4 | 0.25 | 8 | 0 | 17 | 0.0588 |
| 16_3797794 | NA | NA | 2 | 0.5 | 3 | 0 | 1 | 1 | 6 | 0.3333 | 1 | 0 | 5 | 0.6 | 3 | 0.3333 | 8 | 0.375 | 17 | 0.4118 |
| 16_3802885 | NA | NA | NA | NA | 2 | 0.5 | 1 | 0 | 3 | 0.3333 | 1 | 0 | 4 | 0 | 3 | 0 | 7 | 0.2857 | 15 | 0.1333 |
| 16_3802931 | NA | NA | NA | NA | 2 | 1 | NA | NA | 2 | 1 | 1 | 1 | 3 | 0.6667 | 2 | 1 | 5 | 0.4 | 11 | 0.6364 |
| 16_3803045 | NA | NA | NA | NA | 2 | 1 | 1 | 0 | 3 | 0.6667 | 1 | 0 | 4 | 0.75 | 4 | 1 | 8 | 0.875 | 17 | 0.8235 |
| 16_3803181 | NA | NA | NA | NA | 2 | 0 | 1 | 1 | 3 | 0.3333 | 1 | 0 | 5 | 0 | 3 | 0 | 8 | 0.125 | 17 | 0.0588 |
| 16_3803259 | NA | NA | NA | NA | 2 | 1 | 1 | 1 | 3 | 1 | 1 | 1 | 3 | 0.6667 | 3 | 1 | 7 | 0.7143 | 14 | 0.7857 |
| 16_3803381 | NA | NA | NA | NA | 2 | 1 | 1 | 1 | 3 | 1 | NA | NA | 5 | 0.8 | 4 | 1 | 5 | 0.6 | 14 | 0.7857 |
| 16_3803548 | NA | NA | NA | NA | 2 | 0.5 | 1 | 0 | 3 | 0.3333 | 1 | 0 | 4 | 0.5 | 4 | 0.5 | 6 | 0.6667 | 15 | 0.5333 |
| 16_3803664 | NA | NA | NA | NA | 1 | 1 | 1 | 1 | 2 | 1 | 1 | 0 | 3 | 0.3333 | 4 | 0.5 | 4 | 0.5 | 12 | 0.4167 |
| 16_3803834 | NA | NA | NA | NA | 1 | 0 | 1 | 1 | 2 | 0.5 | 1 | 0 | 3 | 0 | 3 | 1 | 4 | 0.5 | 11 | 0.4545 |
| 16_3803931 | NA | NA | NA | NA | 1 | 0 | 1 | 1 | 2 | 0.5 | NA | NA | 2 | 1 | 2 | 1 | 3 | 0.6667 | 7 | 0.8571 |
| 16_3804404 | NA | NA | NA | NA | 1 | 1 | 1 | 1 | 2 | 1 | NA | NA | 1 | 1 | 2 | 1 | 1 | 1 | 4 | 1 |
| 16_3804599 | NA | NA | NA | NA | NA | NA | 1 | 1 | 1 | 1 | 1 | 1 | NA | NA | 4 | 0.75 | 2 | 0 | 7 | 0.5714 |
| 16_3804677 | NA | NA | NA | NA | 1 | 0 | 1 | 1 | 2 | 0.5 | 1 | 1 | 1 | 0 | 4 | 0.25 | 2 | 0.5 | 8 | 0.375 |
| 16_3804951 | NA | NA | NA | NA | 1 | 1 | 1 | 1 | 2 | 1 | 1 | 1 | 1 | 1 | 4 | 0.75 | 2 | 1 | 8 | 0.875 |
| 16_3857902 | 5 | 0.4 | 10 | 0.1 | 4 | 0.25 | 3 | 0.3333 | 22 | 0.2273 | 2 | 0.5 | 9 | 0 | 4 | 0 | 4 | 0 | 19 | 0.0526 |
| 16_3857915 | 9 | 0.2222 | 13 | 0.4615 | 8 | 0.625 | 7 | 0.4286 | 37 | 0.4324 | 4 | 1 | 14 | 0.3571 | 6 | 0.5 | 6 | 1 | 30 | 0.6 |
| 16_3857934 | 3 | 0 | 8 | 0 | 7 | 0 | 4 | 0.5 | 22 | 0.0909 | 2 | 0.5 | 9 | 0 | 3 | 0 | 3 | 0.3333 | 17 | 0.1176 |
| 16_3857987 | 6 | 0.1667 | 9 | 0.1111 | 3 | 0 | 4 | 0 | 22 | 0.0909 | 2 | 0 | 7 | 0 | 3 | 0 | 4 | 0.25 | 16 | 0.0625 |
| 16_3899432 | 7 | 1 | 5 | 1 | 1 | 1 | 2 | 1 | 15 | 1 | 3 | 0.6667 | 4 | 1 | 3 | 1 | 5 | 0.8 | 15 | 0.8667 |
| 16_3899484 | 7 | 0.8571 | 5 | 0.4 | 1 | 1 | 2 | 1 | 15 | 0.7333 | 3 | 0.6667 | 5 | 0.8 | 3 | 0.6667 | 6 | 1 | 17 | 0.8235 |
| 16_3899568 | 6 | 0.5 | 4 | 0 | 1 | 0 | 2 | 0 | 13 | 0.2308 | 3 | 0 | 5 | 0.4 | 1 | 0 | 4 | 0.25 | 13 | 0.2308 |
| 16_3900735 | 4 | 0.25 | 4 | 0.5 | 3 | 0.3333 | 2 | 0 | 13 | 0.3077 | 4 | 0 | 5 | 0.2 | 4 | 0.25 | 4 | 0.25 | 17 | 0.1765 |
| 16_3900855 | 8 | 0.125 | 9 | 0 | 6 | 0.1667 | 6 | 0 | 29 | 0.069 | 6 | 0.1667 | 9 | 0.1111 | 7 | 0.1429 | 7 | 0.2857 | 29 | 0.1724 |
| 16_3901085 | 9 | 0.2222 | 8 | 0.125 | 8 | 0 | 5 | 0.2 | 30 | 0.1333 | 7 | 0.2857 | 9 | 0.1111 | 6 | 0 | 6 | 0.1667 | 28 | 0.1429 |
| 16_3901196 | 6 | 0 | 7 | 0.1429 | 8 | 0.25 | 4 | 0 | 25 | 0.12 | 7 | 0.2857 | 10 | 0.1 | 8 | 0.125 | 6 | 0.5 | 31 | 0.2258 |
| 16_3901241 | 7 | 0.8571 | 6 | 0.8333 | 4 | 1 | 1 | 1 | 18 | 0.8889 | 6 | 0.6667 | 8 | 0.875 | 4 | 0.5 | 5 | 1 | 23 | 0.7826 |
| 16_3901320 | 8 | 0.5 | 8 | 0.375 | 6 | 0.6667 | 4 | 0.25 | 26 | 0.4615 | 7 | 0.4286 | 8 | 0.375 | 6 | 0.1667 | 5 | 0.4 | 26 | 0.3462 |
| 16_3901367 | 8 | 0.375 | 8 | 0 | 6 | 0 | 6 | 0.1667 | 28 | 0.1429 | 7 | 0.1429 | 8 | 0.25 | 8 | 0 | 6 | 0.1667 | 29 | 0.1379 |
| 16_3901434 | 8 | 0.125 | 7 | 0.1429 | 6 | 0 | 3 | 0 | 24 | 0.0833 | 4 | 0 | 8 | 0.125 | 8 | 0.125 | 4 | 0 | 24 | 0.0833 |
| 16_3901457 | 6 | 0.1667 | 6 | 0.1667 | 4 | 0.5 | 5 | 0.4 | 21 | 0.2857 | 5 | 0.4 | 8 | 0.5 | 6 | 0.1667 | 4 | 1 | 23 | 0.4783 |
| 16_3901623 | 8 | 0.125 | 6 | 0 | 5 | 0.2 | 5 | 0.2 | 24 | 0.125 | 6 | 0.5 | 8 | 0.125 | 8 | 0.25 | 4 | 0.25 | 26 | 0.2692 |
| 16_3901651 | 7 | 0.4286 | 6 | 0.5 | 5 | 0.2 | 4 | 0.5 | 22 | 0.4091 | 6 | 0.6667 | 8 | 0.375 | 7 | 0.2857 | 5 | 0.6 | 26 | 0.4615 |
| 16_3901706 | 5 | 0.6 | 5 | 0.2 | 5 | 0.2 | 3 | 1 | 18 | 0.4444 | 6 | 0.5 | 8 | 0.5 | 4 | 0.25 | 5 | 0.8 | 23 | 0.5217 |
| 16_3901972 | 3 | 1 | 4 | 0.5 | 1 | 1 | 2 | 1 | 10 | 0.8 | 3 | 1 | 5 | 1 | 3 | 1 | 5 | 1 | 16 | 1 |
| 16_3902029 | 3 | 0 | 5 | 0.2 | 3 | 0 | 2 | 0 | 13 | 0.0769 | 2 | 1 | 2 | 0.5 | 4 | 0.25 | 3 | 0.6667 | 11 | 0.5455 |
| 16_3902047 | 4 | 1 | 5 | 0.8 | 3 | 1 | 3 | 1 | 15 | 0.9333 | 6 | 1 | 7 | 0.8571 | 5 | 0.6 | 5 | 1 | 23 | 0.8696 |
| 16_3902083 | 2 | 1 | 3 | 0.6667 | 3 | 1 | 1 | 1 | 9 | 0.8889 | 3 | 1 | 3 | 1 | 3 | 0.6667 | 5 | 1 | 14 | 0.9286 |
| 16_3902103 | 3 | 0.3333 | 5 | 0.4 | 3 | 0.6667 | 3 | 1 | 14 | 0.5714 | 4 | 0.75 | 6 | 1 | 4 | 1 | 5 | 0.6 | 19 | 0.8421 |

| | | | | | | | | | | | | | | | | | | | | |
|---|---|---|---|---|---|---|---|---|---|---|---|---|---|---|---|---|---|---|---|---|
| 16_3902272 | 2 | 0.5 | 4 | 0.5 | 1 | 0 | 3 | 1 | 10 | 0.6 | 4 | 0.75 | 4 | 1 | 3 | 0.6667 | 4 | 1 | 15 | 0.8667 |
| 16_3902434 | 3 | 1 | 4 | 1 | 2 | 1 | NA | NA | 9 | 1 | 5 | 1 | 4 | 1 | 4 | 1 | 4 | 1 | 17 | 1 |
| 16_3902530 | 3 | 1 | 4 | 0.5 | 3 | 0.6667 | 3 | 0.3333 | 13 | 0.6154 | 6 | 0.8333 | 5 | 0.8 | 4 | 1 | 5 | 0.4 | 20 | 0.75 |
| 16_3902557 | 2 | 0.5 | 4 | 1 | 3 | 1 | 2 | 1 | 11 | 0.9091 | 6 | 0.6667 | 5 | 0.8 | 4 | 1 | 5 | 1 | 20 | 0.85 |
| 16_3902684 | 2 | 1 | 2 | 0 | 2 | 0 | 2 | 0 | 8 | 0.25 | 4 | 0 | 3 | 0 | 4 | 0 | 4 | 0.25 | 15 | 0.0667 |
| 16_3902688 | 3 | 0 | 2 | 0 | 3 | 0 | 2 | 0.5 | 10 | 0.1 | 5 | 0 | 3 | 0 | 4 | 0 | 5 | 0.2 | 17 | 0.0588 |
| 16_3902704 | 2 | 0 | 3 | 0.3333 | 3 | 0.3333 | 3 | 1 | 11 | 0.4545 | 6 | 0.6667 | 3 | 0.3333 | 3 | 0.6667 | 5 | 0.2 | 17 | 0.4706 |
| 16_3902711 | 3 | 0 | 3 | 0.6667 | 3 | 0.3333 | 3 | 0 | 12 | 0.25 | 5 | 0.2 | 3 | 0 | 4 | 0.25 | 3 | 0.3333 | 15 | 0.2 |
| 16_3902731 | 3 | 0.3333 | 3 | 0.3333 | 2 | 0.5 | 2 | 0 | 10 | 0.3 | 6 | 0.8333 | 3 | 0.6667 | 3 | 0.6667 | 5 | 0.8 | 17 | 0.7647 |
| 16_3902813 | 3 | 0.3333 | 2 | 0 | 3 | 0 | 2 | 0 | 10 | 0.1 | 6 | 0.1667 | 3 | 0 | 4 | 0 | 5 | 0.2 | 18 | 0.1111 |
| 16_3902875 | 3 | 1 | 2 | 1 | 3 | 1 | 2 | 1 | 10 | 1 | 5 | 1 | 3 | 1 | 3 | 1 | 5 | 1 | 16 | 1 |
| 16_3903018 | 3 | 0.3333 | 2 | 0.5 | 2 | 1 | 2 | 0.5 | 9 | 0.5556 | 5 | 0.6 | 3 | 0.3333 | 3 | 0.6667 | 4 | 0.75 | 15 | 0.6 |
| 16_3903249 | 3 | 1 | 2 | 0 | 2 | 0.5 | 2 | 1 | 9 | 0.6667 | 4 | 1 | 1 | 0 | 2 | 1 | 4 | 1 | 11 | 0.9091 |
| 16_3903456 | 3 | 1 | 2 | 1 | 2 | 1 | 3 | 1 | 10 | 1 | 3 | 1 | 2 | 1 | 3 | 1 | 3 | 1 | 11 | 1 |
| 16_3903561 | 3 | 1 | 2 | 0 | 2 | 1 | 2 | 1 | 9 | 0.7778 | 2 | 0.5 | 2 | 0.5 | 4 | 0.75 | 3 | 1 | 11 | 0.7273 |
| 16_3903617 | 2 | 0.5 | 1 | 0 | 1 | 0 | 2 | 0 | 6 | 0.1667 | NA | NA | 1 | 0 | 2 | 0.5 | 1 | 0 | 4 | 0.25 |
| 16_3903801 | 3 | 0.3333 | 2 | 0.5 | 1 | 0 | 2 | 0 | 8 | 0.25 | 1 | 0 | 2 | 0 | 3 | 0.3333 | 3 | 0.6667 | 9 | 0.3333 |
| 16_3904061 | 3 | 0.3333 | 2 | 1 | 2 | 0.5 | 2 | 0 | 9 | 0.4444 | 1 | 1 | 1 | 0 | 3 | 0 | 3 | 0 | 8 | 0.125 |
| 16_4103372 | 9 | 0.4444 | 4 | 0.5 | 13 | 0.3077 | 8 | 0.375 | 34 | 0.3824 | 4 | 0 | 10 | 0.3 | 11 | 0.6364 | 5 | 0 | 30 | 0.3333 |
| 16_4103519 | 8 | 0 | 5 | 0.4 | 15 | 0.0667 | 7 | 0.1429 | 35 | 0.1143 | 7 | 0.1429 | 9 | 0.1111 | 9 | 0.3333 | 6 | 0.5 | 31 | 0.2581 |
| 16_4104067 | 8 | 0.25 | 3 | 0 | 10 | 0.2 | 8 | 0.5 | 29 | 0.2759 | 7 | 0 | 8 | 0.375 | 10 | 0.3 | 5 | 0 | 30 | 0.2 |
| 16_4104086 | 8 | 0.25 | 3 | 0 | 10 | 0 | 7 | 0 | 28 | 0.0714 | 6 | 0 | 7 | 0.1429 | 7 | 0.1429 | 4 | 0.25 | 24 | 0.125 |
| 16_4104126 | 8 | 0.125 | 3 | 0 | 9 | 0.2222 | 8 | 0 | 28 | 0.1071 | 7 | 0.1429 | 7 | 0.1429 | 8 | 0 | 5 | 0 | 27 | 0.0741 |
| 16_4104186 | 7 | 0.4286 | 3 | 0.3333 | 11 | 0.3636 | 7 | 0.4286 | 28 | 0.3929 | 6 | 0.1667 | 5 | 0.2 | 10 | 0.5 | 5 | 0.8 | 26 | 0.4231 |
| 16_4104520 | 5 | 0.4 | 2 | 1 | 6 | 0.3333 | 6 | 0.1667 | 19 | 0.3684 | 5 | 1 | 4 | 0.25 | 7 | 0.4286 | 3 | 0.3333 | 19 | 0.5263 |
| 16_4104555 | 4 | 0.25 | 1 | 0 | 9 | 0.4444 | 4 | 0.25 | 18 | 0.3333 | 5 | 0 | 5 | 0.4 | 8 | 0.375 | 3 | 0.6667 | 21 | 0.3333 |
| 16_4104604 | 5 | 0.8 | 2 | 1 | 9 | 1 | 6 | 1 | 22 | 0.9545 | 6 | 1 | 4 | 1 | 8 | 0.875 | 3 | 1 | 21 | 0.9524 |
| 16_4104769 | 5 | 0.4 | 1 | 0 | 9 | 0.2222 | 5 | 0.2 | 20 | 0.25 | 6 | 0.5 | 4 | 0.5 | 8 | 0.625 | 3 | 0.6667 | 21 | 0.5714 |
| 16_4104851 | 3 | 0 | 1 | 0 | 9 | 0 | 5 | 0.2 | 18 | 0.0556 | 3 | 0.3333 | 3 | 0 | 8 | 0 | 3 | 0 | 17 | 0.0588 |
| 16_4105419 | 3 | 1 | 1 | 1 | 5 | 0.8 | 5 | 1 | 14 | 0.9286 | 2 | 0.5 | 2 | 0.5 | 4 | 0.5 | 1 | 1 | 9 | 0.5556 |
| 16_4105795 | 3 | 1 | 1 | 1 | 4 | 1 | 5 | 0.4 | 13 | 0.7692 | 2 | 1 | 2 | 1 | 5 | 0.8 | 1 | 1 | 10 | 0.9 |
| 16_4105832 | 1 | 1 | 1 | 1 | 3 | 0.6667 | 4 | 1 | 9 | 0.8889 | 2 | 1 | 3 | 1 | 6 | 1 | NA | NA | 11 | 1 |
| 16_4105958 | 4 | 1 | 1 | 1 | 5 | 0.8 | 4 | 0.5 | 14 | 0.7857 | 2 | 1 | 2 | 0.5 | 5 | 0.8 | 1 | 1 | 10 | 0.8 |
| 16_4106042 | 2 | 0 | 1 | 1 | 5 | 0 | 4 | 0 | 12 | 0.0833 | 2 | 1 | 2 | 0 | 5 | 0.2 | 1 | 0 | 10 | 0.3 |
| 16_4106084 | 4 | 1 | 1 | 1 | 5 | 0.8 | 5 | 1 | 15 | 0.9333 | 2 | 1 | 2 | 1 | 6 | 1 | 1 | 1 | 11 | 1 |
| 16_4106142 | 4 | 0.75 | 1 | 0 | 5 | 0.6 | 5 | 1 | 15 | 0.7333 | NA | NA | 2 | 1 | 5 | 1 | NA | NA | 7 | 1 |
| 16_4106153 | 3 | 0.3333 | NA | NA | 3 | 0.3333 | 4 | 0.25 | 10 | 0.3 | 2 | 0 | 2 | 0.5 | 3 | 0 | NA | NA | 7 | 0.1429 |
| 16_4106159 | 3 | 0.6667 | 1 | 1 | 4 | 0.75 | 4 | 0.75 | 12 | 0.75 | 1 | 0 | 2 | 0 | 5 | 0.6 | NA | NA | 8 | 0.375 |
| 16_4106231 | 3 | 1 | 2 | 1 | 4 | 0.25 | 5 | 0.2 | 14 | 0.5 | 2 | 0.5 | 1 | 1 | 4 | 0 | NA | NA | 7 | 0.2857 |
| 16_4106258 | 4 | 0 | 2 | 0.5 | 4 | 0 | 4 | 0 | 14 | 0.0714 | 2 | 0 | 1 | 0 | 5 | 0.2 | NA | NA | 8 | 0.125 |
| 16_4106314 | 3 | 0.3333 | 2 | 0.5 | 4 | 0 | 5 | 0.6 | 14 | 0.3571 | NA | NA | NA | NA | 4 | 0.5 | NA | NA | 4 | 0.5 |
| 16_4106525 | 3 | 1 | 1 | 1 | 5 | 0.8 | 4 | 1 | 13 | 0.9231 | 2 | 1 | 1 | 1 | 4 | 1 | NA | NA | 7 | 1 |
| 16_4106603 | 2 | 1 | 1 | 1 | 1 | 1 | 3 | 0.6667 | 7 | 0.8571 | NA | NA | NA | NA | 2 | 1 | NA | NA | 2 | 1 |
| 16_4106679 | 3 | 0.3333 | 1 | 0 | 5 | 0 | 4 | 0 | 13 | 0.0769 | 1 | 0 | 1 | 0 | 3 | 0.3333 | NA | NA | 5 | 0.2 |
| 16_4106747 | 3 | 0 | 1 | 0 | 5 | 0.2 | 4 | 0 | 13 | 0.0769 | 1 | 0 | 1 | 0 | 3 | 0.3333 | NA | NA | 5 | 0.2 |
| 16_4235866 | 3 | 1 | NA | NA | 2 | 1 | 2 | 1 | 7 | 1 | 1 | 1 | 1 | 1 | NA | NA | NA | NA | 2 | 1 |
| 16_4235930 | 2 | 1 | NA | NA | 2 | 1 | 2 | 0.5 | 6 | 0.8333 | 1 | 1 | 1 | 1 | NA | NA | NA | NA | 2 | 1 |
| 16_4235956 | 2 | 1 | NA | NA | 2 | 0.5 | 2 | 0.5 | 6 | 0.6667 | 1 | 1 | 1 | 1 | NA | NA | NA | NA | 2 | 1 |
| 16_4236082 | 3 | 1 | NA | NA | 2 | 1 | 2 | 1 | 7 | 1 | 1 | 1 | 1 | 1 | NA | NA | NA | NA | 2 | 1 |
| 16_4236214 | 1 | 0 | NA | NA | 2 | 0.5 | NA | NA | 3 | 0.3333 | NA | NA | 1 | 1 | NA | NA | NA | NA | 1 | 1 |
| 16_4236468 | 3 | 1 | NA | NA | 1 | 1 | 1 | 1 | 5 | 1 | 1 | 1 | 1 | 1 | NA | NA | NA | NA | 2 | 1 |
| 16_4236610 | 2 | 0.5 | NA | NA | 2 | 1 | 1 | 1 | 5 | 0.8 | 1 | 1 | 1 | 0 | NA | NA | NA | NA | 2 | 0.5 |
| 16_4236766 | 2 | 1 | NA | NA | 2 | 1 | 1 | 1 | 5 | 1 | 1 | 1 | 1 | 1 | NA | NA | NA | NA | 2 | 1 |
| 16_4236824 | 2 | 0.5 | NA | NA | 2 | 0.5 | NA | NA | 4 | 0.5 | 1 | 1 | 1 | 1 | NA | NA | NA | NA | 2 | 1 |
| 16_4236887 | 1 | 0 | NA | NA | 2 | 0.5 | 1 | 0 | 4 | 0.25 | 1 | 1 | 1 | 1 | NA | NA | NA | NA | 2 | 1 |
| 16_4237234 | 1 | 1 | NA | NA | 2 | 1 | NA | NA | 3 | 1 | 1 | 1 | NA | NA | NA | NA | NA | NA | 1 | 1 |
| 16_4298212 | 1 | 1 | 1 | 0 | NA | NA | 1 | 0 | 3 | 0.3333 | NA | NA | 1 | 1 | NA | NA | 1 | 0 | 2 | 0.5 |
| 16_4300551 | NA | NA | NA | NA | NA | NA | 1 | 1 | 1 | 1 | NA | NA | 1 | 1 | NA | NA | NA | NA | 1 | 1 |
| 16_4300650 | NA | NA | NA | NA | NA | NA | 1 | 1 | 1 | 1 | NA | NA | 1 | 1 | NA | NA | NA | NA | 1 | 1 |
| 16_4300664 | NA | NA | NA | NA | NA | NA | 1 | 1 | 1 | 1 | NA | NA | 1 | 1 | NA | NA | NA | NA | 1 | 1 |
| 16_4300692 | NA | NA | NA | NA | NA | NA | 1 | 1 | 1 | 1 | NA | NA | 1 | 1 | NA | NA | NA | NA | 1 | 1 |
| 16_4300869 | NA | NA | NA | NA | NA | NA | 1 | 1 | 1 | 1 | NA | NA | 1 | 1 | NA | NA | NA | NA | 1 | 1 |
| 16_4443212 | NA | NA | NA | NA | 1 | 0 | 1 | 1 | 2 | 0.5 | 1 | 0 | 1 | 1 | 1 | 0 | NA | NA | 3 | 0.3333 |
| 16_4443236 | NA | NA | NA | NA | 1 | 1 | NA | NA | 1 | 1 | 1 | 1 | 1 | 0 | 1 | 0 | NA | NA | 3 | 0.3333 |
| 16_4443511 | NA | NA | NA | NA | 1 | 1 | 2 | 1 | 3 | 1 | 1 | 0 | 1 | 1 | 1 | 1 | NA | NA | 3 | 0.6667 |

| | | | | | | | | | | | | | | | | | | | | |
|---|---|---|---|---|---|---|---|---|---|---|---|---|---|---|---|---|---|---|---|---|
| 16_4443557 | NA | NA | NA | NA | 1 | 1 | 2 | 1 | 3 | 1 | 1 | 1 | 1 | 1 | NA | NA | NA | NA | 2 | 1 |
| 16_4443615 | NA | NA | NA | NA | 1 | 1 | 2 | 0.5 | 3 | 0.6667 | 1 | 1 | 1 | 1 | 1 | 0 | NA | NA | 3 | 0.6667 |
| 16_4443730 | NA | NA | NA | NA | 1 | 1 | 2 | 1 | 3 | 1 | 1 | 1 | 1 | 1 | 1 | 1 | NA | NA | 3 | 1 |
| 16_4443806 | NA | NA | NA | NA | 1 | 1 | 2 | 0 | 3 | 0.3333 | 1 | 1 | NA | NA | 1 | 0 | NA | NA | 2 | 0.5 |
| 16_4445162 | 9 | 0.6667 | 5 | 0.4 | 6 | 1 | 8 | 0.75 | 28 | 0.7143 | 7 | 0.8571 | 9 | 0.7778 | 6 | 1 | 7 | 0.8571 | 29 | 0.8621 |
| 16_4445267 | 7 | 0 | 5 | 0.2 | 4 | 0.5 | 6 | 0.1667 | 22 | 0.1818 | 5 | 0.2 | 9 | 0.1111 | 7 | 0.1429 | 5 | 0 | 26 | 0.1154 |
| 16_4466160 | NA | NA | NA | NA | NA | NA | 1 | 1 | 1 | 1 | 1 | 1 | NA | NA | NA | NA | 1 | 1 | 2 | 1 |
| 16_4466302 | 1 | 1 | NA | NA | NA | NA | 1 | 1 | 2 | 1 | 1 | 1 | NA | NA | NA | NA | 1 | 1 | 2 | 1 |
| 16_4466374 | 1 | 1 | NA | NA | NA | NA | 1 | 1 | 2 | 1 | 1 | 1 | NA | NA | NA | NA | 1 | 1 | 2 | 1 |
| 16_4466379 | 1 | 0 | NA | NA | NA | NA | 1 | 1 | 2 | 0.5 | 1 | 1 | NA | NA | NA | NA | 1 | 0 | 2 | 0.5 |
| 16_4467004 | 1 | 0 | NA | NA | NA | NA | 1 | 1 | 2 | 0.5 | 1 | 1 | NA | NA | NA | NA | 2 | 1 | 3 | 1 |
| 16_4467280 | 1 | 1 | NA | NA | NA | NA | 1 | 1 | 2 | 1 | NA | NA | NA | NA | 1 | 1 | 1 | 1 | 2 | 1 |
| 16_4467424 | 1 | 0 | NA | NA | 1 | 1 | 1 | 1 | 3 | 0.6667 | 1 | 1 | NA | NA | 1 | 1 | 1 | 0 | 3 | 0.6667 |
| 16_4467684 | 1 | 1 | NA | NA | 1 | 0 | 1 | 1 | 3 | 0.6667 | 1 | 1 | 1 | 1 | 1 | 1 | 1 | 1 | 4 | 1 |
| 16_4467886 | 1 | 1 | NA | NA | 1 | 1 | 1 | 1 | 3 | 1 | 1 | 1 | 1 | 1 | 1 | 1 | 2 | 1 | 5 | 1 |
| 16_4467909 | 1 | 1 | NA | NA | 1 | 0 | 1 | 0 | 3 | 0.3333 | 1 | 1 | 1 | 1 | NA | NA | 2 | 0.5 | 4 | 0.75 |
| 16_4468077 | NA | NA | NA | NA | 1 | 1 | 1 | 0 | 2 | 0.5 | 1 | 0 | 1 | 1 | 1 | 1 | NA | NA | 3 | 0.6667 |
| 16_4468237 | 2 | 1 | NA | NA | NA | NA | 1 | 1 | 3 | 1 | 1 | 1 | 1 | 1 | 1 | 1 | 1 | 1 | 4 | 1 |
| 16_4468585 | 2 | 0.5 | NA | NA | 1 | 1 | 1 | 0 | 4 | 0.5 | 1 | 0 | 1 | 0 | 1 | 1 | 2 | 0.5 | 5 | 0.4 |
| 16_4468602 | 1 | 0 | NA | NA | 1 | 1 | 1 | 1 | 3 | 0.6667 | 1 | 0 | 1 | 0 | 1 | 1 | 2 | 0 | 5 | 0.2 |
| 16_4524616 | 7 | 0 | 5 | 0 | 9 | 0.2222 | 5 | 0 | 26 | 0.0769 | 6 | 0 | 9 | 0.1111 | 10 | 0.1 | 6 | 0.1667 | 31 | 0.0968 |
| 16_4524635 | 6 | 0.3333 | 7 | 0 | 7 | 0.4286 | 6 | 0.3333 | 26 | 0.2692 | 6 | 0.3333 | 7 | 0.1429 | 11 | 0.5455 | 6 | 0.3333 | 30 | 0.3667 |
| 16_4524712 | 7 | 1 | 7 | 1 | 7 | 0.8571 | 7 | 1 | 28 | 0.9643 | 7 | 1 | 10 | 1 | 11 | 0.9091 | 7 | 1 | 35 | 0.9714 |
| 16_4525088 | 13 | 0.1538 | 9 | 0.2222 | 8 | 0.125 | 9 | 0.1111 | 39 | 0.1538 | 7 | 0.2857 | 11 | 0.0909 | 11 | 0.1818 | 11 | 0.0909 | 40 | 0.15 |
| 16_4529304 | 4 | 0.25 | 3 | 1 | 5 | 0.6 | 3 | 1 | 15 | 0.6667 | 2 | 1 | 4 | 0.5 | 7 | 1 | 2 | 1 | 15 | 0.8667 |
| 16_4529376 | 3 | 0 | 3 | 0.3333 | 4 | 0 | 4 | 0.25 | 14 | 0.1429 | 2 | 0 | 3 | 0 | 9 | 0.1111 | 1 | 0 | 15 | 0.0667 |
| 16_4536933 | 1 | 0 | 1 | 0 | 3 | 0.3333 | 3 | 0 | 8 | 0.125 | 2 | 0 | 3 | 0 | 3 | 0.3333 | 2 | 0 | 10 | 0.1 |
| 16_4573877 | NA | NA | 1 | 1 | NA | NA | NA | NA | 1 | 1 | NA | NA | 1 | 1 | NA | NA | NA | NA | 1 | 1 |
| 16_4573894 | NA | NA | 1 | 1 | NA | NA | NA | NA | 1 | 1 | NA | NA | 1 | 1 | NA | NA | NA | NA | 1 | 1 |
| 16_4574180 | NA | NA | 1 | 1 | NA | NA | NA | NA | 1 | 1 | 1 | 0 | 1 | 1 | NA | NA | NA | NA | 2 | 0.5 |
| 16_4574325 | NA | NA | 1 | 1 | NA | NA | NA | NA | 1 | 1 | NA | NA | 1 | 1 | NA | NA | NA | NA | 1 | 1 |
| 16_4574476 | NA | NA | 1 | 1 | NA | NA | NA | NA | 1 | 1 | NA | NA | 1 | 1 | NA | NA | NA | NA | 1 | 1 |
| 16_4580291 | 1 | 0 | 3 | 0.6667 | 1 | 1 | 2 | 0 | 7 | 0.4286 | 3 | 0.6667 | NA | NA | 3 | 0.6667 | 1 | 1 | 7 | 0.7143 |
| 16_4580307 | 1 | 0 | 3 | 0.3333 | 1 | 0 | 2 | 0.5 | 7 | 0.2857 | 3 | 0.6667 | 1 | 1 | 3 | 0.6667 | 1 | 0 | 8 | 0.625 |
| 16_4582600 | 12 | 0.6667 | 20 | 0.6 | 15 | 0.4667 | 15 | 0.4 | 62 | 0.5323 | 17 | 0.6471 | 15 | 0.6 | 16 | 0.5 | 9 | 0.7778 | 57 | 0.614 |
| 16_4582624 | 9 | 0.1111 | 19 | 0.0526 | 12 | 0.0833 | 13 | 0.0769 | 53 | 0.0755 | 12 | 0.0833 | 12 | 0.1667 | 15 | 0.0667 | 9 | 0.1111 | 48 | 0.1042 |
| 16_4582744 | 11 | 0.7273 | 19 | 0.3158 | 15 | 0.7333 | 15 | 0.3333 | 60 | 0.5 | 13 | 0.6154 | 14 | 0.3571 | 13 | 0.6154 | 9 | 0.5556 | 49 | 0.5306 |
| 16_4582815 | 13 | 0.1538 | 21 | 0.2857 | 16 | 0.3125 | 17 | 0.2353 | 67 | 0.2537 | 15 | 0.6667 | 21 | 0.3333 | 15 | 0.5333 | 11 | 0.3636 | 62 | 0.4677 |
| 16_4582843 | 12 | 0.25 | 19 | 0.4737 | 16 | 0.625 | 15 | 0.4 | 62 | 0.4516 | 14 | 0.6429 | 20 | 0.45 | 13 | 0.3846 | 10 | 0.8 | 57 | 0.5439 |
| 16_4583435 | 2 | 0 | 2 | 0.5 | NA | NA | 2 | 0 | 6 | 0.1667 | 2 | 0.5 | 1 | 1 | 3 | 0.3333 | 1 | 0 | 7 | 0.4286 |
| 16_4583448 | 1 | 0 | 2 | 0.5 | 1 | 0 | 2 | 0.5 | 6 | 0.3333 | 2 | 0.5 | 1 | 1 | 2 | 0.5 | 1 | 0 | 6 | 0.5 |
| 16_4583519 | 2 | 0 | 2 | 0 | 1 | 1 | 2 | 0 | 7 | 0.1429 | 2 | 0.5 | 2 | 0.5 | 3 | 0.3333 | 1 | 0 | 8 | 0.375 |
| 16_4583945 | 11 | 0.4545 | 19 | 0.1053 | 20 | 0.15 | 20 | 0.1 | 70 | 0.1714 | 13 | 0.2308 | 22 | 0.0909 | 16 | 0.125 | 11 | 0.2727 | 62 | 0.1613 |
| 16_4583973 | 13 | 0.0769 | 23 | 0.087 | 22 | 0.1364 | 18 | 0 | 76 | 0.0789 | 14 | 0.0714 | 22 | 0.0455 | 18 | 0 | 12 | 0.25 | 66 | 0.0758 |
| 16_4584515 | 1 | 0 | 1 | 0 | 3 | 0.3333 | 3 | 0 | 8 | 0.125 | 2 | 0.5 | 2 | 0 | 1 | 1 | 1 | 0 | 6 | 0.3333 |
| 16_4584748 | 5 | 0 | 10 | 0.4 | 7 | 0.4286 | 7 | 0.2857 | 29 | 0.3103 | 4 | 0.25 | 12 | 0.25 | 7 | 0.1429 | 6 | 0 | 29 | 0.1724 |
| 16_4584814 | 6 | 0.1667 | 8 | 0.25 | 7 | 0.1429 | 5 | 0 | 26 | 0.1538 | NA | NA | 10 | 0 | 4 | 0.25 | 3 | 0 | 17 | 0.0588 |
| 16_4584886 | 3 | 0 | 7 | 0.1429 | 9 | 0.1111 | 8 | 0.125 | 27 | 0.1111 | 2 | 0.5 | 8 | 0.125 | 7 | 0.1429 | 5 | 0.2 | 22 | 0.1818 |
| 16_4585643 | 29 | 0.2759 | 34 | 0.4706 | 30 | 0.3667 | 26 | 0.0769 | 119 | 0.3109 | 26 | 0.2692 | 39 | 0.2821 | 31 | 0.2581 | 25 | 0.4 | 121 | 0.2975 |
| 16_4585735 | 31 | 0.0645 | 38 | 0.1316 | 31 | 0.0968 | 27 | 0.1481 | 127 | 0.1102 | 29 | 0.069 | 41 | 0.0976 | 38 | 0.0526 | 23 | 0.087 | 131 | 0.0763 |
| 16_4585843 | 26 | 0.0769 | 34 | 0.0294 | 26 | 0.1538 | 26 | 0.0769 | 112 | 0.0804 | 26 | 0.1154 | 40 | 0.075 | 28 | 0 | 23 | 0.1304 | 117 | 0.0769 |
| 16_4586156 | 25 | 0.32 | 25 | 0.52 | 23 | 0.5217 | 20 | 0.25 | 93 | 0.4086 | 20 | 0.55 | 37 | 0.4865 | 26 | 0.3077 | 15 | 0.6667 | 98 | 0.4796 |
| 16_4586386 | 28 | 0.8214 | 31 | 0.871 | 22 | 0.6818 | 19 | 0.6842 | 100 | 0.78 | 23 | 0.6522 | 33 | 0.7576 | 25 | 0.88 | 16 | 0.875 | 97 | 0.7835 |
| 16_4586392 | 29 | 0.069 | 32 | 0.0938 | 25 | 0.04 | 22 | 0.0455 | 108 | 0.0648 | 25 | 0.08 | 40 | 0.05 | 33 | 0.0606 | 18 | 0 | 116 | 0.0517 |
| 16_4586642 | 28 | 0.1429 | 35 | 0.1429 | 22 | 0.0455 | 20 | 0.15 | 105 | 0.1238 | 20 | 0.15 | 36 | 0.1944 | 27 | 0.1111 | 18 | 0.1667 | 101 | 0.1584 |
| 16_4692768 | 16 | 0.3125 | 34 | 0.3529 | 22 | 0.5 | 15 | 0.4667 | 87 | 0.4023 | 23 | 0.3043 | 39 | 0.4359 | 31 | 0.3871 | 29 | 0.4483 | 122 | 0.4016 |
| 16_4692897 | 16 | 0.625 | 30 | 0.4333 | 16 | 0.625 | 17 | 0.6471 | 79 | 0.557 | 22 | 0.6818 | 34 | 0.7647 | 26 | 0.6923 | 25 | 0.56 | 107 | 0.6822 |
| 16_4693255 | 11 | 0.6364 | 21 | 0.7143 | 17 | 0.7647 | 15 | 0.8 | 64 | 0.7344 | 19 | 0.8947 | 27 | 0.8148 | 19 | 0.7368 | 20 | 0.8 | 85 | 0.8118 |
| 16_4693260 | 13 | 0.3077 | 29 | 0.3448 | 21 | 0.4286 | 16 | 0.5625 | 79 | 0.4051 | 18 | 0.5 | 35 | 0.4571 | 26 | 0.5769 | 27 | 0.5185 | 106 | 0.5094 |
| 16_4693351 | 9 | 0 | 22 | 0.0909 | 19 | 0.1053 | 15 | 0 | 65 | 0.0615 | 18 | 0.0556 | 28 | 0.1429 | 21 | 0.0952 | 24 | 0.0417 | 91 | 0.0879 |
| 16_4693456 | 10 | 0.1 | 24 | 0.0833 | 21 | 0.1905 | 14 | 0.2143 | 69 | 0.1449 | 18 | 0.0556 | 29 | 0.1034 | 28 | 0.2143 | 21 | 0.2857 | 96 | 0.1667 |
| 16_4693484 | 10 | 0.2 | 25 | 0.24 | 19 | 0.3158 | 15 | 0.2667 | 69 | 0.2609 | 19 | 0.3158 | 30 | 0.3667 | 26 | 0.3077 | 22 | 0.2273 | 97 | 0.3093 |
| 16_4697910 | 3 | 0 | 2 | 0 | 1 | 0 | 2 | 0.5 | 8 | 0.125 | 1 | 1 | 4 | 0 | 3 | 0.3333 | 4 | 0.25 | 12 | 0.25 |
| 16_4754938 | 17 | 0.1176 | 20 | 0.1 | 13 | 0 | 15 | 0.2 | 65 | 0.1077 | 12 | 0.1667 | 23 | 0.087 | 24 | 0.0833 | 10 | 0 | 69 | 0.087 |
| 16_4755010 | 14 | 0.2857 | 19 | 0.5263 | 12 | 0.5833 | 20 | 0.5 | 65 | 0.4769 | 15 | 0.7333 | 21 | 0.4762 | 23 | 0.6957 | 10 | 0.7 | 69 | 0.6377 |

| | | | | | | | | | | | | | | | | | | | | |
|---|---|---|---|---|---|---|---|---|---|---|---|---|---|---|---|---|---|---|---|---|
| 16_4755098 | 16 | 0.5 | 20 | 0.4 | 11 | 0.2727 | 21 | 0.5714 | 68 | 0.4559 | 13 | 0.4615 | 19 | 0.6842 | 24 | 0.5833 | 11 | 0.3636 | 67 | 0.5522 |
| 16_4755135 | 17 | 0.8235 | 21 | 0.9048 | 13 | 0.8462 | 21 | 0.8095 | 72 | 0.8472 | 14 | 0.8571 | 22 | 0.9091 | 20 | 0.85 | 11 | 1 | 67 | 0.8955 |
| 16_4755437 | 19 | 0.0526 | 26 | 0.1923 | 14 | 0 | 25 | 0.08 | 84 | 0.0952 | 16 | 0 | 28 | 0.3929 | 24 | 0.125 | 11 | 0.1818 | 79 | 0.2025 |
| 16_4757489 | 4 | 0 | 10 | 0.1 | 10 | 0 | 12 | 0.0833 | 36 | 0.0556 | 7 | 0 | 9 | 0 | 14 | 0.0714 | 8 | 0.125 | 38 | 0.0526 |
| 16_4758389 | 10 | 0 | 10 | 0 | 14 | 0.0714 | 12 | 0.1667 | 46 | 0.0652 | 8 | 0.125 | 11 | 0.0909 | 13 | 0.0769 | 10 | 0 | 42 | 0.0714 |
| 16_4758399 | 9 | 0.1111 | 11 | 0.0909 | 15 | 0.0667 | 13 | 0 | 48 | 0.0625 | 8 | 0.375 | 11 | 0 | 10 | 0.2 | 12 | 0 | 41 | 0.122 |
| 16_4758445 | 9 | 0.2222 | 11 | 0.1818 | 17 | 0.0588 | 16 | 0.125 | 53 | 0.1321 | 9 | 0.1111 | 12 | 0 | 15 | 0.1333 | 13 | 0.0769 | 49 | 0.0816 |
| 16_4758625 | 6 | 1 | 5 | 1 | 12 | 1 | 13 | 0.9231 | 36 | 0.9722 | 3 | 1 | 11 | 0.7273 | 10 | 1 | 12 | 1 | 36 | 0.9167 |
| 16_4758771 | 10 | 0.4 | 11 | 0.4545 | 18 | 0.2778 | 13 | 0.3846 | 52 | 0.3654 | 6 | 0.3333 | 12 | 0.4167 | 15 | 0.2667 | 13 | 0.5385 | 46 | 0.3913 |
| 16_4759531 | 8 | 0.25 | 5 | 0.4 | 6 | 0.1667 | 3 | 0.3333 | 22 | 0.2727 | 7 | 0.1429 | 1 | 0 | 6 | 0 | 1 | 1 | 15 | 0.1333 |
| 16_4784293 | NA | NA | NA | NA | 1 | 1 | NA | NA | 1 | 1 | 1 | 1 | NA | NA | NA | NA | NA | NA | 1 | 1 |
| 16_4786641 | NA | NA | NA | NA | 2 | 1 | NA | NA | 2 | 1 | 1 | 1 | NA | NA | NA | NA | NA | NA | 1 | 1 |
| 16_4795570 | NA | NA | NA | NA | 2 | 0.5 | NA | NA | 2 | 0.5 | 1 | 1 | NA | NA | NA | NA | NA | NA | 1 | 1 |
| 16_4826811 | 18 | 0.7222 | 31 | 0.7742 | 24 | 0.625 | 19 | 0.7368 | 92 | 0.7174 | 18 | 0.8333 | 12 | 0.75 | 16 | 0.875 | 25 | 0.76 | 71 | 0.8028 |
| 16_4826830 | 22 | 0.0455 | 26 | 0.1154 | 21 | 0.0952 | 16 | 0 | 85 | 0.0706 | 23 | 0.0435 | 13 | 0.1538 | 8 | 0 | 22 | 0.0455 | 66 | 0.0606 |
| 16_4832171 | 18 | 0.5556 | 24 | 0.4583 | 14 | 0.2857 | 14 | 0.4286 | 70 | 0.4429 | 15 | 0.4 | 10 | 0 | 13 | 0.3846 | 16 | 0.375 | 54 | 0.3148 |
| 16_4832186 | 14 | 0.1429 | 23 | 0.0435 | 12 | 0.4167 | 8 | 0.25 | 57 | 0.1754 | 8 | 0.125 | 10 | 0.2 | 11 | 0.0909 | 14 | 0.2143 | 43 | 0.1628 |
| 16_4832251 | 18 | 0.8333 | 28 | 0.6429 | 18 | 0.6111 | 13 | 0.6923 | 77 | 0.6883 | 12 | 0.9167 | 13 | 0.4615 | 14 | 0.5 | 19 | 0.5789 | 58 | 0.6034 |
| 16_4832280 | 15 | 0.0667 | 18 | 0.0556 | 9 | 0 | 8 | 0.125 | 50 | 0.06 | 7 | 0.1429 | 6 | 0 | 9 | 0 | 12 | 0.0833 | 34 | 0.0588 |
| 16_4832395 | 16 | 0.875 | 23 | 0.8261 | 14 | 0.7857 | 12 | 0.8333 | 65 | 0.8308 | 13 | 0.8462 | 13 | 0.6154 | 13 | 0.4615 | 17 | 0.7059 | 56 | 0.6607 |
| 16_4832437 | 18 | 0.1111 | 24 | 0.1667 | 19 | 0.0526 | 16 | 0.125 | 77 | 0.1169 | 15 | 0.2 | 13 | 0.0769 | 13 | 0 | 19 | 0.1053 | 60 | 0.1 |
| 16_4832780 | 14 | 0 | 16 | 0 | 16 | 0.125 | 14 | 0.0714 | 60 | 0.05 | 8 | 0 | 9 | 0 | 13 | 0.1538 | 13 | 0.0769 | 43 | 0.0698 |
| 16_4832798 | 15 | 0.0667 | 21 | 0.0952 | 20 | 0.05 | 14 | 0.1429 | 70 | 0.0857 | 10 | 0 | 13 | 0.0769 | 13 | 0.0769 | 14 | 0.1429 | 50 | 0.08 |
| 16_4832809 | 12 | 0.6667 | 22 | 0.8636 | 12 | 0.6667 | 13 | 0.6154 | 59 | 0.7288 | 11 | 0.7273 | 11 | 0.6364 | 11 | 0.7273 | 11 | 0.7273 | 44 | 0.7045 |
| 16_4832849 | 15 | 0 | 22 | 0.2727 | 16 | 0.25 | 11 | 0.2727 | 64 | 0.2031 | 13 | 0.2308 | 11 | 0.3636 | 14 | 0.0714 | 12 | 0.3333 | 50 | 0.24 |
| 16_4832908 | 10 | 0.3 | 18 | 0.2222 | 16 | 0.375 | 8 | 0 | 52 | 0.25 | 10 | 0.2 | 12 | 0.3333 | 10 | 0.2 | 15 | 0.3333 | 47 | 0.2766 |
| 16_4832957 | 12 | 1 | 15 | 0.9333 | 9 | 1 | 8 | 1 | 44 | 0.9773 | 8 | 1 | 9 | 0.8889 | 7 | 1 | 11 | 0.9091 | 35 | 0.9429 |
| 16_4832963 | 15 | 0.6 | 18 | 0.8333 | 14 | 0.6429 | 10 | 0.9 | 57 | 0.7368 | 9 | 0.7778 | 10 | 0.4 | 12 | 0.5 | 12 | 0.5 | 43 | 0.5349 |
| 16_4833007 | 12 | 0.6667 | 19 | 0.6842 | 15 | 0.6 | 10 | 0.8 | 56 | 0.6786 | 9 | 0.8889 | 10 | 0.7 | 13 | 0.6154 | 15 | 0.6667 | 47 | 0.7021 |
| 16_4833016 | 15 | 0.6 | 24 | 0.6667 | 18 | 0.7222 | 11 | 0.3636 | 68 | 0.6176 | 12 | 0.6667 | 13 | 0.6154 | 13 | 0.6923 | 16 | 0.5 | 54 | 0.6111 |
| 16_4833064 | 14 | 0.7143 | 21 | 0.4286 | 12 | 0.6667 | 9 | 0.3333 | 56 | 0.5357 | 9 | 0.6667 | 12 | 0.3333 | 10 | 0.5 | 14 | 0.6429 | 45 | 0.5333 |
| 16_4833153 | 16 | 0.125 | 21 | 0.1905 | 14 | 0.2143 | 9 | 0.2222 | 60 | 0.1833 | 11 | 0.0909 | 11 | 0.1818 | 12 | 0 | 14 | 0.0714 | 48 | 0.0833 |
| 16_4833268 | 14 | 0.1429 | 18 | 0.1111 | 16 | 0 | 9 | 0.3333 | 57 | 0.1228 | 9 | 0.1111 | 10 | 0.1 | 10 | 0.1 | 15 | 0.0667 | 44 | 0.0909 |
| 16_4833276 | 14 | 0.7143 | 20 | 0.95 | 16 | 0.8125 | 10 | 0.8 | 60 | 0.8333 | 9 | 0.6667 | 10 | 0.8 | 13 | 0.7692 | 14 | 0.9286 | 46 | 0.8043 |
| 16_4833309 | 15 | 0.2 | 17 | 0.1765 | 17 | 0.0588 | 9 | 0.3333 | 58 | 0.1724 | 8 | 0 | 10 | 0.1 | 9 | 0.2222 | 13 | 0.2308 | 40 | 0.15 |
| 16_4833320 | 12 | 0.6667 | 18 | 0.9444 | 15 | 0.7333 | 10 | 0.8 | 55 | 0.8 | 10 | 0.9 | 7 | 0.7143 | 9 | 0.6667 | 11 | 0.5455 | 37 | 0.7027 |
| 16_4867089 | NA | NA | NA | NA | 1 | 0 | 1 | 1 | 2 | 0.5 | NA | NA | NA | NA | 1 | 1 | NA | NA | 1 | 1 |
| 16_4867278 | NA | NA | NA | NA | NA | NA | 1 | 1 | 1 | 1 | NA | NA | 1 | 0 | 1 | 1 | NA | NA | 2 | 0.5 |
| 16_4867382 | NA | NA | NA | NA | 1 | 0 | 1 | 1 | 2 | 0.5 | NA | NA | 1 | 1 | 1 | 0 | NA | NA | 2 | 0.5 |
| 16_4867410 | NA | NA | NA | NA | 1 | 0 | 1 | 1 | 2 | 0.5 | NA | NA | 1 | 1 | 1 | 1 | NA | NA | 2 | 1 |
| 16_4882439 | NA | NA | NA | NA | 1 | 1 | 5 | 0.4 | 6 | 0.5 | NA | NA | NA | NA | 1 | 1 | NA | NA | 1 | 1 |
| 16_4882466 | NA | NA | NA | NA | 1 | 1 | 5 | 0.8 | 6 | 0.8333 | NA | NA | NA | NA | 1 | 1 | NA | NA | 1 | 1 |
| 16_4882484 | NA | NA | NA | NA | 1 | 1 | 6 | 0.6667 | 7 | 0.7143 | NA | NA | NA | NA | 1 | 1 | NA | NA | 1 | 1 |
| 16_4894907 | NA | NA | 1 | 0 | 2 | 1 | 5 | 0.6 | 8 | 0.625 | 1 | 1 | 1 | 1 | 2 | 0.5 | NA | NA | 4 | 0.75 |
| 16_4894931 | NA | NA | 1 | 1 | 1 | 0 | 6 | 0.1667 | 8 | 0.25 | NA | NA | 1 | 1 | 2 | 0 | NA | NA | 3 | 0.3333 |
| 16_4895105 | NA | NA | 1 | 1 | 2 | 1 | 5 | 1 | 8 | 1 | 1 | 1 | 1 | 1 | 2 | 0.5 | NA | NA | 4 | 0.75 |
| 16_4895331 | NA | NA | 1 | 0 | 2 | 0.5 | 6 | 0.3333 | 9 | 0.3333 | 1 | 1 | 2 | 0.5 | 2 | 1 | NA | NA | 5 | 0.8 |
| 16_4895494 | NA | NA | 1 | 0 | 3 | 0.3333 | 6 | 0.5 | 10 | 0.4 | 1 | 1 | 2 | 1 | 1 | 0 | NA | NA | 4 | 0.75 |
| 16_4895787 | NA | NA | 1 | 1 | 3 | 0.6667 | 5 | 0.8 | 9 | 0.7778 | 1 | 1 | 3 | 1 | 2 | 0.5 | NA | NA | 6 | 0.8333 |
| 16_4895899 | NA | NA | 1 | 1 | 4 | 0.75 | 8 | 0.25 | 13 | 0.4615 | 1 | 0 | 3 | 0.3333 | 3 | 1 | NA | NA | 7 | 0.5714 |
| 16_4899382 | 1 | 0 | NA | NA | 5 | 0.8 | 6 | 0.6667 | 12 | 0.6667 | 2 | 0.5 | 2 | 0.5 | 2 | 1 | NA | NA | 6 | 0.6667 |
| 16_4901987 | 1 | 0 | 1 | 1 | 4 | 1 | 10 | 0.8 | 16 | 0.8125 | 1 | 1 | 3 | 0.3333 | 4 | 0.75 | NA | NA | 8 | 0.625 |
| 16_4902040 | 1 | 0 | 1 | 1 | 4 | 1 | 7 | 0.7143 | 13 | 0.7692 | 1 | 1 | 3 | 0.6667 | 2 | 0.5 | NA | NA | 6 | 0.6667 |
| 16_4902170 | 1 | 0 | NA | NA | 5 | 0.4 | 7 | 0.2857 | 13 | 0.3077 | 2 | 0 | 3 | 0.3333 | 4 | 0.25 | NA | NA | 9 | 0.2222 |
| 16_4902246 | 1 | 1 | 1 | 1 | 6 | 0.8333 | 8 | 0.625 | 16 | 0.75 | 2 | 1 | 3 | 1 | 4 | 0.75 | NA | NA | 9 | 0.8889 |
| 16_4902433 | 1 | 0 | NA | NA | 5 | 0.2 | 9 | 0.5556 | 15 | 0.4 | 3 | 1 | 3 | 0.6667 | 5 | 0.6 | NA | NA | 11 | 0.7273 |
| 16_4902464 | 1 | 0 | 1 | 0 | 6 | 0.3333 | 9 | 0.2222 | 17 | 0.2353 | 3 | 0.3333 | 3 | 0.3333 | 5 | 0.2 | 1 | 1 | 12 | 0.3333 |
| 16_4902483 | 1 | 0 | 1 | 0 | 4 | 0 | 8 | 0.125 | 14 | 0.0714 | 2 | 0.5 | 3 | 0 | 4 | 0.25 | 1 | 0 | 10 | 0.2 |
| 16_4902651 | 2 | 1 | NA | NA | 7 | 0.2857 | 10 | 0.7 | 19 | 0.5789 | 3 | 1 | 2 | 1 | 5 | 0.6 | NA | NA | 10 | 0.8 |
| 16_4903238 | 2 | 0.5 | 2 | 1 | 7 | 0.2857 | 10 | 0.2 | 21 | 0.3333 | 3 | 0 | 4 | 0.25 | 5 | 0.2 | 1 | 0 | 13 | 0.1538 |
| 16_4903531 | 2 | 0 | 2 | 0 | 7 | 0.1429 | 10 | 0.1 | 21 | 0.0952 | 3 | 0 | 4 | 0 | 4 | 0.25 | 1 | 0 | 12 | 0.0833 |
| 16_5001808 | 3 | 0.3333 | 1 | 0 | NA | NA | 1 | 1 | 5 | 0.4 | 1 | 0 | 1 | 1 | 1 | 1 | 1 | 0 | 4 | 0.5 |
| 16_5001816 | 3 | 0.3333 | 1 | 1 | NA | NA | 1 | 0 | 5 | 0.4 | 1 | 0 | 1 | 1 | 1 | 1 | NA | NA | 3 | 0.6667 |
| 16_5001830 | 3 | 0.6667 | 1 | 1 | NA | NA | 1 | 0 | 5 | 0.6 | NA | NA | 1 | 1 | NA | NA | NA | NA | 1 | 1 |
| 16_5002075 | 2 | 1 | NA | NA | NA | NA | NA | NA | 2 | 1 | 1 | 1 | 1 | 0 | 1 | 0 | NA | NA | 3 | 0.3333 |

| | | | | | | | | | | | | | | | | | | | | |
|---|---|---|---|---|---|---|---|---|---|---|---|---|---|---|---|---|---|---|---|---|
| 16_500217( | 2 | 0.5 | 1 | 1 | NA | NA | 1 | 1 | 4 | 0.75 | 1 | 1 | 1 | 0 | 1 | 1 | NA | NA | 3 | 0.6667 |
| 16_500222 | 3 | 1 | 1 | 1 | NA | NA | NA | NA | 4 | 1 | 1 | 1 | 1 | 1 | 1 | 1 | NA | NA | 3 | 1 |
| 16_500229( | 3 | 0.3333 | 1 | 1 | NA | NA | NA | NA | 4 | 0.5 | 1 | 0 | 1 | 0 | 1 | 1 | 1 | 0 | 4 | 0.25 |
| 16_500270 | 3 | 0 | 1 | 0 | NA | NA | 1 | 1 | 5 | 0.2 | 1 | 1 | 1 | 0 | 1 | 0 | NA | NA | 3 | 0.3333 |
| 16_500272 | 3 | 0 | 1 | 1 | NA | NA | 1 | 1 | 5 | 0.4 | 1 | 0 | 1 | 1 | 1 | 0 | NA | NA | 3 | 0.3333 |
| 16_500317 | 3 | 0.3333 | 1 | 1 | NA | NA | 1 | 0 | 5 | 0.4 | 1 | 1 | 1 | 0 | 1 | 0 | 1 | 0 | 4 | 0.25 |
| 16_500347 | 3 | 0.3333 | 1 | 0 | NA | NA | 2 | 0.5 | 6 | 0.3333 | 2 | 0.5 | 1 | 1 | 1 | 0 | 1 | 1 | 5 | 0.6 |
| 16_500489 | 5 | 0.8 | 2 | 0 | 2 | 0.5 | 2 | 0 | 11 | 0.4545 | 2 | 1 | 1 | 0 | NA | NA | 2 | 0 | 5 | 0.4 |
| 16_500501( | 5 | 1 | 3 | 0.6667 | 2 | 1 | 2 | 1 | 12 | 0.9167 | 1 | 1 | 1 | 0 | 1 | 0 | 2 | 1 | 5 | 0.6 |
| 16_500502 | 5 | 0 | 3 | 0 | 2 | 0 | 2 | 0.5 | 12 | 0.0833 | 2 | 0 | 1 | 1 | 1 | 1 | 1 | 0 | 5 | 0.4 |
| 16_501343( | NA | NA | 2 | 1 | 4 | 1 | NA | NA | 6 | 1 | 1 | 1 | 1 | 1 | 1 | 0 | 1 | 1 | 4 | 0.75 |
| 16_501346 | NA | NA | 2 | 0 | 4 | 0.5 | NA | NA | 6 | 0.3333 | 1 | 0 | 1 | 1 | 1 | 0 | 1 | 0 | 4 | 0.25 |
| 16_501365 | NA | NA | 2 | 1 | 2 | 1 | NA | NA | 4 | 1 | 1 | 0 | NA | NA | 1 | 1 | NA | NA | 2 | 0.5 |
| 16_501371( | NA | NA | 2 | 1 | 3 | 1 | NA | NA | 5 | 1 | 1 | 1 | 1 | 1 | 1 | 1 | 1 | 1 | 4 | 1 |
| 16_506251 | 6 | 0.1667 | 4 | 0.5 | 8 | 0.125 | 6 | 0 | 24 | 0.1667 | 8 | 0.125 | 7 | 0.1429 | 4 | 0 | 7 | 0.1429 | 26 | 0.1154 |
| 16_506263 | 7 | 0.5714 | 3 | 0.3333 | 5 | 0 | 3 | 0 | 18 | 0.2778 | 9 | 0.3333 | 6 | 0.6667 | 4 | 0.5 | 6 | 0.3333 | 25 | 0.44 |
| 16_513309 | 4 | 0.25 | 3 | 0.3333 | NA | NA | 1 | 0 | 8 | 0.25 | 2 | 0.5 | 3 | 0.3333 | 3 | 0 | 1 | 1 | 9 | 0.3333 |
| 16_513314 | 2 | 0.5 | 2 | 0 | NA | NA | 1 | 0 | 5 | 0.2 | 2 | 0 | 2 | 0.5 | 3 | 0.3333 | 1 | 0 | 8 | 0.25 |
| 16_513316 | 3 | 0.3333 | NA | NA | NA | NA | 1 | 0 | 4 | 0.25 | 1 | 0 | 3 | 0.3333 | 1 | 1 | 1 | 0 | 6 | 0.3333 |
| 16_722772 | 14 | 0.1429 | 18 | 0.0556 | 10 | 0.1 | 21 | 0.1429 | 63 | 0.1111 | 6 | 0.1667 | 20 | 0.2 | 21 | 0.0476 | 18 | 0.2222 | 65 | 0.1538 |
| 16_722937( | 32 | 0.0938 | 40 | 0.075 | 36 | 0.0556 | 44 | 0.0909 | 152 | 0.0789 | 20 | 0.25 | 41 | 0.0976 | 50 | 0.1 | 34 | 0.1471 | 145 | 0.131 |
| 16_722981 | 37 | 0.2162 | 38 | 0.1842 | 31 | 0.129 | 44 | 0.2273 | 150 | 0.1933 | 16 | 0.125 | 33 | 0.2121 | 53 | 0.1887 | 34 | 0.2647 | 136 | 0.2059 |
| 16_722993 | 29 | 0.3448 | 34 | 0.4706 | 28 | 0.1429 | 33 | 0.2121 | 124 | 0.2984 | 15 | 0.2 | 25 | 0.44 | 52 | 0.3462 | 31 | 0.3548 | 123 | 0.3496 |
| 16_829587 | NA | NA | 6 | 0.1667 | NA | NA | 3 | 0 | 9 | 0.1111 | NA | NA | NA | NA | 1 | 0 | 4 | 0.25 | 5 | 0.2 |
| 16_843648( | 27 | 0.0741 | 15 | 0 | 17 | 0.1176 | 16 | 0.0625 | 75 | 0.0667 | 16 | 0 | 26 | 0.0769 | 19 | 0 | 24 | 0.125 | 85 | 0.0588 |
| 16_843659 | 19 | 0.5789 | 13 | 0.4615 | 11 | 0.3636 | 12 | 0.25 | 55 | 0.4364 | 13 | 0.5385 | 20 | 0.45 | 19 | 0.2632 | 19 | 0.4211 | 71 | 0.4085 |
| 16_843666 | 26 | 0.0769 | 15 | 0 | 18 | 0.1111 | 14 | 0.0714 | 73 | 0.0685 | 17 | 0.2941 | 28 | 0.0714 | 20 | 0.05 | 23 | 0.2174 | 88 | 0.1477 |
| 16_843674( | 27 | 0.1111 | 12 | 0.3333 | 12 | 0.1667 | 19 | 0.1579 | 70 | 0.1714 | 11 | 0.1818 | 20 | 0.2 | 20 | 0.15 | 19 | 0.2105 | 70 | 0.1857 |
| 16_843678 | 24 | 0.125 | 16 | 0.125 | 18 | 0.1667 | 17 | 0.1765 | 75 | 0.1467 | 15 | 0.2 | 26 | 0.1154 | 18 | 0 | 22 | 0.1364 | 81 | 0.1111 |
| 16_843693 | 28 | 0.0714 | 16 | 0.0625 | 18 | 0.1111 | 24 | 0.0833 | 86 | 0.0814 | 16 | 0.1875 | 31 | 0.129 | 23 | 0.0435 | 22 | 0.2727 | 92 | 0.1522 |
| 16_843699 | 28 | 0.4643 | 14 | 0.2857 | 22 | 0.1818 | 26 | 0.2308 | 90 | 0.3 | 20 | 0.35 | 28 | 0.3571 | 24 | 0.2917 | 26 | 0.3846 | 98 | 0.3469 |
| 16_843708 | 31 | 0.0323 | 13 | 0 | 24 | 0.125 | 26 | 0.1154 | 94 | 0.0745 | 20 | 0.1 | 31 | 0.0968 | 25 | 0.12 | 26 | 0 | 102 | 0.0784 |
| 16_843716 | 31 | 0.4839 | 14 | 0.5714 | 21 | 0.5238 | 25 | 0.44 | 91 | 0.4945 | 17 | 0.2941 | 27 | 0.3704 | 22 | 0.4091 | 24 | 0.4167 | 90 | 0.3778 |
| 16_843728 | 34 | 0.0882 | 19 | 0.1579 | 28 | 0.0714 | 25 | 0.12 | 106 | 0.1038 | 23 | 0.0435 | 32 | 0.125 | 26 | 0 | 29 | 0.0345 | 110 | 0.0545 |
| 16_843735 | 31 | 0.2258 | 15 | 0.1333 | 26 | 0.1538 | 29 | 0.069 | 101 | 0.1485 | 22 | 0.1364 | 34 | 0.0882 | 27 | 0.1852 | 30 | 0.2 | 113 | 0.1504 |
| 16_843743( | 35 | 0.1429 | 18 | 0.2778 | 29 | 0.069 | 32 | 0.125 | 114 | 0.1404 | 24 | 0.0833 | 38 | 0.1053 | 27 | 0.0741 | 27 | 0.3704 | 116 | 0.1552 |
| 16_843750( | 31 | 0.2903 | 19 | 0.3684 | 27 | 0.3704 | 29 | 0.3448 | 106 | 0.3396 | 20 | 0.35 | 37 | 0.2432 | 25 | 0.64 | 31 | 0.4194 | 113 | 0.3982 |
| 16_843759 | 36 | 0.1389 | 17 | 0.1765 | 30 | 0.1 | 26 | 0.1154 | 109 | 0.1284 | 19 | 0.3158 | 38 | 0.1842 | 28 | 0.1786 | 28 | 0.1071 | 113 | 0.1858 |
| 16_843770 | 36 | 0.1111 | 20 | 0.1 | 31 | 0.0968 | 29 | 0.1034 | 116 | 0.1034 | 22 | 0.1818 | 44 | 0.1364 | 33 | 0.0909 | 27 | 0.1481 | 126 | 0.1349 |
| 16_843803 | 43 | 0.4884 | 20 | 0.4 | 29 | 0.4138 | 35 | 0.3143 | 127 | 0.4094 | 24 | 0.3333 | 45 | 0.2889 | 35 | 0.3429 | 32 | 0.4375 | 136 | 0.3456 |
| 16_843819 | 43 | 0.0698 | 22 | 0.0909 | 31 | 0.0323 | 37 | 0.1081 | 133 | 0.0752 | 24 | 0.2083 | 43 | 0.0233 | 32 | 0.125 | 30 | 0.1333 | 129 | 0.1085 |
| 16_843858 | 48 | 0.3333 | 23 | 0.3043 | 36 | 0.1944 | 38 | 0.3158 | 145 | 0.2897 | 25 | 0.12 | 46 | 0.1739 | 31 | 0.129 | 33 | 0.2424 | 135 | 0.1704 |
| 16_843872 | 51 | 0.0392 | 24 | 0.0833 | 31 | 0.1613 | 38 | 0.1579 | 144 | 0.1042 | 26 | 0.1923 | 49 | 0.1224 | 40 | 0.075 | 34 | 0.1176 | 149 | 0.1208 |
| 16_843889( | 51 | 0.1765 | 27 | 0.1481 | 34 | 0.1765 | 43 | 0.093 | 155 | 0.1484 | 28 | 0.2143 | 54 | 0.1111 | 44 | 0.1818 | 37 | 0.1351 | 163 | 0.1534 |
| 16_843898 | 55 | 0.4727 | 26 | 0.4615 | 38 | 0.3684 | 42 | 0.3571 | 161 | 0.4161 | 30 | 0.2333 | 60 | 0.4667 | 46 | 0.2391 | 37 | 0.3784 | 173 | 0.3468 |
| 16_843917 | 52 | 0.0962 | 25 | 0.24 | 34 | 0.0588 | 42 | 0.0952 | 153 | 0.1111 | 31 | 0.0323 | 54 | 0.1852 | 45 | 0.1556 | 41 | 0.1951 | 171 | 0.152 |
| 16_843933( | 59 | 0.1356 | 27 | 0.1481 | 45 | 0.0667 | 47 | 0.0851 | 178 | 0.1067 | 38 | 0.0263 | 60 | 0.1667 | 54 | 0.1296 | 45 | 0.1556 | 197 | 0.1269 |
| 16_845270 | NA | NA | 2 | 0.5 | 1 | 0 | NA | NA | 3 | 0.3333 | 3 | 0.6667 | 3 | 0.3333 | 4 | 0.25 | 4 | 0.5 | 14 | 0.4286 |
| 16_845274 | NA | NA | 2 | 0 | 2 | 0.5 | NA | NA | 4 | 0.25 | 3 | 0.3333 | 3 | 0 | 3 | 0.3333 | 3 | 0 | 12 | 0.1667 |
| 16_845287 | NA | NA | 1 | 1 | 1 | 0 | NA | NA | 2 | 0.5 | 3 | 0 | 4 | 0.25 | 4 | 0 | 3 | 0 | 14 | 0.0714 |
| 16_845332 | NA | NA | 2 | 0.5 | 1 | 0 | NA | NA | 3 | 0.3333 | 2 | 0 | 3 | 0.3333 | 4 | 0 | 2 | 0 | 11 | 0.0909 |
| 16_845341( | NA | NA | 2 | 0.5 | 1 | 0 | NA | NA | 3 | 0.3333 | 1 | 0 | 4 | 0 | 4 | 0 | 3 | 0.3333 | 12 | 0.0833 |
| 16_845353 | NA | NA | 2 | 0.5 | 1 | 0 | NA | NA | 3 | 0.3333 | 2 | 0.5 | 1 | 0 | 3 | 0 | 2 | 0.5 | 8 | 0.25 |
| 16_845363 | NA | NA | 2 | 1 | NA | NA | NA | NA | 2 | 1 | 2 | 0.5 | 3 | 1 | 3 | 1 | 2 | 1 | 10 | 0.9 |
| 16_845366 | NA | NA | 2 | 0.5 | 1 | 0 | NA | NA | 3 | 0.3333 | 2 | 1 | 4 | 0.25 | 3 | 0.3333 | 2 | 0.5 | 11 | 0.4545 |
| 16_845370 | NA | NA | 2 | 0 | 1 | 1 | NA | NA | 3 | 0.3333 | 2 | 0 | 3 | 0 | 3 | 0.3333 | 2 | 0 | 10 | 0.1 |
| 16_845378 | NA | NA | 2 | 0 | 1 | 1 | NA | NA | 3 | 0.3333 | 2 | 0 | 4 | 0.75 | 3 | 0.6667 | 2 | 0.5 | 11 | 0.5455 |
| 16_845380 | NA | NA | 1 | 1 | 2 | 0.5 | NA | NA | 3 | 0.6667 | 2 | 1 | 4 | 0.25 | 3 | 1 | 2 | 1 | 11 | 0.7273 |
| 16_845388 | NA | NA | 2 | 0.5 | 2 | 1 | NA | NA | 4 | 0.75 | 2 | 1 | 2 | 1 | 3 | 1 | 2 | 1 | 9 | 1 |
| 16_845396 | NA | NA | 2 | 0.5 | 1 | 0 | NA | NA | 3 | 0.3333 | 2 | 0 | 3 | 0.3333 | 2 | 0.5 | 1 | 0 | 8 | 0.25 |
| 16_845401( | NA | NA | 2 | 1 | 2 | 1 | NA | NA | 4 | 1 | 2 | 1 | 3 | 1 | 2 | 1 | 1 | 1 | 8 | 1 |
| 16_845419 | NA | NA | 2 | 1 | 2 | 1 | NA | NA | 4 | 1 | 1 | 1 | 3 | 1 | 2 | 0.5 | 1 | 1 | 7 | 0.8571 |
| 16_847471 | 2 | 0 | 4 | 0.5 | 3 | 0 | 2 | 0 | 11 | 0.1818 | NA | NA | 3 | 0 | 1 | 0 | 1 | 1 | 5 | 0.2 |
| 16_847515( | 2 | 0 | 4 | 0.25 | 3 | 0.3333 | 3 | 0 | 12 | 0.1667 | NA | NA | 3 | 0.3333 | 1 | 0 | 1 | 0 | 5 | 0.2 |
| 16_847528 | 3 | 1 | 3 | 0.3333 | 3 | 0 | 4 | 0 | 13 | 0.3077 | NA | NA | 4 | 0.25 | 1 | 0 | 1 | 0 | 6 | 0.1667 |

| | | | | | | | | | | | | | | | | | | | | |
|---|---|---|---|---|---|---|---|---|---|---|---|---|---|---|---|---|---|---|---|---|
| 16_8478254 | 10 | 0.3 | 8 | 0.375 | 8 | 0.125 | 8 | 0.125 | 34 | 0.2353 | 4 | 0.75 | 4 | 0.75 | 9 | 0.4444 | 8 | 0.125 | 25 | 0.44 |
| 16_8478296 | 11 | 0.2727 | 8 | 0.375 | 10 | 0.1 | 10 | 0.2 | 39 | 0.2308 | 4 | 0.75 | 5 | 0.4 | 9 | 0.5556 | 9 | 0.5556 | 27 | 0.5556 |
| 16_8478354 | 13 | 0.5385 | 9 | 0.3333 | 7 | 0.4286 | 10 | 0.2 | 39 | 0.3846 | 3 | 0.6667 | 5 | 0.4 | 9 | 0.5556 | 9 | 0.4444 | 26 | 0.5 |
| 16_8478451 | 10 | 1 | 7 | 0.5714 | 7 | 0.7143 | 9 | 0.8889 | 33 | 0.8182 | 2 | 1 | 4 | 1 | 10 | 0.8 | 9 | 0.8889 | 25 | 0.88 |
| 16_8478729 | 11 | 0.7273 | 5 | 0.4 | 11 | 0.5455 | 9 | 0.4444 | 36 | 0.5556 | 3 | 0.6667 | 6 | 0.5 | 9 | 0.3333 | 10 | 0.3 | 28 | 0.3929 |
| 16_8478767 | 13 | 0.6154 | 5 | 0.4 | 8 | 0.625 | 10 | 0.6 | 36 | 0.5833 | 3 | 0.6667 | 7 | 0.8571 | 9 | 0.7778 | 9 | 0.8889 | 28 | 0.8214 |
| 16_8478802 | 12 | 0.1667 | 5 | 0.4 | 8 | 0.25 | 8 | 0.375 | 33 | 0.2727 | 2 | 0 | 5 | 0.6 | 9 | 0.3333 | 9 | 0.1111 | 25 | 0.28 |
| 16_8506711 | 1 | 0 | 1 | 0 | 3 | 0 | 6 | 0.5 | 11 | 0.2727 | NA | NA | 1 | 0 | 1 | 1 | 2 | 0.5 | 4 | 0.5 |
| 16_8507074 | 1 | 0 | NA | NA | 2 | 0.5 | 5 | 0.2 | 8 | 0.25 | NA | NA | 1 | 0 | 1 | 0 | 2 | 0.5 | 4 | 0.25 |
| 16_8507693 | 11 | 0.1818 | 13 | 0.0769 | 8 | 0.25 | 16 | 0.125 | 48 | 0.1458 | 9 | 0 | 21 | 0.1905 | 10 | 0.3 | 6 | 0 | 46 | 0.1522 |
| 16_8507995 | 9 | 0 | 13 | 0.1538 | 8 | 0 | 12 | 0.0833 | 42 | 0.0714 | 6 | 0 | 16 | 0.0625 | 10 | 0 | 6 | 0.1667 | 38 | 0.0526 |
| 16_8508476 | 10 | 0.5 | 9 | 0.5556 | 7 | 0.4286 | 13 | 0.3077 | 39 | 0.4359 | 8 | 0.125 | 13 | 0.3846 | 12 | 0.4167 | 7 | 0.7143 | 40 | 0.4 |
| 16_8508617 | 7 | 0.2857 | 9 | 0.5556 | 3 | 0 | 11 | 0.1818 | 30 | 0.3 | 5 | 0.4 | 10 | 0.3 | 6 | 0.1667 | 5 | 0.8 | 26 | 0.3846 |
| 16_8514888 | NA | NA | NA | NA | 1 | 0 | 3 | 0.3333 | 4 | 0.25 | NA | NA | 1 | 1 | NA | NA | 1 | 0 | 2 | 0.5 |
| 16_8515277 | NA | NA | NA | NA | NA | NA | 2 | 0.5 | 2 | 0.5 | NA | NA | 1 | 1 | NA | NA | NA | NA | 1 | 1 |
| 16_8515551 | 7 | 0.1429 | 9 | 0.3333 | 2 | 0.5 | 10 | 0.1 | 28 | 0.2143 | 3 | 0 | 8 | 0.25 | 6 | 0 | 2 | 0.5 | 19 | 0.1579 |
| 16_8648161 | 1 | 1 | 1 | 0 | 1 | 1 | 2 | 1 | 5 | 0.8 | 4 | 0.75 | NA | NA | 1 | 0 | 1 | 0 | 6 | 0.5 |
| 16_8660933 | 4 | 0 | 8 | 0.125 | 8 | 0.125 | 8 | 0.375 | 28 | 0.1786 | 6 | 0.1667 | 2 | 0 | 1 | 0 | 2 | 0 | 11 | 0.0909 |
| 16_8660991 | 4 | 0 | 7 | 0 | 6 | 0.1667 | 8 | 0.125 | 25 | 0.08 | 5 | 0.2 | 1 | 0 | 3 | 0 | 2 | 0 | 11 | 0.0909 |
| 16_8661010 | 3 | 0.3333 | 4 | 0 | 5 | 0.2 | 8 | 0.75 | 20 | 0.4 | 4 | 0.75 | 2 | 0.5 | 1 | 1 | 1 | 1 | 8 | 0.75 |
| 16_8661014 | 4 | 0.25 | 8 | 0.5 | 6 | 0.3333 | 7 | 0.1429 | 25 | 0.32 | 4 | 0.5 | 2 | 1 | 2 | 0 | 2 | 0.5 | 10 | 0.5 |
| 16_8661068 | 5 | 0.6 | 5 | 0.8 | 9 | 0.6667 | 9 | 0.7778 | 28 | 0.7143 | 5 | 0.8 | 3 | 1 | 3 | 0.6667 | 1 | 1 | 12 | 0.8333 |
| 16_8661160 | 2 | 0 | 1 | 1 | 2 | 0.5 | 5 | 0.2 | 10 | 0.3 | 3 | 0.3333 | 2 | 0 | NA | NA | NA | NA | 5 | 0.2 |
| 16_8661846 | 1 | 0 | 2 | 0 | 3 | 0.3333 | 4 | 0 | 10 | 0.1 | 3 | 0.3333 | 2 | 0 | NA | NA | NA | NA | 5 | 0.2 |
| 16_8663647 | 2 | 0.5 | 3 | 0 | 1 | 0 | 4 | 0 | 10 | 0.1 | NA | NA | NA | NA | 2 | 0.5 | 1 | 0 | 3 | 0.3333 |
| 16_8669663 | 3 | 0.3333 | 1 | 1 | 1 | 0 | 3 | 1 | 8 | 0.625 | 2 | 0 | NA | NA | 2 | 1 | NA | NA | 4 | 0.5 |
| 16_8669864 | 3 | 0 | 1 | 0 | NA | NA | 3 | 0.3333 | 7 | 0.1429 | 2 | 0 | NA | NA | 2 | 0.5 | 1 | 0 | 5 | 0.2 |
| 16_8669962 | 3 | 0 | 2 | 0.5 | 1 | 0 | 3 | 0.3333 | 9 | 0.2222 | 2 | 0 | NA | NA | 2 | 1 | 1 | 0 | 5 | 0.4 |
| 16_8670116 | 3 | 0.3333 | 2 | 0.5 | 1 | 0 | 3 | 1 | 9 | 0.5556 | 2 | 1 | NA | NA | 2 | 1 | 1 | 0 | 5 | 0.8 |
| 16_8670215 | 4 | 0.25 | 2 | 0.5 | 1 | 1 | 3 | 0 | 10 | 0.3 | 2 | 0.5 | NA | NA | 1 | 0 | 1 | 0 | 4 | 0.25 |
| 16_8673731 | 3 | 0.3333 | 5 | 0.4 | 5 | 0 | 5 | 0 | 18 | 0.1667 | 3 | 0.3333 | 3 | 0.3333 | 2 | 0.5 | 3 | 0 | 11 | 0.2727 |
| 16_8673758 | 3 | 1 | 6 | 1 | 5 | 1 | 5 | 1 | 19 | 1 | 3 | 1 | 3 | 1 | 3 | 1 | 3 | 1 | 12 | 1 |
| 16_8673778 | 3 | 0.6667 | 5 | 0.8 | 5 | 0.8 | 4 | 0 | 17 | 0.5882 | 3 | 0.3333 | 2 | 0 | 2 | 0.5 | 2 | 1 | 9 | 0.4444 |
| 16_8673814 | 4 | 0.75 | 6 | 0.8333 | 6 | 0.8333 | 3 | 1 | 19 | 0.8421 | 3 | 0.6667 | 3 | 1 | 3 | 1 | 3 | 1 | 12 | 0.9167 |
| 16_8673825 | 4 | 0 | 6 | 0.1667 | 5 | 0.4 | 4 | 0.5 | 19 | 0.2632 | 3 | 0.3333 | 2 | 0.5 | 2 | 0 | 2 | 0 | 9 | 0.2222 |
| 16_8673868 | 3 | 0.6667 | 6 | 1 | 5 | 1 | 4 | 1 | 18 | 0.9444 | 3 | 1 | 3 | 0.6667 | 2 | 1 | 2 | 1 | 10 | 0.9 |
| 16_8673903 | 2 | 0 | 7 | 0.1429 | 5 | 0 | 5 | 0 | 19 | 0.0526 | 3 | 0.3333 | 2 | 0.5 | 2 | 0 | 2 | 0 | 9 | 0.2222 |
| 16_8673910 | 2 | 0 | 6 | 0.3333 | 6 | 0.5 | 4 | 0.25 | 18 | 0.3333 | 2 | 0.5 | 3 | 0.3333 | 2 | 0.5 | 1 | 0 | 8 | 0.375 |
| 16_8674096 | 4 | 0.25 | 7 | 0 | 6 | 0 | 3 | 0 | 20 | 0.05 | 2 | 0 | 3 | 0.3333 | 2 | 0 | 3 | 0 | 10 | 0.1 |
| 16_8674115 | 4 | 0.25 | 7 | 0.4286 | 6 | 0.6667 | 5 | 0.6 | 22 | 0.5 | 3 | 0.6667 | 4 | 0.5 | 2 | 1 | 3 | 1 | 12 | 0.75 |
| 16_8674121 | 5 | 0.2 | 7 | 0.4286 | 6 | 0.1667 | 5 | 0 | 23 | 0.2174 | 2 | 0 | 4 | 0.5 | 3 | 0 | 2 | 1 | 11 | 0.3636 |
| 16_8674190 | 4 | 0 | 7 | 0.1429 | 5 | 0 | 4 | 0 | 20 | 0.05 | 3 | 0 | 4 | 0 | 2 | 0.5 | 2 | 0 | 11 | 0.0909 |
| 16_8674555 | 3 | 0 | 8 | 0.125 | 4 | 0.25 | 3 | 0 | 18 | 0.1111 | 2 | 0 | 2 | 0 | 3 | 0.3333 | 3 | 0 | 10 | 0.1 |
| 16_8674699 | 5 | 0.4 | 6 | 0.1667 | 6 | 0.1667 | 5 | 0 | 22 | 0.1818 | 3 | 0.3333 | 3 | 0.3333 | 2 | 0 | 2 | 0 | 10 | 0.2 |
| 16_8675010 | 6 | 0.3333 | 6 | 0.5 | 4 | 0 | 5 | 0.2 | 21 | 0.2857 | 3 | 0.3333 | 3 | 0 | 3 | 0.6667 | 2 | 0 | 11 | 0.2727 |
| 16_8675811 | 8 | 0.5 | 7 | 0.7143 | 6 | 0.5 | 7 | 0.4286 | 28 | 0.5357 | 4 | 0.75 | 5 | 0.4 | 3 | 0 | 2 | 0 | 14 | 0.3571 |
| 16_8675818 | 7 | 0.1429 | 6 | 0.1667 | 7 | 0 | 6 | 0 | 26 | 0.0769 | 4 | 0 | 4 | 0 | 5 | 0.2 | 3 | 0 | 16 | 0.0625 |
| 16_8676660 | 2 | 0 | 2 | 0 | 3 | 0 | 1 | 1 | 8 | 0.125 | 2 | 0 | 2 | 0 | 2 | 0.5 | NA | NA | 6 | 0.1667 |
| 16_8680492 | NA | NA | 1 | 1 | 1 | 1 | 1 | 1 | 3 | 1 | NA | NA | 1 | 1 | 2 | 1 | NA | NA | 3 | 1 |
| 16_9386049 | 17 | 0.4118 | 14 | 0.5 | 13 | 0.3077 | 14 | 0.4286 | 58 | 0.4138 | 5 | 0.4 | 10 | 0.3 | 9 | 0.3333 | 10 | 0.5 | 34 | 0.3824 |
| 16_9386190 | 17 | 0.2941 | 12 | 0.1667 | 11 | 0.4545 | 12 | 0.25 | 52 | 0.2885 | 6 | 0 | 10 | 0.3 | 8 | 0.375 | 11 | 0.4545 | 35 | 0.3143 |
| 16_9387153 | 13 | 0.0769 | 11 | 0.0909 | 6 | 0 | 8 | 0 | 38 | 0.0526 | 4 | 0 | 8 | 0 | 7 | 0.1429 | 9 | 0.1111 | 28 | 0.0714 |
| 16_9387541 | 13 | 0.2308 | 6 | 0.3333 | 6 | 0.1667 | 7 | 0 | 32 | 0.1875 | 3 | 0.3333 | 5 | 0.4 | 7 | 0.5714 | 7 | 0.5714 | 22 | 0.5 |
| 16_9394845 | 1 | 1 | NA | NA | 2 | 0.5 | NA | NA | 3 | 0.6667 | NA | NA | NA | NA | 1 | 1 | NA | NA | 1 | 1 |
| 16_9395842 | 1 | 1 | NA | NA | 3 | 1 | NA | NA | 4 | 1 | NA | NA | NA | NA | 1 | 1 | NA | NA | 1 | 1 |
| 16_9395893 | 2 | 0 | NA | NA | 3 | 0.3333 | NA | NA | 5 | 0.2 | NA | NA | NA | NA | 1 | 1 | NA | NA | 1 | 1 |
| 16_9395922 | 2 | 1 | NA | NA | 3 | 0.6667 | NA | NA | 5 | 0.8 | NA | NA | NA | NA | 1 | 1 | NA | NA | 1 | 1 |
| 16_9395932 | 2 | 1 | NA | NA | 2 | 1 | NA | NA | 4 | 1 | NA | NA | NA | NA | 1 | 1 | NA | NA | 1 | 1 |
| 16_9396007 | 2 | 0.5 | NA | NA | 3 | 0 | NA | NA | 5 | 0.2 | NA | NA | NA | NA | 1 | 1 | NA | NA | 1 | 1 |
| 16_9396037 | NA | NA | NA | NA | 3 | 1 | NA | NA | 3 | 1 | NA | NA | NA | NA | 1 | 1 | NA | NA | 1 | 1 |
| 16_9396081 | 1 | 1 | NA | NA | 2 | 0 | NA | NA | 3 | 0.3333 | NA | NA | NA | NA | 1 | 1 | NA | NA | 1 | 1 |
| 16_1010029 | 1 | 0 | 2 | 0.5 | NA | NA | 2 | 0.5 | 5 | 0.4 | 2 | 1 | 1 | 1 | 2 | 0.5 | 5 | 0.4 | 10 | 0.6 |
| 16_1010040 | 1 | 0 | 3 | 0.3333 | NA | NA | 3 | 0 | 7 | 0.1429 | 2 | 0.5 | 1 | 0 | 2 | 0 | 4 | 0.25 | 9 | 0.2222 |
| 16_1010127 | 1 | 0 | 3 | 0.6667 | NA | NA | 1 | 0 | 5 | 0.4 | 1 | 0 | 1 | 0 | 1 | 1 | 4 | 0.25 | 7 | 0.2857 |
| 16_1010176 | 1 | 1 | 2 | 0 | NA | NA | 2 | 0.5 | 5 | 0.4 | 2 | 1 | 1 | 1 | 2 | 0 | 5 | 0.8 | 10 | 0.7 |

| | | | | | | | | | | | | | | | | | | | | |
|---|---|---|---|---|---|---|---|---|---|---|---|---|---|---|---|---|---|---|---|---|
| 16_1010185 | 1 | 0 | 2 | 0.5 | NA | NA | 2 | 0 | 5 | 0.2 | 2 | 0 | NA | NA | 2 | 0.5 | 6 | 0.1667 | 10 | 0.2 |
| 16_1017614 | 3 | 0 | 3 | 0.3333 | 1 | 0 | 4 | 0.5 | 11 | 0.2727 | 4 | 0.25 | NA | NA | 4 | 0.5 | 4 | 0.75 | 12 | 0.5 |
| 16_1021280 | 3 | 0 | 1 | 0 | 1 | 1 | 3 | 0.3333 | 8 | 0.25 | 4 | 0 | NA | NA | 3 | 0.3333 | 2 | 0 | 9 | 0.1111 |
| 16_1021286 | 2 | 0.5 | 1 | 0 | 1 | 0 | 4 | 0 | 8 | 0.125 | 3 | 0 | NA | NA | 2 | 0 | 2 | 0.5 | 7 | 0.1429 |
| 16_1021287 | 3 | 0 | 1 | 0 | 1 | 0 | 4 | 0.25 | 9 | 0.1111 | 4 | 0.25 | NA | NA | 3 | 0 | 2 | 0 | 9 | 0.1111 |
| 16_1021290 | 3 | 0.6667 | 1 | 0 | 1 | 0 | 3 | 0.3333 | 8 | 0.375 | 4 | 0.25 | NA | NA | 3 | 0 | 1 | 0 | 8 | 0.125 |
| 16_1021300 | 3 | 0.3333 | 1 | 0 | 1 | 0 | 3 | 0.3333 | 8 | 0.25 | 3 | 0.3333 | NA | NA | 3 | 0.3333 | 2 | 0.5 | 8 | 0.375 |
| 16_1023869 | 3 | 0 | 4 | 0 | 1 | 0 | 4 | 0.25 | 12 | 0.0833 | 1 | 0 | 4 | 0 | 1 | 1 | 2 | 0 | 8 | 0.125 |
| 16_1023885 | 4 | 0.75 | 3 | 1 | 1 | 1 | 3 | 1 | 11 | 0.9091 | NA | NA | 6 | 0.5 | 1 | 1 | 2 | 1 | 9 | 0.6667 |
| 16_1023893 | 5 | 0 | 2 | 0 | 1 | 1 | 4 | 0 | 12 | 0.0833 | NA | NA | 3 | 0.3333 | NA | NA | 1 | 0 | 4 | 0.25 |
| 16_1023911 | 5 | 1 | 3 | 1 | 1 | 1 | 4 | 1 | 13 | 1 | NA | NA | 4 | 1 | 1 | 1 | 2 | 1 | 7 | 1 |
| 16_1023920 | 3 | 0.6667 | 3 | 0.3333 | 1 | 1 | 4 | 1 | 11 | 0.7273 | NA | NA | 5 | 0.4 | 1 | 1 | 2 | 1 | 8 | 0.625 |
| 16_1023937 | 4 | 0.5 | 2 | 0.5 | 1 | 1 | 3 | 0.6667 | 10 | 0.6 | NA | NA | 3 | 0.3333 | NA | NA | 1 | 1 | 4 | 0.5 |
| 16_1023944 | 4 | 0.75 | 3 | 0.6667 | 1 | 1 | 4 | 0.75 | 12 | 0.75 | NA | NA | 5 | 0.2 | 1 | 0 | 2 | 1 | 8 | 0.375 |
| 16_1023954 | 4 | 0.75 | 3 | 1 | 1 | 1 | 3 | 0.6667 | 11 | 0.8182 | NA | NA | 4 | 1 | 1 | 1 | 2 | 0.5 | 7 | 0.8571 |
| 16_1023972 | 2 | 0 | 2 | 0.5 | 1 | 0 | 4 | 0.25 | 9 | 0.2222 | NA | NA | 3 | 0 | 1 | 0 | 2 | 0.5 | 6 | 0.1667 |
| 16_1023972 | 4 | 0.75 | 3 | 1 | 1 | 1 | 4 | 1 | 12 | 0.9167 | NA | NA | 4 | 1 | 1 | 1 | 2 | 1 | 7 | 1 |
| 16_1023983 | 3 | 0.6667 | 3 | 0.6667 | 1 | 1 | 4 | 0.75 | 11 | 0.7273 | NA | NA | 4 | 0.5 | 1 | 1 | 2 | 1 | 7 | 0.7143 |
| 16_1024017 | 2 | 0 | 2 | 0 | 1 | 1 | 3 | 0 | 8 | 0.125 | NA | NA | 3 | 0 | 1 | 0 | 2 | 0.5 | 6 | 0.1667 |
| 16_1024040 | 2 | 1 | 2 | 0.5 | 1 | 0 | 4 | 0.75 | 9 | 0.6667 | NA | NA | 3 | 1 | 1 | 1 | 2 | 0.5 | 6 | 0.8333 |
| 16_1024040 | 4 | 0.25 | 1 | 0 | 1 | 0 | 3 | 0 | 9 | 0.1111 | NA | NA | 2 | 0.5 | 1 | 0 | 2 | 0 | 5 | 0.2 |
| 16_1024049 | 3 | 0.6667 | 2 | 1 | 1 | 1 | 4 | 0.75 | 10 | 0.8 | NA | NA | 3 | 1 | 1 | 1 | 1 | 1 | 5 | 1 |
| 16_1024059 | 1 | 1 | 2 | 0.5 | NA | NA | 3 | 1 | 6 | 0.8333 | NA | NA | 2 | 0.5 | 1 | 1 | 2 | 1 | 5 | 0.8 |
| 16_1024067 | 3 | 0 | 2 | 0.5 | 1 | 0 | 4 | 0.5 | 10 | 0.3 | NA | NA | 2 | 0.5 | NA | NA | 2 | 1 | 4 | 0.75 |
| 16_1024070 | 2 | 1 | 2 | 0.5 | 1 | 0 | 4 | 0.75 | 9 | 0.6667 | NA | NA | 3 | 0.6667 | 1 | 0 | 2 | 1 | 6 | 0.6667 |
| 16_1024129 | 3 | 0.3333 | 1 | 0 | 6 | 0.1667 | NA | NA | 10 | 0.2 | 1 | 0 | 3 | 0.6667 | 3 | 0.3333 | 2 | 0 | 9 | 0.3333 |
| 16_1024142 | 1 | 0 | 1 | 0 | 1 | 1 | NA | NA | 3 | 0.3333 | NA | NA | 3 | 0.6667 | 2 | 0.5 | 2 | 0.5 | 7 | 0.5714 |
| 16_1024146 | 1 | 1 | NA | NA | 1 | 1 | NA | NA | 2 | 1 | NA | NA | 3 | 1 | 2 | 0.5 | 1 | 0 | 6 | 0.6667 |
| 16_1024195 | 1 | 1 | 2 | 1 | 1 | 0 | NA | NA | 4 | 0.75 | NA | NA | 3 | 0 | 3 | 0 | 2 | 0.5 | 8 | 0.125 |
| 16_1024197 | 1 | 0 | 1 | 0 | 1 | 1 | NA | NA | 3 | 0.3333 | NA | NA | 3 | 0.6667 | 2 | 0.5 | 2 | 0 | 7 | 0.4286 |
| 16_1024216 | 1 | 0 | 2 | 0.5 | 1 | 1 | NA | NA | 4 | 0.5 | NA | NA | 3 | 0.3333 | 3 | 0.3333 | 1 | 0 | 7 | 0.2857 |
| 16_1024225 | 1 | 0 | 4 | 0.25 | 1 | 1 | 1 | 0 | 7 | 0.2857 | NA | NA | 3 | 0.3333 | 3 | 0 | 2 | 0 | 8 | 0.125 |
| 16_1024298 | NA | NA | 3 | 0.3333 | 1 | 0 | 2 | 1 | 6 | 0.5 | NA | NA | 1 | 1 | NA | NA | NA | NA | 1 | 1 |
| 16_1032131 | NA | NA | 1 | 1 | 2 | 1 | NA | NA | 3 | 1 | NA | NA | 1 | 0 | 1 | 1 | 1 | 0 | 3 | 0.3333 |
| 16_1032162 | NA | NA | NA | NA | NA | NA | 1 | 1 | 1 | 1 | NA | NA | 1 | 1 | 1 | 1 | 1 | 1 | 3 | 1 |
| 16_1032163 | NA | NA | 1 | 1 | 2 | 0.5 | 1 | 1 | 4 | 0.75 | NA | NA | 1 | 1 | 1 | 1 | NA | NA | 2 | 1 |
| 16_1032171 | NA | NA | 1 | 1 | 2 | 1 | 1 | 1 | 4 | 1 | NA | NA | 1 | 1 | 1 | 1 | 1 | 1 | 3 | 1 |
| 16_1034823 | 8 | 0.625 | 11 | 0.7273 | 8 | 0.625 | 5 | 0.8 | 32 | 0.6875 | 9 | 0.7778 | 4 | 1 | 8 | 0.75 | 14 | 0.2857 | 35 | 0.6 |
| 16_1034832 | 9 | 0.8889 | 8 | 0.75 | 3 | 1 | 6 | 0.8333 | 26 | 0.8462 | 6 | 0.8333 | 3 | 0.3333 | 7 | 0.5714 | 11 | 0.9091 | 27 | 0.7407 |
| 16_1034833 | 9 | 0.2222 | 10 | 0.4 | 8 | 0.125 | 7 | 0.1429 | 34 | 0.2353 | 8 | 0.125 | 2 | 0 | 9 | 0.2222 | 12 | 0.25 | 31 | 0.1935 |
| 16_1034840 | 9 | 0 | 10 | 0.2 | 8 | 0.625 | 7 | 0.1429 | 34 | 0.2353 | 6 | 0 | 3 | 0.3333 | 8 | 0.25 | 11 | 0.5455 | 28 | 0.3214 |
| 16_1034842 | 10 | 0.8 | 11 | 0.5455 | 5 | 0.6 | 7 | 0.2857 | 33 | 0.5758 | 5 | 0.2 | 2 | 0.5 | 6 | 0.1667 | 12 | 0.6667 | 25 | 0.44 |
| 16_1034852 | 7 | 0.1429 | 10 | 0.1 | 8 | 0 | 7 | 0 | 32 | 0.0625 | 6 | 0.1667 | 3 | 0 | 7 | 0.1429 | 11 | 0.0909 | 27 | 0.1111 |
| 16_1034861 | 4 | 0 | 7 | 0.1429 | 5 | 0.4 | 4 | 0.25 | 20 | 0.2 | 6 | 0.1667 | 2 | 0 | 7 | 0.1429 | 9 | 0.1111 | 24 | 0.125 |
| 16_1036015 | 7 | 0 | 9 | 0.1111 | 5 | 0 | 6 | 0.1667 | 27 | 0.0741 | 5 | 0 | 3 | 0 | 7 | 0 | 11 | 0.1818 | 26 | 0.0769 |
| 16_1036016 | 2 | 0 | 5 | 0.2 | 4 | 0.5 | 3 | 0 | 14 | 0.2143 | 4 | 0.25 | 1 | 0 | 3 | 0 | 7 | 0.1429 | 15 | 0.1333 |
| 16_1036033 | 7 | 0.2857 | 7 | 0.4286 | 6 | 0.3333 | 6 | 0.6667 | 26 | 0.4231 | 6 | 0 | NA | NA | 7 | 0.1429 | 11 | 0.0909 | 24 | 0.0833 |
| 16_1036034 | 6 | 0.3333 | 8 | 0.625 | 5 | 0.4 | 4 | 0.25 | 23 | 0.4348 | 5 | 0.6 | NA | NA | 6 | 0.3333 | 11 | 0.5455 | 22 | 0.5 |
| 16_1036036 | 6 | 0 | 7 | 0.2857 | 4 | 0.5 | 4 | 0 | 21 | 0.1905 | 4 | 0 | NA | NA | 5 | 0 | 10 | 0.1 | 19 | 0.0526 |
| 16_1036037 | 5 | 0.8 | 3 | 1 | 1 | 1 | 2 | 1 | 11 | 0.9091 | 4 | 1 | NA | NA | 3 | 0.6667 | 8 | 0.875 | 15 | 0.8667 |
| 16_1044956 | 1 | 0 | 1 | 0 | 4 | 0.25 | NA | NA | 6 | 0.1667 | 2 | 0.5 | 3 | 0 | 7 | 0.5714 | 1 | 0 | 13 | 0.3846 |
| 16_1044957 | 1 | 1 | 2 | 1 | 4 | 1 | NA | NA | 7 | 1 | 3 | 0.6667 | 3 | 1 | 4 | 1 | NA | NA | 10 | 0.9 |
| 16_1044958 | 1 | 1 | 4 | 1 | 2 | 1 | NA | NA | 7 | 1 | 3 | 1 | 3 | 1 | 7 | 1 | 1 | 1 | 14 | 1 |
| 16_1044965 | 1 | 0 | 3 | 0 | 4 | 0.5 | NA | NA | 8 | 0.25 | 2 | 0 | 3 | 0 | 6 | 0.3333 | 1 | 0 | 12 | 0.1667 |
| 16_1044971 | 1 | 1 | 4 | 1 | 4 | 1 | NA | NA | 9 | 1 | 2 | 0.5 | 3 | 0.6667 | 6 | 1 | 1 | 1 | 12 | 0.8333 |
| 16_1044982 | 2 | 1 | 2 | 1 | 2 | 0 | NA | NA | 6 | 0.6667 | 3 | 0.6667 | 2 | 0.5 | 6 | 0.3333 | NA | NA | 11 | 0.4545 |
| 16_1044983 | 1 | 1 | 4 | 1 | 2 | 1 | NA | NA | 7 | 1 | 3 | 1 | 3 | 1 | 7 | 1 | NA | NA | 13 | 1 |
| 16_1044999 | 2 | 1 | 4 | 1 | 4 | 1 | NA | NA | 10 | 1 | 3 | 0.6667 | 3 | 0.3333 | 8 | 0.875 | 3 | 0.6667 | 17 | 0.7059 |
| 16_1045013 | 2 | 1 | 3 | 1 | 4 | 1 | NA | NA | 9 | 1 | 2 | 0.5 | 3 | 0.6667 | 7 | 1 | 3 | 0.6667 | 15 | 0.8 |
| 16_1045035 | 2 | 1 | 4 | 0.75 | 4 | 1 | NA | NA | 10 | 0.9 | 3 | 0.6667 | 3 | 1 | 7 | 1 | 3 | 1 | 16 | 0.9375 |
| 16_1045039 | 2 | 1 | 4 | 0.75 | 5 | 0.6 | NA | NA | 11 | 0.7273 | 2 | 0 | 3 | 0.6667 | 7 | 0.5714 | 3 | 0.6667 | 15 | 0.5333 |
| 16_1045042 | 3 | 1 | 4 | 1 | 6 | 1 | NA | NA | 13 | 1 | 3 | 0.6667 | 4 | 1 | 7 | 0.8571 | 3 | 1 | 17 | 0.8824 |
| 16_1045044 | 3 | 0.6667 | 4 | 1 | 6 | 0.8333 | NA | NA | 13 | 0.8462 | 3 | 1 | 4 | 1 | 5 | 1 | 3 | 0.6667 | 15 | 0.9333 |
| 16_1045048 | 2 | 0 | 3 | 1 | 5 | 1 | NA | NA | 10 | 0.8 | 3 | 1 | 3 | 0.6667 | 6 | 1 | 3 | 0.6667 | 15 | 0.8667 |
| 16_1045049 | 3 | 0.3333 | 4 | 1 | 6 | 0.6667 | NA | NA | 13 | 0.6923 | 3 | 0.6667 | 4 | 0.5 | 5 | 0.6 | 3 | 0.3333 | 15 | 0.5333 |

| | | | | | | | | | | | | | | | | | | | | |
|---|---|---|---|---|---|---|---|---|---|---|---|---|---|---|---|---|---|---|---|---|
| 16_105596 | 2 | 1 | 2 | 1 | 3 | 0.6667 | 1 | 1 | 8 | 0.875 | NA | NA | 1 | 1 | 1 | 1 | 2 | 1 | 4 | 1 |
| 16_105596 | 2 | 0.5 | 2 | 0.5 | 3 | 0 | 1 | 0 | 8 | 0.25 | NA | NA | 1 | 0 | 1 | 1 | 2 | 1 | 4 | 0.75 |
| 16_105598 | 2 | 0 | 2 | 0.5 | 4 | 0.5 | 1 | 0 | 9 | 0.3333 | NA | NA | 1 | 1 | 1 | 1 | 2 | 0.5 | 4 | 0.75 |
| 16_105598 | 2 | 0 | 4 | 0 | 4 | 0.75 | 1 | 1 | 11 | 0.3636 | NA | NA | 1 | 0 | 1 | 0 | 2 | 1 | 4 | 0.5 |
| 16_105599 | 1 | 0 | 3 | 0 | 4 | 0 | 1 | 1 | 9 | 0.1111 | NA | NA | NA | NA | 1 | 1 | 1 | 0 | 2 | 0.5 |
| 16_105599 | 1 | 0 | 2 | 1 | 2 | 0 | 1 | 1 | 6 | 0.5 | NA | NA | 1 | 0 | 1 | 1 | 2 | 0 | 4 | 0.25 |
| 16_105599 | 2 | 0.5 | 2 | 0.5 | 3 | 0.3333 | 1 | 1 | 8 | 0.5 | NA | NA | 1 | 1 | 1 | 0 | 1 | 0 | 3 | 0.3333 |
| 16_105601 | 3 | 0 | 3 | 0.3333 | 4 | 0 | 2 | 0 | 12 | 0.0833 | NA | NA | 1 | 0 | 1 | 1 | 2 | 0 | 4 | 0.25 |
| 16_105601 | 3 | 1 | 4 | 0.75 | 4 | 1 | 2 | 1 | 13 | 0.9231 | NA | NA | 1 | 1 | 1 | 1 | 2 | 1 | 4 | 1 |
| 16_105603 | 3 | 0.6667 | 2 | 1 | 4 | 0.5 | 2 | 1 | 11 | 0.7273 | NA | NA | 1 | 1 | 1 | 1 | 3 | 0.6667 | 5 | 0.8 |
| 16_105606 | 2 | 0 | 4 | 0.25 | 3 | 0.3333 | 1 | 0 | 10 | 0.2 | NA | NA | 1 | 0 | 2 | 0.5 | 2 | 0.5 | 5 | 0.4 |
| 16_105606 | 3 | 0 | 3 | 0 | 5 | 0.2 | NA | NA | 11 | 0.0909 | NA | NA | 1 | 0 | 2 | 0 | 3 | 0.3333 | 6 | 0.1667 |
| 16_105611 | 2 | 0.5 | 3 | 0.6667 | 6 | 0.5 | 4 | 0 | 15 | 0.4 | NA | NA | 1 | 0 | 3 | 1 | 3 | 0.6667 | 7 | 0.7143 |
| 16_105615 | 3 | 0 | 3 | 0 | 6 | 0.1667 | 6 | 0.3333 | 18 | 0.1667 | NA | NA | 1 | 0 | 2 | 0.5 | 4 | 0.5 | 7 | 0.4286 |
| 16_105618 | 3 | 1 | 3 | 1 | 5 | 1 | 5 | 0.8 | 16 | 0.9375 | NA | NA | 3 | 1 | 2 | 0.5 | 4 | 1 | 9 | 0.8889 |
| 16_105618 | 3 | 0.6667 | 4 | 1 | 6 | 0.8333 | 4 | 0.5 | 17 | 0.7647 | 1 | 1 | 3 | 0.6667 | 1 | 1 | 4 | 0.5 | 9 | 0.6667 |
| 16_105619 | 3 | 0.3333 | 3 | 1 | 6 | 0.5 | 4 | 0.25 | 16 | 0.5 | 1 | 1 | 3 | 0 | 3 | 0.6667 | 5 | 0.2 | 12 | 0.3333 |
| 16_105619 | 3 | 1 | 3 | 0.6667 | 4 | 0.75 | 5 | 0.4 | 15 | 0.6667 | 2 | 0.5 | 3 | 0.6667 | 3 | 0.3333 | 3 | 0.3333 | 11 | 0.4545 |
| 16_105620 | 3 | 0.3333 | 4 | 0 | 6 | 0 | 6 | 0 | 19 | 0.0526 | 1 | 0 | 3 | 0.3333 | 3 | 0 | 5 | 0.2 | 12 | 0.1667 |
| 16_105620 | 3 | 0 | 3 | 0.3333 | 4 | 0.25 | 5 | 0.6 | 15 | 0.3333 | 1 | 1 | 1 | 1 | 3 | 0.3333 | 4 | 0.25 | 9 | 0.4444 |
| 16_105626 | 3 | 0 | 3 | 0 | 7 | 0.2857 | 5 | 0.2 | 18 | 0.1667 | 3 | 0 | 4 | 0.25 | 3 | 0 | 3 | 0 | 13 | 0.0769 |
| 16_105626 | 3 | 0 | 6 | 0.1667 | 7 | 0.1429 | 4 | 0.25 | 20 | 0.15 | 3 | 0 | 6 | 0.1667 | 3 | 0 | 4 | 0.25 | 16 | 0.125 |
| 16_105626 | 3 | 0 | 6 | 0 | 8 | 0.25 | 4 | 0 | 21 | 0.0952 | 3 | 0 | 5 | 0.4 | 2 | 0.5 | 4 | 0.5 | 14 | 0.3571 |
| 16_105626 | 2 | 0 | 6 | 0.1667 | 8 | 0 | 5 | 0.2 | 21 | 0.0952 | 3 | 0 | 6 | 0.3333 | 3 | 0.3333 | 4 | 0.25 | 16 | 0.25 |
| 16_107784 | 4 | 1 | 1 | 0 | 4 | 0.75 | 3 | 1 | 12 | 0.8333 | 3 | 1 | 3 | 0.6667 | 4 | 0.5 | 3 | 0.6667 | 13 | 0.6923 |
| 16_107784 | 4 | 0 | 1 | 0 | 4 | 0.25 | 3 | 0 | 12 | 0.0833 | 3 | 0 | 3 | 0.3333 | 4 | 0.25 | 3 | 0 | 13 | 0.1538 |
| 16_107785 | 4 | 1 | NA | NA | 3 | 0.6667 | 2 | 0 | 9 | 0.6667 | 1 | 0 | NA | NA | 2 | 1 | 1 | 0 | 4 | 0.5 |
| 16_107786 | 4 | 1 | 1 | 1 | 4 | 1 | 2 | 1 | 11 | 1 | 3 | 1 | 1 | 1 | 3 | 0.6667 | 3 | 1 | 10 | 0.9 |
| 16_107786 | 4 | 0.25 | 1 | 1 | 4 | 0.5 | 1 | 0 | 10 | 0.4 | 3 | 1 | 1 | 1 | 4 | 0 | 2 | 1 | 10 | 0.6 |
| 16_107788 | 4 | 1 | 1 | 1 | 4 | 1 | 2 | 1 | 11 | 1 | 3 | 0.6667 | NA | NA | 3 | 0.6667 | 3 | 1 | 9 | 0.7778 |
| 16_108791 | NA | NA | NA | NA | 1 | 1 | 1 | 1 | 2 | 1 | NA | NA | NA | NA | NA | NA | 1 | 1 | 1 | 1 |
| 16_108792 | NA | NA | NA | NA | 1 | 1 | 1 | 1 | 2 | 1 | NA | NA | NA | NA | NA | NA | 1 | 1 | 1 | 1 |
| 16_108900 | 8 | 0.125 | 9 | 0 | 10 | 0.2 | 12 | 0 | 39 | 0.0769 | 2 | 0 | 12 | 0.25 | 7 | 0.2857 | 8 | 0.25 | 29 | 0.2414 |
| 16_108901 | 8 | 0 | 14 | 0 | 10 | 0 | 12 | 0.3333 | 44 | 0.0909 | 2 | 0 | 11 | 0.0909 | 8 | 0.125 | 12 | 0 | 33 | 0.0606 |
| 16_108901 | 9 | 1 | 12 | 0.9167 | 11 | 0.8182 | 6 | 1 | 38 | 0.9211 | 1 | 1 | 10 | 1 | 6 | 1 | 4 | 1 | 21 | 1 |
| 16_108905 | 10 | 0.2 | 14 | 0.1429 | 12 | 0 | 12 | 0.25 | 48 | 0.1458 | 5 | 0.4 | 14 | 0.1429 | 10 | 0.1 | 12 | 0.3333 | 41 | 0.2195 |
| 16_108905 | 11 | 0.4545 | 14 | 0.4286 | 9 | 0.3333 | 13 | 0.3077 | 47 | 0.383 | 4 | 0.25 | 15 | 0.4667 | 8 | 0.125 | 8 | 0.5 | 35 | 0.3714 |
| 16_108907 | 10 | 0.4 | 11 | 0.5455 | 13 | 0.3846 | 10 | 0.7 | 44 | 0.5 | 5 | 0.6 | 13 | 0.7692 | 8 | 0.75 | 10 | 0.5 | 36 | 0.6667 |
| 16_108907 | 9 | 0.8889 | 10 | 0.9 | 8 | 1 | 9 | 0.8889 | 36 | 0.9167 | 6 | 1 | 11 | 1 | 4 | 1 | 8 | 0.875 | 29 | 0.9655 |
| 16_108908 | 14 | 0.8571 | 19 | 1 | 15 | 1 | 14 | 1 | 62 | 0.9677 | 6 | 0.8333 | 16 | 0.9375 | 8 | 0.75 | 11 | 1 | 41 | 0.9024 |
| 16_108908 | 13 | 0.9231 | 19 | 0.9474 | 14 | 1 | 15 | 0.9333 | 61 | 0.9508 | 4 | 1 | 18 | 1 | 9 | 0.8889 | 10 | 1 | 41 | 0.9756 |
| 16_108909 | 9 | 1 | 16 | 0.9375 | 10 | 1 | 13 | 1 | 48 | 0.9792 | 3 | 1 | 12 | 0.9167 | 9 | 1 | 9 | 1 | 33 | 0.9697 |
| 16_108911 | 16 | 0.125 | 18 | 0 | 17 | 0.2353 | 14 | 0 | 65 | 0.0923 | 10 | 0.1 | 21 | 0.1905 | 10 | 0.1 | 11 | 0.1818 | 52 | 0.1538 |
| 16_108912 | 13 | 0.2308 | 19 | 0.3158 | 19 | 0.3158 | 13 | 0.5385 | 64 | 0.3438 | 11 | 0.1818 | 18 | 0.1667 | 10 | 0.4 | 9 | 0.2222 | 48 | 0.2292 |
| 16_108919 | 20 | 0.2 | 21 | 0 | 17 | 0.1765 | 19 | 0.1053 | 77 | 0.1169 | 13 | 0.0769 | 25 | 0.32 | 13 | 0 | 19 | 0 | 70 | 0.1286 |
| 16_108924 | 21 | 0.1429 | 27 | 0.2593 | 26 | 0.1923 | 29 | 0.069 | 103 | 0.165 | 15 | 0.0667 | 27 | 0.1111 | 18 | 0.2778 | 22 | 0.1818 | 82 | 0.1585 |
| 16_108926 | 20 | 0.15 | 28 | 0.25 | 23 | 0.2609 | 28 | 0.3214 | 99 | 0.2525 | 15 | 0.3333 | 29 | 0.3103 | 22 | 0.0909 | 25 | 0.2 | 91 | 0.2308 |
| 16_108926 | 25 | 0.12 | 28 | 0.0714 | 22 | 0.1364 | 26 | 0.1538 | 101 | 0.1188 | 16 | 0.0625 | 30 | 0.1333 | 21 | 0.0476 | 22 | 0.0455 | 89 | 0.0787 |
| 16_108926 | 29 | 0.6897 | 30 | 0.7667 | 25 | 0.72 | 27 | 0.7037 | 111 | 0.7207 | 15 | 0.6667 | 30 | 0.7 | 21 | 0.5714 | 24 | 0.5417 | 90 | 0.6222 |
| 16_108928 | 2 | 0.5 | 3 | 0.6667 | 3 | 0 | 2 | 0 | 10 | 0.3 | NA | NA | 1 | 1 | NA | NA | 6 | 0.6667 | 7 | 0.7143 |
| 16_108928 | 2 | 1 | 3 | 0 | 3 | 0.6667 | 2 | 1 | 10 | 0.6 | NA | NA | 2 | 1 | NA | NA | 6 | 1 | 8 | 1 |
| 16_108931 | 2 | 0.5 | 3 | 0.3333 | 3 | 0.3333 | 2 | 0 | 10 | 0.3 | NA | NA | 3 | 1 | NA | NA | 6 | 0 | 9 | 0.3333 |
| 16_108932 | 2 | 0.5 | 3 | 0 | 3 | 1 | 2 | 0.5 | 10 | 0.5 | NA | NA | 3 | 0 | NA | NA | 7 | 0.1429 | 10 | 0.1 |
| 16_108933 | 2 | 0 | 3 | 0.6667 | 1 | 0 | 2 | 1 | 8 | 0.5 | NA | NA | 2 | 0 | NA | NA | 5 | 0.4 | 7 | 0.2857 |
| 16_108934 | 2 | 0.5 | 4 | 0 | 3 | 0 | 2 | 0 | 11 | 0.0909 | NA | NA | 3 | 0 | NA | NA | 7 | 0.1429 | 10 | 0.1 |
| 16_109051 | 2 | 0.5 | NA | NA | 1 | 0 | NA | NA | 3 | 0.3333 | NA | NA | 1 | 1 | NA | NA | NA | NA | 1 | 1 |
| 16_109057 | 4 | 0.75 | 3 | 1 | 2 | 1 | 1 | 1 | 10 | 0.9 | NA | NA | 1 | 1 | NA | NA | 2 | 1 | 3 | 1 |
| 16_109058 | 3 | 0 | 2 | 0 | 3 | 0.3333 | NA | NA | 8 | 0.125 | NA | NA | 2 | 0.5 | NA | NA | 1 | 0 | 3 | 0.3333 |
| 16_109060 | 4 | 0.5 | 2 | 1 | 3 | 0.6667 | 1 | 1 | 10 | 0.7 | NA | NA | 2 | 0.5 | NA | NA | 3 | 1 | 5 | 0.8 |
| 16_109061 | 1 | 1 | 2 | 0.5 | 3 | 0.3333 | NA | NA | 6 | 0.5 | NA | NA | 2 | 0.5 | NA | NA | 3 | 1 | 5 | 0.8 |
| 16_109061 | 3 | 0.6667 | 3 | 0.3333 | 3 | 0 | NA | NA | 9 | 0.3333 | NA | NA | 2 | 1 | NA | NA | 1 | 0 | 3 | 0.6667 |
| 16_109062 | 4 | 0.75 | 3 | 1 | 3 | 0.6667 | NA | NA | 10 | 0.8 | NA | NA | 2 | 1 | NA | NA | 3 | 1 | 5 | 1 |
| 16_109549 | 1 | 1 | 4 | 0.25 | 4 | 0 | 3 | 0.6667 | 12 | 0.3333 | NA | NA | 7 | 0.1429 | 3 | 0 | 1 | 0 | 11 | 0.0909 |
| 16_109552 | 1 | 1 | 2 | 1 | 3 | 0 | 2 | 0.5 | 8 | 0.5 | NA | NA | 6 | 0.3333 | 2 | 0 | 1 | 0 | 9 | 0.2222 |
| 16_110110 | 5 | 0.4 | 10 | 0.5 | 14 | 0.3571 | 10 | 0.4 | 39 | 0.4103 | 5 | 0.6 | 6 | 0.1667 | 10 | 0.2 | 6 | 0.1667 | 27 | 0.2593 |

| | | | | | | | | | | | | | | | | | | | | |
|---|---|---|---|---|---|---|---|---|---|---|---|---|---|---|---|---|---|---|---|---|
| 16_110110 | 6 | 0.6667 | 11 | 0.6364 | 9 | 0.5556 | 10 | 0.6 | 36 | 0.6111 | 5 | 0.8 | 3 | 0.6667 | 7 | 0.5714 | 5 | 0.6 | 20 | 0.65 |
| 16_110113 | 4 | 0 | 8 | 0.375 | 13 | 0.3846 | 10 | 0.3 | 35 | 0.3143 | 5 | 0.2 | 5 | 0.2 | 8 | 0.375 | 5 | 0.2 | 23 | 0.2609 |
| 16_110113 | 4 | 0.25 | 7 | 0.1429 | 11 | 0.5455 | 9 | 0.6667 | 31 | 0.4516 | 5 | 0.2 | 5 | 0.6 | 7 | 0.2857 | 5 | 0.4 | 22 | 0.3636 |
| 16_110115 | 4 | 0 | 7 | 0.4286 | 11 | 0.3636 | 10 | 0.3 | 32 | 0.3125 | 5 | 0 | 6 | 0.3333 | 8 | 0.5 | 5 | 0.8 | 24 | 0.4167 |
| 16_110343 | 3 | 0 | 9 | 0 | 4 | 0.25 | 2 | 0 | 18 | 0.0556 | 2 | 0 | 7 | 0 | 5 | 0 | 6 | 0.1667 | 20 | 0.05 |
| 16_110346 | 7 | 0.1429 | 10 | 0 | 4 | 0 | 7 | 0.1429 | 28 | 0.0714 | 3 | 0.6667 | 14 | 0.0714 | 8 | 0.125 | 9 | 0.1111 | 34 | 0.1471 |
| 16_110347 | 5 | 0 | 9 | 0.1111 | 6 | 0.3333 | 7 | 0.1429 | 27 | 0.1481 | 3 | 0.6667 | 14 | 0.0714 | 9 | 0.1111 | 9 | 0 | 35 | 0.1143 |
| 16_110349 | 6 | 0.3333 | 8 | 0.625 | 5 | 0.2 | 6 | 1 | 25 | 0.56 | 3 | 1 | 13 | 0.3846 | 9 | 0.5556 | 8 | 0.5 | 33 | 0.5152 |
| 16_110357 | 4 | 0.25 | 9 | 0 | 5 | 0.2 | 4 | 0.25 | 22 | 0.1364 | 2 | 0.5 | 10 | 0 | 5 | 0.2 | 7 | 0.2857 | 24 | 0.1667 |
| 16_110358 | 3 | 0.6667 | 7 | 0.4286 | 4 | 0.5 | 3 | 0.6667 | 17 | 0.5294 | 2 | 0.5 | 10 | 0.5 | 4 | 1 | 7 | 0.5714 | 23 | 0.6087 |
| 16_110359 | 4 | 0.25 | 7 | 0.1429 | 3 | 0.3333 | 4 | 0.5 | 18 | 0.2778 | 3 | 0.6667 | 10 | 0.4 | 6 | 0.1667 | 7 | 0.2857 | 26 | 0.3462 |
| 16_110359 | 4 | 0.75 | 8 | 0.375 | 4 | 0.5 | 3 | 0.6667 | 19 | 0.5263 | 3 | 1 | 8 | 0.125 | 6 | 0.5 | 7 | 0.7143 | 24 | 0.5 |
| 16_110360 | 3 | 0.3333 | 6 | 0 | 2 | 0 | 3 | 0.6667 | 14 | 0.2143 | 2 | 1 | 9 | 0.3333 | 4 | 0.75 | 7 | 0.5714 | 22 | 0.5455 |
| 16_110360 | 4 | 0.75 | 9 | 0.7778 | 2 | 0.5 | 2 | 0 | 17 | 0.6471 | 3 | 1 | 10 | 0.5 | 5 | 0.8 | 6 | 0.6667 | 24 | 0.6667 |
| 16_110375 | 3 | 0 | 9 | 0 | 3 | 0.3333 | 1 | 0 | 16 | 0.0625 | 2 | 0 | 6 | 0.1667 | 5 | 0 | 3 | 0 | 16 | 0.0625 |
| 16_110376 | 2 | 0.5 | 9 | 0.3333 | 4 | 0 | 1 | 0 | 16 | 0.25 | 2 | 0 | 8 | 0.25 | 4 | 0.25 | 4 | 0.5 | 18 | 0.2778 |
| 16_110376 | 3 | 0 | 7 | 0.2857 | 5 | 0.4 | 2 | 0.5 | 17 | 0.2941 | 1 | 0 | 8 | 0.375 | 4 | 0.25 | 4 | 0.25 | 17 | 0.2941 |
| 16_110378 | 3 | 0.3333 | 6 | 0.1667 | 5 | 0 | 2 | 0 | 16 | 0.125 | 2 | 0 | 8 | 0.125 | 3 | 0 | 5 | 0 | 18 | 0.0556 |
| 16_110384 | 10 | 0.5 | 11 | 0.6364 | 7 | 0.1429 | 6 | 0.8333 | 34 | 0.5294 | 6 | 0.3333 | 9 | 0.5556 | 7 | 0.5714 | 7 | 0.4286 | 29 | 0.4828 |
| 16_110384 | 11 | 0.4545 | 13 | 0.3077 | 8 | 0.625 | 6 | 0.6667 | 38 | 0.4737 | 5 | 0.8 | 12 | 0.6667 | 5 | 0.6 | 5 | 0.6 | 27 | 0.6667 |
| 16_110386 | 8 | 0.625 | 12 | 0.5 | 6 | 0.6667 | 6 | 0.3333 | 32 | 0.5312 | 5 | 0 | 11 | 0.3636 | 5 | 0.4 | 8 | 0.375 | 29 | 0.3103 |
| 16_110386 | 9 | 0.3333 | 11 | 0.2727 | 7 | 0.4286 | 6 | 0 | 33 | 0.2727 | 7 | 0.1429 | 10 | 0.2 | 7 | 0.2857 | 6 | 0.3333 | 30 | 0.2333 |
| 16_110387 | 9 | 0.1111 | 11 | 0 | 7 | 0.1429 | 7 | 0.1429 | 34 | 0.0882 | 7 | 0 | 11 | 0.1818 | 6 | 0 | 6 | 0 | 30 | 0.0667 |
| 16_110717 | 3 | 0 | 3 | 0.3333 | 5 | 0.2 | 3 | 0 | 14 | 0.1429 | 5 | 0.2 | 5 | 0.2 | 2 | 0 | 3 | 0 | 15 | 0.1333 |
| 16_110719 | 2 | 0.5 | 3 | 0 | 3 | 0.3333 | 1 | 0 | 9 | 0.2222 | 3 | 0.3333 | 5 | 0.4 | 3 | 0.3333 | 2 | 0 | 13 | 0.3077 |
| 16_110720 | 2 | 0 | 2 | 0.5 | 3 | 0.6667 | 1 | 0 | 8 | 0.375 | 2 | 0.5 | 5 | 0.8 | 2 | 1 | 3 | 0.6667 | 12 | 0.75 |
| 16_115760 | 2 | 1 | NA | NA | 1 | 1 | NA | NA | 3 | 1 | 1 | 1 | 1 | 0 | 3 | 0.3333 | 1 | 0 | 6 | 0.3333 |
| 16_115766 | 2 | 0.5 | NA | NA | 1 | 0 | 2 | 0 | 5 | 0.2 | 2 | 0.5 | 1 | 0 | 2 | 0.5 | 1 | 0 | 6 | 0.3333 |
| 16_115766 | 2 | 0 | NA | NA | NA | NA | 1 | 1 | 3 | 0.3333 | 1 | 0 | 1 | 0 | 3 | 0.3333 | 1 | 0 | 6 | 0.1667 |
| 16_115986 | 1 | 1 | 3 | 1 | 2 | 1 | NA | NA | 6 | 1 | 1 | 0 | 2 | 1 | 3 | 1 | 1 | 1 | 7 | 0.8571 |
| 16_115987 | 3 | 0 | 3 | 0.3333 | 3 | 0.3333 | NA | NA | 9 | 0.2222 | 1 | 0 | 3 | 0.3333 | 4 | 0.25 | 1 | 0 | 9 | 0.2222 |
| 16_115988 | 3 | 0 | 3 | 0.3333 | 3 | 0.3333 | NA | NA | 9 | 0.2222 | 1 | 0 | 3 | 0.3333 | 4 | 0 | 1 | 0 | 9 | 0.1111 |
| 16_115989 | 3 | 0.3333 | 3 | 0 | 2 | 0.5 | NA | NA | 8 | 0.25 | 1 | 0 | 3 | 1 | 4 | 0.5 | 1 | 1 | 9 | 0.6667 |
| 16_115990 | 3 | 0.3333 | 2 | 0 | 3 | 0 | NA | NA | 8 | 0.125 | 1 | 0 | 3 | 0.3333 | 4 | 0 | 1 | 1 | 9 | 0.2222 |
| 16_116221 | 5 | 0.2 | 7 | 0 | 4 | 0.25 | 7 | 0.1429 | 23 | 0.1304 | 5 | 0.2 | 3 | 0 | 2 | 0 | 4 | 0 | 14 | 0.0714 |
| 16_116224 | 5 | 1 | 4 | 1 | 3 | 0.6667 | 4 | 0.75 | 16 | 0.875 | 2 | 1 | 2 | 1 | 2 | 1 | 4 | 0.75 | 10 | 0.9 |
| 16_116225 | 3 | 1 | 2 | 1 | 1 | 1 | 4 | 1 | 10 | 1 | 1 | 1 | NA | NA | 3 | 1 | 1 | 1 | 5 | 1 |
| 16_116226 | 6 | 0.5 | 5 | 0.8 | 6 | 0.5 | 7 | 1 | 24 | 0.7083 | 4 | 0.5 | 4 | 0.25 | 3 | 0.3333 | 3 | 1 | 14 | 0.5 |
| 16_116227 | 4 | 0.25 | 3 | 0 | 2 | 0 | 4 | 0.25 | 13 | 0.1538 | 2 | 0.5 | 2 | 0.5 | 2 | 0 | 1 | 1 | 7 | 0.4286 |
| 16_116227 | 7 | 1 | 5 | 1 | 6 | 1 | 7 | 1 | 25 | 1 | 4 | 1 | 3 | 1 | 3 | 1 | 4 | 1 | 14 | 1 |
| 16_116227 | 6 | 0.8333 | 4 | 1 | 5 | 0.6 | 7 | 0.7143 | 22 | 0.7727 | 4 | 0.75 | 2 | 0.5 | 3 | 0.6667 | 4 | 0.75 | 13 | 0.6923 |
| 16_116229 | 5 | 0.2 | 5 | 0.6 | 4 | 0.5 | 7 | 0.5714 | 21 | 0.4762 | 3 | 0 | 2 | 1 | 4 | 0 | 4 | 0 | 13 | 0.1538 |
| 16_116230 | 7 | 0.5714 | 3 | 0 | 5 | 0 | 7 | 0.5714 | 22 | 0.3636 | 2 | 0.5 | 2 | 0 | 3 | 0 | 2 | 0 | 9 | 0.1111 |
| 16_116230 | 6 | 0.5 | 5 | 0.6 | 4 | 0.25 | 6 | 0.6667 | 21 | 0.5238 | 2 | 1 | 2 | 1 | 2 | 0.5 | 4 | 0.75 | 10 | 0.8 |
| 16_116230 | 4 | 1 | 4 | 1 | 3 | 1 | 3 | 1 | 14 | 1 | 1 | 1 | NA | NA | 2 | 1 | 2 | 1 | 5 | 1 |
| 16_116230 | 4 | 1 | 2 | 1 | 3 | 0.3333 | 3 | 0.6667 | 12 | 0.75 | 1 | 1 | 1 | 1 | 2 | 1 | 3 | 0.6667 | 7 | 0.8571 |
| 16_116231 | 5 | 0 | 2 | 0.5 | 1 | 0 | 6 | 0 | 14 | 0.0714 | 1 | 0 | 2 | 0 | 3 | 0.3333 | 3 | 0 | 9 | 0.1111 |
| 16_116232 | 5 | 0.4 | 4 | 0.5 | 4 | 0 | 5 | 0 | 18 | 0.2222 | 3 | 0 | NA | NA | 3 | 0.3333 | 3 | 0.6667 | 9 | 0.3333 |
| 16_116232 | 6 | 0.5 | 4 | 0 | 4 | 0 | 6 | 0.3333 | 20 | 0.25 | 3 | 0 | 3 | 0 | 3 | 0.3333 | 4 | 0.25 | 13 | 0.1538 |
| 16_116463 | 5 | 0 | 4 | 0.5 | 3 | 0 | 4 | 0 | 16 | 0.125 | 3 | 0 | 2 | 0 | 2 | 0 | 3 | 0.3333 | 10 | 0.1 |
| 16_116463 | 5 | 0.2 | 4 | 0.25 | 5 | 0.2 | 3 | 0 | 17 | 0.1765 | 3 | 0 | NA | NA | 2 | 0.5 | 2 | 0 | 7 | 0.1429 |
| 16_116464 | 6 | 0 | 4 | 0 | 5 | 0 | 3 | 0.3333 | 18 | 0.0556 | 2 | 0 | 1 | 0 | 2 | 0.5 | 3 | 0 | 8 | 0.125 |
| 16_117030 | NA | NA | 1 | 1 | 1 | 1 | NA | NA | 2 | 1 | 1 | 1 | NA | NA | NA | NA | NA | NA | 1 | 1 |
| 16_118027 | 3 | 0.6667 | 1 | 1 | 2 | 0.5 | 2 | 0.5 | 8 | 0.625 | NA | NA | 5 | 1 | 1 | 1 | 3 | 1 | 9 | 1 |
| 16_118027 | 3 | 0 | 1 | 0 | 2 | 0.5 | 2 | 0 | 8 | 0.125 | 1 | 0 | 6 | 0 | 1 | 0 | 3 | 0.3333 | 11 | 0.0909 |
| 16_118028 | 3 | 0.6667 | NA | NA | 3 | 0.3333 | 2 | 0 | 8 | 0.375 | 1 | 0 | 5 | 0 | 1 | 1 | 3 | 0.3333 | 10 | 0.2 |
| 16_120854 | 5 | 0.4 | 2 | 0 | 4 | 0.75 | 7 | 0.5714 | 18 | 0.5 | 2 | 0.5 | 4 | 0.75 | 2 | 1 | 2 | 0.5 | 10 | 0.7 |
| 16_120855 | 5 | 0.2 | 2 | 0 | 3 | 0.3333 | 8 | 0 | 18 | 0.1111 | 1 | 0 | 5 | 0.2 | 2 | 0 | 2 | 0 | 10 | 0.1 |
| 16_120855 | 5 | 0.2 | 3 | 0.3333 | 4 | 0.5 | 9 | 0.2222 | 21 | 0.2857 | 1 | 0 | 5 | 0.2 | 2 | 0.5 | 3 | 0.3333 | 11 | 0.2727 |
| 16_120855 | 5 | 0.8 | 3 | 1 | 4 | 1 | 11 | 1 | 23 | 0.9565 | 2 | 1 | 5 | 1 | 3 | 1 | 3 | 1 | 13 | 1 |
| 16_120856 | 5 | 0.8 | 2 | 1 | 4 | 1 | 11 | 1 | 22 | 0.9545 | 2 | 1 | 5 | 1 | 3 | 1 | 3 | 1 | 13 | 1 |
| 16_120857 | 6 | 0 | 2 | 0.5 | 4 | 0 | 11 | 0.0909 | 23 | 0.087 | 2 | 0.5 | 5 | 0.4 | 3 | 0 | 2 | 0.5 | 12 | 0.3333 |
| 16_120858 | 2 | 0.5 | 3 | 0.3333 | 3 | 0 | 10 | 0.5 | 18 | 0.3889 | 2 | 0.5 | 5 | 0.8 | 2 | 0 | 3 | 0.3333 | 12 | 0.5 |
| 16_120858 | 6 | 1 | 3 | 1 | 5 | 1 | 10 | 1 | 24 | 1 | 2 | 1 | 4 | 1 | 3 | 1 | 3 | 1 | 12 | 1 |
| 16_120859 | 3 | 0.3333 | 3 | 0 | 5 | 0.2 | 11 | 0.1818 | 22 | 0.1818 | 2 | 0 | 5 | 0.2 | 3 | 0 | 3 | 0.3333 | 13 | 0.1538 |

| | | | | | | | | | | | | | | | | | | | | |
|---|---|---|---|---|---|---|---|---|---|---|---|---|---|---|---|---|---|---|---|---|
| 16_120859 | 4 | 0.75 | 3 | 1 | 5 | 0.6 | 8 | 0.375 | 20 | 0.6 | 1 | 0 | 3 | 0.6667 | 3 | 0.6667 | 3 | 1 | 10 | 0.7 |
| 16_120860 | 6 | 0 | 3 | 0.6667 | 5 | 0 | 11 | 0.1818 | 25 | 0.16 | 1 | 1 | 4 | 0 | 4 | 0 | 2 | 0 | 11 | 0.0909 |
| 16_120862 | 6 | 0.8333 | 3 | 1 | 5 | 0.8 | 12 | 0.9167 | 26 | 0.8846 | 2 | 1 | 5 | 1 | 4 | 1 | 4 | 1 | 15 | 1 |
| 16_120862 | 5 | 0.8 | 3 | 1 | 6 | 0.6667 | 7 | 0.8571 | 21 | 0.8095 | 2 | 0.5 | 4 | 0.75 | 4 | 0.5 | 3 | 0.6667 | 13 | 0.6154 |
| 16_120863 | 5 | 0.8 | 3 | 0 | 6 | 0 | 10 | 0 | 24 | 0.1667 | 2 | 0.5 | 5 | 0 | 2 | 0.5 | 3 | 0.3333 | 12 | 0.25 |
| 16_120863 | 4 | 0.25 | 3 | 0 | 7 | 0 | 8 | 0.25 | 22 | 0.1364 | 3 | 0.3333 | 5 | 0.2 | 4 | 0.25 | 4 | 0 | 16 | 0.1875 |
| 16_120863 | 5 | 0.4 | 1 | 1 | 8 | 0.875 | 11 | 0.5455 | 25 | 0.64 | 4 | 0.5 | 5 | 0.6 | 4 | 0.75 | 4 | 0.75 | 17 | 0.6471 |
| 16_120864 | 5 | 0 | 3 | 0.6667 | 8 | 0.25 | 12 | 0.25 | 28 | 0.25 | 3 | 0.3333 | 5 | 0.6 | 2 | 0 | 4 | 0.25 | 14 | 0.3571 |
| 16_120864 | 3 | 0.6667 | 2 | 0.5 | 5 | 0.2 | 9 | 0.4444 | 19 | 0.4211 | 2 | 1 | 5 | 0.6 | 3 | 0 | 1 | 1 | 11 | 0.5455 |
| 16_120864 | 5 | 0.8 | 3 | 1 | 8 | 0.875 | 12 | 1 | 28 | 0.9286 | 4 | 1 | 5 | 1 | 5 | 0.8 | 4 | 1 | 18 | 0.9444 |
| 16_120864 | 6 | 0 | 4 | 0.25 | 6 | 0.1667 | 12 | 0 | 28 | 0.0714 | 3 | 0 | 2 | 0 | 4 | 0.25 | 4 | 0 | 13 | 0.0769 |
| 16_120865 | 5 | 0.4 | 2 | 1 | 5 | 0.8 | 7 | 0.5714 | 19 | 0.6316 | 4 | 1 | 5 | 0.8 | 4 | 1 | 4 | 0.75 | 17 | 0.8824 |
| 16_120865 | 5 | 0.2 | 3 | 0.6667 | 8 | 0.125 | 12 | 0.0833 | 28 | 0.1786 | 4 | 0 | 5 | 0.2 | 5 | 0.6 | 3 | 0 | 17 | 0.2353 |
| 16_120865 | 6 | 0.8333 | 2 | 1 | 4 | 1 | 9 | 1 | 21 | 0.9524 | 3 | 0.6667 | 4 | 1 | 2 | 1 | 2 | 1 | 11 | 0.9091 |
| 16_120869 | 7 | 0.1429 | 3 | 0.3333 | 6 | 0.3333 | 9 | 0.2222 | 25 | 0.24 | 4 | 0.25 | 4 | 0 | 3 | 0.6667 | 5 | 0.4 | 16 | 0.3125 |
| 16_129481 | 1 | 0 | NA | NA | 2 | 0.5 | 1 | 1 | 4 | 0.5 | NA | NA | 1 | 1 | 1 | 1 | 2 | 1 | 4 | 1 |
| 16_129481 | 1 | 1 | 1 | 0 | 3 | 1 | 1 | 0 | 6 | 0.6667 | NA | NA | 1 | 1 | NA | NA | 2 | 1 | 3 | 1 |
| 16_129483 | 2 | 0.5 | 1 | 0 | 5 | 0.2 | 1 | 0 | 9 | 0.2222 | 1 | 0 | 2 | 0 | 1 | 0 | 1 | 1 | 5 | 0.2 |
| 16_129483 | 2 | 0 | 1 | 0 | 5 | 0.2 | 1 | 0 | 9 | 0.1111 | 1 | 0 | 2 | 0.5 | 1 | 0 | 1 | 0 | 5 | 0.2 |
| 16_129655 | 1 | 1 | 3 | 1 | 6 | 1 | 2 | 1 | 12 | 1 | 2 | 1 | 2 | 1 | 1 | 1 | 3 | 1 | 8 | 1 |
| 16_129658 | 2 | 0.5 | 3 | 0.3333 | 8 | 0.375 | 2 | 1 | 15 | 0.4667 | 3 | 0.3333 | 2 | 1 | NA | NA | 3 | 0.3333 | 8 | 0.5 |
| 16_129658 | 2 | 0.5 | 2 | 1 | 9 | 1 | 2 | 1 | 15 | 0.9333 | 3 | 1 | 2 | 1 | 1 | 1 | 2 | 1 | 8 | 1 |
| 16_129659 | 1 | 1 | 3 | 0.6667 | 9 | 0.7778 | 1 | 1 | 14 | 0.7857 | 3 | 1 | 2 | 1 | NA | NA | 3 | 0.6667 | 8 | 0.875 |
| 16_129659 | NA | NA | 1 | 1 | 7 | 1 | 1 | 1 | 9 | 1 | 3 | 1 | 2 | 1 | NA | NA | 1 | 1 | 6 | 1 |
| 16_129659 | 2 | 0 | 3 | 0 | 8 | 0.125 | 1 | 0 | 14 | 0.0714 | 4 | 0 | 2 | 0.5 | 1 | 0 | 2 | 0 | 9 | 0.1111 |
| 16_129660 | 2 | 0.5 | 3 | 0.3333 | 8 | 0.375 | 1 | 0 | 14 | 0.3571 | 4 | 0.5 | 2 | 0 | 1 | 1 | 2 | 0.5 | 9 | 0.4444 |
| 16_129661 | NA | NA | 3 | 1 | 8 | 1 | 2 | 1 | 13 | 1 | 4 | 1 | 1 | 1 | 1 | 1 | 2 | 1 | 8 | 1 |
| 16_129663 | 5 | 0.6 | 2 | 0.5 | 9 | 0.6667 | 1 | 0 | 17 | 0.5882 | 4 | 0.5 | 2 | 0 | NA | NA | 3 | 0.6667 | 9 | 0.4444 |
| 16_129664 | 5 | 0.6 | 3 | 1 | 7 | 1 | 3 | 1 | 18 | 0.8889 | 5 | 1 | 2 | 1 | 1 | 1 | 3 | 1 | 11 | 1 |
| 16_129665 | 3 | 0 | 2 | 0 | 6 | 0.1667 | 4 | 0.25 | 15 | 0.1333 | 4 | 0 | 2 | 0.5 | NA | NA | 2 | 0 | 8 | 0.125 |
| 16_131763 | 1 | 1 | 1 | 1 | NA | NA | 2 | 1 | 4 | 1 | NA | NA | NA | NA | 2 | 0.5 | NA | NA | 2 | 0.5 |
| 16_132186 | 1 | 1 | 1 | 0 | 2 | 0 | 3 | 0 | 7 | 0.1429 | 2 | 0 | 3 | 0.3333 | 4 | 0 | 2 | 0 | 11 | 0.0909 |
| 16_132187 | 1 | 0 | 1 | 1 | 2 | 0.5 | 3 | 0.6667 | 7 | 0.5714 | 2 | 1 | 2 | 0 | 3 | 0.6667 | 1 | 0 | 8 | 0.5 |
| 16_132188 | 1 | 0 | 1 | 1 | 2 | 0.5 | 2 | 0 | 6 | 0.3333 | 2 | 0.5 | 3 | 0.3333 | 4 | 0.25 | 2 | 0.5 | 11 | 0.3636 |
| 16_132189 | 1 | 1 | NA | NA | 2 | 0 | 3 | 0.6667 | 6 | 0.5 | 2 | 0 | 2 | 0 | 2 | 0 | 2 | 0.5 | 8 | 0.125 |
| 16_132193 | NA | NA | 2 | 1 | 2 | 1 | 3 | 1 | 7 | 1 | 2 | 1 | 3 | 1 | 4 | 1 | 3 | 1 | 12 | 1 |
| 16_132194 | NA | NA | 1 | 0 | 2 | 0.5 | 3 | 0 | 6 | 0.1667 | 2 | 0 | 3 | 0.6667 | 5 | 0.2 | 3 | 0.6667 | 13 | 0.3846 |
| 16_132301 | 2 | 0 | 4 | 0 | 4 | 0.25 | 3 | 0 | 13 | 0.0769 | 3 | 0 | 6 | 0.1667 | 9 | 0.3333 | 4 | 0 | 22 | 0.1818 |
| 16_132301 | 2 | 1 | 3 | 0.6667 | 3 | 1 | 3 | 1 | 11 | 0.9091 | NA | NA | 5 | 0.8 | 7 | 0.5714 | 3 | 0.6667 | 15 | 0.6667 |
| 16_132301 | 2 | 0 | 5 | 0.2 | 5 | 0 | 3 | 0.3333 | 15 | 0.1333 | 3 | 0 | 7 | 0.1429 | 7 | 0.1429 | 4 | 0.25 | 21 | 0.1429 |
| 16_132302 | 3 | 0.3333 | 4 | 0.25 | 3 | 0 | 3 | 0.3333 | 13 | 0.2308 | 2 | 0 | 7 | 0 | 8 | 0.125 | 3 | 0 | 20 | 0.05 |
| 16_132303 | 3 | 0.3333 | 7 | 0.5714 | 4 | 1 | 3 | 0.6667 | 17 | 0.6471 | 5 | 0.4 | 7 | 0.5714 | 9 | 0.5556 | 4 | 0.75 | 25 | 0.56 |
| 16_132304 | 3 | 0.3333 | 7 | 0.4286 | 4 | 0.25 | 2 | 1 | 16 | 0.4375 | 5 | 0.2 | 8 | 0.75 | 8 | 0.25 | 3 | 0 | 24 | 0.375 |
| 16_132304 | 2 | 1 | 7 | 0.5714 | 4 | 0.5 | 3 | 1 | 16 | 0.6875 | 4 | 0.75 | 8 | 0.75 | 9 | 0.5556 | 4 | 0.75 | 25 | 0.68 |
| 16_132304 | 2 | 0.5 | 7 | 0.8571 | 3 | 0.3333 | 3 | 0.6667 | 15 | 0.6667 | 5 | 0.8 | 7 | 0.7143 | 8 | 1 | 4 | 0.75 | 24 | 0.8333 |
| 16_132305 | 2 | 0.5 | 7 | 0.5714 | 2 | 0.5 | 3 | 0.3333 | 14 | 0.5 | 2 | 0.5 | 7 | 1 | 9 | 0.4444 | 4 | 0.5 | 22 | 0.6364 |
| 16_132305 | 2 | 1 | 7 | 0.4286 | 4 | 0.25 | 3 | 0.6667 | 16 | 0.5 | 4 | 0.5 | 6 | 0.6667 | 9 | 0.3333 | 4 | 0.75 | 23 | 0.5217 |
| 16_132305 | 2 | 1 | 5 | 1 | 4 | 1 | 3 | 1 | 14 | 1 | 4 | 1 | 8 | 1 | 7 | 1 | 3 | 1 | 22 | 1 |
| 16_132306 | 3 | 0.3333 | 6 | 0.1667 | 4 | 0.25 | 3 | 0.3333 | 16 | 0.25 | 3 | 0 | 9 | 0 | 9 | 0.3333 | 4 | 0 | 25 | 0.12 |
| 16_132307 | 3 | 0.6667 | 7 | 1 | 5 | 0.8 | 3 | 1 | 18 | 0.8889 | 5 | 1 | 9 | 1 | 9 | 1 | 4 | 1 | 27 | 1 |
| 16_132307 | 3 | 1 | 6 | 1 | 5 | 1 | 3 | 1 | 17 | 1 | 4 | 1 | 9 | 0.8889 | 8 | 1 | 4 | 1 | 25 | 0.96 |
| 16_132310 | 3 | 0.3333 | 5 | 0.8 | 3 | 0 | 3 | 0.6667 | 14 | 0.5 | 3 | 0.6667 | 8 | 0.375 | 7 | 0.4286 | 5 | 0.4 | 23 | 0.4348 |
| 16_132312 | 5 | 0.6 | 6 | 0.5 | 5 | 0.4 | 4 | 0.25 | 20 | 0.45 | 5 | 0.2 | 7 | 0.8571 | 10 | 0.7 | 3 | 0.6667 | 25 | 0.64 |
| 16_132312 | 4 | 0 | 9 | 0.1111 | 4 | 0 | 3 | 0 | 20 | 0.05 | 4 | 0 | 9 | 0 | 10 | 0.2 | 5 | 0 | 28 | 0.0714 |
| 16_132313 | 5 | 0.2 | 7 | 0.5714 | 6 | 0.5 | 5 | 0.2 | 23 | 0.3913 | 5 | 0 | 9 | 0.1111 | 13 | 0.2308 | 4 | 0.25 | 31 | 0.1613 |
| 16_132313 | 5 | 0.8 | 7 | 0.8571 | 4 | 0.75 | 5 | 0.8 | 21 | 0.8095 | 4 | 1 | 6 | 0.8333 | 13 | 0.8462 | 5 | 1 | 28 | 0.8929 |
| 16_132315 | 4 | 0.25 | 6 | 0.1667 | 5 | 0 | 5 | 0 | 20 | 0.1 | 2 | 0.5 | 6 | 0.3333 | 11 | 0.4545 | 3 | 0.6667 | 22 | 0.4545 |
| 16_132347 | 20 | 0.05 | 34 | 0.1176 | 19 | 0 | 29 | 0.069 | 102 | 0.0686 | 13 | 0.0769 | 27 | 0.1111 | 31 | 0.0968 | 26 | 0.1154 | 97 | 0.1031 |
| 16_132349 | 22 | 0.3182 | 34 | 0.3235 | 19 | 0.1579 | 28 | 0.1786 | 103 | 0.2524 | 13 | 0.1538 | 25 | 0.2 | 33 | 0.1212 | 24 | 0.2083 | 95 | 0.1684 |
| 16_132350 | 22 | 0.2727 | 36 | 0.1667 | 20 | 0.15 | 21 | 0.3333 | 99 | 0.2222 | 13 | 0.0769 | 25 | 0.12 | 29 | 0.2069 | 26 | 0.1154 | 93 | 0.1398 |
| 16_133565 | 2 | 0 | 4 | 0.25 | 2 | 0 | 5 | 0 | 13 | 0.0769 | NA | NA | 4 | 0.25 | 2 | 0 | 1 | 0 | 7 | 0.1429 |
| 16_135031 | 1 | 1 | 1 | 1 | 1 | 1 | 2 | 0 | 5 | 0.6 | 2 | 0.5 | 2 | 0 | 1 | 0 | 2 | 0 | 7 | 0.1429 |
| 16_135152 | 2 | 0 | 3 | 0.3333 | 1 | 0 | 3 | 0 | 9 | 0.1111 | 2 | 0 | 4 | 0.25 | 1 | 0 | 2 | 0 | 9 | 0.1111 |
| 16_135168 | 2 | 0.5 | 2 | 0.5 | 1 | 0 | 2 | 0.5 | 7 | 0.4286 | 2 | 0 | 3 | 0.3333 | 1 | 0 | 1 | 1 | 7 | 0.2857 |
| 16_135170 | 1 | 0 | 1 | 1 | NA | NA | NA | NA | 2 | 0.5 | 1 | 0 | 2 | 0.5 | NA | NA | NA | NA | 3 | 0.3333 |

| | | | | | | | | | | | | | | | | | | | | |
|---|---|---|---|---|---|---|---|---|---|---|---|---|---|---|---|---|---|---|---|---|
| 16_1352017 | 1 | 0 | 2 | 0 | 2 | 0.5 | 3 | 0.3333 | 8 | 0.25 | 2 | 0.5 | 1 | 0 | NA | NA | 3 | 0.6667 | 6 | 0.5 |
| 16_1352018 | 1 | 0 | 3 | 0 | 2 | 0.5 | 3 | 0.3333 | 9 | 0.2222 | 2 | 0 | 2 | 0.5 | 1 | 0 | 3 | 0 | 8 | 0.125 |
| 16_1352019 | NA | NA | 3 | 1 | 2 | 1 | 3 | 1 | 8 | 1 | 2 | 1 | 2 | 1 | 1 | 1 | 3 | 1 | 8 | 1 |
| 16_1352028 | 1 | 0 | 3 | 0 | 2 | 0 | 3 | 0.3333 | 9 | 0.1111 | 2 | 0.5 | 2 | 0 | NA | NA | 2 | 0.5 | 6 | 0.3333 |
| 16_1352031 | 1 | 0 | 3 | 0.6667 | 2 | 1 | 3 | 1 | 9 | 0.7778 | 2 | 0.5 | 2 | 1 | 1 | 1 | 2 | 1 | 7 | 0.8571 |
| 16_1352034 | 1 | 0 | 3 | 0.3333 | 2 | 1 | 3 | 1 | 9 | 0.6667 | 2 | 0.5 | 2 | 0.5 | 1 | 0 | 3 | 0.6667 | 8 | 0.5 |
| 16_1352109 | 1 | 1 | 3 | 0.6667 | 1 | 1 | 3 | 0.3333 | 8 | 0.625 | 1 | 0 | 3 | 0.3333 | NA | NA | 2 | 0.5 | 6 | 0.3333 |
| 16_1352133 | 1 | 0 | 6 | 0.1667 | 2 | 0 | 3 | 0.3333 | 12 | 0.1667 | 1 | 0 | 4 | 0 | 2 | 0 | 2 | 0.5 | 9 | 0.1111 |
| 16_1363097 | 3 | 0 | 4 | 0 | 6 | 0.3333 | 7 | 0 | 20 | 0.1 | 1 | 0 | 7 | 0 | 5 | 0 | 7 | 0.1429 | 20 | 0.05 |
| 16_1363101 | 3 | 0 | 4 | 0 | 7 | 0.1429 | 6 | 0.3333 | 20 | 0.15 | 1 | 0 | 9 | 0 | 5 | 0.2 | 7 | 0.2857 | 22 | 0.1364 |
| 16_1363119 | 1 | 1 | 6 | 0.6667 | 6 | 0.6667 | 7 | 1 | 20 | 0.8 | 1 | 1 | 9 | 0.8889 | 3 | 0.3333 | 6 | 1 | 19 | 0.8421 |
| 16_1363123 | 2 | 0.5 | 5 | 0 | 6 | 0.1667 | 3 | 0.3333 | 16 | 0.1875 | 1 | 0 | 7 | 0.1429 | 5 | 0.2 | 7 | 0.1429 | 20 | 0.15 |
| 16_1363130 | 2 | 0 | 4 | 0 | 8 | 0.125 | 7 | 0.2857 | 21 | 0.1429 | 1 | 0 | 6 | 0 | 2 | 0.5 | 7 | 0.1429 | 16 | 0.125 |
| 16_1363133 | 3 | 0 | 4 | 0.25 | 6 | 0.3333 | 7 | 0.4286 | 20 | 0.3 | 1 | 0 | 9 | 0.1111 | 4 | 0.75 | 6 | 0.5 | 20 | 0.35 |
| 16_1363150 | 4 | 0 | 6 | 0 | 8 | 0.125 | 5 | 0.2 | 23 | 0.087 | NA | NA | 8 | 0.125 | 5 | 0.4 | 7 | 0.1429 | 20 | 0.2 |
| 16_1363159 | 4 | 0.25 | 6 | 0.1667 | 8 | 0.25 | 6 | 0.1667 | 24 | 0.2083 | 3 | 0 | 9 | 0.3333 | 4 | 0 | 6 | 0 | 22 | 0.1364 |
| 16_1363177 | 3 | 0.3333 | 5 | 0 | 9 | 0.1111 | 6 | 0 | 23 | 0.087 | 3 | 0 | 9 | 0.3333 | 5 | 0 | 8 | 0.125 | 25 | 0.16 |
| 16_1363199 | 4 | 0 | 9 | 0.1111 | 10 | 0.2 | 10 | 0.2 | 33 | 0.1515 | 4 | 0 | 11 | 0.1818 | 4 | 0.5 | 10 | 0.4 | 29 | 0.2759 |
| 16_1363239 | 4 | 0.25 | 10 | 0.1 | 9 | 0 | 10 | 0.3 | 33 | 0.1515 | 5 | 0.2 | 12 | 0.1667 | 5 | 0 | 8 | 0.125 | 30 | 0.1333 |
| 16_1365251 | 7 | 1 | 8 | 1 | 8 | 0.875 | 4 | 1 | 27 | 0.963 | 2 | 1 | 7 | 0.8571 | 7 | 0.8571 | 6 | 1 | 22 | 0.9091 |
| 16_1365253 | 6 | 0 | 8 | 0.125 | 8 | 0.125 | 4 | 0.5 | 26 | 0.1538 | 2 | 0.5 | 7 | 0.2857 | 4 | 0 | 6 | 0.3333 | 19 | 0.2632 |
| 16_1365373 | 2 | 0.5 | NA | NA | 1 | 1 | 1 | 1 | 4 | 0.75 | 1 | 1 | 1 | 0 | 1 | 1 | NA | NA | 3 | 0.6667 |
| 16_1365386 | 2 | 0 | NA | NA | 1 | 0 | 1 | 1 | 4 | 0.25 | 1 | 1 | 1 | 0 | NA | NA | NA | NA | 2 | 0.5 |
| 16_1365401 | 2 | 0 | NA | NA | 1 | 1 | 1 | 0 | 4 | 0.25 | 1 | 0 | 2 | 0.5 | NA | NA | NA | NA | 3 | 0.3333 |
| 16_1365415 | 1 | 0 | NA | NA | 1 | 1 | 2 | 0 | 4 | 0.25 | 1 | 1 | 2 | 0.5 | NA | NA | NA | NA | 3 | 0.6667 |
| 16_1365419 | 2 | 0 | NA | NA | 1 | 0 | 2 | 0.5 | 5 | 0.2 | 1 | 0 | 2 | 0.5 | NA | NA | NA | NA | 3 | 0.3333 |
| 16_1375884 | 2 | 0.5 | 1 | 0 | 6 | 0.3333 | 2 | 0.5 | 11 | 0.3636 | 1 | 0 | 6 | 0.1667 | 4 | 0.5 | 3 | 0 | 14 | 0.2143 |
| 16_1376466 | 5 | 0.6 | 4 | 0.5 | 14 | 0.8571 | 6 | 0.8333 | 29 | 0.7586 | 5 | 0.6 | 10 | 0.7 | 13 | 0.6923 | 9 | 0.8889 | 37 | 0.7297 |
| 16_1376479 | 5 | 1 | 3 | 0.3333 | 12 | 0.75 | 7 | 0.2857 | 27 | 0.6296 | 3 | 0.3333 | 10 | 0.7 | 11 | 0.8182 | 8 | 0.75 | 32 | 0.7188 |
| 16_1376481 | 6 | 0.8333 | 4 | 0.75 | 14 | 0.7143 | 7 | 0.7143 | 31 | 0.7419 | 5 | 0.6 | 10 | 0.9 | 11 | 0.6364 | 8 | 0.625 | 34 | 0.7059 |
| 16_1376489 | 6 | 1 | 5 | 1 | 14 | 0.9286 | 7 | 1 | 32 | 0.9688 | 5 | 1 | 11 | 0.8182 | 13 | 0.9231 | 10 | 1 | 39 | 0.9231 |
| 16_1376504 | 5 | 0 | 6 | 0.1667 | 14 | 0.2143 | 8 | 0 | 33 | 0.1212 | 6 | 0 | 10 | 0 | 16 | 0 | 9 | 0.3333 | 41 | 0.0732 |
| 16_1376590 | 7 | 0 | 4 | 0.5 | 15 | 0.2667 | 10 | 0.3 | 36 | 0.25 | 4 | 0 | 15 | 0.0667 | 16 | 0.1875 | 10 | 0.1 | 45 | 0.1111 |
| 16_1376691 | 10 | 0.1 | 3 | 0 | 7 | 0.5714 | 4 | 0.25 | 24 | 0.25 | 2 | 0 | 9 | 0.2222 | 7 | 0.1429 | 8 | 0 | 26 | 0.1154 |
| 16_1376702 | 5 | 0.2 | 3 | 0 | 7 | 0.2857 | 2 | 1 | 17 | 0.2941 | 2 | 0.5 | 7 | 0.1429 | 6 | 0.3333 | 6 | 0.1667 | 21 | 0.2381 |
| 16_1376716 | 8 | 0 | 4 | 0 | 9 | 0.1111 | 5 | 0.2 | 26 | 0.0769 | 3 | 0.3333 | 8 | 0.375 | 8 | 0.25 | 9 | 0.1111 | 28 | 0.25 |
| 16_1376725 | 6 | 0 | 3 | 0 | 5 | 0.2 | 4 | 0 | 18 | 0.0556 | 1 | 0 | 6 | 0.5 | 7 | 0.1429 | 2 | 0 | 16 | 0.25 |
| 16_1376731 | 6 | 0.3333 | 3 | 0.6667 | 5 | 0.6 | 5 | 0.4 | 19 | 0.4737 | 1 | 0 | 5 | 0.6 | 8 | 0.5 | 2 | 0 | 16 | 0.4375 |
| 16_1376732 | 6 | 0 | 1 | 0 | 4 | 0 | 5 | 0.2 | 16 | 0.0625 | 1 | 0 | 6 | 0.1667 | 8 | 0.25 | 2 | 0 | 17 | 0.1765 |
| 16_1376733 | 6 | 0 | 2 | 0 | 5 | 0.4 | 5 | 0.4 | 18 | 0.2222 | 1 | 0 | 5 | 0.8 | 8 | 0.5 | 2 | 0.5 | 16 | 0.5625 |
| 16_1376735 | 5 | 0 | 3 | 0.3333 | 5 | 0.2 | 5 | 0 | 18 | 0.1111 | 1 | 0 | 6 | 0.1667 | 8 | 0.125 | 2 | 0.5 | 17 | 0.1765 |
| 16_1392910 | 6 | 0 | 8 | 0 | 7 | 0.2857 | 7 | 0 | 28 | 0.0714 | 4 | 0 | 8 | 0.125 | 8 | 0.25 | 4 | 0.25 | 24 | 0.1667 |
| 16_1392914 | 7 | 0 | 6 | 0.1667 | 8 | 0 | 8 | 0.375 | 29 | 0.1379 | 5 | 0 | 8 | 0 | 6 | 0.5 | 4 | 0 | 23 | 0.1304 |
| 16_1392934 | 5 | 0.4 | 5 | 0.4 | 7 | 0.1429 | 7 | 0.1429 | 24 | 0.25 | 4 | 0 | 6 | 0.5 | 8 | 0.375 | 5 | 0.2 | 23 | 0.3043 |
| 16_1392935 | 6 | 0.1667 | 7 | 0 | 8 | 0 | 5 | 0.2 | 26 | 0.0769 | 3 | 0.3333 | 7 | 0.2857 | 7 | 0.4286 | 3 | 0 | 20 | 0.3 |
| 16_1392938 | 7 | 0 | 7 | 0 | 8 | 0 | 7 | 0.4286 | 29 | 0.1034 | 3 | 0 | 7 | 0.1429 | 5 | 0 | 3 | 0 | 18 | 0.0556 |
| 16_1394388 | 5 | 0 | 5 | 0 | 5 | 0 | 4 | 0.25 | 19 | 0.0526 | 2 | 0 | 3 | 0.3333 | 5 | 0 | 4 | 0 | 14 | 0.0714 |
| 16_1396037 | 2 | 0.5 | 3 | 0.3333 | 1 | 1 | 2 | 0.5 | 8 | 0.5 | 2 | 0 | 1 | 1 | 2 | 0 | 3 | 0.6667 | 8 | 0.375 |
| 16_1397019 | 2 | 0 | 3 | 0.3333 | NA | NA | 2 | 1 | 7 | 0.4286 | 2 | 0.5 | 1 | 0 | 2 | 0.5 | 3 | 0.3333 | 8 | 0.375 |
| 16_1397029 | 1 | 0 | 3 | 1 | NA | NA | 2 | 1 | 6 | 0.8333 | 2 | 0.5 | 1 | 1 | 2 | 1 | 2 | 1 | 7 | 0.8571 |
| 16_1397039 | 1 | 1 | 3 | 0 | NA | NA | 2 | 0.5 | 6 | 0.3333 | 1 | 1 | 1 | 1 | 1 | 0 | 3 | 0.3333 | 6 | 0.5 |
| 16_1400968 | 3 | 0.3333 | 5 | 0.2 | 5 | 0.6 | 2 | 0.5 | 15 | 0.4 | 3 | 0.3333 | NA | NA | 3 | 0.3333 | 3 | 0.3333 | 9 | 0.3333 |
| 16_1401002 | 3 | 0 | 5 | 0.2 | 5 | 0 | 2 | 0.5 | 15 | 0.1333 | 2 | 0 | NA | NA | 2 | 0.5 | 2 | 0 | 6 | 0.1667 |
| 16_1401007 | 5 | 0.6 | 6 | 0.3333 | 4 | 1 | 3 | 0.6667 | 18 | 0.6111 | 3 | 1 | NA | NA | 2 | 1 | 3 | 1 | 8 | 1 |
| 16_1401010 | 4 | 0.25 | 6 | 0.1667 | 6 | 0 | 3 | 0 | 19 | 0.1053 | 2 | 0 | NA | NA | 2 | 0.5 | 3 | 0.3333 | 7 | 0.2857 |
| 16_1401024 | 5 | 0 | 7 | 0.1429 | 5 | 0 | 3 | 0 | 20 | 0.05 | 3 | 0 | 1 | 0 | 2 | 0.5 | 4 | 0.25 | 10 | 0.2 |
| 16_1401025 | 5 | 0 | 5 | 0.2 | 6 | 0.1667 | 3 | 0.3333 | 19 | 0.1579 | 2 | 0.5 | NA | NA | 3 | 0.6667 | 3 | 0 | 8 | 0.375 |
| 16_1401052 | 5 | 0 | 7 | 0.1429 | 6 | 0.3333 | 3 | 0.3333 | 21 | 0.1905 | 3 | 0 | 3 | 0 | 2 | 0.5 | 4 | 0.25 | 12 | 0.1667 |
| 16_1401056 | 5 | 0.6 | 6 | 0.5 | 6 | 0.6667 | 3 | 0 | 20 | 0.5 | 3 | 0.3333 | 3 | 0.3333 | 3 | 0.6667 | 4 | 0.5 | 13 | 0.4615 |
| 16_1401057 | 5 | 0.4 | 5 | 0 | 3 | 0 | 2 | 1 | 15 | 0.2667 | 3 | 0.6667 | 4 | 0.75 | 3 | 1 | 4 | 0.5 | 14 | 0.7143 |
| 16_1401059 | 5 | 0.4 | 7 | 0 | 5 | 0.2 | 2 | 0 | 19 | 0.1579 | 3 | 0.3333 | 4 | 0 | 2 | 0.5 | 3 | 0 | 12 | 0.1667 |
| 16_1401060 | 4 | 1 | 7 | 0.8571 | 7 | 0.5714 | 3 | 0.6667 | 21 | 0.7619 | 3 | 0.6667 | 4 | 0.5 | 3 | 0.6667 | 4 | 1 | 14 | 0.7143 |
| 16_1401285 | NA | NA | 2 | 0.5 | NA | NA | NA | NA | 2 | 0.5 | NA | NA | 1 | 1 | 1 | 1 | NA | NA | 2 | 1 |
| 16_1411724 | 11 | 0.4545 | 11 | 0 | 7 | 0.1429 | 8 | 0.125 | 37 | 0.1892 | 7 | 0.2857 | 13 | 0.0769 | 14 | 0.0714 | 7 | 0.2857 | 41 | 0.1463 |
| 16_1411755 | 11 | 0.2727 | 12 | 0.1667 | 6 | 0 | 11 | 0.1818 | 40 | 0.175 | 7 | 0 | 11 | 0.0909 | 14 | 0.0714 | 10 | 0.1 | 42 | 0.0714 |

| | | | | | | | | | | | | | | | | | | | | |
|---|---|---|---|---|---|---|---|---|---|---|---|---|---|---|---|---|---|---|---|---|
| 16_1427893 | NA | NA | 2 | 0.5 | NA | NA | 1 | 0 | 3 | 0.3333 | 1 | 1 | NA | NA | 1 | 0 | NA | NA | 2 | 0.5 |
| 16_1429145 | NA | NA | 2 | 0.5 | NA | NA | 1 | 1 | 3 | 0.6667 | 1 | 1 | 1 | 1 | 1 | 1 | NA | NA | 3 | 1 |
| 16_1429152 | NA | NA | 2 | 0.5 | NA | NA | 1 | 0 | 3 | 0.3333 | 1 | 0 | 1 | 0 | 3 | 0.3333 | NA | NA | 5 | 0.2 |
| 16_1429203 | NA | NA | 2 | 0 | NA | NA | 1 | 1 | 3 | 0.3333 | 2 | 1 | 1 | 1 | 3 | 0.3333 | NA | NA | 6 | 0.6667 |
| 16_1429211 | NA | NA | 2 | 1 | NA | NA | 1 | 1 | 3 | 1 | 2 | 1 | NA | NA | 3 | 1 | NA | NA | 5 | 1 |
| 16_1429268 | 1 | 0 | 2 | 0 | 1 | 1 | 1 | 0 | 5 | 0.2 | 3 | 0.3333 | 1 | 0 | 3 | 0 | 1 | 0 | 8 | 0.125 |
| 16_1452707 | NA | NA | NA | NA | NA | NA | 1 | 1 | 1 | 1 | NA | NA | 1 | 1 | NA | NA | NA | NA | 1 | 1 |
| 16_1537391 | 7 | 0 | 10 | 0.3 | 4 | 0 | 11 | 0 | 32 | 0.0938 | 5 | 0.2 | 6 | 0 | 16 | 0 | 20 | 0.1 | 47 | 0.0638 |
| 16_1537395 | 7 | 0.1429 | 14 | 0.1429 | 6 | 0 | 12 | 0.0833 | 39 | 0.1026 | 5 | 0.4 | 9 | 0.2222 | 17 | 0.0588 | 17 | 0.1765 | 48 | 0.1667 |
| 16_1537402 | 8 | 0 | 15 | 0.0667 | 5 | 0.2 | 14 | 0.1429 | 42 | 0.0952 | 5 | 0 | 8 | 0.375 | 17 | 0.0588 | 19 | 0.4211 | 49 | 0.2449 |
| 16_1537413 | 9 | 0.7778 | 15 | 1 | 6 | 0.8333 | 13 | 0.9231 | 43 | 0.907 | 6 | 0.8333 | 8 | 0.75 | 18 | 0.8889 | 18 | 0.9444 | 50 | 0.88 |
| 16_1537424 | 7 | 0.2857 | 14 | 0 | 6 | 0.1667 | 14 | 0 | 41 | 0.0732 | 6 | 0.1667 | 10 | 0.1 | 18 | 0.1667 | 19 | 0.0526 | 53 | 0.1132 |
| 16_1537427 | 9 | 0.1111 | 13 | 0.4615 | 6 | 0.3333 | 13 | 0.5385 | 41 | 0.3902 | 5 | 0.2 | 9 | 0.5556 | 16 | 0.375 | 18 | 0.2222 | 48 | 0.3333 |
| 16_1565975 | 3 | 0.6667 | 3 | 0 | 2 | 0.5 | 3 | 0 | 11 | 0.2727 | 2 | 0.5 | 3 | 0.3333 | 3 | 0 | 3 | 0.3333 | 11 | 0.2727 |
| 16_1565977 | 3 | 0 | 3 | 0.3333 | 1 | 1 | 3 | 0 | 10 | 0.2 | 3 | 0 | 3 | 0.3333 | 3 | 0 | 3 | 0.3333 | 12 | 0.1667 |
| 16_1565981 | 2 | 1 | 2 | 0.5 | 2 | 0.5 | 2 | 0.5 | 8 | 0.625 | 2 | 1 | 2 | 0 | 3 | 0.6667 | 3 | 0.3333 | 10 | 0.5 |
| 16_1566000 | 1 | 0 | 3 | 0.3333 | 3 | 0.3333 | 3 | 0.3333 | 10 | 0.3 | 2 | 0 | 2 | 0 | 3 | 0.3333 | 3 | 0 | 10 | 0.1 |
| 16_1568030 | 7 | 0.1429 | 3 | 0 | 4 | 0 | 2 | 0 | 16 | 0.0625 | 6 | 0.1667 | 3 | 0 | 6 | 0 | 2 | 0 | 17 | 0.0588 |
| 16_1568039 | 7 | 0 | 3 | 0.3333 | 4 | 0 | 2 | 0 | 16 | 0.0625 | 6 | 0.1667 | 3 | 0 | 7 | 0.1429 | 2 | 0 | 18 | 0.1111 |
| 16_1568082 | 6 | 0.3333 | 1 | 0 | 2 | 0.5 | 1 | 1 | 10 | 0.4 | 3 | 0.3333 | 2 | 0 | 7 | 0.5714 | 1 | 1 | 13 | 0.4615 |
| 16_1570528 | 4 | 0.25 | 2 | 0 | 2 | 0 | 3 | 0 | 11 | 0.0909 | 2 | 0 | 2 | 0 | 1 | 1 | 1 | 0 | 6 | 0.1667 |
| 16_1570542 | 4 | 0.25 | 2 | 0 | 2 | 0.5 | 4 | 0.5 | 12 | 0.3333 | 2 | 0.5 | 3 | 0.3333 | 2 | 0.5 | 1 | 0 | 8 | 0.375 |
| 16_1570591 | 3 | 0.3333 | 3 | 0.3333 | 2 | 0.5 | 5 | 0.8 | 13 | 0.5385 | 4 | 0.5 | 4 | 0 | 2 | 0 | 2 | 0.5 | 12 | 0.25 |
| 16_1570601 | 3 | 1 | 4 | 1 | 2 | 1 | 6 | 1 | 15 | 1 | 4 | 1 | 4 | 1 | 3 | 1 | 2 | 0.5 | 13 | 0.9231 |
| 16_1570608 | 4 | 0 | 3 | 0 | 2 | 0 | 6 | 0.3333 | 15 | 0.1333 | 3 | 0.3333 | 4 | 0.5 | 3 | 0.6667 | 2 | 0 | 12 | 0.4167 |
| 16_1570612 | 5 | 0.6 | 4 | 0.75 | 2 | 1 | 4 | 1 | 15 | 0.8 | 3 | 1 | 2 | 0.5 | 3 | 1 | 2 | 0 | 10 | 0.7 |
| 16_1570623 | 3 | 0.6667 | 3 | 1 | 2 | 1 | 6 | 0.1667 | 14 | 0.5714 | 3 | 0.3333 | 4 | 0.75 | 3 | 0.3333 | 2 | 0 | 12 | 0.4167 |
| 16_1570624 | 3 | 1 | 4 | 0.75 | 2 | 0 | 5 | 0.8 | 14 | 0.7143 | 2 | 0.5 | 4 | 0.75 | 3 | 0.6667 | 2 | 0.5 | 11 | 0.6364 |
| 16_1570626 | 2 | 0.5 | 3 | 0.6667 | 2 | 1 | 3 | 1 | 10 | 0.8 | 3 | 0.6667 | 3 | 0.3333 | 3 | 0.6667 | 2 | 0.5 | 11 | 0.5455 |
| 16_1609009 | 6 | 0.1667 | NA | NA | NA | NA | 2 | 0 | 8 | 0.125 | NA | NA | 5 | 0.6 | 4 | 0.25 | 2 | 1 | 11 | 0.5455 |
| 16_1609021 | 6 | 0 | NA | NA | NA | NA | 3 | 0.3333 | 9 | 0.1111 | NA | NA | 3 | 0.6667 | 3 | 0 | 3 | 0 | 9 | 0.2222 |
| 16_1612101 | 1 | 1 | 1 | 0 | 1 | 0 | 2 | 1 | 5 | 0.6 | NA | NA | 2 | 0.5 | NA | NA | NA | NA | 2 | 0.5 |
| 16_1612120 | NA | NA | 1 | 0 | 1 | 0 | 2 | 0.5 | 4 | 0.25 | 1 | 0 | 5 | 0.2 | 2 | 0 | 1 | 0 | 9 | 0.1111 |
| 16_1612727 | 2 | 0.5 | 3 | 0 | 1 | 0 | 2 | 0 | 8 | 0.125 | 2 | 0.5 | 4 | 0.25 | 4 | 0.25 | 3 | 0 | 13 | 0.2308 |
| 16_1612729 | 2 | 1 | 4 | 1 | 1 | 1 | 1 | 1 | 8 | 1 | 2 | 1 | 5 | 1 | 2 | 1 | 3 | 1 | 12 | 1 |
| 16_1613090 | 25 | 0.12 | 10 | 0 | 18 | 0.2222 | 14 | 0 | 67 | 0.1045 | 8 | 0.125 | 16 | 0.125 | 13 | 0.0769 | 17 | 0.0588 | 54 | 0.0926 |
| 16_1613132 | 25 | 0.16 | 14 | 0.0714 | 18 | 0.0556 | 15 | 0.2 | 72 | 0.125 | 9 | 0.1111 | 16 | 0.1875 | 16 | 0.125 | 18 | 0.1667 | 59 | 0.1525 |
| 16_1613147 | 19 | 0.1053 | 9 | 0.1111 | 18 | 0.0556 | 14 | 0 | 60 | 0.0667 | 7 | 0.1429 | 16 | 0.0625 | 14 | 0 | 20 | 0.05 | 57 | 0.0526 |
| 16_1613151 | 21 | 0.0476 | 13 | 0.0769 | 18 | 0.1111 | 14 | 0.0714 | 66 | 0.0758 | 8 | 0 | 16 | 0 | 14 | 0.3571 | 20 | 0.25 | 58 | 0.1724 |
| 16_1613163 | 20 | 0.05 | 13 | 0.1538 | 15 | 0.0667 | 16 | 0.1875 | 64 | 0.1094 | 6 | 0.3333 | 16 | 0.0625 | 16 | 0.125 | 18 | 0.0556 | 56 | 0.1071 |
| 16_1613174 | 22 | 0.2273 | 13 | 0.0769 | 19 | 0.1579 | 16 | 0.125 | 70 | 0.1571 | 3 | 0 | 16 | 0.125 | 16 | 0.125 | 21 | 0.0952 | 56 | 0.1071 |
| 16_1613178 | 21 | 0.8571 | 12 | 0.9167 | 19 | 0.8947 | 14 | 0.7857 | 66 | 0.8636 | 5 | 1 | 16 | 0.6875 | 16 | 0.9375 | 19 | 0.8947 | 56 | 0.8571 |
| 16_1613184 | 16 | 0.4375 | 11 | 0.5455 | 18 | 0.6111 | 15 | 0.3333 | 60 | 0.4833 | 4 | 0.75 | 16 | 0.6875 | 12 | 0.5 | 19 | 0.3684 | 51 | 0.5294 |
| 16_1613186 | 20 | 0.1 | 12 | 0.3333 | 19 | 0.1579 | 15 | 0.1333 | 66 | 0.1667 | 5 | 0 | 15 | 0.3333 | 16 | 0.25 | 21 | 0.1905 | 57 | 0.2281 |
| 16_1623512 | 2 | 0.5 | 1 | 0 | 3 | 0.3333 | NA | NA | 6 | 0.3333 | 2 | 0 | NA | NA | 4 | 0 | 4 | 0.25 | 10 | 0.1 |
| 16_1623578 | 1 | 1 | 1 | 0 | 3 | 0 | NA | NA | 5 | 0.2 | 3 | 0.3333 | NA | NA | 3 | 0 | 4 | 0 | 10 | 0.1 |
| 16_1623671 | 1 | 0 | 1 | 0 | 3 | 0.3333 | NA | NA | 5 | 0.2 | 3 | 0 | NA | NA | 1 | 1 | 3 | 0 | 7 | 0.1429 |
| 16_1623925 | 1 | 1 | 1 | 1 | NA | NA | NA | NA | 2 | 1 | NA | NA | NA | NA | 2 | 0.5 | 2 | 1 | 4 | 0.75 |
| 16_1624030 | 1 | 1 | 1 | 1 | NA | NA | NA | NA | 2 | 1 | NA | NA | NA | NA | 2 | 1 | 2 | 1 | 4 | 1 |
| 16_1624045 | 1 | 1 | 1 | 0 | NA | NA | NA | NA | 2 | 0.5 | NA | NA | NA | NA | 1 | 0 | 2 | 1 | 3 | 0.6667 |
| 16_1624055 | NA | NA | 1 | 1 | NA | NA | NA | NA | 1 | 1 | NA | NA | NA | NA | NA | NA | 2 | 1 | 2 | 1 |
| 16_1657151 | 3 | 0 | 1 | 0 | 1 | 1 | 2 | 0.5 | 7 | 0.2857 | 1 | 0 | NA | NA | NA | NA | 2 | 0.5 | 3 | 0.3333 |
| 16_1670932 | 2 | 0 | 1 | 0 | NA | NA | 7 | 0.1429 | 10 | 0.1 | NA | NA | 2 | 0.5 | 3 | 0 | 1 | 0 | 6 | 0.1667 |
| 16_1671072 | 2 | 0 | 3 | 0 | 4 | 0 | 10 | 0.2 | 19 | 0.1053 | 1 | 0 | 2 | 0 | 6 | 0.1667 | 2 | 0.5 | 11 | 0.1818 |
| 16_1674157 | 3 | 0.3333 | 3 | 0.3333 | 1 | 0 | NA | NA | 7 | 0.2857 | 2 | 0.5 | 3 | 0 | 1 | 0 | 2 | 0.5 | 8 | 0.25 |
| 16_1674191 | 3 | 0 | 3 | 0.3333 | 1 | 1 | NA | NA | 7 | 0.2857 | 2 | 0 | 2 | 0.5 | 2 | 1 | 2 | 0 | 8 | 0.375 |
| 16_1674205 | 4 | 0 | 3 | 0.3333 | NA | NA | NA | NA | 7 | 0.1429 | 3 | 0.3333 | 3 | 0 | 2 | 0 | 2 | 0.5 | 10 | 0.2 |
| 16_1674214 | 4 | 0.25 | 2 | 0 | 1 | 0 | NA | NA | 7 | 0.1429 | 3 | 0.3333 | 3 | 0 | 2 | 0 | 2 | 0.5 | 10 | 0.2 |
| 16_1674296 | 4 | 0 | 4 | 0.5 | 3 | 0 | 1 | 0 | 12 | 0.1667 | 1 | 0 | 4 | 0 | 1 | 1 | NA | NA | 6 | 0.1667 |
| 16_1674421 | 3 | 0 | 6 | 0.3333 | 3 | 0 | 2 | 0 | 14 | 0.1429 | 6 | 0 | 6 | 0 | 5 | 0.2 | 3 | 0 | 20 | 0.05 |
| 16_1674502 | 7 | 0.5714 | 8 | 0.25 | 6 | 0.3333 | 4 | 0 | 25 | 0.32 | 6 | 0.5 | 10 | 0.1 | 5 | 0.8 | 7 | 0 | 28 | 0.2857 |
| 16_1685631 | 2 | 0 | NA | NA | 2 | 0.5 | 2 | 0.5 | 6 | 0.3333 | 5 | 0 | 4 | 0.5 | 1 | 1 | 3 | 0.6667 | 13 | 0.3846 |
| 16_1685648 | 2 | 0 | NA | NA | 2 | 0.5 | 2 | 1 | 6 | 0.5 | 5 | 0 | 3 | 0.3333 | 1 | 1 | 2 | 0 | 11 | 0.1818 |
| 16_1685654 | 1 | 1 | NA | NA | 2 | 1 | 3 | 0.3333 | 6 | 0.6667 | 4 | 0.75 | 5 | 0.8 | 1 | 0 | 3 | 1 | 13 | 0.7692 |
| 16_1685667 | 2 | 0.5 | NA | NA | 1 | 0 | 3 | 0 | 6 | 0.1667 | 5 | 0 | 6 | 0.3333 | 3 | 0 | 2 | 0 | 16 | 0.125 |

| | | | | | | | | | | | | | | | | | | | | |
|---|---|---|---|---|---|---|---|---|---|---|---|---|---|---|---|---|---|---|---|---|
| 16_1686179 | 46 | 0.1522 | 41 | 0.122 | 37 | 0.3514 | 49 | 0.1633 | 173 | 0.1908 | 35 | 0.2571 | 43 | 0.3721 | 52 | 0.2692 | 41 | 0.1707 | 171 | 0.269 |
| 16_1686257 | 63 | 0.0635 | 58 | 0.0345 | 50 | 0.12 | 68 | 0.0588 | 239 | 0.0669 | 45 | 0.0667 | 70 | 0.0571 | 69 | 0.087 | 64 | 0.0469 | 248 | 0.0645 |
| 16_1686306 | 78 | 0.0513 | 66 | 0.0758 | 60 | 0.0667 | 76 | 0.0921 | 280 | 0.0714 | 53 | 0.0566 | 74 | 0.0811 | 74 | 0.0541 | 69 | 0.1739 | 270 | 0.0926 |
| 16_1686627 | 1 | 1 | NA | NA | NA | NA | 1 | 1 | 2 | 1 | NA | NA | 1 | 1 | NA | NA | 1 | 0 | 2 | 0.5 |
| 16_1686716 | 1 | 0 | NA | NA | NA | NA | 1 | 1 | 2 | 0.5 | NA | NA | 1 | 1 | NA | NA | 1 | 0 | 2 | 0.5 |
| 16_1690243 | 1 | 1 | 2 | 0.5 | 1 | 1 | 1 | 1 | 5 | 0.8 | NA | NA | 2 | 0.5 | 2 | 0.5 | 2 | 0.5 | 6 | 0.5 |
| 16_1690254 | 2 | 0 | 3 | 0.3333 | 2 | 0.5 | 1 | 0 | 8 | 0.25 | 2 | 0 | 4 | 0.5 | 1 | 0 | 2 | 0 | 9 | 0.2222 |
| 16_1690255 | 2 | 1 | 2 | 0.5 | 2 | 0.5 | 1 | 0 | 7 | 0.5714 | 1 | 0 | 4 | 0.5 | 2 | 0.5 | NA | NA | 7 | 0.4286 |
| 16_1690376 | 1 | 0 | 2 | 0.5 | 1 | 0 | 1 | 0 | 5 | 0.2 | 2 | 0.5 | 2 | 0.5 | 1 | 0 | 4 | 0.25 | 9 | 0.3333 |
| 16_1690387 | NA | NA | 2 | 0.5 | NA | NA | NA | NA | 2 | 0.5 | 2 | 1 | 2 | 1 | 1 | 0 | 4 | 0.25 | 9 | 0.5556 |
| 16_1690405 | 1 | 0 | 2 | 0.5 | NA | NA | NA | NA | 3 | 0.3333 | 2 | 1 | 4 | 0.5 | 1 | 0 | 3 | 0 | 10 | 0.4 |
| 16_1690455 | 4 | 0.5 | 7 | 0.7143 | 5 | 0.6 | 9 | 0.8889 | 25 | 0.72 | 2 | 1 | 5 | 0.4 | 9 | 0.7778 | 8 | 0.625 | 24 | 0.6667 |
| 16_1690458 | 3 | 0 | 6 | 0.1667 | 4 | 0 | 9 | 0.1111 | 22 | 0.0909 | 2 | 0.5 | 2 | 0 | 7 | 0 | 7 | 0 | 18 | 0.0556 |
| 16_1690467 | 4 | 0.25 | 6 | 0.3333 | 4 | 0.25 | 8 | 0.125 | 22 | 0.2273 | 2 | 0 | 6 | 0.1667 | 7 | 0.1429 | 8 | 0 | 23 | 0.087 |
| 16_1690469 | 2 | 0.5 | 1 | 0 | NA | NA | NA | NA | 3 | 0.3333 | 1 | 0 | NA | NA | 1 | 0 | 1 | 1 | 3 | 0.3333 |
| 16_1690472 | 2 | 0.5 | 1 | 0 | NA | NA | NA | NA | 3 | 0.3333 | 1 | 0 | NA | NA | 1 | 1 | 1 | 0 | 3 | 0.3333 |
| 16_1690648 | 1 | 1 | NA | NA | 2 | 0.5 | 4 | 0.25 | 7 | 0.4286 | NA | NA | 1 | 1 | 2 | 0.5 | 2 | 0 | 5 | 0.4 |
| 16_1690665 | 4 | 0.25 | 8 | 0.125 | 6 | 0.1667 | 15 | 0.0667 | 33 | 0.1212 | 4 | 0 | 6 | 0 | 9 | 0.1111 | 9 | 0.1111 | 28 | 0.0714 |
| 16_1694831 | 6 | 0.3333 | 5 | 0 | 1 | 1 | 4 | 0.5 | 16 | 0.3125 | 12 | 0.4167 | 3 | 0 | 6 | 0.5 | 1 | 0 | 22 | 0.3636 |
| 16_1694837 | 5 | 0.6 | 5 | 0.8 | NA | NA | 3 | 1 | 13 | 0.7692 | 9 | 0.7778 | 3 | 0.6667 | 5 | 0.8 | NA | NA | 17 | 0.7647 |
| 16_1694850 | 4 | 0.25 | 4 | 0 | 1 | 0 | 3 | 0.3333 | 12 | 0.1667 | 7 | 0.2857 | 1 | 0 | 5 | 0.4 | 1 | 0 | 14 | 0.2857 |
| 16_1696494 | 3 | 0 | 3 | 0.6667 | 2 | 0 | 8 | 0 | 16 | 0.125 | 1 | 0 | 4 | 0 | 3 | 0.3333 | 3 | 0.6667 | 11 | 0.2727 |
| 16_1696602 | 3 | 0 | 3 | 0.3333 | 2 | 0.5 | 8 | 0.25 | 16 | 0.25 | 2 | 0 | 4 | 0.5 | 5 | 0.6 | 4 | 0.25 | 15 | 0.4 |
| 16_1696613 | 4 | 0.25 | 3 | 0.6667 | 3 | 0 | 7 | 0 | 17 | 0.1765 | 1 | 0 | 4 | 0.25 | 6 | 0.1667 | 5 | 0 | 16 | 0.125 |
| 16_1696631 | 4 | 0 | 3 | 0 | 3 | 0 | 8 | 0.125 | 18 | 0.0556 | 1 | 0 | 3 | 0 | 6 | 0 | 5 | 0.2 | 15 | 0.0667 |
| 16_1696636 | 3 | 0.3333 | 1 | 0 | 2 | 0 | 6 | 0.1667 | 12 | 0.1667 | 1 | 0 | 2 | 0 | 4 | 0 | 4 | 0.25 | 11 | 0.0909 |
| 16_1696639 | 4 | 0.25 | 2 | 0 | 3 | 0 | 8 | 0 | 17 | 0.0588 | 1 | 0 | 4 | 0.25 | 6 | 0 | 3 | 0 | 14 | 0.0714 |
| 16_1696641 | 3 | 0.6667 | 3 | 1 | 3 | 1 | 7 | 0.7143 | 16 | 0.8125 | NA | NA | 4 | 1 | 5 | 0.4 | 3 | 0 | 12 | 0.5 |
| 16_1696642 | 4 | 0.25 | 2 | 0 | 3 | 0 | 7 | 0.1429 | 16 | 0.125 | 1 | 0 | 3 | 0.3333 | 5 | 0.2 | 3 | 0 | 12 | 0.1667 |
| 16_1696659 | 3 | 0.6667 | 3 | 0 | 3 | 0 | 7 | 0 | 16 | 0.125 | 3 | 0.6667 | 3 | 0 | 5 | 0.2 | 5 | 0 | 16 | 0.1875 |
| 16_1696660 | 4 | 0.75 | 3 | 0.3333 | 3 | 0.3333 | 8 | 0.125 | 18 | 0.3333 | 3 | 0.3333 | 2 | 0 | 5 | 0.6 | 5 | 0.4 | 15 | 0.4 |
| 16_1697017 | 15 | 0.0667 | 27 | 0.2593 | 15 | 0.0667 | 17 | 0.2941 | 74 | 0.1892 | 10 | 0.3 | 15 | 0.2667 | 15 | 0.0667 | 24 | 0.2083 | 64 | 0.2031 |
| 16_1697046 | 14 | 0.1429 | 24 | 0.0417 | 15 | 0.0667 | 17 | 0 | 70 | 0.0571 | 11 | 0.0909 | 15 | 0.0667 | 17 | 0.1765 | 20 | 0.05 | 63 | 0.0952 |
| 16_1697061 | 15 | 0.2 | 29 | 0.3448 | 16 | 0.125 | 15 | 0.2667 | 75 | 0.2533 | 12 | 0.25 | 18 | 0.4444 | 16 | 0.25 | 20 | 0.25 | 66 | 0.303 |
| 16_1697078 | 10 | 0.2 | 24 | 0.1667 | 12 | 0.3333 | 14 | 0.3571 | 60 | 0.25 | 9 | 0.2222 | 14 | 0.2857 | 14 | 0.1429 | 19 | 0.2632 | 56 | 0.2321 |
| 16_1697083 | 13 | 0.4615 | 23 | 0.3043 | 10 | 0.1 | 15 | 0.2 | 61 | 0.2787 | 10 | 0.2 | 13 | 0.3077 | 12 | 0.4167 | 17 | 0.4118 | 52 | 0.3462 |
| 16_1697096 | 11 | 0.3636 | 25 | 0.24 | 13 | 0.1538 | 16 | 0.125 | 65 | 0.2154 | 13 | 0.6154 | 17 | 0.5294 | 12 | 0.1667 | 15 | 0.6 | 57 | 0.4912 |
| 16_1697100 | 10 | 0 | 21 | 0.0476 | 11 | 0.2727 | 12 | 0 | 54 | 0.0741 | 10 | 0.1 | 14 | 0.1429 | 11 | 0 | 17 | 0 | 52 | 0.0577 |
| 16_1697124 | 12 | 0.25 | 23 | 0.3913 | 15 | 0.2667 | 16 | 0.125 | 66 | 0.2727 | 12 | 0.3333 | 16 | 0.375 | 13 | 0.5385 | 19 | 0.5789 | 60 | 0.4667 |
| 16_1697186 | 11 | 0.2727 | 26 | 0.3077 | 11 | 0.5455 | 19 | 0.2632 | 67 | 0.3284 | 13 | 0.3846 | 19 | 0.2105 | 17 | 0.1765 | 24 | 0.2083 | 73 | 0.2329 |
| 16_1697191 | 8 | 0.875 | 24 | 0.875 | 11 | 0.9091 | 16 | 0.875 | 59 | 0.8814 | 13 | 0.9231 | 19 | 0.8947 | 16 | 0.8125 | 24 | 0.9167 | 72 | 0.8889 |
| 16_1697199 | 9 | 1 | 20 | 0.85 | 15 | 0.8667 | 19 | 0.8947 | 63 | 0.8889 | 15 | 0.7333 | 19 | 0.7368 | 20 | 1 | 23 | 0.9565 | 77 | 0.8701 |
| 16_1697206 | 12 | 0.9167 | 27 | 0.4074 | 14 | 0.2857 | 19 | 0.5263 | 72 | 0.5 | 16 | 0.5625 | 18 | 0.4444 | 15 | 0.4 | 24 | 0.5833 | 73 | 0.5068 |
| 16_1702859 | 1 | 1 | NA | NA | NA | NA | NA | NA | 1 | 1 | NA | NA | 1 | 1 | NA | NA | NA | NA | 1 | 1 |
| 16_1702885 | 1 | 1 | NA | NA | NA | NA | NA | NA | 1 | 1 | NA | NA | 1 | 1 | NA | NA | NA | NA | 1 | 1 |
| 16_1702895 | 1 | 1 | NA | NA | NA | NA | NA | NA | 1 | 1 | NA | NA | 1 | 1 | NA | NA | NA | NA | 1 | 1 |
| 16_1702987 | 1 | 1 | NA | NA | NA | NA | NA | NA | 1 | 1 | NA | NA | 1 | 1 | NA | NA | NA | NA | 1 | 1 |
| 16_1703071 | 1 | 1 | NA | NA | NA | NA | NA | NA | 1 | 1 | NA | NA | 1 | 1 | NA | NA | NA | NA | 1 | 1 |
| 16_1703099 | 1 | 1 | NA | NA | NA | NA | NA | NA | 1 | 1 | NA | NA | 1 | 1 | NA | NA | NA | NA | 1 | 1 |
| 16_1703119 | 1 | 1 | NA | NA | NA | NA | NA | NA | 1 | 1 | NA | NA | 1 | 1 | NA | NA | NA | NA | 1 | 1 |
| 16_1703121 | 1 | 1 | NA | NA | NA | NA | NA | NA | 1 | 1 | NA | NA | 1 | 1 | NA | NA | NA | NA | 1 | 1 |
| 16_1703158 | 1 | 1 | NA | NA | NA | NA | NA | NA | 1 | 1 | NA | NA | 1 | 1 | NA | NA | NA | NA | 1 | 1 |
| 16_1709546 | 2 | 1 | NA | NA | 1 | 0 | 1 | 1 | 4 | 0.75 | 1 | 1 | NA | NA | NA | NA | 1 | 1 | 2 | 1 |
| 16_1709556 | 13 | 0.0769 | 4 | 0 | 11 | 0 | 6 | 0.1667 | 34 | 0.0588 | 8 | 0.25 | 8 | 0 | 6 | 0 | 11 | 0.1818 | 33 | 0.1212 |
| 16_1709569 | 5 | 0.2 | 1 | 0 | 2 | 0.5 | 2 | 0.5 | 10 | 0.3 | 3 | 0.3333 | 5 | 0 | 1 | 0 | 3 | 0 | 12 | 0.0833 |
| 16_1709622 | 8 | 0 | 6 | 0.3333 | 8 | 0.375 | 5 | 0.4 | 27 | 0.2593 | 8 | 0.5 | 7 | 0 | 7 | 0 | 10 | 0.3 | 32 | 0.2188 |
| 16_1709830 | 23 | 0.0435 | 23 | 0 | 19 | 0.2105 | 16 | 0 | 81 | 0.0617 | 9 | 0 | 27 | 0.0741 | 20 | 0.1 | 18 | 0.2222 | 74 | 0.1081 |
| 16_1709834 | 22 | 0.0909 | 27 | 0 | 19 | 0.1053 | 16 | 0.125 | 84 | 0.0714 | 11 | 0 | 25 | 0.16 | 19 | 0.1579 | 21 | 0.2857 | 76 | 0.1711 |
| 16_1710084 | 3 | 1 | NA | NA | 4 | 0.75 | 3 | 1 | 10 | 0.9 | 3 | 0.3333 | 2 | 1 | 1 | 1 | 1 | 1 | 7 | 0.7143 |
| 16_1710090 | 5 | 0.6 | NA | NA | 5 | 0.8 | 4 | 0.5 | 14 | 0.6429 | 4 | 0.25 | 3 | 0.6667 | 1 | 0 | 2 | 0 | 10 | 0.3 |
| 16_1724724 | 1 | 1 | 2 | 0 | NA | NA | NA | NA | 3 | 0.3333 | NA | NA | NA | NA | 2 | 0 | 3 | 0.3333 | 5 | 0.2 |
| 16_1724833 | 2 | 0 | 6 | 0.1667 | NA | NA | 4 | 0.5 | 12 | 0.25 | 1 | 1 | NA | NA | 3 | 0.3333 | 4 | 0.25 | 8 | 0.375 |
| 16_1727039 | 3 | 0.3333 | 4 | 0 | 2 | 0.5 | 4 | 0 | 13 | 0.1538 | 2 | 0 | 2 | 0.5 | 3 | 1 | 1 | 0 | 8 | 0.5 |
| 16_1727041 | 3 | 0 | 4 | 0 | 2 | 0 | 5 | 0.2 | 14 | 0.0714 | 2 | 0.5 | 3 | 0 | 4 | 0 | 1 | 0 | 10 | 0.1 |
| 16_1727046 | 4 | 0.5 | 3 | 1 | 2 | 1 | 5 | 0.6 | 14 | 0.7143 | 2 | 1 | 3 | 0.6667 | 3 | 0.3333 | 1 | 1 | 9 | 0.6667 |

| | | | | | | | | | | | | | | | | | | | | |
|---|---|---|---|---|---|---|---|---|---|---|---|---|---|---|---|---|---|---|---|---|
| 16_1727048 | 4 | 0.75 | 4 | 1 | 2 | 1 | 4 | 0.75 | 14 | 0.8571 | 2 | 0.5 | 3 | 0.6667 | 3 | 0.6667 | 1 | 1 | 9 | 0.6667 |
| 16_1728683 | 3 | 1 | 4 | 0.75 | 2 | 0.5 | 5 | 1 | 14 | 0.8571 | 1 | 1 | 3 | 0.6667 | 4 | 0.75 | 1 | 1 | 9 | 0.7778 |
| 16_1728703 | 5 | 0 | 4 | 0.25 | NA | NA | 2 | 0.5 | 11 | 0.1818 | 2 | 0 | 4 | 0.25 | 4 | 0 | 1 | 0 | 11 | 0.0909 |
| 16_1728705 | 4 | 0 | 4 | 0 | 2 | 0.5 | 4 | 0 | 14 | 0.0714 | 3 | 0 | 4 | 0 | 4 | 0 | 1 | 1 | 12 | 0.0833 |
| 16_1728715 | 4 | 0.75 | 4 | 1 | 2 | 1 | 5 | 0.8 | 15 | 0.8667 | 3 | 0.6667 | 4 | 1 | 4 | 0.75 | NA | NA | 11 | 0.8182 |
| 16_1730164 | 5 | 0.2 | 4 | 0 | 2 | 0 | 7 | 0 | 18 | 0.0556 | 3 | 0.3333 | 3 | 0 | 4 | 0.25 | 1 | 0 | 11 | 0.1818 |
| 16_1730188 | 2 | 1 | 2 | 1 | NA | NA | 4 | 1 | 8 | 1 | 2 | 1 | 3 | 1 | 3 | 1 | NA | NA | 8 | 1 |
| 16_1730192 | 5 | 1 | 4 | 1 | 2 | 1 | 6 | 1 | 17 | 1 | 3 | 1 | 3 | 0.6667 | 5 | 1 | 2 | 1 | 13 | 0.9231 |
| 16_1730192 | 6 | 0.3333 | 4 | 0 | 2 | 0.5 | 5 | 0 | 17 | 0.1765 | 3 | 0.6667 | 4 | 0.25 | 4 | 0 | 2 | 0 | 13 | 0.2308 |
| 16_1730206 | 4 | 1 | 1 | 0 | 2 | 0.5 | 4 | 0 | 11 | 0.4545 | 3 | 0.3333 | 5 | 1 | 3 | 0 | 3 | 0.6667 | 14 | 0.5714 |
| 16_1730211 | 5 | 0 | 4 | 0.25 | 2 | 0 | 6 | 0 | 17 | 0.0588 | 3 | 0 | 5 | 0 | 5 | 0.4 | 3 | 0 | 16 | 0.125 |
| 16_1730212 | 6 | 0 | 4 | 0.5 | 2 | 0.5 | 5 | 0 | 17 | 0.1765 | 2 | 0.5 | 5 | 0 | 4 | 0 | 3 | 0 | 14 | 0.0714 |
| 16_1730213 | 4 | 1 | 4 | 0.75 | 2 | 1 | 4 | 0.75 | 14 | 0.8571 | 2 | 0.5 | 5 | 1 | 5 | 0.8 | 3 | 1 | 15 | 0.8667 |
| 16_1730219 | 6 | 0.5 | 4 | 0 | 2 | 1 | 5 | 0.4 | 17 | 0.4118 | 3 | 0 | 5 | 0.2 | 4 | 0.75 | 2 | 1 | 14 | 0.4286 |
| 16_1730222 | 5 | 0.4 | 4 | 0.5 | 2 | 0.5 | 5 | 0 | 16 | 0.3125 | 2 | 0 | 5 | 0.4 | 5 | 0 | 2 | 0 | 14 | 0.1429 |
| 16_1730226 | 5 | 0.6 | 4 | 1 | 3 | 0.3333 | 4 | 0.25 | 16 | 0.5625 | 2 | 1 | 4 | 0.25 | 3 | 0.3333 | 3 | 0.6667 | 12 | 0.5 |
| 16_1730280 | 7 | 0.2857 | 4 | 0 | 4 | 0.25 | 4 | 0 | 19 | 0.1579 | 2 | 0 | 8 | 0.125 | 5 | 0 | 3 | 0 | 18 | 0.0556 |
| 16_1730300 | 9 | 0.4444 | 4 | 0 | 4 | 0.25 | 6 | 0 | 23 | 0.2174 | 4 | 0.25 | 7 | 0 | 4 | 0.25 | 3 | 0.3333 | 18 | 0.1667 |
| 16_1730354 | 8 | 0.125 | 5 | 0.4 | 3 | 0 | 7 | 0 | 23 | 0.1304 | 3 | 0 | 7 | 0.1429 | 4 | 0 | 3 | 0 | 17 | 0.0588 |
| 16_1730410 | 10 | 0.1 | 4 | 0 | 4 | 0 | 8 | 0.125 | 26 | 0.0769 | 4 | 0 | 7 | 0.1429 | 5 | 0.2 | 4 | 0 | 20 | 0.1 |
| 16_1730457 | 10 | 0.5 | 5 | 0.4 | 3 | 0.6667 | 9 | 0.3333 | 27 | 0.4444 | 4 | 0.75 | 9 | 0.4444 | 6 | 0.5 | 3 | 0.6667 | 22 | 0.5455 |
| 16_1730482 | 11 | 0.1818 | 6 | 0.3333 | 2 | 0.5 | 8 | 0.125 | 27 | 0.2222 | 5 | 0.4 | 7 | 0.2857 | 7 | 0.1429 | 4 | 0 | 23 | 0.2174 |
| 16_1730484 | 9 | 0.5556 | 5 | 1 | 5 | 0.6 | 5 | 0.8 | 24 | 0.7083 | 4 | 0.75 | 7 | 0.5714 | 6 | 0.5 | 4 | 1 | 21 | 0.6667 |
| 16_1730493 | 10 | 0.8 | 6 | 0.5 | 6 | 0.6667 | 7 | 0.7143 | 29 | 0.6897 | 4 | 1 | 6 | 0.8333 | 5 | 0.6 | 4 | 0.25 | 19 | 0.6842 |
| 16_1730495 | 11 | 0.7273 | 6 | 0.6667 | 6 | 0.6667 | 9 | 0.3333 | 32 | 0.5938 | 5 | 0.4 | 9 | 0.3333 | 8 | 0.125 | 4 | 0.5 | 26 | 0.3077 |
| 16_1730500 | 11 | 0.1818 | 6 | 0 | 6 | 0.3333 | 7 | 0 | 30 | 0.1333 | 5 | 0 | 7 | 0.1429 | 7 | 0 | 4 | 0.25 | 23 | 0.087 |
| 16_1732284 | 1 | 0 | 2 | 0 | 6 | 0.1667 | 4 | 0 | 13 | 0.0769 | 4 | 0 | NA | NA | 1 | 1 | 2 | 0 | 7 | 0.1429 |
| 16_1732504 | 3 | 0 | 1 | 0 | 7 | 0.2857 | 4 | 0.25 | 15 | 0.2 | 5 | 0.8 | 3 | 0.3333 | 3 | 0.6667 | 2 | 1 | 13 | 0.6923 |
| 16_1732525 | 3 | 0.3333 | 3 | 0.6667 | 7 | 0 | 5 | 0 | 18 | 0.1667 | 6 | 0.3333 | 2 | 0 | 3 | 0 | 4 | 0.25 | 15 | 0.2 |
| 16_1734231 | 8 | 0.25 | 17 | 0.2353 | 9 | 0.1111 | 12 | 0.4167 | 46 | 0.2609 | 5 | 0.2 | 23 | 0.3043 | 8 | 0 | 14 | 0.2857 | 50 | 0.24 |
| 16_1734348 | 1 | 1 | 2 | 1 | NA | NA | 1 | 0 | 4 | 0.75 | NA | NA | 3 | 0.3333 | 2 | 0.5 | 2 | 1 | 7 | 0.5714 |
| 16_1734359 | 8 | 0.125 | 12 | 0.0833 | 4 | 0 | 8 | 0.125 | 32 | 0.0938 | 3 | 0 | 13 | 0.3077 | 4 | 0 | 12 | 0.0833 | 32 | 0.1562 |
| 16_1734359 | 10 | 0.6 | 19 | 0.6316 | 10 | 0.7 | 13 | 0.3846 | 52 | 0.5769 | 6 | 0.5 | 22 | 0.5455 | 9 | 0.4444 | 13 | 0.6154 | 50 | 0.54 |
| 16_1734380 | 10 | 0.2 | 20 | 0.15 | 8 | 0.25 | 11 | 0.3636 | 49 | 0.2245 | 6 | 0 | 22 | 0.2727 | 12 | 0.3333 | 15 | 0.0667 | 55 | 0.2 |
| 16_1734409 | 14 | 0 | 26 | 0.0385 | 13 | 0.2308 | 15 | 0.0667 | 68 | 0.0735 | 6 | 0 | 25 | 0.16 | 13 | 0.1538 | 18 | 0.0556 | 62 | 0.1129 |
| 16_1734594 | NA | NA | 2 | 1 | 1 | 1 | NA | NA | 3 | 1 | NA | NA | NA | NA | 1 | 1 | 1 | 1 | 2 | 1 |
| 16_1734599 | 5 | 0.6 | 2 | 0.5 | 11 | 0.7273 | 4 | 1 | 22 | 0.7273 | 3 | 0.6667 | 5 | 0.4 | 4 | 0.25 | 6 | 0.6667 | 18 | 0.5 |
| 16_1734600 | 5 | 1 | 3 | 1 | 12 | 0.8333 | 5 | 1 | 25 | 0.92 | 4 | 1 | 9 | 1 | 5 | 1 | 4 | 0.75 | 22 | 0.9545 |
| 16_1734604 | 9 | 0.8889 | 2 | 1 | 11 | 0.9091 | 4 | 0.5 | 26 | 0.8462 | 2 | 1 | 9 | 1 | 6 | 1 | 3 | 1 | 20 | 1 |
| 16_1734608 | 10 | 1 | 2 | 1 | 11 | 0.7273 | 4 | 1 | 27 | 0.8889 | 2 | 0.5 | 8 | 1 | 4 | 1 | 6 | 1 | 20 | 0.95 |
| 16_1734609 | 10 | 0.8 | 3 | 1 | 10 | 0.8 | 3 | 1 | 26 | 0.8462 | 3 | 0.3333 | 10 | 0.9 | 6 | 0.8333 | 5 | 0.4 | 24 | 0.7083 |
| 16_1734610 | 5 | 0.4 | 3 | 0.3333 | 11 | 0.3636 | 4 | 0.25 | 23 | 0.3478 | 3 | 1 | 9 | 0.5556 | 5 | 0.2 | 4 | 0.25 | 21 | 0.4762 |
| 16_1734622 | 6 | 0.6667 | NA | NA | 8 | 0.5 | 4 | 0.75 | 18 | 0.6111 | 3 | 0.6667 | 7 | 0.4286 | 4 | 0.5 | 4 | 0.5 | 18 | 0.5 |
| 16_1734647 | 7 | 0 | 2 | 0 | 9 | 0.3333 | 3 | 0.3333 | 21 | 0.1905 | 3 | 0 | 8 | 0.25 | 1 | 0 | 6 | 0.1667 | 18 | 0.1667 |
| 16_1738706 | 1 | 1 | NA | NA | NA | NA | 1 | 0 | 2 | 0.5 | 1 | 0 | 2 | 0 | 1 | 1 | 3 | 0 | 7 | 0.1429 |
| 16_1738811 | 2 | 0.5 | NA | NA | NA | NA | 1 | 0 | 3 | 0.3333 | NA | NA | 2 | 0.5 | NA | NA | 2 | 0 | 4 | 0.25 |
| 16_1738815 | 1 | 1 | NA | NA | NA | NA | NA | NA | 1 | 1 | NA | NA | 1 | 1 | NA | NA | 1 | 1 | 2 | 1 |
| 16_1738832 | 2 | 0.5 | NA | NA | NA | NA | NA | NA | 2 | 0.5 | NA | NA | 2 | 1 | NA | NA | 2 | 0.5 | 4 | 0.75 |
| 16_1738836 | 2 | 0.5 | NA | NA | NA | NA | NA | NA | 2 | 0.5 | NA | NA | 2 | 1 | NA | NA | 2 | 1 | 4 | 1 |
| 16_1738858 | 2 | 0.5 | NA | NA | NA | NA | NA | NA | 2 | 0.5 | NA | NA | 2 | 0.5 | NA | NA | 2 | 0.5 | 4 | 0.5 |
| 16_1738961 | 1 | 1 | NA | NA | NA | NA | NA | NA | 1 | 1 | NA | NA | 1 | 1 | NA | NA | NA | NA | 1 | 1 |
| 16_1739071 | NA | NA | 3 | 0 | 4 | 0.5 | NA | NA | 7 | 0.2857 | 2 | 1 | 1 | 1 | NA | NA | 3 | 0.6667 | 6 | 0.8333 |
| 16_1739091 | 8 | 0.25 | 8 | 0.25 | 4 | 0.5 | 16 | 0.375 | 36 | 0.3333 | 10 | 0.6 | 6 | 0.5 | 11 | 0.3636 | 8 | 0.375 | 35 | 0.4571 |
| 16_1739094 | 13 | 1 | 10 | 0.7 | 9 | 0.8889 | 17 | 0.8824 | 49 | 0.8776 | 12 | 0.8333 | 5 | 1 | 12 | 1 | 10 | 0.7 | 39 | 0.8718 |
| 16_1739097 | 13 | 0.9231 | 9 | 0.8889 | 8 | 0.875 | 17 | 1 | 47 | 0.9362 | 13 | 0.8462 | 6 | 0.8333 | 15 | 0.8667 | 12 | 1 | 46 | 0.8913 |
| 16_1739120 | 13 | 0 | 9 | 0 | 8 | 0.25 | 14 | 0.0714 | 44 | 0.0682 | 12 | 0 | 6 | 0.3333 | 12 | 0 | 11 | 0.0909 | 41 | 0.0732 |
| 16_1739131 | 13 | 0.1538 | 9 | 0.1111 | 7 | 0.1429 | 13 | 0.1538 | 42 | 0.1429 | 13 | 0.1538 | 6 | 0.1667 | 14 | 0.0714 | 11 | 0.2727 | 44 | 0.1591 |
| 16_1739193 | 13 | 0.0769 | 8 | 0.125 | 6 | 0.1667 | 14 | 0 | 41 | 0.0732 | 11 | 0 | 6 | 0 | 10 | 0 | 11 | 0.1818 | 38 | 0.0526 |
| 16_1739208 | 12 | 0.0833 | 8 | 0 | 5 | 0.2 | 14 | 0.1429 | 39 | 0.1026 | 11 | 0.0909 | 5 | 0 | 10 | 0.1 | 13 | 0.0769 | 39 | 0.0769 |
| 16_1739211 | 12 | 0.6667 | 8 | 0.5 | 6 | 0.8333 | 14 | 0.5714 | 40 | 0.625 | 10 | 0.8 | 6 | 0.6667 | 8 | 0.75 | 12 | 0.75 | 36 | 0.75 |
| 16_1739212 | 8 | 0.125 | 6 | 0.5 | 7 | 0.4286 | 13 | 0.3077 | 34 | 0.3235 | 7 | 0.2857 | 6 | 0.3333 | 7 | 0 | 10 | 0.2 | 30 | 0.2 |
| 16_1739227 | 10 | 0.2 | 8 | 0.125 | 4 | 0.25 | 10 | 0.1 | 32 | 0.1562 | 9 | 0.2222 | 4 | 0 | 7 | 0 | 10 | 0.2 | 30 | 0.1333 |
| 16_1739234 | 5 | 0.2 | 4 | 0.25 | 4 | 0.5 | 9 | 0.5556 | 22 | 0.4091 | 8 | 0.25 | 4 | 0 | 9 | 0 | 7 | 0.2857 | 28 | 0.1429 |
| 16_1739301 | 5 | 0 | 5 | 0 | 4 | 0 | 6 | 0.3333 | 20 | 0.1 | 8 | 0.125 | 3 | 0 | 7 | 0.1429 | 4 | 0.25 | 22 | 0.1364 |
| 16_1739360 | 5 | 0.6 | 6 | 0.6667 | 3 | 0.6667 | 6 | 0.6667 | 20 | 0.65 | 6 | 0.6667 | NA | NA | 4 | 1 | 7 | 0.7143 | 17 | 0.7647 |

| | | | | | | | | | | | | | | | | | | | | |
|---|---|---|---|---|---|---|---|---|---|---|---|---|---|---|---|---|---|---|---|---|
| 16_173936 | 4 | 0.5 | 2 | 0.5 | 2 | 0.5 | 4 | 0.5 | 12 | 0.5 | 6 | 0.6667 | NA | NA | 1 | 0 | 6 | 0.6667 | 13 | 0.6154 |
| 16_173936 | 4 | 1 | 4 | 1 | 1 | 1 | 4 | 1 | 13 | 1 | 5 | 0.8 | NA | NA | 4 | 1 | 7 | 1 | 16 | 0.9375 |
| 16_173937 | 5 | 0 | 6 | 0.1667 | 2 | 0 | 5 | 0 | 18 | 0.0556 | 7 | 0 | NA | NA | 2 | 0 | 6 | 0.1667 | 15 | 0.0667 |
| 16_173937 | 5 | 0.4 | 6 | 0.3333 | NA | NA | 5 | 0 | 16 | 0.25 | 7 | 0.2857 | NA | NA | 2 | 0 | 6 | 0.1667 | 15 | 0.2 |
| 16_174600 | NA | NA | NA | NA | NA | NA | 1 | 1 | 1 | 1 | 1 | 1 | NA | NA | NA | NA | 2 | 0 | 3 | 0.3333 |
| 16_174602 | NA | NA | NA | NA | NA | NA | 1 | 1 | 1 | 1 | 1 | 1 | NA | NA | NA | NA | 2 | 0 | 3 | 0.3333 |
| 16_174603 | NA | NA | NA | NA | NA | NA | 1 | 1 | 1 | 1 | 1 | 1 | NA | NA | NA | NA | 2 | 1 | 3 | 1 |
| 16_174605 | 3 | 0.3333 | 5 | 0 | 2 | 0 | 3 | 0 | 13 | 0.0769 | 3 | 0 | 2 | 0 | 3 | 0.3333 | 2 | 0.5 | 10 | 0.2 |
| 16_174606 | 3 | 0.3333 | 6 | 0.3333 | 3 | 0.3333 | 3 | 0.3333 | 15 | 0.3333 | 3 | 0 | 2 | 0 | 3 | 0 | 2 | 0.5 | 10 | 0.1 |
| 16_174609 | NA | NA | NA | NA | NA | NA | 1 | 1 | 1 | 1 | NA | NA | NA | NA | NA | NA | 1 | 1 | 1 | 1 |
| 16_174610 | 3 | 0.6667 | 6 | 0 | 3 | 0 | 3 | 0.3333 | 15 | 0.2 | 4 | 0.25 | 2 | 0 | 4 | 0.25 | 2 | 0.5 | 12 | 0.25 |
| 16_174612 | 3 | 0.3333 | 5 | 0 | 2 | 0 | 3 | 0.3333 | 13 | 0.1538 | 3 | 0.3333 | 2 | 0.5 | 4 | 0 | 2 | 1 | 11 | 0.3636 |
| 16_174614 | NA | NA | 1 | 1 | NA | NA | 1 | 0 | 2 | 0.5 | NA | NA | NA | NA | NA | NA | 1 | 1 | 1 | 1 |
| 16_174673 | 2 | 1 | NA | NA | NA | NA | NA | NA | 2 | 1 | NA | NA | 2 | 0.5 | 1 | 0 | NA | NA | 3 | 0.3333 |
| 16_174678 | 3 | 1 | 2 | 0.5 | NA | NA | NA | NA | 5 | 0.8 | NA | NA | 2 | 0.5 | 1 | 1 | NA | NA | 3 | 0.6667 |
| 16_174682 | 3 | 0.3333 | 1 | 0 | NA | NA | NA | NA | 4 | 0.25 | NA | NA | 2 | 0.5 | 2 | 0 | NA | NA | 4 | 0.25 |
| 16_174684 | 5 | 0 | 5 | 0.2 | 1 | 1 | NA | NA | 11 | 0.1818 | 1 | 0 | 4 | 0.5 | 3 | 0.3333 | 1 | 0 | 9 | 0.3333 |
| 16_174692 | 4 | 0 | 5 | 0.2 | 4 | 0 | 6 | 0 | 19 | 0.0526 | 6 | 0 | 8 | 0.375 | 8 | 0 | 8 | 0 | 30 | 0.1 |
| 16_174698 | 2 | 0 | 6 | 0 | 3 | 0 | 6 | 0.1667 | 17 | 0.0588 | 4 | 0.25 | 9 | 0 | 5 | 0 | 7 | 0.1429 | 25 | 0.08 |
| 16_174699 | 1 | 0 | 4 | 0.25 | 3 | 0 | 6 | 0 | 14 | 0.0714 | 4 | 0 | 10 | 0.2 | 6 | 0 | 5 | 0.4 | 25 | 0.16 |
| 16_174771 | 1 | 1 | NA | NA | NA | NA | NA | NA | 1 | 1 | 1 | 1 | NA | NA | NA | NA | NA | NA | 1 | 1 |
| 16_174781 | 1 | 1 | NA | NA | NA | NA | NA | NA | 1 | 1 | 1 | 1 | NA | NA | NA | NA | NA | NA | 1 | 1 |
| 16_175944 | 3 | 0.3333 | 3 | 0.6667 | 4 | 0 | 3 | 0.3333 | 13 | 0.3077 | 4 | 0.75 | 4 | 0.75 | 3 | 0.3333 | 3 | 0.6667 | 14 | 0.6429 |
| 16_175944 | 3 | 0.6667 | 5 | 0.6 | 6 | 0.5 | 3 | 0.3333 | 17 | 0.5294 | 3 | 0.3333 | 3 | 0.6667 | 4 | 0.5 | 2 | 0 | 12 | 0.4167 |
| 16_175945 | 3 | 0.6667 | 5 | 0.6 | 6 | 0.6667 | 4 | 0.25 | 18 | 0.5556 | 4 | 0.75 | 4 | 0.5 | 4 | 0.5 | 2 | 0.5 | 14 | 0.5714 |
| 16_175945 | 3 | 1 | 6 | 1 | 7 | 0.5714 | 4 | 0.5 | 20 | 0.75 | 4 | 1 | 4 | 1 | 4 | 1 | 3 | 1 | 15 | 1 |
| 16_175946 | 3 | 1 | 5 | 0.6 | 6 | 0.3333 | 4 | 0.5 | 18 | 0.5556 | 4 | 1 | 4 | 0.25 | 3 | 0.6667 | 2 | 0 | 13 | 0.5385 |
| 16_175947 | 3 | 0 | 5 | 0.2 | 5 | 0.2 | 4 | 0.25 | 17 | 0.1765 | 3 | 0.3333 | 4 | 0 | 3 | 0.3333 | 1 | 0 | 11 | 0.1818 |
| 16_175947 | 3 | 0.6667 | 5 | 0.8 | 6 | 1 | 4 | 0.75 | 18 | 0.8333 | 4 | 0.5 | 4 | 1 | 2 | 1 | 3 | 1 | 13 | 0.8462 |
| 16_175949 | 1 | 1 | 4 | 0.25 | 4 | 0.25 | 4 | 0.75 | 13 | 0.4615 | 5 | 0.8 | 4 | 0.25 | 4 | 0.5 | 2 | 0 | 15 | 0.4667 |
| 16_176104 | 4 | 0.5 | 7 | 0.2857 | 7 | 0.5714 | 3 | 0 | 21 | 0.381 | 3 | 0 | 3 | 1 | 3 | 0 | 4 | 0 | 13 | 0.2308 |
| 16_176105 | 6 | 0.3333 | 12 | 0.0833 | 12 | 0.25 | 6 | 0 | 36 | 0.1667 | 5 | 0.4 | 6 | 0.3333 | 6 | 0 | 5 | 0.2 | 22 | 0.2273 |
| 16_176248 | 1 | 1 | NA | NA | NA | NA | NA | NA | 1 | 1 | 1 | 0 | 2 | 0.5 | NA | NA | NA | NA | 3 | 0.3333 |
| 16_176250 | 1 | 1 | NA | NA | NA | NA | NA | NA | 1 | 1 | 1 | 0 | 2 | 0.5 | NA | NA | 1 | 0 | 4 | 0.25 |
| 16_176250 | 1 | 1 | NA | NA | NA | NA | NA | NA | 1 | 1 | 1 | 1 | 2 | 1 | NA | NA | 1 | 1 | 4 | 1 |
| 16_176252 | 1 | 1 | NA | NA | NA | NA | NA | NA | 1 | 1 | 1 | 1 | 2 | 0 | NA | NA | 1 | 0 | 4 | 0.25 |
| 16_176256 | 1 | 1 | 1 | 1 | NA | NA | NA | NA | 2 | 1 | 1 | 1 | 1 | 0 | NA | NA | 1 | 1 | 3 | 0.6667 |
| 16_176535 | NA | NA | NA | NA | 2 | 0 | 1 | 1 | 3 | 0.3333 | 1 | 1 | NA | NA | 1 | 0 | NA | NA | 2 | 0.5 |
| 16_176557 | NA | NA | 3 | 1 | NA | NA | NA | NA | 3 | 1 | 1 | 1 | 1 | 0 | NA | NA | NA | NA | 2 | 0.5 |
| 16_176557 | NA | NA | 2 | 0.5 | 1 | 1 | 1 | 1 | 4 | 0.75 | 1 | 1 | NA | NA | 1 | 1 | NA | NA | 2 | 1 |
| 16_176559 | NA | NA | 2 | 0.5 | 1 | 1 | 1 | 1 | 4 | 0.75 | 1 | 1 | 1 | 0 | 1 | 1 | NA | NA | 3 | 0.6667 |
| 16_176559 | NA | NA | 2 | 1 | 1 | 1 | 1 | 1 | 4 | 1 | 1 | 1 | 1 | 1 | 1 | 0 | NA | NA | 3 | 0.6667 |
| 16_176564 | NA | NA | NA | NA | NA | NA | 1 | 1 | 1 | 1 | 1 | 1 | 1 | 0 | NA | NA | 1 | 1 | 3 | 0.6667 |
| 16_176584 | 10 | 0.1 | 6 | 0 | 12 | 0.1667 | 9 | 0.2222 | 37 | 0.1351 | 12 | 0.5 | 5 | 0 | 11 | 0.4545 | 8 | 0.5 | 36 | 0.4167 |
| 16_176585 | 10 | 0 | 6 | 0.3333 | 10 | 0 | 7 | 0.2857 | 33 | 0.1212 | 12 | 0.1667 | 6 | 0.1667 | 12 | 0.0833 | 10 | 0 | 40 | 0.1 |
| 16_176586 | 11 | 0.3636 | 7 | 0.2857 | 9 | 0.3333 | 7 | 0.5714 | 34 | 0.3824 | 12 | 0.5833 | 5 | 0.6 | 11 | 0.3636 | 10 | 0.5 | 38 | 0.5 |
| 16_176587 | 11 | 0.0909 | 7 | 0.2857 | 10 | 0.1 | 9 | 0.4444 | 37 | 0.2162 | 10 | 0.2 | 5 | 0.2 | 13 | 0.4615 | 8 | 0.25 | 36 | 0.3056 |
| 16_176587 | 10 | 0.3 | 6 | 0 | 9 | 0.1111 | 7 | 0.1429 | 32 | 0.1562 | 10 | 0.3 | 5 | 0.2 | 13 | 0.2308 | 10 | 0.4 | 38 | 0.2895 |
| 16_176587 | 11 | 0.1818 | 7 | 0 | 10 | 0 | 8 | 0 | 36 | 0.0556 | 12 | 0.1667 | 6 | 0 | 12 | 0 | 8 | 0 | 38 | 0.0526 |
| 16_176589 | 8 | 0.25 | 7 | 0.1429 | 9 | 0.1111 | 9 | 0.3333 | 33 | 0.2121 | 12 | 0.1667 | 4 | 0 | 11 | 0.0909 | 8 | 0.125 | 35 | 0.1143 |
| 16_176594 | 6 | 0.5 | 6 | 0.6667 | 8 | 0.625 | 7 | 1 | 27 | 0.7037 | 9 | 0.6667 | 4 | 0 | 10 | 0.6 | 6 | 0.8333 | 29 | 0.5862 |
| 16_176597 | 9 | 0.3333 | 5 | 0.4 | 7 | 0.4286 | 9 | 0.6667 | 30 | 0.4667 | 7 | 0.4286 | 5 | 0.2 | 11 | 0.0909 | 5 | 0.6 | 28 | 0.2857 |
| 16_176598 | 8 | 0.25 | 7 | 0.5714 | 8 | 0.375 | 8 | 0.5 | 31 | 0.4194 | 9 | 0.5556 | 5 | 0.4 | 13 | 0.2308 | 6 | 0.6667 | 33 | 0.4242 |
| 16_176599 | 5 | 0.4 | 6 | 0.8333 | 8 | 0.25 | 8 | 0.625 | 27 | 0.5185 | 8 | 0.75 | 5 | 0.6 | 11 | 0.8182 | 5 | 0.6 | 29 | 0.7241 |
| 16_176602 | 2 | 1 | 3 | 1 | 6 | 1 | 6 | 1 | 17 | 1 | 7 | 0.8571 | 4 | 0.75 | 8 | 0.875 | 4 | 0.75 | 23 | 0.8261 |
| 16_176602 | 6 | 0.5 | 4 | 0 | 6 | 0.5 | 6 | 0.8333 | 22 | 0.5 | 8 | 0.375 | 4 | 0.25 | 8 | 0.25 | 5 | 0.6 | 25 | 0.36 |
| 16_176603 | 4 | 0.75 | 3 | 1 | 3 | 0.3333 | 4 | 1 | 14 | 0.7857 | 5 | 1 | NA | NA | 3 | 0.3333 | 6 | 0.8333 | 14 | 0.7857 |
| 16_176603 | 6 | 0.3333 | 5 | 0.2 | 6 | 0 | 5 | 0.4 | 22 | 0.2273 | 8 | 0.5 | 3 | 0.3333 | 7 | 0.2857 | 3 | 0 | 21 | 0.3333 |
| 16_176605 | 6 | 0.1667 | 3 | 1 | 7 | 0 | 6 | 0.5 | 22 | 0.3182 | 7 | 0.4286 | 4 | 0.25 | 5 | 0.4 | 4 | 0.5 | 20 | 0.4 |
| 16_176605 | 6 | 0 | 4 | 0 | 7 | 0.1429 | 5 | 0.2 | 22 | 0.0909 | 7 | 0.2857 | 3 | 0 | 6 | 0.3333 | 6 | 0.1667 | 22 | 0.2273 |
| 16_176605 | 6 | 0 | 4 | 0.25 | 7 | 0.2857 | 6 | 0.3333 | 23 | 0.2174 | 9 | 0.4444 | 4 | 0.75 | 7 | 0.2857 | 6 | 0.6667 | 26 | 0.5 |
| 16_176619 | 4 | 0 | 3 | 0 | 5 | 0.4 | 5 | 0.2 | 17 | 0.1765 | 8 | 0.375 | 5 | 0.4 | 5 | 0.2 | 5 | 0.6 | 23 | 0.3913 |
| 16_177186 | 1 | 1 | 1 | 1 | 4 | 0.75 | 3 | 0.6667 | 9 | 0.7778 | 1 | 1 | 2 | 1 | 2 | 1 | 2 | 1 | 7 | 1 |
| 16_177188 | 2 | 0.5 | 1 | 0 | 4 | 0 | 3 | 0 | 10 | 0.1 | 2 | 0 | 2 | 0 | 2 | 0.5 | 2 | 0 | 8 | 0.125 |
| 16_177189 | 2 | 0.5 | 1 | 1 | 4 | 1 | 3 | 0.6667 | 10 | 0.8 | 2 | 1 | 2 | 1 | 2 | 1 | 1 | 1 | 7 | 1 |

| | | | | | | | | | | | | | | | | | | | | |
|---|---|---|---|---|---|---|---|---|---|---|---|---|---|---|---|---|---|---|---|---|
| 16_1771908 | 1 | 1 | NA | NA | 3 | 1 | 2 | 0.5 | 6 | 0.8333 | 2 | 0.5 | 2 | 0 | 2 | 1 | 1 | 1 | 7 | 0.5714 |
| 16_1771916 | 2 | 0.5 | 1 | 0 | 3 | 0 | 3 | 0 | 9 | 0.1111 | 2 | 0 | 1 | 0 | 2 | 0 | 2 | 0.5 | 7 | 0.1429 |
| 16_1771928 | 2 | 1 | 1 | 1 | 4 | 1 | 3 | 1 | 10 | 1 | 2 | 1 | 2 | 1 | 2 | 1 | 2 | 1 | 8 | 1 |
| 16_1771936 | 2 | 0 | 1 | 0 | 3 | 0 | 3 | 0.3333 | 9 | 0.1111 | 2 | 0 | 2 | 0.5 | 2 | 0 | 1 | 0 | 7 | 0.1429 |
| 16_1771937 | 1 | 1 | 1 | 0 | 4 | 0.5 | 3 | 0.6667 | 9 | 0.5556 | 2 | 1 | 2 | 0 | 2 | 0.5 | 2 | 0.5 | 8 | 0.5 |
| 16_1771939 | 2 | 1 | 1 | 1 | 4 | 0 | 3 | 0.6667 | 10 | 0.5 | 2 | 0.5 | 2 | 0.5 | 2 | 0 | 2 | 0.5 | 8 | 0.375 |
| 16_1771950 | 2 | 1 | 1 | 1 | 4 | 0.25 | 3 | 0 | 10 | 0.4 | 2 | 1 | 2 | 0 | 2 | 0.5 | 2 | 0.5 | 8 | 0.5 |
| 16_1771963 | 1 | 0 | 3 | 0.6667 | 5 | 1 | 3 | 1 | 12 | 0.8333 | 2 | 1 | 5 | 1 | 3 | 1 | 5 | 0.6 | 15 | 0.8667 |
| 16_1771964 | 4 | 0.5 | 2 | 0 | 4 | 0.25 | 3 | 0.3333 | 13 | 0.3077 | 1 | 0 | 2 | 0.5 | 4 | 0.25 | 3 | 0.3333 | 10 | 0.3 |
| 16_1771973 | 3 | 0.6667 | 3 | 1 | 3 | 0.6667 | 2 | 1 | 11 | 0.8182 | 1 | 1 | 4 | 0.75 | 4 | 1 | 3 | 0.6667 | 12 | 0.8333 |
| 16_1771980 | 5 | 0.2 | 3 | 0.3333 | 6 | 0.1667 | 3 | 0 | 17 | 0.1765 | 2 | 0 | 5 | 0.2 | 2 | 0 | 5 | 0.2 | 14 | 0.1429 |
| 16_1771987 | 5 | 0.4 | 3 | 0.6667 | 4 | 0.75 | 4 | 0.75 | 16 | 0.625 | 1 | 1 | 4 | 0.75 | 2 | 0.5 | 5 | 0.4 | 12 | 0.5833 |
| 16_1771989 | 5 | 0.6 | 4 | 0.5 | 5 | 0.4 | 4 | 1 | 18 | 0.6111 | 1 | 1 | 5 | 0.8 | 2 | 1 | 5 | 1 | 13 | 0.9231 |
| 16_1771989 | 5 | 0.4 | 4 | 0 | 5 | 0 | 3 | 0 | 17 | 0.1176 | 1 | 0 | 5 | 0.4 | 2 | 0 | 4 | 0.25 | 12 | 0.25 |
| 16_1771991 | 3 | 0.3333 | 3 | 0.6667 | 6 | 0.6667 | 4 | 0.5 | 16 | 0.5625 | 1 | 0 | 4 | 0.75 | 2 | 0.5 | 2 | 0 | 9 | 0.4444 |
| 16_1771997 | 5 | 0 | 4 | 0.25 | 6 | 0 | 4 | 0.25 | 19 | 0.1053 | 1 | 0 | 5 | 0.2 | 2 | 0 | 5 | 0 | 13 | 0.0769 |
| 16_1788469 | 12 | 0.4167 | 4 | 0.5 | 10 | 0.2 | 10 | 0.4 | 36 | 0.3611 | 7 | 0.1429 | 12 | 0 | 8 | 0 | 9 | 0.3333 | 36 | 0.1111 |
| 16_1788514 | NA | NA | NA | NA | 1 | 1 | NA | NA | 1 | 1 | 1 | 1 | NA | NA | NA | NA | 1 | 0 | 2 | 0.5 |
| 16_1788888 | 9 | 0.4444 | 6 | 0.1667 | 12 | 0.3333 | 10 | 0.4 | 37 | 0.3514 | 4 | 0.25 | 10 | 0.6 | 9 | 0.1111 | 8 | 0.5 | 31 | 0.3871 |
| 16_1788920 | 12 | 0.0833 | 5 | 0.2 | 11 | 0 | 12 | 0 | 40 | 0.05 | 7 | 0.1429 | 11 | 0.0909 | 11 | 0.0909 | 10 | 0.2 | 39 | 0.1282 |
| 16_1788963 | 5 | 0 | 5 | 0 | 6 | 0.1667 | 1 | 0 | 17 | 0.0588 | 3 | 0 | 7 | 0 | 6 | 0 | 4 | 0.25 | 20 | 0.05 |
| 16_1788980 | 6 | 0 | 6 | 0.1667 | 5 | 0.2 | 1 | 0 | 18 | 0.1111 | 2 | 0 | 7 | 0.1429 | 5 | 0 | 5 | 0 | 19 | 0.0526 |
| 16_1788986 | 6 | 0 | 6 | 0.1667 | 5 | 0.2 | 2 | 0 | 19 | 0.1053 | 2 | 0.5 | 6 | 0.1667 | 5 | 0 | 3 | 0 | 16 | 0.125 |
| 16_1790220 | 1 | 1 | NA | NA | NA | NA | NA | NA | 1 | 1 | NA | NA | 1 | 1 | 1 | 1 | 1 | 0 | 3 | 0.6667 |
| 16_1790232 | 2 | 0.5 | NA | NA | NA | NA | NA | NA | 2 | 0.5 | NA | NA | 1 | 0 | 1 | 1 | 1 | 0 | 3 | 0.3333 |
| 16_1790246 | 4 | 0.25 | 4 | 0 | 4 | 0 | 3 | 0.3333 | 15 | 0.1333 | NA | NA | 2 | 0 | 4 | 0.25 | 4 | 0.25 | 10 | 0.2 |
| 16_1790686 | 3 | 0 | 3 | 0 | 3 | 0.6667 | 2 | 0 | 11 | 0.1818 | NA | NA | 2 | 0 | 3 | 0.3333 | 4 | 0 | 9 | 0.1111 |
| 16_1796893 | 3 | 0.3333 | 3 | 0 | 4 | 0.75 | 1 | 1 | 11 | 0.4545 | 6 | 0.5 | 1 | 1 | NA | NA | 1 | 0 | 8 | 0.5 |
| 16_1796902 | 3 | 1 | 4 | 0.75 | 3 | 0.6667 | 1 | 0 | 11 | 0.7273 | 6 | 0.6667 | 1 | 1 | NA | NA | 1 | 0 | 8 | 0.625 |
| 16_1796920 | 2 | 0 | 4 | 0.25 | 4 | 0 | 1 | 0 | 11 | 0.0909 | 6 | 0.1667 | 1 | 0 | NA | NA | 1 | 0 | 8 | 0.125 |
| 16_1796924 | 3 | 1 | 4 | 1 | 5 | 0.6 | 1 | 1 | 13 | 0.8462 | 7 | 0.8571 | 1 | 1 | NA | NA | 1 | 1 | 9 | 0.8889 |
| 16_1796936 | 3 | 0 | 6 | 0.3333 | 5 | 0.6 | 1 | 0 | 15 | 0.3333 | 6 | 0.5 | 1 | 1 | NA | NA | 1 | 1 | 8 | 0.625 |
| 16_1796951 | 4 | 0.5 | 8 | 0.625 | 4 | 0.25 | 1 | 0 | 17 | 0.4706 | 7 | 0.7143 | 1 | 1 | NA | NA | 1 | 1 | 9 | 0.7778 |
| 16_1796967 | 3 | 0 | 7 | 0.1429 | 6 | 0 | 1 | 0 | 17 | 0.0588 | 7 | 0.2857 | 1 | 0 | NA | NA | NA | NA | 8 | 0.25 |
| 16_1796971 | 3 | 1 | 7 | 1 | 5 | 1 | 1 | 1 | 16 | 1 | 5 | 1 | 1 | 1 | NA | NA | 1 | 0 | 7 | 0.8571 |
| 16_1796972 | 2 | 1 | 3 | 1 | 3 | 1 | NA | NA | 8 | 1 | 5 | 1 | 1 | 1 | NA | NA | NA | NA | 6 | 1 |
| 16_1796975 | 1 | 1 | 5 | 0.8 | 4 | 0.75 | 1 | 1 | 11 | 0.8182 | 5 | 0.8 | 1 | 0 | NA | NA | 2 | 0 | 8 | 0.5 |
| 16_1796979 | 4 | 1 | 8 | 0.875 | 5 | 1 | 1 | 1 | 18 | 0.9444 | 6 | 1 | NA | NA | 1 | 1 | 2 | 1 | 9 | 1 |
| 16_1796989 | 4 | 0.5 | 5 | 0.2 | 3 | 0 | 1 | 1 | 13 | 0.3077 | 7 | 0.1429 | 1 | 1 | NA | NA | 1 | 0 | 9 | 0.2222 |
| 16_1796997 | 1 | 1 | 5 | 0.8 | 3 | 0.6667 | NA | NA | 9 | 0.7778 | 5 | 1 | NA | NA | 1 | 1 | 1 | 0 | 7 | 0.8571 |
| 16_1797006 | 5 | 0 | 7 | 0.2857 | 6 | 0 | 1 | 0 | 19 | 0.1053 | 7 | 0.4286 | 1 | 0 | NA | NA | 2 | 0 | 10 | 0.3 |
| 16_1797009 | 3 | 0 | 6 | 0.1667 | 5 | 0.2 | 1 | 1 | 15 | 0.2 | 7 | 0.1429 | 1 | 0 | 1 | 0 | 2 | 0 | 11 | 0.0909 |
| 16_1803878 | 6 | 0 | 10 | 0 | 5 | 0.4 | 12 | 0 | 33 | 0.0606 | 9 | 0.1111 | 1 | 0 | 3 | 0 | 7 | 0.1429 | 20 | 0.1 |
| 16_1803921 | 5 | 0.2 | 7 | 0.7143 | 2 | 0 | 8 | 0.25 | 22 | 0.3636 | 6 | 0.5 | 3 | 0.6667 | 1 | 0 | 5 | 0.6 | 15 | 0.5333 |
| 16_1803932 | 3 | 0.6667 | 2 | 1 | 2 | 1 | 2 | 0.5 | 9 | 0.7778 | 3 | 1 | 1 | 1 | NA | NA | 3 | 1 | 7 | 1 |
| 16_1803935 | 5 | 1 | 6 | 1 | 3 | 1 | 8 | 0.75 | 22 | 0.9091 | 8 | 0.875 | 3 | 1 | 1 | 1 | 5 | 1 | 17 | 0.9412 |
| 16_1803941 | 4 | 0.25 | 8 | 0 | 3 | 0.3333 | 9 | 0 | 24 | 0.0833 | 7 | 0.2857 | 3 | 0.3333 | 2 | 0 | 4 | 0.25 | 16 | 0.25 |
| 16_1803951 | 4 | 0.25 | 8 | 0.375 | 4 | 0 | 8 | 0.125 | 24 | 0.2083 | 8 | 0.375 | 3 | 0 | 2 | 0.5 | 5 | 0.2 | 18 | 0.2778 |
| 16_1804058 | 2 | 1 | 2 | 1 | 2 | 1 | 2 | 1 | 8 | 1 | 3 | 1 | 5 | 0.6 | 7 | 0.8571 | 3 | 0.6667 | 18 | 0.7778 |
| 16_1804067 | 2 | 0.5 | 2 | 0 | 3 | 0 | 1 | 0 | 8 | 0.125 | 3 | 0 | 5 | 0 | 6 | 0.1667 | 3 | 0 | 17 | 0.0588 |
| 16_1804082 | 2 | 1 | 2 | 1 | 4 | 0.5 | 1 | 1 | 9 | 0.7778 | 4 | 0.75 | 4 | 0.75 | 7 | 0.7143 | 3 | 1 | 18 | 0.7778 |
| 16_1804093 | 2 | 0.5 | 2 | 0.5 | 3 | 0.3333 | 1 | 0 | 8 | 0.375 | 4 | 0.25 | 4 | 0 | 8 | 0.25 | 3 | 0 | 19 | 0.1579 |
| 16_1804095 | 2 | 1 | 2 | 1 | 3 | 0.6667 | 1 | 1 | 8 | 0.875 | 4 | 0.75 | 4 | 0.25 | 7 | 0.8571 | 2 | 0.5 | 17 | 0.6471 |
| 16_1804106 | 1 | 1 | 1 | 0 | 2 | 0.5 | NA | NA | 4 | 0.5 | 2 | 0 | 3 | 0.3333 | 5 | 0.2 | 3 | 0 | 13 | 0.1538 |
| 16_1804109 | 2 | 0.5 | 1 | 0 | 2 | 1 | 1 | 0 | 6 | 0.5 | 4 | 0 | 2 | 0.5 | 6 | 0.8333 | 3 | 0.6667 | 15 | 0.5333 |
| 16_1804223 | 7 | 0.4286 | 9 | 0.3333 | 6 | 0 | 7 | 0.1429 | 29 | 0.2414 | 9 | 0.2222 | 4 | 0.5 | 7 | 0.2857 | 6 | 0 | 26 | 0.2308 |
| 16_1804228 | 7 | 0.7143 | 9 | 0.7778 | 6 | 0.8333 | 8 | 0.625 | 30 | 0.7333 | 12 | 0.6667 | 4 | 0.5 | 5 | 0.8 | 7 | 0.2857 | 28 | 0.5714 |
| 16_1804249 | 5 | 0.2 | 3 | 0 | 5 | 0 | 6 | 0.5 | 19 | 0.2105 | 3 | 0.6667 | 2 | 0 | 4 | 0.25 | 6 | 0.1667 | 15 | 0.2667 |
| 16_1804250 | 3 | 0.6667 | 6 | 0.6667 | 4 | 0.75 | 4 | 0.25 | 17 | 0.5882 | 8 | 0.25 | 3 | 1 | 6 | 0.6667 | 4 | 0.5 | 21 | 0.5238 |
| 16_1804253 | 6 | 0.3333 | 7 | 0.5714 | 5 | 0.6 | 5 | 0.4 | 23 | 0.4783 | 9 | 0.4444 | 3 | 1 | 6 | 0.6667 | 6 | 0.3333 | 24 | 0.5417 |
| 16_1804256 | 5 | 0 | 7 | 0.2857 | 5 | 0.4 | 6 | 0.3333 | 23 | 0.2609 | 7 | 0.2857 | 2 | 0.5 | 6 | 0.3333 | 5 | 0.2 | 20 | 0.3 |
| 16_1804609 | 5 | 0.2 | 1 | 0 | 1 | 0 | 4 | 0.25 | 11 | 0.1818 | 8 | 0.125 | 1 | 1 | 5 | 0.2 | 4 | 0 | 18 | 0.1667 |
| 16_1805785 | 10 | 0.1 | 13 | 0 | 18 | 0.2778 | 14 | 0.1429 | 55 | 0.1455 | 15 | 0.1333 | 9 | 0.2222 | 16 | 0.125 | 19 | 0.2105 | 59 | 0.1695 |
| 16_1805786 | 9 | 0.1111 | 14 | 0.2857 | 17 | 0.2353 | 13 | 0.4615 | 53 | 0.283 | 11 | 0.5455 | 9 | 0.3333 | 17 | 0.3529 | 19 | 0.1053 | 56 | 0.3036 |
| 16_1805935 | NA | NA | 1 | 0 | NA | NA | 1 | 1 | 2 | 0.5 | NA | NA | NA | NA | 1 | 1 | NA | NA | 1 | 1 |

| | | | | | | | | | | | | | | | | | | | | |
|---|---|---|---|---|---|---|---|---|---|---|---|---|---|---|---|---|---|---|---|---|
| 16_180676 | 1 | 1 | 1 | 0 | NA | NA | 3 | 0.6667 | 5 | 0.6 | 1 | 1 | 2 | 0.5 | 2 | 0.5 | 1 | 0 | 6 | 0.5 |
| 16_180676 | NA | NA | NA | NA | NA | NA | 2 | 1 | 2 | 1 | 1 | 1 | 2 | 1 | 2 | 0 | 2 | 1 | 7 | 0.7143 |
| 16_180676 | 1 | 1 | NA | NA | NA | NA | 3 | 0 | 4 | 0.25 | 1 | 0 | 3 | 0 | 2 | 0.5 | 3 | 0 | 9 | 0.1111 |
| 16_180678 | 1 | 0 | 1 | 0 | NA | NA | 2 | 0.5 | 4 | 0.25 | 1 | 1 | 3 | 0 | 1 | 0 | 3 | 0 | 8 | 0.125 |
| 16_180712 | 1 | 0 | 7 | 0.1429 | NA | NA | 3 | 0 | 11 | 0.0909 | 2 | 0.5 | 5 | 0 | 3 | 0.3333 | 4 | 0 | 14 | 0.1429 |
| 16_180725 | 3 | 0 | 2 | 0 | 1 | 0 | 4 | 0.75 | 10 | 0.3 | 3 | 0.3333 | 3 | 0.3333 | 3 | 0.3333 | 6 | 0.1667 | 15 | 0.2667 |
| 16_180729 | 6 | 0.3333 | 3 | 0 | NA | NA | 6 | 0.1667 | 15 | 0.2 | 4 | 0 | 4 | 0.25 | 3 | 0.3333 | 5 | 0.2 | 16 | 0.1875 |
| 16_180731 | 5 | 0.4 | 2 | 0 | 1 | 1 | 6 | 0.3333 | 14 | 0.3571 | 4 | 0.5 | 4 | 0 | 4 | 0 | 6 | 0.5 | 18 | 0.2778 |
| 16_180731 | NA | NA | 3 | 0 | 1 | 1 | 5 | 0.6 | 9 | 0.4444 | 4 | 0.5 | 3 | 0.3333 | 4 | 0.75 | 5 | 0.2 | 16 | 0.4375 |
| 16_180732 | 2 | 0.5 | 2 | 0 | 1 | 0 | 5 | 0.4 | 10 | 0.3 | 3 | 0.3333 | 3 | 0.3333 | 3 | 0 | 5 | 0.2 | 14 | 0.2143 |
| 16_180732 | 3 | 0 | 3 | 0 | NA | NA | 6 | 0.5 | 12 | 0.25 | 4 | 0.25 | 3 | 0 | 4 | 0 | 5 | 0 | 16 | 0.0625 |
| 16_180732 | 2 | 0 | 3 | 0 | 1 | 0 | 6 | 0.6667 | 12 | 0.3333 | 4 | 0.5 | 3 | 0 | 3 | 0.3333 | 4 | 1 | 14 | 0.5 |
| 16_180963 | 2 | 0 | 3 | 0 | 1 | 0 | 5 | 0.2 | 11 | 0.0909 | 3 | 0.3333 | 2 | 0 | 3 | 0 | 4 | 0 | 12 | 0.0833 |
| 16_181020 | 1 | 1 | 2 | 1 | NA | NA | 1 | 1 | 4 | 1 | 1 | 1 | 1 | 1 | NA | NA | 2 | 1 | 4 | 1 |
| 16_181020 | 2 | 0 | 2 | 0 | 1 | 0 | 3 | 0.6667 | 8 | 0.25 | 1 | 1 | 1 | 0 | 1 | 0 | 3 | 0.3333 | 6 | 0.3333 |
| 16_181021 | 3 | 0.3333 | NA | NA | NA | NA | 2 | 1 | 5 | 0.6 | 1 | 1 | 1 | 1 | 1 | 1 | 3 | 0.6667 | 6 | 0.8333 |
| 16_181021 | 2 | 0.5 | 1 | 1 | NA | NA | 2 | 0.5 | 5 | 0.6 | NA | NA | 1 | 0 | NA | NA | 1 | 1 | 2 | 0.5 |
| 16_181022 | 3 | 0 | 2 | 0.5 | 1 | 0 | 2 | 0 | 8 | 0.125 | 2 | 0.5 | 1 | 0 | NA | NA | 3 | 0 | 6 | 0.1667 |
| 16_181023 | 2 | 0.5 | 2 | 0.5 | NA | NA | 1 | 0 | 5 | 0.4 | 1 | 1 | 1 | 1 | NA | NA | 4 | 0.75 | 6 | 0.8333 |
| 16_181023 | 3 | 1 | 2 | 1 | 1 | 1 | 2 | 0.5 | 8 | 0.875 | 1 | 1 | 1 | 1 | NA | NA | 3 | 0.6667 | 5 | 0.8 |
| 16_181188 | 9 | 0.3333 | 5 | 0 | 19 | 0.1579 | 12 | 0.4167 | 45 | 0.2444 | 8 | 0 | 5 | 0.6 | 5 | 0.2 | 8 | 0.375 | 26 | 0.2692 |
| 16_181189 | 10 | 0.1 | 5 | 0.2 | 19 | 0.1579 | 10 | 0.1 | 44 | 0.1364 | 6 | 0 | 7 | 0.1429 | 5 | 0.2 | 7 | 0.4286 | 25 | 0.2 |
| 16_181189 | 10 | 0.4 | 5 | 0.8 | 18 | 0.4444 | 7 | 0.4286 | 40 | 0.475 | 8 | 0.25 | 6 | 0.1667 | 4 | 0.25 | 8 | 0.5 | 26 | 0.3077 |
| 16_181190 | 6 | 0.3333 | 5 | 0.4 | 14 | 0.5 | 9 | 0.2222 | 34 | 0.3824 | 8 | 0.25 | 5 | 0.4 | 4 | 0.25 | 7 | 0.2857 | 24 | 0.2917 |
| 16_181191 | 10 | 0 | 4 | 0.25 | 16 | 0.0625 | 10 | 0.1 | 40 | 0.075 | 6 | 0 | 6 | 0.1667 | 5 | 0.2 | 6 | 0.3333 | 23 | 0.1739 |
| 16_181192 | 7 | 0.2857 | 3 | 0 | 12 | 0.25 | 6 | 0 | 28 | 0.1786 | 6 | 0 | 7 | 0.1429 | 4 | 0 | 6 | 0.1667 | 23 | 0.087 |
| 16_182152 | NA | NA | 1 | 0 | 1 | 1 | 2 | 0 | 4 | 0.25 | 2 | 0.5 | NA | NA | NA | NA | NA | NA | 2 | 0.5 |
| 16_182167 | NA | NA | 1 | 0 | 1 | 0 | 2 | 0.5 | 4 | 0.25 | 2 | 1 | NA | NA | NA | NA | NA | NA | 2 | 1 |
| 16_182246 | 3 | 1 | 2 | 0.5 | 2 | 0 | 4 | 0 | 11 | 0.3636 | 2 | 0 | 2 | 0.5 | 3 | 0.3333 | 1 | 1 | 8 | 0.375 |
| 16_182246 | 3 | 0.3333 | 3 | 0 | 3 | 0 | 4 | 0 | 13 | 0.0769 | 2 | 0 | 2 | 0 | 3 | 0.3333 | 1 | 0 | 8 | 0.125 |
| 16_182250 | 2 | 0 | 2 | 0.5 | 4 | 0 | 5 | 0.2 | 13 | 0.1538 | 2 | 0 | 4 | 0.5 | 2 | 0 | 1 | 1 | 9 | 0.3333 |
| 16_182250 | 2 | 0 | 3 | 0 | 2 | 0 | 4 | 0.25 | 11 | 0.0909 | 2 | 0 | 2 | 0.5 | 3 | 0 | 1 | 0 | 8 | 0.125 |
| 16_182250 | 3 | 0 | 2 | 0.5 | 3 | 0.3333 | 3 | 0.3333 | 11 | 0.2727 | 1 | 1 | 2 | 0 | 3 | 0 | 1 | 1 | 7 | 0.2857 |
| 16_182251 | 2 | 0 | 2 | 0.5 | 1 | 1 | 1 | 0 | 6 | 0.3333 | NA | NA | 2 | 0.5 | NA | NA | NA | NA | 2 | 0.5 |
| 16_182255 | 1 | 1 | 2 | 0.5 | 1 | 1 | 1 | 1 | 5 | 0.8 | NA | NA | 2 | 1 | NA | NA | NA | NA | 2 | 1 |
| 16_182258 | 1 | 0 | 4 | 0.5 | 3 | 0.3333 | 6 | 0.3333 | 14 | 0.3571 | 4 | 0.25 | 5 | 0.4 | 2 | 0.5 | 4 | 0.25 | 15 | 0.3333 |
| 16_182262 | 17 | 0.6471 | 13 | 0.6154 | 8 | 0.5 | 10 | 0.2 | 48 | 0.5208 | 10 | 0.7 | 10 | 0.4 | 14 | 0.2857 | 15 | 0.5333 | 49 | 0.4694 |
| 16_183199 | NA | NA | 1 | 1 | 1 | 1 | NA | NA | 2 | 1 | NA | NA | 2 | 1 | NA | NA | 1 | 1 | 3 | 1 |
| 16_183200 | NA | NA | 1 | 1 | NA | NA | 1 | 1 | 2 | 1 | NA | NA | 2 | 1 | NA | NA | 1 | 1 | 3 | 1 |
| 16_183201 | NA | NA | NA | NA | 1 | 1 | NA | NA | 1 | 1 | NA | NA | 2 | 1 | NA | NA | 1 | 1 | 3 | 1 |
| 16_183202 | NA | NA | 1 | 1 | 1 | 1 | 1 | 1 | 3 | 1 | NA | NA | 2 | 1 | NA | NA | NA | NA | 2 | 1 |
| 16_183202 | NA | NA | 1 | 0 | 1 | 1 | 1 | 1 | 3 | 0.6667 | NA | NA | 2 | 1 | NA | NA | 1 | 1 | 3 | 1 |
| 16_183202 | NA | NA | 1 | 1 | 1 | 1 | 1 | 1 | 3 | 1 | NA | NA | 2 | 1 | NA | NA | 1 | 1 | 3 | 1 |
| 16_183203 | NA | NA | NA | NA | 1 | 1 | 1 | 1 | 2 | 1 | NA | NA | 2 | 1 | NA | NA | NA | NA | 2 | 1 |
| 16_183203 | NA | NA | 1 | 1 | 1 | 1 | 1 | 1 | 3 | 1 | NA | NA | 1 | 1 | NA | NA | 1 | 1 | 2 | 1 |
| 16_183218 | NA | NA | 1 | 0 | 1 | 1 | 2 | 0 | 4 | 0.25 | NA | NA | 1 | 1 | NA | NA | 1 | 1 | 2 | 1 |
| 16_184391 | NA | NA | 2 | 1 | 1 | 1 | 1 | 1 | 4 | 1 | NA | NA | 2 | 1 | 1 | 1 | 1 | 1 | 4 | 1 |
| 16_184393 | 2 | 0.5 | 4 | 0 | 2 | 0 | 3 | 0.3333 | 11 | 0.1818 | 1 | 0 | 4 | 0.25 | NA | NA | 6 | 0.3333 | 11 | 0.2727 |
| 16_184394 | 2 | 1 | 5 | 0.2 | 4 | 0.75 | 3 | 0.3333 | 14 | 0.5 | 1 | 1 | 4 | 0.5 | NA | NA | 5 | 0.6 | 10 | 0.6 |
| 16_184394 | 2 | 0 | 3 | 0 | 3 | 0.3333 | 4 | 0 | 12 | 0.0833 | 1 | 0 | 5 | 0.4 | NA | NA | 6 | 0.1667 | 12 | 0.25 |
| 16_184394 | 2 | 1 | 5 | 0.6 | 3 | 0.3333 | 4 | 0.25 | 14 | 0.5 | 1 | 0 | 4 | 1 | NA | NA | 5 | 0.8 | 10 | 0.8 |
| 16_184394 | 2 | 1 | 4 | 0.75 | 4 | 1 | 4 | 1 | 14 | 0.9286 | 1 | 1 | 5 | 0.6 | NA | NA | 6 | 0.8333 | 12 | 0.75 |
| 16_184399 | 1 | 1 | 4 | 1 | 4 | 1 | 3 | 0.3333 | 12 | 0.8333 | 1 | 1 | 6 | 0.8333 | NA | NA | 4 | 0.75 | 11 | 0.8182 |
| 16_184401 | 2 | 0 | 4 | 0.25 | 4 | 0 | 3 | 0 | 13 | 0.0769 | 1 | 1 | 4 | 0.25 | NA | NA | 4 | 0 | 9 | 0.2222 |
| 16_184403 | 2 | 0 | 3 | 0.3333 | 3 | 0 | 3 | 0 | 11 | 0.0909 | 1 | 0 | 5 | 0.4 | NA | NA | 3 | 0.3333 | 9 | 0.3333 |
| 16_184407 | 77 | 0.2857 | 90 | 0.2556 | 80 | 0.3125 | 55 | 0.3455 | 302 | 0.2947 | 67 | 0.3284 | 73 | 0.3562 | 63 | 0.2698 | 65 | 0.2923 | 268 | 0.3134 |
| 16_184413 | 71 | 0.8592 | 95 | 0.8 | 77 | 0.7662 | 44 | 0.6818 | 287 | 0.7875 | 73 | 0.8219 | 66 | 0.6818 | 63 | 0.7937 | 69 | 0.7536 | 271 | 0.7638 |
| 16_184413 | 87 | 0.2529 | 104 | 0.2212 | 83 | 0.1566 | 53 | 0.2453 | 327 | 0.2171 | 78 | 0.2308 | 71 | 0.2254 | 75 | 0.28 | 82 | 0.2439 | 306 | 0.2451 |
| 16_184414 | 84 | 0.0833 | 96 | 0.0521 | 84 | 0.0952 | 52 | 0.0385 | 316 | 0.0696 | 75 | 0.1067 | 72 | 0.2083 | 68 | 0.1471 | 77 | 0.0649 | 292 | 0.1301 |
| 16_184449 | 37 | 0.027 | 47 | 0.0851 | 36 | 0.0833 | 15 | 0 | 135 | 0.0593 | 36 | 0.0556 | 23 | 0.1304 | 34 | 0.0882 | 37 | 0.0811 | 130 | 0.0846 |
| 16_184451 | 35 | 0.0571 | 43 | 0.0465 | 36 | 0.0556 | 16 | 0.0625 | 130 | 0.0538 | 35 | 0.0571 | 25 | 0.04 | 34 | 0.0294 | 35 | 0.1143 | 129 | 0.062 |
| 16_185958 | 8 | 0.125 | 14 | 0.2143 | 7 | 0.1429 | 3 | 0 | 32 | 0.1562 | 4 | 0.25 | 3 | 0 | 2 | 0 | 3 | 0.3333 | 12 | 0.1667 |
| 16_185958 | 8 | 0 | 8 | 0.25 | 5 | 0.2 | 3 | 0.3333 | 24 | 0.1667 | 3 | 0 | 3 | 0.3333 | 2 | 0 | 4 | 0 | 12 | 0.0833 |
| 16_185963 | 7 | 1 | 12 | 0.8333 | 5 | 1 | 3 | 1 | 27 | 0.9259 | 3 | 0.6667 | 2 | 1 | 1 | 1 | 4 | 1 | 10 | 0.9 |
| 16_185964 | 7 | 0.1429 | 14 | 0 | 5 | 0.2 | 4 | 0.5 | 30 | 0.1333 | 5 | 0 | 3 | 0 | 1 | 1 | 4 | 0 | 13 | 0.0769 |

| | | | | | | | | | | | | | | | | | | | | |
|---|---|---|---|---|---|---|---|---|---|---|---|---|---|---|---|---|---|---|---|---|
| 16_1859653 | 8 | 0.125 | 12 | 0.25 | 6 | 0 | 4 | 0 | 30 | 0.1333 | 5 | 0 | 3 | 0 | 1 | 0 | 4 | 0.25 | 13 | 0.0769 |
| 16_1859658 | 8 | 0 | 12 | 0.0833 | 6 | 0 | 4 | 0.25 | 30 | 0.0667 | 3 | 0.3333 | 4 | 0.25 | 2 | 0 | 3 | 0 | 12 | 0.1667 |
| 16_1859658 | 8 | 0.5 | 13 | 0.1538 | 7 | 0.4286 | 2 | 0.5 | 30 | 0.3333 | 5 | 0.2 | 3 | 0.3333 | 2 | 0.5 | 2 | 0.5 | 12 | 0.3333 |
| 16_1859660 | 8 | 1 | 13 | 0.7692 | 7 | 0.8571 | 4 | 0.75 | 32 | 0.8438 | 4 | 1 | 2 | 1 | 1 | 1 | 3 | 1 | 10 | 1 |
| 16_1859669 | 6 | 0.5 | 8 | 0.625 | 4 | 0.5 | 2 | 1 | 20 | 0.6 | 1 | 1 | 3 | 0.6667 | 2 | 0.5 | 3 | 0.3333 | 9 | 0.5556 |
| 16_1859685 | 9 | 1 | 13 | 1 | 6 | 1 | 3 | 1 | 31 | 1 | 4 | 0.75 | 2 | 1 | 1 | 1 | 3 | 1 | 10 | 0.9 |
| 16_1859686 | 9 | 0.2222 | 15 | 0.6667 | 7 | 0.4286 | 3 | 0.6667 | 34 | 0.5 | 4 | 0 | 3 | 0.3333 | 2 | 0 | 5 | 0.6 | 14 | 0.2857 |
| 16_1865302 | 15 | 0.1333 | 12 | 0.1667 | 9 | 0.1111 | 9 | 0 | 45 | 0.1111 | 10 | 0.1 | 7 | 0.1429 | 5 | 0 | 12 | 0.3333 | 34 | 0.1765 |
| 16_1865306 | 14 | 0.2143 | 13 | 0.3077 | 10 | 0.1 | 9 | 0.2222 | 46 | 0.2174 | 8 | 0.25 | 11 | 0.2727 | 9 | 0 | 16 | 0.1875 | 44 | 0.1818 |
| 16_1865307 | 18 | 0.0556 | 12 | 0.1667 | 11 | 0.2727 | 9 | 0.2222 | 50 | 0.16 | 10 | 0.1 | 11 | 0 | 9 | 0.1111 | 16 | 0.0625 | 46 | 0.0652 |
| 16_1865318 | 9 | 0.5556 | 12 | 0.5 | 7 | 0.2857 | 7 | 0.2857 | 35 | 0.4286 | 5 | 0.6 | 8 | 0.25 | 7 | 0.5714 | 11 | 0.4545 | 31 | 0.4516 |
| 16_1865321 | 11 | 0.5455 | 12 | 0.25 | 6 | 0.1667 | 9 | 0.3333 | 38 | 0.3421 | 8 | 0.5 | 8 | 0.125 | 5 | 0 | 9 | 0.3333 | 30 | 0.2667 |
| 16_1865345 | 12 | 0.25 | 12 | 0.4167 | 9 | 0.3333 | 9 | 0.4444 | 42 | 0.3571 | 6 | 0.5 | 8 | 0.25 | 5 | 0.4 | 7 | 0.2857 | 26 | 0.3462 |
| 16_1865367 | 11 | 0.5455 | 5 | 0.4 | 7 | 0.1429 | 4 | 0 | 27 | 0.3333 | 5 | 0.2 | 7 | 0.1429 | 5 | 0.2 | 8 | 0.25 | 25 | 0.2 |
| 16_1865792 | 1 | 1 | 1 | 1 | NA | NA | 1 | 1 | 3 | 1 | 3 | 0.3333 | 3 | 1 | 2 | 1 | 1 | 1 | 9 | 0.7778 |
| 16_1865793 | 1 | 0 | 1 | 1 | NA | NA | 1 | 0 | 3 | 0.3333 | 2 | 0 | 3 | 0.3333 | 1 | 0 | 1 | 0 | 7 | 0.1429 |
| 16_1865799 | 1 | 1 | 1 | 1 | NA | NA | 1 | 1 | 3 | 1 | 3 | 0.6667 | 3 | 0.6667 | 2 | 0.5 | 1 | 1 | 9 | 0.6667 |
| 16_1865800 | 2 | 0.5 | 1 | 1 | NA | NA | NA | NA | 3 | 0.6667 | 3 | 0.3333 | 3 | 0.6667 | 2 | 0.5 | 1 | 1 | 9 | 0.5556 |
| 16_1865802 | 2 | 0.5 | 1 | 0 | NA | NA | 1 | 0 | 4 | 0.25 | 3 | 0 | 4 | 0 | 1 | 0 | 1 | 1 | 9 | 0.1111 |
| 16_1865810 | 3 | 0.3333 | 1 | 0 | NA | NA | NA | NA | 4 | 0.25 | 3 | 0.6667 | 4 | 0.5 | 2 | 0 | 1 | 1 | 10 | 0.5 |
| 16_1865835 | 3 | 1 | 1 | 1 | NA | NA | 1 | 1 | 5 | 1 | 3 | 1 | 5 | 1 | 2 | 1 | 2 | 1 | 12 | 1 |
| 16_1865879 | 3 | 1 | NA | NA | NA | NA | 1 | 1 | 4 | 1 | 5 | 0.8 | 5 | 0.8 | 2 | 1 | 2 | 1 | 14 | 0.8571 |
| 16_1869087 | 5 | 0.2 | 2 | 0 | 4 | 0 | 4 | 0 | 15 | 0.0667 | 4 | 0 | 5 | 0 | 7 | 0.1429 | 1 | 1 | 17 | 0.1176 |
| 16_1869090 | 3 | 0.3333 | 2 | 0 | 3 | 0 | 4 | 0 | 12 | 0.0833 | 4 | 0 | 4 | 0.25 | 6 | 0 | 1 | 1 | 15 | 0.1333 |
| 16_1869106 | 5 | 0 | 6 | 0.1667 | 6 | 0 | 5 | 0.2 | 22 | 0.0909 | 7 | 0.1429 | 5 | 0 | 9 | 0.1111 | 7 | 0 | 28 | 0.0714 |
| 16_1869108 | 5 | 0.8 | 7 | 0.8571 | 8 | 0.875 | 5 | 0.8 | 25 | 0.84 | 6 | 1 | 7 | 0.8571 | 9 | 1 | 7 | 1 | 29 | 0.9655 |
| 16_1874626 | 1 | 0 | 5 | 0.2 | 3 | 0.3333 | 2 | 0.5 | 11 | 0.2727 | NA | NA | 3 | 0.6667 | 1 | 0 | 2 | 0 | 6 | 0.3333 |
| 16_1878841 | 1 | 1 | 3 | 0 | NA | NA | 2 | 0 | 6 | 0.1667 | 4 | 0.25 | 2 | 0.5 | 2 | 0.5 | 2 | 0.5 | 10 | 0.4 |
| 16_1878846 | 1 | 0 | 2 | 0 | NA | NA | 2 | 0.5 | 5 | 0.2 | 3 | 0.6667 | 2 | 0.5 | 1 | 0 | 2 | 0.5 | 8 | 0.5 |
| 16_1878847 | 1 | 1 | 3 | 1 | NA | NA | 1 | 1 | 5 | 1 | 4 | 1 | 2 | 1 | 1 | 1 | 3 | 1 | 10 | 1 |
| 16_1878854 | NA | NA | 1 | 1 | NA | NA | 1 | 0 | 2 | 0.5 | 2 | 1 | NA | NA | 1 | 0 | 2 | 0.5 | 5 | 0.6 |
| 16_1878855 | 1 | 1 | 3 | 0 | NA | NA | 2 | 0 | 6 | 0.1667 | 4 | 0.25 | 2 | 0 | 2 | 0.5 | 2 | 0.5 | 10 | 0.3 |
| 16_1878872 | 1 | 0 | 3 | 0.3333 | NA | NA | 1 | 0 | 5 | 0.2 | 4 | 0 | 2 | 0 | 2 | 0.5 | 3 | 0 | 11 | 0.0909 |
| 16_1878876 | 1 | 1 | 3 | 0.6667 | 1 | 1 | 2 | 0.5 | 7 | 0.7143 | 4 | 0.75 | 1 | 1 | 1 | 1 | 3 | 0.3333 | 9 | 0.6667 |
| 16_1878884 | NA | NA | 3 | 0.6667 | 1 | 1 | 1 | 0 | 5 | 0.6 | 3 | 0.6667 | 2 | 1 | 1 | 1 | NA | NA | 6 | 0.8333 |
| 16_1992556 | 4 | 1 | 2 | 1 | NA | NA | NA | NA | 6 | 1 | 3 | 1 | 1 | 1 | 6 | 0.8333 | 1 | 1 | 11 | 0.9091 |
| 16_1992595 | 6 | 0.1667 | 3 | 0.3333 | NA | NA | NA | NA | 9 | 0.2222 | 2 | 0 | 1 | 0 | 7 | 0.1429 | 1 | 1 | 11 | 0.1818 |
| 16_1992605 | 4 | 1 | 3 | 1 | NA | NA | NA | NA | 7 | 1 | 3 | 1 | 1 | 1 | 7 | 0.8571 | 2 | 1 | 13 | 0.9231 |
| 16_1992610 | 5 | 1 | 3 | 1 | NA | NA | NA | NA | 8 | 1 | 3 | 0.6667 | 1 | 1 | 5 | 1 | 2 | 1 | 11 | 0.9091 |
| 16_1992613 | 5 | 1 | 3 | 1 | NA | NA | NA | NA | 8 | 1 | 1 | 1 | NA | NA | 6 | 1 | 2 | 1 | 9 | 1 |
| 16_1992627 | 5 | 0.2 | 3 | 0.3333 | NA | NA | NA | NA | 8 | 0.25 | 2 | 1 | 1 | 0 | 4 | 0 | 2 | 0 | 9 | 0.2222 |
| 16_1992628 | 6 | 0.3333 | 3 | 0.3333 | NA | NA | NA | NA | 9 | 0.3333 | 3 | 0.3333 | 1 | 0 | 7 | 0 | 1 | 1 | 12 | 0.1667 |
| 16_1994181 | 7 | 0.2857 | 3 | 1 | NA | NA | NA | NA | 10 | 0.5 | 2 | 0.5 | 1 | 0 | 6 | 0.6667 | 2 | 0.5 | 11 | 0.5455 |
| 16_1994193 | 6 | 0.6667 | 3 | 0.6667 | NA | NA | NA | NA | 9 | 0.6667 | 2 | 1 | 1 | 1 | 7 | 0.8571 | 2 | 0.5 | 12 | 0.8333 |
| 16_1994197 | 7 | 0 | 4 | 0.25 | NA | NA | NA | NA | 11 | 0.0909 | 3 | 0 | 1 | 0 | 8 | 0.25 | 2 | 0 | 14 | 0.1429 |
| 16_1994208 | 7 | 0.2857 | 4 | 0.5 | 1 | 0 | NA | NA | 12 | 0.3333 | 3 | 0.6667 | 1 | 0 | 8 | 0.5 | 1 | 0 | 13 | 0.4615 |
| 16_1994253 | 7 | 0 | 4 | 0.25 | 2 | 0 | NA | NA | 13 | 0.0769 | 3 | 0 | 1 | 0 | 8 | 0 | 4 | 0.5 | 16 | 0.125 |
| 16_1994260 | 7 | 0 | 5 | 0 | 2 | 0.5 | NA | NA | 14 | 0.0714 | 3 | 0 | 1 | 0 | 8 | 0 | 4 | 0.5 | 16 | 0.125 |
| 16_1994503 | 3 | 0.3333 | 1 | 0 | 5 | 0 | 4 | 0.25 | 13 | 0.1538 | 2 | 0.5 | 3 | 0 | 5 | 0 | 3 | 0 | 13 | 0.0769 |
| 16_1994541 | 3 | 0.3333 | 3 | 0.6667 | 5 | 0.4 | 5 | 0.2 | 16 | 0.375 | 1 | 0 | 3 | 0.6667 | 4 | 0.5 | 3 | 1 | 11 | 0.6364 |
| 16_1994550 | 3 | 0.6667 | 3 | 0.3333 | 5 | 0.6 | 2 | 0.5 | 13 | 0.5385 | 3 | 0.6667 | 3 | 0.6667 | 5 | 0.2 | 3 | 1 | 14 | 0.5714 |
| 16_1994550 | 3 | 0 | 2 | 0 | 4 | 0.25 | 3 | 0 | 12 | 0.0833 | 3 | 0.6667 | 2 | 0 | 4 | 0.25 | 3 | 0.6667 | 12 | 0.4167 |
| 16_1994583 | 4 | 0.25 | 4 | 0 | 5 | 0.2 | 4 | 0.25 | 17 | 0.1765 | 3 | 0.3333 | 2 | 0.5 | 5 | 0 | 3 | 0.3333 | 13 | 0.2308 |
| 16_1994589 | 4 | 0.25 | 4 | 0 | 4 | 0.25 | 5 | 0.2 | 17 | 0.1765 | 4 | 0 | 3 | 0.3333 | 5 | 0 | 4 | 0 | 16 | 0.0625 |
| 16_1994600 | 5 | 0 | 4 | 0 | 6 | 0.1667 | 4 | 0.25 | 19 | 0.1053 | 4 | 0.25 | 3 | 0.3333 | 5 | 0.2 | 3 | 0.3333 | 15 | 0.2667 |
| 16_1994605 | 4 | 0.75 | 3 | 0.6667 | 5 | 0.4 | 2 | 1 | 14 | 0.6429 | 4 | 0.5 | 3 | 0.6667 | 5 | 0.2 | 4 | 1 | 16 | 0.5625 |
| 16_1994607 | 4 | 0.5 | 4 | 0.25 | 6 | 0 | 5 | 0 | 19 | 0.1579 | 4 | 0.25 | 2 | 0 | 4 | 0 | 4 | 0 | 14 | 0.0714 |
| 16_1999502 | 7 | 0 | 5 | 0 | 5 | 0.2 | 7 | 0.1429 | 24 | 0.0833 | 4 | 0 | 3 | 0 | 8 | 0.125 | 6 | 0.1667 | 21 | 0.0952 |
| 16_2004959 | NA | NA | 2 | 0 | NA | NA | 1 | 1 | 3 | 0.3333 | 1 | 0 | NA | NA | 2 | 0.5 | NA | NA | 3 | 0.3333 |
| 16_2004960 | NA | NA | 2 | 0.5 | NA | NA | 1 | 0 | 3 | 0.3333 | 2 | 0 | NA | NA | 2 | 0.5 | 1 | 0 | 5 | 0.2 |
| 16_2004967 | NA | NA | 2 | 0.5 | NA | NA | 1 | 0 | 3 | 0.3333 | 2 | 1 | NA | NA | 2 | 0 | 1 | 0 | 5 | 0.4 |
| 16_2004980 | NA | NA | 2 | 1 | NA | NA | 1 | 1 | 3 | 1 | 1 | 0 | NA | NA | 3 | 1 | 1 | 1 | 5 | 0.8 |
| 16_2004984 | NA | NA | 2 | 1 | NA | NA | 1 | 1 | 3 | 1 | 1 | 1 | NA | NA | 3 | 1 | 1 | 1 | 5 | 1 |
| 16_2005001 | NA | NA | 2 | 1 | NA | NA | 1 | 0 | 3 | 0.6667 | 2 | 1 | NA | NA | 3 | 0.6667 | 1 | 0 | 6 | 0.6667 |
| 16_2005014 | NA | NA | 2 | 0.5 | 1 | 0 | 1 | 0 | 4 | 0.25 | 1 | 0 | NA | NA | 3 | 0.3333 | 1 | 0 | 5 | 0.2 |

| | | | | | | | | | | | | | | | | | | | | |
|---|---|---|---|---|---|---|---|---|---|---|---|---|---|---|---|---|---|---|---|---|
| 16_200502 | NA | NA | 2 | 1 | 2 | 1 | 1 | 1 | 5 | 1 | 2 | 1 | NA | NA | 3 | 0.6667 | 1 | 0 | 6 | 0.6667 |
| 16_200504 | NA | NA | 1 | 1 | 2 | 0.5 | 1 | 0 | 4 | 0.5 | 3 | 0.6667 | NA | NA | 2 | 1 | 2 | 1 | 7 | 0.8571 |
| 16_200506 | NA | NA | 2 | 0 | 1 | 1 | 2 | 0 | 5 | 0.2 | 3 | 0.3333 | NA | NA | 3 | 0 | 2 | 0 | 8 | 0.125 |
| 16_200511 | NA | NA | 1 | 1 | 3 | 0.3333 | 2 | 0.5 | 6 | 0.5 | 3 | 0.3333 | NA | NA | 3 | 0 | 1 | 0 | 7 | 0.1429 |
| 16_200522 | 7 | 0.1429 | 8 | 0 | 3 | 0.3333 | 9 | 0.2222 | 27 | 0.1481 | 3 | 0.6667 | 5 | 0.2 | 7 | 0.1429 | 5 | 0 | 20 | 0.2 |
| 16_200536 | 3 | 0 | 2 | 0 | 2 | 0 | 6 | 0.3333 | 13 | 0.1538 | 1 | 0 | 1 | 0 | 7 | 0.1429 | 3 | 0 | 12 | 0.0833 |
| 16_200544 | 4 | 0 | 4 | 0.5 | 2 | 0.5 | 6 | 0.3333 | 16 | 0.3125 | 2 | 0 | 2 | 0 | 7 | 0.1429 | 4 | 0.25 | 15 | 0.1333 |
| 16_200544 | 2 | 0.5 | 4 | 0 | 2 | 0.5 | 6 | 0.1667 | 14 | 0.2143 | 2 | 0.5 | 2 | 0 | 6 | 0.1667 | 4 | 0 | 14 | 0.1429 |
| 16_200545 | 3 | 0 | 3 | 0.3333 | 2 | 1 | 6 | 0.6667 | 14 | 0.5 | 2 | 1 | 2 | 0.5 | 7 | 0.7143 | 4 | 0.5 | 15 | 0.6667 |
| 16_200545 | 2 | 1 | 1 | 0 | 2 | 0 | 6 | 0.6667 | 11 | 0.5455 | 2 | 0 | 2 | 0 | 6 | 0 | 4 | 0.75 | 14 | 0.2143 |
| 16_200545 | 3 | 0.6667 | 3 | 1 | 2 | 1 | 4 | 0.75 | 12 | 0.8333 | 2 | 0 | 2 | 0.5 | 7 | 1 | 4 | 0.75 | 15 | 0.7333 |
| 16_200547 | 3 | 0 | 3 | 0 | 2 | 0.5 | 6 | 0.5 | 14 | 0.2857 | 2 | 0.5 | 1 | 0 | 6 | 0.5 | 3 | 0.3333 | 12 | 0.4167 |
| 16_200987 | 9 | 0 | 8 | 0.375 | 2 | 0 | 5 | 0 | 24 | 0.125 | 6 | 0.3333 | 5 | 0 | 4 | 0 | 6 | 0 | 21 | 0.0952 |
| 16_202114 | 2 | 0 | 4 | 0.25 | 4 | 0 | 1 | 0 | 11 | 0.0909 | 1 | 0 | 3 | 0 | 2 | 0.5 | 2 | 0 | 8 | 0.125 |
| 16_202114 | 2 | 0 | 4 | 0.75 | 4 | 1 | 1 | 1 | 11 | 0.7273 | NA | NA | 3 | 0.6667 | 2 | 1 | 2 | 1 | 7 | 0.8571 |
| 16_202186 | NA | NA | 5 | 0.2 | 3 | 0 | NA | NA | 8 | 0.125 | 1 | 0 | 3 | 0.3333 | 1 | 0 | 1 | 0 | 6 | 0.1667 |
| 16_202197 | 2 | 0.5 | 4 | 0.25 | 3 | 0 | 3 | 0.6667 | 12 | 0.3333 | 1 | 1 | 4 | 0.25 | 5 | 0.4 | NA | NA | 10 | 0.4 |
| 16_202198 | 3 | 0.3333 | 4 | 0.5 | 2 | 0.5 | 3 | 0.3333 | 12 | 0.4167 | 1 | 1 | 4 | 0.5 | 3 | 0.6667 | NA | NA | 8 | 0.625 |
| 16_202200 | 2 | 0.5 | 2 | 0 | 1 | 0 | 2 | 0 | 7 | 0.1429 | NA | NA | 4 | 0 | 3 | 0.3333 | NA | NA | 7 | 0.1429 |
| 16_202202 | 2 | 0 | 1 | 0 | 1 | 1 | 2 | 1 | 6 | 0.5 | NA | NA | 4 | 0.5 | 2 | 0.5 | NA | NA | 6 | 0.5 |
| 16_202229 | NA | NA | NA | NA | 1 | 0 | 1 | 1 | 2 | 0.5 | NA | NA | 2 | 1 | 2 | 0 | NA | NA | 4 | 0.5 |
| 16_202231 | 1 | 0 | NA | NA | 1 | 0 | 1 | 1 | 3 | 0.3333 | NA | NA | 3 | 0.3333 | 1 | 0 | NA | NA | 4 | 0.25 |
| 16_202242 | 2 | 0.5 | 1 | 0 | 1 | 0 | 1 | 1 | 5 | 0.4 | 1 | 0 | 3 | 0.3333 | 2 | 0 | 2 | 0 | 8 | 0.125 |
| 16_203207 | 1 | 0 | 1 | 0 | 7 | 0 | 5 | 0.4 | 14 | 0.1429 | 5 | 0 | 6 | 0.1667 | 6 | 0.1667 | 3 | 0 | 20 | 0.1 |
| 16_203279 | 3 | 0.3333 | 7 | 0.1429 | 9 | 0.1111 | 6 | 0.8333 | 25 | 0.32 | 8 | 0.25 | 7 | 0.1429 | 8 | 0.125 | 6 | 0 | 29 | 0.1379 |
| 16_203498 | 3 | 0 | NA | NA | 1 | 0 | 2 | 0.5 | 6 | 0.1667 | 1 | 0 | NA | NA | 3 | 0.3333 | NA | NA | 4 | 0.25 |
| 16_203501 | 3 | 0.6667 | 1 | 0 | 1 | 0 | 2 | 1 | 7 | 0.5714 | 3 | 0.3333 | NA | NA | 2 | 0.5 | 1 | 1 | 6 | 0.5 |
| 16_203511 | 2 | 0.5 | 1 | 0 | NA | NA | 5 | 0.4 | 8 | 0.375 | 3 | 0 | NA | NA | 4 | 0.25 | 2 | 0 | 9 | 0.1111 |
| 16_203518 | 4 | 0 | 3 | 0.3333 | 1 | 0 | 6 | 0 | 14 | 0.0714 | 4 | 0.25 | 2 | 0 | 4 | 0 | 2 | 0 | 12 | 0.0833 |
| 16_203627 | 13 | 0.1538 | 24 | 0.0417 | 17 | 0.3529 | 21 | 0.1905 | 75 | 0.1733 | 15 | 0.3333 | 27 | 0.2963 | 17 | 0.3529 | 23 | 0.3043 | 82 | 0.3171 |
| 16_203792 | 9 | 0.6667 | 10 | 0.8 | 8 | 0.625 | 4 | 0.5 | 31 | 0.6774 | 3 | 0.3333 | 4 | 0.25 | 5 | 0.4 | 6 | 0.6667 | 18 | 0.4444 |
| 16_203793 | 9 | 0.7778 | 8 | 0.5 | 5 | 0.8 | 5 | 0.6 | 27 | 0.6667 | 3 | 1 | 1 | 1 | 5 | 0.2 | 5 | 0.6 | 14 | 0.5714 |
| 16_203793 | 6 | 0.5 | 9 | 0.6667 | 6 | 0.6667 | 4 | 0.25 | 25 | 0.56 | 1 | 1 | 4 | 0.5 | 4 | 1 | 6 | 0.5 | 15 | 0.6667 |
| 16_203793 | 8 | 0 | 9 | 0.1111 | 9 | 0.1111 | 4 | 0 | 30 | 0.0667 | 3 | 0 | 3 | 0.3333 | 6 | 0.1667 | 7 | 0.4286 | 19 | 0.2632 |
| 16_203796 | NA | NA | 3 | 0.3333 | NA | NA | NA | NA | 3 | 0.3333 | NA | NA | 1 | 0 | NA | NA | 1 | 1 | 2 | 0.5 |
| 16_203801 | NA | NA | 3 | 0.3333 | NA | NA | NA | NA | 3 | 0.3333 | NA | NA | 1 | 0 | NA | NA | 1 | 1 | 2 | 0.5 |
| 16_203803 | NA | NA | 1 | 1 | NA | NA | NA | NA | 1 | 1 | NA | NA | NA | NA | NA | NA | 1 | 1 | 1 | 1 |
| 16_204207 | 1 | 1 | 1 | 1 | 3 | 0.6667 | 3 | 0.6667 | 8 | 0.75 | 1 | 0 | NA | NA | NA | NA | 2 | 0.5 | 3 | 0.3333 |
| 16_204235 | 3 | 0.3333 | 3 | 0.6667 | 3 | 0 | 3 | 0.3333 | 12 | 0.3333 | 2 | 0.5 | 1 | 0 | NA | NA | 1 | 0 | 4 | 0.25 |
| 16_204235 | 3 | 0 | 4 | 0 | 3 | 0.3333 | 4 | 0.25 | 14 | 0.1429 | 2 | 0.5 | 1 | 0 | NA | NA | 2 | 0 | 5 | 0.2 |
| 16_204236 | 3 | 0.6667 | 3 | 0.3333 | 3 | 0.6667 | 3 | 0.6667 | 12 | 0.5833 | 2 | 1 | 1 | 1 | NA | NA | 2 | 0.5 | 5 | 0.8 |
| 16_204237 | 3 | 0.6667 | 4 | 0.75 | 3 | 0.3333 | 4 | 0.75 | 14 | 0.6429 | 2 | 1 | 1 | 1 | NA | NA | 2 | 0.5 | 5 | 0.8 |
| 16_204237 | 2 | 0.5 | 4 | 0.75 | 3 | 0 | 3 | 0.6667 | 12 | 0.5 | 2 | 1 | 1 | 1 | NA | NA | 2 | 1 | 5 | 1 |
| 16_204242 | 1 | 1 | 4 | 0.75 | 2 | 1 | 2 | 1 | 9 | 0.8889 | 1 | 0 | 6 | 1 | 2 | 0.5 | 1 | 1 | 10 | 0.8 |
| 16_204374 | 1 | 1 | NA | NA | NA | NA | NA | NA | 1 | 1 | 1 | 0 | NA | NA | 1 | 1 | 1 | 0 | 3 | 0.3333 |
| 16_204381 | 20 | 0.45 | 40 | 0.425 | 27 | 0.3333 | 15 | 0.2 | 102 | 0.3725 | 43 | 0.2326 | 15 | 0.2 | 15 | 0.2667 | 19 | 0.2632 | 92 | 0.2391 |
| 16_204381 | 23 | 0.0435 | 37 | 0.0811 | 21 | 0.0476 | 15 | 0 | 96 | 0.0521 | 37 | 0.027 | 16 | 0.125 | 20 | 0.05 | 20 | 0.15 | 93 | 0.0753 |
| 16_204382 | 20 | 0.05 | 35 | 0.0857 | 23 | 0 | 14 | 0.0714 | 92 | 0.0543 | 37 | 0.0541 | 15 | 0 | 19 | 0.0526 | 18 | 0.1111 | 89 | 0.0562 |
| 16_204383 | 19 | 0.6316 | 38 | 0.5526 | 24 | 0.625 | 11 | 0.8182 | 92 | 0.6196 | 38 | 0.6053 | 15 | 0.4667 | 19 | 0.8421 | 19 | 0.6316 | 91 | 0.6374 |
| 16_204385 | 18 | 0.8333 | 32 | 0.6875 | 21 | 0.7143 | 11 | 0.5455 | 82 | 0.7073 | 28 | 0.7857 | 16 | 0.6875 | 18 | 0.8889 | 15 | 0.8667 | 77 | 0.8052 |
| 16_204386 | 20 | 0.1 | 36 | 0.0833 | 21 | 0.1429 | 12 | 0 | 89 | 0.0899 | 36 | 0.0833 | 16 | 0 | 18 | 0.1111 | 19 | 0.1053 | 89 | 0.0787 |
| 16_204642 | 2 | 0 | 3 | 0.6667 | 2 | 1 | 3 | 0.3333 | 10 | 0.5 | 1 | 1 | 1 | 0 | 3 | 0.3333 | NA | NA | 5 | 0.4 |
| 16_204820 | 40 | 0.175 | 29 | 0.1034 | 25 | 0.24 | 36 | 0.1667 | 130 | 0.1692 | 28 | 0.1071 | 22 | 0.2273 | 30 | 0.2 | 27 | 0.1111 | 107 | 0.1589 |
| 16_204981 | 2 | 1 | 1 | 0 | 1 | 1 | 5 | 0.6 | 9 | 0.6667 | 2 | 1 | 1 | 1 | 4 | 0.75 | 4 | 0.75 | 11 | 0.8182 |
| 16_204981 | 2 | 0.5 | 1 | 0 | 4 | 0 | 5 | 0.2 | 12 | 0.1667 | 4 | 0 | 2 | 0 | 5 | 0 | 6 | 0.1667 | 17 | 0.0588 |
| 16_204982 | 3 | 0 | 1 | 0 | 5 | 0.2 | 5 | 0.4 | 14 | 0.2143 | 4 | 0.25 | 3 | 0 | 2 | 0 | 7 | 0.1429 | 16 | 0.125 |
| 16_204983 | 3 | 0.6667 | 2 | 0 | 5 | 0 | 4 | 0.5 | 14 | 0.2857 | 3 | 0.6667 | 2 | 0.5 | 5 | 0.2 | 4 | 0 | 14 | 0.2857 |
| 16_204984 | 2 | 0.5 | 3 | 0 | 4 | 0.75 | 4 | 0 | 13 | 0.3077 | 5 | 0.4 | 2 | 0.5 | 3 | 0.6667 | 7 | 0.2857 | 17 | 0.4118 |
| 16_205112 | 16 | 0.75 | 18 | 0.8333 | 15 | 0.8 | 10 | 0.5 | 59 | 0.7458 | 15 | 0.6667 | 11 | 0.7273 | 13 | 0.5385 | 17 | 0.7059 | 56 | 0.6607 |
| 16_205112 | 13 | 0.9231 | 16 | 0.875 | 14 | 0.8571 | 11 | 1 | 54 | 0.9074 | 14 | 0.8571 | 9 | 1 | 8 | 1 | 19 | 0.7895 | 50 | 0.88 |
| 16_205115 | 14 | 0.2857 | 22 | 0.3636 | 9 | 0.2222 | 11 | 0.5455 | 56 | 0.3571 | 11 | 0.2727 | 11 | 0.2727 | 12 | 0.25 | 19 | 0.3158 | 53 | 0.283 |
| 16_205202 | 24 | 0.1667 | 28 | 0.3571 | 15 | 0 | 18 | 0.2222 | 85 | 0.2118 | 19 | 0.1053 | 14 | 0.1429 | 19 | 0.2105 | 12 | 0.3333 | 64 | 0.1875 |
| 16_205204 | 23 | 0.4783 | 26 | 0.2308 | 16 | 0.375 | 17 | 0.5294 | 82 | 0.3902 | 20 | 0.35 | 14 | 0.3571 | 20 | 0.35 | 11 | 0.3636 | 65 | 0.3538 |
| 16_205205 | 24 | 1 | 26 | 0.9231 | 16 | 1 | 17 | 1 | 83 | 0.9759 | 20 | 0.95 | 12 | 1 | 15 | 0.9333 | 12 | 1 | 59 | 0.9661 |
| 16_205205 | 23 | 0.1304 | 32 | 0.125 | 18 | 0.1667 | 15 | 0 | 88 | 0.1136 | 20 | 0.15 | 13 | 0.1538 | 18 | 0.0556 | 13 | 0.1538 | 64 | 0.125 |

| | | | | | | | | | | | | | | | | | | | | |
|---|---|---|---|---|---|---|---|---|---|---|---|---|---|---|---|---|---|---|---|---|
| 16_205206 | 26 | 0.6923 | 28 | 0.6786 | 20 | 0.45 | 17 | 0.5882 | 91 | 0.6154 | 19 | 0.4737 | 15 | 0.3333 | 21 | 0.5238 | 12 | 0.75 | 67 | 0.5075 |
| 16_205209 | 27 | 0.0741 | 29 | 0.0345 | 22 | 0.0455 | 18 | 0.1667 | 96 | 0.0729 | 22 | 0.0909 | 16 | 0 | 18 | 0.1111 | 13 | 0.1538 | 69 | 0.087 |
| 16_205210 | 24 | 0.2083 | 34 | 0.4118 | 23 | 0.3043 | 18 | 0.2222 | 99 | 0.303 | 20 | 0.2 | 16 | 0.0625 | 21 | 0.3333 | 11 | 0.3636 | 68 | 0.2353 |
| 16_205213 | 22 | 0.6364 | 28 | 0.5714 | 16 | 0.625 | 17 | 0.7059 | 83 | 0.6265 | 15 | 0.6 | 11 | 0.4545 | 18 | 0.8333 | 12 | 0.6667 | 56 | 0.6607 |
| 16_205213 | 18 | 0.3889 | 27 | 0.3333 | 21 | 0.4286 | 14 | 0.2857 | 80 | 0.3625 | 17 | 0.3529 | 12 | 0.3333 | 19 | 0.1579 | 12 | 0.75 | 60 | 0.3667 |
| 16_205214 | 15 | 0.4 | 15 | 0.2 | 16 | 0.375 | 11 | 0.5455 | 57 | 0.3684 | 6 | 0.3333 | 10 | 0.2 | 13 | 0.2308 | 7 | 0.2857 | 36 | 0.25 |
| 16_205216 | 22 | 0.5909 | 27 | 0.4074 | 18 | 0.4444 | 16 | 0.5 | 83 | 0.4819 | 16 | 0.3125 | 14 | 0.6429 | 18 | 0.5 | 12 | 0.5833 | 60 | 0.5 |
| 16_205221 | 4 | 0 | 7 | 0.1429 | 2 | 0 | 4 | 0 | 17 | 0.0588 | 3 | 0 | 6 | 0 | 4 | 0.25 | 3 | 0 | 16 | 0.0625 |
| 16_205408 | 3 | 1 | 5 | 0.8 | 4 | 0.75 | 6 | 0.6667 | 18 | 0.7778 | 9 | 0.6667 | 8 | 1 | 4 | 1 | 3 | 1 | 24 | 0.875 |
| 16_205408 | 3 | 0 | 7 | 0.4286 | 3 | 0 | 5 | 0 | 18 | 0.1667 | 11 | 0.1818 | 8 | 0 | 3 | 0 | 3 | 0 | 25 | 0.08 |
| 16_205443 | 1 | 1 | NA | NA | NA | NA | NA | NA | 1 | 1 | NA | NA | 1 | 1 | NA | NA | 1 | 1 | 2 | 1 |
| 16_205443 | 1 | 1 | NA | NA | NA | NA | NA | NA | 1 | 1 | NA | NA | 2 | 1 | NA | NA | 1 | 1 | 3 | 1 |
| 16_205444 | 1 | 1 | NA | NA | NA | NA | NA | NA | 1 | 1 | NA | NA | 2 | 1 | NA | NA | 1 | 1 | 3 | 1 |
| 16_205444 | 1 | 1 | NA | NA | NA | NA | NA | NA | 1 | 1 | NA | NA | 2 | 0.5 | NA | NA | 1 | 0 | 3 | 0.3333 |
| 16_205445 | 1 | 1 | NA | NA | NA | NA | NA | NA | 1 | 1 | NA | NA | 2 | 0.5 | NA | NA | 1 | 1 | 3 | 0.6667 |
| 16_205446 | 1 | 1 | NA | NA | NA | NA | NA | NA | 1 | 1 | NA | NA | 2 | 0.5 | NA | NA | 1 | 1 | 3 | 0.6667 |
| 16_205447 | 1 | 1 | NA | NA | NA | NA | NA | NA | 1 | 1 | NA | NA | 1 | 1 | NA | NA | 1 | 0 | 2 | 0.5 |
| 16_205608 | 2 | 0.5 | 1 | 1 | 4 | 0.25 | NA | NA | 7 | 0.4286 | 3 | 0.3333 | 2 | 1 | 3 | 0.6667 | 2 | 0.5 | 10 | 0.6 |
| 16_205609 | 1 | 1 | 1 | 0 | 2 | 0 | NA | NA | 4 | 0.25 | 3 | 0.3333 | 1 | 1 | 3 | 0.6667 | 2 | 1 | 9 | 0.6667 |
| 16_210332 | 2 | 1 | 1 | 1 | 1 | 0 | NA | NA | 4 | 0.75 | NA | NA | NA | NA | NA | NA | 2 | 1 | 2 | 1 |
| 16_210413 | 2 | 0.5 | 1 | 1 | 1 | 1 | NA | NA | 4 | 0.75 | NA | NA | NA | NA | 2 | 0.5 | 2 | 1 | 4 | 0.75 |
| 16_210413 | 2 | 1 | 1 | 1 | 1 | 0 | NA | NA | 4 | 0.75 | NA | NA | NA | NA | 1 | 1 | 2 | 1 | 3 | 1 |
| 16_210414 | 2 | 0 | 1 | 0 | 1 | 1 | NA | NA | 4 | 0.25 | NA | NA | NA | NA | 2 | 0 | 2 | 0.5 | 4 | 0.25 |
| 16_210416 | 1 | 0 | 1 | 0 | 1 | 1 | NA | NA | 3 | 0.3333 | NA | NA | NA | NA | 2 | 0 | 2 | 0.5 | 4 | 0.25 |
| 16_210417 | 2 | 0.5 | 1 | 1 | 1 | 1 | NA | NA | 4 | 0.75 | NA | NA | NA | NA | 2 | 0.5 | 2 | 0.5 | 4 | 0.5 |
| 16_210418 | 3 | 1 | NA | NA | 2 | 1 | NA | NA | 5 | 1 | 1 | 1 | NA | NA | 2 | 0.5 | 2 | 0 | 5 | 0.4 |
| 16_211513 | NA | NA | 1 | 1 | NA | NA | NA | NA | 1 | 1 | NA | NA | 1 | 1 | 1 | 1 | NA | NA | 2 | 1 |
| 16_211516 | NA | NA | 1 | 1 | NA | NA | NA | NA | 1 | 1 | NA | NA | 1 | 1 | 1 | 1 | NA | NA | 2 | 1 |
| 16_213190 | 3 | 0 | 3 | 0 | 2 | 0 | 2 | 0.5 | 10 | 0.1 | NA | NA | 3 | 0.3333 | 3 | 0.3333 | 1 | 1 | 7 | 0.4286 |
| 16_214630 | 5 | 0.2 | 6 | 0.1667 | 4 | 0.5 | 3 | 0.3333 | 18 | 0.2778 | 2 | 0.5 | 6 | 0 | 5 | 0 | 2 | 0 | 15 | 0.0667 |
| 16_214681 | 8 | 0.5 | 6 | 0.1667 | 4 | 0 | 11 | 0.1818 | 29 | 0.2414 | 6 | 0.1667 | 9 | 0.2222 | 11 | 0.2727 | 7 | 0.2857 | 33 | 0.2424 |
| 16_214682 | 9 | 0.3333 | 7 | 0.1429 | 7 | 0.2857 | 9 | 0.1111 | 32 | 0.2188 | 5 | 0.2 | 10 | 0.1 | 8 | 0.125 | 6 | 0.1667 | 29 | 0.1379 |
| 16_214688 | 11 | 0.1818 | 5 | 0 | 8 | 0 | 8 | 0 | 32 | 0.0625 | 4 | 0 | 10 | 0.1 | 10 | 0.1 | 4 | 0 | 28 | 0.0714 |
| 16_214689 | 6 | 0 | 5 | 0.4 | 7 | 0.1429 | 7 | 0.1429 | 25 | 0.16 | 3 | 0 | 8 | 0.125 | 6 | 0.1667 | 3 | 0.3333 | 20 | 0.15 |
| 16_214695 | 8 | 0.125 | 2 | 0 | 6 | 0.3333 | 7 | 0.1429 | 23 | 0.1739 | 3 | 0 | 10 | 0.2 | 6 | 0.1667 | 4 | 0 | 23 | 0.1304 |
| 16_214706 | 5 | 0 | 1 | 0 | 5 | 0.2 | 4 | 0.25 | 15 | 0.1333 | 3 | 0.3333 | 4 | 0.25 | 6 | 0 | 3 | 0.3333 | 16 | 0.1875 |
| 16_214709 | 5 | 0.4 | 2 | 0.5 | 4 | 0.25 | 5 | 0.2 | 16 | 0.3125 | 3 | 0.3333 | 5 | 0 | 6 | 0.1667 | 2 | 0.5 | 16 | 0.1875 |
| 16_214713 | 5 | 0.6 | 1 | 0 | 3 | 0.3333 | 3 | 0 | 12 | 0.3333 | 2 | 0 | 3 | 0 | 5 | 0.2 | 1 | 0 | 11 | 0.0909 |
| 16_214713 | 4 | 0.5 | 2 | 0 | 3 | 0 | 3 | 0.3333 | 12 | 0.25 | 2 | 0 | 4 | 0 | 3 | 0.3333 | 2 | 1 | 11 | 0.2727 |
| 16_214714 | 5 | 0.6 | 2 | 0 | 3 | 0.6667 | 4 | 0.25 | 14 | 0.4286 | 3 | 0 | 4 | 0.25 | 4 | 0.5 | 2 | 0 | 13 | 0.2308 |
| 16_214714 | 5 | 0.4 | 2 | 0 | 1 | 0 | 3 | 0.6667 | 11 | 0.3636 | 3 | 0.3333 | 4 | 0 | 3 | 0.3333 | 2 | 0.5 | 12 | 0.25 |
| 16_214717 | 4 | 1 | 1 | 1 | 1 | 1 | 2 | 1 | 8 | 1 | 2 | 1 | 3 | 1 | 3 | 1 | 1 | 1 | 9 | 1 |
| 16_214719 | 3 | 1 | NA | NA | 1 | 1 | 3 | 0.6667 | 7 | 0.8571 | NA | NA | 1 | 0 | 4 | 0.5 | 1 | 1 | 6 | 0.5 |
| 16_214720 | 2 | 1 | NA | NA | 3 | 1 | 2 | 1 | 7 | 1 | NA | NA | 2 | 0 | 4 | 0.75 | 1 | 1 | 7 | 0.5714 |
| 16_214721 | 3 | 1 | NA | NA | 2 | 1 | 2 | 1 | 7 | 1 | NA | NA | 3 | 1 | 3 | 1 | 1 | 1 | 7 | 1 |
| 16_214721 | 2 | 0.5 | NA | NA | 3 | 0.3333 | 2 | 0.5 | 7 | 0.4286 | NA | NA | 3 | 0.6667 | 5 | 0.4 | 1 | 1 | 9 | 0.5556 |
| 16_214723 | 4 | 0.25 | NA | NA | 2 | 0.5 | 2 | 0 | 8 | 0.25 | NA | NA | 2 | 0.5 | 2 | 0 | 1 | 1 | 5 | 0.4 |
| 16_214723 | 3 | 1 | NA | NA | 2 | 0 | 1 | 0 | 6 | 0.5 | NA | NA | 2 | 0.5 | 4 | 0.5 | 1 | 0 | 7 | 0.4286 |
| 16_215802 | 2 | 0 | 1 | 0 | 1 | 0 | 2 | 0.5 | 6 | 0.1667 | NA | NA | 1 | 0 | 2 | 1 | NA | NA | 3 | 0.6667 |
| 16_215804 | 2 | 1 | 1 | 1 | 1 | 1 | 1 | 1 | 5 | 1 | NA | NA | NA | NA | 1 | 1 | NA | NA | 1 | 1 |
| 16_215805 | 2 | 0.5 | 1 | 0 | 1 | 1 | 1 | 0 | 5 | 0.4 | NA | NA | NA | NA | 2 | 0.5 | NA | NA | 2 | 0.5 |
| 16_215818 | 2 | 1 | 1 | 1 | 1 | 0 | NA | NA | 4 | 0.75 | NA | NA | NA | NA | 2 | 1 | NA | NA | 2 | 1 |
| 16_215852 | 1 | 1 | 1 | 1 | NA | NA | NA | NA | 2 | 1 | NA | NA | NA | NA | 2 | 1 | NA | NA | 2 | 1 |
| 16_216248 | 2 | 0.5 | NA | NA | 2 | 0 | 1 | 0 | 5 | 0.2 | 1 | 1 | NA | NA | NA | NA | NA | NA | 1 | 1 |
| 16_216274 | NA | NA | 1 | 1 | 2 | 1 | 1 | 0 | 4 | 0.75 | 1 | 1 | NA | NA | NA | NA | 1 | 1 | 2 | 1 |
| 16_216275 | NA | NA | 1 | 1 | 2 | 1 | 1 | 1 | 4 | 1 | NA | NA | NA | NA | NA | NA | 1 | 1 | 1 | 1 |
| 16_216276 | NA | NA | 1 | 0 | 2 | 0.5 | 1 | 0 | 4 | 0.25 | NA | NA | NA | NA | NA | NA | 1 | 1 | 1 | 1 |
| 16_217107 | NA | NA | 1 | 1 | NA | NA | NA | NA | 1 | 1 | 1 | 1 | 1 | 1 | NA | NA | NA | NA | 2 | 1 |
| 16_217108 | 1 | 0 | 1 | 1 | NA | NA | NA | NA | 2 | 0.5 | 1 | 0 | 1 | 1 | NA | NA | NA | NA | 2 | 0.5 |
| 16_217111 | 1 | 0 | 1 | 1 | NA | NA | NA | NA | 2 | 0.5 | 1 | 1 | 1 | 1 | NA | NA | NA | NA | 2 | 1 |
| 16_217406 | 2 | 0.5 | 1 | 1 | NA | NA | NA | NA | 3 | 0.6667 | 1 | 1 | NA | NA | NA | NA | 1 | 1 | 2 | 1 |
| 16_217406 | 3 | 1 | 1 | 1 | NA | NA | NA | NA | 4 | 1 | 1 | 1 | 1 | 1 | NA | NA | 1 | 1 | 3 | 1 |
| 16_217407 | 3 | 0.3333 | 1 | 1 | NA | NA | NA | NA | 4 | 0.5 | 1 | 1 | 1 | 1 | NA | NA | 1 | 0 | 3 | 0.6667 |
| 16_217408 | 2 | 0.5 | 1 | 0 | NA | NA | NA | NA | 3 | 0.3333 | 1 | 1 | 1 | 1 | NA | NA | NA | NA | 2 | 1 |
| 16_217408 | 2 | 1 | 1 | 1 | NA | NA | NA | NA | 3 | 1 | 1 | 1 | 1 | 1 | NA | NA | 1 | 1 | 3 | 1 |

| | | | | | | | | | | | | | | | | | | | | |
|---|---|---|---|---|---|---|---|---|---|---|---|---|---|---|---|---|---|---|---|---|
| 16_217408 | 3 | 0.3333 | 1 | 0 | NA | NA | NA | NA | 4 | 0.25 | 1 | 0 | 1 | 1 | NA | NA | 1 | 1 | 3 | 0.6667 |
| 16_217467 | 2 | 0.5 | NA | NA | NA | NA | NA | NA | 2 | 0.5 | 1 | 1 | 1 | 1 | NA | NA | 2 | 0.5 | 4 | 0.75 |
| 16_217468 | 3 | 0.3333 | 1 | 1 | NA | NA | NA | NA | 4 | 0.5 | 1 | 1 | 1 | 1 | NA | NA | 2 | 1 | 4 | 1 |
| 16_217469 | 3 | 1 | 1 | 1 | NA | NA | NA | NA | 4 | 1 | 1 | 0 | 1 | 0 | NA | NA | 2 | 1 | 4 | 0.5 |
| 16_217470 | 2 | 1 | 1 | 1 | NA | NA | NA | NA | 3 | 1 | 1 | 1 | 1 | 1 | NA | NA | 2 | 1 | 4 | 1 |
| 16_217470 | 2 | 1 | 1 | 1 | NA | NA | NA | NA | 3 | 1 | NA | NA | 1 | 1 | 1 | 1 | 1 | 1 | 3 | 1 |
| 16_217470 | 3 | 0.3333 | 1 | 0 | NA | NA | NA | NA | 4 | 0.25 | 1 | 0 | 1 | 1 | 2 | 0.5 | 2 | 0 | 6 | 0.3333 |
| 16_217471 | 3 | 1 | 1 | 1 | NA | NA | NA | NA | 4 | 1 | 1 | 0 | 1 | 1 | 2 | 1 | 2 | 1 | 6 | 0.8333 |
| 16_217484 | 2 | 0.5 | 1 | 0 | NA | NA | NA | NA | 3 | 0.3333 | 1 | 0 | 1 | 0 | 2 | 0.5 | 1 | 1 | 5 | 0.4 |
| 16_217487 | 3 | 1 | 1 | 0 | NA | NA | NA | NA | 4 | 0.75 | NA | NA | 4 | 0.5 | 3 | 0 | 1 | 1 | 8 | 0.375 |
| 16_217489 | 3 | 0.6667 | 1 | 0 | NA | NA | NA | NA | 4 | 0.5 | 1 | 0 | 4 | 0.5 | 3 | 0.3333 | 2 | 0.5 | 10 | 0.4 |
| 16_217490 | 3 | 1 | NA | NA | NA | NA | NA | NA | 3 | 1 | 1 | 1 | 4 | 1 | 3 | 1 | 2 | 1 | 10 | 1 |
| 16_217490 | 3 | 1 | 1 | 0 | NA | NA | NA | NA | 4 | 0.75 | 1 | 0 | 4 | 0.75 | 2 | 0.5 | 2 | 1 | 9 | 0.6667 |
| 16_217491 | 3 | 0.3333 | 1 | 1 | NA | NA | NA | NA | 4 | 0.5 | 1 | 0 | 4 | 0.5 | 3 | 0.6667 | 2 | 0 | 10 | 0.4 |
| 16_217491 | 2 | 0.5 | 2 | 0 | NA | NA | NA | NA | 4 | 0.25 | 1 | 0 | 4 | 0.75 | 3 | 0.6667 | 2 | 0.5 | 10 | 0.6 |
| 16_217496 | 2 | 0.5 | 1 | 0 | NA | NA | NA | NA | 3 | 0.3333 | 1 | 1 | 5 | 0.8 | 3 | 1 | 2 | 0.5 | 11 | 0.8182 |
| 16_217499 | 5 | 0 | 3 | 0.3333 | NA | NA | NA | NA | 8 | 0.125 | 1 | 0 | 5 | 0 | 4 | 0.25 | 2 | 0 | 12 | 0.0833 |
| 16_217507 | 3 | 0 | 3 | 0.3333 | 1 | 0 | NA | NA | 7 | 0.1429 | 1 | 0 | 6 | 0.1667 | 3 | 0.3333 | 2 | 0 | 12 | 0.1667 |
| 16_217507 | 4 | 0 | 3 | 0.3333 | 1 | 0 | NA | NA | 8 | 0.125 | 1 | 0 | 5 | 0.2 | 4 | 0.5 | 2 | 0 | 12 | 0.25 |
| 16_217509 | 3 | 0.6667 | 2 | 0.5 | 2 | 0.5 | NA | NA | 7 | 0.5714 | 1 | 1 | 3 | 0.6667 | 6 | 0.5 | 3 | 0.3333 | 13 | 0.5385 |
| 16_217509 | 4 | 0.25 | 3 | 0.3333 | 2 | 0 | NA | NA | 9 | 0.2222 | 1 | 1 | 6 | 0.1667 | 6 | 0.1667 | 3 | 0 | 16 | 0.1875 |
| 16_217511 | 3 | 1 | 3 | 1 | 2 | 0.5 | NA | NA | 8 | 0.875 | 1 | 1 | 6 | 1 | 6 | 1 | 3 | 1 | 16 | 1 |
| 16_217512 | 3 | 1 | 2 | 1 | 2 | 0 | NA | NA | 7 | 0.7143 | 1 | 0 | 8 | 0.5 | 5 | 0.6 | 3 | 0 | 17 | 0.4118 |
| 16_217513 | 4 | 0.5 | 2 | 0.5 | 3 | 0.3333 | NA | NA | 9 | 0.4444 | 1 | 1 | 7 | 0.1429 | 5 | 0.4 | 3 | 0 | 16 | 0.25 |
| 16_217515 | 5 | 0.2 | 2 | 0 | NA | NA | NA | NA | 7 | 0.1429 | 2 | 0.5 | 5 | 0.6 | 2 | 0.5 | 2 | 0.5 | 11 | 0.5455 |
| 16_217515 | 4 | 0.25 | 3 | 0.6667 | 1 | 0 | NA | NA | 8 | 0.375 | 1 | 0 | 8 | 0.375 | 6 | 0 | 3 | 0 | 18 | 0.1667 |
| 16_217515 | 2 | 0 | 3 | 0.3333 | 3 | 0.3333 | NA | NA | 8 | 0.25 | 2 | 0 | 7 | 0.1429 | 6 | 0.6667 | 3 | 0 | 18 | 0.2778 |
| 16_217532 | 4 | 1 | 6 | 0.5 | 6 | 0.3333 | 3 | 0.6667 | 19 | 0.5789 | 4 | 0.75 | 3 | 0 | 9 | 0.4444 | 2 | 1 | 18 | 0.5 |
| 16_217533 | 4 | 0.5 | 6 | 0.3333 | 5 | 0.6 | 2 | 0 | 17 | 0.4118 | 4 | 0.25 | 3 | 0.6667 | 10 | 0.4 | 2 | 0.5 | 19 | 0.4211 |
| 16_217533 | 5 | 0.6 | 6 | 0 | 8 | 0.375 | 3 | 0.3333 | 22 | 0.3182 | 5 | 0.6 | 3 | 0.6667 | 9 | 0.4444 | 1 | 1 | 18 | 0.5556 |
| 16_217534 | 4 | 0.75 | 6 | 0.8333 | 7 | 0.5714 | 3 | 0.6667 | 20 | 0.7 | 5 | 0.6 | 3 | 1 | 11 | 1 | 2 | 1 | 21 | 0.9048 |
| 16_217534 | 5 | 1 | 5 | 1 | 6 | 1 | 4 | 1 | 20 | 1 | 4 | 1 | 3 | 1 | 9 | 1 | 2 | 1 | 18 | 1 |
| 16_217535 | 4 | 1 | 6 | 1 | 8 | 0.875 | 4 | 0.5 | 22 | 0.8636 | 5 | 1 | 3 | 1 | 10 | 0.9 | 2 | 1 | 20 | 0.95 |
| 16_217537 | 4 | 0.5 | 6 | 0.1667 | 6 | 0 | 4 | 0 | 20 | 0.15 | 4 | 0.75 | 3 | 0.3333 | 10 | 0.1 | 1 | 0 | 18 | 0.2778 |
| 16_218656 | 10 | 0.7 | 7 | 0.7143 | 5 | 1 | 8 | 0.875 | 30 | 0.8 | 7 | 0.5714 | 10 | 0.6 | 8 | 0.875 | 10 | 0.9 | 35 | 0.7429 |
| 16_218658 | 9 | 0.2222 | 9 | 0.4444 | 5 | 0.4 | 8 | 0.625 | 31 | 0.4194 | 8 | 0.25 | 8 | 0.5 | 7 | 0.5714 | 10 | 0.7 | 33 | 0.5152 |
| 16_218661 | 11 | 0.1818 | 12 | 0.25 | 8 | 0.125 | 8 | 0.5 | 39 | 0.2564 | 13 | 0.1538 | 11 | 0.0909 | 9 | 0.3333 | 10 | 0.4 | 43 | 0.2326 |
| 16_218668 | 4 | 0 | 2 | 0 | 4 | 0 | 5 | 0.2 | 15 | 0.0667 | 4 | 0 | 3 | 0.3333 | 2 | 0 | 4 | 0 | 13 | 0.0769 |
| 16_218668 | 5 | 0.4 | 1 | 0 | 3 | 0 | 4 | 0 | 13 | 0.1538 | 4 | 0.25 | 4 | 0.75 | 2 | 0 | 4 | 0.25 | 14 | 0.3571 |
| 16_218678 | 6 | 0 | 2 | 0 | 6 | 0.5 | 6 | 0.1667 | 20 | 0.2 | 4 | 0.25 | 3 | 0.3333 | 3 | 0 | 6 | 0.1667 | 16 | 0.1875 |
| 16_218780 | NA | NA | 1 | 0 | NA | NA | 1 | 1 | 2 | 0.5 | 1 | 1 | NA | NA | NA | NA | NA | NA | 1 | 1 |
| 16_218783 | NA | NA | 1 | 1 | NA | NA | NA | NA | 1 | 1 | 1 | 1 | NA | NA | NA | NA | NA | NA | 1 | 1 |
| 16_220646 | 18 | 0.3889 | 15 | 0.1333 | 12 | 0.25 | 19 | 0.2632 | 64 | 0.2656 | 6 | 0.1667 | 18 | 0.1667 | 14 | 0.2143 | 13 | 0.1538 | 51 | 0.1765 |
| 16_220648 | 16 | 0.75 | 16 | 0.375 | 15 | 0.7333 | 17 | 0.5294 | 64 | 0.5938 | 8 | 0.125 | 14 | 0.5714 | 14 | 0.6429 | 15 | 0.5333 | 51 | 0.5098 |
| 16_220655 | 10 | 0 | 14 | 0.0714 | 12 | 0 | 12 | 0.1667 | 48 | 0.0625 | 8 | 0 | 15 | 0.0667 | 11 | 0.0909 | 16 | 0.0625 | 50 | 0.06 |
| 16_220655 | 11 | 0.0909 | 11 | 0.0909 | 15 | 0.2 | 15 | 0.2 | 52 | 0.1538 | 7 | 0.4286 | 16 | 0.375 | 13 | 0.2308 | 16 | 0.3125 | 52 | 0.3269 |
| 16_222002 | 10 | 0.4 | 10 | 0.2 | 6 | 0.1667 | 10 | 0.4 | 36 | 0.3056 | 9 | 0.4444 | 10 | 0.8 | 13 | 0.5385 | 11 | 0.1818 | 43 | 0.4884 |
| 16_222002 | 8 | 0.125 | 8 | 0 | 5 | 0.4 | 11 | 0.3636 | 32 | 0.2188 | 7 | 0.2857 | 9 | 0 | 13 | 0.3077 | 13 | 0.0769 | 42 | 0.1667 |
| 16_222005 | 9 | 0.3333 | 8 | 0.125 | 9 | 0.4444 | 10 | 0.3 | 36 | 0.3056 | 7 | 0.4286 | 10 | 0.5 | 10 | 0.5 | 12 | 0.5 | 39 | 0.4872 |
| 16_222006 | 9 | 0.1111 | 7 | 0.1429 | 7 | 0 | 8 | 0.25 | 31 | 0.129 | 7 | 0.1429 | 10 | 0.1 | 10 | 0.2 | 10 | 0 | 37 | 0.1081 |
| 16_222006 | 8 | 0 | 10 | 0 | 7 | 0.2857 | 8 | 0.25 | 33 | 0.1212 | 7 | 0 | 9 | 0.1111 | 11 | 0.1818 | 12 | 0.0833 | 39 | 0.1026 |
| 16_222010 | 4 | 0 | 7 | 0.1429 | 5 | 0 | 6 | 0.1667 | 22 | 0.0909 | 3 | 0 | 7 | 0.2857 | 6 | 0 | 11 | 0.1818 | 27 | 0.1481 |
| 16_222015 | 7 | 0 | 8 | 0.625 | 7 | 0 | 8 | 0.375 | 30 | 0.2667 | 3 | 0 | 8 | 0.125 | 6 | 0 | 8 | 0.125 | 25 | 0.08 |
| 16_222017 | 3 | 0.3333 | 6 | 0.3333 | 6 | 0.1667 | 6 | 0.1667 | 21 | 0.2381 | 2 | 0.5 | 6 | 0 | 3 | 0.3333 | 8 | 0 | 19 | 0.1053 |
| 16_222017 | 4 | 0.25 | 8 | 0.625 | 7 | 0.7143 | 6 | 0.3333 | 25 | 0.52 | 2 | 0.5 | 9 | 0.5556 | 6 | 0.5 | 8 | 0.25 | 25 | 0.44 |
| 16_222018 | 3 | 1 | 4 | 0.75 | 4 | 0.75 | 8 | 0.75 | 19 | 0.7895 | 4 | 0.75 | 5 | 0.8 | 4 | 1 | 5 | 1 | 18 | 0.8889 |
| 16_222018 | 4 | 0.25 | 4 | 0.5 | 6 | 0.3333 | 6 | 0.5 | 20 | 0.4 | 3 | 0.3333 | 8 | 0.25 | 4 | 0.25 | 6 | 0.1667 | 21 | 0.2381 |
| 16_222018 | 5 | 0.4 | 7 | 0.4286 | 6 | 0.1667 | 8 | 0.25 | 26 | 0.3077 | 4 | 0.25 | 9 | 0 | 6 | 0 | 8 | 0.375 | 27 | 0.1481 |
| 16_222018 | 4 | 1 | 7 | 1 | 7 | 0.8571 | 6 | 1 | 24 | 0.9583 | 4 | 1 | 8 | 1 | 5 | 1 | 8 | 1 | 25 | 1 |
| 16_222019 | 4 | 0 | 8 | 0.5 | 5 | 0.4 | 7 | 0.2857 | 24 | 0.3333 | 3 | 0.3333 | 8 | 0.25 | 6 | 0.1667 | 7 | 0 | 24 | 0.1667 |
| 16_222020 | 5 | 0.6 | 8 | 0.375 | 5 | 0.4 | 8 | 0.5 | 26 | 0.4615 | 3 | 0 | 8 | 0.125 | 6 | 0.1667 | 9 | 0.1111 | 26 | 0.1154 |
| 16_222020 | 4 | 0.25 | 7 | 0.1429 | 4 | 0.25 | 6 | 0.1667 | 21 | 0.1905 | 2 | 1 | 7 | 0.1429 | 4 | 0.25 | 7 | 0 | 20 | 0.2 |
| 16_222020 | 3 | 0 | 8 | 0.25 | 5 | 0 | 7 | 0.1429 | 23 | 0.1304 | 2 | 0 | 8 | 0 | 6 | 0.1667 | 6 | 0.1667 | 22 | 0.0909 |
| 16_222022 | 4 | 0.5 | 3 | 0 | 3 | 0.3333 | 4 | 0 | 14 | 0.2143 | 3 | 0 | 7 | 0.1429 | 4 | 0.25 | 7 | 0.7143 | 21 | 0.3333 |
| 16_222023 | 5 | 0 | 5 | 0.4 | 4 | 0.25 | 3 | 0.3333 | 17 | 0.2353 | 3 | 0.3333 | 6 | 0.1667 | 5 | 0.8 | 7 | 0.1429 | 21 | 0.3333 |

| | | | | | | | | | | | | | | | | | | | | |
|---|---|---|---|---|---|---|---|---|---|---|---|---|---|---|---|---|---|---|---|---|
| 16_222855 | 4 | 0 | 10 | 0.5 | 4 | 0.5 | 4 | 0.25 | 22 | 0.3636 | 5 | 0 | 11 | 0.3636 | 10 | 0.3 | 12 | 0.3333 | 38 | 0.2895 |
| 16_222857 | 5 | 1 | 13 | 1 | 6 | 1 | 6 | 1 | 30 | 1 | 9 | 1 | 10 | 1 | 12 | 1 | 12 | 0.9167 | 43 | 0.9767 |
| 16_222857 | 3 | 0.6667 | 12 | 0.3333 | 6 | 0.5 | 6 | 0.6667 | 27 | 0.4815 | 9 | 0.6667 | 9 | 0.6667 | 11 | 0.5455 | 14 | 0.3571 | 43 | 0.5349 |
| 16_222858 | 5 | 0.4 | 12 | 0.6667 | 4 | 0.75 | 6 | 0.8333 | 27 | 0.6667 | 9 | 0.5556 | 10 | 0.5 | 11 | 0.6364 | 12 | 0.9167 | 42 | 0.6667 |
| 16_222858 | 5 | 0 | 12 | 0.1667 | 5 | 0.2 | 5 | 0.4 | 27 | 0.1852 | 9 | 0 | 8 | 0.25 | 12 | 0.1667 | 13 | 0.2308 | 42 | 0.1667 |
| 16_222860 | 4 | 0.5 | 12 | 0.5833 | 5 | 0.2 | 5 | 0.4 | 26 | 0.4615 | 7 | 0.4286 | 8 | 0.5 | 8 | 0.25 | 10 | 0.5 | 33 | 0.4242 |
| 16_222861 | 4 | 0.75 | 12 | 0.75 | 4 | 0.75 | 5 | 0.6 | 25 | 0.72 | 7 | 1 | 9 | 0.7778 | 8 | 0.625 | 13 | 1 | 37 | 0.8649 |
| 16_222863 | 5 | 0.2 | 12 | 0.4167 | 4 | 0.75 | 4 | 0.5 | 25 | 0.44 | 6 | 0.5 | 10 | 0.2 | 8 | 0.625 | 14 | 0.2857 | 38 | 0.3684 |
| 16_222864 | 4 | 0 | 11 | 0.2727 | 4 | 0.75 | 4 | 0.25 | 23 | 0.3043 | 5 | 0.6 | 4 | 0.5 | 6 | 0.1667 | 10 | 0.3 | 25 | 0.36 |
| 16_222864 | 4 | 0.25 | 14 | 0.0714 | 5 | 0.2 | 6 | 0 | 29 | 0.1034 | 6 | 0 | 9 | 0.1111 | 7 | 0.1429 | 11 | 0.2727 | 33 | 0.1515 |
| 16_222866 | 4 | 1 | 12 | 1 | 3 | 1 | 6 | 0.6667 | 25 | 0.92 | 4 | 1 | 9 | 1 | 6 | 0.8333 | 12 | 0.75 | 31 | 0.871 |
| 16_222867 | 4 | 1 | 12 | 0.8333 | 4 | 0.75 | 6 | 1 | 26 | 0.8846 | 5 | 0.8 | 8 | 0.875 | 6 | 1 | 11 | 0.7273 | 30 | 0.8333 |
| 16_222867 | 4 | 0.5 | 10 | 0.5 | 3 | 0.3333 | 5 | 0.4 | 22 | 0.4545 | 5 | 0.2 | 9 | 0.4444 | 6 | 0.3333 | 13 | 0.3077 | 33 | 0.3333 |
| 16_222867 | 3 | 0 | 10 | 0.2 | 3 | 0.3333 | 2 | 0 | 18 | 0.1667 | 4 | 0.75 | 7 | 0.2857 | 7 | 0.1429 | 9 | 0.5556 | 27 | 0.4074 |
| 16_222868 | 3 | 0.3333 | 12 | 0.0833 | 4 | 0.5 | 5 | 0.2 | 24 | 0.2083 | 4 | 0 | 9 | 0.3333 | 6 | 0.3333 | 13 | 0.1538 | 32 | 0.2188 |
| 16_222868 | 4 | 0 | 13 | 0.1538 | 4 | 0.25 | 5 | 0 | 26 | 0.1154 | 5 | 0 | 9 | 0.2222 | 7 | 0 | 12 | 0.1667 | 33 | 0.1212 |
| 16_222869 | 4 | 1 | 13 | 1 | 4 | 0.75 | 4 | 1 | 25 | 0.96 | 4 | 1 | 9 | 1 | 7 | 0.7143 | 13 | 1 | 33 | 0.9394 |
| 16_222870 | 4 | 0.5 | 13 | 0.3846 | 4 | 0.5 | 4 | 0.75 | 25 | 0.48 | 5 | 1 | 9 | 0.7778 | 8 | 0.75 | 13 | 0.4615 | 35 | 0.6857 |
| 16_222870 | 4 | 0.5 | 12 | 0.75 | 4 | 0.5 | 3 | 0.6667 | 23 | 0.6522 | 5 | 0.6 | 8 | 0.5 | 8 | 0.75 | 12 | 0.75 | 33 | 0.6667 |
| 16_222870 | 4 | 0 | 12 | 0.3333 | 4 | 0 | 5 | 0.4 | 25 | 0.24 | 4 | 0.25 | 8 | 0.375 | 8 | 0.125 | 12 | 0.5 | 32 | 0.3438 |
| 16_222871 | NA | NA | 3 | 0.3333 | NA | NA | NA | NA | 3 | 0.3333 | NA | NA | 4 | 0.25 | 2 | 0.5 | 3 | 0 | 9 | 0.2222 |
| 16_222872 | 4 | 1 | 12 | 1 | 3 | 1 | 5 | 1 | 24 | 1 | 4 | 1 | 7 | 1 | 7 | 0.8571 | 12 | 1 | 30 | 0.9667 |
| 16_222872 | 4 | 0.5 | 10 | 0.6 | 2 | 1 | 4 | 1 | 20 | 0.7 | 4 | 0.75 | 6 | 1 | 3 | 0.6667 | 10 | 0.7 | 23 | 0.7826 |
| 16_222875 | 4 | 0.25 | 10 | 0.2 | 3 | 0 | 5 | 0.2 | 22 | 0.1818 | 2 | 0 | 7 | 0 | 6 | 0 | 12 | 0.1667 | 27 | 0.0741 |
| 16_222877 | 3 | 0.6667 | 9 | 0.4444 | 3 | 1 | 4 | 0.5 | 19 | 0.5789 | 3 | 0.3333 | 7 | 0.8571 | 5 | 0.6 | 11 | 1 | 26 | 0.8077 |
| 16_222877 | 2 | 1 | 6 | 1 | 2 | 1 | 5 | 1 | 15 | 1 | 3 | 1 | 6 | 1 | 5 | 1 | 10 | 1 | 24 | 1 |
| 16_222880 | 3 | 0.6667 | 9 | 0.3333 | 3 | 0.6667 | 5 | 0.4 | 20 | 0.45 | 2 | 0 | 5 | 0 | 4 | 0 | 10 | 0.3 | 21 | 0.1429 |
| 16_226371 | 1 | 1 | NA | NA | NA | NA | NA | NA | 1 | 1 | NA | NA | 1 | 1 | 1 | 1 | NA | NA | 2 | 1 |
| 16_226410 | 6 | 0.5 | 5 | 0.8 | 1 | 0 | NA | NA | 12 | 0.5833 | 1 | 1 | 3 | 0.6667 | 1 | 1 | NA | NA | 5 | 0.8 |
| 16_226410 | 5 | 0.8 | 5 | 0.2 | 1 | 1 | NA | NA | 11 | 0.5455 | 1 | 0 | 4 | 0.25 | NA | NA | NA | NA | 5 | 0.2 |
| 16_226411 | 6 | 0.3333 | 5 | 0.6 | 1 | 1 | 1 | 1 | 13 | 0.5385 | 1 | 1 | 4 | 0.75 | 1 | 1 | NA | NA | 6 | 0.8333 |
| 16_226411 | 5 | 0.2 | 4 | 0 | 1 | 0 | 1 | 1 | 11 | 0.1818 | 1 | 1 | 3 | 1 | 1 | 1 | NA | NA | 5 | 1 |
| 16_226412 | 5 | 0.4 | 5 | 0.2 | 1 | 0 | 1 | 1 | 12 | 0.3333 | 1 | 0 | 3 | 0.3333 | 1 | 0 | NA | NA | 5 | 0.2 |
| 16_226446 | 5 | 1 | 2 | 0.5 | 1 | 1 | 1 | 1 | 9 | 0.8889 | NA | NA | 2 | 1 | 1 | 1 | NA | NA | 3 | 1 |
| 16_226605 | 5 | 0.4 | 4 | 0 | 1 | 0 | 1 | 0 | 11 | 0.1818 | 1 | 0 | 4 | 0.25 | 1 | 0 | NA | NA | 6 | 0.1667 |
| 16_226606 | 2 | 0.5 | 2 | 1 | NA | NA | 1 | 1 | 5 | 0.8 | 1 | 0 | 3 | 0.6667 | 1 | 1 | NA | NA | 5 | 0.6 |
| 16_226904 | 8 | 0.5 | 7 | 0.4286 | 6 | 0.1667 | 7 | 0.1429 | 28 | 0.3214 | 8 | 0.125 | 4 | 0 | 10 | 0.1 | 9 | 0 | 31 | 0.0645 |
| 16_226905 | 7 | 0.5714 | 8 | 0.75 | 6 | 0.8333 | 8 | 0.375 | 29 | 0.6207 | 10 | 0.5 | 6 | 0.3333 | 11 | 0.4545 | 9 | 0.7778 | 36 | 0.5278 |
| 16_226906 | 9 | 0.4444 | 8 | 0.125 | 8 | 0.125 | 9 | 0.1111 | 34 | 0.2059 | 11 | 0.3636 | 5 | 0.2 | 10 | 0 | 9 | 0.3333 | 35 | 0.2286 |
| 16_226907 | 11 | 0.4545 | 8 | 0.75 | 7 | 0.7143 | 7 | 0.4286 | 33 | 0.5758 | 11 | 0.3636 | 5 | 0.2 | 9 | 0.8889 | 10 | 0.6 | 35 | 0.5429 |
| 16_226910 | 8 | 0.75 | 9 | 0.7778 | 4 | 0.75 | 3 | 0 | 24 | 0.6667 | 4 | 0.75 | 3 | 0.6667 | 4 | 0.25 | 5 | 0.8 | 16 | 0.625 |
| 16_229319 | 75 | 0.1067 | 77 | 0.0649 | 52 | 0.1154 | 74 | 0.0811 | 278 | 0.0899 | 46 | 0.1304 | 71 | 0.0563 | 66 | 0.1212 | 54 | 0.0741 | 237 | 0.0928 |
| 16_229320 | 85 | 0.2824 | 81 | 0.2346 | 57 | 0.1404 | 84 | 0.2143 | 307 | 0.2248 | 49 | 0.2449 | 77 | 0.1818 | 74 | 0.1757 | 61 | 0.3443 | 261 | 0.2299 |
| 16_229322 | 92 | 0.4022 | 81 | 0.5432 | 61 | 0.4426 | 86 | 0.4884 | 320 | 0.4688 | 52 | 0.5192 | 81 | 0.4938 | 75 | 0.4933 | 62 | 0.5484 | 270 | 0.5111 |
| 16_229323 | 73 | 0.1781 | 68 | 0.2353 | 60 | 0.2 | 72 | 0.2778 | 273 | 0.2234 | 45 | 0.2444 | 73 | 0.2466 | 65 | 0.1846 | 67 | 0.2985 | 250 | 0.244 |
| 16_231401 | 2 | 1 | 2 | 1 | NA | NA | NA | NA | 4 | 1 | 2 | 1 | 1 | 0 | NA | NA | NA | NA | 3 | 0.6667 |
| 16_231401 | 3 | 0 | 3 | 0.6667 | NA | NA | NA | NA | 6 | 0.3333 | 2 | 1 | 1 | 0 | NA | NA | NA | NA | 3 | 0.6667 |
| 16_231766 | 3 | 0.6667 | 3 | 1 | NA | NA | NA | NA | 6 | 0.8333 | 2 | 1 | 1 | 1 | NA | NA | NA | NA | 3 | 1 |
| 16_231769 | 1 | 0 | 2 | 0.5 | 1 | 0 | 1 | 1 | 5 | 0.4 | NA | NA | 1 | 1 | NA | NA | NA | NA | 1 | 1 |
| 16_231786 | 3 | 0.3333 | 4 | 0 | 1 | 1 | 1 | 0 | 9 | 0.2222 | 2 | 0 | 2 | 0.5 | NA | NA | NA | NA | 4 | 0.25 |
| 16_231788 | 4 | 0.25 | 6 | 0.1667 | 1 | 0 | 1 | 0 | 12 | 0.1667 | 2 | 0 | 2 | 0.5 | NA | NA | NA | NA | 4 | 0.25 |
| 16_232875 | 1 | 1 | 1 | 1 | NA | NA | NA | NA | 2 | 1 | 1 | 1 | NA | NA | NA | NA | 1 | 1 | 2 | 1 |
| 16_232876 | 1 | 1 | 1 | 0 | NA | NA | NA | NA | 2 | 0.5 | 1 | 1 | NA | NA | NA | NA | 1 | 1 | 2 | 1 |
| 16_232876 | 1 | 0 | 1 | 1 | NA | NA | NA | NA | 2 | 0.5 | 1 | 1 | NA | NA | NA | NA | 1 | 1 | 2 | 1 |
| 16_232880 | 1 | 1 | 1 | 1 | NA | NA | NA | NA | 2 | 1 | 1 | 1 | NA | NA | NA | NA | 1 | 1 | 2 | 1 |
| 16_232985 | NA | NA | NA | NA | NA | NA | 1 | 1 | 1 | 1 | NA | NA | NA | NA | NA | NA | 1 | 1 | 1 | 1 |
| 16_232985 | NA | NA | 1 | 1 | NA | NA | 1 | 0 | 2 | 0.5 | NA | NA | NA | NA | 1 | 1 | 1 | 1 | 2 | 1 |
| 16_234318 | 2 | 1 | NA | NA | 1 | 1 | NA | NA | 3 | 1 | 2 | 1 | NA | NA | NA | NA | NA | NA | 2 | 1 |
| 16_234319 | 3 | 1 | NA | NA | 2 | 1 | NA | NA | 5 | 1 | 2 | 1 | NA | NA | NA | NA | NA | NA | 2 | 1 |
| 16_234319 | 3 | 0.3333 | NA | NA | 2 | 0 | NA | NA | 5 | 0.2 | 1 | 1 | NA | NA | NA | NA | NA | NA | 1 | 1 |
| 16_234321 | 3 | 0.6667 | NA | NA | 1 | 1 | NA | NA | 4 | 0.75 | 1 | 1 | NA | NA | NA | NA | NA | NA | 1 | 1 |
| 16_234322 | 3 | 0.3333 | NA | NA | 2 | 0 | NA | NA | 5 | 0.2 | 2 | 1 | NA | NA | NA | NA | NA | NA | 2 | 1 |
| 16_237083 | 86 | 0.0814 | 111 | 0.0991 | 141 | 0.0709 | 88 | 0.0682 | 426 | 0.0798 | 110 | 0.0727 | 85 | 0.0353 | 90 | 0.0778 | 86 | 0.1512 | 371 | 0.0836 |
| 16_237839 | 6 | 0.1667 | 16 | 0.25 | 6 | 0.5 | 2 | 0.5 | 30 | 0.3 | 2 | 0.5 | 8 | 0.5 | 9 | 0.2222 | 2 | 0 | 21 | 0.3333 |
| 16_237842 | 4 | 0 | 14 | 0.2857 | 5 | 0 | 2 | 0.5 | 25 | 0.2 | 1 | 0 | 8 | 0 | 9 | 0.3333 | 3 | 0 | 21 | 0.1429 |

| | | | | | | | | | | | | | | | | | | | | |
|---|---|---|---|---|---|---|---|---|---|---|---|---|---|---|---|---|---|---|---|---|
| 16_237910 | 3 | 0.3333 | 9 | 0.4444 | 3 | 0.3333 | 1 | 1 | 16 | 0.4375 | NA | NA | 6 | 0.5 | 5 | 0 | 1 | 0 | 12 | 0.25 |
| 16_237915 | 3 | 0 | 7 | 0.1429 | 3 | 0.3333 | 1 | 0 | 14 | 0.1429 | NA | NA | 3 | 0.3333 | 5 | 0.4 | 2 | 0 | 10 | 0.3 |
| 16_237936 | 3 | 0.6667 | 6 | 0 | 1 | 0 | NA | NA | 10 | 0.2 | NA | NA | 2 | 0 | 4 | 0.5 | 2 | 0 | 8 | 0.25 |
| 16_242113 | 1 | 0 | 1 | 0 | 1 | 0 | 2 | 0.5 | 5 | 0.2 | 1 | 0 | 2 | 0.5 | NA | NA | NA | NA | 3 | 0.3333 |
| 16_242114 | 4 | 0 | 1 | 0 | 4 | 0.25 | 2 | 0 | 11 | 0.0909 | 1 | 0 | 2 | 0.5 | 2 | 0 | 2 | 0.5 | 7 | 0.2857 |
| 16_242114 | 4 | 0 | 2 | 0.5 | 4 | 0.25 | 2 | 0 | 12 | 0.1667 | 1 | 0 | 2 | 0 | 2 | 0 | 1 | 1 | 6 | 0.1667 |
| 16_242115 | 4 | 0 | 2 | 0 | 5 | 0 | 2 | 0.5 | 13 | 0.0769 | 1 | 0 | 1 | 0 | 2 | 0 | 2 | 0.5 | 6 | 0.1667 |
| 16_242115 | 4 | 0.5 | 2 | 0.5 | 5 | 0.4 | 2 | 0.5 | 13 | 0.4615 | 1 | 0 | NA | NA | 2 | 0.5 | 2 | 0 | 5 | 0.2 |
| 16_242116 | 4 | 0 | 2 | 0.5 | 3 | 0 | 2 | 0.5 | 11 | 0.1818 | 1 | 0 | 2 | 0 | 2 | 0.5 | 2 | 0 | 7 | 0.1429 |
| 16_243493 | 4 | 0 | 5 | 0.2 | 4 | 0 | 1 | 0 | 14 | 0.0714 | 1 | 0 | 9 | 0.1111 | 5 | 0 | 3 | 0.3333 | 18 | 0.1111 |
| 16_261775 | NA | NA | NA | NA | 1 | 1 | NA | NA | 1 | 1 | 1 | 1 | NA | NA | NA | NA | 1 | 1 | 2 | 1 |
| 16_261777 | NA | NA | NA | NA | 1 | 1 | NA | NA | 1 | 1 | NA | NA | NA | NA | NA | NA | 2 | 1 | 2 | 1 |
| 16_265336 | 1 | 1 | 1 | 1 | 1 | 1 | 2 | 0 | 5 | 0.6 | 1 | 1 | 1 | 1 | 2 | 0.5 | NA | NA | 4 | 0.75 |
| 16_265339 | 1 | 1 | 2 | 1 | 2 | 1 | 2 | 0.5 | 7 | 0.8571 | 1 | 1 | 1 | 0 | 2 | 0.5 | NA | NA | 4 | 0.5 |
| 16_265340 | 1 | 0 | 2 | 0 | 2 | 0 | 2 | 0.5 | 7 | 0.1429 | 1 | 1 | 1 | 0 | 1 | 0 | NA | NA | 3 | 0.3333 |
| 16_265411 | NA | NA | NA | NA | 1 | 0 | 1 | 1 | 2 | 0.5 | 2 | 0.5 | NA | NA | 1 | 0 | 1 | 1 | 4 | 0.5 |
| 16_265412 | NA | NA | NA | NA | 1 | 1 | 1 | 1 | 2 | 1 | 2 | 0 | NA | NA | 1 | 0 | 1 | 1 | 4 | 0.25 |
| 16_265415 | NA | NA | NA | NA | 1 | 1 | NA | NA | 1 | 1 | 2 | 1 | NA | NA | 1 | 0 | NA | NA | 3 | 0.6667 |
| 16_265417 | NA | NA | NA | NA | 2 | 0.5 | NA | NA | 2 | 0.5 | 2 | 0 | NA | NA | 1 | 0 | 1 | 1 | 4 | 0.25 |
| 16_265418 | NA | NA | NA | NA | 1 | 1 | NA | NA | 1 | 1 | 1 | 1 | NA | NA | 2 | 0.5 | 1 | 1 | 4 | 0.75 |
| 16_265418 | NA | NA | NA | NA | 2 | 1 | NA | NA | 2 | 1 | 2 | 1 | NA | NA | 2 | 0.5 | 1 | 1 | 5 | 0.8 |
| 16_265419 | NA | NA | NA | NA | 2 | 1 | NA | NA | 2 | 1 | 1 | 1 | NA | NA | 2 | 1 | 1 | 1 | 4 | 1 |
| 16_265422 | NA | NA | NA | NA | 2 | 0.5 | NA | NA | 2 | 0.5 | 2 | 1 | NA | NA | 1 | 0 | 2 | 0.5 | 5 | 0.6 |
| 16_265468 | 1 | 1 | NA | NA | NA | NA | 1 | 1 | 2 | 1 | NA | NA | NA | NA | NA | NA | 1 | 1 | 1 | 1 |
| 16_265468 | 1 | 1 | NA | NA | NA | NA | 1 | 1 | 2 | 1 | NA | NA | NA | NA | NA | NA | 1 | 1 | 1 | 1 |
| 16_265469 | 1 | 1 | NA | NA | NA | NA | 1 | 1 | 2 | 1 | NA | NA | NA | NA | NA | NA | 1 | 1 | 1 | 1 |
| 16_265472 | 1 | 0 | NA | NA | NA | NA | 1 | 1 | 2 | 0.5 | NA | NA | NA | NA | NA | NA | 1 | 1 | 1 | 1 |
| 16_265476 | 1 | 0 | 1 | 1 | NA | NA | 1 | 0 | 3 | 0.3333 | NA | NA | NA | NA | NA | NA | 1 | 1 | 1 | 1 |
| 16_265481 | 1 | 1 | NA | NA | NA | NA | 1 | 1 | 2 | 1 | NA | NA | NA | NA | NA | NA | 1 | 1 | 1 | 1 |
| 16_272650 | 1 | 1 | 1 | 0 | 2 | 0.5 | 4 | 0.5 | 8 | 0.5 | 3 | 1 | 2 | 0.5 | 3 | 1 | NA | NA | 8 | 0.875 |
| 16_272704 | 10 | 0.1 | 5 | 0.2 | 5 | 0.6 | 6 | 0.1667 | 26 | 0.2308 | 4 | 0.25 | 4 | 0.5 | 8 | 0.25 | 3 | 0 | 19 | 0.2632 |
| 16_272705 | 9 | 0.3333 | 6 | 0.5 | 6 | 0.5 | 5 | 0 | 26 | 0.3462 | 3 | 0.3333 | 4 | 0 | 9 | 0.3333 | 2 | 0.5 | 18 | 0.2778 |
| 16_272705 | 10 | 0.4 | 6 | 0.3333 | 6 | 0 | 6 | 0 | 28 | 0.2143 | 4 | 0.25 | 4 | 0 | 8 | 0 | 2 | 0 | 18 | 0.0556 |
| 16_272708 | 8 | 0 | 6 | 0.1667 | 6 | 0 | 3 | 0.3333 | 23 | 0.087 | 4 | 0 | 5 | 0.2 | 8 | 0 | 2 | 0 | 19 | 0.0526 |
| 16_279809 | 10 | 0.8 | 18 | 0.7778 | 11 | 0.7273 | 13 | 0.7692 | 52 | 0.7692 | 9 | 0.5556 | 8 | 0.5 | 7 | 0.7143 | 16 | 0.75 | 40 | 0.65 |
| 16_279810 | 13 | 0.9231 | 21 | 0.6667 | 17 | 0.7059 | 12 | 0.75 | 63 | 0.746 | 10 | 0.9 | 8 | 0.625 | 7 | 0.5714 | 18 | 0.6667 | 43 | 0.6977 |
| 16_279811 | 8 | 0.5 | 22 | 0.4545 | 15 | 0.4 | 13 | 0.6154 | 58 | 0.4828 | 10 | 0.5 | 7 | 0.4286 | 7 | 0.2857 | 14 | 0.5714 | 38 | 0.4737 |
| 16_282639 | 1 | 0 | 2 | 0.5 | 1 | 0 | 1 | 0 | 5 | 0.2 | 1 | 0 | 2 | 0 | 2 | 0 | 2 | 0.5 | 7 | 0.1429 |
| 16_282641 | 1 | 1 | 2 | 0.5 | NA | NA | 1 | 0 | 4 | 0.5 | 1 | 0 | 1 | 1 | 1 | 1 | 2 | 0.5 | 5 | 0.6 |
| 16_282641 | 1 | 1 | 2 | 0.5 | NA | NA | 1 | 1 | 4 | 0.75 | 1 | 1 | 2 | 1 | 2 | 0.5 | 2 | 0.5 | 7 | 0.7143 |
| 16_282641 | 1 | 1 | 1 | 0 | 1 | 0 | NA | NA | 3 | 0.3333 | NA | NA | 2 | 0.5 | 2 | 0 | 2 | 0 | 6 | 0.1667 |
| 16_282643 | 1 | 0 | 1 | 1 | NA | NA | 1 | 1 | 3 | 0.6667 | 1 | 1 | 2 | 1 | NA | NA | 1 | 0 | 4 | 0.75 |
| 16_282643 | 1 | 1 | 1 | 1 | 1 | 1 | 1 | 1 | 4 | 1 | 1 | 1 | 1 | 1 | NA | NA | 1 | 1 | 3 | 1 |
| 16_282644 | 1 | 1 | 2 | 1 | NA | NA | 1 | 1 | 4 | 1 | NA | NA | 2 | 0.5 | 1 | 0 | 1 | 1 | 4 | 0.5 |
| 16_282645 | 1 | 0 | 2 | 0.5 | 1 | 0 | 1 | 0 | 5 | 0.2 | 1 | 1 | 2 | 0.5 | NA | NA | 1 | 0 | 4 | 0.5 |
| 16_282645 | 1 | 0 | 2 | 0 | NA | NA | 1 | 1 | 4 | 0.25 | 1 | 0 | 2 | 1 | NA | NA | 1 | 1 | 4 | 0.75 |
| 16_282648 | 1 | 0 | 1 | 1 | NA | NA | 1 | 1 | 3 | 0.6667 | 1 | 1 | 1 | 0 | NA | NA | NA | NA | 2 | 0.5 |
| 16_282650 | 1 | 0 | 1 | 1 | 1 | 0 | 1 | 0 | 4 | 0.25 | 1 | 1 | 1 | 1 | NA | NA | NA | NA | 2 | 1 |
| 16_283899 | 1 | 1 | NA | NA | NA | NA | NA | NA | 1 | 1 | NA | NA | NA | NA | NA | NA | 1 | 1 | 1 | 1 |
| 16_286450 | 3 | 0.6667 | 4 | 0.25 | 2 | 0.5 | 4 | 0 | 13 | 0.3077 | 4 | 0.5 | 4 | 0.25 | 6 | 0 | 3 | 0.6667 | 17 | 0.2941 |
| 16_286451 | NA | NA | 3 | 0 | 3 | 0.3333 | 4 | 0 | 10 | 0.1 | 3 | 0.3333 | 2 | 0.5 | 5 | 0.2 | 2 | 0 | 12 | 0.25 |
| 16_286452 | 2 | 0 | 3 | 0 | 4 | 0.25 | 4 | 0.25 | 13 | 0.1538 | 4 | 0.5 | 4 | 0 | 4 | 0.25 | 3 | 0 | 15 | 0.2 |
| 16_286459 | 2 | 0 | 1 | 0 | 2 | 0.5 | 2 | 0 | 7 | 0.1429 | 2 | 0 | 4 | 0 | 3 | 0.3333 | 3 | 0 | 12 | 0.0833 |
| 16_286460 | 2 | 0 | 1 | 0 | 2 | 0.5 | 3 | 0 | 8 | 0.125 | 2 | 0 | 4 | 0 | 2 | 0.5 | 3 | 0.3333 | 11 | 0.1818 |
| 16_286462 | 2 | 0.5 | NA | NA | 2 | 0 | 1 | 1 | 5 | 0.4 | 3 | 0 | 4 | 0.25 | 1 | 0 | 1 | 0 | 9 | 0.1111 |
| 16_286463 | 2 | 0.5 | 1 | 0 | 2 | 0 | 2 | 0 | 7 | 0.1429 | 3 | 0 | 3 | 0.3333 | 2 | 0 | 2 | 0 | 10 | 0.1 |
| 16_286463 | 2 | 1 | 1 | 1 | 2 | 1 | 1 | 1 | 6 | 1 | 2 | 1 | 3 | 1 | 2 | 1 | 2 | 0.5 | 9 | 0.8889 |
| 16_286464 | 1 | 1 | 1 | 1 | 1 | 1 | 3 | 0.6667 | 6 | 0.8333 | 2 | 1 | 4 | 1 | 2 | 1 | 1 | 1 | 9 | 1 |
| 16_286464 | 2 | 0.5 | 1 | 1 | 2 | 0 | 1 | 1 | 6 | 0.5 | NA | NA | 2 | 0.5 | 2 | 1 | 2 | 1 | 6 | 0.8333 |
| 16_286465 | 2 | 1 | 1 | 1 | 1 | 1 | 3 | 0.6667 | 7 | 0.8571 | 2 | 1 | 3 | 1 | 1 | 1 | NA | NA | 6 | 1 |
| 16_286466 | 1 | 1 | 1 | 1 | 1 | 0 | NA | NA | 3 | 0.6667 | 2 | 1 | 2 | 0.5 | NA | NA | 1 | 0 | 5 | 0.6 |
| 16_286466 | 2 | 0 | 1 | 1 | 2 | 0 | 2 | 0.5 | 7 | 0.2857 | 2 | 0 | 3 | 0 | 1 | 0 | 1 | 1 | 7 | 0.1429 |
| 16_286469 | 2 | 1 | 1 | 1 | 1 | 1 | 1 | 1 | 5 | 1 | 3 | 1 | 2 | 1 | 1 | 1 | NA | NA | 6 | 1 |
| 16_286471 | 2 | 1 | 1 | 1 | 1 | 1 | 1 | 1 | 5 | 1 | 3 | 0.6667 | 2 | 1 | NA | NA | NA | NA | 5 | 0.8 |
| 16_286475 | 2 | 1 | 1 | 1 | 2 | 1 | 1 | 1 | 6 | 1 | NA | NA | 2 | 1 | NA | NA | NA | NA | 2 | 1 |

| | | | | | | | | | | | | | | | | | | | | |
|---|---|---|---|---|---|---|---|---|---|---|---|---|---|---|---|---|---|---|---|---|
| 16_2864762 | 1 | 1 | 1 | 1 | 1 | 1 | 1 | 1 | 4 | 1 | 2 | 1 | 1 | 1 | 1 | 1 | NA | NA | 4 | 1 |
| 16_2864767 | 2 | 0 | 1 | 1 | 2 | 0 | 2 | 0.5 | 7 | 0.2857 | 3 | 0.6667 | 1 | 0 | 1 | 1 | NA | NA | 5 | 0.6 |
| 16_2904696 | 4 | 0.25 | 1 | 0 | 3 | 0 | 2 | 0 | 10 | 0.1 | 1 | 0 | 2 | 0 | 3 | 0 | 3 | 0.3333 | 9 | 0.1111 |
| 16_2904710 | 4 | 1 | 1 | 1 | 3 | 0.6667 | 2 | 1 | 10 | 0.9 | 2 | 0.5 | 2 | 1 | 2 | 0.5 | 3 | 0.6667 | 9 | 0.6667 |
| 16_2904729 | 4 | 0.75 | NA | NA | 2 | 0.5 | 2 | 0.5 | 8 | 0.625 | 1 | 1 | 1 | 1 | 2 | 0.5 | 4 | 0.5 | 8 | 0.625 |
| 16_2904734 | 4 | 1 | 1 | 1 | 2 | 1 | 2 | 0.5 | 9 | 0.8889 | 1 | 1 | 1 | 1 | 2 | 1 | 3 | 0.3333 | 7 | 0.7143 |
| 16_2904754 | 4 | 0.25 | 1 | 0 | 2 | 0 | 2 | 0 | 9 | 0.1111 | 1 | 0 | NA | NA | 2 | 0.5 | 4 | 0 | 7 | 0.1429 |
| 16_2904781 | 3 | 0.3333 | 1 | 1 | 2 | 0 | 1 | 0 | 7 | 0.2857 | NA | NA | 2 | 1 | 2 | 1 | 3 | 0.3333 | 7 | 0.7143 |
| 16_2904810 | 4 | 0 | 2 | 0.5 | 2 | 0 | 2 | 0 | 10 | 0.1 | 1 | 0 | 3 | 0.3333 | 2 | 0.5 | 4 | 0.25 | 10 | 0.3 |
| 16_2904880 | 5 | 0.2 | 3 | 0 | 2 | 0.5 | 4 | 0 | 14 | 0.1429 | 1 | 0 | 4 | 0.25 | 3 | 0.3333 | 5 | 0 | 13 | 0.1538 |
| 16_2904885 | 5 | 0.4 | 3 | 0.3333 | 2 | 1 | 4 | 0.25 | 14 | 0.4286 | 1 | 0 | 4 | 0.25 | 2 | 0 | 5 | 0 | 12 | 0.0833 |
| 16_2921511 | NA | NA | 1 | 0 | 1 | 1 | 1 | 0 | 3 | 0.3333 | 1 | 0 | 1 | 1 | 1 | 1 | 1 | 1 | 4 | 0.75 |
| 16_2921516 | NA | NA | 1 | 0 | 1 | 1 | 1 | 0 | 3 | 0.3333 | 1 | 1 | 1 | 0 | 1 | 0 | 1 | 0 | 4 | 0.25 |
| 16_2940502 | 2 | 0.5 | 1 | 0 | 2 | 0 | 5 | 0 | 10 | 0.1 | 2 | 0 | 2 | 0 | 5 | 0.2 | 1 | 1 | 10 | 0.2 |
| 16_2940506 | 1 | 0 | 3 | 0.6667 | 2 | 0 | 5 | 0 | 11 | 0.1818 | 3 | 0 | 2 | 0 | 6 | 0.3333 | 1 | 0 | 12 | 0.1667 |
| 16_2947079 | 7 | 0.1429 | 9 | 0.1111 | 7 | 0.1429 | 8 | 0.25 | 31 | 0.1613 | 4 | 0.25 | 5 | 0 | 11 | 0.0909 | 13 | 0.2308 | 33 | 0.1515 |
| 16_2947103 | 9 | 0.4444 | 10 | 0.4 | 8 | 0.625 | 10 | 0.2 | 37 | 0.4054 | 7 | 0.1429 | 6 | 0.1667 | 13 | 0.4615 | 14 | 0.3571 | 40 | 0.325 |
| 16_2947117 | 10 | 0.1 | 11 | 0.0909 | 7 | 0.1429 | 8 | 0 | 36 | 0.0833 | 6 | 0 | 5 | 0 | 13 | 0 | 14 | 0.1429 | 38 | 0.0526 |
| 16_2947254 | 11 | 0.2727 | 9 | 0 | 10 | 0.4 | 14 | 0.3571 | 44 | 0.2727 | 2 | 0 | 8 | 0.25 | 19 | 0.1579 | 8 | 0.25 | 37 | 0.1892 |
| 16_2947329 | 14 | 0.1429 | 19 | 0.0526 | 22 | 0.0909 | 19 | 0.0526 | 74 | 0.0811 | 11 | 0.2727 | 10 | 0.1 | 21 | 0.1905 | 18 | 0.2222 | 60 | 0.2 |
| 16_2988616 | 3 | 0 | 5 | 0.2 | 2 | 0.5 | NA | NA | 10 | 0.2 | 2 | 0.5 | 2 | 0.5 | 1 | 0 | NA | NA | 5 | 0.4 |
| 16_2988621 | 2 | 0.5 | 5 | 0 | 2 | 0.5 | NA | NA | 9 | 0.2222 | 1 | 1 | 1 | 1 | 1 | 0 | NA | NA | 3 | 0.6667 |
| 16_2988624 | 3 | 1 | 4 | 0.75 | 2 | 0.5 | NA | NA | 9 | 0.7778 | 2 | 0.5 | 2 | 0.5 | NA | NA | NA | NA | 4 | 0.5 |
| 16_2988630 | 3 | 1 | 6 | 1 | 2 | 1 | NA | NA | 11 | 1 | 2 | 1 | 2 | 1 | 1 | 1 | NA | NA | 5 | 1 |
| 16_2988631 | 3 | 0.6667 | 7 | 0.5714 | 2 | 0.5 | NA | NA | 12 | 0.5833 | 1 | 1 | 2 | 1 | 1 | 0 | NA | NA | 4 | 0.75 |
| 16_2988633 | 2 | 1 | 7 | 0.5714 | 2 | 1 | NA | NA | 11 | 0.7273 | 2 | 1 | 2 | 1 | 1 | 1 | NA | NA | 5 | 1 |
| 16_3013386 | 1 | 0 | 3 | 0 | 3 | 0.3333 | 1 | 0 | 8 | 0.125 | 2 | 0 | 3 | 0.3333 | 2 | 0 | 2 | 0 | 9 | 0.1111 |
| 16_3033205 | NA | NA | 2 | 0.5 | 1 | 0 | 1 | 0 | 4 | 0.25 | NA | NA | NA | NA | 2 | 0.5 | 1 | 0 | 3 | 0.3333 |
| 16_3033238 | NA | NA | NA | NA | 2 | 0 | 2 | 0.5 | 4 | 0.25 | 1 | 0 | NA | NA | 1 | 1 | NA | NA | 2 | 0.5 |
| 16_3033242 | NA | NA | 2 | 1 | 2 | 1 | 2 | 1 | 6 | 1 | 1 | 1 | NA | NA | 1 | 1 | 1 | 0 | 3 | 0.6667 |
| 16_3033253 | 1 | 0 | 5 | 0.2 | 4 | 0 | 1 | 1 | 11 | 0.1818 | 1 | 0 | NA | NA | 2 | 0.5 | 2 | 0.5 | 5 | 0.4 |
| 16_3033266 | 1 | 1 | 4 | 0.75 | 4 | 0.5 | 1 | 0 | 10 | 0.6 | NA | NA | NA | NA | 3 | 0.3333 | 2 | 1 | 5 | 0.6 |
| 16_3033271 | 1 | 0 | 2 | 0.5 | 5 | 0.6 | 2 | 0 | 10 | 0.4 | NA | NA | NA | NA | 2 | 0.5 | 2 | 0.5 | 4 | 0.5 |
| 16_3033272 | 2 | 1 | 5 | 1 | 4 | 0.25 | 2 | 1 | 13 | 0.7692 | 1 | 0 | NA | NA | 1 | 1 | 1 | 1 | 3 | 0.6667 |
| 16_3033290 | 3 | 0.6667 | 5 | 1 | 7 | 0.5714 | 3 | 1 | 18 | 0.7778 | 3 | 0.3333 | 4 | 0.5 | 6 | 0.8333 | 3 | 0.6667 | 16 | 0.625 |
| 16_3033293 | 4 | 0.5 | 4 | 1 | 8 | 0.625 | 3 | 0.6667 | 19 | 0.6842 | 3 | 1 | 5 | 0.8 | 5 | 0.8 | 3 | 0 | 16 | 0.6875 |
| 16_3033302 | 3 | 1 | 4 | 1 | 7 | 0.5714 | 3 | 1 | 17 | 0.8235 | 1 | 0 | 5 | 0.8 | 5 | 0 | 3 | 0.3333 | 14 | 0.3571 |
| 16_3033305 | 3 | 0.6667 | 4 | 0.75 | 7 | 0 | 3 | 0.3333 | 17 | 0.3529 | 1 | 0 | 5 | 0.2 | 4 | 0.25 | 2 | 0.5 | 12 | 0.25 |
| 16_3033309 | 3 | 0.6667 | 5 | 1 | 5 | 0.8 | 3 | 0.6667 | 16 | 0.8125 | 2 | 0 | 3 | 0.3333 | 4 | 0.75 | 3 | 0.6667 | 12 | 0.5 |
| 16_3033316 | 2 | 0 | 1 | 0 | 5 | 0.2 | 2 | 0 | 10 | 0.1 | 2 | 0 | 3 | 0 | 2 | 0.5 | 3 | 0.3333 | 10 | 0.2 |
| 16_3033318 | 4 | 0.25 | 5 | 0.2 | 6 | 0.5 | 3 | 0.3333 | 18 | 0.3333 | 1 | 0 | 4 | 0.25 | 3 | 0 | 3 | 0.3333 | 11 | 0.1818 |
| 16_3033320 | 3 | 0.6667 | 4 | 1 | 4 | 1 | 1 | 1 | 12 | 0.9167 | 1 | 1 | 4 | 0.25 | 2 | 1 | 2 | 1 | 9 | 0.6667 |
| 16_3033324 | 4 | 0 | 2 | 0 | 6 | 0.1667 | 3 | 0 | 15 | 0.0667 | 2 | 0 | 3 | 0 | 3 | 0.3333 | 3 | 0 | 11 | 0.0909 |
| 16_3033326 | 4 | 0 | 2 | 1 | 6 | 0.1667 | 3 | 0 | 15 | 0.2 | 2 | 0 | 3 | 0 | 3 | 0.3333 | 3 | 0.3333 | 11 | 0.1818 |
| 16_3033333 | 3 | 0.6667 | 3 | 1 | 5 | 0.6 | 2 | 1 | 13 | 0.7692 | 2 | 0.5 | 3 | 0.6667 | 3 | 1 | 3 | 1 | 11 | 0.8182 |
| 16_3033334 | 3 | 1 | 5 | 0.8 | 6 | 0.6667 | 3 | 1 | 17 | 0.8235 | 2 | 1 | 4 | 0.75 | 3 | 1 | 3 | 0.6667 | 12 | 0.8333 |
| 16_3033346 | 3 | 0 | 4 | 0.25 | 6 | 0.1667 | 3 | 0.3333 | 16 | 0.1875 | 1 | 1 | 3 | 0.3333 | 2 | 1 | 3 | 0 | 9 | 0.4444 |
| 16_3033347 | 3 | 0.3333 | 4 | 0.25 | 5 | 0.2 | 3 | 0.3333 | 15 | 0.2667 | 2 | 0.5 | 4 | 0.5 | 3 | 0.3333 | 3 | 0.6667 | 12 | 0.5 |
| 16_3033350 | 1 | 1 | 4 | 0.25 | 5 | 0.4 | 2 | 0.5 | 12 | 0.4167 | 2 | 0 | 4 | 0.5 | 4 | 0.25 | 3 | 0.3333 | 13 | 0.3077 |
| 16_3033353 | 3 | 0 | 5 | 0 | 6 | 0.1667 | 3 | 0 | 17 | 0.0588 | 2 | 0.5 | 4 | 0 | 3 | 0 | 3 | 0 | 12 | 0.0833 |
| 16_3036840 | 3 | 0 | 2 | 0 | 1 | 1 | 2 | 0 | 8 | 0.125 | NA | NA | 1 | 1 | NA | NA | NA | NA | 1 | 1 |
| 16_3036878 | 3 | 0.6667 | 2 | 0.5 | 1 | 1 | 1 | 0 | 7 | 0.5714 | NA | NA | 2 | 0.5 | NA | NA | NA | NA | 2 | 0.5 |
| 16_3038022 | 2 | 0 | 4 | 1 | NA | NA | 1 | 1 | 7 | 0.7143 | NA | NA | 1 | 1 | 1 | 1 | 4 | 0.25 | 6 | 0.5 |
| 16_3038039 | 1 | 1 | 4 | 1 | NA | NA | 2 | 1 | 7 | 1 | NA | NA | 1 | 1 | 4 | 1 | 4 | 1 | 9 | 1 |
| 16_3038045 | 3 | 0.3333 | 3 | 0 | NA | NA | 2 | 0.5 | 8 | 0.25 | NA | NA | 1 | 0 | 4 | 0.25 | 4 | 0.5 | 9 | 0.3333 |
| 16_3038049 | 3 | 0 | 2 | 0.5 | NA | NA | 2 | 0 | 7 | 0.1429 | NA | NA | NA | NA | 4 | 0 | 2 | 0.5 | 6 | 0.1667 |
| 16_3039000 | 2 | 0.5 | 3 | 0 | NA | NA | 2 | 0 | 7 | 0.1429 | NA | NA | 1 | 0 | NA | NA | 3 | 0.3333 | 4 | 0.25 |
| 16_3039004 | 2 | 0.5 | 3 | 0.3333 | NA | NA | NA | NA | 5 | 0.4 | NA | NA | 1 | 1 | NA | NA | 2 | 0.5 | 3 | 0.6667 |
| 16_3039021 | 2 | 0 | 4 | 0.5 | NA | NA | 2 | 0.5 | 8 | 0.375 | NA | NA | 1 | 0 | NA | NA | 3 | 0.3333 | 4 | 0.25 |
| 16_3039026 | 2 | 0 | 4 | 0.5 | NA | NA | 2 | 0 | 8 | 0.25 | NA | NA | 1 | 0 | NA | NA | 4 | 0.25 | 5 | 0.2 |
| 16_3041878 | 1 | 1 | NA | NA | 1 | 0 | 1 | 0 | 3 | 0.3333 | NA | NA | 1 | 0 | 1 | 1 | NA | NA | 2 | 0.5 |
| 16_3041882 | 1 | 1 | NA | NA | 1 | 1 | 1 | 1 | 3 | 1 | NA | NA | 1 | 1 | 1 | 1 | NA | NA | 2 | 1 |
| 16_3041885 | 1 | 1 | NA | NA | 1 | 1 | 1 | 1 | 3 | 1 | NA | NA | NA | NA | 1 | 1 | NA | NA | 1 | 1 |
| 16_3041896 | 1 | 1 | NA | NA | 1 | 0 | 1 | 1 | 3 | 0.6667 | NA | NA | 1 | 1 | 1 | 0 | NA | NA | 2 | 0.5 |
| 16_3041912 | 1 | 0 | NA | NA | 2 | 0.5 | 1 | 0 | 4 | 0.25 | NA | NA | 1 | 0 | 1 | 1 | NA | NA | 2 | 0.5 |

| | | | | | | | | | | | | | | | | | | | | |
|---|---|---|---|---|---|---|---|---|---|---|---|---|---|---|---|---|---|---|---|---|
| 16_304197 | 1 | 1 | NA | NA | 1 | 1 | 1 | 1 | 3 | 1 | NA | NA | 1 | 1 | 1 | 1 | NA | NA | 2 | 1 |
| 16_304201 | 1 | 1 | 2 | 0.5 | 2 | 0.5 | 1 | 1 | 6 | 0.6667 | NA | NA | 1 | 1 | 1 | 1 | NA | NA | 2 | 1 |
| 16_304202 | NA | NA | 2 | 1 | NA | NA | 1 | 0 | 3 | 0.6667 | NA | NA | NA | NA | 1 | 1 | NA | NA | 1 | 1 |
| 16_304203 | 2 | 1 | 2 | 1 | 2 | 1 | 1 | 1 | 7 | 1 | 1 | 1 | 1 | 1 | 1 | 1 | NA | NA | 3 | 1 |
| 16_304203 | 1 | 1 | 1 | 0 | 2 | 1 | NA | NA | 4 | 0.75 | 1 | 0 | 1 | 1 | 1 | 1 | NA | NA | 3 | 0.6667 |
| 16_304204 | 2 | 1 | 1 | 1 | 1 | 1 | 2 | 1 | 6 | 1 | 1 | 1 | 1 | 1 | 2 | 0.5 | NA | NA | 4 | 0.75 |
| 16_304204 | 1 | 0 | 2 | 0 | 2 | 0 | 2 | 1 | 7 | 0.2857 | 1 | 1 | 1 | 0 | 2 | 0 | NA | NA | 4 | 0.25 |
| 16_304204 | 2 | 0 | 2 | 0 | 1 | 1 | 1 | 0 | 6 | 0.1667 | 1 | 0 | 1 | 0 | 2 | 0.5 | NA | NA | 4 | 0.25 |
| 16_304205 | 2 | 1 | 2 | 1 | 2 | 0.5 | 2 | 0 | 8 | 0.625 | 1 | 0 | 1 | 0 | 2 | 0.5 | NA | NA | 4 | 0.25 |
| 16_304207 | 2 | 0.5 | 2 | 0.5 | 2 | 0.5 | 1 | 1 | 7 | 0.5714 | 1 | 1 | 1 | 0 | 2 | 0.5 | NA | NA | 4 | 0.5 |
| 16_304214 | 6 | 0 | 4 | 0.25 | 5 | 0 | 1 | 0 | 16 | 0.0625 | 1 | 1 | 1 | 0 | 2 | 0 | NA | NA | 4 | 0.25 |
| 16_307746 | 3 | 0 | 4 | 0.25 | 4 | 0.5 | 4 | 0.25 | 15 | 0.2667 | 3 | 0.3333 | 5 | 0.6 | 2 | 0 | 3 | 0.6667 | 13 | 0.4615 |
| 16_307746 | 3 | 0 | 4 | 0.25 | 4 | 0 | 6 | 0.3333 | 17 | 0.1765 | 4 | 0.5 | 3 | 0.6667 | 3 | 0 | 3 | 0.6667 | 13 | 0.4615 |
| 16_307751 | 2 | 0.5 | 3 | 0 | 4 | 0.5 | 5 | 0.4 | 14 | 0.3571 | 3 | 0.6667 | 4 | 0.75 | 2 | 0 | 3 | 0 | 12 | 0.4167 |
| 16_307751 | 3 | 0.6667 | 2 | 0.5 | 4 | 0.5 | 5 | 0.8 | 14 | 0.6429 | 4 | 0.5 | 4 | 0.5 | 2 | 0 | 2 | 0.5 | 12 | 0.4167 |
| 16_307751 | 3 | 0.3333 | 3 | 0 | 4 | 0 | 6 | 0.1667 | 16 | 0.125 | 4 | 0.5 | 5 | 0.2 | 2 | 0 | 4 | 0 | 15 | 0.2 |
| 16_307754 | 2 | 0.5 | 3 | 0.3333 | 4 | 0.5 | 6 | 0.8333 | 15 | 0.6 | 3 | 0.3333 | 5 | 1 | 2 | 0.5 | 3 | 1 | 13 | 0.7692 |
| 16_307756 | 2 | 1 | 3 | 1 | 4 | 1 | 5 | 1 | 14 | 1 | 3 | 1 | 4 | 1 | 2 | 1 | 1 | 1 | 10 | 1 |
| 16_307758 | 2 | 1 | NA | NA | 3 | 1 | 5 | 1 | 10 | 1 | 2 | 1 | 3 | 1 | 2 | 1 | NA | NA | 7 | 1 |
| 16_307759 | 2 | 1 | 2 | 1 | 2 | 1 | 5 | 1 | 11 | 1 | 3 | 1 | 4 | 1 | 2 | 1 | 1 | 1 | 10 | 1 |
| 16_307760 | 2 | 1 | 2 | 1 | 3 | 1 | 5 | 1 | 12 | 1 | 3 | 1 | 4 | 1 | 2 | 1 | NA | NA | 9 | 1 |
| 16_307761 | 1 | 1 | 2 | 1 | 2 | 1 | 3 | 1 | 8 | 1 | 3 | 0.6667 | 3 | 1 | 2 | 1 | NA | NA | 8 | 0.875 |
| 16_307761 | NA | NA | NA | NA | 1 | 1 | 1 | 1 | 2 | 1 | NA | NA | 1 | 1 | NA | NA | NA | NA | 1 | 1 |
| 16_307761 | 2 | 1 | 2 | 0.5 | NA | NA | 3 | 1 | 7 | 0.8571 | 3 | 1 | 4 | 0.75 | 2 | 1 | NA | NA | 9 | 0.8889 |
| 16_307761 | 1 | 1 | 2 | 0.5 | 3 | 0.3333 | 4 | 1 | 10 | 0.7 | 3 | 0.3333 | 4 | 0.75 | 2 | 1 | NA | NA | 9 | 0.6667 |
| 16_307762 | 1 | 1 | 2 | 1 | 3 | 0.6667 | 4 | 0.5 | 10 | 0.7 | 2 | 1 | 3 | 0.6667 | 2 | 1 | NA | NA | 7 | 0.8571 |
| 16_307762 | 1 | 1 | NA | NA | 3 | 1 | 4 | 0.75 | 8 | 0.875 | 3 | 1 | 3 | 0.6667 | 1 | 1 | NA | NA | 7 | 0.8571 |
| 16_307763 | 1 | 1 | 1 | 1 | 1 | 0 | 3 | 1 | 6 | 0.8333 | 3 | 0.3333 | 4 | 1 | NA | NA | NA | NA | 7 | 0.7143 |
| 16_307763 | 1 | 1 | 2 | 1 | 3 | 0.6667 | 5 | 1 | 11 | 0.9091 | 3 | 1 | 4 | 0.75 | 2 | 1 | NA | NA | 9 | 0.8889 |
| 16_307763 | NA | NA | 2 | 0 | 1 | 0 | 5 | 0.4 | 8 | 0.25 | 3 | 0 | 4 | 0.25 | 2 | 0 | NA | NA | 9 | 0.1111 |
| 16_307764 | 1 | 1 | 1 | 1 | 3 | 0.3333 | 5 | 0.2 | 10 | 0.4 | 3 | 0 | 4 | 0.5 | 1 | 0 | NA | NA | 8 | 0.25 |
| 16_309120 | 13 | 0 | 17 | 0.0588 | 9 | 0.2222 | 10 | 0.1 | 49 | 0.0816 | 3 | 0 | 8 | 0.25 | 14 | 0.0714 | 13 | 0.1538 | 38 | 0.1316 |
| 16_309151 | 3 | 0 | 6 | 0 | 4 | 0.25 | 4 | 0 | 17 | 0.0588 | NA | NA | 2 | 0 | 2 | 0 | 4 | 0.25 | 8 | 0.125 |
| 16_309183 | 1 | 0 | 2 | 0.5 | 1 | 0 | NA | NA | 4 | 0.25 | NA | NA | 4 | 0 | 3 | 0.3333 | NA | NA | 7 | 0.1429 |
| 16_309206 | 1 | 0 | 2 | 0 | 1 | 0 | 1 | 1 | 5 | 0.2 | 1 | 1 | 1 | 0 | 2 | 0.5 | 1 | 0 | 5 | 0.4 |
| 16_309206 | 1 | 0 | 2 | 0.5 | 1 | 0 | 1 | 1 | 5 | 0.4 | 1 | 0 | 2 | 0.5 | 2 | 1 | 1 | 0 | 6 | 0.5 |
| 16_310723 | 94 | 0.0957 | 60 | 0.0833 | 63 | 0.0635 | 112 | 0.0804 | 329 | 0.0821 | 35 | 0.0286 | 99 | 0.1414 | 98 | 0.0816 | 56 | 0.0893 | 288 | 0.0972 |
| 16_310723 | 89 | 0.0674 | 64 | 0.125 | 56 | 0.125 | 103 | 0.0583 | 312 | 0.0865 | 37 | 0.1081 | 103 | 0.1456 | 92 | 0.1196 | 52 | 0.2308 | 284 | 0.1479 |
| 16_310725 | 87 | 0.1494 | 54 | 0.1111 | 58 | 0.1207 | 95 | 0.1474 | 294 | 0.1361 | 37 | 0.2162 | 91 | 0.1648 | 95 | 0.1158 | 50 | 0.22 | 273 | 0.1648 |
| 16_310733 | 101 | 0.4158 | 70 | 0.3714 | 79 | 0.4684 | 105 | 0.4857 | 355 | 0.4394 | 40 | 0.425 | 114 | 0.4035 | 110 | 0.4818 | 68 | 0.5588 | 332 | 0.4639 |
| 16_310770 | 7 | 0.8571 | 7 | 0.4286 | 4 | 0.5 | 15 | 0.4 | 33 | 0.5152 | 6 | 0.8333 | 8 | 0.5 | 10 | 0.5 | 12 | 0.6667 | 36 | 0.6111 |
| 16_310771 | 6 | 0 | 7 | 0.4286 | 2 | 0 | 15 | 0.0667 | 30 | 0.1333 | 6 | 0.1667 | 9 | 0.1111 | 11 | 0.1818 | 10 | 0.2 | 36 | 0.1667 |
| 16_310772 | 90 | 0.3333 | 60 | 0.3667 | 60 | 0.2167 | 97 | 0.4433 | 307 | 0.3518 | 35 | 0.4 | 105 | 0.419 | 93 | 0.4194 | 67 | 0.4179 | 300 | 0.4167 |
| 16_311161 | 75 | 0.3333 | 65 | 0.3846 | 45 | 0.2444 | 40 | 0.3 | 225 | 0.3244 | 35 | 0.5143 | 18 | 0.5556 | 24 | 0.4583 | 27 | 0.4444 | 104 | 0.4904 |
| 16_311161 | 76 | 0.1316 | 70 | 0.2 | 48 | 0.125 | 35 | 0.1429 | 229 | 0.1528 | 39 | 0.1282 | 21 | 0.0476 | 24 | 0.125 | 26 | 0.1538 | 110 | 0.1182 |
| 16_311162 | 82 | 0.0732 | 76 | 0.0789 | 52 | 0.0769 | 48 | 0.0625 | 258 | 0.0736 | 41 | 0.0488 | 24 | 0.0833 | 30 | 0.1 | 29 | 0.0345 | 124 | 0.0645 |
| 16_312755 | 5 | 0.8 | 7 | 1 | 2 | 1 | 7 | 1 | 21 | 0.9524 | 6 | 1 | 10 | 0.9 | 7 | 1 | 3 | 1 | 26 | 0.9615 |
| 16_312756 | 7 | 1 | 6 | 1 | 2 | 1 | 9 | 1 | 24 | 1 | 7 | 1 | 9 | 1 | 8 | 1 | 3 | 1 | 27 | 1 |
| 16_312758 | 4 | 1 | 8 | 1 | 2 | 1 | 7 | 1 | 21 | 1 | 8 | 0.875 | 9 | 1 | 7 | 1 | 3 | 1 | 27 | 0.963 |
| 16_312758 | NA | NA | NA | NA | NA | NA | 3 | 1 | 3 | 1 | NA | NA | 1 | 1 | NA | NA | NA | NA | 1 | 1 |
| 16_312759 | 5 | 0 | 9 | 0.3333 | 2 | 0 | 10 | 0 | 26 | 0.1154 | 7 | 0 | 9 | 0.2222 | 8 | 0 | 4 | 0 | 28 | 0.0714 |
| 16_312765 | 1 | 1 | 2 | 0 | 1 | 0 | 2 | 0 | 6 | 0.1667 | NA | NA | 3 | 0.6667 | 2 | 0 | 1 | 0 | 6 | 0.3333 |
| 16_312766 | 1 | 1 | 1 | 0 | 2 | 0 | 3 | 0.6667 | 7 | 0.4286 | NA | NA | 3 | 0.3333 | 2 | 0.5 | 1 | 1 | 6 | 0.5 |
| 16_312771 | 1 | 1 | 3 | 0.3333 | 2 | 0.5 | 1 | 1 | 7 | 0.5714 | NA | NA | 4 | 0.5 | NA | NA | 3 | 0.3333 | 7 | 0.4286 |
| 16_312772 | 1 | 1 | 1 | 0 | 3 | 0.6667 | 1 | 1 | 6 | 0.6667 | NA | NA | 3 | 1 | 1 | 1 | 3 | 0.6667 | 7 | 0.8571 |
| 16_312774 | NA | NA | 2 | 0.5 | 2 | 0.5 | 1 | 1 | 5 | 0.6 | NA | NA | 1 | 0 | 1 | 0 | 2 | 0.5 | 4 | 0.25 |
| 16_314096 | NA | NA | NA | NA | NA | NA | 1 | 1 | 1 | 1 | NA | NA | 1 | 1 | NA | NA | NA | NA | 1 | 1 |
| 16_314106 | NA | NA | NA | NA | NA | NA | 1 | 1 | 1 | 1 | NA | NA | 1 | 1 | NA | NA | NA | NA | 1 | 1 |
| 16_315146 | 2 | 0 | NA | NA | NA | NA | 2 | 0.5 | 4 | 0.25 | 1 | 1 | 3 | 0 | NA | NA | NA | NA | 4 | 0.25 |
| 16_316916 | 17 | 0.1765 | 21 | 0 | 11 | 0 | 27 | 0.037 | 76 | 0.0526 | 11 | 0.0909 | 23 | 0.0435 | 20 | 0.1 | 15 | 0.0667 | 69 | 0.0725 |
| 16_316920 | 20 | 0.25 | 26 | 0.1538 | 18 | 0.1667 | 28 | 0.0714 | 92 | 0.1522 | 15 | 0.0667 | 27 | 0.0741 | 25 | 0.12 | 17 | 0.3529 | 84 | 0.1429 |
| 16_316926 | NA | NA | NA | NA | NA | NA | 1 | 1 | 1 | 1 | NA | NA | 2 | 0 | 1 | 1 | NA | NA | 3 | 0.3333 |
| 16_316930 | NA | NA | NA | NA | NA | NA | 1 | 1 | 1 | 1 | NA | NA | 2 | 0 | 2 | 0.5 | NA | NA | 4 | 0.25 |
| 16_316931 | NA | NA | NA | NA | NA | NA | 1 | 1 | 1 | 1 | NA | NA | 2 | 0 | 2 | 0.5 | NA | NA | 4 | 0.25 |
| 16_316935 | NA | NA | NA | NA | NA | NA | 1 | 1 | 1 | 1 | NA | NA | 1 | 1 | 3 | 0.6667 | NA | NA | 4 | 0.75 |

| | | | | | | | | | | | | | | | | | | | | |
|---|---|---|---|---|---|---|---|---|---|---|---|---|---|---|---|---|---|---|---|---|
| 16_316935 | NA | NA | NA | NA | NA | NA | 1 | 1 | 1 | 1 | NA | NA | 2 | 0 | 3 | 0.6667 | NA | NA | 5 | 0.4 |
| 16_316938 | NA | NA | NA | NA | NA | NA | 1 | 1 | 1 | 1 | NA | NA | 1 | 1 | 3 | 0 | NA | NA | 4 | 0.25 |
| 16_317632 | 1 | 1 | 1 | 1 | 1 | 1 | NA | NA | 3 | 1 | 1 | 1 | NA | NA | 1 | 1 | NA | NA | 2 | 1 |
| 16_317634 | 1 | 0 | NA | NA | 1 | 1 | NA | NA | 2 | 0.5 | 1 | 1 | NA | NA | 1 | 1 | NA | NA | 2 | 1 |
| 16_317636 | 1 | 1 | 1 | 1 | 1 | 1 | NA | NA | 3 | 1 | NA | NA | NA | NA | 1 | 1 | NA | NA | 1 | 1 |
| 16_317636 | 2 | 0.5 | 1 | 1 | 1 | 1 | NA | NA | 4 | 0.75 | 1 | 1 | NA | NA | 1 | 1 | NA | NA | 2 | 1 |
| 16_317644 | 1 | 1 | 1 | 1 | 2 | 1 | NA | NA | 4 | 1 | 1 | 1 | NA | NA | NA | NA | NA | NA | 1 | 1 |
| 16_317646 | 2 | 1 | 1 | 1 | 2 | 0.5 | NA | NA | 5 | 0.8 | 1 | 1 | NA | NA | NA | NA | NA | NA | 1 | 1 |
| 16_317659 | 2 | 0.5 | 1 | 1 | 2 | 0.5 | 1 | 1 | 6 | 0.6667 | NA | NA | 1 | 0 | 2 | 1 | 1 | 1 | 4 | 0.75 |
| 16_317662 | 1 | 0 | 1 | 1 | 2 | 0.5 | 1 | 0 | 5 | 0.4 | NA | NA | 1 | 0 | 1 | 1 | 1 | 1 | 3 | 0.6667 |
| 16_317663 | 1 | 0 | NA | NA | 2 | 0 | 2 | 1 | 5 | 0.4 | NA | NA | 1 | 0 | 2 | 1 | 1 | 0 | 4 | 0.5 |
| 16_317664 | 2 | 0 | NA | NA | 2 | 0 | 2 | 1 | 6 | 0.3333 | NA | NA | 1 | 0 | 2 | 0.5 | 1 | 1 | 4 | 0.5 |
| 16_317665 | 2 | 0.5 | NA | NA | 1 | 1 | 2 | 0.5 | 5 | 0.6 | NA | NA | 1 | 1 | NA | NA | 1 | 1 | 2 | 1 |
| 16_317667 | 15 | 0.6667 | 17 | 0.4118 | 16 | 0.5 | 16 | 0.375 | 64 | 0.4844 | 12 | 0.4167 | 16 | 0.5625 | 9 | 0.3333 | 7 | 0.7143 | 44 | 0.5 |
| 16_317667 | 14 | 1 | 14 | 1 | 13 | 1 | 16 | 1 | 57 | 1 | 8 | 1 | 17 | 1 | 10 | 1 | 8 | 1 | 43 | 1 |
| 16_317668 | 13 | 0.8462 | 15 | 0.9333 | 11 | 0.9091 | 17 | 0.8824 | 56 | 0.8929 | 9 | 1 | 17 | 0.8824 | 8 | 1 | 8 | 1 | 42 | 0.9524 |
| 16_317669 | 9 | 0.2222 | 14 | 0.2143 | 10 | 0 | 15 | 0.1333 | 48 | 0.1458 | 6 | 0 | 12 | 0.3333 | 6 | 0.5 | 6 | 0.3333 | 30 | 0.3 |
| 16_317669 | 12 | 0.8333 | 17 | 0.5294 | 11 | 0.7273 | 17 | 0.6471 | 57 | 0.6667 | 8 | 0.25 | 17 | 0.5882 | 8 | 0.625 | 7 | 1 | 40 | 0.6 |
| 16_317670 | 13 | 0 | 15 | 0.2 | 11 | 0.1818 | 16 | 0.375 | 55 | 0.2 | 8 | 0.25 | 19 | 0.4211 | 9 | 0.1111 | 7 | 0.1429 | 43 | 0.2791 |
| 16_317758 | 4 | 0 | 8 | 0.125 | 6 | 0 | 7 | 0.2857 | 25 | 0.12 | 4 | 0.25 | 7 | 0 | 7 | 0 | 2 | 0 | 20 | 0.05 |
| 16_317759 | 4 | 0.25 | 8 | 0 | 7 | 0.1429 | 8 | 0.125 | 27 | 0.1111 | 4 | 0 | 8 | 0.125 | 10 | 0.2 | 4 | 0 | 26 | 0.1154 |
| 16_317761 | 4 | 0.5 | 9 | 0.1111 | 5 | 0 | 10 | 0 | 28 | 0.1071 | 3 | 0.3333 | 7 | 0 | 9 | 0.1111 | 3 | 0 | 22 | 0.0909 |
| 16_317761 | 5 | 0.2 | 9 | 0.5556 | 4 | 0.5 | 10 | 0.1 | 28 | 0.3214 | 3 | 0.6667 | 7 | 0 | 9 | 0.3333 | 3 | 0.3333 | 22 | 0.2727 |
| 16_317761 | 5 | 0 | 9 | 0.1111 | 6 | 0.1667 | 10 | 0.1 | 30 | 0.1 | 3 | 0.6667 | 8 | 0 | 9 | 0 | 3 | 0 | 23 | 0.087 |
| 16_317767 | 4 | 0 | 9 | 0.2222 | 8 | 0 | 12 | 0.0833 | 33 | 0.0909 | 3 | 0 | 12 | 0.0833 | 8 | 0.125 | 4 | 0.25 | 27 | 0.1111 |
| 16_317767 | 5 | 0 | 11 | 0.5455 | 7 | 0.4286 | 13 | 0.1538 | 36 | 0.3056 | 2 | 0 | 12 | 0.0833 | 10 | 0.2 | 3 | 0.3333 | 27 | 0.1481 |
| 16_317822 | 1 | 0 | 1 | 0 | 1 | 1 | 5 | 0.2 | 8 | 0.25 | 1 | 0 | 5 | 0.2 | 1 | 1 | 2 | 0.5 | 9 | 0.3333 |
| 16_317822 | 1 | 1 | 1 | 1 | NA | NA | 4 | 0.5 | 6 | 0.6667 | 1 | 0 | 3 | 0.6667 | NA | NA | 2 | 0.5 | 6 | 0.5 |
| 16_318058 | NA | NA | NA | NA | 1 | 1 | NA | NA | 1 | 1 | 1 | 1 | NA | NA | NA | NA | NA | NA | 1 | 1 |
| 16_318079 | NA | NA | 1 | 0 | 1 | 0 | 4 | 0.25 | 6 | 0.1667 | NA | NA | 2 | 0 | NA | NA | 2 | 0.5 | 4 | 0.25 |
| 16_318080 | NA | NA | 1 | 0 | 1 | 1 | 4 | 0 | 6 | 0.1667 | NA | NA | 2 | 0.5 | NA | NA | 1 | 0 | 3 | 0.3333 |
| 16_318080 | NA | NA | 1 | 1 | 1 | 0 | 4 | 0.75 | 6 | 0.6667 | NA | NA | 2 | 1 | NA | NA | 1 | 1 | 3 | 1 |
| 16_318083 | NA | NA | NA | NA | NA | NA | 1 | 1 | 1 | 1 | NA | NA | 1 | 1 | NA | NA | NA | NA | 1 | 1 |
| 16_318084 | NA | NA | NA | NA | 1 | 1 | 2 | 0.5 | 3 | 0.6667 | NA | NA | 2 | 0.5 | NA | NA | NA | NA | 2 | 0.5 |
| 16_318085 | NA | NA | NA | NA | 1 | 1 | 3 | 0 | 4 | 0.25 | NA | NA | 2 | 0.5 | NA | NA | NA | NA | 2 | 0.5 |
| 16_318089 | NA | NA | NA | NA | NA | NA | 4 | 0.25 | 4 | 0.25 | NA | NA | 2 | 0.5 | NA | NA | NA | NA | 2 | 0.5 |
| 16_318090 | NA | NA | NA | NA | 1 | 1 | 2 | 1 | 3 | 1 | NA | NA | 1 | 1 | NA | NA | NA | NA | 1 | 1 |
| 16_318368 | 4 | 0.25 | 5 | 0.4 | 4 | 0.25 | 10 | 0.6 | 23 | 0.4348 | 2 | 0.5 | 3 | 0.3333 | 4 | 0.25 | 5 | 0.8 | 14 | 0.5 |
| 16_318369 | 4 | 0.25 | 5 | 0 | 5 | 0 | 9 | 0.1111 | 23 | 0.087 | 3 | 0 | 1 | 0 | 4 | 0 | 4 | 0.25 | 12 | 0.0833 |
| 16_318372 | 3 | 1 | 5 | 0.4 | 3 | 0.3333 | 7 | 0.2857 | 18 | 0.4444 | 4 | 0.5 | 1 | 1 | 2 | 0.5 | 5 | 0.8 | 12 | 0.6667 |
| 16_318379 | 3 | 0.6667 | 5 | 0.2 | 6 | 0 | 9 | 0.1111 | 23 | 0.1739 | 4 | 0 | 2 | 0 | 4 | 0.5 | 4 | 0 | 14 | 0.1429 |
| 16_318382 | 2 | 0 | 5 | 0 | 5 | 0 | 8 | 0.125 | 20 | 0.05 | 2 | 0.5 | 2 | 0 | 5 | 0 | 4 | 0.25 | 13 | 0.1538 |
| 16_318387 | 2 | 0.5 | 4 | 0 | 4 | 0 | 7 | 0.5714 | 17 | 0.2941 | 3 | 0.3333 | 1 | 1 | 4 | 0.25 | 3 | 0.6667 | 11 | 0.4545 |
| 16_318387 | 2 | 1 | 5 | 1 | 4 | 1 | 6 | 0.8333 | 17 | 0.9412 | 2 | 1 | 2 | 0.5 | 4 | 1 | 3 | 0.6667 | 11 | 0.8182 |
| 16_318388 | 2 | 0 | 4 | 0.75 | 5 | 0.8 | 8 | 0.625 | 19 | 0.6316 | 2 | 1 | 1 | 0 | 4 | 0.5 | 3 | 1 | 10 | 0.7 |
| 16_318388 | NA | NA | 1 | 0 | 1 | 1 | 1 | 1 | 3 | 0.6667 | 2 | 0.5 | 2 | 0 | 1 | 1 | 2 | 0.5 | 7 | 0.4286 |
| 16_318389 | 2 | 0.5 | 5 | 0.8 | 4 | 1 | 7 | 1 | 18 | 0.8889 | 2 | 1 | 1 | 1 | 3 | 1 | 3 | 1 | 9 | 1 |
| 16_319260 | 12 | 0.4167 | 10 | 0.3 | 11 | 0.4545 | 7 | 0.2857 | 40 | 0.375 | 10 | 0.2 | 8 | 0.5 | 11 | 0.3636 | 4 | 0.25 | 33 | 0.3333 |
| 16_319260 | 12 | 0.0833 | 8 | 0 | 13 | 0.0769 | 6 | 0 | 39 | 0.0513 | 10 | 0 | 9 | 0.2222 | 9 | 0.1111 | 2 | 0.5 | 30 | 0.1333 |
| 16_319261 | 13 | 0.2308 | 7 | 0 | 13 | 0 | 6 | 0.1667 | 39 | 0.1026 | 10 | 0.1 | 8 | 0.125 | 11 | 0.0909 | 5 | 0 | 34 | 0.0882 |
| 16_319261 | 13 | 0.9231 | 9 | 1 | 15 | 0.9333 | 7 | 1 | 44 | 0.9545 | 12 | 0.8333 | 10 | 1 | 15 | 0.9333 | 5 | 1 | 42 | 0.9286 |
| 16_319262 | 10 | 0.5 | 10 | 0.3 | 13 | 0.0769 | 7 | 0.2857 | 40 | 0.275 | 12 | 0.1667 | 10 | 0.3 | 15 | 0.0667 | 5 | 0.2 | 42 | 0.1667 |
| 16_319262 | 13 | 1 | 10 | 1 | 15 | 1 | 8 | 1 | 46 | 1 | 12 | 0.9167 | 11 | 0.9091 | 15 | 1 | 5 | 1 | 43 | 0.9535 |
| 16_319265 | 13 | 0.9231 | 10 | 0.9 | 17 | 0.9412 | 10 | 0.8 | 50 | 0.9 | 11 | 0.5455 | 12 | 0.5833 | 15 | 0.6667 | 6 | 0.8333 | 44 | 0.6364 |
| 16_319265 | 12 | 0.5 | 9 | 0.4444 | 16 | 0.5 | 8 | 0.75 | 45 | 0.5333 | 11 | 0.3636 | 10 | 0.3 | 10 | 0.2 | 6 | 0.5 | 37 | 0.3243 |
| 16_319617 | NA | NA | NA | NA | 1 | 0 | 1 | 1 | 2 | 0.5 | 2 | 0.5 | 3 | 0.6667 | NA | NA | 2 | 0.5 | 7 | 0.5714 |
| 16_319618 | NA | NA | NA | NA | NA | NA | 1 | 1 | 1 | 1 | 3 | 0.6667 | 3 | 1 | 1 | 1 | NA | NA | 7 | 0.8571 |
| 16_319620 | NA | NA | NA | NA | 1 | 1 | 1 | 1 | 2 | 1 | 3 | 1 | 4 | 0.5 | 1 | 1 | 2 | 1 | 10 | 0.8 |
| 16_319620 | NA | NA | NA | NA | 1 | 1 | 1 | 0 | 2 | 0.5 | 3 | 0.3333 | 3 | 0.6667 | 1 | 1 | 2 | 0.5 | 9 | 0.5556 |
| 16_319621 | NA | NA | NA | NA | 1 | 1 | NA | NA | 1 | 1 | 3 | 0.6667 | 1 | 0 | NA | NA | 2 | 0 | 6 | 0.3333 |
| 16_319622 | NA | NA | NA | NA | 1 | 0 | 1 | 1 | 2 | 0.5 | 3 | 0.3333 | NA | NA | NA | NA | 2 | 0.5 | 5 | 0.4 |
| 16_319622 | NA | NA | NA | NA | 1 | 1 | 1 | 1 | 2 | 1 | 3 | 1 | 4 | 1 | 1 | 1 | 2 | 1 | 10 | 1 |
| 16_319623 | NA | NA | NA | NA | 1 | 1 | 1 | 0 | 2 | 0.5 | 3 | 0.6667 | 2 | 1 | 1 | 1 | 2 | 0.5 | 8 | 0.75 |
| 16_319624 | NA | NA | NA | NA | 1 | 0 | 1 | 1 | 2 | 0.5 | 2 | 1 | 3 | 0 | 1 | 0 | 1 | 1 | 7 | 0.4286 |
| 16_319624 | NA | NA | NA | NA | 1 | 0 | 1 | 1 | 2 | 0.5 | 3 | 0.6667 | 4 | 0.5 | 1 | 1 | 2 | 1 | 10 | 0.7 |

| | | | | | | | | | | | | | | | | | | | | |
|---|---|---|---|---|---|---|---|---|---|---|---|---|---|---|---|---|---|---|---|---|
| 16_319628 | NA | NA | NA | NA | NA | NA | 1 | 1 | 1 | 1 | 1 | 0 | 2 | 0.5 | 1 | 1 | 2 | 0 | 6 | 0.3333 |
| 16_319628 | NA | NA | NA | NA | 1 | 0 | 1 | 1 | 2 | 0.5 | 3 | 0.3333 | 2 | 0 | 1 | 0 | 2 | 0 | 8 | 0.125 |
| 16_319629 | NA | NA | NA | NA | 1 | 1 | NA | NA | 1 | 1 | 2 | 0.5 | 1 | 0 | 1 | 1 | 2 | 1 | 6 | 0.6667 |
| 16_319631 | NA | NA | NA | NA | 1 | 1 | 1 | 1 | 2 | 1 | 2 | 0.5 | 2 | 0.5 | 1 | 0 | 2 | 0 | 7 | 0.2857 |
| 16_319632 | NA | NA | NA | NA | NA | NA | 1 | 1 | 1 | 1 | 2 | 1 | 2 | 0.5 | 1 | 1 | 2 | 1 | 7 | 0.8571 |
| 16_319635 | NA | NA | NA | NA | 1 | 1 | 1 | 1 | 2 | 1 | 2 | 0.5 | 2 | 1 | 1 | 1 | 2 | 1 | 7 | 0.8571 |
| 16_319635 | NA | NA | NA | NA | 1 | 0 | 1 | 1 | 2 | 0.5 | 2 | 0 | 2 | 0 | 1 | 1 | 2 | 0 | 7 | 0.1429 |
| 16_320498 | 4 | 0.25 | 3 | 0 | 2 | 0 | 4 | 0 | 13 | 0.0769 | 5 | 0.2 | NA | NA | 3 | 0 | 3 | 0 | 11 | 0.0909 |
| 16_320500 | 3 | 0.3333 | 4 | 0 | 4 | 0.25 | 3 | 0.3333 | 14 | 0.2143 | 4 | 0 | NA | NA | 2 | 0 | 3 | 0.6667 | 9 | 0.2222 |
| 16_320506 | 6 | 0.1667 | 8 | 0.125 | 3 | 0 | 9 | 0 | 26 | 0.0769 | 9 | 0.1111 | 4 | 0 | 4 | 0 | 10 | 0.1 | 27 | 0.0741 |
| 16_320507 | 7 | 0.5714 | 6 | 0.3333 | 2 | 0.5 | 5 | 0.6 | 20 | 0.5 | 7 | 0.2857 | 4 | 0.25 | 4 | 0.5 | 10 | 0.3 | 25 | 0.32 |
| 16_320509 | 6 | 0 | 8 | 0.125 | 3 | 0 | 3 | 0 | 20 | 0.05 | 8 | 0 | 1 | 0 | 2 | 0.5 | 9 | 0 | 20 | 0.05 |
| 16_320514 | 6 | 0.1667 | 3 | 0 | NA | NA | 3 | 0.6667 | 12 | 0.25 | 5 | 0 | NA | NA | 3 | 0.6667 | 7 | 0.1429 | 15 | 0.2 |
| 16_320517 | 6 | 0.6667 | 1 | 1 | NA | NA | 4 | 0.5 | 11 | 0.6364 | 5 | 0.4 | 2 | 1 | 1 | 0 | 5 | 1 | 13 | 0.6923 |
| 16_320518 | 7 | 0 | 2 | 0 | 1 | 0 | 5 | 0.2 | 15 | 0.0667 | 3 | 0 | 2 | 0 | 3 | 0 | 7 | 0.1429 | 15 | 0.0667 |
| 16_320518 | 6 | 1 | 2 | 0.5 | 1 | 1 | 6 | 1 | 15 | 0.9333 | 4 | 0.75 | 2 | 0.5 | 2 | 1 | 5 | 1 | 13 | 0.8462 |
| 16_320519 | 6 | 0.1667 | 1 | 0 | 1 | 0 | 6 | 0.1667 | 14 | 0.1429 | 4 | 0.5 | 2 | 0 | 3 | 0 | 7 | 0 | 16 | 0.125 |
| 16_320519 | 6 | 1 | 2 | 1 | NA | NA | 5 | 1 | 13 | 1 | 4 | 1 | 2 | 1 | 3 | 1 | 7 | 1 | 16 | 1 |
| 16_320519 | 6 | 1 | 1 | 1 | 1 | 1 | 5 | 0.8 | 13 | 0.9231 | 4 | 1 | 2 | 1 | 3 | 1 | 5 | 1 | 14 | 1 |
| 16_320520 | 6 | 0.8333 | 1 | 1 | NA | NA | 4 | 1 | 11 | 0.9091 | 4 | 1 | 2 | 0.5 | 2 | 1 | 7 | 0.7143 | 15 | 0.8 |
| 16_320572 | 5 | 0.4 | 2 | 1 | 1 | 1 | 5 | 0.6 | 13 | 0.6154 | 4 | 0.5 | NA | NA | 3 | 0.3333 | 6 | 0.6667 | 13 | 0.5385 |
| 16_320574 | 5 | 0.4 | 2 | 1 | NA | NA | 5 | 0.6 | 12 | 0.5833 | 2 | 0.5 | 1 | 1 | 3 | 0.3333 | 6 | 0.6667 | 12 | 0.5833 |
| 16_320706 | NA | NA | NA | NA | NA | NA | 2 | 0.5 | 2 | 0.5 | NA | NA | 1 | 0 | 3 | 0 | 3 | 0.3333 | 7 | 0.1429 |
| 16_320707 | NA | NA | NA | NA | NA | NA | 2 | 0.5 | 2 | 0.5 | NA | NA | 1 | 1 | 3 | 1 | 3 | 0.6667 | 7 | 0.8571 |
| 16_320709 | NA | NA | NA | NA | NA | NA | 2 | 1 | 2 | 1 | NA | NA | 1 | 0 | 3 | 0.3333 | 3 | 1 | 7 | 0.5714 |
| 16_320710 | NA | NA | NA | NA | NA | NA | 2 | 1 | 2 | 1 | NA | NA | 1 | 1 | 2 | 1 | 3 | 1 | 6 | 1 |
| 16_320710 | NA | NA | NA | NA | NA | NA | 2 | 0.5 | 2 | 0.5 | NA | NA | 1 | 1 | 2 | 0 | 3 | 0.6667 | 6 | 0.5 |
| 16_320754 | NA | NA | 1 | 0 | NA | NA | 2 | 0.5 | 3 | 0.3333 | NA | NA | 1 | 0 | 1 | 1 | 3 | 0.3333 | 5 | 0.4 |
| 16_320757 | 1 | 1 | 1 | 1 | NA | NA | 3 | 1 | 5 | 1 | 1 | 1 | NA | NA | 3 | 1 | 1 | 1 | 5 | 1 |
| 16_320967 | 2 | 1 | NA | NA | NA | NA | 1 | 1 | 3 | 1 | 3 | 1 | 1 | 0 | NA | NA | NA | NA | 4 | 0.75 |
| 16_320971 | 3 | 0.6667 | 1 | 0 | NA | NA | 1 | 1 | 5 | 0.6 | 2 | 0.5 | 1 | 0 | NA | NA | NA | NA | 3 | 0.3333 |
| 16_321174 | 2 | 1 | 2 | 1 | NA | NA | 1 | 1 | 5 | 1 | 4 | 1 | 1 | 1 | 3 | 1 | NA | NA | 8 | 1 |
| 16_321177 | 2 | 1 | 2 | 0.5 | NA | NA | NA | NA | 4 | 0.75 | 3 | 0.6667 | 2 | 1 | 2 | 0.5 | NA | NA | 7 | 0.7143 |
| 16_321177 | 3 | 0.3333 | 1 | 0 | NA | NA | 1 | 0 | 5 | 0.2 | 4 | 0.5 | 2 | 0.5 | 1 | 0 | NA | NA | 7 | 0.4286 |
| 16_321177 | 4 | 0.75 | 2 | 0.5 | NA | NA | NA | NA | 6 | 0.6667 | 4 | 0.25 | 2 | 0 | 3 | 0.6667 | NA | NA | 9 | 0.3333 |
| 16_321177 | 2 | 0.5 | 2 | 0.5 | NA | NA | 1 | 0 | 5 | 0.4 | 4 | 0.5 | 2 | 0.5 | 3 | 1 | NA | NA | 9 | 0.6667 |
| 16_321179 | 4 | 0.5 | 2 | 0 | NA | NA | 1 | 0 | 7 | 0.2857 | 3 | 0.3333 | 2 | 0 | 2 | 0 | NA | NA | 7 | 0.1429 |
| 16_321179 | 4 | 0.25 | 2 | 0 | NA | NA | 1 | 0 | 7 | 0.1429 | 3 | 0.3333 | 2 | 0.5 | 3 | 0.3333 | NA | NA | 8 | 0.375 |
| 16_321185 | 4 | 0.25 | 1 | 0 | NA | NA | 1 | 0 | 6 | 0.1667 | 4 | 0.25 | 1 | 0 | 3 | 0 | NA | NA | 8 | 0.125 |
| 16_321198 | 1 | 0 | NA | NA | 1 | 1 | NA | NA | 2 | 0.5 | 1 | 0 | 1 | 1 | NA | NA | NA | NA | 2 | 0.5 |
| 16_322003 | 1 | 1 | NA | NA | NA | NA | NA | NA | 1 | 1 | NA | NA | 1 | 1 | NA | NA | 1 | 1 | 2 | 1 |
| 16_322036 | 1 | 1 | NA | NA | NA | NA | NA | NA | 1 | 1 | NA | NA | 2 | 1 | NA | NA | 1 | 1 | 3 | 1 |
| 16_322037 | 1 | 1 | NA | NA | NA | NA | NA | NA | 1 | 1 | NA | NA | 1 | 1 | NA | NA | 1 | 1 | 2 | 1 |
| 16_322038 | 1 | 1 | NA | NA | NA | NA | NA | NA | 1 | 1 | NA | NA | 2 | 1 | NA | NA | 1 | 1 | 3 | 1 |
| 16_322903 | 1 | 1 | 3 | 0.3333 | 1 | 1 | 1 | 1 | 6 | 0.6667 | NA | NA | 1 | 1 | 1 | 0 | NA | NA | 2 | 0.5 |
| 16_322904 | NA | NA | 3 | 0.3333 | 1 | 1 | 1 | 1 | 5 | 0.6 | NA | NA | 1 | 1 | 1 | 0 | NA | NA | 2 | 0.5 |
| 16_322904 | 1 | 1 | 3 | 0.6667 | 1 | 0 | NA | NA | 5 | 0.6 | NA | NA | 2 | 0.5 | 1 | 1 | NA | NA | 3 | 0.6667 |
| 16_322904 | 1 | 0 | 3 | 0.3333 | 1 | 0 | 1 | 0 | 6 | 0.1667 | NA | NA | 2 | 0.5 | 1 | 0 | NA | NA | 3 | 0.3333 |
| 16_322905 | 1 | 1 | 3 | 0.3333 | 1 | 1 | 1 | 1 | 6 | 0.6667 | NA | NA | 1 | 1 | 1 | 0 | NA | NA | 2 | 0.5 |
| 16_322908 | 1 | 1 | 3 | 1 | 1 | 1 | 2 | 1 | 7 | 1 | NA | NA | 1 | 1 | 1 | 1 | NA | NA | 2 | 1 |
| 16_322911 | 1 | 1 | 3 | 0 | 1 | 0 | 1 | 1 | 6 | 0.3333 | NA | NA | 2 | 0.5 | NA | NA | NA | NA | 2 | 0.5 |
| 16_322928 | 2 | 0.5 | 4 | 0 | 3 | 0 | 4 | 0 | 13 | 0.0769 | 1 | 0 | 5 | 0.2 | 1 | 0 | 2 | 0 | 9 | 0.1111 |
| 16_324490 | NA | NA | 3 | 1 | 3 | 1 | 2 | 1 | 8 | 1 | 4 | 0.5 | 2 | 1 | 3 | 1 | 4 | 0.75 | 13 | 0.7692 |
| 16_324889 | NA | NA | 3 | 0 | 5 | 0 | 7 | 0.1429 | 15 | 0.0667 | 3 | 0 | 3 | 0 | 3 | 0.3333 | NA | NA | 9 | 0.1111 |
| 16_324902 | 1 | 0 | 2 | 0 | 4 | 0.25 | 6 | 0 | 13 | 0.0769 | 4 | 0 | 4 | 0 | 4 | 0.25 | NA | NA | 12 | 0.0833 |
| 16_324985 | 1 | 1 | 4 | 0.5 | 11 | 0.2727 | 9 | 0.3333 | 25 | 0.36 | 4 | 0.5 | 4 | 0.5 | 6 | 0.5 | 3 | 0.3333 | 17 | 0.4706 |
| 16_324986 | NA | NA | 2 | 0.5 | 10 | 0.5 | 9 | 0.5556 | 21 | 0.5238 | 3 | 0.3333 | 5 | 0.4 | 9 | 0.5556 | 2 | 0 | 19 | 0.4211 |
| 16_324986 | 1 | 0 | 6 | 0.1667 | 11 | 0.0909 | 13 | 0.3077 | 31 | 0.1935 | 5 | 0.2 | 7 | 0 | 9 | 0.1111 | 3 | 0.3333 | 24 | 0.125 |
| 16_324988 | 2 | 1 | 6 | 0.5 | 12 | 0.5 | 13 | 0.6154 | 33 | 0.5758 | 5 | 0.6 | 6 | 0.3333 | 10 | 0.6 | 3 | 0.6667 | 24 | 0.5417 |
| 16_324990 | 3 | 0 | 4 | 0.25 | 12 | 0.25 | 9 | 0.1111 | 28 | 0.1786 | 7 | 0.4286 | 6 | 0 | 10 | 0 | 3 | 0 | 26 | 0.1154 |
| 16_324990 | 3 | 0 | 5 | 0.8 | 13 | 0.3846 | 13 | 0.1538 | 34 | 0.3235 | 5 | 0.4 | 9 | 0.3333 | 10 | 0.2 | 2 | 0.5 | 26 | 0.3077 |
| 16_324991 | 3 | 0 | 6 | 0.3333 | 16 | 0.1875 | 14 | 0.2857 | 39 | 0.2308 | 7 | 0.1429 | 11 | 0.0909 | 10 | 0.1 | 3 | 0 | 31 | 0.0968 |
| 16_324991 | 3 | 0 | 8 | 0.25 | 15 | 0.3333 | 14 | 0 | 40 | 0.175 | 10 | 0.1 | 10 | 0.1 | 11 | 0.1818 | 3 | 0 | 34 | 0.1176 |
| 16_325018 | 12 | 0.3333 | 15 | 0.1333 | 18 | 0.1667 | 15 | 0.1333 | 60 | 0.1833 | 11 | 0.1818 | 21 | 0.1429 | 19 | 0.1053 | 10 | 0.1 | 61 | 0.1311 |
| 16_325018 | 10 | 0.4 | 13 | 0.4615 | 13 | 0.3077 | 14 | 0.5 | 50 | 0.42 | 13 | 0.2308 | 19 | 0.4737 | 13 | 0.3077 | 7 | 0.4286 | 52 | 0.3654 |

| | | | | | | | | | | | | | | | | | | | | |
|---|---|---|---|---|---|---|---|---|---|---|---|---|---|---|---|---|---|---|---|---|
| 16_3250200 | 8 | 0.25 | 13 | 0.3077 | 16 | 0.25 | 19 | 0.3158 | 56 | 0.2857 | 8 | 0.25 | 16 | 0.1875 | 20 | 0.15 | 5 | 0.2 | 49 | 0.1837 |
| 16_3250203 | 10 | 0.3 | 15 | 0.6667 | 19 | 0.5789 | 15 | 0.4667 | 59 | 0.5254 | 11 | 0.8182 | 18 | 0.6111 | 16 | 0.3125 | 8 | 0.5 | 53 | 0.5472 |
| 16_3250206 | 12 | 0.3333 | 17 | 0.2353 | 22 | 0.2727 | 18 | 0.2222 | 69 | 0.2609 | 17 | 0.0588 | 21 | 0.2381 | 21 | 0.0476 | 11 | 0.0909 | 70 | 0.1143 |
| 16_3264277 | 5 | 0.6 | 12 | 0.1667 | 9 | 0.2222 | 6 | 0.3333 | 32 | 0.2812 | 5 | 0.2 | 4 | 0.25 | 11 | 0.2727 | 6 | 0.1667 | 26 | 0.2308 |
| 16_3264393 | 7 | 0.1429 | 8 | 0.125 | 8 | 0 | 7 | 0.2857 | 30 | 0.1333 | 3 | 0.6667 | 5 | 0.4 | 8 | 0.25 | 3 | 0 | 19 | 0.3158 |
| 16_3264412 | 6 | 0.1667 | 8 | 0.125 | 8 | 0.5 | 6 | 0.3333 | 28 | 0.2857 | 4 | 0.25 | 7 | 0.1429 | 6 | 0.1667 | 2 | 0 | 19 | 0.1579 |
| 16_3264428 | 5 | 1 | 4 | 0.75 | 3 | 0.6667 | 4 | 0.75 | 16 | 0.8125 | 4 | 0.75 | 6 | 0.8333 | 4 | 0.5 | 2 | 1 | 16 | 0.75 |
| 16_3266362 | 1 | 1 | 3 | 0.3333 | 1 | 1 | NA | NA | 5 | 0.6 | 1 | 0 | NA | NA | 2 | 1 | NA | NA | 3 | 0.6667 |
| 16_3272598 | 11 | 0.3636 | 5 | 0.4 | 4 | 0.25 | 11 | 0.2727 | 31 | 0.3226 | 3 | 0 | 7 | 0.5714 | 8 | 0 | 8 | 0.5 | 26 | 0.3077 |
| 16_3272601 | 9 | 0.7778 | 6 | 0.6667 | 4 | 0.5 | 11 | 0.9091 | 30 | 0.7667 | 3 | 0.6667 | 6 | 0.6667 | 6 | 0.3333 | 8 | 0.5 | 23 | 0.5217 |
| 16_3272602 | 11 | 0.5455 | 4 | 0.75 | 4 | 0.75 | 10 | 0.6 | 29 | 0.6207 | 2 | 1 | 9 | 0.8889 | 6 | 0.5 | 8 | 0.625 | 25 | 0.72 |
| 16_3272608 | 12 | 0.0833 | 6 | 0.1667 | 3 | 0 | 12 | 0.1667 | 33 | 0.1212 | 3 | 0.3333 | 9 | 0.2222 | 6 | 0.5 | 9 | 0.1111 | 27 | 0.2593 |
| 16_3272611 | 11 | 0.2727 | 4 | 0 | 4 | 0.5 | 12 | 0.3333 | 31 | 0.2903 | 3 | 0.3333 | 8 | 0.125 | 8 | 0.25 | 7 | 0 | 26 | 0.1538 |
| 16_3272624 | 11 | 0.1818 | 5 | 0 | 4 | 0 | 10 | 0.1 | 30 | 0.1 | 3 | 0 | 8 | 0.125 | 7 | 0.1429 | 10 | 0.1 | 28 | 0.1071 |
| 16_3272629 | 10 | 0.2 | 6 | 0.1667 | 4 | 0.5 | 11 | 0.2727 | 31 | 0.2581 | 4 | 0.25 | 8 | 0.5 | 9 | 0.2222 | 10 | 0.3 | 31 | 0.3226 |
| 16_3272644 | 11 | 0.1818 | 6 | 0.1667 | 5 | 0 | 9 | 0 | 31 | 0.0968 | 3 | 0 | 10 | 0.2 | 8 | 0.125 | 10 | 0.3 | 31 | 0.1935 |
| 16_3272676 | 11 | 0.0909 | 7 | 0.1429 | 4 | 0 | 12 | 0.25 | 34 | 0.1471 | 2 | 0 | 10 | 0.5 | 8 | 0.25 | 9 | 0.4444 | 29 | 0.3793 |
| 16_3272677 | 12 | 0.1667 | 7 | 0.1429 | 4 | 0 | 9 | 0.3333 | 32 | 0.1875 | 3 | 0 | 10 | 0.3 | 9 | 0.2222 | 9 | 0 | 31 | 0.1613 |
| 16_3272763 | 15 | 0.1333 | 8 | 0 | 6 | 0 | 7 | 0 | 36 | 0.0556 | 3 | 0 | 9 | 0 | 5 | 0.2 | 10 | 0.3 | 27 | 0.1481 |
| 16_3272772 | 14 | 0.0714 | 11 | 0 | 4 | 0.25 | 11 | 0.0909 | 40 | 0.075 | 3 | 0.6667 | 10 | 0 | 7 | 0 | 11 | 0.0909 | 31 | 0.0968 |
| 16_3283058 | NA | NA | NA | NA | NA | NA | 1 | 1 | 1 | 1 | NA | NA | 1 | 1 | 3 | 0.3333 | NA | NA | 4 | 0.5 |
| 16_3283089 | NA | NA | NA | NA | NA | NA | 2 | 0.5 | 2 | 0.5 | NA | NA | 1 | 0 | 2 | 0.5 | NA | NA | 3 | 0.3333 |
| 16_3283286 | 2 | 1 | 4 | 0.75 | 4 | 0.5 | 4 | 0.5 | 14 | 0.6429 | 2 | 1 | 3 | 1 | 2 | 0 | 2 | 0.5 | 9 | 0.6667 |
| 16_3283292 | 2 | 0.5 | 4 | 0.75 | 4 | 0.25 | 5 | 0.4 | 15 | 0.4667 | 2 | 0.5 | 3 | 0.3333 | 2 | 1 | 2 | 0 | 9 | 0.4444 |
| 16_3283309 | 1 | 1 | 4 | 0.75 | 3 | 0.6667 | 6 | 0.1667 | 14 | 0.5 | 1 | 0 | 2 | 0 | 2 | 0 | 2 | 0.5 | 7 | 0.1429 |
| 16_3283312 | 2 | 0 | 4 | 0.5 | 3 | 0 | 6 | 0.1667 | 15 | 0.2 | 2 | 0 | 3 | 0.3333 | 2 | 0.5 | 1 | 0 | 8 | 0.25 |
| 16_3283321 | 2 | 1 | 3 | 0.3333 | 3 | 1 | 6 | 0.8333 | 14 | 0.7857 | 2 | 1 | 2 | 0.5 | 1 | 1 | 2 | 1 | 7 | 0.8571 |
| 16_3283321 | 2 | 0 | 3 | 0.3333 | 4 | 0 | 6 | 0 | 15 | 0.0667 | 2 | 1 | 3 | 0.3333 | 2 | 0 | 1 | 0 | 8 | 0.375 |
| 16_3283327 | 1 | 1 | 1 | 1 | 3 | 1 | 4 | 1 | 9 | 1 | 1 | 1 | 2 | 1 | 2 | 0.5 | 1 | 1 | 6 | 0.8333 |
| 16_3283335 | 2 | 1 | 4 | 1 | 3 | 0.6667 | 6 | 0.8333 | 15 | 0.8667 | 1 | 1 | 3 | 0.6667 | 1 | 1 | 1 | 1 | 6 | 0.8333 |
| 16_3283347 | 2 | 0 | 3 | 0.3333 | 3 | 0.3333 | 5 | 0.6 | 13 | 0.3846 | 2 | 0.5 | 3 | 0.3333 | 2 | 1 | 2 | 0.5 | 9 | 0.5556 |
| 16_3283347 | 2 | 1 | 3 | 1 | 4 | 1 | 5 | 1 | 14 | 1 | 2 | 1 | 2 | 1 | 2 | 1 | 2 | 0.5 | 8 | 0.875 |
| 16_3283360 | 3 | 0 | 5 | 0.6 | 4 | 0.25 | 6 | 0.5 | 18 | 0.3889 | 2 | 0 | 3 | 0.3333 | 2 | 0 | 2 | 0 | 9 | 0.1111 |
| 16_3283363 | 1 | 0 | 1 | 0 | 3 | 0.3333 | 6 | 0.1667 | 11 | 0.1818 | 2 | 0.5 | 3 | 0 | 1 | 0 | 2 | 0 | 8 | 0.125 |
| 16_3283363 | 1 | 0 | 5 | 0.6 | 3 | 0 | 6 | 0 | 15 | 0.2 | 2 | 0.5 | 3 | 0.3333 | 2 | 0.5 | 2 | 0 | 9 | 0.3333 |
| 16_3283371 | 3 | 0.6667 | 4 | 0.25 | 3 | 0.3333 | 5 | 0 | 15 | 0.2667 | 2 | 0 | 3 | 0 | 2 | 0.5 | 2 | 0 | 9 | 0.1111 |
| 16_3283374 | 3 | 0.3333 | 5 | 0.4 | 4 | 0.25 | 6 | 0 | 18 | 0.2222 | 2 | 0 | 2 | 0 | 2 | 0.5 | 2 | 0 | 8 | 0.125 |
| 16_3283375 | 3 | 0 | 5 | 0.2 | 4 | 0.25 | 6 | 0 | 18 | 0.1111 | 2 | 1 | 3 | 0 | 2 | 0.5 | 2 | 0 | 9 | 0.3333 |
| 16_3283377 | 3 | 0.6667 | 3 | 1 | 4 | 1 | 5 | 1 | 15 | 0.9333 | 2 | 1 | 3 | 1 | 2 | 1 | 2 | 0 | 9 | 0.7778 |
| 16_3283379 | 2 | 1 | 4 | 1 | 4 | 0.75 | 4 | 0.75 | 14 | 0.8571 | 1 | 1 | 3 | 1 | 1 | 1 | 1 | 1 | 6 | 1 |
| 16_3283392 | 3 | 0.6667 | 5 | 0 | 4 | 0 | 5 | 0 | 17 | 0.1176 | 2 | 0 | 3 | 0.3333 | 2 | 0 | 2 | 0 | 9 | 0.1111 |
| 16_3283400 | 3 | 1 | 5 | 1 | 4 | 0.75 | 5 | 0.8 | 17 | 0.8824 | NA | NA | 3 | 1 | 1 | 1 | 3 | 1 | 7 | 1 |
| 16_3283418 | 3 | 0.3333 | 5 | 0 | 5 | 0.2 | 5 | 0.2 | 18 | 0.1667 | 2 | 0 | 2 | 0.5 | 2 | 0 | 3 | 0.6667 | 9 | 0.3333 |
| 16_3283433 | 2 | 0 | 6 | 0.3333 | 4 | 0.25 | 5 | 0.6 | 17 | 0.3529 | 3 | 0.3333 | 1 | 0 | 2 | 0.5 | 3 | 0 | 9 | 0.2222 |
| 16_3283514 | 7 | 0.1429 | 4 | 0.25 | 8 | 0.375 | 8 | 0 | 27 | 0.1852 | 4 | 0.25 | 4 | 0 | 3 | 0.3333 | 3 | 0 | 14 | 0.1429 |
| 16_3283552 | 6 | 0.5 | 4 | 0 | 8 | 0.25 | 8 | 0.25 | 26 | 0.2692 | 3 | 0 | 3 | 0.3333 | 3 | 0.3333 | 4 | 0.25 | 13 | 0.2308 |
| 16_3283554 | 6 | 0.1667 | 5 | 0 | 7 | 0 | 7 | 0.2857 | 25 | 0.12 | 3 | 0 | 4 | 0 | 3 | 0.3333 | 2 | 0 | 12 | 0.0833 |
| 16_3283570 | 7 | 0.2857 | 5 | 0.6 | 7 | 0.5714 | 8 | 0.25 | 27 | 0.4074 | 4 | 0.25 | 3 | 1 | 3 | 0.6667 | 4 | 0.5 | 14 | 0.5714 |
| 16_3294405 | NA | NA | NA | NA | 1 | 1 | 1 | 0 | 2 | 0.5 | 1 | 0 | NA | NA | 3 | 0.3333 | NA | NA | 4 | 0.25 |
| 16_3294412 | NA | NA | NA | NA | 1 | 1 | 1 | 1 | 2 | 1 | 1 | 1 | NA | NA | 2 | 1 | NA | NA | 3 | 1 |
| 16_3294413 | NA | NA | NA | NA | 1 | 1 | NA | NA | 1 | 1 | 1 | 1 | NA | NA | 3 | 0.3333 | NA | NA | 4 | 0.5 |
| 16_3294420 | NA | NA | NA | NA | 1 | 1 | 1 | 1 | 2 | 1 | 1 | 1 | NA | NA | 3 | 1 | NA | NA | 4 | 1 |
| 16_3294451 | NA | NA | NA | NA | 1 | 0 | 1 | 1 | 2 | 0.5 | 1 | 1 | NA | NA | 1 | 1 | NA | NA | 2 | 1 |
| 16_3294452 | NA | NA | NA | NA | 1 | 1 | 1 | 1 | 2 | 1 | 1 | 1 | NA | NA | 1 | 1 | NA | NA | 2 | 1 |
| 16_3294483 | NA | NA | NA | NA | 1 | 1 | 2 | 1 | 3 | 1 | 1 | 1 | NA | NA | 2 | 1 | NA | NA | 3 | 1 |
| 16_3294498 | NA | NA | NA | NA | 1 | 1 | 2 | 0.5 | 3 | 0.6667 | 1 | 1 | NA | NA | 2 | 1 | NA | NA | 3 | 1 |
| 16_3294520 | NA | NA | NA | NA | 1 | 0 | 2 | 0.5 | 3 | 0.3333 | 1 | 0 | NA | NA | 2 | 0.5 | NA | NA | 3 | 0.3333 |
| 16_3294552 | NA | NA | NA | NA | 1 | 0 | 2 | 0.5 | 3 | 0.3333 | 1 | 0 | NA | NA | 2 | 0.5 | NA | NA | 3 | 0.3333 |
| 16_3300600 | NA | NA | 1 | 1 | NA | NA | 1 | 0 | 2 | 0.5 | 1 | 1 | NA | NA | 1 | 0 | NA | NA | 2 | 0.5 |
| 16_3300617 | NA | NA | 1 | 1 | NA | NA | 1 | 1 | 2 | 1 | 1 | 1 | NA | NA | 1 | 1 | NA | NA | 2 | 1 |
| 16_3304733 | 1 | 1 | 1 | 0 | NA | NA | 3 | 0 | 5 | 0.2 | 3 | 0.3333 | 1 | 0 | 2 | 0 | 1 | 0 | 7 | 0.1429 |
| 16_3304756 | 1 | 1 | NA | NA | NA | NA | 3 | 0.6667 | 4 | 0.75 | 2 | 0.5 | 1 | 1 | 2 | 1 | 1 | 1 | 6 | 0.8333 |
| 16_3304767 | 2 | 0.5 | 1 | 1 | NA | NA | 2 | 0 | 5 | 0.4 | 1 | 1 | 1 | 0 | NA | NA | 1 | 0 | 3 | 0.3333 |
| 16_3306218 | 3 | 1 | 1 | 0 | NA | NA | 2 | 1 | 6 | 0.8333 | 2 | 1 | 1 | 0 | NA | NA | 2 | 1 | 5 | 0.8 |
| 16_3306218 | 2 | 1 | NA | NA | NA | NA | 1 | 1 | 3 | 1 | 3 | 0.3333 | NA | NA | 2 | 0 | NA | NA | 5 | 0.2 |

| | | | | | | | | | | | | | | | | | | | | |
|---|---|---|---|---|---|---|---|---|---|---|---|---|---|---|---|---|---|---|---|---|
| 16_3306225 | 3 | 0.6667 | 1 | 0 | NA | NA | 3 | 0.3333 | 7 | 0.4286 | 2 | 0.5 | 1 | 0 | 3 | 0.3333 | 1 | 1 | 7 | 0.4286 |
| 16_3306226 | 3 | 0.6667 | 1 | 1 | NA | NA | 3 | 1 | 7 | 0.8571 | 3 | 0.3333 | NA | NA | 3 | 0.3333 | 2 | 0.5 | 8 | 0.375 |
| 16_3306233 | 1 | 1 | NA | NA | NA | NA | 2 | 0.5 | 3 | 0.6667 | 3 | 1 | 1 | 1 | 3 | 1 | 1 | 1 | 8 | 1 |
| 16_3306246 | 3 | 0.3333 | 1 | 0 | NA | NA | 3 | 0 | 7 | 0.1429 | 2 | 0 | 1 | 0 | 2 | 0 | 2 | 0.5 | 7 | 0.1429 |
| 16_3306247 | 3 | 1 | 1 | 0 | NA | NA | 1 | 1 | 5 | 0.8 | 3 | 1 | NA | NA | 3 | 0.3333 | 2 | 1 | 8 | 0.75 |
| 16_3306248 | 2 | 1 | 1 | 1 | NA | NA | 3 | 0.6667 | 6 | 0.8333 | 1 | 1 | NA | NA | 2 | 1 | 1 | 1 | 4 | 1 |
| 16_3306272 | 2 | 1 | 1 | 1 | NA | NA | 3 | 1 | 6 | 1 | 2 | 1 | 2 | 1 | 2 | 1 | NA | NA | 6 | 1 |
| 16_3306278 | 1 | 0 | 1 | 0 | NA | NA | 2 | 0.5 | 4 | 0.25 | 4 | 0.5 | 2 | 1 | 1 | 1 | 1 | 1 | 8 | 0.75 |
| 16_3306283 | 3 | 0.3333 | 1 | 0 | NA | NA | 4 | 0.25 | 8 | 0.25 | 4 | 0.25 | 2 | 0 | 3 | 0.6667 | 2 | 0 | 11 | 0.2727 |
| 16_3306287 | 3 | 0.6667 | NA | NA | NA | NA | 3 | 0.3333 | 6 | 0.5 | 4 | 0.5 | 2 | 0.5 | 3 | 0.6667 | 2 | 0.5 | 11 | 0.5455 |
| 16_3306293 | 3 | 0.3333 | 1 | 1 | NA | NA | 4 | 0.25 | 8 | 0.375 | 4 | 0 | 2 | 0.5 | 3 | 0 | 2 | 0.5 | 11 | 0.1818 |
| 16_3306318 | 2 | 0.5 | NA | NA | NA | NA | 5 | 0.4 | 7 | 0.4286 | 5 | 0.2 | 3 | 0.3333 | 2 | 0 | 2 | 0 | 12 | 0.1667 |
| 16_3311904 | 8 | 0.625 | 1 | 0 | 5 | 0 | 1 | 1 | 15 | 0.4 | 4 | 0.25 | 4 | 0.25 | NA | NA | 7 | 0.1429 | 15 | 0.2 |
| 16_3325513 | 1 | 0 | 5 | 0 | 3 | 0 | 5 | 0.2 | 14 | 0.0714 | 1 | 0 | 2 | 0.5 | 2 | 0.5 | 3 | 0.3333 | 8 | 0.375 |
| 16_3325524 | 1 | 0 | 5 | 0 | 2 | 0.5 | 3 | 0.3333 | 11 | 0.1818 | 1 | 1 | 2 | 0.5 | 3 | 0.3333 | 4 | 0.75 | 10 | 0.6 |
| 16_3325532 | 1 | 0 | 5 | 0 | 3 | 0 | 6 | 0.1667 | 15 | 0.0667 | 1 | 0 | 2 | 0.5 | 2 | 0 | 5 | 0 | 10 | 0.1 |
| 16_3331612 | 2 | 0 | 7 | 0.1429 | 4 | 0.25 | 4 | 0 | 17 | 0.1176 | 1 | 0 | 2 | 0 | 3 | 0.3333 | 5 | 0.2 | 11 | 0.1818 |
| 16_3331618 | 1 | 0 | 3 | 0.3333 | 2 | 0.5 | 4 | 1 | 10 | 0.6 | NA | NA | 2 | 0 | 3 | 0.3333 | 4 | 0 | 9 | 0.1111 |
| 16_3331628 | 2 | 0.5 | 6 | 0.5 | 5 | 0.2 | 7 | 0 | 20 | 0.25 | NA | NA | 2 | 0.5 | 2 | 0.5 | 5 | 0 | 9 | 0.2222 |
| 16_3331631 | 2 | 0 | 6 | 0 | 4 | 0 | 8 | 0.125 | 20 | 0.05 | 1 | 0 | 2 | 0 | 3 | 0.3333 | 4 | 0.25 | 10 | 0.2 |
| 16_3331633 | 2 | 0 | 4 | 0.25 | 4 | 0.25 | 8 | 0.125 | 18 | 0.1667 | 1 | 0 | 2 | 0.5 | 3 | 0.3333 | 6 | 0.3333 | 12 | 0.3333 |
| 16_3331663 | 3 | 1 | 8 | 0.75 | 4 | 1 | 7 | 1 | 22 | 0.9091 | 1 | 1 | 2 | 1 | 3 | 1 | 8 | 1 | 14 | 1 |
| 16_3331695 | 3 | 0.3333 | 8 | 0.25 | 4 | 0 | 9 | 0.2222 | 24 | 0.2083 | 1 | 0 | 4 | 0 | 5 | 0.6 | 7 | 0.1429 | 17 | 0.2353 |
| 16_3331699 | 4 | 0.75 | 5 | 0.8 | 3 | 1 | 6 | 1 | 18 | 0.8889 | 2 | 1 | 3 | 1 | 4 | 0.75 | 7 | 0.8571 | 16 | 0.875 |
| 16_3331703 | 4 | 0.75 | 7 | 0.2857 | 4 | 0.5 | 9 | 0.4444 | 24 | 0.4583 | 2 | 1 | 4 | 0.5 | 5 | 0.4 | 8 | 0.625 | 19 | 0.5789 |
| 16_3331717 | 4 | 1 | 8 | 0.625 | 3 | 1 | 6 | 1 | 21 | 0.8571 | 2 | 1 | 3 | 1 | 5 | 1 | 8 | 1 | 18 | 1 |
| 16_3331717 | 3 | 0.3333 | 5 | 0.4 | 4 | 0.5 | 5 | 0.2 | 17 | 0.3529 | 2 | 0.5 | 4 | 0.75 | 4 | 0 | 7 | 0.4286 | 17 | 0.4118 |
| 16_3331721 | 3 | 0.3333 | 7 | 0 | 5 | 0.2 | 8 | 0.125 | 23 | 0.1304 | 2 | 0 | 3 | 0 | 5 | 0.2 | 7 | 0 | 17 | 0.0588 |
| 16_3331722 | 3 | 1 | 8 | 0.75 | 4 | 1 | 7 | 1 | 22 | 0.9091 | 2 | 1 | 2 | 1 | 3 | 1 | 6 | 1 | 13 | 1 |
| 16_3331733 | 4 | 0.25 | 7 | 0.4286 | 3 | 0 | 7 | 0.1429 | 21 | 0.2381 | 2 | 0 | 4 | 0.75 | 5 | 0.2 | 5 | 0 | 16 | 0.25 |
| 16_3331736 | 4 | 1 | 8 | 1 | 4 | 1 | 6 | 1 | 22 | 1 | 2 | 1 | 4 | 1 | 5 | 1 | 6 | 1 | 17 | 1 |
| 16_3331737 | 3 | 0.6667 | 8 | 0.625 | 4 | 1 | 6 | 0.5 | 21 | 0.6667 | 2 | 0.5 | 3 | 1 | 5 | 1 | 7 | 0.8571 | 17 | 0.8824 |
| 16_3331744 | 3 | 1 | 5 | 1 | 4 | 1 | 8 | 0.75 | 20 | 0.9 | 2 | 0.5 | 4 | 1 | 4 | 0.75 | 8 | 0.625 | 18 | 0.7222 |
| 16_3331758 | 4 | 0.75 | 8 | 1 | 5 | 1 | 8 | 1 | 25 | 0.96 | 2 | 1 | 3 | 1 | 4 | 1 | 5 | 1 | 14 | 1 |
| 16_3331759 | 3 | 0.6667 | 7 | 0.7143 | 1 | 1 | 9 | 0.4444 | 20 | 0.6 | 1 | 1 | 2 | 1 | 3 | 0.6667 | 6 | 0.6667 | 12 | 0.75 |
| 16_3331760 | 4 | 0.5 | 8 | 0.125 | 4 | 0.5 | 8 | 0.25 | 24 | 0.2917 | 2 | 0 | 4 | 0.25 | 5 | 0.2 | 8 | 0.25 | 19 | 0.2105 |
| 16_3331761 | 4 | 1 | 8 | 1 | 5 | 1 | 9 | 0.8889 | 26 | 0.9615 | 2 | 1 | 3 | 0.6667 | 5 | 0.8 | 8 | 1 | 18 | 0.8889 |
| 16_3331770 | 4 | 0.5 | 6 | 0.5 | 5 | 1 | 8 | 0.625 | 23 | 0.6522 | 2 | 1 | 4 | 0.75 | 5 | 0.6 | 9 | 0.7778 | 20 | 0.75 |
| 16_3331775 | 3 | 0.3333 | 7 | 0 | 3 | 0.3333 | 8 | 0.375 | 21 | 0.2381 | 2 | 0.5 | 3 | 0 | 5 | 0.4 | 8 | 0.25 | 18 | 0.2778 |
| 16_3331785 | 3 | 0 | 5 | 0 | 4 | 0 | 6 | 0.1667 | 18 | 0.0556 | NA | NA | 4 | 0.25 | 2 | 0 | 5 | 0.2 | 11 | 0.1818 |
| 16_3331789 | 3 | 0.3333 | 6 | 0.5 | 6 | 0.8333 | 8 | 0.75 | 23 | 0.6522 | 1 | 0 | 3 | 0.6667 | 4 | 1 | 7 | 0.7143 | 15 | 0.7333 |
| 16_3331798 | 3 | 1 | 8 | 0.75 | 7 | 0.8571 | 9 | 0.8889 | 27 | 0.8519 | 3 | 0.6667 | 4 | 1 | 4 | 1 | 7 | 0.7143 | 18 | 0.8333 |
| 16_3331813 | 5 | 0 | 9 | 0.1111 | 8 | 0.25 | 9 | 0.4444 | 31 | 0.2258 | 2 | 0 | 4 | 0.25 | 5 | 0.2 | 8 | 0.125 | 19 | 0.1579 |
| 16_3331875 | 2 | 0.5 | 6 | 0 | 7 | 0.2857 | 8 | 0.5 | 23 | 0.3043 | 2 | 0 | 5 | 0 | 4 | 0.5 | 3 | 0 | 14 | 0.1429 |
| 16_3331879 | 1 | 1 | 7 | 0.8571 | 5 | 0.6 | 5 | 0.4 | 18 | 0.6667 | 2 | 0.5 | 4 | 0.75 | 4 | 1 | 2 | 1 | 12 | 0.8333 |
| 16_3332121 | 3 | 0.3333 | 5 | 0.4 | 5 | 0.2 | 5 | 0 | 18 | 0.2222 | 5 | 0.2 | 4 | 0.25 | 4 | 0.25 | 3 | 0 | 16 | 0.1875 |
| 16_3332236 | 4 | 0.5 | 7 | 0.1429 | 7 | 0.1429 | 7 | 0 | 25 | 0.16 | 5 | 0 | 4 | 0 | 5 | 0.2 | 4 | 0.25 | 18 | 0.1111 |
| 16_3332245 | 4 | 0.5 | 7 | 0 | 7 | 0.2857 | 7 | 0.1429 | 25 | 0.2 | 4 | 0 | 4 | 0.25 | 5 | 0 | 3 | 0.3333 | 16 | 0.125 |
| 16_3332335 | 4 | 0.25 | 7 | 0.1429 | 4 | 0 | 6 | 0.1667 | 21 | 0.1429 | 4 | 0 | 2 | 0 | 3 | 0.6667 | 2 | 0 | 11 | 0.1818 |
| 16_3332403 | 2 | 0 | 5 | 0.4 | 6 | 0.5 | 1 | 0 | 14 | 0.3571 | 3 | 0.3333 | 4 | 0 | 1 | 0 | 1 | 0 | 9 | 0.1111 |
| 16_3354010 | NA | NA | NA | NA | NA | NA | 1 | 1 | 1 | 1 | NA | NA | 1 | 1 | 1 | 0 | NA | NA | 2 | 0.5 |
| 16_3354126 | 1 | 0 | 1 | 1 | NA | NA | NA | NA | 2 | 0.5 | NA | NA | 1 | 1 | 1 | 0 | NA | NA | 2 | 0.5 |
| 16_3354685 | 1 | 1 | 2 | 0.5 | 1 | 1 | 1 | 1 | 5 | 0.8 | NA | NA | 1 | 1 | 1 | 0 | NA | NA | 2 | 0.5 |
| 16_3354703 | 1 | 1 | 1 | 1 | 1 | 1 | 1 | 1 | 4 | 1 | 1 | 0 | 1 | 1 | 1 | 0 | NA | NA | 3 | 0.3333 |
| 16_3354741 | 1 | 1 | 1 | 1 | 1 | 1 | 2 | 0.5 | 5 | 0.8 | 1 | 1 | 1 | 1 | 2 | 1 | NA | NA | 4 | 1 |
| 16_3358706 | NA | NA | 2 | 0 | 1 | 1 | 2 | 1 | 5 | 0.6 | 1 | 1 | 1 | 0 | 2 | 0.5 | 1 | 1 | 5 | 0.6 |
| 16_3358709 | 1 | 0 | 1 | 1 | 1 | 0 | 1 | 1 | 4 | 0.5 | 1 | 1 | 1 | 1 | 1 | 0 | 1 | 1 | 4 | 0.75 |
| 16_3358710 | 1 | 0 | 2 | 0 | 1 | 1 | 2 | 0.5 | 6 | 0.3333 | 1 | 0 | 1 | 0 | 2 | 0.5 | 1 | 0 | 5 | 0.2 |
| 16_3358712 | 1 | 0 | 3 | 1 | 1 | 1 | 1 | 0 | 6 | 0.6667 | 1 | 1 | 1 | 1 | 2 | 0 | 1 | 1 | 5 | 0.6 |
| 16_3358728 | 1 | 1 | 3 | 0.6667 | 1 | 1 | 1 | 1 | 6 | 0.8333 | 1 | 1 | 1 | 1 | 1 | 1 | 1 | 1 | 4 | 1 |
| 16_3358734 | 1 | 1 | 3 | 0.3333 | 1 | 1 | 1 | 1 | 6 | 0.6667 | 1 | 0 | 1 | 0 | 2 | 0.5 | 1 | 0 | 5 | 0.2 |
| 16_3358748 | 1 | 1 | 4 | 0.5 | 1 | 1 | 1 | 0 | 7 | 0.5714 | 1 | 0 | 1 | 1 | 2 | 0 | 1 | 1 | 5 | 0.4 |
| 16_3358753 | 1 | 0 | 3 | 0.3333 | 1 | 0 | 1 | 0 | 6 | 0.1667 | 1 | 1 | 1 | 0 | 2 | 0.5 | 1 | 0 | 5 | 0.4 |
| 16_3358818 | 2 | 1 | 3 | 0 | 1 | 1 | 1 | 1 | 7 | 0.5714 | 1 | 0 | 1 | 0 | 2 | 0.5 | 1 | 0 | 5 | 0.2 |
| 16_3358827 | 2 | 0.5 | 1 | 1 | 1 | 0 | 2 | 0 | 6 | 0.3333 | 2 | 0.5 | 1 | 0 | 2 | 0.5 | 1 | 1 | 6 | 0.5 |

| | | | | | | | | | | | | | | | | | | | | |
|---|---|---|---|---|---|---|---|---|---|---|---|---|---|---|---|---|---|---|---|---|
| 16_3358836 | 2 | 1 | 3 | 0 | 1 | 0 | 1 | 0 | 7 | 0.2857 | 2 | 0 | 1 | 0 | 2 | 0 | 1 | 1 | 6 | 0.1667 |
| 16_3358849 | 2 | 0 | 3 | 0 | 2 | 0.5 | 2 | 0 | 9 | 0.1111 | 1 | 0 | 1 | 1 | 2 | 0.5 | 1 | 0 | 5 | 0.4 |
| 16_3358862 | 1 | 0 | 3 | 0 | 2 | 0.5 | 2 | 1 | 8 | 0.375 | 2 | 1 | 1 | 1 | 2 | 1 | 1 | 0 | 6 | 0.8333 |
| 16_3374024 | 6 | 0.6667 | 2 | 0.5 | 5 | 0.4 | 6 | 0.8333 | 19 | 0.6316 | 3 | 0.6667 | 2 | 0 | NA | NA | 2 | 1 | 7 | 0.5714 |
| 16_3374040 | 6 | 0 | 3 | 0 | 6 | 0 | 7 | 0.4286 | 22 | 0.1364 | 5 | 0.2 | 2 | 0.5 | 1 | 0 | 4 | 0.25 | 12 | 0.25 |
| 16_3374052 | 6 | 0.1667 | 3 | 0 | 6 | 0.1667 | 7 | 0 | 22 | 0.0909 | 4 | 0.25 | 1 | 0 | 1 | 0 | 4 | 0 | 10 | 0.1 |
| 16_3376911 | 8 | 0.75 | 7 | 0.7143 | 5 | 0.6 | 6 | 0.6667 | 26 | 0.6923 | 6 | 0.6667 | 3 | 0.3333 | 4 | 0.5 | 6 | 1 | 19 | 0.6842 |
| 16_3376914 | 7 | 0.2857 | 7 | 0 | 7 | 0.2857 | 6 | 0 | 27 | 0.1481 | 7 | 0.1429 | 3 | 0.3333 | 3 | 0 | 5 | 0.4 | 18 | 0.2222 |
| 16_3376915 | 9 | 0.1111 | 9 | 0 | 7 | 0.1429 | 6 | 0 | 31 | 0.0645 | 7 | 0.1429 | 4 | 0 | 4 | 0 | 5 | 0 | 20 | 0.05 |
| 16_3376917 | 9 | 0.8889 | 7 | 1 | 7 | 1 | 7 | 0.8571 | 30 | 0.9333 | 6 | 1 | 4 | 0.25 | 4 | 0.5 | 5 | 0.8 | 19 | 0.6842 |
| 16_3376918 | 8 | 1 | 8 | 0.375 | 8 | 0.75 | 7 | 0.2857 | 31 | 0.6129 | 6 | 0.3333 | 2 | 0 | 3 | 0.3333 | 5 | 0.4 | 16 | 0.3125 |
| 16_3376925 | 8 | 0.625 | 8 | 0.75 | 8 | 0.625 | 7 | 0.7143 | 31 | 0.6774 | 7 | 0.8571 | 4 | 0.75 | 4 | 0.25 | 5 | 0.8 | 20 | 0.7 |
| 16_3376931 | 9 | 0.3333 | 8 | 0.375 | 7 | 0.4286 | 7 | 0.2857 | 31 | 0.3548 | 7 | 0.5714 | 5 | 0.2 | 4 | 0.25 | 6 | 0.5 | 22 | 0.4091 |
| 16_3376937 | 8 | 0.25 | 8 | 0 | 6 | 0.1667 | 7 | 0 | 29 | 0.1034 | 7 | 0 | 5 | 0 | 4 | 0 | 4 | 0.25 | 20 | 0.05 |
| 16_3377695 | 1 | 0 | NA | NA | 1 | 1 | 1 | 1 | 3 | 0.6667 | 1 | 0 | 3 | 0.3333 | NA | NA | 1 | 0 | 5 | 0.2 |
| 16_3377704 | 4 | 1 | 2 | 1 | 2 | 1 | 2 | 1 | 10 | 1 | 2 | 1 | 3 | 1 | NA | NA | 1 | 1 | 6 | 1 |
| 16_3377704 | NA | NA | NA | NA | 1 | 1 | NA | NA | 1 | 1 | 1 | 1 | NA | NA | NA | NA | 1 | 1 | 2 | 1 |
| 16_3377708 | 4 | 0.25 | 2 | 0.5 | 3 | 0.3333 | 2 | 0 | 11 | 0.2727 | 3 | 0.3333 | 4 | 0 | NA | NA | 2 | 0.5 | 9 | 0.2222 |
| 16_3377712 | 4 | 0.5 | 2 | 0.5 | 2 | 0 | NA | NA | 8 | 0.375 | 2 | 0.5 | 3 | 0 | NA | NA | 1 | 0 | 6 | 0.1667 |
| 16_3378202 | 3 | 0.3333 | 1 | 1 | 1 | 0 | 1 | 0 | 6 | 0.3333 | 1 | 1 | 2 | 0 | NA | NA | 2 | 0 | 5 | 0.2 |
| 16_3378207 | 3 | 0.6667 | 2 | 0 | 2 | 0.5 | 1 | 0 | 8 | 0.375 | 1 | 0 | 3 | 0.6667 | NA | NA | 2 | 0.5 | 6 | 0.5 |
| 16_3378209 | 4 | 0.5 | 2 | 1 | 2 | 1 | 2 | 0 | 10 | 0.6 | 1 | 1 | 3 | 0.3333 | NA | NA | 1 | 0 | 5 | 0.4 |
| 16_3378221 | 3 | 0.3333 | 2 | 0 | 2 | 0.5 | 2 | 0 | 9 | 0.2222 | 1 | 0 | 3 | 0.3333 | NA | NA | 2 | 0.5 | 6 | 0.3333 |
| 16_3378224 | 4 | 1 | 2 | 1 | 2 | 0.5 | 2 | 1 | 10 | 0.9 | NA | NA | 3 | 0.3333 | NA | NA | 1 | 1 | 4 | 0.5 |
| 16_3378243 | 4 | 1 | 2 | 0.5 | 1 | 1 | 2 | 0.5 | 9 | 0.7778 | 1 | 1 | 3 | 1 | NA | NA | 1 | 0 | 5 | 0.8 |
| 16_3378246 | 4 | 1 | 1 | 1 | 2 | 0 | 1 | 0 | 8 | 0.625 | 1 | 0 | 1 | 1 | NA | NA | 1 | 1 | 3 | 0.6667 |
| 16_3379013 | 2 | 0.5 | 5 | 0.4 | 8 | 0 | 5 | 0.2 | 20 | 0.2 | 2 | 0 | 1 | 0 | 1 | 0 | 4 | 0.25 | 8 | 0.125 |
| 16_3379025 | 3 | 0.3333 | 6 | 0 | 7 | 0.1429 | 5 | 0.2 | 21 | 0.1429 | 3 | 0 | 1 | 1 | 1 | 0 | 4 | 0 | 9 | 0.1111 |
| 16_3379027 | 3 | 0.3333 | 7 | 0.1429 | 7 | 0 | 5 | 0.2 | 22 | 0.1364 | 3 | 0.3333 | 1 | 0 | 1 | 1 | 4 | 0 | 9 | 0.2222 |
| 16_3379058 | 3 | 0 | 7 | 0.2857 | 6 | 0 | 3 | 0 | 19 | 0.1053 | 3 | 0 | 2 | 0 | 2 | 0 | 4 | 0.25 | 11 | 0.0909 |
| 16_3379059 | 3 | 1 | 6 | 0.8333 | 6 | 0.3333 | 6 | 0.6667 | 21 | 0.6667 | 3 | 0.6667 | 2 | 0 | 2 | 1 | 6 | 1 | 13 | 0.7692 |
| 16_3379077 | 2 | 0.5 | 5 | 0.8 | 5 | 0.2 | 6 | 0.5 | 18 | 0.5 | 2 | 0.5 | 2 | 0 | 2 | 1 | 6 | 0.3333 | 12 | 0.4167 |
| 16_3379113 | 2 | 0 | 4 | 0 | 5 | 0.2 | 6 | 0.1667 | 17 | 0.1176 | 1 | 0 | 2 | 0 | 2 | 0.5 | 4 | 0 | 9 | 0.1111 |
| 16_3379466 | 3 | 0.3333 | 4 | 0.25 | 5 | 0 | 5 | 0.4 | 17 | 0.2353 | 4 | 0.25 | 3 | 0.6667 | 2 | 0.5 | 4 | 0.25 | 13 | 0.3846 |
| 16_3379466 | 3 | 0.6667 | 3 | 0.3333 | 4 | 0 | 3 | 0.6667 | 13 | 0.3846 | 3 | 0.3333 | 3 | 0 | 1 | 0 | 4 | 0.5 | 11 | 0.2727 |
| 16_3379506 | 3 | 0 | 5 | 0 | 4 | 0 | 4 | 0.25 | 16 | 0.0625 | 1 | 0 | 2 | 0.5 | 2 | 0 | 3 | 0 | 8 | 0.125 |
| 16_3379579 | 1 | 1 | 3 | 1 | 4 | 1 | 4 | 1 | 12 | 1 | 2 | 1 | 3 | 1 | 1 | 1 | 3 | 1 | 9 | 1 |
| 16_3379582 | 2 | 0.5 | 3 | 0.6667 | 4 | 0.75 | 1 | 0 | 10 | 0.6 | 2 | 0 | 2 | 0.5 | 1 | 0 | 2 | 1 | 7 | 0.4286 |
| 16_3379584 | 2 | 1 | 3 | 1 | 3 | 1 | 2 | 1 | 10 | 1 | 2 | 1 | 2 | 1 | NA | NA | 3 | 1 | 7 | 1 |
| 16_3379586 | 1 | 0 | 4 | 0 | 5 | 0 | 3 | 0.3333 | 13 | 0.0769 | 2 | 0 | 2 | 0.5 | 1 | 0 | 3 | 0 | 8 | 0.125 |
| 16_3379590 | 2 | 1 | 2 | 1 | 5 | 0.8 | 3 | 1 | 12 | 0.9167 | 2 | 1 | 3 | 1 | 1 | 1 | 2 | 1 | 8 | 1 |
| 16_3379601 | 2 | 0.5 | 4 | 0.5 | 5 | 0.8 | 2 | 0.5 | 13 | 0.6154 | 1 | 1 | 3 | 1 | 1 | 1 | 2 | 0.5 | 7 | 0.8571 |
| 16_3379617 | 1 | 1 | 4 | 1 | 3 | 1 | 2 | 1 | 10 | 1 | 1 | 1 | 3 | 1 | 1 | 1 | 2 | 1 | 7 | 1 |
| 16_3379625 | 2 | 0 | 4 | 0 | 4 | 0.25 | 1 | 0 | 11 | 0.0909 | 1 | 1 | 3 | 0.3333 | 1 | 0 | 2 | 1 | 7 | 0.5714 |
| 16_3379628 | 1 | 0 | 2 | 1 | 3 | 0.3333 | 1 | 1 | 7 | 0.5714 | 1 | 0 | 3 | 0 | 1 | 0 | 1 | 1 | 6 | 0.1667 |
| 16_3383396 | 3 | 0.3333 | 3 | 0 | NA | NA | 1 | 1 | 7 | 0.2857 | NA | NA | 3 | 0.6667 | 4 | 0 | NA | NA | 7 | 0.2857 |
| 16_3383398 | 5 | 0 | 2 | 0 | NA | NA | 1 | 1 | 8 | 0.125 | NA | NA | 3 | 0 | 4 | 0.25 | NA | NA | 7 | 0.1429 |
| 16_3397252 | 15 | 0.3333 | 42 | 0.3095 | 23 | 0.3043 | 18 | 0.3333 | 98 | 0.3163 | 21 | 0.3333 | 14 | 0.2143 | 24 | 0.2917 | 18 | 0.4444 | 77 | 0.3247 |
| 16_3397371 | 5 | 0.4 | 24 | 0.4167 | 17 | 0.5882 | 13 | 0.5385 | 59 | 0.4915 | 14 | 0.2857 | 8 | 0 | 16 | 0.4375 | 11 | 0.4545 | 49 | 0.3265 |
| 16_3397374 | 6 | 0.1667 | 25 | 0.16 | 15 | 0 | 11 | 0.0909 | 57 | 0.1053 | 16 | 0.25 | 9 | 0 | 16 | 0.125 | 13 | 0.0769 | 54 | 0.1296 |
| 16_3450349 | NA | NA | 1 | 1 | NA | NA | NA | NA | 1 | 1 | NA | NA | NA | NA | 1 | 1 | NA | NA | 1 | 1 |
| 16_3482094 | NA | NA | 5 | 0.8 | NA | NA | 1 | 1 | 6 | 0.8333 | 2 | 0.5 | 2 | 1 | 1 | 1 | 2 | 1 | 7 | 0.8571 |
| 16_3482155 | NA | NA | 5 | 0.4 | NA | NA | 2 | 1 | 7 | 0.5714 | 4 | 0.5 | 4 | 1 | 1 | 1 | 3 | 0.6667 | 12 | 0.75 |
| 16_3482160 | NA | NA | 5 | 0.2 | NA | NA | 2 | 0.5 | 7 | 0.2857 | 3 | 0 | 3 | 1 | NA | NA | 3 | 0.6667 | 9 | 0.5556 |
| 16_3482189 | NA | NA | 4 | 0.25 | NA | NA | 2 | 0.5 | 6 | 0.3333 | 4 | 0.5 | 3 | 0.3333 | 1 | 1 | 3 | 0 | 11 | 0.3636 |
| 16_3482191 | NA | NA | 5 | 0.4 | NA | NA | 2 | 0.5 | 7 | 0.4286 | 4 | 0 | 3 | 1 | 1 | 1 | 3 | 0 | 11 | 0.3636 |
| 16_3482268 | NA | NA | 5 | 0 | NA | NA | 3 | 0.3333 | 8 | 0.125 | 4 | 0.25 | 4 | 0.75 | 1 | 0 | 2 | 0 | 11 | 0.3636 |
| 16_3492679 | 4 | 0.25 | 12 | 0 | 7 | 0.1429 | 4 | 0 | 27 | 0.0741 | 5 | 0.4 | 5 | 0.2 | 6 | 0.3333 | 5 | 0.2 | 21 | 0.2857 |
| 16_3492681 | 5 | 0.2 | 13 | 0.0769 | 9 | 0 | 4 | 0 | 31 | 0.0645 | 5 | 0 | 4 | 0 | 7 | 0 | 6 | 0.3333 | 22 | 0.0909 |
| 16_3492684 | 5 | 0 | 14 | 0.2857 | 7 | 0.2857 | 4 | 0.75 | 30 | 0.3 | 4 | 0.25 | 5 | 0.2 | 9 | 0.1111 | 5 | 0 | 23 | 0.1304 |
| 16_3492892 | 4 | 0 | 4 | 0.25 | 5 | 0.2 | 1 | 0 | 14 | 0.1429 | 1 | 0 | NA | NA | 4 | 0.25 | 2 | 0.5 | 7 | 0.2857 |
| 16_3492895 | 5 | 0.2 | 4 | 0 | 5 | 0 | 3 | 0 | 17 | 0.0588 | 1 | 0 | NA | NA | 6 | 0.1667 | 2 | 0 | 9 | 0.1111 |
| 16_3497304 | 1 | 1 | NA | NA | NA | NA | NA | NA | 1 | 1 | NA | NA | NA | NA | 1 | 1 | NA | NA | 1 | 1 |
| 16_3497340 | 1 | 1 | NA | NA | NA | NA | NA | NA | 1 | 1 | NA | NA | NA | NA | 1 | 1 | NA | NA | 1 | 1 |
| 16_3497462 | 1 | 1 | NA | NA | NA | NA | NA | NA | 1 | 1 | NA | NA | NA | NA | 1 | 1 | NA | NA | 1 | 1 |

| | | | | | | | | | | | | | | | | | | | | |
|---|---|---|---|---|---|---|---|---|---|---|---|---|---|---|---|---|---|---|---|---|
| 16_3497554 | 4 | 0 | 2 | 0 | 1 | 0 | 4 | 0.25 | 11 | 0.0909 | 4 | 0.25 | 2 | 0 | 6 | 0 | 2 | 0 | 14 | 0.0714 |
| 16_3497558 | 4 | 0.25 | 2 | 0 | 1 | 0 | 3 | 0 | 10 | 0.1 | 5 | 0.2 | 2 | 0.5 | 5 | 0 | 5 | 0 | 17 | 0.1176 |
| 16_3497562 | 2 | 0 | 2 | 0 | NA | NA | 3 | 0.3333 | 7 | 0.1429 | 4 | 0.25 | 2 | 0 | 6 | 0.1667 | 4 | 0.25 | 16 | 0.1875 |
| 16_3497578 | 4 | 0.5 | 1 | 0 | 2 | 1 | 4 | 0.5 | 11 | 0.5455 | 5 | 0.6 | 3 | 0 | 6 | 0.1667 | 5 | 0.6 | 19 | 0.3684 |
| 16_3497605 | 8 | 0.125 | 3 | 0 | 4 | 0 | 3 | 0.3333 | 18 | 0.1111 | 6 | 0.3333 | 3 | 0.3333 | 10 | 0 | 4 | 0 | 23 | 0.1304 |
| 16_3497608 | 8 | 0.125 | 3 | 0 | 4 | 0 | 7 | 0.5714 | 22 | 0.2273 | 5 | 0.6 | 5 | 0.2 | 11 | 0.0909 | 5 | 0.6 | 26 | 0.3077 |
| 16_3497614 | 7 | 0.4286 | 4 | 0.25 | 3 | 0.3333 | 5 | 0 | 19 | 0.2632 | 5 | 0 | 7 | 0 | 14 | 0.0714 | 4 | 0.25 | 30 | 0.0667 |
| 16_3497621 | 8 | 0 | 3 | 0 | 5 | 0.2 | 6 | 0.1667 | 22 | 0.0909 | 6 | 0.1667 | 8 | 0.125 | 13 | 0.0769 | 3 | 0 | 30 | 0.1 |
| 16_3497622 | 8 | 0.625 | 2 | 0.5 | 5 | 1 | 5 | 0.8 | 20 | 0.75 | 5 | 0.8 | 8 | 0.625 | 13 | 0.6154 | 5 | 1 | 31 | 0.7097 |
| 16_3497623 | 7 | 1 | 4 | 0.75 | 3 | 0.3333 | 6 | 0.8333 | 20 | 0.8 | 6 | 0.8333 | 8 | 0.75 | 13 | 0.8462 | 5 | 1 | 32 | 0.8438 |
| 16_3497700 | 9 | 0.6667 | 7 | 0.7143 | 9 | 0.5556 | 3 | 0.6667 | 28 | 0.6429 | 6 | 0.5 | 13 | 0.3077 | 17 | 0.5294 | 7 | 0.5714 | 43 | 0.4651 |
| 16_3497706 | 10 | 0.4 | 7 | 0.7143 | 8 | 0.5 | 7 | 0.5714 | 32 | 0.5312 | 7 | 0.7143 | 15 | 0.4 | 21 | 0.4762 | 8 | 0.375 | 51 | 0.4706 |
| 16_3497714 | 11 | 0.0909 | 7 | 0.1429 | 11 | 0.1818 | 8 | 0.125 | 37 | 0.1351 | 6 | 0.3333 | 18 | 0.1111 | 23 | 0.3043 | 7 | 0.1429 | 54 | 0.2222 |
| 16_3497735 | 12 | 0.25 | 9 | 0.2222 | 13 | 0.2308 | 10 | 0.6 | 44 | 0.3182 | 7 | 0.4286 | 14 | 0.4286 | 21 | 0.3333 | 10 | 0.2 | 52 | 0.3462 |
| 16_3497746 | 14 | 0.5 | 12 | 0.5 | 13 | 0.7692 | 13 | 0.6154 | 52 | 0.5962 | 11 | 0.7273 | 21 | 0.5714 | 24 | 0.3333 | 11 | 0.2727 | 67 | 0.4627 |
| 16_3512407 | 48 | 0.25 | 35 | 0.2 | 58 | 0.2241 | 44 | 0.25 | 185 | 0.2324 | 27 | 0.1481 | 60 | 0.1667 | 72 | 0.25 | 47 | 0.1702 | 206 | 0.1942 |
| 16_3512415 | 48 | 0.2917 | 31 | 0.2258 | 53 | 0.3208 | 43 | 0.3488 | 175 | 0.3029 | 23 | 0.2174 | 61 | 0.3115 | 67 | 0.2985 | 40 | 0.35 | 191 | 0.3037 |
| 16_3512423 | 43 | 0.7907 | 31 | 0.8387 | 49 | 0.6327 | 42 | 0.6667 | 165 | 0.7212 | 27 | 0.8148 | 57 | 0.7719 | 67 | 0.7463 | 45 | 0.7778 | 196 | 0.7704 |
| 16_3512434 | 47 | 0.5745 | 37 | 0.2973 | 56 | 0.3571 | 51 | 0.3725 | 191 | 0.4031 | 29 | 0.4138 | 62 | 0.4355 | 76 | 0.4474 | 44 | 0.4545 | 211 | 0.4408 |
| 16_3512435 | 54 | 0.3333 | 37 | 0.4054 | 58 | 0.3103 | 49 | 0.2857 | 198 | 0.3283 | 31 | 0.2903 | 71 | 0.3239 | 74 | 0.2973 | 48 | 0.3958 | 224 | 0.3259 |
| 16_3512436 | 56 | 0.125 | 36 | 0.0833 | 58 | 0.1207 | 53 | 0.1887 | 203 | 0.133 | 30 | 0.1333 | 66 | 0.1515 | 76 | 0.2368 | 51 | 0.1176 | 223 | 0.1704 |
| 16_3512452 | 55 | 0.6727 | 34 | 0.6471 | 55 | 0.5636 | 40 | 0.55 | 184 | 0.6087 | 25 | 0.68 | 63 | 0.5714 | 76 | 0.6711 | 39 | 0.7179 | 203 | 0.6502 |
| 16_3512458 | 47 | 0.1277 | 35 | 0.0571 | 50 | 0.06 | 36 | 0.0556 | 168 | 0.0774 | 21 | 0 | 55 | 0.0909 | 62 | 0.0806 | 39 | 0.0513 | 177 | 0.0678 |
| 16_3512458 | 55 | 0.3273 | 36 | 0.2778 | 60 | 0.2667 | 54 | 0.1667 | 205 | 0.2585 | 28 | 0.2143 | 75 | 0.24 | 81 | 0.358 | 45 | 0.2444 | 229 | 0.2795 |
| 16_3512464 | 57 | 0.5088 | 36 | 0.4167 | 53 | 0.3208 | 48 | 0.3333 | 194 | 0.3969 | 26 | 0.3077 | 68 | 0.3971 | 75 | 0.2933 | 34 | 0.4412 | 203 | 0.3547 |
| 16_3512466 | 58 | 0.8966 | 39 | 0.8718 | 59 | 0.8814 | 52 | 0.8077 | 208 | 0.8654 | 29 | 0.8966 | 80 | 0.875 | 91 | 0.8901 | 52 | 0.8654 | 252 | 0.881 |
| 16_3512469 | 55 | 0.3818 | 38 | 0.2368 | 59 | 0.2034 | 49 | 0.102 | 201 | 0.2338 | 26 | 0.1538 | 71 | 0.1831 | 73 | 0.2055 | 45 | 0.2222 | 215 | 0.1953 |
| 16_3512477 | 37 | 0.2432 | 27 | 0.2593 | 41 | 0.0976 | 30 | 0 | 135 | 0.1481 | 20 | 0.25 | 50 | 0.1 | 50 | 0.14 | 33 | 0.2121 | 153 | 0.1569 |
| 16_3513392 | 2 | 0.5 | 1 | 1 | NA | NA | 1 | 0 | 4 | 0.5 | 2 | 0 | 3 | 0 | 7 | 0 | 4 | 0.25 | 16 | 0.0625 |
| 16_3513404 | 2 | 0.5 | 1 | 0 | 2 | 0.5 | 1 | 1 | 6 | 0.5 | 2 | 1 | 4 | 0.75 | 8 | 0.75 | 4 | 1 | 18 | 0.8333 |
| 16_3513408 | 2 | 0.5 | 1 | 0 | 1 | 0 | 3 | 0 | 7 | 0.1429 | 2 | 0 | 5 | 0.2 | 8 | 0.125 | 5 | 0.2 | 20 | 0.15 |
| 16_3513420 | 2 | 0 | 1 | 0 | 2 | 0.5 | 2 | 0 | 7 | 0.1429 | 2 | 0.5 | 5 | 0 | 8 | 0.125 | 4 | 0 | 19 | 0.1053 |
| 16_3513427 | 1 | 1 | 1 | 0 | 1 | 0 | 1 | 0 | 4 | 0.25 | 2 | 0.5 | 4 | 0 | 5 | 0 | 3 | 0.3333 | 14 | 0.1429 |
| 16_3513440 | 2 | 0.5 | 1 | 1 | 2 | 1 | 3 | 0.6667 | 8 | 0.75 | 2 | 1 | 5 | 1 | 7 | 1 | 4 | 1 | 18 | 1 |
| 16_3513453 | 1 | 1 | 1 | 1 | 1 | 1 | 3 | 1 | 6 | 1 | 1 | 1 | 4 | 1 | 8 | 1 | 4 | 1 | 17 | 1 |
| 16_3513456 | 1 | 1 | NA | NA | 1 | 1 | 3 | 1 | 5 | 1 | 2 | 1 | 4 | 1 | 5 | 0.8 | 4 | 1 | 15 | 0.9333 |
| 16_3548362 | 2 | 0.5 | 2 | 0 | 2 | 0.5 | 4 | 0.75 | 10 | 0.5 | 3 | 0.6667 | 2 | 0.5 | 4 | 0.5 | 1 | 0 | 10 | 0.5 |
| 16_3548366 | 2 | 1 | NA | NA | 2 | 0.5 | 6 | 0.6667 | 10 | 0.7 | 2 | 1 | 2 | 1 | 4 | 1 | 1 | 1 | 9 | 1 |
| 16_3548370 | 2 | 0.5 | 3 | 0.3333 | 2 | 1 | 6 | 0.8333 | 13 | 0.6923 | 2 | 0.5 | 2 | 1 | 6 | 0.8333 | 3 | 0.6667 | 13 | 0.7692 |
| 16_3548374 | 1 | 1 | 2 | 0.5 | 3 | 1 | 6 | 0.5 | 12 | 0.6667 | 3 | 0.6667 | 2 | 0.5 | 6 | 0.8333 | 3 | 0.6667 | 14 | 0.7143 |
| 16_3548380 | 1 | 1 | 2 | 0.5 | 2 | 1 | 5 | 1 | 10 | 0.9 | 3 | 1 | 2 | 1 | 5 | 0.6 | 2 | 1 | 12 | 0.8333 |
| 16_3548392 | 3 | 1 | 3 | 1 | 3 | 1 | 6 | 0.8333 | 15 | 0.9333 | 2 | 1 | 2 | 0.5 | 6 | 1 | 2 | 1 | 12 | 0.9167 |
| 16_3548394 | 3 | 1 | 3 | 0.3333 | 3 | 0.3333 | 6 | 0.6667 | 15 | 0.6 | 3 | 1 | 2 | 1 | 5 | 0.4 | 3 | 0 | 13 | 0.5385 |
| 16_3548399 | 3 | 0.6667 | 4 | 0.25 | 4 | 0.25 | 4 | 0.75 | 15 | 0.4667 | 3 | 0 | 2 | 0.5 | 6 | 0.5 | 2 | 0 | 13 | 0.3077 |
| 16_3548417 | 1 | 0 | 3 | 0 | 2 | 1 | 5 | 0.4 | 11 | 0.3636 | 3 | 0.3333 | 2 | 0 | 6 | 0.3333 | 3 | 0.3333 | 14 | 0.2857 |
| 16_3548426 | 3 | 0.6667 | 2 | 1 | 4 | 1 | 6 | 0.5 | 15 | 0.7333 | 3 | 1 | 2 | 1 | 5 | 1 | 3 | 1 | 13 | 1 |
| 16_3548427 | 1 | 0 | 4 | 0.75 | 3 | 1 | 6 | 0.8333 | 14 | 0.7857 | 3 | 1 | 2 | 1 | 6 | 1 | 3 | 1 | 14 | 1 |
| 16_3548430 | 3 | 0.3333 | 3 | 0.6667 | 4 | 0.5 | 6 | 0.8333 | 16 | 0.625 | 2 | 0.5 | 2 | 0.5 | 4 | 0.75 | 4 | 0.5 | 12 | 0.5833 |
| 16_3548436 | 3 | 0 | 4 | 0.25 | 4 | 0 | 5 | 0.2 | 16 | 0.125 | 3 | 0 | 2 | 0 | 6 | 0 | 4 | 0.25 | 15 | 0.0667 |
| 16_3548440 | 4 | 0.5 | 4 | 0.25 | 4 | 0.25 | 6 | 0.6667 | 18 | 0.4444 | 3 | 0.3333 | 2 | 0.5 | 6 | 0.6667 | 4 | 0.5 | 15 | 0.5333 |
| 16_3548444 | 4 | 0.25 | 4 | 0.25 | 4 | 0 | 5 | 0.8 | 17 | 0.3529 | 3 | 0.3333 | 2 | 0.5 | 5 | 0.4 | 4 | 0.5 | 14 | 0.4286 |
| 16_3548463 | 1 | 1 | NA | NA | 1 | 0 | NA | NA | 2 | 0.5 | NA | NA | NA | NA | 1 | 1 | NA | NA | 1 | 1 |
| 16_3551755 | 11 | 0.0909 | 9 | 0.1111 | 6 | 0.1667 | 2 | 0 | 28 | 0.1071 | 8 | 0 | 2 | 0.5 | 6 | 0.5 | 3 | 0.6667 | 19 | 0.3158 |
| 16_3551784 | 18 | 0.0556 | 16 | 0.1875 | 11 | 0 | 7 | 0.4286 | 52 | 0.1346 | 13 | 0.1538 | 7 | 0.2857 | 11 | 0.2727 | 9 | 0 | 40 | 0.175 |
| 16_3551788 | 17 | 0.2353 | 18 | 0.5556 | 10 | 0.4 | 8 | 0.5 | 53 | 0.4151 | 13 | 0.3077 | 7 | 0.4286 | 11 | 0.5455 | 9 | 0.5556 | 40 | 0.45 |
| 16_3551795 | 12 | 0.0833 | 15 | 0.0667 | 11 | 0 | 7 | 0.2857 | 45 | 0.0889 | 15 | 0.2 | 5 | 0 | 11 | 0.0909 | 8 | 0.125 | 39 | 0.1282 |
| 16_3551807 | 13 | 0.6154 | 13 | 0.4615 | 8 | 0.375 | 6 | 0.6667 | 40 | 0.525 | 11 | 0.6364 | 4 | 0 | 9 | 0.5556 | 5 | 0.6 | 29 | 0.5172 |
| 16_3551808 | 12 | 0.0833 | 11 | 0.1818 | 8 | 0.375 | 7 | 0 | 38 | 0.1579 | 14 | 0.5 | 6 | 0.3333 | 7 | 0.1429 | 7 | 0.1429 | 34 | 0.3235 |
| 16_3551813 | 15 | 0.4 | 15 | 0.2667 | 10 | 0.4 | 6 | 0.3333 | 46 | 0.3478 | 12 | 0.3333 | 7 | 0.2857 | 8 | 0.25 | 8 | 0.625 | 35 | 0.3714 |
| 16_3551824 | 13 | 0.6923 | 17 | 0.8235 | 9 | 0.7778 | 7 | 0.8571 | 46 | 0.7826 | 9 | 0.7778 | 7 | 0.5714 | 9 | 0.7778 | 9 | 0.8889 | 34 | 0.7647 |
| 16_3551825 | 13 | 0.8462 | 13 | 0.7692 | 9 | 0.6667 | 6 | 1 | 41 | 0.8049 | 11 | 0.9091 | 8 | 0.875 | 9 | 0.8889 | 7 | 1 | 35 | 0.9143 |
| 16_3551826 | 13 | 0 | 15 | 0.2 | 11 | 0 | 6 | 0.3333 | 45 | 0.1111 | 14 | 0.1429 | 8 | 0 | 9 | 0 | 7 | 0.2857 | 38 | 0.1053 |
| 16_3551830 | 14 | 0.2857 | 12 | 0.25 | 10 | 0.1 | 8 | 0.375 | 44 | 0.25 | 11 | 0.2727 | 7 | 0.2857 | 7 | 0.5714 | 8 | 0.5 | 33 | 0.3939 |
| 16_3564529 | 1 | 1 | 5 | 0.4 | NA | NA | 2 | 0 | 8 | 0.375 | 1 | 0 | 1 | 1 | NA | NA | NA | NA | 2 | 0.5 |
| 16_3564556 | 2 | 1 | 7 | 1 | 1 | 1 | 3 | 1 | 13 | 1 | NA | NA | 1 | 1 | NA | NA | NA | NA | 1 | 1 |

| | | | | | | | | | | | | | | | | | | | | |
|---|---|---|---|---|---|---|---|---|---|---|---|---|---|---|---|---|---|---|---|---|
| 16_3564557 | 2 | 1 | 6 | 0.8333 | 1 | 0 | 1 | 0 | 10 | 0.7 | NA | NA | 1 | 1 | NA | NA | NA | NA | 1 | 1 |
| 16_3564559 | 2 | 0 | 8 | 0.25 | 1 | 0 | 3 | 0.3333 | 14 | 0.2143 | 1 | 0 | 1 | 1 | NA | NA | NA | NA | 2 | 0.5 |
| 16_3564568 | 2 | 0.5 | 8 | 0.125 | 1 | 1 | 3 | 0.3333 | 14 | 0.2857 | 1 | 0 | 1 | 1 | NA | NA | NA | NA | 2 | 0.5 |
| 16_3564569 | 2 | 1 | 7 | 1 | 1 | 1 | 3 | 1 | 13 | 1 | 1 | 1 | 1 | 1 | NA | NA | NA | NA | 2 | 1 |
| 16_3564577 | 2 | 0 | 6 | 0.3333 | 1 | 0 | 2 | 0.5 | 11 | 0.2727 | 1 | 1 | 1 | 0 | NA | NA | NA | NA | 2 | 0.5 |
| 16_3564829 | NA | NA | 1 | 1 | NA | NA | NA | NA | 1 | 1 | 1 | 1 | NA | NA | NA | NA | NA | NA | 1 | 1 |
| 16_3575290 | NA | NA | NA | NA | 1 | 1 | NA | NA | 1 | 1 | NA | NA | NA | NA | 1 | 1 | NA | NA | 1 | 1 |
| 16_3575300 | NA | NA | NA | NA | 1 | 1 | NA | NA | 1 | 1 | NA | NA | NA | NA | 1 | 1 | NA | NA | 1 | 1 |
| 16_3575324 | NA | NA | NA | NA | 1 | 1 | NA | NA | 1 | 1 | NA | NA | NA | NA | 1 | 1 | NA | NA | 1 | 1 |
| 16_3575331 | NA | NA | NA | NA | 1 | 1 | NA | NA | 1 | 1 | NA | NA | NA | NA | 1 | 1 | NA | NA | 1 | 1 |
| 16_3585417 | 10 | 0.2 | 15 | 0.2 | 8 | 0.125 | 9 | 0.1111 | 42 | 0.1667 | 4 | 0 | 9 | 0 | 14 | 0.0714 | 10 | 0.1 | 37 | 0.0541 |
| 16_3585419 | 9 | 0.1111 | 17 | 0.1765 | 7 | 0.4286 | 8 | 0 | 41 | 0.1707 | 5 | 0.2 | 10 | 0.2 | 14 | 0.2143 | 9 | 0.1111 | 38 | 0.1842 |
| 16_3585431 | 11 | 0 | 18 | 0.1667 | 8 | 0.125 | 9 | 0.1111 | 46 | 0.1087 | 5 | 0.2 | 11 | 0 | 15 | 0.0667 | 10 | 0.1 | 41 | 0.0732 |
| 16_3585461 | 11 | 0.8182 | 18 | 0.9444 | 9 | 0.8889 | 7 | 0.8571 | 45 | 0.8889 | 3 | 1 | 9 | 0.8889 | 11 | 0.9091 | 11 | 0.8182 | 34 | 0.8824 |
| 16_3585464 | 12 | 0.4167 | 14 | 0.4286 | 9 | 0.5556 | 7 | 0.5714 | 42 | 0.4762 | 4 | 0.75 | 13 | 0.5385 | 15 | 0.6 | 9 | 0.4444 | 41 | 0.561 |
| 16_3585465 | 12 | 0 | 20 | 0.25 | 8 | 0.25 | 9 | 0.1111 | 49 | 0.1633 | 4 | 0.25 | 12 | 0.3333 | 15 | 0.1333 | 11 | 0.2727 | 42 | 0.2381 |
| 16_3585466 | 13 | 0.0769 | 19 | 0.0526 | 8 | 0.25 | 9 | 0 | 49 | 0.0816 | 1 | 0 | 11 | 0 | 12 | 0.25 | 11 | 0 | 35 | 0.0857 |
| 16_3585470 | 11 | 0.4545 | 17 | 0.4118 | 8 | 0.625 | 7 | 0.4286 | 43 | 0.4651 | 2 | 1 | 10 | 0.3 | 12 | 0.4167 | 10 | 0.4 | 34 | 0.4118 |
| 16_3585494 | 12 | 0.1667 | 20 | 0.2 | 10 | 0.4 | 10 | 0.4 | 52 | 0.2692 | 5 | 0.8 | 13 | 0.3077 | 14 | 0.2857 | 7 | 0.4286 | 39 | 0.3846 |
| 16_3585512 | 11 | 0.0909 | 21 | 0.0952 | 10 | 0.1 | 10 | 0.5 | 52 | 0.1731 | 6 | 0.5 | 14 | 0.1429 | 16 | 0.0625 | 10 | 0.2 | 46 | 0.1739 |
| 16_3585554 | 13 | 0.3077 | 20 | 0.25 | 10 | 0.2 | 10 | 0.4 | 53 | 0.283 | 6 | 0.5 | 15 | 0.0667 | 18 | 0.1111 | 9 | 0.1111 | 48 | 0.1458 |
| 16_3585559 | 12 | 0.5 | 19 | 0.5263 | 10 | 0 | 10 | 0.3 | 51 | 0.3725 | 6 | 0.8333 | 12 | 0.3333 | 16 | 0.5625 | 9 | 0.4444 | 43 | 0.5116 |
| 16_3585564 | 13 | 0.3077 | 21 | 0.2857 | 9 | 0.1111 | 12 | 0.1667 | 55 | 0.2364 | 7 | 0.2857 | 15 | 0.3333 | 15 | 0.4 | 11 | 0.2727 | 48 | 0.3333 |
| 16_3585566 | 14 | 0.1429 | 21 | 0.0952 | 9 | 0 | 9 | 0.1111 | 53 | 0.0943 | 7 | 0.5714 | 14 | 0.1429 | 18 | 0.0556 | 11 | 0.4545 | 50 | 0.24 |
| 16_3585570 | 12 | 0.4167 | 19 | 0.3684 | 10 | 0.2 | 10 | 0.5 | 51 | 0.3725 | 6 | 0.5 | 16 | 0.3125 | 16 | 0.25 | 8 | 0.5 | 46 | 0.3478 |
| 16_3585603 | 15 | 0.1333 | 22 | 0.0909 | 12 | 0 | 11 | 0 | 60 | 0.0667 | 7 | 0.4286 | 18 | 0 | 20 | 0.1 | 13 | 0 | 58 | 0.0862 |
| 16_3585604 | 13 | 0.8462 | 21 | 0.6667 | 12 | 0.9167 | 12 | 1 | 58 | 0.8276 | 7 | 0.7143 | 16 | 0.625 | 18 | 0.7222 | 13 | 0.6923 | 54 | 0.6852 |
| 16_3585624 | 16 | 0.0625 | 23 | 0.087 | 11 | 0.0909 | 13 | 0.2308 | 63 | 0.1111 | 7 | 0 | 18 | 0.0556 | 22 | 0.1364 | 14 | 0.2143 | 61 | 0.1148 |
| 16_3585629 | 14 | 0.2143 | 21 | 0.2381 | 11 | 0 | 11 | 0.2727 | 57 | 0.193 | 7 | 0.2857 | 18 | 0.1111 | 19 | 0.2105 | 13 | 0.0769 | 57 | 0.1579 |
| 16_3585657 | 9 | 0 | 16 | 0.0625 | 10 | 0 | 10 | 0.2 | 45 | 0.0667 | 5 | 0 | 14 | 0.1429 | 15 | 0 | 8 | 0.125 | 42 | 0.0714 |
| 16_3585706 | 12 | 0.3333 | 20 | 0.35 | 10 | 0.3 | 12 | 0.75 | 54 | 0.4259 | 8 | 0.5 | 13 | 0.3846 | 17 | 0.4706 | 10 | 0.3 | 48 | 0.4167 |
| 16_3585718 | 12 | 0.0833 | 19 | 0.1053 | 12 | 0.1667 | 11 | 0.1818 | 54 | 0.1296 | 8 | 0 | 14 | 0.1429 | 16 | 0.0625 | 11 | 0.1818 | 49 | 0.102 |
| 16_3585729 | 8 | 0.25 | 18 | 0.5556 | 10 | 0.7 | 11 | 0.7273 | 47 | 0.5745 | 9 | 0.5556 | 10 | 0.6 | 15 | 0.6667 | 12 | 0.6667 | 46 | 0.6304 |
| 16_3586230 | NA | NA | 1 | 1 | 2 | 1 | NA | NA | 3 | 1 | NA | NA | NA | NA | NA | NA | 1 | 1 | 1 | 1 |
| 16_3586237 | NA | NA | NA | NA | 2 | 0.5 | NA | NA | 2 | 0.5 | NA | NA | NA | NA | NA | NA | 1 | 1 | 1 | 1 |
| 16_3586248 | NA | NA | 1 | 1 | 2 | 1 | NA | NA | 3 | 1 | NA | NA | NA | NA | NA | NA | 1 | 1 | 1 | 1 |
| 16_3586256 | NA | NA | NA | NA | 1 | 1 | NA | NA | 1 | 1 | NA | NA | NA | NA | NA | NA | 1 | 1 | 1 | 1 |
| 16_3586259 | NA | NA | NA | NA | 1 | 1 | NA | NA | 1 | 1 | NA | NA | NA | NA | NA | NA | 1 | 1 | 1 | 1 |
| 16_3586270 | NA | NA | 1 | 1 | 2 | 1 | NA | NA | 3 | 1 | NA | NA | NA | NA | NA | NA | 1 | 1 | 1 | 1 |
| 16_3586284 | NA | NA | NA | NA | 1 | 1 | NA | NA | 1 | 1 | NA | NA | NA | NA | NA | NA | 1 | 1 | 1 | 1 |
| 16_3586286 | NA | NA | 1 | 1 | 2 | 1 | NA | NA | 3 | 1 | NA | NA | NA | NA | NA | NA | 1 | 1 | 1 | 1 |
| 16_3586295 | NA | NA | 1 | 0 | 1 | 1 | NA | NA | 2 | 0.5 | NA | NA | NA | NA | NA | NA | 1 | 1 | 1 | 1 |
| 16_3586307 | NA | NA | 1 | 0 | 2 | 1 | NA | NA | 3 | 0.6667 | NA | NA | NA | NA | NA | NA | 1 | 1 | 1 | 1 |
| 16_3586412 | 1 | 0 | NA | NA | 2 | 0.5 | NA | NA | 3 | 0.3333 | NA | NA | NA | NA | NA | NA | 1 | 1 | 1 | 1 |
| 16_3586430 | 19 | 0.1053 | 15 | 0.2 | 9 | 0 | 12 | 0.25 | 55 | 0.1455 | 15 | 0 | 17 | 0.1765 | 13 | 0.0769 | 13 | 0.1538 | 58 | 0.1034 |
| 16_3659020 | NA | NA | 1 | 1 | 1 | 0 | 1 | 0 | 3 | 0.3333 | 3 | 0.3333 | NA | NA | 1 | 0 | NA | NA | 4 | 0.25 |
| 16_3659235 | 4 | 0.5 | 2 | 0 | 1 | 1 | 1 | 0 | 8 | 0.375 | 3 | 0.6667 | 3 | 0 | 2 | 0 | 1 | 0 | 9 | 0.2222 |
| 16_3673261 | NA | NA | NA | NA | 1 | 1 | NA | NA | 1 | 1 | NA | NA | NA | NA | 1 | 1 | NA | NA | 1 | 1 |
| 16_3673298 | NA | NA | NA | NA | 1 | 1 | NA | NA | 1 | 1 | NA | NA | NA | NA | 1 | 1 | NA | NA | 1 | 1 |
| 16_3673322 | NA | NA | NA | NA | 1 | 1 | NA | NA | 1 | 1 | NA | NA | NA | NA | 2 | 0.5 | NA | NA | 2 | 0.5 |
| 16_3673720 | 1 | 1 | NA | NA | NA | NA | 1 | 1 | 2 | 1 | 1 | 0 | NA | NA | 4 | 0.5 | NA | NA | 5 | 0.4 |
| 16_3673848 | NA | NA | NA | NA | 1 | 1 | 1 | 0 | 2 | 0.5 | NA | NA | NA | NA | 4 | 0.5 | NA | NA | 4 | 0.5 |
| 16_3673888 | 1 | 1 | NA | NA | NA | NA | 1 | 0 | 2 | 0.5 | 1 | 0 | 1 | 1 | 5 | 0.2 | NA | NA | 7 | 0.2857 |
| 16_3673892 | 1 | 0 | NA | NA | 1 | 0 | 1 | 1 | 3 | 0.3333 | 1 | 0 | 1 | 0 | 5 | 0.4 | NA | NA | 7 | 0.2857 |
| 16_3673914 | 1 | 1 | NA | NA | 1 | 1 | 1 | 1 | 3 | 1 | 1 | 1 | 2 | 1 | 5 | 0.8 | NA | NA | 8 | 0.875 |
| 16_3673971 | 1 | 1 | 3 | 0.3333 | 1 | 1 | 2 | 0.5 | 7 | 0.5714 | 2 | 0.5 | 1 | 1 | 5 | 0.6 | 1 | 1 | 9 | 0.6667 |
| 16_3673980 | 1 | 1 | 2 | 1 | 1 | 1 | 3 | 0.3333 | 7 | 0.7143 | 3 | 1 | 3 | 1 | 4 | 0.25 | 1 | 1 | 11 | 0.7273 |
| 16_3673989 | 1 | 0 | 3 | 0.6667 | 1 | 0 | 3 | 0.6667 | 8 | 0.5 | 2 | 0 | 2 | 1 | 2 | 0 | 1 | 0 | 7 | 0.2857 |
| 16_3675232 | 3 | 0 | 5 | 0.4 | 2 | 0 | 5 | 0.2 | 15 | 0.2 | 3 | 0.6667 | 7 | 0.4286 | 6 | 0.3333 | 6 | 0.3333 | 22 | 0.4091 |
| 16_3675233 | 2 | 0 | 4 | 0 | NA | NA | 5 | 0.4 | 11 | 0.1818 | 3 | 0.6667 | 7 | 0.2857 | 7 | 0.4286 | 5 | 0.4 | 22 | 0.4091 |
| 16_3675234 | 2 | 1 | 6 | 0.5 | 2 | 0 | 5 | 0.8 | 15 | 0.6 | 3 | 0 | 7 | 0.2857 | 7 | 0.4286 | 3 | 0 | 20 | 0.25 |
| 16_3675246 | 2 | 0 | 5 | 0 | 3 | 0 | 4 | 0.25 | 14 | 0.0714 | 3 | 0.3333 | 6 | 0.1667 | 6 | 0.1667 | 3 | 0 | 18 | 0.1667 |
| 16_3675275 | 2 | 0.5 | 4 | 0.5 | 1 | 1 | 4 | 0.5 | 11 | 0.5455 | 2 | 0.5 | 2 | 1 | 5 | 0.4 | 4 | 1 | 13 | 0.6923 |
| 16_3675287 | 3 | 0.6667 | 6 | 0.5 | 3 | 0.6667 | 6 | 0.6667 | 18 | 0.6111 | 1 | 1 | 7 | 0.7143 | 7 | 0.7143 | 6 | 0.5 | 21 | 0.6667 |
| 16_3675288 | 4 | 0.5 | 5 | 0.2 | 3 | 0 | 4 | 0.25 | 16 | 0.25 | NA | NA | 7 | 0.2857 | 5 | 0 | 5 | 0.4 | 17 | 0.2353 |

| | | | | | | | | | | | | | | | | | | | | |
|---|---|---|---|---|---|---|---|---|---|---|---|---|---|---|---|---|---|---|---|---|
| 16_367529 | 4 | 0 | 6 | 0.5 | 5 | 0.6 | 5 | 0.2 | 20 | 0.35 | 2 | 0.5 | 7 | 0.1429 | 7 | 0.7143 | 6 | 0.3333 | 22 | 0.4091 |
| 16_367529 | 4 | 0.75 | 5 | 0.6 | 6 | 0.8333 | 7 | 0.7143 | 22 | 0.7273 | 2 | 1 | 6 | 0.5 | 8 | 0.5 | 5 | 0.8 | 21 | 0.619 |
| 16_367529 | 3 | 0 | 4 | 0.25 | 5 | 0 | 4 | 0.25 | 16 | 0.125 | 2 | 0.5 | 7 | 0.1429 | 7 | 0.1429 | 6 | 0.1667 | 22 | 0.1818 |
| 16_367529 | 4 | 1 | 6 | 0.8333 | 7 | 1 | 5 | 0.8 | 22 | 0.9091 | 2 | 0.5 | 7 | 1 | 6 | 0.6667 | 4 | 0.75 | 19 | 0.7895 |
| 16_367530 | 4 | 0.75 | 6 | 0.6667 | 7 | 0.5714 | 7 | 0.8571 | 24 | 0.7083 | 3 | 0 | 7 | 0.7143 | 6 | 0.5 | 5 | 0.8 | 21 | 0.5714 |
| 16_367530 | 4 | 0.75 | 5 | 0.8 | 4 | 0.25 | 3 | 1 | 16 | 0.6875 | NA | NA | 3 | 1 | 3 | 1 | 4 | 0.5 | 10 | 0.8 |
| 16_367531 | 4 | 0.25 | 6 | 0.1667 | 7 | 0.2857 | 6 | 0.5 | 23 | 0.3043 | 2 | 0 | 7 | 0 | 6 | 0.3333 | 5 | 0.2 | 20 | 0.15 |
| 16_367532 | 4 | 1 | 4 | 0.75 | 8 | 0.75 | 7 | 0.7143 | 23 | 0.7826 | 3 | 0.6667 | 7 | 0.8571 | 8 | 0.875 | 6 | 0.8333 | 24 | 0.8333 |
| 16_367533 | 4 | 0.5 | 6 | 0.1667 | 7 | 0.1429 | 8 | 0.25 | 25 | 0.24 | 3 | 0.3333 | 8 | 0.25 | 8 | 0.375 | 5 | 0.2 | 24 | 0.2917 |
| 16_367533 | 3 | 1 | 5 | 0.8 | 6 | 0.6667 | 7 | 0.8571 | 21 | 0.8095 | 2 | 1 | 7 | 0.5714 | 8 | 0.375 | 6 | 0.5 | 23 | 0.5217 |
| 16_367835 | NA | NA | 1 | 1 | NA | NA | NA | NA | 1 | 1 | NA | NA | NA | NA | NA | NA | 2 | 0.5 | 2 | 0.5 |
| 16_369139 | NA | NA | 1 | 0 | NA | NA | 1 | 1 | 2 | 0.5 | NA | NA | 1 | 0 | 1 | 1 | NA | NA | 2 | 0.5 |
| 16_369144 | NA | NA | 1 | 0 | NA | NA | 1 | 1 | 2 | 0.5 | NA | NA | 1 | 0 | 1 | 1 | NA | NA | 2 | 0.5 |
| 16_369278 | 5 | 0.2 | 10 | 0 | 9 | 0.1111 | 8 | 0 | 32 | 0.0625 | 2 | 0 | 4 | 0.5 | 6 | 0 | 9 | 0 | 21 | 0.0952 |
| 16_369281 | 6 | 0 | 7 | 0.2857 | 10 | 0.4 | 7 | 0.4286 | 30 | 0.3 | 2 | 0 | 3 | 0.3333 | 6 | 0.1667 | 9 | 0.5556 | 20 | 0.35 |
| 16_369285 | 2 | 0 | 6 | 0.1667 | 8 | 0 | 5 | 0.2 | 21 | 0.0952 | 2 | 0 | 3 | 0.3333 | 6 | 0 | 6 | 0 | 17 | 0.0588 |
| 16_369290 | 4 | 0 | 5 | 0 | 7 | 0.1429 | 4 | 0 | 20 | 0.05 | 1 | 0 | 2 | 0 | 5 | 0.2 | 6 | 0 | 14 | 0.0714 |
| 16_369298 | 3 | 0 | 4 | 0.25 | 4 | 0.25 | 3 | 0.3333 | 14 | 0.2143 | NA | NA | 3 | 0.3333 | 4 | 0 | 5 | 0.2 | 12 | 0.1667 |
| 16_369303 | 5 | 0 | 13 | 0 | 8 | 0.125 | 6 | 0.1667 | 32 | 0.0625 | 3 | 0.6667 | 5 | 0 | 6 | 0 | 7 | 0.2857 | 21 | 0.1905 |
| 16_369305 | 5 | 0 | 7 | 0 | 4 | 0.25 | 3 | 0 | 19 | 0.0526 | 2 | 0 | 1 | 0 | 3 | 0 | 3 | 0.3333 | 9 | 0.1111 |
| 16_369322 | 4 | 0 | 8 | 0 | 5 | 0 | 2 | 0.5 | 19 | 0.0526 | 2 | 0 | 2 | 0.5 | 3 | 0 | 3 | 0 | 10 | 0.1 |
| 16_369323 | 4 | 0.25 | 9 | 0.3333 | 6 | 0 | 3 | 0.3333 | 22 | 0.2273 | 3 | 0.3333 | 2 | 0.5 | 4 | 0.25 | 3 | 1 | 12 | 0.5 |
| 16_369324 | 4 | 0.5 | 8 | 0.125 | 5 | 0.2 | 1 | 1 | 18 | 0.2778 | 3 | 0.3333 | 2 | 0.5 | 3 | 0.3333 | 3 | 0.6667 | 11 | 0.4545 |
| 16_369324 | 2 | 0 | 8 | 0.5 | 5 | 0.2 | 2 | 0 | 17 | 0.2941 | 3 | 0.6667 | 2 | 1 | 3 | 1 | 2 | 1 | 10 | 0.9 |
| 16_369325 | 4 | 0.25 | 9 | 0.5556 | 5 | 0.6 | 2 | 0.5 | 20 | 0.5 | 3 | 0.6667 | 2 | 0.5 | 3 | 0.3333 | 3 | 0.6667 | 11 | 0.5455 |
| 16_369326 | 3 | 0 | 8 | 0.375 | 4 | 0 | 3 | 0.3333 | 18 | 0.2222 | 3 | 0.3333 | 2 | 0.5 | 3 | 0 | 2 | 0.5 | 10 | 0.3 |
| 16_369332 | 4 | 0.25 | 6 | 0 | 2 | 0 | 3 | 0 | 15 | 0.0667 | 3 | 0 | 2 | 0 | 3 | 0 | 4 | 0.25 | 12 | 0.0833 |
| 16_369345 | 3 | 0.6667 | 3 | 0.3333 | 2 | 1 | 3 | 0.6667 | 11 | 0.6364 | 3 | 1 | 1 | 1 | 2 | 1 | 3 | 0.3333 | 9 | 0.7778 |
| 16_369345 | 4 | 0 | 3 | 0.3333 | 2 | 0 | 2 | 0 | 11 | 0.0909 | 1 | 0 | NA | NA | 2 | 0.5 | 3 | 0 | 6 | 0.1667 |
| 16_369346 | 3 | 0.3333 | 3 | 0 | 2 | 0.5 | 1 | 0 | 9 | 0.2222 | 2 | 0 | 1 | 0 | 2 | 0.5 | 2 | 0.5 | 7 | 0.2857 |
| 16_369356 | 3 | 0.3333 | 2 | 0 | 1 | 0 | 2 | 0 | 8 | 0.125 | 3 | 0 | 1 | 0 | 1 | 0 | 3 | 0.3333 | 8 | 0.125 |
| 16_369357 | 4 | 1 | 1 | 0 | 1 | 1 | 3 | 1 | 9 | 0.8889 | 2 | 1 | 1 | 1 | 2 | 1 | 3 | 0.6667 | 8 | 0.875 |
| 16_369358 | 4 | 0.25 | 3 | 0.6667 | 1 | 0 | 3 | 0 | 11 | 0.2727 | 3 | 0.6667 | NA | NA | 1 | 0 | 3 | 0.3333 | 7 | 0.4286 |
| 16_369358 | 3 | 1 | 3 | 0.6667 | 1 | 1 | 3 | 0.3333 | 10 | 0.7 | 2 | 1 | 1 | 0 | 1 | 1 | 3 | 0.3333 | 7 | 0.5714 |
| 16_369359 | 3 | 0.6667 | 3 | 1 | 1 | 1 | 2 | 1 | 9 | 0.8889 | 2 | 1 | 1 | 0 | 2 | 1 | 3 | 0.6667 | 8 | 0.75 |
| 16_369548 | 2 | 0 | 3 | 0.3333 | 1 | 1 | 3 | 0 | 9 | 0.2222 | 1 | 0 | 1 | 0 | 1 | 0 | 2 | 0.5 | 5 | 0.2 |
| 16_373303 | 5 | 0.8 | NA | NA | 2 | 0.5 | 3 | 0.3333 | 10 | 0.6 | NA | NA | 1 | 1 | 2 | 0.5 | 3 | 0.3333 | 6 | 0.5 |
| 16_373306 | 5 | 0 | 1 | 1 | 2 | 0 | 2 | 0 | 10 | 0.1 | NA | NA | 2 | 0.5 | 2 | 0 | 3 | 0 | 7 | 0.1429 |
| 16_373306 | 7 | 0.1429 | 1 | 1 | 2 | 0 | 2 | 0 | 12 | 0.1667 | NA | NA | 2 | 0 | 2 | 0.5 | 3 | 0 | 7 | 0.1429 |
| 16_373307 | 6 | 0.8333 | 2 | 1 | 2 | 1 | 2 | 1 | 12 | 0.9167 | NA | NA | 2 | 1 | 2 | 1 | 3 | 1 | 7 | 1 |
| 16_373307 | 6 | 0.1667 | 3 | 0 | 2 | 0 | 2 | 0 | 13 | 0.0769 | NA | NA | 2 | 0.5 | 1 | 0 | 3 | 0 | 6 | 0.1667 |
| 16_373309 | 6 | 0.5 | 2 | 0.5 | 1 | 0 | 2 | 0.5 | 11 | 0.4545 | NA | NA | 2 | 0 | 1 | 0 | 2 | 0.5 | 5 | 0.2 |
| 16_373310 | 6 | 0.6667 | 2 | 1 | 1 | 1 | 1 | 1 | 10 | 0.8 | NA | NA | 2 | 1 | NA | NA | 2 | 1 | 4 | 1 |
| 16_373312 | 4 | 0.25 | 2 | 0.5 | 1 | 1 | 2 | 0.5 | 9 | 0.4444 | NA | NA | 2 | 0.5 | 1 | 0 | 2 | 0 | 5 | 0.2 |
| 16_373313 | 5 | 0.8 | 2 | 0.5 | 1 | 0 | 2 | 0 | 10 | 0.5 | NA | NA | 2 | 0 | 1 | 0 | 2 | 0.5 | 5 | 0.2 |
| 16_373320 | 4 | 1 | 2 | 1 | 1 | 1 | 1 | 1 | 8 | 1 | NA | NA | 2 | 1 | NA | NA | 1 | 1 | 3 | 1 |
| 16_373362 | 3 | 0 | 2 | 0.5 | 1 | 0 | NA | NA | 6 | 0.1667 | NA | NA | 1 | 1 | NA | NA | 2 | 1 | 3 | 1 |
| 16_373562 | 3 | 0.6667 | 1 | 0 | 1 | 1 | 1 | 1 | 6 | 0.6667 | NA | NA | NA | NA | NA | NA | 1 | 1 | 1 | 1 |
| 16_373562 | 3 | 1 | 1 | 1 | NA | NA | 1 | 1 | 5 | 1 | NA | NA | 1 | 0 | NA | NA | 1 | 1 | 2 | 0.5 |
| 16_373563 | 3 | 0 | 1 | 1 | NA | NA | 1 | 1 | 5 | 0.4 | NA | NA | 1 | 1 | NA | NA | 1 | 0 | 2 | 0.5 |
| 16_373564 | 3 | 0.3333 | 1 | 1 | NA | NA | 1 | 1 | 5 | 0.6 | NA | NA | 1 | 1 | NA | NA | 1 | 0 | 2 | 0.5 |
| 16_373915 | 4 | 0.5 | 1 | 0 | NA | NA | 2 | 1 | 7 | 0.5714 | 1 | 1 | 2 | 0.5 | 1 | 1 | NA | NA | 4 | 0.75 |
| 16_373917 | 4 | 0.25 | NA | NA | 1 | 1 | 2 | 0.5 | 7 | 0.4286 | 1 | 1 | 2 | 0.5 | 1 | 1 | NA | NA | 4 | 0.75 |
| 16_373917 | 4 | 0.25 | 1 | 0 | 1 | 0 | 2 | 0 | 8 | 0.125 | 1 | 1 | 2 | 0 | 1 | 0 | NA | NA | 4 | 0.25 |
| 16_373920 | 4 | 0 | 1 | 1 | 1 | 1 | 3 | 0.3333 | 9 | 0.3333 | 1 | 0 | 3 | 0.3333 | 1 | 0 | NA | NA | 5 | 0.2 |
| 16_374677 | 1 | 1 | NA | NA | NA | NA | NA | NA | 1 | 1 | NA | NA | 1 | 1 | NA | NA | 1 | 1 | 2 | 1 |
| 16_376545 | 3 | 0 | 4 | 0.25 | 4 | 0 | 2 | 0 | 13 | 0.0769 | 3 | 0.3333 | 1 | 1 | 3 | 0.6667 | 1 | 0 | 8 | 0.5 |
| 16_376549 | 4 | 0 | 3 | 0 | 4 | 0 | 4 | 0.25 | 15 | 0.0667 | 3 | 0 | 1 | 0 | 5 | 0 | 2 | 0.5 | 11 | 0.0909 |
| 16_376566 | 5 | 0.2 | 4 | 0.5 | 6 | 0.1667 | 5 | 0.8 | 20 | 0.4 | 4 | 0 | 5 | 0 | 7 | 0.8571 | 4 | 0.25 | 20 | 0.35 |
| 16_376988 | 1 | 0 | 2 | 0.5 | 4 | 0.25 | 3 | 0 | 10 | 0.2 | 1 | 0 | 2 | 0.5 | 4 | 0.25 | 4 | 0.5 | 11 | 0.3636 |
| 16_377016 | 1 | 0 | 2 | 0.5 | 4 | 0.25 | 3 | 0.3333 | 10 | 0.3 | 1 | 1 | 2 | 0.5 | 4 | 0.25 | 4 | 0.75 | 11 | 0.5455 |
| 16_377016 | 1 | 1 | 2 | 1 | 3 | 1 | 3 | 1 | 9 | 1 | 2 | 1 | 2 | 1 | 3 | 1 | 4 | 1 | 11 | 1 |
| 16_377017 | 1 | 0 | 3 | 0 | 4 | 0 | 2 | 0.5 | 10 | 0.1 | 2 | 0 | 2 | 0 | 4 | 0.25 | 5 | 0.6 | 13 | 0.3077 |
| 16_377018 | NA | NA | 2 | 0.5 | 3 | 0.3333 | 3 | 0.3333 | 8 | 0.375 | 3 | 0.3333 | 1 | 0 | 4 | 0.5 | 4 | 0.25 | 12 | 0.3333 |
| 16_377018 | 1 | 1 | 1 | 0 | 3 | 1 | 3 | 0.6667 | 8 | 0.75 | 3 | 0.6667 | 1 | 1 | 3 | 0.6667 | 4 | 0.5 | 11 | 0.6364 |

| | | | | | | | | | | | | | | | | | | | | |
|---|---|---|---|---|---|---|---|---|---|---|---|---|---|---|---|---|---|---|---|---|
| 16_377018 | 1 | 1 | 2 | 1 | 2 | 1 | 1 | 1 | 6 | 1 | 2 | 1 | 1 | 0 | 4 | 1 | 4 | 1 | 11 | 0.9091 |
| 16_377019 | 1 | 0 | 3 | 0.3333 | 1 | 0 | 3 | 0 | 8 | 0.125 | 2 | 0 | 2 | 0 | 3 | 0 | 5 | 0.6 | 12 | 0.25 |
| 16_377020 | 1 | 0 | 3 | 0 | 2 | 0 | 3 | 0.3333 | 9 | 0.1111 | 2 | 0 | 1 | 1 | 4 | 0 | 5 | 0.2 | 12 | 0.1667 |
| 16_377021 | 2 | 0 | 3 | 0 | 2 | 0.5 | 3 | 0.3333 | 10 | 0.2 | 3 | 0.6667 | 2 | 0.5 | 3 | 0.6667 | 4 | 0.75 | 12 | 0.6667 |
| 16_377021 | 1 | 0 | 2 | 0.5 | 3 | 0.3333 | 3 | 1 | 9 | 0.5556 | 3 | 0 | 2 | 0 | 4 | 0 | 5 | 0.4 | 14 | 0.1429 |
| 16_377022 | 2 | 0.5 | 2 | 0 | 1 | 1 | 3 | 0.3333 | 8 | 0.375 | 2 | 0.5 | 2 | 1 | 2 | 0.5 | 3 | 0.6667 | 9 | 0.6667 |
| 16_377022 | 1 | 1 | 2 | 1 | 2 | 0.5 | 2 | 1 | 7 | 0.8571 | 2 | 0.5 | 1 | 1 | 4 | 0.75 | 4 | 0.75 | 11 | 0.7273 |
| 16_377023 | 1 | 0 | 3 | 0 | 4 | 0.25 | 3 | 0 | 11 | 0.0909 | 3 | 0.3333 | 1 | 1 | 3 | 0 | 5 | 0.6 | 12 | 0.4167 |
| 16_377023 | 2 | 0.5 | 3 | 1 | 3 | 0 | 3 | 0.3333 | 11 | 0.4545 | 3 | 0.6667 | 2 | 0 | 4 | 0.5 | 5 | 0.6 | 14 | 0.5 |
| 16_377025 | 2 | 0 | 3 | 0 | 2 | 0.5 | 2 | 1 | 9 | 0.3333 | 3 | 0.3333 | 2 | 0 | 4 | 0.5 | 5 | 0.4 | 14 | 0.3571 |
| 16_377025 | 2 | 0 | 3 | 0 | 2 | 0.5 | 1 | 0 | 8 | 0.125 | 2 | 0.5 | 2 | 0 | 4 | 0 | 5 | 0 | 13 | 0.0769 |
| 16_377027 | 2 | 0 | 4 | 0 | 3 | 0.3333 | 3 | 0.3333 | 12 | 0.1667 | 3 | 0.3333 | 2 | 0 | 5 | 0.2 | 5 | 0.2 | 15 | 0.2 |
| 16_377028 | 2 | 0 | 3 | 0 | 3 | 0.6667 | 3 | 0.6667 | 11 | 0.3636 | 3 | 0 | 1 | 0 | 3 | 0 | 6 | 0.6667 | 13 | 0.3077 |
| 16_377030 | 2 | 0.5 | 4 | 0.25 | 3 | 0.6667 | 4 | 0.75 | 13 | 0.5385 | 3 | 1 | 2 | 1 | 4 | 0.5 | 6 | 0.8333 | 15 | 0.8 |
| 16_377032 | 2 | 0 | 5 | 0 | 3 | 0.3333 | 4 | 0.25 | 14 | 0.1429 | 2 | 0 | 2 | 0 | 5 | 0.2 | 6 | 0.1667 | 15 | 0.1333 |
| 16_377033 | 2 | 0.5 | 5 | 1 | 3 | 1 | 4 | 1 | 14 | 0.9286 | 3 | 1 | 2 | 1 | 5 | 1 | 7 | 1 | 17 | 1 |
| 16_377034 | 2 | 0 | 4 | 0 | 3 | 0.3333 | 4 | 0.25 | 13 | 0.1538 | 3 | 0.3333 | 2 | 0 | 3 | 0.3333 | 7 | 0.1429 | 15 | 0.2 |
| 16_377034 | 2 | 0 | 7 | 0 | 3 | 0 | 4 | 0.25 | 16 | 0.0625 | 3 | 0 | 2 | 0 | 5 | 0.2 | 6 | 0 | 16 | 0.0625 |
| 16_377035 | 2 | 0 | 6 | 0.1667 | 2 | 0 | 3 | 0.3333 | 13 | 0.1538 | 3 | 0 | 2 | 0 | 5 | 0 | 7 | 0.2857 | 17 | 0.1176 |
| 16_377040 | 1 | 1 | 6 | 1 | 4 | 0.75 | 4 | 1 | 15 | 0.9333 | 2 | 1 | 2 | 0 | 8 | 0.875 | 8 | 0.875 | 20 | 0.8 |
| 16_377040 | 1 | 0 | 3 | 0.3333 | 2 | 0 | 2 | 0 | 8 | 0.125 | 3 | 0 | 2 | 0 | 7 | 0 | 7 | 0.1429 | 19 | 0.0526 |
| 16_377040 | 1 | 0 | 8 | 0.125 | 3 | 0.3333 | 4 | 0 | 16 | 0.125 | 3 | 0.3333 | 2 | 0 | 8 | 0.25 | 7 | 0.2857 | 20 | 0.25 |
| 16_377041 | 2 | 0.5 | 7 | 0.1429 | 4 | 0 | 5 | 0 | 18 | 0.1111 | 4 | 0.25 | 3 | 0 | 6 | 0 | 8 | 0.125 | 21 | 0.0952 |
| 16_377042 | 2 | 1 | 8 | 0.875 | 4 | 0.75 | 5 | 0.8 | 19 | 0.8421 | 4 | 0.75 | 2 | 1 | 8 | 0.75 | 7 | 0.7143 | 21 | 0.7619 |
| 16_377045 | 4 | 0.25 | 7 | 0 | 5 | 0 | 6 | 0.1667 | 22 | 0.0909 | 4 | 0 | 4 | 0 | 11 | 0.2727 | 9 | 0.2222 | 28 | 0.1786 |
| 16_377046 | 3 | 0.3333 | 5 | 0.4 | 3 | 0 | 5 | 0.2 | 16 | 0.25 | 4 | 0 | 2 | 0 | 9 | 0.2222 | 8 | 0.25 | 23 | 0.1739 |
| 16_377047 | 4 | 0.5 | 8 | 0.375 | 4 | 0.25 | 5 | 0.6 | 21 | 0.4286 | 5 | 0.8 | 4 | 0.25 | 11 | 0.3636 | 10 | 0.3 | 30 | 0.4 |
| 16_377062 | 8 | 0.125 | 13 | 0.1538 | 10 | 0.1 | 11 | 0 | 42 | 0.0952 | 7 | 0 | 4 | 0 | 16 | 0.0625 | 12 | 0.1667 | 39 | 0.0769 |
| 16_379100 | 1 | 0 | NA | NA | 2 | 0 | 3 | 0.3333 | 6 | 0.1667 | 1 | 0 | NA | NA | 2 | 0 | 3 | 0.6667 | 6 | 0.3333 |
| 16_379104 | 1 | 1 | 1 | 1 | 3 | 0.6667 | 4 | 0.75 | 9 | 0.7778 | 1 | 1 | 1 | 0 | 1 | 1 | 3 | 0.3333 | 6 | 0.5 |
| 16_379107 | 2 | 1 | 1 | 1 | 5 | 1 | 4 | 0.75 | 12 | 0.9167 | 3 | 0.6667 | 1 | 0 | 2 | 0.5 | 4 | 0.25 | 10 | 0.4 |
| 16_380637 | 12 | 0.0833 | 8 | 0.125 | 16 | 0.0625 | 14 | 0 | 50 | 0.06 | 7 | 0.1429 | 7 | 0.2857 | 8 | 0 | 12 | 0 | 34 | 0.0882 |
| 16_380655 | 23 | 0.4348 | 14 | 0.3571 | 28 | 0.2857 | 23 | 0.3478 | 88 | 0.3523 | 17 | 0.4118 | 21 | 0.5238 | 20 | 0.45 | 26 | 0.3462 | 84 | 0.4286 |
| 16_380659 | 7 | 0.5714 | 2 | 0 | 1 | 0 | 4 | 0.25 | 14 | 0.3571 | 6 | 0.6667 | 2 | 0 | 2 | 0.5 | 4 | 0.75 | 14 | 0.5714 |
| 16_380660 | 7 | 0.5714 | 2 | 0 | 1 | 0 | 4 | 0.5 | 14 | 0.4286 | 7 | 0.5714 | 1 | 0 | 2 | 0 | 7 | 0.5714 | 17 | 0.4706 |
| 16_380661 | 6 | 0.3333 | 1 | 0 | 1 | 1 | 4 | 0 | 12 | 0.25 | 7 | 0.1429 | 2 | 0.5 | 2 | 0.5 | 6 | 0.3333 | 17 | 0.2941 |
| 16_380662 | 4 | 0 | 1 | 0 | NA | NA | 2 | 1 | 7 | 0.2857 | 4 | 0.25 | 1 | 1 | NA | NA | 2 | 0.5 | 7 | 0.4286 |
| 16_381982 | 1 | 1 | NA | NA | NA | NA | NA | NA | 1 | 1 | NA | NA | 1 | 1 | NA | NA | NA | NA | 1 | 1 |
| 16_382662 | 17 | 0.7059 | 17 | 0.8824 | 12 | 0.8333 | 10 | 0.6 | 56 | 0.7679 | 16 | 0.5625 | 19 | 0.5789 | 10 | 0.7 | 16 | 0.625 | 61 | 0.6066 |
| 16_382663 | 18 | 0.5556 | 15 | 0.5333 | 14 | 0.2857 | 10 | 0.4 | 57 | 0.4561 | 15 | 0.2667 | 19 | 0.3684 | 13 | 0.6923 | 14 | 0.4286 | 61 | 0.4262 |
| 16_382705 | 14 | 0.0714 | 15 | 0.1333 | 12 | 0 | 5 | 0.2 | 46 | 0.087 | 11 | 0.0909 | 14 | 0.1429 | 11 | 0.0909 | 11 | 0.0909 | 47 | 0.1064 |
| 16_383182 | NA | NA | 4 | 0.25 | NA | NA | 1 | 0 | 5 | 0.2 | 2 | 0.5 | NA | NA | 2 | 0.5 | NA | NA | 4 | 0.5 |
| 16_383183 | 1 | 1 | 2 | 1 | 1 | 1 | 2 | 1 | 6 | 1 | 1 | 1 | NA | NA | 1 | 1 | NA | NA | 2 | 1 |
| 16_383185 | 1 | 0 | 5 | 0.4 | 1 | 0 | 2 | 0 | 9 | 0.2222 | 2 | 0.5 | NA | NA | 2 | 0 | NA | NA | 4 | 0.25 |
| 16_383190 | 5 | 0.2 | 8 | 0.125 | 4 | 0 | 3 | 0 | 20 | 0.1 | 3 | 0 | 4 | 0 | 4 | 0.25 | 3 | 0 | 14 | 0.0714 |
| 16_383190 | 3 | 0.3333 | 8 | 0.125 | 4 | 0 | 4 | 0 | 19 | 0.1053 | 3 | 0 | 4 | 0.25 | 4 | 0 | 1 | 0 | 12 | 0.0833 |
| 16_383192 | 3 | 0.3333 | 4 | 0.25 | 4 | 0.5 | 3 | 0.6667 | 14 | 0.4286 | 3 | 0 | 2 | 0 | 2 | 0.5 | 2 | 0.5 | 9 | 0.2222 |
| 16_383192 | 4 | 0.25 | 7 | 0.4286 | 5 | 0.4 | 4 | 0.5 | 20 | 0.4 | 3 | 0.3333 | 3 | 0 | 4 | 0.5 | 2 | 1 | 12 | 0.4167 |
| 16_383454 | 4 | 0 | 1 | 1 | 1 | 0 | 2 | 0 | 8 | 0.125 | 1 | 0 | NA | NA | 1 | 0 | 3 | 0.3333 | 5 | 0.2 |
| 16_383456 | 10 | 0.2 | 7 | 0.5714 | 5 | 0.6 | 4 | 0.5 | 26 | 0.4231 | 1 | 1 | 7 | 0.2857 | 5 | 0.8 | 6 | 0.1667 | 19 | 0.4211 |
| 16_383457 | 8 | 0.5 | 6 | 0.8333 | 5 | 0.6 | 2 | 0 | 21 | 0.5714 | 2 | 0.5 | 9 | 0.5556 | 4 | 0.5 | 4 | 0.75 | 19 | 0.5789 |
| 16_383457 | 9 | 0.2222 | 5 | 0 | 5 | 0 | 3 | 0 | 22 | 0.0909 | 2 | 0 | 12 | 0.0833 | 5 | 0 | 6 | 0.1667 | 25 | 0.08 |
| 16_383458 | 7 | 1 | 6 | 0.5 | 4 | 1 | 1 | 0 | 18 | 0.7778 | 1 | 1 | 6 | 0.6667 | 2 | 0.5 | 4 | 0.75 | 13 | 0.6923 |
| 16_383460 | 7 | 0.2857 | 5 | 0.2 | 4 | 0 | 2 | 0.5 | 18 | 0.2222 | 1 | 0 | 7 | 0.1429 | 2 | 0 | 4 | 0.5 | 14 | 0.2143 |
| 16_383461 | 10 | 1 | 9 | 1 | 7 | 1 | 3 | 1 | 29 | 1 | 2 | 1 | 9 | 1 | 5 | 1 | 7 | 1 | 23 | 1 |
| 16_383462 | 10 | 1 | 8 | 0.875 | 5 | 0.8 | 2 | 0.5 | 25 | 0.88 | NA | NA | 11 | 0.8182 | 4 | 0.75 | 4 | 0.75 | 19 | 0.7895 |
| 16_383464 | 8 | 0.5 | 9 | 0.2222 | 6 | 0.3333 | 3 | 0.6667 | 26 | 0.3846 | 2 | 0.5 | 7 | 0.1429 | 4 | 0.75 | 5 | 0.4 | 18 | 0.3889 |
| 16_383464 | 8 | 0.75 | 8 | 0.75 | 4 | 0.75 | 2 | 1 | 22 | 0.7727 | 1 | 1 | 6 | 0.3333 | 4 | 0.25 | 6 | 0.6667 | 17 | 0.4706 |
| 16_383466 | 6 | 0.5 | 8 | 0.5 | 6 | 0.6667 | 3 | 0.3333 | 23 | 0.5217 | 2 | 1 | 9 | 0.4444 | 4 | 0.5 | 6 | 0.8333 | 21 | 0.619 |
| 16_383466 | 7 | 1 | 9 | 1 | 5 | 0.8 | 3 | 1 | 24 | 0.9583 | 2 | 1 | 10 | 1 | 5 | 1 | 6 | 1 | 23 | 1 |
| 16_383467 | 7 | 1 | 9 | 1 | 6 | 0.8333 | 2 | 0.5 | 24 | 0.9167 | 1 | 1 | 10 | 1 | 3 | 1 | 6 | 1 | 20 | 1 |
| 16_383468 | 8 | 1 | 9 | 0.8889 | 5 | 0.8 | 3 | 1 | 25 | 0.92 | 2 | 0.5 | 11 | 0.9091 | 4 | 1 | 6 | 1 | 23 | 0.913 |
| 16_383468 | 8 | 0.375 | 6 | 0.8333 | 3 | 0.6667 | 2 | 0.5 | 19 | 0.5789 | 1 | 0 | 8 | 0.25 | 3 | 0.3333 | 4 | 0.75 | 16 | 0.375 |
| 16_383469 | 6 | 0.6667 | 6 | 0.5 | 5 | 0.6 | 3 | 1 | 20 | 0.65 | 1 | 0 | 8 | 0.625 | 2 | 1 | 6 | 0.3333 | 17 | 0.5294 |
| 16_383469 | 7 | 0 | 9 | 0.2222 | 4 | 0 | 3 | 0 | 23 | 0.087 | 2 | 0 | 11 | 0.1818 | 4 | 0.25 | 5 | 0.2 | 22 | 0.1818 |

| | | | | | | | | | | | | | | | | | | | | |
|---|---|---|---|---|---|---|---|---|---|---|---|---|---|---|---|---|---|---|---|---|
| 16_3834699 | 5 | 1 | 8 | 1 | 3 | 0.6667 | 2 | 1 | 18 | 0.9444 | 2 | 1 | 6 | 1 | 3 | 1 | 3 | 1 | 14 | 1 |
| 16_3834699 | 4 | 0.5 | 7 | 0.2857 | 4 | 0.5 | 1 | 0 | 16 | 0.375 | NA | NA | 10 | 0.5 | 3 | 0 | 4 | 0.75 | 17 | 0.4706 |
| 16_3834707 | 8 | 0.25 | 7 | 0.4286 | 3 | 0 | 3 | 0 | 21 | 0.2381 | 2 | 0 | 9 | 0.2222 | 4 | 0 | 4 | 0 | 19 | 0.1053 |
| 16_3834707 | 7 | 0.4286 | 8 | 0.125 | 4 | 0.25 | 3 | 0.3333 | 22 | 0.2727 | 2 | 0.5 | 9 | 0.2222 | 4 | 0.5 | 5 | 0.2 | 20 | 0.3 |
| 16_3841149 | NA | NA | 2 | 0.5 | 2 | 0.5 | 4 | 0.5 | 8 | 0.5 | 2 | 0 | 1 | 0 | 1 | 1 | 1 | 0 | 5 | 0.2 |
| 16_3841878 | 3 | 0 | 4 | 0.5 | 5 | 0.4 | 2 | 0 | 14 | 0.2857 | NA | NA | 1 | 1 | 1 | 0 | 2 | 0.5 | 4 | 0.5 |
| 16_3841914 | 3 | 0.6667 | 4 | 0.25 | 3 | 0.6667 | 2 | 0 | 12 | 0.4167 | 1 | 0 | 1 | 0 | 1 | 1 | 2 | 0.5 | 5 | 0.4 |
| 16_3841942 | 3 | 0 | 3 | 0 | 3 | 0.3333 | 2 | 0.5 | 11 | 0.1818 | NA | NA | NA | NA | 1 | 0 | 2 | 0.5 | 3 | 0.3333 |
| 16_3841977 | 3 | 1 | 3 | 1 | 4 | 0.75 | 2 | 1 | 12 | 0.9167 | NA | NA | NA | NA | NA | NA | 1 | 1 | 1 | 1 |
| 16_3842012 | 3 | 0 | 3 | 1 | 4 | 0.25 | 1 | 1 | 11 | 0.4545 | NA | NA | NA | NA | NA | NA | 1 | 1 | 1 | 1 |
| 16_3842047 | 2 | 0.5 | 3 | 1 | 2 | 1 | 2 | 1 | 9 | 0.8889 | NA | NA | NA | NA | NA | NA | 1 | 1 | 1 | 1 |
| 16_3842063 | 2 | 0.5 | 3 | 0.3333 | 2 | 0.5 | 1 | 1 | 8 | 0.5 | NA | NA | NA | NA | NA | NA | 1 | 1 | 1 | 1 |
| 16_3857250 | NA | NA | NA | NA | 1 | 1 | 1 | 0 | 2 | 0.5 | 1 | 1 | 2 | 0 | NA | NA | NA | NA | 3 | 0.3333 |
| 16_3857264 | 2 | 1 | NA | NA | 2 | 0.5 | 1 | 1 | 5 | 0.8 | 1 | 1 | 5 | 0.8 | 2 | 0.5 | 1 | 1 | 9 | 0.7778 |
| 16_3858838 | 2 | 1 | 3 | 0.6667 | 2 | 0.5 | 4 | 0.25 | 11 | 0.5455 | 1 | 0 | 3 | 0.3333 | 2 | 1 | 2 | 0 | 8 | 0.375 |
| 16_3858841 | 3 | 0.6667 | 3 | 0.3333 | 3 | 0.3333 | 5 | 0.6 | 14 | 0.5 | 2 | 0.5 | 9 | 0.5556 | 2 | 0 | 2 | 1 | 15 | 0.5333 |
| 16_3858843 | 3 | 0.6667 | 2 | 0.5 | 4 | 0.75 | 5 | 0.4 | 14 | 0.5714 | 2 | 0.5 | 8 | 0.375 | 2 | 0 | 1 | 1 | 13 | 0.3846 |
| 16_3858845 | 3 | 0.3333 | 3 | 0.6667 | 3 | 1 | 5 | 0.6 | 14 | 0.6429 | 2 | 1 | 10 | 0.5 | 2 | 0.5 | 2 | 1 | 16 | 0.625 |
| 16_3858847 | 3 | 0.3333 | 3 | 0.6667 | 3 | 0.6667 | 5 | 0.6 | 14 | 0.5714 | 2 | 0.5 | 10 | 0.7 | 2 | 0 | 2 | 1 | 16 | 0.625 |
| 16_3858862 | 3 | 1 | 3 | 1 | 5 | 0.6 | 4 | 0.25 | 15 | 0.6667 | 2 | 1 | 10 | 0.6 | 1 | 1 | 2 | 1 | 15 | 0.7333 |
| 16_3858891 | 3 | 0.6667 | 3 | 1 | 3 | 0.6667 | 3 | 0.6667 | 12 | 0.75 | 2 | 0 | 10 | 0.5 | 2 | 0.5 | 2 | 1 | 16 | 0.5 |
| 16_3858892 | 3 | 0.6667 | 3 | 0 | 4 | 0.25 | 4 | 0.25 | 14 | 0.2857 | NA | NA | 9 | 0.3333 | 2 | 0.5 | 2 | 0 | 13 | 0.3077 |
| 16_3858907 | 3 | 0 | 2 | 0 | 4 | 0.25 | 5 | 0 | 14 | 0.0714 | 2 | 0 | 8 | 0.25 | 2 | 0 | 2 | 0 | 14 | 0.1429 |
| 16_3884520 | 3 | 0.3333 | 3 | 0.3333 | 3 | 0 | 6 | 0 | 15 | 0.1333 | 3 | 0.3333 | 3 | 0.3333 | 3 | 0 | 1 | 0 | 10 | 0.2 |
| 16_3884528 | 3 | 1 | 1 | 1 | 3 | 1 | 3 | 1 | 10 | 1 | 2 | 1 | 1 | 1 | 2 | 1 | 1 | 1 | 6 | 1 |
| 16_3884537 | 1 | 1 | 3 | 1 | 2 | 1 | 6 | 1 | 12 | 1 | 3 | 1 | 3 | 1 | 3 | 1 | 1 | 0 | 10 | 0.9 |
| 16_3884542 | 3 | 0 | 3 | 0.6667 | 3 | 0 | 4 | 0.5 | 13 | 0.3077 | 3 | 0.3333 | 3 | 0.3333 | 3 | 0 | NA | NA | 9 | 0.2222 |
| 16_3884556 | 3 | 0 | 4 | 0.5 | 3 | 0.3333 | 6 | 0.1667 | 16 | 0.25 | 3 | 0.3333 | 3 | 0.6667 | 3 | 0.3333 | 1 | 1 | 10 | 0.5 |
| 16_3884559 | 3 | 1 | 4 | 1 | 2 | 1 | 4 | 1 | 13 | 1 | 3 | 0.6667 | 3 | 1 | 3 | 1 | 1 | 1 | 10 | 0.9 |
| 16_3884578 | 3 | 0.3333 | 3 | 0.3333 | 4 | 0 | 7 | 0.2857 | 17 | 0.2353 | 2 | 0.5 | 4 | 0.25 | 3 | 0 | 1 | 1 | 10 | 0.3 |
| 16_3884645 | 3 | 0.3333 | 9 | 0.1111 | 4 | 0 | 8 | 0.125 | 24 | 0.125 | 4 | 0.5 | 4 | 0 | 6 | 0 | 2 | 0 | 16 | 0.125 |
| 16_3884686 | 3 | 0.6667 | 10 | 0.4 | 5 | 0 | 8 | 0.25 | 26 | 0.3077 | 4 | 0.25 | 5 | 0 | 6 | 0 | 2 | 0 | 17 | 0.0588 |
| 16_3884693 | 2 | 0 | 10 | 0.3 | 6 | 0.1667 | 6 | 0.3333 | 24 | 0.25 | 3 | 0 | 2 | 0 | 6 | 0.1667 | 2 | 0 | 13 | 0.0769 |
| 16_3884705 | 1 | 1 | 7 | 0.2857 | 5 | 0 | 6 | 0.5 | 19 | 0.3158 | 1 | 0 | 4 | 0.75 | 5 | 0 | 2 | 0 | 12 | 0.25 |
| 16_3884726 | 1 | 1 | 9 | 0.6667 | 2 | 0.5 | 7 | 0.5714 | 19 | 0.6316 | 4 | 0 | 4 | 0.25 | 7 | 0.2857 | 2 | 0 | 17 | 0.1765 |
| 16_3884726 | NA | NA | 1 | 1 | NA | NA | NA | NA | 1 | 1 | 1 | 1 | 1 | 0 | NA | NA | NA | NA | 2 | 0.5 |
| 16_4199486 | 1 | 1 | 3 | 0 | 1 | 0 | 1 | 0 | 6 | 0.1667 | 5 | 0 | 4 | 0.25 | 2 | 0 | 4 | 0.25 | 15 | 0.1333 |
| 16_4199489 | 1 | 1 | 3 | 0 | 1 | 0 | 1 | 1 | 6 | 0.3333 | 5 | 0.4 | 4 | 0.25 | 1 | 1 | 4 | 0 | 14 | 0.2857 |
| 16_4199491 | 1 | 1 | 3 | 0 | 1 | 0 | 1 | 0 | 6 | 0.1667 | 6 | 0.3333 | 4 | 0.25 | 2 | 0 | 4 | 0 | 16 | 0.1875 |
| 16_4199507 | NA | NA | 3 | 0 | 1 | 0 | 2 | 0.5 | 6 | 0.1667 | 4 | 0.75 | 3 | 0 | 1 | 0 | 3 | 0.3333 | 11 | 0.3636 |
| 16_4199966 | NA | NA | 4 | 0.25 | 1 | 0 | NA | NA | 5 | 0.2 | 1 | 1 | 7 | 0.1429 | 2 | 0 | 2 | 0 | 12 | 0.1667 |
| 16_4200075 | NA | NA | 9 | 0.1111 | 2 | 0 | 1 | 0 | 12 | 0.0833 | 1 | 0 | 12 | 0.0833 | 6 | 0.3333 | 4 | 0 | 23 | 0.1304 |
| 16_4200119 | NA | NA | 8 | 0.625 | 3 | 0.3333 | NA | NA | 11 | 0.5455 | 1 | 0 | 6 | 0 | 5 | 0.4 | 4 | 0.75 | 16 | 0.3125 |
| 16_4200144 | NA | NA | 9 | 0.3333 | 3 | 0.3333 | 1 | 0 | 13 | 0.3077 | 1 | 0 | 11 | 0 | 5 | 0.2 | 3 | 0.3333 | 20 | 0.1 |
| 16_4200175 | NA | NA | 9 | 0.3333 | 6 | 0.3333 | 1 | 1 | 16 | 0.375 | 2 | 0.5 | 10 | 0.1 | 5 | 0.2 | 5 | 0.4 | 22 | 0.2273 |
| 16_4206939 | 30 | 0 | 56 | 0.125 | 42 | 0.119 | 47 | 0.0851 | 175 | 0.0914 | 42 | 0.1429 | 42 | 0.1429 | 46 | 0.0217 | 38 | 0.1579 | 168 | 0.1131 |
| 16_4211262 | 31 | 0.1935 | 49 | 0.3265 | 39 | 0.2821 | 47 | 0.1915 | 166 | 0.253 | 37 | 0.2162 | 41 | 0.1951 | 49 | 0.1633 | 36 | 0.1389 | 163 | 0.1779 |
| 16_4345749 | NA | NA | 1 | 1 | NA | NA | NA | NA | 1 | 1 | 1 | 0 | 2 | 0 | 2 | 0.5 | 1 | 0 | 6 | 0.1667 |
| 16_4345860 | NA | NA | 3 | 0.6667 | 1 | 1 | NA | NA | 4 | 0.75 | 2 | 0 | 2 | 0.5 | 2 | 1 | 1 | 1 | 7 | 0.5714 |
| 16_4345918 | 1 | 0 | 5 | 0 | 2 | 0.5 | NA | NA | 8 | 0.125 | 2 | 0 | 4 | 0 | 2 | 0.5 | 1 | 0 | 9 | 0.1111 |
| 16_4345934 | 1 | 0 | 5 | 0 | 2 | 0.5 | NA | NA | 8 | 0.125 | 2 | 0 | 4 | 0 | 2 | 0.5 | NA | NA | 8 | 0.125 |
| 16_4346119 | 5 | 0.8 | 5 | 0.8 | 5 | 0.6 | 4 | 1 | 19 | 0.7895 | 4 | 0.75 | 6 | 0.8333 | 6 | 1 | 2 | 1 | 18 | 0.8889 |
| 16_4346132 | 4 | 0.25 | 5 | 0.2 | 6 | 0.5 | 2 | 0 | 17 | 0.2941 | 4 | 0.75 | 8 | 0.375 | 7 | 0.2857 | 4 | 0.5 | 23 | 0.4348 |
| 16_4346153 | 7 | 0.1429 | 7 | 0.1429 | 7 | 0 | 6 | 0.1667 | 27 | 0.1111 | 4 | 0.25 | 11 | 0 | 8 | 0.25 | 4 | 0.25 | 27 | 0.1481 |
| 16_4346155 | 6 | 0.1667 | 5 | 0.6 | 6 | 0.3333 | 4 | 0.5 | 21 | 0.381 | 3 | 0.6667 | 7 | 0.5714 | 9 | 0.4444 | 4 | 0.5 | 23 | 0.5217 |
| 16_4346157 | 7 | 0.1429 | 7 | 0.2857 | 8 | 0.125 | 7 | 0.1429 | 29 | 0.1724 | 4 | 0.25 | 10 | 0.2 | 8 | 0.375 | 2 | 0 | 24 | 0.25 |
| 16_4346179 | 7 | 0.2857 | 6 | 0.3333 | 8 | 0 | 8 | 0.125 | 29 | 0.1724 | 5 | 0 | 11 | 0.1818 | 9 | 0 | 5 | 0 | 30 | 0.0667 |
| 16_4346183 | 8 | 0.125 | 7 | 0.1429 | 8 | 0.125 | 8 | 0 | 31 | 0.0968 | 5 | 0.2 | 11 | 0.0909 | 10 | 0.1 | 5 | 0.2 | 31 | 0.129 |
| 16_4346251 | 10 | 0.1 | 10 | 0.2 | 9 | 0 | 12 | 0.1667 | 41 | 0.122 | 7 | 0.1429 | 13 | 0.0769 | 12 | 0.0833 | 7 | 0.2857 | 39 | 0.1282 |
| 16_4364176 | 1 | 1 | NA | NA | NA | NA | 2 | 0.5 | 3 | 0.6667 | 1 | 1 | 3 | 0.3333 | 1 | 1 | 1 | 1 | 6 | 0.6667 |
| 16_4364177 | 1 | 1 | NA | NA | NA | NA | 2 | 0 | 3 | 0.3333 | 1 | 1 | 3 | 0.6667 | 1 | 1 | 1 | 0 | 6 | 0.6667 |
| 16_4364304 | 1 | 0 | NA | NA | NA | NA | 2 | 0.5 | 3 | 0.3333 | 1 | 1 | 3 | 1 | 1 | 0 | 1 | 0 | 6 | 0.6667 |
| 16_4364312 | 1 | 1 | NA | NA | NA | NA | 2 | 0.5 | 3 | 0.6667 | 1 | 0 | 2 | 0.5 | 1 | 1 | 1 | 1 | 5 | 0.6 |
| 16_4364313 | 1 | 1 | NA | NA | NA | NA | 1 | 1 | 2 | 1 | 1 | 0 | 2 | 0 | 1 | 1 | 1 | 0 | 5 | 0.2 |
| 16_4364321 | 1 | 1 | NA | NA | NA | NA | 2 | 1 | 3 | 1 | NA | NA | 4 | 1 | 1 | 1 | 1 | 1 | 6 | 1 |

| | | | | | | | | | | | | | | | | | | | | |
|---|---|---|---|---|---|---|---|---|---|---|---|---|---|---|---|---|---|---|---|---|
| 16_436432 | 1 | 0 | NA | NA | NA | NA | 2 | 0.5 | 3 | 0.3333 | 1 | 0 | 3 | 0.3333 | 1 | 0 | 1 | 0 | 6 | 0.1667 |
| 16_436434 | 1 | 1 | NA | NA | NA | NA | 1 | 0 | 2 | 0.5 | 1 | 1 | 2 | 1 | 1 | 1 | 1 | 1 | 5 | 1 |
| 16_436436 | 1 | 0 | NA | NA | NA | NA | 2 | 0.5 | 3 | 0.3333 | 1 | 0 | 3 | 0.3333 | 1 | 0 | 1 | 0 | 6 | 0.1667 |
| 16_436436 | 1 | 0 | NA | NA | NA | NA | 2 | 0.5 | 3 | 0.3333 | 1 | 0 | 4 | 0.25 | NA | NA | 1 | 0 | 6 | 0.1667 |
| 16_436438 | 1 | 1 | NA | NA | NA | NA | 2 | 0.5 | 3 | 0.6667 | NA | NA | 4 | 0.75 | 1 | 0 | 1 | 0 | 6 | 0.5 |
| 16_436440 | 1 | 1 | 1 | 0 | NA | NA | 2 | 0.5 | 4 | 0.5 | 1 | 0 | 3 | 0.3333 | 1 | 1 | 1 | 0 | 6 | 0.3333 |
| 16_436451 | 1 | 1 | 1 | 0 | NA | NA | NA | NA | 2 | 0.5 | 1 | 0 | 4 | 0.5 | 3 | 0 | 2 | 0 | 10 | 0.2 |
| 16_436827 | NA | NA | 2 | 0 | 2 | 0 | 5 | 0.2 | 9 | 0.1111 | NA | NA | 2 | 0 | 2 | 0 | 2 | 0.5 | 6 | 0.1667 |
| 16_436832 | NA | NA | NA | NA | NA | NA | 3 | 1 | 3 | 1 | NA | NA | 1 | 1 | 2 | 0.5 | 2 | 1 | 5 | 0.8 |
| 16_436832 | NA | NA | NA | NA | NA | NA | 2 | 1 | 2 | 1 | NA | NA | 2 | 1 | 2 | 1 | 1 | 1 | 5 | 1 |
| 16_436833 | NA | NA | 1 | 1 | 1 | 0 | 3 | 0 | 5 | 0.2 | NA | NA | 2 | 0.5 | 2 | 0 | NA | NA | 4 | 0.25 |
| 16_436834 | NA | NA | 1 | 1 | NA | NA | 2 | 1 | 3 | 1 | NA | NA | 2 | 1 | 2 | 1 | 1 | 1 | 5 | 1 |
| 16_436834 | NA | NA | 1 | 0 | 1 | 0 | 3 | 0.6667 | 5 | 0.4 | NA | NA | 2 | 0.5 | 2 | 0 | NA | NA | 4 | 0.25 |
| 16_436834 | NA | NA | 1 | 1 | 1 | 1 | 2 | 1 | 4 | 1 | NA | NA | 2 | 1 | 2 | 1 | 1 | 1 | 5 | 1 |
| 16_437856 | 2 | 0 | 2 | 0 | NA | NA | 4 | 0.25 | 8 | 0.125 | 5 | 0.2 | 5 | 0 | 2 | 0 | 6 | 0.3333 | 18 | 0.1667 |
| 16_437860 | 1 | 1 | 2 | 0.5 | NA | NA | 4 | 0.25 | 7 | 0.4286 | 4 | 0 | 3 | 0 | 1 | 0 | 4 | 0.25 | 12 | 0.0833 |
| 16_437861 | 1 | 0 | 1 | 1 | NA | NA | 4 | 0.25 | 6 | 0.3333 | 5 | 0.6 | 5 | 0.2 | 1 | 0 | 6 | 0.5 | 17 | 0.4118 |
| 16_437861 | 1 | 0 | 2 | 0.5 | NA | NA | 4 | 0.5 | 7 | 0.4286 | 5 | 0.8 | 4 | 0.25 | 1 | 0 | 4 | 0.25 | 14 | 0.4286 |
| 16_437865 | 1 | 0 | 2 | 0 | NA | NA | 3 | 0.3333 | 6 | 0.1667 | 5 | 0 | 4 | 0.25 | 1 | 1 | 4 | 0 | 14 | 0.1429 |
| 16_437873 | 1 | 1 | 1 | 0 | NA | NA | 3 | 0.3333 | 5 | 0.4 | 4 | 0.25 | 2 | 0 | NA | NA | 4 | 0.5 | 10 | 0.3 |
| 16_437875 | 1 | 0 | 1 | 0 | NA | NA | 3 | 0.3333 | 5 | 0.2 | 4 | 0.25 | 3 | 0 | 1 | 1 | 3 | 0.3333 | 11 | 0.2727 |
| 16_437879 | 1 | 0 | 1 | 1 | NA | NA | 3 | 0.3333 | 5 | 0.4 | 1 | 0 | 3 | 0.3333 | 1 | 0 | 2 | 0 | 7 | 0.1429 |
| 16_437882 | 1 | 0 | 1 | 1 | NA | NA | NA | NA | 2 | 0.5 | 3 | 0 | 2 | 0 | 1 | 1 | 3 | 0.6667 | 9 | 0.3333 |
| 16_437883 | 1 | 0 | 1 | 0 | NA | NA | 1 | 1 | 3 | 0.3333 | 2 | 0 | 3 | 0.3333 | 1 | 0 | 2 | 0 | 8 | 0.125 |
| 16_437883 | 1 | 1 | 1 | 1 | NA | NA | 1 | 1 | 3 | 1 | 3 | 1 | 3 | 1 | 1 | 1 | 2 | 1 | 9 | 1 |
| 16_437884 | 1 | 1 | 1 | 1 | NA | NA | NA | NA | 2 | 1 | 2 | 1 | 3 | 1 | 1 | 1 | 3 | 1 | 9 | 1 |
| 16_437885 | 1 | 0 | NA | NA | NA | NA | 1 | 1 | 2 | 0.5 | 3 | 0.3333 | 3 | 0 | 1 | 0 | 3 | 0 | 10 | 0.1 |
| 16_437885 | 1 | 1 | 1 | 1 | NA | NA | NA | NA | 2 | 1 | 2 | 1 | 2 | 1 | NA | NA | 1 | 1 | 5 | 1 |
| 16_437886 | 1 | 0 | 1 | 1 | NA | NA | 1 | 0 | 3 | 0.3333 | 2 | 0.5 | 3 | 1 | 1 | 1 | 3 | 1 | 9 | 0.8889 |
| 16_437886 | 1 | 1 | 1 | 1 | 1 | 1 | 1 | 1 | 4 | 1 | 3 | 0.3333 | 3 | 1 | 1 | 1 | 3 | 1 | 10 | 0.8 |
| 16_437888 | 1 | 1 | 1 | 1 | 1 | 0 | NA | NA | 3 | 0.6667 | 2 | 0.5 | 3 | 0.6667 | 1 | 1 | 2 | 1 | 8 | 0.75 |
| 16_437889 | 1 | 1 | 1 | 0 | 1 | 0 | NA | NA | 3 | 0.3333 | 2 | 1 | 3 | 1 | 1 | 1 | 2 | 1 | 8 | 1 |
| 16_437889 | 1 | 1 | 1 | 1 | NA | NA | NA | NA | 2 | 1 | 3 | 1 | 2 | 0.5 | 1 | 1 | 2 | 1 | 8 | 0.875 |
| 16_437889 | 1 | 1 | 1 | 1 | NA | NA | NA | NA | 2 | 1 | 3 | 1 | 3 | 0 | 1 | 1 | 2 | 0.5 | 9 | 0.5556 |
| 16_437892 | 1 | 0 | 1 | 1 | NA | NA | NA | NA | 2 | 0.5 | 3 | 0 | 3 | 0 | 1 | 0 | 1 | 1 | 8 | 0.125 |
| 16_437893 | 1 | 1 | 1 | 0 | NA | NA | NA | NA | 2 | 0.5 | 1 | 1 | 3 | 0 | NA | NA | 1 | 0 | 5 | 0.2 |
| 16_437893 | 1 | 1 | 1 | 1 | NA | NA | NA | NA | 2 | 1 | 2 | 1 | 3 | 1 | 1 | 1 | NA | NA | 6 | 1 |
| 16_437894 | 1 | 1 | 1 | 1 | NA | NA | NA | NA | 2 | 1 | 1 | 1 | 2 | 1 | 1 | 0 | 1 | 1 | 5 | 0.8 |
| 16_437894 | 1 | 1 | 1 | 1 | NA | NA | NA | NA | 2 | 1 | 2 | 1 | 3 | 1 | NA | NA | 1 | 1 | 6 | 1 |
| 16_437896 | 1 | 1 | 1 | 0 | NA | NA | NA | NA | 2 | 0.5 | 2 | 0.5 | 2 | 0 | 1 | 0 | 1 | 0 | 6 | 0.1667 |
| 16_437938 | 1 | 1 | 1 | 0 | NA | NA | NA | NA | 2 | 0.5 | 2 | 0.5 | 3 | 0.3333 | NA | NA | 1 | 0 | 6 | 0.3333 |
| 16_437942 | 1 | 1 | 1 | 0 | NA | NA | NA | NA | 2 | 0.5 | 2 | 0.5 | 1 | 0 | NA | NA | 1 | 1 | 4 | 0.5 |
| 16_439061 | 60 | 0.15 | 56 | 0.1607 | 47 | 0.1915 | 50 | 0.2 | 213 | 0.1737 | 33 | 0.3333 | 44 | 0.2955 | 43 | 0.1628 | 44 | 0.2273 | 164 | 0.25 |
| 16_439064 | 43 | 0.186 | 47 | 0.0851 | 38 | 0.1842 | 48 | 0.125 | 176 | 0.142 | 20 | 0.1 | 29 | 0.0345 | 39 | 0.1795 | 32 | 0.1875 | 120 | 0.1333 |
| 16_439077 | 40 | 0.175 | 33 | 0.1818 | 27 | 0.2222 | 33 | 0.2727 | 133 | 0.2105 | 21 | 0.1429 | 26 | 0.0769 | 30 | 0.3 | 29 | 0.2069 | 106 | 0.1887 |
| 16_439800 | 6 | 1 | 8 | 1 | 3 | 0.3333 | 5 | 1 | 22 | 0.9091 | 3 | 1 | 6 | 0.8333 | 4 | 1 | 8 | 1 | 21 | 0.9524 |
| 16_439801 | 5 | 0.8 | 9 | 0.6667 | 5 | 1 | 6 | 1 | 25 | 0.84 | 2 | 1 | 7 | 1 | 4 | 0.75 | 9 | 0.8889 | 22 | 0.9091 |
| 16_439804 | 5 | 0.2 | 9 | 0 | 6 | 0 | 5 | 0.2 | 25 | 0.08 | 3 | 0 | 9 | 0 | 4 | 0.5 | 9 | 0.1111 | 25 | 0.12 |
| 16_439806 | 6 | 0 | 9 | 0.1111 | 4 | 0.25 | 5 | 0 | 24 | 0.0833 | 5 | 0.2 | 7 | 0 | 7 | 0.1429 | 8 | 0.125 | 27 | 0.1111 |
| 16_439809 | 8 | 0.25 | 11 | 0 | 6 | 0.1667 | 5 | 0.6 | 30 | 0.2 | 5 | 0.2 | 12 | 0.1667 | 6 | 0.1667 | 8 | 0.125 | 31 | 0.1613 |
| 16_439813 | 6 | 0.1667 | 7 | 0.2857 | 7 | 0 | 5 | 0 | 25 | 0.12 | 6 | 0 | 10 | 0.1 | 6 | 0.3333 | 6 | 0.1667 | 28 | 0.1429 |
| 16_439816 | 4 | 0 | 3 | 0.3333 | 4 | 0 | 1 | 0 | 12 | 0.0833 | 1 | 0 | 4 | 0.25 | 2 | 0 | 3 | 0 | 10 | 0.1 |
| 16_439835 | 1 | 0 | 2 | 0 | 4 | 0 | 3 | 0.3333 | 10 | 0.1 | 5 | 0.2 | 3 | 0 | NA | NA | 3 | 0.3333 | 11 | 0.1818 |
| 16_440408 | NA | NA | 1 | 1 | NA | NA | NA | NA | 1 | 1 | 1 | 1 | 2 | 1 | NA | NA | 2 | 1 | 5 | 1 |
| 16_440410 | NA | NA | 1 | 1 | NA | NA | NA | NA | 1 | 1 | 1 | 1 | 1 | 1 | 1 | 1 | 3 | 1 | 6 | 1 |
| 16_440414 | NA | NA | 1 | 1 | NA | NA | NA | NA | 1 | 1 | 1 | 1 | 1 | 0 | 1 | 1 | 2 | 1 | 5 | 0.8 |
| 16_440417 | 1 | 1 | 1 | 1 | NA | NA | 1 | 1 | 3 | 1 | 1 | 1 | 1 | 1 | 1 | 1 | 3 | 1 | 6 | 1 |
| 16_440418 | 1 | 1 | 1 | 1 | NA | NA | 1 | 1 | 3 | 1 | 1 | 1 | 2 | 0.5 | 1 | 1 | 2 | 1 | 6 | 0.8333 |
| 16_440418 | NA | NA | 1 | 1 | NA | NA | 1 | 1 | 2 | 1 | 2 | 1 | NA | NA | 1 | 1 | 3 | 1 | 6 | 1 |
| 16_440419 | 1 | 0 | 1 | 1 | NA | NA | 1 | 0 | 3 | 0.3333 | 2 | 1 | 2 | 0.5 | 1 | 1 | 2 | 0.5 | 7 | 0.7143 |
| 16_440421 | 1 | 0 | 1 | 1 | NA | NA | 1 | 1 | 3 | 0.6667 | 2 | 1 | 2 | 0.5 | 1 | 1 | 2 | 1 | 7 | 0.8571 |
| 16_440421 | 1 | 1 | 1 | 0 | NA | NA | NA | NA | 2 | 0.5 | 2 | 1 | 2 | 0 | 1 | 1 | 3 | 0.6667 | 8 | 0.625 |
| 16_440422 | NA | NA | NA | NA | NA | NA | 1 | 1 | 1 | 1 | 2 | 0.5 | 1 | 1 | NA | NA | 1 | 1 | 4 | 0.75 |
| 16_440422 | NA | NA | 1 | 1 | NA | NA | 1 | 1 | 2 | 1 | 2 | 0 | 1 | 0 | 1 | 0 | 3 | 1 | 7 | 0.4286 |
| 16_440427 | 1 | 0 | 1 | 1 | NA | NA | 1 | 0 | 3 | 0.3333 | 2 | 0 | 3 | 0 | 3 | 0 | 3 | 0.3333 | 11 | 0.0909 |

| | | | | | | | | | | | | | | | | | | | | |
|---|---|---|---|---|---|---|---|---|---|---|---|---|---|---|---|---|---|---|---|---|
| 16_4404627 | 3 | 0 | 7 | 0 | 5 | 0.2 | 4 | 0 | 19 | 0.0526 | 4 | 0 | 4 | 0 | 3 | 0 | 4 | 0.25 | 15 | 0.0667 |
| 16_4404653 | 4 | 0.25 | 8 | 0.125 | 5 | 0.2 | 4 | 0.25 | 21 | 0.1905 | 4 | 0.25 | 4 | 0 | 3 | 1 | 3 | 0.3333 | 14 | 0.3571 |
| 16_4404659 | 3 | 0 | 8 | 0 | 5 | 0.2 | 3 | 0.3333 | 19 | 0.1053 | 4 | 0 | 4 | 0 | 3 | 0.3333 | 4 | 0 | 15 | 0.0667 |
| 16_4404755 | 6 | 0 | 9 | 0.1111 | 9 | 0.1111 | 6 | 0 | 30 | 0.0667 | 5 | 0 | 5 | 0.2 | 4 | 0 | 4 | 0 | 18 | 0.0556 |
| 16_4406064 | NA | NA | 8 | 0.625 | 5 | 0.2 | 1 | 1 | 14 | 0.5 | 6 | 0.1667 | 1 | 0 | 1 | 1 | 2 | 0 | 10 | 0.2 |
| 16_4406064 | 2 | 0 | 2 | 0.5 | NA | NA | NA | NA | 4 | 0.25 | NA | NA | NA | NA | 1 | 1 | 1 | 0 | 2 | 0.5 |
| 16_4406075 | 2 | 0.5 | 9 | 0.7778 | 5 | 1 | NA | NA | 16 | 0.8125 | 4 | 0.75 | 1 | 1 | 2 | 1 | 4 | 0.75 | 11 | 0.8182 |
| 16_4406093 | NA | NA | 9 | 0 | 5 | 0.2 | 1 | 0 | 15 | 0.0667 | 7 | 0 | 1 | 0 | 2 | 0 | 4 | 0.25 | 14 | 0.0714 |
| 16_4406099 | 1 | 0 | 5 | 0.4 | 3 | 0 | 1 | 0 | 10 | 0.2 | 6 | 0.5 | 1 | 0 | 2 | 0.5 | 3 | 0.6667 | 12 | 0.5 |
| 16_4406105 | 1 | 0 | 7 | 0 | 2 | 1 | 1 | 1 | 11 | 0.2727 | 7 | 0.4286 | 1 | 1 | 2 | 0.5 | 4 | 0.5 | 14 | 0.5 |
| 16_4406117 | NA | NA | 8 | 1 | 4 | 1 | 1 | 1 | 13 | 1 | 7 | 1 | 1 | 1 | NA | NA | 2 | 1 | 10 | 1 |
| 16_4406119 | 1 | 0 | 9 | 0.1111 | 4 | 0 | 1 | 0 | 15 | 0.0667 | 6 | 0.1667 | 1 | 0 | 2 | 0.5 | 3 | 0 | 12 | 0.1667 |
| 16_4406121 | 1 | 0 | 6 | 1 | 2 | 1 | NA | NA | 9 | 0.8889 | 5 | 0.8 | NA | NA | 1 | 1 | 2 | 1 | 8 | 0.875 |
| 16_4406126 | 1 | 1 | 7 | 0.7143 | 4 | 1 | 1 | 1 | 13 | 0.8462 | 5 | 0.6 | 1 | 1 | 1 | 1 | 1 | 1 | 8 | 0.75 |
| 16_4406128 | 1 | 0 | 7 | 0.5714 | 4 | 0.75 | 1 | 0 | 13 | 0.5385 | 7 | 0.7143 | 1 | 1 | 2 | 1 | 3 | 0.3333 | 13 | 0.6923 |
| 16_4406133 | 1 | 1 | 9 | 0.5556 | 4 | 0.75 | 1 | 1 | 15 | 0.6667 | 6 | 0.6667 | 1 | 0 | 2 | 0.5 | 3 | 0.6667 | 12 | 0.5833 |
| 16_4406141 | 1 | 1 | 9 | 0.4444 | 4 | 0.25 | 1 | 0 | 15 | 0.4 | 7 | 0.1429 | 1 | 0 | 2 | 0 | 3 | 0.3333 | 13 | 0.1538 |
| 16_4406150 | 1 | 1 | 9 | 0.7778 | 4 | 0.5 | 1 | 1 | 15 | 0.7333 | 7 | 0.8571 | 1 | 1 | 2 | 0 | 3 | 0.6667 | 13 | 0.6923 |
| 16_4406150 | 1 | 0 | 9 | 0.1111 | 4 | 0.25 | 1 | 0 | 15 | 0.1333 | 7 | 0 | 1 | 0 | 2 | 0.5 | 2 | 0.5 | 12 | 0.1667 |
| 16_4406152 | 1 | 0 | 8 | 0.25 | 4 | 0 | 1 | 0 | 14 | 0.1429 | 4 | 0 | 1 | 1 | 2 | 0 | 3 | 0 | 10 | 0.1 |
| 16_4406155 | 1 | 1 | 5 | 0.6 | 4 | 0.25 | NA | NA | 10 | 0.5 | 3 | 0.3333 | 1 | 1 | 1 | 0 | 3 | 0.6667 | 8 | 0.5 |
| 16_4406156 | 1 | 0 | 9 | 0.3333 | 4 | 0.25 | 1 | 0 | 15 | 0.2667 | 6 | 0.1667 | 1 | 1 | 2 | 0 | 3 | 0.3333 | 12 | 0.25 |
| 16_4406179 | NA | NA | 8 | 0.75 | 4 | 0.75 | 1 | 1 | 13 | 0.7692 | 6 | 0.8333 | 1 | 1 | 2 | 0 | 2 | 0.5 | 11 | 0.6364 |
| 16_4406184 | 1 | 1 | 6 | 0.1667 | 2 | 0 | 1 | 0 | 10 | 0.2 | 6 | 0.1667 | 1 | 0 | 1 | 1 | 1 | 0 | 9 | 0.2222 |
| 16_4406184 | NA | NA | 7 | 1 | 4 | 0.75 | 1 | 1 | 12 | 0.9167 | 5 | 1 | 1 | 1 | NA | NA | 3 | 1 | 9 | 1 |
| 16_4415288 | 1 | 0 | 2 | 0.5 | 1 | 1 | NA | NA | 4 | 0.5 | 2 | 1 | NA | NA | NA | NA | NA | NA | 2 | 1 |
| 16_4419956 | NA | NA | NA | NA | 2 | 1 | 1 | 1 | 3 | 1 | NA | NA | 1 | 1 | NA | NA | 1 | 0 | 2 | 0.5 |
| 16_4430566 | 1 | 1 | NA | NA | 1 | 0 | 3 | 0.6667 | 5 | 0.6 | 2 | 0 | NA | NA | NA | NA | 1 | 1 | 3 | 0.3333 |
| 16_4455718 | 2 | 0.5 | 1 | 0 | 3 | 0.3333 | 3 | 0.6667 | 9 | 0.4444 | 1 | 1 | 1 | 1 | 2 | 1 | 2 | 0.5 | 6 | 0.8333 |
| 16_4455724 | 3 | 0 | 1 | 0 | 2 | 0.5 | 2 | 0 | 8 | 0.125 | 1 | 0 | 1 | 1 | 2 | 0 | 2 | 0 | 6 | 0.1667 |
| 16_4455852 | 2 | 0.5 | 1 | 1 | 4 | 0.25 | 2 | 0.5 | 9 | 0.4444 | 1 | 1 | NA | NA | 1 | 0 | NA | NA | 2 | 0.5 |
| 16_4455942 | NA | NA | NA | NA | 1 | 1 | NA | NA | 1 | 1 | NA | NA | 1 | 1 | NA | NA | 2 | 0 | 3 | 0.3333 |
| 16_4455971 | NA | NA | NA | NA | 1 | 1 | NA | NA | 1 | 1 | NA | NA | 2 | 1 | NA | NA | 1 | 1 | 3 | 1 |
| 16_4456649 | 2 | 0 | 1 | 1 | 2 | 0.5 | 2 | 0 | 7 | 0.2857 | 1 | 1 | 2 | 1 | 1 | 0 | NA | NA | 4 | 0.75 |
| 16_4456649 | 1 | 0 | 1 | 0 | 2 | 0.5 | 2 | 0 | 6 | 0.1667 | 1 | 0 | 1 | 1 | NA | NA | NA | NA | 2 | 0.5 |
| 16_4456651 | 2 | 0 | NA | NA | 2 | 0.5 | 1 | 0 | 5 | 0.2 | 1 | 0 | 2 | 0.5 | 1 | 0 | NA | NA | 4 | 0.25 |
| 16_4456663 | 2 | 0 | 1 | 0 | 2 | 0 | 1 | 1 | 6 | 0.1667 | NA | NA | 1 | 1 | 1 | 0 | NA | NA | 2 | 0.5 |
| 16_4491596 | 1 | 1 | NA | NA | NA | NA | NA | NA | 1 | 1 | 1 | 1 | NA | NA | NA | NA | NA | NA | 1 | 1 |
| 16_4494787 | 2 | 0.5 | 1 | 1 | NA | NA | NA | NA | 3 | 0.6667 | 1 | 1 | NA | NA | NA | NA | 1 | 1 | 2 | 1 |
| 16_4496228 | 8 | 0.25 | 4 | 0.5 | 3 | 0.3333 | 5 | 0.4 | 20 | 0.35 | 2 | 1 | 11 | 0.3636 | 2 | 0.5 | 6 | 0.3333 | 21 | 0.4286 |
| 16_4496231 | 6 | 0.1667 | 2 | 0 | 3 | 0 | 5 | 0.2 | 16 | 0.125 | 1 | 0 | 10 | 0.2 | 1 | 0 | 4 | 0.25 | 16 | 0.1875 |
| 16_4496236 | 6 | 0 | 4 | 0 | 4 | 0 | 5 | 0.2 | 19 | 0.0526 | 1 | 0 | 10 | 0.2 | 2 | 0 | 5 | 0.2 | 18 | 0.1667 |
| 16_4496247 | 7 | 0.1429 | 4 | 0.5 | 4 | 0.25 | 5 | 0.8 | 20 | 0.4 | 2 | 0 | 8 | 0.125 | 1 | 0 | 6 | 0.3333 | 17 | 0.1765 |
| 16_4496255 | 8 | 0.125 | 4 | 0 | 4 | 0.25 | 5 | 0 | 21 | 0.0952 | 2 | 0 | 11 | 0.0909 | 2 | 0 | 5 | 0 | 20 | 0.05 |
| 16_4496425 | 6 | 0.5 | 3 | 0 | 2 | 0.5 | 4 | 0.25 | 15 | 0.3333 | 1 | 1 | 7 | 0.2857 | 1 | 1 | 6 | 0.3333 | 15 | 0.4 |
| 16_4496430 | 7 | 0.8571 | 2 | 1 | 1 | 1 | 5 | 1 | 15 | 0.9333 | 2 | 1 | 8 | 1 | 2 | 1 | 6 | 1 | 18 | 1 |
| 16_4496442 | 3 | 0.3333 | 2 | 0 | NA | NA | 4 | 0.5 | 9 | 0.3333 | NA | NA | 4 | 0.75 | 2 | 0 | 3 | 0.6667 | 9 | 0.5556 |
| 16_4496454 | 4 | 0 | 4 | 0 | 2 | 0 | 4 | 0.25 | 14 | 0.0714 | 2 | 0 | 9 | 0.1111 | 1 | 0 | 4 | 0 | 16 | 0.0625 |
| 16_4496456 | 5 | 0.4 | 4 | 0.5 | 1 | 1 | 5 | 0.8 | 15 | 0.6 | 2 | 0 | 8 | 0.75 | 2 | 0.5 | 6 | 0.8333 | 18 | 0.6667 |
| 16_4496458 | 4 | 1 | 2 | 0.5 | 3 | 0.6667 | 3 | 1 | 12 | 0.8333 | NA | NA | 8 | 0.75 | 2 | 1 | 6 | 0.8333 | 16 | 0.8125 |
| 16_4496460 | 5 | 1 | 3 | 1 | 3 | 0.6667 | 5 | 1 | 16 | 0.9375 | 2 | 0.5 | 8 | 1 | 2 | 1 | 6 | 1 | 18 | 0.9444 |
| 16_4502516 | NA | NA | NA | NA | 1 | 1 | 1 | 0 | 2 | 0.5 | NA | NA | NA | NA | 1 | 1 | 3 | 0 | 4 | 0.25 |
| 16_4502557 | NA | NA | NA | NA | 1 | 1 | NA | NA | 1 | 1 | NA | NA | 1 | 0 | 1 | 1 | 2 | 0.5 | 4 | 0.5 |
| 16_4502573 | NA | NA | NA | NA | NA | NA | 1 | 1 | 1 | 1 | NA | NA | 1 | 1 | 1 | 1 | 2 | 1 | 4 | 1 |
| 16_4502574 | NA | NA | NA | NA | 1 | 1 | 1 | 0 | 2 | 0.5 | NA | NA | 1 | 0 | 2 | 0 | 3 | 0.3333 | 6 | 0.1667 |
| 16_4502591 | NA | NA | NA | NA | 1 | 1 | 1 | 1 | 2 | 1 | NA | NA | 1 | 0 | 2 | 1 | 3 | 0.6667 | 6 | 0.6667 |
| 16_4502596 | NA | NA | NA | NA | 1 | 1 | 1 | 1 | 2 | 1 | NA | NA | NA | NA | 1 | 1 | NA | NA | 1 | 1 |
| 16_4503670 | NA | NA | 1 | 0 | 1 | 1 | 1 | 0 | 3 | 0.3333 | NA | NA | 1 | 0 | 1 | 1 | 1 | 0 | 3 | 0.3333 |
| 16_4503681 | NA | NA | 1 | 1 | 1 | 0 | 1 | 1 | 3 | 0.6667 | NA | NA | 1 | 1 | 1 | 1 | 2 | 0 | 4 | 0.5 |
| 16_4503681 | NA | NA | 1 | 1 | 1 | 0 | 1 | 0 | 3 | 0.3333 | NA | NA | 1 | 1 | 1 | 1 | 2 | 0.5 | 4 | 0.75 |
| 16_4503682 | NA | NA | 1 | 1 | 1 | 1 | 1 | 1 | 3 | 1 | NA | NA | 1 | 1 | 1 | 1 | 2 | 1 | 4 | 1 |
| 16_4503684 | NA | NA | NA | NA | 1 | 1 | 1 | 0 | 2 | 0.5 | NA | NA | 1 | 1 | 1 | 1 | 2 | 0.5 | 4 | 0.75 |
| 16_4503685 | NA | NA | NA | NA | 1 | 1 | 1 | 0 | 2 | 0.5 | NA | NA | 1 | 0 | 1 | 1 | 1 | 1 | 3 | 0.6667 |
| 16_4520342 | 10 | 0.3 | 15 | 0.2 | 11 | 0.3636 | 9 | 0.5556 | 45 | 0.3333 | 8 | 0.5 | 10 | 0.1 | 15 | 0.2 | 15 | 0.2 | 48 | 0.2292 |
| 16_4520348 | 9 | 0.1111 | 12 | 0 | 9 | 0.2222 | 9 | 0 | 39 | 0.0769 | 6 | 0 | 8 | 0.125 | 10 | 0 | 14 | 0.0714 | 38 | 0.0526 |

| | | | | | | | | | | | | | | | | | | | | |
|---|---|---|---|---|---|---|---|---|---|---|---|---|---|---|---|---|---|---|---|---|
| 16_452035 | 10 | 0.2 | 17 | 0.0588 | 9 | 0.2222 | 9 | 0.1111 | 45 | 0.1333 | 8 | 0 | 10 | 0 | 16 | 0.1875 | 14 | 0.2857 | 48 | 0.1458 |
| 16_452036 | 9 | 1 | 15 | 0.8667 | 7 | 0.8571 | 8 | 1 | 39 | 0.9231 | 7 | 0.8571 | 8 | 1 | 14 | 0.9286 | 13 | 1 | 42 | 0.9524 |
| 16_452036 | 9 | 0.3333 | 14 | 0.2143 | 7 | 0.5714 | 7 | 0.2857 | 37 | 0.3243 | 7 | 0.5714 | 8 | 0.5 | 14 | 0.6429 | 10 | 0.7 | 39 | 0.6154 |
| 16_452037 | 9 | 0 | 16 | 0.0625 | 8 | 0.125 | 8 | 0.25 | 41 | 0.0976 | 7 | 0.1429 | 9 | 0.3333 | 16 | 0.1875 | 13 | 0.2308 | 45 | 0.2222 |
| 16_452037 | 9 | 0.3333 | 16 | 0.0625 | 7 | 0.4286 | 7 | 0.4286 | 39 | 0.2564 | 7 | 0.2857 | 8 | 0.375 | 16 | 0.3125 | 12 | 0.5 | 43 | 0.3721 |
| 16_452150 | 8 | 0 | 11 | 0.3636 | 7 | 0.2857 | 6 | 0.3333 | 32 | 0.25 | 4 | 0 | 8 | 0.25 | 10 | 0.3 | 9 | 0.3333 | 31 | 0.2581 |
| 16_452151 | 7 | 0.2857 | 13 | 0.0769 | 8 | 0 | 8 | 0 | 36 | 0.0833 | 7 | 0.1429 | 8 | 0.125 | 16 | 0.0625 | 11 | 0 | 42 | 0.0714 |
| 16_452174 | 8 | 0.25 | 11 | 0.2727 | 6 | 0.3333 | 7 | 0.1429 | 32 | 0.25 | 6 | 0.1667 | 7 | 0.4286 | 13 | 0.3077 | 9 | 0.4444 | 35 | 0.3429 |
| 16_455317 | 43 | 0.186 | 63 | 0.2063 | 42 | 0.119 | 35 | 0.2 | 183 | 0.1803 | 43 | 0.1628 | 36 | 0.0833 | 33 | 0.1212 | 36 | 0.1667 | 148 | 0.1351 |
| 16_455317 | 49 | 0.1224 | 60 | 0.1 | 42 | 0.0238 | 29 | 0.1379 | 180 | 0.0944 | 41 | 0.1463 | 32 | 0.1875 | 32 | 0.0938 | 35 | 0.0857 | 140 | 0.1286 |
| 16_455449 | 3 | 0 | 5 | 0.4 | 3 | 0 | 1 | 0 | 12 | 0.1667 | 1 | 1 | 1 | 0 | NA | NA | NA | NA | 2 | 0.5 |
| 16_455460 | NA | NA | NA | NA | 1 | 0 | 1 | 1 | 2 | 0.5 | 1 | 1 | NA | NA | NA | NA | NA | NA | 1 | 1 |
| 16_455520 | 3 | 0 | 1 | 0 | 2 | 0 | 5 | 0.2 | 11 | 0.0909 | 2 | 0 | 3 | 0.3333 | 3 | 0 | 3 | 0.3333 | 11 | 0.1818 |
| 16_455520 | 1 | 1 | NA | NA | 2 | 0.5 | 4 | 0.5 | 7 | 0.5714 | 1 | 1 | 3 | 0.6667 | 1 | 1 | 3 | 0.6667 | 8 | 0.75 |
| 16_455521 | 3 | 0.6667 | 3 | 0 | 2 | 0 | 5 | 0 | 13 | 0.1538 | 2 | 0.5 | 2 | 0.5 | 3 | 0.3333 | 3 | 0.3333 | 10 | 0.4 |
| 16_455522 | 3 | 1 | 3 | 0 | 1 | 0 | 5 | 0.6 | 12 | 0.5 | 2 | 0.5 | 3 | 0 | 2 | 0 | 3 | 0.6667 | 10 | 0.3 |
| 16_457805 | 3 | 0 | 2 | 0 | 3 | 0 | 2 | 0.5 | 10 | 0.1 | 4 | 0.25 | 2 | 0.5 | NA | NA | 1 | 0 | 7 | 0.2857 |
| 16_457810 | 3 | 0.3333 | 2 | 1 | 2 | 0.5 | 1 | 1 | 8 | 0.625 | 3 | 0 | 2 | 0.5 | NA | NA | 1 | 0 | 6 | 0.1667 |
| 16_457829 | 1 | 0 | NA | NA | 2 | 0.5 | 2 | 0 | 5 | 0.2 | 1 | 1 | 1 | 1 | NA | NA | 1 | 0 | 3 | 0.6667 |
| 16_457840 | NA | NA | NA | NA | 2 | 1 | 2 | 0.5 | 4 | 0.75 | 1 | 1 | 1 | 1 | NA | NA | NA | NA | 2 | 1 |
| 16_457842 | NA | NA | 1 | 0 | 2 | 1 | 1 | 1 | 4 | 0.75 | 1 | 1 | 1 | 1 | NA | NA | NA | NA | 2 | 1 |
| 16_457843 | NA | NA | NA | NA | 1 | 1 | 2 | 1 | 3 | 1 | 1 | 1 | 1 | 1 | NA | NA | NA | NA | 2 | 1 |
| 16_457843 | NA | NA | NA | NA | 2 | 0.5 | 2 | 0 | 4 | 0.25 | 1 | 0 | 1 | 1 | NA | NA | NA | NA | 2 | 0.5 |
| 16_457844 | NA | NA | NA | NA | 1 | 1 | 2 | 1 | 3 | 1 | 1 | 1 | 1 | 0 | NA | NA | NA | NA | 2 | 0.5 |
| 16_457846 | NA | NA | NA | NA | 2 | 0.5 | 2 | 0 | 4 | 0.25 | NA | NA | 1 | 1 | NA | NA | NA | NA | 1 | 1 |
| 16_457851 | NA | NA | NA | NA | 2 | 0 | 2 | 0.5 | 4 | 0.25 | 1 | 0 | 1 | 1 | NA | NA | NA | NA | 2 | 0.5 |
| 16_457851 | NA | NA | NA | NA | 1 | 0 | 2 | 1 | 3 | 0.6667 | 1 | 1 | 1 | 1 | NA | NA | NA | NA | 2 | 1 |
| 16_457852 | NA | NA | NA | NA | 2 | 1 | 2 | 0.5 | 4 | 0.75 | NA | NA | 1 | 1 | NA | NA | NA | NA | 1 | 1 |
| 16_457854 | NA | NA | NA | NA | 2 | 1 | 1 | 1 | 3 | 1 | NA | NA | 1 | 1 | NA | NA | NA | NA | 1 | 1 |
| 16_462154 | 1 | 1 | NA | NA | 1 | 1 | NA | NA | 2 | 1 | NA | NA | 2 | 0.5 | NA | NA | NA | NA | 2 | 0.5 |
| 16_462154 | 1 | 1 | NA | NA | 1 | 0 | NA | NA | 2 | 0.5 | NA | NA | 2 | 0.5 | NA | NA | NA | NA | 2 | 0.5 |
| 16_462632 | NA | NA | NA | NA | 2 | 0.5 | 1 | 1 | 3 | 0.6667 | NA | NA | 1 | 0 | 1 | 1 | 1 | 0 | 3 | 0.3333 |
| 16_462688 | 1 | 0 | NA | NA | 1 | 1 | 1 | 0 | 3 | 0.3333 | 1 | 1 | 3 | 0.6667 | 1 | 0 | NA | NA | 5 | 0.6 |
| 16_462689 | 1 | 0 | NA | NA | 1 | 0 | 1 | 1 | 3 | 0.3333 | 1 | 0 | 2 | 0.5 | NA | NA | NA | NA | 3 | 0.3333 |
| 16_462689 | 1 | 1 | NA | NA | 1 | 1 | NA | NA | 2 | 1 | 1 | 0 | 2 | 1 | 1 | 0 | NA | NA | 4 | 0.5 |
| 16_462689 | 1 | 1 | NA | NA | 1 | 1 | 1 | 1 | 3 | 1 | 1 | 1 | 3 | 1 | 1 | 0 | NA | NA | 5 | 0.8 |
| 16_462689 | NA | NA | NA | NA | 1 | 1 | 1 | 1 | 2 | 1 | 1 | 1 | 3 | 1 | 1 | 1 | NA | NA | 5 | 1 |
| 16_487447 | 5 | 0 | 7 | 0.1429 | 6 | 0.3333 | 18 | 0.2778 | 36 | 0.2222 | 3 | 0.3333 | 5 | 0.4 | 7 | 0.1429 | 5 | 0 | 20 | 0.2 |
| 16_487447 | 5 | 0 | 8 | 0 | 7 | 0.1429 | 17 | 0.0588 | 37 | 0.0541 | 2 | 0 | 4 | 0 | 7 | 0.1429 | 7 | 0.1429 | 20 | 0.1 |
| 16_487458 | 1 | 0 | 6 | 0.6667 | 4 | 0.25 | 14 | 0.2143 | 25 | 0.32 | NA | NA | 1 | 1 | 5 | 0.6 | 4 | 0.25 | 10 | 0.5 |
| 16_487461 | 1 | 0 | 4 | 0.25 | 3 | 0 | 10 | 0 | 18 | 0.0556 | NA | NA | 1 | 0 | 4 | 0.25 | 3 | 0 | 8 | 0.125 |
| 16_487462 | 1 | 0 | 4 | 0.25 | 4 | 0.25 | 12 | 0 | 21 | 0.0952 | NA | NA | NA | NA | 3 | 0.6667 | 4 | 0.25 | 7 | 0.4286 |
| 16_495841 | 6 | 0 | 9 | 0.4444 | 7 | 0.2857 | 6 | 0.3333 | 28 | 0.2857 | 3 | 0.3333 | 2 | 0 | 7 | 0.2857 | 6 | 0 | 18 | 0.1667 |
| 16_495842 | 8 | 0.125 | 9 | 0.1111 | 8 | 0.25 | 8 | 0.25 | 33 | 0.1818 | 4 | 0.25 | 3 | 0 | 7 | 0.2857 | 7 | 0 | 21 | 0.1429 |
| 16_496881 | 9 | 0.2222 | 16 | 0.0625 | 9 | 0 | 8 | 0.125 | 42 | 0.0952 | 7 | 0 | 12 | 0.25 | 11 | 0 | 11 | 0.0909 | 41 | 0.0976 |
| 16_496883 | 8 | 0.125 | 16 | 0.375 | 10 | 0.4 | 8 | 0.25 | 42 | 0.3095 | 7 | 0.4286 | 12 | 0.3333 | 10 | 0.2 | 11 | 0.7273 | 40 | 0.425 |
| 16_496883 | 9 | 0.4444 | 17 | 0.2941 | 8 | 0.375 | 7 | 0.2857 | 41 | 0.3415 | 6 | 0.1667 | 11 | 0.1818 | 9 | 0.4444 | 9 | 0.7778 | 35 | 0.4 |
| 16_497312 | 13 | 0.1538 | 21 | 0.0952 | 12 | 0.1667 | 10 | 0.1 | 56 | 0.125 | 11 | 0 | 13 | 0.0769 | 14 | 0.2143 | 17 | 0.1765 | 55 | 0.1273 |
| 16_497317 | 6 | 0 | 15 | 0.0667 | 7 | 0 | 6 | 0.1667 | 34 | 0.0588 | 6 | 0 | 8 | 0 | 7 | 0.1429 | 14 | 0.1429 | 35 | 0.0857 |
| 16_497321 | 5 | 0.4 | 14 | 0.0714 | 6 | 0 | 7 | 0.2857 | 32 | 0.1562 | 7 | 0.1429 | 11 | 0 | 7 | 0 | 15 | 0.1333 | 40 | 0.075 |
| 16_497321 | 5 | 0 | 11 | 0.0909 | 8 | 0.25 | 7 | 0.2857 | 31 | 0.1613 | 6 | 0.1667 | 10 | 0.1 | 8 | 0 | 12 | 0.1667 | 36 | 0.1111 |
| 16_497326 | 6 | 0.1667 | 14 | 0.0714 | 8 | 0.125 | 8 | 0 | 36 | 0.0833 | 6 | 0 | 11 | 0.0909 | 9 | 0.2222 | 14 | 0.0714 | 40 | 0.1 |
| 16_497327 | 7 | 0 | 16 | 0 | 7 | 0.1429 | 8 | 0.125 | 38 | 0.0526 | 7 | 0 | 11 | 0 | 6 | 0.1667 | 15 | 0.0667 | 39 | 0.0513 |
| 16_500134 | 3 | 1 | 2 | 1 | NA | NA | 1 | 1 | 6 | 1 | NA | NA | 2 | 1 | 1 | 1 | NA | NA | 3 | 1 |
| 16_500135 | 3 | 0.3333 | 2 | 0.5 | 1 | 0 | 1 | 0 | 7 | 0.2857 | 1 | 0 | 2 | 1 | 1 | 0 | NA | NA | 4 | 0.5 |
| 16_500135 | 2 | 1 | 2 | 0.5 | NA | NA | 1 | 1 | 5 | 0.8 | 1 | 0 | 1 | 1 | 1 | 1 | NA | NA | 3 | 0.6667 |
| 16_500137 | 2 | 0 | 2 | 0.5 | NA | NA | 1 | 0 | 5 | 0.2 | 1 | 0 | 2 | 1 | 1 | 0 | NA | NA | 4 | 0.5 |
| 16_500168 | 1 | 1 | NA | NA | NA | NA | NA | NA | 1 | 1 | NA | NA | NA | NA | NA | NA | 1 | 1 | 1 | 1 |
| 16_505400 | 2 | 1 | 1 | 1 | NA | NA | 2 | 1 | 5 | 1 | 1 | 0 | NA | NA | 1 | 0 | 3 | 0.6667 | 5 | 0.4 |
| 16_505403 | 1 | 1 | 2 | 1 | NA | NA | 2 | 1 | 5 | 1 | 1 | 1 | NA | NA | 1 | 1 | 1 | 1 | 3 | 1 |
| 16_505403 | 1 | 0 | 2 | 0 | NA | NA | 1 | 1 | 4 | 0.25 | 1 | 0 | NA | NA | 1 | 1 | 1 | 0 | 3 | 0.3333 |
| 16_505405 | 1 | 0 | 2 | 0 | NA | NA | 2 | 0.5 | 5 | 0.2 | 1 | 0 | NA | NA | 1 | 0 | 1 | 1 | 3 | 0.3333 |
| 16_505405 | 1 | 1 | 2 | 1 | NA | NA | 1 | 1 | 4 | 1 | 1 | 1 | NA | NA | NA | NA | 1 | 1 | 2 | 1 |
| 16_505407 | 1 | 0 | 1 | 1 | NA | NA | 2 | 0.5 | 4 | 0.5 | NA | NA | NA | NA | 1 | 1 | 1 | 0 | 2 | 0.5 |
| 16_505408 | 1 | 0 | 2 | 0.5 | NA | NA | 2 | 0 | 5 | 0.2 | 1 | 0 | NA | NA | 1 | 1 | 1 | 0 | 3 | 0.3333 |

| | | | | | | | | | | | | | | | | | | | | |
|---|---|---|---|---|---|---|---|---|---|---|---|---|---|---|---|---|---|---|---|---|
| 16_505408 | 1 | 0 | 2 | 0.5 | NA | NA | 2 | 0.5 | 5 | 0.4 | NA | NA | NA | NA | 1 | 0 | 1 | 1 | 2 | 0.5 |
| 16_505410 | 1 | 0 | 1 | 1 | NA | NA | 2 | 0.5 | 4 | 0.5 | 1 | 1 | NA | NA | 1 | 1 | 1 | 1 | 3 | 1 |
| 16_505414 | NA | NA | NA | NA | NA | NA | 2 | 0.5 | 2 | 0.5 | 1 | 0 | NA | NA | 1 | 0 | 1 | 1 | 3 | 0.3333 |
| 16_505468 | NA | NA | NA | NA | NA | NA | 1 | 1 | 1 | 1 | 1 | 1 | NA | NA | NA | NA | NA | NA | 1 | 1 |
| 16_505470 | NA | NA | NA | NA | NA | NA | 1 | 1 | 1 | 1 | 1 | 1 | NA | NA | NA | NA | NA | NA | 1 | 1 |
| 16_505472 | NA | NA | NA | NA | NA | NA | 1 | 1 | 1 | 1 | 1 | 1 | NA | NA | NA | NA | NA | NA | 1 | 1 |
| 16_505475 | NA | NA | NA | NA | NA | NA | 1 | 1 | 1 | 1 | 1 | 1 | NA | NA | NA | NA | NA | NA | 1 | 1 |
| 16_505482 | NA | NA | NA | NA | NA | NA | 1 | 1 | 1 | 1 | 1 | 1 | NA | NA | 1 | 1 | NA | NA | 2 | 1 |
| 16_505485 | NA | NA | 1 | 1 | NA | NA | 1 | 0 | 2 | 0.5 | 1 | 0 | NA | NA | 1 | 1 | NA | NA | 2 | 0.5 |
| 16_520248 | NA | NA | NA | NA | 1 | 1 | NA | NA | 1 | 1 | NA | NA | NA | NA | 1 | 1 | NA | NA | 1 | 1 |
| 16_520249 | NA | NA | NA | NA | 1 | 1 | NA | NA | 1 | 1 | NA | NA | NA | NA | 1 | 1 | NA | NA | 1 | 1 |
| 16_520251 | NA | NA | NA | NA | 1 | 1 | NA | NA | 1 | 1 | NA | NA | NA | NA | 1 | 1 | NA | NA | 1 | 1 |
| 16_520281 | 1 | 1 | NA | NA | 1 | 1 | 1 | 0 | 3 | 0.6667 | 2 | 1 | 3 | 1 | NA | NA | 1 | 0 | 6 | 0.8333 |
| 16_520696 | 30 | 0.9667 | 25 | 0.96 | 31 | 1 | 40 | 0.95 | 126 | 0.9683 | 16 | 1 | 35 | 0.9714 | 48 | 1 | 31 | 1 | 130 | 0.9923 |
| 16_520697 | 31 | 0.3548 | 27 | 0.4444 | 32 | 0.1875 | 42 | 0.4048 | 132 | 0.3485 | 18 | 0.5556 | 30 | 0.4333 | 44 | 0.3409 | 26 | 0.5385 | 118 | 0.4407 |
| 16_520697 | 35 | 0.0857 | 28 | 0.1429 | 34 | 0.1471 | 45 | 0.0889 | 142 | 0.1127 | 17 | 0 | 39 | 0.1282 | 53 | 0.1321 | 34 | 0.1176 | 143 | 0.1119 |
| 16_520698 | 29 | 0.2069 | 24 | 0.0417 | 35 | 0.1143 | 39 | 0.0769 | 127 | 0.1102 | 17 | 0.1176 | 35 | 0.0571 | 49 | 0.1633 | 28 | 0.1429 | 129 | 0.124 |
| 16_520705 | 21 | 0 | 15 | 0.0667 | 24 | 0.125 | 30 | 0.1 | 90 | 0.0778 | 13 | 0.0769 | 22 | 0.0455 | 31 | 0.0645 | 23 | 0.087 | 89 | 0.0674 |
| 16_520706 | 24 | 0.125 | 13 | 0.3846 | 21 | 0.1429 | 29 | 0.1034 | 87 | 0.1609 | 16 | 0.125 | 20 | 0.1 | 32 | 0.0938 | 20 | 0.15 | 88 | 0.1136 |
| 16_520719 | 20 | 0.1 | 12 | 0.25 | 17 | 0.2941 | 25 | 0.24 | 74 | 0.2162 | 12 | 0.0833 | 26 | 0.1154 | 32 | 0.1562 | 14 | 0.0714 | 84 | 0.119 |
| 16_520719 | 19 | 0.1053 | 11 | 0.0909 | 18 | 0.2222 | 23 | 0.087 | 71 | 0.1268 | 10 | 0 | 24 | 0.1667 | 33 | 0.2121 | 17 | 0.1176 | 84 | 0.1548 |
| 16_520943 | 10 | 0 | 8 | 0 | 12 | 0 | 17 | 0.1765 | 47 | 0.0638 | 10 | 0.2 | 18 | 0.1111 | 19 | 0 | 12 | 0.1667 | 59 | 0.1017 |
| 16_521260 | 8 | 0.25 | 6 | 0 | 8 | 0.125 | 10 | 0 | 32 | 0.0938 | 9 | 0.1111 | 11 | 0.0909 | 16 | 0 | 8 | 0.125 | 44 | 0.0682 |
| 16_522728 | 6 | 0.8333 | 4 | 0.25 | 5 | 0.2 | 11 | 0.4545 | 26 | 0.4615 | 6 | 0.6667 | 9 | 0.5556 | 15 | 0.6 | 7 | 0.8571 | 37 | 0.6486 |
| 16_522729 | 6 | 0 | 6 | 0 | 6 | 0 | 10 | 0.2 | 28 | 0.0714 | 6 | 0.1667 | 9 | 0 | 12 | 0.0833 | 7 | 0 | 34 | 0.0588 |
| 16_522730 | 5 | 1 | 5 | 0.4 | 5 | 0.6 | 7 | 0.7143 | 22 | 0.6818 | 4 | 0.5 | 6 | 1 | 7 | 0.8571 | 7 | 0.5714 | 24 | 0.75 |
| 16_544397 | NA | NA | 2 | 1 | NA | NA | 2 | 1 | 4 | 1 | 1 | 0 | 2 | 0.5 | 1 | 1 | 1 | 1 | 5 | 0.6 |
| 16_544397 | NA | NA | 1 | 1 | NA | NA | 2 | 1 | 3 | 1 | 1 | 1 | 1 | 1 | 1 | 1 | 1 | 1 | 4 | 1 |
| 16_556637 | 4 | 0 | 2 | 0 | 3 | 0.3333 | 4 | 0 | 13 | 0.0769 | 3 | 0 | 3 | 0 | NA | NA | 8 | 0.125 | 14 | 0.0714 |
| 16_556637 | 4 | 0.5 | 5 | 0.6 | 3 | 0.3333 | 4 | 0.25 | 16 | 0.4375 | 2 | 0.5 | 3 | 0.3333 | 1 | 1 | 9 | 0.2222 | 15 | 0.3333 |
| 16_556638 | 3 | 0.6667 | 3 | 0.3333 | 3 | 1 | 4 | 0 | 13 | 0.4615 | 2 | 0.5 | 3 | 0.3333 | NA | NA | 9 | 0.3333 | 14 | 0.3571 |
| 16_556640 | 3 | 0 | 2 | 0 | 3 | 0.6667 | 3 | 0 | 11 | 0.1818 | 3 | 0.3333 | 2 | 0.5 | NA | NA | 7 | 0.4286 | 12 | 0.4167 |
| 16_556640 | 2 | 1 | 1 | 1 | 3 | 0.3333 | 3 | 0.3333 | 9 | 0.5556 | 3 | 0.3333 | 1 | 0 | NA | NA | 7 | 0.4286 | 11 | 0.3636 |
| 16_556641 | 2 | 0 | 2 | 0 | 2 | 0.5 | 3 | 0 | 9 | 0.1111 | 3 | 0 | 2 | 0.5 | NA | NA | 7 | 0.1429 | 12 | 0.1667 |
| 16_556642 | 1 | 0 | 1 | 0 | 3 | 0 | 3 | 0.6667 | 8 | 0.25 | 1 | 0 | 2 | 0 | NA | NA | 5 | 0.4 | 8 | 0.25 |
| 16_556643 | 2 | 1 | 2 | 0.5 | 2 | 1 | 2 | 1 | 8 | 0.875 | 1 | 1 | 2 | 1 | NA | NA | 6 | 1 | 9 | 1 |
| 16_556644 | 2 | 0 | 2 | 0.5 | 2 | 0.5 | 2 | 0 | 8 | 0.25 | NA | NA | 2 | 0 | NA | NA | 7 | 0.1429 | 9 | 0.1111 |
| 16_556645 | 2 | 0 | 1 | 0 | 1 | 0 | 2 | 0.5 | 6 | 0.1667 | 1 | 0 | 2 | 0 | NA | NA | 6 | 0.3333 | 9 | 0.2222 |
| 16_556645 | 2 | 0.5 | 2 | 0 | 1 | 1 | 2 | 0 | 7 | 0.2857 | 1 | 0 | NA | NA | NA | NA | 7 | 0.1429 | 8 | 0.125 |
| 16_556645 | NA | NA | 1 | 0 | 2 | 0.5 | 2 | 1 | 5 | 0.6 | 1 | 0 | 2 | 0.5 | NA | NA | 6 | 1 | 9 | 0.7778 |
| 16_556646 | 1 | 0 | 2 | 1 | NA | NA | 1 | 1 | 4 | 0.75 | NA | NA | 1 | 1 | NA | NA | 3 | 0.3333 | 4 | 0.5 |
| 16_556647 | 1 | 1 | NA | NA | 1 | 1 | 1 | 1 | 3 | 1 | NA | NA | 1 | 1 | NA | NA | 5 | 0.8 | 6 | 0.8333 |
| 16_556647 | 1 | 1 | 1 | 1 | 1 | 1 | 1 | 1 | 4 | 1 | 1 | 1 | 1 | 0 | NA | NA | 3 | 1 | 5 | 0.8 |
| 16_556647 | 1 | 1 | 1 | 1 | 1 | 1 | 1 | 0 | 4 | 0.75 | 1 | 1 | 1 | 1 | NA | NA | 4 | 0.25 | 6 | 0.5 |
| 16_556647 | 1 | 1 | 1 | 0 | 1 | 1 | 1 | 1 | 4 | 0.75 | 1 | 1 | 1 | 1 | NA | NA | 3 | 0.6667 | 5 | 0.8 |
| 16_556649 | 1 | 1 | 2 | 1 | 1 | 1 | 1 | 1 | 5 | 1 | NA | NA | 1 | 1 | NA | NA | 1 | 1 | 2 | 1 |
| 16_556865 | NA | NA | 1 | 1 | NA | NA | 1 | 1 | 2 | 1 | 1 | 1 | NA | NA | NA | NA | 4 | 0.25 | 5 | 0.4 |
| 16_556866 | NA | NA | 1 | 1 | NA | NA | 1 | 1 | 2 | 1 | NA | NA | NA | NA | NA | NA | 3 | 1 | 3 | 1 |
| 16_556867 | NA | NA | 1 | 1 | NA | NA | 1 | 0 | 2 | 0.5 | NA | NA | NA | NA | NA | NA | 4 | 0.5 | 4 | 0.5 |
| 16_557206 | 1 | 0 | 1 | 1 | 1 | 0 | 2 | 1 | 5 | 0.6 | 1 | 0 | 2 | 0.5 | 2 | 0 | 2 | 1 | 7 | 0.4286 |
| 16_557938 | 2 | 0 | 1 | 0 | 3 | 0.3333 | NA | NA | 6 | 0.1667 | 1 | 0 | NA | NA | 2 | 1 | 3 | 0.6667 | 6 | 0.6667 |
| 16_557940 | 2 | 1 | NA | NA | 3 | 1 | NA | NA | 5 | 1 | 1 | 1 | NA | NA | 2 | 1 | 2 | 1 | 5 | 1 |
| 16_557942 | 2 | 0.5 | 1 | 0 | 2 | 0.5 | NA | NA | 5 | 0.4 | 1 | 0 | NA | NA | 3 | 0.3333 | 2 | 0.5 | 6 | 0.3333 |
| 16_557945 | 1 | 0 | 1 | 1 | 1 | 0 | NA | NA | 3 | 0.3333 | 1 | 0 | NA | NA | 2 | 0.5 | 2 | 0 | 5 | 0.2 |
| 16_557947 | 2 | 1 | NA | NA | 2 | 0.5 | NA | NA | 4 | 0.75 | NA | NA | NA | NA | 3 | 1 | 2 | 1 | 5 | 1 |
| 16_557948 | 2 | 0.5 | 1 | 0 | 2 | 0.5 | NA | NA | 5 | 0.4 | 1 | 1 | NA | NA | 3 | 0 | 2 | 0.5 | 6 | 0.3333 |
| 16_558106 | 2 | 0.5 | 2 | 0.5 | 1 | 0 | 1 | 0 | 6 | 0.3333 | 4 | 0 | 1 | 0 | 5 | 0.2 | 1 | 0 | 11 | 0.0909 |
| 16_558108 | 2 | 1 | 3 | 1 | 1 | 1 | 1 | 0 | 7 | 0.8571 | 4 | 0.25 | 3 | 0 | 5 | 0.4 | 2 | 1 | 14 | 0.3571 |
| 16_558109 | 2 | 0.5 | 3 | 0 | 1 | 0 | 1 | 1 | 7 | 0.2857 | 4 | 0.75 | 2 | 0.5 | 5 | 0 | 2 | 0 | 13 | 0.3077 |
| 16_558110 | 2 | 0 | 3 | 0.3333 | 1 | 1 | 1 | 0 | 7 | 0.2857 | 4 | 1 | 3 | 0 | 5 | 0 | 2 | 0.5 | 14 | 0.3571 |
| 16_558111 | 2 | 0.5 | 3 | 0.6667 | 1 | 1 | 2 | 0.5 | 8 | 0.625 | 4 | 0.25 | 3 | 0.6667 | 5 | 0.4 | 2 | 0.5 | 14 | 0.4286 |
| 16_558112 | 2 | 1 | 3 | 0.6667 | NA | NA | 2 | 1 | 7 | 0.8571 | 4 | 0.5 | 3 | 0.6667 | 4 | 0.75 | 2 | 1 | 13 | 0.6923 |
| 16_558112 | 2 | 0.5 | 3 | 0.6667 | 1 | 0 | 3 | 0.6667 | 9 | 0.5556 | 4 | 0.5 | 4 | 0.5 | 5 | 0.4 | 2 | 1 | 15 | 0.5333 |
| 16_558113 | 2 | 0.5 | 3 | 0.3333 | 1 | 0 | 1 | 0 | 7 | 0.2857 | 4 | 0.25 | 4 | 0.25 | 4 | 0 | 1 | 0 | 13 | 0.1538 |
| 16_558113 | 2 | 0 | 3 | 0 | 1 | 0 | 3 | 0.3333 | 9 | 0.1111 | 4 | 0.25 | 3 | 0.3333 | 4 | 0 | 2 | 0.5 | 13 | 0.2308 |

| | | | | | | | | | | | | | | | | | | | | |
|---|---|---|---|---|---|---|---|---|---|---|---|---|---|---|---|---|---|---|---|---|
| 16_5582724 | 4 | 0 | 5 | 0 | 2 | 0.5 | 3 | 0 | 14 | 0.0714 | 3 | 0 | 4 | 0.25 | 7 | 0 | 2 | 0 | 16 | 0.0625 |
| 16_5582730 | 1 | 0 | 5 | 0.6 | 1 | 1 | 3 | 0.3333 | 10 | 0.5 | 5 | 0.6 | 4 | 0.5 | 5 | 0.2 | 5 | 0.2 | 19 | 0.3684 |
| 16_5582732 | 4 | 0.25 | 4 | 0 | 2 | 0 | 3 | 0.3333 | 13 | 0.1538 | 6 | 0 | 4 | 0.25 | 7 | 0.7143 | 5 | 0.2 | 22 | 0.3182 |
| 16_5582734 | 4 | 1 | 5 | 1 | 2 | 1 | 3 | 1 | 14 | 1 | 6 | 0.8333 | 5 | 1 | 7 | 1 | 6 | 1 | 24 | 0.9583 |
| 16_5582737 | 4 | 0.25 | 5 | 0.8 | 2 | 1 | 3 | 1 | 14 | 0.7143 | 6 | 0.8333 | 5 | 0.6 | 7 | 1 | 6 | 0.8333 | 24 | 0.8333 |
| 16_5582740 | 5 | 0.4 | 4 | 0.5 | 2 | 0.5 | 3 | 0.3333 | 14 | 0.4286 | 5 | 0.4 | 4 | 0.25 | 7 | 0 | 6 | 0.5 | 22 | 0.2727 |
| 16_5582747 | 6 | 0.6667 | 4 | 0.5 | 2 | 1 | 2 | 1 | 14 | 0.7143 | 6 | 0.5 | 3 | 0.6667 | 4 | 0.75 | 6 | 0.3333 | 19 | 0.5263 |
| 16_5582747 | 5 | 1 | 4 | 1 | 3 | 1 | 3 | 0.6667 | 15 | 0.9333 | 7 | 1 | 5 | 1 | 6 | 1 | 7 | 1 | 25 | 1 |
| 16_5582750 | 6 | 0 | 5 | 0.4 | 3 | 0.6667 | 3 | 0 | 17 | 0.2353 | 7 | 0 | 5 | 0.4 | 6 | 0 | 6 | 0.1667 | 24 | 0.125 |
| 16_5582758 | 3 | 0.3333 | 3 | 0 | 3 | 0 | 2 | 0 | 11 | 0.0909 | 6 | 0 | 4 | 0.25 | 6 | 0.3333 | 2 | 0 | 18 | 0.1667 |
| 16_5582764 | 6 | 0.8333 | 4 | 1 | 2 | 1 | 3 | 1 | 15 | 0.9333 | 5 | 1 | 5 | 1 | 6 | 1 | 6 | 1 | 22 | 1 |
| 16_5582777 | 6 | 1 | 5 | 1 | 3 | 1 | 4 | 1 | 18 | 1 | 6 | 1 | 3 | 1 | 5 | 1 | 6 | 1 | 20 | 1 |
| 16_5582792 | 5 | 0 | 4 | 0 | 3 | 0.3333 | 3 | 0 | 15 | 0.0667 | 7 | 0.1429 | 4 | 0.25 | 6 | 0 | 7 | 0 | 24 | 0.0833 |
| 16_5582842 | 6 | 0 | 5 | 0 | 2 | 0 | 4 | 0.25 | 17 | 0.0588 | 5 | 0 | 4 | 0 | 7 | 0.1429 | 3 | 0 | 19 | 0.0526 |
| 16_5583684 | 24 | 0.1667 | 29 | 0.1034 | 17 | 0.1176 | 21 | 0.0476 | 91 | 0.1099 | 10 | 0 | 20 | 0 | 24 | 0.0833 | 25 | 0.08 | 79 | 0.0506 |
| 16_5583707 | 25 | 0.04 | 26 | 0.0385 | 14 | 0 | 21 | 0.1905 | 86 | 0.0698 | 9 | 0 | 13 | 0.0769 | 23 | 0.087 | 22 | 0.0455 | 67 | 0.0597 |
| 16_5583753 | 15 | 0.1333 | 20 | 0.1 | 12 | 0.25 | 18 | 0.2222 | 65 | 0.1692 | 6 | 0 | 17 | 0.1176 | 12 | 0.25 | 21 | 0.1429 | 56 | 0.1429 |
| 16_5584476 | 8 | 0.875 | 13 | 0.5385 | 3 | 0.3333 | 8 | 0.625 | 32 | 0.625 | 7 | 0.5714 | 8 | 0.5 | 2 | 0.5 | 8 | 0.5 | 25 | 0.52 |
| 16_5584492 | 4 | 1 | 4 | 1 | 1 | 1 | 3 | 0.6667 | 12 | 0.9167 | 2 | 1 | 2 | 1 | 3 | 0.6667 | 2 | 1 | 9 | 0.8889 |
| 16_5585419 | 2 | 0 | 3 | 0 | 3 | 0 | 2 | 0.5 | 10 | 0.1 | 1 | 0 | 2 | 0 | 3 | 0.3333 | NA | NA | 6 | 0.1667 |
| 16_5585444 | 2 | 1 | 2 | 1 | 3 | 1 | 1 | 1 | 8 | 1 | 1 | 1 | 2 | 1 | 3 | 1 | NA | NA | 6 | 1 |
| 16_5585455 | 2 | 0.5 | 1 | 1 | 1 | 0 | 1 | 0 | 5 | 0.4 | 1 | 0 | 2 | 0.5 | 2 | 0 | NA | NA | 5 | 0.2 |
| 16_5585479 | 2 | 1 | 3 | 1 | 3 | 1 | 2 | 1 | 10 | 1 | 1 | 1 | 1 | 1 | 3 | 1 | NA | NA | 5 | 1 |
| 16_5585484 | 2 | 0.5 | 3 | 0.3333 | 3 | 1 | 2 | 1 | 10 | 0.7 | 1 | 0 | 1 | 1 | 3 | 1 | NA | NA | 5 | 0.8 |
| 16_5585499 | 1 | 1 | 2 | 1 | 2 | 1 | 2 | 1 | 7 | 1 | 1 | 0 | 1 | 1 | 2 | 1 | NA | NA | 4 | 0.75 |
| 16_5585502 | 2 | 1 | 3 | 0 | 2 | 0 | 2 | 0 | 9 | 0.2222 | 1 | 0 | 1 | 0 | 3 | 0.3333 | NA | NA | 5 | 0.2 |
| 16_5585504 | 2 | 0.5 | 3 | 0.6667 | 3 | 1 | 2 | 0.5 | 10 | 0.7 | 1 | 1 | 1 | 1 | 3 | 1 | NA | NA | 5 | 1 |
| 16_5585514 | 2 | 0 | 3 | 0.6667 | 3 | 0 | 2 | 0.5 | 10 | 0.3 | 1 | 0 | 1 | 0 | 3 | 0.3333 | NA | NA | 5 | 0.2 |
| 16_5585548 | 2 | 1 | 2 | 1 | NA | NA | 2 | 1 | 6 | 1 | NA | NA | 1 | 1 | 2 | 1 | NA | NA | 3 | 1 |
| 16_5585562 | 2 | 0 | 1 | 1 | 3 | 0.6667 | 2 | 1 | 8 | 0.625 | 1 | 0 | 1 | 0 | 3 | 0.3333 | NA | NA | 5 | 0.2 |
| 16_5587001 | NA | NA | NA | NA | 1 | 1 | NA | NA | 1 | 1 | NA | NA | NA | NA | 1 | 1 | NA | NA | 1 | 1 |
| 16_5587111 | NA | NA | NA | NA | 1 | 1 | 1 | 1 | 2 | 1 | NA | NA | 1 | 1 | 1 | 1 | NA | NA | 2 | 1 |
| 16_5587119 | 1 | 1 | NA | NA | 1 | 1 | 1 | 1 | 3 | 1 | 1 | 1 | NA | NA | 1 | 1 | NA | NA | 2 | 1 |
| 16_5587154 | 1 | 1 | NA | NA | 1 | 1 | 1 | 0 | 3 | 0.6667 | 1 | 1 | 1 | 1 | 3 | 0.3333 | NA | NA | 5 | 0.6 |
| 16_5587202 | 2 | 0 | 1 | 0 | 1 | 1 | 1 | 0 | 5 | 0.2 | 1 | 0 | 1 | 1 | 1 | 0 | NA | NA | 3 | 0.3333 |
| 16_5587228 | 2 | 0.5 | 1 | 0 | 1 | 0 | 1 | 1 | 5 | 0.4 | 1 | 0 | 1 | 1 | 3 | 0.3333 | NA | NA | 5 | 0.4 |
| 16_5587276 | 3 | 0 | 1 | 0 | 1 | 0 | 1 | 1 | 6 | 0.1667 | 1 | 0 | 1 | 0 | 2 | 0.5 | NA | NA | 4 | 0.25 |
| 16_5597382 | 3 | 0 | 5 | 0.2 | NA | NA | 1 | 0 | 9 | 0.1111 | 2 | 0 | 1 | 1 | 3 | 0.3333 | 1 | 0 | 7 | 0.2857 |
| 16_5600861 | 4 | 0.25 | 4 | 0.25 | 1 | 0 | 4 | 0.25 | 13 | 0.2308 | 2 | 0 | 2 | 0 | 4 | 0.25 | 3 | 0 | 11 | 0.0909 |
| 16_5600866 | 5 | 0.4 | 6 | 0.1667 | 2 | 0 | 4 | 0.75 | 17 | 0.3529 | 2 | 0 | 3 | 0 | 6 | 0.1667 | 3 | 0 | 14 | 0.0714 |
| 16_5600869 | 5 | 0.2 | 6 | 0.1667 | 2 | 0 | 5 | 0 | 18 | 0.1111 | 2 | 0 | NA | NA | 5 | 0.2 | 3 | 0 | 10 | 0.1 |
| 16_5600890 | 3 | 0.3333 | 5 | 0.2 | 2 | 0.5 | 4 | 0 | 14 | 0.2143 | 2 | 0.5 | 3 | 0.3333 | 5 | 0.2 | 2 | 0 | 12 | 0.25 |
| 16_5600998 | 5 | 0 | 6 | 0.1667 | 4 | 0.25 | 5 | 0 | 20 | 0.1 | 2 | 0 | 5 | 0 | 7 | 0 | 4 | 0.25 | 18 | 0.0556 |
| 16_5601021 | 5 | 0 | 7 | 0.1429 | 4 | 0 | 5 | 0.2 | 21 | 0.0952 | 1 | 1 | 5 | 0.4 | 6 | 0 | 4 | 0.25 | 16 | 0.25 |
| 16_5601026 | 7 | 0 | 7 | 0.1429 | 4 | 0 | 6 | 0.1667 | 24 | 0.0833 | 2 | 0 | 7 | 0 | 6 | 0 | 5 | 0.2 | 20 | 0.05 |
| 16_5650938 | 2 | 0 | 4 | 0 | 5 | 0.2 | NA | NA | 11 | 0.0909 | 2 | 0 | NA | NA | NA | NA | 2 | 0.5 | 4 | 0.25 |
| 16_5651108 | 11 | 0.1818 | 11 | 0.2727 | 9 | 0.2222 | 9 | 0.2222 | 40 | 0.225 | 5 | 0.2 | 15 | 0.0667 | 7 | 0.5714 | 13 | 0.1538 | 40 | 0.2 |
| 16_5651114 | 12 | 0.1667 | 9 | 0.2222 | 7 | 0 | 7 | 0.1429 | 35 | 0.1429 | 5 | 0.2 | 12 | 0.25 | 6 | 0.1667 | 12 | 0.1667 | 35 | 0.2 |
| 16_5694230 | 7 | 0.2857 | 4 | 0 | 5 | 0 | 8 | 0.375 | 24 | 0.2083 | 6 | 0.1667 | 7 | 0.2857 | 19 | 0.2632 | 7 | 0.1429 | 39 | 0.2308 |
| 16_5697311 | 14 | 0.7143 | 11 | 0.6364 | 14 | 0.8571 | 15 | 0.8 | 54 | 0.7593 | 5 | 0.8 | 21 | 0.619 | 26 | 0.7308 | 12 | 0.8333 | 64 | 0.7188 |
| 16_5697311 | 15 | 0.4 | 14 | 0.3571 | 12 | 0.0833 | 12 | 0.4167 | 53 | 0.3208 | 5 | 0.2 | 22 | 0.4545 | 33 | 0.3333 | 15 | 0.3333 | 75 | 0.36 |
| 16_5697316 | 18 | 0 | 17 | 0.1176 | 19 | 0.1579 | 23 | 0.087 | 77 | 0.0909 | 9 | 0.1111 | 27 | 0.0741 | 41 | 0.0976 | 23 | 0.0435 | 100 | 0.08 |
| 16_5697319 | 19 | 0.1053 | 16 | 0.25 | 18 | 0.0556 | 21 | 0.1905 | 74 | 0.1486 | 9 | 0.2222 | 28 | 0.1429 | 41 | 0.122 | 22 | 0.2727 | 100 | 0.17 |
| 16_5697361 | 20 | 0.3 | 19 | 0.1053 | 19 | 0.2105 | 22 | 0.2727 | 80 | 0.225 | 9 | 0.2222 | 25 | 0.16 | 47 | 0.0851 | 19 | 0.1053 | 100 | 0.12 |
| 16_5697372 | 22 | 0 | 14 | 0.0714 | 16 | 0.0625 | 21 | 0.0952 | 73 | 0.0548 | 7 | 0 | 23 | 0 | 43 | 0.0465 | 19 | 0.1579 | 92 | 0.0543 |
| 16_5697608 | 4 | 0.25 | 2 | 0 | 5 | 0 | 4 | 0.25 | 15 | 0.1333 | NA | NA | 4 | 0 | 4 | 0.25 | 4 | 0 | 12 | 0.0833 |
| 16_5697622 | 5 | 0 | 2 | 0 | 7 | 0 | 4 | 0.25 | 18 | 0.0556 | NA | NA | 5 | 0.2 | 7 | 0.1429 | 5 | 0 | 17 | 0.1176 |
| 16_5699063 | 4 | 0.25 | 2 | 1 | 1 | 0 | 4 | 0.5 | 11 | 0.4545 | NA | NA | 1 | 1 | 2 | 0 | 2 | 0.5 | 5 | 0.4 |
| 16_5702741 | NA | NA | NA | NA | NA | NA | 2 | 1 | 2 | 1 | NA | NA | 1 | 1 | NA | NA | NA | NA | 1 | 1 |
| 16_5739064 | NA | NA | NA | NA | NA | NA | 1 | 1 | 1 | 1 | NA | NA | NA | NA | 1 | 1 | NA | NA | 1 | 1 |
| 16_5739066 | NA | NA | NA | NA | NA | NA | 1 | 1 | 1 | 1 | NA | NA | NA | NA | 1 | 1 | NA | NA | 1 | 1 |
| 16_5739231 | NA | NA | NA | NA | NA | NA | 1 | 1 | 1 | 1 | NA | NA | NA | NA | 1 | 1 | NA | NA | 1 | 1 |
| 16_5739239 | NA | NA | NA | NA | NA | NA | 1 | 1 | 1 | 1 | NA | NA | NA | NA | 1 | 1 | NA | NA | 1 | 1 |
| 16_5739285 | NA | NA | NA | NA | NA | NA | 1 | 1 | 1 | 1 | NA | NA | NA | NA | 1 | 1 | NA | NA | 1 | 1 |
| 16_5739295 | NA | NA | NA | NA | NA | NA | 1 | 1 | 1 | 1 | NA | NA | NA | NA | 1 | 1 | NA | NA | 1 | 1 |

| | | | | | | | | | | | | | | | | | | | | |
|---|---|---|---|---|---|---|---|---|---|---|---|---|---|---|---|---|---|---|---|---|
| 16_582858 | NA | NA | NA | NA | 2 | 1 | NA | NA | 2 | 1 | NA | NA | NA | NA | 1 | 1 | NA | NA | 1 | 1 |
| 16_582859 | NA | NA | 1 | 1 | 3 | 1 | 1 | 1 | 5 | 1 | NA | NA | NA | NA | 1 | 1 | NA | NA | 1 | 1 |
| 16_582860 | NA | NA | 1 | 1 | 2 | 0.5 | 1 | 1 | 4 | 0.75 | NA | NA | NA | NA | 1 | 1 | NA | NA | 1 | 1 |
| 16_582877 | NA | NA | 2 | 0.5 | 4 | 0.25 | 2 | 0.5 | 8 | 0.375 | NA | NA | NA | NA | 2 | 0.5 | NA | NA | 2 | 0.5 |
| 16_582878 | NA | NA | 3 | 0 | 4 | 0 | 1 | 1 | 8 | 0.125 | NA | NA | NA | NA | 2 | 1 | NA | NA | 2 | 1 |
| 16_582906 | 1 | 1 | NA | NA | NA | NA | NA | NA | 1 | 1 | 1 | 0 | 3 | 0.3333 | NA | NA | 2 | 0 | 6 | 0.1667 |
| 16_582909 | 1 | 1 | NA | NA | NA | NA | NA | NA | 1 | 1 | 3 | 0.3333 | 3 | 0.3333 | NA | NA | 2 | 0.5 | 8 | 0.375 |
| 16_584908 | 3 | 0 | 3 | 0 | 1 | 0 | 3 | 0.3333 | 10 | 0.1 | 1 | 1 | 3 | 0 | 2 | 0 | 2 | 0 | 8 | 0.125 |
| 16_584991 | 4 | 0.5 | 5 | 0.6 | 4 | 0.75 | 4 | 0 | 17 | 0.4706 | 3 | 1 | 3 | 0.6667 | 3 | 1 | 3 | 0 | 12 | 0.6667 |
| 16_584991 | 5 | 0 | 4 | 0.25 | 3 | 0 | 4 | 0 | 16 | 0.0625 | 5 | 0.2 | 3 | 0 | 2 | 0.5 | 4 | 0.5 | 14 | 0.2857 |
| 16_585499 | 8 | 0.25 | 11 | 0.5455 | 7 | 0.5714 | 8 | 0.25 | 34 | 0.4118 | 6 | 0.5 | 7 | 0.2857 | 7 | 0.4286 | 4 | 0.75 | 24 | 0.4583 |
| 16_585499 | 8 | 0.125 | 11 | 0 | 7 | 0.4286 | 8 | 0 | 34 | 0.1176 | 5 | 0.4 | 5 | 0.6 | 7 | 0.1429 | 4 | 0.25 | 21 | 0.3333 |
| 16_585536 | 9 | 0.3333 | 13 | 0.1538 | 8 | 0.125 | 9 | 0.1111 | 39 | 0.1795 | 6 | 0.3333 | 10 | 0.3 | 7 | 0.1429 | 6 | 0.1667 | 29 | 0.2414 |
| 16_592960 | 1 | 1 | 3 | 0.6667 | 2 | 0.5 | 3 | 0.6667 | 9 | 0.6667 | NA | NA | 1 | 1 | 2 | 0.5 | NA | NA | 3 | 0.6667 |
| 16_592961 | 1 | 0 | 3 | 0.3333 | 2 | 0.5 | 3 | 0.6667 | 9 | 0.4444 | NA | NA | 2 | 0.5 | 2 | 0.5 | NA | NA | 4 | 0.5 |
| 16_593124 | 3 | 0 | 3 | 0.3333 | 2 | 0 | 3 | 0 | 11 | 0.0909 | NA | NA | 2 | 0.5 | 2 | 0 | NA | NA | 4 | 0.25 |
| 16_593124 | 3 | 0.3333 | 3 | 0.3333 | 3 | 0.3333 | 3 | 0.3333 | 12 | 0.3333 | NA | NA | 2 | 0 | 2 | 0.5 | NA | NA | 4 | 0.25 |
| 16_593130 | 1 | 1 | 2 | 0 | 1 | 1 | 2 | 0.5 | 6 | 0.5 | 4 | 0.75 | NA | NA | NA | NA | 3 | 1 | 7 | 0.8571 |
| 16_593135 | 1 | 1 | 2 | 0.5 | 1 | 0 | 2 | 0 | 6 | 0.3333 | 4 | 1 | NA | NA | NA | NA | 3 | 0.3333 | 7 | 0.7143 |
| 16_593136 | 1 | 1 | 2 | 0 | 1 | 0 | 2 | 0.5 | 6 | 0.3333 | 3 | 0 | NA | NA | NA | NA | 3 | 0.3333 | 6 | 0.1667 |
| 16_593136 | 1 | 1 | 1 | 1 | 1 | 1 | 2 | 0.5 | 5 | 0.8 | 4 | 0.5 | NA | NA | NA | NA | 3 | 1 | 7 | 0.7143 |
| 16_593137 | 1 | 1 | 1 | 1 | 1 | 1 | 1 | 0 | 4 | 0.75 | 1 | 1 | NA | NA | NA | NA | 2 | 1 | 3 | 1 |
| 16_593138 | 1 | 1 | 2 | 1 | 1 | 1 | 1 | 1 | 5 | 1 | 3 | 1 | NA | NA | NA | NA | 2 | 1 | 5 | 1 |
| 16_593139 | 1 | 1 | 1 | 0 | NA | NA | 1 | 1 | 3 | 0.6667 | 3 | 0.6667 | NA | NA | NA | NA | 2 | 1 | 5 | 0.8 |
| 16_593151 | 1 | 1 | 1 | 0 | NA | NA | NA | NA | 2 | 0.5 | 4 | 0.25 | NA | NA | NA | NA | 3 | 0 | 7 | 0.1429 |
| 16_593751 | NA | NA | 1 | 1 | NA | NA | NA | NA | 1 | 1 | NA | NA | NA | NA | NA | NA | 1 | 1 | 1 | 1 |
| 16_593755 | NA | NA | 1 | 1 | NA | NA | NA | NA | 1 | 1 | 1 | 0 | NA | NA | NA | NA | 1 | 1 | 2 | 0.5 |
| 16_594340 | 8 | 0.25 | 3 | 0.3333 | 9 | 0.3333 | 5 | 0.2 | 25 | 0.28 | 5 | 0 | 11 | 0.1818 | 9 | 0.3333 | 2 | 0 | 27 | 0.1852 |
| 16_594340 | 7 | 0.7143 | 3 | 0.6667 | 6 | 0.1667 | 5 | 0.4 | 21 | 0.4762 | 3 | 0.6667 | 10 | 0.7 | 7 | 0.7143 | 3 | 0.6667 | 23 | 0.6957 |
| 16_594627 | 4 | 0 | 4 | 0.25 | 8 | 0 | 3 | 0.3333 | 19 | 0.1053 | 5 | 0 | 3 | 0 | 4 | 0 | 6 | 0.1667 | 18 | 0.0556 |
| 16_594630 | 2 | 0 | 5 | 0.4 | 6 | 0.3333 | 3 | 0.6667 | 16 | 0.375 | 3 | 0.3333 | 4 | 0.25 | 5 | 0.4 | 6 | 0.1667 | 18 | 0.2778 |
| 16_594634 | 1 | 0 | 4 | 0.25 | 6 | 0.1667 | 3 | 0.6667 | 14 | 0.2857 | 3 | 0.3333 | 3 | 0 | 4 | 0.5 | 6 | 0 | 16 | 0.1875 |
| 16_594634 | 1 | 0 | 2 | 0.5 | 2 | 0 | NA | NA | 5 | 0.2 | 2 | 0 | 1 | 0 | 2 | 0.5 | 5 | 0 | 10 | 0.1 |
| 16_594656 | 1 | 0 | 2 | 0 | 2 | 0 | 2 | 0.5 | 7 | 0.1429 | 3 | 0 | 1 | 1 | 2 | 0 | 1 | 0 | 7 | 0.1429 |
| 16_594735 | 1 | 0 | 2 | 0.5 | 1 | 0 | 1 | 0 | 5 | 0.2 | 1 | 0 | NA | NA | 1 | 0 | 1 | 1 | 3 | 0.3333 |
| 16_594737 | NA | NA | 1 | 1 | 1 | 0 | NA | NA | 2 | 0.5 | NA | NA | NA | NA | 1 | 1 | NA | NA | 1 | 1 |
| 16_594738 | 1 | 0 | 2 | 0.5 | NA | NA | NA | NA | 3 | 0.3333 | 1 | 0 | NA | NA | 2 | 0.5 | NA | NA | 3 | 0.3333 |
| 16_594740 | 1 | 1 | 2 | 1 | 1 | 1 | NA | NA | 4 | 1 | 1 | 1 | NA | NA | 2 | 1 | 1 | 0 | 4 | 0.75 |
| 16_594741 | 1 | 0 | 2 | 1 | 1 | 0 | NA | NA | 4 | 0.5 | 1 | 1 | NA | NA | 2 | 1 | 1 | 0 | 4 | 0.75 |
| 16_594741 | 1 | 1 | 2 | 0.5 | NA | NA | NA | NA | 3 | 0.6667 | 1 | 0 | NA | NA | 2 | 0.5 | 1 | 1 | 4 | 0.5 |
| 16_594741 | 1 | 0 | 2 | 0.5 | 1 | 1 | NA | NA | 4 | 0.5 | NA | NA | NA | NA | 2 | 0.5 | NA | NA | 2 | 0.5 |
| 16_594744 | NA | NA | 3 | 1 | NA | NA | NA | NA | 3 | 1 | 1 | 1 | NA | NA | 1 | 0 | NA | NA | 2 | 0.5 |
| 16_600260 | NA | NA | 2 | 0.5 | NA | NA | NA | NA | 2 | 0.5 | 1 | 0 | NA | NA | NA | NA | 1 | 1 | 2 | 0.5 |
| 16_625389 | 2 | 0.5 | 2 | 0.5 | 2 | 0 | 1 | 1 | 7 | 0.4286 | 2 | 0 | NA | NA | NA | NA | 1 | 1 | 3 | 0.3333 |
| 16_625390 | 2 | 0 | 1 | 0 | 1 | 0 | 2 | 0.5 | 6 | 0.1667 | 3 | 0.3333 | 1 | 1 | NA | NA | NA | NA | 4 | 0.5 |
| 16_625529 | 1 | 0 | 1 | 1 | 2 | 0.5 | 1 | 0 | 5 | 0.4 | 2 | 0 | 2 | 0.5 | NA | NA | 2 | 0 | 6 | 0.1667 |
| 16_625553 | NA | NA | NA | NA | 1 | 1 | NA | NA | 1 | 1 | NA | NA | NA | NA | NA | NA | 1 | 1 | 1 | 1 |
| 16_625556 | 2 | 0.5 | NA | NA | 1 | 1 | NA | NA | 3 | 0.6667 | 1 | 1 | NA | NA | NA | NA | 1 | 1 | 2 | 1 |
| 16_626156 | NA | NA | 1 | 1 | 1 | 1 | NA | NA | 2 | 1 | NA | NA | 3 | 1 | 3 | 1 | NA | NA | 6 | 1 |
| 16_626157 | NA | NA | 1 | 1 | 1 | 1 | NA | NA | 2 | 1 | NA | NA | 3 | 0.6667 | 4 | 0.75 | NA | NA | 7 | 0.7143 |
| 16_626158 | NA | NA | 1 | 1 | 1 | 0 | NA | NA | 2 | 0.5 | NA | NA | 2 | 0 | 3 | 0.3333 | NA | NA | 5 | 0.2 |
| 16_626159 | NA | NA | 1 | 1 | 1 | 0 | NA | NA | 2 | 0.5 | NA | NA | 2 | 0.5 | 3 | 0 | NA | NA | 5 | 0.2 |
| 16_626159 | NA | NA | 1 | 1 | 1 | 1 | NA | NA | 2 | 1 | NA | NA | 2 | 0.5 | 3 | 1 | NA | NA | 5 | 0.8 |
| 16_626160 | NA | NA | 1 | 1 | 1 | 0 | NA | NA | 2 | 0.5 | NA | NA | 3 | 0 | 3 | 0.3333 | NA | NA | 6 | 0.1667 |
| 16_627485 | 2 | 0 | 1 | 1 | 1 | 1 | NA | NA | 4 | 0.5 | 2 | 1 | 2 | 1 | 1 | 1 | 1 | 1 | 6 | 1 |
| 16_645865 | 4 | 1 | 3 | 1 | 4 | 0.5 | 1 | 0 | 12 | 0.75 | 3 | 0.6667 | 6 | 0.6667 | 2 | 0.5 | 4 | 0.75 | 15 | 0.6667 |
| 16_645865 | 5 | 0.8 | 4 | 0.5 | 4 | 0.75 | 1 | 0 | 14 | 0.6429 | NA | NA | 4 | 0.75 | 2 | 0.5 | 6 | 0.6667 | 12 | 0.6667 |
| 16_645865 | 5 | 1 | 3 | 1 | 4 | 1 | 1 | 1 | 13 | 1 | 2 | 0.5 | 2 | 1 | 2 | 1 | 5 | 1 | 11 | 0.9091 |
| 16_645865 | 6 | 0.3333 | 4 | 0.25 | 5 | 0.4 | 1 | 0 | 16 | 0.3125 | 3 | 0.3333 | 5 | 0 | 1 | 1 | 5 | 0.4 | 14 | 0.2857 |
| 16_645866 | 5 | 0.2 | 3 | 0 | 4 | 0 | 1 | 0 | 13 | 0.0769 | 2 | 0.5 | 5 | 0.2 | 2 | 0 | 5 | 0 | 14 | 0.1429 |
| 16_645867 | 5 | 0.4 | 1 | 0 | 3 | 0.3333 | NA | NA | 9 | 0.3333 | 2 | 0 | 2 | 0.5 | 2 | 0.5 | 6 | 0.1667 | 12 | 0.25 |
| 16_645867 | 6 | 0 | 3 | 0 | 5 | 0.2 | 1 | 0 | 15 | 0.0667 | 2 | 0 | 5 | 0.2 | 2 | 0 | 6 | 0.1667 | 15 | 0.1333 |
| 16_645867 | 6 | 0.5 | 3 | 0 | 5 | 0.2 | 1 | 0 | 15 | 0.2667 | 3 | 0.3333 | 5 | 0.2 | 2 | 0.5 | 5 | 0.4 | 15 | 0.3333 |
| 16_645868 | 6 | 0.1667 | 2 | 0 | 3 | 0 | 1 | 0 | 12 | 0.0833 | 3 | 0 | 4 | 0 | 2 | 0 | 3 | 0.3333 | 12 | 0.0833 |
| 16_645869 | 3 | 1 | 3 | 1 | 3 | 1 | 1 | 1 | 10 | 1 | 1 | 1 | 4 | 1 | 1 | 1 | 5 | 1 | 11 | 1 |

| | | | | | | | | | | | | | | | | | | | | |
|---|---|---|---|---|---|---|---|---|---|---|---|---|---|---|---|---|---|---|---|---|
| 16_645870 | 6 | 0.8333 | 3 | 1 | 4 | 1 | 1 | 1 | 14 | 0.9286 | 2 | 1 | 5 | 1 | 2 | 1 | 6 | 1 | 15 | 1 |
| 16_645870 | 6 | 0.8333 | 3 | 1 | 5 | 1 | 1 | 1 | 15 | 0.9333 | 3 | 1 | 4 | 1 | 2 | 1 | 5 | 1 | 14 | 1 |
| 16_645871 | 4 | 0.75 | 2 | 0 | 4 | 0.5 | NA | NA | 10 | 0.5 | 2 | 0.5 | 5 | 0.8 | 2 | 0.5 | 3 | 0.6667 | 12 | 0.6667 |
| 16_645871 | 5 | 0 | NA | NA | 3 | 0.3333 | NA | NA | 8 | 0.125 | 1 | 0 | 5 | 0.4 | 2 | 0 | 3 | 0.3333 | 11 | 0.2727 |
| 16_645872 | 6 | 0.3333 | 3 | 0 | 4 | 0.25 | 1 | 0 | 14 | 0.2143 | 3 | 0.3333 | 4 | 0 | 2 | 0 | 6 | 0 | 15 | 0.0667 |
| 16_645872 | 6 | 0.6667 | 3 | 1 | 4 | 1 | 1 | 0 | 14 | 0.7857 | 2 | 1 | 4 | 1 | 2 | 1 | 5 | 0.8 | 13 | 0.9231 |
| 16_646011 | NA | NA | 1 | 0 | 4 | 0.75 | NA | NA | 5 | 0.6 | NA | NA | 5 | 0.4 | 1 | 0 | NA | NA | 6 | 0.3333 |
| 16_646012 | NA | NA | 1 | 0 | 3 | 0.3333 | NA | NA | 4 | 0.25 | NA | NA | 4 | 0 | 2 | 1 | 1 | 0 | 7 | 0.2857 |
| 16_646034 | 1 | 0 | 1 | 0 | 4 | 0.25 | 1 | 0 | 7 | 0.1429 | 1 | 0 | 4 | 0.5 | 1 | 0 | 1 | 1 | 7 | 0.4286 |
| 16_646034 | 1 | 1 | 1 | 0 | 2 | 0.5 | 1 | 1 | 5 | 0.6 | 1 | 0 | 4 | 0.75 | NA | NA | 1 | 0 | 6 | 0.5 |
| 16_646051 | NA | NA | 1 | 0 | 2 | 1 | 1 | 1 | 4 | 0.75 | 1 | 0 | 2 | 1 | 1 | 0 | 1 | 1 | 5 | 0.6 |
| 16_646052 | NA | NA | 1 | 1 | 3 | 0.3333 | 1 | 1 | 5 | 0.6 | 1 | 0 | 3 | 0.3333 | 1 | 0 | 1 | 0 | 6 | 0.1667 |
| 16_646053 | NA | NA | 1 | 1 | 3 | 1 | 1 | 1 | 5 | 1 | 1 | 0 | 4 | 0.75 | 1 | 0 | 1 | 1 | 7 | 0.5714 |
| 16_646053 | NA | NA | NA | NA | 3 | 0.3333 | 1 | 0 | 4 | 0.25 | 1 | 1 | 4 | 0.25 | 1 | 0 | 1 | 1 | 7 | 0.4286 |
| 16_646054 | NA | NA | 1 | 1 | 2 | 1 | 1 | 1 | 4 | 1 | 1 | 0 | 3 | 1 | 1 | 1 | 1 | 1 | 6 | 0.8333 |
| 16_646054 | NA | NA | 1 | 0 | 3 | 0.6667 | 1 | 0 | 5 | 0.4 | 1 | 0 | 4 | 0.5 | 1 | 0 | 1 | 1 | 7 | 0.4286 |
| 16_646054 | NA | NA | NA | NA | 2 | 0 | 1 | 1 | 3 | 0.3333 | 1 | 0 | 4 | 0.25 | 1 | 0 | 1 | 0 | 7 | 0.1429 |
| 16_646055 | NA | NA | 1 | 0 | 2 | 0.5 | 1 | 0 | 4 | 0.25 | 1 | 0 | 4 | 0.75 | 1 | 0 | 1 | 0 | 7 | 0.4286 |
| 16_646058 | NA | NA | 1 | 1 | 2 | 0.5 | 1 | 1 | 4 | 0.75 | 1 | 1 | 1 | 0 | 1 | 0 | 1 | 0 | 4 | 0.25 |
| 16_646059 | NA | NA | 1 | 1 | 2 | 1 | 1 | 1 | 4 | 1 | 1 | 1 | 3 | 1 | 1 | 1 | 1 | 1 | 6 | 1 |
| 16_646060 | NA | NA | NA | NA | 2 | 0.5 | 1 | 1 | 3 | 0.6667 | NA | NA | 2 | 1 | 1 | 1 | NA | NA | 3 | 1 |
| 16_646062 | NA | NA | 1 | 1 | 1 | 1 | 1 | 1 | 3 | 1 | 1 | 1 | 2 | 0.5 | 1 | 1 | NA | NA | 4 | 0.75 |
| 16_646064 | NA | NA | 1 | 1 | 1 | 1 | 1 | 1 | 3 | 1 | 1 | 1 | 2 | 1 | 1 | 1 | NA | NA | 4 | 1 |
| 16_646066 | NA | NA | 1 | 0 | 1 | 0 | 1 | 1 | 3 | 0.3333 | 1 | 1 | 2 | 0 | 1 | 1 | NA | NA | 4 | 0.5 |
| 16_646068 | NA | NA | 1 | 0 | 1 | 0 | 1 | 1 | 3 | 0.3333 | 1 | 0 | 2 | 1 | 1 | 1 | NA | NA | 4 | 0.75 |
| 16_646732 | 6 | 0.5 | 4 | 0.25 | 3 | 0.3333 | 7 | 0.1429 | 20 | 0.3 | NA | NA | 2 | 0.5 | 5 | 0 | 7 | 0.4286 | 14 | 0.2857 |
| 16_646733 | 7 | 0 | 4 | 0 | 3 | 0.3333 | 7 | 0.1429 | 21 | 0.0952 | 1 | 0 | 3 | 0.3333 | 2 | 0 | 7 | 0 | 13 | 0.0769 |
| 16_646759 | 6 | 0.1667 | 4 | 0 | 3 | 0.3333 | 6 | 0.1667 | 19 | 0.1579 | 1 | 1 | 3 | 0 | 5 | 0 | 7 | 0 | 16 | 0.0625 |
| 16_646759 | 6 | 1 | 3 | 1 | 3 | 1 | 6 | 0.8333 | 18 | 0.9444 | 1 | 1 | 3 | 1 | 5 | 1 | 7 | 0.8571 | 16 | 0.9375 |
| 16_646760 | 8 | 0.5 | 1 | 0 | 3 | 0.6667 | 3 | 0.3333 | 15 | 0.4667 | NA | NA | 3 | 0.3333 | 3 | 0.6667 | 6 | 0.5 | 12 | 0.5 |
| 16_646761 | 8 | 0.25 | 4 | 0.5 | 3 | 0.3333 | 7 | 0.1429 | 22 | 0.2727 | 1 | 0 | 3 | 0.3333 | 4 | 0 | 6 | 0 | 14 | 0.0714 |
| 16_646761 | 7 | 0.8571 | 5 | 1 | 2 | 1 | 6 | 1 | 20 | 0.95 | NA | NA | 3 | 1 | 4 | 1 | 5 | 1 | 12 | 1 |
| 16_646761 | 8 | 0.125 | 5 | 0.6 | 3 | 0 | 6 | 0 | 22 | 0.1818 | 2 | 0 | 3 | 0.3333 | 5 | 0.2 | 6 | 0.1667 | 16 | 0.1875 |
| 16_646762 | 7 | 1 | 5 | 1 | 3 | 1 | 7 | 1 | 22 | 1 | 2 | 1 | 3 | 1 | 5 | 1 | 7 | 1 | 17 | 1 |
| 16_646764 | 7 | 0.1429 | 5 | 0 | 2 | 0 | 7 | 0.1429 | 21 | 0.0952 | 3 | 0 | 3 | 0.6667 | 5 | 0 | 6 | 0 | 17 | 0.1176 |
| 16_646766 | 7 | 0 | 4 | 0 | 2 | 0 | 7 | 0.1429 | 20 | 0.05 | 3 | 0 | 2 | 0.5 | 5 | 0 | 6 | 0 | 16 | 0.0625 |
| 16_646768 | 8 | 0.625 | 5 | 0.2 | 3 | 0.6667 | 7 | 0 | 23 | 0.3478 | 3 | 0.6667 | 3 | 0 | 5 | 0.2 | 7 | 0.2857 | 18 | 0.2778 |
| 16_646769 | 8 | 0.75 | 4 | 0.5 | 3 | 0.6667 | 6 | 0.8333 | 21 | 0.7143 | 1 | 0 | 3 | 0.6667 | 5 | 0.6 | 5 | 0 | 14 | 0.3571 |
| 16_646783 | 6 | 0 | 2 | 0 | 5 | 0.4 | 7 | 0.1429 | 20 | 0.15 | 2 | 0 | 1 | 0 | 4 | 0 | 4 | 0.25 | 11 | 0.0909 |
| 16_646786 | 11 | 0 | 5 | 0.2 | 8 | 0.125 | 8 | 0 | 32 | 0.0625 | 5 | 0.2 | 2 | 0 | 7 | 0 | 8 | 0.25 | 22 | 0.1364 |
| 16_647423 | NA | NA | NA | NA | NA | NA | 1 | 1 | 1 | 1 | NA | NA | NA | NA | 1 | 0 | 1 | 1 | 2 | 0.5 |
| 16_647465 | NA | NA | NA | NA | NA | NA | 1 | 1 | 1 | 1 | 1 | 1 | 2 | 1 | NA | NA | NA | NA | 3 | 1 |
| 16_647466 | NA | NA | NA | NA | NA | NA | 1 | 1 | 1 | 1 | 1 | 1 | 1 | 1 | NA | NA | NA | NA | 2 | 1 |
| 16_647467 | NA | NA | NA | NA | NA | NA | 1 | 1 | 1 | 1 | 1 | 1 | 2 | 0.5 | NA | NA | NA | NA | 3 | 0.6667 |
| 16_647468 | NA | NA | NA | NA | NA | NA | 1 | 1 | 1 | 1 | 1 | 1 | 2 | 1 | NA | NA | NA | NA | 3 | 1 |
| 16_647470 | NA | NA | NA | NA | NA | NA | 1 | 1 | 1 | 1 | 1 | 0 | 2 | 1 | NA | NA | NA | NA | 3 | 0.6667 |
| 16_647470 | NA | NA | NA | NA | NA | NA | 1 | 1 | 1 | 1 | 1 | 1 | 2 | 1 | NA | NA | NA | NA | 3 | 1 |
| 16_653361 | 15 | 0.3333 | 13 | 0.0769 | 7 | 0.2857 | 7 | 0 | 42 | 0.1905 | 9 | 0 | 6 | 0.1667 | 7 | 0.1429 | 14 | 0.2143 | 36 | 0.1389 |
| 16_653361 | 14 | 0 | 12 | 0.1667 | 4 | 0.25 | 8 | 0 | 38 | 0.0789 | 8 | 0 | 7 | 0.1429 | 6 | 0.1667 | 14 | 0.0714 | 35 | 0.0857 |
| 16_653363 | 13 | 0 | 9 | 0.1111 | 3 | 0 | 9 | 0.1111 | 34 | 0.0588 | 4 | 0 | 7 | 0.1429 | 5 | 0.2 | 13 | 0.1538 | 29 | 0.1379 |
| 16_653366 | 11 | 0.0909 | 8 | 0.125 | 2 | 0 | 8 | 0 | 29 | 0.069 | 8 | 0 | 7 | 0 | 6 | 0 | 13 | 0.1538 | 34 | 0.0588 |
| 16_653368 | 6 | 0 | 6 | 0 | 2 | 0 | 6 | 0.1667 | 20 | 0.05 | 5 | 0 | 6 | 0 | 3 | 0 | 13 | 0.1538 | 27 | 0.0741 |
| 16_653369 | 9 | 0.1111 | 6 | 0 | 2 | 0 | 5 | 0.2 | 22 | 0.0909 | 4 | 0 | 5 | 0 | 5 | 0 | 13 | 0.1538 | 27 | 0.0741 |
| 16_653598 | NA | NA | 1 | 1 | NA | NA | NA | NA | 1 | 1 | NA | NA | NA | NA | 2 | 1 | NA | NA | 2 | 1 |
| 16_653599 | NA | NA | 1 | 1 | NA | NA | NA | NA | 1 | 1 | NA | NA | NA | NA | 2 | 0.5 | NA | NA | 2 | 0.5 |
| 16_653599 | NA | NA | 1 | 1 | NA | NA | NA | NA | 1 | 1 | NA | NA | NA | NA | 2 | 0.5 | NA | NA | 2 | 0.5 |
| 16_653599 | NA | NA | 1 | 1 | NA | NA | NA | NA | 1 | 1 | NA | NA | NA | NA | 2 | 0.5 | NA | NA | 2 | 0.5 |
| 16_664530 | 7 | 0.2857 | 9 | 0.1111 | 17 | 0.1765 | 12 | 0.25 | 45 | 0.2 | 6 | 0 | 6 | 0 | 14 | 0.2143 | 6 | 0.1667 | 32 | 0.125 |
| 16_664533 | 8 | 0 | 11 | 0.0909 | 16 | 0.125 | 13 | 0.0769 | 48 | 0.0833 | 3 | 0 | 13 | 0.0769 | 14 | 0.1429 | 5 | 0.4 | 35 | 0.1429 |
| 16_664535 | 9 | 0.2222 | 14 | 0.2143 | 18 | 0.1667 | 13 | 0.4615 | 54 | 0.2593 | 4 | 0.25 | 15 | 0.1333 | 15 | 0.2667 | 7 | 0.1429 | 41 | 0.1951 |
| 16_664536 | 7 | 0 | 13 | 0.0769 | 19 | 0.0526 | 13 | 0.0769 | 52 | 0.0577 | 3 | 0 | 15 | 0 | 17 | 0.1176 | 5 | 0.2 | 40 | 0.075 |
| 16_664537 | 7 | 0.1429 | 13 | 0.3077 | 18 | 0.0556 | 11 | 0.0909 | 49 | 0.1429 | 3 | 0.3333 | 14 | 0.1429 | 15 | 0.0667 | 5 | 0.2 | 37 | 0.1351 |
| 16_664541 | 7 | 0.4286 | 13 | 0.5385 | 12 | 0.5833 | 8 | 0.125 | 40 | 0.45 | 2 | 1 | 11 | 0.3636 | 12 | 0.4167 | 5 | 0.6 | 30 | 0.4667 |
| 16_664565 | 3 | 0 | 5 | 0 | 7 | 0 | 5 | 0.2 | 20 | 0.05 | 1 | 0 | 5 | 0.2 | 7 | 0.1429 | 3 | 0 | 16 | 0.125 |
| 16_664581 | 2 | 0.5 | 9 | 0.1111 | 10 | 0.2 | 7 | 0.2857 | 28 | 0.2143 | 3 | 0.3333 | 5 | 0 | 9 | 0.2222 | 9 | 0.1111 | 26 | 0.1538 |

| | | | | | | | | | | | | | | | | | | | | |
|---|---|---|---|---|---|---|---|---|---|---|---|---|---|---|---|---|---|---|---|---|
| 16_664582 | 2 | 0 | 8 | 0.25 | 9 | 0.1111 | 6 | 0.1667 | 25 | 0.16 | 3 | 0 | 5 | 0.4 | 7 | 0 | 8 | 0 | 23 | 0.087 |
| 16_664595 | 7 | 0.1429 | 11 | 0.3636 | 10 | 0.2 | 3 | 0.3333 | 31 | 0.2581 | 8 | 0.375 | 5 | 0.4 | 6 | 0.5 | 9 | 0.5556 | 28 | 0.4643 |
| 16_664596 | 7 | 0.1429 | 15 | 0.4 | 13 | 0.2308 | 6 | 0.3333 | 41 | 0.2927 | 9 | 0.3333 | 7 | 0.2857 | 7 | 0.1429 | 9 | 0.3333 | 32 | 0.2812 |
| 16_664597 | 7 | 0.8571 | 11 | 0.9091 | 12 | 0.75 | 5 | 0.8 | 35 | 0.8286 | 10 | 0.7 | 6 | 0.8333 | 6 | 1 | 10 | 0.9 | 32 | 0.8438 |
| 16_664599 | 6 | 0.1667 | 10 | 0.2 | 12 | 0.1667 | 4 | 0.75 | 32 | 0.25 | 8 | 0.375 | 5 | 0.2 | 5 | 0.2 | 9 | 0.2222 | 27 | 0.2593 |
| 16_664599 | 7 | 0.2857 | 13 | 0.4615 | 13 | 0.1538 | 5 | 0.4 | 38 | 0.3158 | 10 | 0.4 | 6 | 0.1667 | 5 | 0.2 | 10 | 0.3 | 31 | 0.2903 |
| 16_728013 | 4 | 0 | 1 | 0 | NA | NA | 3 | 0.3333 | 8 | 0.125 | 2 | 1 | 1 | 1 | 1 | 0 | 3 | 0.6667 | 7 | 0.7143 |
| 16_728212 | 4 | 0.25 | 2 | 0 | NA | NA | 8 | 0.125 | 14 | 0.1429 | 2 | 0.5 | 4 | 0.25 | 4 | 0.25 | 5 | 0 | 15 | 0.2 |
| 16_728212 | 3 | 0 | 2 | 0.5 | NA | NA | 9 | 0 | 14 | 0.0714 | 1 | 0 | 4 | 0.25 | 3 | 0.3333 | 5 | 0 | 13 | 0.1538 |
| 16_728326 | 5 | 0 | 3 | 0.3333 | 1 | 0 | 8 | 0 | 17 | 0.0588 | 2 | 0 | 4 | 0 | 4 | 0.25 | 5 | 0 | 15 | 0.0667 |
| 16_728413 | 5 | 1 | 3 | 1 | NA | NA | 9 | 0.6667 | 17 | 0.8235 | 2 | 1 | 3 | 1 | 5 | 0.8 | 5 | 1 | 15 | 0.9333 |
| 16_728414 | 7 | 0.2857 | 3 | 0 | 1 | 0 | 7 | 0.1429 | 18 | 0.1667 | 2 | 0 | 3 | 0 | 3 | 0.3333 | 4 | 0.75 | 12 | 0.3333 |
| 16_728415 | 7 | 0 | 2 | 0 | 1 | 0 | 10 | 0.1 | 20 | 0.05 | 2 | 0 | 2 | 0 | 3 | 0.3333 | 5 | 0.2 | 12 | 0.1667 |
| 16_728416 | 5 | 0.2 | 3 | 0 | 1 | 0 | 10 | 0.3 | 19 | 0.2105 | 2 | 0 | 3 | 0.3333 | 4 | 0 | 5 | 0.4 | 14 | 0.2143 |
| 16_728417 | 7 | 0.1429 | 4 | 0.25 | 1 | 0 | 9 | 0 | 21 | 0.0952 | 1 | 0 | 5 | 0 | 5 | 0 | 5 | 0.2 | 16 | 0.0625 |
| 16_728419 | 6 | 0.3333 | 2 | 0.5 | 1 | 0 | 10 | 0.2 | 19 | 0.2632 | 1 | 0 | 5 | 0.2 | 7 | 0 | 6 | 0.3333 | 19 | 0.1579 |
| 16_728423 | 7 | 0 | 5 | 0.2 | 1 | 1 | 10 | 0.2 | 23 | 0.1739 | 4 | 0.25 | 6 | 0.5 | 7 | 0 | 8 | 0.375 | 25 | 0.28 |
| 16_728424 | 6 | 0.5 | 4 | 0 | 1 | 0 | 10 | 0.9 | 21 | 0.5714 | 3 | 0.3333 | 6 | 0.5 | 5 | 0.6 | 8 | 0.875 | 22 | 0.6364 |
| 16_728425 | 9 | 0 | 5 | 0 | 1 | 0 | 11 | 0.1818 | 26 | 0.0769 | 4 | 0 | 6 | 0.1667 | 8 | 0.125 | 8 | 0.125 | 26 | 0.1154 |
| 16_728426 | 9 | 0.3333 | 4 | 0.25 | 1 | 0 | 11 | 0.1818 | 25 | 0.24 | 2 | 0 | 4 | 0.25 | 7 | 0.2857 | 8 | 0.125 | 21 | 0.1905 |
| 16_736897 | NA | NA | 4 | 0 | 1 | 1 | 2 | 0.5 | 7 | 0.2857 | NA | NA | 2 | 0.5 | 3 | 0 | 1 | 0 | 6 | 0.1667 |
| 16_736913 | 5 | 0 | 10 | 0.4 | 7 | 0.1429 | 10 | 0.3 | 32 | 0.25 | 4 | 0.75 | 2 | 0.5 | 8 | 0.5 | 9 | 0.1111 | 23 | 0.3913 |
| 16_736925 | 1 | 0 | 10 | 0.1 | 6 | 0.1667 | 8 | 0.125 | 25 | 0.12 | 2 | 0 | 3 | 0 | 5 | 0.2 | 6 | 0 | 16 | 0.0625 |
| 16_736926 | NA | NA | 4 | 0.25 | 1 | 1 | 3 | 0.3333 | 8 | 0.375 | 1 | 0 | 3 | 0 | 2 | 0.5 | 3 | 0 | 9 | 0.1111 |
| 16_736926 | 1 | 0 | 8 | 0.625 | 3 | 0.6667 | 5 | 0.6 | 17 | 0.5882 | 2 | 0.5 | 2 | 0.5 | 5 | 0.4 | 5 | 0.6 | 14 | 0.5 |
| 16_736927 | 1 | 0 | 7 | 0.1429 | 4 | 0 | 8 | 0.125 | 20 | 0.1 | 2 | 0 | 3 | 0 | 4 | 0.25 | 5 | 0 | 14 | 0.0714 |
| 16_753976 | 1 | 0 | NA | NA | 2 | 0.5 | 1 | 1 | 4 | 0.5 | 1 | 0 | 1 | 0 | NA | NA | 1 | 1 | 3 | 0.3333 |
| 16_753977 | 1 | 1 | NA | NA | 1 | 0 | 1 | 1 | 3 | 0.6667 | 1 | 1 | NA | NA | NA | NA | 1 | 1 | 2 | 1 |
| 16_755522 | 8 | 0.25 | 6 | 0.1667 | 5 | 0 | 11 | 0.0909 | 30 | 0.1333 | 6 | 0.3333 | 7 | 0.1429 | 7 | 0.4286 | 3 | 0.3333 | 23 | 0.3043 |
| 16_755522 | 6 | 0 | 6 | 0.3333 | 3 | 0 | 10 | 0.1 | 25 | 0.12 | 5 | 0.2 | 3 | 0 | 5 | 0 | 5 | 0.2 | 18 | 0.1111 |
| 16_755523 | 7 | 0 | 7 | 0.2857 | 6 | 0.1667 | 8 | 0 | 28 | 0.1071 | 6 | 0 | 7 | 0.1429 | 5 | 0.2 | 5 | 0 | 23 | 0.087 |
| 16_755531 | 9 | 0.1111 | 8 | 0.375 | 5 | 0.2 | 7 | 0.1429 | 29 | 0.2069 | 5 | 0.6 | 5 | 0.2 | 6 | 0.1667 | 7 | 0.4286 | 23 | 0.3478 |
| 16_755535 | 8 | 0 | 10 | 0.1 | 3 | 0.3333 | 6 | 0 | 27 | 0.0741 | 5 | 0 | 4 | 0.25 | 4 | 0.25 | 6 | 0 | 19 | 0.1053 |
| 16_755542 | 6 | 0.3333 | 9 | 0.3333 | 5 | 0.2 | 5 | 0.2 | 25 | 0.28 | 5 | 0.4 | 2 | 0 | 4 | 0.25 | 3 | 1 | 14 | 0.4286 |
| 16_755542 | 6 | 0.1667 | 8 | 0.25 | 4 | 0.75 | 5 | 0.2 | 23 | 0.3043 | 5 | 1 | 3 | 0.3333 | 4 | 0.5 | 3 | 0.6667 | 15 | 0.6667 |
| 16_755543 | 5 | 0.2 | 8 | 0.25 | 5 | 0.2 | 4 | 0.25 | 22 | 0.2273 | 4 | 0 | 2 | 0 | 4 | 0 | 3 | 0.3333 | 13 | 0.0769 |
| 16_755545 | 6 | 0.5 | 6 | 0.8333 | 5 | 0.8 | 5 | 0.6 | 22 | 0.6818 | 5 | 0.8 | 3 | 1 | 5 | 0 | 3 | 0.6667 | 16 | 0.5625 |
| 16_755545 | 7 | 1 | 5 | 1 | 4 | 1 | 4 | 1 | 20 | 1 | 4 | 1 | 3 | 0.6667 | 4 | 1 | 2 | 1 | 13 | 0.9231 |
| 16_755546 | 7 | 0.4286 | 7 | 0 | 4 | 0.25 | 4 | 0 | 22 | 0.1818 | 5 | 0.4 | 3 | 0.3333 | 4 | 0 | 3 | 0.3333 | 15 | 0.2667 |
| 16_755546 | 4 | 0.5 | 2 | 0.5 | 2 | 1 | 2 | 0.5 | 10 | 0.6 | 2 | 0.5 | 3 | 0.3333 | 1 | 0 | 1 | 0 | 7 | 0.2857 |
| 16_755546 | 7 | 1 | 5 | 1 | 3 | 1 | 3 | 1 | 18 | 1 | 5 | 1 | 2 | 1 | 3 | 1 | 3 | 1 | 13 | 1 |
| 16_755547 | 7 | 0.5714 | 7 | 0.2857 | 4 | 0.5 | 4 | 0.25 | 22 | 0.4091 | 5 | 0.4 | 3 | 0 | 5 | 0.2 | 2 | 0 | 15 | 0.2 |
| 16_755548 | 6 | 0.6667 | 8 | 0.875 | 2 | 1 | 3 | 1 | 19 | 0.8421 | 4 | 1 | 3 | 1 | 4 | 0.75 | 3 | 1 | 14 | 0.9286 |
| 16_755549 | 6 | 0.1667 | 8 | 0.125 | 3 | 0 | 4 | 0 | 21 | 0.0952 | 3 | 0.3333 | 3 | 0 | 5 | 0 | 2 | 0 | 13 | 0.0769 |
| 16_755549 | 5 | 0.6 | 6 | 0.3333 | 4 | 0.75 | 4 | 0.5 | 19 | 0.5263 | 3 | 0.3333 | 3 | 1 | 5 | 0.2 | 1 | 1 | 12 | 0.5 |
| 16_755550 | 6 | 0 | 8 | 0 | 4 | 0.25 | 4 | 0.25 | 22 | 0.0909 | 4 | 0 | 2 | 0 | 5 | 0.2 | 2 | 0 | 13 | 0.0769 |
| 16_755550 | 6 | 0.3333 | 5 | 0.6 | 4 | 0.75 | 3 | 0 | 18 | 0.4444 | 4 | 0.75 | 2 | 0 | 4 | 0.25 | 2 | 0 | 12 | 0.3333 |
| 16_755551 | 3 | 1 | 3 | 1 | 2 | 1 | 3 | 1 | 11 | 1 | 1 | 1 | 1 | 1 | 2 | 1 | 1 | 1 | 5 | 1 |
| 16_755551 | 6 | 0.8333 | 5 | 0.8 | 4 | 0.75 | 3 | 0.6667 | 18 | 0.7778 | 3 | 0.6667 | 2 | 0.5 | 2 | 0.5 | 2 | 1 | 9 | 0.6667 |
| 16_755551 | 5 | 0.4 | 5 | 0.2 | 4 | 0.25 | 3 | 0.3333 | 17 | 0.2941 | 4 | 0.25 | 2 | 0.5 | 3 | 0.3333 | 2 | 0 | 11 | 0.2727 |
| 16_755551 | 6 | 1 | 5 | 1 | 4 | 0.5 | 4 | 1 | 19 | 0.8947 | 3 | 0.6667 | 2 | 1 | 3 | 0.6667 | 2 | 1 | 10 | 0.8 |
| 16_755551 | 6 | 0.3333 | 6 | 0 | 2 | 0 | 3 | 0 | 17 | 0.1176 | 2 | 0 | 2 | 0.5 | 2 | 0 | 1 | 0 | 7 | 0.1429 |
| 16_755552 | 5 | 0.4 | 6 | 0.1667 | 3 | 0 | 3 | 0.3333 | 17 | 0.2353 | 3 | 0 | 2 | 0 | 3 | 0 | 2 | 0.5 | 10 | 0.1 |
| 16_755620 | 5 | 0 | 4 | 0 | 4 | 0 | 2 | 0.5 | 15 | 0.0667 | 3 | 0 | 2 | 0 | 3 | 0 | 2 | 0.5 | 10 | 0.1 |
| 16_760849 | 2 | 1 | 3 | 1 | 4 | 0.75 | 3 | 1 | 12 | 0.9167 | 1 | 0 | 6 | 1 | 3 | 0.3333 | 8 | 0.625 | 18 | 0.6667 |
| 16_760850 | 3 | 0.3333 | 3 | 1 | 4 | 0.75 | 4 | 0.75 | 14 | 0.7143 | 4 | 0.25 | 7 | 1 | 3 | 0.6667 | 7 | 0.8571 | 21 | 0.7619 |
| 16_760851 | 3 | 0 | 4 | 0.5 | 3 | 0 | 3 | 0 | 13 | 0.1538 | 4 | 0 | 8 | 0.25 | 3 | 0.3333 | 9 | 0 | 24 | 0.125 |
| 16_760855 | 1 | 0 | 6 | 0.1667 | 2 | 0 | 3 | 0 | 12 | 0.0833 | 3 | 0.3333 | 6 | 0.3333 | 1 | 0 | 8 | 0.125 | 18 | 0.2222 |
| 16_760861 | 1 | 0 | 4 | 1 | 1 | 1 | 2 | 1 | 8 | 0.875 | 3 | 0.6667 | 3 | 1 | 1 | 0 | 8 | 0.5 | 15 | 0.6 |
| 16_760864 | 1 | 0 | 6 | 0 | 1 | 1 | 1 | 0 | 9 | 0.1111 | 3 | 0.3333 | 3 | 0 | 1 | 0 | 9 | 0.1111 | 16 | 0.125 |
| 16_760864 | 1 | 0 | 6 | 0.3333 | 1 | 0 | 2 | 0 | 10 | 0.2 | 3 | 0 | 3 | 0 | 1 | 0 | 8 | 0.125 | 15 | 0.0667 |
| 16_760870 | 1 | 0 | 2 | 0.5 | 1 | 0 | 2 | 0 | 6 | 0.1667 | 3 | 0.3333 | 3 | 0 | 1 | 0 | 7 | 0 | 14 | 0.0714 |
| 16_760878 | 1 | 0 | 2 | 0.5 | 1 | 0 | 1 | 0 | 5 | 0.2 | 3 | 0 | 2 | 0.5 | 1 | 0 | 4 | 0 | 10 | 0.1 |
| 16_760880 | 1 | 1 | 3 | 1 | NA | NA | 1 | 1 | 5 | 1 | 3 | 1 | 3 | 1 | 1 | 1 | 5 | 1 | 12 | 1 |
| 16_760881 | 1 | 1 | 4 | 1 | 1 | 1 | 2 | 1 | 8 | 1 | 2 | 1 | 3 | 1 | 1 | 1 | 5 | 1 | 11 | 1 |

| | | | | | | | | | | | | | | | | | | | | |
|---|---|---|---|---|---|---|---|---|---|---|---|---|---|---|---|---|---|---|---|---|
| 16_760882 | 1 | 1 | 2 | 0.5 | NA | NA | 1 | 0 | 4 | 0.5 | 1 | 0 | 3 | 0.6667 | 1 | 1 | 4 | 0.5 | 9 | 0.5556 |
| 16_760882 | 1 | 0 | 3 | 0 | NA | NA | 1 | 1 | 5 | 0.2 | 2 | 0 | 3 | 0.6667 | 1 | 1 | 4 | 0.75 | 10 | 0.6 |
| 16_760882 | 1 | 0 | 3 | 0.6667 | 1 | 1 | 2 | 0.5 | 7 | 0.5714 | 2 | 0 | 3 | 0.3333 | 1 | 0 | 5 | 0.2 | 11 | 0.1818 |
| 16_760883 | 1 | 0 | 4 | 0 | 1 | 1 | 2 | 0.5 | 8 | 0.25 | 2 | 1 | 2 | 0 | 1 | 1 | 3 | 0.3333 | 8 | 0.5 |
| 16_760883 | 1 | 1 | 3 | 1 | NA | NA | 2 | 1 | 6 | 1 | 2 | 1 | 2 | 1 | 1 | 1 | 2 | 1 | 7 | 1 |
| 16_760883 | NA | NA | 2 | 1 | 1 | 1 | 2 | 1 | 5 | 1 | 3 | 1 | 3 | 1 | 1 | 1 | 4 | 1 | 11 | 1 |
| 16_760884 | 1 | 0 | 3 | 0.3333 | 1 | 1 | 2 | 0.5 | 7 | 0.4286 | 3 | 0 | 3 | 0.3333 | 1 | 0 | 4 | 0 | 11 | 0.0909 |
| 16_760885 | 1 | 0 | 4 | 0.5 | 1 | 0 | 2 | 0 | 8 | 0.25 | 3 | 0.3333 | 3 | 0.3333 | 1 | 1 | 3 | 0 | 10 | 0.3 |
| 16_760885 | 1 | 1 | 4 | 1 | 1 | 1 | 2 | 1 | 8 | 1 | 2 | 1 | 3 | 1 | 1 | 1 | 3 | 1 | 9 | 1 |
| 16_760886 | 1 | 1 | 3 | 1 | NA | NA | 2 | 1 | 6 | 1 | 3 | 1 | 1 | 1 | 1 | 1 | 3 | 1 | 8 | 1 |
| 16_760886 | 1 | 0 | 2 | 1 | 1 | 1 | 2 | 0.5 | 6 | 0.6667 | 2 | 0.5 | 3 | 0.6667 | 1 | 1 | 3 | 0.6667 | 9 | 0.6667 |
| 16_760886 | 1 | 1 | 4 | 0.5 | NA | NA | 2 | 0.5 | 7 | 0.5714 | 2 | 1 | 3 | 0.6667 | 1 | 1 | 3 | 0.6667 | 9 | 0.7778 |
| 16_760887 | NA | NA | 2 | 1 | 1 | 1 | 2 | 0.5 | 5 | 0.8 | 3 | 0.3333 | 2 | 1 | NA | NA | 3 | 1 | 8 | 0.75 |
| 16_760890 | NA | NA | 4 | 0 | 1 | 1 | 2 | 0 | 7 | 0.1429 | 2 | 0.5 | NA | NA | NA | NA | 3 | 0 | 5 | 0.2 |
| 16_760891 | 1 | 1 | 3 | 1 | 1 | 1 | 2 | 1 | 7 | 1 | 3 | 0.6667 | 1 | 1 | NA | NA | 2 | 1 | 6 | 0.8333 |
| 16_760893 | 1 | 0 | 3 | 0 | 1 | 1 | NA | NA | 5 | 0.2 | 2 | 0.5 | NA | NA | NA | NA | NA | NA | 2 | 0.5 |
| 16_760894 | NA | NA | 4 | 1 | NA | NA | 2 | 0.5 | 6 | 0.8333 | 3 | 1 | 2 | 0.5 | NA | NA | 2 | 1 | 7 | 0.8571 |
| 16_760894 | 1 | 1 | 4 | 0.5 | 1 | 0 | 2 | 0.5 | 8 | 0.5 | 3 | 0 | 2 | 0.5 | NA | NA | 2 | 0 | 7 | 0.1429 |
| 16_760896 | 1 | 1 | 3 | 0.3333 | NA | NA | 1 | 0 | 5 | 0.4 | 2 | 0.5 | 2 | 0 | NA | NA | 1 | 0 | 5 | 0.2 |
| 16_760897 | 1 | 0 | 3 | 0.3333 | 1 | 1 | 2 | 0 | 7 | 0.2857 | 3 | 0.6667 | 2 | 0 | NA | NA | 2 | 0 | 7 | 0.2857 |
| 16_760899 | 1 | 1 | 2 | 1 | 1 | 1 | 2 | 0.5 | 6 | 0.8333 | 3 | 0.6667 | 2 | 1 | NA | NA | 1 | 1 | 6 | 0.8333 |
| 16_760899 | NA | NA | 2 | 0 | 1 | 1 | 1 | 1 | 4 | 0.5 | 3 | 0.3333 | NA | NA | NA | NA | 2 | 0.5 | 5 | 0.4 |
| 16_760899 | 1 | 0 | 3 | 0.3333 | 1 | 0 | 2 | 0 | 7 | 0.1429 | 3 | 0 | 1 | 0 | NA | NA | 2 | 0.5 | 6 | 0.1667 |
| 16_760900 | 1 | 0 | 2 | 0 | 1 | 0 | 2 | 1 | 6 | 0.3333 | 3 | 0 | 2 | 0 | NA | NA | 2 | 0.5 | 7 | 0.1429 |
| 16_760902 | 1 | 1 | 3 | 0.6667 | 1 | 0 | 2 | 0 | 7 | 0.4286 | 3 | 0 | 2 | 0 | NA | NA | 2 | 0.5 | 7 | 0.1429 |
| 16_760906 | 1 | 0 | 1 | 1 | 1 | 1 | 1 | 0 | 4 | 0.5 | 1 | 0 | 2 | 0 | NA | NA | 1 | 1 | 4 | 0.25 |
| 16_760909 | 1 | 0 | 2 | 1 | 1 | 1 | 1 | 0 | 5 | 0.6 | 1 | 1 | 1 | 1 | NA | NA | 1 | 0 | 3 | 0.6667 |
| 16_760911 | NA | NA | NA | NA | 1 | 0 | 1 | 1 | 2 | 0.5 | 1 | 1 | NA | NA | NA | NA | 1 | 1 | 2 | 1 |
| 16_760912 | NA | NA | NA | NA | 1 | 1 | 1 | 1 | 2 | 1 | 1 | 1 | NA | NA | NA | NA | 1 | 1 | 2 | 1 |
| 16_760914 | NA | NA | 1 | 0 | 1 | 1 | NA | NA | 2 | 0.5 | 1 | 1 | NA | NA | NA | NA | 1 | 0 | 2 | 0.5 |
| 16_768806 | 2 | 0.5 | 2 | 0 | 2 | 0 | 1 | 0 | 7 | 0.1429 | 2 | 0 | 1 | 0 | 2 | 0.5 | 1 | 1 | 6 | 0.3333 |
| 16_768807 | 2 | 0.5 | 2 | 0 | 2 | 0 | 2 | 0 | 8 | 0.125 | 2 | 0 | 1 | 1 | 2 | 0 | 2 | 0 | 7 | 0.1429 |
| 16_768906 | 2 | 0 | 2 | 0 | 2 | 0 | 2 | 0.5 | 8 | 0.125 | 2 | 0 | 1 | 1 | 2 | 0 | 2 | 0 | 7 | 0.1429 |
| 16_769123 | 2 | 0 | 2 | 0.5 | 3 | 0 | 2 | 0 | 9 | 0.1111 | 4 | 0.25 | 1 | 0 | 2 | 0 | 2 | 0 | 9 | 0.1111 |
| 16_769123 | 1 | 1 | 3 | 0.6667 | 3 | 0.6667 | 2 | 1 | 9 | 0.7778 | 4 | 0.5 | 2 | 1 | 2 | 0.5 | 2 | 1 | 10 | 0.7 |
| 16_769124 | 2 | 0.5 | 2 | 0 | 3 | 0.3333 | 2 | 1 | 9 | 0.4444 | 3 | 0.3333 | 2 | 0.5 | 2 | 1 | 2 | 0 | 9 | 0.4444 |
| 16_769124 | 2 | 0.5 | 2 | 1 | 2 | 0.5 | 1 | 1 | 7 | 0.7143 | 4 | 0.75 | 1 | 1 | 2 | 1 | 1 | 1 | 8 | 0.875 |
| 16_769125 | 3 | 0.3333 | 4 | 0.25 | 3 | 0.3333 | 2 | 0 | 12 | 0.25 | 4 | 0.5 | 2 | 0 | 2 | 0.5 | 2 | 0.5 | 10 | 0.4 |
| 16_769125 | 3 | 0.3333 | 3 | 1 | 3 | 0.3333 | 2 | 0 | 11 | 0.4545 | 3 | 0.3333 | 2 | 0.5 | 2 | 0.5 | 2 | 0 | 9 | 0.3333 |
| 16_769125 | 3 | 0 | 4 | 0.25 | 3 | 0 | 2 | 0.5 | 12 | 0.1667 | 4 | 0.25 | 2 | 0.5 | 2 | 0 | 1 | 0 | 9 | 0.2222 |
| 16_769125 | 2 | 0.5 | 4 | 1 | 2 | 0.5 | 2 | 0.5 | 10 | 0.7 | 4 | 0.5 | 2 | 1 | 2 | 0.5 | 2 | 1 | 10 | 0.7 |
| 16_769127 | 2 | 0 | 5 | 0.4 | 3 | 0.3333 | 2 | 0 | 12 | 0.25 | 4 | 0.25 | 3 | 0 | 2 | 0 | 2 | 0 | 11 | 0.0909 |
| 16_769130 | 3 | 0 | 6 | 0.3333 | 3 | 0.3333 | 3 | 0 | 15 | 0.2 | 3 | 0.3333 | 2 | 0 | 2 | 0 | 2 | 0 | 9 | 0.1111 |
| 16_781366 | 3 | 0 | 1 | 0 | 5 | 0.2 | 4 | 0 | 13 | 0.0769 | 1 | 0 | 4 | 0.25 | 2 | 0 | 5 | 0 | 12 | 0.0833 |
| 16_781569 | 3 | 0 | 5 | 0.4 | 6 | 0.1667 | 6 | 0.3333 | 20 | 0.25 | 4 | 0 | 5 | 0.2 | 13 | 0.3077 | 4 | 0.5 | 26 | 0.2692 |
| 16_787473 | NA | NA | NA | NA | NA | NA | 1 | 1 | 1 | 1 | NA | NA | NA | NA | 1 | 1 | NA | NA | 1 | 1 |
| 16_787474 | NA | NA | NA | NA | NA | NA | 1 | 1 | 1 | 1 | NA | NA | NA | NA | 2 | 1 | NA | NA | 2 | 1 |
| 16_787475 | NA | NA | NA | NA | NA | NA | 1 | 1 | 1 | 1 | NA | NA | NA | NA | 1 | 1 | NA | NA | 1 | 1 |
| 16_814202 | 2 | 0 | 3 | 0.3333 | 1 | 1 | 4 | 0.25 | 10 | 0.3 | NA | NA | 1 | 1 | 6 | 0.1667 | 7 | 0.1429 | 14 | 0.2143 |
| 16_814203 | 2 | 0.5 | 4 | 0.25 | 1 | 1 | 4 | 0.5 | 11 | 0.4545 | NA | NA | 2 | 0 | 5 | 0.4 | 9 | 0.3333 | 16 | 0.3125 |
| 16_814204 | 2 | 0.5 | 4 | 0.25 | 1 | 0 | 3 | 0.3333 | 10 | 0.3 | NA | NA | 1 | 1 | 6 | 0.3333 | 7 | 0.5714 | 14 | 0.5 |
| 16_814204 | 2 | 0 | 4 | 0.5 | 2 | 0 | 4 | 0 | 12 | 0.1667 | NA | NA | 1 | 0 | 5 | 0.4 | 8 | 0.125 | 14 | 0.2143 |
| 16_814204 | 1 | 0 | 3 | 0 | 1 | 1 | 4 | 0.25 | 9 | 0.2222 | NA | NA | 1 | 0 | 5 | 0 | 6 | 0.1667 | 12 | 0.0833 |
| 16_814205 | 2 | 1 | 4 | 0.5 | 2 | 0 | 4 | 0.5 | 12 | 0.5 | NA | NA | 2 | 0 | 6 | 0.1667 | 9 | 0.3333 | 17 | 0.2353 |
| 16_814235 | 1 | 0 | 3 | 0 | 1 | 0 | 6 | 0.1667 | 11 | 0.0909 | 2 | 0 | 1 | 0 | 9 | 0.1111 | 7 | 0 | 19 | 0.0526 |
| 16_845713 | 2 | 0.5 | 6 | 0 | 3 | 0 | 4 | 0 | 15 | 0.0667 | 5 | 0.2 | NA | NA | 5 | 0 | 1 | 0 | 11 | 0.0909 |
| 16_846198 | 1 | 0 | 5 | 0 | 3 | 0.3333 | 2 | 0 | 11 | 0.0909 | 4 | 0.25 | 1 | 0 | 4 | 0 | 2 | 0 | 11 | 0.0909 |
| 16_846199 | 1 | 0 | 6 | 0 | 4 | 0.25 | 4 | 0 | 15 | 0.0667 | 4 | 0.25 | 2 | 0 | 4 | 0.25 | 2 | 0.5 | 12 | 0.25 |
| 16_846199 | 1 | 0 | 4 | 0.5 | 4 | 0.5 | 4 | 0.75 | 13 | 0.5385 | 4 | 0.75 | 2 | 0 | 4 | 0.75 | 2 | 0.5 | 12 | 0.5833 |
| 16_846201 | 1 | 0 | 6 | 0.1667 | 2 | 0.5 | 3 | 0 | 12 | 0.1667 | 3 | 0.3333 | 2 | 0 | 3 | 0 | 1 | 0 | 9 | 0.1111 |
| 16_846202 | 1 | 0 | 4 | 0.75 | 2 | 1 | 4 | 0.75 | 11 | 0.7273 | 4 | 0.75 | 1 | 1 | 4 | 0.75 | 2 | 0.5 | 11 | 0.7273 |
| 16_846203 | 1 | 0 | 6 | 0 | 2 | 0 | 3 | 0.6667 | 12 | 0.1667 | 3 | 0.3333 | 2 | 0.5 | 5 | 0.4 | 2 | 0 | 12 | 0.3333 |
| 16_846205 | 1 | 0 | 5 | 0.2 | 3 | 0 | 3 | 0 | 12 | 0.0833 | 5 | 0 | 3 | 0 | 6 | 0 | 2 | 0.5 | 16 | 0.0625 |
| 16_846218 | 3 | 0.3333 | 5 | 0 | 3 | 0.3333 | 3 | 0.3333 | 14 | 0.2143 | 4 | 0.25 | 4 | 0 | 5 | 0 | 2 | 0 | 15 | 0.0667 |
| 16_846220 | 2 | 0 | 5 | 0 | 6 | 0.1667 | 3 | 0.6667 | 16 | 0.1875 | 4 | 0 | 3 | 0.3333 | 4 | 0.25 | 2 | 0 | 13 | 0.1538 |

| | | | | | | | | | | | | | | | | | | | | |
|---|---|---|---|---|---|---|---|---|---|---|---|---|---|---|---|---|---|---|---|---|
| 16_846224 | 1 | 0 | 5 | 0.2 | 6 | 0.6667 | 2 | 0 | 14 | 0.3571 | 2 | 0 | 4 | 0 | 3 | 0.3333 | 2 | 0 | 11 | 0.0909 |
| 16_846225 | 2 | 0.5 | 4 | 0.5 | 6 | 0.3333 | 2 | 0 | 14 | 0.3571 | 4 | 0.25 | 2 | 0.5 | 4 | 0.75 | 2 | 0 | 12 | 0.4167 |
| 16_846226 | 3 | 0.3333 | 6 | 0 | 6 | 0.1667 | 3 | 0 | 18 | 0.1111 | 4 | 0 | 4 | 0.25 | 6 | 0 | 2 | 0 | 16 | 0.0625 |
| 16_846543 | 1 | 1 | NA | NA | 2 | 0 | 4 | 0.25 | 7 | 0.2857 | 1 | 0 | 4 | 0.25 | 2 | 0.5 | 1 | 1 | 8 | 0.375 |
| 16_846572 | 1 | 1 | 3 | 0 | 2 | 1 | 4 | 0.25 | 10 | 0.4 | 1 | 0 | 4 | 0.75 | 3 | 0.3333 | 3 | 0 | 11 | 0.3636 |
| 16_846573 | 1 | 1 | 3 | 0.3333 | 2 | 1 | 4 | 0.25 | 10 | 0.5 | 1 | 0 | 4 | 0.25 | 3 | 0 | 3 | 0 | 11 | 0.0909 |
| 16_846573 | 1 | 1 | 3 | 0.3333 | 2 | 1 | 4 | 0.5 | 10 | 0.6 | 1 | 0 | 4 | 0.75 | 3 | 0.6667 | 2 | 0.5 | 10 | 0.6 |
| 16_846573 | 1 | 0 | 3 | 0.3333 | 1 | 0 | 5 | 0.4 | 10 | 0.3 | 1 | 1 | 3 | 0.3333 | 3 | 0.6667 | 3 | 0.6667 | 10 | 0.6 |
| 16_846573 | 1 | 0 | 3 | 0.3333 | 2 | 0 | 5 | 0 | 11 | 0.0909 | 1 | 1 | 3 | 0 | 3 | 0.3333 | 3 | 0 | 10 | 0.2 |
| 16_846574 | 1 | 0 | 3 | 0.3333 | 2 | 0.5 | 4 | 0 | 10 | 0.2 | 1 | 0 | 4 | 0 | 3 | 0.3333 | 3 | 0.6667 | 11 | 0.2727 |
| 16_846574 | 1 | 0 | 4 | 0.25 | 2 | 0 | 5 | 0 | 12 | 0.0833 | 1 | 0 | 4 | 0.25 | 3 | 0 | 3 | 0 | 11 | 0.0909 |
| 16_846575 | 1 | 0 | 4 | 0.25 | 1 | 0 | 3 | 0.3333 | 9 | 0.2222 | 1 | 1 | 4 | 0.5 | 3 | 0.3333 | 3 | 0.6667 | 11 | 0.5455 |
| 16_846579 | 1 | 0 | 4 | 0 | 1 | 1 | 4 | 0.25 | 10 | 0.2 | 1 | 0 | 5 | 0.4 | 3 | 0.6667 | 3 | 0.3333 | 12 | 0.4167 |
| 16_846605 | NA | NA | 4 | 0.25 | 2 | 0 | 9 | 0.1111 | 15 | 0.1333 | 2 | 0 | 5 | 0.2 | 4 | 0 | 3 | 0 | 14 | 0.0714 |
| 16_847495 | 3 | 0.3333 | 4 | 0 | 2 | 0 | 3 | 0.3333 | 12 | 0.1667 | 2 | 0 | 3 | 0.3333 | 5 | 0.2 | 3 | 0.3333 | 13 | 0.2308 |
| 16_847497 | 10 | 0 | 9 | 0.2222 | 8 | 0.25 | 4 | 0 | 31 | 0.129 | 10 | 0.2 | 15 | 0.2667 | 9 | 0.1111 | 5 | 0.2 | 39 | 0.2051 |
| 16_847497 | 9 | 1 | 8 | 1 | 7 | 0.8571 | 3 | 1 | 27 | 0.963 | 10 | 0.9 | 14 | 1 | 9 | 0.7778 | 6 | 1 | 39 | 0.9231 |
| 16_847498 | 10 | 0 | 8 | 0 | 7 | 0.1429 | 4 | 0.25 | 29 | 0.069 | 9 | 0.1111 | 14 | 0.2857 | 5 | 0.2 | 4 | 0 | 32 | 0.1875 |
| 16_847498 | 6 | 0.1667 | 8 | 0.5 | 4 | 0.25 | 3 | 0.6667 | 21 | 0.381 | 7 | 0.4286 | 10 | 0.5 | 5 | 0.8 | 3 | 0.3333 | 25 | 0.52 |
| 16_847500 | 8 | 0.125 | 8 | 0 | 7 | 0.4286 | 4 | 0 | 27 | 0.1481 | 10 | 0 | 12 | 0 | 7 | 0.2857 | 3 | 0 | 32 | 0.0625 |
| 16_847501 | 6 | 0.5 | 7 | 0.1429 | 4 | 0.25 | 3 | 0 | 20 | 0.25 | 8 | 0.625 | 7 | 0.1429 | 6 | 0.3333 | 5 | 0.4 | 26 | 0.3846 |
| 16_847503 | 5 | 0 | 9 | 0.3333 | 5 | 0.2 | 3 | 0.6667 | 22 | 0.2727 | 8 | 0.375 | 9 | 0.5556 | 8 | 0.375 | 4 | 0.25 | 29 | 0.4138 |
| 16_847518 | 3 | 0 | 2 | 0.5 | 3 | 0 | 4 | 0 | 12 | 0.0833 | 5 | 0 | 4 | 0 | 5 | 0 | 3 | 0.3333 | 17 | 0.0588 |
| 16_847520 | 107 | 0.4673 | 135 | 0.5111 | 114 | 0.5351 | 131 | 0.3664 | 487 | 0.4682 | 106 | 0.5189 | 117 | 0.4017 | 121 | 0.5785 | 112 | 0.5 | 456 | 0.5 |
| 16_847521 | 97 | 0.2062 | 139 | 0.259 | 117 | 0.2393 | 118 | 0.3305 | 471 | 0.2611 | 97 | 0.3196 | 108 | 0.3611 | 107 | 0.2617 | 101 | 0.2574 | 413 | 0.3002 |
| 16_847521 | 116 | 0.1379 | 146 | 0.0959 | 124 | 0.121 | 135 | 0.1407 | 521 | 0.1228 | 107 | 0.1589 | 113 | 0.1327 | 120 | 0.1583 | 120 | 0.1583 | 460 | 0.1522 |
| 16_847522 | 82 | 0.1951 | 104 | 0.1154 | 93 | 0.1613 | 91 | 0.0989 | 370 | 0.1405 | 85 | 0.1529 | 83 | 0.1325 | 82 | 0.1463 | 82 | 0.0976 | 332 | 0.1325 |
| 16_855925 | NA | NA | NA | NA | NA | NA | 1 | 1 | 1 | 1 | 1 | 1 | NA | NA | NA | NA | NA | NA | 1 | 1 |
| 16_855925 | NA | NA | 1 | 1 | NA | NA | 1 | 1 | 2 | 1 | 1 | 1 | NA | NA | NA | NA | NA | NA | 1 | 1 |
| 16_871645 | NA | NA | 2 | 0.5 | 2 | 1 | 1 | 0 | 5 | 0.6 | 2 | 0 | 1 | 1 | 3 | 1 | 1 | 1 | 7 | 0.7143 |
| 16_871651 | NA | NA | 2 | 0 | 3 | 0.6667 | 1 | 0 | 6 | 0.3333 | 2 | 0.5 | 1 | 0 | 3 | 0 | 1 | 0 | 7 | 0.1429 |
| 16_871752 | 1 | 0 | 5 | 0.2 | 3 | 0.6667 | 4 | 0.25 | 13 | 0.3077 | 1 | 0 | 6 | 0.5 | 5 | 0 | 2 | 0 | 14 | 0.2143 |
| 16_871752 | 1 | 1 | 5 | 0.4 | 5 | 0.4 | 5 | 0.2 | 16 | 0.375 | 1 | 1 | 7 | 0.2857 | 4 | 0 | 1 | 0 | 13 | 0.2308 |
| 16_871754 | 1 | 0 | 5 | 0 | 5 | 0.2 | 5 | 0.2 | 16 | 0.125 | 1 | 0 | 7 | 0.1429 | 4 | 0 | 2 | 0 | 14 | 0.0714 |
| 16_871755 | 1 | 1 | 4 | 0.5 | 5 | 0.2 | 4 | 0.25 | 14 | 0.3571 | 1 | 0 | 5 | 0.2 | 3 | 0 | 2 | 0 | 11 | 0.0909 |
| 16_871756 | NA | NA | 5 | 0.6 | 3 | 0.6667 | 5 | 1 | 13 | 0.7692 | 1 | 0 | 5 | 0.8 | 4 | 1 | 3 | 1 | 13 | 0.8462 |
| 16_871756 | 1 | 0 | 5 | 0.2 | 4 | 0 | 5 | 0 | 15 | 0.0667 | 1 | 0 | 6 | 0.1667 | 4 | 0 | 3 | 0 | 14 | 0.0714 |
| 16_871758 | 1 | 0 | 5 | 0 | 4 | 0.25 | 5 | 0 | 15 | 0.0667 | 1 | 0 | 6 | 0.1667 | 3 | 0 | 3 | 0 | 13 | 0.0769 |
| 16_871776 | NA | NA | 5 | 0 | 4 | 0 | 3 | 0.3333 | 12 | 0.0833 | 1 | 0 | 5 | 0 | 3 | 0 | 2 | 0.5 | 11 | 0.0909 |
| 16_872125 | NA | NA | 3 | 0.3333 | 3 | 0.3333 | 2 | 0.5 | 8 | 0.375 | 1 | 0 | 3 | 0.6667 | 1 | 0 | 4 | 0.25 | 9 | 0.3333 |
| 16_872127 | NA | NA | 3 | 0.6667 | 2 | 1 | 2 | 1 | 7 | 0.8571 | 1 | 1 | 2 | 0 | NA | NA | 3 | 0.6667 | 6 | 0.5 |
| 16_872128 | NA | NA | 2 | 0.5 | 3 | 0.3333 | 2 | 0.5 | 7 | 0.4286 | NA | NA | 2 | 0.5 | 1 | 1 | 3 | 0.3333 | 6 | 0.5 |
| 16_872129 | NA | NA | 2 | 0.5 | 3 | 0.6667 | 2 | 0 | 7 | 0.4286 | NA | NA | 2 | 0 | NA | NA | 2 | 0.5 | 4 | 0.25 |
| 16_872130 | NA | NA | 3 | 0.3333 | 2 | 0 | 2 | 0.5 | 7 | 0.2857 | NA | NA | 1 | 1 | 1 | 0 | 3 | 0 | 5 | 0.2 |
| 16_872131 | NA | NA | 3 | 0 | 3 | 0 | 1 | 1 | 7 | 0.1429 | NA | NA | 2 | 0.5 | NA | NA | 3 | 0 | 5 | 0.2 |
| 16_872132 | NA | NA | 3 | 0.3333 | 1 | 0 | 2 | 0.5 | 6 | 0.3333 | NA | NA | 2 | 0 | 1 | 1 | 3 | 0.3333 | 6 | 0.3333 |
| 16_872312 | NA | NA | 1 | 0 | 2 | 1 | 2 | 0 | 5 | 0.4 | 1 | 0 | 3 | 0.3333 | 2 | 0 | NA | NA | 6 | 0.1667 |
| 16_872312 | NA | NA | 1 | 0 | 2 | 1 | 2 | 0.5 | 5 | 0.6 | NA | NA | 5 | 0.8 | 3 | 0 | NA | NA | 8 | 0.5 |
| 16_872313 | NA | NA | 1 | 1 | 2 | 1 | 2 | 0 | 5 | 0.6 | 1 | 1 | 5 | 1 | 3 | 1 | NA | NA | 9 | 1 |
| 16_872337 | NA | NA | 1 | 0 | 2 | 0 | 1 | 1 | 4 | 0.25 | 1 | 0 | 5 | 0.2 | 2 | 0 | NA | NA | 8 | 0.125 |
| 16_872762 | NA | NA | 2 | 1 | 4 | 0.75 | 3 | 0.3333 | 9 | 0.6667 | NA | NA | 5 | 0.6 | 4 | 0.75 | 2 | 0.5 | 11 | 0.6364 |
| 16_872763 | NA | NA | 2 | 0 | 5 | 0 | 3 | 0.3333 | 10 | 0.1 | NA | NA | 5 | 0.2 | 4 | 0 | 2 | 0 | 11 | 0.0909 |
| 16_872763 | NA | NA | 2 | 0.5 | 3 | 0.6667 | 2 | 0.5 | 7 | 0.5714 | NA | NA | 4 | 0.75 | 2 | 1 | 2 | 1 | 8 | 0.875 |
| 16_872763 | NA | NA | 2 | 1 | 3 | 0.3333 | 2 | 0.5 | 7 | 0.5714 | NA | NA | 4 | 0.5 | 4 | 0.75 | 1 | 0 | 9 | 0.5556 |
| 16_872764 | NA | NA | 1 | 1 | 5 | 0.4 | 1 | 0 | 7 | 0.4286 | NA | NA | 5 | 0.2 | 4 | 0.25 | 2 | 0.5 | 11 | 0.2727 |
| 16_872765 | NA | NA | 2 | 1 | 5 | 0 | 3 | 0 | 10 | 0.2 | NA | NA | 5 | 0.4 | 4 | 0.25 | 3 | 0 | 12 | 0.25 |
| 16_872765 | NA | NA | 2 | 1 | 5 | 0.6 | 3 | 0.6667 | 10 | 0.7 | NA | NA | 5 | 0.8 | 4 | 1 | 4 | 0.75 | 13 | 0.8462 |
| 16_872765 | NA | NA | 1 | 1 | 3 | 0 | 3 | 0 | 7 | 0.1429 | NA | NA | 5 | 0 | 4 | 0.25 | 4 | 0 | 13 | 0.0769 |
| 16_872766 | NA | NA | 2 | 0 | 4 | 0.25 | 3 | 0.3333 | 9 | 0.2222 | NA | NA | 3 | 0 | 3 | 0 | 3 | 0.6667 | 9 | 0.2222 |
| 16_872800 | 1 | 0 | 1 | 0 | 5 | 0 | 3 | 0.3333 | 10 | 0.1 | NA | NA | 4 | 0 | 3 | 0.3333 | 3 | 0 | 10 | 0.1 |
| 16_875264 | NA | NA | 1 | 1 | NA | NA | NA | NA | 1 | 1 | 1 | 1 | NA | NA | NA | NA | NA | NA | 1 | 1 |
| 16_875266 | NA | NA | 1 | 1 | NA | NA | NA | NA | 1 | 1 | 1 | 1 | NA | NA | NA | NA | NA | NA | 1 | 1 |
| 16_875267 | NA | NA | 1 | 1 | NA | NA | NA | NA | 1 | 1 | 1 | 1 | NA | NA | NA | NA | NA | NA | 1 | 1 |
| 16_875271 | NA | NA | 1 | 1 | NA | NA | NA | NA | 1 | 1 | 1 | 1 | NA | NA | NA | NA | NA | NA | 1 | 1 |
| 16_875272 | NA | NA | 1 | 1 | NA | NA | NA | NA | 1 | 1 | 1 | 1 | NA | NA | NA | NA | NA | NA | 1 | 1 |

| | | | | | | | | | | | | | | | | | | | | |
|---|---|---|---|---|---|---|---|---|---|---|---|---|---|---|---|---|---|---|---|---|
| 16_875273 | NA | NA | 1 | 1 | NA | NA | NA | NA | 1 | 1 | 1 | 1 | NA | NA | NA | NA | NA | NA | 1 | 1 |
| 16_875274 | NA | NA | 1 | 1 | NA | NA | NA | NA | 1 | 1 | 1 | 1 | NA | NA | NA | NA | NA | NA | 1 | 1 |
| 16_875274 | NA | NA | 1 | 1 | NA | NA | NA | NA | 1 | 1 | 1 | 1 | NA | NA | NA | NA | NA | NA | 1 | 1 |
| 16_875289 | 3 | 0.3333 | 1 | 0 | 2 | 0.5 | 1 | 0 | 7 | 0.2857 | 3 | 0.3333 | 2 | 0 | NA | NA | 1 | 0 | 6 | 0.1667 |
| 16_875293 | 2 | 0.5 | 1 | 0 | 2 | 0 | 2 | 0.5 | 7 | 0.2857 | 5 | 0.2 | 3 | 0 | NA | NA | 1 | 0 | 9 | 0.1111 |
| 16_875294 | 3 | 0.3333 | 2 | 0 | 3 | 0 | 3 | 1 | 11 | 0.3636 | 5 | 0.6 | 4 | 0.5 | NA | NA | 1 | 0 | 10 | 0.5 |
| 16_875298 | 2 | 0 | 3 | 0 | 4 | 0.25 | 3 | 0.3333 | 12 | 0.1667 | 5 | 0 | 2 | 0 | NA | NA | 2 | 0.5 | 9 | 0.1111 |
| 16_875299 | 2 | 0.5 | 3 | 0.6667 | 4 | 0.75 | 3 | 1 | 12 | 0.75 | 5 | 0.8 | 4 | 1 | NA | NA | 2 | 1 | 11 | 0.9091 |
| 16_876929 | 2 | 0.5 | 1 | 1 | 1 | 0 | 2 | 0 | 6 | 0.3333 | NA | NA | 1 | 0 | 4 | 0.25 | NA | NA | 5 | 0.2 |
| 16_876930 | 1 | 1 | NA | NA | 1 | 1 | 2 | 0 | 4 | 0.5 | NA | NA | 1 | 0 | 4 | 0.25 | NA | NA | 5 | 0.2 |
| 16_876930 | 2 | 1 | NA | NA | 1 | 1 | 2 | 1 | 5 | 1 | NA | NA | NA | NA | 4 | 1 | NA | NA | 4 | 1 |
| 16_876930 | 1 | 1 | 1 | 1 | NA | NA | 3 | 1 | 5 | 1 | NA | NA | 1 | 1 | 4 | 1 | NA | NA | 5 | 1 |
| 16_876931 | 2 | 0 | 1 | 1 | 1 | 1 | 2 | 0.5 | 6 | 0.5 | NA | NA | 1 | 1 | 4 | 0.5 | NA | NA | 5 | 0.6 |
| 16_877111 | 1 | 0 | NA | NA | 1 | 1 | 1 | 0 | 3 | 0.3333 | NA | NA | NA | NA | 3 | 0.3333 | NA | NA | 3 | 0.3333 |
| 16_877447 | 1 | 1 | NA | NA | 2 | 0.5 | NA | NA | 3 | 0.6667 | NA | NA | NA | NA | 2 | 0.5 | NA | NA | 2 | 0.5 |
| 16_895851 | 6 | 0.5 | 13 | 0.1538 | 10 | 0.2 | 13 | 0.2308 | 42 | 0.2381 | 11 | 0.1818 | 19 | 0.2105 | 11 | 0.0909 | 15 | 0.2667 | 56 | 0.1964 |
| 16_895851 | 7 | 0.1429 | 13 | 0.0769 | 8 | 0 | 14 | 0.0714 | 42 | 0.0714 | 11 | 0.1818 | 21 | 0.0952 | 14 | 0.0714 | 16 | 0.125 | 62 | 0.1129 |
| 16_895855 | 6 | 0 | 12 | 0.25 | 8 | 0 | 14 | 0.1429 | 40 | 0.125 | 9 | 0.1111 | 15 | 0.0667 | 12 | 0 | 16 | 0.1875 | 52 | 0.0962 |
| 16_895856 | 5 | 0.6 | 10 | 0.5 | 8 | 0.75 | 11 | 0.4545 | 34 | 0.5588 | 7 | 0.7143 | 15 | 0.6667 | 10 | 0.4 | 10 | 0.5 | 42 | 0.5714 |
| 16_895857 | 6 | 0 | 12 | 0.1667 | 8 | 0.125 | 14 | 0 | 40 | 0.075 | 10 | 0.1 | 17 | 0.1765 | 12 | 0.3333 | 14 | 0.1429 | 53 | 0.1887 |
| 16_895858 | 5 | 0.2 | 12 | 0.0833 | 7 | 0.1429 | 12 | 0.1667 | 36 | 0.1389 | 10 | 0.1 | 18 | 0.1111 | 11 | 0.2727 | 15 | 0.0667 | 54 | 0.1296 |
| 16_895858 | 4 | 0.25 | 13 | 0 | 5 | 0.4 | 13 | 0.3846 | 35 | 0.2286 | 10 | 0.3 | 17 | 0.4706 | 11 | 0.0909 | 16 | 0.5 | 54 | 0.3704 |
| 16_895860 | 3 | 0 | 13 | 0.2308 | 6 | 0.6667 | 12 | 0.25 | 34 | 0.2941 | 10 | 0.3 | 15 | 0.4 | 11 | 0.7273 | 15 | 0.3333 | 51 | 0.4314 |
| 16_895860 | 3 | 0.3333 | 11 | 0.1818 | 7 | 0.2857 | 10 | 0.1 | 31 | 0.1935 | 9 | 0.3333 | 16 | 0.1875 | 10 | 0.2 | 15 | 0.2 | 50 | 0.22 |
| 16_895860 | 4 | 0.75 | 11 | 0.4545 | 4 | 0.25 | 11 | 0.7273 | 30 | 0.5667 | 8 | 0.5 | 13 | 0.4615 | 11 | 0.5455 | 11 | 0.7273 | 43 | 0.5581 |
| 16_895860 | 4 | 0.75 | 13 | 0.6923 | 7 | 0.5714 | 13 | 0.6923 | 37 | 0.6757 | 9 | 0.7778 | 16 | 0.6875 | 10 | 0.7 | 12 | 0.5833 | 47 | 0.6809 |
| 16_895861 | 4 | 0.25 | 13 | 0.0769 | 6 | 0.3333 | 13 | 0.1538 | 36 | 0.1667 | 9 | 0 | 18 | 0.1667 | 11 | 0.1818 | 14 | 0.0714 | 52 | 0.1154 |
| 16_895862 | 4 | 0.5 | 11 | 0.5455 | 4 | 0.5 | 12 | 0.5833 | 31 | 0.5484 | 8 | 0.625 | 17 | 0.6471 | 8 | 0.625 | 11 | 0.6364 | 44 | 0.6364 |
| 16_895862 | 3 | 1 | 5 | 0.8 | 6 | 1 | 10 | 0.8 | 24 | 0.875 | 7 | 1 | 14 | 0.7857 | 8 | 0.875 | 10 | 0.7 | 39 | 0.8205 |
| 16_895862 | 4 | 0 | 11 | 0.1818 | 6 | 0 | 11 | 0 | 32 | 0.0625 | 10 | 0 | 16 | 0.125 | 11 | 0 | 13 | 0.1538 | 50 | 0.08 |
| 16_895862 | 4 | 0.25 | 12 | 0 | 6 | 0 | 11 | 0.1818 | 33 | 0.0909 | 9 | 0 | 14 | 0 | 9 | 0.2222 | 13 | 0.0769 | 45 | 0.0667 |
| 16_895863 | 3 | 1 | 9 | 0.3333 | 4 | 0.5 | 5 | 0.4 | 21 | 0.4762 | 7 | 0.4286 | 14 | 0.5 | 7 | 0.4286 | 10 | 0.6 | 38 | 0.5 |
| 16_895863 | 4 | 0.25 | 10 | 0.1 | 5 | 0.4 | 10 | 0.2 | 29 | 0.2069 | 6 | 0.1667 | 15 | 0.2 | 8 | 0.5 | 12 | 0.3333 | 41 | 0.2927 |
| 16_896816 | NA | NA | 2 | 0 | NA | NA | 4 | 0.25 | 6 | 0.1667 | 3 | 0.3333 | 8 | 0 | 2 | 0 | 1 | 0 | 14 | 0.0714 |
| 16_896818 | NA | NA | 2 | 0.5 | NA | NA | 4 | 0 | 6 | 0.1667 | 4 | 0.25 | 7 | 0.1429 | 2 | 0 | 1 | 1 | 14 | 0.2143 |
| 16_896818 | NA | NA | 2 | 0 | NA | NA | 4 | 0.5 | 6 | 0.3333 | 2 | 0 | 7 | 0.1429 | 2 | 0 | NA | NA | 11 | 0.0909 |
| 16_896946 | NA | NA | 2 | 0.5 | NA | NA | 2 | 0 | 4 | 0.25 | 2 | 0.5 | 5 | 0.6 | 2 | 0.5 | 1 | 1 | 10 | 0.6 |
| 16_896947 | NA | NA | 1 | 1 | NA | NA | 2 | 0 | 3 | 0.3333 | 2 | 0.5 | 5 | 0.2 | 2 | 0 | 1 | 0 | 10 | 0.2 |
| 16_896947 | NA | NA | 2 | 1 | NA | NA | 1 | 0 | 3 | 0.6667 | 1 | 1 | 3 | 0.6667 | 2 | 0.5 | 1 | 1 | 7 | 0.7143 |
| 16_896947 | 1 | 1 | NA | NA | NA | NA | 1 | 1 | 2 | 1 | 1 | 1 | NA | NA | 1 | 1 | 2 | 0.5 | 4 | 0.75 |
| 16_896948 | NA | NA | 2 | 1 | NA | NA | 2 | 1 | 4 | 1 | 2 | 0.5 | 4 | 0.25 | 2 | 0.5 | 1 | 0 | 9 | 0.3333 |
| 16_896950 | 1 | 1 | NA | NA | NA | NA | 1 | 1 | 2 | 1 | 1 | 1 | NA | NA | 1 | 1 | 2 | 1 | 4 | 1 |
| 16_896952 | 2 | 1 | NA | NA | NA | NA | 1 | 1 | 3 | 1 | 1 | 1 | NA | NA | 1 | 1 | 2 | 1 | 4 | 1 |
| 16_896953 | NA | NA | 1 | 0 | NA | NA | 2 | 0.5 | 3 | 0.3333 | 2 | 1 | 3 | 0.3333 | 1 | 1 | 1 | 0 | 7 | 0.5714 |
| 16_900166 | NA | NA | 1 | 1 | NA | NA | NA | NA | 1 | 1 | NA | NA | NA | NA | 1 | 1 | NA | NA | 1 | 1 |
| 16_900232 | 5 | 0.6 | NA | NA | 1 | 0 | 2 | 0.5 | 8 | 0.5 | 1 | 1 | 1 | 0 | NA | NA | 1 | 0 | 3 | 0.3333 |
| 16_900258 | NA | NA | 4 | 0 | 2 | 0.5 | 3 | 0 | 9 | 0.1111 | 3 | 0.3333 | 3 | 0 | 4 | 0 | 4 | 0 | 14 | 0.0714 |
| 16_900261 | 1 | 0 | 5 | 0.8 | 1 | 0 | 3 | 0.6667 | 10 | 0.6 | 4 | 0.5 | 2 | 1 | 5 | 0.2 | 4 | 0.25 | 15 | 0.4 |
| 16_900261 | 1 | 0 | 4 | 0 | 2 | 0.5 | 1 | 0 | 8 | 0.125 | 2 | 0 | 2 | 0 | 4 | 0.5 | 4 | 0.25 | 12 | 0.25 |
| 16_900261 | 1 | 1 | 5 | 0.8 | 2 | 0.5 | 3 | 1 | 11 | 0.8182 | 4 | 0.75 | 3 | 1 | 4 | 1 | 4 | 1 | 15 | 0.9333 |
| 16_900262 | 1 | 0 | 3 | 0.6667 | 1 | 0 | 3 | 0 | 8 | 0.25 | 3 | 0 | 3 | 0 | 4 | 0.25 | 2 | 0.5 | 12 | 0.1667 |
| 16_900263 | NA | NA | 3 | 0.3333 | 1 | 1 | 3 | 0.3333 | 7 | 0.4286 | 3 | 0.6667 | 3 | 0 | 2 | 0.5 | 3 | 0.3333 | 11 | 0.3636 |
| 16_900263 | 1 | 1 | 4 | 0.5 | 1 | 1 | 3 | 0.3333 | 9 | 0.5556 | 4 | 0.75 | 2 | 0.5 | 5 | 0.8 | 3 | 0.6667 | 14 | 0.7143 |
| 16_900264 | NA | NA | 4 | 0.25 | 1 | 1 | 2 | 0.5 | 7 | 0.4286 | 2 | 0.5 | 1 | 1 | 3 | 0 | 2 | 0.5 | 8 | 0.375 |
| 16_900264 | 1 | 0 | 5 | 0 | 2 | 1 | 3 | 0.6667 | 11 | 0.3636 | 3 | 0.6667 | 3 | 0.3333 | 4 | 0.75 | 4 | 0.75 | 14 | 0.6429 |
| 16_900265 | 1 | 0 | 5 | 0 | 2 | 0.5 | 2 | 0.5 | 10 | 0.2 | 2 | 0.5 | 3 | 0.6667 | 2 | 0 | 1 | 1 | 8 | 0.5 |
| 16_900266 | 1 | 0 | 4 | 0.25 | 2 | 0 | 3 | 0.3333 | 10 | 0.2 | 3 | 0 | 3 | 0 | 5 | 0.4 | 4 | 0 | 15 | 0.1333 |
| 16_900269 | NA | NA | 2 | 0 | 1 | 1 | 1 | 1 | 4 | 0.5 | NA | NA | 3 | 0.6667 | 2 | 0.5 | 2 | 0.5 | 7 | 0.5714 |
| 16_900359 | 1 | 0 | 1 | 1 | 2 | 1 | 1 | 0 | 5 | 0.6 | NA | NA | 1 | 1 | NA | NA | NA | NA | 1 | 1 |
| 16_900366 | 1 | 0 | 1 | 0 | 2 | 0 | 1 | 1 | 5 | 0.2 | NA | NA | 2 | 0.5 | 1 | 0 | NA | NA | 3 | 0.3333 |
| 16_900379 | 1 | 0 | 1 | 0 | 1 | 1 | NA | NA | 3 | 0.3333 | NA | NA | 1 | 1 | NA | NA | NA | NA | 1 | 1 |
| 16_902931 | NA | NA | 3 | 0.3333 | 1 | 0 | NA | NA | 4 | 0.25 | 2 | 0 | 2 | 0.5 | NA | NA | NA | NA | 4 | 0.25 |
| 16_902932 | NA | NA | 1 | 1 | 1 | 1 | 1 | 1 | 3 | 1 | 2 | 1 | 2 | 1 | NA | NA | NA | NA | 4 | 1 |
| 16_902932 | NA | NA | 2 | 1 | 1 | 1 | NA | NA | 3 | 1 | 2 | 1 | 2 | 1 | NA | NA | NA | NA | 4 | 1 |
| 16_902934 | NA | NA | 3 | 1 | 1 | 1 | 1 | 1 | 5 | 1 | 1 | 1 | 1 | 1 | NA | NA | NA | NA | 2 | 1 |

| | | | | | | | | | | | | | | | | | | | | |
|---|---|---|---|---|---|---|---|---|---|---|---|---|---|---|---|---|---|---|---|---|
| 16_902936 | NA | NA | 2 | 0.5 | NA | NA | 1 | 1 | 3 | 0.6667 | 1 | 1 | 1 | 1 | NA | NA | NA | NA | 2 | 1 |
| 16_902937 | NA | NA | 2 | 1 | 1 | 1 | 1 | 1 | 4 | 1 | 2 | 1 | 1 | 1 | NA | NA | NA | NA | 3 | 1 |
| 16_902938 | NA | NA | 3 | 0.3333 | 1 | 0 | 1 | 1 | 5 | 0.4 | 2 | 0 | 2 | 1 | NA | NA | NA | NA | 4 | 0.5 |
| 16_902938 | NA | NA | 2 | 0.5 | NA | NA | 1 | 0 | 3 | 0.3333 | 2 | 0 | 2 | 0.5 | NA | NA | NA | NA | 4 | 0.25 |
| 16_902940 | NA | NA | 2 | 0.5 | 1 | 0 | 1 | 1 | 4 | 0.5 | 2 | 0.5 | 1 | 1 | NA | NA | NA | NA | 3 | 0.6667 |
| 16_902943 | NA | NA | 3 | 0.6667 | 1 | 0 | NA | NA | 4 | 0.5 | 2 | 1 | 2 | 0 | NA | NA | NA | NA | 4 | 0.5 |
| 16_902947 | NA | NA | 4 | 0.25 | NA | NA | 2 | 0 | 6 | 0.1667 | 2 | 0.5 | 2 | 0 | NA | NA | NA | NA | 4 | 0.25 |
| 16_902950 | NA | NA | 2 | 0.5 | 1 | 0 | 2 | 0 | 5 | 0.2 | 2 | 0.5 | 2 | 0 | NA | NA | NA | NA | 4 | 0.25 |
| 16_902951 | NA | NA | 3 | 0.3333 | 1 | 0 | 2 | 0 | 6 | 0.1667 | 2 | 0.5 | 2 | 0.5 | NA | NA | NA | NA | 4 | 0.5 |
| 16_902951 | NA | NA | 5 | 0.2 | 1 | 0 | 2 | 0 | 8 | 0.125 | 2 | 0.5 | 2 | 0 | NA | NA | NA | NA | 4 | 0.25 |
| 16_902951 | NA | NA | 3 | 0.3333 | 1 | 0 | 1 | 0 | 5 | 0.2 | 2 | 0.5 | 1 | 0 | NA | NA | NA | NA | 3 | 0.3333 |
| 16_902954 | NA | NA | 4 | 0.5 | 1 | 0 | 2 | 0 | 7 | 0.2857 | 2 | 0 | 2 | 0.5 | NA | NA | NA | NA | 4 | 0.25 |
| 16_902955 | NA | NA | 4 | 1 | 1 | 1 | 2 | 0.5 | 7 | 0.8571 | 3 | 1 | 2 | 1 | NA | NA | NA | NA | 5 | 1 |
| 16_902955 | NA | NA | 4 | 0.5 | 1 | 0 | 2 | 0 | 7 | 0.2857 | 2 | 0.5 | 1 | 1 | NA | NA | NA | NA | 3 | 0.6667 |
| 16_902955 | NA | NA | 4 | 0.25 | 1 | 0 | 2 | 0.5 | 7 | 0.2857 | 3 | 1 | 2 | 0 | NA | NA | NA | NA | 5 | 0.6 |
| 16_902956 | NA | NA | 5 | 1 | 1 | 1 | 2 | 0.5 | 8 | 0.875 | 3 | 1 | 2 | 1 | NA | NA | NA | NA | 5 | 1 |
| 16_902956 | NA | NA | 1 | 1 | 1 | 1 | 1 | 1 | 3 | 1 | 3 | 1 | 1 | 1 | NA | NA | NA | NA | 4 | 1 |
| 16_902959 | NA | NA | 7 | 0.8571 | 1 | 1 | 1 | 1 | 9 | 0.8889 | 3 | 1 | 2 | 1 | NA | NA | NA | NA | 5 | 1 |
| 16_902960 | NA | NA | 6 | 0.5 | 1 | 1 | 2 | 0.5 | 9 | 0.5556 | 3 | 0 | 2 | 0.5 | NA | NA | NA | NA | 5 | 0.2 |
| 16_902963 | NA | NA | 4 | 0.5 | 1 | 0 | 2 | 0 | 7 | 0.2857 | 3 | 0.3333 | 4 | 0.25 | NA | NA | NA | NA | 7 | 0.2857 |
| 16_905487 | NA | NA | 2 | 0.5 | 2 | 0 | 1 | 0 | 5 | 0.2 | 1 | 1 | 1 | 0 | 2 | 0 | 1 | 0 | 5 | 0.2 |
| 16_905487 | NA | NA | 1 | 1 | 2 | 1 | NA | NA | 3 | 1 | 1 | 1 | NA | NA | 1 | 1 | 1 | 1 | 3 | 1 |
| 16_905488 | NA | NA | 2 | 0.5 | 2 | 0.5 | 1 | 0 | 5 | 0.4 | NA | NA | 2 | 0 | 1 | 1 | 1 | 0 | 4 | 0.25 |
| 16_905489 | NA | NA | 2 | 1 | 1 | 1 | 1 | 0 | 4 | 0.75 | NA | NA | 2 | 0.5 | NA | NA | 1 | 0 | 3 | 0.3333 |
| 16_905504 | NA | NA | 2 | 1 | 2 | 1 | 1 | 0 | 5 | 0.8 | NA | NA | 1 | 1 | 1 | 0 | 1 | 1 | 3 | 0.6667 |
| 16_905506 | NA | NA | 2 | 0.5 | 2 | 0.5 | 1 | 0 | 5 | 0.4 | NA | NA | 1 | 0 | NA | NA | 1 | 1 | 2 | 0.5 |
| 16_907010 | 2 | 1 | NA | NA | NA | NA | NA | NA | 2 | 1 | NA | NA | NA | NA | 1 | 1 | NA | NA | 1 | 1 |
| 16_907012 | 2 | 1 | NA | NA | NA | NA | NA | NA | 2 | 1 | NA | NA | NA | NA | 1 | 1 | NA | NA | 1 | 1 |
| 16_907013 | 1 | 1 | NA | NA | NA | NA | NA | NA | 1 | 1 | NA | NA | NA | NA | 1 | 1 | NA | NA | 1 | 1 |
| 16_907229 | 11 | 0.0909 | 5 | 0.2 | 4 | 0.25 | 6 | 0 | 26 | 0.1154 | 3 | 0 | 11 | 0.3636 | 6 | 0 | 7 | 0.2857 | 27 | 0.2222 |
| 16_907331 | 34 | 0.1471 | 44 | 0.0682 | 25 | 0.24 | 44 | 0.1136 | 147 | 0.1293 | 25 | 0.16 | 31 | 0.0968 | 25 | 0.16 | 23 | 0 | 104 | 0.1058 |
| 16_907332 | 40 | 0.25 | 45 | 0.2 | 26 | 0.2308 | 41 | 0.1707 | 152 | 0.2105 | 25 | 0.32 | 31 | 0.1613 | 28 | 0.3214 | 21 | 0.1905 | 105 | 0.2476 |
| 16_907343 | 17 | 0.1176 | 29 | 0.2759 | 15 | 0.2667 | 26 | 0.1154 | 87 | 0.1954 | 11 | 0.0909 | 20 | 0.1 | 15 | 0.1333 | 9 | 0 | 55 | 0.0909 |
| 16_907344 | 24 | 0.125 | 29 | 0.1379 | 15 | 0.0667 | 30 | 0.1333 | 98 | 0.1224 | 15 | 0.1333 | 22 | 0.0455 | 16 | 0 | 13 | 0.1538 | 66 | 0.0758 |
| 16_907347 | 22 | 0.2273 | 31 | 0.129 | 18 | 0.0556 | 27 | 0.1111 | 98 | 0.1327 | 11 | 0 | 24 | 0.0417 | 16 | 0.125 | 15 | 0.0667 | 66 | 0.0606 |
| 16_907351 | 16 | 1 | 26 | 1 | 16 | 1 | 26 | 0.9615 | 84 | 0.9881 | 8 | 1 | 19 | 0.9474 | 12 | 0.9167 | 11 | 1 | 50 | 0.96 |
| 16_907351 | 17 | 0.5294 | 29 | 0.3793 | 17 | 0.5294 | 26 | 0.5385 | 89 | 0.4831 | 11 | 0.8182 | 18 | 0.3889 | 11 | 0.8182 | 13 | 0.4615 | 53 | 0.5849 |
| 16_907352 | 14 | 0.9286 | 22 | 0.7273 | 13 | 0.6923 | 26 | 0.7692 | 75 | 0.7733 | 9 | 0.6667 | 16 | 1 | 5 | 0.8 | 12 | 0.6667 | 42 | 0.8095 |
| 16_907353 | 18 | 0.8889 | 25 | 1 | 12 | 1 | 27 | 0.963 | 82 | 0.9634 | 10 | 1 | 17 | 0.9412 | 8 | 1 | 13 | 0.9231 | 48 | 0.9583 |
| 16_907354 | 20 | 0.95 | 24 | 0.9583 | 13 | 1 | 25 | 1 | 82 | 0.9756 | 6 | 1 | 18 | 1 | 9 | 1 | 12 | 0.9167 | 45 | 0.9778 |
| 16_907356 | 13 | 1 | 16 | 0.9375 | 14 | 0.9286 | 16 | 1 | 59 | 0.9661 | 7 | 1 | 14 | 1 | 4 | 1 | 7 | 1 | 32 | 1 |
| 16_907356 | 11 | 0.9091 | 17 | 0.6471 | 10 | 0.9 | 19 | 0.5263 | 57 | 0.7018 | 8 | 0.75 | 10 | 0.4 | 5 | 0.6 | 9 | 0.6667 | 32 | 0.5938 |
| 16_907356 | 14 | 0.3571 | 24 | 0.3333 | 9 | 0.4444 | 23 | 0.2609 | 70 | 0.3286 | 7 | 0.2857 | 16 | 0.25 | 7 | 0.1429 | 12 | 0.3333 | 42 | 0.2619 |
| 16_907357 | 17 | 0.1176 | 25 | 0.2 | 12 | 0.3333 | 24 | 0.0833 | 78 | 0.1667 | 9 | 0.2222 | 17 | 0.0588 | 8 | 0 | 10 | 0.1 | 44 | 0.0909 |
| 16_908109 | 2 | 0 | 2 | 0.5 | 5 | 0 | 3 | 0 | 12 | 0.0833 | 1 | 0 | 3 | 0.3333 | 3 | 0 | 2 | 0 | 9 | 0.1111 |
| 16_908113 | 2 | 1 | 1 | 0 | 5 | 0.6 | 2 | 1 | 10 | 0.7 | 1 | 1 | 2 | 0.5 | 3 | 0.6667 | 2 | 0 | 8 | 0.5 |
| 16_908113 | 2 | 0 | 1 | 0 | 5 | 0.2 | 3 | 0 | 11 | 0.0909 | 2 | 0.5 | 2 | 0 | 4 | 0.25 | 1 | 0 | 9 | 0.2222 |
| 16_908113 | 1 | 0 | NA | NA | 5 | 0.2 | 3 | 0 | 9 | 0.1111 | 2 | 0.5 | 1 | 0 | 4 | 0 | 2 | 0 | 9 | 0.1111 |
| 16_908138 | 5 | 0.2 | 9 | 0 | 8 | 0.125 | 7 | 0 | 29 | 0.069 | 4 | 0.25 | 3 | 0.3333 | 9 | 0.1111 | 4 | 0 | 20 | 0.15 |
| 16_908311 | NA | NA | 4 | 0 | 2 | 0 | 6 | 0.1667 | 12 | 0.0833 | 1 | 0 | 1 | 0 | 1 | 1 | 3 | 0 | 6 | 0.1667 |
| 16_908314 | NA | NA | 4 | 1 | NA | NA | 3 | 1 | 7 | 1 | 1 | 1 | NA | NA | NA | NA | 2 | 1 | 3 | 1 |
| 16_908317 | NA | NA | 4 | 0.5 | 2 | 0.5 | 5 | 0.2 | 11 | 0.3636 | 1 | 0 | NA | NA | 1 | 0 | 4 | 0.5 | 6 | 0.3333 |
| 16_910232 | 1 | 1 | 2 | 0.5 | NA | NA | 2 | 0.5 | 5 | 0.6 | 3 | 0.6667 | 2 | 1 | 2 | 1 | 2 | 1 | 9 | 0.8889 |
| 16_910233 | 1 | 1 | 2 | 1 | NA | NA | 3 | 0.6667 | 6 | 0.8333 | 2 | 1 | 2 | 0.5 | 2 | 1 | 2 | 1 | 8 | 0.875 |
| 16_910234 | 1 | 1 | 3 | 0 | NA | NA | 3 | 1 | 7 | 0.5714 | 3 | 1 | 2 | 0 | 1 | 1 | 1 | 1 | 7 | 0.7143 |
| 16_910234 | 1 | 0 | 2 | 0.5 | NA | NA | 1 | 0 | 4 | 0.25 | 1 | 0 | 2 | 0.5 | 1 | 0 | 1 | 0 | 5 | 0.2 |
| 16_910243 | 2 | 1 | 4 | 0.75 | 1 | 0 | 3 | 0.6667 | 10 | 0.7 | 2 | 0.5 | 2 | 1 | 2 | 0.5 | 1 | 1 | 7 | 0.7143 |
| 16_910243 | 2 | 1 | 4 | 1 | 1 | 1 | 3 | 1 | 10 | 1 | 2 | 1 | 2 | 1 | 1 | 1 | 2 | 1 | 7 | 1 |
| 16_910244 | 2 | 0 | 5 | 0 | 1 | 0 | 3 | 0.3333 | 11 | 0.0909 | 3 | 0.3333 | 1 | 0 | 2 | 0 | 1 | 1 | 7 | 0.2857 |
| 16_910244 | 2 | 1 | 5 | 0.4 | 1 | 0 | 4 | 0 | 12 | 0.3333 | 3 | 0.3333 | 2 | 0 | 3 | 0 | 1 | 0 | 9 | 0.1111 |
| 16_910244 | 2 | 0.5 | 5 | 1 | 1 | 1 | 3 | 1 | 11 | 0.9091 | 2 | 1 | 1 | 1 | 2 | 1 | 2 | 1 | 7 | 1 |
| 16_910244 | 1 | 1 | 3 | 0.6667 | 1 | 0 | 1 | 0 | 6 | 0.5 | 1 | 0 | 2 | 1 | 3 | 0.6667 | 1 | 0 | 7 | 0.5714 |
| 16_910245 | 2 | 0.5 | 4 | 0.5 | 1 | 0 | 3 | 0.3333 | 10 | 0.4 | 2 | 0 | 2 | 0 | 2 | 0.5 | NA | NA | 6 | 0.1667 |
| 16_910245 | 2 | 0.5 | 5 | 0.4 | 1 | 1 | 3 | 0.6667 | 11 | 0.5455 | 3 | 0.6667 | NA | NA | 3 | 0 | 2 | 0.5 | 8 | 0.375 |
| 16_910245 | 2 | 0.5 | 3 | 0.6667 | NA | NA | 3 | 0.6667 | 8 | 0.625 | 1 | 1 | 1 | 0 | 2 | 1 | 1 | 0 | 5 | 0.6 |

| | | | | | | | | | | | | | | | | | | | | |
|---|---|---|---|---|---|---|---|---|---|---|---|---|---|---|---|---|---|---|---|---|
| 16_910246( | 2 | 0 | 3 | 0.6667 | 1 | 0 | 3 | 0 | 9 | 0.2222 | 3 | 0.6667 | 1 | 1 | 2 | 0.5 | 2 | 0.5 | 8 | 0.625 |
| 16_910246! | 1 | 0 | 4 | 0.25 | 1 | 0 | 2 | 0 | 8 | 0.125 | 2 | 0 | 2 | 0.5 | 2 | 0.5 | 1 | 1 | 7 | 0.4286 |
| 16_910246{ | 2 | 0 | 5 | 0.2 | NA | NA | 4 | 0.25 | 11 | 0.1818 | 3 | 0.3333 | 2 | 0.5 | 3 | 0 | 2 | 0.5 | 10 | 0.3 |
| 16_910246! | 1 | 0 | 3 | 0 | 1 | 0 | 4 | 0.25 | 9 | 0.1111 | 3 | 0.3333 | 2 | 0 | 3 | 0 | 2 | 0 | 10 | 0.1 |
| 16_910669{ | 11 | 0.1818 | 14 | 0.1429 | 11 | 0 | 4 | 0.25 | 40 | 0.125 | 11 | 0.1818 | 8 | 0.125 | 3 | 0 | 8 | 0.25 | 30 | 0.1667 |
| 16_910670! | 15 | 0.6667 | 16 | 0.5 | 12 | 0.5 | 7 | 0.7143 | 50 | 0.58 | 11 | 0.4545 | 10 | 0.5 | 3 | 0.6667 | 11 | 0.5455 | 35 | 0.5143 |
| 16_910670! | 9 | 0.5556 | 10 | 0.6 | 10 | 0.4 | 7 | 0.5714 | 36 | 0.5278 | 7 | 0.2857 | 10 | 0.4 | 3 | 0.3333 | 10 | 0.8 | 30 | 0.5 |
| 16_910673! | 15 | 0.6 | 15 | 0.3333 | 15 | 0.4 | 7 | 0.5714 | 52 | 0.4615 | 11 | 0.5455 | 12 | 0.5 | 3 | 0.3333 | 12 | 0.6667 | 38 | 0.5526 |
| 16_910676 | 16 | 0.9375 | 18 | 0.7778 | 16 | 1 | 8 | 0.875 | 58 | 0.8966 | 13 | 1 | 14 | 0.7857 | 5 | 1 | 13 | 0.9231 | 45 | 0.9111 |
| 16_910676! | 13 | 0.3077 | 16 | 0.25 | 11 | 0.3636 | 8 | 0.25 | 48 | 0.2917 | 12 | 0.25 | 13 | 0.0769 | 4 | 0.25 | 12 | 0.1667 | 41 | 0.1707 |
| 16_910676 | 14 | 1 | 17 | 0.7647 | 13 | 0.7692 | 9 | 0.7778 | 53 | 0.8302 | 11 | 0.5455 | 14 | 0.9286 | 5 | 0.8 | 12 | 0.9167 | 42 | 0.8095 |
| 16_910684 | 20 | 0.7 | 20 | 0.95 | 18 | 0.7778 | 10 | 0.8 | 68 | 0.8088 | 15 | 0.8667 | 18 | 0.6667 | 8 | 0.875 | 14 | 0.6429 | 55 | 0.7455 |
| 16_910685! | 15 | 0.4667 | 20 | 0.35 | 14 | 0.4286 | 10 | 0.2 | 59 | 0.3729 | 14 | 0.2857 | 13 | 0.4615 | 7 | 0.1429 | 14 | 0.2143 | 48 | 0.2917 |
| 16_910686 | 18 | 0.2222 | 22 | 0 | 19 | 0.0526 | 12 | 0 | 71 | 0.0704 | 15 | 0.0667 | 19 | 0.1579 | 10 | 0.1 | 15 | 0.1333 | 59 | 0.1186 |
| 16_910687 | 19 | 0.0526 | 22 | 0.1818 | 14 | 0.2857 | 12 | 0.1667 | 67 | 0.1642 | 15 | 0.0667 | 18 | 0.0556 | 8 | 0 | 14 | 0.0714 | 55 | 0.0545 |
| 16_911906 | 2 | 0 | 6 | 0.1667 | 3 | 0.6667 | 3 | 0.6667 | 14 | 0.3571 | 6 | 0.1667 | 2 | 0.5 | 1 | 0 | 3 | 0 | 12 | 0.1667 |
| 16_912006 | NA | NA | 2 | 1 | 2 | 0 | NA | NA | 4 | 0.5 | NA | NA | NA | NA | 1 | 1 | 1 | 0 | 2 | 0.5 |
| 16_912007 | NA | NA | 3 | 1 | 2 | 0.5 | NA | NA | 5 | 0.8 | NA | NA | NA | NA | 1 | 1 | 1 | 1 | 2 | 1 |
| 16_912009 | NA | NA | 2 | 1 | 2 | 0.5 | NA | NA | 4 | 0.75 | NA | NA | NA | NA | 2 | 1 | 1 | 1 | 3 | 1 |
| 16_912011 | NA | NA | 2 | 0.5 | 2 | 1 | NA | NA | 4 | 0.75 | NA | NA | NA | NA | 2 | 0.5 | 1 | 1 | 3 | 0.6667 |
| 16_912012 | NA | NA | 3 | 1 | 2 | 1 | NA | NA | 5 | 1 | NA | NA | 1 | 1 | 2 | 1 | 1 | 1 | 4 | 1 |
| 16_912013 | NA | NA | 2 | 0.5 | 2 | 1 | NA | NA | 4 | 0.75 | NA | NA | 1 | 1 | 2 | 1 | 1 | 1 | 4 | 1 |
| 16_912014( | NA | NA | 3 | 0.6667 | 2 | 1 | NA | NA | 5 | 0.8 | NA | NA | 1 | 1 | 2 | 1 | NA | NA | 3 | 1 |
| 16_912014( | NA | NA | 3 | 0.3333 | 2 | 0.5 | NA | NA | 5 | 0.4 | NA | NA | 1 | 1 | 2 | 1 | 1 | 1 | 4 | 1 |
| 16_912014! | NA | NA | 1 | 1 | 2 | 0.5 | NA | NA | 3 | 0.6667 | NA | NA | 1 | 1 | 2 | 0.5 | 1 | 0 | 4 | 0.5 |
| 16_912016 | NA | NA | 4 | 1 | 2 | 1 | NA | NA | 6 | 1 | NA | NA | 1 | 1 | 2 | 0.5 | NA | NA | 3 | 0.6667 |
| 16_912020( | NA | NA | 3 | 0 | 1 | 1 | NA | NA | 4 | 0.25 | NA | NA | 1 | 0 | 2 | 0.5 | 1 | 1 | 4 | 0.5 |
| 16_912021 | NA | NA | 3 | 0.3333 | 1 | 1 | NA | NA | 4 | 0.5 | NA | NA | 1 | 1 | 1 | 1 | 1 | 1 | 3 | 1 |
| 16_912021! | NA | NA | 2 | 1 | 2 | 0.5 | NA | NA | 4 | 0.75 | NA | NA | 1 | 0 | 2 | 1 | 1 | 1 | 4 | 0.75 |
| 16_912022 | NA | NA | 1 | 1 | 2 | 1 | NA | NA | 3 | 1 | NA | NA | 1 | 0 | 2 | 1 | 1 | 1 | 4 | 0.75 |
| 16_912220 | 2 | 0 | 2 | 0.5 | NA | NA | 2 | 0 | 6 | 0.1667 | 1 | 0 | NA | NA | 2 | 0.5 | 1 | 0 | 4 | 0.25 |
| 16_912221 | 1 | 1 | 2 | 0.5 | NA | NA | 2 | 0 | 5 | 0.4 | 1 | 0 | 1 | 0 | 3 | 0.6667 | 1 | 1 | 6 | 0.5 |
| 16_912221 | 1 | 0 | 1 | 0 | NA | NA | 1 | 1 | 3 | 0.3333 | 1 | 0 | 1 | 0 | 2 | 0.5 | 1 | 0 | 5 | 0.2 |
| 16_912223 | 2 | 0.5 | 3 | 0 | NA | NA | 2 | 0 | 7 | 0.1429 | 2 | 0 | 3 | 0.3333 | 3 | 0 | 1 | 0 | 9 | 0.1111 |
| 16_912225 | 2 | 0.5 | 2 | 0 | NA | NA | 2 | 0.5 | 6 | 0.3333 | 3 | 0.3333 | 2 | 0.5 | 4 | 0.5 | NA | NA | 9 | 0.4444 |
| 16_913021( | 3 | 0.6667 | 2 | 0.5 | 2 | 0 | 2 | 0.5 | 9 | 0.4444 | NA | NA | 5 | 0.6 | 3 | 1 | 4 | 0.75 | 12 | 0.75 |
| 16_913022 | 4 | 0.75 | 2 | 0 | 2 | 0 | 3 | 1 | 11 | 0.5455 | 1 | 0 | 5 | 0.4 | 1 | 1 | 4 | 0 | 11 | 0.2727 |
| 16_913022 | 4 | 0.5 | 3 | 0.6667 | 2 | 0.5 | 3 | 0.3333 | 12 | 0.5 | 1 | 1 | 5 | 0.6 | 3 | 0.3333 | 4 | 0.75 | 13 | 0.6154 |
| 16_913022 | 3 | 1 | 3 | 1 | 3 | 0.6667 | 3 | 1 | 12 | 0.9167 | 1 | 1 | 5 | 0.6 | 3 | 0.6667 | 4 | 1 | 13 | 0.7692 |
| 16_913023 | 4 | 0.25 | 3 | 0 | 1 | 0 | 2 | 0 | 10 | 0.1 | 1 | 0 | 5 | 0.2 | 2 | 0.5 | 3 | 0.6667 | 11 | 0.3636 |
| 16_913023 | 2 | 1 | 3 | 1 | 2 | 1 | 2 | 1 | 9 | 1 | NA | NA | 4 | 1 | 1 | 1 | 2 | 1 | 7 | 1 |
| 16_913024( | 4 | 0.25 | 2 | 0 | 4 | 0.5 | 3 | 0.3333 | 13 | 0.3077 | 1 | 0 | 5 | 0.2 | NA | NA | 3 | 0 | 9 | 0.1111 |
| 16_913024 | 4 | 0.5 | 3 | 0.6667 | 4 | 0.75 | 3 | 0.6667 | 14 | 0.6429 | 1 | 0 | 5 | 0.6 | 3 | 1 | 3 | 1 | 12 | 0.75 |
| 16_913026 | 4 | 0.75 | 3 | 0.6667 | 2 | 1 | 3 | 0 | 12 | 0.5833 | NA | NA | 3 | 0.6667 | 3 | 0.3333 | 2 | 0.5 | 8 | 0.5 |
| 16_913026 | 4 | 0.5 | 3 | 0.3333 | 3 | 0.6667 | 3 | 0 | 13 | 0.3846 | 1 | 1 | 3 | 0.3333 | 3 | 0.6667 | 1 | 0 | 8 | 0.5 |
| 16_913027( | 3 | 0.3333 | 4 | 0.25 | 3 | 0.3333 | 2 | 0 | 12 | 0.25 | NA | NA | 4 | 0.25 | 3 | 0.6667 | 3 | 0.3333 | 10 | 0.4 |
| 16_913027 | 4 | 0.25 | 3 | 0 | 3 | 0 | 3 | 0 | 13 | 0.0769 | 1 | 1 | 4 | 0.25 | 2 | 0 | 2 | 0 | 9 | 0.2222 |
| 16_913030 | 6 | 0.1667 | 5 | 0 | 3 | 0 | 3 | 0 | 17 | 0.0588 | 2 | 0 | 3 | 0.3333 | 3 | 0 | 1 | 0 | 9 | 0.1111 |
| 16_913032 | 6 | 0.6667 | 4 | 1 | 2 | 0 | 3 | 0 | 15 | 0.5333 | 2 | 0.5 | 4 | 0.75 | 3 | 0.3333 | 2 | 0.5 | 11 | 0.5455 |
| 16_913032! | 6 | 0 | 5 | 0 | 3 | 0.3333 | 3 | 0 | 17 | 0.0588 | 2 | 0 | 5 | 0.2 | 2 | 0 | 2 | 0.5 | 11 | 0.1818 |
| 16_913034 | 6 | 0.6667 | 6 | 1 | 1 | 1 | 4 | 0.75 | 17 | 0.8235 | 2 | 1 | 5 | 0.8 | 1 | 1 | 3 | 1 | 11 | 0.9091 |
| 16_913036 | 5 | 0.2 | 5 | 0 | 4 | 0 | 4 | 0 | 18 | 0.0556 | 2 | 1 | 6 | 0.3333 | 2 | 1 | 4 | 0 | 14 | 0.4286 |
| 16_913038 | 5 | 0 | 6 | 0.3333 | 4 | 0.75 | 4 | 0 | 19 | 0.2632 | 2 | 0 | 4 | 0.5 | 2 | 0 | 3 | 0 | 11 | 0.1818 |
| 16_913040 | 6 | 0.3333 | 6 | 0.3333 | 5 | 0.4 | 4 | 0 | 21 | 0.2857 | 4 | 0 | 6 | 0 | 4 | 0.25 | 4 | 0.25 | 18 | 0.1111 |
| 16_913040 | 6 | 0.3333 | 6 | 0.5 | 4 | 0.75 | 4 | 1 | 20 | 0.6 | 4 | 0.5 | 7 | 0.8571 | 3 | 0.3333 | 4 | 0.5 | 18 | 0.6111 |
| 16_913040 | 6 | 0.8333 | 5 | 0.6 | 5 | 0.6 | 4 | 0.75 | 20 | 0.7 | 3 | 0.6667 | 7 | 0.5714 | 4 | 0.75 | 4 | 0.5 | 18 | 0.6111 |
| 16_913062 | 2 | 0 | 1 | 1 | 1 | 0 | 1 | 0 | 5 | 0.2 | NA | NA | 1 | 1 | NA | NA | NA | NA | 1 | 1 |
| 16_913073 | 3 | 0 | 1 | 1 | NA | NA | 1 | 0 | 5 | 0.2 | NA | NA | 1 | 1 | NA | NA | NA | NA | 1 | 1 |
| 16_913598 | 4 | 0.25 | 6 | 0 | 8 | 0.25 | 5 | 0 | 23 | 0.1304 | 7 | 0 | 5 | 0.2 | 3 | 0.3333 | 7 | 0 | 22 | 0.0909 |
| 16_913599 | 4 | 1 | 7 | 0.7143 | 7 | 0.8571 | 4 | 0.5 | 22 | 0.7727 | 5 | 0.8 | 5 | 0.8 | 3 | 0.3333 | 8 | 0.875 | 21 | 0.7619 |
| 16_913600 | 3 | 0 | 7 | 0 | 9 | 0.1111 | 5 | 0.2 | 24 | 0.0833 | 6 | 0 | 3 | 0 | 4 | 0 | 7 | 0.1429 | 20 | 0.05 |
| 16_913601( | 5 | 0.4 | 6 | 0 | 5 | 0.2 | 5 | 0.6 | 21 | 0.2857 | 6 | 0.1667 | 4 | 0.25 | 4 | 0.75 | 7 | 0.4286 | 21 | 0.381 |
| 16_913601 | 6 | 0.1667 | 7 | 0.1429 | 7 | 0.1429 | 4 | 0.5 | 24 | 0.2083 | 7 | 0.4286 | 5 | 0.2 | 4 | 0.25 | 9 | 0.3333 | 25 | 0.32 |
| 16_913603( | 5 | 0 | 4 | 0 | 6 | 0 | 3 | 0.3333 | 18 | 0.0556 | 5 | 0 | 4 | 0 | 3 | 0 | 6 | 0.1667 | 18 | 0.0556 |
| 16_913725( | 16 | 0.0625 | 15 | 0.0667 | 14 | 0 | 12 | 0.0833 | 57 | 0.0526 | 5 | 0.6 | 8 | 0.375 | 12 | 0.0833 | 17 | 0.0588 | 42 | 0.1905 |

| | | | | | | | | | | | | | | | | | | | | |
|---|---|---|---|---|---|---|---|---|---|---|---|---|---|---|---|---|---|---|---|---|
| 16_9137254 | 12 | 0.25 | 15 | 0.0667 | 13 | 0.0769 | 13 | 0.3077 | 53 | 0.1698 | 5 | 0.2 | 8 | 0.25 | 12 | 0.1667 | 11 | 0.3636 | 36 | 0.25 |
| 16_9141127 | NA | NA | 1 | 1 | 3 | 0 | 1 | 1 | 5 | 0.4 | 2 | 0.5 | NA | NA | 5 | 0 | 1 | 0 | 8 | 0.125 |
| 16_9141154 | NA | NA | 1 | 1 | 3 | 0.3333 | 1 | 1 | 5 | 0.6 | 2 | 1 | NA | NA | 4 | 0.75 | 1 | 1 | 7 | 0.8571 |
| 16_9141226 | NA | NA | 1 | 1 | 2 | 0 | 1 | 0 | 4 | 0.25 | 1 | 0 | NA | NA | 5 | 0.6 | NA | NA | 6 | 0.5 |
| 16_9141232 | NA | NA | 1 | 0 | 3 | 1 | 1 | 0 | 5 | 0.6 | 1 | 0 | NA | NA | 4 | 0.5 | 1 | 1 | 6 | 0.5 |
| 16_9141271 | NA | NA | NA | NA | 2 | 1 | 1 | 1 | 3 | 1 | 2 | 1 | NA | NA | 4 | 1 | 1 | 1 | 7 | 1 |
| 16_9141282 | NA | NA | NA | NA | 2 | 1 | 1 | 1 | 3 | 1 | 1 | 1 | NA | NA | 3 | 1 | 1 | 1 | 5 | 1 |
| 16_9141292 | NA | NA | NA | NA | 2 | 1 | 1 | 0 | 3 | 0.6667 | 2 | 0.5 | NA | NA | 4 | 0.5 | NA | NA | 6 | 0.5 |
| 16_9141295 | NA | NA | NA | NA | 2 | 1 | 1 | 1 | 3 | 1 | 1 | 1 | NA | NA | 1 | 1 | 1 | 1 | 3 | 1 |
| 16_9141300 | NA | NA | NA | NA | 1 | 1 | 1 | 1 | 2 | 1 | 2 | 1 | NA | NA | 3 | 1 | 1 | 1 | 6 | 1 |
| 16_9141308 | NA | NA | NA | NA | 2 | 1 | 1 | 1 | 3 | 1 | 2 | 1 | NA | NA | 2 | 1 | 1 | 1 | 5 | 1 |
| 16_9141344 | NA | NA | NA | NA | 1 | 0 | 1 | 1 | 2 | 0.5 | 2 | 0.5 | NA | NA | 2 | 0.5 | 1 | 0 | 5 | 0.4 |
| 16_9141352 | NA | NA | NA | NA | 2 | 0.5 | 1 | 0 | 3 | 0.3333 | 1 | 1 | NA | NA | 2 | 1 | 1 | 1 | 4 | 1 |
| 16_9141359 | NA | NA | NA | NA | 2 | 1 | 1 | 1 | 3 | 1 | 2 | 1 | NA | NA | 2 | 0.5 | 1 | 1 | 5 | 0.8 |
| 16_9141398 | NA | NA | NA | NA | 2 | 0.5 | 1 | 0 | 3 | 0.3333 | 1 | 0 | NA | NA | 2 | 1 | 1 | 1 | 4 | 0.75 |
| 16_9141420 | NA | NA | NA | NA | 2 | 0.5 | 1 | 1 | 3 | 0.6667 | 1 | 1 | NA | NA | 1 | 0 | 1 | 0 | 3 | 0.3333 |
| 16_9141833 | 3 | 1 | 1 | 1 | 2 | 1 | 8 | 0.875 | 14 | 0.9286 | 3 | 0.6667 | 4 | 0.75 | NA | NA | 4 | 0.75 | 11 | 0.7273 |
| 16_9143069 | 3 | 0 | 4 | 0.25 | 1 | 0 | NA | NA | 8 | 0.125 | 2 | 0 | 3 | 0.3333 | 2 | 0 | NA | NA | 7 | 0.1429 |
| 16_9143072 | 1 | 0 | 2 | 0.5 | 1 | 0 | NA | NA | 4 | 0.25 | 1 | 1 | 2 | 0.5 | 3 | 0 | NA | NA | 6 | 0.3333 |
| 16_9143080 | 4 | 0.25 | 5 | 0.2 | 2 | 0.5 | 2 | 0 | 13 | 0.2308 | 5 | 0.4 | 4 | 0.75 | 3 | 0.6667 | NA | NA | 12 | 0.5833 |
| 16_9143084 | 2 | 0.5 | 4 | 0.75 | 2 | 0 | 1 | 0 | 9 | 0.4444 | 3 | 0 | 5 | 0.6 | 1 | 1 | NA | NA | 9 | 0.4444 |
| 16_9143096 | 3 | 0.3333 | 4 | 0.25 | 3 | 0 | 3 | 0 | 13 | 0.1538 | 5 | 0.2 | 4 | 0.25 | 3 | 0 | NA | NA | 12 | 0.1667 |
| 16_9145173 | 1 | 1 | 1 | 1 | NA | NA | NA | NA | 2 | 1 | 1 | 1 | NA | NA | 1 | 0 | NA | NA | 2 | 0.5 |
| 16_9145185 | 1 | 1 | 1 | 1 | NA | NA | NA | NA | 2 | 1 | 1 | 1 | NA | NA | 1 | 1 | NA | NA | 2 | 1 |
| 16_9145188 | 1 | 1 | 1 | 1 | NA | NA | NA | NA | 2 | 1 | 1 | 1 | NA | NA | 1 | 1 | NA | NA | 2 | 1 |
| 16_9145210 | 1 | 1 | 1 | 0 | NA | NA | NA | NA | 2 | 0.5 | 1 | 0 | NA | NA | 3 | 0.3333 | NA | NA | 4 | 0.25 |
| 16_9145217 | 1 | 0 | 1 | 1 | NA | NA | NA | NA | 2 | 0.5 | 1 | 1 | NA | NA | 3 | 0.3333 | NA | NA | 4 | 0.5 |
| 16_9145293 | 1 | 1 | 1 | 0 | NA | NA | NA | NA | 2 | 0.5 | 2 | 0.5 | 1 | 1 | 4 | 0.25 | NA | NA | 7 | 0.4286 |
| 16_9145333 | 1 | 0 | 3 | 0.3333 | NA | NA | NA | NA | 4 | 0.25 | 2 | 0 | 2 | 0.5 | 6 | 0.3333 | 1 | 0 | 11 | 0.2727 |
| 16_9145335 | 1 | 1 | 3 | 0.3333 | NA | NA | NA | NA | 4 | 0.5 | 2 | 1 | 1 | 1 | 6 | 0.6667 | 1 | 0 | 10 | 0.7 |
| 16_9145343 | 1 | 1 | 3 | 1 | NA | NA | NA | NA | 4 | 1 | 2 | 1 | 2 | 1 | 5 | 1 | 1 | 1 | 10 | 1 |
| 16_9145349 | 1 | 1 | 3 | 1 | NA | NA | NA | NA | 4 | 1 | 2 | 1 | 2 | 0.5 | 6 | 1 | 1 | 1 | 11 | 0.9091 |
| 16_9145352 | 1 | 1 | 3 | 1 | NA | NA | NA | NA | 4 | 1 | 2 | 0.5 | 1 | 1 | 6 | 1 | 1 | 1 | 10 | 0.9 |
| 16_9145355 | 1 | 1 | 3 | 1 | NA | NA | NA | NA | 4 | 1 | 2 | 1 | 2 | 1 | 5 | 0.8 | 1 | 1 | 10 | 0.9 |
| 16_9145358 | 1 | 1 | 3 | 0.6667 | NA | NA | NA | NA | 4 | 0.75 | 2 | 0.5 | 2 | 0.5 | 6 | 0.8333 | 1 | 1 | 11 | 0.7273 |
| 16_9145361 | 1 | 1 | 3 | 0.6667 | NA | NA | NA | NA | 4 | 0.75 | 2 | 0.5 | 2 | 1 | 6 | 0.6667 | 1 | 1 | 11 | 0.7273 |
| 16_9145364 | 1 | 1 | 3 | 1 | NA | NA | NA | NA | 4 | 1 | 2 | 1 | 2 | 0.5 | 6 | 0.8333 | 1 | 1 | 11 | 0.8182 |
| 16_9145367 | 1 | 1 | 3 | 0.3333 | NA | NA | NA | NA | 4 | 0.5 | 2 | 0 | 2 | 0.5 | 6 | 0.6667 | 1 | 1 | 11 | 0.5455 |
| 16_9145373 | 1 | 0 | 4 | 0.5 | 1 | 0 | NA | NA | 6 | 0.3333 | 2 | 0.5 | 2 | 0.5 | 5 | 0.6 | 1 | 1 | 10 | 0.6 |
| 16_9145376 | 2 | 0.5 | 4 | 0.5 | 2 | 1 | NA | NA | 8 | 0.625 | NA | NA | 2 | 0 | 6 | 0.5 | 1 | 0 | 9 | 0.3333 |
| 16_9145379 | 2 | 0.5 | 3 | 0.6667 | 3 | 0 | NA | NA | 8 | 0.375 | 2 | 0.5 | 2 | 0 | 3 | 0.6667 | 1 | 0 | 8 | 0.375 |
| 16_9145382 | 2 | 0 | 3 | 0.6667 | 2 | 0 | NA | NA | 7 | 0.2857 | 1 | 0 | 2 | 0.5 | 5 | 0.4 | 2 | 0.5 | 10 | 0.4 |
| 16_9145411 | 4 | 0 | 4 | 0.5 | 3 | 0 | NA | NA | 11 | 0.1818 | 1 | 0 | 3 | 0 | 6 | 0 | 3 | 0.3333 | 13 | 0.0769 |
| 16_9145473 | 6 | 0.1667 | 6 | 0 | 6 | 0 | 2 | 0 | 20 | 0.05 | 2 | 0 | 5 | 0 | 9 | 0 | 3 | 0.6667 | 19 | 0.1053 |
| 16_9145483 | 6 | 0.5 | 6 | 0.3333 | 6 | 0.3333 | 2 | 0.5 | 20 | 0.4 | 1 | 0 | 5 | 0.6 | 8 | 0.375 | 3 | 0.3333 | 17 | 0.4118 |
| 16_9145494 | 5 | 0.2 | 6 | 0.1667 | 5 | 0.4 | 3 | 0 | 19 | 0.2105 | 3 | 0.3333 | 5 | 0.2 | 9 | 0.3333 | 3 | 0.6667 | 20 | 0.35 |
| 16_9145497 | 5 | 0.8 | 5 | 0.2 | 7 | 0.4286 | 2 | 0.5 | 19 | 0.4737 | 3 | 0.6667 | 6 | 0.6667 | 9 | 0.8889 | 2 | 1 | 20 | 0.8 |
| 16_9145506 | 7 | 0.4286 | 6 | 0.5 | 7 | 0.2857 | 3 | 0.6667 | 23 | 0.4348 | 3 | 0 | 6 | 0.1667 | 10 | 0.2 | 3 | 0.6667 | 22 | 0.2273 |
| 16_9145514 | 6 | 0.6667 | 6 | 0.3333 | 6 | 0.3333 | 3 | 0.3333 | 21 | 0.4286 | 3 | 0.3333 | 4 | 0.25 | 10 | 0.6 | 2 | 0 | 19 | 0.4211 |
| 16_9145518 | 7 | 0 | 7 | 0.2857 | 5 | 0.2 | 2 | 0 | 21 | 0.1429 | 3 | 0 | 6 | 0.5 | 9 | 0.2222 | 3 | 0 | 21 | 0.2381 |
| 16_9145527 | 6 | 0.3333 | 6 | 0.3333 | 6 | 0.3333 | 2 | 0 | 20 | 0.3 | 4 | 0.5 | 6 | 0.1667 | 10 | 0.2 | 2 | 0 | 22 | 0.2273 |
| 16_9145538 | 4 | 0.75 | 6 | 0.1667 | 7 | 0.1429 | 3 | 0 | 20 | 0.25 | 4 | 0.25 | 6 | 0 | 8 | 0.25 | 5 | 0.2 | 23 | 0.1739 |
| 16_9145547 | 5 | 1 | 7 | 0.7143 | 6 | 1 | 1 | 1 | 19 | 0.8947 | 4 | 0.75 | 6 | 0.6667 | 11 | 0.9091 | 5 | 0.8 | 26 | 0.8077 |
| 16_9145564 | 6 | 0 | 8 | 0.125 | 7 | 0 | 3 | 0.3333 | 24 | 0.0833 | 3 | 0 | 8 | 0 | 10 | 0.2 | 4 | 0 | 25 | 0.08 |
| 16_9145576 | 7 | 1 | 5 | 1 | 7 | 0.8571 | 4 | 0.75 | 23 | 0.913 | 3 | 0.6667 | 8 | 0.75 | 10 | 1 | 5 | 1 | 26 | 0.8846 |
| 16_9145585 | 5 | 0.2 | 8 | 0.375 | 7 | 0.2857 | 4 | 0.25 | 24 | 0.2917 | 5 | 0.6 | 10 | 0.3 | 12 | 0.5 | 6 | 0.3333 | 33 | 0.4242 |
| 16_9145589 | 6 | 0.5 | 8 | 0.75 | 9 | 0.6667 | 5 | 0.4 | 28 | 0.6071 | 5 | 0.6 | 10 | 0.8 | 12 | 0.6667 | 6 | 0.8333 | 33 | 0.7273 |
| 16_9145590 | 7 | 0.4286 | 8 | 0.625 | 8 | 0.5 | 4 | 0.5 | 27 | 0.5185 | 5 | 0.2 | 10 | 0.5 | 11 | 0.5455 | 6 | 0.8333 | 32 | 0.5312 |
| 16_9145605 | 7 | 0.1429 | 8 | 0.25 | 8 | 0.125 | 5 | 0 | 28 | 0.1429 | 5 | 0 | 10 | 0.3 | 10 | 0.1 | 6 | 0.1667 | 31 | 0.1613 |
| 16_9145615 | 6 | 0.3333 | 7 | 0.1429 | 6 | 0 | 3 | 0.6667 | 22 | 0.2273 | 5 | 0 | 9 | 0.2222 | 11 | 0.2727 | 7 | 0.1429 | 32 | 0.1875 |
| 16_9145624 | 5 | 0 | 6 | 0.1667 | 5 | 0 | 4 | 0 | 20 | 0.05 | 4 | 0 | 8 | 0 | 10 | 0.2 | 7 | 0.1429 | 29 | 0.1034 |
| 16_9145630 | 6 | 0 | 7 | 0.4286 | 9 | 0.1111 | 7 | 0.1429 | 29 | 0.1724 | 5 | 0 | 12 | 0.5 | 10 | 0.3 | 8 | 0.375 | 35 | 0.3429 |
| 16_9145632 | 7 | 0.5714 | 7 | 0.7143 | 8 | 0.5 | 5 | 0.8 | 27 | 0.6296 | 4 | 0.5 | 10 | 0.6 | 10 | 0.7 | 8 | 0.625 | 32 | 0.625 |
| 16_9145637 | 7 | 0.1429 | 8 | 0.25 | 8 | 0 | 7 | 0.1429 | 30 | 0.1333 | 5 | 0.2 | 12 | 0.3333 | 11 | 0.4545 | 8 | 0.125 | 36 | 0.3056 |
| 16_9145655 | 6 | 0.3333 | 7 | 0.4286 | 6 | 0.3333 | 6 | 0.3333 | 25 | 0.36 | 5 | 0.4 | 12 | 0.3333 | 12 | 0.5833 | 8 | 0.5 | 37 | 0.4595 |

| | | | | | | | | | | | | | | | | | | | | |
|---|---|---|---|---|---|---|---|---|---|---|---|---|---|---|---|---|---|---|---|---|
| 16_914565 | 7 | 0.7143 | 6 | 0.8333 | 8 | 0.75 | 8 | 1 | 29 | 0.8276 | 5 | 0.8 | 13 | 0.8462 | 12 | 0.8333 | 8 | 1 | 38 | 0.8684 |
| 16_914567 | 7 | 0.4286 | 8 | 0.5 | 8 | 0.625 | 8 | 0.375 | 31 | 0.4839 | 4 | 0 | 12 | 0.25 | 12 | 0.4167 | 8 | 0.25 | 36 | 0.2778 |
| 16_914567 | 5 | 0.6 | 8 | 0.5 | 8 | 0.375 | 7 | 0.5714 | 28 | 0.5 | 2 | 0.5 | 14 | 0.3571 | 13 | 0.6154 | 7 | 0.8571 | 36 | 0.5556 |
| 16_914593 | 2 | 0 | 4 | 0.25 | 5 | 0 | 2 | 0 | 13 | 0.0769 | 1 | 0 | 6 | 0 | 3 | 0.3333 | 1 | 0 | 11 | 0.0909 |
| 16_914595 | 3 | 0.6667 | 4 | 0.25 | 5 | 0.8 | 3 | 0.6667 | 15 | 0.6 | 1 | 0 | 6 | 0.6667 | 4 | 0.5 | NA | NA | 11 | 0.5455 |
| 16_914595 | 3 | 0.6667 | 3 | 0.6667 | 5 | 0.8 | 2 | 1 | 13 | 0.7692 | 1 | 1 | 6 | 0.3333 | 5 | 0.8 | 1 | 1 | 13 | 0.6154 |
| 16_914596 | 3 | 1 | 4 | 1 | 5 | 1 | 3 | 1 | 15 | 1 | 1 | 1 | 6 | 0.8333 | 5 | 0.8 | NA | NA | 12 | 0.8333 |
| 16_914601 | 5 | 0.2 | 4 | 0.25 | 4 | 0.25 | 2 | 0 | 15 | 0.2 | 2 | 0 | 6 | 0 | 7 | 0.1429 | 2 | 0.5 | 17 | 0.1176 |
| 16_914601 | 5 | 1 | 4 | 0.75 | 4 | 1 | 2 | 1 | 15 | 0.9333 | 2 | 0.5 | 7 | 0.8571 | 5 | 0.8 | 2 | 0.5 | 16 | 0.75 |
| 16_914603 | 3 | 0.3333 | 2 | 0 | 2 | 0 | 2 | 0 | 9 | 0.1111 | 1 | 0 | 7 | 0 | 7 | 0.1429 | 3 | 0 | 18 | 0.0556 |
| 16_914605 | 4 | 0 | NA | NA | 2 | 0.5 | 2 | 0.5 | 8 | 0.25 | NA | NA | 7 | 0.1429 | 6 | 0 | 1 | 1 | 14 | 0.1429 |
| 16_914609 | 4 | 0 | 2 | 0.5 | 2 | 0 | 2 | 0 | 10 | 0.1 | 1 | 0 | 8 | 0.125 | 6 | 0 | NA | NA | 15 | 0.0667 |
| 16_914609 | 3 | 0.3333 | 2 | 0 | 3 | 0 | 1 | 0 | 9 | 0.1111 | 1 | 0 | 8 | 0 | 6 | 0 | 2 | 0.5 | 17 | 0.0588 |
| 16_914700 | NA | NA | 2 | 0.5 | NA | NA | 2 | 0 | 4 | 0.25 | 1 | 0 | 2 | 0.5 | 3 | 0 | NA | NA | 6 | 0.1667 |
| 16_914714 | NA | NA | 1 | 0 | NA | NA | 3 | 0.3333 | 4 | 0.25 | 3 | 0 | 2 | 0 | 7 | 0.1429 | 3 | 0.3333 | 15 | 0.1333 |
| 16_914728 | NA | NA | NA | NA | 1 | 1 | NA | NA | 1 | 1 | NA | NA | 2 | 0.5 | NA | NA | NA | NA | 2 | 0.5 |
| 16_914732 | NA | NA | NA | NA | 2 | 1 | NA | NA | 2 | 1 | NA | NA | 2 | 1 | NA | NA | NA | NA | 2 | 1 |
| 16_914751 | 10 | 0.3 | 7 | 0.7143 | 7 | 0.2857 | 20 | 0.3 | 44 | 0.3636 | 14 | 0.4286 | 18 | 0.3889 | 21 | 0.381 | 15 | 0.4 | 68 | 0.3971 |
| 16_914757 | NA | NA | NA | NA | 2 | 0.5 | NA | NA | 2 | 0.5 | NA | NA | 2 | 0.5 | 1 | 1 | NA | NA | 3 | 0.6667 |
| 16_914768 | 3 | 0.3333 | 5 | 0.2 | 5 | 0.2 | 2 | 0 | 15 | 0.2 | 3 | 0.3333 | 3 | 0 | 3 | 0 | 3 | 0.3333 | 12 | 0.1667 |
| 16_914768 | 3 | 0.6667 | 4 | 0.25 | 6 | 0.3333 | 2 | 0 | 15 | 0.3333 | 2 | 0 | 4 | 0.5 | 3 | 0.6667 | 2 | 0 | 11 | 0.3636 |
| 16_914780 | 3 | 0.6667 | 4 | 0.25 | 3 | 0 | 1 | 0 | 11 | 0.2727 | 3 | 0.3333 | 4 | 0 | 1 | 0 | 1 | 0 | 9 | 0.1111 |
| 16_914780 | 3 | 0 | 5 | 0 | 6 | 0.1667 | 2 | 0 | 16 | 0.0625 | 3 | 0.3333 | 4 | 0.25 | 3 | 0 | 2 | 0 | 12 | 0.1667 |
| 16_914781 | 3 | 0.3333 | 5 | 0.2 | 6 | 0.1667 | 2 | 0 | 16 | 0.1875 | 3 | 0.3333 | 4 | 0 | 2 | 0 | 1 | 0 | 10 | 0.1 |
| 16_914791 | 3 | 0.6667 | 5 | 0 | 3 | 0.6667 | 3 | 0.3333 | 14 | 0.3571 | 3 | 0 | 1 | 1 | 2 | 0.5 | 2 | 0.5 | 8 | 0.375 |
| 16_914810 | NA | NA | 4 | 0 | NA | NA | 2 | 0.5 | 6 | 0.1667 | 1 | 1 | NA | NA | NA | NA | 2 | 1 | 3 | 1 |
| 16_914850 | NA | NA | 2 | 1 | NA | NA | 1 | 1 | 3 | 1 | NA | NA | NA | NA | NA | NA | 1 | 1 | 1 | 1 |
| 16_914851 | NA | NA | 1 | 1 | NA | NA | 1 | 0 | 2 | 0.5 | NA | NA | NA | NA | NA | NA | 1 | 1 | 1 | 1 |
| 16_914851 | NA | NA | 1 | 1 | NA | NA | 1 | 0 | 2 | 0.5 | 1 | 1 | NA | NA | NA | NA | 1 | 1 | 2 | 1 |
| 16_914860 | NA | NA | 1 | 1 | NA | NA | NA | NA | 1 | 1 | 1 | 1 | NA | NA | NA | NA | NA | NA | 1 | 1 |
| 16_914864 | 5 | 0.4 | 9 | 0.5556 | 6 | 0.5 | 3 | 1 | 23 | 0.5652 | 6 | 0.3333 | 5 | 0.2 | 7 | 0.7143 | 2 | 0.5 | 20 | 0.45 |
| 16_914913 | 4 | 0 | 4 | 0 | 5 | 0 | 6 | 0.1667 | 19 | 0.0526 | 7 | 0.1429 | 5 | 0 | 4 | 0 | 2 | 0 | 18 | 0.0556 |
| 16_914914 | 3 | 0.6667 | 4 | 0 | 4 | 0.25 | 4 | 0.75 | 15 | 0.4 | 7 | 0.4286 | 4 | 0.25 | 4 | 0.25 | 1 | 0 | 16 | 0.3125 |
| 16_914916 | 3 | 1 | 4 | 0.25 | 5 | 0.6 | 6 | 0.5 | 18 | 0.5556 | 6 | 0.6667 | 4 | 0 | 4 | 0.25 | 2 | 0 | 16 | 0.3125 |
| 16_914921 | 3 | 0 | 2 | 0.5 | 4 | 0 | 4 | 0.25 | 13 | 0.1538 | 4 | 0.25 | 5 | 0 | 3 | 0 | 1 | 1 | 13 | 0.1538 |
| 16_916666 | 7 | 0 | 10 | 0.2 | 6 | 0 | 4 | 0.25 | 27 | 0.1111 | 4 | 0 | 5 | 0 | 10 | 0.1 | 6 | 0.1667 | 25 | 0.08 |
| 16_916672 | 2 | 0 | 5 | 0.2 | 2 | 0 | 1 | 0 | 10 | 0.1 | 2 | 0.5 | 1 | 0 | 6 | 0.1667 | 2 | 0.5 | 11 | 0.2727 |
| 16_916681 | 2 | 1 | 2 | 0.5 | 2 | 0 | 1 | 0 | 7 | 0.4286 | 2 | 0 | 3 | 0.3333 | 4 | 0 | 3 | 0 | 12 | 0.0833 |
| 16_917022 | 6 | 0.1667 | 5 | 0 | 5 | 0 | 4 | 0.25 | 20 | 0.1 | 2 | 0 | 5 | 0 | 4 | 0 | 4 | 0.5 | 15 | 0.1333 |
| 16_917087 | 2 | 0 | 6 | 0.1667 | 3 | 0 | 2 | 0 | 13 | 0.0769 | 3 | 0 | 4 | 0 | 5 | 0.2 | 4 | 0 | 16 | 0.0625 |
| 16_917089 | 6 | 0.1667 | 7 | 0.1429 | 7 | 0 | 4 | 0 | 24 | 0.0833 | 3 | 0 | 5 | 0.2 | 6 | 0 | 5 | 0.2 | 19 | 0.1053 |
| 16_917092 | 4 | 0.25 | 3 | 0.3333 | 2 | 1 | 2 | 0.5 | 11 | 0.4545 | 2 | 0.5 | 2 | 0.5 | 4 | 0.5 | 3 | 0.6667 | 11 | 0.5455 |
| 16_917165 | 4 | 0 | 8 | 0.125 | 1 | 0 | 1 | 0 | 14 | 0.0714 | 1 | 0 | 3 | 0 | 3 | 0.3333 | 1 | 0 | 8 | 0.125 |
| 16_917167 | 4 | 0 | 8 | 0 | 3 | 0.3333 | 1 | 0 | 16 | 0.0625 | 2 | 0.5 | 2 | 0 | 3 | 0.6667 | 1 | 0 | 8 | 0.375 |
| 16_917170 | 3 | 0 | 6 | 0 | 3 | 0.3333 | 1 | 0 | 13 | 0.0769 | 1 | 0 | 2 | 0.5 | 3 | 0.3333 | 2 | 0 | 8 | 0.25 |
| 16_917170 | 4 | 0 | 6 | 0 | 3 | 0.3333 | 1 | 0 | 14 | 0.0714 | 1 | 0 | 3 | 0 | 2 | 0 | 2 | 0.5 | 8 | 0.125 |
| 16_917221 | 74 | 0.2432 | 79 | 0.1899 | 76 | 0.3158 | 64 | 0.1875 | 293 | 0.2355 | 75 | 0.2533 | 77 | 0.1688 | 85 | 0.2 | 60 | 0.2833 | 297 | 0.2222 |
| 16_918744 | NA | NA | NA | NA | 1 | 0 | 2 | 0.5 | 3 | 0.3333 | NA | NA | NA | NA | NA | NA | 1 | 1 | 1 | 1 |
| 16_918744 | NA | NA | NA | NA | 1 | 1 | 2 | 1 | 3 | 1 | NA | NA | NA | NA | NA | NA | 1 | 1 | 1 | 1 |
| 16_918748 | NA | NA | NA | NA | 1 | 1 | 2 | 0 | 3 | 0.3333 | NA | NA | NA | NA | NA | NA | 1 | 1 | 1 | 1 |
| 16_918750 | NA | NA | NA | NA | 1 | 1 | 2 | 1 | 3 | 1 | NA | NA | NA | NA | NA | NA | 1 | 1 | 1 | 1 |
| 16_918756 | NA | NA | NA | NA | 1 | 1 | 1 | 0 | 2 | 0.5 | NA | NA | NA | NA | NA | NA | 1 | 1 | 1 | 1 |
| 16_918757 | NA | NA | NA | NA | 1 | 1 | 2 | 1 | 3 | 1 | NA | NA | NA | NA | NA | NA | 1 | 1 | 1 | 1 |
| 16_918759 | NA | NA | NA | NA | 1 | 1 | 2 | 1 | 3 | 1 | NA | NA | NA | NA | NA | NA | 1 | 1 | 1 | 1 |
| 16_918759 | NA | NA | NA | NA | 1 | 1 | 1 | 1 | 2 | 1 | NA | NA | NA | NA | NA | NA | 2 | 1 | 2 | 1 |
| 16_918760 | NA | NA | NA | NA | 1 | 1 | 2 | 0.5 | 3 | 0.6667 | NA | NA | NA | NA | NA | NA | 2 | 1 | 2 | 1 |
| 16_918760 | NA | NA | NA | NA | NA | NA | 2 | 1 | 2 | 1 | NA | NA | NA | NA | NA | NA | 2 | 1 | 2 | 1 |
| 16_918761 | NA | NA | NA | NA | 1 | 1 | 2 | 1 | 3 | 1 | NA | NA | NA | NA | NA | NA | 2 | 0.5 | 2 | 0.5 |
| 16_918762 | NA | NA | NA | NA | 1 | 1 | 2 | 1 | 3 | 1 | NA | NA | NA | NA | NA | NA | 2 | 1 | 2 | 1 |
| 16_918765 | NA | NA | NA | NA | 1 | 1 | 2 | 0 | 3 | 0.3333 | NA | NA | NA | NA | NA | NA | 2 | 0.5 | 2 | 0.5 |
| 16_918806 | NA | NA | 1 | 0 | 1 | 0 | 5 | 0.4 | 7 | 0.2857 | 1 | 0 | 1 | 0 | 1 | 1 | 6 | 0.3333 | 9 | 0.3333 |
| 16_918840 | 3 | 0.3333 | 1 | 1 | 2 | 0 | 11 | 0.0909 | 17 | 0.1765 | 1 | 0 | 7 | 0 | 4 | 0 | 8 | 0.125 | 20 | 0.05 |
| 16_919087 | 10 | 0.5 | 8 | 0.5 | 2 | 0.5 | 8 | 0 | 28 | 0.3571 | 7 | 0.2857 | 13 | 0.3077 | 8 | 0.375 | 6 | 0.6667 | 34 | 0.3824 |
| 16_919089 | 11 | 0.2727 | 7 | 0.1429 | 5 | 0.4 | 11 | 0.3636 | 34 | 0.2941 | 9 | 0.7778 | 14 | 0.2857 | 10 | 0.2 | 6 | 0.5 | 39 | 0.4103 |
| 16_919090 | 7 | 0.1429 | 8 | 0.125 | 4 | 0 | 10 | 0.1 | 29 | 0.1034 | 11 | 0.1818 | 13 | 0.0769 | 9 | 0 | 6 | 0.1667 | 39 | 0.1026 |

| | | | | | | | | | | | | | | | | | | | | |
|---|---|---|---|---|---|---|---|---|---|---|---|---|---|---|---|---|---|---|---|---|
| 16_921891 | 6 | 0.1667 | 10 | 0 | 5 | 0.2 | 7 | 0 | 28 | 0.0714 | 9 | 0.2222 | 7 | 0.1429 | 14 | 0.0714 | 7 | 0.2857 | 37 | 0.1622 |
| 16_921894 | 4 | 0 | 9 | 0.1111 | 6 | 0 | 5 | 0.4 | 24 | 0.125 | 9 | 0.2222 | 8 | 0.125 | 11 | 0.0909 | 8 | 0 | 36 | 0.1111 |
| 16_921896 | 4 | 0.25 | 7 | 0.2857 | 5 | 0.2 | 7 | 0.1429 | 23 | 0.2174 | 6 | 0.1667 | 7 | 0 | 13 | 0.2308 | 7 | 0 | 33 | 0.1212 |
| 16_921897 | 1 | 0 | 8 | 0.5 | 3 | 0.3333 | 5 | 0 | 17 | 0.2941 | 9 | 0.2222 | 8 | 0 | 11 | 0.3636 | 7 | 0.4286 | 35 | 0.2571 |
| 16_921900 | 4 | 0.25 | 6 | 0.3333 | 4 | 0.5 | 8 | 0.375 | 22 | 0.3636 | 6 | 0 | 9 | 0.1111 | 11 | 0.2727 | 6 | 0 | 32 | 0.125 |
| 16_921900 | 4 | 0.25 | 8 | 0.125 | 4 | 0.25 | 8 | 0.125 | 24 | 0.1667 | 7 | 0.1429 | 8 | 0.25 | 12 | 0.0833 | 9 | 0.3333 | 36 | 0.1944 |
| 16_921900 | 3 | 0.3333 | 10 | 0.5 | 3 | 0.3333 | 6 | 0.3333 | 22 | 0.4091 | 7 | 0.5714 | 9 | 0.6667 | 10 | 0.6 | 8 | 0.5 | 34 | 0.5882 |
| 16_921900 | 2 | 1 | 8 | 1 | 4 | 1 | 7 | 1 | 21 | 1 | 8 | 0.875 | 8 | 1 | 12 | 0.9167 | 8 | 1 | 36 | 0.9444 |
| 16_921901 | 2 | 1 | 8 | 1 | 4 | 1 | 7 | 1 | 21 | 1 | 7 | 1 | 8 | 1 | 12 | 0.8333 | 9 | 1 | 36 | 0.9444 |
| 16_921903 | 3 | 0.3333 | 7 | 0.5714 | 2 | 0 | 6 | 0.5 | 18 | 0.4444 | 8 | 0.5 | 8 | 0.375 | 7 | 0.4286 | 7 | 0.2857 | 30 | 0.4 |
| 16_921905 | 4 | 0 | 9 | 0 | 4 | 0.5 | 9 | 0.1111 | 26 | 0.1154 | 8 | 0.125 | 8 | 0 | 11 | 0.0909 | 6 | 0.1667 | 33 | 0.0909 |
| 16_923364 | 3 | 0.3333 | 1 | 0 | 1 | 0 | NA | NA | 5 | 0.2 | 1 | 0 | 5 | 0 | 2 | 0 | 1 | 1 | 9 | 0.1111 |
| 16_923364 | 4 | 0.25 | 1 | 0 | 2 | 0.5 | 1 | 0 | 8 | 0.25 | 1 | 0 | 5 | 0.2 | 3 | 0.3333 | 1 | 0 | 10 | 0.2 |
| 16_923365 | 6 | 0.3333 | 1 | 0 | 2 | 1 | 1 | 0 | 10 | 0.4 | 2 | 0.5 | 6 | 0.5 | 3 | 0.3333 | 1 | 1 | 12 | 0.5 |
| 16_923373 | 6 | 0.3333 | 2 | 0 | 2 | 0 | 1 | 0 | 11 | 0.1818 | 3 | 0 | 6 | 0.3333 | 4 | 0.25 | 2 | 0.5 | 15 | 0.2667 |
| 16_923373 | 6 | 0.3333 | 2 | 0 | 2 | 0 | 1 | 0 | 11 | 0.1818 | 3 | 0 | 8 | 0.375 | 4 | 0.25 | 2 | 0 | 17 | 0.2353 |
| 16_934069 | 17 | 0.0588 | 11 | 0.0909 | 9 | 0.1111 | 19 | 0.1053 | 56 | 0.0893 | 10 | 0.1 | 15 | 0.1333 | 9 | 0 | 8 | 0.125 | 42 | 0.0952 |
| 16_934069 | 15 | 0.3333 | 11 | 0.2727 | 12 | 0.1667 | 15 | 0.4667 | 53 | 0.3208 | 10 | 0.3 | 15 | 0.2 | 10 | 0.1 | 8 | 0.375 | 43 | 0.2326 |
| 16_934069 | 12 | 0 | 12 | 0 | 15 | 0.1333 | 17 | 0.1765 | 56 | 0.0893 | 10 | 0.2 | 14 | 0 | 10 | 0.1 | 7 | 0 | 41 | 0.0732 |
| 16_934069 | 15 | 0.4 | 13 | 0.0769 | 14 | 0.2857 | 19 | 0.0526 | 61 | 0.1967 | 10 | 0.1 | 16 | 0.1875 | 9 | 0.1111 | 9 | 0.1111 | 44 | 0.1364 |
| 16_934070 | 17 | 0.1765 | 12 | 0.25 | 15 | 0.0667 | 17 | 0.2941 | 61 | 0.1967 | 10 | 0.1 | 16 | 0.125 | 10 | 0.1 | 8 | 0.25 | 44 | 0.1364 |
| 16_934071 | 14 | 0.2857 | 12 | 0.0833 | 9 | 0.1111 | 15 | 0.3333 | 50 | 0.22 | 6 | 0.1667 | 16 | 0.0625 | 8 | 0.25 | 7 | 0 | 37 | 0.1081 |
| 16_934071 | 15 | 0.2 | 11 | 0.0909 | 10 | 0.1 | 16 | 0.1875 | 52 | 0.1538 | 9 | 0 | 15 | 0.0667 | 9 | 0.1111 | 6 | 0 | 39 | 0.0513 |
| 16_934071 | 15 | 0.7333 | 12 | 0.5 | 10 | 0.8 | 14 | 0.8571 | 51 | 0.7255 | 9 | 0.4444 | 15 | 0.6667 | 9 | 0.7778 | 6 | 0.6667 | 39 | 0.641 |
| 16_934804 | 10 | 0.3 | 5 | 0.2 | 2 | 0.5 | 12 | 0.0833 | 29 | 0.2069 | 3 | 0 | 9 | 0 | 6 | 0 | 5 | 0.4 | 23 | 0.087 |
| 16_934874 | 11 | 0.4545 | 5 | 0.4 | NA | NA | 14 | 0.6429 | 30 | 0.5333 | 3 | 0.6667 | 9 | 0.6667 | 10 | 0.3 | 5 | 0.6 | 27 | 0.5185 |
| 16_934875 | 12 | 0.1667 | 6 | 0 | 2 | 0.5 | 15 | 0.0667 | 35 | 0.1143 | 4 | 0 | 9 | 0.3333 | 11 | 0.0909 | 5 | 0.2 | 29 | 0.1724 |
| 16_934875 | 10 | 0.5 | 7 | 0.5714 | 2 | 1 | 11 | 0.2727 | 30 | 0.4667 | 3 | 0 | 9 | 0.1111 | 9 | 0.4444 | 5 | 0.4 | 26 | 0.2692 |
| 16_934876 | 11 | 0.0909 | 7 | 0 | 2 | 0 | 13 | 0.1538 | 33 | 0.0909 | 3 | 0 | 7 | 0.1429 | 10 | 0.1 | 6 | 0 | 26 | 0.0769 |
| 16_934876 | 11 | 0.4545 | 7 | 0.1429 | 2 | 0 | 16 | 0.1875 | 36 | 0.25 | 3 | 0 | 10 | 0.3 | 10 | 0.2 | 5 | 0.6 | 28 | 0.2857 |
| 16_934877 | 10 | 0.1 | 7 | 0 | 1 | 0 | 14 | 0.1429 | 32 | 0.0938 | 3 | 0 | 9 | 0.4444 | 9 | 0.2222 | 6 | 0 | 27 | 0.2222 |
| 16_934877 | 10 | 0.3 | 7 | 0.1429 | 1 | 0 | 9 | 0.3333 | 27 | 0.2593 | 3 | 0.6667 | 6 | 0.5 | 7 | 0.1429 | 5 | 0.4 | 21 | 0.381 |
| 16_935593 | NA | NA | 1 | 1 | 3 | 0.6667 | 2 | 1 | 6 | 0.8333 | 2 | 1 | NA | NA | NA | NA | NA | NA | 2 | 1 |
| 16_935595 | 1 | 1 | 1 | 1 | 2 | 0 | 2 | 0.5 | 6 | 0.5 | 2 | 1 | NA | NA | 1 | 1 | NA | NA | 3 | 1 |
| 16_935665 | 1 | 0 | 1 | 0 | 3 | 0 | 2 | 0.5 | 7 | 0.1429 | 2 | 0.5 | NA | NA | 1 | 0 | NA | NA | 3 | 0.3333 |
| 16_935666 | 1 | 1 | 1 | 0 | 3 | 0.3333 | 2 | 0 | 7 | 0.2857 | 1 | 1 | NA | NA | 1 | 0 | NA | NA | 2 | 0.5 |
| 16_935667 | 1 | 0 | 1 | 0 | 5 | 0.6 | 2 | 0.5 | 9 | 0.4444 | 2 | 0.5 | NA | NA | 1 | 0 | NA | NA | 3 | 0.3333 |
| 16_935671 | 1 | 1 | NA | NA | 2 | 0.5 | 2 | 1 | 5 | 0.8 | 2 | 1 | NA | NA | 1 | 1 | 1 | 0 | 4 | 0.75 |
| 16_935672 | 2 | 0 | 1 | 0 | 5 | 0.4 | 3 | 0.3333 | 11 | 0.2727 | 2 | 0 | NA | NA | 1 | 0 | 1 | 1 | 4 | 0.25 |
| 16_935674 | 1 | 0 | NA | NA | 3 | 0.3333 | 2 | 0.5 | 6 | 0.3333 | 2 | 0 | NA | NA | 1 | 0 | 1 | 1 | 4 | 0.25 |
| 16_935675 | 2 | 0 | 1 | 1 | 5 | 0.6 | 3 | 0 | 11 | 0.3636 | 2 | 0 | NA | NA | 1 | 1 | 1 | 0 | 4 | 0.25 |
| 16_935678 | 2 | 0.5 | 1 | 1 | 5 | 0.8 | 2 | 1 | 10 | 0.8 | 2 | 0.5 | NA | NA | 1 | 1 | 3 | 0 | 6 | 0.3333 |
| 16_936070 | 4 | 0.25 | 7 | 0.2857 | 12 | 0.25 | 5 | 0.2 | 28 | 0.25 | 3 | 0 | 3 | 0.6667 | 8 | 0.375 | 4 | 0 | 18 | 0.2778 |
| 16_936072 | 5 | 0.6 | 8 | 0.25 | 14 | 0.5 | 8 | 0.375 | 35 | 0.4286 | 5 | 0.8 | 4 | 1 | 8 | 0.375 | 6 | 0.5 | 23 | 0.6087 |
| 16_936072 | 4 | 0.25 | 8 | 0 | 13 | 0 | 5 | 0.2 | 30 | 0.0667 | 4 | 0 | 4 | 0 | 7 | 0.1429 | 5 | 0 | 20 | 0.05 |
| 16_936073 | 3 | 0 | 8 | 0.5 | 14 | 0.2143 | 5 | 0.4 | 30 | 0.3 | 5 | 0.6 | 3 | 0.6667 | 7 | 0.2857 | 5 | 0.6 | 20 | 0.5 |
| 16_936672 | 2 | 0.5 | 1 | 0 | NA | NA | 3 | 0 | 6 | 0.1667 | 1 | 0 | 1 | 0 | 1 | 1 | 2 | 0 | 5 | 0.2 |
| 16_936677 | 1 | 1 | 1 | 1 | NA | NA | 2 | 1 | 4 | 1 | 1 | 1 | 1 | 1 | 1 | 1 | 2 | 1 | 5 | 1 |
| 16_936677 | 1 | 0 | 1 | 1 | NA | NA | 3 | 0.6667 | 5 | 0.6 | 2 | 1 | 1 | 1 | 1 | 1 | 2 | 1 | 6 | 1 |
| 16_936678 | 2 | 1 | 1 | 1 | NA | NA | 3 | 0.6667 | 6 | 0.8333 | 2 | 0.5 | 1 | 1 | 1 | 0 | 2 | 1 | 6 | 0.6667 |
| 16_936678 | 2 | 0 | 1 | 1 | NA | NA | 4 | 0 | 7 | 0.1429 | 1 | 0 | 1 | 0 | 1 | 1 | 2 | 0 | 5 | 0.2 |
| 16_936679 | 1 | 0 | 1 | 1 | NA | NA | 4 | 0.25 | 6 | 0.3333 | 2 | 0.5 | 1 | 0 | NA | NA | 2 | 0.5 | 5 | 0.4 |
| 16_936718 | 2 | 1 | 3 | 0 | 1 | 0 | 5 | 1 | 11 | 0.6364 | 2 | 1 | 1 | 0 | 3 | 0.3333 | 2 | 0 | 8 | 0.375 |
| 16_936719 | 2 | 0 | 3 | 0 | 1 | 0 | 4 | 0.25 | 10 | 0.1 | 2 | 0 | 1 | 0 | 3 | 0.3333 | 2 | 0.5 | 8 | 0.25 |
| 16_936719 | NA | NA | 3 | 0.6667 | 1 | 1 | 4 | 1 | 8 | 0.875 | 1 | 1 | 1 | 0 | 1 | 1 | 1 | 1 | 4 | 0.75 |
| 16_936720 | 1 | 0 | 3 | 0 | 1 | 1 | 5 | 0.4 | 10 | 0.3 | 2 | 0.5 | 1 | 0 | 3 | 0.3333 | 2 | 0 | 8 | 0.25 |
| 16_939304 | 1 | 1 | 2 | 1 | NA | NA | 1 | 1 | 4 | 1 | 2 | 1 | 1 | 1 | NA | NA | NA | NA | 3 | 1 |
| 16_939305 | 1 | 1 | 1 | 1 | NA | NA | 1 | 1 | 3 | 1 | 2 | 1 | 1 | 0 | NA | NA | NA | NA | 3 | 0.6667 |
| 16_939309 | 1 | 0 | 2 | 1 | NA | NA | 1 | 0 | 4 | 0.5 | 2 | 0.5 | 1 | 0 | NA | NA | NA | NA | 3 | 0.3333 |
| 16_939317 | 1 | 1 | 1 | 0 | 1 | 0 | 1 | 1 | 4 | 0.5 | 1 | 1 | 1 | 0 | NA | NA | NA | NA | 2 | 0.5 |
| 16_939320 | 1 | 0 | 1 | 1 | 1 | 1 | 1 | 1 | 4 | 0.75 | 1 | 0 | 1 | 1 | NA | NA | NA | NA | 2 | 0.5 |
| 16_940683 | 1 | 1 | 1 | 1 | NA | NA | NA | NA | 2 | 1 | 1 | 1 | NA | NA | NA | NA | NA | NA | 1 | 1 |
| 16_940685 | 1 | 1 | 1 | 1 | NA | NA | NA | NA | 2 | 1 | 1 | 1 | NA | NA | NA | NA | NA | NA | 1 | 1 |
| 16_941655 | 11 | 0.0909 | 9 | 0.1111 | 14 | 0.4286 | 8 | 0.375 | 42 | 0.2619 | 1 | 0 | 8 | 0.375 | 8 | 0.5 | 7 | 0.1429 | 24 | 0.3333 |
| 16_942304 | 8 | 0.625 | 24 | 0.625 | 15 | 0.5333 | 18 | 0.3333 | 65 | 0.5231 | 8 | 0.5 | 12 | 0.5833 | 16 | 0.5 | 18 | 0.5 | 54 | 0.5185 |

| | | | | | | | | | | | | | | | | | | | | |
|---|---|---|---|---|---|---|---|---|---|---|---|---|---|---|---|---|---|---|---|---|
| 16_942435 | 12 | 0 | 25 | 0.08 | 20 | 0.15 | 27 | 0.2593 | 84 | 0.1429 | 10 | 0.2 | 24 | 0.0833 | 23 | 0.087 | 25 | 0.04 | 82 | 0.0854 |
| 16_942436 | 14 | 0.0714 | 29 | 0.0345 | 21 | 0.1429 | 30 | 0.0333 | 94 | 0.0638 | 13 | 0 | 24 | 0.0833 | 23 | 0.087 | 27 | 0.1111 | 87 | 0.0805 |
| 16_942437 | 12 | 0.9167 | 27 | 0.963 | 23 | 0.9565 | 26 | 1 | 88 | 0.9659 | 12 | 0.9167 | 24 | 1 | 26 | 1 | 27 | 1 | 89 | 0.9888 |
| 16_942437 | 13 | 0 | 25 | 0.28 | 22 | 0.0909 | 25 | 0.08 | 85 | 0.1294 | 10 | 0 | 19 | 0.1053 | 23 | 0.1304 | 25 | 0.2 | 77 | 0.1299 |
| 16_942437 | 15 | 0.1333 | 28 | 0.0714 | 22 | 0.0909 | 29 | 0.1379 | 94 | 0.1064 | 12 | 0.25 | 25 | 0.08 | 26 | 0.1538 | 31 | 0.2258 | 94 | 0.1702 |
| 16_942607 | 22 | 0.4091 | 42 | 0.5476 | 38 | 0.4474 | 37 | 0.3784 | 139 | 0.4532 | 26 | 0.3462 | 25 | 0.32 | 40 | 0.4 | 46 | 0.413 | 137 | 0.3796 |
| 16_942607 | 24 | 0.0833 | 42 | 0.0714 | 43 | 0.0233 | 36 | 0.0833 | 145 | 0.0621 | 21 | 0.0476 | 35 | 0.1714 | 36 | 0.0278 | 44 | 0.0227 | 136 | 0.0662 |
| 16_942691 | 39 | 0.2821 | 64 | 0.125 | 59 | 0.1356 | 57 | 0.193 | 219 | 0.1735 | 42 | 0.2857 | 49 | 0.2041 | 53 | 0.0943 | 59 | 0.1695 | 203 | 0.1823 |
| 16_942694 | 43 | 0.186 | 59 | 0.1356 | 59 | 0.1525 | 59 | 0.2034 | 220 | 0.1682 | 47 | 0.234 | 43 | 0.2326 | 52 | 0.3269 | 58 | 0.2414 | 200 | 0.26 |
| 16_942898 | NA | NA | NA | NA | 2 | 1 | NA | NA | 2 | 1 | 1 | 1 | NA | NA | NA | NA | NA | NA | 1 | 1 |
| 16_942900 | NA | NA | NA | NA | 2 | 0.5 | NA | NA | 2 | 0.5 | 1 | 1 | NA | NA | NA | NA | NA | NA | 1 | 1 |
| 16_942901 | NA | NA | NA | NA | 2 | 1 | NA | NA | 2 | 1 | 1 | 1 | NA | NA | NA | NA | NA | NA | 1 | 1 |
| 16_942988 | 3 | 0 | 4 | 0.25 | 5 | 0 | 4 | 0 | 16 | 0.0625 | 2 | 0.5 | 3 | 0 | 2 | 0.5 | 2 | 0 | 9 | 0.2222 |
| 16_942988 | 3 | 0 | 4 | 0 | 5 | 0.2 | 3 | 0 | 15 | 0.0667 | 2 | 0 | 3 | 0 | 2 | 0.5 | 2 | 0 | 9 | 0.1111 |
| 16_942988 | 2 | 0 | 2 | 0.5 | 5 | 0.2 | 2 | 1 | 11 | 0.3636 | NA | NA | 2 | 0.5 | 2 | 0 | 3 | 0 | 7 | 0.1429 |
| 16_942991 | 2 | 0.5 | 3 | 0.6667 | 5 | 0.4 | 4 | 0.5 | 14 | 0.5 | 2 | 1 | 4 | 0.25 | 2 | 1 | 2 | 0.5 | 10 | 0.6 |
| 16_942992 | 3 | 0.3333 | 3 | 0.6667 | 5 | 0 | 3 | 0.3333 | 14 | 0.2857 | 2 | 0 | 4 | 0.5 | 2 | 0.5 | 3 | 0 | 11 | 0.2727 |
| 16_942992 | 2 | 0.5 | 3 | 1 | 2 | 0 | 3 | 0.6667 | 10 | 0.6 | 1 | 1 | 3 | 1 | 2 | 0.5 | 1 | 1 | 7 | 0.8571 |
| 16_942993 | 3 | 0 | 2 | 0 | 5 | 0.4 | 4 | 0 | 14 | 0.1429 | 2 | 0 | 5 | 0.2 | 3 | 0 | 1 | 0 | 11 | 0.0909 |
| 16_942994 | 2 | 1 | 3 | 1 | 4 | 0.75 | 2 | 0 | 11 | 0.7273 | 1 | 1 | 4 | 1 | 2 | 1 | 2 | 1 | 9 | 1 |
| 16_942994 | 3 | 0.6667 | 3 | 0.6667 | 4 | 0.5 | 3 | 1 | 13 | 0.6923 | 2 | 1 | 5 | 0.4 | 2 | 0 | 2 | 1 | 11 | 0.5455 |
| 16_942995 | 2 | 0 | 3 | 0.3333 | 4 | 0.5 | 3 | 0.6667 | 12 | 0.4167 | 1 | 1 | 4 | 0.5 | 1 | 1 | 2 | 0.5 | 8 | 0.625 |
| 16_942998 | 3 | 0.3333 | 4 | 0 | 4 | 0 | 4 | 0 | 15 | 0.0667 | 2 | 0 | 1 | 0 | 3 | 0.3333 | 1 | 0 | 7 | 0.1429 |
| 16_942998 | 3 | 1 | 3 | 1 | 5 | 1 | 4 | 0.75 | 15 | 0.9333 | 2 | 1 | 2 | 1 | 3 | 1 | 1 | 1 | 8 | 1 |
| 16_942998 | 2 | 0.5 | 4 | 0 | 4 | 0.5 | 4 | 0 | 14 | 0.2143 | 1 | 1 | 2 | 1 | 3 | 1 | NA | NA | 6 | 1 |
| 16_942998 | 3 | 0 | 4 | 0.25 | 4 | 0.25 | 4 | 0.75 | 15 | 0.3333 | 2 | 0 | 1 | 1 | 3 | 0.3333 | 1 | 1 | 7 | 0.4286 |
| 16_943280 | 1 | 0 | 1 | 1 | 1 | 1 | NA | NA | 3 | 0.6667 | NA | NA | 1 | 1 | 1 | 0 | 1 | 0 | 3 | 0.3333 |
| 16_943281 | 1 | 1 | 1 | 0 | 1 | 0 | NA | NA | 3 | 0.3333 | NA | NA | 1 | 0 | 1 | 1 | 1 | 0 | 3 | 0.3333 |
| 16_943711 | 1 | 0 | NA | NA | 2 | 1 | NA | NA | 3 | 0.6667 | NA | NA | 2 | 0.5 | NA | NA | 2 | 0 | 4 | 0.25 |
| 16_944926 | 3 | 1 | NA | NA | 3 | 1 | 2 | 0.5 | 8 | 0.875 | 3 | 1 | 2 | 1 | 3 | 0.6667 | 5 | 0.4 | 13 | 0.6923 |
| 16_944928 | 3 | 0.3333 | NA | NA | 5 | 0 | 2 | 0 | 10 | 0.1 | 3 | 0 | 4 | 0 | 4 | 0 | 7 | 0.1429 | 18 | 0.0556 |
| 16_944928 | 3 | 1 | NA | NA | 3 | 1 | 2 | 1 | 8 | 1 | 2 | 1 | 3 | 1 | 4 | 1 | 5 | 1 | 14 | 1 |
| 16_944928 | 4 | 1 | 1 | 1 | 3 | 1 | 2 | 1 | 10 | 1 | 2 | 0.5 | 3 | 1 | 4 | 0.5 | 3 | 1 | 12 | 0.75 |
| 16_944929 | 5 | 1 | 1 | 1 | 4 | 1 | 3 | 1 | 13 | 1 | 5 | 1 | 4 | 1 | 5 | 1 | 7 | 1 | 21 | 1 |
| 16_944929 | 4 | 1 | 1 | 1 | 4 | 0.75 | 2 | 1 | 11 | 0.9091 | 5 | 0.8 | 4 | 0.5 | 5 | 1 | 7 | 0.7143 | 21 | 0.7619 |
| 16_944929 | 3 | 0.3333 | 1 | 0 | 4 | 0.25 | 3 | 0.3333 | 11 | 0.2727 | 3 | 1 | 2 | 0 | 4 | 0.25 | 7 | 0.4286 | 16 | 0.4375 |
| 16_944929 | 5 | 0.8 | 1 | 1 | 3 | 1 | 3 | 1 | 12 | 0.9167 | 5 | 1 | 3 | 0.6667 | 5 | 1 | 6 | 0.8333 | 19 | 0.8947 |
| 16_944929 | 5 | 0.2 | 1 | 1 | 4 | 0.25 | 3 | 0 | 13 | 0.2308 | 5 | 0.2 | 3 | 0 | 5 | 0 | 6 | 0.5 | 19 | 0.2105 |
| 16_944930 | 4 | 0.5 | 1 | 0 | 3 | 0.3333 | 3 | 0.6667 | 11 | 0.4545 | 5 | 0.6 | 3 | 1 | 5 | 0.4 | 6 | 0.1667 | 19 | 0.4737 |
| 16_944930 | 3 | 0.3333 | 1 | 0 | 3 | 0.3333 | 2 | 0 | 9 | 0.2222 | 5 | 0.2 | 4 | 0.25 | 4 | 0.5 | 7 | 0.7143 | 20 | 0.45 |
| 16_944931 | 3 | 0.3333 | 1 | 0 | 4 | 0.25 | 2 | 0 | 10 | 0.2 | 4 | 0 | 3 | 0 | 5 | 0 | 5 | 0.2 | 17 | 0.0588 |
| 16_944931 | 5 | 0.6 | 2 | 1 | 4 | 0.75 | 2 | 0.5 | 13 | 0.6923 | 4 | 0.5 | 4 | 0.5 | 5 | 0.4 | 6 | 0.8333 | 19 | 0.5789 |
| 16_944932 | 6 | 1 | 2 | 1 | 4 | 1 | 3 | 1 | 15 | 1 | 4 | 1 | 4 | 1 | 5 | 1 | 8 | 0.75 | 21 | 0.9048 |
| 16_944936 | 4 | 0.25 | 2 | 0 | 3 | 0.3333 | 1 | 1 | 10 | 0.3 | 5 | 0 | 8 | 0 | 4 | 0 | 10 | 0.4 | 27 | 0.1481 |
| 16_944936 | 6 | 0.6667 | 3 | 0.6667 | 3 | 1 | 2 | 1 | 14 | 0.7857 | 4 | 1 | 8 | 0.875 | 4 | 1 | 10 | 0.9 | 26 | 0.9231 |
| 16_944937 | 6 | 0.3333 | 3 | 0 | 5 | 0.4 | 2 | 1 | 16 | 0.375 | 2 | 0.5 | 8 | 0.375 | 3 | 0.3333 | 10 | 0.5 | 23 | 0.4348 |
| 16_944939 | 6 | 1 | 3 | 1 | 4 | 0.75 | 3 | 1 | 16 | 0.9375 | 3 | 0.6667 | 7 | 1 | 2 | 1 | 9 | 0.8889 | 21 | 0.9048 |
| 16_944939 | 6 | 0.6667 | 3 | 0.6667 | 5 | 1 | 3 | 0.6667 | 17 | 0.7647 | 5 | 0.6 | 7 | 1 | 4 | 0.5 | 9 | 0.7778 | 25 | 0.76 |
| 16_944939 | 7 | 0 | 3 | 0 | 3 | 0.3333 | 3 | 0.6667 | 16 | 0.1875 | 3 | 0 | 8 | 0.125 | 4 | 0 | 9 | 0.4444 | 24 | 0.2083 |
| 16_944940 | 5 | 1 | 4 | 1 | 4 | 0.75 | 3 | 1 | 16 | 0.9375 | 4 | 0.5 | 9 | 1 | 4 | 1 | 8 | 0.875 | 25 | 0.88 |
| 16_944940 | 7 | 0.1429 | 4 | 0 | 6 | 0 | 3 | 0 | 20 | 0.05 | 5 | 0 | 9 | 0.2222 | 4 | 0 | 8 | 0.25 | 26 | 0.1538 |
| 16_944942 | 3 | 0 | 4 | 0 | 4 | 0.25 | 3 | 0.3333 | 14 | 0.1429 | 4 | 0 | 7 | 0 | 3 | 0.3333 | 4 | 0 | 18 | 0.0556 |
| 16_944942 | 4 | 0.5 | 5 | 0 | 6 | 0.3333 | 3 | 0.6667 | 18 | 0.3333 | 5 | 0.2 | 7 | 0 | 5 | 0 | 9 | 0.4444 | 26 | 0.1923 |
| 16_944944 | 7 | 0.1429 | 6 | 0 | 6 | 0.1667 | 2 | 0.5 | 21 | 0.1429 | 3 | 0.6667 | 10 | 0.6 | 5 | 0.8 | 7 | 0.2857 | 25 | 0.56 |
| 16_944945 | 7 | 0 | 6 | 0 | 4 | 0.25 | 2 | 0 | 19 | 0.0526 | 5 | 0.2 | 8 | 0 | 5 | 0.2 | 10 | 0 | 28 | 0.0714 |
| 16_944955 | 8 | 0.125 | 11 | 0.0909 | 9 | 0.2222 | 3 | 0.3333 | 31 | 0.1613 | 10 | 0.1 | 12 | 0 | 8 | 0 | 14 | 0.2857 | 44 | 0.1136 |
| 16_944958 | 5 | 0.2 | 5 | 0 | NA | NA | NA | NA | 10 | 0.1 | 6 | 0.1667 | 4 | 0.25 | 1 | 0 | 4 | 0 | 15 | 0.1333 |
| 16_944958 | 5 | 0.2 | 5 | 0.2 | NA | NA | NA | NA | 10 | 0.2 | 6 | 0 | 4 | 0.25 | 1 | 0 | 4 | 0 | 15 | 0.0667 |
| 16_945464 | 6 | 0.1667 | 5 | 0 | 2 | 0 | NA | NA | 13 | 0.0769 | 2 | 0.5 | 2 | 0 | 2 | 0 | 5 | 0 | 11 | 0.0909 |
| 16_945496 | NA | NA | 3 | 0 | 1 | 1 | NA | NA | 4 | 0.25 | 2 | 0 | NA | NA | 1 | 1 | NA | NA | 3 | 0.3333 |
| 16_945496 | NA | NA | 4 | 0.5 | 1 | 1 | NA | NA | 5 | 0.6 | 2 | 0.5 | NA | NA | 1 | 1 | NA | NA | 3 | 0.6667 |
| 16_945497 | NA | NA | 4 | 0.5 | 1 | 0 | NA | NA | 5 | 0.4 | 2 | 0 | NA | NA | 1 | 1 | NA | NA | 3 | 0.3333 |
| 16_945497 | 1 | 0 | 4 | 0.75 | 1 | 0 | NA | NA | 6 | 0.5 | 2 | 0.5 | NA | NA | 1 | 0 | NA | NA | 3 | 0.3333 |
| 16_945497 | 1 | 0 | 3 | 0.3333 | 1 | 0 | NA | NA | 5 | 0.2 | 2 | 0.5 | NA | NA | 2 | 0.5 | NA | NA | 4 | 0.5 |
| 16_945498 | 1 | 0 | 3 | 1 | 1 | 0 | NA | NA | 5 | 0.6 | 2 | 0 | NA | NA | 2 | 0.5 | NA | NA | 4 | 0.25 |

| | | | | | | | | | | | | | | | | | | | | |
|---|---|---|---|---|---|---|---|---|---|---|---|---|---|---|---|---|---|---|---|---|
| 16_945498 | NA | NA | 2 | 1 | 1 | 1 | NA | NA | 3 | 1 | 2 | 0.5 | NA | NA | 2 | 1 | NA | NA | 4 | 0.75 |
| 16_945499 | NA | NA | 3 | 0.6667 | NA | NA | NA | NA | 3 | 0.6667 | 2 | 0.5 | NA | NA | 1 | 0 | NA | NA | 3 | 0.3333 |
| 16_945500 | NA | NA | 3 | 1 | NA | NA | NA | NA | 3 | 1 | 2 | 0.5 | NA | NA | 1 | 1 | NA | NA | 3 | 0.6667 |
| 16_945500 | NA | NA | 3 | 1 | NA | NA | NA | NA | 3 | 1 | 2 | 0 | NA | NA | 1 | 1 | NA | NA | 3 | 0.3333 |
| 16_946334 | NA | NA | 2 | 1 | NA | NA | NA | NA | 2 | 1 | NA | NA | NA | NA | 1 | 1 | NA | NA | 1 | 1 |
| 16_946336 | NA | NA | 2 | 1 | NA | NA | NA | NA | 2 | 1 | NA | NA | NA | NA | 1 | 1 | NA | NA | 1 | 1 |
| 16_946337 | NA | NA | 2 | 1 | NA | NA | NA | NA | 2 | 1 | NA | NA | NA | NA | 1 | 1 | NA | NA | 1 | 1 |
| 16_955172 | 10 | 0.2 | 8 | 0 | 11 | 0 | 9 | 0 | 38 | 0.0526 | 3 | 0 | 11 | 0 | 11 | 0.0909 | 6 | 0.3333 | 31 | 0.0968 |
| 16_955200 | 9 | 0.2222 | 8 | 0.375 | 14 | 0 | 9 | 0 | 40 | 0.125 | 3 | 0 | 15 | 0.1333 | 14 | 0.1429 | 9 | 0.1111 | 41 | 0.122 |
| 16_955201 | 11 | 0.1818 | 8 | 0.25 | 14 | 0.5714 | 9 | 0.2222 | 42 | 0.3333 | 3 | 0.6667 | 14 | 0.1429 | 11 | 0.2727 | 9 | 0.2222 | 37 | 0.2432 |
| 16_955201 | 9 | 0.8889 | 6 | 0.3333 | 14 | 0.5714 | 8 | 0.75 | 37 | 0.6486 | 4 | 0.75 | 13 | 0.7692 | 11 | 0.7273 | 9 | 0.8889 | 37 | 0.7838 |
| 16_955201 | 9 | 1 | 6 | 0.6667 | 14 | 0.7143 | 7 | 0.8571 | 36 | 0.8056 | 2 | 1 | 14 | 0.8571 | 12 | 0.8333 | 6 | 0.8333 | 34 | 0.8529 |
| 16_955201 | 9 | 0.1111 | 10 | 0.1 | 15 | 0.0667 | 9 | 0.1111 | 43 | 0.093 | 4 | 0.25 | 14 | 0.0714 | 14 | 0.1429 | 9 | 0 | 41 | 0.0976 |
| 16_955202 | 10 | 1 | 9 | 1 | 14 | 1 | 10 | 1 | 43 | 1 | 4 | 1 | 14 | 1 | 12 | 0.9167 | 8 | 1 | 38 | 0.9737 |
| 16_955202 | 10 | 0.9 | 7 | 0.5714 | 16 | 0.875 | 5 | 0.6 | 38 | 0.7895 | 2 | 1 | 8 | 1 | 8 | 0.625 | 7 | 0.5714 | 25 | 0.76 |
| 16_955204 | 11 | 1 | 8 | 0.875 | 16 | 0.9375 | 10 | 0.9 | 45 | 0.9333 | 2 | 1 | 17 | 0.9412 | 11 | 1 | 7 | 1 | 37 | 0.973 |
| 16_955205 | 11 | 0.9091 | 10 | 0.6 | 15 | 0.8 | 11 | 0.9091 | 47 | 0.8085 | 2 | 0.5 | 16 | 0.8125 | 11 | 0.7273 | 7 | 1 | 36 | 0.8056 |
| 16_955209 | 12 | 0.4167 | 12 | 0.25 | 14 | 0.3571 | 11 | 0.2727 | 49 | 0.3265 | 5 | 0.2 | 16 | 0.375 | 15 | 0.3333 | 9 | 0.7778 | 45 | 0.4222 |
| 16_955210 | 9 | 0.7778 | 9 | 0.5556 | 14 | 0.7857 | 11 | 0.7273 | 43 | 0.7209 | 1 | 0 | 13 | 0.5385 | 11 | 0.7273 | 9 | 0.7778 | 34 | 0.6471 |
| 16_955215 | 15 | 0.2667 | 19 | 0.0526 | 17 | 0 | 14 | 0 | 65 | 0.0769 | 7 | 0 | 20 | 0.15 | 18 | 0.1111 | 12 | 0.1667 | 57 | 0.1228 |
| 16_955215 | 15 | 0.2 | 17 | 0.2353 | 14 | 0.0714 | 15 | 0.2667 | 61 | 0.1967 | 5 | 0.2 | 17 | 0 | 14 | 0.3571 | 11 | 0.1818 | 47 | 0.1702 |
| 16_955216 | 11 | 0.4545 | 18 | 0.3889 | 16 | 0.375 | 13 | 0.4615 | 58 | 0.4138 | 7 | 0 | 20 | 0.45 | 16 | 0.5625 | 12 | 0.5 | 55 | 0.4364 |
| 16_955216 | 17 | 0.4706 | 20 | 0.5 | 15 | 0.2667 | 16 | 0.25 | 68 | 0.3824 | 8 | 0.25 | 22 | 0.4545 | 19 | 0.5263 | 14 | 0.2143 | 63 | 0.3968 |
| 16_955217 | 18 | 0.1111 | 19 | 0.1579 | 16 | 0 | 17 | 0.2353 | 70 | 0.1286 | 8 | 0.25 | 18 | 0.0556 | 17 | 0.2353 | 12 | 0 | 55 | 0.1273 |
| 16_955217 | 19 | 0.4737 | 20 | 0.5 | 18 | 0.4444 | 17 | 0.1176 | 74 | 0.3919 | 10 | 0.3 | 19 | 0.4211 | 22 | 0.3182 | 13 | 0.6154 | 64 | 0.4062 |
| 16_955218 | 20 | 0.7 | 19 | 0.6842 | 18 | 0.5 | 15 | 0.5333 | 72 | 0.6111 | 9 | 0.5556 | 22 | 0.5909 | 20 | 0.5 | 14 | 0.7143 | 65 | 0.5846 |
| 16_955218 | 16 | 0.125 | 20 | 0.55 | 17 | 0.4706 | 18 | 0.4444 | 71 | 0.4085 | 9 | 0.4444 | 14 | 0.2857 | 18 | 0.5556 | 11 | 0.5455 | 52 | 0.4615 |
| 16_955218 | 18 | 0.0556 | 20 | 0.05 | 19 | 0.1579 | 17 | 0 | 74 | 0.0676 | 11 | 0.0909 | 19 | 0.0526 | 19 | 0 | 10 | 0.2 | 59 | 0.0678 |
| 16_955218 | 17 | 0.9412 | 20 | 0.95 | 18 | 0.7778 | 19 | 0.7895 | 74 | 0.8649 | 11 | 0.9091 | 18 | 0.7222 | 17 | 0.7647 | 7 | 1 | 53 | 0.8113 |
| 16_955219 | 18 | 0 | 20 | 0.1 | 22 | 0.0455 | 21 | 0.0952 | 81 | 0.0617 | 9 | 0 | 22 | 0.1364 | 22 | 0.0455 | 11 | 0.1818 | 64 | 0.0938 |
| 16_957939 | 4 | 0.75 | 4 | 0.25 | 4 | 0.75 | 4 | 0.75 | 16 | 0.625 | 1 | 0 | 6 | 0.6667 | 5 | 0.8 | 6 | 0.5 | 18 | 0.6111 |
| 16_957943 | 3 | 0 | 4 | 0 | 4 | 0 | 4 | 0.25 | 15 | 0.0667 | 1 | 1 | 5 | 0 | 5 | 0.4 | 5 | 0.2 | 16 | 0.25 |
| 16_957943 | 3 | 0 | 3 | 0 | 3 | 0.3333 | 4 | 0 | 13 | 0.0769 | 1 | 0 | 5 | 0 | 5 | 0 | 6 | 0.1667 | 17 | 0.0588 |
| 16_957944 | 3 | 0.3333 | 4 | 0.25 | 3 | 0.3333 | 4 | 0 | 14 | 0.2143 | 1 | 0 | 5 | 0.2 | 5 | 0 | 6 | 0 | 17 | 0.0588 |
| 16_957947 | 2 | 0 | 4 | 0.25 | 2 | 0 | 2 | 0 | 10 | 0.1 | 1 | 0 | 5 | 0.2 | 3 | 0.3333 | 4 | 0 | 13 | 0.1538 |
| 16_958035 | 2 | 1 | NA | NA | 1 | 1 | 2 | 1 | 5 | 1 | NA | NA | 1 | 1 | NA | NA | NA | NA | 1 | 1 |
| 16_958037 | 2 | 1 | 2 | 1 | 1 | 1 | 2 | 0.5 | 7 | 0.8571 | 1 | 1 | 2 | 0.5 | 1 | 1 | 3 | 1 | 7 | 0.8571 |
| 16_958037 | 3 | 0.3333 | 2 | 0.5 | 1 | 1 | 3 | 1 | 9 | 0.6667 | 1 | 1 | 2 | 1 | 1 | 0 | 3 | 0.6667 | 7 | 0.7143 |
| 16_958037 | 2 | 1 | 2 | 0.5 | 1 | 0 | 2 | 1 | 7 | 0.7143 | 1 | 0 | 2 | 0 | 2 | 1 | 3 | 0.6667 | 8 | 0.5 |
| 16_958037 | 2 | 0 | 2 | 0.5 | 1 | 0 | 3 | 0.3333 | 8 | 0.25 | 1 | 0 | 2 | 1 | 1 | 0 | 2 | 0.5 | 6 | 0.5 |
| 16_958038 | 2 | 1 | 2 | 1 | NA | NA | 2 | 0 | 6 | 0.6667 | NA | NA | 1 | 0 | 2 | 1 | 3 | 0 | 6 | 0.3333 |
| 16_958039 | 2 | 0.5 | 2 | 0.5 | 1 | 0 | 3 | 0.6667 | 8 | 0.5 | 1 | 1 | 1 | 0 | 1 | 1 | 3 | 0.6667 | 6 | 0.6667 |
| 16_958039 | 2 | 0.5 | 2 | 0 | 1 | 1 | 3 | 0.3333 | 8 | 0.375 | 1 | 1 | 1 | 1 | 1 | 0 | 3 | 1 | 6 | 0.8333 |
| 16_958040 | 1 | 0 | 1 | 0 | 1 | 0 | 2 | 1 | 5 | 0.4 | 1 | 0 | NA | NA | 1 | 1 | 3 | 0.3333 | 5 | 0.4 |
| 16_958041 | 2 | 0 | 1 | 0 | 1 | 0 | 3 | 0.3333 | 7 | 0.1429 | 1 | 0 | 1 | 1 | NA | NA | 3 | 0.6667 | 5 | 0.6 |
| 16_958042 | 2 | 0 | 2 | 0.5 | NA | NA | 2 | 0 | 6 | 0.1667 | 1 | 0 | NA | NA | 1 | 0 | 1 | 1 | 3 | 0.3333 |
| 16_958042 | 2 | 0 | 1 | 1 | 1 | 0 | 3 | 0 | 7 | 0.1429 | 1 | 0 | 1 | 0 | 1 | 0 | 3 | 0.3333 | 6 | 0.1667 |
| 16_958042 | 2 | 0.5 | 2 | 1 | 1 | 1 | 2 | 1 | 7 | 0.8571 | 1 | 0 | 1 | 1 | 1 | 1 | 3 | 1 | 6 | 0.8333 |
| 16_958052 | 2 | 0 | 1 | 0 | 1 | 1 | 3 | 0 | 7 | 0.1429 | 1 | 0 | 1 | 1 | 1 | 0 | 3 | 0.6667 | 6 | 0.5 |
| 16_958055 | 2 | 0 | 1 | 1 | 1 | 0 | 3 | 0 | 7 | 0.1429 | NA | NA | 1 | 0 | 1 | 0 | 2 | 0.5 | 4 | 0.25 |
| 16_958056 | 2 | 0.5 | 1 | 0 | 1 | 0 | 3 | 0 | 7 | 0.1429 | NA | NA | 1 | 0 | 2 | 0 | 2 | 0.5 | 5 | 0.2 |
| 16_958091 | 2 | 0 | 1 | 0 | NA | NA | 3 | 0.3333 | 6 | 0.1667 | NA | NA | 1 | 0 | 1 | 1 | 1 | 0 | 3 | 0.3333 |
| 16_958103 | 2 | 0.5 | 1 | 1 | NA | NA | 2 | 0 | 5 | 0.4 | NA | NA | 1 | 0 | 2 | 0.5 | 1 | 1 | 4 | 0.5 |
| 16_958682 | 6 | 0 | 9 | 0.2222 | 8 | 0 | 11 | 0 | 34 | 0.0588 | 3 | 0 | 12 | 0.1667 | 12 | 0.0833 | 5 | 0.2 | 32 | 0.125 |
| 16_958682 | 2 | 0 | 3 | 0.6667 | 6 | 0 | 3 | 0 | 14 | 0.1429 | 2 | 0 | 7 | 0.1429 | 9 | 0.3333 | 5 | 0 | 23 | 0.1739 |
| 16_958683 | 5 | 0 | 7 | 0.5714 | 7 | 0.1429 | 8 | 0.125 | 27 | 0.2222 | 3 | 0 | 10 | 0.2 | 11 | 0.1818 | 4 | 0.25 | 28 | 0.1786 |
| 16_958687 | 6 | 0 | 7 | 0.1429 | 8 | 0.125 | 9 | 0.1111 | 30 | 0.1 | 2 | 0 | 11 | 0.0909 | 11 | 0.1818 | 4 | 0 | 28 | 0.1071 |
| 16_958687 | 4 | 0 | 9 | 0.3333 | 8 | 0.125 | 10 | 0.2 | 31 | 0.1935 | 3 | 0 | 10 | 0.2 | 10 | 0 | 5 | 0 | 28 | 0.0714 |
| 16_958695 | 5 | 0.8 | 8 | 0.875 | 4 | 1 | 6 | 0.6667 | 23 | 0.8261 | 1 | 1 | 9 | 0.8889 | 9 | 1 | 3 | 1 | 22 | 0.9545 |
| 16_958696 | 4 | 0.25 | 5 | 0 | 1 | 0 | 3 | 0 | 13 | 0.0769 | 1 | 0 | 8 | 0 | 7 | 0.1429 | 2 | 0 | 18 | 0.0556 |
| 16_958836 | NA | NA | NA | NA | NA | NA | 1 | 1 | 1 | 1 | 1 | 1 | NA | NA | NA | NA | NA | NA | 1 | 1 |
| 16_958843 | NA | NA | NA | NA | NA | NA | 1 | 1 | 1 | 1 | 1 | 1 | NA | NA | NA | NA | NA | NA | 1 | 1 |
| 16_959572 | 10 | 0.9 | 9 | 1 | 5 | 0.8 | 7 | 1 | 31 | 0.9355 | 7 | 0.8571 | 11 | 0.8182 | 10 | 0.8 | 12 | 0.8333 | 40 | 0.825 |
| 16_959574 | 10 | 0 | 8 | 0.125 | 7 | 0.2857 | 11 | 0.0909 | 36 | 0.1111 | 8 | 0.375 | 14 | 0.2857 | 10 | 0.2 | 14 | 0.2857 | 46 | 0.2826 |
| 16_959586 | 13 | 0.1538 | 11 | 0.4545 | 12 | 0.25 | 14 | 0.1429 | 50 | 0.24 | 10 | 0.2 | 16 | 0.4375 | 14 | 0.0714 | 18 | 0.2778 | 58 | 0.2586 |

| | | | | | | | | | | | | | | | | | | | | |
|---|---|---|---|---|---|---|---|---|---|---|---|---|---|---|---|---|---|---|---|---|
| 16_9595893 | 15 | 0 | 8 | 0.125 | 15 | 0.2667 | 15 | 0 | 53 | 0.0943 | 10 | 0 | 15 | 0.1333 | 15 | 0 | 16 | 0.0625 | 56 | 0.0536 |
| 16_9595981 | NA | NA | 1 | 1 | NA | NA | NA | NA | 1 | 1 | NA | NA | 1 | 1 | NA | NA | NA | NA | 1 | 1 |
| 16_9595999 | NA | NA | 1 | 1 | NA | NA | NA | NA | 1 | 1 | NA | NA | 1 | 1 | NA | NA | NA | NA | 1 | 1 |
| 16_9611651 | 1 | 1 | 2 | 1 | 2 | 1 | 1 | 1 | 6 | 1 | NA | NA | 1 | 0 | 1 | 1 | 1 | 0 | 3 | 0.3333 |
| 16_9611673 | 1 | 0 | 1 | 0 | 2 | 1 | 1 | 1 | 5 | 0.6 | 1 | 1 | 3 | 1 | 3 | 1 | 1 | 1 | 8 | 1 |
| 16_9611680 | 1 | 0 | 2 | 0.5 | 2 | 0 | 1 | 0 | 6 | 0.1667 | 1 | 0 | 2 | 0 | 2 | 0.5 | 1 | 0 | 6 | 0.1667 |
| 16_9611682 | 1 | 1 | 2 | 1 | 2 | 1 | 1 | 1 | 6 | 1 | NA | NA | 4 | 1 | 3 | 1 | 1 | 1 | 8 | 1 |
| 16_9611686 | 1 | 1 | 2 | 1 | 2 | 1 | 1 | 1 | 6 | 1 | 1 | 1 | 4 | 1 | 2 | 1 | 1 | 1 | 8 | 1 |
| 16_9611698 | 1 | 1 | 2 | 1 | 2 | 1 | 1 | 1 | 6 | 1 | 1 | 1 | 5 | 1 | 3 | 1 | 1 | 1 | 10 | 1 |
| 16_9611715 | 1 | 1 | 2 | 0.5 | 3 | 1 | 1 | 1 | 7 | 0.8571 | 1 | 1 | 4 | 0.75 | 1 | 1 | 1 | 1 | 7 | 0.8571 |
| 16_9611723 | 1 | 1 | 2 | 1 | 3 | 0.6667 | 1 | 0 | 7 | 0.7143 | 1 | 1 | 5 | 1 | 3 | 0.3333 | 1 | 0 | 10 | 0.7 |
| 16_9611726 | NA | NA | 2 | 0.5 | 3 | 1 | 1 | 1 | 6 | 0.8333 | 1 | 1 | 5 | 0.8 | 3 | 0.3333 | 1 | 1 | 10 | 0.7 |
| 16_9611727 | 1 | 0 | 2 | 0.5 | 3 | 0.3333 | NA | NA | 6 | 0.3333 | NA | NA | 2 | 0 | 2 | 1 | 1 | 0 | 5 | 0.4 |
| 16_9611727 | 1 | 1 | 1 | 1 | 3 | 1 | 1 | 1 | 6 | 1 | NA | NA | 4 | 1 | 3 | 1 | 1 | 1 | 8 | 1 |
| 16_9611730 | 1 | 1 | 2 | 1 | 3 | 1 | NA | NA | 6 | 1 | 1 | 1 | 5 | 1 | 3 | 0.6667 | NA | NA | 9 | 0.8889 |
| 16_9611731 | 1 | 1 | 2 | 0.5 | 3 | 1 | 1 | 1 | 7 | 0.8571 | 1 | 1 | 5 | 0.6 | 3 | 0.6667 | 1 | 1 | 10 | 0.7 |
| 16_9611732 | 1 | 0 | 2 | 0 | 2 | 0 | 1 | 1 | 6 | 0.1667 | NA | NA | 4 | 0.25 | 3 | 0.3333 | 1 | 0 | 8 | 0.25 |
| 16_9611737 | 1 | 1 | 1 | 1 | 3 | 1 | 1 | 1 | 6 | 1 | 1 | 1 | 2 | 1 | 2 | 1 | 1 | 1 | 6 | 1 |
| 16_9611745 | 1 | 0 | 1 | 1 | 3 | 0.6667 | 1 | 1 | 6 | 0.6667 | 1 | 0 | 5 | 0.6 | 3 | 0.6667 | 1 | 1 | 10 | 0.6 |
| 16_9611753 | 1 | 0 | 2 | 0.5 | 3 | 0.3333 | 1 | 0 | 7 | 0.2857 | 1 | 1 | 5 | 0.2 | 3 | 0.6667 | 1 | 1 | 10 | 0.5 |
| 16_9611762 | 1 | 0 | 2 | 1 | 1 | 1 | NA | NA | 4 | 0.75 | 1 | 1 | 5 | 1 | 2 | 1 | 2 | 1 | 10 | 1 |
| 16_9611805 | 1 | 0 | 6 | 0 | 5 | 0.2 | 1 | 0 | 13 | 0.0769 | 1 | 0 | 7 | 0 | 3 | 0.3333 | 4 | 0.25 | 15 | 0.1333 |
| 16_9611812 | 1 | 0 | 5 | 0 | 5 | 0 | 1 | 1 | 12 | 0.0833 | 1 | 0 | 6 | 0 | 2 | 0 | 4 | 0.25 | 13 | 0.0769 |
| 16_9611854 | 2 | 1 | 8 | 1 | 5 | 0.2 | 1 | 1 | 16 | 0.75 | 1 | 1 | 9 | 0.6667 | 3 | 0.3333 | 4 | 0.75 | 17 | 0.6471 |
| 16_9611858 | 2 | 0.5 | 5 | 0.2 | 5 | 0 | 1 | 0 | 13 | 0.1538 | 1 | 0 | 4 | 0 | 3 | 0.3333 | 4 | 0.25 | 12 | 0.1667 |
| 16_9611875 | 2 | 1 | 8 | 0.125 | 5 | 0.4 | 1 | 1 | 16 | 0.375 | 2 | 1 | 8 | 0.375 | 3 | 0.3333 | 5 | 0 | 18 | 0.3333 |
| 16_9611882 | 1 | 1 | 9 | 1 | 5 | 0.6 | 1 | 1 | 16 | 0.875 | 2 | 1 | 9 | 1 | 3 | 1 | 6 | 1 | 20 | 1 |
| 16_9611905 | 2 | 0.5 | 9 | 0 | 6 | 0.1667 | 2 | 0 | 19 | 0.1053 | 2 | 0 | 12 | 0.0833 | 4 | 0.25 | 5 | 0 | 23 | 0.087 |
| 16_9611923 | 2 | 0 | 11 | 0.0909 | 7 | 0.1429 | 1 | 0 | 21 | 0.0952 | 2 | 0 | 10 | 0.1 | 4 | 0.25 | 6 | 0.1667 | 22 | 0.1364 |
| 16_9611924 | 2 | 0 | 10 | 0 | 7 | 0.2857 | 1 | 0 | 20 | 0.1 | 1 | 0 | 12 | 0.1667 | 3 | 0.3333 | 6 | 0.1667 | 22 | 0.1818 |
| 16_9612052 | 3 | 0.6667 | 16 | 0.625 | 8 | 0.625 | 4 | 0.25 | 31 | 0.5806 | 7 | 0.7143 | 14 | 0.2143 | 5 | 0.4 | 7 | 0.2857 | 33 | 0.3636 |
| 16_9612059 | 3 | 1 | 13 | 0.7692 | 8 | 0.75 | 4 | 0.75 | 28 | 0.7857 | 6 | 0.8333 | 10 | 0.9 | 3 | 1 | 8 | 0.875 | 27 | 0.8889 |
| 16_9612081 | 3 | 0.3333 | 17 | 0.5882 | 7 | 0.5714 | 5 | 0.2 | 32 | 0.5 | 8 | 0.625 | 15 | 0.4667 | 6 | 0.8333 | 8 | 0.625 | 37 | 0.5946 |
| 16_9612082 | 4 | 0.25 | 13 | 0.0769 | 7 | 0 | 5 | 0 | 29 | 0.069 | 5 | 0.2 | 12 | 0.1667 | 4 | 0 | 8 | 0.125 | 29 | 0.1379 |
| 16_9612083 | 4 | 0 | 16 | 0.0625 | 6 | 0.1667 | 5 | 0.2 | 31 | 0.0968 | 8 | 0.375 | 13 | 0.0769 | 6 | 0.3333 | 8 | 0 | 35 | 0.1714 |
| 16_9612084 | 4 | 0.5 | 12 | 0 | 6 | 0.3333 | 4 | 0 | 26 | 0.1538 | 5 | 0 | 12 | 0 | 4 | 0.25 | 7 | 0.2857 | 28 | 0.1071 |
| 16_9612085 | 5 | 0.4 | 14 | 0.4286 | 7 | 0.1429 | 5 | 0.4 | 31 | 0.3548 | 8 | 0.375 | 13 | 0.3846 | 6 | 0.3333 | 8 | 0.375 | 35 | 0.3714 |
| 16_9612088 | 5 | 1 | 17 | 0.8824 | 8 | 0.625 | 5 | 0.6 | 35 | 0.8 | 8 | 1 | 14 | 0.9286 | 5 | 1 | 8 | 0.75 | 35 | 0.9143 |
| 16_9617414 | NA | NA | NA | NA | NA | NA | 1 | 1 | 1 | 1 | NA | NA | NA | NA | NA | NA | 1 | 1 | 1 | 1 |
| 16_9620455 | NA | NA | NA | NA | NA | NA | 1 | 1 | 1 | 1 | NA | NA | NA | NA | NA | NA | 1 | 1 | 1 | 1 |
| 16_9620456 | NA | NA | NA | NA | NA | NA | 1 | 1 | 1 | 1 | NA | NA | NA | NA | NA | NA | 1 | 1 | 1 | 1 |
| 16_9620494 | NA | NA | NA | NA | NA | NA | 1 | 1 | 1 | 1 | NA | NA | NA | NA | NA | NA | 1 | 1 | 1 | 1 |
| 16_9639383 | 1 | 0 | 1 | 1 | 2 | 0 | 2 | 0 | 6 | 0.1667 | 2 | 0.5 | 1 | 0 | NA | NA | 1 | 1 | 4 | 0.5 |
| 16_9639394 | 1 | 1 | 1 | 1 | 2 | 1 | 3 | 1 | 7 | 1 | 2 | 1 | NA | NA | 1 | 1 | 1 | 1 | 4 | 1 |
| 16_9639408 | 1 | 1 | NA | NA | 2 | 1 | 3 | 1 | 6 | 1 | 1 | 1 | NA | NA | 1 | 1 | NA | NA | 2 | 1 |
| 16_9639424 | NA | NA | 1 | 1 | 2 | 1 | 3 | 0.6667 | 6 | 0.8333 | 2 | 1 | NA | NA | NA | NA | 1 | 0 | 3 | 0.6667 |
| 16_9639431 | NA | NA | 1 | 1 | 2 | 0.5 | 3 | 1 | 6 | 0.8333 | 2 | 1 | NA | NA | 1 | 1 | 1 | 1 | 4 | 1 |
| 16_9639435 | NA | NA | 1 | 1 | 1 | 1 | 3 | 1 | 5 | 1 | 2 | 1 | NA | NA | 1 | 1 | 1 | 1 | 4 | 1 |
| 16_9639442 | NA | NA | NA | NA | 2 | 1 | 3 | 0.6667 | 5 | 0.8 | 1 | 1 | NA | NA | 1 | 1 | 1 | 0 | 3 | 0.6667 |
| 16_9639449 | NA | NA | 1 | 0 | 2 | 0.5 | 3 | 0.3333 | 6 | 0.3333 | 2 | 0.5 | NA | NA | NA | NA | 1 | 1 | 3 | 0.6667 |
| 16_9639452 | NA | NA | 1 | 1 | 2 | 0.5 | 3 | 0 | 6 | 0.3333 | 2 | 0.5 | NA | NA | NA | NA | 1 | 0 | 3 | 0.3333 |
| 16_9639453 | NA | NA | 1 | 1 | 2 | 0.5 | 3 | 1 | 6 | 0.8333 | 1 | 1 | NA | NA | NA | NA | 1 | 1 | 2 | 1 |
| 16_9639457 | NA | NA | 1 | 1 | 2 | 1 | 3 | 0.6667 | 6 | 0.8333 | 2 | 1 | NA | NA | NA | NA | 1 | 1 | 3 | 1 |
| 16_9640935 | NA | NA | 1 | 0 | NA | NA | 3 | 0.6667 | 4 | 0.5 | 2 | 1 | NA | NA | 1 | 1 | 1 | 1 | 4 | 1 |
| 16_9644505 | NA | NA | 1 | 1 | 1 | 1 | 1 | 0 | 3 | 0.6667 | NA | NA | NA | NA | NA | NA | 1 | 1 | 1 | 1 |
| 16_9759377 | 1 | 0 | 1 | 0 | 1 | 1 | 1 | 0 | 4 | 0.25 | 2 | 0 | 2 | 0 | 3 | 0.3333 | 3 | 0.3333 | 10 | 0.2 |
| 16_9759378 | 1 | 0 | NA | NA | 2 | 1 | 1 | 0 | 4 | 0.5 | 2 | 0 | 2 | 0.5 | 2 | 0 | 3 | 0.3333 | 9 | 0.2222 |
| 16_9759387 | 1 | 1 | 1 | 1 | 2 | 1 | 1 | 1 | 5 | 1 | 2 | 1 | 3 | 0.6667 | 2 | 1 | 3 | 1 | 10 | 0.9 |
| 16_9759396 | 1 | 1 | 1 | 1 | 1 | 1 | 1 | 1 | 4 | 1 | 1 | 1 | 3 | 1 | 2 | 1 | 3 | 0.3333 | 9 | 0.7778 |
| 16_9759418 | 1 | 0 | 1 | 1 | 1 | 1 | NA | NA | 3 | 0.6667 | NA | NA | 2 | 0.5 | 1 | 1 | 3 | 0.3333 | 6 | 0.5 |
| 16_9759419 | 1 | 1 | NA | NA | 2 | 0.5 | 1 | 1 | 4 | 0.75 | 1 | 0 | 3 | 0.6667 | 2 | 1 | 3 | 0 | 9 | 0.4444 |
| 16_9759424 | 1 | 0 | 1 | 0 | 2 | 0.5 | 1 | 0 | 5 | 0.2 | 1 | 0 | 3 | 0.3333 | 2 | 0.5 | 3 | 0 | 9 | 0.2222 |
| 16_9759462 | 1 | 1 | 1 | 0 | 2 | 0.5 | NA | NA | 4 | 0.5 | 2 | 1 | 2 | 1 | 1 | 0 | 3 | 0.6667 | 8 | 0.75 |
| 16_9759484 | 1 | 0 | 1 | 0 | 1 | 1 | 1 | 1 | 4 | 0.5 | 1 | 1 | 1 | 0 | 1 | 1 | 2 | 0.5 | 5 | 0.6 |
| 16_9759504 | 1 | 1 | 1 | 1 | 2 | 0.5 | NA | NA | 4 | 0.75 | 1 | 1 | 1 | 0 | 1 | 0 | 3 | 0.3333 | 6 | 0.3333 |

| | | | | | | | | | | | | | | | | | | | | |
|---|---|---|---|---|---|---|---|---|---|---|---|---|---|---|---|---|---|---|---|---|
| 16_9759567 | NA | NA | 1 | 0 | 2 | 0.5 | 1 | 0 | 4 | 0.25 | 1 | 0 | 1 | 1 | 1 | 0 | NA | NA | 3 | 0.3333 |
| 16_9759567 | NA | NA | 1 | 1 | 2 | 1 | 1 | 1 | 4 | 1 | 1 | 1 | 1 | 0 | 1 | 1 | 1 | 0 | 4 | 0.5 |
| 16_9759572 | NA | NA | 1 | 1 | 2 | 1 | 1 | 1 | 4 | 1 | 1 | 1 | NA | NA | 1 | 1 | 1 | 0 | 3 | 0.6667 |
| 16_9759576 | NA | NA | 1 | 0 | 2 | 1 | 1 | 1 | 4 | 0.75 | 1 | 0 | NA | NA | 1 | 1 | NA | NA | 2 | 0.5 |
| 16_9766016 | 2 | 1 | NA | NA | NA | NA | 3 | 0.6667 | 5 | 0.8 | NA | NA | NA | NA | 1 | 1 | NA | NA | 1 | 1 |
| 16_9766024 | 1 | 1 | NA | NA | NA | NA | 2 | 1 | 3 | 1 | NA | NA | NA | NA | 1 | 1 | NA | NA | 1 | 1 |
| 16_9766028 | 2 | 1 | NA | NA | 1 | 1 | 2 | 1 | 5 | 1 | NA | NA | NA | NA | 1 | 1 | NA | NA | 1 | 1 |
| 16_9766037 | 1 | 1 | NA | NA | NA | NA | 2 | 1 | 3 | 1 | NA | NA | NA | NA | 1 | 1 | NA | NA | 1 | 1 |
| 16_9775086 | 1 | 0 | 2 | 1 | NA | NA | NA | NA | 3 | 0.6667 | 1 | 1 | NA | NA | 1 | 1 | 2 | 0.5 | 4 | 0.75 |
| 16_9775090 | 1 | 0 | 2 | 0.5 | NA | NA | NA | NA | 3 | 0.3333 | NA | NA | NA | NA | 1 | 0 | 1 | 1 | 2 | 0.5 |
| 16_9775101 | 1 | 0 | 4 | 0.25 | 1 | 0 | 1 | 0 | 7 | 0.1429 | 1 | 0 | 1 | 0 | 2 | 0.5 | 3 | 0.3333 | 7 | 0.2857 |
| 16_9775103 | 1 | 0 | 4 | 0.75 | 1 | 1 | 1 | 1 | 7 | 0.7143 | 2 | 1 | 2 | 0.5 | 2 | 0.5 | 3 | 0.6667 | 9 | 0.6667 |
| 16_9775107 | 1 | 1 | 4 | 1 | 1 | 0 | 1 | 1 | 7 | 0.8571 | 1 | 1 | 2 | 1 | 2 | 1 | 3 | 1 | 8 | 1 |
| 16_9775113 | 1 | 1 | 4 | 1 | 1 | 1 | 1 | 1 | 7 | 1 | 2 | 1 | 2 | 1 | 2 | 1 | 3 | 1 | 9 | 1 |
| 16_9775115 | 1 | 1 | 3 | 0.6667 | 1 | 1 | NA | NA | 5 | 0.8 | 1 | 1 | 1 | 1 | 1 | 1 | 1 | 1 | 4 | 1 |
| 16_9775134 | NA | NA | 3 | 1 | 1 | 1 | 1 | 1 | 5 | 1 | 1 | 1 | 2 | 0 | 1 | 1 | 1 | 1 | 5 | 0.6 |
| 16_9775168 | 1 | 1 | 4 | 0.5 | 1 | 0 | 1 | 1 | 7 | 0.5714 | 1 | 1 | 2 | 0.5 | 2 | 0 | 2 | 0.5 | 7 | 0.4286 |
| 16_9775171 | 1 | 1 | 2 | 1 | 1 | 1 | 1 | 1 | 5 | 1 | 2 | 1 | 2 | 1 | 1 | 1 | 2 | 1 | 7 | 1 |
| 16_9775179 | 1 | 1 | 3 | 1 | 1 | 1 | NA | NA | 5 | 1 | 1 | 1 | 2 | 0.5 | 1 | 1 | 2 | 0.5 | 6 | 0.6667 |
| 16_9775183 | 1 | 0 | 3 | 0.6667 | 1 | 0 | 1 | 0 | 6 | 0.3333 | NA | NA | 1 | 0 | 1 | 0 | 2 | 0.5 | 4 | 0.25 |
| 16_9775185 | 1 | 1 | 4 | 0 | NA | NA | 1 | 0 | 6 | 0.1667 | 1 | 0 | 2 | 0.5 | 2 | 0.5 | 3 | 0.3333 | 8 | 0.375 |
| 16_9775195 | 1 | 0 | 2 | 0.5 | 1 | 0 | 1 | 1 | 5 | 0.4 | 1 | 1 | 2 | 0 | 2 | 0 | 2 | 0.5 | 7 | 0.2857 |
| 16_9775197 | 1 | 1 | 3 | 0.3333 | 1 | 1 | 1 | 1 | 6 | 0.6667 | 2 | 0.5 | 2 | 1 | 1 | 0 | 1 | 0 | 6 | 0.5 |
| 16_9775274 | 1 | 1 | 2 | 0.5 | 1 | 1 | 1 | 1 | 5 | 0.8 | 1 | 1 | NA | NA | 2 | 1 | 2 | 1 | 5 | 1 |
| 16_9775300 | 1 | 1 | 2 | 0 | 1 | 0 | 1 | 0 | 5 | 0.2 | 2 | 0.5 | 1 | 1 | 2 | 0 | 2 | 0 | 7 | 0.2857 |
| 16_9775324 | NA | NA | 1 | 1 | 1 | 1 | 1 | 1 | 3 | 1 | 1 | 0 | 2 | 0.5 | 2 | 0.5 | 2 | 1 | 7 | 0.5714 |
| 16_9775336 | 1 | 0 | 1 | 1 | 1 | 0 | 1 | 0 | 4 | 0.25 | 1 | 1 | 2 | 0.5 | 1 | 0 | 1 | 0 | 5 | 0.4 |
| 16_9775342 | 1 | 1 | 1 | 1 | 1 | 1 | 1 | 0 | 4 | 0.75 | NA | NA | 2 | 1 | 1 | 1 | 1 | 1 | 4 | 1 |
| 16_9775353 | NA | NA | 1 | 0 | 1 | 1 | 1 | 1 | 3 | 0.6667 | NA | NA | 2 | 0.5 | 1 | 1 | 1 | 0 | 4 | 0.5 |
| 16_9775365 | 1 | 1 | NA | NA | NA | NA | NA | NA | 1 | 1 | NA | NA | 2 | 0.5 | 1 | 1 | 1 | 0 | 4 | 0.5 |
| 16_9775402 | 1 | 0 | NA | NA | 1 | 1 | 1 | 1 | 3 | 0.6667 | NA | NA | 1 | 1 | 1 | 1 | NA | NA | 2 | 1 |
| 16_9775406 | 1 | 1 | NA | NA | 1 | 0 | 1 | 1 | 3 | 0.6667 | NA | NA | 1 | 1 | 1 | 1 | 1 | 1 | 3 | 1 |
| 16_9775408 | 1 | 1 | NA | NA | NA | NA | 1 | 1 | 2 | 1 | NA | NA | 1 | 1 | NA | NA | 1 | 1 | 2 | 1 |
| 16_9775420 | 1 | 0 | 2 | 0 | 1 | 0 | 1 | 1 | 5 | 0.2 | NA | NA | 1 | 0 | 1 | 1 | 1 | 0 | 3 | 0.3333 |
| 16_9781137 | 1 | 0 | NA | NA | 2 | 0.5 | 1 | 1 | 4 | 0.5 | NA | NA | 1 | 1 | NA | NA | NA | NA | 1 | 1 |
| 16_9781257 | 1 | 0 | 1 | 0 | 2 | 0 | 1 | 1 | 5 | 0.2 | NA | NA | 1 | 1 | NA | NA | NA | NA | 1 | 1 |
| 16_9781731 | NA | NA | NA | NA | NA | NA | 1 | 1 | 1 | 1 | NA | NA | 2 | 0.5 | NA | NA | NA | NA | 2 | 0.5 |
| 17_3113077 | 2 | 0.5 | 1 | 0 | 1 | 0 | 1 | 0 | 5 | 0.2 | NA | NA | NA | NA | 1 | 0 | 2 | 0.5 | 3 | 0.3333 |
| 17_3114456 | 1 | 1 | 1 | 0 | 1 | 1 | 1 | 1 | 4 | 0.75 | NA | NA | NA | NA | 1 | 1 | NA | NA | 1 | 1 |
| 17_3114491 | 1 | 1 | 1 | 1 | 1 | 1 | 1 | 1 | 4 | 1 | NA | NA | NA | NA | NA | NA | 1 | 1 | 1 | 1 |
| 17_3114508 | 1 | 1 | NA | NA | NA | NA | NA | NA | 1 | 1 | NA | NA | NA | NA | NA | NA | 1 | 1 | 1 | 1 |
| 17_3114524 | 1 | 0 | NA | NA | 1 | 1 | 1 | 0 | 3 | 0.3333 | NA | NA | NA | NA | NA | NA | 1 | 1 | 1 | 1 |
| 17_3114581 | 1 | 1 | NA | NA | 1 | 1 | 1 | 0 | 3 | 0.6667 | NA | NA | NA | NA | NA | NA | 1 | 1 | 1 | 1 |
| 17_3162305 | NA | NA | NA | NA | NA | NA | 1 | 1 | 1 | 1 | 3 | 1 | NA | NA | 1 | 1 | 1 | 0 | 5 | 0.8 |
| 17_3162561 | NA | NA | NA | NA | NA | NA | 1 | 1 | 1 | 1 | 3 | 1 | NA | NA | 2 | 1 | 1 | 1 | 6 | 1 |
| 17_3162572 | NA | NA | NA | NA | NA | NA | 1 | 1 | 1 | 1 | 3 | 0.6667 | NA | NA | 2 | 0.5 | 1 | 0 | 6 | 0.5 |
| 17_3162602 | NA | NA | NA | NA | NA | NA | 1 | 1 | 1 | 1 | 3 | 0.6667 | NA | NA | 2 | 1 | 1 | 1 | 6 | 0.8333 |
| 17_3162658 | NA | NA | NA | NA | NA | NA | 1 | 1 | 1 | 1 | 2 | 1 | NA | NA | 1 | 1 | 1 | 0 | 4 | 0.75 |
| 17_3162777 | NA | NA | NA | NA | NA | NA | 1 | 1 | 1 | 1 | 3 | 0.6667 | NA | NA | 2 | 1 | NA | NA | 5 | 0.8 |
| 17_3247123 | 5 | 0.4 | 2 | 0 | 5 | 0.2 | 5 | 0.4 | 17 | 0.2941 | 2 | 0.5 | 5 | 0.2 | 2 | 0 | 3 | 0.3333 | 12 | 0.25 |
| 17_3247224 | 3 | 0.6667 | NA | NA | 2 | 0.5 | 4 | 0.75 | 9 | 0.6667 | 1 | 1 | NA | NA | 1 | 0 | 2 | 0 | 4 | 0.25 |
| 17_3247273 | 5 | 0.8 | 3 | 1 | 5 | 1 | 5 | 1 | 18 | 0.9444 | 2 | 1 | 2 | 1 | 2 | 1 | 4 | 1 | 10 | 1 |
| 17_3247315 | 4 | 1 | 2 | 1 | 6 | 0.8333 | 5 | 1 | 17 | 0.9412 | 2 | 1 | 3 | 1 | 2 | 1 | 4 | 0.75 | 11 | 0.9091 |
| 17_3247324 | 3 | 1 | 3 | 0.6667 | 5 | 0.4 | 5 | 1 | 16 | 0.75 | 1 | 1 | 5 | 0.8 | 2 | 0.5 | 4 | 0.75 | 12 | 0.75 |
| 17_3247357 | 5 | 1 | 3 | 1 | 6 | 1 | 4 | 1 | 18 | 1 | 2 | 1 | 2 | 1 | 2 | 1 | 4 | 0.75 | 10 | 0.9 |
| 17_3247384 | 4 | 0.75 | 3 | 0.3333 | 6 | 0.5 | 5 | 0.2 | 18 | 0.4444 | 2 | 0 | 4 | 0.75 | 2 | 0.5 | 4 | 0.5 | 12 | 0.5 |
| 17_3247446 | 5 | 0.4 | 1 | 0 | 6 | 0.5 | 5 | 0.6 | 17 | 0.4706 | 2 | 0.5 | 4 | 0.5 | 2 | 0.5 | 2 | 0.5 | 10 | 0.5 |
| 17_3247459 | 4 | 1 | 1 | 1 | 5 | 1 | 5 | 0.8 | 15 | 0.9333 | 2 | 1 | 4 | 1 | 2 | 1 | 4 | 1 | 12 | 1 |
| 17_3247576 | 3 | 0.3333 | 3 | 0.3333 | 5 | 0.2 | 4 | 0.75 | 15 | 0.4 | 1 | 1 | 5 | 0 | NA | NA | 3 | 0 | 9 | 0.1111 |
| 17_3247647 | 5 | 1 | 3 | 1 | 6 | 0.8333 | 5 | 0.4 | 19 | 0.7895 | 2 | 1 | 7 | 0.5714 | 2 | 0.5 | 4 | 0.5 | 15 | 0.6 |
| 17_3247689 | 5 | 0 | 3 | 0.3333 | 6 | 0.1667 | 5 | 0 | 19 | 0.1053 | 2 | 0 | 6 | 0.3333 | 1 | 0 | 4 | 0 | 13 | 0.1538 |
| 17_3247759 | 5 | 0 | 3 | 0.3333 | 6 | 0 | 5 | 0 | 19 | 0.0526 | 3 | 0.3333 | 4 | 0.25 | 2 | 0 | 4 | 0 | 13 | 0.1538 |
| 17_3247869 | 4 | 0.75 | 3 | 1 | 5 | 1 | 4 | 0.75 | 16 | 0.875 | 3 | 1 | 5 | 0.8 | 2 | 0.5 | 3 | 1 | 13 | 0.8462 |
| 17_3248082 | 5 | 0 | 1 | 0 | 4 | 0.25 | 5 | 0 | 15 | 0.0667 | 1 | 0 | 5 | 0.2 | 1 | 0 | 1 | 0 | 8 | 0.125 |
| 17_3248131 | 4 | 0.75 | 2 | 0 | 6 | 0.6667 | 6 | 0.5 | 18 | 0.5556 | 3 | 1 | 5 | 0.4 | 2 | 1 | 3 | 0.6667 | 13 | 0.6923 |

| | | | | | | | | | | | | | | | | | | | | |
|---|---|---|---|---|---|---|---|---|---|---|---|---|---|---|---|---|---|---|---|---|
| 17_324822 | 5 | 0.6 | 3 | 0.6667 | 6 | 0.5 | 5 | 0.4 | 19 | 0.5263 | 2 | 0.5 | 5 | 0.6 | 1 | 0 | 4 | 0.75 | 12 | 0.5833 |
| 17_324833 | 5 | 0.6 | 4 | 0.5 | 6 | 0.5 | 6 | 0.3333 | 21 | 0.4762 | 2 | 0.5 | 6 | 0.5 | 2 | 1 | 4 | 0.5 | 14 | 0.5714 |
| 17_324836 | 5 | 0.6 | 2 | 1 | 6 | 0.6667 | 6 | 0.8333 | 19 | 0.7368 | 3 | 1 | 6 | 0.3333 | 3 | 0.6667 | 3 | 0.6667 | 15 | 0.6 |
| 17_324837 | 5 | 0.8 | 4 | 1 | 6 | 0.8333 | 4 | 0.75 | 19 | 0.8421 | 3 | 0.6667 | 6 | 0.6667 | 2 | 1 | 2 | 0.5 | 13 | 0.6923 |
| 17_324838 | 4 | 0.5 | 4 | 0.25 | 5 | 0.6 | 5 | 0.4 | 18 | 0.4444 | 2 | 0 | 6 | 0.1667 | 3 | 0.3333 | 3 | 0.3333 | 14 | 0.2143 |
| 17_324847 | 5 | 1 | 4 | 1 | 6 | 0.8333 | 6 | 0.8333 | 21 | 0.9048 | 3 | 1 | 6 | 0.8333 | 3 | 1 | 4 | 1 | 16 | 0.9375 |
| 17_324848 | 5 | 0.6 | 4 | 0.25 | 5 | 0.8 | 5 | 0.4 | 19 | 0.5263 | 3 | 1 | 8 | 0.25 | 3 | 0.6667 | 4 | 0.75 | 18 | 0.5556 |
| 17_324855 | 4 | 0.25 | 3 | 0.6667 | 4 | 0.25 | 4 | 0.5 | 15 | 0.4 | 1 | 1 | 7 | 0.1429 | 3 | 0 | 2 | 0.5 | 13 | 0.2308 |
| 17_324858 | 4 | 0 | 3 | 0 | 4 | 0 | 5 | 0.4 | 16 | 0.125 | 3 | 0 | 5 | 0.4 | 3 | 0 | 1 | 0 | 12 | 0.1667 |
| 17_324863 | 3 | 0 | 3 | 0 | 4 | 0 | 5 | 0.4 | 15 | 0.1333 | 3 | 0.3333 | 5 | 0 | 2 | 0.5 | 2 | 0 | 12 | 0.1667 |
| 17_324864 | 3 | 0.3333 | 3 | 0 | 5 | 0.2 | 3 | 0.3333 | 14 | 0.2143 | 2 | 0.5 | 6 | 0.1667 | 2 | 0 | 2 | 0.5 | 12 | 0.25 |
| 17_324889 | 4 | 0 | 3 | 0 | 5 | 0.2 | 3 | 0 | 15 | 0.0667 | 2 | 0 | 6 | 0.1667 | 4 | 0 | 2 | 0 | 14 | 0.0714 |
| 17_324891 | 4 | 1 | 3 | 0.6667 | 5 | 0.8 | 5 | 0.6 | 17 | 0.7647 | 3 | 1 | 7 | 0.7143 | 4 | 0.5 | 2 | 1 | 16 | 0.75 |
| 17_330841 | 4 | 0.25 | 1 | 0 | NA | NA | 1 | 0 | 6 | 0.1667 | NA | NA | NA | NA | NA | NA | 1 | 1 | 1 | 1 |
| 17_330880 | 4 | 0.25 | 1 | 0 | 2 | 0 | 1 | 0 | 8 | 0.125 | 1 | 1 | 1 | 0 | NA | NA | 3 | 0 | 5 | 0.2 |
| 17_330914 | 3 | 0 | 1 | 0 | 2 | 0.5 | 1 | 0 | 7 | 0.1429 | 1 | 0 | 1 | 0 | NA | NA | 1 | 1 | 3 | 0.3333 |
| 17_353285 | 6 | 0 | 1 | 0 | 4 | 0 | 3 | 0.3333 | 14 | 0.0714 | 4 | 0.25 | 4 | 0 | 1 | 0 | 3 | 0 | 12 | 0.0833 |
| 17_354086 | NA | NA | NA | NA | 2 | 0.5 | NA | NA | 2 | 0.5 | 1 | 0 | NA | NA | 2 | 0.5 | 3 | 0.3333 | 6 | 0.3333 |
| 17_354087 | NA | NA | NA | NA | 2 | 0.5 | 1 | 0 | 3 | 0.3333 | 1 | 0 | NA | NA | 1 | 0 | 3 | 0.3333 | 5 | 0.2 |
| 17_356846 | 12 | 0.9167 | 8 | 0.25 | 8 | 0.75 | 13 | 0.6923 | 41 | 0.6829 | 13 | 0.7692 | 10 | 0.7 | 13 | 0.8462 | 12 | 0.75 | 48 | 0.7708 |
| 17_356853 | 17 | 0.1765 | 9 | 0.4444 | 11 | 0.1818 | 16 | 0.125 | 53 | 0.2075 | 14 | 0.0714 | 14 | 0.0714 | 14 | 0.2857 | 13 | 0 | 55 | 0.1091 |
| 17_356863 | 15 | 0.5333 | 7 | 0.7143 | 10 | 0.6 | 15 | 0.8667 | 47 | 0.6809 | 14 | 0.9286 | 10 | 0.8 | 11 | 0.8182 | 15 | 0.6667 | 50 | 0.8 |
| 17_356867 | 10 | 1 | 6 | 0.6667 | 10 | 1 | 16 | 0.9375 | 42 | 0.9286 | 12 | 1 | 8 | 1 | 15 | 0.9333 | 11 | 1 | 46 | 0.9783 |
| 17_356870 | 10 | 0.9 | 9 | 0.8889 | 10 | 0.9 | 17 | 0.8824 | 46 | 0.8913 | 15 | 0.9333 | 12 | 0.9167 | 13 | 0.9231 | 12 | 1 | 52 | 0.9423 |
| 17_356883 | 16 | 0.875 | 9 | 0.8889 | 11 | 0.8182 | 14 | 0.8571 | 50 | 0.86 | 14 | 0.7143 | 13 | 0.9231 | 13 | 0.9231 | 12 | 0.9167 | 52 | 0.8654 |
| 17_356885 | 14 | 0.0714 | 11 | 0.2727 | 11 | 0.1818 | 12 | 0 | 48 | 0.125 | 14 | 0 | 12 | 0.0833 | 12 | 0.25 | 15 | 0.1333 | 53 | 0.1132 |
| 17_356887 | 14 | 0.2857 | 11 | 0.5455 | 7 | 0.4286 | 14 | 0.2857 | 46 | 0.3696 | 13 | 0.3077 | 11 | 0.2727 | 11 | 0.5455 | 13 | 0.2308 | 48 | 0.3333 |
| 17_356889 | 17 | 0.4118 | 9 | 0.3333 | 10 | 0.2 | 16 | 0.4375 | 52 | 0.3654 | 14 | 0.2857 | 13 | 0.1538 | 11 | 0.0909 | 13 | 0.3077 | 51 | 0.2157 |
| 17_356971 | 3 | 0.6667 | 1 | 1 | NA | NA | NA | NA | 4 | 0.75 | 2 | 0.5 | NA | NA | 3 | 0.6667 | 1 | 0 | 6 | 0.5 |
| 17_356973 | 3 | 0.3333 | 1 | 1 | NA | NA | NA | NA | 4 | 0.5 | 2 | 0 | NA | NA | 3 | 0.3333 | 1 | 0 | 6 | 0.1667 |
| 17_356978 | 3 | 1 | 1 | 1 | NA | NA | NA | NA | 4 | 1 | 2 | 0.5 | NA | NA | 3 | 1 | 1 | 1 | 6 | 0.8333 |
| 17_357023 | NA | NA | 1 | 0 | 1 | 1 | NA | NA | 2 | 0.5 | NA | NA | NA | NA | 1 | 0 | 1 | 1 | 2 | 0.5 |
| 17_538693 | 3 | 0.3333 | NA | NA | 1 | 1 | 4 | 0.25 | 8 | 0.375 | NA | NA | 4 | 0.25 | 1 | 0 | 6 | 0.5 | 11 | 0.3636 |
| 17_538695 | 3 | 1 | NA | NA | 1 | 0 | 4 | 0.5 | 8 | 0.625 | NA | NA | 4 | 0.75 | 1 | 1 | 5 | 0.4 | 10 | 0.6 |
| 17_538707 | 4 | 0.25 | 1 | 0 | 2 | 0.5 | 4 | 0.25 | 11 | 0.2727 | NA | NA | 3 | 0.3333 | 1 | 0 | 6 | 0.3333 | 10 | 0.3 |
| 17_538717 | 2 | 0.5 | 1 | 0 | 2 | 0 | 3 | 0.3333 | 8 | 0.25 | NA | NA | 4 | 0 | 1 | 0 | 5 | 0.2 | 10 | 0.1 |
| 17_539186 | 5 | 0.2 | 2 | 0 | 3 | 0 | 3 | 0 | 13 | 0.0769 | 1 | 0 | 6 | 0 | 1 | 0 | 5 | 0.2 | 13 | 0.0769 |
| 17_539193 | 5 | 0.2 | 2 | 0 | 3 | 0 | 4 | 0.25 | 14 | 0.1429 | 1 | 0 | 5 | 0.4 | 2 | 0.5 | 6 | 0.1667 | 14 | 0.2857 |
| 17_539199 | 5 | 0.4 | 2 | 0 | 3 | 0.3333 | 4 | 0 | 14 | 0.2143 | 1 | 0 | 3 | 0.3333 | NA | NA | 5 | 0.2 | 9 | 0.2222 |
| 17_539201 | 5 | 0.4 | 2 | 0 | 3 | 0 | 4 | 0 | 14 | 0.1429 | 1 | 0 | 5 | 0.2 | 2 | 0.5 | 6 | 0.3333 | 14 | 0.2857 |
| 17_539210 | 5 | 0.2 | 2 | 0 | 3 | 0 | 5 | 0 | 15 | 0.0667 | 2 | 0 | 5 | 0 | 2 | 0 | 6 | 0.3333 | 15 | 0.1333 |
| 17_539229 | 5 | 0.2 | 2 | 0 | 3 | 0 | 7 | 0 | 17 | 0.0588 | 1 | 0 | 6 | 0 | 3 | 0 | 6 | 0.3333 | 16 | 0.125 |
| 17_539242 | 5 | 1 | 2 | 1 | 4 | 1 | 6 | 1 | 17 | 1 | 2 | 1 | 6 | 0.8333 | 2 | 1 | 7 | 1 | 17 | 0.9412 |
| 17_539261 | 4 | 0.75 | 4 | 0.75 | 4 | 1 | 8 | 1 | 20 | 0.9 | 1 | 1 | 5 | 0.8 | 3 | 1 | 8 | 0.75 | 17 | 0.8235 |
| 17_539265 | 4 | 1 | 5 | 0.8 | 4 | 1 | 7 | 1 | 20 | 0.95 | 3 | 0.6667 | 6 | 1 | 3 | 1 | 7 | 1 | 19 | 0.9474 |
| 17_539270 | 3 | 0.6667 | 3 | 1 | 2 | 1 | 4 | 1 | 12 | 0.9167 | 1 | 0 | 3 | 0.6667 | 3 | 0.6667 | 6 | 0.6667 | 13 | 0.6154 |
| 17_539271 | 5 | 0.6 | 5 | 0.2 | 5 | 0.6 | 7 | 0.2857 | 22 | 0.4091 | 2 | 1 | 5 | 0.6 | 2 | 0 | 6 | 0.6667 | 15 | 0.6 |
| 17_539280 | 5 | 0.4 | 5 | 0.6 | 5 | 0.8 | 8 | 0.375 | 23 | 0.5217 | 2 | 0.5 | 6 | 0.5 | 3 | 0.3333 | 9 | 0.4444 | 20 | 0.45 |
| 17_539285 | 5 | 1 | 5 | 1 | 5 | 1 | 8 | 0.875 | 23 | 0.9565 | 3 | 0.6667 | 6 | 1 | 3 | 1 | 10 | 0.8 | 22 | 0.8636 |
| 17_539286 | 5 | 0 | 5 | 0 | 5 | 0.2 | 8 | 0.125 | 23 | 0.087 | 3 | 0 | 6 | 0 | 3 | 0 | 9 | 0.2222 | 21 | 0.0952 |
| 17_539287 | 3 | 0.3333 | 5 | 0 | 3 | 0.3333 | 7 | 0.2857 | 18 | 0.2222 | 2 | 0 | 4 | 0.75 | 3 | 0 | 8 | 0.25 | 17 | 0.2941 |
| 17_539289 | 5 | 0.6 | 4 | 0.5 | 4 | 1 | 7 | 0.8571 | 20 | 0.75 | 3 | 0.6667 | 6 | 1 | 3 | 1 | 9 | 0.7778 | 21 | 0.8571 |
| 17_539290 | 5 | 0.2 | 5 | 0.4 | 5 | 0.4 | 8 | 0.375 | 23 | 0.3478 | 2 | 0 | 5 | 0.2 | 3 | 0 | 10 | 0.5 | 20 | 0.3 |
| 17_539302 | 5 | 0.4 | 4 | 0.75 | 4 | 0.5 | 6 | 0.5 | 19 | 0.5263 | 2 | 0.5 | 6 | 0.5 | 3 | 0.6667 | 8 | 0.625 | 19 | 0.5789 |
| 17_539303 | 6 | 0.8333 | 4 | 1 | 5 | 0.8 | 6 | 0.6667 | 21 | 0.8095 | 3 | 1 | 7 | 1 | 3 | 0.6667 | 9 | 0.8889 | 22 | 0.9091 |
| 17_539308 | 4 | 0.5 | 5 | 0.4 | 3 | 0 | 7 | 0.1429 | 19 | 0.2632 | 2 | 1 | 7 | 0.4286 | 4 | 0.25 | 6 | 0.1667 | 19 | 0.3684 |
| 17_539319 | 5 | 0.4 | 4 | 0 | 4 | 0.25 | 6 | 0.5 | 19 | 0.3158 | 1 | 1 | 7 | 0.2857 | 4 | 0.5 | 7 | 0.4286 | 19 | 0.4211 |
| 17_539330 | 5 | 1 | 6 | 1 | 5 | 1 | 7 | 0.8571 | 23 | 0.9565 | 3 | 1 | 6 | 0.8333 | 4 | 1 | 9 | 0.8889 | 22 | 0.9091 |
| 17_539339 | 4 | 0.75 | 5 | 1 | 5 | 0.6 | 7 | 0.5714 | 21 | 0.7143 | 3 | 0.3333 | 7 | 0.5714 | 4 | 1 | 7 | 0.8571 | 21 | 0.7143 |
| 17_539341 | 2 | 0 | 5 | 0 | 4 | 0.25 | 3 | 0 | 14 | 0.0714 | 2 | 0 | 7 | 0.1429 | 4 | 0.25 | 6 | 0.1667 | 19 | 0.1579 |
| 17_539371 | 3 | 0.6667 | 5 | 0.4 | 7 | 0.2857 | 3 | 0 | 18 | 0.3333 | 1 | 0 | 6 | 0.1667 | 3 | 0.3333 | 5 | 0.2 | 15 | 0.2 |
| 17_539453 | 6 | 1 | 5 | 1 | 11 | 1 | 5 | 0.4 | 27 | 0.8889 | 2 | 1 | 8 | 0.75 | 6 | 0.8333 | 7 | 0.8571 | 23 | 0.8261 |
| 17_546130 | 21 | 0.3333 | 15 | 0.2667 | 15 | 0.2667 | 18 | 0 | 69 | 0.2174 | 15 | 0.2 | 7 | 0.4286 | 17 | 0.3529 | 14 | 0.1429 | 53 | 0.2642 |
| 17_546140 | 16 | 0.0625 | 12 | 0.3333 | 13 | 0.0769 | 13 | 0.3077 | 54 | 0.1852 | 11 | 0.0909 | 7 | 0.1429 | 10 | 0.2 | 10 | 0.2 | 38 | 0.1579 |
| 17_546142 | 16 | 0.5 | 15 | 0.4 | 14 | 0.4286 | 17 | 0.5882 | 62 | 0.4839 | 12 | 0.3333 | 8 | 0.125 | 17 | 0.4706 | 13 | 0.3077 | 50 | 0.34 |

| | | | | | | | | | | | | | | | | | | | | |
|---|---|---|---|---|---|---|---|---|---|---|---|---|---|---|---|---|---|---|---|---|
| 17_546146 | 20 | 0.1 | 15 | 0.0667 | 15 | 0.0667 | 14 | 0.3571 | 64 | 0.1406 | 15 | 0.2 | 8 | 0 | 15 | 0.0667 | 14 | 0 | 52 | 0.0769 |
| 17_546157 | 20 | 0.9 | 11 | 0.8182 | 12 | 0.5833 | 17 | 0.5882 | 60 | 0.7333 | 14 | 0.6429 | 8 | 0.625 | 15 | 0.7333 | 13 | 0.7692 | 50 | 0.7 |
| 17_553934 | NA | NA | NA | NA | 1 | 1 | NA | NA | 1 | 1 | NA | NA | 1 | 1 | NA | NA | NA | NA | 1 | 1 |
| 17_554305 | 2 | 0.5 | 1 | 1 | 2 | 0 | NA | NA | 5 | 0.4 | 3 | 0.3333 | 2 | 0.5 | 3 | 0.3333 | 2 | 1 | 10 | 0.5 |
| 17_554348 | 2 | 0 | 1 | 0 | 3 | 0.3333 | 2 | 0 | 8 | 0.125 | 4 | 0.25 | 2 | 0.5 | 2 | 0.5 | 3 | 0 | 11 | 0.2727 |
| 17_554366 | 2 | 0 | NA | NA | 4 | 0.25 | 2 | 0.5 | 8 | 0.25 | 4 | 0 | 3 | 0 | 3 | 0.3333 | 3 | 0 | 13 | 0.0769 |
| 17_554375 | 3 | 0.3333 | 1 | 1 | 4 | 0.5 | 2 | 0 | 10 | 0.4 | 4 | 1 | 3 | 0.6667 | 4 | 0.5 | 3 | 0.3333 | 14 | 0.6429 |
| 17_554394 | 3 | 0.3333 | 2 | 0 | 3 | 0 | 1 | 1 | 9 | 0.2222 | 3 | 0 | 2 | 0.5 | 4 | 0 | 3 | 0.3333 | 12 | 0.1667 |
| 17_580317 | 3 | 0.6667 | 6 | 0.6667 | 6 | 0.8333 | 7 | 1 | 22 | 0.8182 | 7 | 0.7143 | 6 | 0.5 | 7 | 0.7143 | 7 | 0.5714 | 27 | 0.6296 |
| 17_580331 | 5 | 0.8 | 6 | 0.3333 | 6 | 0.8333 | 9 | 0.5556 | 26 | 0.6154 | 7 | 0.4286 | 6 | 0.8333 | 9 | 0.7778 | 7 | 0.8571 | 29 | 0.7241 |
| 17_580339 | 4 | 0 | 6 | 0.3333 | 5 | 0 | 9 | 0.4444 | 24 | 0.25 | 5 | 0.4 | 6 | 0.1667 | 8 | 0.25 | 8 | 0.25 | 27 | 0.2593 |
| 17_580342 | 4 | 1 | 7 | 1 | 6 | 1 | 8 | 0.875 | 25 | 0.96 | 6 | 0.8333 | 5 | 0.8 | 7 | 0.8571 | 8 | 0.875 | 26 | 0.8462 |
| 17_580346 | 5 | 0 | 6 | 0.5 | 3 | 0.3333 | 6 | 0 | 20 | 0.2 | 6 | 0.1667 | 6 | 0 | 7 | 0.2857 | 8 | 0.125 | 27 | 0.1481 |
| 17_580355 | 6 | 0.8333 | 7 | 1 | 5 | 1 | 7 | 1 | 25 | 0.96 | 5 | 0.8 | 6 | 0.8333 | 5 | 1 | 7 | 0.7143 | 23 | 0.8261 |
| 17_580359 | 6 | 0.5 | 6 | 0.6667 | 6 | 0.3333 | 9 | 0.3333 | 27 | 0.4444 | 5 | 0.6 | 7 | 0.7143 | 9 | 0.5556 | 8 | 0.625 | 29 | 0.6207 |
| 17_580360 | 6 | 0.8333 | 8 | 0.875 | 6 | 1 | 8 | 1 | 28 | 0.9286 | 6 | 1 | 8 | 1 | 9 | 0.8889 | 8 | 1 | 31 | 0.9677 |
| 17_580381 | 8 | 0.125 | 8 | 0.25 | 5 | 0.6 | 7 | 0.2857 | 28 | 0.2857 | 6 | 0.1667 | 8 | 0.25 | 6 | 0.1667 | 8 | 0.375 | 28 | 0.25 |
| 17_580381 | 3 | 0 | 3 | 0.3333 | 3 | 0 | 4 | 0 | 13 | 0.0769 | 5 | 0 | 5 | 0 | 4 | 0 | 5 | 0.2 | 19 | 0.0526 |
| 17_580385 | 7 | 0.8571 | 5 | 1 | 5 | 1 | 8 | 1 | 25 | 0.96 | 5 | 0.8 | 7 | 0.8571 | 7 | 0.8571 | 7 | 0.8571 | 26 | 0.8462 |
| 17_580394 | 7 | 0.8571 | 7 | 1 | 5 | 1 | 8 | 0.625 | 27 | 0.8519 | 6 | 1 | 7 | 0.4286 | 6 | 0.8333 | 7 | 0.8571 | 26 | 0.7692 |
| 17_580395 | 6 | 0.3333 | 6 | 0.5 | 5 | 0.6 | 6 | 0.8333 | 23 | 0.5652 | 5 | 0.6 | 9 | 0.7778 | 6 | 0.5 | 8 | 0.25 | 28 | 0.5357 |
| 17_580404 | 5 | 1 | 6 | 0.8333 | 5 | 1 | 7 | 1 | 23 | 0.9565 | 4 | 1 | 8 | 0.75 | 4 | 1 | 6 | 0.8333 | 22 | 0.8636 |
| 17_580406 | 6 | 0.1667 | 5 | 0.6 | 5 | 0 | 7 | 0.4286 | 23 | 0.3043 | 7 | 0.2857 | 8 | 0.125 | 8 | 0.25 | 8 | 0.25 | 31 | 0.2258 |
| 17_586670 | 13 | 0.0769 | 8 | 0 | 12 | 0.0833 | 4 | 0 | 37 | 0.0541 | 6 | 0 | 1 | 0 | 5 | 0.2 | 5 | 0 | 17 | 0.0588 |
| 17_608303 | 2 | 0 | 4 | 0.25 | NA | NA | NA | NA | 6 | 0.1667 | 3 | 0 | 3 | 0.3333 | 2 | 0 | 3 | 0.3333 | 11 | 0.1818 |
| 17_608410 | 3 | 0 | 6 | 0.1667 | 1 | 0 | 1 | 0 | 11 | 0.0909 | 3 | 0 | 2 | 0 | 2 | 0 | 4 | 0.25 | 11 | 0.0909 |
| 17_608772 | 3 | 0.3333 | 7 | 0.1429 | 2 | 0 | 1 | 1 | 13 | 0.2308 | 4 | 0 | 4 | 0.5 | 2 | 0.5 | 4 | 0.25 | 14 | 0.2857 |
| 17_608778 | 3 | 0.3333 | 1 | 0 | 1 | 1 | NA | NA | 5 | 0.4 | 2 | 0.5 | 4 | 0.5 | 1 | 1 | 1 | 1 | 8 | 0.625 |
| 17_608811 | 3 | 0.6667 | 5 | 0.4 | 3 | 0.6667 | NA | NA | 11 | 0.5455 | 4 | 0.5 | 3 | 0.3333 | 3 | 0.6667 | 3 | 0.6667 | 13 | 0.5385 |
| 17_608819 | 3 | 1 | 5 | 0.8 | 3 | 0.6667 | 1 | 1 | 12 | 0.8333 | 5 | 0.8 | 4 | 0.75 | 1 | 1 | 2 | 0.5 | 12 | 0.75 |
| 17_608824 | 4 | 0 | 7 | 0.1429 | 3 | 0 | 1 | 0 | 15 | 0.0667 | 6 | 0 | 5 | 0.2 | 3 | 0 | 5 | 0.2 | 19 | 0.1053 |
| 17_608825 | 4 | 1 | 6 | 1 | 3 | 1 | 1 | 1 | 14 | 1 | 5 | 0.8 | 5 | 1 | 1 | 0 | 5 | 1 | 16 | 0.875 |
| 17_608831 | 4 | 1 | 6 | 1 | 1 | 1 | 1 | 1 | 12 | 1 | 5 | 1 | 4 | 1 | 3 | 1 | 5 | 1 | 17 | 1 |
| 17_608833 | 2 | 1 | 4 | 1 | 2 | 1 | 1 | 1 | 9 | 1 | 4 | 0.5 | 3 | 0.6667 | 2 | 1 | 4 | 1 | 13 | 0.7692 |
| 17_608834 | 4 | 0 | 7 | 0.1429 | 3 | 0 | 1 | 0 | 15 | 0.0667 | 6 | 0 | 5 | 0.2 | 3 | 0 | 5 | 0 | 19 | 0.0526 |
| 17_608836 | 4 | 1 | 7 | 0.5714 | 3 | 0.3333 | 1 | 0 | 15 | 0.6 | 4 | 0.25 | 5 | 0.6 | 3 | 0 | 5 | 0.8 | 17 | 0.4706 |
| 17_608837 | 4 | 1 | 7 | 1 | 3 | 1 | NA | NA | 14 | 1 | 5 | 0.8 | 4 | 1 | 2 | 1 | 5 | 1 | 16 | 0.9375 |
| 17_608855 | 3 | 1 | 6 | 1 | 4 | 1 | 2 | 1 | 15 | 1 | 6 | 1 | 5 | 1 | 2 | 0.5 | 6 | 1 | 19 | 0.9474 |
| 17_608880 | 2 | 0 | 8 | 0.125 | 2 | 0 | 2 | 0.5 | 14 | 0.1429 | 6 | 0.1667 | 5 | 0 | 2 | 0.5 | 6 | 0 | 19 | 0.1053 |
| 17_608882 | 3 | 0.6667 | 5 | 0.6 | 3 | 1 | 2 | 0.5 | 13 | 0.6923 | 6 | 0.5 | 6 | 1 | 2 | 0 | 4 | 0.5 | 18 | 0.6111 |
| 17_608891 | 3 | 0.3333 | 6 | 0.6667 | 3 | 0.3333 | 1 | 1 | 13 | 0.5385 | 5 | 0.6 | 5 | 0.4 | 2 | 1 | 5 | 0.6 | 17 | 0.5882 |
| 17_608899 | 3 | 0 | 6 | 0.1667 | 3 | 0.3333 | 2 | 0 | 14 | 0.1429 | 4 | 0 | 5 | 0.2 | 2 | 0 | 5 | 0.2 | 16 | 0.125 |
| 17_608909 | 3 | 0.3333 | 5 | 0 | 2 | 1 | 2 | 0 | 12 | 0.25 | 6 | 0.5 | 4 | 0.75 | 2 | 0.5 | 5 | 0.8 | 17 | 0.6471 |
| 17_608912 | 2 | 0.5 | 5 | 0.2 | 3 | 0.3333 | 2 | 0.5 | 12 | 0.3333 | 5 | 0.4 | 6 | 0.3333 | 2 | 0 | 4 | 0 | 17 | 0.2353 |
| 17_608919 | 2 | 0.5 | 6 | 0.1667 | 2 | 0.5 | 2 | 0 | 12 | 0.25 | 5 | 0.2 | 6 | 0.3333 | 2 | 0 | 5 | 0.2 | 18 | 0.2222 |
| 17_608930 | 3 | 1 | 5 | 0.8 | 4 | 1 | 2 | 0.5 | 14 | 0.8571 | 6 | 0.8333 | 6 | 1 | 2 | 1 | 5 | 1 | 19 | 0.9474 |
| 17_608931 | 3 | 0.3333 | 5 | 0.2 | 4 | 0.5 | 2 | 0 | 14 | 0.2857 | 4 | 0 | 5 | 0.4 | 2 | 0.5 | 5 | 0.2 | 16 | 0.25 |
| 17_608944 | 3 | 0.3333 | 6 | 0.1667 | 3 | 0.3333 | 1 | 0 | 13 | 0.2308 | 4 | 0 | 3 | 0.3333 | 2 | 0 | 4 | 0.25 | 13 | 0.1538 |
| 17_608960 | 3 | 0.3333 | 2 | 0.5 | 3 | 0 | 1 | 0 | 9 | 0.2222 | 3 | 1 | 3 | 0.6667 | 2 | 0.5 | 2 | 0 | 10 | 0.6 |
| 17_609112 | 1 | 0 | NA | NA | 4 | 0.25 | 2 | 0.5 | 7 | 0.2857 | 2 | 1 | 3 | 0 | 1 | 1 | 2 | 0 | 8 | 0.375 |
| 17_609113 | NA | NA | NA | NA | 3 | 0.6667 | 1 | 0 | 4 | 0.5 | NA | NA | 2 | 0.5 | NA | NA | 2 | 1 | 4 | 0.75 |
| 17_609121 | 1 | 0 | NA | NA | 3 | 0.3333 | 2 | 0 | 6 | 0.1667 | 2 | 0.5 | 3 | 0 | 1 | 1 | 2 | 0.5 | 8 | 0.375 |
| 17_609142 | NA | NA | NA | NA | 3 | 0.3333 | 2 | 0 | 5 | 0.2 | 2 | 0 | 3 | 0 | 1 | 1 | 2 | 0 | 8 | 0.125 |
| 17_609148 | 1 | 0 | NA | NA | 4 | 0 | 2 | 0.5 | 7 | 0.1429 | 2 | 0.5 | 3 | 0 | 1 | 0 | 1 | 0 | 7 | 0.1429 |
| 17_609150 | 1 | 0 | NA | NA | 4 | 0.75 | 2 | 1 | 7 | 0.7143 | 2 | 1 | 3 | 0.3333 | 1 | 1 | 2 | 0.5 | 8 | 0.625 |
| 17_609271 | NA | NA | NA | NA | 3 | 0.3333 | 1 | 0 | 4 | 0.25 | 2 | 0.5 | NA | NA | NA | NA | 1 | 1 | 3 | 0.6667 |
| 17_609279 | NA | NA | NA | NA | 3 | 0 | 2 | 0.5 | 5 | 0.2 | 2 | 0.5 | 2 | 0.5 | 1 | 0 | 1 | 0 | 6 | 0.3333 |
| 17_609317 | NA | NA | NA | NA | 3 | 0.3333 | 2 | 0 | 5 | 0.2 | 2 | 0.5 | 2 | 0 | 1 | 0 | 1 | 0 | 6 | 0.1667 |
| 17_609407 | NA | NA | NA | NA | 2 | 1 | 1 | 0 | 3 | 0.6667 | 2 | 1 | 1 | 1 | 1 | 1 | 1 | 1 | 5 | 1 |
| 17_609412 | NA | NA | NA | NA | 2 | 1 | 1 | 1 | 3 | 1 | 2 | 1 | 1 | 1 | 1 | 1 | 1 | 1 | 5 | 1 |
| 17_609419 | NA | NA | NA | NA | 2 | 0.5 | 1 | 0 | 3 | 0.3333 | 2 | 1 | 1 | 1 | 1 | 1 | 1 | 1 | 5 | 1 |
| 17_609439 | NA | NA | NA | NA | 1 | 1 | NA | NA | 1 | 1 | 2 | 0.5 | 1 | 1 | NA | NA | 1 | 0 | 4 | 0.5 |
| 17_613482 | 9 | 0 | 23 | 0.087 | 11 | 0.2727 | 14 | 0.0714 | 57 | 0.1053 | 17 | 0.2353 | 11 | 0.0909 | 13 | 0.1538 | 14 | 0.2143 | 55 | 0.1818 |
| 17_613488 | 9 | 0.1111 | 16 | 0.25 | 13 | 0.1538 | 15 | 0.1333 | 53 | 0.1698 | 18 | 0.1667 | 9 | 0.2222 | 14 | 0.0714 | 14 | 0.4286 | 55 | 0.2182 |
| 17_613489 | 9 | 0.2222 | 19 | 0.5263 | 13 | 0.6923 | 13 | 0.7692 | 54 | 0.5741 | 18 | 0.5 | 10 | 0.5 | 10 | 0.5 | 14 | 0.5 | 52 | 0.5 |

| | | | | | | | | | | | | | | | | | | | | |
|---|---|---|---|---|---|---|---|---|---|---|---|---|---|---|---|---|---|---|---|---|
| 17_613496 | 8 | 0 | 21 | 0.0952 | 15 | 0 | 15 | 0.1333 | 59 | 0.0678 | 18 | 0.1111 | 11 | 0.1818 | 13 | 0 | 13 | 0 | 55 | 0.0727 |
| 17_613508 | 8 | 0.25 | 22 | 0.1364 | 14 | 0.0714 | 17 | 0.1765 | 61 | 0.1475 | 20 | 0.15 | 10 | 0.1 | 13 | 0.1538 | 16 | 0.4375 | 59 | 0.2203 |
| 17_613511 | 3 | 0 | 13 | 0.2308 | 10 | 0.3 | 13 | 0.4615 | 39 | 0.3077 | 18 | 0.2222 | 9 | 0.2222 | 9 | 0 | 12 | 0.3333 | 48 | 0.2083 |
| 17_613526 | 2 | 0.5 | 13 | 0.1538 | 12 | 0.0833 | 8 | 0.125 | 35 | 0.1429 | 15 | 0 | 7 | 0.1429 | 8 | 0.25 | 10 | 0.5 | 40 | 0.2 |
| 17_613657 | NA | NA | 3 | 0.3333 | 5 | 0.2 | 1 | 0 | 9 | 0.2222 | 1 | 1 | 2 | 0 | 1 | 1 | 2 | 0.5 | 6 | 0.5 |
| 17_613666 | 1 | 0 | 2 | 0.5 | 4 | 0.25 | 2 | 0 | 9 | 0.2222 | 2 | 0.5 | 3 | 0 | 1 | 1 | 2 | 0 | 8 | 0.25 |
| 17_613689 | 1 | 0 | 2 | 0 | 3 | 0.6667 | 2 | 0 | 8 | 0.25 | 1 | 0 | 2 | 0 | 1 | 1 | 2 | 0.5 | 6 | 0.3333 |
| 17_617479 | NA | NA | 1 | 1 | NA | NA | 1 | 0 | 2 | 0.5 | NA | NA | NA | NA | 1 | 1 | NA | NA | 1 | 1 |
| 17_617498 | NA | NA | 1 | 1 | NA | NA | NA | NA | 1 | 1 | NA | NA | NA | NA | 1 | 1 | NA | NA | 1 | 1 |
| 17_618856 | NA | NA | 1 | 1 | NA | NA | 1 | 0 | 2 | 0.5 | NA | NA | NA | NA | 1 | 1 | NA | NA | 1 | 1 |
| 17_618894 | NA | NA | 1 | 1 | NA | NA | 1 | 1 | 2 | 1 | NA | NA | NA | NA | 1 | 1 | NA | NA | 1 | 1 |
| 17_618912 | NA | NA | 1 | 1 | NA | NA | NA | NA | 1 | 1 | NA | NA | NA | NA | 1 | 1 | NA | NA | 1 | 1 |
| 17_628223 | 1 | 1 | 1 | 1 | NA | NA | 1 | 1 | 3 | 1 | NA | NA | 1 | 0 | 1 | 1 | 1 | 1 | 3 | 0.6667 |
| 17_628250 | 1 | 1 | 1 | 1 | NA | NA | NA | NA | 2 | 1 | 1 | 0 | 1 | 1 | 2 | 0.5 | 1 | 0 | 5 | 0.4 |
| 17_628269 | 1 | 1 | 1 | 1 | NA | NA | 2 | 0.5 | 4 | 0.75 | NA | NA | 1 | 1 | 2 | 0.5 | 2 | 1 | 5 | 0.8 |
| 17_628276 | 1 | 1 | 1 | 1 | NA | NA | 2 | 1 | 4 | 1 | NA | NA | 1 | 1 | 2 | 1 | 1 | 1 | 4 | 1 |
| 17_628322 | 1 | 1 | 1 | 0 | NA | NA | 2 | 0 | 4 | 0.25 | NA | NA | 1 | 0 | NA | NA | 2 | 0.5 | 3 | 0.3333 |
| 17_628339 | 1 | 1 | 2 | 0.5 | NA | NA | 1 | 1 | 4 | 0.75 | NA | NA | NA | NA | 2 | 0.5 | 2 | 1 | 4 | 0.75 |
| 17_628391 | 1 | 1 | 2 | 0.5 | 1 | 1 | 3 | 0.6667 | 7 | 0.7143 | NA | NA | 2 | 1 | 1 | 1 | 3 | 0.6667 | 6 | 0.8333 |
| 17_628407 | 1 | 1 | 2 | 1 | 1 | 1 | 3 | 1 | 7 | 1 | NA | NA | 2 | 1 | 2 | 0.5 | 4 | 1 | 8 | 0.875 |
| 17_628408 | 1 | 1 | 2 | 1 | 1 | 1 | 3 | 1 | 7 | 1 | 1 | 1 | NA | NA | 1 | 1 | 3 | 1 | 5 | 1 |
| 17_628660 | NA | NA | 3 | 0.3333 | 1 | 0 | 4 | 0.5 | 8 | 0.375 | 2 | 0.5 | 2 | 0.5 | 3 | 1 | 4 | 0.25 | 11 | 0.5455 |
| 17_628662 | 2 | 0.5 | 3 | 0.6667 | NA | NA | 3 | 0.3333 | 8 | 0.5 | 1 | 1 | 2 | 0.5 | 3 | 0.3333 | 4 | 1 | 10 | 0.7 |
| 17_628705 | 2 | 0 | 3 | 0.3333 | 1 | 0 | 5 | 0 | 11 | 0.0909 | 1 | 0 | 3 | 0 | 3 | 0 | 3 | 0.3333 | 10 | 0.1 |
| 17_628721 | 2 | 0 | 1 | 1 | 1 | 0 | 5 | 0.4 | 9 | 0.3333 | 3 | 0.3333 | 3 | 0.3333 | 2 | 0.5 | 5 | 0.4 | 13 | 0.3846 |
| 17_628722 | 2 | 0 | 2 | 0 | 1 | 0 | 4 | 0.5 | 9 | 0.2222 | 2 | 0 | 4 | 0 | 3 | 0.3333 | 3 | 0.3333 | 12 | 0.1667 |
| 17_628782 | 2 | 0 | 4 | 0.5 | 2 | 0.5 | 4 | 0 | 12 | 0.25 | 2 | 0 | 5 | 0 | 3 | 0.3333 | 6 | 0.3333 | 16 | 0.1875 |
| 17_628790 | 2 | 0 | 3 | 0.3333 | 2 | 0.5 | 4 | 0 | 11 | 0.1818 | 2 | 0 | 4 | 0.25 | 3 | 0 | 4 | 0.25 | 13 | 0.1538 |
| 17_628819 | 1 | 0 | 3 | 0.3333 | 1 | 0 | 4 | 0 | 9 | 0.1111 | 4 | 0 | 6 | 0 | 3 | 0 | 6 | 0.1667 | 19 | 0.0526 |
| 17_629014 | 7 | 0.4286 | 6 | 0.3333 | 5 | 0.6 | 10 | 0.2 | 28 | 0.3571 | 9 | 0.2222 | 8 | 0.625 | 8 | 0.125 | 9 | 0.3333 | 34 | 0.3235 |
| 17_629016 | 7 | 0 | 8 | 0.25 | 5 | 0.2 | 10 | 0.1 | 30 | 0.1333 | 8 | 0.375 | 10 | 0.2 | 8 | 0.125 | 10 | 0.4 | 36 | 0.2778 |
| 17_629021 | 7 | 0.1429 | 9 | 0.1111 | 5 | 0 | 12 | 0 | 33 | 0.0606 | 9 | 0.2222 | 9 | 0 | 9 | 0 | 9 | 0.1111 | 36 | 0.0833 |
| 17_635564 | 1 | 0 | 7 | 0.1429 | 2 | 0 | 5 | 0 | 15 | 0.0667 | 2 | 0 | 6 | 0 | 7 | 0 | 5 | 0.2 | 20 | 0.05 |
| 17_635571 | 1 | 1 | 4 | 0.25 | 1 | 0 | 3 | 0 | 9 | 0.2222 | 2 | 0 | 5 | 0.6 | 3 | 0.3333 | 4 | 0.75 | 14 | 0.5 |
| 17_635571 | 1 | 0 | 7 | 0 | 1 | 0 | 5 | 0.2 | 14 | 0.0714 | 1 | 0 | 6 | 0.1667 | 6 | 0.3333 | 4 | 0 | 17 | 0.1765 |
| 17_635574 | 1 | 0 | 5 | 0.2 | 1 | 1 | 3 | 0 | 10 | 0.2 | 2 | 0 | 5 | 0.4 | 3 | 0 | 4 | 0 | 14 | 0.1429 |
| 17_635580 | 1 | 1 | 5 | 0.2 | 2 | 1 | 5 | 0.6 | 13 | 0.5385 | 2 | 1 | 5 | 0.8 | 4 | 0.5 | 5 | 1 | 16 | 0.8125 |
| 17_635626 | 1 | 0 | 3 | 0.6667 | 2 | 1 | 5 | 0.6 | 11 | 0.6364 | 2 | 0 | 5 | 0.4 | 5 | 0.2 | 3 | 0.6667 | 15 | 0.3333 |
| 17_635664 | 1 | 1 | 4 | 0.25 | 3 | 0.6667 | 6 | 0.6667 | 14 | 0.5714 | 2 | 1 | 5 | 0.8 | 6 | 0.5 | 3 | 0.6667 | 16 | 0.6875 |
| 17_635669 | 1 | 0 | 4 | 0.5 | 2 | 0.5 | 6 | 0 | 13 | 0.2308 | 2 | 0 | 3 | 0.3333 | 5 | 0.2 | 2 | 0.5 | 12 | 0.25 |
| 17_635671 | 1 | 0 | 2 | 0 | 1 | 1 | 6 | 0.3333 | 10 | 0.3 | 1 | 0 | 5 | 0.4 | 6 | 0 | 2 | 0 | 14 | 0.1429 |
| 17_635700 | NA | NA | NA | NA | NA | NA | 1 | 1 | 1 | 1 | NA | NA | NA | NA | NA | NA | 1 | 1 | 1 | 1 |
| 17_635711 | 3 | 0 | 4 | 0.25 | 2 | 0 | 6 | 0 | 15 | 0.0667 | 2 | 0 | 4 | 0 | 6 | 0 | 3 | 0.3333 | 15 | 0.0667 |
| 17_635712 | 3 | 1 | 2 | 1 | 1 | 1 | 5 | 0.8 | 11 | 0.9091 | 1 | 1 | 3 | 1 | 6 | 0.5 | 1 | 1 | 11 | 0.7273 |
| 17_635713 | 3 | 0 | 3 | 0 | 2 | 0 | 7 | 0.1429 | 15 | 0.0667 | 2 | 0 | 5 | 0.2 | 7 | 0.4286 | 3 | 0 | 17 | 0.2353 |
| 17_635740 | 3 | 0 | 4 | 0 | 4 | 0.25 | 8 | 0 | 19 | 0.0526 | 2 | 0 | 5 | 0 | 8 | 0 | 3 | 0.3333 | 18 | 0.0556 |
| 17_635741 | 1 | 0 | 4 | 0.25 | 3 | 0.3333 | 8 | 0 | 16 | 0.125 | 2 | 0.5 | 5 | 0.2 | 8 | 0 | 3 | 0.3333 | 18 | 0.1667 |
| 17_635742 | 2 | 0.5 | 3 | 0 | 1 | 0 | 6 | 0.1667 | 12 | 0.1667 | 2 | 0 | 3 | 0.3333 | 5 | 0 | 2 | 0 | 12 | 0.0833 |
| 17_635763 | 3 | 0.3333 | 4 | 0 | 3 | 0.3333 | 8 | 0 | 18 | 0.1111 | 1 | 0 | 4 | 0 | 8 | 0.25 | 3 | 0.3333 | 16 | 0.1875 |
| 17_635765 | 3 | 0 | 4 | 0 | 4 | 0.25 | 8 | 0 | 19 | 0.0526 | 2 | 0 | 5 | 0.2 | 8 | 0 | 3 | 0.3333 | 18 | 0.1111 |
| 17_635770 | NA | NA | NA | NA | NA | NA | 1 | 1 | 1 | 1 | NA | NA | NA | NA | NA | NA | 1 | 1 | 1 | 1 |
| 17_635857 | NA | NA | NA | NA | NA | NA | 1 | 1 | 1 | 1 | NA | NA | NA | NA | NA | NA | 1 | 1 | 1 | 1 |
| 17_635864 | NA | NA | NA | NA | NA | NA | 1 | 1 | 1 | 1 | NA | NA | NA | NA | NA | NA | 1 | 1 | 1 | 1 |
| 17_635893 | NA | NA | NA | NA | NA | NA | 1 | 1 | 1 | 1 | NA | NA | NA | NA | NA | NA | 1 | 1 | 1 | 1 |
| 17_636102 | 2 | 0 | NA | NA | 1 | 1 | 4 | 0.25 | 7 | 0.2857 | NA | NA | 1 | 1 | 2 | 0 | 2 | 0.5 | 5 | 0.4 |
| 17_636116 | 2 | 0 | NA | NA | NA | NA | 4 | 0.25 | 6 | 0.1667 | NA | NA | 1 | 1 | 3 | 0.3333 | 1 | 0 | 5 | 0.4 |
| 17_670335 | 4 | 0 | 5 | 0 | 6 | 0 | 2 | 0.5 | 17 | 0.0588 | 4 | 0.25 | 4 | 0 | 2 | 0 | 2 | 0 | 12 | 0.0833 |
| 17_670710 | 3 | 0 | 9 | 0.1111 | 4 | 0 | 3 | 0 | 19 | 0.0526 | 7 | 0 | 8 | 0.25 | 12 | 0.1667 | 5 | 0 | 32 | 0.125 |
| 17_670738 | 3 | 0.3333 | 6 | 0 | 3 | 0 | 3 | 0 | 15 | 0.0667 | 7 | 0.1429 | 7 | 0.1429 | 9 | 0.3333 | 4 | 0.25 | 27 | 0.2222 |
| 17_670739 | 3 | 1 | 5 | 0.6 | 1 | 1 | 3 | 1 | 12 | 0.8333 | 5 | 1 | 7 | 1 | 6 | 0.8333 | 4 | 0.75 | 22 | 0.9091 |
| 17_670740 | 1 | 0 | 5 | 0 | 2 | 0 | 3 | 0.3333 | 11 | 0.0909 | 5 | 0 | 6 | 0.1667 | 11 | 0.0909 | 3 | 0 | 25 | 0.08 |
| 17_670764 | 2 | 0 | 5 | 0 | 2 | 0 | 2 | 0.5 | 11 | 0.0909 | 5 | 0.2 | 5 | 0.4 | 7 | 0.1429 | 3 | 0.3333 | 20 | 0.25 |
| 17_670778 | 3 | 1 | 5 | 1 | 1 | 1 | 4 | 1 | 13 | 1 | 7 | 1 | 8 | 0.875 | 10 | 0.9 | 2 | 1 | 27 | 0.9259 |
| 17_670780 | 3 | 0 | 6 | 0.1667 | 3 | 0.3333 | 3 | 0.3333 | 15 | 0.2 | 6 | 0.3333 | 7 | 0.1429 | 10 | 0.4 | 2 | 0 | 25 | 0.28 |
| 17_670782 | 3 | 1 | 7 | 0.5714 | 2 | 0 | 3 | 1 | 15 | 0.6667 | 7 | 0.5714 | 8 | 0.375 | 10 | 0.5 | 2 | 0.5 | 27 | 0.4815 |

| | | | | | | | | | | | | | | | | | | | | |
|---|---|---|---|---|---|---|---|---|---|---|---|---|---|---|---|---|---|---|---|---|
| 17_670785 | 3 | 0.6667 | 7 | 0.4286 | 3 | 0 | 3 | 0.3333 | 16 | 0.375 | 6 | 0.1667 | 7 | 0.2857 | 8 | 0.125 | 2 | 0 | 23 | 0.1739 |
| 17_670787 | 3 | 1 | 8 | 0.75 | 2 | 1 | 3 | 1 | 16 | 0.875 | 6 | 0.5 | 8 | 0.875 | 9 | 1 | 3 | 1 | 26 | 0.8462 |
| 17_670826 | 3 | 0.6667 | 4 | 0.25 | 2 | 1 | 3 | 1 | 12 | 0.6667 | 5 | 0.4 | 7 | 0.8571 | 8 | 0.5 | 2 | 1 | 22 | 0.6364 |
| 17_670840 | 3 | 0.3333 | 5 | 0 | 1 | 0 | 3 | 0 | 12 | 0.0833 | 5 | 0 | 7 | 0 | 7 | 0.4286 | 2 | 0 | 21 | 0.1429 |
| 17_670871 | 4 | 1 | 6 | 0.6667 | 5 | 1 | 3 | 0.3333 | 18 | 0.7778 | 4 | 0.5 | 6 | 0.6667 | 6 | 0.6667 | 4 | 0.75 | 20 | 0.65 |
| 17_670880 | 5 | 0.2 | 6 | 0 | 4 | 0.25 | 3 | 0 | 18 | 0.1111 | 4 | 0.25 | 7 | 0 | 7 | 0.2857 | 2 | 0 | 20 | 0.15 |
| 17_670880 | 2 | 0 | 3 | 0.3333 | 5 | 0.2 | 2 | 1 | 12 | 0.3333 | 5 | 0.2 | 4 | 0.5 | 6 | 0.5 | 3 | 0.3333 | 18 | 0.3889 |
| 17_700615 | 11 | 0.1818 | 14 | 0.2143 | 9 | 0.4444 | 8 | 0.75 | 42 | 0.3571 | 18 | 0.2222 | 4 | 0.25 | 9 | 0.1111 | 9 | 0.4444 | 40 | 0.25 |
| 17_700619 | 11 | 0 | 15 | 0.1333 | 8 | 0.5 | 9 | 0.1111 | 43 | 0.1628 | 22 | 0.2727 | 7 | 0.1429 | 9 | 0.3333 | 10 | 0 | 48 | 0.2083 |
| 17_700668 | 11 | 0.2727 | 9 | 0.3333 | 8 | 0.125 | 5 | 0 | 33 | 0.2121 | 13 | 0.1538 | 5 | 0.2 | 4 | 0.75 | 6 | 0.1667 | 28 | 0.25 |
| 17_706587 | NA | NA | NA | NA | NA | NA | 1 | 1 | 1 | 1 | 1 | 1 | 1 | 1 | NA | NA | NA | NA | 2 | 1 |
| 17_706594 | NA | NA | NA | NA | NA | NA | 1 | 1 | 1 | 1 | NA | NA | 1 | 1 | NA | NA | NA | NA | 1 | 1 |
| 17_717221 | NA | NA | 5 | 0 | 3 | 0.3333 | 1 | 0 | 9 | 0.1111 | 2 | 0.5 | 2 | 0 | 1 | 0 | 3 | 0 | 8 | 0.125 |
| 17_719026 | 1 | 1 | NA | NA | NA | NA | 1 | 0 | 2 | 0.5 | NA | NA | NA | NA | 1 | 1 | NA | NA | 1 | 1 |
| 17_719057 | 1 | 1 | NA | NA | NA | NA | 1 | 1 | 2 | 1 | NA | NA | NA | NA | 1 | 1 | NA | NA | 1 | 1 |
| 17_719087 | 1 | 1 | NA | NA | NA | NA | 1 | 1 | 2 | 1 | NA | NA | NA | NA | 1 | 1 | NA | NA | 1 | 1 |
| 17_719104 | 1 | 0 | NA | NA | NA | NA | 1 | 1 | 2 | 0.5 | NA | NA | NA | NA | 1 | 1 | NA | NA | 1 | 1 |
| 17_722272 | 3 | 0.3333 | 7 | 0.1429 | 3 | 0.6667 | 9 | 0.3333 | 22 | 0.3182 | 5 | 0 | 1 | 1 | 7 | 0 | 4 | 0 | 17 | 0.0588 |
| 17_722273 | 4 | 0.5 | 6 | 0.3333 | 3 | 0.6667 | 10 | 0.1 | 23 | 0.3043 | 3 | 0.6667 | 1 | 0 | 7 | 0 | 3 | 0.3333 | 14 | 0.2143 |
| 17_722309 | 3 | 1 | 4 | 1 | 2 | 0.5 | 6 | 1 | 15 | 0.9333 | NA | NA | 1 | 1 | 4 | 1 | 3 | 1 | 8 | 1 |
| 17_722312 | 2 | 1 | 5 | 1 | 3 | 1 | 12 | 1 | 22 | 1 | 4 | 1 | 1 | 1 | 4 | 1 | 4 | 1 | 13 | 1 |
| 17_722325 | 4 | 1 | 6 | 1 | 3 | 1 | 12 | 1 | 25 | 1 | 3 | 1 | 1 | 1 | 4 | 1 | 4 | 1 | 12 | 1 |
| 17_722328 | 4 | 1 | 3 | 1 | 2 | 1 | 10 | 0.9 | 19 | 0.9474 | 2 | 1 | 1 | 1 | 6 | 1 | 2 | 1 | 11 | 1 |
| 17_722332 | 3 | 0 | 5 | 1 | 2 | 0.5 | 10 | 0.8 | 20 | 0.7 | 2 | 0 | 1 | 1 | 6 | 0.6667 | 4 | 1 | 13 | 0.6923 |
| 17_722334 | 4 | 1 | 4 | 1 | 3 | 0.6667 | 11 | 0.8182 | 22 | 0.8636 | 2 | 0.5 | 1 | 1 | 5 | 0.8 | 3 | 0.3333 | 11 | 0.6364 |
| 17_722335 | 4 | 0.5 | 6 | 0.6667 | 3 | 0.3333 | 10 | 0.7 | 23 | 0.6087 | 2 | 0.5 | 1 | 1 | 3 | 0.3333 | 2 | 0 | 8 | 0.375 |
| 17_722336 | 4 | 1 | 6 | 0.3333 | 2 | 0.5 | 11 | 0.6364 | 23 | 0.6087 | 2 | 0.5 | 1 | 0 | 5 | 0.8 | 4 | 1 | 12 | 0.75 |
| 17_722338 | 4 | 0.75 | 4 | 0.5 | 2 | 1 | 11 | 0.7273 | 21 | 0.7143 | 1 | 1 | 1 | 0 | 6 | 0.5 | 3 | 0.6667 | 11 | 0.5455 |
| 17_722339 | 4 | 0.75 | 4 | 0.75 | 3 | 0.6667 | 11 | 0.6364 | 22 | 0.6818 | 2 | 1 | 1 | 0 | 4 | 1 | 4 | 0.5 | 11 | 0.7273 |
| 17_722348 | 4 | 0 | 6 | 0.1667 | 3 | 0 | 12 | 0.0833 | 25 | 0.08 | 2 | 0 | 1 | 1 | 5 | 0.2 | 3 | 0 | 11 | 0.1818 |
| 17_722350 | 3 | 1 | 6 | 1 | 2 | 1 | 10 | 0.9 | 21 | 0.9524 | 1 | 1 | 1 | 1 | 5 | 0.8 | 3 | 0.3333 | 10 | 0.7 |
| 17_722351 | 3 | 0 | 4 | 0.25 | 3 | 0 | 10 | 0.1 | 20 | 0.1 | 1 | 0 | NA | NA | 4 | 0.25 | 4 | 0 | 9 | 0.1111 |
| 17_722370 | 2 | 0.5 | 5 | 1 | 3 | 1 | 7 | 0.8571 | 17 | 0.8824 | 2 | 1 | 1 | 1 | 5 | 1 | 4 | 0.75 | 12 | 0.9167 |
| 17_722371 | 3 | 0 | 5 | 0.2 | 2 | 0 | 11 | 0.2727 | 21 | 0.1905 | 2 | 0 | 1 | 1 | 3 | 0 | 4 | 0.25 | 10 | 0.2 |
| 17_722373 | 3 | 1 | 5 | 1 | 3 | 1 | 11 | 1 | 22 | 1 | 2 | 1 | 1 | 1 | 5 | 1 | 2 | 1 | 10 | 1 |
| 17_722384 | 3 | 0 | 5 | 0 | 3 | 0 | 9 | 0.1111 | 20 | 0.05 | 2 | 0 | 1 | 0 | 4 | 0 | 4 | 0.25 | 11 | 0.0909 |
| 17_722384 | 2 | 1 | 5 | 0.8 | 3 | 0.3333 | 7 | 0.7143 | 17 | 0.7059 | 1 | 0 | 1 | 1 | 3 | 0.3333 | 4 | 0.5 | 9 | 0.4444 |
| 17_722396 | 3 | 0.6667 | 5 | 0.6 | 3 | 1 | 10 | 1 | 21 | 0.8571 | 2 | 0.5 | 1 | 0 | 4 | 0.75 | 4 | 0.75 | 11 | 0.6364 |
| 17_722399 | 2 | 0 | 5 | 0.2 | 3 | 0.3333 | 9 | 0.5556 | 19 | 0.3684 | 2 | 0.5 | 1 | 1 | 5 | 0.2 | 4 | 0.25 | 12 | 0.3333 |
| 17_722402 | 3 | 0 | 5 | 0 | 3 | 0.3333 | 10 | 0.3 | 21 | 0.1905 | 2 | 0 | 1 | 0 | 5 | 0.6 | 4 | 0 | 12 | 0.25 |
| 17_722403 | 3 | 0 | 5 | 0.2 | 1 | 1 | 9 | 0.2222 | 18 | 0.2222 | 2 | 0.5 | NA | NA | 3 | 0 | 3 | 0 | 8 | 0.125 |
| 17_722412 | 3 | 0.3333 | 3 | 0 | 3 | 0.3333 | 9 | 0.2222 | 18 | 0.2222 | 2 | 0 | 1 | 1 | 5 | 0.2 | 4 | 0 | 12 | 0.1667 |
| 17_722421 | NA | NA | NA | NA | NA | NA | 1 | 1 | 1 | 1 | 1 | 1 | NA | NA | NA | NA | NA | NA | 1 | 1 |
| 17_722422 | 3 | 0.6667 | 4 | 0.5 | 3 | 0 | 10 | 0.5 | 20 | 0.45 | 2 | 0.5 | 1 | 1 | 4 | 0.5 | 3 | 0.3333 | 10 | 0.5 |
| 17_722425 | NA | NA | NA | NA | NA | NA | 1 | 1 | 1 | 1 | 1 | 1 | NA | NA | NA | NA | NA | NA | 1 | 1 |
| 17_725158 | 5 | 0.2 | 7 | 0.1429 | 5 | 0 | 2 | 0 | 19 | 0.1053 | NA | NA | 2 | 0 | 3 | 0.3333 | 5 | 0 | 10 | 0.1 |
| 17_756765 | 3 | 0 | 5 | 0 | 7 | 0.1429 | 5 | 0 | 20 | 0.05 | 4 | 0 | 3 | 0.3333 | 1 | 0 | 5 | 0.4 | 13 | 0.2308 |
| 17_756771 | 4 | 0.5 | 5 | 0.8 | 7 | 0.7143 | 4 | 0.75 | 20 | 0.7 | 3 | 1 | 3 | 0.3333 | 2 | 1 | 5 | 0.4 | 13 | 0.6154 |
| 17_756779 | 3 | 0 | 4 | 0.25 | 7 | 0.2857 | 5 | 0.6 | 19 | 0.3158 | 3 | 0.3333 | 3 | 0.3333 | 2 | 0.5 | 5 | 0.2 | 13 | 0.3077 |
| 17_756780 | 4 | 0.25 | 5 | 0.4 | 8 | 0.25 | 5 | 0 | 22 | 0.2273 | 4 | 0 | 3 | 0.3333 | 1 | 0 | 4 | 0.25 | 12 | 0.1667 |
| 17_756797 | 5 | 0 | 5 | 0.2 | 7 | 0 | 5 | 0.2 | 22 | 0.0909 | 4 | 0 | 3 | 0 | 2 | 0 | 4 | 0.25 | 13 | 0.0769 |
| 17_756814 | 5 | 0 | 5 | 0.2 | 8 | 0.125 | 8 | 0.125 | 26 | 0.1154 | 4 | 0 | 4 | 0.25 | 2 | 0 | 4 | 0 | 14 | 0.0714 |
| 17_756831 | 5 | 0.2 | 4 | 0.5 | 8 | 0.375 | 8 | 0.25 | 25 | 0.32 | 5 | 0 | 5 | 0 | 2 | 0.5 | 4 | 0.5 | 16 | 0.1875 |
| 17_756847 | NA | NA | 6 | 0.5 | 5 | 1 | 12 | 0.3333 | 23 | 0.5217 | 3 | 0.3333 | 3 | 0 | 2 | 0.5 | 5 | 0.6 | 13 | 0.3846 |
| 17_756862 | 5 | 0 | 2 | 0 | 8 | 0.125 | 11 | 0.0909 | 26 | 0.0769 | 5 | 0 | 4 | 0.25 | 1 | 1 | 6 | 0 | 16 | 0.125 |
| 17_756868 | 5 | 0 | 5 | 0 | 8 | 0.375 | 13 | 0.3077 | 31 | 0.2258 | 5 | 0.2 | 5 | 0.2 | 2 | 0.5 | 5 | 0.4 | 17 | 0.2941 |
| 17_756880 | 5 | 0 | 5 | 0.2 | 7 | 0.1429 | 14 | 0.2857 | 31 | 0.1935 | 5 | 0.2 | 5 | 0.2 | 3 | 0 | 6 | 0 | 19 | 0.1053 |
| 17_756884 | 5 | 0 | 6 | 0 | 8 | 0 | 14 | 0.1429 | 33 | 0.0606 | 5 | 0 | 5 | 0.2 | 3 | 0 | 6 | 0 | 19 | 0.0526 |
| 17_756889 | 5 | 0.2 | 5 | 0 | 6 | 0 | 14 | 0.2143 | 30 | 0.1333 | 4 | 0 | 5 | 0.2 | 3 | 0.6667 | 7 | 0 | 19 | 0.1579 |
| 17_756894 | 5 | 0 | 4 | 0.25 | 7 | 0.5714 | 13 | 0.2308 | 29 | 0.2759 | 5 | 0.6 | 4 | 0.75 | 2 | 0.5 | 7 | 0.2857 | 18 | 0.5 |
| 17_756911 | 5 | 0.2 | 4 | 0.25 | 8 | 0 | 11 | 0 | 28 | 0.0714 | 4 | 0 | 4 | 0.25 | 3 | 0 | 7 | 0 | 18 | 0.0556 |
| 17_756919 | 3 | 0.6667 | 4 | 1 | 7 | 0.4286 | 9 | 0.8889 | 23 | 0.7391 | 4 | 0.5 | 5 | 0.4 | 3 | 1 | 6 | 0.6667 | 18 | 0.6111 |
| 17_756923 | 4 | 0 | 5 | 0.4 | 8 | 0.375 | 14 | 0.5 | 31 | 0.3871 | 5 | 0.4 | 4 | 0.25 | 2 | 0.5 | 6 | 0.5 | 17 | 0.4118 |
| 17_756991 | 6 | 0.1667 | 6 | 0 | 9 | 0 | 12 | 0.0833 | 33 | 0.0606 | 7 | 0.2857 | 6 | 0.1667 | 5 | 0 | 10 | 0.2 | 28 | 0.1786 |
| 17_756996 | 5 | 0 | 5 | 0 | 10 | 0.1 | 14 | 0.0714 | 34 | 0.0588 | 6 | 0.1667 | 7 | 0.1429 | 5 | 0.2 | 10 | 0 | 28 | 0.1071 |

| | | | | | | | | | | | | | | | | | | | | |
|---|---|---|---|---|---|---|---|---|---|---|---|---|---|---|---|---|---|---|---|---|
| 17_757002( | 7 | 0.5714 | 6 | 0.8333 | 9 | 0.6667 | 14 | 0.7143 | 36 | 0.6944 | 7 | 0.5714 | 5 | 0.4 | 5 | 0.6 | 9 | 0.6667 | 26 | 0.5769 |
| 17_757009' | 6 | 0 | 3 | 0.6667 | 6 | 0.3333 | 12 | 0.0833 | 27 | 0.1852 | 7 | 0 | 2 | 0 | 4 | 0 | 10 | 0.3 | 23 | 0.1304 |
| 17_757038! | 9 | 0.2222 | 7 | 0.4286 | 9 | 0.3333 | 14 | 0.5 | 39 | 0.3846 | 9 | 0.3333 | 6 | 0.1667 | 3 | 0.3333 | 8 | 0.5 | 26 | 0.3462 |
| 17_757043 | 12 | 0.8333 | 7 | 0.8571 | 10 | 1 | 12 | 0.9167 | 41 | 0.9024 | 8 | 0.5 | 5 | 1 | 5 | 0.8 | 7 | 1 | 25 | 0.8 |
| 17_757048! | 12 | 0.4167 | 8 | 0.375 | 8 | 0.25 | 13 | 0.2308 | 41 | 0.3171 | 5 | 0.2 | 6 | 0.1667 | 4 | 0.25 | 9 | 0 | 24 | 0.125 |
| 17_795755 | NA | NA | 2 | 1 | NA | NA | 2 | 1 | 4 | 1 | NA | NA | NA | NA | 4 | 1 | NA | NA | 4 | 1 |
| 17_795775 | NA | NA | 2 | 0.5 | NA | NA | 2 | 0.5 | 4 | 0.5 | NA | NA | NA | NA | 2 | 1 | NA | NA | 2 | 1 |
| 17_795778 | NA | NA | 2 | 1 | NA | NA | 2 | 0.5 | 4 | 0.75 | NA | NA | NA | NA | 3 | 1 | NA | NA | 3 | 1 |
| 17_795786( | NA | NA | 1 | 1 | NA | NA | 2 | 1 | 3 | 1 | NA | NA | NA | NA | 2 | 1 | NA | NA | 2 | 1 |
| 17_795796 | NA | NA | 2 | 0.5 | NA | NA | 1 | 1 | 3 | 0.6667 | NA | NA | NA | NA | 2 | 1 | NA | NA | 2 | 1 |
| 17_795812 | NA | NA | 1 | 0 | NA | NA | 2 | 0.5 | 3 | 0.3333 | NA | NA | NA | NA | 3 | 0.6667 | NA | NA | 3 | 0.6667 |
| 17_797980 | NA | NA | 1 | 1 | NA | NA | 1 | 1 | 2 | 1 | NA | NA | NA | NA | 1 | 1 | NA | NA | 1 | 1 |
| 17_798024 | NA | NA | 1 | 1 | NA | NA | NA | NA | 1 | 1 | NA | NA | NA | NA | 1 | 1 | NA | NA | 1 | 1 |
| 17_798306! | NA | NA | 1 | 1 | NA | NA | NA | NA | 1 | 1 | NA | NA | NA | NA | 1 | 1 | NA | NA | 1 | 1 |
| 17_799173 | NA | NA | NA | NA | NA | NA | 2 | 0.5 | 2 | 0.5 | NA | NA | NA | NA | 1 | 1 | NA | NA | 1 | 1 |
| 17_799247( | NA | NA | NA | NA | NA | NA | 2 | 0.5 | 2 | 0.5 | NA | NA | NA | NA | 1 | 1 | NA | NA | 1 | 1 |
| 17_810002 | 1 | 1 | NA | NA | 2 | 1 | 2 | 1 | 5 | 1 | 1 | 1 | NA | NA | 1 | 0 | 1 | 1 | 3 | 0.6667 |
| 17_810016 | NA | NA | NA | NA | 1 | 0 | 2 | 1 | 3 | 0.6667 | 1 | 1 | NA | NA | NA | NA | NA | NA | 1 | 1 |
| 17_810017 | 1 | 1 | NA | NA | 1 | 0 | 2 | 1 | 4 | 0.75 | 1 | 1 | NA | NA | 1 | 1 | 1 | 1 | 3 | 1 |
| 17_810018 | 1 | 1 | NA | NA | 1 | 1 | 1 | 0 | 3 | 0.6667 | NA | NA | NA | NA | 1 | 0 | 1 | 1 | 2 | 0.5 |
| 17_810034 | 1 | 1 | NA | NA | 2 | 0 | 1 | 0 | 4 | 0.25 | 1 | 1 | NA | NA | NA | NA | 1 | 0 | 2 | 0.5 |
| 17_810079 | 1 | 1 | NA | NA | 2 | 0.5 | 1 | 1 | 4 | 0.75 | 1 | 1 | NA | NA | NA | NA | 1 | 1 | 2 | 1 |
| 17_810088 | 1 | 1 | NA | NA | 2 | 1 | 1 | 1 | 4 | 1 | 1 | 1 | NA | NA | NA | NA | 1 | 1 | 2 | 1 |
| 17_810111 | 1 | 1 | NA | NA | NA | NA | 1 | 1 | 2 | 1 | 1 | 1 | NA | NA | NA | NA | NA | NA | 1 | 1 |
| 17_810129( | NA | NA | NA | NA | 1 | 1 | 1 | 1 | 2 | 1 | 1 | 1 | NA | NA | NA | NA | 1 | 0 | 2 | 0.5 |
| 17_810140 | 1 | 1 | NA | NA | 1 | 0 | 1 | 1 | 3 | 0.6667 | 1 | 1 | NA | NA | NA | NA | NA | NA | 1 | 1 |
| 17_819825 | 2 | 0 | 2 | 0.5 | 3 | 0.3333 | 7 | 0.2857 | 14 | 0.2857 | 5 | 0.2 | NA | NA | 3 | 0.3333 | 3 | 0 | 11 | 0.1818 |
| 17_820000 | 1 | 0 | NA | NA | 2 | 0.5 | NA | NA | 3 | 0.3333 | NA | NA | 1 | 1 | NA | NA | NA | NA | 1 | 1 |
| 17_820027 | 1 | 0 | NA | NA | 1 | 1 | NA | NA | 2 | 0.5 | NA | NA | 1 | 1 | NA | NA | NA | NA | 1 | 1 |
| 17_821207 | NA | NA | NA | NA | NA | NA | 2 | 0.5 | 2 | 0.5 | NA | NA | NA | NA | NA | NA | 1 | 1 | 1 | 1 |
| 17_833564 | NA | NA | NA | NA | NA | NA | 1 | 1 | 1 | 1 | NA | NA | 1 | 1 | NA | NA | NA | NA | 1 | 1 |
| 17_841354 | 1 | 1 | 3 | 1 | 1 | 1 | 2 | 1 | 7 | 1 | 3 | 1 | 1 | 0 | 2 | 1 | 5 | 1 | 11 | 0.9091 |
| 17_841356 | 1 | 0 | 3 | 0.6667 | 1 | 0 | 1 | 0 | 6 | 0.3333 | 4 | 0.25 | 1 | 0 | 3 | 0 | 4 | 0.25 | 12 | 0.1667 |
| 17_841388 | 1 | 0 | 2 | 0.5 | 2 | 0 | 2 | 0 | 7 | 0.1429 | 4 | 0.25 | 1 | 0 | 3 | 0.3333 | 5 | 0.2 | 13 | 0.2308 |
| 17_841444 | 1 | 0 | 3 | 0.3333 | 2 | 0.5 | 2 | 0 | 8 | 0.25 | 4 | 0.25 | 1 | 0 | 2 | 0.5 | 3 | 0 | 10 | 0.2 |
| 17_841496 | 1 | 1 | 1 | 0 | 1 | 1 | 2 | 0 | 5 | 0.4 | 2 | 0 | NA | NA | 2 | 0.5 | NA | NA | 4 | 0.25 |
| 17_841501( | NA | NA | 2 | 0 | 2 | 0.5 | 2 | 0 | 6 | 0.1667 | 3 | 0.3333 | 1 | 0 | 2 | 0 | NA | NA | 6 | 0.1667 |
| 17_841504( | 1 | 1 | 2 | 1 | 2 | 1 | 2 | 0 | 7 | 0.7143 | 3 | 0 | 1 | 0 | 1 | 1 | 1 | 1 | 6 | 0.3333 |
| 17_854593 | 7 | 0.1429 | 5 | 0 | 6 | 0 | 9 | 0.1111 | 27 | 0.0741 | 2 | 0 | 7 | 0 | 4 | 0.25 | NA | NA | 13 | 0.0769 |
| 17_855932( | 1 | 1 | NA | NA | 1 | 0 | NA | NA | 2 | 0.5 | 1 | 1 | NA | NA | NA | NA | 1 | 1 | 2 | 1 |
| 17_855937 | 2 | 0.5 | NA | NA | 2 | 0 | NA | NA | 4 | 0.25 | 2 | 0.5 | NA | NA | 1 | 0 | 1 | 1 | 4 | 0.5 |
| 17_855947 | 2 | 1 | NA | NA | 2 | 0 | NA | NA | 4 | 0.5 | 2 | 0.5 | NA | NA | 1 | 0 | NA | NA | 3 | 0.3333 |
| 17_855984( | 2 | 0.5 | 2 | 0 | 2 | 0.5 | 2 | 0 | 8 | 0.25 | 2 | 0.5 | 1 | 0 | 3 | 0 | 2 | 0 | 8 | 0.125 |
| 17_856017( | 4 | 1 | 2 | 0 | 2 | 0 | 3 | 0 | 11 | 0.3636 | 3 | 0 | 1 | 1 | 3 | 0 | 3 | 0.3333 | 10 | 0.2 |
| 17_856024 | 4 | 0.5 | 2 | 1 | NA | NA | 4 | 0.75 | 10 | 0.7 | 2 | 0.5 | 1 | 1 | 2 | 1 | 3 | 0.6667 | 8 | 0.75 |
| 17_856042 | 4 | 0.5 | 2 | 0 | 2 | 0 | 3 | 0.3333 | 11 | 0.2727 | 3 | 0 | 1 | 0 | 2 | 0.5 | 3 | 0.3333 | 9 | 0.2222 |
| 17_856074 | 5 | 1 | 2 | 1 | 3 | 1 | 6 | 1 | 16 | 1 | 3 | 0.6667 | 1 | 1 | 3 | 1 | 3 | 1 | 10 | 0.9 |
| 17_856078! | 5 | 0.6 | 1 | 0 | 3 | 0.6667 | 5 | 0.2 | 14 | 0.4286 | 2 | 1 | 1 | 0 | 2 | 0 | 3 | 0.6667 | 8 | 0.5 |
| 17_856085 | 6 | 0 | 2 | 0 | 3 | 0.3333 | 6 | 0.1667 | 17 | 0.1176 | 3 | 0 | 1 | 0 | 2 | 0.5 | 3 | 0 | 9 | 0.1111 |
| 17_856087 | 4 | 0.75 | 1 | 1 | 2 | 0.5 | 6 | 0.8333 | 13 | 0.7692 | 3 | 0.6667 | 1 | 1 | 3 | 0.6667 | 3 | 1 | 10 | 0.8 |
| 17_856087 | 5 | 1 | 2 | 1 | 2 | 1 | 6 | 1 | 15 | 1 | 3 | 1 | 1 | 1 | 3 | 1 | 2 | 1 | 9 | 1 |
| 17_856092( | 7 | 0.5714 | 2 | 0 | 3 | 0.6667 | 6 | 0.3333 | 18 | 0.4444 | 2 | 0 | NA | NA | 2 | 0.5 | 3 | 0 | 7 | 0.1429 |
| 17_856097! | 8 | 0.375 | 2 | 0.5 | 3 | 0.3333 | 5 | 0.6 | 18 | 0.4444 | 4 | 0.75 | 1 | 0 | 3 | 0.6667 | 3 | 0 | 11 | 0.4545 |
| 17_856098 | 7 | 0.7143 | 2 | 0.5 | 3 | 1 | 4 | 0.75 | 16 | 0.75 | 2 | 1 | 1 | 1 | 3 | 0.6667 | 3 | 1 | 9 | 0.8889 |
| 17_856099 | 6 | 0.5 | 3 | 0.6667 | 2 | 0 | 6 | 0.1667 | 17 | 0.3529 | 2 | 0 | 1 | 0 | 3 | 0 | 3 | 0.3333 | 9 | 0.1111 |
| 17_856106 | 9 | 0.5556 | 2 | 0.5 | 2 | 1 | 6 | 0.3333 | 19 | 0.5263 | 4 | 0.25 | 1 | 0 | 3 | 0.3333 | 3 | 0.3333 | 11 | 0.2727 |
| 17_856110! | 6 | 0.3333 | 3 | 0.3333 | 2 | 0 | 5 | 0.6 | 16 | 0.375 | 5 | 0.2 | 1 | 0 | 3 | 0.3333 | 3 | 0.3333 | 12 | 0.25 |
| 17_856114 | 9 | 0.7778 | 2 | 1 | 3 | 1 | 4 | 0.75 | 18 | 0.8333 | 5 | 0.8 | 1 | 0 | 3 | 0.3333 | 2 | 1 | 11 | 0.6364 |
| 17_856286 | 6 | 0.1667 | 4 | 0.25 | 6 | 0.1667 | 3 | 0 | 19 | 0.1579 | 4 | 0 | 4 | 0.5 | NA | NA | 3 | 0 | 11 | 0.1818 |
| 17_856287 | 5 | 0.6 | 4 | 0.5 | 4 | 0.75 | 3 | 1 | 16 | 0.6875 | 3 | 1 | 4 | 0.75 | NA | NA | 3 | 1 | 10 | 0.9 |
| 17_860800 | NA | NA | 1 | 1 | NA | NA | NA | NA | 1 | 1 | NA | NA | NA | NA | NA | NA | 1 | 1 | 1 | 1 |
| 17_916181 | 4 | 0.5 | 3 | 0 | 3 | 0 | 6 | 0.3333 | 16 | 0.25 | NA | NA | 8 | 0.25 | 7 | 0.1429 | 3 | 0 | 18 | 0.1667 |
| 17_920048 | 16 | 0.125 | 8 | 0.125 | 10 | 0.1 | 23 | 0.3043 | 57 | 0.193 | 8 | 0 | 20 | 0.1 | 19 | 0.1579 | 11 | 0.0909 | 58 | 0.1034 |
| 17_920054 | 12 | 0.75 | 7 | 1 | 9 | 0.5556 | 20 | 0.75 | 48 | 0.75 | 7 | 0.8571 | 23 | 0.6087 | 15 | 0.6667 | 16 | 0.625 | 61 | 0.6557 |
| 17_920064( | 14 | 0.9286 | 8 | 1 | 9 | 1 | 19 | 0.8947 | 50 | 0.94 | 5 | 1 | 21 | 0.9524 | 16 | 1 | 15 | 1 | 57 | 0.9825 |

| | | | | | | | | | | | | | | | | | | | | |
|---|---|---|---|---|---|---|---|---|---|---|---|---|---|---|---|---|---|---|---|---|
| 17_920076 | 13 | 0.2308 | 5 | 0.8 | 8 | 0.5 | 17 | 0.3529 | 43 | 0.3953 | 7 | 0.2857 | 19 | 0.3684 | 21 | 0.381 | 13 | 0.3077 | 60 | 0.35 |
| 17_920086 | 18 | 1 | 8 | 0.875 | 12 | 1 | 20 | 0.9 | 58 | 0.9483 | 8 | 0.75 | 24 | 0.9583 | 22 | 0.8182 | 16 | 1 | 70 | 0.9 |
| 17_920089 | 18 | 0.1111 | 8 | 0.125 | 13 | 0.0769 | 23 | 0.3043 | 62 | 0.1774 | 9 | 0 | 23 | 0.087 | 21 | 0.0952 | 15 | 0.1333 | 68 | 0.0882 |
| 17_920090 | 14 | 0.1429 | 6 | 0.1667 | 3 | 0.6667 | 16 | 0.0625 | 39 | 0.1538 | 5 | 0 | 18 | 0.3333 | 13 | 0.4615 | 9 | 0.2222 | 45 | 0.3111 |
| 17_920092 | 18 | 0.9444 | 7 | 1 | 13 | 0.9231 | 24 | 0.75 | 62 | 0.871 | 10 | 0.8 | 22 | 0.9091 | 22 | 0.8636 | 12 | 1 | 66 | 0.8939 |
| 17_920097 | 20 | 0.55 | 6 | 0.8333 | 12 | 0.75 | 27 | 0.3333 | 65 | 0.5231 | 8 | 0.5 | 23 | 0.6087 | 25 | 0.4 | 16 | 0.5625 | 72 | 0.5139 |
| 17_920108 | 19 | 0.3158 | 7 | 0.5714 | 12 | 0.4167 | 29 | 0.2069 | 67 | 0.3134 | 11 | 0.4545 | 21 | 0.3333 | 21 | 0.2857 | 17 | 0.4706 | 70 | 0.3714 |
| 17_920110 | 20 | 0.4 | 9 | 0.2222 | 14 | 0.2143 | 31 | 0.3226 | 74 | 0.3108 | 10 | 0.3 | 26 | 0.4231 | 25 | 0.36 | 18 | 0.2778 | 79 | 0.3544 |
| 17_920154 | 24 | 0.1667 | 8 | 0.125 | 13 | 0 | 36 | 0.0833 | 81 | 0.0988 | 10 | 0.1 | 23 | 0.1739 | 27 | 0.037 | 22 | 0.2727 | 82 | 0.1463 |
| 17_920169 | 26 | 0.3462 | 9 | 0.4444 | 13 | 0.3846 | 37 | 0.1622 | 85 | 0.2824 | 11 | 0.3636 | 26 | 0.3846 | 30 | 0.2 | 23 | 0.4348 | 90 | 0.3333 |
| 17_920187 | 27 | 0.1481 | 8 | 0 | 12 | 0.0833 | 34 | 0.0588 | 81 | 0.0864 | 12 | 0.0833 | 20 | 0.05 | 29 | 0 | 18 | 0.1111 | 79 | 0.0506 |
| 17_920241 | 33 | 0.5758 | 10 | 0.7 | 18 | 0.5 | 45 | 0.5556 | 106 | 0.566 | 16 | 0.6875 | 27 | 0.7778 | 41 | 0.561 | 28 | 0.5357 | 112 | 0.625 |
| 17_920244 | 35 | 0.9143 | 8 | 1 | 19 | 0.9474 | 44 | 0.8864 | 106 | 0.9151 | 15 | 1 | 27 | 1 | 40 | 0.975 | 28 | 0.8571 | 110 | 0.9545 |
| 17_920264 | 29 | 0.1034 | 10 | 0.1 | 17 | 0.1176 | 42 | 0.0714 | 98 | 0.0918 | 15 | 0.4667 | 30 | 0.0333 | 43 | 0.1395 | 31 | 0.0645 | 119 | 0.1345 |
| 17_920340 | 30 | 0.1667 | 13 | 0.3077 | 22 | 0.3182 | 46 | 0.1087 | 111 | 0.1892 | 13 | 0.2308 | 44 | 0.0455 | 51 | 0.1373 | 26 | 0.3462 | 134 | 0.1567 |
| 17_920446 | 57 | 0.193 | 24 | 0.2083 | 38 | 0.1316 | 68 | 0.2647 | 187 | 0.2086 | 21 | 0.381 | 59 | 0.1695 | 75 | 0.2 | 47 | 0.1702 | 202 | 0.203 |
| 17_920454 | 60 | 0.05 | 25 | 0.08 | 39 | 0.0769 | 67 | 0.0448 | 191 | 0.0576 | 21 | 0 | 60 | 0.0333 | 75 | 0.0667 | 53 | 0.0943 | 209 | 0.0574 |
| 17_920470 | 57 | 0.0877 | 22 | 0.0909 | 39 | 0.0256 | 58 | 0.1552 | 176 | 0.0966 | 23 | 0.2174 | 60 | 0.1833 | 73 | 0.0959 | 52 | 0.1731 | 208 | 0.1538 |
| 17_920472 | 1 | 1 | NA | NA | NA | NA | NA | NA | 1 | 1 | NA | NA | 1 | 1 | 1 | 1 | NA | NA | 2 | 1 |
| 17_920484 | 63 | 0.4444 | 28 | 0.3214 | 42 | 0.4762 | 65 | 0.4615 | 198 | 0.4394 | 27 | 0.2963 | 69 | 0.5797 | 81 | 0.4691 | 58 | 0.4828 | 235 | 0.4851 |
| 17_104262 | 4 | 0 | 14 | 0.2857 | 3 | 0.3333 | 9 | 0.1111 | 30 | 0.2 | 5 | 0.2 | 6 | 0.1667 | 6 | 0 | 6 | 0 | 23 | 0.087 |
| 17_104281 | 3 | 1 | 8 | 0.75 | 5 | 0.2 | 5 | 0.6 | 21 | 0.619 | 2 | 0.5 | 5 | 0.6 | 1 | 0 | 5 | 0.6 | 13 | 0.5385 |
| 17_104283 | 4 | 0.25 | 8 | 0.25 | 5 | 0.2 | 5 | 0 | 22 | 0.1818 | 2 | 0 | 5 | 0 | 3 | 0.3333 | 5 | 0 | 15 | 0.0667 |
| 17_104283 | 4 | 0.5 | 6 | 0.3333 | 3 | 0 | 4 | 0 | 17 | 0.2353 | 2 | 0.5 | 5 | 0.4 | 2 | 0 | 5 | 0.4 | 14 | 0.3571 |
| 17_104285 | 4 | 0 | 5 | 0.2 | 4 | 0.5 | 5 | 0.2 | 18 | 0.2222 | 2 | 0 | 4 | 0.25 | 3 | 0.3333 | 4 | 0 | 13 | 0.1538 |
| 17_104296 | 32 | 0.2812 | 53 | 0.3585 | 44 | 0.2955 | 41 | 0.1707 | 170 | 0.2824 | 32 | 0.2188 | 31 | 0.0968 | 26 | 0.0385 | 22 | 0.2273 | 111 | 0.1441 |
| 17_104325 | 16 | 0.125 | 25 | 0.2 | 20 | 0.25 | 14 | 0 | 75 | 0.16 | 19 | 0.1053 | 8 | 0.25 | 12 | 0.3333 | 14 | 0.0714 | 53 | 0.1698 |
| 17_104325 | 18 | 0.0556 | 22 | 0.0455 | 22 | 0 | 14 | 0.1429 | 76 | 0.0526 | 17 | 0.1765 | 13 | 0.1538 | 13 | 0.0769 | 15 | 0 | 58 | 0.1034 |
| 17_104340 | 7 | 0.4286 | 2 | 1 | 6 | 0.3333 | 4 | 0.5 | 19 | 0.4737 | 7 | 0.5714 | 6 | 0.3333 | 7 | 0.5714 | 3 | 0.3333 | 23 | 0.4783 |
| 17_104341 | 8 | 0.5 | 2 | 1 | 5 | 0.6 | 4 | 0.75 | 19 | 0.6316 | 6 | 0.3333 | 6 | 0.6667 | 6 | 0.6667 | 4 | 0.25 | 22 | 0.5 |
| 17_104342 | 7 | 0.1429 | 2 | 0 | 5 | 0.2 | 4 | 0 | 18 | 0.1111 | 6 | 0.1667 | 4 | 0 | 6 | 0.1667 | 5 | 0 | 21 | 0.0952 |
| 17_104342 | 2 | 1 | 1 | 1 | 4 | 0.5 | 3 | 0.6667 | 10 | 0.7 | 2 | 0.5 | 4 | 1 | 2 | 1 | 1 | 1 | 9 | 0.8889 |
| 17_104342 | NA | NA | NA | NA | 1 | 1 | 1 | 1 | 2 | 1 | 2 | 0 | 1 | 1 | 1 | 1 | 2 | 0.5 | 6 | 0.5 |
| 17_104342 | NA | NA | 1 | 1 | NA | NA | NA | NA | 1 | 1 | NA | NA | 1 | 1 | 1 | 0 | 1 | 1 | 3 | 0.6667 |
| 17_104343 | 7 | 0.7143 | 3 | 0.3333 | 6 | 0.3333 | 4 | 0.5 | 20 | 0.5 | 5 | 0.4 | 7 | 0.4286 | 6 | 0.6667 | 6 | 0.3333 | 24 | 0.4583 |
| 17_106221 | 4 | 0.5 | 7 | 0.7143 | 1 | 1 | 1 | 1 | 13 | 0.6923 | NA | NA | 2 | 1 | 3 | 0.6667 | 2 | 0.5 | 7 | 0.7143 |
| 17_106221 | 3 | 0 | 7 | 0.1429 | 1 | 0 | 1 | 0 | 12 | 0.0833 | NA | NA | 2 | 0.5 | 4 | 0.25 | 2 | 0 | 8 | 0.25 |
| 17_106222 | 3 | 0 | 6 | 0.1667 | 1 | 0 | NA | NA | 10 | 0.1 | NA | NA | 2 | 0.5 | 4 | 0.25 | 2 | 0 | 8 | 0.25 |
| 17_117756 | NA | NA | NA | NA | NA | NA | 2 | 1 | 2 | 1 | NA | NA | NA | NA | 1 | 0 | 2 | 1 | 3 | 0.6667 |
| 17_117756 | NA | NA | NA | NA | NA | NA | 2 | 1 | 2 | 1 | 1 | 1 | NA | NA | 1 | 1 | 2 | 0.5 | 4 | 0.75 |
| 17_122819 | 1 | 1 | 1 | 1 | NA | NA | 1 | 0 | 3 | 0.6667 | 1 | 1 | NA | NA | NA | NA | 1 | 0 | 2 | 0.5 |
| 17_123706 | 4 | 0.25 | 7 | 0.2857 | 5 | 0 | 6 | 0 | 22 | 0.1364 | 3 | 0 | 2 | 0 | 2 | 0 | 5 | 0.2 | 12 | 0.0833 |
| 17_124379 | 7 | 1 | 13 | 0.8462 | 7 | 1 | 8 | 1 | 35 | 0.9429 | 6 | 1 | 3 | 1 | 4 | 1 | 8 | 1 | 21 | 1 |
| 17_124379 | 5 | 0 | 13 | 0.3846 | 7 | 0.2857 | 7 | 0.1429 | 32 | 0.25 | 4 | 0 | 3 | 0.6667 | 3 | 0 | 5 | 0.2 | 15 | 0.2 |
| 17_124380 | 6 | 0.3333 | 9 | 0.4444 | 5 | 0.6 | 6 | 0.5 | 26 | 0.4615 | 5 | 0.6 | 3 | 1 | 3 | 0.6667 | 7 | 0.5714 | 18 | 0.6667 |
| 17_124380 | 7 | 0.7143 | 13 | 0.3846 | 7 | 0.4286 | 8 | 0.25 | 35 | 0.4286 | 6 | 0.5 | 4 | 0.75 | 3 | 0.3333 | 7 | 0.5714 | 20 | 0.55 |
| 17_124466 | 2 | 0 | 4 | 0.5 | 3 | 0.3333 | 7 | 0 | 16 | 0.1875 | 3 | 0.3333 | 8 | 0.25 | 2 | 0.5 | 5 | 0 | 18 | 0.2222 |
| 17_124466 | 1 | 0 | 3 | 0 | 3 | 0 | 5 | 0.4 | 12 | 0.1667 | 3 | 0.3333 | 7 | 0.2857 | 3 | 0.3333 | 4 | 0.25 | 17 | 0.2941 |
| 17_124467 | 2 | 0.5 | 4 | 0.5 | 3 | 0.6667 | 8 | 0.375 | 17 | 0.4706 | 3 | 0.3333 | 7 | 1 | 3 | 0 | 4 | 1 | 17 | 0.7059 |
| 17_124468 | 1 | 0 | 4 | 0.5 | 2 | 0 | 6 | 0.6667 | 13 | 0.4615 | 2 | 0 | 5 | 0.6 | 2 | 0 | 5 | 0.2 | 14 | 0.2857 |
| 17_124468 | 2 | 0 | 2 | 0.5 | 2 | 0 | 6 | 0 | 12 | 0.0833 | 1 | 0 | 3 | 0 | 2 | 1 | 3 | 0 | 9 | 0.2222 |
| 17_124468 | 2 | 0.5 | 3 | 1 | 2 | 0.5 | 4 | 0 | 11 | 0.4545 | 3 | 1 | 5 | 0.8 | 3 | 0.6667 | 6 | 0.8333 | 17 | 0.8235 |
| 17_124469 | 1 | 1 | 4 | 0.75 | 3 | 0.3333 | 4 | 1 | 12 | 0.75 | 1 | 1 | 7 | 1 | 1 | 1 | 5 | 1 | 14 | 1 |
| 17_124469 | 2 | 0.5 | 4 | 1 | 1 | 0 | 7 | 0.2857 | 14 | 0.5 | 3 | 0.3333 | 4 | 0.25 | 1 | 0 | 5 | 0.4 | 13 | 0.3077 |
| 17_124898 | NA | NA | 1 | 1 | 2 | 0 | 1 | 0 | 4 | 0.25 | NA | NA | NA | NA | 1 | 0 | 2 | 0.5 | 3 | 0.3333 |
| 17_124898 | NA | NA | 1 | 0 | 1 | 1 | 1 | 1 | 3 | 0.6667 | NA | NA | 1 | 0 | 1 | 1 | 3 | 0.3333 | 5 | 0.4 |
| 17_124898 | NA | NA | 1 | 1 | 1 | 0 | 1 | 1 | 3 | 0.6667 | NA | NA | 1 | 0 | 1 | 0 | 3 | 0.3333 | 5 | 0.2 |
| 17_124899 | NA | NA | 1 | 1 | 1 | 0 | 1 | 1 | 3 | 0.6667 | NA | NA | 1 | 1 | 1 | 1 | 2 | 1 | 4 | 1 |
| 17_124899 | NA | NA | 1 | 0 | 1 | 0 | 1 | 1 | 3 | 0.3333 | NA | NA | 1 | 1 | 1 | 0 | 3 | 0.3333 | 5 | 0.4 |
| 17_124900 | NA | NA | 1 | 0 | 1 | 1 | 1 | 0 | 3 | 0.3333 | NA | NA | NA | NA | 1 | 0 | 3 | 1 | 4 | 0.75 |
| 17_124902 | NA | NA | 1 | 1 | 1 | 1 | 1 | 0 | 3 | 0.6667 | NA | NA | 1 | 1 | NA | NA | 2 | 0.5 | 3 | 0.6667 |
| 17_124903 | NA | NA | 1 | 1 | 1 | 0 | 1 | 1 | 3 | 0.6667 | NA | NA | 1 | 1 | 1 | 1 | 2 | 1 | 4 | 1 |
| 17_124906 | NA | NA | 1 | 0 | 2 | 0.5 | 1 | 0 | 4 | 0.25 | NA | NA | 1 | 1 | 1 | 0 | 1 | 0 | 3 | 0.3333 |
| 17_124908 | NA | NA | 1 | 0 | 1 | 1 | 1 | 0 | 3 | 0.3333 | NA | NA | 1 | 1 | 1 | 0 | 1 | 1 | 3 | 0.6667 |
| 17_124909 | NA | NA | 1 | 1 | NA | NA | 1 | 1 | 2 | 1 | NA | NA | 1 | 1 | 1 | 1 | 1 | 1 | 3 | 1 |

| | | | | | | | | | | | | | | | | | | | | |
|---|---|---|---|---|---|---|---|---|---|---|---|---|---|---|---|---|---|---|---|---|
| 17_1253708 | NA | NA | 1 | 1 | NA | NA | NA | NA | 1 | 1 | 1 | 0 | NA | NA | NA | NA | 1 | 1 | 2 | 0.5 |
| 17_1253779 | NA | NA | 2 | 0.5 | NA | NA | NA | NA | 2 | 0.5 | 1 | 0 | NA | NA | NA | NA | 1 | 1 | 2 | 0.5 |
| 17_1253784 | 1 | 0 | 2 | 1 | NA | NA | NA | NA | 3 | 0.6667 | 2 | 0.5 | NA | NA | NA | NA | 1 | 1 | 3 | 0.6667 |
| 17_1253799 | 1 | 0 | 1 | 1 | NA | NA | NA | NA | 2 | 0.5 | 2 | 0.5 | NA | NA | NA | NA | 1 | 1 | 3 | 0.6667 |
| 17_1254347 | 1 | 1 | NA | NA | NA | NA | NA | NA | 1 | 1 | 1 | 1 | NA | NA | NA | NA | NA | NA | 1 | 1 |
| 17_1263932 | 1 | 1 | NA | NA | 2 | 1 | 1 | 0 | 4 | 0.75 | NA | NA | 2 | 1 | 1 | 0 | 1 | 1 | 4 | 0.75 |
| 17_1263937 | 1 | 1 | NA | NA | 2 | 1 | 1 | 1 | 4 | 1 | NA | NA | 2 | 1 | 1 | 1 | 1 | 1 | 4 | 1 |
| 17_1263937 | 1 | 1 | NA | NA | 2 | 0.5 | 1 | 1 | 4 | 0.75 | NA | NA | 2 | 1 | 2 | 1 | NA | NA | 4 | 1 |
| 17_1263952 | 1 | 0 | NA | NA | 2 | 0.5 | 1 | 1 | 4 | 0.5 | NA | NA | 1 | 0 | 2 | 0 | 1 | 1 | 4 | 0.25 |
| 17_1263954 | 1 | 1 | NA | NA | 1 | 1 | 1 | 1 | 3 | 1 | NA | NA | 2 | 0.5 | 2 | 1 | 1 | 1 | 5 | 0.8 |
| 17_1263956 | 1 | 1 | NA | NA | 2 | 1 | 1 | 0 | 4 | 0.75 | NA | NA | 2 | 0.5 | 2 | 0.5 | 1 | 1 | 5 | 0.6 |
| 17_1263977 | 1 | 0 | NA | NA | 2 | 0.5 | 1 | 0 | 4 | 0.25 | NA | NA | 2 | 0.5 | 2 | 0.5 | 1 | 0 | 5 | 0.4 |
| 17_1263991 | 1 | 1 | NA | NA | 1 | 1 | 1 | 1 | 3 | 1 | NA | NA | 2 | 1 | 2 | 1 | 1 | 1 | 5 | 1 |
| 17_1264003 | 1 | 1 | NA | NA | 2 | 0 | 1 | 1 | 4 | 0.5 | NA | NA | NA | NA | 2 | 0 | 1 | 1 | 3 | 0.3333 |
| 17_1264015 | 1 | 1 | NA | NA | 2 | 1 | 1 | 1 | 4 | 1 | NA | NA | 2 | 0.5 | 2 | 1 | 1 | 1 | 5 | 0.8 |
| 17_1264016 | 1 | 1 | NA | NA | 2 | 0 | 1 | 0 | 4 | 0.25 | NA | NA | 1 | 1 | 2 | 0.5 | 1 | 1 | 4 | 0.75 |
| 17_1264023 | 1 | 0 | NA | NA | 2 | 1 | NA | NA | 3 | 0.6667 | NA | NA | 2 | 0.5 | 2 | 0.5 | 1 | 0 | 5 | 0.4 |
| 17_1290205 | 5 | 0 | 4 | 0 | 1 | 1 | 4 | 0 | 14 | 0.0714 | 2 | 0 | 1 | 1 | NA | NA | 2 | 0 | 5 | 0.2 |
| 17_1290218 | 5 | 0.4 | 5 | 0.6 | 1 | 0 | 2 | 0 | 13 | 0.3846 | 2 | 0.5 | 2 | 0.5 | NA | NA | 2 | 0 | 6 | 0.3333 |
| 17_1290249 | 5 | 0.8 | 3 | 0.3333 | 1 | 1 | 3 | 0.3333 | 12 | 0.5833 | NA | NA | NA | NA | NA | NA | 3 | 0.6667 | 3 | 0.6667 |
| 17_1290253 | 5 | 0.8 | 5 | 1 | 1 | 1 | 3 | 0.6667 | 14 | 0.8571 | NA | NA | NA | NA | NA | NA | 3 | 1 | 3 | 1 |
| 17_1290256 | 5 | 1 | 5 | 0.8 | 1 | 1 | 3 | 1 | 14 | 0.9286 | NA | NA | NA | NA | NA | NA | 3 | 0.6667 | 3 | 0.6667 |
| 17_1290267 | 4 | 0.75 | 5 | 0 | 1 | 0 | 3 | 0.3333 | 13 | 0.3077 | NA | NA | NA | NA | NA | NA | 2 | 0.5 | 2 | 0.5 |
| 17_1290285 | 5 | 1 | 4 | 0.75 | 1 | 1 | 3 | 1 | 13 | 0.9231 | NA | NA | NA | NA | NA | NA | 3 | 0.6667 | 3 | 0.6667 |
| 17_1306013 | 2 | 0.5 | 6 | 0.5 | 2 | 0.5 | 1 | 0 | 11 | 0.4545 | 9 | 0.6667 | 2 | 0 | 4 | 0.25 | 1 | 1 | 16 | 0.5 |
| 17_1306014 | 4 | 0.25 | 5 | 0 | 3 | 0.3333 | 1 | 0 | 13 | 0.1538 | 9 | 0.3333 | 2 | 0.5 | 4 | 0.25 | 1 | 1 | 16 | 0.375 |
| 17_1306019 | 3 | 1 | 6 | 1 | 4 | 0.5 | 1 | 1 | 14 | 0.8571 | 6 | 0.8333 | 1 | 1 | 4 | 1 | 2 | 0.5 | 13 | 0.8462 |
| 17_1306019 | 2 | 0 | 5 | 0.4 | 3 | 0.3333 | 1 | 0 | 11 | 0.2727 | 3 | 0.3333 | 1 | 1 | 4 | 0 | 1 | 0 | 9 | 0.2222 |
| 17_1306024 | 3 | 1 | 5 | 1 | 4 | 0.75 | NA | NA | 12 | 0.9167 | 6 | 1 | 2 | 0.5 | 4 | 1 | 2 | 0.5 | 14 | 0.8571 |
| 17_1306030 | 3 | 1 | 6 | 0.8333 | 4 | 0.25 | 1 | 0 | 14 | 0.6429 | 5 | 0.4 | 1 | 1 | 4 | 0.5 | 2 | 0.5 | 12 | 0.5 |
| 17_1306035 | 3 | 0 | 4 | 0.25 | 2 | 0 | 1 | 0 | 10 | 0.1 | 4 | 0.25 | 2 | 0 | 3 | 0 | 1 | 0 | 10 | 0.1 |
| 17_1306042 | 3 | 1 | 5 | 1 | 3 | 1 | NA | NA | 11 | 1 | 5 | 1 | 2 | 1 | 2 | 1 | 2 | 1 | 11 | 1 |
| 17_1306044 | 3 | 1 | 6 | 0.8333 | 4 | 0.75 | NA | NA | 13 | 0.8462 | 4 | 1 | 2 | 1 | 3 | 1 | 2 | 1 | 11 | 1 |
| 17_1306045 | 3 | 1 | 6 | 1 | 3 | 1 | 1 | 1 | 13 | 1 | 6 | 0.8333 | 1 | 1 | 3 | 1 | 2 | 0.5 | 12 | 0.8333 |
| 17_1306064 | 3 | 0 | 5 | 0.2 | 3 | 0 | 1 | 0 | 12 | 0.0833 | 5 | 0.2 | 1 | 0 | 3 | 0.3333 | 2 | 0 | 11 | 0.1818 |
| 17_1306087 | 3 | 0 | 5 | 0 | 3 | 0.6667 | 1 | 0 | 12 | 0.1667 | 3 | 0 | 1 | 0 | 3 | 0.6667 | 2 | 0 | 9 | 0.2222 |
| 17_1306109 | 2 | 1 | 4 | 1 | 1 | 1 | 1 | 1 | 8 | 1 | 2 | 1 | 1 | 1 | 3 | 1 | 2 | 0.5 | 8 | 0.875 |
| 17_1306110 | 3 | 0 | 5 | 0.6 | 2 | 0.5 | 1 | 0 | 11 | 0.3636 | 3 | 0.3333 | 1 | 1 | 3 | 0.6667 | 2 | 0 | 9 | 0.4444 |
| 17_1306113 | 1 | 1 | 5 | 1 | 2 | 1 | NA | NA | 8 | 1 | 4 | 1 | 1 | 1 | 3 | 1 | 2 | 1 | 10 | 1 |
| 17_1306114 | 2 | 0.5 | 5 | 0.4 | 1 | 1 | 1 | 0 | 9 | 0.4444 | 1 | 1 | 1 | 0 | 2 | 1 | 2 | 0.5 | 6 | 0.6667 |
| 17_1306115 | 3 | 0.6667 | 4 | 0.75 | 2 | 0 | 1 | 0 | 10 | 0.5 | 3 | 1 | 1 | 1 | 2 | 1 | 2 | 0.5 | 8 | 0.875 |
| 17_1306141 | 3 | 0 | 5 | 0.2 | 3 | 0.3333 | NA | NA | 11 | 0.1818 | 3 | 0.3333 | 1 | 0 | 3 | 0.3333 | 2 | 0 | 9 | 0.2222 |
| 17_1313023 | 7 | 0.1429 | 6 | 0 | 5 | 0.2 | 10 | 0.1 | 28 | 0.1071 | 1 | 0 | 8 | 0.25 | 6 | 0.3333 | 6 | 0.1667 | 21 | 0.2381 |
| 17_1313030 | 14 | 0 | 15 | 0.1333 | 9 | 0 | 11 | 0.0909 | 49 | 0.0612 | 10 | 0.1 | 13 | 0.0769 | 9 | 0.1111 | 11 | 0 | 43 | 0.0698 |
| 17_1313039 | 12 | 0.5833 | 16 | 0.4375 | 15 | 0.6667 | 12 | 0.5833 | 55 | 0.5636 | 10 | 0.6 | 13 | 0.3846 | 12 | 0.5833 | 15 | 0.4667 | 50 | 0.5 |
| 17_1313042 | 14 | 0.0714 | 19 | 0.1579 | 13 | 0.2308 | 15 | 0.2667 | 61 | 0.1803 | 10 | 0.1 | 14 | 0.2143 | 13 | 0.1538 | 15 | 0.1333 | 52 | 0.1538 |
| 17_1314335 | 37 | 0.3784 | 32 | 0.4375 | 43 | 0.4186 | 31 | 0.3548 | 143 | 0.3986 | 31 | 0.4194 | 40 | 0.425 | 35 | 0.4286 | 27 | 0.4815 | 133 | 0.4361 |
| 17_1314351 | 7 | 0.8571 | 3 | 0.6667 | 5 | 0.6 | 3 | 0.3333 | 18 | 0.6667 | 5 | 0.8 | 4 | 0.75 | 4 | 0.5 | NA | NA | 13 | 0.6923 |
| 17_1314352 | NA | NA | 2 | 0 | 1 | 1 | 1 | 1 | 4 | 0.5 | NA | NA | 2 | 0.5 | 1 | 1 | NA | NA | 3 | 0.6667 |
| 17_1314375 | 8 | 0 | 5 | 0.2 | 6 | 0.1667 | 3 | 0 | 22 | 0.0909 | 7 | 0 | 6 | 0.1667 | 6 | 0.5 | NA | NA | 19 | 0.2105 |
| 17_1314389 | 5 | 0 | 2 | 0.5 | 2 | 0 | 1 | 0 | 10 | 0.1 | 3 | 0 | 4 | 0.25 | 3 | 0 | NA | NA | 10 | 0.1 |
| 17_1316196 | 8 | 0 | 7 | 0.1429 | 7 | 0.1429 | 4 | 0 | 26 | 0.0769 | 7 | 0.1429 | 15 | 0.0667 | 4 | 0.25 | 8 | 0 | 34 | 0.0882 |
| 17_1316267 | 5 | 0 | 5 | 0.2 | 4 | 0 | 3 | 0 | 17 | 0.0588 | 7 | 0 | 12 | 0.1667 | 3 | 0 | 6 | 0 | 28 | 0.0714 |
| 17_1316272 | 4 | 0 | 5 | 0.2 | 4 | 0.25 | 2 | 1 | 15 | 0.2667 | 5 | 0.8 | 7 | 0.2857 | 2 | 0 | 3 | 1 | 17 | 0.5294 |
| 17_1316274 | 4 | 0.25 | 7 | 0 | 4 | 0 | 2 | 0 | 17 | 0.0588 | 5 | 0 | 13 | 0 | 3 | 0 | 6 | 0.3333 | 27 | 0.0741 |
| 17_1318612 | 6 | 0.3333 | 3 | 0 | 1 | 1 | 6 | 0 | 16 | 0.1875 | 2 | 0 | 5 | 0.4 | 7 | 0.1429 | 4 | 0.25 | 18 | 0.2222 |
| 17_1318614 | 3 | 0 | 3 | 0 | 2 | 0 | 4 | 0.25 | 12 | 0.0833 | 2 | 0 | 4 | 0.25 | 7 | 0 | 3 | 0 | 16 | 0.0625 |
| 17_1318638 | 2 | 0.5 | 1 | 0 | 2 | 0 | 4 | 0.25 | 9 | 0.2222 | 1 | 1 | 3 | 0.3333 | 6 | 0.6667 | 2 | 0.5 | 12 | 0.5833 |
| 17_1318642 | NA | NA | 1 | 0 | 1 | 0 | 4 | 1 | 6 | 0.6667 | 2 | 1 | 2 | 1 | 7 | 0.8571 | 2 | 1 | 13 | 0.9231 |
| 17_1318647 | 2 | 0.5 | 1 | 0 | 1 | 0 | 6 | 0.1667 | 10 | 0.2 | 2 | 0.5 | 3 | 0 | 8 | 0.25 | 2 | 0 | 15 | 0.2 |
| 17_1318650 | 2 | 0 | 1 | 0 | 2 | 0.5 | 4 | 0.25 | 9 | 0.2222 | 2 | 0.5 | 5 | 0.2 | 9 | 0.1111 | 1 | 1 | 17 | 0.2353 |
| 17_1318658 | 2 | 1 | 1 | 0 | 1 | 1 | 6 | 0.6667 | 10 | 0.7 | 1 | 1 | 3 | 0.6667 | 9 | 0.8889 | 2 | 0.5 | 15 | 0.8 |
| 17_1318664 | 2 | 0 | 1 | 1 | 2 | 0 | 5 | 0.2 | 10 | 0.2 | 2 | 0.5 | 4 | 0 | 9 | 0.2222 | 2 | 0.5 | 17 | 0.2353 |
| 17_1318671 | 1 | 0 | 1 | 0 | 2 | 0.5 | 5 | 0.8 | 9 | 0.5556 | 2 | 0.5 | 3 | 0.3333 | 9 | 0.3333 | 2 | 0 | 16 | 0.3125 |
| 17_1318674 | 1 | 1 | 1 | 1 | 1 | 0 | 4 | 1 | 7 | 0.8571 | NA | NA | 4 | 0.75 | 7 | 0.4286 | 1 | 0 | 12 | 0.5 |

| | | | | | | | | | | | | | | | | | | | | |
|---|---|---|---|---|---|---|---|---|---|---|---|---|---|---|---|---|---|---|---|---|
| 17_1318678 | 2 | 0.5 | 1 | 0 | 1 | 0 | 5 | 0.4 | 9 | 0.3333 | 1 | 0 | 4 | 0.25 | 9 | 0.3333 | 2 | 0 | 16 | 0.25 |
| 17_1319341 | 9 | 0.6667 | 4 | 0.25 | 6 | 0.5 | 4 | 0.75 | 23 | 0.5652 | 4 | 0.75 | 6 | 0.5 | 5 | 0.8 | NA | NA | 15 | 0.6667 |
| 17_1319360 | 9 | 0.8889 | 6 | 1 | 7 | 0.8571 | 5 | 1 | 27 | 0.9259 | 3 | 1 | 7 | 0.8571 | 6 | 1 | 3 | 1 | 19 | 0.9474 |
| 17_1319364 | 8 | 0.25 | 5 | 0.4 | 5 | 0.4 | 3 | 0.3333 | 21 | 0.3333 | 5 | 0.4 | 7 | 0.4286 | 6 | 0 | 2 | 0.5 | 20 | 0.3 |
| 17_1319366 | 9 | 0.4444 | 5 | 0.8 | 7 | 0.7143 | 4 | 1 | 25 | 0.68 | 5 | 0.6 | 7 | 0.7143 | 6 | 0.6667 | 2 | 0 | 20 | 0.6 |
| 17_1319367 | 10 | 0.6 | 6 | 0.3333 | 6 | 0.8333 | 3 | 1 | 25 | 0.64 | 4 | 0.25 | 6 | 0.5 | 6 | 0.5 | 3 | 0 | 19 | 0.3684 |
| 17_1319369 | 10 | 0.5 | 6 | 0.5 | 7 | 0.2857 | 5 | 0.4 | 28 | 0.4286 | 5 | 0.4 | 8 | 0.5 | 6 | 0.6667 | 4 | 0.75 | 23 | 0.5652 |
| 17_1319370 | 10 | 0.4 | 6 | 0 | 6 | 0.1667 | 5 | 0.4 | 27 | 0.2593 | 5 | 0.2 | 7 | 0.2857 | 6 | 0 | 3 | 0 | 21 | 0.1429 |
| 17_1319377 | 6 | 0.3333 | 6 | 0 | 4 | 0 | 4 | 0 | 20 | 0.1 | 4 | 0.25 | 7 | 0.1429 | 6 | 0.3333 | 3 | 0 | 20 | 0.2 |
| 17_1319385 | 7 | 0 | 5 | 0 | 3 | 0 | 3 | 0.3333 | 18 | 0.0556 | 5 | 0 | 7 | 0 | 3 | 0 | 4 | 0.25 | 19 | 0.0526 |
| 17_1319389 | 9 | 0.1111 | 6 | 0 | 6 | 0.1667 | 4 | 0.5 | 25 | 0.16 | 5 | 0.4 | 7 | 0.1429 | 6 | 0.3333 | 4 | 0 | 22 | 0.2273 |
| 17_1319390 | 8 | 0.75 | 5 | 0.6 | 5 | 0.6 | 2 | 1 | 20 | 0.7 | 5 | 0.6 | 6 | 0.8333 | 4 | 0.25 | 4 | 0.25 | 19 | 0.5263 |
| 17_1319394 | 9 | 0.5556 | 6 | 0.8333 | 6 | 0.5 | 5 | 0.6 | 26 | 0.6154 | 5 | 0.4 | 6 | 0.5 | 6 | 0.6667 | 4 | 0.5 | 21 | 0.5238 |
| 17_1319400 | 8 | 0.375 | 6 | 0.3333 | 7 | 0.2857 | 4 | 0.25 | 25 | 0.32 | 5 | 0.2 | 6 | 0.5 | 6 | 0.3333 | 4 | 0 | 21 | 0.2857 |
| 17_1319408 | 6 | 0.8333 | 3 | 0.6667 | 6 | 1 | 2 | 0.5 | 17 | 0.8235 | 4 | 0.5 | 5 | 0.8 | 5 | 0.8 | 4 | 1 | 18 | 0.7778 |
| 17_1319408 | 6 | 0.5 | 1 | 1 | 2 | 0.5 | 2 | 1 | 11 | 0.6364 | 3 | 0.3333 | 4 | 0.25 | 4 | 0.75 | 3 | 0.6667 | 14 | 0.5 |
| 17_1319411 | 6 | 0 | 5 | 0.4 | 6 | 0.1667 | 3 | 0.3333 | 20 | 0.2 | 1 | 0 | 6 | 0.1667 | 6 | 0 | 3 | 0 | 16 | 0.0625 |
| 17_1319415 | 8 | 0.25 | 5 | 0.2 | 6 | 0.3333 | 4 | 0.25 | 23 | 0.2609 | 5 | 0.4 | 6 | 0.5 | 6 | 0.1667 | 4 | 0.25 | 21 | 0.3333 |
| 17_1319421 | 5 | 0.8 | 6 | 0.8333 | 5 | 1 | 8 | 0.875 | 24 | 0.875 | 5 | 1 | 6 | 0.8333 | 8 | 0.75 | 3 | 1 | 22 | 0.8636 |
| 17_1319423 | 8 | 0.375 | 3 | 0 | 7 | 0.2857 | 7 | 0.1429 | 25 | 0.24 | 4 | 0.25 | 9 | 0.1111 | 10 | 0.1 | 4 | 0.5 | 27 | 0.1852 |
| 17_1319423 | 9 | 0.1111 | 6 | 0.3333 | 7 | 0.1429 | 7 | 0 | 29 | 0.1379 | 5 | 0 | 10 | 0.1 | 10 | 0.1 | 3 | 0 | 28 | 0.0714 |
| 17_1319424 | 9 | 0.1111 | 5 | 0.2 | 6 | 0.3333 | 7 | 0.1429 | 27 | 0.1852 | 4 | 0.75 | 9 | 0.1111 | 10 | 0 | 5 | 0.4 | 28 | 0.2143 |
| 17_1319425 | 9 | 0.4444 | 6 | 0.3333 | 7 | 0.5714 | 6 | 0.3333 | 28 | 0.4286 | 4 | 0.5 | 9 | 0.6667 | 7 | 0.2857 | 3 | 0.3333 | 23 | 0.4783 |
| 17_1323426 | 9 | 0.2222 | 18 | 0.2222 | 4 | 0 | 9 | 0.2222 | 40 | 0.2 | 4 | 0 | 9 | 0.3333 | 4 | 0 | 10 | 0.1 | 27 | 0.1481 |
| 17_1323492 | 8 | 0.125 | 17 | 0.1765 | 5 | 0 | 11 | 0.0909 | 41 | 0.122 | 5 | 0.2 | 12 | 0.0833 | 4 | 0.25 | 12 | 0 | 33 | 0.0909 |
| 17_1325410 | NA | NA | 1 | 1 | NA | NA | NA | NA | 1 | 1 | 1 | 1 | NA | NA | NA | NA | NA | NA | 1 | 1 |
| 17_1325416 | 1 | 0 | 2 | 0.5 | NA | NA | NA | NA | 3 | 0.3333 | 1 | 1 | NA | NA | NA | NA | NA | NA | 1 | 1 |
| 17_1325594 | 1 | 1 | 1 | 0 | NA | NA | NA | NA | 2 | 0.5 | 1 | 1 | NA | NA | NA | NA | NA | NA | 1 | 1 |
| 17_1325654 | 1 | 0 | 2 | 0.5 | 1 | 0 | NA | NA | 4 | 0.25 | 1 | 1 | NA | NA | 1 | 1 | NA | NA | 2 | 1 |
| 17_1325686 | NA | NA | 1 | 0 | 1 | 0 | 1 | 1 | 3 | 0.3333 | 1 | 1 | NA | NA | 1 | 0 | NA | NA | 2 | 0.5 |
| 17_1325743 | 1 | 1 | 1 | 1 | 2 | 0.5 | 1 | 1 | 5 | 0.8 | 1 | 1 | NA | NA | NA | NA | 1 | 1 | 2 | 1 |
| 17_1325832 | 1 | 1 | 1 | 0 | 2 | 0 | 1 | 0 | 5 | 0.2 | 1 | 0 | NA | NA | 1 | 1 | 1 | 0 | 3 | 0.3333 |
| 17_1325862 | 2 | 0 | 2 | 1 | 3 | 0.3333 | 1 | 0 | 8 | 0.375 | NA | NA | NA | NA | 1 | 1 | 2 | 0 | 3 | 0.3333 |
| 17_1325867 | 2 | 0.5 | 1 | 1 | 3 | 0.6667 | 1 | 1 | 7 | 0.7143 | NA | NA | NA | NA | 1 | 1 | 2 | 0.5 | 3 | 0.6667 |
| 17_1370994 | 1 | 1 | NA | NA | NA | NA | NA | NA | 1 | 1 | 1 | 1 | NA | NA | NA | NA | NA | NA | 1 | 1 |
| 17_1371075 | 1 | 1 | NA | NA | NA | NA | NA | NA | 1 | 1 | 1 | 1 | NA | NA | NA | NA | NA | NA | 1 | 1 |
| 17_1410851 | 1 | 0 | 2 | 0.5 | NA | NA | 2 | 0.5 | 5 | 0.4 | NA | NA | 3 | 0.6667 | 3 | 0.3333 | 2 | 0.5 | 8 | 0.5 |
| 17_1410852 | 5 | 0.2 | 1 | 0 | 1 | 0 | 3 | 0 | 10 | 0.1 | 1 | 0 | 2 | 0 | 4 | 0 | 3 | 0.6667 | 10 | 0.2 |
| 17_1410877 | 6 | 0.1667 | 3 | 0 | NA | NA | 5 | 0.4 | 14 | 0.2143 | 2 | 0.5 | 4 | 0.25 | 5 | 0.2 | 3 | 0 | 14 | 0.2143 |
| 17_1410885 | 4 | 0 | 3 | 0 | NA | NA | 5 | 0.2 | 12 | 0.0833 | 2 | 0 | 3 | 0 | 5 | 0 | 1 | 1 | 11 | 0.0909 |
| 17_1410902 | 6 | 0.6667 | 2 | 1 | 2 | 1 | 4 | 1 | 14 | 0.8571 | 1 | 1 | 5 | 0.2 | 4 | 0.75 | 4 | 0.5 | 14 | 0.5 |
| 17_1410907 | 6 | 0.1667 | 2 | 0 | 2 | 0.5 | 5 | 0.2 | 15 | 0.2 | 2 | 0 | 3 | 0.3333 | 5 | 0.2 | 4 | 0 | 14 | 0.1429 |
| 17_1410913 | 6 | 0 | 3 | 0 | 2 | 0.5 | 5 | 0 | 16 | 0.0625 | 2 | 0 | 4 | 0 | 5 | 0.4 | 4 | 0.25 | 15 | 0.2 |
| 17_1410919 | 7 | 0 | 3 | 0.3333 | 2 | 0 | 2 | 0 | 14 | 0.0714 | 2 | 0 | 4 | 0 | 4 | 0 | 3 | 0.3333 | 13 | 0.0769 |
| 17_1411122 | 6 | 0 | 2 | 0 | 2 | 0.5 | 2 | 0 | 12 | 0.0833 | 1 | 0 | 5 | 0 | 5 | 0 | 3 | 0.3333 | 14 | 0.0714 |
| 17_1411656 | NA | NA | 1 | 1 | 1 | 0 | 1 | 0 | 3 | 0.3333 | 4 | 0.25 | NA | NA | 2 | 0 | 1 | 0 | 7 | 0.1429 |
| 17_1411662 | NA | NA | 2 | 1 | 1 | 1 | 1 | 1 | 4 | 1 | 4 | 1 | 1 | 0 | 2 | 1 | NA | NA | 7 | 0.8571 |
| 17_1411663 | NA | NA | 1 | 1 | 1 | 1 | NA | NA | 2 | 1 | 4 | 0.75 | 1 | 0 | 2 | 0.5 | 1 | 0 | 8 | 0.5 |
| 17_1411667 | NA | NA | 2 | 1 | NA | NA | 1 | 1 | 3 | 1 | 4 | 1 | 1 | 1 | 1 | 1 | NA | NA | 6 | 1 |
| 17_1411700 | 1 | 1 | 2 | 0.5 | NA | NA | 1 | 1 | 4 | 0.75 | 4 | 0.75 | 1 | 1 | NA | NA | 1 | 1 | 6 | 0.8333 |
| 17_1411718 | 1 | 0 | 2 | 0.5 | 1 | 1 | 1 | 0 | 5 | 0.4 | 4 | 0.25 | 1 | 0 | 2 | 0.5 | 1 | 0 | 8 | 0.25 |
| 17_1411753 | 2 | 0.5 | 2 | 0 | 1 | 0 | 1 | 0 | 6 | 0.1667 | 4 | 0 | 1 | 0 | 2 | 0 | 2 | 0.5 | 9 | 0.1111 |
| 17_1411796 | 2 | 0 | 2 | 0 | 1 | 1 | 1 | 0 | 6 | 0.1667 | 4 | 0 | 1 | 0 | 2 | 0 | 1 | 1 | 8 | 0.125 |
| 17_1411933 | 8 | 0.125 | 2 | 0 | 3 | 0.3333 | 3 | 0 | 16 | 0.125 | 2 | 0.5 | 5 | 0.4 | 7 | 0.1429 | 3 | 0.3333 | 17 | 0.2941 |
| 17_1412416 | 5 | 0 | 2 | 0.5 | 3 | 0 | 4 | 0 | 14 | 0.0714 | 3 | 0 | 7 | 0 | 6 | 0.1667 | 6 | 0.1667 | 22 | 0.0909 |
| 17_1412425 | 5 | 0.8 | 2 | 1 | 4 | 1 | 4 | 0.5 | 15 | 0.8 | 3 | 1 | 6 | 1 | 7 | 0.5714 | 6 | 0.6667 | 22 | 0.7727 |
| 17_1412432 | 6 | 0 | 2 | 0 | 4 | 0.25 | 4 | 0.25 | 16 | 0.125 | 4 | 0.25 | 7 | 0.1429 | 8 | 0 | 6 | 0 | 25 | 0.08 |
| 17_1412440 | 8 | 0 | 2 | 0 | 5 | 0.4 | 5 | 0 | 20 | 0.1 | 4 | 0 | 7 | 0.1429 | 9 | 0.1111 | 6 | 0 | 26 | 0.0769 |
| 17_1412450 | 5 | 0.2 | NA | NA | 3 | 1 | 4 | 0.75 | 12 | 0.5833 | 4 | 0.5 | 5 | 0.4 | 7 | 0.4286 | 4 | 0 | 20 | 0.35 |
| 17_1412520 | 10 | 0.2 | 4 | 0 | 5 | 0 | 6 | 0.1667 | 25 | 0.12 | 8 | 0 | 11 | 0.0909 | 8 | 0 | 8 | 0.125 | 35 | 0.0571 |
| 17_1412580 | 10 | 0.1 | 3 | 0.3333 | 4 | 0.25 | 5 | 0 | 22 | 0.1364 | 4 | 0.25 | 5 | 0.2 | 6 | 0.1667 | 5 | 0 | 20 | 0.15 |
| 17_1439102 | 1 | 0 | NA | NA | NA | NA | 1 | 1 | 2 | 0.5 | NA | NA | NA | NA | 1 | 1 | NA | NA | 1 | 1 |
| 17_1441559 | 6 | 0 | 2 | 0 | 5 | 0.2 | 4 | 0 | 17 | 0.0588 | 1 | 0 | 9 | 0 | 4 | 0.25 | 4 | 0 | 18 | 0.0556 |
| 17_1441560 | 5 | 0.4 | 1 | 0 | 5 | 0.6 | 4 | 0.75 | 15 | 0.5333 | 2 | 0.5 | 7 | 0.2857 | 3 | 0.3333 | 4 | 0.25 | 16 | 0.3125 |
| 17_1441567 | 6 | 0 | 1 | 0 | 4 | 0.75 | 4 | 0 | 15 | 0.2 | 2 | 0 | 6 | 0 | 3 | 0.3333 | 4 | 0 | 15 | 0.0667 |

| | | | | | | | | | | | | | | | | | | | | |
|---|---|---|---|---|---|---|---|---|---|---|---|---|---|---|---|---|---|---|---|---|
| 17_1441589 | 5 | 0 | 1 | 1 | 4 | 0.25 | 4 | 0.25 | 14 | 0.2143 | 1 | 0 | 7 | 0.1429 | 3 | 0.6667 | 3 | 0 | 14 | 0.2143 |
| 17_1441593 | 5 | 0.2 | 1 | 0 | 4 | 0 | 4 | 0 | 14 | 0.0714 | 1 | 0 | 5 | 0.2 | 3 | 0 | 3 | 0 | 12 | 0.0833 |
| 17_1441601 | 5 | 0.2 | 1 | 1 | 4 | 0.25 | 4 | 0.75 | 14 | 0.4286 | 1 | 0 | 6 | 0.3333 | 4 | 0.5 | 3 | 0.6667 | 14 | 0.4286 |
| 17_1441626 | 5 | 0.4 | 1 | 0 | 4 | 0.25 | 3 | 0 | 13 | 0.2308 | 1 | 1 | 6 | 0.1667 | 3 | 0 | 3 | 0 | 13 | 0.1538 |
| 17_1441688 | 3 | 0.6667 | 1 | 1 | 4 | 1 | 4 | 0.75 | 12 | 0.8333 | NA | NA | 4 | 0.75 | 4 | 0.5 | 3 | 0.3333 | 11 | 0.5455 |
| 17_1441690 | 3 | 0.6667 | 1 | 1 | 4 | 1 | 4 | 1 | 12 | 0.9167 | NA | NA | 5 | 1 | 3 | 1 | 3 | 0.3333 | 11 | 0.8182 |
| 17_1441695 | 3 | 0 | 1 | 0 | 4 | 0.25 | 4 | 0 | 12 | 0.0833 | NA | NA | 5 | 0 | 4 | 0.25 | 3 | 0.3333 | 12 | 0.1667 |
| 17_1441705 | 3 | 1 | 1 | 1 | 4 | 1 | 4 | 1 | 12 | 1 | NA | NA | 5 | 1 | 4 | 1 | 3 | 0.6667 | 12 | 0.9167 |
| 17_1441707 | 2 | 0.5 | 1 | 1 | 3 | 0 | 4 | 0 | 10 | 0.2 | NA | NA | 3 | 0 | 4 | 0 | 3 | 0.3333 | 10 | 0.1 |
| 17_1441708 | 3 | 1 | 1 | 0 | 4 | 1 | 3 | 1 | 11 | 0.9091 | NA | NA | 4 | 1 | 4 | 1 | 3 | 1 | 11 | 1 |
| 17_1441728 | 3 | 1 | 1 | 1 | 2 | 1 | 3 | 1 | 9 | 1 | NA | NA | 5 | 0.8 | 4 | 1 | 3 | 1 | 12 | 0.9167 |
| 17_1441729 | 3 | 0 | 1 | 1 | 3 | 0.3333 | 3 | 0 | 10 | 0.2 | NA | NA | 4 | 0 | 3 | 0 | 3 | 0.3333 | 10 | 0.1 |
| 17_1441734 | 3 | 0 | 1 | 0 | 3 | 0.3333 | 2 | 0 | 9 | 0.1111 | NA | NA | 3 | 0.3333 | 4 | 0 | 3 | 0 | 10 | 0.1 |
| 17_1516403 | NA | NA | 2 | 1 | 2 | 0 | 2 | 1 | 6 | 0.6667 | 1 | 1 | 1 | 1 | 1 | 1 | 4 | 0.75 | 7 | 0.8571 |
| 17_1516408 | NA | NA | 2 | 1 | 2 | 1 | 2 | 0.5 | 6 | 0.8333 | 3 | 1 | 1 | 1 | 1 | 0 | 4 | 1 | 9 | 0.8889 |
| 17_1516410 | NA | NA | 2 | 1 | 1 | 0 | 2 | 1 | 5 | 0.8 | 1 | 1 | 1 | 0 | 1 | 0 | 4 | 0.75 | 7 | 0.5714 |
| 17_1516421 | NA | NA | 2 | 1 | 2 | 1 | 3 | 1 | 7 | 1 | 2 | 1 | 1 | 1 | 1 | 1 | 4 | 1 | 8 | 1 |
| 17_1516426 | NA | NA | 1 | 1 | 2 | 1 | 2 | 1 | 5 | 1 | 3 | 0.6667 | 1 | 1 | NA | NA | 4 | 0.75 | 8 | 0.75 |
| 17_1516426 | NA | NA | 1 | 1 | 1 | 1 | 2 | 1 | 4 | 1 | 2 | 1 | 1 | 1 | NA | NA | 2 | 1 | 5 | 1 |
| 17_1516440 | NA | NA | 2 | 1 | 2 | 1 | 2 | 1 | 6 | 1 | 3 | 0.6667 | 1 | 1 | 1 | 1 | 4 | 0.75 | 9 | 0.7778 |
| 17_1516449 | NA | NA | 2 | 0.5 | 2 | 0 | 3 | 0.3333 | 7 | 0.2857 | 3 | 0.3333 | 1 | 0 | 1 | 0 | 4 | 0.5 | 9 | 0.3333 |
| 17_1516451 | NA | NA | 2 | 0.5 | 2 | 0.5 | 3 | 0 | 7 | 0.2857 | 4 | 0.75 | 1 | 1 | 1 | 0 | 4 | 0.5 | 10 | 0.6 |
| 17_1516475 | NA | NA | 1 | 0 | 2 | 1 | 2 | 0 | 5 | 0.4 | 4 | 0.75 | 1 | 0 | 1 | 0 | 3 | 0 | 9 | 0.3333 |
| 17_1516477 | NA | NA | 1 | 1 | 2 | 0.5 | 3 | 0.3333 | 6 | 0.5 | 2 | 0 | 1 | 0 | 1 | 0 | 4 | 0.5 | 8 | 0.25 |
| 17_1516493 | 1 | 0 | 1 | 1 | 3 | 0 | 2 | 0 | 7 | 0.1429 | 3 | 0.6667 | 1 | 0 | 1 | 0 | 4 | 0 | 9 | 0.2222 |
| 17_1516520 | 1 | 1 | 1 | 0 | 2 | 0 | 2 | 0 | 6 | 0.1667 | 2 | 0.5 | 1 | 0 | 1 | 0 | 3 | 0 | 7 | 0.1429 |
| 17_1516557 | 1 | 0 | NA | NA | 2 | 0.5 | NA | NA | 3 | 0.3333 | 2 | 1 | 1 | 0 | 1 | 0 | 3 | 0.3333 | 7 | 0.4286 |
| 17_1516592 | 1 | 0 | 1 | 0 | 2 | 0.5 | 2 | 0 | 6 | 0.1667 | 2 | 0.5 | 1 | 0 | 1 | 0 | 2 | 0.5 | 6 | 0.3333 |
| 17_1516594 | NA | NA | 1 | 0 | 1 | 1 | 1 | 0 | 3 | 0.3333 | 2 | 0.5 | 1 | 0 | 1 | 1 | 3 | 0.6667 | 7 | 0.5714 |
| 17_1516813 | 3 | 0 | 3 | 0.3333 | 3 | 0 | 3 | 0 | 12 | 0.0833 | 1 | 0 | 5 | 0 | 3 | 0.3333 | 4 | 0 | 13 | 0.0769 |
| 17_1516980 | 2 | 0 | 4 | 0 | 1 | 0 | 3 | 0.3333 | 10 | 0.1 | 1 | 0 | 3 | 0.3333 | NA | NA | 2 | 0 | 6 | 0.1667 |
| 17_1521247 | 1 | 1 | NA | NA | NA | NA | NA | NA | 1 | 1 | NA | NA | NA | NA | NA | NA | 1 | 1 | 1 | 1 |
| 17_1521265 | 1 | 1 | 3 | 0.3333 | NA | NA | 1 | 1 | 5 | 0.6 | 1 | 0 | NA | NA | 1 | 0 | 2 | 0.5 | 4 | 0.25 |
| 17_1521282 | 1 | 1 | 3 | 0.6667 | NA | NA | 1 | 1 | 5 | 0.8 | NA | NA | NA | NA | NA | NA | 1 | 1 | 1 | 1 |
| 17_1521289 | 1 | 0 | 2 | 1 | NA | NA | 1 | 0 | 4 | 0.5 | NA | NA | NA | NA | NA | NA | 1 | 1 | 1 | 1 |
| 17_1523511 | NA | NA | 1 | 1 | NA | NA | NA | NA | 1 | 1 | NA | NA | NA | NA | NA | NA | 1 | 1 | 1 | 1 |
| 17_1523528 | NA | NA | 2 | 1 | NA | NA | NA | NA | 2 | 1 | NA | NA | NA | NA | NA | NA | 1 | 1 | 1 | 1 |
| 17_1523532 | NA | NA | 2 | 0.5 | NA | NA | NA | NA | 2 | 0.5 | NA | NA | NA | NA | NA | NA | 1 | 1 | 1 | 1 |
| 17_1523967 | 4 | 0.25 | 2 | 0.5 | NA | NA | 1 | 0 | 7 | 0.2857 | 3 | 0 | NA | NA | 1 | 0 | 1 | 1 | 5 | 0.2 |
| 17_1524233 | NA | NA | 1 | 1 | 1 | 0 | 3 | 0 | 5 | 0.2 | NA | NA | 1 | 0 | 1 | 1 | NA | NA | 2 | 0.5 |
| 17_1524234 | NA | NA | 1 | 0 | 1 | 0 | 3 | 0.3333 | 5 | 0.2 | NA | NA | 1 | 1 | 1 | 0 | NA | NA | 2 | 0.5 |
| 17_1524383 | 1 | 1 | NA | NA | 1 | 1 | 2 | 1 | 4 | 1 | 1 | 1 | 1 | 1 | 3 | 0.6667 | 3 | 1 | 8 | 0.875 |
| 17_1524401 | 6 | 0.3333 | NA | NA | 1 | 0 | 5 | 0 | 12 | 0.1667 | 3 | 0 | 2 | 0 | 5 | 0.4 | 3 | 0.3333 | 13 | 0.2308 |
| 17_1524402 | 6 | 0.6667 | 1 | 1 | 1 | 1 | 5 | 0.8 | 13 | 0.7692 | 3 | 0 | 2 | 0.5 | 4 | 0.75 | 4 | 0.25 | 13 | 0.3846 |
| 17_1524419 | 4 | 0.5 | NA | NA | 1 | 1 | 3 | 0.6667 | 8 | 0.625 | 2 | 0.5 | 1 | 1 | 1 | 1 | 1 | 1 | 5 | 0.8 |
| 17_1524420 | 4 | 0.5 | NA | NA | 1 | 1 | 4 | 0.5 | 9 | 0.5556 | 2 | 1 | 1 | 1 | 1 | 0 | 1 | 1 | 5 | 0.8 |
| 17_1528395 | 1 | 1 | 1 | 1 | 2 | 1 | 1 | 0 | 5 | 0.8 | NA | NA | 1 | 1 | 2 | 0.5 | 2 | 1 | 5 | 0.8 |
| 17_1528440 | 1 | 0 | 1 | 1 | 2 | 0.5 | 1 | 1 | 5 | 0.6 | NA | NA | 1 | 0 | 2 | 0 | 2 | 0.5 | 5 | 0.2 |
| 17_1529447 | NA | NA | NA | NA | 1 | 1 | 1 | 0 | 2 | 0.5 | NA | NA | NA | NA | 1 | 1 | NA | NA | 1 | 1 |
| 17_1558802 | NA | NA | 1 | 0 | 1 | 1 | NA | NA | 2 | 0.5 | NA | NA | 2 | 1 | 2 | 1 | 2 | 0.5 | 6 | 0.8333 |
| 17_1558806 | NA | NA | 1 | 1 | 1 | 1 | 1 | 1 | 3 | 1 | NA | NA | 2 | 1 | 2 | 1 | 1 | 1 | 5 | 1 |
| 17_1558809 | NA | NA | 1 | 1 | 1 | 1 | 1 | 1 | 3 | 1 | NA | NA | 1 | 0 | 3 | 0.6667 | 1 | 1 | 5 | 0.6 |
| 17_1558875 | NA | NA | 1 | 1 | 1 | 0 | 1 | 1 | 3 | 0.6667 | NA | NA | 2 | 0.5 | 2 | 1 | 2 | 1 | 6 | 0.8333 |
| 17_1558880 | NA | NA | 1 | 0 | 1 | 1 | 1 | 0 | 3 | 0.3333 | NA | NA | 2 | 0 | 1 | 0 | 2 | 0.5 | 5 | 0.2 |
| 17_1558881 | NA | NA | NA | NA | 1 | 1 | 1 | 0 | 2 | 0.5 | NA | NA | 2 | 0 | 2 | 0.5 | 2 | 1 | 6 | 0.5 |
| 17_1558891 | NA | NA | 1 | 1 | 1 | 0 | 1 | 0 | 3 | 0.3333 | NA | NA | 2 | 0 | 1 | 0 | 2 | 0.5 | 5 | 0.2 |
| 17_1558929 | NA | NA | 1 | 0 | 1 | 1 | 1 | 1 | 3 | 0.6667 | NA | NA | 2 | 0 | 1 | 1 | NA | NA | 3 | 0.3333 |
| 17_1559598 | 1 | 1 | NA | NA | NA | NA | NA | NA | 1 | 1 | NA | NA | NA | NA | NA | NA | 1 | 1 | 1 | 1 |
| 17_1562233 | 1 | 1 | 4 | 0.5 | 4 | 0.25 | 5 | 0.6 | 14 | 0.5 | 1 | 0 | 3 | 0.3333 | 1 | 0 | 5 | 0.4 | 10 | 0.3 |
| 17_1562246 | 1 | 0 | 4 | 0.25 | 4 | 0 | 5 | 0.6 | 14 | 0.2857 | 2 | 0 | 5 | 0.4 | 2 | 0 | 4 | 0 | 13 | 0.1538 |
| 17_1562247 | 1 | 1 | 4 | 0.75 | 5 | 0.2 | 5 | 0.8 | 15 | 0.6 | 2 | 1 | 5 | 1 | 2 | 0.5 | 5 | 1 | 14 | 0.9286 |
| 17_1562250 | 1 | 1 | 4 | 0.25 | 3 | 0.3333 | 4 | 0.25 | 12 | 0.3333 | 2 | 0 | 5 | 0 | 1 | 0 | 4 | 0.25 | 12 | 0.0833 |
| 17_1562255 | 1 | 0 | 4 | 0.25 | 4 | 0.25 | 5 | 0 | 14 | 0.1429 | 2 | 0.5 | 5 | 0 | 2 | 0 | 4 | 0.5 | 13 | 0.2308 |
| 17_1562256 | 1 | 0 | 5 | 0.6 | 4 | 0 | 4 | 0.25 | 14 | 0.2857 | 2 | 0.5 | 5 | 0.6 | 1 | 0 | 5 | 0.4 | 13 | 0.4615 |
| 17_1562258 | 1 | 0 | 5 | 0 | 4 | 0.75 | 4 | 0.25 | 14 | 0.2857 | 2 | 0 | 5 | 0 | 2 | 0 | 4 | 0.25 | 13 | 0.0769 |

| | | | | | | | | | | | | | | | | | | | | |
|---|---|---|---|---|---|---|---|---|---|---|---|---|---|---|---|---|---|---|---|---|
| 17_1562271 | 1 | 0 | 4 | 0 | 5 | 0 | 5 | 0.2 | 15 | 0.0667 | 1 | 0 | 6 | 0.3333 | 3 | 0 | 5 | 0 | 15 | 0.1333 |
| 17_1562282 | 1 | 0 | 5 | 0.8 | 5 | 0 | 6 | 0.8333 | 17 | 0.5294 | 1 | 1 | 6 | 0.8333 | 3 | 0 | 5 | 0.6 | 15 | 0.6 |
| 17_1562299 | 1 | 0 | 5 | 0 | 5 | 0 | 6 | 0.3333 | 17 | 0.1176 | 2 | 0.5 | 7 | 0 | 3 | 0.3333 | 4 | 0.25 | 16 | 0.1875 |
| 17_1562319 | 2 | 0.5 | 6 | 0.1667 | 4 | 0.5 | 6 | 0.1667 | 18 | 0.2778 | 3 | 0 | 8 | 0.375 | 4 | 0 | 5 | 0 | 20 | 0.15 |
| 17_1562327 | 2 | 1 | 5 | 0.2 | 5 | 0.8 | 6 | 1 | 18 | 0.7222 | 3 | 1 | 8 | 0.625 | 4 | 1 | 4 | 0.75 | 19 | 0.7895 |
| 17_1562339 | NA | NA | 1 | 1 | 1 | 0 | 2 | 1 | 4 | 0.75 | NA | NA | 1 | 1 | NA | NA | NA | NA | 1 | 1 |
| 17_1562360 | NA | NA | 1 | 1 | 1 | 0 | 2 | 0.5 | 4 | 0.5 | NA | NA | 2 | 0.5 | 1 | 0 | NA | NA | 3 | 0.3333 |
| 17_1562366 | NA | NA | 1 | 0 | 1 | 0 | 2 | 0.5 | 4 | 0.25 | NA | NA | 2 | 0 | 1 | 1 | NA | NA | 3 | 0.3333 |
| 17_1562419 | NA | NA | 1 | 0 | 1 | 1 | 2 | 0 | 4 | 0.25 | NA | NA | 2 | 0.5 | 1 | 0 | NA | NA | 3 | 0.3333 |
| 17_1565208 | 5 | 0 | 5 | 0.2 | 7 | 0.1429 | 4 | 0.25 | 21 | 0.1429 | 2 | 0 | 5 | 0.2 | 5 | 0 | 4 | 0 | 16 | 0.0625 |
| 17_1565220 | 3 | 0.3333 | 7 | 0.1429 | 10 | 0.2 | 6 | 0.1667 | 26 | 0.1923 | 3 | 0 | 4 | 0 | 6 | 0 | 4 | 0.25 | 17 | 0.0588 |
| 17_1565231 | 2 | 0.5 | 3 | 1 | 3 | 0 | NA | NA | 8 | 0.5 | NA | NA | 2 | 0.5 | 1 | 0 | 4 | 0.25 | 7 | 0.2857 |
| 17_1569628 | 41 | 0.4146 | 36 | 0.3889 | 44 | 0.4773 | 42 | 0.3333 | 163 | 0.4049 | 27 | 0.4074 | 48 | 0.375 | 31 | 0.3871 | 44 | 0.4091 | 150 | 0.3933 |
| 17_1572459 | 2 | 0.5 | 2 | 0 | NA | NA | 3 | 0 | 7 | 0.1429 | 3 | 0.3333 | 4 | 0 | 1 | 0 | NA | NA | 8 | 0.125 |
| 17_1572484 | 2 | 0.5 | 2 | 0 | NA | NA | 4 | 0 | 8 | 0.125 | 3 | 0.3333 | 5 | 0 | 1 | 0 | 1 | 0 | 10 | 0.1 |
| 17_1573713 | 3 | 0 | 3 | 0.3333 | 2 | 0.5 | 2 | 0.5 | 10 | 0.3 | 4 | 0.75 | 3 | 0 | 2 | 0.5 | NA | NA | 9 | 0.4444 |
| 17_1573736 | 4 | 0.25 | 2 | 0.5 | 2 | 0 | 3 | 0.6667 | 11 | 0.3636 | 4 | 1 | 5 | 0.6 | 3 | 0.6667 | 1 | 0 | 13 | 0.6923 |
| 17_1573751 | 4 | 0 | 4 | 0.5 | 4 | 0.25 | 3 | 0 | 15 | 0.2 | 4 | 0.75 | 4 | 0 | 3 | 0 | 1 | 0 | 12 | 0.25 |
| 17_1599032 | NA | NA | NA | NA | 2 | 1 | 2 | 0.5 | 4 | 0.75 | 1 | 0 | 2 | 1 | 1 | 1 | 2 | 0.5 | 6 | 0.6667 |
| 17_1599035 | NA | NA | NA | NA | 1 | 1 | 1 | 0 | 2 | 0.5 | NA | NA | 2 | 0 | 1 | 1 | 2 | 0 | 5 | 0.2 |
| 17_1599045 | NA | NA | NA | NA | 2 | 0.5 | 2 | 0.5 | 4 | 0.5 | NA | NA | NA | NA | 1 | 1 | 1 | 0 | 2 | 0.5 |
| 17_1599052 | NA | NA | NA | NA | 2 | 0.5 | 2 | 1 | 4 | 0.75 | NA | NA | 2 | 0.5 | 1 | 1 | 2 | 0.5 | 5 | 0.6 |
| 17_1599061 | NA | NA | NA | NA | 2 | 1 | 2 | 1 | 4 | 1 | 1 | 1 | 2 | 1 | 1 | 1 | 2 | 1 | 6 | 1 |
| 17_1599066 | NA | NA | NA | NA | 2 | 0.5 | 2 | 0.5 | 4 | 0.5 | 1 | 0 | 2 | 0.5 | 1 | 1 | 2 | 0 | 6 | 0.3333 |
| 17_1599085 | NA | NA | NA | NA | 2 | 0 | 2 | 0.5 | 4 | 0.25 | 1 | 1 | 2 | 1 | 1 | 1 | 2 | 1 | 6 | 1 |
| 17_1599088 | NA | NA | NA | NA | 2 | 1 | 1 | 0 | 3 | 0.6667 | NA | NA | 2 | 0.5 | NA | NA | 1 | 0 | 3 | 0.3333 |
| 17_1599094 | NA | NA | NA | NA | 1 | 1 | 1 | 1 | 2 | 1 | NA | NA | 2 | 1 | 1 | 1 | 1 | 1 | 4 | 1 |
| 17_1599102 | NA | NA | NA | NA | 2 | 1 | 2 | 1 | 4 | 1 | 1 | 1 | 2 | 1 | 1 | 1 | 2 | 1 | 6 | 1 |
| 17_1599186 | 1 | 1 | NA | NA | NA | NA | 1 | 1 | 2 | 1 | NA | NA | NA | NA | 1 | 0 | 1 | 1 | 2 | 0.5 |
| 17_1602575 | 7 | 0.2857 | 9 | 0.2222 | 4 | 0 | 6 | 0.1667 | 26 | 0.1923 | 8 | 0.125 | 11 | 0.0909 | 11 | 0.2727 | 11 | 0.1818 | 41 | 0.1707 |
| 17_1602597 | 10 | 0.5 | 9 | 0.5556 | 3 | 0.6667 | 6 | 0.8333 | 28 | 0.6071 | 8 | 0.5 | 11 | 0.8182 | 12 | 0.5 | 10 | 0.5 | 41 | 0.5854 |
| 17_1602651 | 8 | 0.125 | 9 | 0 | 5 | 0.4 | 5 | 0.2 | 27 | 0.1481 | 5 | 0 | 12 | 0 | 9 | 0.1111 | 9 | 0.1111 | 35 | 0.0571 |
| 17_1602744 | 5 | 1 | 3 | 1 | 3 | 1 | 5 | 1 | 16 | 1 | 6 | 1 | 11 | 1 | 7 | 1 | 7 | 1 | 31 | 1 |
| 17_1602761 | 6 | 0.8333 | 5 | 0.8 | 3 | 0.6667 | 4 | 0.75 | 18 | 0.7778 | 5 | 0.6 | 10 | 0.9 | 9 | 0.8889 | 8 | 0.875 | 32 | 0.8438 |
| 17_1602765 | 5 | 1 | 5 | 1 | 2 | 1 | 3 | 1 | 15 | 1 | 4 | 0.75 | 10 | 0.9 | 6 | 1 | 8 | 0.5 | 28 | 0.7857 |
| 17_1602773 | 6 | 1 | 5 | 1 | 2 | 1 | 4 | 1 | 17 | 1 | 5 | 1 | 10 | 1 | 9 | 1 | 7 | 1 | 31 | 1 |
| 17_1602778 | 6 | 1 | 5 | 1 | 3 | 1 | 3 | 1 | 17 | 1 | 5 | 1 | 10 | 1 | 7 | 1 | 8 | 1 | 30 | 1 |
| 17_1602781 | 6 | 0.6667 | 4 | 0.75 | 3 | 0.6667 | 3 | 0.6667 | 16 | 0.6875 | 5 | 0.4 | 10 | 0.6 | 8 | 0.75 | 6 | 0.3333 | 29 | 0.5517 |
| 17_1602800 | 5 | 0 | 5 | 0.2 | 2 | 0 | 4 | 0 | 16 | 0.0625 | 5 | 0.2 | 9 | 0.1111 | 9 | 0 | 6 | 0 | 29 | 0.069 |
| 17_1604100 | 6 | 0.3333 | 4 | 0.25 | 2 | 0 | 4 | 0 | 16 | 0.1875 | 5 | 0.2 | 10 | 0.1 | 8 | 0.5 | 6 | 0.5 | 29 | 0.3103 |
| 17_1604106 | 5 | 1 | 5 | 1 | 2 | 0.5 | 2 | 1 | 14 | 0.9286 | 4 | 0.75 | 7 | 0.8571 | 5 | 0.6 | 2 | 1 | 18 | 0.7778 |
| 17_1604110 | 5 | 0.4 | 4 | 0.75 | 2 | 0 | 4 | 0.25 | 15 | 0.4 | 3 | 1 | 10 | 0.4 | 6 | 0.3333 | 6 | 0.3333 | 25 | 0.44 |
| 17_1604111 | 5 | 0.2 | 4 | 0.25 | 2 | 0 | 4 | 0 | 15 | 0.1333 | 6 | 0.3333 | 9 | 0.1111 | 8 | 0 | 8 | 0.375 | 31 | 0.1935 |
| 17_1604113 | NA | NA | 1 | 0 | NA | NA | 1 | 1 | 2 | 0.5 | NA | NA | 3 | 1 | 3 | 0.6667 | 1 | 1 | 7 | 0.8571 |
| 17_1604117 | 5 | 0 | 5 | 0.4 | 2 | 0 | 4 | 0 | 16 | 0.125 | 5 | 0 | 10 | 0.1 | 8 | 0.125 | 7 | 0 | 30 | 0.0667 |
| 17_1604120 | 5 | 0.6 | 5 | 0.6 | NA | NA | 4 | 0.75 | 14 | 0.6429 | 6 | 0.8333 | 10 | 0.5 | 8 | 0.625 | 7 | 0.7143 | 31 | 0.6452 |
| 17_1604121 | 3 | 1 | 3 | 1 | NA | NA | 4 | 1 | 10 | 1 | 2 | 1 | 9 | 1 | 6 | 1 | 8 | 0.75 | 25 | 0.92 |
| 17_1604129 | 3 | 1 | 1 | 0 | NA | NA | 1 | 0 | 5 | 0.6 | 3 | 0.6667 | 8 | 0.625 | 4 | 0.25 | 5 | 0.4 | 20 | 0.5 |
| 17_1604134 | 4 | 0.25 | 5 | 0.4 | NA | NA | 4 | 0.25 | 13 | 0.3077 | 5 | 0.2 | 8 | 0.625 | 8 | 0.25 | 6 | 0.5 | 27 | 0.4074 |
| 17_1604136 | 3 | 0 | 5 | 0.2 | NA | NA | 2 | 0 | 10 | 0.1 | 5 | 0.2 | 7 | 0.4286 | 5 | 0.2 | 5 | 0.2 | 22 | 0.2727 |
| 17_1719301 | 1 | 0 | NA | NA | 1 | 1 | NA | NA | 2 | 0.5 | NA | NA | NA | NA | 2 | 0.5 | NA | NA | 2 | 0.5 |
| 17_1719317 | 1 | 1 | NA | NA | NA | NA | NA | NA | 1 | 1 | NA | NA | 1 | 1 | 4 | 1 | NA | NA | 5 | 1 |
| 17_1719318 | 1 | 1 | NA | NA | NA | NA | NA | NA | 1 | 1 | NA | NA | 1 | 1 | 3 | 0.3333 | NA | NA | 4 | 0.5 |
| 17_1719320 | NA | NA | NA | NA | 1 | 1 | NA | NA | 1 | 1 | NA | NA | 1 | 1 | 3 | 1 | NA | NA | 4 | 1 |
| 17_1719360 | 1 | 1 | NA | NA | 1 | 1 | NA | NA | 2 | 1 | NA | NA | NA | NA | 2 | 1 | NA | NA | 2 | 1 |
| 17_1719378 | 1 | 1 | NA | NA | 1 | 1 | NA | NA | 2 | 1 | NA | NA | 1 | 1 | 4 | 1 | NA | NA | 5 | 1 |
| 17_1719402 | 1 | 1 | NA | NA | 2 | 0.5 | 1 | 1 | 4 | 0.75 | NA | NA | 2 | 1 | 4 | 1 | NA | NA | 6 | 1 |
| 17_1719412 | 1 | 1 | NA | NA | 2 | 0 | 1 | 0 | 4 | 0.25 | NA | NA | 1 | 0 | 5 | 0.6 | NA | NA | 6 | 0.5 |
| 17_1726010 | 1 | 1 | NA | NA | NA | NA | NA | NA | 1 | 1 | NA | NA | 1 | 1 | 1 | 1 | NA | NA | 2 | 1 |
| 17_1726014 | 1 | 1 | NA | NA | NA | NA | NA | NA | 1 | 1 | NA | NA | 1 | 0 | 1 | 1 | NA | NA | 2 | 0.5 |
| 17_1726018 | 1 | 1 | NA | NA | NA | NA | NA | NA | 1 | 1 | NA | NA | 1 | 1 | 1 | 0 | NA | NA | 2 | 0.5 |
| 17_1726082 | 1 | 1 | NA | NA | NA | NA | NA | NA | 1 | 1 | NA | NA | 1 | 1 | 1 | 1 | NA | NA | 2 | 1 |
| 17_1730786 | NA | NA | 1 | 1 | NA | NA | 1 | 0 | 2 | 0.5 | NA | NA | NA | NA | 1 | 0 | 2 | 0.5 | 3 | 0.3333 |
| 17_1730812 | NA | NA | 1 | 1 | NA | NA | 1 | 1 | 2 | 1 | NA | NA | NA | NA | 1 | 0 | 2 | 0.5 | 3 | 0.3333 |
| 17_1742499 | NA | NA | 1 | 1 | 1 | 0 | NA | NA | 2 | 0.5 | NA | NA | 1 | 0 | 2 | 0.5 | NA | NA | 3 | 0.3333 |

| | | | | | | | | | | | | | | | | | | | | |
|---|---|---|---|---|---|---|---|---|---|---|---|---|---|---|---|---|---|---|---|---|
| 17_1742524 | NA | NA | 1 | 1 | 1 | 0 | NA | NA | 2 | 0.5 | NA | NA | 1 | 0 | 3 | 0.3333 | NA | NA | 4 | 0.25 |
| 17_1742520 | NA | NA | 1 | 0 | 1 | 1 | NA | NA | 2 | 0.5 | NA | NA | 1 | 0 | 2 | 0.5 | NA | NA | 3 | 0.3333 |
| 17_1742572 | NA | NA | 1 | 0 | 1 | 1 | NA | NA | 2 | 0.5 | NA | NA | 1 | 1 | 2 | 1 | NA | NA | 3 | 1 |
| 17_1742600 | NA | NA | 1 | 1 | 1 | 0 | NA | NA | 2 | 0.5 | NA | NA | 1 | 0 | 2 | 1 | NA | NA | 3 | 0.6667 |
| 17_1742609 | NA | NA | 2 | 0.5 | 1 | 1 | NA | NA | 3 | 0.6667 | NA | NA | 1 | 1 | 3 | 1 | NA | NA | 4 | 1 |
| 17_1742617 | NA | NA | 2 | 0 | 1 | 1 | NA | NA | 3 | 0.3333 | NA | NA | 1 | 0 | 3 | 0.3333 | NA | NA | 4 | 0.25 |
| 17_1742619 | NA | NA | 2 | 0 | 2 | 0.5 | NA | NA | 4 | 0.25 | NA | NA | 1 | 0 | 3 | 0.6667 | NA | NA | 4 | 0.5 |
| 17_1742641 | NA | NA | 2 | 1 | 2 | 1 | NA | NA | 4 | 1 | NA | NA | 1 | 1 | 3 | 1 | NA | NA | 4 | 1 |
| 17_1742644 | NA | NA | 2 | 0.5 | 2 | 0.5 | NA | NA | 4 | 0.5 | NA | NA | NA | NA | 1 | 1 | NA | NA | 1 | 1 |
| 17_1748630 | NA | NA | 1 | 1 | NA | NA | NA | NA | 1 | 1 | NA | NA | 1 | 0 | NA | NA | 1 | 1 | 2 | 0.5 |
| 17_1753872 | NA | NA | NA | NA | 1 | 1 | NA | NA | 1 | 1 | NA | NA | NA | NA | 1 | 1 | 1 | 0 | 2 | 0.5 |
| 17_1753899 | NA | NA | NA | NA | 1 | 1 | NA | NA | 1 | 1 | NA | NA | NA | NA | 1 | 1 | 1 | 0 | 2 | 0.5 |
| 17_1753923 | NA | NA | NA | NA | 1 | 1 | NA | NA | 1 | 1 | NA | NA | NA | NA | 1 | 1 | NA | NA | 1 | 1 |
| 17_1753942 | 1 | 0 | NA | NA | 1 | 1 | NA | NA | 2 | 0.5 | NA | NA | NA | NA | 1 | 1 | 1 | 0 | 2 | 0.5 |
| 17_1753959 | 1 | 0 | NA | NA | 1 | 1 | NA | NA | 2 | 0.5 | NA | NA | NA | NA | 1 | 1 | 1 | 0 | 2 | 0.5 |
| 17_1754042 | NA | NA | NA | NA | 1 | 1 | NA | NA | 1 | 1 | NA | NA | NA | NA | 1 | 0 | 1 | 1 | 2 | 0.5 |
| 17_1754214 | NA | NA | NA | NA | 1 | 1 | NA | NA | 1 | 1 | NA | NA | NA | NA | 1 | 1 | NA | NA | 1 | 1 |
| 17_1754234 | NA | NA | NA | NA | 1 | 1 | NA | NA | 1 | 1 | NA | NA | NA | NA | 1 | 1 | NA | NA | 1 | 1 |
| 17_1754240 | NA | NA | NA | NA | 1 | 1 | NA | NA | 1 | 1 | NA | NA | NA | NA | 1 | 1 | 1 | 1 | 2 | 1 |
| 17_1754261 | NA | NA | NA | NA | 1 | 1 | NA | NA | 1 | 1 | NA | NA | NA | NA | NA | NA | 1 | 1 | 1 | 1 |
| 17_1756572 | 2 | 0 | 2 | 0.5 | 2 | 0 | 5 | 0 | 11 | 0.0909 | 2 | 0 | 2 | 0.5 | 5 | 0 | NA | NA | 9 | 0.1111 |
| 17_1760739 | 1 | 1 | 2 | 0.5 | 1 | 0 | 5 | 0.2 | 9 | 0.3333 | 2 | 0.5 | 1 | 0 | 3 | 0.3333 | 2 | 0.5 | 8 | 0.375 |
| 17_1760744 | 2 | 1 | 2 | 0.5 | 1 | 1 | 7 | 0.7143 | 12 | 0.75 | 2 | 1 | NA | NA | 3 | 0.6667 | 2 | 1 | 7 | 0.8571 |
| 17_1760772 | 1 | 0 | 2 | 0 | 1 | 0 | 7 | 0.1429 | 11 | 0.0909 | 2 | 0 | 1 | 1 | 2 | 0 | 1 | 0 | 6 | 0.1667 |
| 17_1760776 | 2 | 0 | 2 | 0.5 | 1 | 1 | 7 | 1 | 12 | 0.75 | 2 | 0.5 | 1 | 1 | 3 | 0.6667 | 2 | 0.5 | 8 | 0.625 |
| 17_1761412 | 1 | 0 | 2 | 0.5 | NA | NA | 3 | 0 | 6 | 0.1667 | 3 | 0.6667 | 2 | 0 | 4 | 0 | 2 | 0 | 11 | 0.1818 |
| 17_1767743 | 2 | 0.5 | 6 | 0.1667 | 4 | 0 | 4 | 0.25 | 16 | 0.1875 | 2 | 0.5 | 7 | 0.1429 | NA | NA | 8 | 0.125 | 17 | 0.1765 |
| 17_1767752 | 3 | 0.3333 | 6 | 0 | 5 | 0 | 6 | 0.1667 | 20 | 0.1 | 4 | 0 | 8 | 0 | NA | NA | 10 | 0.2 | 22 | 0.0909 |
| 17_1805860 | NA | NA | 2 | 0 | NA | NA | 2 | 0.5 | 4 | 0.25 | NA | NA | NA | NA | NA | NA | 2 | 0.5 | 2 | 0.5 |
| 17_1808650 | NA | NA | NA | NA | 1 | 1 | 1 | 1 | 2 | 1 | NA | NA | NA | NA | 1 | 1 | NA | NA | 1 | 1 |
| 17_1808651 | NA | NA | NA | NA | 1 | 1 | 1 | 1 | 2 | 1 | NA | NA | 1 | 1 | 1 | 1 | NA | NA | 2 | 1 |
| 17_1808670 | NA | NA | NA | NA | 1 | 1 | 1 | 1 | 2 | 1 | NA | NA | NA | NA | 1 | 1 | NA | NA | 1 | 1 |
| 17_1808739 | NA | NA | NA | NA | 1 | 1 | 1 | 1 | 2 | 1 | NA | NA | NA | NA | 1 | 1 | NA | NA | 1 | 1 |
| 17_1808760 | NA | NA | NA | NA | 1 | 1 | 1 | 1 | 2 | 1 | NA | NA | NA | NA | 1 | 1 | NA | NA | 1 | 1 |
| 17_2118582 | 93 | 0.0753 | 110 | 0.0545 | 86 | 0.1047 | 99 | 0.101 | 388 | 0.0825 | 62 | 0.0323 | 89 | 0.0899 | 57 | 0.1053 | 78 | 0.0641 | 286 | 0.0734 |
| 17_2124570 | NA | NA | NA | NA | 1 | 1 | NA | NA | 1 | 1 | NA | NA | NA | NA | 1 | 1 | NA | NA | 1 | 1 |
| 17_2124585 | NA | NA | 1 | 1 | 1 | 1 | NA | NA | 2 | 1 | NA | NA | NA | NA | NA | NA | 1 | 1 | 1 | 1 |
| 17_2124590 | NA | NA | NA | NA | 1 | 1 | 1 | 1 | 2 | 1 | NA | NA | NA | NA | 1 | 1 | 1 | 1 | 2 | 1 |
| 17_2124621 | NA | NA | 2 | 0 | 2 | 0.5 | NA | NA | 4 | 0.25 | NA | NA | NA | NA | 1 | 0 | 2 | 0.5 | 3 | 0.3333 |
| 17_2124628 | NA | NA | 2 | 1 | 1 | 0 | 1 | 0 | 4 | 0.5 | NA | NA | NA | NA | NA | NA | 1 | 1 | 1 | 1 |
| 17_2124669 | NA | NA | 2 | 0.5 | 2 | 0 | NA | NA | 4 | 0.25 | NA | NA | NA | NA | 1 | 0 | 2 | 0.5 | 3 | 0.3333 |
| 17_2124690 | NA | NA | 1 | 1 | 2 | 0.5 | 1 | 1 | 4 | 0.75 | NA | NA | NA | NA | 1 | 1 | 2 | 1 | 3 | 1 |
| 17_2124714 | NA | NA | 5 | 0.4 | 2 | 0 | 1 | 0 | 8 | 0.25 | NA | NA | NA | NA | 1 | 0 | 2 | 1 | 3 | 0.6667 |
| 17_2124721 | NA | NA | 5 | 0.8 | 2 | 1 | NA | NA | 7 | 0.8571 | NA | NA | NA | NA | 1 | 1 | 2 | 1 | 3 | 1 |
| 17_2124722 | NA | NA | 3 | 0.3333 | 2 | 0 | 1 | 0 | 6 | 0.1667 | NA | NA | NA | NA | 1 | 1 | 2 | 0 | 3 | 0.3333 |
| 17_2124737 | NA | NA | 5 | 0.8 | 2 | 1 | 1 | 1 | 8 | 0.875 | NA | NA | NA | NA | 2 | 0.5 | 2 | 1 | 4 | 0.75 |
| 17_2124760 | NA | NA | 5 | 0.2 | 2 | 0 | 1 | 1 | 8 | 0.25 | NA | NA | NA | NA | 2 | 0.5 | 2 | 0 | 4 | 0.25 |
| 17_2124777 | NA | NA | 3 | 0.3333 | 1 | 0 | 1 | 1 | 5 | 0.4 | 1 | 1 | NA | NA | 2 | 0 | 1 | 0 | 4 | 0.25 |
| 17_2124857 | NA | NA | 5 | 0.2 | 2 | 0.5 | 1 | 0 | 8 | 0.25 | NA | NA | NA | NA | 3 | 0 | 2 | 0.5 | 5 | 0.2 |
| 17_2124888 | NA | NA | 4 | 0 | 3 | 0.3333 | 2 | 0 | 9 | 0.1111 | NA | NA | 1 | 0 | 3 | 0.3333 | 2 | 0 | 6 | 0.1667 |
| 17_2150441 | 1 | 0 | 3 | 0.6667 | 1 | 0 | 3 | 0.3333 | 8 | 0.375 | 3 | 0.3333 | 1 | 1 | 3 | 0 | 2 | 0 | 9 | 0.2222 |
| 17_2161739 | 1 | 1 | NA | NA | NA | NA | 1 | 1 | 2 | 1 | NA | NA | NA | NA | NA | NA | 1 | 1 | 1 | 1 |
| 17_2161746 | NA | NA | NA | NA | NA | NA | 1 | 1 | 1 | 1 | NA | NA | NA | NA | NA | NA | 1 | 1 | 1 | 1 |
| 17_2161789 | 1 | 0 | NA | NA | NA | NA | 1 | 1 | 2 | 0.5 | NA | NA | NA | NA | NA | NA | 1 | 1 | 1 | 1 |
| 17_2161839 | 1 | 1 | NA | NA | NA | NA | 1 | 0 | 2 | 0.5 | NA | NA | NA | NA | NA | NA | 2 | 1 | 2 | 1 |
| 17_2161878 | 1 | 1 | NA | NA | NA | NA | 1 | 1 | 2 | 1 | NA | NA | NA | NA | NA | NA | 1 | 1 | 1 | 1 |
| 17_2161886 | 1 | 1 | NA | NA | NA | NA | 1 | 1 | 2 | 1 | NA | NA | NA | NA | NA | NA | 2 | 1 | 2 | 1 |
| 17_2161914 | NA | NA | NA | NA | NA | NA | 1 | 1 | 1 | 1 | NA | NA | NA | NA | NA | NA | 2 | 1 | 2 | 1 |
| 17_2161920 | 1 | 1 | NA | NA | NA | NA | 1 | 1 | 2 | 1 | NA | NA | NA | NA | NA | NA | 2 | 1 | 2 | 1 |
| 17_2161924 | 1 | 1 | NA | NA | NA | NA | 1 | 0 | 2 | 0.5 | NA | NA | NA | NA | NA | NA | 2 | 1 | 2 | 1 |
| 17_2168370 | NA | NA | 1 | 0 | 1 | 1 | NA | NA | 2 | 0.5 | NA | NA | NA | NA | NA | NA | 1 | 1 | 1 | 1 |
| 17_2168387 | NA | NA | 1 | 1 | 1 | 1 | NA | NA | 2 | 1 | NA | NA | NA | NA | NA | NA | 1 | 1 | 1 | 1 |
| 17_2168392 | NA | NA | 1 | 1 | 1 | 1 | NA | NA | 2 | 1 | NA | NA | NA | NA | NA | NA | 1 | 1 | 1 | 1 |
| 17_2168538 | 1 | 1 | 1 | 1 | 1 | 1 | NA | NA | 3 | 1 | NA | NA | 1 | 1 | NA | NA | 1 | 1 | 2 | 1 |
| 17_2168566 | 1 | 1 | 1 | 0 | 1 | 1 | NA | NA | 3 | 0.6667 | NA | NA | 1 | 1 | NA | NA | 2 | 1 | 3 | 1 |

| | | | | | | | | | | | | | | | | | | | | |
|---|---|---|---|---|---|---|---|---|---|---|---|---|---|---|---|---|---|---|---|---|
| 17_216857 | 1 | 1 | 1 | 0 | 1 | 1 | NA | NA | 3 | 0.6667 | NA | NA | 1 | 0 | NA | NA | 2 | 0.5 | 3 | 0.3333 |
| 17_216961 | NA | NA | 1 | 1 | NA | NA | NA | NA | 1 | 1 | NA | NA | NA | NA | 1 | 1 | 1 | 1 | 2 | 1 |
| 17_216962 | NA | NA | 1 | 1 | NA | NA | NA | NA | 1 | 1 | NA | NA | NA | NA | 1 | 1 | 1 | 0 | 2 | 0.5 |
| 17_216963 | NA | NA | 1 | 1 | NA | NA | NA | NA | 1 | 1 | NA | NA | NA | NA | 1 | 0 | 1 | 1 | 2 | 0.5 |
| 17_216964 | NA | NA | 1 | 1 | NA | NA | NA | NA | 1 | 1 | NA | NA | NA | NA | NA | NA | 1 | 1 | 1 | 1 |
| 17_216965 | NA | NA | 1 | 1 | NA | NA | NA | NA | 1 | 1 | NA | NA | NA | NA | 1 | 1 | 1 | 0 | 2 | 0.5 |
| 17_216971 | NA | NA | 1 | 1 | NA | NA | NA | NA | 1 | 1 | NA | NA | NA | NA | 1 | 1 | 1 | 1 | 2 | 1 |
| 17_216972 | NA | NA | 1 | 1 | NA | NA | NA | NA | 1 | 1 | NA | NA | NA | NA | 1 | 1 | 1 | 0 | 2 | 0.5 |
| 17_216979 | NA | NA | 1 | 1 | NA | NA | NA | NA | 1 | 1 | NA | NA | NA | NA | 1 | 1 | 1 | 0 | 2 | 0.5 |
| 17_216980 | NA | NA | 1 | 1 | NA | NA | NA | NA | 1 | 1 | NA | NA | NA | NA | 1 | 0 | 1 | 1 | 2 | 0.5 |
| 17_216980 | NA | NA | 1 | 1 | NA | NA | NA | NA | 1 | 1 | NA | NA | NA | NA | 1 | 1 | NA | NA | 1 | 1 |
| 17_217284 | 3 | 0.6667 | 1 | 0 | NA | NA | NA | NA | 4 | 0.5 | NA | NA | 1 | 1 | NA | NA | 1 | 1 | 2 | 1 |
| 17_217284 | 3 | 0.3333 | 1 | 0 | NA | NA | NA | NA | 4 | 0.25 | NA | NA | 2 | 0.5 | NA | NA | 1 | 0 | 3 | 0.3333 |
| 17_217284 | 3 | 0.6667 | 1 | 0 | NA | NA | NA | NA | 4 | 0.5 | NA | NA | 2 | 0.5 | NA | NA | 1 | 0 | 3 | 0.3333 |
| 17_217284 | 3 | 1 | 1 | 0 | NA | NA | NA | NA | 4 | 0.75 | NA | NA | 2 | 0.5 | NA | NA | 1 | 1 | 3 | 0.6667 |
| 17_217285 | 3 | 1 | 1 | 1 | NA | NA | NA | NA | 4 | 1 | NA | NA | 2 | 1 | NA | NA | 1 | 1 | 3 | 1 |
| 17_217287 | 3 | 1 | 1 | 0 | NA | NA | NA | NA | 4 | 0.75 | NA | NA | 2 | 1 | NA | NA | 1 | 0 | 3 | 0.6667 |
| 17_217289 | 3 | 1 | 1 | 0 | NA | NA | NA | NA | 4 | 0.75 | NA | NA | 2 | 1 | 1 | 1 | NA | NA | 3 | 1 |
| 17_217291 | 3 | 0.3333 | 1 | 0 | NA | NA | NA | NA | 4 | 0.25 | NA | NA | 2 | 0.5 | 1 | 0 | 1 | 0 | 4 | 0.25 |
| 17_217296 | 1 | 1 | 1 | 1 | NA | NA | 1 | 1 | 3 | 1 | NA | NA | 2 | 0.5 | 1 | 1 | 1 | 1 | 4 | 0.75 |
| 17_217297 | 3 | 0 | 1 | 1 | 1 | 0 | 1 | 0 | 6 | 0.1667 | NA | NA | 1 | 0 | 1 | 1 | 1 | 0 | 3 | 0.3333 |
| 17_217298 | 3 | 1 | 1 | 1 | 1 | 1 | 1 | 1 | 6 | 1 | NA | NA | 1 | 1 | 1 | 1 | 1 | 1 | 3 | 1 |
| 17_217506 | NA | NA | 1 | 1 | NA | NA | NA | NA | 1 | 1 | NA | NA | NA | NA | 1 | 1 | NA | NA | 1 | 1 |
| 17_217507 | NA | NA | 1 | 1 | NA | NA | NA | NA | 1 | 1 | NA | NA | NA | NA | 1 | 1 | NA | NA | 1 | 1 |
| 17_218071 | NA | NA | NA | NA | 2 | 0.5 | 1 | 1 | 3 | 0.6667 | NA | NA | NA | NA | 1 | 1 | NA | NA | 1 | 1 |
| 17_218072 | NA | NA | NA | NA | 2 | 1 | 1 | 1 | 3 | 1 | NA | NA | NA | NA | 1 | 1 | NA | NA | 1 | 1 |
| 17_218080 | NA | NA | NA | NA | 2 | 1 | 1 | 1 | 3 | 1 | NA | NA | NA | NA | 1 | 1 | NA | NA | 1 | 1 |
| 17_218081 | NA | NA | NA | NA | 2 | 0.5 | 1 | 1 | 3 | 0.6667 | NA | NA | NA | NA | 1 | 1 | NA | NA | 1 | 1 |
| 17_218086 | 1 | 1 | NA | NA | 1 | 0 | 1 | 1 | 3 | 0.6667 | NA | NA | NA | NA | 1 | 0 | 1 | 1 | 2 | 0.5 |
| 17_218087 | 1 | 1 | NA | NA | 2 | 1 | NA | NA | 3 | 1 | NA | NA | NA | NA | NA | NA | 2 | 1 | 2 | 1 |
| 17_218803 | 1 | 0 | 2 | 0.5 | 1 | 0 | 2 | 0 | 6 | 0.1667 | NA | NA | 2 | 0.5 | NA | NA | 1 | 0 | 3 | 0.3333 |
| 17_218804 | NA | NA | 2 | 1 | 1 | 1 | 2 | 1 | 5 | 1 | NA | NA | 2 | 0.5 | NA | NA | 1 | 0 | 3 | 0.3333 |
| 17_218805 | 1 | 1 | 1 | 1 | 1 | 1 | 2 | 0.5 | 5 | 0.8 | NA | NA | 1 | 1 | NA | NA | 1 | 1 | 2 | 1 |
| 17_218807 | 1 | 1 | 2 | 1 | 2 | 1 | 2 | 0.5 | 7 | 0.8571 | NA | NA | 2 | 1 | NA | NA | NA | NA | 2 | 1 |
| 17_218807 | 1 | 0 | 2 | 0 | 1 | 0 | 2 | 0.5 | 6 | 0.1667 | NA | NA | 2 | 0.5 | NA | NA | NA | NA | 2 | 0.5 |
| 17_218809 | NA | NA | 2 | 1 | 2 | 1 | 2 | 1 | 6 | 1 | NA | NA | 2 | 1 | NA | NA | NA | NA | 2 | 1 |
| 17_218809 | 1 | 1 | 2 | 0.5 | 2 | 1 | 2 | 0.5 | 7 | 0.7143 | NA | NA | 2 | 1 | NA | NA | NA | NA | 2 | 1 |
| 17_218810 | NA | NA | 2 | 0.5 | 2 | 1 | 2 | 1 | 6 | 0.8333 | NA | NA | 2 | 1 | NA | NA | NA | NA | 2 | 1 |
| 17_218811 | 1 | 1 | 2 | 0.5 | 2 | 1 | 2 | 0 | 7 | 0.5714 | NA | NA | 1 | 1 | NA | NA | NA | NA | 1 | 1 |
| 17_218814 | 1 | 0 | 2 | 0 | 2 | 0.5 | 2 | 0.5 | 7 | 0.2857 | NA | NA | 2 | 0.5 | NA | NA | NA | NA | 2 | 0.5 |
| 17_218814 | 1 | 1 | 2 | 0 | 2 | 0 | 2 | 0 | 7 | 0.1429 | NA | NA | 2 | 0.5 | NA | NA | NA | NA | 2 | 0.5 |
| 17_218814 | 1 | 1 | 2 | 1 | 2 | 1 | 2 | 1 | 7 | 1 | NA | NA | 2 | 1 | NA | NA | NA | NA | 2 | 1 |
| 17_218816 | 1 | 1 | 2 | 0 | 1 | 0 | 2 | 0 | 6 | 0.1667 | NA | NA | 2 | 0.5 | NA | NA | NA | NA | 2 | 0.5 |
| 17_218835 | 1 | 1 | 1 | 0 | 2 | 1 | NA | NA | 4 | 0.75 | 1 | 1 | 1 | 1 | 1 | 1 | NA | NA | 3 | 1 |
| 17_218836 | 2 | 0.5 | 1 | 0 | 1 | 1 | 1 | 1 | 5 | 0.6 | 1 | 0 | 1 | 1 | 1 | 1 | NA | NA | 3 | 0.6667 |
| 17_218837 | 1 | 1 | 1 | 1 | 1 | 1 | 1 | 1 | 4 | 1 | 1 | 1 | 1 | 1 | 1 | 1 | NA | NA | 3 | 1 |
| 17_218837 | 2 | 0 | 1 | 1 | 2 | 0 | 1 | 0 | 6 | 0.1667 | 1 | 1 | 1 | 1 | 1 | 0 | NA | NA | 3 | 0.6667 |
| 17_219091 | NA | NA | NA | NA | 2 | 0.5 | 1 | 0 | 3 | 0.3333 | NA | NA | 2 | 0.5 | NA | NA | NA | NA | 2 | 0.5 |
| 17_219092 | NA | NA | NA | NA | 2 | 0.5 | 1 | 0 | 3 | 0.3333 | NA | NA | 2 | 0.5 | NA | NA | NA | NA | 2 | 0.5 |
| 17_219096 | NA | NA | NA | NA | 2 | 0.5 | 1 | 1 | 3 | 0.6667 | NA | NA | 1 | 1 | NA | NA | NA | NA | 1 | 1 |
| 17_219098 | NA | NA | NA | NA | 2 | 0.5 | 1 | 0 | 3 | 0.3333 | NA | NA | 2 | 0.5 | NA | NA | NA | NA | 2 | 0.5 |
| 17_219098 | NA | NA | NA | NA | 2 | 0.5 | 1 | 1 | 3 | 0.6667 | NA | NA | 2 | 0.5 | NA | NA | NA | NA | 2 | 0.5 |
| 17_219099 | NA | NA | NA | NA | 2 | 0 | 1 | 1 | 3 | 0.3333 | NA | NA | 2 | 0.5 | NA | NA | NA | NA | 2 | 0.5 |
| 17_219102 | NA | NA | NA | NA | 2 | 0.5 | 1 | 0 | 3 | 0.3333 | NA | NA | 2 | 0.5 | NA | NA | NA | NA | 2 | 0.5 |
| 17_219104 | NA | NA | NA | NA | 2 | 1 | 1 | 1 | 3 | 1 | NA | NA | 2 | 0.5 | NA | NA | NA | NA | 2 | 0.5 |
| 17_219105 | NA | NA | NA | NA | 2 | 1 | 1 | 1 | 3 | 1 | NA | NA | 1 | 1 | NA | NA | NA | NA | 1 | 1 |
| 17_219106 | NA | NA | NA | NA | 2 | 0.5 | 1 | 0 | 3 | 0.3333 | NA | NA | 2 | 0.5 | NA | NA | NA | NA | 2 | 0.5 |
| 17_219107 | NA | NA | NA | NA | 3 | 1 | 1 | 1 | 4 | 1 | NA | NA | 2 | 1 | NA | NA | NA | NA | 2 | 1 |
| 17_219107 | NA | NA | NA | NA | 2 | 1 | 1 | 1 | 3 | 1 | NA | NA | 2 | 1 | NA | NA | NA | NA | 2 | 1 |
| 17_219110 | NA | NA | NA | NA | 3 | 1 | 1 | 1 | 4 | 1 | NA | NA | 2 | 1 | NA | NA | NA | NA | 2 | 1 |
| 17_219411 | NA | NA | NA | NA | 1 | 1 | 1 | 1 | 2 | 1 | NA | NA | NA | NA | NA | NA | 1 | 1 | 1 | 1 |
| 17_219411 | NA | NA | NA | NA | 2 | 0 | 2 | 1 | 4 | 0.5 | NA | NA | NA | NA | NA | NA | 1 | 1 | 1 | 1 |
| 17_219412 | NA | NA | NA | NA | 2 | 1 | 1 | 1 | 3 | 1 | NA | NA | NA | NA | NA | NA | 1 | 1 | 1 | 1 |
| 17_219412 | NA | NA | NA | NA | 2 | 0.5 | 2 | 0 | 4 | 0.25 | NA | NA | NA | NA | NA | NA | 1 | 1 | 1 | 1 |
| 17_219413 | NA | NA | NA | NA | 1 | 1 | 2 | 0.5 | 3 | 0.6667 | NA | NA | NA | NA | NA | NA | 1 | 1 | 1 | 1 |

| | | | | | | | | | | | | | | | | | | | | |
|---|---|---|---|---|---|---|---|---|---|---|---|---|---|---|---|---|---|---|---|---|
| 17_219413 | NA | NA | NA | NA | 2 | 1 | 2 | 1 | 4 | 1 | NA | NA | NA | NA | NA | NA | 1 | 1 | 1 | 1 |
| 17_219414 | NA | NA | NA | NA | 1 | 1 | 2 | 1 | 3 | 1 | NA | NA | NA | NA | NA | NA | 1 | 1 | 1 | 1 |
| 17_219415 | NA | NA | NA | NA | 2 | 0.5 | 2 | 1 | 4 | 0.75 | NA | NA | NA | NA | NA | NA | 1 | 1 | 1 | 1 |
| 17_219416 | NA | NA | NA | NA | 2 | 0.5 | 2 | 0 | 4 | 0.25 | NA | NA | NA | NA | NA | NA | 1 | 1 | 1 | 1 |
| 17_219417 | NA | NA | NA | NA | 1 | 1 | 2 | 1 | 3 | 1 | NA | NA | NA | NA | NA | NA | 1 | 1 | 1 | 1 |
| 17_219423 | NA | NA | NA | NA | 1 | 1 | 2 | 1 | 3 | 1 | NA | NA | NA | NA | NA | NA | 1 | 1 | 1 | 1 |
| 17_219424 | NA | NA | NA | NA | 2 | 0.5 | 2 | 0 | 4 | 0.25 | NA | NA | NA | NA | NA | NA | 1 | 1 | 1 | 1 |
| 17_219425 | NA | NA | NA | NA | 1 | 1 | 2 | 1 | 3 | 1 | NA | NA | NA | NA | NA | NA | 1 | 1 | 1 | 1 |
| 17_219428 | NA | NA | 1 | 0 | 1 | 1 | 2 | 1 | 4 | 0.75 | NA | NA | NA | NA | 1 | 0 | 1 | 1 | 2 | 0.5 |
| 17_219428 | NA | NA | 1 | 0 | 2 | 0.5 | 2 | 0.5 | 5 | 0.4 | NA | NA | NA | NA | 1 | 0 | 1 | 1 | 2 | 0.5 |
| 17_219437 | NA | NA | 1 | 0 | 2 | 0 | 2 | 0.5 | 5 | 0.2 | NA | NA | NA | NA | 1 | 0 | 1 | 1 | 2 | 0.5 |
| 17_220982 | NA | NA | 2 | 0.5 | 1 | 0 | NA | NA | 3 | 0.3333 | NA | NA | NA | NA | 1 | 1 | NA | NA | 1 | 1 |
| 17_220985 | NA | NA | 2 | 0.5 | 1 | 1 | NA | NA | 3 | 0.6667 | NA | NA | NA | NA | 1 | 1 | NA | NA | 1 | 1 |
| 17_221461 | 2 | 0 | 1 | 0 | 3 | 0 | 2 | 0.5 | 8 | 0.125 | NA | NA | 4 | 0.25 | 3 | 0.3333 | 4 | 0 | 11 | 0.1818 |
| 17_221462 | 1 | 0 | 2 | 0 | 3 | 0.6667 | 2 | 0 | 8 | 0.25 | NA | NA | 3 | 0.3333 | 3 | 0 | 5 | 0 | 11 | 0.0909 |
| 17_221464 | 2 | 0 | 2 | 0.5 | 4 | 0.25 | 2 | 0 | 10 | 0.2 | NA | NA | 3 | 0 | 3 | 0.3333 | 5 | 0.2 | 11 | 0.1818 |
| 17_221465 | 2 | 1 | 2 | 0 | 4 | 0.25 | 2 | 0.5 | 10 | 0.4 | NA | NA | 2 | 0.5 | 2 | 0.5 | 2 | 0 | 6 | 0.3333 |
| 17_221481 | 1 | 0 | 2 | 1 | 3 | 0.6667 | 1 | 0 | 7 | 0.5714 | NA | NA | 3 | 0.6667 | 2 | 0.5 | 4 | 0.5 | 9 | 0.5556 |
| 17_221481 | 1 | 0 | 2 | 1 | 3 | 0.6667 | 1 | 1 | 7 | 0.7143 | NA | NA | 2 | 1 | 2 | 1 | 2 | 1 | 6 | 1 |
| 17_221481 | 1 | 1 | 2 | 0.5 | 3 | 0.3333 | 1 | 1 | 7 | 0.5714 | NA | NA | 3 | 0 | 2 | 1 | 4 | 1 | 9 | 0.6667 |
| 17_221482 | 1 | 0 | 2 | 0 | 3 | 0.3333 | 1 | 0 | 7 | 0.1429 | NA | NA | 3 | 0 | 2 | 0 | 3 | 0.3333 | 8 | 0.125 |
| 17_221482 | 1 | 1 | 2 | 0 | 3 | 0 | 1 | 0 | 7 | 0.1429 | NA | NA | 3 | 0.3333 | 2 | 0 | 4 | 0 | 9 | 0.1111 |
| 17_222111 | NA | NA | NA | NA | 2 | 1 | 1 | 0 | 3 | 0.6667 | 1 | 0 | 2 | 1 | NA | NA | 2 | 1 | 5 | 0.8 |
| 17_222111 | NA | NA | NA | NA | 1 | 0 | 2 | 0.5 | 3 | 0.3333 | 2 | 0.5 | 1 | 0 | NA | NA | 2 | 0 | 5 | 0.2 |
| 17_222113 | NA | NA | 1 | 0 | 2 | 0.5 | 2 | 0.5 | 5 | 0.4 | 2 | 0.5 | 1 | 0 | NA | NA | 1 | 1 | 4 | 0.5 |
| 17_222113 | NA | NA | 1 | 0 | 2 | 1 | 2 | 0 | 5 | 0.4 | 2 | 0 | 1 | 0 | NA | NA | 2 | 0.5 | 5 | 0.2 |
| 17_222114 | NA | NA | 1 | 1 | 2 | 0.5 | 2 | 0.5 | 5 | 0.6 | 2 | 0 | 2 | 1 | NA | NA | 2 | 0.5 | 6 | 0.5 |
| 17_222130 | 1 | 0 | 2 | 0 | 1 | 0 | 3 | 0.3333 | 7 | 0.1429 | 3 | 0 | 5 | 0.2 | 1 | 0 | 3 | 0 | 12 | 0.0833 |
| 17_222858 | 1 | 0 | 1 | 0 | 5 | 0.4 | 4 | 0.25 | 11 | 0.2727 | 3 | 0 | 3 | 0.3333 | 3 | 0.3333 | 1 | 0 | 10 | 0.2 |
| 17_222858 | 1 | 0 | 1 | 0 | 5 | 0.2 | 5 | 0.2 | 12 | 0.1667 | 3 | 0 | 4 | 0 | 3 | 0.3333 | 1 | 0 | 11 | 0.0909 |
| 17_222859 | 1 | 0 | 1 | 0 | 5 | 0.4 | 4 | 0 | 11 | 0.1818 | 3 | 0.6667 | 3 | 0.3333 | 3 | 0.3333 | 1 | 0 | 10 | 0.4 |
| 17_222859 | 1 | 0 | 1 | 1 | 5 | 0.4 | 4 | 0.25 | 11 | 0.3636 | 3 | 0.3333 | 4 | 0 | 3 | 0.3333 | 1 | 1 | 11 | 0.2727 |
| 17_222875 | NA | NA | 1 | 1 | 2 | 0 | 4 | 0.5 | 7 | 0.4286 | NA | NA | 1 | 0 | 3 | 0.6667 | NA | NA | 4 | 0.5 |
| 17_222876 | NA | NA | NA | NA | 2 | 1 | 4 | 0.5 | 6 | 0.6667 | 1 | 1 | 1 | 0 | 2 | 1 | 1 | 0 | 5 | 0.6 |
| 17_225050 | 2 | 0 | 1 | 0 | 3 | 0.3333 | 3 | 0.6667 | 9 | 0.3333 | NA | NA | 1 | 0 | 4 | 0.5 | 1 | 1 | 6 | 0.5 |
| 17_225051 | 1 | 0 | 1 | 0 | 2 | 0 | 4 | 0.25 | 8 | 0.125 | NA | NA | NA | NA | 3 | 0.6667 | 1 | 0 | 4 | 0.5 |
| 17_225051 | 2 | 0.5 | 1 | 1 | 3 | 0.3333 | 4 | 0.75 | 10 | 0.6 | NA | NA | NA | NA | 3 | 0.6667 | 1 | 1 | 4 | 0.75 |
| 17_225052 | 2 | 0.5 | 1 | 0 | 3 | 0.6667 | 4 | 0 | 10 | 0.3 | NA | NA | NA | NA | 4 | 0.25 | 1 | 0 | 5 | 0.2 |
| 17_225053 | 1 | 0 | 1 | 0 | 3 | 0 | 4 | 0.25 | 9 | 0.1111 | NA | NA | 1 | 1 | 4 | 0.5 | 1 | 1 | 6 | 0.6667 |
| 17_225056 | 1 | 0 | 1 | 0 | 2 | 0.5 | 4 | 0.25 | 8 | 0.25 | NA | NA | 1 | 1 | 3 | 0 | 1 | 0 | 5 | 0.2 |
| 17_225061 | 2 | 1 | NA | NA | 1 | 0 | 4 | 0.25 | 7 | 0.4286 | NA | NA | NA | NA | 3 | 1 | 1 | 0 | 4 | 0.75 |
| 17_225062 | 2 | 0.5 | 1 | 0 | 2 | 0 | 4 | 0 | 9 | 0.1111 | NA | NA | NA | NA | 3 | 0.3333 | 1 | 0 | 4 | 0.25 |
| 17_225075 | 1 | 0 | NA | NA | NA | NA | 2 | 0.5 | 3 | 0.3333 | NA | NA | NA | NA | 1 | 1 | 1 | 1 | 2 | 1 |
| 17_225076 | 1 | 1 | NA | NA | NA | NA | 3 | 1 | 4 | 1 | NA | NA | NA | NA | 1 | 1 | 1 | 0 | 2 | 0.5 |
| 17_225076 | 1 | 0 | NA | NA | NA | NA | 2 | 0.5 | 3 | 0.3333 | NA | NA | NA | NA | 2 | 0 | 1 | 1 | 3 | 0.3333 |
| 17_225572 | 1 | 0 | 1 | 1 | 1 | 0 | 2 | 0 | 5 | 0.2 | NA | NA | 1 | 1 | 1 | 1 | 2 | 0 | 4 | 0.5 |
| 17_225574 | 1 | 0 | NA | NA | 3 | 0.6667 | 1 | 1 | 5 | 0.6 | NA | NA | NA | NA | 1 | 0 | 2 | 0.5 | 3 | 0.3333 |
| 17_225587 | NA | NA | NA | NA | 1 | 1 | 1 | 1 | 2 | 1 | NA | NA | 1 | 1 | NA | NA | 1 | 0 | 2 | 0.5 |
| 17_225588 | NA | NA | NA | NA | 1 | 1 | 1 | 0 | 2 | 0.5 | 1 | 0 | 1 | 0 | NA | NA | 2 | 0.5 | 4 | 0.25 |
| 17_225616 | NA | NA | NA | NA | 1 | 1 | NA | NA | 1 | 1 | 1 | 1 | NA | NA | 1 | 0 | 1 | 0 | 3 | 0.3333 |
| 17_226736 | NA | NA | NA | NA | 1 | 1 | 2 | 1 | 3 | 1 | 1 | 1 | 3 | 0.6667 | 2 | 0.5 | NA | NA | 6 | 0.6667 |
| 17_226751 | 2 | 0 | 2 | 0 | 3 | 0 | 3 | 0.3333 | 10 | 0.1 | 1 | 1 | 3 | 0 | 4 | 0 | 1 | 0 | 9 | 0.1111 |
| 17_226752 | 3 | 0.3333 | 2 | 0.5 | 3 | 0.3333 | 3 | 0.3333 | 11 | 0.3636 | 1 | 1 | 2 | 0.5 | 4 | 0 | 1 | 0 | 8 | 0.25 |
| 17_229296 | NA | NA | NA | NA | 1 | 1 | 1 | 0 | 2 | 0.5 | NA | NA | NA | NA | 2 | 0.5 | 1 | 1 | 3 | 0.6667 |
| 17_229296 | NA | NA | NA | NA | NA | NA | 1 | 1 | 1 | 1 | NA | NA | NA | NA | 2 | 0.5 | 1 | 0 | 3 | 0.3333 |
| 17_229298 | NA | NA | NA | NA | NA | NA | 1 | 1 | 1 | 1 | NA | NA | NA | NA | 1 | 1 | 1 | 1 | 2 | 1 |
| 17_229318 | NA | NA | 1 | 1 | NA | NA | 2 | 0.5 | 3 | 0.6667 | NA | NA | NA | NA | 2 | 0.5 | 1 | 0 | 3 | 0.3333 |
| 17_229318 | NA | NA | 1 | 1 | 1 | 1 | NA | NA | 2 | 1 | NA | NA | NA | NA | 2 | 1 | NA | NA | 2 | 1 |
| 17_229324 | 1 | 0 | 1 | 1 | 1 | 0 | 1 | 1 | 4 | 0.5 | NA | NA | NA | NA | 1 | 1 | 1 | 0 | 2 | 0.5 |
| 17_229325 | NA | NA | 1 | 1 | NA | NA | 1 | 1 | 2 | 1 | NA | NA | NA | NA | 1 | 1 | 1 | 1 | 2 | 1 |
| 17_229326 | 1 | 1 | 1 | 0 | 1 | 1 | 1 | 0 | 4 | 0.5 | NA | NA | NA | NA | 1 | 1 | 1 | 1 | 2 | 1 |
| 17_229327 | 1 | 1 | 1 | 1 | 1 | 0 | 1 | 1 | 4 | 0.75 | NA | NA | NA | NA | 1 | 1 | 1 | 0 | 2 | 0.5 |
| 17_229329 | 1 | 1 | 1 | 1 | 1 | 1 | 1 | 1 | 4 | 1 | NA | NA | NA | NA | 1 | 1 | 1 | 1 | 2 | 1 |
| 17_229331 | 1 | 0 | 1 | 0 | 1 | 0 | 1 | 1 | 4 | 0.25 | NA | NA | NA | NA | 1 | 1 | 1 | 0 | 2 | 0.5 |
| 17_229337 | 1 | 0 | NA | NA | 1 | 1 | 1 | 1 | 3 | 0.6667 | NA | NA | NA | NA | 1 | 0 | 1 | 1 | 2 | 0.5 |

| | | | | | | | | | | | | | | | | | | | | |
|---|---|---|---|---|---|---|---|---|---|---|---|---|---|---|---|---|---|---|---|---|
| 17_229344! | 1 | 0 | NA | NA | 1 | 1 | 1 | 1 | 3 | 0.6667 | NA | NA | NA | NA | NA | NA | 1 | 1 | 1 | 1 |
| 17_229346! | NA | NA | NA | NA | NA | NA | 1 | 1 | 1 | 1 | NA | NA | NA | NA | NA | NA | 1 | 1 | 1 | 1 |
| 17_229347( | 1 | 1 | NA | NA | 1 | 1 | 1 | 1 | 3 | 1 | NA | NA | NA | NA | NA | NA | 1 | 1 | 1 | 1 |
| 17_230688 | 1 | 1 | 1 | 0 | 1 | 0 | NA | NA | 3 | 0.3333 | NA | NA | 2 | 0.5 | NA | NA | 1 | 1 | 3 | 0.6667 |
| 17_230689( | 2 | 0.5 | 1 | 0 | 1 | 0 | NA | NA | 4 | 0.25 | NA | NA | 1 | 1 | NA | NA | 1 | 0 | 2 | 0.5 |
| 17_230698 | 1 | 1 | 1 | 0 | 1 | 0 | NA | NA | 3 | 0.3333 | NA | NA | 1 | 1 | NA | NA | 1 | 0 | 2 | 0.5 |
| 17_230700 | 1 | 1 | 1 | 0 | 1 | 0 | NA | NA | 3 | 0.3333 | NA | NA | 1 | 1 | NA | NA | 1 | 1 | 2 | 1 |
| 17_230700! | 1 | 1 | 1 | 0 | 1 | 0 | NA | NA | 3 | 0.3333 | NA | NA | 1 | 0 | NA | NA | 1 | 1 | 2 | 0.5 |
| 17_230703! | 1 | 1 | NA | NA | 1 | 0 | NA | NA | 2 | 0.5 | NA | NA | 1 | 1 | NA | NA | 1 | 1 | 2 | 1 |
| 17_230708 | 1 | 1 | NA | NA | 1 | 0 | NA | NA | 2 | 0.5 | NA | NA | 1 | 1 | NA | NA | 1 | 0 | 2 | 0.5 |
| 17_230713! | NA | NA | 1 | 1 | 1 | 0 | NA | NA | 2 | 0.5 | NA | NA | 1 | 1 | NA | NA | 1 | 0 | 2 | 0.5 |
| 17_230715( | 1 | 0 | 1 | 1 | 1 | 1 | NA | NA | 3 | 0.6667 | 1 | 1 | 4 | 1 | 1 | 1 | 1 | 1 | 7 | 1 |
| 17_230715 | 3 | 0.3333 | NA | NA | 1 | 0 | NA | NA | 4 | 0.25 | 1 | 0 | 4 | 0.25 | NA | NA | 1 | 0 | 6 | 0.1667 |
| 17_230715 | 3 | 0 | 1 | 1 | 1 | 0 | NA | NA | 5 | 0.2 | 1 | 0 | 4 | 0.25 | 1 | 0 | 1 | 0 | 7 | 0.1429 |
| 17_233936 | 1 | 0 | NA | NA | 1 | 1 | NA | NA | 2 | 0.5 | NA | NA | 2 | 1 | 2 | 1 | 1 | 0 | 5 | 0.8 |
| 17_233938 | 1 | 1 | NA | NA | 2 | 0 | NA | NA | 3 | 0.3333 | NA | NA | 1 | 0 | 3 | 0.6667 | 1 | 0 | 5 | 0.4 |
| 17_233939 | 1 | 1 | NA | NA | 1 | 1 | NA | NA | 2 | 1 | NA | NA | 2 | 1 | 4 | 0.25 | 1 | 0 | 7 | 0.4286 |
| 17_233940 | 1 | 1 | NA | NA | 1 | 1 | NA | NA | 2 | 1 | NA | NA | 2 | 0.5 | 3 | 1 | 1 | 1 | 6 | 0.8333 |
| 17_233940! | 1 | 0 | NA | NA | 1 | 1 | NA | NA | 2 | 0.5 | NA | NA | 2 | 0 | 3 | 0.3333 | 1 | 0 | 6 | 0.1667 |
| 17_233944 | 1 | 1 | NA | NA | 1 | 1 | NA | NA | 2 | 1 | NA | NA | 2 | 1 | 3 | 0.3333 | 1 | 1 | 6 | 0.6667 |
| 17_233945( | 1 | 1 | NA | NA | NA | NA | NA | NA | 1 | 1 | NA | NA | 2 | 1 | 3 | 1 | NA | NA | 5 | 1 |
| 17_233948 | 1 | 1 | NA | NA | 1 | 1 | NA | NA | 2 | 1 | NA | NA | 2 | 1 | 3 | 1 | 1 | 1 | 6 | 1 |
| 17_233950( | NA | NA | NA | NA | 1 | 1 | NA | NA | 1 | 1 | NA | NA | 1 | 1 | 3 | 1 | 1 | 1 | 5 | 1 |
| 17_233954 | NA | NA | NA | NA | 1 | 1 | NA | NA | 1 | 1 | NA | NA | 2 | 0.5 | 3 | 0.6667 | NA | NA | 5 | 0.6 |
| 17_233958! | NA | NA | NA | NA | 1 | 1 | NA | NA | 1 | 1 | NA | NA | 2 | 0.5 | 3 | 0.6667 | NA | NA | 5 | 0.6 |
| 17_233959! | NA | NA | NA | NA | 1 | 1 | NA | NA | 1 | 1 | NA | NA | 1 | 0 | 3 | 0.3333 | NA | NA | 4 | 0.25 |
| 17_233959( | NA | NA | NA | NA | 1 | 1 | NA | NA | 1 | 1 | NA | NA | 2 | 1 | 3 | 0.6667 | NA | NA | 5 | 0.8 |
| 17_237758( | 1 | 1 | 1 | 0 | NA | NA | 1 | 1 | 3 | 0.6667 | 2 | 0.5 | 1 | 1 | NA | NA | 1 | 1 | 4 | 0.75 |
| 17_237758 | 1 | 1 | NA | NA | 1 | 1 | 1 | 1 | 3 | 1 | 1 | 1 | NA | NA | NA | NA | 1 | 1 | 2 | 1 |
| 17_237761 | 1 | 1 | 1 | 1 | 1 | 1 | 1 | 1 | 4 | 1 | 1 | 1 | 1 | 1 | NA | NA | 1 | 1 | 3 | 1 |
| 17_237762 | 1 | 1 | 2 | 1 | 1 | 1 | 1 | 1 | 5 | 1 | 1 | 1 | 1 | 1 | NA | NA | 1 | 1 | 3 | 1 |
| 17_237762 | NA | NA | 2 | 1 | NA | NA | 1 | 1 | 3 | 1 | 1 | 1 | 1 | 1 | NA | NA | NA | NA | 2 | 1 |
| 17_237763 | 1 | 1 | 2 | 1 | 1 | 1 | 1 | 1 | 5 | 1 | 1 | 1 | 1 | 1 | NA | NA | 1 | 1 | 3 | 1 |
| 17_237764 | 1 | 1 | 2 | 1 | 1 | 1 | NA | NA | 4 | 1 | 1 | 1 | 1 | 1 | NA | NA | 1 | 1 | 3 | 1 |
| 17_237765( | NA | NA | 1 | 1 | 1 | 1 | NA | NA | 2 | 1 | 1 | 1 | 1 | 1 | NA | NA | NA | NA | 2 | 1 |
| 17_237765 | 1 | 1 | 2 | 0.5 | 1 | 1 | 1 | 0 | 5 | 0.6 | 1 | 1 | NA | NA | NA | NA | 1 | 1 | 2 | 1 |
| 17_237766 | 1 | 0 | 1 | 1 | 1 | 1 | 1 | 1 | 4 | 0.75 | 1 | 0 | NA | NA | NA | NA | 1 | 1 | 2 | 0.5 |
| 17_237770 | 1 | 1 | NA | NA | NA | NA | NA | NA | 1 | 1 | NA | NA | 1 | 1 | NA | NA | 1 | 1 | 2 | 1 |
| 17_237804( | 1 | 1 | 1 | 1 | NA | NA | NA | NA | 2 | 1 | 1 | 1 | 1 | 1 | NA | NA | NA | NA | 2 | 1 |
| 17_237949 | 3 | 0 | 3 | 0.3333 | 3 | 1 | 2 | 1 | 11 | 0.5455 | NA | NA | NA | NA | 2 | 1 | 2 | 0 | 4 | 0.5 |
| 17_237949 | 3 | 0 | 2 | 0.5 | 3 | 0 | 2 | 0.5 | 10 | 0.2 | 1 | 0 | NA | NA | 2 | 0.5 | 2 | 1 | 5 | 0.6 |
| 17_237951( | 2 | 1 | 2 | 0.5 | 3 | 1 | 1 | 1 | 8 | 0.875 | 1 | 1 | NA | NA | 2 | 1 | 2 | 0.5 | 5 | 0.8 |
| 17_237954 | 4 | 0.25 | 3 | 0.6667 | 2 | 0.5 | 2 | 0 | 11 | 0.3636 | 1 | 0 | NA | NA | 2 | 0.5 | 1 | 0 | 4 | 0.25 |
| 17_237956 | 4 | 1 | 3 | 1 | 2 | 1 | 2 | 1 | 11 | 1 | 1 | 1 | NA | NA | 1 | 1 | 1 | 1 | 3 | 1 |
| 17_237959 | 4 | 0.5 | 2 | 0.5 | 2 | 1 | 1 | 1 | 9 | 0.6667 | 1 | 1 | NA | NA | NA | NA | NA | NA | 1 | 1 |
| 17_237960 | 3 | 0.6667 | 2 | 0.5 | NA | NA | 2 | 1 | 7 | 0.7143 | 1 | 0 | NA | NA | 1 | 1 | NA | NA | 2 | 0.5 |
| 17_237960 | 4 | 0 | 2 | 0 | 2 | 0.5 | 2 | 0 | 10 | 0.1 | 1 | 0 | NA | NA | 1 | 0 | 1 | 1 | 3 | 0.3333 |
| 17_238940 | 33 | 0.2424 | 19 | 0.1579 | 30 | 0.1 | 23 | 0.1304 | 105 | 0.1619 | 33 | 0.1818 | 22 | 0.0909 | 23 | 0.087 | 39 | 0.1282 | 117 | 0.1282 |
| 17_238941 | 33 | 0.2424 | 19 | 0.0526 | 28 | 0.0714 | 24 | 0.125 | 104 | 0.1346 | 28 | 0.1071 | 22 | 0.0909 | 23 | 0.2174 | 39 | 0.2051 | 112 | 0.1607 |
| 17_238948 | 1 | 1 | 1 | 0 | NA | NA | NA | NA | 2 | 0.5 | NA | NA | NA | NA | 2 | 0.5 | NA | NA | 2 | 0.5 |
| 17_239552( | 8 | 0.125 | 10 | 0 | 8 | 0.25 | 7 | 0 | 33 | 0.0909 | 3 | 0 | 5 | 0 | 3 | 0.3333 | 2 | 0 | 13 | 0.0769 |
| 17_239552! | 8 | 0.125 | 7 | 0.1429 | 8 | 0.25 | 5 | 0 | 28 | 0.1429 | 3 | 0.3333 | 5 | 0 | 5 | 0 | 5 | 0.2 | 18 | 0.1111 |
| 17_239554 | 8 | 0.25 | 6 | 0.3333 | 9 | 0.2222 | 6 | 0 | 29 | 0.2069 | 3 | 0 | 7 | 0.2857 | 5 | 0.4 | 5 | 0 | 20 | 0.2 |
| 17_239556! | 8 | 0.25 | 7 | 0.1429 | 8 | 0.375 | 6 | 0.1667 | 29 | 0.2414 | 3 | 0.3333 | 6 | 0.3333 | 4 | 0 | 3 | 0.3333 | 16 | 0.25 |
| 17_239557 | 13 | 0.7692 | 10 | 0.5 | 7 | 0.2857 | 9 | 0.7778 | 39 | 0.6154 | 8 | 0.375 | 11 | 0.5455 | 8 | 0.75 | 8 | 0.75 | 35 | 0.6 |
| 17_239557 | 12 | 0.6667 | 8 | 0.5 | 10 | 0.2 | 10 | 0.6 | 40 | 0.5 | 6 | 0.6667 | 10 | 0.6 | 7 | 0.7143 | 6 | 0.3333 | 29 | 0.5862 |
| 17_239558 | 15 | 1 | 10 | 1 | 13 | 1 | 12 | 1 | 50 | 1 | 8 | 1 | 12 | 1 | 7 | 1 | 5 | 1 | 32 | 1 |
| 17_239558 | 13 | 0.8462 | 8 | 1 | 13 | 0.9231 | 12 | 0.9167 | 46 | 0.913 | 7 | 0.8571 | 11 | 0.9091 | 8 | 0.75 | 7 | 0.8571 | 33 | 0.8485 |
| 17_239558! | 12 | 0.1667 | 7 | 0.2857 | 12 | 0.0833 | 9 | 0.2222 | 40 | 0.175 | 5 | 0.2 | 9 | 0 | 7 | 0.1429 | 6 | 0.3333 | 27 | 0.1481 |
| 17_239559! | 14 | 0.4286 | 9 | 0.4444 | 10 | 0.6 | 13 | 0.6154 | 46 | 0.5217 | 6 | 0.5 | 8 | 0.875 | 6 | 0.5 | 4 | 0.75 | 24 | 0.6667 |
| 17_239560( | 14 | 0.9286 | 10 | 0.7 | 11 | 0.9091 | 13 | 0.7692 | 48 | 0.8333 | 7 | 1 | 11 | 0.9091 | 7 | 0.7143 | 7 | 0.8571 | 32 | 0.875 |
| 17_239560 | 14 | 0.5 | 9 | 0.3333 | 9 | 0 | 13 | 0.3077 | 45 | 0.3111 | 5 | 0.2 | 9 | 0.3333 | 7 | 0.1429 | 7 | 0.5714 | 28 | 0.3214 |
| 17_239562 | 10 | 0.1 | 6 | 0.1667 | 11 | 0 | 8 | 0.25 | 35 | 0.1143 | 5 | 0.2 | 10 | 0.1 | 4 | 0.25 | 5 | 0.2 | 24 | 0.1667 |
| 17_239564 | 10 | 1 | 6 | 0.8333 | 9 | 1 | 10 | 1 | 35 | 0.9714 | 3 | 1 | 7 | 1 | 6 | 0.6667 | 5 | 1 | 21 | 0.9048 |
| 17_239564! | 12 | 0.6667 | 8 | 0.875 | 12 | 0.5 | 7 | 0.8571 | 39 | 0.6923 | 4 | 1 | 8 | 0.375 | 5 | 0.4 | 6 | 0.5 | 23 | 0.5217 |

| | | | | | | | | | | | | | | | | | | | | |
|---|---|---|---|---|---|---|---|---|---|---|---|---|---|---|---|---|---|---|---|---|
| 17_2395652 | 11 | 0 | 6 | 0.1667 | 12 | 0.0833 | 8 | 0 | 37 | 0.0541 | 4 | 0 | 6 | 0.3333 | 5 | 0 | 5 | 0 | 20 | 0.1 |
| 17_2395654 | 11 | 0.7273 | 7 | 0.4286 | 11 | 0.4545 | 9 | 0.5556 | 38 | 0.5526 | 4 | 0.75 | 7 | 0.2857 | 5 | 0.8 | 6 | 0.5 | 22 | 0.5455 |
| 17_2395896 | 2 | 0.5 | 5 | 0.2 | 3 | 0.6667 | 3 | 0.6667 | 13 | 0.4615 | 2 | 0 | 3 | 0.3333 | 3 | 0 | 4 | 0 | 12 | 0.0833 |
| 17_2397200 | 32 | 0.5938 | 36 | 0.6944 | 38 | 0.5 | 34 | 0.7647 | 140 | 0.6357 | 23 | 0.4348 | 34 | 0.4412 | 29 | 0.5517 | 23 | 0.5652 | 109 | 0.4954 |
| 17_2397220 | 33 | 0.3333 | 36 | 0.2778 | 39 | 0.2051 | 31 | 0.2903 | 139 | 0.2734 | 26 | 0.2308 | 34 | 0.2059 | 32 | 0.1875 | 21 | 0.2381 | 113 | 0.2124 |
| 17_2397223 | 25 | 0.16 | 26 | 0.1154 | 27 | 0.037 | 29 | 0.1379 | 107 | 0.1121 | 20 | 0.1 | 29 | 0.0345 | 20 | 0.1 | 18 | 0.0556 | 87 | 0.069 |
| 17_2397225 | 29 | 0.069 | 30 | 0.3 | 37 | 0.2703 | 30 | 0.4 | 126 | 0.2619 | 22 | 0.2727 | 30 | 0.2 | 28 | 0.25 | 21 | 0.2857 | 101 | 0.2475 |
| 17_2397236 | 23 | 0.1304 | 30 | 0.1667 | 28 | 0.2143 | 25 | 0.24 | 106 | 0.1887 | 19 | 0.1053 | 25 | 0.12 | 21 | 0.1429 | 19 | 0.1053 | 84 | 0.119 |
| 17_2397238 | 30 | 0.6 | 34 | 0.4706 | 33 | 0.6364 | 32 | 0.5 | 129 | 0.5504 | 26 | 0.3846 | 26 | 0.5769 | 29 | 0.4828 | 21 | 0.5238 | 102 | 0.4902 |
| 17_2397255 | 27 | 0.8889 | 31 | 0.9677 | 32 | 0.9375 | 29 | 0.931 | 119 | 0.9328 | 23 | 0.8261 | 26 | 0.9615 | 25 | 0.92 | 20 | 0.95 | 94 | 0.9149 |
| 17_2397256 | 25 | 0.16 | 30 | 0.0667 | 26 | 0.1154 | 31 | 0 | 112 | 0.0804 | 20 | 0.05 | 30 | 0.1 | 24 | 0.1667 | 20 | 0.05 | 94 | 0.0957 |
| 17_2397263 | 24 | 1 | 28 | 1 | 28 | 0.9643 | 26 | 0.9615 | 106 | 0.9811 | 18 | 1 | 26 | 0.8846 | 29 | 0.8621 | 16 | 0.9375 | 89 | 0.9101 |
| 17_2397268 | 27 | 0.8148 | 28 | 0.8571 | 29 | 0.8966 | 29 | 0.7931 | 113 | 0.8407 | 22 | 0.7273 | 27 | 0.7407 | 27 | 0.963 | 19 | 0.9474 | 95 | 0.8421 |
| 17_2397972 | 11 | 0.6364 | 13 | 0.3077 | 10 | 0.3 | 16 | 0.4375 | 50 | 0.42 | 11 | 0.2727 | 14 | 0.3571 | 10 | 0.3 | 13 | 0.5385 | 48 | 0.375 |
| 17_2397978 | 15 | 0.8 | 16 | 0.4375 | 13 | 0.7692 | 15 | 0.5333 | 59 | 0.6271 | 11 | 0.8182 | 18 | 0.7222 | 14 | 0.7857 | 12 | 0.4167 | 55 | 0.6909 |
| 17_2397980 | 14 | 0.1429 | 19 | 0.2632 | 13 | 0.1538 | 14 | 0.2143 | 60 | 0.2 | 11 | 0.0909 | 17 | 0.0588 | 14 | 0.2857 | 14 | 0.2857 | 56 | 0.1786 |
| 17_2398134 | 11 | 0 | 11 | 0.2727 | 8 | 0.125 | 10 | 0.1 | 40 | 0.125 | 8 | 0.125 | 13 | 0.1538 | 11 | 0.3636 | 7 | 0 | 39 | 0.1795 |
| 17_2398138 | 12 | 0 | 10 | 0.1 | 6 | 0 | 8 | 0.25 | 36 | 0.0833 | 6 | 0.5 | 13 | 0.3077 | 8 | 0.25 | 6 | 0 | 33 | 0.2727 |
| 17_2398217 | 9 | 0.4444 | 6 | 0.6667 | 5 | 0.8 | 8 | 0.625 | 28 | 0.6071 | 8 | 0.5 | 12 | 0.5 | 8 | 0.5 | 5 | 0.8 | 33 | 0.5455 |
| 17_2403233 | NA | NA | 1 | 1 | NA | NA | NA | NA | 1 | 1 | 1 | 1 | NA | NA | NA | NA | NA | NA | 1 | 1 |
| 17_2403414 | 3 | 0 | 2 | 0 | 1 | 0 | 1 | 1 | 7 | 0.1429 | 5 | 0.2 | 1 | 0 | 1 | 0 | NA | NA | 7 | 0.1429 |
| 17_2403560 | 3 | 0.3333 | 6 | 0.1667 | 3 | 0.3333 | 1 | 0 | 13 | 0.2308 | 8 | 0.25 | 2 | 0 | 4 | 0.25 | 3 | 0.6667 | 17 | 0.2941 |
| 17_2403564 | 3 | 0.3333 | 6 | 0 | 3 | 0 | 2 | 0 | 14 | 0.0714 | 8 | 0 | 2 | 0 | 4 | 0.25 | 3 | 0 | 17 | 0.0588 |
| 17_2403582 | 4 | 0 | 9 | 0 | 3 | 0.3333 | 2 | 0 | 18 | 0.0556 | 9 | 0.1111 | 2 | 0 | 3 | 0 | 3 | 0 | 17 | 0.0588 |
| 17_2403587 | 5 | 0 | 7 | 0.1429 | 2 | 0 | 2 | 0 | 16 | 0.0625 | 9 | 0.2222 | 2 | 0 | 4 | 0.25 | 3 | 0 | 18 | 0.1667 |
| 17_2403615 | 4 | 0.25 | 9 | 0.2222 | 3 | 0.6667 | 2 | 0.5 | 18 | 0.3333 | 9 | 0.5556 | 2 | 0 | 4 | 0.75 | 3 | 1 | 18 | 0.6111 |
| 17_2403641 | 2 | 0 | 9 | 0.4444 | 3 | 0 | 2 | 1 | 16 | 0.375 | 8 | 0.625 | 2 | 0 | 6 | 0.3333 | 3 | 0.3333 | 19 | 0.4211 |
| 17_2403646 | 5 | 0.6 | 8 | 0.5 | 4 | 1 | 2 | 0.5 | 19 | 0.6316 | 9 | 0.6667 | 1 | 1 | 5 | 1 | 2 | 1 | 17 | 0.8235 |
| 17_2403662 | 5 | 0.2 | 10 | 0.1 | 4 | 0 | 2 | 0 | 21 | 0.0952 | 9 | 0.1111 | 2 | 0 | 6 | 0.1667 | 3 | 0 | 20 | 0.1 |
| 17_2403666 | 5 | 0 | 10 | 0.2 | 4 | 0 | 2 | 0 | 21 | 0.0952 | 8 | 0.25 | 2 | 0 | 5 | 0 | 1 | 0 | 16 | 0.125 |
| 17_2403689 | 3 | 0.3333 | 10 | 0.7 | 3 | 0.6667 | 1 | 1 | 17 | 0.6471 | 5 | 0.4 | 2 | 1 | 5 | 0.8 | 2 | 0.5 | 14 | 0.6429 |
| 17_2403704 | 5 | 0.4 | 11 | 0.1818 | 3 | 0 | 2 | 0 | 21 | 0.1905 | 9 | 0.2222 | 2 | 0 | 5 | 0.2 | 3 | 0.3333 | 19 | 0.2105 |
| 17_2403716 | 5 | 0.4 | 11 | 0.2727 | 4 | 0.5 | 3 | 0 | 23 | 0.3043 | 8 | 0 | 2 | 0 | 6 | 0.6667 | 5 | 0.4 | 21 | 0.2857 |
| 17_2403721 | 5 | 0.2 | 11 | 0.1818 | 4 | 0.5 | 3 | 0.3333 | 23 | 0.2609 | 8 | 0.125 | 2 | 0 | 5 | 0 | 5 | 0.4 | 20 | 0.15 |
| 17_2403727 | 5 | 0.8 | 11 | 0.8182 | 4 | 0.5 | 2 | 1 | 22 | 0.7727 | 8 | 1 | 2 | 0 | 6 | 0.8333 | 3 | 0.6667 | 19 | 0.7895 |
| 17_2403729 | 6 | 0.6667 | 11 | 0.2727 | 4 | 0.25 | 4 | 0.5 | 25 | 0.4 | 7 | 0.4286 | 2 | 0.5 | 6 | 0.5 | 5 | 0.4 | 20 | 0.45 |
| 17_2403737 | 6 | 0.1667 | 12 | 0.1667 | 4 | 0 | 4 | 0.25 | 26 | 0.1538 | 8 | 0 | 2 | 0 | 6 | 0.1667 | 5 | 0.2 | 21 | 0.0952 |
| 17_2403761 | 6 | 0.1667 | 16 | 0.0625 | 4 | 0 | 5 | 0 | 31 | 0.0645 | 9 | 0.1111 | 2 | 0 | 6 | 0.1667 | 6 | 0 | 23 | 0.087 |
| 17_2403764 | 6 | 0.1667 | 16 | 0.0625 | 2 | 0.5 | 3 | 0 | 27 | 0.1111 | 9 | 0.1111 | 2 | 0 | 5 | 0 | 4 | 0 | 20 | 0.05 |
| 17_2403764 | 5 | 0 | 15 | 0.1333 | 3 | 0 | 2 | 0.5 | 25 | 0.12 | 6 | 0.1667 | 2 | 0 | 6 | 0 | 2 | 0 | 16 | 0.0625 |
| 17_2403791 | 5 | 0.2 | 13 | 0 | 3 | 0.3333 | 2 | 0 | 23 | 0.087 | 6 | 0 | 2 | 0 | 6 | 0.1667 | 5 | 0 | 19 | 0.0526 |
| 17_2403793 | 1 | 0 | 12 | 0.1667 | 4 | 0.5 | 2 | 0 | 19 | 0.2105 | 5 | 0.2 | 2 | 0 | 5 | 0 | 2 | 0 | 14 | 0.0714 |
| 17_2403824 | 5 | 0.2 | 17 | 0.1176 | 3 | 0.3333 | 2 | 0 | 27 | 0.1481 | 10 | 0.1 | 3 | 0 | 9 | 0.2222 | 3 | 0.3333 | 25 | 0.16 |
| 17_2403923 | 9 | 0.1111 | 24 | 0.25 | 5 | 0.2 | 7 | 0.2857 | 45 | 0.2222 | 9 | 0.2222 | 8 | 0.375 | 8 | 0.25 | 9 | 0.2222 | 34 | 0.2647 |
| 17_2403964 | 9 | 0.4444 | 22 | 0.5 | 6 | 0.3333 | 5 | 0.6 | 42 | 0.4762 | 9 | 0.6667 | 7 | 0.4286 | 7 | 0.7143 | 10 | 0.4 | 33 | 0.5455 |
| 17_2403966 | 10 | 0.2 | 26 | 0.1154 | 6 | 0.1667 | 6 | 0.1667 | 48 | 0.1458 | 11 | 0.0909 | 9 | 0.1111 | 9 | 0.2222 | 10 | 0.1 | 39 | 0.1282 |
| 17_2403968 | 10 | 0.1 | 29 | 0.069 | 6 | 0.1667 | 7 | 0.1429 | 52 | 0.0962 | 10 | 0.3 | 7 | 0.1429 | 8 | 0.125 | 10 | 0 | 35 | 0.1429 |
| 17_2403969 | 10 | 0.2 | 27 | 0.0741 | 9 | 0.1111 | 5 | 0 | 51 | 0.098 | 10 | 0.1 | 9 | 0.1111 | 9 | 0 | 10 | 0 | 38 | 0.0526 |
| 17_2403973 | 5 | 0.8 | 27 | 0.8148 | 7 | 0.8571 | 6 | 0.6667 | 45 | 0.8 | 10 | 0.8 | 5 | 1 | 9 | 0.7778 | 8 | 0.875 | 32 | 0.8438 |
| 17_2403974 | 6 | 0 | 20 | 0.05 | 7 | 0.1429 | 6 | 0 | 39 | 0.0513 | 11 | 0.1818 | 4 | 0.25 | 6 | 0 | 10 | 0.1 | 31 | 0.129 |
| 17_2403977 | 9 | 0.2222 | 26 | 0.1538 | 9 | 0.2222 | 7 | 0 | 51 | 0.1569 | 11 | 0.0909 | 9 | 0.1111 | 9 | 0 | 9 | 0.1111 | 38 | 0.0789 |
| 17_2403982 | 5 | 0 | 17 | 0.1176 | 4 | 0 | 5 | 0 | 31 | 0.0645 | 9 | 0.2222 | 6 | 0.1667 | 6 | 0 | 4 | 0 | 25 | 0.12 |
| 17_2403987 | 10 | 0.9 | 29 | 0.7586 | 8 | 0.875 | 4 | 1 | 51 | 0.8235 | 10 | 0.8 | 8 | 0.75 | 9 | 1 | 9 | 0.8889 | 36 | 0.8611 |
| 17_2404011 | 10 | 0 | 27 | 0.037 | 9 | 0.1111 | 8 | 0.25 | 54 | 0.0741 | 11 | 0.0909 | 9 | 0 | 10 | 0 | 10 | 0.1 | 40 | 0.05 |
| 17_2404015 | 7 | 0.2857 | 25 | 0.04 | 6 | 0.1667 | 5 | 0 | 43 | 0.093 | 11 | 0.0909 | 7 | 0 | 8 | 0.25 | 9 | 0.1111 | 35 | 0.1143 |
| 17_2404030 | 11 | 0.2727 | 33 | 0.1515 | 9 | 0.1111 | 9 | 0 | 62 | 0.1452 | 12 | 0.25 | 10 | 0.1 | 10 | 0.3 | 11 | 0.5455 | 43 | 0.3023 |
| 17_2404036 | 8 | 0.625 | 29 | 0.5517 | 10 | 0.7 | 12 | 0.4167 | 59 | 0.5593 | 11 | 0.5455 | 9 | 0.7778 | 8 | 0.625 | 10 | 0.4 | 38 | 0.5789 |
| 17_2404091 | NA | NA | 6 | 0.1667 | 1 | 1 | 5 | 0.4 | 12 | 0.3333 | 3 | 0.3333 | 2 | 1 | 3 | 0.3333 | 2 | 0.5 | 10 | 0.5 |
| 17_2404114 | NA | NA | 8 | 0 | 1 | 0 | 3 | 0.3333 | 12 | 0.0833 | 2 | 0 | 2 | 0.5 | 3 | 0.6667 | 3 | 0.3333 | 10 | 0.4 |
| 17_2404230 | 1 | 1 | 20 | 0.4 | 4 | 0.75 | 3 | 0 | 28 | 0.4286 | 4 | 0.5 | 3 | 0.6667 | 2 | 1 | 2 | 0.5 | 11 | 0.6364 |
| 17_2404234 | 1 | 1 | 22 | 0.3636 | 3 | 0.3333 | 3 | 0 | 29 | 0.3448 | 3 | 0 | 2 | 0 | 3 | 0.3333 | 2 | 0 | 10 | 0.1 |
| 17_2404235 | 1 | 1 | 22 | 0.2273 | 4 | 0.5 | 4 | 0 | 31 | 0.2581 | 4 | 0.25 | 3 | 0.6667 | 3 | 0.3333 | 1 | 1 | 11 | 0.4545 |
| 17_2404238 | 1 | 1 | 22 | 0.1818 | 2 | 0.5 | 4 | 0 | 29 | 0.2069 | 4 | 0.5 | 4 | 0.5 | 3 | 0.3333 | 2 | 0.5 | 13 | 0.4615 |
| 17_2404254 | 1 | 1 | 22 | 0.2273 | 3 | 0.3333 | 4 | 0 | 30 | 0.2333 | 4 | 0.25 | 4 | 0.5 | 3 | 0.3333 | 2 | 0 | 13 | 0.3077 |
| 17_2404329 | 17 | 0.1176 | 42 | 0.0952 | 16 | 0.125 | 24 | 0.0833 | 99 | 0.101 | 12 | 0.0833 | 17 | 0 | 12 | 0.1667 | 15 | 0.0667 | 56 | 0.0714 |

| | | | | | | | | | | | | | | | | | | | | |
|---|---|---|---|---|---|---|---|---|---|---|---|---|---|---|---|---|---|---|---|---|
| 17_2404352 | 19 | 0.0526 | 52 | 0.0962 | 20 | 0.2 | 28 | 0.0714 | 119 | 0.1008 | 19 | 0.1579 | 20 | 0.05 | 15 | 0.2667 | 18 | 0.2222 | 72 | 0.1667 |
| 17_2426690 | 6 | 0 | 2 | 0 | 4 | 0.25 | 3 | 0.3333 | 15 | 0.1333 | 7 | 0.1429 | 6 | 0 | 5 | 0.2 | 1 | 0 | 19 | 0.1053 |
| 17_2426703 | 6 | 0.1667 | 2 | 0.5 | 2 | 0 | 2 | 0 | 12 | 0.1667 | 6 | 0 | 6 | 0 | 5 | 0.2 | 1 | 0 | 18 | 0.0556 |
| 17_2426706 | 5 | 0 | 1 | 0 | 3 | 0.3333 | 1 | 0 | 10 | 0.1 | 7 | 0 | 4 | 0.25 | 4 | 0 | 1 | 0 | 16 | 0.0625 |
| 17_2426717 | 6 | 0 | 1 | 1 | 3 | 0.3333 | 2 | 0 | 12 | 0.1667 | 4 | 0 | 3 | 0 | 3 | 0 | 1 | 1 | 11 | 0.0909 |
| 17_2426727 | 6 | 0 | 2 | 0 | 3 | 0.3333 | 4 | 0 | 15 | 0.0667 | 7 | 0.2857 | 6 | 0 | 4 | 0 | 1 | 1 | 18 | 0.1667 |
| 17_2426736 | 6 | 0.3333 | 2 | 1 | 3 | 0.6667 | 4 | 0.75 | 15 | 0.6 | 7 | 0.7143 | 5 | 0.4 | 4 | 0.5 | 1 | 1 | 17 | 0.5882 |
| 17_2426772 | 8 | 0.875 | 3 | 1 | 3 | 1 | 3 | 0.6667 | 17 | 0.8824 | 7 | 0.8571 | 6 | 0.8333 | 4 | 0.75 | 2 | 0.5 | 19 | 0.7895 |
| 17_2426781 | 7 | 1 | 2 | 1 | 3 | 1 | 3 | 1 | 15 | 1 | 6 | 1 | 3 | 1 | 3 | 0.6667 | 2 | 0.5 | 14 | 0.8571 |
| 17_2426795 | 8 | 0.25 | 2 | 0.5 | 4 | 0.25 | 4 | 0.25 | 18 | 0.2778 | 6 | 0.3333 | 6 | 0 | 3 | 0.3333 | 3 | 0.3333 | 18 | 0.2222 |
| 17_2426803 | 8 | 0.5 | 2 | 0.5 | 3 | 1 | 4 | 0.5 | 17 | 0.5882 | 6 | 0.3333 | 6 | 0.6667 | 1 | 0 | 1 | 0 | 14 | 0.4286 |
| 17_2426808 | 7 | 0.2857 | 2 | 1 | 4 | 0.75 | 3 | 0.6667 | 16 | 0.5625 | 7 | 0.5714 | 5 | 0.4 | 4 | 0.75 | 3 | 0.6667 | 19 | 0.5789 |
| 17_2426813 | 6 | 0.8333 | 2 | 1 | 3 | 1 | NA | NA | 11 | 0.9091 | 5 | 1 | 4 | 0.75 | 2 | 1 | 1 | 0 | 12 | 0.8333 |
| 17_2429417 | 9 | 0 | 9 | 0 | 9 | 0 | 10 | 0.2 | 37 | 0.0541 | 5 | 0 | 14 | 0.1429 | 8 | 0 | 9 | 0.1111 | 36 | 0.0833 |
| 17_2429422 | 8 | 0 | 10 | 0.1 | 9 | 0 | 10 | 0.1 | 37 | 0.0541 | 4 | 0 | 12 | 0.1667 | 8 | 0 | 9 | 0.1111 | 33 | 0.0909 |
| 17_2429451 | 6 | 0.1667 | 11 | 0.1818 | 8 | 0.25 | 10 | 0.1 | 35 | 0.1714 | 5 | 0.2 | 10 | 0.3 | 8 | 0.25 | 11 | 0.0909 | 34 | 0.2059 |
| 17_2429677 | 4 | 0 | 4 | 0.25 | 6 | 0.1667 | 7 | 0.1429 | 21 | 0.1429 | 1 | 0 | 5 | 0 | 3 | 0 | 4 | 0.25 | 13 | 0.0769 |
| 17_2429743 | 3 | 0 | 4 | 0 | 5 | 0.2 | 7 | 0.1429 | 19 | 0.1053 | 1 | 0 | 5 | 0 | 2 | 0.5 | 3 | 0.3333 | 11 | 0.1818 |
| 17_2429759 | 3 | 0 | 3 | 0 | 4 | 0 | 4 | 0.25 | 14 | 0.0714 | 1 | 0 | 4 | 0.25 | 2 | 0 | 3 | 0.3333 | 10 | 0.2 |
| 17_2429764 | 3 | 0 | 3 | 0.6667 | 4 | 0.5 | 6 | 0.8333 | 16 | 0.5625 | 1 | 0 | 2 | 0 | NA | NA | 4 | 0.75 | 7 | 0.4286 |
| 17_2429782 | 1 | 0 | 1 | 0 | 3 | 0.3333 | 5 | 0 | 10 | 0.1 | 1 | 0 | 3 | 0.6667 | 1 | 0 | 3 | 0.3333 | 8 | 0.375 |
| 17_2429783 | 3 | 0.3333 | 2 | 0.5 | 3 | 0.3333 | 4 | 0.25 | 12 | 0.3333 | 1 | 0 | 2 | 0.5 | 1 | 0 | 2 | 0 | 6 | 0.1667 |
| 17_2437494 | 2 | 1 | 5 | 0.2 | 5 | 0.2 | 2 | 0.5 | 14 | 0.3571 | 1 | 0 | 5 | 0.2 | 1 | 0 | 1 | 0 | 8 | 0.125 |
| 17_2438180 | NA | NA | NA | NA | NA | NA | 1 | 1 | 1 | 1 | 1 | 1 | 1 | 1 | NA | NA | NA | NA | 2 | 1 |
| 17_2438298 | 214 | 0.2243 | 221 | 0.2036 | 201 | 0.2289 | 189 | 0.1958 | 825 | 0.2133 | 187 | 0.1872 | 191 | 0.2094 | 173 | 0.1965 | 189 | 0.2328 | 740 | 0.2068 |
| 17_2438309 | 220 | 0.0818 | 241 | 0.0996 | 202 | 0.104 | 193 | 0.0674 | 856 | 0.0888 | 206 | 0.0874 | 204 | 0.0882 | 185 | 0.0919 | 192 | 0.1146 | 787 | 0.0953 |
| 17_2438320 | 194 | 0.2216 | 214 | 0.1869 | 186 | 0.2204 | 172 | 0.2093 | 766 | 0.2089 | 167 | 0.1796 | 192 | 0.2135 | 163 | 0.1656 | 158 | 0.1456 | 680 | 0.1779 |
| 17_2438331 | 166 | 0.0301 | 189 | 0.0688 | 154 | 0.0779 | 147 | 0.0476 | 656 | 0.0564 | 170 | 0.0529 | 158 | 0.0759 | 140 | 0.05 | 146 | 0.0685 | 614 | 0.0619 |
| 17_2438854 | 3 | 1 | 4 | 0.5 | 2 | 0 | 1 | 1 | 10 | 0.6 | 1 | 1 | NA | NA | NA | NA | NA | NA | 1 | 1 |
| 17_2439449 | 4 | 0.5 | 9 | 0.6667 | 3 | 0.6667 | 8 | 0.625 | 24 | 0.625 | 4 | 0.75 | 5 | 0.4 | 4 | 0.25 | 2 | 0.5 | 15 | 0.4667 |
| 17_2439480 | 3 | 0 | 8 | 0.125 | 6 | 0.3333 | 8 | 0 | 25 | 0.12 | 2 | 0 | 5 | 0.2 | 4 | 0 | 2 | 0 | 13 | 0.0769 |
| 17_2439761 | 1 | 1 | 2 | 0 | 2 | 0 | 5 | 0 | 10 | 0.1 | 1 | 1 | 3 | 0 | 1 | 0 | NA | NA | 5 | 0.2 |
| 17_2439808 | 1 | 1 | 3 | 1 | 3 | 1 | 5 | 0.6 | 12 | 0.8333 | 1 | 1 | 3 | 0.6667 | 2 | 0.5 | NA | NA | 6 | 0.6667 |
| 17_2439814 | NA | NA | 3 | 0.6667 | 2 | 0.5 | 6 | 0.5 | 11 | 0.5455 | 1 | 0 | 3 | 0.6667 | 2 | 0.5 | NA | NA | 6 | 0.5 |
| 17_2439815 | 1 | 1 | 3 | 1 | 3 | 0.3333 | 5 | 0.4 | 12 | 0.5833 | 1 | 0 | 3 | 0.3333 | 2 | 1 | NA | NA | 6 | 0.5 |
| 17_2439821 | 1 | 1 | 3 | 1 | 3 | 1 | 6 | 0.8333 | 13 | 0.9231 | NA | NA | 2 | 1 | 2 | 0.5 | NA | NA | 4 | 0.75 |
| 17_2439826 | NA | NA | NA | NA | NA | NA | 1 | 1 | 1 | 1 | NA | NA | NA | NA | 1 | 1 | NA | NA | 1 | 1 |
| 17_2439830 | 1 | 1 | 2 | 1 | 3 | 1 | 5 | 0.6 | 11 | 0.8182 | 1 | 0 | 1 | 0 | 1 | 1 | NA | NA | 3 | 0.3333 |
| 17_2439838 | 1 | 1 | 2 | 1 | 3 | 0.6667 | 4 | 0.75 | 10 | 0.8 | 1 | 0 | 3 | 0.6667 | NA | NA | NA | NA | 4 | 0.5 |
| 17_2439843 | 1 | 1 | 2 | 0.5 | 4 | 0.75 | 3 | 0.6667 | 10 | 0.7 | 1 | 1 | 2 | 1 | NA | NA | NA | NA | 3 | 1 |
| 17_2439845 | 1 | 0 | 1 | 0 | 4 | 0.25 | 2 | 0 | 8 | 0.125 | 1 | 0 | 2 | 0.5 | NA | NA | NA | NA | 3 | 0.3333 |
| 17_2439861 | 1 | 1 | 3 | 0.6667 | 3 | 0.3333 | 2 | 0 | 9 | 0.4444 | 1 | 1 | 2 | 0.5 | NA | NA | NA | NA | 3 | 0.6667 |
| 17_2439865 | NA | NA | 3 | 1 | 3 | 0.3333 | 3 | 0.3333 | 9 | 0.5556 | 1 | 1 | 2 | 1 | NA | NA | NA | NA | 3 | 1 |
| 17_2439868 | 1 | 1 | 3 | 1 | 2 | 1 | 3 | 1 | 9 | 1 | NA | NA | 2 | 1 | NA | NA | NA | NA | 2 | 1 |
| 17_2439885 | 1 | 1 | 1 | 1 | 3 | 1 | 2 | 1 | 7 | 1 | NA | NA | 1 | 1 | NA | NA | NA | NA | 1 | 1 |
| 17_2439887 | 1 | 1 | 3 | 0.6667 | 4 | 0.75 | 3 | 0.3333 | 11 | 0.6364 | 1 | 1 | 1 | 1 | NA | NA | NA | NA | 2 | 1 |
| 17_2439900 | NA | NA | 2 | 1 | 3 | 1 | 1 | 1 | 6 | 1 | 1 | 1 | 1 | 1 | NA | NA | NA | NA | 2 | 1 |
| 17_2439930 | NA | NA | 2 | 0.5 | 4 | 0.5 | 1 | 0 | 7 | 0.4286 | NA | NA | 1 | 1 | NA | NA | NA | NA | 1 | 1 |
| 17_2439947 | 1 | 1 | 2 | 1 | 3 | 0.6667 | 1 | 1 | 7 | 0.8571 | NA | NA | 1 | 1 | NA | NA | NA | NA | 1 | 1 |
| 17_2439952 | 1 | 1 | 2 | 0 | 3 | 0.6667 | 1 | 1 | 7 | 0.5714 | 1 | 1 | 1 | 1 | NA | NA | NA | NA | 2 | 1 |
| 17_2439967 | 1 | 0 | 3 | 0.3333 | NA | NA | NA | NA | 4 | 0.25 | 1 | 0 | 1 | 1 | NA | NA | NA | NA | 2 | 0.5 |
| 17_2440431 | 1 | 0 | 3 | 0.3333 | 2 | 0.5 | 1 | 0 | 7 | 0.2857 | 1 | 1 | 1 | 1 | NA | NA | NA | NA | 2 | 1 |
| 17_2440455 | 1 | 1 | 2 | 1 | 1 | 1 | NA | NA | 4 | 1 | 1 | 1 | NA | NA | NA | NA | NA | NA | 1 | 1 |
| 17_2440459 | 1 | 1 | 3 | 0.3333 | 1 | 0 | NA | NA | 5 | 0.4 | 1 | 1 | NA | NA | NA | NA | NA | NA | 1 | 1 |
| 17_2440491 | 1 | 1 | 2 | 0 | 2 | 1 | NA | NA | 5 | 0.6 | 1 | 1 | NA | NA | NA | NA | NA | NA | 1 | 1 |
| 17_2440496 | 1 | 0 | 2 | 0.5 | 1 | 0 | NA | NA | 4 | 0.25 | 1 | 1 | NA | NA | NA | NA | NA | NA | 1 | 1 |
| 17_2443478 | 1 | 1 | 2 | 0.5 | 1 | 1 | NA | NA | 4 | 0.75 | 1 | 1 | 3 | 0.6667 | NA | NA | NA | NA | 4 | 0.75 |
| 17_2444280 | NA | NA | 1 | 0 | NA | NA | 1 | 1 | 2 | 0.5 | NA | NA | NA | NA | NA | NA | 1 | 1 | 1 | 1 |
| 17_2444289 | NA | NA | 1 | 1 | NA | NA | NA | NA | 1 | 1 | NA | NA | NA | NA | NA | NA | 1 | 1 | 1 | 1 |
| 17_2444313 | NA | NA | 1 | 1 | NA | NA | 1 | 1 | 2 | 1 | NA | NA | NA | NA | NA | NA | 1 | 1 | 1 | 1 |
| 17_2462849 | 10 | 0.7 | 5 | 0.6 | 9 | 0.6667 | 5 | 0.8 | 29 | 0.6897 | 5 | 1 | 7 | 0.8571 | 9 | 0.6667 | 7 | 0.8571 | 28 | 0.8214 |
| 17_2462852 | 12 | 0.0833 | 2 | 0 | 9 | 0.2222 | 4 | 0.25 | 27 | 0.1481 | 4 | 0.5 | 7 | 0.1429 | 7 | 0.2857 | 7 | 0.1429 | 25 | 0.24 |
| 17_2462871 | 10 | 0.1 | 1 | 0 | 11 | 0.0909 | 1 | 0 | 23 | 0.087 | 2 | 0.5 | 7 | 0 | 6 | 0.1667 | 9 | 0.3333 | 24 | 0.2083 |
| 17_2462872 | 13 | 0.2308 | 5 | 0.4 | 12 | 0.75 | 4 | 0.25 | 34 | 0.4412 | 3 | 0.3333 | 8 | 0.5 | 8 | 0.25 | 10 | 0.3 | 29 | 0.3448 |
| 17_2462879 | 10 | 0.9 | 3 | 0.3333 | 4 | 1 | 1 | 1 | 18 | 0.8333 | 3 | 1 | 5 | 1 | 3 | 0.6667 | 4 | 0.75 | 15 | 0.8667 |

| | | | | | | | | | | | | | | | | | | | | |
|---|---|---|---|---|---|---|---|---|---|---|---|---|---|---|---|---|---|---|---|---|
| 17_2462894 | 12 | 0.75 | 4 | 1 | 5 | 1 | 4 | 1 | 25 | 0.88 | 3 | 1 | 7 | 1 | 8 | 0.75 | 6 | 1 | 24 | 0.9167 |
| 17_2462894 | 12 | 0.1667 | 5 | 0 | 10 | 0 | 5 | 0 | 32 | 0.0625 | 5 | 0.2 | 8 | 0 | 8 | 0.125 | 8 | 0 | 29 | 0.069 |
| 17_2464416 | 20 | 0.2 | 18 | 0.0556 | 11 | 0.0909 | 14 | 0.2143 | 63 | 0.1429 | 20 | 0.3 | 12 | 0.25 | 15 | 0.3333 | 13 | 0.0769 | 60 | 0.25 |
| 17_2464449 | 22 | 0.0455 | 17 | 0.1176 | 13 | 0.0769 | 15 | 0 | 67 | 0.0597 | 19 | 0.0526 | 13 | 0.1538 | 17 | 0.0588 | 14 | 0 | 63 | 0.0635 |
| 17_2465809 | 4 | 0.5 | 4 | 0 | 2 | 0 | 3 | 0.3333 | 13 | 0.2308 | 5 | 0.6 | 3 | 0.6667 | 4 | 0.25 | 4 | 0.25 | 16 | 0.4375 |
| 17_2465974 | NA | NA | 3 | 1 | NA | NA | NA | NA | 3 | 1 | 2 | 0.5 | NA | NA | 1 | 1 | 3 | 1 | 6 | 0.8333 |
| 17_2465988 | NA | NA | 3 | 1 | NA | NA | NA | NA | 3 | 1 | 2 | 1 | NA | NA | 1 | 1 | 3 | 0.3333 | 6 | 0.6667 |
| 17_2465989 | NA | NA | 3 | 0.3333 | NA | NA | NA | NA | 3 | 0.3333 | 2 | 0 | NA | NA | 1 | 0 | 3 | 0.3333 | 6 | 0.1667 |
| 17_2465990 | NA | NA | 3 | 0.6667 | NA | NA | NA | NA | 3 | 0.6667 | 2 | 0.5 | NA | NA | 1 | 1 | 3 | 1 | 6 | 0.8333 |
| 17_2465993 | NA | NA | 3 | 0.6667 | NA | NA | NA | NA | 3 | 0.6667 | 2 | 0.5 | NA | NA | 1 | 0 | 3 | 0.3333 | 6 | 0.3333 |
| 17_2465998 | NA | NA | 2 | 1 | NA | NA | NA | NA | 2 | 1 | 2 | 1 | NA | NA | NA | NA | 2 | 0 | 4 | 0.5 |
| 17_2466281 | 2 | 0 | 1 | 0 | 2 | 0.5 | 3 | 0 | 8 | 0.125 | 2 | 0.5 | NA | NA | 3 | 0 | 2 | 0 | 7 | 0.1429 |
| 17_2466292 | 2 | 0.5 | 3 | 0.6667 | 2 | 0.5 | 3 | 0 | 10 | 0.4 | 3 | 0.3333 | NA | NA | 5 | 0.2 | 4 | 0 | 12 | 0.1667 |
| 17_2466303 | 2 | 1 | 2 | 0.5 | 2 | 1 | 3 | 0.6667 | 9 | 0.7778 | 3 | 0 | NA | NA | 4 | 0.75 | 3 | 0.3333 | 10 | 0.4 |
| 17_2466305 | 2 | 0 | 3 | 0 | 1 | 1 | 3 | 0 | 9 | 0.1111 | 3 | 0.3333 | NA | NA | 4 | 0.25 | 4 | 0 | 11 | 0.1818 |
| 17_2468956 | 11 | 0.2727 | 15 | 0 | 5 | 0 | 7 | 0.1429 | 38 | 0.1053 | 11 | 0.0909 | 8 | 0.125 | 8 | 0.25 | 9 | 0 | 36 | 0.1111 |
| 17_2468983 | 10 | 0.2 | 13 | 0.2308 | 9 | 0.3333 | 10 | 0.3 | 42 | 0.2619 | 11 | 0.3636 | 10 | 0.1 | 5 | 0.4 | 8 | 0.25 | 34 | 0.2647 |
| 17_2469008 | 8 | 1 | 17 | 0.8235 | 9 | 0.7778 | 10 | 0.8 | 44 | 0.8409 | 14 | 0.7857 | 9 | 0.6667 | 7 | 0.8571 | 8 | 0.75 | 38 | 0.7632 |
| 17_2469026 | 10 | 0.7 | 12 | 0.25 | 6 | 0.1667 | 9 | 0.4444 | 37 | 0.4054 | 12 | 0.3333 | 9 | 0.1111 | 8 | 0.25 | 9 | 0.1111 | 38 | 0.2105 |
| 17_2469036 | 9 | 0.3333 | 15 | 0.2667 | 8 | 0.125 | 9 | 0.2222 | 41 | 0.2439 | 12 | 0.3333 | 9 | 0.1111 | 9 | 0.2222 | 10 | 0 | 40 | 0.175 |
| 17_2469046 | 11 | 0.8182 | 16 | 0.5625 | 8 | 0.75 | 9 | 0.8889 | 44 | 0.7273 | 16 | 0.75 | 10 | 0.7 | 8 | 0.75 | 9 | 0.5556 | 43 | 0.6977 |
| 17_2469049 | 12 | 0.25 | 19 | 0.1579 | 11 | 0.0909 | 10 | 0.2 | 52 | 0.1731 | 14 | 0.0714 | 9 | 0.2222 | 9 | 0 | 11 | 0.2727 | 43 | 0.1395 |
| 17_2469081 | 3 | 0.6667 | 4 | 0.5 | 6 | 0.6667 | 2 | 0.5 | 15 | 0.6 | 3 | 0.6667 | 5 | 0.6 | 2 | 1 | 5 | 0.6 | 15 | 0.6667 |
| 17_2469090 | 3 | 0 | 4 | 0 | 6 | 0.1667 | 3 | 0 | 16 | 0.0625 | 3 | 0 | 5 | 0 | 3 | 0 | 7 | 0.1429 | 18 | 0.0556 |
| 17_2469091 | 3 | 0 | 5 | 0 | 5 | 0 | 2 | 0.5 | 15 | 0.0667 | 3 | 0 | 5 | 0.2 | 3 | 0 | 6 | 0 | 17 | 0.0588 |
| 17_2469113 | 3 | 0 | 5 | 0 | 5 | 0 | 3 | 0.3333 | 16 | 0.0625 | 3 | 0 | 5 | 0.2 | 4 | 0.25 | 7 | 0 | 19 | 0.1053 |
| 17_2469114 | 2 | 0.5 | 4 | 0.25 | 7 | 0.5714 | 3 | 0.3333 | 16 | 0.4375 | 2 | 0 | 5 | 0.6 | 3 | 0.3333 | 6 | 0.6667 | 16 | 0.5 |
| 17_2469121 | 3 | 0 | 2 | 0.5 | 5 | 0.4 | 4 | 1 | 14 | 0.5 | 2 | 0 | 2 | 1 | 3 | 0 | 6 | 0.1667 | 13 | 0.2308 |
| 17_2469150 | 4 | 0.25 | 4 | 0.25 | 6 | 0.6667 | 3 | 0.6667 | 17 | 0.4706 | 2 | 0 | 5 | 0.8 | 3 | 0.6667 | 6 | 0 | 16 | 0.375 |
| 17_2469153 | 3 | 0 | 3 | 0.3333 | 5 | 0.2 | 3 | 0.3333 | 14 | 0.2143 | NA | NA | 6 | 0 | 3 | 0 | 6 | 0.3333 | 15 | 0.1333 |
| 17_2469161 | 4 | 0.5 | 3 | 0 | 6 | 0 | 4 | 0.25 | 17 | 0.1765 | 2 | 0 | 6 | 0 | 3 | 0.3333 | 7 | 0.1429 | 18 | 0.1111 |
| 17_2469163 | 3 | 0.3333 | 3 | 0 | 5 | 0.2 | 3 | 0 | 14 | 0.1429 | 2 | 0 | 6 | 0.1667 | 4 | 0.25 | 6 | 0 | 18 | 0.1111 |
| 17_2469168 | 4 | 0.75 | 5 | 0.6 | 6 | 0.5 | 4 | 0.75 | 19 | 0.6316 | 1 | 1 | 5 | 0.4 | 4 | 0.75 | 7 | 0.7143 | 17 | 0.6471 |
| 17_2469179 | 2 | 0 | 5 | 0 | 8 | 0.375 | 5 | 0.2 | 20 | 0.2 | 3 | 0.3333 | 6 | 0 | 3 | 0.3333 | 7 | 0.2857 | 19 | 0.2105 |
| 17_2469191 | 4 | 0 | 4 | 0.25 | 7 | 0 | 5 | 0 | 20 | 0.05 | 2 | 0 | 3 | 0 | 3 | 0.3333 | 6 | 0 | 14 | 0.0714 |
| 17_2469259 | 3 | 0.3333 | 7 | 0 | 3 | 0 | 7 | 0 | 20 | 0.05 | 5 | 0.2 | 6 | 0.1667 | 2 | 0 | 5 | 0 | 18 | 0.1111 |
| 17_2469268 | 3 | 0 | 7 | 0.1429 | 3 | 0 | 5 | 0 | 18 | 0.0556 | 4 | 0.25 | 4 | 0 | 2 | 0 | 5 | 0 | 15 | 0.0667 |
| 17_2469270 | 2 | 0.5 | 6 | 0.3333 | 3 | 0.6667 | 7 | 0.8571 | 18 | 0.6111 | 5 | 1 | 6 | 0.1667 | 2 | 1 | 5 | 0.8 | 18 | 0.6667 |
| 17_2469294 | 4 | 0.25 | 6 | 0 | NA | NA | 5 | 0.2 | 15 | 0.1333 | 3 | 0.3333 | 2 | 0 | 2 | 0 | 3 | 0 | 10 | 0.1 |
| 17_2469296 | 3 | 0.3333 | 7 | 0.1429 | 2 | 0.5 | 7 | 0.1429 | 19 | 0.2105 | 4 | 0 | 6 | 0.1667 | 2 | 0.5 | 2 | 1 | 14 | 0.2857 |
| 17_2469305 | 4 | 0 | 6 | 0 | 3 | 0.3333 | 8 | 0.5 | 21 | 0.2381 | 4 | 0.75 | 6 | 0.1667 | 2 | 0 | 2 | 0.5 | 14 | 0.3571 |
| 17_2469328 | 4 | 0.25 | 6 | 0 | 2 | 0 | 6 | 0.1667 | 18 | 0.1111 | 4 | 0.25 | 4 | 0 | 2 | 0 | 3 | 0 | 13 | 0.0769 |
| 17_2469377 | 3 | 0.3333 | 7 | 0.1429 | 1 | 0 | 7 | 0 | 18 | 0.1111 | 2 | 0.5 | 5 | 0.2 | 2 | 0 | 3 | 0 | 12 | 0.1667 |
| 17_2469378 | 3 | 0 | 5 | 0.4 | 2 | 0 | 4 | 0 | 14 | 0.1429 | 2 | 0 | 5 | 0.8 | 2 | 0 | 2 | 0.5 | 11 | 0.4545 |
| 17_2469395 | 2 | 0 | 7 | 0.2857 | 2 | 0 | 5 | 0.2 | 16 | 0.1875 | 2 | 0.5 | 5 | 0 | 2 | 0 | 3 | 0 | 12 | 0.0833 |
| 17_2469451 | 3 | 0.6667 | 4 | 0.5 | NA | NA | 4 | 0.75 | 11 | 0.6364 | 1 | 0 | 4 | 0.25 | 1 | 0 | 4 | 0.75 | 10 | 0.4 |
| 17_2469452 | 3 | 0 | 6 | 0.1667 | 2 | 0 | 3 | 0.3333 | 14 | 0.1429 | 1 | 1 | 5 | 0 | 1 | 1 | 4 | 0 | 11 | 0.1818 |
| 17_2469454 | 3 | 0 | 5 | 0.2 | 1 | 0 | 3 | 0.3333 | 12 | 0.1667 | 1 | 0 | 5 | 0 | 1 | 0 | 4 | 0.25 | 11 | 0.0909 |
| 17_2469456 | 2 | 0 | 5 | 0.6 | 2 | 0.5 | 4 | 0.75 | 13 | 0.5385 | NA | NA | 2 | 0 | 1 | 1 | 3 | 0.6667 | 6 | 0.5 |
| 17_2469462 | 3 | 0.3333 | 5 | 0.2 | 2 | 0 | 4 | 0.75 | 14 | 0.3571 | 1 | 0 | 4 | 0.25 | 1 | 1 | 4 | 0.25 | 10 | 0.3 |
| 17_2469464 | 3 | 0.6667 | 6 | 0.1667 | 2 | 0 | 3 | 0.6667 | 14 | 0.3571 | 1 | 1 | 4 | 0.25 | 1 | 1 | 4 | 0 | 10 | 0.3 |
| 17_2469467 | 3 | 1 | 4 | 0.5 | 2 | 1 | 4 | 0.75 | 13 | 0.7692 | 1 | 1 | 5 | 1 | 1 | 1 | 3 | 1 | 10 | 1 |
| 17_2469477 | 3 | 0.6667 | 6 | 0.5 | 2 | 1 | 4 | 0.75 | 15 | 0.6667 | 1 | 1 | 4 | 0.75 | 1 | 1 | 3 | 0.6667 | 9 | 0.7778 |
| 17_2469488 | 3 | 0.3333 | 5 | 0 | 1 | 0 | 4 | 0.5 | 13 | 0.2308 | 1 | 0 | 4 | 0.25 | 1 | 1 | 3 | 0.3333 | 9 | 0.3333 |
| 17_2472105 | NA | NA | 1 | 1 | NA | NA | 1 | 0 | 2 | 0.5 | NA | NA | NA | NA | 1 | 1 | 1 | 1 | 2 | 1 |
| 17_2472299 | 1 | 0 | 2 | 0 | 4 | 0 | 4 | 0.5 | 11 | 0.1818 | 2 | 0.5 | 2 | 0 | 5 | 0.2 | 4 | 0 | 13 | 0.1538 |
| 17_2472318 | 2 | 0.5 | 4 | 0.25 | 4 | 0.25 | 4 | 0.75 | 14 | 0.4286 | 3 | 0.3333 | 1 | 0 | 5 | 0.6 | 7 | 0.7143 | 16 | 0.5625 |
| 17_2472319 | 1 | 0 | 4 | 0.25 | 4 | 0.25 | 3 | 0.3333 | 12 | 0.25 | 3 | 0.3333 | 1 | 1 | 4 | 0 | 5 | 0.4 | 13 | 0.3077 |
| 17_2472329 | 2 | 0 | 5 | 0 | 4 | 0.25 | 4 | 0 | 15 | 0.0667 | 2 | 0.5 | 3 | 0 | 4 | 0.25 | 4 | 0.5 | 13 | 0.3077 |
| 17_2472360 | 2 | 0 | 6 | 0 | 4 | 0 | 4 | 0.25 | 16 | 0.0625 | 3 | 0.3333 | 3 | 0 | 5 | 0 | 7 | 0 | 18 | 0.0556 |
| 17_2472376 | 2 | 0 | 4 | 0.5 | 4 | 0.5 | 5 | 0.6 | 15 | 0.4667 | 3 | 0 | 3 | 0.3333 | 4 | 1 | 5 | 0.4 | 15 | 0.4667 |
| 17_2472382 | 2 | 0.5 | 5 | 0.4 | 4 | 0.25 | 5 | 0.6 | 16 | 0.4375 | 3 | 0.3333 | 3 | 0 | 5 | 0.4 | 5 | 0.6 | 16 | 0.375 |
| 17_2472393 | 3 | 0 | 6 | 0.1667 | 5 | 0 | 5 | 0 | 19 | 0.0526 | 6 | 0.3333 | 3 | 0.6667 | 4 | 0.25 | 7 | 0.5714 | 20 | 0.45 |
| 17_2472399 | 3 | 0.6667 | 6 | 0.8333 | 5 | 0.8 | 4 | 1 | 18 | 0.8333 | 6 | 0.1667 | 3 | 1 | 5 | 0.6 | 7 | 0.7143 | 21 | 0.5714 |
| 17_2472420 | 2 | 0.5 | 5 | 0.8 | 5 | 1 | 4 | 0.75 | 16 | 0.8125 | 4 | 0.75 | 2 | 1 | 6 | 1 | 7 | 1 | 19 | 0.9474 |

| | | | | | | | | | | | | | | | | | | | | |
|---|---|---|---|---|---|---|---|---|---|---|---|---|---|---|---|---|---|---|---|---|
| 17_247257 | 5 | 0 | 11 | 0.0909 | 7 | 0.2857 | 6 | 0.3333 | 29 | 0.1724 | 7 | 0 | 4 | 0.25 | 7 | 0.1429 | 10 | 0 | 28 | 0.0714 |
| 17_247258 | 6 | 0 | 11 | 0.0909 | 6 | 0.1667 | 6 | 0.1667 | 29 | 0.1034 | 7 | 0.2857 | 5 | 0.2 | 7 | 0 | 11 | 0.0909 | 30 | 0.1333 |
| 17_247265 | 6 | 0.1667 | 13 | 0.0769 | 7 | 0.1429 | 7 | 0 | 33 | 0.0909 | 4 | 0.25 | 4 | 0.25 | 7 | 0.1429 | 11 | 0.0909 | 26 | 0.1538 |
| 17_247265 | 3 | 0.3333 | 6 | 0 | 4 | 0.75 | 5 | 0.2 | 18 | 0.2778 | 5 | 0.6 | 3 | 0.3333 | 3 | 0.3333 | 7 | 0.4286 | 18 | 0.4444 |
| 17_247266 | 5 | 0.4 | 8 | 0.25 | 5 | 0.2 | 7 | 0.1429 | 25 | 0.24 | 5 | 0.6 | 4 | 0.5 | 6 | 0.5 | 10 | 0.1 | 25 | 0.36 |
| 17_247271 | 4 | 0 | 10 | 0.1 | 4 | 0.25 | 4 | 0 | 22 | 0.0909 | 7 | 0.5714 | 3 | 0.6667 | 9 | 0.1111 | 13 | 0.3846 | 32 | 0.375 |
| 17_247273 | 7 | 0.1429 | 14 | 0.2857 | 5 | 0.2 | 4 | 0.25 | 30 | 0.2333 | 6 | 0.1667 | 5 | 0.4 | 8 | 0.375 | 11 | 0.4545 | 30 | 0.3667 |
| 17_247278 | 4 | 0.25 | 5 | 0 | 8 | 0 | 7 | 0.1429 | 24 | 0.0833 | 3 | 0 | 2 | 0 | 5 | 0.2 | 3 | 0 | 13 | 0.0769 |
| 17_247280 | 5 | 0.2 | 6 | 0 | 8 | 0.125 | 7 | 0 | 26 | 0.0769 | 3 | 0.3333 | 1 | 0 | 6 | 0 | 4 | 0.25 | 14 | 0.1429 |
| 17_247280 | 5 | 0.8 | 6 | 0.1667 | 6 | 0.1667 | 4 | 0.75 | 21 | 0.4286 | 2 | 0.5 | 1 | 0 | 5 | 0 | 3 | 0.6667 | 11 | 0.2727 |
| 17_247293 | 4 | 0 | 4 | 0.25 | 6 | 0 | 4 | 0 | 18 | 0.0556 | 3 | 0 | 1 | 0 | 5 | 0.2 | 4 | 0 | 13 | 0.0769 |
| 17_247482 | 3 | 0.6667 | 7 | 0.5714 | 1 | 1 | 2 | 0.5 | 13 | 0.6154 | 3 | 0.6667 | 3 | 0.6667 | 3 | 0 | 2 | 0 | 11 | 0.3636 |
| 17_247484 | 4 | 0.5 | 6 | 0.8333 | 2 | 1 | 1 | 1 | 13 | 0.7692 | 3 | 1 | 4 | 0.5 | 3 | 0.3333 | 4 | 0.5 | 14 | 0.5714 |
| 17_247944 | 1 | 1 | NA | NA | NA | NA | NA | NA | 1 | 1 | NA | NA | NA | NA | 1 | 1 | NA | NA | 1 | 1 |
| 17_247945 | NA | NA | 1 | 1 | NA | NA | NA | NA | 1 | 1 | NA | NA | NA | NA | 1 | 1 | NA | NA | 1 | 1 |
| 17_247946 | 1 | 1 | 1 | 1 | NA | NA | NA | NA | 2 | 1 | NA | NA | NA | NA | 1 | 1 | NA | NA | 1 | 1 |
| 17_247946 | 1 | 0 | 1 | 1 | NA | NA | NA | NA | 2 | 0.5 | NA | NA | NA | NA | 1 | 1 | NA | NA | 1 | 1 |
| 17_247950 | 1 | 1 | 1 | 1 | NA | NA | NA | NA | 2 | 1 | NA | NA | NA | NA | 1 | 1 | NA | NA | 1 | 1 |
| 17_247993 | 1 | 1 | 1 | 1 | NA | NA | 2 | 0.5 | 4 | 0.75 | 1 | 0 | NA | NA | 2 | 1 | NA | NA | 3 | 0.6667 |
| 17_248099 | 6 | 0 | 1 | 0 | 2 | 0 | 3 | 0.3333 | 12 | 0.0833 | 2 | 0 | 2 | 0 | 1 | 0 | 3 | 0.3333 | 8 | 0.125 |
| 17_248143 | 7 | 0.5714 | 7 | 0.8571 | 5 | 0.8 | 6 | 0.8333 | 25 | 0.76 | 5 | 1 | 5 | 0.8 | 4 | 1 | 3 | 0.6667 | 17 | 0.8824 |
| 17_248144 | 7 | 0.4286 | 6 | 0.6667 | 6 | 0.8333 | 6 | 0.8333 | 25 | 0.68 | 4 | 0.25 | 6 | 0.6667 | 5 | 1 | 4 | 0.5 | 19 | 0.6316 |
| 17_248145 | 7 | 0.1429 | 7 | 0.1429 | 6 | 0.3333 | 7 | 0 | 27 | 0.1481 | 5 | 0.2 | 6 | 0 | 4 | 0 | 4 | 0 | 19 | 0.0526 |
| 17_248147 | 7 | 0.1429 | 7 | 0 | 6 | 0.3333 | 8 | 0.125 | 28 | 0.1429 | 6 | 0 | 6 | 0 | 5 | 0.4 | 5 | 0.2 | 22 | 0.1364 |
| 17_248147 | 8 | 0 | 6 | 0.1667 | 5 | 0.2 | 8 | 0.25 | 27 | 0.1481 | 5 | 0.4 | 6 | 0 | 4 | 0 | 4 | 0.5 | 19 | 0.2105 |
| 17_248151 | 5 | 0 | 3 | 0 | 5 | 0.2 | 5 | 0 | 18 | 0.0556 | 3 | 0.3333 | 7 | 0.1429 | 3 | 0 | 2 | 0 | 15 | 0.1333 |
| 17_248152 | 11 | 0.6364 | 6 | 0.1667 | 6 | 0.5 | 8 | 0.375 | 31 | 0.4516 | 5 | 0.2 | 9 | 0.5556 | 2 | 0.5 | 8 | 0.5 | 24 | 0.4583 |
| 17_248152 | 8 | 0.125 | 6 | 0 | 6 | 0.1667 | 6 | 0.1667 | 26 | 0.1154 | 6 | 0.3333 | 6 | 0 | 3 | 0.6667 | 7 | 0.1429 | 22 | 0.2273 |
| 17_248155 | 12 | 0.1667 | 11 | 0.2727 | 6 | 0 | 5 | 0.2 | 34 | 0.1765 | 4 | 0 | 10 | 0.1 | 10 | 0.1 | 4 | 0.25 | 28 | 0.1071 |
| 17_248155 | 16 | 0.375 | 18 | 0.1111 | 9 | 0.2222 | 8 | 0.25 | 51 | 0.2353 | 5 | 0 | 12 | 0.1667 | 16 | 0.0625 | 3 | 0 | 36 | 0.0833 |
| 17_248171 | 11 | 0.0909 | 14 | 0.0714 | 8 | 0 | 4 | 0 | 37 | 0.0541 | 4 | 0.25 | 7 | 0 | 10 | 0.2 | 3 | 0 | 24 | 0.125 |
| 17_248172 | 10 | 0 | 15 | 0.0667 | 8 | 0 | 6 | 0.1667 | 39 | 0.0513 | 5 | 0.2 | 8 | 0.125 | 13 | 0.0769 | 3 | 0 | 29 | 0.1034 |
| 17_248188 | 10 | 0 | 12 | 0.1667 | 4 | 0 | 6 | 0.1667 | 32 | 0.0938 | 4 | 0.25 | 6 | 0 | 12 | 0 | 3 | 0.3333 | 25 | 0.08 |
| 17_248189 | 9 | 0.3333 | 8 | 0.125 | 4 | 0 | 6 | 0.3333 | 27 | 0.2222 | 2 | 0 | 5 | 0.2 | 11 | 0 | 3 | 0.3333 | 21 | 0.0952 |
| 17_248587 | 6 | 0.3333 | 7 | 0.8571 | 5 | 0.6 | 4 | 1 | 22 | 0.6818 | 9 | 0.5556 | 9 | 0.6667 | 8 | 0.75 | 8 | 0.5 | 34 | 0.6176 |
| 17_248591 | 5 | 0.2 | 6 | 0.1667 | 6 | 0.3333 | 6 | 0.5 | 23 | 0.3043 | 8 | 0.5 | 9 | 0.3333 | 6 | 0.3333 | 6 | 0.3333 | 29 | 0.3793 |
| 17_248592 | 10 | 0 | 6 | 0.1667 | 6 | 0.5 | 6 | 0.3333 | 28 | 0.2143 | 9 | 0.4444 | 9 | 0.1111 | 6 | 0 | 7 | 0 | 31 | 0.1613 |
| 17_248986 | 4 | 0.25 | 7 | 0.1429 | 1 | 1 | 9 | 0.2222 | 21 | 0.2381 | 2 | 0.5 | 6 | 0.3333 | 3 | 0.6667 | 3 | 0 | 14 | 0.3571 |
| 17_248991 | 5 | 0.8 | 3 | 0.3333 | 1 | 1 | 8 | 0.875 | 17 | 0.7647 | 2 | 0.5 | 3 | 1 | 1 | 1 | 4 | 0.75 | 10 | 0.8 |
| 17_248991 | 5 | 0.8 | 6 | 0 | 2 | 0 | 8 | 0 | 21 | 0.1905 | 2 | 0.5 | 4 | 0.25 | NA | NA | 6 | 0.3333 | 12 | 0.3333 |
| 17_248999 | 7 | 0.2857 | 8 | 0.125 | 2 | 0 | 12 | 0.1667 | 29 | 0.1724 | 2 | 0 | 5 | 0.4 | 2 | 1 | 7 | 0.1429 | 16 | 0.3125 |
| 17_249002 | 7 | 0.4286 | 9 | 0.1111 | 3 | 0.6667 | 12 | 0 | 31 | 0.1935 | 2 | 0.5 | 6 | 0.3333 | 4 | 0 | 5 | 0 | 17 | 0.1765 |
| 17_249003 | 6 | 0.8333 | 9 | 0.6667 | 2 | 1 | 11 | 0.6364 | 28 | 0.7143 | 2 | 1 | 6 | 0.8333 | 4 | 1 | 5 | 0.8 | 17 | 0.8824 |
| 17_249003 | 8 | 0.125 | 8 | 0 | 4 | 0 | 12 | 0.0833 | 32 | 0.0625 | 2 | 0 | 5 | 0 | 5 | 0 | 8 | 0.125 | 20 | 0.05 |
| 17_249005 | 8 | 0.25 | 9 | 0.7778 | 4 | 0.5 | 12 | 0.5 | 33 | 0.5152 | 2 | 0.5 | 8 | 0.25 | 4 | 0.75 | 8 | 0.125 | 22 | 0.3182 |
| 17_249007 | 7 | 0.4286 | 8 | 0 | 4 | 0.25 | 12 | 0.1667 | 31 | 0.1935 | 2 | 0 | 6 | 0.3333 | 4 | 0.25 | 6 | 0.1667 | 18 | 0.2222 |
| 17_249009 | 7 | 0.2857 | 8 | 0 | 3 | 0 | 12 | 0 | 30 | 0.0667 | 2 | 0.5 | 8 | 0 | 7 | 0 | 6 | 0.1667 | 23 | 0.087 |
| 17_249049 | 11 | 0.0909 | 24 | 0.0417 | 17 | 0.0588 | 15 | 0.2 | 67 | 0.0896 | 5 | 0.2 | 11 | 0.0909 | 8 | 0 | 14 | 0 | 38 | 0.0526 |
| 17_249050 | 18 | 0.1111 | 30 | 0.0333 | 19 | 0.1053 | 21 | 0.1905 | 88 | 0.1023 | 10 | 0 | 19 | 0.1579 | 14 | 0.0714 | 14 | 0.1429 | 57 | 0.1053 |
| 17_249051 | 18 | 0.0556 | 30 | 0.1333 | 16 | 0.125 | 19 | 0.1579 | 83 | 0.1205 | 9 | 0.2222 | 21 | 0.0476 | 14 | 0.1429 | 16 | 0.1875 | 60 | 0.1333 |
| 17_249121 | NA | NA | 4 | 0.5 | 4 | 0.5 | NA | NA | 8 | 0.5 | 1 | 0 | 4 | 0.25 | NA | NA | 2 | 0.5 | 7 | 0.2857 |
| 17_249121 | 1 | 0 | 5 | 0.2 | 4 | 0.5 | NA | NA | 10 | 0.3 | 2 | 0 | 4 | 0.25 | 2 | 1 | 2 | 0 | 10 | 0.3 |
| 17_249123 | 1 | 1 | 4 | 1 | 1 | 1 | NA | NA | 6 | 1 | 2 | 1 | 2 | 1 | 3 | 1 | 2 | 1 | 9 | 1 |
| 17_249124 | 2 | 1 | 5 | 0.8 | 2 | 1 | NA | NA | 9 | 0.8889 | 2 | 1 | 2 | 0.5 | 2 | 1 | 2 | 1 | 8 | 0.875 |
| 17_249124 | 2 | 1 | 4 | 0.75 | 2 | 1 | NA | NA | 8 | 0.875 | 1 | 1 | 4 | 0.5 | 2 | 1 | 1 | 1 | 8 | 0.75 |
| 17_249125 | 3 | 0.6667 | 5 | 0.4 | 3 | 1 | NA | NA | 11 | 0.6364 | 2 | 0.5 | 4 | 0.25 | 2 | 0.5 | 2 | 0 | 10 | 0.3 |
| 17_249125 | 2 | 1 | 4 | 0.25 | 1 | 1 | NA | NA | 7 | 0.5714 | NA | NA | 3 | 0 | 2 | 1 | 2 | 1 | 7 | 0.5714 |
| 17_249126 | 3 | 1 | 3 | 1 | 3 | 1 | NA | NA | 9 | 1 | 1 | 1 | 3 | 0.6667 | 1 | 1 | 2 | 1 | 7 | 0.8571 |
| 17_249127 | 3 | 0.3333 | 3 | 0 | 1 | 0 | NA | NA | 7 | 0.1429 | 2 | 0.5 | 3 | 0 | NA | NA | 2 | 0 | 7 | 0.1429 |
| 17_249128 | 2 | 0.5 | 3 | 0.3333 | 2 | 0 | NA | NA | 7 | 0.2857 | 1 | 0 | 4 | 0.75 | 1 | 0 | 1 | 0 | 7 | 0.4286 |
| 17_249130 | 1 | 1 | 2 | 1 | 2 | 1 | NA | NA | 5 | 1 | 2 | 0.5 | 3 | 1 | 2 | 1 | 2 | 1 | 9 | 0.8889 |
| 17_249131 | 3 | 1 | 3 | 1 | 3 | 0.6667 | NA | NA | 9 | 0.8889 | 1 | 1 | 2 | 0.5 | 1 | 1 | 2 | 1 | 6 | 0.8333 |
| 17_249144 | 5 | 0.2 | 4 | 0.25 | 2 | 0 | NA | NA | 11 | 0.1818 | 2 | 0.5 | 4 | 0.25 | 1 | 0 | 4 | 0.75 | 11 | 0.4545 |
| 17_249148 | 4 | 0.25 | 2 | 0 | 2 | 0 | NA | NA | 8 | 0.125 | 1 | 0 | 3 | 0 | 1 | 0 | 4 | 0.5 | 9 | 0.2222 |
| 17_249197 | 3 | 0 | 5 | 0 | 2 | 0 | 6 | 0.1667 | 16 | 0.0625 | 3 | 0.3333 | 1 | 0 | 4 | 0 | 4 | 0.25 | 12 | 0.1667 |

| | | | | | | | | | | | | | | | | | | | | |
|---|---|---|---|---|---|---|---|---|---|---|---|---|---|---|---|---|---|---|---|---|
| 17_2493982 | 1 | 0 | 1 | 0 | 1 | 1 | 2 | 0.5 | 5 | 0.4 | 1 | 0 | 1 | 0 | 2 | 0.5 | 2 | 0 | 6 | 0.1667 |
| 17_2493996 | 70 | 0.0857 | 77 | 0.0649 | 79 | 0.1392 | 107 | 0.1308 | 333 | 0.1081 | 58 | 0.1034 | 72 | 0.1667 | 113 | 0.1327 | 83 | 0.1325 | 326 | 0.135 |
| 17_2494163 | 52 | 0.0962 | 35 | 0.1143 | 44 | 0.0682 | 37 | 0.0811 | 168 | 0.0893 | 32 | 0.0625 | 42 | 0.119 | 54 | 0.1111 | 39 | 0 | 167 | 0.0778 |
| 17_2496129 | 5 | 0.2 | 2 | 0 | 9 | 0.2222 | 5 | 0.2 | 21 | 0.1905 | 9 | 0.1111 | 3 | 0 | 2 | 0.5 | 6 | 0.1667 | 20 | 0.15 |
| 17_2496144 | 4 | 0.25 | 2 | 0 | 8 | 0 | 5 | 0.2 | 19 | 0.1053 | 9 | 0.1111 | 1 | 1 | 1 | 1 | 6 | 0.3333 | 17 | 0.2941 |
| 17_2496157 | 5 | 0.2 | 4 | 0 | 9 | 0.2222 | 6 | 0.1667 | 24 | 0.1667 | 9 | 0.1111 | 3 | 0.3333 | 2 | 0 | 6 | 0.3333 | 20 | 0.2 |
| 17_2506787 | 1 | 0 | 6 | 0.1667 | 4 | 0 | 2 | 0.5 | 13 | 0.1538 | 3 | 0.3333 | NA | NA | 3 | 0 | 2 | 0 | 8 | 0.125 |
| 17_2506810 | 2 | 0 | 6 | 0.1667 | 5 | 0 | 3 | 0 | 16 | 0.0625 | 3 | 0.3333 | NA | NA | 5 | 0.4 | 4 | 0.25 | 12 | 0.3333 |
| 17_2506814 | 2 | 1 | 5 | 0.8 | 5 | 0.6 | 4 | 0.75 | 16 | 0.75 | 4 | 0.75 | NA | NA | 5 | 0.8 | 5 | 0.8 | 14 | 0.7857 |
| 17_2506815 | 1 | 1 | 3 | 0.6667 | 4 | 0.5 | 5 | 0.4 | 13 | 0.5385 | 1 | 1 | NA | NA | 3 | 0.3333 | 3 | 0.3333 | 7 | 0.4286 |
| 17_2506825 | 2 | 1 | 3 | 0.6667 | 5 | 1 | 4 | 0.75 | 14 | 0.8571 | 4 | 1 | NA | NA | 5 | 1 | 4 | 1 | 13 | 1 |
| 17_2506826 | 2 | 0 | 4 | 0.25 | 5 | 0 | 4 | 0 | 15 | 0.0667 | 4 | 0 | NA | NA | 5 | 0 | 5 | 0.2 | 14 | 0.0714 |
| 17_2506849 | 1 | 1 | 4 | 0.75 | 2 | 0.5 | 4 | 0.5 | 11 | 0.6364 | 2 | 0.5 | NA | NA | 4 | 1 | 3 | 1 | 9 | 0.8889 |
| 17_2506862 | 2 | 1 | 4 | 1 | 3 | 1 | 4 | 1 | 13 | 1 | 3 | 1 | NA | NA | 4 | 0.75 | 2 | 1 | 9 | 0.8889 |
| 17_2506870 | 2 | 0.5 | 4 | 0.75 | 3 | 0.3333 | 4 | 0 | 13 | 0.3846 | 3 | 0.3333 | NA | NA | 4 | 0.5 | 2 | 0.5 | 9 | 0.4444 |
| 17_2506876 | 2 | 0.5 | 3 | 1 | 3 | 0.6667 | 4 | 0.5 | 12 | 0.6667 | 3 | 1 | NA | NA | 3 | 0.6667 | 3 | 0.3333 | 9 | 0.6667 |
| 17_2506879 | 2 | 1 | 4 | 1 | 3 | 1 | 4 | 0.75 | 13 | 0.9231 | 3 | 0.6667 | NA | NA | 3 | 0.6667 | 3 | 0.6667 | 9 | 0.6667 |
| 17_2506899 | 2 | 0 | 4 | 0 | 3 | 0 | 3 | 0.3333 | 12 | 0.0833 | 3 | 0 | NA | NA | 2 | 0 | 2 | 0.5 | 7 | 0.1429 |
| 17_2506930 | 1 | 0 | NA | NA | 2 | 0.5 | NA | NA | 3 | 0.3333 | 1 | 0 | 3 | 0.3333 | NA | NA | NA | NA | 4 | 0.25 |
| 17_2506939 | 2 | 0 | 1 | 0 | 2 | 0.5 | NA | NA | 5 | 0.2 | 1 | 0 | 2 | 0.5 | NA | NA | NA | NA | 3 | 0.3333 |
| 17_2508329 | 26 | 0.0769 | 17 | 0.1765 | 17 | 0.2353 | 20 | 0 | 80 | 0.1125 | 18 | 0.1111 | 16 | 0.0625 | 14 | 0.2143 | 14 | 0.2143 | 62 | 0.1452 |
| 17_2510172 | 7 | 0.2857 | 6 | 0.1667 | 8 | 0.25 | 6 | 0 | 27 | 0.1852 | 3 | 0.6667 | 4 | 0 | 6 | 0.1667 | 8 | 0.375 | 21 | 0.2857 |
| 17_2510203 | 7 | 0.2857 | 5 | 0.6 | 8 | 0.75 | 5 | 0.6 | 25 | 0.56 | 4 | 0.75 | 4 | 0.25 | 4 | 0.25 | 8 | 0.5 | 20 | 0.45 |
| 17_2510207 | 4 | 0 | 5 | 0.2 | 8 | 0.375 | 6 | 0 | 23 | 0.1739 | 3 | 0.3333 | 4 | 0 | 7 | 0.2857 | 7 | 0 | 21 | 0.1429 |
| 17_2510208 | 7 | 0.4286 | 6 | 0.5 | 7 | 0.8571 | 5 | 0.2 | 25 | 0.52 | 4 | 0 | 4 | 1 | 7 | 0.4286 | 7 | 0.7143 | 22 | 0.5455 |
| 17_2510210 | 7 | 0.1429 | 6 | 0.1667 | 9 | 0 | 7 | 0.2857 | 29 | 0.1379 | 4 | 0.25 | 5 | 0 | 7 | 0.1429 | 6 | 0.1667 | 22 | 0.1364 |
| 17_2511049 | 6 | 0.5 | 5 | 0.2 | 2 | 0 | 5 | 0 | 18 | 0.2222 | 3 | 0.6667 | 12 | 0.0833 | 7 | 0.1429 | 4 | 0 | 26 | 0.1538 |
| 17_2511057 | 5 | 0.8 | 5 | 1 | 4 | 1 | 5 | 0.6 | 19 | 0.8421 | 4 | 1 | 9 | 0.8889 | 6 | 1 | 4 | 1 | 23 | 0.9565 |
| 17_2511074 | 5 | 0.8 | 5 | 1 | 3 | 1 | 7 | 0.5714 | 20 | 0.8 | 4 | 0.25 | 9 | 0.4444 | 6 | 0.6667 | 6 | 1 | 25 | 0.6 |
| 17_2511078 | 6 | 0.3333 | 5 | 0.8 | 4 | 0.5 | 6 | 0 | 21 | 0.381 | 5 | 0.4 | 11 | 0.5455 | 7 | 0.7143 | 7 | 0.2857 | 30 | 0.5 |
| 17_2511088 | 7 | 0.5714 | 5 | 0.6 | 4 | 0.75 | 6 | 0.6667 | 22 | 0.6364 | 6 | 0.5 | 10 | 0.6 | 6 | 0.8333 | 6 | 0.5 | 28 | 0.6071 |
| 17_2511090 | 6 | 0 | 6 | 0 | 4 | 0.5 | 7 | 0 | 23 | 0.087 | 5 | 0.2 | 10 | 0 | 8 | 0 | 6 | 0.5 | 29 | 0.1379 |
| 17_2511114 | 1 | 0 | 5 | 0.2 | 4 | 0.25 | NA | NA | 10 | 0.2 | NA | NA | 2 | 0.5 | 1 | 1 | NA | NA | 3 | 0.6667 |
| 17_2511115 | 1 | 0 | 5 | 0.4 | 3 | 0.6667 | NA | NA | 9 | 0.4444 | NA | NA | 2 | 0 | 1 | 1 | NA | NA | 3 | 0.3333 |
| 17_2511135 | 1 | 0 | 3 | 1 | 4 | 0.75 | NA | NA | 8 | 0.75 | NA | NA | 2 | 1 | 1 | 1 | NA | NA | 3 | 1 |
| 17_2511136 | 1 | 1 | 5 | 1 | 4 | 0.75 | NA | NA | 10 | 0.9 | NA | NA | 2 | 1 | 1 | 1 | NA | NA | 3 | 1 |
| 17_2511144 | 1 | 0 | 5 | 0.2 | 3 | 0.3333 | NA | NA | 9 | 0.2222 | NA | NA | 2 | 0.5 | 1 | 1 | NA | NA | 3 | 0.6667 |
| 17_2511150 | 1 | 1 | 5 | 0.4 | 4 | 0.25 | NA | NA | 10 | 0.4 | NA | NA | 2 | 0 | 1 | 1 | NA | NA | 3 | 0.3333 |
| 17_2511164 | 1 | 0 | 3 | 0 | 2 | 0.5 | NA | NA | 6 | 0.1667 | NA | NA | 1 | 1 | 1 | 0 | NA | NA | 2 | 0.5 |
| 17_2511452 | 3 | 1 | 2 | 1 | 4 | 1 | 3 | 0.6667 | 12 | 0.9167 | 1 | 1 | 2 | 1 | 1 | 1 | 1 | 1 | 5 | 1 |
| 17_2511464 | 7 | 0.1429 | 8 | 0 | 12 | 0.1667 | 12 | 0.0833 | 39 | 0.1026 | 4 | 0 | 7 | 0.2857 | 7 | 0.1429 | 4 | 0 | 22 | 0.1364 |
| 17_2511498 | 8 | 0.375 | 10 | 0.5 | 17 | 0.6471 | 14 | 0.5 | 49 | 0.5306 | 7 | 0.2857 | 8 | 0.75 | 11 | 0.4545 | 8 | 0.375 | 34 | 0.4706 |
| 17_2511500 | 9 | 0.5556 | 10 | 0.3 | 12 | 0.4167 | 16 | 0.5 | 47 | 0.4468 | 3 | 0 | 7 | 0.2857 | 10 | 0.6 | 4 | 0.5 | 24 | 0.4167 |
| 17_2511506 | 8 | 0.5 | 7 | 0 | 16 | 0.1875 | 13 | 0.2308 | 44 | 0.2273 | 6 | 0 | 8 | 0.125 | 11 | 0.3636 | 4 | 0 | 29 | 0.1724 |
| 17_2511510 | 10 | 0.6 | 10 | 0.6 | 15 | 0.6667 | 14 | 0.5 | 49 | 0.5918 | 7 | 0.2857 | 8 | 0.625 | 12 | 0.4167 | 8 | 0.875 | 35 | 0.5429 |
| 17_2511627 | 3 | 0.3333 | 10 | 0 | 4 | 0 | 10 | 0.1 | 27 | 0.0741 | 7 | 0 | 8 | 0 | 7 | 0.1429 | 9 | 0.1111 | 31 | 0.0645 |
| 17_2511688 | 31 | 0.0645 | 28 | 0.0714 | 29 | 0.0345 | 21 | 0.0952 | 109 | 0.0642 | 22 | 0.0455 | 28 | 0.1071 | 14 | 0 | 24 | 0.125 | 88 | 0.0795 |
| 17_2511720 | 30 | 0.1 | 26 | 0.1154 | 24 | 0.0833 | 26 | 0.0769 | 106 | 0.0943 | 18 | 0.0556 | 22 | 0.0909 | 18 | 0.2222 | 20 | 0.05 | 78 | 0.1026 |
| 17_2511762 | 26 | 0.1154 | 29 | 0.0345 | 24 | 0.0833 | 23 | 0.2174 | 102 | 0.1078 | 22 | 0.1364 | 23 | 0.1304 | 20 | 0.05 | 21 | 0.0952 | 86 | 0.1047 |
| 17_2511788 | 21 | 0.2381 | 24 | 0.0833 | 18 | 0.1111 | 20 | 0 | 83 | 0.1084 | 18 | 0.1667 | 20 | 0.1 | 16 | 0.1875 | 17 | 0.1765 | 71 | 0.1549 |
| 17_2511993 | 15 | 0 | 17 | 0.0588 | 14 | 0.0714 | 13 | 0.0769 | 59 | 0.0508 | 11 | 0 | 13 | 0.1538 | 8 | 0.125 | 13 | 0 | 45 | 0.0667 |
| 17_2512415 | 8 | 0.125 | 10 | 0.1 | 10 | 0.1 | 8 | 0 | 36 | 0.0833 | 7 | 0.1429 | 7 | 0.1429 | 3 | 0 | 3 | 0.3333 | 20 | 0.15 |
| 17_2518042 | 5 | 0 | 5 | 0 | 1 | 1 | 1 | 0 | 12 | 0.0833 | 3 | 0 | 3 | 0 | 2 | 0 | 3 | 0.3333 | 11 | 0.0909 |
| 17_2518088 | 4 | 0.25 | 3 | 0 | 1 | 0 | 1 | 0 | 9 | 0.1111 | 3 | 0.3333 | 2 | 0 | 1 | 1 | 1 | 0 | 7 | 0.2857 |
| 17_2518109 | 4 | 0 | 4 | 0.25 | 1 | 0 | 1 | 0 | 10 | 0.1 | 2 | 0 | 2 | 0 | 1 | 1 | 2 | 0 | 7 | 0.1429 |
| 17_2518130 | 4 | 0.25 | 4 | 0.25 | NA | NA | 1 | 0 | 9 | 0.2222 | 2 | 0.5 | 2 | 0.5 | NA | NA | 1 | 0 | 5 | 0.4 |
| 17_2518139 | 3 | 0.3333 | 4 | 0.25 | 1 | 0 | 1 | 0 | 9 | 0.2222 | 3 | 0.3333 | 2 | 0 | NA | NA | 2 | 0.5 | 7 | 0.2857 |
| 17_2518175 | 3 | 0.3333 | 4 | 0 | 1 | 0 | 1 | 0 | 9 | 0.1111 | 2 | 0 | 2 | 0 | NA | NA | 2 | 0.5 | 6 | 0.1667 |
| 17_2518355 | 2 | 0.5 | 2 | 1 | 1 | 0 | 1 | 0 | 6 | 0.5 | 3 | 0.3333 | 1 | 0 | NA | NA | NA | NA | 4 | 0.25 |
| 17_2518359 | 2 | 1 | 2 | 1 | 1 | 0 | 1 | 0 | 6 | 0.6667 | 3 | 0.6667 | 1 | 1 | NA | NA | NA | NA | 4 | 0.75 |
| 17_2518374 | 3 | 0.3333 | 2 | 0.5 | NA | NA | 1 | 0 | 6 | 0.3333 | 3 | 0.3333 | NA | NA | NA | NA | NA | NA | 3 | 0.3333 |
| 17_2518375 | 2 | 1 | 1 | 1 | 1 | 1 | NA | NA | 4 | 1 | 2 | 0.5 | NA | NA | NA | NA | NA | NA | 2 | 0.5 |
| 17_2520181 | 1 | 1 | NA | NA | NA | NA | 1 | 1 | 2 | 1 | 2 | 1 | NA | NA | NA | NA | NA | NA | 2 | 1 |
| 17_2520193 | 1 | 0 | NA | NA | NA | NA | 1 | 1 | 2 | 0.5 | 2 | 0.5 | NA | NA | NA | NA | NA | NA | 2 | 0.5 |
| 17_2520194 | 1 | 1 | NA | NA | NA | NA | 1 | 1 | 2 | 1 | 3 | 1 | NA | NA | NA | NA | NA | NA | 3 | 1 |

| | | | | | | | | | | | | | | | | | | | | |
|---|---|---|---|---|---|---|---|---|---|---|---|---|---|---|---|---|---|---|---|---|
| 17_2520217 | 1 | 1 | NA | NA | NA | NA | 1 | 1 | 2 | 1 | 2 | 1 | NA | NA | NA | NA | NA | NA | 2 | 1 |
| 17_2527683 | 4 | 0.25 | 3 | 0 | 3 | 0 | 1 | 0 | 11 | 0.0909 | 1 | 1 | 1 | 0 | 4 | 0 | 1 | 0 | 7 | 0.1429 |
| 17_2527718 | 4 | 1 | 5 | 0.8 | 3 | 0.6667 | NA | NA | 12 | 0.8333 | NA | NA | 1 | 0 | 4 | 0.75 | 1 | 1 | 6 | 0.6667 |
| 17_2527722 | 4 | 0.5 | 4 | 0.5 | 3 | 0.6667 | 1 | 0 | 12 | 0.5 | NA | NA | 1 | 0 | 4 | 0.5 | 1 | 1 | 6 | 0.5 |
| 17_2527729 | 3 | 0 | 5 | 0 | 2 | 0 | 1 | 1 | 11 | 0.0909 | 1 | 0 | 1 | 0 | 4 | 0.25 | NA | NA | 6 | 0.1667 |
| 17_2529930 | 4 | 0 | 3 | 0.6667 | 3 | 0.3333 | NA | NA | 10 | 0.3 | NA | NA | 1 | 0 | 3 | 0.3333 | NA | NA | 4 | 0.25 |
| 17_2529931 | 4 | 0.25 | 3 | 0 | 3 | 0 | 1 | 0 | 11 | 0.0909 | 1 | 0 | 1 | 0 | 4 | 0.25 | NA | NA | 6 | 0.1667 |
| 17_2529933 | 3 | 1 | 4 | 0.75 | 3 | 0.6667 | 1 | 1 | 11 | 0.8182 | 1 | 1 | 1 | 0 | 2 | 0.5 | NA | NA | 4 | 0.5 |
| 17_2529936 | 2 | 0.5 | 4 | 0.25 | 1 | 1 | 1 | 0 | 8 | 0.375 | 1 | 0 | 1 | 0 | 3 | 0.3333 | NA | NA | 5 | 0.2 |
| 17_2529955 | 4 | 0.25 | 4 | 0.25 | 2 | 0.5 | 1 | 0 | 11 | 0.2727 | 1 | 1 | NA | NA | 3 | 0 | NA | NA | 4 | 0.25 |
| 17_2529955 | 4 | 0.75 | 4 | 0.75 | 3 | 0.3333 | 1 | 0 | 12 | 0.5833 | 1 | 1 | NA | NA | 3 | 1 | NA | NA | 4 | 1 |
| 17_2529969 | 2 | 0 | 3 | 0.3333 | 3 | 0.3333 | NA | NA | 8 | 0.25 | NA | NA | NA | NA | 2 | 0.5 | NA | NA | 2 | 0.5 |
| 17_2529987 | 2 | 0 | 4 | 0.25 | 2 | 0 | NA | NA | 8 | 0.125 | NA | NA | NA | NA | 1 | 1 | NA | NA | 1 | 1 |
| 17_2531778 | NA | NA | 3 | 0.3333 | 1 | 0 | 5 | 0.2 | 9 | 0.2222 | 3 | 0.6667 | 2 | 1 | 1 | 0 | 2 | 1 | 8 | 0.75 |
| 17_2531789 | 1 | 1 | 3 | 0 | 1 | 0 | 4 | 0.25 | 9 | 0.2222 | 2 | 0.5 | 2 | 0.5 | 3 | 1 | 2 | 0.5 | 9 | 0.6667 |
| 17_2531822 | 1 | 0 | 1 | 1 | 1 | 0 | 4 | 0.75 | 7 | 0.5714 | 2 | 0.5 | 1 | 0 | 3 | 0.6667 | 2 | 1 | 8 | 0.625 |
| 17_2531826 | 1 | 0 | 2 | 0 | 2 | 0.5 | 5 | 0.6 | 10 | 0.4 | 4 | 0.25 | 2 | 1 | 4 | 0 | 2 | 0 | 12 | 0.25 |
| 17_2531837 | 1 | 0 | 2 | 0 | 2 | 0 | 6 | 0.6667 | 11 | 0.3636 | 4 | 1 | 2 | 0 | 4 | 0 | 1 | 0 | 11 | 0.3636 |
| 17_2531869 | 1 | 0 | 3 | 0 | NA | NA | 1 | 1 | 5 | 0.2 | NA | NA | 2 | 0.5 | 1 | 1 | 3 | 0 | 6 | 0.3333 |
| 17_2531902 | NA | NA | 2 | 0.5 | NA | NA | 2 | 0 | 4 | 0.25 | NA | NA | 2 | 0.5 | 2 | 0 | 1 | 1 | 5 | 0.4 |
| 17_2538001 | 4 | 0 | 6 | 0.1667 | 3 | 0.3333 | 3 | 0 | 16 | 0.125 | 2 | 0 | 3 | 0 | 7 | 0.1429 | 4 | 0.25 | 16 | 0.125 |
| 17_2538005 | 5 | 0.4 | 6 | 0.5 | 2 | 0 | 3 | 0 | 16 | 0.3125 | 3 | 0.3333 | 3 | 0.3333 | 7 | 0.2857 | 4 | 0.75 | 17 | 0.4118 |
| 17_2538036 | 6 | 0 | 6 | 0.1667 | 3 | 0 | 3 | 0.3333 | 18 | 0.1111 | 3 | 0 | 3 | 0 | 8 | 0 | 4 | 0.25 | 18 | 0.0556 |
| 17_2538070 | 5 | 0.2 | 6 | 0 | 4 | 0.25 | 3 | 0 | 18 | 0.1111 | 3 | 0.3333 | 4 | 0 | 7 | 0 | 3 | 0 | 17 | 0.0588 |
| 17_2538091 | 6 | 0.1667 | 5 | 0 | 3 | 0.3333 | 3 | 0 | 17 | 0.1176 | 4 | 0.25 | 5 | 0 | 8 | 0.25 | 4 | 0 | 21 | 0.1429 |
| 17_2538094 | 7 | 0.1429 | 6 | 0.3333 | 4 | 0 | 3 | 0.3333 | 20 | 0.2 | 4 | 0.5 | 5 | 0 | 9 | 0.2222 | 4 | 0.5 | 22 | 0.2727 |
| 17_2540385 | 4 | 0.25 | 7 | 0.7143 | 3 | 0.6667 | 4 | 0.25 | 18 | 0.5 | 5 | 0 | 4 | 0.5 | 4 | 0.25 | 6 | 0.3333 | 19 | 0.2632 |
| 17_2540397 | 5 | 1 | 7 | 1 | 2 | 1 | 4 | 1 | 18 | 1 | 6 | 1 | 4 | 1 | 5 | 1 | 7 | 1 | 22 | 1 |
| 17_2540409 | 5 | 1 | 9 | 1 | 3 | 1 | 5 | 1 | 22 | 1 | 5 | 0.8 | 5 | 1 | 5 | 1 | 7 | 1 | 22 | 0.9545 |
| 17_2540413 | 4 | 1 | 8 | 1 | 3 | 1 | 4 | 0.75 | 19 | 0.9474 | 3 | 1 | 2 | 1 | 4 | 1 | 6 | 0.8333 | 15 | 0.9333 |
| 17_2540417 | 5 | 1 | 9 | 1 | 3 | 1 | 4 | 1 | 21 | 1 | 6 | 1 | 6 | 1 | 5 | 1 | 7 | 1 | 24 | 1 |
| 17_2540420 | 5 | 0.2 | 6 | 0.3333 | 2 | 0 | 4 | 0 | 17 | 0.1765 | 5 | 0.6 | 5 | 0 | 4 | 0.25 | 6 | 0.1667 | 20 | 0.25 |
| 17_2540424 | 4 | 0.25 | 7 | 0.1429 | 4 | 0.25 | 3 | 0.3333 | 18 | 0.2222 | 6 | 0 | 5 | 0.2 | 5 | 0.2 | 5 | 0.4 | 21 | 0.1905 |
| 17_2540427 | 4 | 0.75 | 6 | 1 | 3 | 1 | 4 | 0.75 | 17 | 0.8824 | 4 | 1 | 4 | 0.75 | 5 | 0.8 | 6 | 1 | 19 | 0.8947 |
| 17_2540441 | 5 | 0.4 | 10 | 0.3 | 5 | 0.4 | 4 | 0.25 | 24 | 0.3333 | 4 | 0.25 | 6 | 0.6667 | 6 | 0.5 | 7 | 0.1429 | 23 | 0.3913 |
| 17_2540441 | 5 | 0 | 8 | 0.125 | 5 | 0.2 | 4 | 0.25 | 22 | 0.1364 | 5 | 0.4 | 5 | 0 | 5 | 0 | 7 | 0.2857 | 22 | 0.1818 |
| 17_2540447 | 4 | 0.25 | 9 | 0.1111 | 4 | 0.25 | 4 | 0.25 | 21 | 0.1905 | 5 | 0 | 6 | 0 | 5 | 0.4 | 6 | 0 | 22 | 0.0909 |
| 17_2540451 | 3 | 1 | 3 | 0.6667 | 4 | 0.75 | 2 | 1 | 12 | 0.8333 | 1 | 1 | 2 | 0.5 | 2 | 0.5 | 1 | 1 | 6 | 0.6667 |
| 17_2540460 | 2 | 1 | 1 | 0 | 5 | 0 | 1 | 1 | 9 | 0.3333 | 1 | 0 | NA | NA | 2 | 0 | 1 | 1 | 4 | 0.25 |
| 17_2540472 | 2 | 0 | 1 | 1 | 5 | 0 | 1 | 1 | 9 | 0.2222 | 1 | 1 | 1 | 0 | 3 | 0 | 1 | 0 | 6 | 0.1667 |
| 17_2540508 | 3 | 0.3333 | 1 | 0 | 6 | 0 | NA | NA | 10 | 0.1 | 2 | 0.5 | 1 | 0 | 1 | 0 | NA | NA | 4 | 0.25 |
| 17_2540617 | 1 | 0 | NA | NA | 1 | 1 | NA | NA | 2 | 0.5 | 1 | 1 | 1 | 0 | 2 | 0 | NA | NA | 4 | 0.25 |
| 17_2544881 | NA | NA | 1 | 1 | NA | NA | NA | NA | 1 | 1 | NA | NA | 1 | 1 | NA | NA | NA | NA | 1 | 1 |
| 17_2544886 | NA | NA | 1 | 1 | NA | NA | NA | NA | 1 | 1 | NA | NA | 1 | 1 | NA | NA | NA | NA | 1 | 1 |
| 17_2544903 | NA | NA | 1 | 1 | NA | NA | NA | NA | 1 | 1 | NA | NA | 1 | 1 | NA | NA | NA | NA | 1 | 1 |
| 17_2545084 | 3 | 0 | 6 | 0 | 3 | 0 | 7 | 0.1429 | 19 | 0.0526 | 3 | 0 | 4 | 0.25 | 3 | 0 | 7 | 0 | 17 | 0.0588 |
| 17_2545120 | 1 | 0 | 2 | 0.5 | NA | NA | NA | NA | 3 | 0.3333 | 1 | 0 | 1 | 1 | 1 | 0 | NA | NA | 3 | 0.3333 |
| 17_2545133 | 3 | 0 | 4 | 0 | 2 | 0 | 8 | 0.125 | 17 | 0.0588 | 1 | 0 | 4 | 0 | 3 | 0.3333 | 6 | 0 | 14 | 0.0714 |
| 17_2545164 | 4 | 0.5 | 6 | 0.8333 | 3 | 0.6667 | 7 | 0.5714 | 20 | 0.65 | 3 | 1 | 3 | 1 | 3 | 0.6667 | 8 | 0.625 | 17 | 0.7647 |
| 17_2545166 | 3 | 0 | 6 | 0.1667 | 3 | 0 | 6 | 0 | 18 | 0.0556 | 2 | 0 | 2 | 0 | 3 | 0.3333 | 4 | 0 | 11 | 0.0909 |
| 17_2545168 | 1 | 1 | 2 | 0.5 | 1 | 0 | 1 | 1 | 5 | 0.6 | 1 | 1 | 1 | 0 | 1 | 1 | NA | NA | 3 | 0.6667 |
| 17_2545168 | 4 | 0.25 | 6 | 0.3333 | 3 | 0 | 8 | 0.5 | 21 | 0.3333 | 4 | 0.25 | 4 | 0 | 4 | 0 | 7 | 0.1429 | 19 | 0.1053 |
| 17_2545170 | 1 | 1 | 1 | 1 | 1 | 1 | 1 | 1 | 4 | 1 | 1 | 1 | 1 | 1 | 1 | 1 | NA | NA | 3 | 1 |
| 17_2545181 | 4 | 0 | 5 | 0.2 | 3 | 0 | 8 | 0.125 | 20 | 0.1 | 3 | 0 | 5 | 0.2 | 4 | 0 | 8 | 0 | 20 | 0.05 |
| 17_2545182 | 4 | 0.5 | 6 | 0.3333 | 3 | 0.6667 | 6 | 0.5 | 19 | 0.4737 | 3 | 0 | 5 | 0.4 | 4 | 0.5 | 8 | 0.25 | 20 | 0.3 |
| 17_2545183 | 1 | 0 | 2 | 1 | 1 | 1 | 1 | 0 | 5 | 0.6 | 1 | 1 | 1 | 0 | 1 | 1 | NA | NA | 3 | 0.6667 |
| 17_2545189 | NA | NA | 2 | 0.5 | 1 | 0 | 1 | 1 | 4 | 0.5 | 1 | 1 | NA | NA | 1 | 1 | NA | NA | 2 | 1 |
| 17_2545189 | 4 | 1 | 3 | 0.6667 | 3 | 1 | 7 | 1 | 17 | 0.9412 | 4 | 0.75 | 4 | 1 | 4 | 1 | 6 | 1 | 18 | 0.9444 |
| 17_2545190 | 1 | 0 | 2 | 1 | 1 | 0 | 1 | 1 | 5 | 0.6 | 1 | 1 | 1 | 0 | 1 | 1 | NA | NA | 3 | 0.6667 |
| 17_2545196 | 4 | 0.25 | 5 | 0.4 | 2 | 0 | 8 | 0.25 | 19 | 0.2632 | 4 | 0.75 | 4 | 0.75 | 2 | 0 | 9 | 0.3333 | 19 | 0.4737 |
| 17_2545198 | NA | NA | 1 | 0 | 1 | 1 | 1 | 1 | 3 | 0.6667 | 1 | 1 | NA | NA | 1 | 1 | NA | NA | 2 | 1 |
| 17_2545206 | 4 | 1 | 5 | 1 | 3 | 1 | 7 | 0.8571 | 19 | 0.9474 | 3 | 1 | 4 | 1 | 4 | 1 | 9 | 0.7778 | 20 | 0.9 |
| 17_2545210 | 4 | 0.5 | 5 | 0.8 | 4 | 0.75 | 6 | 0.6667 | 19 | 0.6842 | 2 | 0.5 | 4 | 1 | 4 | 1 | 8 | 0.875 | 18 | 0.8889 |
| 17_2545211 | 1 | 1 | 5 | 1 | 4 | 1 | 4 | 1 | 14 | 1 | 1 | 1 | 1 | 1 | 4 | 1 | 5 | 1 | 11 | 1 |
| 17_2545229 | 4 | 1 | 3 | 1 | 3 | 0.6667 | 7 | 0.2857 | 17 | 0.6471 | 2 | 1 | 5 | 1 | 4 | 0.75 | 8 | 1 | 19 | 0.9474 |

| | | | | | | | | | | | | | | | | | | | | |
|---|---|---|---|---|---|---|---|---|---|---|---|---|---|---|---|---|---|---|---|---|
| 17_2545237 | 4 | 0 | 8 | 0 | 4 | 0 | 9 | 0.2222 | 25 | 0.08 | 4 | 0.25 | 5 | 0 | 4 | 0.25 | 9 | 0 | 22 | 0.0909 |
| 17_2545258 | 3 | 0 | 6 | 0.1667 | 4 | 0.25 | 9 | 0.2222 | 22 | 0.1818 | 5 | 0 | 6 | 0.1667 | 4 | 0 | 6 | 0.1667 | 21 | 0.0952 |
| 17_2545346 | 11 | 0.2727 | 18 | 0.2778 | 13 | 0.3077 | 16 | 0.25 | 58 | 0.2759 | 17 | 0.2353 | 21 | 0.2857 | 19 | 0.2105 | 21 | 0.3333 | 78 | 0.2692 |
| 17_2545348 | 11 | 0.0909 | 16 | 0.0625 | 14 | 0.0714 | 17 | 0 | 58 | 0.0517 | 16 | 0 | 24 | 0.0833 | 19 | 0.1053 | 19 | 0.0526 | 78 | 0.0641 |
| 17_2545353 | 10 | 0.3 | 14 | 0.4286 | 13 | 0.1538 | 13 | 0.3846 | 50 | 0.32 | 15 | 0.2667 | 22 | 0.2727 | 12 | 0.3333 | 18 | 0.3889 | 67 | 0.3134 |
| 17_2545353 | NA | NA | 1 | 1 | NA | NA | 1 | 0 | 2 | 0.5 | NA | NA | 1 | 1 | NA | NA | NA | NA | 1 | 1 |
| 17_2546114 | 7 | 0.1429 | 7 | 0.2857 | 2 | 0 | 4 | 0.5 | 20 | 0.25 | 2 | 0.5 | 2 | 0 | 4 | 0.25 | 2 | 1 | 10 | 0.4 |
| 17_2546158 | 10 | 0.1 | 9 | 0.2222 | 3 | 0.3333 | 4 | 0.25 | 26 | 0.1923 | 3 | 0 | 5 | 0.2 | 7 | 0.1429 | 3 | 0.3333 | 18 | 0.1667 |
| 17_2546163 | 10 | 0.7 | 10 | 0.6 | 3 | 0.6667 | 4 | 0.5 | 27 | 0.6296 | 3 | 0.6667 | 5 | 0.2 | 4 | 0.5 | 3 | 0.3333 | 15 | 0.4 |
| 17_2546169 | 10 | 0.8 | 7 | 0.5714 | 3 | 1 | 4 | 0.75 | 24 | 0.75 | 3 | 0.6667 | 5 | 0.4 | 5 | 1 | 2 | 1 | 15 | 0.7333 |
| 17_2546171 | 10 | 0.6 | 10 | 0.3 | 3 | 0 | 4 | 0.5 | 27 | 0.4074 | 3 | 0.3333 | 6 | 0 | 7 | 0.4286 | 3 | 0.3333 | 19 | 0.2632 |
| 17_2546204 | 2 | 0 | 1 | 0 | 4 | 0 | 6 | 0.1667 | 13 | 0.0769 | 2 | 0 | 3 | 0.3333 | 2 | 0 | 4 | 0 | 11 | 0.0909 |
| 17_2546225 | 1 | 0 | 2 | 0 | 4 | 0 | 5 | 0.2 | 12 | 0.0833 | 2 | 0 | 3 | 0.3333 | 2 | 0 | 5 | 0 | 12 | 0.0833 |
| 17_2546229 | 2 | 0.5 | 2 | 1 | 4 | 0.75 | 5 | 0.4 | 13 | 0.6154 | 2 | 1 | 3 | 0.3333 | 2 | 0.5 | 5 | 0.4 | 12 | 0.5 |
| 17_2546235 | 2 | 0 | 2 | 0.5 | 3 | 0.3333 | 5 | 0.8 | 12 | 0.5 | 1 | 1 | 1 | 0 | 2 | 0 | 5 | 0.2 | 9 | 0.2222 |
| 17_2546237 | 1 | 0 | 1 | 0 | 3 | 0 | 6 | 0.3333 | 11 | 0.1818 | 2 | 0 | 3 | 0.3333 | 2 | 0 | 5 | 0 | 12 | 0.0833 |
| 17_2546249 | 2 | 0 | 2 | 0 | 4 | 0.5 | 4 | 0.25 | 12 | 0.25 | 2 | 0 | 3 | 0.3333 | 2 | 0 | 5 | 0.2 | 12 | 0.1667 |
| 17_2546256 | 2 | 0 | 2 | 0 | 4 | 0.5 | 4 | 0 | 12 | 0.1667 | 2 | 1 | 3 | 0.3333 | 2 | 0 | 6 | 0.3333 | 13 | 0.3846 |
| 17_2546261 | 2 | 0.5 | 2 | 0 | 4 | 0 | 6 | 0 | 14 | 0.0714 | 2 | 0.5 | 3 | 0 | 2 | 0 | 6 | 0 | 13 | 0.0769 |
| 17_2546291 | NA | NA | 2 | 0 | 3 | 0 | 5 | 0.6 | 10 | 0.3 | 2 | 1 | 2 | 0.5 | 1 | 0 | 4 | 0.25 | 9 | 0.4444 |
| 17_2546294 | 1 | 0 | 2 | 0 | 4 | 0.25 | 2 | 0 | 9 | 0.1111 | 2 | 0 | 2 | 0.5 | 2 | 0.5 | 4 | 0.5 | 10 | 0.4 |
| 17_2547989 | 23 | 0.2609 | 9 | 0.6667 | 21 | 0.2857 | 19 | 0.3158 | 72 | 0.3333 | 22 | 0.2727 | 19 | 0.4211 | 13 | 0.5385 | 11 | 0.3636 | 65 | 0.3846 |
| 17_2548005 | 25 | 0.08 | 14 | 0.1429 | 25 | 0.08 | 24 | 0.1667 | 88 | 0.1136 | 25 | 0.12 | 20 | 0.15 | 19 | 0 | 17 | 0.2353 | 81 | 0.1235 |
| 17_2548011 | 26 | 0.1154 | 14 | 0.0714 | 26 | 0.1923 | 28 | 0.1786 | 94 | 0.1489 | 28 | 0.1429 | 21 | 0.2381 | 22 | 0.2273 | 16 | 0.3125 | 87 | 0.2184 |
| 17_2548014 | 22 | 0.2727 | 9 | 0.3333 | 22 | 0.2727 | 28 | 0.1429 | 81 | 0.2346 | 20 | 0.2 | 22 | 0.0455 | 19 | 0.1053 | 14 | 0.5 | 75 | 0.1867 |
| 17_2548016 | 24 | 0.0417 | 14 | 0.2857 | 24 | 0.125 | 27 | 0.1111 | 89 | 0.1236 | 25 | 0.04 | 15 | 0 | 19 | 0 | 16 | 0.1875 | 75 | 0.0533 |
| 17_2548051 | 23 | 0.0435 | 10 | 0.1 | 17 | 0.0588 | 21 | 0.0952 | 71 | 0.0704 | 21 | 0.0952 | 14 | 0.0714 | 16 | 0.125 | 13 | 0.0769 | 64 | 0.0938 |
| 17_2548052 | 21 | 0.2857 | 13 | 0.3846 | 24 | 0.5 | 29 | 0.5172 | 87 | 0.4368 | 24 | 0.4167 | 19 | 0.2632 | 21 | 0.1905 | 16 | 0.5625 | 80 | 0.35 |
| 17_2548056 | 21 | 0.5238 | 12 | 0.5833 | 20 | 0.8 | 24 | 0.7083 | 77 | 0.6623 | 19 | 0.5263 | 18 | 0.5 | 19 | 0.4737 | 13 | 0.8462 | 69 | 0.5652 |
| 17_2548069 | 21 | 0.7619 | 11 | 0.8182 | 24 | 0.8333 | 24 | 0.7083 | 80 | 0.775 | 21 | 0.8095 | 21 | 0.8095 | 18 | 0.6111 | 14 | 0.9286 | 74 | 0.7838 |
| 17_2548090 | 27 | 0.1111 | 19 | 0.0526 | 30 | 0.0667 | 31 | 0.1613 | 107 | 0.1028 | 30 | 0.1667 | 31 | 0.0645 | 30 | 0.0667 | 25 | 0.08 | 116 | 0.0948 |
| 17_2548135 | 33 | 0.1515 | 18 | 0.1667 | 29 | 0.2759 | 28 | 0.2143 | 108 | 0.2037 | 27 | 0.1111 | 33 | 0.1212 | 26 | 0.1923 | 25 | 0.16 | 111 | 0.1441 |
| 17_2549182 | 2 | 0.5 | NA | NA | NA | NA | NA | NA | 2 | 0.5 | NA | NA | 1 | 0 | 1 | 0 | 1 | 1 | 3 | 0.3333 |
| 17_2549314 | 1 | 0 | 1 | 1 | 1 | 0 | NA | NA | 3 | 0.3333 | 1 | 0 | 2 | 0.5 | NA | NA | NA | NA | 3 | 0.3333 |
| 17_2559352 | 3 | 0 | 6 | 0 | 8 | 0.125 | 3 | 0 | 20 | 0.05 | 5 | 0 | 10 | 0.2 | 5 | 0.2 | 4 | 0 | 24 | 0.125 |
| 17_2559399 | 3 | 0 | 4 | 0.25 | 6 | 0.3333 | 1 | 0 | 14 | 0.2143 | 5 | 0.2 | 9 | 0.5556 | 4 | 0.25 | 4 | 0 | 22 | 0.3182 |
| 17_2559407 | 3 | 0 | 4 | 0.25 | 6 | 0.3333 | 2 | 0.5 | 15 | 0.2667 | 5 | 0 | 10 | 0.2 | 4 | 0 | 4 | 0 | 23 | 0.087 |
| 17_2559417 | 3 | 1 | 4 | 1 | 6 | 1 | 2 | 1 | 15 | 1 | 4 | 0.75 | 7 | 0.8571 | 3 | 0.6667 | 4 | 1 | 18 | 0.8333 |
| 17_2559421 | 1 | 0 | 1 | 1 | 4 | 0.25 | 1 | 1 | 7 | 0.4286 | 3 | 0.6667 | 2 | 0.5 | 1 | 0 | 1 | 0 | 7 | 0.4286 |
| 17_2559421 | 2 | 0.5 | 4 | 0.75 | 4 | 0.5 | 1 | 1 | 11 | 0.6364 | 3 | 0.6667 | 7 | 0.8571 | 3 | 0.6667 | 3 | 0.6667 | 16 | 0.75 |
| 17_2559428 | 1 | 1 | 4 | 1 | 6 | 1 | 2 | 1 | 13 | 1 | 5 | 1 | 8 | 0.75 | 5 | 1 | 3 | 1 | 21 | 0.9048 |
| 17_2559430 | 4 | 0.5 | 5 | 0.6 | 6 | 0.6667 | 2 | 0.5 | 17 | 0.5882 | 5 | 0.6 | 8 | 0.75 | 5 | 0.6 | 3 | 1 | 21 | 0.7143 |
| 17_2559431 | 2 | 1 | 5 | 0.2 | 4 | 0.5 | 1 | 1 | 12 | 0.5 | 5 | 0.2 | 7 | 0.5714 | 3 | 0.6667 | 3 | 0 | 18 | 0.3889 |
| 17_2559432 | 2 | 0.5 | 3 | 0.6667 | 6 | 0.5 | 1 | 0 | 12 | 0.5 | 4 | 1 | 6 | 0.6667 | 5 | 0.8 | 3 | 1 | 18 | 0.8333 |
| 17_2559435 | 3 | 0.3333 | 2 | 0.5 | 3 | 1 | 1 | 0 | 9 | 0.5556 | 3 | 0.3333 | 3 | 1 | 5 | 0.6 | 2 | 1 | 13 | 0.6923 |
| 17_2559440 | 3 | 0.3333 | 5 | 0.4 | 6 | 0.1667 | 2 | 0.5 | 16 | 0.3125 | 5 | 0.2 | 8 | 0.375 | 6 | 0 | 3 | 0 | 22 | 0.1818 |
| 17_2559444 | 3 | 0.6667 | 4 | 0.75 | 4 | 1 | 2 | 1 | 13 | 0.8462 | 5 | 0.8 | 6 | 0.6667 | 5 | 0.8 | 3 | 1 | 19 | 0.7895 |
| 17_2559447 | 3 | 0 | 4 | 0.25 | 6 | 0 | 2 | 0 | 15 | 0.0667 | 5 | 0 | 6 | 0 | 5 | 0 | 3 | 0.3333 | 19 | 0.0526 |
| 17_2559455 | 3 | 0.3333 | 4 | 0 | 6 | 0 | 2 | 0 | 15 | 0.0667 | 5 | 0 | 6 | 0 | 5 | 0.2 | 3 | 0 | 19 | 0.0526 |
| 17_2559467 | NA | NA | 2 | 0 | 5 | 1 | 1 | 1 | 8 | 0.75 | 2 | 0.5 | 4 | 0.75 | 2 | 0.5 | 3 | 0.6667 | 11 | 0.6364 |
| 17_2559471 | 1 | 1 | 4 | 0.5 | 5 | 0.6 | 1 | 1 | 11 | 0.6364 | 3 | 0.3333 | 6 | 0.5 | 2 | 1 | 3 | 0.3333 | 14 | 0.5 |
| 17_2559475 | 2 | 0.5 | 4 | 1 | 5 | 0.4 | 1 | 0 | 12 | 0.5833 | 3 | 0.3333 | 6 | 0.6667 | 4 | 0 | 3 | 1 | 16 | 0.5 |
| 17_2569120 | NA | NA | 1 | 0 | NA | NA | 1 | 1 | 2 | 0.5 | 2 | 1 | 1 | 1 | 2 | 1 | 1 | 1 | 6 | 1 |
| 17_2578577 | 8 | 0.375 | 1 | 0 | 6 | 0.5 | 6 | 0.6667 | 21 | 0.4762 | 7 | 0.2857 | 5 | 0.4 | 6 | 0.3333 | 9 | 0.4444 | 27 | 0.3704 |
| 17_2578584 | 9 | 1 | 3 | 1 | 6 | 1 | 8 | 1 | 26 | 1 | 8 | 0.875 | 4 | 0.5 | 5 | 1 | 9 | 0.6667 | 26 | 0.7692 |
| 17_2578585 | 8 | 1 | 2 | 1 | 6 | 1 | 7 | 1 | 23 | 1 | 6 | 1 | 5 | 1 | 4 | 1 | 8 | 1 | 23 | 1 |
| 17_2578589 | 8 | 1 | 2 | 0.5 | 5 | 1 | 8 | 1 | 23 | 0.9565 | 8 | 1 | 6 | 1 | 6 | 0.8333 | 9 | 1 | 29 | 0.9655 |
| 17_2578593 | 8 | 0.375 | 3 | 0.6667 | 5 | 0.6 | 7 | 0.2857 | 23 | 0.4348 | 8 | 0.375 | 5 | 0.6 | 5 | 0.4 | 9 | 0.6667 | 27 | 0.5185 |
| 17_2578598 | 7 | 1 | 1 | 1 | 4 | 1 | 7 | 1 | 19 | 1 | 7 | 0.5714 | 5 | 1 | 4 | 1 | 8 | 0.875 | 24 | 0.8333 |
| 17_2578607 | 6 | 0.1667 | 3 | 0 | 3 | 0 | 5 | 0 | 17 | 0.0588 | 6 | 0.1667 | 5 | 0.2 | 4 | 0.25 | 6 | 0.1667 | 21 | 0.1905 |
| 17_2578609 | 5 | 1 | 3 | 1 | 5 | 1 | 7 | 1 | 20 | 1 | 8 | 1 | 4 | 1 | 6 | 1 | 8 | 1 | 26 | 1 |
| 17_2578613 | 8 | 1 | 3 | 1 | 5 | 0.8 | 5 | 0.8 | 21 | 0.9048 | 7 | 1 | 5 | 1 | 6 | 1 | 8 | 0.875 | 26 | 0.9615 |
| 17_2578614 | 3 | 0.6667 | 3 | 0 | 5 | 0.2 | 6 | 0.1667 | 17 | 0.2353 | 5 | 0.6 | 3 | 0 | 5 | 0 | 8 | 0.25 | 21 | 0.2381 |
| 17_2578634 | 7 | 0.1429 | 2 | 0.5 | 5 | 0.2 | 6 | 0.1667 | 20 | 0.2 | 8 | 0.5 | 5 | 0.2 | 6 | 0.8333 | 9 | 0.4444 | 28 | 0.5 |
| 17_2578658 | 5 | 0.4 | 2 | 0.5 | 5 | 0 | 5 | 0.2 | 17 | 0.2353 | 7 | 0.4286 | 4 | 0.25 | 4 | 0 | 9 | 0.1111 | 24 | 0.2083 |

| | | | | | | | | | | | | | | | | | | | | |
|---|---|---|---|---|---|---|---|---|---|---|---|---|---|---|---|---|---|---|---|---|
| 17_2578667 | 7 | 0.5714 | 2 | 0 | 5 | 0.2 | 6 | 0.1667 | 20 | 0.3 | 6 | 0.3333 | 5 | 0.2 | 6 | 0.5 | 9 | 0.1111 | 26 | 0.2692 |
| 17_2578684 | 7 | 0.1429 | 2 | 0 | 5 | 0.2 | 5 | 0.4 | 19 | 0.2105 | 7 | 0.1429 | 5 | 0.6 | 4 | 0 | 7 | 0.1429 | 23 | 0.2174 |
| 17_2578691 | 7 | 0.4286 | 2 | 1 | 4 | 0.25 | 4 | 1 | 17 | 0.5882 | 7 | 0.1429 | 3 | 0 | 4 | 1 | 5 | 0.6 | 19 | 0.4211 |
| 17_2578693 | 6 | 0.1667 | 2 | 0 | 4 | 0 | 3 | 0 | 15 | 0.0667 | 6 | 0 | 2 | 0 | 2 | 0.5 | 6 | 0 | 16 | 0.0625 |
| 17_2578694 | 7 | 0.1429 | 2 | 0 | 5 | 0.2 | 4 | 0.25 | 18 | 0.1667 | 7 | 0.2857 | 4 | 0.25 | 5 | 0.2 | 7 | 0.1429 | 23 | 0.2174 |
| 17_2578926 | 3 | 0.3333 | 1 | 1 | 3 | 0.3333 | 5 | 0.4 | 12 | 0.4167 | 3 | 0 | 3 | 0.3333 | 2 | 0 | 4 | 0.25 | 12 | 0.1667 |
| 17_2578941 | 4 | 0.5 | 2 | 0.5 | 3 | 0.3333 | 5 | 0.6 | 14 | 0.5 | 6 | 0.5 | 2 | 0.5 | NA | NA | 5 | 0.4 | 13 | 0.4615 |
| 17_2578948 | 3 | 0.6667 | NA | NA | 2 | 0.5 | 3 | 0.6667 | 8 | 0.625 | 4 | 0.5 | 2 | 0.5 | NA | NA | 3 | 0 | 9 | 0.3333 |
| 17_2579310 | 5 | 0.8 | 3 | 0.6667 | 3 | 0.6667 | 5 | 0.6 | 16 | 0.6875 | 1 | 0 | 4 | 1 | 4 | 0.75 | 4 | 1 | 13 | 0.8462 |
| 17_2579312 | 5 | 0 | 3 | 0 | 3 | 0.3333 | 4 | 0 | 15 | 0.0667 | NA | NA | 6 | 0.1667 | 4 | 0.25 | 2 | 0 | 12 | 0.1667 |
| 17_2579321 | 5 | 0.2 | 2 | 0.5 | 3 | 0.3333 | 4 | 0 | 14 | 0.2143 | NA | NA | 4 | 0.75 | 4 | 0 | 4 | 0.25 | 12 | 0.3333 |
| 17_2579325 | 5 | 0.6 | 3 | 0 | 2 | 0 | 5 | 0.4 | 15 | 0.3333 | 1 | 0 | 5 | 0.2 | 3 | 0.6667 | 2 | 0 | 11 | 0.2727 |
| 17_2579330 | 5 | 0.4 | 2 | 0.5 | 3 | 0.3333 | 4 | 0.25 | 14 | 0.3571 | 1 | 0 | 3 | 1 | 1 | 1 | 4 | 0.5 | 9 | 0.6667 |
| 17_2579335 | 5 | 0 | 3 | 0 | 4 | 0.5 | 5 | 0.2 | 17 | 0.1765 | NA | NA | 6 | 0.5 | 4 | 0 | 4 | 0 | 14 | 0.2143 |
| 17_2579358 | 4 | 0.25 | 3 | 0.3333 | 2 | 0.5 | 5 | 0.2 | 14 | 0.2857 | NA | NA | 4 | 0 | 3 | 0.3333 | 5 | 0.6 | 12 | 0.3333 |
| 17_2579359 | 5 | 0.4 | 3 | 0.3333 | 2 | 0 | 4 | 0.75 | 14 | 0.4286 | NA | NA | 3 | 1 | 3 | 0.3333 | 4 | 0.75 | 10 | 0.7 |
| 17_2579363 | 5 | 0.8 | 5 | 0.8 | 2 | 1 | 4 | 1 | 16 | 0.875 | NA | NA | 6 | 1 | 3 | 1 | 4 | 1 | 13 | 1 |
| 17_2579372 | 6 | 0.1667 | 4 | 0.5 | 4 | 0.25 | 4 | 0.25 | 18 | 0.2778 | NA | NA | 5 | 0.2 | 4 | 0.25 | 6 | 0.3333 | 15 | 0.2667 |
| 17_2579375 | 5 | 0.8 | 5 | 0.6 | 3 | 1 | 5 | 0.8 | 18 | 0.7778 | 1 | 1 | 6 | 1 | 1 | 1 | 6 | 1 | 14 | 1 |
| 17_2579384 | 4 | 0.25 | 5 | 0.6 | 4 | 0.5 | 5 | 0.8 | 18 | 0.5556 | 1 | 1 | 4 | 1 | 3 | 0.6667 | 4 | 0.75 | 12 | 0.8333 |
| 17_2579388 | 5 | 0.2 | 5 | 0.2 | 4 | 0.25 | 5 | 0.2 | 19 | 0.2105 | 1 | 0 | 7 | 0.1429 | 4 | 0 | 5 | 0.2 | 17 | 0.1176 |
| 17_2580467 | 1 | 0 | 2 | 0.5 | 3 | 0.3333 | NA | NA | 6 | 0.3333 | 1 | 0 | 1 | 1 | NA | NA | NA | NA | 2 | 0.5 |
| 17_2580469 | 2 | 0 | 1 | 1 | 2 | 0.5 | NA | NA | 5 | 0.4 | 1 | 0 | 1 | 0 | 1 | 0 | 1 | 1 | 4 | 0.25 |
| 17_2588117 | NA | NA | 1 | 1 | NA | NA | NA | NA | 1 | 1 | NA | NA | NA | NA | 1 | 1 | 1 | 1 | 2 | 1 |
| 17_2588163 | 2 | 0 | 4 | 0.25 | 4 | 0.25 | 1 | 0 | 11 | 0.1818 | NA | NA | 3 | 0.3333 | 3 | 0.6667 | 2 | 0 | 8 | 0.375 |
| 17_2588168 | 2 | 0 | 4 | 0 | 5 | 0.2 | 1 | 0 | 12 | 0.0833 | 1 | 0 | 3 | 0 | 5 | 0.2 | 2 | 0 | 11 | 0.0909 |
| 17_2588169 | 1 | 0 | 4 | 0.25 | 4 | 0 | 1 | 0 | 10 | 0.1 | 1 | 1 | 3 | 0.3333 | 5 | 0.2 | 2 | 0 | 11 | 0.2727 |
| 17_2593804 | 17 | 0.1765 | 22 | 0.0455 | 15 | 0.0667 | 25 | 0.12 | 79 | 0.1013 | 18 | 0.0556 | 17 | 0.0588 | 12 | 0.1667 | 12 | 0.0833 | 59 | 0.0847 |
| 17_2593807 | 1 | 0 | NA | NA | NA | NA | 1 | 1 | 2 | 0.5 | NA | NA | NA | NA | NA | NA | 1 | 1 | 1 | 1 |
| 17_2594742 | 1 | 0 | 1 | 1 | NA | NA | NA | NA | 2 | 0.5 | NA | NA | NA | NA | 1 | 1 | NA | NA | 1 | 1 |
| 17_2594742 | 1 | 0 | 1 | 1 | NA | NA | NA | NA | 2 | 0.5 | NA | NA | NA | NA | 1 | 1 | NA | NA | 1 | 1 |
| 17_2594959 | 5 | 1 | 5 | 1 | 10 | 1 | 4 | 1 | 24 | 1 | 4 | 0.5 | 5 | 1 | 14 | 0.8571 | 8 | 1 | 31 | 0.871 |
| 17_2594968 | 10 | 0.6 | 6 | 0.5 | 10 | 0.5 | 8 | 0.75 | 34 | 0.5882 | 6 | 0 | 6 | 0.3333 | 10 | 0.6 | 11 | 0.3636 | 33 | 0.3636 |
| 17_2594973 | 8 | 0.875 | 6 | 1 | 12 | 1 | 7 | 1 | 33 | 0.9697 | 7 | 0.8571 | 7 | 1 | 15 | 1 | 10 | 0.9 | 39 | 0.9487 |
| 17_2594974 | 9 | 0.3333 | 5 | 0.6 | 9 | 0.4444 | 5 | 0 | 28 | 0.3571 | 6 | 0.3333 | 5 | 0.2 | 14 | 0.3571 | 10 | 0.3 | 35 | 0.3143 |
| 17_2594981 | 8 | 0 | 7 | 0.2857 | 12 | 0.1667 | 8 | 0.25 | 35 | 0.1714 | 8 | 0.125 | 3 | 0 | 11 | 0.1818 | 10 | 0.2 | 32 | 0.1562 |
| 17_2595001 | 9 | 0.1111 | 6 | 0.1667 | 12 | 0.0833 | 9 | 0.2222 | 36 | 0.1389 | 6 | 0.1667 | 7 | 0.5714 | 13 | 0.2308 | 11 | 0.1818 | 37 | 0.2703 |
| 17_2595003 | 8 | 0.75 | 6 | 0.8333 | 11 | 0.8182 | 8 | 0.875 | 33 | 0.8182 | 7 | 0.7143 | 6 | 1 | 15 | 0.8667 | 10 | 0.9 | 38 | 0.8684 |
| 17_2595010 | 8 | 0.625 | 6 | 0.6667 | 10 | 0.7 | 8 | 0.625 | 32 | 0.6562 | 10 | 0.7 | 6 | 0.3333 | 13 | 0.4615 | 11 | 0.4545 | 40 | 0.5 |
| 17_2599939 | 1 | 1 | 3 | 0 | 1 | 0 | 2 | 0 | 7 | 0.1429 | 3 | 0 | 3 | 0.3333 | 1 | 0 | 2 | 0.5 | 9 | 0.2222 |
| 17_2600106 | 2 | 0 | 4 | 0.25 | 2 | 0 | 3 | 0 | 11 | 0.0909 | 4 | 0 | 2 | 0.5 | 2 | 0 | 2 | 0 | 10 | 0.1 |
| 17_2600114 | 1 | 0 | 3 | 0.3333 | 2 | 0.5 | 4 | 0.5 | 10 | 0.4 | 2 | 0.5 | 1 | 1 | NA | NA | 2 | 1 | 5 | 0.8 |
| 17_2600126 | 1 | 0 | 3 | 0 | 2 | 0.5 | 3 | 0 | 9 | 0.1111 | 3 | 0 | 2 | 0.5 | 2 | 0 | 2 | 0 | 9 | 0.1111 |
| 17_2600128 | 2 | 0 | 3 | 0.6667 | 2 | 1 | 3 | 0.3333 | 10 | 0.5 | 3 | 0.6667 | 2 | 1 | 1 | 0 | 2 | 1 | 8 | 0.75 |
| 17_2600128 | 1 | 1 | 4 | 0.75 | 2 | 1 | 4 | 1 | 11 | 0.9091 | 4 | 1 | 2 | 1 | 2 | 1 | 1 | 1 | 9 | 1 |
| 17_2600136 | 3 | 1 | 3 | 0.6667 | 2 | 0.5 | 5 | 0.8 | 13 | 0.7692 | 4 | 0.75 | 3 | 1 | 2 | 1 | 3 | 1 | 12 | 0.9167 |
| 17_2600150 | 5 | 0 | 3 | 0.3333 | 3 | 0.3333 | 4 | 0.25 | 15 | 0.2 | 3 | 0.3333 | 3 | 0 | 2 | 0 | 3 | 0.3333 | 11 | 0.1818 |
| 17_2600153 | 5 | 0 | 3 | 0 | 3 | 0.3333 | 3 | 0.6667 | 14 | 0.2143 | 4 | 0.75 | 3 | 0.6667 | 2 | 0 | 2 | 0.5 | 11 | 0.5455 |
| 17_2600165 | 5 | 0 | 4 | 0.25 | 4 | 0 | 4 | 0.5 | 17 | 0.1765 | 4 | 0.25 | 3 | 0 | 2 | 0.5 | 2 | 0.5 | 11 | 0.2727 |
| 17_2600166 | 5 | 0.2 | 3 | 0.6667 | 4 | 0.75 | 2 | 1 | 14 | 0.5714 | 3 | 1 | 1 | 1 | 2 | 0 | 2 | 1 | 8 | 0.75 |
| 17_2600182 | 7 | 0.7143 | 10 | 0.7 | 11 | 0.5455 | 3 | 1 | 31 | 0.6774 | 4 | 1 | 11 | 0.8182 | 9 | 0.6667 | 11 | 0.5455 | 35 | 0.7143 |
| 17_2600184 | 8 | 0.25 | 12 | 0.4167 | 13 | 0.2308 | 3 | 0 | 36 | 0.2778 | 4 | 0.25 | 10 | 0.2 | 11 | 0.1818 | 12 | 0.0833 | 37 | 0.1622 |
| 17_2600194 | 7 | 0.1429 | 11 | 0.1818 | 12 | 0.25 | 3 | 0 | 33 | 0.1818 | 5 | 0 | 10 | 0.1 | 8 | 0.125 | 13 | 0.1538 | 36 | 0.1111 |
| 17_2600198 | 4 | 0.5 | 8 | 0.5 | 5 | 0.6 | 3 | 0.6667 | 20 | 0.55 | 4 | 0.25 | 9 | 0.8889 | 8 | 0.5 | 9 | 0.4444 | 30 | 0.5667 |
| 17_2600214 | NA | NA | NA | NA | NA | NA | 1 | 1 | 1 | 1 | 1 | 1 | NA | NA | NA | NA | 1 | 1 | 2 | 1 |
| 17_2600235 | NA | NA | 2 | 0 | 2 | 0.5 | 1 | 0 | 5 | 0.2 | 1 | 0 | NA | NA | 2 | 0 | 2 | 0.5 | 5 | 0.2 |
| 17_2600949 | 13 | 0.5385 | 18 | 0.3333 | 12 | 0.25 | 12 | 0.4167 | 55 | 0.3818 | 7 | 0.2857 | 12 | 0.4167 | 15 | 0.4 | 10 | 0.1 | 44 | 0.3182 |
| 17_2601101 | 5 | 0.4 | 3 | 0.3333 | NA | NA | 3 | 0.3333 | 11 | 0.3636 | NA | NA | 2 | 0 | 3 | 0.3333 | 3 | 0.6667 | 8 | 0.375 |
| 17_2601120 | 8 | 0.125 | 18 | 0.2778 | 7 | 0.1429 | 11 | 0.3636 | 44 | 0.25 | 6 | 0.3333 | 8 | 0.375 | 13 | 0.3077 | 11 | 0.6364 | 38 | 0.4211 |
| 17_2601121 | 8 | 0.25 | 18 | 0.1111 | 7 | 0 | 11 | 0 | 44 | 0.0909 | 6 | 0 | 12 | 0.1667 | 14 | 0.0714 | 12 | 0.3333 | 44 | 0.1591 |
| 17_2601292 | NA | NA | NA | NA | NA | NA | 1 | 1 | 1 | 1 | 1 | 1 | NA | NA | NA | NA | 2 | 0.5 | 3 | 0.6667 |
| 17_2601299 | NA | NA | 1 | 1 | NA | NA | 1 | 1 | 2 | 1 | NA | NA | NA | NA | NA | NA | 2 | 0.5 | 2 | 0.5 |
| 17_2601316 | NA | NA | 2 | 0.5 | NA | NA | 1 | 0 | 3 | 0.3333 | 1 | 1 | NA | NA | NA | NA | 1 | 0 | 2 | 0.5 |
| 17_2601327 | NA | NA | 2 | 1 | NA | NA | 1 | 0 | 3 | 0.6667 | 1 | 1 | NA | NA | NA | NA | NA | NA | 1 | 1 |
| 17_2601329 | NA | NA | 2 | 0.5 | NA | NA | 1 | 1 | 3 | 0.6667 | 1 | 1 | NA | NA | NA | NA | NA | NA | 1 | 1 |

| | | | | | | | | | | | | | | | | | | | | |
|---|---|---|---|---|---|---|---|---|---|---|---|---|---|---|---|---|---|---|---|---|
| 17_260136 | 2 | 0 | 4 | 0 | 3 | 0 | 2 | 0.5 | 11 | 0.0909 | 1 | 0 | 2 | 0.5 | 2 | 0 | 2 | 0 | 7 | 0.1429 |
| 17_260138 | 12 | 0.25 | 5 | 0.2 | 6 | 0 | 5 | 0.2 | 28 | 0.1786 | 9 | 0.2222 | 2 | 0 | 5 | 0.2 | 7 | 0 | 23 | 0.1304 |
| 17_260139 | 15 | 0.8667 | 7 | 0.8571 | 7 | 1 | 3 | 1 | 32 | 0.9062 | 10 | 0.4 | 3 | 0.6667 | 5 | 0.4 | 7 | 0.8571 | 25 | 0.56 |
| 17_260139 | 14 | 0.7857 | 7 | 0.8571 | 8 | 1 | 3 | 1 | 32 | 0.875 | 9 | 0.8889 | 3 | 1 | 5 | 1 | 6 | 1 | 23 | 0.9565 |
| 17_260153 | 11 | 0.0909 | 3 | 0 | 3 | 0 | 1 | 0 | 18 | 0.0556 | 3 | 0 | 3 | 0.3333 | 2 | 0 | 1 | 0 | 9 | 0.1111 |
| 17_260155 | 10 | 0.4 | 5 | 0.2 | 6 | 0 | 1 | 0 | 22 | 0.2273 | 5 | 0.4 | 2 | 0.5 | 4 | 0 | 4 | 0.25 | 15 | 0.2667 |
| 17_260356 | 51 | 0.2353 | 34 | 0.0294 | 33 | 0.0909 | 49 | 0.1224 | 167 | 0.1317 | 42 | 0.1667 | 65 | 0.1231 | 55 | 0.1273 | 36 | 0.0833 | 198 | 0.1263 |
| 17_260358 | 81 | 0.0617 | 62 | 0.0484 | 75 | 0.08 | 76 | 0.0658 | 294 | 0.0646 | 64 | 0.1406 | 108 | 0.0833 | 98 | 0.0408 | 64 | 0.0938 | 334 | 0.0838 |
| 17_260361 | NA | NA | 1 | 1 | NA | NA | NA | NA | 1 | 1 | NA | NA | 1 | 1 | NA | NA | 1 | 0 | 2 | 0.5 |
| 17_260367 | 86 | 0.2674 | 65 | 0.4154 | 72 | 0.25 | 84 | 0.25 | 307 | 0.2899 | 66 | 0.2727 | 110 | 0.2455 | 109 | 0.3211 | 64 | 0.3438 | 349 | 0.2923 |
| 17_260368 | 69 | 0.5942 | 53 | 0.5849 | 60 | 0.4333 | 70 | 0.5714 | 252 | 0.5476 | 57 | 0.6491 | 85 | 0.6 | 84 | 0.5595 | 57 | 0.5965 | 283 | 0.5972 |
| 17_260387 | 97 | 0.3918 | 72 | 0.3472 | 95 | 0.3895 | 97 | 0.3402 | 361 | 0.3684 | 78 | 0.3974 | 134 | 0.3358 | 127 | 0.2835 | 91 | 0.3956 | 430 | 0.3442 |
| 17_260389 | 109 | 0.4679 | 78 | 0.3718 | 107 | 0.4579 | 119 | 0.4538 | 413 | 0.4431 | 84 | 0.4286 | 145 | 0.4345 | 125 | 0.464 | 103 | 0.4563 | 457 | 0.4464 |
| 17_260392 | 110 | 0.1182 | 86 | 0.093 | 108 | 0.1019 | 99 | 0.101 | 403 | 0.1042 | 81 | 0.1235 | 137 | 0.1168 | 114 | 0.1754 | 95 | 0.0737 | 427 | 0.1241 |
| 17_260392 | 102 | 0.1373 | 75 | 0.1333 | 102 | 0.1275 | 102 | 0.1373 | 381 | 0.1339 | 77 | 0.1299 | 132 | 0.1667 | 129 | 0.1938 | 103 | 0.0971 | 441 | 0.1519 |
| 17_260400 | 157 | 0.0637 | 122 | 0.082 | 147 | 0.0748 | 148 | 0.0676 | 574 | 0.0714 | 122 | 0.082 | 192 | 0.0781 | 170 | 0.0824 | 122 | 0.0984 | 606 | 0.0842 |
| 17_260430 | 1 | 0 | 1 | 0 | 1 | 1 | NA | NA | 3 | 0.3333 | 2 | 0.5 | NA | NA | NA | NA | NA | NA | 2 | 0.5 |
| 17_260431 | 1 | 1 | 1 | 1 | 1 | 0 | NA | NA | 3 | 0.6667 | 2 | 0.5 | NA | NA | NA | NA | NA | NA | 2 | 0.5 |
| 17_260436 | 2 | 1 | 1 | 1 | 1 | 0 | NA | NA | 4 | 0.75 | 3 | 0.6667 | 1 | 0 | NA | NA | NA | NA | 4 | 0.5 |
| 17_260448 | 2 | 0.5 | 1 | 1 | 1 | 0 | NA | NA | 4 | 0.5 | 3 | 0.3333 | 1 | 1 | NA | NA | NA | NA | 4 | 0.5 |
| 17_260449 | 2 | 1 | 1 | 0 | 1 | 0 | NA | NA | 4 | 0.5 | 3 | 0.3333 | NA | NA | NA | NA | NA | NA | 3 | 0.3333 |
| 17_260452 | 2 | 0.5 | 1 | 0 | 1 | 1 | 2 | 0.5 | 6 | 0.5 | 3 | 0.3333 | 1 | 1 | NA | NA | NA | NA | 4 | 0.5 |
| 17_260453 | 2 | 1 | 1 | 1 | 1 | 0 | NA | NA | 4 | 0.75 | 3 | 0.3333 | 2 | 0 | NA | NA | NA | NA | 5 | 0.2 |
| 17_260454 | 2 | 0.5 | 1 | 0 | 1 | 0 | 1 | 0 | 5 | 0.2 | 3 | 0.3333 | 2 | 0.5 | NA | NA | NA | NA | 5 | 0.4 |
| 17_260467 | 3 | 0 | 1 | 0 | 2 | 0.5 | 1 | 0 | 7 | 0.1429 | 3 | 0 | 2 | 0.5 | 1 | 0 | NA | NA | 6 | 0.1667 |
| 17_260467 | 3 | 0.3333 | 1 | 0 | 2 | 0 | 1 | 0 | 7 | 0.1429 | 3 | 0.3333 | 2 | 0 | 1 | 1 | NA | NA | 6 | 0.3333 |
| 17_260496 | 1 | 0 | 2 | 0.5 | 1 | 1 | 1 | 1 | 5 | 0.6 | 1 | 0 | 3 | 0.3333 | 3 | 1 | 1 | 1 | 8 | 0.625 |
| 17_260497 | 20 | 0.05 | 13 | 0.1538 | 24 | 0.125 | 22 | 0.1818 | 79 | 0.1266 | 26 | 0.4615 | 19 | 0.0526 | 22 | 0.2727 | 20 | 0.15 | 87 | 0.2529 |
| 17_260497 | 19 | 0.4737 | 14 | 0.6429 | 24 | 0.375 | 23 | 0.4783 | 80 | 0.475 | 25 | 0.32 | 26 | 0.5 | 25 | 0.24 | 20 | 0.5 | 96 | 0.3854 |
| 17_260498 | 1 | 0 | 2 | 0 | 2 | 0 | 1 | 1 | 6 | 0.1667 | 1 | 0 | 3 | 0 | 3 | 0.3333 | 1 | 1 | 8 | 0.25 |
| 17_260498 | 24 | 0.5 | 16 | 0.375 | 27 | 0.2963 | 30 | 0.4 | 97 | 0.3918 | 27 | 0.3333 | 31 | 0.5161 | 28 | 0.4643 | 24 | 0.5 | 110 | 0.4545 |
| 17_260505 | 5 | 0.2 | 1 | 1 | 2 | 0 | 1 | 0 | 9 | 0.2222 | 1 | 0 | 3 | 1 | 5 | 0.8 | 4 | 0.5 | 13 | 0.6923 |
| 17_260515 | NA | NA | 1 | 1 | 2 | 0 | 2 | 0.5 | 5 | 0.4 | NA | NA | NA | NA | 2 | 0.5 | NA | NA | 2 | 0.5 |
| 17_260524 | 1 | 1 | NA | NA | 3 | 1 | NA | NA | 4 | 1 | NA | NA | NA | NA | NA | NA | 1 | 1 | 1 | 1 |
| 17_260525 | 1 | 1 | NA | NA | 3 | 1 | NA | NA | 4 | 1 | NA | NA | NA | NA | NA | NA | 1 | 1 | 1 | 1 |
| 17_260526 | 1 | 1 | 1 | 0 | NA | NA | NA | NA | 2 | 0.5 | NA | NA | NA | NA | 1 | 1 | NA | NA | 1 | 1 |
| 17_260527 | 1 | 0 | 1 | 1 | NA | NA | NA | NA | 2 | 0.5 | NA | NA | NA | NA | 1 | 1 | NA | NA | 1 | 1 |
| 17_260528 | 1 | 1 | 1 | 1 | NA | NA | NA | NA | 2 | 1 | NA | NA | NA | NA | 1 | 1 | NA | NA | 1 | 1 |
| 17_260532 | 1 | 1 | 1 | 1 | NA | NA | NA | NA | 2 | 1 | NA | NA | NA | NA | 1 | 1 | NA | NA | 1 | 1 |
| 17_260532 | 1 | 0 | 2 | 1 | NA | NA | NA | NA | 3 | 0.6667 | NA | NA | NA | NA | 1 | 1 | NA | NA | 1 | 1 |
| 17_260533 | 1 | 0 | 2 | 0.5 | NA | NA | NA | NA | 3 | 0.3333 | NA | NA | NA | NA | 1 | 1 | NA | NA | 1 | 1 |
| 17_260534 | 7 | 0.2857 | 10 | 0.3 | 9 | 0.1111 | 9 | 0.2222 | 35 | 0.2286 | 9 | 0.2222 | 11 | 0.3636 | 10 | 0.1 | 7 | 0.2857 | 37 | 0.2432 |
| 17_260535 | 5 | 0 | 12 | 0.1667 | 9 | 0.1111 | 7 | 0 | 33 | 0.0909 | 6 | 0.1667 | 8 | 0.25 | 7 | 0 | 7 | 0.2857 | 28 | 0.1786 |
| 17_260536 | 6 | 1 | 9 | 0.7778 | 8 | 1 | 8 | 0.875 | 31 | 0.9032 | 6 | 1 | 9 | 0.8889 | 4 | 0.75 | 4 | 1 | 23 | 0.913 |
| 17_260538 | 3 | 0 | 11 | 0.0909 | 9 | 0.1111 | 6 | 0.1667 | 29 | 0.1034 | 4 | 0 | 6 | 0.3333 | 7 | 0.1429 | 5 | 0.2 | 22 | 0.1818 |
| 17_260575 | 3 | 0 | 2 | 0.5 | 3 | 0 | 3 | 0 | 11 | 0.0909 | 4 | 0 | 3 | 0 | 5 | 0.2 | 3 | 0 | 15 | 0.0667 |
| 17_260578 | 8 | 0.125 | 9 | 0 | 8 | 0.125 | 12 | 0 | 37 | 0.0541 | 8 | 0 | 13 | 0.0769 | 12 | 0.1667 | 13 | 0 | 46 | 0.0652 |
| 17_260581 | 9 | 0 | 9 | 0.2222 | 9 | 0 | 12 | 0.0833 | 39 | 0.0769 | 9 | 0.2222 | 12 | 0 | 12 | 0 | 14 | 0.0714 | 47 | 0.0638 |
| 17_260582 | 7 | 0.7143 | 7 | 0.7143 | 7 | 0.8571 | 10 | 0.8 | 31 | 0.7742 | 7 | 0.8571 | 9 | 0.7778 | 10 | 0.7 | 12 | 0.9167 | 38 | 0.8158 |
| 17_260583 | 8 | 0.125 | 11 | 0 | 9 | 0.2222 | 12 | 0.0833 | 40 | 0.1 | 9 | 0.1111 | 11 | 0.0909 | 13 | 0.0769 | 13 | 0 | 46 | 0.0652 |
| 17_260638 | 1 | 0 | 2 | 0.5 | 1 | 0 | NA | NA | 4 | 0.25 | NA | NA | 1 | 0 | 1 | 0 | 1 | 1 | 3 | 0.3333 |
| 17_260640 | NA | NA | 2 | 0.5 | 1 | 0 | NA | NA | 3 | 0.3333 | NA | NA | 1 | 1 | 1 | 0 | 2 | 0.5 | 4 | 0.5 |
| 17_260828 | 5 | 0.8 | 1 | 0 | 4 | 0.75 | 4 | 0.75 | 14 | 0.7143 | 2 | 1 | 4 | 1 | 4 | 0.75 | 2 | 1 | 12 | 0.9167 |
| 17_260955 | 7 | 0.2857 | 9 | 0.2222 | 3 | 0.3333 | 5 | 0.2 | 24 | 0.25 | 3 | 0.3333 | 1 | 0 | NA | NA | 2 | 1 | 6 | 0.5 |
| 17_260956 | 5 | 0.2 | 5 | 0 | 3 | 0 | 3 | 0 | 16 | 0.0625 | 1 | 0 | 1 | 1 | NA | NA | 2 | 0 | 4 | 0.25 |
| 17_260959 | 8 | 0.125 | 10 | 0.3 | 5 | 0.2 | 6 | 0.1667 | 29 | 0.2069 | 5 | 0.2 | 1 | 0 | NA | NA | 3 | 0 | 9 | 0.1111 |
| 17_261011 | 9 | 0 | 8 | 0.25 | 7 | 0.4286 | 13 | 0.0769 | 37 | 0.1622 | 3 | 0.3333 | 3 | 0 | 13 | 0.2308 | 12 | 0.25 | 31 | 0.2258 |
| 17_261011 | 10 | 0 | 10 | 0.2 | 9 | 0 | 14 | 0.0714 | 43 | 0.0698 | 2 | 0 | 3 | 0 | 12 | 0.0833 | 11 | 0.3636 | 28 | 0.1786 |
| 17_261012 | 8 | 0.125 | 10 | 0.5 | 9 | 0.6667 | 12 | 0.5833 | 39 | 0.4872 | 2 | 0.5 | 4 | 0.5 | 13 | 0.4615 | 11 | 0.5455 | 30 | 0.5 |
| 17_261016 | 10 | 0.6 | 9 | 0.6667 | 7 | 0.8571 | 12 | 0.9167 | 38 | 0.7632 | 3 | 1 | 2 | 1 | 11 | 0.8182 | 10 | 0.8 | 26 | 0.8462 |
| 17_261017 | 9 | 0.3333 | 6 | 0 | 7 | 0.1429 | 11 | 0.0909 | 33 | 0.1515 | 3 | 0 | 3 | 0 | 12 | 0.0833 | 10 | 0.2 | 28 | 0.1071 |
| 17_261023 | 7 | 0 | 8 | 0.25 | 6 | 0.5 | 11 | 0.1818 | 32 | 0.2188 | 2 | 0 | 1 | 0 | 7 | 0.2857 | 9 | 0.2222 | 19 | 0.2105 |
| 17_261023 | 6 | 0.1667 | 8 | 0.125 | 7 | 0.5714 | 11 | 0.2727 | 32 | 0.2812 | 3 | 0.3333 | 2 | 0.5 | 7 | 0.7143 | 8 | 0.625 | 20 | 0.6 |
| 17_261023 | 6 | 0.1667 | 6 | 0.1667 | 7 | 0 | 11 | 0.2727 | 30 | 0.1667 | 2 | 0 | 1 | 1 | 7 | 0.4286 | 9 | 0.7778 | 19 | 0.5789 |
| 17_261025 | 7 | 0.5714 | 8 | 0 | 7 | 0.4286 | 11 | 0.0909 | 33 | 0.2424 | 3 | 0.3333 | 2 | 0 | 8 | 0.125 | 9 | 0.3333 | 22 | 0.2273 |

| | | | | | | | | | | | | | | | | | | | | |
|---|---|---|---|---|---|---|---|---|---|---|---|---|---|---|---|---|---|---|---|---|
| 17_2610257 | 6 | 0.8333 | 7 | 0.7143 | 7 | 0.8571 | 11 | 0.7273 | 31 | 0.7742 | 2 | 1 | 2 | 0.5 | 8 | 1 | 8 | 1 | 20 | 0.95 |
| 17_2610259 | 6 | 0.3333 | 6 | 0.1667 | 5 | 0.4 | 10 | 0.5 | 27 | 0.3704 | 2 | 0 | 2 | 0 | 7 | 0.2857 | 9 | 0.2222 | 20 | 0.2 |
| 17_2610263 | 2 | 0 | 3 | 0.6667 | 4 | 0.75 | 7 | 0.4286 | 16 | 0.5 | NA | NA | 2 | 1 | 2 | 0.5 | 5 | 0.4 | 9 | 0.5556 |
| 17_2614553 | 10 | 0.1 | 7 | 0.1429 | 18 | 0.2222 | 8 | 0.25 | 43 | 0.186 | 6 | 0 | 10 | 0.1 | 13 | 0.3077 | 9 | 0.4444 | 38 | 0.2368 |
| 17_2614641 | 9 | 0.2222 | 6 | 0.3333 | 13 | 0 | 9 | 0 | 37 | 0.1081 | 5 | 0.4 | 9 | 0 | 12 | 0 | 7 | 0 | 33 | 0.0606 |
| 17_2614651 | 8 | 0.25 | 5 | 0 | 14 | 0.1429 | 7 | 0 | 34 | 0.1176 | 6 | 0.1667 | 8 | 0.25 | 11 | 0 | 7 | 0.1429 | 32 | 0.125 |
| 17_2614656 | 8 | 0 | 5 | 0.2 | 13 | 0.0769 | 7 | 0.1429 | 33 | 0.0909 | 6 | 0 | 7 | 0 | 13 | 0.0769 | 7 | 0.1429 | 33 | 0.0606 |
| 17_2614658 | 8 | 0.375 | 7 | 0 | 14 | 0.2857 | 6 | 0.5 | 35 | 0.2857 | 6 | 0.3333 | 7 | 0.2857 | 11 | 0.4545 | 8 | 0.25 | 32 | 0.3438 |
| 17_2615073 | 3 | 0.6667 | 5 | 0 | 12 | 0.4167 | 4 | 0.25 | 24 | 0.3333 | NA | NA | 3 | 0 | 6 | 0.1667 | 5 | 0 | 14 | 0.0714 |
| 17_2615074 | 6 | 0.1667 | 4 | 0.5 | 11 | 0.2727 | 4 | 0 | 25 | 0.24 | 4 | 0.25 | 3 | 0.3333 | 8 | 0 | 6 | 0 | 21 | 0.0952 |
| 17_2615587 | 6 | 0.5 | 5 | 0.4 | 9 | 0.3333 | 5 | 0.4 | 25 | 0.4 | 2 | 0.5 | 2 | 0.5 | 6 | 0.1667 | 5 | 0.8 | 15 | 0.4667 |
| 17_2615607 | 3 | 0.3333 | 3 | 1 | 5 | 1 | 3 | 0 | 14 | 0.6429 | 1 | 1 | 3 | 0.6667 | 4 | 0.25 | 5 | 0.2 | 13 | 0.3846 |
| 17_2615611 | 4 | 0.5 | 4 | 0.5 | 6 | 0.3333 | 3 | 0 | 17 | 0.3529 | 2 | 0.5 | 3 | 1 | 5 | 0.4 | 4 | 0.25 | 14 | 0.5 |
| 17_2615623 | 2 | 0.5 | 2 | 0 | 7 | 0.2857 | 4 | 0.25 | 15 | 0.2667 | 1 | 0 | 2 | 0 | 4 | 0 | 5 | 0.2 | 12 | 0.0833 |
| 17_2615634 | 3 | 0.6667 | 2 | 0 | 5 | 0 | 1 | 0 | 11 | 0.1818 | 1 | 0 | 2 | 0 | 5 | 0.2 | 5 | 0.4 | 13 | 0.2308 |
| 17_2615659 | NA | NA | 1 | 1 | NA | NA | NA | NA | 1 | 1 | NA | NA | 1 | 1 | NA | NA | NA | NA | 1 | 1 |
| 17_2617744 | 5 | 0.6 | 7 | 0.4286 | 7 | 0.1429 | 8 | 0.625 | 27 | 0.4444 | 5 | 0.6 | 6 | 0.1667 | 5 | 0.4 | 3 | 0.3333 | 19 | 0.3684 |
| 17_2617774 | 5 | 0 | 7 | 0 | 6 | 0.1667 | 6 | 0.5 | 24 | 0.1667 | 5 | 0.2 | 5 | 0.2 | 3 | 0 | 2 | 0 | 15 | 0.1333 |
| 17_2617796 | 4 | 0.25 | 3 | 0.6667 | 3 | 0.3333 | 4 | 0 | 14 | 0.2857 | 4 | 0.5 | NA | NA | 1 | 0 | 1 | 0 | 6 | 0.3333 |
| 17_2617825 | 5 | 0.2 | 5 | 0.2 | 5 | 0.4 | 5 | 0.2 | 20 | 0.25 | 5 | 0 | 1 | 0 | 1 | 0 | 2 | 0.5 | 9 | 0.1111 |
| 17_2617853 | 5 | 0.2 | 6 | 0 | 4 | 0 | 5 | 0 | 20 | 0.05 | 4 | 0.25 | 1 | 1 | 1 | 0 | 2 | 0.5 | 8 | 0.375 |
| 17_2617860 | 3 | 0 | 5 | 0.2 | 5 | 0.2 | 5 | 0.2 | 18 | 0.1667 | 3 | 0.3333 | 1 | 0 | 1 | 1 | 2 | 0 | 7 | 0.2857 |
| 17_2617940 | 4 | 0.75 | 5 | 0 | 3 | 0.3333 | 5 | 0.2 | 17 | 0.2941 | 2 | 0.5 | 1 | 0 | NA | NA | 2 | 0 | 5 | 0.2 |
| 17_2618347 | 2 | 1 | 2 | 1 | 2 | 1 | 3 | 1 | 9 | 1 | 2 | 0.5 | 1 | 0 | 1 | 1 | 1 | 0 | 5 | 0.4 |
| 17_2618394 | 2 | 0.5 | 2 | 0.5 | 1 | 1 | 2 | 0.5 | 7 | 0.5714 | NA | NA | 1 | 1 | 1 | 0 | NA | NA | 2 | 0.5 |
| 17_2618395 | 3 | 0 | 2 | 0 | 2 | 0 | 3 | 0.3333 | 10 | 0.1 | 1 | 1 | 1 | 0 | 1 | 0 | NA | NA | 3 | 0.3333 |
| 17_2618415 | 3 | 1 | 2 | 1 | 2 | 1 | 3 | 1 | 10 | 1 | 2 | 1 | NA | NA | NA | NA | NA | NA | 2 | 1 |
| 17_2618431 | 3 | 1 | NA | NA | 1 | 1 | 2 | 1 | 6 | 1 | 2 | 1 | 1 | 1 | 1 | 1 | NA | NA | 4 | 1 |
| 17_2620884 | 3 | 0 | 9 | 0.2222 | 3 | 0.3333 | 10 | 0.3 | 25 | 0.24 | 5 | 0.2 | 6 | 0.1667 | 9 | 0 | 9 | 0.1111 | 29 | 0.1034 |
| 17_2620886 | 4 | 0 | 13 | 0 | 6 | 0.1667 | 12 | 0.0833 | 35 | 0.0571 | 6 | 0 | 9 | 0.1111 | 7 | 0 | 11 | 0.0909 | 33 | 0.0606 |
| 17_2620890 | 3 | 0 | 11 | 0.0909 | 6 | 0.1667 | 9 | 0 | 29 | 0.069 | 6 | 0.1667 | 9 | 0 | 7 | 0.1429 | 11 | 0.0909 | 33 | 0.0909 |
| 17_2620892 | 4 | 0 | 12 | 0.1667 | 5 | 0.6 | 9 | 0.2222 | 30 | 0.2333 | 5 | 0 | 7 | 0 | 8 | 0.125 | 9 | 0.2222 | 29 | 0.1034 |
| 17_2620899 | 4 | 1 | 12 | 0.9167 | 5 | 1 | 10 | 1 | 31 | 0.9677 | 4 | 0.75 | 5 | 1 | 7 | 0.7143 | 9 | 0.7778 | 25 | 0.8 |
| 17_2620901 | 4 | 0.25 | 12 | 0.25 | 5 | 1 | 5 | 0.6 | 26 | 0.4615 | 3 | 0.6667 | 6 | 0.3333 | 6 | 0.5 | 9 | 0.4444 | 24 | 0.4583 |
| 17_2620912 | 2 | 0.5 | 12 | 0.1667 | 4 | 0 | 5 | 0.2 | 23 | 0.1739 | 4 | 0.5 | 6 | 0 | 9 | 0.1111 | 8 | 0.25 | 27 | 0.1852 |
| 17_2620925 | 1 | 1 | 10 | 0.2 | 6 | 0.5 | 7 | 0.1429 | 24 | 0.2917 | 3 | 0.6667 | 8 | 0.375 | 7 | 0.5714 | 8 | 0.5 | 26 | 0.5 |
| 17_2620932 | 2 | 0 | 9 | 0.1111 | 6 | 0.1667 | 8 | 0 | 25 | 0.08 | 5 | 0 | 6 | 0 | 8 | 0.125 | 7 | 0.1429 | 26 | 0.0769 |
| 17_2620952 | 2 | 0.5 | 8 | 0.125 | 5 | 0.4 | 7 | 0 | 22 | 0.1818 | 5 | 0 | 9 | 0.1111 | 7 | 0.2857 | 7 | 0.1429 | 28 | 0.1429 |
| 17_2620971 | 2 | 1 | 9 | 1 | 5 | 0.8 | 6 | 0.8333 | 22 | 0.9091 | 6 | 1 | 9 | 1 | 10 | 0.9 | 8 | 1 | 33 | 0.9697 |
| 17_2620981 | 1 | 0 | 4 | 0 | 4 | 0.25 | 7 | 0.1429 | 16 | 0.125 | 3 | 0 | 7 | 0.2857 | 7 | 0.1429 | 6 | 0.1667 | 23 | 0.1739 |
| 17_2628662 | 1 | 1 | NA | NA | 1 | 1 | 3 | 0.6667 | 5 | 0.8 | NA | NA | 2 | 0.5 | 2 | 0 | 1 | 0 | 5 | 0.2 |
| 17_2628667 | 2 | 0.5 | 1 | 1 | 1 | 0 | 4 | 1 | 8 | 0.75 | NA | NA | 1 | 1 | 1 | 0 | NA | NA | 2 | 0.5 |
| 17_2628670 | 2 | 0 | 3 | 0 | 1 | 0 | 4 | 0.25 | 10 | 0.1 | NA | NA | 2 | 0.5 | 3 | 0.3333 | 2 | 0 | 7 | 0.2857 |
| 17_2628671 | 1 | 1 | 3 | 0.3333 | 1 | 1 | 3 | 0.6667 | 8 | 0.625 | NA | NA | 1 | 0 | 3 | 1 | 2 | 0.5 | 6 | 0.6667 |
| 17_2628679 | 2 | 0.5 | 2 | 1 | 1 | 0 | 2 | 1 | 7 | 0.7143 | NA | NA | 2 | 1 | NA | NA | 1 | 0 | 3 | 0.6667 |
| 17_2628693 | 1 | 0 | 1 | 0 | 1 | 1 | 3 | 0.3333 | 6 | 0.3333 | NA | NA | 2 | 0 | NA | NA | 2 | 0.5 | 4 | 0.25 |
| 17_2628701 | 2 | 0.5 | 2 | 0 | 1 | 0 | 4 | 0 | 9 | 0.1111 | NA | NA | 1 | 1 | 2 | 0 | 2 | 0 | 5 | 0.2 |
| 17_2628719 | 2 | 0 | 1 | 1 | 1 | 0 | 4 | 0 | 8 | 0.125 | NA | NA | 1 | 1 | 2 | 1 | NA | NA | 3 | 1 |
| 17_2628751 | 2 | 0 | 2 | 0.5 | 1 | 1 | 4 | 0 | 9 | 0.2222 | NA | NA | 1 | 0 | 2 | 0.5 | 2 | 0 | 5 | 0.2 |
| 17_2628752 | 2 | 0 | 2 | 0 | 1 | 0 | 4 | 0.5 | 9 | 0.2222 | NA | NA | 1 | 0 | 2 | 1 | 1 | 0 | 4 | 0.5 |
| 17_2628799 | 1 | 0 | 2 | 0 | NA | NA | 2 | 0.5 | 5 | 0.2 | NA | NA | 1 | 0 | 1 | 0 | 2 | 0.5 | 4 | 0.25 |
| 17_2630040 | 5 | 1 | 2 | 1 | 4 | 0.75 | 4 | 0.5 | 15 | 0.8 | 8 | 0.875 | 3 | 0.6667 | 7 | 0.7143 | 6 | 0.6667 | 24 | 0.75 |
| 17_2630050 | 4 | 0.25 | 2 | 0 | 3 | 0 | 3 | 0.3333 | 12 | 0.1667 | 7 | 0 | 2 | 0 | 9 | 0.1111 | 6 | 0.1667 | 24 | 0.0833 |
| 17_2630070 | 5 | 0.2 | 2 | 0 | 2 | 0 | 3 | 0.6667 | 12 | 0.25 | 6 | 0.5 | 3 | 0 | 5 | 0.2 | 4 | 0.5 | 18 | 0.3333 |
| 17_2630082 | 5 | 0 | 2 | 0 | 2 | 0 | 4 | 0.25 | 13 | 0.0769 | 6 | 0.1667 | 4 | 0.25 | 8 | 0.25 | 6 | 0 | 24 | 0.1667 |
| 17_2630093 | 5 | 0.6 | 1 | 1 | 2 | 1 | 2 | 0 | 10 | 0.6 | 4 | 0 | 4 | 0.25 | 7 | 0.2857 | 3 | 0.3333 | 18 | 0.2222 |
| 17_2630103 | 4 | 0.25 | 2 | 0.5 | 2 | 0.5 | 1 | 0 | 9 | 0.3333 | 4 | 0.25 | 2 | 0 | 8 | 0.5 | 6 | 0 | 20 | 0.25 |
| 17_2630913 | 2 | 0 | 1 | 0 | NA | NA | 2 | 0.5 | 5 | 0.2 | 2 | 0.5 | 2 | 0.5 | 3 | 0.6667 | 2 | 0.5 | 9 | 0.5556 |
| 17_2630941 | NA | NA | NA | NA | NA | NA | 1 | 1 | 1 | 1 | 1 | 1 | NA | NA | NA | NA | NA | NA | 1 | 1 |
| 17_2634093 | 2 | 0.5 | NA | NA | 1 | 1 | NA | NA | 3 | 0.6667 | NA | NA | 3 | 0 | 2 | 0 | 2 | 0.5 | 7 | 0.1429 |
| 17_2634103 | 2 | 0.5 | 1 | 0 | 1 | 1 | NA | NA | 4 | 0.5 | NA | NA | 3 | 0 | 1 | 1 | 2 | 0 | 6 | 0.1667 |
| 17_2634122 | 2 | 0.5 | 1 | 1 | 1 | 1 | NA | NA | 4 | 0.75 | NA | NA | 3 | 1 | 2 | 1 | 2 | 0.5 | 7 | 0.8571 |
| 17_2634123 | 2 | 1 | 1 | 1 | 1 | 1 | NA | NA | 4 | 1 | NA | NA | 3 | 0.6667 | 2 | 0.5 | 2 | 0.5 | 7 | 0.5714 |
| 17_2634543 | 10 | 0.2 | 12 | 0.1667 | 9 | 0 | 17 | 0.1765 | 48 | 0.1458 | 10 | 0.3 | 7 | 0 | 11 | 0.1818 | 11 | 0.1818 | 39 | 0.1795 |
| 17_2634546 | 14 | 0 | 12 | 0.0833 | 10 | 0.1 | 19 | 0.0526 | 55 | 0.0545 | 10 | 0.1 | 7 | 0.1429 | 12 | 0 | 13 | 0.1538 | 42 | 0.0952 |

| | | | | | | | | | | | | | | | | | | | | |
|---|---|---|---|---|---|---|---|---|---|---|---|---|---|---|---|---|---|---|---|---|
| 17_263617( | NA | NA | NA | NA | 1 | 1 | NA | NA | 1 | 1 | NA | NA | 1 | 1 | 2 | 0 | NA | NA | 3 | 0.3333 |
| 17_263619! | NA | NA | NA | NA | 1 | 1 | NA | NA | 1 | 1 | NA | NA | 1 | 1 | 1 | 1 | NA | NA | 2 | 1 |
| 17_263620' | NA | NA | NA | NA | 1 | 1 | NA | NA | 1 | 1 | NA | NA | 1 | 1 | 2 | 0.5 | NA | NA | 3 | 0.6667 |
| 17_263621! | NA | NA | NA | NA | 1 | 1 | NA | NA | 1 | 1 | NA | NA | 1 | 1 | 2 | 1 | NA | NA | 3 | 1 |
| 17_263623( | NA | NA | NA | NA | 1 | 1 | NA | NA | 1 | 1 | NA | NA | 1 | 1 | 2 | 1 | NA | NA | 3 | 1 |
| 17_264067! | NA | NA | 1 | 1 | NA | NA | NA | NA | 1 | 1 | NA | NA | 1 | 0 | 2 | 1 | NA | NA | 3 | 0.6667 |
| 17_264069: | NA | NA | 1 | 1 | 1 | 0 | NA | NA | 2 | 0.5 | NA | NA | 1 | 0 | 2 | 0.5 | NA | NA | 3 | 0.3333 |
| 17_264144: | 1 | 1 | 2 | 0 | 3 | 0 | NA | NA | 6 | 0.1667 | NA | NA | 1 | 0 | 2 | 0.5 | NA | NA | 3 | 0.3333 |
| 17_264183( | 12 | 0.5833 | 13 | 0.6923 | 11 | 0.8182 | 8 | 0.625 | 44 | 0.6818 | 14 | 0.7143 | 13 | 1 | 9 | 0.7778 | 10 | 0.7 | 46 | 0.8043 |
| 17_264183' | 11 | 0.2727 | 13 | 0.3846 | 11 | 0.2727 | 8 | 0 | 43 | 0.2558 | 13 | 0.4615 | 12 | 0.1667 | 8 | 0.375 | 12 | 0.25 | 45 | 0.3111 |
| 17_264298: | 1 | 0 | 3 | 0.3333 | 2 | 0 | 4 | 0.5 | 10 | 0.3 | NA | NA | 1 | 0 | 5 | 0.2 | 2 | 0.5 | 8 | 0.25 |
| 17_264319 | 2 | 0 | 1 | 0 | NA | NA | 1 | 1 | 4 | 0.25 | 3 | 0.3333 | 1 | 0 | 2 | 0 | NA | NA | 6 | 0.1667 |
| 17_265006! | 2 | 0 | 7 | 0.1429 | 4 | 0 | 5 | 0.2 | 18 | 0.1111 | 5 | 0 | 6 | 0 | 2 | 0.5 | 1 | 0 | 14 | 0.0714 |
| 17_265017! | 1 | 0 | 4 | 0 | 6 | 0.1667 | 6 | 0 | 17 | 0.0588 | 1 | 0 | 4 | 0 | 1 | 0 | 2 | 0.5 | 8 | 0.125 |
| 17_265023 | 1 | 0 | 5 | 0 | 3 | 0 | 7 | 0.1429 | 16 | 0.0625 | 5 | 0.2 | 5 | 0 | 1 | 0 | 2 | 0 | 13 | 0.0769 |
| 17_265035! | 7 | 0 | 9 | 0.2222 | 6 | 0.1667 | 11 | 0.0909 | 33 | 0.1212 | 6 | 0.1667 | 9 | 0.4444 | 6 | 0.1667 | 5 | 0 | 26 | 0.2308 |
| 17_265040: | 6 | 0.8333 | 8 | 1 | 6 | 0.8333 | 11 | 0.8182 | 31 | 0.871 | 6 | 0.8333 | 9 | 0.7778 | 6 | 0.8333 | 3 | 1 | 24 | 0.8333 |
| 17_265044: | 8 | 0.5 | 11 | 0.5455 | 7 | 0.4286 | 13 | 0.6923 | 39 | 0.5641 | 8 | 0.5 | 10 | 0.2 | 7 | 0.2857 | 7 | 0.5714 | 32 | 0.375 |
| 17_266606: | 1 | 0 | 1 | 0 | NA | NA | 2 | 0.5 | 4 | 0.25 | 1 | 1 | NA | NA | 1 | 1 | NA | NA | 2 | 1 |
| 17_266609( | 1 | 1 | 1 | 1 | NA | NA | 2 | 1 | 4 | 1 | NA | NA | NA | NA | 1 | 1 | NA | NA | 1 | 1 |
| 17_266643! | 4 | 0.5 | 8 | 0.25 | 4 | 0 | 11 | 0 | 27 | 0.1481 | 1 | 1 | 4 | 0.25 | 5 | 0 | 10 | 0.2 | 20 | 0.2 |
| 17_266651: | 8 | 0 | 9 | 0.1111 | 4 | 0 | 14 | 0.0714 | 35 | 0.0571 | 3 | 0.3333 | 5 | 0.2 | 6 | 0.3333 | 10 | 0.1 | 24 | 0.2083 |
| 17_266651 | 6 | 0 | 10 | 0.2 | 4 | 0 | 12 | 0.0833 | 32 | 0.0938 | 3 | 0 | 5 | 0.4 | 6 | 0.5 | 11 | 0.1818 | 25 | 0.28 |
| 17_267249: | 4 | 0 | 10 | 0.2 | 4 | 0 | 4 | 0 | 22 | 0.0909 | 4 | 0.25 | 8 | 0 | 7 | 0 | 8 | 0.25 | 27 | 0.1111 |
| 17_267249! | 5 | 0.6 | 10 | 1 | 7 | 1 | 3 | 0.6667 | 25 | 0.88 | 4 | 1 | 8 | 0.875 | 8 | 1 | 8 | 0.875 | 28 | 0.9286 |
| 17_267250: | 5 | 0 | 8 | 0.25 | 6 | 0.1667 | 4 | 0 | 23 | 0.1304 | 5 | 0.4 | 8 | 0.25 | 7 | 0.4286 | 8 | 0.125 | 28 | 0.2857 |
| 17_267250! | 4 | 0.25 | 8 | 0.125 | 5 | 0.2 | 4 | 0.25 | 21 | 0.1905 | 4 | 0.25 | 7 | 0.1429 | 4 | 0.25 | 6 | 0 | 21 | 0.1429 |
| 17_267250( | 4 | 0.25 | 10 | 0.2 | 6 | 0.1667 | 4 | 0 | 24 | 0.1667 | 4 | 0 | 7 | 0.1429 | 7 | 0.4286 | 4 | 0.25 | 22 | 0.2273 |
| 17_267250 | 5 | 0 | 9 | 0.3333 | 6 | 0.5 | 3 | 0 | 23 | 0.2609 | 4 | 0 | 5 | 0.2 | 6 | 0.3333 | 6 | 0.5 | 21 | 0.2857 |
| 17_267251! | 1 | 1 | 4 | 0.75 | 1 | 0 | 3 | 1 | 9 | 0.7778 | 2 | 1 | 3 | 0.6667 | NA | NA | 2 | 1 | 7 | 0.8571 |
| 17_267252! | 5 | 0.2 | 8 | 0.375 | 6 | 0.3333 | 4 | 0 | 23 | 0.2609 | 4 | 0.5 | 7 | 0.1429 | 5 | 0.4 | 5 | 0.4 | 21 | 0.3333 |
| 17_267253( | 1 | 1 | 5 | 0.6 | 4 | 1 | 2 | 0.5 | 12 | 0.75 | 2 | 1 | 5 | 0.4 | 4 | 1 | 4 | 0.75 | 15 | 0.7333 |
| 17_267254( | 3 | 0 | 7 | 0 | 5 | 0.2 | 3 | 0 | 18 | 0.0556 | 4 | 0 | 5 | 0 | 5 | 0.2 | 7 | 0.2857 | 21 | 0.1429 |
| 17_267255: | 3 | 0.3333 | 7 | 0.4286 | 4 | 0.5 | 3 | 0.6667 | 17 | 0.4706 | 3 | 0.6667 | 6 | 0.5 | 5 | 0.6 | 6 | 0.3333 | 20 | 0.5 |
| 17_267270! | 3 | 0 | 6 | 0.1667 | 4 | 0.25 | 2 | 0 | 15 | 0.1333 | 2 | 0 | 5 | 0.4 | 4 | 0.25 | 7 | 0.1429 | 18 | 0.2222 |
| 17_268745( | 15 | 0.3333 | 9 | 0.1111 | 15 | 0.2 | 11 | 0 | 50 | 0.18 | 6 | 0.1667 | 6 | 0.1667 | 5 | 0 | 11 | 0.1818 | 28 | 0.1429 |
| 17_269184! | 6 | 0.3333 | 8 | 0 | 5 | 0.2 | 11 | 0.2727 | 30 | 0.2 | 8 | 0.25 | 7 | 0.1429 | 14 | 0.0714 | 9 | 0.2222 | 38 | 0.1579 |
| 17_269184! | 5 | 1 | 6 | 0.5 | 4 | 0.75 | 8 | 0.625 | 23 | 0.6957 | 6 | 0.6667 | 4 | 0.75 | 10 | 0.6 | 5 | 0.6 | 25 | 0.64 |
| 17_269610! | 1 | 0 | 1 | 0 | 1 | 0 | 3 | 0.3333 | 6 | 0.1667 | NA | NA | NA | NA | 2 | 0.5 | 1 | 0 | 3 | 0.3333 |
| 17_269612( | 1 | 0 | NA | NA | 1 | 0 | 3 | 0.6667 | 5 | 0.4 | NA | NA | NA | NA | 2 | 1 | 2 | 0.5 | 4 | 0.75 |
| 17_269612 | 1 | 0 | 1 | 1 | 1 | 1 | 3 | 0.3333 | 6 | 0.5 | NA | NA | NA | NA | 2 | 0.5 | 3 | 0.3333 | 5 | 0.4 |
| 17_269613' | 1 | 1 | 1 | 1 | 2 | 1 | 2 | 1 | 6 | 1 | NA | NA | NA | NA | 2 | 1 | 3 | 1 | 5 | 1 |
| 17_269614 | 1 | 1 | 1 | 1 | 2 | 1 | 4 | 0.75 | 8 | 0.875 | NA | NA | NA | NA | 2 | 1 | 3 | 0.6667 | 5 | 0.8 |
| 17_269615! | 1 | 1 | 1 | 1 | 2 | 0.5 | 4 | 0.75 | 8 | 0.75 | NA | NA | NA | NA | 2 | 0.5 | 2 | 0.5 | 4 | 0.5 |
| 17_269617( | 1 | 1 | 1 | 1 | 2 | 0.5 | 4 | 0.25 | 8 | 0.5 | NA | NA | NA | NA | 2 | 1 | 2 | 0.5 | 4 | 0.75 |
| 17_269619! | NA | NA | 2 | 0 | 2 | 0.5 | 3 | 0.3333 | 7 | 0.2857 | NA | NA | NA | NA | 2 | 0.5 | 3 | 0.3333 | 5 | 0.4 |
| 17_269903! | 2 | 0.5 | 4 | 0.25 | 3 | 0 | 4 | 0.25 | 13 | 0.2308 | NA | NA | 1 | 0 | 3 | 0 | 5 | 0.2 | 9 | 0.1111 |
| 17_269930! | 3 | 0.3333 | 6 | 0.3333 | 7 | 0.1429 | 6 | 0 | 22 | 0.1818 | 1 | 0 | 1 | 0 | 7 | 0.1429 | 7 | 0 | 16 | 0.0625 |
| 17_269933! | 4 | 0.5 | 6 | 0.3333 | 5 | 0.2 | 5 | 0 | 20 | 0.25 | NA | NA | 1 | 0 | 8 | 0 | 8 | 0.375 | 17 | 0.1765 |
| 17_269944( | 1 | 0 | 3 | 0.3333 | 5 | 0.2 | 2 | 0 | 11 | 0.1818 | 1 | 0 | 2 | 0 | 7 | 0.2857 | 6 | 0.1667 | 16 | 0.1875 |
| 17_269950' | 3 | 0.3333 | 4 | 0 | 6 | 0 | 4 | 0 | 17 | 0.0588 | 1 | 0 | 3 | 0.3333 | 8 | 0.125 | 7 | 0.1429 | 19 | 0.1579 |
| 17_270110: | 2 | 0 | 3 | 0 | 9 | 0.1111 | 6 | 0.3333 | 20 | 0.15 | 4 | 0 | 3 | 0 | 2 | 0.5 | 3 | 0 | 12 | 0.0833 |
| 17_270111 | 2 | 1 | 3 | 0.6667 | 9 | 0.8889 | 4 | 1 | 18 | 0.8889 | 3 | 1 | 4 | 1 | 3 | 1 | 3 | 0.6667 | 13 | 0.9231 |
| 17_270111! | 2 | 0.5 | 3 | 0 | 8 | 0.125 | 4 | 0.25 | 17 | 0.1765 | 3 | 0 | 4 | 0.5 | 3 | 0 | 3 | 0.3333 | 13 | 0.2308 |
| 17_270111! | 2 | 1 | 3 | 1 | 6 | 1 | 6 | 1 | 17 | 1 | 1 | 1 | 3 | 1 | 1 | 1 | 2 | 1 | 7 | 1 |
| 17_270112: | 2 | 0.5 | 3 | 0 | 8 | 0.375 | 5 | 0 | 18 | 0.2222 | 4 | 0.25 | 4 | 0.5 | 3 | 0.3333 | 3 | 0.6667 | 14 | 0.4286 |
| 17_270113 | 2 | 1 | 3 | 1 | 9 | 0.7778 | 6 | 1 | 20 | 0.9 | 3 | 1 | 4 | 1 | 2 | 1 | 3 | 1 | 12 | 1 |
| 17_270113! | 2 | 1 | 3 | 0.6667 | 7 | 0.5714 | 3 | 0.6667 | 15 | 0.6667 | 2 | 1 | 4 | 0.5 | 2 | 0.5 | 2 | 1 | 10 | 0.7 |
| 17_271431 | NA | NA | 1 | 1 | NA | NA | NA | NA | 1 | 1 | NA | NA | NA | NA | 1 | 1 | NA | NA | 1 | 1 |
| 17_271432 | NA | NA | 1 | 1 | NA | NA | NA | NA | 1 | 1 | NA | NA | NA | NA | 1 | 1 | NA | NA | 1 | 1 |
| 17_271435! | NA | NA | 1 | 1 | NA | NA | NA | NA | 1 | 1 | NA | NA | NA | NA | 1 | 1 | NA | NA | 1 | 1 |
| 17_271564( | 3 | 0.3333 | 5 | 0 | 5 | 0.2 | 2 | 0 | 15 | 0.1333 | 6 | 0.1667 | 4 | 0.25 | 3 | 0 | 3 | 0 | 16 | 0.125 |
| 17_271567 | 2 | 0 | 5 | 0.2 | 6 | 0 | 1 | 0 | 14 | 0.0714 | 5 | 0 | 4 | 0 | 2 | 0 | 3 | 0.3333 | 14 | 0.0714 |
| 17_271576: | 1 | 1 | 2 | 0 | 2 | 0 | 1 | 0 | 6 | 0.1667 | 1 | 0 | NA | NA | 1 | 1 | 1 | 1 | 3 | 0.6667 |
| 17_271760 | NA | NA | 4 | 0.25 | 1 | 0 | 4 | 0 | 9 | 0.1111 | 3 | 0 | 3 | 0 | 1 | 0 | 3 | 0.3333 | 10 | 0.1 |

| | | | | | | | | | | | | | | | | | | | | |
|---|---|---|---|---|---|---|---|---|---|---|---|---|---|---|---|---|---|---|---|---|
| 17_2717618 | 2 | 0.5 | 3 | 0 | 2 | 0 | 5 | 0.2 | 12 | 0.1667 | 6 | 0.1667 | 4 | 0 | 3 | 0 | 4 | 0 | 17 | 0.0588 |
| 17_2717620 | 2 | 0 | 4 | 0.25 | 1 | 0 | 5 | 0 | 12 | 0.0833 | 6 | 0.6667 | 4 | 0.25 | 1 | 0 | 2 | 0.5 | 13 | 0.4615 |
| 17_2717623 | 1 | 1 | 5 | 1 | 2 | 1 | 5 | 0.6 | 13 | 0.8462 | 5 | 0.6 | 3 | 0.3333 | 3 | 0.6667 | 3 | 1 | 14 | 0.6429 |
| 17_2717639 | 2 | 0 | 5 | 0.2 | 3 | 0.3333 | 5 | 0.4 | 15 | 0.2667 | 5 | 0 | 5 | 0.2 | 3 | 0 | 4 | 0.25 | 17 | 0.1176 |
| 17_2717670 | NA | NA | 6 | 0.1667 | 3 | 0 | 6 | 0 | 15 | 0.0667 | 5 | 0 | 6 | 0 | 5 | 0.2 | 4 | 0 | 20 | 0.05 |
| 17_2717865 | 2 | 0 | 7 | 0 | 3 | 0 | 6 | 0.1667 | 18 | 0.0556 | 5 | 0 | 6 | 0.1667 | 8 | 0.125 | 3 | 0 | 22 | 0.0909 |
| 17_2718041 | 3 | 0.3333 | 6 | 0.5 | 4 | 0.5 | 7 | 0.2857 | 20 | 0.4 | 9 | 0.5556 | 7 | 0.4286 | 9 | 0.1111 | 5 | 0.2 | 30 | 0.3333 |
| 17_2718048 | 5 | 0.2 | 8 | 0.75 | 3 | 0.6667 | 5 | 0.6 | 21 | 0.5714 | 8 | 0.625 | 7 | 0.8571 | 9 | 0.5556 | 6 | 1 | 30 | 0.7333 |
| 17_2718055 | 5 | 0 | 8 | 0.25 | 4 | 0.25 | 7 | 0.1429 | 24 | 0.1667 | 8 | 0.5 | 6 | 0.3333 | 8 | 0.375 | 5 | 0.2 | 27 | 0.3704 |
| 17_2718060 | 3 | 0 | 8 | 0.125 | 2 | 0 | 4 | 0 | 17 | 0.0588 | 8 | 0.5 | 5 | 0 | 7 | 0.4286 | 5 | 0.2 | 25 | 0.32 |
| 17_2718067 | 5 | 1 | 8 | 1 | 5 | 1 | 6 | 0.8333 | 24 | 0.9583 | 9 | 0.8889 | 8 | 1 | 8 | 0.75 | 5 | 0.6 | 30 | 0.8333 |
| 17_2718085 | 6 | 0.1667 | 8 | 0.375 | 5 | 0.2 | 5 | 0.4 | 24 | 0.2917 | 6 | 0.5 | 7 | 0.1429 | 8 | 0.125 | 4 | 0 | 25 | 0.2 |
| 17_2718099 | 6 | 0.8333 | 9 | 0.8889 | 6 | 0.8333 | 8 | 1 | 29 | 0.8966 | 7 | 0.8571 | 8 | 1 | 7 | 1 | 6 | 1 | 28 | 0.9643 |
| 17_2718103 | 6 | 0.6667 | 4 | 0.75 | 3 | 0.6667 | 6 | 0.6667 | 19 | 0.6842 | 5 | 0.6 | 8 | 0.5 | 7 | 0.2857 | 2 | 0 | 22 | 0.4091 |
| 17_2718110 | 5 | 0.2 | 11 | 0.7273 | 5 | 0.4 | 8 | 0.75 | 29 | 0.5862 | 10 | 0.4 | 8 | 0.5 | 8 | 0.875 | 5 | 0.8 | 31 | 0.6129 |
| 17_2718115 | 5 | 0.6 | 11 | 0.6364 | 6 | 0.6667 | 7 | 0.7143 | 29 | 0.6552 | 9 | 0.6667 | 9 | 0.5556 | 9 | 0.5556 | 4 | 0.5 | 31 | 0.5806 |
| 17_2718129 | 7 | 0 | 12 | 0.0833 | 6 | 0 | 8 | 0.125 | 33 | 0.0606 | 8 | 0 | 8 | 0.375 | 8 | 0.25 | 7 | 0 | 31 | 0.1613 |
| 17_2718972 | 7 | 0.2857 | 21 | 0.381 | 11 | 0.3636 | 14 | 0.2857 | 53 | 0.3396 | 13 | 0.3077 | 14 | 0.4286 | 13 | 0.3846 | 8 | 0.5 | 48 | 0.3958 |
| 17_2718974 | 11 | 0.3636 | 22 | 0.4091 | 10 | 0.5 | 14 | 0.3571 | 57 | 0.4035 | 12 | 0.4167 | 12 | 0.25 | 11 | 0.7273 | 9 | 0.3333 | 44 | 0.4318 |
| 17_2719090 | 10 | 0.7 | 29 | 0.2759 | 14 | 0.5 | 17 | 0.4118 | 70 | 0.4143 | 17 | 0.3529 | 23 | 0.4348 | 20 | 0.5 | 17 | 0.2353 | 77 | 0.3896 |
| 17_2719260 | NA | NA | 1 | 1 | 2 | 0 | NA | NA | 3 | 0.3333 | NA | NA | 1 | 0 | 1 | 1 | NA | NA | 2 | 0.5 |
| 17_2719277 | NA | NA | 1 | 0 | 2 | 0.5 | NA | NA | 3 | 0.3333 | NA | NA | 1 | 0 | 1 | 1 | NA | NA | 2 | 0.5 |
| 17_2719285 | NA | NA | NA | NA | 2 | 1 | NA | NA | 2 | 1 | NA | NA | 1 | 1 | 1 | 1 | NA | NA | 2 | 1 |
| 17_2719289 | NA | NA | 1 | 1 | 3 | 1 | NA | NA | 4 | 1 | NA | NA | 1 | 1 | 1 | 0 | NA | NA | 2 | 0.5 |
| 17_2719327 | NA | NA | 1 | 1 | 2 | 0.5 | NA | NA | 3 | 0.6667 | NA | NA | 1 | 0 | 1 | 1 | NA | NA | 2 | 0.5 |
| 17_2719350 | NA | NA | 1 | 0 | 3 | 0.6667 | NA | NA | 4 | 0.5 | 1 | 1 | 1 | 1 | 1 | 0 | NA | NA | 3 | 0.6667 |
| 17_2719363 | NA | NA | 1 | 0 | 2 | 1 | NA | NA | 3 | 0.6667 | 1 | 1 | 2 | 0.5 | 1 | 0 | NA | NA | 4 | 0.5 |
| 17_2719364 | NA | NA | 1 | 1 | 1 | 0 | NA | NA | 2 | 0.5 | NA | NA | 2 | 0.5 | 1 | 0 | NA | NA | 3 | 0.3333 |
| 17_2719368 | NA | NA | 1 | 1 | 3 | 0.6667 | NA | NA | 4 | 0.75 | 1 | 1 | 2 | 0.5 | 2 | 0.5 | NA | NA | 5 | 0.6 |
| 17_2719392 | NA | NA | 1 | 1 | 3 | 1 | NA | NA | 4 | 1 | 1 | 1 | 1 | 1 | 2 | 1 | NA | NA | 4 | 1 |
| 17_2719396 | NA | NA | 1 | 1 | 3 | 1 | NA | NA | 4 | 1 | 1 | 1 | 3 | 1 | 1 | 1 | NA | NA | 5 | 1 |
| 17_2719397 | NA | NA | 1 | 0 | 2 | 1 | NA | NA | 3 | 0.6667 | 1 | 1 | 3 | 0.3333 | 2 | 1 | NA | NA | 6 | 0.6667 |
| 17_2719400 | NA | NA | 1 | 1 | 1 | 1 | NA | NA | 2 | 1 | 1 | 0 | 3 | 1 | 2 | 0.5 | NA | NA | 6 | 0.6667 |
| 17_2719403 | NA | NA | 1 | 1 | 3 | 1 | NA | NA | 4 | 1 | 1 | 1 | 3 | 1 | 2 | 1 | NA | NA | 6 | 1 |
| 17_2719421 | NA | NA | 1 | 0 | 2 | 0.5 | NA | NA | 3 | 0.3333 | 1 | 0 | 3 | 0.6667 | 2 | 0 | NA | NA | 6 | 0.3333 |
| 17_2719492 | NA | NA | 3 | 0 | 3 | 0.3333 | NA | NA | 6 | 0.1667 | 3 | 0 | 3 | 0.3333 | 2 | 0 | 1 | 1 | 9 | 0.2222 |
| 17_2719493 | NA | NA | 3 | 0 | 2 | 0.5 | NA | NA | 5 | 0.2 | 1 | 0 | 3 | 0.3333 | 1 | 0 | 1 | 1 | 6 | 0.3333 |
| 17_2719501 | NA | NA | 3 | 0.6667 | 3 | 0.6667 | NA | NA | 6 | 0.6667 | 3 | 0.6667 | 3 | 0.3333 | 2 | 0 | 1 | 0 | 9 | 0.3333 |
| 17_2719506 | NA | NA | 3 | 0 | 3 | 0.6667 | NA | NA | 6 | 0.3333 | 3 | 0.6667 | 2 | 0.5 | 1 | 1 | 1 | 1 | 7 | 0.7143 |
| 17_2719515 | NA | NA | 3 | 0.3333 | 3 | 0 | NA | NA | 6 | 0.1667 | 3 | 0.6667 | 3 | 0 | 2 | 0 | 1 | 0 | 9 | 0.2222 |
| 17_2719518 | NA | NA | 1 | 1 | NA | NA | NA | NA | 1 | 1 | 1 | 1 | NA | NA | NA | NA | NA | NA | 1 | 1 |
| 17_2719530 | NA | NA | 1 | 1 | NA | NA | NA | NA | 1 | 1 | 1 | 1 | NA | NA | NA | NA | NA | NA | 1 | 1 |
| 17_2723968 | NA | NA | 2 | 0.5 | NA | NA | 1 | 0 | 3 | 0.3333 | 1 | 0 | 1 | 0 | 2 | 0.5 | NA | NA | 4 | 0.25 |
| 17_2736384 | NA | NA | NA | NA | NA | NA | 1 | 1 | 1 | 1 | NA | NA | 1 | 1 | 1 | 0 | NA | NA | 2 | 0.5 |
| 17_2740860 | 3 | 0 | 2 | 0 | 2 | 0 | 7 | 0.1429 | 14 | 0.0714 | 2 | 0 | 3 | 0.3333 | 5 | 0 | 8 | 0.125 | 18 | 0.1111 |
| 17_2740865 | 1 | 0 | 2 | 0 | 1 | 0 | 3 | 0.3333 | 7 | 0.1429 | 2 | 0.5 | 2 | 0.5 | 4 | 0.25 | 3 | 0.3333 | 11 | 0.3636 |
| 17_2740892 | 4 | 0 | 3 | 0 | 5 | 0 | 9 | 0.2222 | 21 | 0.0952 | 4 | 0 | 4 | 0.25 | 9 | 0 | 10 | 0.1 | 27 | 0.0741 |
| 17_2740950 | 2 | 0.5 | 2 | 0 | 2 | 0 | 4 | 0.25 | 10 | 0.2 | 2 | 0 | 3 | 0.3333 | 7 | 0.1429 | 6 | 0.1667 | 18 | 0.1667 |
| 17_2740955 | 2 | 1 | 3 | 0 | 4 | 0 | 6 | 0 | 15 | 0.1333 | 2 | 0 | 3 | 0.6667 | 6 | 0.1667 | 7 | 0.2857 | 18 | 0.2778 |
| 17_2740956 | 3 | 0.3333 | 3 | 0 | 4 | 0.5 | 6 | 0.5 | 16 | 0.375 | 1 | 1 | 3 | 0.6667 | 6 | 0.3333 | 7 | 0.5714 | 17 | 0.5294 |
| 17_2740964 | 2 | 0.5 | 3 | 1 | 4 | 0.75 | 5 | 1 | 14 | 0.8571 | 1 | 1 | 3 | 1 | 6 | 0.5 | 5 | 0.4 | 15 | 0.6 |
| 17_2742263 | NA | NA | 1 | 0 | 4 | 0 | 4 | 0.25 | 9 | 0.1111 | 1 | 0 | 2 | 0 | 3 | 0.3333 | 4 | 0.25 | 10 | 0.2 |
| 17_2742291 | NA | NA | 1 | 0 | 4 | 0.25 | 2 | 0 | 7 | 0.1429 | NA | NA | NA | NA | NA | NA | 3 | 0.3333 | 3 | 0.3333 |
| 17_2742390 | 1 | 1 | NA | NA | NA | NA | NA | NA | 1 | 1 | NA | NA | NA | NA | 2 | 0 | 1 | 1 | 3 | 0.3333 |
| 17_2742398 | 1 | 1 | NA | NA | NA | NA | NA | NA | 1 | 1 | NA | NA | NA | NA | 1 | 1 | NA | NA | 1 | 1 |
| 17_2742406 | 1 | 1 | NA | NA | NA | NA | NA | NA | 1 | 1 | NA | NA | NA | NA | 2 | 1 | 1 | 1 | 3 | 1 |
| 17_2764562 | 5 | 0.8 | 6 | 1 | 1 | 1 | 6 | 1 | 18 | 0.9444 | 4 | 0.75 | 11 | 0.9091 | 9 | 0.8889 | 4 | 1 | 28 | 0.8929 |
| 17_2764574 | 5 | 0 | 4 | 0.25 | 2 | 0 | 6 | 0.1667 | 17 | 0.1176 | 3 | 0 | 13 | 0.3077 | 8 | 0.375 | 5 | 0.4 | 29 | 0.3103 |
| 17_2765361 | 6 | 0.1667 | 5 | 0 | 2 | 0 | 4 | 0.75 | 17 | 0.2353 | 3 | 0.3333 | 11 | 0.0909 | 8 | 0.125 | 6 | 0.1667 | 28 | 0.1429 |
| 17_2765362 | 5 | 0.6 | 5 | 0 | 3 | 0.6667 | 3 | 0 | 16 | 0.3125 | 3 | 0.3333 | 12 | 0.3333 | 8 | 0.125 | 6 | 0.3333 | 29 | 0.2759 |
| 17_2765370 | 4 | 0.75 | 5 | 0.4 | 3 | 1 | 4 | 0.75 | 16 | 0.6875 | 3 | 0.6667 | 11 | 0.6364 | 7 | 0.8571 | 7 | 0.8571 | 28 | 0.75 |
| 17_2765374 | 5 | 1 | 5 | 1 | 2 | 1 | 4 | 1 | 16 | 1 | 2 | 1 | 11 | 1 | 6 | 1 | 4 | 1 | 23 | 1 |
| 17_2765375 | 6 | 0 | 5 | 0.2 | 2 | 0.5 | 4 | 0.25 | 17 | 0.1765 | 3 | 0.6667 | 12 | 0.3333 | 7 | 0.2857 | 7 | 0.1429 | 29 | 0.3103 |
| 17_2765376 | 4 | 1 | 4 | 1 | 2 | 1 | 4 | 1 | 14 | 1 | 3 | 1 | 12 | 1 | 7 | 0.8571 | 6 | 1 | 28 | 0.9643 |
| 17_2765378 | 3 | 0.6667 | 5 | 0.4 | 1 | 1 | 2 | 0 | 11 | 0.4545 | 3 | 0.3333 | 10 | 0.7 | 6 | 0 | 5 | 0.6 | 24 | 0.4583 |

| | | | | | | | | | | | | | | | | | | | | |
|---|---|---|---|---|---|---|---|---|---|---|---|---|---|---|---|---|---|---|---|---|
| 17_276539 | 6 | 0.5 | 4 | 1 | 2 | 1 | 3 | 0.6667 | 15 | 0.7333 | 3 | 1 | 9 | 0.8889 | 8 | 0.625 | 5 | 0.6 | 25 | 0.76 |
| 17_276541 | 4 | 1 | 2 | 1 | 1 | 1 | 3 | 1 | 10 | 1 | 3 | 0.6667 | 10 | 1 | 8 | 1 | 5 | 1 | 26 | 0.9615 |
| 17_276542 | 5 | 0.2 | 4 | 0.25 | 2 | 0 | 2 | 0 | 13 | 0.1538 | 3 | 0 | 9 | 0.2222 | 9 | 0.2222 | 4 | 0.5 | 25 | 0.24 |
| 17_276542 | 4 | 0.75 | 4 | 0.5 | 2 | 0 | 2 | 0.5 | 12 | 0.5 | 3 | 1 | 8 | 0.875 | 9 | 0.7778 | 4 | 0.75 | 24 | 0.8333 |
| 17_276572 | 3 | 0 | 1 | 0 | 1 | 0 | 3 | 0.3333 | 8 | 0.125 | 2 | 0 | 7 | 0.1429 | 6 | 0 | 5 | 0 | 20 | 0.05 |
| 17_277218 | 2 | 0.5 | NA | NA | 1 | 1 | 2 | 1 | 5 | 0.8 | 1 | 1 | 3 | 1 | 4 | 0.75 | 3 | 0.6667 | 11 | 0.8182 |
| 17_277218 | 1 | 0 | NA | NA | 1 | 0 | 2 | 1 | 4 | 0.5 | 1 | 0 | 5 | 0 | 3 | 0 | 3 | 0.3333 | 12 | 0.0833 |
| 17_277220 | 2 | 0.5 | NA | NA | NA | NA | 1 | 1 | 3 | 0.6667 | 1 | 1 | 1 | 1 | 2 | 1 | 3 | 1 | 7 | 1 |
| 17_277222 | 2 | 0.5 | 1 | 1 | 1 | 0 | 1 | 1 | 5 | 0.6 | 1 | 1 | 3 | 1 | 6 | 0.3333 | 2 | 0.5 | 12 | 0.5833 |
| 17_277223 | 2 | 0 | 1 | 0 | 1 | 0 | 2 | 0.5 | 6 | 0.1667 | 1 | 0 | 3 | 0 | 5 | 0 | 3 | 0.3333 | 12 | 0.0833 |
| 17_277223 | 2 | 1 | 1 | 0 | 1 | 0 | 2 | 0.5 | 6 | 0.5 | NA | NA | 3 | 0.3333 | 5 | 0.4 | 3 | 0.3333 | 11 | 0.3636 |
| 17_277986 | 43 | 0.2558 | 46 | 0.3261 | 39 | 0.1795 | 36 | 0.1667 | 164 | 0.2378 | 32 | 0.3438 | 44 | 0.2955 | 30 | 0.2 | 36 | 0.3611 | 142 | 0.3028 |
| 17_277989 | 41 | 0.0244 | 40 | 0.15 | 39 | 0.1282 | 35 | 0 | 155 | 0.0774 | 26 | 0.0769 | 39 | 0.0513 | 34 | 0.0588 | 39 | 0.1026 | 138 | 0.0725 |
| 17_277990 | 42 | 0.0952 | 40 | 0.075 | 36 | 0.0278 | 36 | 0.0556 | 154 | 0.0649 | 25 | 0.12 | 34 | 0.0588 | 28 | 0.0714 | 37 | 0.0811 | 124 | 0.0806 |
| 17_277996 | 30 | 0.0333 | 29 | 0.0345 | 28 | 0.1429 | 28 | 0.1429 | 115 | 0.087 | 22 | 0.1818 | 24 | 0.0417 | 26 | 0.0769 | 23 | 0.1739 | 95 | 0.1158 |
| 17_277997 | 30 | 0.0333 | 32 | 0.1562 | 31 | 0.0323 | 29 | 0.1724 | 122 | 0.0984 | 22 | 0.0455 | 23 | 0.087 | 24 | 0.125 | 25 | 0.16 | 94 | 0.1064 |
| 17_278422 | 17 | 0 | 19 | 0.1579 | 17 | 0.0588 | 11 | 0.0909 | 64 | 0.0781 | 13 | 0 | 12 | 0 | 8 | 0.125 | 12 | 0.25 | 45 | 0.0889 |
| 17_279276 | 16 | 0.25 | 24 | 0.0833 | 15 | 0.2 | 8 | 0.125 | 63 | 0.1587 | 11 | 0.0909 | 15 | 0.1333 | 15 | 0.0667 | 10 | 0.3 | 51 | 0.1373 |
| 17_279277 | 18 | 0.3333 | 26 | 0.4615 | 17 | 0.4706 | 10 | 0.5 | 71 | 0.4366 | 12 | 0.4167 | 12 | 0.75 | 17 | 0.3529 | 12 | 0.5833 | 53 | 0.5094 |
| 17_279288 | 20 | 0.05 | 29 | 0.1724 | 25 | 0.08 | 14 | 0.1429 | 88 | 0.1136 | 14 | 0 | 15 | 0.0667 | 18 | 0.0556 | 18 | 0.1111 | 65 | 0.0615 |
| 17_279704 | 30 | 0.0667 | 20 | 0.05 | 19 | 0.0526 | 22 | 0.1818 | 91 | 0.0879 | 14 | 0.1429 | 20 | 0 | 24 | 0.0833 | 19 | 0 | 77 | 0.0519 |
| 17_279705 | 22 | 0.2273 | 17 | 0.1176 | 21 | 0.0476 | 20 | 0.3 | 80 | 0.175 | 12 | 0.1667 | 19 | 0.2632 | 21 | 0.2381 | 14 | 0.4286 | 66 | 0.2727 |
| 17_279722 | 8 | 0.625 | 3 | 0.6667 | 8 | 0.5 | 5 | 0.4 | 24 | 0.5417 | 5 | 0.4 | 7 | 0.8571 | 8 | 0.125 | 4 | 0.75 | 24 | 0.5 |
| 17_279724 | 5 | 0.8 | 2 | 0 | 5 | 0.2 | 2 | 0 | 14 | 0.3571 | 3 | 0.3333 | 5 | 0.4 | 6 | 0.3333 | 3 | 0 | 17 | 0.2941 |
| 17_279725 | 5 | 0.2 | 3 | 0.3333 | 5 | 0 | 4 | 0 | 17 | 0.1176 | 3 | 0.3333 | 6 | 0.3333 | 6 | 0.1667 | 3 | 0 | 18 | 0.2222 |
| 17_279727 | 5 | 0.6 | 2 | 0.5 | 4 | 0.25 | 5 | 0 | 16 | 0.3125 | 1 | 0 | 5 | 0.6 | 7 | 0.4286 | 4 | 0.25 | 17 | 0.4118 |
| 17_279727 | 5 | 1 | 2 | 0 | 6 | 0.8333 | 4 | 0.5 | 17 | 0.7059 | 1 | 0 | 6 | 0.8333 | 7 | 0.8571 | 3 | 0.6667 | 17 | 0.7647 |
| 17_279727 | 5 | 0.6 | 1 | 1 | 6 | 0.6667 | 4 | 0.75 | 16 | 0.6875 | 2 | 0 | 7 | 0.5714 | 7 | 0.7143 | 4 | 0 | 20 | 0.45 |
| 17_279728 | 6 | 0 | 1 | 0 | 6 | 0.3333 | 4 | 0 | 17 | 0.1176 | 1 | 0 | 6 | 0 | 7 | 0.2857 | 4 | 0.25 | 18 | 0.1667 |
| 17_279728 | 3 | 1 | 1 | 1 | 2 | 0.5 | 1 | 1 | 7 | 0.8571 | 2 | 1 | 4 | 1 | 5 | 0.8 | 2 | 1 | 13 | 0.9231 |
| 17_279869 | 3 | 0.6667 | 1 | 1 | 2 | 0.5 | 2 | 1 | 8 | 0.75 | NA | NA | 4 | 1 | 5 | 0.8 | 3 | 1 | 12 | 0.9167 |
| 17_279870 | 3 | 1 | 1 | 1 | 2 | 0.5 | 2 | 1 | 8 | 0.875 | NA | NA | 2 | 1 | 4 | 0.25 | 3 | 1 | 9 | 0.6667 |
| 17_279871 | 2 | 0 | 2 | 0.5 | 4 | 0.25 | 3 | 0.6667 | 11 | 0.3636 | NA | NA | 6 | 0.6667 | 6 | 0.1667 | 4 | 0.25 | 16 | 0.375 |
| 17_281054 | NA | NA | 1 | 1 | NA | NA | NA | NA | 1 | 1 | NA | NA | 1 | 1 | NA | NA | 1 | 1 | 2 | 1 |
| 17_281055 | NA | NA | 1 | 1 | NA | NA | NA | NA | 1 | 1 | NA | NA | 1 | 1 | NA | NA | 1 | 1 | 2 | 1 |
| 17_281061 | NA | NA | 1 | 1 | NA | NA | NA | NA | 1 | 1 | NA | NA | 1 | 0 | NA | NA | 1 | 1 | 2 | 0.5 |
| 17_281061 | 1 | 0 | 1 | 1 | NA | NA | NA | NA | 2 | 0.5 | NA | NA | 1 | 1 | NA | NA | 1 | 1 | 2 | 1 |
| 17_281064 | 1 | 0 | 1 | 1 | NA | NA | NA | NA | 2 | 0.5 | NA | NA | 1 | 1 | NA | NA | 1 | 0 | 2 | 0.5 |
| 17_281149 | 1 | 1 | 1 | 1 | NA | NA | NA | NA | 2 | 1 | NA | NA | NA | NA | NA | NA | 1 | 1 | 1 | 1 |
| 17_281150 | 1 | 1 | 1 | 1 | NA | NA | NA | NA | 2 | 1 | NA | NA | 1 | 1 | NA | NA | 2 | 1 | 3 | 1 |
| 17_281150 | NA | NA | 1 | 1 | NA | NA | NA | NA | 1 | 1 | NA | NA | 1 | 1 | NA | NA | NA | NA | 1 | 1 |
| 17_281151 | 1 | 1 | NA | NA | NA | NA | NA | NA | 1 | 1 | NA | NA | 1 | 1 | NA | NA | 1 | 0 | 2 | 0.5 |
| 17_281154 | 1 | 0 | 1 | 1 | NA | NA | NA | NA | 2 | 0.5 | NA | NA | 1 | 0 | NA | NA | 2 | 1 | 3 | 0.6667 |
| 17_281189 | 3 | 0.3333 | 3 | 0 | NA | NA | 1 | 0 | 7 | 0.1429 | 1 | 0 | 1 | 0 | NA | NA | 3 | 0.3333 | 5 | 0.2 |
| 17_281189 | 3 | 0.3333 | NA | NA | 1 | 0 | NA | NA | 4 | 0.25 | 1 | 0 | 1 | 1 | NA | NA | 3 | 0.3333 | 5 | 0.4 |
| 17_281203 | 2 | 0 | 5 | 0.4 | 7 | 0 | 5 | 0 | 19 | 0.1053 | 5 | 0 | 5 | 0 | 8 | 0.25 | 3 | 0 | 21 | 0.0952 |
| 17_281204 | 3 | 0.6667 | 7 | 0.7143 | 8 | 0.75 | 5 | 0.6 | 23 | 0.6957 | 6 | 0.6667 | 6 | 0.5 | 8 | 0.625 | 4 | 1 | 24 | 0.6667 |
| 17_281204 | 2 | 0.5 | 7 | 0.5714 | 9 | 0.5556 | 4 | 1 | 22 | 0.6364 | 6 | 0.3333 | 7 | 0.2857 | 7 | 0.2857 | 4 | 0 | 24 | 0.25 |
| 17_281204 | 3 | 1 | 8 | 1 | 9 | 0.8889 | 5 | 1 | 25 | 0.96 | 6 | 1 | 7 | 0.7143 | 9 | 0.8889 | 4 | 1 | 26 | 0.8846 |
| 17_281205 | 3 | 0 | 10 | 0.2 | 11 | 0.0909 | 5 | 0.2 | 29 | 0.1379 | 6 | 0.1667 | 6 | 0 | 9 | 0.1111 | 4 | 0.25 | 25 | 0.12 |
| 17_281206 | 2 | 0.5 | 11 | 0.5455 | 9 | 0.2222 | 5 | 0.2 | 27 | 0.3704 | 6 | 0.1667 | 7 | 0 | 9 | 0.5556 | 4 | 0.5 | 26 | 0.3077 |
| 17_281266 | 2 | 0 | 7 | 0.4286 | 6 | 0.1667 | 3 | 0.6667 | 18 | 0.3333 | 5 | 0.2 | 3 | 0 | 3 | 0 | 4 | 0 | 15 | 0.0667 |
| 17_281271 | NA | NA | 2 | 1 | 1 | 1 | 1 | 1 | 4 | 1 | 1 | 1 | NA | NA | NA | NA | NA | NA | 1 | 1 |
| 17_282271 | 2 | 0.5 | NA | NA | NA | NA | NA | NA | 2 | 0.5 | NA | NA | NA | NA | NA | NA | 2 | 1 | 2 | 1 |
| 17_282785 | NA | NA | NA | NA | NA | NA | 1 | 1 | 1 | 1 | NA | NA | NA | NA | NA | NA | 1 | 1 | 1 | 1 |
| 17_282788 | NA | NA | NA | NA | NA | NA | 1 | 1 | 1 | 1 | NA | NA | NA | NA | NA | NA | 1 | 1 | 1 | 1 |
| 17_282790 | NA | NA | NA | NA | NA | NA | 2 | 0.5 | 2 | 0.5 | NA | NA | NA | NA | NA | NA | 1 | 1 | 1 | 1 |
| 17_282794 | 1 | 0 | NA | NA | NA | NA | 2 | 0.5 | 3 | 0.3333 | NA | NA | NA | NA | 1 | 1 | 1 | 1 | 2 | 1 |
| 17_282796 | 2 | 0.5 | NA | NA | 1 | 0 | 2 | 0 | 5 | 0.2 | NA | NA | NA | NA | 1 | 0 | 1 | 1 | 2 | 0.5 |
| 17_282798 | 4 | 0.25 | NA | NA | 1 | 1 | 2 | 0.5 | 7 | 0.4286 | NA | NA | NA | NA | 2 | 1 | 2 | 0.5 | 4 | 0.75 |
| 17_282801 | 2 | 0.5 | NA | NA | NA | NA | 2 | 0 | 4 | 0.25 | NA | NA | NA | NA | 4 | 0.5 | 1 | 0 | 5 | 0.4 |
| 17_282811 | 4 | 0.5 | NA | NA | NA | NA | 1 | 1 | 5 | 0.6 | NA | NA | 1 | 1 | 3 | 0.3333 | NA | NA | 4 | 0.5 |
| 17_282813 | 4 | 1 | NA | NA | 1 | 1 | 1 | 1 | 6 | 1 | NA | NA | 1 | 1 | 5 | 1 | 2 | 1 | 8 | 1 |
| 17_282813 | 4 | 0 | NA | NA | 1 | 1 | 2 | 0 | 7 | 0.1429 | NA | NA | NA | NA | 2 | 0 | 1 | 1 | 3 | 0.3333 |
| 17_282814 | 4 | 0.5 | NA | NA | 1 | 0 | 2 | 0.5 | 7 | 0.4286 | NA | NA | 1 | 0 | 4 | 0.25 | 1 | 1 | 6 | 0.3333 |

| | | | | | | | | | | | | | | | | | | | | |
|---|---|---|---|---|---|---|---|---|---|---|---|---|---|---|---|---|---|---|---|---|
| 17_2839039 | 1 | 1 | NA | NA | 1 | 0 | NA | NA | 2 | 0.5 | NA | NA | NA | NA | 2 | 0.5 | 1 | 1 | 3 | 0.6667 |
| 17_2839066 | 1 | 1 | NA | NA | NA | NA | NA | NA | 1 | 1 | NA | NA | NA | NA | 1 | 1 | 1 | 1 | 2 | 1 |
| 17_2839085 | 1 | 1 | NA | NA | NA | NA | NA | NA | 1 | 1 | NA | NA | NA | NA | 2 | 0.5 | NA | NA | 2 | 0.5 |
| 17_2839151 | 1 | 1 | NA | NA | NA | NA | NA | NA | 1 | 1 | NA | NA | NA | NA | 2 | 1 | 3 | 0.6667 | 5 | 0.8 |
| 17_2839161 | 1 | 1 | NA | NA | NA | NA | NA | NA | 1 | 1 | NA | NA | 1 | 0 | 2 | 0 | 3 | 0.3333 | 6 | 0.1667 |
| 17_2839167 | 1 | 1 | NA | NA | NA | NA | NA | NA | 1 | 1 | NA | NA | 1 | 1 | 2 | 1 | 3 | 1 | 6 | 1 |
| 17_2839183 | 1 | 1 | NA | NA | NA | NA | NA | NA | 1 | 1 | NA | NA | NA | NA | 2 | 1 | 2 | 1 | 4 | 1 |
| 17_2839269 | NA | NA | 1 | 1 | NA | NA | NA | NA | 1 | 1 | NA | NA | 1 | 0 | 3 | 0.3333 | 2 | 0 | 6 | 0.1667 |
| 17_2839287 | NA | NA | 1 | 1 | NA | NA | NA | NA | 1 | 1 | NA | NA | 1 | 1 | 3 | 0.6667 | 3 | 0.6667 | 7 | 0.7143 |
| 17_2839334 | NA | NA | NA | NA | NA | NA | 1 | 1 | 1 | 1 | NA | NA | NA | NA | 3 | 0.6667 | 3 | 0.3333 | 6 | 0.5 |
| 17_2839335 | NA | NA | 1 | 1 | NA | NA | 1 | 0 | 2 | 0.5 | NA | NA | 1 | 1 | 3 | 0.3333 | 2 | 0.5 | 6 | 0.5 |
| 17_2839339 | NA | NA | 1 | 1 | NA | NA | 1 | 0 | 2 | 0.5 | NA | NA | 1 | 0 | 3 | 0.3333 | 3 | 0 | 7 | 0.1429 |
| 17_2839346 | NA | NA | 1 | 1 | NA | NA | 1 | 1 | 2 | 1 | NA | NA | 1 | 1 | 3 | 1 | 2 | 1 | 6 | 1 |
| 17_2839353 | NA | NA | 1 | 1 | NA | NA | 1 | 0 | 2 | 0.5 | NA | NA | 1 | 1 | 2 | 0 | 2 | 0 | 5 | 0.2 |
| 17_2839357 | NA | NA | 1 | 1 | NA | NA | 1 | 1 | 2 | 1 | NA | NA | 1 | 1 | 2 | 1 | 3 | 1 | 6 | 1 |
| 17_2842449 | 11 | 0.3636 | 14 | 0.7143 | 11 | 0.4545 | 13 | 0.3077 | 49 | 0.4694 | 15 | 0.6 | 16 | 0.625 | 14 | 0.5 | 12 | 0.5833 | 57 | 0.5789 |
| 17_2842462 | 8 | 0.375 | 16 | 0.625 | 11 | 0.7273 | 13 | 0.4615 | 48 | 0.5625 | 14 | 0.6429 | 15 | 0.6 | 13 | 0.5385 | 10 | 0.7 | 52 | 0.6154 |
| 17_2842473 | 12 | 0.1667 | 16 | 0.375 | 13 | 0.2308 | 16 | 0.3125 | 57 | 0.2807 | 17 | 0.0588 | 15 | 0.1333 | 16 | 0.125 | 12 | 0.3333 | 60 | 0.15 |
| 17_2842474 | 11 | 0 | 15 | 0.0667 | 12 | 0.1667 | 13 | 0.0769 | 51 | 0.0784 | 12 | 0.0833 | 15 | 0.0667 | 13 | 0 | 11 | 0.0909 | 51 | 0.0588 |
| 17_2851749 | NA | NA | 1 | 1 | NA | NA | NA | NA | 1 | 1 | NA | NA | 1 | 1 | NA | NA | NA | NA | 1 | 1 |
| 17_2851881 | NA | NA | 3 | 1 | 1 | 1 | NA | NA | 4 | 1 | 4 | 1 | 2 | 0 | NA | NA | 2 | 0.5 | 8 | 0.625 |
| 17_2851887 | NA | NA | 3 | 0.6667 | 1 | 1 | NA | NA | 4 | 0.75 | 2 | 1 | 2 | 1 | 1 | 1 | 4 | 1 | 9 | 1 |
| 17_2851924 | NA | NA | 3 | 0 | 1 | 1 | NA | NA | 4 | 0.25 | 4 | 0.25 | 2 | 0.5 | 1 | 0 | 5 | 0.2 | 12 | 0.25 |
| 17_2851987 | NA | NA | 4 | 0 | 2 | 0.5 | NA | NA | 6 | 0.1667 | 4 | 0 | 1 | 0 | 1 | 0 | 5 | 0.2 | 11 | 0.0909 |
| 17_2852040 | 1 | 1 | 5 | 0 | 1 | 1 | 1 | 0 | 8 | 0.25 | 5 | 0 | 2 | 0.5 | 3 | 0 | 4 | 0.25 | 14 | 0.1429 |
| 17_2853481 | NA | NA | 1 | 1 | 5 | 1 | 1 | 1 | 7 | 1 | NA | NA | 2 | 1 | NA | NA | 1 | 1 | 3 | 1 |
| 17_2853489 | NA | NA | NA | NA | 5 | 0.2 | 1 | 0 | 6 | 0.1667 | 1 | 0 | 2 | 0 | 1 | 0 | 1 | 1 | 5 | 0.2 |
| 17_2854524 | 1 | 1 | 1 | 0 | 5 | 0.4 | 1 | 1 | 8 | 0.5 | 1 | 1 | 3 | 0.3333 | 1 | 0 | 1 | 0 | 6 | 0.3333 |
| 17_2861863 | 16 | 0.1875 | 15 | 0.0667 | 8 | 0.125 | 15 | 0.0667 | 54 | 0.1111 | 11 | 0.1818 | 9 | 0 | 8 | 0.25 | 4 | 0.5 | 32 | 0.1875 |
| 17_2861871 | 15 | 0 | 15 | 0.0667 | 6 | 0.1667 | 12 | 0.0833 | 48 | 0.0625 | 9 | 0 | 8 | 0.125 | 8 | 0 | 3 | 0.3333 | 28 | 0.0714 |
| 17_2861876 | 16 | 0.125 | 13 | 0 | 7 | 0.1429 | 10 | 0.1 | 46 | 0.087 | 8 | 0.25 | 10 | 0 | 7 | 0 | 4 | 0.25 | 29 | 0.1034 |
| 17_2861879 | 11 | 0.1818 | 10 | 0.1 | 7 | 0.1429 | 10 | 0.2 | 38 | 0.1579 | 9 | 0.1111 | 8 | 0.125 | 8 | 0.125 | 2 | 0.5 | 27 | 0.1481 |
| 17_2861958 | 10 | 0 | 10 | 0 | 6 | 0 | 8 | 0.25 | 34 | 0.0588 | 7 | 0.2857 | 6 | 0.1667 | 5 | 0.2 | 2 | 0.5 | 20 | 0.25 |
| 17_2861994 | 5 | 0.2 | 4 | 0 | 3 | 0 | 5 | 0 | 17 | 0.0588 | 5 | 0.2 | 4 | 0 | 4 | 0.25 | 1 | 1 | 14 | 0.2143 |
| 17_2861995 | 8 | 1 | 7 | 0.8571 | 3 | 0.6667 | 7 | 1 | 25 | 0.92 | 5 | 1 | 5 | 0.8 | 6 | 0.8333 | 1 | 1 | 17 | 0.8824 |
| 17_2861998 | 7 | 0.1429 | 7 | 0.5714 | 2 | 0.5 | 5 | 0.8 | 21 | 0.4762 | 4 | 0.75 | 4 | 0.5 | 3 | 0 | 1 | 1 | 12 | 0.5 |
| 17_2879489 | 1 | 0 | 2 | 0.5 | 1 | 0 | 1 | 1 | 5 | 0.4 | NA | NA | NA | NA | 1 | 0 | 1 | 1 | 2 | 0.5 |
| 17_2879522 | 1 | 1 | 2 | 1 | 1 | 1 | 1 | 1 | 5 | 1 | NA | NA | NA | NA | 1 | 1 | 1 | 1 | 2 | 1 |
| 17_2880241 | 1 | 0 | 2 | 0 | 1 | 0 | 1 | 1 | 5 | 0.2 | NA | NA | 1 | 0 | 1 | 1 | 1 | 0 | 3 | 0.3333 |
| 17_2880674 | 5 | 0.2 | 4 | 0.25 | 1 | 0 | 4 | 0.75 | 14 | 0.3571 | 5 | 0.4 | NA | NA | 5 | 0.4 | 5 | 0.2 | 15 | 0.3333 |
| 17_2880695 | 4 | 0 | 5 | 0 | 1 | 0 | 4 | 0.25 | 14 | 0.0714 | 3 | 0 | NA | NA | 4 | 0.5 | 2 | 0.5 | 9 | 0.3333 |
| 17_2880726 | 2 | 0.5 | 4 | 0 | 1 | 0 | 3 | 0 | 10 | 0.1 | 2 | 0.5 | NA | NA | 4 | 0.5 | 4 | 0.25 | 10 | 0.4 |
| 17_2880759 | 2 | 0.5 | 4 | 0 | 1 | 0 | 3 | 0 | 10 | 0.1 | 4 | 0.25 | NA | NA | 4 | 0 | 4 | 0.25 | 12 | 0.1667 |
| 17_2880800 | 2 | 1 | 3 | 0.3333 | 1 | 1 | 3 | 0.6667 | 9 | 0.6667 | 3 | 0.6667 | NA | NA | 4 | 1 | 2 | 0.5 | 9 | 0.7778 |
| 17_2880860 | 4 | 0 | 7 | 0.2857 | 3 | 0.6667 | 6 | 0.5 | 20 | 0.35 | 5 | 0.6 | 4 | 0 | 3 | 0.6667 | 6 | 0.5 | 18 | 0.4444 |
| 17_2880869 | 1 | 0 | 6 | 0.3333 | 2 | 0.5 | 5 | 0.6 | 14 | 0.4286 | 4 | 0.25 | 3 | 0.3333 | 1 | 0 | 4 | 0.5 | 12 | 0.3333 |
| 17_2880879 | 1 | 0 | 3 | 1 | 1 | 0 | 2 | 1 | 7 | 0.7143 | 1 | 1 | NA | NA | 1 | 0 | 1 | 0 | 3 | 0.3333 |
| 17_2880880 | 3 | 0.6667 | 7 | 0.5714 | 1 | 1 | 6 | 0.8333 | 17 | 0.7059 | 3 | 0.6667 | 3 | 0.3333 | 2 | 1 | 6 | 0.6667 | 14 | 0.6429 |
| 17_2880899 | 3 | 0.6667 | 6 | 1 | 2 | 1 | 5 | 1 | 16 | 0.9375 | 3 | 1 | 3 | 0.6667 | 3 | 0.6667 | 7 | 0.7143 | 16 | 0.75 |
| 17_2881860 | NA | NA | 4 | 0.75 | 1 | 0 | 1 | 1 | 6 | 0.6667 | NA | NA | 3 | 0.3333 | 1 | 0 | 3 | 0.3333 | 7 | 0.2857 |
| 17_2881863 | NA | NA | 5 | 0 | NA | NA | 5 | 0.2 | 10 | 0.1 | NA | NA | 3 | 0.3333 | 1 | 0 | 4 | 0.25 | 8 | 0.25 |
| 17_2881868 | NA | NA | 4 | 1 | NA | NA | 5 | 1 | 9 | 1 | NA | NA | 2 | 1 | 1 | 1 | 5 | 0.8 | 8 | 0.875 |
| 17_2896537 | 11 | 0.3636 | 3 | 0 | 10 | 0 | 5 | 0.4 | 29 | 0.2069 | 2 | 0 | 8 | 0.375 | 5 | 0.2 | 3 | 0.3333 | 18 | 0.2778 |
| 17_2896621 | 9 | 0.6667 | 3 | 0.3333 | 7 | 0.1429 | 5 | 0.6 | 24 | 0.4583 | 5 | 0.8 | 8 | 0.75 | 4 | 0.25 | 4 | 0.75 | 21 | 0.6667 |
| 17_2896631 | 8 | 0.25 | 4 | 0.25 | 8 | 0.375 | 7 | 0.1429 | 27 | 0.2593 | 4 | 0 | 7 | 0 | 5 | 0.2 | 7 | 0.1429 | 23 | 0.087 |
| 17_2896635 | 11 | 0.8182 | 3 | 1 | 11 | 0.9091 | 7 | 0.8571 | 32 | 0.875 | 5 | 1 | 7 | 1 | 5 | 1 | 6 | 1 | 23 | 1 |
| 17_2896636 | 11 | 0.1818 | 5 | 0 | 10 | 0 | 9 | 0 | 35 | 0.0571 | 5 | 0 | 8 | 0.25 | 7 | 0.1429 | 7 | 0 | 27 | 0.1111 |
| 17_2896637 | 10 | 0.2 | 5 | 0 | 10 | 0.1 | 8 | 0.125 | 33 | 0.1212 | 5 | 0 | 10 | 0.3 | 8 | 0.25 | 7 | 0 | 30 | 0.1667 |
| 17_2896642 | 12 | 0.6667 | 6 | 0.6667 | 9 | 0.5556 | 9 | 0.3333 | 36 | 0.5556 | 5 | 0.6 | 11 | 0.6364 | 5 | 0.8 | 7 | 0.7143 | 28 | 0.6786 |
| 17_2896678 | 13 | 0.7692 | 6 | 0.5 | 10 | 0.5 | 9 | 0.6667 | 38 | 0.6316 | 6 | 0.5 | 15 | 0.8 | 8 | 0.5 | 8 | 0.5 | 37 | 0.6216 |
| 17_2896683 | 14 | 0.4286 | 6 | 0.3333 | 10 | 0.5 | 11 | 0.3636 | 41 | 0.4146 | 6 | 0 | 15 | 0.5333 | 10 | 0.3 | 9 | 0.2222 | 40 | 0.325 |
| 17_2896696 | 13 | 0.0769 | 8 | 0 | 9 | 0.1111 | 7 | 0.1429 | 37 | 0.0811 | 7 | 0.1429 | 16 | 0.0625 | 10 | 0.1 | 9 | 0.1111 | 42 | 0.0952 |
| 17_2896700 | 14 | 0.0714 | 7 | 0.2857 | 11 | 0 | 11 | 0.2727 | 43 | 0.1395 | 7 | 0.1429 | 16 | 0.25 | 10 | 0 | 9 | 0 | 42 | 0.119 |
| 17_2896718 | 11 | 0.7273 | 9 | 1 | 8 | 0.375 | 9 | 0.6667 | 37 | 0.7027 | 7 | 0.4286 | 16 | 0.6875 | 10 | 0.7 | 7 | 0.5714 | 40 | 0.625 |
| 17_2896718 | 10 | 0.9 | 7 | 0.4286 | 8 | 0.625 | 8 | 0.625 | 33 | 0.6667 | 4 | 1 | 11 | 0.9091 | 7 | 0.5714 | 7 | 1 | 29 | 0.8621 |

| | | | | | | | | | | | | | | | | | | | | |
|---|---|---|---|---|---|---|---|---|---|---|---|---|---|---|---|---|---|---|---|---|
| 17_289672 | 13 | 0 | 5 | 0 | 10 | 0.2 | 7 | 0.1429 | 35 | 0.0857 | 5 | 0.2 | 16 | 0.125 | 10 | 0.3 | 8 | 0 | 39 | 0.1538 |
| 17_290259 | NA | NA | NA | NA | NA | NA | 2 | 1 | 2 | 1 | NA | NA | NA | NA | 2 | 0.5 | NA | NA | 2 | 0.5 |
| 17_290260 | 1 | 1 | NA | NA | NA | NA | 1 | 1 | 2 | 1 | NA | NA | NA | NA | 2 | 1 | 1 | 1 | 3 | 1 |
| 17_290402 | 1 | 0 | 1 | 0 | NA | NA | 3 | 0.6667 | 5 | 0.4 | 2 | 0.5 | 1 | 1 | 3 | 1 | 2 | 0.5 | 8 | 0.75 |
| 17_290402 | 1 | 1 | 1 | 1 | NA | NA | 4 | 1 | 6 | 1 | 2 | 1 | 1 | 0 | 3 | 1 | 2 | 1 | 8 | 0.875 |
| 17_290404 | 1 | 0 | 1 | 0 | NA | NA | 3 | 0.3333 | 5 | 0.2 | 1 | 0 | 1 | 0 | 2 | 0 | 2 | 0.5 | 6 | 0.1667 |
| 17_290555 | 1 | 0 | 1 | 0 | NA | NA | 5 | 0.2 | 7 | 0.1429 | NA | NA | 2 | 0 | 6 | 0 | 3 | 0.3333 | 11 | 0.0909 |
| 17_290557 | 1 | 0 | 1 | 0 | NA | NA | 6 | 0.3333 | 8 | 0.25 | 1 | 0 | 1 | 0 | 7 | 0.2857 | 2 | 1 | 11 | 0.3636 |
| 17_290557 | 1 | 0 | 1 | 1 | NA | NA | 6 | 0 | 8 | 0.125 | 1 | 0 | 3 | 0.3333 | 4 | 0 | 2 | 0 | 10 | 0.1 |
| 17_290557 | 1 | 1 | 1 | 1 | NA | NA | 5 | 0.6 | 7 | 0.7143 | 1 | 1 | 2 | 0.5 | 7 | 0.5714 | 3 | 0.6667 | 13 | 0.6154 |
| 17_290558 | 1 | 1 | 1 | 0 | NA | NA | 7 | 0.2857 | 9 | 0.3333 | NA | NA | 3 | 0 | 7 | 0.1429 | 3 | 0 | 13 | 0.0769 |
| 17_290569 | 2 | 0.5 | 3 | 0 | NA | NA | 7 | 0.2857 | 12 | 0.25 | 3 | 0 | 4 | 0 | 8 | 0.25 | 2 | 1 | 17 | 0.2353 |
| 17_290571 | 1 | 1 | 3 | 0 | 1 | 1 | 7 | 0.1429 | 12 | 0.25 | 5 | 0.4 | 5 | 0.6 | 8 | 0.625 | 4 | 0.75 | 22 | 0.5909 |
| 17_290573 | 2 | 0 | 4 | 0.25 | 2 | 0 | 7 | 0 | 15 | 0.0667 | 5 | 0.2 | 5 | 0 | 6 | 0 | 4 | 0 | 20 | 0.05 |
| 17_291830 | 1 | 0 | 3 | 0 | 3 | 0 | 4 | 0.25 | 11 | 0.0909 | 1 | 0 | 4 | 0 | 2 | 0.5 | 3 | 0 | 10 | 0.1 |
| 17_291830 | 2 | 0 | 4 | 0 | 3 | 0 | 4 | 0.25 | 13 | 0.0769 | 1 | 0 | 4 | 0 | 2 | 0.5 | 3 | 0 | 10 | 0.1 |
| 17_291858 | 3 | 1 | 3 | 1 | 2 | 1 | 3 | 1 | 11 | 1 | 1 | 1 | 5 | 1 | 1 | 1 | 5 | 1 | 12 | 1 |
| 17_291859 | 3 | 1 | 4 | 0.5 | 3 | 1 | 5 | 1 | 15 | 0.8667 | 1 | 1 | 3 | 1 | 2 | 1 | 5 | 1 | 11 | 1 |
| 17_291859 | 3 | 0.6667 | 4 | 0.75 | 4 | 0.5 | 4 | 0.75 | 15 | 0.6667 | 1 | 0 | 4 | 0.25 | 1 | 1 | 3 | 0.6667 | 9 | 0.4444 |
| 17_291862 | 2 | 1 | 4 | 0.5 | 2 | 1 | 4 | 1 | 12 | 0.8333 | 2 | 1 | 5 | 0.4 | 3 | 1 | 4 | 0.75 | 14 | 0.7143 |
| 17_291862 | 4 | 0 | 3 | 0 | 3 | 0 | 5 | 0.2 | 15 | 0.0667 | 2 | 0 | 7 | 0.1429 | 4 | 0.25 | 5 | 0.2 | 18 | 0.1667 |
| 17_291862 | 4 | 0.75 | 4 | 0.25 | 3 | 0.3333 | 3 | 0.3333 | 14 | 0.4286 | 2 | 0 | 7 | 0.4286 | 4 | 0 | 5 | 0.2 | 18 | 0.2222 |
| 17_291903 | 3 | 0.3333 | 2 | 0.5 | 5 | 0.2 | 1 | 0 | 11 | 0.2727 | 3 | 0.3333 | NA | NA | 2 | 0.5 | 2 | 0.5 | 7 | 0.4286 |
| 17_291907 | 3 | 0.3333 | 2 | 1 | 3 | 0.6667 | 1 | 0 | 9 | 0.5556 | 3 | 1 | NA | NA | 2 | 0.5 | 3 | 0 | 8 | 0.5 |
| 17_291907 | 4 | 0.5 | 2 | 1 | 4 | 0.5 | 2 | 0 | 12 | 0.5 | 4 | 0.75 | NA | NA | 2 | 1 | 3 | 0.3333 | 9 | 0.6667 |
| 17_291908 | 4 | 0.25 | 2 | 0.5 | 2 | 0.5 | 2 | 0.5 | 10 | 0.4 | 2 | 0.5 | NA | NA | 1 | 0 | 2 | 0 | 5 | 0.2 |
| 17_291909 | 4 | 0.5 | 2 | 0.5 | 4 | 0.25 | 2 | 0.5 | 12 | 0.4167 | 4 | 0.25 | NA | NA | 2 | 0.5 | 2 | 0.5 | 8 | 0.375 |
| 17_291909 | 4 | 0.25 | 2 | 0.5 | 4 | 0.5 | 2 | 0 | 12 | 0.3333 | 3 | 0.3333 | NA | NA | 2 | 0 | 2 | 0.5 | 7 | 0.2857 |
| 17_291910 | 3 | 0.3333 | 2 | 0 | NA | NA | 1 | 0 | 6 | 0.1667 | 2 | 0.5 | NA | NA | 2 | 0 | 2 | 0.5 | 6 | 0.3333 |
| 17_291911 | 3 | 1 | 1 | 1 | NA | NA | 2 | 1 | 6 | 1 | 4 | 1 | NA | NA | 1 | 1 | 2 | 1 | 7 | 1 |
| 17_291913 | 4 | 0 | 2 | 0 | 2 | 0.5 | 2 | 0 | 10 | 0.1 | 4 | 0 | NA | NA | 2 | 0 | 2 | 0.5 | 8 | 0.125 |
| 17_291913 | 3 | 0.3333 | 2 | 0.5 | 2 | 0.5 | 2 | 0.5 | 9 | 0.4444 | 4 | 0.5 | NA | NA | 2 | 0.5 | 2 | 0 | 8 | 0.375 |
| 17_291914 | 2 | 0.5 | 2 | 0.5 | 2 | 0.5 | 1 | 1 | 7 | 0.5714 | 4 | 0.75 | NA | NA | 1 | 1 | 2 | 1 | 7 | 0.8571 |
| 17_291916 | 3 | 0 | 2 | 0.5 | 2 | 0.5 | 2 | 0.5 | 9 | 0.3333 | 3 | 0.3333 | NA | NA | 2 | 0 | 2 | 0 | 7 | 0.1429 |
| 17_292606 | 35 | 0.2 | 32 | 0.0625 | 22 | 0.1364 | 23 | 0.3043 | 112 | 0.1696 | 13 | 0.2308 | 31 | 0.0323 | 32 | 0.125 | 32 | 0.1562 | 108 | 0.1204 |
| 17_292621 | NA | NA | 3 | 0 | 3 | 0 | 4 | 0.25 | 10 | 0.1 | 2 | 0 | 2 | 0 | 3 | 0.3333 | 5 | 0.2 | 12 | 0.1667 |
| 17_293174 | 7 | 0.2857 | 3 | 0 | 5 | 0.6 | 4 | 0.25 | 19 | 0.3158 | 7 | 0.5714 | 5 | 0 | 7 | 0.1429 | 4 | 0 | 23 | 0.2174 |
| 17_293175 | 10 | 0.2 | 4 | 0.5 | 6 | 0.5 | 6 | 0.1667 | 26 | 0.3077 | 7 | 0 | 8 | 0.25 | 7 | 0 | 4 | 0.5 | 26 | 0.1538 |
| 17_293176 | 13 | 0.1538 | 4 | 0 | 5 | 0 | 6 | 0 | 28 | 0.0714 | 7 | 0.1429 | 7 | 0 | 7 | 0.1429 | 5 | 0 | 26 | 0.0769 |
| 17_293191 | 17 | 0.0588 | 5 | 0 | 11 | 0.0909 | 7 | 0 | 40 | 0.05 | 10 | 0.3 | 11 | 0 | 11 | 0.1818 | 9 | 0.1111 | 41 | 0.1463 |
| 17_293195 | 18 | 0.3889 | 7 | 0.1429 | 11 | 0 | 5 | 0.4 | 41 | 0.2439 | 9 | 0.3333 | 12 | 0.1667 | 15 | 0.4667 | 9 | 0.4444 | 45 | 0.3556 |
| 17_293778 | 26 | 0.1923 | 9 | 0.2222 | 20 | 0.05 | 29 | 0.069 | 84 | 0.119 | 12 | 0.3333 | 29 | 0.2759 | 32 | 0.2188 | 25 | 0.16 | 98 | 0.2347 |
| 17_294701 | 6 | 0 | 3 | 0 | 2 | 0 | 3 | 0.3333 | 14 | 0.0714 | 2 | 0 | 7 | 0 | 3 | 0 | 4 | 0.25 | 16 | 0.0625 |
| 17_294703 | 6 | 0.8333 | 4 | 1 | 1 | 1 | 6 | 0.6667 | 17 | 0.8235 | 3 | 1 | 4 | 0.5 | 3 | 1 | 4 | 1 | 14 | 0.8571 |
| 17_294705 | 5 | 0 | 2 | 0 | 2 | 0 | 4 | 0.5 | 13 | 0.1538 | 2 | 0 | 7 | 0.1429 | 3 | 0 | 4 | 0 | 16 | 0.0625 |
| 17_295011 | 11 | 0.1818 | 12 | 0.25 | 11 | 0.3636 | 7 | 0.2857 | 41 | 0.2683 | 4 | 0.5 | 6 | 0.3333 | 9 | 0.3333 | 5 | 0.2 | 24 | 0.3333 |
| 17_295206 | 1 | 1 | NA | NA | 1 | 0 | NA | NA | 2 | 0.5 | 1 | 0 | 1 | 1 | 1 | 1 | 1 | 0 | 4 | 0.5 |
| 17_295206 | 1 | 1 | NA | NA | 2 | 1 | NA | NA | 3 | 1 | NA | NA | 1 | 1 | 1 | 1 | 1 | 1 | 3 | 1 |
| 17_295208 | 1 | 0 | NA | NA | 2 | 0 | 1 | 1 | 4 | 0.25 | NA | NA | 1 | 1 | 1 | 1 | 1 | 0 | 3 | 0.6667 |
| 17_295404 | NA | NA | 1 | 1 | NA | NA | NA | NA | 1 | 1 | NA | NA | NA | NA | 1 | 1 | NA | NA | 1 | 1 |
| 17_295405 | NA | NA | 1 | 1 | NA | NA | NA | NA | 1 | 1 | NA | NA | NA | NA | 1 | 1 | NA | NA | 1 | 1 |
| 17_295476 | 1 | 1 | NA | NA | NA | NA | NA | NA | 1 | 1 | 1 | 1 | NA | NA | NA | NA | NA | NA | 1 | 1 |
| 17_295515 | 53 | 0.0566 | 58 | 0.069 | 48 | 0.0417 | 57 | 0.0877 | 216 | 0.0648 | 56 | 0.0357 | 47 | 0.0426 | 55 | 0.0909 | 46 | 0.0652 | 204 | 0.0588 |
| 17_295981 | NA | NA | 1 | 1 | 1 | 1 | 1 | 1 | 3 | 1 | NA | NA | 1 | 1 | NA | NA | 3 | 0.6667 | 4 | 0.75 |
| 17_295983 | NA | NA | 2 | 0.5 | 1 | 0 | 1 | 0 | 4 | 0.25 | NA | NA | 1 | 0 | NA | NA | 3 | 0.6667 | 4 | 0.5 |
| 17_297874 | 4 | 0.5 | 1 | 0 | 2 | 0 | 2 | 0.5 | 9 | 0.3333 | 4 | 0.25 | 1 | 1 | 3 | 0.3333 | 2 | 0 | 10 | 0.3 |
| 17_297874 | 4 | 0.25 | 1 | 0 | 4 | 0 | 3 | 0 | 12 | 0.0833 | 5 | 0.2 | 4 | 0 | 4 | 0 | 1 | 0 | 14 | 0.0714 |
| 17_297875 | 3 | 0.6667 | NA | NA | 4 | 0 | 4 | 0 | 11 | 0.1818 | 5 | 0 | 4 | 0.25 | 4 | 0.25 | 2 | 0.5 | 15 | 0.2 |
| 17_298239 | 5 | 0.6 | 2 | 0 | 5 | 0.6 | 3 | 1 | 15 | 0.6 | 5 | 1 | 4 | 0.75 | 5 | 1 | 3 | 0.6667 | 17 | 0.8824 |
| 17_298239 | 4 | 0.5 | 2 | 0 | 6 | 0.1667 | 4 | 0.25 | 16 | 0.25 | 5 | 0.2 | 4 | 0.25 | 6 | 0.6667 | 3 | 1 | 18 | 0.5 |
| 17_298239 | 4 | 0.25 | 2 | 0 | 5 | 0.2 | 4 | 0.75 | 15 | 0.3333 | 5 | 1 | 4 | 0.5 | 3 | 0.3333 | 2 | 0.5 | 14 | 0.6429 |
| 17_298240 | 5 | 0.2 | 2 | 0 | 5 | 0 | 1 | 0 | 13 | 0.0769 | 5 | 0.4 | 4 | 0 | 6 | 0.3333 | 2 | 0.5 | 17 | 0.2941 |
| 17_298242 | 5 | 0.2 | 2 | 0 | 8 | 0 | 4 | 0 | 19 | 0.0526 | 3 | 0 | 4 | 0 | 7 | 0.1429 | 3 | 0.3333 | 17 | 0.1176 |
| 17_298242 | 3 | 1 | 1 | 0 | 3 | 0.6667 | 3 | 0.6667 | 10 | 0.7 | 4 | 1 | 2 | 1 | 5 | 0.6 | 2 | 0.5 | 13 | 0.7692 |
| 17_298243 | 4 | 0 | NA | NA | 7 | 0 | 4 | 0.25 | 15 | 0.0667 | 6 | 0.1667 | 1 | 0 | 7 | 0 | 2 | 0 | 16 | 0.0625 |

| | | | | | | | | | | | | | | | | | | | | |
|---|---|---|---|---|---|---|---|---|---|---|---|---|---|---|---|---|---|---|---|---|
| 17_2982444 | 5 | 0.2 | 1 | 0 | 9 | 0.7778 | 5 | 1 | 20 | 0.65 | 6 | 0.6667 | 3 | 0.6667 | 7 | 0.1429 | 5 | 0.4 | 21 | 0.4286 |
| 17_2982549 | 7 | 0.1429 | 2 | 0 | 12 | 0 | 8 | 0.125 | 29 | 0.069 | 5 | 0 | 4 | 0 | 7 | 0.1429 | 4 | 0.25 | 20 | 0.1 |
| 17_2982576 | 9 | 0.1111 | 3 | 0 | 14 | 0.1429 | 7 | 0.1429 | 33 | 0.1212 | 6 | 0 | 5 | 0.2 | 9 | 0.3333 | 6 | 0.5 | 26 | 0.2692 |
| 17_2984569 | 3 | 0.3333 | 3 | 0.6667 | 1 | 0 | 1 | 0 | 8 | 0.375 | 2 | 0 | 2 | 0 | 6 | 0.5 | NA | NA | 10 | 0.3 |
| 17_2984592 | 2 | 0 | 3 | 1 | 1 | 1 | 1 | 0 | 7 | 0.5714 | 4 | 0.5 | 2 | 1 | 7 | 0.4286 | NA | NA | 13 | 0.5385 |
| 17_2984598 | 2 | 1 | 3 | 1 | 1 | 1 | 1 | 1 | 7 | 1 | 5 | 1 | 2 | 1 | 6 | 0.6667 | NA | NA | 13 | 0.8462 |
| 17_2985488 | 2 | 0 | 4 | 0.25 | 1 | 0 | 2 | 0 | 9 | 0.1111 | 4 | 0.75 | NA | NA | 5 | 0.6 | NA | NA | 9 | 0.6667 |
| 17_2986019 | 7 | 0.1429 | 4 | 1 | 1 | 0 | 3 | 0.3333 | 15 | 0.4 | 5 | 0.2 | 3 | 0 | 8 | 0 | 1 | 1 | 17 | 0.1176 |
| 17_2986022 | 6 | 0 | 4 | 0 | 1 | 0 | 3 | 0.3333 | 14 | 0.0714 | 4 | 0 | 4 | 0 | 9 | 0.1111 | 1 | 0 | 18 | 0.0556 |
| 17_2986023 | 7 | 0 | 4 | 1 | 1 | 0 | 3 | 0.3333 | 15 | 0.3333 | 5 | 0.6 | 4 | 0.25 | 8 | 0.25 | 1 | 0 | 18 | 0.3333 |
| 17_2986032 | 7 | 0.1429 | 4 | 0.5 | NA | NA | 3 | 0.6667 | 14 | 0.3571 | 5 | 0.4 | 4 | 0.75 | 9 | 0.6667 | 1 | 1 | 19 | 0.6316 |
| 17_2986040 | 7 | 0 | 4 | 0 | 1 | 0 | 3 | 0.3333 | 15 | 0.0667 | 5 | 0 | 4 | 0.25 | 10 | 0.1 | 1 | 0 | 20 | 0.1 |
| 17_2986041 | 8 | 0.625 | 3 | 0.6667 | 1 | 0 | 3 | 1 | 15 | 0.6667 | 4 | 0.5 | 4 | 0.5 | 8 | 0.5 | 1 | 0 | 17 | 0.4706 |
| 17_2986042 | 5 | 0.2 | 3 | 0 | NA | NA | 3 | 0.3333 | 11 | 0.1818 | 5 | 0 | 3 | 0 | 8 | 0.125 | 1 | 0 | 17 | 0.0588 |
| 17_2986050 | 9 | 0.5556 | 3 | 0 | 1 | 0 | 2 | 0.5 | 15 | 0.4 | 5 | 0.2 | 3 | 0.3333 | 6 | 0.1667 | NA | NA | 14 | 0.2143 |
| 17_2992046 | 7 | 0 | 5 | 0 | 3 | 0 | 2 | 0.5 | 17 | 0.0588 | 4 | 0 | 7 | 0 | 3 | 0 | 6 | 0.1667 | 20 | 0.05 |
| 17_2992134 | 8 | 0.5 | 8 | 0.75 | 3 | 0.3333 | 1 | 0 | 20 | 0.55 | 4 | 0.5 | 7 | 0.2857 | 3 | 0 | 6 | 0.5 | 20 | 0.35 |
| 17_2992135 | 8 | 0.875 | 7 | 0.8571 | 5 | 0.6 | 2 | 0.5 | 22 | 0.7727 | 3 | 1 | 5 | 1 | 3 | 1 | 5 | 0.8 | 16 | 0.9375 |
| 17_2992142 | 9 | 0 | 8 | 0.25 | 5 | 0.2 | 4 | 0.25 | 26 | 0.1538 | 5 | 0.2 | 8 | 0 | 4 | 0 | 8 | 0.5 | 25 | 0.2 |
| 17_2992143 | 9 | 0 | 7 | 0 | 5 | 0.4 | 4 | 0 | 25 | 0.08 | 5 | 0 | 6 | 0.1667 | 4 | 0 | 8 | 0.25 | 23 | 0.1304 |
| 17_2992145 | 9 | 0.5556 | 10 | 0.6 | 5 | 0.6 | 4 | 0.5 | 28 | 0.5714 | 4 | 0.5 | 6 | 0.5 | 3 | 0.6667 | 7 | 0.8571 | 20 | 0.65 |
| 17_2992150 | 7 | 0.7143 | 8 | 0.75 | 4 | 1 | 4 | 1 | 23 | 0.8261 | 4 | 0.75 | 6 | 0.6667 | 4 | 0.75 | 7 | 0.7143 | 21 | 0.7143 |
| 17_2992185 | 6 | 0.1667 | 7 | 0 | 3 | 0 | 4 | 0 | 20 | 0.05 | 6 | 0.1667 | 7 | 0 | 5 | 0.2 | 9 | 0.1111 | 27 | 0.1111 |
| 17_2992334 | 1 | 0 | 1 | 0 | 1 | 1 | NA | NA | 3 | 0.3333 | 2 | 0.5 | 2 | 0.5 | NA | NA | 4 | 0.5 | 8 | 0.5 |
| 17_2992351 | 1 | 0 | 1 | 1 | 3 | 1 | 1 | 1 | 6 | 0.8333 | 3 | 0.6667 | 2 | 1 | NA | NA | 3 | 1 | 8 | 0.875 |
| 17_2992382 | 1 | 0 | 1 | 1 | 4 | 0.75 | 1 | 0 | 7 | 0.5714 | 3 | 0.6667 | 2 | 1 | NA | NA | 3 | 0.3333 | 8 | 0.625 |
| 17_2992423 | 2 | 0 | 3 | 0 | 5 | 0.2 | 1 | 0 | 11 | 0.0909 | 4 | 0 | 2 | 0 | NA | NA | 3 | 0.3333 | 9 | 0.1111 |
| 17_2992460 | 1 | 0 | 2 | 0 | 5 | 0.4 | 1 | 0 | 9 | 0.2222 | 3 | 0 | 3 | 0 | 1 | 0 | 4 | 0.25 | 11 | 0.0909 |
| 17_3004751 | 1 | 0 | 3 | 0.3333 | 2 | 0 | 1 | 0 | 7 | 0.1429 | 1 | 0 | NA | NA | 2 | 0 | 5 | 0.2 | 8 | 0.125 |
| 17_3004755 | 1 | 1 | 3 | 0.3333 | 2 | 0 | 1 | 0 | 7 | 0.2857 | 1 | 0 | 1 | 0 | 2 | 0.5 | 5 | 0.2 | 9 | 0.2222 |
| 17_3004756 | 1 | 0 | 3 | 0.3333 | 2 | 0 | 1 | 0 | 7 | 0.1429 | 1 | 0 | 1 | 1 | 2 | 0.5 | 5 | 0.4 | 9 | 0.4444 |
| 17_3004780 | 1 | 1 | 3 | 1 | 2 | 0.5 | 1 | 0 | 7 | 0.7143 | 1 | 1 | 1 | 1 | 1 | 0 | 5 | 0.8 | 8 | 0.75 |
| 17_3004781 | NA | NA | 2 | 1 | 2 | 1 | 1 | 1 | 5 | 1 | 1 | 1 | 1 | 0 | NA | NA | 3 | 1 | 5 | 0.8 |
| 17_3004796 | 1 | 1 | 3 | 0.6667 | 1 | 1 | 1 | 1 | 6 | 0.8333 | 1 | 1 | 1 | 1 | 2 | 1 | 5 | 0.8 | 9 | 0.8889 |
| 17_3004798 | 1 | 1 | 3 | 1 | 1 | 1 | 1 | 1 | 6 | 1 | 1 | 1 | 1 | 1 | 2 | 1 | 5 | 1 | 9 | 1 |
| 17_3004800 | 1 | 0 | 3 | 0.3333 | 2 | 0 | 1 | 1 | 7 | 0.2857 | 1 | 0 | 1 | 0 | 2 | 0 | 5 | 0.4 | 9 | 0.2222 |
| 17_3004804 | 1 | 0 | 2 | 0 | 2 | 0.5 | NA | NA | 5 | 0.2 | 1 | 0 | 1 | 0 | 2 | 0.5 | 5 | 0 | 9 | 0.1111 |
| 17_3004812 | NA | NA | 1 | 1 | 2 | 1 | 1 | 1 | 4 | 1 | 1 | 1 | NA | NA | NA | NA | 4 | 1 | 5 | 1 |
| 17_3004816 | 1 | 1 | 2 | 0 | 2 | 0 | 1 | 0 | 6 | 0.1667 | 1 | 0 | NA | NA | 2 | 0.5 | 5 | 0 | 8 | 0.125 |
| 17_3004824 | 1 | 1 | 1 | 1 | NA | NA | 1 | 1 | 3 | 1 | 1 | 1 | NA | NA | 1 | 0 | 4 | 0.5 | 6 | 0.5 |
| 17_3005059 | 1 | 1 | NA | NA | 2 | 0.5 | NA | NA | 3 | 0.6667 | NA | NA | NA | NA | NA | NA | 3 | 1 | 3 | 1 |
| 17_3005067 | 1 | 1 | 1 | 1 | 2 | 1 | NA | NA | 4 | 1 | 1 | 1 | NA | NA | NA | NA | 2 | 0.5 | 3 | 0.6667 |
| 17_3005070 | 1 | 0 | 1 | 1 | 2 | 0.5 | NA | NA | 4 | 0.5 | 1 | 0 | NA | NA | NA | NA | 2 | 1 | 3 | 0.6667 |
| 17_3005077 | 1 | 1 | 1 | 1 | 2 | 0.5 | NA | NA | 4 | 0.75 | 1 | 0 | NA | NA | NA | NA | 2 | 1 | 3 | 0.6667 |
| 17_3005082 | 1 | 0 | 1 | 1 | 1 | 1 | NA | NA | 3 | 0.6667 | 1 | 1 | NA | NA | NA | NA | 2 | 1 | 3 | 1 |
| 17_3005088 | 1 | 0 | 1 | 1 | 2 | 1 | NA | NA | 4 | 0.75 | 1 | 1 | NA | NA | NA | NA | 2 | 0.5 | 3 | 0.6667 |
| 17_3005094 | 1 | 1 | 1 | 1 | 2 | 0.5 | NA | NA | 4 | 0.75 | 1 | 1 | NA | NA | NA | NA | 2 | 1 | 3 | 1 |
| 17_3005096 | 1 | 1 | 1 | 1 | 2 | 0.5 | NA | NA | 4 | 0.75 | 1 | 1 | NA | NA | NA | NA | 2 | 1 | 3 | 1 |
| 17_3005099 | 1 | 1 | 1 | 1 | 2 | 1 | NA | NA | 4 | 1 | 1 | 1 | NA | NA | NA | NA | 1 | 1 | 2 | 1 |
| 17_3042709 | 7 | 0 | 7 | 0 | 5 | 0.2 | 13 | 0.2308 | 32 | 0.125 | 8 | 0 | 6 | 0.1667 | 5 | 0 | 4 | 0.25 | 23 | 0.087 |
| 17_3042720 | 9 | 0.1111 | 8 | 0 | 8 | 0.375 | 15 | 0.1333 | 40 | 0.15 | 9 | 0.3333 | 7 | 0.1429 | 5 | 0.6 | 5 | 0.2 | 26 | 0.3077 |
| 17_3042744 | 7 | 0.2857 | 8 | 0.125 | 7 | 0.4286 | 14 | 0.2143 | 36 | 0.25 | 9 | 0.2222 | 3 | 0.3333 | 5 | 0.2 | 6 | 0.3333 | 23 | 0.2609 |
| 17_3042753 | 8 | 0.625 | 9 | 0.2222 | 7 | 0.5714 | 15 | 0.2 | 39 | 0.359 | 7 | 0.2857 | 3 | 0.3333 | 6 | 0.3333 | 5 | 0.2 | 21 | 0.2857 |
| 17_3042775 | 8 | 0 | 9 | 0.2222 | 8 | 0 | 15 | 0 | 40 | 0.05 | 8 | 0.125 | 3 | 0 | 6 | 0.1667 | 6 | 0.1667 | 23 | 0.1304 |
| 17_3042776 | 6 | 0.1667 | 8 | 0.125 | 7 | 0.4286 | 12 | 0 | 33 | 0.1515 | 6 | 0.1667 | 2 | 0 | 6 | 0.1667 | 4 | 0.25 | 18 | 0.1667 |
| 17_3042804 | 11 | 0 | 11 | 0 | 8 | 0.25 | 16 | 0.0625 | 46 | 0.0652 | 9 | 0 | 4 | 0.25 | 7 | 0 | 5 | 0.2 | 25 | 0.08 |
| 17_3042809 | 11 | 0.4545 | 11 | 0.4545 | 8 | 0.25 | 16 | 0.1875 | 46 | 0.3261 | 9 | 0.1111 | 5 | 0.8 | 7 | 0.2857 | 6 | 0.8333 | 27 | 0.4444 |
| 17_3042834 | 10 | 0.2 | 12 | 0.0833 | 9 | 0 | 17 | 0.1765 | 48 | 0.125 | 12 | 0 | 6 | 0 | 9 | 0.2222 | 7 | 0 | 34 | 0.0588 |
| 17_3042872 | 12 | 0 | 13 | 0 | 10 | 0 | 19 | 0.2105 | 54 | 0.0741 | 11 | 0.1818 | 6 | 0.1667 | 10 | 0.1 | 7 | 0 | 34 | 0.1176 |
| 17_3042899 | NA | NA | 2 | 0 | NA | NA | 3 | 0.6667 | 5 | 0.4 | 1 | 1 | 2 | 0.5 | 2 | 0 | 1 | 1 | 6 | 0.5 |
| 17_3043618 | 1 | 1 | 1 | 0 | NA | NA | 3 | 0 | 5 | 0.2 | NA | NA | 3 | 0.3333 | 2 | 0 | 1 | 0 | 6 | 0.1667 |
| 17_3043643 | 1 | 1 | NA | NA | 1 | 0 | 4 | 0.25 | 6 | 0.3333 | NA | NA | 3 | 0.3333 | 2 | 0.5 | 1 | 0 | 6 | 0.3333 |
| 17_3043673 | 2 | 0.5 | 2 | 0 | 5 | 0.4 | 5 | 0.6 | 14 | 0.4286 | 5 | 0.4 | 2 | 0.5 | 6 | 0.5 | 1 | 1 | 14 | 0.5 |
| 17_3043675 | 4 | 0 | 2 | 0 | 4 | 0 | 3 | 0.3333 | 13 | 0.0769 | 5 | 0.2 | 1 | 1 | 7 | 0 | 1 | 0 | 14 | 0.1429 |
| 17_3043676 | 4 | 0.25 | 2 | 0.5 | 3 | 1 | 7 | 0.8571 | 16 | 0.6875 | 3 | 1 | 1 | 1 | 7 | 1 | 1 | 1 | 12 | 1 |

| | | | | | | | | | | | | | | | | | | | | |
|---|---|---|---|---|---|---|---|---|---|---|---|---|---|---|---|---|---|---|---|---|
| 17_304369 | 4 | 1 | 3 | 0.3333 | 3 | 0.6667 | 2 | 1 | 12 | 0.75 | 5 | 0.8 | NA | NA | 7 | 1 | 2 | 0.5 | 14 | 0.8571 |
| 17_304370 | 4 | 0.5 | 3 | 0.3333 | 3 | 0.3333 | 3 | 0.6667 | 13 | 0.4615 | 4 | 0.5 | 1 | 0 | 7 | 0.4286 | 2 | 0.5 | 14 | 0.4286 |
| 17_304370 | 5 | 0.4 | 2 | 0 | 4 | 1 | 4 | 1 | 15 | 0.6667 | 5 | 0.4 | 1 | 1 | 6 | 0.6667 | 2 | 0.5 | 14 | 0.5714 |
| 17_304370 | 5 | 0.2 | 2 | 0 | 4 | 0.25 | 3 | 0.3333 | 14 | 0.2143 | 4 | 0.25 | 1 | 0 | 6 | 0.5 | 2 | 0.5 | 13 | 0.3846 |
| 17_304370 | 4 | 0.25 | 2 | 0 | 3 | 0.3333 | 4 | 0 | 13 | 0.1538 | 5 | 0 | 1 | 0 | 7 | 0.1429 | 2 | 0 | 15 | 0.0667 |
| 17_304370 | 4 | 0.5 | 2 | 0.5 | 3 | 1 | 3 | 0.6667 | 12 | 0.6667 | 4 | 0.25 | 1 | 1 | 6 | 1 | 2 | 0 | 13 | 0.6154 |
| 17_304371 | 4 | 0.75 | 2 | 0 | 4 | 0.75 | 3 | 1 | 13 | 0.6923 | 4 | 0.5 | 1 | 0 | 5 | 1 | 2 | 1 | 12 | 0.75 |
| 17_304371 | 4 | 0.75 | 2 | 0.5 | 4 | 0.5 | 3 | 0.3333 | 13 | 0.5385 | 3 | 0.6667 | 1 | 1 | 5 | 1 | 1 | 0 | 10 | 0.8 |
| 17_304372 | 4 | 0 | 2 | 0 | 3 | 0 | 4 | 0.25 | 13 | 0.0769 | 4 | 0 | 1 | 0 | 6 | 0.5 | 2 | 0 | 13 | 0.2308 |
| 17_304375 | 5 | 1 | 2 | 1 | 2 | 0.5 | 3 | 1 | 12 | 0.9167 | 5 | 1 | 1 | 1 | 5 | 0.6 | 1 | 1 | 12 | 0.8333 |
| 17_304376 | 5 | 0.6 | 3 | 0.3333 | 1 | 0 | 3 | 0 | 12 | 0.3333 | 5 | 0.8 | 1 | 1 | 5 | 0.2 | 1 | 1 | 12 | 0.5833 |
| 17_304382 | 4 | 0.25 | 3 | 0 | 1 | 0 | 2 | 0.5 | 10 | 0.2 | 2 | 0.5 | 1 | 0 | 5 | 0 | NA | NA | 8 | 0.125 |
| 17_304383 | 3 | 0.6667 | 2 | 0.5 | 1 | 0 | 2 | 1 | 8 | 0.625 | 1 | 0 | NA | NA | 4 | 0.25 | NA | NA | 5 | 0.2 |
| 17_304384 | 3 | 0.3333 | 3 | 0 | 1 | 0 | 2 | 0 | 9 | 0.1111 | 3 | 0.3333 | NA | NA | 4 | 0 | NA | NA | 7 | 0.1429 |
| 17_304384 | 3 | 0.6667 | 3 | 1 | 2 | 1 | 2 | 1 | 10 | 0.9 | 2 | 0.5 | 1 | 1 | 4 | 1 | NA | NA | 7 | 0.8571 |
| 17_304385 | 3 | 0.6667 | 1 | 0 | 2 | 0.5 | 1 | 1 | 7 | 0.5714 | 2 | 0.5 | 1 | 1 | 4 | 1 | NA | NA | 7 | 0.8571 |
| 17_304387 | 3 | 0.6667 | 2 | 0.5 | 2 | 0.5 | 1 | 1 | 8 | 0.625 | 1 | 1 | 1 | 1 | 3 | 1 | NA | NA | 5 | 1 |
| 17_304387 | 3 | 1 | 3 | 1 | 2 | 1 | 1 | 1 | 9 | 1 | 3 | 0.6667 | 1 | 1 | 4 | 0.5 | NA | NA | 8 | 0.625 |
| 17_304392 | 1 | 0 | 1 | 0 | 2 | 0.5 | NA | NA | 4 | 0.25 | 2 | 0.5 | NA | NA | 2 | 0.5 | NA | NA | 4 | 0.5 |
| 17_304392 | NA | NA | 2 | 0.5 | 2 | 0 | 1 | 0 | 5 | 0.2 | 2 | 0.5 | NA | NA | 3 | 0.6667 | NA | NA | 5 | 0.6 |
| 17_307464 | 2 | 0.5 | 1 | 1 | 1 | 0 | 1 | 0 | 5 | 0.4 | 2 | 0.5 | NA | NA | 1 | 1 | NA | NA | 3 | 0.6667 |
| 17_307464 | 2 | 0.5 | NA | NA | 1 | 1 | 1 | 0 | 4 | 0.5 | 1 | 1 | NA | NA | NA | NA | NA | NA | 1 | 1 |
| 17_307961 | 1 | 1 | NA | NA | NA | NA | NA | NA | 1 | 1 | NA | NA | NA | NA | 1 | 1 | NA | NA | 1 | 1 |
| 17_308109 | 4 | 0 | 3 | 0 | 5 | 0 | 3 | 0.3333 | 15 | 0.0667 | 1 | 0 | NA | NA | 2 | 0.5 | 1 | 0 | 4 | 0.25 |
| 17_308114 | 3 | 0 | NA | NA | 4 | 0.25 | 1 | 0 | 8 | 0.125 | 1 | 0 | 1 | 1 | 2 | 0 | 1 | 0 | 5 | 0.2 |
| 17_308120 | 16 | 0.0625 | 10 | 0.2 | 11 | 0.0909 | 13 | 0.1538 | 50 | 0.12 | 15 | 0 | 14 | 0.1429 | 8 | 0.125 | 13 | 0.0769 | 50 | 0.08 |
| 17_308124 | 15 | 0.2667 | 10 | 0.3 | 8 | 0.5 | 15 | 0.3333 | 48 | 0.3333 | 12 | 0.25 | 10 | 0.3 | 6 | 0.1667 | 12 | 0.25 | 40 | 0.25 |
| 17_308125 | 15 | 0.1333 | 13 | 0.2308 | 12 | 0.1667 | 15 | 0.0667 | 55 | 0.1455 | 16 | 0.0625 | 14 | 0.0714 | 10 | 0.2 | 13 | 0.1538 | 53 | 0.1132 |
| 17_308127 | 8 | 0.25 | 13 | 0.1538 | 13 | 0.3846 | 13 | 0.2308 | 47 | 0.2553 | 16 | 0.25 | 13 | 0.2308 | 8 | 0.375 | 13 | 0.0769 | 50 | 0.22 |
| 17_308128 | 8 | 0 | 16 | 0.25 | 13 | 0.3077 | 12 | 0.0833 | 49 | 0.1837 | 15 | 0.1333 | 16 | 0.1875 | 8 | 0.125 | 11 | 0.1818 | 50 | 0.16 |
| 17_308130 | 11 | 0.5455 | 16 | 0.75 | 10 | 0.7 | 12 | 0.8333 | 49 | 0.7143 | 14 | 0.7143 | 16 | 0.875 | 9 | 0.8889 | 11 | 0.7273 | 50 | 0.8 |
| 17_308130 | 7 | 0.1429 | 14 | 0.0714 | 10 | 0 | 8 | 0.25 | 39 | 0.1026 | 12 | 0 | 15 | 0.2667 | 8 | 0.125 | 11 | 0.2727 | 46 | 0.1739 |
| 17_308131 | 7 | 0 | 16 | 0.0625 | 13 | 0.0769 | 11 | 0.0909 | 47 | 0.0638 | 14 | 0.1429 | 14 | 0 | 12 | 0.1667 | 11 | 0.1818 | 51 | 0.1176 |
| 17_311554 | 1 | 1 | NA | NA | 1 | 0 | 1 | 0 | 3 | 0.3333 | NA | NA | NA | NA | NA | NA | 1 | 1 | 1 | 1 |
| 17_311556 | 1 | 1 | NA | NA | NA | NA | 1 | 1 | 2 | 1 | NA | NA | NA | NA | NA | NA | 1 | 1 | 1 | 1 |
| 17_311557 | 1 | 1 | NA | NA | 1 | 1 | NA | NA | 2 | 1 | NA | NA | NA | NA | NA | NA | 1 | 1 | 1 | 1 |
| 17_311557 | 1 | 1 | 1 | 0 | NA | NA | NA | NA | 2 | 0.5 | NA | NA | NA | NA | NA | NA | 1 | 1 | 1 | 1 |
| 17_311563 | 1 | 1 | 1 | 1 | 1 | 1 | 1 | 1 | 4 | 1 | NA | NA | NA | NA | NA | NA | 1 | 1 | 1 | 1 |
| 17_311573 | 1 | 0 | 1 | 0 | NA | NA | 1 | 1 | 3 | 0.3333 | NA | NA | NA | NA | NA | NA | 1 | 1 | 1 | 1 |
| 17_311579 | 1 | 1 | NA | NA | 1 | 1 | 1 | 1 | 3 | 1 | NA | NA | NA | NA | NA | NA | 1 | 1 | 1 | 1 |
| 17_311580 | 1 | 1 | NA | NA | 1 | 1 | 1 | 1 | 3 | 1 | NA | NA | NA | NA | NA | NA | 1 | 1 | 1 | 1 |
| 17_311583 | 1 | 0 | 1 | 0 | 1 | 1 | 1 | 1 | 4 | 0.5 | NA | NA | NA | NA | NA | NA | 1 | 1 | 1 | 1 |
| 17_311588 | NA | NA | 1 | 1 | 1 | 0 | 1 | 0 | 3 | 0.3333 | 1 | 0 | NA | NA | NA | NA | 1 | 1 | 2 | 0.5 |
| 17_311590 | 1 | 0 | 1 | 1 | NA | NA | 1 | 0 | 3 | 0.3333 | 1 | 0 | NA | NA | NA | NA | 1 | 1 | 2 | 0.5 |
| 17_313332 | 6 | 1 | 3 | 0.6667 | 5 | 1 | 6 | 0.8333 | 20 | 0.9 | 2 | 1 | 7 | 1 | 10 | 0.9 | 4 | 1 | 23 | 0.9565 |
| 17_313333 | 6 | 0.1667 | 4 | 0 | 5 | 0.4 | 5 | 0.2 | 20 | 0.2 | 1 | 0 | 6 | 0 | 11 | 0.0909 | 5 | 0.2 | 23 | 0.087 |
| 17_313333 | 4 | 1 | 2 | 0.5 | 4 | 1 | 6 | 1 | 16 | 0.9375 | 4 | 1 | 5 | 0.8 | 11 | 1 | 6 | 1 | 26 | 0.9615 |
| 17_313334 | 6 | 1 | 2 | 0 | 4 | 0.75 | 6 | 0.6667 | 18 | 0.7222 | 3 | 1 | 7 | 0.5714 | 10 | 0.7 | 5 | 1 | 25 | 0.76 |
| 17_313335 | 5 | 0.4 | 5 | 0.6 | 5 | 1 | 6 | 0.8333 | 21 | 0.7143 | 3 | 0 | 6 | 0.1667 | 13 | 0.2308 | 5 | 0.2 | 27 | 0.1852 |
| 17_313336 | 10 | 0.3 | 4 | 0.25 | 7 | 0.4286 | 7 | 0.4286 | 28 | 0.3571 | 4 | 0.5 | 6 | 0.1667 | 11 | 0.2727 | 6 | 0.1667 | 27 | 0.2593 |
| 17_313338 | 10 | 0.8 | 4 | 0.25 | 5 | 0.6 | 10 | 0.8 | 29 | 0.6897 | 4 | 1 | 6 | 0.5 | 10 | 0.8 | 7 | 0.7143 | 27 | 0.7407 |
| 17_313339 | 10 | 0.4 | 5 | 0.4 | 8 | 0.625 | 11 | 0.8182 | 34 | 0.5882 | 4 | 0.25 | 9 | 0.5556 | 11 | 0.8182 | 6 | 0.8333 | 30 | 0.6667 |
| 17_313341 | 11 | 0.1818 | 9 | 0 | 7 | 0 | 11 | 0.2727 | 38 | 0.1316 | 4 | 0.25 | 11 | 0 | 12 | 0.0833 | 5 | 0.2 | 32 | 0.0938 |
| 17_313345 | 11 | 0.1818 | 12 | 0.1667 | 11 | 0.2727 | 12 | 0.3333 | 46 | 0.2391 | 6 | 0.1667 | 12 | 0.0833 | 13 | 0.2308 | 7 | 0.2857 | 38 | 0.1842 |
| 17_313347 | 1 | 0 | 1 | 0 | 1 | 0 | 2 | 0.5 | 5 | 0.2 | 1 | 0 | 2 | 0 | 4 | 0 | 2 | 1 | 9 | 0.2222 |
| 17_313360 | 2 | 0 | 1 | 0 | 4 | 0.25 | 3 | 0 | 10 | 0.1 | 2 | 0 | 3 | 0 | 4 | 0 | 1 | 1 | 10 | 0.1 |
| 17_313368 | 1 | 1 | NA | NA | 4 | 1 | 2 | 1 | 7 | 1 | 2 | 0.5 | 2 | 1 | 1 | 1 | 3 | 0.6667 | 8 | 0.75 |
| 17_313368 | 2 | 0 | 2 | 0 | 5 | 0.2 | 3 | 0 | 12 | 0.0833 | 3 | 0.3333 | 4 | 0.5 | 3 | 0.6667 | 3 | 0.3333 | 13 | 0.4615 |
| 17_315157 | NA | NA | NA | NA | 1 | 1 | NA | NA | 1 | 1 | NA | NA | NA | NA | NA | NA | 1 | 1 | 1 | 1 |
| 17_315688 | NA | NA | 1 | 1 | NA | NA | 2 | 0 | 3 | 0.3333 | NA | NA | NA | NA | NA | NA | 2 | 1 | 2 | 1 |
| 17_315690 | NA | NA | NA | NA | 1 | 1 | 1 | 0 | 2 | 0.5 | NA | NA | NA | NA | 1 | 0 | 1 | 1 | 2 | 0.5 |
| 17_315690 | NA | NA | 1 | 1 | 1 | 1 | 2 | 0.5 | 4 | 0.75 | NA | NA | NA | NA | 1 | 0 | 2 | 0.5 | 3 | 0.3333 |
| 17_315692 | NA | NA | NA | NA | NA | NA | 2 | 1 | 2 | 1 | 1 | 0 | NA | NA | 1 | 1 | 2 | 1 | 4 | 0.75 |
| 17_315694 | NA | NA | 1 | 0 | 2 | 0.5 | 2 | 0 | 5 | 0.2 | 1 | 0 | NA | NA | 1 | 1 | 2 | 0.5 | 4 | 0.5 |
| 17_316539 | 1 | 1 | NA | NA | NA | NA | NA | NA | 1 | 1 | NA | NA | 1 | 1 | NA | NA | NA | NA | 1 | 1 |

| | | | | | | | | | | | | | | | | | | | | |
|---|---|---|---|---|---|---|---|---|---|---|---|---|---|---|---|---|---|---|---|---|
| 17_3169518 | NA | NA | 1 | 1 | 1 | 0 | 1 | 0 | 3 | 0.3333 | 1 | 1 | 1 | 0 | NA | NA | NA | NA | 2 | 0.5 |
| 17_3174634 | 3 | 0.3333 | 2 | 0.5 | 2 | 0 | NA | NA | 7 | 0.2857 | 1 | 0 | 1 | 1 | 3 | 0.3333 | 2 | 0 | 7 | 0.2857 |
| 17_3174646 | 4 | 1 | 4 | 0.75 | 1 | 1 | NA | NA | 9 | 0.8889 | 2 | 1 | NA | NA | 3 | 0.6667 | 2 | 1 | 7 | 0.8571 |
| 17_3174652 | 5 | 0.6 | 3 | 1 | 2 | 0.5 | NA | NA | 10 | 0.7 | 2 | 0.5 | 1 | 1 | 2 | 0 | 3 | 0.6667 | 8 | 0.5 |
| 17_3174662 | 3 | 0 | 2 | 0 | 2 | 0.5 | NA | NA | 7 | 0.1429 | 3 | 0.3333 | 1 | 0 | 2 | 1 | 1 | 0 | 7 | 0.4286 |
| 17_3174724 | 5 | 0 | 5 | 0.2 | 2 | 0 | NA | NA | 12 | 0.0833 | 3 | 0 | 1 | 0 | 3 | 0 | 4 | 0.25 | 11 | 0.0909 |
| 17_3182397 | 1 | 1 | 1 | 1 | NA | NA | NA | NA | 2 | 1 | NA | NA | 1 | 1 | NA | NA | 2 | 1 | 3 | 1 |
| 17_3183184 | 3 | 0.3333 | 3 | 0 | 4 | 0 | 4 | 0.5 | 14 | 0.2143 | 3 | 0 | 2 | 0 | 1 | 0 | 4 | 0.25 | 10 | 0.1 |
| 17_3183186 | 3 | 0.3333 | 4 | 0.25 | 3 | 0.3333 | 6 | 0.3333 | 16 | 0.3125 | 4 | 0.25 | 3 | 0.6667 | 1 | 0 | 2 | 0 | 10 | 0.3 |
| 17_3183197 | 3 | 0.3333 | 6 | 0.1667 | 6 | 0 | 7 | 0 | 22 | 0.0909 | 4 | 0 | 4 | 0.25 | 2 | 0 | 5 | 0 | 15 | 0.0667 |
| 17_3185205 | 1 | 0 | NA | NA | 2 | 0.5 | 1 | 1 | 4 | 0.5 | 2 | 0.5 | 1 | 0 | 1 | 0 | 2 | 0 | 6 | 0.1667 |
| 17_3206341 | 2 | 0.5 | 3 | 0 | 6 | 0.5 | 1 | 1 | 12 | 0.4167 | 4 | 0.5 | 4 | 0.5 | NA | NA | 2 | 0.5 | 10 | 0.5 |
| 17_3206344 | 2 | 1 | 4 | 0.5 | 5 | 0.8 | 1 | 1 | 12 | 0.75 | 4 | 1 | 3 | 0.6667 | NA | NA | 2 | 1 | 9 | 0.8889 |
| 17_3206344 | 2 | 0 | 4 | 0.75 | 4 | 0 | 1 | 1 | 11 | 0.3636 | 3 | 1 | 4 | 0.75 | NA | NA | 2 | 0.5 | 9 | 0.7778 |
| 17_3206357 | 1 | 1 | 3 | 0.6667 | 5 | 1 | 1 | 1 | 10 | 0.9 | 3 | 0.3333 | 4 | 0.25 | NA | NA | 1 | 0 | 8 | 0.25 |
| 17_3206378 | 1 | 0 | 2 | 0 | 2 | 0.5 | 1 | 0 | 6 | 0.1667 | 3 | 0 | 4 | 0.25 | NA | NA | 1 | 0 | 8 | 0.125 |
| 17_3206380 | NA | NA | 2 | 1 | 3 | 1 | 1 | 1 | 6 | 1 | 4 | 0.75 | 2 | 0.5 | NA | NA | 1 | 1 | 7 | 0.7143 |
| 17_3206382 | 1 | 1 | 3 | 0.6667 | 2 | 0 | 1 | 1 | 7 | 0.5714 | 4 | 0.25 | 4 | 0.25 | NA | NA | 1 | 0 | 9 | 0.2222 |
| 17_3206430 | 1 | 0 | NA | NA | 3 | 0.3333 | 1 | 0 | 5 | 0.2 | 4 | 0.25 | 4 | 0 | NA | NA | 1 | 0 | 9 | 0.1111 |
| 17_3206444 | 1 | 0 | NA | NA | 2 | 0 | 1 | 1 | 4 | 0.25 | 4 | 0.5 | 3 | 0 | NA | NA | 1 | 0 | 8 | 0.25 |
| 17_3206451 | 1 | 0 | 1 | 1 | 2 | 0.5 | NA | NA | 4 | 0.5 | 4 | 0.5 | 4 | 0.25 | NA | NA | 1 | 0 | 9 | 0.3333 |
| 17_3206512 | 2 | 1 | NA | NA | 1 | 1 | NA | NA | 3 | 1 | 3 | 1 | 1 | 0 | NA | NA | NA | NA | 4 | 0.75 |
| 17_3206514 | 2 | 1 | NA | NA | 1 | 1 | NA | NA | 3 | 1 | 3 | 1 | 1 | 1 | NA | NA | NA | NA | 4 | 1 |
| 17_3206518 | 1 | 1 | NA | NA | 1 | 0 | NA | NA | 2 | 0.5 | 3 | 1 | 1 | 1 | NA | NA | NA | NA | 4 | 1 |
| 17_3206527 | 2 | 0.5 | NA | NA | NA | NA | NA | NA | 2 | 0.5 | 3 | 0.6667 | 1 | 1 | NA | NA | NA | NA | 4 | 0.75 |
| 17_3206530 | 2 | 0.5 | 1 | 1 | NA | NA | NA | NA | 3 | 0.6667 | 1 | 1 | NA | NA | NA | NA | NA | NA | 1 | 1 |
| 17_3206531 | 2 | 1 | 1 | 1 | NA | NA | NA | NA | 3 | 1 | 3 | 1 | 1 | 1 | NA | NA | NA | NA | 4 | 1 |
| 17_3226494 | 1 | 0 | NA | NA | 2 | 0.5 | NA | NA | 3 | 0.3333 | NA | NA | 1 | 1 | NA | NA | NA | NA | 1 | 1 |
| 17_3227572 | 1 | 0 | 2 | 0.5 | 2 | 1 | NA | NA | 5 | 0.6 | NA | NA | 2 | 0 | NA | NA | 1 | 1 | 3 | 0.3333 |
| 17_3227587 | NA | NA | 5 | 0.6 | 1 | 0 | NA | NA | 6 | 0.5 | NA | NA | 1 | 1 | NA | NA | 1 | 1 | 2 | 1 |
| 17_3227950 | 2 | 0.5 | 4 | 0.75 | 3 | 1 | 1 | 1 | 10 | 0.8 | 1 | 1 | 3 | 1 | NA | NA | 1 | 1 | 5 | 1 |
| 17_3227957 | 1 | 0 | 5 | 0.2 | 2 | 0 | 1 | 1 | 9 | 0.2222 | 1 | 1 | 2 | 0.5 | NA | NA | 2 | 0 | 5 | 0.4 |
| 17_3227962 | NA | NA | 2 | 0 | 1 | 1 | NA | NA | 3 | 0.3333 | 1 | 0 | 2 | 0.5 | NA | NA | 1 | 1 | 4 | 0.5 |
| 17_3227970 | 1 | 1 | 3 | 0.6667 | 2 | 1 | NA | NA | 6 | 0.8333 | 1 | 1 | 3 | 1 | NA | NA | NA | NA | 4 | 1 |
| 17_3228052 | 2 | 0 | 3 | 0.3333 | 2 | 0 | NA | NA | 7 | 0.1429 | 1 | 1 | 3 | 0.6667 | 1 | 1 | 2 | 0.5 | 7 | 0.7143 |
| 17_3228073 | 1 | 1 | 4 | 0.5 | 1 | 0 | NA | NA | 6 | 0.5 | 1 | 0 | 3 | 0.3333 | 1 | 0 | 1 | 0 | 6 | 0.1667 |
| 17_3236059 | 1 | 1 | NA | NA | NA | NA | NA | NA | 1 | 1 | 1 | 1 | NA | NA | NA | NA | NA | NA | 1 | 1 |
| 17_3241581 | 9 | 0.1111 | 8 | 0.5 | 5 | 0 | 9 | 0.2222 | 31 | 0.2258 | 6 | 0 | 8 | 0.25 | 7 | 0.1429 | 8 | 0.125 | 29 | 0.1379 |
| 17_3241605 | 5 | 0 | 3 | 0.3333 | NA | NA | 4 | 0.25 | 12 | 0.1667 | 4 | 0.25 | 5 | 0.2 | 4 | 0 | 5 | 0.4 | 18 | 0.2222 |
| 17_3241618 | 5 | 0.6 | 6 | 0.1667 | 2 | 0.5 | 7 | 0.2857 | 20 | 0.35 | 4 | 0.25 | 6 | 0.5 | 5 | 0.2 | 5 | 0.4 | 20 | 0.35 |
| 17_3241620 | 6 | 0.5 | 6 | 0.5 | 1 | 1 | 8 | 0.625 | 21 | 0.5714 | 3 | 0.6667 | 7 | 0.4286 | 6 | 0.3333 | 6 | 0.8333 | 22 | 0.5455 |
| 17_3241624 | 6 | 1 | 6 | 1 | 3 | 0.3333 | 7 | 0.8571 | 22 | 0.8636 | 4 | 1 | 7 | 1 | 7 | 0.8571 | 7 | 1 | 25 | 0.96 |
| 17_3241633 | 4 | 0 | 3 | 0 | 1 | 0 | 5 | 0.2 | 13 | 0.0769 | 3 | 0.3333 | 4 | 0 | 4 | 0 | 4 | 0.25 | 15 | 0.1333 |
| 17_3241635 | 6 | 1 | 5 | 0.4 | 3 | 0.6667 | 8 | 0.625 | 22 | 0.6818 | 5 | 0.8 | 5 | 1 | 6 | 0.5 | 6 | 0.6667 | 22 | 0.7273 |
| 17_3241637 | 6 | 0.6667 | 6 | 0.8333 | 2 | 1 | 8 | 1 | 22 | 0.8636 | 5 | 0.8 | 5 | 1 | 7 | 0.8571 | 7 | 0.8571 | 24 | 0.875 |
| 17_3241639 | 5 | 0 | 6 | 0.1667 | 3 | 0 | 8 | 0.125 | 22 | 0.0909 | 3 | 0 | 4 | 0 | 6 | 0.1667 | 6 | 0 | 19 | 0.0526 |
| 17_3241645 | 6 | 0.8333 | 6 | 0.5 | 2 | 1 | 7 | 0.7143 | 21 | 0.7143 | 5 | 0.8 | 6 | 0.6667 | 7 | 0.5714 | 7 | 0.8571 | 25 | 0.72 |
| 17_3241653 | 6 | 0.1667 | 3 | 0.3333 | 3 | 0.6667 | 7 | 0.4286 | 19 | 0.3684 | 4 | 0.75 | 4 | 0.75 | 4 | 0.25 | 6 | 0.1667 | 18 | 0.4444 |
| 17_3241668 | 5 | 0.6 | 2 | 0 | 3 | 0.3333 | 7 | 0.5714 | 17 | 0.4706 | 3 | 0.6667 | 3 | 0.6667 | 4 | 0.25 | 5 | 0.8 | 15 | 0.6 |
| 17_3241670 | 4 | 0 | 4 | 0.5 | 2 | 0.5 | 8 | 0.5 | 18 | 0.3889 | 4 | 0.75 | 5 | 0.8 | 7 | 0.2857 | 5 | 0.4 | 21 | 0.5238 |
| 17_3241671 | 6 | 0.5 | 5 | 0.2 | 2 | 0 | 8 | 0.125 | 21 | 0.2381 | 3 | 0 | 5 | 0.2 | 7 | 0.4286 | 5 | 0.4 | 20 | 0.3 |
| 17_3252338 | 2 | 0 | 1 | 0 | 4 | 0 | 5 | 0.2 | 12 | 0.0833 | 3 | 0.3333 | 7 | 0 | 8 | 0.125 | 7 | 0 | 25 | 0.08 |
| 17_3252343 | 4 | 0 | 1 | 0 | 5 | 0.2 | 6 | 0 | 16 | 0.0625 | 3 | 0 | 9 | 0 | 7 | 0 | 7 | 0.2857 | 26 | 0.0769 |
| 17_3252426 | 1 | 0 | 1 | 1 | 4 | 0.25 | 5 | 0.4 | 11 | 0.3636 | 3 | 0.3333 | 7 | 0.8571 | 5 | 0.6 | 5 | 0.8 | 20 | 0.7 |
| 17_3252447 | 1 | 1 | 1 | 0 | 4 | 0.5 | 6 | 0.5 | 12 | 0.5 | 3 | 0.3333 | 6 | 0.3333 | 6 | 0.5 | 7 | 0.7143 | 22 | 0.5 |
| 17_3253550 | NA | NA | NA | NA | 1 | 0 | 4 | 0.75 | 5 | 0.6 | 1 | 0 | 1 | 0 | 1 | 1 | 3 | 0 | 6 | 0.1667 |
| 17_3253553 | 1 | 0 | NA | NA | 1 | 0 | 4 | 0.5 | 6 | 0.3333 | 1 | 1 | NA | NA | 1 | 1 | 3 | 0.6667 | 5 | 0.8 |
| 17_3254060 | 6 | 0.3333 | 12 | 0.25 | 12 | 0.1667 | 10 | 0.1 | 40 | 0.2 | 9 | 0.3333 | 11 | 0.2727 | 11 | 0.1818 | 12 | 0.25 | 43 | 0.2558 |
| 17_3255222 | 4 | 0 | 5 | 0.4 | 4 | 0 | 4 | 0.25 | 17 | 0.1765 | NA | NA | 7 | 0.2857 | 8 | 0.25 | 3 | 0 | 18 | 0.2222 |
| 17_3255228 | 5 | 0.4 | 4 | 0.5 | 4 | 1 | 4 | 0.75 | 17 | 0.6471 | NA | NA | 8 | 0.5 | 8 | 0.25 | 2 | 0 | 18 | 0.3333 |
| 17_3255252 | 4 | 0.5 | 5 | 0.6 | 4 | 0.75 | 4 | 0.75 | 17 | 0.6471 | NA | NA | 10 | 0.7 | 8 | 0.625 | 3 | 0.3333 | 21 | 0.619 |
| 17_3255269 | 3 | 0 | 5 | 0 | 2 | 0.5 | 3 | 0.6667 | 13 | 0.2308 | NA | NA | 9 | 0.5556 | 7 | 0.4286 | 3 | 0.3333 | 19 | 0.4737 |
| 17_3255277 | 4 | 0 | 5 | 0.2 | 3 | 0.3333 | 4 | 0.5 | 16 | 0.25 | NA | NA | 9 | 0.1111 | 6 | 0 | 3 | 0.3333 | 18 | 0.1111 |
| 17_3255296 | 3 | 0 | 3 | 0.3333 | 1 | 1 | 5 | 0.2 | 12 | 0.25 | 1 | 0 | 7 | 0.2857 | 5 | 0.2 | 3 | 0.3333 | 16 | 0.25 |
| 17_3255298 | 4 | 0.25 | 4 | 0.5 | 3 | 0.6667 | 3 | 0.3333 | 14 | 0.4286 | 1 | 0 | 9 | 0.7778 | 5 | 0.6 | 3 | 0.6667 | 18 | 0.6667 |

| | | | | | | | | | | | | | | | | | | | | |
|---|---|---|---|---|---|---|---|---|---|---|---|---|---|---|---|---|---|---|---|---|
| 17_325532 | 4 | 0.5 | 5 | 0.8 | 2 | 0.5 | 4 | 1 | 15 | 0.7333 | 1 | 1 | 8 | 1 | 8 | 0.625 | 3 | 0.6667 | 20 | 0.8 |
| 17_325534 | 3 | 0 | 5 | 0.4 | 2 | 0 | 3 | 0 | 13 | 0.1538 | 1 | 0 | 9 | 0.4444 | 3 | 0 | 2 | 0.5 | 15 | 0.3333 |
| 17_325738 | 4 | 0.25 | 1 | 1 | 5 | 0 | 3 | 0 | 13 | 0.1538 | 5 | 0 | 2 | 0.5 | 6 | 0.3333 | NA | NA | 13 | 0.2308 |
| 17_325764 | 3 | 0 | 5 | 0.2 | 4 | 0.5 | 1 | 1 | 13 | 0.3077 | 2 | 0 | 4 | 0.25 | 5 | 0.2 | NA | NA | 11 | 0.1818 |
| 17_325764 | 3 | 0 | 4 | 0.25 | 3 | 0.3333 | 2 | 0 | 12 | 0.1667 | 1 | 0 | 3 | 0.3333 | 3 | 0 | NA | NA | 7 | 0.1429 |
| 17_325768 | 2 | 0.5 | 4 | 0.25 | 3 | 0 | 1 | 0 | 10 | 0.2 | 1 | 0 | 4 | 0 | 4 | 0.25 | NA | NA | 9 | 0.1111 |
| 17_325782 | 1 | 0 | 2 | 0.5 | 2 | 0 | 2 | 0.5 | 7 | 0.2857 | 1 | 1 | 2 | 0 | 3 | 0 | NA | NA | 6 | 0.1667 |
| 17_325805 | 1 | 1 | 1 | 1 | 3 | 0.3333 | 1 | 1 | 6 | 0.6667 | 1 | 0 | 3 | 0.3333 | 3 | 0.3333 | NA | NA | 7 | 0.2857 |
| 17_327703 | NA | NA | 1 | 1 | 2 | 0 | NA | NA | 3 | 0.3333 | NA | NA | NA | NA | NA | NA | 1 | 1 | 1 | 1 |
| 17_329921 | NA | NA | NA | NA | 2 | 1 | 2 | 1 | 4 | 1 | 2 | 0.5 | 2 | 0.5 | NA | NA | 3 | 0.6667 | 7 | 0.5714 |
| 17_329923 | NA | NA | NA | NA | 3 | 0.3333 | 2 | 0.5 | 5 | 0.4 | 3 | 1 | 4 | 1 | NA | NA | 2 | 0 | 9 | 0.7778 |
| 17_329925 | NA | NA | NA | NA | 3 | 0.6667 | 2 | 1 | 5 | 0.8 | 3 | 0.6667 | 3 | 0.6667 | NA | NA | 3 | 0.6667 | 9 | 0.6667 |
| 17_329925 | NA | NA | NA | NA | 3 | 1 | 2 | 1 | 5 | 1 | 2 | 1 | 2 | 1 | NA | NA | 4 | 1 | 8 | 1 |
| 17_329926 | NA | NA | NA | NA | 3 | 0.3333 | 2 | 0.5 | 5 | 0.4 | 1 | 1 | 3 | 0.6667 | NA | NA | 3 | 0.6667 | 7 | 0.7143 |
| 17_329926 | NA | NA | NA | NA | 3 | 1 | 2 | 0.5 | 5 | 0.8 | 2 | 1 | 4 | 1 | NA | NA | 3 | 0.3333 | 9 | 0.7778 |
| 17_329927 | NA | NA | NA | NA | 3 | 1 | 2 | 1 | 5 | 1 | 2 | 1 | 4 | 1 | NA | NA | 1 | 1 | 7 | 1 |
| 17_329927 | NA | NA | NA | NA | 2 | 0.5 | 1 | 1 | 3 | 0.6667 | 2 | 0.5 | 2 | 0 | NA | NA | 2 | 0.5 | 6 | 0.3333 |
| 17_329927 | NA | NA | NA | NA | 3 | 0.6667 | 2 | 1 | 5 | 0.8 | 1 | 1 | 4 | 0.5 | NA | NA | 2 | 0.5 | 7 | 0.5714 |
| 17_329927 | NA | NA | NA | NA | 2 | 0.5 | NA | NA | 2 | 0.5 | 1 | 0 | 2 | 0.5 | NA | NA | 2 | 0 | 5 | 0.2 |
| 17_329928 | NA | NA | NA | NA | 3 | 1 | 2 | 1 | 5 | 1 | 1 | 1 | 2 | 1 | NA | NA | 4 | 1 | 7 | 1 |
| 17_329928 | NA | NA | NA | NA | 3 | 0 | 2 | 1 | 5 | 0.4 | 1 | 1 | 4 | 0.25 | NA | NA | 3 | 1 | 8 | 0.625 |
| 17_329928 | NA | NA | NA | NA | 3 | 0.3333 | 1 | 1 | 4 | 0.5 | 2 | 0.5 | 2 | 0 | NA | NA | 4 | 0.75 | 8 | 0.5 |
| 17_329930 | NA | NA | NA | NA | 3 | 0.6667 | 1 | 1 | 4 | 0.75 | 2 | 1 | 3 | 1 | NA | NA | 2 | 1 | 7 | 1 |
| 17_329930 | NA | NA | NA | NA | 2 | 0.5 | 1 | 0 | 3 | 0.3333 | 2 | 1 | 2 | 0.5 | NA | NA | 4 | 0.25 | 8 | 0.5 |
| 17_329930 | NA | NA | NA | NA | 3 | 0.3333 | 2 | 0.5 | 5 | 0.4 | 2 | 0.5 | 3 | 0.6667 | NA | NA | 3 | 0.3333 | 8 | 0.5 |
| 17_329932 | NA | NA | NA | NA | 3 | 0.6667 | 1 | 1 | 4 | 0.75 | 2 | 1 | 2 | 1 | NA | NA | 3 | 1 | 7 | 1 |
| 17_329933 | NA | NA | NA | NA | 2 | 0.5 | 1 | 0 | 3 | 0.3333 | 2 | 0.5 | 2 | 0 | NA | NA | 2 | 1 | 6 | 0.5 |
| 17_329933 | NA | NA | NA | NA | 3 | 0.3333 | NA | NA | 3 | 0.3333 | 2 | 0 | 1 | 0 | NA | NA | 3 | 0.3333 | 6 | 0.1667 |
| 17_329935 | NA | NA | NA | NA | 3 | 1 | NA | NA | 3 | 1 | 1 | 1 | 1 | 1 | NA | NA | 3 | 0.6667 | 5 | 0.8 |
| 17_329940 | NA | NA | NA | NA | 3 | 0.3333 | NA | NA | 3 | 0.3333 | 2 | 1 | 2 | 0 | NA | NA | 1 | 0 | 5 | 0.4 |
| 17_329943 | NA | NA | NA | NA | 2 | 0.5 | NA | NA | 2 | 0.5 | 2 | 0.5 | 1 | 1 | NA | NA | 2 | 0.5 | 5 | 0.6 |
| 17_329943 | NA | NA | NA | NA | 1 | 1 | NA | NA | 1 | 1 | 2 | 1 | 1 | 1 | NA | NA | 1 | 1 | 4 | 1 |
| 17_329946 | NA | NA | NA | NA | 2 | 0.5 | NA | NA | 2 | 0.5 | 2 | 1 | NA | NA | NA | NA | 2 | 1 | 4 | 1 |
| 17_329946 | NA | NA | NA | NA | 2 | 0.5 | NA | NA | 2 | 0.5 | 2 | 0.5 | 1 | 0 | NA | NA | 2 | 0 | 5 | 0.2 |
| 17_329947 | NA | NA | NA | NA | 2 | 1 | NA | NA | 2 | 1 | 2 | 0.5 | 1 | 1 | NA | NA | 1 | 1 | 4 | 0.75 |
| 17_329949 | NA | NA | NA | NA | 2 | 1 | NA | NA | 2 | 1 | 2 | 0 | 1 | 1 | NA | NA | 2 | 0.5 | 5 | 0.4 |
| 17_330073 | 1 | 1 | 1 | 1 | NA | NA | NA | NA | 2 | 1 | NA | NA | 1 | 1 | NA | NA | NA | NA | 1 | 1 |
| 17_330078 | 1 | 1 | 1 | 1 | NA | NA | NA | NA | 2 | 1 | NA | NA | 1 | 1 | NA | NA | NA | NA | 1 | 1 |
| 17_330078 | 1 | 1 | 1 | 1 | NA | NA | NA | NA | 2 | 1 | NA | NA | 1 | 1 | NA | NA | NA | NA | 1 | 1 |
| 17_330079 | 1 | 0 | 1 | 1 | NA | NA | NA | NA | 2 | 0.5 | NA | NA | 1 | 1 | NA | NA | NA | NA | 1 | 1 |
| 17_330082 | NA | NA | 1 | 1 | NA | NA | NA | NA | 1 | 1 | NA | NA | 1 | 1 | NA | NA | NA | NA | 1 | 1 |
| 17_330084 | 1 | 1 | 1 | 1 | NA | NA | NA | NA | 2 | 1 | NA | NA | 1 | 1 | NA | NA | NA | NA | 1 | 1 |
| 17_330163 | NA | NA | 1 | 1 | NA | NA | NA | NA | 1 | 1 | 1 | 1 | NA | NA | NA | NA | NA | NA | 1 | 1 |
| 17_330371 | NA | NA | 4 | 0 | 3 | 0.6667 | 2 | 0 | 9 | 0.2222 | NA | NA | 1 | 1 | NA | NA | 2 | 0 | 3 | 0.3333 |
| 17_330376 | NA | NA | 6 | 0.1667 | 1 | 0 | NA | NA | 7 | 0.1429 | 1 | 0 | 1 | 0 | 1 | 0 | 3 | 0.3333 | 6 | 0.1667 |
| 17_330376 | NA | NA | 5 | 0.4 | 2 | 0 | 1 | 1 | 8 | 0.375 | 1 | 1 | 2 | 0 | NA | NA | 3 | 0.3333 | 6 | 0.3333 |
| 17_330378 | NA | NA | 4 | 0.5 | 2 | 0.5 | 1 | 1 | 7 | 0.5714 | NA | NA | 1 | 0 | 1 | 0 | 2 | 0.5 | 4 | 0.25 |
| 17_330378 | NA | NA | 4 | 0.5 | 2 | 1 | 1 | 1 | 7 | 0.7143 | NA | NA | 1 | 1 | 1 | 0 | 3 | 0.6667 | 5 | 0.6 |
| 17_330379 | NA | NA | 4 | 1 | 2 | 1 | 1 | 1 | 7 | 1 | NA | NA | 2 | 1 | 1 | 0 | 2 | 1 | 5 | 0.8 |
| 17_330379 | NA | NA | 2 | 0.5 | 1 | 0 | 1 | 0 | 4 | 0.25 | 1 | 0 | 1 | 0 | 1 | 0 | 3 | 0.3333 | 6 | 0.1667 |
| 17_330379 | NA | NA | 4 | 1 | 2 | 0.5 | 1 | 1 | 7 | 0.8571 | 1 | 1 | 2 | 1 | 1 | 0 | 3 | 0.6667 | 7 | 0.7143 |
| 17_330381 | NA | NA | 4 | 1 | 2 | 1 | NA | NA | 6 | 1 | NA | NA | 2 | 1 | 1 | 0 | 3 | 1 | 6 | 0.8333 |
| 17_330382 | NA | NA | 4 | 0.25 | 2 | 0 | 1 | 0 | 7 | 0.1429 | 1 | 0 | 2 | 0.5 | 1 | 0 | 3 | 0 | 7 | 0.1429 |
| 17_330382 | NA | NA | 3 | 1 | 2 | 1 | 1 | 1 | 6 | 1 | 1 | 1 | 2 | 1 | 1 | 1 | 3 | 1 | 7 | 1 |
| 17_330383 | NA | NA | 4 | 0.75 | 2 | 1 | 1 | 0 | 7 | 0.7143 | NA | NA | 2 | 0.5 | 1 | 0 | 3 | 1 | 6 | 0.6667 |
| 17_330383 | NA | NA | 2 | 0.5 | 2 | 1 | NA | NA | 4 | 0.75 | NA | NA | 2 | 0.5 | 1 | 1 | 1 | 1 | 4 | 0.75 |
| 17_330383 | NA | NA | 3 | 0.6667 | 2 | 0.5 | 1 | 1 | 6 | 0.6667 | 1 | 0 | 2 | 0.5 | 1 | 1 | 1 | 0 | 5 | 0.4 |
| 17_330386 | NA | NA | 4 | 0 | 2 | 1 | 1 | 0 | 7 | 0.2857 | 1 | 0 | 1 | 1 | 1 | 0 | 1 | 1 | 4 | 0.5 |
| 17_330386 | NA | NA | 3 | 0 | 2 | 0.5 | 1 | 1 | 6 | 0.3333 | NA | NA | 2 | 0.5 | 1 | 0 | 1 | 0 | 4 | 0.25 |
| 17_330387 | NA | NA | 3 | 1 | 2 | 1 | 1 | 1 | 6 | 1 | 1 | 1 | 2 | 0.5 | NA | NA | 2 | 1 | 5 | 0.8 |
| 17_330388 | NA | NA | 1 | 1 | 2 | 1 | 1 | 1 | 4 | 1 | NA | NA | NA | NA | 1 | 1 | 2 | 1 | 3 | 1 |
| 17_330390 | NA | NA | 2 | 0.5 | 2 | 0 | 1 | 0 | 5 | 0.2 | 1 | 0 | 2 | 0 | 1 | 0 | 2 | 0.5 | 6 | 0.1667 |
| 17_330390 | NA | NA | 2 | 1 | 2 | 1 | 1 | 1 | 5 | 1 | NA | NA | 2 | 1 | NA | NA | 1 | 1 | 3 | 1 |
| 17_330392 | NA | NA | 2 | 0.5 | 1 | 0 | 1 | 0 | 4 | 0.25 | 1 | 0 | 2 | 0.5 | 1 | 0 | NA | NA | 4 | 0.25 |
| 17_330392 | NA | NA | NA | NA | 2 | 1 | 1 | 1 | 3 | 1 | NA | NA | 2 | 1 | 1 | 1 | 2 | 1 | 5 | 1 |

| | | | | | | | | | | | | | | | | | | | | |
|---|---|---|---|---|---|---|---|---|---|---|---|---|---|---|---|---|---|---|---|---|
| 17_330396 | NA | NA | 3 | 0 | 2 | 0.5 | 1 | 0 | 6 | 0.1667 | NA | NA | 2 | 0 | 1 | 0 | 2 | 0.5 | 5 | 0.2 |
| 17_330398 | NA | NA | 1 | 1 | 2 | 0 | NA | NA | 3 | 0.3333 | NA | NA | 2 | 0 | NA | NA | 1 | 1 | 3 | 0.3333 |
| 17_330398 | NA | NA | NA | NA | 2 | 1 | NA | NA | 2 | 1 | 1 | 1 | 1 | 1 | 1 | 1 | 1 | 1 | 4 | 1 |
| 17_330398 | NA | NA | 2 | 0.5 | 1 | 1 | NA | NA | 3 | 0.6667 | NA | NA | 2 | 1 | 1 | 1 | 1 | 0 | 4 | 0.75 |
| 17_331011 | NA | NA | NA | NA | NA | NA | 1 | 1 | 1 | 1 | NA | NA | 1 | 1 | NA | NA | 1 | 1 | 2 | 1 |
| 17_331016 | NA | NA | NA | NA | NA | NA | 1 | 1 | 1 | 1 | NA | NA | 1 | 1 | NA | NA | NA | NA | 1 | 1 |
| 17_331017 | NA | NA | NA | NA | NA | NA | 1 | 1 | 1 | 1 | NA | NA | 1 | 0 | NA | NA | 1 | 1 | 2 | 0.5 |
| 17_331020 | NA | NA | NA | NA | NA | NA | 1 | 1 | 1 | 1 | NA | NA | 1 | 1 | NA | NA | 1 | 1 | 2 | 1 |
| 17_331023 | NA | NA | NA | NA | NA | NA | 1 | 1 | 1 | 1 | NA | NA | NA | NA | NA | NA | 1 | 1 | 1 | 1 |
| 17_331027 | NA | NA | NA | NA | NA | NA | 1 | 1 | 1 | 1 | NA | NA | 1 | 1 | NA | NA | 1 | 0 | 2 | 0.5 |
| 17_331243 | 1 | 0 | NA | NA | 1 | 1 | 2 | 0 | 4 | 0.25 | 4 | 0.75 | 1 | 1 | 1 | 1 | 1 | 0 | 7 | 0.7143 |
| 17_331244 | 1 | 0 | NA | NA | 1 | 0 | 2 | 0.5 | 4 | 0.25 | 4 | 0.75 | 2 | 0 | 1 | 0 | 1 | 0 | 8 | 0.375 |
| 17_331963 | NA | NA | 1 | 1 | NA | NA | 1 | 1 | 2 | 1 | NA | NA | 1 | 1 | NA | NA | 2 | 0.5 | 3 | 0.6667 |
| 17_331963 | NA | NA | NA | NA | NA | NA | 1 | 1 | 1 | 1 | NA | NA | NA | NA | NA | NA | 2 | 1 | 2 | 1 |
| 17_331967 | NA | NA | 1 | 1 | NA | NA | 1 | 1 | 2 | 1 | NA | NA | 1 | 1 | NA | NA | 2 | 1 | 3 | 1 |
| 17_331968 | NA | NA | 1 | 1 | NA | NA | 1 | 1 | 2 | 1 | 1 | 1 | 1 | 1 | NA | NA | 2 | 1 | 4 | 1 |
| 17_331971 | NA | NA | 1 | 0 | NA | NA | 1 | 1 | 2 | 0.5 | 1 | 0 | 1 | 0 | NA | NA | 2 | 0.5 | 4 | 0.25 |
| 17_331971 | NA | NA | 1 | 1 | NA | NA | 1 | 0 | 2 | 0.5 | 1 | 0 | 1 | 0 | NA | NA | 2 | 0.5 | 4 | 0.25 |
| 17_331972 | NA | NA | 1 | 0 | NA | NA | 1 | 1 | 2 | 0.5 | 1 | 0 | 1 | 1 | NA | NA | 1 | 0 | 3 | 0.3333 |
| 17_331973 | NA | NA | 1 | 1 | NA | NA | NA | NA | 1 | 1 | 1 | 1 | 1 | 1 | NA | NA | 2 | 1 | 4 | 1 |
| 17_331973 | NA | NA | 1 | 1 | NA | NA | NA | NA | 1 | 1 | 1 | 0 | 1 | 1 | NA | NA | 2 | 0.5 | 4 | 0.5 |
| 17_331974 | NA | NA | 1 | 1 | NA | NA | 1 | 0 | 2 | 0.5 | 1 | 0 | 1 | 0 | NA | NA | 2 | 1 | 4 | 0.5 |
| 17_331975 | NA | NA | 1 | 1 | NA | NA | NA | NA | 1 | 1 | 1 | 1 | NA | NA | NA | NA | 2 | 1 | 3 | 1 |
| 17_332218 | 3 | 0 | 1 | 0 | 2 | 0.5 | NA | NA | 6 | 0.1667 | 2 | 0.5 | 4 | 0.5 | 1 | 1 | 2 | 0.5 | 9 | 0.5556 |
| 17_332218 | 2 | 0.5 | 1 | 1 | 1 | 0 | NA | NA | 4 | 0.5 | 1 | 1 | 4 | 0.25 | 2 | 0 | NA | NA | 7 | 0.2857 |
| 17_332218 | 3 | 1 | 1 | 1 | 2 | 1 | NA | NA | 6 | 1 | 2 | 1 | 4 | 1 | 2 | 1 | 2 | 1 | 10 | 1 |
| 17_332219 | 2 | 0.5 | 1 | 1 | 2 | 0 | NA | NA | 5 | 0.4 | 1 | 1 | 4 | 0.25 | NA | NA | 2 | 0 | 7 | 0.2857 |
| 17_332220 | 2 | 0.5 | 1 | 0 | 2 | 1 | NA | NA | 5 | 0.6 | 2 | 0.5 | 4 | 0.5 | 2 | 0.5 | 2 | 1 | 10 | 0.6 |
| 17_332221 | 3 | 0.3333 | 1 | 1 | 2 | 0 | NA | NA | 6 | 0.3333 | 2 | 0.5 | 3 | 0.3333 | 2 | 0.5 | 2 | 0 | 9 | 0.3333 |
| 17_332221 | 3 | 0.3333 | NA | NA | 2 | 0 | NA | NA | 5 | 0.2 | 3 | 0 | 4 | 0 | 2 | 1 | 2 | 1 | 11 | 0.3636 |
| 17_332223 | 5 | 0.2 | 1 | 0 | 2 | 0 | NA | NA | 8 | 0.125 | 2 | 0 | 3 | 0.3333 | 1 | 0 | 2 | 0.5 | 8 | 0.25 |
| 17_332223 | 3 | 0 | 1 | 1 | 2 | 0 | NA | NA | 6 | 0.1667 | 3 | 0.6667 | 4 | 0.25 | 2 | 0.5 | 2 | 0 | 11 | 0.3636 |
| 17_332224 | 5 | 0.2 | 1 | 0 | 2 | 0.5 | NA | NA | 8 | 0.25 | 2 | 0 | 3 | 0 | 2 | 0.5 | 2 | 0 | 9 | 0.1111 |
| 17_332225 | 5 | 0.4 | 1 | 0 | 2 | 0 | NA | NA | 8 | 0.25 | 3 | 0.3333 | 4 | 0.5 | 2 | 0 | 2 | 0 | 11 | 0.2727 |
| 17_332226 | 5 | 0 | 1 | 0 | 2 | 0.5 | NA | NA | 8 | 0.125 | 2 | 0 | 6 | 0 | 2 | 0 | 2 | 0.5 | 12 | 0.0833 |
| 17_332227 | 4 | 0 | 1 | 1 | 2 | 0 | NA | NA | 7 | 0.1429 | 3 | 0.3333 | 6 | 0.6667 | 2 | 0.5 | 1 | 0 | 12 | 0.5 |
| 17_332230 | 5 | 0.4 | 1 | 1 | 2 | 0.5 | NA | NA | 8 | 0.5 | 3 | 0.6667 | 4 | 0.5 | 2 | 0.5 | 2 | 0.5 | 11 | 0.5455 |
| 17_332231 | 6 | 0.3333 | 1 | 1 | 2 | 0 | NA | NA | 9 | 0.3333 | 3 | 0.6667 | 7 | 0.4286 | 2 | 0 | 2 | 0 | 14 | 0.3571 |
| 17_332236 | 6 | 0.3333 | 2 | 0 | 2 | 0 | NA | NA | 10 | 0.2 | 3 | 0 | 7 | 0.1429 | 2 | 0 | 2 | 0 | 14 | 0.0714 |
| 17_332238 | 4 | 0.25 | 1 | 0 | 1 | 0 | NA | NA | 6 | 0.1667 | 4 | 0.5 | 3 | 0 | 2 | 0.5 | 1 | 1 | 10 | 0.4 |
| 17_332238 | 6 | 0.1667 | 1 | 1 | 1 | 1 | NA | NA | 8 | 0.375 | 3 | 0.3333 | 6 | 0.1667 | 2 | 0 | 2 | 0.5 | 13 | 0.2308 |
| 17_332239 | 5 | 0.4 | 2 | 0 | 1 | 1 | NA | NA | 8 | 0.375 | 4 | 0 | 7 | 0.8571 | 2 | 1 | 2 | 0.5 | 15 | 0.6 |
| 17_332240 | 6 | 0.6667 | 1 | 0 | 2 | 0 | NA | NA | 9 | 0.4444 | 4 | 0.25 | 7 | 0.4286 | 2 | 0.5 | 2 | 0 | 15 | 0.3333 |
| 17_332367 | 2 | 1 | 2 | 1 | 3 | 1 | 1 | 0 | 8 | 0.875 | 5 | 0.6 | 2 | 1 | 1 | 1 | 1 | 1 | 9 | 0.7778 |
| 17_332371 | 3 | 0 | 1 | 0 | 3 | 0.3333 | 2 | 0 | 9 | 0.1111 | 6 | 0.1667 | 1 | 1 | 1 | 0 | 3 | 0 | 11 | 0.1818 |
| 17_332371 | 3 | 0 | 1 | 0 | 2 | 0.5 | 2 | 0 | 8 | 0.125 | 5 | 0.4 | 2 | 0.5 | 1 | 0 | 3 | 0 | 11 | 0.2727 |
| 17_332372 | 2 | 1 | 1 | 1 | 2 | 1 | 2 | 1 | 7 | 1 | 4 | 1 | 1 | 1 | 1 | 1 | 2 | 1 | 8 | 1 |
| 17_332373 | 2 | 1 | 1 | 1 | 3 | 1 | 2 | 1 | 8 | 1 | 4 | 1 | 2 | 1 | 1 | 0 | 2 | 1 | 9 | 0.8889 |
| 17_332373 | 3 | 0.3333 | 1 | 0 | 2 | 0 | 2 | 0 | 8 | 0.125 | 4 | 0.25 | 2 | 0 | 1 | 0 | 1 | 0 | 8 | 0.125 |
| 17_332373 | 2 | 1 | 1 | 1 | 2 | 0.5 | 2 | 0.5 | 7 | 0.7143 | 4 | 1 | 2 | 1 | 2 | 0.5 | 2 | 1 | 10 | 0.9 |
| 17_332375 | 2 | 1 | NA | NA | 1 | 0 | 2 | 1 | 5 | 0.8 | 3 | 1 | 2 | 1 | 1 | 1 | 2 | 1 | 8 | 1 |
| 17_332375 | 2 | 0.5 | NA | NA | 3 | 0 | 1 | 0 | 6 | 0.1667 | 4 | 0.25 | 2 | 0 | 2 | 0 | 2 | 0 | 10 | 0.1 |
| 17_332376 | 3 | 0.3333 | 1 | 0 | 3 | 0 | 2 | 0 | 9 | 0.1111 | 3 | 0.3333 | 2 | 0 | 2 | 0 | 2 | 0.5 | 9 | 0.2222 |
| 17_332376 | 1 | 1 | 1 | 1 | 2 | 0.5 | 1 | 1 | 5 | 0.8 | 4 | 1 | 2 | 1 | 1 | 1 | 1 | 1 | 8 | 1 |
| 17_332378 | 3 | 0 | 1 | 0 | 3 | 0.6667 | 2 | 0 | 9 | 0.2222 | 4 | 0.5 | 2 | 0 | 2 | 0 | 2 | 0 | 10 | 0.2 |
| 17_332379 | 3 | 1 | 1 | 1 | 2 | 1 | 2 | 1 | 8 | 1 | 4 | 1 | 2 | 1 | 1 | 1 | 2 | 1 | 9 | 1 |
| 17_332379 | 2 | 1 | 1 | 1 | 2 | 0 | 1 | 1 | 6 | 0.6667 | 2 | 1 | 2 | 1 | 1 | 1 | 2 | 1 | 7 | 1 |
| 17_332380 | 2 | 0 | 1 | 1 | 3 | 0 | 1 | 0 | 7 | 0.1429 | 4 | 0.25 | 2 | 0 | 1 | 0 | 2 | 0 | 9 | 0.1111 |
| 17_332382 | 2 | 0.5 | 1 | 0 | 3 | 0 | 1 | 0 | 7 | 0.1429 | 4 | 0 | 2 | 0 | 1 | 1 | 2 | 0 | 9 | 0.1111 |
| 17_332382 | 2 | 1 | 1 | 1 | 2 | 1 | 1 | 1 | 6 | 1 | 3 | 1 | 2 | 1 | 1 | 1 | 2 | 1 | 8 | 1 |
| 17_332384 | 1 | 0 | 1 | 0 | 2 | 0.5 | NA | NA | 4 | 0.25 | 3 | 0.6667 | 2 | 0 | 1 | 0 | 2 | 0 | 8 | 0.25 |
| 17_332385 | 1 | 1 | 1 | 0 | 3 | 0.3333 | 1 | 0 | 6 | 0.3333 | 4 | 0.5 | 2 | 0.5 | 1 | 1 | 2 | 0 | 9 | 0.4444 |
| 17_332387 | 2 | 0 | 1 | 0 | 3 | 0.6667 | 1 | 0 | 7 | 0.2857 | 4 | 0 | 2 | 1 | NA | NA | 1 | 0 | 7 | 0.2857 |
| 17_332387 | 2 | 0.5 | 1 | 1 | 3 | 0.3333 | 1 | 0 | 7 | 0.4286 | 2 | 0.5 | 2 | 0.5 | 1 | 0 | 2 | 0.5 | 7 | 0.4286 |
| 17_332387 | 2 | 0 | 1 | 0 | 3 | 0.3333 | 1 | 0 | 7 | 0.1429 | 4 | 0.25 | 2 | 0 | 1 | 1 | 2 | 0 | 9 | 0.2222 |

| | | | | | | | | | | | | | | | | | | | | |
|---|---|---|---|---|---|---|---|---|---|---|---|---|---|---|---|---|---|---|---|---|
| 17_3332368 | 1 | 0 | NA | NA | 5 | 0.2 | 2 | 0 | 8 | 0.125 | 1 | 0 | 1 | 0 | NA | NA | 4 | 0.25 | 6 | 0.1667 |
| 17_3332373 | 1 | 1 | NA | NA | 5 | 0.2 | 1 | 0 | 7 | 0.2857 | 1 | 0 | 1 | 1 | 1 | 0 | 4 | 0.5 | 7 | 0.4286 |
| 17_3332378 | 1 | 1 | NA | NA | 5 | 1 | 2 | 1 | 8 | 1 | 1 | 1 | 1 | 0 | 1 | 1 | 3 | 1 | 6 | 0.8333 |
| 17_3332381 | NA | NA | NA | NA | 3 | 0.6667 | 2 | 1 | 5 | 0.8 | NA | NA | 1 | 1 | 1 | 1 | 4 | 0.75 | 6 | 0.8333 |
| 17_3332390 | 1 | 0 | NA | NA | 5 | 0.2 | 2 | 0.5 | 8 | 0.25 | 1 | 1 | 1 | 1 | 1 | 1 | 4 | 0 | 7 | 0.4286 |
| 17_3332397 | 1 | 1 | NA | NA | 5 | 1 | 2 | 1 | 8 | 1 | 1 | 1 | 1 | 1 | 1 | 1 | 3 | 1 | 6 | 1 |
| 17_3332422 | 1 | 1 | NA | NA | 5 | 1 | 1 | 1 | 7 | 1 | 1 | 1 | 1 | 1 | 1 | 1 | 4 | 1 | 7 | 1 |
| 17_3332424 | NA | NA | NA | NA | 5 | 0.8 | 2 | 1 | 7 | 0.8571 | 1 | 1 | 1 | 1 | 1 | 1 | 4 | 0.5 | 7 | 0.7143 |
| 17_3332434 | 1 | 1 | NA | NA | 3 | 1 | 1 | 0 | 5 | 0.8 | 1 | 1 | 1 | 1 | 1 | 1 | 3 | 0.6667 | 6 | 0.8333 |
| 17_3332450 | 1 | 1 | NA | NA | 4 | 1 | 1 | 1 | 6 | 1 | 1 | 1 | 1 | 1 | 1 | 1 | 4 | 1 | 7 | 1 |
| 17_3332462 | 1 | 1 | NA | NA | 3 | 0.6667 | 2 | 1 | 6 | 0.8333 | 1 | 1 | 1 | 0 | NA | NA | 4 | 1 | 6 | 0.8333 |
| 17_3332463 | 1 | 1 | NA | NA | 4 | 0.75 | 2 | 0.5 | 7 | 0.7143 | 1 | 1 | 1 | 1 | 1 | 0 | 4 | 0.75 | 7 | 0.7143 |
| 17_3332473 | 1 | 1 | NA | NA | 5 | 0.2 | 2 | 0 | 8 | 0.25 | 1 | 1 | 1 | 0 | 1 | 0 | 4 | 0 | 7 | 0.1429 |
| 17_3332477 | 1 | 1 | NA | NA | 5 | 1 | 2 | 0.5 | 8 | 0.875 | 1 | 1 | 1 | 1 | 1 | 1 | 2 | 1 | 5 | 1 |
| 17_3332493 | 1 | 1 | NA | NA | 5 | 0.2 | 2 | 0 | 8 | 0.25 | 1 | 0 | 1 | 1 | 1 | 0 | 3 | 0.6667 | 6 | 0.5 |
| 17_3332541 | 1 | 0 | NA | NA | 5 | 1 | 2 | 0.5 | 8 | 0.75 | 1 | 1 | 1 | 1 | NA | NA | 2 | 1 | 4 | 1 |
| 17_3332552 | NA | NA | NA | NA | 5 | 0.6 | 1 | 1 | 6 | 0.6667 | NA | NA | 1 | 1 | NA | NA | 1 | 1 | 2 | 1 |
| 17_3332670 | 1 | 0 | NA | NA | 2 | 0.5 | 2 | 0 | 5 | 0.2 | 1 | 0 | 1 | 1 | NA | NA | 2 | 1 | 4 | 0.75 |
| 17_3345940 | NA | NA | 1 | 1 | NA | NA | NA | NA | 1 | 1 | NA | NA | 2 | 0.5 | 2 | 0.5 | NA | NA | 4 | 0.5 |
| 17_3352095 | 2 | 0.5 | 2 | 0 | 1 | 0 | 2 | 1 | 7 | 0.4286 | NA | NA | 1 | 0 | 1 | 0 | 2 | 0.5 | 4 | 0.25 |
| 17_3352106 | 2 | 0.5 | 3 | 0.3333 | 1 | 0 | 2 | 0 | 8 | 0.25 | NA | NA | 1 | 1 | 1 | 0 | 1 | 1 | 3 | 0.6667 |
| 17_3352108 | 2 | 1 | 3 | 1 | 1 | 1 | 2 | 1 | 8 | 1 | NA | NA | 1 | 1 | 1 | 1 | 2 | 1 | 4 | 1 |
| 17_3352131 | 1 | 1 | 2 | 0 | 1 | 1 | 2 | 0.5 | 6 | 0.5 | NA | NA | 2 | 0.5 | 1 | 0 | 2 | 1 | 5 | 0.6 |
| 17_3352132 | 2 | 0.5 | 3 | 0.3333 | 1 | 1 | 2 | 1 | 8 | 0.625 | NA | NA | 2 | 0 | 1 | 0 | 2 | 0.5 | 5 | 0.2 |
| 17_3352136 | 2 | 0.5 | 3 | 0.6667 | 1 | 1 | 2 | 0.5 | 8 | 0.625 | NA | NA | 2 | 0.5 | 1 | 0 | 2 | 0.5 | 5 | 0.4 |
| 17_3352141 | 2 | 1 | 3 | 0.3333 | 1 | 1 | 1 | 1 | 7 | 0.7143 | NA | NA | 2 | 0.5 | 1 | 1 | 2 | 0 | 5 | 0.4 |
| 17_3352142 | 2 | 0 | 3 | 0.6667 | 1 | 0 | 2 | 0 | 8 | 0.25 | NA | NA | 2 | 0.5 | 1 | 0 | 1 | 0 | 4 | 0.25 |
| 17_3352146 | 2 | 0 | 3 | 0 | 1 | 0 | 2 | 0.5 | 8 | 0.125 | NA | NA | 1 | 1 | 1 | 0 | 1 | 0 | 3 | 0.3333 |
| 17_3352190 | 2 | 0.5 | 3 | 0.3333 | 1 | 1 | 2 | 0.5 | 8 | 0.5 | NA | NA | 1 | 0 | 1 | 0 | 2 | 0.5 | 4 | 0.25 |
| 17_3352245 | 1 | 0 | 3 | 0 | 2 | 0.5 | 2 | 0.5 | 8 | 0.25 | NA | NA | 1 | 0 | 1 | 1 | 1 | 1 | 3 | 0.6667 |
| 17_3352267 | 2 | 0 | 3 | 0 | 2 | 0.5 | 2 | 1 | 9 | 0.3333 | NA | NA | 1 | 1 | 1 | 1 | 1 | 0 | 3 | 0.6667 |
| 17_3352325 | 2 | 1 | 2 | 1 | 2 | 1 | NA | NA | 6 | 1 | NA | NA | 2 | 1 | 1 | 1 | 1 | 1 | 4 | 1 |
| 17_3352331 | 2 | 0 | 2 | 1 | 1 | 0 | 1 | 1 | 6 | 0.5 | NA | NA | 2 | 0 | 1 | 1 | 2 | 0.5 | 5 | 0.4 |
| 17_3352335 | 2 | 0.5 | 3 | 1 | 3 | 1 | 1 | 1 | 9 | 0.8889 | NA | NA | 1 | 1 | 1 | 1 | 1 | 1 | 3 | 1 |
| 17_3352340 | 2 | 0.5 | 3 | 0 | 1 | 0 | 1 | 0 | 7 | 0.1429 | NA | NA | 2 | 0 | 1 | 0 | 2 | 0.5 | 5 | 0.2 |
| 17_3352409 | 2 | 0 | 2 | 0.5 | 1 | 0 | 1 | 0 | 6 | 0.1667 | NA | NA | 4 | 0.5 | 2 | 0 | NA | NA | 6 | 0.3333 |
| 17_3352427 | 1 | 1 | 2 | 1 | 1 | 1 | 1 | 1 | 5 | 1 | NA | NA | 3 | 0.6667 | 2 | 0 | NA | NA | 5 | 0.4 |
| 17_3352443 | 2 | 0.5 | 2 | 1 | 2 | 1 | 1 | 1 | 7 | 0.8571 | NA | NA | 4 | 1 | 1 | 0 | 1 | 1 | 6 | 0.8333 |
| 17_3352444 | 2 | 1 | 2 | 1 | 2 | 1 | 1 | 1 | 7 | 1 | NA | NA | 5 | 0.8 | 2 | 1 | 1 | 1 | 8 | 0.875 |
| 17_3352452 | 1 | 0 | 2 | 1 | 1 | 0 | NA | NA | 4 | 0.5 | NA | NA | 4 | 0.5 | 1 | 0 | NA | NA | 5 | 0.4 |
| 17_3352453 | 2 | 1 | 2 | 0.5 | 2 | 1 | 1 | 1 | 7 | 0.8571 | NA | NA | 5 | 0.8 | 2 | 1 | NA | NA | 7 | 0.8571 |
| 17_3352476 | 2 | 0.5 | 2 | 0 | 1 | 0 | 2 | 0.5 | 7 | 0.2857 | NA | NA | 4 | 0.25 | 1 | 0 | NA | NA | 5 | 0.2 |
| 17_3352496 | 2 | 0 | 2 | 0.5 | 2 | 0.5 | 2 | 0 | 8 | 0.25 | NA | NA | 6 | 0.3333 | 2 | 0 | 1 | 0 | 9 | 0.2222 |
| 17_3352512 | 1 | 0 | 2 | 0 | 2 | 0.5 | 2 | 0 | 7 | 0.1429 | 1 | 1 | 6 | 0.1667 | 2 | 0 | 1 | 0 | 10 | 0.2 |
| 17_3352550 | 2 | 1 | 2 | 1 | 1 | 0 | 2 | 0.5 | 7 | 0.7143 | 2 | 1 | 5 | 0.8 | 2 | 1 | 1 | 1 | 10 | 0.9 |
| 17_3352567 | 2 | 0 | NA | NA | 2 | 0.5 | 2 | 0 | 6 | 0.1667 | 2 | 1 | 4 | 0.25 | 1 | 0 | 2 | 0 | 9 | 0.3333 |
| 17_3352583 | 2 | 1 | NA | NA | 2 | 0.5 | 2 | 1 | 6 | 0.8333 | 2 | 1 | 1 | 1 | 1 | 1 | 1 | 1 | 5 | 1 |
| 17_3352585 | 2 | 0 | NA | NA | 2 | 0 | 2 | 0.5 | 6 | 0.1667 | 2 | 0.5 | 3 | 0.3333 | 1 | 0 | 1 | 0 | 7 | 0.2857 |
| 17_3352614 | 1 | 0 | 1 | 0 | 2 | 0.5 | 2 | 0 | 6 | 0.1667 | 2 | 0.5 | 4 | 0 | 2 | 0 | 1 | 1 | 9 | 0.2222 |
| 17_3353455 | 1 | 1 | NA | NA | NA | NA | NA | NA | 1 | 1 | NA | NA | NA | NA | 1 | 1 | NA | NA | 1 | 1 |
| 17_3353469 | 1 | 1 | NA | NA | NA | NA | NA | NA | 1 | 1 | NA | NA | NA | NA | 1 | 1 | NA | NA | 1 | 1 |
| 17_3353530 | 1 | 1 | NA | NA | NA | NA | NA | NA | 1 | 1 | NA | NA | NA | NA | 1 | 1 | NA | NA | 1 | 1 |
| 17_3353533 | 1 | 1 | NA | NA | NA | NA | NA | NA | 1 | 1 | NA | NA | NA | NA | 2 | 1 | NA | NA | 2 | 1 |
| 17_3353540 | 1 | 1 | NA | NA | NA | NA | NA | NA | 1 | 1 | NA | NA | NA | NA | 2 | 0.5 | NA | NA | 2 | 0.5 |
| 17_3354854 | 2 | 0.5 | 2 | 0 | 4 | 0 | 4 | 0 | 12 | 0.0833 | 2 | 0.5 | NA | NA | 2 | 0 | 1 | 1 | 5 | 0.4 |
| 17_3355252 | NA | NA | 1 | 0 | 1 | 0 | 2 | 0.5 | 4 | 0.25 | 1 | 1 | NA | NA | NA | NA | NA | NA | 1 | 1 |
| 17_3359943 | 1 | 0 | 1 | 1 | 1 | 0 | 1 | 0 | 4 | 0.25 | NA | NA | NA | NA | 1 | 0 | 1 | 1 | 2 | 0.5 |
| 17_3359984 | 1 | 0 | NA | NA | 1 | 0 | 1 | 1 | 3 | 0.3333 | NA | NA | NA | NA | 1 | 0 | 1 | 1 | 2 | 0.5 |
| 17_3359991 | NA | NA | NA | NA | 1 | 1 | 1 | 1 | 2 | 1 | NA | NA | NA | NA | NA | NA | 1 | 1 | 1 | 1 |
| 17_3360001 | 1 | 1 | 1 | 1 | 1 | 1 | 1 | 1 | 4 | 1 | NA | NA | NA | NA | NA | NA | 1 | 1 | 1 | 1 |
| 17_3360011 | 1 | 0 | 1 | 1 | 1 | 1 | NA | NA | 3 | 0.6667 | NA | NA | NA | NA | NA | NA | 1 | 1 | 1 | 1 |
| 17_3360015 | NA | NA | 1 | 1 | 1 | 1 | 1 | 1 | 3 | 1 | NA | NA | NA | NA | NA | NA | 1 | 1 | 1 | 1 |
| 17_3360038 | 1 | 1 | 1 | 1 | 1 | 1 | NA | NA | 3 | 1 | NA | NA | NA | NA | NA | NA | 1 | 1 | 1 | 1 |
| 17_3360055 | 1 | 0 | NA | NA | 1 | 1 | NA | NA | 2 | 0.5 | NA | NA | NA | NA | NA | NA | 1 | 1 | 1 | 1 |
| 17_3360117 | NA | NA | 1 | 1 | 1 | 1 | NA | NA | 2 | 1 | NA | NA | NA | NA | NA | NA | 1 | 1 | 1 | 1 |

| | | | | | | | | | | | | | | | | | | | | |
|---|---|---|---|---|---|---|---|---|---|---|---|---|---|---|---|---|---|---|---|---|
| 17_338489 | 2 | 0 | 3 | 0.3333 | 4 | 0.25 | 3 | 0 | 12 | 0.1667 | 2 | 0 | 5 | 0 | 3 | 0.3333 | 1 | 0 | 11 | 0.0909 |
| 17_338657 | NA | NA | NA | NA | 1 | 0 | 1 | 1 | 2 | 0.5 | NA | NA | 1 | 0 | 2 | 0.5 | 1 | 0 | 4 | 0.25 |
| 17_338692 | 9 | 0.5556 | 11 | 0.7273 | 14 | 0.4286 | 13 | 0.6923 | 47 | 0.5957 | 10 | 0.6 | 5 | 0.4 | 12 | 0.5833 | 14 | 0.7143 | 41 | 0.6098 |
| 17_339045 | NA | NA | 1 | 1 | 2 | 0.5 | NA | NA | 3 | 0.6667 | NA | NA | NA | NA | 1 | 1 | NA | NA | 1 | 1 |
| 17_339048 | NA | NA | NA | NA | 1 | 1 | NA | NA | 1 | 1 | NA | NA | 1 | 1 | NA | NA | NA | NA | 1 | 1 |
| 17_339053 | NA | NA | NA | NA | 1 | 1 | NA | NA | 1 | 1 | NA | NA | 1 | 1 | NA | NA | NA | NA | 1 | 1 |
| 17_339065 | NA | NA | NA | NA | 2 | 0.5 | 1 | 0 | 3 | 0.3333 | NA | NA | 1 | 1 | 1 | 0 | 1 | 1 | 3 | 0.6667 |
| 17_339106 | 6 | 0.5 | 4 | 0.75 | 9 | 0.5556 | 10 | 0.7 | 29 | 0.6207 | 10 | 0.9 | 9 | 0.8889 | 4 | 0.5 | 3 | 1 | 26 | 0.8462 |
| 17_339107 | 13 | 0.7692 | 5 | 0.8 | 10 | 0.7 | 12 | 0.6667 | 40 | 0.725 | 10 | 0.8 | 12 | 0.8333 | 7 | 1 | 8 | 0.75 | 37 | 0.8378 |
| 17_339108 | 6 | 0.3333 | 3 | 0 | 10 | 0.2 | 8 | 0.125 | 27 | 0.1852 | 7 | 0 | 7 | 0.1429 | 6 | 0.3333 | 5 | 0 | 25 | 0.12 |
| 17_339108 | 13 | 0.3846 | 4 | 0 | 11 | 0.3636 | 13 | 0.3846 | 41 | 0.3415 | 9 | 0.4444 | 13 | 0.2308 | 7 | 0.1429 | 8 | 0.125 | 37 | 0.2432 |
| 17_339109 | 13 | 0.1538 | 5 | 0 | 10 | 0.2 | 12 | 0 | 40 | 0.1 | 10 | 0.1 | 13 | 0.0769 | 7 | 0 | 8 | 0.25 | 38 | 0.1053 |
| 17_339109 | 11 | 0 | 3 | 0.3333 | 8 | 0.125 | 12 | 0.0833 | 34 | 0.0882 | 9 | 0.1111 | 9 | 0.1111 | 7 | 0.1429 | 6 | 0.1667 | 31 | 0.129 |
| 17_339110 | 12 | 0.8333 | 5 | 1 | 9 | 0.7778 | 13 | 0.7692 | 39 | 0.8205 | 8 | 0.875 | 12 | 0.5833 | 6 | 1 | 8 | 0.75 | 34 | 0.7647 |
| 17_339150 | 10 | 0.2 | 5 | 0.2 | 10 | 0.1 | 10 | 0.2 | 35 | 0.1714 | 4 | 0 | 10 | 0.3 | 4 | 0 | 5 | 0 | 23 | 0.1304 |
| 17_339288 | 10 | 0.2 | 4 | 0 | 9 | 0 | 12 | 0.0833 | 35 | 0.0857 | 8 | 0.125 | 10 | 0.2 | 6 | 0 | 4 | 0 | 28 | 0.1071 |
| 17_339656 | 24 | 0.1667 | 35 | 0.2571 | 28 | 0.1429 | 25 | 0.16 | 112 | 0.1875 | 17 | 0.1765 | 25 | 0.16 | 21 | 0.1429 | 30 | 0.0667 | 93 | 0.129 |
| 17_339659 | 37 | 0.8919 | 36 | 0.8889 | 37 | 0.8649 | 27 | 0.7778 | 137 | 0.8613 | 33 | 0.7576 | 27 | 0.8148 | 23 | 0.7826 | 32 | 0.8438 | 115 | 0.8 |
| 17_339661 | 32 | 0.0938 | 34 | 0.2647 | 35 | 0.2 | 29 | 0.2759 | 130 | 0.2077 | 32 | 0.2188 | 29 | 0.3103 | 17 | 0.3529 | 31 | 0.2258 | 109 | 0.2661 |
| 17_339665 | 26 | 0.4231 | 34 | 0.4412 | 26 | 0.3846 | 22 | 0.4545 | 108 | 0.4259 | 28 | 0.3929 | 25 | 0.32 | 20 | 0.25 | 25 | 0.52 | 98 | 0.3776 |
| 17_339665 | 32 | 0.375 | 36 | 0.4444 | 30 | 0.4333 | 27 | 0.4074 | 125 | 0.416 | 31 | 0.3871 | 34 | 0.4706 | 26 | 0.3846 | 27 | 0.4815 | 118 | 0.4322 |
| 17_339666 | 29 | 0.2414 | 34 | 0.2941 | 34 | 0.1765 | 25 | 0 | 122 | 0.1885 | 28 | 0.0714 | 30 | 0.1333 | 23 | 0.0435 | 26 | 0.1538 | 107 | 0.1028 |
| 17_339794 | NA | NA | 1 | 1 | 1 | 0 | NA | NA | 2 | 0.5 | NA | NA | 1 | 1 | 1 | 0 | NA | NA | 2 | 0.5 |
| 17_339795 | NA | NA | 1 | 1 | 1 | 0 | NA | NA | 2 | 0.5 | NA | NA | 1 | 1 | 1 | 0 | NA | NA | 2 | 0.5 |
| 17_339806 | NA | NA | 1 | 1 | 1 | 0 | 1 | 0 | 3 | 0.3333 | NA | NA | NA | NA | 1 | 1 | NA | NA | 1 | 1 |
| 17_339806 | NA | NA | 1 | 1 | 1 | 1 | 1 | 0 | 3 | 0.6667 | NA | NA | 1 | 1 | 1 | 0 | 1 | 0 | 3 | 0.3333 |
| 17_339940 | 7 | 0 | 3 | 0.3333 | 1 | 0 | NA | NA | 11 | 0.0909 | 3 | 0 | NA | NA | 1 | 0 | 1 | 1 | 5 | 0.2 |
| 17_339940 | 7 | 0.7143 | 3 | 0.3333 | NA | NA | NA | NA | 10 | 0.6 | 2 | 1 | NA | NA | 2 | 0.5 | 1 | 0 | 5 | 0.6 |
| 17_339940 | 6 | 0.1667 | 3 | 0 | 2 | 0.5 | NA | NA | 11 | 0.1818 | 3 | 0.3333 | NA | NA | 2 | 0 | 1 | 0 | 6 | 0.1667 |
| 17_339942 | 2 | 0.5 | 2 | 1 | 2 | 1 | NA | NA | 6 | 0.8333 | 3 | 1 | NA | NA | 1 | 1 | 1 | 1 | 5 | 1 |
| 17_339942 | 4 | 0 | 3 | 0.3333 | 1 | 0 | NA | NA | 8 | 0.125 | 2 | 0 | NA | NA | 2 | 0.5 | NA | NA | 4 | 0.25 |
| 17_339943 | 5 | 0.8 | 3 | 1 | 2 | 0.5 | NA | NA | 10 | 0.8 | 3 | 1 | NA | NA | 2 | 1 | NA | NA | 5 | 1 |
| 17_339944 | 5 | 0.6 | 3 | 0 | 1 | 1 | NA | NA | 9 | 0.4444 | 1 | 0 | NA | NA | 2 | 0.5 | 1 | 1 | 4 | 0.5 |
| 17_340002 | 2 | 0.5 | 1 | 0 | 1 | 1 | NA | NA | 4 | 0.5 | 1 | 0 | NA | NA | 2 | 0.5 | 1 | 0 | 4 | 0.25 |
| 17_340370 | 2 | 0 | 1 | 1 | 1 | 1 | 2 | 0.5 | 6 | 0.5 | 2 | 0.5 | NA | NA | 1 | 0 | NA | NA | 3 | 0.3333 |
| 17_340371 | 1 | 0 | 1 | 1 | NA | NA | 1 | 1 | 3 | 0.6667 | 2 | 0.5 | NA | NA | NA | NA | 1 | 1 | 3 | 0.6667 |
| 17_340371 | 3 | 1 | 1 | 0 | 1 | 0 | 1 | 1 | 6 | 0.6667 | 2 | 0.5 | NA | NA | 1 | 1 | 1 | 0 | 4 | 0.5 |
| 17_340372 | 3 | 1 | NA | NA | 1 | 1 | 3 | 1 | 7 | 1 | 1 | 0 | NA | NA | 1 | 1 | 1 | 1 | 3 | 0.6667 |
| 17_340380 | 3 | 0 | 3 | 0.6667 | 1 | 1 | 2 | 0.5 | 9 | 0.4444 | 2 | 0.5 | NA | NA | 1 | 0 | 1 | 0 | 4 | 0.25 |
| 17_340383 | NA | NA | 1 | 1 | NA | NA | 2 | 0.5 | 3 | 0.6667 | 1 | 1 | NA | NA | NA | NA | NA | NA | 1 | 1 |
| 17_340384 | NA | NA | 1 | 1 | NA | NA | 2 | 0 | 3 | 0.3333 | 1 | 1 | NA | NA | NA | NA | NA | NA | 1 | 1 |
| 17_340666 | 5 | 0 | 1 | 1 | 5 | 0.2 | 2 | 0 | 13 | 0.1538 | 3 | 0 | 3 | 0.3333 | 6 | 0.3333 | 3 | 0.3333 | 15 | 0.2667 |
| 17_340667 | NA | NA | 1 | 1 | NA | NA | NA | NA | 1 | 1 | NA | NA | 1 | 1 | NA | NA | NA | NA | 1 | 1 |
| 17_340668 | NA | NA | 1 | 1 | NA | NA | NA | NA | 1 | 1 | NA | NA | 1 | 1 | NA | NA | NA | NA | 1 | 1 |
| 17_340669 | 6 | 0 | 1 | 1 | 5 | 0 | 3 | 0 | 15 | 0.0667 | 2 | 0 | 3 | 0.3333 | 6 | 0 | 3 | 0 | 14 | 0.0714 |
| 17_340671 | 5 | 0 | 1 | 0 | 5 | 0.2 | 2 | 0 | 13 | 0.0769 | 3 | 0 | 3 | 0.3333 | 6 | 0.1667 | 2 | 0 | 14 | 0.1429 |
| 17_340671 | NA | NA | 1 | 1 | NA | NA | 1 | 1 | 2 | 1 | NA | NA | 1 | 1 | NA | NA | NA | NA | 1 | 1 |
| 17_340674 | 6 | 0.3333 | 2 | 0.5 | 4 | 0.25 | 3 | 0.6667 | 15 | 0.4 | 4 | 0.25 | 3 | 0.3333 | 5 | 0.4 | 3 | 0.3333 | 15 | 0.3333 |
| 17_340674 | 5 | 0.2 | 2 | 0.5 | 5 | 0.2 | 3 | 0.3333 | 15 | 0.2667 | 4 | 0.25 | 4 | 0.25 | 5 | 0.4 | 4 | 0 | 17 | 0.2353 |
| 17_340675 | 6 | 0.6667 | 2 | 1 | 5 | 1 | 3 | 1 | 16 | 0.875 | 2 | 1 | 4 | 1 | 7 | 0.7143 | 4 | 1 | 17 | 0.8824 |
| 17_340675 | 6 | 1 | 2 | 0.5 | 5 | 0.8 | 4 | 0.75 | 17 | 0.8235 | 3 | 0.3333 | 5 | 0.8 | 7 | 0.7143 | 4 | 1 | 19 | 0.7368 |
| 17_340676 | 3 | 0.6667 | 1 | 1 | 5 | 0.4 | 3 | 1 | 12 | 0.6667 | 2 | 0.5 | 5 | 0.8 | 6 | 0.5 | 3 | 1 | 16 | 0.6875 |
| 17_340676 | 5 | 0.2 | 1 | 0 | 5 | 0.4 | 4 | 0.5 | 15 | 0.3333 | 4 | 0.25 | 5 | 0 | 5 | 0.4 | 3 | 0 | 17 | 0.1765 |
| 17_340893 | 1 | 1 | NA | NA | NA | NA | 1 | 1 | 2 | 1 | NA | NA | 1 | 1 | NA | NA | NA | NA | 1 | 1 |
| 17_340894 | 1 | 1 | NA | NA | NA | NA | 1 | 0 | 2 | 0.5 | NA | NA | 1 | 1 | NA | NA | NA | NA | 1 | 1 |
| 17_340897 | 1 | 0 | NA | NA | NA | NA | 1 | 1 | 2 | 0.5 | NA | NA | 1 | 1 | NA | NA | NA | NA | 1 | 1 |
| 17_340910 | 1 | 1 | NA | NA | NA | NA | 1 | 1 | 2 | 1 | NA | NA | 1 | 1 | NA | NA | NA | NA | 1 | 1 |
| 17_340910 | NA | NA | NA | NA | NA | NA | 1 | 1 | 1 | 1 | NA | NA | 1 | 1 | NA | NA | NA | NA | 1 | 1 |
| 17_340923 | 2 | 1 | NA | NA | NA | NA | 1 | 0 | 3 | 0.6667 | 1 | 0 | NA | NA | NA | NA | 2 | 0.5 | 3 | 0.3333 |
| 17_340924 | 1 | 1 | NA | NA | NA | NA | 1 | 1 | 2 | 1 | 1 | 0 | 1 | 0 | NA | NA | 1 | 1 | 3 | 0.3333 |
| 17_340924 | 1 | 1 | NA | NA | NA | NA | 1 | 1 | 2 | 1 | NA | NA | 1 | 1 | NA | NA | 2 | 1 | 3 | 1 |
| 17_340925 | 2 | 0 | NA | NA | NA | NA | 1 | 1 | 3 | 0.3333 | NA | NA | 1 | 1 | NA | NA | 2 | 1 | 3 | 1 |
| 17_340925 | 2 | 0 | NA | NA | NA | NA | 1 | 1 | 3 | 0.3333 | 1 | 0 | 1 | 0 | NA | NA | 2 | 1 | 4 | 0.5 |
| 17_341247 | NA | NA | NA | NA | 1 | 1 | NA | NA | 1 | 1 | NA | NA | NA | NA | 1 | 1 | NA | NA | 1 | 1 |
| 17_341272 | 2 | 0 | 2 | 1 | 2 | 1 | 3 | 1 | 9 | 0.7778 | NA | NA | 2 | 1 | 2 | 1 | NA | NA | 4 | 1 |

| | | | | | | | | | | | | | | | | | | | | |
|---|---|---|---|---|---|---|---|---|---|---|---|---|---|---|---|---|---|---|---|---|
| 17_3412735 | 2 | 0.5 | 2 | 0 | 2 | 0 | 3 | 0.3333 | 9 | 0.2222 | NA | NA | 1 | 0 | 3 | 0.3333 | NA | NA | 4 | 0.25 |
| 17_3413022 | 4 | 0.5 | 2 | 0 | 1 | 0 | 2 | 0 | 9 | 0.2222 | 1 | 0 | 1 | 0 | 2 | 1 | 3 | 0.3333 | 7 | 0.4286 |
| 17_3413027 | 6 | 0 | 3 | 0.3333 | 1 | 0 | 3 | 0 | 13 | 0.0769 | 1 | 0 | 2 | 0.5 | 1 | 0 | 2 | 0 | 6 | 0.1667 |
| 17_3413043 | 4 | 0.25 | 4 | 0.5 | 1 | 1 | 3 | 0 | 12 | 0.3333 | NA | NA | 2 | 0 | 2 | 0.5 | 3 | 0.6667 | 7 | 0.4286 |
| 17_3413050 | 6 | 0 | 4 | 0.25 | 1 | 0 | 3 | 0.3333 | 14 | 0.1429 | 1 | 0 | 2 | 0 | 3 | 0 | 3 | 0.3333 | 9 | 0.1111 |
| 17_3413051 | 6 | 0.8333 | 4 | 0.5 | 1 | 0 | 2 | 0.5 | 13 | 0.6154 | 1 | 0 | 3 | 0.6667 | 3 | 0 | 2 | 0 | 9 | 0.2222 |
| 17_3413054 | 6 | 1 | 3 | 0.3333 | 1 | 1 | 3 | 0.6667 | 13 | 0.7692 | 1 | 1 | 3 | 1 | 3 | 0.3333 | 2 | 1 | 9 | 0.7778 |
| 17_3413063 | 6 | 0.5 | 4 | 0.5 | 1 | 1 | 3 | 0.3333 | 14 | 0.5 | 1 | 1 | 3 | 0.3333 | 4 | 0.25 | 3 | 0.6667 | 11 | 0.4545 |
| 17_3413073 | 6 | 0.8333 | 4 | 1 | 2 | 1 | 4 | 0.75 | 16 | 0.875 | 1 | 1 | 2 | 1 | 5 | 0.8 | 3 | 1 | 11 | 0.9091 |
| 17_3413084 | 6 | 0.3333 | 4 | 0.25 | 2 | 0 | 4 | 0.5 | 16 | 0.3125 | 1 | 0 | 2 | 0.5 | 4 | 0.25 | 3 | 0 | 10 | 0.2 |
| 17_3413103 | 6 | 0.3333 | 3 | 0 | 2 | 0 | 3 | 0 | 14 | 0.1429 | 1 | 0 | 3 | 0.3333 | 4 | 0 | 2 | 0 | 10 | 0.1 |
| 17_3413161 | 7 | 0.1429 | 4 | 0 | 3 | 0 | 6 | 0 | 20 | 0.05 | 1 | 0 | 3 | 0.3333 | 4 | 0.5 | 2 | 0 | 10 | 0.3 |
| 17_3413235 | 2 | 0 | 8 | 0.375 | 3 | 0.6667 | 6 | 0.5 | 19 | 0.4211 | NA | NA | 4 | 0.5 | 5 | 0.4 | 3 | 1 | 12 | 0.5833 |
| 17_3413251 | 6 | 0.3333 | 8 | 0.25 | 3 | 0.6667 | 6 | 0.1667 | 23 | 0.3043 | 1 | 1 | 4 | 0.25 | 4 | 0.5 | 3 | 0.6667 | 12 | 0.5 |
| 17_3413268 | 5 | 0.2 | 5 | 0.2 | 2 | 0 | 8 | 0.25 | 20 | 0.2 | 1 | 0 | 4 | 0.5 | 3 | 0 | 1 | 0 | 9 | 0.2222 |
| 17_3413436 | 7 | 0 | 7 | 0.2857 | 3 | 0 | 7 | 0.1429 | 24 | 0.125 | NA | NA | 5 | 0 | 5 | 0 | 3 | 0.3333 | 13 | 0.0769 |
| 17_3413613 | 4 | 0.5 | 1 | 0 | 1 | 0 | 1 | 0 | 7 | 0.2857 | 1 | 0 | 2 | 0.5 | NA | NA | NA | NA | 3 | 0.3333 |
| 17_3413615 | 4 | 0.75 | 2 | 0.5 | 1 | 0 | 1 | 1 | 8 | 0.625 | 1 | 1 | 1 | 0 | NA | NA | NA | NA | 2 | 0.5 |
| 17_3413617 | 4 | 1 | 2 | 1 | 1 | 1 | 1 | 1 | 8 | 1 | 1 | 1 | 2 | 1 | NA | NA | NA | NA | 3 | 1 |
| 17_3413622 | 3 | 1 | 2 | 1 | 1 | 1 | 1 | 1 | 7 | 1 | 1 | 1 | 2 | 1 | NA | NA | NA | NA | 3 | 1 |
| 17_3413656 | 3 | 0.3333 | 2 | 0 | 1 | 1 | 1 | 0 | 7 | 0.2857 | 1 | 0 | 1 | 1 | 1 | 0 | 1 | 0 | 4 | 0.25 |
| 17_3413674 | 3 | 0 | 1 | 0 | 2 | 0.5 | 2 | 1 | 8 | 0.375 | 1 | 1 | 4 | 0.75 | 1 | 1 | NA | NA | 6 | 0.8333 |
| 17_3413685 | 5 | 0.2 | 3 | 0.3333 | 2 | 1 | 2 | 0.5 | 12 | 0.4167 | 1 | 0 | 4 | 0.25 | 1 | 0 | 3 | 0 | 9 | 0.1111 |
| 17_3413694 | 6 | 0.8333 | 3 | 1 | 1 | 1 | 2 | 1 | 12 | 0.9167 | 1 | 1 | 3 | 1 | 1 | 1 | 4 | 1 | 9 | 1 |
| 17_3413696 | 5 | 0 | 3 | 0.3333 | 2 | 0.5 | 2 | 0.5 | 12 | 0.25 | 1 | 0 | 3 | 0.3333 | 1 | 0 | 4 | 0 | 9 | 0.1111 |
| 17_3413700 | 5 | 1 | 1 | 1 | 1 | 1 | NA | NA | 7 | 1 | 1 | 1 | 3 | 1 | 1 | 1 | 4 | 1 | 9 | 1 |
| 17_3413705 | 6 | 0.1667 | 3 | 0 | 2 | 0 | 2 | 0 | 13 | 0.0769 | 1 | 0 | 4 | 0.25 | 1 | 0 | 4 | 0 | 10 | 0.1 |
| 17_3413706 | 4 | 0.5 | 3 | 0.6667 | 2 | 1 | 3 | 0 | 12 | 0.5 | 1 | 1 | 4 | 0 | 1 | 1 | 4 | 0.5 | 10 | 0.4 |
| 17_3413718 | 5 | 0.4 | 3 | 0.6667 | 1 | 0 | 2 | 0.5 | 11 | 0.4545 | 1 | 0 | 2 | 0 | 1 | 0 | 4 | 0.5 | 8 | 0.25 |
| 17_3413721 | 6 | 0.1667 | 3 | 0.3333 | 2 | 0.5 | 2 | 1 | 13 | 0.3846 | 1 | 1 | 3 | 1 | 1 | 1 | 3 | 0.3333 | 8 | 0.75 |
| 17_3413727 | 5 | 0.6 | 4 | 0.75 | 2 | 0.5 | 3 | 1 | 14 | 0.7143 | 2 | 0.5 | 3 | 0.6667 | 1 | 1 | 3 | 0.6667 | 9 | 0.6667 |
| 17_3413734 | 4 | 1 | 4 | 0.25 | 2 | 0.5 | 3 | 0.3333 | 13 | 0.5385 | 2 | 1 | 4 | 0.25 | 1 | 0 | 3 | 0.6667 | 10 | 0.5 |
| 17_3413738 | 5 | 0.6 | 4 | 0.25 | 2 | 0 | 3 | 0 | 14 | 0.2857 | 2 | 0.5 | 4 | 0 | 1 | 0 | 2 | 0 | 9 | 0.1111 |
| 17_3413750 | 6 | 0.8333 | 4 | 1 | 2 | 0.5 | 2 | 0.5 | 14 | 0.7857 | 1 | 1 | 2 | 1 | 2 | 1 | 4 | 1 | 9 | 1 |
| 17_3413753 | 6 | 0.6667 | 5 | 1 | 2 | 1 | 3 | 0.6667 | 16 | 0.8125 | 2 | 1 | 5 | 0.6 | 2 | 0.5 | 4 | 1 | 13 | 0.7692 |
| 17_3413759 | 4 | 0.75 | 5 | 1 | 1 | 1 | 4 | 0.5 | 14 | 0.7857 | 2 | 0.5 | 4 | 1 | 2 | 0.5 | 4 | 0.75 | 12 | 0.75 |
| 17_3413760 | 6 | 0.1667 | 6 | 0.1667 | NA | NA | 4 | 0 | 16 | 0.125 | 1 | 0 | 3 | 0.3333 | 2 | 0 | 4 | 0.25 | 10 | 0.2 |
| 17_3413785 | 6 | 0.6667 | 6 | 0.8333 | 1 | 1 | 4 | 0.5 | 17 | 0.7059 | 2 | 0.5 | 4 | 1 | 2 | 0.5 | 4 | 0.75 | 12 | 0.75 |
| 17_3413794 | 6 | 0.1667 | 6 | 0 | NA | NA | 4 | 0 | 16 | 0.0625 | 2 | 0 | 6 | 0.1667 | 3 | 0 | 3 | 0 | 14 | 0.0714 |
| 17_3413802 | 9 | 0.5556 | 6 | 0.5 | 1 | 0 | 4 | 0 | 20 | 0.4 | 2 | 0.5 | 5 | 0.4 | 4 | 0 | 5 | 0.4 | 16 | 0.3125 |
| 17_3414466 | 4 | 0.25 | 2 | 0 | 3 | 0.6667 | 4 | 0.25 | 13 | 0.3077 | 2 | 0.5 | 5 | 0.4 | 4 | 0.5 | 7 | 0.8571 | 18 | 0.6111 |
| 17_3414467 | 5 | 0.2 | 1 | 0 | 2 | 0 | 2 | 0 | 10 | 0.1 | 2 | 0 | 4 | 0 | 3 | 0.3333 | 5 | 0 | 14 | 0.0714 |
| 17_3414504 | 5 | 0 | 3 | 0 | 2 | 0.5 | 4 | 0 | 14 | 0.0714 | 2 | 0 | 7 | 0.1429 | 5 | 0 | 4 | 0 | 18 | 0.0556 |
| 17_3414749 | 5 | 0 | 2 | 0 | 5 | 0.2 | 5 | 0 | 17 | 0.0588 | 5 | 0 | 6 | 0 | 3 | 0 | 6 | 0.1667 | 20 | 0.05 |
| 17_3414759 | 4 | 0 | 1 | 1 | 3 | 0 | 5 | 0.4 | 13 | 0.2308 | 4 | 0 | 5 | 0.6 | 3 | 0 | 7 | 0.2857 | 19 | 0.2632 |
| 17_3414795 | 8 | 0 | 4 | 0.75 | 5 | 0 | 5 | 0.2 | 22 | 0.1818 | 5 | 0 | 5 | 0.2 | 5 | 0 | 7 | 0.1429 | 22 | 0.0909 |
| 17_3414809 | 9 | 0 | 2 | 0 | 8 | 0.25 | 6 | 0 | 25 | 0.08 | 5 | 0.2 | 5 | 0.2 | 5 | 0.2 | 6 | 0 | 21 | 0.1429 |
| 17_3415474 | 2 | 0.5 | 2 | 1 | 4 | 0.75 | NA | NA | 8 | 0.75 | 1 | 0 | 3 | 0.3333 | 2 | 0 | 3 | 0.6667 | 9 | 0.3333 |
| 17_3415633 | 2 | 0.5 | 1 | 0 | 4 | 0 | 2 | 0 | 9 | 0.1111 | 1 | 0 | 6 | 0 | 5 | 0.2 | 3 | 1 | 15 | 0.2667 |
| 17_3415636 | 2 | 1 | 3 | 1 | 5 | 0.8 | 2 | 0.5 | 12 | 0.8333 | 1 | 1 | 6 | 1 | 5 | 0.8 | 5 | 0.8 | 17 | 0.8824 |
| 17_3415642 | 2 | 0.5 | 2 | 0.5 | 5 | 0.6 | 2 | 0.5 | 11 | 0.5455 | 1 | 1 | 5 | 0.8 | 3 | 0.3333 | 3 | 0.6667 | 12 | 0.6667 |
| 17_3415658 | 2 | 0.5 | 2 | 0 | 5 | 0 | 2 | 0 | 11 | 0.0909 | 2 | 0 | 4 | 0.25 | 4 | 0 | 6 | 0 | 16 | 0.0625 |
| 17_3415738 | 1 | 0 | 2 | 0 | 2 | 0.5 | NA | NA | 5 | 0.2 | NA | NA | 1 | 0 | NA | NA | 2 | 0.5 | 3 | 0.3333 |
| 17_3416731 | NA | NA | 5 | 0.4 | 1 | 1 | 2 | 0 | 8 | 0.375 | NA | NA | 1 | 0 | 3 | 0.3333 | NA | NA | 4 | 0.25 |
| 17_3417501 | 2 | 0.5 | 1 | 0 | NA | NA | NA | NA | 3 | 0.3333 | NA | NA | NA | NA | 1 | 1 | 2 | 0.5 | 3 | 0.6667 |
| 17_3417706 | 2 | 0 | 3 | 0 | 2 | 0 | 2 | 0.5 | 9 | 0.1111 | 1 | 1 | 3 | 0 | 2 | 0 | 4 | 0 | 10 | 0.1 |
| 17_3418520 | 8 | 0.5 | 8 | 0.625 | 9 | 0.4444 | 5 | 0.2 | 30 | 0.4667 | 7 | 0.2857 | 10 | 0.2 | 7 | 0.2857 | 6 | 0.3333 | 30 | 0.2667 |
| 17_3420091 | NA | NA | NA | NA | 1 | 1 | NA | NA | 1 | 1 | 1 | 0 | NA | NA | 2 | 0.5 | 3 | 0 | 6 | 0.1667 |
| 17_3420099 | NA | NA | NA | NA | 1 | 1 | NA | NA | 1 | 1 | 1 | 0 | NA | NA | 2 | 1 | 3 | 0 | 6 | 0.3333 |
| 17_3420148 | NA | NA | NA | NA | 1 | 1 | NA | NA | 1 | 1 | 1 | 1 | NA | NA | 2 | 1 | 2 | 1 | 5 | 1 |
| 17_3420160 | NA | NA | NA | NA | 1 | 1 | NA | NA | 1 | 1 | 1 | 1 | NA | NA | 2 | 0.5 | 3 | 0.6667 | 6 | 0.6667 |
| 17_3420193 | NA | NA | NA | NA | 1 | 1 | NA | NA | 1 | 1 | 1 | 1 | NA | NA | NA | NA | 2 | 1 | 3 | 1 |
| 17_3420198 | NA | NA | NA | NA | 1 | 1 | NA | NA | 1 | 1 | 1 | 0 | NA | NA | 2 | 1 | 3 | 1 | 6 | 0.8333 |
| 17_3420201 | NA | NA | NA | NA | 1 | 1 | NA | NA | 1 | 1 | 1 | 1 | NA | NA | 2 | 1 | 1 | 1 | 4 | 1 |
| 17_3420209 | NA | NA | NA | NA | 1 | 1 | NA | NA | 1 | 1 | NA | NA | NA | NA | 2 | 0.5 | 2 | 0.5 | 4 | 0.5 |

| | | | | | | | | | | | | | | | | | | | | |
|---|---|---|---|---|---|---|---|---|---|---|---|---|---|---|---|---|---|---|---|---|
| 17_342152 | 12 | 0.5 | 15 | 0.2 | 6 | 0.1667 | 6 | 0.5 | 39 | 0.3333 | 10 | 0.4 | 2 | 0.5 | 4 | 0.5 | 3 | 0.3333 | 19 | 0.4211 |
| 17_342153 | 8 | 0.25 | 13 | 0.2308 | 5 | 0.2 | 5 | 0.2 | 31 | 0.2258 | 8 | 0.125 | 3 | 0.6667 | 3 | 0.3333 | 2 | 0 | 16 | 0.25 |
| 17_342153 | 11 | 0.6364 | 14 | 0.7143 | 6 | 0.3333 | 6 | 0.8333 | 37 | 0.6486 | 9 | 0.3333 | 3 | 0.6667 | 4 | 0.5 | 3 | 0.3333 | 19 | 0.4211 |
| 17_342392 | NA | NA | 1 | 1 | NA | NA | NA | NA | 1 | 1 | NA | NA | NA | NA | NA | NA | 1 | 1 | 1 | 1 |
| 17_342401 | 4 | 0 | 7 | 0.1429 | 4 | 0 | 4 | 0 | 19 | 0.0526 | 3 | 0.3333 | 4 | 0 | 6 | 0 | 4 | 0 | 17 | 0.0588 |
| 17_342401 | 5 | 0.4 | 9 | 0.1111 | 4 | 0 | 4 | 0 | 22 | 0.1364 | 3 | 0 | 4 | 0 | 5 | 0.4 | 3 | 0 | 15 | 0.1333 |
| 17_342401 | 3 | 0.6667 | 6 | 0.8333 | 4 | 0.75 | 2 | 0.5 | 15 | 0.7333 | 1 | 0 | 4 | 1 | 6 | 0.8333 | 3 | 1 | 14 | 0.8571 |
| 17_342402 | 3 | 0.6667 | 8 | 0.25 | 3 | 0.3333 | 3 | 0.6667 | 17 | 0.4118 | 3 | 0.3333 | 1 | 1 | 4 | 0.25 | 4 | 0.5 | 12 | 0.4167 |
| 17_342402 | 4 | 0.25 | 8 | 0.5 | 4 | 0 | 3 | 0 | 19 | 0.2632 | 3 | 0 | 2 | 0.5 | 6 | 0.1667 | 4 | 0.25 | 15 | 0.2 |
| 17_342403 | 5 | 0.6 | 8 | 0.125 | 4 | 0.25 | 3 | 0.6667 | 20 | 0.35 | 2 | 0.5 | 2 | 0 | 5 | 0 | 4 | 0.5 | 13 | 0.2308 |
| 17_342407 | 3 | 1 | 7 | 0.7143 | 3 | 0.3333 | 2 | 1 | 15 | 0.7333 | 1 | 1 | 3 | 0.6667 | 5 | 0.6 | 3 | 1 | 12 | 0.75 |
| 17_342414 | 2 | 1 | 6 | 0.5 | 3 | 0 | 2 | 0.5 | 13 | 0.4615 | 1 | 0 | 2 | 0 | 2 | 0.5 | 3 | 0.6667 | 8 | 0.375 |
| 17_342422 | 2 | 0.5 | 5 | 0.8 | NA | NA | 1 | 1 | 8 | 0.75 | 2 | 0.5 | 1 | 1 | 1 | 0 | 2 | 1 | 6 | 0.6667 |
| 17_342451 | 6 | 0.1667 | 4 | 0.5 | 8 | 0.5 | 5 | 0.2 | 23 | 0.3478 | 3 | 0.3333 | 6 | 0.6667 | 4 | 0.25 | 5 | 0 | 18 | 0.3333 |
| 17_342464 | 5 | 0 | 3 | 0 | 8 | 0 | 4 | 0.5 | 20 | 0.1 | 2 | 0 | 5 | 0.2 | 2 | 0 | 4 | 0.75 | 13 | 0.3077 |
| 17_342474 | 12 | 0.5833 | 14 | 0.8571 | 12 | 0.6667 | 13 | 0.6923 | 51 | 0.7059 | 11 | 0.9091 | 11 | 0.7273 | 8 | 0.875 | 13 | 0.7692 | 43 | 0.814 |
| 17_342474 | 9 | 0.1111 | 14 | 0.0714 | 10 | 0.2 | 13 | 0.0769 | 46 | 0.1087 | 12 | 0.0833 | 10 | 0.2 | 8 | 0.25 | 9 | 0.3333 | 39 | 0.2051 |
| 17_342475 | 9 | 0 | 15 | 0.0667 | 12 | 0.1667 | 14 | 0.2857 | 50 | 0.14 | 9 | 0 | 10 | 0 | 7 | 0.1429 | 11 | 0.0909 | 37 | 0.0541 |
| 17_342475 | 6 | 0 | 14 | 0.1429 | 10 | 0.2 | 11 | 0.0909 | 41 | 0.122 | 7 | 0.1429 | 7 | 0 | 8 | 0.125 | 6 | 0.1667 | 28 | 0.1071 |
| 17_342476 | 11 | 0.1818 | 11 | 0.1818 | 10 | 0.2 | 13 | 0.0769 | 45 | 0.1556 | 6 | 0.3333 | 8 | 0.375 | 8 | 0.125 | 11 | 0.1818 | 33 | 0.2424 |
| 17_342479 | 7 | 0.5714 | 8 | 0.25 | 8 | 0.375 | 11 | 0.6364 | 34 | 0.4706 | 2 | 0 | 7 | 0.5714 | 6 | 0 | 6 | 0.3333 | 21 | 0.2857 |
| 17_342498 | 5 | 0.6 | 4 | 0.25 | 3 | 0.3333 | 8 | 0.375 | 20 | 0.4 | 6 | 0.1667 | 4 | 0.25 | 3 | 0.3333 | 6 | 0.5 | 19 | 0.3158 |
| 17_342498 | 5 | 0.6 | 3 | 1 | 4 | 1 | 8 | 0.375 | 20 | 0.65 | 8 | 0.5 | 4 | 0.5 | 2 | 0.5 | 4 | 0.5 | 18 | 0.5 |
| 17_342499 | 3 | 1 | 4 | 0.75 | 4 | 1 | 7 | 0.8571 | 18 | 0.8889 | 5 | 1 | 5 | 0.8 | 4 | 1 | 5 | 1 | 19 | 0.9474 |
| 17_342501 | 4 | 0.25 | 3 | 0 | 4 | 0.5 | 8 | 0.375 | 19 | 0.3158 | 7 | 0.4286 | 3 | 0.3333 | 2 | 0 | 5 | 0 | 17 | 0.2353 |
| 17_342501 | 2 | 0 | 3 | 0 | 3 | 0.3333 | 7 | 0.1429 | 15 | 0.1333 | 5 | 0 | 5 | 0.2 | 2 | 0 | 4 | 0 | 16 | 0.0625 |
| 17_342501 | 3 | 0.3333 | 2 | 0 | 3 | 0.3333 | 7 | 0.2857 | 15 | 0.2667 | 5 | 0.4 | 4 | 0.5 | 3 | 0.3333 | 5 | 0.4 | 17 | 0.4118 |
| 17_342503 | 2 | 0 | 2 | 0.5 | 2 | 0.5 | 3 | 0 | 9 | 0.2222 | 3 | 0.3333 | 2 | 0 | NA | NA | 2 | 0.5 | 7 | 0.2857 |
| 17_342504 | 2 | 0 | 2 | 0.5 | 2 | 0 | 4 | 0 | 10 | 0.1 | 4 | 0.25 | 2 | 0 | 2 | 0 | 2 | 0 | 10 | 0.1 |
| 17_342565 | 7 | 0.7143 | 8 | 0.625 | 6 | 0.6667 | 7 | 0.4286 | 28 | 0.6071 | 4 | 1 | 9 | 0.8889 | 9 | 0.6667 | 6 | 0.6667 | 28 | 0.7857 |
| 17_342565 | 6 | 0.1667 | 7 | 0.2857 | 9 | 0.1111 | 9 | 0.1111 | 31 | 0.1613 | 3 | 0 | 10 | 0.5 | 14 | 0.2143 | 6 | 0.5 | 33 | 0.3333 |
| 17_342565 | 9 | 0.2222 | 8 | 0.125 | 9 | 0.2222 | 8 | 0.5 | 34 | 0.2647 | 4 | 0.5 | 9 | 0 | 14 | 0.2857 | 7 | 0.7143 | 34 | 0.3235 |
| 17_342566 | 9 | 0.5556 | 8 | 1 | 8 | 0.75 | 8 | 0.875 | 33 | 0.7879 | 4 | 1 | 11 | 0.9091 | 15 | 0.6667 | 8 | 0.875 | 38 | 0.8158 |
| 17_342568 | 5 | 0 | 9 | 0.3333 | 8 | 0 | 9 | 0.1111 | 31 | 0.129 | 3 | 0.3333 | 8 | 0.25 | 7 | 0.1429 | 6 | 0.5 | 24 | 0.2917 |
| 17_342569 | 9 | 0.2222 | 12 | 0.1667 | 11 | 0.0909 | 8 | 0.25 | 40 | 0.175 | 5 | 0.4 | 8 | 0.125 | 11 | 0.1818 | 8 | 0.125 | 32 | 0.1875 |
| 17_342569 | 9 | 0.2222 | 10 | 0 | 11 | 0.0909 | 7 | 0.1429 | 37 | 0.1081 | 4 | 0 | 10 | 0 | 13 | 0.0769 | 8 | 0.125 | 35 | 0.0571 |
| 17_342570 | 9 | 0.2222 | 12 | 0.1667 | 12 | 0.0833 | 9 | 0.1111 | 42 | 0.1429 | 4 | 0.25 | 11 | 0.2727 | 14 | 0 | 9 | 0.4444 | 38 | 0.2105 |
| 17_342572 | 7 | 0.1429 | 9 | 0.3333 | 9 | 0.2222 | 7 | 0.4286 | 32 | 0.2812 | 4 | 0 | 8 | 0.125 | 9 | 0.1111 | 7 | 0.1429 | 28 | 0.1071 |
| 17_343314 | 13 | 0.4615 | 14 | 0.4286 | 13 | 0.4615 | 16 | 0.4375 | 56 | 0.4464 | 12 | 0.5 | 13 | 0.8462 | 21 | 0.5238 | 11 | 0.6364 | 57 | 0.614 |
| 17_343314 | 19 | 0.2105 | 19 | 0.0526 | 17 | 0.1176 | 17 | 0 | 72 | 0.0972 | 11 | 0.0909 | 14 | 0.1429 | 24 | 0.0833 | 15 | 0.1333 | 64 | 0.1094 |
| 17_343315 | 15 | 0.0667 | 13 | 0.0769 | 16 | 0.25 | 17 | 0 | 61 | 0.0984 | 11 | 0.0909 | 11 | 0.2727 | 18 | 0 | 9 | 0 | 49 | 0.0816 |
| 17_343325 | 11 | 0 | 14 | 0 | 12 | 0.1667 | 15 | 0.1333 | 52 | 0.0769 | 10 | 0 | 13 | 0.0769 | 20 | 0.05 | 11 | 0.0909 | 54 | 0.0556 |
| 17_343338 | 11 | 0.0909 | 14 | 0.1429 | 10 | 0.1 | 12 | 0 | 47 | 0.0851 | 6 | 0 | 8 | 0.375 | 14 | 0.0714 | 9 | 0.2222 | 37 | 0.1622 |
| 17_343339 | 11 | 0.0909 | 14 | 0.0714 | 10 | 0.1 | 13 | 0.0769 | 48 | 0.0833 | 8 | 0 | 10 | 0.1 | 14 | 0.1429 | 11 | 0 | 43 | 0.0698 |
| 17_343340 | 9 | 0.1111 | 13 | 0.2308 | 9 | 0.2222 | 11 | 0.0909 | 42 | 0.1667 | 8 | 0.125 | 9 | 0.2222 | 12 | 0.25 | 11 | 0.2727 | 40 | 0.225 |
| 17_343376 | 8 | 0.25 | 8 | 0.625 | 9 | 0.3333 | 10 | 0.3 | 35 | 0.3714 | 5 | 0.2 | 7 | 0.1429 | 9 | 0.4444 | 7 | 0.5714 | 28 | 0.3571 |
| 17_343378 | 7 | 0.2857 | 6 | 0.3333 | 6 | 0.3333 | 7 | 0.4286 | 26 | 0.3462 | 5 | 0.6 | 5 | 0.6 | 5 | 0.2 | 5 | 0.4 | 20 | 0.45 |
| 17_343378 | 7 | 0.7143 | 6 | 0.6667 | 6 | 0.8333 | 9 | 0.6667 | 28 | 0.7143 | 5 | 0.4 | 5 | 0.8 | 8 | 0.375 | 5 | 0.2 | 23 | 0.4348 |
| 17_343381 | 7 | 0.1429 | 5 | 0 | 7 | 0.1429 | 6 | 0 | 25 | 0.08 | 6 | 0 | 4 | 0.25 | 9 | 0.1111 | 5 | 0 | 24 | 0.0833 |
| 17_343381 | 7 | 0.2857 | 4 | 0.75 | 6 | 0.8333 | 5 | 0.6 | 22 | 0.5909 | 5 | 0.4 | 5 | 0.2 | 8 | 0.5 | 5 | 0.4 | 23 | 0.3913 |
| 17_343381 | 7 | 0.4286 | 4 | 0.5 | 5 | 0.8 | 6 | 0.5 | 22 | 0.5455 | 5 | 0.4 | 4 | 0.5 | 9 | 0.6667 | 5 | 0.6 | 23 | 0.5652 |
| 17_343384 | 3 | 1 | 2 | 0.5 | 3 | 0.6667 | 4 | 0.5 | 12 | 0.6667 | 1 | 0 | 2 | 0.5 | 3 | 1 | 2 | 0.5 | 8 | 0.625 |
| 17_343385 | 2 | 1 | 2 | 0.5 | 3 | 1 | 4 | 1 | 11 | 0.9091 | 3 | 0.3333 | 3 | 1 | 4 | 1 | 2 | 0 | 12 | 0.6667 |
| 17_343385 | 2 | 0.5 | 2 | 0.5 | 3 | 0.3333 | 4 | 0.25 | 11 | 0.3636 | 2 | 0 | 3 | 0 | 4 | 0.25 | 2 | 0 | 11 | 0.0909 |
| 17_343386 | 1 | 0 | 2 | 0.5 | 2 | 0.5 | 4 | 0 | 9 | 0.2222 | 2 | 0.5 | 2 | 0.5 | 2 | 0 | NA | NA | 6 | 0.3333 |
| 17_343542 | 1 | 0 | 1 | 1 | NA | NA | 1 | 0 | 3 | 0.3333 | 1 | 1 | 1 | 0 | 1 | 0 | NA | NA | 3 | 0.3333 |
| 17_343543 | 1 | 0 | NA | NA | NA | NA | 1 | 1 | 2 | 0.5 | 1 | 1 | 1 | 1 | 1 | 1 | NA | NA | 3 | 1 |
| 17_343544 | 1 | 0 | 1 | 1 | NA | NA | 1 | 1 | 3 | 0.6667 | NA | NA | 1 | 1 | 2 | 0.5 | NA | NA | 3 | 0.6667 |
| 17_343586 | NA | NA | NA | NA | NA | NA | 1 | 1 | 1 | 1 | NA | NA | NA | NA | 1 | 1 | NA | NA | 1 | 1 |
| 17_344351 | 1 | 1 | NA | NA | 1 | 1 | NA | NA | 2 | 1 | NA | NA | 1 | 1 | NA | NA | NA | NA | 1 | 1 |
| 17_344864 | NA | NA | 3 | 0.3333 | NA | NA | 1 | 0 | 4 | 0.25 | NA | NA | NA | NA | 1 | 1 | 1 | 0 | 2 | 0.5 |
| 17_348066 | 3 | 1 | 2 | 1 | NA | NA | NA | NA | 5 | 1 | 2 | 0.5 | 1 | 0 | 1 | 1 | NA | NA | 4 | 0.5 |
| 17_348070 | 2 | 1 | 2 | 1 | NA | NA | 1 | 1 | 5 | 1 | 2 | 1 | 1 | 1 | 1 | 1 | 1 | 1 | 5 | 1 |
| 17_348071 | 2 | 0.5 | 1 | 1 | NA | NA | 2 | 1 | 5 | 0.8 | 2 | 1 | 1 | 1 | NA | NA | 1 | 1 | 4 | 1 |
| 17_348073 | 3 | 0.6667 | 2 | 1 | NA | NA | 2 | 0.5 | 7 | 0.7143 | 2 | 0 | 1 | 1 | 1 | 1 | NA | NA | 4 | 0.5 |

| | | | | | | | | | | | | | | | | | | | | |
|---|---|---|---|---|---|---|---|---|---|---|---|---|---|---|---|---|---|---|---|---|
| 17_3481494 | 6 | 0.6667 | 8 | 1 | 6 | 0.8333 | 5 | 0.6 | 25 | 0.8 | 9 | 0.6667 | 7 | 0.7143 | 9 | 1 | 12 | 0.8333 | 37 | 0.8108 |
| 17_3481496 | 7 | 0 | 8 | 0.25 | 6 | 0 | 6 | 0.1667 | 27 | 0.1111 | 9 | 0.1111 | 10 | 0 | 9 | 0.2222 | 11 | 0 | 39 | 0.0769 |
| 17_3481517 | 5 | 0.6 | 9 | 0.4444 | 5 | 0.4 | 5 | 0.6 | 24 | 0.5 | 9 | 0.6667 | 9 | 0.5556 | 9 | 0.6667 | 12 | 0.8333 | 39 | 0.6923 |
| 17_3481526 | 6 | 0.3333 | 9 | 0.5556 | 4 | 1 | 6 | 0.1667 | 25 | 0.48 | 8 | 0.125 | 6 | 0.1667 | 11 | 0.1818 | 10 | 0.8 | 35 | 0.3429 |
| 17_3481527 | 4 | 0.25 | 4 | 0.25 | 4 | 0.25 | 5 | 0.2 | 17 | 0.2353 | 5 | 0.2 | 6 | 0.3333 | 9 | 0.2222 | 6 | 0.5 | 26 | 0.3077 |
| 17_3481546 | 7 | 0.2857 | 8 | 0.5 | 5 | 0 | 5 | 0.2 | 25 | 0.28 | 8 | 0.5 | 8 | 0.375 | 10 | 0.4 | 10 | 0.4 | 36 | 0.4167 |
| 17_3481623 | 2 | 0.5 | 3 | 0 | 2 | 0.5 | 3 | 0 | 10 | 0.2 | 3 | 0 | 3 | 0 | 7 | 0.1429 | 2 | 0 | 15 | 0.0667 |
| 17_3482021 | 15 | 0.0667 | 17 | 0.1176 | 11 | 0.0909 | 20 | 0.05 | 63 | 0.0794 | 13 | 0.1538 | 14 | 0 | 8 | 0 | 19 | 0.1053 | 54 | 0.0741 |
| 17_3482031 | 14 | 0 | 13 | 0.2308 | 10 | 0.2 | 11 | 0.3636 | 48 | 0.1875 | 12 | 0.3333 | 12 | 0.0833 | 9 | 0.2222 | 17 | 0.1176 | 50 | 0.18 |
| 17_3482046 | 8 | 0.125 | 11 | 0.0909 | 6 | 0.5 | 15 | 0.3333 | 40 | 0.25 | 13 | 0.0769 | 8 | 0.125 | 3 | 0 | 14 | 0.2143 | 38 | 0.1316 |
| 17_3483156 | 32 | 0.9062 | 11 | 1 | 15 | 0.8 | 13 | 1 | 71 | 0.9155 | 5 | 1 | 23 | 0.9565 | 12 | 1 | 16 | 1 | 56 | 0.9821 |
| 17_3483162 | 41 | 0.7317 | 18 | 0.5 | 21 | 0.619 | 21 | 0.381 | 101 | 0.5941 | 7 | 0.7143 | 24 | 0.8333 | 18 | 0.7222 | 22 | 0.7273 | 71 | 0.7606 |
| 17_3483174 | 32 | 0.7812 | 13 | 0.6923 | 16 | 0.625 | 17 | 0.6471 | 78 | 0.7051 | 9 | 0.7778 | 24 | 0.6667 | 16 | 0.5625 | 17 | 0.8824 | 66 | 0.7121 |
| 17_3483174 | 35 | 0.9429 | 18 | 0.9444 | 22 | 0.9545 | 19 | 1 | 94 | 0.9574 | 9 | 1 | 28 | 0.9286 | 15 | 0.8667 | 18 | 0.9444 | 70 | 0.9286 |
| 17_3483182 | 32 | 0.3438 | 18 | 0.3889 | 24 | 0.5 | 24 | 0.2917 | 98 | 0.3776 | 10 | 0.3 | 28 | 0.2143 | 16 | 0.375 | 21 | 0.2857 | 75 | 0.28 |
| 17_3483188 | 41 | 0.4878 | 18 | 0.7222 | 24 | 0.7083 | 26 | 0.6154 | 109 | 0.6055 | 9 | 0.6667 | 30 | 0.5 | 18 | 0.6111 | 22 | 0.4091 | 79 | 0.519 |
| 17_3483595 | 5 | 0.6 | 5 | 0.8 | 3 | 0.3333 | 6 | 0.5 | 19 | 0.5789 | 6 | 0.1667 | 6 | 0.5 | 11 | 0.5455 | 7 | 0.2857 | 30 | 0.4 |
| 17_3483643 | 4 | 0 | 5 | 0.2 | 1 | 0 | 5 | 0 | 15 | 0.0667 | 6 | 0.1667 | 2 | 0 | 8 | 0 | 4 | 0 | 20 | 0.05 |
| 17_3484485 | 2 | 0 | 1 | 0 | NA | NA | 5 | 0.2 | 8 | 0.125 | 3 | 0 | 3 | 0 | 7 | 0 | 3 | 0.3333 | 16 | 0.0625 |
| 17_3484704 | NA | NA | NA | NA | NA | NA | 1 | 1 | 1 | 1 | 1 | 1 | 1 | 1 | NA | NA | NA | NA | 2 | 1 |
| 17_3484708 | NA | NA | NA | NA | NA | NA | 1 | 1 | 1 | 1 | 1 | 1 | 2 | 1 | NA | NA | NA | NA | 3 | 1 |
| 17_3484710 | 2 | 0 | 2 | 0.5 | NA | NA | 1 | 0 | 5 | 0.2 | 1 | 0 | 2 | 0.5 | 1 | 0 | NA | NA | 4 | 0.25 |
| 17_3484713 | NA | NA | NA | NA | NA | NA | 1 | 1 | 1 | 1 | 1 | 1 | 1 | 1 | NA | NA | NA | NA | 2 | 1 |
| 17_3484734 | NA | NA | NA | NA | NA | NA | 1 | 1 | 1 | 1 | 1 | 1 | 2 | 1 | NA | NA | 1 | 1 | 4 | 1 |
| 17_3484736 | NA | NA | NA | NA | NA | NA | 1 | 1 | 1 | 1 | 1 | 0 | 2 | 1 | NA | NA | NA | NA | 3 | 0.6667 |
| 17_3485136 | 21 | 0.1429 | 27 | 0.1111 | 9 | 0.1111 | 17 | 0.0588 | 74 | 0.1081 | 10 | 0.1 | 24 | 0.0833 | 24 | 0.0417 | 23 | 0.087 | 81 | 0.0741 |
| 17_3485170 | 25 | 0.72 | 32 | 0.8438 | 12 | 0.9167 | 17 | 0.9412 | 86 | 0.8372 | 12 | 0.75 | 28 | 0.7857 | 27 | 0.8148 | 22 | 0.8182 | 89 | 0.7978 |
| 17_3485173 | 22 | 0.8182 | 29 | 0.7586 | 9 | 1 | 14 | 0.7857 | 74 | 0.8108 | 13 | 0.8462 | 22 | 0.7727 | 23 | 0.7826 | 17 | 0.8824 | 75 | 0.8133 |
| 17_3485184 | 22 | 0.1364 | 24 | 0.0833 | 11 | 0.0909 | 19 | 0.0526 | 76 | 0.0921 | 15 | 0.0667 | 24 | 0.125 | 25 | 0.2 | 17 | 0 | 81 | 0.1111 |
| 17_3485206 | 24 | 0.1667 | 25 | 0.08 | 9 | 0.1111 | 18 | 0.1111 | 76 | 0.1184 | 19 | 0.1053 | 24 | 0.0833 | 27 | 0 | 20 | 0.05 | 90 | 0.0556 |
| 17_3486441 | 1 | 1 | 1 | 1 | 1 | 0 | NA | NA | 3 | 0.6667 | NA | NA | 2 | 1 | 2 | 1 | NA | NA | 4 | 1 |
| 17_3486445 | 1 | 0 | 1 | 0 | 1 | 1 | NA | NA | 3 | 0.3333 | NA | NA | 2 | 0 | 2 | 0.5 | NA | NA | 4 | 0.25 |
| 17_3486465 | 1 | 1 | 1 | 1 | NA | NA | 1 | 1 | 3 | 1 | NA | NA | 1 | 0 | 2 | 1 | NA | NA | 3 | 0.6667 |
| 17_3486465 | 1 | 1 | 1 | 0 | 1 | 1 | NA | NA | 3 | 0.6667 | NA | NA | 2 | 0.5 | 2 | 0 | 1 | 0 | 5 | 0.2 |
| 17_3486467 | 1 | 1 | NA | NA | 1 | 1 | 1 | 1 | 3 | 1 | NA | NA | 2 | 1 | 2 | 1 | 1 | 1 | 5 | 1 |
| 17_3486481 | 1 | 0 | 2 | 1 | NA | NA | 1 | 0 | 4 | 0.5 | NA | NA | 2 | 0.5 | 2 | 1 | 1 | 1 | 5 | 0.8 |
| 17_3486488 | NA | NA | 2 | 1 | 2 | 1 | 2 | 1 | 6 | 1 | NA | NA | 2 | 1 | 2 | 0.5 | 1 | 1 | 5 | 0.8 |
| 17_3486498 | 1 | 0 | 2 | 0.5 | 2 | 0.5 | 1 | 0 | 6 | 0.3333 | NA | NA | 1 | 1 | 2 | 1 | 1 | 1 | 4 | 1 |
| 17_3486512 | 1 | 1 | 3 | 0.3333 | 2 | 0.5 | 1 | 0 | 7 | 0.4286 | NA | NA | 2 | 1 | 2 | 1 | 1 | 1 | 5 | 1 |
| 17_3486525 | 1 | 0 | 3 | 0.3333 | 2 | 0 | 2 | 0 | 8 | 0.125 | NA | NA | NA | NA | 3 | 0.6667 | 1 | 0 | 4 | 0.5 |
| 17_3487351 | 5 | 0 | 9 | 0 | 8 | 0.25 | 3 | 0 | 25 | 0.08 | 2 | 0 | 8 | 0.125 | 4 | 0 | 3 | 0 | 17 | 0.0588 |
| 17_3494745 | 2 | 0.5 | 8 | 0.125 | 3 | 0 | 4 | 0 | 17 | 0.1176 | 8 | 0.125 | 2 | 0 | 4 | 0 | 3 | 0.3333 | 17 | 0.1176 |
| 17_3505637 | 3 | 0 | 3 | 0 | 5 | 0.2 | NA | NA | 11 | 0.0909 | 4 | 0.25 | 4 | 0 | 3 | 0 | NA | NA | 11 | 0.0909 |
| 17_3505639 | 2 | 0 | 3 | 0.3333 | 5 | 0 | NA | NA | 10 | 0.1 | 5 | 0.2 | 5 | 0.4 | 1 | 0 | NA | NA | 11 | 0.2727 |
| 17_3505643 | 4 | 0 | NA | NA | 5 | 0.4 | NA | NA | 9 | 0.2222 | 3 | 0 | 4 | 0.25 | 1 | 0 | NA | NA | 8 | 0.125 |
| 17_3505646 | 4 | 0 | 3 | 0 | 5 | 0.2 | NA | NA | 12 | 0.0833 | 5 | 0 | 4 | 0.25 | 3 | 0 | NA | NA | 12 | 0.0833 |
| 17_3505648 | 4 | 0.5 | 3 | 0.3333 | 5 | 0.2 | NA | NA | 12 | 0.3333 | 5 | 0 | 3 | 0.3333 | 3 | 0 | NA | NA | 11 | 0.0909 |
| 17_3505651 | 3 | 0.3333 | 3 | 0.3333 | 4 | 0.5 | NA | NA | 10 | 0.4 | 4 | 0.25 | 2 | 0.5 | 3 | 0.3333 | NA | NA | 9 | 0.3333 |
| 17_3505661 | 5 | 0.4 | 3 | 1 | 5 | 0 | NA | NA | 13 | 0.3846 | 5 | 0 | 4 | 0.25 | 3 | 0 | NA | NA | 12 | 0.0833 |
| 17_3505681 | 1 | 1 | 1 | 1 | NA | NA | NA | NA | 2 | 1 | NA | NA | 1 | 1 | NA | NA | NA | NA | 1 | 1 |
| 17_3505699 | 3 | 1 | 3 | 1 | 6 | 0.6667 | 1 | 1 | 13 | 0.8462 | 5 | 0.8 | 5 | 0.8 | 4 | 0.25 | NA | NA | 14 | 0.6429 |
| 17_3505731 | 4 | 0.25 | 3 | 0 | 5 | 0 | 1 | 0 | 13 | 0.0769 | 4 | 0.25 | 5 | 0.2 | 4 | 0 | NA | NA | 13 | 0.1538 |
| 17_3505909 | 4 | 0 | 9 | 0.3333 | 3 | 0 | 5 | 0.2 | 21 | 0.1905 | 3 | 0 | 4 | 0 | 6 | 0 | 3 | 0.3333 | 16 | 0.0625 |
| 17_3511452 | NA | NA | NA | NA | NA | NA | 1 | 1 | 1 | 1 | NA | NA | NA | NA | 1 | 1 | NA | NA | 1 | 1 |
| 17_3513279 | 16 | 0.75 | 25 | 0.68 | 27 | 0.6296 | 21 | 0.8095 | 89 | 0.7079 | 15 | 0.7333 | 24 | 0.8333 | 19 | 0.4211 | 18 | 0.6667 | 76 | 0.6711 |
| 17_3513283 | 22 | 0.2273 | 19 | 0.1579 | 29 | 0.1724 | 23 | 0.1304 | 93 | 0.172 | 15 | 0.2 | 27 | 0.1481 | 18 | 0.2778 | 17 | 0.1765 | 77 | 0.1948 |
| 17_3513291 | 18 | 0.2222 | 25 | 0.16 | 29 | 0.1724 | 23 | 0.2174 | 95 | 0.1895 | 14 | 0.2143 | 25 | 0.12 | 19 | 0.1579 | 16 | 0.1875 | 74 | 0.1622 |
| 17_3515382 | 6 | 0.3333 | 10 | 0.4 | 3 | 0.6667 | 3 | 0.6667 | 22 | 0.4545 | 10 | 0.5 | 3 | 0.3333 | 3 | 0.3333 | 5 | 0.8 | 21 | 0.5238 |
| 17_3515384 | 7 | 0 | 10 | 0.3 | 3 | 0.3333 | 3 | 0 | 23 | 0.1739 | 9 | 0.2222 | 4 | 0.25 | 3 | 0 | 6 | 0 | 22 | 0.1364 |
| 17_3515386 | 5 | 0.6 | 10 | 0.4 | 2 | 1 | 2 | 1 | 19 | 0.5789 | 8 | 0.375 | 2 | 0.5 | 3 | 0.6667 | 4 | 0.75 | 17 | 0.5294 |
| 17_3515390 | 6 | 1 | 9 | 0.8889 | 3 | 1 | 3 | 1 | 21 | 0.9524 | 9 | 1 | 1 | 1 | 2 | 1 | 6 | 0.8333 | 18 | 0.9444 |
| 17_3515395 | 6 | 0.1667 | 9 | 0.6667 | 3 | 0.3333 | 3 | 0.6667 | 21 | 0.4762 | 10 | 0.5 | 5 | 0.6 | 4 | 1 | 7 | 0.7143 | 26 | 0.6538 |
| 17_3515405 | 7 | 0.2857 | 11 | 0.1818 | 4 | 0.5 | 4 | 0.25 | 26 | 0.2692 | 11 | 0.5455 | 5 | 0.4 | 3 | 0.6667 | 9 | 0.3333 | 28 | 0.4643 |
| 17_3515407 | 3 | 1 | 6 | 0.8333 | 2 | 1 | 1 | 1 | 12 | 0.9167 | 6 | 0.8333 | 2 | 1 | 2 | 0.5 | 5 | 1 | 15 | 0.8667 |
| 17_3515411 | 6 | 0.1667 | 8 | 0.625 | 4 | 0.5 | 3 | 0.6667 | 21 | 0.4762 | 9 | 0.3333 | 4 | 0.25 | 1 | 1 | 5 | 0.4 | 19 | 0.3684 |

| | | | | | | | | | | | | | | | | | | | | |
|---|---|---|---|---|---|---|---|---|---|---|---|---|---|---|---|---|---|---|---|---|
| 17_3515412 | 5 | 1 | 7 | 1 | 2 | 0 | 3 | 1 | 17 | 0.8824 | 6 | 1 | 4 | 1 | 2 | 1 | 6 | 1 | 18 | 1 |
| 17_3515412 | 6 | 0.6667 | 8 | 0.75 | 5 | 0.4 | 4 | 0.75 | 23 | 0.6522 | 11 | 0.5455 | 5 | 0.4 | 3 | 1 | 4 | 1 | 23 | 0.6522 |
| 17_3515453 | 3 | 0.6667 | 3 | 0.6667 | 2 | 0 | 3 | 0.6667 | 11 | 0.5455 | 3 | 1 | 3 | 0 | 1 | 1 | 2 | 0.5 | 9 | 0.5556 |
| 17_3515465 | 2 | 1 | 4 | 1 | 2 | 0 | 1 | 1 | 9 | 0.7778 | 3 | 1 | 3 | 1 | 2 | 1 | 4 | 1 | 12 | 1 |
| 17_3515470 | 4 | 1 | 4 | 1 | 2 | 0.5 | 2 | 1 | 12 | 0.9167 | 3 | 0.6667 | 3 | 1 | 4 | 1 | 4 | 1 | 14 | 0.9286 |
| 17_3517590 | 4 | 0.5 | 1 | 1 | 4 | 0.25 | NA | NA | 9 | 0.4444 | 4 | 0.5 | 2 | 0.5 | 1 | 0 | 2 | 0.5 | 9 | 0.4444 |
| 17_3517601 | 4 | 1 | 1 | 1 | 2 | 1 | 1 | 1 | 8 | 1 | 4 | 0.75 | 2 | 1 | NA | NA | 2 | 1 | 8 | 0.875 |
| 17_3517607 | 3 | 0.6667 | 1 | 1 | 3 | 0.3333 | 1 | 0 | 8 | 0.5 | 7 | 0.4286 | 2 | 0 | 1 | 0 | NA | NA | 10 | 0.3 |
| 17_3517623 | 4 | 1 | NA | NA | 2 | 1 | 1 | 1 | 7 | 1 | 6 | 1 | 2 | 1 | NA | NA | 3 | 1 | 11 | 1 |
| 17_3517626 | NA | NA | NA | NA | 2 | 1 | NA | NA | 2 | 1 | 3 | 0 | 1 | 0 | NA | NA | 2 | 0.5 | 6 | 0.1667 |
| 17_3517668 | 2 | 0.5 | 1 | 0 | 2 | 0 | 1 | 0 | 6 | 0.1667 | 7 | 0.5714 | 1 | 0 | NA | NA | 3 | 0.6667 | 11 | 0.5455 |
| 17_3517683 | 3 | 0.3333 | 1 | 1 | 2 | 0 | NA | NA | 6 | 0.3333 | 7 | 0.5714 | 2 | 0.5 | NA | NA | 1 | 0 | 10 | 0.5 |
| 17_3517692 | 3 | 0 | 1 | 1 | 3 | 0.3333 | 1 | 0 | 8 | 0.25 | 7 | 0 | 2 | 0 | NA | NA | 3 | 0.3333 | 12 | 0.0833 |
| 17_3517700 | 3 | 0.6667 | 1 | 1 | 2 | 0 | 1 | 0 | 7 | 0.4286 | 5 | 0 | 2 | 0.5 | NA | NA | 3 | 0 | 10 | 0.1 |
| 17_3517722 | 3 | 0.6667 | 1 | 1 | 1 | 0 | 1 | 0 | 6 | 0.5 | 5 | 0.2 | 2 | 0 | NA | NA | 3 | 0.3333 | 10 | 0.2 |
| 17_3518846 | NA | NA | 2 | 0.5 | 1 | 1 | 1 | 1 | 4 | 0.75 | 2 | 1 | NA | NA | 1 | 1 | 4 | 0.75 | 7 | 0.8571 |
| 17_3518851 | NA | NA | 2 | 0.5 | NA | NA | 1 | 0 | 3 | 0.3333 | 2 | 0.5 | NA | NA | 1 | 0 | 4 | 0.25 | 7 | 0.2857 |
| 17_3518859 | NA | NA | NA | NA | 1 | 1 | 1 | 1 | 2 | 1 | 2 | 1 | NA | NA | 1 | 1 | 4 | 1 | 7 | 1 |
| 17_3518862 | NA | NA | 1 | 0 | NA | NA | 1 | 1 | 2 | 0.5 | 1 | 1 | NA | NA | NA | NA | 4 | 0.5 | 5 | 0.6 |
| 17_3518864 | NA | NA | 2 | 0 | 1 | 1 | 1 | 0 | 4 | 0.25 | 2 | 0.5 | NA | NA | 1 | 0 | 4 | 0.25 | 7 | 0.2857 |
| 17_3518884 | 2 | 0 | 1 | 1 | 1 | 1 | NA | NA | 4 | 0.5 | 3 | 0.3333 | 1 | 0 | 4 | 0.5 | 3 | 0 | 11 | 0.2727 |
| 17_3518886 | 2 | 0 | 2 | 0 | 1 | 0 | 2 | 0.5 | 7 | 0.1429 | 3 | 0 | 2 | 1 | 4 | 0 | 3 | 0 | 12 | 0.1667 |
| 17_3518892 | 1 | 1 | 1 | 1 | 1 | 1 | 2 | 1 | 5 | 1 | 3 | 1 | 3 | 1 | 3 | 1 | 3 | 1 | 12 | 1 |
| 17_3518894 | 1 | 1 | 2 | 0.5 | 1 | 0 | 2 | 0 | 6 | 0.3333 | 3 | 0 | 3 | 0.6667 | 4 | 0.25 | 4 | 0 | 14 | 0.2143 |
| 17_3518900 | NA | NA | 2 | 0.5 | 1 | 1 | 2 | 0 | 5 | 0.4 | 3 | 0.6667 | 3 | 0.3333 | 4 | 0.75 | 3 | 0.3333 | 13 | 0.5385 |
| 17_3518915 | 1 | 1 | 2 | 1 | 1 | 1 | 2 | 1 | 6 | 1 | 3 | 1 | 2 | 1 | 4 | 0.75 | 3 | 1 | 12 | 0.9167 |
| 17_3518952 | 1 | 1 | 2 | 1 | NA | NA | 1 | 0 | 4 | 0.75 | 2 | 1 | 2 | 0.5 | 4 | 0.5 | 3 | 0.3333 | 11 | 0.5455 |
| 17_3518984 | 1 | 0 | 2 | 0 | NA | NA | 1 | 1 | 4 | 0.25 | 3 | 0.3333 | 2 | 0 | 3 | 0.3333 | 3 | 0.3333 | 11 | 0.2727 |
| 17_3518992 | 1 | 0 | 2 | 0.5 | NA | NA | 2 | 0.5 | 5 | 0.4 | 3 | 0.3333 | 2 | 0 | 4 | 0.25 | 2 | 0.5 | 11 | 0.2727 |
| 17_3519593 | 1 | 1 | NA | NA | NA | NA | 1 | 1 | 2 | 1 | 1 | 1 | 1 | 1 | NA | NA | NA | NA | 2 | 1 |
| 17_3519622 | NA | NA | NA | NA | 1 | 1 | 1 | 1 | 2 | 1 | 1 | 1 | 1 | 1 | NA | NA | NA | NA | 2 | 1 |
| 17_3519628 | 1 | 1 | NA | NA | 1 | 1 | 1 | 1 | 3 | 1 | 1 | 1 | 1 | 1 | NA | NA | NA | NA | 2 | 1 |
| 17_3519638 | 1 | 0 | NA | NA | 1 | 0 | 1 | 1 | 3 | 0.3333 | NA | NA | 1 | 1 | NA | NA | NA | NA | 1 | 1 |
| 17_3519692 | 1 | 1 | NA | NA | 1 | 1 | NA | NA | 2 | 1 | 1 | 1 | 1 | 1 | NA | NA | NA | NA | 2 | 1 |
| 17_3519709 | 1 | 1 | NA | NA | NA | NA | 1 | 1 | 2 | 1 | 1 | 1 | 1 | 1 | NA | NA | NA | NA | 2 | 1 |
| 17_3519766 | NA | NA | NA | NA | NA | NA | 1 | 1 | 1 | 1 | 1 | 1 | NA | NA | NA | NA | NA | NA | 1 | 1 |
| 17_3524479 | 1 | 1 | 1 | 0 | NA | NA | NA | NA | 2 | 0.5 | NA | NA | NA | NA | NA | NA | 1 | 1 | 1 | 1 |
| 17_3524540 | 1 | 1 | 1 | 0 | NA | NA | NA | NA | 2 | 0.5 | NA | NA | NA | NA | NA | NA | 1 | 1 | 1 | 1 |
| 17_3527754 | 4 | 0.75 | 1 | 1 | 1 | 0 | 1 | 1 | 7 | 0.7143 | 8 | 0.75 | 2 | 1 | 2 | 1 | 3 | 0.3333 | 15 | 0.7333 |
| 17_3529715 | 1 | 1 | 2 | 0 | 1 | 0 | NA | NA | 4 | 0.25 | 2 | 0.5 | 2 | 0 | NA | NA | 5 | 0.2 | 9 | 0.2222 |
| 17_3529720 | 1 | 1 | 3 | 0 | 1 | 0 | NA | NA | 5 | 0.2 | 2 | 0 | 2 | 0.5 | NA | NA | 4 | 0.25 | 8 | 0.25 |
| 17_3532179 | 14 | 0.1429 | 15 | 0.1333 | 15 | 0.0667 | 12 | 0.1667 | 56 | 0.125 | 9 | 0.1111 | 9 | 0 | 13 | 0.1538 | 7 | 0 | 38 | 0.0789 |
| 17_3534236 | 5 | 0.6 | 1 | 0 | 2 | 0 | 1 | 0 | 9 | 0.3333 | 3 | 0 | 3 | 0.3333 | 2 | 0 | 2 | 0.5 | 10 | 0.2 |
| 17_3534243 | 7 | 1 | 1 | 1 | 2 | 1 | 2 | 1 | 12 | 1 | 5 | 0.8 | 3 | 0.6667 | 3 | 1 | 2 | 1 | 13 | 0.8462 |
| 17_3534322 | 9 | 0 | 1 | 1 | 4 | 0 | 3 | 0.3333 | 17 | 0.1176 | 5 | 0 | 3 | 0.3333 | 3 | 0 | 2 | 0 | 13 | 0.0769 |
| 17_3534361 | 9 | 0.8889 | 3 | 0.6667 | 4 | 0.75 | 5 | 1 | 21 | 0.8571 | 5 | 1 | 3 | 0.3333 | 3 | 1 | 2 | 0.5 | 13 | 0.7692 |
| 17_3534365 | 8 | 0.75 | 2 | 1 | 1 | 1 | 4 | 0.5 | 15 | 0.7333 | 4 | 0.5 | 3 | 1 | 3 | 0.6667 | 2 | 0.5 | 12 | 0.6667 |
| 17_3534367 | 7 | 0.1429 | 2 | 0.5 | 4 | 0 | 5 | 0.2 | 18 | 0.1667 | 4 | 0.25 | 2 | 0 | 2 | 0.5 | 2 | 0 | 10 | 0.2 |
| 17_3534369 | 9 | 0.5556 | 2 | 0.5 | 3 | 0 | 4 | 0.5 | 18 | 0.4444 | 5 | 0.2 | 3 | 0.3333 | 2 | 1 | 2 | 0.5 | 12 | 0.4167 |
| 17_3534370 | 8 | 0.75 | 3 | 0.3333 | 3 | 0.6667 | 4 | 0 | 18 | 0.5 | 5 | 0.4 | 3 | 0.3333 | 2 | 0.5 | 2 | 0.5 | 12 | 0.4167 |
| 17_3534596 | 1 | 0 | NA | NA | 1 | 1 | 1 | 0 | 3 | 0.3333 | 1 | 1 | NA | NA | NA | NA | NA | NA | 1 | 1 |
| 17_3534622 | 2 | 1 | 1 | 1 | 1 | 1 | 1 | 1 | 5 | 1 | 1 | 1 | NA | NA | NA | NA | NA | NA | 1 | 1 |
| 17_3534623 | 2 | 1 | 1 | 1 | 1 | 1 | 1 | 1 | 5 | 1 | 1 | 1 | NA | NA | NA | NA | NA | NA | 1 | 1 |
| 17_3534637 | 2 | 0 | NA | NA | 1 | 1 | 1 | 1 | 4 | 0.5 | 1 | 1 | NA | NA | NA | NA | NA | NA | 1 | 1 |
| 17_3534672 | 4 | 0.75 | 1 | 1 | 2 | 1 | 2 | 1 | 9 | 0.8889 | 1 | 1 | NA | NA | NA | NA | NA | NA | 1 | 1 |
| 17_3534697 | 3 | 0 | 1 | 0 | 1 | 1 | 3 | 0.6667 | 8 | 0.375 | 1 | 1 | NA | NA | NA | NA | NA | NA | 1 | 1 |
| 17_3534714 | 3 | 0.6667 | 2 | 0.5 | 1 | 1 | 2 | 1 | 8 | 0.75 | 1 | 0 | NA | NA | NA | NA | 1 | 1 | 2 | 0.5 |
| 17_3534757 | 4 | 0 | 3 | 0.3333 | 2 | 0 | 4 | 0 | 13 | 0.0769 | 1 | 0 | 2 | 0.5 | 1 | 0 | 2 | 0 | 6 | 0.1667 |
| 17_3534770 | 4 | 0.25 | 3 | 0 | 1 | 0 | 4 | 0.25 | 12 | 0.1667 | 1 | 0 | 3 | 0 | 1 | 1 | 2 | 0 | 7 | 0.1429 |
| 17_3537444 | 15 | 0.1333 | 13 | 0.1538 | 6 | 0.1667 | 9 | 0.1111 | 43 | 0.1395 | 10 | 0.4 | 15 | 0.1333 | 12 | 0.3333 | 6 | 0 | 43 | 0.2326 |
| 17_3537457 | 17 | 0.2353 | 15 | 0.1333 | 7 | 0.1429 | 13 | 0.2308 | 52 | 0.1923 | 15 | 0.2 | 16 | 0.0625 | 11 | 0.0909 | 7 | 0.4286 | 49 | 0.1633 |
| 17_3537460 | 14 | 1 | 17 | 0.9412 | 6 | 1 | 15 | 1 | 52 | 0.9808 | 14 | 0.8571 | 15 | 0.9333 | 13 | 1 | 8 | 0.75 | 50 | 0.9 |
| 17_3537462 | 16 | 0.0625 | 15 | 0.0667 | 7 | 0.1429 | 13 | 0.0769 | 51 | 0.0784 | 15 | 0.2 | 19 | 0.0526 | 16 | 0.125 | 9 | 0 | 59 | 0.1017 |
| 17_3537464 | 16 | 0.0625 | 15 | 0.0667 | 6 | 0.1667 | 15 | 0 | 52 | 0.0577 | 15 | 0.0667 | 17 | 0.1176 | 16 | 0 | 9 | 0.1111 | 57 | 0.0702 |
| 17_3537469 | 17 | 0.1176 | 15 | 0.1333 | 6 | 0 | 15 | 0.2 | 53 | 0.1321 | 14 | 0.2857 | 16 | 0.125 | 16 | 0.1875 | 9 | 0.1111 | 55 | 0.1818 |

| | | | | | | | | | | | | | | | | | | | | |
|---|---|---|---|---|---|---|---|---|---|---|---|---|---|---|---|---|---|---|---|---|
| 17_3537472 | 17 | 0.7059 | 12 | 0.9167 | 4 | 1 | 10 | 0.7 | 43 | 0.7907 | 12 | 0.6667 | 16 | 0.6875 | 11 | 0.6364 | 6 | 0.8333 | 45 | 0.6889 |
| 17_3537479 | 15 | 0.0667 | 16 | 0 | 6 | 0 | 12 | 0.1667 | 49 | 0.0612 | 12 | 0.0833 | 17 | 0.1765 | 16 | 0.0625 | 9 | 0.1111 | 54 | 0.1111 |
| 17_3537480 | 15 | 0.5333 | 13 | 0.4615 | 7 | 0.5714 | 14 | 0.5 | 49 | 0.5102 | 11 | 0.5455 | 17 | 0.4118 | 16 | 0.1875 | 9 | 0.2222 | 53 | 0.3396 |
| 17_3537490 | 15 | 0.1333 | 14 | 0.1429 | 7 | 0.1429 | 12 | 0.0833 | 48 | 0.125 | 10 | 0.4 | 18 | 0.0556 | 14 | 0.1429 | 6 | 0 | 48 | 0.1458 |
| 17_3537503 | 16 | 0.375 | 10 | 0.5 | 6 | 0.6667 | 11 | 0 | 43 | 0.3488 | 11 | 0.1818 | 16 | 0.375 | 13 | 0.6154 | 6 | 0.1667 | 46 | 0.3696 |
| 17_3537532 | 15 | 0.4667 | 12 | 0.6667 | 4 | 0.5 | 11 | 1 | 42 | 0.6667 | 12 | 0.4167 | 13 | 0.5385 | 14 | 0.6429 | 6 | 0.8333 | 45 | 0.5778 |
| 17_3537553 | 13 | 1 | 10 | 0.7 | 3 | 1 | 8 | 0.75 | 34 | 0.8529 | 11 | 0.9091 | 16 | 1 | 12 | 0.75 | 4 | 0.75 | 43 | 0.8837 |
| 17_3537562 | 16 | 0.9375 | 10 | 0.9 | 5 | 1 | 10 | 1 | 41 | 0.9512 | 12 | 0.75 | 15 | 1 | 14 | 1 | 6 | 1 | 47 | 0.9362 |
| 17_3537569 | 13 | 0.1538 | 8 | 0 | 4 | 0 | 7 | 0 | 32 | 0.0625 | 8 | 0 | 12 | 0.3333 | 10 | 0.2 | 5 | 0.4 | 35 | 0.2286 |
| 17_3537588 | 13 | 0.3077 | 10 | 0.1 | 5 | 0.4 | 5 | 0.4 | 33 | 0.2727 | 11 | 0.2727 | 13 | 0.3077 | 9 | 0.3333 | 4 | 0 | 37 | 0.2703 |
| 17_3537688 | 7 | 0.4286 | 9 | 0 | 3 | 0 | 4 | 0 | 23 | 0.1304 | 7 | 0.1429 | 9 | 0.2222 | 6 | 0 | 1 | 0 | 23 | 0.1304 |
| 17_3537695 | 11 | 0.0909 | 8 | 0.125 | 4 | 0.5 | 5 | 0.4 | 28 | 0.2143 | 9 | 0.1111 | 10 | 0.3 | 7 | 0.2857 | 3 | 0.3333 | 29 | 0.2414 |
| 17_3537821 | 10 | 0.1 | 8 | 0 | 4 | 0.25 | 4 | 0 | 26 | 0.0769 | 5 | 0 | 11 | 0.3636 | 8 | 0.125 | 4 | 0 | 28 | 0.1786 |
| 17_3537876 | 8 | 0.125 | 7 | 0.2857 | 4 | 0 | 4 | 0.25 | 23 | 0.1739 | 3 | 0 | 8 | 0.25 | 6 | 0.1667 | 4 | 0.5 | 21 | 0.2381 |
| 17_3537878 | 6 | 0.1667 | 5 | 0 | 4 | 0.25 | 3 | 0 | 18 | 0.1111 | 7 | 0 | 6 | 0 | 7 | 0.1429 | 4 | 0.25 | 24 | 0.0833 |
| 17_3537886 | 7 | 0.2857 | 6 | 0 | 4 | 0 | 5 | 0 | 22 | 0.0909 | 4 | 0.25 | 7 | 0.1429 | 6 | 0.1667 | 3 | 0 | 20 | 0.15 |
| 17_3543916 | 3 | 0 | 2 | 0 | 2 | 1 | 1 | 0 | 8 | 0.25 | NA | NA | 1 | 0 | 2 | 0.5 | 2 | 0 | 5 | 0.2 |
| 17_3543918 | NA | NA | 2 | 0.5 | 2 | 0.5 | 1 | 0 | 5 | 0.4 | NA | NA | 1 | 1 | 2 | 1 | 2 | 1 | 5 | 1 |
| 17_3543928 | 2 | 0.5 | 1 | 1 | 2 | 1 | 1 | 0 | 6 | 0.6667 | NA | NA | 1 | 0 | 2 | 0.5 | 2 | 0 | 5 | 0.2 |
| 17_3543934 | 2 | 0.5 | 2 | 0 | 2 | 0 | 1 | 0 | 7 | 0.1429 | NA | NA | 1 | 0 | 2 | 0.5 | 1 | 0 | 4 | 0.25 |
| 17_3543970 | 2 | 1 | 1 | 1 | 1 | 1 | 1 | 1 | 5 | 1 | NA | NA | 1 | 1 | 1 | 1 | 2 | 1 | 4 | 1 |
| 17_3543974 | 2 | 1 | NA | NA | 1 | 1 | 1 | 0 | 4 | 0.75 | NA | NA | 1 | 1 | 2 | 1 | 2 | 1 | 5 | 1 |
| 17_3545679 | 66 | 0.0909 | 69 | 0.1739 | 58 | 0.1379 | 52 | 0.1538 | 245 | 0.1388 | 45 | 0.1111 | 58 | 0.2069 | 56 | 0.1786 | 50 | 0.16 | 209 | 0.1675 |
| 17_3545680 | 77 | 0.1169 | 69 | 0.1014 | 61 | 0.0328 | 58 | 0.1034 | 265 | 0.0906 | 46 | 0.1087 | 60 | 0.1167 | 60 | 0.1 | 56 | 0.125 | 222 | 0.1126 |
| 17_3545696 | 81 | 0.0247 | 71 | 0.0845 | 61 | 0.0492 | 59 | 0.0508 | 272 | 0.0515 | 53 | 0.0189 | 62 | 0.129 | 63 | 0.0952 | 58 | 0.0517 | 236 | 0.0763 |
| 17_3545698 | 77 | 0.4675 | 68 | 0.4706 | 63 | 0.4444 | 54 | 0.4074 | 262 | 0.4504 | 49 | 0.5306 | 64 | 0.5156 | 59 | 0.5085 | 48 | 0.4792 | 220 | 0.5091 |
| 17_3545708 | 72 | 0.125 | 70 | 0.2 | 56 | 0.1964 | 54 | 0.1111 | 252 | 0.1587 | 48 | 0.1667 | 54 | 0.1667 | 56 | 0.2143 | 50 | 0.2 | 208 | 0.1875 |
| 17_3545711 | 82 | 0.122 | 74 | 0.1216 | 65 | 0.1538 | 55 | 0.2727 | 276 | 0.1594 | 52 | 0.0769 | 60 | 0.2 | 65 | 0.2308 | 58 | 0.1379 | 235 | 0.166 |
| 17_3545713 | 57 | 0.0702 | 55 | 0.0364 | 48 | 0.0833 | 51 | 0.0784 | 211 | 0.0664 | 40 | 0.075 | 55 | 0.0727 | 58 | 0.1034 | 42 | 0.119 | 195 | 0.0923 |
| 17_3545719 | 75 | 0.3733 | 73 | 0.3014 | 63 | 0.3651 | 59 | 0.3898 | 270 | 0.3556 | 48 | 0.3542 | 60 | 0.3667 | 61 | 0.4426 | 53 | 0.3585 | 222 | 0.3829 |
| 17_3545724 | 69 | 0.5072 | 69 | 0.3478 | 58 | 0.431 | 52 | 0.4231 | 248 | 0.4274 | 50 | 0.44 | 51 | 0.5686 | 51 | 0.4706 | 56 | 0.4286 | 208 | 0.476 |
| 17_3545738 | 76 | 0.1711 | 75 | 0.1733 | 62 | 0.2097 | 59 | 0.1864 | 272 | 0.1838 | 53 | 0.1698 | 55 | 0.3273 | 57 | 0.193 | 59 | 0.1864 | 224 | 0.2188 |
| 17_3545742 | 87 | 0.1034 | 87 | 0.092 | 69 | 0.1014 | 61 | 0.0492 | 304 | 0.0888 | 56 | 0.0179 | 65 | 0.0923 | 66 | 0.0909 | 60 | 0.05 | 247 | 0.0648 |
| 17_3545748 | 74 | 0.0541 | 61 | 0.0656 | 55 | 0.0545 | 54 | 0.037 | 244 | 0.0533 | 44 | 0.0909 | 56 | 0.1071 | 52 | 0.1538 | 53 | 0.1132 | 205 | 0.1171 |
| 17_3546820 | 1 | 0 | 5 | 0.2 | 1 | 0 | 4 | 0.25 | 11 | 0.1818 | 2 | 0 | 5 | 0 | 3 | 0.3333 | 3 | 0 | 13 | 0.0769 |
| 17_3546873 | 1 | 0 | 6 | 0.1667 | 1 | 0 | 4 | 0 | 12 | 0.0833 | 3 | 0 | 6 | 0.1667 | 6 | 0 | 4 | 0.25 | 19 | 0.1053 |
| 17_3546888 | 1 | 0 | 4 | 0.5 | 1 | 1 | 4 | 0.5 | 10 | 0.5 | 5 | 0.4 | 6 | 0.5 | 5 | 0.6 | 5 | 0.4 | 21 | 0.4762 |
| 17_3546894 | 1 | 0 | 6 | 0.5 | 2 | 0.5 | 3 | 1 | 12 | 0.5833 | 4 | 0.25 | 7 | 0.4286 | 4 | 0.75 | 5 | 0 | 20 | 0.35 |
| 17_3546904 | 1 | 0 | 7 | 0.2857 | 2 | 1 | 4 | 0.5 | 14 | 0.4286 | 4 | 0.25 | 6 | 0.8333 | 6 | 0.3333 | 4 | 0.5 | 20 | 0.5 |
| 17_3546984 | 1 | 0 | 10 | 0.1 | 3 | 0 | 5 | 0.2 | 19 | 0.1053 | 6 | 0 | 11 | 0 | 7 | 0 | 7 | 0.4286 | 31 | 0.0968 |
| 17_3548286 | 7 | 0.1429 | 16 | 0.125 | 3 | 0.6667 | 7 | 0.1429 | 33 | 0.1818 | 8 | 0.125 | 4 | 0.25 | 2 | 0 | 5 | 0 | 19 | 0.1053 |
| 17_3548586 | 7 | 0 | 15 | 0.0667 | 3 | 0.6667 | 10 | 0.1 | 35 | 0.1143 | 10 | 0.3 | 2 | 0.5 | 4 | 0 | 8 | 0 | 24 | 0.1667 |
| 17_3548597 | 9 | 0.6667 | 17 | 0.1765 | 3 | 0.6667 | 12 | 0.0833 | 41 | 0.2927 | 12 | 0.5 | 6 | 0.5 | 5 | 0.4 | 11 | 0.4545 | 34 | 0.4706 |
| 17_3548600 | 9 | 0.4444 | 17 | 0.1176 | 3 | 0.3333 | 10 | 0.2 | 39 | 0.2308 | 10 | 0.1 | 6 | 0.1667 | 5 | 0.2 | 10 | 0.2 | 31 | 0.1613 |
| 17_3548607 | 9 | 0.1111 | 18 | 0.0556 | 3 | 0 | 10 | 0.3 | 40 | 0.125 | 11 | 0.3636 | 6 | 0.3333 | 3 | 0.6667 | 8 | 0.5 | 28 | 0.4286 |
| 17_3548618 | 9 | 0.3333 | 21 | 0.0476 | 3 | 0 | 11 | 0.0909 | 44 | 0.1136 | 12 | 0.0833 | 5 | 0.6 | 5 | 0.2 | 11 | 0.3636 | 33 | 0.2727 |
| 17_3564850 | 1 | 1 | 1 | 1 | NA | NA | NA | NA | 2 | 1 | NA | NA | NA | NA | 1 | 1 | NA | NA | 1 | 1 |
| 17_3574242 | NA | NA | 1 | 1 | NA | NA | NA | NA | 1 | 1 | NA | NA | NA | NA | 1 | 1 | NA | NA | 1 | 1 |
| 17_3574243 | NA | NA | 1 | 1 | NA | NA | NA | NA | 1 | 1 | NA | NA | NA | NA | 1 | 1 | NA | NA | 1 | 1 |
| 17_3574269 | NA | NA | 1 | 1 | NA | NA | NA | NA | 1 | 1 | NA | NA | NA | NA | 1 | 1 | NA | NA | 1 | 1 |
| 17_3582705 | NA | NA | 2 | 1 | NA | NA | 1 | 1 | 3 | 1 | NA | NA | NA | NA | NA | NA | 1 | 1 | 1 | 1 |
| 17_3582710 | NA | NA | 1 | 1 | NA | NA | NA | NA | 1 | 1 | NA | NA | NA | NA | NA | NA | 1 | 1 | 1 | 1 |
| 17_3582723 | NA | NA | 2 | 0.5 | NA | NA | NA | NA | 2 | 0.5 | NA | NA | NA | NA | NA | NA | 1 | 1 | 1 | 1 |
| 17_3596676 | 1 | 1 | 1 | 1 | NA | NA | NA | NA | 2 | 1 | 1 | 0 | 1 | 1 | NA | NA | NA | NA | 2 | 0.5 |
| 17_3596706 | 1 | 1 | 1 | 1 | NA | NA | 2 | 1 | 4 | 1 | NA | NA | 2 | 1 | NA | NA | NA | NA | 2 | 1 |
| 17_3596718 | 2 | 1 | 1 | 1 | NA | NA | NA | NA | 3 | 1 | 1 | 1 | 2 | 1 | NA | NA | 1 | 1 | 4 | 1 |
| 17_3596737 | 2 | 1 | 2 | 0.5 | 1 | 0 | 1 | 1 | 6 | 0.6667 | 2 | 1 | 3 | 0 | NA | NA | 2 | 0.5 | 7 | 0.4286 |
| 17_3596738 | 2 | 1 | 2 | 1 | 2 | 1 | 2 | 0.5 | 8 | 0.875 | 2 | 0.5 | 4 | 0.75 | NA | NA | 2 | 0 | 8 | 0.5 |
| 17_3597886 | 2 | 0 | 5 | 0.2 | 1 | 1 | 5 | 0.2 | 13 | 0.2308 | NA | NA | 3 | 0.3333 | NA | NA | NA | NA | 3 | 0.3333 |
| 17_3599254 | 9 | 0 | 4 | 0.25 | 11 | 0.0909 | 8 | 0.125 | 32 | 0.0938 | 4 | 0 | 10 | 0.4 | 5 | 0.2 | 3 | 0 | 22 | 0.2273 |
| 17_3599272 | 7 | 0.2857 | 2 | 1 | 7 | 0.5714 | 9 | 0.2222 | 25 | 0.4 | 6 | 0.5 | 8 | 0.375 | 2 | 0 | 3 | 0.3333 | 19 | 0.3684 |
| 17_3599279 | 6 | 0.6667 | 2 | 1 | 8 | 0.625 | 10 | 0.8 | 26 | 0.7308 | 5 | 0.6 | 11 | 1 | 4 | 1 | 1 | 1 | 21 | 0.9048 |
| 17_3599286 | 8 | 0.25 | 2 | 0 | 7 | 0 | 10 | 0.1 | 27 | 0.1111 | 7 | 0.1429 | 10 | 0 | 4 | 0 | 3 | 0.3333 | 24 | 0.0833 |
| 17_3599295 | 8 | 1 | 1 | 1 | 8 | 1 | 10 | 0.9 | 27 | 0.963 | 6 | 0.8333 | 10 | 0.9 | 4 | 0.75 | 3 | 1 | 23 | 0.8696 |
| 17_3599297 | 8 | 0.875 | 2 | 0.5 | 5 | 1 | 9 | 1 | 24 | 0.9167 | 5 | 0.4 | 10 | 0.6 | 4 | 0.5 | 3 | 0.6667 | 22 | 0.5455 |

| | | | | | | | | | | | | | | | | | | | | |
|---|---|---|---|---|---|---|---|---|---|---|---|---|---|---|---|---|---|---|---|---|
| 17_3599302 | 8 | 0.375 | 2 | 0.5 | 9 | 0.4444 | 10 | 0.3 | 29 | 0.3793 | 5 | 0.6 | 8 | 0.375 | 5 | 0.4 | 3 | 0.3333 | 21 | 0.4286 |
| 17_3599321 | 8 | 1 | 2 | 0.5 | 9 | 1 | 8 | 0.875 | 27 | 0.9259 | 5 | 1 | 8 | 0.875 | 5 | 0.6 | 3 | 1 | 21 | 0.8571 |
| 17_3599326 | 8 | 0.125 | 2 | 0 | 8 | 0 | 7 | 0.1429 | 25 | 0.08 | 5 | 0.2 | 7 | 0.1429 | 4 | 0 | 3 | 0 | 19 | 0.1053 |
| 17_3613315 | 1 | 1 | NA | NA | NA | NA | 2 | 0 | 3 | 0.3333 | NA | NA | 2 | 0.5 | 1 | 1 | 3 | 0.6667 | 6 | 0.6667 |
| 17_3613327 | 2 | 0 | 2 | 0 | 1 | 0 | 2 | 0.5 | 7 | 0.1429 | NA | NA | 3 | 0.3333 | 1 | 0 | 3 | 0 | 7 | 0.1429 |
| 17_3613330 | 2 | 0 | 2 | 1 | NA | NA | 1 | 0 | 5 | 0.4 | NA | NA | 2 | 0.5 | 1 | 0 | 3 | 0.3333 | 6 | 0.3333 |
| 17_3613335 | 2 | 0 | 2 | 0 | 1 | 0 | 2 | 1 | 7 | 0.2857 | NA | NA | 3 | 0.6667 | 1 | 1 | 3 | 0.6667 | 7 | 0.7143 |
| 17_3613336 | 2 | 1 | 1 | 1 | 1 | 1 | 2 | 1 | 6 | 1 | NA | NA | 3 | 1 | 1 | 1 | 3 | 1 | 7 | 1 |
| 17_3614344 | 7 | 0.1429 | 10 | 0.1 | 3 | 0.3333 | 11 | 0.1818 | 31 | 0.1613 | 13 | 0.1538 | 10 | 0.2 | 7 | 0.1429 | 6 | 0.1667 | 36 | 0.1667 |
| 17_3614536 | 8 | 0.25 | 9 | 0.2222 | 9 | 0 | 8 | 0.25 | 34 | 0.1765 | 6 | 0.1667 | 11 | 0.0909 | 7 | 0.4286 | 10 | 0.1 | 34 | 0.1765 |
| 17_3614554 | 7 | 0 | 13 | 0 | 10 | 0.1 | 5 | 0.2 | 35 | 0.0571 | 5 | 0 | 12 | 0.0833 | 7 | 0 | 7 | 0.2857 | 31 | 0.0968 |
| 17_3614631 | 51 | 0.1569 | 86 | 0.1395 | 85 | 0.1059 | 77 | 0.1169 | 299 | 0.1271 | 62 | 0.0484 | 68 | 0.1029 | 49 | 0.0408 | 54 | 0.1481 | 233 | 0.0858 |
| 17_3614634 | 33 | 0.6364 | 58 | 0.3966 | 66 | 0.2879 | 52 | 0.3846 | 209 | 0.3971 | 37 | 0.5405 | 40 | 0.575 | 33 | 0.4848 | 38 | 0.5263 | 148 | 0.5338 |
| 17_3614686 | 40 | 0.125 | 70 | 0.1571 | 70 | 0.1571 | 65 | 0.1692 | 245 | 0.1551 | 51 | 0.0784 | 57 | 0.2456 | 42 | 0.2143 | 53 | 0.1887 | 203 | 0.1823 |
| 17_3620341 | 2 | 0 | 5 | 0.2 | 3 | 0 | 2 | 0 | 12 | 0.0833 | 2 | 0.5 | 4 | 0 | 3 | 0 | 1 | 0 | 10 | 0.1 |
| 17_3620345 | 2 | 0.5 | 4 | 0.5 | 3 | 0 | 2 | 1 | 11 | 0.4545 | 1 | 1 | 3 | 0.6667 | 3 | 0.3333 | 1 | 0 | 8 | 0.5 |
| 17_3620353 | NA | NA | 2 | 0.5 | 1 | 0 | 2 | 0.5 | 5 | 0.4 | 2 | 0 | 2 | 0 | 1 | 1 | NA | NA | 5 | 0.2 |
| 17_3620371 | 18 | 0.1111 | 7 | 0.2857 | 12 | 0.25 | 7 | 0.2857 | 44 | 0.2045 | 12 | 0.3333 | 15 | 0.2667 | 10 | 0.2 | 8 | 0.125 | 45 | 0.2444 |
| 17_3620766 | 1 | 0 | 2 | 0.5 | 1 | 0 | 2 | 0.5 | 6 | 0.3333 | 1 | 0 | 2 | 0.5 | NA | NA | 3 | 0.3333 | 6 | 0.3333 |
| 17_3620784 | NA | NA | 2 | 0.5 | 1 | 0 | 2 | 0 | 5 | 0.2 | 1 | 0 | 2 | 0.5 | NA | NA | 3 | 0 | 6 | 0.1667 |
| 17_3622135 | 7 | 0.4286 | 5 | 0.4 | 2 | 0 | 3 | 0.3333 | 17 | 0.3529 | 3 | 0.3333 | 6 | 0.1667 | 3 | 0.3333 | NA | NA | 12 | 0.25 |
| 17_3622137 | 4 | 0.5 | 6 | 0.3333 | 3 | 0.6667 | 3 | 0.3333 | 16 | 0.4375 | 2 | 0.5 | 6 | 0.3333 | 6 | 0.1667 | 1 | 0 | 15 | 0.2667 |
| 17_3622140 | 7 | 0.1429 | 6 | 0.3333 | 3 | 0 | 3 | 0 | 19 | 0.1579 | 2 | 0 | 6 | 0.1667 | 6 | 0 | 1 | 0 | 15 | 0.0667 |
| 17_3622145 | 6 | 0.3333 | 6 | 0.3333 | 2 | 0 | 3 | 0 | 17 | 0.2353 | 3 | 0.3333 | 6 | 0.1667 | 6 | 0.1667 | 1 | 0 | 16 | 0.1875 |
| 17_3622155 | 6 | 0.5 | 5 | 0.2 | 3 | 0.6667 | 3 | 0.3333 | 17 | 0.4118 | 3 | 0.3333 | 5 | 0.2 | 6 | 0.3333 | 1 | 0 | 15 | 0.2667 |
| 17_3622827 | 1 | 0 | 2 | 0.5 | 1 | 1 | NA | NA | 4 | 0.5 | 2 | 1 | 2 | 0.5 | 2 | 0.5 | 2 | 0.5 | 8 | 0.625 |
| 17_3622829 | 1 | 0 | 2 | 0 | 1 | 1 | NA | NA | 4 | 0.25 | 1 | 0 | 2 | 0 | 2 | 0 | 2 | 0.5 | 7 | 0.1429 |
| 17_3622839 | 1 | 1 | 2 | 0 | 1 | 0 | NA | NA | 4 | 0.25 | 2 | 0 | 2 | 0 | 2 | 0 | 2 | 0.5 | 8 | 0.125 |
| 17_3622847 | 1 | 1 | 2 | 1 | 1 | 1 | NA | NA | 4 | 1 | 1 | 0 | 2 | 0.5 | 2 | 0 | 2 | 1 | 7 | 0.4286 |
| 17_3622850 | 1 | 0 | 1 | 1 | 1 | 0 | NA | NA | 3 | 0.3333 | 2 | 0 | 1 | 0 | 2 | 0 | 3 | 0.3333 | 8 | 0.125 |
| 17_3624151 | 5 | 0.8 | 4 | 0.75 | 2 | 0 | 5 | 0.4 | 16 | 0.5625 | 4 | 0.5 | 4 | 0 | 5 | 0 | 9 | 0.2222 | 22 | 0.1818 |
| 17_3624156 | 4 | 0.75 | 4 | 0.25 | 4 | 0.5 | 4 | 0.75 | 16 | 0.5625 | 4 | 0.75 | 3 | 0 | 4 | 0.75 | 8 | 0.625 | 19 | 0.5789 |
| 17_3624163 | 4 | 0 | 4 | 0 | 4 | 0.25 | 5 | 0 | 17 | 0.0588 | 3 | 0.3333 | 4 | 0 | 5 | 0 | 7 | 0 | 19 | 0.0526 |
| 17_3624165 | 4 | 0.25 | 5 | 0.4 | 5 | 0 | 4 | 0 | 18 | 0.1667 | 3 | 0.3333 | 4 | 0 | 6 | 0.1667 | 9 | 0.3333 | 22 | 0.2273 |
| 17_3624217 | 4 | 1 | 7 | 0.7143 | 4 | 1 | 4 | 0.75 | 19 | 0.8421 | 4 | 0.75 | 3 | 0.6667 | 7 | 1 | 11 | 0.9091 | 25 | 0.88 |
| 17_3624235 | 5 | 0.6 | 7 | 0.8571 | 5 | 1 | 5 | 0.6 | 22 | 0.7727 | 5 | 0.8 | 6 | 0.8333 | 5 | 0.8 | 9 | 0.6667 | 25 | 0.76 |
| 17_3624235 | 4 | 0 | 6 | 0.6667 | 3 | 0.6667 | 4 | 1 | 17 | 0.5882 | 4 | 0.25 | 2 | 0.5 | 7 | 0.4286 | 9 | 0.5556 | 22 | 0.4545 |
| 17_3624237 | 5 | 1 | 7 | 0.8571 | 3 | 0.6667 | 3 | 0.6667 | 18 | 0.8333 | 4 | 0.5 | 5 | 0.8 | 7 | 0.5714 | 8 | 0.75 | 24 | 0.6667 |
| 17_3624250 | 5 | 0.4 | 6 | 0.3333 | 4 | 0.75 | 5 | 0.4 | 20 | 0.45 | 5 | 0.2 | 5 | 0.4 | 5 | 0.2 | 7 | 0 | 22 | 0.1818 |
| 17_3624271 | 4 | 0.5 | 8 | 0.25 | 5 | 0.4 | 4 | 0.5 | 21 | 0.381 | 5 | 0.2 | 5 | 0 | 7 | 0 | 10 | 0.2 | 27 | 0.1111 |
| 17_3624285 | 5 | 0.4 | 7 | 0.2857 | 6 | 0.1667 | 7 | 0.1429 | 25 | 0.24 | 5 | 0.4 | 6 | 0.1667 | 9 | 0 | 11 | 0.0909 | 31 | 0.129 |
| 17_3626775 | 1 | 1 | 5 | 0.4 | 3 | 0.6667 | 9 | 0.3333 | 18 | 0.4444 | 2 | 0 | 2 | 0.5 | 1 | 0 | 5 | 0.2 | 10 | 0.2 |
| 17_3626776 | 5 | 0 | 10 | 0 | 5 | 0 | 12 | 0.1667 | 32 | 0.0625 | 3 | 0 | 2 | 0.5 | 7 | 0.2857 | 12 | 0.0833 | 24 | 0.1667 |
| 17_3626778 | 5 | 0.2 | 12 | 0.4167 | 6 | 0.1667 | 12 | 0.1667 | 35 | 0.2571 | 2 | 0.5 | 3 | 0.3333 | 9 | 0.3333 | 11 | 0.1818 | 25 | 0.28 |
| 17_3626783 | 4 | 1 | 8 | 1 | 5 | 1 | 11 | 0.6364 | 28 | 0.8571 | 3 | 0.6667 | 2 | 1 | 10 | 0.9 | 11 | 0.7273 | 26 | 0.8077 |
| 17_3626789 | 3 | 0.6667 | 8 | 0.5 | 7 | 0.7143 | 13 | 0.6154 | 31 | 0.6129 | 3 | 0.6667 | 3 | 0.6667 | 10 | 0.7 | 9 | 0.4444 | 25 | 0.6 |
| 17_3626792 | 4 | 0.75 | 6 | 0.6667 | 5 | 0.6 | 8 | 0.625 | 23 | 0.6522 | 2 | 0.5 | NA | NA | 5 | 0.2 | 8 | 0.75 | 15 | 0.5333 |
| 17_3626794 | 5 | 0 | 12 | 0 | 7 | 0 | 14 | 0.2857 | 38 | 0.1053 | 4 | 0 | 3 | 0 | 13 | 0.2308 | 11 | 0.0909 | 31 | 0.129 |
| 17_3626800 | 3 | 0 | 10 | 0 | 6 | 0.1667 | 14 | 0.0714 | 33 | 0.0606 | 4 | 0 | 3 | 0 | 12 | 0.0833 | 10 | 0.2 | 29 | 0.1034 |
| 17_3626806 | 5 | 0.8 | 9 | 0.5556 | 7 | 0.8571 | 10 | 1 | 31 | 0.8065 | 4 | 1 | 1 | 1 | 8 | 0.875 | 9 | 0.6667 | 22 | 0.8182 |
| 17_3627404 | 2 | 0 | 7 | 0.1429 | 2 | 0 | 5 | 0 | 16 | 0.0625 | 2 | 0 | NA | NA | 4 | 0 | 3 | 0.6667 | 9 | 0.2222 |
| 17_3628349 | 1 | 0 | 4 | 0.25 | 4 | 0 | 4 | 0 | 13 | 0.0769 | 6 | 0.1667 | 4 | 0.25 | 6 | 0 | 2 | 0 | 18 | 0.1111 |
| 17_3628364 | 3 | 0 | 3 | 0.6667 | 5 | 0 | 4 | 0.25 | 15 | 0.2 | 6 | 0 | 4 | 0 | 4 | 0 | 1 | 1 | 15 | 0.0667 |
| 17_3628368 | 3 | 0 | 4 | 0.25 | 6 | 0.5 | 4 | 0.5 | 17 | 0.3529 | 6 | 0.5 | 4 | 0.25 | 4 | 0 | 1 | 0 | 15 | 0.2667 |
| 17_3628378 | 3 | 0 | 3 | 0.3333 | 6 | 0 | 4 | 0 | 16 | 0.0625 | 5 | 0.2 | 3 | 0.3333 | 6 | 0 | NA | NA | 14 | 0.1429 |
| 17_3628382 | 2 | 1 | 4 | 0.25 | 5 | 0.6 | 5 | 0.6 | 16 | 0.5625 | 7 | 0.2857 | 4 | 0.75 | 2 | 0 | 2 | 0.5 | 15 | 0.4 |
| 17_3628384 | 3 | 0 | 4 | 0.5 | 4 | 0.25 | 5 | 0.4 | 16 | 0.3125 | 7 | 0.4286 | 3 | 0 | 5 | 0 | 1 | 0 | 16 | 0.1875 |
| 17_3628400 | 3 | 0 | 4 | 0.25 | 3 | 0 | 5 | 0.2 | 15 | 0.1333 | 5 | 0.2 | 4 | 0 | 4 | 0 | 3 | 0 | 16 | 0.0625 |
| 17_3628410 | 2 | 0.5 | 3 | 0.6667 | 2 | 0 | 5 | 1 | 12 | 0.6667 | 7 | 0.7143 | 4 | 0.25 | 5 | 0.4 | 3 | 0.6667 | 19 | 0.5263 |
| 17_3628411 | 2 | 0.5 | 4 | 1 | 2 | 1 | 3 | 1 | 11 | 0.9091 | 6 | 0.5 | 4 | 0.5 | 5 | 0.6 | 3 | 0.6667 | 18 | 0.5556 |
| 17_3628431 | 3 | 1 | 4 | 1 | NA | NA | 5 | 1 | 12 | 1 | 4 | 1 | 4 | 1 | 4 | 1 | 3 | 0.6667 | 15 | 0.9333 |
| 17_3628437 | 3 | 0 | 4 | 0.25 | 2 | 0 | 5 | 0 | 14 | 0.0714 | 6 | 0.3333 | 4 | 0 | 5 | 0.2 | 3 | 0.3333 | 18 | 0.2222 |
| 17_3628438 | 2 | 0.5 | 4 | 0.5 | 2 | 1 | 5 | 0.8 | 13 | 0.6923 | 6 | 0.8333 | 4 | 0.5 | 5 | 1 | 3 | 0 | 18 | 0.6667 |
| 17_3628458 | 2 | 0.5 | 4 | 0.75 | 2 | 0.5 | 4 | 1 | 12 | 0.75 | 5 | 0.8 | 3 | 0 | 5 | 0.6 | 3 | 0.3333 | 16 | 0.5 |
| 17_3628460 | 2 | 0 | 3 | 0.3333 | 2 | 0 | 4 | 0.25 | 11 | 0.1818 | 4 | 0.25 | 2 | 0 | 5 | 0 | 3 | 0 | 14 | 0.0714 |

| | | | | | | | | | | | | | | | | | | | | |
|---|---|---|---|---|---|---|---|---|---|---|---|---|---|---|---|---|---|---|---|---|
| 17_362904 | 1 | 1 | NA | NA | 1 | 0 | NA | NA | 2 | 0.5 | NA | NA | 1 | 1 | NA | NA | NA | NA | 1 | 1 |
| 17_362999 | 17 | 0.4118 | 8 | 0.5 | 6 | 0 | 8 | 0.5 | 39 | 0.3846 | 10 | 0.4 | 12 | 0.5 | 9 | 0.1111 | 8 | 0.125 | 39 | 0.3077 |
| 17_363000 | 20 | 0.05 | 13 | 0.0769 | 12 | 0.3333 | 10 | 0.1 | 55 | 0.1273 | 11 | 0 | 17 | 0.1176 | 11 | 0 | 9 | 0.2222 | 48 | 0.0833 |
| 17_363000 | 14 | 0.2857 | 11 | 0.0909 | 8 | 0.125 | 7 | 0 | 40 | 0.15 | 11 | 0.1818 | 14 | 0.2857 | 10 | 0.2 | 12 | 0.25 | 47 | 0.234 |
| 17_363001 | 22 | 0.8636 | 12 | 0.8333 | 14 | 0.7143 | 11 | 1 | 59 | 0.8475 | 11 | 0.8182 | 21 | 0.7619 | 10 | 0.7 | 12 | 0.75 | 54 | 0.7593 |
| 17_363001 | 24 | 0.4583 | 12 | 0.4167 | 15 | 0.4 | 11 | 0.3636 | 62 | 0.4194 | 11 | 0.4545 | 20 | 0.4 | 10 | 0.6 | 9 | 0.4444 | 50 | 0.46 |
| 17_363001 | 22 | 0.8636 | 12 | 0.9167 | 15 | 0.8 | 9 | 1 | 58 | 0.8793 | 11 | 0.8182 | 18 | 0.7222 | 9 | 1 | 9 | 0.6667 | 47 | 0.7872 |
| 17_363002 | 18 | 0.1111 | 12 | 0 | 18 | 0.1111 | 9 | 0 | 57 | 0.0702 | 11 | 0.0909 | 22 | 0.0455 | 11 | 0 | 11 | 0.0909 | 55 | 0.0545 |
| 17_363243 | 5 | 0 | 8 | 0.125 | 3 | 0.3333 | 5 | 0 | 21 | 0.0952 | 4 | 0 | 3 | 0 | 2 | 0 | 4 | 0.25 | 13 | 0.0769 |
| 17_363484 | 2 | 1 | NA | NA | 2 | 1 | 1 | 1 | 5 | 1 | NA | NA | 1 | 1 | NA | NA | 1 | 1 | 2 | 1 |
| 17_363486 | 2 | 0.5 | NA | NA | 2 | 0 | 1 | 1 | 5 | 0.4 | NA | NA | 1 | 1 | NA | NA | 1 | 0 | 2 | 0.5 |
| 17_363486 | 2 | 1 | NA | NA | 2 | 1 | NA | NA | 4 | 1 | NA | NA | 1 | 1 | NA | NA | 1 | 0 | 2 | 0.5 |
| 17_363493 | 2 | 0 | 3 | 0.3333 | 1 | 1 | 3 | 0.3333 | 9 | 0.3333 | 4 | 0.5 | NA | NA | 1 | 0 | 1 | 0 | 6 | 0.3333 |
| 17_363493 | 2 | 0 | 3 | 0.6667 | NA | NA | 3 | 0.6667 | 8 | 0.5 | 3 | 0.6667 | NA | NA | 1 | 1 | 1 | 0 | 5 | 0.6 |
| 17_363494 | 3 | 0.3333 | 3 | 1 | 2 | 1 | 3 | 0.6667 | 11 | 0.7273 | 4 | 0.75 | NA | NA | 1 | 1 | 2 | 1 | 7 | 0.8571 |
| 17_363495 | 2 | 0 | 2 | 0 | 2 | 1 | 3 | 0 | 9 | 0.2222 | 4 | 0 | NA | NA | 2 | 0.5 | 1 | 0 | 7 | 0.1429 |
| 17_363496 | 3 | 0 | 3 | 0.3333 | 3 | 0 | 3 | 0 | 12 | 0.0833 | 4 | 0.5 | NA | NA | 2 | 0 | 2 | 0 | 8 | 0.25 |
| 17_363515 | 1 | 1 | 2 | 0 | 1 | 0 | 2 | 0 | 6 | 0.1667 | 3 | 0 | NA | NA | 2 | 0.5 | 1 | 0 | 6 | 0.1667 |
| 17_365698 | NA | NA | 7 | 0.5714 | 2 | 1 | 6 | 0.5 | 15 | 0.6 | 1 | 1 | 2 | 0.5 | 3 | 0.6667 | NA | NA | 6 | 0.6667 |
| 17_365711 | NA | NA | 2 | 0 | 1 | 0 | 5 | 0.2 | 8 | 0.125 | 1 | 1 | 2 | 0.5 | 2 | 0 | NA | NA | 5 | 0.4 |
| 17_365713 | NA | NA | 3 | 0.6667 | 1 | 0 | 3 | 0.3333 | 7 | 0.4286 | NA | NA | 2 | 1 | 1 | 1 | NA | NA | 3 | 1 |
| 17_365714 | NA | NA | 2 | 0 | 1 | 0 | 4 | 0.25 | 7 | 0.1429 | 1 | 0 | 2 | 0.5 | 2 | 0.5 | NA | NA | 5 | 0.4 |
| 17_365715 | NA | NA | 3 | 0.3333 | 1 | 1 | 5 | 0.4 | 9 | 0.4444 | 1 | 1 | 2 | 0 | 2 | 0.5 | NA | NA | 5 | 0.4 |
| 17_365715 | NA | NA | 2 | 1 | 1 | 1 | 4 | 0.75 | 7 | 0.8571 | 1 | 1 | 1 | 1 | 2 | 1 | NA | NA | 4 | 1 |
| 17_365715 | NA | NA | 3 | 0.3333 | 1 | 0 | 5 | 0 | 9 | 0.1111 | 1 | 0 | 1 | 0 | 2 | 0.5 | NA | NA | 4 | 0.25 |
| 17_365715 | NA | NA | 3 | 0.3333 | 1 | 0 | 5 | 0.6 | 9 | 0.4444 | 1 | 0 | 1 | 0 | 2 | 0.5 | NA | NA | 4 | 0.25 |
| 17_365715 | NA | NA | 2 | 1 | 1 | 1 | 4 | 0.75 | 7 | 0.8571 | 1 | 1 | 1 | 1 | 2 | 1 | NA | NA | 4 | 1 |
| 17_365797 | NA | NA | 3 | 0.6667 | 1 | 1 | 2 | 0.5 | 6 | 0.6667 | NA | NA | 1 | 1 | NA | NA | NA | NA | 1 | 1 |
| 17_365832 | NA | NA | NA | NA | NA | NA | 1 | 1 | 1 | 1 | NA | NA | NA | NA | NA | NA | 1 | 1 | 1 | 1 |
| 17_366284 | NA | NA | 1 | 1 | 2 | 0 | 1 | 0 | 4 | 0.25 | NA | NA | NA | NA | 2 | 0.5 | 2 | 0 | 4 | 0.25 |
| 17_366285 | NA | NA | 1 | 1 | 2 | 0.5 | 1 | 0 | 4 | 0.5 | 1 | 0 | NA | NA | 2 | 0.5 | 2 | 0 | 5 | 0.2 |
| 17_366291 | NA | NA | 1 | 1 | NA | NA | 2 | 0 | 3 | 0.3333 | 1 | 0 | NA | NA | 2 | 0.5 | 1 | 0 | 4 | 0.25 |
| 17_371683 | 1 | 1 | 1 | 0 | 1 | 1 | NA | NA | 3 | 0.6667 | 1 | 1 | 2 | 1 | NA | NA | 1 | 1 | 4 | 1 |
| 17_371686 | 1 | 1 | 2 | 0 | 1 | 1 | 1 | 1 | 5 | 0.6 | 1 | 0 | 3 | 0.6667 | 1 | 0 | 1 | 1 | 6 | 0.5 |
| 17_371688 | 1 | 0 | 2 | 1 | 1 | 1 | 1 | 1 | 5 | 0.8 | 1 | 1 | 3 | 0.3333 | NA | NA | 1 | 0 | 5 | 0.4 |
| 17_371688 | 1 | 1 | 2 | 0 | 1 | 0 | 1 | 0 | 5 | 0.2 | 1 | 1 | 3 | 0.3333 | 1 | 0 | 1 | 1 | 6 | 0.5 |
| 17_371689 | 1 | 0 | 2 | 0.5 | 1 | 1 | 1 | 0 | 5 | 0.4 | 1 | 0 | 2 | 1 | 1 | 1 | 1 | 0 | 5 | 0.6 |
| 17_371690 | 1 | 1 | 1 | 1 | 1 | 1 | NA | NA | 3 | 1 | 1 | 1 | NA | NA | 1 | 1 | 1 | 1 | 3 | 1 |
| 17_371690 | 1 | 1 | 2 | 0 | 1 | 1 | 1 | 1 | 5 | 0.6 | 1 | 1 | 3 | 0.3333 | 1 | 1 | 1 | 1 | 6 | 0.6667 |
| 17_371690 | 1 | 0 | 2 | 0 | 1 | 1 | 1 | 0 | 5 | 0.2 | 1 | 1 | 3 | 0.3333 | NA | NA | 1 | 0 | 5 | 0.4 |
| 17_371691 | 1 | 0 | 2 | 0.5 | 1 | 0 | 1 | 1 | 5 | 0.4 | 1 | 0 | 3 | 0.6667 | 1 | 1 | 1 | 1 | 6 | 0.6667 |
| 17_371692 | 2 | 0.5 | 2 | 0 | 1 | 0 | 1 | 0 | 6 | 0.1667 | 1 | 1 | 3 | 0.3333 | NA | NA | NA | NA | 4 | 0.5 |
| 17_371693 | 2 | 0.5 | 2 | 0 | 1 | 0 | 1 | 0 | 6 | 0.1667 | 1 | 0 | 3 | 1 | 1 | 1 | 1 | 0 | 6 | 0.6667 |
| 17_371697 | 2 | 0.5 | NA | NA | 1 | 0 | 1 | 1 | 4 | 0.5 | 1 | 0 | 3 | 0 | 1 | 1 | 1 | 0 | 6 | 0.1667 |
| 17_371698 | 2 | 0.5 | 1 | 0 | 1 | 0 | 1 | 1 | 5 | 0.4 | 1 | 1 | 3 | 0.6667 | 1 | 1 | 1 | 0 | 6 | 0.6667 |
| 17_371700 | 2 | 0.5 | 1 | 1 | 1 | 1 | NA | NA | 4 | 0.75 | 1 | 1 | 3 | 0.3333 | 1 | 1 | 1 | 1 | 6 | 0.6667 |
| 17_371701 | 2 | 0.5 | 1 | 0 | 1 | 0 | 1 | 0 | 5 | 0.2 | 1 | 1 | 3 | 0 | 1 | 0 | 1 | 0 | 6 | 0.1667 |
| 17_372600 | 2 | 1 | 1 | 0 | 5 | 0 | 6 | 0.3333 | 14 | 0.2857 | 4 | 0.5 | 3 | 0 | 1 | 0 | 5 | 0.2 | 13 | 0.2308 |
| 17_372601 | 15 | 0.4 | 15 | 0.1333 | 18 | 0.3889 | 6 | 0.3333 | 54 | 0.3148 | 15 | 0.2 | 16 | 0.375 | 19 | 0.2105 | 20 | 0.35 | 70 | 0.2857 |
| 17_372602 | 23 | 0.4783 | 21 | 0.4286 | 20 | 0.5 | 12 | 0.5 | 76 | 0.4737 | 16 | 0.25 | 18 | 0.2778 | 21 | 0.3333 | 18 | 0.3333 | 73 | 0.3014 |
| 17_372758 | 1 | 1 | NA | NA | 1 | 1 | NA | NA | 2 | 1 | 1 | 1 | 1 | 1 | NA | NA | 1 | 1 | 3 | 1 |
| 17_372759 | 2 | 0 | NA | NA | 1 | 1 | 1 | 0 | 4 | 0.25 | 1 | 1 | 1 | 0 | 1 | 0 | 1 | 1 | 4 | 0.5 |
| 17_372764 | 2 | 0 | NA | NA | 1 | 1 | 1 | 1 | 4 | 0.5 | 1 | 1 | 1 | 1 | 1 | 0 | 1 | 1 | 4 | 0.75 |
| 17_372764 | 1 | 0 | NA | NA | 1 | 1 | 1 | 0 | 3 | 0.3333 | NA | NA | 1 | 1 | 1 | 0 | 1 | 0 | 3 | 0.3333 |
| 17_373218 | 5 | 0 | 10 | 0.5 | 7 | 0.1429 | 9 | 0.5556 | 31 | 0.3548 | 7 | 0.2857 | 9 | 0.2222 | 6 | 0.3333 | 3 | 0.3333 | 25 | 0.28 |
| 17_373218 | 6 | 0.1667 | 10 | 0.4 | 9 | 0.1111 | 9 | 0.3333 | 34 | 0.2647 | 9 | 0.3333 | 8 | 0.375 | 6 | 0 | 4 | 0.5 | 27 | 0.2963 |
| 17_373220 | 5 | 0.6 | 6 | 0.1667 | 8 | 0.125 | 6 | 0 | 25 | 0.2 | 7 | 0.1429 | 7 | 0.1429 | 5 | 0 | 1 | 0 | 20 | 0.1 |
| 17_373220 | 3 | 0.6667 | 4 | 0.5 | 2 | 1 | 3 | 0.3333 | 12 | 0.5833 | 6 | 0.1667 | 4 | 0.75 | 2 | 0 | 1 | 0 | 13 | 0.3077 |
| 17_373221 | 4 | 0 | 4 | 0 | 6 | 0.1667 | 6 | 0.1667 | 20 | 0.1 | 7 | 0.1429 | 6 | 0.1667 | 5 | 0 | 2 | 0 | 20 | 0.1 |
| 17_373222 | 3 | 0 | 5 | 0.2 | 5 | 0.2 | 6 | 0.3333 | 19 | 0.2105 | 6 | 0.1667 | 6 | 0.1667 | 5 | 0 | 1 | 0 | 18 | 0.1111 |
| 17_373223 | 2 | 0.5 | 5 | 0.2 | 6 | 0.1667 | 3 | 0 | 16 | 0.1875 | 6 | 0 | 5 | 0.8 | 4 | 0.25 | 1 | 0 | 16 | 0.3125 |
| 17_373313 | 2 | 0.5 | 3 | 0.3333 | 4 | 0 | 5 | 0.4 | 14 | 0.2857 | 4 | 0.5 | 4 | 0.5 | 4 | 0.5 | NA | NA | 12 | 0.5 |
| 17_373735 | 33 | 0.1818 | 27 | 0.2963 | 25 | 0.4 | 40 | 0.325 | 125 | 0.296 | 23 | 0.4348 | 32 | 0.4688 | 23 | 0.3043 | 26 | 0.3462 | 104 | 0.3942 |
| 17_373736 | 24 | 0.2083 | 19 | 0.1579 | 21 | 0.1905 | 32 | 0.2188 | 96 | 0.1979 | 17 | 0.1176 | 27 | 0.2593 | 20 | 0.25 | 18 | 0.1111 | 82 | 0.1951 |
| 17_373850 | 60 | 0.3833 | 65 | 0.3385 | 62 | 0.2742 | 80 | 0.275 | 267 | 0.3146 | 52 | 0.3462 | 79 | 0.3924 | 45 | 0.4 | 61 | 0.377 | 237 | 0.3797 |

| | | | | | | | | | | | | | | | | | | | | |
|---|---|---|---|---|---|---|---|---|---|---|---|---|---|---|---|---|---|---|---|---|
| 17_3738509 | 50 | 0.58 | 54 | 0.6481 | 49 | 0.6531 | 59 | 0.6271 | 212 | 0.6274 | 39 | 0.6154 | 68 | 0.5735 | 40 | 0.7 | 47 | 0.7234 | 194 | 0.6443 |
| 17_3738521 | 1 | 1 | 1 | 0 | NA | NA | 1 | 1 | 3 | 0.6667 | 2 | 0 | NA | NA | 1 | 0 | 3 | 0.3333 | 6 | 0.1667 |
| 17_3738551 | 1 | 0 | 3 | 0.3333 | NA | NA | NA | NA | 4 | 0.25 | 4 | 0.25 | NA | NA | 1 | 0 | 2 | 0 | 7 | 0.1429 |
| 17_3738574 | 1 | 1 | 4 | 1 | 1 | 1 | NA | NA | 6 | 1 | 3 | 0.6667 | NA | NA | NA | NA | 1 | 0 | 4 | 0.5 |
| 17_3738643 | NA | NA | 1 | 1 | NA | NA | 2 | 0 | 3 | 0.3333 | NA | NA | NA | NA | NA | NA | 3 | 0.3333 | 3 | 0.3333 |
| 17_4015905 | 12 | 0.25 | 15 | 0.1333 | 19 | 0.2105 | 11 | 0 | 57 | 0.1579 | 17 | 0.2353 | 10 | 0.3 | 8 | 0.25 | 15 | 0.3333 | 50 | 0.28 |
| 17_4015907 | 28 | 0.2143 | 32 | 0.1562 | 42 | 0.1429 | 30 | 0.1 | 132 | 0.1515 | 31 | 0.2258 | 28 | 0.1786 | 34 | 0.0882 | 28 | 0.25 | 121 | 0.1818 |
| 17_4124814 | 2 | 0 | 6 | 0.6667 | 2 | 0 | 1 | 1 | 11 | 0.4545 | 2 | 1 | 2 | 1 | 3 | 0.6667 | 3 | 0.3333 | 10 | 0.7 |
| 17_4124820 | 1 | 0 | 6 | 0.6667 | 2 | 1 | 1 | 1 | 10 | 0.7 | 1 | 1 | 1 | 1 | 3 | 0.6667 | 3 | 0.3333 | 8 | 0.625 |
| 17_4124825 | 2 | 0.5 | 7 | 0.2857 | 3 | 0 | 1 | 0 | 13 | 0.2308 | 2 | 0 | 2 | 0.5 | 4 | 0 | 2 | 0 | 10 | 0.1 |
| 17_4124828 | 2 | 0 | 6 | 0 | 3 | 0.3333 | NA | NA | 11 | 0.0909 | NA | NA | 2 | 0.5 | 4 | 0.25 | 3 | 0 | 9 | 0.2222 |
| 17_4124834 | 2 | 0 | 5 | 0.2 | 3 | 0.3333 | 1 | 0 | 11 | 0.1818 | 2 | 0 | 2 | 0 | 4 | 0.5 | 3 | 0 | 11 | 0.1818 |
| 17_4125799 | 2 | 0.5 | 6 | 0.8333 | 3 | 0 | 2 | 0.5 | 13 | 0.5385 | 2 | 0 | 2 | 0.5 | 3 | 0.6667 | 4 | 0.75 | 11 | 0.5455 |
| 17_4127160 | 5 | 0.4 | 6 | 0.3333 | 3 | 0 | 4 | 0 | 18 | 0.2222 | 3 | 0 | 2 | 0 | 4 | 0.25 | 4 | 0.25 | 13 | 0.1538 |
| 17_4127241 | 6 | 0.6667 | 6 | 0.3333 | 5 | 0.2 | 4 | 0.25 | 21 | 0.381 | 1 | 1 | 7 | 0.2857 | 1 | 0 | 6 | 0.1667 | 15 | 0.2667 |
| 17_4283198 | 2 | 1 | NA | NA | 1 | 0 | 2 | 0.5 | 5 | 0.6 | NA | NA | 1 | 1 | 1 | 1 | NA | NA | 2 | 1 |
| 17_4283202 | 1 | 1 | NA | NA | 1 | 0 | 1 | 1 | 3 | 0.6667 | NA | NA | 1 | 1 | NA | NA | 1 | 0 | 2 | 0.5 |
| 17_4310430 | 2 | 1 | 5 | 0.2 | 5 | 0 | 2 | 0.5 | 14 | 0.2857 | 4 | 0.5 | 5 | 0.2 | 2 | 0 | 4 | 0 | 15 | 0.2 |
| 17_4310712 | 1 | 0 | 2 | 0 | 3 | 0.3333 | 2 | 0 | 8 | 0.125 | 1 | 1 | 4 | 0 | 1 | 1 | 2 | 0 | 8 | 0.25 |
| 17_4310717 | 1 | 0 | 2 | 0 | 3 | 0.3333 | 2 | 0.5 | 8 | 0.25 | 1 | 0 | 4 | 0.25 | 1 | 0 | 2 | 0.5 | 8 | 0.25 |
| 17_4310718 | 1 | 1 | 3 | 1 | 3 | 0.6667 | 2 | 0.5 | 9 | 0.7778 | 1 | 0 | 4 | 0.5 | 1 | 1 | 2 | 1 | 8 | 0.625 |
| 17_4334852 | 6 | 0 | 2 | 0 | 2 | 0 | 5 | 0.4 | 15 | 0.1333 | 1 | 0 | 7 | 0.2857 | 2 | 0 | 2 | 1 | 12 | 0.3333 |
| 17_4334867 | 7 | 0.2857 | 2 | 0 | 2 | 0 | 5 | 0.2 | 16 | 0.1875 | 2 | 0 | 7 | 0.2857 | 1 | 0 | 2 | 0 | 12 | 0.1667 |
| 17_4334871 | 6 | 1 | 1 | 1 | 2 | 1 | 5 | 0.8 | 14 | 0.9286 | 2 | 1 | 6 | 0.8333 | 2 | 1 | 2 | 0.5 | 12 | 0.8333 |
| 17_4334876 | 7 | 0.1429 | 2 | 0.5 | 3 | 0 | 6 | 0 | 18 | 0.1111 | 2 | 0 | 7 | 0 | 2 | 0 | 2 | 0.5 | 13 | 0.0769 |
| 17_4334887 | 4 | 0.25 | 3 | 1 | 3 | 1 | 6 | 0.6667 | 16 | 0.6875 | 2 | 0.5 | 6 | 0.8333 | 2 | 1 | 2 | 0.5 | 12 | 0.75 |
| 17_4334896 | 5 | 1 | 4 | 0.75 | 2 | 1 | 6 | 1 | 17 | 0.9412 | 1 | 1 | 4 | 0.75 | 2 | 1 | 1 | 1 | 8 | 0.875 |
| 17_4334899 | 3 | 0.3333 | 4 | 0.5 | 3 | 0 | 8 | 0.375 | 18 | 0.3333 | NA | NA | 8 | 0.125 | 2 | 0 | 3 | 0.6667 | 13 | 0.2308 |
| 17_4335062 | 3 | 0.3333 | 4 | 0.25 | 3 | 0 | 6 | 0 | 16 | 0.125 | 2 | 0 | 8 | 0.125 | 1 | 0 | 4 | 0 | 15 | 0.0667 |
| 17_4335065 | 5 | 0.6 | 5 | 0.6 | 3 | 0.6667 | 6 | 0.3333 | 19 | 0.5263 | 2 | 0.5 | 10 | 0.3 | 1 | 1 | 4 | 0.25 | 17 | 0.3529 |
| 17_4335070 | 6 | 0.3333 | 4 | 0.25 | 3 | 0.6667 | 7 | 0.5714 | 20 | 0.45 | 2 | 0.5 | 9 | 0.4444 | 3 | 0.3333 | 4 | 0.5 | 18 | 0.4444 |
| 17_4335096 | 8 | 1 | 6 | 0.6667 | 2 | 1 | 9 | 0.8889 | 25 | 0.88 | 1 | 1 | 11 | 1 | 2 | 0 | 4 | 0.75 | 18 | 0.8333 |
| 17_4337601 | 10 | 0.4 | 4 | 0 | 1 | 0 | 7 | 0.5714 | 22 | 0.3636 | 3 | 0 | 3 | 0 | 2 | 0 | 3 | 0.3333 | 11 | 0.0909 |
| 17_4339867 | 10 | 0.1 | 8 | 0 | 3 | 0.3333 | 11 | 0 | 32 | 0.0625 | 5 | 0 | 12 | 0.0833 | 4 | 0 | 5 | 0.2 | 26 | 0.0769 |
| 17_4339882 | 10 | 0.9 | 8 | 1 | 4 | 0.75 | 12 | 0.9167 | 34 | 0.9118 | 5 | 1 | 12 | 0.9167 | 5 | 1 | 8 | 1 | 30 | 0.9667 |
| 17_4339886 | 8 | 0.375 | 8 | 0.375 | 4 | 0.25 | 10 | 0.2 | 30 | 0.3 | 4 | 0.25 | 7 | 0.1429 | 3 | 0.6667 | 7 | 0.1429 | 21 | 0.2381 |
| 17_4339941 | 11 | 0 | 9 | 0 | 7 | 0.1429 | 12 | 0.0833 | 39 | 0.0513 | 4 | 0 | 11 | 0.0909 | 5 | 0 | 9 | 0.1111 | 29 | 0.069 |
| 17_4339961 | 14 | 0.3571 | 9 | 0 | 7 | 0.5714 | 11 | 0.4545 | 41 | 0.3415 | 6 | 0.1667 | 10 | 0.6 | 5 | 0.8 | 8 | 0.5 | 29 | 0.5172 |
| 17_4339965 | 12 | 0.4167 | 6 | 0.5 | 4 | 0.25 | 7 | 0.4286 | 29 | 0.4138 | 4 | 0.25 | 10 | 0.5 | 3 | 0.3333 | 7 | 0 | 24 | 0.2917 |
| 17_4339967 | 12 | 0.1667 | 11 | 0 | 7 | 0.2857 | 13 | 0.2308 | 43 | 0.1628 | 6 | 0 | 10 | 0.2 | 5 | 0.4 | 11 | 0.0909 | 32 | 0.1562 |
| 17_4339989 | 15 | 0.4667 | 11 | 0.3636 | 7 | 0.7143 | 16 | 0.4375 | 49 | 0.4694 | 6 | 0.1667 | 12 | 0.75 | 6 | 0.3333 | 11 | 0.1818 | 35 | 0.4 |
| 17_4340001 | 15 | 0.0667 | 13 | 0.0769 | 7 | 0.1429 | 14 | 0 | 49 | 0.0612 | 6 | 0 | 12 | 0.25 | 8 | 0.125 | 11 | 0 | 37 | 0.1081 |
| 17_4346223 | NA | NA | 1 | 1 | NA | NA | 1 | 1 | 2 | 1 | NA | NA | 1 | 0 | 2 | 0.5 | NA | NA | 3 | 0.3333 |
| 17_4346238 | NA | NA | 1 | 1 | NA | NA | 1 | 0 | 2 | 0.5 | NA | NA | 1 | 1 | 1 | 1 | NA | NA | 2 | 1 |
| 17_4376092 | 1 | 1 | 3 | 0.3333 | 5 | 0.8 | 2 | 1 | 11 | 0.7273 | NA | NA | NA | NA | 2 | 0.5 | 1 | 1 | 3 | 0.6667 |
| 17_4376105 | 1 | 0 | 3 | 0.6667 | 5 | 0.2 | 2 | 0 | 11 | 0.2727 | NA | NA | NA | NA | 2 | 0.5 | 1 | 1 | 3 | 0.6667 |
| 17_4376112 | 1 | 1 | 3 | 0.3333 | 5 | 0.6 | 1 | 1 | 10 | 0.6 | NA | NA | NA | NA | 2 | 0.5 | NA | NA | 2 | 0.5 |
| 17_4376120 | 1 | 1 | 3 | 0.6667 | 4 | 1 | 1 | 1 | 9 | 0.8889 | NA | NA | NA | NA | 2 | 0.5 | 1 | 1 | 3 | 0.6667 |
| 17_4376144 | 1 | 0 | 3 | 0.6667 | 2 | 0.5 | 2 | 1 | 8 | 0.625 | 1 | 1 | NA | NA | 1 | 0 | 1 | 1 | 3 | 0.6667 |
| 17_4376181 | 1 | 0 | 3 | 0.3333 | 7 | 0.4286 | 2 | 0 | 13 | 0.3077 | 1 | 1 | 1 | 1 | 1 | 1 | 2 | 0.5 | 5 | 0.8 |
| 17_4376192 | 2 | 0.5 | 3 | 0 | 5 | 0 | 2 | 1 | 12 | 0.25 | 1 | 0 | 1 | 0 | 3 | 0 | 2 | 0.5 | 7 | 0.1429 |
| 17_4376635 | 2 | 1 | 3 | 0.6667 | 8 | 0.5 | 2 | 1 | 15 | 0.6667 | 2 | 0.5 | 2 | 1 | 3 | 0.6667 | 2 | 1 | 9 | 0.7778 |
| 17_4376638 | 2 | 0 | 3 | 0.3333 | 7 | 0.2857 | 2 | 0 | 14 | 0.2143 | 2 | 0 | 2 | 0 | 2 | 1 | 2 | 0 | 8 | 0.25 |
| 17_4387958 | 15 | 0 | 30 | 0.0667 | 19 | 0 | 14 | 0.1429 | 78 | 0.0513 | 12 | 0.0833 | 14 | 0.2143 | 17 | 0 | 10 | 0 | 53 | 0.0755 |
| 17_4395245 | 1 | 0 | NA | NA | 2 | 1 | 1 | 0 | 4 | 0.5 | NA | NA | NA | NA | 1 | 1 | NA | NA | 1 | 1 |
| 17_4395322 | 4 | 0 | 2 | 0 | 5 | 0.2 | 4 | 0.25 | 15 | 0.1333 | NA | NA | 1 | 0 | 4 | 0.25 | 2 | 0 | 7 | 0.1429 |
| 17_4395328 | 5 | 0 | 3 | 0 | 6 | 0.1667 | 4 | 0 | 18 | 0.0556 | 2 | 0 | 4 | 0 | 4 | 0.25 | 4 | 0 | 14 | 0.0714 |
| 17_4395346 | 5 | 0 | 4 | 0 | 6 | 0.1667 | 5 | 0.2 | 20 | 0.1 | 2 | 0 | 3 | 0.3333 | 4 | 0.25 | 4 | 0.5 | 13 | 0.3077 |
| 17_4395347 | 5 | 0.2 | 3 | 0 | 6 | 0 | 5 | 0.4 | 19 | 0.1579 | 2 | 1 | 2 | 1 | 4 | 0.5 | 3 | 0.3333 | 11 | 0.6364 |
| 17_4395366 | 5 | 0.2 | 3 | 0 | 5 | 0.8 | 4 | 0.25 | 17 | 0.3529 | 1 | 1 | 2 | 0.5 | 4 | 0.25 | 4 | 0.75 | 11 | 0.5455 |
| 17_4395373 | 5 | 0 | 2 | 0 | 5 | 0 | 5 | 0.2 | 17 | 0.0588 | 1 | 0 | 2 | 1 | 3 | 0 | 3 | 0 | 9 | 0.2222 |
| 17_4395375 | 5 | 0.2 | 3 | 0 | 4 | 0.25 | 5 | 0 | 17 | 0.1176 | 2 | 0 | 3 | 0.3333 | 4 | 0.5 | 4 | 0.5 | 13 | 0.3846 |
| 17_4395397 | 1 | 0 | 2 | 0.5 | 3 | 0 | 6 | 0 | 12 | 0.0833 | 2 | 0 | 2 | 0.5 | 4 | 0 | 4 | 0 | 12 | 0.0833 |
| 17_4395411 | 5 | 0 | 2 | 0.5 | 4 | 0.5 | 6 | 0.3333 | 17 | 0.2941 | 2 | 0.5 | 3 | 0 | 5 | 0 | 4 | 0 | 14 | 0.0714 |
| 17_4395415 | 4 | 0.5 | 3 | 0.3333 | 3 | 0.3333 | 6 | 0.5 | 16 | 0.4375 | 1 | 0 | 1 | 1 | 5 | 0.4 | 4 | 0 | 11 | 0.2727 |

| | | | | | | | | | | | | | | | | | | | | |
|---|---|---|---|---|---|---|---|---|---|---|---|---|---|---|---|---|---|---|---|---|
| 17_4395423 | 4 | 0.5 | 2 | 0.5 | 3 | 0.3333 | 6 | 0.3333 | 15 | 0.4 | 2 | 1 | 2 | 0.5 | 5 | 0.8 | 3 | 1 | 12 | 0.8333 |
| 17_4395424 | 4 | 0.25 | 1 | 1 | 4 | 0.25 | 6 | 0.1667 | 15 | 0.2667 | 2 | 0 | 2 | 0 | 4 | 0.5 | 3 | 0 | 11 | 0.1818 |
| 17_4395427 | 5 | 0.2 | 2 | 0 | 3 | 0.3333 | 4 | 0.25 | 14 | 0.2143 | 2 | 0.5 | 3 | 0.3333 | 4 | 0.25 | 4 | 0.75 | 13 | 0.4615 |
| 17_4395433 | 4 | 0.5 | 3 | 1 | 4 | 0.25 | 5 | 0.4 | 16 | 0.5 | 2 | 0.5 | 2 | 0.5 | 5 | 0.4 | 1 | 0 | 10 | 0.4 |
| 17_4395439 | 5 | 1 | 2 | 0.5 | 3 | 1 | 6 | 1 | 16 | 0.9375 | 2 | 1 | 4 | 1 | 1 | 1 | 4 | 1 | 11 | 1 |
| 17_4397507 | 1 | 0 | 1 | 0 | NA | NA | 1 | 1 | 3 | 0.3333 | 2 | 0.5 | 2 | 0 | 1 | 1 | 1 | 0 | 6 | 0.3333 |
| 17_4406122 | NA | NA | NA | NA | 1 | 1 | NA | NA | 1 | 1 | NA | NA | NA | NA | 1 | 1 | NA | NA | 1 | 1 |
| 17_4414723 | 12 | 0 | 6 | 0 | 12 | 0 | 16 | 0.1875 | 46 | 0.0652 | 7 | 0.1429 | 18 | 0.0556 | 15 | 0.2 | 12 | 0 | 52 | 0.0962 |
| 17_4414726 | 11 | 0.1818 | 8 | 0 | 13 | 0.0769 | 17 | 0.2353 | 49 | 0.1429 | 8 | 0.25 | 17 | 0.2941 | 16 | 0.25 | 14 | 0.3571 | 55 | 0.2909 |
| 17_4426436 | 8 | 0.25 | 17 | 0.2941 | 14 | 0.1429 | 9 | 0.1111 | 48 | 0.2083 | 11 | 0.3636 | 6 | 0.1667 | 9 | 0.5556 | 10 | 0.1 | 36 | 0.3056 |
| 17_4434795 | 25 | 0.08 | 36 | 0.1111 | 31 | 0.0323 | 30 | 0.0667 | 122 | 0.0738 | 20 | 0.05 | 24 | 0 | 24 | 0.0417 | 22 | 0.1364 | 90 | 0.0556 |
| 17_4434816 | 24 | 0.0417 | 31 | 0.0323 | 27 | 0.1111 | 26 | 0.1538 | 108 | 0.0833 | 16 | 0.125 | 21 | 0.1429 | 23 | 0.087 | 28 | 0.1429 | 88 | 0.125 |
| 17_4434941 | 28 | 0.3214 | 32 | 0.2188 | 34 | 0.3824 | 27 | 0.2963 | 121 | 0.3058 | 21 | 0.2857 | 21 | 0.381 | 26 | 0.3077 | 28 | 0.4286 | 96 | 0.3542 |
| 17_4434962 | 24 | 0.0833 | 31 | 0.0323 | 24 | 0.25 | 24 | 0.0417 | 103 | 0.0971 | 19 | 0.1053 | 22 | 0.0455 | 26 | 0.0769 | 23 | 0.1304 | 90 | 0.0889 |
| 17_4435024 | 33 | 0.2424 | 37 | 0.2162 | 34 | 0.2353 | 32 | 0.2188 | 136 | 0.2279 | 26 | 0.1923 | 24 | 0.2083 | 32 | 0.125 | 29 | 0.1379 | 111 | 0.1622 |
| 17_4439175 | 22 | 0.3182 | 12 | 0.0833 | 17 | 0.2353 | 21 | 0.1429 | 72 | 0.2083 | 20 | 0.25 | 8 | 0 | 13 | 0.2308 | 21 | 0.0952 | 62 | 0.1613 |
| 17_4439178 | 19 | 0.8421 | 13 | 0.4615 | 11 | 0.8182 | 22 | 0.7273 | 65 | 0.7231 | 18 | 0.6111 | 6 | 0.6667 | 12 | 0.8333 | 20 | 0.85 | 56 | 0.75 |
| 17_4439185 | 28 | 0.1786 | 14 | 0.0714 | 16 | 0.3125 | 23 | 0.087 | 81 | 0.1605 | 21 | 0.2381 | 10 | 0.3 | 12 | 0.25 | 24 | 0.2917 | 67 | 0.2687 |
| 17_4439187 | 19 | 0.0526 | 9 | 0 | 13 | 0.0769 | 15 | 0.1333 | 56 | 0.0714 | 16 | 0.0625 | 6 | 0 | 10 | 0.2 | 14 | 0.0714 | 46 | 0.087 |
| 17_4439195 | 24 | 0.2917 | 13 | 0.2308 | 13 | 0.2308 | 13 | 0.0769 | 63 | 0.2222 | 17 | 0.1765 | 5 | 0 | 10 | 0 | 19 | 0.2105 | 51 | 0.1373 |
| 17_4439206 | 27 | 0.1481 | 17 | 0.0588 | 19 | 0.1579 | 23 | 0 | 86 | 0.093 | 18 | 0.2222 | 10 | 0.1 | 15 | 0.2 | 24 | 0.1667 | 67 | 0.1791 |
| 17_4439213 | 28 | 0.0714 | 16 | 0.125 | 16 | 0.0625 | 23 | 0.0435 | 83 | 0.0723 | 19 | 0.1579 | 8 | 0 | 15 | 0 | 20 | 0.05 | 62 | 0.0645 |
| 17_4439216 | 24 | 0.9583 | 15 | 0.9333 | 15 | 1 | 17 | 0.9412 | 71 | 0.9577 | 19 | 0.9474 | 7 | 1 | 11 | 0.9091 | 21 | 0.9048 | 58 | 0.931 |
| 17_4439232 | 32 | 0.5938 | 16 | 0.4375 | 17 | 0.5882 | 25 | 0.64 | 90 | 0.5778 | 15 | 0.5333 | 11 | 0.6364 | 13 | 0.1538 | 25 | 0.48 | 64 | 0.4531 |
| 17_4439402 | 14 | 0.3571 | 6 | 0.5 | 11 | 0.2727 | 14 | 0.2143 | 45 | 0.3111 | 10 | 0.5 | 9 | 0.2222 | 10 | 0.5 | 7 | 0.2857 | 36 | 0.3889 |
| 17_4439470 | 9 | 0.1111 | 2 | 0 | 5 | 0 | 10 | 0.1 | 26 | 0.0769 | 3 | 0 | 2 | 0 | 7 | 0.1429 | NA | NA | 12 | 0.0833 |
| 17_4439488 | 8 | 0.25 | NA | NA | 5 | 0.4 | 5 | 0 | 18 | 0.2222 | 4 | 0 | 3 | 0.3333 | 5 | 0 | 2 | 0 | 14 | 0.0714 |
| 17_4439619 | 25 | 0.32 | 10 | 0.5 | 14 | 0.4286 | 27 | 0.2593 | 76 | 0.3421 | 17 | 0.3529 | 15 | 0.0667 | 16 | 0.5 | 17 | 0.4118 | 65 | 0.3385 |
| 17_4439627 | 29 | 0.3793 | 16 | 0.25 | 17 | 0.4118 | 30 | 0.4 | 92 | 0.3696 | 21 | 0.3333 | 15 | 0.1333 | 20 | 0.35 | 20 | 0.4 | 76 | 0.3158 |
| 17_4439635 | 27 | 0 | 15 | 0.1333 | 18 | 0.0556 | 26 | 0.1538 | 86 | 0.0814 | 22 | 0 | 15 | 0 | 20 | 0.15 | 22 | 0.0909 | 79 | 0.0633 |
| 17_4439713 | 27 | 0.2222 | 17 | 0.1765 | 21 | 0.381 | 27 | 0.1852 | 92 | 0.2391 | 23 | 0.3043 | 19 | 0.2632 | 23 | 0.2174 | 20 | 0.35 | 85 | 0.2824 |
| 17_4440793 | 6 | 0.1667 | 1 | 0 | 7 | 0.1429 | 4 | 0 | 18 | 0.1111 | 6 | 0 | 5 | 0.2 | 2 | 0 | 2 | 0 | 15 | 0.0667 |
| 17_4440823 | 5 | 0.2 | 1 | 1 | 7 | 0 | 4 | 0 | 17 | 0.1176 | 4 | 0.5 | 3 | 0 | 2 | 0 | 1 | 1 | 10 | 0.3 |
| 17_4440827 | 5 | 0.4 | 1 | 0 | 6 | 0 | 5 | 0 | 17 | 0.1176 | 4 | 0 | 3 | 0 | 2 | 0 | 1 | 1 | 10 | 0.1 |
| 17_4440853 | 3 | 1 | 1 | 1 | 4 | 1 | 4 | 0.75 | 12 | 0.9167 | 4 | 1 | 2 | 1 | 2 | 0.5 | NA | NA | 8 | 0.875 |
| 17_4440857 | 4 | 0.5 | 1 | 0 | 7 | 0.2857 | 3 | 0 | 15 | 0.2667 | 4 | 0.5 | 3 | 0 | 1 | 0 | 1 | 0 | 9 | 0.2222 |
| 17_4440865 | 4 | 0 | 1 | 0 | 6 | 0 | 3 | 0.6667 | 14 | 0.1429 | 4 | 0.75 | 3 | 0.3333 | 2 | 0.5 | 1 | 0 | 10 | 0.5 |
| 17_4440871 | 4 | 0.5 | 1 | 0 | 3 | 1 | 3 | 0.3333 | 11 | 0.5455 | 4 | 0.25 | 1 | 1 | 2 | 0 | 1 | 1 | 8 | 0.375 |
| 17_4440883 | 4 | 0.25 | 1 | 0 | 5 | 0.2 | 4 | 0.25 | 14 | 0.2143 | 3 | 0 | 3 | 0.6667 | 2 | 0 | 1 | 0 | 9 | 0.2222 |
| 17_4440897 | 3 | 0.3333 | 1 | 0 | 5 | 0.6 | 2 | 1 | 11 | 0.5455 | 4 | 0.5 | 1 | 0 | 2 | 0 | NA | NA | 7 | 0.2857 |
| 17_4440913 | 3 | 0.6667 | 2 | 0.5 | 6 | 0.6667 | 3 | 0.6667 | 14 | 0.6429 | 3 | 1 | 2 | 0.5 | 2 | 0 | 1 | 1 | 8 | 0.625 |
| 17_4440919 | 3 | 0 | 2 | 0.5 | 6 | 0.1667 | 2 | 0 | 13 | 0.1538 | 3 | 0.3333 | 3 | 0 | 2 | 0 | 1 | 0 | 9 | 0.1111 |
| 17_4441012 | 2 | 0.5 | 2 | 0 | 8 | 0.375 | 2 | 0 | 14 | 0.2857 | 2 | 0.5 | 3 | 0 | 2 | 0 | NA | NA | 7 | 0.1429 |
| 17_4441022 | 2 | 0 | 2 | 0.5 | 8 | 0 | 2 | 0 | 14 | 0.0714 | 2 | 0 | 3 | 0.3333 | 2 | 0 | NA | NA | 7 | 0.1429 |
| 17_4441037 | 2 | 1 | 1 | 1 | 7 | 0.7143 | 2 | 1 | 12 | 0.8333 | 2 | 0.5 | 3 | 0.6667 | 2 | 0.5 | NA | NA | 7 | 0.5714 |
| 17_4441040 | 1 | 1 | NA | NA | 6 | 0.3333 | NA | NA | 7 | 0.4286 | NA | NA | 3 | 0.6667 | 1 | 1 | NA | NA | 4 | 0.75 |
| 17_4441042 | 2 | 1 | 1 | 1 | 7 | 1 | 2 | 1 | 12 | 1 | 2 | 1 | 1 | 1 | 2 | 1 | NA | NA | 5 | 1 |
| 17_4441050 | 1 | 0 | 1 | 1 | 7 | 0.5714 | 1 | 0 | 10 | 0.5 | 2 | 0.5 | 3 | 0.6667 | 2 | 0 | NA | NA | 7 | 0.4286 |
| 17_4441278 | 1 | 0 | 1 | 1 | 3 | 1 | 2 | 0.5 | 7 | 0.7143 | 2 | 0 | 2 | 0.5 | 2 | 0.5 | NA | NA | 6 | 0.3333 |
| 17_4441284 | 1 | 0 | NA | NA | 5 | 1 | 1 | 1 | 7 | 0.8571 | 2 | 0 | 3 | 1 | 1 | 1 | NA | NA | 6 | 0.6667 |
| 17_4441284 | 1 | 0 | 1 | 0 | 6 | 0.8333 | 2 | 1 | 10 | 0.7 | 2 | 1 | 3 | 1 | 2 | 0.5 | NA | NA | 7 | 0.8571 |
| 17_4441303 | 1 | 0 | 1 | 1 | 4 | 0.5 | 2 | 1 | 8 | 0.625 | 2 | 1 | 3 | 1 | 2 | 1 | NA | NA | 7 | 1 |
| 17_4441306 | 1 | 1 | 1 | 1 | 5 | 1 | 2 | 1 | 9 | 1 | 2 | 1 | 3 | 1 | 2 | 1 | NA | NA | 7 | 1 |
| 17_4441322 | 1 | 1 | 1 | 1 | 4 | 0.75 | 1 | 0 | 7 | 0.7143 | 1 | 0 | 2 | 0 | 2 | 0.5 | NA | NA | 5 | 0.2 |
| 17_4441331 | 1 | 0 | 1 | 1 | 3 | 0 | NA | NA | 5 | 0.2 | 2 | 0.5 | 2 | 0 | 1 | 0 | NA | NA | 5 | 0.2 |
| 17_4441334 | 1 | 0 | 1 | 0 | 3 | 0 | 2 | 0.5 | 7 | 0.1429 | 2 | 0 | 3 | 0.3333 | 2 | 0 | NA | NA | 7 | 0.1429 |
| 17_4441346 | NA | NA | NA | NA | 3 | 0.3333 | 1 | 0 | 4 | 0.25 | 2 | 0 | 3 | 0 | 2 | 0.5 | NA | NA | 7 | 0.1429 |
| 17_4441349 | 1 | 1 | NA | NA | 1 | 1 | 2 | 1 | 4 | 1 | 2 | 0 | 3 | 0.6667 | 1 | 1 | NA | NA | 6 | 0.5 |
| 17_4491823 | NA | NA | NA | NA | 1 | 1 | NA | NA | 1 | 1 | NA | NA | NA | NA | 1 | 1 | NA | NA | 1 | 1 |
| 17_4491905 | NA | NA | NA | NA | 1 | 1 | NA | NA | 1 | 1 | NA | NA | NA | NA | 1 | 1 | NA | NA | 1 | 1 |
| 17_4504618 | 1 | 1 | NA | NA | NA | NA | NA | NA | 1 | 1 | 1 | 1 | NA | NA | NA | NA | NA | NA | 1 | 1 |
| 17_4504685 | 1 | 0 | NA | NA | 1 | 1 | NA | NA | 2 | 0.5 | 1 | 0 | NA | NA | NA | NA | 1 | 1 | 2 | 0.5 |
| 17_4504773 | NA | NA | NA | NA | 1 | 1 | NA | NA | 1 | 1 | 1 | 1 | NA | NA | NA | NA | NA | NA | 1 | 1 |
| 17_4570296 | 7 | 0.8571 | 13 | 1 | 2 | 1 | 2 | 0.5 | 24 | 0.9167 | 3 | 1 | 3 | 0.3333 | NA | NA | 4 | 1 | 10 | 0.8 |
| 17_4570297 | 7 | 0.2857 | 17 | 0.3529 | NA | NA | 2 | 0 | 26 | 0.3077 | 5 | 0 | 3 | 0.3333 | NA | NA | 6 | 0.1667 | 14 | 0.1429 |

| | | | | | | | | | | | | | | | | | | | | |
|---|---|---|---|---|---|---|---|---|---|---|---|---|---|---|---|---|---|---|---|---|
| 17_457030 | 7 | 0.7143 | 16 | 0.5625 | 4 | 0.75 | 3 | 0.3333 | 30 | 0.6 | 6 | 0.1667 | 3 | 0.6667 | NA | NA | 7 | 0.5714 | 16 | 0.4375 |
| 17_457031 | 8 | 1 | 20 | 0.8 | 3 | 1 | 2 | 1 | 33 | 0.8788 | 4 | 1 | 3 | 1 | NA | NA | 5 | 1 | 12 | 1 |
| 17_457040 | 5 | 0 | 18 | 0.2222 | 4 | 0.5 | 3 | 0 | 30 | 0.2 | 6 | 0 | 3 | 0.3333 | NA | NA | 7 | 0 | 16 | 0.0625 |
| 17_457501 | NA | NA | 1 | 1 | NA | NA | NA | NA | 1 | 1 | NA | NA | NA | NA | 2 | 1 | NA | NA | 2 | 1 |
| 17_457502 | NA | NA | 1 | 1 | NA | NA | NA | NA | 1 | 1 | NA | NA | NA | NA | 2 | 0.5 | NA | NA | 2 | 0.5 |
| 17_457505 | NA | NA | 2 | 1 | NA | NA | NA | NA | 2 | 1 | NA | NA | NA | NA | 2 | 0.5 | NA | NA | 2 | 0.5 |
| 17_457508 | NA | NA | 2 | 0.5 | NA | NA | NA | NA | 2 | 0.5 | NA | NA | NA | NA | 2 | 0.5 | NA | NA | 2 | 0.5 |
| 17_457509 | NA | NA | 2 | 0.5 | NA | NA | NA | NA | 2 | 0.5 | NA | NA | NA | NA | 1 | 1 | NA | NA | 1 | 1 |
| 17_457513 | NA | NA | 2 | 0.5 | NA | NA | NA | NA | 2 | 0.5 | NA | NA | NA | NA | 2 | 0.5 | 1 | 0 | 3 | 0.3333 |
| 17_457515 | NA | NA | 2 | 1 | NA | NA | NA | NA | 2 | 1 | NA | NA | NA | NA | 2 | 1 | 1 | 1 | 3 | 1 |
| 17_457516 | NA | NA | 2 | 1 | NA | NA | NA | NA | 2 | 1 | NA | NA | NA | NA | 1 | 0 | 1 | 1 | 2 | 0.5 |
| 17_457517 | NA | NA | 2 | 1 | NA | NA | 1 | 1 | 3 | 1 | NA | NA | NA | NA | 2 | 1 | 1 | 1 | 3 | 1 |
| 17_457518 | NA | NA | 2 | 0.5 | NA | NA | 1 | 0 | 3 | 0.3333 | NA | NA | NA | NA | 2 | 0.5 | 1 | 0 | 3 | 0.3333 |
| 17_457519 | NA | NA | 2 | 0 | NA | NA | 1 | 1 | 3 | 0.3333 | NA | NA | NA | NA | 1 | 0 | 1 | 1 | 2 | 0.5 |
| 17_457519 | NA | NA | 2 | 1 | NA | NA | 1 | 1 | 3 | 1 | NA | NA | NA | NA | 2 | 1 | 1 | 1 | 3 | 1 |
| 17_457520 | NA | NA | 2 | 1 | NA | NA | 1 | 1 | 3 | 1 | NA | NA | NA | NA | 1 | 1 | 1 | 1 | 2 | 1 |
| 17_457526 | NA | NA | 2 | 0.5 | 1 | 0 | 1 | 1 | 4 | 0.5 | 1 | 1 | 1 | 1 | 3 | 0.3333 | 2 | 0.5 | 7 | 0.5714 |
| 17_458313 | 1 | 0 | 2 | 0 | 1 | 0 | 5 | 0.6 | 9 | 0.3333 | 2 | 0.5 | 3 | 0 | 1 | 0 | 3 | 0.3333 | 9 | 0.2222 |
| 17_458315 | 3 | 1 | 2 | 0 | 3 | 0.6667 | 5 | 0.8 | 13 | 0.6923 | 3 | 1 | 4 | 0.75 | 4 | 0.5 | 5 | 1 | 16 | 0.8125 |
| 17_458316 | 3 | 1 | 2 | 0.5 | 3 | 0.3333 | 5 | 0.6 | 13 | 0.6154 | 3 | 0.3333 | 3 | 0.3333 | 4 | 1 | 4 | 0.75 | 14 | 0.6429 |
| 17_458316 | 3 | 0 | 1 | 1 | 3 | 0.3333 | 5 | 0.6 | 12 | 0.4167 | 3 | 0 | 3 | 0 | 4 | 0.25 | 4 | 0.5 | 14 | 0.2143 |
| 17_458316 | 3 | 1 | 1 | 1 | 2 | 0.5 | 4 | 0.25 | 10 | 0.6 | 3 | 1 | 2 | 1 | 3 | 1 | 3 | 1 | 11 | 1 |
| 17_458530 | 11 | 0.1818 | 23 | 0 | 18 | 0.0556 | 9 | 0.1111 | 61 | 0.0656 | 15 | 0.1333 | 4 | 0.25 | 4 | 0.25 | 14 | 0 | 37 | 0.1081 |
| 17_458530 | 11 | 0.3636 | 19 | 0.2105 | 17 | 0.1176 | 9 | 0.2222 | 56 | 0.2143 | 15 | 0.1333 | 5 | 0 | 4 | 0 | 13 | 0.3077 | 37 | 0.1622 |
| 17_458548 | 6 | 0 | 9 | 0 | 2 | 0 | 2 | 0.5 | 19 | 0.0526 | 8 | 0.125 | 1 | 0 | 1 | 0 | 4 | 0 | 14 | 0.0714 |
| 17_458550 | 3 | 0 | 6 | 0.5 | 3 | 0.3333 | 1 | 0 | 13 | 0.3077 | 7 | 0.2857 | NA | NA | NA | NA | 2 | 0.5 | 9 | 0.3333 |
| 17_458553 | 3 | 0.3333 | 7 | 0.2857 | NA | NA | 1 | 0 | 11 | 0.2727 | 6 | 0.1667 | 1 | 0 | NA | NA | NA | NA | 7 | 0.1429 |
| 17_458554 | 2 | 0.5 | 8 | 0.25 | NA | NA | 1 | 0 | 11 | 0.2727 | 6 | 0.5 | 1 | 0 | NA | NA | 2 | 0.5 | 9 | 0.4444 |
| 17_458555 | 3 | 0.6667 | 6 | 0 | NA | NA | 1 | 0 | 10 | 0.2 | 7 | 0.5714 | 1 | 0 | NA | NA | 1 | 1 | 9 | 0.5556 |
| 17_458559 | 3 | 0.3333 | 4 | 0.25 | NA | NA | 1 | 0 | 8 | 0.25 | 6 | 0.3333 | 1 | 0 | NA | NA | 1 | 0 | 8 | 0.25 |
| 17_458562 | 3 | 1 | 6 | 1 | NA | NA | 1 | 1 | 10 | 1 | 6 | 1 | 1 | 1 | NA | NA | 1 | 1 | 8 | 1 |
| 17_458563 | 3 | 1 | 3 | 1 | 1 | 1 | 1 | 1 | 8 | 1 | 3 | 1 | NA | NA | NA | NA | NA | NA | 3 | 1 |
| 17_458563 | 2 | 0.5 | 5 | 1 | 1 | 1 | NA | NA | 8 | 0.875 | 4 | 1 | 1 | 1 | NA | NA | 1 | 1 | 6 | 1 |
| 17_458564 | 3 | 0.3333 | 5 | 0.6 | 1 | 0 | 1 | 0 | 10 | 0.4 | 5 | 0.2 | 1 | 1 | NA | NA | NA | NA | 6 | 0.3333 |
| 17_458566 | 2 | 1 | 4 | 1 | 1 | 1 | 1 | 1 | 8 | 1 | 6 | 1 | 1 | 1 | NA | NA | 1 | 1 | 8 | 1 |
| 17_458577 | 3 | 0 | 5 | 0.2 | 1 | 0 | NA | NA | 9 | 0.1111 | 5 | 0.2 | 1 | 0 | NA | NA | 1 | 0 | 7 | 0.1429 |
| 17_458773 | 1 | 1 | 6 | 0.1667 | 1 | 0 | 3 | 0.3333 | 11 | 0.2727 | 3 | 0.3333 | 3 | 1 | 2 | 0.5 | 2 | 0.5 | 10 | 0.6 |
| 17_458773 | 1 | 0 | 6 | 0.1667 | 2 | 0 | 4 | 0.25 | 13 | 0.1538 | 2 | 0.5 | 3 | 0.3333 | 3 | 0 | 2 | 0 | 10 | 0.2 |
| 17_458775 | 1 | 1 | 5 | 0.8 | 2 | 1 | 4 | 1 | 12 | 0.9167 | 2 | 0.5 | 3 | 1 | 2 | 0 | 1 | 1 | 8 | 0.625 |
| 17_458775 | 1 | 1 | 5 | 0.6 | 2 | 1 | 3 | 0.6667 | 11 | 0.7273 | 4 | 0.5 | 2 | 1 | 3 | 1 | 1 | 0 | 10 | 0.7 |
| 17_458778 | 2 | 0 | 3 | 0.3333 | 2 | 0.5 | 2 | 0.5 | 9 | 0.3333 | 1 | 0 | 2 | 0.5 | 3 | 0.3333 | 1 | 0 | 7 | 0.2857 |
| 17_458779 | 2 | 1 | 5 | 0.8 | NA | NA | 3 | 1 | 10 | 0.9 | 1 | 1 | 4 | 1 | 3 | 1 | 3 | 0.6667 | 11 | 0.9091 |
| 17_458780 | 2 | 0.5 | 7 | 0.2857 | 2 | 0 | 3 | 0.3333 | 14 | 0.2857 | 2 | 0 | 3 | 0.3333 | 4 | 0 | 3 | 0.3333 | 12 | 0.1667 |
| 17_458785 | 2 | 0.5 | 6 | 0.1667 | 1 | 0 | 2 | 1 | 11 | 0.3636 | 3 | 0.3333 | 4 | 0 | 1 | 1 | 3 | 0.6667 | 11 | 0.3636 |
| 17_458792 | 196 | 0.0408 | 219 | 0.0411 | 178 | 0.0562 | 187 | 0.0695 | 780 | 0.0513 | 184 | 0.087 | 200 | 0.085 | 221 | 0.0905 | 187 | 0.0802 | 792 | 0.0859 |
| 17_458806 | 126 | 0.1746 | 150 | 0.0933 | 110 | 0.2182 | 128 | 0.1719 | 514 | 0.1595 | 115 | 0.1826 | 125 | 0.168 | 145 | 0.2069 | 126 | 0.1667 | 511 | 0.182 |
| 17_458963 | 4 | 0.75 | 6 | 0.3333 | 5 | 0.6 | 6 | 0.3333 | 21 | 0.4762 | 4 | 0.5 | 3 | 0.6667 | 2 | 1 | 4 | 1 | 13 | 0.7692 |
| 17_458963 | 4 | 0.5 | 6 | 0 | 4 | 0 | 6 | 0 | 20 | 0.1 | 4 | 0.25 | 1 | 0 | 1 | 1 | 3 | 0 | 9 | 0.2222 |
| 17_458964 | 4 | 0.75 | 5 | 0.6 | 4 | 1 | 4 | 0.5 | 17 | 0.7059 | 3 | 1 | 3 | 1 | 2 | 0.5 | 3 | 1 | 11 | 0.9091 |
| 17_458964 | 3 | 0.6667 | 1 | 1 | 3 | 0.6667 | 4 | 0.75 | 11 | 0.7273 | 2 | 0.5 | 4 | 0.25 | 2 | 1 | 3 | 0.6667 | 11 | 0.5455 |
| 17_458965 | 3 | 0.6667 | 5 | 0 | 3 | 0.3333 | 4 | 0 | 15 | 0.2 | 4 | 0.25 | 3 | 0 | 2 | 0 | 4 | 0 | 13 | 0.0769 |
| 17_459712 | 6 | 0.1667 | 10 | 0.3 | 7 | 0.2857 | 8 | 0.25 | 31 | 0.2581 | 7 | 0.1429 | 10 | 0.4 | 11 | 0.3636 | 7 | 0.1429 | 35 | 0.2857 |
| 17_459716 | 3 | 0 | 5 | 0.2 | 8 | 0.125 | 5 | 0 | 21 | 0.0952 | 6 | 0 | 8 | 0.5 | 9 | 0 | 6 | 0.1667 | 29 | 0.1724 |
| 17_459716 | 4 | 0.75 | 13 | 0.6923 | 12 | 0.9167 | 6 | 0.6667 | 35 | 0.7714 | 8 | 0.875 | 11 | 0.6364 | 12 | 0.75 | 10 | 0.9 | 41 | 0.7805 |
| 17_459904 | 1 | 0 | 2 | 0 | 3 | 0 | 3 | 0.3333 | 9 | 0.1111 | 2 | 0 | 6 | 0.1667 | 5 | 0 | 3 | 0 | 16 | 0.0625 |
| 17_460092 | 3 | 0.3333 | 4 | 0.75 | 5 | 0.6 | 4 | 0.5 | 16 | 0.5625 | 9 | 0.3333 | 2 | 0 | 8 | 0 | 3 | 0 | 22 | 0.1364 |
| 17_460092 | 2 | 1 | 3 | 0.3333 | 7 | 0.1429 | 5 | 0.2 | 17 | 0.2941 | 11 | 0.3636 | 2 | 1 | 6 | 0 | 2 | 0.5 | 21 | 0.3333 |
| 17_460092 | 3 | 1 | 5 | 1 | 8 | 1 | 5 | 1 | 21 | 1 | 13 | 1 | 4 | 1 | 6 | 1 | 3 | 1 | 26 | 1 |
| 17_460093 | 1 | 0 | 5 | 0.6 | 8 | 0.5 | 4 | 0.75 | 18 | 0.5556 | 13 | 0.7692 | 3 | 1 | 8 | 0.875 | 3 | 0.6667 | 27 | 0.8148 |
| 17_463286 | 1 | 0 | 6 | 0.1667 | 7 | 0.2857 | 7 | 0 | 21 | 0.1429 | 5 | 0 | 6 | 0 | 4 | 0.25 | 1 | 0 | 16 | 0.0625 |
| 17_463287 | 1 | 0 | 8 | 0.375 | 9 | 0.3333 | 7 | 0.2857 | 25 | 0.32 | 6 | 0.3333 | 5 | 0.2 | 3 | 0 | 1 | 1 | 15 | 0.2667 |
| 17_463287 | 1 | 0 | 8 | 0.375 | 8 | 0.125 | 7 | 0 | 24 | 0.1667 | 4 | 0.25 | 6 | 0 | 4 | 0.25 | 3 | 0 | 17 | 0.1176 |
| 17_463288 | 1 | 0 | 8 | 0 | 9 | 0.1111 | 5 | 0.2 | 23 | 0.087 | 6 | 0.5 | 6 | 0.1667 | 4 | 0 | 3 | 0 | 19 | 0.2105 |
| 17_463288 | 1 | 0 | 8 | 0.5 | 9 | 0.1111 | 6 | 0.6667 | 24 | 0.375 | 4 | 0.25 | 6 | 0.5 | 4 | 0.5 | 2 | 0 | 16 | 0.375 |
| 17_463290 | 1 | 1 | 8 | 0.875 | 9 | 0.5556 | 6 | 0.8333 | 24 | 0.75 | 5 | 0.6 | 6 | 0.6667 | 3 | 1 | 3 | 0.6667 | 17 | 0.7059 |

| | | | | | | | | | | | | | | | | | | | | |
|---|---|---|---|---|---|---|---|---|---|---|---|---|---|---|---|---|---|---|---|---|
| 17_4632912 | NA | NA | 6 | 1 | 8 | 0.75 | 6 | 0.6667 | 20 | 0.8 | 5 | 0.6 | 6 | 0.6667 | 3 | 1 | 3 | 1 | 17 | 0.7647 |
| 17_4632912 | 1 | 0 | 7 | 1 | 7 | 0.5714 | 4 | 0.5 | 19 | 0.6842 | 6 | 0.5 | 5 | 0.8 | 4 | 0.75 | 3 | 0.6667 | 18 | 0.6667 |
| 17_4632915 | 1 | 0 | 6 | 0.1667 | 7 | 0 | 3 | 0 | 17 | 0.0588 | 5 | 0 | 5 | 0 | 4 | 0.25 | 2 | 0 | 16 | 0.0625 |
| 17_4632916 | 1 | 0 | 7 | 0.2857 | 8 | 0 | 4 | 0.25 | 20 | 0.15 | 5 | 0 | 5 | 0 | 4 | 0 | 2 | 0.5 | 16 | 0.0625 |
| 17_4632927 | NA | NA | 7 | 0.2857 | 6 | 0.1667 | 4 | 0.25 | 17 | 0.2353 | 4 | 0.25 | 5 | 0.2 | 4 | 0.25 | 2 | 0.5 | 15 | 0.2667 |
| 17_4632932 | NA | NA | 7 | 1 | 6 | 1 | 4 | 1 | 17 | 1 | 5 | 1 | 3 | 1 | 2 | 1 | 2 | 1 | 12 | 1 |
| 17_4632933 | 1 | 0 | 7 | 0.4286 | 6 | 0.6667 | 2 | 0 | 16 | 0.4375 | 4 | 0.5 | 3 | 0.3333 | 3 | 0.3333 | 2 | 0.5 | 12 | 0.4167 |
| 17_4632937 | 1 | 0 | 5 | 0.6 | 8 | 0.375 | 3 | 0.3333 | 17 | 0.4118 | 3 | 0.3333 | 4 | 0 | 3 | 0.3333 | 1 | 0 | 11 | 0.1818 |
| 17_4632954 | 1 | 1 | 5 | 1 | 8 | 1 | 5 | 0.6 | 19 | 0.8947 | 3 | 1 | 4 | 0.5 | 4 | 1 | 2 | 0 | 13 | 0.6923 |
| 17_4632957 | 1 | 0 | 2 | 0 | 5 | 0 | 3 | 0.3333 | 11 | 0.0909 | 3 | 0.3333 | 1 | 0 | 4 | 0 | 1 | 0 | 9 | 0.1111 |
| 17_4632959 | 1 | 0 | 5 | 0 | 8 | 0.125 | 4 | 0 | 18 | 0.0556 | 3 | 0.3333 | 4 | 0 | 3 | 0 | 2 | 0 | 12 | 0.0833 |
| 17_4632963 | 1 | 1 | 5 | 0.4 | 7 | 0 | 3 | 0 | 16 | 0.1875 | 3 | 0 | 4 | 0 | 4 | 0.25 | 2 | 0 | 13 | 0.0769 |
| 17_4632967 | 1 | 1 | 4 | 0 | 5 | 0 | 3 | 0.3333 | 13 | 0.1538 | 1 | 0 | 2 | 0.5 | 3 | 0 | 1 | 0 | 7 | 0.1429 |
| 17_4634227 | NA | NA | 3 | 1 | 4 | 0.75 | 3 | 0.6667 | 10 | 0.8 | 1 | 1 | 2 | 0 | 3 | 1 | NA | NA | 6 | 0.6667 |
| 17_4634280 | 1 | 0 | 1 | 1 | 2 | 0.5 | 1 | 0 | 5 | 0.4 | NA | NA | 2 | 0 | 2 | 0.5 | NA | NA | 4 | 0.25 |
| 17_4634289 | 1 | 0 | 1 | 1 | NA | NA | NA | NA | 2 | 0.5 | NA | NA | 1 | 1 | 2 | 0.5 | NA | NA | 3 | 0.6667 |
| 17_4643953 | 16 | 0.1875 | 20 | 0.15 | 18 | 0.0556 | 11 | 0 | 65 | 0.1077 | 11 | 0.2727 | 11 | 0.0909 | 13 | 0.1538 | 9 | 0.3333 | 44 | 0.2045 |
| 17_4643958 | 16 | 0.5 | 20 | 0.3 | 18 | 0.5556 | 9 | 0.3333 | 63 | 0.4286 | 14 | 0.4286 | 11 | 0.4545 | 9 | 0.6667 | 7 | 0.1429 | 41 | 0.439 |
| 17_4644028 | 3 | 0 | 3 | 0.3333 | 4 | 0.25 | 2 | 0.5 | 12 | 0.25 | 2 | 1 | 2 | 0.5 | 1 | 0 | 3 | 0 | 8 | 0.375 |
| 17_4644032 | 2 | 0.5 | 3 | 0 | 3 | 0 | 2 | 0 | 10 | 0.1 | 1 | 0 | 2 | 0 | 1 | 0 | 3 | 0.6667 | 7 | 0.2857 |
| 17_4645501 | 4 | 0.25 | 3 | 0 | 2 | 0 | 1 | 0 | 10 | 0.1 | 2 | 0 | 2 | 0 | 1 | 0 | 2 | 0.5 | 7 | 0.1429 |
| 17_4645839 | 1 | 1 | NA | NA | 4 | 0.75 | 2 | 1 | 7 | 0.8571 | 3 | 0.6667 | NA | NA | 3 | 1 | 1 | 1 | 7 | 0.8571 |
| 17_4645857 | 1 | 1 | NA | NA | 4 | 0.75 | 2 | 1 | 7 | 0.8571 | 4 | 1 | NA | NA | 3 | 0.6667 | 1 | 1 | 8 | 0.875 |
| 17_4645868 | 1 | 1 | NA | NA | 4 | 1 | 3 | 1 | 8 | 1 | 4 | 1 | NA | NA | 4 | 1 | 2 | 1 | 10 | 1 |
| 17_4645872 | 1 | 1 | NA | NA | 4 | 0.5 | 3 | 0 | 8 | 0.375 | 4 | 0.5 | NA | NA | 4 | 1 | 2 | 0 | 10 | 0.6 |
| 17_4645879 | NA | NA | NA | NA | 3 | 0 | 3 | 0.3333 | 6 | 0.1667 | 3 | 0.3333 | NA | NA | 3 | 0.3333 | 1 | 0 | 7 | 0.2857 |
| 17_4645880 | NA | NA | NA | NA | 3 | 0 | 2 | 0.5 | 5 | 0.2 | 3 | 0 | NA | NA | 3 | 0 | 2 | 1 | 8 | 0.25 |
| 17_4645883 | NA | NA | NA | NA | 2 | 0.5 | 2 | 0 | 4 | 0.25 | 2 | 0.5 | NA | NA | 3 | 0 | 1 | 1 | 6 | 0.3333 |
| 17_4645886 | 1 | 1 | NA | NA | 4 | 0.5 | 2 | 0 | 7 | 0.4286 | 4 | 0.25 | NA | NA | 4 | 0.75 | 2 | 0.5 | 10 | 0.5 |
| 17_4645891 | 1 | 0 | NA | NA | 2 | 0 | 3 | 0.3333 | 6 | 0.1667 | 4 | 0.25 | NA | NA | NA | NA | 1 | 1 | 5 | 0.4 |
| 17_4645891 | 1 | 1 | NA | NA | 3 | 1 | 2 | 1 | 6 | 1 | 1 | 1 | NA | NA | 3 | 1 | 1 | 1 | 5 | 1 |
| 17_4645902 | NA | NA | NA | NA | 1 | 1 | 1 | 1 | 2 | 1 | 4 | 1 | NA | NA | 2 | 1 | 1 | 1 | 7 | 1 |
| 17_4645909 | 1 | 0 | NA | NA | 2 | 1 | 3 | 1 | 6 | 0.8333 | 4 | 1 | NA | NA | 4 | 0.5 | 2 | 0.5 | 10 | 0.7 |
| 17_4646391 | NA | NA | NA | NA | 3 | 0.3333 | 1 | 0 | 4 | 0.25 | 4 | 0 | NA | NA | 4 | 0.25 | 2 | 0 | 10 | 0.1 |
| 17_4647929 | 3 | 1 | 1 | 1 | 2 | 1 | 1 | 0 | 7 | 0.8571 | 2 | 0.5 | 1 | 1 | 6 | 0.5 | 5 | 0.6 | 14 | 0.5714 |
| 17_4647952 | 5 | 0.4 | 2 | 0 | 2 | 0 | 1 | 1 | 10 | 0.3 | 1 | 0 | 2 | 0.5 | 4 | 0.25 | 5 | 0.4 | 12 | 0.3333 |
| 17_4647955 | 7 | 0.4286 | 1 | 0 | 3 | 0 | 1 | 0 | 12 | 0.25 | 2 | 0.5 | 2 | 0 | 5 | 0 | 7 | 0.2857 | 16 | 0.1875 |
| 17_4647956 | 5 | 0.4 | 2 | 0 | 3 | 0 | 2 | 0 | 12 | 0.1667 | 2 | 0.5 | 2 | 0 | 7 | 0.2857 | 8 | 0 | 19 | 0.1579 |
| 17_4647968 | 7 | 0.7143 | 1 | 1 | 3 | 0.3333 | 2 | 0 | 13 | 0.5385 | 2 | 1 | 2 | 0.5 | 7 | 0.5714 | 8 | 0.75 | 19 | 0.6842 |
| 17_4647969 | 7 | 0.5714 | 2 | 1 | 4 | 1 | 2 | 0.5 | 15 | 0.7333 | 2 | 0.5 | 2 | 1 | 7 | 0.5714 | 7 | 0.7143 | 18 | 0.6667 |
| 17_4647971 | 5 | 0.6 | 2 | 0 | 3 | 0 | 2 | 0 | 12 | 0.25 | 2 | 0 | 1 | 0 | 6 | 0.1667 | 7 | 0.2857 | 16 | 0.1875 |
| 17_4647972 | 5 | 1 | 2 | 1 | 3 | 1 | 2 | 1 | 12 | 1 | 3 | 0.6667 | 2 | 1 | 9 | 0.6667 | 8 | 0.75 | 22 | 0.7273 |
| 17_4647975 | 6 | 1 | 2 | 1 | 4 | 1 | 2 | 1 | 14 | 1 | 3 | 1 | 2 | 1 | 9 | 0.7778 | 8 | 1 | 22 | 0.9091 |
| 17_4647983 | 5 | 0.6 | 2 | 0 | 2 | 0 | 2 | 0 | 11 | 0.2727 | 3 | 0.3333 | 1 | 0 | 8 | 0.125 | 8 | 0.5 | 20 | 0.3 |
| 17_4647999 | 7 | 0 | 1 | 0 | 4 | 0 | 2 | 0.5 | 14 | 0.0714 | 4 | 0 | 2 | 0.5 | 11 | 0.3636 | 7 | 0.2857 | 24 | 0.2917 |
| 17_4648281 | NA | NA | 2 | 1 | NA | NA | NA | NA | 2 | 1 | NA | NA | NA | NA | 1 | 1 | NA | NA | 1 | 1 |
| 17_4648294 | 1 | 0 | 2 | 0.5 | NA | NA | 1 | 1 | 4 | 0.5 | NA | NA | 1 | 1 | 1 | 1 | NA | NA | 2 | 1 |
| 17_4648296 | 1 | 0 | 1 | 0 | NA | NA | 1 | 1 | 3 | 0.3333 | NA | NA | 1 | 1 | 1 | 0 | NA | NA | 2 | 0.5 |
| 17_4648305 | 1 | 1 | 2 | 1 | NA | NA | 1 | 1 | 4 | 1 | NA | NA | 1 | 1 | 1 | 1 | NA | NA | 2 | 1 |
| 17_4648311 | 1 | 0 | 1 | 0 | NA | NA | 1 | 1 | 3 | 0.3333 | NA | NA | 1 | 1 | 1 | 1 | NA | NA | 2 | 1 |
| 17_4648336 | 1 | 1 | 2 | 1 | NA | NA | 1 | 1 | 4 | 1 | NA | NA | 1 | 1 | 1 | 1 | NA | NA | 2 | 1 |
| 17_4648338 | 1 | 1 | 2 | 1 | NA | NA | 1 | 1 | 4 | 1 | NA | NA | 1 | 1 | 1 | 0 | NA | NA | 2 | 0.5 |
| 17_4648342 | 1 | 1 | 2 | 0 | NA | NA | 1 | 1 | 4 | 0.5 | NA | NA | 1 | 1 | NA | NA | NA | NA | 1 | 1 |
| 17_4648357 | NA | NA | 1 | 1 | NA | NA | NA | NA | 1 | 1 | NA | NA | 1 | 1 | 1 | 0 | NA | NA | 2 | 0.5 |
| 17_4648378 | 1 | 1 | 1 | 1 | NA | NA | 1 | 1 | 3 | 1 | NA | NA | 1 | 1 | 1 | 1 | NA | NA | 2 | 1 |
| 17_4655315 | 5 | 0.6 | 2 | 1 | 3 | 1 | 1 | 0 | 11 | 0.7273 | 1 | 1 | 3 | 0.3333 | 4 | 1 | 3 | 1 | 11 | 0.8182 |
| 17_4655317 | 3 | 0.3333 | 6 | 0.5 | 4 | 0.25 | 2 | 0.5 | 15 | 0.4 | 3 | 0.6667 | 3 | 0.3333 | 3 | 0.6667 | 4 | 0.75 | 13 | 0.6154 |
| 17_4655319 | 5 | 0.2 | 6 | 0.5 | 4 | 0 | 1 | 0 | 16 | 0.25 | 3 | 0 | 4 | 0.25 | 5 | 0 | 4 | 0.25 | 16 | 0.125 |
| 17_4655339 | 4 | 0.25 | 5 | 0.6 | 4 | 0.25 | 1 | 0 | 14 | 0.3571 | 3 | 0.3333 | 3 | 0.3333 | 5 | 0.6 | 3 | 0 | 14 | 0.3571 |
| 17_4655352 | 4 | 0.75 | 5 | 0.8 | 2 | 0.5 | 1 | 1 | 12 | 0.75 | 3 | 1 | 2 | 1 | 4 | 1 | 3 | 0.3333 | 12 | 0.8333 |
| 17_4655358 | 3 | 0.3333 | 4 | 0.5 | 3 | 1 | 1 | 0 | 11 | 0.5455 | 4 | 0.5 | NA | NA | 3 | 0.3333 | 3 | 0.3333 | 10 | 0.4 |
| 17_4655400 | 5 | 0 | 6 | 0 | 4 | 0 | 2 | 0.5 | 17 | 0.0588 | 5 | 0.2 | 3 | 0 | 6 | 0.3333 | 4 | 0 | 18 | 0.1667 |
| 17_4655503 | 4 | 0.25 | 7 | 0.1429 | 4 | 0 | 7 | 0.1429 | 22 | 0.1364 | 4 | 0.5 | 9 | 0.1111 | 3 | 0.6667 | 5 | 0 | 21 | 0.2381 |
| 17_4656270 | 8 | 0.875 | 9 | 0.6667 | 9 | 0.7778 | 14 | 0.7857 | 40 | 0.775 | 11 | 0.7273 | 7 | 0.8571 | 11 | 0.7273 | 16 | 0.5 | 45 | 0.6667 |
| 17_4656280 | 8 | 0.375 | 8 | 0.375 | 10 | 0.3 | 12 | 0.1667 | 38 | 0.2895 | 16 | 0.375 | 5 | 0.6 | 10 | 0.4 | 17 | 0.0588 | 48 | 0.2917 |

| | | | | | | | | | | | | | | | | | | | | |
|---|---|---|---|---|---|---|---|---|---|---|---|---|---|---|---|---|---|---|---|---|
| 17_465628 | 9 | 0.3333 | 8 | 0.375 | 8 | 0.5 | 13 | 0.6923 | 38 | 0.5 | 15 | 0.3333 | 9 | 0.1111 | 9 | 0.4444 | 17 | 0.5882 | 50 | 0.4 |
| 17_465629 | 9 | 0 | 9 | 0.3333 | 10 | 0.2 | 12 | 0.3333 | 40 | 0.225 | 15 | 0.1333 | 9 | 0.3333 | 12 | 0.0833 | 18 | 0.2222 | 54 | 0.1852 |
| 17_465634 | 2 | 0.5 | NA | NA | NA | NA | 1 | 0 | 3 | 0.3333 | 1 | 1 | NA | NA | 1 | 1 | 1 | 0 | 3 | 0.6667 |
| 17_465652 | 8 | 0 | 3 | 0 | 1 | 1 | 1 | 0 | 13 | 0.0769 | 4 | 0 | 5 | 0.2 | 5 | 0 | 7 | 0.1429 | 21 | 0.0952 |
| 17_465680 | 10 | 0 | 12 | 0.0833 | 4 | 0.25 | 6 | 0.3333 | 32 | 0.125 | 10 | 0.3 | 9 | 0.3333 | 7 | 0.1429 | 7 | 0.2857 | 33 | 0.2727 |
| 17_465681 | 8 | 0.125 | 13 | 0.1538 | 4 | 0 | 6 | 0 | 31 | 0.0968 | 10 | 0 | 6 | 0 | 8 | 0.25 | 10 | 0.1 | 34 | 0.0882 |
| 17_465688 | 4 | 0.75 | 2 | 1 | 2 | 0.5 | 2 | 0 | 10 | 0.6 | 2 | 0.5 | 1 | 0 | 1 | 0 | 2 | 0.5 | 6 | 0.3333 |
| 17_465689 | 3 | 0.3333 | 1 | 0 | 3 | 1 | 2 | 1 | 9 | 0.6667 | 4 | 1 | 1 | 0 | NA | NA | 2 | 1 | 7 | 0.8571 |
| 17_465692 | 3 | 0.3333 | 1 | 0 | 3 | 0 | 3 | 0.3333 | 10 | 0.2 | 3 | 0.3333 | 1 | 1 | 1 | 0 | 2 | 0.5 | 7 | 0.4286 |
| 17_465692 | 3 | 0.6667 | 1 | 0 | 2 | 1 | 2 | 0.5 | 8 | 0.625 | 3 | 1 | NA | NA | 1 | 0 | 2 | 0.5 | 6 | 0.6667 |
| 17_465693 | 2 | 1 | NA | NA | NA | NA | 2 | 1 | 4 | 1 | 1 | 1 | NA | NA | 1 | 0 | NA | NA | 2 | 0.5 |
| 17_465695 | 4 | 0.5 | 1 | 0 | 2 | 0.5 | 2 | 0.5 | 9 | 0.4444 | 3 | 0.3333 | NA | NA | 1 | 0 | 2 | 0 | 6 | 0.1667 |
| 17_465695 | 4 | 0 | 1 | 0 | 2 | 0.5 | 1 | 1 | 8 | 0.25 | 3 | 0.3333 | 1 | 0 | 1 | 0 | 2 | 0.5 | 7 | 0.2857 |
| 17_465696 | 4 | 0.75 | NA | NA | 2 | 0 | 2 | 0.5 | 8 | 0.5 | 3 | 0.3333 | 1 | 0 | 1 | 0 | 2 | 1 | 7 | 0.4286 |
| 17_465696 | 4 | 0.5 | 1 | 0 | 1 | 0 | 1 | 0 | 7 | 0.2857 | 3 | 0.3333 | 1 | 0 | 1 | 1 | 2 | 0.5 | 7 | 0.4286 |
| 17_465697 | 4 | 0.75 | 1 | 0 | 2 | 0 | 2 | 0.5 | 9 | 0.4444 | 3 | 0.3333 | NA | NA | NA | NA | 2 | 0 | 5 | 0.2 |
| 17_465697 | 2 | 0.5 | 1 | 0 | 1 | 0 | 1 | 0 | 5 | 0.2 | 3 | 0.6667 | 1 | 0 | 1 | 0 | 3 | 0 | 8 | 0.25 |
| 17_465699 | 4 | 0.25 | 1 | 1 | 2 | 0.5 | 2 | 0 | 9 | 0.3333 | 3 | 0.3333 | 1 | 1 | 1 | 0 | 3 | 0 | 8 | 0.25 |
| 17_465701 | 3 | 1 | 1 | 0 | 2 | 1 | 1 | 1 | 7 | 0.8571 | 2 | 1 | NA | NA | 1 | 1 | 1 | 1 | 4 | 1 |
| 17_466135 | 2 | 0 | 3 | 0.6667 | 2 | 0 | 2 | 0 | 9 | 0.2222 | 2 | 0 | 3 | 0.6667 | 5 | 0 | 2 | 0.5 | 12 | 0.25 |
| 17_466135 | 2 | 0.5 | 2 | 0.5 | 3 | 0.3333 | 2 | 0.5 | 9 | 0.4444 | 2 | 0.5 | 3 | 0.3333 | 4 | 0 | 2 | 0 | 11 | 0.1818 |
| 17_466135 | 3 | 0.6667 | 2 | 0.5 | 3 | 0 | 1 | 1 | 9 | 0.4444 | 1 | 0 | 3 | 0.3333 | 3 | 0.6667 | 3 | 0.6667 | 10 | 0.5 |
| 17_466136 | 4 | 0.5 | 3 | 0.3333 | 3 | 1 | 2 | 0.5 | 12 | 0.5833 | 2 | 1 | 3 | 0.3333 | 5 | 0.4 | 3 | 0.6667 | 13 | 0.5385 |
| 17_466136 | 4 | 0.5 | 2 | 1 | 3 | 1 | 1 | 1 | 10 | 0.8 | 3 | 0.6667 | 3 | 0.6667 | 3 | 1 | 3 | 1 | 12 | 0.8333 |
| 17_466138 | 4 | 0.25 | 2 | 0.5 | 3 | 0 | 2 | 0 | 11 | 0.1818 | 3 | 0.3333 | 1 | 0 | 5 | 0.4 | 4 | 0 | 13 | 0.2308 |
| 17_466140 | 5 | 0 | 3 | 0.3333 | 3 | 0.3333 | 4 | 1 | 15 | 0.4 | 1 | 1 | 2 | 0.5 | 6 | 0.3333 | 3 | 0.6667 | 12 | 0.5 |
| 17_466344 | 1 | 1 | NA | NA | NA | NA | 1 | 1 | 2 | 1 | 1 | 1 | NA | NA | NA | NA | NA | NA | 1 | 1 |
| 17_466348 | NA | NA | NA | NA | NA | NA | 1 | 1 | 1 | 1 | 1 | 1 | NA | NA | NA | NA | NA | NA | 1 | 1 |
| 17_466349 | 1 | 1 | NA | NA | NA | NA | 1 | 0 | 2 | 0.5 | 1 | 1 | NA | NA | NA | NA | NA | NA | 1 | 1 |
| 17_466349 | 1 | 1 | NA | NA | NA | NA | 1 | 1 | 2 | 1 | 1 | 1 | NA | NA | NA | NA | NA | NA | 1 | 1 |
| 17_466351 | 1 | 0 | NA | NA | NA | NA | 1 | 1 | 2 | 0.5 | 1 | 1 | NA | NA | NA | NA | NA | NA | 1 | 1 |
| 17_466352 | 1 | 1 | NA | NA | NA | NA | 1 | 1 | 2 | 1 | 1 | 1 | NA | NA | NA | NA | NA | NA | 1 | 1 |
| 17_466354 | 1 | 1 | NA | NA | NA | NA | 1 | 1 | 2 | 1 | 1 | 1 | NA | NA | NA | NA | NA | NA | 1 | 1 |
| 17_466355 | 1 | 1 | NA | NA | NA | NA | 1 | 1 | 2 | 1 | 1 | 1 | NA | NA | NA | NA | NA | NA | 1 | 1 |
| 17_466356 | 1 | 1 | NA | NA | NA | NA | 1 | 1 | 2 | 1 | 1 | 1 | NA | NA | NA | NA | NA | NA | 1 | 1 |
| 17_467078 | NA | NA | NA | NA | NA | NA | 3 | 0.6667 | 3 | 0.6667 | NA | NA | NA | NA | NA | NA | 1 | 1 | 1 | 1 |
| 17_467079 | NA | NA | NA | NA | NA | NA | 3 | 1 | 3 | 1 | NA | NA | NA | NA | NA | NA | 1 | 1 | 1 | 1 |
| 17_467096 | NA | NA | NA | NA | NA | NA | 3 | 0.3333 | 3 | 0.3333 | NA | NA | 1 | 0 | NA | NA | 1 | 1 | 2 | 0.5 |
| 17_467096 | NA | NA | NA | NA | NA | NA | 3 | 1 | 3 | 1 | NA | NA | 1 | 1 | NA | NA | 1 | 1 | 2 | 1 |
| 17_467098 | NA | NA | NA | NA | NA | NA | 3 | 0.6667 | 3 | 0.6667 | NA | NA | 1 | 1 | NA | NA | 1 | 1 | 2 | 1 |
| 17_467102 | NA | NA | NA | NA | NA | NA | 3 | 0.3333 | 3 | 0.3333 | NA | NA | 2 | 1 | NA | NA | 1 | 0 | 3 | 0.6667 |
| 17_467103 | NA | NA | NA | NA | NA | NA | 3 | 0.3333 | 3 | 0.3333 | NA | NA | 2 | 0.5 | NA | NA | 1 | 0 | 3 | 0.3333 |
| 17_467105 | NA | NA | NA | NA | NA | NA | 2 | 1 | 2 | 1 | NA | NA | 2 | 0.5 | NA | NA | NA | NA | 2 | 0.5 |
| 17_467106 | NA | NA | NA | NA | NA | NA | 2 | 1 | 2 | 1 | NA | NA | 2 | 0.5 | NA | NA | 1 | 0 | 3 | 0.3333 |
| 17_467108 | NA | NA | NA | NA | NA | NA | 3 | 0.3333 | 3 | 0.3333 | NA | NA | 2 | 0.5 | NA | NA | NA | NA | 2 | 0.5 |
| 17_467227 | NA | NA | 2 | 0 | 3 | 0.6667 | NA | NA | 5 | 0.4 | NA | NA | 2 | 0.5 | 1 | 0 | NA | NA | 3 | 0.3333 |
| 17_467231 | NA | NA | 3 | 0.3333 | 2 | 1 | NA | NA | 5 | 0.6 | NA | NA | 2 | 1 | 1 | 1 | NA | NA | 3 | 1 |
| 17_467234 | 1 | 0 | 3 | 0.3333 | 2 | 0.5 | NA | NA | 6 | 0.3333 | NA | NA | 2 | 0.5 | 1 | 0 | NA | NA | 3 | 0.3333 |
| 17_467235 | 1 | 1 | 2 | 1 | 3 | 1 | NA | NA | 6 | 1 | NA | NA | 2 | 0.5 | 1 | 0 | NA | NA | 3 | 0.3333 |
| 17_467236 | 1 | 1 | 2 | 1 | 3 | 0.3333 | NA | NA | 6 | 0.6667 | NA | NA | 2 | 1 | 1 | 0 | NA | NA | 3 | 0.6667 |
| 17_467237 | 1 | 0 | 2 | 0 | 2 | 0.5 | NA | NA | 5 | 0.2 | NA | NA | 1 | 0 | 2 | 0.5 | NA | NA | 3 | 0.3333 |
| 17_467238 | 1 | 0 | 2 | 0.5 | 2 | 0.5 | NA | NA | 5 | 0.4 | NA | NA | 2 | 0 | 2 | 0.5 | NA | NA | 4 | 0.25 |
| 17_467238 | 1 | 0 | 3 | 0.3333 | 2 | 0.5 | NA | NA | 6 | 0.3333 | NA | NA | 2 | 1 | 2 | 0 | NA | NA | 4 | 0.5 |
| 17_467238 | 1 | 0 | 1 | 0 | 3 | 0.6667 | NA | NA | 5 | 0.4 | NA | NA | 2 | 0.5 | 2 | 0.5 | NA | NA | 4 | 0.5 |
| 17_467238 | 2 | 0 | 2 | 0.5 | 3 | 0 | NA | NA | 7 | 0.1429 | NA | NA | 1 | 0 | 2 | 0.5 | NA | NA | 3 | 0.3333 |
| 17_467239 | 2 | 0.5 | 2 | 0 | 2 | 0 | NA | NA | 6 | 0.1667 | NA | NA | 2 | 0.5 | 2 | 0 | NA | NA | 4 | 0.25 |
| 17_467240 | 1 | 1 | 2 | 1 | 3 | 1 | NA | NA | 6 | 1 | NA | NA | 2 | 0.5 | 2 | 0 | NA | NA | 4 | 0.25 |
| 17_467240 | 1 | 1 | 2 | 0 | 3 | 1 | NA | NA | 6 | 0.6667 | NA | NA | 2 | 0.5 | 2 | 1 | NA | NA | 4 | 0.75 |
| 17_467241 | NA | NA | 2 | 0.5 | 3 | 0.3333 | NA | NA | 5 | 0.4 | NA | NA | 2 | 0.5 | 2 | 0.5 | NA | NA | 4 | 0.5 |
| 17_467241 | NA | NA | 2 | 1 | 3 | 0.6667 | NA | NA | 5 | 0.8 | NA | NA | 1 | 1 | 2 | 1 | NA | NA | 3 | 1 |
| 17_467242 | 2 | 0 | 2 | 0 | 3 | 0.3333 | NA | NA | 7 | 0.1429 | NA | NA | 1 | 1 | 3 | 0.3333 | NA | NA | 4 | 0.5 |
| 17_467243 | NA | NA | 3 | 0 | 3 | 0.6667 | NA | NA | 6 | 0.3333 | NA | NA | 2 | 0.5 | 3 | 0.6667 | NA | NA | 5 | 0.6 |
| 17_467243 | 1 | 1 | 2 | 1 | 3 | 1 | NA | NA | 6 | 1 | NA | NA | 2 | 1 | 3 | 1 | NA | NA | 5 | 1 |
| 17_467245 | 1 | 0 | 3 | 0.3333 | 3 | 0.3333 | NA | NA | 7 | 0.2857 | NA | NA | 2 | 0 | 3 | 0.6667 | NA | NA | 5 | 0.4 |
| 17_467245 | 2 | 1 | 3 | 0.6667 | 2 | 0.5 | NA | NA | 7 | 0.7143 | NA | NA | 2 | 0 | 3 | 0.3333 | NA | NA | 5 | 0.2 |

| | | | | | | | | | | | | | | | | | | | | |
|---|---|---|---|---|---|---|---|---|---|---|---|---|---|---|---|---|---|---|---|---|
| 17_467245( | 1 | 0 | 2 | 0 | 2 | 0.5 | NA | NA | 5 | 0.2 | NA | NA | 2 | 0.5 | 1 | 0 | NA | NA | 3 | 0.3333 |
| 17_467246! | 2 | 0.5 | 1 | 1 | 3 | 1 | NA | NA | 6 | 0.8333 | NA | NA | 2 | 1 | 2 | 1 | NA | NA | 4 | 1 |
| 17_467247‹ | 2 | 0 | 2 | 0.5 | 3 | 1 | NA | NA | 7 | 0.5714 | NA | NA | 2 | 0.5 | 3 | 0.6667 | NA | NA | 5 | 0.6 |
| 17_467252 | 2 | 0.5 | 2 | 1 | 3 | 1 | NA | NA | 7 | 0.8571 | NA | NA | 2 | 1 | 3 | 1 | 1 | 1 | 6 | 1 |
| 17_467253 | 1 | 1 | 1 | 1 | 3 | 1 | 1 | 1 | 6 | 1 | NA | NA | 2 | 0.5 | 3 | 1 | 1 | 1 | 6 | 0.8333 |
| 17_467253‹ | 1 | 0 | 3 | 0 | 4 | 0.25 | 1 | 0 | 9 | 0.1111 | NA | NA | 2 | 0 | 3 | 0.6667 | 1 | 1 | 6 | 0.5 |
| 17_467253! | 1 | 0 | 4 | 0.25 | 4 | 0.25 | 1 | 0 | 10 | 0.2 | NA | NA | 2 | 1 | 3 | 0.3333 | 2 | 0.5 | 7 | 0.5714 |
| 17_467253ε | 1 | 0 | 2 | 0.5 | 3 | 1 | 1 | 0 | 7 | 0.5714 | NA | NA | 2 | 0.5 | 3 | 0 | 1 | 0 | 6 | 0.1667 |
| 17_467255 | 2 | 0 | 3 | 0.3333 | 4 | 0 | 1 | 0 | 10 | 0.1 | NA | NA | 2 | 0.5 | 3 | 0.6667 | 2 | 1 | 7 | 0.7143 |
| 17_467255‹ | 1 | 0 | 3 | 0.6667 | 4 | 0.5 | 1 | 1 | 9 | 0.5556 | NA | NA | 2 | 0.5 | 2 | 0.5 | 1 | 1 | 5 | 0.6 |
| 17_467316 | 1 | 0 | NA | NA | NA | NA | 3 | 0.3333 | 4 | 0.25 | NA | NA | 1 | 1 | NA | NA | 4 | 0.75 | 5 | 0.8 |
| 17_467317 | 1 | 0 | NA | NA | NA | NA | 4 | 0.25 | 5 | 0.2 | NA | NA | 1 | 1 | NA | NA | 3 | 0 | 4 | 0.25 |
| 17_467524 | NA | NA | 2 | 0.5 | 3 | 0 | 3 | 0.3333 | 8 | 0.25 | 1 | 0 | 1 | 0 | 3 | 0.3333 | 1 | 0 | 6 | 0.1667 |
| 17_467536 | 4 | 0 | 4 | 0 | 5 | 0 | 5 | 0.2 | 18 | 0.0556 | 5 | 0 | 4 | 0.25 | 7 | 0.1429 | 5 | 0 | 21 | 0.0952 |
| 17_467537 | 5 | 0.2 | 5 | 0.2 | 5 | 0 | 5 | 0.2 | 20 | 0.15 | 4 | 0 | 5 | 0 | 9 | 0.2222 | 5 | 0 | 23 | 0.087 |
| 17_467545 | 3 | 0.3333 | 6 | 0.1667 | 2 | 0.5 | 4 | 0.5 | 15 | 0.3333 | 5 | 0 | 2 | 0.5 | 5 | 0.2 | 3 | 0.3333 | 15 | 0.2 |
| 17_467546 | 3 | 0.3333 | 4 | 0 | 2 | 0.5 | 3 | 0.3333 | 12 | 0.25 | 4 | 0.5 | 2 | 0 | 4 | 0 | 3 | 0.3333 | 13 | 0.2308 |
| 17_467547‹ | 4 | 0 | 6 | 0 | 3 | 0.3333 | 4 | 0.25 | 17 | 0.1176 | 5 | 0.2 | 3 | 0 | 7 | 0 | 2 | 0.5 | 17 | 0.1176 |
| 17_467547 | 4 | 0.25 | 6 | 0.1667 | 2 | 0 | 3 | 0.6667 | 15 | 0.2667 | 4 | 0 | 3 | 0 | 6 | 0.1667 | 1 | 0 | 14 | 0.0714 |
| 17_467557 | 4 | 0 | 3 | 0.3333 | 2 | 0 | 2 | 1 | 11 | 0.2727 | 2 | 0 | 3 | 0 | 6 | 0.3333 | NA | NA | 11 | 0.1818 |
| 17_467558 | 3 | 0.6667 | 4 | 0 | 2 | 0.5 | 2 | 0 | 11 | 0.2727 | 4 | 0.25 | 2 | 0 | 4 | 0.5 | NA | NA | 10 | 0.3 |
| 17_467560 | 3 | 0 | 3 | 0 | 1 | 0 | 2 | 0.5 | 9 | 0.1111 | 3 | 0 | 2 | 0.5 | 4 | 0.25 | NA | NA | 9 | 0.2222 |
| 17_467561 | 4 | 0.5 | 2 | 0.5 | 2 | 0 | 1 | 0 | 9 | 0.3333 | 4 | 0.25 | 3 | 0 | 4 | 0.5 | NA | NA | 11 | 0.2727 |
| 17_467561 | 2 | 1 | 2 | 1 | 2 | 0.5 | 1 | 1 | 7 | 0.8571 | 4 | 1 | 2 | 1 | 2 | 1 | NA | NA | 8 | 1 |
| 17_467572 | 4 | 0.25 | 2 | 1 | 1 | 0 | 2 | 0.5 | 9 | 0.4444 | 4 | 0.5 | 3 | 0.3333 | 2 | 0.5 | NA | NA | 9 | 0.4444 |
| 17_467573 | 3 | 0 | 2 | 0.5 | 1 | 0 | 2 | 1 | 8 | 0.375 | 3 | 0 | 3 | 1 | 1 | 1 | NA | NA | 7 | 0.5714 |
| 17_467575 | 4 | 0.25 | 1 | 0 | 1 | 1 | 2 | 0 | 8 | 0.25 | 3 | 0 | 2 | 0.5 | 2 | 0 | NA | NA | 7 | 0.1429 |
| 17_467575 | 4 | 0.5 | 2 | 0.5 | 1 | 0 | 2 | 0.5 | 9 | 0.4444 | 3 | 0.6667 | 1 | 0 | 3 | 1 | NA | NA | 7 | 0.7143 |
| 17_467576 | 4 | 1 | 2 | 0.5 | 1 | 0 | 1 | 1 | 8 | 0.75 | 3 | 0.6667 | 2 | 1 | 3 | 0.6667 | NA | NA | 8 | 0.75 |
| 17_467576 | 3 | 1 | 2 | 1 | 1 | 1 | 2 | 0.5 | 8 | 0.875 | 3 | 1 | 2 | 1 | 2 | 1 | NA | NA | 7 | 1 |
| 17_467578 | 2 | 1 | 1 | 0 | NA | NA | NA | NA | 3 | 0.6667 | 1 | 0 | 1 | 1 | 2 | 0.5 | NA | NA | 4 | 0.5 |
| 17_467578 | 3 | 1 | 2 | 1 | 1 | 1 | 2 | 1 | 8 | 1 | 3 | 1 | 2 | 1 | 3 | 1 | NA | NA | 8 | 1 |
| 17_467579 | 3 | 0.6667 | 2 | 0.5 | 1 | 0 | 2 | 1 | 8 | 0.625 | 2 | 1 | 2 | 0.5 | 4 | 0.25 | NA | NA | 8 | 0.5 |
| 17_467580 | 3 | 0.6667 | NA | NA | 1 | 1 | 1 | 1 | 5 | 0.8 | 3 | 1 | 1 | 1 | 1 | 1 | NA | NA | 5 | 1 |
| 17_467581 | 2 | 0.5 | 1 | 0 | 1 | 0 | 2 | 0 | 6 | 0.1667 | 2 | 0.5 | 2 | 0.5 | 1 | 0 | NA | NA | 5 | 0.4 |
| 17_467585 | 2 | 0 | 2 | 0 | 1 | 0 | 1 | 1 | 6 | 0.1667 | 3 | 0.6667 | 2 | 0.5 | 2 | 0 | NA | NA | 7 | 0.4286 |
| 17_467714 | 1 | 0 | 3 | 0.3333 | NA | NA | 1 | 0 | 5 | 0.2 | NA | NA | 1 | 0 | 4 | 0.25 | NA | NA | 5 | 0.2 |
| 17_467717 | 1 | 1 | 5 | 0.8 | NA | NA | 1 | 1 | 7 | 0.8571 | NA | NA | 1 | 1 | 4 | 1 | NA | NA | 5 | 1 |
| 17_467719 | 1 | 0 | 5 | 0.2 | NA | NA | 1 | 0 | 7 | 0.1429 | NA | NA | NA | NA | 3 | 0.3333 | NA | NA | 3 | 0.3333 |
| 17_467781 | 3 | 0.3333 | 6 | 0 | 1 | 0 | 2 | 0 | 12 | 0.0833 | 3 | 0.3333 | 2 | 0 | 5 | 0 | NA | NA | 10 | 0.1 |
| 17_467781 | 3 | 0.6667 | 6 | 0.1667 | 1 | 1 | 2 | 1 | 12 | 0.5 | 2 | 0 | 2 | 0 | 6 | 0.3333 | NA | NA | 10 | 0.2 |
| 17_467782 | 2 | 0 | 5 | 0.4 | 1 | 0 | 2 | 1 | 10 | 0.4 | 3 | 0 | 2 | 0 | 6 | 0.1667 | NA | NA | 11 | 0.0909 |
| 17_467782 | 2 | 0.5 | 5 | 0.6 | 1 | 1 | 1 | 1 | 9 | 0.6667 | 3 | 0.6667 | 1 | 0 | 5 | 0.6 | NA | NA | 9 | 0.5556 |
| 17_467783 | 2 | 1 | 5 | 1 | 1 | 1 | 1 | 1 | 9 | 1 | 2 | 1 | 2 | 1 | 4 | 0.75 | NA | NA | 8 | 0.875 |
| 17_467783 | 3 | 0.3333 | 6 | 0 | 1 | 0 | 1 | 0 | 11 | 0.0909 | 3 | 0 | 2 | 0 | 5 | 0.2 | NA | NA | 10 | 0.1 |
| 17_467784 | 3 | 0 | 6 | 0 | 1 | 1 | 1 | 0 | 11 | 0.0909 | 3 | 0.3333 | 1 | 0 | 5 | 0.4 | NA | NA | 9 | 0.3333 |
| 17_467785 | 3 | 0.6667 | 3 | 0 | 1 | 0 | 1 | 0 | 8 | 0.25 | 2 | 0 | 1 | 1 | 4 | 1 | NA | NA | 7 | 0.7143 |
| 17_468116 | 5 | 0.8 | 5 | 0.8 | 3 | 0.6667 | 5 | 0.8 | 18 | 0.7778 | 2 | 1 | 6 | 0.6667 | 5 | 0.6 | 2 | 1 | 15 | 0.7333 |
| 17_468116 | 5 | 0.2 | 5 | 0 | 5 | 0.2 | 7 | 0.1429 | 22 | 0.1364 | 2 | 0 | 7 | 0.1429 | 7 | 0 | 3 | 0.3333 | 19 | 0.1053 |
| 17_468117 | 8 | 0.125 | 7 | 0.2857 | 4 | 0 | 7 | 0 | 26 | 0.1154 | 2 | 0.5 | 9 | 0.1111 | 8 | 0.125 | 4 | 0.25 | 23 | 0.1739 |
| 17_468305 | 1 | 0 | NA | NA | 1 | 0 | 2 | 0.5 | 4 | 0.25 | 1 | 0 | NA | NA | 1 | 0 | 3 | 0.3333 | 5 | 0.2 |
| 17_468308 | 3 | 0 | 1 | 0 | 2 | 0 | 3 | 0.3333 | 9 | 0.1111 | 2 | 0 | 6 | 0.1667 | 2 | 0.5 | 4 | 0.75 | 14 | 0.3571 |
| 17_468488 | NA | NA | NA | NA | NA | NA | 1 | 1 | 1 | 1 | NA | NA | 1 | 0 | 1 | 1 | NA | NA | 2 | 0.5 |
| 17_468488 | NA | NA | NA | NA | NA | NA | 1 | 1 | 1 | 1 | NA | NA | 1 | 0 | 1 | 1 | NA | NA | 2 | 0.5 |
| 17_468492 | 1 | 0 | NA | NA | 1 | 0 | 3 | 0.3333 | 5 | 0.2 | 1 | 0 | 4 | 0 | 2 | 0.5 | 1 | 0 | 8 | 0.125 |
| 17_468494 | 1 | 0 | NA | NA | 2 | 0 | 3 | 0.6667 | 6 | 0.3333 | 1 | 0 | 4 | 0.75 | 2 | 0.5 | 1 | 1 | 8 | 0.625 |
| 17_468542 | NA | NA | NA | NA | 2 | 0 | 3 | 0.6667 | 5 | 0.4 | NA | NA | 3 | 0.3333 | 1 | 0 | 1 | 1 | 5 | 0.4 |
| 17_468543 | NA | NA | NA | NA | 2 | 0 | 3 | 0.6667 | 5 | 0.4 | NA | NA | 2 | 0.5 | 1 | 0 | 1 | 1 | 4 | 0.5 |
| 17_468544 | NA | NA | NA | NA | 1 | 0 | 2 | 0.5 | 3 | 0.3333 | NA | NA | 3 | 0.3333 | 1 | 0 | 1 | 1 | 5 | 0.4 |
| 17_468591 | 4 | 0.25 | 1 | 0 | 2 | 0 | 1 | 0 | 8 | 0.125 | 5 | 0.2 | 1 | 0 | 6 | 0.1667 | 1 | 0 | 13 | 0.1538 |
| 17_468593 | 7 | 0 | 4 | 0.5 | 4 | 0 | 4 | 0 | 19 | 0.1053 | 4 | 0 | 5 | 0 | 9 | 0.1111 | 2 | 0 | 20 | 0.05 |
| 17_468594 | 7 | 0.2857 | 2 | 0.5 | 4 | 0 | 6 | 0.5 | 19 | 0.3158 | 3 | 0.3333 | 5 | 0.2 | 6 | 0 | 2 | 0 | 16 | 0.125 |
| 17_468596 | 6 | 0 | 4 | 0.5 | 3 | 0 | 3 | 0 | 16 | 0.125 | 3 | 0.6667 | 4 | 0.25 | 8 | 0.25 | 3 | 0.3333 | 18 | 0.3333 |
| 17_468597 | 8 | 0.5 | 4 | 0.75 | 3 | 0.6667 | 4 | 0.25 | 19 | 0.5263 | 4 | 0.5 | 4 | 0.75 | 8 | 0.75 | 2 | 0.5 | 18 | 0.6667 |
| 17_468598 | 8 | 1 | 4 | 0.75 | 3 | 0.6667 | 4 | 0.75 | 19 | 0.8421 | 3 | 1 | 5 | 0.4 | 6 | 0.6667 | 2 | 1 | 16 | 0.6875 |

| | | | | | | | | | | | | | | | | | | | | |
|---|---|---|---|---|---|---|---|---|---|---|---|---|---|---|---|---|---|---|---|---|
| 17_4685992 | 5 | 0.4 | 2 | 0 | 1 | 0 | NA | NA | 8 | 0.25 | 4 | 0 | 2 | 0.5 | 5 | 0.4 | 2 | 0 | 13 | 0.2308 |
| 17_4686253 | 5 | 0.2 | 3 | 0.3333 | NA | NA | 3 | 0.3333 | 11 | 0.2727 | 4 | 0.25 | 5 | 0 | 6 | 0 | 2 | 0 | 17 | 0.0588 |
| 17_4686614 | NA | NA | NA | NA | 1 | 0 | 1 | 1 | 2 | 0.5 | NA | NA | 2 | 0 | NA | NA | 1 | 1 | 3 | 0.3333 |
| 17_4686651 | 1 | 1 | 1 | 1 | 1 | 1 | 1 | 0 | 4 | 0.75 | NA | NA | 1 | 0 | NA | NA | 1 | 1 | 2 | 0.5 |
| 17_4686664 | 1 | 1 | 2 | 0 | NA | NA | NA | NA | 3 | 0.3333 | 1 | 0 | 3 | 0.6667 | 3 | 0.3333 | 1 | 0 | 8 | 0.375 |
| 17_4686705 | 1 | 0 | 1 | 0 | 1 | 1 | 2 | 0 | 5 | 0.2 | NA | NA | 2 | 1 | NA | NA | NA | NA | 2 | 1 |
| 17_4686707 | 2 | 0.5 | 1 | 0 | 1 | 0 | 2 | 0 | 6 | 0.1667 | NA | NA | 2 | 0.5 | 1 | 0 | 1 | 0 | 4 | 0.25 |
| 17_4686724 | 1 | 1 | 1 | 1 | 1 | 1 | 2 | 1 | 5 | 1 | NA | NA | 2 | 0.5 | 2 | 1 | 1 | 0 | 5 | 0.6 |
| 17_4686725 | 1 | 1 | 1 | 1 | 1 | 0 | 2 | 1 | 5 | 0.8 | NA | NA | 2 | 0.5 | 2 | 0 | NA | NA | 4 | 0.25 |
| 17_4686726 | 1 | 1 | NA | NA | 1 | 1 | 2 | 0.5 | 4 | 0.75 | NA | NA | 1 | 0 | 2 | 1 | 1 | 0 | 4 | 0.5 |
| 17_4686729 | 1 | 0 | 1 | 1 | 1 | 0 | 3 | 0 | 6 | 0.1667 | NA | NA | 2 | 0.5 | 2 | 0 | 1 | 0 | 5 | 0.2 |
| 17_4686733 | 1 | 1 | NA | NA | 1 | 1 | 3 | 0.6667 | 5 | 0.8 | NA | NA | 2 | 0.5 | 2 | 0.5 | 1 | 0 | 5 | 0.4 |
| 17_4686741 | 1 | 1 | 1 | 1 | NA | NA | 3 | 1 | 5 | 1 | NA | NA | 1 | 0 | 2 | 0.5 | 1 | 0 | 4 | 0.25 |
| 17_4686771 | 1 | 1 | 1 | 0 | 1 | 0 | 3 | 0.3333 | 6 | 0.3333 | NA | NA | 3 | 0 | 2 | 0 | 1 | 1 | 6 | 0.1667 |
| 17_4686814 | 1 | 0 | 1 | 0 | 1 | 1 | 4 | 0 | 7 | 0.1429 | NA | NA | 3 | 0.6667 | 4 | 0 | 1 | 0 | 8 | 0.25 |
| 17_4686818 | 2 | 0.5 | 2 | 0 | 1 | 0 | 4 | 0 | 9 | 0.1111 | NA | NA | 3 | 0 | 4 | 0.25 | 1 | 0 | 8 | 0.125 |
| 17_4687604 | 1 | 0 | NA | NA | 2 | 0.5 | 2 | 0.5 | 5 | 0.4 | NA | NA | 1 | 1 | 1 | 1 | NA | NA | 2 | 1 |
| 17_4696103 | 8 | 0.375 | 10 | 0.4 | 10 | 0.2 | 3 | 0.3333 | 31 | 0.3226 | 1 | 0 | 6 | 0.3333 | 11 | 0.1818 | 6 | 0.1667 | 24 | 0.2083 |
| 17_4696537 | 1 | 1 | NA | NA | NA | NA | NA | NA | 1 | 1 | NA | NA | 1 | 1 | NA | NA | 1 | 0 | 2 | 0.5 |
| 17_4697839 | 2 | 1 | 6 | 0.6667 | 2 | 1 | 2 | 1 | 12 | 0.8333 | 3 | 1 | 3 | 1 | 8 | 0.75 | 2 | 1 | 16 | 0.875 |
| 17_4697840 | 2 | 1 | 6 | 0.6667 | 1 | 1 | 2 | 0.5 | 11 | 0.7273 | 4 | 0.75 | 3 | 0.3333 | 6 | 0.6667 | 2 | 0.5 | 15 | 0.6 |
| 17_4697847 | 2 | 0 | 6 | 0.3333 | 1 | 1 | 1 | 1 | 10 | 0.4 | 3 | 0.3333 | 3 | 0.6667 | 7 | 0.2857 | 2 | 1 | 15 | 0.4667 |
| 17_4697859 | 1 | 1 | 6 | 1 | 2 | 1 | 1 | 1 | 10 | 1 | 3 | 1 | 3 | 1 | 6 | 0.8333 | 2 | 0.5 | 14 | 0.8571 |
| 17_4697863 | 2 | 1 | 5 | 0.6 | 2 | 1 | 2 | 0.5 | 11 | 0.7273 | 2 | 1 | 2 | 0.5 | 7 | 0.4286 | 1 | 0 | 12 | 0.5 |
| 17_4697874 | 2 | 0.5 | 3 | 1 | 2 | 1 | 1 | 1 | 8 | 0.875 | 2 | 1 | 1 | 1 | 6 | 1 | 2 | 1 | 11 | 1 |
| 17_4697885 | 2 | 0 | 5 | 0.2 | 1 | 0 | 2 | 0 | 10 | 0.1 | 4 | 0.25 | 2 | 0 | 8 | 0 | 1 | 0 | 15 | 0.0667 |
| 17_4697887 | 2 | 1 | 5 | 0.8 | 2 | 1 | 2 | 0.5 | 11 | 0.8182 | 2 | 0.5 | 3 | 0.3333 | 7 | 1 | 2 | 1 | 14 | 0.7857 |
| 17_4697890 | 2 | 0.5 | 6 | 0 | 2 | 0 | 1 | 1 | 11 | 0.1818 | 4 | 0 | 3 | 0 | 8 | 0.125 | 2 | 0 | 17 | 0.0588 |
| 17_4698522 | 1 | 1 | 1 | 0 | NA | NA | NA | NA | 2 | 0.5 | 2 | 0 | 1 | 0 | 1 | 1 | 1 | 0 | 5 | 0.2 |
| 17_4698550 | 21 | 0.1905 | 33 | 0.2424 | 26 | 0.0769 | 26 | 0.1154 | 106 | 0.1604 | 31 | 0.0645 | 20 | 0.1 | 25 | 0.12 | 25 | 0.04 | 101 | 0.0792 |
| 17_4698571 | 17 | 0.3529 | 19 | 0.3158 | 21 | 0.2857 | 20 | 0.25 | 77 | 0.2987 | 20 | 0.2 | 13 | 0.1538 | 20 | 0.2 | 17 | 0.0588 | 70 | 0.1571 |
| 17_4698589 | 22 | 0.8182 | 28 | 0.8929 | 31 | 0.7419 | 27 | 0.9259 | 108 | 0.8426 | 30 | 0.9 | 20 | 0.8 | 25 | 1 | 24 | 0.875 | 99 | 0.899 |
| 17_4698592 | 18 | 0.6111 | 19 | 0.5789 | 23 | 0.7391 | 22 | 0.6818 | 82 | 0.6585 | 22 | 0.6818 | 14 | 0.9286 | 21 | 0.5714 | 19 | 0.5789 | 76 | 0.6711 |
| 17_4698593 | 24 | 0.1667 | 24 | 0.1667 | 29 | 0.2069 | 26 | 0.3462 | 103 | 0.2233 | 28 | 0.1786 | 18 | 0.0556 | 25 | 0.24 | 24 | 0.2083 | 95 | 0.1789 |
| 17_4699385 | 13 | 0.3077 | 21 | 0.2381 | 15 | 0.2667 | 16 | 0.3125 | 65 | 0.2769 | 16 | 0.1875 | 13 | 0.1538 | 19 | 0.1579 | 14 | 0.5 | 62 | 0.2419 |
| 17_4699395 | 11 | 0 | 18 | 0.0556 | 18 | 0 | 13 | 0.1538 | 60 | 0.05 | 14 | 0 | 14 | 0.0714 | 17 | 0.0588 | 10 | 0.1 | 55 | 0.0545 |
| 17_4699423 | 12 | 0.25 | 16 | 0.0625 | 14 | 0.0714 | 15 | 0.1333 | 57 | 0.1228 | 10 | 0.1 | 13 | 0.3077 | 25 | 0.08 | 11 | 0.1818 | 59 | 0.1525 |
| 17_4699425 | 8 | 0 | 8 | 0 | 11 | 0.0909 | 11 | 0.0909 | 38 | 0.0526 | 8 | 0 | 9 | 0.2222 | 20 | 0.05 | 10 | 0 | 47 | 0.0638 |
| 17_4699433 | 13 | 0 | 15 | 0.0667 | 11 | 0.0909 | 12 | 0.0833 | 51 | 0.0588 | 10 | 0 | 12 | 0.0833 | 21 | 0 | 8 | 0.25 | 51 | 0.0588 |
| 17_4699434 | 13 | 0.3846 | 17 | 0.4706 | 12 | 0.75 | 13 | 0.4615 | 55 | 0.5091 | 9 | 0.3333 | 14 | 0.4286 | 22 | 0.5909 | 10 | 0.4 | 55 | 0.4727 |
| 17_4699436 | 10 | 0 | 14 | 0.2143 | 14 | 0.1429 | 15 | 0.1333 | 53 | 0.1321 | 10 | 0 | 16 | 0.0625 | 22 | 0.0909 | 11 | 0.1818 | 59 | 0.0847 |
| 17_4699452 | 12 | 0.0833 | 17 | 0.2353 | 14 | 0.6429 | 15 | 0.4667 | 58 | 0.3621 | 8 | 0.375 | 14 | 0.3571 | 22 | 0.4091 | 9 | 0.3333 | 53 | 0.3774 |
| 17_4699462 | 12 | 0.5833 | 17 | 0.5882 | 14 | 0.5714 | 14 | 0.4286 | 57 | 0.5439 | 9 | 0.5556 | 13 | 0.3846 | 22 | 0.4091 | 11 | 0.3636 | 55 | 0.4182 |
| 17_4702289 | 6 | 0 | 2 | 0.5 | 1 | 0 | 2 | 0 | 11 | 0.0909 | 3 | 0 | 3 | 0.3333 | 2 | 0.5 | 2 | 0 | 10 | 0.2 |
| 17_4702302 | 6 | 0 | 3 | 0.3333 | 1 | 1 | 2 | 0 | 12 | 0.1667 | 3 | 1 | 4 | 0.5 | 2 | 0 | 2 | 0.5 | 11 | 0.5455 |
| 17_4702307 | 6 | 0.8333 | 2 | 1 | 1 | 1 | 4 | 1 | 13 | 0.9231 | 4 | 1 | 4 | 1 | 2 | 1 | 3 | 1 | 13 | 1 |
| 17_4702318 | 7 | 0.5714 | 3 | 0 | 2 | 0.5 | 4 | 0.25 | 16 | 0.375 | 4 | 0.25 | 4 | 0 | 3 | 0.3333 | 2 | 0 | 13 | 0.1538 |
| 17_4702319 | 6 | 0.1667 | 3 | 0 | 2 | 0 | 2 | 0 | 13 | 0.0769 | 4 | 0.25 | 3 | 0 | 3 | 0 | 3 | 0 | 13 | 0.0769 |
| 17_4702323 | 6 | 0.3333 | 2 | 0 | 2 | 0 | 5 | 0 | 15 | 0.1333 | 5 | 0.2 | 4 | 0.25 | 3 | 0.3333 | 3 | 0 | 15 | 0.2 |
| 17_4704698 | 3 | 0 | 7 | 0 | 8 | 0.125 | 8 | 0.125 | 26 | 0.0769 | 4 | 0.25 | 7 | 0 | 4 | 0 | 5 | 0 | 20 | 0.05 |
| 17_4704710 | 4 | 1 | 7 | 1 | 9 | 1 | 6 | 0.8333 | 26 | 0.9615 | 3 | 1 | 6 | 1 | 7 | 1 | 6 | 1 | 22 | 1 |
| 17_4704715 | 3 | 0 | 5 | 0.2 | 6 | 0 | 5 | 0.4 | 19 | 0.1579 | 3 | 0 | 6 | 0 | 8 | 0.25 | 3 | 0 | 20 | 0.1 |
| 17_4704716 | 4 | 1 | 6 | 0.8333 | 7 | 0.8571 | 6 | 1 | 23 | 0.913 | 3 | 0.6667 | 4 | 1 | 5 | 0.8 | 5 | 0.8 | 17 | 0.8235 |
| 17_4704717 | 3 | 1 | 6 | 0.8333 | 9 | 0.8889 | 5 | 0.6 | 23 | 0.8261 | 3 | 0.3333 | 6 | 0.8333 | 5 | 0.8 | 5 | 0.8 | 19 | 0.7368 |
| 17_4704725 | 3 | 1 | 8 | 0.75 | 8 | 0.875 | 8 | 0.875 | 27 | 0.8519 | 4 | 1 | 6 | 1 | 7 | 1 | 6 | 1 | 23 | 1 |
| 17_4704753 | 4 | 1 | 6 | 0.6667 | 8 | 0.25 | 6 | 0.6667 | 24 | 0.5833 | 4 | 0.5 | 6 | 0.6667 | 6 | 0.8333 | 6 | 0.6667 | 22 | 0.6818 |
| 17_4704756 | 3 | 0.6667 | 5 | 0.6 | 8 | 1 | 4 | 0.5 | 20 | 0.75 | 3 | 0.6667 | 4 | 1 | 5 | 0.8 | 4 | 1 | 16 | 0.875 |
| 17_4704758 | 4 | 0 | 6 | 0 | 7 | 0.4286 | 6 | 0.1667 | 23 | 0.1739 | 4 | 0 | 4 | 0.25 | 5 | 0 | 6 | 0.3333 | 19 | 0.1579 |
| 17_4704771 | 3 | 0.6667 | 4 | 1 | 6 | 0.6667 | 4 | 0.75 | 17 | 0.7647 | 1 | 1 | 4 | 0.5 | 6 | 0.3333 | 4 | 1 | 15 | 0.6 |
| 17_4708411 | 1 | 1 | NA | NA | 1 | 0 | NA | NA | 2 | 0.5 | 1 | 0 | 1 | 0 | 1 | 0 | 4 | 0.25 | 7 | 0.1429 |
| 17_4708412 | 2 | 0.5 | NA | NA | 1 | 1 | NA | NA | 3 | 0.6667 | 1 | 0 | 1 | 1 | 1 | 0 | 4 | 0.5 | 7 | 0.4286 |
| 17_4708413 | 2 | 0.5 | NA | NA | 1 | 1 | NA | NA | 3 | 0.6667 | NA | NA | NA | NA | 1 | 1 | 4 | 1 | 5 | 1 |
| 17_4708415 | 1 | 1 | NA | NA | 1 | 1 | NA | NA | 2 | 1 | 1 | 1 | 1 | 1 | 1 | 1 | 4 | 1 | 7 | 1 |
| 17_4708427 | 2 | 0 | NA | NA | 1 | 1 | NA | NA | 3 | 0.3333 | 1 | 0 | 1 | 0 | 1 | 0 | 5 | 0.2 | 8 | 0.125 |
| 17_4708507 | 3 | 0.3333 | NA | NA | NA | NA | 1 | 1 | 4 | 0.5 | 1 | 1 | 1 | 1 | 2 | 0 | 3 | 0.6667 | 7 | 0.5714 |

| | | | | | | | | | | | | | | | | | | | | |
|---|---|---|---|---|---|---|---|---|---|---|---|---|---|---|---|---|---|---|---|---|
| 17_4708522 | 2 | 0.5 | 1 | 1 | NA | NA | NA | NA | 3 | 0.6667 | 1 | 0 | 1 | 1 | 2 | 1 | 3 | 0.3333 | 7 | 0.5714 |
| 17_4708560 | 3 | 0.6667 | 5 | 0.2 | 1 | 0 | 3 | 0.3333 | 12 | 0.3333 | 4 | 0.25 | 1 | 0 | 4 | 0.5 | 4 | 0.25 | 13 | 0.3077 |
| 17_4708577 | 3 | 0.3333 | 7 | 0 | 1 | 0 | 3 | 0.3333 | 14 | 0.1429 | 4 | 0.25 | 2 | 0 | 6 | 0.3333 | 3 | 0 | 15 | 0.2 |
| 17_4708587 | 2 | 0.5 | 4 | 0.25 | 1 | 0 | 2 | 0 | 9 | 0.2222 | 3 | 0 | 1 | 0 | 3 | 0.3333 | 2 | 1 | 9 | 0.3333 |
| 17_4708607 | 5 | 0 | 11 | 0 | 4 | 0.25 | 3 | 0.6667 | 23 | 0.1304 | 5 | 0.2 | 8 | 0 | 9 | 0.1111 | 3 | 0 | 25 | 0.08 |
| 17_4708631 | 4 | 0 | 5 | 0 | 4 | 0.25 | NA | NA | 13 | 0.0769 | 2 | 0 | 2 | 0.5 | 7 | 0 | 1 | 0 | 12 | 0.0833 |
| 17_4708635 | 7 | 0.8571 | 8 | 1 | 3 | 1 | 3 | 1 | 21 | 0.9524 | 5 | 0.8 | 9 | 0.7778 | 8 | 0.875 | 2 | 0.5 | 24 | 0.7917 |
| 17_4708652 | 10 | 0 | 12 | 0.3333 | 5 | 0.2 | 3 | 0.3333 | 30 | 0.2 | 5 | 0.2 | 9 | 0.1111 | 8 | 0.25 | 5 | 0 | 27 | 0.1481 |
| 17_4708657 | 8 | 0.25 | 10 | 0.1 | 4 | 0 | 4 | 0.5 | 26 | 0.1923 | 4 | 0 | 9 | 0 | 8 | 0.125 | 4 | 0.25 | 25 | 0.08 |
| 17_4710937 | 2 | 0 | 3 | 0 | 7 | 0.2857 | 5 | 0 | 17 | 0.1176 | 3 | 0 | 2 | 1 | 4 | 0 | 5 | 0.2 | 14 | 0.2143 |
| 17_4710942 | 2 | 0.5 | 3 | 0.3333 | 4 | 0.75 | 4 | 0.75 | 13 | 0.6154 | 2 | 0.5 | 1 | 1 | 3 | 0.3333 | 5 | 0.4 | 11 | 0.4545 |
| 17_4711022 | 2 | 0 | 2 | 0.5 | 5 | 0.2 | 3 | 0.3333 | 12 | 0.25 | 3 | 0.3333 | 1 | 0 | 4 | 0.75 | 4 | 0.25 | 12 | 0.4167 |
| 17_4711100 | 1 | 1 | 2 | 0.5 | 3 | 0 | 3 | 0.3333 | 9 | 0.3333 | 2 | 0 | 1 | 1 | 3 | 0.3333 | 3 | 0 | 9 | 0.2222 |
| 17_4711156 | 1 | 1 | 1 | 1 | 3 | 1 | 3 | 0.6667 | 8 | 0.875 | 2 | 1 | 1 | 1 | 2 | 0.5 | 3 | 1 | 8 | 0.875 |
| 17_4711157 | 1 | 0 | 1 | 1 | 5 | 0.8 | 3 | 0.6667 | 10 | 0.7 | 1 | 1 | 1 | 1 | 2 | 0.5 | 3 | 1 | 7 | 0.8571 |
| 17_4711160 | 1 | 1 | 1 | 1 | 2 | 0.5 | 2 | 0.5 | 6 | 0.6667 | 1 | 1 | 1 | 1 | 2 | 0.5 | 3 | 1 | 7 | 0.8571 |
| 17_4711194 | 1 | 0 | 1 | 0 | 3 | 0.3333 | 2 | 0.5 | 7 | 0.2857 | 2 | 0.5 | 1 | 0 | 1 | 1 | 3 | 0 | 7 | 0.2857 |
| 17_4711201 | 1 | 1 | 1 | 1 | 4 | 1 | 2 | 1 | 8 | 1 | 2 | 1 | 1 | 1 | 1 | 1 | 3 | 1 | 7 | 1 |
| 17_4711210 | 1 | 1 | 1 | 1 | 5 | 0.8 | 2 | 0.5 | 9 | 0.7778 | 2 | 0.5 | 1 | 1 | 1 | 1 | 1 | 1 | 5 | 0.8 |
| 17_4711211 | 1 | 0 | 1 | 0 | 4 | 0 | 2 | 0.5 | 8 | 0.125 | 2 | 0.5 | 1 | 0 | 1 | 0 | 1 | 0 | 5 | 0.2 |
| 17_4711216 | 1 | 1 | NA | NA | 2 | 0.5 | 3 | 1 | 6 | 0.8333 | 1 | 0 | 1 | 0 | 1 | 1 | 2 | 0.5 | 5 | 0.4 |
| 17_4711217 | 1 | 0 | 1 | 0 | 3 | 0 | 3 | 0.3333 | 8 | 0.125 | 2 | 0.5 | 1 | 0 | 1 | 0 | 2 | 0 | 6 | 0.1667 |
| 17_4711224 | 1 | 1 | 1 | 1 | 2 | 0.5 | 3 | 0.3333 | 7 | 0.5714 | 2 | 0.5 | 1 | 0 | 1 | 1 | 2 | 1 | 6 | 0.6667 |
| 17_4711229 | 1 | 1 | 1 | 1 | 4 | 0.5 | 3 | 0.3333 | 9 | 0.5556 | 2 | 0.5 | 1 | 0 | 1 | 1 | 2 | 0.5 | 6 | 0.5 |
| 17_4711243 | 1 | 0 | NA | NA | 3 | 0 | 2 | 0.5 | 6 | 0.1667 | 2 | 0.5 | 1 | 0 | NA | NA | 2 | 0.5 | 5 | 0.4 |
| 17_4711245 | NA | NA | 1 | 1 | 3 | 0.6667 | 3 | 0.3333 | 7 | 0.5714 | 2 | 1 | 1 | 1 | NA | NA | 1 | 1 | 4 | 1 |
| 17_4711249 | 1 | 0 | NA | NA | 4 | 0.5 | 3 | 0.6667 | 8 | 0.5 | NA | NA | NA | NA | NA | NA | 2 | 1 | 2 | 1 |
| 17_4711253 | 1 | 0 | 1 | 0 | 2 | 0.5 | 3 | 0.6667 | 7 | 0.4286 | 1 | 1 | NA | NA | NA | NA | 2 | 0.5 | 3 | 0.6667 |
| 17_4711262 | 1 | 1 | 1 | 1 | 3 | 0.3333 | 3 | 0.3333 | 8 | 0.5 | 1 | 0 | NA | NA | NA | NA | 1 | 1 | 2 | 0.5 |
| 17_4716718 | 1 | 0 | 2 | 0 | 1 | 0 | 3 | 0.3333 | 7 | 0.1429 | 1 | 0 | 3 | 0.3333 | 1 | 0 | 2 | 0 | 7 | 0.1429 |
| 17_4720225 | 5 | 0.8 | 3 | 0.3333 | 6 | 0.6667 | 4 | 0.75 | 18 | 0.6667 | 5 | 1 | 3 | 1 | 3 | 0.3333 | 5 | 1 | 16 | 0.875 |
| 17_4720240 | 7 | 0.2857 | 4 | 0.25 | 6 | 0.3333 | 5 | 0.2 | 22 | 0.2727 | 7 | 0 | 4 | 0 | 4 | 0.25 | 4 | 0.25 | 19 | 0.1053 |
| 17_4720244 | 6 | 0.6667 | 2 | 1 | 6 | 0.5 | 5 | 0.4 | 19 | 0.5789 | 6 | 1 | 5 | 0.6 | 4 | 0.75 | 3 | 1 | 18 | 0.8333 |
| 17_4720250 | 8 | 1 | 4 | 0.75 | 5 | 1 | 7 | 1 | 24 | 0.9583 | 7 | 1 | 7 | 0.8571 | 4 | 1 | 5 | 1 | 23 | 0.9565 |
| 17_4720251 | 8 | 0.5 | 4 | 0.75 | 6 | 0.8333 | 7 | 0.8571 | 25 | 0.72 | 7 | 0.8571 | 6 | 0.6667 | 4 | 0.75 | 5 | 1 | 22 | 0.8182 |
| 17_4720254 | 5 | 0.8 | 2 | 0.5 | 3 | 1 | 2 | 1 | 12 | 0.8333 | 1 | 0 | 4 | 0.5 | 1 | 1 | NA | NA | 6 | 0.5 |
| 17_4720254 | 8 | 0.875 | 4 | 0.75 | 6 | 0.8333 | 8 | 0.875 | 26 | 0.8462 | 7 | 1 | 6 | 0.8333 | 4 | 0.75 | 5 | 0.8 | 22 | 0.8636 |
| 17_4720259 | 8 | 0.375 | 3 | 0.3333 | 6 | 0.3333 | 8 | 0.25 | 25 | 0.32 | 9 | 0.2222 | 8 | 0.125 | 3 | 0 | 5 | 0.4 | 25 | 0.2 |
| 17_4720262 | 7 | 0.4286 | 3 | 0.3333 | 4 | 0.75 | 8 | 0.625 | 22 | 0.5455 | 9 | 0.6667 | 7 | 0.5714 | 4 | 1 | 5 | 0.2 | 25 | 0.6 |
| 17_4720277 | 1 | 1 | 1 | 1 | 1 | 1 | NA | NA | 3 | 1 | 3 | 1 | 3 | 1 | 4 | 1 | 2 | 1 | 12 | 1 |
| 17_4723964 | 7 | 0.1429 | 5 | 0 | 3 | 0.6667 | 11 | 0.1818 | 26 | 0.1923 | 6 | 0 | 8 | 0 | 6 | 0.1667 | 8 | 0.375 | 28 | 0.1429 |
| 17_4723988 | 7 | 0 | 5 | 0 | 2 | 0 | 13 | 0.3846 | 27 | 0.1852 | 5 | 0.2 | 8 | 0 | 5 | 0.4 | 10 | 0 | 28 | 0.1071 |
| 17_4723992 | 7 | 0.2857 | 6 | 0.5 | 3 | 0.6667 | 12 | 0.5 | 28 | 0.4643 | 5 | 0.6 | 8 | 0.375 | 6 | 0.3333 | 9 | 0.4444 | 28 | 0.4286 |
| 17_4724025 | 7 | 0.1429 | 5 | 0.4 | 2 | 0.5 | 10 | 0.5 | 24 | 0.375 | 4 | 0.25 | 8 | 0.125 | 4 | 0.25 | 8 | 0.5 | 24 | 0.2917 |
| 17_4724031 | 7 | 0.1429 | 5 | 0.6 | 2 | 0 | 11 | 0.5455 | 25 | 0.4 | 3 | 0.6667 | 6 | 0.5 | 4 | 0.5 | 9 | 0.4444 | 22 | 0.5 |
| 17_4724067 | 5 | 0.4 | 3 | 0 | 1 | 1 | 11 | 0.1818 | 20 | 0.25 | 3 | 0.6667 | 7 | 0 | 5 | 0.4 | 8 | 0.25 | 23 | 0.2609 |
| 17_4724069 | 6 | 0.5 | 5 | 0.6 | 2 | 0.5 | 9 | 0.2222 | 22 | 0.4091 | 1 | 1 | 7 | 0.4286 | 5 | 0.8 | 9 | 0.6667 | 22 | 0.6364 |
| 17_4724074 | 5 | 0.4 | 5 | 0.6 | 2 | 0.5 | 10 | 0.6 | 22 | 0.5455 | 3 | 1 | 7 | 0.5714 | 4 | 0.25 | 10 | 0.4 | 24 | 0.5 |
| 17_4724079 | 5 | 0.6 | 4 | 1 | 2 | 0 | 10 | 0.8 | 21 | 0.7143 | 3 | 0.6667 | 6 | 0.8333 | 4 | 1 | 8 | 0.5 | 21 | 0.7143 |
| 17_4724082 | 5 | 0.6 | 4 | 0.75 | 2 | 1 | 11 | 0.8182 | 22 | 0.7727 | 2 | 0 | 5 | 0.6 | 3 | 0.6667 | 8 | 0.75 | 18 | 0.6111 |
| 17_4724087 | 5 | 0 | 5 | 0 | 2 | 0.5 | 10 | 0.6 | 22 | 0.3182 | 3 | 0.6667 | 6 | 0.3333 | 3 | 0 | 7 | 0.4286 | 19 | 0.3684 |
| 17_4724089 | 3 | 1 | 4 | 0 | 2 | 1 | 10 | 0.6 | 19 | 0.5789 | 3 | 0.3333 | 7 | 0.2857 | 3 | 0.3333 | 10 | 0.3 | 23 | 0.3043 |
| 17_4724092 | 5 | 0.2 | 5 | 0.2 | 1 | 0 | 9 | 0.3333 | 20 | 0.25 | 3 | 0 | 7 | 0.1429 | 4 | 0.5 | 8 | 0.25 | 22 | 0.2273 |
| 17_4724113 | 5 | 0.6 | 2 | 0.5 | 2 | 0.5 | 10 | 0.6 | 19 | 0.5789 | 2 | 1 | 7 | 0.8571 | 4 | 0.75 | 7 | 1 | 20 | 0.9 |
| 17_4724122 | 5 | 0 | 2 | 0 | 2 | 0 | 10 | 0.3 | 19 | 0.1579 | 2 | 0 | 6 | 0 | 4 | 0.25 | 8 | 0 | 20 | 0.05 |
| 17_4724126 | 5 | 0.4 | 2 | 0 | 1 | 0 | 9 | 0.1111 | 17 | 0.1765 | 2 | 0 | 6 | 0.1667 | 4 | 0 | 7 | 0 | 19 | 0.0526 |
| 17_4724126 | 4 | 0 | 2 | 0.5 | 1 | 0 | 8 | 0.25 | 15 | 0.2 | 1 | 0 | 6 | 0.5 | 4 | 0.75 | 5 | 0.8 | 16 | 0.625 |
| 17_4724130 | 4 | 0.25 | 1 | 0 | 2 | 0 | 9 | 0.4444 | 16 | 0.3125 | 2 | 0.5 | 6 | 0.1667 | 4 | 0 | 7 | 0.1429 | 19 | 0.1579 |
| 17_4762542 | NA | NA | NA | NA | 1 | 1 | NA | NA | 1 | 1 | NA | NA | 1 | 1 | 1 | 1 | NA | NA | 2 | 1 |
| 17_4762563 | NA | NA | 1 | 1 | 1 | 0 | 1 | 1 | 3 | 0.6667 | 1 | 0 | NA | NA | 1 | 1 | NA | NA | 2 | 0.5 |
| 17_4762569 | 1 | 1 | 1 | 0 | 1 | 1 | 1 | 0 | 4 | 0.5 | 1 | 0 | 1 | 1 | 1 | 0 | NA | NA | 3 | 0.3333 |
| 17_4762583 | 1 | 1 | 1 | 0 | NA | NA | 1 | 1 | 3 | 0.6667 | 1 | 0 | 1 | 0 | 1 | 1 | NA | NA | 3 | 0.3333 |
| 17_4762604 | 1 | 0 | 1 | 1 | 1 | 1 | 1 | 0 | 4 | 0.5 | 1 | 1 | 1 | 0 | 1 | 1 | NA | NA | 3 | 0.6667 |
| 17_4762626 | NA | NA | 1 | 0 | 1 | 0 | 3 | 0.3333 | 5 | 0.2 | 1 | 1 | 1 | 0 | 1 | 0 | NA | NA | 3 | 0.3333 |
| 17_4762631 | NA | NA | 1 | 1 | 1 | 0 | 2 | 0 | 4 | 0.25 | 1 | 0 | 1 | 1 | 1 | 0 | NA | NA | 3 | 0.3333 |

| | | | | | | | | | | | | | | | | | | | | |
|---|---|---|---|---|---|---|---|---|---|---|---|---|---|---|---|---|---|---|---|---|
| 17_476344 | 2 | 0.5 | NA | NA | 1 | 0 | 2 | 0.5 | 5 | 0.4 | 1 | 1 | 1 | 1 | NA | NA | NA | NA | 2 | 1 |
| 17_476699 | 4 | 0.5 | 4 | 0.75 | 3 | 1 | 8 | 0.5 | 19 | 0.6316 | 3 | 0.6667 | 2 | 1 | 1 | 0 | 1 | 1 | 7 | 0.7143 |
| 17_476699 | 5 | 0.4 | 6 | 0.1667 | 2 | 0 | 7 | 0.2857 | 20 | 0.25 | 4 | 0.25 | 2 | 0 | 3 | 0 | 1 | 0 | 10 | 0.1 |
| 17_476700 | 4 | 0.25 | 5 | 0.6 | 3 | 0.6667 | 6 | 0.3333 | 18 | 0.4444 | 3 | 0 | 2 | 0 | 2 | 0 | 1 | 1 | 8 | 0.125 |
| 17_476701 | 3 | 1 | 5 | 1 | 3 | 1 | 7 | 1 | 18 | 1 | 4 | 1 | 1 | 1 | 3 | 1 | 1 | 1 | 9 | 1 |
| 17_476701 | 5 | 0.4 | 6 | 0.5 | 3 | 1 | 8 | 0.5 | 22 | 0.5455 | 4 | 0.75 | 2 | 1 | 3 | 0.6667 | 1 | 0 | 10 | 0.7 |
| 17_476702 | 4 | 1 | 5 | 0.8 | 2 | 1 | 8 | 0.875 | 19 | 0.8947 | 4 | 0.5 | 2 | 1 | 2 | 1 | 1 | 1 | 9 | 0.7778 |
| 17_476718 | 8 | 0.125 | 7 | 0 | 3 | 0 | 8 | 0.25 | 26 | 0.1154 | 1 | 0 | 4 | 0 | 2 | 0.5 | 1 | 0 | 8 | 0.125 |
| 17_476721 | 8 | 0.25 | 8 | 0 | 4 | 0.25 | 9 | 0.1111 | 29 | 0.1379 | 4 | 0.25 | 5 | 0 | 3 | 0.6667 | 1 | 0 | 13 | 0.2308 |
| 17_476726 | 8 | 0 | 5 | 0 | 3 | 0.6667 | 4 | 0.25 | 20 | 0.15 | 4 | 0 | 2 | 0 | 4 | 0.5 | 1 | 1 | 11 | 0.2727 |
| 17_477225 | NA | NA | 1 | 1 | 2 | 0.5 | 7 | 0.2857 | 10 | 0.4 | 1 | 0 | 7 | 0.2857 | 2 | 0 | 3 | 0.6667 | 13 | 0.3077 |
| 17_477993 | NA | NA | 2 | 0.5 | NA | NA | 2 | 1 | 4 | 0.75 | 2 | 0 | 3 | 0 | 2 | 0.5 | 1 | 0 | 8 | 0.125 |
| 17_477994 | NA | NA | 2 | 0.5 | 2 | 0 | 2 | 0 | 6 | 0.1667 | 2 | 0 | 4 | 0.25 | 2 | 0 | 2 | 0.5 | 10 | 0.2 |
| 17_477995 | NA | NA | 2 | 0.5 | 2 | 0 | 2 | 1 | 6 | 0.5 | 2 | 0.5 | 4 | 0.5 | 2 | 0 | 1 | 0 | 9 | 0.3333 |
| 17_477995 | NA | NA | 2 | 0 | 2 | 0.5 | 2 | 1 | 6 | 0.5 | 2 | 0 | 4 | 0.75 | 2 | 0 | 2 | 0.5 | 10 | 0.4 |
| 17_477996 | NA | NA | 2 | 1 | 2 | 1 | 1 | 1 | 5 | 1 | 2 | 1 | 4 | 1 | 2 | 1 | 2 | 1 | 10 | 1 |
| 17_477996 | NA | NA | 2 | 0.5 | 2 | 0 | 2 | 0.5 | 6 | 0.3333 | 2 | 0.5 | 4 | 0 | 2 | 0 | 2 | 0.5 | 10 | 0.2 |
| 17_477996 | NA | NA | 2 | 0.5 | 2 | 0 | 2 | 0 | 6 | 0.1667 | 1 | 0 | 3 | 0.3333 | 2 | 0.5 | 2 | 1 | 8 | 0.5 |
| 17_477999 | NA | NA | 2 | 1 | 1 | 0 | 2 | 1 | 5 | 0.8 | 1 | 0 | 4 | 0.75 | 2 | 0.5 | 2 | 1 | 9 | 0.6667 |
| 17_478001 | NA | NA | 2 | 1 | 1 | 1 | 2 | 1 | 5 | 1 | 2 | 1 | 4 | 1 | 2 | 1 | 2 | 1 | 10 | 1 |
| 17_478001 | NA | NA | 2 | 1 | NA | NA | 2 | 1 | 4 | 1 | 1 | 1 | 3 | 0.6667 | 2 | 0.5 | 2 | 1 | 8 | 0.75 |
| 17_478002 | NA | NA | 2 | 1 | NA | NA | NA | NA | 2 | 1 | 1 | 0 | 4 | 0.75 | 2 | 0.5 | 1 | 0 | 8 | 0.5 |
| 17_478003 | NA | NA | 2 | 1 | 1 | 1 | 1 | 1 | 4 | 1 | 2 | 1 | 3 | 0.6667 | 1 | 1 | 2 | 0.5 | 8 | 0.75 |
| 17_478005 | NA | NA | 1 | 1 | NA | NA | 1 | 1 | 2 | 1 | 2 | 1 | 1 | 1 | 2 | 1 | 1 | 1 | 6 | 1 |
| 17_478005 | NA | NA | 2 | 0.5 | 1 | 1 | 1 | 1 | 4 | 0.75 | 2 | 0.5 | 2 | 0 | 1 | 1 | 1 | 0 | 6 | 0.3333 |
| 17_478006 | NA | NA | 2 | 0.5 | 1 | 0 | 1 | 0 | 4 | 0.25 | 2 | 0 | 3 | 0.3333 | 2 | 0 | 2 | 0.5 | 9 | 0.2222 |
| 17_478039 | NA | NA | 1 | 1 | NA | NA | NA | NA | 1 | 1 | NA | NA | 1 | 1 | NA | NA | NA | NA | 1 | 1 |
| 17_479097 | 6 | 0 | 5 | 0.2 | 1 | 0 | 2 | 0 | 14 | 0.0714 | 2 | 0.5 | 3 | 0 | 3 | 0 | 1 | 0 | 9 | 0.1111 |
| 17_479118 | 1 | 0 | 2 | 1 | NA | NA | 2 | 0.5 | 5 | 0.6 | 1 | 0 | NA | NA | 2 | 0.5 | NA | NA | 3 | 0.3333 |
| 17_479123 | 3 | 1 | 6 | 1 | 4 | 1 | 4 | 0.75 | 17 | 0.9412 | 3 | 1 | 3 | 1 | 5 | 1 | 1 | 1 | 12 | 1 |
| 17_479123 | 2 | 0.5 | 4 | 0 | 3 | 0 | 4 | 0.5 | 13 | 0.2308 | 3 | 0.3333 | 3 | 0.3333 | 4 | 0 | NA | NA | 10 | 0.2 |
| 17_479124 | 3 | 0.3333 | 5 | 0.2 | 3 | 0 | 4 | 0.5 | 15 | 0.2667 | 4 | 0.25 | 4 | 0.5 | 4 | 0.75 | 1 | 0 | 13 | 0.4615 |
| 17_479124 | 3 | 0.6667 | 5 | 0.2 | 2 | 0 | 4 | 0 | 14 | 0.2143 | 5 | 0.6 | 4 | 0.25 | 5 | 0.8 | 1 | 0 | 15 | 0.5333 |
| 17_479124 | 2 | 0.5 | 4 | 0.25 | 2 | 0 | 3 | 0.3333 | 11 | 0.2727 | 3 | 0.3333 | 3 | 0 | 4 | 0.75 | 1 | 0 | 11 | 0.3636 |
| 17_479124 | 3 | 0.3333 | 1 | 1 | 2 | 1 | 3 | 0.3333 | 9 | 0.5556 | 3 | 0.6667 | 1 | 1 | 2 | 1 | NA | NA | 6 | 0.8333 |
| 17_479126 | 3 | 0.3333 | 4 | 0.25 | 3 | 0.3333 | 2 | 0 | 12 | 0.25 | 4 | 0.5 | 2 | 0 | 5 | 0.2 | 1 | 0 | 12 | 0.25 |
| 17_479223 | 1 | 1 | 1 | 0 | 1 | 0 | NA | NA | 3 | 0.3333 | 1 | 1 | NA | NA | 1 | 0 | 1 | 0 | 3 | 0.3333 |
| 17_479338 | 6 | 0.1667 | 1 | 0 | 3 | 0.3333 | 1 | 0 | 11 | 0.1818 | 3 | 0 | 2 | 0.5 | 3 | 0 | 3 | 0.3333 | 11 | 0.1818 |
| 17_479339 | 6 | 0.1667 | 1 | 0 | 3 | 0 | 1 | 0 | 11 | 0.0909 | 2 | 0 | 2 | 0 | 2 | 0.5 | 3 | 0 | 9 | 0.1111 |
| 17_479339 | 6 | 0.1667 | 1 | 0 | 4 | 0 | 1 | 0 | 12 | 0.0833 | 3 | 0 | 2 | 0 | 3 | 0.3333 | 3 | 0 | 11 | 0.0909 |
| 17_479339 | 5 | 0.6 | 1 | 0 | 4 | 0.5 | 2 | 1 | 12 | 0.5833 | 3 | 1 | 2 | 0.5 | 3 | 1 | 3 | 0.6667 | 11 | 0.8182 |
| 17_479340 | 6 | 0.8333 | 1 | 0 | 4 | 0.5 | 2 | 0.5 | 13 | 0.6154 | 2 | 0 | 1 | 1 | 3 | 0.3333 | 3 | 0.6667 | 9 | 0.4444 |
| 17_479340 | 4 | 1 | NA | NA | 2 | 1 | 2 | 1 | 8 | 1 | 2 | 0.5 | 2 | 1 | 3 | 1 | 3 | 1 | 10 | 0.9 |
| 17_479341 | 3 | 1 | 1 | 1 | 4 | 0.75 | 2 | 1 | 10 | 0.9 | 2 | 1 | 3 | 1 | 3 | 1 | 1 | 1 | 9 | 1 |
| 17_479344 | 5 | 0.4 | 1 | 0 | 4 | 0.75 | 2 | 0 | 12 | 0.4167 | 1 | 0 | 3 | 0.3333 | 3 | 1 | 3 | 0 | 10 | 0.4 |
| 17_479344 | 6 | 0.3333 | 1 | 0 | 3 | 0.3333 | 2 | 0.5 | 12 | 0.3333 | 1 | 0 | 3 | 0.3333 | 3 | 0.3333 | 2 | 0.5 | 9 | 0.3333 |
| 17_479345 | 6 | 0.3333 | 1 | 0 | 3 | 0 | 2 | 0.5 | 12 | 0.25 | 1 | 0 | 3 | 0.6667 | 3 | 0.6667 | 3 | 0.3333 | 10 | 0.5 |
| 17_479347 | 5 | 0.2 | 1 | 0 | 3 | 0 | 2 | 0 | 11 | 0.0909 | 2 | 0 | 3 | 0 | 4 | 0.25 | 3 | 0 | 12 | 0.0833 |
| 17_479348 | 4 | 0 | NA | NA | 4 | 0.25 | 2 | 0 | 10 | 0.1 | 2 | 0 | 3 | 0.3333 | 4 | 0.25 | 3 | 0.3333 | 12 | 0.25 |
| 17_479952 | NA | NA | 1 | 1 | NA | NA | 2 | 1 | 3 | 1 | NA | NA | NA | NA | NA | NA | 1 | 1 | 1 | 1 |
| 17_479956 | NA | NA | 1 | 0 | NA | NA | 2 | 1 | 3 | 0.6667 | NA | NA | NA | NA | 1 | 1 | 1 | 0 | 2 | 0.5 |
| 17_479990 | 4 | 1 | 6 | 0.6667 | 3 | 0.6667 | 1 | 1 | 14 | 0.7857 | 1 | 0 | 1 | 1 | 1 | 1 | 2 | 0.5 | 5 | 0.6 |
| 17_479991 | 4 | 0.25 | 3 | 0.3333 | 1 | 0 | 2 | 0.5 | 10 | 0.3 | 2 | 0 | NA | NA | 1 | 0 | 3 | 0.3333 | 6 | 0.1667 |
| 17_479991 | 4 | 0 | 4 | 0 | 2 | 0.5 | 2 | 0 | 12 | 0.0833 | 2 | 0 | NA | NA | NA | NA | 3 | 0.3333 | 5 | 0.2 |
| 17_479992 | 4 | 0.75 | 5 | 1 | 3 | 1 | 1 | 1 | 13 | 0.9231 | 2 | 1 | 1 | 1 | 1 | 1 | 2 | 1 | 6 | 1 |
| 17_479993 | 4 | 0.75 | 4 | 0.5 | 3 | 0.6667 | 1 | 0 | 12 | 0.5833 | 2 | 0.5 | 1 | 0 | NA | NA | 3 | 0 | 6 | 0.1667 |
| 17_479993 | 4 | 0.25 | 3 | 0 | 3 | 0 | 2 | 0 | 12 | 0.0833 | 2 | 0 | 1 | 0 | 1 | 0 | 3 | 0.3333 | 7 | 0.1429 |
| 17_479994 | 1 | 0 | 3 | 0.6667 | 3 | 0.3333 | 1 | 0 | 8 | 0.375 | 2 | 0.5 | 1 | 0 | 1 | 1 | 3 | 0 | 7 | 0.2857 |
| 17_479998 | NA | NA | 2 | 1 | 1 | 0 | NA | NA | 3 | 0.6667 | 1 | 1 | NA | NA | 1 | 0 | 3 | 0.6667 | 5 | 0.6 |
| 17_480141 | 4 | 0.75 | 4 | 0.25 | 5 | 0.4 | 2 | 0 | 15 | 0.4 | 2 | 0 | 3 | 0.6667 | 1 | 0 | 2 | 0 | 8 | 0.25 |
| 17_480142 | 2 | 0.5 | 3 | 1 | 3 | 0.6667 | 2 | 0.5 | 10 | 0.7 | 1 | 0 | 1 | 1 | 1 | 1 | NA | NA | 3 | 0.6667 |
| 17_480143 | 1 | 1 | 1 | 0 | 2 | 1 | 2 | 1 | 6 | 0.8333 | 1 | 0 | 1 | 1 | 1 | 1 | NA | NA | 3 | 0.6667 |
| 17_480146 | 2 | 1 | 4 | 0.5 | 6 | 0.6667 | 2 | 0.5 | 14 | 0.6429 | 2 | 0.5 | 3 | 0.6667 | 1 | 0 | 1 | 0 | 7 | 0.4286 |
| 17_480148 | 3 | 0.6667 | 4 | 0.5 | 5 | 1 | 3 | 1 | 15 | 0.8 | 1 | 1 | 2 | 1 | 1 | 1 | 2 | 0.5 | 6 | 0.8333 |
| 17_480149 | 4 | 0 | 3 | 0.3333 | 6 | 0.1667 | 4 | 0 | 17 | 0.1176 | 3 | 0.3333 | 1 | 0 | 1 | 0 | 1 | 0 | 6 | 0.1667 |

| | | | | | | | | | | | | | | | | | | | | |
|---|---|---|---|---|---|---|---|---|---|---|---|---|---|---|---|---|---|---|---|---|
| 17_4801498 | 3 | 0.3333 | 3 | 0 | 5 | 0.4 | 3 | 0 | 14 | 0.2143 | 3 | 0.3333 | 3 | 0 | NA | NA | 2 | 0.5 | 8 | 0.25 |
| 17_4810260 | NA | NA | 1 | 1 | 1 | 1 | 1 | 1 | 3 | 1 | 5 | 1 | 2 | 1 | NA | NA | 3 | 1 | 10 | 1 |
| 17_4810289 | NA | NA | 1 | 1 | 1 | 1 | NA | NA | 2 | 1 | 5 | 1 | 1 | 0 | NA | NA | 3 | 1 | 9 | 0.8889 |
| 17_4810307 | NA | NA | 1 | 1 | 1 | 1 | NA | NA | 2 | 1 | 5 | 0.6 | 2 | 0 | NA | NA | 2 | 1 | 9 | 0.5556 |
| 17_4810314 | NA | NA | 1 | 1 | 1 | 0 | 1 | 0 | 3 | 0.3333 | 5 | 0.4 | 2 | 0.5 | NA | NA | 3 | 0.6667 | 10 | 0.5 |
| 17_4812685 | 1 | 1 | NA | NA | NA | NA | NA | NA | 1 | 1 | 1 | 1 | NA | NA | NA | NA | NA | NA | 1 | 1 |
| 17_4817889 | 5 | 0.2 | 6 | 0.1667 | 3 | 0 | 2 | 0 | 16 | 0.125 | 1 | 0 | 5 | 0.2 | 3 | 0.3333 | 5 | 0.4 | 14 | 0.2857 |
| 17_4817935 | 5 | 0.4 | 6 | 0.1667 | 6 | 0.1667 | NA | NA | 17 | 0.2353 | 2 | 0 | 6 | 0.3333 | 2 | 0.5 | 3 | 0 | 13 | 0.2308 |
| 17_4817937 | 3 | 0.3333 | 5 | 0.4 | 4 | 0.5 | 2 | 1 | 14 | 0.5 | 3 | 0.6667 | 4 | 0.25 | 3 | 0.6667 | 4 | 0.5 | 14 | 0.5 |
| 17_4817947 | 2 | 1 | 6 | 0.1667 | 2 | 1 | 2 | 0.5 | 12 | 0.5 | 3 | 0.3333 | 5 | 1 | 4 | 0.25 | 4 | 0.25 | 16 | 0.5 |
| 17_4817959 | 3 | 0 | 1 | 1 | 3 | 0 | 2 | 0.5 | 9 | 0.2222 | 1 | 1 | 4 | 0.25 | 4 | 0.5 | 5 | 0 | 14 | 0.2857 |
| 17_4817960 | 2 | 0 | 5 | 0.2 | 5 | 0 | 2 | 0.5 | 14 | 0.1429 | 3 | 0 | 4 | 0.25 | 4 | 0.25 | 5 | 0 | 16 | 0.125 |
| 17_4818220 | 3 | 0 | 6 | 0 | 4 | 0 | 2 | 0.5 | 15 | 0.0667 | NA | NA | 2 | 0.5 | 3 | 0.3333 | 3 | 0 | 8 | 0.25 |
| 17_4865927 | NA | NA | 3 | 0.3333 | 3 | 0.3333 | 1 | 1 | 7 | 0.4286 | 2 | 1 | 1 | 0 | 2 | 0.5 | 2 | 0.5 | 7 | 0.5714 |
| 17_4869404 | 4 | 0.25 | 2 | 0 | 5 | 0 | 1 | 0 | 12 | 0.0833 | 1 | 0 | 4 | 0 | 1 | 1 | 2 | 0.5 | 8 | 0.25 |
| 17_4869475 | 5 | 0 | 2 | 0 | 3 | 0 | 2 | 0.5 | 12 | 0.0833 | NA | NA | 3 | 0.3333 | 2 | 0 | 2 | 0 | 7 | 0.1429 |
| 17_4869554 | 2 | 0.5 | 2 | 0 | 4 | 0.25 | 2 | 0 | 10 | 0.2 | NA | NA | 2 | 0 | 1 | 0 | 1 | 1 | 4 | 0.25 |
| 17_4869560 | 3 | 0 | 2 | 0.5 | 4 | 0.25 | 2 | 0.5 | 11 | 0.2727 | NA | NA | 3 | 0.6667 | 1 | 0 | 1 | 0 | 5 | 0.4 |
| 17_4869566 | 3 | 0.3333 | 1 | 1 | 2 | 0 | 2 | 0 | 8 | 0.25 | NA | NA | 4 | 0.25 | 1 | 0 | 1 | 0 | 6 | 0.1667 |
| 17_4869569 | 3 | 0 | 2 | 0.5 | 3 | 0.6667 | 2 | 0 | 10 | 0.3 | NA | NA | 3 | 0.3333 | 1 | 0 | 1 | 1 | 5 | 0.4 |
| 17_4869571 | 2 | 0.5 | 2 | 0 | 2 | 0 | 2 | 0.5 | 8 | 0.25 | NA | NA | 3 | 1 | 1 | 1 | 1 | 1 | 5 | 1 |
| 17_4869583 | 3 | 0.6667 | 1 | 0 | 3 | 0.3333 | 2 | 0.5 | 9 | 0.4444 | NA | NA | 2 | 0 | 1 | 0 | 1 | 1 | 4 | 0.25 |
| 17_4869587 | 3 | 0.3333 | 1 | 0 | 2 | 0 | 2 | 0 | 8 | 0.125 | NA | NA | 3 | 1 | 1 | 0 | 1 | 0 | 5 | 0.6 |
| 17_4869610 | 3 | 1 | 1 | 1 | 2 | 0.5 | 1 | 1 | 7 | 0.8571 | NA | NA | 3 | 1 | 1 | 1 | 1 | 1 | 5 | 1 |
| 17_4869611 | 3 | 0.3333 | 1 | 1 | 2 | 1 | 1 | 1 | 7 | 0.7143 | NA | NA | 2 | 1 | 1 | 1 | 1 | 1 | 4 | 1 |
| 17_4869616 | 2 | 1 | 1 | 0 | 2 | 0 | 1 | 0 | 6 | 0.3333 | NA | NA | 3 | 1 | 1 | 0 | 1 | 1 | 5 | 0.8 |
| 17_4869621 | 3 | 0.6667 | NA | NA | 2 | 1 | 1 | 1 | 6 | 0.8333 | NA | NA | 3 | 0.6667 | 1 | 0 | 1 | 1 | 5 | 0.6 |
| 17_4869625 | 2 | 1 | 1 | 1 | 2 | 1 | 1 | 1 | 6 | 1 | NA | NA | 3 | 1 | 1 | 1 | 1 | 1 | 5 | 1 |
| 17_4871833 | NA | NA | 4 | 0.25 | 3 | 0.3333 | NA | NA | 7 | 0.2857 | 4 | 0 | 2 | 0 | 3 | 0.3333 | NA | NA | 9 | 0.1111 |
| 17_4872387 | NA | NA | 5 | 0.4 | 3 | 0.6667 | 1 | 1 | 9 | 0.5556 | 3 | 0.6667 | 1 | 0 | 3 | 0.3333 | 1 | 1 | 8 | 0.5 |
| 17_4872388 | NA | NA | 6 | 0 | 4 | 0.25 | 1 | 0 | 11 | 0.0909 | 3 | 0 | 2 | 0 | 3 | 0.3333 | 1 | 0 | 9 | 0.1111 |
| 17_4872390 | NA | NA | 6 | 0.6667 | 2 | 1 | 1 | 1 | 9 | 0.7778 | 4 | 0.75 | 2 | 0.5 | 1 | 1 | NA | NA | 7 | 0.7143 |
| 17_4923942 | 1 | 0 | NA | NA | NA | NA | 1 | 1 | 2 | 0.5 | 2 | 1 | NA | NA | NA | NA | NA | NA | 2 | 1 |
| 17_4937713 | 1 | 0 | 2 | 0.5 | NA | NA | 1 | 1 | 4 | 0.5 | 4 | 0.75 | NA | NA | NA | NA | 2 | 0.5 | 6 | 0.6667 |
| 17_4937721 | 1 | 1 | 4 | 0.5 | NA | NA | 1 | 0 | 6 | 0.5 | 3 | 0.6667 | NA | NA | NA | NA | 2 | 0.5 | 5 | 0.6 |
| 17_4937727 | 1 | 1 | 2 | 0.5 | NA | NA | 1 | 0 | 4 | 0.5 | 4 | 0.25 | NA | NA | NA | NA | 2 | 1 | 6 | 0.5 |
| 17_4937731 | 1 | 1 | 4 | 0.25 | NA | NA | NA | NA | 5 | 0.4 | 3 | 0.6667 | NA | NA | NA | NA | 2 | 1 | 5 | 0.8 |
| 17_4937738 | 1 | 0 | 4 | 0.5 | NA | NA | 1 | 0 | 6 | 0.3333 | 4 | 0.25 | NA | NA | NA | NA | 2 | 0.5 | 6 | 0.3333 |
| 17_4937740 | 1 | 0 | 3 | 1 | NA | NA | NA | NA | 4 | 0.75 | 4 | 0.5 | NA | NA | NA | NA | 2 | 1 | 6 | 0.6667 |
| 17_4937746 | 1 | 0 | 4 | 0.25 | NA | NA | 1 | 0 | 6 | 0.1667 | 5 | 0.4 | NA | NA | NA | NA | 2 | 0 | 7 | 0.2857 |
| 17_4937747 | 1 | 1 | 3 | 0 | NA | NA | NA | NA | 4 | 0.25 | 5 | 0.6 | NA | NA | NA | NA | 2 | 1 | 7 | 0.7143 |
| 17_4937751 | 1 | 1 | 3 | 0.3333 | NA | NA | 1 | 1 | 5 | 0.6 | 4 | 1 | NA | NA | NA | NA | 2 | 0 | 6 | 0.6667 |
| 17_4937756 | 1 | 0 | 5 | 0.2 | NA | NA | 1 | 0 | 7 | 0.1429 | 4 | 0.5 | NA | NA | NA | NA | 1 | 0 | 5 | 0.4 |
| 17_4937770 | 1 | 0 | 5 | 0.2 | NA | NA | 1 | 0 | 7 | 0.1429 | 5 | 0.2 | NA | NA | NA | NA | 2 | 0 | 7 | 0.1429 |
| 17_4937786 | NA | NA | 4 | 0.25 | NA | NA | 1 | 0 | 5 | 0.2 | 3 | 0.3333 | NA | NA | NA | NA | 2 | 0.5 | 5 | 0.4 |
| 17_4937791 | 1 | 0 | 4 | 1 | NA | NA | 1 | 1 | 6 | 0.8333 | 4 | 0.5 | NA | NA | NA | NA | 1 | 0 | 5 | 0.4 |
| 17_4937792 | NA | NA | 6 | 1 | NA | NA | 1 | 1 | 7 | 1 | 5 | 1 | NA | NA | NA | NA | 2 | 1 | 7 | 1 |
| 17_4937795 | 1 | 1 | 6 | 1 | NA | NA | 2 | 1 | 9 | 1 | 5 | 1 | NA | NA | NA | NA | 2 | 1 | 7 | 1 |
| 17_4937797 | 1 | 1 | 5 | 1 | NA | NA | 2 | 1 | 8 | 1 | 5 | 1 | NA | NA | NA | NA | 2 | 1 | 7 | 1 |
| 17_4937798 | 1 | 1 | 4 | 1 | NA | NA | 2 | 0.5 | 7 | 0.8571 | 5 | 0.6 | NA | NA | NA | NA | 2 | 1 | 7 | 0.7143 |
| 17_4937811 | 1 | 1 | 6 | 0.6667 | NA | NA | 2 | 1 | 9 | 0.7778 | 5 | 0.8 | NA | NA | 1 | 1 | 1 | 1 | 7 | 0.8571 |
| 17_4940361 | 1 | 1 | 6 | 0.1667 | NA | NA | 3 | 0 | 10 | 0.2 | 5 | 0.2 | NA | NA | 1 | 0 | 2 | 0 | 8 | 0.125 |
| 17_4940371 | 1 | 1 | 6 | 1 | 1 | 1 | 2 | 1 | 10 | 1 | 6 | 1 | NA | NA | 1 | 1 | 1 | 1 | 8 | 1 |
| 17_4940374 | 1 | 1 | 6 | 0.8333 | 1 | 1 | 1 | 1 | 9 | 0.8889 | 4 | 1 | NA | NA | 1 | 1 | 2 | 1 | 7 | 1 |
| 17_4940387 | 1 | 1 | 8 | 0.375 | 1 | 1 | 3 | 0.6667 | 13 | 0.5385 | 6 | 0.3333 | NA | NA | 1 | 1 | 2 | 1 | 9 | 0.5556 |
| 17_4940389 | 1 | 1 | 6 | 0.3333 | 1 | 0 | 3 | 0.6667 | 11 | 0.4545 | 6 | 0.3333 | NA | NA | 1 | 0 | 2 | 0.5 | 9 | 0.3333 |
| 17_4940391 | 1 | 1 | 7 | 0.7143 | 1 | 0 | 3 | 1 | 12 | 0.75 | 6 | 0.8333 | NA | NA | 1 | 1 | 2 | 1 | 9 | 0.8889 |
| 17_4940393 | 1 | 1 | 7 | 0.8571 | 1 | 1 | 3 | 0.6667 | 12 | 0.8333 | 6 | 0.8333 | NA | NA | 1 | 1 | 2 | 1 | 9 | 0.8889 |
| 17_4940405 | NA | NA | 5 | 0.4 | 1 | 1 | 1 | 1 | 7 | 0.5714 | 5 | 0.2 | NA | NA | NA | NA | 1 | 0 | 6 | 0.1667 |
| 17_4940406 | NA | NA | 5 | 1 | 1 | 1 | 2 | 1 | 8 | 1 | 4 | 1 | NA | NA | 1 | 1 | 2 | 1 | 7 | 1 |
| 17_4940418 | 1 | 1 | 7 | 0.5714 | 1 | 1 | 1 | 0 | 10 | 0.6 | 4 | 0.5 | NA | NA | 2 | 1 | 2 | 1 | 8 | 0.75 |
| 17_4940420 | NA | NA | 7 | 0.8571 | 1 | 1 | 3 | 1 | 11 | 0.9091 | 6 | 1 | NA | NA | 1 | 1 | 2 | 0.5 | 9 | 0.8889 |
| 17_4940435 | 1 | 1 | 6 | 1 | 1 | 0 | 3 | 1 | 11 | 0.9091 | 6 | 1 | NA | NA | 2 | 1 | 3 | 1 | 11 | 1 |
| 17_4940447 | 1 | 1 | 7 | 1 | 2 | 1 | 3 | 1 | 13 | 1 | 5 | 1 | NA | NA | 2 | 1 | 3 | 1 | 10 | 1 |
| 17_4940448 | 1 | 0 | 6 | 0.3333 | 2 | 0.5 | 2 | 1 | 11 | 0.4545 | 3 | 0 | NA | NA | 1 | 0 | 1 | 1 | 5 | 0.2 |

| | | | | | | | | | | | | | | | | | | | | |
|---|---|---|---|---|---|---|---|---|---|---|---|---|---|---|---|---|---|---|---|---|
| 17_4976124 | 1 | 0 | NA | NA | NA | NA | 1 | 1 | 2 | 0.5 | 1 | 0 | 1 | 1 | 2 | 0 | 3 | 0 | 7 | 0.1429 |
| 17_4976198 | 1 | 0 | NA | NA | NA | NA | 1 | 1 | 2 | 0.5 | 1 | 0 | 2 | 0.5 | 2 | 0 | 3 | 0 | 8 | 0.125 |
| 17_4976330 | 6 | 0 | 13 | 0.1538 | 14 | 0.2143 | 7 | 0 | 40 | 0.125 | 5 | 0.4 | 7 | 0 | 10 | 0.1 | 9 | 0.3333 | 31 | 0.1935 |
| 17_4976352 | 7 | 0.1429 | 11 | 0.0909 | 13 | 0.0769 | 4 | 0 | 35 | 0.0857 | 6 | 0 | 6 | 0.1667 | 11 | 0 | 7 | 0.2857 | 30 | 0.1 |
| 17_4976354 | 4 | 0.75 | 12 | 0.6667 | 14 | 0.8571 | 7 | 0.7143 | 37 | 0.7568 | 5 | 0.8 | 7 | 0.7143 | 12 | 0.8333 | 5 | 0.8 | 29 | 0.7931 |
| 17_4976354 | 6 | 0.5 | 12 | 0.25 | 11 | 0.3636 | 8 | 0.625 | 37 | 0.4054 | 3 | 0.6667 | 5 | 0.2 | 12 | 0.5 | 6 | 0.3333 | 26 | 0.4231 |
| 17_4976370 | 6 | 0.3333 | 9 | 0.2222 | 11 | 0.2727 | 7 | 0.4286 | 33 | 0.303 | 6 | 0.1667 | 6 | 0.1667 | 11 | 0.1818 | 7 | 0 | 30 | 0.1333 |
| 17_4976377 | 5 | 0 | 10 | 0 | 12 | 0.1667 | 7 | 0 | 34 | 0.0588 | 5 | 0.2 | 5 | 0 | 11 | 0.1818 | 8 | 0 | 29 | 0.1034 |
| 17_4976379 | 5 | 0.8 | 10 | 0.9 | 12 | 1 | 7 | 1 | 34 | 0.9412 | 7 | 0.2857 | 6 | 0.8333 | 12 | 0.9167 | 4 | 0.75 | 29 | 0.7241 |
| 17_4976380 | 6 | 0.1667 | 12 | 0 | 13 | 0.0769 | 5 | 0.2 | 36 | 0.0833 | 8 | 0.125 | 5 | 0 | 12 | 0.0833 | 7 | 0 | 32 | 0.0625 |
| 17_4976386 | 4 | 0 | 10 | 0.3 | 10 | 0.6 | 7 | 0.4286 | 31 | 0.3871 | 5 | 0 | 5 | 0 | 9 | 0.1111 | 7 | 0.1429 | 26 | 0.0769 |
| 17_4976399 | 5 | 0.2 | 12 | 0.4167 | 12 | 0.5 | 6 | 0.5 | 35 | 0.4286 | 7 | 0.5714 | 6 | 0.3333 | 12 | 0.3333 | 8 | 0.125 | 33 | 0.3333 |
| 17_4976418 | 6 | 0.1667 | 10 | 0 | 8 | 0.125 | 5 | 0.2 | 29 | 0.1034 | 7 | 0.2857 | 5 | 0.2 | 11 | 0.2727 | 5 | 0 | 28 | 0.2143 |
| 17_4976553 | 4 | 0.25 | 7 | 0.5714 | 5 | 1 | 4 | 0.75 | 20 | 0.65 | 6 | 0.6667 | 5 | 0.8 | 7 | 0.5714 | 2 | 0.5 | 20 | 0.65 |
| 17_4976554 | 4 | 0.25 | 7 | 0.5714 | 6 | 0.1667 | 5 | 0.4 | 22 | 0.3636 | 6 | 0.3333 | 5 | 0.4 | 8 | 0.5 | 4 | 0.25 | 23 | 0.3913 |
| 17_4976555 | 3 | 0.6667 | 4 | 0.5 | 6 | 0.5 | 5 | 0.4 | 18 | 0.5 | 3 | 0.6667 | 6 | 0.3333 | 7 | 0.2857 | 3 | 0.6667 | 19 | 0.4211 |
| 17_4976564 | 3 | 0.3333 | 7 | 0.4286 | 5 | 0.6 | 4 | 0 | 19 | 0.3684 | 6 | 0.3333 | 6 | 0 | 7 | 0.2857 | 3 | 0 | 22 | 0.1818 |
| 17_4978169 | NA | NA | NA | NA | 1 | 1 | 2 | 0 | 3 | 0.3333 | NA | NA | 3 | 0 | 1 | 0 | 1 | 1 | 5 | 0.2 |
| 17_5030078 | 3 | 0.6667 | 1 | 0 | 1 | 1 | 2 | 0.5 | 7 | 0.5714 | 1 | 0 | NA | NA | 1 | 1 | NA | NA | 2 | 0.5 |
| 17_5030078 | 3 | 0 | 1 | 0 | 1 | 0 | 2 | 0.5 | 7 | 0.1429 | 1 | 1 | NA | NA | 1 | 0 | NA | NA | 2 | 0.5 |
| 17_5030085 | 2 | 0.5 | 1 | 0 | 1 | 1 | 2 | 1 | 6 | 0.6667 | 1 | 0 | NA | NA | 1 | 1 | NA | NA | 2 | 0.5 |
| 17_5030088 | 3 | 1 | 1 | 1 | 1 | 1 | 2 | 1 | 7 | 1 | 1 | 1 | NA | NA | 1 | 1 | NA | NA | 2 | 1 |
| 17_5030126 | 2 | 0 | 1 | 0 | 1 | 1 | 2 | 0.5 | 6 | 0.3333 | 1 | 0 | NA | NA | 1 | 1 | NA | NA | 2 | 0.5 |
| 17_5058716 | 1 | 0 | NA | NA | 1 | 0 | 1 | 1 | 3 | 0.3333 | NA | NA | NA | NA | 3 | 0.6667 | 2 | 0 | 5 | 0.4 |
| 17_5070267 | 1 | 1 | NA | NA | NA | NA | NA | NA | 1 | 1 | NA | NA | 1 | 1 | NA | NA | NA | NA | 1 | 1 |
| 17_5070283 | 1 | 1 | NA | NA | NA | NA | NA | NA | 1 | 1 | NA | NA | 1 | 1 | NA | NA | NA | NA | 1 | 1 |
| 17_5070305 | 1 | 1 | NA | NA | NA | NA | NA | NA | 1 | 1 | NA | NA | 1 | 1 | NA | NA | NA | NA | 1 | 1 |
| 17_5070345 | 1 | 1 | NA | NA | NA | NA | NA | NA | 1 | 1 | NA | NA | 1 | 1 | NA | NA | NA | NA | 1 | 1 |
| 17_5073023 | NA | NA | NA | NA | 1 | 1 | NA | NA | 1 | 1 | NA | NA | 1 | 1 | 1 | 0 | NA | NA | 2 | 0.5 |
| 17_5073357 | NA | NA | NA | NA | 1 | 1 | NA | NA | 1 | 1 | NA | NA | 1 | 1 | NA | NA | 1 | 0 | 2 | 0.5 |
| 17_5091340 | 3 | 0.6667 | 1 | 1 | 2 | 0.5 | 2 | 0.5 | 8 | 0.625 | NA | NA | 3 | 0.3333 | 3 | 0.6667 | 6 | 0.3333 | 12 | 0.4167 |
| 17_5091343 | 2 | 1 | 1 | 0 | 2 | 0.5 | 2 | 0 | 7 | 0.4286 | NA | NA | 3 | 0.3333 | 3 | 1 | 5 | 0.8 | 11 | 0.7273 |
| 17_5091373 | 3 | 1 | 1 | 1 | 2 | 0.5 | 2 | 1 | 8 | 0.875 | NA | NA | 3 | 1 | 3 | 1 | 7 | 0.8571 | 13 | 0.9231 |
| 17_5091380 | 3 | 1 | 1 | 1 | 3 | 0.3333 | 2 | 1 | 9 | 0.7778 | 1 | 1 | 3 | 1 | 3 | 1 | 9 | 1 | 16 | 1 |
| 17_5091385 | 3 | 1 | NA | NA | 2 | 1 | 2 | 0.5 | 7 | 0.8571 | 2 | 0.5 | 3 | 0.6667 | 3 | 0.6667 | 8 | 0.75 | 16 | 0.6875 |
| 17_5091395 | 4 | 0.5 | 1 | 1 | 1 | 0 | 1 | 1 | 7 | 0.5714 | 2 | 0.5 | 2 | 0.5 | 3 | 0.6667 | 7 | 0.2857 | 14 | 0.4286 |
| 17_5091400 | 3 | 1 | 1 | 1 | 2 | 1 | 1 | 1 | 7 | 1 | 1 | 1 | 3 | 0.6667 | 4 | 0.75 | 9 | 0.8889 | 17 | 0.8235 |
| 17_5091408 | 3 | 0.3333 | 2 | 0.5 | 4 | 0.75 | 2 | 0 | 11 | 0.4545 | 2 | 0.5 | 4 | 0.75 | 4 | 0 | 9 | 0.6667 | 19 | 0.5263 |
| 17_5091430 | 4 | 0.5 | 2 | 0.5 | 3 | 0 | 2 | 0 | 11 | 0.2727 | 1 | 1 | 4 | 1 | 3 | 0.3333 | 6 | 0.5 | 14 | 0.6429 |
| 17_5091484 | 5 | 0.6 | 3 | 0.3333 | 3 | 1 | 2 | 1 | 13 | 0.6923 | 2 | 1 | 6 | 0.6667 | 5 | 0.6 | 7 | 0.4286 | 20 | 0.6 |
| 17_5091498 | 6 | 1 | 5 | 1 | 4 | 1 | 3 | 0.6667 | 18 | 0.9444 | 3 | 1 | 5 | 0.6 | 5 | 0.8 | 9 | 0.8889 | 22 | 0.8182 |
| 17_5091504 | 7 | 0 | 5 | 0.2 | 5 | 0.6 | 3 | 0 | 20 | 0.2 | 3 | 0.3333 | 5 | 0 | 5 | 0 | 8 | 0.25 | 21 | 0.1429 |
| 17_5091510 | 7 | 0.2857 | 5 | 0.4 | 3 | 0.3333 | 4 | 0 | 19 | 0.2632 | 4 | 0 | 6 | 0.5 | 4 | 0 | 5 | 0.2 | 19 | 0.2105 |
| 17_5091513 | 3 | 0 | 5 | 1 | 2 | 0 | 3 | 1 | 13 | 0.6154 | 4 | 1 | 5 | 0.4 | 5 | 0.4 | 10 | 0.4 | 24 | 0.5 |
| 17_5091516 | 6 | 0.3333 | 4 | 0.5 | 4 | 0.25 | 4 | 0.25 | 18 | 0.3333 | 4 | 0.75 | 6 | 0.3333 | 5 | 0.4 | 10 | 0.6 | 25 | 0.52 |
| 17_5091523 | 7 | 0.5714 | 5 | 0.4 | 2 | 1 | 4 | 0.75 | 18 | 0.6111 | 4 | 1 | 5 | 0.8 | 4 | 1 | 11 | 0.5455 | 24 | 0.75 |
| 17_5091528 | 6 | 0.5 | 4 | 0.25 | 4 | 0 | 4 | 0.75 | 18 | 0.3889 | 5 | 0.4 | 5 | 0.6 | 3 | 0.3333 | 9 | 0.7778 | 22 | 0.5909 |
| 17_5091533 | 7 | 0.2857 | 5 | 0.4 | 5 | 0.6 | 4 | 0.25 | 21 | 0.381 | 4 | 0.5 | 5 | 0.2 | 5 | 0.2 | 10 | 0.3 | 24 | 0.2917 |
| 17_5091539 | 6 | 0.8333 | 4 | 1 | 4 | 0.5 | 4 | 1 | 18 | 0.8333 | 5 | 0.4 | 6 | 0.5 | 5 | 0.4 | 10 | 0.6 | 26 | 0.5 |
| 17_5091540 | 6 | 0 | 4 | 0.25 | 6 | 0 | 3 | 0 | 19 | 0.0526 | 4 | 0.25 | 5 | 0 | 6 | 0 | 7 | 0.1429 | 22 | 0.0909 |
| 17_5091546 | 7 | 0.1429 | 3 | 0.3333 | 4 | 0 | 2 | 0 | 16 | 0.125 | 5 | 0.6 | 4 | 0.5 | 5 | 0.4 | 6 | 0.1667 | 20 | 0.4 |
| 17_5091558 | 8 | 0.25 | 5 | 0 | 7 | 0 | 3 | 0 | 23 | 0.087 | 6 | 0 | 6 | 0 | 6 | 0.1667 | 10 | 0.1 | 28 | 0.0714 |
| 17_5091564 | 7 | 0 | 4 | 0.25 | 2 | 0 | 2 | 0 | 15 | 0.0667 | 6 | 0 | 4 | 0 | 4 | 0 | 6 | 0.3333 | 20 | 0.1 |
| 17_5091572 | 7 | 0.7143 | 5 | 0.4 | 7 | 0.4286 | 3 | 0.3333 | 22 | 0.5 | 3 | 0.3333 | 5 | 0.2 | 5 | 0.4 | 9 | 0.3333 | 22 | 0.3182 |
| 17_5091573 | 4 | 0.25 | 3 | 0.3333 | 5 | 0.2 | 3 | 0.3333 | 15 | 0.2667 | 4 | 0 | 5 | 0.6 | 4 | 0.5 | 8 | 0.125 | 21 | 0.2857 |
| 17_5091579 | 8 | 0 | 5 | 0 | 8 | 0 | 4 | 0.5 | 25 | 0.08 | 7 | 0.1429 | 7 | 0.1429 | 5 | 0 | 11 | 0.2727 | 30 | 0.1667 |
| 17_5099482 | 9 | 0.4444 | 7 | 0.5714 | 7 | 0.4286 | 3 | 0.3333 | 26 | 0.4615 | 6 | 0.8333 | 8 | 0.75 | 7 | 0.5714 | 10 | 0.7 | 31 | 0.7097 |
| 17_5099491 | 9 | 0.3333 | 7 | 0.5714 | 9 | 0.2222 | 5 | 0.4 | 30 | 0.3667 | 9 | 0.3333 | 10 | 0.2 | 8 | 0.25 | 13 | 0.1538 | 40 | 0.225 |
| 17_5099494 | 8 | 0.875 | 6 | 1 | 9 | 1 | 3 | 0.6667 | 26 | 0.9231 | 8 | 0.75 | 10 | 0.9 | 7 | 0.8571 | 10 | 1 | 35 | 0.8857 |
| 17_5099508 | 6 | 0.6667 | 6 | 0.6667 | 11 | 0.5455 | 5 | 0.6 | 28 | 0.6071 | 7 | 0.4286 | 10 | 0.6 | 8 | 0.5 | 10 | 0.5 | 35 | 0.5143 |
| 17_5099510 | 7 | 0.7143 | 6 | 0.8333 | 7 | 0.7143 | 4 | 0.75 | 24 | 0.75 | 7 | 0.7143 | 6 | 0.8333 | 6 | 0.5 | 12 | 0.75 | 31 | 0.7097 |
| 17_5099521 | 3 | 0 | 4 | 0.25 | 2 | 0 | 2 | 0 | 11 | 0.0909 | 5 | 0.2 | 7 | 0 | 4 | 0 | 8 | 0.125 | 24 | 0.0833 |
| 17_5099535 | 6 | 0 | 6 | 0.3333 | 3 | 0 | 3 | 0.3333 | 18 | 0.1667 | 5 | 0 | 7 | 0.1429 | 4 | 0.5 | 7 | 0.2857 | 23 | 0.2174 |
| 17_5099538 | 5 | 0.2 | 5 | 0.2 | 3 | 0 | 2 | 0.5 | 15 | 0.2 | 5 | 0 | 5 | 0 | 3 | 0 | 9 | 0.2222 | 22 | 0.0909 |
| 17_5100480 | NA | NA | 1 | 1 | NA | NA | NA | NA | 1 | 1 | NA | NA | NA | NA | NA | NA | 1 | 1 | 1 | 1 |

| | | | | | | | | | | | | | | | | | | | | |
|---|---|---|---|---|---|---|---|---|---|---|---|---|---|---|---|---|---|---|---|---|
| 17_510058 | NA | NA | 1 | 1 | 2 | 0.5 | NA | NA | 3 | 0.6667 | NA | NA | NA | NA | NA | NA | 2 | 0.5 | 2 | 0.5 |
| 17_510062 | NA | NA | 1 | 1 | 3 | 0.3333 | NA | NA | 4 | 0.5 | NA | NA | NA | NA | NA | NA | 2 | 0.5 | 2 | 0.5 |
| 17_510063 | NA | NA | 1 | 1 | 2 | 0 | NA | NA | 3 | 0.3333 | NA | NA | NA | NA | NA | NA | 2 | 0.5 | 2 | 0.5 |
| 17_510065 | 1 | 0 | 1 | 0 | 3 | 0.3333 | NA | NA | 5 | 0.2 | NA | NA | NA | NA | NA | NA | 3 | 0.3333 | 3 | 0.3333 |
| 17_510067 | 1 | 0 | NA | NA | 2 | 0.5 | NA | NA | 3 | 0.3333 | NA | NA | NA | NA | NA | NA | 4 | 0.5 | 4 | 0.5 |
| 17_510401 | 1 | 0 | 1 | 1 | 4 | 0.25 | NA | NA | 6 | 0.3333 | 2 | 1 | 2 | 0.5 | 1 | 0 | NA | NA | 5 | 0.6 |
| 17_510404 | NA | NA | NA | NA | 5 | 0.6 | NA | NA | 5 | 0.6 | 2 | 0.5 | 2 | 0.5 | NA | NA | NA | NA | 4 | 0.5 |
| 17_510406 | 2 | 0 | NA | NA | 5 | 0.2 | NA | NA | 7 | 0.1429 | 1 | 0 | 1 | 1 | NA | NA | NA | NA | 2 | 0.5 |
| 17_510411 | 6 | 0.6667 | 4 | 0.75 | 7 | 0.5714 | 1 | 1 | 18 | 0.6667 | 1 | 0 | 4 | 0.5 | 2 | 0.5 | 1 | 1 | 8 | 0.5 |
| 17_510413 | 6 | 0.6667 | 3 | 1 | 6 | 0.5 | 1 | 1 | 16 | 0.6875 | 1 | 1 | 5 | 0.8 | 1 | 1 | 2 | 1 | 9 | 0.8889 |
| 17_510419 | 3 | 0 | 4 | 0 | 3 | 0.3333 | 1 | 0 | 11 | 0.0909 | NA | NA | 4 | 0.25 | 1 | 0 | 1 | 0 | 6 | 0.1667 |
| 17_510424 | 4 | 0.75 | 3 | 0.6667 | 5 | 0.6 | 1 | 0 | 13 | 0.6154 | NA | NA | 5 | 0.4 | 2 | 0.5 | 2 | 0.5 | 9 | 0.4444 |
| 17_510428 | 2 | 1 | 3 | 1 | 2 | 1 | NA | NA | 7 | 1 | NA | NA | 3 | 1 | 1 | 1 | 1 | 1 | 5 | 1 |
| 17_510429 | 4 | 0.75 | 4 | 0.75 | 5 | 0.6 | 1 | 0 | 14 | 0.6429 | NA | NA | 3 | 1 | 1 | 1 | 1 | 1 | 5 | 1 |
| 17_510429 | 3 | 1 | 4 | 0.75 | 5 | 0.8 | NA | NA | 12 | 0.8333 | NA | NA | 3 | 1 | 1 | 1 | 1 | 0 | 5 | 0.8 |
| 17_520435 | NA | NA | 2 | 0.5 | 2 | 1 | NA | NA | 4 | 0.75 | 1 | 1 | NA | NA | NA | NA | NA | NA | 1 | 1 |
| 17_520464 | 2 | 0.5 | 2 | 1 | 2 | 1 | 2 | 1 | 8 | 0.875 | 2 | 1 | 3 | 1 | NA | NA | 2 | 1 | 7 | 1 |
| 17_520464 | 1 | 0 | 2 | 0 | 2 | 0.5 | 1 | 0 | 6 | 0.1667 | 3 | 0 | 3 | 0.6667 | 1 | 0 | 2 | 0 | 9 | 0.2222 |
| 17_520465 | 2 | 0.5 | 4 | 0 | 1 | 0 | 3 | 0 | 10 | 0.1 | 2 | 0 | 3 | 0.6667 | 1 | 0 | 3 | 0.6667 | 9 | 0.4444 |
| 17_520467 | 2 | 1 | 2 | 0.5 | 2 | 1 | 3 | 1 | 9 | 0.8889 | 3 | 1 | 1 | 1 | 1 | 1 | 2 | 0.5 | 7 | 0.8571 |
| 17_520467 | 1 | 1 | 2 | 1 | 2 | 0 | 2 | 1 | 7 | 0.7143 | 3 | 0.3333 | 1 | 0 | 1 | 1 | NA | NA | 5 | 0.4 |
| 17_520467 | 2 | 1 | 2 | 1 | 2 | 1 | 2 | 0.5 | 8 | 0.875 | 3 | 0.3333 | 1 | 0 | 1 | 0 | 2 | 0.5 | 7 | 0.2857 |
| 17_520467 | 2 | 0.5 | 3 | 1 | NA | NA | 2 | 1 | 7 | 0.8571 | 3 | 1 | 1 | 0 | 1 | 1 | 2 | 0.5 | 7 | 0.7143 |
| 17_520468 | 2 | 0 | 2 | 0.5 | 2 | 1 | 1 | 0 | 7 | 0.4286 | 3 | 0.6667 | 1 | 0 | NA | NA | 1 | 0 | 5 | 0.4 |
| 17_520468 | 2 | 0.5 | 2 | 0.5 | 2 | 0.5 | 3 | 0.3333 | 9 | 0.4444 | 3 | 0.3333 | 1 | 0 | NA | NA | 1 | 0 | 5 | 0.2 |
| 17_520469 | 1 | 1 | 2 | 1 | 2 | 1 | 3 | 1 | 8 | 1 | 1 | 1 | 1 | 1 | 1 | 1 | 1 | 1 | 4 | 1 |
| 17_520469 | 2 | 1 | 2 | 1 | 2 | 1 | 3 | 1 | 9 | 1 | 3 | 1 | NA | NA | NA | NA | NA | NA | 3 | 1 |
| 17_520470 | 1 | 1 | 2 | 0.5 | 2 | 1 | 2 | 1 | 7 | 0.8571 | 2 | 1 | 1 | 1 | 1 | 1 | 1 | 1 | 5 | 1 |
| 17_520471 | NA | NA | 3 | 0.3333 | 1 | 1 | 1 | 1 | 5 | 0.6 | 3 | 0 | 1 | 0 | 1 | 1 | 1 | 1 | 6 | 0.3333 |
| 17_520471 | 1 | 1 | 2 | 0.5 | 2 | 0 | 1 | 1 | 6 | 0.5 | 2 | 0 | 1 | 1 | 1 | 1 | 1 | 0 | 5 | 0.4 |
| 17_520472 | 1 | 0 | 2 | 0.5 | 2 | 0.5 | 1 | 0 | 6 | 0.3333 | 1 | 0 | 1 | 0 | 1 | 1 | 1 | 0 | 4 | 0.25 |
| 17_520898 | 1 | 0 | 1 | 1 | NA | NA | 1 | 1 | 3 | 0.6667 | 3 | 1 | NA | NA | 1 | 1 | 1 | 0 | 5 | 0.8 |
| 17_520898 | 1 | 0 | 1 | 1 | NA | NA | 1 | 0 | 3 | 0.3333 | 1 | 0 | NA | NA | 1 | 1 | 1 | 0 | 3 | 0.3333 |
| 17_520899 | 1 | 0 | 1 | 1 | NA | NA | 1 | 0 | 3 | 0.3333 | 3 | 0.3333 | NA | NA | 1 | 1 | 1 | 1 | 5 | 0.6 |
| 17_521188 | NA | NA | 1 | 0 | NA | NA | 1 | 1 | 2 | 0.5 | 2 | 0 | NA | NA | 1 | 1 | NA | NA | 3 | 0.3333 |
| 17_537909 | 12 | 0.5 | 18 | 0.7222 | 20 | 0.45 | 14 | 0.6429 | 64 | 0.5781 | 5 | 0.2 | 20 | 0.6 | 13 | 0.5385 | 15 | 0.8 | 53 | 0.6038 |
| 17_537909 | 11 | 0.4545 | 18 | 0.3333 | 19 | 0.3158 | 12 | 0.3333 | 60 | 0.35 | 5 | 0.4 | 19 | 0.3684 | 10 | 0.3 | 14 | 0.5 | 48 | 0.3958 |
| 17_538134 | 13 | 0.1538 | 19 | 0 | 20 | 0.05 | 17 | 0.0588 | 69 | 0.058 | 7 | 0 | 18 | 0.1111 | 10 | 0.1 | 11 | 0 | 46 | 0.0652 |
| 17_538134 | 17 | 0.1765 | 20 | 0.2 | 21 | 0.0952 | 18 | 0 | 76 | 0.1184 | 10 | 0 | 21 | 0.1429 | 11 | 0.1818 | 16 | 0 | 58 | 0.0862 |
| 17_538135 | 16 | 0.6875 | 22 | 0.8636 | 23 | 0.6522 | 20 | 0.95 | 81 | 0.7901 | 9 | 0.8889 | 24 | 0.8333 | 15 | 0.8667 | 17 | 0.9412 | 65 | 0.8769 |
| 17_538135 | 17 | 0.1176 | 22 | 0 | 23 | 0.1304 | 22 | 0 | 84 | 0.0595 | 9 | 0.1111 | 25 | 0.08 | 16 | 0.0625 | 16 | 0 | 66 | 0.0606 |
| 17_538136 | 19 | 0.5263 | 23 | 0.3913 | 24 | 0.5417 | 19 | 0.5263 | 85 | 0.4941 | 9 | 0.5556 | 22 | 0.4091 | 16 | 0.375 | 17 | 0.2941 | 64 | 0.3906 |
| 17_539777 | 1 | 1 | 2 | 0 | 4 | 0.25 | 2 | 0.5 | 9 | 0.3333 | 2 | 0.5 | NA | NA | 3 | 0 | 3 | 0.3333 | 8 | 0.25 |
| 17_539780 | 1 | 0 | 3 | 0.3333 | 4 | 0 | 2 | 0.5 | 10 | 0.2 | 2 | 0.5 | NA | NA | 3 | 0.3333 | 3 | 0 | 8 | 0.25 |
| 17_539794 | 1 | 0 | 2 | 0 | 4 | 0.25 | 5 | 0.4 | 12 | 0.25 | 3 | 0.3333 | 1 | 0 | 5 | 0.4 | 4 | 0 | 13 | 0.2308 |
| 17_545815 | 2 | 0 | 2 | 0 | 3 | 0 | 5 | 0.2 | 12 | 0.0833 | NA | NA | 4 | 0.25 | 2 | 0 | 1 | 0 | 7 | 0.1429 |
| 17_545816 | 1 | 0 | 2 | 0.5 | 3 | 0 | 5 | 0.4 | 11 | 0.2727 | NA | NA | 3 | 0.6667 | 2 | 0 | 1 | 0 | 6 | 0.3333 |
| 17_545817 | 2 | 0 | 2 | 0.5 | 2 | 1 | 2 | 0.5 | 8 | 0.5 | NA | NA | 2 | 1 | 1 | 1 | 1 | 1 | 4 | 1 |
| 17_545822 | 1 | 0 | 2 | 0 | NA | NA | 5 | 0.6 | 8 | 0.375 | NA | NA | 1 | 1 | 2 | 0 | 1 | 0 | 4 | 0.25 |
| 17_545823 | 2 | 0 | 1 | 0 | 1 | 0 | 4 | 0.25 | 8 | 0.125 | NA | NA | 2 | 0.5 | 3 | 0 | 1 | 0 | 6 | 0.1667 |
| 17_545835 | 2 | 0.5 | 2 | 0.5 | 2 | 0.5 | 3 | 0.6667 | 9 | 0.5556 | NA | NA | 2 | 1 | 3 | 0 | 2 | 0 | 7 | 0.2857 |
| 17_545836 | 1 | 1 | 1 | 1 | 2 | 1 | 3 | 0.6667 | 7 | 0.8571 | NA | NA | NA | NA | 2 | 0 | 2 | 0.5 | 4 | 0.25 |
| 17_545837 | 1 | 1 | 2 | 0 | 2 | 0.5 | 3 | 0.6667 | 8 | 0.5 | NA | NA | 2 | 1 | 3 | 0.3333 | 2 | 0.5 | 7 | 0.5714 |
| 17_545838 | NA | NA | NA | NA | 2 | 0.5 | 1 | 1 | 3 | 0.6667 | NA | NA | 1 | 0 | 1 | 0 | 2 | 0.5 | 4 | 0.25 |
| 17_545839 | NA | NA | NA | NA | 2 | 1 | 2 | 1 | 4 | 1 | NA | NA | 1 | 1 | 2 | 0.5 | 2 | 1 | 5 | 0.8 |
| 17_546057 | NA | NA | NA | NA | NA | NA | 1 | 1 | 1 | 1 | NA | NA | 1 | 1 | NA | NA | NA | NA | 1 | 1 |
| 17_557891 | 1 | 1 | 2 | 0.5 | NA | NA | 1 | 0 | 4 | 0.5 | 1 | 0 | 3 | 0.6667 | 1 | 0 | 3 | 0 | 8 | 0.25 |
| 17_557892 | 1 | 1 | 2 | 1 | NA | NA | 1 | 1 | 4 | 1 | 1 | 1 | 2 | 1 | 1 | 1 | NA | NA | 4 | 1 |
| 17_557892 | 1 | 1 | 1 | 1 | NA | NA | 1 | 0 | 3 | 0.6667 | 1 | 1 | 3 | 0.6667 | NA | NA | 2 | 0.5 | 6 | 0.6667 |
| 17_557892 | 1 | 0 | 2 | 0.5 | NA | NA | 1 | 0 | 4 | 0.25 | 1 | 0 | 2 | 1 | 2 | 0 | 4 | 0.25 | 9 | 0.3333 |
| 17_557892 | 1 | 0 | 2 | 0 | NA | NA | 1 | 1 | 4 | 0.25 | 1 | 1 | 3 | 0.3333 | 2 | 1 | 4 | 0.5 | 10 | 0.6 |
| 17_557894 | 1 | 1 | 2 | 0 | NA | NA | 1 | 1 | 4 | 0.5 | 1 | 0 | 3 | 0 | 1 | 1 | 3 | 0.6667 | 8 | 0.375 |
| 17_557894 | 1 | 1 | 2 | 1 | NA | NA | 1 | 1 | 4 | 1 | 1 | 0 | 3 | 1 | 2 | 1 | 4 | 0.75 | 10 | 0.8 |
| 17_557898 | 1 | 0 | 1 | 1 | NA | NA | 1 | 1 | 3 | 0.6667 | 1 | 1 | 1 | 0 | 2 | 0 | 3 | 0.3333 | 7 | 0.2857 |
| 17_557997 | 2 | 1 | 2 | 1 | NA | NA | 1 | 1 | 5 | 1 | NA | NA | 1 | 1 | 3 | 1 | 1 | 1 | 5 | 1 |

| | | | | | | | | | | | | | | | | | | | | |
|---|---|---|---|---|---|---|---|---|---|---|---|---|---|---|---|---|---|---|---|---|
| 17_558053 | 2 | 0 | 2 | 0.5 | 1 | 0 | 2 | 0.5 | 7 | 0.2857 | 1 | 1 | 3 | 0 | 1 | 0 | 3 | 0 | 8 | 0.125 |
| 17_558173 | 1 | 0 | 2 | 1 | 1 | 1 | 1 | 1 | 5 | 0.8 | 1 | 0 | 3 | 0.6667 | 3 | 0.6667 | 3 | 0.6667 | 10 | 0.6 |
| 17_558174 | 1 | 1 | 2 | 1 | 1 | 1 | 1 | 1 | 5 | 1 | NA | NA | 2 | 1 | 1 | 1 | 2 | 1 | 5 | 1 |
| 17_558174 | 1 | 0 | 2 | 1 | 1 | 1 | 1 | 1 | 5 | 0.8 | 1 | 1 | 3 | 0.3333 | 2 | 0.5 | 3 | 1 | 9 | 0.6667 |
| 17_558182 | 1 | 0 | 3 | 0 | 1 | 1 | 1 | 0 | 6 | 0.1667 | 1 | 0 | 4 | 0 | 3 | 0.6667 | 3 | 0.3333 | 11 | 0.2727 |
| 17_558184 | 1 | 0 | 3 | 0 | 1 | 0 | 1 | 1 | 6 | 0.1667 | 1 | 0 | 4 | 0.25 | 2 | 0.5 | 2 | 0 | 9 | 0.2222 |
| 17_558187 | 1 | 0 | 2 | 0 | 1 | 1 | 2 | 0.5 | 6 | 0.3333 | 1 | 0 | 4 | 0.25 | 3 | 0.6667 | 1 | 0 | 9 | 0.3333 |
| 17_558196 | 1 | 0 | 3 | 1 | 1 | 1 | 2 | 0 | 7 | 0.5714 | 1 | 1 | 4 | 0.5 | 2 | 0.5 | 3 | 0 | 10 | 0.4 |
| 17_558200 | 1 | 0 | 3 | 0 | 2 | 0 | 2 | 0.5 | 8 | 0.125 | 1 | 0 | 4 | 0.25 | 2 | 0 | 3 | 0 | 10 | 0.1 |
| 17_559591 | NA | NA | NA | NA | NA | NA | 1 | 1 | 1 | 1 | NA | NA | NA | NA | NA | NA | 1 | 1 | 1 | 1 |
| 17_559593 | NA | NA | NA | NA | NA | NA | 1 | 1 | 1 | 1 | NA | NA | NA | NA | NA | NA | 1 | 1 | 1 | 1 |
| 17_561724 | 2 | 1 | 1 | 1 | 1 | 0 | NA | NA | 4 | 0.75 | NA | NA | 1 | 1 | NA | NA | NA | NA | 1 | 1 |
| 17_561727 | 2 | 1 | 2 | 1 | 2 | 1 | NA | NA | 6 | 1 | 1 | 1 | 2 | 1 | NA | NA | NA | NA | 3 | 1 |
| 17_561731 | 2 | 1 | 1 | 1 | 1 | 1 | NA | NA | 4 | 1 | 1 | 1 | 1 | 1 | NA | NA | NA | NA | 2 | 1 |
| 17_561732 | 2 | 1 | 2 | 0.5 | 2 | 0.5 | NA | NA | 6 | 0.6667 | 1 | 1 | 2 | 0 | NA | NA | NA | NA | 3 | 0.3333 |
| 17_561734 | 2 | 1 | 2 | 0.5 | 2 | 0.5 | NA | NA | 6 | 0.6667 | 1 | 0 | 2 | 0.5 | NA | NA | NA | NA | 3 | 0.3333 |
| 17_561734 | 2 | 0 | 2 | 0.5 | 2 | 0.5 | NA | NA | 6 | 0.3333 | 1 | 0 | 2 | 0.5 | NA | NA | NA | NA | 3 | 0.3333 |
| 17_562043 | NA | NA | 1 | 1 | 2 | 0 | 1 | 1 | 4 | 0.5 | 2 | 0.5 | 2 | 0.5 | NA | NA | 1 | 1 | 5 | 0.6 |
| 17_562044 | NA | NA | 1 | 1 | 2 | 1 | NA | NA | 3 | 1 | 2 | 0.5 | 2 | 1 | NA | NA | 1 | 1 | 5 | 0.8 |
| 17_562044 | NA | NA | 1 | 0 | 2 | 0 | 1 | 1 | 4 | 0.25 | 2 | 0.5 | 2 | 0 | NA | NA | 1 | 0 | 5 | 0.2 |
| 17_562045 | NA | NA | 1 | 1 | 2 | 0 | 1 | 0 | 4 | 0.25 | 1 | 0 | 2 | 0.5 | NA | NA | 1 | 1 | 4 | 0.5 |
| 17_562047 | NA | NA | 1 | 0 | 2 | 0.5 | 1 | 0 | 4 | 0.25 | 2 | 0 | 2 | 0 | NA | NA | 1 | 1 | 5 | 0.2 |
| 17_562047 | NA | NA | 1 | 1 | 1 | 1 | 1 | 1 | 3 | 1 | 2 | 1 | 2 | 1 | NA | NA | 1 | 1 | 5 | 1 |
| 17_562747 | 2 | 0.5 | NA | NA | NA | NA | NA | NA | 2 | 0.5 | NA | NA | 1 | 0 | 1 | 1 | 2 | 0 | 4 | 0.25 |
| 17_562748 | 3 | 0.3333 | NA | NA | NA | NA | NA | NA | 3 | 0.3333 | 1 | 0 | 1 | 0 | 1 | 0 | 2 | 0.5 | 5 | 0.2 |
| 17_562752 | 2 | 1 | NA | NA | NA | NA | NA | NA | 2 | 1 | 1 | 1 | 1 | 1 | 3 | 0.6667 | 2 | 1 | 7 | 0.8571 |
| 17_562784 | 1 | 0 | 4 | 0.25 | 4 | 0.25 | 3 | 0 | 12 | 0.1667 | NA | NA | NA | NA | 2 | 0.5 | 1 | 0 | 3 | 0.3333 |
| 17_562784 | 1 | 0 | 4 | 0.25 | 3 | 0 | 2 | 0 | 10 | 0.1 | NA | NA | 1 | 0 | 2 | 0 | 1 | 1 | 4 | 0.25 |
| 17_562833 | 2 | 0 | 4 | 0.25 | 4 | 0.25 | 3 | 0.3333 | 13 | 0.2308 | NA | NA | NA | NA | 3 | 0.6667 | 1 | 0 | 4 | 0.5 |
| 17_562834 | 2 | 0.5 | 3 | 0.3333 | 5 | 0.6 | 3 | 0.6667 | 13 | 0.5385 | NA | NA | 1 | 0 | 3 | 0.6667 | 1 | 0 | 5 | 0.4 |
| 17_563238 | 2 | 1 | 5 | 0.8 | 3 | 0.3333 | 2 | 0.5 | 12 | 0.6667 | 2 | 0.5 | 2 | 0.5 | 4 | 0.75 | 3 | 0.3333 | 11 | 0.5455 |
| 17_563240 | 4 | 0.75 | 6 | 0.6667 | 2 | 1 | 5 | 0.8 | 17 | 0.7647 | 2 | 0.5 | 5 | 0.8 | 4 | 0.75 | 2 | 1 | 13 | 0.7692 |
| 17_563241 | 3 | 1 | 7 | 0.8571 | 4 | 1 | 5 | 0.8 | 19 | 0.8947 | 2 | 1 | 6 | 0.6667 | 6 | 0.5 | 2 | 0.5 | 16 | 0.625 |
| 17_563241 | 3 | 0.3333 | 7 | 0.1429 | 4 | 0 | 3 | 0.3333 | 17 | 0.1765 | 2 | 0 | 6 | 0 | 6 | 0.1667 | 3 | 0.3333 | 17 | 0.1176 |
| 17_563242 | 4 | 0.25 | 6 | 0 | 4 | 0.25 | 5 | 0 | 19 | 0.1053 | 2 | 0 | 6 | 0.1667 | 6 | 0.1667 | 2 | 0 | 16 | 0.125 |
| 17_563244 | 2 | 0.5 | 5 | 0.4 | 4 | 0.25 | 5 | 0.4 | 16 | 0.375 | 2 | 0 | 6 | 0.1667 | 6 | 0.5 | 3 | 0.3333 | 17 | 0.2941 |
| 17_563245 | 3 | 0.3333 | 5 | 0.2 | 3 | 0.3333 | 4 | 0 | 15 | 0.2 | 2 | 0.5 | 5 | 0.2 | 6 | 0.5 | 3 | 0.3333 | 16 | 0.375 |
| 17_563258 | 3 | 0 | 5 | 0 | 3 | 0 | 4 | 0.25 | 15 | 0.0667 | 1 | 0 | 5 | 0 | 6 | 0.1667 | 2 | 0 | 14 | 0.0714 |
| 17_563754 | 6 | 0.6667 | 7 | 0.4286 | 12 | 0.0833 | 7 | 0.1429 | 32 | 0.2812 | 5 | 0 | 8 | 0.25 | 8 | 0.125 | 9 | 0.1111 | 30 | 0.1333 |
| 17_563767 | 1 | 1 | NA | NA | NA | NA | NA | NA | 1 | 1 | 1 | 1 | NA | NA | NA | NA | NA | NA | 1 | 1 |
| 17_563867 | 5 | 0 | 4 | 0.25 | 3 | 0 | 5 | 0 | 17 | 0.0588 | 4 | 0 | 7 | 0.1429 | 2 | 0 | 5 | 0 | 18 | 0.0556 |
| 17_564035 | NA | NA | 1 | 1 | 1 | 1 | 2 | 0.5 | 4 | 0.75 | NA | NA | 1 | 0 | NA | NA | 1 | 1 | 2 | 0.5 |
| 17_564078 | 2 | 0 | 3 | 0.3333 | 6 | 0.1667 | 3 | 0.6667 | 14 | 0.2857 | 3 | 0 | 3 | 0 | 3 | 0 | 3 | 0.6667 | 12 | 0.1667 |
| 17_564083 | 2 | 0.5 | 2 | 0.5 | 6 | 0.6667 | 3 | 0 | 13 | 0.4615 | NA | NA | 3 | 0.6667 | 3 | 1 | 3 | 0.6667 | 9 | 0.7778 |
| 17_564086 | 3 | 0.3333 | 3 | 0 | 4 | 0 | 3 | 0.3333 | 13 | 0.1538 | NA | NA | 2 | 0 | 2 | 1 | 3 | 0 | 7 | 0.2857 |
| 17_564086 | 3 | 0.3333 | 3 | 0.3333 | 4 | 0 | 2 | 0 | 12 | 0.1667 | NA | NA | 3 | 0.3333 | 3 | 0.3333 | 3 | 0 | 9 | 0.2222 |
| 17_564088 | 3 | 0.3333 | 3 | 0 | 6 | 0 | 4 | 0 | 16 | 0.0625 | NA | NA | 2 | 0 | 3 | 0 | 2 | 0.5 | 7 | 0.1429 |
| 17_564090 | 3 | 0.6667 | 1 | 0 | 7 | 0.5714 | 2 | 0.5 | 13 | 0.5385 | NA | NA | 2 | 1 | 2 | 1 | 1 | 0 | 5 | 0.8 |
| 17_564106 | 3 | 0.3333 | 1 | 0 | 4 | 0 | 1 | 0 | 9 | 0.1111 | NA | NA | 1 | 0 | 2 | 0.5 | NA | NA | 3 | 0.3333 |
| 17_564108 | 3 | 0.3333 | 1 | 0 | 4 | 0.25 | NA | NA | 8 | 0.25 | NA | NA | 1 | 1 | 2 | 0.5 | 2 | 0.5 | 5 | 0.6 |
| 17_564108 | 1 | 1 | 1 | 1 | 4 | 1 | 1 | 1 | 7 | 1 | NA | NA | 1 | 1 | 1 | 1 | 2 | 1 | 4 | 1 |
| 17_564109 | 3 | 0 | 1 | 0 | 4 | 0.5 | 1 | 0 | 9 | 0.2222 | NA | NA | 1 | 0 | 1 | 0 | 2 | 0.5 | 4 | 0.25 |
| 17_564109 | 3 | 0.6667 | 1 | 0 | 4 | 0.75 | 1 | 1 | 9 | 0.6667 | NA | NA | 1 | 0 | 1 | 0 | 2 | 0.5 | 4 | 0.25 |
| 17_564110 | 2 | 1 | 1 | 0 | 3 | 1 | 1 | 1 | 7 | 0.8571 | NA | NA | 1 | 1 | 1 | 1 | 1 | 1 | 3 | 1 |
| 17_564110 | 2 | 1 | NA | NA | 4 | 0.5 | NA | NA | 6 | 0.6667 | NA | NA | 1 | 1 | 1 | 1 | 2 | 1 | 4 | 1 |
| 17_564111 | 3 | 0.3333 | 1 | 0 | 4 | 0.25 | 1 | 1 | 9 | 0.3333 | NA | NA | NA | NA | 1 | 1 | 2 | 0 | 3 | 0.3333 |
| 17_564112 | 3 | 0.6667 | 1 | 0 | 5 | 0.8 | 1 | 1 | 10 | 0.7 | NA | NA | 1 | 1 | 1 | 1 | 1 | 1 | 3 | 1 |
| 17_564112 | 3 | 0.3333 | 1 | 0 | 5 | 0.4 | 1 | 0 | 10 | 0.3 | NA | NA | 1 | 1 | 1 | 0 | 1 | 0 | 3 | 0.3333 |
| 17_564113 | 3 | 0.6667 | 1 | 1 | 3 | 0.6667 | 1 | 1 | 8 | 0.75 | NA | NA | 1 | 1 | 1 | 0 | NA | NA | 2 | 0.5 |
| 17_564113 | 3 | 0.3333 | 1 | 0 | 4 | 0 | 1 | 0 | 9 | 0.1111 | NA | NA | NA | NA | 1 | 1 | 1 | 0 | 2 | 0.5 |
| 17_564114 | 3 | 0.6667 | 1 | 1 | 3 | 1 | NA | NA | 7 | 0.8571 | NA | NA | 1 | 1 | 1 | 0 | 1 | 1 | 3 | 0.6667 |
| 17_564114 | 3 | 0.3333 | 1 | 1 | 4 | 0.75 | 1 | 1 | 9 | 0.6667 | NA | NA | 1 | 1 | NA | NA | 1 | 0 | 2 | 0.5 |
| 17_564114 | 1 | 0 | 1 | 0 | 4 | 0.25 | 1 | 0 | 7 | 0.1429 | NA | NA | NA | NA | 1 | 1 | NA | NA | 1 | 1 |
| 17_564115 | 3 | 0.6667 | 1 | 0 | 3 | 0.3333 | 1 | 1 | 8 | 0.5 | NA | NA | 1 | 0 | 1 | 0 | 1 | 1 | 3 | 0.3333 |
| 17_564116 | 2 | 1 | 1 | 0 | 3 | 0.6667 | 1 | 1 | 7 | 0.7143 | NA | NA | 1 | 1 | 1 | 1 | 1 | 1 | 3 | 1 |

| | | | | | | | | | | | | | | | | | | | | |
|---|---|---|---|---|---|---|---|---|---|---|---|---|---|---|---|---|---|---|---|---|
| 17_564117 | 3 | 0.6667 | NA | NA | 3 | 0.3333 | 1 | 0 | 7 | 0.4286 | NA | NA | 1 | 0 | NA | NA | 1 | 1 | 2 | 0.5 |
| 17_564118 | 2 | 0.5 | NA | NA | 2 | 0.5 | 1 | 0 | 5 | 0.4 | NA | NA | 1 | 1 | NA | NA | 1 | 1 | 2 | 1 |
| 17_564118 | 3 | 0.6667 | 1 | 0 | 4 | 0.25 | 1 | 0 | 9 | 0.3333 | NA | NA | 1 | 0 | NA | NA | 1 | 1 | 2 | 0.5 |
| 17_564118 | 3 | 1 | 1 | 0 | 3 | 0.6667 | NA | NA | 7 | 0.7143 | NA | NA | 1 | 1 | NA | NA | 1 | 1 | 2 | 1 |
| 17_564119 | 3 | 1 | 1 | 1 | 4 | 1 | 1 | 1 | 9 | 1 | NA | NA | 1 | 0 | NA | NA | 1 | 1 | 2 | 0.5 |
| 17_564119 | 2 | 0 | 1 | 0 | 4 | 0.25 | 1 | 0 | 8 | 0.125 | NA | NA | 1 | 1 | NA | NA | 1 | 0 | 2 | 0.5 |
| 17_564120 | 3 | 0.6667 | 1 | 1 | 4 | 0.75 | 1 | 1 | 9 | 0.7778 | NA | NA | 1 | 1 | NA | NA | 1 | 1 | 2 | 1 |
| 17_564121 | 3 | 0.6667 | 1 | 1 | 3 | 0.3333 | 1 | 0 | 8 | 0.5 | NA | NA | 1 | 0 | NA | NA | 1 | 1 | 2 | 0.5 |
| 17_564122 | 3 | 0.6667 | 1 | 1 | 4 | 0.25 | 1 | 1 | 9 | 0.5556 | NA | NA | 1 | 1 | NA | NA | 1 | 1 | 2 | 1 |
| 17_564123 | 2 | 1 | 1 | 0 | 3 | 0.3333 | 1 | 1 | 7 | 0.5714 | NA | NA | 1 | 1 | NA | NA | 1 | 1 | 2 | 1 |
| 17_564124 | 3 | 1 | 1 | 1 | 4 | 1 | 1 | 1 | 9 | 1 | NA | NA | 1 | 1 | NA | NA | NA | NA | 1 | 1 |
| 17_564124 | 3 | 0.3333 | 1 | 0 | 3 | 0.3333 | 1 | 1 | 8 | 0.375 | NA | NA | 1 | 0 | NA | NA | 1 | 1 | 2 | 0.5 |
| 17_564126 | 3 | 0.6667 | 1 | 1 | 4 | 0.75 | 1 | 1 | 9 | 0.7778 | NA | NA | NA | NA | NA | NA | 1 | 1 | 1 | 1 |
| 17_564128 | 2 | 0.5 | 1 | 0 | 4 | 0.5 | 1 | 1 | 8 | 0.5 | NA | NA | 1 | 0 | NA | NA | 1 | 1 | 2 | 0.5 |
| 17_564129 | 2 | 0.5 | NA | NA | 3 | 0.6667 | 1 | 1 | 6 | 0.6667 | NA | NA | 1 | 1 | NA | NA | NA | NA | 1 | 1 |
| 17_564130 | 2 | 1 | 1 | 1 | 3 | 0.6667 | 1 | 1 | 7 | 0.8571 | NA | NA | 1 | 0 | NA | NA | 1 | 1 | 2 | 0.5 |
| 17_564130 | 2 | 0.5 | 1 | 1 | 4 | 0 | 1 | 0 | 8 | 0.25 | NA | NA | 1 | 1 | NA | NA | 1 | 1 | 2 | 1 |
| 17_564131 | 2 | 0.5 | 1 | 0 | 4 | 0.25 | 1 | 0 | 8 | 0.25 | NA | NA | 1 | 1 | NA | NA | 1 | 0 | 2 | 0.5 |
| 17_564131 | 2 | 0 | 1 | 0 | 4 | 0.5 | 1 | 1 | 8 | 0.375 | NA | NA | 1 | 1 | NA | NA | 1 | 0 | 2 | 0.5 |
| 17_564132 | 1 | 1 | NA | NA | 3 | 0.3333 | 1 | 0 | 5 | 0.4 | NA | NA | 1 | 0 | NA | NA | 1 | 1 | 2 | 0.5 |
| 17_564134 | 2 | 0.5 | 1 | 1 | 3 | 0.6667 | 1 | 1 | 7 | 0.7143 | NA | NA | 1 | 1 | NA | NA | NA | NA | 1 | 1 |
| 17_564135 | 2 | 1 | 1 | 0 | 3 | 1 | 1 | 1 | 7 | 0.8571 | NA | NA | 1 | 1 | NA | NA | NA | NA | 1 | 1 |
| 17_564137 | 2 | 0 | 1 | 1 | 2 | 1 | 1 | 0 | 6 | 0.5 | NA | NA | 1 | 1 | NA | NA | NA | NA | 1 | 1 |
| 17_564268 | 1 | 1 | 2 | 1 | NA | NA | NA | NA | 3 | 1 | 1 | 1 | NA | NA | NA | NA | 2 | 1 | 3 | 1 |
| 17_564271 | 1 | 1 | 1 | 0 | 1 | 1 | NA | NA | 3 | 0.6667 | 1 | 1 | NA | NA | NA | NA | 1 | 1 | 2 | 1 |
| 17_564274 | 1 | 0 | 3 | 0.6667 | NA | NA | NA | NA | 4 | 0.5 | 1 | 1 | NA | NA | NA | NA | 2 | 1 | 3 | 1 |
| 17_564307 | 10 | 0.7 | 10 | 0.6 | 10 | 0.8 | 12 | 0.5 | 42 | 0.6429 | 11 | 0.6364 | 9 | 0.5556 | 11 | 0.5455 | 10 | 0.8 | 41 | 0.6341 |
| 17_564308 | 9 | 0.8889 | 8 | 0.75 | 8 | 0.875 | 13 | 0.5385 | 38 | 0.7368 | 10 | 0.5 | 9 | 0.8889 | 7 | 0.7143 | 7 | 0.8571 | 33 | 0.7273 |
| 17_564308 | 9 | 0.1111 | 10 | 0 | 8 | 0.125 | 11 | 0 | 38 | 0.0526 | 9 | 0 | 8 | 0.25 | 9 | 0 | 9 | 0 | 35 | 0.0571 |
| 17_564310 | 9 | 1 | 8 | 1 | 10 | 0.9 | 10 | 1 | 37 | 0.973 | 10 | 0.9 | 8 | 1 | 10 | 0.9 | 10 | 1 | 38 | 0.9474 |
| 17_564310 | 11 | 1 | 4 | 0.75 | 8 | 0.875 | 12 | 1 | 35 | 0.9429 | 10 | 1 | 8 | 1 | 10 | 1 | 10 | 1 | 38 | 1 |
| 17_564312 | 9 | 0.7778 | 6 | 0.6667 | 9 | 0.6667 | 9 | 0.5556 | 33 | 0.6667 | 9 | 0.8889 | 6 | 0.8333 | 10 | 0.9 | 6 | 0.8333 | 31 | 0.871 |
| 17_564312 | 6 | 0.5 | 6 | 0 | 4 | 0.5 | 12 | 0.6667 | 28 | 0.4643 | 7 | 0.5714 | 6 | 0.3333 | 9 | 0.4444 | 8 | 0.375 | 30 | 0.4333 |
| 17_564314 | 7 | 0.1429 | 5 | 0 | 6 | 0 | 11 | 0.2727 | 29 | 0.1379 | 9 | 0.2222 | 8 | 0.125 | 9 | 0.1111 | 8 | 0.125 | 34 | 0.1471 |
| 17_564315 | 3 | 0.6667 | 3 | 0.6667 | 3 | 0.3333 | 6 | 0.6667 | 15 | 0.6 | 7 | 0.5714 | 5 | 0.8 | 2 | 0 | 1 | 1 | 15 | 0.6 |
| 17_564315 | 7 | 0.8571 | 5 | 1 | 7 | 0.8571 | 9 | 0.7778 | 28 | 0.8571 | 9 | 0.7778 | 6 | 1 | 7 | 0.7143 | 7 | 1 | 29 | 0.8621 |
| 17_564316 | 2 | 0 | 6 | 0.1667 | 3 | 0.3333 | 7 | 0.1429 | 18 | 0.1667 | 4 | 0.25 | 3 | 0 | 3 | 0 | 5 | 0 | 15 | 0.0667 |
| 17_564317 | 8 | 0 | 4 | 0 | 8 | 0.25 | 13 | 0.1538 | 33 | 0.1212 | 5 | 0.4 | 7 | 0.2857 | 8 | 0.25 | 8 | 0.125 | 28 | 0.25 |
| 17_564788 | 1 | 1 | 1 | 0 | NA | NA | NA | NA | 2 | 0.5 | 1 | 1 | 1 | 1 | NA | NA | NA | NA | 2 | 1 |
| 17_564788 | 1 | 1 | 1 | 1 | NA | NA | NA | NA | 2 | 1 | 1 | 1 | 1 | 1 | NA | NA | NA | NA | 2 | 1 |
| 17_564790 | 1 | 1 | 1 | 1 | NA | NA | NA | NA | 2 | 1 | 1 | 1 | 1 | 1 | NA | NA | NA | NA | 2 | 1 |
| 17_564793 | 1 | 1 | 2 | 0.5 | NA | NA | NA | NA | 3 | 0.6667 | 1 | 1 | 1 | 0 | NA | NA | NA | NA | 2 | 0.5 |
| 17_564795 | 1 | 1 | 2 | 1 | NA | NA | NA | NA | 3 | 1 | 1 | 1 | 1 | 1 | NA | NA | NA | NA | 2 | 1 |
| 17_564796 | 1 | 1 | 2 | 0 | NA | NA | NA | NA | 3 | 0.3333 | 1 | 0 | 1 | 0 | NA | NA | 1 | 1 | 3 | 0.3333 |
| 17_564797 | 1 | 1 | 1 | 1 | NA | NA | NA | NA | 2 | 1 | 1 | 1 | 1 | 1 | NA | NA | 1 | 1 | 3 | 1 |
| 17_564798 | 1 | 1 | 2 | 0 | NA | NA | NA | NA | 3 | 0.3333 | 1 | 0 | 1 | 1 | 1 | 1 | 1 | 1 | 4 | 0.75 |
| 17_564798 | 1 | 1 | 2 | 0.5 | NA | NA | NA | NA | 3 | 0.6667 | 1 | 1 | 1 | 1 | 1 | 1 | 1 | 1 | 4 | 1 |
| 17_564799 | NA | NA | 2 | 1 | NA | NA | NA | NA | 2 | 1 | 1 | 1 | 1 | 0 | 1 | 1 | 1 | 1 | 4 | 0.75 |
| 17_564799 | 1 | 1 | NA | NA | NA | NA | NA | NA | 1 | 1 | 1 | 1 | 1 | 0 | 1 | 0 | 1 | 0 | 4 | 0.25 |
| 17_564835 | 8 | 0 | 3 | 0 | 4 | 0 | 6 | 0.3333 | 21 | 0.0952 | 6 | 0.1667 | 6 | 0.1667 | 1 | 0 | 7 | 0 | 20 | 0.1 |
| 17_564836 | 7 | 0.1429 | 5 | 0 | 5 | 0 | 3 | 0.3333 | 20 | 0.1 | 6 | 0 | 4 | 0 | 2 | 0 | 6 | 0.3333 | 18 | 0.1111 |
| 17_564837 | 10 | 0.1 | 10 | 0.2 | 7 | 0.2857 | 9 | 0.3333 | 36 | 0.2222 | 9 | 0.1111 | 6 | 0.1667 | 7 | 0.2857 | 10 | 0.3 | 32 | 0.2188 |
| 17_564837 | 5 | 0 | 5 | 0.4 | 3 | 0 | 3 | 0 | 16 | 0.125 | 6 | 0.3333 | 6 | 0.3333 | 8 | 0.25 | 5 | 0.2 | 25 | 0.28 |
| 17_564838 | 10 | 0.3 | 4 | 0 | 6 | 0.8333 | 8 | 0.25 | 28 | 0.3571 | 9 | 0.2222 | 6 | 0.1667 | 6 | 0.1667 | 8 | 0.125 | 29 | 0.1724 |
| 17_564889 | NA | NA | 1 | 1 | NA | NA | NA | NA | 1 | 1 | NA | NA | 1 | 1 | NA | NA | NA | NA | 1 | 1 |
| 17_564940 | 4 | 0 | 3 | 0 | 8 | 0.125 | 10 | 0.3 | 25 | 0.16 | 4 | 0 | 5 | 0.2 | 9 | 0 | 4 | 0.25 | 22 | 0.0909 |
| 17_564940 | 5 | 0.4 | 6 | 0.3333 | 10 | 0 | 10 | 0 | 31 | 0.129 | 3 | 0 | 6 | 0.3333 | 7 | 0 | 4 | 0 | 20 | 0.1 |
| 17_564941 | 5 | 0.2 | 5 | 0 | 11 | 0.3636 | 10 | 0.3 | 31 | 0.2581 | 3 | 0.3333 | 7 | 0 | 9 | 0.2222 | 5 | 0 | 24 | 0.125 |
| 17_564942 | 5 | 0 | 5 | 0.4 | 9 | 0 | 8 | 0.375 | 27 | 0.1852 | 4 | 0.5 | 4 | 0 | 6 | 0.6667 | 5 | 0.4 | 19 | 0.4211 |
| 17_565060 | 4 | 0 | 2 | 0.5 | 5 | 0.2 | 4 | 0 | 15 | 0.1333 | 4 | 0 | 2 | 0 | 4 | 0.25 | 4 | 0 | 14 | 0.0714 |
| 17_565060 | 3 | 0 | 3 | 0 | 5 | 0 | 5 | 0.2 | 16 | 0.0625 | 4 | 0 | 2 | 0.5 | 5 | 0 | 4 | 0 | 15 | 0.0667 |
| 17_565248 | 1 | 0 | NA | NA | 1 | 1 | 3 | 0.3333 | 5 | 0.4 | 1 | 1 | 1 | 0 | 2 | 0.5 | NA | NA | 4 | 0.5 |
| 17_565643 | 5 | 0.2 | 2 | 0 | 3 | 0 | 2 | 0 | 12 | 0.0833 | NA | NA | 1 | 0 | 3 | 0.3333 | 3 | 0.3333 | 7 | 0.2857 |
| 17_565648 | 3 | 0.6667 | 3 | 0 | 3 | 0 | 2 | 0.5 | 11 | 0.2727 | NA | NA | 3 | 0 | 3 | 0.3333 | 5 | 0.2 | 11 | 0.1818 |
| 17_565649 | 4 | 0.25 | 2 | 0 | 3 | 0 | 2 | 0.5 | 11 | 0.1818 | NA | NA | 3 | 0.6667 | 3 | 0 | 4 | 0.25 | 10 | 0.3 |

| | | | | | | | | | | | | | | | | | | | | |
|---|---|---|---|---|---|---|---|---|---|---|---|---|---|---|---|---|---|---|---|---|
| 17_5656502 | 3 | 0 | 1 | 0 | 3 | 0.3333 | 1 | 0 | 8 | 0.125 | NA | NA | 3 | 0.3333 | 3 | 0 | 5 | 0.2 | 11 | 0.1818 |
| 17_5656506 | 3 | 0.3333 | 2 | 0.5 | 4 | 0.75 | 3 | 1 | 12 | 0.6667 | NA | NA | 3 | 1 | 4 | 0.5 | 6 | 0.5 | 13 | 0.6154 |
| 17_5656511 | 5 | 0.6 | 3 | 1 | 3 | 0.6667 | 2 | 0.5 | 13 | 0.6923 | NA | NA | 2 | 0.5 | 4 | 0.5 | 6 | 0.3333 | 12 | 0.4167 |
| 17_5656520 | 5 | 0.6 | 5 | 0.4 | 4 | 0.5 | 3 | 0 | 17 | 0.4118 | NA | NA | 3 | 0.3333 | 3 | 0.3333 | 7 | 0.4286 | 13 | 0.3846 |
| 17_5656542 | 4 | 0.5 | 4 | 0 | 4 | 0 | 2 | 0 | 14 | 0.1429 | NA | NA | 2 | 0.5 | 2 | 0 | 6 | 0 | 10 | 0.1 |
| 17_5656549 | 4 | 0.25 | 4 | 0 | 4 | 0 | 2 | 0.5 | 14 | 0.1429 | NA | NA | 4 | 0.25 | 4 | 0 | 6 | 0 | 14 | 0.0714 |
| 17_5656556 | 4 | 0.75 | 5 | 0.8 | 3 | 1 | 3 | 1 | 15 | 0.8667 | 1 | 1 | 4 | 0.5 | 4 | 0.75 | 5 | 1 | 14 | 0.7857 |
| 17_5656570 | 4 | 1 | 4 | 0.75 | 3 | 1 | 3 | 1 | 14 | 0.9286 | 1 | 1 | 2 | 1 | 4 | 1 | 10 | 0.9 | 17 | 0.9412 |
| 17_5656572 | 5 | 1 | 5 | 0.6 | 4 | 1 | 4 | 1 | 18 | 0.8889 | 2 | 0.5 | 5 | 1 | 4 | 0.75 | 9 | 0.8889 | 20 | 0.85 |
| 17_5656577 | 3 | 0.6667 | 3 | 0.6667 | 3 | 0 | 3 | 0 | 12 | 0.3333 | 2 | 0.5 | 5 | 0.4 | 2 | 0 | 8 | 0.25 | 17 | 0.2941 |
| 17_5656580 | 3 | 1 | 4 | 0.5 | 4 | 0.75 | 4 | 1 | 15 | 0.8 | 2 | 0.5 | 4 | 0.5 | 4 | 0.75 | 7 | 0.7143 | 17 | 0.6471 |
| 17_5656595 | 6 | 0.1667 | 5 | 0 | 4 | 0 | 3 | 0 | 18 | 0.0556 | 2 | 0 | 6 | 0.3333 | 4 | 0.25 | 10 | 0.3 | 22 | 0.2727 |
| 17_5656625 | 6 | 1 | 6 | 1 | 3 | 0.6667 | 4 | 1 | 19 | 0.9474 | 2 | 1 | 6 | 1 | 4 | 1 | 10 | 1 | 22 | 1 |
| 17_5656627 | 6 | 1 | 4 | 1 | 2 | 1 | 4 | 1 | 16 | 1 | 2 | 1 | 5 | 1 | 4 | 0.75 | 9 | 1 | 20 | 0.95 |
| 17_5656632 | 5 | 1 | 3 | 1 | 3 | 1 | 3 | 1 | 14 | 1 | 1 | 1 | 2 | 1 | 4 | 1 | 9 | 1 | 16 | 1 |
| 17_5656639 | 6 | 1 | 6 | 1 | 1 | 1 | 3 | 1 | 16 | 1 | 2 | 1 | 6 | 0.8333 | 4 | 0.75 | 11 | 0.9091 | 23 | 0.8696 |
| 17_5656647 | 6 | 1 | 2 | 1 | 3 | 0.6667 | 3 | 1 | 14 | 0.9286 | 2 | 1 | 5 | 1 | 3 | 1 | 10 | 1 | 20 | 1 |
| 17_5656648 | 4 | 0 | 2 | 1 | 3 | 0.3333 | 3 | 0 | 12 | 0.25 | 1 | 0 | 3 | 0.6667 | 3 | 0 | 6 | 0.3333 | 13 | 0.3077 |
| 17_5656649 | 5 | 0.2 | 4 | 0 | 4 | 0 | 3 | 0.3333 | 16 | 0.125 | 2 | 0.5 | 6 | 0.3333 | 2 | 0 | 7 | 0.4286 | 17 | 0.3529 |
| 17_5656650 | 4 | 1 | 3 | 1 | 2 | 0.5 | 1 | 1 | 10 | 0.9 | 2 | 1 | 3 | 1 | 3 | 1 | 5 | 1 | 13 | 1 |
| 17_5656659 | 5 | 0.6 | 5 | 0.2 | 4 | 0.25 | 3 | 0.6667 | 17 | 0.4118 | NA | NA | 7 | 0.5714 | 3 | 0.3333 | 10 | 0.7 | 20 | 0.6 |
| 17_5656663 | 5 | 0.6 | 3 | 0.6667 | 3 | 1 | 2 | 1 | 13 | 0.7692 | 2 | 0.5 | 5 | 0.4 | 3 | 0.3333 | 5 | 0.4 | 15 | 0.4 |
| 17_5656670 | 6 | 0.6667 | 5 | 0.8 | 4 | 0.75 | 4 | 0.75 | 19 | 0.7368 | 3 | 0.6667 | 5 | 0.6 | 4 | 0.75 | 11 | 0.8182 | 23 | 0.7391 |
| 17_5656673 | 5 | 0 | 4 | 0 | 4 | 0.25 | 3 | 0 | 16 | 0.0625 | 3 | 0 | 7 | 0 | 4 | 0.25 | 12 | 0.0833 | 26 | 0.0769 |
| 17_5656677 | 6 | 0.1667 | 3 | 0 | 2 | 0 | 3 | 0.6667 | 14 | 0.2143 | 2 | 0 | 6 | 0 | 5 | 0.2 | 11 | 0.0909 | 24 | 0.0833 |
| 17_5657665 | 1 | 1 | 1 | 1 | NA | NA | 3 | 0.3333 | 5 | 0.6 | 2 | 1 | NA | NA | 2 | 1 | 1 | 0 | 5 | 0.8 |
| 17_5657666 | 2 | 0.5 | 1 | 1 | NA | NA | 2 | 0.5 | 5 | 0.6 | 1 | 1 | 1 | 1 | 1 | 1 | 1 | 0 | 4 | 0.75 |
| 17_5657678 | 2 | 0 | 1 | 1 | NA | NA | 3 | 0.6667 | 6 | 0.5 | 2 | 1 | 1 | 0 | 2 | 0 | NA | NA | 5 | 0.4 |
| 17_5657685 | 1 | 1 | 1 | 1 | NA | NA | 2 | 0.5 | 4 | 0.75 | 1 | 1 | 1 | 1 | 2 | 1 | 1 | 0 | 5 | 0.8 |
| 17_5657696 | 2 | 0.5 | 1 | 1 | NA | NA | 1 | 0 | 4 | 0.5 | 2 | 1 | 1 | 1 | 1 | 1 | 1 | 1 | 5 | 1 |
| 17_5657706 | 2 | 0 | 1 | 1 | NA | NA | 3 | 0 | 6 | 0.1667 | 2 | 0 | 1 | 0 | 2 | 0 | 1 | 1 | 6 | 0.1667 |
| 17_5657729 | 2 | 0.5 | 1 | 1 | NA | NA | 3 | 0.6667 | 6 | 0.6667 | 2 | 1 | 1 | 1 | 2 | 0 | 1 | 0 | 6 | 0.5 |
| 17_5657730 | 2 | 0.5 | 1 | 1 | NA | NA | 2 | 0.5 | 5 | 0.6 | 1 | 0 | 1 | 1 | 1 | 0 | NA | NA | 3 | 0.3333 |
| 17_5657733 | 2 | 1 | 1 | 1 | NA | NA | 3 | 1 | 6 | 1 | 2 | 1 | 1 | 1 | 2 | 0.5 | NA | NA | 5 | 0.8 |
| 17_5657754 | 1 | 1 | 1 | 0 | NA | NA | 1 | 0 | 3 | 0.3333 | 2 | 0.5 | 1 | 1 | 2 | 0.5 | 1 | 0 | 6 | 0.5 |
| 17_5657757 | 1 | 1 | 1 | 1 | NA | NA | 3 | 0.6667 | 5 | 0.8 | 1 | 1 | 1 | 1 | 2 | 0 | 1 | 0 | 5 | 0.4 |
| 17_5657763 | 2 | 0.5 | 1 | 1 | NA | NA | 3 | 1 | 6 | 0.8333 | 2 | 1 | 1 | 1 | 1 | 1 | 1 | 1 | 5 | 1 |
| 17_5657771 | 2 | 0.5 | NA | NA | NA | NA | 2 | 0.5 | 4 | 0.5 | 1 | 0 | 1 | 0 | 2 | 1 | 1 | 1 | 5 | 0.6 |
| 17_5657787 | 1 | 0 | 1 | 0 | NA | NA | 2 | 1 | 4 | 0.5 | 2 | 0.5 | 1 | 0 | 1 | 0 | 1 | 1 | 5 | 0.4 |
| 17_5657793 | 2 | 1 | 1 | 0 | NA | NA | 1 | 0 | 4 | 0.5 | 2 | 1 | 1 | 1 | 2 | 0.5 | 1 | 1 | 6 | 0.8333 |
| 17_5657839 | 2 | 1 | 1 | 1 | NA | NA | NA | NA | 3 | 1 | 2 | 1 | 1 | 1 | 2 | 1 | NA | NA | 5 | 1 |
| 17_5657842 | 2 | 0 | 1 | 0 | NA | NA | 1 | 1 | 4 | 0.25 | 1 | 1 | NA | NA | 2 | 0 | NA | NA | 3 | 0.3333 |
| 17_5657852 | 2 | 0.5 | 1 | 1 | NA | NA | 1 | 0 | 4 | 0.5 | 2 | 1 | 1 | 1 | 2 | 1 | NA | NA | 5 | 1 |
| 17_5657872 | 2 | 0.5 | 1 | 1 | NA | NA | 1 | 0 | 4 | 0.5 | 1 | 1 | 1 | 1 | 2 | 1 | NA | NA | 4 | 1 |
| 17_5657875 | 2 | 0.5 | 1 | 1 | NA | NA | 1 | 1 | 4 | 0.75 | 2 | 1 | 1 | 1 | 1 | 1 | NA | NA | 4 | 1 |
| 17_5657889 | NA | NA | 1 | 1 | NA | NA | 1 | 0 | 2 | 0.5 | 1 | 1 | 1 | 1 | 1 | 0 | NA | NA | 3 | 0.6667 |
| 17_5657903 | 2 | 1 | 1 | 1 | NA | NA | 1 | 1 | 4 | 1 | 1 | 1 | 1 | 1 | 1 | 1 | NA | NA | 3 | 1 |
| 17_5658618 | 2 | 0 | 3 | 0.6667 | 1 | 0 | 5 | 0.2 | 11 | 0.2727 | 5 | 0 | 1 | 0 | 5 | 0.2 | 3 | 0 | 14 | 0.0714 |
| 17_5658621 | 2 | 0 | 3 | 0.6667 | 2 | 0 | 5 | 0 | 12 | 0.1667 | 5 | 0.2 | 1 | 0 | 4 | 0 | 3 | 0 | 13 | 0.0769 |
| 17_5670864 | 6 | 0.3333 | 4 | 0 | 6 | 0.3333 | 4 | 0.5 | 20 | 0.3 | 4 | 0.25 | 5 | 0 | 5 | 0.2 | 3 | 0.6667 | 17 | 0.2353 |
| 17_5670875 | 5 | 0.2 | 3 | 0 | 7 | 0 | 4 | 0 | 19 | 0.0526 | 5 | 0.2 | 3 | 0 | 4 | 0 | 1 | 0 | 13 | 0.0769 |
| 17_5670877 | 9 | 0 | 6 | 0.3333 | 7 | 0 | 5 | 0 | 27 | 0.0741 | 7 | 0 | 5 | 0.2 | 4 | 0 | 5 | 0.2 | 21 | 0.0952 |
| 17_5670879 | 7 | 0.1429 | 5 | 0.4 | 7 | 0.2857 | 3 | 0 | 22 | 0.2273 | 6 | 0.5 | 5 | 0.2 | 5 | 0 | 3 | 0.3333 | 19 | 0.2632 |
| 17_5670882 | 7 | 0.7143 | 2 | 0.5 | 4 | 0.75 | 3 | 0.3333 | 16 | 0.625 | 5 | 0.4 | 5 | 0.4 | 4 | 0.25 | 3 | 1 | 17 | 0.4706 |
| 17_5670883 | 7 | 0.2857 | 4 | 0.5 | 6 | 0.1667 | 6 | 0.3333 | 23 | 0.3043 | 7 | 0.2857 | 5 | 0 | 5 | 0.2 | 4 | 0.5 | 21 | 0.2381 |
| 17_5670891 | 4 | 1 | 2 | 1 | 6 | 1 | 3 | 1 | 15 | 1 | 2 | 1 | 3 | 1 | 5 | 0.6 | 2 | 1 | 12 | 0.8333 |
| 17_5670892 | 1 | 1 | NA | NA | NA | NA | NA | NA | 1 | 1 | NA | NA | NA | NA | NA | NA | 1 | 1 | 1 | 1 |
| 17_5670896 | 8 | 0.5 | 5 | 0.4 | 7 | 0.8571 | 4 | 0 | 24 | 0.5 | 3 | 0.6667 | 5 | 0.6 | 6 | 0.5 | 5 | 0.8 | 19 | 0.6316 |
| 17_5670909 | 5 | 0.4 | 6 | 0 | 8 | 0.625 | 3 | 0 | 22 | 0.3182 | 7 | 0.2857 | 4 | 0 | 4 | 0.25 | 5 | 0.2 | 20 | 0.2 |
| 17_5670927 | 8 | 0.25 | 1 | 0 | 6 | 0 | 3 | 0 | 18 | 0.1111 | 4 | 0.25 | 5 | 0.4 | 3 | 0 | 4 | 0 | 16 | 0.1875 |
| 17_5670935 | 9 | 0 | 6 | 0.1667 | 6 | 0.1667 | 5 | 0 | 26 | 0.0769 | 8 | 0.125 | 6 | 0 | 8 | 0.25 | 5 | 0.2 | 27 | 0.1481 |
| 17_5670948 | 9 | 1 | 5 | 1 | 8 | 0.875 | 6 | 0.8333 | 28 | 0.9286 | 7 | 0.7143 | 5 | 0.6 | 8 | 0.875 | 7 | 1 | 27 | 0.8148 |
| 17_5670949 | 7 | 1 | 1 | 1 | 5 | 0.8 | 3 | 0.6667 | 16 | 0.875 | 5 | 0.8 | 6 | 0.8333 | 5 | 1 | 4 | 1 | 20 | 0.9 |
| 17_5670953 | 9 | 0.1111 | 2 | 0 | 7 | 0.2857 | 4 | 0.5 | 22 | 0.2273 | 6 | 0 | 5 | 0 | 7 | 0.1429 | 7 | 0.2857 | 25 | 0.12 |
| 17_5670961 | 6 | 0.5 | 3 | 0.3333 | 7 | 0.2857 | 6 | 0.3333 | 22 | 0.3636 | 7 | 0.1429 | 5 | 0.6 | 9 | 0.3333 | 6 | 0.8333 | 27 | 0.4444 |

| | | | | | | | | | | | | | | | | | | | | |
|---|---|---|---|---|---|---|---|---|---|---|---|---|---|---|---|---|---|---|---|---|
| 17_5670978 | 9 | 0.6667 | 6 | 0.5 | 8 | 0.75 | 6 | 0.1667 | 29 | 0.5517 | 5 | 0.4 | 7 | 0.5714 | 5 | 0.2 | 3 | 0.6667 | 20 | 0.45 |
| 17_5671958 | 33 | 0.303 | 24 | 0.4167 | 34 | 0.3235 | 28 | 0.2143 | 119 | 0.3109 | 20 | 0.3 | 26 | 0.3462 | 29 | 0.3793 | 24 | 0.375 | 99 | 0.3535 |
| 17_5671959 | 39 | 0.3846 | 37 | 0.4595 | 36 | 0.4444 | 35 | 0.4571 | 147 | 0.4354 | 30 | 0.5333 | 35 | 0.3429 | 42 | 0.5 | 28 | 0.3929 | 135 | 0.4444 |
| 17_5671968 | 41 | 0 | 40 | 0.075 | 45 | 0.0222 | 38 | 0.1579 | 164 | 0.061 | 34 | 0.0588 | 42 | 0.0714 | 41 | 0.0488 | 34 | 0.1176 | 151 | 0.0728 |
| 17_5671970 | 44 | 0.1591 | 41 | 0.0488 | 42 | 0.119 | 39 | 0.0513 | 166 | 0.0964 | 36 | 0.0833 | 47 | 0.0851 | 45 | 0.1111 | 35 | 0.0857 | 163 | 0.092 |
| 17_5674312 | 8 | 0.375 | 10 | 0.3 | 6 | 0.3333 | 6 | 0.5 | 30 | 0.3667 | 4 | 0.5 | 5 | 0.8 | 11 | 0.4545 | NA | NA | 20 | 0.55 |
| 17_5674317 | 9 | 0.5556 | 8 | 0.625 | 3 | 0.6667 | 3 | 0.3333 | 23 | 0.5652 | 4 | 0.75 | 4 | 0.5 | 12 | 0.5 | NA | NA | 20 | 0.55 |
| 17_5686999 | 1 | 0 | 4 | 0.25 | 7 | 0.4286 | NA | NA | 12 | 0.3333 | 5 | 0 | 3 | 0.3333 | 7 | 0.2857 | 2 | 0 | 17 | 0.1765 |
| 17_5687005 | 2 | 0.5 | 5 | 0.6 | 10 | 0.2 | NA | NA | 17 | 0.3529 | 6 | 0 | 7 | 0.4286 | 9 | 0.5556 | 3 | 0 | 25 | 0.32 |
| 17_5687009 | 2 | 0 | 4 | 0 | 10 | 0.1 | NA | NA | 16 | 0.0625 | 5 | 0 | 5 | 0 | 6 | 0.5 | 3 | 0 | 19 | 0.1579 |
| 17_5687018 | 2 | 0 | 3 | 0 | 7 | 0.1429 | NA | NA | 12 | 0.0833 | 3 | 0 | 6 | 0.1667 | 8 | 0.25 | 2 | 0 | 19 | 0.1579 |
| 17_5687021 | 2 | 0 | 5 | 0.6 | 6 | 0.5 | NA | NA | 13 | 0.4615 | 2 | 0.5 | 4 | 0.5 | 4 | 1 | 2 | 0.5 | 12 | 0.6667 |
| 17_5687797 | NA | NA | NA | NA | 1 | 1 | NA | NA | 1 | 1 | NA | NA | 2 | 0.5 | 1 | 1 | 1 | 1 | 4 | 0.75 |
| 17_5688254 | NA | NA | NA | NA | 2 | 0.5 | NA | NA | 2 | 0.5 | NA | NA | NA | NA | 2 | 0.5 | 2 | 0.5 | 4 | 0.5 |
| 17_5688258 | NA | NA | NA | NA | 3 | 1 | NA | NA | 3 | 1 | NA | NA | NA | NA | 2 | 1 | 2 | 1 | 4 | 1 |
| 17_5688341 | 1 | 1 | NA | NA | 2 | 1 | NA | NA | 3 | 1 | 1 | 1 | NA | NA | NA | NA | NA | NA | 1 | 1 |
| 17_5690099 | 1 | 0 | 2 | 0 | 1 | 0 | 6 | 0.1667 | 10 | 0.1 | 2 | 0 | 2 | 0 | 2 | 0 | 3 | 0.6667 | 9 | 0.2222 |
| 17_5690599 | 4 | 0.5 | 3 | 0.6667 | 1 | 0 | 4 | 0 | 12 | 0.3333 | 3 | 0 | 4 | 0.5 | 5 | 0.6 | 5 | 0.6 | 17 | 0.4706 |
| 17_5690602 | 4 | 0.25 | NA | NA | 1 | 0 | 6 | 0.1667 | 11 | 0.1818 | 3 | 0 | 3 | 0 | 5 | 0.2 | 4 | 0 | 15 | 0.0667 |
| 17_5690913 | NA | NA | 1 | 1 | NA | NA | NA | NA | 1 | 1 | NA | NA | 2 | 0.5 | NA | NA | NA | NA | 2 | 0.5 |
| 17_5690920 | NA | NA | 2 | 0.5 | NA | NA | NA | NA | 2 | 0.5 | NA | NA | 2 | 1 | NA | NA | NA | NA | 2 | 1 |
| 17_5691305 | 3 | 0 | 9 | 0.1111 | 4 | 0 | 1 | 0 | 17 | 0.0588 | 3 | 0.3333 | 1 | 0 | 8 | 0.125 | 7 | 0.1429 | 19 | 0.1579 |
| 17_5691463 | 30 | 0.4 | 28 | 0.3571 | 21 | 0.381 | 27 | 0.4074 | 106 | 0.3868 | 23 | 0.5217 | 20 | 0.55 | 23 | 0.2174 | 13 | 0.4615 | 79 | 0.4304 |
| 17_5691728 | NA | NA | 2 | 1 | NA | NA | 3 | 1 | 5 | 1 | 2 | 1 | 1 | 1 | 1 | 1 | NA | NA | 4 | 1 |
| 17_5691756 | NA | NA | 1 | 1 | NA | NA | 3 | 1 | 4 | 1 | 2 | 1 | 2 | 1 | 1 | 1 | NA | NA | 5 | 1 |
| 17_5691763 | NA | NA | 1 | 0 | NA | NA | 4 | 0.25 | 5 | 0.2 | 2 | 1 | 2 | 0 | 1 | 1 | NA | NA | 5 | 0.6 |
| 17_5691767 | NA | NA | 2 | 0 | NA | NA | 3 | 0.6667 | 5 | 0.4 | 2 | 1 | 2 | 0 | 1 | 1 | NA | NA | 5 | 0.6 |
| 17_5691770 | NA | NA | 1 | 0 | NA | NA | 1 | 1 | 2 | 0.5 | 2 | 0.5 | 2 | 0.5 | 1 | 1 | NA | NA | 5 | 0.6 |
| 17_5691789 | NA | NA | 2 | 0 | NA | NA | 3 | 0.3333 | 5 | 0.2 | 2 | 0 | 1 | 0 | 2 | 0.5 | NA | NA | 5 | 0.2 |
| 17_5692082 | 5 | 0.2 | 4 | 0.5 | 3 | 0 | 6 | 0 | 18 | 0.1667 | NA | NA | 1 | 0 | 4 | 0.25 | 5 | 0.4 | 10 | 0.3 |
| 17_5692093 | 66 | 0.0758 | 63 | 0.1111 | 57 | 0.1228 | 63 | 0.1905 | 249 | 0.1245 | 37 | 0.0811 | 38 | 0.0526 | 38 | 0.0526 | 40 | 0.225 | 153 | 0.1046 |
| 17_5692136 | 10 | 0.1 | 8 | 0.25 | 11 | 0.0909 | 10 | 0.1 | 39 | 0.1282 | 7 | 0.1429 | 8 | 0.375 | 10 | 0.2 | 5 | 0.2 | 30 | 0.2333 |
| 17_5692144 | 6 | 0.1667 | 8 | 0.125 | 10 | 0.1 | 8 | 0 | 32 | 0.0938 | 8 | 0.125 | 5 | 0.2 | 8 | 0.375 | 5 | 0.4 | 26 | 0.2692 |
| 17_5692225 | 5 | 0.2 | 3 | 0.3333 | 3 | 0 | 2 | 0 | 13 | 0.1538 | 4 | 0.5 | 3 | 0.3333 | 7 | 0 | 2 | 0 | 16 | 0.1875 |
| 17_5692567 | 1 | 0 | 2 | 0 | NA | NA | 2 | 0.5 | 5 | 0.2 | 1 | 0 | NA | NA | 1 | 1 | NA | NA | 2 | 0.5 |
| 17_5692619 | 4 | 0.25 | 3 | 0 | 2 | 0 | 3 | 0 | 12 | 0.0833 | 4 | 0.25 | 3 | 0 | 3 | 0.3333 | 2 | 0 | 12 | 0.1667 |
| 17_5701805 | 14 | 1 | 11 | 0.8182 | 6 | 1 | 10 | 1 | 41 | 0.9512 | 14 | 0.8571 | 3 | 0.6667 | 6 | 1 | 6 | 0.8333 | 29 | 0.8621 |
| 17_5701812 | 12 | 0.4167 | 12 | 0.1667 | 8 | 0 | 11 | 0.1818 | 43 | 0.2093 | 13 | 0.3077 | 7 | 0.2857 | 9 | 0.2222 | 8 | 0.25 | 37 | 0.2703 |
| 17_5701818 | 10 | 0.5 | 11 | 0.5455 | 5 | 0.4 | 11 | 0.6364 | 37 | 0.5405 | 11 | 0.6364 | 4 | 0.25 | 9 | 0.7778 | 7 | 0.7143 | 31 | 0.6452 |
| 17_5701820 | 13 | 0 | 13 | 0.0769 | 8 | 0.125 | 14 | 0.0714 | 48 | 0.0625 | 12 | 0 | 6 | 0.1667 | 12 | 0 | 10 | 0.1 | 40 | 0.05 |
| 17_5701832 | 14 | 0.3571 | 14 | 0.4286 | 10 | 0.3 | 14 | 0.5714 | 52 | 0.4231 | 13 | 0.5385 | 7 | 0.1429 | 11 | 0.2727 | 10 | 0.4 | 41 | 0.3659 |
| 17_5701839 | 16 | 0.5 | 13 | 0.5385 | 8 | 0.375 | 12 | 0.5 | 49 | 0.4898 | 13 | 0.6154 | 5 | 0.6 | 11 | 0.6364 | 9 | 0.6667 | 38 | 0.6316 |
| 17_5701848 | 17 | 0.1176 | 13 | 0.0769 | 9 | 0.2222 | 13 | 0.4615 | 52 | 0.2115 | 14 | 0.2857 | 6 | 0.1667 | 10 | 0.3 | 11 | 0.4545 | 41 | 0.3171 |
| 17_5701857 | 17 | 0.9412 | 15 | 1 | 8 | 0.875 | 14 | 0.9286 | 54 | 0.9444 | 16 | 0.9375 | 5 | 0.8 | 8 | 1 | 10 | 1 | 39 | 0.9487 |
| 17_5701861 | 14 | 0.6429 | 15 | 0.7333 | 9 | 0.7778 | 14 | 0.7857 | 52 | 0.7308 | 16 | 0.75 | 4 | 0.75 | 11 | 0.7273 | 9 | 0.5556 | 40 | 0.7 |
| 17_5701871 | 15 | 0.2 | 13 | 0.3077 | 9 | 0.3333 | 14 | 0.2143 | 51 | 0.2549 | 15 | 0.2 | 5 | 0 | 10 | 0 | 11 | 0.2727 | 41 | 0.1463 |
| 17_5701874 | 12 | 0.4167 | 10 | 0.5 | 8 | 0.5 | 14 | 0.6429 | 44 | 0.5227 | 11 | 0.5455 | 5 | 0.2 | 10 | 0.4 | 9 | 0.5556 | 35 | 0.4571 |
| 17_5702105 | 1 | 0 | NA | NA | 2 | 0.5 | 4 | 0.25 | 7 | 0.2857 | 1 | 0 | 3 | 0.3333 | 1 | 0 | 1 | 0 | 6 | 0.1667 |
| 17_5702241 | 2 | 1 | 2 | 1 | 2 | 1 | 2 | 1 | 8 | 1 | NA | NA | 2 | 0.5 | 1 | 1 | NA | NA | 3 | 0.6667 |
| 17_5702263 | 2 | 1 | 2 | 1 | 2 | 1 | 3 | 0.6667 | 9 | 0.8889 | NA | NA | 2 | 1 | 1 | 0 | NA | NA | 3 | 0.6667 |
| 17_5705833 | NA | NA | 1 | 0 | 1 | 0 | 2 | 0.5 | 4 | 0.25 | 2 | 0.5 | NA | NA | 1 | 0 | NA | NA | 3 | 0.3333 |
| 17_5705835 | NA | NA | 1 | 1 | 1 | 0 | 2 | 0.5 | 4 | 0.5 | 2 | 1 | 1 | 1 | 1 | 0 | NA | NA | 4 | 0.75 |
| 17_5707559 | 2 | 0 | 2 | 0 | 2 | 0.5 | 2 | 0 | 8 | 0.125 | 1 | 0 | 2 | 0 | 2 | 0.5 | 3 | 0 | 8 | 0.125 |
| 17_5707681 | 5 | 0.2 | 6 | 0.6667 | 3 | 0 | 5 | 0.4 | 19 | 0.3684 | 3 | 0 | 8 | 0.25 | 7 | 0.5714 | 9 | 0.3333 | 27 | 0.3333 |
| 17_5707690 | 4 | 0 | 5 | 0.2 | 3 | 0 | 2 | 0 | 14 | 0.0714 | 3 | 0.3333 | 6 | 0.1667 | 5 | 0.2 | 4 | 0 | 18 | 0.1667 |
| 17_5707698 | 4 | 0.25 | 5 | 0 | 3 | 0 | 6 | 0.3333 | 18 | 0.1667 | 2 | 0 | 6 | 0.3333 | 6 | 0 | 6 | 0.1667 | 20 | 0.15 |
| 17_5707727 | NA | NA | NA | NA | NA | NA | 1 | 1 | 1 | 1 | NA | NA | NA | NA | 1 | 1 | NA | NA | 1 | 1 |
| 17_5707847 | NA | NA | NA | NA | NA | NA | 1 | 1 | 1 | 1 | NA | NA | NA | NA | 1 | 1 | NA | NA | 1 | 1 |
| 17_5707868 | NA | NA | NA | NA | NA | NA | 1 | 1 | 1 | 1 | NA | NA | NA | NA | 1 | 1 | NA | NA | 1 | 1 |
| 17_5707887 | NA | NA | NA | NA | NA | NA | 1 | 1 | 1 | 1 | NA | NA | NA | NA | 1 | 1 | NA | NA | 1 | 1 |
| 17_5719991 | 6 | 0.1667 | 10 | 0 | 10 | 0.2 | 15 | 0 | 41 | 0.0732 | 4 | 0 | 6 | 0 | 17 | 0.1765 | 8 | 0.375 | 35 | 0.1714 |
| 17_5720077 | 5 | 0.2 | 8 | 0.5 | 10 | 0.3 | 9 | 0 | 32 | 0.25 | 4 | 0.25 | 6 | 0.1667 | 12 | 0.5 | 6 | 0.3333 | 28 | 0.3571 |
| 17_5720084 | 5 | 0 | 9 | 0.2222 | 10 | 0 | 10 | 0.1 | 34 | 0.0882 | 4 | 0 | 4 | 0 | 11 | 0.1818 | 5 | 0 | 24 | 0.0833 |
| 17_5720127 | 5 | 0 | 7 | 0.1429 | 6 | 0 | 10 | 0.3 | 28 | 0.1429 | 3 | 0.3333 | 4 | 0.25 | 11 | 0.1818 | 5 | 0 | 23 | 0.1739 |
| 17_5720694 | 5 | 0 | 2 | 0 | 5 | 0.2 | 9 | 0.1111 | 21 | 0.0952 | 2 | 0 | 5 | 0 | 5 | 0 | 5 | 0.2 | 17 | 0.0588 |

| | | | | | | | | | | | | | | | | | | | | |
|---|---|---|---|---|---|---|---|---|---|---|---|---|---|---|---|---|---|---|---|---|
| 17_5720699 | 5 | 0.8 | 2 | 0.5 | 5 | 0.6 | 9 | 0.8889 | 21 | 0.7619 | 2 | 1 | 6 | 0.5 | 4 | 0.5 | 4 | 1 | 16 | 0.6875 |
| 17_5720722 | 3 | 1 | 1 | 1 | 4 | 0.75 | 8 | 0.25 | 16 | 0.5625 | 1 | 0 | 3 | 0 | 5 | 0.6 | 5 | 0.4 | 14 | 0.3571 |
| 17_5729742 | 2 | 0.5 | 9 | 0.2222 | 8 | 0.25 | 4 | 0.25 | 23 | 0.2609 | 9 | 0 | 9 | 0.1111 | 14 | 0.2143 | 7 | 0.2857 | 39 | 0.1538 |
| 17_5730306 | 6 | 0.3333 | 15 | 0.2 | 15 | 0.1333 | 12 | 0.0833 | 48 | 0.1667 | 11 | 0.0909 | 14 | 0.2143 | 16 | 0.1875 | 11 | 0.3636 | 52 | 0.2115 |
| 17_5730313 | 4 | 0 | 13 | 0.3846 | 11 | 0.1818 | 13 | 0 | 41 | 0.1707 | 11 | 0.3636 | 13 | 0.2308 | 18 | 0.2222 | 8 | 0.125 | 50 | 0.24 |
| 17_5730314 | 6 | 0.1667 | 15 | 0.3333 | 16 | 0.375 | 11 | 0.5455 | 48 | 0.375 | 13 | 0.3077 | 12 | 0.5833 | 19 | 0.3684 | 10 | 0.1 | 54 | 0.3519 |
| 17_5730676 | NA | NA | 1 | 1 | NA | NA | NA | NA | 1 | 1 | NA | NA | NA | NA | 1 | 1 | NA | NA | 1 | 1 |
| 17_5730704 | NA | NA | 1 | 1 | NA | NA | NA | NA | 1 | 1 | NA | NA | NA | NA | 1 | 1 | NA | NA | 1 | 1 |
| 17_5732818 | 12 | 0 | 23 | 0.087 | 21 | 0.0952 | 15 | 0 | 71 | 0.0563 | 11 | 0.0909 | 20 | 0.05 | 18 | 0.0556 | 16 | 0.125 | 65 | 0.0769 |
| 17_5732882 | 10 | 0 | 17 | 0.1765 | 15 | 0.1333 | 15 | 0.2 | 57 | 0.1404 | 11 | 0 | 13 | 0.2308 | 14 | 0 | 10 | 0.1 | 48 | 0.0833 |
| 17_5732949 | 8 | 0.25 | 14 | 0.3571 | 11 | 0.4545 | 11 | 0.2727 | 44 | 0.3409 | 9 | 0.4444 | 11 | 0.3636 | 10 | 0.6 | 9 | 0.5556 | 39 | 0.4872 |
| 17_5732952 | 6 | 0.8333 | 9 | 1 | 9 | 0.7778 | 6 | 1 | 30 | 0.9 | 9 | 1 | 11 | 0.8182 | 5 | 1 | 7 | 0.8571 | 32 | 0.9062 |
| 17_5732953 | 8 | 0.25 | 11 | 0.4545 | 11 | 0.2727 | 10 | 0.4 | 40 | 0.35 | 9 | 0.3333 | 9 | 0.4444 | 7 | 0.4286 | 5 | 0.6 | 30 | 0.4333 |
| 17_5732979 | 9 | 0.1111 | 9 | 0.2222 | 11 | 0.0909 | 6 | 0 | 35 | 0.1143 | 6 | 0.3333 | 11 | 0.0909 | 7 | 0 | 6 | 0.1667 | 30 | 0.1333 |
| 17_5732981 | 8 | 0.25 | 11 | 0.2727 | 8 | 0.25 | 7 | 0.2857 | 34 | 0.2647 | 5 | 0.2 | 12 | 0.3333 | 8 | 0.125 | 5 | 0.2 | 30 | 0.2333 |
| 17_5732986 | 7 | 0 | 13 | 0.1538 | 11 | 0.1818 | 7 | 0.4286 | 38 | 0.1842 | 8 | 0.125 | 11 | 0.3636 | 9 | 0.3333 | 6 | 0.3333 | 34 | 0.2941 |
| 17_5735083 | 23 | 0.3478 | 39 | 0.2821 | 34 | 0.4118 | 27 | 0.2222 | 123 | 0.3171 | 20 | 0.45 | 26 | 0.2692 | 35 | 0.2857 | 19 | 0.4211 | 100 | 0.34 |
| 17_5735188 | 14 | 0.2857 | 37 | 0.1081 | 34 | 0.0882 | 24 | 0.125 | 109 | 0.1284 | 18 | 0.0556 | 25 | 0.12 | 31 | 0.0645 | 19 | 0.1579 | 93 | 0.0968 |
| 17_5735226 | 13 | 0.6154 | 37 | 0.4595 | 27 | 0.4815 | 17 | 0.5882 | 94 | 0.5106 | 19 | 0.4211 | 21 | 0.4286 | 30 | 0.6333 | 21 | 0.4286 | 91 | 0.4945 |
| 17_5735235 | 14 | 0.2143 | 35 | 0.1143 | 28 | 0.0357 | 18 | 0.0556 | 95 | 0.0947 | 21 | 0 | 24 | 0.2083 | 29 | 0.1034 | 17 | 0.1765 | 91 | 0.1209 |
| 17_5735240 | 13 | 0.9231 | 34 | 0.9412 | 24 | 0.875 | 18 | 1 | 89 | 0.9326 | 19 | 0.9474 | 21 | 0.9048 | 28 | 0.9286 | 18 | 0.9444 | 86 | 0.9302 |
| 17_5735247 | 8 | 0.25 | 10 | 0.1 | 7 | 0.1429 | 5 | 0 | 30 | 0.1333 | 5 | 0.2 | 7 | 0 | 12 | 0.25 | 9 | 0.1111 | 33 | 0.1515 |
| 17_5738759 | 76 | 0.1316 | 140 | 0.0929 | 109 | 0.0642 | 68 | 0.1618 | 393 | 0.1043 | 100 | 0.13 | 65 | 0.0154 | 62 | 0.0645 | 74 | 0.0811 | 301 | 0.0797 |
| 17_5738809 | 62 | 0.1613 | 121 | 0.2314 | 82 | 0.2683 | 56 | 0.2857 | 321 | 0.2368 | 79 | 0.1392 | 57 | 0.2632 | 55 | 0.2545 | 64 | 0.3594 | 255 | 0.2471 |
| 17_5738813 | 53 | 0.283 | 108 | 0.1574 | 71 | 0.2113 | 52 | 0.1731 | 284 | 0.1972 | 69 | 0.1594 | 51 | 0.098 | 53 | 0.0943 | 58 | 0.1724 | 231 | 0.1342 |
| 17_5738835 | 62 | 0.2903 | 103 | 0.2427 | 75 | 0.28 | 51 | 0.3137 | 291 | 0.2749 | 67 | 0.2388 | 51 | 0.2549 | 53 | 0.1887 | 52 | 0.3077 | 223 | 0.2466 |
| 17_5738843 | 59 | 0.0508 | 106 | 0.0755 | 72 | 0.0417 | 51 | 0.0196 | 288 | 0.0521 | 73 | 0.0411 | 49 | 0.0408 | 52 | 0.0769 | 56 | 0.0714 | 230 | 0.0565 |
| 17_5738844 | 58 | 0.3276 | 96 | 0.1875 | 74 | 0.2162 | 49 | 0.3265 | 277 | 0.2491 | 69 | 0.2174 | 47 | 0.2128 | 49 | 0.1429 | 52 | 0.2692 | 217 | 0.212 |
| 17_5738858 | 56 | 0.3393 | 103 | 0.2913 | 72 | 0.3056 | 55 | 0.3636 | 286 | 0.3182 | 70 | 0.3286 | 46 | 0.4348 | 48 | 0.2292 | 57 | 0.386 | 221 | 0.3439 |
| 17_5754193 | 2 | 0 | 2 | 0 | 4 | 0.25 | 1 | 0 | 9 | 0.1111 | 3 | 0 | 3 | 0 | 1 | 1 | 1 | 0 | 8 | 0.125 |
| 17_5754198 | 3 | 0.3333 | 5 | 0 | 4 | 0 | 2 | 0 | 14 | 0.0714 | 3 | 0 | 5 | 0.2 | 3 | 0 | 4 | 0 | 15 | 0.0667 |
| 17_5754208 | 5 | 0.2 | 5 | 0.4 | 5 | 0 | 1 | 0 | 16 | 0.1875 | 3 | 0 | 5 | 0.2 | 3 | 0.3333 | 5 | 0.4 | 16 | 0.25 |
| 17_5754210 | 5 | 0.8 | 5 | 1 | 4 | 1 | 1 | 1 | 15 | 0.9333 | 2 | 0.5 | 6 | 0.5 | NA | NA | 4 | 0.25 | 12 | 0.4167 |
| 17_5754235 | 5 | 0 | 4 | 0.25 | 4 | 0 | 1 | 0 | 14 | 0.0714 | 3 | 0.3333 | 6 | 0 | 3 | 0 | 3 | 0 | 15 | 0.0667 |
| 17_5754393 | 5 | 0.2 | 5 | 0 | 4 | 0 | NA | NA | 14 | 0.0714 | 3 | 0 | 5 | 0 | 2 | 0 | 4 | 0.25 | 14 | 0.0714 |
| 17_5872459 | NA | NA | 3 | 0.3333 | 1 | 0 | 1 | 0 | 5 | 0.2 | 3 | 0 | 1 | 0 | 1 | 1 | 3 | 0 | 8 | 0.125 |
| 17_5930895 | NA | NA | 1 | 1 | NA | NA | NA | NA | 1 | 1 | 1 | 1 | NA | NA | NA | NA | NA | NA | 1 | 1 |
| 17_6291014 | NA | NA | 2 | 1 | 1 | 0 | 3 | 1 | 6 | 0.8333 | 2 | 1 | 2 | 0.5 | NA | NA | 2 | 0.5 | 6 | 0.6667 |
| 17_6291072 | NA | NA | 2 | 0.5 | 1 | 0 | 2 | 0 | 5 | 0.2 | NA | NA | 2 | 1 | NA | NA | 3 | 0 | 5 | 0.4 |
| 17_6291100 | NA | NA | 2 | 0 | 1 | 0 | 2 | 0.5 | 5 | 0.2 | NA | NA | 1 | 0 | NA | NA | 3 | 0.3333 | 4 | 0.25 |
| 17_6291102 | NA | NA | 2 | 1 | 1 | 1 | 2 | 1 | 5 | 1 | NA | NA | 2 | 0 | NA | NA | 4 | 0.75 | 6 | 0.5 |
| 17_6291203 | NA | NA | 1 | 0 | 1 | 0 | 3 | 0.6667 | 5 | 0.4 | 1 | 0 | 4 | 0.25 | NA | NA | 3 | 0 | 8 | 0.125 |
| 17_6291215 | NA | NA | 1 | 0 | 1 | 1 | 5 | 0.4 | 7 | 0.4286 | 1 | 0 | 4 | 0.5 | NA | NA | 2 | 0.5 | 7 | 0.4286 |
| 17_6291221 | NA | NA | NA | NA | 1 | 0 | 4 | 0.25 | 5 | 0.2 | 1 | 0 | 4 | 0.25 | NA | NA | 2 | 0 | 7 | 0.1429 |
| 17_6291237 | NA | NA | 1 | 1 | 1 | 0 | 2 | 0.5 | 4 | 0.5 | NA | NA | 4 | 1 | NA | NA | 2 | 1 | 6 | 1 |
| 17_6291246 | NA | NA | 1 | 1 | 1 | 0 | 2 | 0.5 | 4 | 0.5 | 1 | 1 | 4 | 0.5 | NA | NA | 2 | 1 | 7 | 0.7143 |
| 17_6291425 | NA | NA | NA | NA | 1 | 1 | 1 | 1 | 2 | 1 | NA | NA | 1 | 1 | NA | NA | NA | NA | 1 | 1 |
| 17_6291434 | NA | NA | NA | NA | 1 | 1 | 2 | 0 | 3 | 0.3333 | NA | NA | 1 | 1 | NA | NA | NA | NA | 1 | 1 |
| 17_6335944 | 2 | 0.5 | 1 | 1 | 1 | 0 | NA | NA | 4 | 0.5 | 2 | 0.5 | NA | NA | 2 | 0.5 | 3 | 0 | 7 | 0.2857 |
| 17_6336009 | 2 | 0.5 | NA | NA | 1 | 0 | NA | NA | 3 | 0.3333 | 2 | 1 | NA | NA | 2 | 0.5 | 2 | 0 | 6 | 0.5 |
| 17_6336478 | NA | NA | NA | NA | 1 | 1 | NA | NA | 1 | 1 | 1 | 1 | 2 | 1 | NA | NA | 1 | 1 | 4 | 1 |
| 17_6336498 | NA | NA | NA | NA | 3 | 1 | NA | NA | 3 | 1 | 1 | 0 | 3 | 0.3333 | NA | NA | NA | NA | 4 | 0.25 |
| 17_6336507 | NA | NA | NA | NA | 3 | 0.3333 | NA | NA | 3 | 0.3333 | 1 | 1 | 1 | 1 | NA | NA | 1 | 1 | 3 | 1 |
| 17_6336516 | NA | NA | NA | NA | 2 | 1 | NA | NA | 2 | 1 | NA | NA | 1 | 1 | NA | NA | 1 | 1 | 2 | 1 |
| 17_6336527 | NA | NA | NA | NA | 2 | 0.5 | NA | NA | 2 | 0.5 | 1 | 1 | 1 | 1 | 1 | 0 | 1 | 1 | 4 | 0.75 |
| 17_6336698 | 2 | 1 | 3 | 0.6667 | 6 | 0.6667 | 2 | 0.5 | 13 | 0.6923 | 2 | 0 | 10 | 0.5 | 4 | 0.5 | 5 | 0.2 | 21 | 0.381 |
| 17_6336704 | 2 | 0.5 | 4 | 0.25 | 6 | 0.1667 | 4 | 0.25 | 16 | 0.25 | 4 | 0.5 | 9 | 0.2222 | 5 | 0 | 5 | 0 | 23 | 0.1739 |
| 17_6336726 | 2 | 0 | 3 | 0 | 5 | 0.2 | 3 | 0 | 13 | 0.0769 | 2 | 0 | 9 | 0.1111 | 4 | 0 | 2 | 0 | 17 | 0.0588 |
| 17_6336733 | 1 | 0 | 4 | 0 | 5 | 0 | 3 | 0.3333 | 13 | 0.0769 | 3 | 0 | 8 | 0.25 | 4 | 0 | 2 | 0 | 17 | 0.1176 |
| 17_6379484 | 2 | 0.5 | 1 | 1 | 4 | 0.5 | 2 | 0 | 9 | 0.4444 | 3 | 0.6667 | 6 | 0.3333 | 3 | 0.6667 | 1 | 1 | 13 | 0.5385 |
| 17_6380611 | 2 | 0.5 | 1 | 0 | 4 | 0.25 | 3 | 0.6667 | 10 | 0.4 | NA | NA | 7 | 0.2857 | 1 | 0 | 3 | 0 | 11 | 0.1818 |
| 17_6380650 | 1 | 0 | 1 | 1 | 3 | 0.3333 | 2 | 0.5 | 7 | 0.4286 | NA | NA | 4 | 0.5 | 1 | 0 | NA | NA | 5 | 0.4 |
| 17_6380667 | 1 | 0 | 1 | 0 | 2 | 0 | 3 | 0.3333 | 7 | 0.1429 | NA | NA | 5 | 0.6 | 2 | 0 | 1 | 0 | 8 | 0.375 |
| 17_6380685 | 1 | 1 | 1 | 0 | 2 | 0.5 | 2 | 1 | 6 | 0.6667 | NA | NA | 4 | 0.75 | NA | NA | NA | NA | 4 | 0.75 |
| 17_6380728 | 1 | 0 | 1 | 1 | 1 | 0 | 2 | 0 | 5 | 0.2 | NA | NA | 3 | 0.3333 | 1 | 0 | NA | NA | 4 | 0.25 |

| | | | | | | | | | | | | | | | | | | | | |
|---|---|---|---|---|---|---|---|---|---|---|---|---|---|---|---|---|---|---|---|---|
| 17_6380732 | 2 | 1 | NA | NA | 1 | 1 | NA | NA | 3 | 1 | 1 | 0 | 2 | 1 | 1 | 0 | 1 | 1 | 5 | 0.6 |
| 17_6380747 | 2 | 1 | NA | NA | 2 | 0 | NA | NA | 4 | 0.5 | 1 | 0 | 3 | 0.3333 | 2 | 0.5 | 2 | 0 | 8 | 0.25 |
| 17_6380772 | 2 | 0 | NA | NA | 2 | 0.5 | NA | NA | 4 | 0.25 | 1 | 0 | 3 | 0 | 2 | 0 | 2 | 0.5 | 8 | 0.125 |
| 17_6380781 | 2 | 0 | NA | NA | 2 | 0.5 | NA | NA | 4 | 0.25 | 1 | 0 | 3 | 0.3333 | 2 | 0 | 2 | 0 | 8 | 0.125 |
| 17_6380803 | 2 | 0.5 | NA | NA | 2 | 0 | NA | NA | 4 | 0.25 | 1 | 0 | 2 | 0.5 | 2 | 0 | 2 | 0.5 | 7 | 0.2857 |
| 17_6380846 | 2 | 0 | NA | NA | 5 | 0.2 | NA | NA | 7 | 0.1429 | 2 | 0 | 3 | 0 | 4 | 0.25 | 1 | 0 | 10 | 0.1 |
| 17_6380851 | 2 | 0 | NA | NA | 5 | 0.2 | NA | NA | 7 | 0.1429 | 3 | 0.3333 | 5 | 0 | 4 | 0.25 | 2 | 0 | 14 | 0.1429 |
| 17_6380876 | NA | NA | NA | NA | 5 | 0.2 | NA | NA | 5 | 0.2 | 3 | 0.3333 | 4 | 0.5 | 3 | 0.3333 | 2 | 0.5 | 12 | 0.4167 |
| 17_6380898 | 2 | 0 | NA | NA | 5 | 0.6 | NA | NA | 7 | 0.4286 | 3 | 0.6667 | 3 | 0.3333 | 3 | 0 | 2 | 0 | 11 | 0.2727 |
| 17_6380903 | 2 | 1 | NA | NA | 5 | 1 | NA | NA | 7 | 1 | 2 | 1 | 5 | 1 | 1 | 1 | 1 | 1 | 9 | 1 |
| 17_6444604 | NA | NA | 1 | 1 | NA | NA | 1 | 1 | 2 | 1 | NA | NA | 2 | 1 | 1 | 0 | 3 | 0.6667 | 6 | 0.6667 |
| 17_6458990 | 44 | 0.0455 | 46 | 0.0652 | 32 | 0.1562 | 52 | 0.1731 | 174 | 0.1092 | 30 | 0.0667 | 31 | 0.0323 | 43 | 0.2093 | 49 | 0.0816 | 153 | 0.1046 |
| 17_6459001 | 38 | 0.0526 | 45 | 0.0889 | 34 | 0.0882 | 46 | 0.1087 | 163 | 0.0859 | 31 | 0.0323 | 34 | 0.1471 | 40 | 0.15 | 49 | 0.1633 | 154 | 0.1299 |
| 17_6459006 | 43 | 0.0233 | 45 | 0.0222 | 35 | 0.1143 | 53 | 0.1509 | 176 | 0.0795 | 33 | 0.1212 | 34 | 0.1765 | 45 | 0.0444 | 49 | 0.1224 | 161 | 0.1118 |
| 17_6459025 | 34 | 0.0294 | 40 | 0.05 | 33 | 0.0909 | 45 | 0.0889 | 152 | 0.0658 | 29 | 0 | 31 | 0.1613 | 42 | 0.119 | 42 | 0.0714 | 144 | 0.0903 |
| 17_6461567 | 27 | 0.2593 | 36 | 0.0833 | 26 | 0.3846 | 35 | 0.2286 | 124 | 0.2258 | 26 | 0.3077 | 19 | 0.4737 | 42 | 0.3095 | 34 | 0.2941 | 121 | 0.3306 |
| 17_6461592 | 27 | 0.1111 | 35 | 0.0857 | 23 | 0 | 34 | 0.1471 | 119 | 0.0924 | 24 | 0.0417 | 20 | 0.05 | 35 | 0.0571 | 32 | 0.125 | 111 | 0.0721 |
| 17_6461603 | 25 | 0.4 | 32 | 0.5625 | 21 | 0.2381 | 30 | 0.3667 | 108 | 0.4074 | 17 | 0.5882 | 15 | 0.3333 | 29 | 0.4138 | 24 | 0.625 | 85 | 0.4941 |
| 17_6461639 | 22 | 0.8636 | 29 | 0.7586 | 22 | 0.6818 | 30 | 0.7667 | 103 | 0.767 | 15 | 0.7333 | 15 | 0.9333 | 26 | 0.8462 | 28 | 0.8929 | 84 | 0.8571 |
| 17_6505984 | 2 | 0.5 | NA | NA | NA | NA | NA | NA | 2 | 0.5 | 2 | 0.5 | NA | NA | NA | NA | 1 | 0 | 3 | 0.3333 |
| 17_6506075 | 3 | 0.3333 | 1 | 1 | 1 | 0 | 1 | 1 | 6 | 0.5 | 3 | 0.6667 | 1 | 0 | NA | NA | 2 | 0.5 | 6 | 0.5 |
| 17_6588746 | 34 | 0.0882 | 34 | 0.1176 | 34 | 0.1176 | 37 | 0.0811 | 139 | 0.1007 | 28 | 0.1071 | 31 | 0.0323 | 23 | 0.087 | 28 | 0.0714 | 110 | 0.0727 |
| 17_6588748 | 31 | 0.129 | 35 | 0.1714 | 34 | 0.2353 | 34 | 0.2941 | 134 | 0.209 | 30 | 0.2 | 31 | 0.0968 | 25 | 0.2 | 27 | 0.2222 | 113 | 0.177 |
| 17_6595893 | 9 | 0.8889 | 15 | 0.6667 | 10 | 0.7 | 4 | 0.75 | 38 | 0.7368 | 7 | 0.8571 | 6 | 0.8333 | 7 | 0.8571 | 4 | 1 | 24 | 0.875 |
| 17_6595895 | 10 | 0.4 | 14 | 0.2857 | 12 | 0.25 | 3 | 0 | 39 | 0.2821 | 8 | 0.125 | 6 | 0.5 | 9 | 0.5556 | 6 | 0.1667 | 29 | 0.3448 |
| 17_6595909 | 8 | 0.875 | 13 | 0.7692 | 13 | 0.9231 | 4 | 1 | 38 | 0.8684 | 6 | 1 | 6 | 1 | 8 | 1 | 6 | 0.8333 | 26 | 0.9615 |
| 17_6595930 | 8 | 0.25 | 12 | 0 | 13 | 0.1538 | 2 | 0 | 35 | 0.1143 | 8 | 0.25 | 4 | 0.5 | 6 | 0.1667 | 5 | 0.2 | 23 | 0.2609 |
| 17_6595984 | 5 | 0.4 | 9 | 0.2222 | 12 | 0.4167 | 2 | 0.5 | 28 | 0.3571 | 5 | 0.2 | 4 | 0.5 | 6 | 0.3333 | 2 | 0.5 | 17 | 0.3529 |
| 17_6596008 | 8 | 0 | 10 | 0.3 | 12 | 0.3333 | 3 | 0.3333 | 33 | 0.2424 | 6 | 0.1667 | 4 | 0.25 | 7 | 0 | 5 | 0.2 | 22 | 0.1364 |
| 17_6596078 | 6 | 1 | 8 | 0.625 | 13 | 0.7692 | 3 | 0.6667 | 30 | 0.7667 | 7 | 0.8571 | 2 | 1 | 7 | 0.5714 | 3 | 0.6667 | 19 | 0.7368 |
| 17_6596085 | 6 | 0.5 | 8 | 0 | 9 | 0.1111 | 3 | 0.3333 | 26 | 0.1923 | 7 | 0 | 1 | 0 | 7 | 0.1429 | 2 | 0 | 17 | 0.0588 |
| 17_6596086 | 6 | 0 | 7 | 0.1429 | 12 | 0.0833 | 3 | 0 | 28 | 0.0714 | 7 | 0 | 3 | 0 | 7 | 0.1429 | 3 | 0 | 20 | 0.05 |
| 17_6596121 | 7 | 0.4286 | 5 | 0.2 | 7 | 0.2857 | 2 | 0 | 21 | 0.2857 | 6 | 0 | NA | NA | 5 | 0.2 | 3 | 0 | 14 | 0.0714 |
| 17_6596127 | 5 | 0.2 | 6 | 0.5 | 8 | 0.5 | 1 | 1 | 20 | 0.45 | 7 | 0.5714 | 2 | 1 | 6 | 0.3333 | 3 | 1 | 18 | 0.6111 |
| 17_6613141 | 1 | 1 | 3 | 0.6667 | 2 | 1 | 2 | 1 | 8 | 0.875 | 1 | 1 | 3 | 0.6667 | 1 | 1 | NA | NA | 5 | 0.8 |
| 17_6613156 | NA | NA | 3 | 0.3333 | 2 | 0.5 | 1 | 0 | 6 | 0.3333 | NA | NA | 4 | 0.75 | 1 | 0 | NA | NA | 5 | 0.6 |
| 17_6613159 | NA | NA | 2 | 0.5 | 2 | 1 | 2 | 1 | 6 | 0.8333 | 1 | 1 | 3 | 0.3333 | 1 | 0 | 1 | 1 | 6 | 0.5 |
| 17_6613164 | NA | NA | 3 | 0.3333 | 1 | 0 | 2 | 0.5 | 6 | 0.3333 | NA | NA | 4 | 0.25 | 1 | 0 | 1 | 1 | 6 | 0.3333 |
| 17_6614792 | 1 | 0 | 3 | 0.3333 | 2 | 0 | NA | NA | 6 | 0.1667 | NA | NA | 5 | 0 | 3 | 0.3333 | NA | NA | 8 | 0.125 |
| 17_6614807 | 1 | 1 | 1 | 1 | 1 | 1 | 1 | 1 | 4 | 1 | NA | NA | 4 | 0.75 | 2 | 0.5 | 1 | 0 | 7 | 0.5714 |
| 17_6614819 | 1 | 0 | 2 | 0.5 | 2 | 0 | 1 | 1 | 6 | 0.3333 | NA | NA | 6 | 0.5 | 3 | 0.3333 | 1 | 0 | 10 | 0.4 |
| 17_6614826 | NA | NA | 2 | 0.5 | 2 | 0 | 1 | 0 | 5 | 0.2 | 1 | 0 | 5 | 0.2 | 3 | 0 | 1 | 0 | 10 | 0.1 |
| 17_6614836 | 1 | 1 | 2 | 0.5 | 2 | 1 | NA | NA | 5 | 0.8 | 1 | 1 | 7 | 0.4286 | 3 | 0.6667 | 1 | 1 | 12 | 0.5833 |
| 17_6614840 | 1 | 0 | 2 | 1 | 2 | 0.5 | NA | NA | 5 | 0.6 | 1 | 0 | 4 | 0.25 | 3 | 0 | 1 | 0 | 9 | 0.1111 |
| 17_6614857 | 1 | 0 | 2 | 0 | 1 | 1 | 1 | 0 | 5 | 0.2 | 1 | 0 | 5 | 0.4 | 2 | 0 | 1 | 1 | 9 | 0.3333 |
| 17_6614879 | 1 | 0 | 4 | 0.25 | 2 | 0 | NA | NA | 7 | 0.1429 | 3 | 0.3333 | 6 | 0.1667 | 4 | 0.5 | 1 | 0 | 14 | 0.2857 |
| 17_6615567 | 11 | 0 | 7 | 0 | 9 | 0.1111 | 13 | 0.0769 | 40 | 0.05 | 11 | 0.0909 | 5 | 0.2 | 6 | 0.1667 | 6 | 0.1667 | 28 | 0.1429 |
| 17_6615589 | 10 | 0.1 | 9 | 0.2222 | 8 | 0 | 10 | 0 | 37 | 0.0811 | 9 | 0 | 6 | 0.3333 | 5 | 0.2 | 6 | 0.1667 | 26 | 0.1538 |
| 17_6615599 | 5 | 0.4 | 6 | 0.5 | 4 | 0.5 | 7 | 0.2857 | 22 | 0.4091 | 9 | 0.4444 | 3 | 0 | 5 | 0.2 | 3 | 0.3333 | 20 | 0.3 |
| 17_6615602 | 9 | 0.1111 | 9 | 0.7778 | 4 | 0.25 | 9 | 0.5556 | 31 | 0.4516 | 7 | 0.4286 | 3 | 0.3333 | 3 | 0.6667 | 5 | 0 | 18 | 0.3333 |
| 17_6615606 | 10 | 0.8 | 9 | 1 | 7 | 0.8571 | 9 | 1 | 35 | 0.9143 | 10 | 0.9 | 6 | 0.8333 | 6 | 1 | 6 | 0.8333 | 28 | 0.8929 |
| 17_6615617 | 8 | 0.125 | 6 | 0.1667 | 4 | 0.25 | 8 | 0.375 | 26 | 0.2308 | 9 | 0.4444 | 5 | 0 | 5 | 0.2 | 5 | 0.2 | 24 | 0.25 |
| 17_6615646 | 7 | 0.1429 | 4 | 0 | 6 | 0.1667 | 8 | 0 | 25 | 0.08 | 8 | 0.125 | 3 | 0.3333 | 3 | 0 | 4 | 0.25 | 18 | 0.1667 |
| 17_6615679 | 5 | 0.8 | 6 | 1 | 6 | 1 | 9 | 0.8889 | 26 | 0.9231 | 9 | 0.7778 | 5 | 1 | 7 | 1 | 3 | 1 | 24 | 0.9167 |
| 17_6615688 | 8 | 0.75 | 8 | 1 | 6 | 1 | 9 | 1 | 31 | 0.9355 | 10 | 0.8 | 5 | 0.8 | 6 | 1 | 5 | 0.8 | 26 | 0.8462 |
| 17_6616473 | 7 | 0.2857 | 4 | 0 | 7 | 0.1429 | 6 | 0.1667 | 24 | 0.1667 | 8 | 0 | 4 | 0 | 5 | 0 | 5 | 0.6 | 22 | 0.1364 |
| 17_6616837 | 4 | 0.25 | 4 | 0 | 6 | 0 | 4 | 0 | 18 | 0.0556 | 6 | 0.1667 | 3 | 0 | 4 | 0.25 | 2 | 0 | 15 | 0.1333 |
| 17_6617131 | 5 | 0 | 1 | 0 | 6 | 0.1667 | 3 | 0.3333 | 15 | 0.1333 | 7 | 0 | 3 | 0 | 4 | 0.25 | 4 | 0.25 | 18 | 0.1111 |
| 17_6617151 | 5 | 0.8 | 1 | 1 | 6 | 0.8333 | 3 | 0.6667 | 15 | 0.8 | 6 | 0.8333 | 3 | 0.6667 | 3 | 0.6667 | 5 | 0.8 | 17 | 0.7647 |
| 17_6623019 | 2 | 0 | 4 | 0 | 3 | 0 | 5 | 0.4 | 14 | 0.1429 | 4 | 0.25 | 2 | 0 | 7 | 0.1429 | 6 | 0.1667 | 19 | 0.1579 |
| 17_6623234 | 1 | 0 | 1 | 1 | 1 | 0 | 2 | 0 | 5 | 0.2 | 1 | 0 | 1 | 1 | 1 | 0 | 6 | 0 | 9 | 0.1111 |
| 17_6623329 | 1 | 1 | 1 | 1 | 2 | 0.5 | 3 | 0.6667 | 7 | 0.7143 | NA | NA | NA | NA | 1 | 1 | 4 | 0.5 | 5 | 0.6 |
| 17_6623332 | 1 | 1 | 1 | 0 | 2 | 0 | 2 | 1 | 6 | 0.5 | 1 | 0 | NA | NA | 1 | 0 | 5 | 0.8 | 7 | 0.5714 |
| 17_6623661 | 1 | 0 | 1 | 1 | 1 | 0 | 1 | 0 | 4 | 0.25 | 1 | 1 | NA | NA | 1 | 0 | 5 | 0.2 | 7 | 0.2857 |
| 17_6627289 | 1 | 0 | 2 | 1 | 2 | 0 | 4 | 0 | 9 | 0.2222 | 3 | 0.3333 | NA | NA | 2 | 0 | 3 | 0 | 8 | 0.125 |

| | | | | | | | | | | | | | | | | | | | | |
|---|---|---|---|---|---|---|---|---|---|---|---|---|---|---|---|---|---|---|---|---|
| 17_6627474 | 4 | 1 | 3 | 1 | 5 | 1 | 14 | 1 | 26 | 1 | 4 | 1 | 3 | 1 | 8 | 1 | 8 | 1 | 23 | 1 |
| 17_6627476 | 4 | 1 | 3 | 1 | 6 | 1 | 14 | 0.9286 | 27 | 0.963 | 3 | 1 | 3 | 1 | 7 | 1 | 6 | 1 | 19 | 1 |
| 17_6627485 | 4 | 0.75 | 3 | 0 | 5 | 0.8 | 14 | 0.7857 | 26 | 0.6923 | 3 | 0.6667 | 4 | 1 | 8 | 0.875 | 6 | 0.8333 | 21 | 0.8571 |
| 17_6627499 | 4 | 0 | 3 | 0 | 5 | 0.4 | 12 | 0.25 | 24 | 0.2083 | 3 | 0.3333 | 4 | 0.5 | 5 | 0 | 6 | 0.3333 | 18 | 0.2778 |
| 17_6627501 | 4 | 0.25 | 3 | 0 | 5 | 0 | 13 | 0.4615 | 25 | 0.28 | 3 | 0.3333 | 3 | 0 | 7 | 0.1429 | 6 | 0.3333 | 19 | 0.2105 |
| 17_6627502 | 3 | 0 | 3 | 0 | 4 | 0 | 13 | 0.1538 | 23 | 0.087 | 2 | 0 | 4 | 0 | 6 | 0 | 6 | 0.1667 | 18 | 0.0556 |
| 17_6627503 | 2 | 1 | 3 | 0.6667 | 3 | 0.6667 | 12 | 0.5 | 20 | 0.6 | 3 | 0.6667 | 3 | 0 | 5 | 0.4 | 4 | 0.75 | 15 | 0.4667 |
| 17_6627634 | NA | NA | 2 | 0 | 4 | 0 | 11 | 0.1818 | 17 | 0.1176 | 1 | 0 | 3 | 0 | 4 | 0 | 2 | 0.5 | 10 | 0.1 |
| 17_6627716 | 1 | 1 | 2 | 0.5 | 3 | 0.3333 | 10 | 0.2 | 16 | 0.3125 | NA | NA | 1 | 0 | 5 | 0.2 | 2 | 0 | 8 | 0.125 |
| 17_6629017 | 1 | 0 | NA | NA | 3 | 0.6667 | 5 | 0 | 9 | 0.2222 | NA | NA | 1 | 1 | 4 | 0.25 | 1 | 1 | 6 | 0.5 |
| 17_6629028 | NA | NA | NA | NA | 3 | 0.3333 | 4 | 0.5 | 7 | 0.4286 | NA | NA | 1 | 1 | 3 | 0.3333 | 1 | 0 | 5 | 0.4 |
| 17_6629031 | 1 | 1 | NA | NA | 3 | 0 | 4 | 0.25 | 8 | 0.25 | NA | NA | 1 | 1 | 4 | 0 | 1 | 0 | 6 | 0.1667 |
| 17_6629049 | NA | NA | NA | NA | 3 | 0 | 5 | 0.2 | 8 | 0.125 | NA | NA | 1 | 0 | 2 | 0 | 1 | 1 | 4 | 0.25 |
| 17_6629067 | 1 | 0 | NA | NA | 2 | 0.5 | 6 | 0.1667 | 9 | 0.2222 | NA | NA | 1 | 1 | 3 | 0 | 1 | 0 | 5 | 0.2 |
| 17_6629072 | NA | NA | NA | NA | 2 | 1 | 5 | 1 | 7 | 1 | NA | NA | NA | NA | 4 | 1 | 1 | 1 | 5 | 1 |
| 17_6629087 | 1 | 1 | NA | NA | 1 | 0 | 4 | 0.25 | 6 | 0.3333 | NA | NA | 1 | 0 | 4 | 0.25 | 1 | 0 | 6 | 0.1667 |
| 17_6629098 | NA | NA | NA | NA | 3 | 0 | 4 | 0.25 | 7 | 0.1429 | NA | NA | 1 | 1 | 3 | 0 | 1 | 0 | 5 | 0.2 |
| 17_6629105 | 1 | 0 | NA | NA | 3 | 0 | 5 | 0.2 | 9 | 0.1111 | NA | NA | 1 | 0 | 4 | 0.25 | 1 | 0 | 6 | 0.1667 |
| 17_6629107 | 1 | 0 | NA | NA | 3 | 0.3333 | 5 | 0.4 | 9 | 0.3333 | NA | NA | 1 | 0 | 3 | 0.6667 | 1 | 0 | 5 | 0.4 |
| 17_6629110 | 1 | 0 | NA | NA | 3 | 0.6667 | 4 | 0.75 | 8 | 0.625 | NA | NA | NA | NA | 4 | 0.75 | NA | NA | 4 | 0.75 |
| 17_6629119 | 1 | 1 | NA | NA | 3 | 0 | 5 | 0.6 | 9 | 0.4444 | NA | NA | 1 | 0 | 3 | 0.3333 | 1 | 0 | 5 | 0.2 |
| 17_6629140 | 1 | 0 | NA | NA | 3 | 0 | 4 | 0.25 | 8 | 0.125 | NA | NA | 1 | 0 | 2 | 0.5 | 1 | 0 | 4 | 0.25 |
| 17_6629164 | 1 | 0 | NA | NA | 3 | 0.3333 | 2 | 1 | 6 | 0.5 | NA | NA | 1 | 0 | 2 | 0 | 1 | 1 | 4 | 0.25 |
| 17_6629207 | 1 | 0 | NA | NA | 2 | 0 | 2 | 1 | 5 | 0.4 | NA | NA | NA | NA | 2 | 0.5 | 1 | 0 | 3 | 0.3333 |
| 17_6629278 | NA | NA | NA | NA | 2 | 0.5 | 2 | 1 | 4 | 0.75 | NA | NA | NA | NA | 2 | 0.5 | 1 | 1 | 3 | 0.6667 |
| 17_6629342 | NA | NA | NA | NA | 1 | 1 | 2 | 0 | 3 | 0.3333 | NA | NA | NA | NA | 2 | 0.5 | 1 | 0 | 3 | 0.3333 |
| 17_6632564 | 1 | 1 | NA | NA | NA | NA | NA | NA | 1 | 1 | NA | NA | NA | NA | 1 | 1 | 1 | 0 | 2 | 0.5 |
| 17_6632609 | 1 | 1 | NA | NA | NA | NA | NA | NA | 1 | 1 | NA | NA | NA | NA | 1 | 1 | 1 | 1 | 2 | 1 |
| 17_6642611 | 3 | 0 | 3 | 0.3333 | 7 | 0.5714 | 6 | 0.5 | 19 | 0.4211 | 4 | 0.25 | 3 | 0.3333 | 5 | 0.2 | 3 | 0.3333 | 15 | 0.2667 |
| 17_6642628 | 1 | 1 | 3 | 1 | 7 | 0.8571 | 6 | 1 | 17 | 0.9412 | 3 | 1 | 3 | 1 | 3 | 1 | 5 | 1 | 14 | 1 |
| 17_6642637 | 4 | 0.5 | 2 | 0.5 | 4 | 0.75 | 3 | 0.3333 | 13 | 0.5385 | 4 | 0.25 | 2 | 0.5 | 3 | 0.3333 | 4 | 0.5 | 13 | 0.3846 |
| 17_6642649 | 2 | 1 | NA | NA | 1 | 0 | 1 | 0 | 4 | 0.5 | 1 | 0 | 1 | 0 | 1 | 1 | 1 | 0 | 4 | 0.25 |
| 17_6642661 | 2 | 0.5 | NA | NA | 2 | 1 | 1 | 0 | 5 | 0.6 | 1 | 0 | 1 | 1 | 1 | 1 | 2 | 0 | 5 | 0.4 |
| 17_6642662 | 2 | 0.5 | NA | NA | 2 | 0 | 1 | 0 | 5 | 0.2 | 2 | 0.5 | 1 | 0 | 1 | 1 | 2 | 0.5 | 6 | 0.5 |
| 17_6642667 | 2 | 0.5 | NA | NA | 2 | 1 | 1 | 1 | 5 | 0.8 | 2 | 0.5 | 1 | 0 | 1 | 1 | 2 | 0.5 | 6 | 0.5 |
| 17_6642669 | 2 | 0 | NA | NA | 2 | 0.5 | 1 | 0 | 5 | 0.2 | 2 | 0.5 | 1 | 1 | 1 | 0 | 2 | 0 | 6 | 0.3333 |
| 17_6642680 | 2 | 1 | NA | NA | 2 | 0 | 1 | 0 | 5 | 0.4 | 2 | 0.5 | 1 | 0 | 1 | 0 | 2 | 0.5 | 6 | 0.3333 |
| 17_6642685 | 2 | 0 | NA | NA | 2 | 1 | 1 | 0 | 5 | 0.4 | 2 | 0 | 1 | 0 | NA | NA | 1 | 1 | 4 | 0.25 |
| 17_6642707 | 2 | 0.5 | NA | NA | 2 | 0 | 2 | 0 | 6 | 0.1667 | 2 | 0 | 1 | 0 | 2 | 0.5 | 2 | 0 | 7 | 0.1429 |
| 17_6664434 | 6 | 0.3333 | 9 | 0.4444 | 6 | 0.3333 | 9 | 0.5556 | 30 | 0.4333 | 3 | 0 | 6 | 0.5 | 9 | 0.5556 | 4 | 0.5 | 22 | 0.4545 |
| 17_6664434 | 6 | 0.5 | 6 | 0.5 | 6 | 0.1667 | 10 | 0.7 | 28 | 0.5 | 2 | 0.5 | 8 | 0.625 | 8 | 0.125 | 3 | 0.3333 | 21 | 0.381 |
| 17_6664445 | 6 | 0.3333 | 8 | 0.625 | 6 | 0.3333 | 8 | 0.75 | 28 | 0.5357 | 2 | 0.5 | 6 | 0.8333 | 8 | 0.25 | 3 | 0.3333 | 19 | 0.4737 |
| 17_6664446 | 7 | 0.4286 | 7 | 0.5714 | 6 | 0 | 8 | 0.5 | 28 | 0.3929 | 2 | 0.5 | 7 | 0.2857 | 8 | 0.625 | 2 | 1 | 19 | 0.5263 |
| 17_6664472 | 5 | 0 | 6 | 0.3333 | 6 | 0 | 9 | 0.1111 | 26 | 0.1154 | 2 | 0 | 5 | 0.2 | 6 | 0.1667 | 4 | 0.5 | 17 | 0.2353 |
| 17_6664476 | 5 | 0 | 5 | 0.4 | 6 | 0 | 8 | 0.125 | 24 | 0.125 | 3 | 0 | 4 | 0 | 6 | 0 | 5 | 0.2 | 18 | 0.0556 |
| 17_6664478 | 5 | 0.8 | 4 | 0.5 | 5 | 0.6 | 3 | 0.3333 | 17 | 0.5882 | 1 | 1 | 3 | 1 | 7 | 0.4286 | 5 | 0.4 | 16 | 0.5625 |
| 17_6664481 | 5 | 0.2 | 4 | 0.25 | 6 | 0.1667 | 8 | 0 | 23 | 0.1304 | 1 | 0 | 5 | 0.4 | 8 | 0.125 | 4 | 0.25 | 18 | 0.2222 |
| 17_6664501 | 5 | 0 | 6 | 0.3333 | 4 | 0 | 7 | 0 | 22 | 0.0909 | 2 | 0 | 6 | 0 | 8 | 0.125 | 3 | 0 | 19 | 0.0526 |
| 17_6664999 | 3 | 1 | 5 | 0.8 | 3 | 1 | 5 | 0.8 | 16 | 0.875 | 2 | 1 | 4 | 1 | 8 | 1 | 4 | 1 | 18 | 1 |
| 17_6665000 | 3 | 0 | 6 | 0.3333 | 3 | 0 | 7 | 0 | 19 | 0.1053 | 3 | 0 | 4 | 0 | 8 | 0 | 3 | 0.6667 | 18 | 0.1111 |
| 17_6665029 | 2 | 0.5 | 5 | 0.2 | 3 | 0.6667 | 5 | 0 | 15 | 0.2667 | NA | NA | 1 | 1 | 6 | 0.5 | 4 | 1 | 11 | 0.7273 |
| 17_6665034 | 3 | 0 | 7 | 0 | 3 | 0.3333 | 6 | 0 | 19 | 0.0526 | 3 | 0 | 2 | 0.5 | 5 | 0 | 4 | 0 | 14 | 0.0714 |
| 17_6665042 | 3 | 0 | 5 | 0.2 | 3 | 0.3333 | 4 | 0 | 15 | 0.1333 | 3 | 0.3333 | 3 | 0 | 5 | 0 | 4 | 0 | 15 | 0.0667 |
| 17_6665057 | 2 | 0.5 | 6 | 0.8333 | 3 | 1 | 5 | 0.4 | 16 | 0.6875 | 3 | 0.6667 | 2 | 0.5 | 5 | 0.4 | 2 | 0.5 | 12 | 0.5 |
| 17_6665068 | 2 | 0.5 | 5 | 0.4 | 3 | 0 | 4 | 0.25 | 14 | 0.2857 | 2 | 0.5 | 1 | 0 | 4 | 0.5 | 3 | 0.3333 | 10 | 0.4 |
| 17_6665074 | 2 | 0 | 6 | 0.1667 | 3 | 0 | 6 | 0 | 17 | 0.0588 | 2 | 0 | 1 | 0 | 5 | 0 | 4 | 0.25 | 12 | 0.0833 |
| 17_6665078 | 2 | 1 | 3 | 1 | 3 | 1 | 4 | 1 | 12 | 1 | 1 | 1 | NA | NA | 5 | 1 | 3 | 1 | 9 | 1 |
| 17_6665082 | 3 | 0.3333 | 4 | 0.25 | 2 | 0.5 | 5 | 0.6 | 14 | 0.4286 | 2 | 0.5 | 2 | 1 | 4 | 0.25 | 4 | 0.5 | 12 | 0.5 |
| 17_6665089 | 2 | 0 | 6 | 0.6667 | 2 | 0 | 5 | 1 | 15 | 0.6 | 2 | 0.5 | 2 | 0 | 5 | 0.4 | 4 | 0.25 | 13 | 0.3077 |
| 17_6665093 | 3 | 0.3333 | 5 | 0.2 | 1 | 0 | 4 | 0 | 13 | 0.1538 | 2 | 0.5 | 1 | 0 | 5 | 0 | 1 | 0 | 9 | 0.1111 |
| 17_6665100 | 2 | 0 | 7 | 0.5714 | 2 | 0 | 4 | 0.5 | 15 | 0.4 | 1 | 1 | 2 | 0.5 | 5 | 0.4 | 4 | 0.5 | 12 | 0.5 |
| 17_6665114 | 3 | 1 | 6 | 1 | 2 | 1 | 4 | 0.25 | 15 | 0.8 | 1 | 1 | 2 | 1 | 5 | 0.8 | 3 | 1 | 11 | 0.9091 |
| 17_6665525 | 2 | 1 | 4 | 0 | 1 | 0 | 4 | 0.25 | 11 | 0.2727 | 1 | 0 | 2 | 0 | 4 | 0.25 | 2 | 0 | 9 | 0.1111 |
| 17_6668617 | NA | NA | 2 | 0 | NA | NA | 1 | 1 | 3 | 0.3333 | 2 | 0 | NA | NA | 3 | 0 | 2 | 0.5 | 7 | 0.1429 |
| 17_6668640 | NA | NA | 2 | 1 | NA | NA | NA | NA | 2 | 1 | 1 | 1 | NA | NA | 3 | 1 | 2 | 1 | 6 | 1 |

| | | | | | | | | | | | | | | | | | | | | |
|---|---|---|---|---|---|---|---|---|---|---|---|---|---|---|---|---|---|---|---|---|
| 17_666864! | NA | NA | 2 | 1 | NA | NA | 1 | 1 | 3 | 1 | 1 | 1 | NA | NA | 3 | 0.6667 | 2 | 1 | 6 | 0.8333 |
| 17_666866! | NA | NA | 1 | 1 | NA | NA | NA | NA | 1 | 1 | 2 | 1 | NA | NA | 1 | 0 | 1 | 0 | 4 | 0.5 |
| 17_666873: | NA | NA | 1 | 1 | NA | NA | NA | NA | 1 | 1 | 1 | 0 | NA | NA | 2 | 0.5 | 1 | 0 | 4 | 0.25 |
| 17_667560: | NA | NA | 1 | 1 | NA | NA | NA | NA | 1 | 1 | NA | NA | NA | NA | 1 | 1 | NA | NA | 1 | 1 |
| 17_668016 | 16 | 0.0625 | 13 | 0.2308 | 10 | 0 | 23 | 0.2609 | 62 | 0.1613 | 10 | 0.1 | 21 | 0.3333 | 24 | 0.125 | 13 | 0.1538 | 68 | 0.1912 |
| 17_668017! | 16 | 0.375 | 13 | 0.3077 | 10 | 0.3 | 24 | 0.2917 | 63 | 0.3175 | 10 | 0.4 | 22 | 0.4545 | 22 | 0.3636 | 17 | 0.2941 | 71 | 0.3803 |
| 17_668017! | 14 | 0.0714 | 13 | 0 | 8 | 0.125 | 18 | 0.0556 | 53 | 0.0566 | 10 | 0.1 | 15 | 0.1333 | 18 | 0.0556 | 13 | 0 | 56 | 0.0714 |
| 17_668019: | 14 | 0 | 15 | 0.0667 | 9 | 0 | 22 | 0.0909 | 60 | 0.05 | 7 | 0 | 20 | 0 | 21 | 0.1429 | 13 | 0.0769 | 61 | 0.0656 |
| 17_668021( | 16 | 0.5 | 14 | 0.2857 | 10 | 0.4 | 22 | 0.3636 | 62 | 0.3871 | 9 | 0.1111 | 17 | 0.5294 | 21 | 0.4286 | 16 | 0.5 | 63 | 0.4286 |
| 17_668021: | 15 | 0.6667 | 12 | 1 | 7 | 0.7143 | 20 | 0.85 | 54 | 0.8148 | 5 | 1 | 14 | 0.9286 | 19 | 0.9474 | 10 | 0.8 | 48 | 0.9167 |
| 17_668021: | 16 | 0.3125 | 13 | 0.4615 | 8 | 0.375 | 24 | 0.4583 | 61 | 0.4098 | 8 | 0.5 | 17 | 0.4706 | 23 | 0.4348 | 14 | 0.5 | 62 | 0.4677 |
| 17_668022! | 14 | 0.1429 | 13 | 0.2308 | 9 | 0.1111 | 20 | 0.15 | 56 | 0.1607 | 5 | 0 | 13 | 0.3077 | 18 | 0.2222 | 12 | 0.0833 | 48 | 0.1875 |
| 17_668023: | 15 | 0 | 15 | 0.1333 | 7 | 0.1429 | 21 | 0.0476 | 58 | 0.069 | 8 | 0.125 | 18 | 0.0556 | 23 | 0.2609 | 14 | 0.2857 | 63 | 0.1905 |
| 17_668024: | 12 | 0.25 | 14 | 0.2857 | 8 | 0.25 | 20 | 0.3 | 54 | 0.2778 | 7 | 0.1429 | 16 | 0.5 | 23 | 0.1739 | 12 | 0.25 | 58 | 0.2759 |
| 17_668025( | 11 | 0.2727 | 13 | 0.3077 | 4 | 0.5 | 18 | 0.4444 | 46 | 0.3696 | 6 | 0.1667 | 15 | 0.4 | 21 | 0.2857 | 13 | 0.6923 | 55 | 0.4 |
| 17_668026' | 10 | 0.3 | 14 | 0.2143 | 6 | 0.6667 | 19 | 0.1579 | 49 | 0.2653 | 7 | 0.1429 | 17 | 0.2353 | 22 | 0.2727 | 13 | 0.1538 | 59 | 0.2203 |
| 17_669744( | 3 | 0.3333 | 3 | 0.3333 | 3 | 0.6667 | 5 | 0.4 | 14 | 0.4286 | NA | NA | 2 | 1 | 3 | 0 | 1 | 0 | 6 | 0.3333 |
| 17_669744' | 4 | 0 | 4 | 0 | 3 | 0 | 6 | 0.1667 | 17 | 0.0588 | NA | NA | 2 | 0.5 | 3 | 0 | 3 | 0.3333 | 8 | 0.25 |
| 17_669745· | 1 | 0 | 4 | 0.5 | 3 | 0.6667 | 5 | 0.6 | 13 | 0.5385 | NA | NA | 2 | 0 | 3 | 0.6667 | 3 | 0.6667 | 8 | 0.5 |
| 17_691367! | NA | NA | NA | NA | NA | NA | 1 | 1 | 1 | 1 | NA | NA | NA | NA | 1 | 1 | NA | NA | 1 | 1 |
| 17_691369! | NA | NA | NA | NA | NA | NA | 1 | 1 | 1 | 1 | NA | NA | NA | NA | 1 | 1 | NA | NA | 1 | 1 |
| 17_691370( | NA | NA | NA | NA | NA | NA | 1 | 1 | 1 | 1 | NA | NA | NA | NA | 1 | 1 | NA | NA | 1 | 1 |
| 17_691371( | NA | NA | NA | NA | NA | NA | 1 | 1 | 1 | 1 | NA | NA | NA | NA | 1 | 1 | NA | NA | 1 | 1 |
| 17_691371! | NA | NA | NA | NA | NA | NA | 1 | 1 | 1 | 1 | NA | NA | NA | NA | 1 | 1 | NA | NA | 1 | 1 |
| 17_691475! | NA | NA | 1 | 1 | NA | NA | NA | NA | 1 | 1 | NA | NA | NA | NA | NA | NA | 1 | 1 | 1 | 1 |
| 17_691480' | NA | NA | 1 | 1 | 1 | 0 | NA | NA | 2 | 0.5 | NA | NA | NA | NA | NA | NA | 2 | 1 | 2 | 1 |
| 17_691480· | NA | NA | 1 | 1 | 1 | 1 | NA | NA | 2 | 1 | NA | NA | NA | NA | NA | NA | 2 | 1 | 2 | 1 |
| 17_691483: | NA | NA | 1 | 1 | 1 | 1 | NA | NA | 2 | 1 | NA | NA | NA | NA | NA | NA | 1 | 1 | 1 | 1 |
| 17_691484! | NA | NA | 1 | 1 | 1 | 1 | NA | NA | 2 | 1 | NA | NA | NA | NA | 1 | 0 | 1 | 1 | 2 | 0.5 |
| 17_691486: | NA | NA | 1 | 1 | 1 | 1 | NA | NA | 2 | 1 | NA | NA | NA | NA | 1 | 1 | 2 | 1 | 3 | 1 |
| 17_691486· | NA | NA | 1 | 1 | 1 | 1 | NA | NA | 2 | 1 | NA | NA | NA | NA | 1 | 1 | 2 | 1 | 3 | 1 |
| 17_691487' | NA | NA | 1 | 1 | 1 | 1 | NA | NA | 2 | 1 | NA | NA | NA | NA | 1 | 1 | 1 | 1 | 2 | 1 |
| 17_691487: | NA | NA | 1 | 0 | 1 | 1 | NA | NA | 2 | 0.5 | NA | NA | NA | NA | 1 | 0 | 2 | 1 | 3 | 0.6667 |
| 17_691488! | NA | NA | 1 | 1 | NA | NA | NA | NA | 1 | 1 | NA | NA | NA | NA | 1 | 1 | 2 | 1 | 3 | 1 |
| 17_691489! | NA | NA | 1 | 1 | 1 | 1 | NA | NA | 2 | 1 | NA | NA | NA | NA | 1 | 0 | 1 | 1 | 2 | 0.5 |
| 17_691490' | NA | NA | 2 | 0.5 | 1 | 1 | NA | NA | 3 | 0.6667 | NA | NA | NA | NA | 1 | 0 | 2 | 1 | 3 | 0.6667 |
| 17_691491· | NA | NA | 3 | 0.6667 | 1 | 1 | NA | NA | 4 | 0.75 | NA | NA | NA | NA | 1 | 1 | 2 | 1 | 3 | 1 |
| 17_691492( | NA | NA | 3 | 0.6667 | 1 | 0 | NA | NA | 4 | 0.5 | NA | NA | NA | NA | 1 | 0 | 2 | 1 | 3 | 0.6667 |
| 17_691493: | NA | NA | 2 | 0.5 | 1 | 1 | NA | NA | 3 | 0.6667 | NA | NA | NA | NA | 1 | 1 | 2 | 1 | 3 | 1 |
| 17_691494: | NA | NA | 1 | 1 | 1 | 0 | NA | NA | 2 | 0.5 | NA | NA | 1 | 0 | 1 | 0 | 2 | 1 | 4 | 0.5 |
| 17_691496: | NA | NA | 2 | 0.5 | 1 | 0 | NA | NA | 3 | 0.3333 | NA | NA | 1 | 0 | 1 | 1 | 2 | 0.5 | 4 | 0.5 |
| 17_691496( | NA | NA | 2 | 0.5 | 1 | 0 | NA | NA | 3 | 0.3333 | NA | NA | 1 | 0 | 1 | 0 | 2 | 0.5 | 4 | 0.25 |
| 17_691497: | NA | NA | 1 | 1 | 1 | 1 | NA | NA | 2 | 1 | NA | NA | 1 | 0 | 1 | 0 | 2 | 1 | 4 | 0.5 |
| 17_691498: | NA | NA | 3 | 1 | 1 | 0 | NA | NA | 4 | 0.75 | NA | NA | 1 | 0 | NA | NA | 1 | 1 | 2 | 0.5 |
| 17_691500' | NA | NA | 3 | 0.3333 | 1 | 0 | NA | NA | 4 | 0.25 | NA | NA | 1 | 0 | NA | NA | 2 | 0.5 | 3 | 0.3333 |
| 17_695172( | 1 | 0 | 4 | 0.25 | 8 | 0 | 4 | 0 | 17 | 0.0588 | 5 | 0 | 2 | 0 | 2 | 0 | 6 | 0.1667 | 15 | 0.0667 |
| 17_695897( | NA | NA | 2 | 0.5 | NA | NA | NA | NA | 2 | 0.5 | 2 | 1 | NA | NA | NA | NA | NA | NA | 2 | 1 |
| 17_695900( | NA | NA | 2 | 0.5 | 1 | 1 | NA | NA | 3 | 0.6667 | 2 | 0.5 | 1 | 0 | NA | NA | NA | NA | 3 | 0.3333 |
| 17_695958 | 16 | 0.0625 | 16 | 0 | 18 | 0.0556 | 13 | 0.1538 | 63 | 0.0635 | 11 | 0 | 13 | 0.1538 | 6 | 0 | 15 | 0.1333 | 45 | 0.0889 |
| 17_695960! | 10 | 0.1 | 9 | 0.4444 | 13 | 0.3077 | 7 | 0 | 39 | 0.2308 | 6 | 0 | 5 | 0 | 7 | 0.4286 | 8 | 0.25 | 26 | 0.1923 |
| 17_695961( | 16 | 0.5 | 16 | 0.25 | 19 | 0.4211 | 12 | 0.5833 | 63 | 0.4286 | 11 | 0.3636 | 12 | 0.5833 | 10 | 0.2 | 15 | 0.4 | 48 | 0.3958 |
| 17_695961( | 15 | 0.2 | 16 | 0.125 | 15 | 0.2 | 9 | 0.1111 | 55 | 0.1636 | 12 | 0.0833 | 10 | 0.1 | 9 | 0.2222 | 10 | 0.2 | 41 | 0.1463 |
| 17_695962: | 13 | 0.3846 | 16 | 0.625 | 16 | 0.3125 | 10 | 0.3 | 55 | 0.4182 | 13 | 0.4615 | 9 | 0.1111 | 8 | 0.5 | 11 | 0.4545 | 41 | 0.3902 |
| 17_695964· | 15 | 0.1333 | 14 | 0.1429 | 21 | 0.0476 | 11 | 0.0909 | 61 | 0.0984 | 14 | 0.0714 | 12 | 0.1667 | 9 | 0 | 15 | 0.0667 | 50 | 0.08 |
| 17_696910! | 1 | 1 | NA | NA | NA | NA | 1 | 1 | 2 | 1 | 2 | 0 | NA | NA | 3 | 0.6667 | 2 | 0 | 7 | 0.2857 |
| 17_696946( | 3 | 1 | NA | NA | 1 | 1 | 2 | 1 | 6 | 1 | 2 | 0.5 | NA | NA | 4 | 0.75 | 5 | 0.8 | 11 | 0.7273 |
| 17_696949· | 1 | 1 | 1 | 1 | 1 | 1 | 3 | 1 | 6 | 1 | 2 | 1 | 1 | 1 | 4 | 1 | 6 | 1 | 13 | 1 |
| 17_696950( | 4 | 0.5 | 1 | 0 | 1 | 0 | 3 | 0 | 9 | 0.2222 | 2 | 0 | 1 | 0 | 6 | 0.3333 | 6 | 0 | 15 | 0.1333 |
| 17_696950! | 2 | 0.5 | NA | NA | 1 | 0 | 2 | 0 | 5 | 0.2 | 2 | 0.5 | NA | NA | 3 | 0.3333 | 3 | 0 | 8 | 0.25 |
| 17_696952( | 3 | 0.6667 | NA | NA | 2 | 1 | 2 | 1 | 7 | 0.8571 | 3 | 0.6667 | 1 | 1 | 4 | 0.75 | 6 | 0.8333 | 14 | 0.7857 |
| 17_696953! | 3 | 0.3333 | 1 | 1 | NA | NA | 2 | 0.5 | 6 | 0.5 | 3 | 1 | 1 | 0 | 4 | 0.75 | 5 | 0.8 | 13 | 0.7692 |
| 17_696954: | 3 | 0.3333 | 1 | 0 | 2 | 0 | 2 | 1 | 8 | 0.375 | 3 | 0.3333 | 1 | 1 | 5 | 0.4 | 7 | 0.4286 | 16 | 0.4375 |
| 17_696954· | 3 | 0.3333 | 1 | 0 | 2 | 0 | 2 | 0 | 8 | 0.125 | 2 | 0 | NA | NA | 6 | 0 | 7 | 0.4286 | 15 | 0.2 |
| 17_696955' | 4 | 0 | 1 | 0 | 2 | 0 | 3 | 0.3333 | 10 | 0.1 | 3 | 0 | 1 | 0 | 6 | 0.1667 | 8 | 0.125 | 18 | 0.1111 |
| 17_696955: | 3 | 0.6667 | 1 | 1 | 1 | 1 | 2 | 1 | 7 | 0.8571 | 3 | 1 | 1 | 1 | 5 | 1 | 6 | 0.8333 | 15 | 0.9333 |

| | | | | | | | | | | | | | | | | | | | | |
|---|---|---|---|---|---|---|---|---|---|---|---|---|---|---|---|---|---|---|---|---|
| 17_696955 | 4 | 0 | 1 | 0 | 2 | 0.5 | 3 | 0.3333 | 10 | 0.2 | 2 | 0.5 | 1 | 1 | 6 | 0.3333 | 6 | 0.3333 | 15 | 0.4 |
| 17_696955 | 2 | 1 | NA | NA | NA | NA | 2 | 0.5 | 4 | 0.75 | 2 | 0.5 | NA | NA | 2 | 0.5 | 4 | 1 | 8 | 0.75 |
| 17_696956 | 4 | 0.75 | 1 | 1 | 2 | 1 | 3 | 0.6667 | 10 | 0.8 | 3 | 0.6667 | 1 | 1 | 6 | 0.6667 | 7 | 0.8571 | 17 | 0.7647 |
| 17_696956 | 2 | 1 | NA | NA | 1 | 1 | 2 | 1 | 5 | 1 | 3 | 1 | 1 | 1 | 5 | 1 | 8 | 1 | 17 | 1 |
| 17_696957 | 3 | 0 | NA | NA | 2 | 0 | 2 | 0.5 | 7 | 0.1429 | 3 | 0 | 1 | 0 | 4 | 0.25 | 7 | 0.2857 | 15 | 0.2 |
| 17_696957 | 3 | 0.6667 | 1 | 1 | 2 | 1 | 3 | 0.6667 | 9 | 0.7778 | 2 | 0.5 | 1 | 1 | 4 | 1 | 8 | 0.875 | 15 | 0.8667 |
| 17_696960 | 4 | 0 | 1 | 1 | 2 | 0 | 3 | 0 | 10 | 0.1 | 1 | 0 | 1 | 0 | 6 | 0.1667 | 7 | 0.5714 | 15 | 0.3333 |
| 17_696961 | 2 | 0 | 1 | 0 | 2 | 0 | 2 | 0.5 | 7 | 0.1429 | 2 | 0 | 1 | 0 | 6 | 0 | 6 | 0.1667 | 15 | 0.0667 |
| 17_696961 | 2 | 0 | 1 | 0 | 2 | 0.5 | 3 | 0.3333 | 8 | 0.25 | 2 | 0.5 | 1 | 0 | 6 | 0.1667 | 5 | 0.4 | 14 | 0.2857 |
| 17_696962 | 4 | 0 | 1 | 0 | 3 | 0.3333 | 2 | 0.5 | 10 | 0.2 | 1 | 1 | 1 | 0 | 4 | 0 | 6 | 0.1667 | 12 | 0.1667 |
| 17_696964 | 3 | 0.3333 | 1 | 1 | 3 | 0.6667 | 2 | 0.5 | 9 | 0.5556 | 2 | 0.5 | 1 | 1 | 5 | 0.6 | 7 | 0.5714 | 15 | 0.6 |
| 17_696965 | 4 | 0 | 1 | 0 | 3 | 0.3333 | 3 | 0 | 11 | 0.0909 | 2 | 0 | 2 | 0 | 7 | 0.1429 | 6 | 0 | 17 | 0.0588 |
| 17_696967 | 2 | 0 | NA | NA | 3 | 0 | 1 | 1 | 6 | 0.1667 | 2 | 0.5 | 2 | 0 | 5 | 0.2 | 4 | 0.25 | 13 | 0.2308 |
| 17_696968 | 4 | 0 | NA | NA | 4 | 0 | 3 | 0.6667 | 11 | 0.1818 | NA | NA | 1 | 0 | 6 | 0.3333 | 6 | 0.1667 | 13 | 0.2308 |
| 17_696969 | 2 | 0 | NA | NA | 4 | 0.5 | 1 | 1 | 7 | 0.4286 | 2 | 0.5 | 3 | 0.3333 | 6 | 0.1667 | 8 | 0.375 | 19 | 0.3158 |
| 17_696974 | 2 | 0.5 | 1 | 1 | 3 | 0.3333 | 2 | 0 | 8 | 0.375 | NA | NA | 3 | 0 | 5 | 0.2 | 4 | 0.5 | 12 | 0.25 |
| 17_697954 | 2 | 0.5 | 1 | 1 | NA | NA | 1 | 0 | 4 | 0.5 | 1 | 0 | 1 | 1 | NA | NA | 1 | 0 | 3 | 0.3333 |
| 17_709159 | 1 | 1 | NA | NA | NA | NA | NA | NA | 1 | 1 | NA | NA | 1 | 1 | NA | NA | NA | NA | 1 | 1 |
| 17_709167 | 1 | 1 | NA | NA | NA | NA | NA | NA | 1 | 1 | NA | NA | 1 | 1 | NA | NA | NA | NA | 1 | 1 |
| 17_709173 | 1 | 1 | NA | NA | NA | NA | NA | NA | 1 | 1 | NA | NA | 1 | 1 | NA | NA | NA | NA | 1 | 1 |
| 17_709183 | 1 | 1 | NA | NA | NA | NA | NA | NA | 1 | 1 | NA | NA | 1 | 1 | NA | NA | NA | NA | 1 | 1 |
| 17_709184 | 1 | 1 | NA | NA | NA | NA | NA | NA | 1 | 1 | NA | NA | 1 | 1 | NA | NA | NA | NA | 1 | 1 |
| 17_709184 | 1 | 1 | NA | NA | NA | NA | NA | NA | 1 | 1 | NA | NA | 1 | 1 | NA | NA | NA | NA | 1 | 1 |
| 17_709189 | 1 | 1 | NA | NA | NA | NA | NA | NA | 1 | 1 | NA | NA | 1 | 1 | NA | NA | NA | NA | 1 | 1 |
| 17_710682 | 4 | 0 | 7 | 0.1429 | 3 | 0 | 1 | 0 | 15 | 0.0667 | 5 | 0.2 | 1 | 0 | NA | NA | 1 | 0 | 7 | 0.1429 |
| 17_710938 | 6 | 0.1667 | 8 | 0.5 | 6 | 0.1667 | 3 | 0 | 23 | 0.2609 | 6 | 0.3333 | 2 | 0.5 | 3 | 0 | 2 | 0.5 | 13 | 0.3077 |
| 17_710938 | 6 | 0.8333 | 8 | 0.875 | 6 | 0.6667 | 2 | 0.5 | 22 | 0.7727 | 6 | 0.8333 | 2 | 0.5 | 3 | 1 | 1 | 1 | 12 | 0.8333 |
| 17_710940 | 9 | 0 | 9 | 0.2222 | 7 | 0 | 4 | 0.25 | 29 | 0.1034 | 7 | 0.1429 | 2 | 0.5 | 3 | 0.3333 | 4 | 0 | 16 | 0.1875 |
| 17_710940 | 8 | 0.125 | 8 | 0.375 | 6 | 0.1667 | 3 | 0.3333 | 25 | 0.24 | 8 | 0.375 | 2 | 0 | 3 | 0.3333 | 2 | 0.5 | 15 | 0.3333 |
| 17_711222 | 8 | 0.125 | 10 | 0.1 | 9 | 0.1111 | 3 | 0 | 30 | 0.1 | 7 | 0 | 3 | 0 | 2 | 0.5 | 4 | 0.5 | 16 | 0.1875 |
| 17_711222 | 8 | 0 | 11 | 0.0909 | 9 | 0 | 4 | 0.25 | 32 | 0.0625 | 9 | 0.1111 | 2 | 0 | 3 | 0 | 2 | 0.5 | 16 | 0.125 |
| 17_711222 | 7 | 0.2857 | 10 | 0.5 | 9 | 0.2222 | 4 | 0 | 30 | 0.3 | 9 | 0.1111 | 2 | 0 | 2 | 0 | 2 | 1 | 15 | 0.2 |
| 17_711223 | 1 | 0 | 4 | 0.25 | NA | NA | 1 | 0 | 6 | 0.1667 | 1 | 0 | 1 | 0 | 1 | 1 | 3 | 0.3333 | 6 | 0.3333 |
| 17_711223 | NA | NA | 3 | 1 | NA | NA | 1 | 1 | 4 | 1 | 1 | 1 | NA | NA | NA | NA | 2 | 0.5 | 3 | 0.6667 |
| 17_711252 | 7 | 0 | 7 | 0.1429 | 10 | 0.1 | 3 | 0.3333 | 27 | 0.1111 | 7 | 0 | 1 | 0 | 3 | 0.3333 | 3 | 0 | 14 | 0.0714 |
| 17_711253 | 6 | 0.5 | 7 | 0 | 9 | 0.4444 | 3 | 0 | 25 | 0.28 | 10 | 0.4 | 1 | 0 | 3 | 1 | 3 | 0 | 17 | 0.4118 |
| 17_711254 | 8 | 0.125 | 7 | 0.1429 | 10 | 0 | 3 | 0 | 28 | 0.0714 | 12 | 0 | 2 | 0 | 3 | 0.6667 | 5 | 0 | 22 | 0.0909 |
| 17_711255 | 8 | 0.75 | 9 | 0.4444 | 11 | 0.5455 | 3 | 0.6667 | 31 | 0.5806 | 13 | 0.6154 | 1 | 1 | 2 | 1 | 5 | 0.6 | 21 | 0.6667 |
| 17_711257 | 7 | 0 | 7 | 0 | 9 | 0.2222 | 2 | 0 | 25 | 0.08 | 11 | 0.0909 | 2 | 0 | 2 | 0 | 3 | 0 | 18 | 0.0556 |
| 17_711260 | 10 | 0 | 8 | 0.125 | 14 | 0.2143 | 4 | 0 | 36 | 0.1111 | 14 | 0 | 2 | 0 | 3 | 0.3333 | 5 | 0.2 | 24 | 0.0833 |
| 17_711262 | 9 | 0.2222 | 13 | 0.3846 | 14 | 0.5 | 6 | 0.1667 | 42 | 0.3571 | 15 | 0.2667 | 3 | 0 | 3 | 0.3333 | 5 | 0.2 | 26 | 0.2308 |
| 17_711267 | 11 | 0.0909 | 20 | 0.1 | 17 | 0.0588 | 6 | 0 | 54 | 0.0741 | 20 | 0.1 | 5 | 0 | 7 | 0.1429 | 9 | 0.1111 | 41 | 0.0976 |
| 17_712805 | NA | NA | 1 | 0 | 1 | 1 | NA | NA | 2 | 0.5 | NA | NA | NA | NA | 2 | 0 | 1 | 1 | 3 | 0.3333 |
| 17_712810 | 48 | 0.0625 | 41 | 0.0244 | 38 | 0.0789 | 32 | 0.0938 | 159 | 0.0629 | 33 | 0.0303 | 40 | 0.025 | 38 | 0.0789 | 33 | 0.0909 | 144 | 0.0556 |
| 17_712811 | 80 | 0.0625 | 75 | 0.1067 | 53 | 0.1509 | 60 | 0.0167 | 268 | 0.0821 | 53 | 0.1132 | 63 | 0.0476 | 56 | 0.0714 | 55 | 0.0727 | 227 | 0.0749 |
| 17_712812 | 67 | 0.1045 | 66 | 0.0606 | 54 | 0.0926 | 58 | 0.1034 | 245 | 0.0898 | 44 | 0.1364 | 63 | 0.1429 | 51 | 0.1176 | 48 | 0.125 | 206 | 0.1311 |
| 17_713014 | 16 | 0.1875 | 8 | 0.125 | 7 | 0.1429 | 9 | 0 | 40 | 0.125 | 9 | 0.1111 | 8 | 0.125 | 6 | 0.3333 | 8 | 0.125 | 31 | 0.1613 |
| 17_715539 | 1 | 0 | NA | NA | 1 | 1 | 4 | 0 | 6 | 0.1667 | 1 | 0 | 1 | 0 | 3 | 0 | 3 | 0.3333 | 8 | 0.125 |
| 17_715540 | 1 | 0 | 1 | 1 | 1 | 0 | 1 | 0 | 4 | 0.25 | 2 | 0 | 2 | 0 | 2 | 0 | 2 | 0.5 | 8 | 0.125 |
| 17_715540 | 2 | 0 | 1 | 1 | 1 | 0 | 5 | 0.2 | 9 | 0.2222 | 2 | 0 | 2 | 0 | 4 | 0.5 | 3 | 0 | 11 | 0.1818 |
| 17_715540 | 2 | 1 | 1 | 1 | 1 | 0 | 4 | 0.75 | 8 | 0.75 | 2 | 0.5 | 1 | 1 | 4 | 0.25 | 3 | 0.6667 | 10 | 0.5 |
| 17_715541 | 3 | 1 | NA | NA | 1 | 1 | 5 | 1 | 9 | 1 | 2 | 1 | 3 | 0.6667 | 3 | 1 | 3 | 1 | 11 | 0.9091 |
| 17_715542 | 3 | 1 | 1 | 1 | 1 | 1 | 4 | 1 | 9 | 1 | 1 | 1 | 2 | 1 | 3 | 0.6667 | 1 | 1 | 7 | 0.8571 |
| 17_715542 | 3 | 0.3333 | 1 | 0 | 1 | 0 | 6 | 0.3333 | 11 | 0.2727 | 2 | 0 | 3 | 0.3333 | 3 | 0.3333 | 3 | 0.6667 | 11 | 0.3636 |
| 17_715544 | 2 | 1 | NA | NA | 1 | 1 | 2 | 1 | 5 | 1 | NA | NA | 1 | 1 | 1 | 0 | 1 | 0 | 3 | 0.3333 |
| 17_715544 | 3 | 0.3333 | 1 | 1 | 1 | 0 | 5 | 1 | 10 | 0.7 | 2 | 0.5 | 3 | 1 | 4 | 1 | 3 | 1 | 12 | 0.9167 |
| 17_715545 | 3 | 0 | 1 | 1 | 1 | 1 | 5 | 0 | 10 | 0.2 | 1 | 0 | 3 | 0.3333 | 4 | 0.25 | 4 | 0.75 | 12 | 0.4167 |
| 17_715546 | 3 | 0.3333 | 1 | 1 | 1 | 1 | 4 | 0.5 | 9 | 0.5556 | 1 | 0 | 3 | 0.3333 | 5 | 0.4 | 4 | 0.75 | 13 | 0.4615 |
| 17_715547 | 3 | 0 | 1 | 0 | 1 | 0 | 5 | 0.6 | 10 | 0.3 | 2 | 0 | 4 | 0.5 | 3 | 1 | 3 | 0.3333 | 12 | 0.5 |
| 17_715547 | 2 | 0.5 | NA | NA | NA | NA | 4 | 0.75 | 6 | 0.6667 | 1 | 0 | 4 | 0 | 4 | 0.5 | 3 | 0.6667 | 12 | 0.3333 |
| 17_715547 | 2 | 0.5 | 2 | 0 | 1 | 0 | 3 | 0 | 8 | 0.125 | 2 | 0 | 1 | 1 | 3 | 0.6667 | 4 | 0.5 | 10 | 0.5 |
| 17_715549 | 3 | 0.3333 | 2 | 0.5 | 1 | 0 | 4 | 0.25 | 10 | 0.3 | 3 | 0.6667 | 3 | 0.6667 | 3 | 0.6667 | 2 | 0 | 11 | 0.5455 |
| 17_715551 | 3 | 0 | 3 | 0 | 1 | 0 | 7 | 0.2857 | 14 | 0.1429 | 3 | 0 | 5 | 0.2 | 5 | 0 | 4 | 0 | 17 | 0.0588 |
| 17_715551 | 3 | 0 | 3 | 0.6667 | 1 | 0 | 8 | 0.75 | 15 | 0.5333 | 3 | 0.3333 | 5 | 0.4 | 5 | 0.4 | 4 | 0.25 | 17 | 0.3529 |
| 17_715553 | 3 | 0.6667 | 3 | 0.6667 | 1 | 0 | 8 | 0.375 | 15 | 0.4667 | 3 | 0.3333 | 4 | 0.5 | 5 | 0.4 | 5 | 0.8 | 17 | 0.5294 |

| | | | | | | | | | | | | | | | | | | | | |
|---|---|---|---|---|---|---|---|---|---|---|---|---|---|---|---|---|---|---|---|---|
| 17_7155534 | 3 | 0 | 3 | 0.3333 | 1 | 0 | 5 | 0.2 | 12 | 0.1667 | 3 | 0.3333 | 6 | 0.3333 | 4 | 0.5 | 3 | 0 | 16 | 0.3125 |
| 17_7155552 | 3 | 0.6667 | 3 | 0.6667 | 1 | 0 | 6 | 0.3333 | 13 | 0.4615 | 3 | 0 | 6 | 0.1667 | 4 | 0.25 | 4 | 0.5 | 17 | 0.2353 |
| 17_7155571 | 3 | 0 | 2 | 0 | 1 | 0 | 9 | 0.1111 | 15 | 0.0667 | 3 | 0.3333 | 5 | 0 | 5 | 0 | 4 | 0 | 17 | 0.0588 |
| 17_7155572 | 2 | 0.5 | 3 | 0.3333 | 1 | 0 | 6 | 0.3333 | 12 | 0.3333 | 2 | 0 | 4 | 0.25 | 4 | 0 | 3 | 0.3333 | 13 | 0.1538 |
| 17_7155608 | 3 | 0.6667 | 5 | 0.4 | 2 | 0.5 | 8 | 0.75 | 18 | 0.6111 | 3 | 0.3333 | 6 | 0.1667 | 7 | 0.1429 | 3 | 0 | 19 | 0.1579 |
| 17_7155650 | 3 | 0 | 4 | 0 | 2 | 0 | 8 | 0.125 | 17 | 0.0588 | 4 | 0.25 | 4 | 0 | 7 | 0 | 4 | 0 | 19 | 0.0526 |
| 17_7155925 | 2 | 1 | 1 | 1 | NA | NA | NA | NA | 3 | 1 | 1 | 1 | NA | NA | NA | NA | NA | NA | 1 | 1 |
| 17_7155936 | 2 | 1 | 1 | 1 | NA | NA | NA | NA | 3 | 1 | 1 | 1 | NA | NA | NA | NA | NA | NA | 1 | 1 |
| 17_7155956 | 2 | 0.5 | 1 | 0 | NA | NA | NA | NA | 3 | 0.3333 | 1 | 1 | NA | NA | NA | NA | NA | NA | 1 | 1 |
| 17_7155968 | 2 | 0.5 | 1 | 1 | NA | NA | NA | NA | 3 | 0.6667 | 1 | 1 | NA | NA | NA | NA | NA | NA | 1 | 1 |
| 17_7155981 | NA | NA | 1 | 1 | NA | NA | NA | NA | 1 | 1 | 1 | 1 | NA | NA | NA | NA | NA | NA | 1 | 1 |
| 17_7155984 | 2 | 0.5 | 1 | 1 | NA | NA | NA | NA | 3 | 0.6667 | 1 | 1 | NA | NA | NA | NA | NA | NA | 1 | 1 |
| 17_7165182 | 2 | 0.5 | 7 | 0 | 1 | 0 | 3 | 0 | 13 | 0.0769 | 1 | 0 | 7 | 0.5714 | 7 | 0 | 6 | 0 | 21 | 0.1905 |
| 17_7165203 | 2 | 1 | 7 | 0.4286 | 1 | 1 | 2 | 1 | 12 | 0.6667 | 1 | 0 | 5 | 0.6 | 4 | 0.25 | 4 | 0 | 14 | 0.2857 |
| 17_7165207 | 1 | 1 | 3 | 1 | 1 | 1 | 1 | 0 | 6 | 0.8333 | 1 | 1 | 7 | 0.4286 | 7 | 0.5714 | 4 | 0.25 | 19 | 0.4737 |
| 17_7189588 | 2 | 0 | 6 | 0.1667 | 1 | 0 | 4 | 0.75 | 13 | 0.3077 | 2 | 0 | 3 | 0.3333 | 1 | 0 | 2 | 0.5 | 8 | 0.25 |
| 17_7189593 | 2 | 0.5 | 4 | 0 | 1 | 0 | 4 | 0.25 | 11 | 0.1818 | 4 | 0 | 4 | 0.25 | 1 | 0 | 2 | 0 | 11 | 0.0909 |
| 17_7190511 | 3 | 1 | 5 | 1 | 1 | 1 | 5 | 1 | 14 | 1 | 4 | 1 | 1 | 1 | NA | NA | NA | NA | 5 | 1 |
| 17_7190521 | 3 | 0 | 6 | 0.6667 | 1 | 0 | 3 | 0.6667 | 13 | 0.4615 | 4 | 0.25 | 1 | 0 | 1 | 0 | 2 | 0 | 8 | 0.125 |
| 17_7190522 | 3 | 1 | 6 | 0.8333 | 1 | 1 | 5 | 0.8 | 15 | 0.8667 | 4 | 1 | 2 | 0.5 | NA | NA | 1 | 1 | 7 | 0.8571 |
| 17_7190531 | 3 | 1 | 5 | 1 | 1 | 1 | 5 | 1 | 14 | 1 | 4 | 1 | 1 | 1 | 1 | 1 | 2 | 1 | 8 | 1 |
| 17_7190537 | 3 | 0.3333 | 5 | 0.2 | 1 | 0 | 4 | 0.5 | 13 | 0.3077 | 4 | 0.25 | 1 | 0 | 1 | 0 | 2 | 0 | 8 | 0.125 |
| 17_7190544 | 3 | 0.3333 | 5 | 0 | 1 | 0 | 5 | 0 | 14 | 0.0714 | 4 | 0 | 1 | 0 | 1 | 0 | 2 | 0.5 | 8 | 0.125 |
| 17_7196458 | 4 | 0.25 | 5 | 0.2 | 2 | 1 | 6 | 0 | 17 | 0.2353 | 1 | 0 | 3 | 0.3333 | 5 | 0.2 | 1 | 0 | 10 | 0.2 |
| 17_7196482 | 4 | 0 | 5 | 0.4 | 2 | 1 | 5 | 0 | 16 | 0.25 | 1 | 0 | 2 | 0.5 | 5 | 0.6 | NA | NA | 8 | 0.5 |
| 17_7196486 | 5 | 0.4 | 5 | 0 | 1 | 0 | 3 | 0.6667 | 14 | 0.2857 | 1 | 0 | 3 | 0.6667 | 3 | 0.3333 | NA | NA | 7 | 0.4286 |
| 17_7196501 | 5 | 0.2 | 4 | 0.25 | 1 | 0 | 5 | 0.2 | 15 | 0.2 | 1 | 0 | 3 | 0.3333 | 2 | 0 | NA | NA | 6 | 0.1667 |
| 17_7196537 | 5 | 0 | 4 | 0.25 | 2 | 0.5 | 5 | 0 | 16 | 0.125 | 1 | 0 | 4 | 0 | 2 | 1 | 1 | 0 | 8 | 0.25 |
| 17_7196580 | 7 | 0.1429 | 5 | 0.4 | 3 | 0.6667 | 5 | 0.4 | 20 | 0.35 | 1 | 0 | 3 | 1 | 2 | 1 | 2 | 0.5 | 8 | 0.75 |
| 17_7196583 | 7 | 0 | 5 | 0.4 | 3 | 0.6667 | 5 | 0.4 | 20 | 0.3 | 1 | 1 | 3 | 0.6667 | 2 | 0.5 | 2 | 0.5 | 8 | 0.625 |
| 17_7196586 | 7 | 0.2857 | 3 | 0 | 2 | 1 | NA | NA | 12 | 0.3333 | NA | NA | 4 | 0.25 | 2 | 0.5 | 1 | 1 | 7 | 0.4286 |
| 17_7203636 | 1 | 0 | 1 | 0 | 1 | 1 | 2 | 0.5 | 5 | 0.4 | 1 | 0 | 2 | 0.5 | 1 | 0 | NA | NA | 4 | 0.25 |
| 17_7203641 | 1 | 1 | 2 | 1 | 2 | 1 | 2 | 0.5 | 7 | 0.8571 | 1 | 1 | 2 | 1 | 1 | 1 | NA | NA | 4 | 1 |
| 17_7208487 | NA | NA | NA | NA | 2 | 0.5 | NA | NA | 2 | 0.5 | NA | NA | NA | NA | 2 | 0.5 | NA | NA | 2 | 0.5 |
| 17_7208503 | NA | NA | NA | NA | 3 | 1 | NA | NA | 3 | 1 | NA | NA | NA | NA | 2 | 1 | 1 | 1 | 3 | 1 |
| 17_7208609 | NA | NA | NA | NA | 1 | 1 | NA | NA | 1 | 1 | NA | NA | 1 | 1 | NA | NA | 1 | 0 | 2 | 0.5 |
| 17_7208670 | NA | NA | NA | NA | 2 | 1 | NA | NA | 2 | 1 | NA | NA | 1 | 1 | NA | NA | 1 | 1 | 2 | 1 |
| 17_7208673 | NA | NA | NA | NA | 2 | 1 | NA | NA | 2 | 1 | NA | NA | 1 | 1 | NA | NA | NA | NA | 1 | 1 |
| 17_7208692 | NA | NA | NA | NA | 2 | 0.5 | NA | NA | 2 | 0.5 | NA | NA | 1 | 0 | NA | NA | 1 | 1 | 2 | 0.5 |
| 17_7208696 | NA | NA | NA | NA | 2 | 0.5 | NA | NA | 2 | 0.5 | NA | NA | 1 | 1 | NA | NA | 1 | 1 | 2 | 1 |
| 17_7208703 | NA | NA | NA | NA | 2 | 0.5 | NA | NA | 2 | 0.5 | NA | NA | 1 | 1 | NA | NA | 1 | 1 | 2 | 1 |
| 17_7209029 | NA | NA | NA | NA | 1 | 1 | NA | NA | 1 | 1 | 1 | 0 | 1 | 1 | NA | NA | 1 | 0 | 3 | 0.3333 |
| 17_7216336 | 6 | 0.8333 | 6 | 1 | 7 | 0.8571 | 6 | 0.6667 | 25 | 0.84 | 2 | 1 | 8 | 1 | 9 | 1 | 6 | 0.6667 | 25 | 0.92 |
| 17_7216339 | 8 | 0.375 | 5 | 0.2 | 9 | 0.3333 | 5 | 0 | 27 | 0.2593 | 2 | 0.5 | 10 | 0.6 | 6 | 0.6667 | 6 | 0.6667 | 24 | 0.625 |
| 17_7216370 | 7 | 0 | 5 | 0.2 | 8 | 0.125 | 7 | 0.1429 | 27 | 0.1111 | 3 | 0.3333 | 10 | 0.5 | 10 | 0.5 | 6 | 0.5 | 29 | 0.4828 |
| 17_7216379 | 10 | 0 | 5 | 0.2 | 9 | 0.2222 | 8 | 0.125 | 32 | 0.125 | 2 | 0.5 | 12 | 0 | 11 | 0.0909 | 7 | 0.1429 | 32 | 0.0938 |
| 17_7217087 | 1 | 1 | NA | NA | NA | NA | NA | NA | 1 | 1 | NA | NA | 1 | 1 | NA | NA | 1 | 1 | 2 | 1 |
| 17_7217595 | 1 | 1 | 1 | 0 | NA | NA | NA | NA | 2 | 0.5 | NA | NA | NA | NA | 1 | 1 | 1 | 1 | 2 | 1 |
| 17_7217596 | 1 | 0 | 1 | 1 | NA | NA | 1 | 0 | 3 | 0.3333 | NA | NA | 1 | 0 | 1 | 1 | 2 | 0.5 | 4 | 0.5 |
| 17_7217634 | 1 | 0 | 1 | 1 | NA | NA | NA | NA | 2 | 0.5 | NA | NA | NA | NA | 1 | 1 | NA | NA | 1 | 1 |
| 17_7217634 | 2 | 0.5 | 1 | 0 | NA | NA | 1 | 0 | 4 | 0.25 | NA | NA | 1 | 0 | 1 | 0 | 1 | 1 | 3 | 0.3333 |
| 17_7217637 | 2 | 1 | 1 | 1 | NA | NA | 1 | 1 | 4 | 1 | NA | NA | NA | NA | 1 | 0 | 1 | 1 | 2 | 0.5 |
| 17_7217648 | 2 | 0.5 | 1 | 1 | NA | NA | NA | NA | 3 | 0.6667 | NA | NA | 1 | 1 | NA | NA | 1 | 0 | 2 | 0.5 |
| 17_7217666 | 2 | 1 | 1 | 0 | NA | NA | NA | NA | 3 | 0.6667 | NA | NA | 1 | 1 | NA | NA | 1 | 1 | 2 | 1 |
| 17_7217672 | 2 | 1 | NA | NA | NA | NA | NA | NA | 2 | 1 | NA | NA | NA | NA | NA | NA | 1 | 1 | 1 | 1 |
| 17_7315565 | 15 | 0.3333 | 22 | 0.4091 | 9 | 0.2222 | 15 | 0.1333 | 61 | 0.2951 | 8 | 0.375 | 22 | 0.3182 | 17 | 0.3529 | 13 | 0.4615 | 60 | 0.3667 |
| 17_7315589 | 10 | 0.8 | 23 | 1 | 9 | 0.8889 | 11 | 0.8182 | 53 | 0.9057 | 8 | 0.75 | 21 | 0.9524 | 9 | 0.7778 | 10 | 1 | 48 | 0.8958 |
| 17_7315593 | 15 | 0.2 | 29 | 0.1379 | 16 | 0.1875 | 15 | 0.3333 | 75 | 0.2 | 8 | 0.125 | 22 | 0.1364 | 18 | 0.1111 | 15 | 0.1333 | 63 | 0.127 |
| 17_7315632 | 19 | 0.5789 | 35 | 0.6 | 23 | 0.7391 | 21 | 0.4762 | 98 | 0.602 | 11 | 0.5455 | 26 | 0.7308 | 25 | 0.68 | 19 | 0.6316 | 81 | 0.6667 |
| 17_7315640 | 17 | 0.8824 | 36 | 0.8056 | 22 | 0.7273 | 19 | 0.6842 | 94 | 0.7766 | 10 | 0.8 | 30 | 0.9 | 24 | 0.8333 | 20 | 0.85 | 84 | 0.8571 |
| 17_7315685 | 11 | 0.0909 | 22 | 0.1364 | 18 | 0.2778 | 13 | 0.3077 | 64 | 0.2031 | 3 | 0 | 16 | 0.25 | 12 | 0.1667 | 16 | 0.25 | 47 | 0.2128 |
| 17_7324643 | 2 | 0.5 | 5 | 0.4 | 2 | 0 | 3 | 0 | 12 | 0.25 | 3 | 0.3333 | 1 | 0 | 1 | 0 | 1 | 0 | 6 | 0.1667 |
| 17_7324648 | 2 | 1 | 4 | 1 | 1 | 1 | 1 | 1 | 8 | 1 | NA | NA | 1 | 1 | 1 | 0 | NA | NA | 2 | 0.5 |
| 17_7324675 | 1 | 0 | 6 | 0 | 2 | 0.5 | NA | NA | 9 | 0.1111 | 2 | 0.5 | 2 | 0.5 | 1 | 0 | 1 | 0 | 6 | 0.3333 |
| 17_7324689 | 2 | 0.5 | 7 | 0 | 3 | 0 | 2 | 0 | 14 | 0.0714 | 3 | 0 | 2 | 0 | NA | NA | 2 | 1 | 7 | 0.2857 |

| | | | | | | | | | | | | | | | | | | | | |
|---|---|---|---|---|---|---|---|---|---|---|---|---|---|---|---|---|---|---|---|---|
| 17_7324789 | 2 | 0 | 8 | 0.125 | 3 | 0 | 4 | 0 | 17 | 0.0588 | 4 | 0.25 | 3 | 0 | 3 | 0.3333 | 2 | 0 | 12 | 0.1667 |
| 17_7324861 | 2 | 0 | 6 | 0 | 3 | 0.3333 | 4 | 0 | 15 | 0.0667 | 4 | 0.25 | 2 | 0 | 2 | 0 | NA | NA | 8 | 0.125 |
| 17_7354694 | 3 | 0 | 3 | 0.3333 | 3 | 0 | 1 | 1 | 10 | 0.2 | 2 | 0.5 | NA | NA | NA | NA | 3 | 0.6667 | 5 | 0.6 |
| 17_7354714 | 4 | 1 | 3 | 1 | 3 | 1 | 1 | 1 | 11 | 1 | 4 | 1 | 1 | 1 | 1 | 1 | 3 | 1 | 9 | 1 |
| 17_7354715 | 4 | 1 | 3 | 0.6667 | 3 | 1 | 1 | 1 | 11 | 0.9091 | 3 | 0.6667 | NA | NA | NA | NA | 3 | 1 | 6 | 0.8333 |
| 17_7354790 | 3 | 0 | 3 | 0.3333 | 5 | 0 | 1 | 0 | 12 | 0.0833 | 3 | 0.3333 | 2 | 0 | 2 | 0 | 3 | 0 | 10 | 0.1 |
| 17_7354905 | 5 | 0 | 4 | 0 | 6 | 0.1667 | NA | NA | 15 | 0.0667 | 4 | 0.25 | 1 | 0 | 3 | 0 | 4 | 0.25 | 12 | 0.1667 |
| 17_7354983 | 7 | 0.1429 | 5 | 0.6 | 8 | 0.125 | 3 | 0.6667 | 23 | 0.3043 | 4 | 0.25 | 2 | 0.5 | 3 | 0 | 4 | 0.5 | 13 | 0.3077 |
| 17_7355013 | 8 | 0.375 | 6 | 0.5 | 8 | 0.625 | 1 | 1 | 23 | 0.5217 | 3 | 0.3333 | 2 | 0 | 3 | 0 | 3 | 0.6667 | 11 | 0.2727 |
| 17_7357183 | NA | NA | NA | NA | 1 | 1 | NA | NA | 1 | 1 | NA | NA | NA | NA | NA | NA | 1 | 1 | 1 | 1 |
| 17_7357233 | NA | NA | 1 | 1 | NA | NA | NA | NA | 1 | 1 | NA | NA | NA | NA | NA | NA | 1 | 1 | 1 | 1 |
| 17_7357238 | NA | NA | 1 | 1 | 1 | 1 | NA | NA | 2 | 1 | NA | NA | NA | NA | NA | NA | 1 | 1 | 1 | 1 |
| 17_7380076 | 1 | 0 | 1 | 0 | NA | NA | 1 | 1 | 3 | 0.3333 | 1 | 0 | NA | NA | 1 | 1 | 1 | 0 | 3 | 0.3333 |
| 17_7380091 | 1 | 1 | 2 | 0.5 | NA | NA | 1 | 0 | 4 | 0.5 | 3 | 0.3333 | 1 | 0 | 1 | 0 | 1 | 0 | 6 | 0.1667 |
| 17_7380103 | 1 | 0 | 2 | 0.5 | NA | NA | 1 | 0 | 4 | 0.25 | 3 | 0 | 1 | 0 | 1 | 1 | NA | NA | 5 | 0.2 |
| 17_7380112 | 1 | 0 | 2 | 0.5 | NA | NA | 1 | 0 | 4 | 0.25 | 3 | 0 | 1 | 0 | 1 | 1 | 1 | 0 | 6 | 0.1667 |
| 17_7380118 | NA | NA | 2 | 1 | NA | NA | NA | NA | 2 | 1 | 3 | 0.6667 | NA | NA | NA | NA | NA | NA | 3 | 0.6667 |
| 17_7411210 | 3 | 0.3333 | 4 | 0.5 | 2 | 0.5 | 4 | 0.75 | 13 | 0.5385 | 3 | 0 | 1 | 0 | 2 | 0.5 | 1 | 0 | 7 | 0.1429 |
| 17_7411210 | 4 | 0.5 | 2 | 0.5 | 3 | 1 | 3 | 0.6667 | 12 | 0.6667 | 2 | 0.5 | 1 | 0 | 3 | 0.6667 | 3 | 0.6667 | 9 | 0.5556 |
| 17_7411230 | 3 | 1 | 2 | 1 | 5 | 0.6 | 5 | 0.2 | 15 | 0.6 | 3 | 0.6667 | 2 | 1 | 4 | 1 | 5 | 0.8 | 14 | 0.8571 |
| 17_7413448 | 4 | 0.25 | 5 | 0 | 6 | 0.6667 | 6 | 0 | 21 | 0.2381 | 5 | 0.2 | 3 | 0 | 5 | 0.4 | 7 | 0.2857 | 20 | 0.25 |
| 17_7413511 | 3 | 0.3333 | 5 | 0 | 6 | 0 | 5 | 0 | 19 | 0.0526 | 4 | 0 | 4 | 0.25 | 5 | 0 | 6 | 0 | 19 | 0.0526 |
| 17_7413693 | 4 | 0.25 | 5 | 0 | 6 | 0 | 7 | 0.1429 | 22 | 0.0909 | 5 | 0.2 | 4 | 0.25 | 5 | 0.2 | 7 | 0.2857 | 21 | 0.2381 |
| 17_7413703 | 4 | 0 | 5 | 0.2 | 6 | 0.1667 | 7 | 0.5714 | 22 | 0.2727 | 5 | 0 | 3 | 0.3333 | 5 | 0.2 | 5 | 0 | 18 | 0.1111 |
| 17_7413708 | 3 | 0 | 4 | 0.5 | 5 | 0.4 | 6 | 0.1667 | 18 | 0.2778 | 5 | 0 | 4 | 0.5 | 6 | 0.1667 | 6 | 0 | 21 | 0.1429 |
| 17_7413716 | 2 | 0 | 5 | 0.2 | 6 | 0 | 7 | 0.2857 | 20 | 0.15 | 5 | 0.2 | 4 | 0.25 | 4 | 0.25 | 4 | 0.25 | 17 | 0.2353 |
| 17_7413785 | 5 | 0 | 7 | 0 | 6 | 0.5 | 7 | 0 | 25 | 0.12 | 5 | 0.2 | 3 | 0.3333 | 6 | 0.1667 | 10 | 0.2 | 24 | 0.2083 |
| 17_7413794 | 5 | 0.2 | 7 | 0.2857 | 7 | 0.4286 | 6 | 0.6667 | 25 | 0.4 | 4 | 0.25 | 4 | 0.75 | 5 | 0.4 | 9 | 0.2222 | 22 | 0.3636 |
| 17_7413798 | 4 | 0 | 5 | 0.2 | 6 | 0.1667 | 8 | 0.125 | 23 | 0.1304 | 5 | 0.2 | 4 | 0.5 | 3 | 0.6667 | 8 | 0.125 | 20 | 0.3 |
| 17_7413801 | 6 | 1 | 6 | 0.8333 | 7 | 1 | 5 | 0.8 | 24 | 0.9167 | 3 | 0.6667 | 4 | 0.75 | 6 | 1 | 9 | 0.7778 | 22 | 0.8182 |
| 17_7413804 | 6 | 0.6667 | 6 | 0.3333 | 6 | 0.1667 | 8 | 0.375 | 26 | 0.3846 | 5 | 0.4 | 4 | 0.25 | 6 | 0.6667 | 10 | 0.1 | 25 | 0.32 |
| 17_7413812 | 8 | 0.625 | 5 | 0.4 | 5 | 0.4 | 5 | 0.8 | 23 | 0.5652 | 5 | 0.6 | 3 | 0.3333 | 3 | 0.3333 | 6 | 0.6667 | 17 | 0.5294 |
| 17_7413832 | 7 | 0.2857 | 7 | 0.1429 | 7 | 0.1429 | 8 | 0.375 | 29 | 0.2414 | 8 | 0 | 7 | 0.4286 | 5 | 0.4 | 10 | 0 | 30 | 0.1667 |
| 17_7413847 | 7 | 0.8571 | 7 | 0.1429 | 5 | 0.6 | 6 | 0.5 | 25 | 0.52 | 7 | 0.7143 | 7 | 0.2857 | 5 | 0.8 | 9 | 0.3333 | 28 | 0.5 |
| 17_7413859 | 6 | 0 | 5 | 0 | 8 | 0.125 | 9 | 0.1111 | 28 | 0.0714 | 10 | 0.2 | 7 | 0.2857 | 6 | 0.3333 | 9 | 0.2222 | 32 | 0.25 |
| 17_7413866 | 8 | 0.75 | 4 | 0.75 | 8 | 0.75 | 5 | 0.8 | 25 | 0.76 | 8 | 0.875 | 7 | 0.7143 | 6 | 1 | 8 | 0.875 | 29 | 0.8621 |
| 17_7413876 | 8 | 1 | 7 | 0.8571 | 6 | 0.8333 | 8 | 1 | 29 | 0.931 | 9 | 0.8889 | 7 | 1 | 6 | 1 | 8 | 0.875 | 30 | 0.9333 |
| 17_7413880 | 6 | 1 | 8 | 0.875 | 8 | 1 | 3 | 1 | 25 | 0.96 | 8 | 1 | 4 | 1 | 5 | 1 | 7 | 1 | 24 | 1 |
| 17_7413880 | 12 | 1 | 7 | 0.5714 | 7 | 0.8571 | 8 | 1 | 34 | 0.8824 | 10 | 0.9 | 7 | 0.8571 | 6 | 0.8333 | 9 | 0.8889 | 32 | 0.875 |
| 17_7413883 | 11 | 0.2727 | 7 | 0.4286 | 8 | 0.5 | 10 | 0.3 | 36 | 0.3611 | 10 | 0.1 | 8 | 0 | 6 | 0.5 | 11 | 0.2727 | 35 | 0.2 |
| 17_7419129 | 1 | 1 | 4 | 1 | 1 | 0 | NA | NA | 6 | 0.8333 | 2 | 1 | NA | NA | 1 | 1 | 2 | 1 | 5 | 1 |
| 17_7419133 | 1 | 0 | 4 | 0.25 | 1 | 0 | NA | NA | 6 | 0.1667 | 1 | 1 | NA | NA | 1 | 0 | 3 | 0.3333 | 5 | 0.4 |
| 17_7450794 | 2 | 0.5 | 1 | 1 | 3 | 0.3333 | 3 | 0.6667 | 9 | 0.5556 | 4 | 0.5 | 6 | 0.1667 | 3 | 0 | 6 | 0.3333 | 19 | 0.2632 |
| 17_7450807 | 2 | 0 | NA | NA | 3 | 0.3333 | 3 | 0.3333 | 8 | 0.25 | 3 | 0.3333 | 3 | 0.3333 | 2 | 0 | 7 | 0.5714 | 15 | 0.4 |
| 17_7450815 | 2 | 0.5 | 1 | 1 | 3 | 0 | 3 | 0.6667 | 9 | 0.4444 | 3 | 0.6667 | 6 | 0.5 | 3 | 0 | 7 | 0.2857 | 19 | 0.3684 |
| 17_7450823 | 1 | 0 | 1 | 0 | 3 | 0.3333 | 4 | 0 | 9 | 0.1111 | 4 | 0.5 | 6 | 0 | 2 | 0 | 6 | 0.6667 | 18 | 0.3333 |
| 17_7460649 | 6 | 0 | 5 | 0.2 | 10 | 0.1 | 8 | 0.25 | 29 | 0.1379 | 3 | 0 | 6 | 0.3333 | 9 | 0.1111 | 9 | 0.1111 | 27 | 0.1481 |
| 17_7469533 | 1 | 1 | 6 | 0 | 5 | 0 | 2 | 0 | 14 | 0.0714 | 7 | 0.1429 | 5 | 0 | 3 | 0 | 4 | 0 | 19 | 0.0526 |
| 17_7469538 | 3 | 0 | 6 | 0.5 | 5 | 0.2 | 2 | 0 | 16 | 0.25 | 8 | 0.375 | 5 | 0.2 | 4 | 0.5 | 6 | 0.1667 | 23 | 0.3043 |
| 17_7469545 | 3 | 0 | 7 | 0.1429 | 5 | 0 | 1 | 0 | 16 | 0.0625 | 9 | 0.2222 | 5 | 0 | 4 | 0 | 5 | 0.2 | 23 | 0.1304 |
| 17_7469564 | 1 | 0 | 3 | 0 | 2 | 1 | 1 | 1 | 7 | 0.4286 | 4 | 0.75 | 2 | 1 | 4 | 0.25 | 4 | 0 | 14 | 0.4286 |
| 17_7469585 | 2 | 0 | 4 | 0.5 | 3 | 0 | 1 | 0 | 10 | 0.2 | 5 | 0 | 2 | 0.5 | 4 | 0 | 4 | 0.25 | 15 | 0.1333 |
| 17_7469648 | 3 | 0.3333 | 4 | 0 | 3 | 0 | 1 | 0 | 11 | 0.0909 | 6 | 0 | 2 | 0.5 | 4 | 0.5 | 3 | 0 | 15 | 0.2 |
| 17_7469654 | 1 | 0 | 3 | 0.3333 | 3 | 0.3333 | 1 | 0 | 8 | 0.25 | 3 | 1 | 1 | 0 | 4 | 0 | 3 | 0 | 11 | 0.2727 |
| 17_7473010 | 1 | 0 | 1 | 0 | NA | NA | 3 | 0.3333 | 5 | 0.2 | NA | NA | NA | NA | 2 | 0 | 3 | 1 | 5 | 0.6 |
| 17_7473033 | 2 | 0 | 1 | 1 | NA | NA | 2 | 0.5 | 5 | 0.4 | NA | NA | NA | NA | 2 | 0.5 | 3 | 0.3333 | 5 | 0.4 |
| 17_7473037 | 1 | 1 | NA | NA | NA | NA | 3 | 1 | 4 | 1 | NA | NA | NA | NA | 1 | 1 | 3 | 1 | 4 | 1 |
| 17_7473040 | 1 | 1 | 1 | 1 | NA | NA | 3 | 1 | 5 | 1 | NA | NA | NA | NA | 2 | 1 | 4 | 1 | 6 | 1 |
| 17_7473051 | 1 | 1 | 1 | 1 | NA | NA | 3 | 0.6667 | 5 | 0.8 | NA | NA | NA | NA | 2 | 1 | 2 | 1 | 4 | 1 |
| 17_7473058 | 1 | 1 | 1 | 1 | NA | NA | 3 | 0.6667 | 5 | 0.8 | NA | NA | 1 | 1 | 1 | 1 | 5 | 0.6 | 7 | 0.7143 |
| 17_7494699 | 2 | 0.5 | 3 | 0.6667 | 2 | 0.5 | NA | NA | 7 | 0.5714 | NA | NA | 2 | 0.5 | 2 | 1 | NA | NA | 4 | 0.75 |
| 17_7495677 | 1 | 0 | 6 | 0.3333 | 2 | 1 | 3 | 0.6667 | 12 | 0.5 | NA | NA | 3 | 0.6667 | 3 | 0.6667 | 3 | 1 | 9 | 0.7778 |
| 17_7495693 | 2 | 0 | 5 | 0 | 3 | 0 | 3 | 0.3333 | 13 | 0.0769 | NA | NA | 1 | 0 | 2 | 0.5 | 3 | 0.3333 | 6 | 0.3333 |
| 17_7495703 | 2 | 0 | 6 | 0.3333 | 2 | 0 | 3 | 0 | 13 | 0.1538 | NA | NA | 3 | 0 | 3 | 0.3333 | 3 | 0.3333 | 9 | 0.2222 |
| 17_7495718 | 2 | 0.5 | 6 | 0 | 3 | 0.6667 | 3 | 0.3333 | 14 | 0.2857 | NA | NA | 2 | 0.5 | 3 | 0.3333 | 3 | 0.6667 | 8 | 0.5 |

| | | | | | | | | | | | | | | | | | | | | |
|---|---|---|---|---|---|---|---|---|---|---|---|---|---|---|---|---|---|---|---|---|
| 17_7495723 | 2 | 0 | 6 | 0 | 2 | 0.5 | 3 | 0 | 13 | 0.0769 | NA | NA | 2 | 0 | 3 | 0.3333 | 4 | 0 | 9 | 0.1111 |
| 17_7496746 | 4 | 0.25 | 7 | 0.1429 | 3 | 0.3333 | 5 | 0 | 19 | 0.1579 | NA | NA | 4 | 0 | 4 | 0 | 4 | 0.25 | 12 | 0.0833 |
| 17_7500936 | 5 | 0.4 | 15 | 0.3333 | 10 | 0.4 | 10 | 0.3 | 40 | 0.35 | 3 | 0 | 8 | 0.125 | 7 | 0.1429 | 9 | 0.3333 | 27 | 0.1852 |
| 17_7500947 | 5 | 0.2 | 16 | 0.125 | 12 | 0.1667 | 9 | 0.2222 | 42 | 0.1667 | 4 | 0 | 8 | 0.125 | 7 | 0 | 9 | 0.2222 | 28 | 0.1071 |
| 17_7501017 | 2 | 0 | 7 | 0.2857 | 6 | 0.3333 | 5 | 0.2 | 20 | 0.25 | 2 | 0.5 | 2 | 0.5 | 1 | 0 | 6 | 0.1667 | 11 | 0.2727 |
| 17_7569773 | NA | NA | 1 | 1 | NA | NA | 1 | 1 | 2 | 1 | NA | NA | NA | NA | 1 | 1 | 1 | 1 | 2 | 1 |
| 17_7581022 | NA | NA | NA | NA | 1 | 1 | NA | NA | 1 | 1 | 2 | 1 | 2 | 0.5 | 2 | 0.5 | 2 | 0.5 | 8 | 0.625 |
| 17_7583399 | NA | NA | NA | NA | 1 | 1 | NA | NA | 1 | 1 | 1 | 1 | 2 | 0 | 2 | 0 | 1 | 1 | 6 | 0.3333 |
| 17_7583402 | NA | NA | NA | NA | 1 | 1 | NA | NA | 1 | 1 | NA | NA | 3 | 0.3333 | 2 | 1 | 1 | 1 | 6 | 0.6667 |
| 17_7583431 | NA | NA | NA | NA | 1 | 1 | NA | NA | 1 | 1 | 1 | 0 | 3 | 0 | 3 | 0.3333 | 2 | 0.5 | 9 | 0.2222 |
| 17_7583577 | 1 | 0 | 2 | 0.5 | NA | NA | NA | NA | 3 | 0.3333 | 3 | 0.3333 | 3 | 1 | 3 | 0.3333 | 2 | 0 | 11 | 0.4545 |
| 17_7584432 | 4 | 1 | 3 | 0.3333 | 4 | 0.5 | 2 | 0 | 13 | 0.5385 | 3 | 0.3333 | 4 | 0.25 | 3 | 0.3333 | NA | NA | 10 | 0.3 |
| 17_7584559 | 2 | 1 | NA | NA | 1 | 1 | 1 | 0 | 4 | 0.75 | 1 | 1 | 1 | 1 | NA | NA | NA | NA | 2 | 1 |
| 17_7584571 | 2 | 1 | NA | NA | 1 | 0 | 1 | 1 | 4 | 0.75 | 1 | 1 | 1 | 1 | NA | NA | NA | NA | 2 | 1 |
| 17_7868188 | 1 | 1 | 1 | 0 | 2 | 0 | 1 | 0 | 5 | 0.2 | NA | NA | 1 | 1 | NA | NA | NA | NA | 1 | 1 |
| 17_7868189 | 2 | 0 | 1 | 0 | 2 | 0.5 | 1 | 0 | 6 | 0.1667 | NA | NA | 1 | 1 | NA | NA | NA | NA | 1 | 1 |
| 17_7868193 | 2 | 0.5 | 1 | 0 | 2 | 0.5 | 1 | 1 | 6 | 0.5 | NA | NA | 1 | 1 | NA | NA | NA | NA | 1 | 1 |
| 17_7868209 | 1 | 1 | 2 | 0 | 2 | 0 | 2 | 1 | 7 | 0.4286 | NA | NA | 1 | 1 | NA | NA | NA | NA | 1 | 1 |
| 17_7868217 | 1 | 1 | 2 | 0 | 2 | 0 | 3 | 0 | 8 | 0.125 | NA | NA | 1 | 1 | NA | NA | NA | NA | 1 | 1 |
| 17_7868218 | 1 | 1 | 2 | 0 | 2 | 0.5 | 3 | 1 | 8 | 0.625 | NA | NA | 1 | 1 | NA | NA | NA | NA | 1 | 1 |
| 17_7868245 | 1 | 1 | 2 | 0 | 2 | 0 | 3 | 1 | 8 | 0.5 | NA | NA | 1 | 1 | 1 | 1 | NA | NA | 2 | 1 |
| 17_7868258 | 2 | 1 | 2 | 0.5 | 2 | 1 | 5 | 1 | 11 | 0.9091 | NA | NA | 1 | 1 | 1 | 0 | NA | NA | 2 | 0.5 |
| 17_7868330 | 6 | 0.1667 | 7 | 0.2857 | 4 | 0.25 | 10 | 0.2 | 27 | 0.2222 | 3 | 0.3333 | 2 | 0 | 2 | 0 | 7 | 0 | 14 | 0.0714 |
| 17_7868377 | 8 | 0.125 | 7 | 0 | 6 | 0 | 12 | 0.0833 | 33 | 0.0606 | 5 | 0.2 | 3 | 0 | 4 | 0.25 | 9 | 0.1111 | 21 | 0.1429 |
| 17_7868553 | 3 | 1 | 7 | 1 | 6 | 0.8333 | 3 | 1 | 19 | 0.9474 | 1 | 1 | 1 | 1 | 6 | 1 | 3 | 1 | 11 | 1 |
| 17_7868569 | 5 | 0.2 | 8 | 0.125 | 8 | 0.125 | 3 | 0 | 24 | 0.125 | 1 | 0 | NA | NA | 7 | 0.2857 | 5 | 0.4 | 13 | 0.3077 |
| 17_7868582 | 4 | 0 | 7 | 0.1429 | 7 | 0.1429 | 3 | 0.6667 | 21 | 0.1905 | NA | NA | NA | NA | 6 | 0.1667 | 4 | 0.25 | 10 | 0.2 |
| 17_7868589 | 4 | 0 | 8 | 0.75 | 8 | 0.5 | 3 | 0.3333 | 23 | 0.4783 | 1 | 1 | NA | NA | 8 | 0.25 | 4 | 0.5 | 13 | 0.3846 |
| 17_7868592 | 4 | 0.25 | 5 | 0.2 | 6 | 0.5 | 3 | 0.3333 | 18 | 0.3333 | 1 | 1 | NA | NA | 9 | 0.6667 | 4 | 0.25 | 14 | 0.5714 |
| 17_7868603 | 4 | 0.25 | 7 | 0.4286 | 8 | 0.75 | 3 | 0.3333 | 22 | 0.5 | NA | NA | NA | NA | 8 | 0.375 | 4 | 0.75 | 12 | 0.5 |
| 17_7870821 | 2 | 0 | 5 | 0.2 | 6 | 0.1667 | 2 | 0.5 | 15 | 0.2 | NA | NA | NA | NA | 7 | 0 | 4 | 0.25 | 11 | 0.0909 |
| 17_7870829 | 3 | 0 | 5 | 0.2 | 6 | 0 | 2 | 0 | 16 | 0.0625 | NA | NA | NA | NA | 8 | 0.125 | 4 | 0 | 12 | 0.0833 |
| 17_7872543 | 1 | 0 | 1 | 0 | 6 | 0.3333 | 2 | 0 | 10 | 0.2 | NA | NA | NA | NA | 5 | 0.2 | 3 | 0.3333 | 8 | 0.25 |
| 17_7895982 | 1 | 0 | 2 | 0.5 | 4 | 0.25 | 5 | 0 | 12 | 0.1667 | 3 | 0.3333 | 4 | 0 | 5 | 0 | 7 | 0 | 19 | 0.0526 |
| 17_7895983 | 2 | 1 | 2 | 1 | 7 | 0.2857 | 6 | 0.1667 | 17 | 0.4118 | 5 | 0.4 | 6 | 0.3333 | 4 | 0 | 6 | 0.5 | 21 | 0.3333 |
| 17_7895992 | 13 | 0.2308 | 13 | 0.0769 | 15 | 0.1333 | 16 | 0.125 | 57 | 0.1404 | 11 | 0.0909 | 22 | 0.0455 | 12 | 0.1667 | 17 | 0.1176 | 62 | 0.0968 |
| 17_7896027 | 14 | 0 | 12 | 0.1667 | 13 | 0.0769 | 16 | 0.1875 | 55 | 0.1091 | 9 | 0.2222 | 20 | 0.15 | 11 | 0.5455 | 17 | 0.0588 | 57 | 0.2105 |
| 17_7896064 | 5 | 0.2 | 9 | 0.2222 | 9 | 0.1111 | 9 | 0.2222 | 32 | 0.1875 | 7 | 0.4286 | 13 | 0.3846 | 7 | 0.7143 | 11 | 0.1818 | 38 | 0.3947 |
| 17_7896067 | 9 | 0.1111 | 13 | 0.1538 | 12 | 0.0833 | 12 | 0.0833 | 46 | 0.1087 | 9 | 0.3333 | 19 | 0.1053 | 12 | 0.3333 | 16 | 0.0625 | 56 | 0.1786 |
| 17_7896077 | 9 | 0.2222 | 12 | 0.25 | 11 | 0.1818 | 12 | 0 | 44 | 0.1591 | 9 | 0 | 19 | 0.1053 | 12 | 0.1667 | 17 | 0 | 57 | 0.0702 |
| 17_7896081 | 7 | 0.1429 | 8 | 0.375 | 8 | 0.5 | 9 | 0.1111 | 32 | 0.2812 | 7 | 0.5714 | 14 | 0.3571 | 11 | 0.2727 | 12 | 0.0833 | 44 | 0.2955 |
| 17_7896222 | 3 | 0 | 4 | 0.25 | 10 | 0 | 10 | 0.1 | 27 | 0.0741 | 8 | 0.125 | 11 | 0.1818 | 7 | 0 | 12 | 0.1667 | 38 | 0.1316 |
| 17_7896262 | 5 | 0.4 | 3 | 0 | 10 | 0.4 | 9 | 0.4444 | 27 | 0.3704 | 8 | 0.375 | 10 | 0.3 | 5 | 0.6 | 9 | 0.3333 | 32 | 0.375 |
| 17_7896270 | 4 | 0.25 | 4 | 0 | 10 | 0.3 | 9 | 0 | 27 | 0.1481 | 8 | 0.75 | 12 | 0.1667 | 6 | 0.8333 | 12 | 0.4167 | 38 | 0.4737 |
| 17_7896279 | 4 | 1 | 4 | 1 | 10 | 0.9 | 8 | 1 | 26 | 0.9615 | 6 | 0.6667 | 10 | 0.7 | 5 | 1 | 11 | 0.9091 | 32 | 0.8125 |
| 17_7896304 | 3 | 0.3333 | 3 | 0 | 9 | 0.1111 | 8 | 0.125 | 23 | 0.1304 | 7 | 0.2857 | 10 | 0.2 | 4 | 0 | 9 | 0.1111 | 30 | 0.1667 |
| 17_7896308 | 4 | 0 | 4 | 0.25 | 9 | 0.2222 | 9 | 0 | 26 | 0.1154 | 7 | 0.2857 | 10 | 0.2 | 6 | 0.1667 | 9 | 0.1111 | 32 | 0.1875 |
| 17_7905686 | 1 | 0 | NA | NA | 1 | 0 | 1 | 1 | 3 | 0.3333 | 2 | 0 | 3 | 0.3333 | 1 | 1 | 3 | 0 | 9 | 0.2222 |
| 17_7905727 | 1 | 1 | 1 | 1 | NA | NA | NA | NA | 2 | 1 | 2 | 0.5 | 2 | 1 | 1 | 1 | 5 | 0.8 | 10 | 0.8 |
| 17_7905731 | 1 | 0 | 1 | 1 | 1 | 0 | NA | NA | 3 | 0.3333 | 2 | 1 | 3 | 0.6667 | 1 | 1 | 5 | 1 | 11 | 0.9091 |
| 17_7905734 | NA | NA | 1 | 1 | 1 | 0 | NA | NA | 2 | 0.5 | 1 | 0 | 1 | 1 | 1 | 1 | 2 | 1 | 5 | 0.8 |
| 17_7905739 | 1 | 0 | 1 | 1 | 1 | 0 | NA | NA | 3 | 0.3333 | 2 | 1 | 2 | 1 | 1 | 1 | 4 | 0.75 | 9 | 0.8889 |
| 17_7905743 | NA | NA | 1 | 1 | 1 | 1 | NA | NA | 2 | 1 | 2 | 1 | 2 | 1 | 1 | 1 | 3 | 1 | 8 | 1 |
| 17_7905758 | NA | NA | NA | NA | 1 | 1 | NA | NA | 1 | 1 | 1 | 1 | 2 | 1 | 1 | 1 | 2 | 1 | 6 | 1 |
| 17_7905760 | 1 | 1 | NA | NA | 1 | 0 | NA | NA | 2 | 0.5 | 2 | 1 | 3 | 0.6667 | 1 | 1 | 3 | 0.6667 | 9 | 0.7778 |
| 17_7905762 | 1 | 0 | 1 | 1 | 1 | 0 | NA | NA | 3 | 0.3333 | 2 | 0 | 2 | 0.5 | NA | NA | 3 | 0.6667 | 7 | 0.4286 |
| 17_7905779 | 1 | 1 | 1 | 0 | 1 | 0 | NA | NA | 3 | 0.3333 | 1 | 0 | 2 | 0.5 | NA | NA | 3 | 0.3333 | 6 | 0.3333 |
| 17_7905784 | 1 | 0 | 1 | 1 | 1 | 0 | NA | NA | 3 | 0.3333 | 2 | 0.5 | 1 | 1 | NA | NA | 2 | 0.5 | 5 | 0.6 |
| 17_7905795 | 1 | 0 | 1 | 1 | 1 | 0 | NA | NA | 3 | 0.3333 | 2 | 0 | 1 | 0 | NA | NA | 3 | 0.3333 | 6 | 0.1667 |
| 17_7905801 | 1 | 1 | 1 | 1 | 1 | 0 | NA | NA | 3 | 0.6667 | 2 | 0.5 | 2 | 0 | NA | NA | 3 | 0.3333 | 7 | 0.2857 |
| 17_7905874 | 1 | 0 | 1 | 0 | 1 | 1 | NA | NA | 3 | 0.3333 | 2 | 1 | 2 | 0 | NA | NA | 3 | 0.3333 | 7 | 0.4286 |
| 17_7905876 | 1 | 0 | 1 | 1 | 1 | 1 | NA | NA | 3 | 0.6667 | 1 | 1 | 2 | 1 | NA | NA | 3 | 0.6667 | 6 | 0.8333 |
| 17_7905911 | NA | NA | 1 | 1 | NA | NA | NA | NA | 1 | 1 | 2 | 1 | 2 | 0 | NA | NA | 2 | 0.5 | 6 | 0.5 |
| 17_7905923 | 1 | 0 | 1 | 0 | 1 | 1 | NA | NA | 3 | 0.3333 | 2 | 0.5 | 1 | 1 | NA | NA | 3 | 0.3333 | 6 | 0.5 |
| 17_7905930 | 1 | 1 | 1 | 1 | 1 | 1 | NA | NA | 3 | 1 | 2 | 1 | 2 | 1 | NA | NA | 3 | 1 | 7 | 1 |

| | | | | | | | | | | | | | | | | | | | | |
|---|---|---|---|---|---|---|---|---|---|---|---|---|---|---|---|---|---|---|---|---|
| 17_790593 | 1 | 1 | NA | NA | 1 | 1 | NA | NA | 2 | 1 | 1 | 1 | 2 | 0 | NA | NA | 2 | 0.5 | 5 | 0.4 |
| 17_790605 | 6 | 0.1667 | 2 | 0.5 | 1 | 0 | 2 | 0.5 | 11 | 0.2727 | 4 | 0.5 | 2 | 0.5 | NA | NA | 3 | 0 | 9 | 0.3333 |
| 17_790605 | 5 | 0.8 | 4 | 0.25 | 1 | 1 | 2 | 0.5 | 12 | 0.5833 | 3 | 1 | 1 | 1 | 1 | 1 | 2 | 0.5 | 7 | 0.8571 |
| 17_790606 | 6 | 0.5 | 3 | 0.6667 | 2 | 1 | 3 | 1 | 14 | 0.7143 | 5 | 0.8 | 2 | 0.5 | 1 | 0 | 3 | 1 | 11 | 0.7273 |
| 17_791527 | 2 | 0 | 4 | 0 | 1 | 1 | 1 | 0 | 8 | 0.125 | 5 | 0 | 2 | 0 | 5 | 0.2 | 2 | 0 | 14 | 0.0714 |
| 17_791530 | NA | NA | 1 | 0 | 2 | 0.5 | 2 | 0 | 5 | 0.2 | 3 | 0 | NA | NA | 3 | 0.3333 | 1 | 0 | 7 | 0.1429 |
| 17_791531 | NA | NA | 1 | 0 | 2 | 0 | 2 | 0.5 | 5 | 0.2 | 3 | 0.3333 | NA | NA | 3 | 0.3333 | 1 | 1 | 7 | 0.4286 |
| 17_791533 | NA | NA | 1 | 0 | 1 | 1 | 2 | 0.5 | 4 | 0.5 | 2 | 1 | NA | NA | 3 | 0.6667 | 2 | 1 | 7 | 0.8571 |
| 17_791535 | NA | NA | 1 | 0 | 2 | 0 | 2 | 0.5 | 5 | 0.2 | 3 | 0.6667 | NA | NA | 3 | 0.3333 | 2 | 1 | 8 | 0.625 |
| 17_791540 | NA | NA | NA | NA | 2 | 0 | 2 | 0.5 | 4 | 0.25 | 2 | 0.5 | NA | NA | 4 | 0.5 | 3 | 0 | 9 | 0.3333 |
| 17_791540 | NA | NA | NA | NA | 1 | 0 | 3 | 0.3333 | 4 | 0.25 | 2 | 0.5 | NA | NA | 4 | 0.25 | 3 | 0.3333 | 9 | 0.3333 |
| 17_791540 | NA | NA | 1 | 0 | 2 | 0 | 3 | 0.3333 | 6 | 0.1667 | 3 | 0.3333 | NA | NA | 3 | 0 | 3 | 0.3333 | 9 | 0.2222 |
| 17_791541 | NA | NA | 1 | 0 | 2 | 0 | 3 | 0.6667 | 6 | 0.3333 | 3 | 0.3333 | NA | NA | 4 | 0.25 | 3 | 0.3333 | 10 | 0.3 |
| 17_791542 | NA | NA | NA | NA | 2 | 0 | 2 | 0.5 | 4 | 0.25 | 3 | 0.3333 | NA | NA | 4 | 0.75 | 4 | 0.75 | 11 | 0.6364 |
| 17_792271 | 6 | 0.1667 | 1 | 0 | 2 | 0.5 | 9 | 0.1111 | 18 | 0.1667 | 3 | 0 | 3 | 0 | 2 | 0 | 4 | 0.25 | 12 | 0.0833 |
| 17_792272 | 6 | 0 | 1 | 0 | 2 | 0 | 9 | 0.1111 | 18 | 0.0556 | 3 | 0 | 3 | 0 | 4 | 0 | 5 | 0.2 | 15 | 0.0667 |
| 17_792421 | 3 | 1 | 1 | 1 | 1 | 0 | 8 | 0.75 | 13 | 0.7692 | 2 | 1 | 3 | 1 | 1 | 0 | 3 | 1 | 9 | 0.8889 |
| 17_792422 | 2 | 1 | 1 | 1 | 1 | 1 | 5 | 0.8 | 9 | 0.8889 | 3 | 0.6667 | 3 | 1 | 2 | 1 | 3 | 1 | 11 | 0.9091 |
| 17_792422 | 3 | 0.6667 | 1 | 0 | 1 | 1 | 8 | 0.875 | 13 | 0.7692 | 3 | 1 | 4 | 1 | 2 | 1 | 2 | 0.5 | 11 | 0.9091 |
| 17_792424 | 3 | 0.3333 | 1 | 0 | NA | NA | 7 | 0 | 11 | 0.0909 | 2 | 0 | 4 | 0.25 | 1 | 0 | 3 | 0.3333 | 10 | 0.2 |
| 17_792425 | 3 | 0.6667 | 1 | 1 | NA | NA | 7 | 0.4286 | 11 | 0.5455 | 2 | 1 | 4 | 1 | 2 | 0 | 3 | 0.6667 | 11 | 0.7273 |
| 17_792426 | 2 | 0 | 1 | 1 | NA | NA | 7 | 0.4286 | 10 | 0.4 | 2 | 0 | 4 | 0.5 | 2 | 0 | 2 | 1 | 10 | 0.4 |
| 17_792426 | 3 | 1 | 1 | 0 | NA | NA | 6 | 1 | 10 | 0.9 | 2 | 0.5 | 4 | 0.75 | 2 | 1 | 3 | 1 | 11 | 0.8182 |
| 17_792427 | 3 | 0.3333 | NA | NA | NA | NA | 4 | 0 | 7 | 0.1429 | 2 | 0.5 | 3 | 0 | 2 | 0.5 | 2 | 0.5 | 9 | 0.3333 |
| 17_792429 | 3 | 0.3333 | 1 | 1 | NA | NA | 6 | 0.1667 | 10 | 0.3 | 2 | 0.5 | 4 | 0.5 | 2 | 0 | 3 | 0 | 11 | 0.2727 |
| 17_792431 | 2 | 0 | 1 | 0 | NA | NA | 6 | 0.5 | 9 | 0.3333 | 1 | 1 | 3 | 0.3333 | 2 | 0.5 | 3 | 0 | 9 | 0.3333 |
| 17_792432 | 2 | 1 | 1 | 0 | NA | NA | 5 | 0.6 | 8 | 0.625 | 2 | 0.5 | 3 | 1 | 2 | 1 | 2 | 0.5 | 9 | 0.7778 |
| 17_792544 | 4 | 1 | 3 | 1 | 5 | 0.6 | 5 | 0.6 | 17 | 0.7647 | 1 | 1 | 3 | 0.6667 | 6 | 1 | 8 | 0.625 | 18 | 0.7778 |
| 17_792545 | 3 | 1 | 4 | 1 | 6 | 0.6667 | 8 | 0.625 | 21 | 0.7619 | 2 | 0.5 | 3 | 0.6667 | 6 | 0.6667 | 10 | 0.7 | 21 | 0.6667 |
| 17_792545 | 3 | 0.6667 | 4 | 0.25 | 6 | 0.5 | 7 | 0.2857 | 20 | 0.4 | 2 | 0.5 | 4 | 0.5 | 6 | 0.6667 | 8 | 0.25 | 20 | 0.45 |
| 17_792570 | NA | NA | NA | NA | NA | NA | 2 | 1 | 2 | 1 | 1 | 1 | NA | NA | NA | NA | 3 | 1 | 4 | 1 |
| 17_792586 | NA | NA | NA | NA | NA | NA | 3 | 0.3333 | 3 | 0.3333 | NA | NA | NA | NA | NA | NA | 2 | 0.5 | 2 | 0.5 |
| 17_792588 | NA | NA | NA | NA | NA | NA | 3 | 0.6667 | 3 | 0.6667 | 1 | 0 | NA | NA | NA | NA | 3 | 0.3333 | 4 | 0.25 |
| 17_792591 | NA | NA | NA | NA | NA | NA | 3 | 1 | 3 | 1 | 1 | 1 | NA | NA | NA | NA | 3 | 1 | 4 | 1 |
| 17_793201 | NA | NA | NA | NA | NA | NA | 1 | 1 | 1 | 1 | 1 | 0 | NA | NA | NA | NA | 3 | 0.3333 | 4 | 0.25 |
| 17_793202 | NA | NA | NA | NA | NA | NA | 1 | 1 | 1 | 1 | NA | NA | NA | NA | NA | NA | 3 | 1 | 3 | 1 |
| 17_793205 | NA | NA | NA | NA | NA | NA | 1 | 1 | 1 | 1 | NA | NA | NA | NA | NA | NA | 2 | 0.5 | 2 | 0.5 |
| 17_793781 | 5 | 0.4 | 3 | 0 | 1 | 0 | 3 | 0 | 12 | 0.1667 | 4 | 0 | 5 | 0.2 | 3 | 0 | 5 | 0 | 17 | 0.0588 |
| 17_793973 | 3 | 0 | 3 | 0 | 2 | 1 | 5 | 0 | 13 | 0.1538 | 5 | 0.4 | 4 | 0.5 | 3 | 0.3333 | 5 | 0.2 | 17 | 0.3529 |
| 17_796415 | 1 | 1 | 6 | 0.3333 | NA | NA | 3 | 0 | 10 | 0.3 | NA | NA | 5 | 0.4 | 2 | 0 | 3 | 0 | 10 | 0.2 |
| 17_796419 | NA | NA | 6 | 0.1667 | NA | NA | 3 | 0.3333 | 9 | 0.2222 | NA | NA | 3 | 0 | 3 | 0 | 3 | 0.3333 | 9 | 0.1111 |
| 17_796419 | NA | NA | 5 | 0.2 | NA | NA | 2 | 0.5 | 7 | 0.2857 | NA | NA | 3 | 0.3333 | 3 | 0.3333 | 2 | 0 | 8 | 0.25 |
| 17_796420 | 1 | 1 | 4 | 1 | NA | NA | 2 | 1 | 7 | 1 | NA | NA | 3 | 1 | 1 | 1 | 3 | 1 | 7 | 1 |
| 17_796422 | 1 | 1 | 3 | 1 | NA | NA | 1 | 0 | 5 | 0.8 | NA | NA | 3 | 1 | 1 | 1 | 1 | 1 | 5 | 1 |
| 17_796425 | 1 | 1 | 5 | 0.8 | NA | NA | 1 | 1 | 7 | 0.8571 | NA | NA | 2 | 1 | 2 | 1 | 3 | 1 | 7 | 1 |
| 17_796426 | 1 | 1 | 4 | 1 | NA | NA | 1 | 1 | 6 | 1 | NA | NA | 3 | 1 | 2 | 1 | 3 | 1 | 8 | 1 |
| 17_796428 | 1 | 0 | 4 | 0.5 | NA | NA | 1 | 0 | 6 | 0.3333 | NA | NA | 3 | 0.3333 | 1 | 0 | 3 | 0.3333 | 7 | 0.2857 |
| 17_796428 | 1 | 1 | 5 | 0.6 | NA | NA | 1 | 0 | 7 | 0.5714 | NA | NA | 2 | 1 | 2 | 1 | 3 | 1 | 7 | 1 |
| 17_796428 | 1 | 0 | 2 | 1 | NA | NA | 1 | 1 | 4 | 0.75 | NA | NA | 2 | 0.5 | 1 | 0 | 1 | 0 | 4 | 0.25 |
| 17_796429 | 1 | 1 | 4 | 1 | NA | NA | 1 | 1 | 6 | 1 | NA | NA | 3 | 1 | 2 | 1 | 3 | 1 | 8 | 1 |
| 17_796430 | 1 | 0 | 4 | 0.25 | NA | NA | 1 | 0 | 6 | 0.1667 | NA | NA | 2 | 0.5 | 2 | 0.5 | 3 | 0 | 7 | 0.2857 |
| 17_796431 | 1 | 1 | 3 | 0.3333 | NA | NA | 1 | 0 | 5 | 0.4 | NA | NA | 3 | 0.3333 | 2 | 0.5 | 3 | 0.6667 | 8 | 0.5 |
| 17_799290 | 1 | 1 | NA | NA | NA | NA | NA | NA | 1 | 1 | 1 | 1 | NA | NA | NA | NA | NA | NA | 1 | 1 |
| 17_800171 | 4 | 0.5 | 1 | 1 | NA | NA | 2 | 0 | 7 | 0.4286 | NA | NA | NA | NA | NA | NA | 1 | 1 | 1 | 1 |
| 17_801569 | 8 | 0.25 | 8 | 0.125 | 2 | 0 | 10 | 0.4 | 28 | 0.25 | 7 | 0.1429 | 12 | 0 | 7 | 0.1429 | 8 | 0.125 | 34 | 0.0882 |
| 17_801570 | 10 | 0.1 | 9 | 0.1111 | 1 | 0 | 8 | 0.125 | 28 | 0.1071 | 5 | 0.2 | 13 | 0.2308 | 6 | 0 | 8 | 0.125 | 32 | 0.1562 |
| 17_801575 | 7 | 0.2857 | 9 | 0.3333 | NA | NA | 7 | 0.1429 | 23 | 0.2609 | 6 | 0.5 | 9 | 0.3333 | 5 | 0.4 | 8 | 0.375 | 28 | 0.3929 |
| 17_801575 | 5 | 0.8 | 7 | 1 | 1 | 1 | 8 | 0.625 | 21 | 0.8095 | 5 | 0.6 | 11 | 0.7273 | 6 | 0.8333 | 6 | 0.8333 | 28 | 0.75 |
| 17_801575 | 7 | 0.1429 | 9 | 0.1111 | 1 | 0 | 9 | 0.2222 | 26 | 0.1538 | 5 | 0.4 | 12 | 0.0833 | 7 | 0.2857 | 7 | 0.2857 | 31 | 0.2258 |
| 17_801576 | 7 | 0.1429 | 9 | 0.1111 | NA | NA | 9 | 0.1111 | 25 | 0.12 | 5 | 0.2 | 12 | 0.3333 | 8 | 0.375 | 8 | 0.625 | 33 | 0.3939 |
| 17_801580 | 5 | 0.2 | 6 | 0.1667 | NA | NA | 4 | 0.75 | 15 | 0.3333 | 5 | 0.6 | 5 | 0.2 | 1 | 0 | 3 | 0.3333 | 14 | 0.3571 |
| 17_801581 | 5 | 0 | 6 | 0.5 | 1 | 1 | 4 | 0.25 | 16 | 0.3125 | 6 | 0.6667 | 8 | 0.125 | 2 | 0 | 4 | 0.25 | 20 | 0.3 |
| 17_801600 | 6 | 0.1667 | 6 | 0.1667 | 1 | 0 | 3 | 0 | 16 | 0.125 | 5 | 0.4 | 10 | 0.2 | 4 | 0.25 | 6 | 0.5 | 25 | 0.32 |
| 17_801601 | 5 | 0.2 | 5 | 0 | NA | NA | 2 | 0 | 12 | 0.0833 | 4 | 0 | 6 | 0.3333 | 6 | 0.3333 | 5 | 0.2 | 21 | 0.2381 |
| 17_801601 | 6 | 0.8333 | 7 | 0.8571 | NA | NA | 5 | 1 | 18 | 0.8889 | 6 | 0.8333 | 9 | 0.8889 | 6 | 0.8333 | 8 | 1 | 29 | 0.8966 |

| | | | | | | | | | | | | | | | | | | | | |
|---|---|---|---|---|---|---|---|---|---|---|---|---|---|---|---|---|---|---|---|---|
| 17_801602 | 5 | 0.2 | 7 | 0 | 1 | 0 | 5 | 0 | 18 | 0.0556 | 6 | 0 | 11 | 0.2727 | 7 | 0 | 7 | 0 | 31 | 0.0968 |
| 17_801832 | 5 | 0.2 | 5 | 0.2 | 1 | 0 | 2 | 0 | 13 | 0.1538 | 6 | 0 | 7 | 0 | 4 | 0.25 | 6 | 0.1667 | 23 | 0.087 |
| 17_805085 | 8 | 0.125 | 17 | 0.0588 | 16 | 0.1875 | 25 | 0.08 | 66 | 0.1061 | 7 | 0.1429 | 18 | 0.2222 | 17 | 0.1765 | 7 | 0.4286 | 49 | 0.2245 |
| 17_805120 | 5 | 0.2 | 9 | 0.2222 | 5 | 0 | 10 | 0.3 | 29 | 0.2069 | 8 | 0.375 | 7 | 0.1429 | 8 | 0.125 | 3 | 0 | 26 | 0.1923 |
| 17_805353 | 1 | 1 | NA | NA | NA | NA | 1 | 0 | 2 | 0.5 | NA | NA | 1 | 0 | NA | NA | 1 | 1 | 2 | 0.5 |
| 17_805354 | 1 | 1 | NA | NA | NA | NA | 1 | 1 | 2 | 1 | NA | NA | 1 | 1 | NA | NA | 1 | 1 | 2 | 1 |
| 17_805355 | 1 | 1 | NA | NA | NA | NA | 1 | 1 | 2 | 1 | NA | NA | 1 | 1 | NA | NA | 1 | 1 | 2 | 1 |
| 17_805459 | 5 | 0.6 | 3 | 1 | 1 | 1 | 2 | 0 | 11 | 0.6364 | 2 | 0.5 | 5 | 0.8 | 8 | 0.375 | 1 | 0 | 16 | 0.5 |
| 17_805460 | 4 | 0.75 | 3 | 0 | 1 | 0 | 2 | 0 | 10 | 0.3 | 2 | 0 | 4 | 0.5 | 9 | 0.5556 | 1 | 0 | 16 | 0.4375 |
| 17_805460 | 4 | 0.25 | 4 | 0.5 | 1 | 0 | 1 | 0 | 10 | 0.3 | 2 | 0.5 | 5 | 0.2 | 9 | 0.3333 | 1 | 1 | 17 | 0.3529 |
| 17_805460 | 2 | 1 | 1 | 1 | 1 | 1 | 2 | 0.5 | 6 | 0.8333 | 1 | 0 | 4 | 1 | 5 | 0.8 | 1 | 0 | 11 | 0.7273 |
| 17_805461 | 4 | 0 | 2 | 0.5 | 1 | 0 | 2 | 0 | 9 | 0.1111 | 1 | 0 | 4 | 0.5 | 5 | 0.2 | 1 | 0 | 11 | 0.2727 |
| 17_805462 | 2 | 0.5 | 3 | 0.3333 | 1 | 0 | 2 | 0.5 | 8 | 0.375 | 2 | 0 | 1 | 0 | 8 | 0.125 | 1 | 0 | 12 | 0.0833 |
| 17_805587 | 1 | 0 | 1 | 1 | NA | NA | NA | NA | 2 | 0.5 | 1 | 0 | 1 | 0 | 4 | 0.25 | NA | NA | 6 | 0.1667 |
| 17_805823 | NA | NA | 1 | 1 | NA | NA | NA | NA | 1 | 1 | NA | NA | 1 | 1 | 3 | 0.3333 | 1 | 0 | 5 | 0.4 |
| 17_805827 | 1 | 1 | 1 | 1 | NA | NA | NA | NA | 2 | 1 | NA | NA | 1 | 1 | 3 | 1 | 1 | 1 | 5 | 1 |
| 17_805827 | 1 | 1 | 1 | 1 | NA | NA | NA | NA | 2 | 1 | NA | NA | 1 | 1 | 3 | 1 | NA | NA | 4 | 1 |
| 17_805827 | 1 | 1 | 1 | 1 | NA | NA | NA | NA | 2 | 1 | NA | NA | 1 | 1 | 3 | 1 | NA | NA | 4 | 1 |
| 17_806136 | NA | NA | 1 | 1 | NA | NA | NA | NA | 1 | 1 | NA | NA | NA | NA | NA | NA | 1 | 1 | 1 | 1 |
| 17_807018 | 2 | 0 | 4 | 0.5 | 1 | 1 | 2 | 0 | 9 | 0.3333 | 1 | 0 | 6 | 0.1667 | 4 | 0.25 | 2 | 1 | 13 | 0.3077 |
| 17_807027 | 1 | 0 | 8 | 0 | 1 | 1 | 4 | 0 | 14 | 0.0714 | 1 | 0 | 5 | 0.4 | 3 | 0 | 2 | 0 | 11 | 0.1818 |
| 17_807031 | 1 | 0 | 7 | 0 | 1 | 0 | 3 | 0.3333 | 12 | 0.0833 | NA | NA | 4 | 0.25 | 2 | 0 | 2 | 0 | 8 | 0.125 |
| 17_807041 | 2 | 0 | 3 | 0.3333 | 1 | 0 | 2 | 0 | 8 | 0.125 | NA | NA | 5 | 0.2 | 1 | 0 | 3 | 0 | 9 | 0.1111 |
| 17_807041 | 2 | 0 | 3 | 0 | NA | NA | 2 | 0.5 | 7 | 0.1429 | NA | NA | 4 | 0.25 | 2 | 1 | 3 | 0 | 9 | 0.3333 |
| 17_807043 | NA | NA | 2 | 0 | NA | NA | 2 | 0.5 | 4 | 0.25 | NA | NA | 4 | 0.25 | 2 | 0 | 3 | 0 | 9 | 0.1111 |
| 17_807048 | 2 | 0.5 | 6 | 0 | 2 | 0 | 2 | 0 | 12 | 0.0833 | 2 | 0 | 5 | 0 | 5 | 0 | 4 | 0.25 | 16 | 0.0625 |
| 17_807049 | 2 | 0 | 5 | 0.2 | 2 | 0 | 2 | 0.5 | 11 | 0.1818 | 2 | 0 | 3 | 0 | 3 | 0.3333 | 4 | 0 | 12 | 0.0833 |
| 17_807051 | 1 | 0 | 4 | 0.5 | 1 | 1 | 2 | 0.5 | 8 | 0.5 | 1 | 1 | 4 | 0.75 | 3 | 0.6667 | 2 | 0.5 | 10 | 0.7 |
| 17_807053 | 2 | 1 | 5 | 0.8 | 1 | 1 | 2 | 1 | 10 | 0.9 | 1 | 1 | 4 | 0.5 | 5 | 1 | 2 | 0.5 | 12 | 0.75 |
| 17_807054 | 1 | 0 | 4 | 0.5 | 1 | 1 | 2 | 1 | 8 | 0.625 | 1 | 0 | 4 | 0.5 | 5 | 0.4 | 2 | 0.5 | 12 | 0.4167 |
| 17_807054 | 2 | 1 | 4 | 0.5 | 1 | 1 | 2 | 0 | 9 | 0.5556 | 1 | 1 | 5 | 0.4 | 5 | 0.8 | 2 | 0.5 | 13 | 0.6154 |
| 17_807055 | 1 | 0 | 4 | 0.25 | 1 | 1 | 2 | 0 | 8 | 0.25 | 1 | 0 | 5 | 0.8 | 4 | 0.75 | 2 | 0.5 | 12 | 0.6667 |
| 17_807055 | 1 | 0 | 2 | 0.5 | NA | NA | 2 | 0.5 | 5 | 0.4 | 1 | 0 | 4 | 0.25 | 3 | 0 | 1 | 0 | 9 | 0.1111 |
| 17_807056 | 2 | 1 | 3 | 0.6667 | 1 | 1 | 1 | 1 | 7 | 0.8571 | 1 | 1 | 4 | 1 | 3 | 1 | 1 | 1 | 9 | 1 |
| 17_807056 | 1 | 1 | 2 | 0.5 | NA | NA | 2 | 0.5 | 5 | 0.6 | NA | NA | 2 | 1 | 1 | 1 | 2 | 0.5 | 5 | 0.8 |
| 17_807056 | 1 | 1 | NA | NA | NA | NA | NA | NA | 1 | 1 | NA | NA | NA | NA | 1 | 1 | NA | NA | 1 | 1 |
| 17_807057 | 1 | 1 | 2 | 1 | 1 | 1 | 1 | 1 | 5 | 1 | 1 | 1 | 3 | 0.6667 | 3 | 1 | 2 | 0.5 | 9 | 0.7778 |
| 17_807059 | 1 | 1 | 4 | 0.5 | 1 | 1 | 1 | 0 | 7 | 0.5714 | 1 | 0 | 3 | 0.3333 | 3 | 1 | 2 | 0.5 | 9 | 0.5556 |
| 17_807060 | 2 | 1 | 3 | 0.6667 | 1 | 1 | 1 | 0 | 7 | 0.7143 | 1 | 1 | 2 | 1 | 4 | 1 | 2 | 0.5 | 9 | 0.8889 |
| 17_807414 | NA | NA | 1 | 1 | NA | NA | NA | NA | 1 | 1 | NA | NA | NA | NA | 3 | 0.6667 | NA | NA | 3 | 0.6667 |
| 17_807414 | NA | NA | 2 | 0.5 | NA | NA | NA | NA | 2 | 0.5 | NA | NA | NA | NA | 2 | 0.5 | NA | NA | 2 | 0.5 |
| 17_808321 | NA | NA | 2 | 0.5 | 1 | 0 | NA | NA | 3 | 0.3333 | 3 | 1 | NA | NA | 1 | 1 | 1 | 0 | 5 | 0.8 |
| 17_808321 | NA | NA | 2 | 1 | 1 | 1 | 1 | 1 | 4 | 1 | 3 | 0.6667 | NA | NA | 1 | 1 | 1 | 1 | 5 | 0.8 |
| 17_808322 | NA | NA | 2 | 0.5 | 1 | 0 | 1 | 0 | 4 | 0.25 | 3 | 0.3333 | NA | NA | 1 | 0 | 1 | 0 | 5 | 0.2 |
| 17_808324 | NA | NA | 2 | 1 | 2 | 1 | 1 | 1 | 5 | 1 | 3 | 1 | NA | NA | 1 | 1 | 1 | 1 | 5 | 1 |
| 17_808325 | NA | NA | 2 | 1 | NA | NA | NA | NA | 2 | 1 | 3 | 0.6667 | NA | NA | 1 | 1 | 1 | 1 | 5 | 0.8 |
| 17_808326 | NA | NA | 2 | 0.5 | 1 | 0 | 1 | 0 | 4 | 0.25 | 3 | 0.3333 | NA | NA | 1 | 0 | 1 | 0 | 5 | 0.2 |
| 17_808327 | NA | NA | 2 | 1 | 2 | 1 | 1 | 1 | 5 | 1 | 2 | 1 | NA | NA | NA | NA | 1 | 1 | 3 | 1 |
| 17_808891 | 6 | 0.1667 | 6 | 0.1667 | 2 | 0 | 9 | 0.1111 | 23 | 0.1304 | 4 | 0 | 3 | 0.3333 | 3 | 0.3333 | NA | NA | 10 | 0.2 |
| 17_813083 | 5 | 0 | 4 | 0.25 | 4 | 0 | 3 | 0 | 16 | 0.0625 | 3 | 0 | 2 | 0 | 3 | 0 | 3 | 0.3333 | 11 | 0.0909 |
| 17_813083 | 5 | 0.2 | 3 | 0.3333 | 2 | 0.5 | 3 | 0 | 13 | 0.2308 | 4 | 0.25 | 3 | 0.3333 | 2 | 0 | 2 | 0 | 11 | 0.1818 |
| 17_813084 | 6 | 1 | 5 | 1 | 4 | 1 | 3 | 1 | 18 | 1 | 6 | 1 | 3 | 1 | 3 | 1 | 3 | 0.6667 | 15 | 0.9333 |
| 17_813085 | 5 | 1 | 4 | 1 | 4 | 0.75 | 2 | 1 | 15 | 0.9333 | 6 | 1 | 2 | 1 | 3 | 1 | 3 | 1 | 14 | 1 |
| 17_813086 | 4 | 0.25 | 5 | 0.2 | 4 | 0 | 3 | 0 | 16 | 0.125 | 6 | 0.3333 | 3 | 0.6667 | 3 | 0.6667 | 2 | 0 | 14 | 0.4286 |
| 17_813086 | 6 | 0.3333 | 4 | 0.25 | 2 | 0.5 | 3 | 0.3333 | 15 | 0.3333 | 4 | 0.75 | 2 | 0 | 3 | 0.3333 | 3 | 0.6667 | 12 | 0.5 |
| 17_813086 | 6 | 0.5 | 5 | 0.8 | 4 | 0.75 | 3 | 0.3333 | 18 | 0.6111 | 6 | 0.3333 | 2 | 1 | 2 | 1 | 3 | 0.3333 | 13 | 0.5385 |
| 17_813087 | 4 | 0.25 | 3 | 0.6667 | 2 | 0.5 | 3 | 0.3333 | 12 | 0.4167 | 4 | 1 | 1 | 0 | 3 | 0 | NA | NA | 8 | 0.5 |
| 17_813087 | 5 | 1 | 3 | 1 | 4 | 1 | 4 | 1 | 16 | 1 | 6 | 1 | 2 | 1 | 3 | 1 | 3 | 1 | 14 | 1 |
| 17_813089 | 6 | 0.5 | 4 | 0.25 | 3 | 0 | 1 | 1 | 14 | 0.3571 | 5 | 0.6 | 2 | 0 | 3 | 0.6667 | 2 | 1 | 12 | 0.5833 |
| 17_813120 | 1 | 1 | 2 | 1 | 2 | 1 | NA | NA | 5 | 1 | NA | NA | NA | NA | NA | NA | 1 | 1 | 1 | 1 |
| 17_813132 | 1 | 1 | 1 | 0 | 1 | 1 | NA | NA | 3 | 0.6667 | NA | NA | NA | NA | NA | NA | 1 | 1 | 1 | 1 |
| 17_813139 | 1 | 0 | 3 | 0.3333 | 2 | 0 | 1 | 0 | 7 | 0.1429 | NA | NA | NA | NA | NA | NA | 1 | 1 | 1 | 1 |
| 17_813147 | 1 | 0 | 1 | 0 | 4 | 0.5 | 1 | 0 | 7 | 0.2857 | NA | NA | NA | NA | 1 | 1 | 1 | 0 | 2 | 0.5 |
| 17_813338 | 1 | 0 | 1 | 1 | 1 | 1 | 2 | 1 | 5 | 0.8 | 1 | 1 | 2 | 1 | NA | NA | 3 | 1 | 6 | 1 |
| 17_813339 | 1 | 1 | 1 | 1 | 1 | 1 | 2 | 1 | 5 | 1 | 1 | 1 | 2 | 1 | NA | NA | 3 | 1 | 6 | 1 |

| | | | | | | | | | | | | | | | | | | | | |
|---|---|---|---|---|---|---|---|---|---|---|---|---|---|---|---|---|---|---|---|---|
| 17_8133408 | 1 | 1 | 1 | 1 | 1 | 1 | 2 | 1 | 5 | 1 | 1 | 1 | 2 | 1 | NA | NA | 3 | 1 | 6 | 1 |
| 17_8133425 | 1 | 0 | 1 | 1 | NA | NA | 2 | 1 | 4 | 0.75 | 1 | 0 | NA | NA | NA | NA | 3 | 0.3333 | 4 | 0.25 |
| 17_8136160 | NA | NA | 1 | 1 | 1 | 0 | NA | NA | 2 | 0.5 | 1 | 1 | NA | NA | 1 | 1 | 1 | 1 | 3 | 1 |
| 17_8168138 | 5 | 0.8 | 17 | 0.7647 | 12 | 0.8333 | 10 | 0.8 | 44 | 0.7955 | 5 | 0.6 | 5 | 0.8 | 9 | 0.8889 | 4 | 1 | 23 | 0.8261 |
| 17_8168142 | 4 | 0.25 | 14 | 0.2143 | 10 | 0.2 | 11 | 0.2727 | 39 | 0.2308 | 2 | 0 | 4 | 0.25 | 8 | 0.25 | 4 | 0 | 18 | 0.1667 |
| 17_8168170 | 2 | 0 | 13 | 0.1538 | 11 | 0 | 11 | 0.1818 | 37 | 0.1081 | 3 | 0 | 5 | 0.2 | 5 | 0 | 2 | 0 | 15 | 0.0667 |
| 17_8168221 | 4 | 0.5 | 13 | 0.2308 | 13 | 0.2308 | 10 | 0.3 | 40 | 0.275 | 2 | 0 | 5 | 0.2 | 7 | 0.1429 | 2 | 0.5 | 16 | 0.1875 |
| 17_8168226 | 2 | 0 | 14 | 0.1429 | 10 | 0.4 | 9 | 0.4444 | 35 | 0.2857 | 3 | 0.6667 | 5 | 0.4 | 8 | 0 | 1 | 0 | 17 | 0.2353 |
| 17_8168236 | 4 | 1 | 14 | 0.4286 | 12 | 0.5833 | 10 | 0.4 | 40 | 0.525 | 3 | 0.6667 | 4 | 0.25 | 8 | 0.625 | 2 | 0.5 | 17 | 0.5294 |
| 17_8168240 | 3 | 0.6667 | 13 | 0.3846 | 11 | 0.6364 | 8 | 0.625 | 35 | 0.5429 | 3 | 0.3333 | 4 | 0.5 | 7 | 0.2857 | 2 | 1 | 16 | 0.4375 |
| 17_8168257 | 4 | 0.25 | 14 | 0.2143 | 9 | 0.1111 | 9 | 0.4444 | 36 | 0.25 | 3 | 0 | 5 | 0.4 | 7 | 0.2857 | 2 | 0.5 | 17 | 0.2941 |
| 17_8169438 | 2 | 0.5 | 3 | 0 | 3 | 0 | 1 | 0 | 9 | 0.1111 | 1 | 1 | 2 | 0 | 1 | 0 | 3 | 0 | 7 | 0.1429 |
| 17_8169549 | 1 | 0 | 3 | 0.6667 | 2 | 0 | 2 | 0.5 | 8 | 0.375 | 1 | 0 | 2 | 1 | 1 | 0 | 2 | 0.5 | 6 | 0.5 |
| 17_8169560 | NA | NA | 2 | 1 | 2 | 0.5 | 1 | 1 | 5 | 0.8 | NA | NA | 2 | 0.5 | NA | NA | 1 | 1 | 3 | 0.6667 |
| 17_8169603 | NA | NA | 2 | 1 | 2 | 0.5 | 1 | 1 | 5 | 0.8 | 1 | 1 | 1 | 0 | 1 | 0 | NA | NA | 3 | 0.3333 |
| 17_8169603 | NA | NA | 2 | 0.5 | 2 | 0 | 1 | 0 | 5 | 0.2 | NA | NA | 2 | 0.5 | 1 | 1 | NA | NA | 3 | 0.6667 |
| 17_8169604 | NA | NA | 2 | 1 | 2 | 0 | 1 | 1 | 5 | 0.6 | NA | NA | 1 | 1 | NA | NA | NA | NA | 1 | 1 |
| 17_8169611 | NA | NA | 2 | 0.5 | 2 | 1 | 1 | 1 | 5 | 0.8 | NA | NA | 1 | 1 | NA | NA | NA | NA | 1 | 1 |
| 17_8169621 | NA | NA | 2 | 1 | 2 | 1 | 1 | 1 | 5 | 1 | 1 | 0 | 2 | 1 | 1 | 1 | NA | NA | 4 | 0.75 |
| 17_8195664 | NA | NA | NA | NA | 1 | 1 | NA | NA | 1 | 1 | 1 | 1 | 1 | 1 | 1 | 0 | NA | NA | 3 | 0.6667 |
| 17_8195682 | NA | NA | NA | NA | 1 | 1 | NA | NA | 1 | 1 | 1 | 0 | 2 | 1 | NA | NA | NA | NA | 3 | 0.6667 |
| 17_8353182 | 2 | 0.5 | 2 | 0 | 1 | 0 | NA | NA | 5 | 0.2 | 1 | 1 | NA | NA | 1 | 0 | NA | NA | 2 | 0.5 |
| 17_8378519 | 1 | 0 | NA | NA | NA | NA | 1 | 1 | 2 | 0.5 | 1 | 0 | 1 | 1 | 1 | 0 | 1 | 1 | 4 | 0.5 |
| 17_8378533 | 1 | 1 | NA | NA | 1 | 0 | NA | NA | 2 | 0.5 | 1 | 1 | 3 | 0 | 1 | 0 | 1 | 0 | 6 | 0.1667 |
| 17_8378549 | 1 | 0 | NA | NA | 1 | 1 | 1 | 0 | 3 | 0.3333 | 1 | 0 | 3 | 0.3333 | 2 | 0 | 1 | 0 | 7 | 0.1429 |
| 17_8378554 | 1 | 1 | NA | NA | 1 | 1 | 1 | 0 | 3 | 0.6667 | 1 | 0 | 3 | 0.3333 | 2 | 0 | 1 | 0 | 7 | 0.1429 |
| 17_8378571 | 1 | 1 | NA | NA | 1 | 0 | 1 | 1 | 3 | 0.6667 | 1 | 1 | 3 | 0.6667 | 2 | 1 | 1 | 0 | 7 | 0.7143 |
| 17_8378576 | NA | NA | 1 | 1 | 1 | 1 | 1 | 1 | 3 | 1 | 1 | 1 | 3 | 1 | 2 | 1 | 1 | 1 | 7 | 1 |
| 17_8378581 | 1 | 0 | 1 | 1 | 1 | 1 | 1 | 1 | 4 | 0.75 | 2 | 0.5 | 2 | 1 | 2 | 0.5 | 1 | 1 | 7 | 0.7143 |
| 17_8378641 | NA | NA | NA | NA | NA | NA | 1 | 1 | 1 | 1 | 1 | 1 | 2 | 1 | 2 | 1 | 1 | 1 | 6 | 1 |
| 17_8378695 | 3 | 0.3333 | NA | NA | 1 | 0 | 1 | 0 | 5 | 0.2 | 2 | 0 | 3 | 0.3333 | 4 | 0.75 | 2 | 1 | 11 | 0.5455 |
| 17_8378702 | 2 | 1 | NA | NA | 1 | 0 | NA | NA | 3 | 0.6667 | 2 | 1 | 3 | 1 | 4 | 1 | 2 | 1 | 11 | 1 |
| 17_8378707 | 2 | 0 | NA | NA | 1 | 0 | 1 | 1 | 4 | 0.25 | 2 | 1 | 2 | 0 | 4 | 0.25 | 3 | 1 | 11 | 0.5455 |
| 17_8378734 | 3 | 0.3333 | NA | NA | 1 | 0 | 1 | 0 | 5 | 0.2 | 3 | 0.3333 | 3 | 0 | 3 | 0 | 3 | 0 | 12 | 0.0833 |
| 17_8378743 | 3 | 0.3333 | NA | NA | 1 | 1 | 1 | 1 | 5 | 0.6 | 3 | 1 | 2 | 1 | 2 | 1 | 3 | 1 | 10 | 1 |
| 17_8378744 | 3 | 0.6667 | NA | NA | 2 | 1 | 1 | 0 | 6 | 0.6667 | 3 | 0.6667 | 3 | 1 | 3 | 0.3333 | 3 | 0.6667 | 12 | 0.6667 |
| 17_8378752 | 1 | 1 | NA | NA | 2 | 1 | 1 | 1 | 4 | 1 | 2 | 0.5 | 2 | 1 | 3 | 0.3333 | 3 | 1 | 10 | 0.7 |
| 17_8380938 | 40 | 0.05 | 42 | 0.1429 | 37 | 0.2162 | 54 | 0.1852 | 173 | 0.1503 | 30 | 0.1667 | 39 | 0.1282 | 43 | 0.0465 | 40 | 0.1 | 152 | 0.1053 |
| 17_8380950 | 46 | 0.087 | 41 | 0.0488 | 38 | 0.1842 | 55 | 0.1091 | 180 | 0.1056 | 27 | 0.1481 | 36 | 0.0556 | 41 | 0.1463 | 43 | 0.093 | 147 | 0.1088 |
| 17_8380963 | 43 | 0.1163 | 44 | 0.1364 | 40 | 0.25 | 55 | 0.2727 | 182 | 0.1978 | 29 | 0.2069 | 37 | 0.1622 | 44 | 0.25 | 43 | 0.1163 | 153 | 0.183 |
| 17_8380972 | 40 | 0.5 | 41 | 0.6098 | 42 | 0.5476 | 56 | 0.625 | 179 | 0.5754 | 26 | 0.3462 | 39 | 0.5641 | 40 | 0.5 | 47 | 0.4894 | 152 | 0.4868 |
| 17_8380984 | 40 | 0.2 | 39 | 0.2308 | 41 | 0.2195 | 50 | 0.16 | 170 | 0.2 | 22 | 0.2727 | 34 | 0.1176 | 37 | 0.1892 | 37 | 0.1351 | 130 | 0.1692 |
| 17_8380985 | 42 | 0.1429 | 39 | 0.1282 | 41 | 0.1463 | 55 | 0.1273 | 177 | 0.1356 | 23 | 0.087 | 31 | 0.1613 | 35 | 0.0571 | 37 | 0.0811 | 126 | 0.0952 |
| 17_8380994 | 38 | 0.1053 | 40 | 0.025 | 37 | 0.0541 | 49 | 0.0408 | 164 | 0.0549 | 26 | 0.0385 | 34 | 0.0588 | 35 | 0.1143 | 36 | 0 | 131 | 0.0534 |
| 17_8381001 | 37 | 0.9459 | 44 | 0.8864 | 43 | 0.9767 | 53 | 0.9811 | 177 | 0.9492 | 26 | 0.9231 | 34 | 0.9118 | 38 | 0.8947 | 41 | 0.9756 | 139 | 0.9281 |
| 17_8388165 | 1 | 0 | 1 | 0 | 3 | 0 | 6 | 0.3333 | 11 | 0.1818 | NA | NA | 4 | 0 | 3 | 0.3333 | 4 | 0 | 11 | 0.0909 |
| 17_8411216 | NA | NA | 1 | 1 | NA | NA | NA | NA | 1 | 1 | NA | NA | NA | NA | 1 | 1 | 1 | 0 | 2 | 0.5 |
| 17_8411241 | NA | NA | 1 | 1 | NA | NA | NA | NA | 1 | 1 | NA | NA | NA | NA | 1 | 1 | 1 | 0 | 2 | 0.5 |
| 17_8411269 | NA | NA | 1 | 1 | NA | NA | NA | NA | 1 | 1 | NA | NA | NA | NA | 1 | 1 | NA | NA | 1 | 1 |
| 17_8411272 | NA | NA | 1 | 1 | NA | NA | NA | NA | 1 | 1 | NA | NA | NA | NA | NA | NA | 1 | 1 | 1 | 1 |
| 17_8412777 | 2 | 1 | 1 | 1 | 1 | 0 | NA | NA | 4 | 0.75 | 1 | 1 | 3 | 0.6667 | NA | NA | 1 | 1 | 5 | 0.8 |
| 17_8412786 | 3 | 0.3333 | 1 | 0 | 2 | 0 | NA | NA | 6 | 0.1667 | 1 | 0 | 3 | 0.6667 | 1 | 0 | 1 | 0 | 6 | 0.3333 |
| 17_8412789 | 3 | 0.6667 | 1 | 1 | 2 | 0.5 | 1 | 0 | 7 | 0.5714 | 1 | 1 | 4 | 0.5 | 1 | 1 | 1 | 1 | 7 | 0.7143 |
| 17_8412848 | 4 | 0.5 | NA | NA | 3 | 0 | 1 | 0 | 8 | 0.25 | 1 | 0 | 4 | 0 | 1 | 1 | 1 | 0 | 7 | 0.1429 |
| 17_8413030 | 3 | 0 | NA | NA | 4 | 0.25 | 1 | 0 | 8 | 0.125 | NA | NA | 5 | 0 | 4 | 0.25 | 1 | 0 | 10 | 0.1 |
| 17_8449150 | 10 | 0.9 | 7 | 0.4286 | 6 | 0.8333 | 5 | 0.8 | 28 | 0.75 | 3 | 0.6667 | 7 | 1 | 3 | 1 | 1 | 1 | 14 | 0.9286 |
| 17_8449182 | 9 | 0.2222 | 5 | 0 | 6 | 0.1667 | 7 | 0.1429 | 27 | 0.1481 | 2 | 0.5 | 10 | 0.2 | 3 | 0.6667 | 2 | 0 | 17 | 0.2941 |
| 17_8449189 | 10 | 0.1 | 6 | 0 | 6 | 0.1667 | 7 | 0 | 29 | 0.069 | 3 | 0.3333 | 10 | 0 | 4 | 0 | 3 | 0 | 20 | 0.05 |
| 17_8449202 | 7 | 0 | 6 | 0 | 6 | 0.1667 | 6 | 0.1667 | 25 | 0.08 | 3 | 0 | 10 | 0.1 | 3 | 0 | 2 | 0 | 18 | 0.0556 |
| 17_8449268 | 9 | 0.6667 | 5 | 0.6 | 6 | 0.8333 | 6 | 0.5 | 26 | 0.6538 | 3 | 1 | 8 | 0.75 | 1 | 0 | NA | NA | 12 | 0.75 |
| 17_8449275 | 8 | 0.75 | 4 | 0.75 | 4 | 0.75 | 4 | 0.25 | 20 | 0.65 | 3 | 1 | 5 | 1 | NA | NA | 1 | 1 | 9 | 1 |
| 17_8449285 | 7 | 0.2857 | 5 | 0.2 | 6 | 0.1667 | 5 | 0.2 | 23 | 0.2174 | 3 | 0.3333 | 8 | 0.25 | 1 | 0 | 2 | 0 | 14 | 0.2143 |
| 17_8449289 | 4 | 0.25 | 3 | 0.3333 | 6 | 0 | 5 | 0.2 | 18 | 0.1667 | 2 | 0.5 | 6 | 0.1667 | 1 | 0 | 1 | 0 | 10 | 0.2 |
| 17_8449295 | 4 | 1 | 4 | 0.5 | 6 | 1 | 5 | 1 | 19 | 0.8947 | 2 | 0.5 | 7 | 0.8571 | 2 | 1 | 2 | 1 | 13 | 0.8462 |
| 17_8449307 | 4 | 1 | 4 | 0.75 | 4 | 0.75 | 4 | 1 | 16 | 0.875 | 1 | 1 | 4 | 0.5 | 1 | 1 | 1 | 1 | 7 | 0.7143 |

| | | | | | | | | | | | | | | | | | | | | |
|---|---|---|---|---|---|---|---|---|---|---|---|---|---|---|---|---|---|---|---|---|
| 17_844930 | 5 | 1 | 3 | 1 | 6 | 1 | 4 | 1 | 18 | 1 | 1 | 1 | 7 | 1 | 1 | 1 | 1 | 1 | 10 | 1 |
| 17_844931 | 6 | 0.3333 | 4 | 0 | 6 | 0.6667 | 4 | 0.5 | 20 | 0.4 | 3 | 0.3333 | 5 | 0.2 | 2 | 0 | 2 | 0 | 12 | 0.1667 |
| 17_844932 | 6 | 1 | 3 | 1 | 6 | 0.6667 | 3 | 0.6667 | 18 | 0.8333 | 2 | 1 | 6 | 0.8333 | 2 | 1 | 2 | 1 | 12 | 0.9167 |
| 17_844933 | 6 | 0.6667 | 4 | 0.5 | 5 | 0.4 | 3 | 0.6667 | 18 | 0.5556 | 2 | 0.5 | 6 | 0.5 | 1 | 1 | 2 | 1 | 11 | 0.6364 |
| 17_844933 | 7 | 0.8571 | 3 | 1 | 5 | 1 | 3 | 1 | 18 | 0.9444 | 2 | 1 | 5 | 1 | 2 | 1 | 1 | 1 | 10 | 1 |
| 17_844936 | 5 | 1 | 3 | 1 | 5 | 0.6 | 3 | 0.3333 | 16 | 0.75 | 2 | 0.5 | 7 | 0.8571 | 1 | 1 | 1 | 1 | 11 | 0.8182 |
| 17_844939 | 6 | 0.5 | 4 | 0.5 | 6 | 0.6667 | 4 | 0.75 | 20 | 0.6 | 2 | 0.5 | 7 | 0.8571 | 1 | 0 | 1 | 1 | 11 | 0.7273 |
| 17_844940 | 6 | 0 | 3 | 1 | 4 | 0 | 4 | 0.25 | 17 | 0.2353 | 2 | 0 | 6 | 0.3333 | 1 | 0 | 1 | 0 | 10 | 0.2 |
| 17_844942 | 5 | 0.2 | 1 | 0 | 4 | 0.5 | 4 | 0.25 | 14 | 0.2857 | 2 | 0.5 | 5 | 0.4 | 1 | 0 | 1 | 1 | 9 | 0.4444 |
| 17_844945 | 5 | 0 | 3 | 0 | 4 | 0.25 | 3 | 0 | 15 | 0.0667 | 2 | 0.5 | 5 | 0 | 1 | 0 | 1 | 0 | 9 | 0.1111 |
| 17_844958 | NA | NA | NA | NA | 1 | 1 | NA | NA | 1 | 1 | 1 | 1 | NA | NA | NA | NA | NA | NA | 1 | 1 |
| 17_844960 | NA | NA | NA | NA | 1 | 1 | NA | NA | 1 | 1 | 1 | 1 | NA | NA | NA | NA | NA | NA | 1 | 1 |
| 17_844961 | NA | NA | NA | NA | 1 | 1 | NA | NA | 1 | 1 | 1 | 1 | NA | NA | NA | NA | NA | NA | 1 | 1 |
| 17_844984 | 1 | 1 | 1 | 0 | 1 | 1 | 1 | 0 | 4 | 0.5 | NA | NA | 1 | 1 | NA | NA | NA | NA | 1 | 1 |
| 17_844987 | NA | NA | 1 | 1 | 1 | 0 | 1 | 0 | 3 | 0.3333 | NA | NA | 1 | 1 | NA | NA | NA | NA | 1 | 1 |
| 17_844992 | NA | NA | 2 | 0.5 | 1 | 1 | 1 | 1 | 4 | 0.75 | NA | NA | 1 | 1 | NA | NA | 1 | 0 | 2 | 0.5 |
| 17_844993 | NA | NA | 2 | 1 | 1 | 1 | NA | NA | 3 | 1 | NA | NA | 1 | 1 | NA | NA | 1 | 1 | 2 | 1 |
| 17_844994 | NA | NA | 2 | 1 | 1 | 1 | 1 | 1 | 4 | 1 | NA | NA | 1 | 1 | NA | NA | 1 | 1 | 2 | 1 |
| 17_847515 | NA | NA | 1 | 1 | NA | NA | NA | NA | 1 | 1 | NA | NA | 1 | 1 | NA | NA | NA | NA | 1 | 1 |
| 17_847518 | NA | NA | 1 | 1 | NA | NA | NA | NA | 1 | 1 | NA | NA | 1 | 1 | NA | NA | NA | NA | 1 | 1 |
| 17_849627 | 2 | 0 | 1 | 1 | 2 | 0 | 4 | 0.25 | 9 | 0.2222 | NA | NA | 3 | 0.3333 | 2 | 0 | NA | NA | 5 | 0.2 |
| 17_853011 | 5 | 0.2 | 6 | 0 | 9 | 0 | 10 | 0.1 | 30 | 0.0667 | 4 | 0.25 | 4 | 0.25 | 4 | 0.25 | 9 | 0 | 21 | 0.1429 |
| 17_853012 | 5 | 0 | 6 | 0.5 | 7 | 0.1429 | 13 | 0.5385 | 31 | 0.3548 | 5 | 0.4 | 3 | 0.3333 | 5 | 0.4 | 9 | 0.6667 | 22 | 0.5 |
| 17_853013 | 11 | 0 | 7 | 0.1429 | 9 | 0.2222 | 14 | 0.0714 | 41 | 0.0976 | 5 | 0 | 5 | 0.2 | 5 | 0 | 11 | 0.1818 | 26 | 0.1154 |
| 17_853014 | 11 | 0.6364 | 6 | 0.5 | 8 | 0.375 | 12 | 0.4167 | 37 | 0.4865 | 6 | 0.5 | 5 | 0.6 | 4 | 0.75 | 11 | 0.8182 | 26 | 0.6923 |
| 17_853014 | 11 | 0.9091 | 6 | 1 | 9 | 0.8889 | 15 | 0.8 | 41 | 0.878 | 6 | 1 | 5 | 1 | 5 | 0.6 | 9 | 0.8889 | 25 | 0.88 |
| 17_853015 | 10 | 0.2 | 8 | 0.5 | 5 | 0.4 | 11 | 0.2727 | 34 | 0.3235 | 6 | 0 | 3 | 0.3333 | 4 | 0.5 | 9 | 0.4444 | 22 | 0.3182 |
| 17_853015 | 10 | 0.6 | 8 | 0.625 | 7 | 1 | 14 | 0.7857 | 39 | 0.7436 | 6 | 1 | 4 | 1 | 3 | 0.6667 | 9 | 0.5556 | 22 | 0.7727 |
| 17_853016 | 12 | 0.9167 | 8 | 1 | 9 | 0.8889 | 15 | 0.8 | 44 | 0.8864 | 6 | 0.8333 | 5 | 1 | 6 | 0.6667 | 11 | 0.9091 | 28 | 0.8571 |
| 17_853017 | 10 | 1 | 8 | 1 | 9 | 1 | 13 | 1 | 40 | 1 | 5 | 0.6 | 5 | 1 | 6 | 1 | 10 | 1 | 26 | 0.9231 |
| 17_853017 | 10 | 0.2 | 8 | 0.125 | 7 | 0.4286 | 13 | 0.1538 | 38 | 0.2105 | 7 | 0.2857 | 3 | 0 | 4 | 0 | 9 | 0 | 23 | 0.087 |
| 17_853019 | 14 | 0.2143 | 9 | 0.2222 | 10 | 0.1 | 18 | 0.4444 | 51 | 0.2745 | 7 | 0.1429 | 5 | 0.6 | 6 | 0.5 | 11 | 0.2727 | 29 | 0.3448 |
| 17_853211 | 6 | 0 | 8 | 0.125 | 4 | 0.25 | 5 | 0 | 23 | 0.087 | 5 | 0.2 | 5 | 0 | 1 | 1 | 7 | 0.1429 | 18 | 0.1667 |
| 17_853211 | 6 | 0 | 4 | 0.25 | 4 | 0 | 3 | 0.3333 | 17 | 0.1176 | 4 | 0 | 4 | 0.25 | 3 | 0.3333 | 7 | 0.2857 | 18 | 0.2222 |
| 17_853212 | 7 | 0.8571 | 7 | 0.8571 | 4 | 1 | 4 | 0.5 | 22 | 0.8182 | 3 | 1 | 4 | 0.75 | 2 | 1 | 6 | 1 | 15 | 0.9333 |
| 17_853212 | 7 | 0.2857 | 8 | 0.25 | 4 | 0 | 5 | 0.2 | 24 | 0.2083 | 5 | 0 | 5 | 0.4 | 3 | 0 | 5 | 0.4 | 18 | 0.2222 |
| 17_853215 | 8 | 0.125 | 8 | 0.125 | 3 | 0 | 5 | 0.2 | 24 | 0.125 | 4 | 0.25 | 5 | 0.2 | 4 | 0 | 7 | 0.2857 | 20 | 0.2 |
| 17_853230 | 10 | 0.5 | 3 | 1 | 8 | 0.5 | 12 | 0.75 | 33 | 0.6364 | 2 | 1 | 4 | 0.75 | 4 | 1 | 7 | 0.8571 | 17 | 0.8824 |
| 17_853232 | 11 | 0.2727 | 5 | 0 | 9 | 0.2222 | 14 | 0.3571 | 39 | 0.2564 | 2 | 0.5 | 6 | 0.5 | 4 | 0.75 | 8 | 0.375 | 20 | 0.5 |
| 17_853244 | 15 | 0.3333 | 8 | 0 | 14 | 0.3571 | 16 | 0.3125 | 53 | 0.283 | 8 | 0.375 | 7 | 0 | 12 | 0 | 15 | 0.2667 | 42 | 0.1667 |
| 17_853698 | 36 | 0.1389 | 49 | 0.1633 | 47 | 0.1277 | 49 | 0.1633 | 181 | 0.1492 | 43 | 0.186 | 40 | 0.1 | 44 | 0.1364 | 26 | 0.1538 | 153 | 0.1438 |
| 17_853699 | 40 | 0.1 | 53 | 0.0943 | 48 | 0.0625 | 46 | 0 | 187 | 0.0642 | 41 | 0.0732 | 44 | 0.0227 | 44 | 0.1136 | 30 | 0.1 | 159 | 0.0755 |
| 17_853703 | 39 | 0.0769 | 45 | 0.0444 | 44 | 0.0455 | 50 | 0.06 | 178 | 0.0562 | 37 | 0.0811 | 43 | 0.093 | 35 | 0.0857 | 32 | 0.125 | 147 | 0.0952 |
| 17_853706 | 36 | 0.1111 | 39 | 0.0769 | 34 | 0.0294 | 40 | 0.025 | 149 | 0.0604 | 30 | 0.1 | 42 | 0.0714 | 33 | 0.1212 | 27 | 0.1481 | 132 | 0.1061 |
| 17_853707 | 38 | 0.3684 | 37 | 0.5405 | 32 | 0.4375 | 46 | 0.2609 | 153 | 0.3922 | 31 | 0.2258 | 40 | 0.375 | 30 | 0.3 | 28 | 0.3929 | 129 | 0.3256 |
| 17_853710 | 41 | 0.0976 | 42 | 0.1429 | 31 | 0.0323 | 49 | 0.102 | 163 | 0.0982 | 30 | 0.0667 | 41 | 0.122 | 32 | 0.0938 | 29 | 0.1034 | 132 | 0.0985 |
| 17_853710 | 41 | 0.3902 | 41 | 0.5122 | 32 | 0.4062 | 48 | 0.625 | 162 | 0.4938 | 33 | 0.4848 | 44 | 0.3636 | 30 | 0.4333 | 33 | 0.6061 | 140 | 0.4643 |
| 17_853711 | 37 | 0.4865 | 41 | 0.5122 | 28 | 0.4286 | 49 | 0.5714 | 155 | 0.5097 | 32 | 0.4062 | 42 | 0.619 | 32 | 0.4375 | 32 | 0.4688 | 138 | 0.4928 |
| 17_853711 | 36 | 0.7222 | 31 | 0.8065 | 28 | 0.75 | 37 | 0.7568 | 132 | 0.7576 | 21 | 0.8571 | 35 | 0.8 | 26 | 0.7308 | 27 | 0.6667 | 109 | 0.7615 |
| 17_853718 | 28 | 0.3571 | 26 | 0.6154 | 27 | 0.4074 | 34 | 0.7353 | 115 | 0.5391 | 22 | 0.5455 | 24 | 0.5833 | 22 | 0.3636 | 20 | 0.7 | 88 | 0.5455 |
| 17_853720 | 29 | 0.069 | 26 | 0.1538 | 23 | 0.1304 | 32 | 0.1562 | 110 | 0.1273 | 22 | 0.1364 | 28 | 0.1429 | 24 | 0.125 | 26 | 0.1154 | 100 | 0.13 |
| 17_853721 | 33 | 0.1515 | 35 | 0.2286 | 27 | 0.3333 | 38 | 0.0526 | 133 | 0.1805 | 25 | 0.08 | 31 | 0.2903 | 23 | 0.1304 | 24 | 0.1667 | 103 | 0.1748 |
| 17_853909 | NA | NA | NA | NA | 1 | 1 | NA | NA | 1 | 1 | NA | NA | NA | NA | 1 | 1 | 1 | 1 | 2 | 1 |
| 17_853960 | 1 | 1 | 1 | 0 | NA | NA | NA | NA | 2 | 0.5 | 3 | 0.6667 | 1 | 1 | NA | NA | 1 | 0 | 5 | 0.6 |
| 17_853961 | 1 | 1 | 1 | 0 | 1 | 1 | NA | NA | 3 | 0.6667 | 2 | 0.5 | NA | NA | NA | NA | 1 | 1 | 3 | 0.6667 |
| 17_857656 | 3 | 0.3333 | NA | NA | 1 | 0 | 3 | 0.6667 | 7 | 0.4286 | 1 | 0 | NA | NA | 3 | 0.3333 | NA | NA | 4 | 0.25 |
| 17_859092 | NA | NA | NA | NA | 1 | 1 | NA | NA | 1 | 1 | NA | NA | 1 | 1 | NA | NA | NA | NA | 1 | 1 |
| 17_859096 | NA | NA | NA | NA | 1 | 1 | NA | NA | 1 | 1 | NA | NA | 1 | 1 | NA | NA | NA | NA | 1 | 1 |
| 17_859098 | NA | NA | NA | NA | 1 | 1 | NA | NA | 1 | 1 | NA | NA | 1 | 1 | NA | NA | NA | NA | 1 | 1 |
| 17_859099 | NA | NA | NA | NA | 1 | 1 | NA | NA | 1 | 1 | NA | NA | 1 | 1 | NA | NA | NA | NA | 1 | 1 |
| 17_859292 | 4 | 0.75 | 1 | 1 | 5 | 0.8 | 4 | 1 | 14 | 0.8571 | NA | NA | 7 | 0.5714 | 6 | 0.6667 | 3 | 0.6667 | 16 | 0.625 |
| 17_859293 | 1 | 1 | NA | NA | 3 | 0.3333 | NA | NA | 4 | 0.5 | 1 | 1 | 4 | 0.25 | 6 | 0.6667 | 4 | 0.5 | 15 | 0.5333 |
| 17_859293 | 2 | 0.5 | NA | NA | 3 | 0.3333 | 1 | 1 | 6 | 0.5 | 1 | 1 | 6 | 0.5 | 5 | 0.8 | 3 | 0.6667 | 15 | 0.6667 |
| 17_859314 | 5 | 0.8 | 3 | 0.3333 | 6 | 0.5 | 3 | 0 | 17 | 0.4706 | 2 | 0 | 11 | 0.1818 | 6 | 0.3333 | 5 | 0.2 | 24 | 0.2083 |
| 17_859316 | 4 | 0.5 | 2 | 0.5 | 7 | 0.5714 | 3 | 1 | 16 | 0.625 | 1 | 1 | 10 | 0.4 | 7 | 0.1429 | 4 | 0 | 22 | 0.2727 |

| | | | | | | | | | | | | | | | | | | | | |
|---|---|---|---|---|---|---|---|---|---|---|---|---|---|---|---|---|---|---|---|---|
| 17_8593161 | NA | NA | 1 | 1 | 4 | 0.5 | 1 | 1 | 6 | 0.6667 | 1 | 0 | 8 | 0.5 | 6 | 0.3333 | 3 | 0.6667 | 18 | 0.4444 |
| 17_8593166 | 6 | 0.6667 | 3 | 0.3333 | 6 | 0.3333 | 4 | 0.25 | 19 | 0.4211 | 2 | 0 | 9 | 0.3333 | 9 | 0.7778 | 5 | 0.2 | 25 | 0.44 |
| 17_8593168 | 6 | 0.5 | 3 | 0.6667 | 8 | 0.875 | 4 | 0 | 21 | 0.5714 | 2 | 0 | 9 | 0.7778 | 10 | 0.4 | 6 | 0.6667 | 27 | 0.5556 |
| 17_8593171 | 5 | 0.4 | 3 | 0.3333 | 8 | 0.75 | 3 | 0.3333 | 19 | 0.5263 | 2 | 0 | 11 | 0.5455 | 10 | 0.6 | 4 | 0.25 | 27 | 0.4815 |
| 17_8593179 | 6 | 1 | 3 | 1 | 8 | 1 | 5 | 1 | 22 | 1 | 3 | 1 | 10 | 1 | 10 | 1 | 6 | 1 | 29 | 1 |
| 17_8593180 | 6 | 0.1667 | 3 | 0 | 6 | 0.3333 | 5 | 0 | 20 | 0.15 | 3 | 0.3333 | 9 | 0 | 7 | 0.2857 | 5 | 0.2 | 24 | 0.1667 |
| 17_8593181 | 5 | 0 | 4 | 0 | 8 | 0.25 | 5 | 0.4 | 22 | 0.1818 | 2 | 0 | 9 | 0 | 9 | 0.1111 | 7 | 0.1429 | 27 | 0.0741 |
| 17_8593194 | 1 | 0 | 1 | 0 | 5 | 0.2 | 2 | 0 | 9 | 0.1111 | NA | NA | 5 | 0.2 | 5 | 0.2 | 5 | 0 | 15 | 0.1333 |
| 17_8629285 | 1 | 0 | NA | NA | 2 | 1 | NA | NA | 3 | 0.6667 | NA | NA | 1 | 1 | NA | NA | 1 | 1 | 2 | 1 |
| 17_8647546 | 5 | 0.8 | 3 | 1 | 2 | 0.5 | 2 | 0.5 | 12 | 0.75 | 6 | 1 | 4 | 0.5 | 5 | 0.6 | 2 | 0.5 | 17 | 0.7059 |
| 17_8647567 | 6 | 0.3333 | 2 | 0.5 | 2 | 0 | 2 | 0 | 12 | 0.25 | 5 | 0.4 | 6 | 0.1667 | 3 | 0 | 4 | 0.25 | 18 | 0.2222 |
| 17_8647578 | 5 | 0.4 | 3 | 0.6667 | 4 | 0.75 | 3 | 0.3333 | 15 | 0.5333 | 7 | 0.5714 | 7 | 0.4286 | 5 | 0 | 4 | 0.25 | 23 | 0.3478 |
| 17_8647611 | 8 | 0.25 | 5 | 0.4 | 7 | 0.1429 | 4 | 0 | 24 | 0.2083 | 9 | 0 | 9 | 0.2222 | 5 | 0.2 | 7 | 0 | 30 | 0.1 |
| 17_8647613 | 7 | 0.4286 | 5 | 0.2 | 8 | 0.5 | 4 | 0.5 | 24 | 0.4167 | 9 | 0.5556 | 8 | 0.125 | 6 | 0.5 | 7 | 0.2857 | 30 | 0.3667 |
| 17_8672635 | 1 | 1 | NA | NA | NA | NA | NA | NA | 1 | 1 | NA | NA | 1 | 1 | NA | NA | NA | NA | 1 | 1 |
| 17_8672649 | 1 | 1 | NA | NA | NA | NA | NA | NA | 1 | 1 | NA | NA | 1 | 1 | NA | NA | NA | NA | 1 | 1 |
| 17_8672669 | 1 | 1 | NA | NA | NA | NA | NA | NA | 1 | 1 | NA | NA | 1 | 1 | NA | NA | NA | NA | 1 | 1 |
| 17_8680345 | 9 | 0.2222 | 9 | 0.1111 | 6 | 0.3333 | 7 | 0.1429 | 31 | 0.1935 | 5 | 0 | 10 | 0 | 9 | 0.1111 | 6 | 0.3333 | 30 | 0.1 |
| 17_8696200 | 21 | 0.2381 | 16 | 0.1875 | 15 | 0.0667 | 12 | 0.0833 | 64 | 0.1562 | 13 | 0.2308 | 16 | 0.125 | 13 | 0.3846 | 14 | 0.2143 | 56 | 0.2321 |
| 17_8696232 | 16 | 0.4375 | 15 | 0.4667 | 13 | 0.5385 | 10 | 0.3 | 54 | 0.4444 | 14 | 0.3571 | 16 | 0.4375 | 11 | 0.6364 | 12 | 0.5 | 53 | 0.4717 |
| 17_8696246 | 15 | 0.0667 | 12 | 0.0833 | 14 | 0 | 12 | 0.0833 | 53 | 0.0566 | 12 | 0.1667 | 12 | 0 | 13 | 0.0769 | 8 | 0.25 | 45 | 0.1111 |
| 17_8696247 | 20 | 0.75 | 19 | 0.5789 | 13 | 0.7692 | 10 | 0.8 | 62 | 0.7097 | 15 | 0.7333 | 16 | 0.6875 | 14 | 0.8571 | 14 | 0.7143 | 59 | 0.7458 |
| 17_8696249 | 21 | 0.9524 | 19 | 1 | 15 | 1 | 13 | 1 | 68 | 0.9853 | 19 | 0.9474 | 16 | 0.9375 | 13 | 1 | 18 | 0.9444 | 66 | 0.9545 |
| 17_8696276 | 19 | 0.9474 | 26 | 0.9615 | 16 | 1 | 11 | 1 | 72 | 0.9722 | 18 | 0.9444 | 15 | 0.9333 | 14 | 1 | 13 | 1 | 60 | 0.9667 |
| 17_8696289 | 25 | 0.08 | 21 | 0.1905 | 18 | 0.2222 | 13 | 0.3846 | 77 | 0.1948 | 20 | 0.15 | 19 | 0.2105 | 18 | 0.3333 | 15 | 0.2667 | 72 | 0.2361 |
| 17_8696293 | 25 | 0.48 | 23 | 0.5217 | 21 | 0.6667 | 16 | 0.375 | 85 | 0.5176 | 22 | 0.4091 | 20 | 0.5 | 17 | 0.5294 | 16 | 0.6875 | 75 | 0.52 |
| 17_8696294 | 24 | 0.375 | 17 | 0.4118 | 17 | 0.2941 | 12 | 0.25 | 70 | 0.3429 | 18 | 0.3333 | 18 | 0.5 | 17 | 0.4706 | 11 | 0.5455 | 64 | 0.4531 |
| 17_8696301 | 24 | 0.625 | 24 | 0.625 | 21 | 0.4286 | 16 | 0.5625 | 85 | 0.5647 | 22 | 0.5909 | 18 | 0.7222 | 14 | 0.8571 | 18 | 0.6111 | 72 | 0.6806 |
| 17_8696305 | 20 | 0.45 | 23 | 0.4783 | 15 | 0.8 | 14 | 0.7143 | 72 | 0.5833 | 23 | 0.5652 | 18 | 0.5556 | 13 | 0.4615 | 14 | 0.7857 | 68 | 0.5882 |
| 17_8696309 | 26 | 0.3462 | 25 | 0.12 | 21 | 0.1429 | 15 | 0.2 | 87 | 0.2069 | 23 | 0.087 | 19 | 0.2105 | 13 | 0.2308 | 20 | 0.15 | 75 | 0.16 |
| 17_8696313 | 22 | 0.8636 | 23 | 0.8696 | 20 | 0.7 | 16 | 0.875 | 81 | 0.8272 | 18 | 0.7222 | 21 | 0.619 | 18 | 1 | 15 | 0.8667 | 72 | 0.7917 |
| 17_8696318 | 18 | 0.2778 | 17 | 0.2941 | 19 | 0.2632 | 15 | 0.2667 | 69 | 0.2754 | 20 | 0.2 | 17 | 0.2941 | 14 | 0.3571 | 16 | 0.25 | 67 | 0.2687 |
| 17_8696326 | 24 | 0.9167 | 27 | 0.7037 | 24 | 0.75 | 15 | 0.8667 | 90 | 0.8 | 19 | 0.7368 | 20 | 0.75 | 15 | 0.7333 | 17 | 0.9412 | 71 | 0.7887 |
| 17_8696333 | 31 | 0.2258 | 32 | 0.0938 | 24 | 0.0833 | 20 | 0 | 107 | 0.1121 | 25 | 0.16 | 21 | 0.1429 | 18 | 0.1667 | 21 | 0.1429 | 85 | 0.1529 |
| 17_8696339 | 25 | 0.08 | 32 | 0.1875 | 24 | 0.0417 | 21 | 0.0476 | 102 | 0.098 | 24 | 0.0417 | 21 | 0.0952 | 20 | 0.1 | 22 | 0.0909 | 87 | 0.0805 |
| 17_8696340 | 29 | 0.0345 | 31 | 0.0645 | 26 | 0.0385 | 20 | 0.1 | 106 | 0.0566 | 23 | 0.1304 | 23 | 0.0435 | 20 | 0 | 20 | 0.05 | 86 | 0.0581 |
| 17_8696341 | 26 | 1 | 29 | 1 | 25 | 0.92 | 19 | 1 | 99 | 0.9798 | 22 | 1 | 18 | 0.9444 | 18 | 1 | 21 | 0.9524 | 79 | 0.9747 |
| 17_8696352 | 15 | 0.8 | 14 | 0.6429 | 15 | 0.6667 | 14 | 0.7143 | 58 | 0.7069 | 17 | 0.7647 | 12 | 0.4167 | 13 | 0.6154 | 12 | 0.5833 | 54 | 0.6111 |
| 17_8696353 | 21 | 0.0952 | 22 | 0.1364 | 18 | 0.0556 | 17 | 0.0588 | 78 | 0.0897 | 16 | 0.125 | 15 | 0.0667 | 12 | 0 | 18 | 0.0556 | 61 | 0.0656 |
| 17_8696391 | 24 | 0.125 | 20 | 0.1 | 20 | 0.15 | 21 | 0.2381 | 85 | 0.1529 | 23 | 0.0435 | 19 | 0.0526 | 17 | 0.1176 | 16 | 0.1875 | 75 | 0.0933 |
| 17_8696400 | 34 | 0.3824 | 28 | 0.5357 | 29 | 0.3448 | 22 | 0.3182 | 113 | 0.3982 | 20 | 0.4 | 19 | 0.2632 | 20 | 0.4 | 24 | 0.4583 | 83 | 0.3855 |
| 17_8696401 | 36 | 0.25 | 36 | 0.0556 | 32 | 0.1875 | 27 | 0.1111 | 131 | 0.1527 | 29 | 0.1379 | 25 | 0.16 | 27 | 0.1481 | 25 | 0.24 | 106 | 0.1698 |
| 17_8696502 | 46 | 0.5217 | 40 | 0.625 | 42 | 0.7143 | 30 | 0.5333 | 158 | 0.6013 | 36 | 0.5278 | 32 | 0.625 | 29 | 0.7586 | 31 | 0.6774 | 128 | 0.6406 |
| 17_8706156 | 4 | 0 | 5 | 0 | NA | NA | 2 | 0.5 | 11 | 0.0909 | 9 | 0 | 2 | 0 | 2 | 0 | 4 | 0.25 | 17 | 0.0588 |
| 17_8714022 | 18 | 0.0556 | 21 | 0.1429 | 9 | 0.1111 | 25 | 0.24 | 73 | 0.1507 | 22 | 0.0909 | 13 | 0.2308 | 13 | 0.1538 | 18 | 0.1111 | 66 | 0.1364 |
| 17_8714044 | 24 | 0.125 | 25 | 0.08 | 9 | 0.1111 | 28 | 0.0357 | 86 | 0.0814 | 25 | 0.04 | 16 | 0.0625 | 14 | 0.1429 | 21 | 0.0476 | 76 | 0.0658 |
| 17_8730438 | 3 | 1 | 3 | 0.3333 | 2 | 1 | 5 | 0.8 | 13 | 0.7692 | 4 | 0.75 | 3 | 0.3333 | 5 | 0.8 | 2 | 1 | 14 | 0.7143 |
| 17_8730442 | 3 | 0 | 3 | 0.3333 | 3 | 0.3333 | 5 | 0 | 14 | 0.1429 | 4 | 0 | 3 | 0 | 4 | 0.25 | 2 | 0 | 13 | 0.0769 |
| 17_8730443 | 4 | 0 | 2 | 0 | 3 | 0 | 5 | 0.2 | 14 | 0.0714 | 3 | 0.3333 | 3 | 0 | 4 | 0 | 3 | 0 | 13 | 0.0769 |
| 17_8730446 | 4 | 1 | 2 | 1 | 1 | 1 | 5 | 1 | 12 | 1 | 4 | 1 | 2 | 1 | 5 | 1 | 2 | 1 | 13 | 1 |
| 17_8730449 | 4 | 0 | 2 | 0 | 3 | 0.3333 | 5 | 0 | 14 | 0.0714 | 4 | 0 | 3 | 0.3333 | 5 | 0 | 2 | 0 | 14 | 0.0714 |
| 17_8730451 | 4 | 0.75 | 1 | 1 | 2 | 1 | 5 | 0.2 | 12 | 0.5833 | 4 | 0.75 | 2 | 0.5 | 5 | 0.6 | 2 | 0.5 | 13 | 0.6154 |
| 17_8730458 | 3 | 0.3333 | 1 | 0 | 1 | 0 | 4 | 0 | 9 | 0.1111 | 4 | 0.25 | 2 | 0 | 4 | 0.25 | 3 | 0 | 13 | 0.1538 |
| 17_8730466 | 3 | 0 | 1 | 0 | 3 | 0 | 5 | 0.2 | 12 | 0.0833 | 1 | 0 | 3 | 0.3333 | 5 | 0.2 | 2 | 0 | 11 | 0.1818 |
| 17_8730477 | 4 | 0.5 | 2 | 0.5 | 3 | 0.6667 | 5 | 0.6 | 14 | 0.5714 | 3 | 0 | 2 | 0 | 5 | 0.8 | 2 | 0.5 | 12 | 0.4167 |
| 17_8730480 | 4 | 1 | 1 | 1 | 3 | 1 | 3 | 1 | 11 | 1 | 3 | 1 | 2 | 1 | 6 | 1 | 2 | 1 | 13 | 1 |
| 17_8730481 | 4 | 0.5 | 1 | 0 | 3 | 1 | 6 | 0.5 | 14 | 0.5714 | 3 | 0.6667 | 3 | 0.6667 | 6 | 0.5 | 3 | 0.6667 | 15 | 0.6 |
| 17_8730482 | 3 | 0.3333 | 1 | 0 | 3 | 0 | 7 | 0.1429 | 14 | 0.1429 | 4 | 0.25 | 2 | 0 | 5 | 0 | 3 | 0 | 14 | 0.0714 |
| 17_8730488 | 2 | 0 | 2 | 0 | 5 | 0 | 4 | 0.25 | 13 | 0.0769 | 3 | 0 | 4 | 0 | 5 | 0.2 | 5 | 0.2 | 17 | 0.1176 |
| 17_8730514 | 4 | 0 | 2 | 0 | 3 | 0.6667 | 5 | 0.4 | 14 | 0.2857 | 2 | 0 | 2 | 0 | 5 | 0.4 | 1 | 0 | 10 | 0.2 |
| 17_8730547 | 5 | 0.4 | 2 | 0 | 1 | 0 | 6 | 0 | 14 | 0.1429 | 4 | 0.25 | 2 | 0 | 5 | 0 | 2 | 0 | 13 | 0.0769 |
| 17_8730563 | 5 | 0.2 | 2 | 0 | 2 | 0 | 5 | 0 | 14 | 0.0714 | 4 | 0.5 | 3 | 0 | 6 | 0.3333 | 3 | 0 | 16 | 0.25 |
| 17_8730619 | 5 | 0.4 | 2 | 0 | 1 | 0 | 6 | 0.5 | 14 | 0.3571 | 3 | 0.6667 | 2 | 0.5 | 6 | 0 | 1 | 0 | 12 | 0.25 |
| 17_8730620 | 4 | 0 | 2 | 0.5 | 3 | 0 | 4 | 0 | 13 | 0.0769 | 4 | 0.25 | 2 | 0 | 5 | 0.2 | 2 | 0.5 | 13 | 0.2308 |
| 17_8730629 | 5 | 0.2 | 2 | 1 | 3 | 0.6667 | 8 | 0.5 | 18 | 0.5 | 5 | 0.2 | 3 | 0 | 7 | 0.1429 | 4 | 0.5 | 19 | 0.2105 |

| | | | | | | | | | | | | | | | | | | | | |
|---|---|---|---|---|---|---|---|---|---|---|---|---|---|---|---|---|---|---|---|---|
| 17_873063 | 5 | 0.4 | 1 | 0 | 3 | 0.3333 | 7 | 0.4286 | 16 | 0.375 | 4 | 0.25 | 4 | 0 | 8 | 0.375 | 4 | 0.25 | 20 | 0.25 |
| 17_873064 | 4 | 0.25 | 1 | 0 | 4 | 0 | 7 | 0 | 16 | 0.0625 | 2 | 0 | 4 | 0 | 6 | 0.3333 | 3 | 0 | 15 | 0.1333 |
| 17_873065 | 1 | 1 | NA | NA | 3 | 1 | 4 | 1 | 8 | 1 | 1 | 1 | 3 | 1 | 6 | 1 | 2 | 1 | 12 | 1 |
| 17_873067 | 1 | 0 | NA | NA | 3 | 0.3333 | 4 | 0.5 | 8 | 0.375 | 1 | 0 | 2 | 0.5 | 6 | 0 | 2 | 0 | 11 | 0.0909 |
| 17_873067 | 1 | 0 | NA | NA | 4 | 0.25 | 4 | 0 | 9 | 0.1111 | 1 | 0 | 3 | 0.3333 | 4 | 0 | 2 | 0 | 10 | 0.1 |
| 17_873067 | NA | NA | NA | NA | 1 | 1 | NA | NA | 1 | 1 | 1 | 1 | 2 | 1 | 5 | 0.6 | 2 | 0.5 | 10 | 0.7 |
| 17_873069 | 1 | 0 | NA | NA | 4 | 0.75 | 4 | 0 | 9 | 0.3333 | 1 | 0 | 3 | 0 | 4 | 0.25 | 2 | 0 | 10 | 0.1 |
| 17_873070 | 1 | 1 | NA | NA | 3 | 1 | 3 | 0.6667 | 7 | 0.8571 | 1 | 1 | 3 | 0 | 5 | 0.8 | 1 | 0 | 10 | 0.5 |
| 17_873312 | 2 | 0 | 2 | 0 | 4 | 0.25 | 2 | 0 | 10 | 0.1 | 3 | 0 | 2 | 0 | 4 | 0 | 6 | 0.1667 | 15 | 0.0667 |
| 17_873426 | 10 | 0.1 | 10 | 0 | 20 | 0.25 | 10 | 0.1 | 50 | 0.14 | 7 | 0.1429 | 13 | 0.0769 | 9 | 0.1111 | 12 | 0.0833 | 41 | 0.0976 |
| 17_873427 | 14 | 0.0714 | 14 | 0.1429 | 21 | 0.381 | 15 | 0.2 | 64 | 0.2188 | 6 | 0.5 | 12 | 0.4167 | 10 | 0.3 | 13 | 0.1538 | 41 | 0.3171 |
| 17_873427 | 12 | 0.25 | 16 | 0.4375 | 21 | 0.3333 | 16 | 0.1875 | 65 | 0.3077 | 7 | 0.4286 | 15 | 0.1333 | 12 | 0.4167 | 15 | 0.4 | 49 | 0.3265 |
| 17_873428 | 9 | 0 | 11 | 0.0909 | 19 | 0.1579 | 12 | 0 | 51 | 0.0784 | 7 | 0.1429 | 10 | 0.1 | 11 | 0.1818 | 12 | 0.1667 | 40 | 0.15 |
| 17_873430 | 10 | 0.1 | 9 | 0.1111 | 15 | 0.0667 | 10 | 0.4 | 44 | 0.1591 | 6 | 0 | 9 | 0 | 7 | 0.4286 | 13 | 0.2308 | 35 | 0.1714 |
| 17_873875 | 1 | 0 | 1 | 1 | NA | NA | 1 | 1 | 3 | 0.6667 | 2 | 0.5 | 3 | 0.3333 | NA | NA | 1 | 0 | 6 | 0.3333 |
| 17_873876 | 1 | 0 | 1 | 0 | 1 | 0 | 2 | 1 | 5 | 0.4 | 2 | 0.5 | 2 | 0 | NA | NA | 2 | 0 | 6 | 0.1667 |
| 17_874412 | 7 | 0.8571 | 6 | 0.8333 | 3 | 1 | 5 | 0.8 | 21 | 0.8571 | 3 | 1 | 4 | 1 | 11 | 0.9091 | 4 | 1 | 22 | 0.9545 |
| 17_874412 | 7 | 0 | 7 | 0 | 4 | 0.25 | 7 | 0.1429 | 25 | 0.08 | 3 | 0 | 4 | 0 | 10 | 0.2 | 4 | 0 | 21 | 0.0952 |
| 17_874413 | 7 | 0.1429 | 6 | 0.1667 | 3 | 0 | 6 | 0.1667 | 22 | 0.1364 | 2 | 0 | 3 | 0.3333 | 9 | 0.1111 | 3 | 0 | 17 | 0.1176 |
| 17_874413 | 6 | 0 | 5 | 0.2 | 4 | 0.25 | 4 | 0.5 | 19 | 0.2105 | 1 | 1 | 2 | 0 | 9 | 0.5556 | 2 | 0.5 | 14 | 0.5 |
| 17_874413 | 7 | 0.1429 | 4 | 0.5 | 4 | 0.5 | 6 | 0.8333 | 21 | 0.4762 | 3 | 0 | 3 | 0.3333 | 11 | 0.1818 | 4 | 0.25 | 21 | 0.1905 |
| 17_874413 | 7 | 0.2857 | 8 | 0.5 | 4 | 0.75 | 7 | 0.2857 | 26 | 0.4231 | 3 | 0.6667 | 4 | 0.25 | 12 | 0.3333 | 5 | 0.4 | 24 | 0.375 |
| 17_874414 | 2 | 0.5 | 4 | 0.5 | 4 | 0 | 3 | 0.3333 | 13 | 0.3077 | 3 | 0 | 3 | 0.3333 | 8 | 0.375 | 3 | 0 | 17 | 0.2353 |
| 17_874415 | 5 | 0.6 | 8 | 0.375 | 5 | 0.6 | 5 | 0.4 | 23 | 0.4783 | 4 | 0.25 | 4 | 0.25 | 11 | 0.6364 | 4 | 0.25 | 23 | 0.4348 |
| 17_874415 | 5 | 1 | 6 | 1 | 3 | 1 | 6 | 1 | 20 | 1 | 3 | 1 | 3 | 1 | 7 | 0.7143 | 3 | 0.3333 | 16 | 0.75 |
| 17_874416 | 7 | 0.8571 | 8 | 0.625 | 4 | 0.5 | 8 | 0.625 | 27 | 0.6667 | 3 | 0.6667 | 2 | 0.5 | 9 | 0.4444 | 5 | 0.8 | 19 | 0.5789 |
| 17_874416 | 8 | 0.875 | 7 | 0.8571 | 5 | 0.8 | 5 | 0.8 | 25 | 0.84 | 3 | 0.3333 | 3 | 0.6667 | 10 | 0.9 | 5 | 1 | 21 | 0.8095 |
| 17_874419 | 8 | 0.75 | 10 | 0.8 | 5 | 0.6 | 7 | 0.4286 | 30 | 0.6667 | 5 | 0.6 | 6 | 0.5 | 11 | 0.7273 | 4 | 0.25 | 26 | 0.5769 |
| 17_874420 | 9 | 0.1111 | 10 | 0.3 | 3 | 0.3333 | 6 | 0.5 | 28 | 0.2857 | 4 | 0 | 7 | 0.5714 | 11 | 0.5455 | 5 | 0.2 | 27 | 0.4074 |
| 17_874420 | 6 | 0.5 | 8 | 0.625 | 5 | 0.2 | 7 | 0.4286 | 26 | 0.4615 | 5 | 0.8 | 5 | 0.4 | 9 | 0.7778 | 6 | 0.5 | 25 | 0.64 |
| 17_874421 | 8 | 0 | 11 | 0.0909 | 5 | 0.2 | 8 | 0 | 32 | 0.0625 | 6 | 0 | 6 | 0.1667 | 10 | 0.2 | 5 | 0.2 | 27 | 0.1481 |
| 17_874422 | 10 | 1 | 10 | 1 | 5 | 1 | 6 | 1 | 31 | 1 | 6 | 1 | 6 | 1 | 14 | 1 | 7 | 1 | 33 | 1 |
| 17_874422 | 9 | 0.8889 | 9 | 1 | 4 | 1 | 7 | 1 | 29 | 0.9655 | 4 | 0.75 | 5 | 1 | 10 | 1 | 6 | 1 | 25 | 0.96 |
| 17_874422 | 9 | 1 | 11 | 0.9091 | 5 | 0.8 | 7 | 0.8571 | 32 | 0.9062 | 3 | 0.6667 | 7 | 0.8571 | 12 | 0.9167 | 6 | 0.8333 | 28 | 0.8571 |
| 17_874424 | 9 | 1 | 11 | 1 | 4 | 1 | 5 | 1 | 29 | 1 | 5 | 1 | 5 | 1 | 8 | 0.875 | 6 | 1 | 24 | 0.9583 |
| 17_874424 | 10 | 0.3 | 12 | 0.0833 | 4 | 0.5 | 6 | 0.3333 | 32 | 0.25 | 6 | 0.3333 | 7 | 0.2857 | 11 | 0.3636 | 7 | 0.5714 | 31 | 0.3871 |
| 17_874424 | 11 | 0.6364 | 11 | 0.4545 | 6 | 1 | 8 | 0.625 | 36 | 0.6389 | 6 | 1 | 9 | 0.6667 | 13 | 0.6154 | 7 | 0.5714 | 35 | 0.6857 |
| 17_874424 | 9 | 1 | 5 | 1 | 5 | 1 | 6 | 1 | 25 | 1 | 6 | 1 | 7 | 1 | 10 | 1 | 6 | 1 | 29 | 1 |
| 17_874426 | 9 | 0.2222 | 10 | 0.4 | 2 | 0 | 7 | 0.2857 | 28 | 0.2857 | 7 | 0.1429 | 5 | 0.2 | 10 | 0.2 | 6 | 0.3333 | 28 | 0.2143 |
| 17_874427 | 9 | 0.3333 | 10 | 0.6 | 7 | 0.4286 | 7 | 0.2857 | 33 | 0.4242 | 6 | 0.3333 | 6 | 0.3333 | 14 | 0.2857 | 9 | 0.7778 | 35 | 0.4286 |
| 17_874428 | 8 | 0.125 | 7 | 0.2857 | 5 | 0.2 | 6 | 0.1667 | 26 | 0.1923 | 4 | 0 | 5 | 0.2 | 10 | 0.2 | 8 | 0.125 | 27 | 0.1481 |
| 17_874430 | 11 | 0.0909 | 12 | 0.3333 | 6 | 0.8333 | 9 | 0.2222 | 38 | 0.3158 | 6 | 0.1667 | 8 | 0.125 | 16 | 0.125 | 9 | 0.3333 | 39 | 0.1795 |
| 17_874434 | 12 | 0.1667 | 10 | 0.1 | 9 | 0 | 12 | 0 | 43 | 0.0698 | 8 | 0.125 | 7 | 0.2857 | 19 | 0.2105 | 11 | 0.0909 | 45 | 0.1778 |
| 17_874437 | 10 | 0.2 | 12 | 0.0833 | 8 | 0.375 | 11 | 0.2727 | 41 | 0.2195 | 6 | 0.5 | 4 | 0.5 | 19 | 0.1053 | 13 | 0.3077 | 42 | 0.2619 |
| 17_874444 | 14 | 0.0714 | 12 | 0.0833 | 8 | 0.125 | 12 | 0.25 | 46 | 0.1304 | 9 | 0.3333 | 10 | 0.2 | 19 | 0.1053 | 16 | 0.125 | 54 | 0.1667 |
| 17_874444 | 13 | 0.5385 | 11 | 0.7273 | 5 | 1 | 10 | 0.8 | 39 | 0.7179 | 8 | 0.625 | 9 | 0.4444 | 18 | 0.3333 | 15 | 0.4 | 50 | 0.42 |
| 17_874445 | 13 | 0.0769 | 11 | 0 | 7 | 0 | 11 | 0.1818 | 42 | 0.0714 | 9 | 0.2222 | 9 | 0.1111 | 20 | 0.1 | 15 | 0.2667 | 53 | 0.1698 |
| 17_874447 | 15 | 0.0667 | 12 | 0.3333 | 11 | 0.0909 | 13 | 0.1538 | 51 | 0.1569 | 10 | 0 | 14 | 0.0714 | 19 | 0.1053 | 14 | 0.2143 | 57 | 0.1053 |
| 17_874447 | 13 | 0.4615 | 12 | 0.6667 | 8 | 0.375 | 11 | 0.4545 | 44 | 0.5 | 7 | 0.2857 | 12 | 0.3333 | 17 | 0.3529 | 13 | 0.5385 | 49 | 0.3878 |
| 17_874448 | 15 | 0.0667 | 12 | 0.1667 | 11 | 0.0909 | 13 | 0 | 51 | 0.0784 | 10 | 0 | 11 | 0.2727 | 19 | 0.0526 | 13 | 0.2308 | 53 | 0.1321 |
| 17_874448 | 14 | 0.3571 | 12 | 0.1667 | 13 | 0.0769 | 12 | 0.0833 | 51 | 0.1765 | 12 | 0.3333 | 11 | 0.1818 | 18 | 0.1111 | 15 | 0 | 56 | 0.1429 |
| 17_875621 | 15 | 0 | 13 | 0 | 14 | 0.1429 | 12 | 0.0833 | 54 | 0.0556 | 14 | 0.0714 | 13 | 0 | 10 | 0 | 10 | 0.2 | 47 | 0.0638 |
| 17_875622 | 15 | 0.2 | 17 | 0.0588 | 17 | 0.0588 | 18 | 0.2222 | 67 | 0.1343 | 13 | 0 | 13 | 0.3077 | 11 | 0.1818 | 10 | 0.3 | 47 | 0.1915 |
| 17_875626 | 13 | 0.3077 | 16 | 0.25 | 13 | 0.3846 | 13 | 0.1538 | 55 | 0.2727 | 13 | 0.3846 | 8 | 0.5 | 11 | 0.1818 | 10 | 0.5 | 42 | 0.381 |
| 17_875629 | 7 | 0 | 8 | 0.125 | 10 | 0 | 11 | 0.1818 | 36 | 0.0833 | 9 | 0.2222 | 7 | 0.1429 | 7 | 0.1429 | 6 | 0 | 29 | 0.1379 |
| 17_875630 | 9 | 0.3333 | 16 | 0.1875 | 13 | 0.3846 | 13 | 0.6154 | 51 | 0.3725 | 9 | 0.3333 | 7 | 0.5714 | 9 | 0.4444 | 9 | 0.7778 | 34 | 0.5294 |
| 17_875631 | 5 | 0 | 15 | 0 | 11 | 0.1818 | 14 | 0.0714 | 45 | 0.0667 | 10 | 0.3 | 5 | 0.2 | 8 | 0 | 8 | 0.125 | 31 | 0.1613 |
| 17_875632 | 6 | 0 | 16 | 0.3125 | 13 | 0.2308 | 12 | 0.3333 | 47 | 0.2553 | 11 | 0.2727 | 8 | 0.25 | 9 | 0.2222 | 8 | 0.5 | 36 | 0.3056 |
| 17_875633 | 9 | 0.1111 | 15 | 0.0667 | 10 | 0.1 | 12 | 0.0833 | 46 | 0.087 | 11 | 0 | 9 | 0.2222 | 8 | 0.125 | 9 | 0.1111 | 37 | 0.1081 |
| 17_875888 | NA | NA | NA | NA | 1 | 1 | NA | NA | 1 | 1 | NA | NA | NA | NA | NA | NA | 1 | 1 | 1 | 1 |
| 17_875894 | NA | NA | NA | NA | 1 | 1 | NA | NA | 1 | 1 | NA | NA | NA | NA | NA | NA | 1 | 1 | 1 | 1 |
| 17_875899 | NA | NA | NA | NA | 1 | 1 | NA | NA | 1 | 1 | NA | NA | NA | NA | NA | NA | 1 | 1 | 1 | 1 |
| 17_876874 | 1 | 1 | 1 | 1 | NA | NA | NA | NA | 2 | 1 | 1 | 0 | 1 | 1 | 1 | 0 | 2 | 1 | 5 | 0.6 |
| 17_876877 | 1 | 1 | 2 | 1 | 1 | 1 | NA | NA | 4 | 1 | NA | NA | 3 | 0.6667 | 1 | 1 | 2 | 0.5 | 6 | 0.6667 |
| 17_876880 | 1 | 0 | 1 | 1 | 1 | 0 | NA | NA | 3 | 0.3333 | 1 | 0 | 4 | 0.5 | NA | NA | 2 | 0.5 | 7 | 0.4286 |

| | | | | | | | | | | | | | | | | | | | | |
|---|---|---|---|---|---|---|---|---|---|---|---|---|---|---|---|---|---|---|---|---|
| 17_8768805 | 1 | 1 | 1 | 1 | 1 | 1 | NA | NA | 3 | 1 | 1 | 1 | 4 | 0.25 | 1 | 0 | 1 | 0 | 7 | 0.2857 |
| 17_8768817 | 2 | 1 | 2 | 0.5 | 1 | 0 | NA | NA | 5 | 0.6 | 1 | 1 | 5 | 0.4 | 1 | 0 | 2 | 0 | 9 | 0.3333 |
| 17_8768882 | 2 | 0 | 2 | 0.5 | 1 | 0 | NA | NA | 5 | 0.2 | 1 | 0 | 5 | 0.4 | NA | NA | 2 | 0 | 8 | 0.25 |
| 17_8768915 | 3 | 0.3333 | 1 | 0 | 1 | 0 | NA | NA | 5 | 0.2 | 1 | 1 | 5 | 0 | NA | NA | 2 | 0.5 | 8 | 0.25 |
| 17_8774107 | 174 | 0.1034 | 163 | 0.0613 | 126 | 0.0556 | 156 | 0.1218 | 619 | 0.0872 | 121 | 0.1157 | 162 | 0.0864 | 110 | 0.1091 | 153 | 0.0784 | 546 | 0.0952 |
| 17_8774108 | 353 | 0.0878 | 383 | 0.0809 | 303 | 0.0957 | 340 | 0.1029 | 1379 | 0.0914 | 253 | 0.0988 | 334 | 0.0749 | 323 | 0.1053 | 328 | 0.1006 | 1238 | 0.0945 |
| 17_8777176 | NA | NA | NA | NA | NA | NA | 1 | 1 | 1 | 1 | NA | NA | 1 | 1 | NA | NA | 1 | 1 | 2 | 1 |
| 17_8777185 | NA | NA | NA | NA | NA | NA | 1 | 1 | 1 | 1 | NA | NA | NA | NA | NA | NA | 1 | 1 | 1 | 1 |
| 17_8777201 | NA | NA | NA | NA | NA | NA | 1 | 1 | 1 | 1 | NA | NA | 1 | 1 | NA | NA | 1 | 1 | 2 | 1 |
| 17_8777209 | NA | NA | NA | NA | NA | NA | 1 | 1 | 1 | 1 | NA | NA | 1 | 1 | NA | NA | NA | NA | 1 | 1 |
| 17_8802686 | NA | NA | 1 | 1 | 4 | 0.25 | 3 | 0.6667 | 8 | 0.5 | 2 | 0.5 | NA | NA | 1 | 0 | 3 | 0.3333 | 6 | 0.3333 |
| 17_8804944 | NA | NA | 2 | 1 | 3 | 1 | NA | NA | 5 | 1 | NA | NA | 2 | 1 | 1 | 1 | 2 | 1 | 5 | 1 |
| 17_8804949 | NA | NA | 3 | 1 | 3 | 1 | NA | NA | 6 | 1 | NA | NA | 2 | 1 | 1 | 1 | 1 | 1 | 4 | 1 |
| 17_8804975 | NA | NA | 3 | 0.3333 | 3 | 0 | NA | NA | 6 | 0.1667 | NA | NA | 2 | 0 | 1 | 0 | 1 | 1 | 4 | 0.25 |
| 17_8805233 | NA | NA | 1 | 0 | 2 | 0.5 | NA | NA | 3 | 0.3333 | NA | NA | 1 | 0 | NA | NA | 1 | 1 | 2 | 0.5 |
| 17_8805245 | NA | NA | 1 | 1 | 2 | 0.5 | NA | NA | 3 | 0.6667 | NA | NA | 1 | 1 | NA | NA | 1 | 1 | 2 | 1 |
| 17_8805257 | NA | NA | 1 | 1 | 2 | 0 | NA | NA | 3 | 0.3333 | NA | NA | 1 | 1 | NA | NA | 1 | 1 | 2 | 1 |
| 17_8805316 | NA | NA | 1 | 1 | 2 | 1 | NA | NA | 3 | 1 | NA | NA | 1 | 1 | NA | NA | 1 | 1 | 2 | 1 |
| 17_8805322 | NA | NA | 1 | 0 | 2 | 0.5 | NA | NA | 3 | 0.3333 | NA | NA | 1 | 1 | NA | NA | NA | NA | 1 | 1 |
| 17_8805334 | 2 | 0 | 3 | 0.3333 | 4 | 0.25 | NA | NA | 9 | 0.2222 | NA | NA | 2 | 0 | NA | NA | 2 | 0.5 | 4 | 0.25 |
| 17_8805387 | 2 | 0.5 | 3 | 0 | 2 | 0 | NA | NA | 7 | 0.1429 | NA | NA | 1 | 0 | NA | NA | 2 | 0.5 | 3 | 0.3333 |
| 17_8805392 | 2 | 1 | 3 | 1 | 4 | 0.75 | NA | NA | 9 | 0.8889 | NA | NA | 1 | 1 | NA | NA | 2 | 1 | 3 | 1 |
| 17_8805393 | 2 | 0.5 | 3 | 0.3333 | 3 | 0.3333 | NA | NA | 8 | 0.375 | NA | NA | 1 | 0 | NA | NA | 2 | 0.5 | 3 | 0.3333 |
| 17_8805421 | 2 | 0.5 | 3 | 0.3333 | 3 | 1 | NA | NA | 8 | 0.625 | NA | NA | 1 | 1 | NA | NA | 2 | 0.5 | 3 | 0.6667 |
| 17_8810453 | 2 | 0.5 | 3 | 0.3333 | 3 | 0.3333 | NA | NA | 8 | 0.375 | NA | NA | NA | NA | NA | NA | 2 | 0.5 | 2 | 0.5 |
| 17_8810520 | 1 | 1 | 2 | 1 | 1 | 1 | NA | NA | 4 | 1 | NA | NA | 1 | 1 | NA | NA | 1 | 1 | 2 | 1 |
| 17_8810527 | 1 | 1 | 1 | 0 | 2 | 0 | NA | NA | 4 | 0.25 | NA | NA | 1 | 1 | NA | NA | 1 | 0 | 2 | 0.5 |
| 17_8810538 | 1 | 0 | 2 | 0.5 | 1 | 1 | NA | NA | 4 | 0.5 | NA | NA | 1 | 0 | NA | NA | 1 | 1 | 2 | 0.5 |
| 17_8829217 | NA | NA | 2 | 1 | 2 | 1 | NA | NA | 4 | 1 | NA | NA | NA | NA | NA | NA | 1 | 1 | 1 | 1 |
| 17_8829234 | NA | NA | 2 | 0 | 1 | 1 | NA | NA | 3 | 0.3333 | NA | NA | NA | NA | NA | NA | 1 | 1 | 1 | 1 |
| 17_8829288 | 1 | 0 | 3 | 0.3333 | 2 | 1 | 1 | 1 | 7 | 0.5714 | NA | NA | NA | NA | NA | NA | 1 | 1 | 1 | 1 |
| 17_8829292 | 1 | 0 | 2 | 0.5 | 2 | 1 | 1 | 1 | 6 | 0.6667 | NA | NA | NA | NA | NA | NA | 1 | 1 | 1 | 1 |
| 17_8829305 | 1 | 0 | 3 | 0 | 2 | 0.5 | 1 | 1 | 7 | 0.2857 | NA | NA | NA | NA | NA | NA | 1 | 1 | 1 | 1 |
| 17_8829348 | 1 | 0 | 3 | 0.3333 | 3 | 0.3333 | 1 | 0 | 8 | 0.25 | NA | NA | NA | NA | 1 | 0 | 1 | 1 | 2 | 0.5 |
| 17_8829401 | 1 | 1 | 4 | 1 | 4 | 0.5 | 2 | 1 | 11 | 0.8182 | NA | NA | NA | NA | 1 | 0 | 1 | 1 | 2 | 0.5 |
| 17_8829403 | NA | NA | 4 | 0.25 | 4 | 0 | 2 | 0.5 | 10 | 0.2 | NA | NA | NA | NA | 1 | 1 | 1 | 0 | 2 | 0.5 |
| 17_8829425 | NA | NA | 4 | 0 | 4 | 0.25 | 2 | 0.5 | 10 | 0.2 | 1 | 1 | NA | NA | 1 | 1 | 1 | 1 | 3 | 1 |
| 17_8829432 | 1 | 1 | 4 | 0.75 | 4 | 1 | 1 | 1 | 10 | 0.9 | 1 | 1 | NA | NA | 1 | 1 | 1 | 1 | 3 | 1 |
| 17_8829574 | 2 | 0 | 4 | 0.25 | 3 | 0 | 2 | 0 | 11 | 0.0909 | 1 | 0 | NA | NA | 1 | 1 | 1 | 1 | 3 | 0.6667 |
| 17_8829910 | 9 | 1 | 19 | 1 | 16 | 1 | 23 | 1 | 67 | 1 | 5 | 1 | 8 | 1 | 19 | 0.8421 | 15 | 1 | 47 | 0.9362 |
| 17_8829924 | 5 | 0.4 | 16 | 0.5625 | 15 | 0.6667 | 21 | 0.5714 | 57 | 0.5789 | 3 | 0.6667 | 13 | 0.5385 | 18 | 0.6667 | 12 | 0.75 | 46 | 0.6522 |
| 17_8829925 | 9 | 0.8889 | 16 | 1 | 16 | 0.9375 | 23 | 1 | 64 | 0.9688 | 6 | 1 | 10 | 0.8 | 16 | 1 | 14 | 1 | 46 | 0.9565 |
| 17_8829930 | 9 | 0.2222 | 16 | 0.5625 | 12 | 0.4167 | 22 | 0.2273 | 59 | 0.3559 | 2 | 0 | 11 | 0.0909 | 20 | 0.15 | 13 | 0.3077 | 46 | 0.1739 |
| 17_8829934 | 8 | 0.375 | 20 | 0.05 | 15 | 0 | 25 | 0.04 | 68 | 0.0735 | 6 | 0 | 12 | 0.0833 | 19 | 0.0526 | 16 | 0.0625 | 53 | 0.0566 |
| 17_8829936 | 7 | 0.8571 | 17 | 0.9412 | 15 | 0.8667 | 23 | 0.913 | 62 | 0.9032 | 4 | 0.75 | 8 | 1 | 17 | 0.9412 | 15 | 0.8667 | 44 | 0.9091 |
| 17_8829938 | 9 | 0.7778 | 19 | 0.9474 | 16 | 1 | 23 | 0.9565 | 67 | 0.9403 | 5 | 1 | 10 | 1 | 18 | 1 | 14 | 1 | 47 | 1 |
| 17_8829943 | 8 | 0.25 | 19 | 0.0526 | 15 | 0.2 | 22 | 0.1364 | 64 | 0.1406 | 6 | 0 | 11 | 0.0909 | 19 | 0.1579 | 14 | 0.1429 | 50 | 0.12 |
| 17_8829951 | 6 | 0.1667 | 15 | 0.2667 | 12 | 0.25 | 20 | 0.05 | 53 | 0.1698 | 2 | 0 | 11 | 0.1818 | 17 | 0.3529 | 15 | 0.2667 | 45 | 0.2667 |
| 17_8829952 | 8 | 0.25 | 17 | 0.1176 | 13 | 0 | 21 | 0.0952 | 59 | 0.1017 | 5 | 0 | 11 | 0 | 17 | 0.1176 | 15 | 0.1333 | 48 | 0.0833 |
| 17_8837267 | 4 | 0.25 | 5 | 0 | 3 | 0 | 12 | 0.1667 | 24 | 0.125 | 1 | 1 | 5 | 0.2 | 6 | 0.3333 | 2 | 0 | 14 | 0.2857 |
| 17_8879417 | NA | NA | 1 | 1 | NA | NA | 1 | 1 | 2 | 1 | 2 | 0.5 | 2 | 1 | 1 | 1 | 1 | 1 | 6 | 0.8333 |
| 17_8879445 | NA | NA | 1 | 1 | NA | NA | 1 | 1 | 2 | 1 | 2 | 0.5 | 2 | 0 | 1 | 0 | 1 | 1 | 6 | 0.3333 |
| 17_8879472 | NA | NA | 1 | 0 | NA | NA | 2 | 1 | 3 | 0.6667 | 2 | 1 | 2 | 0.5 | 1 | 0 | 1 | 1 | 6 | 0.6667 |
| 17_8879779 | 1 | 0 | 3 | 0 | 2 | 0 | 3 | 0.3333 | 9 | 0.1111 | 2 | 0 | 2 | 0 | 3 | 0.3333 | 4 | 0 | 11 | 0.0909 |
| 17_8885280 | 4 | 0 | 4 | 0.25 | 2 | 0 | 7 | 0.1429 | 17 | 0.1176 | 4 | 0.25 | 8 | 0 | 6 | 0.1667 | 4 | 0 | 22 | 0.0909 |
| 17_8889516 | 8 | 0 | 7 | 0.1429 | 3 | 0.3333 | 8 | 0.375 | 26 | 0.1923 | 5 | 0 | 8 | 0.375 | 7 | 0 | 6 | 0 | 26 | 0.1154 |
| 17_8889524 | 9 | 0.4444 | 8 | 0.25 | 3 | 1 | 8 | 0.25 | 28 | 0.3929 | 5 | 0.2 | 8 | 0.5 | 7 | 0.5714 | 7 | 0.4286 | 27 | 0.4444 |
| 17_8889536 | 8 | 0.375 | 6 | 0.8333 | 1 | 0 | 9 | 0.7778 | 24 | 0.625 | 8 | 0.25 | 9 | 0.4444 | 7 | 0.5714 | 6 | 0.1667 | 30 | 0.3667 |
| 17_9034150 | 27 | 0.2222 | 39 | 0.1282 | 28 | 0.2143 | 21 | 0.1905 | 115 | 0.1826 | 25 | 0.32 | 22 | 0.0909 | 22 | 0.2273 | 28 | 0.2143 | 97 | 0.2165 |
| 17_9034176 | 26 | 0.0385 | 41 | 0.0976 | 32 | 0.0625 | 19 | 0.2632 | 118 | 0.1017 | 26 | 0.3077 | 19 | 0 | 25 | 0.2 | 27 | 0.1481 | 97 | 0.1753 |
| 17_9034362 | 13 | 0.5385 | 21 | 0.5238 | 18 | 0.6111 | 10 | 0.4 | 62 | 0.5323 | 11 | 0.5455 | 9 | 0.6667 | 11 | 0.4545 | 13 | 0.4615 | 44 | 0.5227 |
| 17_9034385 | 13 | 0.1538 | 24 | 0.0833 | 19 | 0.0526 | 10 | 0.1 | 66 | 0.0909 | 11 | 0.0909 | 9 | 0 | 9 | 0.1111 | 12 | 0.0833 | 41 | 0.0732 |
| 17_9034393 | 13 | 0.6154 | 18 | 0.6111 | 16 | 0.75 | 11 | 0.7273 | 58 | 0.6724 | 9 | 0.4444 | 6 | 0.1667 | 8 | 0.125 | 12 | 0.6667 | 35 | 0.4 |
| 17_9034398 | 12 | 0.5833 | 19 | 0.3158 | 14 | 0.2857 | 11 | 0.3636 | 56 | 0.375 | 10 | 0.5 | 8 | 0.375 | 9 | 0.5556 | 12 | 0.3333 | 39 | 0.4359 |
| 17_9034415 | 11 | 0.2727 | 20 | 0.1 | 17 | 0.1176 | 10 | 0.2 | 58 | 0.1552 | 11 | 0.4545 | 9 | 0.2222 | 8 | 0.375 | 10 | 0.2 | 38 | 0.3158 |

| | | | | | | | | | | | | | | | | | | | | |
|---|---|---|---|---|---|---|---|---|---|---|---|---|---|---|---|---|---|---|---|---|
| 17_9034424 | 7 | 0.1429 | 15 | 0.0667 | 12 | 0.0833 | 7 | 0 | 41 | 0.0732 | 6 | 0 | 5 | 0.2 | 6 | 0 | 8 | 0.125 | 25 | 0.08 |
| 17_9034431 | 4 | 0.5 | 7 | 0.2857 | 7 | 0.1429 | 3 | 0.3333 | 21 | 0.2857 | 5 | 0.2 | 3 | 0 | 2 | 0 | 3 | 0.3333 | 13 | 0.1538 |
| 17_9034431 | 1 | 0 | 7 | 0.2857 | 4 | 0.25 | 6 | 0.1667 | 18 | 0.2222 | 4 | 0.25 | 2 | 0 | 3 | 0.3333 | NA | NA | 9 | 0.2222 |
| 17_9034435 | 3 | 0.3333 | 3 | 0 | 6 | 0.1667 | 4 | 0 | 16 | 0.125 | 4 | 0.25 | 2 | 0 | 2 | 0 | 3 | 0.3333 | 11 | 0.1818 |
| 17_9034437 | 8 | 0.375 | 20 | 0.4 | 16 | 0.1875 | 8 | 0.625 | 52 | 0.3654 | 9 | 0.4444 | 4 | 0.5 | 6 | 0.1667 | 9 | 0.5556 | 28 | 0.4286 |
| 17_9034437 | NA | NA | 1 | 1 | NA | NA | NA | NA | 1 | 1 | NA | NA | NA | NA | NA | NA | 2 | 0.5 | 2 | 0.5 |
| 17_9034444 | 5 | 0.8 | 13 | 1 | 11 | 0.9091 | 8 | 0.75 | 37 | 0.8919 | 9 | 0.8889 | 5 | 1 | 6 | 1 | 4 | 1 | 24 | 0.9583 |
| 17_9034445 | 9 | 0.6667 | 18 | 0.3889 | 13 | 0.3077 | 11 | 0.3636 | 51 | 0.4118 | 8 | 0.5 | 6 | 0.3333 | 6 | 0.5 | 7 | 0.4286 | 27 | 0.4444 |
| 17_9034448 | 9 | 1 | 20 | 1 | 14 | 1 | 10 | 1 | 53 | 1 | 10 | 1 | 6 | 1 | 8 | 1 | 11 | 0.9091 | 35 | 0.9714 |
| 17_9076223 | 2 | 0 | 2 | 0.5 | NA | NA | 4 | 0.25 | 8 | 0.25 | 2 | 0.5 | 2 | 0.5 | 1 | 0 | NA | NA | 5 | 0.4 |
| 17_9076252 | 2 | 0.5 | 1 | 1 | NA | NA | 2 | 0.5 | 5 | 0.6 | 2 | 0.5 | 2 | 0.5 | NA | NA | NA | NA | 4 | 0.5 |
| 17_9076263 | NA | NA | 2 | 1 | NA | NA | 3 | 0.3333 | 5 | 0.6 | 1 | 0 | 1 | 1 | NA | NA | NA | NA | 2 | 0.5 |
| 17_9076264 | 2 | 0 | 3 | 0.6667 | NA | NA | 4 | 0.5 | 9 | 0.4444 | 2 | 0.5 | 2 | 0.5 | NA | NA | NA | NA | 4 | 0.5 |
| 17_9076265 | 2 | 1 | 1 | 0 | NA | NA | 3 | 0 | 6 | 0.3333 | 2 | 0 | 2 | 0.5 | NA | NA | NA | NA | 4 | 0.25 |
| 17_9076272 | 1 | 0 | 2 | 0 | NA | NA | 2 | 0.5 | 5 | 0.2 | 2 | 1 | 2 | 0.5 | NA | NA | NA | NA | 4 | 0.75 |
| 17_9076288 | 1 | 0 | 3 | 0 | NA | NA | 2 | 0.5 | 6 | 0.1667 | 1 | 0 | 1 | 1 | NA | NA | NA | NA | 2 | 0.5 |
| 17_9076303 | NA | NA | 2 | 0 | NA | NA | 2 | 1 | 4 | 0.5 | 1 | 1 | 1 | 0 | NA | NA | NA | NA | 2 | 0.5 |
| 17_9076313 | NA | NA | 1 | 0 | NA | NA | 2 | 0.5 | 3 | 0.3333 | 1 | 1 | NA | NA | NA | NA | NA | NA | 1 | 1 |
| 17_9089747 | NA | NA | 4 | 0.5 | 5 | 0.8 | 2 | 0 | 11 | 0.5455 | 3 | 1 | 1 | 1 | 3 | 1 | 1 | 0 | 8 | 0.875 |
| 17_9089751 | NA | NA | 4 | 1 | 6 | 0.6667 | 2 | 0 | 12 | 0.6667 | 3 | 0.6667 | 2 | 0.5 | 3 | 1 | 4 | 0.25 | 12 | 0.5833 |
| 17_9093751 | NA | NA | 4 | 0.25 | 5 | 0 | 2 | 0 | 11 | 0.0909 | 3 | 0 | 2 | 0 | 3 | 0.3333 | 3 | 0.3333 | 11 | 0.1818 |
| 17_9093755 | NA | NA | 3 | 0.6667 | 5 | 0.2 | 2 | 0 | 10 | 0.3 | 3 | 0.3333 | 2 | 0 | 3 | 0.3333 | 3 | 1 | 11 | 0.4545 |
| 17_9095056 | NA | NA | 3 | 0.3333 | 5 | 0.8 | 2 | 0 | 10 | 0.5 | 3 | 0.6667 | 2 | 0.5 | 2 | 0.5 | 3 | 0 | 10 | 0.4 |
| 17_9095072 | NA | NA | 1 | 0 | 5 | 0.2 | 2 | 0 | 8 | 0.125 | 3 | 0 | 2 | 0 | 2 | 0 | 3 | 0.3333 | 10 | 0.1 |
| 17_9095073 | NA | NA | 1 | 0 | 4 | 0.25 | 2 | 0.5 | 7 | 0.2857 | 2 | 1 | 2 | 0 | 1 | 0 | 2 | 0 | 7 | 0.2857 |
| 17_9101170 | NA | NA | 2 | 0.5 | 3 | 0 | NA | NA | 5 | 0.2 | 3 | 0.3333 | NA | NA | 1 | 0 | 1 | 0 | 5 | 0.2 |
| 17_9154445 | NA | NA | NA | NA | 2 | 1 | 1 | 1 | 3 | 1 | 1 | 1 | NA | NA | NA | NA | NA | NA | 1 | 1 |
| 17_9154459 | NA | NA | NA | NA | 2 | 1 | NA | NA | 2 | 1 | 1 | 1 | 1 | 0 | NA | NA | 1 | 0 | 3 | 0.3333 |
| 17_9154471 | NA | NA | NA | NA | 4 | 0.25 | NA | NA | 4 | 0.25 | 1 | 0 | NA | NA | NA | NA | 1 | 1 | 2 | 0.5 |
| 17_9154478 | NA | NA | NA | NA | 4 | 1 | 1 | 1 | 5 | 1 | 1 | 1 | 1 | 1 | NA | NA | 1 | 1 | 3 | 1 |
| 17_9154484 | NA | NA | NA | NA | 4 | 1 | 2 | 1 | 6 | 1 | 1 | 1 | 1 | 1 | NA | NA | 1 | 1 | 3 | 1 |
| 17_9154500 | NA | NA | NA | NA | 3 | 0.6667 | 2 | 1 | 5 | 0.8 | 1 | 1 | 1 | 1 | NA | NA | 1 | 0 | 3 | 0.6667 |
| 17_9154532 | NA | NA | 1 | 1 | 4 | 0.25 | 3 | 1 | 8 | 0.625 | 1 | 1 | 1 | 0 | 1 | 1 | 1 | 1 | 4 | 0.75 |
| 17_9503033 | 4 | 0.25 | NA | NA | 2 | 0 | NA | NA | 6 | 0.1667 | 1 | 0 | NA | NA | 5 | 0.2 | 2 | 0 | 8 | 0.125 |
| 17_9503037 | 2 | 0 | NA | NA | 2 | 0.5 | NA | NA | 4 | 0.25 | 1 | 0 | NA | NA | 4 | 0 | 3 | 1 | 8 | 0.375 |
| 17_9503079 | 2 | 0.5 | NA | NA | 2 | 0.5 | NA | NA | 4 | 0.5 | NA | NA | NA | NA | 5 | 0.4 | 1 | 1 | 6 | 0.5 |
| 17_9503092 | 2 | 0.5 | NA | NA | 2 | 0.5 | NA | NA | 4 | 0.5 | NA | NA | NA | NA | 5 | 0.4 | 1 | 0 | 6 | 0.3333 |
| 17_9503459 | 2 | 1 | NA | NA | 1 | 1 | NA | NA | 3 | 1 | 1 | 1 | NA | NA | 2 | 1 | NA | NA | 3 | 1 |
| 17_9503467 | 2 | 0 | 1 | 0 | 2 | 0.5 | NA | NA | 5 | 0.2 | NA | NA | NA | NA | 2 | 0.5 | NA | NA | 2 | 0.5 |
| 17_9505555 | 1 | 1 | NA | NA | 2 | 0.5 | NA | NA | 3 | 0.6667 | NA | NA | NA | NA | 1 | 1 | NA | NA | 1 | 1 |
| 17_9505556 | 1 | 1 | NA | NA | 2 | 1 | NA | NA | 3 | 1 | NA | NA | NA | NA | 1 | 1 | NA | NA | 1 | 1 |
| 17_9505579 | 1 | 1 | NA | NA | 2 | 0.5 | NA | NA | 3 | 0.6667 | NA | NA | NA | NA | 1 | 1 | NA | NA | 1 | 1 |
| 17_9506422 | 1 | 1 | 1 | 1 | 1 | 1 | NA | NA | 3 | 1 | 1 | 1 | NA | NA | 1 | 1 | NA | NA | 2 | 1 |
| 17_9506544 | 1 | 1 | NA | NA | 1 | 1 | NA | NA | 2 | 1 | 1 | 0 | NA | NA | 1 | 1 | NA | NA | 2 | 0.5 |
| 17_9506563 | 1 | 1 | NA | NA | 1 | 1 | NA | NA | 2 | 1 | 1 | 0 | NA | NA | 1 | 1 | NA | NA | 2 | 0.5 |
| 17_9506755 | 4 | 0.25 | 3 | 0.3333 | 5 | 0.2 | 2 | 0 | 14 | 0.2143 | 3 | 0.3333 | 4 | 0 | 1 | 1 | 1 | 0 | 9 | 0.2222 |
| 17_9506760 | 4 | 0.75 | 3 | 1 | 5 | 0.4 | 3 | 0.6667 | 15 | 0.6667 | 4 | 0.25 | 6 | 0.3333 | 1 | 1 | 1 | 0 | 12 | 0.3333 |
| 17_9506806 | 4 | 0.75 | 3 | 0 | NA | NA | 3 | 0.3333 | 10 | 0.4 | 5 | 0 | 5 | 0.2 | 1 | 0 | 1 | 0 | 12 | 0.0833 |
| 17_9506814 | 6 | 0.3333 | 3 | 0.3333 | 4 | 0.5 | 3 | 0.3333 | 16 | 0.375 | 4 | 0.5 | 4 | 0.5 | 1 | 0 | 1 | 1 | 10 | 0.5 |
| 17_9506894 | 5 | 0.6 | 2 | 0 | 4 | 0.75 | 3 | 0.6667 | 14 | 0.5714 | 4 | 0.25 | 3 | 0.3333 | 2 | 1 | 1 | 1 | 10 | 0.5 |
| 17_9512220 | 3 | 0 | NA | NA | 1 | 1 | 4 | 0 | 8 | 0.125 | NA | NA | 2 | 0 | 3 | 0.3333 | 2 | 0 | 7 | 0.1429 |
| 17_9512254 | 3 | 0.3333 | NA | NA | NA | NA | 3 | 0.6667 | 6 | 0.5 | NA | NA | 1 | 0 | 1 | 0 | 1 | 1 | 3 | 0.3333 |
| 17_9512286 | 2 | 0.5 | NA | NA | NA | NA | 3 | 0 | 5 | 0.2 | NA | NA | 1 | 1 | 2 | 0.5 | NA | NA | 3 | 0.6667 |
| 17_9512291 | 1 | 1 | NA | NA | NA | NA | 3 | 0.3333 | 4 | 0.5 | NA | NA | NA | NA | 2 | 0.5 | 2 | 0 | 4 | 0.25 |
| 17_9512320 | 1 | 1 | NA | NA | NA | NA | 2 | 0.5 | 3 | 0.6667 | NA | NA | 1 | 1 | 2 | 0.5 | 1 | 0 | 4 | 0.5 |
| 17_9512321 | 1 | 0 | NA | NA | NA | NA | 2 | 0.5 | 3 | 0.3333 | NA | NA | 1 | 0 | 2 | 0.5 | 1 | 1 | 4 | 0.5 |
| 17_9512324 | 1 | 1 | NA | NA | NA | NA | 3 | 0 | 4 | 0.25 | NA | NA | 1 | 0 | 2 | 0.5 | 1 | 0 | 4 | 0.25 |
| 17_9512325 | 1 | 0 | NA | NA | NA | NA | 2 | 0.5 | 3 | 0.3333 | NA | NA | 1 | 1 | 2 | 0 | 1 | 1 | 4 | 0.5 |
| 17_9512340 | 1 | 0 | NA | NA | NA | NA | 3 | 0.3333 | 4 | 0.25 | NA | NA | NA | NA | 2 | 0.5 | NA | NA | 2 | 0.5 |
| 17_9512365 | 1 | 1 | NA | NA | NA | NA | 3 | 1 | 4 | 1 | NA | NA | 1 | 1 | 1 | 1 | NA | NA | 2 | 1 |
| 17_9512374 | 1 | 1 | NA | NA | NA | NA | 3 | 1 | 4 | 1 | NA | NA | 1 | 1 | 2 | 1 | NA | NA | 3 | 1 |
| 17_9512376 | 1 | 0 | NA | NA | NA | NA | 3 | 0.3333 | 4 | 0.25 | NA | NA | 1 | 0 | 2 | 0.5 | NA | NA | 3 | 0.3333 |
| 17_9512379 | 1 | 1 | NA | NA | NA | NA | 3 | 1 | 4 | 1 | NA | NA | 1 | 1 | 2 | 1 | NA | NA | 3 | 1 |
| 17_9512409 | 1 | 0 | NA | NA | NA | NA | 3 | 1 | 4 | 0.75 | NA | NA | NA | NA | 2 | 0.5 | NA | NA | 2 | 0.5 |
| 17_9512414 | 1 | 0 | NA | NA | NA | NA | 1 | 1 | 2 | 0.5 | NA | NA | 1 | 1 | 2 | 0.5 | NA | NA | 3 | 0.6667 |

| | | | | | | | | | | | | | | | | | | | | |
|---|---|---|---|---|---|---|---|---|---|---|---|---|---|---|---|---|---|---|---|---|
| 17_951243 | 1 | 0 | NA | NA | NA | NA | 2 | 0.5 | 3 | 0.3333 | NA | NA | 1 | 0 | 2 | 0.5 | NA | NA | 3 | 0.3333 |
| 17_951243 | 1 | 1 | NA | NA | NA | NA | 2 | 1 | 3 | 1 | NA | NA | 1 | 0 | 2 | 0.5 | NA | NA | 3 | 0.3333 |
| 17_951247 | 1 | 1 | NA | NA | NA | NA | 3 | 0.6667 | 4 | 0.75 | NA | NA | 1 | 0 | 2 | 0.5 | NA | NA | 3 | 0.3333 |
| 17_951248 | NA | NA | NA | NA | NA | NA | 3 | 0.6667 | 3 | 0.6667 | NA | NA | 1 | 1 | 2 | 0.5 | NA | NA | 3 | 0.6667 |
| 17_951831 | NA | NA | 1 | 1 | NA | NA | NA | NA | 1 | 1 | NA | NA | NA | NA | NA | NA | 1 | 1 | 1 | 1 |
| 17_951840 | NA | NA | 1 | 0 | NA | NA | 1 | 1 | 2 | 0.5 | NA | NA | NA | NA | NA | NA | 1 | 1 | 1 | 1 |
| 17_951847 | NA | NA | 1 | 1 | NA | NA | 1 | 0 | 2 | 0.5 | NA | NA | NA | NA | NA | NA | 2 | 0.5 | 2 | 0.5 |
| 17_951850 | NA | NA | 1 | 1 | NA | NA | 1 | 1 | 2 | 1 | NA | NA | NA | NA | NA | NA | 2 | 0.5 | 2 | 0.5 |
| 17_951853 | NA | NA | 1 | 0 | NA | NA | 1 | 1 | 2 | 0.5 | NA | NA | NA | NA | NA | NA | 2 | 1 | 2 | 1 |
| 17_951855 | NA | NA | 1 | 1 | NA | NA | 1 | 1 | 2 | 1 | NA | NA | NA | NA | NA | NA | 2 | 1 | 2 | 1 |
| 17_951856 | NA | NA | 1 | 0 | NA | NA | 1 | 1 | 2 | 0.5 | NA | NA | NA | NA | NA | NA | 2 | 0.5 | 2 | 0.5 |
| 17_951856 | NA | NA | NA | NA | NA | NA | 1 | 1 | 1 | 1 | NA | NA | NA | NA | NA | NA | 2 | 1 | 2 | 1 |
| 17_951856 | NA | NA | NA | NA | NA | NA | 1 | 1 | 1 | 1 | NA | NA | NA | NA | NA | NA | 2 | 0.5 | 2 | 0.5 |
| 17_951868 | NA | NA | 1 | 0 | NA | NA | 1 | 1 | 2 | 0.5 | NA | NA | NA | NA | NA | NA | 1 | 1 | 1 | 1 |
| 18_325453 | NA | NA | 1 | 1 | NA | NA | 1 | 0 | 2 | 0.5 | NA | NA | NA | NA | 1 | 1 | NA | NA | 1 | 1 |
| 18_326756 | NA | NA | 3 | 0.3333 | 1 | 1 | 2 | 0.5 | 6 | 0.5 | 2 | 1 | 2 | 0.5 | 2 | 0.5 | 5 | 0.6 | 11 | 0.6364 |
| 18_326761 | 1 | 1 | 6 | 0.3333 | 5 | 0.6 | 2 | 0 | 14 | 0.4286 | 2 | 0.5 | 3 | 0.6667 | 3 | 0.6667 | 3 | 0.3333 | 11 | 0.5455 |
| 18_326790 | NA | NA | 5 | 1 | 4 | 1 | 2 | 1 | 11 | 1 | 2 | 1 | 1 | 1 | 1 | 1 | 4 | 0.75 | 8 | 0.875 |
| 18_326795 | NA | NA | 5 | 0.6 | 4 | 0.75 | 2 | 0 | 11 | 0.5455 | 2 | 1 | 2 | 0.5 | 1 | 1 | 2 | 1 | 7 | 0.8571 |
| 18_326796 | NA | NA | 5 | 0.6 | 3 | 0.6667 | 2 | 0 | 10 | 0.5 | 1 | 1 | 1 | 1 | 2 | 0 | 4 | 0.5 | 8 | 0.5 |
| 18_326796 | NA | NA | 4 | 0.25 | 4 | 1 | 2 | 0.5 | 10 | 0.6 | 2 | 0.5 | 1 | 1 | 2 | 1 | 4 | 0.5 | 9 | 0.6667 |
| 18_326798 | NA | NA | 4 | 0.25 | 3 | 0.3333 | 2 | 0 | 9 | 0.2222 | 1 | 0 | 2 | 1 | 1 | 1 | 4 | 0.75 | 8 | 0.75 |
| 18_326801 | NA | NA | 5 | 0.4 | 4 | 0.5 | 2 | 0.5 | 11 | 0.4545 | 2 | 0.5 | 2 | 0.5 | 2 | 0 | 4 | 0.5 | 10 | 0.4 |
| 18_343491 | 9 | 0.1111 | 6 | 0.1667 | 5 | 0 | 12 | 0 | 32 | 0.0625 | 4 | 0 | 9 | 0 | 12 | 0.3333 | 11 | 0.0909 | 36 | 0.1389 |
| 18_343518 | 11 | 0.3636 | 6 | 0.5 | 3 | 1 | 11 | 0.4545 | 31 | 0.4839 | 4 | 0.75 | 9 | 0.4444 | 9 | 0.7778 | 12 | 0.75 | 34 | 0.6765 |
| 18_351205 | 2 | 1 | 2 | 1 | 1 | 1 | 1 | 1 | 6 | 1 | 1 | 1 | NA | NA | NA | NA | NA | NA | 1 | 1 |
| 18_351210 | 2 | 1 | 3 | 1 | 1 | 1 | 1 | 1 | 7 | 1 | 1 | 1 | NA | NA | 1 | 1 | NA | NA | 2 | 1 |
| 18_351221 | 3 | 1 | 3 | 0.6667 | 1 | 1 | NA | NA | 7 | 0.8571 | NA | NA | NA | NA | 1 | 1 | NA | NA | 1 | 1 |
| 18_351221 | NA | NA | NA | NA | NA | NA | 1 | 1 | 1 | 1 | 1 | 1 | NA | NA | NA | NA | NA | NA | 1 | 1 |
| 18_351225 | 2 | 1 | 3 | 1 | NA | NA | 1 | 1 | 6 | 1 | 1 | 1 | NA | NA | 1 | 1 | NA | NA | 2 | 1 |
| 18_351228 | 3 | 1 | 3 | 0.6667 | 1 | 1 | 1 | 0 | 8 | 0.75 | 1 | 1 | NA | NA | 1 | 1 | NA | NA | 2 | 1 |
| 18_351240 | 3 | 0.3333 | 3 | 0 | 1 | 0 | NA | NA | 7 | 0.1429 | NA | NA | NA | NA | 1 | 1 | NA | NA | 1 | 1 |
| 18_351241 | 3 | 1 | 3 | 1 | 1 | 1 | 1 | 1 | 8 | 1 | 1 | 1 | NA | NA | 1 | 1 | NA | NA | 2 | 1 |
| 18_351248 | 3 | 0 | 3 | 0 | 1 | 0 | 1 | 1 | 8 | 0.125 | 1 | 1 | NA | NA | 1 | 0 | NA | NA | 2 | 0.5 |
| 18_351251 | 3 | 0.6667 | 3 | 0 | 1 | 0 | 1 | 0 | 8 | 0.25 | 1 | 1 | NA | NA | 1 | 1 | NA | NA | 2 | 1 |
| 18_351261 | 2 | 1 | 3 | 1 | 1 | 1 | 1 | 1 | 7 | 1 | 2 | 1 | NA | NA | 1 | 1 | NA | NA | 3 | 1 |
| 18_351266 | 3 | 0.3333 | 3 | 0.6667 | 1 | 1 | NA | NA | 7 | 0.5714 | 2 | 0.5 | NA | NA | 1 | 1 | NA | NA | 3 | 0.6667 |
| 18_438336 | 4 | 0 | 4 | 0 | 3 | 0.3333 | 2 | 0 | 13 | 0.0769 | 3 | 0.3333 | 3 | 0 | 4 | 0 | 5 | 0 | 15 | 0.0667 |
| 18_438709 | 6 | 0.1667 | 4 | 0 | 2 | 0 | 3 | 0 | 15 | 0.0667 | 4 | 0 | 4 | 0 | 3 | 0 | 8 | 0.125 | 19 | 0.0526 |
| 18_439621 | 6 | 0.3333 | 5 | 0.6 | 3 | 0.6667 | 5 | 0.2 | 19 | 0.4211 | 5 | 0 | 6 | 0 | 5 | 0.2 | 7 | 0.4286 | 23 | 0.1739 |
| 18_439622 | 6 | 0.6667 | 5 | 0.8 | 2 | 1 | 5 | 0.8 | 18 | 0.7778 | 4 | 1 | 5 | 0.8 | 3 | 1 | 5 | 1 | 17 | 0.9412 |
| 18_439629 | 6 | 0.5 | 4 | 0.5 | 3 | 0.6667 | 5 | 0.4 | 18 | 0.5 | 3 | 1 | 6 | 0.5 | 4 | 0.5 | 7 | 1 | 20 | 0.75 |
| 18_439641 | 6 | 0 | 5 | 0 | 1 | 1 | 3 | 0.3333 | 15 | 0.1333 | 4 | 0 | 5 | 0.2 | 5 | 0 | 7 | 0.2857 | 21 | 0.1429 |
| 18_439644 | 5 | 0.6 | 4 | 0.5 | 3 | 1 | 4 | 0.75 | 16 | 0.6875 | 5 | 1 | 4 | 0.5 | 3 | 1 | 7 | 0.8571 | 19 | 0.8421 |
| 18_439648 | 6 | 1 | 5 | 0.8 | 2 | 1 | 6 | 1 | 19 | 0.9474 | 4 | 1 | 4 | 1 | 6 | 0.8333 | 5 | 1 | 19 | 0.9474 |
| 18_439649 | 6 | 0.6667 | 3 | 0.6667 | 3 | 0.6667 | 6 | 0.8333 | 18 | 0.7222 | 3 | 1 | 6 | 1 | 5 | 1 | 6 | 1 | 20 | 1 |
| 18_439652 | 6 | 0.3333 | 5 | 0.2 | 2 | 0 | 3 | 0.3333 | 16 | 0.25 | 4 | 0.25 | 4 | 0.25 | 3 | 0 | 6 | 0 | 17 | 0.1176 |
| 18_439664 | 7 | 1 | 2 | 1 | 6 | 0.6667 | 7 | 1 | 22 | 0.9091 | 4 | 1 | 6 | 1 | 5 | 1 | 8 | 0.875 | 23 | 0.9565 |
| 18_464931 | 3 | 0 | 2 | 0 | 1 | 1 | 2 | 0 | 8 | 0.125 | NA | NA | 2 | 0.5 | 5 | 0.2 | 3 | 0 | 10 | 0.2 |
| 18_467262 | 2 | 0.5 | 2 | 0 | 1 | 0 | 2 | 1 | 7 | 0.4286 | NA | NA | 3 | 0.6667 | 5 | 0.2 | 4 | 0 | 12 | 0.25 |
| 18_467263 | 3 | 1 | 2 | 1 | 1 | 1 | 3 | 1 | 9 | 1 | NA | NA | 2 | 0.5 | 1 | 1 | 4 | 0.25 | 7 | 0.4286 |
| 18_467271 | 2 | 0.5 | 2 | 0 | 1 | 0 | 2 | 0 | 7 | 0.1429 | NA | NA | 4 | 0 | 5 | 0.2 | 4 | 0 | 13 | 0.0769 |
| 18_467276 | 4 | 0 | 2 | 0 | 1 | 0 | 3 | 0.3333 | 10 | 0.1 | 1 | 0 | 4 | 0.5 | 7 | 0.5714 | 4 | 0.25 | 16 | 0.4375 |
| 18_467304 | 4 | 0 | 2 | 0 | NA | NA | 4 | 0.25 | 10 | 0.1 | 1 | 0 | 5 | 0 | 8 | 0.125 | 4 | 0 | 18 | 0.0556 |
| 18_467305 | 4 | 0.25 | 2 | 0 | 1 | 0 | 5 | 0 | 12 | 0.0833 | 1 | 0 | 5 | 0 | 8 | 0.125 | 4 | 0.25 | 18 | 0.1111 |
| 18_467311 | 3 | 1 | 2 | 0.5 | 1 | 1 | 5 | 0.4 | 11 | 0.6364 | 1 | 0 | 4 | 0 | 6 | 0.3333 | 3 | 0.3333 | 14 | 0.2143 |
| 18_467318 | 3 | 0.3333 | 3 | 0.6667 | 1 | 1 | 5 | 0.2 | 12 | 0.4167 | 1 | 0 | 4 | 0.5 | 9 | 0.1111 | 4 | 0.5 | 18 | 0.2778 |
| 18_467338 | 4 | 0 | 2 | 0 | 2 | 0.5 | 6 | 0 | 14 | 0.0714 | 1 | 0 | 7 | 0.1429 | 9 | 0.1111 | 4 | 0 | 21 | 0.0952 |
| 18_467365 | 3 | 0.6667 | 2 | 0.5 | 1 | 1 | 5 | 0.2 | 11 | 0.4545 | 1 | 0 | 4 | 1 | 9 | 0.3333 | 4 | 0.75 | 18 | 0.5556 |
| 18_467377 | 3 | 0.3333 | 2 | 0.5 | 2 | 0 | 4 | 0 | 11 | 0.1818 | 1 | 1 | 7 | 0.5714 | 8 | 0.125 | 5 | 0 | 21 | 0.2857 |
| 18_467384 | 4 | 0 | 1 | 0 | 2 | 0 | 6 | 0.1667 | 13 | 0.0769 | 1 | 1 | 6 | 0 | 7 | 0 | 4 | 0 | 18 | 0.0556 |
| 18_467392 | 5 | 0.2 | 2 | 0 | 2 | 0.5 | 7 | 0.1429 | 16 | 0.1875 | 1 | 1 | 6 | 0 | 10 | 0.1 | 5 | 0 | 22 | 0.0909 |
| 18_467404 | 3 | 0.6667 | 3 | 0.6667 | 1 | 1 | 4 | 0.5 | 11 | 0.6364 | 2 | 0.5 | 6 | 1 | 8 | 1 | 5 | 0.4 | 21 | 0.8095 |
| 18_467453 | 6 | 0.8333 | 2 | 1 | 2 | 1 | 6 | 0.8333 | 16 | 0.875 | 2 | 0.5 | 10 | 0.9 | 13 | 0.8462 | 6 | 0.6667 | 31 | 0.8065 |
| 18_467457 | 4 | 0 | 2 | 0 | 2 | 0.5 | 8 | 0.125 | 16 | 0.125 | 1 | 0 | 12 | 0.4167 | 15 | 0.2667 | 6 | 0 | 34 | 0.2647 |

| | | | | | | | | | | | | | | | | | | | | |
|---|---|---|---|---|---|---|---|---|---|---|---|---|---|---|---|---|---|---|---|---|
| 18_4674812 | 6 | 0.8333 | 2 | 0 | 1 | 1 | 11 | 0.7273 | 20 | 0.7 | 2 | 1 | 12 | 0.5833 | 14 | 0.7857 | 6 | 1 | 34 | 0.7647 |
| 18_4675094 | 9 | 0.1111 | 3 | 0.6667 | 2 | 0.5 | 10 | 0.3 | 24 | 0.2917 | 4 | 0 | 12 | 0.0833 | 13 | 0.3846 | 8 | 0.25 | 37 | 0.2162 |
| 18_4675185 | 8 | 0.25 | 3 | 0.3333 | 2 | 0 | 6 | 0.3333 | 19 | 0.2632 | 2 | 0 | 14 | 0.2143 | 14 | 0.2857 | 6 | 0.3333 | 36 | 0.25 |
| 18_4675234 | 8 | 0.75 | 3 | 0.6667 | 1 | 0 | 10 | 0.5 | 22 | 0.5909 | 2 | 0 | 9 | 0.3333 | 15 | 0.6 | 7 | 0.2857 | 33 | 0.4242 |
| 18_4675328 | 7 | 0.1429 | 2 | 0 | 2 | 0 | 10 | 0.7 | 21 | 0.381 | 1 | 0 | 8 | 0.5 | 16 | 0.375 | 8 | 0.625 | 33 | 0.4545 |
| 18_4675356 | 8 | 0.375 | 3 | 0.3333 | 2 | 0.5 | 10 | 0.1 | 23 | 0.2609 | 3 | 0.6667 | 13 | 0.2308 | 13 | 0.2308 | 6 | 0.5 | 35 | 0.3143 |
| 18_4675565 | 10 | 0.7 | 4 | 0.5 | 1 | 1 | 10 | 0.6 | 25 | 0.64 | 4 | 0.5 | 14 | 0.5714 | 15 | 0.6667 | 8 | 0.625 | 41 | 0.6098 |
| 18_4675645 | 11 | 1 | 3 | 1 | 2 | 1 | 8 | 1 | 24 | 1 | 1 | 1 | 10 | 1 | 12 | 1 | 6 | 1 | 29 | 1 |
| 18_4675865 | 11 | 0.8182 | 4 | 0.75 | 3 | 1 | 12 | 0.6667 | 30 | 0.7667 | 2 | 1 | 12 | 0.6667 | 15 | 0.6 | 10 | 0.4 | 39 | 0.5897 |
| 18_4675878 | 9 | 0.7778 | 4 | 1 | 3 | 0.6667 | 6 | 0.3333 | 22 | 0.6818 | 2 | 1 | 13 | 0.6923 | 12 | 0.75 | 7 | 0.8571 | 34 | 0.7647 |
| 18_4675904 | 11 | 0.9091 | 3 | 1 | 3 | 1 | 11 | 0.7273 | 28 | 0.8571 | 4 | 0.75 | 14 | 0.8571 | 17 | 0.7059 | 9 | 0.4444 | 44 | 0.7045 |
| 18_4680343 | 10 | 0.6 | 5 | 0.8 | 3 | 0.6667 | 8 | 0.5 | 26 | 0.6154 | 3 | 0.3333 | 13 | 0.6923 | 15 | 0.4 | 12 | 0.4167 | 43 | 0.4884 |
| 18_4680472 | 10 | 0 | 7 | 0.1429 | 3 | 0 | 10 | 0.1 | 30 | 0.0667 | 5 | 0 | 12 | 0.0833 | 14 | 0.0714 | 11 | 0.0909 | 42 | 0.0714 |
| 18_4680530 | 11 | 0.1818 | 6 | 0.3333 | 3 | 0.3333 | 9 | 0 | 29 | 0.1724 | 5 | 0.2 | 14 | 0.2143 | 12 | 0.1667 | 10 | 0 | 41 | 0.1463 |
| 18_4680705 | 11 | 0.1818 | 6 | 0 | 3 | 0 | 9 | 0.1111 | 29 | 0.1034 | 4 | 0 | 15 | 0.1333 | 17 | 0.2353 | 11 | 0.2727 | 47 | 0.1915 |
| 18_4681236 | 11 | 0.0909 | 7 | 0 | 5 | 0 | 9 | 0.1111 | 32 | 0.0625 | 8 | 0.125 | 15 | 0.1333 | 18 | 0.0556 | 12 | 0 | 53 | 0.0755 |
| 18_4682135 | 16 | 0.125 | 10 | 0 | 7 | 0 | 12 | 0.0833 | 45 | 0.0667 | 12 | 0.1667 | 22 | 0.0455 | 19 | 0.0526 | 10 | 0.1 | 63 | 0.0794 |
| 18_4682634 | 18 | 0.0556 | 13 | 0.0769 | 8 | 0.125 | 16 | 0.0625 | 55 | 0.0727 | 12 | 0.0833 | 24 | 0 | 21 | 0.0952 | 11 | 0.1818 | 68 | 0.0735 |
| 18_4682695 | 20 | 0.65 | 12 | 0.5 | 9 | 0.6667 | 16 | 0.75 | 57 | 0.6491 | 12 | 0.75 | 24 | 0.6667 | 22 | 0.6818 | 12 | 0.5833 | 70 | 0.6714 |
| 18_5118503 | 1 | 0 | NA | NA | 1 | 0 | 4 | 1 | 6 | 0.6667 | 2 | 1 | NA | NA | 3 | 0.6667 | 3 | 0.3333 | 8 | 0.625 |
| 18_5119021 | 2 | 0.5 | NA | NA | 1 | 0 | 4 | 0 | 7 | 0.1429 | 2 | 0.5 | NA | NA | 3 | 0 | 3 | 0.3333 | 8 | 0.25 |
| 18_5210709 | 1 | 1 | NA | NA | 1 | 1 | NA | NA | 2 | 1 | 1 | 0 | 1 | 1 | NA | NA | 1 | 1 | 3 | 0.6667 |
| 18_5210758 | 2 | 1 | NA | NA | 1 | 1 | NA | NA | 3 | 1 | NA | NA | 1 | 1 | NA | NA | 1 | 1 | 2 | 1 |
| 18_5210893 | 2 | 0 | NA | NA | 1 | 1 | NA | NA | 3 | 0.3333 | 1 | 0 | 1 | 1 | NA | NA | 1 | 1 | 3 | 0.6667 |
| 18_5210913 | 2 | 1 | NA | NA | 1 | 1 | NA | NA | 3 | 1 | 1 | 1 | 1 | 1 | NA | NA | 1 | 1 | 3 | 1 |
| 18_5210941 | 1 | 1 | NA | NA | 1 | 1 | NA | NA | 2 | 1 | 1 | 1 | 1 | 1 | NA | NA | NA | NA | 2 | 1 |
| 18_5214009 | 1 | 1 | NA | NA | 1 | 1 | NA | NA | 2 | 1 | 1 | 1 | 1 | 1 | NA | NA | 1 | 1 | 3 | 1 |
| 18_5214173 | 1 | 1 | NA | NA | NA | NA | NA | NA | 1 | 1 | NA | NA | 1 | 1 | NA | NA | 1 | 1 | 2 | 1 |
| 18_5214192 | 1 | 1 | NA | NA | NA | NA | NA | NA | 1 | 1 | NA | NA | 1 | 0 | NA | NA | 1 | 1 | 2 | 0.5 |
| 18_5214295 | 2 | 0.5 | NA | NA | 1 | 1 | NA | NA | 3 | 0.6667 | 1 | 0 | 1 | 0 | NA | NA | 1 | 1 | 3 | 0.3333 |
| 18_5214767 | 1 | 1 | NA | NA | 1 | 1 | NA | NA | 2 | 1 | NA | NA | 1 | 0 | NA | NA | 1 | 1 | 2 | 0.5 |
| 18_5505061 | NA | NA | NA | NA | 1 | 1 | NA | NA | 1 | 1 | NA | NA | 1 | 1 | NA | NA | NA | NA | 1 | 1 |
| 18_5566956 | NA | NA | NA | NA | 4 | 1 | 1 | 0 | 5 | 0.8 | 1 | 1 | NA | NA | NA | NA | NA | NA | 1 | 1 |
| 18_5567207 | 1 | 1 | NA | NA | 5 | 0.8 | 2 | 0.5 | 8 | 0.75 | 1 | 1 | NA | NA | NA | NA | NA | NA | 1 | 1 |
| 18_5567421 | 1 | 1 | NA | NA | 4 | 1 | 1 | 1 | 6 | 1 | 1 | 1 | NA | NA | NA | NA | NA | NA | 1 | 1 |
| 18_5766345 | 1 | 0 | 1 | 0 | 1 | 0 | 2 | 0.5 | 5 | 0.2 | NA | NA | 1 | 1 | NA | NA | 1 | 0 | 2 | 0.5 |
| 18_5766662 | 1 | 0 | 1 | 1 | 1 | 1 | 2 | 0.5 | 5 | 0.6 | NA | NA | 1 | 1 | NA | NA | 1 | 0 | 2 | 0.5 |
| 18_5766721 | 1 | 1 | 1 | 0 | 1 | 0 | 2 | 0.5 | 5 | 0.4 | NA | NA | 1 | 0 | NA | NA | 1 | 1 | 2 | 0.5 |
| 18_5767037 | 1 | 0 | 1 | 1 | 1 | 1 | 3 | 0.3333 | 6 | 0.5 | NA | NA | 1 | 0 | NA | NA | 2 | 0.5 | 3 | 0.3333 |
| 18_5767049 | 1 | 0 | 1 | 1 | 1 | 1 | 3 | 0.3333 | 6 | 0.5 | NA | NA | 1 | 1 | NA | NA | 2 | 0.5 | 3 | 0.6667 |
| 18_5767341 | 1 | 1 | 1 | 1 | 1 | 0 | 1 | 1 | 4 | 0.75 | NA | NA | 1 | 0 | 1 | 1 | 1 | 1 | 3 | 0.6667 |
| 18_5767623 | 1 | 0 | 1 | 1 | 1 | 0 | 3 | 0 | 6 | 0.1667 | NA | NA | 1 | 0 | 1 | 1 | 1 | 0 | 3 | 0.3333 |
| 18_5767697 | 2 | 0 | 1 | 0 | 1 | 0 | 3 | 0.3333 | 7 | 0.1429 | NA | NA | 1 | 1 | 1 | 1 | 1 | 1 | 3 | 1 |
| 18_5767730 | 2 | 1 | 1 | 1 | 2 | 0.5 | 2 | 0.5 | 7 | 0.7143 | NA | NA | 1 | 1 | 1 | 0 | 2 | 0 | 4 | 0.25 |
| 18_5767763 | 2 | 1 | 1 | 1 | 2 | 1 | 2 | 1 | 7 | 1 | NA | NA | 1 | 1 | 1 | 1 | 2 | 1 | 4 | 1 |
| 18_5767778 | 2 | 0 | 1 | 1 | 2 | 1 | 3 | 1 | 8 | 0.75 | NA | NA | 1 | 1 | 1 | 0 | 2 | 1 | 4 | 0.75 |
| 18_5767826 | 2 | 0 | NA | NA | 2 | 1 | 1 | 0 | 5 | 0.4 | NA | NA | 1 | 1 | 1 | 0 | 2 | 0.5 | 4 | 0.5 |
| 18_5767886 | 2 | 1 | NA | NA | 2 | 0.5 | 3 | 0 | 7 | 0.4286 | NA | NA | 1 | 1 | 1 | 1 | 1 | 0 | 3 | 0.6667 |
| 18_5772831 | NA | NA | 2 | 0 | 1 | 0 | 3 | 0.3333 | 6 | 0.1667 | NA | NA | NA | NA | 3 | 0.3333 | 1 | 0 | 4 | 0.25 |
| 18_5772938 | 5 | 1 | 3 | 1 | 5 | 0.8 | 4 | 1 | 17 | 0.9412 | 1 | 1 | 2 | 1 | 5 | 0.8 | 2 | 1 | 10 | 0.9 |
| 18_5773192 | 3 | 0.6667 | 5 | 1 | 5 | 0.8 | 4 | 1 | 17 | 0.8824 | 1 | 1 | 2 | 1 | 4 | 1 | 2 | 1 | 9 | 1 |
| 18_5773224 | 7 | 0.2857 | 6 | 0.3333 | 5 | 0.2 | 8 | 0.375 | 26 | 0.3077 | 1 | 0 | 2 | 0 | 5 | 0.2 | 4 | 0 | 12 | 0.0833 |
| 18_5773267 | 6 | 0 | 5 | 0.2 | 4 | 0 | 5 | 0 | 20 | 0.05 | NA | NA | 2 | 0 | 5 | 0 | 4 | 0.25 | 11 | 0.0909 |
| 18_5773306 | 5 | 0.8 | 6 | 0.8333 | 4 | 0.75 | 7 | 0.4286 | 22 | 0.6818 | 1 | 0 | 2 | 0.5 | 3 | 0.6667 | 4 | 0.75 | 10 | 0.6 |
| 18_5773495 | 8 | 0.25 | 6 | 0.1667 | 4 | 0.25 | 8 | 0.125 | 26 | 0.1923 | 2 | 0 | 3 | 0.3333 | 4 | 0.5 | 4 | 0.5 | 13 | 0.3846 |
| 18_5773588 | 8 | 0.25 | 5 | 0.4 | 5 | 0.2 | 4 | 0 | 22 | 0.2273 | 2 | 0 | 3 | 0 | 4 | 0.5 | 4 | 0.5 | 13 | 0.3077 |
| 18_5773984 | 9 | 0.5556 | 3 | 0 | 4 | 0 | 7 | 0.2857 | 23 | 0.3043 | 3 | 0 | 2 | 0 | 3 | 0.3333 | 4 | 0.25 | 12 | 0.1667 |
| 18_5774054 | 8 | 0.25 | 6 | 0.3333 | 5 | 0.2 | 6 | 0.1667 | 25 | 0.24 | 3 | 0 | 3 | 0 | 3 | 0.6667 | 5 | 0.4 | 14 | 0.2857 |
| 18_5774184 | 10 | 0.5 | 7 | 0.1429 | 4 | 0.5 | 7 | 0.4286 | 28 | 0.3929 | 3 | 0 | 3 | 0 | 3 | 0.3333 | 5 | 0 | 14 | 0.0714 |
| 18_6025934 | 1 | 0 | 5 | 0.2 | 4 | 0 | 9 | 0.1111 | 19 | 0.1053 | 7 | 0.1429 | 3 | 0 | 5 | 0 | 3 | 0.3333 | 18 | 0.1111 |
| 18_6026222 | 1 | 0 | 5 | 0.2 | 4 | 0 | 8 | 0.125 | 18 | 0.1111 | 5 | 0.2 | 4 | 0 | 4 | 0 | 2 | 0 | 15 | 0.0667 |
| 18_6026230 | 1 | 0 | 5 | 0.2 | 4 | 0 | 9 | 0.1111 | 19 | 0.1053 | 4 | 0.5 | 4 | 0.25 | 4 | 0 | 2 | 0 | 14 | 0.2143 |
| 18_6026267 | 1 | 0 | 4 | 0.25 | 3 | 0.3333 | 8 | 0 | 16 | 0.125 | 6 | 0.3333 | 4 | 0 | 4 | 0.25 | 2 | 0.5 | 16 | 0.25 |
| 18_6026312 | 1 | 0 | 3 | 0 | 2 | 0.5 | 6 | 0 | 12 | 0.0833 | 6 | 0.1667 | 4 | 0.25 | 3 | 0 | 2 | 0.5 | 15 | 0.2 |
| 18_6026331 | 1 | 0 | 3 | 0 | 4 | 0.25 | 8 | 0.125 | 16 | 0.125 | 5 | 0.6 | 4 | 0.25 | 3 | 1 | 2 | 0 | 14 | 0.5 |

| | | | | | | | | | | | | | | | | | | | | |
|---|---|---|---|---|---|---|---|---|---|---|---|---|---|---|---|---|---|---|---|---|
| 18_602638 | 1 | 1 | 3 | 1 | 3 | 1 | 6 | 0.6667 | 13 | 0.8462 | 6 | 1 | 3 | 1 | 2 | 1 | 1 | 1 | 12 | 1 |
| 18_602646 | 1 | 1 | 4 | 0.75 | 2 | 1 | 7 | 1 | 14 | 0.9286 | 3 | 1 | 2 | 1 | 2 | 0.5 | 2 | 1 | 9 | 0.8889 |
| 18_602647 | 1 | 0 | 4 | 0.5 | 4 | 0.5 | 7 | 0.2857 | 16 | 0.375 | 3 | 0.6667 | 4 | 0.5 | 1 | 0 | 2 | 0.5 | 10 | 0.5 |
| 18_602654 | 1 | 1 | 4 | 0.5 | 3 | 0 | 8 | 0.125 | 16 | 0.25 | 5 | 0.4 | 2 | 0 | 2 | 0 | NA | NA | 9 | 0.2222 |
| 18_602655 | 1 | 1 | 2 | 0.5 | 2 | 0.5 | 5 | 0 | 10 | 0.3 | 5 | 0.2 | 2 | 0 | 2 | 0 | 1 | 0 | 10 | 0.1 |
| 18_602660 | 1 | 0 | 4 | 0.25 | 3 | 0 | 8 | 0 | 16 | 0.0625 | 6 | 0 | 1 | 0 | 2 | 0 | 2 | 0.5 | 11 | 0.0909 |
| 18_602664 | 1 | 1 | 3 | 0.6667 | 4 | 1 | 6 | 1 | 14 | 0.9286 | 5 | 1 | 3 | 1 | 2 | 1 | 1 | 1 | 11 | 1 |
| 18_611180 | NA | NA | 1 | 0 | 3 | 0.6667 | 1 | 1 | 5 | 0.6 | 1 | 0 | 1 | 1 | 1 | 0 | 1 | 0 | 4 | 0.25 |
| 18_611184 | NA | NA | 3 | 1 | 3 | 0.3333 | 1 | 0 | 7 | 0.5714 | 1 | 1 | 1 | 0 | 1 | 1 | 1 | 0 | 4 | 0.5 |
| 18_611188 | NA | NA | 2 | 1 | 3 | 0.6667 | 1 | 0 | 6 | 0.6667 | 1 | 1 | 1 | 1 | 1 | 1 | 1 | 1 | 4 | 1 |
| 18_611193 | NA | NA | 3 | 0.3333 | 2 | 0 | 1 | 1 | 6 | 0.3333 | 1 | 0 | NA | NA | 1 | 1 | 1 | 1 | 3 | 0.6667 |
| 18_611210 | NA | NA | 3 | 0.3333 | 1 | 0 | 1 | 0 | 5 | 0.2 | 1 | 1 | 1 | 0 | 1 | 0 | 1 | 0 | 4 | 0.25 |
| 18_611211 | NA | NA | 2 | 1 | 3 | 1 | 1 | 0 | 6 | 0.8333 | 1 | 1 | 1 | 1 | 1 | 0 | 1 | 0 | 4 | 0.5 |
| 18_611219 | NA | NA | 3 | 1 | 3 | 0.6667 | 1 | 1 | 7 | 0.8571 | 1 | 1 | 1 | 1 | 1 | 1 | 1 | 1 | 4 | 1 |
| 18_611228 | NA | NA | 2 | 0 | 3 | 0.3333 | 1 | 0 | 6 | 0.1667 | 1 | 1 | 1 | 0 | 1 | 1 | 1 | 0 | 4 | 0.5 |
| 18_620141 | 19 | 0.6316 | 21 | 0.4762 | 22 | 0.6364 | 29 | 0.3448 | 91 | 0.5055 | 17 | 0.1765 | 17 | 0.6471 | 31 | 0.5484 | 25 | 0.48 | 90 | 0.4778 |
| 18_620146 | 16 | 0.0625 | 25 | 0.08 | 21 | 0.0952 | 26 | 0.0385 | 88 | 0.0682 | 15 | 0.0667 | 17 | 0.1176 | 27 | 0.1111 | 23 | 0.1304 | 82 | 0.1098 |
| 18_620146 | 17 | 0.0588 | 23 | 0.0435 | 21 | 0.0476 | 31 | 0.0645 | 92 | 0.0543 | 16 | 0.0625 | 19 | 0.1053 | 28 | 0.0357 | 24 | 0.0833 | 87 | 0.069 |
| 18_620172 | 16 | 0.625 | 19 | 0.6316 | 16 | 0.6875 | 26 | 0.7308 | 77 | 0.6753 | 14 | 0.8571 | 16 | 0.875 | 22 | 0.7273 | 20 | 0.65 | 72 | 0.7639 |
| 18_620343 | 15 | 0.0667 | 16 | 0.1875 | 11 | 0.1818 | 20 | 0.05 | 62 | 0.1129 | 6 | 0.3333 | 11 | 0 | 16 | 0.125 | 18 | 0.1667 | 51 | 0.1373 |
| 18_624120 | 9 | 0.2222 | 6 | 0.1667 | 6 | 0.1667 | 12 | 0.3333 | 33 | 0.2424 | 3 | 0 | 6 | 0.1667 | 9 | 0.3333 | 7 | 0 | 25 | 0.16 |
| 18_624128 | 8 | 0.125 | 3 | 0 | 2 | 1 | 5 | 0.2 | 18 | 0.2222 | 3 | 0 | 4 | 0 | 6 | 0.1667 | 4 | 0.5 | 17 | 0.1765 |
| 18_643609 | 6 | 0 | 6 | 0.1667 | 5 | 0 | 1 | 0 | 18 | 0.0556 | 3 | 0 | 7 | 0 | 11 | 0.0909 | 7 | 0.1429 | 28 | 0.0714 |
| 18_643675 | 3 | 0.3333 | 6 | 0.3333 | 4 | 0 | 1 | 0 | 14 | 0.2143 | 1 | 0 | 5 | 0.4 | 11 | 0 | 6 | 0.3333 | 23 | 0.1739 |
| 18_643696 | 3 | 0 | 5 | 0 | 2 | 0 | 1 | 1 | 11 | 0.0909 | 1 | 0 | 6 | 0.1667 | 9 | 0.1111 | 3 | 0 | 19 | 0.1053 |
| 18_643712 | 2 | 0.5 | 6 | 0.1667 | 3 | 1 | 1 | 1 | 12 | 0.5 | NA | NA | 6 | 0.5 | 9 | 0.5556 | 5 | 0.6 | 20 | 0.55 |
| 18_643720 | 3 | 0 | 5 | 0 | 3 | 0.3333 | NA | NA | 11 | 0.0909 | NA | NA | 4 | 0 | 7 | 0 | 4 | 0.5 | 15 | 0.1333 |
| 18_643721 | 3 | 0.6667 | 5 | 0 | 3 | 0 | 1 | 1 | 12 | 0.25 | 1 | 0 | 6 | 0.5 | 7 | 0 | 5 | 0.2 | 19 | 0.2105 |
| 18_643969 | NA | NA | 4 | 1 | 3 | 1 | 1 | 1 | 8 | 1 | 1 | 1 | 1 | 1 | 5 | 1 | 2 | 1 | 9 | 1 |
| 18_643975 | 1 | 0 | 4 | 0.75 | 3 | 1 | 1 | 0 | 9 | 0.6667 | NA | NA | 2 | 1 | 4 | 1 | 2 | 0 | 8 | 0.75 |
| 18_643981 | 1 | 0 | 4 | 0 | 3 | 0.3333 | 1 | 0 | 9 | 0.1111 | NA | NA | 1 | 1 | 5 | 0.4 | 2 | 0.5 | 8 | 0.5 |
| 18_643984 | 1 | 1 | 3 | 0.3333 | 3 | 0.3333 | 1 | 1 | 8 | 0.5 | 1 | 0 | 2 | 0 | 4 | 0.5 | 3 | 0.6667 | 10 | 0.4 |
| 18_644902 | 1 | 0 | 2 | 0 | 3 | 0.6667 | 1 | 0 | 7 | 0.2857 | 1 | 0 | 1 | 0 | 3 | 1 | 3 | 0 | 8 | 0.375 |
| 18_644905 | 1 | 0 | 2 | 0 | 3 | 0.3333 | 1 | 0 | 7 | 0.1429 | 1 | 0 | 1 | 0 | 3 | 0.3333 | 3 | 0.6667 | 8 | 0.375 |
| 18_644912 | 1 | 1 | 2 | 0.5 | 2 | 0 | 1 | 0 | 6 | 0.3333 | NA | NA | 1 | 0 | 3 | 0 | 3 | 0.3333 | 7 | 0.1429 |
| 18_649071 | NA | NA | 1 | 1 | 1 | 0 | NA | NA | 2 | 0.5 | 1 | 1 | NA | NA | 2 | 1 | 1 | 1 | 4 | 1 |
| 18_649084 | NA | NA | 1 | 1 | 1 | 1 | NA | NA | 2 | 1 | 1 | 1 | NA | NA | 2 | 0.5 | 1 | 0 | 4 | 0.5 |
| 18_649121 | NA | NA | NA | NA | 1 | 1 | NA | NA | 1 | 1 | 1 | 1 | NA | NA | 2 | 0.5 | 1 | 1 | 4 | 0.75 |
| 18_649126 | NA | NA | 1 | 1 | 1 | 0 | NA | NA | 2 | 0.5 | 1 | 1 | NA | NA | 2 | 0 | 1 | 0 | 4 | 0.25 |
| 18_649168 | NA | NA | 1 | 1 | 1 | 0 | NA | NA | 2 | 0.5 | 1 | 1 | NA | NA | 2 | 0 | 1 | 0 | 4 | 0.25 |
| 18_678865 | 3 | 0.6667 | 12 | 0.4167 | 5 | 0.8 | 6 | 0.5 | 26 | 0.5385 | 7 | 0.8571 | 7 | 0.5714 | 9 | 0.7778 | 9 | 0.5556 | 32 | 0.6875 |
| 18_678871 | 2 | 0.5 | 8 | 0.25 | 7 | 0.4286 | 6 | 0.6667 | 23 | 0.4348 | 10 | 0.5 | 8 | 0.25 | 11 | 0.1818 | 8 | 0.25 | 37 | 0.2973 |
| 18_678873 | 3 | 0.3333 | 12 | 0.1667 | 6 | 0.3333 | 4 | 0 | 25 | 0.2 | 8 | 0.25 | 7 | 0.2857 | 10 | 0.2 | 12 | 0.1667 | 37 | 0.2162 |
| 18_679223 | 2 | 1 | NA | NA | NA | NA | NA | NA | 2 | 1 | 1 | 0 | 1 | 0 | 1 | 1 | 2 | 1 | 5 | 0.6 |
| 18_679232 | 2 | 1 | NA | NA | NA | NA | NA | NA | 2 | 1 | 1 | 1 | NA | NA | NA | NA | 2 | 1 | 3 | 1 |
| 18_693551 | 3 | 0.3333 | NA | NA | NA | NA | 1 | 0 | 4 | 0.25 | NA | NA | 1 | 0 | 2 | 0.5 | 1 | 0 | 4 | 0.25 |
| 18_693579 | 3 | 0.6667 | NA | NA | NA | NA | NA | NA | 3 | 0.6667 | NA | NA | 1 | 0 | 2 | 0.5 | 1 | 1 | 4 | 0.5 |
| 18_693592 | 3 | 0.6667 | NA | NA | NA | NA | 1 | 1 | 4 | 0.75 | NA | NA | 1 | 0 | 2 | 1 | 1 | 1 | 4 | 0.75 |
| 18_693650 | 3 | 0.3333 | NA | NA | NA | NA | NA | NA | 3 | 0.3333 | NA | NA | 1 | 0 | 2 | 0 | 1 | 1 | 4 | 0.25 |
| 18_693668 | 2 | 0.5 | NA | NA | NA | NA | NA | NA | 2 | 0.5 | NA | NA | 1 | 1 | 2 | 0.5 | 1 | 1 | 4 | 0.75 |
| 18_693676 | 1 | 1 | NA | NA | NA | NA | NA | NA | 1 | 1 | NA | NA | 1 | 1 | 1 | 1 | 2 | 1 | 4 | 1 |
| 18_693698 | 2 | 0.5 | NA | NA | NA | NA | NA | NA | 2 | 0.5 | NA | NA | 1 | 0 | 1 | 1 | 2 | 0 | 4 | 0.25 |
| 18_735025 | NA | NA | 2 | 0.5 | 1 | 0 | 1 | 0 | 4 | 0.25 | 1 | 1 | 1 | 0 | NA | NA | 2 | 0 | 4 | 0.25 |
| 18_735055 | NA | NA | 3 | 0 | 1 | 0 | 1 | 1 | 5 | 0.2 | 1 | 0 | 1 | 0 | NA | NA | 2 | 0.5 | 4 | 0.25 |
| 18_735100 | NA | NA | 3 | 1 | 1 | 0 | 1 | 1 | 5 | 0.8 | 1 | 1 | 1 | 1 | NA | NA | 1 | 0 | 3 | 0.6667 |
| 18_791059 | 5 | 0.4 | 10 | 0.2 | 4 | 0.25 | 8 | 0.25 | 27 | 0.2593 | 3 | 0.6667 | 4 | 0.25 | 2 | 0 | 3 | 0.3333 | 12 | 0.3333 |
| 18_791060 | 3 | 0 | 9 | 0.3333 | 4 | 0.25 | 6 | 0 | 22 | 0.1818 | 1 | 0 | 6 | 0.5 | 3 | 0.6667 | 3 | 0 | 13 | 0.3846 |
| 18_791062 | 6 | 0.5 | 9 | 0.7778 | 3 | 0.3333 | 6 | 0.6667 | 24 | 0.625 | 3 | 0 | 7 | 0.1429 | 3 | 0.3333 | 4 | 0.5 | 17 | 0.2353 |
| 18_791622 | 6 | 0 | 8 | 0 | 4 | 0.5 | 7 | 0.1429 | 25 | 0.12 | 3 | 0 | 7 | 0.1429 | 3 | 0 | 4 | 0.25 | 17 | 0.1176 |
| 18_791627 | 5 | 0.2 | 10 | 0.1 | 4 | 0.5 | 8 | 0 | 27 | 0.1481 | 2 | 0.5 | 7 | 0.2857 | 3 | 0.6667 | 2 | 0 | 14 | 0.3571 |
| 18_791638 | NA | NA | NA | NA | 1 | 1 | 1 | 0 | 2 | 0.5 | 1 | 1 | NA | NA | NA | NA | NA | NA | 1 | 1 |
| 18_792617 | 6 | 0.3333 | 12 | 0.9167 | 6 | 1 | 9 | 0.4444 | 33 | 0.697 | 3 | 0.3333 | 8 | 0.875 | 5 | 0.6 | 4 | 0.75 | 20 | 0.7 |
| 18_792622 | 8 | 0.375 | 13 | 0.2308 | 5 | 0.2 | 9 | 0 | 35 | 0.2 | 4 | 0.25 | 8 | 0.125 | 6 | 0 | 2 | 0 | 20 | 0.1 |
| 18_792623 | 8 | 0.5 | 10 | 0.6 | 6 | 1 | 10 | 0.7 | 34 | 0.6765 | 4 | 0.75 | 7 | 0.5714 | 5 | 0.6 | 4 | 1 | 20 | 0.7 |
| 18_792625 | 6 | 0.1667 | 13 | 0 | 5 | 0.2 | 8 | 0 | 32 | 0.0625 | 1 | 1 | 7 | 0.2857 | 3 | 0 | 4 | 0.25 | 15 | 0.2667 |

| | | | | | | | | | | | | | | | | | | | | |
|---|---|---|---|---|---|---|---|---|---|---|---|---|---|---|---|---|---|---|---|---|
| 18_792692 | 11 | 0.6364 | 14 | 0.4286 | 11 | 0.9091 | 15 | 0.7333 | 51 | 0.6667 | 3 | 0.6667 | 10 | 0.6 | 8 | 0.5 | 10 | 0.5 | 31 | 0.5484 |
| 18_792699 | 11 | 0 | 13 | 0 | 11 | 0.1818 | 12 | 0.0833 | 47 | 0.0638 | 3 | 0 | 9 | 0.1111 | 9 | 0.2222 | 8 | 0.125 | 29 | 0.1379 |
| 18_792738 | 7 | 0 | 8 | 0 | 10 | 0.1 | 11 | 0.1818 | 36 | 0.0833 | 2 | 0 | 3 | 0 | 3 | 0 | 9 | 0.1111 | 17 | 0.0588 |
| 18_921510 | 1 | 1 | NA | NA | NA | NA | NA | NA | 1 | 1 | 1 | 1 | NA | NA | NA | NA | NA | NA | 1 | 1 |
| 18_931254 | 4 | 0 | 10 | 0.1 | 5 | 0.4 | 6 | 0 | 25 | 0.12 | 5 | 0.2 | 5 | 0.2 | 10 | 0.4 | 4 | 0 | 24 | 0.25 |
| 18_931387 | 2 | 0 | 7 | 0.1429 | 1 | 0 | 5 | 0 | 15 | 0.0667 | 6 | 0.1667 | 2 | 0 | 7 | 0.2857 | 3 | 0.3333 | 18 | 0.2222 |
| 18_931401 | 3 | 0 | 7 | 0.2857 | 1 | 0 | 5 | 0 | 16 | 0.125 | 7 | 0 | 4 | 0.25 | 6 | 0.3333 | 2 | 0 | 19 | 0.1579 |
| 18_931426 | 7 | 0.5714 | 9 | 0.3333 | 6 | 0.3333 | 9 | 0.4444 | 31 | 0.4194 | 7 | 0.4286 | 10 | 0.3 | 9 | 0.4444 | 5 | 0.6 | 31 | 0.4194 |
| 18_931429 | 6 | 0.1667 | 9 | 0.1111 | 6 | 0.1667 | 11 | 0.1818 | 32 | 0.1562 | 8 | 0 | 11 | 0.3636 | 11 | 0.0909 | 4 | 0.5 | 34 | 0.2059 |
| 18_931432 | 6 | 0 | 8 | 0.125 | 5 | 0.2 | 11 | 0.2727 | 30 | 0.1667 | 8 | 0 | 9 | 0.2222 | 11 | 0.2727 | 3 | 0.3333 | 31 | 0.1935 |
| 18_931451 | 9 | 0 | 10 | 0.1 | 8 | 0.125 | 14 | 0.0714 | 41 | 0.0732 | 10 | 0.1 | 15 | 0.0667 | 10 | 0.2 | 4 | 0 | 39 | 0.1026 |
| 18_931609 | 8 | 0.5 | 14 | 0.5 | 4 | 0.75 | 2 | 1 | 28 | 0.5714 | 6 | 0.3333 | 1 | 0 | 9 | 0.4444 | 5 | 0.2 | 21 | 0.3333 |
| 18_931629 | 9 | 0.7778 | 11 | 0.8182 | 4 | 1 | 4 | 0.75 | 28 | 0.8214 | 5 | 1 | NA | NA | 8 | 0.875 | 4 | 0.5 | 17 | 0.8235 |
| 18_931649 | 7 | 1 | 12 | 0.9167 | 3 | 0.6667 | 4 | 1 | 26 | 0.9231 | 5 | 1 | NA | NA | 6 | 1 | 5 | 1 | 16 | 1 |
| 18_931663 | 8 | 0.375 | 13 | 0.0769 | 3 | 0 | 4 | 0 | 28 | 0.1429 | 5 | 0.2 | NA | NA | 7 | 0 | 5 | 0 | 17 | 0.0588 |
| 18_984843 | NA | NA | NA | NA | NA | NA | 1 | 1 | 1 | 1 | NA | NA | 1 | 1 | NA | NA | NA | NA | 1 | 1 |
| 18_984871 | NA | NA | 2 | 0 | NA | NA | 1 | 1 | 3 | 0.3333 | NA | NA | 1 | 1 | NA | NA | NA | NA | 1 | 1 |
| 18_984885 | NA | NA | 2 | 0.5 | NA | NA | 1 | 0 | 3 | 0.3333 | NA | NA | 1 | 1 | NA | NA | NA | NA | 1 | 1 |
| 18_985878 | 1 | 0 | 2 | 1 | 1 | 1 | 1 | 0 | 5 | 0.6 | NA | NA | 1 | 1 | NA | NA | NA | NA | 1 | 1 |
| 18_985882 | 1 | 1 | 2 | 1 | 1 | 0 | 1 | 1 | 5 | 0.8 | NA | NA | 1 | 1 | NA | NA | NA | NA | 1 | 1 |
| 18_985886 | 1 | 0 | 2 | 0.5 | 1 | 1 | 1 | 1 | 5 | 0.6 | NA | NA | 1 | 1 | NA | NA | NA | NA | 1 | 1 |
| 18_985889 | 1 | 1 | 2 | 0.5 | 1 | 1 | 1 | 0 | 5 | 0.6 | NA | NA | 1 | 1 | NA | NA | NA | NA | 1 | 1 |
| 18_987713 | 1 | 0 | 1 | 1 | 1 | 1 | 1 | 1 | 4 | 0.75 | NA | NA | 1 | 1 | NA | NA | NA | NA | 1 | 1 |
| 18_987722 | 1 | 1 | 2 | 1 | 1 | 0 | NA | NA | 4 | 0.75 | NA | NA | 1 | 1 | NA | NA | NA | NA | 1 | 1 |
| 18_999342 | 2 | 0.5 | 7 | 0.7143 | 4 | 0.25 | 4 | 0.5 | 17 | 0.5294 | 2 | 1 | 3 | 0.3333 | 7 | 0.8571 | 6 | 0.5 | 18 | 0.6667 |
| 18_999357 | 3 | 0 | 8 | 0.125 | 4 | 0 | 4 | 0 | 19 | 0.0526 | 3 | 0 | 2 | 0.5 | 6 | 0 | 5 | 0 | 16 | 0.0625 |
| 18_999359 | 3 | 0.6667 | 8 | 0 | 4 | 0 | 5 | 0 | 20 | 0.1 | 3 | 0 | 3 | 0.3333 | 7 | 0 | 5 | 0 | 18 | 0.0556 |
| 18_999592 | 19 | 0.1053 | 21 | 0.0952 | 20 | 0.05 | 21 | 0.1429 | 81 | 0.0988 | 14 | 0.1429 | 22 | 0.2727 | 12 | 0 | 22 | 0 | 70 | 0.1143 |
| 18_999597 | 16 | 0.0625 | 16 | 0.5 | 18 | 0.5556 | 18 | 0.2778 | 68 | 0.3529 | 14 | 0.6429 | 17 | 0.4118 | 6 | 0.3333 | 21 | 0.381 | 58 | 0.4483 |
| 18_999604 | 15 | 0.2 | 16 | 0.3125 | 13 | 0.3846 | 14 | 0.5 | 58 | 0.3448 | 11 | 0.6364 | 20 | 0.1 | 7 | 0.2857 | 16 | 0.3125 | 54 | 0.2963 |
| 18_100655 | 1 | 0 | 5 | 0.4 | 1 | 0 | 3 | 0 | 10 | 0.2 | NA | NA | 1 | 1 | 3 | 0.3333 | 4 | 0.5 | 8 | 0.5 |
| 18_100656 | NA | NA | 5 | 0.4 | 1 | 1 | 2 | 0 | 8 | 0.375 | NA | NA | 1 | 1 | 2 | 0.5 | 3 | 0 | 6 | 0.3333 |
| 18_105604 | NA | NA | NA | NA | 1 | 0 | 3 | 0.6667 | 4 | 0.5 | NA | NA | NA | NA | NA | NA | 3 | 0.6667 | 3 | 0.6667 |
| 18_105605 | NA | NA | NA | NA | 1 | 0 | 2 | 0.5 | 3 | 0.3333 | NA | NA | NA | NA | NA | NA | 3 | 0.3333 | 3 | 0.3333 |
| 18_105606 | NA | NA | NA | NA | 1 | 1 | 3 | 0.3333 | 4 | 0.5 | NA | NA | NA | NA | NA | NA | 3 | 0.3333 | 3 | 0.3333 |
| 18_105606 | NA | NA | NA | NA | 1 | 1 | 3 | 0 | 4 | 0.25 | NA | NA | NA | NA | NA | NA | 3 | 0.3333 | 3 | 0.3333 |
| 18_105608 | NA | NA | NA | NA | 1 | 1 | 3 | 0.3333 | 4 | 0.5 | NA | NA | NA | NA | NA | NA | 3 | 0.3333 | 3 | 0.3333 |
| 18_105614 | NA | NA | NA | NA | 2 | 0.5 | 3 | 0 | 5 | 0.2 | NA | NA | NA | NA | NA | NA | 3 | 0.3333 | 3 | 0.3333 |
| 18_105615 | NA | NA | NA | NA | 2 | 0 | 3 | 0.3333 | 5 | 0.2 | NA | NA | NA | NA | NA | NA | 3 | 0.3333 | 3 | 0.3333 |
| 18_105667 | 5 | 0.4 | 3 | 0 | 2 | 0 | 5 | 0.4 | 15 | 0.2667 | NA | NA | 4 | 0.25 | 1 | 1 | 3 | 0.3333 | 8 | 0.375 |
| 18_105669 | 5 | 0 | 3 | 0 | 2 | 0 | 4 | 0.5 | 14 | 0.1429 | NA | NA | 5 | 0.2 | 1 | 0 | 3 | 0 | 9 | 0.1111 |
| 18_105669 | 5 | 0.4 | 2 | 0.5 | 2 | 0.5 | 4 | 0.25 | 13 | 0.3846 | NA | NA | 5 | 0.4 | 1 | 0 | 4 | 0.5 | 10 | 0.4 |
| 18_105671 | 5 | 0 | 3 | 0.3333 | 1 | 0 | 4 | 0.5 | 13 | 0.2308 | NA | NA | 4 | 0.25 | 1 | 0 | 4 | 0.25 | 9 | 0.2222 |
| 18_105672 | 4 | 0.5 | 3 | 0 | NA | NA | 4 | 0 | 11 | 0.1818 | NA | NA | 4 | 0.25 | 1 | 0 | 4 | 0.5 | 9 | 0.3333 |
| 18_105673 | 3 | 1 | 2 | 1 | 1 | 0 | 3 | 0.6667 | 9 | 0.7778 | NA | NA | 4 | 0.75 | 1 | 1 | 4 | 1 | 9 | 0.8889 |
| 18_105675 | 3 | 1 | 1 | 1 | NA | NA | 3 | 0.3333 | 7 | 0.7143 | NA | NA | 3 | 1 | 1 | 1 | 2 | 1 | 6 | 1 |
| 18_105940 | 2 | 0.5 | 2 | 0.5 | NA | NA | 1 | 1 | 5 | 0.6 | NA | NA | NA | NA | NA | NA | 1 | 1 | 1 | 1 |
| 18_105940 | 1 | 1 | 2 | 0.5 | NA | NA | 2 | 1 | 5 | 0.8 | NA | NA | 1 | 0 | NA | NA | 1 | 1 | 2 | 0.5 |
| 18_105940 | 2 | 1 | 1 | 1 | NA | NA | 1 | 1 | 4 | 1 | NA | NA | 1 | 1 | NA | NA | 1 | 1 | 2 | 1 |
| 18_105941 | 2 | 0 | 2 | 1 | NA | NA | 1 | 1 | 5 | 0.6 | NA | NA | 1 | 1 | NA | NA | NA | NA | 1 | 1 |
| 18_105942 | 2 | 0 | 1 | 0 | NA | NA | 2 | 0.5 | 5 | 0.2 | NA | NA | NA | NA | NA | NA | 1 | 1 | 1 | 1 |
| 18_105944 | 1 | 0 | 1 | 0 | NA | NA | 1 | 1 | 3 | 0.3333 | NA | NA | NA | NA | NA | NA | 1 | 1 | 1 | 1 |
| 18_105950 | 2 | 0.5 | NA | NA | NA | NA | 1 | 1 | 3 | 0.6667 | NA | NA | NA | NA | NA | NA | 1 | 1 | 1 | 1 |
| 18_106450 | 4 | 0.5 | 5 | 0.8 | 6 | 0.8333 | 3 | 1 | 18 | 0.7778 | 8 | 0.875 | 6 | 1 | 9 | 0.8889 | 5 | 0.8 | 28 | 0.8929 |
| 18_106450 | 6 | 0.6667 | 6 | 0.3333 | 5 | 0.4 | 7 | 0.2857 | 24 | 0.4167 | 10 | 0.5 | 7 | 0.1429 | 7 | 0.1429 | 9 | 0.5556 | 33 | 0.3636 |
| 18_106452 | 5 | 0 | 7 | 0 | 7 | 0.2857 | 6 | 0.3333 | 25 | 0.16 | 9 | 0.1111 | 8 | 0.125 | 9 | 0.2222 | 8 | 0.125 | 34 | 0.1471 |
| 18_107046 | 7 | 0.2857 | 6 | 0.3333 | 6 | 0 | 5 | 0 | 24 | 0.1667 | 9 | 0 | 4 | 0.25 | 9 | 0 | 7 | 0.1429 | 29 | 0.069 |
| 18_107263 | 3 | 0.6667 | 1 | 0 | NA | NA | 1 | 1 | 5 | 0.6 | 1 | 0 | NA | NA | 1 | 1 | NA | NA | 2 | 0.5 |
| 18_108124 | 8 | 0.25 | 3 | 0 | NA | NA | 1 | 0 | 12 | 0.1667 | 3 | 0.6667 | 2 | 0 | 2 | 0 | 1 | 0 | 8 | 0.25 |
| 18_108167 | 20 | 0.1 | 15 | 0 | 16 | 0.125 | 15 | 0.1333 | 66 | 0.0909 | 11 | 0 | 14 | 0.1429 | 20 | 0.05 | 15 | 0.1333 | 60 | 0.0833 |
| 18_108180 | 1 | 0 | NA | NA | 2 | 0.5 | 1 | 1 | 4 | 0.5 | NA | NA | 1 | 0 | 3 | 0.3333 | 1 | 0 | 5 | 0.2 |
| 18_119521 | NA | NA | 1 | 1 | 2 | 0 | 1 | 0 | 4 | 0.25 | 1 | 1 | NA | NA | NA | NA | NA | NA | 1 | 1 |
| 18_119550 | NA | NA | NA | NA | 1 | 1 | NA | NA | 1 | 1 | 1 | 1 | NA | NA | NA | NA | NA | NA | 1 | 1 |
| 18_119718 | NA | NA | 1 | 1 | NA | NA | NA | NA | 1 | 1 | NA | NA | 1 | 1 | 1 | 1 | NA | NA | 2 | 1 |
| 18_119719 | NA | NA | 1 | 1 | NA | NA | NA | NA | 1 | 1 | 1 | 1 | 1 | 1 | 1 | 1 | NA | NA | 3 | 1 |

| | | | | | | | | | | | | | | | | | | | | |
|---|---|---|---|---|---|---|---|---|---|---|---|---|---|---|---|---|---|---|---|---|
| 18_1197212 | NA | NA | 1 | 1 | NA | NA | NA | NA | 1 | 1 | 1 | 1 | 1 | 1 | 1 | 0 | NA | NA | 3 | 0.6667 |
| 18_1197304 | NA | NA | 1 | 1 | NA | NA | NA | NA | 1 | 1 | 1 | 1 | 1 | 0 | 1 | 1 | NA | NA | 3 | 0.6667 |
| 18_1207755 | 3 | 0 | 4 | 0.25 | 3 | 0.3333 | 2 | 0 | 12 | 0.1667 | 4 | 0.5 | 7 | 0.7143 | 5 | 1 | 3 | 0.3333 | 19 | 0.6842 |
| 18_1207768 | 3 | 1 | 4 | 0.75 | 3 | 1 | 2 | 0 | 12 | 0.75 | 3 | 0.6667 | 6 | 1 | 5 | 0.8 | 3 | 0.6667 | 17 | 0.8235 |
| 18_1207778 | 3 | 0.3333 | 4 | 0.25 | 3 | 0.6667 | 2 | 0.5 | 12 | 0.4167 | 3 | 0.3333 | 7 | 0.2857 | 6 | 0.3333 | 4 | 0.25 | 20 | 0.3 |
| 18_1207789 | 2 | 1 | 3 | 0.3333 | 2 | 0.5 | 2 | 0 | 9 | 0.4444 | 3 | 0.6667 | 6 | 0.5 | 5 | 0.8 | 4 | 0.25 | 18 | 0.5556 |
| 18_1207795 | 3 | 1 | 4 | 0.75 | 3 | 1 | 3 | 0.6667 | 13 | 0.8462 | 4 | 1 | 6 | 1 | 6 | 1 | 3 | 1 | 19 | 1 |
| 18_1207796 | 3 | 0 | 2 | 0.5 | 3 | 0 | 2 | 0 | 10 | 0.1 | 4 | 0.25 | 5 | 0 | 4 | 0 | 4 | 0 | 17 | 0.0588 |
| 18_1207810 | 4 | 0.5 | 4 | 0.25 | 3 | 1 | 3 | 0 | 14 | 0.4286 | 3 | 1 | 6 | 0.1667 | 6 | 0.1667 | 5 | 0.4 | 20 | 0.35 |
| 18_1207815 | 4 | 0.75 | 4 | 0.25 | 3 | 0.6667 | 2 | 0.5 | 13 | 0.5385 | 3 | 0.6667 | 6 | 0.6667 | 6 | 0.3333 | 5 | 0.6 | 20 | 0.55 |
| 18_1207819 | 2 | 1 | 4 | 0.75 | 2 | 0.5 | 1 | 1 | 9 | 0.7778 | 3 | 0.6667 | 7 | 0.8571 | 5 | 0.6 | 4 | 0.5 | 19 | 0.6842 |
| 18_1207823 | 4 | 1 | 4 | 0.75 | 2 | 1 | 3 | 1 | 13 | 0.9231 | 3 | 0.6667 | 6 | 1 | 4 | 1 | 5 | 1 | 18 | 0.9444 |
| 18_1207854 | 5 | 0.4 | 3 | 0.3333 | 3 | 0 | 3 | 0.6667 | 14 | 0.3571 | 4 | 0.25 | 7 | 0.2857 | 6 | 0.5 | 5 | 0.4 | 22 | 0.3636 |
| 18_1211609 | 1 | 1 | 1 | 0 | NA | NA | 3 | 0.3333 | 5 | 0.4 | 1 | 1 | NA | NA | NA | NA | NA | NA | 1 | 1 |
| 18_1211638 | 1 | 0 | 2 | 1 | NA | NA | 3 | 1 | 6 | 0.8333 | 1 | 1 | NA | NA | NA | NA | NA | NA | 1 | 1 |
| 18_1211643 | 1 | 1 | 1 | 1 | NA | NA | 2 | 1 | 4 | 1 | 1 | 1 | NA | NA | NA | NA | NA | NA | 1 | 1 |
| 18_1211650 | 1 | 0 | 2 | 0 | NA | NA | 3 | 0.6667 | 6 | 0.3333 | NA | NA | 1 | 1 | NA | NA | NA | NA | 1 | 1 |
| 18_1211654 | 2 | 0 | 2 | 1 | NA | NA | 3 | 0 | 7 | 0.2857 | 1 | 0 | 1 | 1 | NA | NA | NA | NA | 2 | 0.5 |
| 18_1228845 | 5 | 1 | 7 | 0.7143 | 3 | 0.6667 | 4 | 0.5 | 19 | 0.7368 | 1 | 1 | 12 | 0.75 | 2 | 1 | 4 | 0.75 | 19 | 0.7895 |
| 18_1228869 | 5 | 0 | 4 | 0.5 | 3 | 0 | 3 | 0 | 15 | 0.1333 | 1 | 0 | 9 | 0.1111 | 3 | 0 | 2 | 0 | 15 | 0.0667 |
| 18_1228888 | 5 | 0.4 | 7 | 0.2857 | 4 | 0.25 | 4 | 0 | 20 | 0.25 | 2 | 1 | 10 | 0.4 | 5 | 0.4 | 4 | 0.5 | 21 | 0.4762 |
| 18_1228930 | 3 | 0.3333 | 5 | 0.4 | 4 | 0 | 2 | 0 | 14 | 0.2143 | 2 | 0.5 | 5 | 0.2 | 6 | 0.5 | 3 | 0 | 16 | 0.3125 |
| 18_1231802 | 2 | 0 | 3 | 0 | 5 | 0 | 5 | 0.2 | 15 | 0.0667 | 3 | 0 | 1 | 0 | 4 | 0.25 | 1 | 0 | 9 | 0.1111 |
| 18_1232189 | 2 | 0 | 6 | 0.1667 | 5 | 0 | 6 | 0 | 19 | 0.0526 | 2 | 0 | 2 | 0.5 | 4 | 0 | 1 | 0 | 9 | 0.1111 |
| 18_1232221 | 3 | 0 | 5 | 0.2 | 4 | 0.25 | 6 | 0 | 18 | 0.1111 | 4 | 0 | 2 | 0 | 5 | 0.2 | 1 | 0 | 12 | 0.0833 |
| 18_1232294 | 6 | 0.1667 | 7 | 0.4286 | 5 | 0.2 | 4 | 0.5 | 22 | 0.3182 | 2 | 0 | 2 | 1 | 11 | 0.1818 | 5 | 0.8 | 20 | 0.4 |
| 18_1232295 | 6 | 0.1667 | 7 | 0 | 7 | 0.4286 | 8 | 0.125 | 28 | 0.1786 | 2 | 0 | 3 | 0.3333 | 10 | 0.3 | 4 | 0.25 | 19 | 0.2632 |
| 18_1232338 | 5 | 0 | 8 | 0 | 5 | 0.2 | 8 | 0.125 | 26 | 0.0769 | 4 | 0.25 | 1 | 0 | 11 | 0.0909 | 4 | 0 | 20 | 0.1 |
| 18_1232339 | 5 | 0.2 | 9 | 0.2222 | 7 | 0.1429 | 8 | 0.375 | 29 | 0.2414 | 3 | 0 | 1 | 0 | 12 | 0.0833 | 4 | 0 | 20 | 0.05 |
| 18_1232361 | 4 | 0 | 9 | 0.4444 | 7 | 0.1429 | 5 | 0.6 | 25 | 0.32 | 3 | 0.3333 | 1 | 1 | 9 | 0.1111 | 4 | 0.25 | 17 | 0.2353 |
| 18_1232390 | 5 | 0.4 | 5 | 0.6 | 4 | 0.5 | 6 | 0.3333 | 20 | 0.45 | 4 | 0.5 | 1 | 1 | 8 | 0.25 | 4 | 0.5 | 17 | 0.4118 |
| 18_1232395 | 5 | 1 | 7 | 0.5714 | 5 | 0.8 | 7 | 0.8571 | 24 | 0.7917 | 4 | 0.75 | 1 | 1 | 8 | 0.875 | 5 | 0.8 | 18 | 0.8333 |
| 18_1232400 | 4 | 0 | 5 | 0 | 5 | 0.6 | 6 | 0 | 20 | 0.15 | 2 | 0.5 | 1 | 0 | 6 | 0.1667 | 5 | 0.6 | 14 | 0.3571 |
| 18_1238556 | NA | NA | 5 | 0.2 | 3 | 0 | NA | NA | 8 | 0.125 | 2 | 0 | 2 | 0 | 3 | 0.3333 | NA | NA | 7 | 0.1429 |
| 18_1238559 | NA | NA | 5 | 0.2 | 4 | 0 | NA | NA | 9 | 0.1111 | 4 | 0.5 | 1 | 0 | 2 | 0 | 1 | 1 | 8 | 0.375 |
| 18_1238747 | NA | NA | 3 | 0.3333 | 1 | 0 | NA | NA | 4 | 0.25 | 5 | 0.4 | 2 | 0 | 3 | 0 | 1 | 0 | 11 | 0.1818 |
| 18_1238767 | NA | NA | 3 | 0.3333 | 1 | 0 | NA | NA | 4 | 0.25 | 5 | 0 | 2 | 0.5 | 2 | 0 | NA | NA | 9 | 0.1111 |
| 18_1239369 | 3 | 0.3333 | 3 | 0 | 1 | 0 | NA | NA | 7 | 0.1429 | NA | NA | 2 | 0.5 | NA | NA | 1 | 0 | 3 | 0.3333 |
| 18_1239392 | 2 | 0.5 | 2 | 0.5 | 1 | 0 | NA | NA | 5 | 0.4 | NA | NA | 3 | 0.6667 | NA | NA | 1 | 0 | 4 | 0.5 |
| 18_1282835 | 6 | 0.1667 | 7 | 0 | 2 | 0 | 2 | 0 | 17 | 0.0588 | 4 | 0 | 4 | 0.25 | 2 | 0 | 2 | 0 | 12 | 0.0833 |
| 18_1282837 | 7 | 0.1429 | 7 | 0.1429 | 3 | 0 | 1 | 1 | 18 | 0.1667 | 4 | 0.25 | 4 | 0.5 | 2 | 0 | 3 | 0.3333 | 13 | 0.3077 |
| 18_1282847 | 5 | 0 | 4 | 0.25 | 3 | 0.6667 | 2 | 0.5 | 14 | 0.2857 | NA | NA | 1 | 1 | 3 | 0.3333 | 2 | 0 | 6 | 0.3333 |
| 18_1286980 | 10 | 0.2 | 8 | 0.125 | 7 | 0 | 3 | 0.3333 | 28 | 0.1429 | 2 | 0.5 | 6 | 0 | 9 | 0 | 3 | 0 | 20 | 0.05 |
| 18_1287133 | 2 | 0.5 | NA | NA | 2 | 0 | 2 | 0.5 | 6 | 0.3333 | 1 | 0 | NA | NA | 3 | 0.3333 | NA | NA | 4 | 0.25 |
| 18_1287146 | 2 | 1 | NA | NA | 2 | 0 | 2 | 0 | 6 | 0.3333 | 1 | 0 | 1 | 1 | 3 | 0.6667 | NA | NA | 5 | 0.6 |
| 18_1287232 | 2 | 0.5 | 1 | 0 | NA | NA | 1 | 0 | 4 | 0.25 | NA | NA | 1 | 1 | 2 | 0.5 | NA | NA | 3 | 0.6667 |
| 18_1288400 | 1 | 1 | 1 | 0 | NA | NA | 1 | 1 | 3 | 0.6667 | NA | NA | NA | NA | 1 | 1 | NA | NA | 1 | 1 |
| 18_1288402 | 1 | 0 | 1 | 1 | NA | NA | 1 | 0 | 3 | 0.3333 | NA | NA | NA | NA | 1 | 1 | NA | NA | 1 | 1 |
| 18_1288408 | NA | NA | 1 | 1 | NA | NA | 1 | 1 | 2 | 1 | NA | NA | NA | NA | 1 | 1 | NA | NA | 1 | 1 |
| 18_1288453 | 1 | 1 | 1 | 1 | NA | NA | NA | NA | 2 | 1 | NA | NA | NA | NA | 1 | 1 | NA | NA | 1 | 1 |
| 18_1288455 | 1 | 1 | 1 | 1 | NA | NA | NA | NA | 2 | 1 | NA | NA | NA | NA | 1 | 1 | NA | NA | 1 | 1 |
| 18_1288495 | 1 | 1 | 1 | 0 | NA | NA | NA | NA | 2 | 0.5 | NA | NA | NA | NA | 1 | 1 | NA | NA | 1 | 1 |
| 18_1288498 | 1 | 1 | 1 | 1 | NA | NA | NA | NA | 2 | 1 | NA | NA | NA | NA | 1 | 1 | NA | NA | 1 | 1 |
| 18_1288710 | 3 | 0.3333 | 4 | 1 | 1 | 1 | NA | NA | 8 | 0.75 | NA | NA | 1 | 1 | NA | NA | NA | NA | 1 | 1 |
| 18_1288929 | 12 | 0.25 | 12 | 0.0833 | 14 | 0.0714 | 12 | 0 | 50 | 0.1 | 6 | 0 | 12 | 0.0833 | 6 | 0 | 8 | 0.125 | 32 | 0.0625 |
| 18_1288932 | 12 | 0.0833 | 12 | 0.1667 | 14 | 0.0714 | 11 | 0.0909 | 49 | 0.102 | 6 | 0 | 15 | 0.0667 | 6 | 0.1667 | 6 | 0.1667 | 33 | 0.0909 |
| 18_1299550 | 2 | 0 | 2 | 0.5 | 4 | 0 | 2 | 0 | 10 | 0.1 | 1 | 0 | 1 | 0 | 2 | 0.5 | 2 | 0.5 | 6 | 0.3333 |
| 18_1305888 | NA | NA | NA | NA | 3 | 0.6667 | 1 | 0 | 4 | 0.5 | NA | NA | NA | NA | 1 | 1 | NA | NA | 1 | 1 |
| 18_1310499 | NA | NA | 2 | 0.5 | NA | NA | NA | NA | 2 | 0.5 | 2 | 0.5 | 1 | 1 | 1 | 0 | NA | NA | 4 | 0.5 |
| 18_1310515 | NA | NA | 2 | 0.5 | NA | NA | NA | NA | 2 | 0.5 | 2 | 0.5 | 1 | 1 | 1 | 0 | NA | NA | 4 | 0.5 |
| 18_1382146 | 2 | 0 | 1 | 0 | 2 | 0.5 | 1 | 1 | 6 | 0.3333 | 1 | 1 | NA | NA | 1 | 0 | 1 | 0 | 3 | 0.3333 |
| 18_1382168 | 1 | 1 | NA | NA | 1 | 0 | 1 | 1 | 3 | 0.6667 | 1 | 1 | NA | NA | NA | NA | 1 | 1 | 2 | 1 |
| 18_1382170 | 1 | 1 | NA | NA | 1 | 0 | 1 | 1 | 3 | 0.6667 | 1 | 1 | NA | NA | NA | NA | NA | NA | 1 | 1 |
| 18_1382179 | 2 | 0.5 | NA | NA | 1 | 0 | 1 | 0 | 4 | 0.25 | 1 | 1 | NA | NA | NA | NA | 1 | 0 | 2 | 0.5 |
| 18_1382202 | 2 | 1 | NA | NA | 1 | 1 | 1 | 1 | 4 | 1 | 1 | 1 | NA | NA | NA | NA | 1 | 1 | 2 | 1 |

| | | | | | | | | | | | | | | | | | | | | |
|---|---|---|---|---|---|---|---|---|---|---|---|---|---|---|---|---|---|---|---|---|
| 18_1397722 | 1 | 1 | NA | NA | NA | NA | 1 | 1 | 2 | 1 | 1 | 1 | NA | NA | NA | NA | 1 | 1 | 2 | 1 |
| 18_1397729 | 1 | 1 | NA | NA | 1 | 1 | NA | NA | 2 | 1 | 1 | 1 | NA | NA | NA | NA | 1 | 0 | 2 | 0.5 |
| 18_1397744 | 1 | 0 | NA | NA | 1 | 1 | 1 | 0 | 3 | 0.3333 | 1 | 0 | NA | NA | NA | NA | 1 | 1 | 2 | 0.5 |
| 18_1397757 | 1 | 0 | NA | NA | 1 | 0 | 1 | 1 | 3 | 0.3333 | 1 | 1 | NA | NA | NA | NA | 1 | 1 | 2 | 1 |
| 18_1397842 | NA | NA | NA | NA | 1 | 1 | 1 | 0 | 2 | 0.5 | 1 | 0 | NA | NA | NA | NA | 1 | 1 | 2 | 0.5 |
| 18_1397854 | 1 | 1 | NA | NA | 1 | 1 | 1 | 0 | 3 | 0.6667 | 1 | 1 | NA | NA | NA | NA | 1 | 1 | 2 | 1 |
| 18_1397876 | 1 | 1 | NA | NA | 1 | 0 | 1 | 0 | 3 | 0.3333 | 1 | 0 | NA | NA | NA | NA | 1 | 1 | 2 | 0.5 |
| 18_1397881 | 1 | 1 | NA | NA | 1 | 0 | NA | NA | 2 | 0.5 | 1 | 1 | NA | NA | NA | NA | 1 | 1 | 2 | 1 |
| 18_1397883 | 1 | 1 | NA | NA | 1 | 1 | 1 | 1 | 3 | 1 | 1 | 1 | NA | NA | NA | NA | NA | NA | 1 | 1 |
| 18_1397888 | 1 | 1 | NA | NA | 1 | 1 | 1 | 1 | 3 | 1 | 1 | 1 | NA | NA | NA | NA | 1 | 0 | 2 | 0.5 |
| 18_1397905 | 1 | 1 | NA | NA | NA | NA | 1 | 1 | 2 | 1 | 1 | 1 | NA | NA | NA | NA | 1 | 0 | 2 | 0.5 |
| 18_1397948 | 1 | 0 | NA | NA | 1 | 1 | 1 | 0 | 3 | 0.3333 | NA | NA | NA | NA | NA | NA | 1 | 1 | 1 | 1 |
| 18_1397982 | 1 | 0 | NA | NA | NA | NA | 1 | 1 | 2 | 0.5 | 1 | 1 | NA | NA | NA | NA | 1 | 0 | 2 | 0.5 |
| 18_1398006 | 1 | 1 | NA | NA | NA | NA | 1 | 0 | 2 | 0.5 | 1 | 1 | NA | NA | NA | NA | NA | NA | 1 | 1 |
| 18_1475968 | 4 | 0 | 2 | 0 | 3 | 0.3333 | 2 | 0 | 11 | 0.0909 | 1 | 0 | 3 | 0 | 3 | 0.3333 | 4 | 0.25 | 11 | 0.1818 |
| 18_1476005 | 4 | 1 | NA | NA | 3 | 1 | 2 | 1 | 9 | 1 | 1 | 1 | 1 | 1 | 2 | 1 | 3 | 0.6667 | 7 | 0.8571 |
| 18_1476007 | 4 | 0.5 | NA | NA | 3 | 0.6667 | 2 | 1 | 9 | 0.6667 | 1 | 1 | 1 | 1 | 2 | 0.5 | 2 | 0 | 6 | 0.5 |
| 18_1476011 | 2 | 0.5 | NA | NA | 2 | 0 | 1 | 0 | 5 | 0.2 | 1 | 0 | 1 | 0 | 1 | 0 | 2 | 0.5 | 5 | 0.2 |
| 18_1476025 | 3 | 0 | NA | NA | 3 | 0 | 2 | 0.5 | 8 | 0.125 | NA | NA | 1 | 0 | 2 | 0.5 | 2 | 0.5 | 5 | 0.4 |
| 18_1476027 | 4 | 0.75 | NA | NA | 3 | 0.6667 | 2 | 0.5 | 9 | 0.6667 | NA | NA | 1 | 1 | 2 | 0 | 2 | 1 | 5 | 0.6 |
| 18_1510198 | 7 | 0 | 12 | 0.25 | 6 | 0.1667 | 13 | 0.3077 | 38 | 0.2105 | 14 | 0.1429 | 21 | 0.1905 | 14 | 0.2143 | 17 | 0.2353 | 66 | 0.197 |
| 18_1510200 | 7 | 0 | 11 | 0.1818 | 5 | 0 | 12 | 0.0833 | 35 | 0.0857 | 13 | 0 | 18 | 0.2222 | 13 | 0 | 13 | 0.2308 | 57 | 0.1228 |
| 18_1510247 | 11 | 0.7273 | 8 | 1 | 10 | 0.7 | 17 | 0.5882 | 46 | 0.7174 | 10 | 0.7 | 23 | 0.6522 | 14 | 0.7143 | 18 | 0.5556 | 65 | 0.6462 |
| 18_1510249 | 11 | 0.0909 | 10 | 0.2 | 11 | 0 | 15 | 0.0667 | 47 | 0.0851 | 11 | 0.2727 | 23 | 0.0435 | 15 | 0.1333 | 16 | 0 | 65 | 0.0923 |
| 18_1510251 | 8 | 0.5 | 9 | 0.8889 | 11 | 0.4545 | 15 | 0.4667 | 43 | 0.5581 | 9 | 0.7778 | 21 | 0.6667 | 12 | 0.6667 | 14 | 0.6429 | 56 | 0.6786 |
| 18_1510253 | 9 | 0.5556 | 10 | 0.8 | 9 | 0.6667 | 13 | 0.8462 | 41 | 0.7317 | 9 | 0.7778 | 20 | 0.8 | 14 | 0.9286 | 16 | 0.8125 | 59 | 0.8305 |
| 18_1510258 | 9 | 0.2222 | 9 | 0.4444 | 9 | 0.1111 | 16 | 0.375 | 43 | 0.3023 | 8 | 0.375 | 23 | 0.3913 | 11 | 0.3636 | 17 | 0.2941 | 59 | 0.3559 |
| 18_1510259 | 6 | 1 | 5 | 1 | 6 | 1 | 12 | 0.9167 | 29 | 0.9655 | 9 | 0.8889 | 23 | 0.913 | 9 | 0.8889 | 18 | 0.8889 | 59 | 0.8983 |
| 18_1510269 | 7 | 0.2857 | 7 | 0.4286 | 6 | 0.6667 | 13 | 0.3077 | 33 | 0.3939 | 7 | 0.5714 | 20 | 0.4 | 10 | 0.3 | 16 | 0.3125 | 53 | 0.3774 |
| 18_1510720 | 7 | 0 | 8 | 0.375 | 7 | 0.1429 | 14 | 0.1429 | 36 | 0.1667 | 9 | 0.1111 | 24 | 0.0833 | 12 | 0.1667 | 16 | 0.1875 | 61 | 0.1311 |
| 18_1510726 | 6 | 0 | 7 | 0 | 8 | 0.25 | 12 | 0 | 33 | 0.0606 | 8 | 0 | 25 | 0.08 | 11 | 0.1818 | 17 | 0.2353 | 61 | 0.1311 |
| 18_1519495 | 2 | 0 | 2 | 0.5 | 1 | 0 | 4 | 0.25 | 9 | 0.2222 | 3 | 0 | 9 | 0.3333 | 5 | 0.2 | 3 | 0 | 20 | 0.2 |
| 18_1519497 | 1 | 1 | 2 | 1 | 1 | 1 | 4 | 1 | 8 | 1 | 3 | 0.6667 | 11 | 0.9091 | 7 | 0.8571 | 6 | 0.8333 | 27 | 0.8519 |
| 18_1519508 | NA | NA | 1 | 0 | NA | NA | 4 | 0.5 | 5 | 0.4 | 1 | 1 | 8 | 0.375 | 6 | 0.6667 | 4 | 0.5 | 19 | 0.5263 |
| 18_1519523 | 1 | 0 | 2 | 0 | NA | NA | 4 | 0.5 | 7 | 0.2857 | 3 | 0 | 9 | 0 | 7 | 0.1429 | 5 | 0.2 | 24 | 0.0833 |
| 18_1519535 | 1 | 1 | 2 | 1 | NA | NA | 3 | 0.6667 | 6 | 0.8333 | 2 | 1 | 4 | 0.75 | 7 | 0.7143 | 4 | 0.75 | 17 | 0.7647 |
| 18_1519538 | 1 | 1 | 2 | 0 | NA | NA | 3 | 0.3333 | 6 | 0.3333 | 3 | 0 | 9 | 0.1111 | 8 | 0 | 4 | 0.25 | 24 | 0.0833 |
| 18_1519540 | NA | NA | NA | NA | NA | NA | 2 | 0.5 | 2 | 0.5 | 2 | 0 | 7 | 0.2857 | 5 | 0.6 | 4 | 0.5 | 18 | 0.3889 |
| 18_1519551 | 1 | 0 | 2 | 1 | NA | NA | 3 | 0.6667 | 6 | 0.6667 | 2 | 1 | 8 | 0.375 | 6 | 0.1667 | 5 | 0.4 | 21 | 0.381 |
| 18_1519564 | 1 | 1 | 2 | 1 | NA | NA | 2 | 0 | 5 | 0.6 | 2 | 0 | 6 | 0.8333 | 8 | 0.625 | 5 | 0.4 | 21 | 0.5714 |
| 18_1519587 | 1 | 1 | 1 | 1 | NA | NA | 2 | 0.5 | 4 | 0.75 | 3 | 0.3333 | 3 | 0.3333 | 6 | 0.5 | 4 | 0.25 | 16 | 0.375 |
| 18_1519626 | NA | NA | 1 | 0 | NA | NA | 2 | 0.5 | 3 | 0.3333 | 1 | 0 | 5 | 0.2 | 1 | 0 | 2 | 0 | 9 | 0.1111 |
| 18_1519633 | 1 | 0 | 1 | 1 | NA | NA | 2 | 1 | 4 | 0.75 | 1 | 0 | 3 | 0 | NA | NA | 2 | 0.5 | 6 | 0.1667 |
| 18_1519655 | NA | NA | NA | NA | NA | NA | 2 | 1 | 2 | 1 | NA | NA | 1 | 1 | NA | NA | 1 | 0 | 2 | 0.5 |
| 18_1519659 | NA | NA | 1 | 0 | NA | NA | 2 | 1 | 3 | 0.6667 | NA | NA | 2 | 0.5 | 1 | 1 | 1 | 0 | 4 | 0.5 |
| 18_1519741 | NA | NA | NA | NA | 1 | 1 | NA | NA | 1 | 1 | NA | NA | 1 | 1 | NA | NA | NA | NA | 1 | 1 |
| 18_1519748 | 1 | 0 | 1 | 1 | NA | NA | 2 | 1 | 4 | 0.75 | NA | NA | NA | NA | NA | NA | 3 | 0.6667 | 3 | 0.6667 |
| 18_1519773 | NA | NA | NA | NA | 1 | 1 | NA | NA | 1 | 1 | NA | NA | 1 | 1 | 1 | 1 | NA | NA | 2 | 1 |
| 18_1519803 | 1 | 1 | NA | NA | 1 | 0 | NA | NA | 2 | 0.5 | NA | NA | 1 | 0 | 1 | 1 | NA | NA | 2 | 0.5 |
| 18_1519880 | 1 | 1 | NA | NA | 2 | 1 | NA | NA | 3 | 1 | NA | NA | NA | NA | 1 | 1 | NA | NA | 1 | 1 |
| 18_1519898 | 1 | 1 | NA | NA | 2 | 0 | NA | NA | 3 | 0.3333 | NA | NA | NA | NA | 1 | 1 | NA | NA | 1 | 1 |
| 18_1519904 | 1 | 0 | NA | NA | 2 | 0.5 | NA | NA | 3 | 0.3333 | NA | NA | 1 | 1 | 1 | 0 | NA | NA | 2 | 0.5 |
| 18_1519938 | 1 | 1 | NA | NA | 2 | 0.5 | NA | NA | 3 | 0.6667 | NA | NA | 1 | 1 | 1 | 1 | NA | NA | 2 | 1 |
| 18_1519942 | 1 | 1 | NA | NA | 2 | 1 | NA | NA | 3 | 1 | NA | NA | 1 | 1 | 1 | 1 | NA | NA | 2 | 1 |
| 18_1519955 | 1 | 1 | NA | NA | 1 | 1 | NA | NA | 2 | 1 | NA | NA | 1 | 1 | NA | NA | NA | NA | 1 | 1 |
| 18_1519960 | 1 | 1 | NA | NA | 2 | 1 | NA | NA | 3 | 1 | NA | NA | 1 | 1 | NA | NA | NA | NA | 1 | 1 |
| 18_1519962 | 1 | 1 | NA | NA | 2 | 0.5 | NA | NA | 3 | 0.6667 | NA | NA | 1 | 1 | 1 | 1 | NA | NA | 2 | 1 |
| 18_1519974 | 1 | 1 | NA | NA | 1 | 0 | NA | NA | 2 | 0.5 | NA | NA | 1 | 1 | NA | NA | NA | NA | 1 | 1 |
| 18_1519975 | 1 | 1 | NA | NA | 2 | 1 | NA | NA | 3 | 1 | NA | NA | NA | NA | 1 | 1 | NA | NA | 1 | 1 |
| 18_1534661 | NA | NA | NA | NA | 1 | 1 | NA | NA | 1 | 1 | 1 | 1 | NA | NA | 1 | 0 | NA | NA | 2 | 0.5 |
| 18_1534671 | NA | NA | NA | NA | 1 | 1 | NA | NA | 1 | 1 | 1 | 1 | NA | NA | 1 | 1 | NA | NA | 2 | 1 |
| 18_1534673 | NA | NA | NA | NA | 1 | 1 | NA | NA | 1 | 1 | 1 | 1 | NA | NA | 1 | 1 | NA | NA | 2 | 1 |
| 18_1534684 | NA | NA | NA | NA | 1 | 1 | NA | NA | 1 | 1 | 1 | 1 | NA | NA | 1 | 1 | NA | NA | 2 | 1 |
| 18_1534706 | NA | NA | NA | NA | 1 | 1 | NA | NA | 1 | 1 | 1 | 1 | NA | NA | 1 | 1 | NA | NA | 2 | 1 |
| 18_1534714 | NA | NA | NA | NA | 1 | 1 | NA | NA | 1 | 1 | 1 | 1 | NA | NA | NA | NA | NA | NA | 1 | 1 |

| | | | | | | | | | | | | | | | | | | | | |
|---|---|---|---|---|---|---|---|---|---|---|---|---|---|---|---|---|---|---|---|---|
| 18_153473 | NA | NA | NA | NA | 1 | 1 | NA | NA | 1 | 1 | 1 | 1 | NA | NA | 1 | 1 | NA | NA | 2 | 1 |
| 18_153474 | NA | NA | NA | NA | 1 | 1 | NA | NA | 1 | 1 | NA | NA | NA | NA | 1 | 1 | NA | NA | 1 | 1 |
| 18_155231 | 9 | 0.3333 | 16 | 0.125 | 8 | 0 | 13 | 0 | 46 | 0.1087 | 10 | 0.1 | 14 | 0.2857 | 6 | 0.3333 | 15 | 0.2 | 45 | 0.2222 |
| 18_155231 | 11 | 0.2727 | 16 | 0.75 | 10 | 0.4 | 13 | 0.3077 | 50 | 0.46 | 11 | 0.6364 | 14 | 0.6429 | 9 | 0.5556 | 15 | 0.6667 | 49 | 0.6327 |
| 18_155231 | 10 | 0.3 | 15 | 0.0667 | 10 | 0 | 12 | 0 | 47 | 0.0851 | 10 | 0 | 13 | 0.1538 | 8 | 0 | 16 | 0.0625 | 47 | 0.0638 |
| 18_155233 | 10 | 0.5 | 10 | 0.6 | 9 | 0.3333 | 10 | 0.1 | 39 | 0.3846 | 10 | 0.4 | 11 | 0.4545 | 9 | 0.3333 | 15 | 0.1333 | 45 | 0.3111 |
| 18_155235 | 10 | 0.2 | 11 | 0 | 7 | 0.1429 | 11 | 0 | 39 | 0.0769 | 8 | 0 | 9 | 0 | 6 | 0 | 15 | 0.1333 | 38 | 0.0526 |
| 18_155236 | 7 | 0.4286 | 12 | 0.0833 | 10 | 0.1 | 10 | 0 | 39 | 0.1282 | 11 | 0.1818 | 9 | 0.1111 | 8 | 0.125 | 15 | 0.0667 | 43 | 0.1163 |
| 18_155236 | 7 | 0.1429 | 14 | 0.0714 | 9 | 0 | 9 | 0 | 39 | 0.0513 | 9 | 0.1111 | 9 | 0.2222 | 8 | 0.125 | 13 | 0.0769 | 39 | 0.1282 |
| 18_155242 | 5 | 0.4 | 12 | 0.0833 | 8 | 0.25 | 10 | 0.1 | 35 | 0.1714 | 9 | 0.2222 | 8 | 0.125 | 7 | 0.1429 | 14 | 0.1429 | 38 | 0.1579 |
| 18_155256 | 3 | 0 | 7 | 0.1429 | 6 | 0.1667 | 7 | 0.1429 | 23 | 0.1304 | 5 | 0.2 | 4 | 0 | 4 | 0 | 12 | 0.0833 | 25 | 0.08 |
| 18_155259 | 3 | 0 | 7 | 0.2857 | 5 | 0.2 | 5 | 0.4 | 20 | 0.25 | 7 | 0.2857 | 4 | 0.5 | 3 | 0.3333 | 9 | 0.2222 | 23 | 0.3043 |
| 18_155261 | 4 | 0.5 | 7 | 0.4286 | 6 | 0.5 | 7 | 0.5714 | 24 | 0.5 | 6 | 0.6667 | 4 | 0.75 | 3 | 0.3333 | 10 | 0.4 | 23 | 0.5217 |
| 18_155261 | 3 | 0.3333 | 7 | 0.8571 | 5 | 0.2 | 6 | 0.5 | 21 | 0.5238 | 4 | 0.75 | 4 | 0.75 | 3 | 0.3333 | 11 | 1 | 22 | 0.8182 |
| 18_155262 | 4 | 0.25 | 6 | 0 | 6 | 0.1667 | 3 | 0 | 19 | 0.1053 | 5 | 0.4 | 4 | 0 | 2 | 0 | 11 | 0.1818 | 22 | 0.1818 |
| 18_155262 | 4 | 0.5 | 4 | 0.25 | 6 | 0.1667 | 5 | 0.2 | 19 | 0.2632 | 5 | 0.6 | 5 | 0.4 | 3 | 0.3333 | 8 | 0.25 | 21 | 0.381 |
| 18_155264 | 4 | 0.5 | 6 | 0.6667 | 6 | 1 | 6 | 0.6667 | 22 | 0.7273 | 3 | 0.3333 | 4 | 1 | 2 | 1 | 9 | 0.7778 | 18 | 0.7778 |
| 18_155264 | 4 | 0.5 | 6 | 0 | 6 | 0.3333 | 7 | 0.4286 | 23 | 0.3043 | 6 | 0.3333 | 4 | 0.5 | 3 | 0.3333 | 12 | 0.5 | 25 | 0.44 |
| 18_155265 | 4 | 0.75 | 6 | 0.6667 | 5 | 0.6 | 4 | 0.5 | 19 | 0.6316 | 2 | 1 | 4 | 0.5 | 1 | 0 | 11 | 0.4545 | 18 | 0.5 |
| 18_155265 | 2 | 0.5 | 5 | 0.4 | 4 | 0.75 | 4 | 0.5 | 15 | 0.5333 | 5 | 0.6 | 3 | 1 | 3 | 1 | 8 | 0.875 | 19 | 0.8421 |
| 18_155265 | 2 | 0 | 4 | 0.25 | 3 | 0.6667 | 5 | 0.2 | 14 | 0.2857 | 4 | 0 | 5 | 0.4 | 2 | 1 | 10 | 0.4 | 21 | 0.381 |
| 18_155267 | 2 | 0 | 5 | 0.4 | 3 | 0.3333 | 6 | 0.1667 | 16 | 0.25 | 4 | 0 | 2 | 0.5 | 2 | 0 | 8 | 0.25 | 16 | 0.1875 |
| 18_155311 | 2 | 0 | 6 | 0.1667 | 4 | 0 | 4 | 0.25 | 16 | 0.125 | 3 | 0 | 2 | 0 | 2 | 0 | 9 | 0.1111 | 16 | 0.0625 |
| 18_155312 | 1 | 0 | 6 | 0 | 4 | 0.25 | 7 | 0.2857 | 18 | 0.1667 | 3 | 0.3333 | 3 | 0 | 2 | 0 | 6 | 0.1667 | 14 | 0.1429 |
| 18_155327 | 2 | 0.5 | 6 | 0.3333 | 3 | 0.6667 | 5 | 0.4 | 16 | 0.4375 | 2 | 1 | 4 | 0 | 2 | 0 | 7 | 0.1429 | 15 | 0.2 |
| 18_155328 | 2 | 1 | 6 | 1 | 3 | 1 | 4 | 1 | 15 | 1 | 1 | 1 | 4 | 1 | 2 | 0.5 | 8 | 0.875 | 15 | 0.8667 |
| 18_155328 | 2 | 1 | 6 | 1 | 3 | 0.3333 | 6 | 1 | 17 | 0.8824 | 2 | 0 | 3 | 1 | 2 | 1 | 7 | 0.7143 | 14 | 0.7143 |
| 18_155329 | NA | NA | 6 | 0.3333 | 4 | 0.25 | 5 | 0 | 15 | 0.2 | 1 | 0 | 3 | 0 | 2 | 0.5 | 5 | 0.2 | 11 | 0.1818 |
| 18_155330 | 2 | 0.5 | 7 | 0.4286 | 4 | 0.25 | 5 | 0.4 | 18 | 0.3889 | 2 | 0 | 4 | 0.25 | NA | NA | 6 | 0.6667 | 12 | 0.4167 |
| 18_167220 | 7 | 0 | 7 | 0 | 9 | 0.1111 | 17 | 0.0588 | 40 | 0.05 | 11 | 0.0909 | 14 | 0.1429 | 12 | 0 | 17 | 0.1765 | 54 | 0.1111 |
| 18_167224 | 8 | 0 | 6 | 0.1667 | 10 | 0 | 15 | 0.2667 | 39 | 0.1282 | 10 | 0.5 | 11 | 0.2727 | 10 | 0.4 | 19 | 0.2632 | 50 | 0.34 |
| 18_167230 | 9 | 0.1111 | 4 | 0 | 9 | 0 | 14 | 0.0714 | 36 | 0.0556 | 10 | 0.1 | 9 | 0 | 10 | 0 | 19 | 0.1053 | 48 | 0.0625 |
| 18_167231 | 8 | 0.375 | 4 | 0 | 6 | 0.3333 | 13 | 0.1538 | 31 | 0.2258 | 8 | 0.125 | 4 | 0 | 7 | 0 | 15 | 0.2667 | 34 | 0.1471 |
| 18_167231 | 7 | 0.8571 | 3 | 1 | 5 | 0.6 | 14 | 0.7143 | 29 | 0.7586 | 9 | 0.7778 | 6 | 0.8333 | 6 | 1 | 15 | 0.8 | 36 | 0.8333 |
| 18_167231 | 8 | 0.375 | 4 | 0.5 | 9 | 0.2222 | 14 | 0.5714 | 35 | 0.4286 | 10 | 0.6 | 8 | 0.375 | 8 | 0.25 | 20 | 0.25 | 46 | 0.3478 |
| 18_167231 | 10 | 0.7 | 4 | 0.5 | 9 | 0.6667 | 15 | 0.6 | 38 | 0.6316 | 12 | 0.75 | 10 | 0.7 | 9 | 0.6667 | 18 | 0.7778 | 49 | 0.7347 |
| 18_167232 | 9 | 0.1111 | 4 | 0 | 9 | 0 | 14 | 0.0714 | 36 | 0.0556 | 10 | 0 | 10 | 0 | 8 | 0.125 | 16 | 0.125 | 44 | 0.0682 |
| 18_167232 | 7 | 0.1429 | 3 | 0.3333 | 9 | 0 | 14 | 0.1429 | 33 | 0.1212 | 10 | 0.1 | 8 | 0.125 | 10 | 0 | 18 | 0.0556 | 46 | 0.0652 |
| 18_167232 | 10 | 0.6 | 4 | 0.5 | 9 | 0.5556 | 15 | 0.4 | 38 | 0.5 | 10 | 0.5 | 9 | 0.6667 | 8 | 0.625 | 20 | 0.75 | 47 | 0.6596 |
| 18_169420 | NA | NA | 1 | 1 | NA | NA | 1 | 0 | 2 | 0.5 | NA | NA | NA | NA | 1 | 1 | NA | NA | 1 | 1 |
| 18_201652 | 1 | 1 | NA | NA | 1 | 1 | NA | NA | 2 | 1 | 1 | 1 | NA | NA | NA | NA | NA | NA | 1 | 1 |
| 18_204163 | NA | NA | NA | NA | 1 | 1 | NA | NA | 1 | 1 | NA | NA | NA | NA | 1 | 1 | 1 | 1 | 2 | 1 |
| 18_208152 | NA | NA | NA | NA | 1 | 1 | NA | NA | 1 | 1 | NA | NA | NA | NA | 1 | 1 | NA | NA | 1 | 1 |
| 18_208161 | NA | NA | NA | NA | 1 | 1 | 1 | 1 | 2 | 1 | NA | NA | NA | NA | 2 | 1 | 1 | 1 | 3 | 1 |
| 18_208205 | 6 | 0 | 7 | 0.1429 | 10 | 0.1 | 8 | 0 | 31 | 0.0645 | 6 | 0.3333 | 10 | 0.2 | 8 | 0 | 7 | 0.1429 | 31 | 0.1613 |
| 18_208207 | 5 | 0.4 | 5 | 0.4 | 10 | 0.2 | 8 | 0.25 | 28 | 0.2857 | 7 | 0.5714 | 9 | 0.5556 | 7 | 0.1429 | 5 | 0.2 | 28 | 0.3929 |
| 18_208208 | 2 | 0.5 | 2 | 0.5 | 8 | 0 | 4 | 0.5 | 16 | 0.25 | 3 | 0.3333 | 7 | 0.1429 | 6 | 0.5 | 4 | 0.25 | 20 | 0.3 |
| 18_208209 | 3 | 0.3333 | 4 | 0.25 | 4 | 0.25 | 8 | 0.5 | 19 | 0.3684 | 4 | 0.75 | 6 | 0.1667 | 6 | 0.8333 | 4 | 0.5 | 20 | 0.55 |
| 18_208211 | 5 | 0.8 | 7 | 0.7143 | 9 | 0.8889 | 8 | 0.875 | 29 | 0.8276 | 7 | 0.8571 | 12 | 0.8333 | 7 | 0.7143 | 7 | 0.8571 | 33 | 0.8182 |
| 18_208211 | 5 | 0 | 7 | 0.2857 | 10 | 0.2 | 9 | 0 | 31 | 0.129 | 7 | 0 | 15 | 0.2 | 8 | 0.375 | 8 | 0 | 38 | 0.1579 |
| 18_208213 | 2 | 1 | 5 | 1 | 8 | 0.875 | 7 | 1 | 22 | 0.9545 | 6 | 1 | 15 | 0.8667 | 6 | 0.8333 | 8 | 0.875 | 35 | 0.8857 |
| 18_208213 | 1 | 1 | 6 | 0.1667 | 8 | 0.375 | 8 | 0.625 | 23 | 0.4348 | 7 | 0.4286 | 11 | 0.3636 | 8 | 0.5 | 7 | 0.2857 | 33 | 0.3939 |
| 18_208214 | 3 | 1 | 6 | 0.8333 | 7 | 0.5714 | 6 | 0.8333 | 22 | 0.7727 | 8 | 0.625 | 12 | 0.8333 | 6 | 0.6667 | 7 | 0.8571 | 33 | 0.7576 |
| 18_208785 | 3 | 0.3333 | 2 | 0 | 3 | 0.6667 | 3 | 0.3333 | 11 | 0.3636 | 3 | 0.3333 | 5 | 0.6 | 4 | 0.25 | 4 | 0 | 16 | 0.3125 |
| 18_208788 | 2 | 0.5 | 1 | 0 | 6 | 0 | 2 | 0.5 | 11 | 0.1818 | 1 | 0 | 3 | 0.3333 | 5 | 0.2 | 2 | 0 | 11 | 0.1818 |
| 18_209509 | 4 | 0.5 | 7 | 0.4286 | 3 | 0.3333 | 2 | 0 | 16 | 0.375 | 1 | 1 | 5 | 0.4 | 5 | 0 | 3 | 0 | 14 | 0.2143 |
| 18_209510 | 3 | 0.6667 | 7 | 0.8571 | 3 | 0.6667 | 2 | 1 | 15 | 0.8 | 1 | 1 | 4 | 0.75 | 5 | 1 | 2 | 1 | 12 | 0.9167 |
| 18_209511 | 4 | 0.75 | 8 | 0.5 | 3 | 0.6667 | 4 | 0.75 | 19 | 0.6316 | 1 | 0 | 6 | 0.5 | 5 | 0.6 | 1 | 0 | 13 | 0.4615 |
| 18_209511 | 5 | 0.4 | 7 | 0.2857 | 2 | 0.5 | 4 | 0.25 | 18 | 0.3333 | 1 | 0 | 5 | 0.2 | 5 | 0.2 | 1 | 1 | 12 | 0.25 |
| 18_209511 | 5 | 0 | 7 | 0 | 2 | 0 | 3 | 0.3333 | 17 | 0.0588 | 1 | 0 | 5 | 0.2 | 4 | 0.25 | NA | NA | 10 | 0.2 |
| 18_209513 | 4 | 0.25 | 7 | 0.2857 | 3 | 0.3333 | 3 | 0 | 17 | 0.2353 | 1 | 0 | 5 | 0 | 4 | 0.25 | 1 | 0 | 11 | 0.0909 |
| 18_209699 | 3 | 0.6667 | 3 | 0 | 2 | 0 | 4 | 0.5 | 12 | 0.3333 | 1 | 0 | 4 | 0 | 2 | 0.5 | NA | NA | 7 | 0.1429 |
| 18_209699 | 5 | 0.6 | 2 | 1 | 2 | 1 | 2 | 1 | 11 | 0.8182 | 1 | 0 | 3 | 0.3333 | 2 | 1 | NA | NA | 6 | 0.5 |
| 18_209700 | 5 | 0.8 | 3 | 0.3333 | 1 | 0 | 4 | 0.25 | 13 | 0.4615 | NA | NA | 4 | 0.25 | 4 | 0 | NA | NA | 8 | 0.125 |
| 18_209701 | 5 | 0.8 | 3 | 0.6667 | 2 | 0.5 | 4 | 0.25 | 14 | 0.5714 | NA | NA | 4 | 0.25 | 4 | 0.75 | NA | NA | 8 | 0.5 |

| | | | | | | | | | | | | | | | | | | | | |
|---|---|---|---|---|---|---|---|---|---|---|---|---|---|---|---|---|---|---|---|---|
| 18_211663 | NA | NA | 1 | 1 | NA | NA | NA | NA | 1 | 1 | NA | NA | NA | NA | NA | NA | 1 | 1 | 1 | 1 |
| 18_212585 | NA | NA | NA | NA | NA | NA | 2 | 1 | 2 | 1 | 2 | 1 | 2 | 1 | NA | NA | 1 | 1 | 5 | 1 |
| 18_212586 | NA | NA | NA | NA | NA | NA | 1 | 1 | 1 | 1 | 2 | 0.5 | 2 | 1 | NA | NA | 1 | 1 | 5 | 0.8 |
| 18_212587 | NA | NA | NA | NA | NA | NA | 2 | 0.5 | 2 | 0.5 | 2 | 0.5 | 1 | 0 | NA | NA | NA | NA | 3 | 0.3333 |
| 18_212590 | NA | NA | NA | NA | NA | NA | 2 | 1 | 2 | 1 | 2 | 1 | 1 | 1 | NA | NA | 1 | 1 | 4 | 1 |
| 18_212591 | NA | NA | NA | NA | NA | NA | 2 | 1 | 2 | 1 | 2 | 0 | 1 | 1 | NA | NA | 1 | 0 | 4 | 0.25 |
| 18_212591 | NA | NA | NA | NA | NA | NA | 2 | 1 | 2 | 1 | 2 | 1 | 1 | 1 | NA | NA | 1 | 1 | 4 | 1 |
| 18_212591 | NA | NA | NA | NA | NA | NA | 2 | 1 | 2 | 1 | 1 | 1 | 1 | 1 | NA | NA | NA | NA | 2 | 1 |
| 18_212593 | NA | NA | NA | NA | NA | NA | 1 | 1 | 1 | 1 | 2 | 1 | 1 | 0 | NA | NA | 1 | 1 | 4 | 0.75 |
| 18_212598 | NA | NA | NA | NA | NA | NA | 2 | 0.5 | 2 | 0.5 | 2 | 0.5 | 1 | 0 | NA | NA | 1 | 0 | 4 | 0.25 |
| 18_212598 | NA | NA | NA | NA | NA | NA | 2 | 1 | 2 | 1 | 2 | 0.5 | 1 | 1 | NA | NA | 1 | 0 | 4 | 0.5 |
| 18_212610 | NA | NA | NA | NA | NA | NA | 1 | 1 | 1 | 1 | 1 | 0 | 1 | 1 | NA | NA | 1 | 0 | 3 | 0.3333 |
| 18_212618 | NA | NA | NA | NA | NA | NA | 1 | 1 | 1 | 1 | 1 | 1 | NA | NA | NA | NA | 1 | 1 | 2 | 1 |
| 18_212621 | 1 | 1 | 1 | 0 | NA | NA | NA | NA | 2 | 0.5 | 1 | 0 | NA | NA | NA | NA | 2 | 0.5 | 3 | 0.3333 |
| 18_212623 | 1 | 1 | 1 | 1 | NA | NA | NA | NA | 2 | 1 | 1 | 1 | NA | NA | NA | NA | 2 | 1 | 3 | 1 |
| 18_212624 | 1 | 1 | NA | NA | NA | NA | NA | NA | 1 | 1 | 1 | 1 | NA | NA | NA | NA | 2 | 1 | 3 | 1 |
| 18_212626 | 1 | 1 | 1 | 1 | NA | NA | NA | NA | 2 | 1 | 1 | 1 | NA | NA | NA | NA | 2 | 0 | 3 | 0.3333 |
| 18_212626 | 1 | 0 | 1 | 1 | NA | NA | NA | NA | 2 | 0.5 | 1 | 1 | NA | NA | NA | NA | 2 | 0.5 | 3 | 0.6667 |
| 18_212628 | NA | NA | 1 | 1 | NA | NA | NA | NA | 1 | 1 | 1 | 1 | NA | NA | NA | NA | 1 | 0 | 2 | 0.5 |
| 18_212630 | NA | NA | 1 | 1 | NA | NA | NA | NA | 1 | 1 | 1 | 0 | NA | NA | NA | NA | 2 | 0.5 | 3 | 0.3333 |
| 18_212811 | NA | NA | 1 | 1 | NA | NA | NA | NA | 1 | 1 | 1 | 1 | NA | NA | NA | NA | 1 | 1 | 2 | 1 |
| 18_212812 | NA | NA | 1 | 1 | NA | NA | NA | NA | 1 | 1 | 1 | 1 | NA | NA | NA | NA | 2 | 0 | 3 | 0.3333 |
| 18_212813 | NA | NA | 1 | 1 | NA | NA | NA | NA | 1 | 1 | 1 | 1 | NA | NA | NA | NA | 2 | 1 | 3 | 1 |
| 18_212818 | NA | NA | 1 | 1 | NA | NA | NA | NA | 1 | 1 | NA | NA | NA | NA | NA | NA | 1 | 1 | 1 | 1 |
| 18_216843 | 1 | 0 | 1 | 0 | 1 | 1 | NA | NA | 3 | 0.3333 | 1 | 1 | NA | NA | NA | NA | NA | NA | 1 | 1 |
| 18_216851 | 1 | 1 | NA | NA | 1 | 1 | NA | NA | 2 | 1 | 1 | 1 | NA | NA | NA | NA | NA | NA | 1 | 1 |
| 18_216852 | 1 | 1 | 1 | 1 | 1 | 0 | NA | NA | 3 | 0.6667 | 1 | 1 | NA | NA | NA | NA | NA | NA | 1 | 1 |
| 18_216852 | 1 | 1 | NA | NA | 1 | 1 | NA | NA | 2 | 1 | 1 | 1 | NA | NA | NA | NA | NA | NA | 1 | 1 |
| 18_216854 | 1 | 1 | 1 | 0 | 1 | 1 | NA | NA | 3 | 0.6667 | 1 | 1 | NA | NA | NA | NA | NA | NA | 1 | 1 |
| 18_216855 | 1 | 1 | 1 | 1 | 1 | 1 | NA | NA | 3 | 1 | 1 | 1 | NA | NA | NA | NA | NA | NA | 1 | 1 |
| 18_217847 | 1 | 1 | NA | NA | 1 | 1 | NA | NA | 2 | 1 | 1 | 1 | NA | NA | NA | NA | NA | NA | 1 | 1 |
| 18_228265 | 7 | 0 | 8 | 0.125 | 4 | 0.25 | 4 | 0.5 | 23 | 0.1739 | 5 | 0 | 4 | 0.25 | 6 | 0.1667 | 8 | 0.125 | 23 | 0.1304 |
| 18_228268 | 6 | 0.3333 | 5 | 0.2 | 4 | 0.5 | 4 | 0.25 | 19 | 0.3158 | 4 | 0.5 | 4 | 0.5 | 4 | 0.25 | 4 | 0.25 | 16 | 0.375 |
| 18_228270 | 7 | 0.4286 | 8 | 0 | 8 | 0 | 6 | 0.3333 | 29 | 0.1724 | 4 | 0.25 | 6 | 0 | 7 | 0.1429 | 8 | 0 | 25 | 0.08 |
| 18_228270 | 7 | 0.2857 | 7 | 0.4286 | 8 | 0.125 | 5 | 0.4 | 27 | 0.2963 | 4 | 0.25 | 6 | 0.3333 | 6 | 0.1667 | 7 | 0.1429 | 23 | 0.2174 |
| 18_228271 | 7 | 0.4286 | 6 | 0.3333 | 6 | 0.3333 | 5 | 0.8 | 24 | 0.4583 | 4 | 0.5 | 6 | 0 | 6 | 0.1667 | 8 | 0.375 | 24 | 0.25 |
| 18_228274 | 7 | 0.2857 | 6 | 0.1667 | 8 | 0.125 | 6 | 0.3333 | 27 | 0.2222 | 3 | 0.6667 | 5 | 0.4 | 7 | 0.2857 | 7 | 0.5714 | 22 | 0.4545 |
| 18_228274 | 7 | 0.4286 | 7 | 0.4286 | 8 | 0.375 | 6 | 0.5 | 28 | 0.4286 | 1 | 0 | 3 | 0.6667 | 5 | 0.4 | 7 | 0.2857 | 16 | 0.375 |
| 18_228274 | 5 | 0 | 5 | 0 | 8 | 0 | 6 | 0.3333 | 24 | 0.0833 | 3 | 0.3333 | 5 | 0 | 6 | 0 | 6 | 0 | 20 | 0.05 |
| 18_228276 | 6 | 0.8333 | 5 | 1 | 5 | 0.8 | 5 | 1 | 21 | 0.9048 | 3 | 1 | 4 | 1 | 2 | 1 | 5 | 0.8 | 14 | 0.9286 |
| 18_228276 | 5 | 0.2 | 4 | 0.75 | 6 | 0.1667 | 4 | 0.75 | 19 | 0.4211 | 3 | 0.3333 | 3 | 0.3333 | 6 | 0.5 | 4 | 0.5 | 16 | 0.4375 |
| 18_228278 | 6 | 1 | 6 | 0.8333 | 7 | 0.8571 | 5 | 1 | 24 | 0.9167 | 3 | 1 | 5 | 1 | 6 | 1 | 5 | 1 | 19 | 1 |
| 18_228278 | 6 | 0.5 | 6 | 0.3333 | 8 | 0.625 | 6 | 0.6667 | 26 | 0.5385 | 3 | 0.6667 | 5 | 0.6 | 6 | 0.6667 | 6 | 0.5 | 20 | 0.6 |
| 18_228279 | 4 | 0.5 | 7 | 0.4286 | 7 | 0.4286 | 3 | 0.6667 | 21 | 0.4762 | 3 | 0.6667 | 5 | 0.8 | 6 | 0.8333 | 5 | 0.6 | 19 | 0.7368 |
| 18_228280 | 2 | 1 | 4 | 1 | 4 | 1 | 5 | 1 | 15 | 1 | 3 | 1 | 3 | 1 | 4 | 1 | 2 | 1 | 12 | 1 |
| 18_228280 | 2 | 1 | 6 | 1 | 5 | 1 | 3 | 0.6667 | 16 | 0.9375 | 2 | 0.5 | 4 | 1 | 5 | 1 | 3 | 1 | 14 | 0.9286 |
| 18_231730 | NA | NA | 1 | 1 | 4 | 1 | 1 | 0 | 6 | 0.8333 | 1 | 1 | 1 | 1 | 4 | 1 | 2 | 0.5 | 8 | 0.875 |
| 18_231730 | NA | NA | 2 | 1 | 4 | 0.75 | 1 | 1 | 7 | 0.8571 | 1 | 0 | 1 | 1 | 4 | 0.5 | 2 | 1 | 8 | 0.625 |
| 18_231730 | NA | NA | 2 | 0 | 4 | 0.25 | 1 | 0 | 7 | 0.1429 | 1 | 1 | 1 | 1 | 3 | 0.3333 | 2 | 1 | 7 | 0.7143 |
| 18_231732 | NA | NA | 2 | 1 | 3 | 0.3333 | 1 | 0 | 6 | 0.5 | NA | NA | 1 | 0 | 3 | 0.3333 | 2 | 0.5 | 6 | 0.3333 |
| 18_231733 | NA | NA | 2 | 1 | 3 | 0.6667 | NA | NA | 5 | 0.8 | NA | NA | 1 | 0 | 2 | 1 | 2 | 0.5 | 5 | 0.6 |
| 18_231734 | NA | NA | 1 | 1 | 1 | 1 | NA | NA | 2 | 1 | NA | NA | 1 | 1 | 1 | 0 | 2 | 0 | 4 | 0.25 |
| 18_231735 | NA | NA | 1 | 0 | 3 | 0.3333 | NA | NA | 4 | 0.25 | NA | NA | 1 | 1 | 1 | 0 | 1 | 0 | 3 | 0.3333 |
| 18_231735 | NA | NA | 1 | 1 | 3 | 0.3333 | NA | NA | 4 | 0.5 | NA | NA | NA | NA | 1 | 1 | 1 | 1 | 2 | 1 |
| 18_231739 | NA | NA | 1 | 1 | NA | NA | NA | NA | 1 | 1 | NA | NA | NA | NA | 2 | 1 | NA | NA | 2 | 1 |
| 18_231741 | NA | NA | 1 | 1 | 1 | 1 | NA | NA | 2 | 1 | NA | NA | NA | NA | 2 | 1 | NA | NA | 2 | 1 |
| 18_231741 | NA | NA | 1 | 1 | 1 | 1 | NA | NA | 2 | 1 | NA | NA | NA | NA | 1 | 1 | NA | NA | 1 | 1 |
| 18_231743 | NA | NA | 1 | 1 | 1 | 0 | NA | NA | 2 | 0.5 | NA | NA | NA | NA | 2 | 1 | NA | NA | 2 | 1 |
| 18_237664 | 1 | 1 | 6 | 0 | 4 | 0 | 8 | 0 | 19 | 0.0526 | 3 | 0.3333 | 3 | 0 | 2 | 0.5 | 4 | 0.25 | 12 | 0.25 |
| 18_237668 | 2 | 0 | 7 | 0.4286 | 6 | 0.1667 | 8 | 0 | 23 | 0.1739 | 3 | 0.3333 | 4 | 0 | 4 | 0.5 | 5 | 0.2 | 16 | 0.25 |
| 18_237676 | NA | NA | 1 | 0 | 3 | 0.3333 | 1 | 0 | 5 | 0.2 | 2 | 0 | 3 | 0.3333 | 3 | 0 | 3 | 0 | 11 | 0.0909 |
| 18_237897 | 3 | 0 | 4 | 0.25 | 2 | 0 | 6 | 0.1667 | 15 | 0.1333 | 5 | 0.2 | 3 | 0.3333 | 7 | 0 | 4 | 0.25 | 19 | 0.1579 |
| 18_237901 | 3 | 0.3333 | 4 | 0 | 2 | 0 | 5 | 0.4 | 14 | 0.2143 | 4 | 0 | 4 | 0.25 | 9 | 0.2222 | 5 | 0 | 22 | 0.1364 |
| 18_237914 | 3 | 0.3333 | 5 | 0 | 2 | 1 | 9 | 0.4444 | 19 | 0.3684 | 7 | 0.2857 | 5 | 0 | 14 | 0.1429 | 6 | 0.1667 | 32 | 0.1562 |
| 18_237915 | 2 | 0 | 8 | 0.375 | 2 | 0.5 | 7 | 0.1429 | 19 | 0.2632 | 5 | 0.2 | 6 | 0.1667 | 8 | 0.375 | 5 | 0 | 24 | 0.2083 |

| | | | | | | | | | | | | | | | | | | | | |
|---|---|---|---|---|---|---|---|---|---|---|---|---|---|---|---|---|---|---|---|---|
| 18_237920 | 4 | 0.25 | 11 | 0 | 6 | 0.1667 | 10 | 0.1 | 31 | 0.0968 | 10 | 0.2 | 8 | 0.125 | 14 | 0 | 8 | 0.25 | 40 | 0.125 |
| 18_237921 | 4 | 0.5 | 11 | 0.1818 | 6 | 0.1667 | 11 | 0.0909 | 32 | 0.1875 | 9 | 0.2222 | 8 | 0 | 15 | 0.2 | 9 | 0.2222 | 41 | 0.1707 |
| 18_237923 | 4 | 0 | 9 | 0 | 7 | 0 | 9 | 0.2222 | 29 | 0.069 | 9 | 0.2222 | 5 | 0.2 | 15 | 0.2667 | 9 | 0.2222 | 38 | 0.2368 |
| 18_237924 | 4 | 0.25 | 13 | 0.0769 | 7 | 0.1429 | 11 | 0.0909 | 35 | 0.1143 | 9 | 0.4444 | 8 | 0.125 | 15 | 0.0667 | 9 | 0 | 41 | 0.1463 |
| 18_240240 | 20 | 0.75 | 13 | 0.8462 | 12 | 0.9167 | 17 | 0.6471 | 62 | 0.7742 | 15 | 1 | 24 | 0.9583 | 18 | 0.7778 | 19 | 0.8421 | 76 | 0.8947 |
| 18_240241 | 18 | 0.5556 | 11 | 0.5455 | 7 | 0.5714 | 14 | 0.3571 | 50 | 0.5 | 14 | 0.4286 | 23 | 0.5217 | 15 | 0.5333 | 15 | 0.4 | 67 | 0.4776 |
| 18_240242 | 21 | 0.4762 | 14 | 0.2143 | 13 | 0.1538 | 21 | 0.1905 | 69 | 0.2754 | 16 | 0.3125 | 24 | 0.1667 | 19 | 0.2105 | 19 | 0.3684 | 78 | 0.2564 |
| 18_240246 | 21 | 0.0952 | 15 | 0.0667 | 15 | 0.0667 | 23 | 0.0435 | 74 | 0.0676 | 17 | 0.2353 | 25 | 0.16 | 18 | 0.2222 | 22 | 0.2273 | 82 | 0.2073 |
| 18_240249 | 23 | 0.1739 | 16 | 0.125 | 11 | 0.0909 | 24 | 0.125 | 74 | 0.1351 | 14 | 0.1429 | 27 | 0.1481 | 20 | 0 | 20 | 0.05 | 81 | 0.0864 |
| 18_240251 | 27 | 0.037 | 16 | 0.0625 | 16 | 0.125 | 22 | 0.1818 | 81 | 0.0988 | 15 | 0 | 25 | 0.04 | 17 | 0.0588 | 18 | 0.1111 | 75 | 0.0533 |
| 18_240260 | 11 | 0.1818 | 15 | 0.1333 | 14 | 0.1429 | 24 | 0.1667 | 64 | 0.1562 | 10 | 0 | 12 | 0.0833 | 14 | 0.1429 | 13 | 0.3077 | 49 | 0.1429 |
| 18_240265 | 18 | 0.1111 | 17 | 0 | 22 | 0.0909 | 33 | 0.0606 | 90 | 0.0667 | 11 | 0.0909 | 18 | 0.1667 | 22 | 0.2273 | 21 | 0.2857 | 72 | 0.2083 |
| 18_240897 | NA | NA | 4 | 1 | NA | NA | 3 | 1 | 7 | 1 | NA | NA | 1 | 1 | 1 | 1 | 2 | 1 | 4 | 1 |
| 18_240897 | NA | NA | 3 | 0.6667 | NA | NA | 4 | 1 | 7 | 0.8571 | NA | NA | 1 | 1 | 1 | 0 | 1 | 1 | 3 | 0.6667 |
| 18_240931 | NA | NA | 5 | 0.6 | NA | NA | 3 | 0.3333 | 8 | 0.5 | NA | NA | 1 | 1 | 1 | 0 | 2 | 0 | 4 | 0.25 |
| 18_240931 | NA | NA | 5 | 0.2 | NA | NA | 4 | 0 | 9 | 0.1111 | NA | NA | 1 | 1 | NA | NA | 2 | 0.5 | 3 | 0.6667 |
| 18_240932 | NA | NA | 5 | 0.4 | NA | NA | 4 | 0.25 | 9 | 0.3333 | NA | NA | 1 | 1 | 1 | 0 | 2 | 0 | 4 | 0.25 |
| 18_240933 | NA | NA | 2 | 0.5 | NA | NA | 2 | 0 | 4 | 0.25 | NA | NA | 1 | 1 | 1 | 0 | 2 | 0.5 | 4 | 0.5 |
| 18_240933 | NA | NA | 4 | 0.25 | NA | NA | 4 | 0 | 8 | 0.125 | NA | NA | 1 | 1 | 1 | 0 | 2 | 0 | 4 | 0.25 |
| 18_240939 | NA | NA | 4 | 0.5 | 1 | 1 | 4 | 0.5 | 9 | 0.5556 | NA | NA | 1 | 1 | 1 | 1 | 1 | 1 | 3 | 1 |
| 18_240944 | NA | NA | 4 | 0.25 | 1 | 1 | 4 | 0 | 9 | 0.2222 | NA | NA | 1 | 1 | 1 | 0 | 1 | 0 | 3 | 0.3333 |
| 18_240953 | NA | NA | 4 | 0.25 | 1 | 1 | 4 | 0.5 | 9 | 0.4444 | NA | NA | 2 | 0.5 | 2 | 0.5 | 1 | 0 | 5 | 0.4 |
| 18_240955 | NA | NA | 4 | 0.25 | 1 | 1 | 3 | 0.6667 | 8 | 0.5 | NA | NA | 1 | 0 | 2 | 0.5 | 1 | 0 | 4 | 0.25 |
| 18_240958 | NA | NA | 3 | 0.3333 | 1 | 1 | 3 | 0.6667 | 7 | 0.5714 | NA | NA | 2 | 0.5 | 1 | 0 | 1 | 0 | 4 | 0.25 |
| 18_240975 | NA | NA | 1 | 0 | 1 | 1 | 2 | 0.5 | 4 | 0.5 | NA | NA | 3 | 0.3333 | 3 | 0.3333 | 1 | 1 | 7 | 0.4286 |
| 18_241358 | 4 | 0.25 | 3 | 0 | 1 | 0 | 3 | 0.3333 | 11 | 0.1818 | 1 | 1 | 5 | 0.4 | 3 | 0 | 5 | 0.4 | 14 | 0.3571 |
| 18_241359 | 4 | 0 | 3 | 0 | 1 | 0 | 3 | 0.3333 | 11 | 0.0909 | 1 | 0 | 5 | 0 | 3 | 0 | 5 | 0.2 | 14 | 0.0714 |
| 18_241359 | 2 | 1 | 3 | 1 | NA | NA | 3 | 1 | 8 | 1 | 1 | 1 | 4 | 1 | 3 | 1 | 3 | 1 | 11 | 1 |
| 18_241360 | 4 | 0 | 3 | 0 | 2 | 0 | 3 | 0.3333 | 12 | 0.0833 | 1 | 0 | 3 | 0.6667 | 3 | 0 | 5 | 0 | 12 | 0.1667 |
| 18_241368 | 5 | 0.2 | 3 | 0.3333 | 2 | 0 | 3 | 0.6667 | 13 | 0.3077 | 1 | 0 | 5 | 0 | 3 | 0 | 4 | 0.25 | 13 | 0.0769 |
| 18_241370 | 5 | 0.2 | 3 | 0.3333 | 2 | 0.5 | 3 | 0 | 13 | 0.2308 | 1 | 0 | 5 | 0 | 2 | 0 | 5 | 0.2 | 13 | 0.0769 |
| 18_241371 | 5 | 0.2 | 3 | 0 | 3 | 0.3333 | 4 | 0 | 15 | 0.1333 | 1 | 1 | 5 | 0.2 | 4 | 0.25 | 5 | 0 | 15 | 0.2 |
| 18_241371 | 5 | 0.2 | 3 | 0.3333 | 3 | 0 | 4 | 0 | 15 | 0.1333 | 1 | 0 | 5 | 0.2 | 4 | 0 | 5 | 0.2 | 15 | 0.1333 |
| 18_241373 | 4 | 0.25 | 3 | 0 | 3 | 0.3333 | 3 | 0 | 13 | 0.1538 | 1 | 0 | 5 | 0.2 | 4 | 0.25 | 7 | 0 | 17 | 0.1176 |
| 18_241374 | 4 | 1 | 2 | 1 | 2 | 1 | 4 | 1 | 12 | 1 | 1 | 1 | 4 | 1 | 3 | 1 | 6 | 0.8333 | 14 | 0.9286 |
| 18_241375 | 4 | 0.25 | 3 | 0 | 1 | 0 | 2 | 0.5 | 10 | 0.2 | 1 | 0 | 5 | 0.2 | 2 | 0.5 | 5 | 0.2 | 13 | 0.2308 |
| 18_241376 | 6 | 0.3333 | 3 | 0.3333 | 2 | 0 | 4 | 0.75 | 15 | 0.4 | 1 | 1 | 4 | 0.25 | 4 | 0.75 | 7 | 0.5714 | 16 | 0.5625 |
| 18_241376 | 6 | 0.1667 | 3 | 0 | 3 | 0 | 5 | 0 | 17 | 0.0588 | 1 | 0 | 5 | 0.2 | 4 | 0 | 6 | 0 | 16 | 0.0625 |
| 18_241429 | 1 | 0 | 2 | 0 | 3 | 0.3333 | NA | NA | 6 | 0.1667 | 3 | 0.6667 | NA | NA | 2 | 0.5 | NA | NA | 5 | 0.6 |
| 18_241430 | 1 | 1 | 2 | 0 | 3 | 0.6667 | 1 | 1 | 7 | 0.5714 | 1 | 0 | NA | NA | 3 | 0.3333 | NA | NA | 4 | 0.25 |
| 18_241459 | 6 | 0 | 4 | 0.25 | 8 | 0.125 | 9 | 0 | 27 | 0.0741 | 6 | 0.1667 | 3 | 0.3333 | 5 | 0 | 8 | 0.125 | 22 | 0.1364 |
| 18_241470 | 4 | 1 | 5 | 1 | 4 | 1 | 5 | 1 | 18 | 1 | 4 | 1 | 2 | 1 | 5 | 1 | 4 | 1 | 15 | 1 |
| 18_241470 | 1 | 1 | NA | NA | 1 | 1 | 2 | 1 | 4 | 1 | 2 | 1 | NA | NA | 3 | 1 | 3 | 1 | 8 | 1 |
| 18_241470 | 1 | 1 | NA | NA | NA | NA | 3 | 1 | 4 | 1 | NA | NA | NA | NA | NA | NA | 1 | 1 | 1 | 1 |
| 18_241471 | 3 | 0.3333 | 4 | 0.5 | 4 | 0.75 | 6 | 0.8333 | 17 | 0.6471 | 3 | 0 | 2 | 0.5 | 6 | 0.6667 | 4 | 0.25 | 15 | 0.4 |
| 18_241471 | 5 | 0.6 | 4 | 1 | 2 | 1 | 7 | 0.2857 | 18 | 0.6111 | 5 | 0.8 | 2 | 0.5 | 5 | 0.8 | 5 | 0.6 | 17 | 0.7059 |
| 18_241471 | 4 | 0 | 3 | 0.6667 | 4 | 0 | 6 | 0.1667 | 17 | 0.1765 | 3 | 0 | 2 | 0.5 | 5 | 0 | 5 | 0.2 | 15 | 0.1333 |
| 18_241471 | 3 | 0 | 5 | 0 | 2 | 0 | 5 | 0.2 | 15 | 0.0667 | 3 | 0.3333 | 1 | 0 | 4 | 0 | 3 | 0 | 11 | 0.0909 |
| 18_241472 | 5 | 1 | 5 | 1 | 4 | 1 | 6 | 1 | 20 | 1 | 5 | 0.8 | 2 | 1 | 6 | 1 | 4 | 1 | 17 | 0.9412 |
| 18_241472 | 4 | 0 | 4 | 0.25 | 4 | 0 | 7 | 0 | 19 | 0.0526 | 4 | 0.25 | 2 | 0 | 6 | 0.5 | 5 | 0.2 | 17 | 0.2941 |
| 18_241473 | 3 | 0.3333 | 5 | 0.4 | 4 | 0 | 7 | 0.1429 | 19 | 0.2105 | 5 | 0.2 | 1 | 0 | 6 | 0.1667 | 5 | 0 | 17 | 0.1176 |
| 18_241473 | 3 | 0.3333 | 4 | 0.25 | 3 | 0 | 4 | 0 | 14 | 0.1429 | 2 | 0.5 | 2 | 0.5 | 5 | 0.2 | 3 | 0 | 12 | 0.25 |
| 18_241473 | 3 | 0.3333 | 5 | 0.2 | 3 | 0 | 4 | 0.25 | 15 | 0.2 | 3 | 0.3333 | 1 | 0 | 6 | 0.1667 | 5 | 0.2 | 15 | 0.2 |
| 18_241475 | 1 | 0 | 3 | 0.3333 | 3 | 0 | 3 | 0.3333 | 10 | 0.2 | 3 | 0.3333 | 2 | 0 | 5 | 0 | 5 | 0 | 15 | 0.0667 |
| 18_241507 | 2 | 0 | 5 | 0.2 | 4 | 0 | 4 | 0 | 15 | 0.0667 | 3 | 0.3333 | 2 | 0 | 6 | 0.1667 | 3 | 0 | 14 | 0.1429 |
| 18_241508 | 3 | 0.6667 | 5 | 0.6 | 3 | 0.6667 | 5 | 0.8 | 16 | 0.6875 | 4 | 0.25 | 2 | 1 | 6 | 0.3333 | 3 | 0.6667 | 15 | 0.4667 |
| 18_241509 | 2 | 0.5 | 5 | 0.8 | NA | NA | 2 | 1 | 9 | 0.7778 | 3 | 0.6667 | 2 | 1 | 5 | 0.8 | 2 | 1 | 12 | 0.8333 |
| 18_241509 | 3 | 0 | 4 | 0.25 | 3 | 0 | 4 | 0 | 14 | 0.0714 | 4 | 0 | 2 | 0 | 5 | 0.2 | 3 | 0.3333 | 14 | 0.1429 |
| 18_242379 | 2 | 0 | 4 | 0.25 | 5 | 0.2 | 5 | 0.2 | 16 | 0.1875 | 3 | 0 | 1 | 1 | 3 | 0.3333 | 4 | 0.25 | 11 | 0.2727 |
| 18_242381 | 1 | 0 | 5 | 0 | 4 | 0 | 4 | 0.25 | 14 | 0.0714 | 3 | 0 | 1 | 0 | 3 | 0 | 3 | 0.3333 | 10 | 0.1 |
| 18_242385 | NA | NA | 4 | 0 | 3 | 0.6667 | 3 | 0 | 10 | 0.2 | 2 | 0.5 | 1 | 0 | 2 | 0 | 3 | 0.6667 | 8 | 0.375 |
| 18_242392 | NA | NA | 2 | 1 | 2 | 1 | 3 | 0.6667 | 7 | 0.8571 | 1 | 1 | NA | NA | NA | NA | 1 | 1 | 2 | 1 |
| 18_242392 | NA | NA | 2 | 1 | 2 | 0.5 | 2 | 1 | 6 | 0.8333 | 2 | 1 | 1 | 1 | 1 | 1 | 2 | 1 | 6 | 1 |
| 18_242393 | NA | NA | 3 | 0.3333 | 2 | 0.5 | 4 | 0 | 9 | 0.2222 | 2 | 0.5 | NA | NA | 2 | 0 | 2 | 0 | 6 | 0.1667 |
| 18_242395 | NA | NA | 3 | 1 | 2 | 1 | 4 | 1 | 9 | 1 | 3 | 1 | 1 | 1 | 2 | 1 | 3 | 1 | 9 | 1 |

| | | | | | | | | | | | | | | | | | | | | |
|---|---|---|---|---|---|---|---|---|---|---|---|---|---|---|---|---|---|---|---|---|
| 18_2423954 | NA | NA | 3 | 0.3333 | 2 | 0 | 4 | 0.25 | 9 | 0.2222 | 3 | 0 | 1 | 1 | 2 | 0 | 1 | 0 | 7 | 0.1429 |
| 18_2441784 | 7 | 0.8571 | 8 | 1 | 2 | 1 | 3 | 1 | 20 | 0.95 | 4 | 1 | 2 | 1 | 1 | 1 | 4 | 1 | 11 | 1 |
| 18_2441791 | 6 | 0 | 7 | 0.5714 | 3 | 0 | 3 | 0 | 19 | 0.2105 | 5 | 0 | 1 | 0 | 2 | 0.5 | 4 | 0.25 | 12 | 0.1667 |
| 18_2441795 | 7 | 0.8571 | 6 | 0.6667 | 3 | 0.6667 | 3 | 1 | 19 | 0.7895 | 5 | 0.2 | NA | NA | 2 | 0.5 | 3 | 0.3333 | 10 | 0.3 |
| 18_2441805 | 4 | 1 | 7 | 1 | 3 | 1 | 2 | 0.5 | 16 | 0.9375 | 5 | 0.8 | 1 | 1 | 1 | 0 | 3 | 0.6667 | 10 | 0.7 |
| 18_2441805 | 5 | 0.8 | 6 | 1 | 4 | 1 | 3 | 1 | 18 | 0.9444 | 4 | 1 | 2 | 0.5 | 2 | 1 | 4 | 1 | 12 | 0.9167 |
| 18_2441812 | 6 | 0.5 | 6 | 1 | 4 | 1 | 3 | 0.6667 | 19 | 0.7895 | 5 | 0.8 | 1 | 1 | 2 | 1 | 4 | 0.75 | 12 | 0.8333 |
| 18_2441837 | 6 | 0.3333 | 6 | 0.3333 | 5 | 0.4 | 3 | 0 | 20 | 0.3 | 4 | 0.25 | 2 | 0 | 2 | 0.5 | 5 | 0.8 | 13 | 0.4615 |
| 18_2441850 | 7 | 0.1429 | 8 | 0.125 | 5 | 0 | 4 | 0 | 24 | 0.0833 | 3 | 0 | 2 | 0 | 2 | 0.5 | 6 | 0 | 13 | 0.0769 |
| 18_2441878 | 8 | 0.375 | 11 | 0.1818 | 5 | 0.2 | 5 | 0.2 | 29 | 0.2414 | 5 | 0.2 | 4 | 0 | 2 | 0 | 7 | 0.1429 | 18 | 0.1111 |
| 18_2441887 | 8 | 0 | 8 | 0.125 | 5 | 0 | 5 | 0.2 | 26 | 0.0769 | 5 | 0 | 4 | 0.25 | 2 | 0 | 5 | 0.2 | 16 | 0.125 |
| 18_2441897 | 7 | 0.4286 | 7 | 0.2857 | 4 | 0.25 | 3 | 0 | 21 | 0.2857 | 2 | 0 | 2 | 0 | 1 | 0 | 5 | 0.4 | 10 | 0.2 |
| 18_2441909 | 6 | 0.5 | 6 | 1 | 5 | 0.8 | 2 | 1 | 19 | 0.7895 | 3 | 0.6667 | 5 | 0.4 | 2 | 1 | 7 | 0.8571 | 17 | 0.7059 |
| 18_2441939 | 4 | 0.25 | 6 | 0.5 | 2 | 0.5 | 3 | 0.3333 | 15 | 0.4 | 4 | 0.5 | 1 | 0 | 1 | 1 | 5 | 0.8 | 11 | 0.6364 |
| 18_2441954 | 5 | 0.6 | 8 | 0.625 | 2 | 0 | 2 | 1 | 17 | 0.5882 | 5 | 0.4 | 4 | 0.5 | 1 | 1 | 6 | 0.6667 | 16 | 0.5625 |
| 18_2461034 | 2 | 0 | NA | NA | 4 | 0.5 | 3 | 0 | 9 | 0.2222 | 1 | 0 | 5 | 0.2 | 4 | 0 | 1 | 0 | 11 | 0.0909 |
| 18_2461038 | 3 | 0 | NA | NA | 4 | 0.25 | 3 | 0 | 10 | 0.1 | 2 | 0 | 4 | 0 | 3 | 0 | 1 | 1 | 10 | 0.1 |
| 18_2461043 | 2 | 0 | NA | NA | 4 | 0.5 | 2 | 0 | 8 | 0.25 | 2 | 0.5 | 5 | 0 | 3 | 0 | 1 | 0 | 11 | 0.0909 |
| 18_2461045 | 3 | 1 | NA | NA | 5 | 1 | 1 | 1 | 9 | 1 | 1 | 1 | 2 | 0.5 | 4 | 1 | 1 | 1 | 8 | 0.875 |
| 18_2461050 | 2 | 1 | NA | NA | 4 | 1 | 3 | 1 | 9 | 1 | 1 | 1 | 3 | 1 | 3 | 1 | NA | NA | 7 | 1 |
| 18_2461066 | 2 | 0 | 1 | 0 | 5 | 1 | 3 | 0.3333 | 11 | 0.5455 | 2 | 0.5 | 3 | 0.6667 | 5 | 0.6 | 1 | 0 | 11 | 0.5455 |
| 18_2461082 | 1 | 0 | 1 | 0 | 3 | 1 | 1 | 1 | 6 | 0.6667 | NA | NA | 2 | 1 | NA | NA | NA | NA | 2 | 1 |
| 18_2461836 | 11 | 0.0909 | 15 | 0.0667 | 12 | 0.1667 | 7 | 0 | 45 | 0.0889 | 4 | 0.25 | 11 | 0 | 10 | 0 | 8 | 0.125 | 33 | 0.0606 |
| 18_2461838 | 10 | 0.2 | 11 | 0.2727 | 12 | 0.4167 | 8 | 0.5 | 41 | 0.3415 | 6 | 0.1667 | 13 | 0.2308 | 9 | 0.1111 | 11 | 0.2727 | 39 | 0.2051 |
| 18_2461970 | 8 | 0.25 | 6 | 0 | 11 | 0.2727 | 7 | 0.4286 | 32 | 0.25 | 4 | 0.25 | 9 | 0.2222 | 8 | 0 | 10 | 0 | 31 | 0.0968 |
| 18_2461977 | 11 | 0.1818 | 6 | 0 | 13 | 0.3077 | 7 | 0.4286 | 37 | 0.2432 | 3 | 0.6667 | 10 | 0.3 | 7 | 0.4286 | 10 | 0.4 | 30 | 0.4 |
| 18_2461988 | 8 | 0.125 | 5 | 0.2 | 9 | 0.1111 | 7 | 0 | 29 | 0.1034 | 2 | 0 | 9 | 0 | 8 | 0.375 | 10 | 0.3 | 29 | 0.2069 |
| 18_2462083 | 8 | 0.25 | 3 | 0 | 8 | 0 | 7 | 0 | 26 | 0.0769 | 3 | 0 | 8 | 0.5 | 8 | 0 | 8 | 0.125 | 27 | 0.1852 |
| 18_2462086 | 8 | 0.25 | 3 | 0 | 8 | 0 | 6 | 0 | 25 | 0.08 | 2 | 0 | 5 | 0.2 | 8 | 0 | 6 | 0.1667 | 21 | 0.0952 |
| 18_2462102 | 7 | 0.2857 | 4 | 0 | 6 | 0.5 | 7 | 0.1429 | 24 | 0.25 | 3 | 0 | 7 | 0.1429 | 7 | 0.1429 | 8 | 0.375 | 25 | 0.2 |
| 18_2462121 | 8 | 0.875 | 3 | 1 | 8 | 0.875 | 5 | 0.8 | 24 | 0.875 | 2 | 1 | 5 | 0.8 | 8 | 1 | 8 | 0.75 | 23 | 0.8696 |
| 18_2462124 | 8 | 0.25 | 3 | 0 | 8 | 0 | 6 | 0.1667 | 25 | 0.12 | 3 | 0 | 7 | 0.1429 | 8 | 0.25 | 7 | 0 | 25 | 0.12 |
| 18_2462125 | 7 | 0 | 4 | 0 | 7 | 0.1429 | 7 | 0.1429 | 25 | 0.08 | 3 | 0.3333 | 7 | 0 | 8 | 0.25 | 7 | 0 | 25 | 0.12 |
| 18_2462134 | 6 | 0 | 4 | 0 | 6 | 0 | 4 | 0.25 | 20 | 0.05 | 3 | 0 | 6 | 0.3333 | 8 | 0.25 | 7 | 0.2857 | 24 | 0.25 |
| 18_2471319 | 2 | 0 | 5 | 0.2 | 4 | 0.25 | 1 | 0 | 12 | 0.1667 | 3 | 0 | 2 | 0.5 | 1 | 0 | 4 | 0 | 10 | 0.1 |
| 18_2473404 | NA | NA | 5 | 0.6 | 2 | 1 | NA | NA | 7 | 0.7143 | NA | NA | 2 | 1 | 1 | 0 | 3 | 1 | 6 | 0.8333 |
| 18_2473405 | NA | NA | 5 | 0.4 | 1 | 1 | NA | NA | 6 | 0.5 | NA | NA | 2 | 0.5 | 1 | 1 | 2 | 0.5 | 5 | 0.6 |
| 18_2473407 | NA | NA | 4 | 0.75 | 2 | 1 | NA | NA | 6 | 0.8333 | NA | NA | 2 | 0.5 | 1 | 0 | 3 | 1 | 6 | 0.6667 |
| 18_2473418 | NA | NA | 4 | 0 | 1 | 1 | NA | NA | 5 | 0.2 | NA | NA | 1 | 0 | 1 | 1 | 2 | 0 | 4 | 0.25 |
| 18_2473421 | NA | NA | 4 | 0.75 | 2 | 0.5 | NA | NA | 6 | 0.6667 | NA | NA | 1 | 1 | 1 | 1 | 1 | 0 | 3 | 0.6667 |
| 18_2473422 | NA | NA | 5 | 0.4 | 2 | 0 | NA | NA | 7 | 0.2857 | NA | NA | 1 | 0 | 1 | 1 | 3 | 0.3333 | 5 | 0.4 |
| 18_2473444 | NA | NA | 4 | 0.25 | 2 | 0 | NA | NA | 6 | 0.1667 | NA | NA | 1 | 0 | 2 | 0 | 3 | 0.6667 | 6 | 0.3333 |
| 18_2473454 | NA | NA | 3 | 0.3333 | 2 | 0 | NA | NA | 5 | 0.2 | NA | NA | 1 | 0 | 2 | 0.5 | 2 | 0 | 5 | 0.2 |
| 18_2473456 | NA | NA | 2 | 0 | 2 | 0.5 | NA | NA | 4 | 0.25 | NA | NA | NA | NA | 2 | 0.5 | 3 | 0.3333 | 5 | 0.4 |
| 18_2473459 | NA | NA | 3 | 0.3333 | 2 | 0 | NA | NA | 5 | 0.2 | NA | NA | 1 | 0 | 2 | 0 | 2 | 0.5 | 5 | 0.2 |
| 18_2484172 | NA | NA | NA | NA | 1 | 1 | NA | NA | 1 | 1 | NA | NA | NA | NA | NA | NA | 1 | 1 | 1 | 1 |
| 18_2522333 | NA | NA | 1 | 1 | 2 | 1 | NA | NA | 3 | 1 | NA | NA | NA | NA | 1 | 1 | 1 | 1 | 2 | 1 |
| 18_2522343 | NA | NA | 2 | 0.5 | 2 | 0 | NA | NA | 4 | 0.25 | NA | NA | NA | NA | 1 | 1 | 1 | 0 | 2 | 0.5 |
| 18_2522347 | NA | NA | 2 | 0.5 | 1 | 1 | NA | NA | 3 | 0.6667 | 2 | 1 | NA | NA | 2 | 0.5 | 1 | 1 | 5 | 0.8 |
| 18_2522354 | NA | NA | 2 | 0.5 | 2 | 0.5 | NA | NA | 4 | 0.5 | 1 | 1 | NA | NA | 1 | 0 | 1 | 0 | 3 | 0.3333 |
| 18_2522367 | NA | NA | 2 | 0.5 | 1 | 1 | NA | NA | 3 | 0.6667 | 2 | 1 | NA | NA | 2 | 0.5 | 1 | 0 | 5 | 0.6 |
| 18_2522385 | 1 | 1 | 3 | 0 | 2 | 0.5 | NA | NA | 6 | 0.3333 | 1 | 0 | NA | NA | 2 | 1 | 1 | 0 | 4 | 0.5 |
| 18_2526605 | NA | NA | 2 | 0 | 4 | 0.5 | NA | NA | 6 | 0.3333 | 3 | 0.3333 | 1 | 0 | 1 | 1 | 1 | 1 | 6 | 0.5 |
| 18_2526612 | 1 | 1 | 3 | 0.3333 | 4 | 0 | NA | NA | 8 | 0.25 | 3 | 0 | NA | NA | 3 | 0.3333 | 1 | 0 | 7 | 0.1429 |
| 18_2526615 | 1 | 1 | 2 | 1 | 4 | 0.5 | NA | NA | 7 | 0.7143 | 3 | 0.6667 | 1 | 1 | 3 | 0.3333 | 1 | 0 | 8 | 0.5 |
| 18_2526639 | 2 | 0 | 3 | 0.3333 | 5 | 0 | 1 | 0 | 11 | 0.0909 | 3 | 0 | 1 | 0 | 3 | 0.3333 | 1 | 0 | 8 | 0.125 |
| 18_2526933 | 1 | 0 | 3 | 1 | 1 | 0 | 2 | 1 | 7 | 0.7143 | 2 | 1 | 3 | 0.3333 | 1 | 0 | 3 | 0.6667 | 9 | 0.5556 |
| 18_2526947 | 3 | 0.6667 | 2 | 0.5 | 1 | 0 | 2 | 0 | 8 | 0.375 | 4 | 0.5 | 4 | 0.5 | 2 | 0 | 4 | 0 | 14 | 0.2857 |
| 18_2526950 | 3 | 0.3333 | 2 | 0 | 1 | 0 | 2 | 0 | 8 | 0.125 | 4 | 0.5 | 3 | 0 | 2 | 0.5 | 4 | 0.25 | 13 | 0.3077 |
| 18_2526963 | 3 | 0.3333 | 2 | 0 | 1 | 0 | 2 | 0 | 8 | 0.125 | 2 | 0 | 4 | 0.5 | 1 | 0 | 3 | 0 | 10 | 0.2 |
| 18_2527177 | 7 | 0 | 15 | 0.1333 | 15 | 0.2 | 8 | 0.25 | 45 | 0.1556 | 13 | 0.1538 | 7 | 0.1429 | 11 | 0.1818 | 15 | 0.2667 | 46 | 0.1957 |
| 18_2527189 | 8 | 0.25 | 17 | 0.1176 | 15 | 0.2667 | 8 | 0.375 | 48 | 0.2292 | 12 | 0.3333 | 7 | 0 | 11 | 0.0909 | 15 | 0.3333 | 45 | 0.2222 |
| 18_2559909 | NA | NA | 1 | 0 | NA | NA | 1 | 1 | 2 | 0.5 | NA | NA | 1 | 0 | NA | NA | 2 | 0.5 | 3 | 0.3333 |
| 18_2559986 | NA | NA | 2 | 0.5 | NA | NA | 1 | 0 | 3 | 0.3333 | NA | NA | 2 | 0.5 | NA | NA | 1 | 0 | 3 | 0.3333 |
| 18_2561114 | 18 | 0.1111 | 63 | 0.1746 | 50 | 0.18 | 24 | 0.1667 | 155 | 0.1677 | 33 | 0.1818 | 45 | 0.1111 | 60 | 0.1833 | 25 | 0.16 | 163 | 0.1595 |

| | | | | | | | | | | | | | | | | | | | | |
|---|---|---|---|---|---|---|---|---|---|---|---|---|---|---|---|---|---|---|---|---|
| 18_2561123 | 20 | 0.4 | 62 | 0.3387 | 47 | 0.2766 | 23 | 0.2609 | 152 | 0.3158 | 32 | 0.1562 | 44 | 0.3182 | 62 | 0.3871 | 22 | 0.3636 | 160 | 0.3188 |
| 18_2561132 | 18 | 0.3889 | 59 | 0.2881 | 47 | 0.3191 | 21 | 0.2381 | 145 | 0.3034 | 31 | 0.3548 | 41 | 0.2927 | 54 | 0.3519 | 18 | 0.1111 | 144 | 0.3056 |
| 18_2561141 | 18 | 0.2222 | 51 | 0.1765 | 45 | 0.1778 | 21 | 0.2857 | 135 | 0.2 | 26 | 0.2692 | 38 | 0.0789 | 58 | 0.1897 | 24 | 0.3333 | 146 | 0.1986 |
| 18_2561142 | 18 | 0.0556 | 54 | 0.1296 | 47 | 0.1489 | 21 | 0.1429 | 140 | 0.1286 | 30 | 0.0667 | 44 | 0.1364 | 53 | 0.3019 | 21 | 0.1905 | 148 | 0.1892 |
| 18_2561144 | 15 | 0.2667 | 52 | 0.1731 | 43 | 0.1628 | 19 | 0.4211 | 129 | 0.2171 | 28 | 0.3571 | 39 | 0.1795 | 53 | 0.2642 | 21 | 0.1429 | 141 | 0.2411 |
| 18_2561159 | 17 | 0.2941 | 53 | 0.6038 | 45 | 0.7111 | 18 | 0.6667 | 133 | 0.609 | 29 | 0.5517 | 38 | 0.5789 | 53 | 0.7358 | 24 | 0.4167 | 144 | 0.6042 |
| 18_2561161 | 16 | 0.3125 | 53 | 0.2453 | 38 | 0.3684 | 18 | 0.1667 | 125 | 0.28 | 29 | 0.4483 | 35 | 0.4 | 51 | 0.3922 | 21 | 0.2381 | 136 | 0.3824 |
| 18_2561165 | 20 | 0 | 56 | 0.0357 | 45 | 0.0889 | 20 | 0.15 | 141 | 0.0638 | 34 | 0.0882 | 41 | 0.0976 | 54 | 0.1852 | 22 | 0 | 151 | 0.1126 |
| 18_2561167 | 21 | 0.619 | 54 | 0.5556 | 45 | 0.5333 | 20 | 0.7 | 140 | 0.5786 | 33 | 0.5455 | 43 | 0.6047 | 54 | 0.6667 | 24 | 0.5 | 154 | 0.5974 |
| 18_2561189 | 18 | 0.3889 | 48 | 0.2083 | 38 | 0.3158 | 16 | 0.3125 | 120 | 0.2833 | 27 | 0.2963 | 36 | 0.2778 | 47 | 0.234 | 20 | 0.25 | 130 | 0.2615 |
| 18_2561191 | 18 | 0.1111 | 43 | 0.2558 | 35 | 0.1714 | 17 | 0.2941 | 113 | 0.2124 | 23 | 0.2609 | 30 | 0.1 | 37 | 0.1351 | 16 | 0.1875 | 106 | 0.1604 |
| 18_2561202 | 18 | 0.1111 | 52 | 0.1154 | 44 | 0.1591 | 18 | 0.1667 | 132 | 0.1364 | 29 | 0.1379 | 39 | 0.1795 | 45 | 0.2889 | 18 | 0 | 131 | 0.1832 |
| 18_2561228 | 20 | 0.55 | 47 | 0.5532 | 42 | 0.5476 | 15 | 0.6 | 124 | 0.5565 | 26 | 0.5769 | 35 | 0.6 | 42 | 0.6905 | 16 | 0.625 | 119 | 0.6303 |
| 18_2561229 | 17 | 0 | 45 | 0.0667 | 40 | 0.075 | 14 | 0.0714 | 116 | 0.0603 | 25 | 0 | 35 | 0.1143 | 43 | 0.0465 | 14 | 0 | 117 | 0.0513 |
| 18_2561263 | 18 | 0.8333 | 42 | 0.7619 | 36 | 0.8611 | 12 | 0.8333 | 108 | 0.8148 | 22 | 0.8636 | 28 | 0.8571 | 32 | 0.8125 | 18 | 0.7778 | 100 | 0.83 |
| 18_2561270 | 14 | 0.9286 | 38 | 0.8421 | 31 | 0.9355 | 13 | 0.8462 | 96 | 0.8854 | 23 | 0.8696 | 28 | 0.75 | 28 | 0.8214 | 12 | 0.9167 | 91 | 0.8242 |
| 18_2588642 | 11 | 0 | 27 | 0.1111 | 21 | 0.1905 | 7 | 0.1429 | 66 | 0.1212 | 9 | 0.1111 | 12 | 0.4167 | 16 | 0 | 11 | 0.1818 | 48 | 0.1667 |
| 18_3040230 | NA | NA | 1 | 0 | 1 | 0 | 3 | 0.6667 | 5 | 0.4 | 1 | 0 | NA | NA | 1 | 1 | NA | NA | 2 | 0.5 |
| 18_3040234 | NA | NA | 2 | 1 | 1 | 1 | 2 | 0.5 | 5 | 0.8 | 1 | 1 | NA | NA | 1 | 1 | NA | NA | 2 | 1 |
| 18_3040240 | NA | NA | 2 | 0.5 | 1 | 0 | 3 | 0.6667 | 6 | 0.5 | 1 | 1 | NA | NA | 1 | 1 | NA | NA | 2 | 1 |
| 18_3040253 | NA | NA | 1 | 0 | 1 | 0 | 3 | 0.3333 | 5 | 0.2 | 1 | 0 | NA | NA | 1 | 1 | NA | NA | 2 | 0.5 |
| 18_3040302 | NA | NA | 2 | 1 | 1 | 1 | 2 | 1 | 5 | 1 | 1 | 1 | NA | NA | 1 | 1 | NA | NA | 2 | 1 |
| 18_3040312 | NA | NA | 1 | 0 | 1 | 1 | 3 | 0.3333 | 5 | 0.4 | 1 | 1 | NA | NA | 1 | 1 | NA | NA | 2 | 1 |
| 18_3040352 | NA | NA | 2 | 1 | 1 | 0 | 3 | 0 | 6 | 0.3333 | 1 | 0 | NA | NA | 1 | 1 | NA | NA | 2 | 0.5 |
| 18_3040354 | NA | NA | 2 | 0.5 | 1 | 1 | 1 | 1 | 4 | 0.75 | 1 | 1 | NA | NA | 1 | 1 | NA | NA | 2 | 1 |
| 18_3040362 | NA | NA | 2 | 0.5 | 1 | 0 | 3 | 0 | 6 | 0.1667 | 1 | 1 | NA | NA | 1 | 1 | NA | NA | 2 | 1 |
| 18_3040416 | 1 | 1 | 3 | 0.3333 | 2 | 0 | 3 | 0.3333 | 9 | 0.3333 | 3 | 0.6667 | NA | NA | 1 | 0 | 1 | 0 | 5 | 0.4 |
| 18_3040417 | NA | NA | 4 | 0.5 | 1 | 0 | 3 | 1 | 8 | 0.625 | 4 | 0.75 | NA | NA | 1 | 1 | 1 | 1 | 6 | 0.8333 |
| 18_3040512 | 1 | 0 | 2 | 0 | 2 | 0.5 | 3 | 0 | 8 | 0.125 | 4 | 0.25 | 1 | 0 | 1 | 0 | 1 | 1 | 7 | 0.2857 |
| 18_3048086 | 11 | 0 | 8 | 0.125 | 7 | 0 | 9 | 0.1111 | 35 | 0.0571 | 7 | 0 | 10 | 0.1 | 10 | 0.1 | 7 | 0.1429 | 34 | 0.0882 |
| 18_3048101 | 12 | 0 | 7 | 0 | 6 | 0.1667 | 7 | 0.2857 | 32 | 0.0938 | 4 | 0.25 | 9 | 0.2222 | 8 | 0 | 6 | 0 | 27 | 0.1111 |
| 18_3048107 | 12 | 0.3333 | 9 | 0.3333 | 5 | 0.6 | 9 | 0.4444 | 35 | 0.4 | 8 | 0.75 | 9 | 0.2222 | 13 | 0.2308 | 6 | 0.3333 | 36 | 0.3611 |
| 18_3106702 | NA | NA | NA | NA | 1 | 1 | NA | NA | 1 | 1 | NA | NA | NA | NA | 1 | 1 | NA | NA | 1 | 1 |
| 18_3106757 | NA | NA | NA | NA | 1 | 1 | NA | NA | 1 | 1 | NA | NA | NA | NA | 1 | 1 | NA | NA | 1 | 1 |
| 18_3157100 | 15 | 0.2 | 18 | 0.1111 | 26 | 0.1923 | 26 | 0.1923 | 85 | 0.1765 | 22 | 0.2273 | 23 | 0.3913 | 29 | 0.3103 | 20 | 0.25 | 94 | 0.2979 |
| 18_3157144 | 11 | 0.5455 | 17 | 0.6471 | 20 | 0.55 | 28 | 0.75 | 76 | 0.6447 | 19 | 0.7895 | 21 | 0.8571 | 27 | 0.7037 | 16 | 0.6875 | 83 | 0.759 |
| 18_3157168 | 11 | 0 | 13 | 0 | 21 | 0.1429 | 23 | 0.087 | 68 | 0.0735 | 16 | 0 | 20 | 0.1 | 25 | 0.12 | 15 | 0.0667 | 76 | 0.0789 |
| 18_3157344 | 8 | 1 | 11 | 0.7273 | 14 | 1 | 20 | 1 | 53 | 0.9434 | 8 | 1 | 10 | 1 | 17 | 0.9412 | 12 | 1 | 47 | 0.9787 |
| 18_3157653 | 8 | 0 | 12 | 0.0833 | 16 | 0.0625 | 19 | 0.0526 | 55 | 0.0545 | 11 | 0.0909 | 11 | 0.0909 | 18 | 0.0556 | 11 | 0.2727 | 51 | 0.1176 |
| 18_3157670 | 7 | 0.2857 | 11 | 0.1818 | 15 | 0.4667 | 19 | 0.2632 | 52 | 0.3077 | 10 | 0.2 | 11 | 0.4545 | 18 | 0.3889 | 11 | 0.5455 | 50 | 0.4 |
| 18_3157676 | 7 | 0.5714 | 12 | 0.0833 | 14 | 0.2857 | 18 | 0.2778 | 51 | 0.2745 | 7 | 0.7143 | 11 | 0.1818 | 17 | 0.4118 | 12 | 0.5 | 47 | 0.4255 |
| 18_3157683 | 7 | 0 | 10 | 0 | 11 | 0.1818 | 13 | 0.0769 | 41 | 0.0732 | 9 | 0 | 9 | 0.2222 | 13 | 0.1538 | 10 | 0.2 | 41 | 0.1463 |
| 18_3157688 | 6 | 0.5 | 12 | 0.8333 | 12 | 0.6667 | 19 | 0.6842 | 49 | 0.6939 | 7 | 0.5714 | 10 | 0.3 | 14 | 0.7857 | 11 | 0.9091 | 42 | 0.6667 |
| 18_3157696 | 7 | 0.1429 | 11 | 0.0909 | 14 | 0.0714 | 18 | 0.0556 | 50 | 0.08 | 9 | 0.2222 | 11 | 0 | 16 | 0.125 | 10 | 0 | 46 | 0.087 |
| 18_3157721 | 7 | 0 | 12 | 0.0833 | 14 | 0 | 18 | 0.1667 | 51 | 0.0784 | 8 | 0.25 | 9 | 0.1111 | 17 | 0.1765 | 8 | 0.25 | 42 | 0.1905 |
| 18_3158042 | 4 | 0.25 | 2 | 0 | 6 | 0 | 8 | 0 | 20 | 0.05 | 2 | 0 | 4 | 0 | 9 | 0.1111 | 5 | 0 | 20 | 0.05 |
| 18_3171479 | 5 | 0 | 9 | 0.2222 | 8 | 0.375 | 11 | 0.4545 | 33 | 0.303 | 4 | 0.25 | 4 | 0 | 10 | 0.6 | 5 | 0.6 | 23 | 0.4348 |
| 18_3171534 | 10 | 0.5 | 14 | 0.3571 | 5 | 0.2 | 9 | 0.1111 | 38 | 0.3158 | 7 | 0.7143 | 14 | 0.5 | 12 | 0.25 | 12 | 0.4167 | 45 | 0.4444 |
| 18_3171548 | 12 | 0.0833 | 19 | 0.3158 | 9 | 0.2222 | 14 | 0.1429 | 54 | 0.2037 | 11 | 0.0909 | 15 | 0.3333 | 12 | 0.1667 | 19 | 0.2632 | 57 | 0.2281 |
| 18_3171554 | 12 | 0.25 | 21 | 0.0476 | 9 | 0.1111 | 13 | 0.0769 | 55 | 0.1091 | 8 | 0.375 | 14 | 0.1429 | 13 | 0 | 17 | 0.1765 | 52 | 0.1538 |
| 18_3171576 | 11 | 0.1818 | 15 | 0.0667 | 7 | 0.1429 | 13 | 0.0769 | 46 | 0.1087 | 7 | 0.1429 | 10 | 0 | 9 | 0.4444 | 16 | 0.125 | 42 | 0.1667 |
| 18_3171619 | 12 | 0.0833 | 19 | 0.0526 | 9 | 0.1111 | 16 | 0 | 56 | 0.0536 | 7 | 0.1429 | 9 | 0 | 9 | 0.1111 | 15 | 0 | 40 | 0.05 |
| 18_3171638 | 12 | 1 | 17 | 0.9412 | 8 | 1 | 12 | 1 | 49 | 0.9796 | 7 | 1 | 10 | 1 | 7 | 1 | 14 | 1 | 38 | 1 |
| 18_3171644 | 10 | 0.5 | 15 | 0.5333 | 7 | 0.2857 | 16 | 0.375 | 48 | 0.4375 | 7 | 0.5714 | 10 | 0.7 | 9 | 0.4444 | 13 | 0.4615 | 39 | 0.5385 |
| 18_3171646 | 10 | 0.2 | 13 | 0.0769 | 7 | 0.2857 | 14 | 0.4286 | 44 | 0.25 | 6 | 0.5 | 10 | 0.1 | 9 | 0.2222 | 12 | 0 | 37 | 0.1622 |
| 18_3171654 | 8 | 0.75 | 13 | 1 | 8 | 1 | 13 | 0.9231 | 42 | 0.9286 | 7 | 1 | 7 | 1 | 7 | 1 | 10 | 1 | 31 | 1 |
| 18_3171655 | 10 | 1 | 14 | 1 | 8 | 1 | 15 | 1 | 47 | 1 | 7 | 1 | 9 | 1 | 9 | 1 | 13 | 0.8462 | 38 | 0.9474 |
| 18_3171662 | 10 | 0.9 | 15 | 0.6667 | 7 | 0.4286 | 16 | 0.5625 | 48 | 0.6458 | 6 | 0.1667 | 9 | 0.5556 | 8 | 0.625 | 11 | 0.7273 | 34 | 0.5588 |
| 18_3185527 | 2 | 0.5 | 2 | 1 | 3 | 0.3333 | 5 | 0.4 | 12 | 0.5 | 2 | 0 | 6 | 0.3333 | 6 | 0.6667 | 3 | 0.6667 | 17 | 0.4706 |
| 18_3185539 | 2 | 0.5 | 2 | 0.5 | 3 | 1 | 6 | 0.6667 | 13 | 0.6923 | 2 | 1 | 6 | 0.3333 | 6 | 0.5 | 3 | 0.6667 | 17 | 0.5294 |
| 18_3185549 | 2 | 1 | 4 | 1 | 3 | 0.6667 | 5 | 1 | 14 | 0.9286 | 2 | 1 | 5 | 0.8 | 6 | 1 | 4 | 0.75 | 17 | 0.8824 |
| 18_3185554 | 2 | 1 | 3 | 0.3333 | 2 | 0.5 | 4 | 0 | 11 | 0.3636 | 2 | 0 | 6 | 0.3333 | 4 | 0.75 | 3 | 0.3333 | 15 | 0.4 |
| 18_3185555 | 2 | 1 | 3 | 1 | 2 | 1 | 6 | 0.8333 | 13 | 0.9231 | 1 | 1 | 3 | 1 | 7 | 1 | 5 | 0.8 | 16 | 0.9375 |
| 18_3185561 | 2 | 1 | 4 | 1 | 3 | 1 | 5 | 0.6 | 14 | 0.8571 | 2 | 1 | 5 | 1 | 7 | 0.8571 | 4 | 0.75 | 18 | 0.8889 |
| 18_3185567 | 1 | 1 | 4 | 1 | 3 | 1 | 4 | 1 | 12 | 1 | 2 | 1 | 3 | 1 | 7 | 1 | 4 | 0.75 | 16 | 0.9375 |

| | | | | | | | | | | | | | | | | | | | | |
|---|---|---|---|---|---|---|---|---|---|---|---|---|---|---|---|---|---|---|---|---|
| 18_318557 | 2 | 1 | 5 | 1 | 3 | 1 | 7 | 0.8571 | 17 | 0.9412 | 2 | 1 | 5 | 1 | 6 | 0.8333 | 3 | 1 | 16 | 0.9375 |
| 18_318557 | 2 | 1 | 5 | 1 | 2 | 1 | 6 | 0.8333 | 15 | 0.9333 | 1 | 1 | 6 | 1 | 6 | 1 | 3 | 1 | 16 | 1 |
| 18_318557 | 2 | 0 | 5 | 0 | 3 | 0.6667 | 6 | 0.1667 | 16 | 0.1875 | 2 | 0.5 | 6 | 0.1667 | 7 | 0.2857 | 5 | 0.2 | 20 | 0.25 |
| 18_318558 | 2 | 0.5 | 5 | 0.2 | 3 | 0.6667 | 8 | 0.25 | 18 | 0.3333 | 2 | 0 | 5 | 0.2 | 7 | 0.4286 | 5 | 0.4 | 19 | 0.3158 |
| 18_318558 | 1 | 0 | 3 | 1 | 1 | 1 | 5 | 1 | 10 | 0.9 | 1 | 0 | 4 | 0.75 | 5 | 1 | 5 | 0.8 | 15 | 0.8 |
| 18_318559 | 3 | 0 | 4 | 0 | 3 | 0.3333 | 7 | 0.1429 | 17 | 0.1176 | 2 | 0.5 | 6 | 0 | 6 | 0 | 3 | 0.3333 | 17 | 0.1176 |
| 18_318559 | 3 | 0.3333 | 5 | 0.6 | 2 | 0 | 7 | 0.2857 | 17 | 0.3529 | 2 | 0 | 5 | 0.6 | 7 | 0.5714 | 5 | 0.2 | 19 | 0.4211 |
| 18_318560 | 1 | 1 | 6 | 1 | 3 | 0.6667 | 7 | 0.8571 | 17 | 0.8824 | 2 | 1 | 5 | 1 | 7 | 0.7143 | 3 | 0.3333 | 17 | 0.7647 |
| 18_318560 | 3 | 0.6667 | 5 | 1 | 2 | 1 | 7 | 0.1429 | 17 | 0.5882 | 1 | 0 | 5 | 0.4 | 7 | 0.4286 | 5 | 0.6 | 18 | 0.4444 |
| 18_318931 | NA | NA | 1 | 0 | 1 | 1 | 1 | 0 | 3 | 0.3333 | NA | NA | 2 | 0.5 | NA | NA | NA | NA | 2 | 0.5 |
| 18_319049 | NA | NA | 3 | 0.6667 | 2 | 0.5 | 1 | 0 | 6 | 0.5 | NA | NA | 2 | 1 | NA | NA | 1 | 1 | 3 | 1 |
| 18_319049 | NA | NA | 3 | 0 | 2 | 1 | 1 | 0 | 6 | 0.3333 | NA | NA | 2 | 1 | NA | NA | 1 | 1 | 3 | 1 |
| 18_319139 | NA | NA | 5 | 0.8 | 2 | 1 | 2 | 0.5 | 9 | 0.7778 | NA | NA | 1 | 1 | NA | NA | 2 | 1 | 3 | 1 |
| 18_319139 | NA | NA | 3 | 0.6667 | 3 | 1 | 2 | 0.5 | 8 | 0.75 | 1 | 1 | 2 | 1 | NA | NA | 2 | 0.5 | 5 | 0.8 |
| 18_319140 | NA | NA | 5 | 0.2 | 3 | 0 | 2 | 0 | 10 | 0.1 | 1 | 1 | 2 | 0.5 | NA | NA | 3 | 0 | 6 | 0.3333 |
| 18_319141 | NA | NA | 5 | 0.8 | 1 | 1 | 2 | 0 | 8 | 0.625 | 1 | 1 | 2 | 1 | NA | NA | 3 | 0.3333 | 6 | 0.6667 |
| 18_319141 | NA | NA | 3 | 1 | 3 | 1 | 2 | 1 | 8 | 1 | 1 | 1 | 2 | 1 | NA | NA | 2 | 1 | 5 | 1 |
| 18_319142 | NA | NA | 4 | 0.25 | 4 | 0 | 2 | 0 | 10 | 0.1 | 1 | 0 | 2 | 0 | 1 | 0 | 3 | 0.3333 | 7 | 0.1429 |
| 18_319142 | NA | NA | 4 | 0 | 3 | 0.3333 | 2 | 0 | 9 | 0.1111 | 1 | 0 | 2 | 0.5 | 1 | 0 | 3 | 0 | 7 | 0.1429 |
| 18_319143 | NA | NA | 3 | 0.3333 | 3 | 0 | 2 | 0 | 8 | 0.125 | NA | NA | 1 | 1 | 1 | 1 | 1 | 0 | 3 | 0.6667 |
| 18_319144 | NA | NA | 4 | 0.25 | 4 | 0 | 2 | 0 | 10 | 0.1 | NA | NA | 2 | 1 | 1 | 0 | 3 | 0.3333 | 6 | 0.5 |
| 18_319144 | NA | NA | 5 | 1 | 3 | 0.6667 | 2 | 1 | 10 | 0.9 | NA | NA | 2 | 1 | 1 | 1 | 2 | 1 | 5 | 1 |
| 18_319146 | NA | NA | 5 | 0 | 2 | 0.5 | 2 | 0 | 9 | 0.1111 | NA | NA | 2 | 0.5 | 1 | 1 | 3 | 0 | 6 | 0.3333 |
| 18_319147 | NA | NA | 4 | 0 | 3 | 0.3333 | 2 | 0 | 9 | 0.1111 | NA | NA | 1 | 0 | 1 | 0 | 3 | 0.3333 | 5 | 0.2 |
| 18_319147 | NA | NA | 4 | 0.25 | 3 | 1 | 3 | 0.3333 | 10 | 0.5 | 1 | 0 | 2 | 1 | 1 | 1 | 3 | 0 | 7 | 0.4286 |
| 18_319148 | NA | NA | 5 | 0.2 | 4 | 0.75 | 3 | 0.3333 | 12 | 0.4167 | 1 | 1 | 2 | 0 | 1 | 0 | 3 | 0.3333 | 7 | 0.2857 |
| 18_319148 | NA | NA | 4 | 0.25 | 4 | 0 | 2 | 0 | 10 | 0.1 | 1 | 0 | 2 | 1 | 1 | 0 | 2 | 0 | 6 | 0.3333 |
| 18_319149 | NA | NA | 4 | 1 | 1 | 0 | 3 | 0.6667 | 8 | 0.75 | 1 | 1 | 1 | 1 | 1 | 1 | 3 | 0.6667 | 6 | 0.8333 |
| 18_319149 | NA | NA | 4 | 0.5 | 3 | 0 | 3 | 0 | 10 | 0.2 | 1 | 1 | 1 | 0 | NA | NA | 3 | 0 | 5 | 0.2 |
| 18_319152 | 1 | 1 | 3 | 1 | 2 | 0.5 | 2 | 1 | 8 | 0.875 | NA | NA | 1 | 1 | 1 | 1 | 3 | 0.6667 | 5 | 0.8 |
| 18_319170 | 3 | 0.3333 | 4 | 0 | 4 | 0 | 4 | 0 | 15 | 0.0667 | NA | NA | 3 | 0.3333 | NA | NA | 3 | 0 | 6 | 0.1667 |
| 18_319292 | 2 | 0 | 4 | 0.75 | 2 | 0.5 | 1 | 0 | 9 | 0.4444 | 5 | 0.2 | 4 | 0.25 | 3 | 0.6667 | 4 | 0.25 | 16 | 0.3125 |
| 18_319294 | 4 | 1 | 5 | 1 | 2 | 1 | 4 | 0.75 | 15 | 0.9333 | 4 | 1 | 4 | 1 | 3 | 0.6667 | 5 | 1 | 16 | 0.9375 |
| 18_319295 | 3 | 0.6667 | 3 | 1 | 2 | 0.5 | 1 | 1 | 9 | 0.7778 | 2 | 0.5 | 4 | 0.75 | 5 | 0.6 | 4 | 1 | 15 | 0.7333 |
| 18_319295 | 2 | 0.5 | 5 | 0 | 2 | 0 | 1 | 0 | 10 | 0.1 | 4 | 0 | 2 | 0 | 4 | 0.25 | 4 | 0 | 14 | 0.0714 |
| 18_319295 | 4 | 0 | 5 | 0.6 | 3 | 0 | 4 | 0.25 | 16 | 0.25 | 5 | 0 | 5 | 0 | 5 | 0.2 | 6 | 0.1667 | 21 | 0.0952 |
| 18_319296 | 3 | 0.3333 | 5 | 0.2 | 3 | 0 | 3 | 0 | 14 | 0.1429 | 4 | 0.25 | 6 | 0.3333 | 5 | 0 | 5 | 0 | 20 | 0.15 |
| 18_319666 | 1 | 0 | 4 | 0 | 2 | 0 | 8 | 0.125 | 15 | 0.0667 | 8 | 0 | 4 | 0 | 3 | 0.3333 | 5 | 0 | 20 | 0.05 |
| 18_319672 | NA | NA | 1 | 1 | 1 | 1 | 2 | 1 | 4 | 1 | 3 | 1 | 1 | 1 | 2 | 1 | 1 | 1 | 7 | 1 |
| 18_319673 | NA | NA | 1 | 0 | 1 | 0 | 1 | 1 | 3 | 0.3333 | 3 | 0 | 2 | 0 | 2 | 0 | 1 | 1 | 8 | 0.125 |
| 18_319673 | NA | NA | 1 | 1 | NA | NA | 1 | 1 | 2 | 1 | 2 | 1 | 2 | 0.5 | 2 | 1 | 1 | 0 | 7 | 0.7143 |
| 18_320211 | 1 | 0 | NA | NA | 1 | 0 | 2 | 0.5 | 4 | 0.25 | 3 | 0.3333 | NA | NA | NA | NA | NA | NA | 3 | 0.3333 |
| 18_320213 | 1 | 1 | 1 | 0 | 1 | 0 | 4 | 0.5 | 7 | 0.4286 | 4 | 0.25 | NA | NA | NA | NA | 1 | 1 | 5 | 0.4 |
| 18_320213 | 2 | 0 | NA | NA | 1 | 0 | 3 | 0.3333 | 6 | 0.1667 | 4 | 0.25 | NA | NA | NA | NA | 1 | 0 | 5 | 0.2 |
| 18_320214 | 1 | 0 | 1 | 0 | 1 | 0 | 3 | 0.3333 | 6 | 0.1667 | 4 | 0.5 | NA | NA | NA | NA | 1 | 1 | 5 | 0.6 |
| 18_320214 | 2 | 0.5 | 1 | 1 | 1 | 1 | 4 | 1 | 8 | 0.875 | 4 | 0.75 | NA | NA | NA | NA | 1 | 1 | 5 | 0.8 |
| 18_320214 | 2 | 0.5 | 1 | 1 | NA | NA | 4 | 0.5 | 7 | 0.5714 | 5 | 0.2 | NA | NA | NA | NA | 1 | 1 | 6 | 0.3333 |
| 18_320215 | 2 | 1 | 1 | 1 | 1 | 1 | 4 | 0.5 | 8 | 0.75 | 5 | 0.8 | NA | NA | NA | NA | 1 | 1 | 6 | 0.8333 |
| 18_320215 | 2 | 0.5 | 1 | 1 | 1 | 0 | 2 | 0 | 6 | 0.3333 | 2 | 0 | NA | NA | NA | NA | 2 | 1 | 4 | 0.5 |
| 18_320217 | 2 | 1 | 1 | 0 | 1 | 0 | 4 | 0.25 | 8 | 0.375 | 5 | 0.2 | NA | NA | 1 | 0 | 2 | 0.5 | 8 | 0.25 |
| 18_320217 | 2 | 0.5 | NA | NA | 1 | 0 | 4 | 0.25 | 7 | 0.2857 | 4 | 0.5 | NA | NA | 1 | 0 | 2 | 0.5 | 7 | 0.4286 |
| 18_320217 | 2 | 1 | NA | NA | 1 | 0 | 3 | 0.6667 | 6 | 0.6667 | 5 | 0.8 | NA | NA | 1 | 1 | NA | NA | 6 | 0.8333 |
| 18_320218 | 2 | 0.5 | 1 | 1 | 1 | 0 | 4 | 0.75 | 8 | 0.625 | 5 | 0.4 | NA | NA | 1 | 1 | 2 | 0.5 | 8 | 0.5 |
| 18_320218 | 2 | 0.5 | 1 | 0 | 1 | 0 | 3 | 0.6667 | 7 | 0.4286 | 4 | 0.25 | NA | NA | NA | NA | 2 | 0.5 | 6 | 0.3333 |
| 18_320219 | 2 | 0.5 | NA | NA | 1 | 0 | 2 | 0.5 | 5 | 0.4 | 5 | 0.4 | NA | NA | 1 | 0 | 2 | 0 | 8 | 0.25 |
| 18_320220 | 1 | 0 | 1 | 1 | 2 | 0 | 4 | 0.5 | 8 | 0.375 | 3 | 0.6667 | NA | NA | NA | NA | 2 | 1 | 5 | 0.8 |
| 18_320220 | 2 | 0 | 1 | 0 | 1 | 0 | 4 | 0.25 | 8 | 0.125 | 5 | 0.6 | NA | NA | NA | NA | 2 | 0.5 | 7 | 0.5714 |
| 18_320297 | 2 | 0.5 | 1 | 1 | 2 | 0 | 3 | 0 | 8 | 0.25 | 5 | 0.4 | NA | NA | 1 | 0 | 2 | 0 | 8 | 0.25 |
| 18_320298 | 3 | 0.3333 | 2 | 0 | 1 | 0 | 3 | 0.3333 | 9 | 0.2222 | 4 | 0 | NA | NA | 1 | 0 | 3 | 0.3333 | 8 | 0.125 |
| 18_320394 | NA | NA | 1 | 1 | 1 | 1 | 1 | 1 | 3 | 1 | NA | NA | 1 | 1 | NA | NA | 1 | 1 | 2 | 1 |
| 18_320395 | NA | NA | 1 | 1 | NA | NA | 1 | 1 | 2 | 1 | NA | NA | 1 | 1 | NA | NA | 1 | 1 | 2 | 1 |
| 18_320397 | NA | NA | NA | NA | 1 | 1 | 1 | 1 | 2 | 1 | NA | NA | 1 | 0 | NA | NA | 1 | 1 | 2 | 0.5 |
| 18_320399 | 5 | 0.2 | 4 | 0.25 | NA | NA | 3 | 0.3333 | 12 | 0.25 | 5 | 0.6 | NA | NA | 2 | 0.5 | 4 | 0.25 | 11 | 0.4545 |
| 18_320399 | 5 | 1 | 3 | 1 | 2 | 1 | 4 | 1 | 14 | 1 | 4 | 1 | NA | NA | 2 | 1 | 4 | 1 | 10 | 1 |
| 18_320400 | NA | NA | 1 | 1 | NA | NA | 1 | 1 | 2 | 1 | NA | NA | 1 | 1 | NA | NA | 1 | 1 | 2 | 1 |

| | | | | | | | | | | | | | | | | | | | | |
|---|---|---|---|---|---|---|---|---|---|---|---|---|---|---|---|---|---|---|---|---|
| 18_320402( | 3 | 1 | 4 | 0.5 | 2 | 0.5 | 4 | 1 | 13 | 0.7692 | 8 | 0.875 | NA | NA | 2 | 0.5 | 2 | 1 | 12 | 0.8333 |
| 18_320402( | 4 | 0.25 | 4 | 0.5 | 3 | 0 | 5 | 0.8 | 16 | 0.4375 | 8 | 0.25 | NA | NA | 2 | 0 | 5 | 0.2 | 15 | 0.2 |
| 18_320403' | NA | NA | 1 | 1 | NA | NA | 1 | 1 | 2 | 1 | NA | NA | 1 | 1 | NA | NA | 1 | 1 | 2 | 1 |
| 18_320403! | NA | NA | NA | NA | NA | NA | 1 | 1 | 1 | 1 | NA | NA | 1 | 1 | NA | NA | 1 | 1 | 2 | 1 |
| 18_320403! | NA | NA | 1 | 1 | NA | NA | NA | NA | 1 | 1 | NA | NA | 1 | 0 | NA | NA | 1 | 1 | 2 | 0.5 |
| 18_320406( | NA | NA | 1 | 1 | NA | NA | 1 | 1 | 2 | 1 | NA | NA | 1 | 1 | NA | NA | 1 | 1 | 2 | 1 |
| 18_320407! | NA | NA | NA | NA | NA | NA | 1 | 1 | 1 | 1 | NA | NA | 1 | 1 | NA | NA | 1 | 0 | 2 | 0.5 |
| 18_320433' | 1 | 0 | 1 | 1 | NA | NA | 1 | 0 | 3 | 0.3333 | 1 | 0 | 2 | 0.5 | 1 | 0 | 2 | 0.5 | 6 | 0.3333 |
| 18_320434' | NA | NA | 1 | 1 | NA | NA | 1 | 1 | 2 | 1 | 1 | 1 | 1 | 1 | NA | NA | 2 | 1 | 4 | 1 |
| 18_320435; | 1 | 1 | 1 | 1 | NA | NA | 1 | 0 | 3 | 0.6667 | 1 | 1 | 2 | 1 | 1 | 0 | 1 | 1 | 5 | 0.8 |
| 18_320435( | 1 | 1 | NA | NA | NA | NA | 1 | 1 | 2 | 1 | 2 | 1 | 1 | 0 | 1 | 1 | 2 | 0.5 | 6 | 0.6667 |
| 18_320436' | 1 | 1 | 1 | 0 | NA | NA | 1 | 0 | 3 | 0.3333 | 2 | 1 | 2 | 0.5 | NA | NA | 2 | 1 | 6 | 0.8333 |
| 18_320436! | 1 | 1 | 1 | 1 | NA | NA | 1 | 1 | 3 | 1 | 2 | 0.5 | 1 | 1 | 1 | 0 | 1 | 1 | 5 | 0.6 |
| 18_320761( | 3 | 0 | 3 | 0 | 8 | 0.125 | 1 | 0 | 15 | 0.0667 | 2 | 0.5 | 3 | 0 | 4 | 0 | 2 | 0 | 11 | 0.0909 |
| 18_320766; | 3 | 0 | 3 | 0.3333 | 6 | 0.5 | 1 | 0 | 13 | 0.3077 | 1 | 0 | 3 | 0.3333 | 2 | 0 | 1 | 0 | 7 | 0.1429 |
| 18_320768; | 2 | 1 | 3 | 0 | 7 | 0.4286 | 1 | 1 | 13 | 0.4615 | 2 | 0.5 | 3 | 0 | 2 | 0.5 | 2 | 0 | 9 | 0.2222 |
| 18_320773; | 1 | 1 | 1 | 0 | 7 | 0.1429 | NA | NA | 9 | 0.2222 | 1 | 1 | 3 | 0 | 3 | 0.3333 | 2 | 0.5 | 9 | 0.3333 |
| 18_320773( | 2 | 0 | 1 | 0 | 6 | 0 | 1 | 1 | 10 | 0.1 | 1 | 0 | 3 | 0.3333 | 3 | 0 | 1 | 0 | 8 | 0.125 |
| 18_320782; | 1 | 0 | 1 | 0 | 3 | 0.3333 | 1 | 0 | 6 | 0.1667 | 1 | 1 | 1 | 1 | 1 | 0 | NA | NA | 3 | 0.6667 |
| 18_320783! | 1 | 0 | 1 | 0 | 3 | 0.6667 | 1 | 0 | 6 | 0.3333 | 1 | 1 | 1 | 1 | NA | NA | NA | NA | 2 | 1 |
| 18_320784! | 1 | 0 | 1 | 0 | 3 | 1 | 1 | 1 | 6 | 0.6667 | 1 | 1 | NA | NA | NA | NA | NA | NA | 1 | 1 |
| 18_320785; | 1 | 0 | 1 | 1 | 3 | 0.6667 | 1 | 0 | 6 | 0.5 | 1 | 1 | NA | NA | NA | NA | NA | NA | 1 | 1 |
| 18_320786( | NA | NA | NA | NA | 2 | 1 | 1 | 1 | 3 | 1 | 1 | 1 | NA | NA | NA | NA | NA | NA | 1 | 1 |
| 18_320786· | NA | NA | 1 | 0 | 3 | 0 | 1 | 1 | 5 | 0.2 | 1 | 1 | NA | NA | NA | NA | NA | NA | 1 | 1 |
| 18_320786! | 1 | 0 | 1 | 0 | 2 | 0.5 | 1 | 0 | 5 | 0.2 | 1 | 1 | NA | NA | NA | NA | NA | NA | 1 | 1 |
| 18_320787; | 1 | 0 | 1 | 0 | 3 | 1 | 1 | 1 | 6 | 0.6667 | 1 | 1 | NA | NA | NA | NA | NA | NA | 1 | 1 |
| 18_320912; | 3 | 0.3333 | 3 | 0 | 2 | 0.5 | 1 | 0 | 9 | 0.2222 | 3 | 0.3333 | 1 | 0 | 3 | 0 | 3 | 0 | 10 | 0.1 |
| 18_322128· | 2 | 0.5 | 1 | 0 | 1 | 1 | 3 | 0.3333 | 7 | 0.4286 | NA | NA | NA | NA | NA | NA | 2 | 1 | 2 | 1 |
| 18_322129( | 2 | 1 | 1 | 0 | 1 | 1 | 4 | 0.5 | 8 | 0.625 | NA | NA | NA | NA | NA | NA | 2 | 0.5 | 2 | 0.5 |
| 18_322130( | 2 | 0 | 1 | 0 | 1 | 1 | 4 | 0 | 8 | 0.125 | NA | NA | NA | NA | NA | NA | 2 | 0.5 | 2 | 0.5 |
| 18_322130! | 1 | 1 | 1 | 0 | 1 | 0 | 4 | 0.5 | 7 | 0.4286 | NA | NA | NA | NA | NA | NA | 2 | 0.5 | 2 | 0.5 |
| 18_322132! | 2 | 0.5 | 1 | 0 | 1 | 1 | 5 | 0.2 | 9 | 0.3333 | NA | NA | NA | NA | NA | NA | 3 | 0.3333 | 3 | 0.3333 |
| 18_322134! | 2 | 0.5 | 2 | 1 | 1 | 1 | 5 | 0.6 | 10 | 0.7 | NA | NA | NA | NA | NA | NA | 4 | 0.5 | 4 | 0.5 |
| 18_322262; | 4 | 0.25 | 3 | 0 | 2 | 0.5 | 5 | 0 | 14 | 0.1429 | NA | NA | 1 | 0 | 1 | 0 | 4 | 0.25 | 6 | 0.1667 |
| 18_322286! | NA | NA | 1 | 1 | NA | NA | NA | NA | 1 | 1 | 1 | 0 | 1 | 1 | NA | NA | NA | NA | 2 | 0.5 |
| 18_322303! | 3 | 1 | 3 | 1 | 2 | 0.5 | 6 | 0.8333 | 14 | 0.8571 | 1 | 1 | 1 | 1 | 1 | 1 | 4 | 1 | 7 | 1 |
| 18_322304( | 3 | 0.6667 | 2 | 0.5 | 2 | 0 | 5 | 0 | 12 | 0.25 | 1 | 1 | 1 | 0 | 1 | 0 | 4 | 0 | 7 | 0.1429 |
| 18_322308( | 2 | 0.5 | NA | NA | NA | NA | 2 | 1 | 4 | 0.75 | 1 | 1 | NA | NA | NA | NA | 1 | 0 | 2 | 0.5 |
| 18_322309' | 1 | 1 | NA | NA | NA | NA | 2 | 0 | 3 | 0.3333 | 1 | 1 | NA | NA | NA | NA | 1 | 1 | 2 | 1 |
| 18_322314! | 2 | 0.5 | NA | NA | NA | NA | 2 | 0.5 | 4 | 0.5 | 1 | 1 | NA | NA | NA | NA | 1 | 1 | 2 | 1 |
| 18_322340! | 2 | 0 | NA | NA | NA | NA | 2 | 0.5 | 4 | 0.25 | NA | NA | NA | NA | 1 | 1 | 2 | 1 | 3 | 1 |
| 18_323677! | 2 | 0 | 6 | 0.3333 | 4 | 0 | 4 | 0.5 | 16 | 0.25 | 3 | 0 | 1 | 0 | 6 | 0.3333 | 1 | 0 | 11 | 0.1818 |
| 18_323678; | 2 | 0 | 4 | 0.25 | 4 | 0.25 | 2 | 0 | 12 | 0.1667 | 2 | 0 | 2 | 0 | 6 | 0.1667 | 1 | 0 | 11 | 0.0909 |
| 18_323691( | 5 | 0 | 9 | 0 | 3 | 0 | 3 | 0.3333 | 20 | 0.05 | 3 | 0 | 2 | 0 | 9 | 0.1111 | 3 | 0 | 17 | 0.0588 |
| 18_323697( | 3 | 0.6667 | 8 | 0.125 | 3 | 0 | 3 | 0 | 17 | 0.1765 | 2 | 0 | 2 | 0.5 | 3 | 0.3333 | 1 | 0 | 8 | 0.25 |
| 18_323697· | 7 | 0 | 11 | 0.1818 | 3 | 0 | 3 | 0 | 24 | 0.0833 | 5 | 0 | 2 | 0.5 | 9 | 0.1111 | 2 | 0 | 18 | 0.1111 |
| 18_323786! | 2 | 0.5 | 2 | 0 | 3 | 0.3333 | 2 | 0 | 9 | 0.2222 | 1 | 1 | 1 | 1 | 1 | 0 | 1 | 0 | 4 | 0.5 |
| 18_324029! | 2 | 0.5 | 5 | 0.2 | 7 | 0.1429 | 3 | 0.6667 | 17 | 0.2941 | 1 | 0 | 2 | 0 | 2 | 0.5 | 4 | 0 | 9 | 0.1111 |
| 18_324030; | 2 | 1 | 3 | 0.6667 | 7 | 0.8571 | 2 | 0.5 | 14 | 0.7857 | 2 | 1 | 2 | 1 | 2 | 1 | 4 | 0.75 | 10 | 0.9 |
| 18_324030! | 1 | 1 | 4 | 0.25 | 5 | 0.2 | 3 | 0 | 13 | 0.2308 | 2 | 0.5 | 2 | 0.5 | 2 | 0 | 3 | 0.6667 | 9 | 0.4444 |
| 18_324051· | 2 | 0.5 | 1 | 0 | 3 | 0.3333 | 1 | 0 | 7 | 0.2857 | NA | NA | NA | NA | 2 | 0 | 3 | 0.3333 | 5 | 0.2 |
| 18_325685( | 24 | 0.25 | 43 | 0.2791 | 19 | 0.3684 | 21 | 0.3333 | 107 | 0.2991 | 22 | 0.3636 | 21 | 0.2381 | 31 | 0.4839 | 19 | 0.4737 | 93 | 0.3978 |
| 18_329989! | 6 | 0.1667 | 4 | 0.25 | NA | NA | 5 | 0.2 | 15 | 0.2 | 3 | 0.3333 | 4 | 0.5 | 2 | 0 | 3 | 0.3333 | 12 | 0.3333 |
| 18_329993( | 8 | 0.625 | 1 | 1 | 1 | 1 | 4 | 0.5 | 14 | 0.6429 | 1 | 1 | 5 | 0.4 | 4 | 0.25 | 3 | 0.6667 | 13 | 0.4615 |
| 18_333179; | 8 | 0.5 | 4 | 0 | 8 | 0.25 | 5 | 0.4 | 25 | 0.32 | NA | NA | 3 | 0 | 9 | 0.4444 | 3 | 0.6667 | 15 | 0.4 |
| 18_333179! | 5 | 0.4 | 1 | 0 | 5 | 0.4 | 4 | 0 | 15 | 0.2667 | NA | NA | 3 | 0 | 6 | 0.1667 | 3 | 0.6667 | 12 | 0.25 |
| 18_333181; | 8 | 0.625 | 4 | 0 | 8 | 0.625 | 5 | 0.4 | 25 | 0.48 | NA | NA | 3 | 1 | 9 | 0.5556 | 3 | 0.3333 | 15 | 0.6 |
| 18_333181! | 7 | 0.2857 | 3 | 0 | 8 | 0.5 | 4 | 0.5 | 22 | 0.3636 | NA | NA | 3 | 0.3333 | 8 | 0.375 | 2 | 0.5 | 13 | 0.3846 |
| 18_333182' | 7 | 0.8571 | 4 | 1 | 8 | 0.875 | 6 | 1 | 25 | 0.92 | NA | NA | 3 | 1 | 10 | 1 | 2 | 1 | 15 | 1 |
| 18_333183( | 7 | 0 | 2 | 0 | 7 | 0 | 3 | 0.3333 | 19 | 0.0526 | NA | NA | 2 | 0 | 7 | 0.1429 | 1 | 0 | 10 | 0.1 |
| 18_333184; | 5 | 0 | 2 | 0 | 8 | 0.25 | 4 | 0 | 19 | 0.1053 | NA | NA | 1 | 0 | 7 | 0.1429 | 1 | 1 | 9 | 0.2222 |
| 18_333185! | 5 | 0.2 | 1 | 0 | 7 | 0.2857 | 2 | 0 | 15 | 0.2 | NA | NA | 2 | 0 | 7 | 0.1429 | 1 | 0 | 10 | 0.1 |
| 18_333186' | 6 | 0 | 2 | 0 | 6 | 0.1667 | 3 | 0 | 17 | 0.0588 | NA | NA | 2 | 0.5 | 7 | 0.1429 | 1 | 0 | 10 | 0.2 |
| 18_333186· | 5 | 0.2 | 2 | 0 | 7 | 0 | 2 | 0 | 16 | 0.0625 | NA | NA | 3 | 0.3333 | 7 | 0.1429 | 1 | 0 | 11 | 0.1818 |
| 18_333274; | 10 | 0 | 3 | 0.3333 | 9 | 0.3333 | 6 | 0 | 28 | 0.1429 | 4 | 0 | 5 | 0.6 | 13 | 0.1538 | 3 | 0.6667 | 25 | 0.28 |

| | | | | | | | | | | | | | | | | | | | | |
|---|---|---|---|---|---|---|---|---|---|---|---|---|---|---|---|---|---|---|---|---|
| 18_3332806 | 10 | 0.2 | 4 | 0 | 12 | 0 | 6 | 0.1667 | 32 | 0.0938 | 5 | 0 | 7 | 0.1429 | 16 | 0.0625 | 6 | 0 | 34 | 0.0588 |
| 18_3332821 | 12 | 0.4167 | 6 | 0 | 14 | 0.0714 | 7 | 0.1429 | 39 | 0.1795 | 6 | 0 | 6 | 0.1667 | 15 | 0.1333 | 7 | 0.1429 | 34 | 0.1176 |
| 18_3332849 | 8 | 0 | 4 | 0.5 | 9 | 0 | 5 | 0.2 | 26 | 0.1154 | 4 | 0 | 6 | 0 | 12 | 0.1667 | 5 | 0 | 27 | 0.0741 |
| 18_3332857 | 11 | 0.8182 | 5 | 0.8 | 10 | 0.3 | 5 | 0.4 | 31 | 0.5806 | 4 | 0.25 | 7 | 0.4286 | 16 | 0.5625 | 5 | 0.6 | 32 | 0.5 |
| 18_3332863 | 15 | 0.5333 | 7 | 0.8571 | 13 | 0.4615 | 7 | 0.2857 | 42 | 0.5238 | 7 | 0.5714 | 10 | 0.7 | 16 | 0.75 | 6 | 0.5 | 39 | 0.6667 |
| 18_3332871 | 17 | 0.2353 | 9 | 0.2222 | 18 | 0.2222 | 9 | 0 | 53 | 0.1887 | 9 | 0.2222 | 9 | 0.1111 | 18 | 0.3333 | 8 | 0.25 | 44 | 0.25 |
| 18_3333265 | 1 | 0 | 3 | 0 | 3 | 0.6667 | 2 | 0 | 9 | 0.2222 | 1 | 0 | 1 | 0 | 3 | 0.3333 | 2 | 0.5 | 7 | 0.2857 |
| 18_3333269 | 1 | 0 | 2 | 0 | 3 | 0.3333 | 1 | 0 | 7 | 0.1429 | 1 | 0 | NA | NA | 2 | 0 | 1 | 1 | 4 | 0.25 |
| 18_3357119 | 1 | 0 | 2 | 0.5 | 5 | 0.6 | 4 | 0.5 | 12 | 0.5 | NA | NA | 1 | 0 | 1 | 1 | 4 | 0.5 | 6 | 0.5 |
| 18_3357145 | 1 | 0 | 2 | 0 | 7 | 0 | 6 | 0.1667 | 16 | 0.0625 | 2 | 0.5 | 3 | 0 | 1 | 0 | 5 | 0 | 11 | 0.0909 |
| 18_3357161 | 1 | 1 | 1 | 0 | 7 | 0.5714 | 5 | 0.4 | 14 | 0.5 | 2 | 1 | NA | NA | 1 | 0 | 4 | 0.75 | 7 | 0.7143 |
| 18_3357164 | 1 | 1 | 1 | 1 | 6 | 0.6667 | 6 | 0.8333 | 14 | 0.7857 | 2 | 1 | 2 | 1 | 1 | 1 | 4 | 0.75 | 9 | 0.8889 |
| 18_3357165 | 1 | 1 | 2 | 0 | 5 | 0 | 5 | 0 | 13 | 0.0769 | 2 | 0.5 | 2 | 0 | NA | NA | 4 | 0 | 8 | 0.125 |
| 18_3443153 | NA | NA | NA | NA | 2 | 0 | 1 | 1 | 3 | 0.3333 | 1 | 0 | NA | NA | NA | NA | 2 | 1 | 3 | 0.6667 |
| 18_3443163 | NA | NA | NA | NA | 2 | 1 | NA | NA | 2 | 1 | 1 | 1 | NA | NA | NA | NA | 3 | 1 | 4 | 1 |
| 18_3444514 | NA | NA | 1 | 1 | 4 | 0.75 | 1 | 1 | 6 | 0.8333 | 1 | 0 | 1 | 1 | NA | NA | 3 | 0.6667 | 5 | 0.6 |
| 18_3444519 | NA | NA | NA | NA | 3 | 0.6667 | 1 | 0 | 4 | 0.5 | 1 | 1 | NA | NA | NA | NA | 2 | 0.5 | 3 | 0.6667 |
| 18_3444529 | NA | NA | 1 | 0 | 4 | 0.5 | 1 | 0 | 6 | 0.3333 | 1 | 0 | 1 | 0 | 1 | 0 | 3 | 0.3333 | 6 | 0.1667 |
| 18_3444546 | NA | NA | 1 | 1 | 4 | 0.25 | 1 | 0 | 6 | 0.3333 | 1 | 0 | 1 | 0 | 1 | 0 | 3 | 0.6667 | 6 | 0.3333 |
| 18_3444549 | NA | NA | 1 | 0 | 4 | 0.25 | 1 | 0 | 6 | 0.1667 | 1 | 0 | 1 | 1 | 1 | 0 | 3 | 0.6667 | 6 | 0.5 |
| 18_3444555 | NA | NA | NA | NA | 4 | 0.75 | 1 | 1 | 5 | 0.8 | 1 | 1 | 1 | 1 | 1 | 1 | 2 | 1 | 5 | 1 |
| 18_3444593 | NA | NA | 3 | 1 | 5 | 0.8 | 1 | 1 | 9 | 0.8889 | NA | NA | 2 | 0 | 1 | 1 | 3 | 0.6667 | 6 | 0.5 |
| 18_3444615 | NA | NA | 4 | 0.5 | 5 | 0.4 | 1 | 1 | 10 | 0.5 | 2 | 1 | 2 | 0.5 | 1 | 0 | 3 | 0.3333 | 8 | 0.5 |
| 18_3444620 | NA | NA | 4 | 0.25 | 4 | 0 | 1 | 0 | 9 | 0.1111 | 2 | 0.5 | 2 | 0 | 1 | 0 | 3 | 0 | 8 | 0.125 |
| 18_3444638 | NA | NA | 3 | 0.6667 | 4 | 0 | 2 | 0.5 | 9 | 0.3333 | 2 | 1 | 1 | 1 | 1 | 1 | 2 | 0 | 6 | 0.6667 |
| 18_3444665 | NA | NA | 4 | 0.25 | 5 | 0.2 | 2 | 0 | 11 | 0.1818 | 2 | 1 | NA | NA | NA | NA | 4 | 0 | 6 | 0.3333 |
| 18_3444688 | 1 | 0 | 4 | 0 | 5 | 0.2 | 1 | 1 | 11 | 0.1818 | 2 | 0.5 | 2 | 0 | 1 | 0 | 3 | 0.3333 | 8 | 0.25 |
| 18_3444722 | 1 | 1 | 4 | 0.5 | 4 | 0.25 | 1 | 0 | 10 | 0.4 | 2 | 0.5 | 1 | 0 | 1 | 0 | 3 | 0.3333 | 7 | 0.2857 |
| 18_3444724 | 1 | 1 | 6 | 0.5 | 6 | 0 | 3 | 0 | 16 | 0.25 | 1 | 0 | 2 | 0.5 | 1 | 0 | 5 | 0.4 | 9 | 0.3333 |
| 18_3444728 | 1 | 0 | 5 | 0 | 6 | 0.1667 | 2 | 0 | 14 | 0.0714 | 2 | 0 | 2 | 0.5 | 1 | 1 | 5 | 0 | 10 | 0.2 |
| 18_3444751 | 1 | 1 | 5 | 0.4 | 6 | 0.3333 | 4 | 0.75 | 16 | 0.5 | NA | NA | 2 | 0.5 | 1 | 0 | 4 | 0.5 | 7 | 0.4286 |
| 18_3444771 | 1 | 1 | 6 | 0 | 4 | 0.25 | 3 | 0.3333 | 14 | 0.2143 | 2 | 0.5 | 2 | 0 | 1 | 0 | 6 | 0 | 11 | 0.0909 |
| 18_3444777 | 1 | 0 | 5 | 0.2 | 6 | 0.1667 | 4 | 0.5 | 16 | 0.25 | 2 | 0.5 | 2 | 0 | 1 | 0 | 5 | 0.2 | 10 | 0.2 |
| 18_3444778 | 1 | 1 | 4 | 1 | 6 | 1 | 4 | 0.75 | 15 | 0.9333 | 2 | 1 | 2 | 0.5 | 1 | 1 | 5 | 1 | 10 | 0.9 |
| 18_3444788 | NA | NA | 6 | 0.5 | 4 | 0.5 | 4 | 0.5 | 14 | 0.5 | 2 | 0.5 | 1 | 0 | 2 | 0.5 | 5 | 0 | 10 | 0.2 |
| 18_3444794 | 1 | 0 | 5 | 0.8 | 5 | 0.4 | 4 | 0 | 15 | 0.4 | 2 | 0 | 1 | 1 | 2 | 0.5 | 4 | 0.5 | 9 | 0.4444 |
| 18_3444803 | NA | NA | 5 | 0.8 | 5 | 1 | 4 | 0.5 | 14 | 0.7857 | 2 | 0.5 | 1 | 1 | 2 | 1 | 7 | 0.7143 | 12 | 0.75 |
| 18_3444804 | NA | NA | 5 | 0.8 | 4 | 1 | 1 | 0 | 10 | 0.8 | 2 | 0.5 | 1 | 1 | 1 | 0 | 5 | 0.4 | 9 | 0.4444 |
| 18_3444834 | 1 | 0 | 7 | 0 | 5 | 0.2 | 5 | 0 | 18 | 0.0556 | 2 | 0 | 1 | 0 | 4 | 0.25 | 7 | 0 | 14 | 0.0714 |
| 18_3444848 | 2 | 0.5 | 7 | 0.4286 | 5 | 0.8 | 5 | 0.4 | 19 | 0.5263 | 2 | 0.5 | 2 | 0.5 | 4 | 0.5 | 7 | 0.7143 | 15 | 0.6 |
| 18_3444850 | 2 | 0 | 7 | 0 | 5 | 0 | 6 | 0.1667 | 20 | 0.05 | 2 | 0 | 2 | 0 | 4 | 0.25 | 6 | 0 | 14 | 0.0714 |
| 18_3444852 | 2 | 0 | 6 | 0.1667 | 5 | 0.2 | 4 | 0.5 | 17 | 0.2353 | 2 | 0.5 | 2 | 0.5 | 4 | 0.25 | 7 | 0.5714 | 15 | 0.4667 |
| 18_3444862 | 1 | 0 | 7 | 0.4286 | 4 | 0.25 | 6 | 0 | 18 | 0.2222 | 2 | 0 | 2 | 0.5 | 3 | 0 | 6 | 0.3333 | 13 | 0.2308 |
| 18_3444867 | 2 | 1 | 5 | 0.6 | 3 | 0 | 6 | 0.1667 | 16 | 0.375 | 3 | 0.3333 | 1 | 0 | 4 | 0.25 | 6 | 0.3333 | 14 | 0.2857 |
| 18_3444891 | 2 | 0.5 | 8 | 0.125 | 5 | 0.4 | 6 | 0 | 21 | 0.1905 | 2 | 0 | 3 | 0 | 3 | 0.3333 | 6 | 0.3333 | 14 | 0.2143 |
| 18_3444894 | 2 | 0.5 | 8 | 0.75 | 6 | 0.1667 | 5 | 0.4 | 21 | 0.4762 | 3 | 0.3333 | 4 | 0.5 | 3 | 0.3333 | 6 | 0.1667 | 16 | 0.3125 |
| 18_3444902 | 2 | 0 | 8 | 0.375 | 6 | 0.1667 | 6 | 0.5 | 22 | 0.3182 | 3 | 0.3333 | 2 | 1 | 3 | 0 | 6 | 0.3333 | 14 | 0.3571 |
| 18_3444920 | 1 | 0 | 9 | 0.4444 | 6 | 0.6667 | 5 | 0.6 | 21 | 0.5238 | 3 | 0.6667 | 4 | 0.25 | 3 | 0 | 6 | 0.5 | 16 | 0.375 |
| 18_3444940 | 3 | 0.6667 | 8 | 0.125 | 6 | 0.1667 | 6 | 0.1667 | 23 | 0.2174 | 3 | 0.3333 | 4 | 0 | 4 | 0.25 | 6 | 0.5 | 17 | 0.2941 |
| 18_3444986 | 4 | 0.75 | 14 | 0.3571 | 8 | 0 | 7 | 0.2857 | 33 | 0.303 | 4 | 0.75 | 5 | 0.2 | 4 | 0.25 | 8 | 0.375 | 21 | 0.381 |
| 18_3444992 | 4 | 0.5 | 13 | 0.2308 | 7 | 0.2857 | 5 | 0.4 | 29 | 0.3103 | 4 | 0.5 | 4 | 0.25 | 4 | 0.25 | 6 | 0.5 | 18 | 0.3889 |
| 18_3444997 | 4 | 0 | 13 | 0 | 8 | 0.125 | 6 | 0.1667 | 31 | 0.0645 | 4 | 0 | 6 | 0.1667 | 5 | 0.2 | 6 | 0.1667 | 21 | 0.1429 |
| 18_3445002 | 5 | 0.8 | 15 | 0.6 | 9 | 0.2222 | 7 | 0.7143 | 36 | 0.5556 | 4 | 0.75 | 6 | 0.3333 | 4 | 0.25 | 10 | 0.2 | 24 | 0.3333 |
| 18_3445012 | 5 | 0.8 | 12 | 1 | 9 | 0.8889 | 7 | 1 | 33 | 0.9394 | 3 | 1 | 6 | 1 | 4 | 1 | 7 | 1 | 20 | 1 |
| 18_3445030 | 5 | 1 | 14 | 0.9286 | 9 | 1 | 7 | 1 | 35 | 0.9714 | 5 | 1 | 6 | 1 | 4 | 1 | 9 | 1 | 24 | 1 |
| 18_3445035 | 5 | 0.6 | 13 | 0.6923 | 8 | 0.75 | 5 | 0.8 | 31 | 0.7097 | 5 | 1 | 8 | 0.5 | 5 | 0.6 | 11 | 0.7273 | 29 | 0.6897 |
| 18_3445047 | 6 | 0 | 13 | 0.0769 | 9 | 0.1111 | 5 | 0 | 33 | 0.0606 | 5 | 0.4 | 7 | 0.1429 | 4 | 0 | 11 | 0.2727 | 27 | 0.2222 |
| 18_3445054 | 6 | 0 | 15 | 0.1333 | 10 | 0.3 | 7 | 0 | 38 | 0.1316 | 6 | 0.1667 | 7 | 0 | 5 | 0 | 12 | 0.1667 | 30 | 0.1 |
| 18_3445071 | 6 | 0.1667 | 15 | 0.1333 | 10 | 0.2 | 7 | 0.4286 | 38 | 0.2105 | 4 | 0.25 | 8 | 0.125 | 4 | 0.25 | 11 | 0.0909 | 27 | 0.1481 |
| 18_3445080 | 6 | 0.3333 | 16 | 0.5 | 11 | 0.6364 | 5 | 0.4 | 38 | 0.5 | 6 | 0.6667 | 9 | 0.4444 | 6 | 0.5 | 10 | 0.5 | 31 | 0.5161 |
| 18_3445090 | 6 | 0 | 18 | 0.1111 | 12 | 0.0833 | 7 | 0.2857 | 43 | 0.1163 | 6 | 0.1667 | 9 | 0.2222 | 7 | 0.1429 | 12 | 0.25 | 34 | 0.2059 |
| 18_3445093 | 4 | 0 | 15 | 0.2667 | 10 | 0.2 | 6 | 0.3333 | 35 | 0.2286 | 3 | 0.3333 | 7 | 0 | 7 | 0.2857 | 12 | 0.3333 | 29 | 0.2414 |
| 18_3445096 | 3 | 0.6667 | 10 | 0.9 | 10 | 0.8 | 7 | 1 | 30 | 0.8667 | 6 | 1 | 7 | 0.8571 | 6 | 0.6667 | 9 | 0.8889 | 28 | 0.8571 |
| 18_3445106 | 5 | 0.2 | 12 | 0.25 | 11 | 0 | 6 | 0 | 34 | 0.1176 | 7 | 0.4286 | 5 | 0 | 5 | 0.4 | 8 | 0.25 | 25 | 0.28 |
| 18_3445109 | 6 | 1 | 14 | 1 | 10 | 1 | 6 | 1 | 36 | 1 | 7 | 1 | 8 | 1 | 4 | 1 | 13 | 1 | 32 | 1 |

| | | | | | | | | | | | | | | | | | | | | |
|---|---|---|---|---|---|---|---|---|---|---|---|---|---|---|---|---|---|---|---|---|
| 18_3445115 | 4 | 0.75 | 14 | 0.5714 | 10 | 0.7 | 9 | 0.8889 | 37 | 0.7027 | 8 | 0.75 | 7 | 0.8571 | 5 | 0.4 | 9 | 0.5556 | 29 | 0.6552 |
| 18_3445120 | 4 | 0.75 | 18 | 0.5 | 12 | 0.5 | 9 | 0.8889 | 43 | 0.6047 | 7 | 0.8571 | 7 | 0.5714 | 4 | 0.5 | 10 | 0.9 | 28 | 0.75 |
| 18_3445121 | 4 | 0 | 16 | 0.125 | 9 | 0 | 7 | 0.1429 | 36 | 0.0833 | 7 | 0 | 7 | 0.1429 | 3 | 0 | 12 | 0.0833 | 29 | 0.069 |
| 18_3445124 | 4 | 0 | 14 | 0.0714 | 11 | 0 | 8 | 0.125 | 37 | 0.0541 | 7 | 0.2857 | 6 | 0.1667 | 7 | 0 | 11 | 0.3636 | 31 | 0.2258 |
| 18_3445129 | 4 | 0.25 | 17 | 0.4118 | 11 | 0.1818 | 7 | 0 | 39 | 0.2564 | 5 | 0 | 8 | 0.25 | 5 | 0.4 | 9 | 0.2222 | 27 | 0.2222 |
| 18_3445130 | 6 | 1 | 17 | 1 | 12 | 0.9167 | 10 | 1 | 45 | 0.9778 | 7 | 1 | 9 | 1 | 6 | 1 | 13 | 0.9231 | 35 | 0.9714 |
| 18_3445132 | 3 | 1 | 14 | 1 | 9 | 0.8889 | 7 | 0.8571 | 33 | 0.9394 | 8 | 0.875 | 6 | 1 | 5 | 1 | 10 | 1 | 29 | 0.9655 |
| 18_3445135 | 5 | 0.6 | 17 | 0.4118 | 12 | 0.8333 | 9 | 0.5556 | 43 | 0.5814 | 8 | 0.875 | 9 | 0.5556 | 6 | 0.6667 | 14 | 0.6429 | 37 | 0.6757 |
| 18_3445142 | 5 | 1 | 19 | 1 | 10 | 1 | 9 | 1 | 43 | 1 | 8 | 1 | 10 | 0.9 | 7 | 1 | 15 | 1 | 40 | 0.975 |
| 18_3445144 | 6 | 0.1667 | 21 | 0.381 | 11 | 0.2727 | 10 | 0.2 | 48 | 0.2917 | 7 | 0.1429 | 9 | 0.1111 | 7 | 0.4286 | 14 | 0.0714 | 37 | 0.1622 |
| 18_3445156 | 6 | 0.3333 | 21 | 0.0952 | 13 | 0.2308 | 10 | 0.1 | 50 | 0.16 | 10 | 0.2 | 9 | 0.2222 | 5 | 0 | 13 | 0 | 37 | 0.1081 |
| 18_3445164 | 7 | 0.7143 | 22 | 0.8636 | 14 | 0.8571 | 12 | 0.8333 | 55 | 0.8364 | 10 | 0.8 | 9 | 0.6667 | 7 | 0.8571 | 15 | 0.6 | 41 | 0.7073 |
| 18_3445169 | 8 | 1 | 22 | 1 | 14 | 0.9286 | 12 | 0.9167 | 56 | 0.9643 | 10 | 1 | 10 | 1 | 7 | 1 | 17 | 1 | 44 | 1 |
| 18_3445170 | 7 | 0.1429 | 21 | 0.0952 | 13 | 0 | 11 | 0.0909 | 52 | 0.0769 | 11 | 0.0909 | 10 | 0 | 7 | 0.1429 | 15 | 0.0667 | 43 | 0.0698 |
| 18_3445242 | 7 | 0.1429 | 21 | 0.4286 | 16 | 0.0625 | 12 | 0.25 | 56 | 0.25 | 10 | 0.3 | 10 | 0.2 | 9 | 0.3333 | 12 | 0.1667 | 41 | 0.2439 |
| 18_3445275 | 2 | 0 | 9 | 0 | 3 | 0 | 6 | 0.1667 | 20 | 0.05 | 4 | 0.25 | 6 | 0 | 2 | 0 | 7 | 0 | 19 | 0.0526 |
| 18_3445432 | 3 | 0 | 18 | 0.0556 | 6 | 0 | 13 | 0.0769 | 40 | 0.05 | 6 | 0.3333 | 7 | 0 | 5 | 0 | 12 | 0.1667 | 30 | 0.1333 |
| 18_3446934 | 9 | 0.2222 | 10 | 0.2 | 11 | 0 | 4 | 0.5 | 34 | 0.1765 | 4 | 0 | 14 | 0.3571 | 11 | 0.1818 | 11 | 0.0909 | 40 | 0.2 |
| 18_3446938 | 9 | 0.2222 | 9 | 0 | 7 | 0.1429 | 4 | 0 | 29 | 0.1034 | 3 | 0 | 9 | 0.1111 | 14 | 0.1429 | 11 | 0.0909 | 37 | 0.1081 |
| 18_3446945 | 8 | 0.125 | 9 | 0.1111 | 12 | 0.0833 | 7 | 0.1429 | 36 | 0.1111 | 4 | 0 | 13 | 0.3077 | 12 | 0 | 9 | 0 | 38 | 0.1053 |
| 18_3446951 | 12 | 0.3333 | 11 | 0.4545 | 10 | 0.3 | 6 | 0.6667 | 39 | 0.4103 | 4 | 0.25 | 11 | 0.4545 | 12 | 0.75 | 10 | 0.5 | 37 | 0.5405 |
| 18_3447805 | 46 | 0.3913 | 52 | 0.3462 | 43 | 0.2326 | 52 | 0.1923 | 193 | 0.2902 | 30 | 0.2667 | 46 | 0.2609 | 39 | 0.2308 | 48 | 0.3333 | 163 | 0.2761 |
| 18_3448011 | 37 | 0.027 | 44 | 0.1136 | 34 | 0.0294 | 49 | 0.0816 | 164 | 0.0671 | 15 | 0.2 | 33 | 0.0606 | 33 | 0.0606 | 37 | 0.1351 | 118 | 0.1017 |
| 18_3457470 | 2 | 0 | 6 | 0.1667 | 7 | 0 | 3 | 0 | 18 | 0.0556 | 5 | 0 | 5 | 0 | 3 | 0 | 3 | 0.3333 | 16 | 0.0625 |
| 18_3457491 | 1 | 0 | 6 | 0.5 | 3 | 0.3333 | 3 | 0 | 13 | 0.3077 | 5 | 0.6 | 5 | 0.4 | 2 | 0.5 | 1 | 0 | 13 | 0.4615 |
| 18_3457502 | 2 | 0 | 5 | 0.8 | 6 | 0.1667 | 4 | 0.25 | 17 | 0.3529 | 4 | 0.5 | 4 | 0 | 3 | 0.3333 | 3 | 0.3333 | 14 | 0.2857 |
| 18_3457518 | 1 | 0 | 6 | 0.3333 | 4 | 0.25 | 3 | 0 | 14 | 0.2143 | 5 | 0.2 | 5 | 0 | 3 | 0.6667 | 3 | 0 | 16 | 0.1875 |
| 18_3457530 | 2 | 0.5 | 7 | 0.5714 | 6 | 0.5 | 2 | 0 | 17 | 0.4706 | 4 | 0.25 | 4 | 0 | 3 | 0.6667 | 3 | 0 | 14 | 0.2143 |
| 18_3457543 | 2 | 0 | 7 | 0.4286 | 8 | 0.25 | 2 | 0.5 | 19 | 0.3158 | 6 | 0.1667 | 5 | 0.6 | 5 | 0.4 | 3 | 0 | 19 | 0.3158 |
| 18_3457545 | 2 | 0 | 7 | 0.2857 | 8 | 0.125 | 4 | 0 | 21 | 0.1429 | 8 | 0.25 | 5 | 0 | 5 | 0.2 | 4 | 0 | 22 | 0.1364 |
| 18_3457564 | 3 | 0 | 5 | 0.2 | 6 | 0.1667 | 3 | 0 | 17 | 0.1176 | 7 | 0 | 5 | 0 | 5 | 0.2 | 3 | 0 | 20 | 0.05 |
| 18_3457571 | 3 | 0.3333 | 8 | 0.25 | 6 | 0.1667 | 4 | 0 | 21 | 0.1905 | 7 | 0.1429 | 5 | 0.2 | 5 | 0.4 | 4 | 0.25 | 21 | 0.2381 |
| 18_3463050 | NA | NA | 2 | 0 | 2 | 0 | 1 | 1 | 5 | 0.2 | 1 | 0 | 2 | 0.5 | 1 | 0 | 2 | 0 | 6 | 0.1667 |
| 18_3473366 | 2 | 0 | 1 | 0 | 1 | 1 | NA | NA | 4 | 0.25 | NA | NA | NA | NA | NA | NA | 1 | 1 | 1 | 1 |
| 18_3474019 | 1 | 0 | 3 | 0 | 2 | 0 | 5 | 0.4 | 11 | 0.1818 | 2 | 0 | 3 | 0 | 2 | 0 | 4 | 0.25 | 11 | 0.0909 |
| 18_3474032 | 1 | 0 | 3 | 0.3333 | 3 | 0.6667 | 5 | 0.2 | 12 | 0.3333 | 2 | 0 | 3 | 0.3333 | 1 | 1 | 3 | 0.6667 | 9 | 0.4444 |
| 18_3474087 | 1 | 1 | 2 | 0 | 2 | 0.5 | 5 | 0 | 10 | 0.2 | 2 | 0 | 2 | 0 | 2 | 0.5 | 3 | 0 | 9 | 0.1111 |
| 18_3474105 | 1 | 1 | 2 | 0 | 2 | 0.5 | 4 | 0 | 9 | 0.2222 | 2 | 0 | 2 | 0 | 2 | 0.5 | 4 | 0.25 | 10 | 0.2 |
| 18_3476413 | 2 | 0 | NA | NA | 4 | 0.25 | NA | NA | 6 | 0.1667 | 3 | 0.3333 | 2 | 0 | 2 | 0 | 2 | 0 | 9 | 0.1111 |
| 18_3476414 | 1 | 1 | NA | NA | 4 | 0.25 | NA | NA | 5 | 0.4 | 3 | 0.6667 | 2 | 0.5 | 2 | 0 | 2 | 0 | 9 | 0.3333 |
| 18_3476427 | 2 | 0.5 | NA | NA | 4 | 0.5 | NA | NA | 6 | 0.5 | 3 | 0 | 1 | 1 | 1 | 0 | 2 | 0 | 7 | 0.1429 |
| 18_3476614 | 5 | 0.2 | 2 | 0 | 5 | 0.2 | 4 | 0.25 | 16 | 0.1875 | 3 | 0.3333 | 1 | 0 | 4 | 0 | 3 | 0 | 11 | 0.0909 |
| 18_3476633 | 5 | 0.4 | 2 | 0.5 | 6 | 0.5 | 5 | 0.8 | 18 | 0.5556 | 2 | 0.5 | 2 | 0.5 | 4 | 1 | 3 | 0.3333 | 11 | 0.6364 |
| 18_3476634 | 7 | 0 | 2 | 0 | 6 | 0.1667 | 3 | 0 | 18 | 0.0556 | 2 | 0.5 | 2 | 0.5 | 4 | 0 | 3 | 0 | 11 | 0.1818 |
| 18_3476642 | 7 | 0.7143 | 1 | 1 | 4 | 1 | 4 | 1 | 16 | 0.875 | 1 | 1 | NA | NA | 3 | 0.3333 | 2 | 0.5 | 6 | 0.5 |
| 18_3476666 | 8 | 0 | 2 | 0 | 5 | 0.4 | 4 | 0 | 19 | 0.1053 | 1 | 0 | 2 | 0.5 | 4 | 0 | 3 | 0.3333 | 10 | 0.2 |
| 18_3476693 | 9 | 0.1111 | 1 | 0 | 4 | 0.25 | 5 | 0.2 | 19 | 0.1579 | 1 | 1 | NA | NA | 3 | 0 | 3 | 0.3333 | 7 | 0.2857 |
| 18_3476696 | 9 | 0.4444 | 1 | 1 | 4 | 0.5 | 5 | 0.2 | 19 | 0.4211 | 1 | 1 | 2 | 0.5 | 4 | 0.5 | 3 | 0.3333 | 10 | 0.5 |
| 18_3490146 | 2 | 1 | 4 | 0.75 | 1 | 0 | 3 | 0.3333 | 10 | 0.6 | 1 | 1 | 2 | 1 | 1 | 1 | 2 | 0 | 6 | 0.6667 |
| 18_3490151 | 1 | 1 | 2 | 1 | 1 | 1 | 2 | 0.5 | 6 | 0.8333 | NA | NA | NA | NA | 1 | 1 | 2 | 1 | 3 | 1 |
| 18_3490198 | 3 | 0 | 3 | 0.3333 | 1 | 0 | 4 | 0 | 11 | 0.0909 | 1 | 0 | 3 | 0 | 2 | 0 | 2 | 1 | 8 | 0.25 |
| 18_3490385 | 2 | 1 | 4 | 0.75 | 2 | 0.5 | 4 | 1 | 12 | 0.8333 | 2 | 0.5 | 3 | 1 | 3 | 1 | 3 | 0.3333 | 11 | 0.7273 |
| 18_3490387 | 4 | 1 | 3 | 1 | 2 | 1 | 4 | 1 | 13 | 1 | 2 | 1 | 3 | 1 | 3 | 1 | 3 | 0.6667 | 11 | 0.9091 |
| 18_3490388 | 4 | 0.5 | 4 | 0.5 | 2 | 0.5 | 4 | 0.25 | 14 | 0.4286 | 2 | 1 | 3 | 0.3333 | 3 | 0.3333 | 3 | 1 | 11 | 0.6364 |
| 18_3490388 | 4 | 0.75 | 4 | 0.25 | 1 | 1 | 4 | 0 | 13 | 0.3846 | 3 | 0 | 3 | 0 | 3 | 0.6667 | 3 | 0.3333 | 12 | 0.25 |
| 18_3490394 | 4 | 1 | 4 | 1 | 2 | 1 | 3 | 1 | 13 | 1 | 2 | 0.5 | 3 | 1 | 2 | 1 | 3 | 1 | 10 | 0.9 |
| 18_3490407 | 4 | 1 | 6 | 0.8333 | 1 | 1 | 4 | 1 | 15 | 0.9333 | 4 | 0.75 | 3 | 1 | 2 | 1 | 4 | 1 | 13 | 0.9231 |
| 18_3490408 | 4 | 0.25 | 4 | 0 | 2 | 0.5 | 5 | 0.2 | 15 | 0.2 | 4 | 0 | 3 | 0.3333 | 3 | 0 | 3 | 0 | 13 | 0.0769 |
| 18_3490492 | 4 | 0 | 7 | 0.1429 | 3 | 0 | 4 | 0 | 18 | 0.0556 | 5 | 0 | 4 | 0 | 6 | 0 | 4 | 0.25 | 19 | 0.0526 |
| 18_3490633 | 6 | 0 | 7 | 0 | 3 | 0.3333 | 4 | 0 | 20 | 0.05 | 5 | 0 | 6 | 0 | 4 | 0.25 | 7 | 0.1429 | 22 | 0.0909 |
| 18_3493681 | NA | NA | NA | NA | 1 | 1 | NA | NA | 1 | 1 | NA | NA | NA | NA | 1 | 1 | NA | NA | 1 | 1 |
| 18_3494168 | NA | NA | 1 | 1 | NA | NA | NA | NA | 1 | 1 | 2 | 0 | 2 | 0 | 1 | 1 | NA | NA | 5 | 0.2 |
| 18_3494218 | NA | NA | 3 | 0.6667 | 1 | 1 | NA | NA | 4 | 0.75 | 2 | 1 | 2 | 1 | 2 | 1 | NA | NA | 6 | 1 |
| 18_3494234 | NA | NA | 3 | 0.6667 | NA | NA | NA | NA | 3 | 0.6667 | 2 | 0.5 | 2 | 1 | 2 | 1 | NA | NA | 6 | 0.8333 |
| 18_3494252 | NA | NA | 3 | 1 | 1 | 1 | NA | NA | 4 | 1 | 2 | 0.5 | 2 | 1 | 2 | 1 | NA | NA | 6 | 0.8333 |

| | | | | | | | | | | | | | | | | | | | | |
|---|---|---|---|---|---|---|---|---|---|---|---|---|---|---|---|---|---|---|---|---|
| 18_349708 | 1 | 0 | 5 | 0.4 | 1 | 0 | 1 | 0 | 8 | 0.25 | 2 | 0.5 | 5 | 0 | 6 | 0.1667 | 2 | 0 | 15 | 0.1333 |
| 18_349709 | NA | NA | 6 | 0.5 | 1 | 0 | 2 | 0 | 9 | 0.3333 | 4 | 0.5 | 4 | 0.5 | 7 | 0.2857 | 1 | 0 | 16 | 0.375 |
| 18_349709 | 1 | 0 | 6 | 0.3333 | NA | NA | 2 | 0 | 9 | 0.2222 | 4 | 0.25 | 6 | 0.6667 | 7 | 0.7143 | 1 | 0 | 18 | 0.5556 |
| 18_349711 | 1 | 0 | 4 | 0.25 | NA | NA | 1 | 0 | 6 | 0.1667 | 3 | 0 | 3 | 0.3333 | 7 | 0 | 1 | 0 | 14 | 0.0714 |
| 18_349711 | 1 | 0 | 7 | 0.1429 | 2 | 0 | 1 | 0 | 11 | 0.0909 | 5 | 0 | 5 | 0.2 | 7 | 0.1429 | 2 | 0 | 19 | 0.1053 |
| 18_349712 | 1 | 1 | 7 | 0 | 1 | 0 | 2 | 0 | 11 | 0.0909 | 5 | 0 | 7 | 0 | 7 | 0.2857 | NA | NA | 19 | 0.1053 |
| 18_349713 | 1 | 1 | 7 | 0.1429 | 2 | 0 | 2 | 0.5 | 12 | 0.25 | 5 | 0 | 4 | 0 | 8 | 0.375 | 2 | 0 | 19 | 0.1579 |
| 18_349713 | 1 | 1 | 6 | 0.6667 | 2 | 0.5 | 2 | 0.5 | 11 | 0.6364 | 3 | 0.6667 | 5 | 0.6 | 3 | 1 | NA | NA | 11 | 0.7273 |
| 18_349715 | 1 | 0 | 8 | 0.25 | 4 | 0.25 | 2 | 1 | 15 | 0.3333 | 6 | 0.3333 | 7 | 0.5714 | 8 | 0.375 | 1 | 0 | 22 | 0.4091 |
| 18_349716 | NA | NA | 6 | 0.1667 | 1 | 0 | 2 | 0 | 9 | 0.1111 | 5 | 0.2 | 7 | 0.2857 | 3 | 0.3333 | 1 | 0 | 16 | 0.25 |
| 18_349716 | 1 | 1 | 6 | 0.5 | 5 | 0.2 | 1 | 0 | 13 | 0.3846 | 7 | 0.7143 | 6 | 0.3333 | 8 | 0.625 | 1 | 1 | 22 | 0.5909 |
| 18_349791 | 5 | 0 | 10 | 0.2 | 16 | 0 | 4 | 0.25 | 35 | 0.0857 | 3 | 0 | 16 | 0.125 | 10 | 0.2 | 9 | 0.1111 | 38 | 0.1316 |
| 18_349791 | 5 | 0.4 | 14 | 0.0714 | 20 | 0.2 | 4 | 0.25 | 43 | 0.186 | 4 | 0 | 13 | 0.2308 | 16 | 0.3125 | 9 | 0.2222 | 42 | 0.2381 |
| 18_349796 | 2 | 0 | 6 | 0 | 9 | 0.1111 | 1 | 0 | 18 | 0.0556 | 3 | 0 | 11 | 0.1818 | 14 | 0.0714 | 4 | 0 | 32 | 0.0938 |
| 18_349797 | 7 | 0.5714 | 16 | 0.5 | 20 | 0.55 | 5 | 0.4 | 48 | 0.5208 | 6 | 1 | 21 | 0.619 | 15 | 0.6667 | 11 | 0.6364 | 53 | 0.6792 |
| 18_349797 | 6 | 0 | 16 | 0.0625 | 18 | 0.1111 | 4 | 0.5 | 44 | 0.1136 | 5 | 0 | 17 | 0.1765 | 13 | 0.0769 | 10 | 0.1 | 45 | 0.1111 |
| 18_349798 | 6 | 0.1667 | 16 | 0.125 | 21 | 0.1429 | 4 | 0 | 47 | 0.1277 | 5 | 0 | 12 | 0.1667 | 18 | 0.1111 | 7 | 0 | 42 | 0.0952 |
| 18_349801 | 8 | 0.25 | 15 | 0.1333 | 24 | 0.2083 | 6 | 0.3333 | 53 | 0.2075 | 9 | 0.1111 | 19 | 0.3684 | 21 | 0.2381 | 11 | 0.2727 | 60 | 0.2667 |
| 18_349801 | 9 | 0.2222 | 21 | 0.2857 | 26 | 0.3077 | 6 | 0.1667 | 62 | 0.2742 | 11 | 0.0909 | 22 | 0.2727 | 23 | 0.5217 | 13 | 0.5385 | 69 | 0.3768 |
| 18_349802 | 10 | 0.5 | 17 | 0.5294 | 27 | 0.7407 | 7 | 0.5714 | 61 | 0.623 | 11 | 0.5455 | 20 | 0.7 | 21 | 0.7143 | 13 | 0.5385 | 65 | 0.6462 |
| 18_349803 | 10 | 0.2 | 18 | 0.1111 | 28 | 0.0714 | 6 | 0 | 62 | 0.0968 | 10 | 0.1 | 22 | 0.2273 | 23 | 0.1304 | 14 | 0.1429 | 69 | 0.1594 |
| 18_349803 | 6 | 0 | 17 | 0.0588 | 24 | 0.0833 | 6 | 0.1667 | 53 | 0.0755 | 8 | 0 | 19 | 0.0526 | 19 | 0.1053 | 14 | 0.0714 | 60 | 0.0667 |
| 18_349804 | 8 | 0.125 | 11 | 0.0909 | 14 | 0.1429 | 2 | 0.5 | 35 | 0.1429 | 7 | 0.2857 | 10 | 0.2 | 10 | 0.2 | 8 | 0.625 | 35 | 0.3143 |
| 18_349946 | 4 | 0.25 | 5 | 0.2 | 6 | 0.5 | 3 | 0.3333 | 18 | 0.3333 | 3 | 0.3333 | 8 | 0.25 | 5 | 0 | 2 | 0 | 18 | 0.1667 |
| 18_349958 | 9 | 1 | 8 | 1 | 9 | 1 | 11 | 1 | 37 | 1 | 4 | 1 | 11 | 1 | 10 | 0.9 | 5 | 0.8 | 30 | 0.9333 |
| 18_349959 | 12 | 0.4167 | 12 | 0.6667 | 14 | 0.2143 | 12 | 0.4167 | 50 | 0.42 | 6 | 0.3333 | 12 | 0.25 | 14 | 0.2857 | 7 | 0.2857 | 39 | 0.2821 |
| 18_349960 | 11 | 0.0909 | 16 | 0.25 | 16 | 0.0625 | 13 | 0 | 56 | 0.1071 | 6 | 0 | 11 | 0.2727 | 17 | 0.2941 | 7 | 0 | 41 | 0.1951 |
| 18_349961 | 13 | 1 | 14 | 1 | 14 | 1 | 12 | 1 | 53 | 1 | 6 | 1 | 15 | 1 | 18 | 1 | 9 | 1 | 48 | 1 |
| 18_349964 | 14 | 0.2143 | 12 | 0.3333 | 15 | 0.3333 | 10 | 0.5 | 51 | 0.3333 | 6 | 0.3333 | 13 | 0.3846 | 16 | 0.3125 | 9 | 0.4444 | 44 | 0.3636 |
| 18_349964 | 12 | 0.5833 | 15 | 0.3333 | 14 | 0.1429 | 9 | 0.3333 | 50 | 0.34 | 8 | 0.5 | 11 | 0.5455 | 16 | 0.5 | 10 | 0.6 | 45 | 0.5333 |
| 18_349964 | 16 | 0.875 | 15 | 1 | 15 | 0.9333 | 12 | 0.9167 | 58 | 0.931 | 6 | 0.8333 | 13 | 1 | 19 | 0.8421 | 10 | 1 | 48 | 0.9167 |
| 18_349964 | 14 | 0.1429 | 11 | 0 | 15 | 0.0667 | 10 | 0.1 | 50 | 0.08 | 6 | 0 | 12 | 0.1667 | 16 | 0.1875 | 9 | 0 | 43 | 0.1163 |
| 18_349967 | 14 | 0.0714 | 19 | 0.0526 | 18 | 0.1667 | 15 | 0 | 66 | 0.0758 | 8 | 0 | 18 | 0.1111 | 23 | 0.087 | 11 | 0.0909 | 60 | 0.0833 |
| 18_349968 | 14 | 1 | 18 | 1 | 17 | 1 | 14 | 1 | 63 | 1 | 8 | 1 | 17 | 0.9412 | 23 | 1 | 11 | 1 | 59 | 0.9831 |
| 18_349968 | 13 | 0.6154 | 15 | 0.8667 | 14 | 0.8571 | 15 | 0.7333 | 57 | 0.7719 | 5 | 0.8 | 12 | 0.75 | 20 | 0.8 | 10 | 0.8 | 47 | 0.7872 |
| 18_349969 | 14 | 0 | 17 | 0.0588 | 18 | 0.1111 | 17 | 0.1176 | 66 | 0.0758 | 8 | 0 | 17 | 0.0588 | 25 | 0.16 | 12 | 0 | 62 | 0.0806 |
| 18_349970 | 14 | 0.3571 | 18 | 0.4444 | 13 | 0.3846 | 17 | 0.1765 | 62 | 0.3387 | 8 | 0.5 | 15 | 0.2667 | 23 | 0.3043 | 11 | 0.2727 | 57 | 0.3158 |
| 18_349970 | 10 | 1 | 11 | 1 | 13 | 1 | 12 | 1 | 46 | 1 | 8 | 1 | 14 | 1 | 21 | 1 | 9 | 1 | 52 | 1 |
| 18_349971 | 19 | 0.1053 | 20 | 0.15 | 19 | 0.0526 | 17 | 0 | 75 | 0.08 | 8 | 0.25 | 18 | 0.0556 | 25 | 0.08 | 12 | 0.0833 | 63 | 0.0952 |
| 18_349973 | 19 | 0.1053 | 20 | 0.2 | 20 | 0.05 | 17 | 0.0588 | 76 | 0.1053 | 12 | 0 | 19 | 0.2105 | 26 | 0.1154 | 12 | 0 | 69 | 0.1014 |
| 18_349976 | 19 | 0 | 21 | 0.1905 | 19 | 0.0526 | 18 | 0 | 77 | 0.0649 | 19 | 0.1053 | 19 | 0.1579 | 24 | 0.25 | 15 | 0.2 | 77 | 0.1818 |
| 18_349977 | 14 | 0.5 | 20 | 0.2 | 16 | 0.25 | 17 | 0.5882 | 67 | 0.3731 | 12 | 0.5 | 18 | 0.3333 | 23 | 0.5652 | 13 | 0.6923 | 66 | 0.5152 |
| 18_350364 | 5 | 0.2 | 4 | 0.5 | 5 | 0 | 6 | 0.1667 | 20 | 0.2 | 6 | 0.1667 | 9 | 0.4444 | 12 | 0.1667 | 17 | 0.1765 | 44 | 0.2273 |
| 18_350368 | 4 | 0.25 | 3 | 0.3333 | 3 | 0 | 3 | 0 | 13 | 0.1538 | 4 | 0.25 | 8 | 0.5 | 9 | 0.5556 | 7 | 0.5714 | 28 | 0.5 |
| 18_350372 | 2 | 0 | 6 | 0.5 | 6 | 0.1667 | 5 | 0.2 | 19 | 0.2632 | 2 | 0 | 5 | 0.4 | 8 | 0.125 | 9 | 0.2222 | 24 | 0.2083 |
| 18_350375 | 1 | 0 | 4 | 0.25 | 5 | 0.8 | 4 | 0.75 | 14 | 0.5714 | 4 | 0.5 | 4 | 0.75 | 8 | 0.375 | 10 | 0.4 | 26 | 0.4615 |
| 18_350378 | 1 | 1 | 8 | 1 | 8 | 1 | 6 | 1 | 23 | 1 | 5 | 1 | 5 | 1 | 8 | 1 | 7 | 1 | 25 | 1 |
| 18_350378 | 1 | 0 | 9 | 0.1111 | 8 | 0.25 | 5 | 0 | 23 | 0.1304 | 4 | 0.25 | 6 | 0.1667 | 6 | 0.1667 | 9 | 0 | 25 | 0.12 |
| 18_350379 | 1 | 0 | 8 | 0.375 | 7 | 0.5714 | 6 | 0.5 | 22 | 0.4545 | 5 | 0.4 | 5 | 0.6 | 7 | 0.4286 | 10 | 0.6 | 27 | 0.5185 |
| 18_350713 | 24 | 0.2083 | 30 | 0.0333 | 24 | 0.125 | 19 | 0.3158 | 97 | 0.1546 | 15 | 0.2 | 16 | 0.125 | 11 | 0.0909 | 19 | 0.2632 | 61 | 0.1803 |
| 18_350713 | 21 | 0.1905 | 28 | 0.25 | 21 | 0 | 15 | 0.1333 | 85 | 0.1529 | 15 | 0.1333 | 13 | 0.0769 | 9 | 0.5556 | 18 | 0 | 55 | 0.1455 |
| 18_353422 | 4 | 0 | 6 | 0.3333 | 6 | 0.3333 | 6 | 0.1667 | 22 | 0.2273 | 5 | 0 | 2 | 0 | 4 | 0 | 6 | 0.3333 | 17 | 0.1176 |
| 18_353422 | 4 | 0 | 5 | 0 | 5 | 0 | 3 | 0.3333 | 17 | 0.0588 | 5 | 0.2 | 2 | 0 | 3 | 0 | 5 | 0 | 15 | 0.0667 |
| 18_353449 | 2 | 0 | 2 | 0.5 | 4 | 0 | 4 | 0 | 12 | 0.0833 | 4 | 0 | 5 | 0 | 3 | 0.3333 | 3 | 0 | 15 | 0.0667 |
| 18_353456 | 5 | 0.2 | 2 | 0.5 | 3 | 0 | 3 | 0 | 13 | 0.1538 | 5 | 0.4 | 6 | 0.3333 | 4 | 0.25 | 4 | 0 | 19 | 0.2632 |
| 18_353456 | 4 | 0.5 | 2 | 0.5 | 3 | 0.3333 | 3 | 0.3333 | 12 | 0.4167 | 5 | 0 | 4 | 0 | 3 | 0 | 4 | 0.25 | 16 | 0.0625 |
| 18_353457 | 4 | 1 | 2 | 1 | 3 | 1 | 2 | 1 | 11 | 1 | 4 | 1 | 6 | 1 | 3 | 1 | 3 | 1 | 16 | 1 |
| 18_353458 | 4 | 1 | 2 | 1 | 3 | 1 | 2 | 0.5 | 11 | 0.9091 | 4 | 0.75 | 5 | 1 | 4 | 1 | 4 | 1 | 17 | 0.9412 |
| 18_353458 | 5 | 0.2 | 2 | 0.5 | 3 | 0.3333 | 2 | 0 | 12 | 0.25 | 5 | 0.2 | 6 | 0.3333 | 4 | 0.5 | 4 | 0 | 19 | 0.2632 |
| 18_353458 | 4 | 1 | 2 | 1 | 3 | 0.6667 | 2 | 1 | 11 | 0.9091 | 4 | 0.5 | 6 | 0.8333 | 4 | 0.75 | 4 | 1 | 18 | 0.7778 |
| 18_353458 | 5 | 0.2 | 2 | 0 | 3 | 0 | 2 | 0 | 12 | 0.0833 | 4 | 0 | 6 | 0.1667 | 4 | 0 | 2 | 0 | 16 | 0.0625 |
| 18_353459 | 4 | 0 | 2 | 0 | 3 | 0.3333 | 1 | 0 | 10 | 0.1 | 5 | 0 | 6 | 0 | 4 | 0 | 4 | 0.25 | 19 | 0.0526 |
| 18_353460 | 4 | 0.75 | 2 | 0.5 | 3 | 0.6667 | 2 | 1 | 11 | 0.7273 | 5 | 0.6 | 4 | 0.75 | 3 | 1 | 4 | 1 | 16 | 0.8125 |
| 18_353460 | 4 | 0.75 | 2 | 0 | 3 | 0 | 2 | 0.5 | 11 | 0.3636 | 5 | 0.8 | 4 | 0 | 4 | 0.5 | 4 | 0 | 17 | 0.3529 |
| 18_353460 | NA | NA | 1 | 0 | 2 | 1 | 2 | 1 | 5 | 0.8 | 4 | 1 | 5 | 0.8 | 4 | 0.75 | 2 | 0.5 | 15 | 0.8 |

| | | | | | | | | | | | | | | | | | | | | |
|---|---|---|---|---|---|---|---|---|---|---|---|---|---|---|---|---|---|---|---|---|
| 18_353464( | 2 | 0.5 | 1 | 0 | 3 | 0.3333 | 1 | 0 | 7 | 0.2857 | 3 | 0 | 4 | 0 | 3 | 0 | 2 | 0.5 | 12 | 0.0833 |
| 18_3534647 | 3 | 0.3333 | 1 | 0 | 2 | 0 | 1 | 0 | 7 | 0.1429 | 4 | 0 | 5 | 0 | 3 | 0 | 3 | 0.3333 | 15 | 0.0667 |
| 18_3534655 | 3 | 0 | 1 | 0 | 1 | 0 | 1 | 1 | 6 | 0.1667 | 3 | 0 | 5 | 0.6 | 2 | 0.5 | 3 | 0 | 13 | 0.3077 |
| 18_3534655 | 3 | 1 | 1 | 1 | 2 | 1 | NA | NA | 6 | 1 | 3 | 1 | 5 | 1 | 2 | 1 | 3 | 1 | 13 | 1 |
| 18_3534668 | 1 | 0 | 1 | 0 | 2 | 1 | 1 | 0 | 5 | 0.4 | 3 | 0.6667 | 4 | 0.25 | 2 | 0.5 | 2 | 0.5 | 11 | 0.4545 |
| 18_353467( | 2 | 1 | 1 | 1 | 1 | 1 | 1 | 0 | 5 | 0.8 | 2 | 1 | 4 | 1 | 2 | 1 | 3 | 0.6667 | 11 | 0.9091 |
| 18_3534679 | 2 | 1 | 1 | 1 | 2 | 0.5 | 1 | 1 | 6 | 0.8333 | 3 | 1 | 4 | 1 | 2 | 1 | 3 | 1 | 12 | 1 |
| 18_3534688 | 2 | 1 | 1 | 1 | 2 | 0.5 | 1 | 0 | 6 | 0.6667 | 3 | 0.6667 | 3 | 0.6667 | 2 | 0.5 | 3 | 0.6667 | 11 | 0.6364 |
| 18_3534698 | 2 | 1 | 1 | 1 | 2 | 1 | 1 | 1 | 6 | 1 | 2 | 1 | 4 | 1 | 2 | 1 | 3 | 1 | 11 | 1 |
| 18_3534778 | 1 | 0 | 1 | 1 | 1 | 0 | 1 | 1 | 4 | 0.5 | 2 | 0.5 | 4 | 0.75 | 2 | 1 | 3 | 1 | 11 | 0.8182 |
| 18_353478( | 1 | 1 | NA | NA | 1 | 0 | NA | NA | 2 | 0.5 | NA | NA | 3 | 0 | 1 | 0 | 3 | 0.6667 | 7 | 0.2857 |
| 18_3538693 | 7 | 0.7143 | 5 | 0.6 | 8 | 0.625 | 9 | 0.5556 | 29 | 0.6207 | 5 | 0.4 | 6 | 0.6667 | 8 | 0.75 | 7 | 0.4286 | 26 | 0.5769 |
| 18_3538705 | 9 | 0.1111 | 6 | 0 | 9 | 0 | 9 | 0.1111 | 33 | 0.0606 | 8 | 0.375 | 6 | 0.1667 | 9 | 0 | 9 | 0.1111 | 32 | 0.1562 |
| 18_3538708 | 8 | 0.125 | 6 | 0.1667 | 8 | 0 | 9 | 0.2222 | 31 | 0.129 | 8 | 0.125 | 6 | 0.5 | 9 | 0.2222 | 8 | 0.5 | 31 | 0.3226 |
| 18_353875( | 7 | 0.8571 | 5 | 1 | 9 | 1 | 8 | 0.875 | 29 | 0.931 | 8 | 0.875 | 7 | 0.8571 | 11 | 0.7273 | 9 | 1 | 35 | 0.8571 |
| 18_353876( | 7 | 0 | 6 | 0 | 7 | 0.1429 | 10 | 0.1 | 30 | 0.0667 | 8 | 0 | 7 | 0.1429 | 12 | 0.1667 | 8 | 0.125 | 35 | 0.1143 |
| 18_353994( | 2 | 0.5 | 9 | 0 | 4 | 0 | 6 | 0.1667 | 21 | 0.0952 | 6 | 0.1667 | 6 | 0 | 3 | 0 | 5 | 0 | 20 | 0.05 |
| 18_3539952 | 2 | 1 | 8 | 0.875 | 5 | 1 | 3 | 0.6667 | 18 | 0.8889 | 6 | 0.8333 | 4 | 1 | 4 | 1 | 6 | 1 | 20 | 0.95 |
| 18_3539953 | 2 | 0.5 | 8 | 0.25 | 4 | 0.5 | 6 | 0.3333 | 20 | 0.35 | 6 | 0.3333 | 6 | 0.1667 | 4 | 0.5 | 4 | 0.25 | 20 | 0.3 |
| 18_3539958 | 1 | 1 | 8 | 0.875 | 5 | 0.8 | 3 | 1 | 17 | 0.8824 | 6 | 1 | 6 | 0.6667 | 4 | 1 | 5 | 1 | 21 | 0.9048 |
| 18_3539961 | 2 | 0 | 6 | 0.3333 | 4 | 0.25 | 5 | 0.4 | 17 | 0.2941 | 6 | 0.3333 | 5 | 0.4 | 4 | 0.5 | 5 | 0.6 | 20 | 0.45 |
| 18_3539964 | 2 | 0 | 7 | 0.2857 | 4 | 0.25 | 2 | 0 | 15 | 0.2 | 5 | 0 | 6 | 0.1667 | 3 | 0.3333 | 4 | 0 | 18 | 0.1111 |
| 18_3539973 | 2 | 0.5 | 9 | 0.3333 | 5 | 0.4 | 4 | 0.25 | 20 | 0.35 | 5 | 0.2 | 6 | 0.5 | 3 | 0 | 3 | 0.6667 | 17 | 0.3529 |
| 18_3539979 | 2 | 0 | 9 | 0.1111 | 5 | 0 | 5 | 0.2 | 21 | 0.0952 | 6 | 0 | 6 | 0 | 3 | 0.3333 | 4 | 0.25 | 19 | 0.1053 |
| 18_3569064 | 1 | 0 | 6 | 0.1667 | 5 | 0 | 1 | 0 | 13 | 0.0769 | 4 | 0.25 | 2 | 0.5 | 3 | 0 | 3 | 0 | 12 | 0.1667 |
| 18_356906( | 1 | 0 | 6 | 0.1667 | 5 | 0 | 1 | 0 | 13 | 0.0769 | 4 | 0.25 | 1 | 0 | 3 | 0 | 3 | 0.3333 | 11 | 0.1818 |
| 18_3570543 | 19 | 0.3158 | 32 | 0.5625 | 21 | 0.5238 | 33 | 0.6061 | 105 | 0.5238 | 5 | 0.2 | 19 | 0.4737 | 25 | 0.64 | 30 | 0.5667 | 79 | 0.5443 |
| 18_3570544 | 20 | 0.45 | 27 | 0.7037 | 21 | 0.6667 | 29 | 0.5517 | 97 | 0.5979 | 4 | 0.5 | 16 | 0.625 | 23 | 0.3913 | 27 | 0.5926 | 70 | 0.5286 |
| 18_357054( | 22 | 0.0455 | 28 | 0 | 19 | 0.0526 | 33 | 0.1212 | 102 | 0.0588 | 4 | 0 | 13 | 0 | 23 | 0.087 | 29 | 0.069 | 69 | 0.058 |
| 18_3570548 | 20 | 0.05 | 25 | 0.08 | 17 | 0.0588 | 30 | 0.0333 | 92 | 0.0543 | 4 | 0 | 12 | 0 | 17 | 0.0588 | 20 | 0.1 | 53 | 0.0566 |
| 18_3570552 | 20 | 0.1 | 26 | 0.1538 | 20 | 0.1 | 33 | 0.0303 | 99 | 0.0909 | 6 | 0 | 21 | 0.1429 | 21 | 0.1429 | 26 | 0 | 74 | 0.0811 |
| 18_3570581 | 20 | 0.1 | 32 | 0.125 | 23 | 0.087 | 33 | 0.0303 | 108 | 0.0833 | 5 | 0 | 21 | 0.0476 | 25 | 0.12 | 30 | 0.0333 | 81 | 0.0617 |
| 18_3570592 | 18 | 0.5 | 20 | 0.45 | 14 | 0.3571 | 23 | 0.4348 | 75 | 0.44 | 4 | 0.75 | 14 | 0.3571 | 23 | 0.4783 | 26 | 0.2692 | 67 | 0.3881 |
| 18_3571775 | 26 | 0.3077 | 46 | 0.2609 | 31 | 0.2903 | 48 | 0.375 | 151 | 0.3113 | 10 | 0.5 | 43 | 0.3488 | 40 | 0.425 | 39 | 0.3846 | 132 | 0.3939 |
| 18_3572371 | 39 | 0.2564 | 49 | 0.3265 | 36 | 0.2222 | 55 | 0.2909 | 179 | 0.2793 | 14 | 0.2857 | 52 | 0.2115 | 44 | 0.2727 | 51 | 0.3922 | 161 | 0.2919 |
| 18_3572391 | 47 | 0.1277 | 63 | 0.0635 | 50 | 0.06 | 64 | 0.0469 | 224 | 0.0714 | 19 | 0.1053 | 60 | 0.1 | 57 | 0.1754 | 59 | 0.1356 | 195 | 0.1333 |
| 18_3572464 | 5 | 0.4 | 6 | 0.3333 | 8 | 0.25 | 8 | 0.875 | 27 | 0.4815 | 4 | 0.5 | 6 | 0.5 | 7 | 0.4286 | 4 | 0.5 | 21 | 0.4762 |
| 18_357494( | 22 | 0.1364 | 24 | 0.2083 | 19 | 0.2105 | 27 | 0.1481 | 92 | 0.1739 | 16 | 0.0625 | 24 | 0.2083 | 27 | 0.1111 | 31 | 0.2581 | 98 | 0.1735 |
| 18_3578562 | 1 | 1 | NA | NA | NA | NA | NA | NA | 1 | 1 | 1 | 1 | NA | NA | NA | NA | NA | NA | 1 | 1 |
| 18_3578621 | 1 | 1 | NA | NA | NA | NA | NA | NA | 1 | 1 | NA | NA | NA | NA | 1 | 1 | NA | NA | 1 | 1 |
| 18_3580422 | 8 | 0 | 9 | 0.3333 | 11 | 0.0909 | 5 | 0.4 | 33 | 0.1818 | 6 | 0 | 9 | 0.2222 | 13 | 0.1538 | 3 | 0.3333 | 31 | 0.1613 |
| 18_3580434 | 7 | 0.2857 | 6 | 0.3333 | 7 | 0.4286 | 3 | 0.3333 | 23 | 0.3478 | 5 | 0.8 | 8 | 0.25 | 10 | 0.5 | 2 | 1 | 25 | 0.52 |
| 18_3580457 | 4 | 0 | 5 | 0.6 | 7 | 0 | 3 | 0.6667 | 19 | 0.2632 | 4 | 0 | 4 | 0.5 | 6 | 0.1667 | NA | NA | 14 | 0.2143 |
| 18_3580465 | 6 | 0.6667 | 8 | 0.625 | 12 | 0.1667 | 3 | 0.6667 | 29 | 0.4483 | 5 | 0.8 | 8 | 0.625 | 10 | 0.6 | 2 | 0 | 25 | 0.6 |
| 18_3580467 | 6 | 1 | 8 | 0.875 | 11 | 0.9091 | 5 | 1 | 30 | 0.9333 | 7 | 1 | 9 | 1 | 8 | 0.875 | 4 | 1 | 28 | 0.9643 |
| 18_358049( | 5 | 0 | 7 | 0.1429 | 9 | 0.1111 | 2 | 0 | 23 | 0.087 | 5 | 0 | 7 | 0.2857 | 8 | 0.375 | 3 | 0 | 23 | 0.2174 |
| 18_3580575 | 3 | 0 | 4 | 0.25 | 7 | 0 | 3 | 0 | 17 | 0.0588 | 2 | 0 | 6 | 0.3333 | 7 | 0.1429 | 4 | 0 | 19 | 0.1579 |
| 18_3580674 | 1 | 0 | 5 | 0.2 | 4 | 0 | 4 | 0.25 | 14 | 0.1429 | 2 | 0.5 | 5 | 0.4 | 6 | 0 | 4 | 0 | 17 | 0.1765 |
| 18_358072( | 2 | 0.5 | 6 | 0 | 3 | 0.3333 | 4 | 0 | 15 | 0.1333 | 1 | 0 | 5 | 0.2 | 7 | 0 | 4 | 0 | 17 | 0.0588 |
| 18_3580774 | 1 | 0 | 3 | 0.6667 | 1 | 0 | 4 | 0.5 | 9 | 0.4444 | 2 | 0.5 | 4 | 0.25 | 7 | 0.2857 | 3 | 0.3333 | 16 | 0.3125 |
| 18_3580821 | 1 | 1 | 3 | 0.3333 | 1 | 1 | 5 | 0.2 | 10 | 0.4 | 2 | 1 | 4 | 0.25 | 6 | 0.3333 | 3 | 0.3333 | 15 | 0.4 |
| 18_3586747 | NA | NA | 3 | 0.6667 | NA | NA | NA | NA | 3 | 0.6667 | NA | NA | 1 | 1 | 1 | 1 | NA | NA | 2 | 1 |
| 18_3586754 | NA | NA | 2 | 1 | NA | NA | NA | NA | 2 | 1 | NA | NA | 1 | 1 | 1 | 0 | NA | NA | 2 | 0.5 |
| 18_3593859 | 42 | 0.0714 | 25 | 0.12 | 21 | 0.0476 | 22 | 0.0455 | 110 | 0.0727 | 39 | 0.0256 | 19 | 0.0526 | 28 | 0.1429 | 27 | 0.1111 | 113 | 0.0796 |
| 18_359386( | 35 | 0.0571 | 24 | 0.0417 | 20 | 0.15 | 20 | 0.05 | 99 | 0.0707 | 35 | 0.0857 | 18 | 0.1667 | 24 | 0.1667 | 25 | 0.2 | 102 | 0.1471 |
| 18_3593869 | 33 | 0.2424 | 22 | 0.4091 | 19 | 0.3684 | 19 | 0.3684 | 93 | 0.3333 | 27 | 0.2963 | 17 | 0.5294 | 21 | 0.1905 | 23 | 0.4348 | 88 | 0.3523 |
| 18_3593888 | 40 | 0.075 | 29 | 0.0345 | 20 | 0.05 | 24 | 0.125 | 113 | 0.0708 | 33 | 0.0909 | 21 | 0.0952 | 26 | 0.2308 | 26 | 0.1538 | 106 | 0.1415 |
| 18_359389( | 40 | 0.2 | 30 | 0.2333 | 21 | 0.1905 | 21 | 0.1905 | 112 | 0.2054 | 35 | 0.1429 | 18 | 0.2222 | 25 | 0.28 | 27 | 0.1481 | 105 | 0.1905 |
| 18_359389( | 34 | 0.0882 | 26 | 0.0385 | 20 | 0.1 | 20 | 0 | 100 | 0.06 | 25 | 0.04 | 16 | 0.0625 | 24 | 0.0417 | 23 | 0.087 | 88 | 0.0568 |
| 18_3593908 | 45 | 0.2222 | 26 | 0.1154 | 23 | 0.0435 | 24 | 0.25 | 118 | 0.1695 | 36 | 0.1944 | 22 | 0.2727 | 26 | 0.2308 | 27 | 0.1852 | 111 | 0.2162 |
| 18_3593917 | 38 | 0.7368 | 31 | 0.6452 | 21 | 0.7143 | 19 | 0.7368 | 109 | 0.7064 | 35 | 0.6571 | 21 | 0.7143 | 22 | 0.5455 | 24 | 0.75 | 102 | 0.6667 |
| 18_3599145 | 4 | 0.25 | 8 | 0 | 4 | 0 | 2 | 0 | 18 | 0.0556 | 7 | 0.1429 | 2 | 0 | 5 | 0 | 2 | 0 | 16 | 0.0625 |
| 18_3599149 | 5 | 0 | 9 | 0.2222 | 4 | 0 | 2 | 0 | 20 | 0.1 | 8 | 0 | 3 | 0 | 5 | 0.2 | 2 | 0 | 18 | 0.0556 |
| 18_3599165 | 4 | 0 | 10 | 0.1 | 4 | 0 | 1 | 0 | 19 | 0.0526 | 3 | 0 | 3 | 0 | 1 | 0 | 2 | 0.5 | 9 | 0.1111 |
| 18_3599169 | 2 | 0.5 | 10 | 0.5 | 3 | 0 | 2 | 0 | 17 | 0.3529 | 6 | 0 | 3 | 0 | 3 | 0.6667 | 1 | 0 | 13 | 0.1538 |

| | | | | | | | | | | | | | | | | | | | | |
|---|---|---|---|---|---|---|---|---|---|---|---|---|---|---|---|---|---|---|---|---|
| 18_359917 | 4 | 0 | 10 | 0.3 | 4 | 0.25 | 2 | 0 | 20 | 0.2 | 6 | 0.3333 | 3 | 0 | 3 | 0 | 1 | 0 | 13 | 0.1538 |
| 18_359917 | 2 | 0 | 11 | 0 | 3 | 0.3333 | 2 | 0 | 18 | 0.0556 | 8 | 0 | 3 | 0.3333 | 5 | 0 | 2 | 0 | 18 | 0.0556 |
| 18_359917 | 4 | 0.25 | 9 | 0 | 4 | 0 | 3 | 0 | 20 | 0.05 | 6 | 0 | 4 | 0 | 5 | 0.2 | 2 | 0.5 | 17 | 0.1176 |
| 18_359919 | 5 | 1 | 9 | 1 | 4 | 1 | 5 | 1 | 23 | 1 | 7 | 1 | 4 | 1 | 6 | 1 | 2 | 1 | 19 | 1 |
| 18_359920 | 5 | 1 | 11 | 0.9091 | 4 | 1 | 5 | 1 | 25 | 0.96 | 9 | 1 | 4 | 0.75 | 7 | 0.8571 | 3 | 1 | 23 | 0.913 |
| 18_359924 | 5 | 1 | 13 | 0.9231 | 3 | 0.6667 | 5 | 0.6 | 26 | 0.8462 | 9 | 1 | 4 | 1 | 8 | 0.875 | 4 | 0.75 | 25 | 0.92 |
| 18_359924 | 5 | 0 | 12 | 0.4167 | 3 | 0.3333 | 4 | 0.25 | 24 | 0.2917 | 7 | 0.4286 | 1 | 0 | 9 | 0.4444 | 4 | 0 | 21 | 0.3333 |
| 18_359927 | 2 | 1 | 3 | 0.6667 | NA | NA | NA | NA | 5 | 0.8 | 4 | 1 | NA | NA | 5 | 1 | 1 | 0 | 10 | 0.9 |
| 18_359928 | 2 | 0.5 | 4 | 0.75 | NA | NA | NA | NA | 6 | 0.6667 | 4 | 0.5 | NA | NA | 5 | 0.4 | 1 | 1 | 10 | 0.5 |
| 18_359928 | 2 | 0 | 4 | 0.75 | 1 | 1 | NA | NA | 7 | 0.5714 | 4 | 0.75 | NA | NA | 5 | 0.6 | 1 | 1 | 10 | 0.7 |
| 18_359931 | 1 | 0 | 4 | 0 | 2 | 0.5 | NA | NA | 7 | 0.1429 | 4 | 0 | NA | NA | 5 | 0.4 | 1 | 0 | 10 | 0.2 |
| 18_359933 | 2 | 0.5 | 3 | 0.6667 | 2 | 0.5 | NA | NA | 7 | 0.5714 | 5 | 0.6 | NA | NA | 6 | 1 | 1 | 0 | 12 | 0.75 |
| 18_359935 | 2 | 1 | 2 | 1 | 1 | 1 | NA | NA | 5 | 1 | 3 | 1 | NA | NA | 3 | 0.6667 | 1 | 1 | 7 | 0.8571 |
| 18_359936 | 2 | 0.5 | 4 | 0.5 | 2 | 0.5 | NA | NA | 8 | 0.5 | 6 | 0.1667 | 1 | 1 | 7 | 0.4286 | 1 | 0 | 15 | 0.3333 |
| 18_359936 | 2 | 0.5 | 2 | 0.5 | 2 | 0.5 | NA | NA | 6 | 0.5 | 2 | 1 | 1 | 0 | 3 | 0.6667 | 1 | 1 | 7 | 0.7143 |
| 18_359936 | NA | NA | 3 | 1 | 1 | 1 | NA | NA | 4 | 1 | 3 | 1 | 1 | 1 | 5 | 1 | NA | NA | 9 | 1 |
| 18_359938 | 7 | 0.1429 | 14 | 0.3571 | 5 | 0.2 | 6 | 0 | 32 | 0.2188 | 12 | 0.25 | 5 | 0 | 9 | 0.3333 | 4 | 0.25 | 30 | 0.2333 |
| 18_359939 | 4 | 0.25 | 11 | 0 | 7 | 0 | 5 | 0.4 | 27 | 0.1111 | 9 | 0.1111 | 6 | 0 | 10 | 0.2 | 3 | 0 | 28 | 0.1071 |
| 18_359940 | 8 | 0.25 | 13 | 0.3846 | 7 | 0.1429 | 6 | 0.1667 | 34 | 0.2647 | 12 | 0.0833 | 6 | 0.1667 | 9 | 0.2222 | 4 | 0.5 | 31 | 0.1935 |
| 18_359940 | 7 | 1 | 13 | 0.9231 | 7 | 1 | 4 | 1 | 31 | 0.9677 | 10 | 1 | 6 | 0.6667 | 9 | 1 | 2 | 1 | 27 | 0.9259 |
| 18_359943 | 6 | 0 | 13 | 0 | 11 | 0.1818 | 5 | 0 | 35 | 0.0571 | 10 | 0.2 | 6 | 0 | 9 | 0.1111 | 4 | 0.25 | 29 | 0.1379 |
| 18_361127 | NA | NA | 1 | 0 | 1 | 1 | 1 | 0 | 3 | 0.3333 | 1 | 0 | 2 | 0.5 | NA | NA | 1 | 1 | 4 | 0.5 |
| 18_361458 | 2 | 0.5 | 5 | 0.4 | 3 | 0.3333 | 2 | 1 | 12 | 0.5 | 7 | 0.2857 | 6 | 0.3333 | 1 | 0 | 2 | 1 | 16 | 0.375 |
| 18_361459 | 2 | 0.5 | 4 | 0.5 | 3 | 0 | 3 | 0.6667 | 12 | 0.4167 | 8 | 0.125 | 6 | 0.1667 | 1 | 1 | 2 | 0.5 | 17 | 0.2353 |
| 18_361460 | 2 | 0 | 3 | 0.3333 | 3 | 0.3333 | 3 | 1 | 11 | 0.4545 | 7 | 0.4286 | 3 | 0.3333 | 1 | 0 | 2 | 0 | 13 | 0.3077 |
| 18_361461 | 2 | 0.5 | 6 | 0.6667 | 3 | 0.3333 | 4 | 0.5 | 15 | 0.5333 | 7 | 0.5714 | 6 | 0.5 | NA | NA | 2 | 0 | 15 | 0.4667 |
| 18_361462 | 2 | 0.5 | 6 | 1 | 4 | 0.75 | 4 | 1 | 16 | 0.875 | 9 | 0.8889 | 5 | 1 | NA | NA | 2 | 1 | 16 | 0.9375 |
| 18_361463 | 1 | 1 | 6 | 0.3333 | 3 | 1 | 3 | 0 | 13 | 0.4615 | 10 | 0.5 | 5 | 0.4 | NA | NA | 1 | 1 | 16 | 0.5 |
| 18_361466 | 2 | 0 | 6 | 0 | 2 | 0.5 | 4 | 0 | 14 | 0.0714 | 8 | 0.125 | 5 | 0 | 3 | 0 | 2 | 0 | 18 | 0.0556 |
| 18_361467 | 3 | 0 | 6 | 0.1667 | 3 | 0 | 4 | 0 | 16 | 0.0625 | 10 | 0 | 7 | 0.2857 | 3 | 0 | 2 | 0 | 22 | 0.0909 |
| 18_361467 | 1 | 1 | 5 | 0.4 | 3 | 1 | 3 | 1 | 12 | 0.75 | 8 | 0.625 | 7 | 0.5714 | 3 | 0 | 2 | 0.5 | 20 | 0.5 |
| 18_361467 | 2 | 1 | 4 | 0.75 | 3 | 1 | 4 | 1 | 13 | 0.9231 | 9 | 0.7778 | 6 | 0.6667 | 3 | 0.6667 | 1 | 1 | 19 | 0.7368 |
| 18_361467 | 3 | 0.6667 | 5 | 0.6 | 1 | 0 | 2 | 1 | 11 | 0.6364 | 5 | 0.2 | 4 | 0.25 | 2 | 0 | NA | NA | 11 | 0.1818 |
| 18_361469 | 3 | 0.3333 | 5 | 0.2 | 3 | 0 | 3 | 0.6667 | 14 | 0.2857 | 8 | 0.375 | 7 | 0.5714 | 4 | 0.5 | 1 | 0 | 20 | 0.45 |
| 18_361481 | 3 | 0.3333 | NA | NA | NA | NA | 1 | 1 | 4 | 0.5 | NA | NA | 1 | 0 | 1 | 0 | 1 | 1 | 3 | 0.3333 |
| 18_361489 | 3 | 0.3333 | NA | NA | NA | NA | NA | NA | 3 | 0.3333 | NA | NA | 1 | 0 | 1 | 0 | 1 | 1 | 3 | 0.3333 |
| 18_361503 | 2 | 0 | 1 | 0 | NA | NA | 1 | 1 | 4 | 0.25 | 1 | 0 | 2 | 0 | 2 | 0.5 | 1 | 0 | 6 | 0.1667 |
| 18_361503 | 2 | 0.5 | 2 | 0 | 1 | 1 | 1 | 1 | 6 | 0.5 | 1 | 0 | 2 | 0.5 | 3 | 0.3333 | 1 | 1 | 7 | 0.4286 |
| 18_361505 | 3 | 0 | 2 | 0.5 | 1 | 0 | NA | NA | 6 | 0.1667 | 1 | 1 | 2 | 0 | 3 | 0 | 2 | 0 | 8 | 0.125 |
| 18_361505 | 3 | 0 | 2 | 0.5 | 1 | 1 | 1 | 1 | 7 | 0.4286 | NA | NA | 2 | 0 | 3 | 0.3333 | 2 | 0 | 7 | 0.1429 |
| 18_361508 | 2 | 0 | 1 | 1 | NA | NA | 2 | 0 | 5 | 0.2 | 1 | 0 | 1 | 1 | 3 | 0 | 1 | 0 | 6 | 0.1667 |
| 18_361509 | 1 | 1 | 1 | 1 | NA | NA | 2 | 1 | 4 | 1 | 1 | 1 | 3 | 1 | NA | NA | 1 | 1 | 5 | 1 |
| 18_361510 | 1 | 1 | NA | NA | NA | NA | NA | NA | 1 | 1 | NA | NA | 2 | 0.5 | 3 | 0.6667 | 1 | 0 | 6 | 0.5 |
| 18_361510 | 3 | 1 | 1 | 1 | NA | NA | 2 | 1 | 6 | 1 | 1 | 1 | 3 | 1 | 3 | 1 | 2 | 1 | 9 | 1 |
| 18_361510 | 3 | 0.6667 | 1 | 1 | NA | NA | 1 | 0 | 5 | 0.6 | NA | NA | 3 | 1 | 2 | 1 | 1 | 1 | 6 | 1 |
| 18_361512 | 3 | 1 | 1 | 1 | NA | NA | 2 | 0.5 | 6 | 0.8333 | 1 | 1 | 3 | 1 | 3 | 1 | 2 | 1 | 9 | 1 |
| 18_361513 | 2 | 0.5 | 1 | 0 | NA | NA | NA | NA | 3 | 0.3333 | 1 | 0 | 3 | 0 | 2 | 0 | 2 | 0.5 | 8 | 0.125 |
| 18_361515 | 3 | 0.6667 | 1 | 1 | NA | NA | 1 | 0 | 5 | 0.6 | 1 | 1 | 3 | 0.3333 | 3 | 0.6667 | 2 | 0.5 | 9 | 0.5556 |
| 18_364204 | 1 | 0 | 3 | 0.3333 | 3 | 0.3333 | 4 | 0.25 | 11 | 0.2727 | 1 | 0 | 3 | 0.3333 | 2 | 0 | 3 | 0.3333 | 9 | 0.2222 |
| 18_364207 | NA | NA | 1 | 0 | 3 | 0.6667 | 3 | 0 | 7 | 0.2857 | NA | NA | 2 | 0.5 | 2 | 0 | 2 | 0.5 | 6 | 0.3333 |
| 18_364208 | NA | NA | 3 | 1 | 2 | 1 | 4 | 0.5 | 9 | 0.7778 | 2 | 1 | 4 | 0.75 | 5 | 1 | 3 | 0.6667 | 14 | 0.8571 |
| 18_364208 | 1 | 0 | 4 | 0.5 | 5 | 0.8 | 5 | 0.8 | 15 | 0.6667 | 1 | 0 | 4 | 1 | 5 | 1 | 4 | 1 | 14 | 0.9286 |
| 18_364210 | 1 | 0 | 4 | 1 | 4 | 1 | 6 | 0.5 | 15 | 0.7333 | 1 | 1 | 6 | 0.8333 | 5 | 0.6 | 3 | 1 | 15 | 0.8 |
| 18_364210 | NA | NA | 3 | 0 | 4 | 0 | 6 | 0.1667 | 13 | 0.0769 | 2 | 0 | 5 | 0.2 | 6 | 0.1667 | 5 | 0 | 18 | 0.1111 |
| 18_364211 | 1 | 0 | 4 | 0.5 | 5 | 0.8 | 5 | 0.4 | 15 | 0.5333 | 2 | 0.5 | 5 | 0.8 | 6 | 0.5 | 4 | 0.75 | 17 | 0.6471 |
| 18_364213 | 1 | 0 | 3 | 0.3333 | 3 | 0.3333 | 5 | 0.2 | 12 | 0.25 | 2 | 0.5 | 4 | 0.25 | 3 | 0 | 2 | 0.5 | 11 | 0.2727 |
| 18_364213 | NA | NA | 4 | 0.25 | 3 | 0.6667 | 5 | 0.6 | 12 | 0.5 | 2 | 0.5 | 6 | 0.3333 | 4 | 0.75 | 4 | 0.5 | 16 | 0.5 |
| 18_364213 | 2 | 0 | 4 | 0.5 | 6 | 0.1667 | 6 | 0 | 18 | 0.1667 | 2 | 0 | 6 | 0.5 | 5 | 0 | 5 | 0.2 | 18 | 0.2222 |
| 18_364214 | NA | NA | 3 | 0.6667 | 3 | 0.3333 | 6 | 0.3333 | 12 | 0.4167 | NA | NA | 2 | 0 | 4 | 1 | 2 | 0 | 8 | 0.5 |
| 18_364214 | 1 | 0 | 3 | 0 | 4 | 0.25 | 6 | 0 | 14 | 0.0714 | 2 | 0 | 4 | 0 | 5 | 0 | 4 | 0.25 | 15 | 0.0667 |
| 18_364214 | 2 | 0 | 2 | 0 | 5 | 0.4 | 5 | 0.2 | 14 | 0.2143 | 2 | 0 | 6 | 0 | 5 | 0.2 | 5 | 0.2 | 18 | 0.1111 |
| 18_364214 | 2 | 0 | 4 | 0.25 | 5 | 0 | 6 | 0.3333 | 17 | 0.1765 | 1 | 0 | 4 | 0 | 6 | 0 | 5 | 0.4 | 16 | 0.125 |
| 18_364214 | 1 | 1 | 3 | 1 | 5 | 1 | 8 | 1 | 17 | 1 | 2 | 1 | 4 | 1 | 6 | 1 | 3 | 1 | 15 | 1 |
| 18_364215 | 1 | 1 | 3 | 1 | 5 | 1 | 7 | 0.8571 | 16 | 0.9375 | 2 | 1 | 6 | 1 | 5 | 1 | 5 | 1 | 18 | 1 |
| 18_364215 | 2 | 0 | 3 | 1 | 6 | 0.8333 | 9 | 0.7778 | 20 | 0.75 | 2 | 1 | 6 | 0.8333 | 6 | 0.8333 | 5 | 1 | 19 | 0.8947 |

| | | | | | | | | | | | | | | | | | | | | |
|---|---|---|---|---|---|---|---|---|---|---|---|---|---|---|---|---|---|---|---|---|
| 18_364216 | 2 | 0 | 3 | 0 | 6 | 0.1667 | 8 | 0.125 | 19 | 0.1053 | 2 | 0 | 6 | 0.1667 | 6 | 0.1667 | 5 | 0.2 | 19 | 0.1579 |
| 18_364219 | 2 | 0 | 3 | 0.6667 | 6 | 0.1667 | 8 | 0.25 | 19 | 0.2632 | 1 | 1 | 7 | 0.2857 | 6 | 0.6667 | 8 | 0.75 | 22 | 0.5909 |
| 18_364229 | NA | NA | 1 | 0 | 2 | 0 | 2 | 0.5 | 5 | 0.2 | 1 | 0 | 3 | 0 | 1 | 1 | 1 | 0 | 6 | 0.1667 |
| 18_364230 | NA | NA | 1 | 1 | 2 | 1 | 2 | 1 | 5 | 1 | 1 | 1 | 3 | 0.3333 | 1 | 1 | 1 | 0 | 6 | 0.5 |
| 18_364326 | 6 | 0.1667 | 15 | 0.2 | 11 | 0 | 11 | 0 | 43 | 0.093 | 8 | 0.25 | 20 | 0.1 | 9 | 0.1111 | 13 | 0.1538 | 50 | 0.14 |
| 18_364332 | 13 | 0.2308 | 21 | 0.0952 | 14 | 0.0714 | 17 | 0 | 65 | 0.0923 | 10 | 0.2 | 24 | 0.0833 | 17 | 0 | 15 | 0.1333 | 66 | 0.0909 |
| 18_364340 | 15 | 0.1333 | 26 | 0 | 16 | 0 | 21 | 0.0952 | 78 | 0.0513 | 12 | 0.0833 | 29 | 0.069 | 23 | 0 | 12 | 0.0833 | 76 | 0.0526 |
| 18_365356 | 14 | 0.1429 | 16 | 0.25 | 8 | 0.125 | 13 | 0.0769 | 51 | 0.1569 | 7 | 0.1429 | 15 | 0.2 | 7 | 0.1429 | 12 | 0.25 | 41 | 0.1951 |
| 18_365372 | 17 | 0.0588 | 16 | 0.3125 | 13 | 0.0769 | 11 | 0.3636 | 57 | 0.193 | 9 | 0.1111 | 13 | 0.1538 | 17 | 0.1176 | 13 | 0.3077 | 52 | 0.1731 |
| 18_365372 | 14 | 0.2143 | 23 | 0.5217 | 14 | 0.2143 | 13 | 0.2308 | 64 | 0.3281 | 11 | 0.3636 | 14 | 0.3571 | 18 | 0.3889 | 12 | 0.4167 | 55 | 0.3818 |
| 18_366385 | 1 | 0 | NA | NA | 3 | 0.3333 | 1 | 0 | 5 | 0.2 | 3 | 0 | 1 | 0 | 4 | 0.25 | 1 | 0 | 9 | 0.1111 |
| 18_366388 | 1 | 0 | NA | NA | 3 | 1 | NA | NA | 4 | 0.75 | 4 | 1 | NA | NA | 4 | 1 | NA | NA | 8 | 1 |
| 18_366389 | 1 | 0 | NA | NA | 2 | 1 | 2 | 1 | 5 | 0.8 | 3 | 0.6667 | 1 | 1 | 4 | 1 | NA | NA | 8 | 0.875 |
| 18_366389 | 1 | 0 | NA | NA | 2 | 0 | 1 | 1 | 4 | 0.25 | 3 | 0.6667 | NA | NA | 4 | 0 | NA | NA | 7 | 0.2857 |
| 18_366389 | 1 | 1 | NA | NA | 2 | 1 | 2 | 0 | 5 | 0.6 | 2 | 0.5 | NA | NA | 3 | 0.6667 | NA | NA | 5 | 0.6 |
| 18_366389 | 2 | 0 | NA | NA | 3 | 0.3333 | 2 | 0.5 | 7 | 0.2857 | 4 | 0.25 | 1 | 1 | 4 | 0.25 | NA | NA | 9 | 0.3333 |
| 18_366390 | 2 | 1 | NA | NA | 3 | 1 | 2 | 1 | 7 | 1 | 4 | 1 | 1 | 1 | 4 | 1 | NA | NA | 9 | 1 |
| 18_366390 | 1 | 0 | NA | NA | 2 | 1 | 2 | 1 | 5 | 0.8 | 2 | 1 | 1 | 1 | 4 | 0.5 | NA | NA | 7 | 0.7143 |
| 18_366391 | 2 | 0.5 | NA | NA | 3 | 0.6667 | 2 | 0.5 | 7 | 0.5714 | 3 | 0.6667 | NA | NA | 4 | 0.25 | NA | NA | 7 | 0.4286 |
| 18_366926 | NA | NA | NA | NA | NA | NA | 1 | 1 | 1 | 1 | 1 | 1 | 2 | 0.5 | 3 | 0.6667 | 3 | 0.6667 | 9 | 0.6667 |
| 18_366926 | NA | NA | NA | NA | 1 | 1 | 1 | 0 | 2 | 0.5 | 1 | 0 | 3 | 0.3333 | 3 | 0 | 4 | 0.25 | 11 | 0.1818 |
| 18_366928 | NA | NA | NA | NA | 1 | 1 | 1 | 1 | 2 | 1 | 1 | 0 | 3 | 1 | 2 | 0.5 | 3 | 0.6667 | 9 | 0.6667 |
| 18_367804 | 2 | 1 | NA | NA | NA | NA | 1 | 0 | 3 | 0.6667 | 1 | 0 | 1 | 1 | 1 | 1 | 1 | 1 | 4 | 0.75 |
| 18_367807 | 2 | 1 | NA | NA | 1 | 1 | 3 | 0.6667 | 6 | 0.8333 | 1 | 1 | 1 | 1 | 1 | 1 | 2 | 1 | 5 | 1 |
| 18_367808 | 3 | 1 | 1 | 0 | 1 | 1 | 3 | 1 | 8 | 0.875 | 1 | 0 | 1 | 1 | NA | NA | 2 | 1 | 4 | 0.75 |
| 18_367809 | 3 | 0.3333 | 1 | 0 | NA | NA | 3 | 0.6667 | 7 | 0.4286 | 1 | 0 | 2 | 0.5 | 1 | 0 | 3 | 0.3333 | 7 | 0.2857 |
| 18_367810 | 3 | 0.6667 | 1 | 0 | 1 | 0 | 3 | 0.3333 | 8 | 0.375 | 1 | 0 | 3 | 0 | 1 | 1 | 6 | 0.3333 | 11 | 0.2727 |
| 18_367812 | 2 | 1 | 1 | 0 | 1 | 1 | 5 | 0.2 | 9 | 0.4444 | 1 | 1 | 2 | 0 | 1 | 0 | 4 | 0.75 | 8 | 0.5 |
| 18_367812 | 3 | 1 | 1 | 0 | NA | NA | 4 | 0.5 | 8 | 0.625 | 1 | 1 | 3 | 1 | 1 | 0 | 5 | 0.4 | 10 | 0.6 |
| 18_367812 | 3 | 0.3333 | 1 | 0 | NA | NA | 5 | 0.2 | 9 | 0.2222 | 1 | 0 | 3 | 0.3333 | 1 | 1 | 4 | 0.5 | 9 | 0.4444 |
| 18_367812 | 3 | 1 | 1 | 1 | 1 | 1 | 5 | 0.8 | 10 | 0.9 | 1 | 1 | 2 | 1 | 1 | 1 | 5 | 1 | 9 | 1 |
| 18_367813 | 3 | 0 | 1 | 0 | 1 | 0 | 4 | 0.25 | 9 | 0.1111 | 1 | 0 | 3 | 0 | NA | NA | 6 | 0.1667 | 10 | 0.1 |
| 18_367814 | 3 | 0.6667 | 1 | 0 | NA | NA | 3 | 1 | 7 | 0.7143 | 1 | 1 | 3 | 0.3333 | NA | NA | 4 | 0.5 | 8 | 0.5 |
| 18_367816 | 3 | 0.3333 | 1 | 0 | 2 | 0.5 | 6 | 0.1667 | 12 | 0.25 | 1 | 0 | 3 | 0.3333 | 1 | 1 | 2 | 0 | 7 | 0.2857 |
| 18_367816 | 3 | 0 | 1 | 0 | 2 | 0 | 5 | 0.4 | 11 | 0.1818 | 1 | 0 | 3 | 0.3333 | 2 | 0.5 | 6 | 0.3333 | 12 | 0.3333 |
| 18_367817 | 2 | 1 | NA | NA | 2 | 0.5 | 6 | 0.6667 | 10 | 0.7 | 1 | 0 | 3 | 0.3333 | 1 | 1 | 6 | 0.5 | 11 | 0.4545 |
| 18_367915 | 3 | 0.3333 | 4 | 0.5 | 2 | 0 | 2 | 0 | 11 | 0.2727 | NA | NA | 3 | 0.3333 | 2 | 0 | 4 | 1 | 9 | 0.5556 |
| 18_368004 | 15 | 0.1333 | 9 | 0.1111 | 9 | 0.3333 | 7 | 0.2857 | 40 | 0.2 | 5 | 0 | 6 | 0.1667 | 8 | 0.25 | 9 | 0.1111 | 28 | 0.1429 |
| 18_368004 | 13 | 0.0769 | 8 | 0.375 | 10 | 0.1 | 9 | 0 | 40 | 0.125 | 6 | 0.1667 | 6 | 0 | 6 | 0 | 7 | 0.2857 | 25 | 0.12 |
| 18_368004 | 13 | 0.3077 | 10 | 0.4 | 9 | 0.4444 | 9 | 0.3333 | 41 | 0.3659 | 7 | 0.2857 | 6 | 0.3333 | 8 | 0.5 | 7 | 0.1429 | 28 | 0.3214 |
| 18_368006 | 11 | 0.0909 | 9 | 0 | 9 | 0.1111 | 9 | 0.1111 | 38 | 0.0789 | 8 | 0 | 5 | 0.2 | 7 | 0.2857 | 9 | 0 | 29 | 0.1034 |
| 18_368042 | 4 | 0 | 3 | 0.3333 | 3 | 0 | 2 | 0 | 12 | 0.0833 | 6 | 0 | 6 | 0 | 2 | 0 | 6 | 0.1667 | 20 | 0.05 |
| 18_368122 | 1 | 0 | NA | NA | NA | NA | 1 | 1 | 2 | 0.5 | 3 | 0 | 4 | 0.5 | 2 | 0 | 3 | 0 | 12 | 0.1667 |
| 18_368148 | 1 | 0 | 3 | 1 | 7 | 0.5714 | NA | NA | 11 | 0.6364 | 1 | 1 | 2 | 0 | 2 | 1 | NA | NA | 5 | 0.6 |
| 18_368149 | 5 | 0.4 | 10 | 0.6 | 8 | 0.25 | 3 | 0.3333 | 26 | 0.4231 | 6 | 0.1667 | 6 | 0.5 | 5 | 0.4 | 2 | 0 | 19 | 0.3158 |
| 18_368149 | 6 | 0 | 11 | 0.0909 | 9 | 0 | 4 | 0.25 | 30 | 0.0667 | 5 | 0 | 3 | 0 | 5 | 0.2 | 1 | 0 | 14 | 0.0714 |
| 18_368149 | 5 | 0.6 | 10 | 0.8 | 7 | 0.7143 | 4 | 1 | 26 | 0.7692 | 5 | 0.6 | 5 | 1 | 5 | 0.8 | 1 | 1 | 16 | 0.8125 |
| 18_368149 | 6 | 0.3333 | 9 | 0.1111 | 8 | 0.125 | 4 | 0.5 | 27 | 0.2222 | 6 | 0 | 7 | 0.1429 | 5 | 0.4 | 2 | 0 | 20 | 0.15 |
| 18_368152 | 7 | 0.1429 | 11 | 0.4545 | 9 | 0.1111 | 4 | 0.75 | 31 | 0.3226 | 6 | 0.3333 | 7 | 0.2857 | 6 | 0.3333 | 3 | 0.6667 | 22 | 0.3636 |
| 18_368155 | 6 | 1 | 8 | 0.625 | 4 | 1 | 3 | 0.6667 | 21 | 0.8095 | 6 | 1 | 7 | 0.8571 | 5 | 0.6 | 2 | 1 | 20 | 0.85 |
| 18_368156 | 8 | 0.625 | 13 | 0.1538 | 9 | 0 | 4 | 0 | 34 | 0.2059 | 7 | 0.1429 | 9 | 0.3333 | 6 | 0 | 4 | 0.25 | 26 | 0.1923 |
| 18_368156 | 4 | 0 | 10 | 0.3 | 7 | 0.1429 | 2 | 0 | 23 | 0.1739 | 3 | 0 | 8 | 0.5 | 4 | 0 | 1 | 1 | 16 | 0.3125 |
| 18_368157 | 7 | 1 | 13 | 1 | 8 | 1 | 4 | 1 | 32 | 1 | 7 | 1 | 8 | 1 | 6 | 1 | 4 | 1 | 25 | 1 |
| 18_368158 | 8 | 0.125 | 13 | 0.0769 | 8 | 0.125 | 3 | 0.3333 | 32 | 0.125 | 8 | 0.125 | 9 | 0.2222 | 4 | 0.5 | 3 | 0.3333 | 24 | 0.25 |
| 18_368158 | 9 | 1 | 11 | 0.9091 | 7 | 0.8571 | 4 | 1 | 31 | 0.9355 | 6 | 1 | 8 | 1 | 4 | 1 | 2 | 1 | 20 | 1 |
| 18_368158 | 8 | 1 | 9 | 1 | 7 | 1 | 3 | 1 | 27 | 1 | 7 | 1 | 6 | 1 | 6 | 1 | 3 | 1 | 22 | 1 |
| 18_368159 | 8 | 0.125 | 9 | 0.4444 | 3 | 0 | 4 | 0 | 24 | 0.2083 | 3 | 0.3333 | 7 | 0.2857 | 4 | 0.25 | 3 | 0 | 17 | 0.2353 |
| 18_368161 | 11 | 0.9091 | 12 | 0.8333 | 8 | 0.875 | 3 | 1 | 34 | 0.8824 | 8 | 1 | 8 | 0.625 | 5 | 0.8 | 4 | 1 | 25 | 0.84 |
| 18_368164 | 7 | 0.5714 | 12 | 0.3333 | 10 | 0.1 | 4 | 0.25 | 33 | 0.303 | 6 | 0.6667 | 8 | 0.25 | 3 | 0.3333 | 3 | 0 | 20 | 0.35 |
| 18_368583 | NA | NA | 1 | 1 | NA | NA | 1 | 1 | 2 | 1 | 1 | 1 | NA | NA | NA | NA | NA | NA | 1 | 1 |
| 18_368583 | NA | NA | 1 | 1 | 1 | 1 | 1 | 0 | 3 | 0.6667 | 1 | 1 | NA | NA | NA | NA | NA | NA | 1 | 1 |
| 18_368584 | NA | NA | 1 | 1 | 1 | 1 | 1 | 1 | 3 | 1 | 1 | 1 | NA | NA | NA | NA | NA | NA | 1 | 1 |
| 18_368755 | 41 | 0.1951 | 39 | 0.2308 | 40 | 0.2 | 23 | 0.2609 | 143 | 0.2168 | 32 | 0.125 | 40 | 0.15 | 34 | 0.1176 | 37 | 0.2432 | 143 | 0.1608 |
| 18_368756 | 40 | 0.25 | 41 | 0.1951 | 45 | 0.1778 | 28 | 0.25 | 154 | 0.2143 | 34 | 0.2647 | 49 | 0.2857 | 40 | 0.25 | 41 | 0.1951 | 164 | 0.25 |
| 18_368962 | NA | NA | 2 | 0.5 | 1 | 1 | NA | NA | 3 | 0.6667 | NA | NA | 3 | 0.3333 | 5 | 0.6 | 3 | 0.6667 | 11 | 0.5455 |

| | | | | | | | | | | | | | | | | | | | | |
|---|---|---|---|---|---|---|---|---|---|---|---|---|---|---|---|---|---|---|---|---|
| 18_3689647 | NA | NA | 1 | 1 | 2 | 0 | 1 | 1 | 4 | 0.5 | NA | NA | 2 | 1 | 3 | 0 | 3 | 1 | 8 | 0.625 |
| 18_3689650 | NA | NA | 2 | 1 | NA | NA | 1 | 1 | 3 | 1 | NA | NA | 2 | 0.5 | 4 | 0.5 | 1 | 1 | 7 | 0.5714 |
| 18_3689652 | NA | NA | 2 | 1 | 2 | 1 | 1 | 1 | 5 | 1 | NA | NA | 2 | 0.5 | 4 | 0.75 | 3 | 0.6667 | 9 | 0.6667 |
| 18_3689662 | NA | NA | 2 | 1 | 3 | 0.6667 | NA | NA | 5 | 0.8 | NA | NA | 3 | 0 | NA | NA | 4 | 0.5 | 7 | 0.2857 |
| 18_3689667 | NA | NA | 3 | 0.3333 | 3 | 0 | NA | NA | 6 | 0.1667 | NA | NA | 2 | 0.5 | 4 | 0 | 3 | 0.3333 | 9 | 0.2222 |
| 18_3689966 | 10 | 0.7 | 23 | 0.7826 | 14 | 0.7857 | 14 | 0.8571 | 61 | 0.7869 | 21 | 0.6667 | 9 | 0.8889 | 19 | 0.8421 | 16 | 0.9375 | 65 | 0.8154 |
| 18_3689969 | 11 | 0.3636 | 22 | 0.2727 | 13 | 0.2308 | 14 | 0.1429 | 60 | 0.25 | 15 | 0.4667 | 6 | 0.5 | 15 | 0.5333 | 13 | 0.2308 | 49 | 0.4286 |
| 18_3689973 | 1 | 0 | 2 | 0 | 3 | 0 | 2 | 0.5 | 8 | 0.125 | 3 | 0 | 4 | 0.25 | 2 | 0.5 | 1 | 0 | 10 | 0.2 |
| 18_3689983 | 9 | 0.4444 | 23 | 0.087 | 11 | 0.2727 | 12 | 0.1667 | 55 | 0.2 | 18 | 0.3333 | 7 | 0.1429 | 13 | 0.1538 | 14 | 0.2857 | 52 | 0.25 |
| 18_3689988 | 8 | 0.25 | 23 | 0.1304 | 7 | 0.1429 | 11 | 0.1818 | 49 | 0.1633 | 15 | 0.0667 | 7 | 0.2857 | 12 | 0.0833 | 10 | 0.2 | 44 | 0.1364 |
| 18_3692423 | NA | NA | 1 | 0 | 1 | 0 | 6 | 0.1667 | 8 | 0.125 | 3 | 0 | 6 | 0.1667 | 2 | 0 | 1 | 0 | 12 | 0.0833 |
| 18_3692424 | 1 | 1 | 2 | 1 | 1 | 0 | 5 | 0.8 | 9 | 0.7778 | 3 | 1 | 6 | 0.8333 | 2 | 0.5 | 1 | 1 | 12 | 0.8333 |
| 18_3692431 | 1 | 0 | 1 | 0 | 1 | 0 | 4 | 0.75 | 7 | 0.4286 | 2 | 0 | 6 | 0.5 | 3 | 0 | 1 | 0 | 12 | 0.25 |
| 18_3692433 | NA | NA | 2 | 1 | 1 | 0 | 5 | 0.8 | 8 | 0.75 | 2 | 1 | 5 | 0.6 | 3 | 0.6667 | 1 | 1 | 11 | 0.7273 |
| 18_3692434 | NA | NA | 1 | 0 | 1 | 1 | 5 | 1 | 7 | 0.8571 | 2 | 1 | 6 | 1 | 2 | 0.5 | NA | NA | 10 | 0.9 |
| 18_3692450 | 1 | 1 | 3 | 1 | 1 | 1 | 6 | 0.8333 | 11 | 0.9091 | 3 | 1 | 5 | 0.6 | 2 | 0.5 | 1 | 1 | 11 | 0.7273 |
| 18_3692459 | 1 | 1 | 3 | 0.6667 | 1 | 0 | 5 | 0.8 | 10 | 0.7 | 3 | 1 | 7 | 0.5714 | 4 | 0.5 | 1 | 0 | 15 | 0.6 |
| 18_3692462 | 1 | 1 | 3 | 1 | 1 | 1 | 4 | 0.5 | 9 | 0.7778 | 3 | 1 | 6 | 0.8333 | 4 | 0.75 | 1 | 1 | 14 | 0.8571 |
| 18_3692471 | NA | NA | 3 | 0 | NA | NA | 6 | 0.1667 | 9 | 0.1111 | 1 | 1 | 5 | 0.2 | 3 | 0 | 1 | 0 | 10 | 0.2 |
| 18_3692472 | 1 | 1 | 3 | 0 | 1 | 0 | 4 | 0.25 | 9 | 0.2222 | 3 | 0.3333 | 7 | 0.2857 | 3 | 0 | 1 | 0 | 14 | 0.2143 |
| 18_3692488 | NA | NA | 2 | 1 | 1 | 1 | 6 | 0.1667 | 9 | 0.4444 | 3 | 0.6667 | 8 | 0.5 | 3 | 0.3333 | NA | NA | 14 | 0.5 |
| 18_3692546 | 1 | 1 | 1 | 1 | NA | NA | 1 | 1 | 3 | 1 | NA | NA | NA | NA | 1 | 1 | NA | NA | 1 | 1 |
| 18_3693099 | 21 | 0.2381 | 24 | 0.4167 | 16 | 0.25 | 22 | 0.2727 | 83 | 0.3012 | 21 | 0.2381 | 19 | 0.4737 | 11 | 0.2727 | 18 | 0.3889 | 69 | 0.3478 |
| 18_3693100 | 23 | 0.0435 | 25 | 0.04 | 16 | 0.125 | 19 | 0.0526 | 83 | 0.0602 | 23 | 0.087 | 18 | 0.1667 | 10 | 0.1 | 18 | 0.1111 | 69 | 0.1159 |
| 18_3693111 | 20 | 0.85 | 27 | 0.9259 | 14 | 0.7857 | 21 | 0.7143 | 82 | 0.8293 | 23 | 0.8261 | 21 | 0.8571 | 13 | 0.9231 | 18 | 0.9444 | 75 | 0.88 |
| 18_3693133 | 20 | 0.15 | 28 | 0.1429 | 21 | 0.0476 | 21 | 0.2381 | 90 | 0.1444 | 24 | 0.2083 | 22 | 0.1364 | 15 | 0.1333 | 21 | 0 | 82 | 0.122 |
| 18_3708811 | NA | NA | NA | NA | 1 | 1 | NA | NA | 1 | 1 | NA | NA | 1 | 1 | 1 | 0 | NA | NA | 2 | 0.5 |
| 18_3709344 | NA | NA | 1 | 1 | 1 | 1 | NA | NA | 2 | 1 | NA | NA | 1 | 1 | NA | NA | NA | NA | 1 | 1 |
| 18_3709442 | NA | NA | 1 | 1 | 2 | 0 | NA | NA | 3 | 0.3333 | NA | NA | 1 | 1 | NA | NA | NA | NA | 1 | 1 |
| 18_3709446 | NA | NA | 1 | 0 | 1 | 1 | NA | NA | 2 | 0.5 | NA | NA | 1 | 1 | NA | NA | NA | NA | 1 | 1 |
| 18_3709474 | NA | NA | 1 | 1 | 2 | 1 | NA | NA | 3 | 1 | NA | NA | 1 | 1 | NA | NA | NA | NA | 1 | 1 |
| 18_3709506 | NA | NA | 1 | 0 | 1 | 1 | NA | NA | 2 | 0.5 | NA | NA | 1 | 1 | NA | NA | NA | NA | 1 | 1 |
| 18_3709514 | NA | NA | 1 | 1 | 2 | 1 | NA | NA | 3 | 1 | NA | NA | 1 | 1 | NA | NA | 1 | 1 | 2 | 1 |
| 18_3709521 | NA | NA | 1 | 1 | 2 | 1 | NA | NA | 3 | 1 | NA | NA | NA | NA | NA | NA | 1 | 1 | 1 | 1 |
| 18_3709540 | NA | NA | 1 | 0 | 1 | 1 | NA | NA | 2 | 0.5 | NA | NA | 1 | 1 | NA | NA | 1 | 1 | 2 | 1 |
| 18_3709555 | NA | NA | 2 | 0.5 | 2 | 0.5 | NA | NA | 4 | 0.5 | NA | NA | 1 | 1 | NA | NA | 1 | 1 | 2 | 1 |
| 18_3709572 | NA | NA | 2 | 0.5 | 2 | 1 | NA | NA | 4 | 0.75 | NA | NA | 1 | 1 | NA | NA | 1 | 1 | 2 | 1 |
| 18_3709582 | NA | NA | 2 | 0 | 2 | 0.5 | NA | NA | 4 | 0.25 | NA | NA | 1 | 1 | NA | NA | 1 | 0 | 2 | 0.5 |
| 18_3710245 | NA | NA | NA | NA | 1 | 1 | NA | NA | 1 | 1 | NA | NA | NA | NA | 1 | 1 | NA | NA | 1 | 1 |
| 18_3710250 | 1 | 1 | NA | NA | 1 | 1 | NA | NA | 2 | 1 | NA | NA | NA | NA | 1 | 1 | NA | NA | 1 | 1 |
| 18_3710254 | 1 | 1 | NA | NA | 1 | 0 | NA | NA | 2 | 0.5 | NA | NA | NA | NA | 1 | 1 | 1 | 1 | 2 | 1 |
| 18_3710276 | NA | NA | NA | NA | 1 | 1 | NA | NA | 1 | 1 | NA | NA | NA | NA | 1 | 0 | 1 | 1 | 2 | 0.5 |
| 18_3710735 | NA | NA | 1 | 1 | NA | NA | NA | NA | 1 | 1 | NA | NA | 1 | 1 | NA | NA | NA | NA | 1 | 1 |
| 18_3710750 | NA | NA | 1 | 1 | NA | NA | NA | NA | 1 | 1 | NA | NA | 1 | 1 | NA | NA | NA | NA | 1 | 1 |
| 18_3710786 | NA | NA | 1 | 1 | NA | NA | NA | NA | 1 | 1 | NA | NA | 1 | 1 | NA | NA | NA | NA | 1 | 1 |
| 18_3710877 | NA | NA | 1 | 1 | NA | NA | NA | NA | 1 | 1 | NA | NA | 1 | 1 | 1 | 1 | 1 | 1 | 3 | 1 |
| 18_3714428 | 2 | 1 | NA | NA | NA | NA | 1 | 1 | 3 | 1 | NA | NA | NA | NA | 1 | 1 | NA | NA | 1 | 1 |
| 18_3714555 | 2 | 1 | NA | NA | NA | NA | 2 | 1 | 4 | 1 | 2 | 0.5 | NA | NA | NA | NA | NA | NA | 2 | 0.5 |
| 18_3714582 | 2 | 1 | NA | NA | NA | NA | 1 | 1 | 3 | 1 | 1 | 1 | NA | NA | NA | NA | NA | NA | 1 | 1 |
| 18_3714584 | 2 | 0.5 | NA | NA | NA | NA | NA | NA | 2 | 0.5 | 1 | 1 | NA | NA | NA | NA | NA | NA | 1 | 1 |
| 18_3714586 | 2 | 0.5 | NA | NA | NA | NA | 1 | 1 | 3 | 0.6667 | 1 | 1 | NA | NA | NA | NA | NA | NA | 1 | 1 |
| 18_3715365 | 1 | 1 | 2 | 0.5 | NA | NA | 1 | 0 | 4 | 0.5 | NA | NA | 2 | 0.5 | NA | NA | NA | NA | 2 | 0.5 |
| 18_3715379 | 1 | 0 | 1 | 1 | NA | NA | NA | NA | 2 | 0.5 | NA | NA | 1 | 1 | NA | NA | NA | NA | 1 | 1 |
| 18_3715385 | NA | NA | 1 | 1 | NA | NA | NA | NA | 1 | 1 | NA | NA | 1 | 1 | NA | NA | NA | NA | 1 | 1 |
| 18_3715405 | 1 | 0 | 2 | 1 | NA | NA | NA | NA | 3 | 0.6667 | NA | NA | 1 | 1 | NA | NA | NA | NA | 1 | 1 |
| 18_3715419 | NA | NA | 1 | 1 | NA | NA | NA | NA | 1 | 1 | NA | NA | 1 | 1 | NA | NA | NA | NA | 1 | 1 |
| 18_3715454 | 1 | 1 | 2 | 1 | NA | NA | 1 | 0 | 4 | 0.75 | NA | NA | 2 | 0.5 | NA | NA | NA | NA | 2 | 0.5 |
| 18_3715493 | 1 | 1 | 2 | 1 | NA | NA | 1 | 1 | 4 | 1 | 1 | 1 | 2 | 1 | NA | NA | 2 | 0.5 | 5 | 0.8 |
| 18_3715496 | 1 | 1 | 2 | 0 | NA | NA | NA | NA | 3 | 0.3333 | 1 | 0 | 2 | 0.5 | NA | NA | 1 | 0 | 4 | 0.25 |
| 18_3715503 | NA | NA | NA | NA | NA | NA | 1 | 1 | 1 | 1 | NA | NA | NA | NA | NA | NA | 1 | 1 | 1 | 1 |
| 18_3715508 | 1 | 1 | 1 | 1 | NA | NA | 2 | 1 | 4 | 1 | 2 | 1 | 2 | 1 | NA | NA | 2 | 1 | 6 | 1 |
| 18_3715511 | NA | NA | 1 | 1 | NA | NA | 1 | 1 | 2 | 1 | NA | NA | NA | NA | NA | NA | 1 | 1 | 1 | 1 |
| 18_3715524 | 1 | 0 | 2 | 0.5 | NA | NA | 3 | 0.6667 | 6 | 0.5 | 2 | 0.5 | 2 | 1 | NA | NA | 2 | 0.5 | 6 | 0.6667 |
| 18_3715536 | NA | NA | 1 | 1 | NA | NA | 2 | 1 | 3 | 1 | 1 | 1 | 1 | 1 | NA | NA | NA | NA | 2 | 1 |
| 18_3715552 | 1 | 1 | NA | NA | NA | NA | NA | NA | 1 | 1 | NA | NA | NA | NA | NA | NA | 1 | 1 | 1 | 1 |

| | | | | | | | | | | | | | | | | | | | | |
|---|---|---|---|---|---|---|---|---|---|---|---|---|---|---|---|---|---|---|---|---|
| 18_371555 | NA | NA | NA | NA | NA | NA | 2 | 0.5 | 2 | 0.5 | 1 | 1 | 1 | 0 | NA | NA | 1 | 0 | 3 | 0.3333 |
| 18_372772 | NA | NA | 2 | 1 | NA | NA | 1 | 0 | 3 | 0.6667 | 1 | 0 | 1 | 1 | 2 | 0 | NA | NA | 4 | 0.25 |
| 18_372772 | NA | NA | 2 | 1 | NA | NA | 1 | 1 | 3 | 1 | 1 | 0 | 1 | 1 | 2 | 1 | NA | NA | 4 | 0.75 |
| 18_372775 | NA | NA | 3 | 1 | NA | NA | 2 | 0 | 5 | 0.6 | 1 | 0 | 1 | 0 | 2 | 0.5 | NA | NA | 4 | 0.25 |
| 18_372778 | NA | NA | 2 | 0 | NA | NA | 2 | 1 | 4 | 0.5 | NA | NA | NA | NA | 1 | 1 | NA | NA | 1 | 1 |
| 18_372779 | NA | NA | 3 | 1 | 1 | 1 | 2 | 1 | 6 | 1 | NA | NA | 1 | 1 | 2 | 1 | NA | NA | 3 | 1 |
| 18_372780 | NA | NA | 3 | 1 | 1 | 1 | 2 | 1 | 6 | 1 | 1 | 0 | 1 | 1 | 2 | 1 | NA | NA | 4 | 0.75 |
| 18_372781 | NA | NA | 3 | 1 | 1 | 1 | 2 | 0.5 | 6 | 0.8333 | 1 | 1 | 1 | 0 | 2 | 0.5 | 1 | 1 | 5 | 0.6 |
| 18_372783 | NA | NA | 2 | 1 | 1 | 1 | 4 | 0.75 | 7 | 0.8571 | 1 | 1 | 1 | 1 | 2 | 1 | 1 | 1 | 5 | 1 |
| 18_372784 | NA | NA | 2 | 1 | 1 | 1 | 3 | 0.6667 | 6 | 0.8333 | 1 | 1 | 1 | 1 | 2 | 1 | NA | NA | 4 | 1 |
| 18_372785 | NA | NA | 2 | 0.5 | NA | NA | 3 | 0 | 5 | 0.2 | 1 | 1 | 1 | 0 | NA | NA | NA | NA | 2 | 0.5 |
| 18_372786 | NA | NA | 2 | 1 | 1 | 0 | 5 | 1 | 8 | 0.875 | 2 | 0.5 | 1 | 0 | 1 | 1 | 1 | 1 | 5 | 0.6 |
| 18_372786 | NA | NA | 3 | 0.3333 | 2 | 0.5 | 6 | 0.6667 | 11 | 0.5455 | 2 | 0.5 | 1 | 1 | 2 | 1 | 1 | 0 | 6 | 0.6667 |
| 18_372788 | NA | NA | 3 | 1 | 1 | 1 | 6 | 1 | 10 | 1 | 2 | 1 | 2 | 1 | 2 | 1 | NA | NA | 6 | 1 |
| 18_372788 | NA | NA | 2 | 1 | 2 | 1 | 5 | 1 | 9 | 1 | 1 | 1 | 2 | 1 | 1 | 1 | 1 | 1 | 5 | 1 |
| 18_372789 | NA | NA | 3 | 1 | 2 | 1 | 6 | 0.5 | 11 | 0.7273 | 3 | 0.6667 | 2 | 0.5 | 2 | 1 | 1 | 1 | 8 | 0.75 |
| 18_372790 | NA | NA | 3 | 0.6667 | 1 | 0 | 6 | 0 | 10 | 0.2 | 3 | 0 | 2 | 0 | 1 | 1 | 1 | 1 | 7 | 0.2857 |
| 18_372790 | NA | NA | 3 | 0.3333 | 2 | 0 | 3 | 0 | 8 | 0.125 | 3 | 0.3333 | 2 | 0.5 | 1 | 0 | 1 | 0 | 7 | 0.2857 |
| 18_372790 | NA | NA | 2 | 1 | 2 | 1 | 6 | 0.5 | 10 | 0.7 | 2 | 1 | 2 | 0.5 | 2 | 1 | NA | NA | 6 | 0.8333 |
| 18_372791 | NA | NA | 3 | 0.3333 | 2 | 1 | 6 | 1 | 11 | 0.8182 | 3 | 0.6667 | 2 | 1 | 2 | 0 | NA | NA | 7 | 0.5714 |
| 18_372791 | NA | NA | 3 | 0.6667 | NA | NA | 6 | 0.5 | 9 | 0.5556 | 2 | 1 | 2 | 0 | 1 | 0 | 1 | 0 | 6 | 0.3333 |
| 18_372793 | NA | NA | 1 | 0 | 2 | 0 | 3 | 0.3333 | 6 | 0.1667 | 2 | 0 | 2 | 0.5 | 2 | 0 | NA | NA | 6 | 0.1667 |
| 18_372794 | NA | NA | 2 | 1 | 2 | 1 | 5 | 1 | 9 | 1 | 4 | 1 | 3 | 0.3333 | 3 | 0.6667 | 1 | 1 | 11 | 0.7273 |
| 18_373180 | 8 | 0.25 | 6 | 0.5 | 8 | 0.375 | 9 | 0.3333 | 31 | 0.3548 | 6 | 0.3333 | 7 | 0.5714 | 6 | 0.3333 | 5 | 0.2 | 24 | 0.375 |
| 18_373180 | 8 | 0.25 | 10 | 0.1 | 6 | 0 | 13 | 0.3077 | 37 | 0.1892 | 8 | 0.375 | 8 | 0.375 | 6 | 0.1667 | 6 | 0.3333 | 28 | 0.3214 |
| 18_373181 | 9 | 0 | 8 | 0.375 | 10 | 0.2 | 9 | 0.1111 | 36 | 0.1667 | 5 | 0.2 | 9 | 0 | 7 | 0.1429 | 6 | 0 | 27 | 0.0741 |
| 18_373182 | 9 | 0.1111 | 10 | 0.2 | 11 | 0.3636 | 8 | 0.5 | 38 | 0.2895 | 7 | 0 | 8 | 0 | 8 | 0 | 5 | 0.4 | 28 | 0.0714 |
| 18_373186 | 9 | 0.1111 | 11 | 0.3636 | 10 | 0.6 | 11 | 0.5455 | 41 | 0.4146 | 10 | 0.1 | 12 | 0.3333 | 9 | 0.2222 | 4 | 0.5 | 35 | 0.2571 |
| 18_373188 | 12 | 0.25 | 13 | 0 | 12 | 0.25 | 15 | 0.2667 | 52 | 0.1923 | 11 | 0.2727 | 12 | 0.25 | 9 | 0.2222 | 7 | 0.5714 | 39 | 0.3077 |
| 18_373188 | 11 | 0.1818 | 12 | 0.0833 | 11 | 0 | 15 | 0.3333 | 49 | 0.1633 | 10 | 0.1 | 13 | 0.0769 | 10 | 0.4 | 7 | 0.2857 | 40 | 0.2 |
| 18_373193 | 11 | 0.0909 | 13 | 0 | 12 | 0 | 16 | 0.125 | 52 | 0.0577 | 8 | 0 | 14 | 0.0714 | 8 | 0 | 6 | 0.1667 | 36 | 0.0556 |
| 18_373195 | 14 | 0.2143 | 16 | 0.1875 | 15 | 0.0667 | 18 | 0.1667 | 63 | 0.1587 | 11 | 0.1818 | 16 | 0 | 14 | 0.2857 | 8 | 0 | 49 | 0.1224 |
| 18_373195 | 13 | 0.3846 | 17 | 0.3529 | 15 | 0.4667 | 17 | 0.5882 | 62 | 0.4516 | 10 | 0.4 | 16 | 0.3125 | 12 | 0.5 | 5 | 0.2 | 43 | 0.3721 |
| 18_373196 | 13 | 0.0769 | 16 | 0 | 11 | 0 | 17 | 0.2941 | 57 | 0.1053 | 11 | 0.1818 | 12 | 0 | 12 | 0.0833 | 8 | 0 | 43 | 0.0698 |
| 18_373201 | 13 | 0.0769 | 18 | 0.0556 | 12 | 0.3333 | 14 | 0.1429 | 57 | 0.1404 | 8 | 0 | 10 | 0 | 10 | 0 | 10 | 0.2 | 38 | 0.0526 |
| 18_374295 | 1 | 1 | 1 | 0 | 1 | 1 | NA | NA | 3 | 0.6667 | 1 | 1 | NA | NA | NA | NA | NA | NA | 1 | 1 |
| 18_374298 | 1 | 1 | 1 | 1 | 1 | 0 | NA | NA | 3 | 0.6667 | 1 | 1 | NA | NA | NA | NA | NA | NA | 1 | 1 |
| 18_374299 | 1 | 1 | 1 | 1 | NA | NA | NA | NA | 2 | 1 | 1 | 1 | NA | NA | NA | NA | NA | NA | 1 | 1 |
| 18_374300 | 1 | 1 | NA | NA | NA | NA | NA | NA | 1 | 1 | 1 | 1 | NA | NA | NA | NA | NA | NA | 1 | 1 |
| 18_374303 | 1 | 0 | 1 | 1 | 1 | 0 | NA | NA | 3 | 0.3333 | 1 | 1 | NA | NA | NA | NA | NA | NA | 1 | 1 |
| 18_374303 | 1 | 1 | 1 | 1 | 1 | 1 | NA | NA | 3 | 1 | 1 | 1 | NA | NA | NA | NA | NA | NA | 1 | 1 |
| 18_374342 | NA | NA | 1 | 1 | NA | NA | 3 | 0.6667 | 4 | 0.75 | NA | NA | 1 | 1 | NA | NA | NA | NA | 1 | 1 |
| 18_374343 | NA | NA | 2 | 0 | 1 | 1 | 3 | 0.3333 | 6 | 0.3333 | NA | NA | 1 | 1 | NA | NA | NA | NA | 1 | 1 |
| 18_374344 | NA | NA | 1 | 1 | 1 | 1 | 3 | 0.6667 | 5 | 0.8 | NA | NA | 2 | 1 | NA | NA | NA | NA | 2 | 1 |
| 18_374345 | NA | NA | 2 | 1 | 1 | 1 | 3 | 0.6667 | 6 | 0.8333 | NA | NA | 2 | 1 | NA | NA | NA | NA | 2 | 1 |
| 18_374348 | NA | NA | 2 | 0.5 | 1 | 0 | 3 | 0 | 6 | 0.1667 | 1 | 0 | 2 | 0.5 | NA | NA | NA | NA | 3 | 0.3333 |
| 18_374349 | NA | NA | 2 | 0 | 1 | 1 | 2 | 0.5 | 5 | 0.4 | 1 | 0 | 1 | 1 | NA | NA | NA | NA | 2 | 0.5 |
| 18_374350 | NA | NA | 2 | 1 | 1 | 0 | 3 | 0.6667 | 6 | 0.6667 | 1 | 1 | 2 | 1 | NA | NA | NA | NA | 3 | 1 |
| 18_374351 | NA | NA | 2 | 0.5 | 1 | 1 | 3 | 0.3333 | 6 | 0.5 | 1 | 1 | 2 | 0.5 | NA | NA | NA | NA | 3 | 0.6667 |
| 18_374351 | NA | NA | 2 | 0.5 | 1 | 0 | 2 | 1 | 5 | 0.6 | 1 | 1 | 1 | 1 | NA | NA | NA | NA | 2 | 1 |
| 18_374352 | NA | NA | 1 | 0 | 1 | 1 | 3 | 0 | 5 | 0.2 | 1 | 1 | 2 | 0.5 | NA | NA | NA | NA | 3 | 0.6667 |
| 18_374352 | NA | NA | 2 | 0 | NA | NA | 2 | 1 | 4 | 0.5 | NA | NA | 1 | 1 | NA | NA | NA | NA | 1 | 1 |
| 18_374354 | NA | NA | 2 | 1 | NA | NA | 3 | 0.6667 | 5 | 0.8 | 1 | 1 | 2 | 1 | NA | NA | NA | NA | 3 | 1 |
| 18_374355 | NA | NA | 1 | 1 | 1 | 1 | 2 | 0.5 | 4 | 0.75 | 1 | 1 | 2 | 1 | NA | NA | NA | NA | 3 | 1 |
| 18_374357 | NA | NA | 2 | 0 | 1 | 1 | 3 | 0 | 6 | 0.1667 | 1 | 0 | 2 | 0.5 | NA | NA | NA | NA | 3 | 0.3333 |
| 18_374358 | NA | NA | 2 | 1 | 1 | 0 | 1 | 0 | 4 | 0.5 | 1 | 0 | 2 | 0.5 | NA | NA | NA | NA | 3 | 0.3333 |
| 18_374358 | NA | NA | 2 | 1 | 1 | 1 | 2 | 0.5 | 5 | 0.8 | 1 | 1 | 1 | 0 | NA | NA | NA | NA | 2 | 0.5 |
| 18_374359 | NA | NA | 1 | 1 | 1 | 1 | 3 | 0.6667 | 5 | 0.8 | NA | NA | 1 | 1 | NA | NA | NA | NA | 1 | 1 |
| 18_374360 | NA | NA | 2 | 1 | 1 | 1 | 3 | 0.6667 | 6 | 0.8333 | NA | NA | 2 | 1 | NA | NA | NA | NA | 2 | 1 |
| 18_374361 | NA | NA | 1 | 1 | 1 | 1 | 3 | 0.6667 | 5 | 0.8 | 1 | 1 | 2 | 1 | NA | NA | NA | NA | 3 | 1 |
| 18_374362 | NA | NA | 2 | 1 | 1 | 1 | 3 | 1 | 6 | 1 | 1 | 1 | 2 | 1 | NA | NA | NA | NA | 3 | 1 |
| 18_374363 | NA | NA | 2 | 1 | 1 | 1 | 3 | 1 | 6 | 1 | 1 | 1 | 2 | 1 | NA | NA | NA | NA | 3 | 1 |
| 18_374366 | NA | NA | 2 | 0.5 | 1 | 1 | 3 | 0.3333 | 6 | 0.5 | 1 | 1 | 1 | 0 | NA | NA | NA | NA | 2 | 0.5 |
| 18_374429 | NA | NA | 2 | 0.5 | 1 | 1 | NA | NA | 3 | 0.6667 | NA | NA | NA | NA | 1 | 1 | NA | NA | 1 | 1 |
| 18_374564 | NA | NA | NA | NA | 1 | 1 | NA | NA | 1 | 1 | NA | NA | NA | NA | NA | NA | 1 | 1 | 1 | 1 |

| | | | | | | | | | | | | | | | | | | | | |
|---|---|---|---|---|---|---|---|---|---|---|---|---|---|---|---|---|---|---|---|---|
| 18_374755( | 1 | 1 | NA | NA | NA | NA | NA | NA | 1 | 1 | NA | NA | 1 | 1 | NA | NA | NA | NA | 1 | 1 |
| 18_374756! | 1 | 1 | NA | NA | NA | NA | NA | NA | 1 | 1 | NA | NA | 1 | 1 | NA | NA | NA | NA | 1 | 1 |
| 18_374765' | 1 | 1 | NA | NA | NA | NA | 1 | 1 | 2 | 1 | NA | NA | 2 | 1 | NA | NA | NA | NA | 2 | 1 |
| 18_374765( | 2 | 0 | NA | NA | NA | NA | 1 | 1 | 3 | 0.3333 | NA | NA | 1 | 1 | NA | NA | 1 | 1 | 2 | 1 |
| 18_374766' | 2 | 0 | NA | NA | NA | NA | 1 | 1 | 3 | 0.3333 | NA | NA | 2 | 0.5 | NA | NA | 1 | 0 | 3 | 0.3333 |
| 18_374766: | 2 | 1 | NA | NA | NA | NA | 1 | 0 | 3 | 0.6667 | NA | NA | 2 | 1 | NA | NA | 1 | 1 | 3 | 1 |
| 18_374767: | 2 | 1 | NA | NA | NA | NA | 1 | 1 | 3 | 1 | NA | NA | 1 | 1 | NA | NA | NA | NA | 1 | 1 |
| 18_374771! | 1 | 1 | NA | NA | 1 | 1 | 1 | 1 | 3 | 1 | NA | NA | 2 | 1 | NA | NA | 1 | 1 | 3 | 1 |
| 18_374772: | 2 | 0.5 | NA | NA | 2 | 1 | 1 | 1 | 5 | 0.8 | NA | NA | 2 | 0.5 | 1 | 1 | NA | NA | 3 | 0.6667 |
| 18_374773' | 2 | 0.5 | NA | NA | 1 | 1 | 1 | 0 | 4 | 0.5 | NA | NA | 1 | 1 | NA | NA | 1 | 0 | 2 | 0.5 |
| 18_374774( | 2 | 0.5 | NA | NA | 1 | 1 | 1 | 0 | 4 | 0.5 | NA | NA | 1 | 1 | 2 | 1 | 1 | 1 | 4 | 1 |
| 18_374882: | 1 | 0 | NA | NA | NA | NA | 2 | 0.5 | 3 | 0.3333 | NA | NA | NA | NA | 2 | 0.5 | NA | NA | 2 | 0.5 |
| 18_3748877 | NA | NA | NA | NA | NA | NA | 1 | 1 | 1 | 1 | NA | NA | NA | NA | 1 | 1 | NA | NA | 1 | 1 |
| 18_374894! | 1 | 0 | NA | NA | NA | NA | 2 | 1 | 3 | 0.6667 | NA | NA | NA | NA | 2 | 0.5 | NA | NA | 2 | 0.5 |
| 18_374894! | 1 | 1 | NA | NA | NA | NA | 2 | 1 | 3 | 1 | NA | NA | NA | NA | 1 | 1 | NA | NA | 1 | 1 |
| 18_374895' | 1 | 1 | NA | NA | NA | NA | 2 | 1 | 3 | 1 | NA | NA | NA | NA | 2 | 1 | NA | NA | 2 | 1 |
| 18_374897! | 1 | 1 | NA | NA | NA | NA | NA | NA | 1 | 1 | NA | NA | NA | NA | 2 | 0.5 | NA | NA | 2 | 0.5 |
| 18_374897! | NA | NA | NA | NA | NA | NA | 1 | 1 | 1 | 1 | NA | NA | NA | NA | 2 | 0.5 | NA | NA | 2 | 0.5 |
| 18_374898: | 1 | 1 | NA | NA | NA | NA | 2 | 1 | 3 | 1 | NA | NA | NA | NA | 1 | 1 | NA | NA | 1 | 1 |
| 18_374899! | 1 | 0 | NA | NA | NA | NA | 2 | 1 | 3 | 0.6667 | NA | NA | NA | NA | 2 | 0.5 | NA | NA | 2 | 0.5 |
| 18_374900! | 1 | 1 | NA | NA | NA | NA | 1 | 1 | 2 | 1 | NA | NA | NA | NA | 1 | 1 | NA | NA | 1 | 1 |
| 18_374902! | 1 | 1 | NA | NA | NA | NA | 2 | 1 | 3 | 1 | NA | NA | NA | NA | 2 | 1 | NA | NA | 2 | 1 |
| 18_374905' | 1 | 1 | NA | NA | NA | NA | 2 | 0.5 | 3 | 0.6667 | NA | NA | NA | NA | 2 | 1 | NA | NA | 2 | 1 |
| 18_374905! | 1 | 0 | NA | NA | NA | NA | 2 | 0.5 | 3 | 0.3333 | NA | NA | NA | NA | 2 | 0.5 | NA | NA | 2 | 0.5 |
| 18_374905! | 1 | 1 | NA | NA | NA | NA | 2 | 1 | 3 | 1 | NA | NA | NA | NA | 1 | 1 | NA | NA | 1 | 1 |
| 18_374906! | 1 | 1 | NA | NA | NA | NA | 2 | 0.5 | 3 | 0.6667 | NA | NA | NA | NA | 2 | 0.5 | NA | NA | 2 | 0.5 |
| 18_375343! | 2 | 1 | NA | NA | 1 | 0 | 1 | 1 | 4 | 0.75 | NA | NA | NA | NA | NA | NA | 1 | 1 | 1 | 1 |
| 18_375344: | NA | NA | NA | NA | 1 | 0 | 1 | 1 | 2 | 0.5 | 1 | 1 | NA | NA | NA | NA | NA | NA | 1 | 1 |
| 18_375344! | 2 | 0.5 | NA | NA | 1 | 1 | 1 | 1 | 4 | 0.75 | 1 | 1 | NA | NA | NA | NA | NA | NA | 1 | 1 |
| 18_375345! | 3 | 1 | NA | NA | 1 | 1 | 1 | 1 | 5 | 1 | 1 | 1 | NA | NA | NA | NA | 1 | 1 | 2 | 1 |
| 18_375347( | 3 | 0.6667 | NA | NA | 1 | 0 | 1 | 0 | 5 | 0.4 | 1 | 0 | 1 | 0 | NA | NA | 1 | 1 | 3 | 0.3333 |
| 18_375347' | 3 | 0.6667 | NA | NA | NA | NA | 1 | 0 | 4 | 0.5 | 1 | 1 | NA | NA | NA | NA | NA | NA | 1 | 1 |
| 18_375348! | 3 | 0.6667 | NA | NA | 1 | 1 | 1 | 1 | 5 | 0.8 | 1 | 0 | 2 | 0.5 | NA | NA | NA | NA | 3 | 0.3333 |
| 18_375350' | 3 | 0 | NA | NA | 1 | 0 | 1 | 1 | 5 | 0.2 | 1 | 1 | 2 | 0 | NA | NA | 1 | 0 | 4 | 0.25 |
| 18_375351: | 3 | 1 | NA | NA | 1 | 1 | 1 | 0 | 5 | 0.8 | 1 | 0 | 2 | 0 | NA | NA | 1 | 1 | 4 | 0.25 |
| 18_375353· | 2 | 0.5 | NA | NA | 1 | 1 | 1 | 1 | 4 | 0.75 | NA | NA | 1 | 1 | NA | NA | 1 | 1 | 2 | 1 |
| 18_375355· | 2 | 1 | NA | NA | 2 | 1 | 1 | 1 | 5 | 1 | 1 | 1 | NA | NA | NA | NA | 1 | 0 | 2 | 0.5 |
| 18_375355· | 3 | 0 | NA | NA | 2 | 0.5 | 1 | 1 | 6 | 0.3333 | 1 | 0 | 1 | 1 | NA | NA | 1 | 1 | 3 | 0.6667 |
| 18_375356( | 3 | 1 | NA | NA | 2 | 0.5 | 1 | 1 | 6 | 0.8333 | 1 | 1 | 1 | 1 | NA | NA | 1 | 1 | 3 | 1 |
| 18_375356: | 2 | 1 | NA | NA | 2 | 1 | 1 | 1 | 5 | 1 | 1 | 1 | 1 | 1 | NA | NA | 1 | 1 | 3 | 1 |
| 18_375357: | 2 | 0 | NA | NA | 2 | 0.5 | 1 | 0 | 5 | 0.2 | 1 | 0 | 2 | 0 | 1 | 1 | 1 | 0 | 5 | 0.2 |
| 18_375357: | 3 | 0.3333 | NA | NA | 2 | 0 | 1 | 1 | 6 | 0.3333 | 1 | 0 | 2 | 0.5 | 1 | 0 | 1 | 1 | 5 | 0.4 |
| 18_375358! | 3 | 0.6667 | NA | NA | 2 | 0.5 | 1 | 1 | 6 | 0.6667 | 1 | 1 | 2 | 0.5 | 1 | 1 | 1 | 0 | 5 | 0.6 |
| 18_375360· | 3 | 1 | NA | NA | 2 | 1 | 1 | 1 | 6 | 1 | 1 | 1 | 1 | 1 | 1 | 1 | NA | NA | 3 | 1 |
| 18_375362! | 2 | 1 | 1 | 1 | 1 | 1 | 1 | 1 | 5 | 1 | NA | NA | 2 | 1 | NA | NA | 1 | 1 | 3 | 1 |
| 18_375363( | 1 | 1 | 1 | 1 | 2 | 1 | 1 | 1 | 5 | 1 | 1 | 1 | 1 | 1 | NA | NA | NA | NA | 2 | 1 |
| 18_375363· | 3 | 0.3333 | 1 | 0 | 2 | 0 | 1 | 1 | 7 | 0.2857 | 1 | 0 | 1 | 0 | 2 | 0.5 | 1 | 1 | 5 | 0.4 |
| 18_375363! | 3 | 1 | 1 | 1 | 2 | 1 | 1 | 1 | 7 | 1 | 1 | 1 | 2 | 1 | 2 | 1 | 1 | 1 | 6 | 1 |
| 18_375367: | 3 | 0.3333 | 2 | 0.5 | 2 | 0 | 1 | 0 | 8 | 0.25 | 1 | 0 | 2 | 0 | 1 | 1 | 2 | 0.5 | 6 | 0.3333 |
| 18_375460( | NA | NA | NA | NA | 1 | 1 | NA | NA | 1 | 1 | 1 | 1 | NA | NA | NA | NA | NA | NA | 1 | 1 |
| 18_375461' | NA | NA | NA | NA | 1 | 1 | NA | NA | 1 | 1 | 1 | 1 | NA | NA | NA | NA | NA | NA | 1 | 1 |
| 18_375461· | NA | NA | NA | NA | 1 | 1 | NA | NA | 1 | 1 | 1 | 1 | NA | NA | NA | NA | NA | NA | 1 | 1 |
| 18_375463! | NA | NA | NA | NA | 1 | 1 | NA | NA | 1 | 1 | 1 | 1 | NA | NA | NA | NA | NA | NA | 1 | 1 |
| 18_375466: | NA | NA | 1 | 1 | NA | NA | NA | NA | 1 | 1 | 1 | 1 | NA | NA | NA | NA | NA | NA | 1 | 1 |
| 18_375467( | NA | NA | 1 | 1 | NA | NA | NA | NA | 1 | 1 | 1 | 1 | NA | NA | NA | NA | NA | NA | 1 | 1 |
| 18_375472( | NA | NA | 1 | 1 | 1 | 1 | NA | NA | 2 | 1 | 1 | 1 | NA | NA | NA | NA | NA | NA | 1 | 1 |
| 18_375484! | NA | NA | NA | NA | 1 | 1 | NA | NA | 1 | 1 | NA | NA | 1 | 1 | NA | NA | NA | NA | 1 | 1 |
| 18_375560! | NA | NA | 1 | 1 | NA | NA | 1 | 1 | 2 | 1 | NA | NA | NA | NA | 1 | 1 | NA | NA | 1 | 1 |
| 18_375563: | NA | NA | 1 | 1 | NA | NA | 1 | 0 | 2 | 0.5 | NA | NA | NA | NA | 1 | 1 | NA | NA | 1 | 1 |
| 18_375569· | NA | NA | 1 | 1 | NA | NA | 1 | 0 | 2 | 0.5 | NA | NA | NA | NA | 1 | 1 | NA | NA | 1 | 1 |
| 18_375569! | NA | NA | 1 | 1 | NA | NA | 1 | 0 | 2 | 0.5 | NA | NA | NA | NA | 1 | 1 | NA | NA | 1 | 1 |
| 18_375581: | NA | NA | 1 | 1 | NA | NA | 1 | 0 | 2 | 0.5 | NA | NA | 1 | 1 | NA | NA | NA | NA | 1 | 1 |
| 18_375688: | 1 | 0 | NA | NA | NA | NA | 1 | 1 | 2 | 0.5 | 2 | 1 | NA | NA | NA | NA | NA | NA | 2 | 1 |
| 18_375690: | 1 | 1 | NA | NA | NA | NA | 1 | 1 | 2 | 1 | 2 | 1 | NA | NA | NA | NA | NA | NA | 2 | 1 |
| 18_375692! | 1 | 1 | NA | NA | NA | NA | 1 | 0 | 2 | 0.5 | 2 | 1 | NA | NA | NA | NA | 1 | 0 | 3 | 0.6667 |

| | | | | | | | | | | | | | | | | | | | | |
|---|---|---|---|---|---|---|---|---|---|---|---|---|---|---|---|---|---|---|---|---|
| 18_375693 | 1 | 1 | NA | NA | NA | NA | NA | NA | 1 | 1 | 1 | 1 | NA | NA | NA | NA | 1 | 1 | 2 | 1 |
| 18_375695 | 1 | 1 | NA | NA | NA | NA | 1 | 0 | 2 | 0.5 | 2 | 0.5 | NA | NA | NA | NA | 1 | 0 | 3 | 0.3333 |
| 18_375696 | 1 | 1 | NA | NA | NA | NA | 1 | 1 | 2 | 1 | 2 | 1 | NA | NA | NA | NA | 1 | 1 | 3 | 1 |
| 18_375699 | 1 | 1 | NA | NA | NA | NA | 1 | 1 | 2 | 1 | 2 | 1 | NA | NA | NA | NA | 1 | 1 | 3 | 1 |
| 18_375699 | 1 | 1 | NA | NA | NA | NA | NA | NA | 1 | 1 | 2 | 1 | NA | NA | NA | NA | 1 | 0 | 3 | 0.6667 |
| 18_375699 | 1 | 0 | NA | NA | NA | NA | 1 | 1 | 2 | 0.5 | 2 | 0.5 | NA | NA | NA | NA | 1 | 0 | 3 | 0.3333 |
| 18_375700 | 1 | 1 | NA | NA | NA | NA | 1 | 0 | 2 | 0.5 | 2 | 0.5 | NA | NA | NA | NA | 1 | 0 | 3 | 0.3333 |
| 18_375702 | NA | NA | NA | NA | NA | NA | 1 | 1 | 1 | 1 | 1 | 1 | NA | NA | NA | NA | NA | NA | 1 | 1 |
| 18_375703 | 2 | 1 | NA | NA | NA | NA | 1 | 1 | 3 | 1 | 1 | 1 | NA | NA | NA | NA | 1 | 1 | 2 | 1 |
| 18_375704 | 2 | 1 | NA | NA | NA | NA | 1 | 0 | 3 | 0.6667 | 2 | 1 | NA | NA | NA | NA | NA | NA | 2 | 1 |
| 18_375704 | 3 | 0.3333 | NA | NA | NA | NA | 1 | 1 | 4 | 0.5 | 2 | 0.5 | NA | NA | NA | NA | 1 | 0 | 3 | 0.3333 |
| 18_375704 | 3 | 1 | NA | NA | NA | NA | 1 | 1 | 4 | 1 | 2 | 1 | NA | NA | NA | NA | 1 | 1 | 3 | 1 |
| 18_375707 | 3 | 1 | NA | NA | NA | NA | NA | NA | 3 | 1 | 2 | 1 | NA | NA | NA | NA | 1 | 1 | 3 | 1 |
| 18_375709 | 3 | 0.3333 | NA | NA | NA | NA | NA | NA | 3 | 0.3333 | 2 | 0.5 | NA | NA | NA | NA | 1 | 0 | 3 | 0.3333 |
| 18_375710 | 3 | 1 | NA | NA | NA | NA | 1 | 1 | 4 | 1 | 2 | 1 | NA | NA | NA | NA | 1 | 1 | 3 | 1 |
| 18_375712 | 3 | 0.6667 | NA | NA | NA | NA | 1 | 0 | 4 | 0.5 | 1 | 0 | NA | NA | NA | NA | 1 | 1 | 2 | 0.5 |
| 18_375717 | 2 | 0 | NA | NA | 1 | 1 | 1 | 0 | 4 | 0.25 | 3 | 0.6667 | NA | NA | NA | NA | 1 | 0 | 4 | 0.5 |
| 18_375718 | 2 | 0 | NA | NA | 1 | 1 | 1 | 1 | 4 | 0.5 | NA | NA | NA | NA | NA | NA | 1 | 1 | 1 | 1 |
| 18_375736 | NA | NA | NA | NA | 1 | 1 | NA | NA | 1 | 1 | NA | NA | NA | NA | NA | NA | 1 | 1 | 1 | 1 |
| 18_375759 | NA | NA | NA | NA | NA | NA | 1 | 1 | 1 | 1 | 1 | 1 | NA | NA | 1 | 0 | NA | NA | 2 | 0.5 |
| 18_375760 | NA | NA | 1 | 1 | NA | NA | 1 | 1 | 2 | 1 | 1 | 1 | NA | NA | 1 | 0 | NA | NA | 2 | 0.5 |
| 18_375762 | 1 | 0 | 1 | 1 | NA | NA | 1 | 0 | 3 | 0.3333 | 1 | 0 | NA | NA | 2 | 0.5 | NA | NA | 3 | 0.3333 |
| 18_375764 | 1 | 0 | 1 | 1 | NA | NA | 1 | 0 | 3 | 0.3333 | 1 | 0 | NA | NA | 2 | 1 | NA | NA | 3 | 0.6667 |
| 18_375764 | 1 | 1 | 1 | 1 | NA | NA | 1 | 1 | 3 | 1 | 1 | 1 | NA | NA | 1 | 1 | NA | NA | 2 | 1 |
| 18_375767 | 1 | 1 | 1 | 1 | NA | NA | 1 | 1 | 3 | 1 | 1 | 1 | NA | NA | 1 | 1 | NA | NA | 2 | 1 |
| 18_375770 | 1 | 0 | 1 | 1 | NA | NA | 1 | 1 | 3 | 0.6667 | NA | NA | NA | NA | 2 | 0.5 | NA | NA | 2 | 0.5 |
| 18_375771 | 1 | 1 | 1 | 1 | NA | NA | 1 | 1 | 3 | 1 | 1 | 1 | NA | NA | 2 | 1 | NA | NA | 3 | 1 |
| 18_375774 | 1 | 1 | 1 | 1 | NA | NA | 1 | 1 | 3 | 1 | 1 | 1 | NA | NA | 3 | 1 | NA | NA | 4 | 1 |
| 18_375775 | 1 | 1 | 1 | 1 | NA | NA | NA | NA | 2 | 1 | 1 | 1 | NA | NA | 1 | 1 | NA | NA | 2 | 1 |
| 18_375776 | 2 | 1 | 1 | 1 | NA | NA | 1 | 1 | 4 | 1 | 1 | 1 | NA | NA | 3 | 1 | NA | NA | 4 | 1 |
| 18_375778 | 2 | 1 | NA | NA | NA | NA | 1 | 1 | 3 | 1 | 1 | 1 | NA | NA | 3 | 1 | NA | NA | 4 | 1 |
| 18_375778 | 2 | 0.5 | 1 | 0 | NA | NA | 1 | 1 | 4 | 0.5 | 1 | 1 | NA | NA | 3 | 0.3333 | NA | NA | 4 | 0.5 |
| 18_375779 | 2 | 1 | 1 | 1 | NA | NA | 1 | 1 | 4 | 1 | 1 | 1 | NA | NA | 3 | 1 | NA | NA | 4 | 1 |
| 18_375788 | 2 | 0 | 2 | 0.5 | NA | NA | NA | NA | 4 | 0.25 | 1 | 1 | 1 | 0 | 3 | 0 | 1 | 0 | 6 | 0.1667 |
| 18_375791 | 3 | 0 | 2 | 0.5 | NA | NA | 1 | 1 | 6 | 0.3333 | 1 | 0 | 1 | 0 | 3 | 0.3333 | 1 | 0 | 6 | 0.1667 |
| 18_375815 | 1 | 1 | NA | NA | NA | NA | NA | NA | 1 | 1 | NA | NA | 1 | 1 | NA | NA | NA | NA | 1 | 1 |
| 18_375818 | 1 | 1 | NA | NA | NA | NA | NA | NA | 1 | 1 | NA | NA | 1 | 1 | NA | NA | NA | NA | 1 | 1 |
| 18_375824 | 1 | 1 | 2 | 1 | NA | NA | NA | NA | 3 | 1 | NA | NA | 1 | 1 | NA | NA | 1 | 1 | 2 | 1 |
| 18_375826 | 1 | 1 | 2 | 0.5 | NA | NA | NA | NA | 3 | 0.6667 | NA | NA | 1 | 1 | NA | NA | 1 | 0 | 2 | 0.5 |
| 18_375827 | 1 | 1 | 1 | 1 | NA | NA | NA | NA | 2 | 1 | NA | NA | 1 | 1 | NA | NA | 1 | 1 | 2 | 1 |
| 18_375828 | 1 | 1 | 2 | 1 | NA | NA | 1 | 1 | 4 | 1 | NA | NA | 1 | 1 | NA | NA | NA | NA | 1 | 1 |
| 18_375830 | 1 | 1 | 2 | 1 | NA | NA | 1 | 1 | 4 | 1 | NA | NA | 1 | 1 | NA | NA | 1 | 1 | 2 | 1 |
| 18_375830 | 1 | 1 | 3 | 1 | NA | NA | 1 | 1 | 5 | 1 | NA | NA | 1 | 1 | NA | NA | 1 | 1 | 2 | 1 |
| 18_375837 | 1 | 1 | 1 | 1 | NA | NA | 1 | 1 | 3 | 1 | NA | NA | 2 | 0.5 | NA | NA | NA | NA | 2 | 0.5 |
| 18_376111 | 1 | 0 | NA | NA | NA | NA | 1 | 1 | 2 | 0.5 | NA | NA | NA | NA | NA | NA | 1 | 1 | 1 | 1 |
| 18_376113 | 1 | 1 | NA | NA | NA | NA | 1 | 0 | 2 | 0.5 | NA | NA | NA | NA | NA | NA | 1 | 1 | 1 | 1 |
| 18_376113 | NA | NA | NA | NA | NA | NA | 1 | 1 | 1 | 1 | NA | NA | NA | NA | NA | NA | 1 | 1 | 1 | 1 |
| 18_376116 | NA | NA | NA | NA | NA | NA | 1 | 1 | 1 | 1 | NA | NA | NA | NA | NA | NA | 1 | 1 | 1 | 1 |
| 18_376118 | NA | NA | NA | NA | NA | NA | 1 | 1 | 1 | 1 | NA | NA | NA | NA | NA | NA | 1 | 1 | 1 | 1 |
| 18_376118 | 1 | 1 | NA | NA | NA | NA | 1 | 1 | 2 | 1 | NA | NA | NA | NA | NA | NA | 1 | 1 | 1 | 1 |
| 18_376119 | 1 | 0 | NA | NA | NA | NA | 1 | 1 | 2 | 0.5 | NA | NA | NA | NA | NA | NA | 1 | 1 | 1 | 1 |
| 18_376124 | 1 | 1 | NA | NA | NA | NA | NA | NA | 1 | 1 | NA | NA | NA | NA | NA | NA | 1 | 1 | 1 | 1 |
| 18_376126 | 1 | 1 | NA | NA | NA | NA | 1 | 1 | 2 | 1 | NA | NA | NA | NA | NA | NA | 1 | 1 | 1 | 1 |
| 18_376127 | 1 | 1 | NA | NA | NA | NA | 1 | 0 | 2 | 0.5 | NA | NA | NA | NA | NA | NA | 1 | 1 | 1 | 1 |
| 18_376128 | NA | NA | NA | NA | NA | NA | 1 | 1 | 1 | 1 | NA | NA | NA | NA | NA | NA | 1 | 1 | 1 | 1 |
| 18_376129 | 1 | 1 | NA | NA | NA | NA | 1 | 1 | 2 | 1 | NA | NA | NA | NA | NA | NA | 1 | 1 | 1 | 1 |
| 18_376130 | NA | NA | NA | NA | NA | NA | 1 | 1 | 1 | 1 | NA | NA | NA | NA | NA | NA | 1 | 1 | 1 | 1 |
| 18_376132 | NA | NA | NA | NA | NA | NA | 1 | 1 | 1 | 1 | NA | NA | NA | NA | NA | NA | 1 | 1 | 1 | 1 |
| 18_376133 | NA | NA | NA | NA | NA | NA | 1 | 1 | 1 | 1 | NA | NA | NA | NA | NA | NA | 1 | 1 | 1 | 1 |
| 18_376133 | NA | NA | NA | NA | NA | NA | 1 | 1 | 1 | 1 | NA | NA | NA | NA | NA | NA | 2 | 1 | 2 | 1 |
| 18_376134 | NA | NA | NA | NA | NA | NA | 1 | 1 | 1 | 1 | NA | NA | NA | NA | NA | NA | 2 | 0.5 | 2 | 0.5 |
| 18_376181 | NA | NA | 3 | 0.6667 | NA | NA | NA | NA | 3 | 0.6667 | NA | NA | NA | NA | NA | NA | 3 | 0.6667 | 3 | 0.6667 |
| 18_376183 | NA | NA | 3 | 0.3333 | NA | NA | NA | NA | 3 | 0.3333 | NA | NA | NA | NA | 1 | 0 | 3 | 0.6667 | 4 | 0.5 |
| 18_376184 | NA | NA | 2 | 1 | NA | NA | NA | NA | 2 | 1 | NA | NA | 1 | 0 | 1 | 0 | 2 | 0.5 | 4 | 0.25 |
| 18_376185 | NA | NA | 3 | 1 | NA | NA | NA | NA | 3 | 1 | NA | NA | NA | NA | 1 | 1 | 3 | 1 | 4 | 1 |

| | | | | | | | | | | | | | | | | | | | | |
|---|---|---|---|---|---|---|---|---|---|---|---|---|---|---|---|---|---|---|---|---|
| 18_376185 | NA | NA | 2 | 0.5 | NA | NA | NA | NA | 2 | 0.5 | NA | NA | 1 | 1 | 1 | 1 | 3 | 1 | 5 | 1 |
| 18_376186 | NA | NA | 3 | 1 | NA | NA | NA | NA | 3 | 1 | NA | NA | 2 | 1 | 1 | 1 | 3 | 1 | 6 | 1 |
| 18_376187 | NA | NA | 3 | 0.3333 | NA | NA | NA | NA | 3 | 0.3333 | NA | NA | 2 | 0 | NA | NA | 3 | 0.3333 | 5 | 0.2 |
| 18_376188 | NA | NA | 3 | 0.6667 | NA | NA | NA | NA | 3 | 0.6667 | NA | NA | 1 | 1 | 1 | 0 | 3 | 0.6667 | 5 | 0.6 |
| 18_376190 | NA | NA | 3 | 1 | NA | NA | NA | NA | 3 | 1 | NA | NA | 1 | 1 | 1 | 0 | 2 | 1 | 4 | 0.75 |
| 18_376191 | 1 | 0 | 3 | 0.6667 | NA | NA | NA | NA | 4 | 0.5 | NA | NA | 2 | 0.5 | 1 | 0 | 3 | 0 | 6 | 0.1667 |
| 18_376194 | NA | NA | 3 | 1 | NA | NA | NA | NA | 3 | 1 | NA | NA | 2 | 0 | 1 | 1 | 1 | 1 | 4 | 0.5 |
| 18_376196 | 1 | 1 | 3 | 1 | NA | NA | NA | NA | 4 | 1 | NA | NA | 2 | 0.5 | NA | NA | 3 | 1 | 5 | 0.8 |
| 18_376197 | 1 | 1 | 3 | 1 | NA | NA | NA | NA | 4 | 1 | 1 | 1 | 2 | 0.5 | NA | NA | 3 | 1 | 6 | 0.8333 |
| 18_376197 | 1 | 1 | 2 | 1 | NA | NA | NA | NA | 3 | 1 | NA | NA | 1 | 1 | 1 | 1 | 3 | 1 | 5 | 1 |
| 18_376198 | 1 | 0 | 2 | 0.5 | NA | NA | NA | NA | 3 | 0.3333 | 1 | 0 | NA | NA | 1 | 0 | 3 | 0.6667 | 5 | 0.4 |
| 18_376199 | 1 | 1 | 1 | 1 | NA | NA | NA | NA | 2 | 1 | NA | NA | 2 | 1 | 1 | 0 | NA | NA | 3 | 0.6667 |
| 18_376200 | 1 | 1 | 1 | 1 | NA | NA | NA | NA | 2 | 1 | NA | NA | 2 | 1 | 1 | 0 | 2 | 1 | 5 | 0.8 |
| 18_376201 | NA | NA | 3 | 1 | NA | NA | NA | NA | 3 | 1 | 1 | 1 | 1 | 1 | 1 | 1 | 3 | 1 | 6 | 1 |
| 18_376207 | 1 | 0 | 3 | 0.3333 | 1 | 0 | NA | NA | 5 | 0.2 | 3 | 0 | 2 | 0 | 1 | 0 | 3 | 0.3333 | 9 | 0.1111 |
| 18_376233 | NA | NA | NA | NA | 1 | 1 | NA | NA | 1 | 1 | 1 | 1 | NA | NA | NA | NA | NA | NA | 1 | 1 |
| 18_376233 | NA | NA | NA | NA | 1 | 0 | 1 | 1 | 2 | 0.5 | 1 | 1 | NA | NA | NA | NA | NA | NA | 1 | 1 |
| 18_376234 | NA | NA | NA | NA | 1 | 1 | NA | NA | 1 | 1 | 1 | 1 | NA | NA | NA | NA | NA | NA | 1 | 1 |
| 18_376237 | NA | NA | NA | NA | 1 | 1 | NA | NA | 1 | 1 | 1 | 1 | NA | NA | NA | NA | NA | NA | 1 | 1 |
| 18_376238 | NA | NA | NA | NA | 1 | 1 | NA | NA | 1 | 1 | 1 | 1 | NA | NA | NA | NA | NA | NA | 1 | 1 |
| 18_376244 | 1 | 1 | NA | NA | NA | NA | 1 | 1 | 2 | 1 | 1 | 1 | NA | NA | NA | NA | NA | NA | 1 | 1 |
| 18_376246 | 1 | 1 | NA | NA | 1 | 1 | 1 | 1 | 3 | 1 | 1 | 1 | NA | NA | NA | NA | NA | NA | 1 | 1 |
| 18_376248 | NA | NA | NA | NA | 1 | 1 | NA | NA | 1 | 1 | 1 | 1 | NA | NA | NA | NA | NA | NA | 1 | 1 |
| 18_376248 | 1 | 0 | NA | NA | 1 | 0 | 1 | 1 | 3 | 0.3333 | 1 | 1 | NA | NA | NA | NA | NA | NA | 1 | 1 |
| 18_376373 | NA | NA | 1 | 1 | 2 | 1 | 1 | 0 | 4 | 0.75 | NA | NA | NA | NA | NA | NA | 1 | 1 | 1 | 1 |
| 18_376374 | NA | NA | 1 | 0 | 2 | 1 | 1 | 0 | 4 | 0.5 | NA | NA | NA | NA | NA | NA | 1 | 1 | 1 | 1 |
| 18_376374 | NA | NA | 1 | 1 | 1 | 1 | 1 | 0 | 3 | 0.6667 | NA | NA | NA | NA | NA | NA | 1 | 1 | 1 | 1 |
| 18_376379 | NA | NA | 1 | 0 | 2 | 0.5 | 1 | 0 | 4 | 0.25 | NA | NA | NA | NA | NA | NA | 1 | 1 | 1 | 1 |
| 18_376381 | NA | NA | 1 | 0 | 2 | 1 | 1 | 0 | 4 | 0.5 | NA | NA | NA | NA | 1 | 1 | 1 | 0 | 2 | 0.5 |
| 18_376381 | NA | NA | 1 | 0 | 2 | 0.5 | 1 | 1 | 4 | 0.5 | NA | NA | NA | NA | 1 | 1 | 1 | 0 | 2 | 0.5 |
| 18_376381 | NA | NA | NA | NA | NA | NA | 1 | 1 | 1 | 1 | NA | NA | NA | NA | 1 | 1 | NA | NA | 1 | 1 |
| 18_376382 | NA | NA | 1 | 0 | 2 | 0 | 1 | 1 | 4 | 0.25 | NA | NA | NA | NA | 1 | 1 | NA | NA | 1 | 1 |
| 18_376385 | NA | NA | 1 | 1 | 2 | 0 | 1 | 1 | 4 | 0.5 | NA | NA | NA | NA | 1 | 1 | 1 | 0 | 2 | 0.5 |
| 18_376385 | NA | NA | 2 | 0.5 | 2 | 0.5 | 1 | 0 | 5 | 0.4 | NA | NA | NA | NA | 1 | 0 | 1 | 1 | 2 | 0.5 |
| 18_376389 | NA | NA | 2 | 1 | 1 | 1 | NA | NA | 3 | 1 | NA | NA | NA | NA | 2 | 1 | NA | NA | 2 | 1 |
| 18_376390 | NA | NA | 2 | 1 | 2 | 0.5 | 1 | 1 | 5 | 0.8 | NA | NA | NA | NA | 1 | 1 | 1 | 1 | 2 | 1 |
| 18_376391 | NA | NA | 2 | 0.5 | 1 | 1 | 1 | 0 | 4 | 0.5 | NA | NA | NA | NA | 2 | 0.5 | 1 | 1 | 3 | 0.6667 |
| 18_376391 | NA | NA | 2 | 0 | 2 | 0.5 | NA | NA | 4 | 0.25 | NA | NA | NA | NA | 2 | 0.5 | 1 | 0 | 3 | 0.3333 |
| 18_376394 | NA | NA | 2 | 1 | 2 | 1 | 1 | 1 | 5 | 1 | NA | NA | NA | NA | 2 | 1 | NA | NA | 2 | 1 |
| 18_376396 | NA | NA | 2 | 1 | 2 | 1 | 1 | 1 | 5 | 1 | NA | NA | NA | NA | 2 | 1 | NA | NA | 2 | 1 |
| 18_376397 | NA | NA | 2 | 0.5 | 1 | 1 | NA | NA | 3 | 0.6667 | NA | NA | NA | NA | 2 | 1 | 2 | 0.5 | 4 | 0.75 |
| 18_376397 | NA | NA | 1 | 0 | 2 | 1 | 1 | 1 | 4 | 0.75 | NA | NA | NA | NA | 1 | 1 | 1 | 0 | 2 | 0.5 |
| 18_376398 | NA | NA | 2 | 1 | 1 | 1 | 1 | 1 | 4 | 1 | NA | NA | NA | NA | 3 | 1 | 1 | 1 | 4 | 1 |
| 18_376468 | NA | NA | NA | NA | 1 | 1 | NA | NA | 1 | 1 | NA | NA | NA | NA | 1 | 1 | NA | NA | 1 | 1 |
| 18_376471 | NA | NA | NA | NA | 1 | 1 | NA | NA | 1 | 1 | NA | NA | NA | NA | 1 | 1 | NA | NA | 1 | 1 |
| 18_376473 | NA | NA | NA | NA | 1 | 1 | NA | NA | 1 | 1 | NA | NA | NA | NA | 1 | 1 | NA | NA | 1 | 1 |
| 18_376476 | NA | NA | NA | NA | 1 | 1 | NA | NA | 1 | 1 | NA | NA | NA | NA | 1 | 1 | NA | NA | 1 | 1 |
| 18_376479 | NA | NA | NA | NA | 1 | 1 | NA | NA | 1 | 1 | NA | NA | NA | NA | 1 | 1 | NA | NA | 1 | 1 |
| 18_376484 | NA | NA | NA | NA | 1 | 1 | NA | NA | 1 | 1 | NA | NA | NA | NA | 1 | 1 | NA | NA | 1 | 1 |
| 18_376484 | NA | NA | NA | NA | 1 | 1 | NA | NA | 1 | 1 | NA | NA | NA | NA | 1 | 1 | NA | NA | 1 | 1 |
| 18_376488 | NA | NA | NA | NA | 1 | 1 | NA | NA | 1 | 1 | NA | NA | NA | NA | 1 | 1 | NA | NA | 1 | 1 |
| 18_376490 | NA | NA | NA | NA | 1 | 1 | NA | NA | 1 | 1 | NA | NA | NA | NA | 1 | 1 | NA | NA | 1 | 1 |
| 18_376490 | NA | NA | NA | NA | 1 | 1 | NA | NA | 1 | 1 | NA | NA | NA | NA | 1 | 1 | NA | NA | 1 | 1 |
| 18_376523 | 1 | 1 | 1 | 1 | 1 | 1 | 1 | 1 | 4 | 1 | 1 | 1 | NA | NA | NA | NA | NA | NA | 1 | 1 |
| 18_376524 | 1 | 1 | 1 | 1 | 1 | 1 | 1 | 1 | 4 | 1 | 1 | 1 | NA | NA | NA | NA | NA | NA | 1 | 1 |
| 18_376526 | 1 | 1 | 1 | 1 | 1 | 1 | 1 | 1 | 4 | 1 | 1 | 1 | NA | NA | NA | NA | NA | NA | 1 | 1 |
| 18_376527 | NA | NA | NA | NA | NA | NA | 1 | 1 | 1 | 1 | 1 | 1 | NA | NA | NA | NA | NA | NA | 1 | 1 |
| 18_376527 | 1 | 0 | 1 | 0 | 1 | 1 | 1 | 1 | 4 | 0.5 | 1 | 1 | NA | NA | NA | NA | NA | NA | 1 | 1 |
| 18_376529 | 1 | 1 | 1 | 1 | 1 | 1 | 1 | 1 | 4 | 1 | 1 | 1 | NA | NA | NA | NA | NA | NA | 1 | 1 |
| 18_376529 | 1 | 1 | NA | NA | 1 | 0 | 1 | 1 | 3 | 0.6667 | 1 | 1 | NA | NA | NA | NA | NA | NA | 1 | 1 |
| 18_376530 | 1 | 1 | 1 | 1 | 1 | 1 | 1 | 1 | 4 | 1 | 1 | 1 | NA | NA | 1 | 1 | NA | NA | 2 | 1 |
| 18_376534 | 1 | 1 | 1 | 1 | 1 | 1 | 1 | 1 | 4 | 1 | 1 | 1 | NA | NA | 1 | 1 | NA | NA | 2 | 1 |
| 18_376537 | 1 | 1 | 2 | 0.5 | 1 | 1 | 1 | 0 | 5 | 0.6 | 1 | 1 | NA | NA | 1 | 1 | NA | NA | 2 | 1 |
| 18_376570 | 1 | 1 | 2 | 0.5 | NA | NA | NA | NA | 3 | 0.6667 | 1 | 1 | NA | NA | NA | NA | NA | NA | 1 | 1 |
| 18_377752 | 3 | 1 | NA | NA | NA | NA | 1 | 1 | 4 | 1 | 3 | 1 | 1 | 1 | 1 | 1 | 2 | 1 | 7 | 1 |

| | | | | | | | | | | | | | | | | | | | | |
|---|---|---|---|---|---|---|---|---|---|---|---|---|---|---|---|---|---|---|---|---|
| 18_3777530 | 1 | 1 | 1 | 1 | NA | NA | 1 | 1 | 3 | 1 | 3 | 1 | 1 | 1 | 1 | 0 | 2 | 0 | 7 | 0.5714 |
| 18_3777540 | 3 | 1 | 1 | 1 | NA | NA | 1 | 1 | 5 | 1 | 3 | 1 | NA | NA | 1 | 1 | 1 | 1 | 5 | 1 |
| 18_3777553 | 2 | 0.5 | 1 | 0 | NA | NA | 1 | 0 | 4 | 0.25 | 3 | 1 | NA | NA | 1 | 1 | 1 | 1 | 5 | 1 |
| 18_3777644 | 1 | 1 | NA | NA | NA | NA | NA | NA | 1 | 1 | 2 | 0.5 | 1 | 1 | NA | NA | 1 | 0 | 4 | 0.5 |
| 18_3779567 | 1 | 1 | NA | NA | 1 | 1 | NA | NA | 2 | 1 | NA | NA | 1 | 1 | 1 | 0 | NA | NA | 2 | 0.5 |
| 18_3779597 | 1 | 1 | NA | NA | 1 | 1 | NA | NA | 2 | 1 | NA | NA | 1 | 1 | 1 | 1 | NA | NA | 2 | 1 |
| 18_3779622 | 1 | 1 | NA | NA | 1 | 0 | NA | NA | 2 | 0.5 | NA | NA | 1 | 1 | NA | NA | NA | NA | 1 | 1 |
| 18_3779636 | 1 | 1 | 1 | 1 | 1 | 1 | NA | NA | 3 | 1 | NA | NA | 1 | 1 | 1 | 1 | NA | NA | 2 | 1 |
| 18_3779645 | 1 | 1 | 1 | 1 | 1 | 0 | NA | NA | 3 | 0.6667 | NA | NA | NA | NA | 1 | 1 | NA | NA | 1 | 1 |
| 18_3779655 | 1 | 1 | 1 | 0 | 1 | 0 | NA | NA | 3 | 0.3333 | NA | NA | 1 | 0 | 1 | 1 | NA | NA | 2 | 0.5 |
| 18_3779666 | 1 | 1 | 1 | 1 | NA | NA | NA | NA | 2 | 1 | NA | NA | 1 | 1 | 1 | 0 | NA | NA | 2 | 0.5 |
| 18_3779678 | 1 | 1 | 1 | 1 | 1 | 0 | NA | NA | 3 | 0.6667 | NA | NA | 1 | 0 | 1 | 1 | NA | NA | 2 | 0.5 |
| 18_3779685 | 1 | 1 | 1 | 1 | 1 | 1 | NA | NA | 3 | 1 | NA | NA | 1 | 1 | 1 | 0 | NA | NA | 2 | 0.5 |
| 18_3779694 | 1 | 1 | 1 | 1 | 1 | 1 | NA | NA | 3 | 1 | NA | NA | 1 | 0 | 1 | 1 | NA | NA | 2 | 0.5 |
| 18_3779719 | 1 | 1 | 1 | 1 | 2 | 1 | NA | NA | 4 | 1 | NA | NA | 1 | 1 | 1 | 1 | NA | NA | 2 | 1 |
| 18_3779725 | 2 | 0.5 | 1 | 0 | 2 | 0 | NA | NA | 5 | 0.2 | NA | NA | 1 | 1 | NA | NA | NA | NA | 1 | 1 |
| 18_3780247 | NA | NA | NA | NA | 1 | 1 | 2 | 1 | 3 | 1 | NA | NA | 1 | 1 | 1 | 1 | 1 | 1 | 3 | 1 |
| 18_3780250 | 1 | 0 | NA | NA | 1 | 0 | 2 | 1 | 4 | 0.5 | NA | NA | 2 | 0.5 | 1 | 0 | 1 | 1 | 4 | 0.5 |
| 18_3780257 | NA | NA | NA | NA | 1 | 1 | 2 | 1 | 3 | 1 | NA | NA | 2 | 0.5 | 1 | 1 | 2 | 0.5 | 5 | 0.6 |
| 18_3780270 | 1 | 1 | NA | NA | 2 | 0 | 2 | 1 | 5 | 0.6 | NA | NA | 1 | 0 | 1 | 1 | 1 | 1 | 3 | 0.6667 |
| 18_3780298 | 1 | 1 | NA | NA | 2 | 1 | 2 | 1 | 5 | 1 | NA | NA | 2 | 1 | 1 | 1 | 1 | 1 | 4 | 1 |
| 18_3780328 | 1 | 0 | NA | NA | 1 | 1 | 2 | 1 | 4 | 0.75 | NA | NA | 2 | 1 | 2 | 1 | 2 | 1 | 6 | 1 |
| 18_3780335 | 1 | 1 | NA | NA | 2 | 1 | 2 | 1 | 5 | 1 | NA | NA | 3 | 0.6667 | 1 | 1 | 2 | 1 | 6 | 0.8333 |
| 18_3780373 | 1 | 0 | 1 | 0 | 2 | 1 | 2 | 0.5 | 6 | 0.5 | NA | NA | 3 | 0.3333 | 2 | 0 | 1 | 1 | 6 | 0.3333 |
| 18_3780381 | 1 | 0 | 1 | 1 | 2 | 0 | 2 | 0.5 | 6 | 0.3333 | NA | NA | 2 | 0.5 | 1 | 0 | 2 | 0.5 | 5 | 0.4 |
| 18_3780394 | 1 | 1 | 1 | 0 | 2 | 1 | 1 | 1 | 5 | 0.8 | NA | NA | 1 | 1 | 2 | 0.5 | 2 | 1 | 5 | 0.8 |
| 18_3780401 | 1 | 1 | NA | NA | 2 | 1 | 2 | 0.5 | 5 | 0.8 | NA | NA | 3 | 1 | 2 | 1 | 1 | 0 | 6 | 0.8333 |
| 18_3780403 | 1 | 1 | NA | NA | 2 | 1 | 2 | 1 | 5 | 1 | NA | NA | 2 | 1 | 1 | 1 | 2 | 1 | 5 | 1 |
| 18_3780405 | NA | NA | 1 | 0 | 2 | 0 | 2 | 1 | 5 | 0.4 | NA | NA | 3 | 0.6667 | 1 | 1 | 2 | 0.5 | 6 | 0.6667 |
| 18_3780410 | 1 | 0 | 1 | 1 | 2 | 0.5 | 1 | 0 | 5 | 0.4 | NA | NA | 3 | 0.3333 | 1 | 0 | 2 | 1 | 6 | 0.5 |
| 18_3780422 | 1 | 1 | 1 | 1 | 1 | 1 | 1 | 1 | 4 | 1 | NA | NA | 3 | 1 | 2 | 0.5 | 2 | 1 | 7 | 0.8571 |
| 18_3780441 | 1 | 0 | 1 | 0 | 2 | 0 | 1 | 1 | 5 | 0.2 | NA | NA | 3 | 0.6667 | 2 | 0.5 | 1 | 1 | 6 | 0.6667 |
| 18_3780451 | 1 | 1 | 1 | 0 | 2 | 0.5 | 1 | 1 | 5 | 0.6 | NA | NA | 3 | 1 | 2 | 1 | 2 | 1 | 7 | 1 |
| 18_3780790 | 2 | 0.5 | NA | NA | NA | NA | 1 | 1 | 3 | 0.6667 | NA | NA | 1 | 0 | 2 | 0.5 | 2 | 0 | 5 | 0.2 |
| 18_3780800 | 1 | 0 | NA | NA | NA | NA | 2 | 0.5 | 3 | 0.3333 | NA | NA | 1 | 1 | 2 | 1 | 1 | 0 | 4 | 0.75 |
| 18_3780804 | 1 | 1 | NA | NA | NA | NA | 2 | 0.5 | 3 | 0.6667 | NA | NA | 1 | 1 | 2 | 1 | 2 | 1 | 5 | 1 |
| 18_3780806 | 2 | 1 | NA | NA | NA | NA | 2 | 1 | 4 | 1 | NA | NA | 1 | 1 | NA | NA | 1 | 1 | 2 | 1 |
| 18_3780873 | 2 | 1 | NA | NA | NA | NA | 3 | 1 | 5 | 1 | NA | NA | NA | NA | 2 | 0.5 | 2 | 0.5 | 4 | 0.5 |
| 18_3780884 | 2 | 0.5 | NA | NA | NA | NA | 3 | 0 | 5 | 0.2 | NA | NA | 1 | 0 | 2 | 0.5 | 2 | 0.5 | 5 | 0.4 |
| 18_3780902 | 2 | 0.5 | NA | NA | NA | NA | 3 | 0 | 5 | 0.2 | NA | NA | 1 | 0 | 2 | 0.5 | 2 | 0.5 | 5 | 0.4 |
| 18_3780910 | 3 | 0 | NA | NA | NA | NA | 2 | 0.5 | 5 | 0.2 | NA | NA | 1 | 0 | 2 | 0.5 | 2 | 0 | 5 | 0.2 |
| 18_3780911 | 2 | 0.5 | NA | NA | NA | NA | 3 | 1 | 5 | 0.8 | NA | NA | 1 | 1 | 1 | 1 | 2 | 0.5 | 4 | 0.75 |
| 18_3780914 | 3 | 0.6667 | NA | NA | NA | NA | 3 | 0.6667 | 6 | 0.6667 | NA | NA | 1 | 1 | 2 | 0.5 | 2 | 0.5 | 5 | 0.6 |
| 18_3780932 | 2 | 1 | NA | NA | NA | NA | 2 | 1 | 4 | 1 | NA | NA | 1 | 1 | 2 | 0.5 | 2 | 0.5 | 5 | 0.6 |
| 18_3780933 | 2 | 0.5 | NA | NA | NA | NA | 2 | 0.5 | 4 | 0.5 | NA | NA | NA | NA | 2 | 0.5 | 2 | 0.5 | 4 | 0.5 |
| 18_3780944 | 3 | 1 | NA | NA | NA | NA | 2 | 1 | 5 | 1 | NA | NA | 1 | 1 | 3 | 1 | 2 | 1 | 6 | 1 |
| 18_3780948 | 3 | 1 | NA | NA | NA | NA | 3 | 0.3333 | 6 | 0.6667 | NA | NA | 1 | 1 | 3 | 1 | NA | NA | 4 | 1 |
| 18_3780956 | 3 | 0 | NA | NA | NA | NA | 3 | 0.3333 | 6 | 0.1667 | NA | NA | 1 | 1 | 3 | 0 | 2 | 0.5 | 6 | 0.3333 |
| 18_3780960 | 2 | 0 | NA | NA | NA | NA | 3 | 0.3333 | 5 | 0.2 | NA | NA | 1 | 0 | 2 | 0.5 | 2 | 0 | 5 | 0.2 |
| 18_3781527 | 1 | 1 | 1 | 0 | 1 | 1 | NA | NA | 3 | 0.6667 | NA | NA | NA | NA | 1 | 1 | NA | NA | 1 | 1 |
| 18_3781538 | NA | NA | 1 | 1 | 1 | 1 | NA | NA | 2 | 1 | NA | NA | NA | NA | 1 | 1 | NA | NA | 1 | 1 |
| 18_3781562 | NA | NA | 1 | 0 | 1 | 1 | NA | NA | 2 | 0.5 | NA | NA | NA | NA | 1 | 1 | NA | NA | 1 | 1 |
| 18_3781571 | 1 | 1 | 1 | 1 | 1 | 0 | NA | NA | 3 | 0.6667 | NA | NA | NA | NA | 1 | 1 | NA | NA | 1 | 1 |
| 18_3781893 | NA | NA | 2 | 1 | NA | NA | NA | NA | 2 | 1 | NA | NA | 1 | 0 | NA | NA | 1 | 1 | 2 | 0.5 |
| 18_3781900 | 1 | 1 | 3 | 0.3333 | NA | NA | 1 | 0 | 5 | 0.4 | NA | NA | 1 | 1 | NA | NA | 1 | 0 | 2 | 0.5 |
| 18_3781934 | 1 | 1 | 3 | 1 | NA | NA | 1 | 0 | 5 | 0.8 | NA | NA | 1 | 0 | NA | NA | 1 | 1 | 2 | 0.5 |
| 18_3781938 | 1 | 0 | 3 | 0.3333 | NA | NA | 1 | 0 | 5 | 0.2 | NA | NA | 1 | 1 | NA | NA | NA | NA | 1 | 1 |
| 18_3781947 | 1 | 1 | 3 | 0.3333 | NA | NA | 1 | 0 | 5 | 0.4 | NA | NA | 1 | 1 | NA | NA | 1 | 1 | 2 | 1 |
| 18_3781967 | 1 | 1 | 3 | 0.6667 | 1 | 1 | 1 | 0 | 6 | 0.6667 | NA | NA | 1 | 1 | NA | NA | 2 | 1 | 3 | 1 |
| 18_3781968 | 1 | 0 | 3 | 0.3333 | 1 | 0 | 1 | 0 | 6 | 0.1667 | NA | NA | 1 | 1 | NA | NA | 1 | 0 | 2 | 0.5 |
| 18_3781975 | 1 | 1 | 2 | 0.5 | 1 | 1 | 1 | 1 | 5 | 0.8 | NA | NA | 1 | 1 | NA | NA | 2 | 1 | 3 | 1 |
| 18_3781993 | 1 | 1 | 1 | 1 | 1 | 0 | 1 | 1 | 4 | 0.75 | NA | NA | 1 | 1 | NA | NA | 1 | 1 | 2 | 1 |
| 18_3781997 | NA | NA | 2 | 0 | 1 | 0 | 1 | 1 | 4 | 0.25 | NA | NA | 1 | 1 | NA | NA | 1 | 1 | 2 | 1 |
| 18_3782002 | 1 | 0 | 2 | 0.5 | 1 | 0 | 1 | 0 | 5 | 0.2 | NA | NA | 1 | 0 | NA | NA | 2 | 0.5 | 3 | 0.3333 |
| 18_3782006 | 1 | 1 | 3 | 1 | 1 | 0 | 1 | 1 | 6 | 0.8333 | NA | NA | 1 | 1 | NA | NA | 2 | 1 | 3 | 1 |

| | | | | | | | | | | | | | | | | | | | | |
|---|---|---|---|---|---|---|---|---|---|---|---|---|---|---|---|---|---|---|---|---|
| 18_378201 | 1 | 1 | 2 | 0.5 | 1 | 0 | 1 | 0 | 5 | 0.4 | NA | NA | 1 | 0 | NA | NA | 2 | 0.5 | 3 | 0.3333 |
| 18_378202 | 1 | 1 | 2 | 0.5 | 1 | 1 | 1 | 1 | 5 | 0.8 | NA | NA | 1 | 1 | NA | NA | 1 | 1 | 2 | 1 |
| 18_378203 | 1 | 1 | 3 | 1 | 1 | 0 | 1 | 1 | 6 | 0.8333 | NA | NA | NA | NA | NA | NA | 2 | 1 | 2 | 1 |
| 18_378203 | 2 | 0.5 | 3 | 0.6667 | 1 | 1 | 2 | 1 | 8 | 0.75 | NA | NA | 1 | 0 | NA | NA | 2 | 0.5 | 3 | 0.3333 |
| 18_378205 | 2 | 0 | 3 | 0.6667 | 1 | 0 | 1 | 0 | 7 | 0.2857 | NA | NA | 1 | 0 | NA | NA | 2 | 0.5 | 3 | 0.3333 |
| 18_378207 | 2 | 0.5 | 3 | 0.6667 | 1 | 0 | 1 | 1 | 7 | 0.5714 | NA | NA | 1 | 0 | NA | NA | 2 | 1 | 3 | 0.6667 |
| 18_378207 | 1 | 0 | 3 | 1 | 1 | 1 | 1 | 0 | 6 | 0.6667 | NA | NA | 1 | 0 | NA | NA | 2 | 0.5 | 3 | 0.3333 |
| 18_378208 | 2 | 0 | 3 | 0.3333 | 1 | 0 | 1 | 0 | 7 | 0.1429 | NA | NA | 1 | 1 | NA | NA | 2 | 0 | 3 | 0.3333 |
| 18_378231 | 1 | 1 | NA | NA | 1 | 1 | NA | NA | 2 | 1 | NA | NA | 1 | 0 | NA | NA | 1 | 1 | 2 | 0.5 |
| 18_378232 | 1 | 1 | NA | NA | 1 | 1 | NA | NA | 2 | 1 | NA | NA | 1 | 1 | NA | NA | 1 | 1 | 2 | 1 |
| 18_378236 | 1 | 1 | NA | NA | 1 | 1 | NA | NA | 2 | 1 | NA | NA | 1 | 0 | NA | NA | 1 | 1 | 2 | 0.5 |
| 18_378241 | 2 | 0.5 | NA | NA | 1 | 1 | NA | NA | 3 | 0.6667 | NA | NA | 1 | 0 | NA | NA | 1 | 1 | 2 | 0.5 |
| 18_378242 | 2 | 0.5 | NA | NA | 1 | 0 | NA | NA | 3 | 0.3333 | NA | NA | 1 | 0 | NA | NA | 1 | 1 | 2 | 0.5 |
| 18_378245 | 2 | 0.5 | NA | NA | NA | NA | NA | NA | 2 | 0.5 | NA | NA | 3 | 1 | 1 | 1 | NA | NA | 4 | 1 |
| 18_378248 | 2 | 1 | NA | NA | 1 | 0 | NA | NA | 3 | 0.6667 | NA | NA | 2 | 1 | 1 | 0 | 1 | 1 | 4 | 0.75 |
| 18_378249 | 2 | 1 | NA | NA | 1 | 0 | NA | NA | 3 | 0.6667 | NA | NA | 2 | 1 | 1 | 1 | 1 | 1 | 4 | 1 |
| 18_378249 | 2 | 1 | NA | NA | 1 | 0 | NA | NA | 3 | 0.6667 | NA | NA | 3 | 0.3333 | 1 | 1 | 2 | 1 | 6 | 0.6667 |
| 18_378251 | 3 | 0.3333 | NA | NA | 1 | 1 | NA | NA | 4 | 0.5 | NA | NA | 3 | 0.6667 | 1 | 0 | 2 | 1 | 6 | 0.6667 |
| 18_378252 | 2 | 1 | 1 | 1 | 1 | 1 | NA | NA | 4 | 1 | NA | NA | 2 | 1 | NA | NA | 2 | 1 | 4 | 1 |
| 18_378252 | 3 | 0.3333 | 1 | 1 | 1 | 1 | NA | NA | 5 | 0.6 | NA | NA | 3 | 0 | 1 | 0 | 2 | 0.5 | 6 | 0.1667 |
| 18_378253 | 3 | 0.6667 | 1 | 0 | NA | NA | NA | NA | 4 | 0.5 | NA | NA | 2 | 0.5 | 1 | 0 | 1 | 0 | 4 | 0.25 |
| 18_378277 | NA | NA | NA | NA | NA | NA | 1 | 1 | 1 | 1 | NA | NA | NA | NA | 1 | 0 | 1 | 1 | 2 | 0.5 |
| 18_378283 | NA | NA | NA | NA | 1 | 1 | 2 | 0.5 | 3 | 0.6667 | NA | NA | NA | NA | 1 | 1 | 2 | 0 | 3 | 0.3333 |
| 18_378284 | NA | NA | NA | NA | 2 | 0.5 | 2 | 0 | 4 | 0.25 | NA | NA | NA | NA | 1 | 0 | 2 | 1 | 3 | 0.6667 |
| 18_378284 | NA | NA | NA | NA | 2 | 0.5 | 1 | 1 | 3 | 0.6667 | NA | NA | NA | NA | 1 | 1 | 2 | 0.5 | 3 | 0.6667 |
| 18_378285 | NA | NA | 1 | 0 | 2 | 0.5 | 2 | 0 | 5 | 0.2 | NA | NA | NA | NA | 1 | 0 | 2 | 0.5 | 3 | 0.3333 |
| 18_378286 | NA | NA | 1 | 1 | 2 | 0.5 | 2 | 0 | 5 | 0.4 | NA | NA | NA | NA | 1 | 1 | 2 | 0.5 | 3 | 0.6667 |
| 18_378286 | NA | NA | 1 | 1 | 2 | 0.5 | 2 | 0 | 5 | 0.4 | NA | NA | NA | NA | 1 | 1 | 2 | 0 | 3 | 0.3333 |
| 18_378287 | NA | NA | 1 | 1 | 2 | 0 | 2 | 0.5 | 5 | 0.4 | NA | NA | 1 | 1 | 1 | 1 | 2 | 0.5 | 4 | 0.75 |
| 18_378291 | NA | NA | NA | NA | 2 | 0.5 | 2 | 0.5 | 4 | 0.5 | NA | NA | 1 | 0 | NA | NA | 2 | 0.5 | 3 | 0.3333 |
| 18_378291 | NA | NA | 1 | 1 | 2 | 1 | 3 | 1 | 6 | 1 | NA | NA | 1 | 1 | 1 | 1 | 2 | 0.5 | 4 | 0.75 |
| 18_378291 | NA | NA | 1 | 0 | 2 | 1 | 3 | 1 | 6 | 0.8333 | NA | NA | 1 | 1 | 1 | 1 | 2 | 0 | 4 | 0.5 |
| 18_378292 | NA | NA | 1 | 0 | 2 | 0.5 | 4 | 0 | 7 | 0.1429 | 1 | 0 | 1 | 1 | 1 | 0 | 2 | 0 | 5 | 0.2 |
| 18_378292 | NA | NA | 1 | 1 | 2 | 0.5 | 4 | 0.25 | 7 | 0.4286 | NA | NA | 1 | 1 | 1 | 1 | 1 | 0 | 3 | 0.6667 |
| 18_378293 | NA | NA | 1 | 1 | 2 | 1 | 2 | 1 | 5 | 1 | 1 | 0 | 1 | 1 | NA | NA | 2 | 1 | 4 | 0.75 |
| 18_378293 | NA | NA | 1 | 1 | 1 | 1 | 1 | 1 | 3 | 1 | 1 | 1 | 1 | 1 | 1 | 1 | 2 | 1 | 5 | 1 |
| 18_378293 | NA | NA | 1 | 1 | 1 | 1 | 3 | 1 | 5 | 1 | 1 | 1 | 1 | 1 | 1 | 1 | 3 | 0.6667 | 6 | 0.8333 |
| 18_378294 | NA | NA | 1 | 0 | 2 | 1 | 4 | 0.75 | 7 | 0.7143 | 1 | 0 | 1 | 0 | 1 | 1 | 3 | 0.3333 | 6 | 0.3333 |
| 18_378295 | NA | NA | 1 | 1 | 2 | 0.5 | 2 | 0.5 | 5 | 0.6 | 1 | 0 | NA | NA | 1 | 1 | 3 | 1 | 5 | 0.8 |
| 18_378297 | NA | NA | 1 | 1 | 2 | 0.5 | 3 | 0.6667 | 6 | 0.6667 | 1 | 1 | 1 | 1 | 1 | 1 | 1 | 1 | 4 | 1 |
| 18_378298 | NA | NA | 1 | 1 | 1 | 1 | 5 | 1 | 7 | 1 | 1 | 0 | 1 | 0 | 1 | 1 | 3 | 1 | 6 | 0.6667 |
| 18_378406 | NA | NA | 1 | 1 | NA | NA | 1 | 1 | 2 | 1 | NA | NA | 1 | 1 | NA | NA | 3 | 1 | 4 | 1 |
| 18_378412 | NA | NA | 2 | 0.5 | NA | NA | 1 | 1 | 3 | 0.6667 | 1 | 1 | 2 | 0 | 1 | 0 | 3 | 0 | 7 | 0.1429 |
| 18_378413 | 1 | 0 | 2 | 0.5 | NA | NA | 1 | 0 | 4 | 0.25 | 1 | 0 | 2 | 0.5 | 1 | 1 | 3 | 0.3333 | 7 | 0.4286 |
| 18_378414 | 1 | 1 | 2 | 1 | NA | NA | 1 | 1 | 4 | 1 | 1 | 1 | 1 | 1 | 1 | 1 | 3 | 1 | 6 | 1 |
| 18_378415 | 1 | 0 | 2 | 0.5 | NA | NA | 1 | 1 | 4 | 0.5 | 1 | 0 | 1 | 0 | NA | NA | 2 | 0.5 | 4 | 0.25 |
| 18_378416 | 1 | 0 | 2 | 0 | 1 | 0 | 1 | 1 | 5 | 0.2 | 1 | 0 | 2 | 0.5 | 1 | 0 | 2 | 0.5 | 6 | 0.3333 |
| 18_378417 | 1 | 0 | 2 | 0 | 1 | 1 | 1 | 0 | 5 | 0.2 | 1 | 0 | 2 | 1 | 1 | 0 | 3 | 0.3333 | 7 | 0.4286 |
| 18_378418 | 2 | 0.5 | 2 | 0 | 1 | 0 | 1 | 1 | 6 | 0.3333 | NA | NA | 2 | 0 | 1 | 1 | 3 | 0.3333 | 6 | 0.3333 |
| 18_378418 | 2 | 1 | 2 | 1 | NA | NA | 1 | 1 | 5 | 1 | NA | NA | 2 | 0.5 | NA | NA | 3 | 1 | 5 | 0.8 |
| 18_378419 | 2 | 0 | 2 | 0.5 | 1 | 0 | 1 | 0 | 6 | 0.1667 | 1 | 0 | 2 | 0.5 | 1 | 1 | 3 | 0.6667 | 7 | 0.5714 |
| 18_378419 | 2 | 0.5 | 2 | 0 | 1 | 0 | 1 | 0 | 6 | 0.1667 | 1 | 0 | 2 | 0.5 | 1 | 0 | 3 | 0 | 7 | 0.1429 |
| 18_378420 | 2 | 0.5 | 2 | 0.5 | NA | NA | 1 | 0 | 5 | 0.4 | NA | NA | 1 | 0 | 1 | 0 | 3 | 0.3333 | 5 | 0.2 |
| 18_378420 | 2 | 1 | 2 | 0.5 | NA | NA | 1 | 0 | 5 | 0.6 | 1 | 1 | 2 | 1 | 1 | 1 | 2 | 0.5 | 6 | 0.8333 |
| 18_378421 | 2 | 1 | 2 | 1 | NA | NA | 1 | 1 | 5 | 1 | 1 | 1 | 1 | 1 | NA | NA | 2 | 1 | 4 | 1 |
| 18_378421 | 1 | 1 | 1 | 1 | 1 | 0 | 1 | 1 | 4 | 0.75 | NA | NA | 2 | 1 | NA | NA | 3 | 1 | 5 | 1 |
| 18_378422 | 2 | 0.5 | 2 | 0.5 | 1 | 1 | 1 | 0 | 6 | 0.5 | NA | NA | 2 | 0.5 | 1 | 0 | 1 | 0 | 4 | 0.25 |
| 18_378425 | 2 | 0.5 | 3 | 0.3333 | 2 | 0 | 1 | 1 | 8 | 0.375 | NA | NA | 2 | 0.5 | 1 | 0 | 2 | 0.5 | 5 | 0.4 |
| 18_378425 | 2 | 0.5 | 2 | 0.5 | 2 | 0.5 | 1 | 0 | 7 | 0.4286 | NA | NA | 3 | 0.6667 | 1 | 1 | 4 | 0.5 | 8 | 0.625 |
| 18_378480 | 1 | 0 | 3 | 0.3333 | NA | NA | 2 | 0 | 6 | 0.1667 | NA | NA | 2 | 0.5 | 1 | 0 | NA | NA | 3 | 0.3333 |
| 18_378481 | 1 | 0 | 2 | 1 | NA | NA | 2 | 0.5 | 5 | 0.6 | NA | NA | 2 | 0 | 1 | 1 | NA | NA | 3 | 0.3333 |
| 18_378484 | 1 | 0 | 3 | 0.6667 | NA | NA | 2 | 0 | 6 | 0.3333 | NA | NA | 1 | 0 | 1 | 1 | NA | NA | 2 | 0.5 |
| 18_378487 | 1 | 0 | 3 | 0.3333 | NA | NA | 2 | 1 | 6 | 0.5 | NA | NA | 2 | 0.5 | 1 | 1 | NA | NA | 3 | 0.6667 |
| 18_378488 | 1 | 1 | 3 | 1 | NA | NA | 2 | 1 | 6 | 1 | 1 | 1 | 2 | 1 | NA | NA | NA | NA | 3 | 1 |
| 18_378490 | 1 | 1 | 3 | 0.3333 | NA | NA | 2 | 0.5 | 6 | 0.5 | 1 | 1 | 2 | 0 | 1 | 1 | NA | NA | 4 | 0.5 |

| | | | | | | | | | | | | | | | | | | | | |
|---|---|---|---|---|---|---|---|---|---|---|---|---|---|---|---|---|---|---|---|---|
| 18_3784909 | NA | NA | 1 | 1 | NA | NA | 2 | 0 | 3 | 0.3333 | 1 | 1 | NA | NA | 1 | 0 | NA | NA | 2 | 0.5 |
| 18_3784943 | 1 | 0 | 4 | 0.25 | 1 | 0 | 2 | 0 | 8 | 0.125 | NA | NA | 2 | 0.5 | 1 | 0 | NA | NA | 3 | 0.3333 |
| 18_3784959 | 2 | 0.5 | 4 | 0.25 | 1 | 1 | 2 | 0.5 | 9 | 0.4444 | NA | NA | 2 | 0.5 | 2 | 0.5 | NA | NA | 4 | 0.5 |
| 18_3784961 | 2 | 0 | 4 | 0.25 | 1 | 1 | 3 | 0.6667 | 10 | 0.4 | NA | NA | 2 | 0 | 3 | 0.3333 | NA | NA | 5 | 0.2 |
| 18_3784964 | 2 | 1 | 3 | 0.6667 | NA | NA | 3 | 0 | 8 | 0.5 | 1 | 1 | 2 | 0.5 | 3 | 0.6667 | NA | NA | 6 | 0.6667 |
| 18_3784966 | 1 | 1 | 4 | 0.75 | 1 | 1 | 3 | 0.6667 | 9 | 0.7778 | 1 | 0 | 2 | 0.5 | 2 | 1 | NA | NA | 5 | 0.6 |
| 18_3784978 | NA | NA | 2 | 1 | 1 | 1 | 2 | 1 | 5 | 1 | 1 | 1 | 2 | 0.5 | 3 | 1 | NA | NA | 6 | 0.8333 |
| 18_3784979 | 2 | 1 | 2 | 1 | 1 | 1 | 3 | 0.6667 | 8 | 0.875 | NA | NA | 2 | 1 | 3 | 1 | NA | NA | 5 | 1 |
| 18_3784990 | 2 | 1 | 5 | 1 | NA | NA | 3 | 1 | 10 | 1 | NA | NA | 2 | 0.5 | 3 | 1 | NA | NA | 5 | 0.8 |
| 18_3784991 | 1 | 0 | 5 | 0.2 | 1 | 0 | 3 | 0 | 10 | 0.1 | 1 | 0 | 1 | 0 | 3 | 0.3333 | NA | NA | 5 | 0.2 |
| 18_3784992 | 2 | 0 | 5 | 0.6 | 1 | 0 | 1 | 1 | 9 | 0.4444 | 1 | 1 | 1 | 1 | 2 | 0.5 | NA | NA | 4 | 0.75 |
| 18_3784994 | 2 | 1 | 5 | 0.4 | 1 | 0 | 3 | 0.6667 | 11 | 0.5455 | 1 | 1 | 2 | 0.5 | 2 | 0.5 | NA | NA | 5 | 0.6 |
| 18_3785003 | 2 | 0 | 5 | 0.2 | 2 | 0 | 3 | 0 | 12 | 0.0833 | 1 | 0 | 2 | 0 | 3 | 0.3333 | NA | NA | 6 | 0.1667 |
| 18_3785004 | 2 | 0.5 | 5 | 0.2 | 2 | 0.5 | 3 | 0.6667 | 12 | 0.4167 | 1 | 1 | 2 | 0 | 2 | 0 | NA | NA | 5 | 0.2 |
| 18_3785028 | 1 | 1 | 5 | 0.6 | 2 | 1 | 1 | 1 | 9 | 0.7778 | 2 | 0.5 | 1 | 1 | 2 | 1 | 2 | 1 | 7 | 0.8571 |
| 18_3785398 | 1 | 1 | NA | NA | NA | NA | NA | NA | 1 | 1 | NA | NA | 1 | 0 | 1 | 1 | NA | NA | 2 | 0.5 |
| 18_3785425 | 1 | 1 | NA | NA | NA | NA | NA | NA | 1 | 1 | NA | NA | 1 | 1 | 1 | 1 | NA | NA | 2 | 1 |
| 18_3785469 | 1 | 1 | NA | NA | NA | NA | NA | NA | 1 | 1 | NA | NA | 1 | 0 | 1 | 1 | NA | NA | 2 | 0.5 |
| 18_3785539 | 1 | 1 | NA | NA | NA | NA | NA | NA | 1 | 1 | NA | NA | 1 | 1 | 1 | 1 | NA | NA | 2 | 1 |
| 18_3785546 | 1 | 1 | NA | NA | NA | NA | NA | NA | 1 | 1 | NA | NA | 1 | 0 | 1 | 1 | NA | NA | 2 | 0.5 |
| 18_3785550 | 1 | 1 | NA | NA | NA | NA | NA | NA | 1 | 1 | NA | NA | 1 | 1 | 1 | 1 | NA | NA | 2 | 1 |
| 18_3785582 | 1 | 1 | NA | NA | NA | NA | NA | NA | 1 | 1 | NA | NA | 1 | 1 | 1 | 1 | NA | NA | 2 | 1 |
| 18_3785986 | NA | NA | NA | NA | 1 | 1 | 1 | 1 | 2 | 1 | NA | NA | 2 | 1 | NA | NA | NA | NA | 2 | 1 |
| 18_3786036 | 1 | 1 | NA | NA | 2 | 1 | NA | NA | 3 | 1 | NA | NA | 3 | 1 | NA | NA | NA | NA | 3 | 1 |
| 18_3786042 | 1 | 0 | NA | NA | 2 | 0.5 | 1 | 0 | 4 | 0.25 | NA | NA | 3 | 0.6667 | NA | NA | NA | NA | 3 | 0.6667 |
| 18_3786060 | 1 | 1 | NA | NA | 3 | 0.3333 | 1 | 0 | 5 | 0.4 | NA | NA | 3 | 0.6667 | NA | NA | NA | NA | 3 | 0.6667 |
| 18_3786073 | 1 | 0 | NA | NA | 3 | 0.3333 | 1 | 1 | 5 | 0.4 | NA | NA | 3 | 0.6667 | NA | NA | NA | NA | 3 | 0.6667 |
| 18_3786084 | 1 | 1 | NA | NA | 3 | 1 | 1 | 0 | 5 | 0.8 | NA | NA | 3 | 0.6667 | NA | NA | NA | NA | 3 | 0.6667 |
| 18_3786088 | 1 | 1 | NA | NA | 2 | 1 | 1 | 1 | 4 | 1 | NA | NA | 2 | 0.5 | NA | NA | NA | NA | 2 | 0.5 |
| 18_3786109 | 1 | 1 | NA | NA | 3 | 1 | 1 | 0 | 5 | 0.8 | NA | NA | 3 | 0.6667 | NA | NA | NA | NA | 3 | 0.6667 |
| 18_3786127 | 1 | 0 | 1 | 0 | 2 | 0.5 | NA | NA | 4 | 0.25 | NA | NA | 3 | 0.6667 | NA | NA | NA | NA | 3 | 0.6667 |
| 18_3786528 | NA | NA | NA | NA | NA | NA | 1 | 1 | 1 | 1 | NA | NA | NA | NA | NA | NA | 1 | 1 | 1 | 1 |
| 18_3786539 | NA | NA | NA | NA | NA | NA | 1 | 1 | 1 | 1 | NA | NA | NA | NA | NA | NA | 1 | 1 | 1 | 1 |
| 18_3786599 | NA | NA | NA | NA | NA | NA | 1 | 1 | 1 | 1 | 1 | 1 | NA | NA | NA | NA | 1 | 1 | 2 | 1 |
| 18_3786647 | NA | NA | NA | NA | NA | NA | 1 | 1 | 1 | 1 | 1 | 0 | NA | NA | NA | NA | 1 | 1 | 2 | 0.5 |
| 18_3786652 | NA | NA | NA | NA | NA | NA | 1 | 1 | 1 | 1 | NA | NA | NA | NA | NA | NA | 1 | 1 | 1 | 1 |
| 18_3787051 | 1 | 1 | NA | NA | NA | NA | NA | NA | 1 | 1 | 2 | 0.5 | 3 | 0 | NA | NA | 1 | 1 | 6 | 0.3333 |
| 18_3787055 | 1 | 1 | NA | NA | NA | NA | NA | NA | 1 | 1 | 2 | 1 | 2 | 0.5 | NA | NA | 1 | 0 | 5 | 0.6 |
| 18_3787077 | 1 | 0 | 1 | 0 | NA | NA | 1 | 1 | 3 | 0.3333 | 2 | 0.5 | 3 | 0.3333 | NA | NA | 2 | 0.5 | 7 | 0.4286 |
| 18_3787109 | NA | NA | 1 | 1 | NA | NA | 1 | 1 | 2 | 1 | 1 | 1 | 2 | 1 | NA | NA | 2 | 1 | 5 | 1 |
| 18_3787115 | 1 | 1 | 1 | 1 | NA | NA | 1 | 0 | 3 | 0.6667 | 2 | 1 | 2 | 0.5 | NA | NA | 2 | 0.5 | 6 | 0.6667 |
| 18_3787126 | 1 | 1 | 1 | 1 | NA | NA | 1 | 1 | 3 | 1 | 2 | 0.5 | 3 | 0.6667 | NA | NA | 2 | 1 | 7 | 0.7143 |
| 18_3787144 | 1 | 1 | 1 | 0 | NA | NA | 1 | 0 | 3 | 0.3333 | 2 | 0.5 | 2 | 0 | NA | NA | 1 | 0 | 5 | 0.2 |
| 18_3787173 | 1 | 1 | 1 | 0 | NA | NA | 2 | 0.5 | 4 | 0.5 | 2 | 0.5 | 2 | 0.5 | NA | NA | 2 | 0 | 6 | 0.3333 |
| 18_3787184 | 1 | 1 | 1 | 0 | NA | NA | 2 | 0 | 4 | 0.25 | 2 | 0 | 3 | 0 | NA | NA | 3 | 0.3333 | 8 | 0.125 |
| 18_3787195 | 1 | 1 | 1 | 0 | NA | NA | 2 | 0 | 4 | 0.25 | 2 | 0.5 | 3 | 0 | NA | NA | 2 | 1 | 7 | 0.4286 |
| 18_3787196 | 1 | 1 | 1 | 1 | NA | NA | 2 | 1 | 4 | 1 | 1 | 1 | 3 | 1 | NA | NA | 3 | 1 | 7 | 1 |
| 18_3787197 | NA | NA | 1 | 1 | NA | NA | 1 | 1 | 2 | 1 | 1 | 1 | 3 | 0.6667 | NA | NA | 1 | 0 | 5 | 0.6 |
| 18_3787202 | 1 | 1 | 1 | 1 | NA | NA | 2 | 1 | 4 | 1 | 1 | 1 | 3 | 1 | NA | NA | 3 | 0.6667 | 7 | 0.8571 |
| 18_3787211 | 1 | 1 | 1 | 0 | NA | NA | 2 | 1 | 4 | 0.75 | 2 | 0.5 | 3 | 0.3333 | NA | NA | 3 | 0.6667 | 8 | 0.5 |
| 18_3787223 | 1 | 0 | 1 | 1 | NA | NA | 2 | 0 | 4 | 0.25 | 1 | 1 | 1 | 1 | NA | NA | 3 | 0.3333 | 5 | 0.6 |
| 18_3787241 | 2 | 1 | 1 | 1 | NA | NA | 2 | 0.5 | 5 | 0.8 | 2 | 0.5 | 3 | 1 | NA | NA | 3 | 0.6667 | 8 | 0.75 |
| 18_3787245 | 1 | 1 | 1 | 1 | NA | NA | 1 | 0 | 3 | 0.6667 | 1 | 1 | 1 | 0 | NA | NA | 2 | 0.5 | 4 | 0.5 |
| 18_3787554 | 3 | 0 | NA | NA | NA | NA | 1 | 1 | 4 | 0.25 | 1 | 0 | 1 | 1 | 1 | 0 | NA | NA | 3 | 0.3333 |
| 18_3787572 | 3 | 0.6667 | NA | NA | 1 | 1 | 1 | 1 | 5 | 0.8 | 1 | 0 | 1 | 1 | 2 | 1 | NA | NA | 4 | 0.75 |
| 18_3787588 | 4 | 0.5 | NA | NA | 1 | 0 | 1 | 0 | 6 | 0.3333 | 1 | 0 | 1 | 0 | 2 | 0.5 | NA | NA | 4 | 0.25 |
| 18_3787593 | 4 | 0.75 | NA | NA | 1 | 0 | 1 | 0 | 6 | 0.5 | 1 | 0 | NA | NA | 2 | 1 | NA | NA | 3 | 0.6667 |
| 18_3787599 | 4 | 0.75 | NA | NA | 1 | 0 | 1 | 1 | 6 | 0.6667 | 1 | 1 | 1 | 1 | 2 | 1 | NA | NA | 4 | 1 |
| 18_3787662 | 4 | 0.25 | NA | NA | NA | NA | 1 | 0 | 5 | 0.2 | 1 | 0 | 1 | 1 | 3 | 0.3333 | 1 | 0 | 6 | 0.3333 |
| 18_3787677 | 4 | 0.5 | NA | NA | 1 | 0 | 2 | 0 | 7 | 0.2857 | 1 | 0 | 1 | 0 | 3 | 0 | 1 | 1 | 6 | 0.1667 |
| 18_3787688 | 3 | 0.3333 | NA | NA | 1 | 0 | 2 | 0 | 6 | 0.1667 | 1 | 0 | 1 | 1 | 3 | 0 | 1 | 0 | 6 | 0.1667 |
| 18_3787708 | 4 | 0.5 | NA | NA | 1 | 0 | 2 | 1 | 7 | 0.5714 | 1 | 0 | 1 | 1 | 2 | 0.5 | 1 | 1 | 5 | 0.6 |
| 18_3787709 | 4 | 0.75 | NA | NA | 1 | 1 | 2 | 1 | 7 | 0.8571 | 1 | 0 | 1 | 1 | 4 | 1 | 1 | 1 | 7 | 0.8571 |
| 18_3787715 | 3 | 0.6667 | NA | NA | 1 | 1 | 1 | 1 | 5 | 0.8 | 1 | 1 | 1 | 1 | 4 | 0.75 | 1 | 1 | 7 | 0.8571 |
| 18_3787724 | 2 | 0.5 | NA | NA | 1 | 1 | NA | NA | 3 | 0.6667 | 1 | 0 | NA | NA | 4 | 0.5 | 1 | 0 | 6 | 0.3333 |

| | | | | | | | | | | | | | | | | | | | | |
|---|---|---|---|---|---|---|---|---|---|---|---|---|---|---|---|---|---|---|---|---|
| 18_378773 | 4 | 0.25 | NA | NA | 1 | 1 | 2 | 0 | 7 | 0.2857 | 1 | 1 | 1 | 0 | 3 | 0.3333 | 1 | 0 | 6 | 0.3333 |
| 18_378807 | NA | NA | 1 | 1 | 2 | 0 | 2 | 0 | 5 | 0.2 | NA | NA | 2 | 0.5 | 1 | 1 | NA | NA | 3 | 0.6667 |
| 18_378813 | NA | NA | 2 | 0 | 2 | 0.5 | 1 | 1 | 5 | 0.4 | 2 | 0.5 | 2 | 0 | 1 | 0 | NA | NA | 5 | 0.2 |
| 18_378816 | NA | NA | 2 | 0 | 2 | 0.5 | 3 | 0.6667 | 7 | 0.4286 | 2 | 0 | 2 | 0 | 2 | 0.5 | NA | NA | 6 | 0.1667 |
| 18_378817 | NA | NA | 2 | 0 | 2 | 1 | 3 | 0 | 7 | 0.2857 | 1 | 1 | 2 | 0 | 2 | 0.5 | NA | NA | 5 | 0.4 |
| 18_378818 | NA | NA | 1 | 1 | 1 | 1 | 1 | 1 | 3 | 1 | 1 | 1 | 1 | 0 | 1 | 1 | NA | NA | 3 | 0.6667 |
| 18_378818 | NA | NA | 2 | 0.5 | 1 | 0 | 2 | 0 | 5 | 0.2 | 2 | 0.5 | 2 | 0 | 2 | 0 | NA | NA | 6 | 0.1667 |
| 18_378818 | NA | NA | 2 | 0 | 1 | 1 | 3 | 0.3333 | 6 | 0.3333 | 2 | 0.5 | 2 | 0 | 1 | 1 | NA | NA | 5 | 0.4 |
| 18_378820 | NA | NA | 3 | 0.3333 | 2 | 1 | 3 | 0.6667 | 8 | 0.625 | 2 | 0.5 | 2 | 0.5 | 2 | 1 | NA | NA | 6 | 0.6667 |
| 18_378820 | NA | NA | 3 | 1 | 2 | 0.5 | 3 | 1 | 8 | 0.875 | 1 | 1 | 2 | 1 | 2 | 1 | NA | NA | 5 | 1 |
| 18_378820 | NA | NA | 1 | 0 | 1 | 0 | 1 | 1 | 3 | 0.3333 | 1 | 0 | 1 | 1 | 2 | 0.5 | NA | NA | 4 | 0.5 |
| 18_378821 | NA | NA | 3 | 1 | 2 | 1 | 3 | 1 | 8 | 1 | 2 | 0.5 | 2 | 0 | 2 | 0.5 | NA | NA | 6 | 0.3333 |
| 18_378822 | NA | NA | 4 | 1 | 2 | 1 | 3 | 1 | 9 | 1 | 1 | 1 | 2 | 1 | 1 | 1 | NA | NA | 4 | 1 |
| 18_378857 | 1 | 1 | NA | NA | NA | NA | NA | NA | 1 | 1 | NA | NA | 1 | 1 | NA | NA | NA | NA | 1 | 1 |
| 18_378859 | 1 | 1 | NA | NA | NA | NA | NA | NA | 1 | 1 | NA | NA | 1 | 1 | 1 | 1 | NA | NA | 2 | 1 |
| 18_378863 | 1 | 1 | NA | NA | NA | NA | NA | NA | 1 | 1 | NA | NA | 1 | 1 | 1 | 1 | NA | NA | 2 | 1 |
| 18_378870 | 1 | 1 | NA | NA | NA | NA | NA | NA | 1 | 1 | NA | NA | 1 | 0 | 1 | 1 | NA | NA | 2 | 0.5 |
| 18_378890 | NA | NA | 1 | 1 | NA | NA | NA | NA | 1 | 1 | 1 | 1 | 1 | 0 | 1 | 0 | NA | NA | 3 | 0.3333 |
| 18_378892 | NA | NA | 1 | 0 | NA | NA | 1 | 1 | 2 | 0.5 | 1 | 1 | 1 | 1 | 1 | 0 | NA | NA | 3 | 0.6667 |
| 18_378893 | NA | NA | 1 | 1 | NA | NA | 1 | 1 | 2 | 1 | 1 | 1 | 1 | 1 | 1 | 1 | NA | NA | 3 | 1 |
| 18_378895 | NA | NA | 1 | 0 | NA | NA | 2 | 1 | 3 | 0.6667 | 1 | 1 | 1 | 1 | NA | NA | NA | NA | 2 | 1 |
| 18_378902 | NA | NA | 2 | 0.5 | NA | NA | 2 | 0.5 | 4 | 0.5 | 2 | 1 | NA | NA | 1 | 1 | NA | NA | 3 | 1 |
| 18_378902 | NA | NA | 2 | 0.5 | NA | NA | 2 | 0 | 4 | 0.25 | 1 | 0 | 1 | 0 | 3 | 1 | NA | NA | 5 | 0.6 |
| 18_378902 | 1 | 1 | 3 | 0.3333 | NA | NA | 3 | 0.3333 | 7 | 0.4286 | 2 | 0.5 | NA | NA | 3 | 0 | NA | NA | 5 | 0.2 |
| 18_378906 | 1 | 0 | 3 | 1 | NA | NA | 3 | 0 | 7 | 0.4286 | 2 | 0 | 1 | 1 | 3 | 0.3333 | NA | NA | 6 | 0.3333 |
| 18_378907 | 1 | 1 | 3 | 0 | NA | NA | 3 | 0.3333 | 7 | 0.2857 | 2 | 1 | 1 | 0 | 3 | 0 | NA | NA | 6 | 0.3333 |
| 18_378907 | 1 | 1 | 2 | 0.5 | NA | NA | 2 | 0.5 | 5 | 0.6 | 1 | 0 | NA | NA | 3 | 0.6667 | NA | NA | 4 | 0.5 |
| 18_378908 | 1 | 0 | 2 | 1 | NA | NA | 3 | 0.6667 | 6 | 0.6667 | 2 | 1 | 1 | 1 | 3 | 1 | NA | NA | 6 | 1 |
| 18_378908 | 1 | 1 | 3 | 1 | NA | NA | 3 | 1 | 7 | 1 | 2 | 1 | 1 | 1 | 2 | 0.5 | NA | NA | 5 | 0.8 |
| 18_378910 | 1 | 1 | 3 | 1 | NA | NA | 3 | 1 | 7 | 1 | 1 | 1 | 1 | 1 | 3 | 1 | NA | NA | 5 | 1 |
| 18_379000 | 1 | 0 | NA | NA | NA | NA | 1 | 1 | 2 | 0.5 | 2 | 0.5 | NA | NA | NA | NA | 1 | 0 | 3 | 0.3333 |
| 18_379000 | 1 | 1 | NA | NA | NA | NA | 1 | 0 | 2 | 0.5 | 2 | 0.5 | NA | NA | NA | NA | 1 | 0 | 3 | 0.3333 |
| 18_379002 | 1 | 1 | NA | NA | NA | NA | 1 | 1 | 2 | 1 | 2 | 1 | NA | NA | NA | NA | 1 | 1 | 3 | 1 |
| 18_379004 | 1 | 1 | NA | NA | NA | NA | 1 | 0 | 2 | 0.5 | 2 | 0.5 | NA | NA | NA | NA | 1 | 0 | 3 | 0.3333 |
| 18_379009 | NA | NA | NA | NA | NA | NA | 1 | 1 | 1 | 1 | 1 | 1 | NA | NA | NA | NA | 1 | 1 | 2 | 1 |
| 18_379010 | 1 | 1 | NA | NA | NA | NA | 2 | 1 | 3 | 1 | 2 | 0.5 | NA | NA | NA | NA | 2 | 1 | 4 | 0.75 |
| 18_379011 | 1 | 0 | NA | NA | NA | NA | 2 | 0.5 | 3 | 0.3333 | 2 | 0 | NA | NA | NA | NA | 2 | 1 | 4 | 0.5 |
| 18_379012 | 1 | 0 | NA | NA | NA | NA | 1 | 1 | 2 | 0.5 | 3 | 0.3333 | NA | NA | NA | NA | 1 | 0 | 4 | 0.25 |
| 18_379013 | 1 | 1 | 1 | 1 | NA | NA | 1 | 0 | 3 | 0.6667 | 3 | 0.3333 | 1 | 0 | NA | NA | 2 | 0.5 | 6 | 0.3333 |
| 18_379014 | 1 | 1 | 1 | 0 | NA | NA | 2 | 1 | 4 | 0.75 | 2 | 1 | 1 | 1 | 1 | 0 | 2 | 1 | 6 | 0.8333 |
| 18_379392 | NA | NA | NA | NA | NA | NA | 1 | 1 | 1 | 1 | NA | NA | NA | NA | NA | NA | 1 | 1 | 1 | 1 |
| 18_379395 | 2 | 0.5 | 3 | 0 | 1 | 0 | 4 | 0 | 10 | 0.1 | 3 | 0 | 1 | 0 | 2 | 0 | 2 | 0.5 | 8 | 0.125 |
| 18_379395 | NA | NA | NA | NA | NA | NA | 1 | 1 | 1 | 1 | NA | NA | NA | NA | NA | NA | 1 | 1 | 1 | 1 |
| 18_379396 | 4 | 0.5 | 3 | 0.6667 | 2 | 0.5 | 6 | 0.3333 | 15 | 0.4667 | 3 | 0.3333 | 4 | 0.25 | 2 | 1 | 3 | 0.6667 | 12 | 0.5 |
| 18_379397 | NA | NA | NA | NA | NA | NA | 1 | 1 | 1 | 1 | NA | NA | NA | NA | NA | NA | 1 | 1 | 1 | 1 |
| 18_379398 | 4 | 0.25 | 4 | 0.5 | 2 | 0.5 | 7 | 0.2857 | 17 | 0.3529 | 5 | 0.4 | 5 | 0.4 | 2 | 0.5 | 3 | 0 | 15 | 0.3333 |
| 18_379398 | 4 | 0.5 | 4 | 0.5 | 2 | 0 | 7 | 0.1429 | 17 | 0.2941 | 4 | 0.25 | 4 | 0.25 | 3 | 0 | 3 | 0.6667 | 14 | 0.2857 |
| 18_379398 | 4 | 0.25 | 3 | 0 | 2 | 0 | 5 | 0.2 | 14 | 0.1429 | 4 | 0 | 5 | 0.2 | 2 | 0.5 | 3 | 0 | 14 | 0.1429 |
| 18_379399 | 4 | 0.25 | 4 | 0.25 | 2 | 0 | 6 | 0.3333 | 16 | 0.25 | 4 | 0 | 5 | 0 | 2 | 0.5 | 3 | 0 | 14 | 0.0714 |
| 18_379401 | 5 | 0.2 | 3 | 0.3333 | 2 | 0 | 7 | 0 | 17 | 0.1176 | 5 | 0.2 | 4 | 0 | 2 | 0 | 4 | 0.25 | 15 | 0.1333 |
| 18_379401 | 5 | 0.4 | 4 | 0.25 | 2 | 0.5 | 7 | 0.5714 | 18 | 0.4444 | 4 | 0.25 | 3 | 0.3333 | 4 | 0.25 | 4 | 0 | 15 | 0.2 |
| 18_379404 | 5 | 0.4 | 4 | 0.5 | 2 | 0 | 7 | 0.4286 | 18 | 0.3889 | 3 | 0.6667 | 5 | 0.6 | 3 | 0.3333 | 4 | 0.75 | 15 | 0.6 |
| 18_379405 | 5 | 0.2 | 4 | 0 | 2 | 0 | 5 | 0 | 16 | 0.0625 | 3 | 0 | 5 | 0.2 | 4 | 0 | 4 | 0.25 | 16 | 0.125 |
| 18_379406 | 5 | 0.4 | 4 | 0.75 | 2 | 1 | 7 | 0.2857 | 18 | 0.5 | 5 | 0 | 5 | 0.2 | 3 | 0 | 5 | 0.4 | 18 | 0.1667 |
| 18_379407 | 4 | 0.75 | 4 | 0.5 | 2 | 1 | 5 | 0.8 | 15 | 0.7333 | 5 | 0.6 | 5 | 1 | 4 | 0.75 | 5 | 0.8 | 19 | 0.7895 |
| 18_379407 | 1 | 1 | 3 | 1 | 1 | 1 | 2 | 0.5 | 7 | 0.8571 | 5 | 1 | 4 | 0.75 | 1 | 1 | 3 | 1 | 13 | 0.9231 |
| 18_379407 | 6 | 0.1667 | 2 | 0 | 3 | 0 | 6 | 0 | 17 | 0.0588 | 5 | 0.4 | 4 | 0 | 3 | 0 | 4 | 0.25 | 16 | 0.1875 |
| 18_379410 | 6 | 0.5 | 6 | 0.3333 | 4 | 0.5 | 7 | 0.4286 | 23 | 0.4348 | 5 | 0.6 | 6 | 0.3333 | 3 | 0.3333 | 6 | 0.6667 | 20 | 0.5 |
| 18_379411 | 6 | 0.1667 | 5 | 0.4 | 4 | 0.25 | 7 | 0.1429 | 22 | 0.2273 | 5 | 0.2 | 6 | 0 | 4 | 0 | 4 | 0 | 19 | 0.0526 |
| 18_379411 | 7 | 1 | 3 | 0.6667 | 3 | 1 | 6 | 0.8333 | 19 | 0.8947 | 4 | 0.75 | 6 | 0.8333 | 3 | 1 | 4 | 0.5 | 17 | 0.7647 |
| 18_379412 | 7 | 0.1429 | 6 | 0.1667 | 2 | 0 | 7 | 0.2857 | 22 | 0.1818 | 3 | 0 | 6 | 0 | 4 | 0.25 | 6 | 0 | 19 | 0.0526 |
| 18_379415 | 4 | 0.75 | 5 | 0.8 | 3 | 0.6667 | 8 | 0.5 | 20 | 0.65 | 5 | 0.4 | 5 | 0.8 | 3 | 0.6667 | 6 | 0.6667 | 19 | 0.6316 |
| 18_379415 | 7 | 0.4286 | 8 | 0.75 | 6 | 0.8333 | 9 | 0.3333 | 30 | 0.5667 | 6 | 0.8333 | 5 | 0.8 | 4 | 0.5 | 6 | 0.3333 | 21 | 0.619 |
| 18_379415 | 4 | 1 | 7 | 0.1429 | 5 | 0.6 | 5 | 0.6 | 21 | 0.5238 | 6 | 0.3333 | 4 | 0 | 4 | 0.75 | 6 | 0.6667 | 20 | 0.45 |
| 18_379416 | 6 | 0.3333 | 4 | 0.5 | 3 | 0 | 7 | 0.2857 | 20 | 0.3 | 7 | 0.1429 | 6 | 0.3333 | 2 | 0 | 6 | 0 | 21 | 0.1429 |

| | | | | | | | | | | | | | | | | | | | | |
|---|---|---|---|---|---|---|---|---|---|---|---|---|---|---|---|---|---|---|---|---|
| 18_379416 | 4 | 0.25 | 6 | 0.1667 | 4 | 0 | 8 | 0 | 22 | 0.0909 | 7 | 0.1429 | 6 | 0 | 2 | 0 | 5 | 0 | 20 | 0.05 |
| 18_379417 | 6 | 0.6667 | 8 | 0.875 | 6 | 1 | 10 | 0.7 | 30 | 0.8 | 6 | 0.6667 | 6 | 0.6667 | 4 | 1 | 7 | 0.5714 | 23 | 0.6957 |
| 18_379418 | 3 | 0.6667 | 6 | 0.8333 | 4 | 0.25 | 8 | 0.5 | 21 | 0.5714 | 7 | 0.2857 | 5 | 0.2 | 3 | 0.6667 | 4 | 0.5 | 19 | 0.3684 |
| 18_379418 | 7 | 1 | 8 | 0.5 | 4 | 0.5 | 9 | 0.8889 | 28 | 0.75 | 8 | 0.5 | 5 | 0.6 | 4 | 0.5 | 6 | 0.8333 | 23 | 0.6087 |
| 18_379487 | NA | NA | 2 | 0 | NA | NA | 1 | 1 | 3 | 0.3333 | 1 | 1 | 2 | 0.5 | NA | NA | 1 | 0 | 4 | 0.5 |
| 18_379488 | NA | NA | 1 | 0 | NA | NA | 1 | 1 | 2 | 0.5 | 1 | 0 | 2 | 0.5 | NA | NA | 1 | 0 | 4 | 0.25 |
| 18_379490 | 1 | 0 | NA | NA | NA | NA | 1 | 1 | 2 | 0.5 | 1 | 0 | 2 | 0 | NA | NA | 2 | 0.5 | 5 | 0.2 |
| 18_379490 | 1 | 0 | 2 | 0 | NA | NA | 2 | 0.5 | 5 | 0.2 | 1 | 0 | 2 | 0 | NA | NA | 2 | 0.5 | 5 | 0.2 |
| 18_379491 | 1 | 1 | 2 | 0 | NA | NA | 2 | 0.5 | 5 | 0.4 | 1 | 0 | 2 | 0.5 | NA | NA | 2 | 0 | 5 | 0.2 |
| 18_379494 | 1 | 0 | 3 | 0.3333 | NA | NA | 2 | 1 | 6 | 0.5 | 1 | 0 | 3 | 0 | NA | NA | 2 | 1 | 6 | 0.3333 |
| 18_379494 | 1 | 0 | 2 | 0 | NA | NA | 2 | 0.5 | 5 | 0.2 | 1 | 0 | 3 | 0.6667 | NA | NA | 2 | 0.5 | 6 | 0.5 |
| 18_379495 | 1 | 1 | 3 | 0.6667 | NA | NA | 2 | 1 | 6 | 0.8333 | 1 | 1 | 3 | 0.6667 | NA | NA | 2 | 1 | 6 | 0.8333 |
| 18_379496 | 1 | 0 | 3 | 0.3333 | NA | NA | 2 | 0 | 6 | 0.1667 | 1 | 0 | 3 | 0 | NA | NA | 2 | 0.5 | 6 | 0.1667 |
| 18_379496 | 1 | 1 | 3 | 0.6667 | NA | NA | 2 | 1 | 6 | 0.8333 | 1 | 1 | 2 | 1 | NA | NA | NA | NA | 3 | 1 |
| 18_379497 | 1 | 1 | 2 | 0.5 | NA | NA | 2 | 0 | 5 | 0.4 | 1 | 1 | 3 | 0 | NA | NA | 2 | 0 | 6 | 0.1667 |
| 18_379498 | 1 | 0 | 3 | 0.6667 | NA | NA | 2 | 0.5 | 6 | 0.5 | 1 | 0 | 3 | 0 | NA | NA | 2 | 0.5 | 6 | 0.1667 |
| 18_379498 | 1 | 1 | 2 | 0 | NA | NA | 2 | 0 | 5 | 0.2 | 1 | 0 | 3 | 0.3333 | NA | NA | 2 | 0 | 6 | 0.1667 |
| 18_379500 | 1 | 0 | 3 | 0.6667 | NA | NA | 2 | 0.5 | 6 | 0.5 | 1 | 1 | 4 | 0.5 | NA | NA | 2 | 0.5 | 7 | 0.5714 |
| 18_379500 | 1 | 1 | 3 | 0.6667 | NA | NA | 2 | 1 | 6 | 0.8333 | 1 | 1 | 4 | 1 | NA | NA | 2 | 1 | 7 | 1 |
| 18_379501 | 1 | 1 | 3 | 0.3333 | NA | NA | 2 | 1 | 6 | 0.6667 | NA | NA | 3 | 0.3333 | NA | NA | 2 | 0 | 5 | 0.2 |
| 18_379501 | NA | NA | 3 | 1 | NA | NA | 1 | 1 | 4 | 1 | 1 | 1 | 3 | 0.6667 | NA | NA | 1 | 0 | 5 | 0.6 |
| 18_379502 | 1 | 1 | 2 | 0.5 | NA | NA | 1 | 1 | 4 | 0.75 | 1 | 1 | 3 | 0.3333 | NA | NA | 2 | 0 | 6 | 0.3333 |
| 18_379502 | 1 | 1 | 3 | 0 | NA | NA | 2 | 0.5 | 6 | 0.3333 | 1 | 0 | 4 | 0.75 | NA | NA | 2 | 0.5 | 7 | 0.5714 |
| 18_379503 | NA | NA | 2 | 0.5 | NA | NA | 2 | 1 | 4 | 0.75 | 1 | 1 | 4 | 1 | NA | NA | 1 | 1 | 6 | 1 |
| 18_379504 | 1 | 0 | 3 | 0.6667 | NA | NA | 2 | 1 | 6 | 0.6667 | 1 | 1 | 4 | 1 | NA | NA | 1 | 0 | 6 | 0.8333 |
| 18_379505 | 1 | 0 | 3 | 0.6667 | NA | NA | 2 | 1 | 6 | 0.6667 | 1 | 1 | 4 | 1 | NA | NA | 1 | 1 | 6 | 1 |
| 18_379505 | 1 | 1 | 3 | 0.6667 | NA | NA | 2 | 0.5 | 6 | 0.6667 | 1 | 1 | 3 | 0.6667 | NA | NA | 2 | 0 | 6 | 0.5 |
| 18_379505 | 1 | 0 | 3 | 0 | NA | NA | 2 | 0.5 | 6 | 0.1667 | 1 | 1 | 4 | 0 | NA | NA | 2 | 0 | 7 | 0.1429 |
| 18_379527 | 4 | 0.5 | 7 | 0.7143 | 4 | 0.25 | 3 | 1 | 18 | 0.6111 | 2 | 0.5 | 3 | 0.3333 | 2 | 1 | 1 | 1 | 8 | 0.625 |
| 18_379528 | 3 | 0.6667 | 3 | 1 | 1 | 1 | 4 | 1 | 11 | 0.9091 | 3 | 0.6667 | 1 | 1 | NA | NA | 1 | 1 | 5 | 0.8 |
| 18_379528 | 3 | 0.3333 | 3 | 1 | NA | NA | 4 | 0.5 | 10 | 0.6 | 2 | 0.5 | 2 | 0 | NA | NA | 1 | 0 | 5 | 0.2 |
| 18_379529 | 3 | 0 | 5 | 0 | NA | NA | 4 | 0.25 | 12 | 0.0833 | 4 | 0.25 | 2 | 0 | 1 | 0 | 3 | 0 | 10 | 0.1 |
| 18_379531 | 3 | 0 | 5 | 0.4 | 1 | 0 | 6 | 0.1667 | 15 | 0.2 | 3 | 0.6667 | 1 | 0 | 1 | 0 | 4 | 0 | 9 | 0.2222 |
| 18_379534 | 3 | 0 | 4 | 0.25 | 1 | 0 | 7 | 0.2857 | 15 | 0.2 | 4 | 0.25 | NA | NA | 2 | 0.5 | 5 | 0 | 11 | 0.1818 |
| 18_379534 | 3 | 0.6667 | 5 | 0.4 | 1 | 1 | 6 | 0.5 | 15 | 0.5333 | 4 | 0.5 | NA | NA | 1 | 0 | 6 | 0.3333 | 11 | 0.3636 |
| 18_379535 | NA | NA | 3 | 0.3333 | 1 | 1 | 6 | 0 | 10 | 0.2 | 1 | 0 | 1 | 1 | 2 | 0 | 6 | 0.1667 | 10 | 0.2 |
| 18_379537 | 2 | 1 | 2 | 1 | 1 | 1 | 7 | 0.5714 | 12 | 0.75 | 4 | 0.5 | NA | NA | 2 | 1 | 6 | 0.6667 | 12 | 0.6667 |
| 18_379538 | 4 | 0.5 | 3 | 0.3333 | NA | NA | 7 | 0.4286 | 14 | 0.4286 | 3 | 0.3333 | 1 | 1 | 1 | 0 | 5 | 0.4 | 10 | 0.4 |
| 18_379538 | 2 | 1 | 5 | 0.4 | 1 | 0 | 6 | 0.3333 | 14 | 0.4286 | 6 | 0.6667 | 1 | 1 | 2 | 1 | 6 | 0.5 | 15 | 0.6667 |
| 18_379539 | 4 | 0.5 | 4 | 0 | 1 | 0 | 8 | 0.25 | 17 | 0.2353 | 5 | 0.2 | 3 | 0 | 2 | 0.5 | 5 | 0.4 | 15 | 0.2667 |
| 18_379540 | NA | NA | 3 | 0.3333 | NA | NA | 3 | 1 | 6 | 0.6667 | 4 | 0.5 | 2 | 0.5 | 2 | 0.5 | 1 | 1 | 9 | 0.5556 |
| 18_379542 | 4 | 1 | 5 | 1 | 2 | 1 | 9 | 1 | 20 | 1 | 8 | 0.75 | 3 | 0 | 3 | 1 | 8 | 0.625 | 22 | 0.6364 |
| 18_379542 | 3 | 0.3333 | 5 | 0 | 2 | 0 | 8 | 0.25 | 18 | 0.1667 | 7 | 0.7143 | 2 | 0.5 | 3 | 1 | 8 | 0.5 | 20 | 0.65 |
| 18_379545 | 6 | 1 | 6 | 1 | 2 | 1 | 11 | 0.9091 | 25 | 0.96 | 10 | 0.8 | 1 | 1 | 3 | 1 | 7 | 0.8571 | 21 | 0.8571 |
| 18_379546 | 7 | 0.5714 | 5 | 0.4 | 1 | 1 | 10 | 0.7 | 23 | 0.6087 | 8 | 0.375 | 2 | 1 | 3 | 0.3333 | 6 | 0.6667 | 19 | 0.5263 |
| 18_379546 | 7 | 0.1429 | 6 | 0 | 2 | 0.5 | 10 | 0 | 25 | 0.08 | 9 | 0.1111 | 1 | 0 | 4 | 0 | 7 | 0.1429 | 21 | 0.0952 |
| 18_379547 | 6 | 0.8333 | 4 | 0.75 | 2 | 1 | 9 | 0.6667 | 21 | 0.7619 | 8 | 1 | 5 | 0.6 | 4 | 1 | 8 | 0.875 | 25 | 0.88 |
| 18_379548 | 2 | 0 | 5 | 0 | 2 | 1 | 9 | 0.2222 | 18 | 0.2222 | 7 | 0.1429 | 4 | 0.25 | 5 | 1 | 6 | 0.1667 | 22 | 0.3636 |
| 18_379549 | 7 | 0.7143 | 6 | 0.6667 | 3 | 0.3333 | 11 | 0.5455 | 27 | 0.5926 | 9 | 0.6667 | 3 | 0.3333 | 6 | 0.8333 | 6 | 0.6667 | 24 | 0.6667 |
| 18_379550 | 9 | 0.8889 | 5 | 1 | 3 | 0.6667 | 13 | 0.8462 | 30 | 0.8667 | 9 | 1 | 4 | 0.75 | 6 | 1 | 9 | 0.8889 | 28 | 0.9286 |
| 18_379550 | 9 | 0.2222 | 6 | 0.5 | 3 | 0 | 14 | 0.5 | 32 | 0.375 | 9 | 0.3333 | 5 | 0.2 | 4 | 0 | 10 | 0.2 | 28 | 0.2143 |
| 18_379550 | 7 | 0.7143 | 8 | 0.875 | 2 | 0.5 | 13 | 0.3846 | 30 | 0.6 | 8 | 0.5 | 5 | 0.6 | 6 | 0.8333 | 9 | 0.5556 | 28 | 0.6071 |
| 18_379770 | 11 | 0.0909 | 6 | 0 | 5 | 1 | 7 | 0.2857 | 29 | 0.2759 | 9 | 0.1111 | 10 | 0.1 | 12 | 0.3333 | 8 | 0.25 | 39 | 0.2051 |
| 18_379771 | 8 | 0 | 5 | 0 | 6 | 0.1667 | 6 | 0.3333 | 25 | 0.12 | 6 | 0.1667 | 9 | 0.1111 | 7 | 0.2857 | 6 | 0 | 28 | 0.1429 |
| 18_379779 | 9 | 0.2222 | 5 | 0.4 | 4 | 0.75 | 6 | 0.5 | 24 | 0.4167 | 6 | 0.3333 | 7 | 0.4286 | 8 | 0.625 | 5 | 0.8 | 26 | 0.5385 |
| 18_379782 | 6 | 0.1667 | 4 | 0 | 2 | 0.5 | 3 | 0 | 15 | 0.1333 | 4 | 0.5 | 5 | 0 | 5 | 0.2 | 2 | 0.5 | 16 | 0.25 |
| 18_379783 | 7 | 0.7143 | 2 | 1 | 3 | 0.3333 | 3 | 1 | 15 | 0.7333 | 5 | 0.4 | 7 | 0.8571 | 4 | 0.25 | 4 | 0.75 | 20 | 0.6 |
| 18_380237 | 7 | 0.2857 | 2 | 0 | 3 | 0 | 2 | 0 | 14 | 0.1429 | 4 | 0 | 3 | 0.6667 | 4 | 0 | 2 | 0 | 13 | 0.1538 |
| 18_380240 | 6 | 0.1667 | 4 | 0 | 2 | 0 | 2 | 0.5 | 14 | 0.1429 | 2 | 0 | 2 | 0 | 5 | 0 | 2 | 0.5 | 11 | 0.0909 |
| 18_380702 | 14 | 0.0714 | 17 | 0.2353 | 13 | 0.2308 | 11 | 0.0909 | 55 | 0.1636 | 4 | 0.25 | 9 | 0 | 16 | 0.0625 | 11 | 0 | 40 | 0.05 |
| 18_380703 | 14 | 0.6429 | 17 | 0.8235 | 9 | 0.7778 | 10 | 0.5 | 50 | 0.7 | 3 | 0.3333 | 9 | 0.8889 | 17 | 0.5882 | 9 | 0.8889 | 38 | 0.7105 |
| 18_380704 | 11 | 0.1818 | 14 | 0.5714 | 9 | 0.2222 | 8 | 0.25 | 42 | 0.3333 | 3 | 0.6667 | 8 | 0.25 | 13 | 0.2308 | 9 | 0.4444 | 33 | 0.3333 |
| 18_380705 | 6 | 0.1667 | 10 | 0 | 9 | 0.1111 | 10 | 0.1 | 35 | 0.0857 | 4 | 0.25 | 5 | 0.2 | 11 | 0 | 6 | 0.3333 | 26 | 0.1538 |
| 18_380705 | 8 | 0.25 | 7 | 0.2857 | 6 | 0 | 5 | 0 | 26 | 0.1538 | 2 | 0 | 7 | 0.5714 | 12 | 0 | 6 | 0.3333 | 27 | 0.2222 |
| 18_380911 | 13 | 0.2308 | 26 | 0.2692 | 14 | 0.2857 | 11 | 0 | 64 | 0.2188 | 14 | 0.1429 | 16 | 0.125 | 16 | 0.4375 | 13 | 0.0769 | 59 | 0.2034 |

| | | | | | | | | | | | | | | | | | | | | |
|---|---|---|---|---|---|---|---|---|---|---|---|---|---|---|---|---|---|---|---|---|
| 18_380912( | 10 | 0.3 | 21 | 0.0952 | 17 | 0.0588 | 8 | 0.25 | 56 | 0.1429 | 11 | 0.0909 | 16 | 0.0625 | 17 | 0.0588 | 11 | 0.0909 | 55 | 0.0727 |
| 18_380912( | 13 | 0.1538 | 29 | 0.1034 | 21 | 0.0476 | 11 | 0.0909 | 74 | 0.0946 | 17 | 0.1176 | 17 | 0.1176 | 19 | 0.3158 | 13 | 0.0769 | 66 | 0.1667 |
| 18_380913 | 15 | 0 | 28 | 0.1071 | 19 | 0.3158 | 12 | 0.0833 | 74 | 0.1351 | 17 | 0.0588 | 13 | 0.1538 | 18 | 0.1667 | 12 | 0.0833 | 60 | 0.1167 |
| 18_380913 | 15 | 0.0667 | 24 | 0.1667 | 20 | 0 | 11 | 0 | 70 | 0.0714 | 16 | 0.125 | 15 | 0.0667 | 14 | 0.2857 | 12 | 0 | 57 | 0.1228 |
| 18_380914( | 13 | 0 | 18 | 0.0556 | 18 | 0.2222 | 13 | 0.2308 | 62 | 0.129 | 15 | 0.3333 | 12 | 0.1667 | 13 | 0.3846 | 9 | 0.3333 | 49 | 0.3061 |
| 18_380914 | 12 | 0.1667 | 24 | 0.1667 | 15 | 0.0667 | 10 | 0 | 61 | 0.1148 | 18 | 0 | 15 | 0.2 | 18 | 0.3333 | 12 | 0.0833 | 63 | 0.1587 |
| 18_380918 | 11 | 0 | 22 | 0.1818 | 11 | 0.1818 | 8 | 0 | 52 | 0.1154 | 18 | 0.1111 | 13 | 0.1538 | 12 | 0.3333 | 9 | 0 | 52 | 0.1538 |
| 18_380921 | 13 | 0.4615 | 26 | 0.2692 | 14 | 0.1429 | 8 | 0 | 61 | 0.2459 | 14 | 0 | 13 | 0 | 24 | 0.2083 | 12 | 0.1667 | 63 | 0.1111 |
| 18_381059 | NA | NA | 2 | 1 | NA | NA | NA | NA | 2 | 1 | NA | NA | 2 | 1 | 3 | 1 | NA | NA | 5 | 1 |
| 18_381060 | NA | NA | 2 | 0.5 | NA | NA | NA | NA | 2 | 0.5 | NA | NA | 2 | 1 | 2 | 1 | NA | NA | 4 | 1 |
| 18_381061 | NA | NA | 2 | 0.5 | NA | NA | NA | NA | 2 | 0.5 | NA | NA | 3 | 0.6667 | 2 | 1 | NA | NA | 5 | 0.8 |
| 18_381063 | NA | NA | 1 | 1 | NA | NA | NA | NA | 1 | 1 | NA | NA | 2 | 0.5 | 1 | 0 | NA | NA | 3 | 0.3333 |
| 18_383216 | 7 | 0.1429 | 3 | 0.3333 | NA | NA | 4 | 0 | 14 | 0.1429 | 7 | 0.2857 | 3 | 0.3333 | 3 | 0 | 2 | 0 | 15 | 0.2 |
| 18_383219 | 4 | 0.5 | 2 | 0.5 | 2 | 0 | 3 | 0.3333 | 11 | 0.3636 | 5 | 0.6 | 3 | 0.3333 | 4 | 0.25 | 3 | 0.6667 | 15 | 0.4667 |
| 18_383221 | 4 | 0 | 1 | 0 | 1 | 0 | 2 | 0.5 | 8 | 0.125 | 3 | 0.3333 | 3 | 0 | 2 | 0 | 3 | 0.3333 | 11 | 0.1818 |
| 18_383221 | 4 | 0.25 | 2 | 0 | 2 | 0 | 4 | 0 | 12 | 0.0833 | 2 | 0 | 3 | 0 | 2 | 0 | 2 | 0.5 | 9 | 0.1111 |
| 18_383223 | 2 | 0.5 | 2 | 1 | 2 | 1 | 3 | 0.6667 | 9 | 0.7778 | 3 | 0.3333 | 3 | 0 | 2 | 0.5 | 2 | 0.5 | 10 | 0.3 |
| 18_383225( | 2 | 1 | 2 | 1 | 2 | 1 | 2 | 0.5 | 8 | 0.875 | 3 | 1 | 3 | 1 | 2 | 1 | 3 | 1 | 11 | 1 |
| 18_383226 | 3 | 1 | 2 | 1 | 1 | 1 | 1 | 1 | 7 | 1 | 3 | 1 | 2 | 1 | 2 | 1 | 3 | 1 | 10 | 1 |
| 18_383227( | 3 | 1 | 2 | 1 | 2 | 0.5 | 1 | 1 | 8 | 0.875 | 3 | 1 | 3 | 1 | 2 | 1 | 3 | 1 | 11 | 1 |
| 18_383228 | 3 | 0.3333 | 1 | 1 | 1 | 0 | 1 | 0 | 6 | 0.3333 | 3 | 0.3333 | 1 | 0 | 1 | 0 | 3 | 1 | 8 | 0.5 |
| 18_383301 | 1 | 0 | 1 | 0 | 1 | 1 | NA | NA | 3 | 0.3333 | NA | NA | NA | NA | 1 | 1 | NA | NA | 1 | 1 |
| 18_383303 | 1 | 0 | 1 | 0 | 1 | 1 | NA | NA | 3 | 0.3333 | NA | NA | 1 | 1 | 1 | 1 | NA | NA | 2 | 1 |
| 18_383304 | 4 | 0.75 | 3 | 0.6667 | 3 | 0.3333 | NA | NA | 10 | 0.6 | 3 | 0.3333 | 4 | 0.75 | 2 | 0.5 | 2 | 1 | 11 | 0.6364 |
| 18_383308( | 3 | 1 | 3 | 0.6667 | 3 | 0.6667 | NA | NA | 9 | 0.7778 | 3 | 0.3333 | 2 | 1 | 1 | 1 | 2 | 1 | 8 | 0.75 |
| 18_383309 | 3 | 0 | 3 | 0.3333 | 3 | 0 | NA | NA | 9 | 0.1111 | 3 | 0.3333 | 2 | 0 | 1 | 0 | 1 | 0 | 7 | 0.1429 |
| 18_383312 | 3 | 0.3333 | 1 | 1 | 2 | 1 | NA | NA | 6 | 0.6667 | 3 | 0 | 1 | 1 | 1 | 0 | 2 | 1 | 7 | 0.4286 |
| 18_383312 | 3 | 0.6667 | 3 | 0.6667 | 2 | 1 | NA | NA | 8 | 0.75 | 3 | 1 | 2 | 1 | 2 | 1 | 2 | 0.5 | 9 | 0.8889 |
| 18_383313 | 3 | 0 | 3 | 0.3333 | 3 | 0 | NA | NA | 9 | 0.1111 | 3 | 0 | 2 | 0 | 2 | 0.5 | 1 | 0 | 8 | 0.125 |
| 18_383315 | 3 | 0.6667 | 2 | 0.5 | 2 | 1 | NA | NA | 7 | 0.7143 | 2 | 1 | 2 | 0.5 | 2 | 1 | 1 | 1 | 7 | 0.8571 |
| 18_383315 | 3 | 0.6667 | 3 | 0.3333 | 2 | 1 | NA | NA | 8 | 0.625 | 2 | 0.5 | 2 | 1 | 2 | 1 | 1 | 1 | 7 | 0.8571 |
| 18_383315 | 3 | 0.3333 | 3 | 0.3333 | 1 | 1 | NA | NA | 7 | 0.4286 | 3 | 0.6667 | 1 | 1 | 1 | 0 | 1 | 0 | 6 | 0.5 |
| 18_383316 | 3 | 0.3333 | 2 | 0.5 | 2 | 0 | NA | NA | 7 | 0.2857 | 3 | 0 | 2 | 0 | NA | NA | 1 | 1 | 6 | 0.1667 |
| 18_383317 | 3 | 0 | 1 | 1 | 2 | 0 | NA | NA | 6 | 0.1667 | 3 | 0.6667 | 1 | 0 | NA | NA | NA | NA | 4 | 0.5 |
| 18_383318 | 2 | 0.5 | 2 | 0.5 | 2 | 0.5 | NA | NA | 6 | 0.5 | 2 | 0 | 1 | 1 | NA | NA | 1 | 1 | 4 | 0.5 |
| 18_383320 | 2 | 0.5 | 2 | 0.5 | 2 | 0.5 | NA | NA | 6 | 0.5 | 2 | 0 | 1 | 1 | NA | NA | 1 | 1 | 4 | 0.5 |
| 18_383323 | 1 | 1 | 2 | 0 | 2 | 0.5 | NA | NA | 5 | 0.4 | 2 | 0.5 | NA | NA | NA | NA | 1 | 0 | 3 | 0.3333 |
| 18_383323 | 2 | 0.5 | 2 | 1 | 2 | 1 | NA | NA | 6 | 0.8333 | 2 | 1 | NA | NA | NA | NA | 1 | 1 | 3 | 1 |
| 18_383324 | NA | NA | 1 | 0 | 2 | 0.5 | NA | NA | 3 | 0.3333 | 1 | 1 | NA | NA | NA | NA | NA | NA | 1 | 1 |
| 18_383359 | 2 | 0.5 | 1 | 0 | 2 | 0.5 | NA | NA | 5 | 0.4 | 2 | 0.5 | NA | NA | NA | NA | NA | NA | 2 | 0.5 |
| 18_383360 | 2 | 1 | 1 | 0 | 2 | 0 | NA | NA | 5 | 0.4 | 2 | 0 | NA | NA | NA | NA | 1 | 1 | 3 | 0.3333 |
| 18_383360 | NA | NA | 1 | 1 | 2 | 1 | NA | NA | 3 | 1 | 2 | 1 | NA | NA | NA | NA | 1 | 1 | 3 | 1 |
| 18_383361 | 2 | 0.5 | NA | NA | 1 | 0 | NA | NA | 3 | 0.3333 | 2 | 0.5 | NA | NA | NA | NA | 1 | 0 | 3 | 0.3333 |
| 18_383362( | 2 | 1 | 1 | 1 | 1 | 0 | NA | NA | 4 | 0.75 | 1 | 1 | NA | NA | NA | NA | NA | NA | 1 | 1 |
| 18_383362 | 2 | 1 | 1 | 1 | 2 | 0.5 | NA | NA | 5 | 0.8 | 1 | 1 | NA | NA | NA | NA | 1 | 0 | 2 | 0.5 |
| 18_383363 | NA | NA | 1 | 1 | 1 | 1 | NA | NA | 2 | 1 | 1 | 0 | NA | NA | NA | NA | 1 | 1 | 2 | 0.5 |
| 18_383830( | NA | NA | NA | NA | NA | NA | 1 | 1 | 1 | 1 | NA | NA | NA | NA | 1 | 1 | NA | NA | 1 | 1 |
| 18_383834 | 3 | 0.3333 | NA | NA | 2 | 0 | 2 | 0.5 | 7 | 0.2857 | 1 | 1 | NA | NA | 3 | 0 | 1 | 0 | 5 | 0.2 |
| 18_383951( | 6 | 0 | 2 | 0 | 5 | 0.6 | 5 | 0.6 | 18 | 0.3333 | 3 | 0 | 7 | 0.2857 | 6 | 0.5 | 5 | 0.6 | 21 | 0.381 |
| 18_383951 | 10 | 0.1 | 2 | 0 | 6 | 0.8333 | 6 | 0.1667 | 24 | 0.2917 | 3 | 0 | 8 | 0.25 | 6 | 0.3333 | 6 | 0.1667 | 23 | 0.2174 |
| 18_383951 | 7 | 0.4286 | 2 | 0.5 | 4 | 0.5 | 3 | 0 | 16 | 0.375 | 3 | 0.6667 | 7 | 0.2857 | 5 | 0.4 | 6 | 0.6667 | 21 | 0.4762 |
| 18_383952 | 9 | 0.8889 | 3 | 0.6667 | 6 | 1 | 4 | 0.75 | 22 | 0.8636 | 3 | 1 | 8 | 1 | 6 | 0.6667 | 6 | 1 | 23 | 0.913 |
| 18_383953 | 7 | 0.1429 | 3 | 0 | 5 | 0.8 | 5 | 0.4 | 20 | 0.35 | 3 | 0.3333 | 7 | 0.4286 | 7 | 0 | 4 | 0.25 | 21 | 0.2381 |
| 18_383955 | 3 | 0 | 2 | 0 | 6 | 0.3333 | 4 | 0 | 15 | 0.1333 | 3 | 0 | 6 | 0 | 5 | 0 | 5 | 0.2 | 19 | 0.0526 |
| 18_384411 | 27 | 0.1481 | 28 | 0.0714 | 26 | 0.1538 | 21 | 0.2381 | 102 | 0.1471 | 23 | 0.087 | 30 | 0.0667 | 19 | 0.1579 | 29 | 0.1724 | 101 | 0.1188 |
| 18_384412 | 24 | 0.125 | 25 | 0.08 | 25 | 0.04 | 21 | 0.0952 | 95 | 0.0842 | 18 | 0.1667 | 27 | 0.1481 | 23 | 0.1739 | 25 | 0.16 | 93 | 0.1613 |
| 18_384412 | 23 | 0.8696 | 25 | 0.52 | 29 | 0.5517 | 24 | 0.75 | 101 | 0.6634 | 20 | 0.6 | 24 | 0.6667 | 20 | 0.6 | 22 | 0.4545 | 86 | 0.5814 |
| 18_384504 | 37 | 0.1081 | 50 | 0.12 | 36 | 0.0556 | 41 | 0.2439 | 164 | 0.1341 | 32 | 0.2188 | 38 | 0.1316 | 37 | 0.2162 | 38 | 0.1842 | 145 | 0.1862 |
| 18_384507 | 37 | 0.0811 | 48 | 0.0625 | 41 | 0.0732 | 37 | 0.027 | 163 | 0.0613 | 38 | 0.0789 | 41 | 0.122 | 35 | 0.0857 | 34 | 0.1176 | 148 | 0.1014 |
| 18_384614 | 4 | 0 | 1 | 1 | 6 | 0.3333 | 2 | 0 | 13 | 0.2308 | 2 | 0 | 2 | 0.5 | 8 | 0 | 2 | 0 | 14 | 0.0714 |
| 18_384616 | 4 | 0.25 | 1 | 0 | 6 | 0.5 | 2 | 0.5 | 13 | 0.3846 | 3 | 0 | 4 | 0 | 7 | 0.2857 | 3 | 0 | 17 | 0.1176 |
| 18_387218 | NA | NA | 3 | 1 | 1 | 1 | NA | NA | 4 | 1 | 3 | 1 | NA | NA | NA | NA | NA | NA | 3 | 1 |
| 18_387220 | NA | NA | 3 | 0.3333 | NA | NA | NA | NA | 3 | 0.3333 | 3 | 0.3333 | NA | NA | NA | NA | NA | NA | 3 | 0.3333 |
| 18_387221 | 1 | 0 | 3 | 1 | 1 | 0 | NA | NA | 5 | 0.6 | 2 | 0.5 | NA | NA | NA | NA | NA | NA | 2 | 0.5 |
| 18_387222 | NA | NA | 3 | 1 | 1 | 1 | NA | NA | 4 | 1 | 2 | 1 | NA | NA | NA | NA | NA | NA | 2 | 1 |

| | | | | | | | | | | | | | | | | | | | | |
|---|---|---|---|---|---|---|---|---|---|---|---|---|---|---|---|---|---|---|---|---|
| 18_3872225 | NA | NA | 2 | 1 | NA | NA | NA | NA | 2 | 1 | 2 | 1 | NA | NA | NA | NA | NA | NA | 2 | 1 |
| 18_3872244 | NA | NA | 3 | 1 | 1 | 1 | NA | NA | 4 | 1 | 2 | 1 | NA | NA | NA | NA | NA | NA | 2 | 1 |
| 18_3872260 | 1 | 0 | 2 | 0.5 | 1 | 1 | NA | NA | 4 | 0.5 | 2 | 0.5 | NA | NA | NA | NA | NA | NA | 2 | 0.5 |
| 18_3872261 | 1 | 0 | 3 | 0.3333 | 1 | 0 | NA | NA | 5 | 0.2 | 2 | 0.5 | NA | NA | NA | NA | NA | NA | 2 | 0.5 |
| 18_3872266 | NA | NA | 3 | 1 | 1 | 1 | NA | NA | 4 | 1 | 2 | 1 | NA | NA | NA | NA | NA | NA | 2 | 1 |
| 18_3872268 | 1 | 1 | 3 | 0.6667 | 1 | 1 | NA | NA | 5 | 0.8 | 2 | 1 | NA | NA | NA | NA | NA | NA | 2 | 1 |
| 18_3872275 | 1 | 0 | 2 | 0.5 | 1 | 0 | NA | NA | 4 | 0.25 | 2 | 1 | NA | NA | NA | NA | NA | NA | 2 | 1 |
| 18_3872333 | 1 | 1 | 1 | 1 | 1 | 1 | NA | NA | 3 | 1 | 1 | 1 | NA | NA | NA | NA | NA | NA | 1 | 1 |
| 18_3872360 | 1 | 1 | NA | NA | NA | NA | NA | NA | 1 | 1 | 1 | 1 | NA | NA | NA | NA | NA | NA | 1 | 1 |
| 18_3897200 | 6 | 0.3333 | 11 | 0 | 6 | 0.1667 | 9 | 0.2222 | 32 | 0.1562 | 4 | 0 | 6 | 0.3333 | 3 | 0 | 6 | 0 | 19 | 0.1053 |
| 18_3897204 | 2 | 0.5 | 4 | 0.5 | 3 | 0.6667 | 8 | 0.625 | 17 | 0.5882 | 1 | 1 | 5 | 1 | 3 | 0.6667 | 2 | 1 | 11 | 0.9091 |
| 18_3897207 | 8 | 0.5 | 9 | 0 | 6 | 0.1667 | 10 | 0.1 | 33 | 0.1818 | 4 | 0.5 | 7 | 0 | 2 | 0 | 6 | 0.1667 | 19 | 0.1579 |
| 18_3897218 | 5 | 0.4 | 8 | 0.5 | 5 | 0.4 | 10 | 0.3 | 28 | 0.3929 | 2 | 0.5 | 6 | 0.5 | 3 | 0.6667 | 6 | 0.5 | 17 | 0.5294 |
| 18_3897239 | 6 | 0.8333 | 10 | 0.6 | 4 | 0.5 | 10 | 0.5 | 30 | 0.6 | 4 | 0.75 | 6 | 0.3333 | 3 | 0.3333 | 5 | 1 | 18 | 0.6111 |
| 18_3897260 | 5 | 0 | 8 | 0.125 | 4 | 0.25 | 10 | 0.2 | 27 | 0.1481 | 4 | 0.5 | 6 | 0.1667 | 3 | 0 | 5 | 0.6 | 18 | 0.3333 |
| 18_3897446 | 5 | 0 | 5 | 0.2 | 3 | 0 | 6 | 0 | 19 | 0.0526 | 3 | 0 | 5 | 0.2 | 2 | 0 | 4 | 0 | 14 | 0.0714 |
| 18_3897471 | 2 | 0.5 | 4 | 0.25 | 3 | 0.6667 | 6 | 0.3333 | 15 | 0.4 | 3 | 0 | 4 | 0.5 | 2 | 0 | 5 | 0.4 | 14 | 0.2857 |
| 18_3897478 | 3 | 0.6667 | 4 | 0 | 3 | 0.6667 | 5 | 0.4 | 15 | 0.4 | 3 | 0 | 5 | 0.2 | 2 | 0 | 5 | 0 | 15 | 0.0667 |
| 18_3897480 | 3 | 0.3333 | 4 | 0.5 | 3 | 0 | 6 | 0.1667 | 16 | 0.25 | 3 | 0 | 5 | 0 | 2 | 0 | 5 | 0.4 | 15 | 0.1333 |
| 18_3897498 | 3 | 0.3333 | 4 | 0 | 2 | 0 | 5 | 0 | 14 | 0.0714 | 3 | 0 | 5 | 0.4 | 2 | 0.5 | 4 | 0 | 14 | 0.2143 |
| 18_3897503 | 3 | 0.6667 | 4 | 0.5 | 2 | 1 | 4 | 1 | 13 | 0.7692 | 2 | 0 | 5 | 1 | 1 | 1 | 3 | 1 | 11 | 0.8182 |
| 18_3910276 | NA | NA | 1 | 1 | NA | NA | NA | NA | 1 | 1 | NA | NA | NA | NA | NA | NA | 1 | 1 | 1 | 1 |
| 18_3950412 | NA | NA | 1 | 1 | NA | NA | NA | NA | 1 | 1 | NA | NA | NA | NA | 2 | 1 | NA | NA | 2 | 1 |
| 18_3950416 | 3 | 0.6667 | 5 | 0.4 | 1 | 1 | 3 | 0.3333 | 12 | 0.5 | NA | NA | 5 | 0.8 | 2 | 0.5 | 3 | 0.6667 | 10 | 0.7 |
| 18_3950417 | 3 | 1 | 4 | 0.25 | 1 | 1 | 3 | 0.3333 | 11 | 0.5455 | NA | NA | 5 | 0.2 | 2 | 0 | 3 | 1 | 10 | 0.4 |
| 18_3950419 | 3 | 0.6667 | 4 | 0.25 | 1 | 0 | 3 | 0 | 11 | 0.2727 | NA | NA | 5 | 0.2 | 2 | 0 | 2 | 0 | 9 | 0.1111 |
| 18_3950441 | 2 | 1 | 4 | 1 | NA | NA | 1 | 1 | 7 | 1 | NA | NA | 4 | 1 | 2 | 1 | 3 | 1 | 9 | 1 |
| 18_3950462 | 2 | 1 | 1 | 1 | 1 | 1 | 3 | 1 | 7 | 1 | NA | NA | 4 | 1 | NA | NA | 2 | 1 | 6 | 1 |
| 18_3950470 | 2 | 1 | 1 | 1 | 1 | 1 | 3 | 1 | 7 | 1 | NA | NA | 4 | 0.75 | 2 | 1 | 2 | 1 | 8 | 0.875 |
| 18_3950517 | 3 | 0.6667 | 6 | 0.6667 | 1 | 1 | 3 | 1 | 13 | 0.7692 | NA | NA | 6 | 0.8333 | 3 | 0.6667 | 5 | 0.8 | 14 | 0.7857 |
| 18_3950531 | 4 | 0.75 | 6 | 0.6667 | 1 | 1 | 3 | 0.3333 | 14 | 0.6429 | NA | NA | 7 | 0.7143 | 3 | 0.6667 | 5 | 0.4 | 15 | 0.6 |
| 18_3950533 | 4 | 0.25 | 6 | 0.1667 | 2 | 0 | 4 | 0.25 | 16 | 0.1875 | NA | NA | 6 | 0.5 | 3 | 0.3333 | 5 | 0.4 | 14 | 0.4286 |
| 18_3950538 | 4 | 1 | 6 | 0.8333 | 2 | 1 | 3 | 1 | 15 | 0.9333 | NA | NA | 7 | 0.8571 | 3 | 1 | 5 | 0.8 | 15 | 0.8667 |
| 18_3950538 | 1 | 1 | 5 | 0.8 | 1 | 1 | 2 | 0 | 9 | 0.6667 | NA | NA | 7 | 0.2857 | 2 | 1 | 4 | 1 | 13 | 0.6154 |
| 18_3950539 | 2 | 1 | 1 | 1 | 1 | 0 | 2 | 0.5 | 6 | 0.6667 | NA | NA | 3 | 1 | 2 | 0.5 | 2 | 0.5 | 7 | 0.7143 |
| 18_3950609 | 5 | 0.2 | 4 | 0.25 | 2 | 0 | 4 | 0 | 15 | 0.1333 | NA | NA | 8 | 0 | 3 | 0.3333 | 4 | 0 | 15 | 0.0667 |
| 18_3950664 | 7 | 0 | 4 | 0 | 5 | 0.2 | 6 | 0.1667 | 22 | 0.0909 | NA | NA | 8 | 0.125 | 4 | 0 | 3 | 0 | 15 | 0.0667 |
| 18_3950803 | 12 | 0.25 | 9 | 0 | 7 | 0 | 7 | 0 | 35 | 0.0857 | 2 | 0 | 12 | 0 | 6 | 0 | 5 | 0.4 | 25 | 0.08 |
| 18_3950906 | 20 | 0.1 | 14 | 0.2143 | 14 | 0.1429 | 10 | 0.3 | 58 | 0.1724 | 4 | 0 | 15 | 0.1333 | 14 | 0.1429 | 7 | 0 | 40 | 0.1 |
| 18_3954417 | 7 | 0 | 8 | 0.25 | 14 | 0 | 8 | 0 | 37 | 0.0541 | 8 | 0 | 5 | 0.2 | 5 | 0.2 | 6 | 0.1667 | 24 | 0.125 |
| 18_3954422 | 8 | 0.125 | 8 | 0.375 | 14 | 0.0714 | 6 | 0 | 36 | 0.1389 | 7 | 0.1429 | 3 | 0 | 5 | 0 | 6 | 0.1667 | 21 | 0.0952 |
| 18_3954471 | 5 | 0 | 6 | 0.1667 | 12 | 0.25 | 8 | 0 | 31 | 0.129 | 5 | 0 | 5 | 0 | 3 | 0 | 3 | 0.3333 | 16 | 0.0625 |
| 18_3954658 | 2 | 0 | 4 | 0 | 10 | 0.1 | 2 | 0 | 18 | 0.0556 | 7 | 0.1429 | 5 | 0 | 2 | 0 | 2 | 0 | 16 | 0.0625 |
| 18_3954668 | 1 | 0 | 3 | 0 | 10 | 0.1 | 4 | 0 | 18 | 0.0556 | 6 | 0.3333 | 5 | 0 | 1 | 0 | 1 | 0 | 13 | 0.1538 |
| 18_3954717 | 2 | 0 | 3 | 0 | 9 | 0.1111 | 2 | 0 | 16 | 0.0625 | 7 | 0.1429 | 4 | 0 | 1 | 0 | 2 | 0.5 | 14 | 0.1429 |
| 18_3954725 | 1 | 0 | 3 | 0.3333 | 10 | 0.1 | 4 | 0.25 | 18 | 0.1667 | 6 | 0.3333 | 4 | 0.5 | 1 | 0 | 1 | 0 | 12 | 0.3333 |
| 18_3954729 | 1 | 0 | 3 | 0 | 9 | 0 | 4 | 0.5 | 17 | 0.1176 | 6 | 0.1667 | 4 | 0 | 1 | 0 | 2 | 0 | 13 | 0.0769 |
| 18_3954733 | 2 | 0 | 3 | 0.3333 | 9 | 0.1111 | 4 | 0 | 18 | 0.1111 | 6 | 0.1667 | 3 | 0 | 1 | 0 | 2 | 0 | 12 | 0.0833 |
| 18_3954734 | 2 | 0 | 2 | 0 | 10 | 0.1 | 4 | 0.25 | 18 | 0.1111 | 4 | 0 | 3 | 0.3333 | 1 | 0 | 2 | 0 | 10 | 0.1 |
| 18_3954738 | 2 | 0 | 3 | 0 | 10 | 0 | 4 | 0.25 | 19 | 0.0526 | 7 | 0.1429 | 4 | 0.25 | 1 | 0 | 2 | 0 | 14 | 0.1429 |
| 18_3954747 | 1 | 0 | 3 | 0 | 7 | 0.1429 | 4 | 0 | 15 | 0.0667 | 7 | 0 | 3 | 0.3333 | 2 | 0 | 1 | 0 | 13 | 0.0769 |
| 18_3954753 | 1 | 1 | 2 | 0 | 8 | 0.25 | 4 | 0.25 | 15 | 0.2667 | 6 | 0.1667 | 3 | 1 | 2 | 0 | 2 | 0 | 13 | 0.3077 |
| 18_3961909 | 1 | 0 | 3 | 0.3333 | 7 | 0.1429 | 3 | 0.3333 | 14 | 0.2143 | 5 | 0.2 | 2 | 1 | 1 | 0 | 2 | 0 | 10 | 0.3 |
| 18_3961914 | 1 | 1 | 3 | 0.3333 | 7 | 0.2857 | 2 | 1 | 13 | 0.4615 | 2 | 0 | 1 | 1 | 1 | 1 | 2 | 0.5 | 6 | 0.5 |
| 18_3961924 | 1 | 1 | 2 | 1 | 7 | 0.8571 | 3 | 1 | 13 | 0.9231 | 4 | 0.75 | 1 | 1 | 1 | 1 | 1 | 1 | 7 | 0.8571 |
| 18_3961931 | 1 | 1 | 2 | 1 | 7 | 0.8571 | 3 | 0.3333 | 13 | 0.7692 | 4 | 0.75 | 2 | 1 | 1 | 1 | 2 | 0.5 | 9 | 0.7778 |
| 18_3961934 | 1 | 1 | 3 | 1 | 7 | 1 | 2 | 1 | 13 | 1 | 3 | 0.6667 | 1 | 1 | 1 | 1 | 1 | 1 | 6 | 0.8333 |
| 18_3961948 | 1 | 1 | 3 | 1 | 7 | 0.7143 | 3 | 0.6667 | 14 | 0.7857 | 3 | 1 | 1 | 0 | NA | NA | 1 | 1 | 5 | 0.8 |
| 18_3961956 | NA | NA | 1 | 0 | 5 | 0.2 | 1 | 1 | 7 | 0.2857 | 2 | 0.5 | NA | NA | 1 | 0 | NA | NA | 3 | 0.3333 |
| 18_3961958 | 1 | 0 | 3 | 0 | 7 | 0.1429 | 3 | 0 | 14 | 0.0714 | 4 | 0.5 | NA | NA | 1 | 0 | 2 | 0.5 | 7 | 0.4286 |
| 18_3961961 | 1 | 0 | 3 | 0.6667 | 7 | 0.5714 | 3 | 0.6667 | 14 | 0.5714 | 3 | 0.3333 | NA | NA | 1 | 0 | 2 | 0.5 | 6 | 0.3333 |
| 18_3961975 | 1 | 0 | 2 | 0.5 | 6 | 0 | 3 | 0 | 12 | 0.0833 | 4 | 0.25 | NA | NA | 1 | 0 | 2 | 0 | 7 | 0.1429 |
| 18_3961987 | 1 | 1 | 2 | 1 | 6 | 0.5 | 3 | 0.3333 | 12 | 0.5833 | 3 | 0.6667 | NA | NA | 1 | 0 | NA | NA | 4 | 0.5 |
| 18_3961993 | 1 | 1 | 2 | 0 | 5 | 0 | 3 | 0.3333 | 11 | 0.1818 | 3 | 0 | NA | NA | 1 | 0 | 2 | 0.5 | 6 | 0.1667 |
| 18_3961995 | 1 | 0 | 2 | 0.5 | 5 | 0.2 | 3 | 0.3333 | 11 | 0.2727 | 3 | 0 | NA | NA | 1 | 1 | 2 | 0 | 6 | 0.1667 |

| | | | | | | | | | | | | | | | | | | | | |
|---|---|---|---|---|---|---|---|---|---|---|---|---|---|---|---|---|---|---|---|---|
| 18_3961997 | 1 | 1 | 2 | 1 | 5 | 1 | 2 | 0.5 | 10 | 0.9 | 1 | 1 | NA | NA | 1 | 1 | 2 | 0.5 | 4 | 0.75 |
| 18_3962020 | NA | NA | 1 | 1 | 5 | 1 | 1 | 1 | 7 | 1 | 2 | 0 | NA | NA | 1 | 1 | NA | NA | 3 | 0.3333 |
| 18_3962028 | NA | NA | 2 | 0.5 | 3 | 0.3333 | 1 | 0 | 6 | 0.3333 | 1 | 0 | NA | NA | 1 | 1 | 2 | 0 | 4 | 0.25 |
| 18_4033713 | NA | NA | NA | NA | 1 | 0 | 1 | 1 | 2 | 0.5 | NA | NA | NA | NA | NA | NA | 2 | 0.5 | 2 | 0.5 |
| 18_4033822 | 2 | 0 | 1 | 0 | 3 | 0 | 2 | 0.5 | 8 | 0.125 | 1 | 0 | 4 | 0 | NA | NA | 4 | 0.25 | 9 | 0.1111 |
| 18_4033864 | 6 | 0.6667 | 10 | 0.9 | 9 | 0.7778 | 8 | 0.75 | 33 | 0.7879 | 3 | 1 | 7 | 0.5714 | 2 | 0.5 | 11 | 0.7273 | 23 | 0.6957 |
| 18_4033877 | 7 | 0.2857 | 11 | 0.1818 | 7 | 0.1429 | 5 | 0 | 30 | 0.1667 | 3 | 0.3333 | 6 | 0 | 2 | 0 | 7 | 0.1429 | 18 | 0.1111 |
| 18_4033913 | 5 | 0.8 | 10 | 0.5 | 9 | 0.5556 | 6 | 0.5 | 30 | 0.5667 | 3 | 0.6667 | 6 | 0.6667 | 2 | 0.5 | 9 | 0.8889 | 20 | 0.75 |
| 18_4033915 | 5 | 1 | 11 | 0.7273 | 8 | 0.875 | 7 | 0.8571 | 31 | 0.8387 | 2 | 1 | 4 | 0.75 | 2 | 0.5 | 10 | 0.9 | 18 | 0.8333 |
| 18_4033922 | 5 | 0 | 11 | 0.1818 | 8 | 0.125 | 7 | 0 | 31 | 0.0968 | 3 | 0 | 5 | 0 | 2 | 0 | 7 | 0.4286 | 17 | 0.1765 |
| 18_4033923 | 4 | 1 | 8 | 1 | 7 | 0.8571 | 3 | 1 | 22 | 0.9545 | 2 | 1 | 4 | 1 | 2 | 1 | 9 | 1 | 17 | 1 |
| 18_4033924 | 4 | 0.75 | 8 | 0.625 | 8 | 0.625 | 7 | 0.4286 | 27 | 0.5926 | 2 | 0 | 5 | 0.6 | 2 | 0.5 | 10 | 0.8 | 19 | 0.6316 |
| 18_4033927 | 4 | 0.75 | 7 | 0.1429 | 7 | 0.7143 | 6 | 0.6667 | 24 | 0.5417 | 1 | 1 | 4 | 0.75 | 1 | 0 | 7 | 0.4286 | 13 | 0.5385 |
| 18_4033929 | 3 | 1 | 9 | 0.6667 | 8 | 0.375 | 7 | 0.7143 | 27 | 0.6296 | 2 | 0.5 | 4 | 0.75 | 1 | 1 | 9 | 0.7778 | 16 | 0.75 |
| 18_4033936 | 4 | 0.25 | 9 | 0.3333 | 8 | 0.625 | 7 | 0.1429 | 28 | 0.3571 | 2 | 0 | 5 | 0.4 | 2 | 0 | 10 | 0.4 | 19 | 0.3158 |
| 18_4066397 | NA | NA | 1 | 1 | NA | NA | NA | NA | 1 | 1 | NA | NA | 3 | 1 | NA | NA | NA | NA | 3 | 1 |
| 18_4066413 | 1 | 1 | 1 | 1 | NA | NA | NA | NA | 2 | 1 | NA | NA | 2 | 1 | NA | NA | NA | NA | 2 | 1 |
| 18_4066454 | 1 | 1 | 1 | 1 | NA | NA | NA | NA | 2 | 1 | NA | NA | 3 | 1 | NA | NA | NA | NA | 3 | 1 |
| 18_4066553 | 1 | 0 | 1 | 0 | 2 | 0.5 | 1 | 0 | 5 | 0.2 | NA | NA | 5 | 0.2 | NA | NA | NA | NA | 5 | 0.2 |
| 18_4066570 | 1 | 0 | 1 | 0 | 2 | 0.5 | NA | NA | 4 | 0.25 | NA | NA | 5 | 0.2 | NA | NA | NA | NA | 5 | 0.2 |
| 18_4066632 | 1 | 0 | 1 | 1 | 2 | 0.5 | NA | NA | 4 | 0.5 | NA | NA | 4 | 0.25 | NA | NA | 1 | 0 | 5 | 0.2 |
| 18_4066969 | 2 | 0.5 | 1 | 0 | 2 | 0 | 2 | 0 | 7 | 0.1429 | NA | NA | 5 | 0.2 | NA | NA | 2 | 0 | 7 | 0.1429 |
| 18_4067515 | 2 | 0 | 2 | 1 | 1 | 0 | 2 | 0 | 7 | 0.2857 | NA | NA | 5 | 0.6 | 2 | 0.5 | 2 | 0.5 | 9 | 0.5556 |
| 18_4067526 | 2 | 0 | 4 | 0.25 | 2 | 0.5 | 1 | 0 | 9 | 0.2222 | 1 | 0 | 7 | 0.1429 | 3 | 0.6667 | 3 | 0.3333 | 14 | 0.2857 |
| 18_4067545 | 2 | 1 | 3 | 1 | 2 | 1 | 2 | 1 | 9 | 1 | 1 | 1 | 5 | 1 | 3 | 0.6667 | 5 | 1 | 14 | 0.9286 |
| 18_4067555 | 2 | 0.5 | 4 | 0.5 | 3 | 0.3333 | 2 | 0 | 11 | 0.3636 | 1 | 0 | 4 | 0.5 | 3 | 0.6667 | 5 | 0.8 | 13 | 0.6154 |
| 18_4067555 | 2 | 0 | 3 | 0 | 4 | 0.75 | 2 | 0 | 11 | 0.2727 | 1 | 0 | 7 | 0.4286 | 3 | 0.3333 | 4 | 0.25 | 15 | 0.3333 |
| 18_4067576 | 4 | 0.25 | 4 | 0 | 3 | 0.6667 | 3 | 0 | 14 | 0.2143 | 1 | 0 | 5 | 0 | 1 | 0 | 5 | 0.4 | 12 | 0.1667 |
| 18_4067582 | 4 | 0 | 6 | 0.1667 | 3 | 0 | 3 | 0 | 16 | 0.0625 | 1 | 0 | 5 | 0.2 | 2 | 0.5 | 5 | 0 | 13 | 0.1538 |
| 18_4067588 | 4 | 0 | 6 | 0 | 3 | 0.3333 | 1 | 0 | 14 | 0.0714 | 1 | 0 | 7 | 0.2857 | 3 | 0 | 6 | 0.1667 | 17 | 0.1765 |
| 18_4067923 | 1 | 0 | 5 | 0.2 | 1 | 0 | 2 | 0 | 9 | 0.1111 | 2 | 0.5 | NA | NA | NA | NA | 2 | 0.5 | 4 | 0.5 |
| 18_4067952 | 1 | 0 | 4 | 0.25 | 2 | 0 | 3 | 0 | 10 | 0.1 | 2 | 0 | NA | NA | NA | NA | 2 | 0.5 | 4 | 0.25 |
| 18_4068026 | 1 | 0 | 6 | 0 | 2 | 0.5 | 3 | 0 | 12 | 0.0833 | 4 | 0.25 | 1 | 0 | 1 | 0 | 2 | 0 | 8 | 0.125 |
| 18_4218686 | NA | NA | NA | NA | NA | NA | 2 | 0.5 | 2 | 0.5 | NA | NA | NA | NA | 2 | 0 | 1 | 1 | 3 | 0.3333 |
| 18_4218704 | NA | NA | NA | NA | NA | NA | 2 | 0.5 | 2 | 0.5 | NA | NA | NA | NA | 2 | 0.5 | 1 | 1 | 3 | 0.6667 |
| 18_4228920 | NA | NA | 1 | 1 | NA | NA | 1 | 1 | 2 | 1 | NA | NA | 2 | 0.5 | 5 | 0.4 | 1 | 1 | 8 | 0.5 |
| 18_4228923 | NA | NA | 1 | 1 | 1 | 0 | 2 | 0 | 4 | 0.25 | NA | NA | 2 | 0.5 | 5 | 0.2 | 2 | 0 | 9 | 0.2222 |
| 18_4228927 | NA | NA | 1 | 1 | 1 | 1 | 2 | 0.5 | 4 | 0.75 | NA | NA | 2 | 0.5 | 5 | 1 | 2 | 1 | 9 | 0.8889 |
| 18_4228931 | NA | NA | 1 | 1 | 1 | 0 | 2 | 0 | 4 | 0.25 | NA | NA | 2 | 0.5 | 5 | 0.2 | 1 | 1 | 8 | 0.375 |
| 18_4228936 | NA | NA | 1 | 1 | 2 | 0.5 | 2 | 0 | 5 | 0.4 | NA | NA | 1 | 1 | 4 | 1 | 2 | 1 | 7 | 1 |
| 18_4228947 | NA | NA | 1 | 1 | 1 | 1 | 2 | 0.5 | 4 | 0.75 | NA | NA | 1 | 1 | 5 | 0.8 | 3 | 1 | 9 | 0.8889 |
| 18_4228948 | NA | NA | 1 | 1 | 2 | 1 | 2 | 0.5 | 5 | 0.8 | NA | NA | NA | NA | 4 | 0.75 | 3 | 1 | 7 | 0.8571 |
| 18_4228951 | NA | NA | 1 | 1 | 2 | 1 | 1 | 1 | 4 | 1 | NA | NA | 1 | 1 | 4 | 1 | 2 | 1 | 7 | 1 |
| 18_4228970 | NA | NA | 1 | 0 | 1 | 1 | 2 | 0.5 | 4 | 0.5 | NA | NA | 1 | 0 | 4 | 0.5 | 1 | 1 | 6 | 0.5 |
| 18_4228980 | NA | NA | 1 | 1 | 1 | 0 | 2 | 0 | 4 | 0.25 | NA | NA | 1 | 0 | 4 | 0.25 | 1 | 0 | 6 | 0.1667 |
| 18_4228991 | NA | NA | 1 | 1 | 1 | 1 | 2 | 0.5 | 4 | 0.75 | NA | NA | 1 | 1 | 5 | 1 | 3 | 1 | 9 | 1 |
| 18_4228994 | NA | NA | 1 | 1 | 1 | 1 | 2 | 1 | 4 | 1 | NA | NA | 1 | 1 | 4 | 1 | 4 | 1 | 9 | 1 |
| 18_4228999 | NA | NA | 1 | 0 | 1 | 0 | 2 | 0.5 | 4 | 0.25 | NA | NA | 1 | 1 | 5 | 0 | 4 | 0 | 10 | 0.1 |
| 18_4229011 | NA | NA | 1 | 1 | 1 | 1 | 2 | 0.5 | 4 | 0.75 | 1 | 1 | 1 | 1 | 4 | 1 | 4 | 1 | 10 | 1 |
| 18_4229026 | NA | NA | 1 | 1 | 2 | 1 | 2 | 0.5 | 5 | 0.8 | 1 | 0 | 1 | 1 | 4 | 0.75 | 4 | 0.5 | 10 | 0.6 |
| 18_4229164 | NA | NA | 1 | 0 | 2 | 0.5 | 2 | 0 | 5 | 0.2 | 1 | 0 | 1 | 0 | 6 | 0.3333 | 1 | 0 | 9 | 0.2222 |
| 18_4229174 | NA | NA | 1 | 0 | 3 | 0.3333 | 2 | 0 | 6 | 0.1667 | 1 | 0 | 4 | 0 | 7 | 0.1429 | 1 | 1 | 13 | 0.1538 |
| 18_4229197 | NA | NA | 1 | 0 | 3 | 0.6667 | 2 | 0 | 6 | 0.3333 | 1 | 0 | 5 | 0 | 7 | 0.4286 | 1 | 1 | 14 | 0.2857 |
| 18_4233550 | 7 | 0.1429 | 7 | 0 | 3 | 0 | 6 | 0.1667 | 23 | 0.087 | 2 | 0 | 2 | 0 | 7 | 0.2857 | 5 | 0 | 16 | 0.125 |
| 18_4247166 | 1 | 1 | 5 | 0.8 | 4 | 0.5 | 1 | 1 | 11 | 0.7273 | 4 | 0.75 | 1 | 1 | 2 | 1 | NA | NA | 7 | 0.8571 |
| 18_4247172 | 1 | 0 | 5 | 0.6 | 2 | 0 | 1 | 1 | 9 | 0.4444 | 2 | 1 | 1 | 0 | 3 | 0 | NA | NA | 6 | 0.3333 |
| 18_4247175 | 1 | 1 | 4 | 0.75 | 3 | 0.3333 | 1 | 0 | 9 | 0.5556 | 2 | 0 | 1 | 0 | 3 | 0.3333 | NA | NA | 6 | 0.1667 |
| 18_4247182 | 1 | 0 | 5 | 0.2 | 4 | 0 | 2 | 0 | 12 | 0.0833 | 4 | 0.25 | 1 | 0 | 3 | 0.3333 | NA | NA | 8 | 0.25 |
| 18_4247298 | 2 | 0 | 5 | 0 | 6 | 0.3333 | 5 | 0.4 | 18 | 0.2222 | 4 | 0 | 3 | 0.3333 | 4 | 0 | 1 | 0 | 12 | 0.0833 |
| 18_4247317 | 2 | 0 | 4 | 0 | 7 | 0.1429 | 5 | 0 | 18 | 0.0556 | 4 | 0.25 | 4 | 0 | 4 | 0 | NA | NA | 12 | 0.0833 |
| 18_4266935 | 2 | 0 | NA | NA | 4 | 0.25 | 3 | 0 | 9 | 0.1111 | 2 | 0 | 3 | 0.3333 | 1 | 0 | 5 | 0.2 | 11 | 0.1818 |
| 18_4270766 | 3 | 1 | 3 | 0.6667 | 7 | 0.5714 | 5 | 0.6 | 18 | 0.6667 | 1 | 1 | 5 | 1 | 3 | 1 | 5 | 0.8 | 14 | 0.9286 |
| 18_4270789 | 2 | 0 | 3 | 0 | 5 | 0.2 | 2 | 0.5 | 12 | 0.1667 | 2 | 0 | 7 | 0.2857 | 3 | 0 | 4 | 0 | 16 | 0.125 |
| 18_4270793 | 4 | 0.25 | 4 | 0.25 | 6 | 0.3333 | 5 | 0.4 | 19 | 0.3158 | 1 | 1 | 6 | 0 | 2 | 0.5 | 5 | 0.6 | 14 | 0.3571 |
| 18_4270797 | 5 | 0 | 5 | 0 | 6 | 0.1667 | 6 | 0.3333 | 22 | 0.1364 | 2 | 0 | 5 | 0.2 | 3 | 0 | 7 | 0.1429 | 17 | 0.1176 |

| | | | | | | | | | | | | | | | | | | | | |
|---|---|---|---|---|---|---|---|---|---|---|---|---|---|---|---|---|---|---|---|---|
| 18_427080( | 5 | 0 | 6 | 0.3333 | 9 | 0.1111 | 6 | 0 | 26 | 0.1154 | 2 | 0 | 7 | 0.1429 | 3 | 0.3333 | 8 | 0 | 20 | 0.1 |
| 18_427082 | 5 | 0.6 | 7 | 0.5714 | 8 | 0.375 | 6 | 0.6667 | 26 | 0.5385 | 2 | 1 | 7 | 0.4286 | 2 | 0.5 | 8 | 0.375 | 19 | 0.4737 |
| 18_427785 | 20 | 0.45 | 30 | 0.6333 | 19 | 0.4737 | 4 | 0.25 | 73 | 0.5205 | 12 | 0.5 | 28 | 0.5714 | 15 | 0.4 | 17 | 0.8824 | 72 | 0.5972 |
| 18_427786 | 19 | 0.3684 | 26 | 0.4231 | 14 | 0.5 | 7 | 0.4286 | 66 | 0.4242 | 10 | 0.3 | 23 | 0.4348 | 15 | 0.5333 | 19 | 0.2105 | 67 | 0.3731 |
| 18_427787( | 20 | 0.3 | 25 | 0.32 | 15 | 0.4667 | 4 | 0 | 64 | 0.3281 | 12 | 0.5 | 27 | 0.3333 | 15 | 0.6667 | 20 | 0.3 | 74 | 0.4189 |
| 18_427788 | 16 | 0.9375 | 24 | 0.9167 | 15 | 1 | 5 | 0.6 | 60 | 0.9167 | 9 | 1 | 21 | 0.9524 | 14 | 1 | 18 | 1 | 62 | 0.9839 |
| 18_428345( | 16 | 0.1875 | 17 | 0.1765 | 13 | 0.0769 | 5 | 0.2 | 51 | 0.1569 | 6 | 0.1667 | 20 | 0.2 | 10 | 0.2 | 14 | 0.1429 | 50 | 0.18 |
| 18_430318( | 11 | 0.1818 | 14 | 0.4286 | 10 | 0.5 | 2 | 0.5 | 37 | 0.3784 | 6 | 0.1667 | 12 | 0.25 | 7 | 0.4286 | 13 | 0.3077 | 38 | 0.2895 |
| 18_430319 | 10 | 0.2 | 13 | 0.3846 | 10 | 0.1 | 3 | 0 | 36 | 0.2222 | 7 | 0.1429 | 13 | 0.3846 | 7 | 0.4286 | 12 | 0.1667 | 39 | 0.2821 |
| 18_430319 | 10 | 0.3 | 16 | 0.25 | 10 | 0.4 | 2 | 0.5 | 38 | 0.3158 | 5 | 0.6 | 10 | 0.6 | 6 | 0.5 | 13 | 0.4615 | 34 | 0.5294 |
| 18_430322( | 3 | 0.3333 | 8 | 0.5 | 2 | 0 | 2 | 1 | 15 | 0.4667 | 2 | 0.5 | 8 | 0.5 | 4 | 0.75 | 7 | 0.7143 | 21 | 0.619 |
| 18_434481 | 2 | 0 | 2 | 0 | NA | NA | 3 | 0.3333 | 7 | 0.1429 | 1 | 0 | 2 | 0 | 5 | 0.2 | 1 | 0 | 9 | 0.1111 |
| 18_434482 | 1 | 0 | 2 | 0.5 | NA | NA | 2 | 0 | 5 | 0.2 | 1 | 1 | 1 | 0 | 4 | 0 | 1 | 1 | 7 | 0.2857 |
| 18_434483 | 2 | 0.5 | 1 | 0 | NA | NA | 1 | 0 | 4 | 0.25 | NA | NA | 2 | 0 | 4 | 0.5 | NA | NA | 6 | 0.3333 |
| 18_434484 | 1 | 1 | 1 | 1 | NA | NA | 2 | 1 | 4 | 1 | 1 | 1 | 1 | 1 | 6 | 1 | 1 | 1 | 9 | 1 |
| 18_434484 | 2 | 0 | 1 | 0 | NA | NA | 1 | 1 | 4 | 0.25 | NA | NA | 2 | 0 | 5 | 0.2 | 1 | 0 | 8 | 0.125 |
| 18_434486 | 2 | 0.5 | 1 | 0 | 1 | 1 | 3 | 1 | 7 | 0.7143 | 1 | 1 | 2 | 0 | 4 | 0.75 | 1 | 1 | 8 | 0.625 |
| 18_434487 | 2 | 0 | 1 | 0 | 1 | 0 | 3 | 0.3333 | 7 | 0.1429 | 1 | 0 | 3 | 0 | 6 | 0.1667 | 1 | 0 | 11 | 0.0909 |
| 18_434491( | 2 | 0 | NA | NA | 2 | 0 | 3 | 0.3333 | 7 | 0.1429 | 1 | 1 | 3 | 0 | 5 | 0.6 | 1 | 0 | 10 | 0.4 |
| 18_434508 | NA | NA | NA | NA | 1 | 1 | NA | NA | 1 | 1 | NA | NA | 1 | 0 | 1 | 1 | NA | NA | 2 | 0.5 |
| 18_434509( | NA | NA | NA | NA | 1 | 1 | NA | NA | 1 | 1 | NA | NA | 1 | 0 | 1 | 1 | NA | NA | 2 | 0.5 |
| 18_434510( | NA | NA | NA | NA | 1 | 1 | NA | NA | 1 | 1 | NA | NA | 1 | 1 | 1 | 1 | NA | NA | 2 | 1 |
| 18_434511 | NA | NA | NA | NA | 1 | 1 | NA | NA | 1 | 1 | NA | NA | 2 | 1 | 1 | 1 | NA | NA | 3 | 1 |
| 18_434542 | 7 | 0.1429 | NA | NA | 4 | 0.25 | 2 | 0 | 13 | 0.1538 | 4 | 0 | 4 | 0.25 | 1 | 0 | 4 | 0 | 13 | 0.0769 |
| 18_434544 | 5 | 0 | NA | NA | 3 | 0.3333 | 1 | 1 | 9 | 0.2222 | 2 | 0.5 | 4 | 0.25 | NA | NA | 2 | 0.5 | 8 | 0.375 |
| 18_434546 | 4 | 0 | NA | NA | 1 | 1 | 3 | 0 | 8 | 0.125 | 4 | 0 | 3 | 0 | NA | NA | 4 | 0.5 | 11 | 0.1818 |
| 18_434552 | 6 | 0.3333 | NA | NA | 3 | 0 | 3 | 0 | 12 | 0.1667 | 4 | 0.5 | 1 | 1 | NA | NA | 2 | 0 | 7 | 0.4286 |
| 18_434553 | 6 | 0.1667 | NA | NA | 3 | 0 | 2 | 0.5 | 11 | 0.1818 | 4 | 0.5 | 1 | 0 | NA | NA | 2 | 0.5 | 7 | 0.4286 |
| 18_434567 | 3 | 0 | NA | NA | 2 | 0.5 | 1 | 0 | 6 | 0.1667 | 2 | 0 | 1 | 0 | 1 | 1 | 1 | 0 | 5 | 0.2 |
| 18_436633( | NA | NA | 1 | 0 | 2 | 0 | 1 | 1 | 4 | 0.25 | 1 | 0 | NA | NA | 3 | 0.3333 | 1 | 0 | 5 | 0.2 |
| 18_436638 | NA | NA | 2 | 0.5 | 1 | 1 | NA | NA | 3 | 0.6667 | 1 | 0 | 1 | 0 | 3 | 0 | 1 | 1 | 6 | 0.1667 |
| 18_436639 | NA | NA | 2 | 1 | 1 | 1 | NA | NA | 3 | 1 | 1 | 1 | 1 | 1 | 3 | 0.3333 | 1 | 1 | 6 | 0.6667 |
| 18_436639 | NA | NA | 2 | 0.5 | 1 | 1 | NA | NA | 3 | 0.6667 | 1 | 0 | 1 | 0 | 3 | 0 | 1 | 1 | 6 | 0.1667 |
| 18_436641( | NA | NA | 2 | 0.5 | 1 | 0 | NA | NA | 3 | 0.3333 | 1 | 0 | 1 | 1 | 3 | 0.3333 | 1 | 0 | 6 | 0.3333 |
| 18_436641 | NA | NA | 1 | 1 | 1 | 1 | NA | NA | 2 | 1 | 1 | 0 | 1 | 1 | 3 | 0.6667 | NA | NA | 5 | 0.6 |
| 18_436642( | NA | NA | 2 | 1 | 1 | 1 | NA | NA | 3 | 1 | 1 | 1 | 1 | 1 | 3 | 0.6667 | 1 | 1 | 6 | 0.8333 |
| 18_436642 | NA | NA | 2 | 1 | NA | NA | NA | NA | 2 | 1 | 1 | 1 | 1 | 1 | 3 | 1 | NA | NA | 5 | 1 |
| 18_436644( | NA | NA | 2 | 0.5 | 1 | 1 | NA | NA | 3 | 0.6667 | 1 | 0 | 1 | 0 | 3 | 0.6667 | 1 | 1 | 6 | 0.5 |
| 18_436644 | 5 | 0.2 | 9 | 0.3333 | 5 | 0.6 | 2 | 0.5 | 21 | 0.381 | 3 | 0.3333 | 2 | 0 | 8 | 0.375 | 1 | 1 | 14 | 0.3571 |
| 18_436645 | 3 | 0 | 7 | 0.1429 | 3 | 0 | 2 | 0 | 15 | 0.0667 | 2 | 1 | 1 | 0 | 5 | 0 | 1 | 0 | 9 | 0.2222 |
| 18_436646 | 3 | 0.3333 | 9 | 0.1111 | 3 | 0.6667 | 1 | 0 | 16 | 0.25 | 3 | 0 | 1 | 0 | 6 | 0.5 | 2 | 0.5 | 12 | 0.3333 |
| 18_436647( | 4 | 0 | 8 | 0.125 | 4 | 0.25 | 4 | 0 | 20 | 0.1 | 3 | 0.6667 | 2 | 0 | 7 | 0.1429 | 1 | 1 | 13 | 0.3077 |
| 18_436647 | 4 | 0 | 6 | 0.3333 | 2 | 0 | 4 | 0.25 | 16 | 0.1875 | 2 | 0 | 1 | 0 | 5 | 0.2 | NA | NA | 8 | 0.125 |
| 18_436647 | 5 | 0.4 | 9 | 0.6667 | 3 | 0.6667 | 4 | 0.5 | 21 | 0.5714 | 3 | 0.6667 | 2 | 0 | 6 | 0.5 | 1 | 1 | 12 | 0.5 |
| 18_436648 | 5 | 0 | 8 | 0 | 3 | 0.3333 | 3 | 0.3333 | 19 | 0.1053 | 3 | 0.3333 | 2 | 0 | 6 | 0.1667 | 2 | 0 | 13 | 0.1538 |
| 18_436648 | 5 | 0 | 9 | 0.2222 | 3 | 0 | 4 | 0 | 21 | 0.0952 | 3 | 0 | 2 | 0.5 | 6 | 0 | 2 | 0 | 13 | 0.0769 |
| 18_436648 | 5 | 0.4 | 8 | 0.375 | 2 | 0.5 | 4 | 0.5 | 19 | 0.4211 | 3 | 0 | 2 | 0 | 6 | 0.5 | 2 | 0.5 | 13 | 0.3077 |
| 18_436649( | 3 | 0.6667 | 7 | 0.5714 | 3 | 0.3333 | 4 | 0.5 | 17 | 0.5294 | 3 | 0 | 2 | 0.5 | 5 | 0.6 | 2 | 0.5 | 12 | 0.4167 |
| 18_437149 | 1 | 0 | 5 | 0.6 | 1 | 0 | 1 | 0 | 8 | 0.375 | 1 | 1 | NA | NA | 2 | 0.5 | NA | NA | 3 | 0.6667 |
| 18_437150 | 1 | 1 | 5 | 0.4 | 2 | 0.5 | 2 | 1 | 10 | 0.6 | 1 | 1 | 1 | 1 | 1 | 1 | NA | NA | 3 | 1 |
| 18_437151 | NA | NA | 5 | 0.8 | 2 | 0.5 | 2 | 1 | 9 | 0.7778 | 1 | 0 | 1 | 1 | 2 | 1 | NA | NA | 4 | 0.75 |
| 18_437151 | NA | NA | 5 | 1 | 2 | 1 | 2 | 1 | 9 | 1 | 1 | 0 | 1 | 1 | 2 | 1 | NA | NA | 4 | 0.75 |
| 18_437152 | NA | NA | 4 | 1 | 2 | 1 | 2 | 0.5 | 8 | 0.875 | 1 | 1 | NA | NA | 2 | 1 | NA | NA | 3 | 1 |
| 18_437237 | NA | NA | 5 | 0.2 | 2 | 1 | 2 | 1 | 9 | 0.5556 | 1 | 0 | 1 | 0 | 2 | 0.5 | NA | NA | 4 | 0.25 |
| 18_437237( | NA | NA | 3 | 0.3333 | 2 | 0.5 | 2 | 0.5 | 7 | 0.4286 | 1 | 0 | 1 | 0 | 1 | 1 | NA | NA | 3 | 0.3333 |
| 18_445506 | NA | NA | 1 | 1 | 1 | 1 | 1 | 0 | 3 | 0.6667 | 1 | 1 | 1 | 1 | NA | NA | NA | NA | 2 | 1 |
| 18_445507 | NA | NA | 1 | 1 | 1 | 1 | 1 | 1 | 3 | 1 | 1 | 1 | NA | NA | NA | NA | NA | NA | 1 | 1 |
| 18_445507 | NA | NA | NA | NA | 1 | 1 | 1 | 1 | 2 | 1 | 1 | 1 | 1 | 1 | NA | NA | NA | NA | 2 | 1 |
| 18_445509 | NA | NA | 1 | 1 | 1 | 0 | 1 | 1 | 3 | 0.6667 | 1 | 1 | NA | NA | NA | NA | NA | NA | 1 | 1 |
| 18_445512( | NA | NA | 1 | 1 | 1 | 1 | 1 | 1 | 3 | 1 | 2 | 1 | 1 | 1 | NA | NA | NA | NA | 3 | 1 |
| 18_445518 | NA | NA | 1 | 1 | 1 | 0 | 1 | 0 | 3 | 0.3333 | 2 | 0.5 | 1 | 0 | NA | NA | NA | NA | 3 | 0.3333 |
| 18_445519( | NA | NA | 1 | 1 | 1 | 0 | 1 | 0 | 3 | 0.3333 | 2 | 0 | 1 | 1 | NA | NA | NA | NA | 3 | 0.3333 |
| 18_445521 | NA | NA | 1 | 1 | 1 | 0 | 1 | 0 | 3 | 0.3333 | 2 | 0.5 | 1 | 0 | NA | NA | NA | NA | 3 | 0.3333 |
| 18_445585( | 4 | 0.25 | 2 | 0 | NA | NA | 1 | 0 | 7 | 0.1429 | 1 | 0 | 2 | 0.5 | NA | NA | 3 | 0 | 6 | 0.1667 |
| 18_445586 | 3 | 0.3333 | 1 | 1 | NA | NA | 1 | 0 | 5 | 0.4 | 1 | 0 | 2 | 0.5 | NA | NA | 1 | 0 | 4 | 0.25 |

| | | | | | | | | | | | | | | | | | | | | |
|---|---|---|---|---|---|---|---|---|---|---|---|---|---|---|---|---|---|---|---|---|
| 18_4455870 | 3 | 1 | NA | NA | NA | NA | 1 | 0 | 4 | 0.75 | 1 | 0 | 2 | 0.5 | NA | NA | 2 | 0.5 | 5 | 0.4 |
| 18_4455889 | 4 | 0.25 | 1 | 0 | NA | NA | NA | NA | 5 | 0.2 | 1 | 0 | 2 | 0 | NA | NA | 3 | 0.3333 | 6 | 0.1667 |
| 18_4456287 | NA | NA | 1 | 1 | NA | NA | NA | NA | 1 | 1 | NA | NA | NA | NA | NA | NA | 1 | 1 | 1 | 1 |
| 18_4502115 | 3 | 0 | 3 | 0 | 3 | 0 | 1 | 1 | 10 | 0.1 | NA | NA | 6 | 0.1667 | 1 | 0 | NA | NA | 7 | 0.1429 |
| 18_4569330 | 5 | 0.4 | 8 | 0.125 | 3 | 0 | 7 | 0.4286 | 23 | 0.2609 | 2 | 0.5 | 3 | 0.6667 | 3 | 0.3333 | 8 | 0.75 | 16 | 0.625 |
| 18_4569346 | 3 | 0 | 5 | 0.6 | 2 | 1 | 3 | 0.6667 | 13 | 0.5385 | 3 | 0.3333 | 4 | 0.5 | 3 | 1 | 8 | 0.625 | 18 | 0.6111 |
| 18_4572508 | 4 | 0.25 | 8 | 0.125 | 2 | 0 | 7 | 0 | 21 | 0.0952 | 3 | 0 | 3 | 0 | 3 | 0 | 6 | 0.1667 | 15 | 0.0667 |
| 18_4572520 | 5 | 0.4 | 7 | 0.7143 | 1 | 1 | 5 | 0.6 | 18 | 0.6111 | 4 | 0.5 | 2 | 1 | 2 | 1 | 6 | 1 | 14 | 0.8571 |
| 18_4572532 | 3 | 0.3333 | 8 | 0.5 | 1 | 0 | 6 | 0.1667 | 18 | 0.3333 | 3 | 0.3333 | 3 | 0.3333 | 1 | 1 | 3 | 0.6667 | 10 | 0.5 |
| 18_4572537 | 3 | 0 | 11 | 0.1818 | 2 | 1 | 8 | 0.125 | 24 | 0.2083 | 2 | 0.5 | 2 | 0.5 | 4 | 0.5 | 8 | 0.375 | 16 | 0.4375 |
| 18_4572540 | 5 | 0 | 10 | 0.3 | 2 | 0.5 | 7 | 0.4286 | 24 | 0.2917 | 4 | 0.25 | 3 | 0 | 4 | 0.25 | 8 | 0.125 | 19 | 0.1579 |
| 18_4572540 | 5 | 0.2 | 10 | 0 | 2 | 0 | 5 | 0.2 | 22 | 0.0909 | 3 | 0.3333 | 4 | 0.5 | 4 | 0.5 | 6 | 0.1667 | 17 | 0.3529 |
| 18_4572542 | 5 | 0 | 10 | 0.2 | 3 | 0.3333 | 8 | 0.5 | 26 | 0.2692 | 3 | 0.3333 | 3 | 0 | 4 | 0.25 | 7 | 0 | 17 | 0.1176 |
| 18_4572547 | 5 | 0.2 | 10 | 0.1 | 2 | 0 | 5 | 0 | 22 | 0.0909 | 3 | 0.3333 | 4 | 0 | 2 | 0.5 | 8 | 0.25 | 17 | 0.2353 |
| 18_4581843 | 6 | 0.6667 | 14 | 0.8571 | 3 | 0.6667 | 11 | 0.9091 | 34 | 0.8235 | 5 | 1 | 5 | 0.8 | 5 | 1 | 9 | 0.7778 | 24 | 0.875 |
| 18_4581850 | 7 | 0.2857 | 16 | 0.0625 | 3 | 0.3333 | 12 | 0.3333 | 38 | 0.2105 | 7 | 0.1429 | 5 | 0.2 | 5 | 0 | 11 | 0.2727 | 28 | 0.1786 |
| 18_4581865 | 5 | 0.6 | 14 | 0.7857 | 3 | 1 | 11 | 0.6364 | 33 | 0.7273 | 6 | 0.8333 | 5 | 1 | 5 | 0.8 | 10 | 0.8 | 26 | 0.8462 |
| 18_4636881 | 1 | 0 | 1 | 0 | 1 | 1 | 1 | 0 | 4 | 0.25 | 2 | 0.5 | 2 | 0 | 2 | 0 | 2 | 0 | 8 | 0.125 |
| 18_4636882 | 2 | 0 | 3 | 0 | 1 | 1 | 1 | 0 | 7 | 0.1429 | 2 | 0.5 | 2 | 0.5 | 2 | 0 | 2 | 0 | 8 | 0.25 |
| 18_4637420 | NA | NA | NA | NA | 1 | 1 | NA | NA | 1 | 1 | 1 | 0 | NA | NA | 2 | 1 | NA | NA | 3 | 0.6667 |
| 18_4637462 | NA | NA | 1 | 1 | NA | NA | NA | NA | 1 | 1 | 1 | 1 | NA | NA | 2 | 0.5 | NA | NA | 3 | 0.6667 |
| 18_4641550 | 2 | 1 | NA | NA | NA | NA | 1 | 0 | 3 | 0.6667 | NA | NA | NA | NA | 2 | 0.5 | NA | NA | 2 | 0.5 |
| 18_4641571 | 2 | 0.5 | 1 | 0 | 2 | 0.5 | 3 | 0.3333 | 8 | 0.375 | NA | NA | 1 | 1 | 1 | 0 | 2 | 0.5 | 4 | 0.5 |
| 18_4642394 | NA | NA | NA | NA | 1 | 0 | 2 | 0.5 | 3 | 0.3333 | NA | NA | 1 | 0 | 3 | 0 | 3 | 0.3333 | 7 | 0.1429 |
| 18_4642396 | NA | NA | 1 | 1 | 1 | 1 | 3 | 1 | 5 | 1 | NA | NA | NA | NA | 2 | 1 | 3 | 0.6667 | 5 | 0.8 |
| 18_4642397 | NA | NA | 1 | 1 | 1 | 0 | 2 | 0 | 4 | 0.25 | NA | NA | 1 | 0 | 3 | 0.3333 | 2 | 0.5 | 6 | 0.3333 |
| 18_4662544 | NA | NA | NA | NA | 1 | 1 | NA | NA | 1 | 1 | NA | NA | NA | NA | NA | NA | 1 | 1 | 1 | 1 |
| 18_4663652 | 1 | 0 | 4 | 0.75 | 1 | 0 | 2 | 0.5 | 8 | 0.5 | 2 | 0 | 2 | 0 | 3 | 0.3333 | 4 | 0.25 | 11 | 0.1818 |
| 18_4663666 | 1 | 0 | 4 | 0.5 | 1 | 0 | 3 | 0 | 9 | 0.2222 | 1 | 0 | 2 | 0 | 1 | 0 | 4 | 0.25 | 8 | 0.125 |
| 18_4663713 | 1 | 1 | 2 | 0.5 | 1 | 0 | 1 | 0 | 5 | 0.4 | 1 | 1 | 1 | 1 | NA | NA | 4 | 0.5 | 6 | 0.6667 |
| 18_4663761 | 2 | 0 | 3 | 1 | 1 | 1 | 3 | 1 | 9 | 0.7778 | 1 | 1 | 1 | 1 | 1 | 1 | 4 | 1 | 7 | 1 |
| 18_4663768 | 2 | 1 | 3 | 1 | 1 | 1 | 2 | 1 | 8 | 1 | 1 | 1 | NA | NA | 1 | 1 | 4 | 0.75 | 6 | 0.8333 |
| 18_4663786 | 2 | 0.5 | 1 | 0 | 1 | 0 | NA | NA | 4 | 0.25 | 1 | 0 | 1 | 1 | NA | NA | 3 | 0 | 5 | 0.2 |
| 18_4663810 | 2 | 0 | 4 | 0.75 | 1 | 0 | NA | NA | 7 | 0.4286 | 1 | 1 | NA | NA | NA | NA | 4 | 0.5 | 5 | 0.6 |
| 18_4663817 | 4 | 1 | 4 | 0.75 | 1 | 1 | 2 | 1 | 11 | 0.9091 | 1 | 1 | NA | NA | NA | NA | 4 | 1 | 5 | 1 |
| 18_4663830 | 4 | 1 | 4 | 0.75 | NA | NA | 1 | 1 | 9 | 0.8889 | 2 | 1 | NA | NA | NA | NA | 2 | 0.5 | 4 | 0.75 |
| 18_4663831 | 2 | 1 | 2 | 1 | NA | NA | NA | NA | 4 | 1 | 1 | 1 | NA | NA | NA | NA | 1 | 0 | 2 | 0.5 |
| 18_4663832 | 3 | 0.6667 | NA | NA | 1 | 1 | 1 | 1 | 5 | 0.8 | 1 | 1 | NA | NA | NA | NA | 1 | 0 | 2 | 0.5 |
| 18_4663837 | 3 | 0 | 4 | 0 | 1 | 0 | 1 | 1 | 9 | 0.1111 | 2 | 0 | NA | NA | NA | NA | 3 | 0.3333 | 5 | 0.2 |
| 18_4663842 | 4 | 1 | 4 | 1 | 1 | 1 | 2 | 1 | 11 | 1 | 1 | 1 | NA | NA | NA | NA | 3 | 1 | 4 | 1 |
| 18_4663871 | 4 | 0 | 3 | 0.3333 | 1 | 1 | 2 | 0.5 | 10 | 0.3 | 2 | 0 | NA | NA | NA | NA | 3 | 0.6667 | 5 | 0.4 |
| 18_4663883 | 1 | 1 | 1 | 1 | 1 | 1 | 1 | 1 | 4 | 1 | 1 | 1 | NA | NA | NA | NA | NA | NA | 1 | 1 |
| 18_4663886 | 2 | 1 | 4 | 1 | 1 | 1 | 1 | 1 | 8 | 1 | NA | NA | NA | NA | NA | NA | 1 | 1 | 1 | 1 |
| 18_4663888 | 4 | 1 | 4 | 1 | 1 | 1 | 2 | 1 | 11 | 1 | 1 | 1 | NA | NA | NA | NA | 2 | 1 | 3 | 1 |
| 18_4663895 | 4 | 0.5 | 4 | 0.75 | NA | NA | 1 | 0 | 9 | 0.5556 | 1 | 0 | NA | NA | NA | NA | 2 | 0.5 | 3 | 0.3333 |
| 18_4663909 | 4 | 0.25 | 3 | 0 | 1 | 0 | 1 | 0 | 9 | 0.1111 | NA | NA | NA | NA | NA | NA | 2 | 0.5 | 2 | 0.5 |
| 18_4663913 | 3 | 0.6667 | 3 | 1 | NA | NA | 1 | 1 | 7 | 0.8571 | 1 | 1 | NA | NA | NA | NA | 1 | 1 | 2 | 1 |
| 18_4663916 | 3 | 1 | 4 | 1 | 1 | 1 | 2 | 1 | 10 | 1 | 1 | 1 | NA | NA | NA | NA | 2 | 1 | 3 | 1 |
| 18_4663918 | 4 | 0.75 | 3 | 0.6667 | NA | NA | 1 | 1 | 8 | 0.75 | NA | NA | NA | NA | NA | NA | 1 | 1 | 1 | 1 |
| 18_4663927 | 4 | 0.25 | 4 | 0 | 1 | 0 | 1 | 0 | 10 | 0.1 | 1 | 1 | NA | NA | NA | NA | 2 | 0 | 3 | 0.3333 |
| 18_4665723 | 3 | 1 | 3 | 1 | 1 | 1 | 1 | 1 | 8 | 1 | 1 | 1 | NA | NA | NA | NA | 2 | 1 | 3 | 1 |
| 18_4665743 | 3 | 0.3333 | 4 | 0.5 | 1 | 0 | 2 | 0 | 10 | 0.3 | 1 | 0 | NA | NA | NA | NA | 2 | 1 | 3 | 0.6667 |
| 18_4665755 | 4 | 1 | 4 | 1 | 1 | 1 | 2 | 1 | 11 | 1 | 1 | 1 | NA | NA | NA | NA | 2 | 1 | 3 | 1 |
| 18_4665770 | 4 | 0 | 2 | 0 | NA | NA | 1 | 1 | 7 | 0.1429 | 1 | 0 | NA | NA | NA | NA | 1 | 1 | 2 | 0.5 |
| 18_4665780 | 4 | 0.75 | 3 | 1 | NA | NA | 1 | 1 | 8 | 0.875 | NA | NA | NA | NA | NA | NA | 2 | 1 | 2 | 1 |
| 18_4665781 | 4 | 0.5 | 4 | 0.25 | 1 | 1 | 2 | 0.5 | 11 | 0.4545 | NA | NA | NA | NA | NA | NA | 2 | 0.5 | 2 | 0.5 |
| 18_4671926 | 30 | 0.4667 | 16 | 0.4375 | 15 | 0.3333 | 19 | 0.3158 | 80 | 0.4 | 14 | 0.2143 | 18 | 0.3333 | 13 | 0.2308 | 17 | 0.4118 | 62 | 0.3065 |
| 18_4672134 | 17 | 0.9412 | 18 | 1 | 16 | 1 | 22 | 0.9545 | 73 | 0.9726 | 11 | 0.9091 | 17 | 1 | 11 | 0.9091 | 15 | 0.8 | 54 | 0.9074 |
| 18_4672137 | 18 | 0.7222 | 18 | 0.5556 | 16 | 0.6875 | 20 | 0.3 | 72 | 0.5556 | 11 | 0.2727 | 20 | 0.35 | 12 | 0.5 | 14 | 0.5714 | 57 | 0.4211 |
| 18_4672145 | 18 | 0.9444 | 16 | 1 | 14 | 0.9286 | 20 | 0.85 | 68 | 0.9265 | 11 | 0.9091 | 18 | 0.9444 | 11 | 1 | 14 | 0.7143 | 54 | 0.8889 |
| 18_4672146 | 15 | 0.8 | 18 | 0.6667 | 14 | 0.9286 | 19 | 0.8947 | 66 | 0.8182 | 11 | 0.7273 | 17 | 0.8824 | 8 | 1 | 12 | 0.75 | 48 | 0.8333 |
| 18_4672170 | 16 | 0.25 | 17 | 0.1176 | 14 | 0.2857 | 18 | 0.1667 | 65 | 0.2 | 10 | 0.3 | 18 | 0.0556 | 9 | 0.3333 | 13 | 0.1538 | 50 | 0.18 |
| 18_4674058 | 5 | 0 | 6 | 0.1667 | 4 | 0.25 | 8 | 0.125 | 23 | 0.1304 | 3 | 0 | 8 | 0.375 | 8 | 0 | 5 | 0.2 | 24 | 0.1667 |
| 18_4674079 | 5 | 0.2 | 7 | 0.2857 | 3 | 0 | 7 | 0 | 22 | 0.1364 | 2 | 0.5 | 8 | 0.25 | 6 | 0.3333 | 5 | 0 | 21 | 0.2381 |
| 18_4674099 | 2 | 1 | 6 | 1 | 4 | 1 | 7 | 1 | 19 | 1 | 3 | 1 | 6 | 1 | 8 | 1 | 4 | 1 | 21 | 1 |

| | | | | | | | | | | | | | | | | | | | | |
|---|---|---|---|---|---|---|---|---|---|---|---|---|---|---|---|---|---|---|---|---|
| 18_467410 | 5 | 0.4 | 6 | 0.3333 | 4 | 0 | 8 | 0.125 | 23 | 0.2174 | 3 | 0.3333 | 9 | 0.2222 | 9 | 0.2222 | 4 | 0.5 | 25 | 0.28 |
| 18_467410 | 6 | 0 | 6 | 0.5 | 5 | 0.2 | 9 | 0.2222 | 26 | 0.2308 | 3 | 1 | 9 | 0.4444 | 9 | 0.4444 | 5 | 0.2 | 26 | 0.4615 |
| 18_467411 | 3 | 1 | 5 | 1 | 5 | 1 | 7 | 1 | 20 | 1 | 1 | 1 | 9 | 1 | 6 | 1 | 5 | 1 | 21 | 1 |
| 18_467411 | 4 | 0.5 | 5 | 0 | 5 | 0.2 | 8 | 0.125 | 22 | 0.1818 | 3 | 0.3333 | 8 | 0.125 | 7 | 0.2857 | 5 | 0.4 | 23 | 0.2609 |
| 18_467412 | 4 | 0.5 | 6 | 0.1667 | 5 | 0.2 | 9 | 0.1111 | 24 | 0.2083 | 2 | 0.5 | 8 | 0.125 | 10 | 0.2 | 4 | 0.25 | 24 | 0.2083 |
| 18_467413 | 6 | 0.5 | 6 | 0.5 | 4 | 0.5 | 8 | 0.5 | 24 | 0.5 | 4 | 1 | 7 | 0.4286 | 8 | 0.625 | 5 | 0.4 | 24 | 0.5833 |
| 18_467413 | 5 | 0.4 | 6 | 0.5 | 6 | 0.5 | 9 | 0.5556 | 26 | 0.5 | 4 | 0.75 | 9 | 0.4444 | 9 | 0.3333 | 5 | 0 | 27 | 0.3704 |
| 18_467414 | 5 | 0.4 | 7 | 0 | 5 | 0.4 | 8 | 0.125 | 25 | 0.2 | 3 | 0.3333 | 8 | 0.125 | 7 | 0.2857 | 4 | 0.25 | 22 | 0.2273 |
| 18_467414 | 4 | 1 | 7 | 1 | 5 | 1 | 8 | 1 | 24 | 1 | 3 | 1 | 8 | 1 | 10 | 1 | 5 | 1 | 26 | 1 |
| 18_467416 | 5 | 0 | 8 | 0.125 | 4 | 0.25 | 9 | 0.2222 | 26 | 0.1538 | 4 | 0 | 7 | 0.1429 | 7 | 0.1429 | 4 | 0.25 | 22 | 0.1364 |
| 18_467416 | 4 | 0.5 | 7 | 0.2857 | 6 | 0.1667 | 9 | 0.6667 | 26 | 0.4231 | 4 | 0.25 | 9 | 0.3333 | 10 | 0.3 | 5 | 0.4 | 28 | 0.3214 |
| 18_468466 | 11 | 0.1818 | 8 | 0.125 | 7 | 0.1429 | 12 | 0.1667 | 38 | 0.1579 | 6 | 0.1667 | 6 | 0.1667 | 11 | 0.1818 | 10 | 0.3 | 33 | 0.2121 |
| 18_468469 | 7 | 0 | 5 | 0 | 6 | 0.3333 | 6 | 0.5 | 24 | 0.2083 | 6 | 0.1667 | 5 | 0 | 5 | 0 | 6 | 0.5 | 22 | 0.1818 |
| 18_468657 | 11 | 0.1818 | 8 | 0.25 | 12 | 0 | 13 | 0.0769 | 44 | 0.1136 | 10 | 0 | 11 | 0.2727 | 6 | 0.1667 | 9 | 0.2222 | 36 | 0.1667 |
| 18_468658 | 11 | 0.0909 | 8 | 0 | 13 | 0.1538 | 14 | 0.2143 | 46 | 0.1304 | 9 | 0.1111 | 11 | 0.1818 | 7 | 0.2857 | 9 | 0 | 36 | 0.1389 |
| 18_468660 | 12 | 0.0833 | 9 | 0 | 15 | 0.2 | 11 | 0.0909 | 47 | 0.1064 | 10 | 0 | 11 | 0 | 9 | 0.3333 | 9 | 0 | 39 | 0.0769 |
| 18_468667 | 11 | 0.0909 | 8 | 0.125 | 13 | 0.2308 | 11 | 0.0909 | 43 | 0.1395 | 10 | 0.1 | 8 | 0.25 | 8 | 0.375 | 9 | 0.2222 | 35 | 0.2286 |
| 18_468667 | 10 | 0.1 | 8 | 0 | 11 | 0.0909 | 11 | 0 | 40 | 0.05 | 9 | 0 | 8 | 0.125 | 9 | 0.1111 | 10 | 0.2 | 36 | 0.1111 |
| 18_468671 | 9 | 0 | 7 | 0.1429 | 13 | 0.2308 | 11 | 0 | 40 | 0.1 | 9 | 0.1111 | 8 | 0.125 | 9 | 0 | 9 | 0 | 35 | 0.0571 |
| 18_468673 | 10 | 0 | 7 | 0.2857 | 14 | 0.2143 | 12 | 0.25 | 43 | 0.186 | 11 | 0.0909 | 10 | 0.2 | 9 | 0.1111 | 9 | 0.1111 | 39 | 0.1282 |
| 18_468674 | 10 | 0 | 7 | 0 | 15 | 0.2 | 12 | 0.0833 | 44 | 0.0909 | 11 | 0.1818 | 10 | 0.1 | 9 | 0 | 10 | 0.1 | 40 | 0.1 |
| 18_468674 | 10 | 0.1 | 7 | 0.1429 | 16 | 0.1875 | 12 | 0.1667 | 45 | 0.1556 | 9 | 0.3333 | 10 | 0.1 | 9 | 0.4444 | 10 | 0.4 | 38 | 0.3158 |
| 18_468674 | 9 | 0.3333 | 7 | 0.4286 | 16 | 0.5625 | 11 | 0.3636 | 43 | 0.4419 | 10 | 0.4 | 9 | 0.3333 | 9 | 0.4444 | 9 | 0.2222 | 37 | 0.3514 |
| 18_468701 | NA | NA | NA | NA | 1 | 0 | 2 | 0.5 | 3 | 0.3333 | NA | NA | 1 | 1 | 1 | 0 | NA | NA | 2 | 0.5 |
| 18_468702 | NA | NA | NA | NA | 1 | 1 | 1 | 0 | 2 | 0.5 | NA | NA | 1 | 1 | 1 | 0 | NA | NA | 2 | 0.5 |
| 18_468702 | NA | NA | NA | NA | 1 | 1 | 2 | 1 | 3 | 1 | NA | NA | 1 | 1 | 1 | 0 | NA | NA | 2 | 0.5 |
| 18_468704 | NA | NA | NA | NA | 1 | 1 | 1 | 1 | 2 | 1 | NA | NA | 1 | 1 | 1 | 1 | NA | NA | 2 | 1 |
| 18_468704 | 8 | 0 | 6 | 0.1667 | 12 | 0 | 10 | 0.1 | 36 | 0.0556 | 7 | 0 | 6 | 0 | 5 | 0 | 8 | 0.25 | 26 | 0.0769 |
| 18_468705 | 8 | 0.375 | 7 | 0.4286 | 13 | 0.4615 | 11 | 0.3636 | 39 | 0.4103 | 9 | 0.1111 | 8 | 0.125 | 7 | 0.5714 | 8 | 0.375 | 32 | 0.2812 |
| 18_469231 | 16 | 0.1875 | 18 | 0.4444 | 17 | 0.2941 | 16 | 0.3125 | 67 | 0.3134 | 13 | 0.3077 | 19 | 0.2105 | 14 | 0.1429 | 15 | 0.4 | 61 | 0.2623 |
| 18_469235 | 8 | 0.25 | 9 | 0.2222 | 13 | 0.2308 | 11 | 0.1818 | 41 | 0.2195 | 6 | 0.1667 | 10 | 0.3 | 8 | 0.125 | 6 | 0.1667 | 30 | 0.2 |
| 18_472202 | 3 | 1 | 4 | 0.5 | 3 | 0 | 7 | 0.5714 | 17 | 0.5294 | 3 | 0 | 5 | 0.6 | 5 | 0.4 | 2 | 1 | 15 | 0.4667 |
| 18_472202 | 5 | 0 | 5 | 0.2 | 4 | 0.25 | 9 | 0.1111 | 23 | 0.1304 | 3 | 0.3333 | 5 | 0.2 | 5 | 0.2 | 2 | 0 | 15 | 0.2 |
| 18_472202 | 5 | 0.4 | 5 | 0.4 | 4 | 0.25 | 6 | 0 | 20 | 0.25 | 1 | 0 | 5 | 0.4 | 5 | 0.4 | 2 | 0 | 13 | 0.3077 |
| 18_473796 | 2 | 0.5 | 5 | 0.4 | NA | NA | 2 | 0 | 9 | 0.3333 | 1 | 0 | NA | NA | 2 | 0.5 | 1 | 0 | 4 | 0.25 |
| 18_473797 | 2 | 1 | 5 | 1 | NA | NA | 2 | 1 | 9 | 1 | 1 | 1 | 1 | 1 | 2 | 1 | 1 | 1 | 5 | 1 |
| 18_473797 | 2 | 0 | 4 | 0 | NA | NA | 2 | 0.5 | 8 | 0.125 | 1 | 1 | 1 | 0 | 2 | 0 | 1 | 0 | 5 | 0.2 |
| 18_473799 | 2 | 0.5 | 4 | 0 | NA | NA | 2 | 0 | 8 | 0.125 | 1 | 0 | 1 | 0 | 2 | 0.5 | 1 | 0 | 5 | 0.2 |
| 18_473803 | 2 | 0.5 | 4 | 0.5 | NA | NA | 2 | 0 | 8 | 0.375 | 1 | 0 | 1 | 0 | 1 | 1 | 1 | 0 | 4 | 0.25 |
| 18_473804 | 2 | 0.5 | 4 | 1 | NA | NA | 2 | 0.5 | 8 | 0.75 | 1 | 1 | 1 | 0 | 1 | 1 | 1 | 0 | 4 | 0.5 |
| 18_473804 | 1 | 1 | 3 | 1 | NA | NA | 2 | 0.5 | 6 | 0.8333 | 1 | 1 | NA | NA | 1 | 1 | 1 | 0 | 3 | 0.6667 |
| 18_473805 | 4 | 0 | 6 | 0.3333 | 4 | 0 | 4 | 0.25 | 18 | 0.1667 | 2 | 0.5 | 2 | 0.5 | 2 | 0 | NA | NA | 6 | 0.3333 |
| 18_473806 | 2 | 0.5 | 6 | 0.8333 | 4 | 0.5 | 3 | 0.6667 | 15 | 0.6667 | 1 | 1 | 1 | 0 | 2 | 0 | 1 | 0 | 5 | 0.2 |
| 18_473806 | 2 | 0 | 5 | 0.2 | 4 | 0 | 2 | 0 | 13 | 0.0769 | 2 | 0.5 | 2 | 0 | 2 | 0 | 1 | 0 | 7 | 0.1429 |
| 18_473808 | 3 | 0.3333 | 5 | 0.4 | 1 | 1 | 4 | 0.25 | 13 | 0.3846 | 2 | 0.5 | 1 | 0 | 2 | 0 | 1 | 1 | 6 | 0.3333 |
| 18_473809 | 3 | 0.3333 | 6 | 0 | 1 | 0 | 4 | 0.5 | 14 | 0.2143 | 2 | 0.5 | 2 | 0 | 2 | 0 | 1 | 0 | 7 | 0.1429 |
| 18_473811 | 2 | 0.5 | 5 | 0.2 | 1 | 1 | 3 | 0.6667 | 11 | 0.4545 | 2 | 0.5 | 2 | 0 | 2 | 0 | 1 | 0 | 7 | 0.1429 |
| 18_473812 | 1 | 0 | 6 | 0.5 | 1 | 1 | 3 | 0.3333 | 11 | 0.4545 | 2 | 1 | 2 | 1 | 2 | 1 | 1 | 1 | 7 | 1 |
| 18_473813 | 2 | 0.5 | 5 | 0.8 | 1 | 1 | 3 | 1 | 11 | 0.8182 | 2 | 1 | 1 | 1 | 1 | 0 | 1 | 1 | 5 | 0.8 |
| 18_473815 | 2 | 1 | 3 | 1 | 1 | 1 | 2 | 1 | 8 | 1 | 2 | 1 | 1 | 1 | 2 | 1 | 1 | 1 | 6 | 1 |
| 18_473816 | 2 | 1 | 5 | 0.8 | 1 | 1 | 2 | 1 | 10 | 0.9 | 1 | 1 | 2 | 1 | 1 | 1 | 1 | 1 | 5 | 1 |
| 18_473817 | 1 | 0 | 5 | 0 | NA | NA | 2 | 0.5 | 8 | 0.125 | 2 | 0.5 | 1 | 0 | 1 | 0 | NA | NA | 4 | 0.25 |
| 18_473817 | 1 | 1 | NA | NA | 1 | 1 | 1 | 1 | 3 | 1 | 1 | 1 | NA | NA | NA | NA | 1 | 1 | 2 | 1 |
| 18_473818 | 2 | 0 | 4 | 0 | 1 | 0 | 2 | 0.5 | 9 | 0.1111 | 2 | 0 | 1 | 0 | 2 | 0.5 | 1 | 0 | 6 | 0.1667 |
| 18_473819 | 2 | 0.5 | 3 | 1 | 1 | 1 | 2 | 1 | 8 | 0.875 | 2 | 1 | 1 | 1 | 2 | 1 | 1 | 1 | 6 | 1 |
| 18_473819 | 2 | 0.5 | 4 | 0.25 | 1 | 1 | 2 | 0 | 9 | 0.3333 | 2 | 0.5 | NA | NA | 2 | 0.5 | 1 | 1 | 5 | 0.6 |
| 18_473821 | 1 | 1 | 2 | 1 | NA | NA | 2 | 1 | 5 | 1 | 2 | 1 | NA | NA | NA | NA | NA | NA | 2 | 1 |
| 18_473821 | 2 | 1 | 4 | 0.75 | 1 | 1 | 2 | 0.5 | 9 | 0.7778 | 2 | 1 | 1 | 1 | 2 | 1 | 1 | 1 | 6 | 1 |
| 18_473823 | 2 | 0.5 | 4 | 0.75 | 1 | 0 | 1 | 0 | 8 | 0.5 | 2 | 1 | 1 | 1 | 1 | 1 | 1 | 1 | 5 | 1 |
| 18_473824 | 2 | 0 | 5 | 1 | NA | NA | 1 | 0 | 8 | 0.625 | 2 | 0.5 | 1 | 0 | 1 | 1 | 1 | 1 | 5 | 0.6 |
| 18_498307 | NA | NA | 2 | 0 | 2 | 0.5 | NA | NA | 4 | 0.25 | NA | NA | 3 | 0.3333 | 1 | 1 | 1 | 1 | 5 | 0.6 |
| 18_498312 | NA | NA | NA | NA | 1 | 1 | NA | NA | 1 | 1 | NA | NA | 1 | 1 | NA | NA | 1 | 1 | 2 | 1 |
| 18_498313 | NA | NA | 1 | 1 | 2 | 1 | NA | NA | 3 | 1 | NA | NA | 3 | 1 | 1 | 1 | 1 | 1 | 5 | 1 |
| 18_498314 | NA | NA | NA | NA | 1 | 1 | NA | NA | 1 | 1 | NA | NA | 2 | 0.5 | NA | NA | 1 | 1 | 3 | 0.6667 |
| 18_500264 | NA | NA | 1 | 1 | 1 | 1 | 1 | 1 | 3 | 1 | NA | NA | NA | NA | NA | NA | 2 | 1 | 2 | 1 |

| | | | | | | | | | | | | | | | | | | | | |
|---|---|---|---|---|---|---|---|---|---|---|---|---|---|---|---|---|---|---|---|---|
| 18_500268 | NA | NA | 1 | 1 | 1 | 1 | 1 | 1 | 3 | 1 | NA | NA | NA | NA | NA | NA | 2 | 1 | 2 | 1 |
| 18_500268 | NA | NA | 2 | 0.5 | 1 | 1 | 1 | 1 | 4 | 0.75 | NA | NA | NA | NA | NA | NA | 1 | 1 | 1 | 1 |
| 18_500269 | NA | NA | 2 | 0.5 | 1 | 1 | 1 | 1 | 4 | 0.75 | NA | NA | NA | NA | 1 | 1 | 2 | 1 | 3 | 1 |
| 18_500270 | NA | NA | 2 | 1 | 1 | 0 | 1 | 1 | 4 | 0.75 | NA | NA | NA | NA | 1 | 1 | 2 | 0.5 | 3 | 0.6667 |
| 18_500359 | NA | NA | 2 | 0 | 1 | 1 | 1 | 0 | 4 | 0.25 | 1 | 1 | NA | NA | 2 | 0 | 2 | 0 | 5 | 0.2 |
| 18_500952 | 1 | 0 | 3 | 0 | 3 | 0 | 2 | 1 | 9 | 0.2222 | 1 | 0 | 2 | 0.5 | 4 | 0 | 5 | 0.2 | 12 | 0.1667 |
| 18_500953 | 1 | 1 | 6 | 0.3333 | 3 | 1 | 3 | 0.6667 | 13 | 0.6154 | 1 | 0 | 2 | 0.5 | 3 | 0.3333 | 4 | 0.5 | 10 | 0.4 |
| 18_500953 | 2 | 0.5 | 6 | 0 | 3 | 0.3333 | 3 | 0 | 14 | 0.1429 | 1 | 0 | 2 | 0 | 5 | 0 | 5 | 0.2 | 13 | 0.0769 |
| 18_500953 | 2 | 0.5 | 6 | 0.1667 | 3 | 0 | 3 | 0.6667 | 14 | 0.2857 | 1 | 1 | 2 | 0 | 5 | 0.2 | 5 | 0 | 13 | 0.1538 |
| 18_500954 | 1 | 1 | 6 | 0.1667 | 1 | 0 | 2 | 0 | 10 | 0.2 | 1 | 1 | 2 | 0 | 1 | 0 | 5 | 0 | 9 | 0.1111 |
| 18_500955 | 2 | 0 | 7 | 0.4286 | 3 | 0.6667 | 3 | 0.3333 | 15 | 0.4 | 1 | 0 | 2 | 0 | 5 | 0.4 | 5 | 0.2 | 13 | 0.2308 |
| 18_500957 | 2 | 0 | 8 | 0.5 | 3 | 0.3333 | 3 | 0.3333 | 16 | 0.375 | 1 | 1 | 4 | 0.5 | 5 | 0.2 | 5 | 0.2 | 15 | 0.3333 |
| 18_500957 | 2 | 0 | 8 | 0.25 | 3 | 0 | 3 | 0 | 16 | 0.125 | 1 | 0 | 4 | 0.5 | 5 | 0.2 | 5 | 0 | 15 | 0.2 |
| 18_500958 | 2 | 0 | 5 | 0.2 | 2 | 0 | 2 | 0.5 | 11 | 0.1818 | NA | NA | 3 | 0.6667 | 3 | 0 | 4 | 0 | 10 | 0.2 |
| 18_500959 | 2 | 0.5 | 8 | 0 | 3 | 0 | 3 | 0.3333 | 16 | 0.125 | 1 | 0 | 4 | 0.5 | 4 | 0.5 | 5 | 0.4 | 14 | 0.4286 |
| 18_500959 | 1 | 1 | 4 | 0.75 | 2 | 0 | 3 | 0.6667 | 10 | 0.6 | 1 | 1 | 3 | 0.6667 | 5 | 0.6 | 4 | 0.5 | 13 | 0.6154 |
| 18_500967 | 3 | 0 | 8 | 0.125 | 3 | 0.3333 | 3 | 0 | 17 | 0.1176 | 1 | 0 | 3 | 0 | 5 | 0.2 | 4 | 0 | 13 | 0.0769 |
| 18_501850 | NA | NA | NA | NA | NA | NA | 1 | 1 | 1 | 1 | 1 | 1 | NA | NA | NA | NA | NA | NA | 1 | 1 |
| 18_501851 | NA | NA | NA | NA | NA | NA | 1 | 1 | 1 | 1 | 1 | 1 | NA | NA | NA | NA | NA | NA | 1 | 1 |
| 18_501853 | NA | NA | NA | NA | NA | NA | 1 | 1 | 1 | 1 | 1 | 1 | NA | NA | NA | NA | 1 | 0 | 2 | 0.5 |
| 18_502237 | 2 | 1 | NA | NA | 1 | 1 | 3 | 1 | 6 | 1 | NA | NA | NA | NA | 2 | 0.5 | 1 | 0 | 3 | 0.3333 |
| 18_502238 | 2 | 1 | NA | NA | 1 | 0 | 3 | 0.3333 | 6 | 0.5 | NA | NA | NA | NA | 3 | 0.6667 | 1 | 0 | 4 | 0.5 |
| 18_502238 | 2 | 0 | 1 | 0 | 1 | 1 | 2 | 0 | 6 | 0.1667 | NA | NA | NA | NA | 3 | 0.3333 | 1 | 0 | 4 | 0.25 |
| 18_502239 | 2 | 0.5 | 1 | 1 | 1 | 1 | 3 | 1 | 7 | 0.8571 | NA | NA | NA | NA | 3 | 0.6667 | 1 | 1 | 4 | 0.75 |
| 18_502239 | 1 | 1 | 1 | 1 | 1 | 1 | 3 | 1 | 6 | 1 | NA | NA | NA | NA | 3 | 1 | 1 | 1 | 4 | 1 |
| 18_502241 | 2 | 0 | 1 | 0 | 1 | 0 | 3 | 0.3333 | 7 | 0.1429 | NA | NA | NA | NA | 3 | 0.3333 | NA | NA | 3 | 0.3333 |
| 18_502243 | 2 | 0 | NA | NA | 1 | 1 | 2 | 0.5 | 5 | 0.4 | NA | NA | NA | NA | 2 | 0.5 | 1 | 1 | 3 | 0.6667 |
| 18_502244 | 1 | 1 | 1 | 0 | 1 | 0 | 3 | 0.6667 | 6 | 0.5 | NA | NA | NA | NA | 2 | 1 | 1 | 1 | 3 | 1 |
| 18_502244 | NA | NA | 1 | 0 | 1 | 1 | 3 | 0 | 5 | 0.2 | NA | NA | NA | NA | 2 | 1 | 1 | 0 | 3 | 0.6667 |
| 18_503289 | 11 | 0 | 10 | 0.3 | 10 | 0 | 9 | 0.3333 | 40 | 0.15 | 4 | 0 | 8 | 0.125 | 10 | 0.2 | 13 | 0.0769 | 35 | 0.1143 |
| 18_503291 | 13 | 0.0769 | 9 | 0.2222 | 11 | 0.0909 | 8 | 0.25 | 41 | 0.1463 | 3 | 0 | 7 | 0.2857 | 11 | 0 | 12 | 0.1667 | 33 | 0.1212 |
| 18_514358 | 2 | 1 | 2 | 0.5 | 1 | 1 | NA | NA | 5 | 0.8 | 1 | 1 | NA | NA | 1 | 1 | NA | NA | 2 | 1 |
| 18_514359 | 2 | 1 | 2 | 0.5 | 1 | 1 | NA | NA | 5 | 0.8 | 1 | 0 | NA | NA | 1 | 1 | NA | NA | 2 | 0.5 |
| 18_514360 | 2 | 0 | 2 | 0 | 1 | 1 | NA | NA | 5 | 0.2 | 1 | 0 | NA | NA | 1 | 1 | NA | NA | 2 | 0.5 |
| 18_514360 | 1 | 1 | 1 | 0 | 1 | 1 | NA | NA | 3 | 0.6667 | NA | NA | NA | NA | 1 | 1 | NA | NA | 1 | 1 |
| 18_514361 | 2 | 1 | 1 | 0 | 1 | 0 | NA | NA | 4 | 0.5 | 1 | 0 | NA | NA | 1 | 1 | NA | NA | 2 | 0.5 |
| 18_514361 | 1 | 0 | 3 | 1 | 1 | 1 | NA | NA | 5 | 0.8 | 1 | 1 | NA | NA | 1 | 1 | NA | NA | 2 | 1 |
| 18_514362 | 2 | 1 | 2 | 1 | NA | NA | 1 | 1 | 5 | 1 | 1 | 1 | NA | NA | 1 | 1 | NA | NA | 2 | 1 |
| 18_514363 | 3 | 0.6667 | 3 | 0.6667 | 1 | 1 | 1 | 1 | 8 | 0.75 | 1 | 1 | NA | NA | 1 | 1 | NA | NA | 2 | 1 |
| 18_514366 | 3 | 0.3333 | 2 | 0 | NA | NA | 1 | 0 | 6 | 0.1667 | 1 | 0 | NA | NA | 1 | 1 | NA | NA | 2 | 0.5 |
| 18_514367 | 3 | 1 | 2 | 1 | 1 | 1 | 1 | 1 | 7 | 1 | 1 | 0 | NA | NA | 1 | 1 | NA | NA | 2 | 0.5 |
| 18_514368 | 3 | 1 | 2 | 1 | 2 | 1 | 1 | 1 | 8 | 1 | 1 | 1 | NA | NA | 1 | 1 | NA | NA | 2 | 1 |
| 18_514369 | 3 | 0.6667 | 1 | 0 | 2 | 1 | 1 | 1 | 7 | 0.7143 | 1 | 1 | NA | NA | 1 | 1 | NA | NA | 2 | 1 |
| 18_514376 | 1 | 0 | 1 | 1 | 2 | 0.5 | NA | NA | 4 | 0.5 | NA | NA | NA | NA | 1 | 1 | NA | NA | 1 | 1 |
| 18_515525 | 1 | 1 | 2 | 1 | NA | NA | NA | NA | 3 | 1 | 1 | 1 | NA | NA | NA | NA | NA | NA | 1 | 1 |
| 18_526214 | 2 | 0.5 | 2 | 0.5 | 1 | 1 | 4 | 0.5 | 9 | 0.5556 | 5 | 0.2 | 2 | 0.5 | 1 | 1 | 2 | 0.5 | 10 | 0.4 |
| 18_526214 | 2 | 1 | 2 | 1 | 2 | 0.5 | 4 | 0.5 | 10 | 0.7 | 5 | 0.4 | 2 | 0.5 | 2 | 0 | 1 | 1 | 10 | 0.4 |
| 18_526214 | 2 | 0.5 | 2 | 0 | 2 | 0.5 | 4 | 0 | 10 | 0.2 | 5 | 0.4 | 2 | 0 | 2 | 0.5 | 1 | 0 | 10 | 0.3 |
| 18_526215 | 2 | 0.5 | 2 | 0.5 | 1 | 1 | 2 | 0.5 | 7 | 0.5714 | 3 | 0 | 1 | 1 | 2 | 0 | 2 | 0.5 | 8 | 0.25 |
| 18_526215 | 2 | 0.5 | 1 | 0 | 1 | 1 | 3 | 0.6667 | 7 | 0.5714 | 2 | 0.5 | 2 | 0.5 | 1 | 0 | 1 | 1 | 6 | 0.5 |
| 18_526216 | 1 | 1 | 2 | 0 | 1 | 1 | 3 | 1 | 7 | 0.7143 | 3 | 1 | 2 | 1 | 3 | 1 | 1 | 1 | 9 | 1 |
| 18_526216 | 2 | 0 | 2 | 0.5 | 3 | 0.3333 | 4 | 0.25 | 11 | 0.2727 | 5 | 0.6 | 2 | 0 | 3 | 0.3333 | 1 | 1 | 11 | 0.4545 |
| 18_526218 | 1 | 1 | 2 | 0.5 | 1 | 1 | 3 | 0.3333 | 7 | 0.5714 | 4 | 0.5 | 2 | 0.5 | 2 | 1 | 1 | 0 | 9 | 0.5556 |
| 18_526218 | 2 | 0 | 1 | 1 | 3 | 0.3333 | 5 | 0.2 | 11 | 0.2727 | 5 | 0.4 | 2 | 0.5 | 3 | 0 | 2 | 0 | 12 | 0.25 |
| 18_533536 | 6 | 0.5 | 11 | 0.3636 | 9 | 0 | 9 | 0 | 35 | 0.2 | 7 | 0 | 14 | 0.3571 | 5 | 0.4 | 6 | 0.1667 | 32 | 0.25 |
| 18_533537 | 6 | 0.1667 | 10 | 0.1 | 8 | 0 | 9 | 0 | 33 | 0.0606 | 7 | 0 | 12 | 0.0833 | 4 | 0 | 7 | 0.1429 | 30 | 0.0667 |
| 18_535227 | 1 | 0 | 6 | 0.1667 | 1 | 0 | 1 | 0 | 9 | 0.1111 | NA | NA | 1 | 1 | 3 | 0 | 1 | 0 | 5 | 0.2 |
| 18_535228 | 1 | 0 | 4 | 0.25 | 2 | 0 | 2 | 1 | 9 | 0.3333 | 1 | 0 | 1 | 0 | 3 | 0.3333 | 1 | 0 | 6 | 0.1667 |
| 18_535228 | 1 | 1 | 6 | 0 | 2 | 0 | 2 | 1 | 11 | 0.2727 | NA | NA | 1 | 0 | 3 | 0.3333 | 1 | 1 | 5 | 0.4 |
| 18_535230 | 1 | 1 | 6 | 0 | 1 | 0 | 2 | 0.5 | 10 | 0.2 | 1 | 0 | 1 | 0 | 3 | 0.3333 | 2 | 0 | 7 | 0.1429 |
| 18_535230 | 1 | 1 | 6 | 0 | 2 | 0.5 | 2 | 0.5 | 11 | 0.2727 | NA | NA | 1 | 0 | 2 | 0.5 | 2 | 0 | 5 | 0.2 |
| 18_535231 | NA | NA | 4 | 0.25 | 2 | 0.5 | 1 | 1 | 7 | 0.4286 | NA | NA | 1 | 0 | 2 | 0.5 | 2 | 0 | 5 | 0.2 |
| 18_535234 | 1 | 0 | 4 | 0.5 | 1 | 1 | 2 | 0.5 | 8 | 0.5 | NA | NA | 1 | 1 | 3 | 0.6667 | 1 | 1 | 5 | 0.8 |
| 18_535234 | NA | NA | 5 | 0.2 | 2 | 0 | 2 | 1 | 9 | 0.3333 | 1 | 1 | NA | NA | 3 | 0.3333 | 1 | 0 | 5 | 0.4 |
| 18_535235 | NA | NA | 6 | 0.6667 | 1 | 1 | 1 | 1 | 8 | 0.75 | 1 | 1 | 1 | 0 | 4 | 0.75 | 1 | 1 | 7 | 0.7143 |

| | | | | | | | | | | | | | | | | | | | | |
|---|---|---|---|---|---|---|---|---|---|---|---|---|---|---|---|---|---|---|---|---|
| 18_5381509 | 6 | 0 | 7 | 0.2857 | 2 | 0 | 5 | 0 | 20 | 0.1 | 2 | 0 | 8 | 0.125 | 4 | 0 | 3 | 0 | 17 | 0.0588 |
| 18_5381556 | 6 | 0.6667 | 11 | 0.5455 | 4 | 0 | 9 | 0.4444 | 30 | 0.4667 | 4 | 0.5 | 9 | 0.5556 | 5 | 0.8 | 3 | 0.3333 | 21 | 0.5714 |
| 18_5381569 | 4 | 0.25 | 7 | 0.1429 | 4 | 0.25 | 8 | 0.375 | 23 | 0.2609 | 3 | 0 | 5 | 0.2 | 5 | 0 | 3 | 0.3333 | 16 | 0.125 |
| 18_5381631 | 4 | 0 | 6 | 0.1667 | 2 | 0 | 7 | 0 | 19 | 0.0526 | 2 | 0 | 5 | 0.2 | 5 | 0.6 | 2 | 0.5 | 14 | 0.3571 |
| 18_5381635 | 2 | 0 | 7 | 0.2857 | 3 | 0 | 6 | 0.3333 | 18 | 0.2222 | 2 | 0 | 6 | 0.1667 | 4 | 0.25 | 2 | 0 | 14 | 0.1429 |
| 18_5381636 | 4 | 0 | 10 | 0.5 | 4 | 0 | 6 | 0 | 24 | 0.2083 | 4 | 0.5 | 6 | 0.1667 | 5 | 0.4 | 1 | 0 | 16 | 0.3125 |
| 18_5381640 | 5 | 0.4 | 9 | 0.6667 | 4 | 0.25 | 7 | 0.4286 | 25 | 0.48 | 4 | 0.25 | 6 | 0.1667 | 5 | 0.2 | 3 | 0.6667 | 18 | 0.2778 |
| 18_5381650 | 5 | 0 | 8 | 0 | 2 | 0 | 5 | 0.2 | 20 | 0.05 | 4 | 0.25 | 7 | 0.1429 | 6 | 0 | 3 | 0 | 20 | 0.1 |
| 18_5385634 | 1 | 0 | 2 | 0.5 | 2 | 0 | 4 | 0 | 9 | 0.1111 | 1 | 0 | 4 | 0.25 | 1 | 0 | 1 | 0 | 7 | 0.1429 |
| 18_5402871 | 7 | 0.5714 | 2 | 0 | 3 | 0.6667 | 2 | 0 | 14 | 0.4286 | 2 | 1 | 5 | 0.8 | 3 | 0.3333 | 1 | 0 | 11 | 0.6364 |
| 18_5402871 | 7 | 0.2857 | 4 | 0.75 | 4 | 0.25 | 3 | 0.3333 | 18 | 0.3889 | 2 | 0 | 3 | 0.6667 | 4 | 0.75 | 1 | 1 | 10 | 0.6 |
| 18_5402884 | 6 | 0 | 4 | 0.25 | 4 | 0.25 | 4 | 0 | 18 | 0.1111 | 2 | 0 | 5 | 0.4 | 4 | 0 | 1 | 0 | 12 | 0.1667 |
| 18_5402886 | 6 | 0.3333 | 4 | 0.5 | 4 | 0.5 | 2 | 0 | 16 | 0.375 | 2 | 0 | 3 | 0.6667 | 2 | 0.5 | 1 | 0 | 8 | 0.375 |
| 18_5402887 | 5 | 1 | 3 | 0.3333 | 4 | 0.75 | 4 | 0.75 | 16 | 0.75 | 2 | 1 | 5 | 0.8 | 4 | 1 | NA | NA | 11 | 0.9091 |
| 18_5402893 | 7 | 0.2857 | 4 | 0.25 | 3 | 0.6667 | 4 | 0.5 | 18 | 0.3889 | 2 | 0.5 | 3 | 0.6667 | 3 | 0.3333 | 1 | 1 | 9 | 0.5556 |
| 18_5402896 | 5 | 0 | 4 | 0 | 4 | 0.25 | 4 | 0 | 17 | 0.0588 | 2 | 0 | 5 | 0 | 4 | 0.25 | 1 | 0 | 12 | 0.0833 |
| 18_5408637 | 9 | 0.2222 | 7 | 0 | 3 | 0 | 6 | 0.1667 | 25 | 0.12 | 4 | 0.25 | 8 | 0.125 | 8 | 0.125 | 5 | 0 | 25 | 0.12 |
| 18_5408637 | 8 | 0 | 7 | 0.1429 | 5 | 0.2 | 6 | 0 | 26 | 0.0769 | 2 | 0 | 3 | 0 | 8 | 0.125 | 5 | 0 | 18 | 0.0556 |
| 18_5408646 | 11 | 0.8182 | 5 | 0.8 | 5 | 1 | 7 | 0.8571 | 28 | 0.8571 | 5 | 1 | 8 | 0.875 | 8 | 0.875 | 6 | 0.8333 | 27 | 0.8889 |
| 18_5408661 | 10 | 0.1 | 7 | 0.4286 | 5 | 0.2 | 6 | 0.3333 | 28 | 0.25 | 5 | 0.4 | 7 | 0.2857 | 11 | 0.6364 | 5 | 0.4 | 28 | 0.4643 |
| 18_5408664 | 10 | 0.4 | 7 | 0.4286 | 4 | 0.25 | 8 | 0.375 | 29 | 0.3793 | 4 | 0.75 | 7 | 0 | 8 | 0.25 | 5 | 0.2 | 24 | 0.25 |
| 18_5408682 | 3 | 0.3333 | 6 | 0.3333 | 4 | 0 | 6 | 0.1667 | 19 | 0.2105 | 3 | 0.3333 | 3 | 0.3333 | 10 | 0.1 | 3 | 0.3333 | 19 | 0.2105 |
| 18_5408686 | 6 | 0.3333 | 5 | 0.2 | 4 | 0 | 4 | 0 | 19 | 0.1579 | 3 | 0 | 4 | 0.25 | 8 | 0.25 | 5 | 0.2 | 20 | 0.2 |
| 18_5408690 | 5 | 0.2 | 4 | 0.25 | 3 | 0.3333 | 5 | 0.4 | 17 | 0.2941 | 3 | 0.3333 | 4 | 0.5 | 6 | 0.3333 | 4 | 0 | 17 | 0.2941 |
| 18_5408826 | 8 | 0.5 | 7 | 0.1429 | 5 | 0.8 | 8 | 0.5 | 28 | 0.4643 | 7 | 0.1429 | 7 | 0.4286 | 10 | 0 | 5 | 0.6 | 29 | 0.2414 |
| 18_5408833 | 6 | 0.1667 | 7 | 0.4286 | 5 | 0.4 | 9 | 0.5556 | 27 | 0.4074 | 5 | 0.2 | 6 | 0.1667 | 9 | 0.4444 | 3 | 0.6667 | 23 | 0.3478 |
| 18_5409614 | NA | NA | 1 | 1 | NA | NA | NA | NA | 1 | 1 | NA | NA | NA | NA | NA | NA | 1 | 1 | 1 | 1 |
| 18_5409629 | NA | NA | 1 | 1 | NA | NA | NA | NA | 1 | 1 | NA | NA | NA | NA | NA | NA | 1 | 1 | 1 | 1 |
| 18_5502845 | 1 | 0 | 2 | 0 | 1 | 1 | NA | NA | 4 | 0.25 | 1 | 1 | NA | NA | NA | NA | NA | NA | 1 | 1 |
| 18_5503028 | 1 | 1 | 1 | 0 | 1 | 1 | NA | NA | 3 | 0.6667 | 2 | 0.5 | 1 | 1 | NA | NA | NA | NA | 3 | 0.6667 |
| 18_5503041 | 1 | 0 | 1 | 0 | 2 | 0.5 | NA | NA | 4 | 0.25 | 2 | 0.5 | NA | NA | NA | NA | NA | NA | 2 | 0.5 |
| 18_5503056 | 1 | 1 | 1 | 1 | 2 | 0.5 | NA | NA | 4 | 0.75 | 2 | 1 | 1 | 1 | NA | NA | NA | NA | 3 | 1 |
| 18_5503060 | 1 | 1 | 2 | 0.5 | 1 | 1 | NA | NA | 4 | 0.75 | 2 | 1 | 1 | 0 | NA | NA | NA | NA | 3 | 0.6667 |
| 18_5503067 | 1 | 0 | 2 | 0.5 | 2 | 1 | NA | NA | 5 | 0.6 | 2 | 0.5 | 1 | 1 | NA | NA | NA | NA | 3 | 0.6667 |
| 18_5503073 | NA | NA | 2 | 0.5 | 2 | 0 | NA | NA | 4 | 0.25 | 2 | 0.5 | 1 | 1 | NA | NA | NA | NA | 3 | 0.6667 |
| 18_5503112 | 1 | 0 | 1 | 1 | 2 | 0.5 | NA | NA | 4 | 0.5 | 2 | 0.5 | 1 | 1 | NA | NA | NA | NA | 3 | 0.6667 |
| 18_5503127 | 1 | 1 | 1 | 1 | 2 | 0.5 | NA | NA | 4 | 0.75 | 2 | 1 | 1 | 1 | NA | NA | NA | NA | 3 | 1 |
| 18_5503137 | 1 | 1 | NA | NA | 2 | 0.5 | NA | NA | 3 | 0.6667 | 2 | 0.5 | 1 | 0 | NA | NA | NA | NA | 3 | 0.3333 |
| 18_5503146 | NA | NA | NA | NA | 2 | 0.5 | NA | NA | 2 | 0.5 | 2 | 1 | 1 | 0 | NA | NA | NA | NA | 3 | 0.6667 |
| 18_5503164 | 1 | 1 | NA | NA | 2 | 0.5 | NA | NA | 3 | 0.6667 | NA | NA | 1 | 1 | NA | NA | NA | NA | 1 | 1 |
| 18_5503165 | 1 | 0 | NA | NA | 2 | 0.5 | NA | NA | 3 | 0.3333 | 1 | 1 | 1 | 0 | NA | NA | NA | NA | 2 | 0.5 |
| 18_5503176 | NA | NA | NA | NA | 2 | 0.5 | NA | NA | 2 | 0.5 | 1 | 1 | 1 | 0 | NA | NA | NA | NA | 2 | 0.5 |
| 18_5503183 | 1 | 1 | NA | NA | 2 | 0.5 | NA | NA | 3 | 0.6667 | 1 | 1 | 1 | 1 | NA | NA | NA | NA | 2 | 1 |
| 18_5503232 | 1 | 0 | NA | NA | 2 | 0.5 | NA | NA | 3 | 0.3333 | 1 | 0 | 1 | 1 | NA | NA | NA | NA | 2 | 0.5 |
| 18_5663655 | 3 | 1 | 1 | 1 | 1 | 1 | NA | NA | 5 | 1 | 3 | 1 | 2 | 1 | NA | NA | NA | NA | 5 | 1 |
| 18_5665868 | 3 | 0 | 1 | 0 | NA | NA | 1 | 1 | 5 | 0.2 | 8 | 0 | 2 | 0.5 | NA | NA | 5 | 0.2 | 15 | 0.1333 |
| 18_5665941 | 2 | 0 | 1 | 0 | NA | NA | 1 | 1 | 4 | 0.25 | 5 | 0.8 | 2 | 0 | NA | NA | 2 | 0.5 | 9 | 0.5556 |
| 18_5665955 | 2 | 0.5 | 1 | 1 | NA | NA | NA | NA | 3 | 0.6667 | 5 | 0.4 | 2 | 1 | NA | NA | 3 | 0 | 10 | 0.4 |
| 18_5665992 | 12 | 0.1667 | 13 | 0.3077 | 8 | 0.375 | 11 | 0.3636 | 44 | 0.2955 | 9 | 0.5556 | 11 | 0.3636 | 6 | 0.1667 | 8 | 0.25 | 34 | 0.3529 |
| 18_5665996 | 12 | 0.0833 | 13 | 0.1538 | 8 | 0 | 10 | 0.2 | 43 | 0.1163 | 10 | 0.2 | 12 | 0.0833 | 6 | 0 | 8 | 0.25 | 36 | 0.1389 |
| 18_5670858 | 4 | 0.75 | 4 | 0.75 | 2 | 0.5 | 5 | 0.4 | 15 | 0.6 | 3 | 1 | 5 | 0.8 | 5 | 0.8 | 4 | 0.75 | 17 | 0.8235 |
| 18_5670860 | 2 | 0.5 | 5 | 0.4 | 2 | 0 | 6 | 0.1667 | 15 | 0.2667 | NA | NA | 4 | 0.25 | 3 | 0.3333 | 4 | 0 | 11 | 0.1818 |
| 18_5670870 | 3 | 0 | 5 | 0.2 | 3 | 0.3333 | 5 | 0.2 | 16 | 0.1875 | 3 | 0.3333 | 4 | 0.5 | 5 | 0.2 | 5 | 0 | 17 | 0.2353 |
| 18_5670880 | 4 | 0.25 | 5 | 0 | 3 | 0.3333 | 6 | 0.1667 | 18 | 0.1667 | 3 | 0 | 5 | 0.2 | 5 | 0 | 3 | 0 | 16 | 0.0625 |
| 18_5670883 | 5 | 0 | 4 | 0 | 1 | 0 | 6 | 0.3333 | 16 | 0.125 | 3 | 0 | 3 | 0 | 4 | 0.25 | 5 | 0 | 15 | 0.0667 |
| 18_5670885 | 5 | 0.8 | 5 | 0.4 | 2 | 1 | 5 | 0.4 | 17 | 0.5882 | 3 | 0.6667 | 5 | 0.4 | 5 | 0.2 | 5 | 0.8 | 18 | 0.5 |
| 18_5670886 | 4 | 0 | 4 | 0.25 | 2 | 0.5 | 6 | 0 | 16 | 0.125 | 3 | 0 | 4 | 0.25 | 4 | 0 | 5 | 0 | 16 | 0.0625 |
| 18_5670888 | 5 | 0.4 | 4 | 0.5 | 4 | 0.25 | 5 | 0.4 | 18 | 0.3889 | 3 | 0.3333 | 5 | 0.4 | 4 | 0 | 5 | 0.6 | 17 | 0.3529 |
| 18_5670902 | 5 | 0.2 | 6 | 0.5 | 5 | 0.2 | 6 | 0 | 22 | 0.2273 | 3 | 0 | 5 | 0.8 | 4 | 0 | 5 | 0.2 | 17 | 0.2941 |
| 18_5670902 | 4 | 0.75 | 6 | 0.8333 | 3 | 0.3333 | 6 | 0.1667 | 19 | 0.5263 | 2 | 0.5 | 4 | 0 | 5 | 0.4 | 5 | 0.4 | 16 | 0.3125 |
| 18_5670905 | 4 | 0 | 6 | 0.3333 | 5 | 0 | 6 | 0.1667 | 21 | 0.1429 | 2 | 0 | 4 | 0 | 3 | 0 | 4 | 0.25 | 13 | 0.0769 |
| 18_5672005 | 6 | 0 | 6 | 0.1667 | 5 | 0 | 7 | 0.2857 | 24 | 0.125 | 2 | 0 | 3 | 0 | 5 | 0 | 6 | 0.1667 | 16 | 0.0625 |
| 18_5672010 | 4 | 0 | 6 | 0.1667 | 4 | 0.5 | 9 | 0.1111 | 23 | 0.1739 | 2 | 0 | 4 | 0.5 | 5 | 0.2 | 8 | 0.125 | 19 | 0.2105 |
| 18_5672014 | 6 | 0.3333 | 7 | 0 | 6 | 0 | 10 | 0.2 | 29 | 0.1379 | 3 | 0.3333 | 6 | 0.1667 | 5 | 0 | 7 | 0 | 21 | 0.0952 |
| 18_5672020 | 6 | 0.6667 | 7 | 0.5714 | 7 | 0.5714 | 10 | 0.4 | 30 | 0.5333 | 3 | 0.6667 | 5 | 0.2 | 4 | 0.5 | 10 | 0.5 | 22 | 0.4545 |

| | | | | | | | | | | | | | | | | | | | | |
|---|---|---|---|---|---|---|---|---|---|---|---|---|---|---|---|---|---|---|---|---|
| 18_5672020 | 6 | 0.1667 | 5 | 0.2 | 5 | 0.4 | 9 | 0.6667 | 25 | 0.4 | 3 | 0.3333 | 6 | 0.3333 | 3 | 1 | 9 | 0.2222 | 21 | 0.381 |
| 18_5672023 | 5 | 0.8 | 6 | 1 | 5 | 0.8 | 9 | 1 | 25 | 0.92 | 2 | 0.5 | 6 | 1 | 3 | 0.6667 | 9 | 0.7778 | 20 | 0.8 |
| 18_5672030 | 4 | 0.25 | 5 | 0 | 2 | 0.5 | 6 | 0 | 17 | 0.1176 | 2 | 0 | 4 | 0.25 | 4 | 0 | 7 | 0.4286 | 17 | 0.2353 |
| 18_5672035 | 3 | 0 | 5 | 0.2 | 2 | 0.5 | 5 | 0 | 15 | 0.1333 | 3 | 0.3333 | 5 | 0 | 4 | 0.25 | 6 | 0 | 18 | 0.1111 |
| 18_5690887 | NA | NA | NA | NA | 1 | 1 | 1 | 0 | 2 | 0.5 | 1 | 0 | NA | NA | 1 | 1 | NA | NA | 2 | 0.5 |
| 18_5690899 | NA | NA | NA | NA | 1 | 1 | 1 | 1 | 2 | 1 | 1 | 1 | NA | NA | 1 | 1 | NA | NA | 2 | 1 |
| 18_5708898 | NA | NA | 1 | 0 | 1 | 1 | 1 | 1 | 3 | 0.6667 | 1 | 1 | NA | NA | NA | NA | NA | NA | 1 | 1 |
| 18_5709030 | 2 | 1 | 4 | 0.5 | 2 | 1 | 2 | 0.5 | 10 | 0.7 | 2 | 0 | 1 | 0 | NA | NA | 4 | 0.75 | 7 | 0.4286 |
| 18_5709036 | 2 | 0.5 | 3 | 0 | 2 | 0.5 | 1 | 0 | 8 | 0.25 | 2 | 0.5 | NA | NA | NA | NA | 4 | 0.5 | 6 | 0.5 |
| 18_5709037 | 1 | 1 | 2 | 0.5 | 2 | 0 | 2 | 0 | 7 | 0.2857 | 2 | 0 | 1 | 0 | NA | NA | 4 | 0.75 | 7 | 0.4286 |
| 18_5709050 | 1 | 1 | 4 | 0.25 | 2 | 0 | 3 | 0.6667 | 10 | 0.4 | 2 | 0.5 | 1 | 0 | NA | NA | 3 | 0 | 6 | 0.1667 |
| 18_5709065 | 1 | 1 | 2 | 1 | 2 | 1 | NA | NA | 5 | 1 | 1 | 1 | 1 | 1 | NA | NA | 2 | 1 | 4 | 1 |
| 18_5709070 | 1 | 1 | 3 | 1 | 2 | 1 | 3 | 1 | 9 | 1 | 2 | 1 | 1 | 1 | NA | NA | 5 | 0.8 | 8 | 0.875 |
| 18_5709075 | NA | NA | 3 | 0.3333 | 2 | 0 | 1 | 0 | 6 | 0.1667 | 2 | 0 | 1 | 1 | NA | NA | 5 | 0.6 | 8 | 0.5 |
| 18_5709081 | 2 | 1 | 3 | 1 | 2 | 1 | 3 | 1 | 10 | 1 | 2 | 1 | 1 | 1 | NA | NA | 3 | 1 | 6 | 1 |
| 18_5709084 | 2 | 0 | 4 | 0 | 1 | 0 | 3 | 0.3333 | 10 | 0.1 | 2 | 0 | 1 | 0 | NA | NA | 5 | 0.2 | 8 | 0.125 |
| 18_5742699 | NA | NA | NA | NA | 3 | 0.3333 | NA | NA | 3 | 0.3333 | 1 | 0 | 1 | 1 | NA | NA | NA | NA | 2 | 0.5 |
| 18_5742703 | NA | NA | NA | NA | 5 | 1 | 1 | 1 | 6 | 1 | 2 | 0 | 1 | 1 | NA | NA | 1 | 1 | 4 | 0.5 |
| 18_5742713 | NA | NA | NA | NA | 4 | 0.75 | 1 | 1 | 5 | 0.8 | 2 | 1 | 1 | 1 | NA | NA | 1 | 0 | 4 | 0.75 |
| 18_5742728 | NA | NA | NA | NA | 5 | 1 | 1 | 0 | 6 | 0.8333 | 2 | 0.5 | 1 | 1 | NA | NA | NA | NA | 3 | 0.6667 |
| 18_5742735 | NA | NA | NA | NA | 2 | 0.5 | 1 | 1 | 3 | 0.6667 | 2 | 1 | 1 | 1 | NA | NA | 1 | 1 | 4 | 1 |
| 18_5742738 | NA | NA | NA | NA | 5 | 0.2 | 1 | 1 | 6 | 0.3333 | 2 | 0 | 1 | 0 | NA | NA | 1 | 1 | 4 | 0.25 |
| 18_5742741 | NA | NA | NA | NA | 4 | 0.75 | NA | NA | 4 | 0.75 | NA | NA | 1 | 1 | NA | NA | 1 | 0 | 2 | 0.5 |
| 18_5742849 | NA | NA | 1 | 0 | 4 | 0.25 | 2 | 0 | 7 | 0.1429 | 3 | 0 | 2 | 0.5 | NA | NA | 3 | 0 | 8 | 0.125 |
| 18_5752239 | NA | NA | NA | NA | 3 | 1 | 3 | 1 | 6 | 1 | 2 | 1 | 1 | 1 | 3 | 0.6667 | 1 | 1 | 7 | 0.8571 |
| 18_5752251 | NA | NA | NA | NA | 4 | 0.75 | 3 | 0 | 7 | 0.4286 | 2 | 0.5 | 3 | 0 | 3 | 0 | 1 | 0 | 9 | 0.1111 |
| 18_5752292 | NA | NA | NA | NA | 3 | 0.3333 | 3 | 0 | 6 | 0.1667 | NA | NA | NA | NA | 1 | 1 | 1 | 0 | 2 | 0.5 |
| 18_5752561 | 2 | 0 | 1 | 1 | 3 | 0.3333 | 5 | 0.6 | 11 | 0.4545 | NA | NA | NA | NA | 1 | 1 | 2 | 0.5 | 3 | 0.6667 |
| 18_5752583 | 2 | 0.5 | 1 | 0 | 3 | 0.3333 | 6 | 0.1667 | 12 | 0.25 | 1 | 0 | NA | NA | 2 | 0.5 | 2 | 0.5 | 5 | 0.4 |
| 18_5791938 | NA | NA | 1 | 1 | NA | NA | NA | NA | 1 | 1 | 1 | 1 | NA | NA | NA | NA | NA | NA | 1 | 1 |
| 18_5791972 | NA | NA | 1 | 1 | NA | NA | NA | NA | 1 | 1 | 1 | 1 | NA | NA | NA | NA | NA | NA | 1 | 1 |
| 18_5801175 | 1 | 0 | NA | NA | 2 | 0.5 | 1 | 1 | 4 | 0.5 | NA | NA | 3 | 0.6667 | NA | NA | 1 | 0 | 4 | 0.5 |
| 18_5801196 | 1 | 0 | NA | NA | 1 | 0 | 1 | 1 | 3 | 0.3333 | NA | NA | 2 | 0.5 | NA | NA | 1 | 0 | 3 | 0.3333 |
| 18_5801229 | 2 | 0.5 | 1 | 0 | 2 | 0 | 2 | 0.5 | 7 | 0.2857 | NA | NA | 3 | 0.6667 | NA | NA | NA | NA | 3 | 0.6667 |
| 18_5801244 | 1 | 0 | 2 | 0.5 | 1 | 0 | 2 | 0.5 | 6 | 0.3333 | 1 | 0 | 5 | 0.6 | NA | NA | 1 | 0 | 7 | 0.4286 |
| 18_5807712 | 3 | 0.6667 | 10 | 0 | 5 | 0.2 | 5 | 0 | 23 | 0.1304 | 5 | 0 | 6 | 0.3333 | 2 | 0.5 | 5 | 0.4 | 18 | 0.2778 |
| 18_5807714 | 2 | 1 | 13 | 0.7692 | 6 | 1 | 7 | 1 | 28 | 0.8929 | 6 | 0.6667 | 7 | 0.7143 | 2 | 1 | 5 | 1 | 20 | 0.8 |
| 18_5807718 | 4 | 0.75 | 9 | 0.5556 | 6 | 1 | 6 | 1 | 25 | 0.8 | 7 | 0.7143 | 7 | 0.7143 | 3 | 1 | 4 | 1 | 21 | 0.8095 |
| 18_5807722 | 4 | 1 | 12 | 0.5833 | 7 | 0.8571 | 8 | 0.625 | 31 | 0.7097 | 8 | 0.375 | 8 | 0.75 | 2 | 1 | 5 | 0.8 | 23 | 0.6522 |
| 18_5807728 | 4 | 0.25 | 14 | 0 | 8 | 0 | 8 | 0.375 | 34 | 0.1176 | 9 | 0 | 7 | 0.1429 | 3 | 0 | 5 | 0.2 | 24 | 0.0833 |
| 18_5807732 | 4 | 1 | 11 | 0.8182 | 8 | 0.625 | 7 | 0.2857 | 30 | 0.6667 | 3 | 0.3333 | 8 | 1 | NA | NA | 3 | 1 | 14 | 0.8571 |
| 18_5807734 | 4 | 0.75 | 10 | 0.8 | 4 | 1 | 9 | 0.7778 | 27 | 0.8148 | 7 | 1 | 6 | 0.8333 | 3 | 0.6667 | 5 | 1 | 21 | 0.9048 |
| 18_5807756 | 4 | 0 | 12 | 0.3333 | 8 | 0.25 | 8 | 0.125 | 32 | 0.2188 | 8 | 0.125 | 6 | 0.5 | 2 | 0 | 6 | 0 | 22 | 0.1818 |
| 18_5880439 | 2 | 0 | NA | NA | 4 | 0.25 | 4 | 0 | 10 | 0.1 | NA | NA | 5 | 0.2 | 1 | 0 | 4 | 0 | 10 | 0.1 |
| 18_5881098 | 2 | 0 | 1 | 0 | 3 | 0.3333 | 6 | 0 | 12 | 0.0833 | 1 | 0 | 4 | 0.25 | 2 | 0 | 4 | 0.25 | 11 | 0.1818 |
| 18_5881102 | 4 | 0.5 | 1 | 0 | 4 | 0.75 | 7 | 0.2857 | 16 | 0.4375 | 1 | 0 | 5 | 0.4 | 2 | 0.5 | 4 | 0.25 | 12 | 0.3333 |
| 18_5881106 | 3 | 0.3333 | 1 | 0 | 2 | 0 | 4 | 0 | 10 | 0.1 | 1 | 0 | 5 | 0.2 | 1 | 0 | 2 | 0 | 9 | 0.1111 |
| 18_5881144 | 4 | 0 | 3 | 0 | 5 | 0.4 | 8 | 0 | 20 | 0.1 | 2 | 0 | 5 | 0.2 | 2 | 0 | 4 | 0 | 13 | 0.0769 |
| 18_5881147 | 3 | 0.6667 | 2 | 0 | 5 | 0.2 | 8 | 0.75 | 18 | 0.5 | 2 | 0 | 4 | 0.75 | 2 | 0 | 4 | 0.5 | 12 | 0.4167 |
| 18_5881281 | 6 | 0 | 3 | 0 | 4 | 0.25 | 8 | 0.25 | 21 | 0.1429 | 2 | 0 | 5 | 0 | 5 | 0 | 4 | 0.25 | 16 | 0.0625 |
| 18_5881326 | 5 | 0 | 2 | 0 | 4 | 0 | 7 | 0.1429 | 18 | 0.0556 | 3 | 0 | 3 | 0 | 4 | 0 | 4 | 0.25 | 14 | 0.0714 |
| 18_5920543 | 2 | 0 | 3 | 0.3333 | NA | NA | 4 | 0.25 | 9 | 0.2222 | NA | NA | 4 | 0.75 | 7 | 0.2857 | 4 | 0.25 | 15 | 0.4 |
| 18_5920548 | 2 | 0.5 | 3 | 0.3333 | 1 | 1 | 3 | 0.6667 | 9 | 0.5556 | NA | NA | 3 | 0.6667 | 8 | 0.375 | 4 | 0.5 | 15 | 0.4667 |
| 18_5920878 | 1 | 1 | 2 | 0.5 | 2 | 1 | 4 | 0.75 | 9 | 0.7778 | 1 | 1 | 7 | 0.5714 | 7 | 0.4286 | 4 | 0.25 | 19 | 0.4737 |
| 18_5920909 | 3 | 0 | 2 | 0.5 | 2 | 0.5 | 5 | 0.2 | 12 | 0.25 | 2 | 0 | 7 | 0.2857 | 6 | 0 | 5 | 0.2 | 20 | 0.15 |
| 18_5930926 | NA | NA | NA | NA | NA | NA | 1 | 1 | 1 | 1 | 1 | 1 | NA | NA | 1 | 1 | NA | NA | 2 | 1 |
| 18_5954248 | NA | NA | NA | NA | 1 | 1 | 1 | 1 | 2 | 1 | 1 | 0 | 2 | 0.5 | NA | NA | NA | NA | 3 | 0.3333 |
| 18_5954270 | NA | NA | NA | NA | 1 | 1 | 1 | 1 | 2 | 1 | 1 | 1 | 2 | 0.5 | NA | NA | NA | NA | 3 | 0.6667 |
| 18_5954299 | NA | NA | NA | NA | 1 | 1 | 1 | 0 | 2 | 0.5 | 2 | 0.5 | 2 | 0 | 2 | 0 | NA | NA | 6 | 0.1667 |
| 18_5954300 | NA | NA | NA | NA | 1 | 0 | 2 | 1 | 3 | 0.6667 | 2 | 0.5 | 2 | 0 | 2 | 0.5 | NA | NA | 6 | 0.3333 |
| 18_5954302 | NA | NA | NA | NA | 1 | 1 | 2 | 1 | 3 | 1 | 1 | 1 | 1 | 1 | 2 | 1 | NA | NA | 4 | 1 |
| 18_5954306 | NA | NA | NA | NA | 1 | 0 | 2 | 1 | 3 | 0.6667 | 2 | 0.5 | 1 | 1 | 1 | 0 | NA | NA | 4 | 0.5 |
| 18_5954308 | NA | NA | NA | NA | 1 | 1 | 2 | 1 | 3 | 1 | 2 | 0.5 | 2 | 1 | 2 | 0.5 | NA | NA | 6 | 0.6667 |
| 18_5954316 | NA | NA | NA | NA | 1 | 1 | 2 | 0 | 3 | 0.3333 | 2 | 0.5 | 2 | 0 | 1 | 1 | NA | NA | 5 | 0.4 |
| 18_5954319 | NA | NA | NA | NA | NA | NA | 2 | 1 | 2 | 1 | 2 | 1 | 2 | 0.5 | 2 | 1 | NA | NA | 6 | 0.8333 |

| | | | | | | | | | | | | | | | | | | | | |
|---|---|---|---|---|---|---|---|---|---|---|---|---|---|---|---|---|---|---|---|---|
| 18_595432( | NA | NA | NA | NA | 1 | 1 | 2 | 1 | 3 | 1 | 2 | 1 | 2 | 0.5 | 2 | 1 | NA | NA | 6 | 0.8333 |
| 18_595433! | NA | NA | NA | NA | 1 | 1 | 2 | 1 | 3 | 1 | 2 | 0.5 | 2 | 0 | 1 | 1 | NA | NA | 5 | 0.4 |
| 18_595434( | 1 | 1 | NA | NA | 1 | 1 | 2 | 1 | 4 | 1 | 1 | 1 | 1 | 1 | 2 | 1 | NA | NA | 4 | 1 |
| 18_595434{ | 1 | 1 | NA | NA | 1 | 1 | 2 | 1 | 4 | 1 | 1 | 1 | 2 | 0 | 1 | 0 | NA | NA | 4 | 0.25 |
| 18_595435( | 1 | 1 | NA | NA | 1 | 1 | NA | NA | 2 | 1 | 2 | 1 | NA | NA | 1 | 1 | NA | NA | 3 | 1 |
| 18_595437! | 1 | 1 | NA | NA | 1 | 1 | NA | NA | 2 | 1 | 2 | 1 | 2 | 0.5 | 2 | 0 | 1 | 1 | 7 | 0.5714 |
| 18_595438: | NA | NA | NA | NA | 1 | 1 | 2 | 1 | 3 | 1 | 2 | 1 | 2 | 1 | 1 | 1 | 1 | 1 | 6 | 1 |
| 18_595438! | 1 | 0 | NA | NA | 1 | 0 | 2 | 1 | 4 | 0.5 | 2 | 1 | 2 | 0 | 1 | 0 | 1 | 0 | 6 | 0.3333 |
| 18_595439( | 1 | 1 | NA | NA | 1 | 0 | 2 | 1 | 4 | 0.75 | 2 | 1 | 2 | 1 | 2 | 1 | 1 | 1 | 7 | 1 |
| 18_595439! | 2 | 1 | NA | NA | 1 | 1 | 2 | 1 | 5 | 1 | 2 | 1 | 2 | 1 | 3 | 1 | 1 | 1 | 8 | 1 |
| 18_595440: | 2 | 1 | NA | NA | 1 | 1 | 2 | 1 | 5 | 1 | 2 | 1 | 2 | 1 | 3 | 1 | 1 | 1 | 8 | 1 |
| 18_595440⁄ | 1 | 0 | NA | NA | 1 | 0 | 2 | 1 | 4 | 0.5 | 2 | 0 | 1 | 1 | 3 | 0 | 1 | 0 | 7 | 0.1429 |
| 18_595441: | 2 | 0.5 | NA | NA | 1 | 0 | 2 | 0.5 | 5 | 0.4 | 1 | 0 | 1 | 0 | 2 | 1 | NA | NA | 4 | 0.5 |
| 18_595442: | 2 | 1 | NA | NA | NA | NA | 2 | 1 | 4 | 1 | 2 | 1 | 2 | 1 | 3 | 1 | 1 | 1 | 8 | 1 |
| 18_606075( | 7 | 0.8571 | 1 | 1 | 5 | 0.6 | 2 | 1 | 15 | 0.8 | 5 | 1 | NA | NA | 4 | 0.5 | 1 | 1 | 10 | 0.8 |
| 18_606076: | 6 | 0.3333 | 1 | 1 | 5 | 0.4 | 2 | 0.5 | 14 | 0.4286 | 3 | 1 | NA | NA | 4 | 0.5 | 1 | 1 | 8 | 0.75 |
| 18_606080( | 5 | 0.2 | 1 | 0 | 5 | 0.2 | 2 | 0.5 | 13 | 0.2308 | 4 | 0.5 | NA | NA | 3 | 0.3333 | NA | NA | 7 | 0.4286 |
| 18_606080! | 5 | 0.6 | 1 | 0 | 4 | 0.5 | 2 | 1 | 12 | 0.5833 | 4 | 1 | NA | NA | 3 | 1 | 1 | 1 | 8 | 1 |
| 18_606081{ | 4 | 0 | 1 | 0 | 6 | 0.3333 | 2 | 0.5 | 13 | 0.2308 | 4 | 0.25 | NA | NA | 4 | 0.5 | 1 | 0 | 9 | 0.3333 |
| 18_606082: | 5 | 0 | 1 | 0 | 6 | 0.3333 | 2 | 0.5 | 14 | 0.2143 | 4 | 0.25 | NA | NA | 4 | 0.25 | NA | NA | 8 | 0.25 |
| 18_606083' | 4 | 0.75 | NA | NA | 4 | 0.5 | 2 | 1 | 10 | 0.7 | 4 | 0.75 | NA | NA | 3 | 0.6667 | 1 | 1 | 8 | 0.75 |
| 18_606083: | 4 | 0.5 | NA | NA | 5 | 0.6 | 2 | 0.5 | 11 | 0.5455 | 4 | 0.25 | NA | NA | 3 | 0.6667 | 1 | 1 | 8 | 0.5 |
| 18_606083{ | 3 | 0.3333 | NA | NA | 5 | 0.2 | 2 | 0 | 10 | 0.2 | 4 | 0.25 | NA | NA | 3 | 0 | 1 | 1 | 8 | 0.25 |
| 18_606083! | 3 | 0.6667 | NA | NA | 5 | 0.8 | 2 | 1 | 10 | 0.8 | 3 | 0.6667 | NA | NA | 2 | 1 | 1 | 1 | 6 | 0.8333 |
| 18_606084( | 4 | 0.75 | NA | NA | 4 | 0.5 | 2 | 0 | 10 | 0.5 | 3 | 0.6667 | NA | NA | 3 | 0.3333 | NA | NA | 6 | 0.5 |
| 18_606084! | 4 | 1 | NA | NA | 5 | 1 | 2 | 1 | 11 | 1 | 3 | 1 | NA | NA | 3 | 1 | 1 | 1 | 7 | 1 |
| 18_606598( | NA | NA | NA | NA | 1 | 1 | NA | NA | 1 | 1 | NA | NA | 1 | 1 | NA | NA | NA | NA | 1 | 1 |
| 18_606599: | NA | NA | NA | NA | 1 | 1 | NA | NA | 1 | 1 | NA | NA | 1 | 1 | NA | NA | NA | NA | 1 | 1 |
| 18_606896! | 7 | 0 | 6 | 0.1667 | 7 | 0.1429 | 4 | 0 | 24 | 0.0833 | 7 | 0.1429 | 9 | 0.1111 | 9 | 0.2222 | 4 | 0.25 | 29 | 0.1724 |
| 18_606897' | 7 | 0 | 6 | 0 | 8 | 0.25 | 5 | 0 | 26 | 0.0769 | 5 | 0.2 | 10 | 0.1 | 10 | 0 | 4 | 0 | 29 | 0.069 |
| 18_606899' | 7 | 0.5714 | 5 | 0.4 | 9 | 0.3333 | 5 | 0.4 | 26 | 0.4231 | 8 | 0.5 | 10 | 0.7 | 12 | 0.3333 | 5 | 0.2 | 35 | 0.4571 |
| 18_606900! | 6 | 0.5 | 6 | 0.1667 | 9 | 1 | 5 | 0.6 | 26 | 0.6154 | 8 | 0.875 | 10 | 0.6 | 11 | 0.6364 | 4 | 0.5 | 33 | 0.6667 |
| 18_606905: | 6 | 0 | 4 | 0 | 8 | 0.25 | 3 | 0.3333 | 21 | 0.1429 | 7 | 0 | 6 | 0.1667 | 10 | 0.1 | 1 | 0 | 24 | 0.0833 |
| 18_606908⁄ | 9 | 0.2222 | 4 | 0 | 9 | 0 | 3 | 0 | 25 | 0.08 | 6 | 0.1667 | 10 | 0.1 | 12 | 0 | 6 | 0.1667 | 34 | 0.0882 |
| 18_607052: | 6 | 0.1667 | 4 | 0.5 | 5 | 0 | 8 | 0.125 | 23 | 0.1739 | 5 | 0.2 | 2 | 0 | 3 | 0 | 8 | 0.125 | 18 | 0.1111 |
| 18_607056: | 8 | 0.25 | 4 | 0.25 | 7 | 0 | 7 | 0.2857 | 26 | 0.1923 | 5 | 0.6 | 3 | 0.3333 | 5 | 0.6 | 8 | 0.25 | 21 | 0.4286 |
| 18_607288( | NA | NA | NA | NA | 1 | 1 | NA | NA | 1 | 1 | NA | NA | 1 | 1 | NA | NA | NA | NA | 1 | 1 |
| 18_607291' | NA | NA | NA | NA | 1 | 1 | NA | NA | 1 | 1 | NA | NA | 1 | 1 | NA | NA | NA | NA | 1 | 1 |
| 18_607338' | 13 | 0 | 13 | 0.2308 | 13 | 0.0769 | 15 | 0.2667 | 54 | 0.1481 | 11 | 0.0909 | 12 | 0.25 | 18 | 0.2222 | 15 | 0.1333 | 56 | 0.1786 |
| 18_607338! | 12 | 0.0833 | 8 | 0.125 | 11 | 0.2727 | 15 | 0.2 | 46 | 0.1739 | 10 | 0.1 | 10 | 0.2 | 15 | 0.2667 | 15 | 0.2667 | 50 | 0.22 |
| 18_607342! | 8 | 0.125 | 10 | 0.1 | 10 | 0 | 13 | 0.2308 | 41 | 0.122 | 8 | 0.125 | 7 | 0.1429 | 16 | 0.3125 | 15 | 0.2 | 46 | 0.2174 |
| 18_607343: | 10 | 0.2 | 10 | 0.1 | 11 | 0.0909 | 13 | 0.0769 | 44 | 0.1136 | 10 | 0.1 | 7 | 0 | 16 | 0.3125 | 13 | 0 | 46 | 0.1304 |
| 18_607344! | 9 | 0.5556 | 11 | 0.0909 | 11 | 0.1818 | 14 | 0.2857 | 45 | 0.2667 | 9 | 0 | 9 | 0.2222 | 14 | 0.0714 | 13 | 0.1538 | 45 | 0.1111 |
| 18_607345' | 10 | 0.1 | 7 | 0 | 6 | 0.1667 | 9 | 0 | 32 | 0.0625 | 3 | 0 | 6 | 0.5 | 13 | 0.2308 | 10 | 0.1 | 32 | 0.2188 |
| 18_607345: | 10 | 0.5 | 10 | 0.5 | 9 | 0.5556 | 10 | 0.9 | 39 | 0.6154 | 8 | 0.875 | 7 | 0.5714 | 14 | 0.4286 | 13 | 0.7692 | 42 | 0.6429 |
| 18_607345⁄ | 7 | 1 | 10 | 1 | 10 | 1 | 12 | 1 | 39 | 1 | 6 | 0.8333 | 9 | 0.8889 | 15 | 0.8667 | 14 | 0.9286 | 44 | 0.8864 |
| 18_607345! | 9 | 1 | 9 | 1 | 11 | 1 | 14 | 1 | 43 | 1 | 7 | 1 | 9 | 1 | 17 | 0.9412 | 13 | 1 | 46 | 0.9783 |
| 18_607346( | 8 | 1 | 9 | 0.6667 | 10 | 0.8 | 12 | 0.9167 | 39 | 0.8462 | 7 | 1 | 9 | 0.7778 | 15 | 0.9333 | 13 | 1 | 44 | 0.9318 |
| 18_607348( | 7 | 0.5714 | 6 | 0.3333 | 7 | 0.5714 | 8 | 0.625 | 28 | 0.5357 | 5 | 0.4 | 4 | 1 | 12 | 0.5 | 7 | 0.8571 | 28 | 0.6429 |
| 18_607348( | 7 | 0.5714 | 6 | 0.3333 | 5 | 0.2 | 7 | 0.1429 | 25 | 0.32 | 4 | 0.5 | 5 | 0.2 | 9 | 0.5556 | 10 | 0.2 | 28 | 0.3571 |
| 18_607348' | 8 | 1 | 9 | 0.8889 | 9 | 1 | 11 | 1 | 37 | 0.973 | 5 | 1 | 8 | 1 | 14 | 1 | 13 | 1 | 40 | 1 |
| 18_607349{ | 9 | 0 | 8 | 0.125 | 7 | 0 | 11 | 0.0909 | 35 | 0.0571 | 6 | 0.3333 | 7 | 0 | 14 | 0 | 12 | 0 | 39 | 0.0513 |
| 18_607350: | 9 | 0.8889 | 8 | 1 | 9 | 1 | 12 | 1 | 38 | 0.9737 | 6 | 1 | 6 | 1 | 14 | 0.9286 | 12 | 1 | 38 | 0.9737 |
| 18_607350: | 9 | 0.1111 | 7 | 0.1429 | 9 | 0 | 12 | 0.0833 | 37 | 0.0811 | 6 | 0.1667 | 8 | 0 | 14 | 0.0714 | 12 | 0.0833 | 40 | 0.075 |
| 18_607350( | 7 | 0.7143 | 7 | 0.7143 | 9 | 1 | 9 | 0.6667 | 32 | 0.7812 | 6 | 0.6667 | 8 | 0.625 | 13 | 0.5385 | 13 | 0.5385 | 40 | 0.575 |
| 18_607353: | 9 | 0.3333 | 8 | 0.25 | 9 | 0.4444 | 8 | 0.375 | 34 | 0.3529 | 3 | 0.3333 | 6 | 0.1667 | 11 | 0.2727 | 11 | 0.2727 | 31 | 0.2581 |
| 18_607354: | 9 | 1 | 7 | 0.7143 | 7 | 0.8571 | 9 | 1 | 32 | 0.9062 | 3 | 1 | 7 | 1 | 11 | 0.7273 | 10 | 0.9 | 31 | 0.871 |
| 18_607355' | 8 | 0.125 | 5 | 0 | 7 | 0 | 8 | 0.125 | 28 | 0.0714 | 4 | 0.25 | 6 | 0 | 13 | 0.1538 | 10 | 0 | 33 | 0.0909 |
| 18_607356⁄ | 8 | 0 | 7 | 0.1429 | 7 | 0 | 7 | 0.1429 | 29 | 0.069 | 4 | 0 | 6 | 0 | 13 | 0 | 11 | 0.2727 | 34 | 0.0882 |
| 18_607358: | 5 | 0.2 | 4 | 0 | 7 | 0.1429 | 4 | 0 | 20 | 0.1 | 2 | 0 | 5 | 0 | 9 | 0.1111 | 9 | 0.1111 | 25 | 0.08 |
| 18_607359: | 7 | 0.1429 | 7 | 0 | 7 | 0.1429 | 8 | 0.25 | 29 | 0.1379 | 4 | 0.25 | 6 | 0.3333 | 11 | 0.3636 | 9 | 0.6667 | 30 | 0.4333 |
| 18_607360: | 7 | 0.7143 | 7 | 0.4286 | 9 | 0.5556 | 8 | 0.125 | 31 | 0.4516 | 4 | 0.5 | 5 | 0.4 | 8 | 0.5 | 8 | 0.25 | 25 | 0.4 |
| 18_607360! | 6 | 0.1667 | 6 | 0 | 9 | 0 | 8 | 0.125 | 29 | 0.069 | 4 | 0 | 7 | 0.1429 | 10 | 0 | 8 | 0.125 | 29 | 0.069 |
| 18_607360( | 5 | 0.4 | 6 | 0.1667 | 7 | 0.2857 | 9 | 0.3333 | 27 | 0.2963 | 4 | 0 | 6 | 0.1667 | 11 | 0.2727 | 9 | 0.2222 | 30 | 0.2 |
| 18_607362( | 5 | 0.8 | 7 | 0.8571 | 7 | 0.8571 | 8 | 0.875 | 27 | 0.8519 | 3 | 0.6667 | 5 | 0.8 | 9 | 0.6667 | 8 | 0.75 | 25 | 0.72 |

| | | | | | | | | | | | | | | | | | | | | |
|---|---|---|---|---|---|---|---|---|---|---|---|---|---|---|---|---|---|---|---|---|
| 18_6073635 | 4 | 0 | 6 | 0.6667 | 5 | 0.2 | 5 | 0.4 | 20 | 0.35 | 2 | 0.5 | 4 | 0.75 | 7 | 0.7143 | 7 | 0.4286 | 20 | 0.6 |
| 18_6073650 | 4 | 0.25 | 6 | 0.1667 | 6 | 0.3333 | 8 | 0.125 | 24 | 0.2083 | 2 | 0 | 6 | 0.3333 | 8 | 0.25 | 7 | 0.1429 | 23 | 0.2174 |
| 18_6073708 | 4 | 0.25 | 5 | 0.2 | 4 | 0 | 7 | 0.2857 | 20 | 0.2 | 1 | 0 | 4 | 0.25 | 6 | 0.1667 | 6 | 0.1667 | 17 | 0.1765 |
| 18_6081844 | 3 | 0 | 4 | 0 | 5 | 0.2 | 6 | 0 | 18 | 0.0556 | 4 | 0 | 4 | 0.25 | 3 | 0 | 3 | 0 | 14 | 0.0714 |
| 18_6081954 | 3 | 0 | 6 | 0.1667 | 5 | 0.2 | 4 | 0.5 | 18 | 0.2222 | 3 | 0.3333 | 6 | 0 | 5 | 0 | 4 | 0.25 | 18 | 0.1111 |
| 18_6082167 | 2 | 0.5 | 4 | 0 | 2 | 0.5 | 1 | 0 | 9 | 0.2222 | 3 | 0.3333 | 3 | 0 | 2 | 0 | 3 | 0.3333 | 11 | 0.1818 |
| 18_6082183 | 2 | 0 | 2 | 0.5 | 2 | 0.5 | 1 | 0 | 7 | 0.2857 | 2 | 0 | 3 | 0 | 2 | 0.5 | 4 | 0.5 | 11 | 0.2727 |
| 18_6082211 | 2 | 0 | 3 | 0.3333 | 2 | 0.5 | 1 | 0 | 8 | 0.25 | 2 | 0 | 3 | 0.6667 | 2 | 0 | 4 | 0.25 | 11 | 0.2727 |
| 18_6082225 | 2 | 1 | 3 | 0.6667 | 1 | 1 | 1 | 0 | 7 | 0.7143 | 2 | 0.5 | 1 | 1 | 2 | 0.5 | 4 | 0 | 9 | 0.3333 |
| 18_6084074 | 1 | 0 | 2 | 0.5 | NA | NA | NA | NA | 3 | 0.3333 | 2 | 1 | 1 | 1 | 2 | 1 | 2 | 0.5 | 7 | 0.8571 |
| 18_6084100 | 1 | 1 | 1 | 1 | NA | NA | NA | NA | 2 | 1 | 1 | 1 | 2 | 1 | 2 | 0 | 2 | 1 | 7 | 0.7143 |
| 18_6084109 | 1 | 1 | 1 | 1 | NA | NA | NA | NA | 2 | 1 | 2 | 0.5 | 2 | 1 | 2 | 0.5 | 2 | 0.5 | 8 | 0.625 |
| 18_6084597 | 1 | 1 | NA | NA | NA | NA | NA | NA | 1 | 1 | NA | NA | 2 | 1 | 1 | 1 | 2 | 0.5 | 5 | 0.8 |
| 18_6084614 | 1 | 1 | 1 | 1 | NA | NA | NA | NA | 2 | 1 | NA | NA | 2 | 1 | 1 | 1 | 2 | 1 | 5 | 1 |
| 18_6084636 | 1 | 1 | 1 | 1 | NA | NA | NA | NA | 2 | 1 | NA | NA | 2 | 1 | 1 | 1 | 1 | 1 | 4 | 1 |
| 18_6084637 | 1 | 1 | 1 | 1 | NA | NA | NA | NA | 2 | 1 | NA | NA | 2 | 1 | 1 | 1 | 2 | 1 | 5 | 1 |
| 18_6084638 | 1 | 1 | NA | NA | NA | NA | NA | NA | 1 | 1 | NA | NA | 2 | 0.5 | 1 | 1 | 2 | 1 | 5 | 0.8 |
| 18_6084649 | 1 | 1 | NA | NA | NA | NA | NA | NA | 1 | 1 | NA | NA | 1 | 1 | 1 | 1 | 2 | 1 | 4 | 1 |
| 18_6084655 | 1 | 1 | 1 | 0 | NA | NA | NA | NA | 2 | 0.5 | NA | NA | 1 | 1 | 1 | 1 | 2 | 0.5 | 4 | 0.75 |
| 18_6094684 | 1 | 0 | 2 | 0 | 1 | 1 | 1 | 0 | 5 | 0.2 | 1 | 1 | 1 | 0 | 2 | 0 | 3 | 0.3333 | 7 | 0.2857 |
| 18_6094685 | 3 | 0 | 3 | 0.3333 | 1 | 0 | 1 | 0 | 8 | 0.125 | 1 | 1 | 1 | 0 | 6 | 0 | 3 | 0 | 11 | 0.0909 |
| 18_6094719 | 3 | 0.6667 | 4 | 0.5 | 2 | 0.5 | 1 | 1 | 10 | 0.6 | 1 | 1 | 1 | 1 | 6 | 0.6667 | 2 | 0 | 10 | 0.6 |
| 18_6099285 | NA | NA | 1 | 1 | NA | NA | NA | NA | 1 | 1 | NA | NA | NA | NA | NA | NA | 1 | 1 | 1 | 1 |
| 18_6111804 | 38 | 0.3947 | 47 | 0.3191 | 28 | 0.1786 | 48 | 0.2708 | 161 | 0.2981 | 46 | 0.2609 | 33 | 0.4545 | 45 | 0.2889 | 31 | 0.2903 | 155 | 0.3161 |
| 18_6111809 | 56 | 0.8929 | 60 | 0.9 | 34 | 0.9118 | 65 | 0.8308 | 215 | 0.8791 | 59 | 0.9831 | 48 | 0.9167 | 54 | 0.9074 | 43 | 0.9302 | 204 | 0.9363 |
| 18_6111813 | 62 | 0.2742 | 59 | 0.2881 | 42 | 0.119 | 64 | 0.1875 | 227 | 0.2247 | 61 | 0.2131 | 53 | 0.2453 | 52 | 0.1731 | 43 | 0.0698 | 209 | 0.1818 |
| 18_6111836 | 67 | 0.806 | 71 | 0.8592 | 48 | 0.8958 | 66 | 0.7576 | 252 | 0.8254 | 72 | 0.8611 | 57 | 0.7368 | 63 | 0.8413 | 52 | 0.8077 | 244 | 0.8156 |
| 18_6111840 | 65 | 0.0308 | 72 | 0.0833 | 48 | 0.1042 | 71 | 0.1268 | 256 | 0.0859 | 69 | 0.087 | 57 | 0.0526 | 64 | 0.1562 | 53 | 0.1132 | 243 | 0.1029 |
| 18_6111841 | 60 | 0.3167 | 56 | 0.1786 | 44 | 0.25 | 66 | 0.3333 | 226 | 0.2743 | 64 | 0.2656 | 49 | 0.2449 | 54 | 0.2963 | 46 | 0.3043 | 213 | 0.277 |
| 18_6111850 | 62 | 0.0645 | 72 | 0.0417 | 45 | 0.0889 | 73 | 0.0822 | 252 | 0.0675 | 62 | 0.0323 | 58 | 0.069 | 64 | 0.0469 | 51 | 0.1176 | 235 | 0.0638 |
| 18_6111863 | NA | NA | 1 | 1 | NA | NA | NA | NA | 1 | 1 | NA | NA | 1 | 1 | NA | NA | NA | NA | 1 | 1 |
| 18_6111863 | 66 | 0.0606 | 70 | 0.0286 | 50 | 0.04 | 72 | 0.0694 | 258 | 0.0504 | 70 | 0.0429 | 58 | 0.069 | 66 | 0.1061 | 47 | 0.0426 | 241 | 0.0664 |
| 18_6111867 | 78 | 0.9103 | 77 | 0.9351 | 53 | 0.9245 | 86 | 0.8953 | 294 | 0.915 | 73 | 0.8356 | 65 | 0.8154 | 69 | 0.8406 | 61 | 0.8852 | 268 | 0.8433 |
| 18_6111954 | 112 | 0.1071 | 100 | 0.18 | 79 | 0.0886 | 96 | 0.125 | 387 | 0.1266 | 98 | 0.1429 | 86 | 0.1047 | 94 | 0.1489 | 91 | 0.1538 | 369 | 0.1382 |
| 18_6111958 | 116 | 0.1897 | 116 | 0.1724 | 87 | 0.1609 | 101 | 0.1485 | 420 | 0.169 | 99 | 0.101 | 81 | 0.0864 | 110 | 0.1364 | 102 | 0.1373 | 392 | 0.1173 |
| 18_6112029 | 146 | 0.0274 | 143 | 0.035 | 115 | 0.0609 | 124 | 0.1129 | 528 | 0.0568 | 123 | 0.0894 | 100 | 0.02 | 139 | 0.0504 | 117 | 0.0684 | 479 | 0.0585 |
| 18_6112111 | 18 | 0 | 17 | 0 | 15 | 0.0667 | 28 | 0.1071 | 78 | 0.0513 | 15 | 0.0667 | 17 | 0.0588 | 21 | 0.0952 | 14 | 0.2143 | 67 | 0.1045 |
| 18_6112866 | 4 | 0.5 | 2 | 0.5 | 6 | 0.3333 | 3 | 0.3333 | 15 | 0.4 | 1 | 1 | 3 | 0 | NA | NA | 2 | 0 | 6 | 0.1667 |
| 18_6112868 | 6 | 0.5 | 1 | 1 | 7 | 0.2857 | 4 | 0.75 | 18 | 0.5 | 1 | 1 | 2 | 0.5 | NA | NA | 2 | 0 | 5 | 0.4 |
| 18_6112950 | 2 | 0.5 | 3 | 0.3333 | 3 | 0 | 3 | 0 | 11 | 0.1818 | NA | NA | NA | NA | NA | NA | 1 | 1 | 1 | 1 |
| 18_6112952 | 4 | 0.75 | 3 | 1 | 3 | 1 | 3 | 1 | 13 | 0.9231 | NA | NA | NA | NA | NA | NA | 1 | 1 | 1 | 1 |
| 18_6112954 | 4 | 0.75 | 2 | 0 | 3 | 0.3333 | 2 | 0 | 11 | 0.3636 | NA | NA | NA | NA | NA | NA | 1 | 1 | 1 | 1 |
| 18_6112968 | 3 | 0 | 2 | 0 | 3 | 0.3333 | 2 | 0.5 | 10 | 0.2 | NA | NA | NA | NA | NA | NA | 1 | 1 | 1 | 1 |
| 18_6121645 | 1 | 0 | 2 | 0.5 | 2 | 0 | 1 | 0 | 6 | 0.1667 | 2 | 0.5 | 3 | 0.3333 | 3 | 0.3333 | NA | NA | 8 | 0.375 |
| 18_6121661 | 1 | 1 | 3 | 0.3333 | 2 | 0.5 | 1 | 0 | 7 | 0.4286 | 2 | 0.5 | 4 | 0.25 | 3 | 0 | NA | NA | 9 | 0.2222 |
| 18_6121664 | 1 | 0 | 2 | 0 | 2 | 0.5 | 1 | 0 | 6 | 0.1667 | 2 | 0 | 4 | 0.5 | 2 | 0 | NA | NA | 8 | 0.25 |
| 18_6121666 | 1 | 1 | 2 | 1 | 1 | 1 | 1 | 1 | 5 | 1 | 2 | 1 | 4 | 0.75 | 3 | 1 | NA | NA | 9 | 0.8889 |
| 18_6121686 | 3 | 0 | 3 | 1 | 2 | 0 | 1 | 0 | 9 | 0.3333 | 2 | 1 | 1 | 1 | 3 | 0.3333 | NA | NA | 6 | 0.6667 |
| 18_6121690 | 2 | 0 | 2 | 1 | 2 | 0 | 1 | 1 | 7 | 0.4286 | 2 | 0.5 | 3 | 0 | 3 | 0 | NA | NA | 8 | 0.125 |
| 18_6121723 | 4 | 0.25 | 3 | 1 | 2 | 0.5 | 1 | 1 | 10 | 0.6 | 2 | 1 | 5 | 0.2 | 3 | 0.3333 | NA | NA | 10 | 0.4 |
| 18_6121724 | 4 | 0.25 | 1 | 0 | 2 | 1 | 1 | 1 | 8 | 0.5 | 2 | 1 | 5 | 0.6 | 3 | 1 | NA | NA | 10 | 0.8 |
| 18_6121731 | 4 | 0.5 | 3 | 1 | 2 | 0.5 | 1 | 0 | 10 | 0.6 | 2 | 0.5 | 5 | 0.4 | 3 | 0.3333 | NA | NA | 10 | 0.4 |
| 18_6121732 | 4 | 1 | 2 | 1 | 2 | 1 | 1 | 1 | 9 | 1 | 1 | 1 | 4 | 0.5 | 3 | 1 | NA | NA | 8 | 0.75 |
| 18_6121769 | 4 | 0.5 | 3 | 1 | 2 | 1 | 1 | 0 | 10 | 0.7 | 2 | 1 | 4 | 0.5 | 2 | 1 | NA | NA | 8 | 0.75 |
| 18_6121772 | 4 | 0.75 | 2 | 0.5 | 2 | 1 | 2 | 1 | 10 | 0.8 | 2 | 1 | 5 | 1 | 3 | 0.6667 | NA | NA | 10 | 0.9 |
| 18_6121781 | 4 | 0.25 | 2 | 0.5 | 2 | 0 | 2 | 0.5 | 10 | 0.3 | 2 | 0 | 5 | 0 | 2 | 0.5 | NA | NA | 9 | 0.1111 |
| 18_6121783 | 3 | 1 | 1 | 1 | 1 | 0 | 1 | 1 | 6 | 0.8333 | 3 | 0.6667 | 3 | 1 | 2 | 1 | NA | NA | 8 | 0.875 |
| 18_6121785 | 2 | 1 | 2 | 0 | 1 | 1 | NA | NA | 5 | 0.6 | 2 | 0 | 4 | 0.25 | NA | NA | NA | NA | 6 | 0.1667 |
| 18_6121789 | 3 | 0.3333 | 2 | 0 | 2 | 0.5 | 1 | 0 | 8 | 0.25 | 2 | 0 | 5 | 0.2 | 1 | 0 | NA | NA | 8 | 0.125 |
| 18_6121801 | 4 | 0.5 | 3 | 0.3333 | 1 | 1 | 2 | 0.5 | 10 | 0.5 | 3 | 0.3333 | 6 | 0 | 3 | 0.3333 | NA | NA | 12 | 0.1667 |
| 18_6124339 | 3 | 0.3333 | 3 | 0.6667 | 7 | 0.2857 | 2 | 0 | 15 | 0.3333 | 6 | 0 | 5 | 0.2 | 4 | 0.5 | 2 | 0.5 | 17 | 0.2353 |
| 18_6126363 | 12 | 0.75 | 7 | 0.7143 | 13 | 0.7692 | 9 | 0.6667 | 41 | 0.7317 | 17 | 0.8824 | 10 | 1 | 10 | 1 | 12 | 0.6667 | 49 | 0.8776 |
| 18_6126367 | 9 | 0.1111 | 7 | 0.1429 | 11 | 0.3636 | 8 | 0.375 | 35 | 0.2571 | 11 | 0.4545 | 11 | 0.4545 | 8 | 0.25 | 9 | 0.2222 | 39 | 0.359 |
| 18_6126386 | 14 | 0.7857 | 7 | 0.5714 | 14 | 0.6429 | 7 | 0.5714 | 42 | 0.6667 | 17 | 0.6471 | 9 | 0.6667 | 8 | 0.625 | 10 | 0.5 | 44 | 0.6136 |
| 18_6126409 | 12 | 0.0833 | 8 | 0.125 | 13 | 0 | 10 | 0.1 | 43 | 0.0698 | 13 | 0.1538 | 9 | 0.2222 | 9 | 0 | 6 | 0 | 37 | 0.1081 |

| | | | | | | | | | | | | | | | | | | | | |
|---|---|---|---|---|---|---|---|---|---|---|---|---|---|---|---|---|---|---|---|---|
| 18_6126417 | 14 | 0.0714 | 8 | 0.125 | 12 | 0 | 11 | 0.0909 | 45 | 0.0667 | 14 | 0.2857 | 8 | 0.125 | 9 | 0.1111 | 8 | 0.25 | 39 | 0.2051 |
| 18_6126468 | 4 | 0.75 | 5 | 1 | 2 | 0.5 | NA | NA | 11 | 0.8182 | 7 | 0.5714 | 1 | 1 | 6 | 0.6667 | 1 | 1 | 15 | 0.6667 |
| 18_6126498 | 4 | 0.75 | 6 | 0.6667 | 2 | 1 | NA | NA | 12 | 0.75 | 7 | 1 | 1 | 0 | 5 | 1 | 1 | 1 | 14 | 0.9286 |
| 18_6126509 | 3 | 1 | 3 | 1 | 1 | 1 | NA | NA | 7 | 1 | 4 | 1 | NA | NA | 1 | 1 | 1 | 1 | 6 | 1 |
| 18_6126510 | 3 | 0.6667 | 5 | 0.8 | 2 | 1 | NA | NA | 10 | 0.8 | 6 | 0.6667 | NA | NA | 3 | 0.6667 | 1 | 1 | 10 | 0.7 |
| 18_6126513 | 2 | 1 | 4 | 1 | 3 | 1 | NA | NA | 9 | 1 | 4 | 0.75 | NA | NA | 4 | 0.75 | NA | NA | 8 | 0.75 |
| 18_6126534 | NA | NA | 5 | 0.2 | 2 | 0 | NA | NA | 7 | 0.1429 | 7 | 0.2857 | 1 | 0 | 3 | 0.3333 | 1 | 0 | 12 | 0.25 |
| 18_6126538 | 1 | 1 | 5 | 1 | 2 | 1 | NA | NA | 8 | 1 | 6 | 1 | NA | NA | 4 | 1 | 1 | 1 | 11 | 1 |
| 18_6126546 | 1 | 1 | 3 | 1 | 3 | 0.3333 | NA | NA | 7 | 0.7143 | 5 | 0.6 | 1 | 0 | 4 | 1 | 1 | 1 | 11 | 0.7273 |
| 18_6126553 | 1 | 0 | 5 | 0.2 | 3 | 0 | NA | NA | 9 | 0.1111 | 7 | 0.2857 | NA | NA | 4 | 0 | 1 | 0 | 12 | 0.1667 |
| 18_6130036 | NA | NA | 2 | 1 | NA | NA | NA | NA | 2 | 1 | 2 | 1 | NA | NA | NA | NA | 1 | 0 | 3 | 0.6667 |
| 18_6132649 | NA | NA | 2 | 0.5 | 1 | 0 | 1 | 0 | 4 | 0.25 | NA | NA | 1 | 1 | NA | NA | 1 | 0 | 2 | 0.5 |
| 18_6133210 | NA | NA | NA | NA | NA | NA | 1 | 1 | 1 | 1 | 1 | 1 | NA | NA | NA | NA | NA | NA | 1 | 1 |
| 18_6139263 | 7 | 0.1429 | 8 | 0 | 8 | 0.125 | 9 | 0 | 32 | 0.0625 | 5 | 0 | 10 | 0.1 | 8 | 0.125 | 5 | 0.6 | 28 | 0.1786 |
| 18_6142502 | 4 | 0.25 | 3 | 0.3333 | 1 | 0 | 6 | 0.3333 | 14 | 0.2857 | 2 | 0 | 8 | 0.125 | 1 | 0 | 4 | 0 | 15 | 0.0667 |
| 18_6142507 | 5 | 0.2 | 4 | 0 | 1 | 1 | 8 | 0 | 18 | 0.1111 | 1 | 0 | 7 | 0.1429 | 1 | 0 | 4 | 0 | 13 | 0.0769 |
| 18_6142539 | 4 | 0 | 1 | 0 | NA | NA | 3 | 0.3333 | 8 | 0.125 | 1 | 0 | 3 | 0 | 1 | 1 | 2 | 0 | 7 | 0.1429 |
| 18_6142769 | 1 | 0 | 2 | 0 | 1 | 1 | 6 | 0 | 10 | 0.1 | 1 | 1 | 6 | 0.1667 | 1 | 0 | NA | NA | 8 | 0.25 |
| 18_6142919 | 1 | 0 | 1 | 0 | 1 | 1 | 2 | 0 | 5 | 0.2 | 1 | 1 | 1 | 0 | 1 | 0 | NA | NA | 3 | 0.3333 |
| 18_6142931 | NA | NA | 1 | 0 | 1 | 1 | 3 | 0 | 5 | 0.2 | 1 | 1 | 1 | 0 | 1 | 0 | NA | NA | 3 | 0.3333 |
| 18_6142940 | NA | NA | 1 | 0 | 1 | 1 | 2 | 0.5 | 4 | 0.5 | 1 | 1 | 1 | 0 | 1 | 0 | NA | NA | 3 | 0.3333 |
| 18_6168838 | 1 | 0 | 1 | 0 | 4 | 0 | 4 | 0.25 | 10 | 0.1 | 2 | 0 | 7 | 0 | 6 | 0.1667 | 5 | 0 | 20 | 0.05 |
| 18_6168891 | 7 | 0.1429 | 12 | 0 | 17 | 0.0588 | 14 | 0.0714 | 50 | 0.06 | 10 | 0.1 | 17 | 0.0588 | 21 | 0.0476 | 15 | 0.0667 | 63 | 0.0635 |
| 18_6171729 | NA | NA | NA | NA | NA | NA | 3 | 1 | 3 | 1 | NA | NA | NA | NA | 1 | 0 | 3 | 1 | 4 | 0.75 |
| 18_6171741 | NA | NA | NA | NA | NA | NA | 3 | 0.3333 | 3 | 0.3333 | 1 | 0 | NA | NA | 3 | 0 | 4 | 0.25 | 8 | 0.125 |
| 18_6171834 | NA | NA | NA | NA | NA | NA | 3 | 1 | 3 | 1 | 1 | 1 | NA | NA | 1 | 0 | 4 | 0.75 | 6 | 0.6667 |
| 18_6171843 | NA | NA | NA | NA | NA | NA | 3 | 1 | 3 | 1 | 1 | 1 | NA | NA | 3 | 0.6667 | 3 | 0.6667 | 7 | 0.7143 |
| 18_6171860 | NA | NA | NA | NA | NA | NA | 2 | 0.5 | 2 | 0.5 | 1 | 1 | NA | NA | 3 | 0.3333 | 4 | 0.75 | 8 | 0.625 |
| 18_6172057 | 10 | 0.4 | 16 | 0.125 | 17 | 0.2941 | 21 | 0.2381 | 64 | 0.25 | 15 | 0.2667 | 19 | 0.2632 | 23 | 0.3043 | 25 | 0.2 | 82 | 0.2561 |
| 18_6172061 | 17 | 0.2353 | 23 | 0.1739 | 24 | 0.125 | 20 | 0.1 | 84 | 0.1548 | 16 | 0.125 | 22 | 0.1364 | 23 | 0.2609 | 23 | 0.1739 | 84 | 0.1786 |
| 18_6172063 | 17 | 0.4706 | 24 | 0.625 | 25 | 0.64 | 21 | 0.381 | 87 | 0.5402 | 14 | 0.5 | 26 | 0.5 | 26 | 0.4615 | 27 | 0.7037 | 93 | 0.5484 |
| 18_6172065 | 18 | 0.2778 | 24 | 0.1667 | 25 | 0.08 | 22 | 0.1364 | 89 | 0.1573 | 15 | 0.2 | 23 | 0.1739 | 25 | 0.16 | 22 | 0.1818 | 85 | 0.1765 |
| 18_6172071 | 15 | 0.0667 | 25 | 0.12 | 25 | 0.08 | 21 | 0.0476 | 86 | 0.0814 | 15 | 0.2667 | 23 | 0.0435 | 25 | 0.12 | 31 | 0.1613 | 94 | 0.1383 |
| 18_6172085 | 17 | 0.0588 | 28 | 0.1786 | 23 | 0.087 | 24 | 0.0833 | 92 | 0.1087 | 14 | 0.1429 | 26 | 0.0769 | 23 | 0.0435 | 24 | 0.0833 | 87 | 0.0805 |
| 18_6172107 | 20 | 0.8 | 25 | 0.76 | 28 | 0.6786 | 25 | 0.6 | 98 | 0.7041 | 20 | 0.8 | 28 | 0.7857 | 29 | 0.6897 | 30 | 0.7667 | 107 | 0.757 |
| 18_6172116 | 18 | 0.3333 | 28 | 0.5 | 31 | 0.5484 | 23 | 0.6087 | 100 | 0.51 | 23 | 0.4783 | 30 | 0.4667 | 29 | 0.5172 | 33 | 0.697 | 115 | 0.5478 |
| 18_6172162 | 10 | 0.1 | 11 | 0.3636 | 21 | 0.1429 | 12 | 0 | 54 | 0.1481 | 10 | 0.1 | 14 | 0.0714 | 20 | 0.15 | 17 | 0.2353 | 61 | 0.1475 |
| 18_6172186 | 10 | 0.2 | 13 | 0.4615 | 22 | 0.2273 | 15 | 0.4667 | 60 | 0.3333 | 11 | 0.3636 | 16 | 0.25 | 18 | 0.3333 | 19 | 0.5789 | 64 | 0.3906 |
| 18_6172271 | 10 | 0 | 20 | 0.05 | 22 | 0.1364 | 18 | 0.1111 | 70 | 0.0857 | 15 | 0.0667 | 16 | 0.0625 | 24 | 0.0833 | 25 | 0.04 | 80 | 0.0625 |
| 18_6172299 | 10 | 0.7 | 22 | 0.6818 | 24 | 0.625 | 19 | 0.5789 | 75 | 0.64 | 13 | 0.4615 | 18 | 0.5556 | 23 | 0.6522 | 21 | 0.6667 | 75 | 0.6 |
| 18_6172304 | 10 | 0.6 | 18 | 0.1111 | 19 | 0.2632 | 12 | 0.4167 | 59 | 0.3051 | 10 | 0.5 | 10 | 0.1 | 21 | 0.0952 | 12 | 0.25 | 53 | 0.2075 |
| 18_6172319 | 10 | 0 | 13 | 0.2308 | 11 | 0.0909 | 10 | 0.3 | 44 | 0.1591 | 3 | 0 | 2 | 0 | 10 | 0.3 | 9 | 0.2222 | 24 | 0.2083 |
| 18_6183030 | 3 | 1 | 6 | 1 | 1 | 1 | 6 | 1 | 16 | 1 | 5 | 0.6 | 3 | 1 | 6 | 1 | 2 | 1 | 16 | 0.875 |
| 18_6183033 | 2 | 0 | 8 | 0.25 | 1 | 0 | 5 | 0 | 16 | 0.125 | 7 | 0 | 4 | 0.25 | 4 | 0 | 2 | 0.5 | 17 | 0.1176 |
| 18_6183036 | 4 | 1 | 9 | 0.7778 | 1 | 1 | 6 | 1 | 20 | 0.9 | 7 | 0.5714 | 4 | 1 | 8 | 0.875 | 2 | 1 | 21 | 0.8095 |
| 18_6183038 | 3 | 0.6667 | 3 | 0.6667 | 1 | 1 | 3 | 1 | 10 | 0.8 | 1 | 1 | 2 | 0.5 | 3 | 1 | 2 | 1 | 8 | 0.875 |
| 18_6183039 | 3 | 0.6667 | 5 | 0.6 | NA | NA | 5 | 0.6 | 13 | 0.6154 | 7 | 0.8571 | 1 | 1 | 5 | 0.8 | 1 | 1 | 14 | 0.8571 |
| 18_6183041 | 3 | 0.6667 | 9 | 0.4444 | 1 | 1 | 5 | 0.6 | 18 | 0.5556 | 7 | 0.4286 | 3 | 0.6667 | 8 | 0.75 | 2 | 1 | 20 | 0.65 |
| 18_6183043 | 4 | 0.5 | 8 | 0.625 | 1 | 1 | 6 | 0.1667 | 19 | 0.4737 | 7 | 0.4286 | 3 | 0 | 8 | 0.5 | 2 | 0 | 20 | 0.35 |
| 18_6183075 | 2 | 0 | 7 | 0.2857 | 1 | 0 | 3 | 0 | 13 | 0.1538 | 5 | 0.2 | 2 | 0 | 3 | 0.3333 | 2 | 0 | 12 | 0.1667 |
| 18_6183078 | 3 | 1 | 7 | 0.8571 | 1 | 1 | 5 | 0.8 | 16 | 0.875 | 6 | 1 | 3 | 0 | 6 | 0.8333 | 2 | 0 | 17 | 0.6471 |
| 18_6183081 | 4 | 0.25 | 8 | 0.25 | 1 | 0 | 5 | 0.4 | 18 | 0.2778 | 5 | 0.4 | 2 | 0 | 7 | 0.4286 | 2 | 0.5 | 16 | 0.375 |
| 18_6183089 | 2 | 1 | 5 | 1 | 1 | 1 | 4 | 0.75 | 12 | 0.9167 | 4 | 0.75 | NA | NA | 5 | 1 | 2 | 1 | 11 | 0.9091 |
| 18_6183095 | 4 | 0 | 8 | 0 | 1 | 0 | 3 | 0.3333 | 16 | 0.0625 | 5 | 0 | 2 | 0 | 6 | 0.1667 | 2 | 0 | 15 | 0.0667 |
| 18_6183669 | 2 | 0.5 | 1 | 0 | 1 | 0 | 2 | 0 | 6 | 0.1667 | 2 | 0 | 2 | 0.5 | 3 | 0.3333 | NA | NA | 7 | 0.2857 |
| 18_6184570 | 3 | 1 | 2 | 0.5 | 1 | 0 | 2 | 1 | 8 | 0.75 | NA | NA | 3 | 0.6667 | NA | NA | 1 | 1 | 4 | 0.75 |
| 18_6184578 | 2 | 0.5 | 2 | 0 | NA | NA | NA | NA | 4 | 0.25 | NA | NA | 3 | 0 | NA | NA | 1 | 1 | 4 | 0.25 |
| 18_6184579 | 3 | 0.3333 | 2 | 0.5 | 1 | 1 | 1 | 1 | 7 | 0.5714 | NA | NA | 3 | 0.3333 | NA | NA | 1 | 0 | 4 | 0.25 |
| 18_6184581 | 3 | 0 | 2 | 0 | 1 | 0 | 1 | 1 | 7 | 0.1429 | NA | NA | 3 | 0.3333 | NA | NA | 1 | 0 | 4 | 0.25 |
| 18_6184582 | 3 | 0.6667 | 1 | 0 | 1 | 0 | 1 | 1 | 6 | 0.5 | NA | NA | 2 | 0 | NA | NA | 1 | 1 | 3 | 0.3333 |
| 18_6184645 | 1 | 1 | 1 | 1 | 1 | 0 | 1 | 1 | 4 | 0.75 | NA | NA | 1 | 0 | NA | NA | 1 | 1 | 2 | 0.5 |
| 18_6184691 | NA | NA | 1 | 1 | 1 | 0 | 1 | 1 | 3 | 0.6667 | NA | NA | 3 | 0 | NA | NA | 1 | 1 | 4 | 0.25 |
| 18_6184798 | NA | NA | 1 | 1 | 1 | 0 | 1 | 0 | 3 | 0.3333 | NA | NA | 2 | 0.5 | NA | NA | NA | NA | 2 | 0.5 |
| 18_6184803 | NA | NA | 1 | 0 | 1 | 0 | 1 | 1 | 3 | 0.3333 | NA | NA | 3 | 0.3333 | NA | NA | NA | NA | 3 | 0.3333 |
| 18_6184813 | 1 | 1 | 5 | 0.6 | 1 | 0 | 1 | 0 | 8 | 0.5 | 2 | 0 | 6 | 0.1667 | 3 | 0.3333 | 3 | 0 | 14 | 0.1429 |

| | | | | | | | | | | | | | | | | | | | | |
|---|---|---|---|---|---|---|---|---|---|---|---|---|---|---|---|---|---|---|---|---|
| 18_618481 | 2 | 0.5 | 4 | 0.75 | 1 | 0 | 1 | 1 | 8 | 0.625 | 2 | 0 | 6 | 0.5 | 3 | 0.6667 | 4 | 1 | 15 | 0.6 |
| 18_618481 | 1 | 1 | 5 | 0.4 | 1 | 0 | 1 | 0 | 8 | 0.375 | 2 | 0.5 | 6 | 0 | 2 | 0.5 | 3 | 0 | 13 | 0.1538 |
| 18_618486 | 1 | 1 | 5 | 0.2 | 1 | 0 | 1 | 0 | 8 | 0.25 | 3 | 0 | 4 | 0.25 | 3 | 0 | 3 | 0.6667 | 13 | 0.2308 |
| 18_618486 | 1 | 0 | 4 | 0.25 | 1 | 0 | NA | NA | 6 | 0.1667 | 2 | 0 | 3 | 0.3333 | 1 | 1 | 3 | 0.6667 | 9 | 0.4444 |
| 18_618488 | 1 | 0 | 4 | 0.75 | 1 | 1 | 1 | 1 | 7 | 0.7143 | 2 | 1 | 4 | 0.75 | 1 | 0 | 2 | 1 | 9 | 0.7778 |
| 18_618489 | NA | NA | 3 | 1 | NA | NA | 1 | 1 | 4 | 1 | 1 | 1 | 3 | 1 | NA | NA | 1 | 1 | 5 | 1 |
| 18_618490 | NA | NA | 2 | 1 | NA | NA | 1 | 1 | 3 | 1 | 2 | 0 | 2 | 1 | 2 | 0.5 | 2 | 0.5 | 8 | 0.5 |
| 18_618635 | 1 | 0 | NA | NA | 2 | 0.5 | 2 | 0 | 5 | 0.2 | NA | NA | 1 | 0 | 4 | 0.25 | 1 | 0 | 6 | 0.1667 |
| 18_618636 | 1 | 1 | NA | NA | 2 | 0.5 | 1 | 1 | 4 | 0.75 | NA | NA | 1 | 1 | 4 | 0.5 | 1 | 0 | 6 | 0.5 |
| 18_618639 | 1 | 1 | NA | NA | 2 | 0.5 | 2 | 0.5 | 5 | 0.6 | NA | NA | 1 | 1 | 4 | 0.75 | NA | NA | 5 | 0.8 |
| 18_618642 | 1 | 1 | NA | NA | 2 | 0 | 1 | 0 | 4 | 0.25 | 1 | 1 | 2 | 0.5 | 3 | 0.3333 | 1 | 0 | 7 | 0.4286 |
| 18_619325 | 2 | 0.5 | 3 | 0 | 2 | 0.5 | NA | NA | 7 | 0.2857 | 2 | 0 | 2 | 0 | 4 | 0 | 3 | 0.3333 | 11 | 0.0909 |
| 18_619326 | 3 | 0 | 4 | 0.5 | 2 | 0 | NA | NA | 9 | 0.2222 | 2 | 0 | 2 | 0 | 4 | 0.5 | 1 | 0 | 9 | 0.2222 |
| 18_619330 | 3 | 0.3333 | 3 | 0.3333 | 3 | 0.3333 | NA | NA | 9 | 0.3333 | 2 | 0 | 2 | 0.5 | 4 | 1 | 2 | 0.5 | 10 | 0.6 |
| 18_619332 | 3 | 0 | 3 | 0.3333 | 2 | 0 | NA | NA | 8 | 0.125 | 2 | 0 | 3 | 0.3333 | 3 | 0.3333 | 2 | 0 | 10 | 0.2 |
| 18_619332 | 3 | 0.3333 | 3 | 0.3333 | 2 | 0 | NA | NA | 8 | 0.25 | 2 | 0 | 2 | 0 | 3 | 1 | 2 | 0 | 9 | 0.3333 |
| 18_619340 | 3 | 0 | 3 | 0.6667 | 1 | 1 | NA | NA | 7 | 0.4286 | 1 | 0 | 2 | 0 | 1 | 1 | 2 | 0.5 | 6 | 0.3333 |
| 18_619342 | 3 | 0 | 2 | 0.5 | 2 | 0.5 | NA | NA | 7 | 0.2857 | 1 | 1 | 3 | 0.6667 | 3 | 0.6667 | 1 | 1 | 8 | 0.75 |
| 18_619343 | 2 | 0 | 3 | 0.3333 | NA | NA | NA | NA | 5 | 0.2 | 1 | 0 | 2 | 0.5 | 3 | 0.3333 | 1 | 0 | 7 | 0.2857 |
| 18_625707 | 1 | 1 | NA | NA | NA | NA | NA | NA | 1 | 1 | NA | NA | NA | NA | 1 | 1 | 1 | 1 | 2 | 1 |
| 18_625707 | 1 | 1 | NA | NA | NA | NA | NA | NA | 1 | 1 | NA | NA | NA | NA | 1 | 1 | 1 | 1 | 2 | 1 |
| 18_625708 | 1 | 1 | NA | NA | NA | NA | NA | NA | 1 | 1 | NA | NA | NA | NA | 1 | 1 | 1 | 0 | 2 | 0.5 |
| 18_625708 | 1 | 1 | NA | NA | NA | NA | NA | NA | 1 | 1 | NA | NA | NA | NA | 1 | 0 | 1 | 1 | 2 | 0.5 |
| 18_625976 | 1 | 1 | NA | NA | NA | NA | NA | NA | 1 | 1 | NA | NA | NA | NA | 1 | 1 | 1 | 1 | 2 | 1 |
| 18_625977 | 1 | 1 | NA | NA | NA | NA | NA | NA | 1 | 1 | NA | NA | NA | NA | 1 | 1 | 1 | 1 | 2 | 1 |
| 18_625983 | 1 | 1 | NA | NA | NA | NA | NA | NA | 1 | 1 | NA | NA | 1 | 1 | 1 | 1 | 1 | 1 | 3 | 1 |
| 18_625984 | 1 | 1 | NA | NA | NA | NA | NA | NA | 1 | 1 | NA | NA | 1 | 1 | 2 | 0.5 | 1 | 1 | 4 | 0.75 |
| 18_625988 | 1 | 1 | NA | NA | NA | NA | NA | NA | 1 | 1 | NA | NA | 1 | 0 | 1 | 1 | 1 | 1 | 3 | 0.6667 |
| 18_625990 | 1 | 1 | NA | NA | 1 | 1 | NA | NA | 2 | 1 | NA | NA | 1 | 1 | 2 | 1 | 1 | 1 | 4 | 1 |
| 18_625990 | NA | NA | NA | NA | 1 | 1 | NA | NA | 1 | 1 | NA | NA | 1 | 1 | 1 | 1 | 1 | 0 | 3 | 0.6667 |
| 18_625990 | 1 | 1 | NA | NA | 1 | 0 | NA | NA | 2 | 0.5 | NA | NA | 1 | 0 | 1 | 1 | 1 | 1 | 3 | 0.6667 |
| 18_625992 | 1 | 1 | NA | NA | 1 | 0 | NA | NA | 2 | 0.5 | NA | NA | 1 | 1 | 2 | 0.5 | 1 | 0 | 4 | 0.5 |
| 18_625995 | 1 | 0 | NA | NA | 1 | 1 | NA | NA | 2 | 0.5 | NA | NA | NA | NA | 2 | 0 | 2 | 0.5 | 4 | 0.25 |
| 18_625996 | 1 | 1 | NA | NA | 1 | 1 | NA | NA | 2 | 1 | NA | NA | 1 | 1 | 2 | 1 | 2 | 1 | 5 | 1 |
| 18_625996 | 1 | 1 | NA | NA | 1 | 0 | NA | NA | 2 | 0.5 | NA | NA | 1 | 1 | 2 | 0.5 | 2 | 0.5 | 5 | 0.6 |
| 18_625997 | 1 | 0 | NA | NA | 1 | 1 | NA | NA | 2 | 0.5 | NA | NA | 1 | 0 | 2 | 0 | 2 | 0.5 | 5 | 0.2 |
| 18_626005 | 1 | 1 | NA | NA | 1 | 0 | NA | NA | 2 | 0.5 | NA | NA | 1 | 1 | 2 | 0.5 | 2 | 0 | 5 | 0.4 |
| 18_626007 | 1 | 1 | NA | NA | 1 | 0 | NA | NA | 2 | 0.5 | NA | NA | 1 | 0 | 2 | 0.5 | 2 | 0 | 5 | 0.2 |
| 18_626351 | 1 | 0 | NA | NA | NA | NA | 2 | 0.5 | 3 | 0.3333 | NA | NA | NA | NA | 1 | 1 | NA | NA | 1 | 1 |
| 18_626352 | 1 | 0 | NA | NA | NA | NA | 2 | 1 | 3 | 0.6667 | NA | NA | NA | NA | 1 | 1 | NA | NA | 1 | 1 |
| 18_626357 | 1 | 0 | NA | NA | 1 | 1 | 3 | 0.3333 | 5 | 0.4 | NA | NA | NA | NA | 1 | 1 | NA | NA | 1 | 1 |
| 18_626377 | 2 | 1 | 6 | 0.1667 | 5 | 0.8 | 7 | 0.5714 | 20 | 0.55 | 1 | 0 | 5 | 0.8 | 3 | 0 | 1 | 1 | 10 | 0.5 |
| 18_626377 | 1 | 0 | 8 | 0.5 | 5 | 0.4 | 5 | 0.6 | 19 | 0.4737 | 1 | 0 | 5 | 0 | 5 | 0.8 | 2 | 0 | 13 | 0.3077 |
| 18_626438 | 2 | 0 | 5 | 0 | 3 | 0 | 7 | 0.1429 | 17 | 0.0588 | 1 | 1 | 3 | 0 | 2 | 0 | 1 | 0 | 7 | 0.1429 |
| 18_626478 | 1 | 0 | 4 | 0.5 | 1 | 1 | 6 | 0.5 | 12 | 0.5 | 1 | 1 | NA | NA | 1 | 1 | 1 | 1 | 3 | 1 |
| 18_626479 | 1 | 0 | 4 | 0 | NA | NA | 6 | 0.1667 | 11 | 0.0909 | 1 | 1 | NA | NA | 1 | 0 | 1 | 0 | 3 | 0.3333 |
| 18_626479 | 1 | 0 | 2 | 0 | 1 | 1 | 5 | 0.4 | 9 | 0.3333 | 1 | 1 | NA | NA | 1 | 1 | 1 | 1 | 3 | 1 |
| 18_626480 | 1 | 1 | 3 | 1 | 1 | 1 | 6 | 0.8333 | 11 | 0.9091 | 1 | 1 | NA | NA | 1 | 1 | 1 | 1 | 3 | 1 |
| 18_626480 | 1 | 1 | 4 | 1 | NA | NA | 7 | 1 | 12 | 1 | 1 | 1 | NA | NA | 1 | 1 | 1 | 1 | 3 | 1 |
| 18_626481 | 1 | 0 | 4 | 0.25 | 1 | 1 | 6 | 0.6667 | 12 | 0.5 | 1 | 1 | 1 | 0 | 1 | 0 | 1 | 1 | 4 | 0.5 |
| 18_626482 | 1 | 1 | 4 | 0.75 | 2 | 0.5 | 4 | 1 | 11 | 0.8182 | 1 | 1 | 1 | 1 | 1 | 1 | 1 | 1 | 4 | 1 |
| 18_626483 | 1 | 0 | 4 | 0.75 | 2 | 0.5 | 6 | 1 | 13 | 0.7692 | 1 | 1 | NA | NA | 1 | 1 | 1 | 1 | 3 | 1 |
| 18_627997 | 1 | 1 | NA | NA | NA | NA | NA | NA | 1 | 1 | NA | NA | NA | NA | NA | NA | 1 | 1 | 1 | 1 |
| 18_630702 | NA | NA | NA | NA | NA | NA | 1 | 1 | 1 | 1 | 2 | 1 | 2 | 1 | 1 | 1 | NA | NA | 5 | 1 |
| 18_630703 | NA | NA | NA | NA | NA | NA | 1 | 1 | 1 | 1 | 2 | 0.5 | 2 | 1 | 1 | 1 | NA | NA | 5 | 0.8 |
| 18_630704 | NA | NA | 2 | 0 | 2 | 0.5 | NA | NA | 4 | 0.25 | 2 | 0.5 | 1 | 0 | NA | NA | NA | NA | 3 | 0.3333 |
| 18_630851 | NA | NA | 5 | 0 | 2 | 0.5 | 1 | 0 | 8 | 0.125 | 4 | 0 | 2 | 0 | 3 | 0 | 1 | 1 | 10 | 0.1 |
| 18_630852 | NA | NA | 5 | 0.8 | 2 | 1 | 1 | 1 | 8 | 0.875 | 3 | 1 | 2 | 1 | 4 | 1 | 1 | 1 | 10 | 1 |
| 18_630853 | NA | NA | 5 | 1 | 2 | 1 | NA | NA | 7 | 1 | 4 | 1 | 2 | 1 | 4 | 1 | 1 | 1 | 11 | 1 |
| 18_630853 | NA | NA | 6 | 0.3333 | 2 | 0 | 1 | 0 | 9 | 0.2222 | 5 | 0 | 2 | 0 | 4 | 0.25 | 1 | 1 | 12 | 0.1667 |
| 18_630853 | NA | NA | 4 | 0.75 | 2 | 1 | 1 | 1 | 7 | 0.8571 | 5 | 0.8 | 2 | 1 | 4 | 0.75 | NA | NA | 11 | 0.8182 |
| 18_630853 | NA | NA | 6 | 0.5 | 2 | 1 | 1 | 1 | 9 | 0.6667 | 4 | 1 | 2 | 1 | 2 | 1 | 1 | 0 | 9 | 0.8889 |
| 18_630854 | NA | NA | 5 | 1 | 2 | 1 | 1 | 1 | 8 | 1 | 5 | 1 | 1 | 1 | 2 | 1 | 1 | 1 | 9 | 1 |
| 18_630855 | NA | NA | 4 | 0.5 | 2 | 0.5 | 1 | 1 | 7 | 0.5714 | 5 | 0.4 | 2 | 0.5 | 5 | 0.8 | 1 | 1 | 13 | 0.6154 |
| 18_630856 | NA | NA | 4 | 1 | 1 | 1 | NA | NA | 5 | 1 | 4 | 0.25 | 2 | 1 | 5 | 1 | 1 | 1 | 12 | 0.75 |

| | | | | | | | | | | | | | | | | | | | | |
|---|---|---|---|---|---|---|---|---|---|---|---|---|---|---|---|---|---|---|---|---|
| 18_6308578 | NA | NA | 5 | 0 | 2 | 0.5 | 1 | 0 | 8 | 0.125 | 3 | 0.3333 | 2 | 0.5 | 6 | 0.5 | 1 | 1 | 12 | 0.5 |
| 18_6308594 | NA | NA | 6 | 0.5 | 1 | 0 | 2 | 0.5 | 9 | 0.4444 | 4 | 0.5 | 2 | 0.5 | 5 | 0.6 | NA | NA | 11 | 0.5455 |
| 18_6308606 | 2 | 0.5 | 6 | 0.8333 | 1 | 0 | 1 | 0 | 10 | 0.6 | 4 | 0.5 | 2 | 0.5 | 6 | 1 | 1 | 1 | 13 | 0.7692 |
| 18_6308632 | 4 | 0 | 5 | 0 | 3 | 0.3333 | 1 | 0 | 13 | 0.0769 | 4 | 0.25 | 3 | 0 | 4 | 0 | NA | NA | 11 | 0.0909 |
| 18_6308663 | 4 | 0.25 | 6 | 0.1667 | 3 | 0.3333 | 1 | 1 | 14 | 0.2857 | 4 | 0 | 2 | 0.5 | 6 | 0.6667 | NA | NA | 12 | 0.4167 |
| 18_6308664 | 4 | 0 | 6 | 0.1667 | 3 | 0 | 3 | 0 | 16 | 0.0625 | 5 | 0 | 2 | 0 | 6 | 0.1667 | 1 | 0 | 14 | 0.0714 |
| 18_6313010 | 11 | 1 | 14 | 0.8571 | 11 | 0.9091 | 13 | 0.7692 | 49 | 0.8776 | 5 | 1 | 10 | 1 | 8 | 1 | 13 | 1 | 36 | 1 |
| 18_6313078 | 10 | 0 | 14 | 0.0714 | 11 | 0.0909 | 13 | 0.0769 | 48 | 0.0625 | 6 | 0.1667 | 10 | 0 | 12 | 0.3333 | 10 | 0.1 | 38 | 0.1579 |
| 18_6313081 | 11 | 0.0909 | 15 | 0.3333 | 11 | 0.2727 | 13 | 0.2308 | 50 | 0.24 | 6 | 0.3333 | 13 | 0.2308 | 12 | 0.5833 | 15 | 0.2667 | 46 | 0.3478 |
| 18_6389372 | 6 | 0 | 2 | 0.5 | NA | NA | 2 | 0 | 10 | 0.1 | 2 | 0 | 3 | 0.3333 | 2 | 0 | 3 | 0 | 10 | 0.1 |
| 18_6391060 | 7 | 0 | 2 | 0.5 | NA | NA | 2 | 0 | 11 | 0.0909 | 2 | 0 | 4 | 0.25 | 4 | 0.25 | 3 | 0 | 13 | 0.1538 |
| 18_6391085 | 6 | 0.5 | 2 | 0.5 | NA | NA | 2 | 0.5 | 10 | 0.5 | 1 | 0 | 4 | 0.25 | 4 | 0.25 | 4 | 0 | 13 | 0.1538 |
| 18_6391086 | 6 | 0.1667 | 2 | 0.5 | 1 | 0 | 1 | 0 | 10 | 0.2 | 2 | 0.5 | 2 | 0 | 4 | 0 | 3 | 0 | 11 | 0.0909 |
| 18_6391087 | 2 | 0.5 | 2 | 0 | 1 | 0 | 1 | 0 | 6 | 0.1667 | 1 | 0 | 3 | 0 | 4 | 0.25 | 4 | 0 | 12 | 0.0833 |
| 18_6391118 | 6 | 0.3333 | 2 | 0 | 1 | 1 | 2 | 0 | 11 | 0.2727 | 2 | 0 | 3 | 0.6667 | 4 | 0.25 | 4 | 0.25 | 13 | 0.3077 |
| 18_6391124 | 6 | 0.1667 | 2 | 0 | 1 | 1 | 2 | 0 | 11 | 0.1818 | 1 | 0 | 3 | 0 | 5 | 0.2 | 6 | 0.1667 | 15 | 0.1333 |
| 18_6391301 | 6 | 0.3333 | 3 | 0 | 1 | 0 | 2 | 0 | 12 | 0.1667 | 2 | 0 | 3 | 0 | 5 | 0.2 | 7 | 0.1429 | 17 | 0.1176 |
| 18_6412036 | 10 | 0.6 | 6 | 0.5 | 4 | 0.25 | 11 | 0.3636 | 31 | 0.4516 | 5 | 0.2 | 5 | 0 | 7 | 0.4286 | 9 | 0.4444 | 26 | 0.3077 |
| 18_6412045 | 10 | 0 | 8 | 0.375 | 3 | 0.6667 | 11 | 0.4545 | 32 | 0.3125 | 6 | 0 | 5 | 0.2 | 9 | 0.4444 | 11 | 0.3636 | 31 | 0.2903 |
| 18_6412050 | 11 | 0.3636 | 9 | 0.1111 | 4 | 0.25 | 10 | 0.4 | 34 | 0.2941 | 5 | 0.4 | 6 | 0.1667 | 8 | 0.875 | 10 | 0.4 | 29 | 0.4828 |
| 18_6412074 | 12 | 0.25 | 7 | 0.4286 | 5 | 0.6 | 11 | 0.4545 | 35 | 0.4 | 2 | 0.5 | 6 | 0.5 | 9 | 0.6667 | 12 | 0.5833 | 29 | 0.5862 |
| 18_6412114 | 13 | 0.1538 | 10 | 0.1 | 6 | 0.1667 | 12 | 0.0833 | 41 | 0.122 | 6 | 0.3333 | 7 | 0 | 10 | 0.3 | 13 | 0.1538 | 36 | 0.1944 |
| 18_6412142 | 16 | 0.125 | 10 | 0 | 8 | 0.25 | 13 | 0.1538 | 47 | 0.1277 | 8 | 0 | 7 | 0 | 13 | 0.0769 | 12 | 0.0833 | 40 | 0.05 |
| 18_6412144 | 16 | 0.0625 | 9 | 0.1111 | 5 | 0.2 | 9 | 0.1111 | 39 | 0.1026 | 7 | 0.1429 | 7 | 0.1429 | 12 | 0 | 12 | 0.0833 | 38 | 0.0789 |
| 18_6412235 | 18 | 0.1667 | 15 | 0.0667 | 11 | 0.1818 | 17 | 0.1765 | 61 | 0.1475 | 10 | 0.2 | 9 | 0.1111 | 18 | 0.3333 | 19 | 0.2105 | 56 | 0.2321 |
| 18_6439850 | 14 | 0 | 9 | 0.3333 | 12 | 0.0833 | 13 | 0.0769 | 48 | 0.1042 | 6 | 0 | 11 | 0.0909 | 21 | 0.2381 | 8 | 0.125 | 46 | 0.1522 |
| 18_6440256 | 18 | 0.1111 | 12 | 0.4167 | 18 | 0.2778 | 18 | 0.2778 | 66 | 0.2576 | 3 | 0 | 17 | 0.2941 | 31 | 0.1935 | 9 | 0.2222 | 60 | 0.2167 |
| 18_6440272 | 19 | 0.4737 | 15 | 0.6 | 17 | 0.5294 | 20 | 0.65 | 71 | 0.5634 | 7 | 0.5714 | 19 | 0.4211 | 36 | 0.5833 | 11 | 0.8182 | 73 | 0.5753 |
| 18_6440287 | 19 | 0.4211 | 13 | 0.6154 | 20 | 0.3 | 21 | 0.3333 | 73 | 0.3973 | 8 | 0.5 | 19 | 0.4211 | 33 | 0.4848 | 11 | 0.4545 | 71 | 0.4648 |
| 18_6440475 | 24 | 0.0833 | 13 | 0.1538 | 22 | 0.3182 | 21 | 0.1429 | 80 | 0.175 | 9 | 0.1111 | 18 | 0.2778 | 29 | 0 | 10 | 0.2 | 66 | 0.1212 |
| 18_6440487 | 22 | 0.9545 | 13 | 0.8462 | 20 | 0.75 | 24 | 0.9167 | 79 | 0.8734 | 8 | 0.75 | 19 | 1 | 37 | 0.8378 | 15 | 0.9333 | 79 | 0.8861 |
| 18_6440494 | 24 | 0.125 | 10 | 0 | 18 | 0.1111 | 25 | 0.08 | 77 | 0.0909 | 9 | 0 | 14 | 0 | 32 | 0.0938 | 15 | 0.1333 | 70 | 0.0714 |
| 18_6440495 | 22 | 0.9091 | 15 | 0.8 | 20 | 0.75 | 24 | 0.7917 | 81 | 0.8148 | 10 | 0.7 | 17 | 0.7647 | 35 | 0.7714 | 15 | 1 | 77 | 0.8052 |
| 18_6440504 | 26 | 0.1154 | 15 | 0.2667 | 20 | 0.15 | 26 | 0.1923 | 87 | 0.1724 | 9 | 0.3333 | 21 | 0.0476 | 36 | 0.1667 | 17 | 0.1176 | 83 | 0.1446 |
| 18_6440504 | 26 | 0.2692 | 13 | 0.5385 | 22 | 0.3636 | 27 | 0.4444 | 88 | 0.3864 | 10 | 0.3 | 21 | 0.1429 | 33 | 0.4242 | 14 | 0.5714 | 78 | 0.359 |
| 18_6440507 | 28 | 0.1786 | 15 | 0.4 | 22 | 0.4545 | 27 | 0.2963 | 92 | 0.3152 | 9 | 0.4444 | 22 | 0.4545 | 33 | 0.2727 | 18 | 0.5 | 82 | 0.3902 |
| 18_6440519 | 14 | 0.7143 | 11 | 0.8182 | 15 | 0.5333 | 25 | 0.56 | 65 | 0.6308 | 7 | 0.7143 | 16 | 0.625 | 25 | 0.56 | 16 | 0.75 | 64 | 0.6406 |
| 18_6440531 | 7 | 0 | 8 | 0.125 | 9 | 0 | 10 | 0.2 | 34 | 0.0882 | 5 | 0.2 | 11 | 0 | 13 | 0 | 11 | 0.1818 | 40 | 0.075 |
| 18_6440552 | 4 | 0.25 | 6 | 0 | 5 | 0.2 | 7 | 0 | 22 | 0.0909 | 5 | 0 | 6 | 0.1667 | 9 | 0 | 10 | 0.1 | 30 | 0.0667 |
| 18_6440583 | 3 | 0 | 7 | 0.1429 | 4 | 0.5 | 7 | 0 | 21 | 0.1429 | 4 | 0.75 | 5 | 0.2 | 9 | 0.2222 | 9 | 0.2222 | 27 | 0.2963 |
| 18_6440709 | 4 | 0.25 | 3 | 0 | 6 | 0.3333 | 11 | 0 | 24 | 0.125 | 4 | 0.25 | 6 | 0 | 8 | 0.25 | 9 | 0.1111 | 27 | 0.1481 |
| 18_6440773 | 6 | 0.1667 | 9 | 0 | 4 | 0 | 7 | 0.1429 | 26 | 0.0769 | 4 | 0 | 6 | 0.1667 | 12 | 0.0833 | 10 | 0 | 32 | 0.0625 |
| 18_6440876 | 9 | 0.3333 | 14 | 0.1429 | 10 | 0.3 | 23 | 0.1739 | 56 | 0.2143 | 7 | 0 | 17 | 0.1176 | 20 | 0.3 | 17 | 0.1765 | 61 | 0.1803 |
| 18_6440890 | 12 | 0.9167 | 16 | 0.9375 | 10 | 0.9 | 19 | 0.7895 | 57 | 0.8772 | 8 | 1 | 18 | 0.8889 | 20 | 0.95 | 17 | 0.8235 | 63 | 0.9048 |
| 18_6440891 | 11 | 0.1818 | 16 | 0.25 | 11 | 0 | 23 | 0.087 | 61 | 0.1311 | 8 | 0.125 | 21 | 0.1905 | 23 | 0.2174 | 19 | 0.2105 | 71 | 0.1972 |
| 18_6440900 | 11 | 0.9091 | 16 | 0.5 | 9 | 1 | 19 | 0.4737 | 55 | 0.6545 | 7 | 0.8571 | 19 | 0.6316 | 15 | 0.5333 | 16 | 0.5 | 57 | 0.5965 |
| 18_6440903 | 13 | 0.8462 | 18 | 0.7778 | 13 | 0.7692 | 24 | 0.625 | 68 | 0.7353 | 9 | 0.6667 | 21 | 0.8571 | 23 | 0.6522 | 17 | 0.7647 | 70 | 0.7429 |
| 18_6440907 | 13 | 0.6923 | 14 | 0.5 | 10 | 0.5 | 15 | 0.4 | 52 | 0.5192 | 8 | 0.75 | 12 | 0.5833 | 17 | 0.5294 | 14 | 0.5 | 51 | 0.5686 |
| 18_6440909 | 13 | 0.3077 | 10 | 0.4 | 11 | 0.1818 | 15 | 0.1333 | 49 | 0.2449 | 7 | 0.2857 | 14 | 0.1429 | 20 | 0.15 | 15 | 0.2 | 56 | 0.1786 |
| 18_6441079 | 1 | 1 | NA | NA | NA | NA | 2 | 0.5 | 3 | 0.6667 | NA | NA | 1 | 1 | 3 | 0 | NA | NA | 4 | 0.25 |
| 18_6441225 | 1 | 0 | NA | NA | NA | NA | 3 | 0.3333 | 4 | 0.25 | 1 | 1 | 2 | 0 | 3 | 0.6667 | 2 | 0.5 | 8 | 0.5 |
| 18_6441228 | 1 | 0 | NA | NA | NA | NA | 1 | 1 | 2 | 0.5 | 1 | 1 | 2 | 0.5 | 3 | 0.6667 | 2 | 1 | 8 | 0.75 |
| 18_6452714 | NA | NA | NA | NA | 1 | 1 | NA | NA | 1 | 1 | NA | NA | NA | NA | 1 | 1 | NA | NA | 1 | 1 |
| 18_6452719 | NA | NA | NA | NA | 1 | 1 | NA | NA | 1 | 1 | NA | NA | NA | NA | 1 | 0 | 1 | 1 | 2 | 0.5 |
| 18_6459025 | 3 | 0 | 7 | 0.2857 | 1 | 1 | 6 | 0 | 17 | 0.1765 | 1 | 0 | 5 | 0 | 4 | 0.25 | 2 | 0.5 | 12 | 0.1667 |
| 18_6459028 | 4 | 0 | 7 | 0.4286 | 1 | 0 | 4 | 0 | 16 | 0.1875 | 1 | 0 | 3 | 0.3333 | 3 | 0 | 2 | 0.5 | 9 | 0.2222 |
| 18_6459085 | 5 | 0 | 12 | 0.25 | 1 | 0 | 9 | 0.2222 | 27 | 0.1852 | 2 | 0.5 | 8 | 0.125 | 5 | 0.4 | 5 | 0.2 | 20 | 0.25 |
| 18_6459090 | 6 | 0 | 13 | 0.0769 | 3 | 0.3333 | 10 | 0.1 | 32 | 0.0938 | 2 | 0 | 7 | 0 | 8 | 0.125 | 5 | 0.2 | 22 | 0.0909 |
| 18_6459105 | 6 | 0.1667 | 10 | 0.4 | 2 | 0.5 | 8 | 0.625 | 26 | 0.4231 | 1 | 0 | 6 | 0.1667 | 3 | 0.3333 | 4 | 0 | 14 | 0.1429 |
| 18_6459108 | 3 | 0 | 7 | 0.1429 | 2 | 0 | 6 | 0.1667 | 18 | 0.1111 | NA | NA | 2 | 0 | 7 | 0.2857 | 5 | 0 | 14 | 0.1429 |
| 18_6459122 | 5 | 0.6 | 11 | 0.2727 | 3 | 0.3333 | 6 | 0.3333 | 25 | 0.36 | 2 | 0 | 6 | 0.1667 | 7 | 0.4286 | 3 | 0.3333 | 18 | 0.2778 |
| 18_6463292 | 18 | 0.2222 | 14 | 0.1429 | 18 | 0.1111 | 17 | 0.1765 | 67 | 0.1642 | 15 | 0.1333 | 15 | 0.2 | 10 | 0.2 | 15 | 0 | 55 | 0.1273 |
| 18_6463347 | 16 | 0.3125 | 8 | 0 | 13 | 0 | 18 | 0.2222 | 55 | 0.1636 | 7 | 0.1429 | 9 | 0.2222 | 6 | 0 | 11 | 0.0909 | 33 | 0.1212 |
| 18_6463354 | 15 | 0.2667 | 7 | 0.1429 | 11 | 0.0909 | 17 | 0.2353 | 50 | 0.2 | 6 | 0.3333 | 8 | 0.375 | 5 | 0.2 | 11 | 0.0909 | 30 | 0.2333 |
| 18_6466392 | NA | NA | 1 | 1 | NA | NA | NA | NA | 1 | 1 | NA | NA | 1 | 1 | NA | NA | NA | NA | 1 | 1 |

| | | | | | | | | | | | | | | | | | | | | |
|---|---|---|---|---|---|---|---|---|---|---|---|---|---|---|---|---|---|---|---|---|
| 18_646641( | NA | NA | 1 | 1 | NA | NA | NA | NA | 1 | 1 | NA | NA | 1 | 1 | NA | NA | NA | NA | 1 | 1 |
| 18_646641ε | NA | NA | 1 | 1 | NA | NA | NA | NA | 1 | 1 | NA | NA | 1 | 1 | NA | NA | NA | NA | 1 | 1 |
| 18_646642( | NA | NA | 1 | 1 | NA | NA | NA | NA | 1 | 1 | NA | NA | 1 | 1 | NA | NA | NA | NA | 1 | 1 |
| 18_646643: | NA | NA | 1 | 1 | NA | NA | NA | NA | 1 | 1 | NA | NA | 1 | 1 | NA | NA | NA | NA | 1 | 1 |
| 18_6532628 | 8 | 0.125 | 4 | 0 | 4 | 0 | 9 | 0.1111 | 25 | 0.08 | 6 | 0.1667 | 14 | 0.1429 | 9 | 0.1111 | 12 | 0.0833 | 41 | 0.122 |
| 18_653460: | 12 | 0.9167 | 10 | 0.8 | 8 | 1 | 11 | 1 | 41 | 0.9268 | 11 | 0.7273 | 21 | 0.9524 | 15 | 0.9333 | 20 | 0.85 | 67 | 0.8806 |
| 18_653462( | 17 | 0.1765 | 8 | 0.125 | 6 | 0 | 14 | 0.2857 | 45 | 0.1778 | 9 | 0.1111 | 17 | 0.1176 | 15 | 0.1333 | 19 | 0.1053 | 60 | 0.1167 |
| 18_653463: | 13 | 0.3077 | 5 | 0.2 | 5 | 0.2 | 12 | 0.1667 | 35 | 0.2286 | 8 | 0 | 14 | 0 | 11 | 0.1818 | 13 | 0.2308 | 46 | 0.1087 |
| 18_653464 | 4 | 0 | 3 | 0 | 2 | 0.5 | 6 | 0.1667 | 15 | 0.1333 | 3 | 0 | 5 | 0 | 4 | 0 | 7 | 0.1429 | 19 | 0.0526 |
| 18_653465 | 6 | 0.8333 | 4 | 1 | 1 | 1 | 6 | 0.3333 | 17 | 0.7059 | 4 | 1 | 7 | 0.5714 | 4 | 0.75 | 9 | 0.6667 | 24 | 0.7083 |
| 18_653466( | 6 | 0.1667 | 3 | 0.3333 | 2 | 0.5 | 7 | 0.2857 | 18 | 0.2778 | 3 | 0.6667 | 5 | 0.2 | 5 | 0.2 | 10 | 0.1 | 23 | 0.2174 |
| 18_653467( | 5 | 1 | 3 | 1 | 2 | 1 | 7 | 1 | 17 | 1 | 2 | 1 | 6 | 0.8333 | 5 | 1 | 11 | 1 | 24 | 0.9583 |
| 18_653467( | 5 | 1 | 3 | 1 | 2 | 1 | 7 | 1 | 17 | 1 | 4 | 1 | 5 | 0.6 | 5 | 1 | 9 | 1 | 23 | 0.913 |
| 18_653468: | 5 | 0.4 | 2 | 1 | 2 | 1 | 6 | 0.8333 | 15 | 0.7333 | 3 | 0.6667 | 5 | 0.6 | 5 | 0.8 | 7 | 1 | 20 | 0.8 |
| 18_653468! | 5 | 1 | 4 | 1 | 2 | 1 | 7 | 0.8571 | 18 | 0.9444 | 4 | 1 | 6 | 1 | 5 | 1 | 11 | 1 | 26 | 1 |
| 18_653468! | 5 | 0.2 | 3 | 0.3333 | 2 | 0 | 6 | 0.1667 | 16 | 0.1875 | 4 | 0 | 6 | 0.1667 | 5 | 0 | 10 | 0.1 | 25 | 0.08 |
| 18_653471( | 5 | 0.6 | 3 | 0.6667 | 2 | 0.5 | 8 | 0.5 | 18 | 0.5556 | 3 | 0.3333 | 5 | 0.4 | 4 | 0.75 | 9 | 0.6667 | 21 | 0.5714 |
| 18_653471: | 3 | 0.3333 | 4 | 0 | 2 | 0 | 8 | 0.25 | 17 | 0.1765 | 3 | 0 | 5 | 0 | 4 | 0.25 | 12 | 0.1667 | 24 | 0.125 |
| 18_653475( | 5 | 0 | 5 | 0.4 | 2 | 0 | 8 | 0.25 | 20 | 0.2 | 3 | 0.6667 | 7 | 0.1429 | 6 | 0.3333 | 12 | 0.5 | 28 | 0.3929 |
| 18_653476! | 4 | 0.25 | 5 | 0.2 | 3 | 0.6667 | 8 | 0.25 | 20 | 0.3 | 2 | 0 | 4 | 0 | 5 | 0.2 | 8 | 0.25 | 19 | 0.1579 |
| 18_653488( | 14 | 0.1429 | 7 | 0.1429 | 5 | 0 | 11 | 0.1818 | 37 | 0.1351 | 4 | 0.25 | 10 | 0 | 7 | 0 | 13 | 0.0769 | 34 | 0.0588 |
| 18_653503( | 16 | 0.0625 | 13 | 0.1538 | 7 | 0.1429 | 16 | 0.5625 | 52 | 0.25 | 8 | 0.125 | 11 | 0.3636 | 14 | 0.2857 | 18 | 0.2778 | 51 | 0.2745 |
| 18_653504! | 17 | 0.1765 | 11 | 0.1818 | 10 | 0 | 16 | 0.0625 | 54 | 0.1111 | 8 | 0.25 | 13 | 0.3077 | 14 | 0.0714 | 18 | 0 | 53 | 0.1321 |
| 18_656092! | 1 | 1 | 1 | 1 | 2 | 0.5 | NA | NA | 4 | 0.75 | NA | NA | 1 | 1 | NA | NA | NA | NA | 1 | 1 |
| 18_656853( | NA | NA | NA | NA | NA | NA | 1 | 1 | 1 | 1 | NA | NA | 1 | 1 | NA | NA | NA | NA | 1 | 1 |
| 18_657564: | NA | NA | NA | NA | 2 | 1 | 1 | 0 | 3 | 0.6667 | 2 | 0.5 | NA | NA | 2 | 1 | 1 | 0 | 5 | 0.6 |
| 18_657565! | NA | NA | NA | NA | 3 | 1 | 1 | 0 | 4 | 0.75 | 1 | 0 | NA | NA | 2 | 0.5 | 1 | 1 | 4 | 0.5 |
| 18_657565! | NA | NA | NA | NA | 3 | 0.6667 | NA | NA | 3 | 0.6667 | 1 | 1 | NA | NA | 1 | 1 | 1 | 0 | 3 | 0.6667 |
| 18_657567! | NA | NA | NA | NA | 3 | 0.3333 | 1 | 0 | 4 | 0.25 | 2 | 0.5 | NA | NA | 2 | 0 | 2 | 0 | 6 | 0.1667 |
| 18_657572: | NA | NA | NA | NA | 3 | 0.3333 | 1 | 1 | 4 | 0.5 | 2 | 0.5 | NA | NA | 2 | 0.5 | 3 | 0 | 7 | 0.2857 |
| 18_657576: | NA | NA | NA | NA | 3 | 1 | 3 | 1 | 6 | 1 | 2 | 1 | NA | NA | 1 | 1 | 3 | 1 | 6 | 1 |
| 18_657576! | NA | NA | NA | NA | 2 | 0.5 | 3 | 0.6667 | 5 | 0.6 | 1 | 0 | NA | NA | NA | NA | 2 | 0.5 | 3 | 0.3333 |
| 18_657580: | NA | NA | NA | NA | 4 | 1 | 3 | 0.6667 | 7 | 0.8571 | 2 | 1 | NA | NA | 2 | 1 | 3 | 0.6667 | 7 | 0.8571 |
| 18_657581: | NA | NA | NA | NA | 4 | 0 | 3 | 0.3333 | 7 | 0.1429 | 3 | 0 | NA | NA | 1 | 1 | 3 | 0.3333 | 7 | 0.2857 |
| 18_657584( | NA | NA | 1 | 1 | 4 | 0.75 | 3 | 0.3333 | 8 | 0.625 | 2 | 0.5 | NA | NA | 1 | 1 | 3 | 0.3333 | 6 | 0.5 |
| 18_658201 | 2 | 0.5 | 3 | 0.3333 | 5 | 0.8 | 2 | 1 | 12 | 0.6667 | 1 | 0 | NA | NA | 2 | 0.5 | 2 | 0.5 | 5 | 0.4 |
| 18_658201 | 3 | 0 | 3 | 0 | 6 | 0.3333 | 3 | 0 | 15 | 0.1333 | 1 | 0 | NA | NA | 3 | 0.3333 | 5 | 0 | 9 | 0.1111 |
| 18_658201! | NA | NA | 1 | 0 | 5 | 0.4 | 1 | 0 | 7 | 0.2857 | 1 | 1 | NA | NA | 2 | 0 | 5 | 0.6 | 8 | 0.5 |
| 18_658204: | 3 | 0.6667 | 3 | 1 | 5 | 1 | NA | NA | 11 | 0.9091 | 1 | 1 | NA | NA | 3 | 0.3333 | 4 | 1 | 8 | 0.75 |
| 18_658204 | 2 | 1 | 4 | 1 | 4 | 1 | 3 | 1 | 13 | 1 | 1 | 1 | NA | NA | 3 | 1 | 5 | 1 | 9 | 1 |
| 18_658205: | 1 | 0 | 3 | 0.6667 | 4 | 0.25 | 2 | 0.5 | 10 | 0.4 | 1 | 1 | NA | NA | 3 | 0 | 4 | 0.5 | 8 | 0.375 |
| 18_658205 | 3 | 0 | 4 | 0.75 | 6 | 0.5 | 3 | 0.3333 | 16 | 0.4375 | 1 | 0 | NA | NA | 3 | 0.3333 | 5 | 0.4 | 9 | 0.3333 |
| 18_658206 | 2 | 0.5 | 3 | 0.3333 | 5 | 0 | 3 | 0.6667 | 13 | 0.3077 | 1 | 0 | NA | NA | 3 | 0.3333 | 7 | 0 | 11 | 0.0909 |
| 18_658218 | 4 | 0.25 | 5 | 0 | 7 | 0.1429 | 6 | 0 | 22 | 0.0909 | 2 | 0 | 2 | 0.5 | 5 | 0.4 | 9 | 0.2222 | 18 | 0.2778 |
| 18_658220 | 3 | 0 | 1 | 0 | 5 | 0.4 | 1 | 0 | 10 | 0.2 | 2 | 0 | NA | NA | 2 | 0 | 7 | 0.1429 | 11 | 0.0909 |
| 18_658220( | 2 | 0.5 | NA | NA | 5 | 0.6 | 1 | 0 | 8 | 0.5 | 2 | 0.5 | NA | NA | NA | NA | 2 | 1 | 4 | 0.75 |
| 18_658222! | 2 | 0.5 | 1 | 0 | 2 | 0 | NA | NA | 5 | 0.2 | 2 | 0 | NA | NA | 2 | 0 | 3 | 1 | 7 | 0.4286 |
| 18_661140 | 8 | 0.75 | 4 | 0.5 | 5 | 0.8 | 8 | 0.625 | 25 | 0.68 | NA | NA | 4 | 0.75 | 7 | 0.7143 | 7 | 1 | 18 | 0.8333 |
| 18_661141! | 6 | 0.5 | 2 | 0.5 | 5 | 0.6 | 2 | 1 | 15 | 0.6 | NA | NA | 2 | 1 | 7 | 0.5714 | 5 | 0.2 | 14 | 0.5 |
| 18_661142( | 6 | 0.3333 | 3 | 0.3333 | 5 | 0.8 | 8 | 0.625 | 22 | 0.5455 | NA | NA | 5 | 0.4 | 7 | 0.4286 | 7 | 0.4286 | 19 | 0.4211 |
| 18_661143! | 6 | 0.1667 | 3 | 0 | 5 | 0 | 7 | 0.4286 | 21 | 0.1905 | NA | NA | 3 | 0.6667 | 8 | 0.125 | 7 | 0.1429 | 18 | 0.2222 |
| 18_661156( | 3 | 0 | 2 | 0 | 4 | 0.25 | 7 | 0 | 16 | 0.0625 | NA | NA | 5 | 0 | 7 | 0 | 4 | 0.5 | 16 | 0.125 |
| 18_669927 | NA | NA | NA | NA | 1 | 1 | NA | NA | 1 | 1 | 1 | 1 | NA | NA | 1 | 1 | 1 | 1 | 3 | 1 |
| 18_669929: | 1 | 1 | NA | NA | NA | NA | NA | NA | 1 | 1 | 1 | 1 | 1 | 1 | 2 | 1 | 1 | 1 | 5 | 1 |
| 18_669929! | 1 | 1 | NA | NA | 1 | 1 | NA | NA | 2 | 1 | 1 | 1 | 1 | 1 | 2 | 1 | 1 | 1 | 5 | 1 |
| 18_669931: | 1 | 1 | NA | NA | 1 | 0 | NA | NA | 2 | 0.5 | 1 | 1 | 1 | 0 | 1 | 1 | 1 | 1 | 4 | 0.75 |
| 18_669934: | 1 | 1 | NA | NA | 1 | 1 | NA | NA | 2 | 1 | 1 | 0 | 1 | 0 | 1 | 1 | 1 | 1 | 4 | 0.5 |
| 18_673386: | 1 | 0 | 5 | 0.6 | 4 | 0.5 | 2 | 0.5 | 12 | 0.5 | 2 | 0 | 3 | 1 | 1 | 0 | 6 | 0.5 | 12 | 0.5 |
| 18_673387( | 3 | 0 | 7 | 0.2857 | 3 | 0.3333 | 1 | 0 | 14 | 0.2143 | 2 | 0 | 3 | 0.3333 | 1 | 0 | 6 | 0.1667 | 12 | 0.1667 |
| 18_673389 | 3 | 0.6667 | 7 | 0.7143 | 3 | 0.6667 | 2 | 1 | 15 | 0.7333 | 3 | 0.3333 | 3 | 1 | 2 | 1 | 6 | 0.5 | 14 | 0.6429 |
| 18_673389 | 1 | 0 | 6 | 0.1667 | 4 | 0 | 2 | 0 | 13 | 0.0769 | 2 | 0 | 3 | 0 | 2 | 0 | 6 | 0.1667 | 13 | 0.0769 |
| 18_673391: | 2 | 1 | 5 | 1 | 4 | 1 | 2 | 1 | 13 | 1 | 2 | 1 | 3 | 1 | 2 | 1 | 3 | 1 | 10 | 1 |
| 18_673391 | 2 | 1 | 8 | 0.5 | 4 | 0.75 | 2 | 0.5 | 16 | 0.625 | 3 | 1 | 3 | 1 | 3 | 0.3333 | 5 | 0.4 | 14 | 0.6429 |
| 18_673393 | 3 | 1 | 5 | 1 | 4 | 1 | NA | NA | 12 | 1 | 2 | 1 | 3 | 1 | 2 | 1 | 5 | 1 | 12 | 1 |
| 18_673394: | 2 | 0.5 | 5 | 0.8 | 3 | 1 | 2 | 1 | 12 | 0.8333 | 2 | 1 | 3 | 1 | 3 | 1 | 6 | 0.5 | 14 | 0.7857 |

| | | | | | | | | | | | | | | | | | | | | |
|---|---|---|---|---|---|---|---|---|---|---|---|---|---|---|---|---|---|---|---|---|
| 18_673394 | 1 | 1 | 6 | 1 | 3 | 1 | 2 | 1 | 12 | 1 | 3 | 1 | 3 | 0.6667 | 3 | 1 | 6 | 0.8333 | 15 | 0.8667 |
| 18_673395 | 1 | 1 | 6 | 1 | 3 | 1 | 2 | 1 | 12 | 1 | 4 | 1 | 3 | 1 | 3 | 1 | 6 | 1 | 16 | 1 |
| 18_673396 | 2 | 1 | 6 | 0.1667 | 1 | 0 | 3 | 0 | 12 | 0.25 | 4 | 0.25 | 3 | 0.3333 | 3 | 0 | 6 | 0.3333 | 16 | 0.25 |
| 18_673405 | NA | NA | 6 | 0 | 2 | 0.5 | 2 | 0 | 10 | 0.1 | 3 | 0.3333 | 1 | 0 | 1 | 0 | 2 | 0 | 7 | 0.1429 |
| 18_673557 | 7 | 0.2857 | 1 | 1 | 3 | 0.3333 | 9 | 0.3333 | 20 | 0.35 | 3 | 0.3333 | 10 | 0.5 | 12 | 0.4167 | 6 | 0.3333 | 31 | 0.4194 |
| 18_673563 | 19 | 0.2632 | 15 | 0.1333 | 19 | 0.1053 | 29 | 0.1034 | 82 | 0.1463 | 17 | 0.1765 | 32 | 0.1562 | 38 | 0.1316 | 28 | 0.0714 | 115 | 0.1304 |
| 18_673583 | 4 | 0 | 1 | 0 | 1 | 0 | 5 | 0.4 | 11 | 0.1818 | NA | NA | 6 | 0.1667 | 4 | 0 | 2 | 0.5 | 12 | 0.1667 |
| 18_673584 | 5 | 1 | 1 | 1 | 1 | 1 | 6 | 0.8333 | 13 | 0.9231 | NA | NA | 5 | 0.4 | 4 | 0.75 | 3 | 0.3333 | 12 | 0.5 |
| 18_673585 | 3 | 0 | 1 | 1 | 1 | 0 | 6 | 0.1667 | 11 | 0.1818 | NA | NA | 6 | 0.1667 | 5 | 0.2 | 3 | 0.3333 | 14 | 0.2143 |
| 18_673586 | 3 | 0.6667 | 1 | 1 | 1 | 1 | 6 | 0.6667 | 11 | 0.7273 | NA | NA | 6 | 0.8333 | 4 | 1 | 1 | 1 | 11 | 0.9091 |
| 18_673587 | 4 | 0.75 | 1 | 0 | 1 | 1 | 5 | 0.4 | 11 | 0.5455 | NA | NA | 5 | 0.8 | 3 | 0.3333 | 3 | 0.6667 | 11 | 0.6364 |
| 18_673587 | 4 | 0.25 | 1 | 0 | 1 | 1 | 7 | 0.1429 | 13 | 0.2308 | NA | NA | 6 | 0.1667 | 5 | 0 | 3 | 0 | 14 | 0.0714 |
| 18_673587 | 4 | 0.75 | 1 | 1 | 1 | 0 | 7 | 0.7143 | 13 | 0.6923 | NA | NA | 6 | 0.5 | 6 | 0.6667 | 2 | 0.5 | 14 | 0.5714 |
| 18_673596 | 44 | 0.0682 | 28 | 0.1429 | 32 | 0.0938 | 47 | 0.0426 | 151 | 0.0795 | 39 | 0.1282 | 38 | 0.0789 | 60 | 0.0833 | 50 | 0.08 | 187 | 0.0909 |
| 18_673705 | 1 | 1 | NA | NA | 1 | 0 | 2 | 0 | 4 | 0.25 | NA | NA | 4 | 0.25 | 3 | 0.6667 | 2 | 0 | 9 | 0.3333 |
| 18_673707 | 2 | 0.5 | NA | NA | 1 | 1 | 2 | 1 | 5 | 0.8 | NA | NA | 4 | 0.5 | 3 | 0.3333 | 2 | 1 | 9 | 0.5556 |
| 18_674516 | NA | NA | 1 | 0 | 2 | 0.5 | NA | NA | 3 | 0.3333 | NA | NA | NA | NA | NA | NA | 2 | 0.5 | 2 | 0.5 |
| 18_675373 | 1 | 0 | NA | NA | 2 | 0.5 | NA | NA | 3 | 0.3333 | 1 | 1 | NA | NA | NA | NA | NA | NA | 1 | 1 |
| 18_675375 | 1 | 1 | NA | NA | 1 | 1 | NA | NA | 2 | 1 | 1 | 1 | NA | NA | NA | NA | NA | NA | 1 | 1 |
| 18_675377 | 1 | 1 | NA | NA | 1 | 1 | NA | NA | 2 | 1 | 1 | 1 | NA | NA | NA | NA | NA | NA | 1 | 1 |
| 18_675378 | 7 | 1 | 8 | 1 | 9 | 1 | 5 | 0.8 | 29 | 0.9655 | 5 | 1 | 7 | 0.8571 | 9 | 1 | 6 | 0.5 | 27 | 0.8519 |
| 18_675379 | 10 | 0 | 7 | 0.1429 | 11 | 0.0909 | 6 | 0 | 34 | 0.0588 | 4 | 0 | 5 | 0 | 12 | 0.25 | 4 | 0 | 25 | 0.12 |
| 18_675379 | 11 | 1 | 9 | 1 | 11 | 1 | 8 | 0.875 | 39 | 0.9744 | 5 | 1 | 8 | 1 | 11 | 1 | 5 | 1 | 29 | 1 |
| 18_675380 | 10 | 0.1 | 9 | 0 | 11 | 0.0909 | 7 | 0 | 37 | 0.0541 | 5 | 0 | 7 | 0 | 9 | 0.1111 | 6 | 0.3333 | 27 | 0.1111 |
| 18_675380 | 10 | 0.6 | 8 | 0.375 | 10 | 0.2 | 7 | 0.4286 | 35 | 0.4 | 5 | 0.6 | 8 | 0.5 | 12 | 0.5 | 6 | 0.5 | 31 | 0.5161 |
| 18_675381 | 11 | 0.1818 | 8 | 0.125 | 11 | 0.1818 | 7 | 0.2857 | 37 | 0.1892 | 5 | 0.2 | 5 | 0 | 12 | 0.4167 | 7 | 0 | 29 | 0.2069 |
| 18_675381 | 7 | 0.5714 | 5 | 0.6 | 8 | 0.25 | 5 | 0.6 | 25 | 0.48 | 3 | 0 | 5 | 0.4 | 9 | 0.1111 | 5 | 0.2 | 22 | 0.1818 |
| 18_675381 | 12 | 0.0833 | 7 | 0.2857 | 11 | 0.1818 | 8 | 0 | 38 | 0.1316 | 4 | 0 | 7 | 0.1429 | 13 | 0.0769 | 5 | 0 | 29 | 0.069 |
| 18_675382 | 8 | 0.25 | 4 | 0.25 | 10 | 0.2 | 4 | 0.25 | 26 | 0.2308 | 3 | 0.3333 | 4 | 0.25 | 7 | 0.2857 | 5 | 0 | 19 | 0.2105 |
| 18_675383 | 13 | 0.4615 | 8 | 0.75 | 10 | 0.4 | 8 | 0.25 | 39 | 0.4615 | 5 | 0.6 | 6 | 0 | 13 | 0.4615 | 6 | 0.6667 | 30 | 0.4333 |
| 18_675384 | 10 | 0.4 | 5 | 0.2 | 7 | 0.2857 | 4 | 0.75 | 26 | 0.3846 | 4 | 0.5 | 6 | 0.6667 | 11 | 0.7273 | 5 | 0.4 | 26 | 0.6154 |
| 18_676100 | 2 | 0.5 | 5 | 0.2 | 4 | 0.5 | 1 | 0 | 12 | 0.3333 | 2 | 0 | 8 | 0.25 | 2 | 0 | 3 | 0.3333 | 15 | 0.2 |
| 18_676105 | 1 | 0 | NA | NA | NA | NA | 1 | 1 | 2 | 0.5 | 1 | 0 | 1 | 1 | NA | NA | NA | NA | 2 | 0.5 |
| 18_676126 | 3 | 0.3333 | 7 | 0.5714 | 9 | 0.7778 | 3 | 0.3333 | 22 | 0.5909 | 3 | 0.6667 | 11 | 0.4545 | 5 | 0.4 | 4 | 0.25 | 23 | 0.4348 |
| 18_676127 | 3 | 0.3333 | 6 | 0.5 | 9 | 0.6667 | 2 | 0.5 | 20 | 0.55 | 3 | 0.6667 | 9 | 0.3333 | 4 | 0.25 | 4 | 0.5 | 20 | 0.4 |
| 18_676127 | 1 | 0 | 5 | 0.6 | 9 | 0.8889 | 2 | 1 | 17 | 0.7647 | 2 | 0.5 | 9 | 0.6667 | 4 | 0.25 | 4 | 0.75 | 19 | 0.5789 |
| 18_676128 | 3 | 0 | 7 | 0.1429 | 8 | 0.125 | 3 | 0 | 21 | 0.0952 | 3 | 0 | 11 | 0.1818 | 4 | 0 | 4 | 0.25 | 22 | 0.1364 |
| 18_676128 | 3 | 1 | 6 | 1 | 8 | 1 | 3 | 1 | 20 | 1 | 2 | 1 | 9 | 0.7778 | 4 | 1 | 4 | 0.75 | 19 | 0.8421 |
| 18_676128 | 3 | 0.3333 | 7 | 0.8571 | 8 | 0.75 | 2 | 0.5 | 20 | 0.7 | 1 | 1 | 8 | 0.75 | 4 | 0.75 | 3 | 0.3333 | 16 | 0.6875 |
| 18_676128 | 3 | 0 | 6 | 0.1667 | 8 | 0.375 | 2 | 0 | 19 | 0.2105 | 2 | 0 | 10 | 0.2 | 3 | 0.3333 | 4 | 0 | 19 | 0.1579 |
| 18_676129 | 3 | 0 | 8 | 0.125 | 8 | 0.25 | 1 | 1 | 20 | 0.2 | 2 | 0 | 11 | 0.1818 | 3 | 0 | 4 | 0 | 20 | 0.1 |
| 18_676129 | 1 | 0 | 5 | 0.4 | 7 | 0 | 2 | 0 | 15 | 0.1333 | 1 | 0 | 6 | 0 | 1 | 1 | 2 | 0 | 10 | 0.1 |
| 18_676215 | 6 | 0 | 8 | 0.125 | 3 | 0 | 2 | 0 | 19 | 0.0526 | 8 | 0.125 | 3 | 0 | 3 | 0.3333 | 2 | 0 | 16 | 0.125 |
| 18_676242 | 9 | 0.7778 | 12 | 0.5833 | 7 | 0.5714 | 7 | 0.4286 | 35 | 0.6 | 8 | 0.625 | 3 | 0.6667 | 7 | 0.8571 | 10 | 0.6 | 28 | 0.6786 |
| 18_676242 | 10 | 0 | 12 | 0.0833 | 8 | 0.125 | 9 | 0 | 39 | 0.0513 | 8 | 0.125 | 5 | 0.2 | 10 | 0 | 13 | 0 | 36 | 0.0556 |
| 18_676243 | 10 | 0.9 | 12 | 1 | 8 | 0.75 | 8 | 0.625 | 38 | 0.8421 | 8 | 0.75 | 6 | 1 | 8 | 1 | 15 | 0.8667 | 37 | 0.8919 |
| 18_676243 | 11 | 0.8182 | 12 | 0.9167 | 7 | 1 | 9 | 0.7778 | 39 | 0.8718 | 8 | 0.625 | 6 | 0.8333 | 6 | 1 | 15 | 0.8 | 35 | 0.8 |
| 18_676245 | 11 | 0.5455 | 11 | 0.8182 | 7 | 0.7143 | 8 | 0.625 | 37 | 0.6757 | 7 | 0.7143 | 6 | 0.6667 | 6 | 0.8333 | 13 | 0.6923 | 32 | 0.7188 |
| 18_676245 | 10 | 0.4 | 9 | 0.3333 | 8 | 0 | 9 | 0.4444 | 36 | 0.3056 | 7 | 0.4286 | 5 | 0.4 | 8 | 0.5 | 12 | 0.1667 | 32 | 0.3438 |
| 18_677581 | NA | NA | 1 | 1 | NA | NA | NA | NA | 1 | 1 | 1 | 0 | NA | NA | 2 | 0.5 | NA | NA | 3 | 0.3333 |
| 18_677987 | 2 | 0 | 6 | 0.1667 | 6 | 0 | 5 | 0 | 19 | 0.0526 | 1 | 0 | 3 | 0.3333 | 2 | 0.5 | 3 | 0.3333 | 9 | 0.3333 |
| 18_677988 | 1 | 1 | 6 | 0.1667 | 5 | 0.2 | 4 | 0 | 16 | 0.1875 | 1 | 1 | 2 | 0 | 1 | 0 | 3 | 0.3333 | 7 | 0.2857 |
| 18_677989 | 1 | 0 | 4 | 0.5 | 5 | 0 | 4 | 0 | 14 | 0.1429 | NA | NA | 3 | 0.3333 | 1 | 0 | 2 | 1 | 6 | 0.5 |
| 18_678054 | 2 | 0 | 2 | 0.5 | 2 | 0 | NA | NA | 6 | 0.1667 | 3 | 0 | NA | NA | 3 | 0.3333 | 2 | 0 | 8 | 0.125 |
| 18_678054 | 2 | 0.5 | 2 | 0.5 | 2 | 0 | NA | NA | 6 | 0.3333 | 3 | 0.6667 | NA | NA | 3 | 0.6667 | 1 | 0 | 7 | 0.5714 |
| 18_678054 | 2 | 1 | 2 | 0.5 | 2 | 0 | NA | NA | 6 | 0.5 | 3 | 0 | NA | NA | 3 | 0.6667 | 1 | 1 | 7 | 0.4286 |
| 18_678055 | 2 | 0 | 2 | 0.5 | 2 | 0 | NA | NA | 6 | 0.1667 | 3 | 0 | NA | NA | 3 | 0 | 1 | 1 | 7 | 0.1429 |
| 18_678056 | 2 | 1 | 2 | 1 | 1 | 1 | NA | NA | 5 | 1 | 2 | 0 | NA | NA | 3 | 0.6667 | 1 | 1 | 6 | 0.5 |
| 18_678057 | 4 | 0.25 | 2 | 0 | 3 | 0 | NA | NA | 9 | 0.1111 | 3 | 0 | 1 | 0 | 3 | 0.3333 | 2 | 0 | 9 | 0.1111 |
| 18_679283 | 11 | 0.0909 | 6 | 0 | 9 | 0 | 11 | 0.1818 | 37 | 0.0811 | 5 | 0.2 | 6 | 0.1667 | 5 | 0 | 8 | 0.375 | 24 | 0.2083 |
| 18_679741 | 1 | 1 | NA | NA | NA | NA | NA | NA | 1 | 1 | 1 | 0 | NA | NA | 1 | 1 | NA | NA | 2 | 0.5 |
| 18_680647 | NA | NA | NA | NA | 1 | 1 | NA | NA | 1 | 1 | 1 | 1 | NA | NA | NA | NA | 1 | 1 | 2 | 1 |
| 18_680655 | NA | NA | NA | NA | 1 | 1 | NA | NA | 1 | 1 | NA | NA | NA | NA | NA | NA | 1 | 1 | 1 | 1 |
| 18_680656 | NA | NA | NA | NA | 1 | 1 | NA | NA | 1 | 1 | 1 | 1 | NA | NA | NA | NA | 1 | 1 | 2 | 1 |
| 18_680656 | NA | NA | NA | NA | 1 | 1 | NA | NA | 1 | 1 | 1 | 1 | NA | NA | NA | NA | 1 | 1 | 2 | 1 |

| | | | | | | | | | | | | | | | | | | | | |
|---|---|---|---|---|---|---|---|---|---|---|---|---|---|---|---|---|---|---|---|---|
| 18_6819749 | NA | NA | 1 | 1 | 2 | 0.5 | 2 | 0.5 | 5 | 0.6 | 2 | 1 | 1 | 0 | 1 | 0 | 2 | 1 | 6 | 0.6667 |
| 18_6819752 | NA | NA | 1 | 1 | 1 | 1 | 3 | 0.6667 | 5 | 0.8 | 2 | 1 | NA | NA | NA | NA | 2 | 0.5 | 4 | 0.75 |
| 18_6819756 | NA | NA | 1 | 0 | 1 | 0 | 3 | 0.3333 | 5 | 0.2 | 2 | 0.5 | NA | NA | 1 | 0 | 1 | 0 | 4 | 0.25 |
| 18_6819761 | NA | NA | 1 | 1 | 2 | 1 | 3 | 0.3333 | 6 | 0.6667 | 2 | 1 | NA | NA | 1 | 0 | 1 | 1 | 4 | 0.75 |
| 18_6819762 | NA | NA | 1 | 0 | 2 | 0.5 | 2 | 0 | 5 | 0.2 | 2 | 0.5 | NA | NA | 1 | 0 | 2 | 1 | 5 | 0.6 |
| 18_6838718 | 1 | 1 | 1 | 0 | 2 | 0.5 | 1 | 1 | 5 | 0.6 | 4 | 0.25 | 2 | 0 | 2 | 0 | 6 | 0.1667 | 14 | 0.1429 |
| 18_6838724 | 1 | 0 | 1 | 0 | 2 | 0.5 | 3 | 0.3333 | 7 | 0.2857 | 4 | 0.25 | 2 | 0 | 2 | 1 | 6 | 0.3333 | 14 | 0.3571 |
| 18_6838728 | 1 | 0 | 1 | 0 | 2 | 0.5 | 3 | 0.6667 | 7 | 0.4286 | 3 | 0 | 2 | 0.5 | 2 | 0.5 | 3 | 0 | 10 | 0.2 |
| 18_6838732 | 1 | 0 | 1 | 0 | 2 | 0 | 2 | 0.5 | 6 | 0.1667 | 3 | 1 | 2 | 0 | 1 | 0 | 5 | 0.2 | 11 | 0.3636 |
| 18_6838733 | 1 | 0 | 1 | 1 | 1 | 1 | 3 | 1 | 6 | 0.8333 | 3 | 1 | 2 | 1 | 1 | 1 | 6 | 0.8333 | 12 | 0.9167 |
| 18_6838734 | 1 | 1 | 1 | 1 | 1 | 1 | 1 | 1 | 4 | 1 | 4 | 1 | 2 | 1 | 2 | 1 | 4 | 0.75 | 12 | 0.9167 |
| 18_6838758 | NA | NA | NA | NA | 2 | 0.5 | 3 | 0.3333 | 5 | 0.4 | 2 | 0 | 2 | 1 | 1 | 1 | 6 | 0.3333 | 11 | 0.4545 |
| 18_6838760 | 1 | 1 | 1 | 0 | 2 | 0 | 3 | 1 | 7 | 0.5714 | 4 | 0.25 | 2 | 0 | 2 | 0 | 6 | 0.3333 | 14 | 0.2143 |
| 18_6838764 | 1 | 0 | 1 | 1 | 2 | 0 | 3 | 0 | 7 | 0.1429 | 4 | 0.25 | 2 | 0 | 3 | 0 | 4 | 0.25 | 13 | 0.1538 |
| 18_6838957 | NA | NA | NA | NA | 2 | 0.5 | 2 | 0.5 | 4 | 0.5 | 1 | 0 | 1 | 1 | 1 | 0 | NA | NA | 3 | 0.3333 |
| 18_6838996 | NA | NA | 3 | 0.3333 | 2 | 0 | 2 | 0.5 | 7 | 0.2857 | 1 | 0 | 1 | 1 | 2 | 0 | NA | NA | 4 | 0.25 |
| 18_6839300 | 3 | 0 | 7 | 0.1429 | 3 | 0 | 2 | 0 | 15 | 0.0667 | 1 | 0 | 2 | 0 | 5 | 0.2 | NA | NA | 8 | 0.125 |
| 18_6839308 | 3 | 1 | 7 | 0.7143 | 5 | 0.6 | 2 | 1 | 17 | 0.7647 | 1 | 0 | 1 | 1 | 2 | 0 | NA | NA | 4 | 0.25 |
| 18_6839313 | 3 | 0.3333 | 7 | 0.7143 | 4 | 0.5 | 2 | 1 | 16 | 0.625 | 2 | 1 | 2 | 0 | 4 | 0.75 | NA | NA | 8 | 0.625 |
| 18_6839335 | 3 | 1 | 2 | 1 | 3 | 1 | 2 | 1 | 10 | 1 | 1 | 0 | 1 | 0 | 3 | 0.6667 | NA | NA | 5 | 0.4 |
| 18_6839564 | 1 | 1 | 2 | 1 | 3 | 0.3333 | 1 | 1 | 7 | 0.7143 | NA | NA | 1 | 0 | 1 | 1 | NA | NA | 2 | 0.5 |
| 18_6839576 | 1 | 1 | 2 | 1 | NA | NA | 1 | 1 | 4 | 1 | NA | NA | 1 | 1 | NA | NA | NA | NA | 1 | 1 |
| 18_6839594 | NA | NA | 1 | 1 | 3 | 0.6667 | 1 | 0 | 5 | 0.6 | NA | NA | 1 | 1 | NA | NA | NA | NA | 1 | 1 |
| 18_6839619 | 1 | 0 | 2 | 0.5 | 3 | 0 | 1 | 0 | 7 | 0.1429 | NA | NA | 1 | 1 | NA | NA | NA | NA | 1 | 1 |
| 18_6839961 | 1 | 0 | 2 | 0.5 | NA | NA | 5 | 0.2 | 8 | 0.25 | 2 | 0 | 3 | 0 | 1 | 0 | 2 | 0.5 | 8 | 0.125 |
| 18_6840030 | 1 | 1 | 1 | 0 | 1 | 0 | 4 | 0.25 | 7 | 0.2857 | NA | NA | 3 | 0.3333 | 1 | 0 | 2 | 0.5 | 6 | 0.3333 |
| 18_6840089 | 1 | 0 | 1 | 1 | 1 | 0 | 5 | 0 | 8 | 0.125 | NA | NA | 3 | 0 | 2 | 0.5 | 2 | 0 | 7 | 0.1429 |
| 18_6840094 | 1 | 0 | 1 | 1 | 1 | 0 | 4 | 0.25 | 7 | 0.2857 | NA | NA | 3 | 0.6667 | 2 | 0.5 | 2 | 0 | 7 | 0.4286 |
| 18_6840104 | 1 | 0 | NA | NA | 1 | 0 | 4 | 0.5 | 6 | 0.3333 | NA | NA | 3 | 0.6667 | 1 | 0 | 1 | 0 | 5 | 0.4 |
| 18_6840118 | 1 | 1 | 1 | 1 | 1 | 0 | 4 | 1 | 7 | 0.8571 | NA | NA | 3 | 0.6667 | 2 | 0.5 | 1 | 1 | 6 | 0.6667 |
| 18_6840135 | 3 | 0.3333 | 1 | 1 | 2 | 0.5 | 5 | 1 | 11 | 0.7273 | NA | NA | 4 | 1 | 3 | 1 | 4 | 1 | 11 | 1 |
| 18_6840137 | 3 | 0 | NA | NA | 2 | 0.5 | 5 | 0.4 | 10 | 0.3 | NA | NA | 2 | 0.5 | 3 | 0.6667 | 3 | 0.3333 | 8 | 0.5 |
| 18_6840141 | 2 | 0 | 1 | 1 | 2 | 0 | 5 | 0.8 | 10 | 0.5 | NA | NA | 4 | 0.5 | 3 | 0.6667 | 4 | 0.5 | 11 | 0.5455 |
| 18_6840142 | 2 | 0.5 | 1 | 1 | 2 | 0 | 4 | 1 | 9 | 0.6667 | NA | NA | 1 | 1 | 2 | 0.5 | 3 | 1 | 6 | 0.8333 |
| 18_6840155 | 3 | 0 | 1 | 0 | 2 | 0 | 5 | 0.4 | 11 | 0.1818 | NA | NA | 4 | 0.25 | 3 | 0 | 4 | 0.75 | 11 | 0.3636 |
| 18_6840173 | 2 | 0 | 1 | 0 | NA | NA | 5 | 0.2 | 8 | 0.125 | NA | NA | 4 | 0.25 | 3 | 0.3333 | 3 | 0 | 10 | 0.2 |
| 18_6840174 | 2 | 1 | 1 | 1 | 1 | 1 | 4 | 1 | 8 | 1 | NA | NA | 4 | 0.75 | 3 | 1 | 3 | 1 | 10 | 0.9 |
| 18_6840185 | 2 | 1 | NA | NA | 2 | 0.5 | 5 | 0.4 | 9 | 0.5556 | NA | NA | 3 | 0.6667 | 2 | 0.5 | 3 | 1 | 8 | 0.75 |
| 18_6840892 | 3 | 0.3333 | 1 | 0 | 2 | 0 | 5 | 0.4 | 11 | 0.2727 | NA | NA | 3 | 0.3333 | 2 | 0 | 3 | 0 | 8 | 0.125 |
| 18_6840896 | 3 | 0.3333 | 1 | 0 | 2 | 0 | 5 | 0.2 | 11 | 0.1818 | NA | NA | 3 | 0.3333 | 3 | 0 | 3 | 0 | 9 | 0.1111 |
| 18_6840904 | 2 | 1 | 1 | 1 | 2 | 1 | 5 | 0.8 | 10 | 0.9 | NA | NA | 3 | 0.3333 | 3 | 1 | 3 | 0.6667 | 9 | 0.6667 |
| 18_6840907 | 2 | 0.5 | 1 | 1 | 2 | 0 | 5 | 0 | 10 | 0.2 | NA | NA | 2 | 0.5 | 3 | 0 | 3 | 0.3333 | 8 | 0.25 |
| 18_6840916 | 1 | 1 | 1 | 1 | 2 | 0.5 | 4 | 0.25 | 8 | 0.5 | NA | NA | 3 | 0.6667 | 3 | 0 | 2 | 0.5 | 8 | 0.375 |
| 18_6840926 | 2 | 0.5 | 1 | 1 | 2 | 0.5 | 5 | 0.8 | 10 | 0.7 | NA | NA | 3 | 0.3333 | 2 | 0.5 | 3 | 0.3333 | 8 | 0.375 |
| 18_6840928 | 2 | 0.5 | 1 | 1 | 2 | 0.5 | 5 | 1 | 10 | 0.8 | NA | NA | 3 | 0.6667 | 3 | 1 | 3 | 1 | 9 | 0.8889 |
| 18_6845220 | NA | NA | 4 | 0 | 3 | 0 | 4 | 0.25 | 11 | 0.0909 | 5 | 0 | 2 | 0 | 3 | 0.3333 | 5 | 0 | 15 | 0.0667 |
| 18_6845454 | 2 | 0 | 5 | 0 | 5 | 0.2 | 4 | 0 | 16 | 0.0625 | 5 | 0.2 | 4 | 0 | 5 | 0 | 5 | 0 | 19 | 0.0526 |
| 18_6845625 | 2 | 1 | 1 | 0 | 2 | 1 | 2 | 1 | 7 | 0.8571 | 4 | 0.5 | 1 | 1 | 2 | 0.5 | 2 | 1 | 9 | 0.6667 |
| 18_6845631 | 2 | 0 | 1 | 0 | 2 | 0 | 2 | 0.5 | 7 | 0.1429 | 4 | 0.25 | 1 | 1 | 3 | 0.3333 | 2 | 0.5 | 10 | 0.4 |
| 18_6845634 | 2 | 0.5 | 1 | 0 | 2 | 0.5 | 1 | 0 | 6 | 0.3333 | 4 | 0.25 | 1 | 0 | 2 | 0 | 1 | 0 | 8 | 0.125 |
| 18_6845645 | 2 | 0.5 | 1 | 0 | 1 | 1 | 2 | 0.5 | 6 | 0.5 | 3 | 0.3333 | NA | NA | 2 | 0.5 | 1 | 0 | 6 | 0.3333 |
| 18_6845649 | 2 | 1 | 1 | 0 | 2 | 1 | 2 | 0.5 | 7 | 0.7143 | 3 | 0.3333 | 1 | 1 | 3 | 0.6667 | 2 | 0.5 | 9 | 0.5556 |
| 18_6845675 | 2 | 0 | 1 | 0 | 3 | 0.3333 | 2 | 0 | 8 | 0.125 | 3 | 0 | 1 | 0 | 2 | 0.5 | 2 | 0 | 8 | 0.125 |
| 18_6847371 | NA | NA | 1 | 1 | NA | NA | NA | NA | 1 | 1 | NA | NA | NA | NA | 1 | 1 | NA | NA | 1 | 1 |
| 18_6981618 | 1 | 0 | 5 | 0 | 5 | 0 | 7 | 0.1429 | 18 | 0.0556 | 3 | 0.3333 | 1 | 0 | 2 | 0.5 | 5 | 0.2 | 11 | 0.2727 |
| 18_6981622 | 1 | 0 | 3 | 0.6667 | 5 | 0.4 | 6 | 0.5 | 15 | 0.4667 | 4 | 0.75 | 1 | 0 | 2 | 0.5 | 6 | 0.5 | 13 | 0.5385 |
| 18_6981632 | 1 | 0 | 5 | 0 | 3 | 0 | 6 | 0.1667 | 15 | 0.0667 | 4 | 0 | 1 | 0 | 3 | 0 | 6 | 0.1667 | 14 | 0.0714 |
| 18_6981633 | 1 | 0 | 5 | 0 | 3 | 0 | 6 | 0.1667 | 15 | 0.0667 | 3 | 0.3333 | 1 | 0 | 2 | 0 | 7 | 0 | 13 | 0.0769 |
| 18_6981645 | 2 | 1 | 8 | 1 | 4 | 0.75 | 6 | 0.8333 | 20 | 0.9 | 4 | 1 | 1 | 1 | 2 | 1 | 7 | 0.8571 | 14 | 0.9286 |
| 18_6981650 | 1 | 0 | 8 | 0.125 | 5 | 0.2 | 6 | 0 | 20 | 0.1 | 4 | 0 | 1 | 0 | 3 | 0.3333 | 7 | 0.1429 | 15 | 0.1333 |
| 18_6981657 | 3 | 0.3333 | 8 | 0.125 | 4 | 0.5 | 6 | 0.3333 | 21 | 0.2857 | 4 | 0 | 1 | 0 | 2 | 0.5 | 7 | 0.1429 | 14 | 0.1429 |
| 18_6981676 | 3 | 0 | 7 | 0.4286 | 3 | 0 | 6 | 0.5 | 19 | 0.3158 | 3 | 0 | 1 | 0 | 3 | 0.3333 | 6 | 0 | 13 | 0.0769 |
| 18_6981677 | 3 | 0 | 6 | 0.3333 | 4 | 0.25 | 6 | 0 | 19 | 0.1579 | 4 | 0 | 2 | 0 | 3 | 0 | 8 | 0.125 | 17 | 0.0588 |
| 18_6981710 | 3 | 0 | 6 | 0.3333 | 2 | 0.5 | 6 | 0.5 | 17 | 0.3529 | 3 | 0 | 2 | 0.5 | 2 | 0.5 | 7 | 0.2857 | 14 | 0.2857 |
| 18_6981779 | 4 | 0 | 9 | 0.3333 | 3 | 0 | 5 | 0 | 21 | 0.1429 | 5 | 0.2 | 3 | 0 | 2 | 0 | 8 | 0.125 | 18 | 0.1111 |

| | | | | | | | | | | | | | | | | | | | | |
|---|---|---|---|---|---|---|---|---|---|---|---|---|---|---|---|---|---|---|---|---|
| 18_698178 | 3 | 0.6667 | 10 | 0.5 | 3 | 0.6667 | 4 | 0.5 | 20 | 0.55 | 5 | 1 | 3 | 1 | 1 | 1 | 8 | 0.75 | 17 | 0.8824 |
| 18_698178 | 4 | 0.25 | 10 | 0 | 3 | 0 | 7 | 0.1429 | 24 | 0.0833 | 6 | 0 | 2 | 0 | 2 | 0 | 9 | 0.1111 | 19 | 0.0526 |
| 18_698180 | 2 | 0 | 9 | 0 | 2 | 0 | 6 | 0.1667 | 19 | 0.0526 | 5 | 0.2 | 1 | 0 | 2 | 0 | 8 | 0 | 16 | 0.0625 |
| 18_698185 | 3 | 0 | 9 | 0.5556 | 2 | 0 | 4 | 0.5 | 18 | 0.3889 | 6 | 0.6667 | 3 | 0.6667 | 2 | 0 | 10 | 0.4 | 21 | 0.4762 |
| 18_698194 | 7 | 0 | 16 | 0.0625 | 3 | 0.3333 | 6 | 0 | 32 | 0.0625 | 8 | 0.125 | 5 | 0 | 2 | 0 | 14 | 0.0714 | 29 | 0.069 |
| 18_698198 | 8 | 0 | 8 | 0.25 | 5 | 0.2 | 9 | 0.1111 | 30 | 0.1333 | 8 | 0 | 4 | 0.25 | 5 | 0.2 | 11 | 0.1818 | 28 | 0.1429 |
| 18_698198 | 13 | 0.2308 | 20 | 0 | 11 | 0.2727 | 9 | 0 | 53 | 0.1132 | 7 | 0.2857 | 8 | 0 | 6 | 0.1667 | 16 | 0.0625 | 37 | 0.1081 |
| 18_698200 | 12 | 0.1667 | 23 | 0.087 | 12 | 0.1667 | 11 | 0.1818 | 58 | 0.1379 | 13 | 0.1538 | 9 | 0.2222 | 7 | 0 | 20 | 0.1 | 49 | 0.1224 |
| 18_698200 | 13 | 0.3846 | 24 | 0.375 | 9 | 0.3333 | 10 | 0.4 | 56 | 0.375 | 12 | 0.3333 | 8 | 0.5 | 9 | 0.5556 | 19 | 0.3684 | 48 | 0.4167 |
| 18_698201 | 10 | 0.7 | 22 | 0.6818 | 13 | 0.7692 | 10 | 0.6 | 55 | 0.6909 | 13 | 0.5385 | 9 | 0.6667 | 8 | 1 | 22 | 0.7727 | 52 | 0.7308 |
| 18_698201 | 13 | 0.0769 | 22 | 0.2727 | 13 | 0.2308 | 11 | 0.1818 | 59 | 0.2034 | 13 | 0.1538 | 9 | 0.2222 | 8 | 0.375 | 22 | 0.1818 | 52 | 0.2115 |
| 18_698201 | 12 | 0.5 | 22 | 0.6364 | 13 | 0.6923 | 9 | 0.5556 | 56 | 0.6071 | 12 | 0.3333 | 8 | 0.375 | 8 | 0.375 | 20 | 0.4 | 48 | 0.375 |
| 18_698203 | 15 | 0.2 | 25 | 0.12 | 12 | 0.3333 | 12 | 0.1667 | 64 | 0.1875 | 14 | 0.1429 | 12 | 0.1667 | 8 | 0.25 | 23 | 0.1304 | 57 | 0.1579 |
| 18_698208 | 17 | 0.0588 | 21 | 0.0952 | 8 | 0.125 | 11 | 0 | 57 | 0.0702 | 11 | 0.0909 | 8 | 0 | 8 | 0.125 | 21 | 0.0476 | 48 | 0.0625 |
| 18_698212 | 2 | 0.5 | 8 | 0.25 | 2 | 0.5 | 2 | 0.5 | 14 | 0.3571 | NA | NA | NA | NA | 1 | 0 | 4 | 0.5 | 5 | 0.4 |
| 18_698213 | 1 | 0 | 6 | 0.1667 | 2 | 0 | 2 | 0 | 11 | 0.0909 | NA | NA | 1 | 1 | 1 | 0 | 4 | 0 | 6 | 0.1667 |
| 18_698214 | 2 | 0.5 | 10 | 0.2 | 2 | 0 | 2 | 0 | 16 | 0.1875 | NA | NA | 1 | 1 | 1 | 0 | 4 | 0.25 | 6 | 0.3333 |
| 18_701128 | 3 | 0 | 1 | 0 | 3 | 0.3333 | 4 | 0 | 11 | 0.0909 | 1 | 1 | 3 | 0 | 2 | 0.5 | 1 | 0 | 7 | 0.2857 |
| 18_701129 | 2 | 0.5 | 1 | 0 | 2 | 0.5 | 4 | 0.25 | 9 | 0.3333 | 2 | 0.5 | 3 | 0.3333 | 1 | 0 | 1 | 0 | 7 | 0.2857 |
| 18_701180 | 3 | 1 | 4 | 1 | 3 | 1 | 4 | 1 | 14 | 1 | 2 | 1 | 1 | 0 | 1 | 1 | 1 | 1 | 5 | 0.8 |
| 18_701183 | 3 | 1 | 4 | 0.5 | 3 | 1 | 4 | 0.5 | 14 | 0.7143 | 2 | 0.5 | 1 | 0 | 1 | 1 | 1 | 1 | 5 | 0.6 |
| 18_701183 | 2 | 1 | 4 | 0.75 | 2 | 1 | 4 | 0.5 | 12 | 0.75 | 2 | 0 | 1 | 1 | 1 | 1 | 1 | 1 | 5 | 0.6 |
| 18_705860 | 2 | 1 | NA | NA | 1 | 1 | 3 | 0.6667 | 6 | 0.8333 | NA | NA | 2 | 1 | 1 | 0 | 3 | 1 | 6 | 0.8333 |
| 18_705862 | 2 | 1 | NA | NA | 1 | 1 | 3 | 0.6667 | 6 | 0.8333 | NA | NA | 2 | 0.5 | 2 | 0.5 | 4 | 0.5 | 8 | 0.5 |
| 18_705863 | 1 | 1 | NA | NA | 1 | 0 | 3 | 0.3333 | 5 | 0.4 | NA | NA | 2 | 0.5 | 2 | 0.5 | 3 | 0 | 7 | 0.2857 |
| 18_706009 | 1 | 0 | NA | NA | 1 | 0 | 2 | 0.5 | 4 | 0.25 | NA | NA | 1 | 0 | 1 | 0 | 1 | 1 | 3 | 0.3333 |
| 18_706010 | 1 | 1 | NA | NA | 1 | 0 | 2 | 0 | 4 | 0.25 | NA | NA | 1 | 0 | 1 | 1 | NA | NA | 2 | 0.5 |
| 18_706011 | 1 | 1 | NA | NA | 1 | 1 | 2 | 0.5 | 4 | 0.75 | NA | NA | 1 | 1 | 1 | 1 | 1 | 1 | 3 | 1 |
| 18_706012 | 1 | 0 | NA | NA | 1 | 0 | 2 | 0.5 | 4 | 0.25 | NA | NA | 1 | 0 | 1 | 1 | 1 | 1 | 3 | 0.6667 |
| 18_706013 | 1 | 1 | NA | NA | 1 | 1 | 2 | 0.5 | 4 | 0.75 | NA | NA | 1 | 0 | 1 | 0 | 1 | 1 | 3 | 0.3333 |
| 18_706420 | 1 | 1 | 1 | 1 | NA | NA | 2 | 1 | 4 | 1 | 2 | 1 | 1 | 1 | 3 | 1 | 6 | 1 | 12 | 1 |
| 18_706420 | 1 | 1 | 2 | 1 | NA | NA | 2 | 0 | 5 | 0.6 | 2 | 0 | 1 | 0 | 2 | 0 | 5 | 0.2 | 10 | 0.1 |
| 18_706421 | 1 | 1 | 2 | 1 | NA | NA | 1 | 1 | 4 | 1 | 2 | 1 | 1 | 1 | 1 | 1 | 5 | 1 | 9 | 1 |
| 18_706421 | 1 | 1 | 2 | 0.5 | NA | NA | 2 | 0.5 | 5 | 0.6 | 2 | 0 | 1 | 0 | 2 | 1 | 5 | 1 | 10 | 0.7 |
| 18_706422 | 1 | 1 | 2 | 1 | NA | NA | 1 | 0 | 4 | 0.75 | 1 | 1 | 1 | 1 | 3 | 0.6667 | 6 | 0.1667 | 11 | 0.4545 |
| 18_706422 | 1 | 1 | 2 | 0.5 | NA | NA | 2 | 0.5 | 5 | 0.6 | 2 | 0.5 | 1 | 0 | 2 | 0 | 5 | 0.4 | 10 | 0.3 |
| 18_706423 | 1 | 1 | 2 | 0 | NA | NA | 2 | 0 | 5 | 0.2 | 2 | 0.5 | 1 | 0 | NA | NA | 6 | 0.3333 | 9 | 0.3333 |
| 18_706423 | 1 | 1 | 2 | 1 | NA | NA | 2 | 1 | 5 | 1 | 2 | 1 | 1 | 1 | 2 | 1 | 6 | 1 | 11 | 1 |
| 18_706424 | 1 | 1 | 1 | 1 | NA | NA | 2 | 1 | 4 | 1 | 1 | 1 | 1 | 1 | 1 | 1 | 4 | 1 | 7 | 1 |
| 18_706426 | 1 | 1 | 1 | 1 | NA | NA | 2 | 1 | 4 | 1 | 2 | 1 | NA | NA | 2 | 1 | 4 | 1 | 8 | 1 |
| 18_706426 | 1 | 1 | 1 | 0 | NA | NA | 2 | 1 | 4 | 0.75 | 2 | 0.5 | 1 | 0 | 2 | 0 | 6 | 0.3333 | 11 | 0.2727 |
| 18_707608 | 4 | 0.25 | 3 | 0.3333 | 5 | 0 | 4 | 0 | 16 | 0.125 | 3 | 0 | 4 | 0.25 | 3 | 0 | 3 | 0 | 13 | 0.0769 |
| 18_707614 | 4 | 0 | 4 | 0.25 | 4 | 0.25 | 4 | 0.25 | 16 | 0.1875 | 3 | 0 | 5 | 0 | 4 | 0 | 3 | 0.3333 | 15 | 0.0667 |
| 18_707619 | 4 | 0 | 4 | 0 | 5 | 0.2 | 4 | 0.25 | 17 | 0.1176 | 3 | 0 | 5 | 0.2 | 5 | 0 | 5 | 0 | 18 | 0.0556 |
| 18_713912 | NA | NA | 1 | 1 | NA | NA | 2 | 0.5 | 3 | 0.6667 | NA | NA | NA | NA | 3 | 0.3333 | NA | NA | 3 | 0.3333 |
| 18_713917 | NA | NA | 1 | 0 | NA | NA | 2 | 0.5 | 3 | 0.3333 | 1 | 1 | NA | NA | 3 | 0.3333 | NA | NA | 4 | 0.5 |
| 18_713918 | NA | NA | 1 | 1 | NA | NA | 2 | 1 | 3 | 1 | 1 | 1 | NA | NA | 3 | 1 | NA | NA | 4 | 1 |
| 18_713919 | NA | NA | 1 | 0 | NA | NA | 2 | 0.5 | 3 | 0.3333 | NA | NA | NA | NA | 2 | 0.5 | NA | NA | 2 | 0.5 |
| 18_713919 | NA | NA | 1 | 1 | NA | NA | 2 | 1 | 3 | 1 | NA | NA | NA | NA | 2 | 1 | NA | NA | 2 | 1 |
| 18_713920 | NA | NA | 1 | 1 | NA | NA | 2 | 0 | 3 | 0.3333 | 1 | 0 | NA | NA | 3 | 0.3333 | NA | NA | 4 | 0.25 |
| 18_713920 | NA | NA | 1 | 1 | NA | NA | 2 | 1 | 3 | 1 | 1 | 1 | NA | NA | 3 | 1 | NA | NA | 4 | 1 |
| 18_737065 | 1 | 0 | 6 | 0.5 | 5 | 0.2 | 2 | 0.5 | 14 | 0.3571 | 1 | 1 | 3 | 0.3333 | 1 | 0 | 3 | 0 | 8 | 0.25 |
| 18_737229 | 2 | 0.5 | 4 | 0.5 | 9 | 0.4444 | 4 | 0.75 | 19 | 0.5263 | NA | NA | 5 | 0.2 | 4 | 0.25 | 3 | 0.6667 | 12 | 0.3333 |
| 18_737229 | 1 | 1 | 4 | 0 | 9 | 0.1111 | 5 | 0.2 | 19 | 0.1579 | NA | NA | 5 | 0.2 | 4 | 0.25 | 4 | 0 | 13 | 0.1538 |
| 18_737230 | 2 | 0.5 | 4 | 0 | 9 | 0 | 5 | 0.2 | 20 | 0.1 | NA | NA | 5 | 0.4 | 4 | 0.25 | 4 | 0.25 | 13 | 0.3077 |
| 18_737232 | 1 | 0 | 2 | 0 | 10 | 0 | 6 | 0.1667 | 19 | 0.0526 | NA | NA | 5 | 0.2 | 3 | 0 | 2 | 0 | 10 | 0.1 |
| 18_737232 | 1 | 0 | 5 | 0.6 | 8 | 0.875 | 6 | 1 | 20 | 0.8 | NA | NA | 5 | 0.8 | 3 | 0.3333 | 5 | 1 | 13 | 0.7692 |
| 18_737233 | NA | NA | 4 | 1 | 8 | 0.75 | 5 | 1 | 17 | 0.8824 | NA | NA | 4 | 1 | 3 | 0.6667 | 3 | 1 | 10 | 0.9 |
| 18_737233 | 1 | 0 | 5 | 0.6 | 8 | 0.75 | 5 | 0.2 | 19 | 0.5263 | NA | NA | 4 | 1 | 4 | 0.5 | 4 | 1 | 12 | 0.8333 |
| 18_737233 | 1 | 1 | 5 | 1 | 8 | 1 | 6 | 1 | 20 | 1 | NA | NA | 5 | 1 | 4 | 0.5 | 5 | 1 | 14 | 0.8571 |
| 18_737234 | 1 | 0 | 4 | 0 | 8 | 0.25 | 5 | 0.2 | 18 | 0.1667 | NA | NA | 4 | 0.5 | 3 | 0.3333 | 5 | 0 | 12 | 0.25 |
| 18_737234 | 1 | 1 | 5 | 1 | 9 | 1 | 5 | 1 | 20 | 1 | NA | NA | 5 | 1 | 3 | 1 | 5 | 1 | 13 | 1 |
| 18_737234 | 1 | 0 | 4 | 0 | 10 | 0.1 | 4 | 0 | 19 | 0.0526 | NA | NA | 5 | 0.2 | 3 | 0 | 4 | 0 | 12 | 0.0833 |
| 18_737234 | 2 | 0.5 | 4 | 0.25 | 7 | 0 | 6 | 0.1667 | 19 | 0.1579 | NA | NA | 4 | 0.25 | 4 | 0 | 4 | 0 | 12 | 0.0833 |
| 18_737235 | NA | NA | 3 | 0 | 9 | 0.2222 | 6 | 0.1667 | 18 | 0.1667 | NA | NA | 3 | 0 | 2 | 0 | 5 | 0.2 | 10 | 0.1 |

| | | | | | | | | | | | | | | | | | | | | |
|---|---|---|---|---|---|---|---|---|---|---|---|---|---|---|---|---|---|---|---|---|
| 18_7372357 | 1 | 0 | 5 | 0 | 10 | 0.1 | 6 | 0.1667 | 22 | 0.0909 | NA | NA | 6 | 0.1667 | 4 | 0.25 | 5 | 0.4 | 15 | 0.2667 |
| 18_7372383 | 2 | 0 | 5 | 0.2 | 10 | 0.3 | 6 | 0.1667 | 23 | 0.2174 | 2 | 0 | 7 | 0 | 4 | 0.25 | 3 | 0.3333 | 16 | 0.125 |
| 18_7372386 | 1 | 0 | 5 | 0 | 9 | 0.1111 | 6 | 0.1667 | 21 | 0.0952 | 2 | 0 | 7 | 0.1429 | 3 | 0.3333 | 3 | 0.3333 | 15 | 0.2 |
| 18_7372390 | 1 | 1 | 4 | 1 | 9 | 1 | 4 | 1 | 18 | 1 | 2 | 1 | 6 | 1 | 3 | 1 | 5 | 1 | 16 | 1 |
| 18_7372396 | 3 | 1 | 4 | 0.75 | 7 | 0.8571 | 4 | 0.75 | 18 | 0.8333 | 2 | 1 | 7 | 0.5714 | 4 | 0.5 | 5 | 1 | 18 | 0.7222 |
| 18_7372435 | 2 | 0 | 5 | 0.2 | 10 | 0 | 5 | 0.2 | 22 | 0.0909 | 2 | 0 | 5 | 0.2 | 7 | 0.2857 | 3 | 0 | 17 | 0.1765 |
| 18_7372443 | 2 | 0 | 6 | 0.1667 | 11 | 0.1818 | 6 | 0.3333 | 25 | 0.2 | 2 | 0 | 5 | 0.4 | 9 | 0.3333 | 2 | 0 | 18 | 0.2778 |
| 18_7372444 | 2 | 0 | 6 | 0.3333 | 9 | 0.2222 | 5 | 0.4 | 22 | 0.2727 | 3 | 0.3333 | 6 | 0.1667 | 7 | 0 | 2 | 0 | 18 | 0.1111 |
| 18_7372463 | 3 | 0 | 4 | 0.25 | 10 | 0.4 | 6 | 0.5 | 23 | 0.3478 | 3 | 0 | 7 | 0 | 9 | 0.3333 | 4 | 0.25 | 23 | 0.1739 |
| 18_7376796 | 6 | 0 | 4 | 0.25 | 5 | 0.4 | 2 | 0 | 17 | 0.1765 | 10 | 0.1 | 5 | 0.4 | 5 | 0.4 | 5 | 0 | 25 | 0.2 |
| 18_7376800 | 3 | 0 | 2 | 0 | 3 | 0.3333 | 2 | 0 | 10 | 0.1 | 7 | 0.1429 | 5 | 0 | 3 | 0.3333 | 4 | 0 | 19 | 0.1053 |
| 18_7376825 | 4 | 0 | 5 | 0 | 5 | 0.2 | 2 | 0 | 16 | 0.0625 | 8 | 0.125 | 4 | 0.25 | 4 | 0 | 6 | 0 | 22 | 0.0909 |
| 18_7376866 | 1 | 1 | 2 | 0 | 1 | 0 | NA | NA | 4 | 0.25 | 1 | 0 | 2 | 0 | 1 | 1 | 1 | 0 | 5 | 0.2 |
| 18_7377192 | 2 | 0.5 | 3 | 0 | 5 | 0.4 | 1 | 1 | 11 | 0.3636 | 3 | 0.3333 | NA | NA | 3 | 0.3333 | 1 | 0 | 7 | 0.2857 |
| 18_7377194 | 3 | 0.3333 | 2 | 0.5 | 8 | 0.125 | 2 | 0.5 | 15 | 0.2667 | 2 | 1 | NA | NA | 2 | 0.5 | 2 | 0 | 6 | 0.5 |
| 18_7377215 | 5 | 0 | 8 | 0.125 | 9 | 0.2222 | 2 | 0 | 24 | 0.125 | 4 | 0 | 1 | 0 | 4 | 0.25 | 6 | 0.1667 | 15 | 0.1333 |
| 18_7377328 | 4 | 0.75 | 7 | 0.8571 | 8 | 0.875 | 1 | 1 | 20 | 0.85 | 2 | 1 | 1 | 1 | 3 | 0.6667 | 5 | 0.8 | 11 | 0.8182 |
| 18_7377333 | 4 | 0 | 8 | 0.25 | 7 | 0.4286 | 1 | 0 | 20 | 0.25 | 1 | 1 | 1 | 0 | 4 | 0.75 | 6 | 0.5 | 12 | 0.5833 |
| 18_7377334 | 3 | 0.6667 | 6 | 0.5 | 7 | 0.1429 | 1 | 0 | 17 | 0.3529 | 3 | 0.3333 | 1 | 0 | 5 | 0.2 | 6 | 0 | 15 | 0.1333 |
| 18_7381090 | 1 | 0 | 5 | 0.2 | 3 | 0 | NA | NA | 9 | 0.1111 | 2 | 0 | 1 | 0 | 3 | 0 | 4 | 0.25 | 10 | 0.1 |
| 18_7381097 | 1 | 1 | 5 | 0.8 | 2 | 1 | NA | NA | 8 | 0.875 | 2 | 1 | 1 | 1 | 3 | 1 | 4 | 0.75 | 10 | 0.9 |
| 18_7389846 | 2 | 0 | 1 | 1 | 1 | 0 | 1 | 1 | 5 | 0.4 | 3 | 0 | 2 | 0.5 | 2 | 0.5 | 1 | 1 | 8 | 0.375 |
| 18_7389900 | 2 | 0 | 1 | 0 | NA | NA | 1 | 1 | 4 | 0.25 | 3 | 0 | 2 | 0.5 | 2 | 0 | 1 | 0 | 8 | 0.125 |
| 18_7390317 | 4 | 0.5 | 3 | 0.3333 | NA | NA | 1 | 1 | 8 | 0.5 | 1 | 1 | 3 | 0.6667 | NA | NA | 1 | 1 | 5 | 0.8 |
| 18_7390366 | 6 | 0 | 3 | 0.3333 | 1 | 0 | 1 | 0 | 11 | 0.0909 | NA | NA | 4 | 0.25 | NA | NA | 1 | 0 | 5 | 0.2 |
| 18_7399625 | NA | NA | NA | NA | 1 | 1 | NA | NA | 1 | 1 | NA | NA | NA | NA | 1 | 1 | NA | NA | 1 | 1 |
| 18_7399652 | NA | NA | NA | NA | 1 | 1 | NA | NA | 1 | 1 | NA | NA | NA | NA | 1 | 1 | NA | NA | 1 | 1 |
| 18_7399653 | NA | NA | NA | NA | 1 | 1 | NA | NA | 1 | 1 | NA | NA | NA | NA | 1 | 1 | NA | NA | 1 | 1 |
| 18_7399718 | NA | NA | NA | NA | 1 | 1 | NA | NA | 1 | 1 | NA | NA | NA | NA | 1 | 1 | NA | NA | 1 | 1 |
| 18_7400333 | NA | NA | 1 | 1 | NA | NA | NA | NA | 1 | 1 | NA | NA | NA | NA | NA | NA | 1 | 1 | 1 | 1 |
| 18_7400477 | NA | NA | 1 | 1 | NA | NA | NA | NA | 1 | 1 | 1 | 0 | NA | NA | NA | NA | 1 | 1 | 2 | 0.5 |
| 18_7400500 | 2 | 0 | 2 | 0.5 | 2 | 0 | 2 | 0 | 8 | 0.125 | 1 | 0 | 2 | 0 | NA | NA | 2 | 0.5 | 5 | 0.2 |
| 18_7400522 | NA | NA | 1 | 1 | NA | NA | NA | NA | 1 | 1 | 1 | 0 | NA | NA | NA | NA | 1 | 1 | 2 | 0.5 |
| 18_7401105 | 4 | 0 | 3 | 0 | 2 | 0 | 2 | 0.5 | 11 | 0.0909 | 3 | 0 | 2 | 0.5 | NA | NA | 4 | 0 | 9 | 0.1111 |
| 18_7401115 | 2 | 0 | 2 | 0.5 | 2 | 1 | 1 | 0 | 7 | 0.4286 | 2 | 0 | 2 | 1 | NA | NA | 2 | 0.5 | 6 | 0.5 |
| 18_7401178 | 2 | 0 | 3 | 0.6667 | 1 | 0 | 2 | 0.5 | 8 | 0.375 | 3 | 0.3333 | 2 | 0 | 1 | 0 | 3 | 0.3333 | 9 | 0.2222 |
| 18_7401196 | 5 | 0.2 | 2 | 0.5 | NA | NA | 2 | 0 | 9 | 0.2222 | 2 | 0.5 | 1 | 1 | 1 | 0 | NA | NA | 4 | 0.5 |
| 18_7406164 | 15 | 0 | 7 | 0.4286 | 12 | 0 | 14 | 0.0714 | 48 | 0.0833 | 9 | 0 | 9 | 0.1111 | 14 | 0.0714 | 9 | 0.2222 | 41 | 0.0976 |
| 18_7406167 | 12 | 0.0833 | 6 | 0 | 11 | 0.0909 | 9 | 0 | 38 | 0.0526 | 7 | 0.1429 | 9 | 0.2222 | 11 | 0.0909 | 8 | 0.25 | 35 | 0.1714 |
| 18_7406168 | 16 | 0.625 | 6 | 0.5 | 13 | 0.4615 | 14 | 0.2143 | 49 | 0.449 | 8 | 0.375 | 10 | 0.6 | 13 | 0.5385 | 8 | 0.25 | 39 | 0.4615 |
| 18_7406186 | 11 | 0.6364 | 8 | 0.75 | 6 | 0.8333 | 14 | 0.5 | 39 | 0.641 | 3 | 1 | 9 | 0.7778 | 10 | 0.7 | 8 | 0.875 | 30 | 0.8 |
| 18_7406309 | 7 | 0.8571 | 7 | 0.7143 | 3 | 0.6667 | 4 | 1 | 21 | 0.8095 | 2 | 0.5 | 4 | 1 | 7 | 0.8571 | 5 | 0.8 | 18 | 0.8333 |
| 18_7406326 | 5 | 0.2 | 3 | 0.3333 | 5 | 0 | 2 | 0.5 | 15 | 0.2 | 2 | 0.5 | 3 | 0 | 6 | 0.3333 | 4 | 0 | 15 | 0.2 |
| 18_7406331 | 8 | 0.125 | 5 | 0.4 | 3 | 0 | 5 | 0 | 21 | 0.1429 | 2 | 0.5 | 4 | 0.5 | 8 | 0.375 | 6 | 0.3333 | 20 | 0.4 |
| 18_7406341 | 8 | 0.625 | 7 | 0.2857 | 5 | 0.8 | 4 | 0.25 | 24 | 0.5 | 3 | 0.3333 | 3 | 0.3333 | 8 | 0.25 | 5 | 0.6 | 19 | 0.3684 |
| 18_7406355 | 7 | 0.7143 | 5 | 0.6 | 4 | 0.5 | 6 | 1 | 22 | 0.7273 | 3 | 0.6667 | 3 | 1 | 6 | 0.6667 | 3 | 1 | 15 | 0.8 |
| 18_7406356 | 7 | 0.7143 | 4 | 0.5 | 5 | 0.8 | 5 | 0.6 | 21 | 0.6667 | 3 | 0.6667 | 3 | 0.3333 | 8 | 1 | 4 | 1 | 18 | 0.8333 |
| 18_7406366 | 5 | 1 | 7 | 0.8571 | 4 | 0.5 | 5 | 0.8 | 21 | 0.8095 | 3 | 0.6667 | 2 | 0.5 | 6 | 0.8333 | 5 | 1 | 16 | 0.8125 |
| 18_7406385 | 9 | 0.8889 | 4 | 1 | 5 | 1 | 4 | 0.75 | 22 | 0.9091 | 3 | 0.6667 | 3 | 1 | 6 | 0.5 | 5 | 0.6 | 17 | 0.6471 |
| 18_7406391 | 6 | 0.5 | 5 | 0.4 | 4 | 0.5 | 3 | 0 | 18 | 0.3889 | 2 | 0.5 | 1 | 1 | 3 | 0.6667 | 4 | 0.75 | 10 | 0.7 |
| 18_7406403 | 7 | 0.8571 | 4 | 1 | 4 | 1 | 6 | 1 | 21 | 0.9524 | 2 | 1 | 2 | 1 | 5 | 1 | 4 | 1 | 13 | 1 |
| 18_7410302 | 4 | 0.5 | 3 | 0 | 4 | 0.5 | 5 | 0.6 | 16 | 0.4375 | 1 | 0 | NA | NA | 3 | 0.3333 | 4 | 0.5 | 8 | 0.375 |
| 18_7410313 | 3 | 0 | 2 | 0 | 4 | 0.75 | 4 | 0.25 | 13 | 0.3077 | 1 | 0 | NA | NA | NA | NA | 3 | 0.3333 | 4 | 0.25 |
| 18_7410314 | 4 | 0.5 | 3 | 0.3333 | 3 | 0 | 4 | 0.5 | 14 | 0.3571 | 1 | 0 | NA | NA | 3 | 0 | 4 | 0.25 | 8 | 0.125 |
| 18_7410318 | 3 | 0.3333 | 2 | 0.5 | 5 | 0.6 | 3 | 0.6667 | 13 | 0.5385 | 1 | 1 | NA | NA | 2 | 1 | 4 | 1 | 7 | 1 |
| 18_7410325 | 3 | 0.3333 | 3 | 0 | 5 | 0.4 | 4 | 0 | 15 | 0.2 | 1 | 0 | NA | NA | 3 | 0.3333 | 4 | 0.5 | 8 | 0.375 |
| 18_7410344 | 3 | 0.3333 | NA | NA | 4 | 0 | 3 | 0.3333 | 10 | 0.2 | 1 | 1 | NA | NA | 2 | 0 | 4 | 1 | 7 | 0.7143 |
| 18_7410349 | 2 | 0.5 | 2 | 0 | 3 | 0.3333 | 4 | 0 | 11 | 0.1818 | 1 | 0 | NA | NA | 1 | 0 | 3 | 0.3333 | 5 | 0.2 |
| 18_7410352 | 2 | 0 | 2 | 0.5 | 3 | 0.6667 | 3 | 0.3333 | 10 | 0.4 | 1 | 1 | NA | NA | 3 | 0.3333 | 4 | 0 | 8 | 0.25 |
| 18_7410355 | 2 | 0.5 | 1 | 0 | 5 | 0.6 | 4 | 0.25 | 12 | 0.4167 | 1 | 0 | NA | NA | 2 | 0.5 | 3 | 0.3333 | 6 | 0.3333 |
| 18_7410356 | 2 | 0 | 1 | 0 | 5 | 0.2 | 4 | 0 | 12 | 0.0833 | 1 | 0 | NA | NA | 3 | 0 | 4 | 0.25 | 8 | 0.125 |
| 18_7410368 | 2 | 0.5 | 1 | 1 | 4 | 1 | 3 | 0.6667 | 10 | 0.8 | 1 | 0 | NA | NA | 2 | 0.5 | 4 | 0.75 | 7 | 0.5714 |
| 18_7410379 | 2 | 0 | 1 | 0 | 4 | 0.25 | 3 | 0 | 10 | 0.1 | 1 | 0 | NA | NA | 2 | 0 | 4 | 0.25 | 7 | 0.1429 |
| 18_7410402 | 1 | 0 | 1 | 0 | 4 | 0.25 | 3 | 0 | 9 | 0.1111 | 1 | 1 | NA | NA | 2 | 0 | 4 | 0.75 | 7 | 0.5714 |
| 18_7410407 | 1 | 1 | 1 | 0 | 5 | 0.6 | 3 | 0.3333 | 10 | 0.5 | NA | NA | NA | NA | 2 | 0 | 4 | 0.5 | 6 | 0.3333 |

| | | | | | | | | | | | | | | | | | | | | |
|---|---|---|---|---|---|---|---|---|---|---|---|---|---|---|---|---|---|---|---|---|
| 18_741040 | 1 | 0 | NA | NA | 5 | 0.6 | 3 | 1 | 9 | 0.6667 | 1 | 1 | NA | NA | 2 | 1 | 4 | 0.25 | 7 | 0.5714 |
| 18_741041 | NA | NA | 1 | 0 | 5 | 0.6 | 3 | 0.3333 | 9 | 0.4444 | 1 | 1 | NA | NA | 2 | 1 | 4 | 0.5 | 7 | 0.7143 |
| 18_741041 | 1 | 0 | 1 | 1 | 5 | 0 | 3 | 0 | 10 | 0.1 | 1 | 0 | NA | NA | 1 | 0 | 4 | 0.25 | 6 | 0.1667 |
| 18_741041 | 1 | 1 | 1 | 0 | 4 | 0.5 | 4 | 0.5 | 10 | 0.5 | 1 | 1 | NA | NA | 2 | 0 | 4 | 0.5 | 7 | 0.4286 |
| 18_741041 | 1 | 0 | 1 | 1 | 5 | 0.4 | 4 | 1 | 11 | 0.6364 | 1 | 0 | NA | NA | 2 | 0 | 4 | 0.25 | 7 | 0.1429 |
| 18_741043 | 1 | 1 | 1 | 0 | 5 | 0.2 | 4 | 0 | 11 | 0.1818 | NA | NA | NA | NA | 2 | 0 | 4 | 0.5 | 6 | 0.3333 |
| 18_741987 | NA | NA | NA | NA | NA | NA | 1 | 1 | 1 | 1 | NA | NA | NA | NA | 1 | 1 | NA | NA | 1 | 1 |
| 18_743526 | 7 | 0 | 4 | 0 | 4 | 0 | 4 | 0.25 | 19 | 0.0526 | 1 | 0 | 5 | 0 | 6 | 0 | 7 | 0.1429 | 19 | 0.0526 |
| 18_744097 | NA | NA | 5 | 0.4 | 4 | 0 | 3 | 0 | 12 | 0.1667 | 2 | 0 | NA | NA | 3 | 0.3333 | 1 | 0 | 6 | 0.1667 |
| 18_744118 | NA | NA | 1 | 0 | 2 | 1 | 3 | 0 | 6 | 0.3333 | 1 | 1 | NA | NA | 4 | 0.25 | 1 | 1 | 6 | 0.5 |
| 18_744121 | NA | NA | 1 | 1 | 3 | 0.6667 | 4 | 0.5 | 8 | 0.625 | 1 | 0 | NA | NA | 1 | 1 | 1 | 0 | 3 | 0.3333 |
| 18_744122 | NA | NA | 1 | 0 | 2 | 0.5 | 4 | 0.25 | 7 | 0.2857 | 1 | 1 | NA | NA | 3 | 1 | NA | NA | 4 | 1 |
| 18_744124 | NA | NA | 1 | 0 | 2 | 0 | 4 | 0.25 | 7 | 0.1429 | 1 | 1 | NA | NA | 3 | 0 | 1 | 0 | 5 | 0.2 |
| 18_744129 | NA | NA | 2 | 0 | 4 | 0 | 3 | 0.3333 | 9 | 0.1111 | 1 | 0 | NA | NA | 4 | 0.25 | NA | NA | 5 | 0.2 |
| 18_744133 | NA | NA | 2 | 0.5 | 4 | 0.25 | 4 | 0.5 | 10 | 0.4 | 1 | 0 | 2 | 0.5 | 4 | 0.5 | 2 | 0.5 | 9 | 0.4444 |
| 18_744133 | NA | NA | 2 | 0 | 4 | 0.5 | 4 | 0.25 | 10 | 0.3 | 1 | 0 | 2 | 0.5 | 4 | 0.25 | 2 | 0.5 | 9 | 0.3333 |
| 18_744135 | NA | NA | 5 | 0.6 | 4 | 0.5 | 4 | 0.5 | 13 | 0.5385 | 1 | 1 | 3 | 0.6667 | 7 | 0.4286 | 2 | 1 | 13 | 0.6154 |
| 18_744136 | NA | NA | 4 | 0 | 4 | 0.25 | 4 | 0 | 12 | 0.0833 | 1 | 1 | 3 | 0.3333 | 6 | 0.1667 | 2 | 0.5 | 12 | 0.3333 |
| 18_744136 | NA | NA | 4 | 0 | 3 | 0.3333 | 4 | 0.5 | 11 | 0.2727 | NA | NA | 3 | 0.3333 | 7 | 0 | 1 | 0 | 11 | 0.0909 |
| 18_744136 | NA | NA | 5 | 1 | 3 | 1 | 4 | 1 | 12 | 1 | 1 | 1 | 3 | 0.6667 | 4 | 1 | 2 | 1 | 10 | 0.9 |
| 18_744138 | NA | NA | 7 | 0.1429 | 4 | 0 | 4 | 0 | 15 | 0.0667 | 1 | 0 | 3 | 0.6667 | 6 | 0 | 2 | 0 | 12 | 0.1667 |
| 18_744140 | NA | NA | 8 | 0.25 | 2 | 0 | 2 | 0 | 12 | 0.1667 | 1 | 0 | 3 | 1 | 5 | 0 | 1 | 0 | 10 | 0.3 |
| 18_744141 | NA | NA | 7 | 1 | 4 | 1 | 4 | 1 | 15 | 1 | 1 | 1 | 3 | 1 | 6 | 1 | 1 | 1 | 11 | 1 |
| 18_744150 | NA | NA | 7 | 0.1429 | 2 | 0 | NA | NA | 9 | 0.1111 | 2 | 0.5 | NA | NA | 1 | 0 | NA | NA | 3 | 0.3333 |
| 18_744152 | NA | NA | 7 | 0.4286 | 4 | 0.25 | NA | NA | 11 | 0.3636 | 2 | 0.5 | NA | NA | 3 | 0.6667 | NA | NA | 5 | 0.6 |
| 18_744154 | NA | NA | 6 | 0.5 | 4 | 0.75 | NA | NA | 10 | 0.6 | 1 | 1 | 1 | 1 | 5 | 0.6 | NA | NA | 7 | 0.7143 |
| 18_749034 | 3 | 1 | 1 | 1 | 5 | 1 | 6 | 0.8333 | 15 | 0.9333 | 3 | 1 | 5 | 0.6 | 9 | 0.5556 | 8 | 0.875 | 25 | 0.72 |
| 18_749036 | 2 | 1 | 1 | 1 | 3 | 0.6667 | 5 | 0.8 | 11 | 0.8182 | 4 | 1 | 6 | 0.8333 | 8 | 0.875 | 7 | 1 | 25 | 0.92 |
| 18_749036 | 3 | 0.3333 | 1 | 0 | 3 | 0.3333 | 6 | 0 | 13 | 0.1538 | 3 | 0 | 5 | 0.6 | 8 | 0.25 | 8 | 0.5 | 24 | 0.375 |
| 18_749037 | 2 | 0 | 1 | 0 | 2 | 0 | 4 | 0.25 | 9 | 0.1111 | 3 | 0.3333 | 4 | 0.25 | 5 | 0.2 | 4 | 0 | 16 | 0.1875 |
| 18_749039 | 2 | 1 | 1 | 1 | 3 | 0.6667 | 2 | 1 | 8 | 0.875 | 1 | 1 | 2 | 1 | 1 | 1 | 3 | 1 | 7 | 1 |
| 18_749039 | 2 | 0 | 1 | 0 | 3 | 0.3333 | 2 | 0 | 8 | 0.125 | 1 | 0 | 2 | 0 | 1 | 1 | 2 | 0 | 6 | 0.1667 |
| 18_749040 | 2 | 0.5 | 2 | 0 | 2 | 0 | 2 | 0.5 | 8 | 0.25 | 1 | 1 | 2 | 1 | 1 | 0 | 4 | 0 | 8 | 0.375 |
| 18_749040 | 1 | 0 | 2 | 0.5 | 2 | 0 | 1 | 0 | 6 | 0.1667 | 1 | 0 | NA | NA | 1 | 0 | 4 | 0.75 | 6 | 0.5 |
| 18_749040 | NA | NA | 2 | 0 | 3 | 0 | 2 | 0.5 | 7 | 0.1429 | 1 | 0 | NA | NA | 1 | 0 | 3 | 0.3333 | 5 | 0.2 |
| 18_749043 | 3 | 0.3333 | 2 | 0.5 | 2 | 0 | 2 | 0 | 9 | 0.2222 | 1 | 0 | 2 | 0.5 | 1 | 0 | 4 | 0 | 8 | 0.125 |
| 18_749055 | 1 | 1 | 1 | 1 | 1 | 1 | NA | NA | 3 | 1 | 1 | 0 | 1 | 0 | NA | NA | 2 | 1 | 4 | 0.5 |
| 18_751424 | 29 | 0.2414 | 16 | 0.0625 | 18 | 0.1667 | 26 | 0.1923 | 89 | 0.1798 | 23 | 0.2609 | 24 | 0.125 | 23 | 0.1304 | 20 | 0.3 | 90 | 0.2 |
| 18_751424 | 38 | 0.7895 | 23 | 0.8261 | 19 | 0.8421 | 29 | 0.8621 | 109 | 0.8257 | 26 | 0.7308 | 27 | 0.5556 | 27 | 0.7407 | 29 | 0.8966 | 109 | 0.7339 |
| 18_751426 | 28 | 0.0714 | 19 | 0 | 16 | 0.0625 | 29 | 0.1379 | 92 | 0.0761 | 20 | 0.1 | 25 | 0.04 | 22 | 0.0455 | 25 | 0.16 | 92 | 0.087 |
| 18_751429 | 29 | 0.1724 | 21 | 0.1905 | 12 | 0.0833 | 26 | 0.1154 | 88 | 0.1477 | 16 | 0.125 | 25 | 0.04 | 23 | 0.1304 | 27 | 0.1111 | 91 | 0.0989 |
| 18_754198 | 7 | 0.2857 | 10 | 0.1 | 9 | 0.4444 | 13 | 0.1538 | 39 | 0.2308 | 5 | 0.4 | 7 | 0.2857 | 7 | 0.1429 | 13 | 0.3846 | 32 | 0.3125 |
| 18_754198 | 7 | 0.5714 | 11 | 0.1818 | 10 | 0.2 | 13 | 0.3077 | 41 | 0.2927 | 5 | 0.8 | 7 | 0.1429 | 8 | 0.25 | 11 | 0.3636 | 31 | 0.3548 |
| 18_754198 | 6 | 0.5 | 10 | 0.5 | 10 | 0.4 | 14 | 0.4286 | 40 | 0.45 | 5 | 0.4 | 7 | 0.2857 | 8 | 0.625 | 13 | 0.3846 | 33 | 0.4242 |
| 18_754199 | 7 | 0 | 8 | 0.625 | 7 | 0.5714 | 9 | 0.2222 | 31 | 0.3548 | 3 | 0.3333 | 7 | 0.5714 | 6 | 0.3333 | 10 | 0.2 | 26 | 0.3462 |
| 18_754199 | 4 | 0 | 8 | 0.25 | 4 | 0.5 | 9 | 0.4444 | 25 | 0.32 | 4 | 0.5 | 7 | 0.2857 | 6 | 0.3333 | 10 | 0.1 | 27 | 0.2593 |
| 18_755006 | NA | NA | 1 | 1 | 2 | 0.5 | 1 | 1 | 4 | 0.75 | NA | NA | 2 | 1 | NA | NA | 1 | 1 | 3 | 1 |
| 18_755010 | 1 | 1 | 2 | 1 | 2 | 1 | 1 | 1 | 6 | 1 | 2 | 1 | 2 | 0.5 | NA | NA | 1 | 1 | 5 | 0.8 |
| 18_755018 | 5 | 0.4 | 1 | 0 | NA | NA | 2 | 0.5 | 8 | 0.375 | 3 | 0 | 2 | 0.5 | 1 | 0 | 1 | 1 | 7 | 0.2857 |
| 18_755020 | 4 | 0.5 | 1 | 1 | 2 | 0.5 | 2 | 1 | 9 | 0.6667 | 2 | 1 | 3 | 0.3333 | 1 | 1 | 1 | 1 | 7 | 0.7143 |
| 18_755023 | 2 | 0.5 | NA | NA | 2 | 0.5 | 2 | 1 | 6 | 0.6667 | 5 | 0.8 | 3 | 0.3333 | 1 | 0 | 1 | 0 | 10 | 0.5 |
| 18_755024 | 4 | 1 | NA | NA | 2 | 1 | NA | NA | 6 | 1 | 3 | 1 | 4 | 1 | 1 | 0 | 1 | 1 | 9 | 0.8889 |
| 18_755270 | 5 | 1 | 1 | 0 | 3 | 0.3333 | 2 | 0.5 | 11 | 0.6364 | 5 | 1 | 4 | 0.5 | 2 | 0.5 | 3 | 0.6667 | 14 | 0.7143 |
| 18_755270 | 5 | 0.6 | 1 | 0 | 3 | 0.3333 | 2 | 0 | 11 | 0.3636 | 5 | 0.4 | 3 | 0.6667 | 1 | 1 | 3 | 0.3333 | 12 | 0.5 |
| 18_755271 | 4 | 0.5 | 2 | 0.5 | 2 | 0.5 | 1 | 0 | 9 | 0.4444 | 5 | 0.2 | 2 | 0.5 | 1 | 0 | 2 | 0.5 | 10 | 0.3 |
| 18_755272 | 4 | 1 | 2 | 1 | 3 | 0.6667 | 1 | 1 | 10 | 0.9 | 6 | 0.6667 | 3 | 1 | 2 | 1 | 3 | 0.6667 | 14 | 0.7857 |
| 18_755272 | 5 | 0.6 | 3 | 0.3333 | 3 | 0 | 2 | 0.5 | 13 | 0.3846 | 4 | 0.5 | 4 | 0.25 | 2 | 0.5 | 2 | 0 | 12 | 0.3333 |
| 18_755274 | 4 | 0.75 | 3 | 0 | 3 | 0.6667 | 2 | 0.5 | 12 | 0.5 | 6 | 0.5 | 2 | 0.5 | 1 | 0 | 3 | 0.3333 | 12 | 0.4167 |
| 18_755274 | 4 | 0.25 | 3 | 0 | 4 | 0 | 1 | 0 | 12 | 0.0833 | 4 | 0 | 4 | 0.25 | 1 | 0 | 3 | 0 | 12 | 0.0833 |
| 18_755279 | 2 | 0.5 | 2 | 0 | 2 | 0.5 | 1 | 1 | 7 | 0.4286 | 4 | 0.25 | 2 | 0 | NA | NA | 3 | 0 | 9 | 0.1111 |
| 18_755280 | 6 | 0 | 5 | 0 | 5 | 0.2 | 2 | 0 | 18 | 0.0556 | 8 | 0.125 | 3 | 0 | 3 | 0 | 4 | 0 | 18 | 0.0556 |
| 18_755286 | 5 | 0 | 4 | 0 | 6 | 0.1667 | 2 | 0 | 17 | 0.0588 | 9 | 0.1111 | 4 | 0 | 3 | 0 | 3 | 0 | 19 | 0.0526 |
| 18_755286 | 6 | 0.6667 | 6 | 0.5 | 5 | 0.8 | 2 | 0 | 19 | 0.5789 | 10 | 0.5 | 4 | 0.25 | 3 | 0.6667 | 3 | 0 | 20 | 0.4 |
| 18_755644 | 5 | 0.6 | 5 | 0.4 | 3 | 0 | 4 | 0.25 | 17 | 0.3529 | 5 | 0 | 3 | 0.3333 | 6 | 0.3333 | 3 | 0 | 17 | 0.1765 |
| 18_755645 | 4 | 0.75 | 5 | 0.2 | 9 | 0.5556 | 7 | 0.2857 | 25 | 0.44 | 6 | 0.6667 | 3 | 0.3333 | 7 | 0.7143 | 7 | 0.5714 | 23 | 0.6087 |

| | | | | | | | | | | | | | | | | | | | | |
|---|---|---|---|---|---|---|---|---|---|---|---|---|---|---|---|---|---|---|---|---|
| 18_7556455 | 2 | 0.5 | 4 | 0 | 6 | 0 | 5 | 0.2 | 17 | 0.1176 | 5 | 0.2 | 2 | 0 | 5 | 0 | 5 | 0.2 | 17 | 0.1176 |
| 18_7556464 | 4 | 0.75 | 5 | 0.6 | 7 | 0.8571 | 6 | 1 | 22 | 0.8182 | 5 | 0.6 | NA | NA | 7 | 0.5714 | 6 | 0.8333 | 18 | 0.6667 |
| 18_7556499 | 3 | 0.3333 | 4 | 0 | 5 | 0 | 5 | 0.2 | 17 | 0.1176 | 1 | 0 | 1 | 0 | 2 | 0.5 | 5 | 0.4 | 9 | 0.3333 |
| 18_7556502 | 4 | 0 | 5 | 0 | 8 | 0.375 | 3 | 0 | 20 | 0.15 | 3 | 0 | 2 | 0 | 5 | 0 | 6 | 0.3333 | 16 | 0.125 |
| 18_7556505 | 4 | 0.75 | 5 | 0.4 | 4 | 0.5 | 6 | 0.6667 | 19 | 0.5789 | 2 | 0.5 | 2 | 0.5 | 6 | 0.6667 | 6 | 0.8333 | 16 | 0.6875 |
| 18_7556525 | 3 | 0 | 3 | 0 | 6 | 0.1667 | 5 | 0 | 17 | 0.0588 | 2 | 1 | 1 | 0 | 5 | 0.2 | 7 | 0 | 15 | 0.2 |
| 18_7556529 | 3 | 0 | 3 | 0 | 3 | 0.3333 | 5 | 0 | 14 | 0.0714 | 4 | 0 | 1 | 0 | 2 | 0 | 7 | 0.1429 | 14 | 0.0714 |
| 18_7556721 | 2 | 1 | NA | NA | 1 | 0 | 4 | 0.5 | 7 | 0.5714 | 1 | 1 | NA | NA | 2 | 0.5 | 2 | 1 | 5 | 0.8 |
| 18_7556780 | 2 | 1 | 2 | 1 | 1 | 0 | 2 | 0 | 7 | 0.5714 | 2 | 1 | 1 | 0 | 2 | 0.5 | 2 | 0.5 | 7 | 0.5714 |
| 18_7556815 | NA | NA | 1 | 0 | 1 | 0 | 1 | 1 | 3 | 0.3333 | 1 | 0 | NA | NA | 1 | 1 | NA | NA | 2 | 0.5 |
| 18_7556818 | 1 | 0 | 1 | 0 | 1 | 1 | 2 | 1 | 5 | 0.6 | 2 | 0.5 | 1 | 0 | 2 | 0.5 | 2 | 0 | 7 | 0.2857 |
| 18_7556819 | NA | NA | 1 | 1 | NA | NA | 2 | 0.5 | 3 | 0.6667 | 2 | 1 | 1 | 1 | 2 | 1 | 2 | 1 | 7 | 1 |
| 18_7556827 | 1 | 0 | 1 | 0 | 1 | 1 | 2 | 0 | 5 | 0.2 | 1 | 0 | 1 | 0 | 2 | 0.5 | 1 | 1 | 5 | 0.4 |
| 18_7556828 | 1 | 0 | 1 | 0 | 1 | 1 | 1 | 0 | 4 | 0.25 | 1 | 0 | 1 | 0 | 2 | 0.5 | 2 | 0 | 6 | 0.1667 |
| 18_7565454 | 1 | 0 | NA | NA | NA | NA | 1 | 1 | 2 | 0.5 | 2 | 0 | NA | NA | 2 | 1 | NA | NA | 4 | 0.5 |
| 18_7627042 | 1 | 1 | NA | NA | NA | NA | 1 | 0 | 2 | 0.5 | 1 | 1 | NA | NA | NA | NA | 1 | 1 | 2 | 1 |
| 18_7627050 | 1 | 1 | NA | NA | 1 | 0 | 1 | 1 | 3 | 0.6667 | 1 | 1 | NA | NA | NA | NA | 1 | 0 | 2 | 0.5 |
| 18_7627053 | 1 | 1 | NA | NA | 1 | 1 | 1 | 0 | 3 | 0.6667 | 1 | 1 | NA | NA | NA | NA | NA | NA | 1 | 1 |
| 18_7627058 | 1 | 1 | NA | NA | 1 | 0 | 1 | 1 | 3 | 0.6667 | 1 | 1 | NA | NA | NA | NA | 1 | 1 | 2 | 1 |
| 18_7627074 | 1 | 1 | NA | NA | NA | NA | 1 | 1 | 2 | 1 | 1 | 1 | NA | NA | NA | NA | 1 | 1 | 2 | 1 |
| 18_7627084 | 1 | 1 | NA | NA | 1 | 1 | 1 | 1 | 3 | 1 | 1 | 1 | NA | NA | NA | NA | 1 | 1 | 2 | 1 |
| 18_7627090 | 1 | 1 | NA | NA | 1 | 0 | 1 | 1 | 3 | 0.6667 | 1 | 1 | NA | NA | NA | NA | NA | NA | 1 | 1 |
| 18_7627948 | 1 | 1 | NA | NA | NA | NA | 1 | 1 | 2 | 1 | 1 | 1 | NA | NA | NA | NA | 1 | 1 | 2 | 1 |
| 18_7627956 | 1 | 1 | NA | NA | 1 | 0 | 1 | 0 | 3 | 0.3333 | 1 | 1 | 1 | 1 | NA | NA | 2 | 0.5 | 4 | 0.75 |
| 18_7627978 | 1 | 0 | NA | NA | 1 | 1 | 1 | 1 | 3 | 0.6667 | 1 | 0 | 1 | 1 | NA | NA | 1 | 1 | 3 | 0.6667 |
| 18_7628015 | 1 | 0 | NA | NA | 1 | 0 | 1 | 1 | 3 | 0.3333 | 1 | 1 | 2 | 0 | NA | NA | 1 | 0 | 4 | 0.25 |
| 18_7628071 | 1 | 1 | NA | NA | 1 | 0 | 2 | 0 | 4 | 0.25 | 1 | 0 | 2 | 0.5 | NA | NA | 3 | 0 | 6 | 0.1667 |
| 18_7628083 | 1 | 0 | NA | NA | 1 | 0 | 2 | 0.5 | 4 | 0.25 | 1 | 0 | 2 | 0 | NA | NA | 2 | 0.5 | 5 | 0.2 |
| 18_7628111 | 1 | 1 | NA | NA | NA | NA | 2 | 1 | 3 | 1 | NA | NA | 2 | 1 | 1 | 1 | NA | NA | 3 | 1 |
| 18_7628112 | NA | NA | NA | NA | 1 | 0 | 2 | 0.5 | 3 | 0.3333 | 1 | 0 | 2 | 0.5 | NA | NA | 2 | 0 | 5 | 0.2 |
| 18_7628113 | 1 | 0 | NA | NA | 1 | 1 | 2 | 1 | 4 | 0.75 | 1 | 1 | 1 | 1 | 1 | 1 | 2 | 1 | 5 | 1 |
| 18_7628153 | 1 | 1 | NA | NA | NA | NA | 1 | 1 | 2 | 1 | NA | NA | NA | NA | 1 | 1 | 2 | 1 | 3 | 1 |
| 18_7639560 | 4 | 0 | 3 | 0 | 1 | 0 | 5 | 0.2 | 13 | 0.0769 | 1 | 1 | 5 | 0.2 | 5 | 0 | 2 | 0 | 13 | 0.1538 |
| 18_7643774 | 4 | 0 | 4 | 0 | 1 | 1 | 5 | 0 | 14 | 0.0714 | 2 | 0 | 5 | 0 | 4 | 0 | 5 | 0.2 | 16 | 0.0625 |
| 18_7643781 | 5 | 0.8 | 4 | 0.75 | 2 | 0.5 | 6 | 0.5 | 17 | 0.6471 | 2 | 1 | 6 | 0.6667 | 4 | 0.5 | 5 | 0.8 | 17 | 0.7059 |
| 18_7643786 | 5 | 1 | 4 | 1 | 2 | 1 | 7 | 1 | 18 | 1 | 2 | 0.5 | 6 | 1 | 4 | 1 | 5 | 1 | 17 | 0.9412 |
| 18_7643800 | 5 | 0.4 | 4 | 0.5 | 2 | 0.5 | 6 | 0.3333 | 17 | 0.4118 | 3 | 0 | 6 | 0.1667 | 6 | 0.5 | 5 | 0.2 | 20 | 0.25 |
| 18_7643801 | 5 | 0.6 | 4 | 0.5 | 1 | 1 | 7 | 0.7143 | 17 | 0.6471 | 3 | 0.6667 | 6 | 0.6667 | 6 | 0.1667 | 5 | 0.8 | 20 | 0.55 |
| 18_7702828 | 15 | 0.2667 | 6 | 0 | 6 | 0.6667 | 12 | 0.4167 | 39 | 0.3333 | 6 | 0.5 | 13 | 0.4615 | 17 | 0.2941 | 10 | 0.5 | 46 | 0.413 |
| 18_7702870 | 5 | 0 | 3 | 0 | 3 | 0 | 9 | 0.1111 | 20 | 0.05 | 1 | 0 | 7 | 0.2857 | 6 | 0 | 10 | 0 | 24 | 0.0833 |
| 18_7702881 | 7 | 0.1429 | 3 | 0 | 3 | 0 | 12 | 0.0833 | 25 | 0.08 | 2 | 0 | 8 | 0.125 | 11 | 0 | 11 | 0.0909 | 32 | 0.0625 |
| 18_7702898 | 7 | 0.1429 | 3 | 0.3333 | 4 | 0 | 12 | 0.1667 | 26 | 0.1538 | 6 | 0.1667 | 11 | 0.1818 | 11 | 0 | 10 | 0.1 | 38 | 0.1053 |
| 18_7704855 | NA | NA | NA | NA | NA | NA | 1 | 1 | 1 | 1 | NA | NA | 1 | 1 | NA | NA | NA | NA | 1 | 1 |
| 18_7704865 | NA | NA | NA | NA | NA | NA | 1 | 1 | 1 | 1 | NA | NA | 1 | 1 | NA | NA | NA | NA | 1 | 1 |
| 18_7704891 | NA | NA | NA | NA | NA | NA | 1 | 1 | 1 | 1 | NA | NA | 1 | 1 | NA | NA | NA | NA | 1 | 1 |
| 18_7705843 | 9 | 1 | 10 | 0.9 | 10 | 0.8 | 12 | 1 | 41 | 0.9268 | 11 | 0.9091 | 11 | 0.7273 | 16 | 1 | 9 | 0.7778 | 47 | 0.8723 |
| 18_7705858 | 9 | 0.4444 | 12 | 0.3333 | 11 | 0.3636 | 10 | 0.5 | 42 | 0.4048 | 11 | 0.4545 | 12 | 0.3333 | 15 | 0.5333 | 11 | 0.4545 | 49 | 0.449 |
| 18_7705873 | 7 | 0.1429 | 12 | 0.1667 | 11 | 0.3636 | 11 | 0.1818 | 41 | 0.2195 | 9 | 0.2222 | 14 | 0.2143 | 11 | 0.3636 | 9 | 0.1111 | 43 | 0.2326 |
| 18_7705874 | 7 | 0.5714 | 13 | 0.2308 | 11 | 0.3636 | 12 | 0.1667 | 43 | 0.3023 | 9 | 0.2222 | 13 | 0.1538 | 19 | 0.2105 | 11 | 0.2727 | 52 | 0.2115 |
| 18_7705893 | 10 | 0.3 | 14 | 0.2143 | 13 | 0.4615 | 11 | 0.2727 | 48 | 0.3125 | 11 | 0.0909 | 14 | 0.2857 | 20 | 0.45 | 10 | 0.3 | 55 | 0.3091 |
| 18_7705973 | 1 | 0 | 2 | 0 | 2 | 0.5 | 1 | 0 | 6 | 0.1667 | NA | NA | NA | NA | 3 | 0 | 1 | 1 | 4 | 0.25 |
| 18_7705976 | 1 | 0 | 2 | 0 | 2 | 1 | 1 | 0 | 6 | 0.3333 | NA | NA | NA | NA | 3 | 1 | 1 | 1 | 4 | 1 |
| 18_7706004 | 1 | 1 | 2 | 0 | 2 | 1 | 1 | 0 | 6 | 0.5 | NA | NA | NA | NA | 3 | 0.6667 | 1 | 0 | 4 | 0.5 |
| 18_7706018 | 1 | 1 | 2 | 0.5 | 2 | 0 | 2 | 0 | 7 | 0.2857 | NA | NA | NA | NA | 3 | 0.3333 | 1 | 0 | 4 | 0.25 |
| 18_7706023 | 1 | 0 | 3 | 0.3333 | 1 | 1 | 2 | 0.5 | 7 | 0.4286 | NA | NA | NA | NA | 3 | 0.3333 | 1 | 0 | 4 | 0.25 |
| 18_7706024 | 1 | 1 | 3 | 0.6667 | 1 | 1 | 2 | 0.5 | 7 | 0.7143 | NA | NA | NA | NA | 2 | 1 | 1 | 0 | 3 | 0.6667 |
| 18_7706028 | 1 | 0 | 3 | 0.6667 | 2 | 0.5 | 2 | 0 | 8 | 0.375 | NA | NA | NA | NA | 2 | 0.5 | 1 | 1 | 3 | 0.6667 |
| 18_7706036 | 1 | 0 | 2 | 0.5 | 2 | 0.5 | 2 | 0 | 7 | 0.2857 | 1 | 0 | NA | NA | 3 | 0.6667 | 1 | 1 | 5 | 0.6 |
| 18_7706112 | 1 | 1 | 3 | 1 | 2 | 1 | 3 | 1 | 9 | 1 | 1 | 1 | NA | NA | 3 | 1 | 1 | 1 | 5 | 1 |
| 18_7706118 | 1 | 0 | 3 | 0.3333 | 1 | 0 | 2 | 0.5 | 7 | 0.2857 | NA | NA | 1 | 0 | 2 | 0.5 | 1 | 0 | 4 | 0.25 |
| 18_7706388 | 2 | 0.5 | NA | NA | 1 | 0 | NA | NA | 3 | 0.3333 | 1 | 1 | NA | NA | NA | NA | NA | NA | 1 | 1 |
| 18_7706498 | 1 | 1 | 1 | 1 | NA | NA | NA | NA | 2 | 1 | 1 | 1 | 2 | 0.5 | 1 | 0 | NA | NA | 4 | 0.5 |
| 18_7706499 | 2 | 1 | 1 | 1 | NA | NA | NA | NA | 3 | 1 | 1 | 1 | 2 | 1 | 1 | 1 | NA | NA | 4 | 1 |
| 18_7706517 | 2 | 0.5 | 1 | 0 | NA | NA | NA | NA | 3 | 0.3333 | 1 | 1 | 2 | 0 | NA | NA | NA | NA | 3 | 0.3333 |
| 18_7718512 | 2 | 0 | 3 | 0.6667 | 5 | 0.8 | 4 | 1 | 14 | 0.7143 | 3 | 0.6667 | 2 | 1 | 2 | 1 | NA | NA | 7 | 0.8571 |

| | | | | | | | | | | | | | | | | | | | | |
|---|---|---|---|---|---|---|---|---|---|---|---|---|---|---|---|---|---|---|---|---|
| 18_7718513 | 2 | 0.5 | 1 | 0 | 3 | 0.6667 | 5 | 1 | 11 | 0.7273 | 3 | 0.6667 | 2 | 1 | 3 | 0.6667 | NA | NA | 8 | 0.75 |
| 18_7718519 | 1 | 0 | 3 | 0.3333 | 4 | 0 | 4 | 0.25 | 12 | 0.1667 | 2 | 0.5 | 3 | 0 | 2 | 0 | NA | NA | 7 | 0.1429 |
| 18_7722350 | 2 | 0.5 | 4 | 0 | 4 | 0.25 | 8 | 0.25 | 18 | 0.2222 | 4 | 0 | 4 | 0.5 | 3 | 0 | 1 | 0 | 12 | 0.1667 |
| 18_7724046 | NA | NA | 2 | 0 | 4 | 0.5 | 10 | 0.2 | 16 | 0.25 | 2 | 0.5 | 4 | 0.25 | 5 | 0.4 | 2 | 0.5 | 13 | 0.3846 |
| 18_7724048 | 1 | 1 | 1 | 1 | 4 | 0.75 | 9 | 1 | 15 | 0.9333 | 3 | 0.6667 | 3 | 1 | 6 | 1 | 3 | 0.3333 | 15 | 0.8 |
| 18_7724049 | 1 | 1 | 2 | 0.5 | 3 | 1 | 8 | 0.875 | 14 | 0.8571 | 2 | 0.5 | 3 | 1 | 6 | 0.6667 | 3 | 0.6667 | 14 | 0.7143 |
| 18_7724052 | 1 | 1 | NA | NA | 2 | 1 | 9 | 0.4444 | 12 | 0.5833 | NA | NA | 2 | 0 | 4 | 1 | 2 | 0 | 8 | 0.5 |
| 18_7724062 | 2 | 0.5 | 1 | 0 | 5 | 0 | 8 | 0.125 | 16 | 0.125 | 3 | 0 | 4 | 0.25 | 4 | 0.25 | 4 | 0 | 15 | 0.1333 |
| 18_7724064 | 3 | 0 | 2 | 0 | 5 | 0.2 | 10 | 0.1 | 20 | 0.1 | 2 | 0 | 4 | 0 | 6 | 0.1667 | 4 | 0.25 | 16 | 0.125 |
| 18_7724066 | 2 | 1 | 2 | 1 | 3 | 1 | 10 | 0.9 | 17 | 0.9412 | 3 | 0.3333 | 4 | 0.75 | 5 | 0.8 | 3 | 0.6667 | 15 | 0.6667 |
| 18_7724072 | 3 | 1 | 2 | 1 | 5 | 1 | 10 | 1 | 20 | 1 | 2 | 1 | 4 | 1 | 6 | 1 | 4 | 1 | 16 | 1 |
| 18_7724081 | 2 | 0 | NA | NA | 2 | 0 | 9 | 0.2222 | 13 | 0.1538 | 4 | 0.25 | 1 | 0 | 3 | 0 | 3 | 0 | 11 | 0.0909 |
| 18_7724084 | 1 | 1 | 2 | 0 | 2 | 1 | 9 | 0.4444 | 14 | 0.5 | 3 | 0.6667 | 1 | 0 | 3 | 0.3333 | 2 | 1 | 9 | 0.5556 |
| 18_7727383 | 2 | 0.5 | 6 | 0.1667 | 2 | 0 | 1 | 1 | 11 | 0.2727 | 5 | 0.2 | NA | NA | 1 | 0 | 5 | 0.4 | 11 | 0.2727 |
| 18_7734237 | 4 | 0 | 10 | 0.3 | 3 | 0.3333 | 3 | 0.6667 | 20 | 0.3 | 9 | 0.1111 | 1 | 1 | 1 | 0 | 8 | 0.125 | 19 | 0.1579 |
| 18_7734258 | 4 | 0.25 | 9 | 0.2222 | 3 | 0.6667 | 3 | 0.3333 | 19 | 0.3158 | 9 | 0.2222 | 1 | 1 | 1 | 0 | 7 | 0.1429 | 18 | 0.2222 |
| 18_7734270 | 4 | 0.25 | 10 | 0.2 | 3 | 0 | 4 | 0 | 21 | 0.1429 | 10 | 0.2 | 1 | 0 | 1 | 0 | 9 | 0.1111 | 21 | 0.1429 |
| 18_7734278 | 5 | 0.4 | 8 | 0.75 | 2 | 1 | 2 | 1 | 17 | 0.7059 | 7 | 0.4286 | NA | NA | 1 | 1 | 8 | 0.875 | 16 | 0.6875 |
| 18_7734279 | 3 | 0.6667 | 9 | 0.6667 | 3 | 0.3333 | 4 | 0.25 | 19 | 0.5263 | 8 | 0.625 | 1 | 1 | 1 | 0 | 7 | 0.4286 | 17 | 0.5294 |
| 18_7734284 | 5 | 1 | 12 | 1 | 2 | 1 | 4 | 0.75 | 23 | 0.9565 | 10 | 1 | NA | NA | 1 | 1 | 9 | 1 | 20 | 1 |
| 18_7734293 | 2 | 1 | 8 | 0.75 | 1 | 1 | 1 | 1 | 12 | 0.8333 | 6 | 0.6667 | 1 | 1 | 1 | 1 | 5 | 0.4 | 13 | 0.6154 |
| 18_7734293 | 1 | 1 | 1 | 0 | NA | NA | 2 | 1 | 4 | 0.75 | NA | NA | NA | NA | NA | NA | 1 | 1 | 1 | 1 |
| 18_7734293 | NA | NA | 1 | 1 | NA | NA | NA | NA | 1 | 1 | 1 | 1 | NA | NA | NA | NA | NA | NA | 1 | 1 |
| 18_7734299 | 5 | 0 | 11 | 0.2727 | 4 | 0 | 2 | 0 | 22 | 0.1364 | 8 | 0 | 1 | 0 | 1 | 0 | 9 | 0.2222 | 19 | 0.1053 |
| 18_7735830 | NA | NA | NA | NA | 1 | 1 | NA | NA | 1 | 1 | 1 | 1 | NA | NA | NA | NA | NA | NA | 1 | 1 |
| 18_7754496 | 1 | 0 | 9 | 0 | 5 | 0 | 3 | 0.3333 | 18 | 0.0556 | NA | NA | 5 | 0 | 6 | 0.1667 | 4 | 0 | 15 | 0.0667 |
| 18_7754649 | NA | NA | 4 | 0.5 | 3 | 0 | 5 | 0.6 | 12 | 0.4167 | 1 | 1 | 4 | 0 | 4 | 0 | 3 | 0.3333 | 12 | 0.1667 |
| 18_7754653 | NA | NA | 4 | 0.5 | 3 | 0.6667 | 7 | 0.4286 | 14 | 0.5 | 1 | 1 | 4 | 0.25 | 5 | 0.4 | 4 | 0.75 | 14 | 0.5 |
| 18_7754659 | NA | NA | 3 | 0.3333 | 2 | 0.5 | 6 | 0.3333 | 11 | 0.3636 | 1 | 0 | 4 | 0.25 | 5 | 0.6 | 3 | 0 | 13 | 0.3077 |
| 18_7754668 | NA | NA | 4 | 0 | 3 | 1 | 6 | 0.5 | 13 | 0.4615 | 1 | 1 | 4 | 0.5 | 3 | 0 | 4 | 0.25 | 12 | 0.3333 |
| 18_7754677 | NA | NA | 4 | 0.5 | 2 | 0 | 7 | 0.2857 | 13 | 0.3077 | 1 | 1 | 4 | 0 | 3 | 0 | 4 | 0.25 | 12 | 0.1667 |
| 18_7754684 | NA | NA | 2 | 0 | 2 | 1 | 7 | 0.5714 | 11 | 0.5455 | 1 | 0 | 4 | 0.5 | 4 | 0.25 | 1 | 0 | 10 | 0.3 |
| 18_7754686 | NA | NA | 3 | 1 | 2 | 1 | 7 | 1 | 12 | 1 | 1 | 1 | 4 | 1 | 5 | 1 | 3 | 1 | 13 | 1 |
| 18_7754689 | NA | NA | 3 | 0 | 2 | 0 | 7 | 0.5714 | 12 | 0.3333 | 1 | 1 | 4 | 0 | 5 | 0.6 | 4 | 0.25 | 14 | 0.3571 |
| 18_7754712 | NA | NA | 1 | 0 | 2 | 0.5 | 4 | 0 | 7 | 0.1429 | NA | NA | 3 | 0 | 2 | 0.5 | 3 | 0.3333 | 8 | 0.25 |
| 18_7754768 | NA | NA | 1 | 1 | 2 | 0.5 | 7 | 0.1429 | 10 | 0.3 | 1 | 0 | 3 | 0.6667 | 2 | 0 | 4 | 0 | 10 | 0.2 |
| 18_7754785 | NA | NA | 1 | 1 | 2 | 0.5 | 5 | 0.8 | 8 | 0.75 | 1 | 0 | 1 | 1 | 2 | 0.5 | 4 | 0.25 | 8 | 0.375 |
| 18_7754792 | NA | NA | 1 | 1 | NA | NA | 6 | 0.8333 | 7 | 0.8571 | NA | NA | 2 | 1 | 4 | 0.75 | 3 | 0.6667 | 9 | 0.7778 |
| 18_7754802 | NA | NA | 1 | 0 | 2 | 0.5 | 6 | 0.6667 | 9 | 0.5556 | 1 | 1 | 1 | 1 | 3 | 0.6667 | 4 | 0.75 | 9 | 0.7778 |
| 18_7754952 | NA | NA | 1 | 0 | 1 | 0 | 6 | 0.1667 | 8 | 0.125 | NA | NA | 1 | 0 | 2 | 0.5 | 2 | 1 | 5 | 0.6 |
| 18_7754954 | NA | NA | 1 | 1 | 2 | 1 | 6 | 0.6667 | 9 | 0.7778 | 1 | 1 | 1 | 1 | 2 | 0.5 | 2 | 0.5 | 6 | 0.6667 |
| 18_7754972 | NA | NA | 1 | 1 | 2 | 1 | 6 | 1 | 9 | 1 | 1 | 1 | NA | NA | 2 | 0.5 | 1 | 1 | 4 | 0.75 |
| 18_7755005 | NA | NA | 1 | 1 | 2 | 0.5 | 6 | 0.6667 | 9 | 0.6667 | 1 | 0 | 1 | 0 | 2 | 0.5 | 1 | 1 | 5 | 0.4 |
| 18_7757207 | NA | NA | 1 | 0 | 1 | 0 | 6 | 0.1667 | 8 | 0.125 | 1 | 0 | 1 | 1 | 2 | 0 | 2 | 0 | 6 | 0.1667 |
| 18_7771944 | 7 | 0.7143 | 7 | 0.5714 | 4 | 0.75 | 3 | 0.6667 | 21 | 0.6667 | 5 | 0.8 | 4 | 0.5 | 7 | 0.7143 | 10 | 0.5 | 26 | 0.6154 |
| 18_7771946 | 7 | 0.7143 | 8 | 0.5 | 7 | 0.5714 | 3 | 1 | 25 | 0.64 | 4 | 0.5 | 5 | 0.6 | 6 | 0.3333 | 8 | 0.75 | 23 | 0.5652 |
| 18_7772292 | 1 | 0 | 4 | 0.75 | 3 | 0.6667 | 1 | 1 | 9 | 0.6667 | 1 | 0 | 4 | 1 | 4 | 0.75 | 2 | 1 | 11 | 0.8182 |
| 18_7775925 | NA | NA | NA | NA | 2 | 1 | NA | NA | 2 | 1 | NA | NA | NA | NA | NA | NA | 2 | 0.5 | 2 | 0.5 |
| 18_7775944 | NA | NA | NA | NA | 1 | 1 | NA | NA | 1 | 1 | NA | NA | NA | NA | NA | NA | 2 | 1 | 2 | 1 |
| 18_7776139 | NA | NA | NA | NA | NA | NA | 1 | 1 | 1 | 1 | NA | NA | 1 | 0 | NA | NA | 2 | 0.5 | 3 | 0.3333 |
| 18_7776167 | 1 | 1 | 3 | 0 | 2 | 0 | 3 | 0 | 9 | 0.1111 | 1 | 0 | 5 | 0.2 | 1 | 0 | 4 | 0.25 | 11 | 0.1818 |
| 18_7776168 | 1 | 0 | 1 | 0 | 1 | 0 | 3 | 0.3333 | 6 | 0.1667 | 1 | 1 | 5 | 0.2 | 2 | 0.5 | 3 | 0 | 11 | 0.2727 |
| 18_7776170 | 1 | 0 | 2 | 1 | 2 | 1 | 3 | 0.6667 | 8 | 0.75 | 1 | 0 | 4 | 0.5 | 2 | 0 | 4 | 1 | 11 | 0.5455 |
| 18_7776204 | 1 | 1 | 2 | 1 | 1 | 1 | 2 | 0.5 | 6 | 0.8333 | 1 | 0 | 3 | 0.3333 | 2 | 1 | 3 | 1 | 9 | 0.6667 |
| 18_7776211 | 1 | 0 | 2 | 0.5 | 2 | 1 | 3 | 0.3333 | 8 | 0.5 | 1 | 1 | 4 | 0.5 | 2 | 0.5 | 4 | 0.5 | 11 | 0.5455 |
| 18_7776212 | 1 | 1 | 2 | 1 | 2 | 0.5 | 1 | 1 | 6 | 0.8333 | 1 | 1 | 4 | 1 | 1 | 1 | 1 | 1 | 7 | 1 |
| 18_7776216 | 1 | 1 | 2 | 0 | 2 | 0 | 2 | 0 | 7 | 0.1429 | 1 | 0 | 4 | 0 | 2 | 0.5 | 4 | 0.25 | 11 | 0.1818 |
| 18_7776220 | 1 | 0 | 2 | 0.5 | 2 | 0.5 | 2 | 0.5 | 7 | 0.4286 | 1 | 1 | 2 | 0.5 | 1 | 0 | 3 | 0.3333 | 7 | 0.4286 |
| 18_7781730 | NA | NA | 1 | 1 | NA | NA | NA | NA | 1 | 1 | NA | NA | 1 | 1 | NA | NA | 1 | 1 | 2 | 1 |
| 18_7781755 | NA | NA | 1 | 1 | 1 | 1 | 1 | 1 | 3 | 1 | NA | NA | 1 | 1 | NA | NA | 1 | 1 | 2 | 1 |
| 18_7781780 | NA | NA | NA | NA | 1 | 1 | NA | NA | 1 | 1 | NA | NA | 1 | 0 | NA | NA | 1 | 1 | 2 | 0.5 |
| 18_7784530 | NA | NA | 2 | 0.5 | NA | NA | 1 | 0 | 3 | 0.3333 | NA | NA | 1 | 1 | NA | NA | NA | NA | 1 | 1 |
| 18_7796997 | 2 | 0 | NA | NA | 3 | 0.3333 | 3 | 1 | 8 | 0.5 | NA | NA | NA | NA | 1 | 1 | 1 | 1 | 2 | 1 |
| 18_7797091 | 1 | 0 | NA | NA | 2 | 0.5 | 3 | 0 | 6 | 0.1667 | NA | NA | NA | NA | 1 | 1 | NA | NA | 1 | 1 |
| 18_7797124 | 2 | 0.5 | NA | NA | 3 | 0.3333 | 3 | 0.6667 | 8 | 0.5 | NA | NA | NA | NA | 1 | 1 | NA | NA | 1 | 1 |

| | | | | | | | | | | | | | | | | | | | | |
|---|---|---|---|---|---|---|---|---|---|---|---|---|---|---|---|---|---|---|---|---|
| 18_780763 | 2 | 0.5 | 2 | 0.5 | 2 | 0.5 | 2 | 0 | 8 | 0.375 | 1 | 1 | 3 | 1 | 3 | 0.6667 | 3 | 0.6667 | 10 | 0.8 |
| 18_780765 | 2 | 0.5 | 2 | 0.5 | 2 | 0.5 | 2 | 0.5 | 8 | 0.5 | 1 | 0 | 2 | 0.5 | 4 | 0.75 | 3 | 0.6667 | 10 | 0.6 |
| 18_780765 | 2 | 0.5 | 3 | 0.6667 | 2 | 0.5 | 3 | 0.6667 | 10 | 0.6 | 2 | 0 | 3 | 0.3333 | 3 | 0.3333 | 2 | 0.5 | 10 | 0.3 |
| 18_780766 | 2 | 1 | 3 | 0.6667 | 2 | 1 | 3 | 0.3333 | 10 | 0.7 | 2 | 0.5 | 3 | 1 | 3 | 0.6667 | 3 | 0.3333 | 11 | 0.6364 |
| 18_780767 | 2 | 1 | 3 | 0.6667 | 2 | 1 | 3 | 0.6667 | 10 | 0.8 | 2 | 1 | 3 | 0.6667 | 3 | 0.6667 | 3 | 1 | 11 | 0.8182 |
| 18_780768 | 2 | 1 | 3 | 0.3333 | 2 | 0.5 | 3 | 1 | 10 | 0.7 | 2 | 0.5 | 3 | 1 | 3 | 1 | 3 | 0.6667 | 11 | 0.8182 |
| 18_780769 | 2 | 0.5 | 2 | 0.5 | 2 | 0 | 4 | 0.5 | 10 | 0.4 | 2 | 0 | 2 | 0 | 3 | 0 | 3 | 0.6667 | 10 | 0.2 |
| 18_780769 | 1 | 0 | 1 | 0 | 2 | 0.5 | 3 | 1 | 7 | 0.5714 | 1 | 0 | 2 | 0.5 | 2 | 0 | 2 | 0.5 | 7 | 0.2857 |
| 18_780769 | 2 | 0 | 2 | 0 | 2 | 0 | 4 | 0.25 | 10 | 0.1 | 3 | 0.3333 | 3 | 0 | 4 | 0 | 3 | 0 | 13 | 0.0769 |
| 18_780770 | 2 | 0 | 3 | 0 | 1 | 0 | 5 | 0.2 | 11 | 0.0909 | 3 | 0 | 3 | 0.3333 | 4 | 0.5 | 2 | 1 | 12 | 0.4167 |
| 18_780770 | 2 | 0 | 3 | 0 | 2 | 0 | 4 | 0.25 | 11 | 0.0909 | 2 | 0 | 3 | 0 | 4 | 0.25 | 3 | 0.3333 | 12 | 0.1667 |
| 18_780771 | 2 | 0 | 3 | 0 | 2 | 0 | 4 | 0.25 | 11 | 0.0909 | 2 | 0 | 3 | 0 | 3 | 0 | 3 | 0.3333 | 11 | 0.0909 |
| 18_780772 | 2 | 0 | 3 | 0 | 2 | 0.5 | 4 | 0.5 | 11 | 0.2727 | 3 | 0 | 2 | 0 | 4 | 0.25 | 3 | 0.6667 | 12 | 0.25 |
| 18_780774 | 2 | 0.5 | 3 | 0.3333 | 2 | 0.5 | 6 | 0.5 | 13 | 0.4615 | 3 | 0.6667 | 3 | 0.3333 | 3 | 0.3333 | 3 | 0.6667 | 12 | 0.5 |
| 18_780774 | 2 | 0.5 | 3 | 0.3333 | 2 | 0.5 | 5 | 1 | 12 | 0.6667 | 3 | 0.6667 | 3 | 0.3333 | 4 | 0.75 | 3 | 1 | 13 | 0.6923 |
| 18_780774 | 2 | 0 | 3 | 0 | 2 | 0.5 | 5 | 0.8 | 12 | 0.4167 | 3 | 0.6667 | 4 | 0.25 | 4 | 0.25 | 3 | 1 | 14 | 0.5 |
| 18_780775 | 2 | 0 | 2 | 0 | 2 | 0.5 | 4 | 0.25 | 10 | 0.2 | 3 | 0 | 3 | 0 | 4 | 0 | 3 | 0.3333 | 13 | 0.0769 |
| 18_780776 | 2 | 0.5 | 2 | 0 | 1 | 0 | 4 | 0 | 9 | 0.1111 | 3 | 0 | 4 | 0 | 4 | 0.25 | 4 | 0.25 | 15 | 0.1333 |
| 18_780778 | 3 | 0.3333 | 2 | 0 | 2 | 1 | 5 | 0.6 | 12 | 0.5 | 3 | 0.6667 | 3 | 0.3333 | 4 | 0.25 | 4 | 0.5 | 14 | 0.4286 |
| 18_780778 | 3 | 0.3333 | 3 | 0 | 2 | 0 | 5 | 0 | 13 | 0.0769 | 3 | 0 | 5 | 0.2 | 3 | 0 | 4 | 0 | 15 | 0.0667 |
| 18_780780 | 3 | 0.3333 | 2 | 0 | 2 | 0 | 4 | 0 | 11 | 0.0909 | 3 | 0 | 5 | 0 | 5 | 0.4 | 3 | 0 | 16 | 0.125 |
| 18_780780 | 3 | 0.3333 | 3 | 0 | 2 | 0 | 6 | 0 | 14 | 0.0714 | 3 | 0 | 4 | 0 | 5 | 0 | 4 | 0.25 | 16 | 0.0625 |
| 18_781435 | 1 | 0 | 2 | 0 | NA | NA | 2 | 0.5 | 5 | 0.2 | 1 | 0 | 1 | 1 | 1 | 0 | 1 | 0 | 4 | 0.25 |
| 18_781438 | 1 | 1 | 2 | 1 | NA | NA | 2 | 1 | 5 | 1 | NA | NA | 1 | 1 | 1 | 1 | 1 | 1 | 3 | 1 |
| 18_781441 | 1 | 0 | 2 | 0.5 | NA | NA | 2 | 0 | 5 | 0.2 | NA | NA | 1 | 0 | 1 | 0 | 1 | 1 | 3 | 0.3333 |
| 18_781454 | NA | NA | 1 | 1 | NA | NA | NA | NA | 1 | 1 | NA | NA | 1 | 0 | 1 | 1 | NA | NA | 2 | 0.5 |
| 18_787938 | NA | NA | NA | NA | 4 | 1 | 1 | 1 | 5 | 1 | 2 | 1 | 1 | 1 | NA | NA | 3 | 0.3333 | 6 | 0.6667 |
| 18_787943 | NA | NA | NA | NA | 3 | 0.6667 | 1 | 1 | 4 | 0.75 | 2 | 1 | 1 | 1 | NA | NA | 3 | 0.6667 | 6 | 0.8333 |
| 18_787951 | NA | NA | NA | NA | 2 | 0.5 | NA | NA | 2 | 0.5 | NA | NA | 1 | 1 | NA | NA | NA | NA | 1 | 1 |
| 18_787956 | NA | NA | NA | NA | 2 | 0.5 | NA | NA | 2 | 0.5 | 1 | 1 | 1 | 1 | NA | NA | NA | NA | 2 | 1 |
| 18_787957 | NA | NA | NA | NA | 2 | 1 | NA | NA | 2 | 1 | 1 | 1 | 1 | 0 | NA | NA | NA | NA | 2 | 0.5 |
| 18_787964 | NA | NA | NA | NA | 1 | 1 | NA | NA | 1 | 1 | 1 | 1 | 1 | 0 | NA | NA | NA | NA | 2 | 0.5 |
| 18_787965 | NA | NA | NA | NA | 1 | 1 | NA | NA | 1 | 1 | 1 | 1 | 1 | 0 | NA | NA | NA | NA | 2 | 0.5 |
| 18_801605 | NA | NA | NA | NA | 2 | 1 | NA | NA | 2 | 1 | NA | NA | NA | NA | 1 | 1 | NA | NA | 1 | 1 |
| 18_801608 | NA | NA | NA | NA | 1 | 1 | NA | NA | 1 | 1 | NA | NA | NA | NA | 1 | 1 | NA | NA | 1 | 1 |
| 18_801610 | 1 | 1 | NA | NA | 2 | 1 | NA | NA | 3 | 1 | NA | NA | NA | NA | 1 | 1 | NA | NA | 1 | 1 |
| 18_801621 | 1 | 1 | NA | NA | 2 | 0.5 | NA | NA | 3 | 0.6667 | 2 | 0 | NA | NA | 1 | 1 | NA | NA | 3 | 0.3333 |
| 18_801622 | NA | NA | NA | NA | 1 | 1 | NA | NA | 1 | 1 | 1 | 1 | NA | NA | 1 | 1 | NA | NA | 2 | 1 |
| 18_801623 | 1 | 1 | NA | NA | 2 | 1 | NA | NA | 3 | 1 | 2 | 0.5 | NA | NA | 1 | 1 | NA | NA | 3 | 0.6667 |
| 18_801625 | NA | NA | NA | NA | 2 | 0.5 | NA | NA | 2 | 0.5 | 2 | 0.5 | NA | NA | 1 | 1 | NA | NA | 3 | 0.6667 |
| 18_801626 | 1 | 1 | NA | NA | 2 | 0.5 | NA | NA | 3 | 0.6667 | 2 | 0.5 | NA | NA | 1 | 1 | NA | NA | 3 | 0.6667 |
| 18_801696 | NA | NA | 2 | 0.5 | NA | NA | 1 | 0 | 3 | 0.3333 | 2 | 1 | 4 | 0.25 | 1 | 1 | 1 | 1 | 8 | 0.625 |
| 18_801704 | 3 | 0 | 5 | 0 | 4 | 0 | 2 | 0.5 | 14 | 0.0714 | 4 | 0 | 7 | 0.1429 | NA | NA | 1 | 0 | 12 | 0.0833 |
| 18_801705 | 3 | 0.6667 | 5 | 0.2 | 4 | 0 | 1 | 0 | 13 | 0.2308 | 4 | 0 | 7 | 0.1429 | NA | NA | 2 | 0 | 13 | 0.0769 |
| 18_801708 | 2 | 1 | 5 | 0.8 | 3 | 0.6667 | 2 | 1 | 12 | 0.8333 | 2 | 1 | 4 | 0.5 | NA | NA | 2 | 0 | 8 | 0.5 |
| 18_801708 | 2 | 0 | 4 | 0.25 | 3 | 0 | 2 | 0 | 11 | 0.0909 | 4 | 0 | 7 | 0.2857 | NA | NA | 2 | 0 | 13 | 0.1538 |
| 18_801709 | NA | NA | 1 | 1 | 3 | 1 | 1 | 1 | 5 | 1 | 3 | 1 | 2 | 1 | NA | NA | 1 | 1 | 6 | 1 |
| 18_801709 | 1 | 1 | 2 | 1 | 2 | 1 | NA | NA | 5 | 1 | 2 | 1 | NA | NA | NA | NA | NA | NA | 2 | 1 |
| 18_801709 | 1 | 1 | 4 | 0 | 3 | 0 | 2 | 0 | 10 | 0.1 | 4 | 0 | 5 | 0.2 | NA | NA | 2 | 0 | 11 | 0.0909 |
| 18_801709 | 1 | 1 | 4 | 0.75 | 3 | 1 | 2 | 1 | 10 | 0.9 | 3 | 1 | 5 | 1 | NA | NA | 2 | 1 | 10 | 1 |
| 18_801710 | 1 | 1 | 1 | 1 | 3 | 0.3333 | 2 | 1 | 7 | 0.7143 | 2 | 1 | 4 | 0.5 | NA | NA | 1 | 1 | 7 | 0.7143 |
| 18_801710 | NA | NA | 1 | 1 | NA | NA | NA | NA | 1 | 1 | NA | NA | 1 | 1 | NA | NA | NA | NA | 1 | 1 |
| 18_801711 | 1 | 1 | 3 | 0 | 3 | 0 | 2 | 0 | 9 | 0.1111 | 3 | 0 | 5 | 0.2 | NA | NA | 2 | 0 | 10 | 0.1 |
| 18_801711 | 1 | 1 | 2 | 0.5 | 3 | 0.6667 | 1 | 1 | 7 | 0.7143 | 2 | 0.5 | 5 | 0.6 | NA | NA | NA | NA | 7 | 0.5714 |
| 18_801711 | 1 | 0 | 3 | 0.3333 | 3 | 0 | 2 | 0.5 | 9 | 0.2222 | 3 | 0.3333 | 6 | 0.1667 | NA | NA | 1 | 0 | 10 | 0.2 |
| 18_801712 | 1 | 1 | 1 | 1 | 3 | 0.6667 | 1 | 1 | 6 | 0.8333 | 3 | 0.3333 | 5 | 0 | NA | NA | 2 | 0.5 | 10 | 0.2 |
| 18_801712 | 1 | 0 | 3 | 0 | 3 | 0.3333 | 2 | 0 | 9 | 0.1111 | 4 | 0.5 | 6 | 0.5 | NA | NA | 2 | 0 | 12 | 0.4167 |
| 18_801714 | 1 | 1 | 1 | 1 | 2 | 1 | 2 | 1 | 6 | 1 | 3 | 1 | 5 | 1 | NA | NA | 1 | 1 | 9 | 1 |
| 18_801714 | 1 | 1 | 2 | 1 | 2 | 0 | 2 | 0 | 7 | 0.4286 | 3 | 0.3333 | 6 | 0.1667 | NA | NA | 1 | 0 | 10 | 0.2 |
| 18_801718 | 1 | 1 | 1 | 0 | 2 | 0.5 | 1 | 0 | 5 | 0.4 | 2 | 0.5 | 5 | 0.2 | NA | NA | NA | NA | 7 | 0.2857 |
| 18_801719 | 1 | 1 | NA | NA | 2 | 1 | 1 | 1 | 4 | 1 | 2 | 0.5 | 4 | 1 | NA | NA | 1 | 1 | 7 | 0.8571 |
| 18_801719 | 1 | 1 | NA | NA | 2 | 1 | 1 | 0 | 4 | 0.75 | 3 | 1 | 5 | 0.8 | NA | NA | NA | NA | 8 | 0.875 |
| 18_801722 | NA | NA | NA | NA | 2 | 1 | 1 | 1 | 3 | 1 | 4 | 1 | 4 | 1 | NA | NA | 1 | 1 | 9 | 1 |
| 18_801725 | NA | NA | NA | NA | 1 | 0 | 1 | 1 | 2 | 0.5 | 3 | 0.6667 | 4 | 0.75 | NA | NA | 1 | 1 | 8 | 0.75 |
| 18_801728 | NA | NA | NA | NA | 1 | 1 | NA | NA | 1 | 1 | 2 | 0.5 | 2 | 1 | NA | NA | 1 | 1 | 5 | 0.8 |

| | | | | | | | | | | | | | | | | | | | | |
|---|---|---|---|---|---|---|---|---|---|---|---|---|---|---|---|---|---|---|---|---|
| 18_801731 | NA | NA | NA | NA | 1 | 1 | 1 | 1 | 2 | 1 | 4 | 0.5 | 1 | 0 | NA | NA | 1 | 0 | 6 | 0.3333 |
| 18_801731 | NA | NA | NA | NA | NA | NA | 1 | 1 | 1 | 1 | 3 | 1 | 2 | 0.5 | NA | NA | 1 | 1 | 6 | 0.8333 |
| 18_801731 | NA | NA | NA | NA | 1 | 1 | 1 | 1 | 2 | 1 | 4 | 1 | 3 | 1 | NA | NA | 1 | 1 | 8 | 1 |
| 18_801732 | NA | NA | NA | NA | 1 | 1 | 1 | 0 | 2 | 0.5 | 2 | 0.5 | 3 | 0 | NA | NA | NA | NA | 5 | 0.2 |
| 18_801734 | NA | NA | NA | NA | 1 | 1 | 1 | 1 | 2 | 1 | 1 | 0 | 3 | 1 | NA | NA | NA | NA | 4 | 0.75 |
| 18_801736 | NA | NA | NA | NA | NA | NA | 1 | 1 | 1 | 1 | 1 | 1 | 3 | 0.6667 | NA | NA | 1 | 1 | 5 | 0.8 |
| 18_801736 | NA | NA | NA | NA | 1 | 1 | 1 | 1 | 2 | 1 | 1 | 1 | 3 | 1 | NA | NA | 1 | 0 | 5 | 0.8 |
| 18_801738 | NA | NA | NA | NA | 1 | 1 | NA | NA | 1 | 1 | 1 | 1 | 2 | 1 | NA | NA | 1 | 1 | 4 | 1 |
| 18_801739 | NA | NA | NA | NA | 1 | 1 | 1 | 1 | 2 | 1 | 1 | 0 | 3 | 0.6667 | NA | NA | 1 | 1 | 5 | 0.6 |
| 18_801951 | NA | NA | 1 | 1 | 2 | 0.5 | NA | NA | 3 | 0.6667 | NA | NA | NA | NA | NA | NA | 1 | 1 | 1 | 1 |
| 18_802331 | NA | NA | 1 | 1 | NA | NA | NA | NA | 1 | 1 | NA | NA | NA | NA | NA | NA | 1 | 1 | 1 | 1 |
| 18_802333 | NA | NA | 1 | 1 | NA | NA | NA | NA | 1 | 1 | NA | NA | NA | NA | NA | NA | 1 | 1 | 1 | 1 |
| 18_802337 | NA | NA | 1 | 1 | NA | NA | NA | NA | 1 | 1 | NA | NA | NA | NA | NA | NA | 1 | 1 | 1 | 1 |
| 18_802337 | NA | NA | 1 | 1 | NA | NA | NA | NA | 1 | 1 | NA | NA | NA | NA | NA | NA | 1 | 1 | 1 | 1 |
| 18_802355 | 6 | 1 | 7 | 0.7143 | 8 | 0.75 | 3 | 0.6667 | 24 | 0.7917 | 4 | 0.75 | 2 | 0 | 4 | 1 | 8 | 0.75 | 18 | 0.7222 |
| 18_802355 | 4 | 0.25 | 7 | 0.2857 | 6 | 0.6667 | 4 | 0.5 | 21 | 0.4286 | 3 | 0.3333 | 3 | 0 | 5 | 0.8 | 6 | 0.5 | 17 | 0.4706 |
| 18_802356 | 6 | 0.1667 | 7 | 0.2857 | 6 | 0.3333 | 4 | 0.5 | 23 | 0.3043 | 3 | 0.3333 | 3 | 0 | 5 | 0.2 | 7 | 0.8571 | 18 | 0.4444 |
| 18_802356 | 4 | 0.75 | 7 | 0.2857 | 6 | 0.6667 | 4 | 0.5 | 21 | 0.5238 | 4 | 0.25 | 3 | 0 | 7 | 0.2857 | 8 | 0.375 | 22 | 0.2727 |
| 18_802358 | 2 | 1 | 5 | 0 | 2 | 0.5 | 4 | 0.25 | 13 | 0.3077 | 3 | 0 | 4 | 0.5 | 7 | 0.1429 | 5 | 0.4 | 19 | 0.2632 |
| 18_802547 | 9 | 0 | 13 | 0.1538 | 7 | 0 | 6 | 0 | 35 | 0.0571 | 9 | 0 | 13 | 0.0769 | 8 | 0.125 | 12 | 0.1667 | 42 | 0.0952 |
| 18_802547 | 16 | 0.125 | 11 | 0 | 9 | 0.5556 | 7 | 0 | 43 | 0.1628 | 8 | 0.375 | 15 | 0.3333 | 8 | 0.375 | 11 | 0 | 42 | 0.2619 |
| 18_802547 | 12 | 0.0833 | 18 | 0.1111 | 9 | 0 | 6 | 0.1667 | 45 | 0.0889 | 11 | 0.2727 | 18 | 0.0556 | 9 | 0.2222 | 13 | 0.0769 | 51 | 0.1373 |
| 18_802548 | 17 | 0.2941 | 18 | 0.3889 | 10 | 0.3 | 6 | 0.1667 | 51 | 0.3137 | 12 | 0.3333 | 19 | 0.3158 | 11 | 0.1818 | 14 | 0.3571 | 56 | 0.3036 |
| 18_802549 | 13 | 0 | 19 | 0.2105 | 8 | 0.125 | 5 | 0.2 | 45 | 0.1333 | 9 | 0 | 16 | 0.25 | 8 | 0 | 12 | 0.25 | 45 | 0.1556 |
| 18_802549 | 14 | 0.5714 | 16 | 0.5 | 8 | 0.5 | 4 | 0.5 | 42 | 0.5238 | 8 | 0.125 | 14 | 0.5 | 11 | 0.3636 | 14 | 0.5 | 47 | 0.4043 |
| 18_802550 | 16 | 0.5 | 16 | 0.3125 | 7 | 0.2857 | 5 | 0.2 | 44 | 0.3636 | 12 | 0.5833 | 17 | 0.5294 | 8 | 0.125 | 13 | 0.3846 | 50 | 0.44 |
| 18_802550 | 11 | 0.3636 | 17 | 0.4118 | 7 | 0 | 5 | 0 | 40 | 0.275 | 7 | 0.5714 | 16 | 0.3125 | 7 | 0.4286 | 10 | 0.8 | 40 | 0.5 |
| 18_802553 | 18 | 0.5 | 17 | 0.4118 | 11 | 0.2727 | 6 | 0.5 | 52 | 0.4231 | 10 | 0.5 | 19 | 0.2105 | 8 | 0.375 | 14 | 0.3571 | 51 | 0.3333 |
| 18_802553 | 20 | 0.45 | 16 | 0.5 | 10 | 0.5 | 6 | 0.6667 | 52 | 0.5 | 7 | 0.7143 | 18 | 0.5556 | 9 | 0.6667 | 11 | 0.4545 | 45 | 0.5778 |
| 18_802553 | 19 | 0.2632 | 15 | 0.2 | 12 | 0.1667 | 4 | 0.25 | 50 | 0.22 | 10 | 0.2 | 17 | 0.2353 | 8 | 0.375 | 11 | 0.1818 | 46 | 0.2391 |
| 18_802554 | 12 | 0.4167 | 12 | 0.5 | 10 | 0.8 | 6 | 0.1667 | 40 | 0.5 | 10 | 0.4 | 14 | 0.7143 | 6 | 0.5 | 9 | 0.4444 | 39 | 0.5385 |
| 18_802554 | 23 | 0.1304 | 21 | 0.1905 | 13 | 0.3846 | 9 | 0.1111 | 66 | 0.197 | 12 | 0.5 | 21 | 0.2381 | 7 | 0.1429 | 15 | 0.2667 | 55 | 0.2909 |
| 18_802555 | 20 | 0.05 | 19 | 0.1579 | 14 | 0.2143 | 9 | 0.1111 | 62 | 0.129 | 11 | 0 | 19 | 0.1053 | 9 | 0.2222 | 16 | 0.0625 | 55 | 0.0909 |
| 18_802555 | 24 | 0.3333 | 22 | 0.1818 | 13 | 0.0769 | 12 | 0 | 71 | 0.1831 | 14 | 0.1429 | 19 | 0 | 9 | 0.4444 | 13 | 0 | 55 | 0.1091 |
| 18_802555 | 21 | 0.7143 | 21 | 0.7143 | 14 | 0.7857 | 10 | 0.9 | 66 | 0.7576 | 15 | 0.7333 | 18 | 0.7778 | 11 | 0.8182 | 12 | 0.9167 | 56 | 0.8036 |
| 18_802556 | 22 | 0.9091 | 19 | 1 | 15 | 1 | 11 | 1 | 67 | 0.9701 | 13 | 1 | 20 | 1 | 10 | 1 | 15 | 0.9333 | 58 | 0.9828 |
| 18_802557 | 14 | 0.1429 | 17 | 0 | 10 | 0.1 | 7 | 0.1429 | 48 | 0.0833 | 9 | 0.1111 | 16 | 0.125 | 9 | 0.1111 | 10 | 0.1 | 44 | 0.1136 |
| 18_802558 | 26 | 0.1154 | 23 | 0.0435 | 15 | 0.0667 | 11 | 0.0909 | 75 | 0.08 | 14 | 0 | 22 | 0.2273 | 10 | 0 | 17 | 0 | 63 | 0.0794 |
| 18_802654 | 7 | 0.1429 | 7 | 0 | 12 | 0.25 | 6 | 0 | 32 | 0.125 | 7 | 0 | 6 | 0.1667 | 10 | 0 | 5 | 0.2 | 28 | 0.0714 |
| 18_802655 | 8 | 0.875 | 7 | 0.8571 | 12 | 1 | 6 | 1 | 33 | 0.9394 | 7 | 0.8571 | 5 | 1 | 10 | 0.8 | 6 | 0.5 | 28 | 0.7857 |
| 18_802655 | 7 | 0.5714 | 7 | 1 | 11 | 1 | 6 | 0.8333 | 31 | 0.871 | 7 | 1 | 6 | 0.6667 | 10 | 0.9 | 5 | 0.6 | 28 | 0.8214 |
| 18_802658 | 1 | 1 | 3 | 0.3333 | NA | NA | 1 | 1 | 5 | 0.6 | 1 | 1 | NA | NA | 2 | 0.5 | 1 | 1 | 4 | 0.75 |
| 18_802666 | 1 | 0 | 3 | 0 | 1 | 0 | 1 | 1 | 6 | 0.1667 | 1 | 1 | NA | NA | 3 | 0.3333 | 1 | 0 | 5 | 0.4 |
| 18_802674 | 1 | 0 | 1 | 0 | 1 | 1 | 1 | 0 | 4 | 0.25 | 1 | 0 | NA | NA | 2 | 0.5 | 1 | 0 | 4 | 0.25 |
| 18_802675 | 1 | 1 | 1 | 1 | 1 | 0 | NA | NA | 3 | 0.6667 | 1 | 0 | NA | NA | 2 | 0.5 | 1 | 0 | 4 | 0.25 |
| 18_802676 | 1 | 1 | NA | NA | 1 | 1 | 1 | 0 | 3 | 0.6667 | 1 | 1 | NA | NA | 2 | 1 | 1 | 1 | 4 | 1 |
| 18_802677 | NA | NA | 1 | 1 | NA | NA | NA | NA | 1 | 1 | NA | NA | NA | NA | 2 | 0 | 1 | 1 | 3 | 0.3333 |
| 18_802678 | 1 | 1 | 1 | 0 | NA | NA | 1 | 1 | 3 | 0.6667 | 1 | 0 | NA | NA | 2 | 0.5 | 1 | 1 | 4 | 0.5 |
| 18_802681 | 1 | 1 | 1 | 0 | 1 | 1 | 1 | 1 | 4 | 0.75 | 1 | 1 | NA | NA | 2 | 1 | 1 | 1 | 4 | 1 |
| 18_802682 | NA | NA | 1 | 1 | 1 | 0 | 1 | 0 | 3 | 0.3333 | 1 | 0 | NA | NA | 2 | 0.5 | NA | NA | 3 | 0.3333 |
| 18_802683 | 1 | 0 | 1 | 1 | NA | NA | 1 | 1 | 3 | 0.6667 | 1 | 1 | NA | NA | 1 | 0 | NA | NA | 2 | 0.5 |
| 18_803000 | 3 | 0.3333 | 1 | 0 | 1 | 0 | 3 | 0.3333 | 8 | 0.25 | NA | NA | 2 | 0.5 | 3 | 0.3333 | 2 | 0.5 | 7 | 0.4286 |
| 18_803002 | 3 | 0.3333 | 1 | 0 | 1 | 0 | 3 | 0.6667 | 8 | 0.375 | NA | NA | 2 | 0.5 | 2 | 0 | 2 | 0.5 | 6 | 0.3333 |
| 18_803002 | 3 | 1 | 1 | 0 | 2 | 0 | 2 | 0.5 | 8 | 0.5 | NA | NA | 2 | 0 | 4 | 0.25 | 2 | 0 | 8 | 0.125 |
| 18_803349 | 3 | 0 | 2 | 0 | 2 | 0 | 2 | 0.5 | 9 | 0.1111 | NA | NA | 2 | 0 | 2 | 0 | 5 | 0.2 | 9 | 0.1111 |
| 18_803350 | 3 | 0.6667 | 2 | 0.5 | 2 | 1 | 3 | 1 | 10 | 0.8 | NA | NA | 2 | 1 | 4 | 0.75 | 4 | 0.5 | 10 | 0.7 |
| 18_803385 | 1 | 1 | NA | NA | 1 | 0 | NA | NA | 2 | 0.5 | 1 | 1 | NA | NA | NA | NA | NA | NA | 1 | 1 |
| 18_803388 | NA | NA | NA | NA | 1 | 1 | 1 | 1 | 2 | 1 | 2 | 0.5 | 4 | 0.75 | 1 | 1 | 2 | 0.5 | 9 | 0.6667 |
| 18_803388 | NA | NA | NA | NA | 2 | 0 | 1 | 1 | 3 | 0.3333 | 1 | 1 | 4 | 0 | 1 | 1 | 2 | 0 | 8 | 0.25 |
| 18_803388 | NA | NA | NA | NA | 2 | 0.5 | 1 | 0 | 3 | 0.3333 | 2 | 0.5 | 4 | 0.75 | 1 | 1 | 2 | 0 | 9 | 0.5556 |
| 18_803389 | NA | NA | NA | NA | 1 | 1 | 1 | 0 | 2 | 0.5 | 1 | 1 | 2 | 1 | 1 | 1 | 2 | 1 | 5 | 1 |
| 18_803389 | NA | NA | NA | NA | 3 | 0.6667 | 1 | 1 | 4 | 0.75 | 2 | 1 | 3 | 0.3333 | 1 | 0 | 2 | 1 | 8 | 0.625 |
| 18_803392 | NA | NA | NA | NA | 2 | 1 | 1 | 0 | 3 | 0.6667 | 2 | 1 | 3 | 1 | 1 | 1 | 2 | 1 | 8 | 1 |
| 18_804514 | 3 | 0.3333 | 2 | 0 | 5 | 0.8 | 4 | 0 | 14 | 0.3571 | 2 | 0 | 2 | 0 | 2 | 0.5 | 2 | 0 | 8 | 0.125 |
| 18_804514 | 3 | 0 | 2 | 0.5 | 5 | 0 | 4 | 0.75 | 14 | 0.2857 | 2 | 0 | 2 | 0 | 2 | 0 | 2 | 0.5 | 8 | 0.125 |

| | | | | | | | | | | | | | | | | | | | | |
|---|---|---|---|---|---|---|---|---|---|---|---|---|---|---|---|---|---|---|---|---|
| 18_804515( | 3 | 0.6667 | 2 | 1 | 5 | 0.2 | 4 | 0.25 | 14 | 0.4286 | 2 | 1 | 3 | 0.3333 | 3 | 0.6667 | 2 | 1 | 10 | 0.7 |
| 18_804515! | 3 | 0 | 2 | 0.5 | 5 | 0.4 | 3 | 0.3333 | 13 | 0.3077 | 2 | 0 | 3 | 0 | 3 | 0.3333 | 2 | 0.5 | 10 | 0.2 |
| 18_804518! | 3 | 0.3333 | 2 | 0.5 | 4 | 0.25 | 3 | 0.3333 | 12 | 0.3333 | 2 | 0.5 | 2 | 1 | 3 | 0 | 2 | 0 | 9 | 0.3333 |
| 18_804925( | 2 | 0.5 | 1 | 0 | 4 | 0.75 | 2 | 0 | 9 | 0.4444 | 1 | 0 | 2 | 1 | 1 | 1 | 1 | 0 | 5 | 0.6 |
| 18_804925! | 2 | 1 | 1 | 0 | 4 | 0.25 | 2 | 0.5 | 9 | 0.4444 | 1 | 1 | 1 | 1 | 2 | 0.5 | 1 | 1 | 5 | 0.8 |
| 18_804927' | 2 | 1 | 1 | 0 | 4 | 0.5 | 1 | 0 | 8 | 0.5 | NA | NA | 2 | 1 | 2 | 0.5 | 1 | 1 | 5 | 0.8 |
| 18_804928 | 1 | 1 | 1 | 1 | 3 | 0.3333 | 1 | 1 | 6 | 0.6667 | NA | NA | 2 | 0 | 1 | 1 | NA | NA | 3 | 0.3333 |
| 18_804929 | 1 | 1 | 1 | 0 | 1 | 0 | 1 | 1 | 4 | 0.5 | NA | NA | 2 | 1 | 1 | 0 | NA | NA | 3 | 0.6667 |
| 18_804930 | 1 | 0 | 1 | 1 | 3 | 0.6667 | NA | NA | 5 | 0.6 | NA | NA | 1 | 1 | 1 | 1 | NA | NA | 2 | 1 |
| 18_804931 | 2 | 1 | 1 | 1 | 3 | 1 | 1 | 1 | 7 | 1 | NA | NA | 2 | 1 | 2 | 1 | NA | NA | 4 | 1 |
| 18_804934( | 2 | 0 | 1 | 1 | 1 | 0 | 1 | 1 | 5 | 0.4 | NA | NA | 1 | 1 | 2 | 0 | NA | NA | 3 | 0.3333 |
| 18_806495( | NA | NA | 3 | 0.3333 | NA | NA | 3 | 0.3333 | 6 | 0.3333 | NA | NA | NA | NA | NA | NA | 2 | 0.5 | 2 | 0.5 |
| 18_806496 | NA | NA | 3 | 0.6667 | NA | NA | 2 | 1 | 5 | 0.8 | NA | NA | NA | NA | 1 | 0 | 1 | 1 | 2 | 0.5 |
| 18_806497 | NA | NA | 2 | 1 | NA | NA | 2 | 1 | 4 | 1 | NA | NA | NA | NA | 1 | 1 | 1 | 1 | 2 | 1 |
| 18_806504! | NA | NA | 3 | 0 | NA | NA | 2 | 0.5 | 5 | 0.2 | NA | NA | NA | NA | 1 | 1 | NA | NA | 1 | 1 |
| 18_806508! | NA | NA | 1 | 1 | NA | NA | 1 | 0 | 2 | 0.5 | NA | NA | NA | NA | 1 | 1 | NA | NA | 1 | 1 |
| 18_806511 | NA | NA | 2 | 1 | NA | NA | 2 | 1 | 4 | 1 | NA | NA | NA | NA | 1 | 1 | NA | NA | 1 | 1 |
| 18_806790 | NA | NA | 2 | 1 | NA | NA | 2 | 1 | 4 | 1 | NA | NA | NA | NA | 1 | 1 | NA | NA | 1 | 1 |
| 18_806928 | 1 | 1 | NA | NA | NA | NA | NA | NA | 1 | 1 | NA | NA | 1 | 1 | NA | NA | NA | NA | 1 | 1 |
| 18_807410 | 1 | 0 | 2 | 1 | NA | NA | NA | NA | 3 | 0.6667 | NA | NA | 1 | 1 | 1 | 1 | NA | NA | 2 | 1 |
| 18_807417 | 1 | 0 | 1 | 1 | NA | NA | NA | NA | 2 | 0.5 | NA | NA | NA | NA | 1 | 1 | NA | NA | 1 | 1 |
| 18_807776 | 1 | 1 | NA | NA | NA | NA | 2 | 0.5 | 3 | 0.6667 | NA | NA | 4 | 1 | NA | NA | NA | NA | 4 | 1 |
| 18_807776 | 2 | 1 | NA | NA | NA | NA | 3 | 1 | 5 | 1 | NA | NA | 4 | 1 | NA | NA | NA | NA | 4 | 1 |
| 18_807776! | 2 | 1 | NA | NA | NA | NA | 2 | 1 | 4 | 1 | NA | NA | 4 | 0.75 | NA | NA | NA | NA | 4 | 0.75 |
| 18_807779! | 2 | 0.5 | NA | NA | 1 | 0 | 3 | 0.6667 | 6 | 0.5 | NA | NA | 2 | 0.5 | NA | NA | NA | NA | 2 | 0.5 |
| 18_807781 | 1 | 1 | NA | NA | NA | NA | 2 | 0.5 | 3 | 0.6667 | NA | NA | 1 | 1 | NA | NA | NA | NA | 1 | 1 |
| 18_807782 | 2 | 1 | NA | NA | NA | NA | 3 | 0.3333 | 5 | 0.6 | NA | NA | 2 | 0.5 | NA | NA | NA | NA | 2 | 0.5 |
| 18_807784 | 4 | 0.25 | 6 | 0.1667 | 3 | 0 | 6 | 0.3333 | 19 | 0.2105 | 6 | 0.1667 | 4 | 0 | 4 | 0 | 6 | 0 | 20 | 0.05 |
| 18_807785 | 8 | 0.75 | 6 | 0.6667 | 3 | 1 | 5 | 0.8 | 22 | 0.7727 | 6 | 0.6667 | 4 | 0 | 2 | 0.5 | 5 | 0.4 | 17 | 0.4118 |
| 18_807787 | 7 | 0.7143 | 5 | 0.4 | 3 | 0.6667 | 2 | 0.5 | 17 | 0.5882 | 4 | 1 | 4 | 0.5 | 3 | 0.6667 | 6 | 0.6667 | 17 | 0.7059 |
| 18_807787! | 6 | 0.8333 | 6 | 0.6667 | 3 | 1 | 3 | 1 | 18 | 0.8333 | 6 | 0.8333 | 4 | 1 | 3 | 1 | 4 | 0.5 | 17 | 0.8235 |
| 18_807788! | 7 | 0.2857 | 6 | 0.1667 | 3 | 0 | 6 | 0 | 22 | 0.1364 | 5 | 0.2 | 2 | 0 | 3 | 0 | 4 | 0.5 | 14 | 0.2143 |
| 18_807789 | 7 | 0.2857 | 4 | 0.5 | 3 | 0.3333 | 5 | 0.8 | 19 | 0.4737 | 5 | 0.4 | 2 | 0.5 | 3 | 0.6667 | 3 | 0 | 13 | 0.3846 |
| 18_807789 | 7 | 0.2857 | 6 | 0.3333 | 3 | 0 | 6 | 0.3333 | 22 | 0.2727 | 6 | 0.3333 | 2 | 0 | 2 | 0 | 5 | 0 | 15 | 0.1333 |
| 18_807790 | 7 | 0.7143 | 4 | 0.5 | 2 | 0 | 4 | 0.75 | 17 | 0.5882 | 3 | 0 | 1 | 1 | 3 | 0.6667 | 4 | 0.75 | 11 | 0.5455 |
| 18_807790 | 8 | 0.375 | 6 | 0.3333 | 3 | 0.3333 | 4 | 0 | 21 | 0.2857 | 6 | 0.1667 | 2 | 0 | 2 | 0.5 | 4 | 0.75 | 14 | 0.3571 |
| 18_807791 | 4 | 0.5 | 4 | 0.25 | NA | NA | 3 | 0.3333 | 11 | 0.3636 | 3 | 0 | 1 | 0 | 2 | 0.5 | 3 | 0 | 9 | 0.1111 |
| 18_807791 | 7 | 0.8571 | 6 | 0.5 | 3 | 1 | 3 | 1 | 19 | 0.7895 | 5 | 0.6 | 1 | 1 | 2 | 1 | 5 | 1 | 13 | 0.8462 |
| 18_807792( | 8 | 1 | 5 | 0.4 | 3 | 0.3333 | 5 | 0.6 | 21 | 0.6667 | 4 | 0.75 | 2 | 1 | 3 | 0.6667 | 5 | 0.6 | 14 | 0.7143 |
| 18_810122( | NA | NA | NA | NA | 1 | 1 | NA | NA | 1 | 1 | NA | NA | NA | NA | 1 | 1 | NA | NA | 1 | 1 |
| 18_810126 | NA | NA | NA | NA | 1 | 1 | NA | NA | 1 | 1 | NA | NA | NA | NA | 1 | 1 | 1 | 1 | 2 | 1 |
| 18_810127( | NA | NA | NA | NA | 1 | 1 | NA | NA | 1 | 1 | NA | NA | NA | NA | 1 | 1 | NA | NA | 1 | 1 |
| 18_810129! | NA | NA | NA | NA | 1 | 1 | NA | NA | 1 | 1 | NA | NA | NA | NA | 1 | 0 | 1 | 1 | 2 | 0.5 |
| 18_825729 | NA | NA | 1 | 1 | 1 | 1 | 1 | 1 | 3 | 1 | NA | NA | 1 | 1 | 1 | 0 | NA | NA | 2 | 0.5 |
| 18_825730( | 1 | 1 | 1 | 1 | 1 | 1 | 1 | 1 | 4 | 1 | NA | NA | 1 | 1 | 2 | 1 | NA | NA | 3 | 1 |
| 18_825731( | 1 | 1 | 1 | 1 | NA | NA | 2 | 1 | 4 | 1 | NA | NA | 1 | 1 | 1 | 1 | NA | NA | 2 | 1 |
| 18_825732! | 1 | 1 | 1 | 0 | 1 | 1 | 2 | 0.5 | 5 | 0.6 | NA | NA | 1 | 0 | 2 | 0.5 | NA | NA | 3 | 0.3333 |
| 18_825762 | 3 | 0.3333 | 2 | 0 | 1 | 0 | 5 | 0.2 | 11 | 0.1818 | 2 | 1 | 2 | 0 | 2 | 0 | 1 | 0 | 7 | 0.2857 |
| 18_825764 | 3 | 0.6667 | 2 | 0 | 1 | 0 | 4 | 0.25 | 10 | 0.3 | 2 | 0.5 | 4 | 0.25 | 3 | 0.3333 | 1 | 0 | 10 | 0.3 |
| 18_826023 | 174 | 0.0632 | 346 | 0.0751 | 334 | 0.0599 | 186 | 0.0968 | 1040 | 0.0721 | 289 | 0.0969 | 182 | 0.0769 | 138 | 0.029 | 156 | 0.0962 | 765 | 0.0797 |
| 18_826024 | 188 | 0.0957 | 370 | 0.0649 | 362 | 0.0497 | 211 | 0.0806 | 1131 | 0.0681 | 317 | 0.0883 | 194 | 0.0567 | 154 | 0.0779 | 163 | 0.1288 | 828 | 0.087 |
| 18_826025 | 184 | 0.1467 | 356 | 0.1994 | 361 | 0.1745 | 185 | 0.2216 | 1086 | 0.186 | 315 | 0.1746 | 181 | 0.1713 | 143 | 0.2098 | 174 | 0.1839 | 813 | 0.182 |
| 18_826026 | 208 | 0.3077 | 399 | 0.3108 | 389 | 0.2699 | 226 | 0.2965 | 1222 | 0.2946 | 348 | 0.2902 | 208 | 0.2788 | 177 | 0.2542 | 182 | 0.2802 | 915 | 0.2787 |
| 18_826029 | 206 | 0.0583 | 412 | 0.0413 | 404 | 0.0644 | 236 | 0.0593 | 1258 | 0.0548 | 374 | 0.0615 | 213 | 0.0657 | 194 | 0.0928 | 191 | 0.0524 | 972 | 0.0669 |
| 18_826030 | 231 | 0.0433 | 442 | 0.0724 | 441 | 0.0862 | 251 | 0.0797 | 1365 | 0.0733 | 395 | 0.0835 | 236 | 0.072 | 197 | 0.0609 | 207 | 0.0531 | 1035 | 0.0705 |
| 18_826030! | 206 | 0.0728 | 394 | 0.0939 | 382 | 0.0942 | 218 | 0.1055 | 1200 | 0.0925 | 355 | 0.0958 | 204 | 0.098 | 173 | 0.1156 | 179 | 0.0894 | 911 | 0.0988 |
| 18_826032 | 234 | 0.0983 | 442 | 0.1357 | 446 | 0.1525 | 257 | 0.1556 | 1379 | 0.1385 | 413 | 0.1453 | 244 | 0.1434 | 203 | 0.1626 | 210 | 0.1476 | 1070 | 0.1486 |
| 18_826120 | 3 | 0.3333 | 2 | 0 | 5 | 0.8 | 4 | 1 | 14 | 0.6429 | NA | NA | 4 | 0.25 | 2 | 0.5 | 1 | 0 | 7 | 0.2857 |
| 18_826121! | 3 | 0 | 2 | 0 | 6 | 0 | 4 | 0.25 | 15 | 0.0667 | NA | NA | 4 | 0 | 2 | 0.5 | 2 | 0 | 8 | 0.125 |
| 18_826122 | 3 | 0.3333 | 1 | 0 | 4 | 0.25 | 4 | 0.25 | 12 | 0.25 | NA | NA | 4 | 0 | 2 | 0.5 | 2 | 0 | 8 | 0.125 |
| 18_826142! | 2 | 0 | 1 | 1 | 4 | 0.25 | 1 | 1 | 8 | 0.375 | 1 | 1 | 2 | 0 | 1 | 0 | 2 | 0.5 | 6 | 0.3333 |
| 18_826145 | 2 | 0 | 2 | 0 | 2 | 0.5 | 1 | 0 | 7 | 0.1429 | 2 | 0 | 5 | 0.6 | 2 | 0.5 | 1 | 0 | 10 | 0.4 |
| 18_826145 | 1 | 1 | 2 | 0.5 | 3 | 0.6667 | 1 | 0 | 7 | 0.5714 | 2 | 0.5 | 4 | 0.25 | 2 | 0.5 | 1 | 0 | 9 | 0.3333 |
| 18_826148( | 3 | 0 | 1 | 0 | 3 | 0.6667 | 1 | 0 | 8 | 0.25 | 1 | 0 | 5 | 0 | 2 | 0.5 | 2 | 0 | 10 | 0.1 |
| 18_826149 | 3 | 0.3333 | 2 | 0 | 2 | 0 | 1 | 0 | 8 | 0.125 | 2 | 0 | 5 | 0 | 2 | 0.5 | 2 | 0 | 11 | 0.0909 |

| | | | | | | | | | | | | | | | | | | | | |
|---|---|---|---|---|---|---|---|---|---|---|---|---|---|---|---|---|---|---|---|---|
| 18_8261494 | 3 | 0 | 2 | 0 | 3 | 0.3333 | 1 | 0 | 9 | 0.1111 | 2 | 0 | 4 | 0 | 2 | 0.5 | 1 | 1 | 9 | 0.2222 |
| 18_8261497 | 3 | 0 | 1 | 0 | 3 | 0.3333 | 1 | 0 | 8 | 0.125 | 2 | 0.5 | 3 | 0 | 2 | 0.5 | 2 | 0.5 | 9 | 0.3333 |
| 18_8261502 | 3 | 0.6667 | 2 | 0 | 3 | 0.3333 | 1 | 0 | 9 | 0.3333 | 1 | 1 | 2 | 0 | 2 | 0.5 | 2 | 0 | 7 | 0.2857 |
| 18_8261508 | 3 | 0.3333 | NA | NA | 1 | 1 | 1 | 0 | 5 | 0.4 | 2 | 1 | 3 | 0.3333 | 1 | 0 | 1 | 0 | 7 | 0.4286 |
| 18_8261509 | 2 | 0.5 | 1 | 0 | 3 | 0 | 1 | 0 | 7 | 0.1429 | 2 | 1 | 3 | 0 | 1 | 1 | 1 | 0 | 7 | 0.4286 |
| 18_8261528 | 3 | 0.3333 | 2 | 0.5 | 2 | 0 | NA | NA | 7 | 0.2857 | 2 | 1 | 3 | 1 | 1 | 1 | 2 | 0.5 | 8 | 0.875 |
| 18_8261545 | 2 | 0 | 2 | 0.5 | 2 | 0 | 1 | 1 | 7 | 0.2857 | 2 | 1 | 4 | 0.5 | 1 | 1 | 2 | 0 | 9 | 0.5556 |
| 18_8261549 | 2 | 1 | 2 | 0.5 | 2 | 1 | NA | NA | 6 | 0.8333 | NA | NA | 4 | 1 | 1 | 1 | NA | NA | 5 | 1 |
| 18_8261561 | 2 | 1 | 2 | 1 | 1 | 1 | NA | NA | 5 | 1 | 1 | 1 | 3 | 1 | 1 | 1 | 1 | 1 | 6 | 1 |
| 18_8261568 | 2 | 1 | 2 | 0 | 2 | 0.5 | NA | NA | 6 | 0.5 | 1 | 1 | 3 | 0.3333 | 1 | 1 | 2 | 0.5 | 7 | 0.5714 |
| 18_8263689 | NA | NA | 1 | 0 | 1 | 1 | NA | NA | 2 | 0.5 | 2 | 0 | 3 | 0.3333 | 1 | 0 | 2 | 0 | 8 | 0.125 |
| 18_8263945 | NA | NA | NA | NA | 1 | 1 | NA | NA | 1 | 1 | 2 | 0.5 | 1 | 0 | NA | NA | 1 | 0 | 4 | 0.25 |
| 18_8271092 | 1 | 0 | 3 | 0 | 1 | 1 | 2 | 0 | 7 | 0.1429 | 1 | 0 | 1 | 0 | 2 | 0.5 | NA | NA | 4 | 0.25 |
| 18_8271119 | NA | NA | 4 | 0.25 | 1 | 1 | NA | NA | 5 | 0.4 | NA | NA | 1 | 1 | 2 | 1 | 1 | 0 | 4 | 0.75 |
| 18_8271121 | 2 | 0.5 | 5 | 0.8 | 1 | 1 | 2 | 1 | 10 | 0.8 | 1 | 1 | 2 | 1 | 1 | 0 | NA | NA | 4 | 0.75 |
| 18_8271134 | 1 | 1 | 5 | 0.8 | 1 | 1 | 2 | 1 | 9 | 0.8889 | 1 | 1 | 2 | 1 | 3 | 0.6667 | 1 | 1 | 7 | 0.8571 |
| 18_8271143 | 2 | 0 | 4 | 0.25 | 1 | 0 | 1 | 0 | 8 | 0.125 | NA | NA | 1 | 0 | 2 | 0.5 | 1 | 0 | 4 | 0.25 |
| 18_8271179 | 2 | 0.5 | 1 | 0 | 1 | 0 | 2 | 0 | 6 | 0.1667 | NA | NA | 1 | 0 | 3 | 0.3333 | 1 | 0 | 5 | 0.2 |
| 18_8271206 | 2 | 0.5 | 4 | 0.25 | 1 | 0 | 2 | 0 | 9 | 0.2222 | 1 | 0 | 2 | 0.5 | 2 | 0.5 | 1 | 0 | 6 | 0.3333 |
| 18_8271266 | 2 | 0.5 | 2 | 0.5 | 1 | 0 | 1 | 1 | 6 | 0.5 | NA | NA | 1 | 1 | NA | NA | NA | NA | 1 | 1 |
| 18_8271431 | 2 | 0 | 2 | 0.5 | 1 | 0 | 2 | 0 | 7 | 0.1429 | 2 | 0 | 1 | 1 | NA | NA | NA | NA | 3 | 0.3333 |
| 18_8271462 | 2 | 0.5 | 2 | 0 | 1 | 1 | 2 | 1 | 7 | 0.5714 | 2 | 0 | 1 | 1 | NA | NA | NA | NA | 3 | 0.3333 |
| 18_8271495 | 2 | 1 | 3 | 1 | NA | NA | 2 | 1 | 7 | 1 | 2 | 1 | 1 | 1 | NA | NA | NA | NA | 3 | 1 |
| 18_8271503 | 2 | 1 | 4 | 0.5 | 1 | 1 | 2 | 1 | 9 | 0.7778 | 2 | 1 | 1 | 1 | NA | NA | NA | NA | 3 | 1 |
| 18_8271525 | 1 | 0 | 3 | 0.3333 | 1 | 1 | 2 | 0.5 | 7 | 0.4286 | 2 | 0.5 | 1 | 0 | NA | NA | NA | NA | 3 | 0.3333 |
| 18_8271557 | 1 | 1 | 3 | 1 | 1 | 1 | 2 | 1 | 7 | 1 | 2 | 1 | 1 | 1 | NA | NA | NA | NA | 3 | 1 |
| 18_8271571 | 2 | 0.5 | 4 | 0.5 | 1 | 1 | 2 | 0.5 | 9 | 0.5556 | 1 | 1 | 1 | 1 | NA | NA | NA | NA | 2 | 1 |
| 18_8271581 | 2 | 1 | 4 | 0.75 | 1 | 1 | 1 | 1 | 8 | 0.875 | 2 | 1 | NA | NA | 1 | 1 | NA | NA | 3 | 1 |
| 18_8271587 | 2 | 0.5 | 3 | 0.3333 | 1 | 0 | 1 | 1 | 7 | 0.4286 | 2 | 0.5 | 1 | 1 | 1 | 0 | NA | NA | 4 | 0.5 |
| 18_8271660 | 2 | 0 | 4 | 0.75 | NA | NA | 4 | 1 | 10 | 0.7 | 2 | 1 | 1 | 0 | 1 | 0 | 1 | 1 | 5 | 0.6 |
| 18_8271710 | 4 | 0.25 | 3 | 0.6667 | 1 | 0 | 5 | 0.6 | 13 | 0.4615 | 2 | 1 | 1 | 0 | 2 | 0 | 1 | 1 | 6 | 0.5 |
| 18_8271712 | 3 | 1 | 4 | 0.75 | 1 | 1 | 4 | 1 | 12 | 0.9167 | 2 | 0.5 | 1 | 1 | 2 | 0.5 | 1 | 1 | 6 | 0.6667 |
| 18_8271720 | 4 | 0.75 | 4 | 1 | 1 | 0 | 4 | 0.25 | 13 | 0.6154 | 2 | 0.5 | 1 | 1 | 2 | 0.5 | 1 | 0 | 6 | 0.5 |
| 18_8271727 | 4 | 0 | 3 | 0 | 1 | 0 | 5 | 0.2 | 13 | 0.0769 | 2 | 0 | 2 | 0 | 1 | 1 | 1 | 0 | 6 | 0.1667 |
| 18_8271745 | 4 | 0 | 4 | 0.5 | 2 | 0 | 4 | 0.5 | 14 | 0.2857 | 2 | 0 | 2 | 0.5 | 2 | 0 | 1 | 0 | 7 | 0.1429 |
| 18_8271751 | 1 | 1 | 3 | 0.6667 | 2 | 0.5 | 5 | 0.6 | 11 | 0.6364 | 2 | 0 | 3 | 0.3333 | 2 | 0 | 1 | 1 | 8 | 0.25 |
| 18_8271775 | 3 | 0.3333 | 5 | 0 | 2 | 0.5 | 4 | 0.25 | 14 | 0.2143 | 2 | 0 | 3 | 0.3333 | 3 | 0.3333 | 1 | 1 | 9 | 0.3333 |
| 18_8271777 | 4 | 0.25 | 5 | 0 | 3 | 0 | 4 | 0.5 | 16 | 0.1875 | 1 | 1 | 3 | 0 | 3 | 0 | 2 | 0 | 9 | 0.1111 |
| 18_8271794 | 3 | 0.3333 | 5 | 0.4 | 4 | 0.25 | 2 | 0 | 14 | 0.2857 | 2 | 1 | 3 | 0 | 2 | 0.5 | 2 | 0 | 9 | 0.3333 |
| 18_8271806 | 4 | 0.75 | 4 | 0.75 | 4 | 1 | 3 | 0 | 15 | 0.6667 | 2 | 1 | 2 | 1 | 3 | 0.6667 | 2 | 0.5 | 9 | 0.7778 |
| 18_8271809 | 4 | 0.5 | 5 | 1 | 3 | 0.6667 | 4 | 0.75 | 16 | 0.75 | 2 | 1 | 2 | 0.5 | 2 | 1 | 2 | 1 | 8 | 0.875 |
| 18_8271811 | 4 | 0 | 4 | 0.5 | 2 | 0.5 | 4 | 0 | 14 | 0.2143 | 2 | 1 | 3 | 0.6667 | 2 | 0 | 2 | 1 | 9 | 0.6667 |
| 18_8271812 | 2 | 0.5 | 5 | 0.4 | 5 | 0.2 | 5 | 0.6 | 17 | 0.4118 | 2 | 0 | 2 | 0.5 | 3 | 0.3333 | 1 | 0 | 8 | 0.25 |
| 18_8271816 | 2 | 0 | 2 | 1 | 3 | 0.3333 | 4 | 0.25 | 11 | 0.3636 | NA | NA | 3 | 0.3333 | 3 | 0.3333 | 1 | 0 | 7 | 0.2857 |
| 18_8271816 | 3 | 1 | 2 | 1 | 4 | 1 | 5 | 0.8 | 14 | 0.9286 | 1 | 1 | 3 | 0.6667 | 2 | 1 | 2 | 1 | 8 | 0.875 |
| 18_8271820 | 4 | 0.75 | 4 | 0 | 4 | 0.25 | 5 | 0.2 | 17 | 0.2941 | 1 | 1 | 3 | 0.3333 | 4 | 0.5 | 1 | 1 | 9 | 0.5556 |
| 18_8271821 | 3 | 0.6667 | 5 | 0.2 | 4 | 0.5 | 2 | 0.5 | 14 | 0.4286 | 1 | 0 | 3 | 0.3333 | 2 | 0 | 2 | 1 | 8 | 0.375 |
| 18_8271830 | 3 | 0.6667 | 5 | 0.2 | 5 | 0.8 | 3 | 0.6667 | 16 | 0.5625 | 1 | 1 | 2 | 1 | 4 | 0.5 | 1 | 1 | 8 | 0.75 |
| 18_8271833 | 4 | 0.5 | 5 | 0.6 | 6 | 0.3333 | 6 | 0.5 | 21 | 0.4762 | 2 | 0.5 | 3 | 0.3333 | 4 | 0.5 | 1 | 1 | 10 | 0.5 |
| 18_8271851 | 4 | 0.25 | 4 | 0.5 | 6 | 0.5 | 7 | 0.2857 | 21 | 0.381 | 2 | 1 | 3 | 0.3333 | 3 | 0 | 2 | 0.5 | 10 | 0.4 |
| 18_8271851 | 4 | 1 | 4 | 1 | 5 | 0.6 | 5 | 0.6 | 18 | 0.7778 | NA | NA | 5 | 0.4 | 1 | 0 | 2 | 0.5 | 8 | 0.375 |
| 18_8271853 | 2 | 0 | 5 | 0 | 4 | 0 | 5 | 0.2 | 16 | 0.0625 | 2 | 0 | 3 | 0 | 3 | 0 | 2 | 0.5 | 10 | 0.1 |
| 18_8271935 | 5 | 0 | 10 | 0.1 | 8 | 0.125 | 7 | 0 | 30 | 0.0667 | 3 | 0 | 6 | 0 | 6 | 0 | 2 | 0.5 | 17 | 0.0588 |
| 18_8300545 | NA | NA | NA | NA | NA | NA | 1 | 1 | 1 | 1 | NA | NA | NA | NA | 1 | 1 | NA | NA | 1 | 1 |
| 18_8300593 | NA | NA | NA | NA | NA | NA | 1 | 1 | 1 | 1 | NA | NA | NA | NA | NA | NA | 1 | 1 | 1 | 1 |
| 18_8300610 | NA | NA | NA | NA | NA | NA | 1 | 1 | 1 | 1 | 1 | 0 | 1 | 1 | 1 | 0 | 1 | 0 | 4 | 0.25 |
| 18_8300620 | NA | NA | NA | NA | NA | NA | 1 | 1 | 1 | 1 | 1 | 0 | 1 | 0 | 1 | 1 | NA | NA | 3 | 0.3333 |
| 18_8301946 | NA | NA | NA | NA | NA | NA | 2 | 0.5 | 2 | 0.5 | 1 | 0 | 2 | 0.5 | 1 | 0 | 1 | 0 | 5 | 0.2 |
| 18_8301949 | NA | NA | NA | NA | NA | NA | 2 | 0.5 | 2 | 0.5 | NA | NA | 2 | 0.5 | 2 | 0 | 1 | 0 | 5 | 0.2 |
| 18_8301958 | NA | NA | NA | NA | NA | NA | 2 | 1 | 2 | 1 | 1 | 0 | 2 | 0.5 | 1 | 0 | 1 | 0 | 5 | 0.2 |
| 18_8301962 | NA | NA | NA | NA | NA | NA | 1 | 1 | 1 | 1 | 1 | 1 | 2 | 1 | 2 | 1 | 1 | 0 | 6 | 0.8333 |
| 18_8301984 | NA | NA | 1 | 1 | NA | NA | 1 | 0 | 2 | 0.5 | 1 | 0 | 2 | 0.5 | 2 | 0.5 | 1 | 0 | 6 | 0.3333 |
| 18_8301991 | NA | NA | 1 | 0 | 1 | 0 | 2 | 0.5 | 4 | 0.25 | 1 | 0 | 2 | 0.5 | 2 | 0 | 1 | 0 | 6 | 0.1667 |
| 18_8302018 | NA | NA | 2 | 0.5 | 1 | 0 | 2 | 0 | 5 | 0.2 | NA | NA | 2 | 0.5 | 2 | 0 | 1 | 0 | 5 | 0.2 |
| 18_8302059 | 1 | 0 | 2 | 0.5 | 1 | 1 | 2 | 0.5 | 6 | 0.5 | 1 | 0 | 2 | 1 | 2 | 0 | 1 | 0 | 6 | 0.3333 |

| | | | | | | | | | | | | | | | | | | | | |
|---|---|---|---|---|---|---|---|---|---|---|---|---|---|---|---|---|---|---|---|---|
| 18_830209 | 1 | 1 | 2 | 0 | 1 | 0 | 2 | 0 | 6 | 0.1667 | 1 | 0 | 2 | 0.5 | 3 | 0 | 1 | 0 | 7 | 0.1429 |
| 18_830219 | 1 | 0 | 1 | 0 | 1 | 0 | 2 | 1 | 5 | 0.4 | 1 | 0 | 1 | 1 | 3 | 0 | NA | NA | 5 | 0.2 |
| 18_830223 | NA | NA | NA | NA | NA | NA | 1 | 1 | 1 | 1 | NA | NA | 1 | 1 | 2 | 0.5 | NA | NA | 3 | 0.6667 |
| 18_830227 | NA | NA | 1 | 0 | 1 | 1 | 2 | 0 | 4 | 0.25 | NA | NA | 1 | 0 | 2 | 1 | 1 | 0 | 4 | 0.5 |
| 18_830442 | NA | NA | NA | NA | 1 | 1 | NA | NA | 1 | 1 | NA | NA | 1 | 1 | 1 | 1 | NA | NA | 2 | 1 |
| 18_830445 | NA | NA | NA | NA | 1 | 1 | NA | NA | 1 | 1 | NA | NA | 1 | 1 | 1 | 1 | NA | NA | 2 | 1 |
| 18_830445 | NA | NA | NA | NA | 1 | 1 | NA | NA | 1 | 1 | NA | NA | 1 | 1 | 1 | 1 | NA | NA | 2 | 1 |
| 18_830446 | NA | NA | NA | NA | 1 | 1 | NA | NA | 1 | 1 | NA | NA | 1 | 1 | 1 | 0 | NA | NA | 2 | 0.5 |
| 18_830447 | NA | NA | NA | NA | 1 | 1 | NA | NA | 1 | 1 | NA | NA | 1 | 1 | 1 | 1 | NA | NA | 2 | 1 |
| 18_830448 | NA | NA | NA | NA | 1 | 1 | NA | NA | 1 | 1 | NA | NA | 1 | 1 | 1 | 1 | NA | NA | 2 | 1 |
| 18_830450 | NA | NA | NA | NA | 1 | 1 | NA | NA | 1 | 1 | NA | NA | 1 | 1 | 1 | 1 | NA | NA | 2 | 1 |
| 18_830453 | NA | NA | NA | NA | 1 | 1 | NA | NA | 1 | 1 | NA | NA | 1 | 1 | 1 | 0 | NA | NA | 2 | 0.5 |
| 18_830461 | NA | NA | NA | NA | 1 | 1 | NA | NA | 1 | 1 | NA | NA | 1 | 1 | 1 | 1 | NA | NA | 2 | 1 |
| 18_830470 | NA | NA | NA | NA | 1 | 1 | NA | NA | 1 | 1 | NA | NA | NA | NA | 1 | 1 | NA | NA | 1 | 1 |
| 18_840113 | 1 | 1 | NA | NA | NA | NA | NA | NA | 1 | 1 | NA | NA | NA | NA | NA | NA | 1 | 1 | 1 | 1 |
| 18_840113 | 1 | 1 | NA | NA | NA | NA | NA | NA | 1 | 1 | NA | NA | NA | NA | NA | NA | 1 | 1 | 1 | 1 |
| 18_840114 | 1 | 1 | NA | NA | NA | NA | NA | NA | 1 | 1 | NA | NA | NA | NA | NA | NA | 1 | 1 | 1 | 1 |
| 18_840115 | 1 | 1 | NA | NA | NA | NA | NA | NA | 1 | 1 | NA | NA | NA | NA | NA | NA | 1 | 1 | 1 | 1 |
| 18_840299 | 3 | 0 | 7 | 0.2857 | 2 | 0.5 | 11 | 0.3636 | 23 | 0.3043 | 2 | 0.5 | 9 | 0.5556 | 5 | 0.8 | 8 | 0.625 | 24 | 0.625 |
| 18_840305 | 2 | 0.5 | 7 | 0.4286 | 1 | 1 | 8 | 0.625 | 18 | 0.5556 | 2 | 1 | 8 | 0.75 | 6 | 0.5 | 5 | 0.8 | 21 | 0.7143 |
| 18_840305 | 2 | 0 | 8 | 0.5 | 1 | 1 | 8 | 0.625 | 19 | 0.5263 | 2 | 0.5 | 8 | 0.5 | 6 | 0.5 | 7 | 0.5714 | 23 | 0.5217 |
| 18_840307 | 2 | 0.5 | 5 | 1 | 1 | 1 | 9 | 0.7778 | 17 | 0.8235 | 2 | 1 | 7 | 1 | 6 | 0.8333 | 6 | 0.6667 | 21 | 0.8571 |
| 18_840308 | 2 | 0 | 7 | 0 | 1 | 0 | 9 | 0.3333 | 19 | 0.1579 | 2 | 0 | 9 | 0.2222 | 6 | 0 | 7 | 0.1429 | 24 | 0.125 |
| 18_840308 | 1 | 0 | 7 | 0.1429 | 1 | 0 | 9 | 0.2222 | 18 | 0.1667 | 2 | 0 | 8 | 0 | 6 | 0.1667 | 7 | 0.1429 | 23 | 0.087 |
| 18_840310 | 2 | 1 | 7 | 1 | 1 | 1 | 7 | 1 | 17 | 1 | 1 | 1 | 6 | 1 | 5 | 1 | 6 | 1 | 18 | 1 |
| 18_840310 | 2 | 0.5 | 7 | 0.2857 | 1 | 1 | 8 | 0.375 | 18 | 0.3889 | 1 | 0 | 6 | 0.3333 | 4 | 0.5 | 7 | 0.5714 | 18 | 0.4444 |
| 18_840311 | 2 | 0 | 6 | 0.1667 | 1 | 0 | 8 | 0 | 17 | 0.0588 | 1 | 0 | 4 | 0 | 3 | 0.3333 | 5 | 0 | 13 | 0.0769 |
| 18_840311 | 2 | 0 | 6 | 0.1667 | 1 | 0 | 8 | 0 | 17 | 0.0588 | 1 | 1 | 7 | 0.7143 | 4 | 0 | 7 | 0 | 19 | 0.3158 |
| 18_840312 | NA | NA | 5 | 0.8 | NA | NA | 6 | 0.3333 | 11 | 0.5455 | NA | NA | 6 | 0.3333 | 2 | 0.5 | 5 | 0.6 | 13 | 0.4615 |
| 18_840312 | 2 | 1 | 7 | 1 | 1 | 1 | 8 | 1 | 18 | 1 | 1 | 1 | 7 | 1 | 3 | 0.6667 | 6 | 1 | 17 | 0.9412 |
| 18_840313 | 1 | 0 | 7 | 1 | NA | NA | 7 | 1 | 15 | 0.9333 | NA | NA | 6 | 1 | 3 | 1 | 5 | 1 | 14 | 1 |
| 18_840313 | 2 | 1 | 7 | 1 | NA | NA | 8 | 0.75 | 17 | 0.8824 | 1 | 1 | 7 | 0.8571 | 4 | 0.75 | 5 | 0.8 | 17 | 0.8235 |
| 18_840314 | 2 | 1 | 7 | 1 | 1 | 1 | 7 | 0.8571 | 17 | 0.9412 | 1 | 1 | 4 | 1 | 2 | 1 | 6 | 1 | 13 | 1 |
| 18_840315 | 1 | 0 | 6 | 0.3333 | 1 | 0 | 7 | 0.4286 | 15 | 0.3333 | 1 | 0 | 6 | 0.3333 | 3 | 0.3333 | 5 | 0.4 | 15 | 0.3333 |
| 18_840315 | 2 | 1 | 3 | 1 | NA | NA | 2 | 0.5 | 7 | 0.8571 | NA | NA | 3 | 1 | 2 | 1 | 4 | 0.25 | 9 | 0.6667 |
| 18_840316 | 2 | 0.5 | 6 | 0.5 | NA | NA | 6 | 0.5 | 14 | 0.5 | 1 | 0 | 6 | 0.6667 | 3 | 0.6667 | 7 | 0.4286 | 17 | 0.5294 |
| 18_840317 | 2 | 0.5 | 6 | 0 | 1 | 1 | 5 | 0 | 14 | 0.1429 | 1 | 0 | 5 | 0 | 3 | 0.3333 | 7 | 0.1429 | 16 | 0.125 |
| 18_840318 | 2 | 0 | 5 | 0.8 | NA | NA | 5 | 1 | 12 | 0.75 | 1 | 1 | 6 | 0.6667 | 3 | 0.6667 | 7 | 0.5714 | 17 | 0.6471 |
| 18_840319 | 1 | 0 | 4 | 0 | NA | NA | 4 | 0.25 | 9 | 0.1111 | 1 | 0 | 4 | 0.5 | 3 | 0.3333 | 6 | 0.1667 | 14 | 0.2857 |
| 18_840319 | 1 | 0 | 4 | 0.25 | NA | NA | 4 | 0.25 | 9 | 0.2222 | 1 | 1 | 5 | 0 | 3 | 0 | 6 | 0 | 15 | 0.0667 |
| 18_840321 | 2 | 0 | 2 | 0.5 | 1 | 0 | 4 | 0.25 | 9 | 0.2222 | 1 | 0 | 5 | 0.2 | 2 | 0 | 4 | 0 | 12 | 0.0833 |
| 18_840321 | 2 | 0 | 4 | 0.5 | NA | NA | 5 | 0.2 | 11 | 0.2727 | 1 | 0 | 6 | 0.3333 | 2 | 0 | 3 | 0.3333 | 12 | 0.25 |
| 18_840322 | NA | NA | 4 | 0 | 1 | 0 | 5 | 0.4 | 10 | 0.2 | NA | NA | 4 | 0.25 | 2 | 0 | 5 | 0 | 11 | 0.0909 |
| 18_840322 | 2 | 0 | 4 | 0.25 | NA | NA | 5 | 0.2 | 11 | 0.1818 | 1 | 0 | 6 | 0 | 3 | 0 | 6 | 0.1667 | 16 | 0.0625 |
| 18_840327 | NA | NA | 3 | 0.3333 | 1 | 0 | 3 | 0.6667 | 7 | 0.4286 | 1 | 1 | 2 | 0 | 2 | 0 | 3 | 0 | 8 | 0.125 |
| 18_840328 | 1 | 0 | 5 | 0 | 1 | 0 | 4 | 0.25 | 11 | 0.0909 | 1 | 1 | 6 | 0.5 | 2 | 0 | 4 | 0 | 13 | 0.3077 |
| 18_840329 | 1 | 0 | 4 | 0.25 | 1 | 0 | 3 | 0.6667 | 9 | 0.3333 | 1 | 0 | 4 | 0.5 | 2 | 1 | 3 | 0 | 10 | 0.4 |
| 18_840332 | 1 | 0 | 3 | 0 | NA | NA | 4 | 0.25 | 8 | 0.125 | 1 | 0 | 3 | 0.3333 | 2 | 0 | 4 | 0.25 | 10 | 0.2 |
| 18_840332 | 2 | 1 | 4 | 0.25 | NA | NA | 4 | 0.25 | 10 | 0.4 | 1 | 0 | 5 | 0.8 | 1 | 1 | 4 | 0.25 | 11 | 0.5455 |
| 18_840334 | 2 | 0 | 3 | 0.3333 | NA | NA | 4 | 0.25 | 9 | 0.2222 | 1 | 1 | 4 | 0.5 | 1 | 0 | 3 | 0 | 9 | 0.3333 |
| 18_840336 | 2 | 0.5 | 2 | 0 | NA | NA | 3 | 0.3333 | 7 | 0.2857 | 1 | 0 | 2 | 0.5 | 1 | 0 | 3 | 0.3333 | 7 | 0.2857 |
| 18_840337 | 2 | 0 | 3 | 0 | NA | NA | 3 | 0.3333 | 8 | 0.125 | 1 | 0 | 4 | 0 | 2 | 0.5 | 1 | 1 | 8 | 0.25 |
| 18_840338 | 1 | 0 | 2 | 0 | NA | NA | 3 | 0.3333 | 6 | 0.1667 | 1 | 1 | 4 | 0 | 1 | 0 | NA | NA | 6 | 0.1667 |
| 18_840340 | 1 | 1 | 2 | 1 | NA | NA | 2 | 1 | 5 | 1 | 1 | 1 | 2 | 1 | 1 | 1 | 1 | 1 | 5 | 1 |
| 18_840340 | 1 | 1 | 3 | 0.6667 | NA | NA | 3 | 0.3333 | 7 | 0.5714 | 1 | 0 | 1 | 0 | 1 | 0 | 1 | 1 | 4 | 0.25 |
| 18_840340 | 1 | 0 | 3 | 0.3333 | NA | NA | 2 | 0 | 6 | 0.1667 | 1 | 1 | 3 | 0 | NA | NA | 2 | 0 | 6 | 0.1667 |
| 18_840342 | 1 | 0 | 3 | 0.3333 | NA | NA | 2 | 0 | 6 | 0.1667 | 1 | 0 | 3 | 0 | 1 | 0 | 2 | 0.5 | 7 | 0.1429 |
| 18_840342 | 1 | 1 | 3 | 0.6667 | NA | NA | 1 | 1 | 5 | 0.8 | 1 | 0 | 1 | 1 | 1 | 0 | 2 | 0.5 | 5 | 0.4 |
| 18_841040 | NA | NA | 1 | 1 | NA | NA | 1 | 0 | 2 | 0.5 | 1 | 0 | 2 | 0.5 | NA | NA | 1 | 0 | 4 | 0.25 |
| 18_841041 | NA | NA | 2 | 0.5 | NA | NA | NA | NA | 2 | 0.5 | NA | NA | 1 | 1 | NA | NA | 1 | 1 | 2 | 1 |
| 18_841133 | NA | NA | 1 | 1 | NA | NA | NA | NA | 1 | 1 | 1 | 0 | 1 | 1 | 1 | 1 | 1 | 1 | 4 | 0.75 |
| 18_841135 | 1 | 1 | 1 | 1 | NA | NA | NA | NA | 2 | 1 | 1 | 1 | 1 | 1 | 1 | 1 | 1 | 1 | 4 | 1 |
| 18_841140 | 1 | 1 | 1 | 0 | NA | NA | NA | NA | 2 | 0.5 | 1 | 1 | NA | NA | 2 | 0.5 | 1 | 1 | 4 | 0.75 |
| 18_841140 | 1 | 0 | 2 | 0.5 | NA | NA | NA | NA | 3 | 0.3333 | 1 | 0 | NA | NA | 2 | 1 | 1 | 1 | 4 | 0.75 |
| 18_841141 | 1 | 1 | 1 | 1 | NA | NA | NA | NA | 2 | 1 | 1 | 0 | NA | NA | 2 | 0.5 | NA | NA | 3 | 0.3333 |

| | | | | | | | | | | | | | | | | | | | | |
|---|---|---|---|---|---|---|---|---|---|---|---|---|---|---|---|---|---|---|---|---|
| 18_8411424 | 1 | 1 | 1 | 0 | 1 | 0 | NA | NA | 3 | 0.3333 | 1 | 1 | NA | NA | 1 | 0 | 1 | 0 | 3 | 0.3333 |
| 18_8411429 | 1 | 1 | 2 | 0 | 1 | 0 | NA | NA | 4 | 0.25 | 1 | 0 | 1 | 0 | 2 | 0 | 1 | 1 | 5 | 0.2 |
| 18_8411500 | 1 | 0 | 3 | 0.3333 | NA | NA | 2 | 0 | 6 | 0.1667 | 1 | 0 | 1 | 0 | 2 | 0 | 2 | 0.5 | 6 | 0.1667 |
| 18_8411542 | 1 | 0 | 3 | 0.3333 | 2 | 0 | 2 | 0.5 | 8 | 0.25 | 2 | 0.5 | 2 | 0 | 2 | 0 | 3 | 0 | 9 | 0.1111 |
| 18_8411543 | 1 | 0 | 2 | 0.5 | 1 | 0 | 1 | 0 | 5 | 0.2 | 2 | 0 | 1 | 1 | 2 | 0 | 2 | 0 | 7 | 0.1429 |
| 18_8411546 | NA | NA | 3 | 0.3333 | 1 | 0 | 2 | 0 | 6 | 0.1667 | 1 | 0 | 2 | 0 | 2 | 0.5 | 1 | 1 | 6 | 0.3333 |
| 18_8422604 | 1 | 1 | 1 | 1 | 2 | 0 | 1 | 0 | 5 | 0.4 | 2 | 0 | 1 | 1 | 2 | 0 | NA | NA | 5 | 0.2 |
| 18_8422642 | NA | NA | 1 | 1 | 1 | 1 | 1 | 0 | 3 | 0.6667 | 1 | 0 | 1 | 1 | 2 | 0.5 | NA | NA | 4 | 0.5 |
| 18_8422665 | NA | NA | NA | NA | 1 | 1 | 1 | 1 | 2 | 1 | 1 | 1 | 2 | 1 | 2 | 0.5 | NA | NA | 5 | 0.8 |
| 18_8422679 | NA | NA | 1 | 1 | 1 | 1 | 1 | 1 | 3 | 1 | 1 | 0 | 1 | 1 | 2 | 1 | NA | NA | 4 | 0.75 |
| 18_8422696 | NA | NA | 1 | 0 | 2 | 1 | NA | NA | 3 | 0.6667 | 1 | 1 | 2 | 1 | 2 | 0.5 | NA | NA | 5 | 0.8 |
| 18_8422703 | NA | NA | 1 | 0 | 2 | 1 | 1 | 1 | 4 | 0.75 | 1 | 1 | 1 | 1 | 2 | 0.5 | NA | NA | 4 | 0.75 |
| 18_8422706 | NA | NA | 1 | 1 | 2 | 1 | 1 | 1 | 4 | 1 | 1 | 0 | 2 | 0.5 | 2 | 0 | NA | NA | 5 | 0.2 |
| 18_8422721 | NA | NA | NA | NA | 1 | 1 | 1 | 1 | 2 | 1 | 1 | 1 | 2 | 1 | 1 | 1 | NA | NA | 4 | 1 |
| 18_8422798 | NA | NA | NA | NA | NA | NA | 1 | 1 | 1 | 1 | 1 | 1 | 1 | 1 | 1 | 1 | NA | NA | 3 | 1 |
| 18_8468582 | 7 | 0.2857 | 9 | 0.2222 | 5 | 0.2 | 3 | 0.3333 | 24 | 0.25 | 6 | 0.3333 | 6 | 0 | 4 | 0.25 | 4 | 0 | 20 | 0.15 |
| 18_8468599 | 8 | 0.125 | 10 | 0.2 | 4 | 0 | 3 | 0.6667 | 25 | 0.2 | 6 | 0 | 6 | 0.5 | 5 | 0.2 | 5 | 0.4 | 22 | 0.2727 |
| 18_8474842 | NA | NA | NA | NA | NA | NA | 1 | 1 | 1 | 1 | NA | NA | 1 | 1 | NA | NA | NA | NA | 1 | 1 |
| 18_8474872 | 1 | 1 | NA | NA | NA | NA | NA | NA | 1 | 1 | NA | NA | 1 | 1 | NA | NA | 1 | 0 | 2 | 0.5 |
| 18_8474874 | NA | NA | NA | NA | NA | NA | 1 | 1 | 1 | 1 | NA | NA | 1 | 1 | NA | NA | 1 | 1 | 2 | 1 |
| 18_8475763 | 1 | 1 | 1 | 1 | NA | NA | NA | NA | 2 | 1 | 1 | 1 | NA | NA | NA | NA | NA | NA | 1 | 1 |
| 18_8475769 | NA | NA | 1 | 1 | NA | NA | 1 | 1 | 2 | 1 | 2 | 1 | NA | NA | NA | NA | NA | NA | 2 | 1 |
| 18_8475776 | 1 | 1 | 1 | 1 | NA | NA | 1 | 0 | 3 | 0.6667 | 2 | 1 | NA | NA | NA | NA | 1 | 1 | 3 | 1 |
| 18_8475786 | 1 | 0 | NA | NA | NA | NA | 1 | 1 | 2 | 0.5 | 2 | 1 | 1 | 1 | NA | NA | NA | NA | 3 | 1 |
| 18_8475799 | 1 | 0 | 1 | 1 | NA | NA | NA | NA | 2 | 0.5 | 1 | 0 | 1 | 1 | NA | NA | 1 | 1 | 3 | 0.6667 |
| 18_8475869 | NA | NA | NA | NA | 1 | 1 | 1 | 1 | 2 | 1 | NA | NA | NA | NA | NA | NA | 1 | 1 | 1 | 1 |
| 18_8475873 | NA | NA | NA | NA | 1 | 1 | NA | NA | 1 | 1 | NA | NA | NA | NA | NA | NA | 1 | 1 | 1 | 1 |
| 18_8475879 | NA | NA | NA | NA | NA | NA | 1 | 1 | 1 | 1 | NA | NA | NA | NA | NA | NA | 1 | 1 | 1 | 1 |
| 18_8475880 | NA | NA | NA | NA | 1 | 1 | NA | NA | 1 | 1 | NA | NA | NA | NA | NA | NA | 1 | 1 | 1 | 1 |
| 18_8475893 | NA | NA | NA | NA | 1 | 0 | 2 | 0.5 | 3 | 0.3333 | NA | NA | NA | NA | NA | NA | 1 | 1 | 1 | 1 |
| 18_8475896 | NA | NA | NA | NA | NA | NA | 2 | 1 | 2 | 1 | NA | NA | NA | NA | NA | NA | 1 | 1 | 1 | 1 |
| 18_8475902 | NA | NA | NA | NA | 1 | 0 | 2 | 0.5 | 3 | 0.3333 | NA | NA | NA | NA | NA | NA | 1 | 1 | 1 | 1 |
| 18_8489774 | 8 | 0.25 | 11 | 0.0909 | 13 | 0.1538 | 13 | 0.1538 | 45 | 0.1556 | 10 | 0.1 | 14 | 0.2143 | 12 | 0 | 13 | 0.0769 | 49 | 0.102 |
| 18_8489786 | 13 | 0.2308 | 16 | 0.3125 | 14 | 0.2857 | 20 | 0.35 | 63 | 0.3016 | 18 | 0.2222 | 16 | 0.25 | 20 | 0.45 | 17 | 0.4118 | 71 | 0.338 |
| 18_8496552 | NA | NA | 1 | 0 | 1 | 0 | 5 | 0.2 | 7 | 0.1429 | 1 | 0 | NA | NA | 5 | 0.4 | 1 | 1 | 7 | 0.4286 |
| 18_8496926 | 1 | 1 | 1 | 1 | NA | NA | 3 | 0.6667 | 5 | 0.8 | NA | NA | NA | NA | 2 | 1 | 2 | 0 | 4 | 0.5 |
| 18_8640416 | 1 | 1 | NA | NA | NA | NA | NA | NA | 1 | 1 | 1 | 1 | NA | NA | NA | NA | NA | NA | 1 | 1 |
| 18_8640460 | 1 | 1 | NA | NA | NA | NA | NA | NA | 1 | 1 | 1 | 1 | NA | NA | NA | NA | NA | NA | 1 | 1 |
| 18_8641319 | 2 | 1 | 4 | 1 | 2 | 0.5 | 4 | 0 | 12 | 0.5833 | 2 | 1 | 5 | 0.6 | 3 | 1 | 3 | 0.3333 | 13 | 0.6923 |
| 18_8641344 | 4 | 0 | 4 | 0.25 | 2 | 0 | 8 | 0.125 | 18 | 0.1111 | 2 | 0 | 7 | 0.4286 | 4 | 0 | 4 | 0.25 | 17 | 0.2353 |
| 18_8641347 | 4 | 0.25 | 5 | 0.2 | 3 | 0.3333 | 9 | 0.1111 | 21 | 0.1905 | 4 | 0 | 6 | 0 | 2 | 0 | 7 | 0.1429 | 19 | 0.0526 |
| 18_8641349 | 4 | 0 | 6 | 0.1667 | 3 | 0 | 10 | 0.3 | 23 | 0.1739 | 4 | 0.25 | 8 | 0.5 | 5 | 0 | 6 | 0.1667 | 23 | 0.2609 |
| 18_8641356 | 4 | 0.25 | 6 | 0 | 3 | 0 | 10 | 0.1 | 23 | 0.087 | 4 | 0 | 6 | 0.1667 | 5 | 0 | 6 | 0.1667 | 21 | 0.0952 |
| 18_8641392 | 5 | 0.8 | 6 | 1 | 2 | 1 | 12 | 0.5833 | 25 | 0.76 | 5 | 0.6 | 10 | 0.7 | 6 | 0.8333 | 5 | 0.6 | 26 | 0.6923 |
| 18_8642278 | 8 | 0.125 | 7 | 0 | 2 | 0.5 | 14 | 0.3571 | 31 | 0.2258 | 6 | 0.6667 | 11 | 0.2727 | 8 | 0.25 | 7 | 0.4286 | 32 | 0.375 |
| 18_8642286 | 7 | 0.4286 | 6 | 0.8333 | 4 | 0.75 | 14 | 0.7143 | 31 | 0.6774 | 6 | 0.5 | 12 | 0.8333 | 8 | 0.375 | 7 | 0.7143 | 33 | 0.6364 |
| 18_8642290 | 8 | 0.25 | 6 | 0.5 | 3 | 0 | 15 | 0.4 | 32 | 0.3438 | 6 | 0.1667 | 11 | 0.0909 | 8 | 0.25 | 9 | 0.5556 | 34 | 0.2647 |
| 18_8651672 | 9 | 0.2222 | 9 | 0 | 4 | 0 | 17 | 0.0588 | 39 | 0.0769 | 5 | 0.2 | 12 | 0.0833 | 8 | 0 | 8 | 0.125 | 33 | 0.0909 |
| 18_8651681 | 10 | 0.9 | 6 | 1 | 5 | 0.8 | 17 | 0.7647 | 38 | 0.8421 | 4 | 1 | 12 | 0.8333 | 9 | 0.7778 | 8 | 0.625 | 33 | 0.7879 |
| 18_8651682 | 9 | 0.2222 | 8 | 0.25 | 4 | 0 | 16 | 0.125 | 37 | 0.1622 | 5 | 0 | 13 | 0.2308 | 6 | 0.1667 | 6 | 0.5 | 30 | 0.2333 |
| 18_8651688 | 10 | 0.7 | 8 | 0.5 | 5 | 0.8 | 15 | 0.3333 | 38 | 0.5263 | 3 | 1 | 15 | 0.6667 | 5 | 0.4 | 7 | 1 | 30 | 0.7333 |
| 18_8651692 | 9 | 0.2222 | 8 | 0.375 | 5 | 0.4 | 17 | 0.4118 | 39 | 0.359 | 6 | 0.3333 | 15 | 0.2667 | 11 | 0.0909 | 9 | 0.5556 | 41 | 0.2927 |
| 18_8651694 | 8 | 0.625 | 5 | 0.4 | 1 | 0 | 13 | 0.6154 | 27 | 0.5556 | 6 | 0.6667 | 10 | 0.6 | 10 | 0.2 | 7 | 0.4286 | 33 | 0.4545 |
| 18_8651741 | 1 | 0 | NA | NA | NA | NA | 1 | 1 | 2 | 0.5 | NA | NA | 1 | 1 | NA | NA | 2 | 0 | 3 | 0.3333 |
| 18_8651906 | 7 | 0.7143 | 4 | 1 | 3 | 1 | 10 | 0.9 | 24 | 0.875 | 4 | 0.5 | 8 | 0.75 | 9 | 0.8889 | 6 | 0.8333 | 27 | 0.7778 |
| 18_8651910 | 10 | 0.7 | 5 | 0.8 | 3 | 0.6667 | 11 | 0.8182 | 29 | 0.7586 | 4 | 0.75 | 8 | 0.625 | 11 | 1 | 8 | 0.875 | 31 | 0.8387 |
| 18_8651915 | 10 | 0.1 | 5 | 0 | 3 | 0.3333 | 12 | 0 | 30 | 0.0667 | 4 | 0 | 11 | 0.1818 | 11 | 0.1818 | 7 | 0.1429 | 33 | 0.1515 |
| 18_8651917 | 9 | 0 | 3 | 0 | 3 | 0.3333 | 10 | 0.2 | 25 | 0.12 | 2 | 0 | 10 | 0 | 9 | 0.1111 | 6 | 0.1667 | 27 | 0.0741 |
| 18_8651927 | 9 | 0.1111 | 4 | 0 | 4 | 0 | 13 | 0.0769 | 30 | 0.0667 | 4 | 0.25 | 11 | 0 | 10 | 0.1 | 7 | 0 | 32 | 0.0625 |
| 18_8651946 | 10 | 0.7 | 4 | 0.75 | 5 | 0.8 | 17 | 0.5294 | 36 | 0.6389 | 4 | 0.5 | 11 | 0.6364 | 12 | 0.5833 | 8 | 0.875 | 35 | 0.6571 |
| 18_8651953 | 13 | 0.2308 | 4 | 0.5 | 5 | 0.2 | 19 | 0.3684 | 41 | 0.3171 | 5 | 0.4 | 12 | 0.4167 | 12 | 0.5 | 8 | 0.625 | 37 | 0.4865 |
| 18_8651956 | 14 | 1 | 5 | 1 | 3 | 1 | 20 | 0.95 | 42 | 0.9762 | 5 | 1 | 13 | 0.9231 | 10 | 0.9 | 6 | 1 | 34 | 0.9412 |
| 18_8651978 | 15 | 0.0667 | 2 | 0 | 4 | 0.25 | 21 | 0.0952 | 42 | 0.0952 | 5 | 0 | 14 | 0.0714 | 11 | 0.0909 | 8 | 0 | 38 | 0.0526 |
| 18_8651982 | 13 | 0.2308 | 4 | 0 | 6 | 0.5 | 20 | 0.35 | 43 | 0.3023 | 5 | 0.2 | 10 | 0.4 | 13 | 0.1538 | 9 | 0 | 37 | 0.1892 |
| 18_8651983 | 15 | 0.1333 | 5 | 0.6 | 6 | 0.3333 | 17 | 0.2353 | 43 | 0.2558 | 3 | 0 | 10 | 0.5 | 13 | 0.3077 | 9 | 0.3333 | 35 | 0.3429 |

| | | | | | | | | | | | | | | | | | | | | |
|---|---|---|---|---|---|---|---|---|---|---|---|---|---|---|---|---|---|---|---|---|
| 18_865225 | 5 | 0 | 7 | 0.1429 | 6 | 0.1667 | 14 | 0.2143 | 32 | 0.1562 | 8 | 0 | 9 | 0 | 12 | 0.25 | 7 | 0.1429 | 36 | 0.1111 |
| 18_865245 | NA | NA | 4 | 0.25 | 3 | 0.3333 | 3 | 0.3333 | 10 | 0.3 | 4 | 0.25 | 5 | 0 | 7 | 0.1429 | 4 | 0.5 | 20 | 0.2 |
| 18_865245 | 1 | 0 | 5 | 0.4 | 4 | 0.25 | 2 | 0 | 12 | 0.25 | 4 | 0 | 4 | 0 | 7 | 0.1429 | 4 | 0.5 | 19 | 0.1579 |
| 18_867349 | NA | NA | 1 | 1 | 2 | 1 | NA | NA | 3 | 1 | NA | NA | 1 | 1 | NA | NA | NA | NA | 1 | 1 |
| 18_867350 | NA | NA | 1 | 1 | 1 | 0 | NA | NA | 2 | 0.5 | NA | NA | 1 | 1 | NA | NA | NA | NA | 1 | 1 |
| 18_867350 | NA | NA | 1 | 0 | 2 | 1 | NA | NA | 3 | 0.6667 | NA | NA | 1 | 1 | NA | NA | NA | NA | 1 | 1 |
| 18_867353 | NA | NA | 2 | 0.5 | 2 | 1 | NA | NA | 4 | 0.75 | 1 | 1 | 1 | 1 | NA | NA | NA | NA | 2 | 1 |
| 18_867354 | NA | NA | 2 | 1 | 1 | 1 | NA | NA | 3 | 1 | 1 | 0 | NA | NA | NA | NA | 1 | 1 | 2 | 0.5 |
| 18_867357 | NA | NA | 1 | 1 | 3 | 1 | NA | NA | 4 | 1 | 1 | 0 | 1 | 1 | NA | NA | 1 | 1 | 3 | 0.6667 |
| 18_867360 | NA | NA | 5 | 0.6 | 3 | 0.3333 | NA | NA | 8 | 0.5 | 1 | 0 | 1 | 0 | NA | NA | 1 | 1 | 3 | 0.3333 |
| 18_867360 | NA | NA | 3 | 0.6667 | 3 | 0.6667 | NA | NA | 6 | 0.6667 | NA | NA | 1 | 1 | NA | NA | 1 | 1 | 2 | 1 |
| 18_867361 | NA | NA | 5 | 0.8 | 3 | 1 | NA | NA | 8 | 0.875 | 1 | 1 | 1 | 1 | NA | NA | 1 | 1 | 3 | 1 |
| 18_867362 | NA | NA | 5 | 0.8 | 3 | 0.3333 | NA | NA | 8 | 0.625 | 1 | 0 | 1 | 1 | NA | NA | 1 | 1 | 3 | 0.6667 |
| 18_867368 | NA | NA | 1 | 1 | 1 | 0 | NA | NA | 2 | 0.5 | NA | NA | 1 | 1 | NA | NA | NA | NA | 1 | 1 |
| 18_888872 | 2 | 1 | 1 | 1 | 2 | 0.5 | 3 | 0.6667 | 8 | 0.75 | NA | NA | 2 | 1 | NA | NA | NA | NA | 2 | 1 |
| 18_888872 | 2 | 0 | 1 | 1 | 1 | 0 | 3 | 0.6667 | 7 | 0.4286 | NA | NA | 2 | 1 | NA | NA | NA | NA | 2 | 1 |
| 18_888873 | 2 | 1 | 1 | 1 | 2 | 1 | 3 | 1 | 8 | 1 | NA | NA | 2 | 1 | NA | NA | NA | NA | 2 | 1 |
| 18_888874 | 2 | 1 | 1 | 1 | 2 | 1 | NA | NA | 5 | 1 | NA | NA | 2 | 1 | NA | NA | NA | NA | 2 | 1 |
| 18_888875 | 2 | 1 | 1 | 1 | 2 | 1 | 3 | 1 | 8 | 1 | NA | NA | 2 | 1 | NA | NA | NA | NA | 2 | 1 |
| 18_888875 | 2 | 1 | 1 | 1 | 1 | 1 | 3 | 0.6667 | 7 | 0.8571 | NA | NA | 1 | 1 | NA | NA | NA | NA | 1 | 1 |
| 18_888876 | 1 | 0 | 1 | 1 | 1 | 0 | 2 | 1 | 5 | 0.6 | NA | NA | 2 | 0.5 | NA | NA | NA | NA | 2 | 0.5 |
| 18_888878 | 1 | 1 | NA | NA | 1 | 1 | 2 | 1 | 4 | 1 | NA | NA | 2 | 1 | NA | NA | NA | NA | 2 | 1 |
| 18_888879 | 2 | 1 | 1 | 1 | 1 | 1 | 4 | 0.75 | 8 | 0.875 | NA | NA | 2 | 1 | NA | NA | NA | NA | 2 | 1 |
| 18_888879 | 1 | 1 | NA | NA | NA | NA | 2 | 1 | 3 | 1 | NA | NA | 1 | 1 | NA | NA | NA | NA | 1 | 1 |
| 18_888880 | 1 | 1 | 1 | 1 | 1 | 0 | 3 | 1 | 6 | 0.8333 | NA | NA | 1 | 1 | NA | NA | NA | NA | 1 | 1 |
| 18_888881 | 2 | 1 | 1 | 1 | 1 | 0 | 3 | 0.3333 | 7 | 0.5714 | NA | NA | 2 | 1 | NA | NA | NA | NA | 2 | 1 |
| 18_888881 | 1 | 0 | 1 | 0 | NA | NA | 3 | 0.6667 | 5 | 0.4 | NA | NA | 2 | 0.5 | NA | NA | NA | NA | 2 | 0.5 |
| 18_888882 | 1 | 1 | 1 | 1 | 1 | 0 | 4 | 0.75 | 7 | 0.7143 | NA | NA | 2 | 0.5 | NA | NA | NA | NA | 2 | 0.5 |
| 18_888882 | 2 | 1 | 1 | 1 | 1 | 1 | 4 | 1 | 8 | 1 | NA | NA | 1 | 1 | NA | NA | NA | NA | 1 | 1 |
| 18_888883 | 1 | 0 | 1 | 0 | 1 | 1 | 3 | 0 | 6 | 0.1667 | NA | NA | 1 | 1 | NA | NA | NA | NA | 1 | 1 |
| 18_893108 | 4 | 0 | 7 | 0 | 6 | 0.3333 | 7 | 0 | 24 | 0.0833 | 2 | 0 | 6 | 0 | 6 | 0 | 3 | 0.3333 | 17 | 0.0588 |
| 18_893192 | 1 | 0 | 1 | 0 | 4 | 0.25 | NA | NA | 6 | 0.1667 | NA | NA | NA | NA | 1 | 1 | NA | NA | 1 | 1 |
| 18_905583 | 4 | 0 | 8 | 0.25 | 3 | 0 | 5 | 0 | 20 | 0.1 | 5 | 0.2 | 6 | 0.5 | 1 | 0 | 10 | 0.2 | 22 | 0.2727 |
| 18_905585 | 5 | 0 | 7 | 0.2857 | 2 | 0 | 5 | 0.6 | 19 | 0.2632 | 6 | 0.5 | 1 | 1 | 1 | 0 | 11 | 0.0909 | 19 | 0.2632 |
| 18_905586 | 4 | 0 | 8 | 0.375 | 2 | 0.5 | 6 | 0.3333 | 20 | 0.3 | 6 | 0.5 | 6 | 0.3333 | 1 | 0 | 11 | 0.0909 | 24 | 0.25 |
| 18_905587 | 5 | 0 | 7 | 0.2857 | 4 | 0 | 6 | 0 | 22 | 0.0909 | 7 | 0.2857 | 8 | 0.25 | 1 | 0 | 13 | 0.3846 | 29 | 0.3103 |
| 18_905589 | 7 | 0.1429 | 9 | 0.1111 | 4 | 0 | 5 | 0.2 | 25 | 0.12 | 6 | 0 | 8 | 0 | 1 | 0 | 13 | 0.1538 | 28 | 0.0714 |
| 18_905595 | 6 | 0.3333 | 10 | 0.2 | 4 | 0 | 8 | 0 | 28 | 0.1429 | 7 | 0.4286 | 9 | 0.2222 | 2 | 0.5 | 12 | 0.1667 | 30 | 0.2667 |
| 18_905607 | 4 | 0 | 4 | 0.25 | 2 | 0 | 6 | 0.1667 | 16 | 0.125 | 3 | 0 | 4 | 0 | 1 | 0 | 10 | 0.5 | 18 | 0.2778 |
| 18_905611 | 6 | 0 | 4 | 0.5 | 2 | 0 | 7 | 0.1429 | 19 | 0.1579 | 3 | 0.3333 | 4 | 0 | 2 | 0 | 11 | 0.3636 | 20 | 0.25 |
| 18_906103 | NA | NA | 5 | 0 | 1 | 0 | 2 | 0.5 | 8 | 0.125 | 2 | 1 | NA | NA | 1 | 0 | NA | NA | 3 | 0.6667 |
| 18_906103 | NA | NA | 4 | 0.25 | 1 | 0 | 1 | 0 | 6 | 0.1667 | 2 | 0.5 | NA | NA | 1 | 0 | NA | NA | 3 | 0.3333 |
| 18_906103 | NA | NA | 4 | 0 | 1 | 0 | 2 | 0.5 | 7 | 0.1429 | 1 | 0 | NA | NA | 1 | 1 | NA | NA | 2 | 0.5 |
| 18_906105 | NA | NA | 4 | 0 | 1 | 0 | 2 | 1 | 7 | 0.2857 | 1 | 0 | NA | NA | 2 | 1 | NA | NA | 3 | 0.6667 |
| 18_906106 | NA | NA | 6 | 0.6667 | 1 | 0 | 2 | 1 | 9 | 0.6667 | 2 | 0.5 | NA | NA | 1 | 1 | NA | NA | 3 | 0.6667 |
| 19_331100 | 1 | 0 | NA | NA | 3 | 0.3333 | 3 | 0.3333 | 7 | 0.2857 | NA | NA | 1 | 1 | 2 | 1 | 2 | 0 | 5 | 0.6 |
| 19_331104 | 1 | 1 | NA | NA | 2 | 0 | 2 | 0.5 | 5 | 0.4 | NA | NA | 1 | 1 | 2 | 0.5 | 2 | 1 | 5 | 0.8 |
| 19_331140 | NA | NA | NA | NA | 2 | 0.5 | 2 | 1 | 4 | 0.75 | NA | NA | 1 | 0 | 2 | 1 | 1 | 0 | 4 | 0.5 |
| 19_331154 | 1 | 1 | NA | NA | 3 | 0 | 2 | 0 | 6 | 0.1667 | NA | NA | 1 | 0 | 2 | 0.5 | 2 | 0.5 | 5 | 0.4 |
| 19_331486 | 1 | 1 | NA | NA | 2 | 0 | 2 | 1 | 5 | 0.6 | NA | NA | 1 | 0 | 1 | 0 | 2 | 1 | 4 | 0.5 |
| 19_331524 | 1 | 1 | NA | NA | 2 | 0.5 | 1 | 1 | 4 | 0.75 | NA | NA | NA | NA | 1 | 0 | 2 | 0.5 | 3 | 0.3333 |
| 19_331526 | 1 | 0 | NA | NA | 1 | 1 | 1 | 1 | 3 | 0.6667 | NA | NA | 1 | 1 | 1 | 1 | 1 | 1 | 3 | 1 |
| 19_331547 | 1 | 1 | NA | NA | NA | NA | NA | NA | 1 | 1 | NA | NA | 1 | 0 | NA | NA | 1 | 1 | 2 | 0.5 |
| 19_343379 | 3 | 0.6667 | 4 | 0.5 | 3 | 0.3333 | 2 | 0.5 | 12 | 0.5 | 2 | 0 | 2 | 0.5 | 1 | 1 | NA | NA | 5 | 0.4 |
| 19_343384 | 3 | 0 | 3 | 0.3333 | 1 | 0 | 1 | 0 | 8 | 0.125 | NA | NA | 2 | 0 | 2 | 1 | NA | NA | 4 | 0.5 |
| 19_343387 | 2 | 1 | 5 | 1 | 3 | 0.6667 | 1 | 1 | 11 | 0.9091 | 2 | 1 | 2 | 0.5 | 2 | 1 | NA | NA | 6 | 0.8333 |
| 19_343391 | 2 | 0 | 4 | 0 | 3 | 0.3333 | 1 | 0 | 10 | 0.1 | 2 | 0.5 | 2 | 0 | 2 | 0.5 | NA | NA | 6 | 0.3333 |
| 19_343416 | 3 | 0 | 5 | 0.2 | 3 | 0.3333 | 2 | 0 | 13 | 0.1538 | 2 | 0 | 2 | 0.5 | 2 | 0 | NA | NA | 6 | 0.1667 |
| 19_343420 | 3 | 0.3333 | 5 | 0.6 | 4 | 0.75 | 2 | 1 | 14 | 0.6429 | 2 | 1 | 2 | 0 | 2 | 1 | NA | NA | 6 | 0.6667 |
| 19_343500 | 5 | 0.2 | 7 | 0.4286 | 3 | 0 | 2 | 0 | 17 | 0.2353 | 2 | 0 | 4 | 0.25 | 2 | 0 | NA | NA | 8 | 0.125 |
| 19_350514 | 6 | 0.3333 | 10 | 0.6 | 1 | 1 | 8 | 0.625 | 25 | 0.56 | 7 | 0.5714 | 8 | 0.625 | 8 | 0.375 | 10 | 0.6 | 33 | 0.5455 |
| 19_350520 | 6 | 0.3333 | 10 | 0.3 | 1 | 1 | 8 | 0.5 | 25 | 0.4 | 9 | 0.4444 | 6 | 0.1667 | 7 | 0.4286 | 7 | 0.4286 | 29 | 0.3793 |
| 19_350525 | 7 | 0.5714 | 12 | 0.6667 | 1 | 1 | 8 | 0.625 | 28 | 0.6429 | 10 | 0.6 | 7 | 0.8571 | 6 | 0.5 | 8 | 0.625 | 31 | 0.6452 |
| 19_350540 | 6 | 0.3333 | 12 | 0.5 | 1 | 0 | 5 | 0.4 | 24 | 0.4167 | 8 | 0.5 | 7 | 0.7143 | 7 | 0.7143 | 9 | 0.6667 | 31 | 0.6452 |
| 19_350547 | 6 | 0.8333 | 7 | 0.8571 | NA | NA | 6 | 0.8333 | 19 | 0.8421 | 7 | 1 | 5 | 0.6 | 5 | 0.8 | 6 | 0.6667 | 23 | 0.7826 |

| | | | | | | | | | | | | | | | | | | | | |
|---|---|---|---|---|---|---|---|---|---|---|---|---|---|---|---|---|---|---|---|---|
| 19_350554( | 5 | 0 | 9 | 0.1111 | 1 | 0 | 3 | 0 | 18 | 0.0556 | 4 | 0.75 | 4 | 0.25 | 5 | 0.4 | 7 | 0 | 20 | 0.3 |
| 19_350683! | 5 | 0.6 | 9 | 0.6667 | 1 | 1 | 6 | 1 | 21 | 0.7619 | 5 | 0.4 | 3 | 0.6667 | 6 | 0.8333 | 6 | 0.8333 | 20 | 0.7 |
| 19_350697( | 5 | 0 | 10 | 0.5 | 1 | 0 | 6 | 0.3333 | 22 | 0.3182 | 6 | 0.1667 | 3 | 0 | 7 | 0.4286 | 6 | 0.3333 | 22 | 0.2727 |
| 19_363517: | 1 | 1 | NA | NA | NA | NA | NA | NA | 1 | 1 | 1 | 0 | 2 | 0.5 | 2 | 1 | 1 | 1 | 6 | 0.6667 |
| 19_376655! | NA | NA | 1 | 1 | 1 | 1 | 3 | 0.6667 | 5 | 0.8 | 2 | 1 | NA | NA | 5 | 0.6 | 2 | 0.5 | 9 | 0.6667 |
| 19_376678 | NA | NA | 1 | 0 | 1 | 0 | 3 | 0.3333 | 5 | 0.2 | 2 | 0 | NA | NA | 6 | 0.1667 | 2 | 0 | 10 | 0.1 |
| 19_376695 | NA | NA | 1 | 1 | 1 | 0 | 4 | 0.25 | 6 | 0.3333 | 4 | 0.5 | NA | NA | 6 | 0.3333 | 2 | 1 | 12 | 0.5 |
| 19_376729 | NA | NA | 2 | 1 | NA | NA | 4 | 0.25 | 6 | 0.5 | 4 | 0.25 | NA | NA | 6 | 0.1667 | 2 | 0 | 12 | 0.1667 |
| 19_376734( | NA | NA | 2 | 1 | 1 | 0 | 4 | 0.75 | 7 | 0.7143 | 4 | 0.75 | NA | NA | 6 | 0.5 | 2 | 0.5 | 12 | 0.5833 |
| 19_376736 | NA | NA | 2 | 0.5 | 1 | 1 | 4 | 0.75 | 7 | 0.7143 | 3 | 0.6667 | NA | NA | 6 | 1 | 1 | 1 | 10 | 0.9 |
| 19_376753 | NA | NA | 2 | 1 | 2 | 1 | 4 | 1 | 8 | 1 | 2 | 1 | NA | NA | 5 | 1 | 1 | 1 | 8 | 1 |
| 19_376755 | NA | NA | 2 | 1 | 2 | 1 | 4 | 1 | 8 | 1 | 4 | 0.5 | NA | NA | 6 | 0.8333 | 2 | 0.5 | 12 | 0.6667 |
| 19_385853 | 1 | 1 | NA | NA | NA | NA | NA | NA | 1 | 1 | NA | NA | NA | NA | NA | NA | 1 | 1 | 1 | 1 |
| 19_386426( | NA | NA | 6 | 0.3333 | NA | NA | 2 | 1 | 8 | 0.5 | NA | NA | NA | NA | 2 | 0.5 | 1 | 0 | 3 | 0.3333 |
| 19_386429 | 1 | 0 | 7 | 0.1429 | NA | NA | 2 | 0 | 10 | 0.1 | NA | NA | NA | NA | 2 | 0 | 3 | 0.3333 | 5 | 0.2 |
| 19_386443 | NA | NA | 8 | 0.625 | NA | NA | 3 | 0.6667 | 11 | 0.6364 | NA | NA | NA | NA | 2 | 1 | 3 | 0.6667 | 5 | 0.8 |
| 19_386446 | 1 | 0 | 7 | 0.2857 | NA | NA | 4 | 0 | 12 | 0.1667 | NA | NA | NA | NA | 2 | 0.5 | 3 | 0 | 5 | 0.2 |
| 19_386457( | NA | NA | 7 | 0.7143 | NA | NA | 1 | 0 | 8 | 0.625 | NA | NA | NA | NA | 1 | 0 | 3 | 0.3333 | 4 | 0.25 |
| 19_386457 | 1 | 0 | 4 | 0.5 | NA | NA | 4 | 0.75 | 9 | 0.5556 | NA | NA | NA | NA | 1 | 1 | 2 | 1 | 3 | 1 |
| 19_386465 | 1 | 0 | 9 | 0.2222 | NA | NA | 3 | 0.3333 | 13 | 0.2308 | NA | NA | NA | NA | 2 | 0 | 3 | 0.6667 | 5 | 0.4 |
| 19_386469 | 1 | 0 | 8 | 0.625 | NA | NA | 2 | 1 | 11 | 0.6364 | 1 | 1 | 1 | 0 | 2 | 0 | 3 | 0.6667 | 7 | 0.4286 |
| 19_386471 | 1 | 0 | 8 | 0.5 | NA | NA | 3 | 0.3333 | 12 | 0.4167 | 1 | 1 | NA | NA | 2 | 1 | 3 | 1 | 6 | 1 |
| 19_386482 | 1 | 1 | 7 | 0.7143 | NA | NA | 2 | 0.5 | 10 | 0.7 | NA | NA | 1 | 1 | 2 | 1 | 1 | 1 | 4 | 1 |
| 19_386493 | 1 | 0 | 10 | 1 | NA | NA | 4 | 1 | 15 | 0.9333 | 1 | 1 | 1 | 1 | 2 | 1 | 5 | 1 | 9 | 1 |
| 19_386499 | 1 | 0 | 9 | 0.3333 | NA | NA | 4 | 0.25 | 14 | 0.2857 | 2 | 0 | NA | NA | 2 | 1 | 4 | 0.25 | 8 | 0.375 |
| 19_386500 | NA | NA | 5 | 0.8 | NA | NA | 3 | 0.6667 | 8 | 0.75 | 2 | 1 | 1 | 1 | NA | NA | 3 | 1 | 6 | 1 |
| 19_386518( | 1 | 0 | 10 | 0.5 | NA | NA | 4 | 0.75 | 15 | 0.5333 | 1 | 1 | 2 | 0 | 3 | 0.3333 | 5 | 0.4 | 11 | 0.3636 |
| 19_386519 | 1 | 0 | 11 | 0.0909 | NA | NA | 4 | 0.25 | 16 | 0.125 | 2 | 0.5 | 2 | 0 | 3 | 0.3333 | 2 | 0 | 9 | 0.2222 |
| 19_386540 | 1 | 1 | 9 | 0.5556 | NA | NA | 3 | 1 | 13 | 0.6923 | 3 | 1 | 3 | 1 | 1 | 1 | 3 | 1 | 10 | 1 |
| 19_386545( | 1 | 1 | 10 | 0.9 | NA | NA | 5 | 1 | 16 | 0.9375 | 3 | 1 | 4 | 1 | 3 | 1 | 5 | 1 | 15 | 1 |
| 19_386546 | 1 | 0 | 11 | 0 | NA | NA | 5 | 0.4 | 17 | 0.1176 | 3 | 0 | 4 | 0.5 | 3 | 0 | 4 | 0.5 | 14 | 0.2857 |
| 19_386547 | 1 | 0 | 9 | 0.3333 | NA | NA | 5 | 0.4 | 15 | 0.3333 | 3 | 0 | 5 | 0.8 | 3 | 0.3333 | 3 | 0.6667 | 14 | 0.5 |
| 19_386552 | 1 | 1 | 11 | 1 | NA | NA | 5 | 1 | 17 | 1 | 3 | 1 | 5 | 1 | 2 | 1 | 3 | 1 | 13 | 1 |
| 19_386554( | 1 | 1 | 9 | 1 | NA | NA | 5 | 0.8 | 15 | 0.9333 | 2 | 1 | 3 | 0.6667 | 2 | 0 | 4 | 1 | 11 | 0.7273 |
| 19_386557 | NA | NA | 8 | 0.875 | NA | NA | 5 | 0.8 | 13 | 0.8462 | 3 | 0.6667 | 4 | 0.5 | 3 | 1 | 4 | 0.75 | 14 | 0.7143 |
| 19_386561 | 1 | 1 | 10 | 0.9 | NA | NA | 4 | 1 | 15 | 0.9333 | 2 | 1 | 4 | 1 | 3 | 1 | 4 | 1 | 13 | 1 |
| 19_386571( | 1 | 0 | 11 | 0.6364 | NA | NA | 5 | 0.8 | 17 | 0.6471 | 2 | 1 | 5 | 0.8 | 3 | 0.6667 | 4 | 1 | 14 | 0.8571 |
| 19_386580 | 1 | 1 | 9 | 0.7778 | NA | NA | 5 | 0.4 | 15 | 0.6667 | 3 | 0.3333 | 4 | 0 | 3 | 0 | 4 | 0.25 | 14 | 0.1429 |
| 19_386583 | 1 | 1 | 12 | 0.6667 | NA | NA | 4 | 0.5 | 17 | 0.6471 | 2 | 0 | 5 | 0.2 | 3 | 0 | 5 | 0.6 | 15 | 0.2667 |
| 19_386590 | 1 | 0 | 12 | 0.3333 | NA | NA | 5 | 0 | 18 | 0.2222 | 3 | 0.3333 | 5 | 0.4 | 2 | 0 | 3 | 0 | 13 | 0.2308 |
| 19_386598 | 1 | 0 | 11 | 0.2727 | NA | NA | 4 | 0.25 | 16 | 0.25 | 3 | 0 | 4 | 0.25 | 2 | 0.5 | 3 | 0 | 12 | 0.1667 |
| 19_386609 | 1 | 0 | 9 | 0.3333 | NA | NA | 4 | 0.25 | 14 | 0.2857 | 2 | 0 | 5 | 0 | 2 | 0.5 | 4 | 0 | 13 | 0.0769 |
| 19_386614 | NA | NA | 5 | 0.8 | NA | NA | 4 | 1 | 9 | 0.8889 | 2 | 1 | 3 | 1 | 1 | 1 | 2 | 1 | 8 | 1 |
| 19_386637 | 1 | 1 | 9 | 0.8889 | NA | NA | 6 | 1 | 16 | 0.9375 | 3 | 0.6667 | 5 | 1 | 3 | 0.3333 | 4 | 1 | 15 | 0.8 |
| 19_386646 | 1 | 0 | 9 | 0 | NA | NA | 6 | 0.1667 | 16 | 0.0625 | 3 | 0.3333 | 6 | 0.1667 | 3 | 0 | 3 | 0 | 15 | 0.1333 |
| 19_386649( | 1 | 0 | 9 | 0.1111 | NA | NA | 5 | 0 | 15 | 0.0667 | 2 | 0.5 | 4 | 0.5 | 2 | 0 | 5 | 0 | 13 | 0.2308 |
| 19_386652 | 1 | 0 | 9 | 0 | NA | NA | 6 | 0.1667 | 16 | 0.0625 | 4 | 0.25 | 6 | 0.1667 | 3 | 0 | 4 | 0 | 17 | 0.1176 |
| 19_386704 | NA | NA | 10 | 0.3 | NA | NA | 5 | 0.2 | 15 | 0.2667 | 3 | 0.3333 | 5 | 0.2 | 2 | 0 | 4 | 0 | 14 | 0.1429 |
| 19_386790 | NA | NA | 11 | 0.9091 | NA | NA | 5 | 1 | 16 | 0.9375 | 4 | 1 | 7 | 0.8571 | 3 | 1 | 6 | 1 | 20 | 0.95 |
| 19_386792( | NA | NA | 5 | 0.2 | 1 | 0 | 3 | 0.3333 | 9 | 0.2222 | 2 | 0 | 3 | 0.6667 | 2 | 0 | 3 | 0.6667 | 10 | 0.4 |
| 19_386799 | NA | NA | 10 | 0.3 | 1 | 0 | 3 | 0.3333 | 14 | 0.2857 | 2 | 1 | 7 | 0.4286 | 3 | 0.3333 | 6 | 0.3333 | 18 | 0.4444 |
| 19_386818 | NA | NA | 12 | 0.6667 | 3 | 0.6667 | 5 | 0.6 | 20 | 0.65 | 3 | 1 | 8 | 0.875 | 4 | 1 | 7 | 0.8571 | 22 | 0.9091 |
| 19_387172 | NA | NA | NA | NA | 1 | 1 | NA | NA | 1 | 1 | 3 | 0 | 1 | 1 | 1 | 0 | NA | NA | 5 | 0.2 |
| 19_390201 | 8 | 0.125 | 1 | 0 | 4 | 0 | 4 | 0 | 17 | 0.0588 | 4 | 0 | 5 | 0.2 | 4 | 0 | 2 | 0 | 15 | 0.0667 |
| 19_390222 | 7 | 0 | 1 | 0 | 3 | 0.3333 | 4 | 0 | 15 | 0.0667 | 4 | 0 | 5 | 0.2 | 7 | 0 | 3 | 0 | 19 | 0.0526 |
| 19_394354 | 15 | 0.2667 | 7 | 0.5714 | 7 | 0.4286 | 7 | 0.5714 | 36 | 0.4167 | 9 | 0.1111 | 9 | 0.1111 | 9 | 0.1111 | 10 | 0.2 | 37 | 0.1351 |
| 19_394360 | 12 | 0.4167 | 9 | 0.3333 | 6 | 0.5 | 7 | 0.7143 | 34 | 0.4706 | 9 | 0.6667 | 10 | 0.6 | 10 | 0.4 | 9 | 0.4444 | 38 | 0.5263 |
| 19_394385 | 14 | 0 | 8 | 0.125 | 8 | 0.125 | 6 | 0.3333 | 36 | 0.1111 | 10 | 0 | 12 | 0.0833 | 8 | 0.125 | 7 | 0 | 37 | 0.0541 |
| 19_394394 | 14 | 0.1429 | 8 | 0.125 | 8 | 0 | 7 | 0.1429 | 37 | 0.1081 | 10 | 0.3 | 10 | 0 | 8 | 0.125 | 7 | 0.1429 | 35 | 0.1429 |
| 19_396375 | 1 | 1 | 1 | 0 | 1 | 1 | NA | NA | 3 | 0.6667 | 2 | 0.5 | NA | NA | NA | NA | NA | NA | 2 | 0.5 |
| 19_396398 | 1 | 1 | 1 | 0 | 1 | 0 | NA | NA | 3 | 0.3333 | 2 | 0.5 | NA | NA | NA | NA | 1 | 0 | 3 | 0.3333 |
| 19_399882 | NA | NA | 5 | 0.2 | 1 | 0 | 1 | 0 | 7 | 0.1429 | 2 | 0 | NA | NA | NA | NA | 3 | 0.3333 | 5 | 0.2 |
| 19_399890( | NA | NA | 4 | 0 | 1 | 0 | 1 | 1 | 6 | 0.1667 | 1 | 0 | 1 | 1 | NA | NA | 2 | 0 | 4 | 0.25 |
| 19_404988 | NA | NA | 2 | 1 | NA | NA | NA | NA | 2 | 1 | NA | NA | 1 | 1 | NA | NA | NA | NA | 1 | 1 |
| 19_405014 | NA | NA | 2 | 0.5 | NA | NA | NA | NA | 2 | 0.5 | NA | NA | 1 | 1 | NA | NA | NA | NA | 1 | 1 |

| | | | | | | | | | | | | | | | | | | | | |
|---|---|---|---|---|---|---|---|---|---|---|---|---|---|---|---|---|---|---|---|---|
| 19_405017 | NA | NA | 2 | 0.5 | NA | NA | NA | NA | 2 | 0.5 | NA | NA | 1 | 1 | NA | NA | NA | NA | 1 | 1 |
| 19_405037 | NA | NA | 2 | 0.5 | NA | NA | NA | NA | 2 | 0.5 | NA | NA | 1 | 1 | NA | NA | NA | NA | 1 | 1 |
| 19_405753 | 20 | 0.05 | 32 | 0.1875 | 20 | 0.1 | 22 | 0.0909 | 94 | 0.117 | 14 | 0.0714 | 26 | 0.1154 | 24 | 0.0417 | 26 | 0.1538 | 90 | 0.1 |
| 19_406018 | 1 | 0 | 1 | 0 | 3 | 0.3333 | 2 | 0.5 | 7 | 0.2857 | NA | NA | 2 | 0 | 1 | 0 | 2 | 0.5 | 5 | 0.2 |
| 19_406026 | 1 | 0 | NA | NA | 3 | 0.3333 | 2 | 0.5 | 6 | 0.3333 | NA | NA | 3 | 0 | 1 | 0 | 2 | 0.5 | 6 | 0.1667 |
| 19_406155 | 2 | 0 | 1 | 1 | NA | NA | 1 | 0 | 4 | 0.25 | 1 | 1 | 1 | 0 | 1 | 0 | 2 | 0.5 | 5 | 0.4 |
| 19_408546 | 36 | 0.2222 | 51 | 0.1373 | 42 | 0.0952 | 38 | 0.2368 | 167 | 0.1677 | 42 | 0.119 | 35 | 0.0857 | 46 | 0.1304 | 34 | 0.0588 | 157 | 0.1019 |
| 19_409034 | NA | NA | NA | NA | 4 | 0.25 | NA | NA | 4 | 0.25 | NA | NA | NA | NA | 1 | 1 | NA | NA | 1 | 1 |
| 19_414873 | 8 | 0.125 | 5 | 0.2 | 6 | 0.1667 | 6 | 0.1667 | 25 | 0.16 | 5 | 0 | 9 | 0 | 13 | 0.3077 | 3 | 0.3333 | 30 | 0.1667 |
| 19_415339 | 7 | 0 | 6 | 0.1667 | 8 | 0.125 | 4 | 0 | 25 | 0.08 | 6 | 0 | 3 | 0 | 4 | 0.5 | 5 | 0.2 | 18 | 0.1667 |
| 19_415487 | 10 | 0.1 | 6 | 0 | 9 | 0.1111 | 5 | 0 | 30 | 0.0667 | 5 | 0 | 4 | 0 | 5 | 0 | 5 | 0.2 | 19 | 0.0526 |
| 19_415504 | 10 | 0 | 6 | 0.5 | 9 | 0.1111 | 5 | 0.2 | 30 | 0.1667 | 6 | 0.1667 | 4 | 0 | 5 | 0.2 | 4 | 0.75 | 19 | 0.2632 |
| 19_415542 | 7 | 0 | 3 | 0 | 6 | 0.1667 | 3 | 0 | 19 | 0.0526 | 1 | 0 | 3 | 0.3333 | 4 | 0.25 | 6 | 0 | 14 | 0.1429 |
| 19_415679 | 9 | 0.2222 | 6 | 0.1667 | 10 | 0.1 | 5 | 0.2 | 30 | 0.1667 | 7 | 0.1429 | 3 | 0 | 4 | 0 | 5 | 0 | 19 | 0.0526 |
| 19_415733 | 1 | 0 | NA | NA | NA | NA | 1 | 1 | 2 | 0.5 | NA | NA | NA | NA | 1 | 0 | 2 | 0.5 | 3 | 0.3333 |
| 19_416360 | 14 | 0.1429 | 11 | 0.1818 | 12 | 0.1667 | 15 | 0.1333 | 52 | 0.1538 | 8 | 0.375 | 11 | 0 | 16 | 0.1875 | 10 | 0.4 | 45 | 0.2222 |
| 19_416373 | 17 | 0.1765 | 15 | 0.2 | 17 | 0.1176 | 13 | 0.3077 | 62 | 0.1935 | 10 | 0.2 | 9 | 0.3333 | 13 | 0.3077 | 11 | 0.3636 | 43 | 0.3023 |
| 19_416378 | 18 | 0.2778 | 13 | 0.0769 | 19 | 0.2105 | 14 | 0.0714 | 64 | 0.1719 | 11 | 0.2727 | 12 | 0.0833 | 19 | 0.2632 | 14 | 0.1429 | 56 | 0.1964 |
| 19_416380 | 19 | 0.6316 | 16 | 0.75 | 15 | 0.6667 | 15 | 0.6667 | 65 | 0.6769 | 11 | 0.2727 | 11 | 0.6364 | 21 | 0.6667 | 12 | 0.75 | 55 | 0.6 |
| 19_418156 | 8 | 0.25 | 8 | 0.375 | 5 | 0.2 | 8 | 0.125 | 29 | 0.2414 | 4 | 0.25 | 9 | 0.2222 | 4 | 0.25 | 8 | 0.125 | 25 | 0.2 |
| 19_418167 | 6 | 0.3333 | 8 | 0.5 | 5 | 1 | 5 | 0.6 | 24 | 0.5833 | 3 | 0.6667 | 4 | 0.75 | 4 | 0.75 | 7 | 0.7143 | 18 | 0.7222 |
| 19_418173 | 2 | 1 | NA | NA | 1 | 1 | NA | NA | 3 | 1 | NA | NA | 2 | 1 | NA | NA | 1 | 0 | 3 | 0.6667 |
| 19_420373 | 7 | 0 | 10 | 0 | 9 | 0.2222 | 10 | 0 | 36 | 0.0556 | 11 | 0 | 14 | 0.2857 | 9 | 0.1111 | 13 | 0 | 47 | 0.1064 |
| 19_420388 | 12 | 0.25 | 13 | 0.0769 | 9 | 0.2222 | 12 | 0.0833 | 46 | 0.1522 | 10 | 0.2 | 10 | 0.3 | 9 | 0.1111 | 15 | 0.2 | 44 | 0.2045 |
| 19_420393 | 11 | 0.1818 | 12 | 0.25 | 9 | 0.5556 | 11 | 0.5455 | 43 | 0.3721 | 12 | 0.4167 | 11 | 0.4545 | 10 | 0.2 | 14 | 0.4286 | 47 | 0.383 |
| 19_420680 | 1 | 1 | 2 | 0.5 | 1 | 0 | 3 | 1 | 7 | 0.7143 | NA | NA | 4 | 0.75 | 1 | 1 | NA | NA | 5 | 0.8 |
| 19_420683 | 1 | 0 | 4 | 0.25 | 2 | 0.5 | 3 | 0.6667 | 10 | 0.4 | NA | NA | 3 | 0.3333 | NA | NA | 1 | 1 | 4 | 0.5 |
| 19_420685 | 1 | 0 | 4 | 0 | 2 | 0.5 | 3 | 0.6667 | 10 | 0.3 | NA | NA | 5 | 0.4 | 1 | 0 | 1 | 1 | 7 | 0.4286 |
| 19_421444 | 1 | 1 | 1 | 0 | 1 | 1 | NA | NA | 3 | 0.6667 | 1 | 0 | NA | NA | 1 | 0 | 1 | 1 | 3 | 0.3333 |
| 19_421508 | 1 | 1 | 1 | 0 | 1 | 1 | NA | NA | 3 | 0.6667 | 1 | 1 | NA | NA | NA | NA | 1 | 1 | 2 | 1 |
| 19_421554 | NA | NA | 1 | 1 | 1 | 1 | NA | NA | 2 | 1 | 1 | 0 | NA | NA | NA | NA | 1 | 1 | 2 | 0.5 |
| 19_421571 | NA | NA | 1 | 1 | 1 | 1 | NA | NA | 2 | 1 | 1 | 1 | NA | NA | NA | NA | 1 | 0 | 2 | 0.5 |
| 19_421590 | NA | NA | 1 | 1 | 1 | 1 | NA | NA | 2 | 1 | 1 | 0 | NA | NA | NA | NA | 1 | 1 | 2 | 0.5 |
| 19_421598 | NA | NA | 1 | 1 | 1 | 1 | NA | NA | 2 | 1 | 1 | 0 | NA | NA | NA | NA | 1 | 1 | 2 | 0.5 |
| 19_422149 | NA | NA | 1 | 1 | NA | NA | NA | NA | 1 | 1 | 1 | 1 | NA | NA | NA | NA | 1 | 1 | 2 | 1 |
| 19_423457 | NA | NA | NA | NA | NA | NA | 1 | 1 | 1 | 1 | NA | NA | 1 | 1 | NA | NA | NA | NA | 1 | 1 |
| 19_423461 | NA | NA | NA | NA | NA | NA | 1 | 1 | 1 | 1 | NA | NA | 1 | 1 | NA | NA | NA | NA | 1 | 1 |
| 19_423481 | NA | NA | NA | NA | NA | NA | 1 | 1 | 1 | 1 | NA | NA | 1 | 1 | NA | NA | NA | NA | 1 | 1 |
| 19_424555 | 1 | 0 | NA | NA | 4 | 1 | 1 | 0 | 6 | 0.6667 | 1 | 0 | NA | NA | 2 | 1 | NA | NA | 3 | 0.6667 |
| 19_424560 | 1 | 0 | NA | NA | 5 | 0.4 | 1 | 0 | 7 | 0.2857 | 1 | 0 | NA | NA | 2 | 0.5 | NA | NA | 3 | 0.3333 |
| 19_424589 | 2 | 0.5 | NA | NA | 5 | 0.6 | 1 | 1 | 8 | 0.625 | 1 | 0 | NA | NA | 2 | 1 | NA | NA | 3 | 0.6667 |
| 19_424594 | 2 | 0 | NA | NA | 5 | 0.2 | 1 | 0 | 8 | 0.125 | NA | NA | 1 | 1 | 1 | 1 | NA | NA | 2 | 1 |
| 19_428340 | 2 | 0 | 7 | 0 | 4 | 0.5 | 3 | 0 | 16 | 0.125 | 2 | 0.5 | 4 | 0 | 4 | 0 | 1 | 0 | 11 | 0.0909 |
| 19_431251 | NA | NA | 4 | 0 | 2 | 0.5 | 4 | 0 | 10 | 0.1 | 5 | 0.4 | 3 | 0.3333 | 1 | 0 | 1 | 0 | 10 | 0.3 |
| 19_431416 | NA | NA | NA | NA | 1 | 1 | NA | NA | 1 | 1 | 1 | 0 | 1 | 0 | 1 | 1 | NA | NA | 3 | 0.3333 |
| 19_431450 | NA | NA | 2 | 0.5 | 4 | 0 | 1 | 1 | 7 | 0.2857 | 5 | 0.2 | 2 | 0 | 1 | 0 | NA | NA | 8 | 0.125 |
| 19_431511 | NA | NA | 2 | 0 | 2 | 0.5 | 1 | 0 | 5 | 0.2 | 3 | 0 | 2 | 0.5 | 1 | 0 | NA | NA | 6 | 0.1667 |
| 19_431539 | NA | NA | 3 | 0.3333 | 3 | 0 | 3 | 0 | 9 | 0.1111 | 5 | 0.2 | 2 | 0 | 1 | 0 | 1 | 0 | 9 | 0.1111 |
| 19_432024 | 5 | 0.2 | 10 | 0.5 | 6 | 0.5 | 5 | 0.4 | 26 | 0.4231 | 10 | 0.1 | 10 | 0.3 | 6 | 0.1667 | 5 | 0.6 | 31 | 0.2581 |
| 19_432027 | 5 | 0.2 | 12 | 0.1667 | 5 | 0.2 | 5 | 0.4 | 27 | 0.2222 | 10 | 0.2 | 11 | 0.2727 | 6 | 0.1667 | 5 | 0.4 | 32 | 0.25 |
| 19_432029 | 3 | 1 | 8 | 0.75 | 5 | 1 | 5 | 0.6 | 21 | 0.8095 | 9 | 0.7778 | 9 | 0.6667 | 4 | 0.75 | 4 | 0.75 | 26 | 0.7308 |
| 19_433645 | 14 | 0.1429 | 21 | 0.2381 | 17 | 0.1765 | 22 | 0.1364 | 74 | 0.1757 | 16 | 0.25 | 22 | 0.1364 | 23 | 0.2174 | 10 | 0.1 | 71 | 0.1831 |
| 19_433647 | 14 | 0.1429 | 20 | 0.1 | 16 | 0.125 | 20 | 0.2 | 70 | 0.1429 | 19 | 0.1579 | 22 | 0.0455 | 23 | 0.1304 | 14 | 0.2143 | 78 | 0.1282 |
| 19_433671 | 11 | 0.2727 | 17 | 0.2941 | 17 | 0.4706 | 17 | 0.1765 | 62 | 0.3065 | 17 | 0.2941 | 17 | 0.3529 | 21 | 0.2381 | 10 | 0.1 | 65 | 0.2615 |
| 19_436636 | 3 | 0.6667 | 1 | 0 | 3 | 1 | 4 | 0 | 11 | 0.4545 | 4 | 0.5 | 9 | 0.6667 | 3 | 0 | 9 | 0.5556 | 25 | 0.52 |
| 19_436686 | 5 | 0.4 | 3 | 0.3333 | 1 | 0 | 5 | 0.2 | 14 | 0.2857 | 4 | 0.25 | 6 | 0 | 3 | 0 | 7 | 0.2857 | 20 | 0.15 |
| 19_436689 | 5 | 0.4 | 3 | 1 | 2 | 0.5 | 4 | 0.75 | 14 | 0.6429 | 4 | 0.5 | 6 | 0.6667 | 3 | 0 | 6 | 0.5 | 19 | 0.4737 |
| 19_436702 | 3 | 0 | 4 | 0.5 | 2 | 0 | 3 | 0.6667 | 12 | 0.3333 | 4 | 0.5 | 6 | 0.3333 | 3 | 0.3333 | 6 | 0.6667 | 19 | 0.4737 |
| 19_436722 | 3 | 0 | 4 | 0 | 2 | 0 | 3 | 0.3333 | 12 | 0.0833 | 2 | 0 | 7 | 0 | 3 | 0 | 6 | 0.3333 | 18 | 0.1111 |
| 19_436727 | 4 | 0 | 4 | 0 | 2 | 0 | 3 | 0.3333 | 13 | 0.0769 | 2 | 0 | 6 | 0.5 | 3 | 0 | 6 | 0 | 17 | 0.1765 |
| 19_436744 | 3 | 0.6667 | 2 | 0 | 2 | 0 | 1 | 0 | 8 | 0.25 | 2 | 0 | 5 | 0.2 | 3 | 0 | 5 | 0.2 | 15 | 0.1333 |
| 19_436752 | 4 | 0.25 | 2 | 0 | 1 | 0 | 2 | 0 | 9 | 0.1111 | NA | NA | 3 | 0.3333 | 2 | 0 | 4 | 0 | 9 | 0.1111 |
| 19_436758 | 4 | 0 | 2 | 0.5 | NA | NA | 2 | 0 | 8 | 0.125 | 1 | 0 | 4 | 0 | 2 | 0 | 3 | 0.3333 | 10 | 0.1 |
| 19_436781 | 2 | 0.5 | 2 | 0.5 | 1 | 0 | 3 | 0.3333 | 8 | 0.375 | 2 | 0 | 6 | 0.5 | 3 | 0 | 5 | 0.4 | 16 | 0.3125 |
| 19_436791 | 3 | 0 | 3 | 0.3333 | NA | NA | 2 | 0 | 8 | 0.125 | 2 | 0 | 6 | 0 | 3 | 0 | 5 | 0.2 | 16 | 0.0625 |

| | | | | | | | | | | | | | | | | | | | | |
|---|---|---|---|---|---|---|---|---|---|---|---|---|---|---|---|---|---|---|---|---|
| 19_4368407 | 2 | 1 | 3 | 0 | NA | NA | 1 | 0 | 6 | 0.3333 | 2 | 0 | 6 | 0.1667 | 3 | 0 | 5 | 0.4 | 16 | 0.1875 |
| 19_4368790 | 3 | 0 | 3 | 0.6667 | NA | NA | 2 | 0 | 8 | 0.25 | 2 | 0 | 6 | 0.1667 | 2 | 0 | 5 | 0.4 | 15 | 0.2 |
| 19_4368966 | 3 | 1 | 3 | 1 | NA | NA | 2 | 1 | 8 | 1 | 2 | 1 | 5 | 1 | 3 | 1 | 3 | 1 | 13 | 1 |
| 19_4369106 | 3 | 0.6667 | 2 | 0.5 | NA | NA | 2 | 0 | 7 | 0.4286 | 1 | 0 | 6 | 0.1667 | 3 | 0.3333 | 2 | 0 | 12 | 0.1667 |
| 19_4369111 | 2 | 0.5 | 2 | 0.5 | NA | NA | 2 | 1 | 6 | 0.6667 | 1 | 0 | 5 | 0.6 | 3 | 0.3333 | 1 | 1 | 10 | 0.5 |
| 19_4369126 | 1 | 0 | 3 | 0 | NA | NA | 1 | 1 | 5 | 0.2 | 1 | 0 | 5 | 0.6 | 2 | 0 | 2 | 0.5 | 10 | 0.4 |
| 19_4369149 | 3 | 1 | 3 | 0.6667 | NA | NA | 1 | 1 | 7 | 0.8571 | NA | NA | 4 | 1 | 2 | 1 | 1 | 1 | 7 | 1 |
| 19_4369189 | 3 | 0.3333 | 2 | 0 | NA | NA | 2 | 0.5 | 7 | 0.2857 | NA | NA | 5 | 0.6 | 2 | 0 | 1 | 1 | 8 | 0.5 |
| 19_4374406 | 2 | 0.5 | NA | NA | NA | NA | 2 | 1 | 4 | 0.75 | 1 | 1 | 4 | 0.75 | 1 | 0 | NA | NA | 6 | 0.6667 |
| 19_4374693 | 2 | 0.5 | NA | NA | NA | NA | 1 | 0 | 3 | 0.3333 | NA | NA | 2 | 0.5 | 1 | 0 | NA | NA | 3 | 0.3333 |
| 19_4374921 | 2 | 0.5 | NA | NA | NA | NA | 2 | 0.5 | 4 | 0.5 | NA | NA | 2 | 0.5 | 1 | 0 | NA | NA | 3 | 0.3333 |
| 19_4393071 | NA | NA | NA | NA | NA | NA | 1 | 1 | 1 | 1 | NA | NA | 1 | 1 | NA | NA | NA | NA | 1 | 1 |
| 19_4495385 | 1 | 1 | 2 | 0.5 | 2 | 0 | 5 | 0.6 | 10 | 0.5 | 2 | 0.5 | 4 | 0 | 3 | 0.3333 | 4 | 0.5 | 13 | 0.3077 |
| 19_4496103 | 1 | 0 | 6 | 0 | 1 | 0 | 2 | 0.5 | 10 | 0.1 | NA | NA | 3 | 0 | 1 | 0 | 6 | 0.1667 | 10 | 0.1 |
| 19_4496220 | NA | NA | 8 | 0.125 | 1 | 0 | 3 | 0 | 12 | 0.0833 | NA | NA | 3 | 0 | NA | NA | 6 | 0.1667 | 9 | 0.1111 |
| 19_4496736 | 1 | 1 | 7 | 0.1429 | 1 | 0 | 2 | 0 | 11 | 0.1818 | NA | NA | 3 | 0 | 2 | 0 | 4 | 0.25 | 9 | 0.1111 |
| 19_4497045 | 1 | 1 | 5 | 0.4 | NA | NA | 1 | 1 | 7 | 0.5714 | NA | NA | 3 | 0.3333 | 2 | 0.5 | 3 | 0.3333 | 8 | 0.375 |
| 19_4497221 | NA | NA | 4 | 0.5 | NA | NA | 2 | 1 | 6 | 0.6667 | NA | NA | 3 | 0.3333 | 2 | 1 | 2 | 0 | 7 | 0.4286 |
| 19_4497356 | 5 | 0.2 | 6 | 0.1667 | 2 | 0 | 4 | 0 | 17 | 0.1176 | 7 | 0.2857 | 5 | 0.6 | 4 | 0 | 6 | 0.3333 | 22 | 0.3182 |
| 19_4497391 | 5 | 0.4 | 7 | 0 | 2 | 0.5 | 4 | 0.75 | 18 | 0.3333 | 6 | 0.5 | 5 | 0.2 | 3 | 0 | 7 | 0.4286 | 21 | 0.3333 |
| 19_4497460 | 6 | 0.6667 | 6 | 0.6667 | 2 | 0 | 4 | 0.75 | 18 | 0.6111 | 8 | 0.25 | 4 | 0.25 | 3 | 0 | 5 | 1 | 20 | 0.4 |
| 19_4497469 | 6 | 0 | 8 | 0.125 | 2 | 0 | 4 | 0 | 20 | 0.05 | 8 | 0.125 | 5 | 0.4 | 4 | 0.25 | 6 | 0 | 23 | 0.1739 |
| 19_4497515 | 5 | 0.2 | 8 | 0.125 | 2 | 0 | 4 | 0.25 | 19 | 0.1579 | 6 | 0 | 5 | 0.2 | 4 | 0 | 6 | 0.1667 | 21 | 0.0952 |
| 19_4497553 | 3 | 0.6667 | 3 | 1 | 2 | 0.5 | 4 | 1 | 12 | 0.8333 | 6 | 0.3333 | 4 | 0.75 | 3 | 0.3333 | 3 | 1 | 16 | 0.5625 |
| 19_4497615 | 3 | 0.3333 | 1 | 0 | 2 | 0 | 1 | 0 | 7 | 0.1429 | 5 | 0.2 | 1 | 1 | 3 | 0.3333 | 3 | 0 | 12 | 0.25 |
| 19_4497719 | 5 | 0 | 6 | 0 | 2 | 0.5 | 3 | 0 | 16 | 0.0625 | 7 | 0 | 5 | 0.2 | 3 | 0 | 5 | 0.2 | 20 | 0.1 |
| 19_4497836 | 4 | 0 | 5 | 0.2 | 2 | 0 | 4 | 0 | 15 | 0.0667 | 6 | 0 | 5 | 0 | 2 | 0 | 5 | 0.2 | 18 | 0.0556 |
| 19_4506662 | 5 | 0.8 | 4 | 0.25 | 1 | 1 | 2 | 1 | 12 | 0.6667 | 6 | 1 | 1 | 1 | 2 | 1 | 4 | 1 | 13 | 1 |
| 19_4506740 | 4 | 0.25 | 4 | 0 | 1 | 0 | 4 | 0 | 13 | 0.0769 | 4 | 0.25 | 1 | 0 | NA | NA | 3 | 0 | 8 | 0.125 |
| 19_4506782 | 5 | 0.8 | 2 | 0.5 | 1 | 1 | 3 | 0.6667 | 11 | 0.7273 | 4 | 0.5 | 1 | 0 | NA | NA | 3 | 0.6667 | 8 | 0.5 |
| 19_4506787 | 4 | 0.25 | 4 | 0 | 1 | 1 | 4 | 0 | 13 | 0.1538 | 2 | 0 | 1 | 0 | NA | NA | 3 | 0.3333 | 6 | 0.1667 |
| 19_4661913 | 3 | 0.6667 | 6 | 0.1667 | 11 | 0.0909 | 3 | 0 | 23 | 0.1739 | 4 | 0.25 | 3 | 0.3333 | 2 | 0.5 | 5 | 0 | 14 | 0.2143 |
| 19_4661955 | 3 | 0 | 5 | 0 | 10 | 0.1 | 2 | 0 | 20 | 0.05 | 5 | 0.4 | 3 | 0 | 3 | 0 | 7 | 0 | 18 | 0.1111 |
| 19_4662256 | 1 | 0 | 7 | 0.4286 | 11 | 0.3636 | 5 | 0 | 24 | 0.2917 | 4 | 0.25 | 3 | 0 | 3 | 0.3333 | 6 | 0 | 16 | 0.125 |
| 19_4662360 | 1 | 0 | 7 | 0.2857 | 9 | 0.4444 | 3 | 0 | 20 | 0.3 | 5 | 0.2 | 3 | 0.3333 | 3 | 0 | 5 | 0.2 | 16 | 0.1875 |
| 19_4662461 | 1 | 1 | 7 | 0.7143 | 9 | 0.3333 | 3 | 0.3333 | 20 | 0.5 | 5 | 0.6 | 3 | 0.6667 | 3 | 1 | 3 | 0.3333 | 14 | 0.6429 |
| 19_4662467 | 1 | 1 | 8 | 0.625 | 11 | 0.9091 | 5 | 0.8 | 25 | 0.8 | 5 | 0.8 | 3 | 1 | 2 | 0.5 | 4 | 0.5 | 14 | 0.7143 |
| 19_4662546 | 1 | 1 | 5 | 1 | 8 | 0.875 | 5 | 0.8 | 19 | 0.8947 | 4 | 1 | NA | NA | 2 | 0.5 | 3 | 0.6667 | 9 | 0.7778 |
| 19_4663329 | 1 | 0 | 6 | 0.8333 | 7 | 0.4286 | 5 | 0.6 | 19 | 0.5789 | 3 | 1 | 3 | 0.6667 | 3 | 0.3333 | 5 | 1 | 14 | 0.7857 |
| 19_4663350 | NA | NA | 5 | 0.8 | 6 | 0.6667 | 4 | 1 | 15 | 0.8 | 6 | 0.6667 | 2 | 1 | 3 | 0.6667 | 5 | 0.6 | 16 | 0.6875 |
| 19_4663370 | 1 | 0 | 7 | 1 | 7 | 0.7143 | 4 | 1 | 19 | 0.8421 | 5 | 1 | 3 | 1 | 3 | 1 | 4 | 1 | 15 | 1 |
| 19_4663530 | 1 | 0 | 6 | 1 | 6 | 1 | 5 | 1 | 18 | 0.9444 | 5 | 1 | 2 | 1 | 3 | 1 | 2 | 1 | 12 | 1 |
| 19_4663577 | 1 | 0 | 7 | 0 | 7 | 0.1429 | 4 | 0.25 | 19 | 0.1053 | 5 | 0 | 3 | 0.3333 | 3 | 0.3333 | 4 | 0 | 15 | 0.1333 |
| 19_4663613 | 1 | 0 | 7 | 0.1429 | 7 | 0 | 5 | 0 | 20 | 0.05 | 5 | 0.2 | 3 | 0 | 3 | 0 | 5 | 0 | 16 | 0.0625 |
| 19_4664003 | 1 | 1 | 6 | 1 | 4 | 1 | 3 | 1 | 14 | 1 | 5 | 0.6 | 3 | 1 | 3 | 1 | 4 | 1 | 15 | 0.8667 |
| 19_4664009 | 1 | 0 | 3 | 0.3333 | 3 | 0.3333 | 5 | 0 | 12 | 0.1667 | 5 | 0.4 | 3 | 0 | 2 | 0 | 3 | 0 | 13 | 0.1538 |
| 19_4664087 | 1 | 0 | 6 | 0.3333 | 4 | 0 | 5 | 0.4 | 16 | 0.25 | 5 | 0.2 | 3 | 0.3333 | 3 | 0.3333 | 4 | 0.25 | 15 | 0.2667 |
| 19_4664252 | NA | NA | 5 | 0.8 | 4 | 1 | 4 | 0.75 | 13 | 0.8462 | 5 | 0.6 | 3 | 1 | 3 | 0.6667 | 3 | 0.6667 | 14 | 0.7143 |
| 19_4664306 | NA | NA | 4 | 1 | 3 | 1 | 5 | 1 | 12 | 1 | 4 | 1 | 3 | 0.6667 | 3 | 1 | 4 | 1 | 14 | 0.9286 |
| 19_4664333 | NA | NA | 4 | 0.25 | 4 | 0 | 4 | 0 | 12 | 0.0833 | 3 | 0.3333 | 2 | 0 | 2 | 0 | 2 | 1 | 9 | 0.3333 |
| 19_4664341 | NA | NA | 3 | 0.3333 | 4 | 0.5 | 5 | 0.4 | 12 | 0.4167 | 3 | 0.3333 | 2 | 0.5 | 3 | 0 | 3 | 0.3333 | 11 | 0.2727 |
| 19_4664377 | NA | NA | 4 | 0.75 | 4 | 0.75 | 4 | 0.25 | 12 | 0.5833 | 5 | 0.6 | 2 | 1 | 3 | 0.3333 | 3 | 0.6667 | 13 | 0.6154 |
| 19_4664459 | NA | NA | 5 | 0.2 | 4 | 0.25 | 3 | 0 | 12 | 0.1667 | 5 | 0.2 | 3 | 0 | 3 | 0 | 4 | 0.25 | 15 | 0.1333 |
| 19_4664614 | NA | NA | NA | NA | 2 | 1 | NA | NA | 2 | 1 | 1 | 0 | 2 | 0.5 | 1 | 0 | 1 | 0 | 5 | 0.2 |
| 19_4664757 | NA | NA | 1 | 1 | 2 | 0 | 1 | 0 | 4 | 0.25 | 2 | 0.5 | 2 | 0 | 1 | 0 | 1 | 0 | 6 | 0.1667 |
| 19_4668266 | 4 | 0.25 | 4 | 0 | NA | NA | 6 | 0 | 14 | 0.0714 | 5 | 0 | 3 | 0 | 3 | 0.3333 | 7 | 0.1429 | 18 | 0.1111 |
| 19_4668383 | 3 | 0.3333 | 2 | 0 | NA | NA | 5 | 0 | 10 | 0.1 | 5 | 0.4 | 3 | 0 | 4 | 0.25 | 6 | 0.1667 | 18 | 0.2222 |
| 19_4669582 | 5 | 0 | 4 | 0.5 | 2 | 0.5 | 8 | 0.125 | 19 | 0.2105 | 7 | 0 | 4 | 0.5 | 4 | 0.25 | 7 | 0 | 22 | 0.1364 |
| 19_4669863 | 4 | 0.25 | 5 | 0 | 2 | 0.5 | 9 | 0 | 20 | 0.1 | 7 | 0.1429 | 4 | 0 | 5 | 0 | 7 | 0.1429 | 23 | 0.087 |
| 19_4671218 | 7 | 0.2857 | 9 | 0.1111 | 5 | 0.4 | 10 | 0.2 | 31 | 0.2258 | 7 | 0.2857 | 10 | 0.5 | 5 | 0.2 | 8 | 0.25 | 30 | 0.3333 |
| 19_4671273 | 9 | 0.4444 | 3 | 0.3333 | 2 | 0.5 | 10 | 0.2 | 24 | 0.3333 | 8 | 0.625 | 9 | 0.3333 | 5 | 0.6 | 8 | 0.375 | 30 | 0.4667 |
| 19_4671449 | 9 | 0.1111 | 8 | 0.5 | 5 | 0 | 11 | 0.0909 | 33 | 0.1818 | 8 | 0.125 | 8 | 0.5 | 4 | 0.25 | 8 | 0.625 | 28 | 0.3929 |
| 19_4672659 | 4 | 0 | 4 | 0.25 | 7 | 0.1429 | 6 | 0 | 21 | 0.0952 | 6 | 0.1667 | 2 | 0 | 5 | 0.2 | 9 | 0 | 22 | 0.0909 |
| 19_4672781 | 3 | 0.6667 | 5 | 0.8 | 7 | 0.5714 | 4 | 0.75 | 19 | 0.6842 | 6 | 0.8333 | 3 | 0.6667 | 5 | 0.4 | 12 | 0.5833 | 26 | 0.6154 |
| 19_4672955 | 1 | 0 | 2 | 0.5 | 5 | 0.4 | 4 | 0.5 | 12 | 0.4167 | 3 | 0 | 2 | 0.5 | 4 | 0 | 8 | 0.625 | 17 | 0.3529 |

| | | | | | | | | | | | | | | | | | | | | |
|---|---|---|---|---|---|---|---|---|---|---|---|---|---|---|---|---|---|---|---|---|
| 19_4673007 | 1 | 1 | 3 | 1 | 4 | 1 | 2 | 0.5 | 10 | 0.9 | 5 | 0.8 | 3 | 0.6667 | 2 | 0.5 | 10 | 0.8 | 20 | 0.75 |
| 19_4673146 | 2 | 0.5 | 3 | 1 | 3 | 0 | 5 | 0.8 | 13 | 0.6154 | 4 | 0.5 | 2 | 1 | 4 | 0.5 | 11 | 0.4545 | 21 | 0.5238 |
| 19_4784279 | 9 | 0.1111 | 6 | 0.1667 | 10 | 0.1 | 5 | 0.2 | 30 | 0.1333 | 4 | 0 | 6 | 0.1667 | 2 | 1 | 6 | 0 | 18 | 0.1667 |
| 19_4784532 | 10 | 0.1 | 6 | 0 | 10 | 0 | 5 | 0.2 | 31 | 0.0645 | 6 | 0 | 6 | 0.1667 | 2 | 0 | 5 | 0 | 19 | 0.0526 |
| 19_4787954 | 16 | 0.0625 | 8 | 0 | 9 | 0.1111 | 8 | 0.125 | 41 | 0.0732 | 8 | 0.125 | 8 | 0.125 | 4 | 0.25 | 8 | 0.125 | 28 | 0.1429 |
| 19_4788249 | 15 | 0.2 | 4 | 0.25 | 8 | 0 | 7 | 0.2857 | 34 | 0.1765 | 9 | 0.1111 | 7 | 0.4286 | 3 | 0 | 10 | 0.1 | 29 | 0.1724 |
| 19_4788278 | 14 | 0.2143 | 6 | 0.1667 | 7 | 0.4286 | 8 | 0.375 | 35 | 0.2857 | 9 | 0.2222 | 11 | 0.1818 | 3 | 0 | 9 | 0.1111 | 32 | 0.1562 |
| 19_4788398 | 18 | 0.5 | 7 | 0.4286 | 11 | 0.2727 | 9 | 0.3333 | 45 | 0.4 | 12 | 0.3333 | 10 | 0.4 | 4 | 0.5 | 13 | 0.3846 | 39 | 0.3846 |
| 19_4796733 | 12 | 0.3333 | 15 | 0.3333 | 16 | 0.1875 | 9 | 0.2222 | 52 | 0.2692 | 12 | 0.3333 | 15 | 0.2667 | 8 | 0.375 | 20 | 0.2 | 55 | 0.2727 |
| 19_4796885 | 20 | 0.25 | 22 | 0.0455 | 21 | 0.0952 | 15 | 0.2667 | 78 | 0.1538 | 18 | 0.0556 | 21 | 0.0476 | 12 | 0.25 | 19 | 0.1053 | 70 | 0.1 |
| 19_4801801 | 37 | 0.0811 | 36 | 0.0556 | 52 | 0.0192 | 25 | 0.08 | 150 | 0.0533 | 33 | 0.0909 | 34 | 0 | 27 | 0.037 | 30 | 0.1 | 124 | 0.0565 |
| 19_4801811 | 37 | 0.7297 | 41 | 0.7317 | 48 | 0.7708 | 28 | 0.7857 | 154 | 0.7532 | 39 | 0.7436 | 41 | 0.6341 | 34 | 0.6471 | 36 | 0.7222 | 150 | 0.6867 |
| 19_4801824 | 35 | 0.0286 | 34 | 0.0588 | 43 | 0.0465 | 28 | 0.0714 | 140 | 0.05 | 35 | 0.1143 | 31 | 0.0323 | 28 | 0.0357 | 29 | 0.1034 | 123 | 0.0732 |
| 19_4801891 | 11 | 0.6364 | 16 | 0.75 | 19 | 0.5789 | 13 | 0.7692 | 59 | 0.678 | 16 | 0.875 | 12 | 0.5833 | 12 | 0.8333 | 13 | 0.5385 | 53 | 0.717 |
| 19_4801904 | 34 | 0.0882 | 38 | 0.0526 | 45 | 0.0444 | 22 | 0.1818 | 139 | 0.0791 | 33 | 0.0606 | 34 | 0.0588 | 28 | 0.1429 | 34 | 0.0882 | 129 | 0.0853 |
| 19_4801960 | 43 | 0.093 | 47 | 0.0638 | 59 | 0.0847 | 30 | 0.0667 | 179 | 0.0782 | 44 | 0.1364 | 42 | 0.0714 | 36 | 0.1389 | 43 | 0.1395 | 165 | 0.1212 |
| 19_4802168 | 44 | 0.0455 | 46 | 0.1087 | 59 | 0.0847 | 35 | 0.0286 | 184 | 0.0707 | 44 | 0.1136 | 43 | 0.0698 | 36 | 0.0556 | 40 | 0.075 | 163 | 0.0798 |
| 19_4802192 | 46 | 0.087 | 46 | 0.0435 | 58 | 0.0345 | 35 | 0.0857 | 185 | 0.0595 | 47 | 0.0638 | 42 | 0.0476 | 34 | 0.0294 | 40 | 0.075 | 163 | 0.0552 |
| 19_4802267 | 38 | 0.3421 | 45 | 0.1778 | 49 | 0.2041 | 34 | 0.1765 | 166 | 0.2229 | 39 | 0.0769 | 36 | 0.0833 | 29 | 0.2414 | 33 | 0.2727 | 137 | 0.1606 |
| 19_4807416 | 4 | 0.5 | 2 | 0.5 | 3 | 0 | 1 | 0 | 10 | 0.3 | NA | NA | 2 | 0.5 | 2 | 0 | 2 | 0.5 | 6 | 0.3333 |
| 19_4807702 | NA | NA | 2 | 1 | NA | NA | 1 | 1 | 3 | 1 | NA | NA | 1 | 1 | NA | NA | NA | NA | 1 | 1 |
| 19_4808345 | 1 | 1 | 1 | 0 | NA | NA | 1 | 1 | 3 | 0.6667 | NA | NA | 1 | 1 | NA | NA | NA | NA | 1 | 1 |
| 19_4812046 | 3 | 0.3333 | 1 | 1 | 1 | 0 | NA | NA | 5 | 0.4 | NA | NA | 1 | 1 | 2 | 0.5 | 2 | 0.5 | 5 | 0.6 |
| 19_4812099 | 3 | 1 | 1 | 1 | 2 | 1 | NA | NA | 6 | 1 | NA | NA | 1 | 1 | 3 | 1 | 2 | 1 | 6 | 1 |
| 19_4812146 | 3 | 1 | 1 | 1 | 1 | 1 | NA | NA | 5 | 1 | NA | NA | 1 | 0 | 3 | 1 | 2 | 1 | 6 | 0.8333 |
| 19_4812168 | 3 | 1 | NA | NA | 2 | 1 | NA | NA | 5 | 1 | NA | NA | 1 | 1 | 2 | 1 | 2 | 1 | 5 | 1 |
| 19_4812248 | 2 | 0 | 1 | 1 | 2 | 1 | NA | NA | 5 | 0.6 | NA | NA | 1 | 0 | 3 | 0.6667 | 1 | 0 | 5 | 0.4 |
| 19_4812411 | 3 | 0.6667 | 1 | 1 | 2 | 1 | NA | NA | 6 | 0.8333 | NA | NA | NA | NA | 3 | 0.6667 | 2 | 0.5 | 5 | 0.6 |
| 19_4812425 | 2 | 1 | 1 | 1 | 2 | 1 | NA | NA | 5 | 1 | NA | NA | 1 | 1 | 3 | 1 | 1 | 1 | 5 | 1 |
| 19_4812491 | 3 | 0.3333 | 1 | 0 | 2 | 0.5 | NA | NA | 6 | 0.3333 | NA | NA | 1 | 0 | 4 | 0.75 | NA | NA | 5 | 0.6 |
| 19_4812641 | 4 | 0.25 | 1 | 0 | 1 | 1 | NA | NA | 6 | 0.3333 | NA | NA | 1 | 0 | 3 | 0 | 2 | 0.5 | 6 | 0.1667 |
| 19_4812669 | 3 | 0 | 1 | 1 | 2 | 0.5 | NA | NA | 6 | 0.3333 | NA | NA | 1 | 1 | 3 | 0 | 2 | 0.5 | 6 | 0.3333 |
| 19_4812764 | NA | NA | NA | NA | 1 | 1 | NA | NA | 1 | 1 | NA | NA | NA | NA | 1 | 1 | NA | NA | 1 | 1 |
| 19_4815770 | 2 | 0 | NA | NA | 2 | 0.5 | NA | NA | 4 | 0.25 | NA | NA | 1 | 1 | 2 | 1 | 1 | 0 | 4 | 0.75 |
| 19_4834660 | NA | NA | 2 | 0.5 | NA | NA | 1 | 1 | 3 | 0.6667 | 2 | 0.5 | 1 | 0 | 1 | 0 | 1 | 1 | 5 | 0.4 |
| 19_4835235 | 3 | 0 | 4 | 0 | 2 | 0 | 6 | 0.3333 | 15 | 0.1333 | 7 | 0 | 4 | 0.25 | 2 | 0 | 4 | 0.25 | 17 | 0.1176 |
| 19_4835478 | 3 | 0 | 4 | 0.75 | 2 | 0 | 6 | 0.5 | 15 | 0.4 | 7 | 0.2857 | 4 | 0 | 2 | 0 | 4 | 0.25 | 17 | 0.1765 |
| 19_4837503 | 1 | 1 | 4 | 0.25 | 1 | 0 | 4 | 0 | 10 | 0.2 | 4 | 0 | 3 | 0 | 2 | 0 | 1 | 1 | 10 | 0.1 |
| 19_4837658 | 1 | 1 | 3 | 0.6667 | NA | NA | 3 | 1 | 7 | 0.8571 | 4 | 0.25 | 3 | 0 | 2 | 0.5 | 2 | 0.5 | 11 | 0.2727 |
| 19_4837710 | 1 | 1 | 4 | 0 | NA | NA | 3 | 0 | 8 | 0.125 | 3 | 0 | 3 | 0.3333 | 1 | 1 | 2 | 0.5 | 9 | 0.3333 |
| 19_4837738 | 1 | 1 | 3 | 0.6667 | NA | NA | 2 | 1 | 6 | 0.8333 | 4 | 1 | 3 | 0.3333 | 2 | 0.5 | 1 | 1 | 10 | 0.7 |
| 19_4837752 | 1 | 1 | 4 | 0.5 | NA | NA | 3 | 0.3333 | 8 | 0.5 | 2 | 0.5 | 3 | 0.6667 | 1 | 0 | 1 | 0 | 7 | 0.4286 |
| 19_4837929 | 1 | 0 | 4 | 0.75 | NA | NA | 3 | 0.6667 | 8 | 0.625 | 4 | 0.5 | 3 | 0.3333 | 1 | 1 | 1 | 1 | 9 | 0.5556 |
| 19_4842515 | 3 | 0 | 3 | 0 | 1 | 0 | 3 | 0.3333 | 10 | 0.1 | 7 | 0.1429 | 2 | 0 | 1 | 0 | 4 | 0 | 14 | 0.0714 |
| 19_4842707 | 3 | 0.3333 | 3 | 0 | NA | NA | 4 | 0.25 | 10 | 0.2 | 5 | 0.2 | NA | NA | 1 | 0 | 3 | 0.3333 | 9 | 0.2222 |
| 19_4842743 | 3 | 0.3333 | 3 | 0 | NA | NA | 5 | 0 | 11 | 0.0909 | 6 | 0.1667 | 2 | 0 | 1 | 0 | 3 | 0.3333 | 12 | 0.1667 |
| 19_4850733 | NA | NA | 1 | 1 | 1 | 1 | NA | NA | 2 | 1 | 1 | 1 | 1 | 1 | NA | NA | 1 | 1 | 3 | 1 |
| 19_4850987 | 2 | 0 | 4 | 0 | 2 | 0 | 8 | 0.25 | 16 | 0.125 | 5 | 0.4 | 5 | 0.2 | 4 | 0 | 5 | 0.6 | 19 | 0.3158 |
| 19_4851065 | 4 | 0 | 5 | 0 | 3 | 0 | 8 | 0.25 | 20 | 0.1 | 4 | 0 | 5 | 0.2 | 5 | 0 | 4 | 0 | 18 | 0.0556 |
| 19_4851139 | 4 | 0.75 | 5 | 0 | 3 | 0 | 7 | 0.1429 | 19 | 0.2105 | 3 | 0 | 5 | 0.4 | 6 | 0 | 5 | 0.4 | 19 | 0.2105 |
| 19_4853594 | 2 | 1 | 3 | 1 | 2 | 1 | 3 | 1 | 10 | 1 | 3 | 1 | 1 | 1 | 2 | 1 | 3 | 1 | 9 | 1 |
| 19_4853866 | 2 | 0.5 | 2 | 0 | 2 | 0 | 3 | 0 | 9 | 0.1111 | 3 | 0.3333 | 1 | 0 | 3 | 0.3333 | 4 | 0.25 | 11 | 0.2727 |
| 19_4853888 | NA | NA | 3 | 0.3333 | 1 | 0 | 1 | 0 | 5 | 0.2 | 2 | 0 | NA | NA | 1 | 0 | 2 | 0.5 | 5 | 0.2 |
| 19_4853893 | 2 | 1 | 3 | 0 | 2 | 0 | 3 | 0.3333 | 10 | 0.3 | 2 | 1 | 1 | 0 | 2 | 0 | 4 | 0.25 | 9 | 0.3333 |
| 19_4853946 | 2 | 0 | 3 | 0.3333 | 2 | 0.5 | 1 | 0 | 8 | 0.25 | 3 | 0.3333 | 1 | 0 | NA | NA | 4 | 0.75 | 8 | 0.5 |
| 19_4875567 | NA | NA | 7 | 0.7143 | NA | NA | 4 | 0.25 | 11 | 0.5455 | 6 | 0.6667 | 3 | 0.3333 | 2 | 0.5 | 4 | 0.5 | 15 | 0.5333 |
| 19_4906153 | 7 | 0.1429 | 5 | 0 | 6 | 0 | 8 | 0.125 | 26 | 0.0769 | 6 | 0 | 2 | 0 | 7 | 0.1429 | 3 | 0 | 18 | 0.0556 |
| 19_4906597 | 6 | 0 | 4 | 0.25 | 6 | 0.1667 | 7 | 0.1429 | 23 | 0.1304 | 7 | 0.1429 | 3 | 0.3333 | 5 | 0.2 | 3 | 0.3333 | 18 | 0.2222 |
| 19_4908320 | 7 | 0 | 6 | 0.6667 | 9 | 0.1111 | 9 | 0.1111 | 31 | 0.1935 | 8 | 0.25 | 3 | 0 | 6 | 0.1667 | 2 | 0 | 19 | 0.1579 |
| 19_4908516 | 7 | 0.1429 | 8 | 0 | 7 | 0.1429 | 8 | 0 | 30 | 0.0667 | 8 | 0 | 4 | 0.5 | 8 | 0.125 | 4 | 0.25 | 24 | 0.1667 |
| 19_4910489 | 6 | 0 | 9 | 0.1111 | 10 | 0.2 | 10 | 0.1 | 35 | 0.1143 | 9 | 0.1111 | 7 | 0.2857 | 14 | 0.2143 | 4 | 0 | 34 | 0.1765 |
| 19_4910510 | 7 | 0 | 10 | 0.2 | 10 | 0.1 | 8 | 0 | 35 | 0.0857 | 10 | 0 | 8 | 0 | 14 | 0.2143 | 5 | 0.2 | 37 | 0.1081 |
| 19_4910606 | 5 | 0.2 | 7 | 0.1429 | 11 | 0.3636 | 10 | 0.3 | 33 | 0.2727 | 10 | 0.1 | 6 | 0.6667 | 9 | 0.1111 | 3 | 0.3333 | 28 | 0.25 |
| 19_4933546 | 2 | 1 | 1 | 1 | 3 | 0.3333 | 1 | 1 | 7 | 0.7143 | NA | NA | 4 | 0.25 | NA | NA | NA | NA | 4 | 0.25 |
| 19_4933616 | 3 | 0.6667 | 1 | 1 | 3 | 0.3333 | 1 | 0 | 8 | 0.5 | 1 | 1 | 4 | 0 | NA | NA | NA | NA | 5 | 0.2 |

| | | | | | | | | | | | | | | | | | | | | |
|---|---|---|---|---|---|---|---|---|---|---|---|---|---|---|---|---|---|---|---|---|
| 19_493379 | 3 | 0.6667 | 1 | 0 | 3 | 0 | 1 | 1 | 8 | 0.375 | 1 | 0 | 4 | 0.25 | NA | NA | NA | NA | 5 | 0.2 |
| 19_493383 | 3 | 0.6667 | 1 | 0 | 3 | 0.3333 | 1 | 1 | 8 | 0.5 | NA | NA | 3 | 0.3333 | NA | NA | NA | NA | 3 | 0.3333 |
| 19_493384 | 3 | 1 | NA | NA | 3 | 0 | 1 | 1 | 7 | 0.5714 | 1 | 1 | 2 | 0 | NA | NA | NA | NA | 3 | 0.3333 |
| 19_493711 | 5 | 0.4 | 1 | 1 | 4 | 0 | 6 | 0.3333 | 16 | 0.3125 | 4 | 0.25 | 3 | 0.3333 | 2 | 0 | 5 | 0.6 | 14 | 0.3571 |
| 19_493730 | 6 | 0.5 | NA | NA | 5 | 0.8 | 3 | 0.6667 | 14 | 0.6429 | 4 | 1 | 3 | 1 | 1 | 1 | 3 | 1 | 11 | 1 |
| 19_493736 | 6 | 0.1667 | NA | NA | 5 | 0.4 | 5 | 0 | 16 | 0.1875 | 4 | 0 | 4 | 0.25 | 2 | 0 | 4 | 0.75 | 14 | 0.2857 |
| 19_493738 | 5 | 0.8 | 1 | 1 | 4 | 1 | 5 | 1 | 15 | 0.9333 | 4 | 0.5 | 4 | 1 | 2 | 1 | 4 | 0.75 | 14 | 0.7857 |
| 19_493746 | 6 | 0.8333 | 1 | 1 | 4 | 1 | 3 | 1 | 14 | 0.9286 | 4 | 1 | 3 | 1 | 2 | 1 | 3 | 0.3333 | 12 | 0.8333 |
| 19_493747 | 5 | 0.4 | 1 | 0 | 4 | 0.25 | 5 | 0.4 | 15 | 0.3333 | 3 | 0.3333 | 3 | 0.3333 | 2 | 0 | 5 | 0.4 | 13 | 0.3077 |
| 19_493748 | 6 | 0.8333 | 1 | 1 | 4 | 0.75 | 5 | 0.8 | 16 | 0.8125 | 3 | 1 | 4 | 0.75 | 2 | 0.5 | 6 | 0.8333 | 15 | 0.8 |
| 19_493761 | 4 | 0.25 | 1 | 0 | 3 | 1 | 4 | 0.75 | 12 | 0.5833 | 4 | 0.5 | 4 | 0.75 | 2 | 0.5 | 6 | 0.6667 | 16 | 0.625 |
| 19_493780 | 5 | 0.6 | 1 | 0 | 2 | 0.5 | 5 | 0.2 | 13 | 0.3846 | 4 | 0.25 | 4 | 0 | 1 | 0 | 6 | 0.1667 | 15 | 0.1333 |
| 19_493789 | 6 | 0.3333 | 1 | 0 | 4 | 0.25 | 5 | 0 | 16 | 0.1875 | 3 | 0 | 3 | 0 | 1 | 0 | 5 | 0.2 | 12 | 0.0833 |
| 19_493878 | 5 | 0 | 1 | 0 | 4 | 0.5 | 2 | 0 | 12 | 0.1667 | 5 | 0.2 | 4 | 0.25 | 2 | 0 | 5 | 0.2 | 16 | 0.1875 |
| 19_493928 | 4 | 0.5 | 1 | 1 | 4 | 0.25 | 3 | 0 | 12 | 0.3333 | 3 | 0.3333 | 3 | 0.3333 | 1 | 0 | 4 | 0 | 11 | 0.1818 |
| 19_493934 | 3 | 0 | 1 | 0 | 2 | 0.5 | 3 | 0 | 9 | 0.1111 | 3 | 0.3333 | 3 | 0.3333 | 1 | 0 | 3 | 0.3333 | 10 | 0.3 |
| 19_493938 | 2 | 0.5 | 1 | 0 | 2 | 0 | 3 | 0 | 8 | 0.125 | 5 | 0.2 | 2 | 0.5 | 1 | 0 | 3 | 0 | 11 | 0.1818 |
| 19_493939 | 4 | 0.25 | 1 | 0 | 4 | 0.75 | 3 | 0 | 12 | 0.3333 | 5 | 0.4 | 3 | 0.3333 | 1 | 1 | 4 | 0.25 | 13 | 0.3846 |
| 19_493940 | 4 | 0.75 | 1 | 0 | 2 | 1 | 2 | 0 | 9 | 0.5556 | 4 | 1 | 3 | 0.6667 | 1 | 1 | 3 | 0.6667 | 11 | 0.8182 |
| 19_493945 | 4 | 0.25 | 1 | 1 | 1 | 0 | 1 | 0 | 7 | 0.2857 | 5 | 0.8 | 1 | 1 | NA | NA | 3 | 0.6667 | 9 | 0.7778 |
| 19_493950 | 3 | 0 | 1 | 1 | 3 | 0 | 3 | 0 | 10 | 0.1 | 4 | 0.25 | 3 | 0 | 1 | 0 | 3 | 0 | 11 | 0.0909 |
| 19_493952 | 4 | 0 | 1 | 1 | 1 | 1 | 3 | 0.3333 | 9 | 0.3333 | 5 | 0 | 3 | 0 | 1 | 0 | 3 | 0.3333 | 12 | 0.0833 |
| 19_493961 | 3 | 0.6667 | NA | NA | 3 | 0.3333 | 2 | 0 | 8 | 0.375 | 4 | 0.25 | 1 | 1 | 1 | 0 | 2 | 1 | 8 | 0.5 |
| 19_493962 | 3 | 1 | 1 | 1 | 3 | 1 | 3 | 0.6667 | 10 | 0.9 | 2 | 1 | 3 | 1 | 1 | 1 | 2 | 1 | 8 | 1 |
| 19_493981 | 3 | 0.3333 | 1 | 0 | 2 | 1 | 3 | 0.3333 | 9 | 0.4444 | 2 | 0.5 | 2 | 0.5 | 1 | 1 | 3 | 0.3333 | 8 | 0.5 |
| 19_493998 | 4 | 0.25 | 1 | 0 | 1 | 0 | 1 | 1 | 7 | 0.2857 | 4 | 0.5 | 2 | 0 | 1 | 0 | 3 | 0.6667 | 10 | 0.4 |
| 19_493999 | 4 | 0.25 | NA | NA | 1 | 1 | 3 | 0 | 8 | 0.25 | 4 | 0.25 | 1 | 0 | 1 | 0 | 3 | 0.6667 | 9 | 0.3333 |
| 19_494054 | 3 | 0.3333 | NA | NA | 1 | 0 | 2 | 0 | 6 | 0.1667 | 2 | 0 | NA | NA | 1 | 0 | 3 | 0.3333 | 6 | 0.1667 |
| 19_494064 | 4 | 0.25 | NA | NA | NA | NA | 2 | 0.5 | 6 | 0.3333 | 2 | 0 | NA | NA | 1 | 0 | 3 | 0.3333 | 6 | 0.1667 |
| 19_495737 | 3 | 0 | 6 | 0.3333 | 6 | 0.3333 | 4 | 0 | 19 | 0.2105 | 3 | 0.3333 | 7 | 0 | 2 | 0 | 1 | 1 | 13 | 0.1538 |
| 19_495754 | 3 | 1 | 7 | 0.2857 | 4 | 0.75 | 3 | 0.3333 | 17 | 0.5294 | 3 | 1 | 7 | 0.8571 | 1 | 1 | 1 | 1 | 12 | 0.9167 |
| 19_498012 | 2 | 0.5 | 3 | 0 | NA | NA | 3 | 0 | 8 | 0.125 | 4 | 0.25 | 3 | 0 | 3 | 0 | NA | NA | 10 | 0.1 |
| 19_498023 | 2 | 1 | 3 | 0.6667 | NA | NA | 2 | 0 | 7 | 0.5714 | 4 | 0.25 | 3 | 0.3333 | 3 | 0 | NA | NA | 10 | 0.2 |
| 19_498029 | 2 | 1 | 2 | 1 | NA | NA | 3 | 0.3333 | 7 | 0.7143 | 3 | 0.6667 | 3 | 0.3333 | 3 | 0.3333 | NA | NA | 9 | 0.4444 |
| 19_498031 | 1 | 0 | 2 | 0 | NA | NA | 2 | 0.5 | 5 | 0.2 | 3 | 0.3333 | 3 | 0.6667 | 2 | 0 | NA | NA | 8 | 0.375 |
| 19_498034 | 2 | 0.5 | 2 | 1 | NA | NA | 3 | 1 | 7 | 0.8571 | 2 | 1 | 3 | 1 | 3 | 1 | NA | NA | 8 | 1 |
| 19_498037 | 1 | 0 | 2 | 0 | NA | NA | 3 | 0.3333 | 6 | 0.1667 | 3 | 0.3333 | 2 | 0 | 2 | 0 | NA | NA | 7 | 0.1429 |
| 19_498042 | NA | NA | 2 | 0.5 | NA | NA | 3 | 1 | 5 | 0.8 | 3 | 1 | 3 | 1 | 2 | 0.5 | NA | NA | 8 | 0.875 |
| 19_498047 | 1 | 1 | 1 | 1 | NA | NA | 2 | 0.5 | 4 | 0.75 | 1 | 1 | 3 | 0.3333 | 3 | 1 | NA | NA | 7 | 0.7143 |
| 19_498198 | 1 | 0 | NA | NA | 3 | 0 | 1 | 1 | 5 | 0.2 | 1 | 1 | 2 | 0.5 | 2 | 0.5 | NA | NA | 5 | 0.6 |
| 19_498212 | 1 | 1 | NA | NA | 3 | 0.3333 | 1 | 1 | 5 | 0.6 | 1 | 1 | 1 | 1 | 2 | 1 | NA | NA | 4 | 1 |
| 19_498224 | 1 | 1 | NA | NA | 2 | 1 | 1 | 1 | 4 | 1 | 1 | 1 | 2 | 1 | 1 | 1 | NA | NA | 4 | 1 |
| 19_498231 | 1 | 1 | NA | NA | 3 | 0.6667 | 1 | 1 | 5 | 0.8 | 1 | 1 | 2 | 1 | 1 | 1 | NA | NA | 4 | 1 |
| 19_498233 | 1 | 1 | NA | NA | 1 | 1 | 1 | 0 | 3 | 0.6667 | 1 | 1 | 2 | 1 | 1 | 1 | NA | NA | 4 | 1 |
| 19_498243 | 1 | 0 | NA | NA | 3 | 0.3333 | 1 | 0 | 5 | 0.2 | 1 | 0 | 1 | 1 | 1 | 0 | NA | NA | 3 | 0.3333 |
| 19_498251 | 1 | 1 | NA | NA | 3 | 1 | 1 | 1 | 5 | 1 | 1 | 0 | 2 | 0.5 | 1 | 1 | NA | NA | 4 | 0.5 |
| 19_498256 | 1 | 1 | NA | NA | 3 | 0.6667 | 1 | 1 | 5 | 0.8 | 1 | 1 | 2 | 1 | NA | NA | NA | NA | 3 | 1 |
| 19_498325 | NA | NA | NA | NA | NA | NA | 4 | 1 | 4 | 1 | NA | NA | 1 | 1 | 1 | 0 | NA | NA | 2 | 0.5 |
| 19_498418 | NA | NA | NA | NA | NA | NA | 3 | 1 | 3 | 1 | NA | NA | NA | NA | 1 | 1 | 1 | 1 | 2 | 1 |
| 19_498448 | 1 | 0 | NA | NA | 3 | 0.3333 | 1 | 1 | 5 | 0.4 | 1 | 0 | 2 | 0.5 | 2 | 0.5 | 1 | 1 | 6 | 0.5 |
| 19_501488 | 8 | 0.5 | 12 | 0.5833 | 5 | 1 | 9 | 0.4444 | 34 | 0.5882 | 7 | 0.1429 | 5 | 0.2 | 7 | 0.4286 | 13 | 0.4615 | 32 | 0.3438 |
| 19_503460 | 2 | 0.5 | 3 | 1 | NA | NA | NA | NA | 5 | 0.8 | 1 | 0 | 1 | 1 | 1 | 0 | 1 | 0 | 4 | 0.25 |
| 19_503463 | 1 | 0 | 3 | 0.6667 | NA | NA | NA | NA | 4 | 0.5 | 1 | 1 | 1 | 1 | 1 | 1 | NA | NA | 3 | 1 |
| 19_503490 | 3 | 1 | 3 | 1 | NA | NA | NA | NA | 6 | 1 | 1 | 0 | 2 | 1 | 2 | 1 | 1 | 1 | 6 | 0.8333 |
| 19_503502 | 2 | 0.5 | 3 | 1 | NA | NA | NA | NA | 5 | 0.8 | 1 | 0 | 1 | 1 | 1 | 0 | 1 | 1 | 4 | 0.5 |
| 19_503589 | 3 | 0.3333 | 2 | 0.5 | NA | NA | NA | NA | 5 | 0.4 | 1 | 1 | 1 | 1 | 1 | 1 | 1 | 0 | 4 | 0.75 |
| 19_503607 | 2 | 1 | 2 | 1 | NA | NA | NA | NA | 4 | 1 | 1 | 1 | 1 | 1 | 1 | 1 | 1 | 1 | 4 | 1 |
| 19_503615 | 3 | 0.6667 | 1 | 1 | NA | NA | NA | NA | 4 | 0.75 | 1 | 0 | 1 | 0 | 1 | 0 | 1 | 1 | 4 | 0.25 |
| 19_503640 | NA | NA | 2 | 0.5 | NA | NA | NA | NA | 2 | 0.5 | 1 | 1 | NA | NA | 1 | 0 | 1 | 1 | 3 | 0.6667 |
| 19_503648 | 3 | 0.6667 | 2 | 1 | NA | NA | NA | NA | 5 | 0.8 | 1 | 0 | 1 | 1 | 1 | 1 | 1 | 0 | 4 | 0.5 |
| 19_503670 | 2 | 1 | 1 | 1 | NA | NA | NA | NA | 3 | 1 | 1 | 1 | NA | NA | NA | NA | 1 | 1 | 2 | 1 |
| 19_503677 | 2 | 0.5 | 1 | 1 | NA | NA | NA | NA | 3 | 0.6667 | NA | NA | NA | NA | 1 | 1 | 1 | 1 | 2 | 1 |
| 19_503683 | 2 | 0.5 | 2 | 0 | NA | NA | NA | NA | 4 | 0.25 | 1 | 0 | 1 | 0 | 1 | 0 | 1 | 1 | 4 | 0.25 |
| 19_503697 | 3 | 0.3333 | 2 | 0.5 | NA | NA | NA | NA | 5 | 0.4 | 1 | 1 | 1 | 1 | 1 | 1 | 1 | 0 | 4 | 0.75 |
| 19_503702 | 3 | 0 | 1 | 1 | NA | NA | NA | NA | 4 | 0.25 | 1 | 0 | 1 | 1 | 1 | 0 | 1 | 0 | 4 | 0.25 |

| | | | | | | | | | | | | | | | | | | | | |
|---|---|---|---|---|---|---|---|---|---|---|---|---|---|---|---|---|---|---|---|---|
| 19_503705 | 2 | 1 | 1 | 0 | NA | NA | NA | NA | 3 | 0.6667 | 1 | 0 | 1 | 1 | 1 | 0 | 1 | 1 | 4 | 0.5 |
| 19_508121 | 1 | 1 | NA | NA | 2 | 0 | 1 | 0 | 4 | 0.25 | 1 | 0 | 1 | 1 | 4 | 0 | 1 | 0 | 7 | 0.1429 |
| 19_508132 | 1 | 1 | NA | NA | 1 | 0 | 1 | 1 | 3 | 0.6667 | 1 | 0 | 2 | 0.5 | 3 | 0.3333 | NA | NA | 6 | 0.3333 |
| 19_508162 | 1 | 0 | NA | NA | 2 | 0.5 | 2 | 0.5 | 5 | 0.4 | 1 | 1 | 2 | 0.5 | 6 | 0.3333 | 1 | 0 | 10 | 0.4 |
| 19_508910 | 8 | 0.125 | 2 | 0 | 6 | 0.1667 | 5 | 0.6 | 21 | 0.2381 | 3 | 0 | 10 | 0.1 | 5 | 0 | 8 | 0.375 | 26 | 0.1538 |
| 19_508927 | 14 | 0.2857 | 8 | 0.125 | 12 | 0.1667 | 9 | 0 | 43 | 0.1628 | 8 | 0 | 16 | 0.25 | 7 | 0.1429 | 15 | 0.3333 | 46 | 0.2174 |
| 19_508931 | 11 | 0.0909 | 8 | 0.125 | 13 | 0 | 8 | 0 | 40 | 0.05 | 8 | 0.125 | 16 | 0.1875 | 7 | 0.1429 | 14 | 0.0714 | 45 | 0.1333 |
| 19_508933 | 15 | 0.5333 | 8 | 0.625 | 10 | 0.3 | 6 | 0.3333 | 39 | 0.4615 | 7 | 0.2857 | 12 | 0.5833 | 9 | 0.6667 | 14 | 0.6429 | 42 | 0.5714 |
| 19_508961 | 14 | 0.6429 | 10 | 0.4 | 12 | 0.5 | 9 | 0.6667 | 45 | 0.5556 | 9 | 0.4444 | 18 | 0.4444 | 10 | 0.8 | 17 | 0.7647 | 54 | 0.6111 |
| 19_508969 | 16 | 1 | 10 | 1 | 12 | 1 | 8 | 0.875 | 46 | 0.9783 | 9 | 1 | 15 | 0.8667 | 7 | 1 | 12 | 0.9167 | 43 | 0.9302 |
| 19_508981 | 11 | 0.2727 | 4 | 0.25 | 9 | 0.2222 | 6 | 0 | 30 | 0.2 | 3 | 0 | 11 | 0.3636 | 8 | 0.25 | 3 | 0.6667 | 25 | 0.32 |
| 19_508994 | 19 | 0.0526 | 12 | 0 | 17 | 0.0588 | 10 | 0.2 | 58 | 0.069 | 10 | 0.2 | 19 | 0 | 15 | 0.1333 | 19 | 0.0526 | 63 | 0.0794 |
| 19_508997 | 20 | 0.15 | 9 | 0.1111 | 18 | 0.1111 | 10 | 0.1 | 57 | 0.1228 | 9 | 0 | 19 | 0.0526 | 14 | 0.3571 | 19 | 0.1579 | 61 | 0.1475 |
| 19_509008 | 1 | 0 | 2 | 1 | NA | NA | NA | NA | 3 | 0.6667 | NA | NA | 1 | 1 | 1 | 1 | 2 | 1 | 4 | 1 |
| 19_509009 | 1 | 0 | 1 | 1 | NA | NA | NA | NA | 2 | 0.5 | NA | NA | NA | NA | 1 | 1 | 1 | 1 | 2 | 1 |
| 19_509018 | 1 | 1 | 2 | 1 | NA | NA | NA | NA | 3 | 1 | NA | NA | 1 | 1 | 2 | 1 | NA | NA | 3 | 1 |
| 19_509019 | 1 | 1 | 2 | 0.5 | NA | NA | NA | NA | 3 | 0.6667 | NA | NA | 1 | 1 | 2 | 0.5 | 2 | 0.5 | 5 | 0.6 |
| 19_509057 | 23 | 0.2174 | 14 | 0 | 17 | 0.1176 | 12 | 0.0833 | 66 | 0.1212 | 12 | 0.25 | 17 | 0 | 17 | 0.1176 | 19 | 0.1579 | 65 | 0.1231 |
| 19_509059 | 19 | 0.0526 | 14 | 0.0714 | 17 | 0.0588 | 9 | 0.1111 | 59 | 0.0678 | 11 | 0 | 17 | 0 | 15 | 0.1333 | 16 | 0.0625 | 59 | 0.0508 |
| 19_509062 | 25 | 0.16 | 15 | 0.2 | 17 | 0.3529 | 7 | 0.5714 | 64 | 0.2656 | 10 | 0.4 | 17 | 0.3529 | 15 | 0.3333 | 18 | 0.3333 | 60 | 0.35 |
| 19_509063 | 21 | 0.9048 | 14 | 0.9286 | 14 | 0.7143 | 9 | 1 | 58 | 0.8793 | 8 | 0.875 | 15 | 0.8667 | 12 | 0.9167 | 11 | 0.9091 | 46 | 0.8913 |
| 19_509064 | 13 | 0.6923 | 11 | 0.3636 | 8 | 0.625 | 5 | 0.8 | 37 | 0.5946 | 4 | 1 | 11 | 0.3636 | 10 | 0.6 | 10 | 0.4 | 35 | 0.5143 |
| 19_509077 | 19 | 0.7895 | 13 | 0.6923 | 12 | 0.75 | 7 | 0.8571 | 51 | 0.7647 | 9 | 0.7778 | 16 | 0.625 | 16 | 0.6875 | 19 | 0.8947 | 60 | 0.75 |
| 19_509958 | 8 | 0.25 | 3 | 0.3333 | 6 | 0.3333 | 10 | 0.4 | 27 | 0.3333 | 1 | 0 | 4 | 0.5 | 5 | 0.4 | 6 | 0.6667 | 16 | 0.5 |
| 19_509964 | 8 | 0.25 | 6 | 0.3333 | 4 | 0.25 | 11 | 0.4545 | 29 | 0.3448 | 1 | 1 | 5 | 0.6 | 7 | 0.2857 | 6 | 0.6667 | 19 | 0.5263 |
| 19_509980 | 8 | 0.625 | 5 | 0.6 | 5 | 0.4 | 8 | 0.625 | 26 | 0.5769 | 1 | 0 | 5 | 0.8 | 6 | 0.6667 | 5 | 0.4 | 17 | 0.5882 |
| 19_509991 | 6 | 0.6667 | 4 | 1 | 5 | 0.8 | 13 | 0.1538 | 28 | 0.5 | 1 | 1 | 4 | 0.75 | 6 | 0.6667 | 5 | 0.8 | 16 | 0.75 |
| 19_510229 | 2 | 0.5 | 2 | 0.5 | 1 | 1 | 1 | 1 | 6 | 0.6667 | NA | NA | 3 | 1 | NA | NA | 3 | 1 | 6 | 1 |
| 19_510240 | 2 | 0.5 | 4 | 0.75 | 2 | 1 | 2 | 1 | 10 | 0.8 | NA | NA | 2 | 1 | 2 | 0.5 | 3 | 1 | 7 | 0.8571 |
| 19_510420 | 1 | 0 | 1 | 0 | 1 | 1 | NA | NA | 3 | 0.3333 | NA | NA | NA | NA | NA | NA | 1 | 1 | 1 | 1 |
| 19_510497 | 7 | 0.1429 | 8 | 0.25 | 2 | 0 | 3 | 0 | 20 | 0.15 | 2 | 0.5 | 5 | 0.4 | 5 | 0.2 | 3 | 0 | 15 | 0.2667 |
| 19_510617 | 6 | 0.1667 | 5 | 0.2 | 5 | 0 | 9 | 0.3333 | 25 | 0.2 | 3 | 0 | 6 | 0.1667 | 6 | 0 | 5 | 0.4 | 20 | 0.15 |
| 19_510619 | 9 | 0.1111 | 7 | 0.1429 | 5 | 0 | 11 | 0.3636 | 32 | 0.1875 | 4 | 0.25 | 8 | 0 | 8 | 0.125 | 9 | 0.4444 | 29 | 0.2069 |
| 19_510657 | 5 | 0 | 5 | 0.6 | 3 | 0 | 8 | 0 | 21 | 0.1429 | 2 | 0 | 7 | 0.5714 | 4 | 0 | 7 | 0.4286 | 20 | 0.35 |
| 19_510688 | 8 | 0.25 | 10 | 0.2 | 4 | 0 | 13 | 0.0769 | 35 | 0.1429 | 5 | 0.4 | 12 | 0.1667 | 8 | 0.25 | 6 | 0 | 31 | 0.1935 |
| 19_511930 | NA | NA | 1 | 1 | NA | NA | NA | NA | 1 | 1 | NA | NA | NA | NA | NA | NA | 1 | 1 | 1 | 1 |
| 19_511936 | NA | NA | 1 | 1 | NA | NA | NA | NA | 1 | 1 | NA | NA | NA | NA | NA | NA | 1 | 1 | 1 | 1 |
| 19_512034 | NA | NA | 1 | 1 | NA | NA | NA | NA | 1 | 1 | NA | NA | NA | NA | NA | NA | 1 | 1 | 1 | 1 |
| 19_512049 | 1 | 1 | 1 | 1 | NA | NA | NA | NA | 2 | 1 | NA | NA | NA | NA | NA | NA | 2 | 1 | 2 | 1 |
| 19_512050 | 1 | 0 | 1 | 1 | NA | NA | NA | NA | 2 | 0.5 | NA | NA | NA | NA | NA | NA | 2 | 1 | 2 | 1 |
| 19_512052 | 1 | 0 | 1 | 1 | NA | NA | NA | NA | 2 | 0.5 | NA | NA | NA | NA | NA | NA | 1 | 1 | 1 | 1 |
| 19_512061 | 1 | 1 | 1 | 1 | NA | NA | NA | NA | 2 | 1 | NA | NA | 1 | 1 | NA | NA | 2 | 0.5 | 3 | 0.6667 |
| 19_513085 | 21 | 0.4762 | 39 | 0.4103 | 28 | 0.6071 | 16 | 0.375 | 104 | 0.4712 | 21 | 0.5714 | 17 | 0.5882 | 18 | 0.3889 | 18 | 0.5556 | 74 | 0.527 |
| 19_513088 | 16 | 0.375 | 25 | 0.24 | 18 | 0.3333 | 12 | 0.25 | 71 | 0.2958 | 15 | 0.5333 | 11 | 0.1818 | 17 | 0.2941 | 15 | 0.0667 | 58 | 0.2759 |
| 19_513098 | 24 | 0.75 | 32 | 0.7188 | 32 | 0.7812 | 18 | 0.8333 | 106 | 0.7642 | 23 | 0.8696 | 13 | 0.8462 | 22 | 0.6364 | 23 | 0.7826 | 81 | 0.7778 |
| 19_513119 | 23 | 0.5217 | 36 | 0.3056 | 28 | 0.5357 | 15 | 0.4667 | 102 | 0.4412 | 21 | 0.2381 | 17 | 0.5294 | 22 | 0.4545 | 20 | 0.45 | 80 | 0.4125 |
| 19_513124 | 24 | 0.0833 | 38 | 0.0263 | 28 | 0 | 16 | 0.1875 | 106 | 0.0566 | 20 | 0 | 18 | 0.1111 | 19 | 0.0526 | 19 | 0.0526 | 76 | 0.0526 |
| 19_513140 | 21 | 0.2381 | 32 | 0.1875 | 25 | 0.16 | 12 | 0.25 | 90 | 0.2 | 15 | 0.2667 | 13 | 0.1538 | 16 | 0.25 | 18 | 0.3333 | 62 | 0.2581 |
| 19_513141 | 18 | 0.3333 | 26 | 0.6154 | 20 | 0.45 | 12 | 0.4167 | 76 | 0.4737 | 14 | 0.5714 | 9 | 0.5556 | 12 | 0.5 | 15 | 0.3333 | 50 | 0.48 |
| 19_513148 | 24 | 0.2917 | 37 | 0.4865 | 28 | 0.3571 | 18 | 0.2778 | 107 | 0.3738 | 17 | 0.2941 | 18 | 0.2778 | 20 | 0.3 | 22 | 0.4545 | 77 | 0.3377 |
| 19_513150 | 15 | 1 | 34 | 0.8235 | 28 | 0.9643 | 14 | 0.9286 | 91 | 0.9121 | 10 | 1 | 14 | 0.8571 | 18 | 0.8889 | 17 | 0.8824 | 59 | 0.8983 |
| 19_513164 | 22 | 0.1818 | 28 | 0.2857 | 25 | 0.32 | 13 | 0.3846 | 88 | 0.2841 | 13 | 0.1538 | 15 | 0.2 | 19 | 0.3158 | 19 | 0.1579 | 66 | 0.2121 |
| 19_513165 | 15 | 1 | 21 | 0.9048 | 25 | 0.96 | 13 | 0.7692 | 74 | 0.9189 | 12 | 1 | 9 | 0.8889 | 10 | 1 | 12 | 1 | 43 | 0.9767 |
| 19_513169 | 21 | 0.1429 | 30 | 0.0667 | 27 | 0.1111 | 15 | 0 | 93 | 0.086 | 15 | 0.2667 | 16 | 0.0625 | 18 | 0.0556 | 20 | 0.1 | 69 | 0.1159 |
| 19_513172 | 19 | 1 | 26 | 0.7308 | 17 | 0.9412 | 16 | 0.875 | 78 | 0.8718 | 13 | 0.9231 | 14 | 0.7143 | 16 | 0.875 | 15 | 1 | 58 | 0.8793 |
| 19_513177 | 15 | 0.0667 | 27 | 0.037 | 22 | 0.0455 | 14 | 0.0714 | 78 | 0.0513 | 14 | 0.0714 | 9 | 0.1111 | 12 | 0.1667 | 17 | 0.1176 | 52 | 0.1154 |
| 19_513190 | 21 | 0.2857 | 27 | 0.3333 | 26 | 0.2692 | 16 | 0.4375 | 90 | 0.3222 | 16 | 0.0625 | 11 | 0.4545 | 17 | 0.2941 | 21 | 0.2381 | 65 | 0.2462 |
| 19_513218 | 19 | 1 | 23 | 1 | 19 | 1 | 15 | 1 | 76 | 1 | 11 | 1 | 12 | 1 | 14 | 1 | 17 | 1 | 54 | 1 |
| 19_513220 | 21 | 0.381 | 21 | 0.4762 | 12 | 0.5 | 13 | 0.3077 | 67 | 0.4179 | 13 | 0.4615 | 6 | 0.6667 | 16 | 0.1875 | 17 | 0.7059 | 52 | 0.4808 |
| 19_513228 | 18 | 0.8889 | 21 | 0.9524 | 15 | 0.9333 | 10 | 0.9 | 64 | 0.9219 | 12 | 0.6667 | 9 | 1 | 12 | 1 | 15 | 0.9333 | 48 | 0.8958 |
| 19_513241 | 19 | 1 | 22 | 1 | 19 | 1 | 15 | 1 | 75 | 1 | 13 | 1 | 12 | 1 | 16 | 0.9375 | 19 | 1 | 60 | 0.9833 |
| 19_513262 | 13 | 0.3846 | 19 | 0.1579 | 17 | 0.3529 | 11 | 0.2727 | 60 | 0.2833 | 11 | 0.1818 | 11 | 0.0909 | 16 | 0.0625 | 15 | 0.1333 | 53 | 0.1132 |
| 19_513265 | 9 | 0.8889 | 11 | 0.8182 | 9 | 0.8889 | 12 | 0.9167 | 41 | 0.878 | 5 | 0.6 | 10 | 0.8 | 7 | 1 | 13 | 0.8462 | 35 | 0.8286 |
| 19_513964 | 1 | 0 | 6 | 0 | 5 | 0 | 2 | 0.5 | 14 | 0.0714 | 1 | 0 | 2 | 0 | 4 | 0 | 3 | 0.3333 | 10 | 0.1 |
| 19_514323 | 51 | 0.0196 | 79 | 0.0759 | 83 | 0.0602 | 52 | 0.0385 | 265 | 0.0528 | 70 | 0.0429 | 76 | 0.0132 | 66 | 0.0758 | 74 | 0.1081 | 286 | 0.0594 |

| | | | | | | | | | | | | | | | | | | | | |
|---|---|---|---|---|---|---|---|---|---|---|---|---|---|---|---|---|---|---|---|---|
| 19_514997 | 16 | 0.125 | 19 | 0.0526 | 21 | 0.0476 | 14 | 0.0714 | 70 | 0.0714 | 14 | 0.0714 | 19 | 0.1053 | 8 | 0 | 15 | 0 | 56 | 0.0536 |
| 19_515022 | 19 | 0.1053 | 29 | 0.1034 | 21 | 0.0952 | 18 | 0.1667 | 87 | 0.1149 | 21 | 0.0952 | 25 | 0.2 | 14 | 0.1429 | 21 | 0.0476 | 81 | 0.1235 |
| 19_515800 | 30 | 0.1333 | 33 | 0 | 19 | 0.0526 | 24 | 0.0833 | 106 | 0.066 | 28 | 0.0357 | 20 | 0.1 | 29 | 0.069 | 32 | 0.0625 | 109 | 0.0642 |
| 19_516832 | 4 | 0 | NA | NA | 2 | 0.5 | NA | NA | 6 | 0.1667 | NA | NA | 1 | 1 | NA | NA | NA | NA | 1 | 1 |
| 19_518193 | NA | NA | 1 | 1 | NA | NA | 1 | 0 | 2 | 0.5 | 1 | 1 | 1 | 0 | 1 | 1 | 2 | 0.5 | 5 | 0.6 |
| 19_518202 | NA | NA | 1 | 1 | NA | NA | 1 | 1 | 2 | 1 | NA | NA | 1 | 0 | 1 | 0 | 1 | 1 | 3 | 0.3333 |
| 19_518328 | NA | NA | 1 | 1 | 2 | 0.5 | NA | NA | 3 | 0.6667 | 1 | 1 | NA | NA | 1 | 1 | 1 | 1 | 3 | 1 |
| 19_518461 | 4 | 0 | 5 | 0.2 | 6 | 0 | 4 | 0 | 19 | 0.0526 | 5 | 0 | 6 | 0 | 6 | 0 | 8 | 0.25 | 25 | 0.08 |
| 19_518474 | 3 | 1 | 4 | 0.75 | 5 | 1 | 2 | 0.5 | 14 | 0.8571 | 3 | 0.3333 | 3 | 0.6667 | 7 | 0.7143 | 9 | 0.5556 | 22 | 0.5909 |
| 19_518480 | 4 | 1 | 5 | 0.6 | 5 | 0.8 | 4 | 1 | 18 | 0.8333 | 5 | 1 | 4 | 0.75 | 6 | 0.8333 | 7 | 0.8571 | 22 | 0.8636 |
| 19_532422 | 19 | 0.1579 | 23 | 0.087 | 17 | 0 | 24 | 0 | 83 | 0.0602 | 16 | 0.0625 | 9 | 0.1111 | 23 | 0 | 19 | 0.1579 | 67 | 0.0746 |
| 19_532423 | 23 | 0.2174 | 29 | 0.1724 | 21 | 0.1905 | 29 | 0.2759 | 102 | 0.2157 | 21 | 0.3333 | 11 | 0.3636 | 23 | 0.2174 | 21 | 0.0952 | 76 | 0.2368 |
| 19_532424 | 23 | 0.2609 | 28 | 0.2143 | 22 | 0.1364 | 28 | 0.2143 | 101 | 0.2079 | 18 | 0.0556 | 11 | 0.4545 | 21 | 0.0476 | 20 | 0.2 | 70 | 0.1571 |
| 19_532440 | 20 | 0.1 | 23 | 0.1304 | 21 | 0 | 30 | 0.1 | 94 | 0.0851 | 19 | 0.1579 | 10 | 0.2 | 23 | 0.0435 | 19 | 0 | 71 | 0.0845 |
| 19_535683 | 6 | 0.1667 | 6 | 0 | 2 | 0 | 3 | 0.3333 | 17 | 0.1176 | 7 | 0 | 5 | 0.2 | 7 | 0 | 1 | 0 | 20 | 0.05 |
| 19_535703 | 6 | 0.1667 | 7 | 0.2857 | 2 | 0 | 3 | 0 | 18 | 0.1667 | 7 | 0.8571 | 6 | 0.1667 | 9 | 0.3333 | 1 | 0 | 23 | 0.4348 |
| 19_535708 | 5 | 0 | 8 | 0.125 | 3 | 0 | 4 | 0 | 20 | 0.05 | 9 | 0.1111 | 7 | 0.1429 | 8 | 0.25 | 1 | 0 | 25 | 0.16 |
| 19_535717 | 5 | 1 | 6 | 0.6667 | 2 | 1 | 5 | 1 | 18 | 0.8889 | 6 | 0.8333 | 5 | 0.8 | 9 | 1 | 1 | 1 | 21 | 0.9048 |
| 19_535739 | 5 | 0.2 | 8 | 0.375 | 2 | 0.5 | 5 | 0.6 | 20 | 0.4 | 9 | 0.5556 | 6 | 0 | 11 | 0.5455 | 2 | 0.5 | 28 | 0.4286 |
| 19_535771 | 6 | 0.8333 | 7 | 1 | 3 | 1 | 2 | 1 | 18 | 0.9444 | 10 | 1 | 5 | 1 | 9 | 1 | 3 | 1 | 27 | 1 |
| 19_535780 | 2 | 0.5 | 5 | 0.4 | 3 | 0 | 6 | 0.5 | 16 | 0.375 | 7 | 0.1429 | 6 | 0.1667 | 9 | 0.5556 | 2 | 0.5 | 24 | 0.3333 |
| 19_535794 | 5 | 0.2 | 8 | 0.125 | 5 | 0.2 | 7 | 0.4286 | 25 | 0.24 | 9 | 0.3333 | 6 | 0.1667 | 12 | 0.5 | 3 | 0 | 30 | 0.3333 |
| 19_535810 | 6 | 0.1667 | 9 | 0.6667 | 7 | 0.4286 | 7 | 0.7143 | 29 | 0.5172 | 8 | 0.5 | 7 | 0.2857 | 12 | 0.6667 | 3 | 0.3333 | 30 | 0.5 |
| 19_535816 | 6 | 0.3333 | 8 | 0.25 | 8 | 0 | 4 | 0.25 | 26 | 0.1923 | 9 | 0.3333 | 6 | 0.5 | 11 | 0.6364 | 2 | 0.5 | 28 | 0.5 |
| 19_535824 | 6 | 0.6667 | 7 | 0.4286 | 5 | 0.6 | 4 | 1 | 22 | 0.6364 | 6 | 0.6667 | 3 | 0.6667 | 9 | 0.7778 | 4 | 0.5 | 22 | 0.6818 |
| 19_535862 | 4 | 0 | 11 | 0.0909 | 11 | 0.0909 | 5 | 0 | 31 | 0.0645 | 9 | 0 | 8 | 0.125 | 15 | 0.1333 | 6 | 0 | 38 | 0.0789 |
| 19_535865 | 5 | 0.2 | 9 | 0 | 11 | 0.0909 | 9 | 0.1111 | 34 | 0.0882 | 9 | 0.2222 | 8 | 0.375 | 11 | 0.0909 | 6 | 0.1667 | 34 | 0.2059 |
| 19_539851 | 2 | 0 | 2 | 0 | 2 | 0 | 2 | 0.5 | 8 | 0.125 | NA | NA | 3 | 0.3333 | 4 | 0 | 1 | 0 | 8 | 0.125 |
| 19_541508 | 6 | 0 | 12 | 0.0833 | 10 | 0.1 | 6 | 0 | 34 | 0.0588 | 9 | 0.1111 | 8 | 0.125 | 12 | 0 | 8 | 0 | 37 | 0.0541 |
| 19_542012 | 9 | 0.3333 | 5 | 0 | 4 | 0.5 | 5 | 0.2 | 23 | 0.2609 | 5 | 0 | 4 | 0.25 | 7 | 0.1429 | 7 | 0.1429 | 23 | 0.1304 |
| 19_542014 | 8 | 0.5 | 5 | 0.6 | 4 | 0.25 | 6 | 0.3333 | 23 | 0.4348 | 5 | 0.2 | 4 | 0.75 | 4 | 0.75 | 6 | 0.6667 | 19 | 0.5789 |
| 19_542017 | 8 | 0.25 | 3 | 0.3333 | 3 | 0 | 5 | 0.2 | 19 | 0.2105 | 2 | 1 | 2 | 0.5 | 3 | 0.3333 | 5 | 0.2 | 12 | 0.4167 |
| 19_542021 | 9 | 0 | 3 | 0.3333 | 4 | 0.25 | 7 | 0.2857 | 23 | 0.1739 | 4 | 0.5 | 5 | 0.2 | 8 | 0 | 8 | 0 | 25 | 0.12 |
| 19_542023 | 2 | 0.5 | 3 | 0.6667 | NA | NA | 3 | 0.3333 | 8 | 0.5 | 2 | 0.5 | 2 | 1 | 3 | 1 | 8 | 0.5 | 15 | 0.6667 |
| 19_542027 | 7 | 0.2857 | 3 | 0.3333 | 2 | 0 | 7 | 0.1429 | 19 | 0.2105 | 4 | 0.75 | 5 | 0.2 | 8 | 0.125 | 6 | 0.1667 | 23 | 0.2609 |
| 19_546546 | NA | NA | NA | NA | 1 | 1 | NA | NA | 1 | 1 | NA | NA | 1 | 1 | NA | NA | NA | NA | 1 | 1 |
| 19_547608 | 8 | 0.875 | 6 | 0.6667 | 5 | 0.8 | 4 | 0.75 | 23 | 0.7826 | 2 | 0.5 | 8 | 0.875 | 1 | 1 | 7 | 1 | 18 | 0.8889 |
| 19_547617 | 8 | 0.25 | 6 | 0.1667 | 5 | 0.2 | 3 | 0.3333 | 22 | 0.2273 | 2 | 0 | 10 | 0.1 | 2 | 0 | 5 | 0.6 | 19 | 0.2105 |
| 19_547620 | 8 | 0.125 | 6 | 0 | 4 | 0.25 | 3 | 0 | 21 | 0.0952 | 3 | 0 | 8 | 0.25 | 2 | 0 | 6 | 0 | 19 | 0.1053 |
| 19_547639 | 4 | 0.5 | 4 | 0.5 | 2 | 0.5 | 2 | 0 | 12 | 0.4167 | 3 | 0.3333 | 4 | 0 | 1 | 0 | 3 | 0.3333 | 11 | 0.1818 |
| 19_547652 | 10 | 0.9 | 10 | 1 | 5 | 1 | 4 | 1 | 29 | 0.9655 | 3 | 1 | 9 | 0.8889 | 3 | 0.6667 | 6 | 1 | 21 | 0.9048 |
| 19_547659 | 12 | 0.0833 | 10 | 0.2 | 5 | 0.2 | 5 | 0 | 32 | 0.125 | 3 | 0 | 9 | 0.2222 | 3 | 0.3333 | 5 | 0 | 20 | 0.15 |
| 19_547670 | 11 | 0.7273 | 11 | 0.9091 | 5 | 0.8 | 5 | 0.8 | 32 | 0.8125 | 4 | 0.75 | 11 | 0.8182 | 5 | 0.8 | 7 | 0.8571 | 27 | 0.8148 |
| 19_547672 | 6 | 1 | 6 | 1 | 3 | 1 | 2 | 1 | 17 | 1 | NA | NA | 4 | 1 | 4 | 0.75 | 4 | 1 | 12 | 0.9167 |
| 19_547673 | 12 | 0.0833 | 10 | 0 | 6 | 0.1667 | 4 | 0 | 32 | 0.0625 | 4 | 0.5 | 10 | 0 | 5 | 0 | 6 | 0.1667 | 25 | 0.12 |
| 19_547682 | 11 | 1 | 8 | 1 | 8 | 1 | 3 | 1 | 30 | 1 | 2 | 1 | 11 | 1 | 4 | 1 | 6 | 1 | 23 | 1 |
| 19_547684 | 11 | 0.4545 | 9 | 0.4444 | 9 | 0.7778 | 4 | 0.5 | 33 | 0.5455 | 3 | 0.3333 | 10 | 0.5 | 6 | 0.6667 | 7 | 0.4286 | 26 | 0.5 |
| 19_547698 | 7 | 0.1429 | 7 | 0 | 8 | 0.125 | 5 | 0 | 27 | 0.0741 | 3 | 0.3333 | 8 | 0 | 6 | 0.1667 | 6 | 0.3333 | 23 | 0.1739 |
| 19_547700 | 14 | 0.8571 | 9 | 0.7778 | 8 | 0.875 | 6 | 0.8333 | 37 | 0.8378 | 4 | 0.75 | 11 | 0.8182 | 9 | 0.8889 | 8 | 0.75 | 32 | 0.8125 |
| 19_547708 | 13 | 0.0769 | 11 | 0.0909 | 9 | 0.3333 | 6 | 0 | 39 | 0.1282 | 5 | 0 | 9 | 0.1111 | 7 | 0.1429 | 8 | 0.125 | 29 | 0.1034 |
| 19_547715 | 15 | 0.9333 | 11 | 1 | 10 | 1 | 6 | 1 | 42 | 0.9762 | 8 | 1 | 11 | 1 | 8 | 1 | 9 | 0.8889 | 36 | 0.9722 |
| 19_547716 | 16 | 0.625 | 9 | 0.4444 | 10 | 0.6 | 7 | 0.4286 | 42 | 0.5476 | 9 | 0.7778 | 11 | 0.3636 | 9 | 0.6667 | 9 | 0.6667 | 38 | 0.6053 |
| 19_547721 | 16 | 0.9375 | 7 | 1 | 8 | 1 | 6 | 1 | 37 | 0.973 | 7 | 0.7143 | 11 | 0.6364 | 5 | 0.8 | 4 | 0.5 | 27 | 0.6667 |
| 19_549993 | NA | NA | 1 | 1 | NA | NA | NA | NA | 1 | 1 | 1 | 0 | NA | NA | 1 | 1 | NA | NA | 2 | 0.5 |
| 19_550338 | 2 | 0 | 1 | 1 | 1 | 0 | NA | NA | 4 | 0.25 | 2 | 0 | 4 | 0.25 | 1 | 0 | 3 | 0 | 10 | 0.1 |
| 19_550466 | 1 | 1 | NA | NA | NA | NA | NA | NA | 1 | 1 | 2 | 0.5 | NA | NA | 1 | 1 | 2 | 1 | 5 | 0.8 |
| 19_550482 | 1 | 1 | NA | NA | NA | NA | NA | NA | 1 | 1 | 2 | 1 | 1 | 1 | 1 | 1 | 1 | 1 | 5 | 1 |
| 19_550485 | 1 | 1 | NA | NA | NA | NA | NA | NA | 1 | 1 | 2 | 1 | 2 | 1 | 1 | 1 | 2 | 1 | 7 | 1 |
| 19_550659 | 17 | 0.1765 | 24 | 0.0417 | 24 | 0.125 | 21 | 0.1429 | 86 | 0.1163 | 23 | 0.1739 | 22 | 0.0455 | 27 | 0.0741 | 22 | 0.0909 | 94 | 0.0957 |
| 19_550662 | 17 | 1 | 24 | 0.9167 | 27 | 1 | 23 | 0.913 | 91 | 0.956 | 25 | 0.96 | 22 | 0.9545 | 26 | 0.9615 | 25 | 0.92 | 98 | 0.949 |
| 19_550670 | 15 | 1 | 18 | 1 | 24 | 1 | 16 | 0.9375 | 73 | 0.9863 | 19 | 1 | 19 | 1 | 18 | 1 | 21 | 1 | 77 | 1 |
| 19_550675 | 20 | 0.95 | 18 | 0.8333 | 21 | 0.9048 | 19 | 0.8421 | 78 | 0.8846 | 23 | 0.6957 | 18 | 0.8889 | 19 | 0.7895 | 18 | 0.9444 | 78 | 0.8205 |
| 19_550682 | 20 | 0.9 | 20 | 0.5 | 18 | 0.6667 | 16 | 0.625 | 74 | 0.6757 | 20 | 0.7 | 17 | 0.8235 | 21 | 0.8571 | 20 | 0.65 | 78 | 0.7564 |
| 19_550709 | 21 | 0.4762 | 17 | 0.5294 | 22 | 0.7727 | 21 | 0.5714 | 81 | 0.5926 | 16 | 0.375 | 18 | 0.5 | 20 | 0.55 | 17 | 0.4706 | 71 | 0.4789 |
| 19_550711 | 17 | 0.3529 | 14 | 0.3571 | 18 | 0.3889 | 13 | 0.5385 | 62 | 0.4032 | 13 | 0.6154 | 14 | 0.5 | 18 | 0.6667 | 20 | 0.5 | 65 | 0.5692 |

| | | | | | | | | | | | | | | | | | | | | |
|---|---|---|---|---|---|---|---|---|---|---|---|---|---|---|---|---|---|---|---|---|
| 19_5507172 | 21 | 0.7143 | 15 | 0.5333 | 20 | 0.75 | 15 | 0.6667 | 71 | 0.6761 | 12 | 0.75 | 15 | 0.8 | 18 | 0.6667 | 17 | 0.7059 | 62 | 0.7258 |
| 19_5531717 | 4 | 0.25 | 2 | 0 | 3 | 0 | 1 | 0 | 10 | 0.1 | NA | NA | 1 | 0 | 4 | 0.5 | 1 | 0 | 6 | 0.3333 |
| 19_5542757 | 138 | 0.1812 | 147 | 0.1361 | 160 | 0.0938 | 125 | 0.176 | 570 | 0.1439 | 162 | 0.1173 | 154 | 0.1753 | 178 | 0.1798 | 132 | 0.1439 | 626 | 0.155 |
| 19_5543881 | 176 | 0.0739 | 179 | 0.1006 | 195 | 0.0667 | 149 | 0.094 | 699 | 0.083 | 205 | 0.0927 | 183 | 0.0874 | 212 | 0.0896 | 162 | 0.0617 | 762 | 0.084 |
| 19_5620681 | 1 | 1 | NA | NA | NA | NA | NA | NA | 1 | 1 | NA | NA | NA | NA | 1 | 1 | NA | NA | 1 | 1 |
| 19_5620863 | 1 | 1 | NA | NA | NA | NA | NA | NA | 1 | 1 | NA | NA | NA | NA | 1 | 1 | NA | NA | 1 | 1 |
| 19_5620938 | 1 | 1 | NA | NA | NA | NA | NA | NA | 1 | 1 | NA | NA | NA | NA | 1 | 1 | NA | NA | 1 | 1 |
| 19_5621150 | 1 | 1 | NA | NA | NA | NA | NA | NA | 1 | 1 | NA | NA | NA | NA | 1 | 1 | NA | NA | 1 | 1 |
| 19_5621228 | 1 | 1 | NA | NA | NA | NA | NA | NA | 1 | 1 | NA | NA | NA | NA | 1 | 1 | NA | NA | 1 | 1 |
| 19_5621287 | 1 | 1 | NA | NA | NA | NA | NA | NA | 1 | 1 | NA | NA | NA | NA | 1 | 1 | NA | NA | 1 | 1 |
| 19_5652874 | 5 | 0.2 | 1 | 0 | 9 | 0.2222 | 7 | 0.2857 | 22 | 0.2273 | 7 | 0.1429 | 2 | 0 | 3 | 0 | 7 | 0.1429 | 19 | 0.1053 |
| 19_5653155 | 13 | 0.1538 | 4 | 0.5 | 6 | 0.1667 | 9 | 0.3333 | 32 | 0.25 | 9 | 0.1111 | 8 | 0.25 | 10 | 0.2 | 2 | 0 | 29 | 0.1724 |
| 19_5653213 | 9 | 0 | 4 | 0.5 | 6 | 0.3333 | 8 | 0.75 | 27 | 0.3704 | 9 | 0.5556 | 6 | 0.1667 | 9 | 0.3333 | 2 | 0 | 26 | 0.3462 |
| 19_5653221 | 8 | 0 | 4 | 0.25 | 6 | 0 | 5 | 0.2 | 23 | 0.087 | 8 | 0 | 7 | 0.1429 | 7 | 0.1429 | 1 | 0 | 23 | 0.087 |
| 19_5653471 | 9 | 0.5556 | 3 | 0.3333 | 4 | 0.75 | 7 | 0.4286 | 23 | 0.5217 | 7 | 0.8571 | 5 | 0.6 | 9 | 0.8889 | 1 | 1 | 22 | 0.8182 |
| 19_5697137 | 3 | 0 | 5 | 0.2 | 1 | 0 | 3 | 0 | 12 | 0.0833 | 2 | 0 | 4 | 0.25 | 3 | 0.6667 | 4 | 0.25 | 13 | 0.3077 |
| 19_5697200 | 3 | 0.3333 | 5 | 0 | 1 | 0 | 3 | 0.3333 | 12 | 0.1667 | 2 | 0 | 4 | 0 | 3 | 0.3333 | 4 | 0.5 | 13 | 0.2308 |
| 19_5697349 | 3 | 0 | 5 | 0.6 | 1 | 0 | 3 | 0.3333 | 12 | 0.3333 | 2 | 0 | 4 | 0.5 | 3 | 0.6667 | 3 | 0.6667 | 12 | 0.5 |
| 19_5697380 | 3 | 0 | 5 | 0.2 | NA | NA | 3 | 0 | 11 | 0.0909 | 2 | 0 | 5 | 0 | 3 | 0 | 3 | 0.3333 | 13 | 0.0769 |
| 19_5697404 | 2 | 1 | 2 | 1 | 1 | 0 | 2 | 0 | 7 | 0.5714 | 1 | 0 | 4 | 1 | 3 | 1 | 3 | 1 | 11 | 0.9091 |
| 19_5697410 | 2 | 1 | 5 | 1 | 1 | 1 | 3 | 1 | 11 | 1 | 2 | 1 | 5 | 1 | 2 | 0.5 | 3 | 1 | 12 | 0.9167 |
| 19_5697647 | 3 | 0.6667 | 6 | 0.5 | 1 | 1 | 3 | 1 | 13 | 0.6923 | 2 | 0.5 | 5 | 0.6 | 3 | 0.6667 | 4 | 0.75 | 14 | 0.6429 |
| 19_5697690 | 3 | 0.3333 | 7 | 0.1429 | 1 | 0 | 3 | 0.6667 | 14 | 0.2857 | 2 | 0 | 4 | 0.25 | 3 | 0 | 4 | 0.25 | 13 | 0.1538 |
| 19_5697741 | 3 | 0.3333 | 7 | 0.1429 | 1 | 1 | 2 | 0 | 13 | 0.2308 | 1 | 0 | 4 | 0 | 2 | 0 | 3 | 0.3333 | 10 | 0.1 |
| 19_5697745 | 3 | 0.6667 | 5 | 0 | 1 | 0 | 3 | 0.6667 | 12 | 0.3333 | 1 | 1 | 5 | 0.2 | 3 | 0.3333 | 2 | 1 | 11 | 0.4545 |
| 19_5698112 | 2 | 0.5 | 6 | 1 | NA | NA | 3 | 0 | 11 | 0.6364 | NA | NA | 2 | 1 | 2 | 0.5 | 4 | 0.75 | 8 | 0.75 |
| 19_5701364 | 2 | 0.5 | 2 | 0.5 | 3 | 0 | 1 | 1 | 8 | 0.375 | 1 | 0 | 1 | 0 | 1 | 1 | 3 | 1 | 6 | 0.6667 |
| 19_5704685 | 1 | 0 | 1 | 1 | 1 | 0 | NA | NA | 3 | 0.3333 | 2 | 0.5 | 1 | 0 | 1 | 0 | 2 | 0.5 | 6 | 0.3333 |
| 19_5704757 | 1 | 1 | 1 | 0 | 1 | 0 | NA | NA | 3 | 0.3333 | 2 | 0.5 | NA | NA | 1 | 0 | 2 | 0 | 5 | 0.2 |
| 19_5704957 | 1 | 0 | 1 | 1 | 1 | 0 | NA | NA | 3 | 0.3333 | 2 | 1 | NA | NA | 1 | 1 | 2 | 0.5 | 5 | 0.8 |
| 19_5714694 | 7 | 0 | 2 | 0 | 4 | 0 | 6 | 0.1667 | 19 | 0.0526 | 3 | 0 | 7 | 0 | 5 | 0.2 | 4 | 0 | 19 | 0.0526 |
| 19_5714775 | 6 | 0.5 | 3 | 0.3333 | 4 | 0.75 | 6 | 0.5 | 19 | 0.5263 | 4 | 0.5 | 8 | 0.625 | 4 | 0.5 | 4 | 0.5 | 20 | 0.55 |
| 19_5714826 | 6 | 0.5 | 2 | 0.5 | 2 | 0.5 | 4 | 0.5 | 14 | 0.5 | 4 | 0 | 7 | 0.5714 | 5 | 0 | 2 | 0.5 | 18 | 0.2778 |
| 19_5714830 | 5 | 0.8 | 2 | 0.5 | 1 | 1 | 5 | 1 | 13 | 0.8462 | 4 | 0.75 | 7 | 0.8571 | 4 | 1 | 4 | 1 | 19 | 0.8947 |
| 19_5714940 | 6 | 0.8333 | 3 | 0.3333 | 4 | 0.75 | 6 | 0.6667 | 19 | 0.6842 | 3 | 0.6667 | 6 | 0.3333 | 2 | 0.5 | 3 | 0.6667 | 14 | 0.5 |
| 19_5715012 | 6 | 0.5 | 2 | 0.5 | 4 | 0.5 | 5 | 0.2 | 17 | 0.4118 | 3 | 1 | 6 | 0.1667 | 4 | 1 | 2 | 1 | 15 | 0.6667 |
| 19_5715020 | 5 | 0.4 | 2 | 1 | NA | NA | 4 | 0.75 | 11 | 0.6364 | 4 | 0 | 6 | 0.5 | 4 | 0.75 | 2 | 0.5 | 16 | 0.4375 |
| 19_5715030 | 7 | 0.1429 | 3 | 0.3333 | 2 | 1 | 6 | 0.6667 | 18 | 0.4444 | 3 | 0.6667 | 8 | 0.25 | 4 | 1 | 2 | 0.5 | 17 | 0.5294 |
| 19_5715044 | 6 | 0.5 | 2 | 0 | 4 | 0 | 5 | 0.6 | 17 | 0.3529 | 4 | 0.5 | 8 | 0.375 | 5 | 0.6 | 2 | 0.5 | 19 | 0.4737 |
| 19_5715208 | 7 | 0.1429 | 2 | 0 | 4 | 0 | 4 | 0 | 17 | 0.0588 | 4 | 0 | 7 | 0.2857 | 5 | 0.2 | 3 | 0.3333 | 19 | 0.2105 |
| 19_5717267 | 6 | 0 | 2 | 0 | 3 | 0 | 4 | 0.25 | 15 | 0.0667 | 2 | 0 | 2 | 0 | 2 | 0.5 | 3 | 0 | 9 | 0.1111 |
| 19_5735393 | 1 | 1 | NA | NA | NA | NA | 1 | 1 | 2 | 1 | NA | NA | 1 | 1 | NA | NA | 2 | 1 | 3 | 1 |
| 19_5735625 | 1 | 1 | NA | NA | NA | NA | NA | NA | 1 | 1 | NA | NA | 1 | 1 | NA | NA | 1 | 1 | 2 | 1 |
| 19_5739874 | 2 | 1 | 1 | 1 | 1 | 0 | 2 | 1 | 6 | 0.8333 | 1 | 0 | 1 | 1 | 1 | 1 | NA | NA | 3 | 0.6667 |
| 19_5739978 | 2 | 0 | 1 | 0 | 1 | 0 | 1 | 1 | 5 | 0.2 | 1 | 0 | 1 | 1 | 1 | 1 | NA | NA | 3 | 0.6667 |
| 19_5739997 | 2 | 1 | 1 | 1 | 1 | 0 | 3 | 0.6667 | 7 | 0.7143 | 1 | 1 | 1 | 0 | 1 | 0 | NA | NA | 3 | 0.3333 |
| 19_5740416 | 2 | 1 | 1 | 0 | 1 | 0 | 3 | 0.6667 | 7 | 0.5714 | 1 | 1 | 1 | 0 | 1 | 0 | 1 | 1 | 4 | 0.5 |
| 19_5740454 | 2 | 0.5 | 1 | 1 | 1 | 1 | 2 | 0.5 | 6 | 0.6667 | NA | NA | 1 | 1 | NA | NA | NA | NA | 1 | 1 |
| 19_5740723 | 2 | 0 | 1 | 0 | 2 | 0 | 3 | 0.3333 | 8 | 0.125 | 1 | 0 | 1 | 0 | 1 | 1 | 1 | 1 | 4 | 0.5 |
| 19_5740735 | 2 | 0.5 | 1 | 1 | 2 | 0 | 3 | 0.6667 | 8 | 0.5 | 1 | 1 | 1 | 0 | 1 | 0 | 1 | 0 | 4 | 0.25 |
| 19_5752238 | 2 | 1 | 3 | 0.3333 | 7 | 0.5714 | 5 | 0.8 | 17 | 0.6471 | 3 | 0.3333 | 2 | 1 | 2 | 1 | 5 | 0.6 | 12 | 0.6667 |
| 19_5752251 | 2 | 0.5 | 3 | 0.3333 | 5 | 0.2 | 4 | 0.75 | 14 | 0.4286 | 2 | 0 | 3 | 0.3333 | 2 | 0.5 | 4 | 0.25 | 11 | 0.2727 |
| 19_5752269 | 1 | 1 | 3 | 0.3333 | 4 | 0.5 | 1 | 1 | 9 | 0.5556 | 1 | 1 | 2 | 0.5 | 2 | 0.5 | 3 | 0.6667 | 8 | 0.625 |
| 19_5752308 | 2 | 0.5 | 3 | 0.6667 | 4 | 1 | 4 | 0.75 | 13 | 0.7692 | 3 | 0.6667 | 3 | 1 | 2 | 0.5 | 3 | 1 | 11 | 0.8182 |
| 19_5752329 | 1 | 0 | 3 | 0 | 7 | 0.1429 | 2 | 0 | 13 | 0.0769 | 3 | 0.3333 | 2 | 0 | 2 | 0 | 4 | 0.5 | 11 | 0.2727 |
| 19_5752477 | 2 | 0.5 | 1 | 0 | 2 | 0.5 | 1 | 1 | 6 | 0.5 | 1 | 1 | 1 | 1 | 2 | 1 | 1 | 1 | 5 | 1 |
| 19_5752501 | 2 | 0.5 | 3 | 0.3333 | 6 | 0.1667 | 6 | 0 | 17 | 0.1765 | 3 | 0.3333 | 2 | 0.5 | 3 | 0 | 3 | 0.3333 | 11 | 0.2727 |
| 19_5752561 | 2 | 0.5 | 2 | 0.5 | 6 | 0.3333 | 4 | 0 | 14 | 0.2857 | 2 | 0 | 3 | 0.3333 | 2 | 0.5 | 4 | 0.75 | 11 | 0.4545 |
| 19_5752570 | 2 | 0.5 | 3 | 0 | 7 | 0 | 6 | 0 | 18 | 0.0556 | 2 | 0 | 3 | 0.3333 | 2 | 0 | 5 | 0 | 12 | 0.0833 |
| 19_5752577 | 1 | 1 | 3 | 0 | 5 | 0 | 3 | 0.3333 | 12 | 0.1667 | 2 | 0 | 3 | 0 | 2 | 0 | 2 | 0.5 | 9 | 0.1111 |
| 19_5752872 | 2 | 0 | 4 | 0 | 6 | 0 | 6 | 0.1667 | 18 | 0.0556 | 2 | 0 | 3 | 0.6667 | 3 | 0 | 4 | 0 | 12 | 0.1667 |
| 19_5758235 | 5 | 0.4 | 2 | 0.5 | 1 | 0 | 2 | 0.5 | 10 | 0.4 | 1 | 0 | 4 | 0.25 | 5 | 0.4 | 3 | 0 | 13 | 0.2308 |
| 19_5758244 | 4 | 0.25 | 2 | 0.5 | 1 | 1 | 2 | 0 | 9 | 0.3333 | NA | NA | 4 | 0.5 | 2 | 1 | 2 | 0 | 8 | 0.5 |
| 19_5766019 | 4 | 0.25 | 1 | 0 | 1 | 0 | 2 | 0.5 | 8 | 0.25 | NA | NA | 2 | 0.5 | 4 | 0.5 | 3 | 0.6667 | 9 | 0.5556 |
| 19_5766185 | 4 | 0.75 | 1 | 0 | 1 | 0 | 1 | 0 | 7 | 0.4286 | NA | NA | 2 | 0 | 3 | 0.3333 | 3 | 0.3333 | 8 | 0.25 |

| | | | | | | | | | | | | | | | | | | | | |
|---|---|---|---|---|---|---|---|---|---|---|---|---|---|---|---|---|---|---|---|---|
| 19_576619 | 4 | 0.5 | 1 | 0 | 1 | 0 | 1 | 1 | 7 | 0.4286 | NA | NA | 2 | 1 | 4 | 0.25 | 3 | 0.6667 | 9 | 0.5556 |
| 19_576622 | 4 | 0.75 | 1 | 1 | 1 | 1 | NA | NA | 6 | 0.8333 | NA | NA | 1 | 1 | 4 | 0.25 | 3 | 0.3333 | 8 | 0.375 |
| 19_576623 | 4 | 0.25 | NA | NA | NA | NA | NA | NA | 4 | 0.25 | NA | NA | 1 | 0 | 4 | 0.25 | 2 | 0.5 | 7 | 0.2857 |
| 19_577834 | 5 | 0.6 | 6 | 0.6667 | 7 | 0.8571 | 9 | 0.6667 | 27 | 0.7037 | 5 | 0.6 | 5 | 0.8 | 6 | 1 | 4 | 0.5 | 20 | 0.75 |
| 19_577838 | 7 | 0.4286 | 9 | 0.3333 | 7 | 0.4286 | 11 | 0.2727 | 34 | 0.3529 | 6 | 0.1667 | 8 | 0.625 | 8 | 0 | 3 | 0 | 25 | 0.24 |
| 19_577867 | 3 | 0 | 11 | 0.0909 | 4 | 0 | 10 | 0.1 | 28 | 0.0714 | 5 | 0 | 8 | 0 | 6 | 0.3333 | 4 | 0 | 23 | 0.087 |
| 19_577876 | 5 | 0 | 8 | 0 | 5 | 0.4 | 10 | 0 | 28 | 0.0714 | 3 | 0 | 5 | 0 | 5 | 0.2 | 2 | 0 | 15 | 0.0667 |
| 19_577882 | 6 | 0 | 10 | 0 | 6 | 0 | 11 | 0.1818 | 33 | 0.0606 | 4 | 0 | 5 | 0.4 | 7 | 0 | 3 | 0 | 19 | 0.1053 |
| 19_577904 | 2 | 0.5 | 1 | 0 | NA | NA | 1 | 1 | 4 | 0.5 | NA | NA | NA | NA | 1 | 1 | NA | NA | 1 | 1 |
| 19_577908 | 2 | 0.5 | 1 | 0 | NA | NA | 1 | 0 | 4 | 0.25 | 1 | 1 | NA | NA | NA | NA | NA | NA | 1 | 1 |
| 19_578047 | 10 | 0 | 9 | 0.1111 | 4 | 0.25 | 12 | 0.1667 | 35 | 0.1143 | 8 | 0 | 4 | 0.25 | 4 | 0 | 13 | 0.2308 | 29 | 0.1379 |
| 19_580822 | 3 | 0.6667 | 5 | 0.6 | 3 | 0.3333 | 2 | 0 | 13 | 0.4615 | 4 | 1 | 4 | 0.75 | 1 | 1 | 1 | 0 | 10 | 0.8 |
| 19_580843 | 3 | 0.3333 | 3 | 0.6667 | 3 | 0.6667 | 1 | 0 | 10 | 0.5 | 4 | 0.75 | 5 | 0.2 | 2 | 0.5 | 2 | 0.5 | 13 | 0.4615 |
| 19_580860 | 8 | 0.5 | 5 | 0.8 | 5 | 0.2 | 11 | 0.4545 | 29 | 0.4828 | 3 | 0.3333 | 7 | 0.1429 | 13 | 0.3077 | 6 | 0.6667 | 29 | 0.3448 |
| 19_582035 | 3 | 0.3333 | 3 | 0.3333 | 2 | 0.5 | 5 | 0.6 | 13 | 0.4615 | NA | NA | 1 | 0 | 3 | 0.3333 | 3 | 0.3333 | 7 | 0.2857 |
| 19_582036 | 2 | 0 | 4 | 0.25 | 3 | 0 | 5 | 0.6 | 14 | 0.2857 | NA | NA | 4 | 0.5 | 2 | 0.5 | 4 | 0 | 10 | 0.3 |
| 19_582038 | 1 | 0 | 3 | 0.3333 | 2 | 0 | 4 | 0.5 | 10 | 0.3 | NA | NA | 2 | 0 | 2 | 0.5 | 3 | 0.6667 | 7 | 0.4286 |
| 19_584863 | 2 | 0 | 5 | 0.2 | 2 | 0 | 1 | 0 | 10 | 0.1 | 2 | 0 | 4 | 0 | 2 | 0 | 7 | 0.1429 | 15 | 0.0667 |
| 19_584878 | 2 | 0.5 | 6 | 0.5 | 2 | 0 | 1 | 0 | 11 | 0.3636 | 2 | 0 | 4 | 0.25 | 2 | 0.5 | 5 | 0.4 | 13 | 0.3077 |
| 19_584884 | 1 | 0 | 6 | 0.1667 | 2 | 0.5 | 1 | 0 | 10 | 0.2 | 2 | 0 | 5 | 0.4 | 1 | 0 | 5 | 0 | 13 | 0.1538 |
| 19_584892 | 2 | 0 | 3 | 0 | 3 | 0.6667 | 1 | 0 | 9 | 0.2222 | 2 | 0 | 4 | 0 | NA | NA | 8 | 0.25 | 14 | 0.1429 |
| 19_584906 | 2 | 1 | 2 | 0.5 | 2 | 0 | 1 | 0 | 7 | 0.4286 | 1 | 0 | 4 | 0.5 | 2 | 0.5 | 2 | 0.5 | 9 | 0.4444 |
| 19_584909 | NA | NA | 2 | 0 | 2 | 0.5 | NA | NA | 4 | 0.25 | 1 | 1 | 2 | 0 | 2 | 0 | 2 | 1 | 7 | 0.4286 |
| 19_584909 | NA | NA | NA | NA | 1 | 1 | NA | NA | 1 | 1 | 1 | 0 | 1 | 1 | NA | NA | 3 | 0.3333 | 5 | 0.4 |
| 19_584977 | NA | NA | 1 | 0 | 1 | 1 | NA | NA | 2 | 0.5 | 1 | 1 | 1 | 0 | NA | NA | 2 | 0.5 | 4 | 0.5 |
| 19_584997 | NA | NA | 1 | 0 | 1 | 1 | NA | NA | 2 | 0.5 | 1 | 1 | 1 | 1 | NA | NA | 2 | 1 | 4 | 1 |
| 19_589262 | 2 | 0.5 | 1 | 1 | 2 | 0.5 | 2 | 0 | 7 | 0.4286 | 4 | 0.5 | 2 | 0.5 | 1 | 1 | 2 | 0.5 | 9 | 0.5556 |
| 19_589337 | 1 | 1 | 5 | 0.4 | 1 | 0 | NA | NA | 7 | 0.4286 | 3 | 1 | 1 | 0 | 1 | 0 | 1 | 0 | 6 | 0.5 |
| 19_589345 | NA | NA | 3 | 1 | 1 | 0 | NA | NA | 4 | 0.75 | 2 | 1 | 1 | 1 | 1 | 1 | 1 | 1 | 5 | 1 |
| 19_589353 | 1 | 1 | 3 | 0 | 1 | 0 | NA | NA | 5 | 0.2 | 2 | 0.5 | NA | NA | 1 | 0 | 1 | 0 | 4 | 0.25 |
| 19_589371 | NA | NA | 1 | 1 | NA | NA | NA | NA | 1 | 1 | NA | NA | 1 | 1 | 1 | 1 | NA | NA | 2 | 1 |
| 19_589386 | NA | NA | 2 | 1 | 1 | 0 | NA | NA | 3 | 0.6667 | 1 | 0 | 1 | 1 | 1 | 0 | NA | NA | 3 | 0.3333 |
| 19_589390 | NA | NA | 3 | 1 | 1 | 1 | NA | NA | 4 | 1 | 1 | 1 | 1 | 1 | 1 | 1 | NA | NA | 3 | 1 |
| 19_589398 | NA | NA | 3 | 0.3333 | 1 | 0 | NA | NA | 4 | 0.25 | NA | NA | 1 | 1 | 1 | 0 | NA | NA | 2 | 0.5 |
| 19_589402 | NA | NA | 3 | 0.3333 | 1 | 1 | NA | NA | 4 | 0.5 | 1 | 1 | 1 | 1 | 1 | 1 | NA | NA | 3 | 1 |
| 19_589405 | NA | NA | 1 | 1 | 1 | 1 | NA | NA | 2 | 1 | 1 | 1 | 1 | 1 | NA | NA | NA | NA | 2 | 1 |
| 19_590129 | 3 | 0.3333 | 1 | 0 | 2 | 0 | NA | NA | 6 | 0.1667 | NA | NA | 1 | 0 | 2 | 1 | 1 | 0 | 4 | 0.5 |
| 19_590186 | 4 | 1 | 4 | 0.75 | 2 | 0 | 1 | 1 | 11 | 0.7273 | 4 | 0.25 | 1 | 0 | 2 | 0 | 1 | 0 | 8 | 0.125 |
| 19_590217 | 2 | 0 | 4 | 0.75 | 3 | 0 | NA | NA | 9 | 0.3333 | 1 | 1 | 1 | 0 | 1 | 1 | NA | NA | 3 | 0.6667 |
| 19_590223 | 2 | 0 | 4 | 0.5 | 2 | 0 | NA | NA | 8 | 0.25 | 2 | 0.5 | 1 | 0 | 1 | 0 | 1 | 0 | 5 | 0.2 |
| 19_590225 | 3 | 1 | 4 | 0.75 | 2 | 0.5 | NA | NA | 9 | 0.7778 | 1 | 1 | 1 | 0 | 1 | 1 | 1 | 1 | 4 | 0.75 |
| 19_590230 | 3 | 0.3333 | 3 | 0 | 2 | 0 | 1 | 1 | 9 | 0.2222 | 4 | 0.25 | 1 | 1 | 2 | 0 | 1 | 1 | 8 | 0.375 |
| 19_590234 | 2 | 1 | 4 | 1 | 2 | 0.5 | NA | NA | 8 | 0.875 | 3 | 1 | 1 | 1 | 1 | 0 | 1 | 0 | 6 | 0.6667 |
| 19_590249 | 1 | 0 | 3 | 0 | 2 | 0.5 | 1 | 0 | 7 | 0.1429 | 3 | 0 | NA | NA | 2 | 1 | 1 | 0 | 6 | 0.3333 |
| 19_590249 | 1 | 1 | 3 | 1 | 1 | 0 | NA | NA | 5 | 0.8 | 1 | 1 | 1 | 1 | NA | NA | 1 | 1 | 3 | 1 |
| 19_590262 | 2 | 0 | 2 | 1 | 2 | 1 | NA | NA | 6 | 0.6667 | 2 | 1 | 1 | 0 | 2 | 0.5 | 1 | 0 | 6 | 0.5 |
| 19_594175 | NA | NA | NA | NA | NA | NA | 1 | 1 | 1 | 1 | NA | NA | 1 | 1 | NA | NA | NA | NA | 1 | 1 |
| 19_594192 | NA | NA | NA | NA | NA | NA | 1 | 1 | 1 | 1 | NA | NA | 1 | 1 | NA | NA | NA | NA | 1 | 1 |
| 19_594218 | NA | NA | NA | NA | NA | NA | 1 | 1 | 1 | 1 | NA | NA | 1 | 1 | NA | NA | NA | NA | 1 | 1 |
| 19_594229 | NA | NA | NA | NA | NA | NA | 1 | 1 | 1 | 1 | NA | NA | 1 | 1 | NA | NA | NA | NA | 1 | 1 |
| 19_594263 | NA | NA | NA | NA | NA | NA | 1 | 1 | 1 | 1 | NA | NA | 1 | 1 | NA | NA | NA | NA | 1 | 1 |
| 19_594295 | NA | NA | NA | NA | NA | NA | 1 | 1 | 1 | 1 | NA | NA | 1 | 1 | NA | NA | NA | NA | 1 | 1 |
| 19_596776 | 3 | 0.6667 | 1 | 1 | 1 | 1 | 1 | 1 | 6 | 0.8333 | 3 | 0.3333 | 1 | 1 | 3 | 1 | 1 | 0 | 8 | 0.625 |
| 19_596783 | 2 | 1 | 1 | 1 | 1 | 1 | 1 | 1 | 5 | 1 | 3 | 1 | 1 | 1 | 3 | 1 | 1 | 1 | 8 | 1 |
| 19_596790 | 3 | 1 | 1 | 1 | 1 | 1 | 1 | 1 | 6 | 1 | 3 | 0.3333 | 1 | 1 | 3 | 1 | 1 | 1 | 8 | 0.75 |
| 19_596794 | 2 | 1 | 1 | 1 | 1 | 1 | NA | NA | 4 | 1 | 3 | 1 | 1 | 1 | 3 | 1 | 1 | 1 | 8 | 1 |
| 19_596805 | 1 | 0 | NA | NA | 1 | 1 | 1 | 0 | 3 | 0.3333 | 3 | 0.3333 | 1 | 0 | NA | NA | 1 | 0 | 5 | 0.2 |
| 19_596813 | 2 | 1 | 1 | 0 | NA | NA | 1 | 0 | 4 | 0.5 | 3 | 0.6667 | 1 | 0 | 1 | 1 | 1 | 0 | 6 | 0.5 |
| 19_596854 | 2 | 0 | NA | NA | 1 | 1 | 1 | 0 | 4 | 0.25 | 2 | 0 | 1 | 0 | 2 | 0.5 | 1 | 0 | 6 | 0.1667 |
| 19_596859 | 1 | 1 | NA | NA | 1 | 1 | 1 | 1 | 3 | 1 | 2 | 1 | 1 | 1 | 2 | 1 | 1 | 1 | 6 | 1 |
| 19_596872 | 1 | 1 | 1 | 1 | 1 | 1 | 1 | 1 | 4 | 1 | 2 | 0.5 | 1 | 1 | 2 | 1 | 1 | 1 | 6 | 0.8333 |
| 19_596873 | 1 | 1 | 1 | 1 | 1 | 1 | 1 | 1 | 4 | 1 | 2 | 1 | NA | NA | 2 | 1 | 1 | 1 | 5 | 1 |
| 19_596879 | 1 | 1 | 1 | 1 | 1 | 1 | 1 | 1 | 4 | 1 | 2 | 1 | 1 | 1 | 1 | 1 | 1 | 1 | 5 | 1 |
| 19_596886 | 1 | 1 | 1 | 0 | NA | NA | 1 | 1 | 3 | 0.6667 | 2 | 0 | 1 | 0 | 2 | 1 | 1 | 0 | 6 | 0.3333 |
| 19_596892 | NA | NA | 1 | 1 | 1 | 1 | 1 | 1 | 3 | 1 | 1 | 1 | 1 | 1 | 2 | 1 | 1 | 1 | 5 | 1 |

| | | | | | | | | | | | | | | | | | | | | |
|---|---|---|---|---|---|---|---|---|---|---|---|---|---|---|---|---|---|---|---|---|
| 19_603914 | 4 | 0.25 | 8 | 0 | 2 | 0 | 4 | 0 | 18 | 0.0556 | 1 | 0 | 4 | 0.25 | NA | NA | 4 | 0 | 9 | 0.1111 |
| 19_608843 | NA | NA | NA | NA | NA | NA | 1 | 1 | 1 | 1 | NA | NA | NA | NA | 1 | 1 | NA | NA | 1 | 1 |
| 19_609863 | 4 | 0.5 | 1 | 0 | 2 | 0 | 4 | 0 | 11 | 0.1818 | 3 | 0.3333 | 1 | 0 | 8 | 0 | 3 | 0 | 15 | 0.0667 |
| 19_610280 | 8 | 1 | 5 | 1 | 3 | 1 | 6 | 1 | 22 | 1 | 6 | 0.8333 | 3 | 1 | 7 | 0.8571 | 8 | 0.75 | 24 | 0.8333 |
| 19_610302 | 7 | 0.7143 | 4 | 0.25 | 4 | 1 | 9 | 0.8889 | 24 | 0.75 | 6 | 0.8333 | 6 | 0.8333 | 8 | 0.625 | 8 | 1 | 28 | 0.8214 |
| 19_610304 | 8 | 0.5 | 5 | 0.2 | 3 | 0.3333 | 8 | 0.375 | 24 | 0.375 | 4 | 0.25 | 4 | 0 | 7 | 0.7143 | 7 | 0.5714 | 22 | 0.4545 |
| 19_610309 | 8 | 0.125 | 5 | 0.2 | 4 | 0 | 8 | 0.125 | 25 | 0.12 | 8 | 0.25 | 6 | 0 | 9 | 0.1111 | 9 | 0.1111 | 32 | 0.125 |
| 19_610310 | 8 | 0.25 | 5 | 0.8 | 3 | 0.6667 | 7 | 0.1429 | 23 | 0.3913 | 8 | 0.375 | 4 | 0.25 | 10 | 0.1 | 8 | 0.375 | 30 | 0.2667 |
| 19_610312 | 9 | 0.4444 | 5 | 0.4 | 3 | 0.3333 | 9 | 0.1111 | 26 | 0.3077 | 7 | 0.4286 | 5 | 0.6 | 9 | 0.5556 | 9 | 0.3333 | 30 | 0.4667 |
| 19_610344 | 8 | 0.875 | 4 | 0.25 | 3 | 0.6667 | 7 | 0.5714 | 22 | 0.6364 | 7 | 0.5714 | 5 | 0.4 | 7 | 0.2857 | 5 | 0.4 | 24 | 0.4167 |
| 19_610354 | 8 | 0.5 | 5 | 0.6 | 3 | 0.6667 | 8 | 0.5 | 24 | 0.5417 | 7 | 0.8571 | 8 | 0.75 | 7 | 0.7143 | 7 | 0.5714 | 29 | 0.7241 |
| 19_610427 | 8 | 0.125 | 7 | 0.1429 | 12 | 0 | 3 | 0 | 30 | 0.0667 | 3 | 0 | 4 | 0.25 | 6 | 0 | 1 | 0 | 14 | 0.0714 |
| 19_610450 | 6 | 0.1667 | 7 | 0 | 11 | 0.0909 | 5 | 0 | 29 | 0.069 | 5 | 0.2 | 6 | 0.1667 | 6 | 0.1667 | 2 | 0 | 19 | 0.1579 |
| 19_610482 | 4 | 0 | 5 | 0.2 | 10 | 0.1 | 2 | 0 | 21 | 0.0952 | 4 | 0 | 2 | 0 | 3 | 0 | 2 | 0.5 | 11 | 0.0909 |
| 19_611154 | 9 | 0.8889 | 5 | 0.6 | 2 | 1 | 5 | 0.8 | 21 | 0.8095 | 2 | 1 | 7 | 0.5714 | 5 | 0.8 | 10 | 0.6 | 24 | 0.6667 |
| 19_611228 | 10 | 0.2 | 3 | 0 | 1 | 0 | 6 | 0.5 | 20 | 0.25 | 2 | 0 | 7 | 0.5714 | 4 | 0.75 | 7 | 0.7143 | 20 | 0.6 |
| 19_611232 | 11 | 0 | 4 | 0.25 | 2 | 0.5 | 5 | 0.2 | 22 | 0.1364 | NA | NA | 6 | 0.5 | 5 | 0.2 | 10 | 0 | 21 | 0.1905 |
| 19_611243 | 11 | 1 | 5 | 1 | 2 | 1 | 3 | 1 | 21 | 1 | 1 | 1 | 7 | 1 | 3 | 1 | 9 | 0.8889 | 20 | 0.95 |
| 19_611244 | 7 | 0.4286 | 3 | 0 | NA | NA | 4 | 0.25 | 14 | 0.2857 | 1 | 0 | 5 | 0.4 | 3 | 0.6667 | 7 | 0.1429 | 16 | 0.3125 |
| 19_611245 | 10 | 0.6 | 1 | 0 | 1 | 1 | 5 | 0.6 | 17 | 0.5882 | NA | NA | 3 | 0.6667 | 4 | 0.5 | 8 | 0.75 | 15 | 0.6667 |
| 19_611803 | 6 | 0 | 4 | 0.25 | 10 | 0.1 | 11 | 0.0909 | 31 | 0.0968 | 9 | 0 | 11 | 0.1818 | 12 | 0 | 5 | 0.2 | 37 | 0.0811 |
| 19_611830 | 11 | 0.8182 | 5 | 0.6 | 8 | 0.625 | 10 | 0.6 | 34 | 0.6765 | 8 | 0.375 | 9 | 0.6667 | 9 | 0.5556 | 5 | 1 | 31 | 0.6129 |
| 19_611911 | 9 | 0 | 5 | 0 | 10 | 0.1 | 9 | 0.1111 | 33 | 0.0606 | 6 | 0 | 7 | 0 | 10 | 0.2 | 3 | 0 | 26 | 0.0769 |
| 19_611990 | 4 | 0 | 2 | 0 | 5 | 0 | 6 | 0.1667 | 17 | 0.0588 | 4 | 0 | 5 | 0 | 5 | 0.2 | 2 | 0 | 16 | 0.0625 |
| 19_612106 | 4 | 0.5 | 2 | 0.5 | 3 | 0 | 6 | 0.1667 | 15 | 0.2667 | 1 | 0 | 4 | 0.25 | 4 | 0 | 2 | 0 | 11 | 0.0909 |
| 19_612625 | NA | NA | NA | NA | NA | NA | 1 | 1 | 1 | 1 | 1 | 0 | 1 | 1 | 1 | 0 | NA | NA | 3 | 0.3333 |
| 19_613080 | 6 | 0.5 | 3 | 0.3333 | 6 | 0.3333 | 4 | 0 | 19 | 0.3158 | NA | NA | 5 | 0.4 | 3 | 0.3333 | 3 | 0.6667 | 11 | 0.4545 |
| 19_613095 | 5 | 0.2 | 4 | 0 | 8 | 0.125 | 4 | 0.25 | 21 | 0.1429 | NA | NA | 5 | 0.4 | 3 | 0 | 4 | 0 | 12 | 0.1667 |
| 19_613108 | 3 | 0 | 2 | 0 | 6 | 0 | 3 | 0.3333 | 14 | 0.0714 | NA | NA | 5 | 0.2 | 2 | 0 | 2 | 0 | 9 | 0.1111 |
| 19_616604 | 3 | 0.6667 | 5 | 0.6 | 2 | 0.5 | 6 | 0.5 | 16 | 0.5625 | 2 | 0.5 | 3 | 1 | 2 | 0.5 | 4 | 0.25 | 11 | 0.5455 |
| 19_616614 | 4 | 0 | 5 | 0 | 5 | 0.2 | 6 | 0 | 20 | 0.05 | 3 | 0 | 3 | 0 | 4 | 0 | 4 | 0.25 | 14 | 0.0714 |
| 19_616615 | 4 | 0 | 5 | 0.2 | 5 | 0 | 6 | 0 | 20 | 0.05 | 2 | 0.5 | 1 | 0 | 2 | 0 | 5 | 0 | 10 | 0.1 |
| 19_616619 | 3 | 0 | 5 | 0.2 | 4 | 0.25 | 6 | 0 | 18 | 0.1111 | 3 | 0.6667 | 1 | 0 | 2 | 0 | 4 | 0.25 | 10 | 0.3 |
| 19_616624 | 4 | 0.75 | 4 | 1 | 5 | 1 | 6 | 0.6667 | 19 | 0.8421 | 2 | 1 | 1 | 1 | 3 | 0.6667 | 5 | 0.8 | 11 | 0.8182 |
| 19_616628 | 4 | 0.25 | 5 | 0.2 | 4 | 0 | 5 | 0.6 | 18 | 0.2778 | 2 | 1 | 2 | 0.5 | 3 | 0.3333 | 5 | 0.6 | 12 | 0.5833 |
| 19_616629 | 3 | 1 | 3 | 1 | 3 | 1 | 5 | 1 | 14 | 1 | 1 | 1 | 2 | 1 | 3 | 0.6667 | 5 | 1 | 11 | 0.9091 |
| 19_616631 | 4 | 0 | 5 | 0.4 | 5 | 0.2 | 6 | 0 | 20 | 0.15 | 2 | 0.5 | 2 | 1 | 3 | 0 | 5 | 0 | 12 | 0.25 |
| 19_616633 | 4 | 0.75 | 5 | 1 | 3 | 0 | 5 | 0.4 | 17 | 0.5882 | 2 | 0.5 | 1 | 0 | 3 | 0.6667 | 4 | 0.75 | 10 | 0.6 |
| 19_616635 | 2 | 0 | 3 | 0.3333 | 3 | 0.3333 | 5 | 0.4 | 13 | 0.3077 | 1 | 1 | NA | NA | 2 | 1 | 2 | 0.5 | 5 | 0.8 |
| 19_616636 | 4 | 1 | 5 | 1 | 4 | 1 | 6 | 1 | 19 | 1 | NA | NA | 2 | 1 | NA | NA | 3 | 1 | 5 | 1 |
| 19_616638 | 3 | 1 | 4 | 0.5 | 4 | 0.5 | 6 | 0.5 | 17 | 0.5882 | 2 | 1 | 1 | 0 | 2 | 0.5 | 3 | 0.6667 | 8 | 0.625 |
| 19_616644 | 4 | 0.5 | 5 | 0 | 2 | 0 | 4 | 0 | 15 | 0.1333 | 1 | 0 | 1 | 0 | 3 | 0 | 4 | 0.5 | 9 | 0.2222 |
| 19_616970 | 9 | 0.1111 | 5 | 0.4 | 6 | 0.3333 | 6 | 0.3333 | 26 | 0.2692 | 9 | 0.3333 | 9 | 0.1111 | 8 | 0.25 | 4 | 0 | 30 | 0.2 |
| 19_616975 | 8 | 0.5 | 6 | 0.8333 | 7 | 0.5714 | 7 | 0.5714 | 28 | 0.6071 | 8 | 0.125 | 8 | 0.375 | 8 | 0.375 | 4 | 0.25 | 28 | 0.2857 |
| 19_616982 | 9 | 0.1111 | 6 | 0.1667 | 6 | 0.3333 | 7 | 0.1429 | 28 | 0.1786 | 9 | 0.3333 | 9 | 0.1111 | 8 | 0.25 | 4 | 0.75 | 30 | 0.3 |
| 19_617405 | 1 | 0 | 1 | 1 | NA | NA | 1 | 0 | 3 | 0.3333 | NA | NA | 1 | 1 | NA | NA | NA | NA | 1 | 1 |
| 19_618449 | 16 | 0.375 | 18 | 0.0556 | 18 | 0.1667 | 13 | 0 | 65 | 0.1538 | 11 | 0.2727 | 19 | 0.1053 | 18 | 0.2778 | 13 | 0.2308 | 61 | 0.2131 |
| 19_618481 | 12 | 0.25 | 16 | 0.125 | 15 | 0.1333 | 11 | 0.0909 | 54 | 0.1481 | 9 | 0.2222 | 17 | 0.1176 | 12 | 0.25 | 14 | 0.1429 | 52 | 0.1731 |
| 19_618754 | 14 | 0.3571 | 19 | 0.4737 | 15 | 0.1333 | 15 | 0.1333 | 63 | 0.2857 | 11 | 0.0909 | 24 | 0.1667 | 13 | 0.1538 | 14 | 0.2857 | 62 | 0.1774 |
| 19_627797 | 29 | 0.069 | 23 | 0.3043 | 20 | 0.3 | 24 | 0.125 | 96 | 0.1875 | 16 | 0.125 | 34 | 0.0882 | 24 | 0.2083 | 12 | 0.1667 | 86 | 0.1395 |
| 19_627807 | 22 | 0.2727 | 20 | 0.4 | 22 | 0.4091 | 21 | 0.3333 | 85 | 0.3529 | 15 | 0.2667 | 29 | 0.2069 | 17 | 0.4706 | 11 | 0.3636 | 72 | 0.3056 |
| 19_627822 | 26 | 0.1154 | 21 | 0.0476 | 17 | 0.2353 | 19 | 0 | 83 | 0.0964 | 16 | 0.0625 | 27 | 0.1481 | 22 | 0.1818 | 8 | 0.125 | 73 | 0.137 |
| 19_627830 | 24 | 0.0833 | 22 | 0.1818 | 23 | 0.1304 | 25 | 0.12 | 94 | 0.1277 | 14 | 0.0714 | 34 | 0.0882 | 20 | 0.3 | 11 | 0.0909 | 79 | 0.1392 |
| 19_628496 | 11 | 0.2727 | 8 | 0.125 | 13 | 0.1538 | 8 | 0.25 | 40 | 0.2 | 7 | 0.2857 | 17 | 0.1176 | 12 | 0.4167 | 9 | 0.2222 | 45 | 0.2444 |
| 19_628500 | 15 | 0.0667 | 10 | 0.2 | 15 | 0.4 | 13 | 0.2308 | 53 | 0.2264 | 9 | 0.5556 | 21 | 0.381 | 14 | 0.2857 | 8 | 0.125 | 52 | 0.3462 |
| 19_628503 | 15 | 0.1333 | 10 | 0 | 13 | 0.0769 | 10 | 0.1 | 48 | 0.0833 | 8 | 0 | 21 | 0.1429 | 12 | 0.25 | 9 | 0.2222 | 50 | 0.16 |
| 19_629785 | 1 | 0 | 1 | 0 | 1 | 1 | 1 | 0 | 4 | 0.25 | 2 | 0 | NA | NA | 1 | 0 | 3 | 0.3333 | 6 | 0.1667 |
| 19_631169 | 3 | 1 | 4 | 0.5 | 5 | 0.4 | 2 | 0 | 14 | 0.5 | 2 | 1 | 4 | 0.5 | 5 | 0.8 | 4 | 0.75 | 15 | 0.7333 |
| 19_631182 | 4 | 0.25 | 1 | 0 | 5 | 0.6 | 2 | 0.5 | 12 | 0.4167 | 2 | 1 | 6 | 0.3333 | 4 | 0.25 | 3 | 0.6667 | 15 | 0.4667 |
| 19_631186 | 5 | 0.8 | 4 | 0.5 | 6 | 0.6667 | 2 | 1 | 17 | 0.7059 | NA | NA | 6 | 0.8333 | 4 | 0.75 | 5 | 1 | 15 | 0.8667 |
| 19_631211 | 5 | 0.8 | 4 | 1 | 7 | 1 | 2 | 1 | 18 | 0.9444 | 4 | 0.5 | 5 | 0.6 | 4 | 0 | 5 | 0.8 | 18 | 0.5 |
| 19_631218 | 5 | 0.8 | 4 | 0.75 | 4 | 1 | 2 | 1 | 15 | 0.8667 | 3 | 0.6667 | 6 | 0.8333 | 4 | 1 | 5 | 0.8 | 18 | 0.8333 |
| 19_631219 | 4 | 1 | 3 | 0.6667 | 3 | 1 | 2 | 1 | 12 | 0.9167 | 3 | 1 | 6 | 0.8333 | 5 | 1 | 5 | 0.8 | 19 | 0.8947 |
| 19_631231 | 3 | 0.6667 | 3 | 0.3333 | 5 | 0.2 | 2 | 0 | 13 | 0.3077 | 3 | 0 | 5 | 0.4 | 4 | 0.5 | 3 | 0.3333 | 15 | 0.3333 |
| 19_634457 | 5 | 0.2 | 7 | 0 | 3 | 0 | 1 | 0 | 16 | 0.0625 | 4 | 0 | 6 | 0.3333 | 3 | 0 | 4 | 0.25 | 17 | 0.1765 |

| | | | | | | | | | | | | | | | | | | | | |
|---|---|---|---|---|---|---|---|---|---|---|---|---|---|---|---|---|---|---|---|---|
| 19_634835 | 6 | 0.6667 | 6 | 0.6667 | 7 | 0.7143 | 1 | 0 | 20 | 0.65 | 3 | 0.6667 | 5 | 0.6 | 3 | 1 | 5 | 0.8 | 16 | 0.75 |
| 19_634849 | 7 | 0.5714 | 7 | 0.4286 | 4 | 0.75 | 1 | 1 | 19 | 0.5789 | 3 | 0.3333 | 2 | 1 | 3 | 0.3333 | 4 | 0.5 | 12 | 0.5 |
| 19_634862 | 6 | 0.8333 | 8 | 1 | 5 | 1 | 1 | 1 | 20 | 0.95 | 6 | 0.6667 | 5 | 1 | 4 | 0.75 | 4 | 1 | 19 | 0.8421 |
| 19_634877 | 5 | 1 | 5 | 1 | 5 | 1 | 1 | 1 | 16 | 1 | 6 | 0.6667 | 5 | 0.8 | 5 | 0.8 | 5 | 1 | 21 | 0.8095 |
| 19_634887 | 4 | 0.25 | 5 | 0.6 | 5 | 0.6 | 1 | 0 | 15 | 0.4667 | 3 | 0.3333 | 5 | 0.4 | 5 | 0.6 | 3 | 0.3333 | 16 | 0.4375 |
| 19_634933 | 4 | 0 | 6 | 0.1667 | 4 | 0 | NA | NA | 14 | 0.0714 | 4 | 0 | 4 | 0.25 | 3 | 0.3333 | 3 | 0 | 14 | 0.1429 |
| 19_638515 | NA | NA | 1 | 0 | 1 | 0 | 1 | 1 | 3 | 0.3333 | NA | NA | 1 | 1 | NA | NA | 1 | 0 | 2 | 0.5 |
| 19_638521 | 1 | 1 | 2 | 0 | NA | NA | NA | NA | 3 | 0.3333 | NA | NA | NA | NA | 2 | 0.5 | 1 | 0 | 3 | 0.3333 |
| 19_638530 | 2 | 0 | 2 | 0.5 | 2 | 0 | 1 | 0 | 7 | 0.1429 | NA | NA | 1 | 0 | 3 | 0.3333 | 1 | 1 | 5 | 0.4 |
| 19_638579 | 2 | 0 | 3 | 0 | 3 | 0.3333 | 4 | 0.25 | 12 | 0.1667 | 1 | 0 | 2 | 0 | 8 | 0.125 | 2 | 0 | 13 | 0.0769 |
| 19_638583 | NA | NA | NA | NA | 1 | 1 | NA | NA | 1 | 1 | 1 | 0 | 1 | 0 | 4 | 0.75 | 2 | 0 | 8 | 0.375 |
| 19_638897 | 3 | 0 | 2 | 0 | 3 | 0 | 6 | 0.1667 | 14 | 0.0714 | 5 | 0.2 | 3 | 0 | 7 | 0 | 2 | 0 | 17 | 0.0588 |
| 19_638986 | 5 | 0.2 | 5 | 0 | 4 | 0.5 | 5 | 0.2 | 19 | 0.2105 | 4 | 0 | 3 | 0.3333 | 7 | 0.1429 | 5 | 0.4 | 19 | 0.2105 |
| 19_638988 | 4 | 0.25 | 4 | 0 | 4 | 0.25 | 6 | 0.1667 | 18 | 0.1667 | 7 | 0.1429 | 3 | 0.6667 | 8 | 0 | 4 | 0.25 | 22 | 0.1818 |
| 19_638992 | 5 | 0.8 | 6 | 0.8333 | 4 | 0.5 | 7 | 0.5714 | 22 | 0.6818 | 3 | 0.3333 | 2 | 1 | 6 | 0.6667 | 4 | 0.5 | 15 | 0.6 |
| 19_639020 | 5 | 1 | 8 | 1 | 4 | 1 | 8 | 1 | 25 | 1 | 7 | 1 | 3 | 1 | 11 | 0.8182 | 6 | 1 | 27 | 0.9259 |
| 19_639028 | 5 | 0.2 | 7 | 0.1429 | 4 | 0.25 | 4 | 0.25 | 20 | 0.2 | 8 | 0.125 | 3 | 0 | 10 | 0.1 | 6 | 0 | 27 | 0.0741 |
| 19_639045 | 6 | 0.5 | 8 | 0.125 | 7 | 0.7143 | 9 | 0.6667 | 30 | 0.5 | 10 | 0.4 | 4 | 0.25 | 10 | 0.4 | 5 | 0.2 | 29 | 0.3448 |
| 19_639055 | 4 | 0.25 | 7 | 0.4286 | 7 | 0.5714 | 8 | 0.625 | 26 | 0.5 | 5 | 0.2 | 3 | 0.3333 | 9 | 0.1111 | 6 | 0.5 | 23 | 0.2609 |
| 19_639063 | 6 | 0.6667 | 9 | 0.5556 | 6 | 0.8333 | 8 | 0.875 | 29 | 0.7241 | 8 | 0.5 | 4 | 0.25 | 12 | 0.4167 | 6 | 0.8333 | 30 | 0.5 |
| 19_639070 | 7 | 0.2857 | 10 | 0.3 | 7 | 0.7143 | 9 | 0.4444 | 33 | 0.4242 | 10 | 0.3 | 5 | 0 | 13 | 0.3077 | 6 | 0.5 | 34 | 0.2941 |
| 19_639077 | 5 | 0 | 8 | 0 | 4 | 0.75 | 5 | 0.8 | 22 | 0.3182 | 7 | 0.4286 | 5 | 0.2 | 7 | 0.2857 | 6 | 0.3333 | 25 | 0.32 |
| 19_640369 | 4 | 0 | 4 | 0.25 | 4 | 0 | 3 | 0.3333 | 15 | 0.1333 | 3 | 0.3333 | 4 | 0.25 | 5 | 0.4 | NA | NA | 12 | 0.3333 |
| 19_640376 | 2 | 0 | 5 | 0.2 | 5 | 0 | 3 | 0.3333 | 15 | 0.1333 | 3 | 0 | 4 | 0.25 | 6 | 0 | 2 | 0.5 | 15 | 0.1333 |
| 19_640377 | 4 | 0 | 4 | 0.25 | 5 | 0.2 | 2 | 0 | 15 | 0.1333 | 3 | 0 | 3 | 0.3333 | 4 | 0 | 2 | 0 | 12 | 0.0833 |
| 19_640473 | 3 | 0.3333 | 3 | 0 | 4 | 0 | 1 | 0 | 11 | 0.0909 | 3 | 0.3333 | 2 | 0.5 | NA | NA | 3 | 0 | 8 | 0.25 |
| 19_640512 | 3 | 0.3333 | 4 | 0.75 | 4 | 0.25 | 3 | 0.6667 | 14 | 0.5 | 3 | 0.3333 | 3 | 0.6667 | NA | NA | 4 | 0 | 10 | 0.3 |
| 19_642592 | 10 | 0.4 | 13 | 0.3846 | 12 | 0.5 | 11 | 0.3636 | 46 | 0.413 | 2 | 0 | 13 | 0.1538 | 13 | 0.5385 | 6 | 0.3333 | 34 | 0.3235 |
| 19_642602 | 12 | 0.3333 | 15 | 0.0667 | 11 | 0.0909 | 10 | 0 | 48 | 0.125 | 2 | 0 | 13 | 0 | 13 | 0.0769 | 6 | 0.1667 | 34 | 0.0588 |
| 19_642611 | 15 | 0.0667 | 14 | 0.1429 | 10 | 0.1 | 10 | 0.1 | 49 | 0.102 | 4 | 0 | 12 | 0 | 9 | 0.2222 | 4 | 0.5 | 29 | 0.1379 |
| 19_642619 | 11 | 0.2727 | 9 | 0.3333 | 7 | 0.4286 | 6 | 0.5 | 33 | 0.3636 | 3 | 0.6667 | 8 | 0.5 | 12 | 0.6667 | 5 | 0.2 | 28 | 0.5357 |
| 19_642637 | 18 | 0.2778 | 19 | 0.3158 | 13 | 0.1538 | 17 | 0.2353 | 67 | 0.2537 | 11 | 0 | 22 | 0.3182 | 19 | 0.3684 | 13 | 0.1538 | 65 | 0.2462 |
| 19_642642 | 16 | 0 | 18 | 0.1111 | 13 | 0.2308 | 17 | 0.0588 | 64 | 0.0938 | 7 | 0 | 20 | 0.2 | 15 | 0.0667 | 11 | 0.0909 | 53 | 0.1132 |
| 19_646503 | 9 | 0.3333 | 7 | 0.1429 | 9 | 0.2222 | 22 | 0.0455 | 47 | 0.1489 | 6 | 0 | 8 | 0.25 | 13 | 0 | 9 | 0 | 36 | 0.0556 |
| 19_647838 | 3 | 0.6667 | 1 | 0 | 1 | 1 | 2 | 0.5 | 7 | 0.5714 | 3 | 0.6667 | 2 | 0 | 3 | 0.6667 | NA | NA | 8 | 0.5 |
| 19_652346 | 4 | 0.25 | 3 | 1 | 1 | 1 | 2 | 0 | 10 | 0.5 | 3 | 0.3333 | 3 | 0.3333 | 3 | 0.6667 | 1 | 1 | 10 | 0.5 |
| 19_652354 | 3 | 0.6667 | 3 | 0.6667 | 2 | 1 | 2 | 1 | 10 | 0.8 | 3 | 1 | 2 | 0.5 | 1 | 0 | 1 | 1 | 7 | 0.7143 |
| 19_652369 | 3 | 0.3333 | 1 | 0 | 2 | 0 | 1 | 1 | 7 | 0.2857 | 3 | 0 | 1 | 0 | 3 | 1 | NA | NA | 7 | 0.4286 |
| 19_652376 | 3 | 0.3333 | 3 | 1 | 2 | 1 | 1 | 1 | 9 | 0.7778 | 3 | 0.3333 | 4 | 1 | 1 | 1 | 1 | 1 | 9 | 0.7778 |
| 19_653125 | 4 | 0.25 | 3 | 0 | 3 | 0 | 2 | 0.5 | 12 | 0.1667 | 3 | 0.3333 | 2 | 0 | 3 | 0.6667 | 1 | 0 | 9 | 0.3333 |
| 19_653128 | 3 | 0 | 2 | 0 | 1 | 1 | 2 | 0 | 8 | 0.125 | 3 | 0 | 4 | 0.25 | 2 | 0 | 1 | 0 | 10 | 0.1 |
| 19_653181 | 3 | 0.3333 | 4 | 1 | 2 | 1 | 2 | 0.5 | 11 | 0.7273 | 4 | 0.25 | 4 | 0.5 | 4 | 0.75 | NA | NA | 12 | 0.5 |
| 19_654070 | 5 | 0.6 | 1 | 1 | 1 | 0 | 2 | 0.5 | 9 | 0.5556 | 4 | 0.75 | 4 | 0.75 | 4 | 0.75 | 1 | 1 | 13 | 0.7692 |
| 19_654088 | 6 | 0.3333 | 2 | 1 | 3 | 0.6667 | 2 | 1 | 13 | 0.6154 | 5 | 0.6 | 4 | 0.5 | 4 | 0.5 | 1 | 1 | 14 | 0.5714 |
| 19_654156 | 7 | 0.1429 | 3 | 0 | 4 | 0 | 2 | 0 | 16 | 0.0625 | 4 | 0.25 | 4 | 0 | 4 | 0.25 | 1 | 0 | 13 | 0.1538 |
| 19_654158 | 7 | 0.1429 | 3 | 0 | 4 | 0 | 2 | 0 | 16 | 0.0625 | 5 | 0 | 3 | 0 | 4 | 0 | 1 | 1 | 13 | 0.0769 |
| 19_654162 | 7 | 0 | 3 | 0 | 3 | 0.3333 | 2 | 0.5 | 15 | 0.1333 | 5 | 0.2 | 4 | 0.25 | 4 | 0.5 | 1 | 0 | 14 | 0.2857 |
| 19_654809 | 7 | 0.2857 | 9 | 0.3333 | 7 | 0.5714 | 5 | 0.4 | 28 | 0.3929 | 12 | 0.4167 | 8 | 0 | 6 | 0.1667 | 4 | 0 | 30 | 0.2 |
| 19_654816 | 6 | 0.6667 | 10 | 0.6 | 5 | 0.8 | 5 | 0.4 | 26 | 0.6154 | 11 | 0.6364 | 7 | 0.4286 | 6 | 0.6667 | 6 | 0.8333 | 30 | 0.6333 |
| 19_658108 | NA | NA | 2 | 1 | 2 | 1 | NA | NA | 4 | 1 | NA | NA | NA | NA | NA | NA | 3 | 1 | 3 | 1 |
| 19_658133 | 15 | 0.2 | 15 | 0.0667 | 14 | 0.2143 | 13 | 0.2308 | 57 | 0.1754 | 19 | 0.0526 | 17 | 0.1176 | 13 | 0.0769 | 8 | 0 | 57 | 0.0702 |
| 19_658158 | 14 | 1 | 16 | 1 | 18 | 0.9444 | 13 | 1 | 61 | 0.9836 | 17 | 1 | 17 | 0.9412 | 15 | 0.9333 | 6 | 1 | 55 | 0.9636 |
| 19_658161 | 15 | 0.7333 | 15 | 0.6 | 19 | 0.6316 | 12 | 0.9167 | 61 | 0.7049 | 20 | 0.7 | 15 | 0.8 | 14 | 0.7143 | 7 | 0.7143 | 56 | 0.7321 |
| 19_658165 | 6 | 0.6667 | 9 | 0.8889 | 9 | 0.7778 | 6 | 0.6667 | 30 | 0.7667 | 7 | 0.7143 | 8 | 0.875 | 7 | 0.7143 | 1 | 1 | 23 | 0.7826 |
| 19_658173 | 12 | 0.4167 | 11 | 0.6364 | 19 | 0.4211 | 13 | 0.3077 | 55 | 0.4364 | 16 | 0.5625 | 12 | 0.4167 | 11 | 0.7273 | 6 | 0.6667 | 45 | 0.5778 |
| 19_658173 | NA | NA | 1 | 1 | NA | NA | 1 | 0 | 2 | 0.5 | 2 | 1 | NA | NA | 1 | 1 | NA | NA | 3 | 1 |
| 19_658174 | 15 | 0.2 | 17 | 0 | 20 | 0.05 | 17 | 0.2353 | 69 | 0.1159 | 17 | 0.1176 | 14 | 0.2857 | 12 | 0.0833 | 8 | 0.125 | 51 | 0.1569 |
| 19_658176 | 9 | 0.1111 | 9 | 0.1111 | 12 | 0.1667 | 9 | 0 | 39 | 0.1026 | 13 | 0.2308 | 10 | 0 | 10 | 0.1 | 3 | 0 | 36 | 0.1111 |
| 19_658205 | 24 | 0.2917 | 26 | 0.2308 | 25 | 0.08 | 18 | 0.1111 | 93 | 0.1828 | 28 | 0.3214 | 25 | 0.24 | 20 | 0.2 | 11 | 0.2727 | 84 | 0.2619 |
| 19_658207 | 20 | 0.25 | 18 | 0.1111 | 22 | 0.2273 | 15 | 0.2667 | 75 | 0.2133 | 26 | 0.1154 | 22 | 0.1364 | 19 | 0.2105 | 8 | 0.125 | 75 | 0.1467 |
| 19_658215 | 29 | 0.2414 | 23 | 0.1304 | 24 | 0.2083 | 17 | 0.2941 | 93 | 0.2151 | 23 | 0.3043 | 20 | 0.35 | 20 | 0.3 | 10 | 0 | 73 | 0.274 |
| 19_658226 | 26 | 0.8077 | 25 | 0.56 | 22 | 0.5909 | 16 | 0.5 | 89 | 0.6292 | 29 | 0.5517 | 25 | 0.6 | 20 | 0.75 | 9 | 0.6667 | 83 | 0.6265 |
| 19_658233 | 24 | 0.0833 | 21 | 0 | 18 | 0.1667 | 16 | 0 | 79 | 0.0633 | 26 | 0.0769 | 26 | 0.0385 | 19 | 0.0526 | 10 | 0.1 | 81 | 0.0617 |
| 19_658288 | 33 | 0.1212 | 31 | 0.0323 | 25 | 0.12 | 25 | 0.08 | 114 | 0.0877 | 43 | 0.1628 | 26 | 0.2692 | 28 | 0.0714 | 16 | 0.25 | 113 | 0.177 |
| 19_658289 | 35 | 0.1429 | 33 | 0.0606 | 26 | 0.0769 | 21 | 0.1429 | 115 | 0.1043 | 46 | 0.087 | 31 | 0.0968 | 24 | 0.0833 | 17 | 0.1765 | 118 | 0.1017 |

| | | | | | | | | | | | | | | | | | | | | |
|---|---|---|---|---|---|---|---|---|---|---|---|---|---|---|---|---|---|---|---|---|
| 19_680681 | 6 | 0 | 7 | 0.1429 | 8 | 0.125 | 4 | 0 | 25 | 0.08 | 6 | 0.3333 | 3 | 0 | 11 | 0.0909 | 5 | 0 | 25 | 0.12 |
| 19_680682 | 4 | 0.25 | 7 | 0 | 5 | 0.2 | 4 | 0 | 20 | 0.1 | 4 | 0 | 3 | 0 | 9 | 0.1111 | 5 | 0.2 | 21 | 0.0952 |
| 19_680685 | 6 | 0.3333 | 7 | 0.2857 | 6 | 0.1667 | 4 | 0.25 | 23 | 0.2609 | 7 | 0.4286 | 1 | 1 | 11 | 0.1818 | 6 | 0 | 25 | 0.24 |
| 19_680687 | 5 | 0.8 | 6 | 0.3333 | 6 | 0.5 | 4 | 0.5 | 21 | 0.5238 | 5 | 0.4 | 2 | 0.5 | 8 | 0.5 | 4 | 0.5 | 19 | 0.4737 |
| 19_680692 | 5 | 0.8 | 5 | 0.6 | 6 | 0.6667 | 3 | 0.6667 | 19 | 0.6842 | 5 | 0.6 | 2 | 0.5 | 10 | 0.4 | 5 | 1 | 22 | 0.5909 |
| 19_688719 | 7 | 0.4286 | 7 | 0.4286 | 6 | 0.5 | 5 | 0.6 | 25 | 0.48 | 8 | 0.5 | 5 | 0.6 | 6 | 0.5 | 7 | 0.5714 | 26 | 0.5385 |
| 19_688735 | 2 | 0 | 1 | 0 | 3 | 0 | 4 | 0.25 | 10 | 0.1 | 6 | 0 | 5 | 0 | 2 | 0 | 4 | 0.25 | 17 | 0.0588 |
| 19_688739 | 2 | 0.5 | 1 | 1 | 1 | 0 | 2 | 0.5 | 6 | 0.5 | NA | NA | 3 | 0 | NA | NA | 3 | 0.3333 | 6 | 0.1667 |
| 19_688743 | NA | NA | 1 | 0 | 1 | 0 | 3 | 0.3333 | 5 | 0.2 | NA | NA | 1 | 0 | NA | NA | 2 | 1 | 3 | 0.6667 |
| 19_688843 | 15 | 0.2667 | 16 | 0.3125 | 15 | 0.4667 | 11 | 0.2727 | 57 | 0.3333 | 7 | 0.2857 | 10 | 0.6 | 8 | 0.5 | 10 | 0.6 | 35 | 0.5143 |
| 19_688912 | 9 | 1 | 5 | 1 | 5 | 0.8 | 6 | 0.8333 | 25 | 0.92 | 2 | 1 | 7 | 0.8571 | 3 | 1 | 7 | 1 | 19 | 0.9474 |
| 19_691563 | 10 | 0.1 | 17 | 0 | 9 | 0.1111 | 9 | 0.1111 | 45 | 0.0667 | 15 | 0.1333 | 7 | 0.1429 | 12 | 0.0833 | 12 | 0.1667 | 46 | 0.1304 |
| 19_691849 | 3 | 0.3333 | 4 | 0 | 3 | 0 | 6 | 0 | 16 | 0.0625 | 7 | 0.1429 | 2 | 0 | 5 | 0 | 6 | 0 | 20 | 0.05 |
| 19_691854 | 4 | 0.5 | 5 | 0.8 | 3 | 0.6667 | 5 | 0.4 | 17 | 0.5882 | 7 | 0.5714 | 3 | 0.6667 | 3 | 0.6667 | 4 | 0.5 | 17 | 0.5882 |
| 19_691856 | 3 | 0.3333 | 3 | 0 | 2 | 0 | 6 | 0.1667 | 14 | 0.1429 | 6 | 0.3333 | 3 | 0.3333 | 4 | 0 | 3 | 0.3333 | 16 | 0.25 |
| 19_691866 | NA | NA | 2 | 0 | 1 | 1 | 2 | 0.5 | 5 | 0.4 | 3 | 0.3333 | NA | NA | 1 | 0 | 2 | 0 | 6 | 0.1667 |
| 19_691871 | 4 | 0.25 | 5 | 0.6 | 2 | 0.5 | 4 | 0 | 15 | 0.3333 | 6 | 0.3333 | 2 | 0 | 3 | 0.3333 | 4 | 0 | 15 | 0.2 |
| 19_691899 | NA | NA | 1 | 1 | 1 | 1 | NA | NA | 2 | 1 | 1 | 1 | NA | NA | NA | NA | NA | NA | 1 | 1 |
| 19_691965 | 1 | 0 | 1 | 1 | NA | NA | 1 | 1 | 3 | 0.6667 | 1 | 1 | 1 | 1 | 1 | 0 | NA | NA | 3 | 0.6667 |
| 19_692876 | 4 | 0 | 5 | 0.2 | 6 | 0 | 5 | 0 | 20 | 0.05 | 5 | 0.2 | 7 | 0 | 8 | 0.125 | 3 | 0 | 23 | 0.087 |
| 19_692891 | 2 | 0 | 8 | 0.375 | 4 | 0.5 | 4 | 0.25 | 18 | 0.3333 | 3 | 0 | 5 | 0.4 | 8 | 0.125 | 3 | 0 | 19 | 0.1579 |
| 19_692915 | 3 | 1 | 8 | 0.875 | 5 | 1 | 5 | 1 | 21 | 0.9524 | 5 | 1 | 4 | 1 | 7 | 0.8571 | 1 | 1 | 17 | 0.9412 |
| 19_695461 | 10 | 0 | 13 | 0.0769 | 3 | 0.3333 | 8 | 0.125 | 34 | 0.0882 | 8 | 0.125 | 11 | 0.1818 | 6 | 0.1667 | 8 | 0.25 | 33 | 0.1818 |
| 19_695519 | 26 | 0.1923 | 30 | 0.0333 | 25 | 0.08 | 22 | 0.0455 | 103 | 0.0874 | 30 | 0.0333 | 28 | 0.1429 | 24 | 0.0417 | 17 | 0.0588 | 99 | 0.0707 |
| 19_695524 | 23 | 0.087 | 27 | 0.1111 | 29 | 0 | 16 | 0 | 95 | 0.0526 | 24 | 0.0833 | 14 | 0 | 20 | 0.1 | 15 | 0 | 73 | 0.0548 |
| 19_695703 | 1 | 0 | NA | NA | 1 | 0 | 3 | 0.3333 | 5 | 0.2 | NA | NA | 1 | 0 | 1 | 1 | NA | NA | 2 | 0.5 |
| 19_695989 | 20 | 0.15 | 26 | 0 | 26 | 0.0769 | 22 | 0.0909 | 94 | 0.0745 | 10 | 0.1 | 20 | 0.15 | 16 | 0.0625 | 16 | 0.0625 | 62 | 0.0968 |
| 19_696003 | 20 | 0.05 | 23 | 0.0435 | 25 | 0.04 | 24 | 0.125 | 92 | 0.0652 | 12 | 0.1667 | 21 | 0.1905 | 19 | 0.0526 | 17 | 0.0588 | 69 | 0.1159 |
| 19_696009 | 18 | 0.3889 | 21 | 0.619 | 22 | 0.3182 | 21 | 0.2857 | 82 | 0.4024 | 11 | 0.1818 | 19 | 0.3158 | 16 | 0.125 | 13 | 0.3077 | 59 | 0.2373 |
| 19_696010 | 21 | 0.0476 | 22 | 0.0455 | 24 | 0.0417 | 21 | 0.0952 | 88 | 0.0568 | 12 | 0.0833 | 21 | 0.1429 | 18 | 0.1111 | 17 | 0.1765 | 68 | 0.1324 |
| 19_697039 | 1 | 0 | 3 | 0.6667 | 2 | 0 | 4 | 0.25 | 10 | 0.3 | 1 | 0 | 1 | 0 | 4 | 0.25 | 1 | 0 | 7 | 0.1429 |
| 19_697055 | 2 | 0.5 | 2 | 0 | 1 | 0 | 4 | 0 | 9 | 0.1111 | NA | NA | 2 | 0 | 3 | 0.3333 | 1 | 0 | 6 | 0.1667 |
| 19_697282 | 1 | 0 | 3 | 0.3333 | 2 | 0.5 | 4 | 0.75 | 10 | 0.5 | 1 | 1 | 1 | 0 | 4 | 0.75 | 1 | 1 | 7 | 0.7143 |
| 19_697629 | 6 | 0.1667 | 3 | 0.3333 | 1 | 0 | 4 | 0.5 | 14 | 0.2857 | 3 | 0.3333 | 4 | 0.5 | 1 | 0 | NA | NA | 8 | 0.375 |
| 19_706605 | 3 | 0 | 1 | 0 | 2 | 0 | 2 | 0.5 | 8 | 0.125 | 1 | 0 | 2 | 1 | 2 | 0 | 4 | 0.5 | 9 | 0.4444 |
| 19_706616 | 3 | 0 | NA | NA | 1 | 0 | 2 | 0.5 | 6 | 0.1667 | 1 | 1 | 2 | 0 | 1 | 1 | 2 | 0 | 6 | 0.3333 |
| 19_706623 | 2 | 0 | 1 | 0 | 3 | 0 | 1 | 1 | 7 | 0.1429 | 1 | 0 | 3 | 0.3333 | 3 | 0 | 3 | 0.3333 | 10 | 0.2 |
| 19_706702 | 1 | 0 | 1 | 1 | 2 | 0.5 | 2 | 0.5 | 6 | 0.5 | 1 | 0 | 3 | 0.6667 | 3 | 0 | 2 | 1 | 9 | 0.4444 |
| 19_706727 | 3 | 0.3333 | NA | NA | 2 | 0 | 2 | 0 | 7 | 0.1429 | 1 | 0 | 3 | 0.3333 | 4 | 0.25 | 1 | 0 | 9 | 0.2222 |
| 19_706731 | 2 | 1 | NA | NA | 3 | 0.3333 | 2 | 0 | 7 | 0.4286 | 1 | 0 | 1 | 1 | 3 | 0.3333 | NA | NA | 5 | 0.4 |
| 19_706737 | 3 | 1 | NA | NA | 3 | 1 | 2 | 1 | 8 | 1 | 1 | 1 | 3 | 1 | 3 | 1 | 1 | 1 | 8 | 1 |
| 19_706741 | 3 | 0 | NA | NA | 3 | 0.3333 | 2 | 0 | 8 | 0.125 | 1 | 0 | 3 | 0 | 3 | 0.3333 | NA | NA | 7 | 0.1429 |
| 19_706765 | 3 | 0.3333 | NA | NA | 2 | 0 | 1 | 0 | 6 | 0.1667 | 1 | 0 | 2 | 0 | 3 | 0.6667 | NA | NA | 6 | 0.3333 |
| 19_706787 | 1 | 1 | NA | NA | 1 | 1 | 2 | 0.5 | 4 | 0.75 | 1 | 1 | 3 | 1 | 3 | 1 | 1 | 1 | 8 | 1 |
| 19_707056 | NA | NA | NA | NA | 1 | 1 | NA | NA | 1 | 1 | NA | NA | 1 | 1 | 2 | 0.5 | 2 | 0.5 | 5 | 0.6 |
| 19_707074 | NA | NA | NA | NA | 1 | 1 | NA | NA | 1 | 1 | NA | NA | 1 | 1 | 1 | 1 | 2 | 1 | 4 | 1 |
| 19_707141 | NA | NA | NA | NA | 1 | 1 | NA | NA | 1 | 1 | 1 | 1 | NA | NA | NA | NA | 3 | 0.3333 | 4 | 0.5 |
| 19_707183 | NA | NA | 1 | 1 | NA | NA | 1 | 1 | 2 | 1 | 2 | 1 | 1 | 1 | NA | NA | 3 | 1 | 6 | 1 |
| 19_707188 | 1 | 1 | NA | NA | NA | NA | 2 | 1 | 3 | 1 | NA | NA | NA | NA | NA | NA | 3 | 0.6667 | 3 | 0.6667 |
| 19_707202 | NA | NA | 1 | 1 | 1 | 1 | 2 | 1 | 4 | 1 | NA | NA | NA | NA | 1 | 1 | 1 | 1 | 2 | 1 |
| 19_707224 | NA | NA | 1 | 0 | 1 | 1 | 1 | 1 | 3 | 0.6667 | NA | NA | NA | NA | 1 | 1 | 1 | 1 | 2 | 1 |
| 19_707228 | NA | NA | 1 | 1 | 1 | 1 | 1 | 1 | 3 | 1 | NA | NA | NA | NA | 1 | 1 | 1 | 1 | 2 | 1 |
| 19_707242 | NA | NA | 1 | 1 | NA | NA | 1 | 0 | 2 | 0.5 | NA | NA | NA | NA | 1 | 0 | 1 | 1 | 2 | 0.5 |
| 19_707253 | NA | NA | 1 | 1 | NA | NA | 1 | 1 | 2 | 1 | NA | NA | NA | NA | 1 | 1 | 1 | 1 | 2 | 1 |
| 19_707269 | NA | NA | 1 | 1 | NA | NA | 1 | 1 | 2 | 1 | NA | NA | NA | NA | 1 | 1 | 1 | 1 | 2 | 1 |
| 19_707296 | NA | NA | 1 | 1 | NA | NA | NA | NA | 1 | 1 | NA | NA | NA | NA | NA | NA | 1 | 1 | 1 | 1 |
| 19_707309 | NA | NA | 1 | 1 | NA | NA | NA | NA | 1 | 1 | NA | NA | NA | NA | 1 | 1 | 1 | 1 | 2 | 1 |
| 19_707329 | NA | NA | 1 | 1 | NA | NA | NA | NA | 1 | 1 | NA | NA | NA | NA | 1 | 1 | 1 | 1 | 2 | 1 |
| 19_707343 | NA | NA | 1 | 1 | NA | NA | NA | NA | 1 | 1 | NA | NA | NA | NA | 1 | 0 | 1 | 1 | 2 | 0.5 |
| 19_707352 | NA | NA | 1 | 1 | NA | NA | NA | NA | 1 | 1 | NA | NA | NA | NA | 1 | 1 | NA | NA | 1 | 1 |
| 19_717593 | 38 | 0.0526 | 42 | 0.119 | 31 | 0.129 | 34 | 0.2059 | 145 | 0.1241 | 32 | 0.1875 | 43 | 0.1163 | 34 | 0.0294 | 21 | 0.1429 | 130 | 0.1154 |
| 19_717640 | 37 | 0.1081 | 46 | 0.087 | 38 | 0.0789 | 28 | 0.1071 | 149 | 0.094 | 35 | 0.0857 | 50 | 0.06 | 37 | 0.1081 | 23 | 0.1304 | 145 | 0.0897 |
| 19_717643 | 29 | 0.1034 | 40 | 0.3 | 32 | 0.3125 | 26 | 0.2692 | 127 | 0.252 | 29 | 0.2414 | 45 | 0.3111 | 33 | 0.1515 | 23 | 0.087 | 130 | 0.2154 |
| 19_719259 | 179 | 0.1229 | 217 | 0.1198 | 210 | 0.119 | 161 | 0.0994 | 767 | 0.116 | 194 | 0.1907 | 178 | 0.2079 | 189 | 0.164 | 182 | 0.1648 | 743 | 0.1817 |
| 19_720353 | 1 | 0 | 3 | 0.3333 | 2 | 0.5 | 1 | 0 | 7 | 0.2857 | 2 | 0.5 | 8 | 0.25 | 2 | 0 | 2 | 0.5 | 14 | 0.2857 |

| | | | | | | | | | | | | | | | | | | | | |
|---|---|---|---|---|---|---|---|---|---|---|---|---|---|---|---|---|---|---|---|---|
| 19_720354 | 1 | 1 | 4 | 0.25 | 2 | 0 | 1 | 0 | 8 | 0.25 | 2 | 0 | 8 | 0.125 | 1 | 0 | 3 | 0 | 14 | 0.0714 |
| 19_720386 | 1 | 0 | 2 | 0.5 | 2 | 0 | 1 | 0 | 6 | 0.1667 | 2 | 0.5 | 6 | 0 | 1 | 0 | 2 | 0.5 | 11 | 0.1818 |
| 19_720401 | 1 | 1 | 3 | 1 | 2 | 0.5 | 1 | 0 | 7 | 0.7143 | 1 | 0 | 8 | 0.75 | 1 | 0 | 2 | 0.5 | 12 | 0.5833 |
| 19_720410 | 1 | 1 | 2 | 0.5 | 1 | 0 | 1 | 0 | 5 | 0.4 | 2 | 0.5 | 7 | 0.4286 | 1 | 0 | 2 | 1 | 12 | 0.5 |
| 19_720416 | NA | NA | 2 | 0.5 | 2 | 0 | 1 | 0 | 5 | 0.2 | 2 | 0.5 | 6 | 0.5 | 1 | 0 | 2 | 0.5 | 11 | 0.4545 |
| 19_720417 | 1 | 1 | 3 | 0 | 2 | 0 | 1 | 0 | 7 | 0.1429 | 2 | 0.5 | 8 | 0.125 | 1 | 0 | 1 | 0 | 12 | 0.1667 |
| 19_720460 | 1 | 0 | 2 | 0 | 2 | 0.5 | 1 | 0 | 6 | 0.1667 | 2 | 0.5 | 7 | 0.2857 | NA | NA | 2 | 0 | 11 | 0.2727 |
| 19_720462 | 1 | 1 | 3 | 0.6667 | 2 | 1 | 1 | 0 | 7 | 0.7143 | 2 | 1 | 7 | 1 | NA | NA | 2 | 1 | 11 | 1 |
| 19_720481 | 1 | 0 | 2 | 0.5 | 2 | 0.5 | 1 | 0 | 6 | 0.3333 | 2 | 0 | 7 | 0.8571 | NA | NA | 2 | 0.5 | 11 | 0.6364 |
| 19_720483 | 1 | 0 | 2 | 0.5 | 2 | 0 | NA | NA | 5 | 0.2 | 2 | 0.5 | 6 | 0.3333 | NA | NA | 2 | 0.5 | 10 | 0.4 |
| 19_720484 | NA | NA | 2 | 0 | 1 | 1 | NA | NA | 3 | 0.3333 | 2 | 0.5 | 6 | 0.8333 | NA | NA | 1 | 1 | 9 | 0.7778 |
| 19_720485 | 1 | 0 | 2 | 0.5 | 2 | 0 | 1 | 0 | 6 | 0.1667 | 2 | 0 | 5 | 0.2 | NA | NA | 2 | 1 | 9 | 0.3333 |
| 19_720488 | 1 | 1 | 2 | 0 | 1 | 1 | 1 | 0 | 5 | 0.4 | 1 | 1 | 5 | 1 | NA | NA | 1 | 1 | 7 | 1 |
| 19_720494 | 1 | 1 | 2 | 1 | 1 | 1 | 1 | 1 | 5 | 1 | 1 | 1 | 6 | 1 | NA | NA | 1 | 1 | 8 | 1 |
| 19_720517 | 1 | 0 | 2 | 0.5 | 2 | 0 | 1 | 0 | 6 | 0.1667 | 2 | 0 | 3 | 0.6667 | NA | NA | 1 | 0 | 6 | 0.3333 |
| 19_720518 | 1 | 1 | 1 | 0 | NA | NA | 1 | 0 | 3 | 0.3333 | 2 | 0.5 | 4 | 0 | NA | NA | 1 | 1 | 7 | 0.2857 |
| 19_720531 | 1 | 1 | 1 | 0 | 2 | 0.5 | 1 | 0 | 5 | 0.4 | 1 | 0 | 3 | 0.6667 | NA | NA | 1 | 0 | 5 | 0.4 |
| 19_720536 | 1 | 0 | 2 | 0 | 2 | 0.5 | 1 | 0 | 6 | 0.1667 | 2 | 0 | 3 | 0.3333 | NA | NA | 1 | 0 | 6 | 0.1667 |
| 19_720540 | 1 | 0 | 2 | 0.5 | 2 | 0 | 1 | 0 | 6 | 0.1667 | 2 | 0 | 3 | 0.6667 | NA | NA | 1 | 0 | 6 | 0.3333 |
| 19_725209 | 3 | 0 | NA | NA | 1 | 1 | NA | NA | 4 | 0.25 | NA | NA | 2 | 0.5 | NA | NA | NA | NA | 2 | 0.5 |
| 19_725406 | 8 | 0.25 | 3 | 0.6667 | 8 | 0.125 | 8 | 0 | 27 | 0.1852 | 3 | 0 | 9 | 0.2222 | 6 | 0.1667 | 12 | 0.25 | 30 | 0.2 |
| 19_725437 | 9 | 0.3333 | 4 | 0.5 | 9 | 0 | 8 | 0.125 | 30 | 0.2 | 6 | 0 | 6 | 0.1667 | 6 | 0.3333 | 12 | 0 | 30 | 0.1 |
| 19_725441 | 6 | 1 | 3 | 1 | 7 | 1 | 7 | 1 | 23 | 1 | 6 | 1 | 8 | 0.875 | 5 | 0.8 | 7 | 0.8571 | 26 | 0.8846 |
| 19_725449 | 8 | 0.625 | 3 | 0.6667 | 5 | 0.2 | 8 | 0.5 | 24 | 0.5 | 5 | 0.4 | 8 | 0.375 | 7 | 0.8571 | 6 | 0.5 | 26 | 0.5385 |
| 19_725459 | 6 | 1 | 2 | 0.5 | 6 | 1 | 9 | 0.8889 | 23 | 0.913 | 5 | 1 | 10 | 1 | 3 | 1 | 11 | 1 | 29 | 1 |
| 19_731918 | NA | NA | NA | NA | 2 | 0.5 | NA | NA | 2 | 0.5 | NA | NA | NA | NA | NA | NA | 1 | 1 | 1 | 1 |
| 19_731946 | NA | NA | NA | NA | 1 | 1 | NA | NA | 1 | 1 | NA | NA | NA | NA | NA | NA | 2 | 1 | 2 | 1 |
| 19_733550 | 3 | 0 | 2 | 0.5 | 1 | 0 | NA | NA | 6 | 0.1667 | 10 | 0.1 | 2 | 0 | 4 | 0.25 | 3 | 0 | 19 | 0.1053 |
| 19_733576 | 2 | 0 | 4 | 0.25 | 1 | 0 | NA | NA | 7 | 0.1429 | 8 | 0.5 | 3 | 0 | 4 | 0 | 5 | 0.4 | 20 | 0.3 |
| 19_735168 | 2 | 0 | 3 | 0.3333 | 1 | 0 | NA | NA | 6 | 0.1667 | 3 | 0 | 1 | 0 | 1 | 0 | 4 | 0.5 | 9 | 0.2222 |
| 19_739484 | 1 | 1 | NA | NA | NA | NA | NA | NA | 1 | 1 | NA | NA | NA | NA | NA | NA | 1 | 1 | 1 | 1 |
| 19_739763 | 1 | 1 | 1 | 1 | 1 | 0 | NA | NA | 3 | 0.6667 | 1 | 0 | NA | NA | NA | NA | 1 | 1 | 2 | 0.5 |
| 19_740020 | NA | NA | 1 | 1 | 2 | 1 | NA | NA | 3 | 1 | NA | NA | NA | NA | 1 | 1 | 3 | 0.6667 | 4 | 0.75 |
| 19_740032 | 2 | 1 | 1 | 1 | 2 | 1 | NA | NA | 5 | 1 | 1 | 1 | NA | NA | 1 | 1 | 1 | 1 | 3 | 1 |
| 19_740034 | 3 | 0 | 1 | 0 | 2 | 0.5 | NA | NA | 6 | 0.1667 | 1 | 1 | NA | NA | 1 | 0 | 3 | 0 | 5 | 0.2 |
| 19_740050 | 3 | 1 | 1 | 1 | 1 | 1 | NA | NA | 5 | 1 | 1 | 1 | NA | NA | 1 | 1 | 4 | 0.75 | 6 | 0.8333 |
| 19_740062 | 3 | 0.3333 | 1 | 0 | 2 | 0 | NA | NA | 6 | 0.1667 | 2 | 0.5 | NA | NA | 2 | 0 | 2 | 0 | 6 | 0.1667 |
| 19_740097 | 3 | 1 | 1 | 1 | 2 | 1 | NA | NA | 6 | 1 | 1 | 1 | NA | NA | 1 | 1 | 2 | 1 | 4 | 1 |
| 19_740102 | 1 | 0 | 1 | 0 | 2 | 0.5 | NA | NA | 4 | 0.25 | 2 | 0.5 | NA | NA | 2 | 0.5 | 3 | 0.3333 | 7 | 0.4286 |
| 19_740109 | 3 | 0.3333 | 1 | 0 | 2 | 0.5 | NA | NA | 6 | 0.3333 | 3 | 0.6667 | NA | NA | 2 | 0.5 | 4 | 0.5 | 9 | 0.5556 |
| 19_740127 | 4 | 0.25 | 1 | 0 | 1 | 0 | NA | NA | 6 | 0.1667 | 2 | 0.5 | NA | NA | 2 | 1 | 4 | 0 | 8 | 0.375 |
| 19_740234 | 5 | 0 | 3 | 0.3333 | 3 | 0 | 1 | 0 | 12 | 0.0833 | 3 | 0 | NA | NA | 6 | 0.5 | 4 | 0.25 | 13 | 0.3077 |
| 19_740341 | 138 | 0.0942 | 156 | 0.1154 | 103 | 0.0194 | 122 | 0.041 | 519 | 0.0732 | 100 | 0.11 | 127 | 0.0472 | 115 | 0.0783 | 127 | 0.1024 | 469 | 0.0832 |
| 19_740369 | 120 | 0.1917 | 132 | 0.2273 | 92 | 0.1848 | 99 | 0.101 | 443 | 0.1806 | 87 | 0.1609 | 115 | 0.2087 | 97 | 0.1546 | 109 | 0.211 | 408 | 0.1863 |
| 19_740399 | 92 | 0.2609 | 118 | 0.2881 | 77 | 0.2208 | 95 | 0.1263 | 382 | 0.2277 | 76 | 0.2368 | 84 | 0.2143 | 82 | 0.1463 | 82 | 0.2073 | 324 | 0.2006 |
| 19_740485 | 103 | 0.0777 | 124 | 0.1371 | 82 | 0.1463 | 92 | 0.0978 | 401 | 0.1147 | 71 | 0.0423 | 105 | 0.0952 | 95 | 0.1474 | 94 | 0.0957 | 365 | 0.0986 |
| 19_740499 | 91 | 0.1099 | 116 | 0.0862 | 73 | 0.1507 | 88 | 0.1023 | 368 | 0.1087 | 66 | 0.0758 | 94 | 0.0638 | 88 | 0.1364 | 87 | 0.1149 | 335 | 0.0985 |
| 19_743368 | 12 | 0.0833 | 14 | 0.3571 | 8 | 0 | 15 | 0.0667 | 49 | 0.1429 | 3 | 0 | 14 | 0.0714 | 9 | 0 | 12 | 0.0833 | 38 | 0.0526 |
| 19_743371 | 10 | 0.4 | 12 | 0.5833 | 5 | 0.6 | 14 | 0.5 | 41 | 0.5122 | 5 | 0.6 | 11 | 0.4545 | 9 | 0.8889 | 12 | 0.6667 | 37 | 0.6486 |
| 19_743375 | 7 | 0.2857 | 11 | 0.2727 | 4 | 0.25 | 9 | 0.4444 | 31 | 0.3226 | 5 | 0.4 | 12 | 0.3333 | 6 | 0.1667 | 10 | 0.3 | 33 | 0.303 |
| 19_743379 | 11 | 0.0909 | 12 | 0.1667 | 6 | 0.3333 | 14 | 0.2857 | 43 | 0.2093 | 4 | 0.25 | 12 | 0.3333 | 9 | 0.1111 | 13 | 0.3846 | 38 | 0.2895 |
| 19_743407 | 11 | 1 | 9 | 1 | 6 | 0.8333 | 11 | 1 | 37 | 0.973 | 5 | 1 | 8 | 1 | 9 | 1 | 11 | 1 | 33 | 1 |
| 19_743409 | 10 | 0.1 | 12 | 0.0833 | 6 | 0 | 9 | 0.1111 | 37 | 0.0811 | 5 | 0 | 10 | 0 | 7 | 0.1429 | 11 | 0.2727 | 33 | 0.1212 |
| 19_743409 | 10 | 0.5 | 10 | 0.8 | 7 | 0.5714 | 11 | 0.4545 | 38 | 0.5789 | 5 | 0.4 | 12 | 0.5833 | 10 | 0.9 | 11 | 0.8182 | 38 | 0.7105 |
| 19_743419 | 10 | 0.6 | 11 | 0.3636 | 7 | 0.1429 | 9 | 0.7778 | 37 | 0.4865 | 6 | 0.6667 | 10 | 0.4 | 9 | 0.3333 | 12 | 0.5833 | 37 | 0.4865 |
| 19_743426 | 6 | 0.8333 | 9 | 0.2222 | 5 | 0.2 | 6 | 0.1667 | 26 | 0.3462 | 4 | 0 | 8 | 0.5 | 6 | 0.1667 | 10 | 0.4 | 28 | 0.3214 |
| 19_743433 | 10 | 0.1 | 11 | 0.0909 | 7 | 0.1429 | 11 | 0 | 39 | 0.0769 | 5 | 0.2 | 11 | 0.0909 | 9 | 0.1111 | 12 | 0.0833 | 37 | 0.1081 |
| 19_743445 | 8 | 0.125 | 7 | 0 | 7 | 0.1429 | 7 | 0.1429 | 29 | 0.1034 | 4 | 0 | 10 | 0.4 | 8 | 0.375 | 9 | 0 | 31 | 0.2258 |
| 19_743447 | 10 | 0.3 | 8 | 0.125 | 7 | 0.4286 | 10 | 0.3 | 35 | 0.2857 | 4 | 0 | 9 | 0.1111 | 8 | 0 | 11 | 0.4545 | 32 | 0.1875 |
| 19_743449 | 10 | 0.5 | 9 | 0.4444 | 5 | 1 | 10 | 0.5 | 34 | 0.5588 | 6 | 0.6667 | 9 | 1 | 8 | 0.625 | 8 | 0.625 | 31 | 0.7419 |
| 19_743455 | 10 | 0.4 | 9 | 0.2222 | 7 | 0.1429 | 10 | 0.2 | 36 | 0.25 | 6 | 0.1667 | 9 | 0.3333 | 9 | 0.2222 | 10 | 0.4 | 34 | 0.2941 |
| 19_743465 | 8 | 0.375 | 9 | 0.2222 | 5 | 0 | 9 | 0.2222 | 31 | 0.2258 | 4 | 0.5 | 8 | 0 | 8 | 0.125 | 9 | 0.3333 | 29 | 0.2069 |
| 19_743478 | 10 | 0.4 | 9 | 0.4444 | 3 | 0.6667 | 11 | 0.5455 | 33 | 0.4848 | 4 | 0.75 | 8 | 0.75 | 8 | 0.75 | 11 | 0.8182 | 31 | 0.7742 |
| 19_743481 | 8 | 0.25 | 8 | 0 | 4 | 0 | 8 | 0.625 | 28 | 0.25 | 4 | 0.5 | 7 | 0.2857 | 4 | 0 | 9 | 0.5556 | 24 | 0.375 |
| 19_743482 | 8 | 0 | 8 | 0.125 | 5 | 0.2 | 8 | 0.25 | 29 | 0.1379 | 4 | 0.25 | 8 | 0 | 8 | 0.25 | 10 | 0.2 | 30 | 0.1667 |

| | | | | | | | | | | | | | | | | | | | | |
|---|---|---|---|---|---|---|---|---|---|---|---|---|---|---|---|---|---|---|---|---|
| 19_743492 | 7 | 0.1429 | 6 | 0.1667 | 5 | 0 | 6 | 0 | 24 | 0.0833 | 4 | 0.25 | 6 | 0.1667 | 8 | 0.125 | 9 | 0.1111 | 27 | 0.1481 |
| 19_743494 | 8 | 0.5 | 5 | 0.4 | 3 | 0.6667 | 6 | 0.6667 | 22 | 0.5455 | 4 | 1 | 6 | 0.6667 | 8 | 0.75 | 9 | 0.4444 | 27 | 0.6667 |
| 19_743500 | 7 | 0.2857 | 6 | 0 | 4 | 0 | 8 | 0.375 | 25 | 0.2 | 4 | 0.5 | 6 | 0.1667 | 7 | 0.1429 | 8 | 0.5 | 25 | 0.32 |
| 19_743500 | 6 | 0.1667 | 4 | 0.25 | 2 | 0 | 3 | 0 | 15 | 0.1333 | 3 | 0 | 3 | 0 | 7 | 0.1429 | 7 | 0 | 20 | 0.05 |
| 19_743509 | 4 | 0.5 | 5 | 0 | 3 | 0 | 8 | 0 | 20 | 0.1 | 3 | 0 | 4 | 0 | 6 | 0.1667 | 7 | 0.2857 | 20 | 0.15 |
| 19_743515 | 5 | 0.2 | 4 | 0 | 4 | 0 | 8 | 0.25 | 21 | 0.1429 | 2 | 0 | 3 | 0.3333 | 5 | 0.2 | 7 | 0.4286 | 17 | 0.2941 |
| 19_743517 | 7 | 0.1429 | 6 | 0 | 4 | 0 | 7 | 0.5714 | 24 | 0.2083 | 3 | 0.3333 | 6 | 0.1667 | 8 | 0.125 | 9 | 0.4444 | 26 | 0.2692 |
| 19_743533 | 5 | 0.2 | 6 | 0.1667 | 4 | 0.25 | 7 | 0.1429 | 22 | 0.1818 | 3 | 0 | 5 | 0 | 7 | 0.2857 | 7 | 0.1429 | 22 | 0.1364 |
| 19_743540 | 6 | 0.1667 | 5 | 0.8 | 4 | 0 | 6 | 0.3333 | 21 | 0.3333 | 3 | 0.3333 | 4 | 0.75 | 7 | 0.4286 | 5 | 0.4 | 19 | 0.4737 |
| 19_743557 | 7 | 0.5714 | 6 | 0.5 | 4 | 0.25 | 8 | 0.625 | 25 | 0.52 | 2 | 0.5 | 5 | 0.4 | 7 | 0.4286 | 7 | 0.1429 | 21 | 0.3333 |
| 19_743559 | 4 | 0.75 | 3 | 0.6667 | 2 | 1 | 3 | 1 | 12 | 0.8333 | 3 | 1 | 5 | 1 | 5 | 1 | 4 | 1 | 17 | 1 |
| 19_743560 | 3 | 0 | 4 | 0.5 | NA | NA | 2 | 0.5 | 9 | 0.3333 | 2 | 0 | 5 | 0.2 | 6 | 0.5 | 3 | 0 | 16 | 0.25 |
| 19_750954 | NA | NA | 1 | 0 | 1 | 1 | NA | NA | 2 | 0.5 | 1 | 1 | NA | NA | NA | NA | 3 | 1 | 4 | 1 |
| 19_750970 | 1 | 1 | 2 | 0.5 | 1 | 1 | NA | NA | 4 | 0.75 | 1 | 1 | NA | NA | NA | NA | 2 | 0.5 | 3 | 0.6667 |
| 19_751151 | 1 | 1 | 1 | 0 | 1 | 0 | NA | NA | 3 | 0.3333 | 1 | 0 | NA | NA | 2 | 0 | 4 | 0.25 | 7 | 0.1429 |
| 19_751153 | NA | NA | 1 | 1 | 1 | 1 | NA | NA | 2 | 1 | NA | NA | NA | NA | NA | NA | 1 | 1 | 1 | 1 |
| 19_751160 | NA | NA | 2 | 0.5 | 1 | 1 | NA | NA | 3 | 0.6667 | 1 | 1 | NA | NA | 1 | 1 | 3 | 1 | 5 | 1 |
| 19_751199 | NA | NA | 1 | 0 | 1 | 1 | NA | NA | 2 | 0.5 | NA | NA | NA | NA | 2 | 1 | 2 | 0.5 | 4 | 0.75 |
| 19_751209 | 1 | 1 | 2 | 0 | 1 | 0 | NA | NA | 4 | 0.25 | 1 | 1 | NA | NA | 2 | 0.5 | 4 | 0.25 | 7 | 0.4286 |
| 19_751503 | 2 | 0.5 | 2 | 0 | 4 | 0 | 3 | 0.3333 | 11 | 0.1818 | 2 | 0 | 2 | 0.5 | 2 | 0 | 1 | 0 | 7 | 0.1429 |
| 19_751516 | 3 | 0 | 2 | 0 | 5 | 0.2 | 2 | 0 | 12 | 0.0833 | 4 | 0 | 4 | 0.25 | 3 | 0 | 2 | 1 | 13 | 0.2308 |
| 19_751654 | 3 | 0 | 2 | 0 | 5 | 0 | 2 | 0.5 | 12 | 0.0833 | 4 | 0 | 4 | 0.25 | 2 | 0 | 1 | 0 | 11 | 0.0909 |
| 19_751666 | 3 | 0 | 2 | 0.5 | 5 | 0 | 2 | 0 | 12 | 0.0833 | 3 | 0 | 4 | 0.25 | 2 | 0 | 1 | 0 | 10 | 0.1 |
| 19_753525 | 1 | 0 | 2 | 0.5 | NA | NA | 1 | 1 | 4 | 0.5 | NA | NA | 1 | 0 | 1 | 0 | 1 | 1 | 3 | 0.3333 |
| 19_753526 | 1 | 0 | 2 | 1 | NA | NA | 1 | 0 | 4 | 0.5 | NA | NA | 1 | 1 | 1 | 1 | 1 | 0 | 3 | 0.6667 |
| 19_755649 | 7 | 0.2857 | 5 | 0 | 8 | 0.125 | 9 | 0.1111 | 29 | 0.1379 | 9 | 0.2222 | 6 | 0.3333 | 9 | 0.2222 | 11 | 0.1818 | 35 | 0.2286 |
| 19_756371 | 3 | 0.3333 | 1 | 0 | 2 | 0 | 3 | 0 | 9 | 0.1111 | 1 | 0 | 2 | 0 | 1 | 0 | 1 | 1 | 5 | 0.2 |
| 19_756586 | 4 | 0 | 1 | 0 | 5 | 0.6 | 4 | 0.5 | 14 | 0.3571 | 1 | 0 | 2 | 0.5 | 1 | 0 | 1 | 0 | 5 | 0.2 |
| 19_764634 | 1 | 1 | 1 | 1 | 2 | 0.5 | 1 | 1 | 5 | 0.8 | NA | NA | NA | NA | NA | NA | 1 | 1 | 1 | 1 |
| 19_764639 | 1 | 0 | 1 | 1 | 2 | 0.5 | 3 | 0.6667 | 7 | 0.5714 | NA | NA | NA | NA | NA | NA | 1 | 1 | 1 | 1 |
| 19_829443 | 1 | 1 | NA | NA | NA | NA | NA | NA | 1 | 1 | NA | NA | NA | NA | 1 | 1 | NA | NA | 1 | 1 |
| 19_858767 | 6 | 0.3333 | 6 | 0.1667 | 4 | 0.5 | NA | NA | 16 | 0.3125 | 3 | 0.3333 | 3 | 1 | NA | NA | 2 | 0.5 | 8 | 0.625 |
| 19_858776 | 7 | 0.2857 | 4 | 0 | 4 | 0 | 1 | 0 | 16 | 0.125 | 3 | 0 | 3 | 0.3333 | NA | NA | 2 | 0 | 8 | 0.125 |
| 19_858902 | 1 | 0 | NA | NA | 1 | 1 | NA | NA | 2 | 0.5 | NA | NA | 2 | 0.5 | NA | NA | 1 | 0 | 3 | 0.3333 |
| 19_860356 | 5 | 0.2 | 1 | 1 | 3 | 0 | 1 | 0 | 10 | 0.2 | 1 | 0 | 1 | 0 | NA | NA | 2 | 0.5 | 4 | 0.25 |
| 19_860366 | 5 | 0.2 | 2 | 0.5 | 3 | 0.3333 | NA | NA | 10 | 0.3 | 1 | 1 | 1 | 0 | NA | NA | 2 | 0 | 4 | 0.25 |
| 19_860374 | 4 | 0 | 1 | 0 | 3 | 0.3333 | 1 | 0 | 9 | 0.1111 | 2 | 0.5 | NA | NA | NA | NA | 3 | 0 | 5 | 0.2 |
| 19_860384 | 4 | 0 | 2 | 0.5 | 2 | 0 | 1 | 0 | 9 | 0.1111 | 2 | 0 | 2 | 0.5 | 1 | 0 | 3 | 0 | 8 | 0.125 |
| 19_860564 | NA | NA | 1 | 1 | NA | NA | NA | NA | 1 | 1 | NA | NA | NA | NA | NA | NA | 1 | 1 | 1 | 1 |
| 19_865530 | 7 | 0 | 4 | 0 | 9 | 0 | 9 | 0.2222 | 29 | 0.069 | 6 | 0 | 11 | 0.1818 | 9 | 0 | 13 | 0 | 39 | 0.0513 |
| 19_865546 | 10 | 0.2 | 9 | 0 | 10 | 0 | 11 | 0 | 40 | 0.05 | 8 | 0 | 12 | 0.1667 | 11 | 0.1818 | 13 | 0.0769 | 44 | 0.1136 |
| 19_865553 | 9 | 0.6667 | 11 | 0.9091 | 9 | 1 | 11 | 0.7273 | 40 | 0.825 | 5 | 0.6 | 13 | 0.8462 | 8 | 0.75 | 13 | 0.8462 | 39 | 0.7949 |
| 19_865561 | 12 | 0.9167 | 12 | 1 | 12 | 1 | 13 | 1 | 49 | 0.9796 | 10 | 0.9 | 15 | 1 | 13 | 1 | 18 | 0.9444 | 56 | 0.9643 |
| 19_865581 | 7 | 1 | 8 | 0.875 | 10 | 0.8 | 10 | 0.9 | 35 | 0.8857 | 5 | 1 | 12 | 0.8333 | 11 | 0.9091 | 15 | 0.9333 | 43 | 0.907 |
| 19_865593 | 14 | 0 | 13 | 0.0769 | 11 | 0.0909 | 13 | 0.0769 | 51 | 0.0588 | 9 | 0.1111 | 15 | 0.0667 | 11 | 0 | 21 | 0.1429 | 56 | 0.0893 |
| 19_865598 | 15 | 0.1333 | 10 | 0.5 | 9 | 0 | 13 | 0.3077 | 47 | 0.234 | 11 | 0.0909 | 18 | 0.1667 | 11 | 0.1818 | 20 | 0 | 60 | 0.1 |
| 19_865609 | 16 | 0.0625 | 13 | 0.1538 | 14 | 0 | 13 | 0.0769 | 56 | 0.0714 | 13 | 0.0769 | 17 | 0.0588 | 15 | 0.1333 | 22 | 0.0455 | 67 | 0.0746 |
| 19_865635 | 12 | 0.1667 | 13 | 0.2308 | 13 | 0.1538 | 12 | 0.1667 | 50 | 0.18 | 12 | 0.0833 | 14 | 0.3571 | 15 | 0.2 | 23 | 0.087 | 64 | 0.1719 |
| 19_865639 | 16 | 0.375 | 14 | 0.3571 | 9 | 0.4444 | 13 | 0.3846 | 52 | 0.3846 | 15 | 0.8667 | 17 | 0.5294 | 17 | 0.7647 | 23 | 0.6087 | 72 | 0.6806 |
| 19_865643 | 13 | 0.3846 | 12 | 0.5 | 10 | 0.4 | 11 | 0.2727 | 46 | 0.3913 | 11 | 0.2727 | 14 | 0.4286 | 16 | 0.4375 | 16 | 0.5625 | 57 | 0.4386 |
| 19_865647 | 16 | 1 | 10 | 1 | 15 | 1 | 13 | 1 | 54 | 1 | 13 | 1 | 12 | 0.9167 | 18 | 1 | 19 | 1 | 62 | 0.9839 |
| 19_865657 | 16 | 1 | 14 | 1 | 17 | 0.9412 | 13 | 1 | 60 | 0.9833 | 15 | 1 | 17 | 0.9412 | 18 | 0.9444 | 24 | 1 | 74 | 0.973 |
| 19_865660 | 14 | 0.1429 | 11 | 0.0909 | 17 | 0.2941 | 10 | 0.1 | 52 | 0.1731 | 14 | 0.2143 | 17 | 0.1765 | 17 | 0.1176 | 21 | 0.2381 | 69 | 0.1884 |
| 19_865678 | 13 | 1 | 16 | 1 | 18 | 1 | 13 | 1 | 60 | 1 | 15 | 1 | 16 | 1 | 19 | 1 | 26 | 0.9615 | 76 | 0.9868 |
| 19_865682 | 14 | 0.7143 | 14 | 0.6429 | 17 | 0.4706 | 12 | 0.5 | 57 | 0.5789 | 11 | 0.5455 | 15 | 0.5333 | 15 | 0.6667 | 21 | 0.4762 | 62 | 0.5484 |
| 19_865683 | 16 | 1 | 18 | 1 | 20 | 1 | 14 | 1 | 68 | 1 | 16 | 1 | 18 | 1 | 20 | 1 | 26 | 1 | 80 | 1 |
| 19_865685 | 17 | 0.2941 | 13 | 0.3077 | 14 | 0.2143 | 12 | 0.1667 | 56 | 0.25 | 13 | 0.2308 | 16 | 0.25 | 17 | 0.3529 | 18 | 0.3333 | 64 | 0.2969 |
| 19_865686 | 18 | 0.2222 | 19 | 0.0526 | 19 | 0 | 13 | 0 | 69 | 0.0725 | 15 | 0.3333 | 18 | 0.1667 | 19 | 0 | 27 | 0 | 79 | 0.1013 |
| 19_865688 | 18 | 0.0556 | 19 | 0.1579 | 16 | 0.0625 | 14 | 0.0714 | 67 | 0.0896 | 13 | 0.0769 | 15 | 0.0667 | 20 | 0.1 | 26 | 0.0769 | 74 | 0.0811 |
| 19_865689 | 13 | 0.5385 | 15 | 0.2667 | 18 | 0.7778 | 13 | 0.6154 | 59 | 0.5593 | 13 | 0.6154 | 13 | 0.6923 | 16 | 0.625 | 21 | 0.7143 | 63 | 0.6667 |
| 19_865716 | 17 | 0 | 14 | 0.0714 | 15 | 0.0667 | 11 | 0.0909 | 57 | 0.0526 | 10 | 0.1 | 18 | 0.0556 | 18 | 0.0556 | 24 | 0.0417 | 70 | 0.0571 |
| 19_865734 | 19 | 0.2105 | 20 | 0.15 | 21 | 0.0476 | 16 | 0.0625 | 76 | 0.1184 | 16 | 0 | 20 | 0.15 | 24 | 0.2083 | 30 | 0.0667 | 90 | 0.1111 |
| 19_865885 | 5 | 0 | 2 | 0 | 1 | 0 | 1 | 1 | 9 | 0.1111 | 1 | 0 | 1 | 0 | 2 | 0.5 | 6 | 0.3333 | 10 | 0.3 |
| 19_865929 | 5 | 0.4 | 1 | 1 | 1 | 1 | 1 | 1 | 8 | 0.625 | NA | NA | 1 | 1 | 3 | 0.3333 | 5 | 0.4 | 9 | 0.4444 |
| 19_865939 | 4 | 0 | NA | NA | NA | NA | 1 | 1 | 5 | 0.2 | 1 | 0 | 2 | 0.5 | 1 | 0 | 5 | 0.8 | 9 | 0.5556 |

| | | | | | | | | | | | | | | | | | | | | |
|---|---|---|---|---|---|---|---|---|---|---|---|---|---|---|---|---|---|---|---|---|
| 19_865941 | 4 | 0 | 1 | 1 | NA | NA | NA | NA | 5 | 0.2 | 1 | 0 | 1 | 0 | 3 | 0.3333 | 5 | 0.6 | 10 | 0.4 |
| 19_865958 | 5 | 0.2 | 1 | 0 | 1 | 1 | 1 | 1 | 8 | 0.375 | 1 | 0 | 2 | 0 | 3 | 0.3333 | 4 | 0.5 | 10 | 0.3 |
| 19_865966 | 5 | 0 | 2 | 0 | 1 | 1 | NA | NA | 8 | 0.125 | 1 | 0 | 2 | 0 | 3 | 0 | 4 | 0.25 | 10 | 0.1 |
| 19_865981 | 5 | 0.2 | 1 | 1 | 1 | 0 | 1 | 0 | 8 | 0.25 | 1 | 1 | 3 | 0.3333 | 2 | 0.5 | 3 | 0.6667 | 9 | 0.5556 |
| 19_865984 | 4 | 0 | 1 | 1 | 1 | 1 | 1 | 1 | 7 | 0.4286 | 1 | 1 | 3 | 0 | 4 | 0.25 | 2 | 0.5 | 10 | 0.3 |
| 19_865988 | 5 | 0.2 | 1 | 0 | 1 | 0 | 1 | 0 | 8 | 0.125 | NA | NA | 4 | 0.25 | 4 | 0.25 | 3 | 0.3333 | 11 | 0.2727 |
| 19_865993 | 4 | 0 | 1 | 0 | 1 | 0 | 1 | 1 | 7 | 0.1429 | NA | NA | 3 | 0 | 4 | 0.25 | 2 | 0 | 9 | 0.1111 |
| 19_866023 | 5 | 0.4 | 2 | 0 | 1 | 0 | 1 | 1 | 9 | 0.3333 | 1 | 1 | 4 | 0.5 | 4 | 0.5 | 3 | 0 | 12 | 0.4167 |
| 19_866051 | 5 | 0 | 2 | 0.5 | 1 | 1 | 1 | 0 | 9 | 0.2222 | 1 | 0 | 4 | 0 | 4 | 0.25 | 4 | 0.5 | 13 | 0.2308 |
| 19_866074 | 5 | 0 | 1 | 1 | NA | NA | 1 | 0 | 7 | 0.1429 | 1 | 0 | 4 | 0.25 | 3 | 0 | 5 | 0.2 | 13 | 0.1538 |
| 19_866087 | 4 | 0.75 | 1 | 0 | NA | NA | 1 | 0 | 6 | 0.5 | NA | NA | 3 | 0.3333 | 3 | 0.3333 | 3 | 0.3333 | 9 | 0.3333 |
| 19_866167 | 5 | 0.2 | 4 | 0.25 | 1 | 0 | 2 | 0.5 | 12 | 0.25 | 1 | 0 | 5 | 0.4 | 5 | 0.2 | 6 | 0 | 17 | 0.1765 |
| 19_866208 | 6 | 0 | 4 | 0.25 | 2 | 0 | 2 | 0 | 14 | 0.0714 | 3 | 0 | 5 | 0.2 | 5 | 0.2 | 7 | 0 | 20 | 0.1 |
| 19_866212 | 5 | 0 | 5 | 0.2 | 3 | 0 | 1 | 0 | 14 | 0.0714 | 3 | 0 | 6 | 0.3333 | 5 | 0.2 | 7 | 0.1429 | 21 | 0.1905 |
| 19_866213 | 6 | 0 | 4 | 0 | 2 | 0 | 2 | 0.5 | 14 | 0.0714 | 3 | 0 | 5 | 0 | 5 | 0 | 7 | 0.1429 | 20 | 0.05 |
| 19_868516 | 20 | 0.15 | 32 | 0.125 | 27 | 0.0741 | 16 | 0.0625 | 95 | 0.1053 | 22 | 0.1364 | 21 | 0 | 16 | 0.1875 | 17 | 0.1176 | 76 | 0.1053 |
| 19_868520 | 20 | 0 | 33 | 0 | 27 | 0.0741 | 19 | 0.2105 | 99 | 0.0606 | 22 | 0.1364 | 19 | 0.1053 | 17 | 0.1765 | 16 | 0.125 | 74 | 0.1351 |
| 19_868531 | 24 | 0.0833 | 30 | 0.0667 | 27 | 0.0741 | 19 | 0.1579 | 100 | 0.09 | 20 | 0.15 | 18 | 0.1111 | 17 | 0.0588 | 13 | 0.0769 | 68 | 0.1029 |
| 19_871782 | 8 | 0.5 | 3 | 0.3333 | 7 | 0.5714 | 7 | 0.5714 | 25 | 0.52 | 7 | 0.4286 | 4 | 0 | 2 | 0 | 2 | 1 | 15 | 0.3333 |
| 19_873248 | 4 | 0 | 7 | 0.1429 | 7 | 0.1429 | 2 | 0 | 20 | 0.1 | 5 | 0.2 | 13 | 0 | 6 | 0.1667 | 4 | 0 | 28 | 0.0714 |
| 19_873255 | 2 | 0.5 | 6 | 1 | 5 | 1 | 3 | 0.6667 | 16 | 0.875 | 6 | 0.8333 | 11 | 0.7273 | 5 | 0.8 | 4 | 0.75 | 26 | 0.7692 |
| 19_873265 | 3 | 0.6667 | 9 | 1 | 7 | 0.8571 | 4 | 0.75 | 23 | 0.8696 | 5 | 1 | 12 | 0.9167 | 6 | 1 | 4 | 0.75 | 27 | 0.9259 |
| 19_873282 | 4 | 0 | 8 | 0.125 | 5 | 0 | 3 | 0 | 20 | 0.05 | 5 | 0 | 14 | 0.0714 | 8 | 0.375 | 4 | 0 | 31 | 0.129 |
| 19_873285 | 3 | 0.6667 | 8 | 0.375 | 4 | 0.25 | 4 | 0.5 | 19 | 0.4211 | 6 | 0.3333 | 15 | 0.5333 | 8 | 0.125 | 6 | 0.8333 | 35 | 0.4571 |
| 19_873285 | 4 | 0.25 | 5 | 0.2 | 3 | 0 | 3 | 0.3333 | 15 | 0.2 | 6 | 0.1667 | 10 | 0 | 6 | 0.1667 | 5 | 0 | 27 | 0.0741 |
| 19_873292 | 4 | 0.5 | 8 | 0.5 | 7 | 0.1429 | 4 | 0.5 | 23 | 0.3913 | 8 | 0.125 | 14 | 0.3571 | 8 | 0 | 5 | 0.2 | 35 | 0.2 |
| 19_873294 | 3 | 0.6667 | 8 | 0.25 | 7 | 0.1429 | 3 | 0 | 21 | 0.2381 | 7 | 0.1429 | 16 | 0.4375 | 9 | 0.6667 | 6 | 0.1667 | 38 | 0.3947 |
| 19_873299 | 2 | 0.5 | 8 | 0 | 7 | 0.1429 | 3 | 0.3333 | 20 | 0.15 | 6 | 0.1667 | 16 | 0.25 | 10 | 0.2 | 6 | 0 | 38 | 0.1842 |
| 19_874806 | 2 | 0 | 6 | 0 | 3 | 0.3333 | 2 | 0 | 13 | 0.0769 | 4 | 0.25 | 3 | 0 | 5 | 0 | 3 | 0 | 15 | 0.0667 |
| 19_874820 | 5 | 0.6 | 6 | 0.5 | 7 | 0.5714 | 7 | 0.2857 | 25 | 0.48 | 8 | 0.125 | 4 | 0.75 | 5 | 0.2 | 3 | 1 | 20 | 0.4 |
| 19_874851 | 6 | 0.6667 | 6 | 0.3333 | 11 | 0.3636 | 3 | 0.6667 | 26 | 0.4615 | 6 | 0.3333 | 6 | 0.1667 | 2 | 0.5 | 7 | 0.4286 | 21 | 0.3333 |
| 19_874963 | 8 | 0 | 8 | 0.25 | 10 | 0.4 | 4 | 0 | 30 | 0.2 | 11 | 0.0909 | 4 | 0.25 | 8 | 0.125 | 7 | 0.1429 | 30 | 0.1333 |
| 19_874964 | 9 | 0.4444 | 9 | 0.8889 | 12 | 0.75 | 4 | 0.75 | 34 | 0.7059 | 10 | 0.9 | 7 | 0.8571 | 8 | 0.625 | 7 | 0.8571 | 32 | 0.8125 |
| 19_874976 | 8 | 0.5 | 8 | 0.5 | 11 | 0.4545 | 3 | 0.6667 | 30 | 0.5 | 9 | 0.4444 | 5 | 0 | 5 | 1 | 11 | 0.2727 | 30 | 0.4 |
| 19_875202 | 1 | 1 | 3 | 0.6667 | 1 | 0 | NA | NA | 5 | 0.6 | 2 | 0 | 1 | 1 | NA | NA | NA | NA | 3 | 0.3333 |
| 19_875223 | 1 | 1 | 4 | 1 | 1 | 1 | 2 | 1 | 8 | 1 | 2 | 0.5 | 1 | 1 | 1 | 1 | 1 | 1 | 5 | 0.8 |
| 19_875228 | 1 | 1 | 4 | 0.25 | 1 | 0 | 1 | 1 | 7 | 0.4286 | 2 | 0.5 | NA | NA | 2 | 0 | 1 | 1 | 5 | 0.4 |
| 19_875239 | 1 | 1 | 5 | 0.4 | 1 | 1 | 1 | 0 | 8 | 0.5 | 2 | 1 | 1 | 0 | 2 | 0.5 | 1 | 0 | 6 | 0.5 |
| 19_875245 | NA | NA | 5 | 0.4 | 1 | 0 | 2 | 0.5 | 8 | 0.375 | 2 | 0.5 | 1 | 0 | 1 | 0 | 1 | 1 | 5 | 0.4 |
| 19_875255 | 1 | 0 | 4 | 0.75 | 1 | 0 | 2 | 0 | 8 | 0.375 | 1 | 1 | 1 | 0 | 1 | 0 | 1 | 0 | 4 | 0.25 |
| 19_877058 | 12 | 0 | 22 | 0.1364 | 13 | 0.0769 | 19 | 0 | 66 | 0.0606 | 20 | 0.15 | 18 | 0 | 14 | 0.2143 | 13 | 0.1538 | 65 | 0.1231 |
| 19_878046 | 1 | 0 | NA | NA | 2 | 1 | NA | NA | 3 | 0.6667 | 1 | 0 | 1 | 1 | NA | NA | 1 | 1 | 3 | 0.6667 |
| 19_878056 | 1 | 1 | NA | NA | 2 | 0 | NA | NA | 3 | 0.3333 | 1 | 1 | 1 | 1 | NA | NA | 1 | 1 | 3 | 1 |
| 19_878057 | 1 | 0 | NA | NA | 2 | 0.5 | NA | NA | 3 | 0.3333 | 1 | 1 | 1 | 1 | NA | NA | 2 | 1 | 4 | 1 |
| 19_878063 | NA | NA | NA | NA | 1 | 1 | NA | NA | 1 | 1 | NA | NA | 1 | 1 | NA | NA | 2 | 1 | 3 | 1 |
| 19_878118 | 1 | 0 | NA | NA | 2 | 0.5 | NA | NA | 3 | 0.3333 | 1 | 1 | 1 | 1 | 1 | 0 | 2 | 0.5 | 5 | 0.6 |
| 19_878127 | 1 | 1 | 1 | 1 | 2 | 1 | NA | NA | 4 | 1 | 1 | 1 | NA | NA | 1 | 1 | 2 | 0.5 | 4 | 0.75 |
| 19_878132 | 1 | 1 | 1 | 0 | 2 | 0 | NA | NA | 4 | 0.25 | 1 | 0 | 1 | 0 | 1 | 0 | 2 | 0.5 | 5 | 0.2 |
| 19_878143 | 1 | 1 | 1 | 1 | 1 | 1 | NA | NA | 3 | 1 | 1 | 1 | 1 | 1 | 1 | 1 | 2 | 1 | 5 | 1 |
| 19_878157 | 1 | 0 | 1 | 1 | 3 | 0 | NA | NA | 5 | 0.2 | NA | NA | 1 | 1 | 1 | 0 | 2 | 0 | 4 | 0.25 |
| 19_878165 | NA | NA | 1 | 0 | 3 | 0.6667 | NA | NA | 4 | 0.5 | 2 | 0.5 | NA | NA | 1 | 0 | 2 | 0.5 | 5 | 0.4 |
| 19_878166 | 1 | 1 | 1 | 1 | 3 | 0.6667 | NA | NA | 5 | 0.8 | 1 | 0 | 1 | 1 | 1 | 1 | 2 | 1 | 5 | 0.8 |
| 19_878176 | 1 | 0 | 1 | 1 | 3 | 0 | NA | NA | 5 | 0.2 | 2 | 0 | 1 | 0 | 1 | 0 | 2 | 0.5 | 6 | 0.1667 |
| 19_878180 | 1 | 1 | 1 | 1 | 3 | 1 | NA | NA | 5 | 1 | 2 | 0.5 | 1 | 1 | 1 | 1 | 2 | 0 | 6 | 0.5 |
| 19_878188 | 1 | 1 | 1 | 1 | 3 | 1 | NA | NA | 5 | 1 | 1 | 1 | 1 | 1 | 1 | 1 | 2 | 1 | 5 | 1 |
| 19_878196 | NA | NA | 1 | 1 | 3 | 1 | NA | NA | 4 | 1 | 2 | 1 | NA | NA | 1 | 1 | 2 | 1 | 5 | 1 |
| 19_878216 | NA | NA | 1 | 1 | 2 | 0 | NA | NA | 3 | 0.3333 | 1 | 0 | 1 | 0 | 1 | 0 | 1 | 1 | 4 | 0.25 |
| 19_878218 | NA | NA | NA | NA | 3 | 0.3333 | NA | NA | 3 | 0.3333 | 1 | 0 | 1 | 1 | 1 | 0 | 1 | 0 | 4 | 0.25 |
| 19_878889 | 1 | 0 | 1 | 1 | 1 | 0 | 1 | 1 | 4 | 0.5 | 1 | 0 | NA | NA | 2 | 0 | 5 | 0.2 | 8 | 0.125 |
| 19_878906 | 1 | 1 | 1 | 1 | 1 | 1 | 1 | 0 | 4 | 0.75 | 3 | 0.3333 | NA | NA | 3 | 0.6667 | 7 | 0.4286 | 13 | 0.4615 |
| 19_878909 | 1 | 1 | 1 | 1 | 1 | 1 | 1 | 1 | 4 | 1 | 3 | 0.6667 | NA | NA | 4 | 1 | 6 | 0.8333 | 13 | 0.8462 |
| 19_879616 | NA | NA | 1 | 1 | 1 | 1 | 1 | 1 | 3 | 1 | 2 | 1 | NA | NA | 4 | 1 | 3 | 1 | 9 | 1 |
| 19_879625 | NA | NA | NA | NA | NA | NA | 1 | 1 | 1 | 1 | 2 | 1 | NA | NA | 2 | 1 | 2 | 1 | 6 | 1 |
| 19_880923 | 1 | 0 | 5 | 0.2 | NA | NA | 1 | 0 | 7 | 0.1429 | 4 | 0 | 5 | 0.4 | 2 | 0 | 2 | 0.5 | 13 | 0.2308 |
| 19_881517 | 4 | 0 | 5 | 0 | 2 | 0 | 4 | 0.25 | 15 | 0.0667 | 5 | 0 | 4 | 0 | 1 | 0 | 9 | 0.1111 | 19 | 0.0526 |

| | | | | | | | | | | | | | | | | | | | | |
|---|---|---|---|---|---|---|---|---|---|---|---|---|---|---|---|---|---|---|---|---|
| 19_881519 | NA | NA | 2 | 0.5 | 2 | 0 | NA | NA | 4 | 0.25 | NA | NA | NA | NA | 2 | 0 | 3 | 0.3333 | 5 | 0.2 |
| 19_881531 | 65 | 0.0769 | 74 | 0.0676 | 59 | 0.0508 | 73 | 0.1096 | 271 | 0.0775 | 60 | 0.05 | 64 | 0.0469 | 62 | 0.0645 | 56 | 0.1071 | 242 | 0.0661 |
| 19_882449 | NA | NA | 1 | 1 | NA | NA | 1 | 0 | 2 | 0.5 | NA | NA | NA | NA | NA | NA | 1 | 1 | 1 | 1 |
| 19_885745 | 3 | 0.3333 | 1 | 1 | 2 | 0 | 2 | 0.5 | 8 | 0.375 | 1 | 0 | 5 | 0.4 | 2 | 0 | 1 | 1 | 9 | 0.3333 |
| 19_885751 | 5 | 0 | 1 | 1 | NA | NA | 2 | 0.5 | 8 | 0.25 | 1 | 0 | 2 | 0.5 | 3 | 0 | 3 | 0 | 9 | 0.1111 |
| 19_885761 | 5 | 0.6 | 1 | 1 | 2 | 1 | 2 | 1 | 10 | 0.8 | NA | NA | 5 | 1 | 3 | 1 | 3 | 1 | 11 | 1 |
| 19_885765 | 11 | 0.6364 | 4 | 0.25 | 3 | 0.6667 | 4 | 0.25 | 22 | 0.5 | 2 | 0.5 | 4 | 0.75 | 5 | 0 | 6 | 0.3333 | 17 | 0.3529 |
| 19_885772 | 7 | 0.8571 | 4 | 1 | 4 | 1 | 5 | 0.8 | 20 | 0.9 | 2 | 1 | 5 | 1 | 3 | 1 | 5 | 0.8 | 15 | 0.9333 |
| 19_885774 | 9 | 0.8889 | 4 | 0.75 | 4 | 1 | 5 | 0.8 | 22 | 0.8636 | 3 | 1 | 6 | 1 | 3 | 0.6667 | 5 | 1 | 17 | 0.9412 |
| 19_885777 | 9 | 0.2222 | 3 | 0 | 4 | 0 | 4 | 0.25 | 20 | 0.15 | 2 | 0 | 5 | 0 | 5 | 0 | 6 | 0.1667 | 18 | 0.0556 |
| 19_885786 | 6 | 0.6667 | 4 | 1 | 2 | 1 | 4 | 1 | 16 | 0.875 | 1 | 1 | 6 | 0.8333 | 3 | 1 | 5 | 1 | 15 | 0.9333 |
| 19_885786 | 7 | 0 | 2 | 0.5 | 2 | 0 | 3 | 0 | 14 | 0.0714 | 2 | 0.5 | 6 | 0 | 4 | 0 | 6 | 0 | 18 | 0.0556 |
| 19_886794 | 3 | 0.3333 | 4 | 1 | 7 | 1 | 5 | 1 | 19 | 0.8947 | 4 | 1 | 4 | 1 | 3 | 0 | 7 | 0.8571 | 18 | 0.7778 |
| 19_886799 | 3 | 0.6667 | 2 | 0 | 4 | 0 | 4 | 0.25 | 13 | 0.2308 | 4 | 0.5 | 4 | 0.75 | 4 | 0.25 | 5 | 0.2 | 17 | 0.4118 |
| 19_886803 | 2 | 1 | 3 | 0 | 3 | 0.6667 | 3 | 0.3333 | 11 | 0.4545 | 4 | 0 | 3 | 0 | 3 | 0.3333 | 3 | 0.3333 | 13 | 0.1538 |
| 19_886804 | NA | NA | 2 | 0 | 1 | 0 | 2 | 0.5 | 5 | 0.2 | 3 | 0.6667 | 3 | 1 | 2 | 0.5 | 6 | 0.6667 | 14 | 0.7143 |
| 19_887249 | 2 | 0 | 2 | 0.5 | NA | NA | 1 | 0 | 5 | 0.2 | NA | NA | NA | NA | 3 | 0.3333 | 1 | 0 | 4 | 0.25 |
| 19_887250 | 2 | 1 | 2 | 0 | NA | NA | NA | NA | 4 | 0.5 | NA | NA | NA | NA | 4 | 1 | 1 | 1 | 5 | 1 |
| 19_887256 | 2 | 1 | 2 | 1 | NA | NA | 1 | 1 | 5 | 1 | NA | NA | NA | NA | 2 | 1 | 1 | 1 | 3 | 1 |
| 19_887261 | 2 | 1 | 2 | 1 | NA | NA | 1 | 1 | 5 | 1 | NA | NA | NA | NA | 4 | 1 | NA | NA | 4 | 1 |
| 19_887271 | 1 | 1 | NA | NA | NA | NA | 1 | 0 | 2 | 0.5 | NA | NA | 1 | 1 | 3 | 0.3333 | 1 | 0 | 5 | 0.4 |
| 19_887278 | 2 | 1 | 2 | 1 | NA | NA | 1 | 1 | 5 | 1 | NA | NA | 1 | 1 | 4 | 1 | 1 | 1 | 6 | 1 |
| 19_887283 | 2 | 0.5 | 1 | 0 | NA | NA | 1 | 1 | 4 | 0.5 | 1 | 0 | 1 | 1 | 4 | 0.5 | 1 | 1 | 7 | 0.5714 |
| 19_887297 | 2 | 0.5 | 2 | 0.5 | NA | NA | 1 | 0 | 5 | 0.4 | NA | NA | 1 | 1 | 4 | 0.75 | 1 | 1 | 6 | 0.8333 |
| 19_887311 | 1 | 0 | 2 | 1 | NA | NA | 1 | 1 | 4 | 0.75 | NA | NA | 1 | 0 | 4 | 0.5 | 1 | 0 | 6 | 0.3333 |
| 19_887312 | 2 | 1 | 2 | 1 | NA | NA | 1 | 1 | 5 | 1 | NA | NA | 1 | 1 | 4 | 1 | 1 | 1 | 6 | 1 |
| 19_887324 | 3 | 1 | 2 | 1 | NA | NA | 1 | 1 | 6 | 1 | NA | NA | 1 | 1 | 2 | 1 | 1 | 1 | 4 | 1 |
| 19_887338 | 4 | 0.25 | 2 | 0 | NA | NA | 1 | 0 | 7 | 0.1429 | NA | NA | 1 | 0 | 4 | 0 | 1 | 1 | 6 | 0.1667 |
| 19_887348 | 2 | 1 | 3 | 1 | NA | NA | 1 | 1 | 6 | 1 | NA | NA | 1 | 1 | 3 | 1 | 1 | 1 | 5 | 1 |
| 19_887350 | 4 | 0.75 | 3 | 0.6667 | NA | NA | NA | NA | 7 | 0.7143 | NA | NA | 1 | 0 | 4 | 0.5 | NA | NA | 5 | 0.4 |
| 19_887358 | 4 | 1 | 2 | 0.5 | NA | NA | 2 | 0.5 | 8 | 0.75 | 1 | 1 | 1 | 1 | 4 | 0.5 | 2 | 1 | 8 | 0.75 |
| 19_887373 | 3 | 1 | 4 | 0.75 | NA | NA | 2 | 1 | 9 | 0.8889 | NA | NA | 1 | 1 | 4 | 1 | 2 | 1 | 7 | 1 |
| 19_887397 | 5 | 0.6 | 3 | 1 | NA | NA | 2 | 0 | 10 | 0.6 | NA | NA | 1 | 1 | 4 | 0.75 | 2 | 1 | 7 | 0.8571 |
| 19_887407 | 4 | 0.5 | 2 | 0 | NA | NA | 2 | 0 | 8 | 0.25 | NA | NA | 1 | 1 | 4 | 0.5 | 2 | 0 | 7 | 0.4286 |
| 19_887414 | 4 | 1 | 2 | 1 | NA | NA | 1 | 1 | 7 | 1 | NA | NA | NA | NA | 2 | 0.5 | 2 | 1 | 4 | 0.75 |
| 19_887422 | 4 | 1 | 2 | 1 | NA | NA | 2 | 1 | 8 | 1 | NA | NA | 1 | 1 | 4 | 1 | NA | NA | 5 | 1 |
| 19_887495 | 4 | 1 | 5 | 1 | 1 | 1 | 2 | 1 | 12 | 1 | NA | NA | NA | NA | 4 | 1 | 2 | 1 | 6 | 1 |
| 19_887509 | 6 | 0.8333 | 6 | 1 | 1 | 1 | 1 | 1 | 14 | 0.9286 | NA | NA | 1 | 1 | 4 | 0.5 | 3 | 0.6667 | 8 | 0.625 |
| 19_887509 | 4 | 0.25 | 5 | 0 | 2 | 0 | 2 | 0.5 | 13 | 0.1538 | NA | NA | 1 | 1 | 4 | 0 | 3 | 0 | 8 | 0.125 |
| 19_887512 | 5 | 0.2 | 6 | 0.1667 | 2 | 0.5 | 3 | 0 | 16 | 0.1875 | NA | NA | 1 | 1 | 4 | 0 | 3 | 0.3333 | 8 | 0.25 |
| 19_889017 | 8 | 0.25 | 9 | 0.1111 | 4 | 0 | 4 | 0 | 25 | 0.12 | 6 | 0 | 1 | 0 | 8 | 0.125 | 5 | 0.4 | 20 | 0.15 |
| 19_889310 | 12 | 0.1667 | 14 | 0.1429 | 6 | 0.1667 | 5 | 0 | 37 | 0.1351 | 6 | 0.1667 | 5 | 0 | 8 | 0.625 | 5 | 0.4 | 24 | 0.3333 |
| 19_889357 | 6 | 0.1667 | 8 | 0.125 | 3 | 0 | 2 | 0 | 19 | 0.1053 | 6 | 0.1667 | 5 | 0 | 5 | 0.2 | 4 | 0 | 20 | 0.1 |
| 19_889377 | 14 | 0.0714 | 18 | 0.1111 | 10 | 0 | 5 | 0.2 | 47 | 0.0851 | 13 | 0 | 6 | 0.1667 | 9 | 0.1111 | 7 | 0 | 35 | 0.0571 |
| 19_889379 | 17 | 0.2353 | 20 | 0.25 | 11 | 0.1818 | 8 | 0 | 56 | 0.1964 | 14 | 0.2857 | 6 | 0.1667 | 13 | 0.3077 | 7 | 0 | 40 | 0.225 |
| 19_890358 | 9 | 0.2222 | 15 | 0.2667 | 8 | 0.5 | 10 | 0.2 | 42 | 0.2857 | 14 | 0.2143 | 15 | 0.7333 | 13 | 0.4615 | 12 | 0.1667 | 54 | 0.4074 |
| 19_890421 | 13 | 0.0769 | 25 | 0.08 | 10 | 0.1 | 11 | 0 | 59 | 0.0678 | 17 | 0.1176 | 16 | 0.25 | 18 | 0.1111 | 12 | 0 | 63 | 0.127 |
| 19_890424 | 13 | 0.2308 | 30 | 0.0333 | 12 | 0.1667 | 10 | 0.4 | 65 | 0.1538 | 18 | 0.2778 | 17 | 0.1765 | 12 | 0.25 | 11 | 0.1818 | 58 | 0.2241 |
| 19_890429 | 15 | 0 | 30 | 0.1 | 12 | 0.0833 | 11 | 0.0909 | 68 | 0.0735 | 21 | 0.1429 | 17 | 0.0588 | 18 | 0.0556 | 12 | 0.0833 | 68 | 0.0882 |
| 19_890444 | 12 | 0.3333 | 27 | 0.1111 | 10 | 0.2 | 11 | 0 | 60 | 0.15 | 20 | 0.4 | 16 | 0.1875 | 20 | 0.25 | 13 | 0.2308 | 69 | 0.2754 |
| 19_890445 | 11 | 0.3636 | 28 | 0.7857 | 11 | 0.8182 | 11 | 0.7273 | 61 | 0.7049 | 20 | 0.7 | 16 | 0.8125 | 19 | 0.8421 | 11 | 0.8182 | 66 | 0.7879 |
| 19_890480 | 2 | 0 | 3 | 0.3333 | 3 | 0 | 1 | 0 | 9 | 0.1111 | 1 | 1 | 2 | 0 | NA | NA | 1 | 0 | 4 | 0.25 |
| 19_890499 | 1 | 0 | 3 | 0.6667 | 2 | 0 | NA | NA | 6 | 0.3333 | NA | NA | 2 | 0.5 | NA | NA | 1 | 0 | 3 | 0.3333 |
| 19_893660 | 2 | 0 | NA | NA | 1 | 0 | 1 | 1 | 4 | 0.25 | 1 | 1 | 3 | 0 | 1 | 1 | NA | NA | 5 | 0.4 |
| 19_893985 | 3 | 1 | NA | NA | 3 | 0.6667 | 3 | 1 | 9 | 0.8889 | 3 | 0.6667 | 3 | 1 | 1 | 1 | 1 | 1 | 8 | 0.875 |
| 19_894255 | 4 | 0.25 | NA | NA | 5 | 0.4 | 3 | 0 | 12 | 0.25 | 3 | 0 | 5 | 0.2 | 1 | 0 | 3 | 0 | 12 | 0.0833 |
| 19_894268 | 2 | 0.5 | 1 | 0 | 5 | 0.2 | 2 | 0 | 10 | 0.2 | 2 | 0 | 5 | 0 | 1 | 1 | 1 | 1 | 9 | 0.2222 |
| 19_894269 | 3 | 0 | 1 | 0 | 5 | 0.2 | 3 | 0.3333 | 12 | 0.1667 | 3 | 0.3333 | 4 | 0.25 | 1 | 0 | 2 | 0 | 10 | 0.2 |
| 19_894287 | 2 | 0.5 | 1 | 1 | 2 | 0 | 1 | 0 | 6 | 0.3333 | 1 | 1 | 3 | 0 | NA | NA | 3 | 0.6667 | 7 | 0.4286 |
| 19_894290 | 4 | 0.5 | 1 | 0 | 4 | 0.25 | 2 | 0 | 11 | 0.2727 | 1 | 0 | 5 | 0 | 1 | 1 | 3 | 0.3333 | 10 | 0.2 |
| 19_894290 | 4 | 0.75 | 1 | 1 | 4 | 0.5 | 3 | 1 | 12 | 0.75 | 3 | 0.6667 | 4 | 0.25 | 1 | 1 | 4 | 0.5 | 12 | 0.5 |
| 19_894301 | 4 | 0 | 1 | 0 | 6 | 0 | 3 | 0.3333 | 14 | 0.0714 | 3 | 0 | 5 | 0.2 | NA | NA | 3 | 0.3333 | 11 | 0.1818 |
| 19_894302 | 4 | 0.75 | 1 | 1 | 5 | 0.6 | 2 | 0.5 | 12 | 0.6667 | 3 | 0.6667 | 5 | 0.6 | 1 | 1 | 5 | 0.8 | 14 | 0.7143 |
| 19_894305 | 2 | 0.5 | 1 | 0 | 2 | 0 | 2 | 0 | 7 | 0.1429 | 2 | 0.5 | 4 | 0.25 | 1 | 0 | 3 | 0 | 10 | 0.2 |
| 19_894334 | 4 | 0.5 | 1 | 1 | 5 | 0.6 | 3 | 0.3333 | 13 | 0.5385 | 3 | 1 | 5 | 0.4 | 1 | 1 | 5 | 0.4 | 14 | 0.5714 |

| | | | | | | | | | | | | | | | | | | | | |
|---|---|---|---|---|---|---|---|---|---|---|---|---|---|---|---|---|---|---|---|---|
| 19_894344 | 3 | 0.3333 | 1 | 0 | 6 | 0.6667 | 1 | 1 | 11 | 0.5455 | 3 | 0.6667 | 2 | 1 | 1 | 1 | 3 | 0.6667 | 9 | 0.7778 |
| 19_895553 | 44 | 0.3409 | 46 | 0.2174 | 46 | 0.3043 | 50 | 0.18 | 186 | 0.2581 | 43 | 0.2326 | 45 | 0.2889 | 50 | 0.24 | 56 | 0.2679 | 194 | 0.2577 |
| 19_895553 | 36 | 0.4722 | 30 | 0.2333 | 32 | 0.25 | 38 | 0.3684 | 136 | 0.3382 | 28 | 0.2857 | 35 | 0.3429 | 38 | 0.3684 | 37 | 0.2973 | 138 | 0.3261 |
| 19_895554 | 2 | 1 | 1 | 1 | 1 | 0 | 2 | 0.5 | 6 | 0.6667 | 1 | 1 | NA | NA | 2 | 1 | NA | NA | 3 | 1 |
| 19_895573 | 2 | 1 | NA | NA | NA | NA | 1 | 1 | 3 | 1 | NA | NA | 1 | 1 | 3 | 0.3333 | NA | NA | 4 | 0.5 |
| 19_899233 | NA | NA | NA | NA | NA | NA | 1 | 1 | 1 | 1 | NA | NA | 1 | 1 | NA | NA | NA | NA | 1 | 1 |
| 19_899250 | NA | NA | NA | NA | NA | NA | 1 | 1 | 1 | 1 | NA | NA | 1 | 1 | NA | NA | NA | NA | 1 | 1 |
| 19_899263 | NA | NA | NA | NA | NA | NA | 1 | 1 | 1 | 1 | NA | NA | 1 | 1 | NA | NA | NA | NA | 1 | 1 |
| 19_899367 | NA | NA | 1 | 0 | NA | NA | 1 | 1 | 2 | 0.5 | NA | NA | 1 | 1 | NA | NA | NA | NA | 1 | 1 |
| 19_899371 | NA | NA | 1 | 0 | NA | NA | 2 | 0.5 | 3 | 0.3333 | NA | NA | 1 | 1 | NA | NA | NA | NA | 1 | 1 |
| 19_899398 | NA | NA | 1 | 0 | NA | NA | 3 | 0.3333 | 4 | 0.25 | NA | NA | 1 | 1 | NA | NA | NA | NA | 1 | 1 |
| 19_899415 | NA | NA | 1 | 0 | NA | NA | 3 | 0.6667 | 4 | 0.5 | NA | NA | 1 | 1 | NA | NA | NA | NA | 1 | 1 |
| 19_899431 | NA | NA | 1 | 0 | NA | NA | 2 | 0.5 | 3 | 0.3333 | NA | NA | 1 | 1 | NA | NA | NA | NA | 1 | 1 |
| 19_899435 | NA | NA | 1 | 0 | NA | NA | 3 | 0.3333 | 4 | 0.25 | NA | NA | 1 | 1 | NA | NA | NA | NA | 1 | 1 |
| 19_899441 | NA | NA | NA | NA | NA | NA | 2 | 1 | 2 | 1 | NA | NA | 1 | 1 | NA | NA | NA | NA | 1 | 1 |
| 19_899457 | NA | NA | NA | NA | NA | NA | 3 | 0.3333 | 3 | 0.3333 | NA | NA | 1 | 1 | NA | NA | NA | NA | 1 | 1 |
| 19_899493 | NA | NA | 1 | 0 | NA | NA | 3 | 0.6667 | 4 | 0.5 | NA | NA | 1 | 1 | NA | NA | NA | NA | 1 | 1 |
| 19_899515 | NA | NA | 1 | 0 | NA | NA | 4 | 0.25 | 5 | 0.2 | NA | NA | 1 | 1 | NA | NA | NA | NA | 1 | 1 |
| 19_899542 | NA | NA | 1 | 1 | 1 | 1 | 4 | 1 | 6 | 1 | NA | NA | 2 | 0.5 | NA | NA | NA | NA | 2 | 0.5 |
| 19_899566 | 1 | 1 | 2 | 0 | 2 | 0 | 4 | 0.25 | 9 | 0.2222 | NA | NA | 1 | 1 | NA | NA | NA | NA | 1 | 1 |
| 19_899585 | 1 | 0 | 2 | 0.5 | 2 | 1 | 5 | 0.6 | 10 | 0.6 | NA | NA | 2 | 0.5 | 1 | 0 | NA | NA | 3 | 0.3333 |
| 19_899619 | 3 | 0.3333 | 5 | 0 | 4 | 0.25 | 9 | 0 | 21 | 0.0952 | 1 | 0 | 4 | 0.5 | 2 | 0 | 2 | 0 | 9 | 0.2222 |
| 19_908893 | 4 | 0.25 | 5 | 0 | 5 | 0 | 4 | 0 | 18 | 0.0556 | 4 | 0.25 | 4 | 0 | 5 | 0 | 1 | 0 | 14 | 0.0714 |
| 19_908907 | 4 | 0 | 7 | 0.2857 | 7 | 0.2857 | 5 | 0.4 | 23 | 0.2609 | 4 | 0.25 | 3 | 0.6667 | 4 | 0.5 | 3 | 0.6667 | 14 | 0.5 |
| 19_908981 | 18 | 0.1111 | 15 | 0.2 | 16 | 0 | 8 | 0.125 | 57 | 0.1053 | 13 | 0 | 4 | 0.5 | 15 | 0.1333 | 7 | 0 | 39 | 0.1026 |
| 19_908993 | 16 | 0 | 11 | 0.2727 | 13 | 0 | 7 | 0 | 47 | 0.0638 | 11 | 0 | 4 | 0.25 | 11 | 0.1818 | 5 | 0 | 31 | 0.0968 |
| 19_909002 | 18 | 0.2222 | 13 | 0.5385 | 13 | 0.3077 | 8 | 0.125 | 52 | 0.3077 | 10 | 0.6 | 5 | 0.6 | 14 | 0.4286 | 7 | 0.5714 | 36 | 0.5278 |
| 19_909029 | 19 | 0.1053 | 11 | 0.0909 | 18 | 0.1667 | 8 | 0.125 | 56 | 0.125 | 10 | 0.1 | 10 | 0.3 | 22 | 0.1364 | 9 | 0.1111 | 51 | 0.1569 |
| 19_909037 | 18 | 0.1667 | 14 | 0.2857 | 17 | 0.3529 | 13 | 0.2308 | 62 | 0.2581 | 13 | 0.2308 | 10 | 0.3 | 23 | 0.2609 | 11 | 0.5455 | 57 | 0.3158 |
| 19_909045 | 20 | 0.1 | 12 | 0.25 | 18 | 0.1111 | 12 | 0 | 62 | 0.1129 | 12 | 0.0833 | 11 | 0.1818 | 19 | 0.0526 | 10 | 0.2 | 52 | 0.1154 |
| 19_909047 | 20 | 0.05 | 13 | 0.0769 | 18 | 0.2222 | 11 | 0 | 62 | 0.0968 | 13 | 0.3077 | 10 | 0.2 | 21 | 0.1905 | 10 | 0.5 | 54 | 0.2778 |
| 19_985004 | 1 | 0 | 3 | 0 | NA | NA | 2 | 0.5 | 6 | 0.1667 | 3 | 0 | 1 | 1 | 1 | 0 | NA | NA | 5 | 0.2 |
| 19_985008 | NA | NA | 4 | 0.5 | NA | NA | 1 | 0 | 5 | 0.4 | 3 | 0 | 1 | 0 | NA | NA | 1 | 1 | 5 | 0.2 |
| 19_985015 | 1 | 0 | 4 | 0.25 | NA | NA | 1 | 0 | 6 | 0.1667 | 3 | 0.3333 | 2 | 0.5 | 1 | 0 | NA | NA | 6 | 0.3333 |
| 19_987087 | NA | NA | 1 | 1 | NA | NA | NA | NA | 1 | 1 | 1 | 1 | NA | NA | NA | NA | NA | NA | 1 | 1 |
| 19_987089 | NA | NA | 1 | 0 | NA | NA | 1 | 1 | 2 | 0.5 | 1 | 1 | NA | NA | NA | NA | NA | NA | 1 | 1 |
| 19_987101 | NA | NA | 1 | 1 | NA | NA | NA | NA | 1 | 1 | 1 | 1 | NA | NA | NA | NA | NA | NA | 1 | 1 |
| 19_996228 | 562 | 0.0391 | 641 | 0.064 | 540 | 0.0667 | 447 | 0.038 | 2190 | 0.053 | 566 | 0.0689 | 399 | 0.0752 | 370 | 0.0486 | 396 | 0.0581 | 1731 | 0.0635 |
| 19_100119 | 2 | 0.5 | NA | NA | 1 | 0 | NA | NA | 3 | 0.3333 | 1 | 1 | 2 | 0.5 | NA | NA | 1 | 0 | 4 | 0.5 |
| 19_100120 | 2 | 0.5 | 2 | 1 | 1 | 1 | 1 | 0 | 6 | 0.6667 | 1 | 1 | 2 | 0.5 | NA | NA | 1 | 1 | 4 | 0.75 |
| 19_100120 | 2 | 1 | 2 | 0.5 | 1 | 1 | 1 | 1 | 6 | 0.8333 | 1 | 1 | 2 | 1 | NA | NA | 1 | 1 | 4 | 1 |
| 19_100120 | 2 | 1 | 2 | 1 | 1 | 1 | 1 | 1 | 6 | 1 | NA | NA | 1 | 0 | NA | NA | 1 | 1 | 2 | 0.5 |
| 19_100122 | 2 | 1 | 2 | 1 | 1 | 1 | 1 | 1 | 6 | 1 | 1 | 1 | 2 | 0 | NA | NA | 1 | 1 | 4 | 0.5 |
| 19_100123 | 2 | 0.5 | 2 | 0.5 | 1 | 0 | 1 | 1 | 6 | 0.5 | 1 | 1 | 2 | 1 | NA | NA | 1 | 1 | 4 | 1 |
| 19_100126 | 2 | 0.5 | 2 | 1 | 1 | 0 | 1 | 0 | 6 | 0.5 | 1 | 1 | 1 | 0 | NA | NA | 1 | 0 | 3 | 0.3333 |
| 19_100355 | 5 | 0.2 | 1 | 1 | 2 | 0.5 | 1 | 1 | 9 | 0.4444 | 1 | 0 | NA | NA | 3 | 0.3333 | 2 | 0 | 6 | 0.1667 |
| 19_100356 | 2 | 0.5 | 1 | 1 | 1 | 1 | 2 | 1 | 6 | 0.8333 | 1 | 1 | NA | NA | 1 | 1 | NA | NA | 2 | 1 |
| 19_100360 | 1 | 1 | 3 | 0 | 1 | 0 | 1 | 1 | 6 | 0.3333 | 1 | 1 | NA | NA | 1 | 1 | NA | NA | 2 | 1 |
| 19_100373 | NA | NA | 2 | 1 | NA | NA | 1 | 0 | 3 | 0.6667 | 1 | 1 | NA | NA | 1 | 0 | NA | NA | 2 | 0.5 |
| 19_101723 | 10 | 0.4 | 19 | 0.3158 | 29 | 0.1034 | 13 | 0.3846 | 71 | 0.2535 | 14 | 0.2143 | 21 | 0.0952 | 17 | 0.0588 | 12 | 0.0833 | 64 | 0.1094 |
| 19_101724 | 15 | 0.2 | 23 | 0.1304 | 30 | 0.0333 | 16 | 0.0625 | 84 | 0.0952 | 22 | 0.1364 | 22 | 0.0909 | 18 | 0.1111 | 17 | 0.1176 | 79 | 0.1139 |
| 19_101728 | 14 | 0.0714 | 18 | 0.1111 | 30 | 0.1 | 18 | 0.0556 | 80 | 0.0875 | 23 | 0.1304 | 26 | 0.2308 | 21 | 0.0476 | 17 | 0.2353 | 87 | 0.1609 |
| 19_101736 | 23 | 0.0435 | 28 | 0 | 37 | 0.1892 | 26 | 0.0769 | 114 | 0.0877 | 29 | 0.1034 | 33 | 0.0606 | 27 | 0.1111 | 23 | 0.087 | 112 | 0.0893 |
| 19_101739 | 20 | 0.1 | 26 | 0.1538 | 31 | 0.129 | 24 | 0.1667 | 101 | 0.1386 | 23 | 0.2174 | 31 | 0.0968 | 25 | 0.2 | 22 | 0.2727 | 101 | 0.1881 |
| 19_101765 | 4 | 0.25 | 1 | 0 | 4 | 0 | 2 | 0.5 | 11 | 0.1818 | 2 | 0 | 4 | 0 | 6 | 0.5 | 1 | 0 | 13 | 0.2308 |
| 19_101767 | 5 | 0.2 | NA | NA | 3 | 0.3333 | 2 | 0 | 10 | 0.2 | 1 | 1 | 3 | 0.3333 | 4 | 0.75 | NA | NA | 8 | 0.625 |
| 19_101768 | 4 | 1 | NA | NA | 4 | 0.75 | 2 | 1 | 10 | 0.9 | 1 | 1 | 2 | 1 | 4 | 1 | 1 | 1 | 8 | 1 |
| 19_101769 | 5 | 0.2 | NA | NA | 3 | 0.3333 | 2 | 1 | 10 | 0.4 | 1 | 0 | 2 | 0 | 6 | 0.3333 | 1 | 0 | 10 | 0.2 |
| 19_101769 | 2 | 0.5 | NA | NA | 3 | 0.3333 | 2 | 0 | 7 | 0.2857 | 2 | 0 | 3 | 0.6667 | 5 | 0.6 | 1 | 1 | 11 | 0.5455 |
| 19_101769 | 5 | 0.4 | NA | NA | 2 | 0 | 2 | 0 | 9 | 0.2222 | 2 | 1 | 3 | 0.3333 | 6 | 0.5 | 1 | 0 | 12 | 0.5 |
| 19_101770 | 5 | 1 | NA | NA | 2 | 1 | 2 | 1 | 9 | 1 | 1 | 1 | 3 | 1 | 5 | 1 | 1 | 1 | 10 | 1 |
| 19_101773 | 2 | 1 | NA | NA | NA | NA | 1 | 1 | 3 | 1 | NA | NA | 2 | 1 | 4 | 1 | 1 | 1 | 7 | 1 |
| 19_101775 | 4 | 0.25 | NA | NA | 2 | 0 | 1 | 0 | 7 | 0.1429 | 1 | 0 | 2 | 0 | 4 | 0.25 | 1 | 0 | 8 | 0.125 |
| 19_101776 | 3 | 0.3333 | NA | NA | 2 | 0 | 2 | 0 | 7 | 0.1429 | NA | NA | 2 | 0 | 4 | 0.25 | 1 | 0 | 7 | 0.1429 |
| 19_101776 | 2 | 0.5 | NA | NA | 2 | 0 | 2 | 0 | 6 | 0.1667 | 1 | 0 | 2 | 0.5 | 3 | 0 | NA | NA | 6 | 0.1667 |

| | | | | | | | | | | | | | | | | | | | | |
|---|---|---|---|---|---|---|---|---|---|---|---|---|---|---|---|---|---|---|---|---|
| 19_101779 | 3 | 1 | NA | NA | 1 | 1 | 1 | 1 | 5 | 1 | NA | NA | 2 | 1 | 3 | 1 | 1 | 1 | 6 | 1 |
| 19_101779 | 3 | 0.6667 | NA | NA | NA | NA | NA | NA | 3 | 0.6667 | NA | NA | 2 | 0.5 | 3 | 0.6667 | 1 | 1 | 6 | 0.6667 |
| 19_101860 | 2 | 0 | 12 | 0.0833 | 3 | 0 | 2 | 0 | 19 | 0.0526 | 4 | 0 | 7 | 0.1429 | 2 | 0 | 5 | 0 | 18 | 0.0556 |
| 19_101864 | 2 | 0 | 10 | 0.4 | 2 | 1 | 1 | 0 | 15 | 0.4 | 3 | 0.3333 | 6 | 0.6667 | 3 | 0.3333 | 2 | 0.5 | 14 | 0.5 |
| 19_101868 | 2 | 0.5 | 13 | 0 | 1 | 0 | 2 | 0 | 18 | 0.0556 | 2 | 0.5 | 4 | 0 | 3 | 0 | 3 | 0.3333 | 12 | 0.1667 |
| 19_101868 | 2 | 0.5 | 10 | 0.1 | 2 | 0 | 2 | 0 | 16 | 0.125 | 2 | 0.5 | 4 | 0 | 2 | 0 | 2 | 0 | 10 | 0.1 |
| 19_101869 | 3 | 0.3333 | 13 | 0.0769 | 2 | 0 | 3 | 0 | 21 | 0.0952 | 3 | 0.3333 | 4 | 0.25 | 3 | 0 | 3 | 0.6667 | 13 | 0.3077 |
| 19_101870 | 3 | 0 | 11 | 0.3636 | 2 | 0 | 3 | 0.3333 | 19 | 0.2632 | 2 | 0 | 4 | 0.5 | 2 | 0 | 2 | 0.5 | 10 | 0.3 |
| 19_101870 | 3 | 0.6667 | 8 | 0 | 2 | 0.5 | 3 | 0 | 16 | 0.1875 | 3 | 0.3333 | 3 | 0.3333 | 3 | 0 | 3 | 0.3333 | 12 | 0.25 |
| 19_101872 | 3 | 0 | 12 | 0.3333 | 2 | 0.5 | 3 | 0 | 20 | 0.25 | 3 | 0 | 4 | 0.25 | 2 | 0.5 | 3 | 0 | 12 | 0.1667 |
| 19_101872 | 3 | 0.6667 | 12 | 0.1667 | 2 | 0.5 | 3 | 0 | 20 | 0.25 | 3 | 0.3333 | 4 | 0 | 3 | 0.3333 | 1 | 0 | 11 | 0.1818 |
| 19_101873 | 2 | 0.5 | 8 | 0 | 1 | 0 | 1 | 0 | 12 | 0.0833 | 3 | 0.3333 | 3 | 0 | 2 | 0 | NA | NA | 8 | 0.125 |
| 19_101873 | 2 | 0 | 9 | 0.3333 | 2 | 0 | 3 | 0 | 16 | 0.1875 | 3 | 0.3333 | 4 | 0.25 | 3 | 0.3333 | 3 | 0.3333 | 13 | 0.3077 |
| 19_101874 | 3 | 0 | 10 | 0.3 | 2 | 0 | 2 | 0 | 17 | 0.1765 | 2 | 0 | 4 | 0 | 2 | 0 | 3 | 0.3333 | 11 | 0.0909 |
| 19_101874 | 1 | 0 | 8 | 0.25 | 1 | 0 | 2 | 0 | 12 | 0.1667 | NA | NA | 3 | 0.3333 | 3 | 0.3333 | 2 | 0.5 | 8 | 0.375 |
| 19_102231 | 6 | 0 | 6 | 0 | 2 | 0 | 6 | 0.1667 | 20 | 0.05 | 5 | 0 | 7 | 0.1429 | 4 | 0 | 4 | 0 | 20 | 0.05 |
| 19_102233 | 5 | 0.2 | 7 | 0 | NA | NA | 5 | 0 | 17 | 0.0588 | 5 | 0.2 | 7 | 0 | 5 | 0 | 6 | 0.1667 | 23 | 0.087 |
| 19_102239 | 4 | 0 | 3 | 0 | NA | NA | 3 | 0.3333 | 10 | 0.1 | 5 | 0.2 | 7 | 0.2857 | 3 | 0.3333 | 4 | 0 | 19 | 0.2105 |
| 19_102242 | 4 | 0 | NA | NA | 1 | 1 | 4 | 0 | 9 | 0.1111 | 5 | 0 | 5 | 0 | 3 | 0 | 5 | 0.2 | 18 | 0.0556 |
| 19_102250 | 4 | 1 | 1 | 1 | 1 | 1 | 3 | 0.6667 | 9 | 0.8889 | 3 | 1 | 4 | 1 | 1 | 1 | 2 | 1 | 10 | 1 |
| 19_102251 | 5 | 0.2 | 1 | 0 | 1 | 0 | 3 | 0 | 10 | 0.1 | 3 | 0.3333 | 4 | 0.25 | 2 | 0 | 4 | 0 | 13 | 0.1538 |
| 19_102251 | 2 | 1 | 1 | 1 | 1 | 1 | 3 | 1 | 7 | 1 | 2 | 1 | 4 | 1 | 2 | 1 | 3 | 0.6667 | 11 | 0.9091 |
| 19_102253 | 3 | 1 | 1 | 1 | 1 | 1 | 3 | 1 | 8 | 1 | 2 | 0.5 | 4 | 1 | 1 | 1 | 3 | 1 | 10 | 0.9 |
| 19_102253 | 5 | 0.4 | 1 | 1 | NA | NA | 3 | 0.6667 | 9 | 0.5556 | 3 | 0.3333 | 4 | 0.75 | 2 | 0 | 4 | 0.75 | 13 | 0.5385 |
| 19_102253 | 5 | 0.8 | 1 | 1 | 1 | 0 | 3 | 0.3333 | 10 | 0.6 | 2 | 0.5 | 5 | 0.4 | 1 | 1 | 2 | 1 | 10 | 0.6 |
| 19_102254 | 5 | 0.2 | 1 | 0 | NA | NA | 3 | 0.3333 | 9 | 0.2222 | 3 | 0 | 4 | 0.5 | 1 | 0 | 2 | 0.5 | 10 | 0.3 |
| 19_102254 | 5 | 0.8 | NA | NA | 1 | 1 | 3 | 0.3333 | 9 | 0.6667 | 3 | 1 | 4 | 0.75 | 2 | 0 | 2 | 1 | 11 | 0.7273 |
| 19_102255 | 4 | 0.25 | 1 | 0 | 1 | 0 | 3 | 0 | 9 | 0.1111 | 2 | 0 | 5 | 0.4 | 2 | 0 | 1 | 0 | 10 | 0.2 |
| 19_102256 | 4 | 1 | 1 | 1 | 1 | 1 | 3 | 0.6667 | 9 | 0.8889 | 3 | 1 | 4 | 0.75 | 1 | 1 | 2 | 1 | 10 | 0.9 |
| 19_102256 | 3 | 0 | NA | NA | 1 | 1 | 3 | 0 | 7 | 0.1429 | 2 | 0.5 | 4 | 0.25 | 2 | 0 | 1 | 0 | 9 | 0.2222 |
| 19_102258 | 1 | 1 | 1 | 0 | 1 | 0 | 3 | 0.3333 | 6 | 0.3333 | 3 | 0.3333 | 4 | 0.25 | 2 | 0 | 1 | 0 | 10 | 0.2 |
| 19_104259 | 6 | 0.5 | 9 | 0.2222 | 6 | 0.1667 | 8 | 0.125 | 29 | 0.2414 | 7 | 0.1429 | 4 | 0 | 6 | 0.1667 | 4 | 0.5 | 21 | 0.1905 |
| 19_104272 | 9 | 0.2222 | 12 | 0.25 | 10 | 0.1 | 10 | 0 | 41 | 0.1463 | 7 | 0.1429 | 10 | 0.1 | 11 | 0.1818 | 8 | 0.25 | 36 | 0.1667 |
| 19_104273 | 5 | 0 | 8 | 0.125 | 7 | 0 | 8 | 0.375 | 28 | 0.1429 | 5 | 0.2 | 9 | 0.3333 | 9 | 0.2222 | 8 | 0.125 | 31 | 0.2258 |
| 19_104288 | 15 | 0.1333 | 20 | 0 | 17 | 0.0588 | 15 | 0.0667 | 67 | 0.0597 | 12 | 0 | 19 | 0.1053 | 20 | 0.1 | 13 | 0.0769 | 64 | 0.0781 |
| 19_104301 | NA | NA | 5 | 0.4 | 3 | 0.3333 | 4 | 0.25 | 12 | 0.3333 | 5 | 0 | 2 | 0 | 1 | 0 | 6 | 0.5 | 14 | 0.2143 |
| 19_104316 | 2 | 0 | 1 | 1 | 2 | 0 | NA | NA | 5 | 0.2 | NA | NA | 6 | 0.1667 | 2 | 1 | 4 | 0.5 | 12 | 0.4167 |
| 19_104318 | 2 | 0 | 2 | 0.5 | 2 | 0 | NA | NA | 6 | 0.1667 | 1 | 0 | 4 | 0 | 3 | 0 | 7 | 0.2857 | 15 | 0.1333 |
| 19_104319 | 2 | 0.5 | 2 | 0 | 1 | 0 | NA | NA | 5 | 0.2 | NA | NA | 3 | 0 | 2 | 0.5 | 6 | 0.1667 | 11 | 0.1818 |
| 19_104320 | 2 | 0 | 2 | 0.5 | 2 | 0 | NA | NA | 6 | 0.1667 | NA | NA | 6 | 0 | 2 | 0 | 6 | 0.3333 | 14 | 0.1429 |
| 19_104324 | 2 | 0.5 | 1 | 1 | 2 | 0 | NA | NA | 5 | 0.4 | NA | NA | 4 | 0.5 | 1 | 1 | 3 | 0.3333 | 8 | 0.5 |
| 19_104328 | 29 | 0.5517 | 11 | 0.7273 | 19 | 0.7368 | 14 | 0.3571 | 73 | 0.589 | 8 | 0.375 | 27 | 0.7037 | 26 | 0.4615 | 14 | 0.4286 | 75 | 0.5333 |
| 19_104328 | 27 | 0.4444 | 20 | 0.3 | 25 | 0.36 | 25 | 0.24 | 97 | 0.3402 | 14 | 0.3571 | 51 | 0.4118 | 40 | 0.275 | 17 | 0.2941 | 122 | 0.3443 |
| 19_104329 | 21 | 0.3333 | 11 | 0.5455 | 10 | 0.3 | 17 | 0.1765 | 59 | 0.322 | 9 | 0.3333 | 27 | 0.1852 | 33 | 0.1212 | 13 | 0.1538 | 82 | 0.1707 |
| 19_104329 | 13 | 0.3077 | 13 | 0.3077 | 12 | 0.0833 | 8 | 0.25 | 46 | 0.2391 | 6 | 0.3333 | 22 | 0.2273 | 20 | 0.05 | 7 | 0 | 55 | 0.1455 |
| 19_104332 | 24 | 0.0833 | 17 | 0.0588 | 23 | 0.1739 | 22 | 0.0455 | 86 | 0.093 | 16 | 0.125 | 50 | 0.06 | 43 | 0.0698 | 18 | 0.0556 | 127 | 0.0709 |
| 19_104334 | 26 | 0.5 | 17 | 0.6471 | 18 | 0.3889 | 23 | 0.3478 | 84 | 0.4643 | 13 | 0.6923 | 45 | 0.3778 | 33 | 0.6364 | 16 | 0.5 | 107 | 0.514 |
| 19_104334 | 21 | 0.2381 | 11 | 0.0909 | 15 | 0.0667 | 14 | 0.0714 | 61 | 0.1311 | 9 | 0 | 31 | 0.129 | 26 | 0.1538 | 15 | 0.2 | 81 | 0.1358 |
| 19_104334 | 21 | 0.4762 | 12 | 0.25 | 16 | 0.4375 | 18 | 0.3889 | 67 | 0.403 | 8 | 0.25 | 38 | 0.2368 | 29 | 0.4483 | 17 | 0.5882 | 92 | 0.3696 |
| 19_104335 | 22 | 0.0909 | 13 | 0.1538 | 18 | 0.2222 | 20 | 0.3 | 73 | 0.1918 | 7 | 0.1429 | 41 | 0.1463 | 30 | 0.1333 | 19 | 0.2105 | 97 | 0.1546 |
| 19_104335 | 23 | 0.087 | 17 | 0.1176 | 19 | 0.1579 | 18 | 0.0556 | 77 | 0.1039 | 12 | 0 | 37 | 0.0811 | 33 | 0.0606 | 15 | 0.0667 | 97 | 0.0619 |
| 19_104336 | 27 | 0.1111 | 13 | 0.0769 | 20 | 0.1 | 19 | 0.1053 | 79 | 0.1013 | 10 | 0 | 40 | 0.075 | 33 | 0.0303 | 15 | 0.2 | 98 | 0.0714 |
| 19_104336 | 17 | 0.7647 | 11 | 0.6364 | 17 | 1 | 15 | 0.8667 | 60 | 0.8333 | 9 | 0.7778 | 27 | 0.9259 | 22 | 0.8636 | 12 | 0.6667 | 70 | 0.8429 |
| 19_104337 | 23 | 1 | 13 | 1 | 18 | 1 | 15 | 1 | 69 | 1 | 10 | 1 | 36 | 1 | 28 | 1 | 15 | 1 | 89 | 1 |
| 19_104338 | 14 | 0.9286 | 12 | 0.9167 | 15 | 1 | 12 | 1 | 53 | 0.9623 | 11 | 0.9091 | 23 | 1 | 19 | 0.9474 | 10 | 1 | 63 | 0.9683 |
| 19_104338 | 19 | 0.5263 | 8 | 0.875 | 14 | 0.6429 | 13 | 0.6923 | 54 | 0.6481 | 7 | 0.4286 | 28 | 0.7143 | 21 | 0.7143 | 13 | 0.3077 | 69 | 0.6087 |
| 19_104338 | 18 | 0.1111 | 8 | 0 | 16 | 0.0625 | 11 | 0.0909 | 53 | 0.0755 | 11 | 0.1818 | 27 | 0.1481 | 23 | 0.087 | 15 | 0.0667 | 76 | 0.1184 |
| 19_104339 | 17 | 0.2353 | 10 | 0.2 | 17 | 0.3529 | 13 | 0.2308 | 57 | 0.2632 | 10 | 0.2 | 32 | 0.2188 | 24 | 0.125 | 14 | 0.1429 | 80 | 0.175 |
| 19_104340 | 9 | 0.4444 | 9 | 0.7778 | 12 | 0.25 | 12 | 0.5 | 42 | 0.4762 | 10 | 0.5 | 27 | 0.5556 | 18 | 0.5556 | 11 | 0.3636 | 66 | 0.5152 |
| 19_104791 | 6 | 0.1667 | 5 | 0.2 | 5 | 0 | 2 | 0.5 | 18 | 0.1667 | 3 | 0 | 7 | 0.2857 | 7 | 0.1429 | 4 | 0 | 21 | 0.1429 |
| 19_104792 | 6 | 0 | 5 | 0.2 | 5 | 0.2 | 2 | 0.5 | 18 | 0.1667 | 3 | 0.3333 | 6 | 0 | 4 | 0 | 3 | 0 | 16 | 0.0625 |
| 19_104797 | 6 | 0.1667 | 5 | 0.2 | 4 | 0 | 1 | 0 | 16 | 0.125 | 3 | 0 | 6 | 0.3333 | 2 | 0 | 3 | 0 | 14 | 0.1429 |
| 19_104798 | 10 | 0.1 | 8 | 0.25 | 8 | 0.125 | 9 | 0.3333 | 35 | 0.2 | 6 | 0.3333 | 10 | 0.3 | 8 | 0.125 | 7 | 0.1429 | 31 | 0.2258 |
| 19_104798 | 8 | 0.5 | 8 | 0.125 | 7 | 0 | 8 | 0.25 | 31 | 0.2258 | 6 | 0.6667 | 9 | 0.3333 | 7 | 0.2857 | 6 | 0.1667 | 28 | 0.3571 |
| 19_104799 | 8 | 0.875 | 6 | 1 | 6 | 0.8333 | 6 | 0.8333 | 26 | 0.8846 | 3 | 1 | 6 | 1 | 3 | 0.6667 | 6 | 1 | 18 | 0.9444 |

| | | | | | | | | | | | | | | | | | | | | |
|---|---|---|---|---|---|---|---|---|---|---|---|---|---|---|---|---|---|---|---|---|
| 19_104800 | 11 | 0.4545 | 8 | 0.25 | 6 | 0.1667 | 9 | 0.6667 | 34 | 0.4118 | 4 | 0.75 | 7 | 0.5714 | 8 | 0.625 | 6 | 0.5 | 25 | 0.6 |
| 19_104800 | 12 | 0.3333 | 8 | 0.25 | 8 | 0.625 | 10 | 0.5 | 38 | 0.4211 | 7 | 0.1429 | 7 | 0.5714 | 7 | 0.2857 | 6 | 0.1667 | 27 | 0.2963 |
| 19_104800 | 12 | 0.5833 | 7 | 0.8571 | 6 | 0.6667 | 7 | 1 | 32 | 0.75 | 4 | 0.25 | 7 | 0.5714 | 6 | 0.3333 | 6 | 1 | 23 | 0.5652 |
| 19_104802 | 11 | 0.6364 | 8 | 0.75 | 6 | 0.3333 | 9 | 0.6667 | 34 | 0.6176 | 6 | 0.5 | 6 | 0.5 | 6 | 0.8333 | 5 | 0.4 | 23 | 0.5652 |
| 19_104802 | 13 | 0.4615 | 8 | 0.5 | 6 | 0.5 | 10 | 0.7 | 37 | 0.5405 | 7 | 0.4286 | 6 | 0.3333 | 8 | 0.25 | 6 | 0.6667 | 27 | 0.4074 |
| 19_104802 | 13 | 1 | 8 | 1 | 7 | 0.8571 | 9 | 0.8889 | 37 | 0.9459 | 7 | 0.7143 | 9 | 0.6667 | 8 | 0.875 | 7 | 0.7143 | 31 | 0.7419 |
| 19_104803 | 8 | 1 | 6 | 0.6667 | 7 | 0.8571 | 8 | 0.5 | 29 | 0.7586 | 6 | 0.8333 | 5 | 0.8 | 7 | 0.7143 | 5 | 0.4 | 23 | 0.6957 |
| 19_104804 | 12 | 0.5 | 8 | 0.375 | 7 | 0.2857 | 9 | 0.4444 | 36 | 0.4167 | 7 | 0.4286 | 8 | 0.625 | 8 | 0.625 | 7 | 0.4286 | 30 | 0.5333 |
| 19_104946 | 8 | 0.25 | 7 | 0.1429 | 6 | 0 | 8 | 0.25 | 29 | 0.1724 | 4 | 0.25 | 5 | 0 | 7 | 0.1429 | 6 | 0.1667 | 22 | 0.1364 |
| 19_105230 | 6 | 0.3333 | 7 | 0.7143 | 9 | 0.5556 | 10 | 0.8 | 32 | 0.625 | 3 | 1 | 9 | 0.5556 | 6 | 0.8333 | 5 | 0.8 | 23 | 0.7391 |
| 19_105231 | 7 | 0.2857 | 6 | 0.1667 | 9 | 0.3333 | 8 | 0.375 | 30 | 0.3 | 3 | 0.6667 | 8 | 0.375 | 7 | 0.4286 | 5 | 1 | 23 | 0.5652 |
| 19_105232 | 7 | 0.2857 | 8 | 0.75 | 9 | 0.5556 | 10 | 0.9 | 34 | 0.6471 | 1 | 1 | 9 | 0.7778 | 5 | 0.8 | 4 | 0.75 | 19 | 0.7895 |
| 19_105233 | 9 | 0.1111 | 8 | 0.125 | 10 | 0.5 | 8 | 0 | 35 | 0.2 | 3 | 0.3333 | 9 | 0 | 7 | 0.1429 | 4 | 0 | 23 | 0.087 |
| 19_105234 | 10 | 0 | 8 | 0.125 | 11 | 0.0909 | 9 | 0 | 38 | 0.0526 | 4 | 0.25 | 11 | 0.2727 | 7 | 0.1429 | 4 | 0 | 26 | 0.1923 |
| 19_105235 | 10 | 0.1 | 8 | 0.125 | 11 | 0 | 9 | 0 | 38 | 0.0526 | 4 | 0 | 10 | 0 | 7 | 0.1429 | 5 | 0.2 | 26 | 0.0769 |
| 19_105237 | 8 | 0.25 | 7 | 0.2857 | 9 | 0.3333 | 10 | 0.5 | 34 | 0.3529 | 3 | 0.3333 | 10 | 0.3 | 7 | 0.5714 | 4 | 0.5 | 24 | 0.4167 |
| 19_105246 | 8 | 0.125 | 8 | 0.25 | 10 | 0.3 | 9 | 0.1111 | 35 | 0.2 | 4 | 0 | 11 | 0.1818 | 8 | 0.25 | 5 | 0 | 28 | 0.1429 |
| 19_105246 | 12 | 0.25 | 9 | 0.4444 | 12 | 0.1667 | 11 | 0.2727 | 44 | 0.2727 | 4 | 0.5 | 13 | 0.3846 | 9 | 0.3333 | 7 | 0.4286 | 33 | 0.3939 |
| 19_105248 | 11 | 0.0909 | 7 | 0 | 11 | 0.0909 | 8 | 0 | 37 | 0.0541 | 3 | 0 | 10 | 0.1 | 11 | 0 | 5 | 0.2 | 29 | 0.069 |
| 19_105280 | 2 | 1 | 2 | 1 | NA | NA | 1 | 0 | 5 | 0.8 | 2 | 0.5 | NA | NA | 2 | 1 | 3 | 0.6667 | 7 | 0.7143 |
| 19_105282 | 1 | 0 | 2 | 0.5 | 1 | 1 | 2 | 0.5 | 6 | 0.5 | 2 | 0.5 | NA | NA | 2 | 0.5 | 3 | 0.3333 | 7 | 0.4286 |
| 19_105282 | 2 | 0.5 | 1 | 1 | NA | NA | 2 | 1 | 5 | 0.8 | 2 | 0.5 | NA | NA | 2 | 1 | 2 | 1 | 6 | 0.8333 |
| 19_105479 | NA | NA | NA | NA | 3 | 0.3333 | 3 | 0 | 6 | 0.1667 | 2 | 0.5 | 2 | 0.5 | 3 | 0.6667 | 2 | 0 | 9 | 0.4444 |
| 19_105481 | NA | NA | 1 | 1 | 3 | 0.3333 | 2 | 0 | 6 | 0.3333 | 1 | 0 | 2 | 0.5 | 2 | 0 | 2 | 0 | 7 | 0.1429 |
| 19_105482 | NA | NA | 1 | 1 | 2 | 0.5 | 2 | 0 | 5 | 0.4 | 2 | 0 | 2 | 0 | 3 | 0 | 2 | 0.5 | 9 | 0.1111 |
| 19_105484 | NA | NA | 1 | 1 | 3 | 1 | 1 | 1 | 5 | 1 | 2 | 1 | 2 | 0.5 | 3 | 0.6667 | 2 | 0.5 | 9 | 0.6667 |
| 19_105484 | NA | NA | 1 | 1 | 3 | 1 | 2 | 0.5 | 6 | 0.8333 | 2 | 1 | 2 | 1 | 3 | 0.6667 | 2 | 0.5 | 9 | 0.7778 |
| 19_105485 | NA | NA | NA | NA | 3 | 0.6667 | 2 | 0 | 5 | 0.4 | 2 | 1 | 1 | 0 | 3 | 1 | 1 | 1 | 7 | 0.8571 |
| 19_105485 | NA | NA | NA | NA | 3 | 0.6667 | 1 | 0 | 4 | 0.5 | 2 | 0.5 | 1 | 0 | 2 | 0 | 1 | 0 | 6 | 0.1667 |
| 19_105488 | NA | NA | 1 | 1 | 2 | 0.5 | 2 | 0.5 | 5 | 0.6 | 2 | 0 | 2 | 0 | 2 | 1 | 1 | 0 | 7 | 0.2857 |
| 19_105550 | 1 | 1 | NA | NA | NA | NA | NA | NA | 1 | 1 | 1 | 1 | NA | NA | 1 | 1 | NA | NA | 2 | 1 |
| 19_105552 | 1 | 1 | NA | NA | NA | NA | NA | NA | 1 | 1 | 2 | 1 | NA | NA | 1 | 1 | NA | NA | 3 | 1 |
| 19_105553 | 1 | 1 | NA | NA | NA | NA | NA | NA | 1 | 1 | 2 | 0.5 | NA | NA | 1 | 1 | NA | NA | 3 | 0.6667 |
| 19_105698 | 3 | 0 | 3 | 0 | NA | NA | 2 | 0.5 | 8 | 0.125 | 1 | 0 | 1 | 0 | 1 | 0 | 3 | 0.3333 | 6 | 0.1667 |
| 19_106067 | 16 | 0 | 29 | 0.069 | 33 | 0.0909 | 28 | 0.1429 | 106 | 0.0849 | 26 | 0.1923 | 22 | 0.0909 | 31 | 0.0645 | 24 | 0.0833 | 103 | 0.1068 |
| 19_106067 | 10 | 0 | 15 | 0.0667 | 21 | 0.0476 | 10 | 0.1 | 56 | 0.0536 | 11 | 0.0909 | 11 | 0.1818 | 17 | 0.0588 | 10 | 0 | 49 | 0.0816 |
| 19_106422 | NA | NA | 1 | 1 | NA | NA | NA | NA | 1 | 1 | NA | NA | NA | NA | NA | NA | 1 | 1 | 1 | 1 |
| 19_106901 | 1 | 0 | NA | NA | 1 | 1 | 2 | 0.5 | 4 | 0.5 | 2 | 0.5 | NA | NA | 1 | 0 | 2 | 0 | 5 | 0.2 |
| 19_106901 | 1 | 0 | NA | NA | 1 | 1 | 1 | 0 | 3 | 0.3333 | 1 | 0 | NA | NA | 1 | 0 | 2 | 0.5 | 4 | 0.25 |
| 19_106902 | 1 | 0 | 1 | 1 | 1 | 1 | 2 | 0 | 5 | 0.4 | 2 | 0.5 | NA | NA | 1 | 0 | 2 | 0 | 5 | 0.2 |
| 19_106905 | 1 | 1 | 1 | 1 | 3 | 0.3333 | 1 | 1 | 6 | 0.6667 | 1 | 1 | NA | NA | 1 | 1 | 2 | 0 | 4 | 0.5 |
| 19_106907 | 1 | 1 | 1 | 1 | 2 | 0.5 | 2 | 0.5 | 6 | 0.6667 | 3 | 0.6667 | NA | NA | 1 | 1 | 2 | 0.5 | 6 | 0.6667 |
| 19_106908 | 1 | 1 | 1 | 0 | 3 | 1 | 1 | 1 | 6 | 0.8333 | 2 | 0.5 | NA | NA | 1 | 0 | 2 | 0.5 | 5 | 0.4 |
| 19_106909 | 1 | 0 | 1 | 0 | 3 | 0.3333 | 2 | 0 | 7 | 0.1429 | 3 | 0.3333 | NA | NA | 1 | 0 | 2 | 0 | 6 | 0.1667 |
| 19_106911 | 1 | 0 | 1 | 1 | 3 | 1 | 2 | 1 | 7 | 0.8571 | 3 | 0.6667 | NA | NA | NA | NA | 2 | 0 | 5 | 0.4 |
| 19_106916 | 1 | 0 | 1 | 0 | 3 | 0.3333 | 3 | 0 | 8 | 0.125 | 2 | 0.5 | 1 | 0 | 2 | 0 | 2 | 0 | 7 | 0.1429 |
| 19_108352 | 20 | 1 | 17 | 1 | 16 | 0.9375 | 19 | 1 | 72 | 0.9861 | 27 | 1 | 21 | 0.9524 | 25 | 0.96 | 10 | 1 | 83 | 0.9759 |
| 19_108354 | 19 | 0.9474 | 16 | 0.6875 | 16 | 0.9375 | 22 | 0.7727 | 73 | 0.8356 | 27 | 0.6667 | 22 | 0.6818 | 24 | 0.875 | 13 | 0.9231 | 86 | 0.7674 |
| 19_108355 | 24 | 0.2917 | 16 | 0.1875 | 19 | 0.2632 | 21 | 0.2381 | 80 | 0.25 | 26 | 0.1923 | 23 | 0.0435 | 25 | 0.08 | 13 | 0.1538 | 87 | 0.1149 |
| 19_108355 | 18 | 0.3333 | 12 | 0.5 | 16 | 0.5625 | 18 | 0.5 | 64 | 0.4688 | 19 | 0.6316 | 19 | 0.5263 | 23 | 0.3913 | 10 | 0.5 | 71 | 0.507 |
| 19_108355 | 22 | 0.2273 | 12 | 0.1667 | 16 | 0.375 | 21 | 0.2381 | 71 | 0.2535 | 24 | 0.2917 | 21 | 0.1429 | 25 | 0.24 | 13 | 0.1538 | 83 | 0.2169 |
| 19_108356 | 17 | 0.4118 | 17 | 0.3529 | 17 | 0.2941 | 20 | 0.2 | 71 | 0.3099 | 24 | 0.375 | 21 | 0.2857 | 23 | 0.4783 | 12 | 0.4167 | 80 | 0.3875 |
| 19_108358 | 16 | 0.1875 | 11 | 0.2727 | 15 | 0.3333 | 18 | 0.6111 | 60 | 0.3667 | 20 | 0.35 | 16 | 0.125 | 22 | 0.3636 | 9 | 0.1111 | 67 | 0.2687 |
| 19_108358 | 13 | 0.8462 | 9 | 0.7778 | 15 | 0.8667 | 21 | 0.8571 | 58 | 0.8448 | 17 | 0.7647 | 15 | 0.8 | 18 | 0.8333 | 9 | 0.7778 | 59 | 0.7966 |
| 19_108358 | 12 | 0.0833 | 8 | 0.125 | 8 | 0.125 | 15 | 0.0667 | 43 | 0.093 | 20 | 0.1 | 11 | 0 | 12 | 0.0833 | 7 | 0 | 50 | 0.06 |
| 19_108358 | 17 | 0.1765 | 13 | 0.3077 | 15 | 0.4 | 20 | 0.15 | 65 | 0.2462 | 20 | 0.2 | 20 | 0.05 | 21 | 0.2857 | 11 | 0.1818 | 72 | 0.1806 |
| 19_108359 | 19 | 0.0526 | 13 | 0.1538 | 14 | 0.0714 | 18 | 0.2778 | 64 | 0.1406 | 21 | 0.0952 | 16 | 0.125 | 18 | 0.0556 | 12 | 0 | 67 | 0.0746 |
| 19_108361 | 13 | 0.3077 | 11 | 0.0909 | 16 | 0.0625 | 19 | 0.0526 | 59 | 0.1186 | 19 | 0 | 16 | 0 | 20 | 0.1 | 11 | 0.1818 | 66 | 0.0606 |
| 19_108484 | 6 | 0.3333 | 6 | 0.1667 | 7 | 0.2857 | 6 | 0.1667 | 25 | 0.24 | 5 | 0.6 | 7 | 0.4286 | 4 | 0.5 | 7 | 0.4286 | 23 | 0.4783 |
| 19_108489 | 7 | 0.1429 | 8 | 0.125 | 5 | 0 | 7 | 0 | 27 | 0.0741 | 6 | 0 | 8 | 0.125 | 3 | 0 | 10 | 0.1 | 27 | 0.0741 |
| 19_108492 | 5 | 0 | 7 | 0.2857 | 6 | 0.1667 | 7 | 0 | 25 | 0.12 | 7 | 0 | 5 | 0.2 | 4 | 0.25 | 8 | 0 | 24 | 0.0833 |
| 19_108826 | 19 | 0.1579 | 23 | 0.0435 | 22 | 0.2273 | 17 | 0.1765 | 81 | 0.1481 | 14 | 0.1429 | 28 | 0.1071 | 29 | 0.069 | 10 | 0.1 | 81 | 0.0988 |
| 19_109199 | 1 | 1 | 1 | 1 | NA | NA | 1 | 1 | 3 | 1 | 1 | 1 | 1 | 1 | NA | NA | NA | NA | 2 | 1 |
| 19_109201 | NA | NA | 2 | 0.5 | NA | NA | 2 | 0 | 4 | 0.25 | 1 | 1 | 1 | 0 | NA | NA | 1 | 0 | 3 | 0.3333 |
| 19_109203 | NA | NA | NA | NA | NA | NA | 2 | 0.5 | 2 | 0.5 | 1 | 0 | 1 | 1 | NA | NA | 2 | 1 | 4 | 0.75 |

| | | | | | | | | | | | | | | | | | | | | |
|---|---|---|---|---|---|---|---|---|---|---|---|---|---|---|---|---|---|---|---|---|
| 19_109203 | 1 | 1 | 1 | 1 | NA | NA | 2 | 0.5 | 4 | 0.75 | 2 | 0.5 | 1 | 1 | NA | NA | 2 | 1 | 5 | 0.8 |
| 19_109204 | NA | NA | 2 | 0 | NA | NA | 2 | 0.5 | 4 | 0.25 | 2 | 0.5 | 1 | 0 | NA | NA | 2 | 0.5 | 5 | 0.4 |
| 19_109205 | 1 | 1 | 2 | 0.5 | NA | NA | 2 | 1 | 5 | 0.8 | 1 | 0 | 1 | 0 | NA | NA | 3 | 1 | 5 | 0.6 |
| 19_109259 | 1 | 1 | 1 | 1 | NA | NA | 2 | 1 | 4 | 1 | 1 | 1 | 1 | 1 | NA | NA | 1 | 0 | 3 | 0.6667 |
| 19_110252 | 1 | 1 | 1 | 1 | 1 | 0 | NA | NA | 3 | 0.6667 | 1 | 0 | NA | NA | NA | NA | 2 | 1 | 3 | 0.6667 |
| 19_117517 | 5 | 0.2 | 4 | 0 | 7 | 0.2857 | 7 | 0 | 23 | 0.1304 | 5 | 0 | 3 | 0 | 6 | 0 | 4 | 0.25 | 18 | 0.0556 |
| 19_117518 | 5 | 0.6 | 4 | 0.75 | 7 | 0.7143 | 8 | 0.375 | 24 | 0.5833 | 6 | 0.6667 | 4 | 0.5 | 5 | 0.4 | 4 | 0.5 | 19 | 0.5263 |
| 19_117518 | 5 | 0.4 | 4 | 0 | 7 | 0.7143 | 6 | 0.6667 | 22 | 0.5 | 4 | 0.75 | 4 | 0.25 | 7 | 0.4286 | 4 | 0.5 | 19 | 0.4737 |
| 19_117518 | 5 | 0.8 | 4 | 0.5 | 7 | 0.7143 | 7 | 0.5714 | 23 | 0.6522 | 5 | 0.8 | 3 | 1 | 4 | 1 | 5 | 1 | 17 | 0.9412 |
| 19_117519 | 5 | 0 | 3 | 0.3333 | 5 | 0.2 | 8 | 0.25 | 21 | 0.1905 | 6 | 0.3333 | 3 | 0 | 6 | 0.3333 | 5 | 0.6 | 20 | 0.35 |
| 19_117521 | 6 | 0.1667 | 6 | 0.1667 | 6 | 0 | 8 | 0.125 | 26 | 0.1154 | 5 | 0 | 5 | 0 | 6 | 0 | 4 | 0.25 | 20 | 0.05 |
| 19_117521 | 3 | 0 | 3 | 0 | 6 | 0.3333 | 7 | 0.2857 | 19 | 0.2105 | 5 | 0.6 | 5 | 0.2 | 6 | 0.3333 | 4 | 0.75 | 20 | 0.45 |
| 19_117522 | NA | NA | 1 | 1 | 2 | 0.5 | 1 | 1 | 4 | 0.75 | 1 | 1 | NA | NA | 1 | 1 | 1 | 0 | 3 | 0.6667 |
| 19_119426 | NA | NA | 1 | 1 | NA | NA | NA | NA | 1 | 1 | NA | NA | 1 | 1 | NA | NA | NA | NA | 1 | 1 |
| 19_119430 | NA | NA | 1 | 1 | NA | NA | NA | NA | 1 | 1 | NA | NA | 1 | 1 | NA | NA | NA | NA | 1 | 1 |
| 19_119435 | 2 | 0 | 1 | 0 | NA | NA | 1 | 1 | 4 | 0.25 | 3 | 0 | 2 | 0 | 5 | 0.2 | 3 | 0 | 13 | 0.0769 |
| 19_119613 | 3 | 0 | 3 | 0 | 1 | 1 | 1 | 0 | 8 | 0.125 | 3 | 0 | 3 | 0 | 7 | 0.2857 | 3 | 0 | 16 | 0.125 |
| 19_119690 | 3 | 0 | 4 | 0.5 | 4 | 0.5 | 3 | 0 | 14 | 0.2857 | 4 | 0.25 | 3 | 0 | 12 | 0.1667 | 4 | 0 | 23 | 0.1304 |
| 19_119693 | 3 | 0.3333 | 4 | 0.25 | 3 | 0.3333 | 4 | 0 | 14 | 0.2143 | 4 | 0 | 4 | 0 | 11 | 0.2727 | 6 | 0.3333 | 25 | 0.2 |
| 19_119693 | 4 | 1 | 3 | 0.6667 | 4 | 1 | 4 | 0.75 | 15 | 0.8667 | 4 | 0.5 | 4 | 1 | 13 | 0.7692 | 6 | 0.8333 | 27 | 0.7778 |
| 19_119694 | 4 | 1 | 4 | 1 | 4 | 0.75 | 4 | 1 | 16 | 0.9375 | 3 | 1 | 4 | 1 | 11 | 0.9091 | 6 | 1 | 24 | 0.9583 |
| 19_119698 | 3 | 0.3333 | 3 | 0.6667 | 3 | 0.3333 | 3 | 0.6667 | 12 | 0.5 | 3 | 0.6667 | 4 | 0.25 | 11 | 0.5455 | 6 | 0.5 | 24 | 0.5 |
| 19_119701 | 5 | 0.4 | 4 | 0.25 | 4 | 0.25 | 2 | 0.5 | 15 | 0.3333 | 4 | 0.75 | 5 | 0.4 | 11 | 0.6364 | 5 | 0.4 | 25 | 0.56 |
| 19_119701 | 4 | 0.75 | 4 | 0.25 | 4 | 0 | 3 | 0 | 15 | 0.2667 | 4 | 0.25 | 6 | 0 | 14 | 0 | 6 | 0.1667 | 30 | 0.0667 |
| 19_119705 | 3 | 0.3333 | 4 | 0.25 | 4 | 0 | 4 | 0.5 | 15 | 0.2667 | 3 | 0 | 6 | 0.1667 | 13 | 0.2308 | 7 | 0.1429 | 29 | 0.1724 |
| 19_119705 | 4 | 0.25 | 5 | 0 | 3 | 0.3333 | 4 | 0 | 16 | 0.125 | 4 | 0 | 5 | 0 | 8 | 0.125 | 8 | 0.125 | 25 | 0.08 |
| 19_124416 | 1 | 0 | 1 | 1 | NA | NA | NA | NA | 2 | 0.5 | 3 | 0.3333 | 1 | 1 | NA | NA | 1 | 1 | 5 | 0.6 |
| 19_124421 | 1 | 0 | 3 | 1 | NA | NA | NA | NA | 4 | 0.75 | 3 | 1 | 1 | 1 | 2 | 0 | 2 | 0.5 | 8 | 0.625 |
| 19_124422 | NA | NA | 2 | 1 | NA | NA | NA | NA | 2 | 1 | 3 | 1 | 1 | 0 | 2 | 1 | 2 | 1 | 8 | 0.875 |
| 19_124422 | 1 | 0 | 1 | 1 | NA | NA | NA | NA | 2 | 0.5 | 3 | 0.6667 | NA | NA | 1 | 0 | 1 | 0 | 5 | 0.4 |
| 19_124424 | 1 | 1 | 3 | 1 | 1 | 1 | NA | NA | 5 | 1 | 4 | 1 | 1 | 1 | 2 | 0.5 | 2 | 1 | 9 | 0.8889 |
| 19_124468 | 1 | 1 | NA | NA | 1 | 1 | NA | NA | 2 | 1 | NA | NA | 2 | 1 | NA | NA | NA | NA | 2 | 1 |
| 19_124468 | NA | NA | 1 | 1 | NA | NA | NA | NA | 1 | 1 | NA | NA | 1 | 1 | NA | NA | NA | NA | 1 | 1 |
| 19_124694 | 2 | 1 | 1 | 1 | NA | NA | NA | NA | 3 | 1 | NA | NA | 1 | 1 | NA | NA | NA | NA | 1 | 1 |
| 19_124696 | 1 | 1 | 1 | 1 | NA | NA | NA | NA | 2 | 1 | NA | NA | 1 | 1 | NA | NA | NA | NA | 1 | 1 |
| 19_127445 | 9 | 0.1111 | 8 | 0.25 | 18 | 0.0556 | 13 | 0.0769 | 48 | 0.1042 | 7 | 0 | 14 | 0.0714 | 14 | 0.1429 | 11 | 0 | 46 | 0.0652 |
| 19_127467 | 5 | 0 | 5 | 0.2 | 7 | 0.1429 | 4 | 0.25 | 21 | 0.1429 | 2 | 0.5 | 4 | 0 | 4 | 0 | 3 | 0 | 13 | 0.0769 |
| 19_127470 | 3 | 0 | 3 | 0 | 7 | 0.1429 | 3 | 0 | 16 | 0.0625 | 2 | 0.5 | 4 | 0 | 5 | 0 | 4 | 0 | 15 | 0.0667 |
| 19_127473 | 3 | 0 | 4 | 0.25 | 6 | 0.1667 | 3 | 0 | 16 | 0.125 | 2 | 0.5 | 3 | 0.3333 | 4 | 0 | 4 | 0 | 13 | 0.1538 |
| 19_127473 | 4 | 0.5 | 6 | 0.6667 | 5 | 0.6 | 3 | 1 | 18 | 0.6667 | 2 | 1 | 5 | 0.8 | 2 | 1 | 5 | 0.6 | 14 | 0.7857 |
| 19_127473 | 4 | 0.75 | 6 | 0.3333 | 7 | 0.8571 | 2 | 0.5 | 19 | 0.6316 | 2 | 0.5 | 2 | 1 | 3 | 0.6667 | 5 | 0.8 | 12 | 0.75 |
| 19_127473 | 3 | 0.3333 | 6 | 0.6667 | 5 | 0.2 | 3 | 0.6667 | 17 | 0.4706 | 1 | 0 | 6 | 0.8333 | 3 | 0.3333 | 4 | 0 | 14 | 0.4286 |
| 19_127473 | 1 | 1 | 3 | 0.6667 | 5 | 0.4 | 2 | 1 | 11 | 0.6364 | 2 | 1 | 3 | 0 | 3 | 0.3333 | 2 | 1 | 10 | 0.5 |
| 19_127474 | 4 | 0.5 | 5 | 0 | 6 | 0 | 2 | 0.5 | 17 | 0.1765 | 2 | 0.5 | 5 | 0 | 2 | 0 | 5 | 0.2 | 14 | 0.1429 |
| 19_127474 | 4 | 1 | 4 | 0.5 | 6 | 0.5 | 2 | 1 | 16 | 0.6875 | 1 | 1 | 5 | 0.8 | 2 | 0.5 | 3 | 1 | 11 | 0.8182 |
| 19_127474 | 4 | 1 | 3 | 0.6667 | 6 | 0.8333 | 2 | 1 | 15 | 0.8667 | 1 | 1 | 3 | 1 | 2 | 1 | 4 | 1 | 10 | 1 |
| 19_127475 | 4 | 1 | 4 | 0.75 | 4 | 1 | 2 | 1 | 14 | 0.9286 | 2 | 1 | 4 | 1 | 1 | 1 | 3 | 1 | 10 | 1 |
| 19_127476 | 2 | 0 | 3 | 0.3333 | 3 | 0 | 1 | 0 | 9 | 0.1111 | 1 | 0 | 5 | 0 | 1 | 0 | 2 | 0.5 | 9 | 0.1111 |
| 19_127477 | 3 | 0.6667 | 4 | 1 | 5 | 1 | NA | NA | 12 | 0.9167 | 1 | 1 | 1 | 1 | 1 | 1 | 2 | 1 | 5 | 1 |
| 19_127478 | 3 | 0.3333 | 5 | 0 | 5 | 0.2 | NA | NA | 13 | 0.1538 | 1 | 1 | 3 | 0.3333 | 1 | 0 | 2 | 0.5 | 7 | 0.4286 |
| 19_144257 | 8 | 0.5 | 5 | 0.2 | 10 | 0.1 | 8 | 0.25 | 31 | 0.2581 | 7 | 0 | 8 | 0.625 | 15 | 0.4 | 10 | 0.6 | 40 | 0.425 |
| 19_144267 | 5 | 0.4 | 2 | 0 | 6 | 0.3333 | 5 | 0 | 18 | 0.2222 | 6 | 0.3333 | 7 | 0 | 8 | 0 | 6 | 0 | 27 | 0.0741 |
| 19_144270 | 5 | 1 | 2 | 1 | 2 | 1 | 4 | 1 | 13 | 1 | 4 | 0.75 | 6 | 0.6667 | 4 | 1 | 5 | 0.6 | 19 | 0.7368 |
| 19_144270 | 6 | 0.3333 | 2 | 0.5 | 2 | 0 | 2 | 0.5 | 12 | 0.3333 | 6 | 0.5 | 7 | 0.8571 | 4 | 0 | 6 | 0.1667 | 23 | 0.4348 |
| 19_158826 | 12 | 0 | 24 | 0.0417 | 18 | 0.1111 | 18 | 0.1667 | 72 | 0.0833 | 19 | 0.1053 | 7 | 0 | 15 | 0.0667 | 10 | 0.1 | 51 | 0.0784 |
| 19_158831 | 10 | 1 | 23 | 1 | 19 | 0.8947 | 17 | 0.9412 | 69 | 0.9565 | 14 | 1 | 7 | 1 | 13 | 0.8462 | 10 | 1 | 44 | 0.9545 |
| 19_158833 | 12 | 0.4167 | 24 | 0.25 | 23 | 0.1304 | 17 | 0.3529 | 76 | 0.2632 | 18 | 0.3333 | 8 | 0 | 16 | 0.0625 | 14 | 0.1429 | 56 | 0.1607 |
| 19_158833 | 11 | 0.1818 | 23 | 0.1304 | 25 | 0.08 | 15 | 0.3333 | 74 | 0.1622 | 18 | 0.1667 | 7 | 0 | 14 | 0.1429 | 16 | 0.125 | 55 | 0.1273 |
| 19_158833 | 12 | 0.1667 | 24 | 0.125 | 24 | 0 | 18 | 0.0556 | 78 | 0.0769 | 17 | 0.1176 | 8 | 0 | 17 | 0.1765 | 15 | 0.0667 | 57 | 0.1053 |
| 19_158833 | 12 | 0.1667 | 23 | 0.0435 | 22 | 0.0909 | 19 | 0.1053 | 76 | 0.0921 | 16 | 0.0625 | 8 | 0 | 15 | 0.0667 | 15 | 0.0667 | 54 | 0.0556 |
| 19_159332 | NA | NA | NA | NA | 1 | 0 | 2 | 1 | 3 | 0.6667 | NA | NA | 1 | 1 | 3 | 0.3333 | NA | NA | 4 | 0.5 |
| 19_159332 | NA | NA | NA | NA | 2 | 0.5 | 2 | 0.5 | 4 | 0.5 | NA | NA | 1 | 1 | 3 | 0.6667 | NA | NA | 4 | 0.75 |
| 19_159333 | NA | NA | NA | NA | 2 | 0 | 2 | 1 | 4 | 0.5 | NA | NA | 1 | 0 | 4 | 1 | NA | NA | 5 | 0.8 |
| 19_159333 | NA | NA | NA | NA | 2 | 1 | 2 | 1 | 4 | 1 | NA | NA | 1 | 1 | 3 | 0.6667 | NA | NA | 4 | 0.75 |
| 19_159334 | NA | NA | NA | NA | 2 | 1 | 2 | 1 | 4 | 1 | NA | NA | NA | NA | 4 | 0.5 | NA | NA | 4 | 0.5 |

| | | | | | | | | | | | | | | | | | | | | |
|---|---|---|---|---|---|---|---|---|---|---|---|---|---|---|---|---|---|---|---|---|
| 19_1593343 | NA | NA | NA | NA | 2 | 0.5 | 1 | 1 | 3 | 0.6667 | NA | NA | 1 | 1 | 4 | 0.75 | NA | NA | 5 | 0.8 |
| 19_1593437 | NA | NA | NA | NA | 1 | 1 | NA | NA | 1 | 1 | 1 | 1 | NA | NA | NA | NA | NA | NA | 1 | 1 |
| 19_1593444 | NA | NA | NA | NA | 1 | 1 | NA | NA | 1 | 1 | 1 | 1 | NA | NA | NA | NA | NA | NA | 1 | 1 |
| 19_1593489 | NA | NA | NA | NA | 1 | 1 | NA | NA | 1 | 1 | 1 | 1 | NA | NA | NA | NA | NA | NA | 1 | 1 |
| 19_1593504 | NA | NA | NA | NA | 1 | 1 | NA | NA | 1 | 1 | 1 | 1 | NA | NA | NA | NA | NA | NA | 1 | 1 |
| 19_1593506 | NA | NA | NA | NA | 1 | 1 | NA | NA | 1 | 1 | 1 | 1 | NA | NA | NA | NA | NA | NA | 1 | 1 |
| 19_1636056 | 2 | 0 | NA | NA | 5 | 0 | 6 | 0.1667 | 13 | 0.0769 | NA | NA | 6 | 0.1667 | 2 | 0.5 | 8 | 0 | 16 | 0.125 |
| 19_1636062 | 2 | 0.5 | NA | NA | 5 | 0 | 5 | 0 | 12 | 0.0833 | 1 | 0 | 4 | 0.25 | 2 | 0 | 7 | 0.1429 | 14 | 0.1429 |
| 19_1636064 | 2 | 0 | NA | NA | 6 | 0 | 7 | 0.1429 | 15 | 0.0667 | 1 | 0 | 7 | 0.1429 | 2 | 0 | 9 | 0 | 19 | 0.0526 |
| 19_1636071 | 2 | 0 | NA | NA | 5 | 0.2 | 7 | 0 | 14 | 0.0714 | 2 | 0 | 7 | 0.2857 | 2 | 0.5 | 10 | 0.1 | 21 | 0.1905 |
| 19_1636073 | 2 | 0 | NA | NA | 6 | 0.1667 | 6 | 0 | 14 | 0.0714 | 1 | 0 | 5 | 0 | 2 | 0 | 10 | 0.1 | 18 | 0.0556 |
| 19_1636077 | 2 | 0 | 1 | 0 | 6 | 0.1667 | 5 | 0 | 14 | 0.0714 | 2 | 0 | 7 | 0.1429 | 1 | 0 | 8 | 0 | 18 | 0.0556 |
| 19_1636081 | 2 | 1 | 1 | 1 | 5 | 1 | 5 | 1 | 13 | 1 | 2 | 1 | 7 | 1 | 2 | 1 | 7 | 1 | 18 | 1 |
| 19_1636087 | 1 | 1 | 1 | 0 | 6 | 0.3333 | 6 | 0.6667 | 14 | 0.5 | 2 | 0.5 | 5 | 0.8 | 2 | 0.5 | 9 | 0.7778 | 18 | 0.7222 |
| 19_1636151 | 2 | 0 | 3 | 0.3333 | 7 | 0 | 9 | 0.1111 | 21 | 0.0952 | 2 | 0 | 9 | 0.1111 | 4 | 0.25 | 10 | 0 | 25 | 0.08 |
| 19_1636229 | 4 | 0.25 | 4 | 0.5 | 4 | 0.5 | 8 | 0.375 | 20 | 0.4 | 1 | 0 | 7 | 0.5714 | 5 | 0.6 | 12 | 0.75 | 25 | 0.64 |
| 19_1636255 | 4 | 0 | 4 | 0 | 5 | 1 | 10 | 0.3 | 23 | 0.3478 | 3 | 0.3333 | 6 | 0.1667 | 6 | 0.1667 | 11 | 0.0909 | 26 | 0.1538 |
| 19_1636260 | 5 | 0.6 | 5 | 1 | 7 | 0.8571 | 8 | 0.875 | 25 | 0.84 | 3 | 0.6667 | 7 | 0.5714 | 7 | 1 | 13 | 0.8462 | 30 | 0.8 |
| 19_1636261 | 5 | 0 | 5 | 0.4 | 7 | 0.4286 | 11 | 0.0909 | 28 | 0.2143 | 3 | 0 | 8 | 0 | 7 | 0.2857 | 14 | 0 | 32 | 0.0625 |
| 19_1636270 | 5 | 0 | 4 | 0 | 6 | 0.3333 | 6 | 0.1667 | 21 | 0.1429 | 3 | 0 | 7 | 0.4286 | 5 | 0.2 | 11 | 0.1818 | 26 | 0.2308 |
| 19_1636439 | 4 | 0.25 | 10 | 0.7 | 12 | 0.4167 | 14 | 0.5714 | 40 | 0.525 | 7 | 0.5714 | 12 | 0.4167 | 8 | 0.5 | 17 | 0.3529 | 44 | 0.4318 |
| 19_1636470 | 8 | 0 | 17 | 0 | 13 | 0.1538 | 20 | 0.15 | 58 | 0.0862 | 11 | 0.1818 | 14 | 0 | 13 | 0 | 24 | 0.1667 | 62 | 0.0968 |
| 19_1636492 | 4 | 0 | 3 | 0.3333 | 3 | 0 | 6 | 0.1667 | 16 | 0.125 | 3 | 0 | 3 | 0.3333 | 5 | 0 | 4 | 0.5 | 15 | 0.2 |
| 19_1659319 | NA | NA | 1 | 1 | 1 | 1 | 1 | 0 | 3 | 0.6667 | NA | NA | NA | NA | 1 | 0 | 2 | 0.5 | 3 | 0.3333 |
| 19_1664202 | 1 | 0 | 1 | 0 | 1 | 1 | NA | NA | 3 | 0.3333 | NA | NA | NA | NA | 1 | 0 | 2 | 0.5 | 3 | 0.3333 |
| 19_1665539 | NA | NA | NA | NA | 1 | 1 | NA | NA | 1 | 1 | 1 | 1 | NA | NA | 1 | 0 | 1 | 1 | 3 | 0.6667 |
| 19_1719373 | 4 | 0 | 4 | 0.25 | 1 | 0 | 2 | 0.5 | 11 | 0.1818 | 2 | 0 | 4 | 0.25 | 3 | 0.3333 | 2 | 0 | 11 | 0.1818 |
| 19_1719819 | 5 | 0.2 | 4 | 0.25 | 2 | 0 | 3 | 0 | 14 | 0.1429 | 3 | 0.3333 | 4 | 0.5 | 3 | 0 | 2 | 0 | 12 | 0.25 |
| 19_1719829 | 6 | 0.3333 | 4 | 0.5 | 2 | 0.5 | 3 | 0.3333 | 15 | 0.4 | 3 | 0 | 4 | 0.25 | 4 | 0 | 3 | 0.3333 | 14 | 0.1429 |
| 19_1719842 | 6 | 0.5 | 4 | 0.5 | 2 | 0.5 | 3 | 0.6667 | 15 | 0.5333 | 2 | 1 | 4 | 0 | 4 | 0.5 | 5 | 1 | 15 | 0.6 |
| 19_1719847 | 6 | 0.1667 | 3 | 0 | 2 | 0 | 3 | 0.3333 | 14 | 0.1429 | 2 | 0 | 2 | 0.5 | 3 | 0 | 5 | 0 | 12 | 0.0833 |
| 19_1719855 | 5 | 0 | 3 | 0 | 2 | 0 | 3 | 0.3333 | 13 | 0.0769 | 3 | 0 | 5 | 0 | 4 | 0 | 5 | 0.2 | 17 | 0.0588 |
| 19_1719862 | 5 | 0.2 | 5 | 0.2 | 1 | 0 | 2 | 0 | 13 | 0.1538 | 1 | 0 | 6 | 0.1667 | 3 | 0 | 5 | 0 | 15 | 0.0667 |
| 19_1719866 | 6 | 0.1667 | 6 | 0 | 2 | 0 | 2 | 0 | 16 | 0.0625 | 3 | 0.3333 | 6 | 0 | 3 | 0 | 4 | 0 | 16 | 0.0625 |
| 19_1719873 | 5 | 0 | 5 | 0.2 | 2 | 0.5 | 3 | 0 | 15 | 0.1333 | 3 | 0 | 6 | 0.1667 | 4 | 0 | 5 | 0.4 | 18 | 0.1667 |
| 19_1719882 | 5 | 0.2 | 5 | 0.4 | 2 | 0.5 | 2 | 0 | 14 | 0.2857 | 3 | 0.3333 | 6 | 0.1667 | 3 | 0.3333 | 5 | 0.2 | 17 | 0.2353 |
| 19_1719923 | 8 | 0.125 | 4 | 0.25 | 2 | 0.5 | 2 | 0 | 16 | 0.1875 | 3 | 0.3333 | 8 | 0.125 | 5 | 0 | 3 | 0.3333 | 19 | 0.1579 |
| 19_1719923 | 5 | 0 | 4 | 0.25 | 2 | 0 | 4 | 0 | 15 | 0.0667 | 2 | 0 | 5 | 0 | 5 | 0 | 5 | 0.2 | 17 | 0.0588 |
| 19_1719948 | 8 | 0 | 6 | 0.1667 | 2 | 0.5 | 4 | 0 | 20 | 0.1 | 3 | 0 | 8 | 0 | 4 | 0 | 4 | 0.25 | 19 | 0.0526 |
| 19_1719958 | 8 | 0.25 | 4 | 0.5 | 2 | 0.5 | 4 | 0.25 | 18 | 0.3333 | 3 | 0.3333 | 7 | 0.4286 | 4 | 0.25 | 3 | 0.3333 | 17 | 0.3529 |
| 19_1720034 | 11 | 0.3636 | 8 | 0 | 2 | 0 | 5 | 0.4 | 26 | 0.2308 | 5 | 0.4 | 9 | 0.1111 | 5 | 0.2 | 7 | 0 | 26 | 0.1538 |
| 19_1720071 | 7 | 0 | 8 | 0.125 | 2 | 0.5 | 5 | 0 | 22 | 0.0909 | 4 | 0.25 | 10 | 0.1 | 3 | 0 | 7 | 0 | 24 | 0.0833 |
| 19_1730620 | NA | NA | 1 | 1 | NA | NA | NA | NA | 1 | 1 | NA | NA | 1 | 1 | NA | NA | NA | NA | 1 | 1 |
| 19_1730624 | NA | NA | 1 | 1 | NA | NA | NA | NA | 1 | 1 | NA | NA | 1 | 1 | NA | NA | NA | NA | 1 | 1 |
| 19_1737712 | 6 | 0.5 | 13 | 0.8462 | 12 | 0.8333 | 6 | 1 | 37 | 0.8108 | 11 | 0.9091 | 9 | 0.8889 | 5 | 0.6 | 3 | 1 | 28 | 0.8571 |
| 19_1737724 | 9 | 0.3333 | 13 | 0.5385 | 13 | 0.3077 | 3 | 0.3333 | 38 | 0.3947 | 8 | 0.25 | 6 | 0.3333 | 5 | 0.4 | 2 | 0.5 | 21 | 0.3333 |
| 19_1737726 | 8 | 0 | 12 | 0.0833 | 12 | 0.0833 | 7 | 0 | 39 | 0.0513 | 5 | 0 | 6 | 0 | 5 | 0.2 | 3 | 0 | 19 | 0.0526 |
| 19_1855854 | 2 | 0.5 | 5 | 0 | 1 | 0 | 2 | 0 | 10 | 0.1 | 6 | 0.1667 | 1 | 0 | 4 | 0 | 1 | 0 | 12 | 0.0833 |
| 19_1868175 | NA | NA | 2 | 0 | 3 | 0.3333 | 2 | 0 | 7 | 0.1429 | 2 | 0.5 | NA | NA | NA | NA | NA | NA | 2 | 0.5 |
| 19_1868436 | NA | NA | 1 | 0 | 2 | 0.5 | 1 | 0 | 4 | 0.25 | NA | NA | NA | NA | 4 | 0.5 | 1 | 0 | 5 | 0.4 |
| 19_1868439 | NA | NA | 1 | 1 | 2 | 0 | NA | NA | 3 | 0.3333 | NA | NA | NA | NA | 4 | 0.5 | NA | NA | 4 | 0.5 |
| 19_1868444 | NA | NA | 1 | 1 | 2 | 0 | 1 | 1 | 4 | 0.5 | NA | NA | NA | NA | 5 | 0.6 | 1 | 0 | 6 | 0.5 |
| 19_1869469 | 3 | 0.6667 | 4 | 0.5 | 3 | 0.6667 | 6 | 0.3333 | 16 | 0.5 | 2 | 0.5 | 10 | 0.8 | 3 | 0.6667 | 5 | 1 | 20 | 0.8 |
| 19_1869470 | 3 | 0.3333 | 4 | 0.25 | 3 | 0 | 4 | 0 | 14 | 0.1429 | 2 | 0.5 | 8 | 0 | 3 | 0.3333 | 5 | 0.2 | 18 | 0.1667 |
| 19_1869472 | 3 | 0.3333 | 3 | 0 | 3 | 0 | 6 | 0 | 15 | 0.0667 | 2 | 0.5 | 10 | 0.1 | 3 | 0.3333 | 5 | 0 | 20 | 0.15 |
| 19_1869481 | 4 | 0.25 | 3 | 0 | 2 | 0 | 6 | 0.1667 | 15 | 0.1333 | 1 | 0 | 6 | 0.3333 | 3 | 0 | 4 | 0.5 | 14 | 0.2857 |
| 19_1869483 | 4 | 0.5 | 3 | 0 | 3 | 0 | 6 | 0.5 | 16 | 0.3125 | 2 | 0 | 9 | 0.2222 | 3 | 0.3333 | 4 | 0.25 | 18 | 0.2222 |
| 19_1869486 | 4 | 0.25 | 2 | 0 | 3 | 0.3333 | 5 | 0.4 | 14 | 0.2857 | 1 | 0 | 9 | 0.1111 | 3 | 0.3333 | 5 | 0 | 18 | 0.1111 |
| 19_1869486 | 3 | 1 | 3 | 0.6667 | 2 | 0.5 | 6 | 0.5 | 14 | 0.6429 | 1 | 0 | 5 | 0.6 | 1 | 0 | 2 | 0.5 | 9 | 0.4444 |
| 19_1869490 | 3 | 0.6667 | 4 | 0.5 | 3 | 0.6667 | 6 | 0.6667 | 16 | 0.625 | 2 | 0 | 9 | 0.5556 | 3 | 0.3333 | 5 | 0.2 | 19 | 0.3684 |
| 19_1869497 | 4 | 0.5 | 4 | 0.25 | 3 | 0 | 6 | 0.1667 | 17 | 0.2353 | 2 | 1 | 7 | 0.1429 | 3 | 0.3333 | 4 | 0.25 | 16 | 0.3125 |
| 19_1869543 | 4 | 0.75 | 5 | 0 | 6 | 0.3333 | 7 | 0 | 22 | 0.2273 | 1 | 0 | 10 | 0.1 | 2 | 0.5 | 5 | 0.2 | 18 | 0.1667 |
| 19_1890977 | 2 | 0 | 6 | 0 | 3 | 0.3333 | 1 | 0 | 12 | 0.0833 | 3 | 0 | 1 | 0 | 1 | 0 | 3 | 0.3333 | 8 | 0.125 |
| 19_1891008 | 1 | 1 | 6 | 0.1667 | 3 | 0.6667 | 2 | 0.5 | 12 | 0.4167 | 2 | 0.5 | 1 | 0 | 1 | 1 | 2 | 0.5 | 6 | 0.5 |
| 19_1891500 | NA | NA | 2 | 1 | NA | NA | NA | NA | 2 | 1 | 1 | 1 | 2 | 0 | 2 | 0 | 1 | 1 | 6 | 0.3333 |

| | | | | | | | | | | | | | | | | | | | | |
|---|---|---|---|---|---|---|---|---|---|---|---|---|---|---|---|---|---|---|---|---|
| 19_1891517 | NA | NA | 2 | 1 | NA | NA | NA | NA | 2 | 1 | 1 | 1 | 3 | 1 | 1 | 1 | 1 | 1 | 6 | 1 |
| 19_1891523 | NA | NA | 3 | 0.3333 | NA | NA | NA | NA | 3 | 0.3333 | 1 | 0 | 2 | 0.5 | 2 | 0.5 | 1 | 1 | 6 | 0.5 |
| 19_1891526 | NA | NA | 2 | 0.5 | NA | NA | NA | NA | 2 | 0.5 | 1 | 0 | 3 | 0.3333 | 2 | 1 | NA | NA | 6 | 0.5 |
| 19_1895507 | NA | NA | 2 | 0.5 | NA | NA | NA | NA | 2 | 0.5 | 1 | 0 | 1 | 0 | 2 | 0.5 | 1 | 1 | 5 | 0.4 |
| 19_1908810 | NA | NA | 1 | 1 | NA | NA | NA | NA | 1 | 1 | 1 | 0 | NA | NA | 1 | 1 | NA | NA | 2 | 0.5 |
| 19_2051201 | 1 | 1 | NA | NA | 1 | 1 | NA | NA | 2 | 1 | 1 | 1 | 2 | 0.5 | NA | NA | 1 | 1 | 4 | 0.75 |
| 19_2051324 | 1 | 1 | 2 | 0 | 2 | 0.5 | NA | NA | 5 | 0.4 | 1 | 0 | 1 | 1 | NA | NA | 2 | 0.5 | 4 | 0.5 |
| 19_2051329 | NA | NA | 2 | 0.5 | 1 | 1 | NA | NA | 3 | 0.6667 | 1 | 0 | 2 | 0.5 | NA | NA | 2 | 1 | 5 | 0.6 |
| 19_2051332 | 1 | 1 | 2 | 1 | 1 | 1 | NA | NA | 4 | 1 | 1 | 1 | 2 | 1 | NA | NA | 2 | 1 | 5 | 1 |
| 19_2051348 | 1 | 1 | 2 | 1 | 1 | 1 | NA | NA | 4 | 1 | 1 | 1 | 2 | 1 | NA | NA | 2 | 1 | 5 | 1 |
| 19_2051349 | 1 | 0 | 2 | 0 | 2 | 0.5 | NA | NA | 5 | 0.2 | 1 | 0 | 1 | 1 | NA | NA | 2 | 0 | 4 | 0.25 |
| 19_2051350 | 1 | 1 | 2 | 1 | NA | NA | NA | NA | 3 | 1 | 1 | 0 | 2 | 1 | NA | NA | 2 | 1 | 5 | 0.8 |
| 19_2051351 | 1 | 1 | 2 | 0 | 1 | 0 | NA | NA | 4 | 0.25 | 1 | 1 | 2 | 0 | NA | NA | 2 | 0.5 | 5 | 0.4 |
| 19_2051356 | 1 | 1 | 2 | 1 | 2 | 1 | NA | NA | 5 | 1 | 1 | 1 | 2 | 1 | NA | NA | 2 | 1 | 5 | 1 |
| 19_2051357 | 1 | 1 | 2 | 1 | 1 | 1 | NA | NA | 4 | 1 | 1 | 1 | 2 | 1 | NA | NA | 2 | 1 | 5 | 1 |
| 19_2051362 | NA | NA | 1 | 1 | 1 | 1 | NA | NA | 2 | 1 | 1 | 1 | 1 | 1 | NA | NA | 1 | 1 | 3 | 1 |
| 19_2051365 | 1 | 1 | 1 | 1 | 2 | 1 | NA | NA | 4 | 1 | NA | NA | 2 | 1 | NA | NA | 2 | 1 | 4 | 1 |
| 19_2051397 | 1 | 1 | 2 | 1 | 2 | 1 | NA | NA | 5 | 1 | NA | NA | 2 | 1 | NA | NA | 2 | 0.5 | 4 | 0.75 |
| 19_2062063 | 10 | 0.1 | 8 | 0 | 10 | 0.1 | 4 | 0.25 | 32 | 0.0938 | 8 | 0 | 5 | 0.2 | 5 | 0 | 6 | 0.3333 | 24 | 0.125 |
| 19_2125782 | 15 | 0.4667 | 28 | 0.25 | 12 | 0.3333 | 15 | 0.4667 | 70 | 0.3571 | 13 | 0.3846 | 16 | 0.3125 | 10 | 0 | 21 | 0.2857 | 60 | 0.2667 |
| 19_2125795 | 18 | 0.7778 | 29 | 0.9655 | 15 | 0.9333 | 16 | 0.875 | 78 | 0.8974 | 13 | 0.8462 | 17 | 1 | 16 | 0.9375 | 20 | 0.8 | 66 | 0.8939 |
| 19_2125796 | 17 | 0.0588 | 30 | 0.1 | 16 | 0.0625 | 19 | 0.1579 | 82 | 0.0976 | 14 | 0.2143 | 19 | 0.1053 | 17 | 0.1176 | 22 | 0.1364 | 72 | 0.1389 |
| 19_2125816 | 16 | 0.5 | 27 | 0.4444 | 18 | 0.1111 | 21 | 0.3333 | 82 | 0.3537 | 13 | 0.3846 | 17 | 0.4118 | 21 | 0.1905 | 20 | 0.35 | 71 | 0.3239 |
| 19_2125835 | 18 | 0.9444 | 32 | 0.9062 | 16 | 0.75 | 21 | 0.7619 | 87 | 0.8506 | 11 | 0.8182 | 20 | 0.85 | 18 | 0.7222 | 21 | 0.7143 | 70 | 0.7714 |
| 19_2125840 | 20 | 0 | 29 | 0.2069 | 15 | 0.0667 | 22 | 0.1818 | 86 | 0.1279 | 14 | 0.2143 | 19 | 0.1053 | 20 | 0.15 | 21 | 0.0952 | 74 | 0.1351 |
| 19_2125849 | 20 | 0.6 | 32 | 0.75 | 18 | 0.3333 | 23 | 0.6957 | 93 | 0.6237 | 13 | 0.8462 | 19 | 0.8421 | 15 | 0.5333 | 21 | 0.4762 | 68 | 0.6618 |
| 19_2125863 | 20 | 0.35 | 32 | 0.5938 | 20 | 0.45 | 19 | 0.4737 | 91 | 0.4835 | 8 | 0.75 | 17 | 0.6471 | 21 | 0.619 | 20 | 0.55 | 66 | 0.6212 |
| 19_2125867 | 22 | 0.0455 | 35 | 0.0286 | 21 | 0.0476 | 25 | 0.16 | 103 | 0.068 | 14 | 0.1429 | 17 | 0 | 22 | 0.2273 | 22 | 0.0455 | 75 | 0.1067 |
| 19_2125874 | 19 | 0.1053 | 35 | 0.1143 | 20 | 0 | 23 | 0.2174 | 97 | 0.1134 | 16 | 0.0625 | 20 | 0.15 | 21 | 0.0476 | 21 | 0.0952 | 78 | 0.0897 |
| 19_2125881 | 21 | 0.1429 | 36 | 0.1667 | 23 | 0.087 | 26 | 0.2692 | 106 | 0.1698 | 16 | 0.1875 | 19 | 0.0526 | 21 | 0 | 24 | 0.25 | 80 | 0.125 |
| 19_2125888 | 22 | 0.0455 | 34 | 0.0882 | 21 | 0.1429 | 24 | 0.125 | 101 | 0.099 | 15 | 0 | 19 | 0.1579 | 21 | 0.0476 | 21 | 0 | 76 | 0.0526 |
| 19_2125894 | 20 | 0.05 | 38 | 0.1842 | 25 | 0.12 | 25 | 0.08 | 108 | 0.1204 | 17 | 0.0588 | 21 | 0 | 23 | 0.087 | 25 | 0.08 | 86 | 0.0581 |
| 19_2126290 | NA | NA | 4 | 0.25 | NA | NA | 1 | 0 | 5 | 0.2 | NA | NA | 2 | 0.5 | NA | NA | NA | NA | 2 | 0.5 |
| 19_2136924 | 12 | 0.0833 | 19 | 0.2105 | 18 | 0.3333 | 18 | 0.0556 | 67 | 0.1791 | 9 | 0.2222 | 29 | 0.1379 | 16 | 0 | 20 | 0.2 | 74 | 0.1351 |
| 19_2136934 | 11 | 0.2727 | 16 | 0.125 | 16 | 0.375 | 15 | 0.2 | 58 | 0.2414 | 7 | 0.1429 | 25 | 0.28 | 15 | 0.3333 | 16 | 0.4375 | 63 | 0.3175 |
| 19_2136937 | 8 | 0 | 17 | 0.1176 | 17 | 0.1176 | 18 | 0.1111 | 60 | 0.1 | 5 | 0.2 | 29 | 0.2414 | 16 | 0.125 | 22 | 0.0909 | 72 | 0.1667 |
| 19_2137001 | 7 | 0 | 10 | 0.1 | 14 | 0.0714 | 17 | 0.1176 | 48 | 0.0833 | 6 | 0.1667 | 24 | 0.0833 | 12 | 0.0833 | 16 | 0.0625 | 58 | 0.0862 |
| 19_2137199 | NA | NA | 1 | 1 | 6 | 0.6667 | 5 | 0.4 | 12 | 0.5833 | 1 | 0 | 9 | 0.7778 | 2 | 0 | 2 | 1 | 14 | 0.6429 |
| 19_2137205 | NA | NA | NA | NA | 5 | 0.6 | 4 | 0.25 | 9 | 0.4444 | NA | NA | 7 | 0.2857 | 2 | 0 | 1 | 0 | 10 | 0.2 |
| 19_2137210 | NA | NA | NA | NA | 6 | 0.6667 | 5 | 0.8 | 11 | 0.7273 | 1 | 1 | 8 | 0.625 | 3 | 0.6667 | 2 | 1 | 14 | 0.7143 |
| 19_2137211 | NA | NA | 1 | 0 | 6 | 0 | 4 | 0.25 | 11 | 0.0909 | 1 | 0 | 9 | 0.1111 | 3 | 0.3333 | 2 | 0.5 | 15 | 0.2 |
| 19_2137215 | NA | NA | 1 | 1 | 6 | 0.8333 | 5 | 0.4 | 12 | 0.6667 | 1 | 0 | 8 | 0.75 | 3 | 0.3333 | 2 | 0.5 | 14 | 0.5714 |
| 19_2137217 | NA | NA | NA | NA | 4 | 0.5 | 3 | 0.3333 | 7 | 0.4286 | 1 | 0 | 9 | 0.4444 | 3 | 0.3333 | 2 | 0.5 | 15 | 0.4 |
| 19_2137219 | NA | NA | 1 | 0 | 6 | 0 | 5 | 0.2 | 12 | 0.0833 | 1 | 0 | 9 | 0.1111 | 3 | 0 | 2 | 0.5 | 15 | 0.1333 |
| 19_2137228 | NA | NA | 1 | 0 | 6 | 0.3333 | 4 | 0 | 11 | 0.1818 | 1 | 0 | 8 | 0 | 2 | 0 | 2 | 0.5 | 13 | 0.0769 |
| 19_2137243 | NA | NA | 1 | 0 | 5 | 0.2 | 5 | 0.2 | 11 | 0.1818 | 1 | 0 | 9 | 0.1111 | 2 | 0 | 2 | 0.5 | 14 | 0.1429 |
| 19_2137248 | NA | NA | 1 | 0 | 3 | 0.6667 | 5 | 0.4 | 9 | 0.4444 | 1 | 0 | 9 | 0.6667 | 2 | 0.5 | 2 | 1 | 14 | 0.6429 |
| 19_2137254 | NA | NA | 1 | 0 | 4 | 0.5 | 4 | 0 | 9 | 0.2222 | 1 | 0 | 7 | 0.1429 | 2 | 0.5 | 2 | 0 | 12 | 0.1667 |
| 19_2157560 | 2 | 0 | 3 | 0 | 8 | 0 | 1 | 1 | 14 | 0.0714 | 2 | 0.5 | 3 | 0 | 9 | 0.3333 | 5 | 0.2 | 19 | 0.2632 |
| 19_2163092 | 2 | 0.5 | 6 | 0.1667 | 4 | 0 | 6 | 0 | 18 | 0.1111 | 3 | 0 | 6 | 0.1667 | 2 | 0 | 1 | 0 | 12 | 0.0833 |
| 19_2165585 | 3 | 1 | 5 | 0.8 | 3 | 0.3333 | 5 | 0.6 | 16 | 0.6875 | 2 | 1 | 6 | 0.3333 | 1 | 1 | 3 | 0.6667 | 12 | 0.5833 |
| 19_2165588 | 4 | 0.5 | 5 | 0.4 | 4 | 0 | 8 | 0.375 | 21 | 0.3333 | 3 | 0.3333 | 7 | 0.1429 | 2 | 0 | 3 | 0.3333 | 15 | 0.2 |
| 19_2165609 | 4 | 0 | 6 | 0.1667 | 6 | 0.1667 | 9 | 0 | 25 | 0.08 | 3 | 0 | 7 | 0.1429 | 1 | 1 | 3 | 0 | 14 | 0.1429 |
| 19_2165828 | 3 | 0.6667 | 5 | 0.2 | 5 | 0.2 | 7 | 0.1429 | 20 | 0.25 | 3 | 0 | 7 | 0 | 3 | 0.3333 | 3 | 0.6667 | 16 | 0.1875 |
| 19_2166151 | 4 | 0.25 | 6 | 0.6667 | 6 | 0.1667 | 9 | 0.4444 | 25 | 0.4 | 5 | 0.4 | 6 | 0.1667 | 3 | 0.6667 | 2 | 0.5 | 16 | 0.375 |
| 19_2166153 | 4 | 0.5 | 5 | 0.4 | 7 | 0.5714 | 11 | 0.2727 | 27 | 0.4074 | 5 | 0.4 | 4 | 0.5 | 3 | 0 | 3 | 0.6667 | 15 | 0.4 |
| 19_2166157 | 3 | 0 | 4 | 0.25 | 5 | 0.4 | 9 | 0.1111 | 21 | 0.1905 | 4 | 0.25 | 6 | 0.3333 | 2 | 0 | 1 | 0 | 13 | 0.2308 |
| 19_2166160 | 4 | 0.5 | 6 | 0.3333 | 6 | 0.1667 | 11 | 0.2727 | 27 | 0.2963 | 5 | 0 | 7 | 0.4286 | 3 | 0.3333 | 3 | 0 | 18 | 0.2222 |
| 19_2166162 | 4 | 1 | 4 | 1 | 6 | 0.8333 | 9 | 1 | 23 | 0.9565 | 4 | 1 | 5 | 1 | 2 | 1 | 2 | 1 | 13 | 1 |
| 19_2166166 | 3 | 0.3333 | 7 | 0.4286 | 6 | 0.3333 | 10 | 0.4 | 26 | 0.3846 | 5 | 0.6 | 6 | 0.5 | 4 | 0.5 | 3 | 0.6667 | 18 | 0.5556 |
| 19_2166170 | 4 | 0 | 7 | 0.1429 | 7 | 0.2857 | 11 | 0.0909 | 29 | 0.1379 | 6 | 0 | 6 | 0 | 4 | 0.5 | 3 | 0.3333 | 19 | 0.1579 |
| 19_2166172 | 4 | 0.25 | 6 | 0.3333 | 4 | 0.5 | 10 | 0.3 | 24 | 0.3333 | 4 | 0.25 | 5 | 0.4 | 4 | 0.5 | 3 | 0 | 16 | 0.3125 |
| 19_2166254 | 5 | 0.4 | 9 | 0.3333 | 8 | 0.375 | 17 | 0.2941 | 39 | 0.3333 | 7 | 0.1429 | 9 | 0.3333 | 5 | 0 | 4 | 0.5 | 25 | 0.24 |
| 19_2177528 | NA | NA | NA | NA | NA | NA | 1 | 1 | 1 | 1 | 1 | 1 | NA | NA | NA | NA | NA | NA | 1 | 1 |
| 19_2177547 | NA | NA | NA | NA | NA | NA | 2 | 0.5 | 2 | 0.5 | 1 | 1 | NA | NA | NA | NA | NA | NA | 1 | 1 |

| | | | | | | | | | | | | | | | | | | | | |
|---|---|---|---|---|---|---|---|---|---|---|---|---|---|---|---|---|---|---|---|---|
| 19_2177554 | NA | NA | NA | NA | NA | NA | 2 | 1 | 2 | 1 | 1 | 1 | NA | NA | NA | NA | NA | NA | 1 | 1 |
| 19_2183367 | 1 | 1 | NA | NA | 1 | 0 | 4 | 0.25 | 6 | 0.3333 | 1 | 0 | 1 | 0 | 2 | 0.5 | 1 | 0 | 5 | 0.2 |
| 19_2183371 | 2 | 1 | NA | NA | 1 | 0 | 4 | 0.75 | 7 | 0.7143 | 1 | 1 | 1 | 0 | 2 | 1 | 1 | 1 | 5 | 0.8 |
| 19_2183405 | 2 | 0.5 | NA | NA | 1 | 0 | 4 | 0.25 | 7 | 0.2857 | 1 | 1 | 1 | 1 | 2 | 0 | 2 | 0.5 | 6 | 0.5 |
| 19_2183446 | 2 | 0 | NA | NA | 1 | 0 | 4 | 0.25 | 7 | 0.1429 | 1 | 0 | 1 | 0 | 2 | 0 | 2 | 0.5 | 6 | 0.1667 |
| 19_2183452 | 2 | 0 | NA | NA | 1 | 0 | 4 | 0.25 | 7 | 0.1429 | 1 | 0 | 1 | 0 | 2 | 0 | 2 | 0.5 | 6 | 0.1667 |
| 19_2183492 | 1 | 1 | 1 | 1 | 1 | 0 | 3 | 1 | 6 | 0.8333 | 1 | 1 | 1 | 1 | 2 | 1 | 2 | 0.5 | 6 | 0.8333 |
| 19_2183499 | 2 | 0.5 | 1 | 1 | 1 | 0 | 3 | 0.3333 | 7 | 0.4286 | 1 | 0 | 1 | 1 | 2 | 0.5 | 2 | 0.5 | 6 | 0.5 |
| 19_2183523 | 2 | 0 | 1 | 0 | NA | NA | 3 | 0.3333 | 6 | 0.1667 | 1 | 0 | 1 | 0 | 1 | 0 | 2 | 0.5 | 5 | 0.2 |
| 19_2191579 | 2 | 0 | 1 | 0 | 1 | 0 | 2 | 0.5 | 6 | 0.1667 | 1 | 0 | 5 | 0.4 | 1 | 0 | 3 | 0.3333 | 10 | 0.3 |
| 19_2296458 | 1 | 1 | NA | NA | 1 | 0 | NA | NA | 2 | 0.5 | 1 | 1 | NA | NA | NA | NA | NA | NA | 1 | 1 |
| 19_2296495 | 2 | 1 | NA | NA | 1 | 1 | NA | NA | 3 | 1 | 1 | 1 | 1 | 0 | NA | NA | NA | NA | 2 | 0.5 |
| 19_2296497 | 2 | 0 | NA | NA | 1 | 1 | NA | NA | 3 | 0.3333 | 1 | 0 | 1 | 1 | NA | NA | NA | NA | 2 | 0.5 |
| 19_2296519 | 1 | 1 | NA | NA | 1 | 0 | NA | NA | 2 | 0.5 | 1 | 1 | 1 | 0 | NA | NA | NA | NA | 2 | 0.5 |
| 19_2296554 | 2 | 0 | NA | NA | 2 | 0.5 | NA | NA | 4 | 0.25 | 1 | 1 | 1 | 0 | NA | NA | NA | NA | 2 | 0.5 |
| 19_2296560 | 2 | 0 | NA | NA | 2 | 1 | NA | NA | 4 | 0.5 | 1 | 0 | 1 | 1 | NA | NA | NA | NA | 2 | 0.5 |
| 19_2296562 | 2 | 1 | NA | NA | 2 | 0.5 | NA | NA | 4 | 0.75 | 1 | 1 | 1 | 1 | NA | NA | NA | NA | 2 | 1 |
| 19_2296579 | 2 | 0.5 | NA | NA | 2 | 0 | NA | NA | 4 | 0.25 | 1 | 0 | 1 | 1 | NA | NA | NA | NA | 2 | 0.5 |
| 19_2296580 | 2 | 0.5 | NA | NA | 2 | 0.5 | NA | NA | 4 | 0.5 | 1 | 1 | 1 | 1 | NA | NA | NA | NA | 2 | 1 |
| 19_2296580 | 1 | 1 | NA | NA | 2 | 0.5 | NA | NA | 3 | 0.6667 | 1 | 0 | 1 | 1 | NA | NA | NA | NA | 2 | 0.5 |
| 19_2296586 | 1 | 0 | NA | NA | 2 | 0.5 | NA | NA | 3 | 0.3333 | 1 | 0 | 1 | 1 | NA | NA | NA | NA | 2 | 0.5 |
| 19_2296614 | 2 | 0.5 | NA | NA | 1 | 1 | 1 | 1 | 4 | 0.75 | 1 | 1 | 3 | 0.6667 | NA | NA | NA | NA | 4 | 0.75 |
| 19_2296635 | 2 | 0.5 | 1 | 1 | 2 | 0.5 | NA | NA | 5 | 0.6 | NA | NA | 2 | 0.5 | 1 | 0 | NA | NA | 3 | 0.3333 |
| 19_2296656 | 2 | 0.5 | 1 | 0 | 2 | 1 | 1 | 1 | 6 | 0.6667 | 1 | 0 | 3 | 0.3333 | 1 | 1 | NA | NA | 5 | 0.4 |
| 19_2296773 | 2 | 0.5 | 1 | 0 | 2 | 0 | 2 | 0.5 | 7 | 0.2857 | 2 | 0 | 1 | 0 | 2 | 0.5 | NA | NA | 5 | 0.2 |
| 19_2296940 | 5 | 0.2 | 1 | 0 | 1 | 1 | 6 | 0.1667 | 13 | 0.2308 | 2 | 0 | 2 | 0 | 4 | 0 | 2 | 0.5 | 10 | 0.1 |
| 19_2297052 | 1 | 0 | 1 | 1 | NA | NA | 2 | 0 | 4 | 0.25 | 1 | 0 | NA | NA | 3 | 0.3333 | NA | NA | 4 | 0.25 |
| 19_2297181 | 1 | 0 | 1 | 1 | NA | NA | 3 | 0.3333 | 5 | 0.4 | 1 | 0 | NA | NA | 4 | 0.25 | NA | NA | 5 | 0.2 |
| 19_2297198 | 1 | 0 | 1 | 0 | NA | NA | 3 | 0.3333 | 5 | 0.2 | 1 | 0 | 1 | 0 | 5 | 0.4 | NA | NA | 7 | 0.2857 |
| 19_2311788 | NA | NA | 1 | 1 | NA | NA | NA | NA | 1 | 1 | NA | NA | 1 | 1 | NA | NA | NA | NA | 1 | 1 |
| 19_2311797 | NA | NA | 1 | 1 | NA | NA | NA | NA | 1 | 1 | NA | NA | 1 | 1 | NA | NA | NA | NA | 1 | 1 |
| 19_2311801 | NA | NA | 1 | 1 | NA | NA | NA | NA | 1 | 1 | NA | NA | 1 | 1 | NA | NA | NA | NA | 1 | 1 |
| 19_2311847 | 1 | 0 | 2 | 0 | 1 | 0 | 7 | 0.1429 | 11 | 0.0909 | 1 | 0 | 3 | 0.3333 | 2 | 0 | 3 | 0 | 9 | 0.1111 |
| 19_2311861 | 3 | 0 | 7 | 0.1429 | 1 | 0 | 10 | 0.1 | 21 | 0.0952 | 3 | 0 | 3 | 0 | 3 | 0.3333 | 4 | 0 | 13 | 0.0769 |
| 19_2311878 | 5 | 0.6 | 8 | 1 | 1 | 1 | 10 | 0.9 | 24 | 0.875 | 4 | 1 | 3 | 1 | 4 | 1 | 5 | 1 | 16 | 1 |
| 19_2311880 | 4 | 0.25 | 7 | 0.1429 | 1 | 0 | 8 | 0.125 | 20 | 0.15 | 3 | 0.3333 | 3 | 0 | 2 | 0 | 5 | 0 | 13 | 0.0769 |
| 19_2311914 | 7 | 0.7143 | 9 | 1 | 2 | 1 | 10 | 0.6 | 28 | 0.7857 | 4 | 1 | 7 | 0.7143 | 7 | 0.4286 | 9 | 0.8889 | 27 | 0.7407 |
| 19_2311925 | 5 | 0.2 | 8 | 0.25 | 2 | 0 | 13 | 0.1538 | 28 | 0.1786 | 6 | 0.1667 | 8 | 0 | 7 | 0 | 7 | 0.2857 | 28 | 0.1071 |
| 19_2311927 | 7 | 0.5714 | 10 | 0.4 | 3 | 0.3333 | 14 | 0.4286 | 34 | 0.4412 | 5 | 0.4 | 9 | 0.3333 | 8 | 0.375 | 9 | 0 | 31 | 0.2581 |
| 19_2311932 | 7 | 0.4286 | 10 | 0.3 | 4 | 0.5 | 12 | 0.4167 | 33 | 0.3939 | 7 | 0.5714 | 9 | 0.4444 | 9 | 0.6667 | 10 | 0.2 | 35 | 0.4571 |
| 19_2314212 | 9 | 0 | 12 | 0.0833 | 5 | 0 | 11 | 0.0909 | 37 | 0.0541 | 6 | 0 | 10 | 0 | 10 | 0.2 | 14 | 0.0714 | 40 | 0.075 |
| 19_2314232 | 10 | 0.7 | 14 | 1 | 4 | 1 | 17 | 0.8824 | 45 | 0.8889 | 9 | 0.7778 | 9 | 0.8889 | 11 | 0.8182 | 14 | 0.9286 | 43 | 0.8605 |
| 19_2314234 | 8 | 0.875 | 9 | 1 | 1 | 1 | 14 | 1 | 32 | 0.9688 | 7 | 1 | 5 | 0.8 | 7 | 1 | 10 | 1 | 29 | 0.9655 |
| 19_2314250 | 9 | 0.1111 | 16 | 0.1875 | 4 | 0 | 16 | 0.0625 | 45 | 0.1111 | 8 | 0.125 | 12 | 0.25 | 11 | 0.0909 | 11 | 0.0909 | 42 | 0.1429 |
| 19_2314251 | 11 | 1 | 15 | 1 | 3 | 1 | 13 | 1 | 42 | 1 | 8 | 1 | 12 | 1 | 11 | 1 | 13 | 1 | 44 | 1 |
| 19_2314261 | 8 | 0.75 | 12 | 0.8333 | 9 | 0.8889 | 18 | 0.6111 | 47 | 0.7447 | 9 | 0.3333 | 11 | 0.6364 | 12 | 0.6667 | 17 | 0.8235 | 49 | 0.6531 |
| 19_2314263 | 10 | 0.1 | 15 | 0.2667 | 8 | 0.25 | 19 | 0 | 52 | 0.1346 | 10 | 0.2 | 17 | 0.1765 | 13 | 0.2308 | 12 | 0.0833 | 52 | 0.1731 |
| 19_2314267 | 13 | 0.7692 | 17 | 1 | 10 | 1 | 22 | 0.8636 | 62 | 0.9032 | 8 | 0.875 | 15 | 1 | 11 | 1 | 15 | 1 | 49 | 0.9796 |
| 19_2314269 | 9 | 0.4444 | 11 | 0.7273 | 9 | 0.5556 | 21 | 0.5238 | 50 | 0.56 | 8 | 1 | 14 | 0.4286 | 13 | 0.5385 | 16 | 0.5625 | 51 | 0.5882 |
| 19_2314276 | 11 | 0.1818 | 14 | 0.0714 | 8 | 0 | 20 | 0.1 | 53 | 0.0943 | 7 | 0 | 12 | 0 | 10 | 0 | 16 | 0.1875 | 45 | 0.0667 |
| 19_2314278 | 11 | 0.8182 | 12 | 1 | 10 | 0.8 | 21 | 0.9524 | 54 | 0.9074 | 8 | 0.875 | 13 | 0.9231 | 13 | 0.9231 | 15 | 0.9333 | 49 | 0.9184 |
| 19_2314280 | 10 | 0.4 | 10 | 0 | 6 | 0.1667 | 16 | 0.0625 | 42 | 0.1429 | 8 | 0.125 | 13 | 0 | 13 | 0.0769 | 15 | 0.1333 | 49 | 0.0816 |
| 19_2314282 | 12 | 0.0833 | 13 | 0.0769 | 8 | 0.125 | 17 | 0.1765 | 50 | 0.12 | 6 | 0 | 10 | 0.4 | 10 | 0.1 | 11 | 0.2727 | 37 | 0.2162 |
| 19_2314283 | 15 | 0.0667 | 16 | 0.125 | 9 | 0.2222 | 20 | 0.15 | 60 | 0.1333 | 9 | 0 | 16 | 0.1875 | 12 | 0.0833 | 15 | 0.0667 | 52 | 0.0962 |
| 19_2314291 | 13 | 0.9231 | 16 | 1 | 10 | 1 | 25 | 0.96 | 64 | 0.9688 | 12 | 0.9167 | 17 | 1 | 16 | 1 | 22 | 0.9545 | 67 | 0.9701 |
| 19_2314299 | 15 | 0.3333 | 16 | 0.125 | 13 | 0.1538 | 25 | 0.24 | 69 | 0.2174 | 11 | 0.1818 | 17 | 0 | 17 | 0.0588 | 21 | 0.0952 | 66 | 0.0758 |
| 19_2314302 | 14 | 0.9286 | 15 | 0.9333 | 13 | 1 | 26 | 0.9615 | 68 | 0.9559 | 12 | 0.9167 | 17 | 0.9412 | 12 | 0.9167 | 17 | 0.9412 | 58 | 0.931 |
| 19_2314306 | 11 | 0.1818 | 15 | 0.0667 | 8 | 0.25 | 18 | 0.3333 | 52 | 0.2115 | 9 | 0.2222 | 11 | 0.5455 | 13 | 0.3846 | 14 | 0.0714 | 47 | 0.2979 |
| 19_2314306 | 12 | 0.5 | 13 | 0.7692 | 11 | 0.7273 | 13 | 0.5385 | 49 | 0.6327 | 8 | 0.875 | 8 | 0.625 | 8 | 0.75 | 16 | 0.5625 | 40 | 0.675 |
| 19_2314319 | 17 | 0.1176 | 17 | 0.1765 | 13 | 0.2308 | 25 | 0.24 | 72 | 0.1944 | 9 | 0.3333 | 14 | 0.2143 | 14 | 0.2857 | 18 | 0.4444 | 55 | 0.3273 |
| 19_2314336 | 15 | 0.0667 | 18 | 0.0556 | 14 | 0.0714 | 25 | 0.08 | 72 | 0.0694 | 12 | 0.0833 | 20 | 0.1 | 16 | 0 | 22 | 0.1364 | 70 | 0.0857 |
| 19_2314336 | 16 | 0 | 20 | 0.05 | 14 | 0.0714 | 27 | 0.0741 | 77 | 0.0519 | 11 | 0.0909 | 19 | 0 | 14 | 0 | 24 | 0.125 | 68 | 0.0588 |
| 19_2314485 | 34 | 0.0588 | 29 | 0.1379 | 25 | 0 | 37 | 0.027 | 125 | 0.056 | 23 | 0.087 | 27 | 0.0741 | 24 | 0.0833 | 35 | 0.0286 | 109 | 0.0642 |
| 19_2318445 | 1 | 0 | 3 | 0.3333 | 3 | 0.3333 | 3 | 0.6667 | 10 | 0.4 | 1 | 1 | 1 | 0 | 2 | 0.5 | 2 | 0.5 | 6 | 0.5 |
| 19_2318573 | NA | NA | 3 | 0 | 2 | 0.5 | 2 | 0 | 7 | 0.1429 | 1 | 0 | NA | NA | 3 | 0 | 2 | 0.5 | 6 | 0.1667 |

| | | | | | | | | | | | | | | | | | | | | |
|---|---|---|---|---|---|---|---|---|---|---|---|---|---|---|---|---|---|---|---|---|
| 19_2318597 | NA | NA | 3 | 0.6667 | 1 | 1 | 1 | 1 | 5 | 0.8 | 1 | 0 | 1 | 0 | 3 | 0.6667 | 3 | 0.6667 | 8 | 0.5 |
| 19_2318601 | NA | NA | 2 | 0 | 1 | 1 | 1 | 1 | 4 | 0.5 | 1 | 0 | 1 | 1 | 3 | 1 | 1 | 1 | 6 | 0.8333 |
| 19_2318602 | NA | NA | 3 | 0.3333 | 1 | 1 | 2 | 1 | 6 | 0.6667 | 2 | 1 | 1 | 1 | 3 | 0.6667 | 3 | 0.6667 | 9 | 0.7778 |
| 19_2369528 | NA | NA | 1 | 0 | 1 | 1 | NA | NA | 2 | 0.5 | NA | NA | 1 | 1 | NA | NA | 1 | 0 | 2 | 0.5 |
| 19_2369752 | NA | NA | 1 | 1 | 1 | 1 | NA | NA | 2 | 1 | NA | NA | 1 | 1 | NA | NA | 1 | 1 | 2 | 1 |
| 19_2369757 | NA | NA | 1 | 1 | 1 | 1 | NA | NA | 2 | 1 | NA | NA | 1 | 1 | NA | NA | 1 | 1 | 2 | 1 |
| 19_2369763 | NA | NA | 1 | 1 | 1 | 0 | NA | NA | 2 | 0.5 | NA | NA | 1 | 1 | NA | NA | 1 | 0 | 2 | 0.5 |
| 19_2369775 | NA | NA | 1 | 1 | 1 | 1 | NA | NA | 2 | 1 | NA | NA | 1 | 1 | NA | NA | 1 | 1 | 2 | 1 |
| 19_2369783 | NA | NA | 1 | 1 | 1 | 1 | NA | NA | 2 | 1 | NA | NA | 1 | 1 | NA | NA | 1 | 1 | 2 | 1 |
| 19_2370533 | 3 | 0 | 4 | 0.5 | 2 | 0.5 | 3 | 0.3333 | 12 | 0.3333 | 2 | 0 | NA | NA | 2 | 0 | 4 | 0.75 | 8 | 0.375 |
| 19_2370601 | 4 | 0.25 | 5 | 0.2 | 4 | 0.5 | 4 | 0.5 | 17 | 0.3529 | 3 | 0.6667 | 1 | 0 | 2 | 0 | 4 | 0.5 | 10 | 0.4 |
| 19_2370638 | 2 | 0 | 1 | 0 | 1 | 0 | 2 | 0.5 | 6 | 0.1667 | NA | NA | NA | NA | 1 | 0 | 3 | 0.3333 | 4 | 0.25 |
| 19_2370844 | 3 | 0.3333 | NA | NA | 2 | 0 | 2 | 0 | 7 | 0.1429 | 2 | 0 | 1 | 1 | 1 | 0 | 4 | 0 | 8 | 0.125 |
| 19_2370876 | 1 | 1 | 2 | 0.5 | 2 | 1 | 5 | 1 | 10 | 0.9 | 4 | 1 | 1 | 1 | 1 | 1 | 4 | 1 | 10 | 1 |
| 19_2387045 | 2 | 0.5 | 3 | 0 | NA | NA | NA | NA | 5 | 0.2 | NA | NA | 2 | 0.5 | 1 | 0 | 2 | 0.5 | 5 | 0.4 |
| 19_2387096 | 3 | 1 | 3 | 0.6667 | 1 | 1 | NA | NA | 7 | 0.8571 | NA | NA | 3 | 1 | 2 | 1 | 3 | 1 | 8 | 1 |
| 19_2387107 | 3 | 0.3333 | 4 | 0 | 1 | 0 | NA | NA | 8 | 0.125 | NA | NA | 2 | 0.5 | 2 | 0 | 3 | 0.6667 | 7 | 0.4286 |
| 19_2387123 | 2 | 0.5 | 4 | 0.75 | 1 | 1 | NA | NA | 7 | 0.7143 | 1 | 0 | 3 | 0.3333 | 2 | 0 | 3 | 0.3333 | 9 | 0.2222 |
| 19_2387127 | 2 | 0.5 | 4 | 0.25 | 1 | 0 | NA | NA | 7 | 0.2857 | 1 | 0 | 2 | 0.5 | 2 | 0 | 3 | 0 | 8 | 0.125 |
| 19_2387135 | 3 | 0.3333 | 3 | 1 | 1 | 0 | NA | NA | 7 | 0.5714 | 1 | 0 | 2 | 0.5 | 2 | 0.5 | 3 | 0 | 8 | 0.25 |
| 19_2392321 | 4 | 0.75 | 7 | 1 | 3 | 0.3333 | 3 | 1 | 17 | 0.8235 | 2 | 0.5 | 4 | 0.25 | 2 | 0 | 5 | 1 | 13 | 0.5385 |
| 19_2392326 | 4 | 0.25 | 7 | 0.1429 | 3 | 0.3333 | 3 | 0.3333 | 17 | 0.2353 | 1 | 0 | 4 | 0.5 | 2 | 0.5 | 3 | 0.3333 | 10 | 0.4 |
| 19_2392332 | 4 | 0 | 7 | 0.1429 | 5 | 0.4 | 3 | 0.3333 | 19 | 0.2105 | 2 | 0 | 4 | 0.25 | 2 | 0 | 5 | 0.2 | 13 | 0.1538 |
| 19_2392500 | 8 | 0.25 | 12 | 0.0833 | 9 | 0 | 5 | 0.2 | 34 | 0.1176 | 5 | 0 | 6 | 0.3333 | 6 | 0 | 7 | 0.2857 | 24 | 0.1667 |
| 19_2392512 | 8 | 0.375 | 11 | 0.0909 | 8 | 0.375 | 6 | 0.3333 | 33 | 0.2727 | 4 | 0 | 7 | 0.2857 | 6 | 0.3333 | 8 | 0.5 | 25 | 0.32 |
| 19_2392532 | 7 | 0.1429 | 13 | 0 | 10 | 0.1 | 7 | 0.1429 | 37 | 0.0811 | 6 | 0 | 6 | 0.1667 | 7 | 0 | 10 | 0.3 | 29 | 0.1379 |
| 19_2392533 | 8 | 0.5 | 10 | 0.2 | 7 | 0.1429 | 6 | 0.6667 | 31 | 0.3548 | 3 | 0 | 5 | 0.2 | 7 | 0 | 9 | 0.6667 | 24 | 0.2917 |
| 19_2392536 | 10 | 0.1 | 13 | 0.0769 | 10 | 0.1 | 7 | 0 | 40 | 0.075 | 5 | 0 | 8 | 0 | 6 | 0.1667 | 10 | 0.1 | 29 | 0.069 |
| 19_2392586 | 8 | 0.125 | 14 | 0.1429 | 7 | 0.1429 | 6 | 0 | 35 | 0.1143 | 5 | 0 | 9 | 0 | 7 | 0 | 10 | 0.2 | 31 | 0.0645 |
| 19_2395078 | 2 | 0.5 | NA | NA | 2 | 0.5 | 2 | 1 | 6 | 0.6667 | 1 | 1 | 2 | 0.5 | 2 | 1 | 1 | 0 | 6 | 0.6667 |
| 19_2395080 | 4 | 0.75 | 1 | 0 | 2 | 0.5 | 4 | 0.75 | 11 | 0.6364 | 2 | 0.5 | 2 | 1 | 3 | 0.6667 | 1 | 1 | 8 | 0.75 |
| 19_2395280 | 3 | 0 | NA | NA | 2 | 0.5 | 5 | 0 | 10 | 0.1 | 1 | 1 | 3 | 0 | 3 | 0.6667 | 2 | 0 | 9 | 0.3333 |
| 19_2395281 | 4 | 0.25 | NA | NA | 1 | 0 | 5 | 0.2 | 10 | 0.2 | 2 | 0 | 2 | 0 | 3 | 0.6667 | 1 | 0 | 8 | 0.25 |
| 19_2395372 | 1 | 1 | NA | NA | NA | NA | NA | NA | 1 | 1 | NA | NA | 1 | 0 | NA | NA | 1 | 1 | 2 | 0.5 |
| 19_2395394 | 2 | 0.5 | NA | NA | NA | NA | NA | NA | 2 | 0.5 | NA | NA | 1 | 1 | NA | NA | 1 | 1 | 2 | 1 |
| 19_2395404 | 2 | 1 | NA | NA | NA | NA | NA | NA | 2 | 1 | NA | NA | NA | NA | NA | NA | 1 | 1 | 1 | 1 |
| 19_2395528 | NA | NA | NA | NA | 1 | 1 | NA | NA | 1 | 1 | NA | NA | NA | NA | 1 | 1 | NA | NA | 1 | 1 |
| 19_2407210 | 3 | 0.3333 | 1 | 1 | 6 | 0.1667 | 3 | 0.3333 | 13 | 0.3077 | 1 | 1 | 5 | 0.4 | 1 | 0 | 2 | 0 | 9 | 0.3333 |
| 19_2407232 | 3 | 0.3333 | 1 | 0 | 7 | 0.1429 | 1 | 0 | 12 | 0.1667 | 1 | 0 | 5 | 0.2 | NA | NA | 1 | 0 | 7 | 0.1429 |
| 19_2407233 | 2 | 1 | NA | NA | 7 | 1 | 1 | 1 | 10 | 1 | NA | NA | 5 | 0.8 | NA | NA | NA | NA | 5 | 0.8 |
| 19_2407244 | 3 | 1 | 1 | 1 | 7 | 0.4286 | 2 | 0.5 | 13 | 0.6154 | 1 | 1 | 5 | 1 | 1 | 1 | 2 | 0.5 | 9 | 0.8889 |
| 19_2407258 | 2 | 0 | 1 | 1 | 7 | 0.1429 | 1 | 0 | 11 | 0.1818 | 1 | 0 | 4 | 0.5 | 1 | 0 | 2 | 0 | 8 | 0.25 |
| 19_2407260 | 1 | 1 | NA | NA | 4 | 0.5 | NA | NA | 5 | 0.6 | 1 | 0 | 4 | 0.5 | NA | NA | 1 | 0 | 6 | 0.3333 |
| 19_2407264 | 2 | 0.5 | 1 | 1 | 5 | 0.4 | 1 | 0 | 9 | 0.4444 | 1 | 0 | 3 | 0 | 1 | 0 | 2 | 1 | 7 | 0.2857 |
| 19_2407266 | 1 | 1 | 1 | 1 | 5 | 1 | 1 | 1 | 8 | 1 | 1 | 1 | 3 | 1 | 1 | 1 | 2 | 1 | 7 | 1 |
| 19_2407277 | 1 | 1 | 1 | 0 | 4 | 0.5 | NA | NA | 6 | 0.5 | NA | NA | 3 | 0.6667 | 1 | 1 | NA | NA | 4 | 0.75 |
| 19_2423890 | 1 | 1 | NA | NA | 1 | 1 | NA | NA | 2 | 1 | NA | NA | 2 | 0.5 | 2 | 0.5 | NA | NA | 4 | 0.5 |
| 19_2423891 | 1 | 0 | NA | NA | 1 | 1 | NA | NA | 2 | 0.5 | NA | NA | 1 | 0 | 2 | 0.5 | NA | NA | 3 | 0.3333 |
| 19_2423908 | 1 | 1 | NA | NA | NA | NA | NA | NA | 1 | 1 | NA | NA | 2 | 0.5 | 2 | 1 | NA | NA | 4 | 0.75 |
| 19_2423924 | 1 | 0 | 1 | 0 | 3 | 0 | 2 | 0.5 | 7 | 0.1429 | 1 | 1 | 2 | 1 | 3 | 0.6667 | 2 | 0.5 | 8 | 0.75 |
| 19_2423930 | 2 | 1 | 1 | 1 | 3 | 1 | 1 | 1 | 7 | 1 | 1 | 1 | 3 | 1 | 4 | 1 | 1 | 1 | 9 | 1 |
| 19_2427262 | 1 | 1 | 1 | 1 | 1 | 1 | 2 | 1 | 5 | 1 | 2 | 0.5 | NA | NA | 1 | 1 | 3 | 0.6667 | 6 | 0.6667 |
| 19_2445363 | 5 | 1 | 5 | 1 | 6 | 1 | 9 | 1 | 25 | 1 | 2 | 1 | 6 | 1 | 3 | 1 | 5 | 1 | 16 | 1 |
| 19_2445373 | 5 | 0.6 | 6 | 0.8333 | 5 | 0.4 | 11 | 0.5455 | 27 | 0.5926 | 2 | 0.5 | 4 | 1 | 6 | 0.6667 | 5 | 0.6 | 17 | 0.7059 |
| 19_2445376 | 4 | 1 | 3 | 1 | 4 | 1 | 5 | 1 | 16 | 1 | 3 | 1 | 1 | 1 | 2 | 1 | 4 | 1 | 10 | 1 |
| 19_2445381 | 3 | 0.3333 | 6 | 0 | 4 | 0.25 | 9 | 0.1111 | 22 | 0.1364 | 1 | 0 | 5 | 0 | 5 | 0 | 4 | 0.25 | 15 | 0.0667 |
| 19_2445382 | 5 | 0.2 | 4 | 0 | 4 | 0 | 4 | 0 | 17 | 0.0588 | 1 | 0 | 4 | 0.25 | 3 | 0 | 4 | 0.5 | 12 | 0.25 |
| 19_2445382 | 6 | 0.5 | 6 | 0.5 | 3 | 0.3333 | 10 | 0.5 | 25 | 0.48 | 3 | 0.6667 | 5 | 0.6 | 6 | 0.5 | 4 | 0.5 | 18 | 0.5556 |
| 19_2445386 | 4 | 1 | 5 | 1 | 5 | 0.8 | 8 | 1 | 22 | 0.9545 | 3 | 1 | 3 | 1 | 3 | 1 | 3 | 1 | 12 | 1 |
| 19_2445388 | 3 | 1 | 5 | 1 | 6 | 1 | 9 | 1 | 23 | 1 | 2 | 1 | 4 | 1 | 4 | 1 | 4 | 1 | 14 | 1 |
| 19_2445410 | 5 | 0.8 | 5 | 0.6 | 4 | 0.25 | 8 | 0.625 | 22 | 0.5909 | 2 | 1 | 4 | 0.5 | 3 | 0.3333 | 4 | 0.25 | 13 | 0.4615 |
| 19_2445413 | 5 | 0.8 | 6 | 0.6667 | 5 | 0.6 | 9 | 0.8889 | 25 | 0.76 | 2 | 0.5 | 5 | 0.4 | 3 | 1 | 4 | 0.5 | 14 | 0.5714 |
| 19_2445414 | 4 | 0.25 | 6 | 0.1667 | 5 | 0 | 8 | 0 | 23 | 0.087 | 2 | 0 | 5 | 0 | 4 | 0.25 | 3 | 0 | 14 | 0.0714 |
| 19_2445422 | 4 | 1 | 5 | 1 | 5 | 1 | 9 | 1 | 23 | 1 | 2 | 1 | 5 | 0.8 | 4 | 1 | 4 | 1 | 15 | 0.9333 |
| 19_2445442 | 4 | 0.25 | 4 | 0 | 4 | 0 | 8 | 0.125 | 20 | 0.1 | 2 | 0 | 5 | 0.2 | 4 | 0.25 | 3 | 0.3333 | 14 | 0.2143 |

| | | | | | | | | | | | | | | | | | | | | |
|---|---|---|---|---|---|---|---|---|---|---|---|---|---|---|---|---|---|---|---|---|
| 19_244544 | 3 | 0.6667 | 5 | 0.6 | 4 | 0 | 9 | 0 | 21 | 0.2381 | 1 | 1 | 5 | 0 | 3 | 0.3333 | 3 | 1 | 12 | 0.4167 |
| 19_246514 | 1 | 0 | NA | NA | NA | NA | 1 | 1 | 2 | 0.5 | 1 | 0 | 2 | 0.5 | 1 | 0 | NA | NA | 4 | 0.25 |
| 19_246551 | 2 | 1 | NA | NA | NA | NA | 1 | 0 | 3 | 0.6667 | 1 | 1 | 2 | 1 | 1 | 1 | NA | NA | 4 | 1 |
| 19_246551 | 2 | 0.5 | NA | NA | NA | NA | 1 | 1 | 3 | 0.6667 | NA | NA | 2 | 0.5 | NA | NA | NA | NA | 2 | 0.5 |
| 19_246551 | 2 | 1 | NA | NA | NA | NA | 1 | 1 | 3 | 1 | NA | NA | 2 | 1 | 1 | 1 | NA | NA | 3 | 1 |
| 19_246554 | 2 | 0.5 | NA | NA | NA | NA | NA | NA | 2 | 0.5 | 1 | 1 | 2 | 0 | 1 | 0 | NA | NA | 4 | 0.25 |
| 19_246554 | 2 | 0 | NA | NA | NA | NA | 1 | 1 | 3 | 0.3333 | NA | NA | 2 | 0 | 1 | 1 | NA | NA | 3 | 0.3333 |
| 19_246556 | 1 | 1 | NA | NA | NA | NA | 1 | 1 | 2 | 1 | 1 | 1 | 2 | 1 | 1 | 1 | NA | NA | 4 | 1 |
| 19_248535 | 2 | 0.5 | 3 | 0.6667 | 1 | 0 | NA | NA | 6 | 0.5 | NA | NA | NA | NA | NA | NA | 2 | 0.5 | 2 | 0.5 |
| 19_248536 | 2 | 0.5 | 2 | 0 | 2 | 0.5 | 1 | 0 | 7 | 0.2857 | 2 | 1 | NA | NA | NA | NA | 1 | 1 | 3 | 1 |
| 19_253876 | NA | NA | 2 | 1 | NA | NA | NA | NA | 2 | 1 | 1 | 1 | NA | NA | NA | NA | NA | NA | 1 | 1 |
| 19_253880 | NA | NA | 1 | 0 | 1 | 1 | NA | NA | 2 | 0.5 | 1 | 1 | NA | NA | NA | NA | NA | NA | 1 | 1 |
| 19_253881 | NA | NA | 2 | 0.5 | 1 | 1 | NA | NA | 3 | 0.6667 | 1 | 1 | NA | NA | NA | NA | NA | NA | 1 | 1 |
| 19_253886 | NA | NA | 2 | 0.5 | 1 | 0 | NA | NA | 3 | 0.3333 | 1 | 1 | NA | NA | NA | NA | NA | NA | 1 | 1 |
| 19_253886 | NA | NA | 2 | 1 | NA | NA | NA | NA | 2 | 1 | 1 | 1 | NA | NA | NA | NA | NA | NA | 1 | 1 |
| 19_253887 | NA | NA | 1 | 1 | 1 | 1 | NA | NA | 2 | 1 | 1 | 1 | NA | NA | NA | NA | NA | NA | 1 | 1 |
| 19_254109 | 1 | 0 | 4 | 0.5 | 1 | 0 | NA | NA | 6 | 0.3333 | 1 | 1 | 1 | 0 | 2 | 0 | NA | NA | 4 | 0.25 |
| 19_254114 | 1 | 0 | 4 | 0.25 | 1 | 1 | NA | NA | 6 | 0.3333 | NA | NA | 1 | 0 | 2 | 0.5 | NA | NA | 3 | 0.3333 |
| 19_266083 | 3 | 0 | 2 | 0 | 2 | 0 | 1 | 1 | 8 | 0.125 | 1 | 0 | 1 | 1 | 2 | 0.5 | 1 | 1 | 5 | 0.6 |
| 19_272155 | 1 | 0 | 2 | 0 | 2 | 0.5 | 2 | 0 | 7 | 0.1429 | NA | NA | 1 | 0 | 1 | 1 | NA | NA | 2 | 0.5 |
| 19_272253 | 3 | 0.6667 | 3 | 0 | 2 | 0.5 | 2 | 1 | 10 | 0.5 | 1 | 0 | 3 | 0 | 3 | 0.3333 | 1 | 0 | 8 | 0.125 |
| 19_272268 | 3 | 0.3333 | 6 | 0 | 2 | 0 | 3 | 0 | 14 | 0.0714 | 1 | 0 | 4 | 0 | 3 | 0 | 2 | 0.5 | 10 | 0.1 |
| 19_272270 | 3 | 0.6667 | 7 | 0.1429 | 2 | 1 | 6 | 0.1667 | 18 | 0.3333 | 1 | 1 | 3 | 0 | 3 | 0.3333 | 2 | 0.5 | 9 | 0.3333 |
| 19_272280 | 3 | 1 | 3 | 0.6667 | 3 | 0.3333 | 3 | 0.3333 | 12 | 0.5833 | 1 | 0 | 3 | 0 | 5 | 0.8 | 4 | 0.75 | 13 | 0.5385 |
| 19_272281 | 3 | 0 | 3 | 0 | 3 | 0.3333 | 3 | 0 | 12 | 0.0833 | NA | NA | 3 | 0.3333 | 5 | 0 | 3 | 0.3333 | 11 | 0.1818 |
| 19_272282 | 2 | 0 | 3 | 1 | 2 | 0 | 3 | 0.6667 | 10 | 0.5 | 1 | 1 | 3 | 0.6667 | 6 | 0.6667 | 4 | 0.75 | 14 | 0.7143 |
| 19_272283 | 3 | 0 | 3 | 0 | 3 | 0.3333 | 3 | 0 | 12 | 0.0833 | 1 | 0 | 3 | 0 | 5 | 0.2 | 4 | 0.25 | 13 | 0.1538 |
| 19_272285 | 2 | 0 | 2 | 0.5 | 2 | 0 | 3 | 0 | 9 | 0.1111 | NA | NA | 2 | 0.5 | 4 | 0 | 3 | 0.3333 | 9 | 0.2222 |
| 19_272296 | 4 | 0.25 | 3 | 0 | 4 | 0 | 3 | 0 | 14 | 0.0714 | 1 | 0 | 4 | 0.25 | 5 | 0.2 | 5 | 0.2 | 15 | 0.2 |
| 19_272296 | 4 | 0 | 4 | 0.25 | 3 | 0 | 3 | 0 | 14 | 0.0714 | 1 | 0 | 4 | 0 | 5 | 0.2 | 5 | 0 | 15 | 0.0667 |
| 19_272296 | 4 | 0.25 | 4 | 0 | 3 | 0 | 3 | 0 | 14 | 0.0714 | 1 | 0 | 4 | 0 | 5 | 0.2 | 5 | 0 | 15 | 0.0667 |
| 19_272309 | 5 | 0 | 5 | 0.6 | 3 | 0 | 3 | 0 | 16 | 0.1875 | 1 | 0 | 4 | 0.25 | 7 | 0 | 5 | 0.2 | 17 | 0.1176 |
| 19_272313 | 7 | 0.7143 | 5 | 0.6 | 6 | 0.8333 | 5 | 0.6 | 23 | 0.6957 | 2 | 0 | 6 | 0.6667 | 9 | 0.6667 | 6 | 0.8333 | 23 | 0.6522 |
| 19_272313 | 6 | 0.1667 | 4 | 0 | 7 | 0 | 3 | 0 | 20 | 0.05 | 1 | 0 | 6 | 0 | 8 | 0 | 4 | 0.25 | 19 | 0.0526 |
| 19_272314 | 9 | 0.2222 | 6 | 0.3333 | 5 | 0.4 | 6 | 0.3333 | 26 | 0.3077 | 2 | 0 | 6 | 0.6667 | 10 | 0.2 | 6 | 0.1667 | 24 | 0.2917 |
| 19_272314 | 8 | 0.125 | 7 | 0 | 6 | 0.1667 | 3 | 0 | 24 | 0.0833 | 2 | 0 | 6 | 0 | 7 | 0 | 5 | 0.2 | 20 | 0.05 |
| 19_273674 | 4 | 0.5 | 2 | 0 | 3 | 1 | 4 | 0.5 | 13 | 0.5385 | 2 | 0 | 4 | 0 | 7 | 0.4286 | 3 | 0.6667 | 16 | 0.3125 |
| 19_273675 | 5 | 0.2 | 2 | 0 | 2 | 0 | 4 | 0 | 13 | 0.0769 | 1 | 0 | 4 | 0 | 5 | 0.2 | 4 | 0 | 14 | 0.0714 |
| 19_273676 | 3 | 0.3333 | 2 | 0.5 | 2 | 0 | 3 | 0.6667 | 10 | 0.4 | 1 | 0 | 4 | 0 | 6 | 0.5 | 3 | 0.3333 | 14 | 0.2857 |
| 19_277395 | 8 | 0.125 | 8 | 0 | 7 | 0.1429 | 5 | 0.2 | 28 | 0.1071 | 4 | 0 | 9 | 0.1111 | 7 | 0.2857 | 3 | 0 | 23 | 0.1304 |
| 19_277399 | 3 | 0 | 3 | 0.3333 | 6 | 0 | 4 | 0.25 | 16 | 0.125 | 3 | 0 | 8 | 0.125 | 6 | 0.3333 | 4 | 0.5 | 21 | 0.2381 |
| 19_277401 | 4 | 0.25 | 5 | 0 | 6 | 0 | 5 | 0 | 20 | 0.05 | 2 | 0 | 8 | 0.25 | 8 | 0.125 | 4 | 0 | 22 | 0.1364 |
| 19_277401 | 4 | 0 | 5 | 0 | 6 | 0 | 5 | 0.2 | 20 | 0.05 | 2 | 0 | 9 | 0 | 6 | 0.1667 | 3 | 0.3333 | 20 | 0.1 |
| 19_277417 | 2 | 0 | 5 | 0 | 3 | 0.3333 | 2 | 0.5 | 12 | 0.1667 | 2 | 0 | 4 | 0.25 | 5 | 0.2 | 1 | 0 | 12 | 0.1667 |
| 19_277418 | 2 | 0.5 | 3 | 0.3333 | 4 | 0.25 | 2 | 0.5 | 11 | 0.3636 | 2 | 0 | 5 | 0.6 | 5 | 0.2 | 1 | 1 | 13 | 0.3846 |
| 19_277419 | 1 | 0 | 4 | 0 | 4 | 0.25 | 2 | 0 | 11 | 0.0909 | 2 | 0 | 5 | 0.2 | 3 | 0 | 1 | 0 | 11 | 0.0909 |
| 19_277420 | 2 | 0 | 5 | 0.2 | 2 | 0 | 2 | 0 | 11 | 0.0909 | 1 | 0 | 5 | 0.2 | 5 | 0 | 1 | 0 | 12 | 0.0833 |
| 19_277443 | 1 | 0 | 3 | 0.3333 | 2 | 0 | NA | NA | 6 | 0.1667 | NA | NA | 2 | 0.5 | 1 | 0 | 1 | 0 | 4 | 0.25 |
| 19_277444 | 1 | 0 | 3 | 0 | 2 | 0.5 | NA | NA | 6 | 0.1667 | NA | NA | 2 | 0.5 | 2 | 0 | 1 | 1 | 5 | 0.4 |
| 19_277446 | 1 | 1 | 3 | 0.6667 | 2 | 0.5 | NA | NA | 6 | 0.6667 | NA | NA | 1 | 1 | 1 | 0 | 1 | 1 | 3 | 0.6667 |
| 19_277447 | 1 | 1 | 3 | 0.6667 | 1 | 1 | NA | NA | 5 | 0.8 | NA | NA | 1 | 1 | 2 | 0 | 1 | 1 | 4 | 0.5 |
| 19_277457 | NA | NA | 3 | 0.6667 | 1 | 0 | NA | NA | 4 | 0.5 | NA | NA | 1 | 1 | NA | NA | NA | NA | 1 | 1 |
| 19_277457 | 1 | 1 | 3 | 1 | 1 | 1 | NA | NA | 5 | 1 | NA | NA | 1 | 0 | 2 | 0.5 | NA | NA | 3 | 0.3333 |
| 19_282380 | NA | NA | 1 | 1 | 1 | 1 | 1 | 1 | 3 | 1 | 1 | 0 | NA | NA | NA | NA | 1 | 1 | 2 | 0.5 |
| 19_282399 | NA | NA | NA | NA | 1 | 1 | 1 | 1 | 2 | 1 | NA | NA | NA | NA | NA | NA | 1 | 1 | 1 | 1 |
| 19_282400 | NA | NA | NA | NA | 1 | 0 | 1 | 1 | 2 | 0.5 | NA | NA | NA | NA | NA | NA | 1 | 1 | 1 | 1 |
| 19_282402 | NA | NA | NA | NA | 1 | 1 | 1 | 1 | 2 | 1 | NA | NA | NA | NA | NA | NA | 1 | 1 | 1 | 1 |
| 19_282402 | NA | NA | NA | NA | 1 | 1 | NA | NA | 1 | 1 | NA | NA | NA | NA | NA | NA | 1 | 1 | 1 | 1 |
| 19_287404 | 2 | 1 | 2 | 0.5 | 3 | 0.6667 | 4 | 0.75 | 11 | 0.7273 | 8 | 0.875 | NA | NA | 4 | 1 | 1 | 0 | 13 | 0.8462 |
| 19_287405 | 2 | 0.5 | 2 | 1 | 2 | 1 | 2 | 1 | 8 | 0.875 | 7 | 0.7143 | 1 | 0 | 3 | 1 | 1 | 1 | 12 | 0.75 |
| 19_287405 | NA | NA | 1 | 1 | 1 | 1 | 4 | 1 | 6 | 1 | 5 | 1 | NA | NA | 2 | 1 | NA | NA | 7 | 1 |
| 19_288892 | 3 | 0.3333 | 6 | 0.3333 | 6 | 0.3333 | 6 | 0.5 | 21 | 0.381 | 3 | 0 | 6 | 0.3333 | 7 | 0.8571 | 2 | 0.5 | 18 | 0.5 |
| 19_288892 | 5 | 0 | 8 | 0.125 | 9 | 0.1111 | 8 | 0 | 30 | 0.0667 | 3 | 0 | 6 | 0 | 8 | 0.125 | 3 | 0 | 20 | 0.05 |
| 19_288892 | 6 | 0.1667 | 7 | 0.2857 | 8 | 0.125 | 8 | 0.25 | 29 | 0.2069 | 3 | 0 | 6 | 0 | 8 | 0 | 2 | 0.5 | 19 | 0.0526 |
| 19_288893 | 6 | 0 | 7 | 0.1429 | 9 | 0.2222 | 8 | 0 | 30 | 0.1 | 4 | 0 | 6 | 0 | 7 | 0.1429 | 3 | 0 | 20 | 0.05 |

| | | | | | | | | | | | | | | | | | | | | |
|---|---|---|---|---|---|---|---|---|---|---|---|---|---|---|---|---|---|---|---|---|
| 19_288893 | 6 | 1 | 6 | 0.8333 | 9 | 1 | 4 | 1 | 25 | 0.96 | 4 | 1 | 6 | 0.8333 | 7 | 1 | 2 | 1 | 19 | 0.9474 |
| 19_288894 | 4 | 0.75 | 6 | 0.3333 | 6 | 0.5 | 4 | 0.5 | 20 | 0.5 | 5 | 0.4 | 2 | 0.5 | 5 | 0.2 | 1 | 1 | 13 | 0.3846 |
| 19_288895 | 6 | 0.5 | 7 | 0.2857 | 9 | 0.5556 | 6 | 0.1667 | 28 | 0.3929 | 3 | 0.3333 | 6 | 0.3333 | 7 | 0.4286 | 2 | 0 | 18 | 0.3333 |
| 19_288897 | 6 | 0.3333 | 8 | 0.25 | 11 | 0.0909 | 6 | 0 | 31 | 0.1613 | 6 | 0.1667 | 7 | 0 | 8 | 0.25 | 3 | 0 | 24 | 0.125 |
| 19_288907 | 11 | 0.2727 | 13 | 0.1538 | 13 | 0 | 11 | 0.0909 | 48 | 0.125 | 6 | 0 | 8 | 0.125 | 12 | 0.25 | 11 | 0.2727 | 37 | 0.1892 |
| 19_288909 | 10 | 0 | 11 | 0.1818 | 12 | 0.0833 | 8 | 0.25 | 41 | 0.122 | 5 | 0 | 9 | 0.1111 | 14 | 0.0714 | 9 | 0 | 37 | 0.0541 |
| 19_288911 | 8 | 0.25 | 14 | 0.3571 | 15 | 0.4 | 10 | 0.2 | 47 | 0.3191 | 6 | 0.3333 | 12 | 0.3333 | 12 | 0.25 | 12 | 0.25 | 42 | 0.2857 |
| 19_289415 | 4 | 0.25 | 4 | 0.5 | 1 | 0 | 4 | 0 | 13 | 0.2308 | 1 | 0 | 2 | 0 | 5 | 0.2 | 2 | 0 | 10 | 0.1 |
| 19_289416 | 7 | 1 | 6 | 0.5 | 5 | 0.8 | 6 | 1 | 24 | 0.8333 | 2 | 1 | 4 | 0.5 | 7 | 1 | 2 | 0.5 | 15 | 0.8 |
| 19_289416 | 9 | 0.1111 | 8 | 0.125 | 4 | 0 | 5 | 0 | 26 | 0.0769 | 2 | 0 | 4 | 0.5 | 6 | 0 | 3 | 0 | 15 | 0.1333 |
| 19_289418 | 8 | 0.125 | 4 | 0 | 5 | 0.2 | 6 | 0 | 23 | 0.087 | 2 | 0 | 4 | 0 | 8 | 0.25 | 5 | 0 | 19 | 0.1053 |
| 19_289419 | 9 | 0.6667 | 8 | 0.75 | 5 | 0.8 | 6 | 0.6667 | 28 | 0.7143 | 3 | 1 | 6 | 0.6667 | 8 | 0.5 | 4 | 0.5 | 21 | 0.619 |
| 19_289419 | 9 | 1 | 8 | 1 | 5 | 1 | 6 | 1 | 28 | 1 | 3 | 1 | 6 | 1 | 6 | 1 | 5 | 1 | 20 | 1 |
| 19_289420 | 5 | 1 | 3 | 1 | 5 | 0.8 | 6 | 0.5 | 19 | 0.7895 | 2 | 1 | 3 | 0.3333 | 7 | 0.7143 | 2 | 1 | 14 | 0.7143 |
| 19_289422 | 9 | 0.4444 | 8 | 0.5 | 4 | 0.5 | 6 | 0 | 27 | 0.3704 | 4 | 0.75 | 5 | 0.4 | 9 | 0.4444 | 5 | 0.4 | 23 | 0.4783 |
| 19_289423 | 8 | 0.375 | 8 | 0.125 | 3 | 0.3333 | 4 | 0.25 | 23 | 0.2609 | 3 | 1 | 5 | 0.2 | 10 | 0.3 | 5 | 0.2 | 23 | 0.3478 |
| 19_289423 | 7 | 0.1429 | 8 | 0 | 5 | 0 | 6 | 0.1667 | 26 | 0.0769 | 4 | 0.25 | 5 | 0.2 | 10 | 0 | 6 | 0.1667 | 25 | 0.12 |
| 19_289424 | 7 | 0.4286 | 7 | 0.1429 | 4 | 0.5 | 5 | 0.2 | 23 | 0.3043 | 3 | 0 | 6 | 0.3333 | 8 | 0 | 6 | 0.1667 | 23 | 0.1304 |
| 19_289424 | 10 | 0 | 6 | 0.1667 | 5 | 0 | 7 | 0.1429 | 28 | 0.0714 | 2 | 0 | 5 | 0 | 10 | 0.3 | 6 | 0 | 23 | 0.1304 |
| 19_289424 | 8 | 0.375 | 5 | 0.8 | 3 | 0.6667 | 7 | 0.2857 | 23 | 0.4783 | 3 | 0.3333 | 3 | 0.3333 | 9 | 0.4444 | 5 | 0.4 | 20 | 0.4 |
| 19_289425 | 5 | 1 | 8 | 0.875 | 4 | 0.5 | 6 | 1 | 23 | 0.8696 | 3 | 1 | 5 | 0.6 | 9 | 1 | 4 | 0.75 | 21 | 0.8571 |
| 19_289428 | 7 | 0 | 8 | 0.5 | 2 | 0.5 | 4 | 0.5 | 21 | 0.3333 | 3 | 0 | 5 | 0.2 | 5 | 0.6 | 5 | 0.6 | 18 | 0.3889 |
| 19_289430 | 8 | 0.125 | 8 | 0.125 | 3 | 0 | 8 | 0.125 | 27 | 0.1111 | 3 | 0 | 4 | 0 | 8 | 0.125 | 4 | 0 | 19 | 0.0526 |
| 19_289433 | 9 | 0.3333 | 9 | 0 | 3 | 0.3333 | 8 | 0 | 29 | 0.1379 | 4 | 0 | 6 | 0 | 9 | 0.1111 | 7 | 0.1429 | 26 | 0.0769 |
| 19_289436 | 9 | 0.4444 | 7 | 0.2857 | 3 | 0.6667 | 8 | 0.25 | 27 | 0.3704 | 4 | 0.25 | 5 | 0.4 | 7 | 0.5714 | 6 | 0.6667 | 22 | 0.5 |
| 19_289436 | 9 | 0.2222 | 9 | 0.3333 | 3 | 0 | 9 | 0.2222 | 30 | 0.2333 | 4 | 0.25 | 5 | 0 | 8 | 0.375 | 6 | 0.1667 | 23 | 0.2174 |
| 19_289436 | 9 | 0.1111 | 10 | 0.2 | 2 | 0 | 9 | 0 | 30 | 0.1 | 4 | 0.25 | 5 | 0.4 | 9 | 0.1111 | 6 | 0.5 | 24 | 0.2917 |
| 19_289439 | 9 | 0.2222 | 10 | 0.3 | 4 | 0.5 | 8 | 0.125 | 31 | 0.2581 | 4 | 0.25 | 6 | 0 | 9 | 0.2222 | 7 | 0.2857 | 26 | 0.1923 |
| 19_289893 | 3 | 0 | 6 | 0 | 12 | 0.1667 | 7 | 0 | 28 | 0.0714 | 5 | 0 | 6 | 0.1667 | 10 | 0 | 7 | 0.1429 | 28 | 0.0714 |
| 19_289921 | 3 | 0.3333 | 2 | 0.5 | 3 | 0.3333 | 2 | 0 | 10 | 0.3 | NA | NA | 3 | 0.3333 | 5 | 0.2 | 1 | 0 | 9 | 0.2222 |
| 19_289929 | 4 | 0.75 | 2 | 1 | 3 | 0 | 2 | 0 | 11 | 0.4545 | 2 | 1 | 2 | 0 | 6 | 0.3333 | 2 | 0 | 12 | 0.3333 |
| 19_289934 | NA | NA | 2 | 1 | 5 | 1 | 5 | 0.8 | 12 | 0.9167 | 4 | 1 | 3 | 1 | 7 | 0.2857 | 6 | 0.5 | 20 | 0.6 |
| 19_289935 | 1 | 1 | 4 | 1 | 7 | 1 | 8 | 0.5 | 20 | 0.8 | 5 | 1 | 3 | 0.6667 | 9 | 0.7778 | 6 | 1 | 23 | 0.8696 |
| 19_289937 | 2 | 0 | 4 | 0.25 | 7 | 0.1429 | 8 | 0.125 | 21 | 0.1429 | 6 | 0 | 1 | 0 | 9 | 0.1111 | 4 | 0.25 | 20 | 0.1 |
| 19_289937 | 2 | 0 | 4 | 0 | 7 | 0.2857 | 8 | 0.125 | 21 | 0.1429 | 5 | 0 | 2 | 0.5 | 9 | 0 | 6 | 0.1667 | 22 | 0.0909 |
| 19_289938 | 2 | 0.5 | 4 | 0.25 | 7 | 0.2857 | 9 | 0.2222 | 22 | 0.2727 | 4 | 0.5 | 2 | 0.5 | 9 | 0.2222 | 6 | 0.1667 | 21 | 0.2857 |
| 19_289948 | 5 | 0.2 | 5 | 0.4 | 7 | 0.4286 | 11 | 0.0909 | 28 | 0.25 | 3 | 0 | 2 | 0 | 7 | 0.1429 | 7 | 0.1429 | 19 | 0.1053 |
| 19_289983 | 19 | 0.1579 | 9 | 0.1111 | 17 | 0.2353 | 14 | 0.0714 | 59 | 0.1525 | 20 | 0.2 | 13 | 0.2308 | 16 | 0.1875 | 13 | 0.0769 | 62 | 0.1774 |
| 19_289993 | 21 | 0.4286 | 10 | 0.5 | 16 | 0.3125 | 11 | 0.4545 | 58 | 0.4138 | 16 | 0.5625 | 13 | 0.2308 | 10 | 0.5 | 13 | 0.2308 | 52 | 0.3846 |
| 19_289994 | 23 | 0.3478 | 8 | 0.375 | 12 | 0.3333 | 11 | 0.1818 | 54 | 0.3148 | 16 | 0.6875 | 12 | 0.4167 | 12 | 0.5833 | 13 | 0.3846 | 53 | 0.5283 |
| 19_289995 | 18 | 0 | 12 | 0.1667 | 16 | 0 | 12 | 0.0833 | 58 | 0.0517 | 15 | 0.0667 | 12 | 0.0833 | 11 | 0 | 12 | 0.0833 | 50 | 0.06 |
| 19_289996 | 20 | 0.3 | 10 | 0.3 | 15 | 0.2 | 12 | 0.25 | 57 | 0.2632 | 13 | 0.3077 | 11 | 0.0909 | 12 | 0.25 | 13 | 0.0769 | 49 | 0.1837 |
| 19_289996 | 18 | 0.1111 | 12 | 0 | 15 | 0 | 12 | 0.0833 | 57 | 0.0526 | 15 | 0.1333 | 12 | 0.1667 | 11 | 0 | 13 | 0 | 51 | 0.0784 |
| 19_290000 | 19 | 0.0526 | 10 | 0 | 13 | 0.3077 | 11 | 0.2727 | 53 | 0.1509 | 14 | 0.1429 | 11 | 0.0909 | 9 | 0.2222 | 11 | 0.0909 | 45 | 0.1333 |
| 19_290001 | 17 | 0.3529 | 11 | 0.6364 | 13 | 0.6154 | 10 | 0.3 | 51 | 0.4706 | 14 | 0.5714 | 10 | 0.4 | 11 | 0.4545 | 11 | 0.3636 | 46 | 0.4565 |
| 19_290006 | NA | NA | 2 | 0.5 | NA | NA | 1 | 1 | 3 | 0.6667 | NA | NA | NA | NA | 1 | 1 | NA | NA | 1 | 1 |
| 19_290016 | 2 | 0 | 2 | 0 | 1 | 1 | 1 | 0 | 6 | 0.1667 | 2 | 0.5 | NA | NA | 2 | 0 | 2 | 0 | 6 | 0.1667 |
| 19_290017 | NA | NA | 2 | 0.5 | NA | NA | NA | NA | 2 | 0.5 | NA | NA | NA | NA | 1 | 1 | NA | NA | 1 | 1 |
| 19_290019 | 1 | 0 | 2 | 0.5 | 1 | 1 | 1 | 0 | 5 | 0.4 | 2 | 0 | NA | NA | 2 | 0.5 | 2 | 0 | 6 | 0.1667 |
| 19_290022 | 3 | 0 | 4 | 0.75 | 2 | 0 | 2 | 0 | 11 | 0.2727 | 1 | 0 | 3 | 0 | 4 | 0.75 | 2 | 0 | 10 | 0.3 |
| 19_290022 | 4 | 0.5 | 1 | 0 | 2 | 0.5 | NA | NA | 7 | 0.4286 | 2 | 0 | 3 | 0.3333 | 3 | 1 | 2 | 1 | 10 | 0.6 |
| 19_290023 | 4 | 0.25 | 4 | 0.25 | 4 | 0 | 2 | 0.5 | 14 | 0.2143 | 3 | 0.3333 | 1 | 0 | 5 | 0.4 | 2 | 0 | 11 | 0.2727 |
| 19_290024 | 4 | 0 | 4 | 0.25 | 4 | 0 | 2 | 0 | 14 | 0.0714 | 4 | 0 | 3 | 0 | 5 | 0 | 2 | 0.5 | 14 | 0.0714 |
| 19_290024 | 4 | 0.25 | 4 | 0 | 4 | 0.25 | 2 | 0 | 14 | 0.1429 | 4 | 0.25 | 3 | 0.3333 | 5 | 0.2 | 2 | 0.5 | 14 | 0.2857 |
| 19_290024 | 3 | 0 | 4 | 0.25 | 4 | 0.25 | 2 | 0 | 13 | 0.1538 | 4 | 0.5 | 2 | 0 | 4 | 0.25 | 2 | 0 | 12 | 0.25 |
| 19_290025 | 4 | 0.5 | 4 | 0.5 | 4 | 0.25 | 2 | 0.5 | 14 | 0.4286 | 4 | 0.25 | 2 | 0 | 4 | 0.25 | 2 | 1 | 12 | 0.3333 |
| 19_290025 | 3 | 0.6667 | 4 | 0.75 | 4 | 0.5 | 2 | 1 | 13 | 0.6923 | 3 | 1 | 3 | 1 | 5 | 0.8 | 2 | 1 | 13 | 0.9231 |
| 19_290027 | 4 | 0.75 | 5 | 0.4 | 4 | 0.75 | 2 | 1 | 15 | 0.6667 | 3 | 0.6667 | 2 | 0.5 | 4 | 0.75 | 2 | 0.5 | 11 | 0.6364 |
| 19_290029 | NA | NA | 1 | 0 | 1 | 1 | NA | NA | 2 | 0.5 | NA | NA | NA | NA | 1 | 0 | 1 | 1 | 2 | 0.5 |
| 19_290030 | 5 | 0.6 | 5 | 0.6 | 4 | 1 | 2 | 1 | 16 | 0.75 | 3 | 0.6667 | 3 | 1 | 6 | 1 | 2 | 1 | 14 | 0.9286 |
| 19_290031 | 5 | 0.8 | 4 | 0.25 | 4 | 0.75 | 3 | 0.6667 | 16 | 0.625 | 3 | 1 | 3 | 1 | 6 | 0.6667 | 2 | 1 | 14 | 0.8571 |
| 19_290032 | 4 | 0.5 | 4 | 1 | 3 | 0.6667 | 3 | 0.3333 | 14 | 0.6429 | 3 | 1 | 2 | 1 | 6 | 1 | 2 | 1 | 13 | 1 |
| 19_290032 | 5 | 0.4 | 4 | 0.25 | 2 | 0.5 | 2 | 0 | 13 | 0.3077 | 3 | 0.3333 | 3 | 0.6667 | 6 | 0.5 | 2 | 0.5 | 14 | 0.5 |
| 19_290033 | NA | NA | 2 | 0 | 1 | 0 | 1 | 1 | 4 | 0.25 | 1 | 0 | NA | NA | 1 | 1 | 1 | 0 | 3 | 0.3333 |
| 19_290034 | 5 | 0.2 | 4 | 0 | 4 | 0.5 | 3 | 0.3333 | 16 | 0.25 | 4 | 0.25 | 2 | 0.5 | 6 | 0.1667 | 1 | 0 | 13 | 0.2308 |

| | | | | | | | | | | | | | | | | | | | | |
|---|---|---|---|---|---|---|---|---|---|---|---|---|---|---|---|---|---|---|---|---|
| 19_290034 | 5 | 0 | 5 | 0.2 | 4 | 0.5 | 4 | 0 | 18 | 0.1667 | 3 | 0.3333 | 2 | 0.5 | 4 | 0.5 | 2 | 0 | 11 | 0.3636 |
| 19_290034 | NA | NA | 1 | 1 | 1 | 1 | 1 | 1 | 3 | 1 | 1 | 1 | NA | NA | 1 | 1 | 1 | 1 | 3 | 1 |
| 19_290035 | NA | NA | 2 | 0.5 | 1 | 0 | 1 | 0 | 4 | 0.25 | 1 | 0 | NA | NA | 1 | 1 | 1 | 1 | 3 | 0.6667 |
| 19_290035 | NA | NA | 1 | 0 | NA | NA | 1 | 1 | 2 | 0.5 | 1 | 0 | NA | NA | 1 | 1 | 1 | 1 | 3 | 0.6667 |
| 19_290035 | 6 | 0 | 6 | 0.1667 | 4 | 0 | 4 | 0 | 20 | 0.05 | 6 | 0 | 2 | 0 | 5 | 0 | 2 | 0.5 | 15 | 0.0667 |
| 19_290036 | 4 | 0.75 | 5 | 0.8 | 2 | 0.5 | 3 | 0.6667 | 14 | 0.7143 | 4 | 1 | 2 | 1 | 6 | 0.8333 | 2 | 0.5 | 14 | 0.8571 |
| 19_290036 | 6 | 0.5 | 5 | 0.6 | 3 | 0 | 4 | 0.5 | 18 | 0.4444 | 4 | 0.5 | 2 | 0.5 | 7 | 0.2857 | 1 | 0 | 14 | 0.3571 |
| 19_290037 | 6 | 0.5 | 7 | 0.1429 | 4 | 0.5 | 4 | 0.25 | 21 | 0.3333 | 4 | 0.5 | 2 | 0.5 | 8 | 0.625 | 2 | 1 | 16 | 0.625 |
| 19_290038 | 3 | 0.3333 | 7 | 0 | 4 | 0.25 | 3 | 0 | 17 | 0.1176 | 2 | 0.5 | 1 | 0 | 4 | 0.25 | NA | NA | 7 | 0.2857 |
| 19_290038 | 6 | 0.5 | 4 | 0.25 | 4 | 0.75 | 4 | 0.75 | 18 | 0.5556 | 6 | 0.8333 | 2 | 1 | 7 | 1 | 2 | 1 | 17 | 0.9412 |
| 19_290040 | 5 | 0 | 6 | 0 | 3 | 0.3333 | 3 | 0 | 17 | 0.0588 | 5 | 0.2 | NA | NA | 5 | 0 | 2 | 0 | 12 | 0.0833 |
| 19_290081 | 2 | 0 | 4 | 0.5 | NA | NA | 1 | 1 | 7 | 0.4286 | NA | NA | 1 | 0 | 5 | 0.2 | NA | NA | 6 | 0.1667 |
| 19_290082 | 3 | 0 | 3 | 0.3333 | NA | NA | 1 | 1 | 7 | 0.2857 | NA | NA | 2 | 0.5 | 6 | 0.3333 | NA | NA | 8 | 0.375 |
| 19_290083 | 2 | 0 | 6 | 0.5 | NA | NA | 1 | 0 | 9 | 0.3333 | NA | NA | 2 | 0.5 | 6 | 0.6667 | NA | NA | 8 | 0.625 |
| 19_290084 | 2 | 0.5 | 6 | 0.5 | NA | NA | 1 | 1 | 9 | 0.5556 | NA | NA | 2 | 0.5 | 5 | 0.4 | NA | NA | 7 | 0.4286 |
| 19_290085 | 3 | 0.3333 | 6 | 0.5 | NA | NA | 1 | 0 | 10 | 0.4 | NA | NA | 2 | 0.5 | 5 | 0.4 | NA | NA | 7 | 0.4286 |
| 19_290085 | 3 | 0 | 6 | 0.1667 | NA | NA | NA | NA | 9 | 0.1111 | NA | NA | 2 | 0 | 5 | 0.2 | NA | NA | 7 | 0.1429 |
| 19_290086 | 5 | 0.8 | 7 | 0.8571 | NA | NA | 1 | 1 | 13 | 0.8462 | NA | NA | 3 | 0.3333 | 5 | 1 | NA | NA | 8 | 0.75 |
| 19_290087 | 4 | 0.5 | 6 | 0.5 | NA | NA | 1 | 1 | 11 | 0.5455 | NA | NA | 3 | 0.3333 | 6 | 0.3333 | NA | NA | 9 | 0.3333 |
| 19_291206 | 3 | 0.3333 | 7 | 0.7143 | 2 | 0.5 | 1 | 0 | 13 | 0.5385 | 3 | 0.6667 | NA | NA | 6 | 0.5 | 2 | 1 | 11 | 0.6364 |
| 19_291206 | 4 | 0.25 | 7 | 0 | 2 | 0.5 | 1 | 0 | 14 | 0.1429 | 3 | 0.3333 | NA | NA | 6 | 0.1667 | 2 | 0 | 11 | 0.1818 |
| 19_291206 | 4 | 0.5 | 7 | 0 | 2 | 0 | 1 | 0 | 14 | 0.1429 | 3 | 0 | NA | NA | 6 | 0 | 2 | 0.5 | 11 | 0.0909 |
| 19_291207 | 4 | 0.25 | 6 | 0.3333 | 1 | 0 | 1 | 0 | 12 | 0.25 | 2 | 0.5 | NA | NA | 6 | 0.1667 | 2 | 1 | 10 | 0.4 |
| 19_291207 | 3 | 1 | 7 | 0.8571 | 1 | 0 | NA | NA | 11 | 0.8182 | 3 | 1 | NA | NA | 6 | 0.5 | 1 | 0 | 10 | 0.6 |
| 19_291207 | 4 | 0 | 7 | 0.2857 | 1 | 0 | 1 | 1 | 13 | 0.2308 | 3 | 0 | NA | NA | 5 | 0.2 | 2 | 0 | 10 | 0.1 |
| 19_291208 | 3 | 1 | 7 | 1 | 2 | 1 | 1 | 1 | 13 | 1 | 3 | 1 | NA | NA | 6 | 1 | 3 | 1 | 12 | 1 |
| 19_291208 | 4 | 0.25 | 6 | 0.3333 | 2 | 0 | NA | NA | 12 | 0.25 | 4 | 0.25 | NA | NA | 6 | 0.3333 | 3 | 0.3333 | 13 | 0.3077 |
| 19_291208 | 3 | 1 | 6 | 0.6667 | 1 | 1 | NA | NA | 10 | 0.8 | 3 | 0.6667 | NA | NA | 3 | 1 | 1 | 1 | 7 | 0.8571 |
| 19_291209 | 5 | 0.6 | 5 | 0.6 | NA | NA | 1 | 0 | 11 | 0.5455 | 3 | 0.6667 | NA | NA | 3 | 0 | 3 | 0.3333 | 9 | 0.3333 |
| 19_291209 | 3 | 0 | 7 | 0.1429 | 2 | 0.5 | 2 | 0.5 | 14 | 0.2143 | 4 | 0.5 | NA | NA | 6 | 0.3333 | 3 | 0.3333 | 13 | 0.3846 |
| 19_291210 | 4 | 0.25 | 5 | 0 | 2 | 0.5 | 2 | 0.5 | 13 | 0.2308 | 5 | 0.2 | NA | NA | 6 | 0.3333 | 2 | 0.5 | 13 | 0.3077 |
| 19_292760 | 4 | 0.25 | 7 | 0 | 2 | 0 | 5 | 0 | 18 | 0.0556 | 2 | 0.5 | 7 | 0 | 2 | 0 | 4 | 0.25 | 15 | 0.1333 |
| 19_292892 | 5 | 0 | 7 | 0.1429 | 1 | 0 | 2 | 0 | 15 | 0.0667 | 4 | 0.25 | 8 | 0.125 | 2 | 0.5 | 4 | 0 | 18 | 0.1667 |
| 19_292892 | 5 | 0.6 | 8 | 0.625 | 2 | 0.5 | 5 | 0.8 | 20 | 0.65 | 4 | 0.75 | 9 | 0.6667 | 4 | 0.25 | 5 | 0.6 | 22 | 0.5909 |
| 19_292892 | 6 | 0.3333 | 9 | 0 | 2 | 0.5 | 4 | 0.25 | 21 | 0.1905 | 5 | 0.6 | 12 | 0.0833 | 4 | 0.25 | 6 | 0.1667 | 27 | 0.2222 |
| 19_292892 | 6 | 0.3333 | 9 | 0.2222 | 2 | 0 | 4 | 0.25 | 21 | 0.2381 | 5 | 0.2 | 12 | 0.3333 | 4 | 0 | 6 | 0.1667 | 27 | 0.2222 |
| 19_292893 | 5 | 0.2 | 9 | 0.5556 | 3 | 1 | 5 | 0.6 | 22 | 0.5455 | 5 | 0.4 | 13 | 0.5385 | 4 | 0.5 | 5 | 0.8 | 27 | 0.5556 |
| 19_292895 | 7 | 0.8571 | 7 | 0.8571 | 1 | 1 | 4 | 0.75 | 19 | 0.8421 | 4 | 1 | 11 | 0.9091 | 4 | 0.75 | 5 | 0.8 | 24 | 0.875 |
| 19_292895 | 5 | 0.2 | 7 | 0 | 2 | 0 | 4 | 0 | 18 | 0.0556 | 4 | 0 | 13 | 0.1538 | 1 | 0 | 6 | 0 | 24 | 0.0833 |
| 19_292898 | 5 | 1 | 6 | 0.8333 | 3 | 0.6667 | 5 | 1 | 19 | 0.8947 | 4 | 0.75 | 13 | 1 | 2 | 0.5 | 2 | 0.5 | 21 | 0.8571 |
| 19_292899 | 5 | 0.2 | 7 | 0 | 3 | 0 | 4 | 0.25 | 19 | 0.1053 | 4 | 0 | 12 | 0.0833 | 2 | 0 | 2 | 0 | 20 | 0.05 |
| 19_292899 | 4 | 0.25 | 6 | 0.3333 | 3 | 0 | 5 | 0.4 | 18 | 0.2778 | 4 | 0.5 | 11 | 0.3636 | 2 | 0 | 2 | 1 | 19 | 0.4211 |
| 19_292900 | 4 | 0.25 | 7 | 0.4286 | 3 | 0.3333 | 4 | 0.25 | 18 | 0.3333 | 3 | 0.3333 | 12 | 0.5 | 1 | 1 | 2 | 1 | 18 | 0.5556 |
| 19_292900 | 5 | 0.2 | 7 | 0.2857 | 5 | 0 | 5 | 0 | 22 | 0.1364 | 4 | 0.25 | 14 | 0.2857 | 2 | 0 | 2 | 0 | 22 | 0.2273 |
| 19_292901 | 4 | 0.25 | 4 | 0 | 3 | 0 | 4 | 0 | 15 | 0.0667 | 4 | 0 | 7 | 0.1429 | 2 | 0 | 1 | 0 | 14 | 0.0714 |
| 19_292901 | 3 | 1 | 3 | 0.6667 | 4 | 0.75 | 3 | 0.3333 | 13 | 0.6923 | 3 | 0.6667 | 9 | 0.6667 | 2 | 0.5 | 1 | 1 | 15 | 0.6667 |
| 19_292902 | 5 | 0.2 | 4 | 0 | 4 | 0 | 3 | 0 | 16 | 0.0625 | 5 | 0 | 8 | 0.125 | 1 | 0 | 1 | 1 | 15 | 0.1333 |
| 19_292987 | 1 | 1 | NA | NA | NA | NA | 2 | 1 | 3 | 1 | NA | NA | 2 | 1 | NA | NA | NA | NA | 2 | 1 |
| 19_293092 | NA | NA | NA | NA | NA | NA | 2 | 1 | 2 | 1 | NA | NA | NA | NA | NA | NA | 1 | 1 | 1 | 1 |
| 19_293261 | 3 | 0.3333 | 4 | 0 | 7 | 0.1429 | 2 | 0 | 16 | 0.125 | 5 | 0.2 | 6 | 0 | 5 | 0 | 3 | 0 | 19 | 0.0526 |
| 19_293263 | 2 | 0 | 3 | 1 | 8 | 0 | 2 | 0.5 | 15 | 0.2667 | 4 | 0.25 | 5 | 0.6 | 3 | 0 | 2 | 0.5 | 14 | 0.3571 |
| 19_293263 | 2 | 0 | 4 | 0 | 9 | 0.1111 | 2 | 0 | 17 | 0.0588 | 3 | 0.3333 | 6 | 0 | 5 | 0.2 | 1 | 0 | 15 | 0.1333 |
| 19_295727 | NA | NA | 1 | 0 | 1 | 0 | 4 | 0.5 | 6 | 0.3333 | NA | NA | NA | NA | 1 | 1 | 1 | 0 | 2 | 0.5 |
| 19_295730 | NA | NA | 3 | 1 | 1 | 1 | 3 | 1 | 7 | 1 | 1 | 1 | NA | NA | 2 | 0.5 | 1 | 0 | 4 | 0.5 |
| 19_295730 | NA | NA | 2 | 0 | 2 | 0 | 4 | 0.25 | 8 | 0.125 | 1 | 1 | NA | NA | 2 | 0 | 1 | 0 | 4 | 0.25 |
| 19_295732 | 1 | 0 | 3 | 0.3333 | 1 | 1 | 4 | 0.25 | 9 | 0.3333 | 1 | 1 | NA | NA | 2 | 0.5 | NA | NA | 3 | 0.6667 |
| 19_295733 | NA | NA | 2 | 0.5 | 2 | 0 | 4 | 0 | 8 | 0.125 | 1 | 1 | NA | NA | 2 | 0 | 1 | 0 | 4 | 0.25 |
| 19_295773 | 1 | 0 | 4 | 0.5 | 2 | 0.5 | 3 | 0 | 10 | 0.3 | NA | NA | NA | NA | 2 | 0 | 1 | 1 | 3 | 0.3333 |
| 19_295773 | 1 | 0 | 4 | 0 | 2 | 0.5 | 4 | 0.25 | 11 | 0.1818 | 1 | 1 | NA | NA | 2 | 0 | 1 | 0 | 4 | 0.25 |
| 19_295813 | 2 | 0.5 | 4 | 0 | 1 | 1 | 4 | 0.25 | 11 | 0.2727 | 1 | 0 | 1 | 1 | 2 | 0.5 | 2 | 0 | 6 | 0.3333 |
| 19_295813 | 3 | 0.6667 | 2 | 1 | 1 | 0 | 7 | 0.2857 | 13 | 0.4615 | 1 | 0 | 1 | 1 | 3 | 0.6667 | 2 | 0 | 7 | 0.4286 |
| 19_295814 | NA | NA | 4 | 0 | NA | NA | 4 | 0.25 | 8 | 0.125 | 1 | 0 | 1 | 1 | 2 | 0.5 | 1 | 0 | 5 | 0.4 |
| 19_295814 | 3 | 0 | 5 | 0 | 2 | 0 | 8 | 0.125 | 18 | 0.0556 | 1 | 1 | 1 | 0 | 3 | 0 | 2 | 0 | 7 | 0.1429 |
| 19_295815 | 3 | 0.6667 | 3 | 1 | 2 | 1 | 8 | 0.5 | 16 | 0.6875 | 1 | 1 | 1 | 1 | 3 | 1 | 1 | 1 | 6 | 1 |
| 19_295816 | 3 | 0 | 5 | 0.2 | 2 | 0 | 8 | 0 | 18 | 0.0556 | 2 | 0 | 1 | 0 | 3 | 0 | 2 | 0.5 | 8 | 0.125 |

| | | | | | | | | | | | | | | | | | | | | |
|---|---|---|---|---|---|---|---|---|---|---|---|---|---|---|---|---|---|---|---|---|
| 19_2958164 | 2 | 0.5 | 4 | 0.75 | 2 | 0 | 5 | 0.6 | 13 | 0.5385 | 2 | 1 | 1 | 0 | 2 | 1 | 2 | 0.5 | 7 | 0.7143 |
| 19_2958165 | 2 | 0.5 | 4 | 0.5 | 2 | 0.5 | 5 | 0.6 | 13 | 0.5385 | 2 | 0 | 1 | 0 | 3 | 0.3333 | 2 | 0.5 | 8 | 0.25 |
| 19_2958168 | 1 | 1 | 4 | 1 | 2 | 1 | 5 | 1 | 12 | 1 | 1 | 1 | 1 | 1 | 1 | 1 | 2 | 1 | 5 | 1 |
| 19_2958205 | 3 | 0 | 3 | 0 | 2 | 0 | 10 | 0.3 | 18 | 0.1667 | 2 | 0 | 1 | 0 | 2 | 1 | 2 | 0 | 7 | 0.2857 |
| 19_2958228 | 3 | 0 | 3 | 0 | 2 | 1 | 12 | 0.25 | 20 | 0.25 | 3 | 0 | 1 | 1 | 3 | 0.3333 | 4 | 0 | 11 | 0.1818 |
| 19_2958241 | 3 | 0 | 4 | 0.25 | 2 | 0.5 | 11 | 0 | 20 | 0.1 | 3 | 0.6667 | 2 | 0 | 3 | 0.3333 | 2 | 0 | 10 | 0.3 |
| 19_2958259 | 1 | 0 | 4 | 0.5 | NA | NA | 10 | 0.5 | 15 | 0.4667 | 3 | 0.3333 | 2 | 0 | 2 | 0 | 3 | 0.3333 | 10 | 0.2 |
| 19_2958783 | 4 | 0 | 7 | 0.1429 | 3 | 0 | 3 | 0 | 17 | 0.0588 | 2 | 0 | 5 | 0 | 3 | 0 | 6 | 0.1667 | 16 | 0.0625 |
| 19_2958844 | 3 | 0 | 7 | 0.2857 | 1 | 0 | 4 | 0.25 | 15 | 0.2 | 2 | 0.5 | 1 | 0 | 4 | 0.25 | 5 | 0.2 | 12 | 0.25 |
| 19_2958866 | 4 | 0.25 | 7 | 0.4286 | 1 | 1 | 4 | 0.25 | 16 | 0.375 | 2 | 0 | 1 | 0 | 4 | 0.25 | 5 | 0.2 | 12 | 0.1667 |
| 19_2958917 | 1 | 0 | 5 | 0.4 | 1 | 0 | 3 | 0 | 10 | 0.2 | 2 | 0 | 1 | 0 | 3 | 0 | 4 | 0.25 | 10 | 0.1 |
| 19_2958965 | 3 | 1 | 4 | 1 | 1 | 1 | 3 | 1 | 11 | 1 | NA | NA | 1 | 1 | 1 | 1 | 2 | 1 | 4 | 1 |
| 19_2958974 | 2 | 1 | 3 | 0.6667 | 1 | 1 | 2 | 0.5 | 8 | 0.75 | 2 | 0.5 | 1 | 1 | NA | NA | 3 | 0.6667 | 6 | 0.6667 |
| 19_2958975 | 2 | 0 | 3 | 0.3333 | NA | NA | 3 | 0.3333 | 8 | 0.25 | 2 | 0.5 | 1 | 0 | NA | NA | 4 | 0 | 7 | 0.1429 |
| 19_2958982 | 1 | 1 | 2 | 1 | 1 | 1 | 1 | 1 | 5 | 1 | 2 | 1 | 1 | 1 | NA | NA | 3 | 0.6667 | 6 | 0.8333 |
| 19_2958988 | 1 | 1 | 2 | 0 | 1 | 1 | 2 | 0.5 | 6 | 0.5 | 1 | 0 | NA | NA | NA | NA | 2 | 0.5 | 3 | 0.3333 |
| 19_2958990 | 3 | 1 | 3 | 1 | NA | NA | 2 | 1 | 8 | 1 | 2 | 1 | 1 | 1 | NA | NA | 4 | 1 | 7 | 1 |
| 19_2959020 | 1 | 0 | 3 | 0.3333 | NA | NA | 1 | 0 | 5 | 0.2 | 1 | 1 | 1 | 0 | NA | NA | 4 | 0.25 | 6 | 0.3333 |
| 19_2960435 | NA | NA | NA | NA | 1 | 1 | NA | NA | 1 | 1 | NA | NA | 1 | 1 | NA | NA | 3 | 0 | 4 | 0.25 |
| 19_2969223 | 1 | 0 | 1 | 0 | 3 | 0 | 1 | 1 | 6 | 0.1667 | NA | NA | 5 | 0.2 | 2 | 0 | NA | NA | 7 | 0.1429 |
| 19_2969276 | 4 | 0.5 | 6 | 0.3333 | 6 | 0.1667 | 2 | 0 | 18 | 0.2778 | 3 | 0.6667 | 9 | 0.2222 | 4 | 0.25 | 3 | 0.3333 | 19 | 0.3158 |
| 19_2969321 | 4 | 0.25 | 5 | 0.2 | 8 | 0.125 | 4 | 0.75 | 21 | 0.2857 | 3 | 0.3333 | 11 | 0 | 3 | 0.3333 | 2 | 0.5 | 19 | 0.1579 |
| 19_2969334 | 4 | 0 | 5 | 0 | 6 | 0.3333 | 4 | 0 | 19 | 0.1053 | 3 | 0.3333 | 11 | 0 | 5 | 0.2 | 2 | 0 | 21 | 0.0952 |
| 19_2969339 | 3 | 0.3333 | 4 | 0.25 | 4 | 0.25 | 2 | 1 | 13 | 0.3846 | 1 | 0 | 7 | 0.8571 | 2 | 0.5 | 1 | 1 | 11 | 0.7273 |
| 19_2969340 | 3 | 0.3333 | 3 | 1 | 4 | 1 | 1 | 1 | 11 | 0.8182 | 3 | 1 | 7 | 0.7143 | 2 | 1 | 1 | 0 | 13 | 0.7692 |
| 19_2969371 | 4 | 0 | 5 | 0.2 | 7 | 0.2857 | 5 | 0.4 | 21 | 0.2381 | 2 | 0 | 8 | 0.125 | 5 | 0.2 | 2 | 0 | 17 | 0.1176 |
| 19_2969382 | 3 | 1 | 3 | 1 | 8 | 1 | 5 | 0.8 | 19 | 0.9474 | 3 | 1 | 8 | 1 | 4 | 1 | 2 | 1 | 17 | 1 |
| 19_2969383 | 3 | 0 | 5 | 0.4 | 7 | 0.1429 | 5 | 0 | 20 | 0.15 | 3 | 0.3333 | 7 | 0.5714 | 5 | 0.2 | 2 | 0 | 17 | 0.3529 |
| 19_2969385 | 3 | 1 | 4 | 0.75 | 7 | 0.7143 | 5 | 1 | 19 | 0.8421 | 3 | 1 | 9 | 0.8889 | 4 | 0.75 | 2 | 1 | 18 | 0.8889 |
| 19_2969388 | 3 | 0 | 5 | 0.2 | 7 | 0.1429 | 5 | 0.2 | 20 | 0.15 | 2 | 0 | 9 | 0.3333 | 5 | 0.4 | 2 | 0.5 | 18 | 0.3333 |
| 19_2969395 | NA | NA | 3 | 0.6667 | 4 | 1 | 3 | 1 | 10 | 0.9 | NA | NA | 4 | 1 | 1 | 1 | 1 | 1 | 6 | 1 |
| 19_2969405 | 4 | 0 | 4 | 0.5 | 7 | 0.5714 | 4 | 0.5 | 19 | 0.4211 | 2 | 1 | 6 | 0.6667 | 5 | 0.8 | 2 | 0 | 15 | 0.6667 |
| 19_2969411 | 4 | 0.25 | 4 | 0.5 | 6 | 0 | 5 | 0.4 | 19 | 0.2632 | 3 | 0 | 7 | 0.4286 | 5 | 0.2 | 2 | 0.5 | 17 | 0.2941 |
| 19_2969434 | 4 | 0.75 | 3 | 1 | 5 | 1 | 4 | 1 | 16 | 0.9375 | 2 | 1 | 8 | 0.75 | 4 | 1 | 2 | 0.5 | 16 | 0.8125 |
| 19_2969438 | 4 | 1 | 3 | 1 | 4 | 0.75 | 5 | 0.8 | 16 | 0.875 | 2 | 1 | 7 | 1 | 2 | 1 | 1 | 0 | 12 | 0.9167 |
| 19_2969449 | 3 | 0.6667 | 4 | 0.75 | 5 | 1 | 5 | 1 | 17 | 0.8824 | 2 | 1 | 6 | 1 | 3 | 0.6667 | 2 | 1 | 13 | 0.9231 |
| 19_2969457 | 3 | 0 | 4 | 0.5 | 6 | 0.5 | 4 | 0.5 | 17 | 0.4118 | 2 | 0 | 6 | 0.1667 | 4 | 0.75 | 2 | 0 | 14 | 0.2857 |
| 19_2969463 | 2 | 1 | 3 | 0.6667 | 3 | 1 | 5 | 1 | 13 | 0.9231 | 2 | 1 | 7 | 1 | 2 | 1 | 1 | 1 | 12 | 1 |
| 19_2969470 | 3 | 0.3333 | 3 | 0.3333 | 6 | 0.5 | 5 | 0.2 | 17 | 0.3529 | 2 | 0.5 | 4 | 0.5 | 3 | 0 | 2 | 0 | 11 | 0.2727 |
| 19_2969478 | 2 | 0.5 | 4 | 0.75 | 4 | 0.5 | 5 | 0.4 | 15 | 0.5333 | 1 | 0 | 5 | 0.8 | 3 | 1 | 2 | 0.5 | 11 | 0.7273 |
| 19_2969498 | 1 | 1 | 3 | 1 | 6 | 1 | 5 | 1 | 15 | 1 | 2 | 1 | 6 | 1 | 2 | 1 | 2 | 0.5 | 12 | 0.9167 |
| 19_2969513 | 1 | 1 | 1 | 1 | 5 | 0.4 | 4 | 0.75 | 11 | 0.6364 | 2 | 0.5 | 4 | 1 | 2 | 1 | 1 | 1 | 9 | 0.8889 |
| 19_2969529 | 1 | 0 | 2 | 0.5 | 4 | 0.5 | 4 | 0.5 | 11 | 0.4545 | 2 | 0.5 | 5 | 0.4 | 2 | 0.5 | 2 | 0.5 | 11 | 0.4545 |
| 19_2969584 | 1 | 0 | 1 | 0 | 6 | 0 | 3 | 0.6667 | 11 | 0.1818 | 1 | 0 | 5 | 0.2 | 2 | 0.5 | 2 | 0.5 | 10 | 0.3 |
| 19_2969586 | 1 | 1 | 1 | 1 | 5 | 0.8 | 4 | 1 | 11 | 0.9091 | 1 | 1 | 6 | 0.8333 | 2 | 1 | 2 | 1 | 11 | 0.9091 |
| 19_2969590 | 1 | 1 | 1 | 1 | 5 | 1 | 4 | 1 | 11 | 1 | NA | NA | 5 | 1 | 1 | 1 | 1 | 1 | 7 | 1 |
| 19_2969590 | 1 | 0 | 1 | 1 | 4 | 0.25 | 2 | 1 | 8 | 0.5 | 1 | 0 | 5 | 0.2 | 1 | 1 | 2 | 0.5 | 9 | 0.3333 |
| 19_2969627 | 1 | 0 | 1 | 1 | 5 | 0 | 4 | 0 | 11 | 0.0909 | 1 | 0 | 6 | 0.1667 | 2 | 0.5 | 2 | 0.5 | 11 | 0.2727 |
| 19_2969641 | 1 | 1 | 1 | 1 | 5 | 1 | 3 | 1 | 10 | 1 | 1 | 1 | 5 | 1 | 2 | 1 | 2 | 1 | 10 | 1 |
| 19_2969646 | NA | NA | 1 | 0 | 5 | 0 | 3 | 0.3333 | 9 | 0.1111 | NA | NA | 6 | 0.1667 | 2 | 0 | 2 | 0 | 10 | 0.1 |
| 19_2969651 | NA | NA | 1 | 1 | 4 | 1 | 3 | 1 | 8 | 1 | 1 | 1 | 6 | 1 | 2 | 1 | 2 | 1 | 11 | 1 |
| 19_2969663 | NA | NA | 1 | 1 | 5 | 0.2 | 3 | 0 | 9 | 0.2222 | 1 | 1 | 5 | 0.4 | 2 | 1 | 2 | 0 | 10 | 0.5 |
| 19_2969668 | NA | NA | 1 | 0 | 5 | 0.8 | 3 | 0.6667 | 9 | 0.6667 | 1 | 0 | 5 | 0.6 | 1 | 0 | 1 | 1 | 8 | 0.5 |
| 19_2969675 | NA | NA | NA | NA | 5 | 0.8 | 3 | 1 | 8 | 0.875 | 1 | 1 | 3 | 0 | 1 | 0 | 1 | 1 | 6 | 0.3333 |
| 19_2969680 | NA | NA | 1 | 0 | 4 | 0.5 | 3 | 0.3333 | 8 | 0.375 | 1 | 0 | 4 | 0.25 | 2 | 0.5 | NA | NA | 7 | 0.2857 |
| 19_2969692 | NA | NA | 1 | 0 | 5 | 0 | 2 | 0.5 | 8 | 0.125 | 1 | 0 | 5 | 0.2 | 2 | 0 | 1 | 0 | 9 | 0.1111 |
| 19_2973124 | NA | NA | 1 | 1 | 2 | 0 | 1 | 1 | 4 | 0.5 | NA | NA | 4 | 0 | 1 | 1 | 1 | 0 | 6 | 0.1667 |
| 19_2973146 | NA | NA | 1 | 0 | 2 | 0.5 | 2 | 1 | 5 | 0.6 | 1 | 1 | 4 | 1 | NA | NA | 1 | 1 | 6 | 1 |
| 19_2973155 | NA | NA | 1 | 1 | NA | NA | 1 | 1 | 2 | 1 | NA | NA | 4 | 1 | 1 | 1 | 1 | 1 | 6 | 1 |
| 19_2973190 | NA | NA | 1 | 0 | 1 | 1 | 1 | 1 | 3 | 0.6667 | 1 | 0 | 3 | 0.6667 | 1 | 1 | 1 | 0 | 6 | 0.5 |
| 19_2973191 | NA | NA | 1 | 0 | NA | NA | 1 | 1 | 2 | 0.5 | 1 | 1 | 3 | 1 | 1 | 1 | 1 | 1 | 6 | 1 |
| 19_2973203 | NA | NA | 1 | 0 | 1 | 1 | 1 | 0 | 3 | 0.3333 | 1 | 0 | 3 | 0.6667 | 1 | 1 | 1 | 0 | 6 | 0.5 |
| 19_2973205 | NA | NA | 1 | 1 | 1 | 1 | 1 | 0 | 3 | 0.6667 | 1 | 0 | 3 | 0.6667 | 1 | 1 | 1 | 0 | 6 | 0.5 |
| 19_2973205 | NA | NA | 1 | 0 | 1 | 1 | 1 | 1 | 3 | 0.6667 | 1 | 1 | 3 | 0.3333 | 1 | 0 | 1 | 1 | 6 | 0.5 |
| 19_2973217 | NA | NA | 1 | 0 | NA | NA | 1 | 1 | 2 | 0.5 | 1 | 0 | 3 | 0.6667 | 1 | 0 | 1 | 1 | 6 | 0.5 |

| | | | | | | | | | | | | | | | | | | | | |
|---|---|---|---|---|---|---|---|---|---|---|---|---|---|---|---|---|---|---|---|---|
| 19_297322 | NA | NA | 1 | 1 | NA | NA | 1 | 1 | 2 | 1 | 1 | 1 | 2 | 1 | 1 | 1 | NA | NA | 4 | 1 |
| 19_297322 | NA | NA | 1 | 0 | NA | NA | 1 | 1 | 2 | 0.5 | 1 | 0 | 3 | 0.3333 | 1 | 0 | 1 | 1 | 6 | 0.3333 |
| 19_297859 | 3 | 0 | 7 | 0 | 9 | 0.1111 | 7 | 0.4286 | 26 | 0.1538 | 2 | 0 | 8 | 0.125 | 10 | 0.2 | 4 | 0 | 24 | 0.125 |
| 19_297859 | 3 | 0.3333 | 6 | 0.5 | 9 | 0.1111 | 8 | 0.125 | 26 | 0.2308 | 2 | 0.5 | 7 | 0.1429 | 11 | 0.1818 | 4 | 0.25 | 24 | 0.2083 |
| 19_297863 | 2 | 0 | 6 | 0.3333 | 6 | 0.1667 | 8 | 0.25 | 22 | 0.2273 | 2 | 0.5 | 5 | 0 | 10 | 0 | 3 | 0 | 20 | 0.05 |
| 19_297864 | 1 | 0 | 6 | 0.3333 | 6 | 0 | 7 | 0.4286 | 20 | 0.25 | 1 | 1 | 6 | 0.3333 | 7 | 0.1429 | 2 | 0 | 16 | 0.25 |
| 19_297865 | 1 | 0 | 4 | 0 | 6 | 0 | 8 | 0.125 | 19 | 0.0526 | 1 | 0 | 6 | 0 | 7 | 0.1429 | 1 | 0 | 15 | 0.0667 |
| 19_297866 | 1 | 0 | 7 | 0 | 6 | 0.3333 | 7 | 0 | 21 | 0.0952 | 1 | 0 | 5 | 0 | 9 | 0.1111 | 3 | 0 | 18 | 0.0556 |
| 19_297867 | 1 | 0 | 6 | 0.3333 | 5 | 0.6 | 7 | 0.2857 | 19 | 0.3684 | 1 | 0 | 6 | 0.3333 | 8 | 0.375 | 3 | 0 | 18 | 0.2778 |
| 19_297868 | 1 | 0 | 5 | 0.2 | 3 | 0.3333 | 6 | 0.5 | 15 | 0.3333 | 1 | 1 | 3 | 0.6667 | 5 | 0.6 | 3 | 0 | 12 | 0.5 |
| 19_297871 | 1 | 1 | 4 | 1 | 1 | 1 | 6 | 1 | 12 | 1 | NA | NA | 4 | 0.75 | 6 | 0.8333 | 3 | 1 | 13 | 0.8462 |
| 19_297871 | 1 | 0 | 5 | 0.4 | 3 | 0.6667 | 6 | 0.8333 | 15 | 0.6 | 1 | 0 | 5 | 0.6 | 8 | 0.5 | 3 | 0.3333 | 17 | 0.4706 |
| 19_297872 | 1 | 1 | 3 | 0 | 3 | 0.3333 | 7 | 0.1429 | 14 | 0.2143 | NA | NA | 4 | 0 | 8 | 0.25 | 3 | 0 | 15 | 0.1333 |
| 19_297872 | 1 | 1 | 4 | 0.75 | 3 | 1 | 5 | 1 | 13 | 0.9231 | 1 | 1 | 5 | 1 | 7 | 0.8571 | 3 | 1 | 16 | 0.9375 |
| 19_297874 | NA | NA | 4 | 1 | 3 | 0.6667 | 4 | 1 | 11 | 0.9091 | 1 | 1 | 5 | 1 | 8 | 1 | 3 | 1 | 17 | 1 |
| 19_297875 | NA | NA | 2 | 1 | 3 | 1 | 2 | 1 | 7 | 1 | 1 | 1 | 5 | 1 | 7 | 1 | 3 | 1 | 16 | 1 |
| 19_297876 | NA | NA | 4 | 1 | 3 | 1 | 5 | 1 | 12 | 1 | 1 | 1 | 5 | 1 | 4 | 1 | 2 | 1 | 12 | 1 |
| 19_297876 | NA | NA | 4 | 0.25 | 3 | 0.6667 | 5 | 0.8 | 12 | 0.5833 | 1 | 1 | 4 | 0.75 | 6 | 0.6667 | 3 | 0.6667 | 14 | 0.7143 |
| 19_297879 | NA | NA | 3 | 0 | 4 | 0.25 | 4 | 0.25 | 11 | 0.1818 | NA | NA | 5 | 0.8 | 6 | 0 | 3 | 0.3333 | 14 | 0.3571 |
| 19_297879 | NA | NA | 1 | 0 | 4 | 0 | 4 | 0.25 | 9 | 0.1111 | 1 | 0 | 3 | 0.3333 | 6 | 0.3333 | 2 | 1 | 12 | 0.4167 |
| 19_297880 | NA | NA | 3 | 1 | 3 | 1 | 5 | 1 | 11 | 1 | 1 | 1 | 5 | 1 | 6 | 1 | 3 | 1 | 15 | 1 |
| 19_297883 | NA | NA | 1 | 1 | 3 | 1 | 5 | 1 | 9 | 1 | 1 | 1 | 3 | 0.6667 | 6 | 0.5 | 2 | 0.5 | 12 | 0.5833 |
| 19_297883 | NA | NA | 1 | 0 | 2 | 0.5 | 4 | 0.25 | 7 | 0.2857 | 1 | 0 | 4 | 0 | 6 | 0.1667 | 3 | 0 | 14 | 0.0714 |
| 19_297883 | NA | NA | 1 | 0 | 3 | 0.3333 | 5 | 0.2 | 9 | 0.2222 | 1 | 0 | 4 | 0 | 6 | 0.1667 | 3 | 0 | 14 | 0.0714 |
| 19_297884 | NA | NA | 1 | 0 | 2 | 0 | 2 | 1 | 5 | 0.4 | 1 | 0 | 2 | 0.5 | 6 | 0.3333 | 3 | 0.6667 | 12 | 0.4167 |
| 19_297884 | NA | NA | 1 | 1 | 2 | 1 | 5 | 1 | 8 | 1 | 1 | 1 | 4 | 1 | 6 | 0.6667 | 3 | 0.3333 | 14 | 0.7143 |
| 19_297885 | NA | NA | 1 | 0 | 3 | 0.3333 | 5 | 0.2 | 9 | 0.2222 | 1 | 0 | 4 | 0.25 | 5 | 0.2 | 3 | 0.3333 | 13 | 0.2308 |
| 19_297885 | NA | NA | 1 | 0 | 3 | 1 | 4 | 0.75 | 8 | 0.75 | 1 | 1 | 3 | 0.6667 | 5 | 0.8 | 3 | 0 | 12 | 0.5833 |
| 19_297886 | NA | NA | 1 | 1 | 3 | 1 | 4 | 1 | 8 | 1 | 1 | 1 | 4 | 1 | 6 | 1 | 3 | 1 | 14 | 1 |
| 19_297886 | NA | NA | 1 | 0 | 3 | 0 | 4 | 0.25 | 8 | 0.125 | 1 | 0 | 4 | 0 | 5 | 0 | 1 | 1 | 11 | 0.0909 |
| 19_297886 | NA | NA | 1 | 0 | 1 | 1 | 5 | 0.6 | 7 | 0.5714 | 1 | 1 | 4 | 0.5 | 6 | 0 | 3 | 0.6667 | 14 | 0.3571 |
| 19_297888 | NA | NA | 1 | 0 | 3 | 0 | 5 | 0.4 | 9 | 0.2222 | 1 | 0 | 3 | 0.3333 | 6 | 0.1667 | 3 | 0 | 13 | 0.1538 |
| 19_297889 | NA | NA | NA | NA | 1 | 0 | 4 | 0.5 | 5 | 0.4 | NA | NA | 4 | 0.5 | 5 | 0 | 2 | 1 | 11 | 0.3636 |
| 19_297889 | NA | NA | 1 | 0 | 3 | 0.3333 | 4 | 0 | 8 | 0.125 | 1 | 0 | 3 | 0 | 6 | 0.5 | 3 | 0 | 13 | 0.2308 |
| 19_297890 | NA | NA | 1 | 1 | 3 | 0.3333 | 5 | 0.6 | 9 | 0.5556 | NA | NA | 3 | 1 | 4 | 0.75 | 3 | 0.6667 | 10 | 0.8 |
| 19_297891 | NA | NA | 1 | 1 | 2 | 0 | 4 | 0 | 7 | 0.1429 | NA | NA | 4 | 0.25 | 6 | 0 | 3 | 0 | 13 | 0.0769 |
| 19_297891 | NA | NA | 1 | 0 | 3 | 0 | 5 | 0.4 | 9 | 0.2222 | NA | NA | 2 | 0.5 | 5 | 0 | 3 | 0 | 10 | 0.1 |
| 19_297891 | NA | NA | 1 | 1 | 3 | 0 | 5 | 0 | 9 | 0.1111 | 1 | 0 | 4 | 0 | 6 | 0.1667 | 1 | 0 | 12 | 0.0833 |
| 19_297892 | NA | NA | 1 | 0 | 3 | 0.3333 | 5 | 0.2 | 9 | 0.2222 | 1 | 0 | 2 | 0 | 6 | 0.3333 | 2 | 0 | 11 | 0.1818 |
| 19_297892 | NA | NA | 1 | 1 | 3 | 1 | 5 | 1 | 9 | 1 | 1 | 1 | 4 | 1 | 6 | 1 | 2 | 1 | 13 | 1 |
| 19_297892 | NA | NA | 1 | 1 | 3 | 1 | 4 | 0.75 | 8 | 0.875 | 1 | 0 | 3 | 0.6667 | 5 | 0.8 | 3 | 0.3333 | 12 | 0.5833 |
| 19_297893 | NA | NA | 1 | 0 | 3 | 0 | 5 | 0.2 | 9 | 0.1111 | 1 | 1 | 3 | 0 | 5 | 0 | 3 | 0 | 12 | 0.0833 |
| 19_297894 | NA | NA | 1 | 1 | 2 | 0.5 | 5 | 0 | 8 | 0.25 | 1 | 0 | 4 | 0.25 | 4 | 0.25 | 3 | 0.3333 | 12 | 0.25 |
| 19_297895 | NA | NA | 1 | 0 | 2 | 0 | 5 | 0.4 | 8 | 0.25 | 1 | 0 | 4 | 0 | 4 | 0.75 | 3 | 0.3333 | 12 | 0.3333 |
| 19_297896 | NA | NA | 1 | 1 | 1 | 1 | 4 | 1 | 6 | 1 | 1 | 1 | 4 | 1 | 3 | 1 | 3 | 1 | 11 | 1 |
| 19_297897 | NA | NA | 1 | 1 | 2 | 0.5 | 4 | 1 | 7 | 0.8571 | NA | NA | 4 | 0.5 | 4 | 1 | 3 | 1 | 11 | 0.8182 |
| 19_297900 | NA | NA | NA | NA | 2 | 1 | 4 | 1 | 6 | 1 | 1 | 1 | 4 | 0.75 | 4 | 1 | 2 | 1 | 11 | 0.9091 |
| 19_299363 | 1 | 1 | 8 | 0.25 | 8 | 0.5 | 7 | 0.1429 | 24 | 0.3333 | 3 | 0.3333 | 5 | 0.2 | 5 | 0.8 | 3 | 0.6667 | 16 | 0.5 |
| 19_299364 | 1 | 0 | 6 | 0.3333 | 8 | 0.25 | 6 | 0 | 21 | 0.1905 | 3 | 0.6667 | 6 | 0.1667 | 6 | 0.5 | 2 | 1 | 17 | 0.4706 |
| 19_299365 | 1 | 0 | 7 | 0.1429 | 7 | 0 | 6 | 0.1667 | 21 | 0.0952 | 4 | 0.25 | 5 | 0 | 6 | 0.1667 | 2 | 0 | 17 | 0.1176 |
| 19_299366 | 1 | 1 | 7 | 0.5714 | 8 | 0.625 | 7 | 0.4286 | 23 | 0.5652 | 3 | 1 | 5 | 0.8 | 6 | 0.8333 | 3 | 0.3333 | 17 | 0.7647 |
| 19_299367 | 1 | 0 | 7 | 0.2857 | 9 | 0.8889 | 6 | 0.5 | 23 | 0.5652 | 4 | 1 | 6 | 0.5 | 4 | 0.5 | 3 | 0.3333 | 17 | 0.5882 |
| 19_300165 | NA | NA | 1 | 0 | 1 | 1 | NA | NA | 2 | 0.5 | 1 | 1 | 4 | 0 | 3 | 0 | 3 | 0 | 11 | 0.0909 |
| 19_300337 | 2 | 0.5 | 3 | 1 | 1 | 0 | NA | NA | 6 | 0.6667 | 1 | 1 | 3 | 0.6667 | 5 | 0.4 | 3 | 1 | 12 | 0.6667 |
| 19_300337 | 2 | 0.5 | 4 | 1 | NA | NA | 1 | 0 | 7 | 0.7143 | NA | NA | 4 | 0.5 | 5 | 0.8 | 2 | 1 | 11 | 0.7273 |
| 19_300338 | 1 | 0 | 3 | 1 | 1 | 0 | 1 | 0 | 6 | 0.5 | 1 | 0 | 3 | 1 | 4 | 0.25 | 3 | 0.3333 | 11 | 0.4545 |
| 19_300703 | 7 | 0.4286 | 3 | 0 | 2 | 0 | 2 | 0.5 | 14 | 0.2857 | 6 | 0.3333 | 4 | 0 | 6 | 0 | 5 | 0.6 | 21 | 0.2381 |
| 19_300703 | 6 | 0.3333 | 6 | 0 | 2 | 0 | 4 | 0.25 | 18 | 0.1667 | 6 | 0 | 5 | 0 | 8 | 0.125 | 4 | 0.25 | 23 | 0.087 |
| 19_300759 | 3 | 0.3333 | 3 | 0.3333 | 3 | 0.3333 | 3 | 0.3333 | 12 | 0.3333 | 6 | 0.3333 | 3 | 0 | 8 | 0 | 8 | 0 | 25 | 0.08 |
| 19_300760 | 2 | 0 | 3 | 0.3333 | 2 | 0 | 1 | 0 | 8 | 0.125 | 6 | 0.1667 | 2 | 0.5 | 7 | 0.4286 | 7 | 0.1429 | 22 | 0.2727 |
| 19_305422 | NA | NA | NA | NA | NA | NA | 1 | 1 | 1 | 1 | 1 | 0 | 3 | 0.6667 | 2 | 1 | 1 | 1 | 7 | 0.7143 |
| 19_305464 | NA | NA | NA | NA | NA | NA | 1 | 1 | 1 | 1 | NA | NA | NA | NA | 2 | 0.5 | 1 | 0 | 3 | 0.3333 |
| 19_305466 | NA | NA | NA | NA | NA | NA | 1 | 1 | 1 | 1 | NA | NA | NA | NA | 2 | 0.5 | 1 | 1 | 3 | 0.6667 |
| 19_310605 | 6 | 0.5 | 1 | 0 | 3 | 0.3333 | 1 | 1 | 11 | 0.4545 | 1 | 1 | 1 | 1 | 1 | 1 | 5 | 0.8 | 8 | 0.875 |
| 19_310606 | 7 | 0.4286 | 1 | 0 | 3 | 0 | 1 | 1 | 12 | 0.3333 | NA | NA | 1 | 0 | 1 | 1 | 5 | 0.4 | 7 | 0.4286 |

| | | | | | | | | | | | | | | | | | | | | |
|---|---|---|---|---|---|---|---|---|---|---|---|---|---|---|---|---|---|---|---|---|
| 19_310609 | 6 | 0.1667 | 1 | 0 | 2 | 0 | NA | NA | 9 | 0.1111 | 1 | 0 | 1 | 0 | 1 | 0 | 1 | 1 | 4 | 0.25 |
| 19_310611 | 8 | 0.375 | 2 | 0 | 4 | 0.75 | 2 | 0.5 | 16 | 0.4375 | 2 | 1 | 1 | 0 | 3 | 0 | 5 | 0.2 | 11 | 0.2727 |
| 19_310612 | 9 | 1 | 2 | 0.5 | 5 | 0.8 | 2 | 1 | 18 | 0.8889 | 2 | 1 | 1 | 1 | 2 | 1 | 5 | 0.8 | 10 | 0.9 |
| 19_310613 | 9 | 0.5556 | 2 | 0.5 | 2 | 0 | 2 | 0.5 | 15 | 0.4667 | 1 | 1 | 1 | 0 | 4 | 0.25 | 4 | 0.5 | 10 | 0.4 |
| 19_310613 | 7 | 0.7143 | 2 | 0.5 | 3 | 0.3333 | 2 | 0 | 14 | 0.5 | 2 | 1 | 1 | 0 | 2 | 1 | 4 | 0.25 | 9 | 0.5556 |
| 19_310614 | 9 | 0.7778 | 2 | 1 | 4 | 0.5 | 2 | 0.5 | 17 | 0.7059 | 1 | 1 | 1 | 1 | 4 | 1 | 4 | 0.5 | 10 | 0.8 |
| 19_310614 | 9 | 0.5556 | 3 | 0 | 5 | 0 | 2 | 0 | 19 | 0.2632 | 2 | 0 | 1 | 0 | 5 | 0.2 | 5 | 0.2 | 13 | 0.1538 |
| 19_310614 | 10 | 0.9 | 3 | 0.6667 | 2 | 0.5 | 1 | 1 | 16 | 0.8125 | 1 | 1 | 1 | 1 | 2 | 1 | 4 | 0.75 | 8 | 0.875 |
| 19_310615 | 10 | 1 | 3 | 1 | 3 | 0.6667 | 1 | 1 | 17 | 0.9412 | 1 | 1 | 1 | 1 | 4 | 1 | 5 | 1 | 11 | 1 |
| 19_310615 | 8 | 0.875 | 3 | 0.6667 | 1 | 1 | NA | NA | 12 | 0.8333 | 1 | 1 | NA | NA | 2 | 1 | 5 | 1 | 8 | 1 |
| 19_310615 | 8 | 0.5 | 2 | 0 | 3 | 0 | NA | NA | 13 | 0.3077 | 1 | 1 | NA | NA | 3 | 0.6667 | 3 | 0.3333 | 7 | 0.5714 |
| 19_310616 | 8 | 0.25 | 3 | 0.3333 | 5 | 0.2 | 1 | 1 | 17 | 0.2941 | 2 | 1 | 1 | 1 | 5 | 0.4 | 5 | 0.2 | 13 | 0.4615 |
| 19_310617 | 9 | 0.1111 | 3 | 0 | 6 | 0.1667 | 2 | 0.5 | 20 | 0.15 | NA | NA | 2 | 0 | 4 | 0.75 | 5 | 0.4 | 11 | 0.4545 |
| 19_310618 | 9 | 0.8889 | 3 | 1 | 6 | 1 | 1 | 1 | 19 | 0.9474 | 1 | 1 | 2 | 1 | 3 | 1 | 6 | 1 | 12 | 1 |
| 19_310618 | 10 | 0.8 | 1 | 1 | 7 | 0.4286 | 2 | 1 | 20 | 0.7 | 2 | 0 | 2 | 1 | 4 | 1 | 4 | 0.25 | 12 | 0.5833 |
| 19_310619 | 9 | 0.3333 | 4 | 0 | 7 | 0.2857 | 3 | 0 | 23 | 0.2174 | 2 | 0.5 | 2 | 0 | 5 | 0.4 | 6 | 0.1667 | 15 | 0.2667 |
| 19_310622 | 10 | 0.2 | 4 | 0.5 | 6 | 0 | 5 | 0 | 25 | 0.16 | 5 | 0.2 | 6 | 0 | 8 | 0 | 9 | 0.2222 | 28 | 0.1071 |
| 19_310627 | 14 | 0.5 | 4 | 0 | 10 | 0.2 | 6 | 0.5 | 34 | 0.3529 | 4 | 0.25 | 7 | 0.1429 | 8 | 0.5 | 12 | 0.4167 | 31 | 0.3548 |
| 19_310631 | 17 | 0.1176 | 10 | 0.1 | 11 | 0 | 7 | 0.4286 | 45 | 0.1333 | 5 | 0.2 | 10 | 0.1 | 7 | 0.2857 | 11 | 0 | 33 | 0.1212 |
| 19_321015 | 6 | 0.5 | 6 | 0.5 | 4 | 0.5 | 10 | 0.3 | 26 | 0.4231 | 7 | 0 | 6 | 0.5 | 4 | 0.25 | 4 | 0.25 | 21 | 0.2381 |
| 19_321015 | 7 | 0.4286 | 6 | 0.3333 | 6 | 0.3333 | 12 | 0.25 | 31 | 0.3226 | 6 | 0.5 | 4 | 0 | 5 | 0 | 2 | 0.5 | 17 | 0.2353 |
| 19_321015 | 5 | 0 | 7 | 0.2857 | 6 | 0.1667 | 12 | 0.3333 | 30 | 0.2333 | 7 | 0.4286 | 4 | 0 | 6 | 0.3333 | 3 | 0.3333 | 20 | 0.3 |
| 19_321016 | 5 | 0.4 | 6 | 0.5 | 6 | 0.8333 | 11 | 0.6364 | 28 | 0.6071 | 6 | 0.6667 | 6 | 0.5 | 6 | 0.3333 | 3 | 0.3333 | 21 | 0.4762 |
| 19_321016 | 6 | 0.3333 | 8 | 0.125 | 5 | 0.4 | 10 | 0.3 | 29 | 0.2759 | 6 | 0 | 6 | 0.1667 | 5 | 0.4 | 4 | 0.25 | 21 | 0.1905 |
| 19_321017 | 3 | 0.6667 | 3 | 0 | 1 | 0 | 5 | 0 | 12 | 0.1667 | NA | NA | 4 | 0.25 | 3 | 0.3333 | 1 | 0 | 8 | 0.25 |
| 19_321369 | 5 | 0.6 | 8 | 0.5 | 6 | 0.3333 | 8 | 0.625 | 27 | 0.5185 | 5 | 0.8 | 6 | 0.6667 | 3 | 0 | 3 | 0.3333 | 17 | 0.5294 |
| 19_321370 | 3 | 1 | 7 | 0.4286 | 4 | 0.5 | 6 | 0.6667 | 20 | 0.6 | 3 | 0.3333 | 6 | 0.8333 | 4 | 0.75 | 3 | 0.6667 | 16 | 0.6875 |
| 19_321371 | 4 | 0.75 | 8 | 0.75 | 6 | 0.6667 | 6 | 0.8333 | 24 | 0.75 | 4 | 0.75 | 6 | 0.5 | 4 | 0.75 | 4 | 0.25 | 18 | 0.5556 |
| 19_321372 | 4 | 0.25 | 7 | 0.1429 | 6 | 0 | 7 | 0.2857 | 24 | 0.1667 | 3 | 0 | 6 | 0 | 4 | 0.25 | 4 | 0.25 | 17 | 0.1176 |
| 19_321372 | 4 | 0.5 | 7 | 0.4286 | 5 | 0.4 | 7 | 0.4286 | 23 | 0.4348 | 3 | 0.3333 | 6 | 0.5 | 5 | 0.6 | 2 | 0 | 16 | 0.4375 |
| 19_321373 | 4 | 0 | 7 | 0.4286 | 6 | 0.5 | 6 | 0.1667 | 23 | 0.3043 | 3 | 0 | 6 | 0 | 5 | 0.2 | 4 | 0 | 18 | 0.0556 |
| 19_321374 | 4 | 0.75 | 8 | 0.875 | 6 | 0.8333 | 7 | 0.8571 | 25 | 0.84 | 3 | 1 | 5 | 0.8 | 5 | 1 | 2 | 1 | 15 | 0.9333 |
| 19_321375 | 3 | 0 | 6 | 0.1667 | 4 | 0.25 | 5 | 0.4 | 18 | 0.2222 | 2 | 0 | 6 | 0.3333 | 4 | 0.25 | 4 | 0.25 | 16 | 0.25 |
| 19_321375 | 3 | 0 | 7 | 0.2857 | 5 | 0.4 | 6 | 0.3333 | 21 | 0.2857 | 3 | 0 | 6 | 0.3333 | 5 | 0.4 | 3 | 0.3333 | 17 | 0.2941 |
| 19_321375 | 2 | 0.5 | 6 | 0.8333 | 3 | 0.3333 | 7 | 0.2857 | 18 | 0.5 | 1 | 1 | 5 | 0.6 | 3 | 0.3333 | 4 | 1 | 13 | 0.6923 |
| 19_321376 | 3 | 1 | 2 | 0.5 | 3 | 1 | 4 | 0.5 | 12 | 0.75 | 3 | 0.6667 | 5 | 0.6 | 3 | 0.3333 | 3 | 0.6667 | 14 | 0.5714 |
| 19_321376 | NA | NA | 1 | 0 | 2 | 0 | 3 | 0.3333 | 6 | 0.1667 | 3 | 0 | 2 | 0 | 1 | 1 | 1 | 0 | 7 | 0.1429 |
| 19_323672 | 2 | 0.5 | 2 | 0.5 | NA | NA | 1 | 1 | 5 | 0.6 | NA | NA | NA | NA | NA | NA | 1 | 1 | 1 | 1 |
| 19_323674 | 2 | 0 | 2 | 1 | NA | NA | 1 | 0 | 5 | 0.4 | NA | NA | NA | NA | 1 | 0 | 1 | 1 | 2 | 0.5 |
| 19_324637 | 6 | 0 | 6 | 0.1667 | 6 | 0.1667 | 3 | 0 | 21 | 0.0952 | 3 | 0 | 4 | 0.25 | 1 | 0 | 3 | 0.6667 | 11 | 0.2727 |
| 19_324638 | 5 | 0 | 4 | 0 | 5 | 0.2 | 1 | 0 | 15 | 0.0667 | 3 | 0 | 2 | 0.5 | 3 | 0 | 5 | 0 | 13 | 0.0769 |
| 19_324639 | 6 | 0 | 4 | 0 | 7 | 0.2857 | 2 | 0 | 19 | 0.1053 | 4 | 0 | 4 | 0.25 | 2 | 0 | 4 | 0 | 14 | 0.0714 |
| 19_324639 | NA | NA | NA | NA | 1 | 1 | NA | NA | 1 | 1 | NA | NA | 1 | 1 | NA | NA | NA | NA | 1 | 1 |
| 19_324640 | NA | NA | NA | NA | 1 | 1 | NA | NA | 1 | 1 | NA | NA | 1 | 1 | NA | NA | NA | NA | 1 | 1 |
| 19_324914 | 5 | 0.2 | 6 | 0.1667 | 6 | 0 | 2 | 0.5 | 19 | 0.1579 | 4 | 0.25 | 3 | 0 | 5 | 0 | 5 | 0 | 17 | 0.0588 |
| 19_324915 | 9 | 0 | 9 | 0.2222 | 10 | 0.1 | 4 | 0 | 32 | 0.0938 | 7 | 0.2857 | 5 | 0 | 5 | 0 | 4 | 0 | 21 | 0.0952 |
| 19_324916 | 11 | 0.0909 | 8 | 0 | 8 | 0.125 | 3 | 0 | 30 | 0.0667 | 9 | 0.1111 | 6 | 0.1667 | 7 | 0.2857 | 5 | 0.2 | 27 | 0.1852 |
| 19_324916 | 9 | 1 | 6 | 0.8333 | 9 | 0.8889 | 3 | 0.6667 | 27 | 0.8889 | 7 | 0.8571 | 6 | 1 | 6 | 1 | 5 | 1 | 24 | 0.9583 |
| 19_324917 | 12 | 0.75 | 8 | 0.75 | 11 | 0.5455 | 5 | 0.6 | 36 | 0.6667 | 9 | 0.6667 | 6 | 1 | 7 | 0.8571 | 3 | 1 | 25 | 0.84 |
| 19_324919 | 11 | 0.8182 | 7 | 0.8571 | 10 | 0.7 | 5 | 0.8 | 33 | 0.7879 | 9 | 0.7778 | 6 | 1 | 7 | 0.7143 | 4 | 0.5 | 26 | 0.7692 |
| 19_324919 | 12 | 0.6667 | 10 | 0.4 | 8 | 0.375 | 4 | 0.5 | 34 | 0.5 | 9 | 0.2222 | 6 | 0.6667 | 6 | 0.1667 | 5 | 0.4 | 26 | 0.3462 |
| 19_324919 | 11 | 0.0909 | 9 | 0.1111 | 10 | 0.3 | 5 | 0.6 | 35 | 0.2286 | 7 | 0.2857 | 6 | 0.1667 | 7 | 0.1429 | 5 | 0.2 | 25 | 0.2 |
| 19_324919 | 12 | 0.3333 | 8 | 0.75 | 8 | 0.25 | 5 | 0.4 | 33 | 0.4242 | 3 | 1 | 5 | 0.4 | 6 | 0.3333 | 4 | 0.25 | 18 | 0.4444 |
| 19_324920 | 11 | 0.0909 | 9 | 0 | 7 | 0.2857 | 3 | 0 | 30 | 0.1 | 5 | 0.2 | 5 | 0 | 6 | 0.1667 | 5 | 0.2 | 21 | 0.1429 |
| 19_324920 | 9 | 0.7778 | 6 | 0.8333 | 9 | 0.6667 | 5 | 0.2 | 29 | 0.6552 | 6 | 0.8333 | 6 | 0.8333 | 4 | 0.75 | 5 | 0.8 | 21 | 0.8095 |
| 19_324921 | 10 | 0.7 | 10 | 0.8 | 13 | 0.3846 | 3 | 0.3333 | 36 | 0.5833 | 9 | 0.4444 | 8 | 0.75 | 6 | 0.8333 | 4 | 0.5 | 27 | 0.6296 |
| 19_324921 | 10 | 1 | 8 | 1 | 11 | 0.9091 | 3 | 0.6667 | 32 | 0.9375 | 7 | 1 | 6 | 1 | 6 | 1 | 5 | 1 | 24 | 1 |
| 19_324922 | 14 | 0.4286 | 9 | 0.4444 | 14 | 0.5 | 4 | 0.5 | 41 | 0.4634 | 9 | 0.5556 | 8 | 0 | 7 | 0.4286 | 5 | 0.6 | 29 | 0.3793 |
| 19_324922 | 13 | 1 | 10 | 0.8 | 10 | 0.7 | 5 | 0.8 | 38 | 0.8421 | 8 | 0.875 | 6 | 0.8333 | 5 | 0.8 | 4 | 0.75 | 23 | 0.8261 |
| 19_324922 | 13 | 0.9231 | 9 | 0.4444 | 13 | 0.7692 | 6 | 1 | 41 | 0.7805 | 8 | 0.625 | 6 | 1 | 6 | 0.8333 | 5 | 0.8 | 25 | 0.8 |
| 19_324923 | 11 | 0.1818 | 10 | 0.4 | 11 | 0.2727 | 6 | 0.1667 | 38 | 0.2632 | 8 | 0.25 | 9 | 0.4444 | 5 | 0.2 | 5 | 0.2 | 27 | 0.2963 |
| 19_324924 | 16 | 0.4375 | 10 | 0.2 | 12 | 0.1667 | 5 | 0.4 | 43 | 0.3023 | 8 | 0.25 | 9 | 0.4444 | 8 | 0 | 5 | 0.4 | 30 | 0.2667 |
| 19_324924 | 12 | 0.5833 | 9 | 0.7778 | 7 | 0.8571 | 3 | 1 | 31 | 0.7419 | 7 | 0.4286 | 6 | 0.6667 | 6 | 0.5 | 4 | 0.5 | 23 | 0.5217 |
| 19_324926 | 15 | 0.2667 | 8 | 0.375 | 11 | 0.1818 | 4 | 0 | 38 | 0.2368 | 7 | 0.4286 | 9 | 0 | 7 | 0.4286 | 3 | 0 | 26 | 0.2308 |
| 19_324927 | 8 | 0.125 | 4 | 0 | 4 | 0 | 3 | 0 | 19 | 0.0526 | 7 | 0 | 7 | 0.2857 | 3 | 0 | 2 | 0 | 19 | 0.1053 |

| | | | | | | | | | | | | | | | | | | | | |
|---|---|---|---|---|---|---|---|---|---|---|---|---|---|---|---|---|---|---|---|---|
| 19_326430( | 2 | 0 | 4 | 0 | 1 | 0 | 6 | 0.1667 | 13 | 0.0769 | NA | NA | 3 | 0 | 5 | 0 | 2 | 0.5 | 10 | 0.1 |
| 19_326431! | 2 | 0 | 4 | 0.5 | 2 | 0 | 6 | 0.1667 | 14 | 0.2143 | 1 | 1 | 3 | 0.3333 | 5 | 0 | 1 | 0 | 10 | 0.2 |
| 19_326434! | 1 | 0 | 4 | 0.25 | 3 | 0 | 8 | 0.5 | 16 | 0.3125 | 2 | 0 | 3 | 0.6667 | 5 | 0.4 | 2 | 0 | 12 | 0.3333 |
| 19_326435! | 2 | 0.5 | 3 | 0 | 3 | 0 | 8 | 0.125 | 16 | 0.125 | 2 | 0 | 3 | 0.6667 | 5 | 0 | 1 | 0 | 11 | 0.1818 |
| 19_327354! | 8 | 0.75 | 11 | 0.5455 | 11 | 0.5455 | 3 | 0.6667 | 33 | 0.6061 | 3 | 0.3333 | 8 | 0.5 | 6 | 0.6667 | 7 | 0.4286 | 24 | 0.5 |
| 19_327359' | 15 | 0.2 | 13 | 0.2308 | 12 | 0.4167 | 9 | 0.4444 | 49 | 0.3061 | 4 | 0.25 | 11 | 0.0909 | 9 | 0.3333 | 9 | 0.1111 | 33 | 0.1818 |
| 19_327773! | 16 | 0.75 | 13 | 0.3077 | 12 | 0.4167 | 9 | 0.5556 | 50 | 0.52 | 3 | 0.6667 | 13 | 0.7692 | 9 | 0.5556 | 9 | 0.7778 | 34 | 0.7059 |
| 19_327973: | 20 | 0.2 | 15 | 0.2 | 14 | 0.2143 | 10 | 0.1 | 59 | 0.1864 | 6 | 0.1667 | 17 | 0.2353 | 11 | 0.2727 | 18 | 0.2778 | 52 | 0.25 |
| 19_327973: | 19 | 0.2105 | 17 | 0.1176 | 14 | 0.2857 | 10 | 0.2 | 60 | 0.2 | 5 | 0.2 | 14 | 0.2857 | 11 | 0.1818 | 16 | 0.25 | 46 | 0.2391 |
| 19_327973( | 17 | 0.5294 | 15 | 0.6 | 13 | 0.6154 | 8 | 0.875 | 53 | 0.6226 | 6 | 0.5 | 16 | 0.5625 | 10 | 0.7 | 13 | 0.4615 | 45 | 0.5556 |
| 19_327973! | 18 | 0.0556 | 16 | 0.25 | 12 | 0.0833 | 8 | 0 | 54 | 0.1111 | 6 | 0.3333 | 16 | 0.0625 | 10 | 0.2 | 15 | 0 | 47 | 0.1064 |
| 19_327974: | 16 | 0.3125 | 16 | 0.375 | 13 | 0 | 10 | 0.3 | 55 | 0.2545 | 6 | 0 | 14 | 0.2143 | 10 | 0.1 | 14 | 0.2857 | 44 | 0.1818 |
| 19_327974! | 14 | 0.5 | 14 | 0.4286 | 14 | 0.4286 | 10 | 0.4 | 52 | 0.4423 | 4 | 0.5 | 14 | 0.5 | 11 | 0.3636 | 13 | 0.5385 | 42 | 0.4762 |
| 19_327974! | 14 | 0.8571 | 18 | 0.8889 | 11 | 1 | 8 | 0.75 | 51 | 0.8824 | 5 | 0.6 | 15 | 0.8 | 12 | 0.8333 | 15 | 0.8667 | 47 | 0.8085 |
| 19_327975: | 17 | 0.2353 | 19 | 0.4211 | 14 | 0.5714 | 10 | 0.4 | 60 | 0.4 | 6 | 0.1667 | 14 | 0.5714 | 11 | 0.2727 | 17 | 0.4118 | 48 | 0.3958 |
| 19_327976! | 17 | 0.4706 | 20 | 0.8 | 13 | 0.6154 | 8 | 0.625 | 58 | 0.6379 | 4 | 0.25 | 18 | 0.7222 | 10 | 0.7 | 18 | 0.7778 | 50 | 0.7 |
| 19_327980! | 12 | 0.0833 | 14 | 0 | 7 | 0.1429 | 6 | 0 | 39 | 0.0513 | 2 | 0.5 | 10 | 0 | 10 | 0.1 | 10 | 0.1 | 32 | 0.0938 |
| 19_328000' | 17 | 0.1176 | 28 | 0.1071 | 20 | 0.1 | 17 | 0 | 82 | 0.0854 | 9 | 0 | 23 | 0.087 | 18 | 0 | 25 | 0.08 | 75 | 0.0533 |
| 19_328001: | 15 | 0.2667 | 22 | 0.3636 | 18 | 0.4444 | 16 | 0.3125 | 71 | 0.3521 | 8 | 0.125 | 21 | 0.3333 | 17 | 0.2941 | 25 | 0.52 | 71 | 0.3662 |
| 19_328018 | 3 | 0 | 7 | 0.1429 | 3 | 0 | 4 | 0.25 | 17 | 0.1176 | 2 | 0.5 | 7 | 0.1429 | 3 | 0 | 2 | 0 | 14 | 0.1429 |
| 19_333698: | 1 | 1 | 1 | 1 | NA | NA | NA | NA | 2 | 1 | NA | NA | NA | NA | NA | NA | 1 | 1 | 1 | 1 |
| 19_333698' | NA | NA | 1 | 1 | NA | NA | NA | NA | 1 | 1 | NA | NA | NA | NA | NA | NA | 1 | 1 | 1 | 1 |
| 19_333699' | 1 | 1 | 1 | 0 | NA | NA | NA | NA | 2 | 0.5 | NA | NA | NA | NA | NA | NA | 1 | 1 | 1 | 1 |
| 19_342175 | 2 | 0 | 3 | 0 | 3 | 0 | 1 | 1 | 9 | 0.1111 | 1 | 0 | 2 | 0.5 | 2 | 0.5 | 2 | 0 | 7 | 0.2857 |
| 19_344713' | NA | NA | 1 | 1 | 4 | 0.75 | 2 | 1 | 7 | 0.8571 | 3 | 1 | NA | NA | 1 | 1 | 4 | 1 | 8 | 1 |
| 19_344714: | NA | NA | 2 | 0.5 | 4 | 0.75 | 2 | 0.5 | 8 | 0.625 | 2 | 0.5 | 1 | 0 | 1 | 0 | 5 | 0.6 | 9 | 0.4444 |
| 19_345645! | 1 | 0 | 3 | 1 | NA | NA | NA | NA | 4 | 0.75 | 2 | 0.5 | 1 | 0 | 2 | 0.5 | NA | NA | 5 | 0.4 |
| 19_345646' | 2 | 0 | 3 | 0.6667 | NA | NA | NA | NA | 5 | 0.4 | 1 | 0 | 1 | 1 | 2 | 0.5 | NA | NA | 4 | 0.5 |
| 19_345648( | 1 | 0 | 3 | 0.3333 | NA | NA | NA | NA | 4 | 0.25 | 2 | 0.5 | 1 | 0 | 2 | 0 | NA | NA | 5 | 0.2 |
| 19_345653! | 2 | 0.5 | 2 | 0.5 | NA | NA | NA | NA | 4 | 0.5 | 3 | 0.3333 | 1 | 1 | 2 | 1 | NA | NA | 6 | 0.6667 |
| 19_345658( | 2 | 0 | 4 | 0.5 | NA | NA | NA | NA | 6 | 0.3333 | 3 | 0.3333 | 1 | 1 | 2 | 1 | NA | NA | 6 | 0.6667 |
| 19_345658' | 2 | 0.5 | 4 | 0.75 | NA | NA | NA | NA | 6 | 0.6667 | 3 | 0 | 1 | 1 | 2 | 1 | NA | NA | 6 | 0.5 |
| 19_345659 | 2 | 0 | 3 | 0.3333 | NA | NA | NA | NA | 5 | 0.2 | 2 | 0.5 | 1 | 1 | 2 | 1 | NA | NA | 5 | 0.8 |
| 19_345659: | 2 | 0.5 | 4 | 0.75 | NA | NA | NA | NA | 6 | 0.6667 | 3 | 1 | 1 | 1 | 2 | 1 | NA | NA | 6 | 1 |
| 19_345889! | 1 | 1 | NA | NA | 1 | 1 | NA | NA | 2 | 1 | NA | NA | NA | NA | NA | NA | 1 | 1 | 1 | 1 |
| 19_345897! | 1 | 1 | 1 | 0 | 1 | 1 | NA | NA | 3 | 0.6667 | NA | NA | NA | NA | NA | NA | 1 | 1 | 1 | 1 |
| 19_345897! | 1 | 0 | NA | NA | 1 | 1 | NA | NA | 2 | 0.5 | NA | NA | NA | NA | NA | NA | 1 | 1 | 1 | 1 |
| 19_345901: | 1 | 0 | 1 | 0 | 1 | 1 | NA | NA | 3 | 0.3333 | 1 | 1 | NA | NA | NA | NA | 2 | 0 | 3 | 0.3333 |
| 19_345901! | 2 | 0.5 | NA | NA | 1 | 0 | NA | NA | 3 | 0.3333 | 1 | 1 | NA | NA | NA | NA | 2 | 1 | 3 | 1 |
| 19_345902( | 1 | 0 | 1 | 1 | 1 | 0 | NA | NA | 3 | 0.3333 | 1 | 0 | NA | NA | NA | NA | 2 | 1 | 3 | 0.6667 |
| 19_345902( | 2 | 1 | 1 | 1 | 1 | 1 | NA | NA | 4 | 1 | 1 | 1 | NA | NA | NA | NA | 2 | 1 | 3 | 1 |
| 19_345903 | 1 | 1 | NA | NA | 1 | 1 | NA | NA | 2 | 1 | 1 | 1 | NA | NA | NA | NA | 1 | 1 | 2 | 1 |
| 19_347844! | NA | NA | 5 | 0.2 | 2 | 0 | 2 | 0 | 9 | 0.1111 | NA | NA | 3 | 0.3333 | 1 | 0 | 1 | 0 | 5 | 0.2 |
| 19_347847! | NA | NA | 3 | 0.6667 | 1 | 0 | NA | NA | 4 | 0.5 | NA | NA | 3 | 0 | 1 | 0 | 2 | 0.5 | 6 | 0.1667 |
| 19_347854: | NA | NA | 2 | 0.5 | NA | NA | 1 | 0 | 3 | 0.3333 | NA | NA | 3 | 0.3333 | 1 | 0 | 1 | 0 | 5 | 0.2 |
| 19_347865( | NA | NA | 1 | 1 | 1 | 0 | NA | NA | 2 | 0.5 | NA | NA | 2 | 0.5 | 1 | 0 | 1 | 0 | 4 | 0.25 |
| 19_347885( | NA | NA | 2 | 0.5 | NA | NA | 1 | 1 | 3 | 0.6667 | 1 | 1 | 3 | 0.3333 | NA | NA | 3 | 0.3333 | 7 | 0.4286 |
| 19_347897( | NA | NA | 2 | 1 | NA | NA | 2 | 0.5 | 4 | 0.75 | NA | NA | 2 | 0.5 | NA | NA | 2 | 0.5 | 4 | 0.5 |
| 19_347897! | NA | NA | 2 | 0 | NA | NA | 1 | 1 | 3 | 0.3333 | NA | NA | 2 | 1 | NA | NA | 2 | 0 | 4 | 0.5 |
| 19_347911: | NA | NA | 2 | 0 | NA | NA | 1 | 1 | 3 | 0.3333 | NA | NA | 2 | 0.5 | NA | NA | 2 | 0 | 4 | 0.25 |
| 19_348047: | NA | NA | 1 | 1 | NA | NA | NA | NA | 1 | 1 | NA | NA | NA | NA | NA | NA | 2 | 1 | 2 | 1 |
| 19_348183: | NA | NA | 1 | 1 | NA | NA | NA | NA | 1 | 1 | NA | NA | 1 | 1 | NA | NA | 2 | 1 | 3 | 1 |
| 19_359363! | NA | NA | 1 | 1 | 1 | 1 | NA | NA | 2 | 1 | 2 | 0.5 | NA | NA | NA | NA | NA | NA | 2 | 0.5 |
| 19_359368' | NA | NA | 1 | 1 | NA | NA | NA | NA | 1 | 1 | 1 | 1 | NA | NA | NA | NA | NA | NA | 1 | 1 |
| 19_359468: | NA | NA | 1 | 1 | 1 | 1 | NA | NA | 2 | 1 | 2 | 0.5 | NA | NA | NA | NA | NA | NA | 2 | 0.5 |
| 19_359469( | NA | NA | 1 | 1 | 1 | 1 | NA | NA | 2 | 1 | 2 | 1 | NA | NA | NA | NA | NA | NA | 2 | 1 |
| 19_359470( | NA | NA | 1 | 1 | 1 | 1 | NA | NA | 2 | 1 | 2 | 1 | NA | NA | NA | NA | NA | NA | 2 | 1 |
| 19_360819' | 2 | 0 | 5 | 0 | NA | NA | 8 | 0.125 | 15 | 0.0667 | 3 | 0.3333 | 1 | 0 | 1 | 0 | 1 | 0 | 6 | 0.1667 |
| 19_364331 | 3 | 0.6667 | 3 | 1 | 1 | 0 | 1 | 1 | 8 | 0.75 | NA | NA | 3 | 1 | 1 | 1 | 5 | 0.8 | 9 | 0.8889 |
| 19_364331! | 4 | 0.75 | 3 | 0.3333 | 1 | 0 | 2 | 0.5 | 10 | 0.5 | 1 | 1 | 3 | 1 | 1 | 0 | 4 | 0 | 9 | 0.4444 |
| 19_364356( | 1 | 0 | NA | NA | 1 | 0 | 2 | 0.5 | 4 | 0.25 | NA | NA | 1 | 0 | 1 | 1 | 3 | 0 | 5 | 0.2 |
| 19_364373! | NA | NA | 2 | 0 | 1 | 0 | 2 | 0.5 | 5 | 0.2 | 1 | 0 | 1 | 1 | 2 | 0.5 | NA | NA | 4 | 0.5 |
| 19_365962: | 1 | 0 | 3 | 1 | 1 | 0 | 2 | 0.5 | 7 | 0.5714 | NA | NA | NA | NA | 3 | 0.3333 | 2 | 0.5 | 5 | 0.4 |
| 19_367094( | 7 | 0.7143 | 4 | 0 | 6 | 0.5 | 6 | 0.3333 | 23 | 0.4348 | 8 | 0.375 | 10 | 0.4 | 5 | 0.6 | 3 | 0 | 26 | 0.3846 |
| 19_367095' | 12 | 0.25 | 7 | 0.2857 | 9 | 0.3333 | 6 | 0.1667 | 34 | 0.2647 | 8 | 0.5 | 11 | 0.3636 | 10 | 0.5 | 4 | 0 | 33 | 0.3939 |

| | | | | | | | | | | | | | | | | | | | | |
|---|---|---|---|---|---|---|---|---|---|---|---|---|---|---|---|---|---|---|---|---|
| 19_3670954 | 11 | 0.8182 | 5 | 0.2 | 6 | 0.5 | 6 | 0.8333 | 28 | 0.6429 | 8 | 0.75 | 11 | 0.7273 | 10 | 0.8 | 3 | 0.3333 | 32 | 0.7188 |
| 19_3670968 | 11 | 0.1818 | 7 | 0 | 9 | 0.1111 | 6 | 0 | 33 | 0.0909 | 8 | 0.125 | 12 | 0.1667 | 9 | 0.1111 | 4 | 0 | 33 | 0.1212 |
| 19_3670969 | 10 | 0.2 | 5 | 0 | 7 | 0.2857 | 4 | 0 | 26 | 0.1538 | 7 | 0.2857 | 11 | 0.1818 | 5 | 0 | 4 | 0 | 27 | 0.1481 |
| 19_3670979 | 12 | 0.1667 | 6 | 0 | 9 | 0.3333 | 4 | 0.25 | 31 | 0.1935 | 8 | 0.125 | 13 | 0.3077 | 9 | 0.4444 | 3 | 0 | 33 | 0.2727 |
| 19_3670980 | 12 | 0.1667 | 5 | 0 | 6 | 0 | 5 | 0 | 28 | 0.0714 | 7 | 0.1429 | 11 | 0 | 10 | 0.1 | 4 | 0 | 32 | 0.0625 |
| 19_3671023 | 8 | 0 | 7 | 0.1429 | 7 | 0.1429 | 4 | 0 | 26 | 0.0769 | 6 | 0.1667 | 8 | 0 | 9 | 0.1111 | 4 | 0 | 27 | 0.0741 |
| 19_3671049 | 6 | 0.8333 | 4 | 0.75 | 7 | 0.8571 | 3 | 1 | 20 | 0.85 | 7 | 0.7143 | 7 | 0.8571 | 9 | 0.7778 | 4 | 0.25 | 27 | 0.7037 |
| 19_3671059 | 6 | 0.1667 | 6 | 0 | 6 | 0.1667 | 3 | 0 | 21 | 0.0952 | 6 | 0.1667 | 6 | 0 | 9 | 0.1111 | 3 | 0 | 24 | 0.0833 |
| 19_3671060 | 7 | 0.5714 | 6 | 0.8333 | 5 | 0.8 | 4 | 0.25 | 22 | 0.6364 | 7 | 0.7143 | 8 | 0.5 | 8 | 0.5 | 4 | 0 | 27 | 0.4815 |
| 19_3671083 | 5 | 0.4 | 6 | 0.5 | 3 | 0 | 2 | 0.5 | 16 | 0.375 | 5 | 0.8 | 4 | 1 | 9 | 0.5556 | 3 | 0.3333 | 21 | 0.6667 |
| 19_3671091 | 7 | 1 | 6 | 1 | 7 | 1 | 3 | 1 | 23 | 1 | 7 | 1 | 6 | 1 | 7 | 0.8571 | 4 | 1 | 24 | 0.9583 |
| 19_3671094 | 6 | 1 | 5 | 1 | 7 | 1 | 4 | 0.75 | 22 | 0.9545 | 6 | 1 | 6 | 1 | 9 | 1 | 4 | 1 | 25 | 1 |
| 19_3671096 | 6 | 0.5 | 7 | 0.1429 | 6 | 0.1667 | 4 | 0.5 | 23 | 0.3043 | 7 | 0.2857 | 6 | 0.1667 | 9 | 0.2222 | 4 | 0.25 | 26 | 0.2308 |
| 19_3671102 | 6 | 0.6667 | 5 | 1 | 6 | 1 | 3 | 0.6667 | 20 | 0.85 | 5 | 1 | 5 | 1 | 7 | 0.8571 | 3 | 0.3333 | 20 | 0.85 |
| 19_3671117 | 7 | 1 | 4 | 1 | 5 | 1 | 4 | 1 | 20 | 1 | 6 | 1 | 6 | 1 | 9 | 1 | 4 | 1 | 25 | 1 |
| 19_3671125 | 7 | 0.5714 | 5 | 0.8 | 5 | 1 | 3 | 0.6667 | 20 | 0.75 | 5 | 0.6 | 6 | 0.3333 | 8 | 0.875 | 3 | 0.6667 | 22 | 0.6364 |
| 19_3671128 | 7 | 0.5714 | 5 | 0.4 | 3 | 0.3333 | 4 | 0.75 | 19 | 0.5263 | 5 | 0.8 | 6 | 0.5 | 7 | 0.4286 | 3 | 0.6667 | 21 | 0.5714 |
| 19_3671130 | 7 | 0.8571 | 5 | 0.8 | 7 | 0.5714 | 3 | 0.6667 | 22 | 0.7273 | 6 | 0.6667 | 5 | 0.2 | 7 | 0.5714 | 4 | 0.5 | 22 | 0.5 |
| 19_3671132 | 6 | 0.1667 | 5 | 0.2 | 7 | 0 | 4 | 0.5 | 22 | 0.1818 | 5 | 0.2 | 6 | 0.5 | 9 | 0.4444 | 4 | 0 | 24 | 0.3333 |
| 19_3671138 | 3 | 0 | 3 | 0.3333 | 5 | 0.2 | 3 | 0 | 14 | 0.1429 | 5 | 0.2 | 5 | 0.4 | 6 | 0 | 3 | 0.6667 | 19 | 0.2632 |
| 19_3671141 | 7 | 0.8571 | 5 | 1 | 7 | 1 | 3 | 1 | 22 | 0.9545 | 6 | 1 | 6 | 1 | 6 | 1 | 3 | 0.6667 | 21 | 0.9524 |
| 19_3671150 | 7 | 1 | 5 | 0.8 | 6 | 1 | 3 | 0.6667 | 21 | 0.9048 | 6 | 1 | 5 | 1 | 8 | 1 | 3 | 1 | 22 | 1 |
| 19_3671170 | 4 | 0 | 4 | 0.25 | 5 | 0.2 | 1 | 0 | 14 | 0.1429 | 2 | 0 | 4 | 0.25 | 4 | 0 | 4 | 0.25 | 14 | 0.1429 |
| 19_3686833 | 11 | 0.4545 | 8 | 0.375 | 11 | 0.1818 | 11 | 0.1818 | 41 | 0.2927 | 5 | 0 | 11 | 0.4545 | 13 | 0.1538 | 7 | 0.4286 | 36 | 0.2778 |
| 19_3686844 | 10 | 0.3 | 4 | 0.25 | 11 | 0.0909 | 10 | 0.3 | 35 | 0.2286 | 6 | 0.5 | 7 | 0.4286 | 9 | 0.1111 | 5 | 0 | 27 | 0.2593 |
| 19_3686851 | 11 | 0.3636 | 8 | 0.375 | 10 | 0.5 | 11 | 0.4545 | 40 | 0.425 | 6 | 0.6667 | 8 | 0.75 | 13 | 0.6154 | 5 | 0.4 | 32 | 0.625 |
| 19_3689535 | 1 | 0 | 1 | 1 | NA | NA | NA | NA | 2 | 0.5 | 2 | 0 | 3 | 0 | 1 | 0 | 1 | 1 | 7 | 0.1429 |
| 19_3689592 | 1 | 0 | 1 | 1 | NA | NA | NA | NA | 2 | 0.5 | 2 | 0 | 3 | 0.3333 | NA | NA | 1 | 1 | 6 | 0.3333 |
| 19_3698880 | 2 | 0.5 | 3 | 0 | 2 | 0 | 5 | 0.4 | 12 | 0.25 | 5 | 0.6 | 2 | 0.5 | 4 | 0 | 4 | 0 | 15 | 0.2667 |
| 19_3698888 | 3 | 1 | 2 | 1 | 2 | 1 | 5 | 0.8 | 12 | 0.9167 | 5 | 0.8 | 1 | 0 | 4 | 0.75 | 6 | 0.8333 | 16 | 0.75 |
| 19_3698961 | 4 | 0 | 4 | 0.25 | 3 | 0 | 5 | 0 | 16 | 0.0625 | 5 | 0 | 2 | 0 | 5 | 0 | 7 | 0.1429 | 19 | 0.0526 |
| 19_3699687 | 8 | 0.625 | 8 | 0.25 | 4 | 0.25 | 9 | 0.4444 | 29 | 0.4138 | 9 | 0.3333 | 14 | 0.2857 | 8 | 0.25 | 4 | 0.25 | 35 | 0.2857 |
| 19_3699702 | 12 | 0.3333 | 10 | 0.2 | 6 | 0 | 8 | 0.125 | 36 | 0.1944 | 8 | 0.25 | 11 | 0.0909 | 11 | 0.1818 | 5 | 0.4 | 35 | 0.2 |
| 19_3699708 | 12 | 0.0833 | 8 | 0.375 | 5 | 0 | 8 | 0 | 33 | 0.1212 | 4 | 0.25 | 8 | 0.125 | 7 | 0.1429 | 5 | 0.6 | 24 | 0.25 |
| 19_3699712 | 11 | 0.5455 | 6 | 0.3333 | 4 | 0.25 | 9 | 0.2222 | 30 | 0.3667 | 7 | 0.5714 | 9 | 0.4444 | 9 | 0.2222 | 4 | 0.5 | 29 | 0.4138 |
| 19_3699715 | 11 | 0.0909 | 10 | 0 | 4 | 0 | 11 | 0.0909 | 36 | 0.0556 | 8 | 0.125 | 11 | 0.0909 | 11 | 0.0909 | 5 | 0 | 35 | 0.0857 |
| 19_3699720 | 11 | 0.0909 | 8 | 0 | 4 | 0 | 9 | 0.1111 | 32 | 0.0625 | 4 | 0 | 11 | 0 | 12 | 0.0833 | 4 | 0.25 | 31 | 0.0645 |
| 19_3699733 | 9 | 0.3333 | 10 | 0.2 | 5 | 0 | 10 | 0.2 | 34 | 0.2059 | 7 | 0.1429 | 11 | 0 | 9 | 0.2222 | 4 | 0.25 | 31 | 0.129 |
| 19_3699873 | 7 | 0 | 7 | 0.1429 | 4 | 0.25 | 6 | 0.1667 | 24 | 0.125 | 7 | 0.1429 | 8 | 0 | 8 | 0.125 | 2 | 0 | 25 | 0.08 |
| 19_3699898 | 5 | 0 | 7 | 0.1429 | 4 | 0.25 | 6 | 0.5 | 22 | 0.2273 | 7 | 0 | 7 | 0.1429 | 10 | 0.1 | 2 | 0.5 | 26 | 0.1154 |
| 19_3699919 | 5 | 0.2 | 5 | 0 | 3 | 0.3333 | 6 | 0 | 19 | 0.1053 | 5 | 0.2 | 8 | 0 | 6 | 0.1667 | 2 | 0 | 21 | 0.0952 |
| 19_3699937 | 3 | 0 | 4 | 0.25 | 4 | 0.25 | 5 | 0 | 16 | 0.125 | 4 | 0.25 | 9 | 0.2222 | 7 | 0.7143 | 3 | 0.3333 | 23 | 0.3913 |
| 19_3699957 | 3 | 0 | 5 | 0.2 | 3 | 0 | 6 | 0 | 17 | 0.0588 | 4 | 0.5 | 8 | 0.25 | 6 | 0 | 3 | 0 | 21 | 0.1905 |
| 19_3699961 | 3 | 0.3333 | 4 | 0.25 | 2 | 0 | 6 | 0.1667 | 15 | 0.2 | 5 | 0.6 | 9 | 0.3333 | 7 | 0.1429 | 3 | 0 | 24 | 0.2917 |
| 19_3699968 | 3 | 0 | 5 | 0.2 | 2 | 0 | 4 | 0 | 14 | 0.0714 | 4 | 0.25 | 7 | 0 | 7 | 0.2857 | 2 | 0 | 20 | 0.15 |
| 19_3699970 | 4 | 0.25 | 5 | 0.4 | 3 | 0.6667 | 5 | 0.6 | 17 | 0.4706 | 4 | 0.25 | 9 | 0.1111 | 7 | 0.2857 | 1 | 0 | 21 | 0.1905 |
| 19_3700209 | NA | NA | 2 | 0 | NA | NA | 2 | 0.5 | 4 | 0.25 | 1 | 1 | 5 | 0 | NA | NA | 1 | 1 | 7 | 0.2857 |
| 19_3719819 | 14 | 0.0714 | 7 | 0 | 7 | 0.2857 | 16 | 0.0625 | 44 | 0.0909 | 11 | 0.0909 | 12 | 0.0833 | 13 | 0.1538 | 15 | 0.0667 | 51 | 0.098 |
| 19_3719891 | 15 | 0.1333 | 7 | 0 | 8 | 0.125 | 16 | 0.125 | 46 | 0.1087 | 14 | 0.1429 | 18 | 0.2778 | 16 | 0.125 | 16 | 0.0625 | 64 | 0.1562 |
| 19_3719898 | 16 | 0.125 | 8 | 0.125 | 11 | 0.0909 | 18 | 0.2778 | 53 | 0.1698 | 17 | 0.1765 | 18 | 0.2778 | 17 | 0.1765 | 17 | 0.0588 | 69 | 0.1739 |
| 19_3719906 | 14 | 0.5714 | 9 | 0.6667 | 11 | 0.5455 | 16 | 0.5625 | 50 | 0.58 | 17 | 0.7059 | 18 | 0.7778 | 15 | 0.5333 | 14 | 0.7143 | 64 | 0.6875 |
| 19_3719907 | 13 | 0.5385 | 8 | 0.625 | 12 | 0.5 | 14 | 0.5 | 47 | 0.5319 | 17 | 0.5882 | 15 | 0.6667 | 16 | 0.375 | 14 | 0.6429 | 62 | 0.5645 |
| 19_3719921 | 12 | 0.0833 | 6 | 0.3333 | 7 | 0.4286 | 15 | 0 | 40 | 0.15 | 15 | 0.3333 | 14 | 0 | 13 | 0.1538 | 15 | 0.1333 | 57 | 0.1579 |
| 19_3719928 | 9 | 0.2222 | 6 | 0.1667 | 8 | 0.125 | 11 | 0.0909 | 34 | 0.1471 | 13 | 0.1538 | 14 | 0.1429 | 13 | 0 | 12 | 0.0833 | 52 | 0.0962 |
| 19_3724622 | 5 | 0.2 | 8 | 0.625 | 3 | 0.3333 | 7 | 0.4286 | 23 | 0.4348 | 4 | 0.5 | 9 | 0.4444 | 5 | 0.6 | 9 | 0.4444 | 27 | 0.4815 |
| 19_3724630 | 4 | 0.25 | 5 | 0.2 | 3 | 0.3333 | 7 | 0 | 19 | 0.1579 | 3 | 0 | 9 | 0.1111 | 6 | 0 | 7 | 0.1429 | 25 | 0.08 |
| 19_3724640 | 5 | 0 | 7 | 0 | 2 | 0.5 | 7 | 0.1429 | 21 | 0.0952 | 3 | 0 | 8 | 0.125 | 5 | 0 | 9 | 0.1111 | 25 | 0.08 |
| 19_3724650 | 3 | 0 | 7 | 0.1429 | 3 | 0 | 6 | 0 | 19 | 0.0526 | 4 | 0 | 9 | 0 | 6 | 0.1667 | 8 | 0.125 | 27 | 0.0741 |
| 19_3724661 | 4 | 1 | NA | NA | 1 | 1 | 6 | 0.8333 | 11 | 0.9091 | 1 | 1 | 5 | 1 | 4 | 0.75 | 5 | 1 | 15 | 0.9333 |
| 19_3724663 | 3 | 0.3333 | 7 | 0.5714 | 2 | 1 | 7 | 0.4286 | 19 | 0.5263 | 3 | 1 | 9 | 0.4444 | 6 | 0.5 | 9 | 0.4444 | 27 | 0.5185 |
| 19_3724670 | 4 | 0 | 7 | 0.2857 | 1 | 0 | 7 | 0.2857 | 19 | 0.2105 | 4 | 0 | 8 | 0 | 4 | 0.5 | 7 | 0.2857 | 23 | 0.1739 |
| 19_3724687 | 2 | 0.5 | 4 | 0.5 | 2 | 0.5 | 2 | 0.5 | 10 | 0.5 | 1 | 1 | 2 | 0 | 3 | 0 | 5 | 0.4 | 11 | 0.2727 |
| 19_3724708 | 4 | 0.25 | 7 | 0 | 3 | 0 | 3 | 0 | 17 | 0.0588 | 2 | 0.5 | 8 | 0.125 | 6 | 0 | 7 | 0 | 23 | 0.087 |
| 19_3724716 | 1 | 0 | 3 | 0 | 1 | 1 | 2 | 0 | 7 | 0.1429 | 2 | 0 | 7 | 0 | 5 | 0 | 4 | 0.25 | 18 | 0.0556 |
| 19_3724718 | 3 | 0.6667 | 6 | 1 | 3 | 1 | 3 | 0.6667 | 15 | 0.8667 | 3 | 0.6667 | 9 | 0.8889 | 6 | 0.5 | 7 | 0.7143 | 25 | 0.72 |

| | | | | | | | | | | | | | | | | | | | | |
|---|---|---|---|---|---|---|---|---|---|---|---|---|---|---|---|---|---|---|---|---|
| 19_372472 | 3 | 0.3333 | 7 | 0.2857 | 3 | 0.6667 | 4 | 0.25 | 17 | 0.3529 | 3 | 0.3333 | 9 | 0.4444 | 6 | 0.1667 | 6 | 0.1667 | 24 | 0.2917 |
| 19_372725 | 3 | 0 | 3 | 0 | 9 | 0.3333 | 3 | 0.3333 | 18 | 0.2222 | 10 | 0.1 | 8 | 0.125 | 7 | 0.1429 | 4 | 0 | 29 | 0.1034 |
| 19_374089 | 2 | 0.5 | NA | NA | NA | NA | NA | NA | 2 | 0.5 | NA | NA | 1 | 1 | NA | NA | NA | NA | 1 | 1 |
| 19_374089 | 2 | 1 | NA | NA | NA | NA | NA | NA | 2 | 1 | NA | NA | 1 | 1 | NA | NA | NA | NA | 1 | 1 |
| 19_374280 | 1 | 0 | 1 | 1 | NA | NA | NA | NA | 2 | 0.5 | 3 | 0 | 1 | 1 | NA | NA | NA | NA | 4 | 0.25 |
| 19_374280 | 1 | 0 | 1 | 1 | NA | NA | NA | NA | 2 | 0.5 | 3 | 0 | 1 | 1 | NA | NA | NA | NA | 4 | 0.25 |
| 19_374282 | 1 | 1 | 1 | 1 | NA | NA | NA | NA | 2 | 1 | 3 | 0.6667 | 2 | 1 | NA | NA | NA | NA | 5 | 0.8 |
| 19_376714 | 3 | 0 | 2 | 0.5 | 3 | 0 | 5 | 0 | 13 | 0.0769 | 1 | 0 | 4 | 0 | 3 | 0.6667 | 1 | 0 | 9 | 0.2222 |
| 19_376715 | 2 | 0 | 2 | 0 | 3 | 0 | 3 | 0.3333 | 10 | 0.1 | 1 | 0 | 5 | 0.4 | 2 | 1 | 1 | 0 | 9 | 0.4444 |
| 19_376715 | 4 | 0.5 | 2 | 0.5 | 4 | 0.25 | 5 | 0.2 | 15 | 0.3333 | 1 | 0 | 6 | 0.1667 | 3 | 0 | 1 | 0 | 11 | 0.0909 |
| 19_376722 | 4 | 0.75 | 2 | 1 | 2 | 0.5 | 2 | 0.5 | 10 | 0.7 | NA | NA | 4 | 0.5 | 2 | 0 | 1 | 1 | 7 | 0.4286 |
| 19_381052 | 4 | 0.25 | 3 | 0.3333 | 2 | 0.5 | 1 | 0 | 10 | 0.3 | 4 | 0.25 | 2 | 0.5 | 1 | 1 | 3 | 0.3333 | 10 | 0.4 |
| 19_381770 | 3 | 0.3333 | NA | NA | 3 | 0 | 2 | 0 | 8 | 0.125 | 1 | 0 | 3 | 0.3333 | 3 | 0 | 2 | 0 | 9 | 0.1111 |
| 19_381770 | 3 | 0.6667 | NA | NA | 3 | 0.6667 | 1 | 1 | 7 | 0.7143 | 1 | 0 | 3 | 0.3333 | 3 | 0.6667 | 2 | 0.5 | 9 | 0.4444 |
| 19_381771 | 3 | 0.3333 | NA | NA | 3 | 0 | 2 | 0 | 8 | 0.125 | 1 | 0 | 2 | 0.5 | 3 | 0 | 2 | 0 | 8 | 0.125 |
| 19_381772 | 3 | 0.3333 | NA | NA | 2 | 0.5 | 2 | 0.5 | 7 | 0.4286 | 1 | 1 | 3 | 0.3333 | 4 | 0.25 | 1 | 0 | 9 | 0.3333 |
| 19_382532 | 2 | 0.5 | 5 | 0 | 7 | 0.1429 | 6 | 0 | 20 | 0.1 | 6 | 0.1667 | 6 | 0.3333 | 5 | 0 | 5 | 0.2 | 22 | 0.1818 |
| 19_382941 | 5 | 0 | 10 | 0.1 | 8 | 0 | 11 | 0.2727 | 34 | 0.1176 | 8 | 0.125 | 10 | 0.1 | 8 | 0.375 | 7 | 0.4286 | 33 | 0.2424 |
| 19_382942 | 6 | 0.3333 | 8 | 0.125 | 7 | 0.2857 | 14 | 0.2857 | 35 | 0.2571 | 11 | 0.0909 | 11 | 0 | 7 | 0 | 8 | 0.25 | 37 | 0.0811 |
| 19_382944 | 6 | 0.1667 | 8 | 0.25 | 6 | 0.3333 | 13 | 0.7692 | 33 | 0.4545 | 9 | 0.4444 | 11 | 0.6364 | 7 | 0.7143 | 8 | 0.375 | 35 | 0.5429 |
| 19_382946 | 9 | 0.3333 | 11 | 0.5455 | 8 | 0.625 | 12 | 0.5 | 40 | 0.5 | 11 | 0.5455 | 11 | 0.5455 | 10 | 0.6 | 9 | 0.5556 | 41 | 0.561 |
| 19_382947 | 4 | 0 | 7 | 0.1429 | 6 | 0.1667 | 6 | 0 | 23 | 0.087 | 7 | 0.4286 | 10 | 0.2 | 7 | 0.2857 | 8 | 0.25 | 32 | 0.2812 |
| 19_382948 | 8 | 0.125 | 11 | 0.2727 | 8 | 0.25 | 14 | 0.2857 | 41 | 0.2439 | 13 | 0.6154 | 13 | 0.3846 | 8 | 0.375 | 8 | 0.375 | 42 | 0.4524 |
| 19_382948 | 7 | 0.4286 | 11 | 0.3636 | 9 | 0.4444 | 13 | 0.6923 | 40 | 0.5 | 12 | 0.5833 | 11 | 0.7273 | 11 | 0.4545 | 8 | 0.5 | 42 | 0.5714 |
| 19_382949 | 8 | 0.125 | 9 | 0 | 9 | 0.2222 | 14 | 0.0714 | 40 | 0.1 | 11 | 0.0909 | 11 | 0.0909 | 11 | 0 | 9 | 0.3333 | 42 | 0.119 |
| 19_382949 | 9 | 0.1111 | 12 | 0 | 8 | 0 | 10 | 0.4 | 39 | 0.1282 | 11 | 0.2727 | 12 | 0.25 | 11 | 0.1818 | 8 | 0 | 42 | 0.1905 |
| 19_382949 | 9 | 0.8889 | 12 | 0.9167 | 8 | 1 | 14 | 1 | 43 | 0.9535 | 10 | 0.9 | 12 | 1 | 11 | 1 | 10 | 1 | 43 | 0.9767 |
| 19_382949 | 8 | 0.5 | 11 | 0.5455 | 7 | 0.5714 | 14 | 0.8571 | 40 | 0.65 | 13 | 0.6923 | 13 | 0.8462 | 11 | 0.9091 | 9 | 0.6667 | 46 | 0.7826 |
| 19_382950 | 7 | 1 | 13 | 0.9231 | 9 | 1 | 15 | 0.9333 | 44 | 0.9545 | 13 | 1 | 13 | 0.9231 | 10 | 1 | 9 | 0.8889 | 45 | 0.9556 |
| 19_382950 | 8 | 0.25 | 7 | 0.7143 | 7 | 0.2857 | 13 | 0.2308 | 35 | 0.3429 | 6 | 0.1667 | 12 | 0.5 | 8 | 0.25 | 8 | 0.75 | 34 | 0.4412 |
| 19_382950 | 9 | 0.2222 | 13 | 0.3077 | 9 | 0.1111 | 15 | 0.0667 | 46 | 0.1739 | 12 | 0 | 13 | 0.0769 | 11 | 0.2727 | 10 | 0.1 | 46 | 0.1087 |
| 19_382951 | 5 | 0 | 11 | 0.4545 | 7 | 0.2857 | 14 | 0.2143 | 37 | 0.2703 | 12 | 0.4167 | 8 | 0.625 | 8 | 0.375 | 7 | 0.5714 | 35 | 0.4857 |
| 19_382954 | 5 | 0.2 | 9 | 0 | 5 | 0.2 | 10 | 0 | 29 | 0.069 | 5 | 0.2 | 4 | 0.25 | 6 | 0 | 9 | 0 | 24 | 0.0833 |
| 19_382956 | 4 | 0 | 9 | 0.2222 | 7 | 0.2857 | 12 | 0.25 | 32 | 0.2188 | 5 | 0.2 | 6 | 0.3333 | 7 | 0.1429 | 9 | 0.1111 | 27 | 0.1852 |
| 19_382957 | 5 | 0 | 9 | 0.5556 | 6 | 0.6667 | 11 | 0.5455 | 31 | 0.4839 | 5 | 0.8 | 5 | 0.4 | 7 | 0.5714 | 8 | 0.5 | 25 | 0.56 |
| 19_382963 | 5 | 0 | 12 | 0.25 | 9 | 0.6667 | 12 | 0.4167 | 38 | 0.3684 | 5 | 0 | 6 | 0.3333 | 8 | 0.25 | 10 | 0 | 29 | 0.1379 |
| 19_382963 | 6 | 0.3333 | 12 | 0.1667 | 8 | 0.375 | 13 | 0.3077 | 39 | 0.2821 | 5 | 0.4 | 6 | 0.3333 | 6 | 0 | 11 | 0.0909 | 28 | 0.1786 |
| 19_382963 | 7 | 0.2857 | 11 | 0.1818 | 6 | 0.1667 | 14 | 0.7143 | 38 | 0.3947 | 5 | 0.8 | 5 | 0.8 | 6 | 0.6667 | 9 | 0.3333 | 25 | 0.6 |
| 19_382965 | 5 | 0.6 | 11 | 0.6364 | 9 | 0.5556 | 14 | 0.4286 | 39 | 0.5385 | 5 | 0.4 | 5 | 0.2 | 7 | 0.5714 | 11 | 0.5455 | 28 | 0.4643 |
| 19_382968 | 3 | 0 | 6 | 0.1667 | 8 | 0.25 | 5 | 0.2 | 22 | 0.1818 | 6 | 0.1667 | 3 | 0.3333 | 3 | 0 | 4 | 0.25 | 16 | 0.1875 |
| 19_382969 | 7 | 0.1429 | 16 | 0.25 | 10 | 0.4 | 17 | 0.1176 | 50 | 0.22 | 6 | 0.3333 | 7 | 0 | 7 | 0 | 14 | 0.4286 | 34 | 0.2353 |
| 19_382969 | 8 | 0.75 | 12 | 0.6667 | 7 | 0.8571 | 15 | 0.6667 | 42 | 0.7143 | 5 | 1 | 5 | 0.8 | 5 | 0.8 | 11 | 0.6364 | 26 | 0.7692 |
| 19_382969 | 7 | 0.4286 | 15 | 0.5333 | 10 | 0.5 | 18 | 0.7222 | 50 | 0.58 | 7 | 0.8571 | 7 | 0.8571 | 8 | 0.5 | 14 | 0.6429 | 36 | 0.6944 |
| 19_382970 | 7 | 0 | 18 | 0.0556 | 10 | 0.2 | 16 | 0 | 51 | 0.0588 | 6 | 0 | 6 | 0.1667 | 8 | 0 | 10 | 0.1 | 30 | 0.0667 |
| 19_382970 | 5 | 0.8 | 11 | 1 | 7 | 1 | 15 | 0.9333 | 38 | 0.9474 | 5 | 1 | 6 | 1 | 6 | 1 | 6 | 1 | 23 | 1 |
| 19_382970 | 5 | 0.4 | 12 | 0.6667 | 6 | 0.5 | 15 | 0.6667 | 38 | 0.6053 | 5 | 0.6 | 6 | 0.8333 | 6 | 0.6667 | 9 | 0.7778 | 26 | 0.7308 |
| 19_382972 | 9 | 0 | 14 | 0.0714 | 8 | 0.125 | 21 | 0.0476 | 52 | 0.0577 | 7 | 0 | 9 | 0.2222 | 8 | 0.125 | 11 | 0.0909 | 35 | 0.1143 |
| 19_382972 | 8 | 0.375 | 15 | 0.2 | 10 | 0.1 | 19 | 0.1579 | 52 | 0.1923 | 8 | 0.125 | 9 | 0.2222 | 7 | 0.1429 | 13 | 0.0769 | 37 | 0.1351 |
| 19_382973 | 6 | 0.6667 | 16 | 0.75 | 10 | 0.6 | 23 | 0.6522 | 55 | 0.6727 | 9 | 0.6667 | 8 | 0.375 | 8 | 0.5 | 13 | 0.7692 | 38 | 0.6053 |
| 19_387713 | 1 | 0 | NA | NA | 1 | 1 | NA | NA | 2 | 0.5 | NA | NA | 1 | 1 | 1 | 0 | NA | NA | 2 | 0.5 |
| 19_387714 | 1 | 1 | NA | NA | 2 | 1 | NA | NA | 3 | 1 | NA | NA | 1 | 1 | 1 | 1 | NA | NA | 2 | 1 |
| 19_387715 | 1 | 0 | NA | NA | 2 | 0.5 | NA | NA | 3 | 0.3333 | NA | NA | 1 | 1 | 1 | 0 | NA | NA | 2 | 0.5 |
| 19_387715 | NA | NA | NA | NA | 2 | 1 | NA | NA | 2 | 1 | NA | NA | 1 | 0 | 1 | 1 | NA | NA | 2 | 0.5 |
| 19_387778 | 1 | 1 | 1 | 0 | 1 | 1 | NA | NA | 3 | 0.6667 | NA | NA | 1 | 1 | 1 | 1 | 2 | 1 | 4 | 1 |
| 19_387779 | 2 | 0.5 | 2 | 1 | 1 | 0 | 2 | 0 | 7 | 0.4286 | NA | NA | 2 | 0.5 | 2 | 0 | 4 | 0 | 8 | 0.125 |
| 19_387779 | 2 | 1 | 2 | 1 | 1 | 1 | 1 | 1 | 6 | 1 | NA | NA | 1 | 1 | 1 | 1 | 4 | 1 | 6 | 1 |
| 19_389051 | 2 | 0.5 | 4 | 0.5 | 1 | 0 | 3 | 0.3333 | 10 | 0.4 | NA | NA | 1 | 1 | 2 | 0.5 | NA | NA | 3 | 0.6667 |
| 19_389051 | 3 | 0.6667 | 3 | 0.6667 | 1 | 1 | 3 | 1 | 10 | 0.8 | 1 | 1 | 2 | 1 | 2 | 1 | NA | NA | 5 | 1 |
| 19_389063 | 3 | 0 | 3 | 0.6667 | 1 | 1 | 3 | 0.6667 | 10 | 0.5 | NA | NA | 2 | 0.5 | 2 | 0.5 | NA | NA | 4 | 0.5 |
| 19_389064 | 3 | 0.6667 | 2 | 0.5 | NA | NA | 2 | 1 | 7 | 0.7143 | NA | NA | 1 | 1 | 2 | 1 | NA | NA | 3 | 1 |
| 19_389065 | 2 | 0.5 | 2 | 1 | 1 | 0 | 3 | 0 | 8 | 0.375 | 1 | 1 | 2 | 1 | 2 | 0.5 | NA | NA | 5 | 0.8 |
| 19_389066 | 3 | 0 | 4 | 0.25 | 1 | 1 | 3 | 0 | 11 | 0.1818 | 1 | 1 | 2 | 0.5 | 3 | 0 | NA | NA | 6 | 0.3333 |
| 19_389069 | 3 | 0 | 4 | 0.5 | 1 | 0 | 4 | 0 | 12 | 0.1667 | NA | NA | 2 | 0 | 3 | 0.3333 | NA | NA | 5 | 0.2 |
| 19_389069 | 2 | 0 | 4 | 0.25 | 1 | 0 | 4 | 0 | 11 | 0.0909 | 1 | 1 | 3 | 0 | 2 | 0 | NA | NA | 6 | 0.1667 |
| 19_402107 | NA | NA | 4 | 0.25 | 1 | 1 | 2 | 0 | 7 | 0.2857 | 1 | 1 | NA | NA | 2 | 0.5 | 1 | 0 | 4 | 0.5 |

| | | | | | | | | | | | | | | | | | | | | |
|---|---|---|---|---|---|---|---|---|---|---|---|---|---|---|---|---|---|---|---|---|
| 19_402109 | NA | NA | 4 | 1 | NA | NA | NA | NA | 4 | 1 | 3 | 0.6667 | NA | NA | 2 | 1 | 1 | 0 | 6 | 0.6667 |
| 19_402810 | 8 | 0 | 14 | 0 | 15 | 0.1333 | 15 | 0.1333 | 52 | 0.0769 | 7 | 0.1429 | 15 | 0.0667 | 8 | 0 | 12 | 0.1667 | 42 | 0.0952 |
| 19_402821 | 6 | 0 | 11 | 0 | 8 | 0.125 | 14 | 0.0714 | 39 | 0.0513 | 3 | 0 | 4 | 0.25 | 3 | 0.3333 | 9 | 0 | 19 | 0.1053 |
| 19_402832 | 5 | 0.2 | 5 | 0 | 5 | 0.4 | 7 | 0.2857 | 22 | 0.2273 | 2 | 0 | 6 | 0.1667 | 2 | 0 | 5 | 0.2 | 15 | 0.1333 |
| 19_402834 | 4 | 1 | 6 | 1 | 7 | 1 | 8 | 0.875 | 25 | 0.96 | 4 | 1 | 7 | 1 | 3 | 1 | 9 | 1 | 23 | 1 |
| 19_402981 | NA | NA | 3 | 0 | 3 | 0 | 2 | 0.5 | 8 | 0.125 | NA | NA | 1 | 0 | 1 | 0 | 5 | 0.2 | 7 | 0.1429 |
| 19_402981 | NA | NA | 3 | 0.6667 | 3 | 0.3333 | 2 | 1 | 8 | 0.625 | NA | NA | 2 | 0.5 | NA | NA | 5 | 0.8 | 7 | 0.7143 |
| 19_402983 | NA | NA | 2 | 1 | 2 | 1 | 3 | 0 | 7 | 0.5714 | NA | NA | 3 | 0.3333 | NA | NA | 3 | 0.6667 | 6 | 0.5 |
| 19_402985 | NA | NA | 1 | 1 | 1 | 0 | 2 | 1 | 4 | 0.75 | NA | NA | 2 | 0.5 | NA | NA | 3 | 0.3333 | 5 | 0.4 |
| 19_402986 | NA | NA | 2 | 0.5 | 3 | 0.6667 | 2 | 0 | 7 | 0.4286 | NA | NA | 3 | 0 | NA | NA | 2 | 0.5 | 5 | 0.2 |
| 19_403139 | 1 | 0 | 1 | 0 | 1 | 1 | 1 | 0 | 4 | 0.25 | NA | NA | NA | NA | NA | NA | 2 | 1 | 2 | 1 |
| 19_403139 | 1 | 0 | 1 | 1 | 1 | 0 | 1 | 0 | 4 | 0.25 | NA | NA | NA | NA | NA | NA | 2 | 0.5 | 2 | 0.5 |
| 19_403140 | 1 | 1 | 1 | 1 | 1 | 1 | 1 | 1 | 4 | 1 | NA | NA | NA | NA | 1 | 1 | 2 | 1 | 3 | 1 |
| 19_403142 | NA | NA | 1 | 1 | NA | NA | 1 | 0 | 2 | 0.5 | NA | NA | NA | NA | NA | NA | 1 | 1 | 1 | 1 |
| 19_403145 | NA | NA | 1 | 1 | 1 | 1 | NA | NA | 2 | 1 | NA | NA | NA | NA | 1 | 1 | 1 | 1 | 2 | 1 |
| 19_403147 | NA | NA | 1 | 1 | 1 | 1 | 1 | 1 | 3 | 1 | NA | NA | NA | NA | 1 | 1 | 1 | 1 | 2 | 1 |
| 19_403148 | NA | NA | 1 | 1 | 1 | 1 | 1 | 1 | 3 | 1 | NA | NA | NA | NA | 1 | 1 | 1 | 1 | 2 | 1 |
| 19_403149 | NA | NA | 1 | 1 | NA | NA | 1 | 1 | 2 | 1 | NA | NA | NA | NA | NA | NA | 1 | 1 | 1 | 1 |
| 19_405387 | 2 | 1 | 4 | 0.5 | 3 | 0.3333 | 2 | 0.5 | 11 | 0.5455 | 5 | 0.2 | 9 | 0.4444 | NA | NA | 1 | 0 | 15 | 0.3333 |
| 19_405387 | 1 | 0 | 2 | 0.5 | 3 | 0.3333 | 2 | 0 | 8 | 0.25 | 5 | 0 | 7 | 0.1429 | NA | NA | NA | NA | 12 | 0.0833 |
| 19_405389 | 1 | 1 | 5 | 0.4 | 3 | 0 | 2 | 0 | 11 | 0.2727 | 5 | 0.4 | 7 | 0.1429 | NA | NA | 1 | 0 | 13 | 0.2308 |
| 19_405390 | 1 | 1 | 4 | 0.5 | 2 | 0.5 | NA | NA | 7 | 0.5714 | 4 | 0.5 | 3 | 0.3333 | NA | NA | NA | NA | 7 | 0.4286 |
| 19_405390 | 1 | 1 | 4 | 0.75 | 3 | 1 | 2 | 1 | 10 | 0.9 | 5 | 0.6 | 6 | 0.8333 | NA | NA | NA | NA | 11 | 0.7273 |
| 19_405390 | 1 | 0 | 3 | 0.3333 | 3 | 1 | 3 | 0.6667 | 10 | 0.6 | 5 | 0.8 | 5 | 0.8 | NA | NA | NA | NA | 10 | 0.8 |
| 19_405392 | 1 | 1 | 3 | 1 | 2 | 1 | 2 | 1 | 8 | 1 | 5 | 1 | 6 | 1 | NA | NA | NA | NA | 11 | 1 |
| 19_405393 | 1 | 0 | 4 | 0.5 | 2 | 0 | 2 | 0.5 | 9 | 0.3333 | 3 | 0 | 7 | 0.1429 | NA | NA | NA | NA | 10 | 0.1 |
| 19_405394 | 1 | 1 | 4 | 0.5 | 1 | 1 | 2 | 1 | 8 | 0.75 | 3 | 0 | 4 | 0.75 | NA | NA | NA | NA | 7 | 0.4286 |
| 19_405395 | 1 | 0 | 4 | 0.5 | 1 | 1 | 1 | 0 | 7 | 0.4286 | 3 | 0 | 7 | 0.1429 | NA | NA | NA | NA | 10 | 0.1 |
| 19_405396 | 1 | 0 | 3 | 0.3333 | 2 | 0 | 2 | 0 | 8 | 0.125 | 1 | 0 | 6 | 0.3333 | NA | NA | NA | NA | 7 | 0.2857 |
| 19_405397 | 1 | 1 | 4 | 0.5 | 2 | 0.5 | 2 | 0 | 9 | 0.4444 | 3 | 0.3333 | 6 | 0.1667 | NA | NA | NA | NA | 9 | 0.2222 |
| 19_405477 | NA | NA | 4 | 0.25 | 3 | 0 | 1 | 0 | 8 | 0.125 | NA | NA | 2 | 0.5 | NA | NA | NA | NA | 2 | 0.5 |
| 19_405487 | NA | NA | 4 | 1 | 3 | 1 | 2 | 0.5 | 9 | 0.8889 | NA | NA | 2 | 1 | NA | NA | 1 | 1 | 3 | 1 |
| 19_405489 | NA | NA | 5 | 0.8 | 3 | 0 | 2 | 0 | 10 | 0.4 | 1 | 0 | 2 | 0.5 | 1 | 0 | 1 | 0 | 5 | 0.2 |
| 19_405489 | NA | NA | 5 | 0.4 | 3 | 0 | 1 | 0 | 9 | 0.2222 | 1 | 1 | 1 | 0 | 1 | 0 | 1 | 0 | 4 | 0.25 |
| 19_405489 | NA | NA | 5 | 0.2 | 3 | 0.6667 | 2 | 0.5 | 10 | 0.4 | NA | NA | 2 | 0.5 | 1 | 0 | 1 | 0 | 4 | 0.25 |
| 19_405498 | NA | NA | 4 | 0.25 | 2 | 0.5 | 2 | 0.5 | 8 | 0.375 | 1 | 1 | 2 | 0.5 | 1 | 0 | 1 | 0 | 5 | 0.4 |
| 19_405498 | NA | NA | 4 | 0.5 | 1 | 1 | 2 | 0 | 7 | 0.4286 | 1 | 1 | 2 | 0.5 | 1 | 1 | 1 | 0 | 5 | 0.6 |
| 19_405849 | 3 | 0.3333 | 1 | 1 | 1 | 0 | 1 | 0 | 6 | 0.3333 | 2 | 0 | 1 | 0 | 3 | 0.3333 | 1 | 0 | 7 | 0.1429 |
| 19_405850 | 4 | 0.25 | 1 | 0 | 1 | 0 | 1 | 0 | 7 | 0.1429 | 3 | 0.6667 | 1 | 0 | 3 | 1 | 1 | 0 | 8 | 0.625 |
| 19_405852 | 3 | 0.6667 | NA | NA | 1 | 0 | 2 | 0.5 | 6 | 0.5 | 2 | 0.5 | 1 | 0 | 3 | 0 | 1 | 0 | 7 | 0.1429 |
| 19_405853 | 3 | 0.6667 | 1 | 0 | 1 | 1 | 1 | 0 | 6 | 0.5 | 3 | 0 | 2 | 0 | 3 | 0.3333 | 1 | 1 | 9 | 0.2222 |
| 19_405854 | 4 | 0.75 | 1 | 1 | 1 | 1 | 2 | 0 | 8 | 0.625 | 3 | 1 | 1 | 0 | 3 | 1 | 1 | 1 | 8 | 0.875 |
| 19_405854 | 2 | 1 | NA | NA | 1 | 1 | 2 | 1 | 5 | 1 | 1 | 1 | 1 | 1 | 3 | 1 | NA | NA | 5 | 1 |
| 19_405855 | 3 | 0.6667 | 1 | 0 | NA | NA | NA | NA | 4 | 0.5 | 3 | 1 | 2 | 0 | 3 | 0.6667 | 1 | 0 | 9 | 0.5556 |
| 19_405856 | 4 | 0.5 | NA | NA | 1 | 0 | 1 | 1 | 6 | 0.5 | 3 | 1 | 2 | 0.5 | 3 | 1 | 1 | 0 | 9 | 0.7778 |
| 19_405857 | 2 | 1 | NA | NA | 1 | 1 | 2 | 0.5 | 5 | 0.8 | 2 | 1 | 1 | 1 | 3 | 0.6667 | 1 | 1 | 7 | 0.8571 |
| 19_405857 | 1 | 1 | NA | NA | 1 | 1 | 2 | 0.5 | 4 | 0.75 | 3 | 1 | 1 | 1 | 3 | 1 | 1 | 1 | 8 | 1 |
| 19_405858 | 3 | 1 | NA | NA | 1 | 1 | 2 | 1 | 6 | 1 | 3 | 0.6667 | 1 | 1 | 3 | 1 | 1 | 1 | 8 | 0.875 |
| 19_405858 | 3 | 0 | NA | NA | 1 | 1 | 2 | 0.5 | 6 | 0.3333 | 3 | 0.6667 | NA | NA | 3 | 0.3333 | 1 | 0 | 7 | 0.4286 |
| 19_405859 | 2 | 0.5 | NA | NA | 1 | 0 | 2 | 0 | 5 | 0.2 | 3 | 0 | 2 | 0.5 | 3 | 0 | 1 | 1 | 9 | 0.2222 |
| 19_406114 | 1 | 1 | 1 | 0 | 2 | 0.5 | 2 | 1 | 6 | 0.6667 | NA | NA | 3 | 0.6667 | NA | NA | NA | NA | 3 | 0.6667 |
| 19_406114 | 2 | 0 | 2 | 0.5 | 3 | 0.3333 | 2 | 0 | 9 | 0.2222 | 2 | 0 | 2 | 0.5 | NA | NA | 1 | 0 | 5 | 0.2 |
| 19_406114 | 2 | 0.5 | 1 | 0 | 4 | 0.75 | 2 | 0.5 | 9 | 0.5556 | 2 | 1 | 3 | 1 | NA | NA | 1 | 1 | 6 | 1 |
| 19_406116 | 2 | 1 | 2 | 1 | 2 | 1 | 4 | 1 | 10 | 1 | 2 | 1 | 2 | 1 | 1 | 1 | 1 | 1 | 6 | 1 |
| 19_406118 | 4 | 0.25 | 2 | 0 | 4 | 0.25 | 4 | 0.25 | 14 | 0.2143 | 2 | 0.5 | 4 | 0.25 | 3 | 0.3333 | 1 | 0 | 10 | 0.3 |
| 19_406124 | 4 | 0 | 6 | 0.3333 | 5 | 0.2 | 5 | 0 | 20 | 0.15 | 3 | 0 | 5 | 0.4 | 6 | 0 | NA | NA | 14 | 0.1429 |
| 19_406126 | 5 | 0 | 6 | 0 | 5 | 0.4 | 3 | 0.3333 | 19 | 0.1579 | 2 | 0 | 7 | 0.1429 | 8 | 0.125 | 1 | 0 | 18 | 0.1111 |
| 19_407146 | NA | NA | 3 | 0 | NA | NA | 1 | 1 | 4 | 0.25 | NA | NA | NA | NA | NA | NA | 1 | 1 | 1 | 1 |
| 19_407272 | NA | NA | 3 | 1 | 1 | 1 | NA | NA | 4 | 1 | 1 | 0 | NA | NA | NA | NA | 3 | 1 | 4 | 0.75 |
| 19_407275 | NA | NA | 3 | 1 | 1 | 1 | NA | NA | 4 | 1 | 1 | 1 | NA | NA | NA | NA | 4 | 1 | 5 | 1 |
| 19_407276 | 1 | 1 | 5 | 1 | 1 | 1 | NA | NA | 7 | 1 | 1 | 1 | NA | NA | NA | NA | 5 | 1 | 6 | 1 |
| 19_407277 | 1 | 1 | 3 | 1 | NA | NA | NA | NA | 4 | 1 | 1 | 1 | NA | NA | NA | NA | 3 | 1 | 4 | 1 |
| 19_407279 | 2 | 0.5 | 4 | 0.5 | 1 | 1 | NA | NA | 7 | 0.5714 | NA | NA | NA | NA | NA | NA | 4 | 0.25 | 4 | 0.25 |
| 19_407289 | 2 | 1 | 3 | 0.6667 | NA | NA | NA | NA | 5 | 0.8 | NA | NA | NA | NA | NA | NA | 2 | 0.5 | 2 | 0.5 |
| 19_407298 | 1 | 0 | 2 | 0 | 1 | 1 | NA | NA | 4 | 0.25 | 1 | 0 | 1 | 0 | NA | NA | 2 | 0.5 | 4 | 0.25 |

| | | | | | | | | | | | | | | | | | | | | |
|---|---|---|---|---|---|---|---|---|---|---|---|---|---|---|---|---|---|---|---|---|
| 19_407300 | 1 | 0 | 2 | 0 | 1 | 1 | NA | NA | 4 | 0.25 | 1 | 0 | 1 | 1 | NA | NA | NA | NA | 2 | 0.5 |
| 19_407301 | 1 | 0 | 2 | 0.5 | 1 | 1 | NA | NA | 4 | 0.5 | 1 | 1 | 1 | 1 | NA | NA | 1 | 0 | 3 | 0.6667 |
| 19_408352 | 1 | 1 | NA | NA | NA | NA | 1 | 1 | 2 | 1 | NA | NA | NA | NA | NA | NA | 1 | 1 | 1 | 1 |
| 19_408354 | 1 | 1 | NA | NA | NA | NA | 1 | 0 | 2 | 0.5 | NA | NA | 1 | 0 | NA | NA | 1 | 1 | 2 | 0.5 |
| 19_409004 | NA | NA | 3 | 1 | NA | NA | 1 | 1 | 4 | 1 | NA | NA | NA | NA | NA | NA | 1 | 1 | 1 | 1 |
| 19_409006 | NA | NA | 3 | 1 | NA | NA | 1 | 1 | 4 | 1 | NA | NA | NA | NA | NA | NA | 1 | 1 | 1 | 1 |
| 19_409009 | NA | NA | 3 | 0 | NA | NA | 1 | 1 | 4 | 0.25 | NA | NA | NA | NA | NA | NA | 1 | 1 | 1 | 1 |
| 19_409010 | NA | NA | 3 | 1 | NA | NA | NA | NA | 3 | 1 | NA | NA | NA | NA | NA | NA | 1 | 1 | 1 | 1 |
| 19_409021 | NA | NA | 1 | 1 | NA | NA | 1 | 0 | 2 | 0.5 | NA | NA | NA | NA | NA | NA | 1 | 1 | 1 | 1 |
| 19_409022 | NA | NA | 3 | 0.3333 | NA | NA | 1 | 0 | 4 | 0.25 | NA | NA | 1 | 1 | NA | NA | 1 | 0 | 2 | 0.5 |
| 19_409025 | NA | NA | 3 | 0.3333 | NA | NA | 1 | 0 | 4 | 0.25 | NA | NA | 1 | 1 | NA | NA | 1 | 1 | 2 | 1 |
| 19_409027 | NA | NA | 3 | 0.3333 | 1 | 1 | 1 | 1 | 5 | 0.6 | NA | NA | 1 | 0 | NA | NA | 1 | 1 | 2 | 0.5 |
| 19_409030 | NA | NA | 3 | 1 | 1 | 1 | 1 | 1 | 5 | 1 | NA | NA | 1 | 1 | NA | NA | 1 | 1 | 2 | 1 |
| 19_409031 | NA | NA | 2 | 1 | 1 | 1 | 1 | 1 | 4 | 1 | NA | NA | 1 | 1 | NA | NA | 1 | 1 | 2 | 1 |
| 19_409031 | NA | NA | 3 | 0.3333 | 1 | 0 | 1 | 0 | 5 | 0.2 | NA | NA | 1 | 0 | NA | NA | 1 | 1 | 2 | 0.5 |
| 19_409034 | NA | NA | 3 | 0.3333 | 1 | 0 | 1 | 0 | 5 | 0.2 | NA | NA | 1 | 0 | NA | NA | 1 | 1 | 2 | 0.5 |
| 19_409036 | NA | NA | 3 | 0.3333 | NA | NA | 1 | 1 | 4 | 0.5 | NA | NA | 1 | 0 | NA | NA | 1 | 1 | 2 | 0.5 |
| 19_409038 | NA | NA | 2 | 1 | 1 | 1 | 1 | 1 | 4 | 1 | NA | NA | 1 | 1 | NA | NA | 1 | 1 | 2 | 1 |
| 19_409038 | NA | NA | 1 | 0 | 1 | 0 | 1 | 1 | 3 | 0.3333 | NA | NA | 1 | 0 | NA | NA | 1 | 1 | 2 | 0.5 |
| 19_409040 | NA | NA | 2 | 0.5 | 1 | 0 | 1 | 1 | 4 | 0.5 | NA | NA | 1 | 0 | NA | NA | 1 | 1 | 2 | 0.5 |
| 19_409040 | NA | NA | 2 | 1 | 1 | 1 | 1 | 1 | 4 | 1 | NA | NA | 1 | 1 | NA | NA | 1 | 1 | 2 | 1 |
| 19_409042 | NA | NA | 2 | 1 | 1 | 1 | 1 | 1 | 4 | 1 | NA | NA | 1 | 1 | NA | NA | 2 | 1 | 3 | 1 |
| 19_409042 | NA | NA | 3 | 0 | 1 | 1 | 1 | 1 | 5 | 0.4 | NA | NA | 1 | 0 | NA | NA | 2 | 1 | 3 | 0.6667 |
| 19_409044 | NA | NA | 3 | 0.6667 | 1 | 1 | 1 | 1 | 5 | 0.8 | NA | NA | 1 | 1 | NA | NA | 2 | 1 | 3 | 1 |
| 19_409044 | NA | NA | 2 | 0.5 | 1 | 0 | 1 | 1 | 4 | 0.5 | NA | NA | 1 | 1 | NA | NA | 1 | 1 | 2 | 1 |
| 19_409045 | NA | NA | 3 | 0 | 1 | 1 | 1 | 1 | 5 | 0.4 | NA | NA | 2 | 0.5 | NA | NA | 2 | 0.5 | 4 | 0.5 |
| 19_409047 | NA | NA | 3 | 0 | 1 | 0 | 2 | 0.5 | 6 | 0.1667 | NA | NA | 2 | 0.5 | NA | NA | 2 | 0.5 | 4 | 0.5 |
| 19_409047 | NA | NA | 2 | 1 | 1 | 0 | 1 | 1 | 4 | 0.75 | NA | NA | 1 | 1 | NA | NA | 1 | 0 | 2 | 0.5 |
| 19_409047 | NA | NA | 3 | 1 | 1 | 1 | 2 | 1 | 6 | 1 | NA | NA | 2 | 0.5 | NA | NA | 1 | 1 | 3 | 0.6667 |
| 19_409048 | NA | NA | 2 | 0.5 | 1 | 0 | 2 | 0 | 5 | 0.2 | NA | NA | 2 | 0 | NA | NA | 1 | 1 | 3 | 0.3333 |
| 19_409049 | NA | NA | 3 | 0.6667 | NA | NA | 2 | 1 | 5 | 0.8 | NA | NA | 2 | 0.5 | NA | NA | 2 | 1 | 4 | 0.75 |
| 19_409049 | NA | NA | 3 | 0.3333 | 1 | 0 | 2 | 0.5 | 6 | 0.3333 | NA | NA | 2 | 1 | NA | NA | 2 | 1 | 4 | 1 |
| 19_409052 | NA | NA | 3 | 0.3333 | 1 | 0 | 2 | 0 | 6 | 0.1667 | 1 | 1 | 2 | 0.5 | NA | NA | 2 | 0 | 5 | 0.4 |
| 19_409055 | NA | NA | 3 | 0.3333 | 1 | 0 | 2 | 0 | 6 | 0.1667 | NA | NA | 2 | 1 | NA | NA | 2 | 0 | 4 | 0.5 |
| 19_409175 | 2 | 0 | 3 | 0.3333 | 3 | 0 | 2 | 0 | 10 | 0.1 | 1 | 0 | NA | NA | 1 | 0 | 2 | 0.5 | 4 | 0.25 |
| 19_409176 | 2 | 1 | 3 | 0 | 2 | 0.5 | 1 | 0 | 8 | 0.375 | 1 | 0 | NA | NA | NA | NA | 2 | 0.5 | 3 | 0.3333 |
| 19_410521 | 3 | 0 | 2 | 0 | 6 | 0.1667 | 2 | 1 | 13 | 0.2308 | 4 | 0 | 3 | 1 | 1 | 0 | NA | NA | 8 | 0.375 |
| 19_410522 | 4 | 0.25 | 2 | 0 | 8 | 0 | 2 | 0 | 16 | 0.0625 | 6 | 0 | 3 | 0.3333 | 2 | 0 | 1 | 0 | 12 | 0.0833 |
| 19_410522 | 4 | 1 | 2 | 1 | 8 | 0.875 | NA | NA | 14 | 0.9286 | 6 | 1 | 2 | 1 | 2 | 1 | 1 | 1 | 11 | 1 |
| 19_411994 | 3 | 0.3333 | 1 | 0 | 5 | 0.2 | 5 | 0.6 | 14 | 0.3571 | 4 | 0.25 | 3 | 0.3333 | 10 | 0.5 | 5 | 0.2 | 22 | 0.3636 |
| 19_412005 | 3 | 0.3333 | 1 | 0 | 5 | 0 | 6 | 0 | 15 | 0.0667 | 5 | 0 | 3 | 0.3333 | 10 | 0 | 7 | 0.1429 | 25 | 0.08 |
| 19_412019 | 4 | 0 | 1 | 0 | 4 | 0 | 5 | 0.2 | 14 | 0.0714 | 3 | 0.3333 | 2 | 0 | 7 | 0 | 3 | 0 | 15 | 0.0667 |
| 19_412030 | 8 | 0.25 | 11 | 0.0909 | 13 | 0.0769 | 14 | 0.2143 | 46 | 0.1522 | 18 | 0.1667 | 12 | 0.0833 | 22 | 0.0455 | 16 | 0.1875 | 68 | 0.1176 |
| 19_412033 | 6 | 0.3333 | 9 | 0.4444 | 15 | 0.5333 | 17 | 0.1176 | 47 | 0.3404 | 13 | 0.3846 | 11 | 0.3636 | 20 | 0.35 | 11 | 0.2727 | 55 | 0.3455 |
| 19_412034 | 6 | 0 | 8 | 0 | 10 | 0.1 | 14 | 0.0714 | 38 | 0.0526 | 9 | 0 | 7 | 0.1429 | 14 | 0 | 9 | 0.1111 | 39 | 0.0513 |
| 19_412034 | 7 | 0 | 9 | 0.1111 | 13 | 0 | 16 | 0.125 | 45 | 0.0667 | 14 | 0 | 10 | 0.1 | 19 | 0.0526 | 12 | 0.3333 | 55 | 0.1091 |
| 19_412528 | NA | NA | NA | NA | 1 | 1 | NA | NA | 1 | 1 | 1 | 1 | NA | NA | NA | NA | NA | NA | 1 | 1 |
| 19_412530 | NA | NA | NA | NA | 1 | 1 | NA | NA | 1 | 1 | 1 | 1 | NA | NA | NA | NA | NA | NA | 1 | 1 |
| 19_412532 | NA | NA | NA | NA | 1 | 1 | NA | NA | 1 | 1 | 1 | 1 | NA | NA | NA | NA | NA | NA | 1 | 1 |
| 19_412532 | NA | NA | NA | NA | 1 | 1 | NA | NA | 1 | 1 | 1 | 1 | NA | NA | NA | NA | NA | NA | 1 | 1 |
| 19_412626 | NA | NA | NA | NA | 1 | 1 | NA | NA | 1 | 1 | NA | NA | NA | NA | NA | NA | 1 | 1 | 1 | 1 |
| 19_415547 | 1 | 0 | NA | NA | 3 | 0 | 4 | 0.25 | 8 | 0.125 | NA | NA | 4 | 0.25 | 2 | 0 | 3 | 0 | 9 | 0.1111 |
| 19_415558 | 2 | 0 | 2 | 0 | 2 | 1 | 5 | 0.2 | 11 | 0.2727 | 1 | 0 | 4 | 0.25 | 2 | 0.5 | 4 | 0.25 | 11 | 0.2727 |
| 19_415819 | 1 | 1 | NA | NA | NA | NA | NA | NA | 1 | 1 | NA | NA | 1 | 1 | 1 | 0 | NA | NA | 2 | 0.5 |
| 19_415840 | 1 | 0 | 2 | 0.5 | NA | NA | NA | NA | 3 | 0.3333 | NA | NA | 2 | 0.5 | 3 | 0.3333 | NA | NA | 5 | 0.4 |
| 19_415843 | NA | NA | 2 | 0.5 | NA | NA | NA | NA | 2 | 0.5 | NA | NA | 2 | 0 | 2 | 0.5 | NA | NA | 4 | 0.25 |
| 19_415865 | 4 | 0.25 | 4 | 0.25 | 2 | 0.5 | 1 | 1 | 11 | 0.3636 | 2 | 0.5 | NA | NA | 1 | 1 | 2 | 1 | 5 | 0.8 |
| 19_415866 | 4 | 0 | 4 | 0.25 | 1 | 0 | 1 | 0 | 10 | 0.1 | 2 | 0.5 | NA | NA | NA | NA | 3 | 0 | 5 | 0.2 |
| 19_415867 | 4 | 0.25 | 3 | 0 | 1 | 0 | NA | NA | 8 | 0.125 | 1 | 0 | NA | NA | 1 | 1 | 2 | 0 | 4 | 0.25 |
| 19_415877 | NA | NA | 2 | 0.5 | NA | NA | NA | NA | 2 | 0.5 | 1 | 0 | NA | NA | 1 | 1 | 2 | 0 | 4 | 0.25 |
| 19_415887 | 1 | 0 | 5 | 1 | NA | NA | NA | NA | 6 | 0.8333 | 7 | 0.7143 | 1 | 1 | 3 | 0.3333 | 4 | 0.75 | 15 | 0.6667 |
| 19_415889 | 1 | 0 | 5 | 0.6 | NA | NA | NA | NA | 6 | 0.5 | 5 | 1 | 1 | 1 | 2 | 0 | 3 | 0.3333 | 11 | 0.6364 |
| 19_415890 | 1 | 0 | 5 | 0.6 | NA | NA | NA | NA | 6 | 0.5 | 6 | 0.8333 | 1 | 1 | 2 | 0.5 | 3 | 0.3333 | 12 | 0.6667 |
| 19_415895 | NA | NA | 4 | 0.5 | NA | NA | NA | NA | 4 | 0.5 | 2 | 1 | NA | NA | NA | NA | 1 | 1 | 3 | 1 |
| 19_415907 | NA | NA | 4 | 0.25 | NA | NA | NA | NA | 4 | 0.25 | 4 | 0.25 | NA | NA | NA | NA | 2 | 0.5 | 6 | 0.3333 |

| | | | | | | | | | | | | | | | | | | | | |
|---|---|---|---|---|---|---|---|---|---|---|---|---|---|---|---|---|---|---|---|---|
| 19_418187( | NA | NA | 1 | 0 | 3 | 0.3333 | 2 | 0 | 6 | 0.1667 | NA | NA | 2 | 0 | 2 | 0.5 | 1 | 1 | 5 | 0.4 |
| 19_418190: | NA | NA | 2 | 0 | 4 | 0.25 | 3 | 0 | 9 | 0.1111 | NA | NA | 2 | 0 | 3 | 0.3333 | 2 | 0 | 7 | 0.1429 |
| 19_418192· | 1 | 0 | 2 | 0 | 5 | 0.2 | 4 | 0.25 | 12 | 0.1667 | NA | NA | 2 | 0.5 | 2 | 0 | 2 | 0 | 6 | 0.1667 |
| 19_418193! | NA | NA | 2 | 0 | 3 | 0.3333 | 4 | 0.25 | 9 | 0.2222 | NA | NA | 2 | 0.5 | 3 | 0 | 2 | 0.5 | 7 | 0.2857 |
| 19_418194( | 1 | 0 | NA | NA | 4 | 1 | 4 | 0.75 | 9 | 0.7778 | NA | NA | 2 | 1 | 3 | 1 | 2 | 0.5 | 7 | 0.8571 |
| 19_418195· | 1 | 1 | 2 | 1 | 4 | 1 | 4 | 1 | 11 | 1 | NA | NA | 2 | 1 | 2 | 1 | 2 | 1 | 6 | 1 |
| 19_418195! | 1 | 0 | 1 | 0 | 4 | 0.5 | 3 | 0 | 9 | 0.2222 | NA | NA | 2 | 0 | 2 | 0.5 | 1 | 0 | 5 | 0.2 |
| 19_418196( | 1 | 1 | 2 | 1 | 3 | 1 | 3 | 1 | 9 | 1 | NA | NA | 3 | 1 | 2 | 1 | 1 | 1 | 6 | 1 |
| 19_418196: | 3 | 1 | 2 | 1 | 5 | 0.8 | 3 | 1 | 13 | 0.9231 | NA | NA | 3 | 1 | 3 | 1 | 1 | 1 | 7 | 1 |
| 19_418197· | 3 | 0.3333 | 2 | 0 | 3 | 0.3333 | 2 | 0.5 | 10 | 0.3 | NA | NA | 2 | 0.5 | 3 | 0.3333 | 3 | 0.3333 | 8 | 0.375 |
| 19_418199( | 2 | 1 | 2 | 1 | 4 | 1 | 3 | 1 | 11 | 1 | 1 | 1 | 2 | 1 | 3 | 1 | 2 | 1 | 8 | 1 |
| 19_418199· | 4 | 0.75 | NA | NA | 2 | 0.5 | 2 | 1 | 8 | 0.75 | 1 | 1 | 1 | 1 | 2 | 0.5 | 2 | 1 | 6 | 0.8333 |
| 19_418199( | 4 | 0.5 | 2 | 0 | 6 | 0.5 | 4 | 0.25 | 16 | 0.375 | 1 | 0 | 1 | 0 | 5 | 0.6 | 2 | 0.5 | 9 | 0.4444 |
| 19_418200: | 3 | 0.6667 | 1 | 0 | 5 | 0.8 | 3 | 0.6667 | 12 | 0.6667 | 1 | 1 | 2 | 0 | 3 | 0.3333 | 2 | 0.5 | 8 | 0.375 |
| 19_418202! | 3 | 0 | 1 | 1 | 1 | 1 | 2 | 0.5 | 7 | 0.4286 | 1 | 0 | 1 | 0 | 2 | 0 | 1 | 1 | 5 | 0.2 |
| 19_418204 | 3 | 0.3333 | 1 | 0 | 2 | 0 | 2 | 0 | 8 | 0.125 | 1 | 0 | 2 | 0 | 4 | 0.25 | 2 | 0 | 9 | 0.1111 |
| 19_418206· | 3 | 0.6667 | 1 | 0 | 1 | 1 | 2 | 0 | 7 | 0.4286 | 1 | 0 | 1 | 0 | 3 | 0.3333 | 1 | 0 | 6 | 0.1667 |
| 19_418208( | 3 | 0.6667 | 1 | 1 | 4 | 0.5 | 1 | 1 | 9 | 0.6667 | 2 | 0.5 | 3 | 0.3333 | 4 | 0.5 | 3 | 0.6667 | 12 | 0.5 |
| 19_418209: | 3 | 0 | 1 | 0 | 5 | 0.6 | 2 | 0 | 11 | 0.2727 | 2 | 0.5 | 3 | 0.3333 | 4 | 0.5 | 3 | 0 | 12 | 0.3333 |
| 19_418209 | 3 | 0.3333 | NA | NA | 2 | 0 | 2 | 0 | 7 | 0.1429 | 1 | 1 | 3 | 0 | 2 | 0 | 2 | 0.5 | 8 | 0.25 |
| 19_418346 | 2 | 0 | 1 | 1 | 2 | 0 | 2 | 1 | 7 | 0.4286 | 2 | 0.5 | NA | NA | 2 | 0 | NA | NA | 4 | 0.25 |
| 19_418354: | 2 | 1 | NA | NA | NA | NA | 1 | 1 | 3 | 1 | 2 | 1 | NA | NA | NA | NA | NA | NA | 2 | 1 |
| 19_418354( | 1 | 1 | 1 | 1 | NA | NA | NA | NA | 2 | 1 | 2 | 1 | NA | NA | NA | NA | NA | NA | 2 | 1 |
| 19_418355( | 2 | 1 | 1 | 1 | NA | NA | 1 | 1 | 4 | 1 | 2 | 1 | NA | NA | NA | NA | NA | NA | 2 | 1 |
| 19_418355! | 2 | 1 | 1 | 1 | NA | NA | 1 | 1 | 4 | 1 | 2 | 0.5 | NA | NA | NA | NA | NA | NA | 2 | 0.5 |
| 19_418356! | 2 | 1 | 1 | 1 | NA | NA | 1 | 1 | 4 | 1 | 2 | 1 | NA | NA | NA | NA | NA | NA | 2 | 1 |
| 19_418356! | 2 | 0.5 | 1 | 0 | NA | NA | 1 | 1 | 4 | 0.5 | 2 | 1 | NA | NA | NA | NA | NA | NA | 2 | 1 |
| 19_418513! | 3 | 0.6667 | 4 | 0 | 2 | 0 | 3 | 0 | 12 | 0.1667 | 1 | 0 | 3 | 0 | 4 | 0.5 | 5 | 0.4 | 13 | 0.3077 |
| 19_418514 | 3 | 0 | 7 | 0.2857 | 2 | 0 | 3 | 0 | 15 | 0.1333 | 3 | 0 | 3 | 0 | 3 | 0.3333 | 5 | 0.2 | 14 | 0.1429 |
| 19_418515· | 3 | 0.6667 | 6 | 1 | 2 | 1 | 1 | 0 | 12 | 0.8333 | 3 | 1 | 3 | 1 | 3 | 1 | 5 | 0.8 | 14 | 0.9286 |
| 19_418515: | 2 | 0.5 | 5 | 0.4 | 2 | 1 | 2 | 0.5 | 11 | 0.5455 | 1 | 1 | 4 | 1 | 3 | 0.6667 | 5 | 0.8 | 13 | 0.8462 |
| 19_418515· | 1 | 0 | 6 | 0.3333 | 2 | 0 | 2 | 0.5 | 11 | 0.2727 | 3 | 1 | 3 | 0.3333 | 1 | 0 | 3 | 0.3333 | 10 | 0.5 |
| 19_419061( | 89 | 0.1124 | 100 | 0.08 | 86 | 0.0814 | 78 | 0.0769 | 353 | 0.0878 | 69 | 0.1159 | 74 | 0.0946 | 86 | 0.0465 | 78 | 0.0641 | 307 | 0.0782 |
| 19_419062! | 97 | 0.0825 | 117 | 0.1026 | 93 | 0.1398 | 89 | 0.1236 | 396 | 0.1111 | 73 | 0.1507 | 80 | 0.1 | 86 | 0.1395 | 83 | 0.1084 | 322 | 0.1242 |
| 19_419064· | 75 | 0.0133 | 84 | 0.0714 | 73 | 0.0685 | 76 | 0.0789 | 308 | 0.0584 | 52 | 0.1346 | 68 | 0.0294 | 73 | 0.0274 | 71 | 0.0845 | 264 | 0.0644 |
| 19_419065! | 90 | 0.0556 | 113 | 0.0619 | 92 | 0.0543 | 83 | 0.1084 | 378 | 0.0688 | 74 | 0.0676 | 74 | 0.0405 | 89 | 0.0787 | 90 | 0.0556 | 327 | 0.0612 |
| 19_419068: | 99 | 0.1414 | 117 | 0.1197 | 108 | 0.1204 | 90 | 0.0889 | 414 | 0.1184 | 76 | 0.1184 | 83 | 0.1325 | 91 | 0.0989 | 88 | 0.2045 | 338 | 0.1391 |
| 19_419121( | 3 | 0 | 4 | 0.25 | 2 | 0 | 2 | 0 | 11 | 0.0909 | 1 | 0 | 2 | 0 | 2 | 0 | 4 | 0.25 | 9 | 0.1111 |
| 19_419121! | 1 | 1 | 4 | 0.25 | 2 | 0.5 | 2 | 0.5 | 9 | 0.4444 | 1 | 0 | 1 | 0 | 2 | 1 | 4 | 1 | 8 | 0.75 |
| 19_419124( | 2 | 0 | 4 | 0.25 | 1 | 0 | 2 | 0 | 9 | 0.1111 | 2 | 0 | 2 | 0.5 | 3 | 0 | 4 | 0.25 | 11 | 0.1818 |
| 19_419321· | 3 | 0.3333 | 3 | 0.6667 | 2 | 0.5 | 2 | 0.5 | 10 | 0.5 | 1 | 0 | 3 | 0.6667 | 3 | 1 | 2 | 0.5 | 9 | 0.6667 |
| 19_419322! | 4 | 0.75 | 5 | 0.8 | 4 | 1 | 1 | 1 | 14 | 0.8571 | 3 | 0.6667 | 5 | 1 | 4 | 0.5 | 3 | 1 | 15 | 0.8 |
| 19_419323( | 6 | 0.1667 | 5 | 0.2 | 4 | 0.25 | 2 | 0 | 17 | 0.1765 | 5 | 0 | 6 | 0.1667 | 4 | 0.25 | 4 | 0 | 19 | 0.1053 |
| 19_419324: | 7 | 0.4286 | 4 | 0.25 | 3 | 0.6667 | 2 | 0.5 | 16 | 0.4375 | 4 | 0.5 | 6 | 0.6667 | 4 | 0.5 | 4 | 0.25 | 18 | 0.5 |
| 19_419324! | 7 | 0.2857 | 5 | 0 | 3 | 0.6667 | 2 | 0.5 | 17 | 0.2941 | 3 | 0.6667 | 7 | 0 | 5 | 0.2 | 4 | 0.5 | 19 | 0.2632 |
| 19_419324! | 6 | 0.3333 | 4 | 0 | 3 | 0.3333 | 2 | 0.5 | 15 | 0.2667 | 4 | 0.25 | 7 | 0.4286 | 3 | 0.3333 | 3 | 0.6667 | 17 | 0.4118 |
| 19_419327( | 5 | 1 | 5 | 0 | 1 | 0 | 3 | 0.6667 | 14 | 0.5 | 1 | 0 | 3 | 0 | 4 | 0.25 | NA | NA | 8 | 0.125 |
| 19_419327! | 7 | 0.2857 | 5 | 0.2 | 2 | 0.5 | 6 | 0 | 20 | 0.2 | 3 | 0 | 5 | 0.2 | 5 | 0.4 | NA | NA | 13 | 0.2308 |
| 19_419328( | 7 | 0.1429 | 5 | 0 | 2 | 0 | 6 | 0.1667 | 20 | 0.1 | 3 | 0 | 5 | 0 | 4 | 0.25 | NA | NA | 12 | 0.0833 |
| 19_419328· | 7 | 0.4286 | 5 | 0.8 | 2 | 0.5 | 6 | 0.5 | 20 | 0.55 | 3 | 1 | 5 | 0.2 | 5 | 0.4 | NA | NA | 13 | 0.4615 |
| 19_420223( | NA | NA | 1 | 1 | 1 | 1 | 2 | 1 | 4 | 1 | NA | NA | 2 | 0.5 | 1 | 1 | 1 | 1 | 4 | 0.75 |
| 19_420224( | NA | NA | 1 | 0 | NA | NA | 2 | 1 | 3 | 0.6667 | NA | NA | 2 | 0.5 | 1 | 1 | 1 | 1 | 4 | 0.75 |
| 19_420224· | NA | NA | 1 | 1 | NA | NA | 2 | 1 | 3 | 1 | NA | NA | 2 | 1 | 1 | 1 | 1 | 1 | 4 | 1 |
| 19_420224: | NA | NA | NA | NA | 1 | 1 | 2 | 1 | 3 | 1 | NA | NA | 2 | 0.5 | 1 | 1 | 1 | 1 | 4 | 0.75 |
| 19_420225 | NA | NA | 1 | 1 | 1 | 1 | 1 | 1 | 3 | 1 | NA | NA | 2 | 1 | 1 | 1 | 1 | 1 | 4 | 1 |
| 19_420228( | NA | NA | NA | NA | NA | NA | 2 | 1 | 2 | 1 | NA | NA | 4 | 0.5 | NA | NA | 1 | 0 | 5 | 0.4 |
| 19_420634: | 9 | 0 | 10 | 0.2 | 10 | 0 | 8 | 0 | 37 | 0.0541 | 3 | 0 | 3 | 0.3333 | 4 | 0.25 | 10 | 0 | 20 | 0.1 |
| 19_420634 | 9 | 0.1111 | 12 | 0.0833 | 13 | 0 | 9 | 0.2222 | 43 | 0.093 | 4 | 0 | 5 | 0 | 5 | 0 | 10 | 0.2 | 24 | 0.0833 |
| 19_420636( | 6 | 0.5 | 8 | 0.375 | 10 | 0.3 | 6 | 0.5 | 30 | 0.4 | 2 | 1 | 4 | 0.75 | 3 | 1 | 4 | 0.75 | 13 | 0.8462 |
| 19_420641: | 6 | 0.1667 | 12 | 0.25 | 12 | 0.0833 | 7 | 0.1429 | 37 | 0.1622 | 3 | 0 | 4 | 0.75 | 5 | 0.2 | 6 | 0.1667 | 18 | 0.2778 |
| 19_420641 | 6 | 0.1667 | 11 | 0.1818 | 9 | 0.3333 | 8 | 0.125 | 34 | 0.2059 | 3 | 0.3333 | 5 | 0 | 5 | 0.2 | 5 | 0 | 18 | 0.1111 |
| 19_420642: | 7 | 1 | 9 | 1 | 11 | 0.9091 | 7 | 1 | 34 | 0.9706 | 3 | 1 | 5 | 1 | 4 | 1 | 6 | 0.8333 | 18 | 0.9444 |
| 19_420642! | 7 | 0.8571 | 11 | 0.5455 | 10 | 0.4 | 7 | 0.2857 | 35 | 0.5143 | 3 | 0.6667 | 5 | 0.6 | 3 | 0 | 5 | 0.4 | 16 | 0.4375 |
| 19_420644! | 7 | 0.2857 | 9 | 0.4444 | 11 | 0.1818 | 7 | 0.2857 | 34 | 0.2941 | 3 | 0.6667 | 6 | 0 | 4 | 0.25 | 6 | 0.3333 | 19 | 0.2632 |
| 19_420793( | 4 | 0.5 | 1 | 0 | 6 | 0.3333 | 4 | 0.25 | 15 | 0.3333 | 2 | 0 | 1 | 0 | 4 | 0.25 | 4 | 0.75 | 11 | 0.3636 |

| | | | | | | | | | | | | | | | | | | | | |
|---|---|---|---|---|---|---|---|---|---|---|---|---|---|---|---|---|---|---|---|---|
| 19_421085 | 3 | 0.3333 | 4 | 0.5 | 5 | 0.4 | 2 | 0 | 14 | 0.3571 | 5 | 0.8 | 3 | 0.6667 | 3 | 0.3333 | 2 | 0 | 13 | 0.5385 |
| 19_421088 | 3 | 1 | 5 | 0.4 | 5 | 0.6 | 2 | 0.5 | 15 | 0.6 | 3 | 1 | 2 | 1 | 1 | 1 | 3 | 0.6667 | 9 | 0.8889 |
| 19_421089 | 7 | 0.7143 | 9 | 0.2222 | 9 | 0.2222 | 2 | 0 | 27 | 0.3333 | 6 | 0.5 | 3 | 0.6667 | 4 | 0.25 | 8 | 0.375 | 21 | 0.4286 |
| 19_421093 | 7 | 0.2857 | 10 | 0.3 | 10 | 0.4 | 4 | 0.25 | 31 | 0.3226 | 6 | 0.1667 | 3 | 0 | 6 | 0.5 | 7 | 0.2857 | 22 | 0.2727 |
| 19_421093 | 7 | 0.2857 | 12 | 0.25 | 10 | 0.4 | 5 | 0.2 | 34 | 0.2941 | 8 | 0.25 | 3 | 0 | 7 | 0.2857 | 8 | 0.25 | 26 | 0.2308 |
| 19_421094 | 6 | 0.5 | 9 | 0.4444 | 4 | 0.5 | 2 | 0.5 | 21 | 0.4762 | 4 | 0.5 | 3 | 1 | 3 | 0 | 3 | 0.6667 | 13 | 0.5385 |
| 19_421095 | 7 | 0.1429 | 12 | 0.0833 | 10 | 0 | 6 | 0.1667 | 35 | 0.0857 | 8 | 0.375 | 3 | 0.3333 | 7 | 0 | 6 | 0 | 24 | 0.1667 |
| 19_421095 | 5 | 0.6 | 8 | 0.375 | 6 | 1 | 5 | 0.8 | 24 | 0.6667 | 4 | 0.5 | 3 | 1 | 6 | 0.8333 | 7 | 1 | 20 | 0.85 |
| 19_421095 | 7 | 0.1429 | 8 | 0.25 | 6 | 0.1667 | 5 | 0.4 | 26 | 0.2308 | 4 | 0.5 | 3 | 0 | 6 | 0 | 6 | 0.3333 | 19 | 0.2105 |
| 19_421096 | 6 | 0.6667 | 12 | 0.3333 | 10 | 0.6 | 6 | 0.5 | 34 | 0.5 | 9 | 0.4444 | 4 | 0.5 | 6 | 0.8333 | 8 | 0.625 | 27 | 0.5926 |
| 19_421097 | 7 | 0.2857 | 11 | 0.2727 | 10 | 0.1 | 6 | 0.3333 | 34 | 0.2353 | 7 | 0.1429 | 4 | 0.5 | 6 | 0.1667 | 8 | 0.375 | 25 | 0.28 |
| 19_421099 | 8 | 0.875 | 13 | 0.9231 | 10 | 1 | 7 | 0.8571 | 38 | 0.9211 | 9 | 1 | 4 | 1 | 7 | 1 | 10 | 1 | 30 | 1 |
| 19_421099 | 8 | 0 | 14 | 0.1429 | 10 | 0.1 | 5 | 0 | 37 | 0.0811 | 6 | 0 | 2 | 0.5 | 5 | 0 | 7 | 0 | 20 | 0.05 |
| 19_421099 | 6 | 0.8333 | 13 | 0.8462 | 11 | 1 | 7 | 0.8571 | 37 | 0.8919 | 10 | 0.9 | 4 | 1 | 7 | 1 | 8 | 1 | 29 | 0.9655 |
| 19_421099 | 9 | 0.4444 | 13 | 0.3846 | 9 | 0.3333 | 7 | 0.1429 | 38 | 0.3421 | 8 | 0.25 | 3 | 0.3333 | 7 | 0.2857 | 9 | 0.5556 | 27 | 0.3704 |
| 19_421101 | 7 | 0.4286 | 15 | 0.4 | 11 | 0.5455 | 9 | 0.3333 | 42 | 0.4286 | 8 | 0.5 | 6 | 0.3333 | 10 | 0.7 | 13 | 0.5385 | 37 | 0.5405 |
| 19_421102 | 8 | 1 | 10 | 0.8 | 12 | 1 | 8 | 0.625 | 38 | 0.8684 | 10 | 1 | 2 | 1 | 12 | 0.9167 | 11 | 1 | 35 | 0.9714 |
| 19_421119 | 9 | 0.7778 | 4 | 0.75 | 8 | 0.625 | 10 | 0.5 | 31 | 0.6452 | 4 | 0.5 | 8 | 0.5 | 8 | 0.375 | 6 | 0.6667 | 26 | 0.5 |
| 19_421120 | 8 | 0.625 | 4 | 0.25 | 9 | 0.5556 | 11 | 0.5455 | 32 | 0.5312 | 4 | 0.25 | 9 | 0.5556 | 11 | 0.3636 | 7 | 0.2857 | 31 | 0.3871 |
| 19_421120 | 10 | 0.7 | 4 | 1 | 9 | 0.7778 | 10 | 0.8 | 33 | 0.7879 | 4 | 0.5 | 10 | 0.5 | 11 | 0.4545 | 4 | 0.75 | 29 | 0.5172 |
| 19_421121 | 6 | 0.3333 | 2 | 0 | 6 | 0 | 11 | 0.1818 | 25 | 0.16 | 3 | 0.3333 | 7 | 0.1429 | 7 | 0.1429 | 2 | 0 | 19 | 0.1579 |
| 19_421367 | NA | NA | 2 | 0.5 | NA | NA | NA | NA | 2 | 0.5 | 2 | 0.5 | 1 | 0 | 1 | 0 | NA | NA | 4 | 0.25 |
| 19_421368 | NA | NA | 2 | 1 | NA | NA | NA | NA | 2 | 1 | 2 | 1 | 1 | 1 | 1 | 1 | NA | NA | 4 | 1 |
| 19_421369 | NA | NA | 2 | 1 | NA | NA | NA | NA | 2 | 1 | 2 | 1 | 1 | 1 | 1 | 1 | NA | NA | 4 | 1 |
| 19_421370 | NA | NA | 2 | 0.5 | NA | NA | NA | NA | 2 | 0.5 | 2 | 0.5 | NA | NA | 1 | 0 | NA | NA | 3 | 0.3333 |
| 19_421376 | NA | NA | 1 | 1 | NA | NA | NA | NA | 1 | 1 | NA | NA | 1 | 1 | 1 | 1 | NA | NA | 2 | 1 |
| 19_421378 | NA | NA | 3 | 0.3333 | NA | NA | NA | NA | 3 | 0.3333 | 2 | 0 | 1 | 1 | 1 | 1 | NA | NA | 4 | 0.5 |
| 19_421379 | 1 | 1 | 3 | 0.6667 | 1 | 1 | NA | NA | 5 | 0.8 | 2 | 0.5 | 1 | 1 | 1 | 0 | NA | NA | 4 | 0.5 |
| 19_421394 | NA | NA | 2 | 1 | NA | NA | 1 | 1 | 3 | 1 | 2 | 1 | 2 | 1 | 1 | 1 | NA | NA | 5 | 1 |
| 19_421398 | 1 | 1 | 3 | 0.6667 | NA | NA | 1 | 0 | 5 | 0.6 | 1 | 0 | 1 | 1 | 1 | 0 | NA | NA | 3 | 0.3333 |
| 19_421399 | NA | NA | 3 | 0.3333 | 1 | 0 | NA | NA | 4 | 0.25 | 1 | 1 | 2 | 0.5 | 1 | 0 | NA | NA | 4 | 0.5 |
| 19_421792 | 1 | 1 | 6 | 0.6667 | 3 | 0.3333 | 2 | 0 | 12 | 0.5 | 1 | 0 | 3 | 0 | 1 | 0 | 4 | 0.25 | 9 | 0.1111 |
| 19_421792 | 1 | 0 | 6 | 0.1667 | 4 | 0 | 3 | 0 | 14 | 0.0714 | 1 | 0 | 3 | 0.3333 | 1 | 0 | 4 | 0.5 | 9 | 0.3333 |
| 19_421798 | 4 | 0 | 5 | 0.2 | 4 | 0 | 5 | 0.4 | 18 | 0.1667 | 2 | 0 | 2 | 0 | 2 | 0 | 4 | 0.25 | 10 | 0.1 |
| 19_421799 | 5 | 0.2 | 7 | 0.2857 | 4 | 0 | 4 | 0.25 | 20 | 0.2 | 2 | 0 | 3 | 0.6667 | 4 | 0.5 | 3 | 0.3333 | 12 | 0.4167 |
| 19_421826 | 1 | 0 | 1 | 0 | 4 | 0.5 | 2 | 1 | 8 | 0.5 | 3 | 0.3333 | 2 | 1 | 1 | 0 | 1 | 1 | 7 | 0.5714 |
| 19_421827 | 1 | 1 | 2 | 1 | 2 | 0.5 | 3 | 0.6667 | 8 | 0.75 | 2 | 1 | 2 | 0 | 1 | 0 | 1 | 0 | 6 | 0.3333 |
| 19_421828 | 2 | 0 | 2 | 0.5 | 3 | 0 | 3 | 0 | 10 | 0.1 | 3 | 0 | 2 | 0 | 2 | 0.5 | 1 | 0 | 8 | 0.125 |
| 19_421834 | 1 | 1 | 1 | 0 | NA | NA | 2 | 0 | 4 | 0.25 | 3 | 0.3333 | 1 | 0 | 2 | 0.5 | NA | NA | 6 | 0.3333 |
| 19_422557 | NA | NA | 1 | 1 | NA | NA | NA | NA | 1 | 1 | NA | NA | NA | NA | 1 | 1 | NA | NA | 1 | 1 |
| 19_422571 | 13 | 0.1538 | 11 | 0.1818 | 8 | 0.125 | 3 | 0 | 35 | 0.1429 | 11 | 0.0909 | 5 | 0.2 | 3 | 0 | 4 | 0.25 | 23 | 0.1304 |
| 19_422572 | 13 | 0.0769 | 17 | 0.0588 | 15 | 0.0667 | 6 | 0.5 | 51 | 0.1176 | 15 | 0.1333 | 10 | 0.2 | 5 | 0 | 10 | 0 | 40 | 0.1 |
| 19_422572 | 12 | 0.6667 | 20 | 0.65 | 16 | 0.75 | 6 | 0.8333 | 54 | 0.7037 | 14 | 0.7143 | 10 | 0.6 | 7 | 0.4286 | 8 | 1 | 39 | 0.6923 |
| 19_422574 | 11 | 0.1818 | 20 | 0.2 | 18 | 0.1667 | 7 | 0.4286 | 56 | 0.2143 | 15 | 0.2 | 9 | 0.5556 | 6 | 0.3333 | 9 | 0.2222 | 39 | 0.3077 |
| 19_422574 | 13 | 0.4615 | 19 | 0.3158 | 15 | 0.4 | 4 | 0.25 | 51 | 0.3725 | 16 | 0.625 | 11 | 0.7273 | 7 | 0.7143 | 8 | 0.375 | 42 | 0.619 |
| 19_422574 | 10 | 0.6 | 15 | 0.4667 | 15 | 0.7333 | 4 | 0.75 | 44 | 0.6136 | 12 | 0.4167 | 9 | 0.8889 | 6 | 0.5 | 6 | 0.5 | 33 | 0.5758 |
| 19_422575 | 15 | 0.6 | 20 | 0.8 | 20 | 0.85 | 7 | 0.5714 | 62 | 0.7419 | 15 | 0.6 | 10 | 0.7 | 7 | 0.7143 | 10 | 0.3 | 42 | 0.5714 |
| 19_422575 | 12 | 0.1667 | 20 | 0.15 | 13 | 0.1538 | 5 | 0.2 | 50 | 0.16 | 15 | 0 | 11 | 0.0909 | 5 | 0 | 7 | 0.1429 | 38 | 0.0526 |
| 19_422583 | 6 | 0.3333 | 12 | 0.4167 | 4 | 0.5 | 4 | 1 | 26 | 0.5 | 8 | 0.125 | 6 | 0.6667 | 1 | 0 | 4 | 0.75 | 19 | 0.4211 |
| 19_422584 | 6 | 0.1667 | 7 | 0.4286 | 1 | 0 | 3 | 0.6667 | 17 | 0.3529 | 5 | 0.4 | 7 | 0.2857 | 1 | 0 | 3 | 0.3333 | 16 | 0.3125 |
| 19_422585 | 9 | 0.1111 | 10 | 0.1 | 6 | 0 | 5 | 0.2 | 30 | 0.1 | 8 | 0.125 | 8 | 0.125 | 1 | 1 | 6 | 0.1667 | 23 | 0.1739 |
| 19_422587 | 6 | 0 | 14 | 0.2143 | 8 | 0.375 | 5 | 0.2 | 33 | 0.2121 | 9 | 0.1111 | 7 | 0 | 1 | 0 | 6 | 0.1667 | 23 | 0.087 |
| 19_422714 | NA | NA | 1 | 1 | NA | NA | NA | NA | 1 | 1 | 2 | 0 | 1 | 1 | NA | NA | 1 | 1 | 4 | 0.5 |
| 19_422715 | 2 | 1 | 17 | 0.7059 | 10 | 0.6 | 2 | 1 | 31 | 0.7097 | 16 | 0.875 | 6 | 0.8333 | 2 | 0.5 | 14 | 0.7143 | 38 | 0.7895 |
| 19_422717 | 2 | 0 | 8 | 0 | 6 | 0 | 1 | 1 | 17 | 0.0588 | 8 | 0.125 | 3 | 0 | 2 | 0 | 11 | 0.3636 | 24 | 0.2083 |
| 19_422718 | 1 | 1 | 14 | 0.7857 | 9 | 0.8889 | 1 | 1 | 25 | 0.84 | 10 | 0.9 | 5 | 0.6 | 1 | 1 | 13 | 0.8462 | 29 | 0.8276 |
| 19_422718 | 1 | 1 | 8 | 0.625 | 8 | 0.5 | 1 | 0 | 18 | 0.5556 | 10 | 0.5 | 4 | 0.5 | 2 | 0.5 | 10 | 0.6 | 26 | 0.5385 |
| 19_422719 | 1 | 0 | 10 | 0 | 8 | 0.125 | NA | NA | 19 | 0.0526 | 8 | 0.125 | 3 | 0 | 1 | 0 | 10 | 0.1 | 22 | 0.0909 |
| 19_422720 | 13 | 0.3077 | 21 | 0.5238 | 18 | 0.5 | 7 | 0.5714 | 59 | 0.4746 | 14 | 0.5714 | 6 | 0.1667 | 4 | 0.5 | 21 | 0.4286 | 45 | 0.4444 |
| 19_422720 | 12 | 0 | 23 | 0.087 | 17 | 0.0588 | 7 | 0.1429 | 59 | 0.0678 | 14 | 0.1429 | 6 | 0 | 3 | 0 | 20 | 0.05 | 43 | 0.0698 |
| 19_422906 | 10 | 0 | 14 | 0.0714 | 10 | 0.1 | 6 | 0 | 40 | 0.05 | 10 | 0.2 | 6 | 0 | 2 | 0 | 18 | 0 | 36 | 0.0556 |
| 19_425514 | NA | NA | NA | NA | NA | NA | 1 | 1 | 1 | 1 | NA | NA | 1 | 1 | 1 | 1 | NA | NA | 2 | 1 |
| 19_425514 | NA | NA | NA | NA | NA | NA | 2 | 0.5 | 2 | 0.5 | 1 | 1 | 1 | 0 | 1 | 0 | NA | NA | 3 | 0.3333 |
| 19_425514 | NA | NA | NA | NA | NA | NA | 2 | 1 | 2 | 1 | 1 | 1 | 1 | 1 | 1 | 1 | NA | NA | 3 | 1 |
| 19_425515 | NA | NA | NA | NA | NA | NA | 2 | 1 | 2 | 1 | 1 | 1 | NA | NA | 1 | 1 | NA | NA | 2 | 1 |

| | | | | | | | | | | | | | | | | | | | | |
|---|---|---|---|---|---|---|---|---|---|---|---|---|---|---|---|---|---|---|---|---|
| 19_4255172 | NA | NA | NA | NA | NA | NA | 2 | 1 | 2 | 1 | 1 | 1 | 1 | 0 | NA | NA | NA | NA | 2 | 0.5 |
| 19_4255178 | NA | NA | NA | NA | NA | NA | 2 | 0.5 | 2 | 0.5 | 1 | 1 | 1 | 0 | 1 | 0 | NA | NA | 3 | 0.3333 |
| 19_4255192 | NA | NA | NA | NA | NA | NA | 2 | 0.5 | 2 | 0.5 | 1 | 0 | 1 | 1 | NA | NA | NA | NA | 2 | 0.5 |
| 19_4255214 | NA | NA | NA | NA | NA | NA | 2 | 1 | 2 | 1 | 1 | 1 | NA | NA | 1 | 1 | NA | NA | 2 | 1 |
| 19_4255247 | NA | NA | NA | NA | NA | NA | 2 | 1 | 2 | 1 | 1 | 1 | 1 | 0 | 1 | 0 | NA | NA | 3 | 0.3333 |
| 19_4255257 | NA | NA | NA | NA | NA | NA | 1 | 1 | 1 | 1 | 1 | 1 | NA | NA | 1 | 0 | NA | NA | 2 | 0.5 |
| 19_4255268 | 1 | 0 | NA | NA | NA | NA | 2 | 0.5 | 3 | 0.3333 | NA | NA | 1 | 0 | 1 | 1 | NA | NA | 2 | 0.5 |
| 19_4257922 | 1 | 1 | 1 | 1 | 2 | 0.5 | 1 | 0 | 5 | 0.6 | 1 | 0 | 1 | 0 | 1 | 1 | 1 | 0 | 4 | 0.25 |
| 19_4257931 | 1 | 1 | 1 | 1 | 2 | 1 | 2 | 1 | 6 | 1 | 1 | 1 | 1 | 1 | NA | NA | 1 | 1 | 3 | 1 |
| 19_4257949 | 1 | 1 | 1 | 1 | 2 | 0.5 | 1 | 0 | 5 | 0.6 | 1 | 1 | 1 | 1 | NA | NA | 1 | 1 | 3 | 1 |
| 19_4257958 | 1 | 0 | 1 | 1 | 2 | 1 | 2 | 1 | 6 | 0.8333 | 1 | 1 | 1 | 1 | 1 | 1 | 1 | 1 | 4 | 1 |
| 19_4257960 | 1 | 0 | 1 | 0 | 2 | 0.5 | 2 | 0 | 6 | 0.1667 | 1 | 0 | 1 | 0 | 1 | 0 | 1 | 1 | 4 | 0.25 |
| 19_4257972 | 1 | 1 | NA | NA | 2 | 0.5 | 1 | 0 | 4 | 0.5 | 1 | 1 | 2 | 1 | NA | NA | 1 | 0 | 4 | 0.75 |
| 19_4257973 | 1 | 0 | NA | NA | 2 | 0.5 | NA | NA | 3 | 0.3333 | 1 | 0 | 3 | 0 | 1 | 1 | 1 | 1 | 6 | 0.3333 |
| 19_4274371 | NA | NA | 1 | 1 | 1 | 0 | 1 | 1 | 3 | 0.6667 | NA | NA | NA | NA | NA | NA | 2 | 0.5 | 2 | 0.5 |
| 19_4274394 | NA | NA | 1 | 0 | NA | NA | 1 | 1 | 2 | 0.5 | 1 | 1 | NA | NA | NA | NA | 2 | 0.5 | 3 | 0.6667 |
| 19_4274402 | NA | NA | 1 | 0 | 1 | 1 | 1 | 1 | 3 | 0.6667 | 1 | 1 | NA | NA | NA | NA | 2 | 0.5 | 3 | 0.6667 |
| 19_4322496 | NA | NA | 2 | 0.5 | 1 | 1 | 1 | 1 | 4 | 0.75 | 1 | 1 | NA | NA | NA | NA | NA | NA | 1 | 1 |
| 19_4322497 | NA | NA | 2 | 1 | 1 | 1 | 1 | 1 | 4 | 1 | 1 | 1 | 1 | 1 | NA | NA | NA | NA | 2 | 1 |
| 19_4322501 | NA | NA | 2 | 1 | 1 | 1 | 1 | 1 | 4 | 1 | 1 | 1 | 1 | 0 | NA | NA | NA | NA | 2 | 0.5 |
| 19_4322503 | NA | NA | 2 | 0.5 | NA | NA | 1 | 1 | 3 | 0.6667 | 1 | 0 | 1 | 1 | NA | NA | NA | NA | 2 | 0.5 |
| 19_4322510 | NA | NA | 2 | 1 | 1 | 1 | 1 | 1 | 4 | 1 | 1 | 1 | 1 | 1 | NA | NA | NA | NA | 2 | 1 |
| 19_4342893 | 2 | 0 | 3 | 0.3333 | 1 | 0 | NA | NA | 6 | 0.1667 | 1 | 1 | 1 | 1 | 1 | 0 | 1 | 1 | 4 | 0.75 |
| 19_4342953 | 3 | 0 | 3 | 0.3333 | 2 | 0 | 1 | 1 | 9 | 0.2222 | 2 | 0.5 | 1 | 0 | 2 | 0 | 1 | 0 | 6 | 0.1667 |
| 19_4342972 | 6 | 0.5 | 2 | 1 | 2 | 0 | 1 | 0 | 11 | 0.4545 | 2 | 0.5 | 1 | 0 | 2 | 0.5 | 1 | 0 | 6 | 0.3333 |
| 19_4342982 | 5 | 0.6 | 2 | 1 | 2 | 0.5 | 1 | 1 | 10 | 0.7 | 2 | 1 | 1 | 1 | 1 | 1 | 1 | 0 | 5 | 0.8 |
| 19_4343002 | 6 | 0.6667 | 4 | 0.75 | 1 | 1 | NA | NA | 11 | 0.7273 | NA | NA | 1 | 1 | 2 | 0.5 | NA | NA | 3 | 0.6667 |
| 19_4343019 | 7 | 0.8571 | 3 | 1 | 2 | 1 | NA | NA | 12 | 0.9167 | 2 | 0.5 | 1 | 1 | 2 | 1 | NA | NA | 5 | 0.8 |
| 19_4343031 | 7 | 0.5714 | 4 | 1 | 2 | 1 | NA | NA | 13 | 0.7692 | 3 | 0.3333 | 1 | 1 | 2 | 1 | 1 | 1 | 7 | 0.7143 |
| 19_4343033 | 6 | 0.5 | 1 | 1 | 2 | 0.5 | 1 | 0 | 10 | 0.5 | 2 | 0 | 1 | 1 | 2 | 1 | 1 | 0 | 6 | 0.5 |
| 19_4343043 | 7 | 0.5714 | 4 | 0.75 | 2 | 0.5 | 1 | 0 | 14 | 0.5714 | 2 | 1 | 1 | 1 | 2 | 0.5 | 1 | 0 | 6 | 0.6667 |
| 19_4348350 | 10 | 0.6 | 4 | 0.5 | 3 | 1 | 2 | 0.5 | 19 | 0.6316 | 4 | 0.75 | 5 | 0.2 | 5 | 0.4 | 2 | 1 | 16 | 0.5 |
| 19_4348357 | 11 | 0.1818 | 6 | 0.1667 | 3 | 0 | 2 | 0 | 22 | 0.1364 | 4 | 0.75 | 5 | 0.4 | 4 | 0 | 3 | 0 | 16 | 0.3125 |
| 19_4348357 | 9 | 0.1111 | 5 | 0 | 3 | 0.6667 | 2 | 0 | 19 | 0.1579 | 3 | 0.3333 | 5 | 0 | 4 | 0.25 | 3 | 0.3333 | 15 | 0.2 |
| 19_4348371 | 5 | 0.4 | 3 | 0.3333 | 1 | 0 | 4 | 0.5 | 13 | 0.3846 | NA | NA | 3 | 0.3333 | 4 | 0.5 | NA | NA | 7 | 0.4286 |
| 19_4348394 | 9 | 0.5556 | 5 | 0.2 | 4 | 0.25 | 4 | 0 | 22 | 0.3182 | 3 | 0.3333 | 4 | 0 | 6 | 0.1667 | 3 | 0.3333 | 16 | 0.1875 |
| 19_4348430 | 11 | 0.2727 | 7 | 0.1429 | 4 | 0 | 3 | 0 | 25 | 0.16 | 4 | 0 | 6 | 0.5 | 6 | 0.3333 | 4 | 0 | 20 | 0.25 |
| 19_4348520 | 14 | 0.3571 | 7 | 0.1429 | 6 | 0.1667 | 8 | 0.25 | 35 | 0.2571 | 7 | 0.1429 | 8 | 0.25 | 6 | 0.3333 | 7 | 0.4286 | 28 | 0.2857 |
| 19_4348830 | 65 | 0.0462 | 78 | 0.1154 | 68 | 0.0588 | 63 | 0.0476 | 274 | 0.0693 | 41 | 0.0732 | 48 | 0.0833 | 62 | 0.0968 | 68 | 0.1029 | 219 | 0.0913 |
| 19_4348837 | 63 | 0.0476 | 71 | 0.0845 | 64 | 0.0781 | 56 | 0.0893 | 254 | 0.0748 | 35 | 0.1714 | 48 | 0.125 | 61 | 0.1475 | 61 | 0.0984 | 205 | 0.1317 |
| 19_4348847 | 61 | 0.7541 | 66 | 0.7424 | 57 | 0.807 | 55 | 0.8182 | 239 | 0.7782 | 32 | 0.7188 | 43 | 0.8372 | 58 | 0.8276 | 55 | 0.6909 | 188 | 0.7713 |
| 19_4348852 | 64 | 0.1562 | 63 | 0.3016 | 66 | 0.2273 | 55 | 0.2545 | 248 | 0.2339 | 38 | 0.2895 | 46 | 0.1739 | 61 | 0.377 | 59 | 0.2373 | 204 | 0.2745 |
| 19_4348862 | 60 | 0.7833 | 66 | 0.803 | 64 | 0.8438 | 53 | 0.8491 | 243 | 0.8189 | 32 | 0.875 | 44 | 0.9091 | 61 | 0.8689 | 57 | 0.7719 | 194 | 0.8505 |
| 19_4365230 | 5 | 0.8 | 5 | 0.8 | 5 | 0.2 | 5 | 0.4 | 20 | 0.55 | 1 | 0 | 3 | 1 | 3 | 1 | 4 | 0.25 | 11 | 0.6364 |
| 19_4365254 | 6 | 0.1667 | 7 | 0.1429 | 7 | 0.1429 | 7 | 0.2857 | 27 | 0.1852 | 3 | 0.3333 | 6 | 0.3333 | 6 | 0 | 6 | 0.3333 | 21 | 0.2381 |
| 19_4365269 | 6 | 0.1667 | 3 | 0.3333 | 3 | 0.3333 | 6 | 0.1667 | 18 | 0.2222 | 2 | 0 | 3 | 0.3333 | 5 | 0.4 | 4 | 0.25 | 14 | 0.2857 |
| 19_4365275 | 4 | 1 | 4 | 0.75 | 5 | 0.6 | 5 | 0.8 | 18 | 0.7778 | 2 | 1 | 5 | 0.8 | 6 | 0.5 | 5 | 0.8 | 18 | 0.7222 |
| 19_4365276 | 5 | 1 | 4 | 1 | 6 | 0.8333 | 4 | 0.75 | 19 | 0.8947 | 2 | 0.5 | 5 | 1 | 4 | 1 | 4 | 0.75 | 15 | 0.8667 |
| 19_4365303 | 6 | 0.3333 | 3 | 1 | 6 | 0.5 | 5 | 0.6 | 20 | 0.55 | 3 | 0 | 3 | 0.3333 | 6 | 0.6667 | 6 | 0.8333 | 18 | 0.5556 |
| 19_4365540 | 2 | 0 | 4 | 0.25 | 4 | 0.5 | 3 | 0.3333 | 13 | 0.3077 | NA | NA | 3 | 0.3333 | 3 | 0 | 2 | 0 | 8 | 0.125 |
| 19_4371379 | NA | NA | NA | NA | 3 | 0.3333 | NA | NA | 3 | 0.3333 | NA | NA | 1 | 1 | 3 | 1 | 1 | 1 | 5 | 1 |
| 19_4371831 | NA | NA | 1 | 1 | 3 | 0.3333 | NA | NA | 4 | 0.5 | NA | NA | 1 | 1 | 3 | 0.6667 | 1 | 0 | 5 | 0.6 |
| 19_4371843 | NA | NA | 1 | 1 | 3 | 0.6667 | NA | NA | 4 | 0.75 | NA | NA | 1 | 0 | 3 | 0.6667 | 1 | 1 | 5 | 0.6 |
| 19_4371845 | NA | NA | NA | NA | 3 | 1 | NA | NA | 3 | 1 | NA | NA | 1 | 1 | NA | NA | NA | NA | 1 | 1 |
| 19_4371853 | NA | NA | 1 | 0 | 3 | 1 | NA | NA | 4 | 0.75 | NA | NA | 1 | 1 | 3 | 1 | 1 | 0 | 5 | 0.8 |
| 19_4371856 | NA | NA | NA | NA | 2 | 1 | NA | NA | 2 | 1 | NA | NA | NA | NA | 3 | 1 | NA | NA | 3 | 1 |
| 19_4371873 | NA | NA | NA | NA | 1 | 1 | NA | NA | 1 | 1 | NA | NA | 1 | 1 | 3 | 1 | 1 | 1 | 5 | 1 |
| 19_4371875 | NA | NA | NA | NA | 2 | 1 | NA | NA | 2 | 1 | NA | NA | 1 | 1 | 3 | 1 | 1 | 1 | 5 | 1 |
| 19_4371883 | NA | NA | 2 | 1 | 3 | 1 | NA | NA | 5 | 1 | NA | NA | 1 | 0 | 3 | 1 | 1 | 1 | 5 | 0.8 |
| 19_4371886 | NA | NA | 2 | 1 | 3 | 0.6667 | NA | NA | 5 | 0.8 | NA | NA | 1 | 1 | 3 | 1 | 1 | 0 | 5 | 0.8 |
| 19_4372212 | NA | NA | 1 | 1 | NA | NA | NA | NA | 1 | 1 | NA | NA | NA | NA | NA | NA | 2 | 0.5 | 2 | 0.5 |
| 19_4372231 | NA | NA | 1 | 1 | NA | NA | NA | NA | 1 | 1 | NA | NA | NA | NA | NA | NA | 2 | 0.5 | 2 | 0.5 |
| 19_4372267 | NA | NA | 1 | 1 | 1 | 1 | NA | NA | 2 | 1 | NA | NA | NA | NA | NA | NA | 2 | 1 | 2 | 1 |
| 19_4372295 | NA | NA | NA | NA | 1 | 1 | NA | NA | 1 | 1 | NA | NA | NA | NA | NA | NA | 1 | 1 | 1 | 1 |
| 19_4372479 | 4 | 0 | 3 | 0.3333 | 5 | 0.2 | 1 | 1 | 13 | 0.2308 | 2 | 0.5 | NA | NA | 8 | 0.25 | NA | NA | 10 | 0.3 |

| | | | | | | | | | | | | | | | | | | | | |
|---|---|---|---|---|---|---|---|---|---|---|---|---|---|---|---|---|---|---|---|---|
| 19_4372485 | 3 | 0.3333 | 2 | 0 | 5 | 0 | 1 | 0 | 11 | 0.0909 | 2 | 0 | NA | NA | 7 | 0.1429 | 1 | 0 | 10 | 0.1 |
| 19_4372490 | 4 | 0 | 3 | 0.3333 | 4 | 0.25 | 1 | 0 | 12 | 0.1667 | 2 | 0 | NA | NA | 6 | 0.1667 | NA | NA | 8 | 0.125 |
| 19_4372496 | 4 | 0 | 2 | 0.5 | 3 | 0 | 1 | 1 | 10 | 0.2 | NA | NA | NA | NA | 5 | 0 | 1 | 1 | 6 | 0.1667 |
| 19_4372500 | 3 | 0.3333 | 3 | 1 | 3 | 1 | 1 | 1 | 10 | 0.8 | 2 | 0.5 | NA | NA | 2 | 1 | 1 | 1 | 5 | 0.8 |
| 19_4372500 | 3 | 0.3333 | 3 | 0.3333 | 5 | 0.4 | 1 | 0 | 12 | 0.3333 | 2 | 0 | NA | NA | 5 | 0.8 | NA | NA | 7 | 0.5714 |
| 19_4372503 | 3 | 1 | 2 | 1 | 4 | 1 | 1 | 1 | 10 | 1 | 2 | 1 | NA | NA | 3 | 0.6667 | NA | NA | 5 | 0.8 |
| 19_4372506 | 4 | 0 | 3 | 0.3333 | 5 | 0 | 1 | 0 | 13 | 0.0769 | 2 | 0 | NA | NA | 5 | 0.2 | 1 | 0 | 8 | 0.125 |
| 19_4372509 | 3 | 0.6667 | 1 | 1 | 4 | 1 | 1 | 1 | 9 | 0.8889 | 2 | 1 | NA | NA | 3 | 0.6667 | 1 | 1 | 6 | 0.8333 |
| 19_4372511 | 4 | 0.25 | 2 | 0 | 5 | 0 | 1 | 0 | 12 | 0.0833 | 2 | 0 | NA | NA | 5 | 0.2 | 1 | 0 | 8 | 0.125 |
| 19_4372513 | 3 | 0.3333 | 3 | 0.3333 | 4 | 0.5 | 1 | 0 | 11 | 0.3636 | 1 | 1 | NA | NA | 3 | 0.3333 | 1 | 1 | 5 | 0.6 |
| 19_4372513 | 4 | 0.75 | 3 | 0.6667 | 5 | 1 | NA | NA | 12 | 0.8333 | 2 | 1 | NA | NA | 4 | 1 | 1 | 1 | 7 | 1 |
| 19_4372518 | 4 | 1 | 3 | 1 | 3 | 0.6667 | 1 | 0 | 11 | 0.8182 | 2 | 0.5 | NA | NA | 3 | 0.6667 | 1 | 0 | 6 | 0.5 |
| 19_4375669 | 1 | 1 | 1 | 1 | 1 | 1 | 1 | 1 | 4 | 1 | 1 | 1 | 1 | 1 | NA | NA | NA | NA | 2 | 1 |
| 19_4383547 | 1 | 1 | 1 | 1 | 1 | 1 | NA | NA | 3 | 1 | 3 | 0.6667 | 1 | 0 | 2 | 1 | 1 | 1 | 7 | 0.7143 |
| 19_4383557 | 1 | 0 | 1 | 1 | 2 | 0.5 | NA | NA | 4 | 0.5 | 3 | 0.6667 | 1 | 1 | 2 | 0.5 | NA | NA | 6 | 0.6667 |
| 19_4383664 | 1 | 1 | NA | NA | 2 | 1 | NA | NA | 3 | 1 | NA | NA | 1 | 0 | 1 | 1 | NA | NA | 2 | 0.5 |
| 19_4383667 | 1 | 1 | NA | NA | NA | NA | NA | NA | 1 | 1 | NA | NA | 1 | 0 | 1 | 1 | NA | NA | 2 | 0.5 |
| 19_4383830 | NA | NA | NA | NA | 2 | 1 | NA | NA | 2 | 1 | NA | NA | 1 | 1 | 1 | 1 | 1 | 1 | 3 | 1 |
| 19_4383848 | 1 | 1 | NA | NA | 2 | 1 | NA | NA | 3 | 1 | NA | NA | 1 | 1 | NA | NA | 1 | 1 | 2 | 1 |
| 19_4383853 | NA | NA | NA | NA | 1 | 1 | NA | NA | 1 | 1 | NA | NA | 1 | 1 | NA | NA | 1 | 1 | 2 | 1 |
| 19_4383857 | NA | NA | NA | NA | 2 | 1 | NA | NA | 2 | 1 | NA | NA | 1 | 1 | NA | NA | NA | NA | 1 | 1 |
| 19_4388957 | 1 | 1 | NA | NA | 1 | 0 | NA | NA | 2 | 0.5 | NA | NA | NA | NA | NA | NA | 1 | 1 | 1 | 1 |
| 19_4388973 | 1 | 1 | NA | NA | 1 | 1 | NA | NA | 2 | 1 | NA | NA | NA | NA | NA | NA | 1 | 1 | 1 | 1 |
| 19_4389002 | 1 | 1 | NA | NA | 1 | 1 | NA | NA | 2 | 1 | NA | NA | NA | NA | NA | NA | 1 | 1 | 1 | 1 |
| 19_4402350 | 2 | 0 | 5 | 0.2 | 4 | 0.25 | 5 | 0.4 | 16 | 0.25 | 4 | 0 | NA | NA | 12 | 0.0833 | 6 | 0.1667 | 22 | 0.0909 |
| 19_4402501 | 2 | 0.5 | 3 | 0.3333 | 3 | 0.3333 | 2 | 0.5 | 10 | 0.4 | 3 | 0 | 2 | 0 | 10 | 0 | 5 | 0.4 | 20 | 0.1 |
| 19_4402529 | 2 | 1 | 3 | 1 | 3 | 1 | 3 | 1 | 11 | 1 | 3 | 1 | 1 | 1 | 9 | 0.8889 | 5 | 0.8 | 18 | 0.8889 |
| 19_4402530 | 2 | 0 | 2 | 0 | 2 | 1 | 1 | 1 | 7 | 0.4286 | 1 | 1 | NA | NA | 4 | 0.5 | 4 | 0.25 | 9 | 0.4444 |
| 19_4402543 | 1 | 0 | 3 | 0.3333 | 3 | 0.3333 | 2 | 0 | 9 | 0.2222 | 2 | 0.5 | 1 | 0 | 8 | 0 | 4 | 0 | 15 | 0.0667 |
| 19_4402545 | 1 | 1 | 3 | 0.3333 | 2 | 1 | 3 | 0 | 9 | 0.4444 | 2 | 1 | 1 | 0 | 9 | 0.6667 | 5 | 1 | 17 | 0.7647 |
| 19_4402548 | 1 | 1 | 3 | 1 | 2 | 0.5 | 3 | 0.6667 | 9 | 0.7778 | 2 | 1 | 1 | 1 | 9 | 0.7778 | 5 | 1 | 17 | 0.8824 |
| 19_4406234 | 1 | 0 | 2 | 0 | 5 | 0.2 | 3 | 0 | 11 | 0.0909 | 5 | 0.2 | 2 | 0 | 5 | 0.4 | 5 | 0.2 | 17 | 0.2353 |
| 19_4406250 | 3 | 0.3333 | 2 | 0.5 | 7 | 0 | 3 | 0 | 15 | 0.1333 | 4 | 0 | 2 | 0.5 | 5 | 0.2 | 5 | 0 | 16 | 0.125 |
| 19_4406281 | 2 | 1 | 2 | 1 | 4 | 0.25 | 3 | 0.3333 | 11 | 0.5455 | 3 | 0.6667 | 1 | 0 | 4 | 0.5 | 3 | 1 | 11 | 0.6364 |
| 19_4406744 | 1 | 0 | NA | NA | 2 | 0.5 | 1 | 0 | 4 | 0.25 | 2 | 0.5 | NA | NA | NA | NA | 3 | 0.3333 | 5 | 0.4 |
| 19_4406752 | 1 | 0 | NA | NA | 2 | 0.5 | 1 | 0 | 4 | 0.25 | 2 | 0.5 | NA | NA | NA | NA | 3 | 0.3333 | 5 | 0.4 |
| 19_4409858 | 1 | 1 | 2 | 1 | 1 | 1 | 2 | 0.5 | 6 | 0.8333 | 1 | 1 | 2 | 0.5 | 1 | 1 | 3 | 0.3333 | 7 | 0.5714 |
| 19_4409860 | 2 | 1 | 2 | 1 | 2 | 1 | 3 | 0 | 9 | 0.6667 | 1 | 0 | 2 | 0.5 | 3 | 0.3333 | 3 | 0.3333 | 9 | 0.3333 |
| 19_4409870 | 2 | 1 | 1 | 0 | 1 | 1 | 1 | 0 | 5 | 0.6 | 1 | 0 | 2 | 1 | 1 | 0 | 2 | 0 | 6 | 0.3333 |
| 19_4409873 | 1 | 1 | 2 | 0.5 | 1 | 0 | 2 | 0.5 | 6 | 0.5 | 2 | 0.5 | 2 | 0.5 | 3 | 0.6667 | 2 | 0.5 | 9 | 0.5556 |
| 19_4409875 | 1 | 0 | 2 | 0.5 | 1 | 1 | 3 | 0.3333 | 7 | 0.4286 | 2 | 0 | 2 | 0 | 3 | 0.6667 | 1 | 0 | 8 | 0.25 |
| 19_4409880 | NA | NA | 1 | 1 | 1 | 1 | 1 | 1 | 3 | 1 | 1 | 0 | 3 | 0.6667 | 2 | 0.5 | 2 | 0.5 | 8 | 0.5 |
| 19_4409883 | 1 | 0 | 2 | 0.5 | 1 | 1 | 2 | 1 | 6 | 0.6667 | 1 | 0 | 3 | 1 | 2 | 1 | 1 | 0 | 7 | 0.7143 |
| 19_4409898 | 1 | 1 | 2 | 1 | 1 | 0 | 3 | 0 | 7 | 0.4286 | 2 | 1 | 4 | 0.5 | 3 | 0.3333 | 2 | 0.5 | 11 | 0.5455 |
| 19_4412787 | 10 | 0.1 | 9 | 0.4444 | 6 | 0 | 11 | 0.1818 | 36 | 0.1944 | 6 | 0.6667 | 10 | 0.2 | 8 | 0.25 | 8 | 0.5 | 32 | 0.375 |
| 19_4412791 | 11 | 0.7273 | 9 | 0.7778 | 3 | 0.6667 | 9 | 0.6667 | 32 | 0.7188 | 6 | 0.3333 | 11 | 0.8182 | 6 | 0.3333 | 7 | 0.4286 | 30 | 0.5333 |
| 19_4412793 | 12 | 0.1667 | 10 | 0.1 | 5 | 0.4 | 9 | 0.1111 | 36 | 0.1667 | 6 | 0 | 10 | 0.3 | 7 | 0.1429 | 8 | 0.375 | 31 | 0.2258 |
| 19_4412798 | 11 | 0.5455 | 5 | 0.4 | 4 | 0.25 | 9 | 0.8889 | 29 | 0.5862 | 4 | 0.25 | 10 | 0.1 | 7 | 0.4286 | 7 | 0.4286 | 28 | 0.2857 |
| 19_4413147 | 11 | 0 | 9 | 0.2222 | 5 | 0.2 | 9 | 0 | 34 | 0.0882 | 7 | 0.1429 | 11 | 0 | 6 | 0.1667 | 7 | 0 | 31 | 0.0645 |
| 19_4423026 | 1 | 1 | NA | NA | NA | NA | NA | NA | 1 | 1 | NA | NA | NA | NA | 1 | 1 | NA | NA | 1 | 1 |
| 19_4428267 | 27 | 0.2593 | 28 | 0.1786 | 27 | 0.2593 | 21 | 0.2381 | 103 | 0.233 | 20 | 0.15 | 29 | 0.1379 | 34 | 0.2353 | 17 | 0.2941 | 100 | 0.2 |
| 19_4428972 | 44 | 0.1364 | 43 | 0.0698 | 48 | 0.125 | 45 | 0.0889 | 180 | 0.1056 | 35 | 0.0571 | 50 | 0.16 | 47 | 0.1489 | 28 | 0.0714 | 160 | 0.1188 |
| 19_4429161 | 49 | 0.6531 | 52 | 0.7115 | 49 | 0.6122 | 48 | 0.7292 | 198 | 0.6768 | 33 | 0.7273 | 54 | 0.6667 | 41 | 0.7561 | 29 | 0.7241 | 157 | 0.7134 |
| 19_4429170 | 40 | 0.05 | 47 | 0.0426 | 51 | 0.098 | 40 | 0.025 | 178 | 0.0562 | 35 | 0.0571 | 51 | 0.0784 | 39 | 0.0769 | 24 | 0.0833 | 149 | 0.0738 |
| 19_4429174 | 56 | 0.3571 | 53 | 0.4717 | 54 | 0.4074 | 48 | 0.4167 | 211 | 0.4123 | 42 | 0.5952 | 55 | 0.3818 | 54 | 0.4444 | 27 | 0.6296 | 178 | 0.4888 |
| 19_4429178 | 50 | 0.36 | 44 | 0.3864 | 43 | 0.4884 | 39 | 0.4359 | 176 | 0.4148 | 34 | 0.5588 | 44 | 0.3864 | 43 | 0.3953 | 21 | 0.6667 | 142 | 0.4718 |
| 19_4429347 | 107 | 0.243 | 114 | 0.2105 | 108 | 0.1667 | 106 | 0.1792 | 435 | 0.2 | 80 | 0.15 | 96 | 0.1562 | 100 | 0.21 | 68 | 0.2353 | 344 | 0.186 |
| 19_4429411 | 1 | 1 | NA | NA | 1 | 1 | NA | NA | 2 | 1 | NA | NA | NA | NA | 2 | 1 | NA | NA | 2 | 1 |
| 19_4429416 | 120 | 0.1417 | 130 | 0.1692 | 133 | 0.1654 | 121 | 0.1074 | 504 | 0.1468 | 92 | 0.1304 | 112 | 0.1964 | 122 | 0.1148 | 94 | 0.1702 | 420 | 0.1524 |
| 19_4429438 | 100 | 0.07 | 102 | 0.0392 | 100 | 0.05 | 81 | 0.0988 | 383 | 0.0627 | 68 | 0.0441 | 91 | 0.0769 | 88 | 0.0455 | 69 | 0.0725 | 316 | 0.0601 |
| 19_4429442 | 120 | 0.0917 | 119 | 0.084 | 127 | 0.0709 | 105 | 0.0286 | 471 | 0.0701 | 89 | 0.0899 | 95 | 0.0632 | 107 | 0.0841 | 94 | 0.1383 | 385 | 0.0935 |
| 19_4429484 | 154 | 0.0714 | 157 | 0.051 | 167 | 0.0659 | 127 | 0.0551 | 605 | 0.0612 | 109 | 0.0459 | 128 | 0.0469 | 136 | 0.0515 | 118 | 0.0763 | 491 | 0.055 |
| 19_4429504 | 157 | 0.0637 | 157 | 0.0573 | 167 | 0.1198 | 147 | 0.0884 | 628 | 0.0828 | 111 | 0.1892 | 128 | 0.0703 | 146 | 0.0479 | 119 | 0.0924 | 504 | 0.0952 |
| 19_4429514 | 162 | 0.1358 | 176 | 0.1364 | 172 | 0.1453 | 155 | 0.1032 | 665 | 0.1308 | 119 | 0.1513 | 134 | 0.1119 | 156 | 0.109 | 124 | 0.1129 | 533 | 0.1201 |
| 19_4438312 | 16 | 0.125 | 19 | 0.1053 | 17 | 0.2353 | 17 | 0.1176 | 69 | 0.1449 | 15 | 0.1333 | 14 | 0.0714 | 13 | 0.0769 | 13 | 0 | 55 | 0.0727 |

| | | | | | | | | | | | | | | | | | | | | |
|---|---|---|---|---|---|---|---|---|---|---|---|---|---|---|---|---|---|---|---|---|
| 19_4438638 | 1 | 0 | 6 | 0.6667 | 5 | 0.8 | 6 | 0.8333 | 18 | 0.7222 | 7 | 0.8571 | 4 | 0.75 | 5 | 0.6 | 3 | 1 | 19 | 0.7895 |
| 19_4438874 | 2 | 0 | 6 | 0.3333 | 7 | 0 | 6 | 0.1667 | 21 | 0.1429 | 5 | 0.2 | 5 | 0 | 5 | 0.4 | 2 | 0 | 17 | 0.1765 |
| 19_4456121 | 1 | 1 | NA | NA | NA | NA | 1 | 0 | 2 | 0.5 | NA | NA | NA | NA | NA | NA | 1 | 1 | 1 | 1 |
| 19_4456458 | 2 | 0 | 6 | 0.1667 | 3 | 0.3333 | 7 | 0.4286 | 18 | 0.2778 | 4 | 0.25 | NA | NA | 5 | 0 | 3 | 0.3333 | 12 | 0.1667 |
| 19_4492344 | 2 | 0.5 | 3 | 0.6667 | NA | NA | NA | NA | 5 | 0.6 | NA | NA | NA | NA | 1 | 0 | 4 | 0.25 | 5 | 0.2 |
| 19_4492403 | 1 | 0 | 2 | 0.5 | NA | NA | 2 | 0.5 | 5 | 0.4 | NA | NA | NA | NA | NA | NA | 4 | 0.75 | 4 | 0.75 |
| 19_4492994 | 2 | 0.5 | 3 | 0.6667 | NA | NA | 2 | 0.5 | 7 | 0.5714 | NA | NA | NA | NA | 1 | 1 | 4 | 0.5 | 5 | 0.6 |
| 19_4493021 | 2 | 1 | 3 | 1 | NA | NA | 2 | 1 | 7 | 1 | NA | NA | NA | NA | NA | NA | 3 | 0.6667 | 3 | 0.6667 |
| 19_4493032 | 2 | 0.5 | 3 | 0.6667 | NA | NA | 1 | 1 | 6 | 0.6667 | NA | NA | NA | NA | 1 | 1 | 4 | 0.5 | 5 | 0.6 |
| 19_4493049 | 2 | 1 | 3 | 1 | NA | NA | 2 | 1 | 7 | 1 | NA | NA | NA | NA | 1 | 0 | 4 | 1 | 5 | 0.8 |
| 19_4493079 | 2 | 0 | 4 | 0.5 | NA | NA | 2 | 0 | 8 | 0.25 | NA | NA | NA | NA | 1 | 1 | 3 | 0 | 4 | 0.25 |
| 19_4496915 | 3 | 0.6667 | 6 | 0 | 1 | 0 | 4 | 0.25 | 14 | 0.2143 | NA | NA | 4 | 0.75 | 5 | 0.4 | 7 | 0.4286 | 16 | 0.5 |
| 19_4496936 | 4 | 0.25 | 5 | 0 | 1 | 1 | 3 | 0.3333 | 13 | 0.2308 | NA | NA | 4 | 0.25 | 5 | 0.2 | 8 | 0 | 17 | 0.1176 |
| 19_4496937 | 4 | 1 | 5 | 0.8 | NA | NA | 4 | 1 | 13 | 0.9231 | NA | NA | 4 | 1 | 4 | 1 | 8 | 0.875 | 16 | 0.9375 |
| 19_4496964 | 4 | 0.5 | 7 | 0 | 2 | 0 | 4 | 0 | 17 | 0.1176 | NA | NA | 4 | 0.25 | 5 | 0.6 | 8 | 0.25 | 17 | 0.3529 |
| 19_4497064 | 6 | 0.3333 | 7 | 0.4286 | 2 | 0 | 3 | 0.6667 | 18 | 0.3889 | NA | NA | 4 | 0.25 | 4 | 0.25 | 8 | 0.25 | 16 | 0.25 |
| 19_4497085 | 5 | 0.2 | 3 | 0 | 2 | 0 | 4 | 0.25 | 14 | 0.1429 | 1 | 1 | 6 | 0 | 5 | 0 | 6 | 0 | 18 | 0.0556 |
| 19_4497129 | 3 | 0.3333 | 1 | 0 | 1 | 1 | 3 | 0.3333 | 8 | 0.375 | 2 | 0.5 | 5 | 0.6 | 5 | 0.4 | 5 | 0.4 | 17 | 0.4706 |
| 19_4497131 | 3 | 0.3333 | 1 | 0 | 1 | 0 | 3 | 0.3333 | 8 | 0.25 | 2 | 0.5 | 3 | 0.3333 | 3 | 0.3333 | 4 | 0.25 | 12 | 0.3333 |
| 19_4498993 | 2 | 0 | NA | NA | 2 | 0.5 | NA | NA | 4 | 0.25 | 3 | 0.3333 | 2 | 0.5 | NA | NA | 1 | 0 | 6 | 0.3333 |
| 19_4499010 | 1 | 0 | 1 | 1 | 2 | 1 | NA | NA | 4 | 0.75 | 3 | 0.6667 | 1 | 1 | 1 | 1 | 1 | 0 | 6 | 0.6667 |
| 19_4499034 | 2 | 0.5 | 1 | 1 | 2 | 1 | NA | NA | 5 | 0.8 | 1 | 0 | 2 | 0.5 | 1 | 0 | 1 | 1 | 5 | 0.4 |
| 19_4499072 | NA | NA | 1 | 1 | 1 | 1 | NA | NA | 2 | 1 | 2 | 0.5 | 2 | 1 | NA | NA | NA | NA | 4 | 0.75 |
| 19_4499144 | NA | NA | 1 | 0 | 1 | 1 | NA | NA | 2 | 0.5 | 2 | 0.5 | 1 | 0 | NA | NA | 1 | 0 | 4 | 0.25 |
| 19_4499156 | 5 | 0.6 | 1 | 1 | 1 | 1 | NA | NA | 7 | 0.7143 | 4 | 1 | 2 | 1 | 1 | 1 | 1 | 1 | 8 | 1 |
| 19_4499357 | 3 | 0 | 3 | 0.6667 | 3 | 0 | 7 | 0.5714 | 16 | 0.375 | 2 | 0 | 1 | 0 | 6 | 0.5 | 3 | 0.3333 | 12 | 0.3333 |
| 19_4499358 | 2 | 0.5 | 4 | 0.75 | 2 | 0 | 5 | 0 | 13 | 0.3077 | 2 | 0 | 2 | 0 | 3 | 0 | 2 | 0.5 | 9 | 0.1111 |
| 19_4499361 | 3 | 0 | 4 | 0 | 4 | 0 | 9 | 0.1111 | 20 | 0.05 | 2 | 0 | 3 | 0 | 3 | 0.3333 | 3 | 0 | 11 | 0.0909 |
| 19_4499363 | 4 | 0 | 5 | 0.2 | 5 | 0 | 8 | 0.125 | 22 | 0.0909 | 2 | 0 | 3 | 0 | 5 | 0.4 | 3 | 0.3333 | 13 | 0.2308 |
| 19_4499383 | 3 | 1 | 2 | 0 | 5 | 0.8 | 9 | 0.6667 | 19 | 0.6842 | 2 | 0.5 | 2 | 0.5 | 3 | 0.3333 | 3 | 1 | 10 | 0.6 |
| 19_4499387 | 8 | 0.625 | 7 | 0.5714 | 8 | 0.375 | 13 | 0.6154 | 36 | 0.5556 | 4 | 0.5 | 10 | 0.6 | 5 | 0.2 | 3 | 0.6667 | 22 | 0.5 |
| 19_4499390 | 8 | 0.875 | 8 | 1 | 10 | 0.7 | 14 | 0.9286 | 40 | 0.875 | 8 | 0.625 | 10 | 0.9 | 6 | 0.8333 | 5 | 0.8 | 29 | 0.7931 |
| 19_4499398 | 7 | 0.1429 | 7 | 0.2857 | 10 | 0.3 | 12 | 0.25 | 36 | 0.25 | 9 | 0.2222 | 9 | 0.2222 | 6 | 0.5 | 5 | 0.4 | 29 | 0.3103 |
| 19_4499401 | 7 | 0.2857 | 5 | 0.4 | 9 | 0 | 12 | 0.5833 | 33 | 0.3333 | 8 | 0.375 | 10 | 0.3 | 6 | 0.3333 | 5 | 0.6 | 29 | 0.3793 |
| 19_4499412 | 8 | 0.625 | 8 | 1 | 10 | 0.9 | 11 | 0.5455 | 37 | 0.7568 | 8 | 0.625 | 10 | 0.8 | 5 | 0.4 | 4 | 1 | 27 | 0.7037 |
| 19_4499413 | 8 | 0.25 | 6 | 0 | 8 | 0 | 11 | 0.0909 | 33 | 0.0909 | 8 | 0 | 9 | 0.1111 | 6 | 0.1667 | 5 | 0 | 28 | 0.0714 |
| 19_4499417 | 7 | 0.2857 | 6 | 0.3333 | 8 | 0.125 | 13 | 0.2308 | 34 | 0.2353 | 6 | 0.1667 | 8 | 0.125 | 6 | 0.3333 | 4 | 0.25 | 24 | 0.2083 |
| 19_4499420 | 8 | 0.875 | 7 | 1 | 7 | 1 | 10 | 1 | 32 | 0.9688 | 8 | 0.75 | 10 | 0.9 | 5 | 0.8 | 5 | 1 | 28 | 0.8571 |
| 19_4499425 | 8 | 0.25 | 8 | 0 | 9 | 0.1111 | 12 | 0.0833 | 37 | 0.1081 | 7 | 0.1429 | 9 | 0.1111 | 5 | 0.2 | 4 | 0.25 | 25 | 0.16 |
| 19_4499589 | NA | NA | NA | NA | 1 | 1 | NA | NA | 1 | 1 | 1 | 1 | NA | NA | NA | NA | NA | NA | 1 | 1 |
| 19_4499643 | NA | NA | 1 | 0 | 2 | 1 | NA | NA | 3 | 0.6667 | 1 | 1 | NA | NA | NA | NA | NA | NA | 1 | 1 |
| 19_4499646 | NA | NA | NA | NA | 3 | 0.6667 | NA | NA | 3 | 0.6667 | 2 | 0.5 | NA | NA | NA | NA | NA | NA | 2 | 0.5 |
| 19_4499652 | NA | NA | 1 | 0 | 3 | 1 | NA | NA | 4 | 0.75 | 2 | 1 | NA | NA | NA | NA | NA | NA | 2 | 1 |
| 19_4499671 | NA | NA | 1 | 1 | 3 | 1 | NA | NA | 4 | 1 | 2 | 1 | NA | NA | NA | NA | NA | NA | 2 | 1 |
| 19_4499678 | NA | NA | 1 | 0 | 3 | 0.3333 | NA | NA | 4 | 0.25 | 2 | 0.5 | NA | NA | NA | NA | NA | NA | 2 | 0.5 |
| 19_4500022 | NA | NA | 1 | 1 | 5 | 1 | NA | NA | 6 | 1 | 2 | 1 | NA | NA | NA | NA | NA | NA | 2 | 1 |
| 19_4500045 | NA | NA | 1 | 0 | 5 | 0.4 | NA | NA | 6 | 0.3333 | 2 | 1 | NA | NA | NA | NA | NA | NA | 2 | 1 |
| 19_4500069 | NA | NA | 1 | 0 | 3 | 0.6667 | NA | NA | 4 | 0.5 | 2 | 0.5 | NA | NA | NA | NA | NA | NA | 2 | 0.5 |
| 19_4500115 | NA | NA | 1 | 1 | 6 | 1 | 1 | 1 | 8 | 1 | 2 | 0.5 | NA | NA | 1 | 0 | NA | NA | 3 | 0.3333 |
| 19_4501222 | 3 | 1 | 6 | 0.6667 | 3 | 0.3333 | NA | NA | 12 | 0.6667 | 3 | 0.6667 | 2 | 0.5 | NA | NA | NA | NA | 5 | 0.6 |
| 19_4501452 | 6 | 0.6667 | 6 | 0.8333 | 3 | 0.6667 | 1 | 1 | 16 | 0.75 | 5 | 1 | 4 | 1 | 1 | 1 | 3 | 1 | 13 | 1 |
| 19_4501465 | 4 | 0.5 | 5 | 0.4 | 3 | 0.6667 | 1 | 0 | 13 | 0.4615 | 4 | 0.25 | 3 | 0.6667 | 1 | 0 | 3 | 0.6667 | 11 | 0.4545 |
| 19_4501469 | 6 | 0.3333 | 6 | 0.6667 | 3 | 0.6667 | 1 | 0 | 16 | 0.5 | 5 | 1 | 3 | 0.6667 | NA | NA | 3 | 1 | 11 | 0.9091 |
| 19_4501473 | 6 | 0 | 5 | 0 | 5 | 0.4 | NA | NA | 16 | 0.125 | 3 | 0 | 4 | 0.25 | NA | NA | 4 | 0.25 | 11 | 0.1818 |
| 19_4501481 | 4 | 1 | 3 | 1 | 7 | 1 | 1 | 1 | 15 | 1 | 5 | 1 | 4 | 1 | 2 | 1 | 3 | 0.6667 | 14 | 0.9286 |
| 19_4501503 | 9 | 0 | 4 | 0.25 | 7 | 0 | NA | NA | 20 | 0.05 | 5 | 0.2 | 2 | 0 | 1 | 0 | 5 | 0 | 13 | 0.0769 |
| 19_4501534 | 6 | 0 | 4 | 0 | 8 | 0.25 | 2 | 1 | 20 | 0.2 | 7 | 0 | 4 | 0.25 | 2 | 0 | 5 | 0 | 18 | 0.0556 |
| 19_4501553 | 3 | 0.6667 | 1 | 1 | 1 | 1 | 1 | 1 | 6 | 0.8333 | 4 | 0.75 | 1 | 1 | NA | NA | NA | NA | 5 | 0.8 |
| 19_4501578 | 1 | 1 | 1 | 0 | 1 | 0 | 1 | 1 | 4 | 0.5 | 1 | 1 | 1 | 0 | 2 | 0 | NA | NA | 4 | 0.25 |
| 19_4501585 | 1 | 0 | 1 | 0 | 1 | 1 | 1 | 0 | 4 | 0.25 | NA | NA | 1 | 1 | 1 | 1 | NA | NA | 2 | 1 |
| 19_4501630 | NA | NA | NA | NA | NA | NA | 1 | 1 | 1 | 1 | NA | NA | 1 | 0 | 2 | 0.5 | NA | NA | 3 | 0.3333 |
| 19_4501916 | NA | NA | 2 | 1 | 1 | 1 | NA | NA | 3 | 1 | NA | NA | NA | NA | 1 | 1 | NA | NA | 1 | 1 |
| 19_4501951 | NA | NA | 2 | 0.5 | 2 | 0 | NA | NA | 4 | 0.25 | NA | NA | NA | NA | 1 | 1 | NA | NA | 1 | 1 |
| 19_4503609 | 2 | 0.5 | 1 | 0 | 1 | 0 | NA | NA | 4 | 0.25 | 1 | 0 | 2 | 0 | 1 | 0 | 3 | 0.3333 | 7 | 0.1429 |
| 19_4504461 | 16 | 0.125 | 14 | 0.3571 | 12 | 0.0833 | 8 | 0 | 50 | 0.16 | 9 | 0 | 10 | 0.3 | 9 | 0.1111 | 17 | 0.1176 | 45 | 0.1333 |

| | | | | | | | | | | | | | | | | | | | | |
|---|---|---|---|---|---|---|---|---|---|---|---|---|---|---|---|---|---|---|---|---|
| 19_450446 | 18 | 0.0556 | 16 | 0.0625 | 13 | 0.0769 | 11 | 0 | 58 | 0.0517 | 11 | 0.0909 | 11 | 0.0909 | 9 | 0.1111 | 16 | 0 | 47 | 0.0638 |
| 19_450676 | 1 | 0 | 1 | 0 | 2 | 0.5 | 2 | 0 | 6 | 0.1667 | 2 | 1 | NA | NA | NA | NA | NA | NA | 2 | 1 |
| 19_452983 | NA | NA | 1 | 1 | NA | NA | NA | NA | 1 | 1 | 1 | 1 | NA | NA | NA | NA | NA | NA | 1 | 1 |
| 19_452997 | NA | NA | 2 | 1 | NA | NA | NA | NA | 2 | 1 | 1 | 1 | NA | NA | NA | NA | 1 | 1 | 2 | 1 |
| 19_452999 | NA | NA | 2 | 0.5 | NA | NA | NA | NA | 2 | 0.5 | 1 | 1 | NA | NA | NA | NA | 1 | 1 | 2 | 1 |
| 19_454962 | 8 | 0.25 | 5 | 0.2 | 2 | 0 | 2 | 0.5 | 17 | 0.2353 | 8 | 0.125 | 6 | 0.1667 | 5 | 0.4 | 8 | 0.125 | 27 | 0.1852 |
| 19_455174 | 12 | 0.0833 | 7 | 0.1429 | 6 | 0 | 7 | 0.1429 | 32 | 0.0938 | 12 | 0.0833 | 12 | 0.1667 | 8 | 0 | 12 | 0 | 44 | 0.0682 |
| 19_455175 | 15 | 0.0667 | 7 | 0.7143 | 7 | 0 | 7 | 0.5714 | 36 | 0.2778 | 13 | 0.2308 | 13 | 0.2308 | 6 | 0.3333 | 10 | 0.3 | 42 | 0.2619 |
| 19_455175 | 12 | 0.25 | 4 | 0.25 | 6 | 0.1667 | 6 | 0.1667 | 28 | 0.2143 | 11 | 0.2727 | 13 | 0.4615 | 8 | 0.5 | 10 | 0.2 | 42 | 0.3571 |
| 19_455176 | 13 | 0.6154 | 7 | 0.4286 | 7 | 0.2857 | 5 | 0.4 | 32 | 0.4688 | 15 | 0.3333 | 15 | 0.7333 | 8 | 0.25 | 12 | 0.4167 | 50 | 0.46 |
| 19_455177 | 9 | 0.1111 | 7 | 0.4286 | 6 | 0.1667 | 6 | 0.8333 | 28 | 0.3571 | 14 | 0.3571 | 14 | 0.4286 | 8 | 0.5 | 13 | 0.2308 | 49 | 0.3673 |
| 19_455177 | 1 | 1 | 1 | 1 | NA | NA | 1 | 1 | 3 | 1 | 3 | 1 | 4 | 0.5 | 1 | 1 | 6 | 0.3333 | 14 | 0.5714 |
| 19_455178 | 2 | 0 | 2 | 0.5 | NA | NA | 2 | 0 | 6 | 0.1667 | 2 | 0 | 3 | 0 | 2 | 0.5 | 7 | 0.2857 | 14 | 0.2143 |
| 19_455180 | 1 | 1 | 2 | 0.5 | NA | NA | 2 | 0 | 5 | 0.4 | 3 | 0 | 6 | 0.3333 | 2 | 0 | 7 | 0.2857 | 18 | 0.2222 |
| 19_455181 | 1 | 1 | 2 | 0.5 | NA | NA | 2 | 1 | 5 | 0.8 | 3 | 1 | 6 | 0.6667 | 1 | 0 | 7 | 0.4286 | 17 | 0.5882 |
| 19_455181 | 2 | 0.5 | 2 | 0.5 | NA | NA | 2 | 0 | 6 | 0.3333 | 3 | 0 | 5 | 0.2 | 2 | 0 | 7 | 0 | 17 | 0.0588 |
| 19_455182 | 2 | 1 | 2 | 1 | 1 | 1 | 1 | 0 | 6 | 0.8333 | 3 | 1 | 7 | 0.5714 | 2 | 0.5 | 7 | 0.8571 | 19 | 0.7368 |
| 19_455183 | 1 | 1 | 2 | 0.5 | 1 | 0 | 2 | 0 | 6 | 0.3333 | 3 | 0 | 7 | 0.1429 | 3 | 0 | 5 | 0.4 | 18 | 0.1667 |
| 19_455185 | 1 | 1 | 2 | 1 | 1 | 1 | 1 | 0 | 5 | 0.8 | 2 | 0.5 | 4 | 0.5 | 2 | 0.5 | 5 | 0.4 | 13 | 0.4615 |
| 19_455188 | 1 | 1 | 2 | 1 | 2 | 1 | 2 | 1 | 7 | 1 | 2 | 1 | 4 | 0.5 | 1 | 1 | 5 | 0.8 | 12 | 0.75 |
| 19_455190 | 1 | 1 | 3 | 1 | 2 | 1 | 3 | 0.3333 | 9 | 0.7778 | 3 | 0.6667 | 5 | 0.6 | 2 | 0.5 | 4 | 0.25 | 14 | 0.5 |
| 19_455191 | 1 | 0 | 4 | 0 | 2 | 0.5 | 3 | 0 | 10 | 0.1 | 2 | 0 | 5 | 0.2 | 2 | 0.5 | 6 | 0.1667 | 15 | 0.2 |
| 19_455193 | 1 | 1 | 3 | 0.3333 | 2 | 0.5 | 2 | 0 | 8 | 0.375 | 2 | 0.5 | 4 | 0 | 3 | 0 | 6 | 0 | 15 | 0.0667 |
| 19_455193 | 1 | 1 | 3 | 1 | 2 | 1 | 2 | 0.5 | 8 | 0.875 | 3 | 0.3333 | 4 | 0.5 | 2 | 0 | 4 | 0.5 | 13 | 0.3846 |
| 19_455193 | 1 | 1 | 3 | 1 | 2 | 1 | 3 | 1 | 9 | 1 | 3 | 0.6667 | 5 | 1 | 2 | 1 | 4 | 0.75 | 14 | 0.8571 |
| 19_455195 | NA | NA | 4 | 0.5 | 2 | 0.5 | 3 | 0.3333 | 9 | 0.4444 | 1 | 0 | 4 | 0.25 | 2 | 1 | 6 | 0 | 13 | 0.2308 |
| 19_455199 | 1 | 0 | 4 | 0 | 2 | 0 | 2 | 0.5 | 9 | 0.1111 | 1 | 0 | 7 | 0.1429 | 4 | 0.25 | 4 | 0 | 16 | 0.125 |
| 19_455212 | 3 | 0 | 9 | 0.2222 | 3 | 0 | 2 | 0 | 17 | 0.1176 | 5 | 0 | 5 | 0 | 5 | 0 | 5 | 0.2 | 20 | 0.05 |
| 19_455213 | 3 | 0 | 9 | 0.1111 | 3 | 0.3333 | 3 | 0.3333 | 18 | 0.1667 | 5 | 0.4 | 6 | 0.1667 | 6 | 0.1667 | 4 | 0 | 21 | 0.1905 |
| 19_455215 | 2 | 1 | 8 | 0.125 | 3 | 0 | 3 | 0.3333 | 16 | 0.25 | 4 | 0.25 | 4 | 0.25 | 5 | 0.2 | 4 | 0.25 | 17 | 0.2353 |
| 19_455409 | NA | NA | 1 | 0 | NA | NA | 1 | 1 | 2 | 0.5 | NA | NA | 3 | 0.6667 | 1 | 1 | 2 | 0.5 | 6 | 0.6667 |
| 19_455412 | NA | NA | 1 | 1 | 1 | 0 | 1 | 0 | 3 | 0.3333 | NA | NA | 2 | 0 | 1 | 1 | 2 | 0.5 | 5 | 0.4 |
| 19_455412 | NA | NA | 1 | 1 | 1 | 1 | 1 | 1 | 3 | 1 | NA | NA | 3 | 1 | 1 | 1 | 1 | 1 | 5 | 1 |
| 19_455412 | 1 | 1 | 1 | 0 | 1 | 0 | 1 | 1 | 4 | 0.5 | NA | NA | 3 | 0.6667 | 1 | 1 | 1 | 0 | 5 | 0.6 |
| 19_455412 | 1 | 1 | 1 | 1 | NA | NA | 1 | 0 | 3 | 0.6667 | NA | NA | 2 | 1 | 1 | 1 | 1 | 1 | 4 | 1 |
| 19_455472 | NA | NA | 1 | 1 | NA | NA | NA | NA | 1 | 1 | NA | NA | 2 | 0.5 | 1 | 1 | NA | NA | 3 | 0.6667 |
| 19_455659 | 12 | 0.4167 | 16 | 0.375 | 9 | 0.3333 | 14 | 0.2857 | 51 | 0.3529 | 11 | 0.2727 | 12 | 0.1667 | 12 | 0.25 | 8 | 0.125 | 43 | 0.2093 |
| 19_455660 | 12 | 0.3333 | 15 | 0.0667 | 8 | 0 | 14 | 0.3571 | 49 | 0.2041 | 10 | 0.2 | 14 | 0.4286 | 11 | 0.1818 | 9 | 0.3333 | 44 | 0.2955 |
| 19_455661 | 9 | 0 | 10 | 0 | 9 | 0.1111 | 12 | 0.1667 | 40 | 0.075 | 10 | 0.2 | 11 | 0.1818 | 9 | 0 | 6 | 0 | 36 | 0.1111 |
| 19_455661 | 12 | 0.0833 | 15 | 0.0667 | 6 | 0 | 12 | 0.0833 | 45 | 0.0667 | 10 | 0.4 | 11 | 0.2727 | 9 | 0.2222 | 8 | 0.25 | 38 | 0.2895 |
| 19_455661 | NA | NA | NA | NA | 1 | 1 | NA | NA | 1 | 1 | 1 | 1 | NA | NA | NA | NA | NA | NA | 1 | 1 |
| 19_455669 | 3 | 0 | 3 | 0.3333 | 2 | 0 | 3 | 0.6667 | 11 | 0.2727 | 2 | 0 | 3 | 0.3333 | 2 | 0 | 3 | 0 | 10 | 0.1 |
| 19_455671 | 2 | 0.5 | 2 | 0 | 1 | 0 | 2 | 0 | 7 | 0.1429 | 2 | 0 | 2 | 0 | 2 | 0 | 2 | 0.5 | 8 | 0.125 |
| 19_455672 | 2 | 0 | 3 | 0 | 2 | 0 | 1 | 1 | 8 | 0.125 | 1 | 0 | 2 | 0 | 2 | 0 | 3 | 0.3333 | 8 | 0.125 |
| 19_457363 | 1 | 0 | 4 | 0.25 | 3 | 0 | NA | NA | 8 | 0.125 | 1 | 1 | 3 | 0.3333 | 5 | 0.6 | 6 | 0 | 15 | 0.3333 |
| 19_457378 | 1 | 0 | 4 | 0.25 | 3 | 0 | NA | NA | 8 | 0.125 | 1 | 0 | NA | NA | 3 | 0 | 3 | 0.3333 | 7 | 0.1429 |
| 19_457387 | 18 | 0.0556 | 16 | 0.0625 | 19 | 0.0526 | 24 | 0.0417 | 77 | 0.0519 | 12 | 0 | 20 | 0.1 | 29 | 0.1034 | 22 | 0.0909 | 83 | 0.0843 |
| 19_457400 | 15 | 0.0667 | 6 | 0 | 14 | 0 | 16 | 0.125 | 51 | 0.0588 | 5 | 0 | 17 | 0.1765 | 21 | 0.0476 | 15 | 0.0667 | 58 | 0.0862 |
| 19_457404 | 14 | 0.7143 | 8 | 0.625 | 15 | 0.8 | 15 | 0.9333 | 52 | 0.7885 | 6 | 0.8333 | 14 | 0.7857 | 15 | 0.8 | 13 | 0.9231 | 48 | 0.8333 |
| 19_457404 | 15 | 0.2 | 8 | 0.125 | 14 | 0.1429 | 21 | 0 | 58 | 0.1034 | 5 | 0.2 | 21 | 0.1905 | 23 | 0.1739 | 14 | 0.0714 | 63 | 0.1587 |
| 19_457405 | 9 | 0.8889 | 5 | 1 | 12 | 0.9167 | 20 | 0.95 | 46 | 0.9348 | 2 | 1 | 15 | 1 | 15 | 1 | 9 | 1 | 41 | 1 |
| 19_457405 | 13 | 0.3077 | 10 | 0.1 | 12 | 0.3333 | 19 | 0.2632 | 54 | 0.2593 | 4 | 0.5 | 18 | 0.2778 | 21 | 0.4762 | 12 | 0.4167 | 55 | 0.4 |
| 19_457431 | 14 | 0 | 10 | 0.2 | 13 | 0.0769 | 16 | 0 | 53 | 0.0566 | 4 | 0.25 | 15 | 0 | 21 | 0 | 12 | 0.25 | 52 | 0.0769 |
| 19_457559 | 4 | 0.75 | 7 | 0.8571 | 7 | 0.5714 | 12 | 0.8333 | 30 | 0.7667 | 3 | 1 | 10 | 0.9 | 11 | 0.6364 | 7 | 0.8571 | 31 | 0.8065 |
| 19_457559 | 6 | 0.3333 | 8 | 0.25 | 7 | 0.2857 | 12 | 0.25 | 33 | 0.2727 | 2 | 0 | 10 | 0.1 | 12 | 0.1667 | 7 | 0.2857 | 31 | 0.1613 |
| 19_457561 | 8 | 0.75 | 6 | 0.6667 | 9 | 0.6667 | 10 | 0.8 | 33 | 0.7273 | 3 | 0.6667 | 10 | 0.7 | 11 | 0.6364 | 6 | 0.6667 | 30 | 0.6667 |
| 19_457561 | 5 | 0.2 | 6 | 0.1667 | 8 | 0.125 | 10 | 0.5 | 29 | 0.2759 | 3 | 0.3333 | 9 | 0.1111 | 11 | 0.0909 | 6 | 0 | 29 | 0.1034 |
| 19_457563 | 4 | 1 | 7 | 0.7143 | 9 | 0.8889 | 11 | 0.6364 | 31 | 0.7742 | 3 | 0.6667 | 10 | 0.7 | 11 | 0.8182 | 7 | 0.5714 | 31 | 0.7097 |
| 19_458066 | 2 | 0 | 2 | 1 | 1 | 0 | 1 | 0 | 6 | 0.3333 | 2 | 0 | 3 | 0 | 2 | 0 | 2 | 0.5 | 9 | 0.1111 |
| 19_458067 | 5 | 0.2 | 2 | 0 | 1 | 0 | 1 | 0 | 9 | 0.1111 | 2 | 0 | 3 | 0.3333 | 1 | 0 | 2 | 0.5 | 8 | 0.25 |
| 19_458069 | 4 | 0 | 2 | 0.5 | 1 | 0 | NA | NA | 7 | 0.1429 | 2 | 0 | 3 | 0.3333 | 1 | 0 | 1 | 0 | 7 | 0.1429 |
| 19_458071 | 4 | 0.5 | 2 | 0 | NA | NA | 1 | 1 | 7 | 0.4286 | 1 | 1 | 2 | 0.5 | 1 | 1 | 2 | 0.5 | 6 | 0.6667 |
| 19_458072 | 3 | 0.6667 | NA | NA | 1 | 1 | 1 | 0 | 5 | 0.6 | 2 | 0.5 | 2 | 0.5 | NA | NA | 2 | 1 | 6 | 0.6667 |
| 19_458073 | 4 | 0.25 | 2 | 0.5 | 1 | 0 | NA | NA | 7 | 0.2857 | 1 | 0 | NA | NA | 1 | 1 | 2 | 0.5 | 4 | 0.5 |
| 19_459926 | NA | NA | 1 | 1 | NA | NA | NA | NA | 1 | 1 | NA | NA | 1 | 1 | NA | NA | NA | NA | 1 | 1 |

| | | | | | | | | | | | | | | | | | | | | |
|---|---|---|---|---|---|---|---|---|---|---|---|---|---|---|---|---|---|---|---|---|
| 19_459926 | NA | NA | 1 | 1 | NA | NA | NA | NA | 1 | 1 | NA | NA | 1 | 1 | NA | NA | NA | NA | 1 | 1 |
| 19_459928 | NA | NA | 1 | 1 | NA | NA | NA | NA | 1 | 1 | NA | NA | 1 | 1 | NA | NA | NA | NA | 1 | 1 |
| 19_459930 | NA | NA | 1 | 1 | NA | NA | NA | NA | 1 | 1 | NA | NA | 1 | 1 | NA | NA | NA | NA | 1 | 1 |
| 19_459930 | NA | NA | 1 | 1 | NA | NA | NA | NA | 1 | 1 | NA | NA | 1 | 1 | NA | NA | NA | NA | 1 | 1 |
| 19_459937 | NA | NA | 1 | 1 | NA | NA | NA | NA | 1 | 1 | NA | NA | NA | NA | 1 | 1 | NA | NA | 1 | 1 |
| 19_459939 | NA | NA | 1 | 1 | NA | NA | NA | NA | 1 | 1 | NA | NA | NA | NA | 1 | 1 | NA | NA | 1 | 1 |
| 19_459941 | NA | NA | 2 | 1 | NA | NA | NA | NA | 2 | 1 | NA | NA | 1 | 1 | 1 | 0 | NA | NA | 2 | 0.5 |
| 19_459942 | NA | NA | 1 | 1 | NA | NA | NA | NA | 1 | 1 | NA | NA | 1 | 1 | 1 | 1 | NA | NA | 2 | 1 |
| 19_459943 | NA | NA | 2 | 1 | NA | NA | NA | NA | 2 | 1 | NA | NA | 1 | 1 | 1 | 1 | NA | NA | 2 | 1 |
| 19_459943 | NA | NA | 2 | 1 | NA | NA | NA | NA | 2 | 1 | NA | NA | 1 | 1 | 1 | 1 | NA | NA | 2 | 1 |
| 19_459943 | NA | NA | 2 | 0.5 | NA | NA | NA | NA | 2 | 0.5 | NA | NA | 1 | 1 | NA | NA | NA | NA | 1 | 1 |
| 19_459945 | NA | NA | 2 | 1 | NA | NA | NA | NA | 2 | 1 | NA | NA | 1 | 1 | 1 | 1 | NA | NA | 2 | 1 |
| 19_459945 | NA | NA | 1 | 1 | NA | NA | NA | NA | 1 | 1 | NA | NA | NA | NA | 1 | 1 | NA | NA | 1 | 1 |
| 19_460199 | 4 | 0 | 2 | 0 | 1 | 0 | 4 | 0.5 | 11 | 0.1818 | 1 | 0 | 3 | 0 | 2 | 0 | 4 | 0.25 | 10 | 0.1 |
| 19_460213 | 16 | 0.8125 | 15 | 0.8667 | 11 | 1 | 12 | 0.9167 | 54 | 0.8889 | 13 | 0.7692 | 13 | 0.7692 | 10 | 0.8 | 12 | 0.8333 | 48 | 0.7917 |
| 19_460214 | 12 | 1 | 8 | 1 | 9 | 1 | 8 | 1 | 37 | 1 | 12 | 0.9167 | 9 | 0.8889 | 8 | 0.875 | 9 | 1 | 38 | 0.9211 |
| 19_460214 | 10 | 0.9 | 11 | 0.9091 | 7 | 0.7143 | 9 | 0.7778 | 37 | 0.8378 | 10 | 1 | 10 | 0.8 | 7 | 0.5714 | 11 | 0.9091 | 38 | 0.8421 |
| 19_460214 | 15 | 0.8667 | 14 | 0.8571 | 10 | 0.9 | 12 | 1 | 51 | 0.902 | 12 | 1 | 12 | 1 | 8 | 1 | 12 | 0.8333 | 44 | 0.9545 |
| 19_460216 | 12 | 0.25 | 14 | 0.1429 | 8 | 0 | 10 | 0.1 | 44 | 0.1364 | 10 | 0.3 | 11 | 0.3636 | 7 | 0 | 9 | 0 | 37 | 0.1892 |
| 19_460218 | 2 | 0 | 6 | 0.1667 | 1 | 0 | 2 | 0 | 11 | 0.0909 | 1 | 1 | 2 | 0 | NA | NA | 1 | 0 | 4 | 0.25 |
| 19_460277 | 1 | 1 | NA | NA | NA | NA | NA | NA | 1 | 1 | 1 | 0 | NA | NA | NA | NA | 1 | 1 | 2 | 0.5 |
| 19_460512 | 1 | 1 | NA | NA | NA | NA | NA | NA | 1 | 1 | 1 | 1 | NA | NA | NA | NA | NA | NA | 1 | 1 |
| 19_460514 | 2 | 1 | NA | NA | 1 | 1 | NA | NA | 3 | 1 | 1 | 1 | NA | NA | NA | NA | NA | NA | 1 | 1 |
| 19_460521 | 2 | 1 | NA | NA | NA | NA | NA | NA | 2 | 1 | 1 | 1 | NA | NA | NA | NA | NA | NA | 1 | 1 |
| 19_460522 | 1 | 0 | NA | NA | 1 | 1 | NA | NA | 2 | 0.5 | 1 | 1 | NA | NA | NA | NA | NA | NA | 1 | 1 |
| 19_460527 | 1 | 1 | NA | NA | 1 | 1 | NA | NA | 2 | 1 | 1 | 1 | NA | NA | NA | NA | NA | NA | 1 | 1 |
| 19_460528 | 1 | 1 | 1 | 1 | 1 | 1 | 1 | 1 | 4 | 1 | 1 | 1 | NA | NA | NA | NA | NA | NA | 1 | 1 |
| 19_460531 | 2 | 0.5 | 1 | 1 | 1 | 1 | NA | NA | 4 | 0.75 | 1 | 1 | NA | NA | NA | NA | NA | NA | 1 | 1 |
| 19_460576 | 3 | 0.3333 | NA | NA | 1 | 1 | NA | NA | 4 | 0.5 | 2 | 1 | NA | NA | 1 | 1 | 1 | 0 | 4 | 0.75 |
| 19_460576 | 2 | 1 | 1 | 1 | NA | NA | 1 | 1 | 4 | 1 | 2 | 0.5 | NA | NA | 1 | 0 | 1 | 0 | 4 | 0.25 |
| 19_460578 | 3 | 0.6667 | 1 | 1 | 2 | 1 | 1 | 0 | 7 | 0.7143 | 3 | 0.3333 | NA | NA | 2 | 1 | 1 | 1 | 6 | 0.6667 |
| 19_460579 | 4 | 1 | 1 | 0 | 2 | 0.5 | 2 | 1 | 9 | 0.7778 | 3 | 0 | NA | NA | 3 | 0.6667 | 1 | 1 | 7 | 0.4286 |
| 19_460611 | 4 | 0 | 1 | 0 | 2 | 1 | 2 | 0 | 9 | 0.2222 | 4 | 0 | 1 | 0 | 5 | 0.6 | 1 | 1 | 11 | 0.3636 |
| 19_460714 | 4 | 0 | 3 | 0 | 4 | 0 | 3 | 0.3333 | 14 | 0.0714 | 2 | 0 | 5 | 0 | 3 | 0 | 4 | 0.25 | 14 | 0.0714 |
| 19_460722 | 3 | 0 | 3 | 0 | 4 | 0 | 4 | 0.25 | 14 | 0.0714 | 2 | 0 | 7 | 0.1429 | 4 | 0 | 5 | 0 | 18 | 0.0556 |
| 19_460726 | 4 | 0.25 | 6 | 0.8333 | 6 | 0.1667 | 3 | 0.3333 | 19 | 0.4211 | 3 | 0.3333 | 9 | 0.6667 | 6 | 0.1667 | 6 | 0.8333 | 24 | 0.5417 |
| 19_460726 | 4 | 0.25 | 5 | 0.8 | 5 | 0.4 | 4 | 0.25 | 18 | 0.4444 | 2 | 0.5 | 9 | 0.5556 | 5 | 0.2 | 6 | 0.6667 | 22 | 0.5 |
| 19_460726 | 4 | 0 | 5 | 0.4 | 6 | 0.1667 | 4 | 0 | 19 | 0.1579 | 3 | 0 | 7 | 0.2857 | 6 | 0.1667 | 7 | 0 | 23 | 0.1304 |
| 19_460726 | 5 | 0 | 6 | 0 | 4 | 0 | 4 | 0.25 | 19 | 0.0526 | 3 | 0.3333 | 9 | 0.1111 | 6 | 0.3333 | 6 | 0.1667 | 24 | 0.2083 |
| 19_460727 | 3 | 1 | 6 | 1 | 5 | 0.6 | 4 | 1 | 18 | 0.8889 | 3 | 0.6667 | 8 | 0.625 | 6 | 0.8333 | 7 | 0.8571 | 24 | 0.75 |
| 19_460727 | 4 | 1 | 6 | 0.6667 | 4 | 1 | 3 | 0.6667 | 17 | 0.8235 | 3 | 0.6667 | 7 | 0.8571 | 6 | 1 | 6 | 1 | 22 | 0.9091 |
| 19_460728 | 4 | 0.25 | 6 | 0.1667 | 5 | 0 | 4 | 0 | 19 | 0.1053 | 3 | 0 | 6 | 0.1667 | 5 | 0 | 6 | 0 | 20 | 0.05 |
| 19_460729 | 5 | 0.4 | 7 | 0 | 4 | 0.25 | 3 | 0 | 19 | 0.1579 | 2 | 0 | 7 | 0 | 6 | 0.3333 | 6 | 0 | 21 | 0.0952 |
| 19_460733 | 4 | 0.25 | 4 | 0 | 6 | 0.5 | 5 | 0 | 19 | 0.2105 | 3 | 0.3333 | 5 | 0.4 | 7 | 0.1429 | 6 | 0.1667 | 21 | 0.2381 |
| 19_460733 | 5 | 0 | 5 | 0 | 5 | 0.2 | 5 | 0 | 20 | 0.05 | 3 | 0 | 7 | 0.2857 | 7 | 0 | 4 | 0 | 21 | 0.0952 |
| 19_460733 | 3 | 0 | 5 | 0.4 | 6 | 0 | 4 | 0 | 18 | 0.1111 | 2 | 0.5 | 5 | 0 | 6 | 0 | 5 | 0 | 18 | 0.0556 |
| 19_460735 | 5 | 0.4 | 5 | 0.4 | 4 | 0.25 | 5 | 0.2 | 19 | 0.3158 | 3 | 0.3333 | 5 | 0.2 | 4 | 0.25 | 5 | 0.4 | 17 | 0.2941 |
| 19_462744 | 9 | 0 | 7 | 0.1429 | 5 | 0 | 10 | 0.2 | 31 | 0.0968 | 6 | 0.1667 | 9 | 0.2222 | 9 | 0.1111 | 9 | 0.1111 | 33 | 0.1515 |
| 19_462745 | 12 | 0.3333 | 8 | 0 | 5 | 0 | 10 | 0.3 | 35 | 0.2 | 6 | 0 | 8 | 0.125 | 9 | 0.1111 | 8 | 0.25 | 31 | 0.129 |
| 19_462748 | 11 | 0.3636 | 9 | 0.3333 | 6 | 0.1667 | 12 | 0.1667 | 38 | 0.2632 | 7 | 0.5714 | 9 | 0.1111 | 7 | 0 | 9 | 0.1111 | 32 | 0.1875 |
| 19_462749 | 6 | 0.5 | 4 | 0.25 | 5 | 0 | 12 | 0.25 | 27 | 0.2593 | 5 | 0.2 | 8 | 0.125 | 5 | 0.4 | 6 | 0 | 24 | 0.1667 |
| 19_463106 | 10 | 0.1 | 7 | 0.2857 | 8 | 0 | 6 | 0 | 31 | 0.0968 | 8 | 0.375 | 7 | 0.1429 | 11 | 0.1818 | 10 | 0.2 | 36 | 0.2222 |
| 19_463106 | 12 | 0.1667 | 6 | 0 | 8 | 0 | 8 | 0 | 34 | 0.0588 | 6 | 0.1667 | 8 | 0.125 | 11 | 0 | 10 | 0.1 | 35 | 0.0857 |
| 19_463117 | 13 | 0 | 5 | 0 | 5 | 0.2 | 8 | 0.125 | 31 | 0.0645 | 7 | 0.5714 | 7 | 0.2857 | 12 | 0.1667 | 10 | 0.2 | 36 | 0.2778 |
| 19_463147 | NA | NA | NA | NA | NA | NA | 1 | 1 | 1 | 1 | NA | NA | NA | NA | 4 | 0.5 | 1 | 0 | 5 | 0.4 |
| 19_463176 | 3 | 0 | 3 | 0 | 6 | 0.1667 | 6 | 0 | 18 | 0.0556 | 4 | 0.5 | 6 | 0 | 3 | 0 | 4 | 0 | 17 | 0.1176 |
| 19_463177 | 3 | 0 | 1 | 0 | 5 | 0.4 | 4 | 0.25 | 13 | 0.2308 | 3 | 0.3333 | 5 | 0.4 | 3 | 0.3333 | 2 | 0.5 | 13 | 0.3846 |
| 19_463179 | 3 | 0 | 4 | 0.5 | 6 | 0 | 10 | 0.4 | 23 | 0.2609 | 2 | 0 | 9 | 0.5556 | 4 | 0.25 | 6 | 0 | 21 | 0.2857 |
| 19_463185 | 3 | 0 | 4 | 0 | 6 | 0 | 8 | 0.25 | 21 | 0.0952 | 4 | 0.25 | 10 | 0.2 | 4 | 0 | 6 | 0 | 24 | 0.125 |
| 19_463186 | 5 | 0.2 | 4 | 0 | 7 | 0.1429 | 8 | 0.25 | 24 | 0.1667 | 4 | 0 | 8 | 0.5 | 4 | 0 | 4 | 0.25 | 20 | 0.25 |
| 19_463187 | 5 | 0.2 | 3 | 0.6667 | 6 | 0.6667 | 8 | 0.625 | 22 | 0.5455 | 4 | 0.25 | 10 | 0.7 | 3 | 0.3333 | 3 | 0.6667 | 20 | 0.55 |
| 19_463188 | 4 | 0.25 | 3 | 0.3333 | 7 | 0.7143 | 7 | 0.5714 | 21 | 0.5238 | 2 | 0.5 | 8 | 0.75 | 4 | 0.5 | 6 | 0.8333 | 20 | 0.7 |
| 19_463631 | 6 | 0.3333 | 2 | 1 | 3 | 0.6667 | 2 | 0 | 13 | 0.4615 | 2 | 0 | 6 | 0.5 | 9 | 0.3333 | 6 | 0.1667 | 23 | 0.3043 |
| 19_463631 | 8 | 0.375 | 3 | 0 | 5 | 0 | 2 | 0.5 | 18 | 0.2222 | 3 | 0.6667 | 9 | 0.3333 | 11 | 0.1818 | 8 | 0.375 | 31 | 0.3226 |
| 19_463632 | 6 | 1 | 2 | 1 | 6 | 0.8333 | 2 | 1 | 16 | 0.9375 | 2 | 1 | 10 | 1 | 9 | 0.8889 | 9 | 0.8889 | 30 | 0.9333 |

| | | | | | | | | | | | | | | | | | | | | |
|---|---|---|---|---|---|---|---|---|---|---|---|---|---|---|---|---|---|---|---|---|
| 19_4636331 | 6 | 0.5 | 3 | 0.3333 | 5 | 0 | 2 | 1 | 16 | 0.375 | 3 | 0.3333 | 10 | 0.4 | 11 | 0.4545 | 3 | 0 | 27 | 0.3704 |
| 19_4636335 | 8 | 0.125 | 3 | 0.3333 | 5 | 0.2 | 2 | 0.5 | 18 | 0.2222 | 3 | 0.3333 | 11 | 0.3636 | 12 | 0.1667 | 8 | 0.25 | 34 | 0.2647 |
| 19_4636339 | 5 | 0.2 | 3 | 0.3333 | 5 | 0 | 3 | 0.3333 | 16 | 0.1875 | 2 | 0 | 9 | 0.5556 | 11 | 0.2727 | 7 | 0.4286 | 29 | 0.3793 |
| 19_4636342 | 7 | 0 | 3 | 0 | 5 | 0.2 | 3 | 0 | 18 | 0.0556 | 3 | 0.3333 | 11 | 0.1818 | 11 | 0 | 7 | 0 | 32 | 0.0938 |
| 19_4636343 | 6 | 0.1667 | 3 | 0 | 5 | 0 | 3 | 0 | 17 | 0.0588 | 3 | 0 | 11 | 0.0909 | 11 | 0.0909 | 7 | 0 | 32 | 0.0625 |
| 19_4636536 | 23 | 0.5652 | 24 | 0.5417 | 18 | 0.3333 | 23 | 0.5217 | 88 | 0.5 | 24 | 0.5833 | 25 | 0.44 | 21 | 0.381 | 18 | 0.3333 | 88 | 0.4432 |
| 19_4636565 | 18 | 0.1667 | 14 | 0.3571 | 16 | 0.3125 | 19 | 0.4737 | 67 | 0.3284 | 24 | 0.125 | 15 | 0.2 | 24 | 0.125 | 13 | 0.3846 | 76 | 0.1842 |
| 19_4636604 | 19 | 0.1053 | 18 | 0 | 20 | 0.05 | 21 | 0.1905 | 78 | 0.0897 | 22 | 0.0455 | 21 | 0 | 20 | 0.1 | 16 | 0.125 | 79 | 0.0633 |
| 19_4636613 | 21 | 0.0476 | 16 | 0 | 19 | 0 | 21 | 0.2381 | 77 | 0.0779 | 20 | 0.25 | 19 | 0.0526 | 17 | 0.0588 | 18 | 0.0556 | 74 | 0.1081 |
| 19_4636648 | 18 | 0.1667 | 13 | 0.0769 | 19 | 0.2105 | 15 | 0.2667 | 65 | 0.1846 | 14 | 0.2143 | 15 | 0.2667 | 17 | 0.2353 | 15 | 0.2 | 61 | 0.2295 |
| 19_4636651 | 18 | 0.1111 | 15 | 0.0667 | 20 | 0.05 | 20 | 0.2 | 73 | 0.1096 | 15 | 0.2 | 21 | 0 | 17 | 0.1765 | 19 | 0.0526 | 72 | 0.0972 |
| 19_4636662 | 12 | 0.1667 | 11 | 0.2727 | 17 | 0 | 10 | 0 | 50 | 0.1 | 9 | 0.3333 | 15 | 0.0667 | 12 | 0.25 | 13 | 0.0769 | 49 | 0.1633 |
| 19_4636663 | 17 | 0.7647 | 13 | 0.7692 | 19 | 0.8421 | 17 | 0.6471 | 66 | 0.7576 | 13 | 0.6923 | 18 | 0.4444 | 15 | 0.6667 | 14 | 0.7143 | 60 | 0.6167 |
| 19_4636672 | 17 | 0.1176 | 13 | 0 | 17 | 0.1176 | 18 | 0.1667 | 65 | 0.1077 | 13 | 0.3077 | 20 | 0.1 | 18 | 0.1111 | 17 | 0.1765 | 68 | 0.1618 |
| 19_4636676 | 12 | 0.0833 | 11 | 0 | 17 | 0.1176 | 13 | 0 | 53 | 0.0566 | 12 | 0.0833 | 18 | 0 | 14 | 0.0714 | 14 | 0.2143 | 58 | 0.0862 |
| 19_4647443 | 2 | 0.5 | 1 | 0 | NA | NA | 2 | 0.5 | 5 | 0.4 | NA | NA | 1 | 1 | NA | NA | NA | NA | 1 | 1 |
| 19_4647444 | 1 | 0 | 2 | 0.5 | NA | NA | 2 | 0 | 5 | 0.2 | 1 | 0 | 1 | 0 | 1 | 1 | 1 | 0 | 4 | 0.25 |
| 19_4647449 | 2 | 0 | 1 | 0 | NA | NA | 2 | 0.5 | 5 | 0.2 | 1 | 0 | 1 | 1 | NA | NA | 1 | 1 | 3 | 0.6667 |
| 19_4647467 | 1 | 1 | 1 | 0 | NA | NA | 1 | 1 | 3 | 0.6667 | 1 | 1 | 1 | 1 | 1 | 1 | 1 | 1 | 4 | 1 |
| 19_4647487 | 1 | 1 | 2 | 1 | NA | NA | 2 | 1 | 5 | 1 | 1 | 1 | 2 | 1 | 1 | 1 | 1 | 1 | 5 | 1 |
| 19_4647489 | NA | NA | 3 | 0.6667 | 1 | 1 | 2 | 0.5 | 6 | 0.6667 | 1 | 1 | 1 | 1 | NA | NA | 1 | 1 | 3 | 1 |
| 19_4647512 | 2 | 0 | 3 | 0 | 1 | 1 | 2 | 0 | 8 | 0.125 | 1 | 0 | 1 | 1 | 1 | 0 | 1 | 0 | 4 | 0.25 |
| 19_4647512 | 2 | 0.5 | 2 | 0.5 | NA | NA | 2 | 0.5 | 6 | 0.5 | 1 | 1 | 2 | 1 | 1 | 0 | 1 | 0 | 5 | 0.6 |
| 19_4647527 | 2 | 0.5 | 3 | 1 | 1 | 0 | 2 | 0 | 8 | 0.5 | 1 | 0 | 2 | 1 | NA | NA | 1 | 0 | 4 | 0.5 |
| 19_4647543 | 1 | 0 | 3 | 0 | 1 | 0 | 2 | 1 | 7 | 0.2857 | 1 | 1 | 2 | 1 | 3 | 0.3333 | NA | NA | 6 | 0.6667 |
| 19_4647569 | 2 | 0.5 | 2 | 0.5 | 1 | 1 | 3 | 0.6667 | 8 | 0.625 | NA | NA | 1 | 1 | 3 | 0.3333 | 1 | 1 | 5 | 0.6 |
| 19_4647588 | 1 | 0 | 2 | 0 | 1 | 0 | 3 | 0.6667 | 7 | 0.2857 | 1 | 0 | 1 | 1 | 3 | 0.6667 | 1 | 0 | 6 | 0.5 |
| 19_4647589 | 1 | 1 | 3 | 0.3333 | 1 | 0 | 3 | 0 | 8 | 0.25 | 1 | 0 | 3 | 0.6667 | 3 | 0.6667 | 1 | 0 | 8 | 0.5 |
| 19_4647659 | 2 | 0.5 | 3 | 0 | 1 | 0 | 3 | 0 | 9 | 0.1111 | 1 | 0 | 2 | 0 | 4 | 0.25 | 1 | 0 | 8 | 0.125 |
| 19_4647664 | 2 | 0 | 3 | 0.3333 | 1 | 0 | 3 | 0 | 9 | 0.1111 | 1 | 0 | 3 | 0 | 4 | 0.25 | 1 | 0 | 9 | 0.1111 |
| 19_4647666 | 1 | 0 | 2 | 0 | 1 | 1 | 3 | 0 | 7 | 0.1429 | 1 | 0 | 2 | 0.5 | 4 | 0.25 | 1 | 1 | 8 | 0.375 |
| 19_4647707 | 2 | 0 | 3 | 0.3333 | 1 | 1 | 3 | 0 | 9 | 0.2222 | NA | NA | 3 | 0.3333 | 4 | 0 | 1 | 0 | 8 | 0.125 |
| 19_4648947 | 2 | 0 | 1 | 0 | 1 | 1 | NA | NA | 4 | 0.25 | 1 | 1 | 1 | 1 | 1 | 1 | NA | NA | 3 | 1 |
| 19_4649018 | 4 | 0 | 4 | 0 | 3 | 1 | NA | NA | 11 | 0.2727 | 5 | 0.4 | 4 | 0.75 | 2 | 1 | 1 | 0 | 12 | 0.5833 |
| 19_4649020 | 2 | 0.5 | 4 | 0.25 | 4 | 0.5 | 1 | 0 | 11 | 0.3636 | 3 | 0.3333 | 2 | 0.5 | 3 | 0.3333 | 2 | 0.5 | 10 | 0.4 |
| 19_4649030 | 5 | 0.2 | 4 | 0 | 9 | 0.1111 | 1 | 0 | 19 | 0.1053 | 3 | 0 | 5 | 0.4 | 3 | 0.3333 | 2 | 0 | 13 | 0.2308 |
| 19_4649036 | 4 | 0.5 | 5 | 0.2 | 6 | 0 | 1 | 0 | 16 | 0.1875 | 5 | 0.6 | 5 | 0 | 3 | 0.3333 | 2 | 0 | 15 | 0.2667 |
| 19_4649038 | 5 | 0.2 | 4 | 0.25 | 8 | 0.125 | NA | NA | 17 | 0.1765 | 5 | 0.4 | 5 | 0.4 | 3 | 0.6667 | 3 | 0.3333 | 16 | 0.4375 |
| 19_4649059 | 3 | 0.3333 | 3 | 1 | 8 | 0.75 | 1 | 1 | 15 | 0.7333 | 5 | 0.8 | 6 | 0.5 | 2 | 1 | 3 | 0.3333 | 16 | 0.625 |
| 19_4649071 | 6 | 0.3333 | 4 | 0.5 | 8 | 0.5 | 1 | 1 | 19 | 0.4737 | 7 | 0.4286 | 6 | 0.5 | 1 | 1 | 6 | 0.3333 | 20 | 0.45 |
| 19_4649075 | 6 | 0.8333 | 6 | 0.6667 | 10 | 0.8 | 1 | 1 | 23 | 0.7826 | 6 | 0.6667 | 5 | 0.6 | 3 | 0.6667 | 4 | 0.5 | 18 | 0.6111 |
| 19_4649084 | 7 | 0.2857 | 6 | 0.1667 | 12 | 0.5833 | 1 | 0 | 26 | 0.3846 | 7 | 0.4286 | 6 | 0.3333 | 3 | 0.3333 | 5 | 0.2 | 21 | 0.3333 |
| 19_4649090 | 7 | 0.7143 | 7 | 0.8571 | 10 | 0.9 | NA | NA | 24 | 0.8333 | 7 | 0.8571 | 7 | 0.7143 | 3 | 1 | 3 | 0.6667 | 20 | 0.8 |
| 19_4649097 | 7 | 0 | 7 | 0.1429 | 7 | 0.2857 | NA | NA | 21 | 0.1429 | 6 | 0.1667 | 2 | 0 | 3 | 0 | 4 | 0 | 15 | 0.0667 |
| 19_4649098 | 6 | 0.1667 | 6 | 0.1667 | 10 | 0.2 | 1 | 0 | 23 | 0.1739 | 6 | 0 | 5 | 0.2 | 3 | 0.3333 | 5 | 0.2 | 19 | 0.1579 |
| 19_4649101 | 7 | 0.7143 | 8 | 0.625 | 12 | 1 | 1 | 1 | 28 | 0.8214 | 4 | 1 | 8 | 0.75 | 3 | 0.3333 | 6 | 1 | 21 | 0.8095 |
| 19_4649141 | 7 | 0.4286 | 8 | 0.625 | 13 | 0.3846 | 1 | 1 | 29 | 0.4828 | 9 | 0.6667 | 8 | 0.875 | 3 | 0.3333 | 9 | 0.4444 | 29 | 0.6207 |
| 19_4650386 | 10 | 0.3 | 11 | 0.7273 | 17 | 0.5294 | 6 | 0.3333 | 44 | 0.5 | 14 | 0.4286 | 11 | 0.5455 | 7 | 0.7143 | 8 | 0.625 | 40 | 0.55 |
| 19_4650393 | 10 | 0.8 | 7 | 1 | 13 | 0.8462 | 3 | 1 | 33 | 0.8788 | 10 | 1 | 12 | 0.6667 | 6 | 0.6667 | 4 | 1 | 32 | 0.8125 |
| 19_4650401 | 11 | 0.0909 | 10 | 0.6 | 20 | 0.7 | 6 | 0.6667 | 47 | 0.5319 | 15 | 0.8667 | 13 | 0.6154 | 7 | 0.4286 | 10 | 0.9 | 45 | 0.7333 |
| 19_4650413 | 11 | 0.3636 | 8 | 0.375 | 17 | 0.4706 | 6 | 0.5 | 42 | 0.4286 | 13 | 0.6154 | 11 | 0.3636 | 9 | 0.3333 | 7 | 0.5714 | 40 | 0.475 |
| 19_4650417 | 12 | 0.0833 | 11 | 0.3636 | 18 | 0.3889 | 7 | 0.1429 | 48 | 0.2708 | 15 | 0.2 | 14 | 0.2143 | 8 | 0.625 | 9 | 0.2222 | 46 | 0.2826 |
| 19_4650436 | 12 | 0.25 | 10 | 0.4 | 18 | 0.2778 | 10 | 0.7 | 50 | 0.38 | 18 | 0.6111 | 15 | 0.4 | 6 | 0.1667 | 6 | 1 | 45 | 0.5333 |
| 19_4650465 | 12 | 0 | 14 | 0.2857 | 19 | 0.1579 | 10 | 0 | 55 | 0.1273 | 22 | 0.0455 | 15 | 0.0667 | 11 | 0.0909 | 9 | 0.1111 | 57 | 0.0702 |
| 19_4650480 | 12 | 0.5833 | 12 | 0.9167 | 16 | 0.75 | 10 | 0.8 | 50 | 0.76 | 20 | 0.85 | 13 | 0.8462 | 10 | 0.8 | 8 | 0.875 | 51 | 0.8431 |
| 19_4651788 | 5 | 0 | 2 | 0 | 2 | 0 | 1 | 1 | 10 | 0.1 | 2 | 0.5 | 2 | 0 | 1 | 0 | 5 | 0.6 | 10 | 0.4 |
| 19_4651795 | 3 | 0 | 1 | 1 | 2 | 0.5 | 2 | 0 | 8 | 0.25 | 2 | 0 | 2 | 0 | 2 | 0 | 4 | 0.25 | 10 | 0.1 |
| 19_4651821 | 5 | 0 | 1 | 0 | 2 | 0.5 | 2 | 0 | 10 | 0.1 | 1 | 0 | 2 | 0 | 2 | 0 | 5 | 0.2 | 10 | 0.1 |
| 19_4651830 | 3 | 0.3333 | 1 | 1 | 1 | 1 | NA | NA | 5 | 0.6 | NA | NA | NA | NA | 1 | 0 | 3 | 0.6667 | 4 | 0.5 |
| 19_4651923 | 4 | 0.25 | NA | NA | 2 | 0 | NA | NA | 6 | 0.1667 | NA | NA | 2 | 0 | 1 | 1 | 4 | 0.25 | 7 | 0.2857 |
| 19_4651937 | 3 | 0.3333 | NA | NA | 2 | 0 | 2 | 0.5 | 7 | 0.2857 | NA | NA | 2 | 0.5 | 2 | 0 | 4 | 0.75 | 8 | 0.5 |
| 19_4664261 | 5 | 0.4 | 1 | 0 | 2 | 0 | 2 | 0.5 | 10 | 0.3 | 3 | 0.3333 | 2 | 0 | 1 | 0 | 1 | 0 | 7 | 0.1429 |
| 19_4664274 | 5 | 0.2 | NA | NA | 1 | 0 | 1 | 0 | 7 | 0.1429 | 3 | 0.3333 | 2 | 0 | 1 | 0 | 1 | 0 | 7 | 0.1429 |
| 19_4664287 | 4 | 0.5 | NA | NA | 2 | 0.5 | 1 | 1 | 7 | 0.5714 | 3 | 0.3333 | 2 | 1 | 2 | 1 | 2 | 0 | 9 | 0.5556 |
| 19_4664297 | 6 | 1 | NA | NA | 3 | 1 | NA | NA | 9 | 1 | 3 | 1 | 1 | 1 | 2 | 0.5 | 2 | 1 | 8 | 0.875 |

| | | | | | | | | | | | | | | | | | | | | |
|---|---|---|---|---|---|---|---|---|---|---|---|---|---|---|---|---|---|---|---|---|
| 19_466429 | 6 | 0.1667 | NA | NA | 3 | 0.6667 | 2 | 0 | 11 | 0.2727 | 2 | 0.5 | 2 | 0 | 2 | 0 | 2 | 0 | 8 | 0.125 |
| 19_466431 | 6 | 1 | 2 | 1 | 3 | 1 | 3 | 1 | 14 | 1 | 3 | 0.6667 | 2 | 1 | 2 | 1 | 2 | 1 | 9 | 0.8889 |
| 19_466431 | 7 | 0.1429 | 1 | 0 | 3 | 0.3333 | 3 | 0 | 14 | 0.1429 | 3 | 0.6667 | 1 | 1 | 2 | 1 | 2 | 0 | 8 | 0.625 |
| 19_466432 | 6 | 1 | 2 | 0 | 3 | 1 | 3 | 0.6667 | 14 | 0.7857 | 3 | 0.6667 | 2 | 0.5 | 2 | 0.5 | 2 | 0.5 | 9 | 0.5556 |
| 19_466432 | 4 | 0.25 | NA | NA | 2 | 0 | 1 | 0 | 7 | 0.1429 | 2 | 0.5 | 1 | 0 | NA | NA | 1 | 0 | 4 | 0.25 |
| 19_466432 | 2 | 0.5 | 1 | 0 | 1 | 1 | NA | NA | 4 | 0.5 | NA | NA | NA | NA | NA | NA | 1 | 1 | 1 | 1 |
| 19_466433 | 7 | 0.2857 | 2 | 0 | 4 | 0.25 | 3 | 0 | 16 | 0.1875 | 2 | 1 | 2 | 0 | 3 | 0.6667 | 2 | 0 | 9 | 0.4444 |
| 19_466434 | 7 | 0.1429 | 3 | 0.3333 | 3 | 0.3333 | 2 | 0 | 15 | 0.2 | 3 | 0.3333 | 2 | 0 | 2 | 0 | 2 | 0 | 9 | 0.1111 |
| 19_466434 | 6 | 1 | 2 | 1 | 5 | 0.8 | 3 | 1 | 16 | 0.9375 | 3 | 1 | 1 | 1 | 2 | 0.5 | 2 | 1 | 8 | 0.875 |
| 19_466434 | 6 | 1 | 2 | 1 | 5 | 1 | 2 | 1 | 15 | 1 | 4 | 0.75 | 2 | 0.5 | 2 | 1 | 1 | 1 | 9 | 0.7778 |
| 19_466436 | 7 | 0.2857 | 2 | 0 | 4 | 0.25 | 3 | 0.6667 | 16 | 0.3125 | 3 | 0 | 2 | 0 | 2 | 0.5 | 2 | 0 | 9 | 0.1111 |
| 19_466436 | 8 | 0.75 | 1 | 1 | 5 | 0.8 | 4 | 0.5 | 18 | 0.7222 | 3 | 1 | 2 | 0 | 3 | 0.6667 | 2 | 1 | 10 | 0.7 |
| 19_466436 | 9 | 0 | 2 | 0 | 5 | 0.2 | 4 | 0 | 20 | 0.05 | 3 | 0 | 2 | 0 | 4 | 0 | 2 | 0.5 | 11 | 0.0909 |
| 19_466437 | 8 | 0.125 | 3 | 0.3333 | 5 | 0.4 | 4 | 0 | 20 | 0.2 | 4 | 0.25 | 2 | 0.5 | 4 | 0.75 | 1 | 1 | 11 | 0.5455 |
| 19_466438 | 6 | 0.8333 | 2 | 1 | 5 | 1 | 3 | 0.6667 | 16 | 0.875 | 4 | 0.5 | 2 | 0.5 | 3 | 0.6667 | 2 | 1 | 11 | 0.6364 |
| 19_466438 | 9 | 0.2222 | 2 | 0 | 4 | 0 | 4 | 0.25 | 19 | 0.1579 | 4 | 0 | 1 | 0 | 3 | 0 | 2 | 0.5 | 10 | 0.1 |
| 19_466441 | 8 | 0.5 | 2 | 0 | 5 | 0.6 | 3 | 0.6667 | 18 | 0.5 | 3 | 1 | 3 | 0.3333 | 3 | 0.6667 | 2 | 0.5 | 11 | 0.6364 |
| 19_466442 | 9 | 0 | 3 | 0 | 3 | 0.3333 | 3 | 0 | 18 | 0.0556 | 4 | 0.25 | 2 | 0.5 | 3 | 0 | 1 | 0 | 10 | 0.2 |
| 19_466442 | 8 | 0.125 | 3 | 0.3333 | 4 | 0.25 | 4 | 0 | 19 | 0.1579 | 4 | 0 | 2 | 0 | 4 | 0.25 | 2 | 0.5 | 12 | 0.1667 |
| 19_466443 | 8 | 0 | 3 | 0.3333 | 4 | 0.25 | 4 | 0 | 19 | 0.1053 | 5 | 0 | 3 | 0.3333 | 4 | 0 | 2 | 0 | 14 | 0.0714 |
| 19_466444 | 9 | 0.2222 | 3 | 0 | 3 | 0 | 4 | 0 | 19 | 0.1053 | 5 | 0.4 | 3 | 0.3333 | 4 | 0.25 | 2 | 0.5 | 14 | 0.3571 |
| 19_466445 | 8 | 0.625 | 3 | 0.3333 | 4 | 1 | 4 | 0.5 | 19 | 0.6316 | 5 | 0.6 | 3 | 0 | 4 | 0.75 | 3 | 0.6667 | 15 | 0.5333 |
| 19_466445 | 7 | 0.2857 | 3 | 0 | 2 | 0 | 2 | 0.5 | 14 | 0.2143 | 5 | 0.4 | 2 | 0.5 | 4 | 0 | 1 | 0 | 12 | 0.25 |
| 19_466450 | 6 | 0 | 5 | 0 | 5 | 0.2 | 4 | 0 | 20 | 0.05 | 6 | 0 | 3 | 0 | 4 | 0.25 | 4 | 0 | 17 | 0.0588 |
| 19_466900 | 3 | 0.6667 | 4 | 0.5 | 2 | 0.5 | 1 | 1 | 10 | 0.6 | 3 | 0.6667 | 2 | 0 | 3 | 0.6667 | 1 | 1 | 9 | 0.5556 |
| 19_466901 | 3 | 0.3333 | 4 | 0.25 | 2 | 0.5 | 1 | 1 | 10 | 0.4 | 1 | 0 | 3 | 0.3333 | 3 | 0.3333 | 1 | 0 | 8 | 0.25 |
| 19_466902 | 3 | 1 | 2 | 0.5 | 2 | 0.5 | 1 | 1 | 8 | 0.75 | 2 | 1 | 3 | 0.6667 | 2 | 1 | 1 | 1 | 8 | 0.875 |
| 19_466902 | 3 | 1 | 2 | 0.5 | 1 | 1 | 1 | 1 | 7 | 0.8571 | 3 | 0.6667 | 3 | 1 | 2 | 0.5 | 1 | 1 | 9 | 0.7778 |
| 19_466902 | 3 | 0.6667 | 4 | 0.75 | NA | NA | 1 | 1 | 8 | 0.75 | 3 | 1 | 2 | 0 | 2 | 1 | 1 | 1 | 8 | 0.75 |
| 19_466913 | 2 | 0 | 2 | 0.5 | 7 | 0.1429 | 2 | 0 | 13 | 0.1538 | 3 | 0 | 3 | 0.3333 | 4 | 0 | NA | NA | 10 | 0.1 |
| 19_466914 | 3 | 0.6667 | 4 | 0.5 | 8 | 0.25 | 2 | 0 | 17 | 0.3529 | 3 | 0 | 3 | 0.6667 | 4 | 0.25 | NA | NA | 10 | 0.3 |
| 19_467290 | 4 | 0 | 2 | 0.5 | 1 | 0 | 5 | 0 | 12 | 0.0833 | 6 | 0 | 2 | 0 | 4 | 0 | 4 | 0.25 | 16 | 0.0625 |
| 19_467290 | 3 | 0 | 2 | 0.5 | 2 | 0.5 | 3 | 1 | 10 | 0.5 | 3 | 0.3333 | 1 | 1 | 2 | 1 | 4 | 0.75 | 10 | 0.7 |
| 19_467291 | 3 | 0 | 2 | 0.5 | 2 | 0.5 | 5 | 0.6 | 12 | 0.4167 | 5 | 0.4 | 2 | 0.5 | 3 | 0.3333 | 3 | 0.3333 | 13 | 0.3846 |
| 19_467483 | NA | NA | NA | NA | 1 | 1 | NA | NA | 1 | 1 | NA | NA | NA | NA | NA | NA | 1 | 1 | 1 | 1 |
| 19_467490 | NA | NA | NA | NA | 1 | 1 | NA | NA | 1 | 1 | NA | NA | NA | NA | NA | NA | 2 | 1 | 2 | 1 |
| 19_467492 | NA | NA | NA | NA | 1 | 1 | NA | NA | 1 | 1 | NA | NA | NA | NA | NA | NA | 2 | 1 | 2 | 1 |
| 19_467495 | NA | NA | NA | NA | 1 | 1 | NA | NA | 1 | 1 | NA | NA | NA | NA | NA | NA | 1 | 1 | 1 | 1 |
| 19_467496 | NA | NA | NA | NA | 1 | 1 | NA | NA | 1 | 1 | NA | NA | NA | NA | NA | NA | 1 | 1 | 1 | 1 |
| 19_467500 | NA | NA | NA | NA | 1 | 1 | NA | NA | 1 | 1 | NA | NA | NA | NA | NA | NA | 1 | 1 | 1 | 1 |
| 19_467501 | NA | NA | NA | NA | 1 | 1 | NA | NA | 1 | 1 | NA | NA | NA | NA | NA | NA | 1 | 1 | 1 | 1 |
| 19_467514 | NA | NA | 1 | 1 | 1 | 1 | 2 | 1 | 4 | 1 | NA | NA | NA | NA | 1 | 1 | NA | NA | 1 | 1 |
| 19_467515 | 1 | 0 | 1 | 1 | 1 | 1 | 2 | 1 | 5 | 0.8 | NA | NA | NA | NA | 1 | 1 | NA | NA | 1 | 1 |
| 19_467516 | 1 | 1 | 1 | 1 | NA | NA | 1 | 1 | 3 | 1 | NA | NA | NA | NA | 2 | 1 | NA | NA | 2 | 1 |
| 19_467517 | NA | NA | 1 | 1 | 1 | 1 | 3 | 0 | 5 | 0.4 | NA | NA | NA | NA | 2 | 0.5 | NA | NA | 2 | 0.5 |
| 19_467517 | 1 | 0 | 1 | 1 | 1 | 0 | 3 | 0 | 6 | 0.1667 | NA | NA | NA | NA | 2 | 1 | NA | NA | 2 | 1 |
| 19_468663 | 3 | 0 | 2 | 0 | 2 | 0 | 4 | 0.25 | 11 | 0.0909 | 1 | 0 | NA | NA | 2 | 0 | 2 | 1 | 5 | 0.4 |
| 19_468663 | 3 | 0.3333 | 2 | 0 | 3 | 0 | 3 | 0 | 11 | 0.0909 | 1 | 1 | NA | NA | 2 | 0 | 2 | 0 | 5 | 0.2 |
| 19_468663 | 3 | 0.6667 | 2 | 1 | 3 | 1 | 4 | 0.5 | 12 | 0.75 | 1 | 1 | NA | NA | 2 | 0.5 | 2 | 1 | 5 | 0.8 |
| 19_468664 | NA | NA | 2 | 0.5 | 1 | 1 | 3 | 0.3333 | 6 | 0.5 | 1 | 1 | NA | NA | NA | NA | NA | NA | 1 | 1 |
| 19_468664 | 2 | 1 | 2 | 1 | 1 | 1 | 2 | 1 | 7 | 1 | NA | NA | NA | NA | 2 | 1 | 2 | 1 | 4 | 1 |
| 19_468665 | 1 | 1 | 2 | 1 | 3 | 1 | 4 | 1 | 10 | 1 | 1 | 1 | NA | NA | 2 | 1 | 2 | 1 | 5 | 1 |
| 19_468665 | 3 | 0 | 2 | 0 | 3 | 0.3333 | 3 | 0 | 11 | 0.0909 | NA | NA | NA | NA | 2 | 0 | 2 | 0.5 | 4 | 0.25 |
| 19_468665 | 3 | 1 | 2 | 1 | 3 | 1 | 3 | 1 | 11 | 1 | 1 | 1 | NA | NA | 2 | 1 | 2 | 1 | 5 | 1 |
| 19_468666 | 3 | 0 | 2 | 0.5 | 3 | 0.3333 | 4 | 0.5 | 12 | 0.3333 | 1 | 1 | NA | NA | 2 | 0.5 | 2 | 0.5 | 5 | 0.6 |
| 19_468666 | 3 | 0.6667 | 1 | 1 | 3 | 0.3333 | 3 | 1 | 10 | 0.7 | 1 | 0 | NA | NA | 2 | 0.5 | 2 | 0.5 | 5 | 0.4 |
| 19_468666 | 3 | 0.3333 | 2 | 0 | 3 | 0.3333 | 4 | 0 | 12 | 0.1667 | 1 | 1 | NA | NA | 2 | 0 | 2 | 0 | 5 | 0.2 |
| 19_468669 | 1 | 1 | 2 | 1 | 2 | 1 | 3 | 0.6667 | 8 | 0.875 | 1 | 1 | NA | NA | 1 | 1 | 1 | 0 | 3 | 0.6667 |
| 19_468709 | NA | NA | 3 | 1 | 2 | 0 | 1 | 1 | 6 | 0.6667 | 1 | 1 | 1 | 0 | NA | NA | 2 | 0.5 | 4 | 0.5 |
| 19_468719 | 1 | 1 | NA | NA | 2 | 0 | 1 | 1 | 4 | 0.5 | NA | NA | 1 | 1 | 2 | 0 | 2 | 0.5 | 5 | 0.4 |
| 19_468722 | NA | NA | NA | NA | 1 | 0 | 1 | 1 | 2 | 0.5 | 1 | 0 | 1 | 1 | 2 | 0 | 1 | 1 | 5 | 0.4 |
| 19_468724 | 1 | 0 | NA | NA | 2 | 0 | 2 | 1 | 5 | 0.4 | 1 | 1 | 1 | 0 | 2 | 0.5 | 2 | 0.5 | 6 | 0.5 |
| 19_468724 | 1 | 0 | NA | NA | 2 | 1 | 2 | 0.5 | 5 | 0.6 | NA | NA | 1 | 1 | 2 | 1 | 1 | 1 | 4 | 1 |
| 19_468725 | 1 | 0 | NA | NA | 1 | 0 | 2 | 0.5 | 4 | 0.25 | 1 | 1 | 1 | 0 | 2 | 0 | 2 | 0.5 | 6 | 0.3333 |
| 19_468726 | NA | NA | NA | NA | 2 | 0 | 2 | 0.5 | 4 | 0.25 | 1 | 1 | 1 | 0 | 1 | 0 | 2 | 0 | 5 | 0.2 |

| | | | | | | | | | | | | | | | | | | | | |
|---|---|---|---|---|---|---|---|---|---|---|---|---|---|---|---|---|---|---|---|---|
| 19_468731 | 1 | 0 | NA | NA | 2 | 1 | 2 | 0 | 5 | 0.4 | 1 | 1 | 1 | 1 | 1 | 1 | 1 | 1 | 4 | 1 |
| 19_468758 | 3 | 0.3333 | 1 | 0 | 2 | 0.5 | 2 | 0 | 8 | 0.25 | NA | NA | 1 | 0 | 2 | 0 | 1 | 1 | 4 | 0.25 |
| 19_468766 | 2 | 0.5 | 1 | 0 | 3 | 0 | 2 | 0 | 8 | 0.125 | 1 | 0 | 1 | 0 | 3 | 0.3333 | 2 | 0 | 7 | 0.1429 |
| 19_468769 | 4 | 0.25 | 2 | 1 | 2 | 0.5 | 3 | 0.3333 | 11 | 0.4545 | 4 | 0.25 | 5 | 0.6 | 5 | 0.6 | 2 | 0.5 | 16 | 0.5 |
| 19_468770 | 4 | 0.5 | 2 | 1 | 3 | 0.3333 | 4 | 0.25 | 13 | 0.4615 | 5 | 0 | 5 | 0.4 | 5 | 0.6 | 3 | 0.3333 | 18 | 0.3333 |
| 19_468771 | 7 | 0 | 5 | 0.2 | 6 | 0 | 8 | 0.25 | 26 | 0.1154 | 9 | 0.1111 | 7 | 0.2857 | 7 | 0.2857 | 4 | 0 | 27 | 0.1852 |
| 19_468772 | 9 | 0.8889 | 4 | 1 | 7 | 1 | 10 | 0.9 | 30 | 0.9333 | 5 | 0.8 | 7 | 0.8571 | 7 | 0.8571 | 5 | 1 | 24 | 0.875 |
| 19_468773 | 7 | 0 | 4 | 0.5 | 7 | 0.1429 | 5 | 0 | 23 | 0.1304 | 7 | 0.2857 | 8 | 0.125 | 3 | 0 | 3 | 0 | 21 | 0.1429 |
| 19_468773 | 6 | 0.5 | 4 | 0.5 | 7 | 0.4286 | 8 | 0.375 | 25 | 0.44 | 7 | 0.2857 | 9 | 0.4444 | 5 | 0.4 | 4 | 1 | 25 | 0.48 |
| 19_470120 | 15 | 0.6 | 17 | 0.7059 | 14 | 0.6429 | 10 | 0.9 | 56 | 0.6964 | 12 | 0.5833 | 11 | 0.7273 | 10 | 0.6 | 16 | 0.5625 | 49 | 0.6122 |
| 19_470122 | 16 | 0.0625 | 21 | 0.0476 | 21 | 0.0952 | 11 | 0.1818 | 69 | 0.087 | 11 | 0.0909 | 12 | 0 | 11 | 0 | 20 | 0.1 | 54 | 0.0556 |
| 19_470123 | 11 | 0.3636 | 22 | 0.1818 | 19 | 0.1579 | 10 | 0.3 | 62 | 0.2258 | 9 | 0.1111 | 10 | 0.3 | 9 | 0 | 16 | 0.1875 | 44 | 0.1591 |
| 19_470124 | 19 | 0.2632 | 22 | 0.1818 | 21 | 0.1429 | 12 | 0.25 | 74 | 0.2027 | 12 | 0.25 | 10 | 0.1 | 13 | 0.0769 | 15 | 0.2 | 50 | 0.16 |
| 19_470124 | 19 | 0.1053 | 24 | 0.0833 | 21 | 0.0476 | 14 | 0.2143 | 78 | 0.1026 | 11 | 0.2727 | 11 | 0.1818 | 11 | 0.0909 | 18 | 0.1111 | 51 | 0.1569 |
| 19_470132 | 25 | 0.16 | 35 | 0.1429 | 26 | 0.1923 | 21 | 0.0952 | 107 | 0.1495 | 21 | 0.0476 | 15 | 0.3333 | 17 | 0.1176 | 21 | 0.0476 | 74 | 0.1216 |
| 19_470133 | 24 | 0.2917 | 34 | 0.0588 | 27 | 0.2593 | 20 | 0.2 | 105 | 0.1905 | 16 | 0.1875 | 17 | 0.1765 | 15 | 0.2 | 21 | 0.2857 | 69 | 0.2174 |
| 19_470134 | 25 | 0.04 | 36 | 0.1389 | 32 | 0.0312 | 25 | 0.08 | 118 | 0.0763 | 19 | 0.0526 | 15 | 0.1333 | 13 | 0.0769 | 22 | 0.0909 | 69 | 0.087 |
| 19_470135 | 24 | 0.9167 | 34 | 0.9118 | 31 | 0.9677 | 27 | 1 | 116 | 0.9483 | 23 | 1 | 18 | 1 | 16 | 0.9375 | 22 | 0.9091 | 79 | 0.962 |
| 19_470136 | 23 | 0.6087 | 37 | 0.5676 | 31 | 0.7742 | 25 | 0.68 | 116 | 0.6552 | 25 | 0.48 | 18 | 0.6667 | 18 | 0.7778 | 22 | 0.7273 | 83 | 0.6506 |
| 19_470136 | 22 | 0.6364 | 32 | 0.8125 | 25 | 0.76 | 23 | 0.6522 | 102 | 0.7255 | 19 | 0.6316 | 16 | 0.5625 | 18 | 0.6667 | 15 | 0.8 | 68 | 0.6618 |
| 19_470136 | 25 | 0.76 | 37 | 0.7297 | 30 | 0.7 | 23 | 0.6957 | 115 | 0.7217 | 24 | 0.5417 | 19 | 0.6316 | 20 | 0.5 | 26 | 0.6538 | 89 | 0.5843 |
| 19_470228 | NA | NA | 2 | 0.5 | 3 | 0.3333 | NA | NA | 5 | 0.4 | 1 | 1 | NA | NA | 1 | 1 | 3 | 0.6667 | 5 | 0.8 |
| 19_470595 | NA | NA | NA | NA | 1 | 1 | NA | NA | 1 | 1 | 2 | 1 | 1 | 1 | NA | NA | NA | NA | 3 | 1 |
| 19_470595 | 1 | 1 | 1 | 1 | 1 | 1 | NA | NA | 3 | 1 | 2 | 1 | 1 | 1 | NA | NA | NA | NA | 3 | 1 |
| 19_470633 | NA | NA | NA | NA | 1 | 1 | NA | NA | 1 | 1 | 1 | 0 | 1 | 1 | NA | NA | NA | NA | 2 | 0.5 |
| 19_470633 | NA | NA | 1 | 1 | 1 | 0 | NA | NA | 2 | 0.5 | 2 | 0 | 1 | 1 | NA | NA | NA | NA | 3 | 0.3333 |
| 19_470709 | 1 | 1 | 1 | 1 | 1 | 0 | NA | NA | 3 | 0.6667 | 1 | 1 | 1 | 1 | 1 | 1 | NA | NA | 3 | 1 |
| 19_470709 | 1 | 1 | 1 | 0 | 1 | 0 | NA | NA | 3 | 0.3333 | 2 | 0.5 | 1 | 0 | 1 | 0 | NA | NA | 4 | 0.25 |
| 19_470711 | 1 | 1 | 1 | 1 | 1 | 1 | NA | NA | 3 | 1 | 2 | 1 | 1 | 0 | 1 | 0 | NA | NA | 4 | 0.5 |
| 19_470712 | NA | NA | 1 | 1 | 1 | 1 | NA | NA | 2 | 1 | 2 | 0.5 | 1 | 0 | 1 | 1 | NA | NA | 4 | 0.5 |
| 19_470713 | 1 | 1 | 1 | 1 | 1 | 1 | NA | NA | 3 | 1 | 2 | 0.5 | 1 | 0 | 1 | 1 | NA | NA | 4 | 0.5 |
| 19_470713 | NA | NA | 1 | 1 | 1 | 0 | NA | NA | 2 | 0.5 | 2 | 0 | 1 | 0 | 1 | 1 | NA | NA | 4 | 0.25 |
| 19_470715 | NA | NA | 1 | 1 | 1 | 1 | NA | NA | 2 | 1 | 1 | 0 | NA | NA | 1 | 1 | NA | NA | 2 | 0.5 |
| 19_471191 | 5 | 0.4 | 2 | 0 | 4 | 0 | 1 | 0 | 12 | 0.1667 | 3 | 0.6667 | NA | NA | 1 | 0 | 1 | 0 | 5 | 0.4 |
| 19_471192 | 5 | 0.6 | 3 | 0 | 4 | 0 | 2 | 0 | 14 | 0.2143 | 3 | 0.3333 | NA | NA | 1 | 1 | 2 | 0 | 6 | 0.3333 |
| 19_471192 | 4 | 0 | 3 | 0.6667 | 4 | 0.25 | 2 | 1 | 13 | 0.3846 | 3 | 0.6667 | NA | NA | 1 | 0 | 3 | 0.3333 | 7 | 0.4286 |
| 19_471193 | 5 | 0.6 | 3 | 0.6667 | 4 | 0.75 | 1 | 1 | 13 | 0.6923 | 4 | 1 | NA | NA | NA | NA | 3 | 0.6667 | 7 | 0.8571 |
| 19_471194 | 3 | 0 | 3 | 0.3333 | 3 | 0 | NA | NA | 9 | 0.1111 | 4 | 0.25 | NA | NA | 1 | 0 | 3 | 0 | 8 | 0.125 |
| 19_471211 | NA | NA | NA | NA | 1 | 1 | NA | NA | 1 | 1 | 1 | 0 | 1 | 1 | NA | NA | 2 | 1 | 4 | 0.75 |
| 19_471211 | NA | NA | NA | NA | 1 | 1 | NA | NA | 1 | 1 | 1 | 1 | NA | NA | NA | NA | 3 | 1 | 4 | 1 |
| 19_471211 | NA | NA | NA | NA | 1 | 1 | NA | NA | 1 | 1 | 1 | 0 | 1 | 1 | NA | NA | 3 | 0 | 5 | 0.2 |
| 19_471566 | NA | NA | NA | NA | 1 | 1 | NA | NA | 1 | 1 | 1 | 1 | NA | NA | NA | NA | NA | NA | 1 | 1 |
| 19_471566 | NA | NA | NA | NA | 1 | 1 | NA | NA | 1 | 1 | 1 | 1 | NA | NA | NA | NA | NA | NA | 1 | 1 |
| 19_471576 | 1 | 1 | NA | NA | NA | NA | NA | NA | 1 | 1 | 1 | 1 | NA | NA | NA | NA | NA | NA | 1 | 1 |
| 19_471674 | 3 | 0 | 4 | 0 | 1 | 1 | 3 | 0 | 11 | 0.0909 | 4 | 0 | 1 | 0 | 1 | 1 | 1 | 0 | 7 | 0.1429 |
| 19_472283 | 11 | 0.1818 | 13 | 0.0769 | 4 | 0.75 | 7 | 0.2857 | 35 | 0.2286 | 13 | 0.3077 | 9 | 0.1111 | 4 | 0.25 | 6 | 0.3333 | 32 | 0.25 |
| 19_472288 | 14 | 0.2143 | 14 | 0.1429 | 6 | 0 | 10 | 0.5 | 44 | 0.2273 | 16 | 0.3125 | 12 | 0.3333 | 6 | 0 | 5 | 0.2 | 39 | 0.2564 |
| 19_472289 | 11 | 0.1818 | 13 | 0 | 6 | 0.1667 | 9 | 0.4444 | 39 | 0.1795 | 10 | 0.1 | 11 | 0.1818 | 4 | 0 | 4 | 0.5 | 29 | 0.1724 |
| 19_472291 | 1 | 1 | NA | NA | 1 | 1 | NA | NA | 2 | 1 | 1 | 1 | NA | NA | NA | NA | 2 | 1 | 3 | 1 |
| 19_472294 | 1 | 0 | NA | NA | 1 | 1 | NA | NA | 2 | 0.5 | NA | NA | NA | NA | NA | NA | 1 | 1 | 1 | 1 |
| 19_472297 | 2 | 0.5 | NA | NA | 1 | 0 | NA | NA | 3 | 0.3333 | 1 | 0 | NA | NA | NA | NA | 2 | 0.5 | 3 | 0.3333 |
| 19_472299 | 2 | 0.5 | NA | NA | 1 | 0 | NA | NA | 3 | 0.3333 | NA | NA | NA | NA | NA | NA | 3 | 0.6667 | 3 | 0.6667 |
| 19_472300 | 2 | 1 | NA | NA | 1 | 1 | NA | NA | 3 | 1 | 1 | 0 | NA | NA | NA | NA | 3 | 0.6667 | 4 | 0.5 |
| 19_472301 | 2 | 0.5 | NA | NA | 1 | 0 | NA | NA | 3 | 0.3333 | 1 | 0 | NA | NA | NA | NA | 3 | 0.3333 | 4 | 0.25 |
| 19_472417 | 14 | 0.2857 | 17 | 0.5882 | 8 | 0.625 | 11 | 0.3636 | 50 | 0.46 | 14 | 0.3571 | 11 | 0.2727 | 6 | 0.3333 | 2 | 1 | 33 | 0.3636 |
| 19_472420 | 13 | 0.0769 | 19 | 0.1053 | 11 | 0.1818 | 11 | 0.3636 | 54 | 0.1667 | 19 | 0.3158 | 14 | 0.4286 | 9 | 0 | 7 | 0.2857 | 49 | 0.2857 |
| 19_472459 | 13 | 0.2308 | 20 | 0.35 | 13 | 0.4615 | 13 | 0.3846 | 59 | 0.3559 | 17 | 0.3529 | 16 | 0.3125 | 14 | 0.4286 | 6 | 0.5 | 53 | 0.3774 |
| 19_472459 | 19 | 0.5789 | 16 | 0.5625 | 14 | 0.8571 | 13 | 0.6154 | 62 | 0.6452 | 16 | 0.625 | 16 | 0.6875 | 14 | 0.6429 | 6 | 0.6667 | 52 | 0.6538 |
| 19_472460 | 17 | 0.2353 | 22 | 0.3182 | 17 | 0.3529 | 17 | 0.2353 | 73 | 0.2877 | 25 | 0.32 | 19 | 0.4211 | 15 | 0.2 | 9 | 0.5556 | 68 | 0.3529 |
| 19_472461 | 14 | 0.7857 | 18 | 0.6111 | 10 | 0.5 | 13 | 0.6154 | 55 | 0.6364 | 21 | 0.5238 | 15 | 0.7333 | 14 | 0.6429 | 7 | 0.8571 | 57 | 0.6491 |
| 19_472461 | 18 | 0.2222 | 20 | 0.2 | 12 | 0.4167 | 18 | 0.1667 | 68 | 0.2353 | 27 | 0.1111 | 16 | 0.25 | 12 | 0.3333 | 6 | 0.8333 | 61 | 0.2623 |
| 19_472464 | 19 | 0.0526 | 23 | 0.1739 | 17 | 0.4118 | 15 | 0.4 | 74 | 0.2432 | 25 | 0.16 | 18 | 0.1667 | 17 | 0.2941 | 10 | 0 | 70 | 0.1714 |
| 19_472466 | 21 | 0.0476 | 23 | 0.0435 | 22 | 0.0455 | 21 | 0.0952 | 87 | 0.0575 | 26 | 0.1538 | 23 | 0.0435 | 17 | 0.0588 | 7 | 0.1429 | 73 | 0.0959 |
| 19_472468 | 19 | 0.2105 | 23 | 0.2174 | 23 | 0.3913 | 18 | 0.4444 | 83 | 0.3133 | 24 | 0.5417 | 21 | 0.4286 | 23 | 0.3478 | 9 | 0.6667 | 77 | 0.4675 |
| 19_472471 | 27 | 0.2222 | 24 | 0.125 | 27 | 0.0741 | 22 | 0.0455 | 100 | 0.12 | 22 | 0.2273 | 23 | 0.1739 | 23 | 0.2174 | 10 | 0.1 | 78 | 0.1923 |

| | | | | | | | | | | | | | | | | | | | | |
|---|---|---|---|---|---|---|---|---|---|---|---|---|---|---|---|---|---|---|---|---|
| 19_472478 | 27 | 0.0741 | 23 | 0.0435 | 22 | 0.0455 | 21 | 0.1429 | 93 | 0.0753 | 29 | 0.069 | 27 | 0.0741 | 20 | 0 | 13 | 0.0769 | 89 | 0.0562 |
| 19_472483 | NA | NA | NA | NA | NA | NA | 3 | 0.3333 | 3 | 0.3333 | 2 | 0 | 1 | 0 | 1 | 0 | 1 | 1 | 5 | 0.2 |
| 19_472487 | 4 | 0.25 | 3 | 0.3333 | 4 | 0 | 6 | 0.3333 | 17 | 0.2353 | 2 | 0 | 4 | 0.25 | 2 | 0 | 4 | 0 | 12 | 0.0833 |
| 19_472489 | 3 | 0.6667 | 4 | 0.25 | 6 | 0.5 | 6 | 0.3333 | 19 | 0.4211 | 1 | 0 | 4 | 0.5 | 3 | 0.6667 | 4 | 0 | 12 | 0.3333 |
| 19_472490 | 2 | 0.5 | 4 | 0 | 3 | 0 | 6 | 0.1667 | 15 | 0.1333 | 2 | 0 | 4 | 0 | 3 | 0 | 2 | 0.5 | 11 | 0.0909 |
| 19_472496 | 1 | 0 | NA | NA | 3 | 0 | 1 | 1 | 5 | 0.2 | 3 | 1 | 1 | 1 | 2 | 1 | 1 | 0 | 7 | 0.8571 |
| 19_472497 | 1 | 0 | NA | NA | 4 | 0.25 | 3 | 0.6667 | 8 | 0.375 | 4 | 0.5 | 1 | 0 | 2 | 0.5 | 1 | 0 | 8 | 0.375 |
| 19_472500 | 1 | 0 | NA | NA | 3 | 0.3333 | 4 | 0.25 | 8 | 0.25 | 2 | 0 | 1 | 0 | 1 | 0 | 1 | 1 | 5 | 0.2 |
| 19_472502 | 1 | 0 | 1 | 1 | 4 | 0 | 4 | 0 | 10 | 0.1 | 2 | 0 | 1 | 0 | 2 | 0.5 | 1 | 0 | 6 | 0.1667 |
| 19_472502 | 1 | 0 | 1 | 1 | 4 | 0.75 | 4 | 1 | 10 | 0.8 | 4 | 0.5 | 1 | 1 | 2 | 1 | 1 | 1 | 8 | 0.75 |
| 19_472504 | 1 | 0 | 1 | 0 | 3 | 0.6667 | 1 | 0 | 6 | 0.3333 | 4 | 0 | 1 | 0 | 2 | 0.5 | 1 | 1 | 8 | 0.25 |
| 19_472513 | 2 | 0 | 1 | 0 | 2 | 0.5 | 2 | 0 | 7 | 0.1429 | 1 | 0 | 3 | 0 | 1 | 1 | 3 | 0 | 8 | 0.125 |
| 19_472518 | 2 | 0 | 2 | 0.5 | 2 | 0 | 1 | 1 | 7 | 0.2857 | NA | NA | 3 | 0.6667 | 1 | 1 | 3 | 0.3333 | 7 | 0.5714 |
| 19_472537 | 3 | 0.6667 | 2 | 0 | 3 | 0.6667 | NA | NA | 8 | 0.5 | 1 | 0 | 3 | 0 | 2 | 0 | 1 | 1 | 7 | 0.1429 |
| 19_472539 | 4 | 0.5 | 2 | 0 | 2 | 0 | NA | NA | 8 | 0.25 | 1 | 0 | 2 | 1 | 1 | 0 | NA | NA | 4 | 0.5 |
| 19_472540 | 2 | 1 | 2 | 1 | 2 | 0.5 | NA | NA | 6 | 0.8333 | NA | NA | 6 | 0.8333 | 2 | 1 | 2 | 0.5 | 10 | 0.8 |
| 19_472540 | 2 | 0.5 | 2 | 0 | 3 | 0.3333 | NA | NA | 7 | 0.2857 | 1 | 1 | 4 | 0.25 | 2 | 0 | 2 | 0.5 | 9 | 0.3333 |
| 19_472540 | 4 | 0.75 | 2 | 0.5 | 3 | 0.3333 | NA | NA | 9 | 0.5556 | 1 | 1 | 4 | 0.5 | 2 | 1 | 2 | 0 | 9 | 0.5556 |
| 19_472540 | 4 | 0 | 2 | 0.5 | 3 | 0 | NA | NA | 9 | 0.1111 | 1 | 0 | 5 | 0.4 | 2 | 0.5 | 2 | 0.5 | 10 | 0.4 |
| 19_472542 | 2 | 1 | 2 | 1 | 3 | 0.6667 | NA | NA | 7 | 0.8571 | 1 | 1 | 5 | 0.8 | 2 | 1 | 2 | 1 | 10 | 0.9 |
| 19_472542 | 3 | 0.3333 | 2 | 0.5 | 2 | 0 | NA | NA | 7 | 0.2857 | NA | NA | 3 | 0.3333 | 2 | 0.5 | 1 | 1 | 6 | 0.5 |
| 19_472542 | 2 | 0 | 2 | 0.5 | 2 | 0 | NA | NA | 6 | 0.1667 | NA | NA | 4 | 0.25 | 2 | 0.5 | 1 | 0 | 7 | 0.2857 |
| 19_472544 | 2 | 0 | 2 | 1 | 3 | 1 | NA | NA | 7 | 0.7143 | NA | NA | 6 | 0.6667 | 2 | 1 | 1 | 1 | 9 | 0.7778 |
| 19_472547 | 2 | 1 | NA | NA | 2 | 1 | NA | NA | 4 | 1 | NA | NA | 2 | 1 | 1 | 1 | 2 | 1 | 5 | 1 |
| 19_472548 | 2 | 1 | NA | NA | 2 | 1 | NA | NA | 4 | 1 | NA | NA | 2 | 1 | 1 | 1 | 2 | 1 | 5 | 1 |
| 19_472551 | 2 | 1 | NA | NA | 2 | 1 | NA | NA | 4 | 1 | NA | NA | 3 | 0.6667 | NA | NA | 2 | 1 | 5 | 0.8 |
| 19_472552 | 1 | 0 | NA | NA | 2 | 0.5 | NA | NA | 3 | 0.3333 | NA | NA | 2 | 0 | NA | NA | 2 | 0.5 | 4 | 0.25 |
| 19_472553 | 2 | 0.5 | NA | NA | 2 | 1 | NA | NA | 4 | 0.75 | NA | NA | 2 | 1 | NA | NA | 1 | 0 | 3 | 0.6667 |
| 19_472553 | 1 | 1 | NA | NA | 2 | 1 | NA | NA | 3 | 1 | NA | NA | 2 | 1 | NA | NA | 1 | 1 | 3 | 1 |
| 19_472553 | 1 | 1 | NA | NA | 2 | 0.5 | NA | NA | 3 | 0.6667 | NA | NA | 2 | 1 | NA | NA | 1 | 1 | 3 | 1 |
| 19_472554 | 1 | 1 | NA | NA | 1 | 1 | NA | NA | 2 | 1 | NA | NA | 2 | 1 | NA | NA | 1 | 1 | 3 | 1 |
| 19_472555 | 1 | 1 | NA | NA | 1 | 0 | NA | NA | 2 | 0.5 | NA | NA | 1 | 0 | NA | NA | 1 | 1 | 2 | 0.5 |
| 19_472558 | 1 | 1 | NA | NA | 1 | 0 | NA | NA | 2 | 0.5 | NA | NA | 2 | 1 | NA | NA | 1 | 0 | 3 | 0.6667 |
| 19_472562 | 1 | 1 | NA | NA | 1 | 0 | NA | NA | 2 | 0.5 | NA | NA | 1 | 1 | NA | NA | NA | NA | 1 | 1 |
| 19_472563 | 1 | 1 | NA | NA | 1 | 0 | NA | NA | 2 | 0.5 | NA | NA | 2 | 0.5 | NA | NA | NA | NA | 2 | 0.5 |
| 19_472564 | 1 | 1 | NA | NA | 1 | 1 | NA | NA | 2 | 1 | NA | NA | 1 | 1 | NA | NA | NA | NA | 1 | 1 |
| 19_474692 | NA | NA | NA | NA | NA | NA | 1 | 1 | 1 | 1 | NA | NA | NA | NA | 1 | 1 | NA | NA | 1 | 1 |
| 19_474694 | NA | NA | NA | NA | NA | NA | 1 | 1 | 1 | 1 | NA | NA | NA | NA | 1 | 1 | NA | NA | 1 | 1 |
| 19_474825 | NA | NA | 1 | 1 | 1 | 1 | 1 | 1 | 3 | 1 | NA | NA | NA | NA | NA | NA | 1 | 1 | 1 | 1 |
| 19_474830 | NA | NA | 1 | 0 | NA | NA | 1 | 1 | 2 | 0.5 | NA | NA | NA | NA | NA | NA | 1 | 1 | 1 | 1 |
| 19_474832 | NA | NA | 1 | 1 | NA | NA | 1 | 1 | 2 | 1 | NA | NA | NA | NA | NA | NA | 1 | 1 | 1 | 1 |
| 19_474833 | NA | NA | 1 | 1 | NA | NA | 1 | 0 | 2 | 0.5 | NA | NA | NA | NA | NA | NA | 1 | 1 | 1 | 1 |
| 19_474834 | NA | NA | 1 | 1 | NA | NA | 1 | 0 | 2 | 0.5 | NA | NA | NA | NA | NA | NA | 1 | 1 | 1 | 1 |
| 19_474835 | NA | NA | 1 | 1 | NA | NA | 1 | 1 | 2 | 1 | NA | NA | NA | NA | NA | NA | 1 | 1 | 1 | 1 |
| 19_474837 | NA | NA | 1 | 1 | NA | NA | 1 | 1 | 2 | 1 | NA | NA | NA | NA | NA | NA | 1 | 1 | 1 | 1 |
| 19_474840 | NA | NA | 1 | 1 | NA | NA | 1 | 1 | 2 | 1 | NA | NA | NA | NA | NA | NA | 1 | 1 | 1 | 1 |
| 19_474895 | NA | NA | NA | NA | 4 | 1 | 1 | 0 | 5 | 0.8 | 1 | 1 | 1 | 0 | 2 | 1 | 2 | 1 | 6 | 0.8333 |
| 19_474897 | 2 | 0 | NA | NA | 4 | 0 | 3 | 0.3333 | 9 | 0.1111 | 2 | 0.5 | 1 | 0 | 3 | 0.6667 | 2 | 0.5 | 8 | 0.5 |
| 19_474898 | 2 | 0.5 | NA | NA | 3 | 0 | 2 | 0 | 7 | 0.1429 | 1 | 0 | 2 | 0 | 2 | 0.5 | 2 | 0 | 7 | 0.1429 |
| 19_474898 | 2 | 0.5 | NA | NA | 4 | 0.75 | 3 | 1 | 9 | 0.7778 | 1 | 1 | 2 | 1 | 3 | 1 | 2 | 1 | 8 | 1 |
| 19_474898 | 2 | 1 | NA | NA | 4 | 1 | 3 | 1 | 9 | 1 | 2 | 0 | 1 | 1 | 3 | 1 | 2 | 1 | 8 | 0.75 |
| 19_474899 | 1 | 1 | NA | NA | 4 | 0.75 | 3 | 1 | 8 | 0.875 | 2 | 0.5 | 2 | 1 | 3 | 1 | 2 | 1 | 9 | 0.8889 |
| 19_474900 | 2 | 0 | NA | NA | 3 | 0.3333 | 3 | 0 | 8 | 0.125 | 2 | 0 | 2 | 0 | 3 | 0.3333 | 1 | 0 | 8 | 0.125 |
| 19_474900 | 2 | 0 | NA | NA | 3 | 0.3333 | 3 | 0 | 8 | 0.125 | 2 | 0 | 2 | 0.5 | 3 | 0 | 1 | 0 | 8 | 0.125 |
| 19_474988 | 1 | 0 | NA | NA | 1 | 1 | 2 | 0.5 | 4 | 0.5 | 1 | 1 | NA | NA | 1 | 1 | NA | NA | 2 | 1 |
| 19_476082 | NA | NA | 2 | 0 | 3 | 0.3333 | 3 | 0.3333 | 8 | 0.25 | NA | NA | NA | NA | 2 | 0 | 1 | 1 | 3 | 0.3333 |
| 19_476082 | NA | NA | 1 | 0 | 2 | 0.5 | 3 | 0 | 6 | 0.1667 | NA | NA | NA | NA | 3 | 0 | 1 | 1 | 4 | 0.25 |
| 19_476085 | NA | NA | 2 | 1 | 2 | 1 | 2 | 1 | 6 | 1 | NA | NA | NA | NA | 3 | 0.6667 | 1 | 1 | 4 | 0.75 |
| 19_476085 | NA | NA | 2 | 0.5 | 3 | 0.3333 | 3 | 0.3333 | 8 | 0.375 | NA | NA | NA | NA | 3 | 0.6667 | 2 | 1 | 5 | 0.8 |
| 19_476090 | NA | NA | 4 | 0.5 | 3 | 0.3333 | 4 | 0.75 | 11 | 0.5455 | 1 | 0 | 1 | 1 | 6 | 0.3333 | 3 | 0.3333 | 11 | 0.3636 |
| 19_476304 | 2 | 1 | 4 | 1 | 11 | 0.5455 | 3 | 1 | 20 | 0.75 | NA | NA | 4 | 0.5 | 6 | 0.6667 | 4 | 1 | 14 | 0.7143 |
| 19_476306 | 1 | 1 | 4 | 0.25 | 5 | 0.8 | 5 | 0.8 | 15 | 0.6667 | NA | NA | 4 | 0.5 | 6 | 0.5 | 5 | 0.6 | 15 | 0.5333 |
| 19_476307 | 1 | 1 | 3 | 1 | 7 | 0.2857 | 5 | 0.2 | 16 | 0.4375 | NA | NA | 4 | 0.5 | 8 | 0.625 | 5 | 0.4 | 17 | 0.5294 |
| 19_476309 | 2 | 0 | 5 | 0.4 | 9 | 0.1111 | 5 | 0.4 | 21 | 0.2381 | NA | NA | 3 | 0 | 7 | 0.4286 | 6 | 0.3333 | 16 | 0.3125 |
| 19_476325 | 5 | 0.2 | 6 | 0 | 10 | 0.2 | 9 | 0.1111 | 30 | 0.1333 | 2 | 0 | 6 | 0 | 9 | 0.1111 | 8 | 0.125 | 25 | 0.08 |

| | | | | | | | | | | | | | | | | | | | | |
|---|---|---|---|---|---|---|---|---|---|---|---|---|---|---|---|---|---|---|---|---|
| 19_476332 | 6 | 0.1667 | 8 | 0.125 | 11 | 0.0909 | 6 | 0.1667 | 31 | 0.129 | 3 | 0.3333 | 4 | 0 | 11 | 0.2727 | 9 | 0.4444 | 27 | 0.2963 |
| 19_476334 | 5 | 0.2 | 6 | 0.5 | 13 | 0.1538 | 9 | 0.1111 | 33 | 0.2121 | 2 | 0 | 6 | 0.1667 | 13 | 0.1538 | 10 | 0.1 | 31 | 0.129 |
| 19_477220 | 10 | 0.2 | 17 | 0.0588 | 13 | 0.1538 | 17 | 0.1176 | 57 | 0.1228 | 5 | 0.2 | 19 | 0.0526 | 16 | 0.125 | 11 | 0.0909 | 51 | 0.098 |
| 19_477221 | 8 | 0.75 | 12 | 0.6667 | 11 | 0.5455 | 12 | 0.5 | 43 | 0.6047 | 5 | 0.8 | 15 | 0.4 | 12 | 0.6667 | 9 | 0.6667 | 41 | 0.5854 |
| 19_477236 | 9 | 0.3333 | 13 | 0.1538 | 12 | 0.3333 | 21 | 0.4286 | 55 | 0.3273 | 3 | 0.3333 | 14 | 0.2857 | 12 | 0.1667 | 7 | 0 | 36 | 0.1944 |
| 19_477239 | 14 | 0.1429 | 22 | 0.1818 | 13 | 0.2308 | 24 | 0.1667 | 73 | 0.1781 | 6 | 0 | 24 | 0.1667 | 22 | 0.1364 | 16 | 0.125 | 68 | 0.1324 |
| 19_481946 | NA | NA | 1 | 1 | NA | NA | 2 | 0.5 | 3 | 0.6667 | NA | NA | 1 | 1 | 1 | 0 | NA | NA | 2 | 0.5 |
| 19_481949 | NA | NA | 1 | 0 | NA | NA | 1 | 1 | 2 | 0.5 | NA | NA | NA | NA | 1 | 1 | NA | NA | 1 | 1 |
| 19_481950 | NA | NA | 1 | 1 | 1 | 1 | 2 | 0 | 4 | 0.5 | NA | NA | 1 | 1 | 1 | 0 | NA | NA | 2 | 0.5 |
| 19_481950 | NA | NA | 1 | 1 | NA | NA | 2 | 0.5 | 3 | 0.6667 | NA | NA | 1 | 1 | 1 | 1 | NA | NA | 2 | 1 |
| 19_481951 | NA | NA | 1 | 1 | 1 | 1 | 2 | 0.5 | 4 | 0.75 | NA | NA | 1 | 1 | NA | NA | NA | NA | 1 | 1 |
| 19_487929 | 3 | 0.3333 | 2 | 0 | 2 | 0.5 | 8 | 0.375 | 15 | 0.3333 | 1 | 0 | 5 | 0.2 | 5 | 0.2 | 1 | 0 | 12 | 0.1667 |
| 19_487929 | 3 | 0.3333 | 2 | 0 | 2 | 0 | 9 | 0.4444 | 16 | 0.3125 | 1 | 0 | 5 | 0.2 | 5 | 0 | 3 | 0.6667 | 14 | 0.2143 |
| 19_487929 | 3 | 0 | 2 | 0 | 2 | 0 | 9 | 0.1111 | 16 | 0.0625 | 1 | 0 | 5 | 0.2 | 5 | 0 | 2 | 0 | 13 | 0.0769 |
| 19_500185 | 2 | 0 | 1 | 1 | NA | NA | 1 | 0 | 4 | 0.25 | 1 | 1 | 1 | 0 | NA | NA | 2 | 0 | 4 | 0.25 |
| 19_501362 | 3 | 0.3333 | 2 | 0.5 | 1 | 0 | 4 | 0 | 10 | 0.2 | NA | NA | 2 | 0.5 | 7 | 0.2857 | 2 | 0 | 11 | 0.2727 |
| 19_501366 | 2 | 0.5 | 1 | 1 | 1 | 1 | 4 | 0.25 | 8 | 0.5 | NA | NA | 2 | 0.5 | 5 | 0.4 | 2 | 0.5 | 9 | 0.4444 |
| 19_501369 | 2 | 0.5 | 1 | 0 | 1 | 0 | 4 | 0 | 8 | 0.125 | NA | NA | 2 | 0 | 4 | 0 | 2 | 0.5 | 8 | 0.125 |
| 19_501379 | 1 | 0 | 1 | 1 | 1 | 0 | 2 | 0 | 5 | 0.2 | NA | NA | 1 | 1 | 5 | 0 | 2 | 0.5 | 8 | 0.25 |
| 19_501380 | NA | NA | 1 | 1 | 1 | 1 | 2 | 0 | 4 | 0.5 | NA | NA | 1 | 1 | 5 | 0.6 | 2 | 0.5 | 8 | 0.625 |
| 19_501393 | NA | NA | 1 | 1 | 1 | 1 | 1 | 0 | 3 | 0.6667 | NA | NA | 2 | 0.5 | 3 | 0.6667 | 3 | 0.6667 | 8 | 0.625 |
| 19_529151 | NA | NA | NA | NA | NA | NA | 1 | 1 | 1 | 1 | 1 | 0 | 1 | 1 | NA | NA | 1 | 0 | 3 | 0.3333 |
| 19_529796 | 2 | 0 | 4 | 0 | 6 | 0.1667 | 3 | 0 | 15 | 0.0667 | 5 | 0.2 | 3 | 0 | 5 | 0 | 4 | 0 | 17 | 0.0588 |
| 19_529796 | 2 | 0 | 8 | 0.25 | 7 | 0.1429 | 4 | 0 | 21 | 0.1429 | 6 | 0 | 3 | 0.3333 | 7 | 0 | 5 | 0.2 | 21 | 0.0952 |
| 19_529797 | NA | NA | 7 | 0.4286 | 5 | 0.4 | 4 | 0.25 | 16 | 0.375 | 3 | 0.6667 | 3 | 0.6667 | 4 | 1 | 3 | 0.3333 | 13 | 0.6923 |
| 19_529798 | 1 | 0 | 4 | 0.25 | 5 | 0 | 1 | 0 | 11 | 0.0909 | 3 | 0 | 3 | 0 | 5 | 0.2 | 3 | 0 | 14 | 0.0714 |
| 19_529799 | 2 | 0 | 7 | 0.1429 | 6 | 0.1667 | 2 | 0 | 17 | 0.1176 | 6 | 0.1667 | 2 | 0 | 6 | 0.1667 | 5 | 0.2 | 19 | 0.1579 |
| 19_529799 | 2 | 0.5 | 7 | 0 | 6 | 0 | 3 | 0 | 18 | 0.0556 | 6 | 0.5 | 3 | 0 | 6 | 0 | 5 | 0.2 | 20 | 0.2 |
| 19_529800 | 2 | 0 | 6 | 0.1667 | 5 | 0 | 2 | 0 | 15 | 0.0667 | 5 | 0 | 2 | 0 | 5 | 0.2 | 4 | 0 | 16 | 0.0625 |
| 19_532336 | 8 | 0.25 | 14 | 0.2143 | 9 | 0.3333 | 6 | 0.3333 | 37 | 0.2703 | 8 | 0.375 | 12 | 0.4167 | 11 | 0.0909 | 10 | 0 | 41 | 0.2195 |
| 19_532338 | 13 | 1 | 12 | 0.8333 | 12 | 0.8333 | 8 | 0.875 | 45 | 0.8889 | 11 | 0.7273 | 14 | 1 | 10 | 0.7 | 8 | 0.875 | 43 | 0.8372 |
| 19_532338 | 13 | 0.0769 | 12 | 0 | 14 | 0.2143 | 7 | 0.1429 | 46 | 0.1087 | 11 | 0 | 15 | 0.2667 | 11 | 0.0909 | 11 | 0.2727 | 48 | 0.1667 |
| 19_532341 | 12 | 0.25 | 13 | 0.1538 | 15 | 0.2 | 6 | 0.3333 | 46 | 0.2174 | 11 | 0.1818 | 13 | 0.0769 | 14 | 0 | 13 | 0.0769 | 51 | 0.0784 |
| 19_532342 | 14 | 0.1429 | 15 | 0 | 14 | 0 | 10 | 0.1 | 53 | 0.0566 | 10 | 0.1 | 12 | 0.0833 | 14 | 0.0714 | 12 | 0 | 48 | 0.0625 |
| 19_532346 | 15 | 0.1333 | 15 | 0.0667 | 16 | 0 | 10 | 0 | 56 | 0.0536 | 13 | 0 | 13 | 0.1538 | 14 | 0.1429 | 16 | 0.125 | 56 | 0.1071 |
| 19_532352 | 12 | 0.0833 | 14 | 0.0714 | 15 | 0.0667 | 12 | 0.0833 | 53 | 0.0755 | 12 | 0.1667 | 16 | 0.125 | 13 | 0.0769 | 19 | 0.1053 | 60 | 0.1167 |
| 19_532992 | 3 | 0.3333 | 4 | 0 | NA | NA | 2 | 0.5 | 9 | 0.2222 | 4 | 0.5 | NA | NA | NA | NA | 2 | 0 | 6 | 0.3333 |
| 19_533601 | 8 | 0.125 | 13 | 0.1538 | 8 | 0.125 | 15 | 0.1333 | 44 | 0.1364 | 17 | 0.1176 | 7 | 0.2857 | 10 | 0.1 | 12 | 0.1667 | 46 | 0.1522 |
| 19_533602 | 7 | 0 | 10 | 0.2 | 8 | 0 | 14 | 0.1429 | 39 | 0.1026 | 15 | 0.2 | 5 | 0 | 10 | 0.1 | 11 | 0.0909 | 41 | 0.122 |
| 19_533602 | 7 | 0.5714 | 10 | 0.4 | 6 | 0.3333 | 14 | 0.2143 | 37 | 0.3514 | 13 | 0.5385 | 4 | 0.75 | 9 | 0.5556 | 11 | 0.4545 | 37 | 0.5405 |
| 19_533602 | 7 | 0.1429 | 7 | 0.2857 | 4 | 0 | 10 | 0 | 28 | 0.1071 | 11 | 0.1818 | 5 | 0 | 9 | 0 | 10 | 0 | 35 | 0.0571 |
| 19_533604 | 10 | 0.3 | 14 | 0.2857 | 8 | 0.25 | 14 | 0.2143 | 46 | 0.2609 | 16 | 0.25 | 7 | 0.5714 | 10 | 0.1 | 13 | 0.3077 | 46 | 0.2826 |
| 19_533607 | 11 | 0.4545 | 19 | 0.4737 | 6 | 0.3333 | 12 | 0.3333 | 48 | 0.4167 | 15 | 0.6667 | 6 | 0.6667 | 11 | 0.4545 | 14 | 0.4286 | 46 | 0.5435 |
| 19_533609 | 9 | 0.2222 | 17 | 0.1176 | 9 | 0.1111 | 16 | 0.375 | 51 | 0.2157 | 18 | 0.1667 | 9 | 0.2222 | 10 | 0.1 | 13 | 0.0769 | 50 | 0.14 |
| 19_533611 | 11 | 0.0909 | 16 | 0 | 7 | 0 | 15 | 0.1333 | 49 | 0.0612 | 18 | 0.0556 | 9 | 0.1111 | 6 | 0.1667 | 12 | 0.0833 | 45 | 0.0889 |
| 19_533615 | 13 | 0.3077 | 18 | 0.3889 | 10 | 0.3 | 14 | 0.3571 | 55 | 0.3455 | 21 | 0.3333 | 9 | 0.4444 | 9 | 0.4444 | 15 | 0.4 | 54 | 0.3889 |
| 19_533686 | 3 | 0 | 5 | 0.4 | 6 | 0.1667 | 6 | 0.5 | 20 | 0.3 | 6 | 0.1667 | 5 | 0.6 | 9 | 0.1111 | 3 | 0.6667 | 23 | 0.3043 |
| 19_533688 | 3 | 0 | 4 | 0.25 | 5 | 0.2 | 5 | 0.2 | 17 | 0.1765 | 5 | 0.4 | 3 | 0.3333 | 8 | 0.25 | 3 | 0 | 19 | 0.2632 |
| 19_533689 | 3 | 0 | 4 | 0 | 6 | 0.1667 | 8 | 0.375 | 21 | 0.1905 | 6 | 0 | 4 | 0 | 7 | 0.1429 | 3 | 0 | 20 | 0.05 |
| 19_535295 | 1 | 1 | 3 | 0.3333 | NA | NA | 1 | 1 | 5 | 0.6 | NA | NA | NA | NA | 2 | 0.5 | 1 | 0 | 3 | 0.3333 |
| 19_535296 | 1 | 1 | 2 | 1 | NA | NA | NA | NA | 3 | 1 | NA | NA | NA | NA | 2 | 1 | NA | NA | 2 | 1 |
| 19_535297 | 2 | 0 | 4 | 0.25 | NA | NA | 1 | 0 | 7 | 0.1429 | NA | NA | 1 | 1 | 1 | 0 | 1 | 0 | 3 | 0.3333 |
| 19_535298 | 1 | 1 | 3 | 1 | NA | NA | NA | NA | 4 | 1 | NA | NA | NA | NA | 2 | 1 | 1 | 1 | 3 | 1 |
| 19_535298 | 2 | 0.5 | 1 | 1 | NA | NA | 1 | 1 | 4 | 0.75 | NA | NA | 1 | 1 | 2 | 1 | 1 | 0 | 4 | 0.75 |
| 19_535299 | 1 | 1 | 4 | 1 | 1 | 1 | 1 | 1 | 7 | 1 | NA | NA | 1 | 1 | 2 | 1 | 1 | 1 | 4 | 1 |
| 19_535300 | 2 | 0.5 | 4 | 0.5 | 1 | 0 | 1 | 0 | 8 | 0.375 | 1 | 0 | NA | NA | 2 | 0.5 | 1 | 0 | 4 | 0.25 |
| 19_535301 | 1 | 1 | 4 | 1 | 1 | 1 | 1 | 1 | 7 | 1 | 1 | 1 | 1 | 1 | 2 | 1 | 1 | 1 | 5 | 1 |
| 19_535302 | 1 | 1 | 3 | 1 | 1 | 1 | 1 | 1 | 6 | 1 | 1 | 1 | 1 | 1 | 2 | 1 | 1 | 1 | 5 | 1 |
| 19_535302 | 2 | 0.5 | 4 | 0.75 | 1 | 1 | 1 | 0 | 8 | 0.625 | 1 | 0 | 1 | 1 | 2 | 0.5 | 1 | 0 | 5 | 0.4 |
| 19_535302 | NA | NA | 4 | 0 | 1 | 0 | 1 | 1 | 6 | 0.1667 | 1 | 0 | 1 | 1 | 2 | 0.5 | 1 | 1 | 5 | 0.6 |
| 19_535303 | 1 | 1 | 2 | 0 | NA | NA | 1 | 0 | 4 | 0.25 | 1 | 1 | 1 | 1 | 2 | 0.5 | 1 | 0 | 5 | 0.6 |
| 19_535304 | 1 | 1 | 4 | 1 | 1 | 1 | 1 | 1 | 7 | 1 | 1 | 1 | 1 | 1 | 2 | 1 | 1 | 1 | 5 | 1 |
| 19_536332 | 8 | 0.25 | 7 | 0 | 8 | 0.5 | 7 | 0 | 30 | 0.2 | 2 | 0.5 | 4 | 0.25 | 7 | 0.1429 | 9 | 0.1111 | 22 | 0.1818 |
| 19_536346 | 1 | 1 | NA | NA | 1 | 0 | 1 | 1 | 3 | 0.6667 | NA | NA | NA | NA | NA | NA | 1 | 1 | 1 | 1 |
| 19_536346 | 1 | 1 | NA | NA | 1 | 1 | 1 | 1 | 3 | 1 | NA | NA | NA | NA | NA | NA | 1 | 1 | 1 | 1 |

| | | | | | | | | | | | | | | | | | | | | |
|---|---|---|---|---|---|---|---|---|---|---|---|---|---|---|---|---|---|---|---|---|
| 19_536346 | 1 | 1 | NA | NA | 1 | 1 | 1 | 1 | 3 | 1 | NA | NA | NA | NA | NA | NA | 1 | 1 | 1 | 1 |
| 19_536349 | 1 | 1 | NA | NA | 1 | 1 | NA | NA | 2 | 1 | NA | NA | NA | NA | NA | NA | 1 | 1 | 1 | 1 |
| 19_536353 | NA | NA | NA | NA | 1 | 1 | 1 | 1 | 2 | 1 | NA | NA | NA | NA | NA | NA | 1 | 1 | 1 | 1 |
| 19_536639 | NA | NA | 1 | 1 | NA | NA | NA | NA | 1 | 1 | 1 | 1 | NA | NA | NA | NA | NA | NA | 1 | 1 |
| 19_536640 | NA | NA | 1 | 1 | NA | NA | NA | NA | 1 | 1 | 1 | 1 | NA | NA | NA | NA | NA | NA | 1 | 1 |
| 19_536642 | NA | NA | 1 | 1 | NA | NA | NA | NA | 1 | 1 | 1 | 1 | NA | NA | NA | NA | NA | NA | 1 | 1 |
| 19_538527 | 1 | 0 | 1 | 1 | 3 | 0.6667 | 1 | 0 | 6 | 0.5 | NA | NA | NA | NA | 1 | 1 | NA | NA | 1 | 1 |
| 19_538553 | 2 | 0 | 1 | 1 | 3 | 0 | 1 | 0 | 7 | 0.1429 | 2 | 0 | NA | NA | 1 | 0 | 1 | 1 | 4 | 0.25 |
| 19_538977 | 6 | 0.1667 | 8 | 0.125 | 8 | 0.125 | 6 | 0.3333 | 28 | 0.1786 | 4 | 0 | 8 | 0.125 | 8 | 0.5 | 9 | 0.1111 | 29 | 0.2069 |
| 19_539179 | 17 | 0 | 16 | 0.25 | 13 | 0.1538 | 11 | 0.2727 | 57 | 0.1579 | 9 | 0 | 13 | 0.1538 | 24 | 0.2083 | 20 | 0.15 | 66 | 0.1515 |
| 19_539189 | NA | NA | 1 | 0 | 1 | 1 | 2 | 0.5 | 4 | 0.5 | 2 | 1 | 3 | 0.3333 | 2 | 0.5 | 2 | 0 | 9 | 0.4444 |
| 19_539197 | 12 | 0.1667 | 8 | 0.125 | 6 | 0 | 12 | 0 | 38 | 0.0789 | 7 | 0.5714 | 13 | 0.1538 | 10 | 0.2 | 8 | 0.125 | 38 | 0.2368 |
| 19_539197 | 8 | 0 | 8 | 0.125 | 7 | 0 | 11 | 0.1818 | 34 | 0.0882 | 7 | 0.1429 | 13 | 0 | 8 | 0.25 | 11 | 0.0909 | 39 | 0.1026 |
| 19_539210 | NA | NA | 2 | 0.5 | NA | NA | 1 | 0 | 3 | 0.3333 | 1 | 1 | NA | NA | NA | NA | NA | NA | 1 | 1 |
| 19_539760 | 5 | 0 | 3 | 0 | 5 | 0.2 | 2 | 0 | 15 | 0.0667 | 3 | 0 | 6 | 0.1667 | 1 | 0 | 7 | 0 | 17 | 0.0588 |
| 19_539761 | 4 | 1 | 3 | 1 | 5 | 0.8 | 2 | 1 | 14 | 0.9286 | 2 | 1 | 5 | 0.8 | 2 | 1 | 6 | 0.6667 | 15 | 0.8 |
| 19_539764 | 2 | 0 | 3 | 0.3333 | 5 | 0.2 | 2 | 0.5 | 12 | 0.25 | 3 | 0.3333 | 4 | 0.5 | 1 | 0 | 6 | 0.3333 | 14 | 0.3571 |
| 19_539764 | 4 | 0.75 | 3 | 0.6667 | 5 | 1 | 2 | 1 | 14 | 0.8571 | 3 | 1 | 6 | 0.6667 | 1 | 1 | 7 | 1 | 17 | 0.8824 |
| 19_539764 | 5 | 0.2 | 3 | 0 | 6 | 0 | 2 | 0.5 | 16 | 0.125 | 3 | 0 | 5 | 0.2 | 2 | 0 | 7 | 0.2857 | 17 | 0.1765 |
| 19_539764 | 5 | 1 | 3 | 1 | 6 | 1 | 2 | 1 | 16 | 1 | 3 | 1 | 5 | 1 | 2 | 1 | 7 | 0.7143 | 17 | 0.8824 |
| 19_539765 | 5 | 0.4 | 3 | 0.3333 | 5 | 0.4 | 2 | 0.5 | 15 | 0.4 | 3 | 0.6667 | 5 | 0.2 | 2 | 0.5 | 5 | 0.2 | 15 | 0.3333 |
| 19_539766 | 3 | 1 | 3 | 1 | 4 | 0.75 | 2 | 1 | 12 | 0.9167 | 2 | 1 | 5 | 0.6 | 2 | 1 | 4 | 0.75 | 13 | 0.7692 |
| 19_539766 | 4 | 0.25 | 2 | 0 | 5 | 0.2 | 2 | 0 | 13 | 0.1538 | 3 | 0.3333 | 7 | 0.1429 | 2 | 1 | 7 | 0 | 19 | 0.2105 |
| 19_539767 | 4 | 0 | 3 | 1 | 3 | 0 | 2 | 0.5 | 12 | 0.3333 | 3 | 0 | 6 | 0.3333 | 2 | 0.5 | 6 | 0.1667 | 17 | 0.2353 |
| 19_540196 | 4 | 0.75 | 4 | 0.5 | 7 | 0.5714 | 4 | 0.75 | 19 | 0.6316 | 3 | 0.6667 | 9 | 0.3333 | 1 | 1 | 7 | 0.7143 | 20 | 0.55 |
| 19_540197 | 5 | 0.6 | 4 | 0.25 | 7 | 0 | 5 | 0.4 | 21 | 0.2857 | 3 | 0 | 10 | 0.3 | 3 | 0.3333 | 8 | 0.5 | 24 | 0.3333 |
| 19_540198 | 5 | 0.6 | 4 | 0.25 | 8 | 0.25 | 4 | 0 | 21 | 0.2857 | 3 | 0 | 10 | 0.1 | 3 | 0.3333 | 6 | 0.6667 | 22 | 0.2727 |
| 19_540209 | 4 | 0.25 | 4 | 0 | 10 | 0 | 4 | 0.25 | 22 | 0.0909 | 4 | 0.25 | 9 | 0.1111 | 7 | 0.2857 | 9 | 0 | 29 | 0.1379 |
| 19_540344 | NA | NA | 2 | 0 | 1 | 1 | NA | NA | 3 | 0.3333 | 2 | 0 | 1 | 1 | NA | NA | 1 | 0 | 4 | 0.25 |
| 19_540345 | NA | NA | 3 | 0.3333 | 1 | 0 | NA | NA | 4 | 0.25 | 2 | 0.5 | 1 | 0 | NA | NA | 2 | 0 | 5 | 0.2 |
| 19_540347 | NA | NA | 3 | 0.6667 | 2 | 0.5 | NA | NA | 5 | 0.6 | 2 | 0.5 | 1 | 1 | NA | NA | 2 | 0.5 | 5 | 0.6 |
| 19_540347 | NA | NA | 3 | 0.3333 | 2 | 0 | NA | NA | 5 | 0.2 | 3 | 0.6667 | 1 | 0 | NA | NA | 3 | 0.6667 | 7 | 0.5714 |
| 19_540349 | NA | NA | 3 | 1 | 1 | 1 | NA | NA | 4 | 1 | 3 | 0.6667 | 1 | 0 | NA | NA | 2 | 1 | 6 | 0.6667 |
| 19_540355 | NA | NA | 3 | 1 | 2 | 1 | NA | NA | 5 | 1 | 6 | 0.8333 | NA | NA | NA | NA | 2 | 1 | 8 | 0.875 |
| 19_540356 | NA | NA | 2 | 1 | 1 | 1 | NA | NA | 3 | 1 | 3 | 0.6667 | 1 | 1 | NA | NA | 4 | 1 | 8 | 0.875 |
| 19_540359 | NA | NA | 6 | 0 | 4 | 0.5 | NA | NA | 10 | 0.2 | 5 | 0.2 | 1 | 0 | NA | NA | 4 | 0 | 10 | 0.1 |
| 19_540360 | NA | NA | 6 | 0.8333 | 4 | 0.75 | NA | NA | 10 | 0.8 | 7 | 0.8571 | 1 | 1 | NA | NA | 2 | 0.5 | 10 | 0.8 |
| 19_540360 | NA | NA | 7 | 0.5714 | 4 | 0.5 | NA | NA | 11 | 0.5455 | 6 | 0.8333 | 1 | 1 | NA | NA | 4 | 0.5 | 11 | 0.7273 |
| 19_540361 | NA | NA | 7 | 0.4286 | 3 | 1 | NA | NA | 10 | 0.6 | 7 | 0.4286 | 1 | 1 | NA | NA | 3 | 1 | 11 | 0.6364 |
| 19_540361 | NA | NA | 7 | 0.4286 | 4 | 0.25 | NA | NA | 11 | 0.3636 | 6 | 0.1667 | 1 | 0 | NA | NA | 4 | 0.25 | 11 | 0.1818 |
| 19_540362 | NA | NA | 7 | 0.8571 | 4 | 0.5 | NA | NA | 11 | 0.7273 | 6 | 0.8333 | 1 | 1 | NA | NA | 4 | 0.5 | 11 | 0.7273 |
| 19_540363 | NA | NA | 6 | 0.1667 | 2 | 0 | NA | NA | 8 | 0.125 | 6 | 0 | 1 | 0 | NA | NA | 4 | 0.25 | 11 | 0.0909 |
| 19_540364 | NA | NA | 7 | 0 | 5 | 0.2 | NA | NA | 12 | 0.0833 | 7 | 0 | 2 | 0.5 | 1 | 0 | 4 | 0 | 14 | 0.0714 |
| 19_540367 | NA | NA | 6 | 0 | 5 | 0.2 | NA | NA | 11 | 0.0909 | 3 | 0.3333 | 2 | 0 | NA | NA | 3 | 0 | 8 | 0.125 |
| 19_540367 | 1 | 0 | 7 | 0.1429 | 6 | 0.1667 | NA | NA | 14 | 0.1429 | 4 | 0.25 | 2 | 0 | 1 | 0 | 5 | 0.2 | 12 | 0.1667 |
| 19_540367 | 1 | 0 | 9 | 0.3333 | 6 | 0.1667 | NA | NA | 16 | 0.25 | 6 | 0.6667 | 2 | 0.5 | NA | NA | 6 | 0.3333 | 14 | 0.5 |
| 19_540368 | 1 | 0 | 7 | 0.1429 | 6 | 0 | NA | NA | 14 | 0.0714 | 5 | 0.2 | 2 | 0 | NA | NA | 5 | 0 | 12 | 0.0833 |
| 19_540369 | NA | NA | 5 | 0.8 | 4 | 0.75 | NA | NA | 9 | 0.7778 | 3 | 0.3333 | 1 | 1 | 1 | 1 | 2 | 1 | 7 | 0.7143 |
| 19_540369 | NA | NA | 8 | 0.5 | 5 | 0 | NA | NA | 13 | 0.3077 | 5 | 0.4 | NA | NA | NA | NA | 2 | 1 | 7 | 0.5714 |
| 19_540372 | 1 | 1 | 11 | 0.1818 | 7 | 0.2857 | NA | NA | 19 | 0.2632 | 6 | 0 | 3 | 0 | 1 | 0 | 3 | 0.3333 | 13 | 0.0769 |
| 19_540372 | 1 | 0 | 13 | 0.1538 | 7 | 0 | NA | NA | 21 | 0.0952 | 6 | 0.3333 | 3 | 0 | 1 | 0 | 5 | 0.2 | 15 | 0.2 |
| 19_550563 | 12 | 0.0833 | 8 | 0 | 10 | 0.1 | 7 | 0 | 37 | 0.0541 | 10 | 0.1 | 10 | 0 | 7 | 0 | 5 | 0.4 | 32 | 0.0938 |
| 19_550584 | 3 | 0 | 3 | 0.3333 | 7 | 0.2857 | 2 | 0.5 | 15 | 0.2667 | 5 | 0.4 | 2 | 0.5 | 2 | 0.5 | 5 | 0.4 | 14 | 0.4286 |
| 19_550590 | 3 | 0 | 2 | 0 | 7 | 0.2857 | 4 | 0 | 16 | 0.125 | 4 | 0 | 2 | 0 | 5 | 0.2 | 3 | 0 | 14 | 0.0714 |
| 19_550590 | 2 | 1 | 2 | 1 | 5 | 0.6 | 4 | 0.5 | 13 | 0.6923 | 5 | 0.8 | 2 | 0.5 | 5 | 0.8 | 3 | 0.6667 | 15 | 0.7333 |
| 19_550592 | 3 | 0 | 2 | 0 | 7 | 0 | 3 | 0.3333 | 15 | 0.0667 | 6 | 0.1667 | 2 | 0.5 | 5 | 0 | 3 | 0 | 16 | 0.125 |
| 19_550593 | 3 | 0 | 2 | 0.5 | 7 | 0 | 3 | 0 | 15 | 0.0667 | 4 | 0.25 | 2 | 0 | 5 | 0 | 3 | 0.3333 | 14 | 0.1429 |
| 19_550594 | 3 | 0.3333 | 2 | 0 | 7 | 0 | 2 | 0.5 | 14 | 0.1429 | 5 | 0 | 1 | 0 | 5 | 0.2 | 2 | 0.5 | 13 | 0.1538 |
| 19_550981 | 1 | 1 | NA | NA | NA | NA | NA | NA | 1 | 1 | NA | NA | 1 | 1 | NA | NA | 2 | 1 | 3 | 1 |
| 19_550982 | 1 | 1 | NA | NA | NA | NA | NA | NA | 1 | 1 | NA | NA | 1 | 1 | NA | NA | 2 | 1 | 3 | 1 |
| 19_550984 | 1 | 1 | NA | NA | NA | NA | NA | NA | 1 | 1 | NA | NA | 1 | 1 | NA | NA | 2 | 1 | 3 | 1 |
| 19_552421 | 1 | 1 | 2 | 0.5 | 1 | 0 | NA | NA | 4 | 0.5 | NA | NA | 1 | 1 | NA | NA | 1 | 0 | 2 | 0.5 |
| 19_552868 | 3 | 1 | 7 | 0.5714 | 1 | 1 | 3 | 0.6667 | 14 | 0.7143 | 2 | 1 | 3 | 0.6667 | 4 | 1 | NA | NA | 9 | 0.8889 |
| 19_552869 | 4 | 0.25 | 7 | 0.2857 | 1 | 1 | 3 | 0 | 15 | 0.2667 | 2 | 0.5 | 3 | 0 | 2 | 0 | NA | NA | 7 | 0.1429 |
| 19_552870 | 4 | 0.5 | 5 | 0.2 | 1 | 1 | 3 | 0 | 13 | 0.3077 | 2 | 0.5 | 1 | 0 | 3 | 0 | NA | NA | 6 | 0.1667 |

| | | | | | | | | | | | | | | | | | | | | |
|---|---|---|---|---|---|---|---|---|---|---|---|---|---|---|---|---|---|---|---|---|
| 19_553028 | 3 | 0 | 6 | 0 | 2 | 0.5 | 2 | 0 | 13 | 0.0769 | 1 | 1 | 1 | 1 | 2 | 0 | NA | NA | 4 | 0.5 |
| 19_553029 | 2 | 1 | 5 | 0.8 | 2 | 1 | 1 | 0 | 10 | 0.8 | 1 | 1 | 1 | 1 | 2 | 0.5 | NA | NA | 4 | 0.75 |
| 19_553030 | NA | NA | NA | NA | 1 | 1 | NA | NA | 1 | 1 | 1 | 1 | 1 | 1 | 1 | 1 | NA | NA | 3 | 1 |
| 19_556123 | 4 | 0 | 5 | 0.4 | 2 | 0 | 5 | 0.4 | 16 | 0.25 | 1 | 0 | 1 | 0 | 3 | 0.3333 | 3 | 0.3333 | 8 | 0.25 |
| 19_556127 | 4 | 0.25 | 5 | 0.4 | 5 | 0 | 2 | 0 | 16 | 0.1875 | 1 | 0 | 1 | 0 | 3 | 0.3333 | 4 | 0 | 9 | 0.1111 |
| 19_556139 | 6 | 0 | 7 | 0.1429 | 5 | 0 | 2 | 0.5 | 20 | 0.1 | 1 | 1 | 2 | 0 | 4 | 0 | 5 | 0.2 | 12 | 0.1667 |
| 19_556143 | 4 | 0.5 | 7 | 0 | 6 | 0 | 2 | 0 | 19 | 0.1053 | 2 | 0 | NA | NA | 4 | 0 | 5 | 0.2 | 11 | 0.0909 |
| 19_556144 | 1 | 0 | 4 | 0.25 | 4 | 0.5 | 1 | 0 | 10 | 0.3 | 2 | 0.5 | NA | NA | 3 | 0.3333 | 4 | 0.75 | 9 | 0.5556 |
| 19_556145 | 5 | 1 | 6 | 1 | 6 | 0.8333 | 1 | 1 | 18 | 0.9444 | 2 | 1 | 1 | 1 | 2 | 1 | 6 | 0.8333 | 11 | 0.9091 |
| 19_556145 | 5 | 0.2 | 6 | 0.1667 | 5 | 0.6 | 2 | 0.5 | 18 | 0.3333 | 2 | 0 | 1 | 0 | 4 | 0.25 | 6 | 0.1667 | 13 | 0.1538 |
| 19_556146 | 5 | 0.2 | 6 | 0.1667 | 6 | 0.1667 | 2 | 1 | 19 | 0.2632 | 1 | 1 | 1 | 1 | 4 | 0.75 | 6 | 0.3333 | 12 | 0.5833 |
| 19_557309 | NA | NA | 1 | 1 | NA | NA | NA | NA | 1 | 1 | NA | NA | 1 | 1 | NA | NA | NA | NA | 1 | 1 |
| 19_559200 | 1 | 0 | NA | NA | 1 | 0 | 2 | 1 | 4 | 0.5 | 3 | 0.3333 | 1 | 0 | 3 | 0 | 1 | 0 | 8 | 0.125 |
| 19_559206 | 3 | 0.6667 | 1 | 1 | 1 | 1 | 3 | 0.3333 | 8 | 0.625 | 2 | 0.5 | 2 | 0 | 4 | 1 | 1 | 0 | 9 | 0.5556 |
| 19_559212 | 2 | 0.5 | 1 | 1 | 1 | 1 | 1 | 1 | 5 | 0.8 | 2 | 0.5 | 2 | 0 | 4 | 0.5 | 1 | 1 | 9 | 0.4444 |
| 19_559214 | 4 | 0.75 | 1 | 1 | 1 | 1 | 2 | 1 | 8 | 0.875 | 3 | 1 | 2 | 0.5 | 2 | 0.5 | 1 | 0 | 8 | 0.625 |
| 19_559214 | 3 | 0 | 1 | 0 | 1 | 0 | 2 | 1 | 7 | 0.2857 | 4 | 0 | 2 | 0 | 3 | 0.3333 | 1 | 0 | 10 | 0.1 |
| 19_559214 | 3 | 0.3333 | 1 | 0 | 1 | 0 | 3 | 0 | 8 | 0.125 | 4 | 0 | 2 | 0 | 4 | 0.25 | 1 | 0 | 11 | 0.0909 |
| 19_564302 | 1 | 0 | NA | NA | NA | NA | 2 | 0.5 | 3 | 0.3333 | NA | NA | 1 | 0 | NA | NA | 1 | 1 | 2 | 0.5 |
| 19_565178 | 1 | 1 | NA | NA | NA | NA | NA | NA | 1 | 1 | NA | NA | 2 | 0 | NA | NA | 1 | 1 | 3 | 0.3333 |
| 19_565325 | 1 | 0 | 1 | 1 | NA | NA | NA | NA | 2 | 0.5 | NA | NA | 2 | 0.5 | NA | NA | NA | NA | 2 | 0.5 |
| 19_565326 | 1 | 0 | 1 | 1 | NA | NA | NA | NA | 2 | 0.5 | NA | NA | 2 | 0.5 | NA | NA | 1 | 1 | 3 | 0.6667 |
| 19_565328 | 1 | 1 | 1 | 1 | NA | NA | NA | NA | 2 | 1 | NA | NA | NA | NA | NA | NA | 1 | 1 | 1 | 1 |
| 19_565335 | 1 | 1 | NA | NA | NA | NA | NA | NA | 1 | 1 | NA | NA | 1 | 1 | NA | NA | 1 | 0 | 2 | 0.5 |
| 19_565353 | 1 | 1 | 1 | 1 | NA | NA | NA | NA | 2 | 1 | NA | NA | 1 | 1 | NA | NA | 1 | 1 | 2 | 1 |
| 19_565355 | 1 | 1 | 1 | 1 | NA | NA | NA | NA | 2 | 1 | NA | NA | 1 | 1 | NA | NA | 1 | 1 | 2 | 1 |
| 19_565355 | 1 | 1 | 1 | 1 | NA | NA | NA | NA | 2 | 1 | NA | NA | 1 | 1 | NA | NA | 1 | 1 | 2 | 1 |
| 19_565359 | 1 | 0 | 1 | 1 | NA | NA | NA | NA | 2 | 0.5 | NA | NA | NA | NA | NA | NA | 1 | 1 | 1 | 1 |
| 19_565359 | 1 | 1 | NA | NA | NA | NA | NA | NA | 1 | 1 | NA | NA | 1 | 1 | NA | NA | 1 | 1 | 2 | 1 |
| 19_565359 | 1 | 1 | 1 | 0 | NA | NA | NA | NA | 2 | 0.5 | NA | NA | 1 | 0 | NA | NA | 1 | 1 | 2 | 0.5 |
| 19_565360 | 1 | 0 | 1 | 1 | NA | NA | NA | NA | 2 | 0.5 | NA | NA | 1 | 1 | NA | NA | 1 | 1 | 2 | 1 |
| 19_565912 | 3 | 0.6667 | 3 | 0 | 3 | 0.3333 | 3 | 0.3333 | 12 | 0.3333 | NA | NA | NA | NA | 1 | 0 | 2 | 1 | 3 | 0.6667 |
| 19_565913 | 2 | 0.5 | 3 | 0.3333 | 3 | 0.3333 | 2 | 0 | 10 | 0.3 | NA | NA | NA | NA | 1 | 0 | 2 | 0.5 | 3 | 0.3333 |
| 19_565915 | 3 | 1 | 2 | 0.5 | 3 | 1 | 3 | 0.6667 | 11 | 0.8182 | NA | NA | NA | NA | 1 | 1 | 2 | 0.5 | 3 | 0.6667 |
| 19_567044 | 1 | 1 | NA | NA | NA | NA | 1 | 1 | 2 | 1 | NA | NA | NA | NA | NA | NA | 1 | 1 | 1 | 1 |
| 19_567046 | NA | NA | NA | NA | NA | NA | 1 | 1 | 1 | 1 | NA | NA | NA | NA | NA | NA | 1 | 1 | 1 | 1 |
| 19_567052 | 1 | 0 | NA | NA | NA | NA | 1 | 1 | 2 | 0.5 | NA | NA | NA | NA | NA | NA | 1 | 1 | 1 | 1 |
| 19_567052 | 1 | 1 | NA | NA | NA | NA | 1 | 1 | 2 | 1 | NA | NA | NA | NA | NA | NA | 1 | 1 | 1 | 1 |
| 19_567094 | 1 | 1 | NA | NA | NA | NA | NA | NA | 1 | 1 | NA | NA | NA | NA | 1 | 1 | NA | NA | 1 | 1 |
| 19_567109 | NA | NA | NA | NA | 1 | 0 | 2 | 0.5 | 3 | 0.3333 | 3 | 0.6667 | 2 | 1 | 3 | 0 | 1 | 0 | 9 | 0.4444 |
| 19_567112 | NA | NA | NA | NA | 2 | 0.5 | 1 | 1 | 3 | 0.6667 | 3 | 1 | 1 | 1 | 3 | 1 | 1 | 1 | 8 | 1 |
| 19_567112 | NA | NA | 1 | 0 | 1 | 1 | 1 | 0 | 3 | 0.3333 | 2 | 0 | 1 | 0 | 3 | 0.3333 | 2 | 0.5 | 8 | 0.25 |
| 19_567116 | 1 | 1 | 1 | 1 | 2 | 1 | 2 | 1 | 6 | 1 | 2 | 1 | 1 | 1 | 5 | 0.8 | 3 | 1 | 11 | 0.9091 |
| 19_567117 | NA | NA | 1 | 0 | 2 | 0.5 | 1 | 1 | 4 | 0.5 | 3 | 1 | 1 | 1 | 5 | 0.8 | 3 | 0.6667 | 12 | 0.8333 |
| 19_567121 | 2 | 0 | 3 | 0 | 1 | 1 | 2 | 0.5 | 8 | 0.25 | 2 | 0.5 | 1 | 0 | 4 | 0.5 | 3 | 0.3333 | 10 | 0.4 |
| 19_567121 | 1 | 1 | 2 | 1 | 2 | 1 | 2 | 1 | 7 | 1 | 1 | 1 | 1 | 1 | 6 | 1 | 4 | 1 | 12 | 1 |
| 19_567122 | 3 | 1 | 3 | 1 | 2 | 0.5 | 3 | 1 | 11 | 0.9091 | 4 | 1 | 1 | 1 | 7 | 1 | 4 | 1 | 16 | 1 |
| 19_567123 | 2 | 0.5 | 4 | 0 | 1 | 0 | 2 | 1 | 9 | 0.3333 | 3 | 0.6667 | 1 | 0 | 6 | 0.1667 | 4 | 0 | 14 | 0.2143 |
| 19_567124 | 3 | 0.6667 | 4 | 0.75 | 2 | 1 | 2 | 0.5 | 11 | 0.7273 | 4 | 1 | NA | NA | 7 | 1 | 4 | 0.25 | 15 | 0.8 |
| 19_567125 | 4 | 0 | 2 | 0 | 2 | 0.5 | 2 | 0 | 10 | 0.1 | 4 | 0 | 2 | 0.5 | 7 | 0.5714 | 4 | 0.25 | 17 | 0.3529 |
| 19_567125 | 4 | 0.5 | 3 | 1 | 2 | 0.5 | 2 | 0.5 | 11 | 0.6364 | 3 | 0.3333 | NA | NA | 7 | 0.8571 | 4 | 0.75 | 14 | 0.7143 |
| 19_567128 | 4 | 0.5 | 3 | 0 | 3 | 0.6667 | 3 | 0 | 13 | 0.3077 | 2 | 0 | 2 | 0.5 | 6 | 0.1667 | 4 | 0.25 | 14 | 0.2143 |
| 19_567129 | 4 | 0.75 | 4 | 1 | 3 | 1 | 4 | 0.75 | 15 | 0.8667 | 4 | 0.75 | 2 | 1 | 8 | 1 | 4 | 1 | 18 | 0.9444 |
| 19_567130 | 3 | 0 | 5 | 0.2 | 3 | 0 | 3 | 0.3333 | 14 | 0.1429 | 4 | 0.25 | 2 | 0.5 | 9 | 0.1111 | 3 | 0 | 18 | 0.1667 |
| 19_567130 | 4 | 0.25 | 4 | 0.25 | 3 | 0 | 3 | 0 | 14 | 0.1429 | 4 | 0.25 | 2 | 0 | 8 | 0.25 | 4 | 0.5 | 18 | 0.2778 |
| 19_567130 | 3 | 0.3333 | 5 | 0.4 | 3 | 0.3333 | 4 | 0 | 15 | 0.2667 | 4 | 0.5 | 1 | 0 | 8 | 0.125 | 5 | 0.2 | 18 | 0.2222 |
| 19_567135 | 1 | 0 | 4 | 0.75 | 1 | 0 | 5 | 0 | 11 | 0.2727 | 4 | 0.25 | 3 | 0.3333 | 8 | 0.125 | 4 | 0.25 | 19 | 0.2105 |
| 19_567761 | NA | NA | 6 | 0.3333 | 3 | 0.3333 | 1 | 0 | 10 | 0.3 | 3 | 0 | 4 | 0.5 | 5 | 0 | 8 | 0.25 | 20 | 0.2 |
| 19_567762 | NA | NA | 5 | 0.2 | 3 | 0.3333 | 1 | 0 | 9 | 0.2222 | 3 | 0 | 4 | 0.5 | 2 | 0 | 6 | 0.3333 | 15 | 0.2667 |
| 19_567765 | NA | NA | 4 | 0.25 | 3 | 0.6667 | 1 | 0 | 8 | 0.375 | 3 | 0.3333 | 4 | 0.5 | 3 | 0.3333 | 6 | 0.1667 | 16 | 0.3125 |
| 19_567768 | NA | NA | 4 | 0.25 | 3 | 0.3333 | NA | NA | 7 | 0.2857 | 3 | 0 | 4 | 0.25 | 3 | 0 | 4 | 0 | 14 | 0.0714 |
| 19_567769 | NA | NA | 5 | 0.2 | 3 | 0.3333 | NA | NA | 8 | 0.25 | 3 | 0 | 4 | 0 | 3 | 0 | 7 | 0.1429 | 17 | 0.0588 |
| 19_567773 | NA | NA | 3 | 0 | 4 | 0.25 | NA | NA | 7 | 0.1429 | 2 | 0 | 4 | 0.25 | 3 | 0 | 5 | 0.2 | 14 | 0.1429 |
| 19_567773 | NA | NA | 3 | 0 | 3 | 0.3333 | 1 | 0 | 7 | 0.1429 | 2 | 0 | 3 | 0.3333 | 3 | 0 | 5 | 0.4 | 13 | 0.2308 |
| 19_567779 | 1 | 1 | 1 | 0 | 4 | 0.75 | 2 | 1 | 8 | 0.75 | 2 | 0 | 4 | 1 | 3 | 0.6667 | 5 | 0.8 | 14 | 0.7143 |

| | | | | | | | | | | | | | | | | | | | | |
|---|---|---|---|---|---|---|---|---|---|---|---|---|---|---|---|---|---|---|---|---|
| 19_567781 | 1 | 0 | 1 | 0 | 4 | 0 | 2 | 0.5 | 8 | 0.125 | 2 | 0 | 4 | 0 | 3 | 0.3333 | 5 | 0 | 14 | 0.0714 |
| 19_567783 | 1 | 0 | 1 | 0 | 3 | 0.3333 | 2 | 0.5 | 7 | 0.2857 | 2 | 1 | 2 | 0.5 | 2 | 0 | 5 | 0.6 | 11 | 0.5455 |
| 19_567794 | NA | NA | NA | NA | 3 | 0.3333 | 1 | 0 | 4 | 0.25 | 1 | 0 | 1 | 0 | 2 | 0 | 1 | 1 | 5 | 0.2 |
| 19_567818 | 1 | 1 | NA | NA | 2 | 0 | 1 | 0 | 4 | 0.25 | NA | NA | 1 | 1 | NA | NA | 2 | 0 | 3 | 0.3333 |
| 19_568015 | 1 | 1 | NA | NA | 2 | 1 | 1 | 1 | 4 | 1 | NA | NA | NA | NA | NA | NA | 1 | 1 | 1 | 1 |
| 19_568015 | 1 | 1 | NA | NA | 1 | 1 | NA | NA | 2 | 1 | NA | NA | NA | NA | NA | NA | 1 | 1 | 1 | 1 |
| 19_568017 | 1 | 1 | NA | NA | 1 | 0 | 1 | 0 | 3 | 0.3333 | NA | NA | NA | NA | NA | NA | 1 | 1 | 1 | 1 |
| 19_568017 | 1 | 0 | NA | NA | NA | NA | 1 | 1 | 2 | 0.5 | NA | NA | NA | NA | NA | NA | 1 | 1 | 1 | 1 |
| 19_568019 | 1 | 1 | NA | NA | 1 | 0 | NA | NA | 2 | 0.5 | NA | NA | NA | NA | NA | NA | 1 | 1 | 1 | 1 |
| 19_569011 | NA | NA | 1 | 0 | NA | NA | 2 | 0.5 | 3 | 0.3333 | NA | NA | 1 | 1 | 1 | 1 | NA | NA | 2 | 1 |
| 19_569012 | NA | NA | 1 | 1 | NA | NA | 2 | 0.5 | 3 | 0.6667 | NA | NA | 1 | 1 | 1 | 0 | NA | NA | 2 | 0.5 |
| 19_569015 | NA | NA | NA | NA | NA | NA | 2 | 1 | 2 | 1 | NA | NA | 2 | 0.5 | 1 | 0 | NA | NA | 3 | 0.3333 |
| 19_569017 | NA | NA | NA | NA | NA | NA | 1 | 1 | 1 | 1 | NA | NA | 2 | 0.5 | 1 | 1 | NA | NA | 3 | 0.6667 |
| 19_569017 | NA | NA | NA | NA | NA | NA | 1 | 1 | 1 | 1 | NA | NA | 2 | 0.5 | 1 | 0 | NA | NA | 3 | 0.3333 |
| 19_570219 | 1 | 0 | 7 | 0.5714 | 8 | 0.5 | 8 | 0.25 | 24 | 0.4167 | 6 | 0 | 4 | 0.25 | 5 | 0.6 | 5 | 0.4 | 20 | 0.3 |
| 19_570219 | 2 | 0.5 | 9 | 0 | 8 | 0.25 | 8 | 0.375 | 27 | 0.2222 | 5 | 0 | 6 | 0.3333 | 5 | 0 | 4 | 0 | 20 | 0.1 |
| 19_570220 | 2 | 1 | 9 | 0.2222 | 8 | 0 | 6 | 0.3333 | 25 | 0.24 | 4 | 0.25 | 5 | 0.4 | 4 | 0.25 | 5 | 0 | 18 | 0.2222 |
| 19_570220 | 3 | 0 | 9 | 0.2222 | 10 | 0.3 | 9 | 0.2222 | 31 | 0.2258 | 6 | 0.3333 | 7 | 0.1429 | 2 | 0.5 | 5 | 0.4 | 20 | 0.3 |
| 19_570222 | 2 | 0 | 8 | 0.5 | 7 | 0.2857 | 5 | 0.4 | 22 | 0.3636 | 4 | 0.5 | 6 | 0.3333 | 2 | 0 | 5 | 0.4 | 17 | 0.3529 |
| 19_570226 | 2 | 0.5 | 5 | 0.4 | 5 | 0.6 | 6 | 0.1667 | 18 | 0.3889 | 4 | 0.25 | 5 | 0.6 | 4 | 0.25 | 3 | 0 | 16 | 0.3125 |
| 19_570229 | 2 | 0.5 | 6 | 0 | 4 | 0 | 5 | 0 | 17 | 0.0588 | 5 | 0 | 5 | 0.2 | 5 | 0 | 2 | 0 | 17 | 0.0588 |
| 19_570234 | 1 | 0 | 5 | 0 | 4 | 0 | 2 | 0.5 | 12 | 0.0833 | 5 | 0 | 5 | 0.8 | 2 | 0.5 | 2 | 0 | 14 | 0.3571 |
| 19_570239 | 1 | 1 | 3 | 0 | NA | NA | 2 | 0.5 | 6 | 0.3333 | 5 | 0.6 | 2 | 0 | 2 | 0.5 | 1 | 1 | 10 | 0.5 |
| 19_570239 | 2 | 0 | 5 | 0.2 | 3 | 0 | 2 | 0 | 12 | 0.0833 | 5 | 0 | 2 | 0 | 2 | 0 | 2 | 0.5 | 11 | 0.0909 |
| 19_570241 | NA | NA | 4 | 0 | 2 | 0 | 2 | 0.5 | 8 | 0.125 | 3 | 0.3333 | 2 | 0.5 | 2 | 0.5 | 2 | 0.5 | 9 | 0.4444 |
| 19_570242 | NA | NA | 4 | 0.5 | 1 | 0 | 2 | 0.5 | 7 | 0.4286 | 2 | 0 | 1 | 0 | 3 | 0.3333 | 2 | 0 | 8 | 0.125 |
| 19_570244 | 1 | 0 | 4 | 0 | 3 | 0.3333 | 2 | 0 | 10 | 0.1 | 3 | 0.3333 | 2 | 0 | 3 | 0 | 1 | 0 | 9 | 0.1111 |
| 19_570246 | 1 | 0 | 4 | 0 | 3 | 0 | 2 | 0.5 | 10 | 0.1 | 3 | 0 | 2 | 0.5 | 3 | 0 | 1 | 0 | 9 | 0.1111 |
| 19_570247 | 1 | 0 | 4 | 0 | 3 | 0 | 2 | 0.5 | 10 | 0.1 | 3 | 0.6667 | 1 | 1 | NA | NA | 1 | 0 | 5 | 0.6 |
| 19_570249 | NA | NA | 4 | 0.25 | 3 | 0.3333 | 2 | 0.5 | 9 | 0.3333 | 3 | 0.6667 | 2 | 0 | 2 | 0 | 1 | 0 | 8 | 0.25 |
| 19_570249 | 1 | 1 | 4 | 0.5 | 3 | 0.3333 | 2 | 0.5 | 10 | 0.5 | 3 | 0.6667 | 2 | 1 | 3 | 0.6667 | 1 | 1 | 9 | 0.7778 |
| 19_570253 | 1 | 1 | 3 | 1 | 3 | 0.6667 | 1 | 1 | 8 | 0.875 | 1 | 1 | 1 | 1 | 1 | 1 | NA | NA | 3 | 1 |
| 19_570254 | NA | NA | 3 | 0.3333 | 3 | 0 | 2 | 0.5 | 8 | 0.25 | 3 | 0.3333 | 2 | 0 | 2 | 1 | NA | NA | 7 | 0.4286 |
| 19_570255 | NA | NA | 2 | 1 | 2 | 0.5 | 2 | 0.5 | 6 | 0.6667 | 2 | 1 | 2 | 1 | 1 | 1 | NA | NA | 5 | 1 |
| 19_570255 | NA | NA | 3 | 1 | 3 | 0.6667 | 2 | 0.5 | 8 | 0.75 | 2 | 0.5 | 2 | 1 | 1 | 1 | NA | NA | 5 | 0.8 |
| 19_570256 | NA | NA | 3 | 1 | 2 | 1 | 2 | 0.5 | 7 | 0.8571 | 1 | 1 | 2 | 1 | NA | NA | NA | NA | 3 | 1 |
| 19_573119 | 3 | 0.6667 | 1 | 0 | NA | NA | 4 | 0.25 | 8 | 0.375 | 1 | 0 | 2 | 0 | NA | NA | 2 | 0.5 | 5 | 0.2 |
| 19_573487 | NA | NA | 1 | 1 | NA | NA | 1 | 1 | 2 | 1 | NA | NA | 1 | 1 | NA | NA | 2 | 0.5 | 3 | 0.6667 |
| 19_573489 | NA | NA | 1 | 1 | NA | NA | 1 | 1 | 2 | 1 | NA | NA | 1 | 1 | NA | NA | 2 | 1 | 3 | 1 |
| 19_573751 | 1 | 1 | 2 | 0.5 | 2 | 0.5 | 1 | 0 | 6 | 0.5 | 3 | 1 | 2 | 1 | 3 | 0.6667 | 3 | 1 | 11 | 0.9091 |
| 19_573752 | 2 | 0.5 | 2 | 0 | 2 | 0.5 | 1 | 0 | 7 | 0.2857 | 5 | 0.4 | 3 | 0 | 2 | 0 | 3 | 0.6667 | 13 | 0.3077 |
| 19_573752 | 1 | 1 | 2 | 1 | 2 | 0.5 | 1 | 0 | 6 | 0.6667 | 3 | 1 | 1 | 1 | 3 | 0.6667 | 3 | 0.3333 | 10 | 0.7 |
| 19_573753 | 2 | 1 | 2 | 1 | 2 | 0 | 1 | 0 | 7 | 0.5714 | 5 | 0 | 3 | 0 | 2 | 0 | 3 | 0.3333 | 13 | 0.0769 |
| 19_573753 | 2 | 1 | 2 | 1 | 1 | 1 | 1 | 1 | 6 | 1 | 5 | 1 | 3 | 1 | 3 | 1 | 3 | 1 | 14 | 1 |
| 19_573753 | 2 | 0.5 | 2 | 0 | 2 | 0 | 1 | 0 | 7 | 0.1429 | 4 | 0 | 1 | 1 | 3 | 0.3333 | 3 | 0.3333 | 11 | 0.2727 |
| 19_573765 | 3 | 0.3333 | 2 | 0 | 3 | 0 | 1 | 0 | 9 | 0.1111 | 5 | 0.2 | 2 | 0 | 1 | 0 | 3 | 0 | 11 | 0.0909 |
| 19_573771 | 6 | 0.1667 | 1 | 0 | 4 | 0.25 | 1 | 0 | 12 | 0.1667 | 4 | 0.25 | 2 | 0.5 | 3 | 0.3333 | 3 | 0 | 12 | 0.25 |
| 19_573779 | 6 | 0.1667 | 3 | 0 | 5 | 0 | 2 | 0 | 16 | 0.0625 | 6 | 0 | 3 | 0.3333 | 4 | 0 | 2 | 0.5 | 15 | 0.1333 |
| 19_573779 | 7 | 0.1429 | 3 | 0 | 7 | 0.1429 | 2 | 0 | 19 | 0.1053 | 7 | 0 | 4 | 0 | 4 | 0.25 | 2 | 0 | 17 | 0.0588 |
| 19_574765 | 4 | 0 | 3 | 0.3333 | 2 | 0 | 5 | 0 | 14 | 0.0714 | 2 | 0 | 1 | 1 | 3 | 0 | NA | NA | 6 | 0.1667 |
| 19_574766 | 3 | 0.3333 | 2 | 0.5 | 2 | 0.5 | 4 | 0.25 | 11 | 0.3636 | 3 | 0.3333 | 2 | 1 | 4 | 0 | NA | NA | 9 | 0.3333 |
| 19_574766 | 5 | 0.2 | 3 | 0.3333 | 1 | 1 | 4 | 0 | 13 | 0.2308 | 3 | 0 | 2 | 0.5 | 3 | 0 | NA | NA | 8 | 0.125 |
| 19_574768 | 2 | 0 | 2 | 0.5 | 2 | 0 | 6 | 0 | 12 | 0.0833 | 3 | 0.3333 | 2 | 0 | 4 | 0 | NA | NA | 9 | 0.1111 |
| 19_574769 | 2 | 0 | 1 | 0 | 2 | 0.5 | 5 | 0.2 | 10 | 0.2 | 4 | 0.75 | 3 | 0.6667 | 3 | 0 | NA | NA | 10 | 0.5 |
| 19_574770 | 4 | 0 | 3 | 0 | 2 | 0 | 6 | 0.1667 | 15 | 0.0667 | 4 | 0.25 | 3 | 0.3333 | 5 | 0 | NA | NA | 12 | 0.1667 |
| 19_574772 | 4 | 0.75 | 2 | 0 | 2 | 1 | 6 | 0.8333 | 14 | 0.7143 | 4 | 0.25 | 3 | 0.6667 | 4 | 0.75 | NA | NA | 11 | 0.5455 |
| 19_578203 | NA | NA | NA | NA | 2 | 1 | NA | NA | 2 | 1 | 2 | 0.5 | NA | NA | NA | NA | NA | NA | 2 | 0.5 |
| 19_578205 | NA | NA | 1 | 1 | 1 | 1 | NA | NA | 2 | 1 | 2 | 1 | NA | NA | 1 | 0 | NA | NA | 3 | 0.6667 |
| 19_581196 | 6 | 0.6667 | 8 | 0.25 | 6 | 0.3333 | 5 | 0.8 | 25 | 0.48 | 2 | 1 | 3 | 0 | 3 | 0.6667 | 5 | 0.6 | 13 | 0.5385 |
| 19_581197 | 6 | 0.1667 | 8 | 0.125 | 5 | 0.4 | 5 | 0.4 | 24 | 0.25 | 2 | 0 | 2 | 1 | 4 | 0.75 | 4 | 0.75 | 12 | 0.6667 |
| 19_581198 | 7 | 1 | 8 | 0.875 | 6 | 1 | 5 | 1 | 26 | 0.9615 | 2 | 1 | 4 | 1 | 5 | 1 | 5 | 1 | 16 | 1 |
| 19_581198 | 4 | 0 | 8 | 0 | 4 | 0 | 4 | 0.25 | 20 | 0.05 | 1 | 0 | 3 | 0.3333 | 3 | 0 | 4 | 0 | 11 | 0.0909 |
| 19_581199 | 5 | 0.6 | 10 | 0.3 | 6 | 0.1667 | 5 | 0.4 | 26 | 0.3462 | 2 | 0 | 4 | 0.25 | 5 | 0.4 | 5 | 0.2 | 16 | 0.25 |
| 19_581200 | 5 | 0.4 | 9 | 0.1111 | 7 | 0 | 5 | 0 | 26 | 0.1154 | 2 | 0 | 4 | 0.25 | 5 | 0.2 | 4 | 0 | 15 | 0.1333 |
| 19_581202 | 3 | 0.6667 | 9 | 0.1111 | 7 | 0.1429 | 5 | 0 | 24 | 0.1667 | 3 | 0 | 4 | 0.75 | 5 | 0 | 4 | 0 | 16 | 0.1875 |

| | | | | | | | | | | | | | | | | | | | | |
|---|---|---|---|---|---|---|---|---|---|---|---|---|---|---|---|---|---|---|---|---|
| 19_5812052 | 5 | 0.6 | 7 | 0.2857 | 7 | 0.1429 | 2 | 0 | 21 | 0.2857 | 2 | 0.5 | 4 | 0.5 | 3 | 0.3333 | 3 | 0 | 12 | 0.3333 |
| 19_5812057 | 4 | 0.25 | 8 | 0.25 | 5 | 0.2 | 5 | 0.4 | 22 | 0.2727 | 3 | 0.6667 | 4 | 0.5 | 5 | 0 | 5 | 0 | 17 | 0.2353 |
| 19_5812064 | 8 | 0.875 | 7 | 0.8571 | 7 | 0.8571 | 5 | 1 | 27 | 0.8889 | 3 | 0.6667 | 4 | 1 | 4 | 1 | 5 | 1 | 16 | 0.9375 |
| 19_5812065 | 6 | 0.3333 | 8 | 0.375 | 7 | 0.4286 | 5 | 0.2 | 26 | 0.3462 | 3 | 0.3333 | 3 | 0.6667 | 5 | 0.6 | 5 | 0.6 | 16 | 0.5625 |
| 19_5812071 | 6 | 1 | 7 | 0.5714 | 6 | 0.3333 | 3 | 1 | 22 | 0.6818 | 3 | 1 | 4 | 0.75 | 3 | 0.6667 | 5 | 0.6 | 15 | 0.7333 |
| 19_5812075 | 4 | 1 | 10 | 0.4 | 6 | 0.6667 | 5 | 1 | 25 | 0.68 | 3 | 0.3333 | 2 | 1 | 5 | 0.2 | 3 | 0.6667 | 13 | 0.4615 |
| 19_5812085 | 9 | 0.6667 | 10 | 0.5 | 6 | 0.8333 | 5 | 0.8 | 30 | 0.6667 | 2 | 1 | 4 | 0.5 | 3 | 1 | 5 | 0.4 | 14 | 0.6429 |
| 19_5812145 | 7 | 0.1429 | 11 | 0.7273 | 7 | 0.2857 | 4 | 0.75 | 29 | 0.4828 | 5 | 0.4 | 2 | 0.5 | 7 | 0.4286 | 2 | 0.5 | 16 | 0.4375 |
| 19_5821991 | 1 | 1 | NA | NA | NA | NA | 3 | 0.6667 | 4 | 0.75 | NA | NA | 1 | 1 | 2 | 1 | NA | NA | 3 | 1 |
| 19_5821993 | 1 | 0 | NA | NA | NA | NA | 3 | 0.3333 | 4 | 0.25 | NA | NA | 1 | 1 | 2 | 0 | NA | NA | 3 | 0.3333 |
| 19_5822404 | 2 | 0 | 1 | 1 | 1 | 1 | 2 | 1 | 6 | 0.6667 | 3 | 0 | 5 | 0.2 | 3 | 0 | 1 | 1 | 12 | 0.1667 |
| 19_5822413 | 4 | 0 | 6 | 0.3333 | 5 | 0.4 | 7 | 0.2857 | 22 | 0.2727 | 5 | 0.6 | 11 | 0.5455 | 6 | 0.5 | 3 | 0.6667 | 25 | 0.56 |
| 19_5822419 | 4 | 0.25 | 4 | 0 | 4 | 0 | 7 | 0.1429 | 19 | 0.1053 | 8 | 0 | 8 | 0.25 | 7 | 0 | 3 | 0 | 26 | 0.0769 |
| 19_5822424 | 4 | 0 | 4 | 0 | 5 | 0.2 | 7 | 0 | 20 | 0.05 | 8 | 0 | 9 | 0.3333 | 7 | 0 | 3 | 0.3333 | 27 | 0.1481 |
| 19_5822427 | 4 | 0.25 | 5 | 0.2 | 5 | 0 | 6 | 0.3333 | 20 | 0.2 | 8 | 0.125 | 11 | 0.3636 | 6 | 0.1667 | 1 | 1 | 26 | 0.2692 |
| 19_5822429 | 3 | 0.3333 | 5 | 0.2 | 4 | 0.25 | 5 | 0.2 | 17 | 0.2353 | 5 | 0.4 | 11 | 0.2727 | 6 | 0.3333 | 2 | 0 | 24 | 0.2917 |
| 19_5822435 | 4 | 1 | 4 | 1 | 6 | 0.5 | 6 | 1 | 20 | 0.85 | 5 | 1 | 9 | 0.8889 | 6 | 0.5 | 3 | 0.6667 | 23 | 0.7826 |
| 19_5822454 | 3 | 0.6667 | 3 | 0.3333 | 4 | 0.5 | 6 | 0.3333 | 16 | 0.4375 | 6 | 0.3333 | 9 | 0.2222 | 5 | 0.2 | 2 | 0.5 | 22 | 0.2727 |
| 19_5822455 | 4 | 0.5 | 3 | 0.3333 | 6 | 0.5 | 5 | 0.4 | 18 | 0.4444 | 6 | 0.3333 | 10 | 0.3 | 6 | 0.1667 | 3 | 0 | 25 | 0.24 |
| 19_5822470 | 2 | 0 | 4 | 0.25 | 6 | 0.1667 | 6 | 0 | 18 | 0.1111 | 6 | 0.1667 | 9 | 0.2222 | 6 | 0 | 3 | 0.3333 | 24 | 0.1667 |
| 19_5822471 | 3 | 0.3333 | 4 | 0.25 | 5 | 0 | 7 | 0.4286 | 19 | 0.2632 | 7 | 0 | 7 | 0.1429 | 6 | 0.1667 | 2 | 0.5 | 22 | 0.1364 |
| 19_5822496 | 3 | 0 | 4 | 0 | 7 | 0.2857 | 6 | 0 | 20 | 0.1 | 4 | 0.5 | 9 | 0.3333 | 8 | 0 | 4 | 0.5 | 25 | 0.28 |
| 19_5822535 | 3 | 0.3333 | 7 | 0.1429 | 6 | 0 | 6 | 0 | 22 | 0.0909 | 7 | 0 | 7 | 0 | 7 | 0.2857 | 2 | 0 | 23 | 0.087 |
| 19_5822568 | 2 | 0 | 6 | 0 | 6 | 0 | 3 | 0.3333 | 17 | 0.0588 | 3 | 0 | 6 | 0 | 6 | 0 | 4 | 0.25 | 19 | 0.0526 |
| 19_5822634 | 2 | 0.5 | 6 | 0.5 | 3 | 0 | 5 | 0.6 | 16 | 0.4375 | 3 | 0.3333 | 8 | 0.625 | 5 | 0.4 | 4 | 0.25 | 20 | 0.45 |
| 19_5822635 | 1 | 1 | 6 | 1 | 5 | 0.8 | 5 | 0.8 | 17 | 0.8824 | 4 | 1 | 6 | 0.8333 | 5 | 0.8 | 3 | 1 | 18 | 0.8889 |
| 19_5822638 | 1 | 0 | 5 | 0.4 | 4 | 0 | 3 | 0 | 13 | 0.1538 | 3 | 0 | 7 | 0.1429 | 5 | 0.2 | 2 | 0 | 17 | 0.1176 |
| 19_5822640 | 1 | 0 | 5 | 0.6 | 5 | 0.2 | 4 | 0 | 15 | 0.2667 | 3 | 0.3333 | 4 | 0.25 | 3 | 0 | 2 | 0 | 12 | 0.1667 |
| 19_5822641 | 2 | 0 | 5 | 0 | 4 | 0 | 5 | 0.2 | 16 | 0.0625 | 3 | 0 | 7 | 0.5714 | 5 | 0.2 | 3 | 0.6667 | 18 | 0.3889 |
| 19_5822649 | 2 | 0.5 | 5 | 0.4 | 5 | 0.4 | 3 | 0 | 15 | 0.3333 | 4 | 0 | 5 | 0.2 | 4 | 0 | 4 | 0.5 | 17 | 0.1765 |
| 19_5822657 | 2 | 1 | 5 | 0.8 | 5 | 1 | 5 | 1 | 17 | 0.9412 | 3 | 1 | 6 | 0.8333 | 4 | 0.75 | 4 | 1 | 17 | 0.8824 |
| 19_5822659 | 2 | 1 | 3 | 0.6667 | 5 | 0.8 | 5 | 0.6 | 15 | 0.7333 | 4 | 1 | 7 | 0.8571 | 2 | 0.5 | 3 | 1 | 16 | 0.875 |
| 19_5822662 | 2 | 1 | 5 | 1 | 2 | 1 | 2 | 1 | 11 | 1 | 1 | 1 | 4 | 1 | 2 | 0.5 | 4 | 1 | 11 | 0.9091 |
| 19_5822665 | 1 | 1 | 5 | 0.8 | 4 | 1 | 4 | 0.75 | 14 | 0.8571 | 2 | 1 | 7 | 0.8571 | 4 | 0.75 | 3 | 0.6667 | 16 | 0.8125 |
| 19_5822670 | 2 | 0.5 | 4 | 0.25 | 5 | 0.4 | 5 | 0.6 | 16 | 0.4375 | 3 | 1 | 4 | 0.5 | 3 | 0 | 3 | 0.6667 | 13 | 0.5385 |
| 19_5822680 | 2 | 0 | 5 | 0.4 | 4 | 0 | 4 | 0.25 | 15 | 0.2 | 3 | 0 | 8 | 0 | 4 | 0 | 4 | 0.5 | 19 | 0.1053 |
| 19_5828876 | 2 | 0 | 1 | 1 | 1 | 0 | 2 | 0.5 | 6 | 0.3333 | 3 | 0 | 5 | 0.2 | 3 | 0 | 1 | 0 | 12 | 0.0833 |
| 19_5878433 | 12 | 0 | 18 | 0.0556 | 17 | 0.1176 | 11 | 0.0909 | 58 | 0.069 | 10 | 0.1 | 9 | 0.1111 | 12 | 0.1667 | 12 | 0 | 43 | 0.093 |
| 19_5878492 | 8 | 0.5 | 9 | 0.4444 | 7 | 0.1429 | 9 | 0.2222 | 33 | 0.3333 | 6 | 0.3333 | 6 | 0.5 | 9 | 0.2222 | 12 | 0.0833 | 33 | 0.2424 |
| 19_5878521 | 9 | 0.4444 | 11 | 0.0909 | 14 | 0.1429 | 10 | 0.2 | 44 | 0.2045 | 6 | 0.3333 | 6 | 0.8333 | 8 | 0 | 12 | 0.0833 | 32 | 0.25 |
| 19_5878531 | 8 | 0.125 | 10 | 0.1 | 13 | 0.1538 | 9 | 0.2222 | 40 | 0.15 | 7 | 0.1429 | 5 | 0.4 | 9 | 0 | 10 | 0.1 | 31 | 0.129 |
| 19_5878536 | 8 | 0.25 | 12 | 0.1667 | 13 | 0.3077 | 9 | 0.3333 | 42 | 0.2619 | 7 | 0.7143 | 7 | 0.4286 | 8 | 0 | 11 | 0.1818 | 33 | 0.303 |
| 19_5878557 | 9 | 1 | 9 | 0.8889 | 13 | 1 | 11 | 1 | 42 | 0.9762 | 6 | 0.8333 | 7 | 1 | 7 | 1 | 11 | 0.8182 | 31 | 0.9032 |
| 19_5878563 | 7 | 1 | 10 | 1 | 14 | 1 | 9 | 1 | 40 | 1 | 6 | 1 | 6 | 1 | 6 | 1 | 10 | 1 | 28 | 1 |
| 19_5878566 | 9 | 0 | 11 | 0 | 12 | 0.0833 | 9 | 0.3333 | 41 | 0.0976 | 5 | 0 | 5 | 0 | 7 | 0.1429 | 11 | 0.0909 | 28 | 0.0714 |
| 19_5878568 | 4 | 0 | 4 | 0.25 | 12 | 0.0833 | 9 | 0 | 29 | 0.069 | 5 | 0.6 | 5 | 0.2 | 5 | 0.2 | 10 | 0.5 | 25 | 0.4 |
| 19_5878569 | 6 | 0 | 8 | 0.125 | 13 | 0.2308 | 10 | 0.1 | 37 | 0.1351 | 8 | 0.125 | 7 | 0.1429 | 7 | 0 | 11 | 0.0909 | 33 | 0.0909 |
| 19_5878570 | 7 | 0.5714 | 8 | 0.375 | 10 | 0.1 | 10 | 0.6 | 35 | 0.4 | 3 | 0.3333 | 4 | 0 | 5 | 0.2 | 8 | 0.375 | 20 | 0.25 |
| 19_5878575 | 6 | 0.6667 | 10 | 0.4 | 12 | 0.5833 | 8 | 0.375 | 36 | 0.5 | 6 | 1 | 7 | 0.7143 | 6 | 0.5 | 11 | 0.2727 | 30 | 0.5667 |
| 19_5878584 | 8 | 0.375 | 11 | 0.0909 | 13 | 0.3077 | 6 | 0.1667 | 38 | 0.2368 | 7 | 0 | 4 | 0 | 6 | 0.1667 | 12 | 0.1667 | 29 | 0.1034 |
| 19_5878586 | 8 | 0.625 | 10 | 0.8 | 11 | 0.9091 | 8 | 0.625 | 37 | 0.7568 | 3 | 0.6667 | 5 | 1 | 7 | 0.8571 | 11 | 0.8182 | 26 | 0.8462 |
| 19_5878588 | 7 | 0.8571 | 11 | 0.8182 | 9 | 0.6667 | 6 | 0.6667 | 33 | 0.7576 | 3 | 1 | 5 | 1 | 6 | 0.3333 | 9 | 0.6667 | 23 | 0.6957 |
| 19_5878591 | 7 | 0.2857 | 8 | 0.125 | 9 | 0 | 5 | 0 | 29 | 0.1034 | 4 | 0.25 | 5 | 0 | 7 | 0.1429 | 8 | 0 | 24 | 0.0833 |
| 19_5878592 | 6 | 0.5 | 6 | 0.5 | 9 | 0.3333 | 7 | 0.2857 | 28 | 0.3929 | 4 | 0.75 | 4 | 0.75 | 5 | 0.4 | 8 | 0.5 | 21 | 0.5714 |
| 19_5878592 | 8 | 0.75 | 10 | 0.6 | 10 | 0.6 | 6 | 0.1667 | 34 | 0.5588 | 5 | 0.8 | 4 | 0.75 | 5 | 0.2 | 8 | 0.5 | 22 | 0.5455 |
| 19_5878594 | 8 | 0.25 | 10 | 0.7 | 8 | 1 | 8 | 0.75 | 34 | 0.6765 | 5 | 0.6 | 5 | 0.6 | 3 | 0.6667 | 11 | 0.4545 | 24 | 0.5417 |
| 19_5878601 | 8 | 0.75 | 8 | 0.875 | 9 | 0.7778 | 5 | 0.6 | 30 | 0.7667 | 6 | 1 | 6 | 1 | 4 | 0.5 | 8 | 0.75 | 24 | 0.8333 |
| 19_5878604 | 7 | 0.1429 | 10 | 0 | 14 | 0.0714 | 8 | 0.125 | 39 | 0.0769 | 5 | 0.2 | 5 | 0.4 | 6 | 0.1667 | 8 | 0.125 | 24 | 0.2083 |
| 19_5878617 | 6 | 0 | 10 | 0 | 8 | 0.125 | 7 | 0.1429 | 31 | 0.0645 | 2 | 0.5 | 3 | 0 | 4 | 0.25 | 8 | 0 | 17 | 0.1176 |
| 19_5878639 | 8 | 0.5 | 9 | 0.3333 | 14 | 0.3571 | 7 | 0.1429 | 38 | 0.3421 | 3 | 0.3333 | 3 | 0.6667 | 6 | 0.3333 | 9 | 0.2222 | 21 | 0.3333 |
| 19_5878732 | 5 | 0 | 9 | 0.1111 | 8 | 0.125 | 6 | 0 | 28 | 0.0714 | 2 | 0 | 3 | 0.3333 | 3 | 0 | 7 | 0 | 15 | 0.0667 |
| 19_5878733 | 5 | 0.4 | 7 | 0.4286 | 8 | 0.375 | 5 | 0.4 | 25 | 0.4 | 1 | 1 | 3 | 0 | 3 | 1 | 5 | 0.2 | 12 | 0.4167 |
| 19_5878751 | 4 | 1 | 6 | 1 | 3 | 1 | 5 | 1 | 18 | 1 | 1 | 1 | 3 | 1 | 2 | 1 | 6 | 0.8333 | 12 | 0.9167 |
| 19_5878765 | 2 | 1 | 7 | 0.7143 | 3 | 1 | 4 | 0.5 | 16 | 0.75 | 1 | 1 | 1 | 1 | 2 | 1 | 3 | 0.6667 | 7 | 0.8571 |
| 19_5878767 | 4 | 0 | 7 | 0.1429 | 4 | 0 | 6 | 0.1667 | 21 | 0.0952 | 2 | 0.5 | 3 | 0.3333 | 1 | 0 | 6 | 0.3333 | 12 | 0.3333 |

| | | | | | | | | | | | | | | | | | | | | |
|---|---|---|---|---|---|---|---|---|---|---|---|---|---|---|---|---|---|---|---|---|
| 19_587877 | 4 | 1 | 6 | 1 | 3 | 1 | 6 | 0.8333 | 19 | 0.9474 | 2 | 1 | 2 | 1 | 1 | 1 | 7 | 0.7143 | 12 | 0.8333 |
| 19_587877 | 3 | 1 | 6 | 0.8333 | 3 | 1 | 6 | 1 | 18 | 0.9444 | 2 | 1 | 2 | 1 | NA | NA | 6 | 1 | 10 | 1 |
| 19_587877 | 3 | 0.6667 | 3 | 1 | 1 | 1 | 3 | 0.6667 | 10 | 0.8 | 1 | 0 | 2 | 1 | NA | NA | 4 | 1 | 7 | 0.8571 |
| 19_587878 | 4 | 1 | 4 | 1 | 3 | 1 | 4 | 1 | 15 | 1 | 1 | 1 | 1 | 1 | NA | NA | 3 | 1 | 5 | 1 |
| 19_587879 | 4 | 0.25 | 6 | 0 | 3 | 0 | 3 | 0 | 16 | 0.0625 | 1 | 1 | 2 | 0 | 1 | 0 | 7 | 0.2857 | 11 | 0.2727 |
| 19_587881 | 3 | 0 | 6 | 0.1667 | 1 | 0 | 3 | 0 | 13 | 0.0769 | 1 | 0 | 1 | 0 | 1 | 0 | 4 | 0.75 | 7 | 0.4286 |
| 19_587882 | 3 | 0.6667 | 5 | 0.6 | 1 | 1 | 4 | 0 | 13 | 0.4615 | 1 | 1 | 2 | 0.5 | 1 | 1 | 6 | 0.6667 | 10 | 0.7 |
| 19_588086 | 1 | 1 | NA | NA | NA | NA | NA | NA | 1 | 1 | NA | NA | NA | NA | 1 | 1 | NA | NA | 1 | 1 |
| 19_588086 | 1 | 1 | NA | NA | NA | NA | NA | NA | 1 | 1 | NA | NA | NA | NA | 1 | 1 | NA | NA | 1 | 1 |
| 19_588087 | 1 | 1 | NA | NA | NA | NA | NA | NA | 1 | 1 | NA | NA | NA | NA | 1 | 1 | NA | NA | 1 | 1 |
| 19_588088 | 1 | 1 | NA | NA | NA | NA | NA | NA | 1 | 1 | NA | NA | NA | NA | 1 | 1 | NA | NA | 1 | 1 |
| 19_589591 | NA | NA | NA | NA | NA | NA | 1 | 1 | 1 | 1 | 1 | 1 | 1 | 1 | NA | NA | NA | NA | 2 | 1 |
| 19_589618 | NA | NA | 4 | 0.5 | 2 | 0.5 | NA | NA | 6 | 0.5 | 1 | 0 | NA | NA | NA | NA | 2 | 1 | 3 | 0.6667 |
| 19_589619 | NA | NA | 4 | 0.25 | 2 | 0 | NA | NA | 6 | 0.1667 | 1 | 1 | NA | NA | NA | NA | 1 | 1 | 2 | 1 |
| 19_589620 | 8 | 0 | 9 | 0.2222 | 5 | 0 | 2 | 0 | 24 | 0.0833 | 4 | 0 | NA | NA | 2 | 0 | 4 | 0.25 | 10 | 0.1 |
| 19_589627 | 4 | 1 | 7 | 0.8571 | 4 | 1 | 1 | 0 | 16 | 0.875 | 3 | 0.6667 | NA | NA | 2 | 0.5 | 3 | 0.6667 | 8 | 0.625 |
| 19_589630 | 4 | 0 | 8 | 0.25 | 5 | 0.2 | NA | NA | 17 | 0.1765 | 2 | 0 | NA | NA | 2 | 0 | 1 | 1 | 5 | 0.2 |
| 19_589631 | 4 | 0 | 7 | 0.4286 | 5 | 0.6 | 1 | 1 | 17 | 0.4118 | 2 | 0 | NA | NA | 1 | 0 | 3 | 0.3333 | 6 | 0.1667 |
| 19_589633 | 4 | 1 | 6 | 1 | 4 | 0.75 | 1 | 1 | 15 | 0.9333 | 1 | 1 | NA | NA | 2 | 1 | 3 | 1 | 6 | 1 |
| 19_589633 | 4 | 0.5 | 7 | 0.8571 | 5 | 1 | 1 | 1 | 17 | 0.8235 | 1 | 1 | NA | NA | 1 | 1 | 3 | 1 | 5 | 1 |
| 19_589634 | 4 | 0.75 | 6 | 0.8333 | 5 | 0.8 | 1 | 1 | 16 | 0.8125 | NA | NA | NA | NA | 1 | 1 | 2 | 1 | 3 | 1 |
| 19_591546 | 2 | 1 | NA | NA | NA | NA | NA | NA | 2 | 1 | NA | NA | NA | NA | 1 | 1 | NA | NA | 1 | 1 |
| 19_591547 | 2 | 1 | NA | NA | NA | NA | NA | NA | 2 | 1 | NA | NA | NA | NA | 2 | 0.5 | NA | NA | 2 | 0.5 |
| 19_591548 | 1 | 1 | NA | NA | NA | NA | NA | NA | 1 | 1 | 1 | 0 | NA | NA | 2 | 0.5 | NA | NA | 3 | 0.3333 |
| 19_591551 | 2 | 0.5 | NA | NA | NA | NA | NA | NA | 2 | 0.5 | 1 | 0 | NA | NA | 2 | 0.5 | NA | NA | 3 | 0.3333 |
| 19_592831 | 1 | 0 | 2 | 0.5 | 1 | 0 | 1 | 0 | 5 | 0.2 | 1 | 0 | NA | NA | 6 | 0 | 4 | 0.25 | 11 | 0.0909 |
| 19_592861 | 3 | 0.6667 | 8 | 0.125 | 3 | 0.3333 | 10 | 0 | 24 | 0.1667 | 4 | 0.25 | 9 | 0.2222 | 10 | 0.1 | 6 | 0 | 29 | 0.1379 |
| 19_592864 | 4 | 0.25 | 7 | 0.1429 | 6 | 0 | 9 | 0 | 26 | 0.0769 | 4 | 0 | 9 | 0.1111 | 10 | 0.2 | 4 | 0.25 | 27 | 0.1481 |
| 19_592867 | 4 | 0.5 | 6 | 0 | 6 | 0 | 6 | 0 | 22 | 0.0909 | 4 | 0 | 10 | 0.2 | 8 | 0.125 | 4 | 0 | 26 | 0.1154 |
| 19_592869 | 5 | 0.8 | 6 | 0.8333 | 6 | 1 | 8 | 0.875 | 25 | 0.88 | 2 | 1 | 7 | 0.8571 | 6 | 0.8333 | 3 | 0.6667 | 18 | 0.8333 |
| 19_592869 | 4 | 0 | 11 | 0.0909 | 6 | 0.1667 | 7 | 0 | 28 | 0.0714 | 6 | 0.1667 | 7 | 0.2857 | 8 | 0.5 | 7 | 0.2857 | 28 | 0.3214 |
| 19_592870 | 5 | 0.2 | 8 | 0.375 | 7 | 0.2857 | 9 | 0.5556 | 29 | 0.3793 | 6 | 0.6667 | 8 | 0.5 | 8 | 0.375 | 4 | 0.25 | 26 | 0.4615 |
| 19_592870 | 5 | 0.6 | 12 | 0.75 | 7 | 0.7143 | 8 | 0.625 | 32 | 0.6875 | 6 | 0.6667 | 9 | 1 | 8 | 0.75 | 7 | 0.8571 | 30 | 0.8333 |
| 19_592870 | 5 | 1 | 11 | 1 | 7 | 1 | 8 | 0.875 | 31 | 0.9677 | 6 | 1 | 8 | 0.875 | 9 | 1 | 5 | 1 | 28 | 0.9643 |
| 19_592870 | 5 | 0.4 | 11 | 0.1818 | 8 | 0.25 | 8 | 0.375 | 32 | 0.2812 | 6 | 0.3333 | 9 | 0.2222 | 8 | 0.375 | 7 | 0.7143 | 30 | 0.4 |
| 19_592870 | 5 | 1 | 11 | 1 | 8 | 1 | 8 | 0.875 | 32 | 0.9688 | 6 | 1 | 10 | 0.9 | 8 | 0.875 | 8 | 1 | 32 | 0.9375 |
| 19_592879 | 2 | 0.5 | 5 | 0.4 | 6 | 0.6667 | 6 | 0.6667 | 19 | 0.5789 | 4 | 0.5 | 8 | 0.5 | 5 | 0.8 | 4 | 0.75 | 21 | 0.619 |
| 19_592879 | 4 | 0.5 | 7 | 0.7143 | 4 | 0.5 | 7 | 0.5714 | 22 | 0.5909 | 6 | 0.3333 | 9 | 0.6667 | 5 | 0.8 | 6 | 0.1667 | 26 | 0.5 |
| 19_592880 | 4 | 1 | 8 | 1 | 5 | 1 | 5 | 1 | 22 | 1 | 6 | 1 | 4 | 1 | 6 | 1 | 4 | 1 | 20 | 1 |
| 19_592880 | 2 | 0 | 5 | 0.2 | 2 | 0 | 5 | 0 | 14 | 0.0714 | 3 | 0 | 8 | 0.625 | 5 | 0.2 | 7 | 0.2857 | 23 | 0.3478 |
| 19_592881 | 3 | 0 | 7 | 0.1429 | 4 | 0.25 | 6 | 0.1667 | 20 | 0.15 | 5 | 0.2 | 9 | 0.3333 | 5 | 0.2 | 6 | 0.1667 | 25 | 0.24 |
| 19_592881 | 4 | 0.5 | 8 | 0.875 | 4 | 0.75 | 7 | 0.7143 | 23 | 0.7391 | 5 | 0.6 | 8 | 0.75 | 7 | 0.8571 | 7 | 0.8571 | 27 | 0.7778 |
| 19_592883 | 4 | 0.25 | 7 | 0.2857 | 5 | 0 | 8 | 0 | 24 | 0.125 | 5 | 0 | 8 | 0.375 | 6 | 0 | 6 | 0.3333 | 25 | 0.2 |
| 19_592884 | 4 | 0 | 5 | 0 | 3 | 0.6667 | 7 | 0.1429 | 19 | 0.1579 | 5 | 0.2 | 8 | 0.125 | 6 | 0.1667 | 7 | 0.1429 | 26 | 0.1538 |
| 19_592884 | 4 | 0 | 8 | 0.125 | 2 | 0 | 6 | 0.1667 | 20 | 0.1 | 5 | 0.2 | 5 | 0 | 6 | 0 | 7 | 0.2857 | 23 | 0.1304 |
| 19_592885 | 4 | 0.25 | 8 | 0.125 | 4 | 0 | 7 | 0.2857 | 23 | 0.1739 | 5 | 0.2 | 8 | 0.125 | 6 | 0.3333 | 7 | 0.2857 | 26 | 0.2308 |
| 19_592885 | 2 | 0 | 5 | 0 | 3 | 0 | 6 | 0.5 | 16 | 0.1875 | 4 | 0.25 | 4 | 0 | 6 | 0.1667 | 5 | 0 | 19 | 0.1053 |
| 19_592887 | 4 | 0.25 | 5 | 0 | 2 | 0.5 | 6 | 0 | 17 | 0.1176 | 2 | 0 | 7 | 0 | 4 | 0.5 | 1 | 0 | 14 | 0.1429 |
| 19_592887 | 4 | 0.5 | 7 | 0.7143 | 3 | 1 | 5 | 0.6 | 19 | 0.6842 | 5 | 0.8 | 7 | 0.8571 | 5 | 0.6 | 4 | 0.75 | 21 | 0.7619 |
| 19_592887 | 4 | 0 | 7 | 0.1429 | 3 | 0.6667 | 6 | 0.1667 | 20 | 0.2 | 5 | 0 | 6 | 0.1667 | 5 | 0.2 | 6 | 0.3333 | 22 | 0.1818 |
| 19_592888 | 4 | 0.75 | 7 | 0.7143 | 4 | 0.75 | 7 | 0.8571 | 22 | 0.7727 | 5 | 0.8 | 8 | 0.625 | 4 | 0.75 | 7 | 0.7143 | 24 | 0.7083 |
| 19_592889 | 3 | 1 | 6 | 1 | 1 | 1 | 5 | 1 | 15 | 1 | 4 | 0.75 | 7 | 0.8571 | 4 | 1 | 6 | 0.6667 | 21 | 0.8095 |
| 19_592889 | 3 | 1 | 7 | 0.5714 | 3 | 1 | 5 | 0.6 | 18 | 0.7222 | 5 | 0.6 | 8 | 0.875 | 5 | 1 | 5 | 1 | 23 | 0.8696 |
| 19_592890 | 4 | 0.5 | 7 | 0.2857 | 4 | 0.25 | 8 | 0.75 | 23 | 0.4783 | 5 | 0.4 | 7 | 0.2857 | 6 | 0.3333 | 6 | 0.5 | 24 | 0.375 |
| 19_592891 | 4 | 0.75 | 5 | 0.8 | 4 | 0.75 | 6 | 0.6667 | 19 | 0.7368 | 5 | 0.8 | 6 | 0.6667 | 6 | 1 | 5 | 1 | 22 | 0.8636 |
| 19_592891 | 2 | 1 | 7 | 1 | 4 | 1 | 5 | 0.6 | 18 | 0.8889 | 5 | 0.4 | 7 | 0.7143 | 6 | 0.8333 | 7 | 0.8571 | 25 | 0.72 |
| 19_592892 | 3 | 0.3333 | 6 | 0 | 4 | 0 | 7 | 0.2857 | 20 | 0.15 | 5 | 0 | 6 | 0.6667 | 6 | 0 | 7 | 0.1429 | 24 | 0.2083 |
| 19_592892 | 3 | 0 | 5 | 0.2 | 4 | 0.25 | 3 | 0.3333 | 15 | 0.2 | 4 | 0.25 | 4 | 0 | 5 | 0.2 | 4 | 0 | 17 | 0.1176 |
| 19_592894 | 2 | 0 | 8 | 0.75 | 4 | 1 | 5 | 1 | 19 | 0.7895 | 5 | 0.6 | 5 | 0.4 | 6 | 0.6667 | 7 | 0.5714 | 23 | 0.5652 |
| 19_592896 | 2 | 0 | 6 | 0.1667 | 3 | 0 | 5 | 0.2 | 16 | 0.125 | 5 | 0 | 7 | 0.1429 | 6 | 0.3333 | 5 | 0 | 23 | 0.1304 |
| 19_592896 | 2 | 0.5 | 6 | 0.8333 | 3 | 0.6667 | 5 | 0.2 | 16 | 0.5625 | 5 | 1 | 5 | 0.6 | 6 | 0.6667 | 7 | 0.7143 | 23 | 0.7391 |
| 19_592897 | 2 | 0 | 7 | 0 | 3 | 0 | 3 | 0.3333 | 15 | 0.0667 | 3 | 0 | 7 | 0.1429 | 3 | 0 | 5 | 0 | 18 | 0.0556 |
| 19_592897 | NA | NA | 2 | 0.5 | 2 | 0 | 3 | 0 | 7 | 0.1429 | 3 | 0.3333 | 5 | 0 | 6 | 0.3333 | 3 | 0.3333 | 17 | 0.2353 |
| 19_592898 | 3 | 0.3333 | 4 | 0.5 | 3 | 0.3333 | 3 | 0.3333 | 13 | 0.3846 | 5 | 0 | 7 | 0.1429 | 4 | 0 | 7 | 0.2857 | 23 | 0.1304 |
| 19_592899 | 2 | 0 | 5 | 0 | 2 | 0 | 5 | 0.4 | 14 | 0.1429 | 4 | 0.25 | 8 | 0.25 | 4 | 0.75 | 7 | 0.4286 | 23 | 0.3913 |

| | | | | | | | | | | | | | | | | | | | | |
|---|---|---|---|---|---|---|---|---|---|---|---|---|---|---|---|---|---|---|---|---|
| 19_5928993 | 3 | 0.6667 | 5 | 0.4 | 2 | 0 | 6 | 0.8333 | 16 | 0.5625 | 4 | 0.25 | 8 | 0 | 6 | 0.6667 | 6 | 0.6667 | 24 | 0.375 |
| 19_5928995 | 3 | 1 | 6 | 0.1667 | 2 | 0.5 | 4 | 1 | 15 | 0.6 | 4 | 1 | 8 | 0.5 | 5 | 0.4 | 6 | 0.6667 | 23 | 0.6087 |
| 19_5929008 | 3 | 0.6667 | 4 | 0 | 2 | 0 | 4 | 0.25 | 13 | 0.2308 | 5 | 0 | 3 | 0 | 4 | 0.5 | 6 | 0 | 18 | 0.1111 |
| 19_5933326 | 4 | 0.25 | 2 | 0 | 2 | 0 | 2 | 0 | 10 | 0.1 | 3 | 0 | 5 | 0 | 5 | 0.4 | 6 | 0.1667 | 19 | 0.1579 |
| 19_5933339 | 4 | 0.25 | 3 | 0.3333 | 2 | 0.5 | 1 | 1 | 10 | 0.4 | 3 | 0.3333 | 4 | 0.5 | 5 | 0.2 | 5 | 0.4 | 17 | 0.3529 |
| 19_5941695 | 1 | 1 | NA | NA | NA | NA | NA | NA | 1 | 1 | 1 | 1 | NA | NA | NA | NA | 1 | 1 | 2 | 1 |
| 19_5941736 | 1 | 1 | NA | NA | NA | NA | NA | NA | 1 | 1 | 1 | 1 | NA | NA | NA | NA | 1 | 0 | 2 | 0.5 |
| 19_5942139 | 1 | 1 | NA | NA | NA | NA | NA | NA | 1 | 1 | 1 | 1 | 1 | 1 | NA | NA | 2 | 0.5 | 4 | 0.75 |
| 19_5944770 | 1 | 1 | 2 | 1 | NA | NA | 1 | 0 | 4 | 0.75 | 1 | 0 | 1 | 0 | 3 | 0.6667 | NA | NA | 5 | 0.4 |
| 19_5944785 | 1 | 0 | 2 | 1 | NA | NA | 3 | 0 | 6 | 0.3333 | NA | NA | 1 | 1 | 3 | 1 | NA | NA | 4 | 1 |
| 19_5945240 | 1 | 1 | 2 | 0.5 | NA | NA | 4 | 0.5 | 7 | 0.5714 | 1 | 0 | 1 | 0 | 4 | 0.25 | NA | NA | 6 | 0.1667 |
| 19_5945250 | NA | NA | 2 | 1 | NA | NA | 3 | 1 | 5 | 1 | 2 | 1 | 1 | 1 | 4 | 0.75 | NA | NA | 7 | 0.8571 |
| 19_5945252 | 1 | 1 | 2 | 0.5 | NA | NA | 4 | 0.5 | 7 | 0.5714 | 2 | 0 | 1 | 0 | 4 | 0.25 | NA | NA | 7 | 0.1429 |
| 19_5945257 | 1 | 1 | 2 | 0.5 | NA | NA | 4 | 0.75 | 7 | 0.7143 | 1 | 1 | 1 | 1 | 3 | 0.6667 | NA | NA | 5 | 0.8 |
| 19_5945263 | 1 | 1 | 3 | 0.6667 | NA | NA | 4 | 0.5 | 8 | 0.625 | 1 | 1 | 1 | 1 | 3 | 0.3333 | NA | NA | 5 | 0.6 |
| 19_5945263 | 1 | 0 | 3 | 0 | NA | NA | 2 | 0.5 | 6 | 0.1667 | 2 | 0.5 | 1 | 1 | 3 | 0.3333 | NA | NA | 6 | 0.5 |
| 19_5945265 | 1 | 0 | 3 | 0.3333 | NA | NA | 3 | 0.3333 | 7 | 0.2857 | 1 | 0 | 1 | 1 | 5 | 0 | NA | NA | 7 | 0.1429 |
| 19_5945269 | 1 | 1 | 2 | 0 | NA | NA | 4 | 0.25 | 7 | 0.2857 | 2 | 0 | NA | NA | 5 | 0.2 | NA | NA | 7 | 0.1429 |
| 19_5945276 | 1 | 0 | 3 | 0.6667 | NA | NA | 4 | 0.25 | 8 | 0.375 | NA | NA | 1 | 0 | 5 | 0.2 | NA | NA | 6 | 0.1667 |
| 19_5945296 | 1 | 1 | 3 | 1 | NA | NA | 3 | 1 | 7 | 1 | 2 | 1 | 1 | 1 | 5 | 1 | NA | NA | 8 | 1 |
| 19_5945334 | NA | NA | 5 | 0.2 | NA | NA | 4 | 0 | 9 | 0.1111 | 1 | 0 | 1 | 0 | 5 | 0.6 | NA | NA | 7 | 0.4286 |
| 19_5945350 | 1 | 1 | 5 | 0 | 1 | 1 | 4 | 0 | 11 | 0.1818 | 1 | 0 | 1 | 1 | 4 | 0.5 | NA | NA | 6 | 0.5 |
| 19_5945354 | 1 | 1 | 5 | 0.8 | NA | NA | 3 | 0.3333 | 9 | 0.6667 | 1 | 0 | 1 | 1 | 4 | 0.5 | NA | NA | 6 | 0.5 |
| 19_5945362 | 1 | 0 | 3 | 0.3333 | 1 | 0 | 4 | 0 | 9 | 0.1111 | 1 | 0 | 1 | 1 | 4 | 0 | 1 | 0 | 7 | 0.1429 |
| 19_5989168 | 4 | 0.25 | 6 | 0 | 2 | 0 | 5 | 0.2 | 17 | 0.1176 | 2 | 0 | 7 | 0.1429 | 5 | 0 | 5 | 0 | 19 | 0.0526 |
| 19_5989192 | 5 | 0 | 4 | 0.25 | 4 | 0 | 6 | 0.3333 | 19 | 0.1579 | 3 | 0 | 6 | 0.1667 | 7 | 0.1429 | 4 | 0.5 | 20 | 0.2 |
| 19_5989255 | 4 | 0.25 | 5 | 0.4 | 4 | 0 | 5 | 0.4 | 18 | 0.2778 | 2 | 0 | 6 | 0 | 5 | 0.2 | 5 | 0.4 | 18 | 0.1667 |
| 19_5989268 | 4 | 0 | 5 | 0.2 | 4 | 0.25 | 6 | 0 | 19 | 0.1053 | 3 | 0 | 6 | 0 | 5 | 0.2 | 5 | 0.4 | 19 | 0.1579 |
| 19_5989300 | 4 | 0 | 4 | 0 | 4 | 0.25 | 6 | 0.1667 | 18 | 0.1111 | 3 | 0 | 4 | 0 | 5 | 0 | 5 | 0.2 | 17 | 0.0588 |
| 19_5989423 | 2 | 0 | 2 | 0.5 | 4 | 0 | 4 | 0 | 12 | 0.0833 | 2 | 0.5 | 3 | 0.6667 | 5 | 0.4 | 4 | 0.25 | 14 | 0.4286 |
| 19_5989464 | 1 | 1 | 1 | 1 | 2 | 0.5 | 4 | 0.25 | 8 | 0.5 | NA | NA | NA | NA | 4 | 0.5 | 4 | 0.25 | 8 | 0.375 |
| 19_5989471 | 1 | 0 | 2 | 0.5 | 3 | 0.6667 | 4 | 0 | 10 | 0.3 | 1 | 0 | 3 | 0 | 2 | 0 | 4 | 0.25 | 10 | 0.1 |
| 19_5989479 | 2 | 0 | 1 | 1 | 4 | 0.5 | 4 | 0 | 11 | 0.2727 | 1 | 0 | 2 | 0 | 4 | 0.25 | 5 | 0.4 | 12 | 0.25 |
| 19_5989491 | 2 | 0 | 2 | 0 | 4 | 0 | 4 | 0.25 | 12 | 0.0833 | 1 | 0 | 2 | 0 | 3 | 0 | 4 | 0.25 | 10 | 0.1 |
| 19_5989506 | 2 | 0.5 | 1 | 1 | 3 | 0.3333 | 3 | 0.3333 | 9 | 0.4444 | 1 | 1 | 2 | 0.5 | 3 | 0.6667 | 5 | 0.8 | 11 | 0.7273 |
| 19_5989512 | 2 | 0.5 | 1 | 1 | 2 | 0.5 | 4 | 0 | 9 | 0.3333 | NA | NA | 2 | 0 | 3 | 0.6667 | 4 | 0.5 | 9 | 0.4444 |
| 19_5989514 | 1 | 1 | 1 | 1 | 2 | 0 | 4 | 0 | 8 | 0.25 | 1 | 0 | 2 | 0.5 | 4 | 0.5 | 5 | 0.4 | 12 | 0.4167 |
| 19_5989515 | 2 | 0.5 | 1 | 0 | 4 | 0 | 4 | 0.25 | 11 | 0.1818 | NA | NA | 2 | 0 | 4 | 0.25 | 4 | 0.25 | 10 | 0.2 |
| 19_5989522 | 2 | 0 | 2 | 0.5 | 2 | 0 | 3 | 0.3333 | 9 | 0.2222 | NA | NA | 2 | 0.5 | 4 | 0.5 | 2 | 0.5 | 8 | 0.5 |
| 19_5989532 | 1 | 0 | 2 | 0 | 1 | 0 | 3 | 0.3333 | 7 | 0.1429 | NA | NA | 2 | 0 | 1 | 0 | 2 | 0.5 | 5 | 0.2 |
| 19_5989533 | 2 | 1 | 2 | 1 | 2 | 1 | 2 | 1 | 8 | 1 | 1 | 1 | 2 | 1 | 4 | 1 | 4 | 1 | 11 | 1 |
| 19_5989545 | NA | NA | 3 | 0.6667 | 2 | 0.5 | 4 | 0.5 | 9 | 0.5556 | 1 | 1 | 3 | 0.6667 | 4 | 0.5 | 4 | 0.25 | 12 | 0.5 |
| 19_5989547 | NA | NA | 2 | 0 | 2 | 0.5 | 4 | 0.5 | 8 | 0.375 | 1 | 0 | 3 | 0.6667 | 4 | 0 | 4 | 0.25 | 12 | 0.25 |
| 19_5989549 | NA | NA | 1 | 0 | 2 | 0.5 | 4 | 0 | 7 | 0.1429 | 1 | 1 | 2 | 0 | 4 | 0 | 3 | 0 | 10 | 0.1 |
| 19_5992411 | NA | NA | 2 | 0.5 | 2 | 0.5 | 1 | 1 | 5 | 0.6 | 1 | 1 | 2 | 0.5 | 4 | 1 | 4 | 0.75 | 11 | 0.8182 |
| 19_5992414 | NA | NA | NA | NA | 1 | 1 | 2 | 0 | 3 | 0.3333 | 1 | 1 | 3 | 0.3333 | 4 | 0.5 | 4 | 0.75 | 12 | 0.5833 |
| 19_5992417 | NA | NA | 1 | 0 | 2 | 0.5 | 2 | 0 | 5 | 0.2 | NA | NA | 2 | 0 | 3 | 0.6667 | 3 | 0.3333 | 8 | 0.375 |
| 19_5992421 | NA | NA | 2 | 1 | 2 | 0 | 2 | 1 | 6 | 0.6667 | NA | NA | 1 | 1 | 4 | 0.75 | 4 | 0.5 | 9 | 0.6667 |
| 19_5992430 | NA | NA | 2 | 1 | 2 | 1 | 2 | 0.5 | 6 | 0.8333 | NA | NA | 3 | 0.3333 | 3 | 1 | 3 | 1 | 9 | 0.7778 |
| 19_5992438 | NA | NA | NA | NA | 1 | 1 | 1 | 1 | 2 | 1 | NA | NA | 1 | 0 | 2 | 0.5 | 4 | 1 | 7 | 0.7143 |
| 19_5992574 | NA | NA | 2 | 0 | 2 | 0 | 2 | 0.5 | 6 | 0.1667 | NA | NA | 1 | 0 | 4 | 0 | 4 | 0.5 | 9 | 0.2222 |
| 19_5992575 | NA | NA | 3 | 0 | 2 | 0.5 | 2 | 0.5 | 7 | 0.2857 | NA | NA | 1 | 0 | 4 | 0.25 | 4 | 0.25 | 9 | 0.2222 |
| 19_5993132 | NA | NA | 2 | 0.5 | 2 | 0.5 | 2 | 0 | 6 | 0.3333 | NA | NA | 1 | 0 | 3 | 0.3333 | 3 | 0 | 7 | 0.1429 |
| 19_5993150 | NA | NA | 2 | 0.5 | 2 | 0.5 | 2 | 0.5 | 6 | 0.5 | NA | NA | NA | NA | 2 | 0.5 | 1 | 1 | 3 | 0.6667 |
| 19_5993161 | NA | NA | 2 | 0 | 1 | 0 | 2 | 1 | 5 | 0.4 | NA | NA | NA | NA | 2 | 0.5 | 3 | 1 | 5 | 0.8 |
| 19_6018776 | 3 | 0 | 7 | 0 | 3 | 0.3333 | 3 | 0 | 16 | 0.0625 | 3 | 0 | 1 | 0 | 5 | 0.2 | 2 | 0 | 11 | 0.0909 |
| 19_6051364 | 5 | 0 | 4 | 0 | 5 | 0 | 4 | 0.25 | 18 | 0.0556 | 4 | 0 | 4 | 0.25 | 5 | 0 | 3 | 0.3333 | 16 | 0.125 |
| 19_6051540 | 3 | 0 | 4 | 0.5 | 4 | 0 | 4 | 0 | 15 | 0.1333 | 2 | 0 | 4 | 0 | 1 | 0 | 1 | 1 | 8 | 0.125 |
| 19_6051595 | 4 | 0 | 7 | 0.4286 | 5 | 0 | 3 | 0.3333 | 19 | 0.2105 | 4 | 0.5 | 3 | 0.6667 | 4 | 0 | 5 | 0.2 | 16 | 0.3125 |
| 19_6051613 | 3 | 0 | 6 | 0 | 6 | 0.3333 | 4 | 0.25 | 19 | 0.1579 | 4 | 0 | 3 | 0.3333 | 4 | 0 | 4 | 0 | 15 | 0.0667 |
| 19_6051662 | 3 | 0.6667 | 8 | 0.875 | 6 | 0.8333 | 4 | 0.75 | 21 | 0.8095 | 2 | 1 | 4 | 1 | 3 | 0.3333 | 7 | 0.7143 | 16 | 0.75 |
| 19_6051678 | 3 | 0 | 6 | 0.1667 | 6 | 0.1667 | 4 | 0.25 | 19 | 0.1579 | 2 | 0 | 4 | 0 | 4 | 0 | 7 | 0.1429 | 17 | 0.0588 |
| 19_6051711 | 1 | 1 | 3 | 1 | 5 | 1 | 3 | 1 | 12 | 1 | 1 | 1 | 3 | 0.6667 | 3 | 0.6667 | 7 | 0.8571 | 14 | 0.7857 |
| 19_6051727 | 3 | 1 | 2 | 1 | 5 | 0.8 | 5 | 0.6 | 15 | 0.8 | 2 | 1 | 3 | 0.6667 | 3 | 0.6667 | 6 | 0.8333 | 14 | 0.7857 |
| 19_6051730 | 3 | 0.6667 | 4 | 0.75 | 5 | 0.8 | 4 | 1 | 16 | 0.8125 | 2 | 1 | 2 | 1 | 3 | 0.6667 | 4 | 0.5 | 11 | 0.7273 |

| | | | | | | | | | | | | | | | | | | | | |
|---|---|---|---|---|---|---|---|---|---|---|---|---|---|---|---|---|---|---|---|---|
| 19_605173 | 2 | 0 | 3 | 0 | 2 | 0 | 4 | 0.25 | 11 | 0.0909 | 1 | 0 | 2 | 0.5 | 3 | 0 | 5 | 0.4 | 11 | 0.2727 |
| 19_605173 | 3 | 0.6667 | 3 | 0.6667 | 5 | 1 | 3 | 0.6667 | 14 | 0.7857 | 1 | 1 | 2 | 0.5 | 3 | 0.6667 | 5 | 0.6 | 11 | 0.6364 |
| 19_605173 | 3 | 1 | 4 | 1 | 5 | 1 | 4 | 1 | 16 | 1 | 2 | 1 | 3 | 1 | 3 | 1 | 6 | 1 | 14 | 1 |
| 19_605174 | 2 | 0.5 | 3 | 0.3333 | 5 | 0.6 | 4 | 0.25 | 14 | 0.4286 | 2 | 0.5 | 2 | 0.5 | 2 | 0 | 6 | 0.3333 | 12 | 0.3333 |
| 19_605174 | 1 | 1 | 4 | 0.25 | 5 | 0.2 | 3 | 0.3333 | 13 | 0.3077 | 2 | 0 | 3 | 0 | 2 | 0.5 | 6 | 0 | 13 | 0.0769 |
| 19_605689 | 3 | 0 | 2 | 0 | 4 | 0.25 | 2 | 0.5 | 11 | 0.1818 | 2 | 0 | 1 | 0 | 1 | 1 | 4 | 0 | 8 | 0.125 |
| 19_607450 | 3 | 0.3333 | 2 | 1 | 1 | 1 | 1 | 1 | 7 | 0.7143 | 2 | 0 | 4 | 0 | 2 | 0 | 1 | 1 | 9 | 0.1111 |
| 19_607454 | 4 | 1 | 4 | 1 | 1 | 1 | 2 | 1 | 11 | 1 | 3 | 1 | 3 | 1 | 2 | 1 | NA | NA | 8 | 1 |
| 19_607458 | 5 | 1 | 4 | 0.5 | NA | NA | 3 | 1 | 12 | 0.8333 | 3 | 1 | 3 | 0.6667 | 3 | 1 | 1 | 1 | 10 | 0.9 |
| 19_607459 | 5 | 0.6 | 4 | 0.25 | 1 | 0 | 4 | 0.25 | 14 | 0.3571 | 3 | 0.3333 | 4 | 0.25 | 3 | 0 | 1 | 0 | 11 | 0.1818 |
| 19_607481 | 2 | 0 | 5 | 0.6 | 1 | 0 | 4 | 0.5 | 12 | 0.4167 | NA | NA | 1 | 1 | 3 | 1 | NA | NA | 4 | 1 |
| 19_607503 | 9 | 0.2222 | 10 | 0.5 | 14 | 0.5714 | 9 | 0.1111 | 42 | 0.381 | 5 | 0 | 8 | 0.375 | 15 | 0.2 | 11 | 0.3636 | 39 | 0.2564 |
| 19_607503 | 9 | 0.8889 | 13 | 0.6154 | 14 | 0.9286 | 8 | 1 | 44 | 0.8409 | 3 | 0.6667 | 8 | 0.75 | 14 | 0.6429 | 15 | 0.8667 | 40 | 0.75 |
| 19_607504 | 9 | 0.2222 | 11 | 0.0909 | 13 | 0.1538 | 9 | 0.2222 | 42 | 0.1667 | 4 | 0 | 9 | 0 | 14 | 0.0714 | 16 | 0.25 | 43 | 0.1163 |
| 19_607505 | 9 | 0.6667 | 10 | 0.3 | 13 | 0.5385 | 10 | 0.2 | 42 | 0.4286 | 4 | 0.5 | 9 | 0.4444 | 11 | 0.6364 | 14 | 0.2143 | 38 | 0.4211 |
| 19_607505 | 8 | 0.25 | 7 | 0.5714 | 14 | 0.1429 | 9 | 0.2222 | 38 | 0.2632 | 3 | 0 | 7 | 0.1429 | 13 | 0.3846 | 16 | 0.375 | 39 | 0.3077 |
| 19_607507 | 6 | 0.1667 | 9 | 0.4444 | 13 | 0.4615 | 9 | 0.4444 | 37 | 0.4054 | 3 | 0 | 8 | 0.625 | 11 | 0.5455 | 13 | 0.2308 | 35 | 0.4 |
| 19_607549 | 6 | 0.1667 | 6 | 0.1667 | 8 | 0.375 | 7 | 0.2857 | 27 | 0.2593 | 1 | 0 | 4 | 0.25 | 12 | 0.25 | 10 | 0.2 | 27 | 0.2222 |
| 19_607550 | 6 | 0.1667 | 6 | 0.1667 | 8 | 0.25 | 6 | 0.5 | 26 | 0.2692 | 1 | 0 | 4 | 0.75 | 12 | 0.0833 | 10 | 0.3 | 27 | 0.2593 |
| 19_607550 | 4 | 0.25 | 6 | 0.3333 | 7 | 0.2857 | 7 | 0.1429 | 24 | 0.25 | 1 | 0 | 3 | 0.6667 | 11 | 0.3636 | 7 | 0.5714 | 22 | 0.4545 |
| 19_607551 | 5 | 0.4 | 6 | 1 | 8 | 1 | 8 | 0.75 | 27 | 0.8148 | 1 | 0 | 4 | 0.5 | 10 | 0.4 | 8 | 0.75 | 23 | 0.5217 |
| 19_607621 | 5 | 0.4 | 4 | 0 | 3 | 0 | 3 | 0 | 15 | 0.1333 | NA | NA | 2 | 0 | 5 | 0.2 | 5 | 0 | 12 | 0.0833 |
| 19_607665 | 4 | 0.25 | 1 | 0 | NA | NA | 2 | 0 | 7 | 0.1429 | NA | NA | 2 | 0.5 | 4 | 0.25 | 3 | 0.3333 | 9 | 0.3333 |
| 19_608235 | NA | NA | 3 | 0 | 3 | 0 | 4 | 0.25 | 10 | 0.1 | 2 | 0.5 | 2 | 0 | 4 | 0 | 3 | 0 | 11 | 0.0909 |
| 19_608272 | 5 | 0.4 | NA | NA | 4 | 0 | 3 | 0.3333 | 12 | 0.25 | 2 | 1 | 2 | 0 | 3 | 0.3333 | 1 | 0 | 8 | 0.375 |
| 19_608525 | 20 | 0.05 | 14 | 0.0714 | 17 | 0.1176 | 18 | 0.1111 | 69 | 0.087 | 17 | 0 | 16 | 0.125 | 10 | 0.1 | 11 | 0 | 54 | 0.0556 |
| 19_610804 | 3 | 0 | 2 | 0 | 2 | 0.5 | 1 | 1 | 8 | 0.25 | NA | NA | 3 | 0.6667 | 2 | 0.5 | 2 | 0.5 | 7 | 0.5714 |
| 19_610805 | 3 | 0.3333 | 1 | 1 | 2 | 0 | 1 | 1 | 7 | 0.4286 | NA | NA | 2 | 0.5 | 2 | 0 | 2 | 1 | 6 | 0.5 |
| 19_610806 | 4 | 1 | 2 | 0.5 | 2 | 0 | 1 | 1 | 9 | 0.6667 | NA | NA | 3 | 1 | 4 | 0.75 | 2 | 1 | 9 | 0.8889 |
| 19_610806 | 3 | 0 | 1 | 0 | NA | NA | 1 | 1 | 5 | 0.2 | NA | NA | 3 | 0.6667 | 4 | 0.5 | NA | NA | 7 | 0.5714 |
| 19_610807 | 4 | 0.25 | 1 | 0 | 2 | 0 | 1 | 0 | 8 | 0.125 | NA | NA | 2 | 0.5 | 4 | 0.25 | 2 | 0.5 | 8 | 0.375 |
| 19_610811 | 1 | 0 | NA | NA | NA | NA | 1 | 1 | 2 | 0.5 | NA | NA | NA | NA | NA | NA | 1 | 1 | 1 | 1 |
| 19_610824 | 2 | 0.5 | NA | NA | NA | NA | 4 | 0.25 | 6 | 0.3333 | NA | NA | NA | NA | NA | NA | 1 | 1 | 1 | 1 |
| 19_611034 | NA | NA | 3 | 0.6667 | 3 | 0 | 1 | 0 | 7 | 0.2857 | NA | NA | 1 | 0 | 1 | 0 | 3 | 0.3333 | 5 | 0.2 |
| 19_611034 | NA | NA | 3 | 1 | 2 | 0 | NA | NA | 5 | 0.6 | NA | NA | 1 | 0 | 1 | 0 | 3 | 1 | 5 | 0.6 |
| 19_611035 | NA | NA | 3 | 0 | 3 | 0.3333 | 1 | 0 | 7 | 0.1429 | NA | NA | 2 | 0 | 1 | 0 | 4 | 0.25 | 7 | 0.1429 |
| 19_611037 | NA | NA | 3 | 0.6667 | 3 | 0 | 1 | 1 | 7 | 0.4286 | NA | NA | 2 | 1 | 1 | 0 | 5 | 0.8 | 8 | 0.75 |
| 19_611038 | NA | NA | 3 | 0.6667 | 3 | 0.6667 | 1 | 1 | 7 | 0.7143 | NA | NA | 2 | 0.5 | 1 | 0 | 5 | 0.6 | 8 | 0.5 |
| 19_611038 | NA | NA | 2 | 0.5 | 2 | 0.5 | 1 | 1 | 5 | 0.6 | NA | NA | 2 | 0.5 | 1 | 0 | 5 | 0.4 | 8 | 0.375 |
| 19_611038 | NA | NA | 3 | 1 | 3 | 0 | 1 | 1 | 7 | 0.5714 | NA | NA | 2 | 0 | 1 | 0 | 4 | 0.5 | 7 | 0.2857 |
| 19_611039 | NA | NA | 2 | 0.5 | 3 | 0.3333 | 1 | 1 | 6 | 0.5 | NA | NA | 2 | 0 | 1 | 0 | 4 | 0.75 | 7 | 0.4286 |
| 19_611042 | NA | NA | 1 | 0 | 3 | 0.6667 | 1 | 1 | 5 | 0.6 | NA | NA | 2 | 0.5 | 1 | 0 | 3 | 0.6667 | 6 | 0.5 |
| 19_611044 | NA | NA | 2 | 1 | 3 | 1 | 1 | 1 | 6 | 1 | NA | NA | 1 | 1 | 1 | 0 | 2 | 0.5 | 4 | 0.5 |
| 19_611048 | NA | NA | 2 | 1 | 3 | 1 | 1 | 1 | 6 | 1 | NA | NA | 2 | 1 | 1 | 0 | 3 | 0.6667 | 6 | 0.6667 |
| 19_611050 | NA | NA | 2 | 1 | 3 | 1 | 1 | 1 | 6 | 1 | NA | NA | 1 | 1 | 1 | 0 | 2 | 1 | 4 | 0.75 |
| 19_611051 | NA | NA | 2 | 0.5 | 3 | 0.6667 | 1 | 0 | 6 | 0.5 | NA | NA | 1 | 0 | 1 | 0 | 2 | 0.5 | 4 | 0.25 |
| 19_611053 | NA | NA | 1 | 0 | 3 | 0.3333 | NA | NA | 4 | 0.25 | NA | NA | 2 | 0 | 1 | 0 | 3 | 0.3333 | 6 | 0.1667 |
| 19_611054 | NA | NA | 2 | 0.5 | 3 | 0.3333 | 1 | 0 | 6 | 0.3333 | NA | NA | 2 | 0.5 | NA | NA | 3 | 0 | 5 | 0.2 |
| 19_611055 | NA | NA | 2 | 0.5 | 3 | 0.6667 | 1 | 1 | 6 | 0.6667 | NA | NA | 2 | 0 | 1 | 0 | 3 | 0.6667 | 6 | 0.3333 |
| 19_611055 | NA | NA | 1 | 1 | 3 | 0.6667 | 1 | 1 | 5 | 0.8 | NA | NA | 2 | 0.5 | 1 | 1 | 3 | 0.6667 | 6 | 0.6667 |
| 19_612076 | NA | NA | 1 | 1 | NA | NA | NA | NA | 1 | 1 | 1 | 1 | NA | NA | NA | NA | 1 | 0 | 2 | 0.5 |
| 19_612130 | 2 | 0.5 | 1 | 0 | 1 | 0 | 4 | 0.5 | 8 | 0.375 | 1 | 0 | 3 | 0 | 1 | 0 | 1 | 1 | 6 | 0.1667 |
| 19_612131 | 2 | 0 | 1 | 0 | 1 | 0 | 6 | 0.1667 | 10 | 0.1 | 2 | 0 | 3 | 0.3333 | 1 | 0 | 1 | 1 | 7 | 0.2857 |
| 19_612133 | 1 | 0 | 1 | 0 | 1 | 0 | 5 | 0.4 | 8 | 0.25 | 2 | 0 | 3 | 0 | 1 | 0 | 1 | 1 | 7 | 0.1429 |
| 19_612134 | 1 | 0 | NA | NA | 1 | 0 | 5 | 0.6 | 7 | 0.4286 | 1 | 0 | 2 | 0 | 1 | 1 | NA | NA | 4 | 0.25 |
| 2_3226036 | 7 | 1 | 8 | 1 | 6 | 1 | 10 | 0.7 | 31 | 0.9032 | 3 | 1 | 10 | 1 | 9 | 0.8889 | 8 | 1 | 30 | 0.9667 |
| 2_3226094 | 8 | 0.25 | 9 | 0.2222 | 8 | 0.375 | 11 | 0.4545 | 36 | 0.3333 | 3 | 0 | 10 | 0.2 | 9 | 0.1111 | 8 | 0.375 | 30 | 0.2 |
| 2_3226222 | 8 | 0.875 | 8 | 0.875 | 8 | 1 | 13 | 1 | 37 | 0.9459 | 3 | 1 | 9 | 1 | 5 | 1 | 8 | 1 | 25 | 1 |
| 2_3226235 | 7 | 0.2857 | 7 | 0.2857 | 6 | 0.6667 | 10 | 0.6 | 30 | 0.4667 | 4 | 0.25 | 6 | 0.1667 | 9 | 0.3333 | 8 | 0.625 | 27 | 0.3704 |
| 2_3226419 | 7 | 0.4286 | 10 | 0 | 7 | 0 | 10 | 0.2 | 34 | 0.1471 | 3 | 0 | 11 | 0 | 6 | 0.1667 | 8 | 0.125 | 28 | 0.0714 |
| 2_3226466 | 8 | 0.25 | 10 | 0.2 | 8 | 0.375 | 14 | 0 | 40 | 0.175 | 3 | 0.3333 | 9 | 0.1111 | 8 | 0.125 | 7 | 0 | 27 | 0.1111 |
| 2_3226518 | 7 | 0.1429 | 11 | 0 | 9 | 0 | 14 | 0.1429 | 41 | 0.0732 | 6 | 0 | 11 | 0.0909 | 8 | 0.25 | 11 | 0.2727 | 36 | 0.1667 |
| 2_3226568 | 8 | 0.5 | 9 | 0.6667 | 9 | 0.5556 | 15 | 0.5333 | 41 | 0.561 | 7 | 0.8571 | 12 | 0.75 | 8 | 0.875 | 9 | 0.4444 | 36 | 0.7222 |
| 2_3226635 | 8 | 0.375 | 9 | 0.2222 | 8 | 0.75 | 15 | 0.5333 | 40 | 0.475 | 8 | 0.625 | 11 | 0.5455 | 9 | 0.3333 | 8 | 0.625 | 36 | 0.5278 |
| 2_3226646 | 8 | 0.125 | 8 | 0.375 | 9 | 0.3333 | 16 | 0.375 | 41 | 0.3171 | 7 | 0.4286 | 12 | 0.1667 | 9 | 0.2222 | 7 | 0.2857 | 35 | 0.2571 |

| | | | | | | | | | | | | | | | | | | | | |
|---|---|---|---|---|---|---|---|---|---|---|---|---|---|---|---|---|---|---|---|---|
| 2_3226913 | 5 | 0.4 | 7 | 0.2857 | 5 | 0.6 | 10 | 0.4 | 27 | 0.4074 | 6 | 0.8333 | 8 | 0.25 | 9 | 0.3333 | 7 | 0.5714 | 30 | 0.4667 |
| 2_3226953 | 6 | 0.8333 | 8 | 0.625 | 5 | 0.8 | 10 | 0.8 | 29 | 0.7586 | 5 | 0.6 | 7 | 0.8571 | 9 | 0.5556 | 7 | 0.5714 | 28 | 0.6429 |
| 2_3227031 | 9 | 0.2222 | 12 | 0.0833 | 9 | 0.1111 | 18 | 0.2778 | 48 | 0.1875 | 8 | 0.125 | 11 | 0.3636 | 11 | 0.4545 | 13 | 0.3077 | 43 | 0.3256 |
| 2_3227090 | 10 | 1 | 13 | 1 | 9 | 1 | 16 | 1 | 48 | 1 | 7 | 1 | 12 | 1 | 10 | 1 | 13 | 1 | 42 | 1 |
| 2_3227112 | 9 | 0.3333 | 7 | 0.5714 | 8 | 0.375 | 13 | 0.3077 | 37 | 0.3784 | 5 | 0.2 | 8 | 0.375 | 10 | 0.3 | 11 | 0.5455 | 34 | 0.3824 |
| 2_3227246 | 11 | 0.5455 | 13 | 0.5385 | 7 | 0.2857 | 16 | 0.3125 | 47 | 0.4255 | 8 | 0.375 | 11 | 0.4545 | 13 | 0.3846 | 12 | 0.4167 | 44 | 0.4091 |
| 2_3227252 | 10 | 0.3 | 10 | 0.3 | 9 | 0.6667 | 17 | 0.2353 | 46 | 0.3478 | 9 | 0.3333 | 12 | 0.5833 | 12 | 0.3333 | 13 | 0.4615 | 46 | 0.4348 |
| 2_3227297 | 9 | 0.7778 | 13 | 0.9231 | 9 | 0.6667 | 15 | 1 | 46 | 0.8696 | 6 | 1 | 12 | 0.9167 | 12 | 0.75 | 14 | 0.8571 | 44 | 0.8636 |
| 2_3227371 | 11 | 0 | 11 | 0 | 10 | 0.3 | 16 | 0.0625 | 48 | 0.0833 | 9 | 0.1111 | 12 | 0 | 11 | 0.0909 | 16 | 0.25 | 48 | 0.125 |
| 2_3227383 | 8 | 0 | 9 | 0.3333 | 9 | 0.2222 | 13 | 0.3846 | 39 | 0.2564 | 7 | 0.1429 | 8 | 0.25 | 9 | 0.4444 | 13 | 0.2308 | 37 | 0.2703 |
| 2_3227692 | 11 | 0.0909 | 15 | 0.1333 | 9 | 0 | 19 | 0.1053 | 54 | 0.0926 | 8 | 0.125 | 11 | 0.1818 | 12 | 0.1667 | 15 | 0.2667 | 46 | 0.1957 |
| 2_3227707 | 10 | 1 | 11 | 0.7273 | 7 | 0.8571 | 12 | 0.8333 | 40 | 0.85 | 5 | 1 | 7 | 1 | 10 | 0.9 | 13 | 0.9231 | 35 | 0.9429 |
| 2_3227711 | 12 | 0 | 12 | 0.25 | 10 | 0.1 | 17 | 0.0588 | 51 | 0.098 | 7 | 0.1429 | 11 | 0 | 10 | 0.1 | 16 | 0.0625 | 44 | 0.0682 |
| 2_3227745 | 11 | 0.1818 | 15 | 0.4 | 10 | 0.2 | 19 | 0.1579 | 55 | 0.2364 | 9 | 0.3333 | 10 | 0.3 | 12 | 0.1667 | 18 | 0.2222 | 49 | 0.2449 |
| 2_3227761 | 11 | 1 | 13 | 1 | 10 | 0.9 | 17 | 0.9412 | 51 | 0.9608 | 8 | 1 | 12 | 0.9167 | 12 | 0.9167 | 15 | 1 | 47 | 0.9574 |
| 2_3227766 | 8 | 0.25 | 14 | 0.1429 | 8 | 0.25 | 17 | 0.2941 | 47 | 0.234 | 6 | 0.1667 | 11 | 0.1818 | 10 | 0.5 | 11 | 0.3636 | 38 | 0.3158 |
| 2_3227880 | 12 | 0.9167 | 14 | 0.5714 | 10 | 0.5 | 17 | 0.5882 | 53 | 0.6415 | 6 | 0.8333 | 11 | 0.5455 | 12 | 0.9167 | 15 | 0.6667 | 44 | 0.7273 |
| 2_3227945 | 11 | 0.7273 | 16 | 0.5 | 11 | 0.4545 | 18 | 0.7222 | 56 | 0.6071 | 10 | 0.6 | 12 | 0.5 | 12 | 0.5833 | 17 | 0.5294 | 51 | 0.549 |
| 2_3228221 | 10 | 0.1 | 12 | 0.0833 | 10 | 0.2 | 16 | 0.4375 | 48 | 0.2292 | 11 | 0.0909 | 7 | 0.1429 | 8 | 0.125 | 16 | 0.25 | 42 | 0.1667 |
| 2_3228270 | 10 | 0.3 | 16 | 0.1875 | 10 | 0.5 | 22 | 0.2273 | 58 | 0.2759 | 14 | 0.1429 | 10 | 0.2 | 10 | 0.1 | 14 | 0.1429 | 48 | 0.1458 |
| 2_3228356 | 2 | 0.5 | 10 | 0.3 | 3 | 0.6667 | 4 | 0 | 19 | 0.3158 | 4 | 0.25 | 7 | 0.2857 | 4 | 0 | 6 | 0.1667 | 21 | 0.1905 |
| 2_3228399 | 8 | 0.375 | 11 | 0.3636 | 7 | 0.4286 | 15 | 0.4 | 41 | 0.3902 | 6 | 0.3333 | 10 | 0.7 | 9 | 0.4444 | 15 | 0.4667 | 40 | 0.5 |
| 2_3228447 | 12 | 0.0833 | 15 | 0.4 | 10 | 0.3 | 19 | 0.4211 | 56 | 0.3214 | 12 | 0.5 | 12 | 0.4167 | 10 | 0.4 | 16 | 0.375 | 50 | 0.42 |
| 2_3228558 | 13 | 0.0769 | 18 | 0.0556 | 11 | 0.1818 | 20 | 0.2 | 62 | 0.129 | 13 | 0.0769 | 15 | 0.3333 | 12 | 0.0833 | 19 | 0.0526 | 59 | 0.1356 |
| 2_3228720 | 13 | 0.2308 | 19 | 0.1579 | 12 | 0 | 17 | 0.2941 | 61 | 0.1803 | 13 | 0.1538 | 15 | 0.2 | 11 | 0.1818 | 20 | 0.1 | 59 | 0.1525 |
| 2_3327113 | 4 | 0 | 9 | 0.1111 | 2 | 0.5 | 7 | 0.1429 | 22 | 0.1364 | 3 | 0.3333 | 7 | 0.2857 | 4 | 0.25 | 3 | 0.3333 | 17 | 0.2941 |
| 2_3327319 | 3 | 0 | 10 | 0 | 3 | 0.3333 | 8 | 0.375 | 24 | 0.1667 | 3 | 0.3333 | 7 | 0 | 4 | 0 | 2 | 0 | 16 | 0.0625 |
| 2_3327378 | 3 | 0.6667 | 10 | 0.3 | 3 | 0.3333 | 7 | 0.2857 | 23 | 0.3478 | 2 | 0 | 4 | 0.25 | 2 | 0.5 | 3 | 0 | 11 | 0.1818 |
| 2_3327764 | 2 | 0 | 8 | 0 | 2 | 0.5 | 8 | 0 | 20 | 0.05 | 3 | 0 | 5 | 0.2 | 5 | 0 | 5 | 0 | 18 | 0.0556 |
| 2_3328488 | 6 | 0 | 10 | 0.1 | 2 | 0 | 7 | 0.2857 | 25 | 0.12 | 1 | 0 | 7 | 0 | 5 | 0 | 6 | 0.1667 | 19 | 0.0526 |
| 2_3330218 | 3 | 1 | 2 | 0.5 | NA | NA | NA | NA | 5 | 0.8 | NA | NA | NA | NA | 2 | 0.5 | NA | NA | 2 | 0.5 |
| 2_3330255 | 1 | 1 | 1 | 1 | NA | NA | 1 | 1 | 3 | 1 | NA | NA | 1 | 1 | 2 | 1 | NA | NA | 3 | 1 |
| 2_3330353 | 2 | 0.5 | 2 | 0.5 | NA | NA | 1 | 1 | 5 | 0.6 | NA | NA | 1 | 1 | 2 | 1 | NA | NA | 3 | 1 |
| 2_3330453 | 2 | 1 | 1 | 1 | NA | NA | 1 | 1 | 4 | 1 | NA | NA | NA | NA | 2 | 1 | NA | NA | 2 | 1 |
| 2_3330532 | 1 | 1 | 2 | 0.5 | NA | NA | 1 | 1 | 4 | 0.75 | NA | NA | NA | NA | 2 | 1 | NA | NA | 2 | 1 |
| 2_3330733 | 2 | 1 | 2 | 1 | NA | NA | 1 | 1 | 5 | 1 | NA | NA | NA | NA | 1 | 1 | NA | NA | 1 | 1 |
| 2_3454006 | NA | NA | NA | NA | 2 | 1 | 1 | 1 | 3 | 1 | NA | NA | 1 | 1 | 2 | 1 | NA | NA | 3 | 1 |
| 2_3454074 | NA | NA | NA | NA | 2 | 1 | 1 | 1 | 3 | 1 | 1 | 0 | 1 | 1 | 1 | 0 | NA | NA | 3 | 0.3333 |
| 2_3454188 | NA | NA | NA | NA | 2 | 1 | 1 | 0 | 3 | 0.6667 | 1 | 1 | 1 | 1 | 2 | 0 | NA | NA | 4 | 0.5 |
| 2_3454249 | NA | NA | 1 | 0 | 2 | 0.5 | 1 | 0 | 4 | 0.25 | 1 | 0 | 1 | 1 | 1 | 1 | NA | NA | 3 | 0.6667 |
| 2_3454271 | NA | NA | NA | NA | 2 | 0 | 1 | 1 | 3 | 0.3333 | 1 | 1 | 1 | 1 | 2 | 0 | NA | NA | 4 | 0.5 |
| 2_3454305 | NA | NA | 1 | 0 | 2 | 0.5 | 1 | 1 | 4 | 0.5 | 1 | 0 | 1 | 0 | 1 | 1 | NA | NA | 3 | 0.3333 |
| 2_3454326 | NA | NA | 1 | 0 | 2 | 0.5 | 1 | 0 | 4 | 0.25 | 1 | 0 | 1 | 1 | 2 | 0 | NA | NA | 4 | 0.25 |
| 2_3454347 | NA | NA | 1 | 1 | 2 | 0.5 | 1 | 1 | 4 | 0.75 | 1 | 0 | 1 | 1 | 2 | 1 | NA | NA | 4 | 0.75 |
| 2_3454431 | NA | NA | NA | NA | 2 | 0 | 1 | 1 | 3 | 0.3333 | 1 | 1 | 1 | 0 | 2 | 0 | NA | NA | 4 | 0.25 |
| 2_3454519 | NA | NA | NA | NA | 1 | 0 | 1 | 1 | 2 | 0.5 | 1 | 1 | 1 | 0 | NA | NA | NA | NA | 2 | 0.5 |
| 2_3454552 | NA | NA | 1 | 1 | 3 | 0 | 1 | 0 | 5 | 0.2 | 1 | 0 | NA | NA | 2 | 0.5 | NA | NA | 3 | 0.3333 |
| 2_3454633 | NA | NA | 1 | 1 | 1 | 1 | 1 | 0 | 3 | 0.6667 | 1 | 0 | 1 | 1 | 2 | 0.5 | NA | NA | 4 | 0.5 |
| 2_3454673 | NA | NA | 1 | 1 | 3 | 0 | 1 | 1 | 5 | 0.4 | 1 | 0 | 1 | 0 | 2 | 0.5 | NA | NA | 4 | 0.25 |
| 2_3454698 | NA | NA | 1 | 1 | 3 | 0.3333 | 1 | 1 | 5 | 0.6 | 1 | 1 | 1 | 1 | 1 | 0 | NA | NA | 3 | 0.6667 |
| 2_3454716 | NA | NA | 1 | 1 | 3 | 0.3333 | 1 | 0 | 5 | 0.4 | 1 | 0 | 1 | 1 | 2 | 0 | NA | NA | 4 | 0.25 |
| 2_3455170 | NA | NA | 1 | 0 | 3 | 0.3333 | 1 | 1 | 5 | 0.4 | 1 | 0 | 1 | 0 | 2 | 1 | NA | NA | 4 | 0.5 |
| 2_3489997 | 3 | 0 | 4 | 0 | 4 | 0.25 | 5 | 0.2 | 16 | 0.125 | 3 | 0.3333 | 6 | 0.1667 | 7 | 0 | 5 | 0 | 21 | 0.0952 |
| 2_3780893 | 2 | 1 | NA | NA | 2 | 1 | NA | NA | 4 | 1 | 1 | 1 | NA | NA | NA | NA | 1 | 1 | 2 | 1 |
| 2_3781008 | 2 | 0 | NA | NA | 1 | 1 | NA | NA | 3 | 0.3333 | 1 | 1 | NA | NA | NA | NA | 1 | 0 | 2 | 0.5 |
| 2_4608825 | 2 | 1 | NA | NA | NA | NA | 3 | 0.6667 | 5 | 0.8 | NA | NA | NA | NA | 1 | 1 | NA | NA | 1 | 1 |
| 2_4616035 | 1 | 0 | NA | NA | 1 | 1 | 4 | 0.5 | 6 | 0.5 | 1 | 1 | 1 | 1 | 2 | 1 | NA | NA | 4 | 1 |
| 2_4616044 | 2 | 1 | NA | NA | 1 | 1 | 4 | 0.75 | 7 | 0.8571 | 1 | 1 | 1 | 0 | 2 | 0.5 | NA | NA | 4 | 0.5 |
| 2_4616162 | 2 | 1 | NA | NA | NA | NA | 4 | 1 | 6 | 1 | 1 | 1 | 1 | 1 | 2 | 1 | NA | NA | 4 | 1 |
| 2_4616379 | 2 | 0 | NA | NA | 1 | 0 | 4 | 0.25 | 7 | 0.1429 | 1 | 0 | 1 | 0 | 2 | 1 | NA | NA | 4 | 0.5 |
| 2_4616438 | 1 | 1 | NA | NA | 1 | 0 | 5 | 0 | 7 | 0.1429 | 1 | 0 | 1 | 1 | 2 | 0.5 | NA | NA | 4 | 0.5 |
| 2_4616454 | 1 | 1 | NA | NA | NA | NA | 2 | 1 | 3 | 1 | NA | NA | NA | NA | 1 | 1 | NA | NA | 1 | 1 |
| 2_4616477 | 2 | 0.5 | NA | NA | 1 | 0 | 4 | 0.25 | 7 | 0.2857 | 1 | 1 | 1 | 0 | 1 | 0 | NA | NA | 3 | 0.3333 |
| 2_4616511 | 2 | 0 | NA | NA | 1 | 0 | 4 | 0.25 | 7 | 0.1429 | 1 | 1 | 1 | 1 | 2 | 0 | NA | NA | 4 | 0.5 |
| 2_4616551 | 1 | 0 | NA | NA | 1 | 0 | 5 | 0.2 | 7 | 0.1429 | 2 | 0.5 | 2 | 0 | 1 | 0 | NA | NA | 5 | 0.2 |

| | | | | | | | | | | | | | | | | | | | | |
|---|---|---|---|---|---|---|---|---|---|---|---|---|---|---|---|---|---|---|---|---|
| 2_4616614 | 2 | 0.5 | NA | NA | 1 | 1 | 5 | 0.8 | 8 | 0.75 | 2 | 1 | 2 | 1 | 2 | 1 | NA | NA | 6 | 1 |
| 2_4616650 | 2 | 0 | NA | NA | 1 | 0 | 4 | 0.25 | 7 | 0.1429 | 2 | 0.5 | 2 | 0 | 2 | 0 | NA | NA | 6 | 0.1667 |
| 2_4616656 | 2 | 1 | NA | NA | 1 | 1 | 5 | 1 | 8 | 1 | 2 | 1 | 2 | 1 | 2 | 1 | NA | NA | 6 | 1 |
| 2_4616671 | 2 | 1 | NA | NA | 1 | 1 | 4 | 1 | 7 | 1 | 2 | 1 | 2 | 1 | 1 | 1 | NA | NA | 5 | 1 |
| 2_4616698 | 2 | 1 | NA | NA | 1 | 1 | 5 | 1 | 8 | 1 | 2 | 1 | 2 | 1 | 1 | 1 | NA | NA | 5 | 1 |
| 2_4616952 | 1 | 0 | NA | NA | 2 | 0 | 2 | 0.5 | 5 | 0.2 | 1 | 1 | 1 | 0 | 1 | 1 | NA | NA | 3 | 0.6667 |
| 2_4617011 | 2 | 0 | NA | NA | 2 | 0.5 | 4 | 0 | 8 | 0.125 | 1 | 0 | 2 | 0.5 | 2 | 0 | NA | NA | 5 | 0.2 |
| 2_4617481 | 2 | 0 | NA | NA | 3 | 0.3333 | 3 | 0 | 8 | 0.125 | 2 | 0.5 | 2 | 0 | 2 | 0 | 1 | 0 | 7 | 0.1429 |
| 2_4617523 | 2 | 0 | NA | NA | 2 | 0.5 | 5 | 0.8 | 9 | 0.5556 | 2 | 1 | 2 | 0.5 | 2 | 0 | 1 | 1 | 7 | 0.5714 |
| 2_4617826 | 2 | 0 | NA | NA | 2 | 0 | 4 | 0.75 | 8 | 0.375 | 2 | 0.5 | 2 | 0.5 | 2 | 1 | 2 | 1 | 8 | 0.75 |
| 2_4618119 | 3 | 0.3333 | NA | NA | 4 | 0 | 6 | 0 | 13 | 0.0769 | 1 | 0 | 2 | 0 | 4 | 0.25 | 2 | 0 | 9 | 0.1111 |
| 2_4618206 | 3 | 0 | NA | NA | 5 | 0.2 | 6 | 0.1667 | 14 | 0.1429 | 2 | 0 | 2 | 0.5 | 4 | 0.25 | 2 | 0 | 10 | 0.2 |
| 2_4618595 | 3 | 0 | 2 | 1 | 5 | 0.4 | 7 | 0.1429 | 17 | 0.2941 | 3 | 0.6667 | 2 | 0 | 4 | 0 | 2 | 0.5 | 11 | 0.2727 |
| 2_4626962 | 1 | 0 | 3 | 0 | 1 | 0 | 3 | 0.3333 | 8 | 0.125 | 2 | 0.5 | 1 | 0 | 4 | 0.25 | 1 | 0 | 8 | 0.25 |
| 2_4627213 | 3 | 0 | 4 | 0.5 | 2 | 0.5 | 3 | 0.3333 | 12 | 0.3333 | 3 | 0.6667 | 6 | 0.3333 | 6 | 0.1667 | 4 | 0.25 | 19 | 0.3158 |
| 2_4627749 | 5 | 0.2 | 5 | 0 | 1 | 1 | 4 | 0.5 | 15 | 0.2667 | NA | NA | 5 | 0.8 | 6 | 0.1667 | 3 | 0.3333 | 14 | 0.4286 |
| 2_4628314 | 4 | 0.25 | 3 | 0 | NA | NA | 2 | 0 | 9 | 0.1111 | NA | NA | 6 | 0.1667 | 6 | 0.1667 | 2 | 0 | 14 | 0.1429 |
| 2_4628332 | 4 | 0.5 | 2 | 0.5 | 1 | 0 | 3 | 0.3333 | 10 | 0.4 | NA | NA | 7 | 0.2857 | 5 | 0.2 | 2 | 0.5 | 14 | 0.2857 |
| 2_4628404 | 4 | 0.25 | 3 | 0.6667 | 1 | 0 | 3 | 0 | 11 | 0.2727 | NA | NA | 6 | 0.5 | 6 | 0 | 2 | 0 | 14 | 0.2143 |
| 2_4628790 | 4 | 0.25 | 3 | 0 | 1 | 0 | 2 | 0.5 | 10 | 0.2 | NA | NA | 6 | 0 | 5 | 0 | 3 | 0.3333 | 14 | 0.0714 |
| 2_4629091 | 4 | 1 | 2 | 1 | 1 | 1 | 3 | 1 | 10 | 1 | NA | NA | 6 | 1 | 5 | 0.8 | 3 | 0.6667 | 14 | 0.8571 |
| 2_4629096 | 3 | 0.6667 | 2 | 1 | 1 | 1 | 3 | 1 | 9 | 0.8889 | NA | NA | 3 | 1 | 5 | 0.8 | 2 | 1 | 10 | 0.9 |
| 2_4629118 | 3 | 0.6667 | 1 | 1 | 1 | 1 | 3 | 1 | 8 | 0.875 | NA | NA | 5 | 1 | 5 | 0.6 | 2 | 1 | 12 | 0.8333 |
| 2_4629127 | 2 | 0 | 2 | 0 | 1 | 0 | 4 | 0.25 | 9 | 0.1111 | NA | NA | 6 | 0.1667 | 5 | 0.4 | 1 | 1 | 12 | 0.3333 |
| 2_4629195 | 3 | 0 | 2 | 0 | 1 | 0 | 3 | 0.3333 | 9 | 0.1111 | NA | NA | 6 | 0.3333 | 5 | 0.4 | 3 | 0 | 14 | 0.2857 |
| 2_4911581 | 8 | 0.25 | 3 | 0 | 4 | 0.25 | 3 | 0.3333 | 18 | 0.2222 | 1 | 0 | 1 | 0 | 2 | 0 | 6 | 0.3333 | 10 | 0.2 |
| 2_4911612 | 7 | 0.5714 | 3 | 0.6667 | 4 | 0.5 | 3 | 0.3333 | 17 | 0.5294 | 1 | 0 | 1 | 0 | 3 | 0.6667 | 6 | 0.3333 | 11 | 0.3636 |
| 2_4911674 | 7 | 1 | 1 | 1 | 3 | 1 | 3 | 1 | 14 | 1 | 1 | 1 | 1 | 1 | 2 | 1 | 4 | 1 | 8 | 1 |
| 2_4911683 | 4 | 0.75 | 1 | 0 | 5 | 0.6 | 2 | 0.5 | 12 | 0.5833 | 1 | 1 | 1 | 0 | 2 | 1 | 5 | 0.6 | 9 | 0.6667 |
| 2_4911717 | 6 | 0.3333 | 2 | 0 | 5 | 0 | 2 | 0 | 15 | 0.1333 | 1 | 1 | 1 | 0 | 3 | 0 | 6 | 0.1667 | 11 | 0.1818 |
| 2_4911788 | 6 | 0.6667 | NA | NA | 5 | 0 | 2 | 0.5 | 13 | 0.3846 | 1 | 1 | 1 | 0 | 2 | 0.5 | 4 | 0.75 | 8 | 0.625 |
| 2_4912017 | 6 | 0 | 3 | 0.3333 | 5 | 0 | 3 | 0.3333 | 17 | 0.1176 | 1 | 0 | 1 | 0 | 2 | 0.5 | 5 | 0 | 9 | 0.1111 |
| 2_4912030 | 7 | 0.2857 | 3 | 0 | 6 | 0 | 4 | 0.25 | 20 | 0.15 | 1 | 0 | 1 | 0 | 1 | 0 | 5 | 0.2 | 8 | 0.125 |
| 2_4912128 | 6 | 0.6667 | 3 | 0.6667 | 6 | 0.3333 | 4 | 0.5 | 19 | 0.5263 | 1 | 0 | 1 | 0 | 2 | 1 | 3 | 0.3333 | 7 | 0.4286 |
| 2_4912242 | 5 | 0.4 | 3 | 0.3333 | 7 | 0 | 4 | 0 | 19 | 0.1579 | 1 | 0 | 1 | 0 | 1 | 1 | 4 | 0.25 | 7 | 0.2857 |
| 2_4912428 | 5 | 0.6 | 2 | 1 | 6 | 0.8333 | 4 | 0.5 | 17 | 0.7059 | 1 | 0 | 1 | 1 | 2 | 0.5 | 4 | 0.5 | 8 | 0.5 |
| 2_4912779 | 6 | 0.3333 | 3 | 0.3333 | 5 | 0 | 4 | 0.25 | 18 | 0.2222 | 1 | 0 | 1 | 0 | 3 | 0.3333 | 4 | 0 | 9 | 0.1111 |
| 2_4912944 | 7 | 0.1429 | 3 | 0 | 5 | 0 | 4 | 0 | 19 | 0.0526 | NA | NA | 1 | 0 | 3 | 0.3333 | 4 | 0 | 8 | 0.125 |
| 2_4913002 | 6 | 0.5 | 2 | 0.5 | 2 | 0.5 | 3 | 0.6667 | 13 | 0.5385 | NA | NA | 1 | 0 | 2 | 0 | 5 | 0.6 | 8 | 0.375 |
| 2_4913028 | 6 | 0.5 | 3 | 0 | 4 | 0 | 4 | 0 | 17 | 0.1765 | 1 | 0 | 1 | 0 | 3 | 0 | 5 | 0.2 | 10 | 0.1 |
| 2_4913087 | 7 | 0.1429 | 3 | 0 | 4 | 0 | 2 | 0 | 16 | 0.0625 | 1 | 1 | 1 | 0 | NA | NA | 3 | 0 | 5 | 0.2 |
| 2_4913156 | 3 | 1 | 3 | 0.3333 | 3 | 1 | 1 | 1 | 10 | 0.8 | 1 | 1 | 1 | 1 | 1 | 1 | 5 | 1 | 8 | 1 |
| 2_4913205 | 7 | 1 | 3 | 0.6667 | 5 | 0.6 | 4 | 1 | 19 | 0.8421 | 1 | 1 | 1 | 0 | 3 | 1 | 4 | 1 | 9 | 0.8889 |
| 2_4913216 | 8 | 1 | 2 | 1 | 3 | 1 | 3 | 1 | 16 | 1 | 1 | 1 | 1 | 1 | 1 | 1 | 3 | 1 | 6 | 1 |
| 2_4913291 | 8 | 0.125 | 3 | 0 | 5 | 0 | 4 | 0 | 20 | 0.05 | 1 | 0 | 1 | 0 | 3 | 0 | 5 | 0.2 | 10 | 0.1 |
| 2_4913308 | 6 | 0 | 1 | 0 | 3 | 0 | 1 | 1 | 11 | 0.0909 | NA | NA | 1 | 0 | 2 | 0 | 3 | 0.3333 | 6 | 0.1667 |
| 2_4913321 | 4 | 1 | NA | NA | 1 | 0 | 1 | 1 | 6 | 0.8333 | NA | NA | NA | NA | 2 | 1 | 1 | 1 | 3 | 1 |
| 2_4943332 | 15 | 0.2 | 17 | 0.2353 | 16 | 0.25 | 15 | 0.2 | 63 | 0.2222 | 15 | 0.2 | 11 | 0.3636 | 8 | 0.375 | 15 | 0.2667 | 49 | 0.2857 |
| 2_4943350 | 16 | 0.25 | 17 | 0 | 17 | 0.1765 | 18 | 0.2778 | 68 | 0.1765 | 14 | 0.2143 | 12 | 0.1667 | 8 | 0 | 18 | 0.1111 | 52 | 0.1346 |
| 2_4943373 | 17 | 0.1176 | 16 | 0.1875 | 15 | 0.0667 | 18 | 0.0556 | 66 | 0.1061 | 15 | 0.0667 | 13 | 0.0769 | 8 | 0 | 14 | 0.0714 | 50 | 0.06 |
| 2_5026175 | 4 | 0.5 | 1 | 0 | NA | NA | 1 | 0 | 6 | 0.3333 | 1 | 1 | 1 | 0 | 4 | 0.5 | 2 | 0.5 | 8 | 0.5 |
| 2_5235605 | 3 | 0.3333 | NA | NA | NA | NA | 1 | 1 | 4 | 0.5 | NA | NA | 1 | 1 | NA | NA | NA | NA | 1 | 1 |
| 2_5299308 | 1 | 1 | 7 | 1 | 2 | 1 | 6 | 1 | 16 | 1 | 4 | 0.75 | 5 | 0.8 | 6 | 0.5 | 5 | 0.6 | 20 | 0.65 |
| 2_5299358 | 6 | 0.8333 | 6 | 0.5 | 4 | 0.75 | 4 | 0.5 | 20 | 0.65 | 9 | 0.6667 | 7 | 0.5714 | 7 | 0.5714 | 6 | 0.8333 | 29 | 0.6552 |
| 2_5299367 | 6 | 1 | 6 | 0.8333 | 5 | 0.8 | 6 | 1 | 23 | 0.913 | 8 | 1 | 6 | 0.8333 | 3 | 0.6667 | 7 | 1 | 24 | 0.9167 |
| 2_5299487 | 5 | 0.4 | 3 | 0.3333 | 6 | 0.1667 | 6 | 0.3333 | 20 | 0.3 | 9 | 0.5556 | 5 | 0.2 | 4 | 0.25 | 5 | 0.6 | 23 | 0.4348 |
| 2_5299499 | 6 | 0.1667 | 7 | 0.1429 | 6 | 0.1667 | 8 | 0.25 | 27 | 0.1852 | 9 | 0.2222 | 8 | 0.125 | 8 | 0.25 | 7 | 0.1429 | 32 | 0.1875 |
| 2_5299584 | 6 | 0.1667 | 6 | 0 | 7 | 0 | 8 | 0.125 | 27 | 0.0741 | 10 | 0 | 6 | 0.1667 | 7 | 0.1429 | 6 | 0.1667 | 29 | 0.1034 |
| 2_5299603 | 6 | 0.1667 | 7 | 0.1429 | 9 | 0 | 7 | 0 | 29 | 0.069 | 9 | 0.1111 | 7 | 0.1429 | 4 | 0 | 6 | 0.1667 | 26 | 0.1154 |
| 2_5299765 | 5 | 0.2 | 5 | 0 | 6 | 0.1667 | 6 | 0 | 22 | 0.0909 | 6 | 0 | 4 | 0 | 4 | 0.25 | 6 | 0.3333 | 20 | 0.15 |
| 2_5300352 | 2 | 0 | 5 | 0.4 | 5 | 0.2 | 5 | 0.2 | 17 | 0.2353 | 5 | 0.6 | 5 | 0 | 4 | 0.25 | 7 | 0.2857 | 21 | 0.2857 |
| 2_5300584 | 2 | 0 | 7 | 0.1429 | 5 | 0.4 | 4 | 0.25 | 18 | 0.2222 | 4 | 0 | 5 | 0 | 3 | 0 | 7 | 0.2857 | 19 | 0.1053 |
| 2_5300689 | 2 | 0 | 6 | 0.1667 | 7 | 0 | 4 | 0 | 19 | 0.0526 | 3 | 0 | 4 | 0 | 4 | 0.25 | 6 | 0 | 17 | 0.0588 |
| 2_5300951 | 2 | 0 | 5 | 0.4 | 7 | 0.2857 | 4 | 0 | 18 | 0.2222 | 4 | 0.5 | 3 | 0 | 3 | 0 | 6 | 0.1667 | 16 | 0.1875 |
| 2_5301641 | 2 | 0 | 4 | 0.5 | 5 | 0 | 3 | 0.3333 | 14 | 0.2143 | 3 | 0 | 3 | 0.3333 | 3 | 0.6667 | 4 | 0 | 13 | 0.2308 |

| | | | | | | | | | | | | | | | | | | | | |
|---|---|---|---|---|---|---|---|---|---|---|---|---|---|---|---|---|---|---|---|---|
| 2_5302248 | 2 | 0.5 | 3 | 0.3333 | 6 | 0.5 | 3 | 0.3333 | 14 | 0.4286 | 2 | 0.5 | 2 | 0.5 | 3 | 0.3333 | 2 | 0 | 9 | 0.3333 |
| 2_5302501 | 2 | 0 | 3 | 0.6667 | 6 | 0.3333 | 3 | 0.3333 | 14 | 0.3571 | 1 | 0 | 1 | 0 | 2 | 0 | 2 | 0.5 | 6 | 0.1667 |
| 2_5302543 | 2 | 0.5 | 3 | 0.3333 | 6 | 0.1667 | 2 | 0.5 | 13 | 0.3077 | 1 | 0 | 1 | 0 | 2 | 0.5 | 2 | 0.5 | 6 | 0.3333 |
| 2_5302552 | 2 | 0 | 1 | 0 | 5 | 0.2 | NA | NA | 8 | 0.125 | 2 | 0 | 1 | 1 | 1 | 0 | 2 | 0.5 | 6 | 0.3333 |
| 2_5302704 | 2 | 0 | 2 | 0 | 5 | 0.2 | 3 | 0 | 12 | 0.0833 | 2 | 0 | 1 | 0 | 2 | 0 | 2 | 0.5 | 7 | 0.1429 |
| 2_5302927 | 2 | 1 | 2 | 1 | 6 | 1 | 2 | 1 | 12 | 1 | 2 | 1 | NA | NA | 1 | 1 | 2 | 1 | 5 | 1 |
| 2_5303070 | 2 | 0 | 2 | 0 | 5 | 0.6 | 3 | 0 | 12 | 0.25 | 1 | 0 | NA | NA | 2 | 0.5 | 1 | 0 | 4 | 0.25 |
| 2_5303530 | 1 | 1 | 2 | 1 | 6 | 1 | 1 | 1 | 10 | 1 | NA | NA | NA | NA | 2 | 1 | 1 | 1 | 3 | 1 |
| 2_5303604 | 1 | 0 | 1 | 1 | 2 | 1 | 1 | 1 | 5 | 0.8 | 1 | 1 | NA | NA | 1 | 0 | 1 | 1 | 3 | 0.6667 |
| 2_5303613 | NA | NA | 1 | 1 | 4 | 1 | 2 | 1 | 7 | 1 | 1 | 1 | NA | NA | 1 | 1 | 1 | 1 | 3 | 1 |
| 2_5303681 | 1 | 1 | 2 | 1 | 5 | 1 | 2 | 0.5 | 10 | 0.9 | NA | NA | NA | NA | 1 | 1 | 2 | 1 | 3 | 1 |
| 2_5799241 | 1 | 0 | 1 | 1 | NA | NA | 2 | 1 | 4 | 0.75 | NA | NA | 1 | 1 | 1 | 1 | 1 | 1 | 3 | 1 |
| 2_5814597 | 1 | 1 | NA | NA | 2 | 1 | 1 | 1 | 4 | 1 | NA | NA | 1 | 1 | 1 | 1 | NA | NA | 2 | 1 |
| 2_5873021 | 4 | 0.5 | 1 | 0 | 2 | 0.5 | 1 | 1 | 8 | 0.5 | 2 | 0.5 | 3 | 0 | 1 | 0 | NA | NA | 6 | 0.1667 |
| 2_5962370 | NA | NA | 2 | 0 | 3 | 0 | 2 | 0.5 | 7 | 0.1429 | NA | NA | 3 | 0.3333 | 1 | 0 | NA | NA | 4 | 0.25 |
| 2_5965916 | NA | NA | 2 | 0 | 2 | 0 | 1 | 1 | 5 | 0.2 | NA | NA | 3 | 0.6667 | 1 | 0 | NA | NA | 4 | 0.5 |
| 2_5965927 | NA | NA | 2 | 0 | 2 | 0.5 | 2 | 0 | 6 | 0.1667 | NA | NA | 2 | 0.5 | 1 | 1 | NA | NA | 3 | 0.6667 |
| 2_5965938 | NA | NA | 2 | 1 | 3 | 0.6667 | 2 | 1 | 7 | 0.8571 | NA | NA | 3 | 1 | 1 | 1 | NA | NA | 4 | 1 |
| 2_5966023 | NA | NA | 1 | 1 | 3 | 0 | NA | NA | 4 | 0.25 | NA | NA | 3 | 0.6667 | NA | NA | NA | NA | 3 | 0.6667 |
| 2_5966037 | NA | NA | 1 | 1 | 1 | 1 | 1 | 1 | 3 | 1 | NA | NA | 2 | 1 | 1 | 1 | NA | NA | 3 | 1 |
| 2_5966168 | 1 | 1 | 1 | 1 | 2 | 0 | 2 | 0.5 | 6 | 0.5 | NA | NA | 3 | 0.6667 | 2 | 0.5 | NA | NA | 5 | 0.6 |
| 2_5966172 | NA | NA | 1 | 1 | 3 | 0.6667 | 1 | 0 | 5 | 0.6 | NA | NA | 3 | 0.6667 | 2 | 0 | NA | NA | 5 | 0.4 |
| 2_5966195 | 1 | 0 | 2 | 0 | 3 | 0 | 1 | 1 | 7 | 0.1429 | NA | NA | 3 | 0.6667 | 2 | 0.5 | NA | NA | 5 | 0.6 |
| 2_5966245 | 1 | 1 | 1 | 1 | 2 | 1 | 1 | 1 | 5 | 1 | NA | NA | 2 | 1 | 2 | 1 | NA | NA | 4 | 1 |
| 2_5966341 | NA | NA | 3 | 0.6667 | 3 | 0.6667 | 2 | 1 | 8 | 0.75 | NA | NA | 3 | 1 | 2 | 1 | NA | NA | 5 | 1 |
| 2_5966487 | 1 | 0 | 3 | 0.3333 | 3 | 0 | 2 | 0 | 9 | 0.1111 | NA | NA | 3 | 0.3333 | 1 | 0 | NA | NA | 4 | 0.25 |
| 2_5966505 | 1 | 1 | 3 | 0 | 3 | 0.3333 | 2 | 0 | 9 | 0.2222 | NA | NA | 3 | 0.3333 | 2 | 0 | NA | NA | 5 | 0.2 |
| 2_6030975 | 1 | 0 | 4 | 0 | 3 | 0 | 4 | 0.25 | 12 | 0.0833 | NA | NA | 4 | 0 | 4 | 0 | 2 | 0.5 | 10 | 0.1 |
| 2_6032123 | NA | NA | 2 | 0 | 3 | 0 | 5 | 0.4 | 10 | 0.2 | NA | NA | 2 | 0.5 | 2 | 0.5 | 1 | 1 | 5 | 0.6 |
| 2_6060338 | 4 | 0.5 | 6 | 0.3333 | 5 | 0 | 8 | 0 | 23 | 0.1739 | NA | NA | 4 | 0.25 | 5 | 0 | 4 | 0 | 13 | 0.0769 |
| 2_6060385 | 4 | 0.75 | 4 | 0.25 | 5 | 0.4 | 8 | 0.625 | 21 | 0.5238 | NA | NA | 2 | 0 | 5 | 0.4 | 4 | 0.75 | 11 | 0.4545 |
| 2_6060540 | 3 | 0.3333 | 6 | 0 | 4 | 0 | 7 | 0 | 20 | 0.05 | NA | NA | 3 | 0 | 5 | 0.2 | 5 | 0 | 13 | 0.0769 |
| 2_6060580 | 4 | 0.5 | 6 | 0.1667 | 3 | 0 | 7 | 0 | 20 | 0.15 | NA | NA | 3 | 0.6667 | 5 | 0.4 | 3 | 0.3333 | 11 | 0.4545 |
| 2_6060597 | 4 | 0.25 | 5 | 0 | 1 | 0 | 4 | 0 | 14 | 0.0714 | NA | NA | 4 | 0.25 | 2 | 0 | 3 | 0.3333 | 9 | 0.2222 |
| 2_6060630 | 4 | 0.75 | 6 | 1 | 5 | 1 | 6 | 0.8333 | 21 | 0.9048 | NA | NA | 4 | 1 | 5 | 1 | 5 | 1 | 14 | 1 |
| 2_6060715 | 3 | 0 | 6 | 0.3333 | 5 | 0.4 | 8 | 0.25 | 22 | 0.2727 | NA | NA | 4 | 0.75 | 6 | 0.6667 | 5 | 0.6 | 15 | 0.6667 |
| 2_6060929 | 2 | 0 | 6 | 0.3333 | 6 | 0.6667 | 8 | 0.125 | 22 | 0.3182 | NA | NA | 4 | 0 | 7 | 0.4286 | 5 | 0.4 | 16 | 0.3125 |
| 2_6060938 | 3 | 1 | 6 | 0.3333 | 4 | 0.5 | 8 | 0.375 | 21 | 0.4762 | NA | NA | 3 | 0.3333 | 5 | 0.8 | 5 | 0.2 | 13 | 0.4615 |
| 2_6061075 | 5 | 0 | 6 | 0.1667 | 4 | 0 | 6 | 0.3333 | 21 | 0.1429 | NA | NA | 4 | 0.25 | 7 | 0 | 5 | 0 | 16 | 0.0625 |
| 2_6445463 | 1 | 1 | 1 | 1 | 1 | 1 | 1 | 1 | 4 | 1 | NA | NA | NA | NA | NA | NA | 1 | 1 | 1 | 1 |
| 2_6445582 | NA | NA | 1 | 1 | 1 | 0 | 1 | 1 | 3 | 0.6667 | NA | NA | NA | NA | NA | NA | 1 | 1 | 1 | 1 |
| 2_6445660 | 1 | 0 | 1 | 1 | 1 | 0 | 1 | 0 | 4 | 0.25 | NA | NA | NA | NA | NA | NA | 1 | 1 | 1 | 1 |
| 2_6446217 | NA | NA | 1 | 1 | 1 | 1 | 1 | 1 | 3 | 1 | NA | NA | NA | NA | NA | NA | 1 | 1 | 1 | 1 |
| 2_6446240 | 1 | 1 | 1 | 1 | 1 | 1 | NA | NA | 3 | 1 | NA | NA | NA | NA | NA | NA | 1 | 1 | 1 | 1 |
| 2_6446314 | 1 | 1 | 1 | 1 | NA | NA | 1 | 1 | 3 | 1 | NA | NA | NA | NA | NA | NA | 1 | 1 | 1 | 1 |
| 2_6446354 | 1 | 0 | 1 | 1 | 1 | 0 | 1 | 1 | 4 | 0.5 | NA | NA | NA | NA | NA | NA | 1 | 1 | 1 | 1 |
| 2_6451056 | 2 | 0.5 | 2 | 0 | 2 | 0 | NA | NA | 6 | 0.1667 | NA | NA | NA | NA | 1 | 1 | NA | NA | 1 | 1 |
| 2_6544522 | 5 | 0.2 | 3 | 0 | 8 | 0.25 | 8 | 0 | 24 | 0.125 | 4 | 0 | 15 | 0.0667 | 7 | 0.2857 | 11 | 0.0909 | 37 | 0.1081 |
| 2_6547080 | 15 | 0.1333 | 28 | 0.0714 | 24 | 0.0417 | 17 | 0.1765 | 84 | 0.0952 | 16 | 0.1875 | 19 | 0.1579 | 25 | 0.12 | 26 | 0.2308 | 86 | 0.1744 |
| 2_6547608 | 12 | 0.0833 | 21 | 0.2381 | 18 | 0.2222 | 20 | 0.25 | 71 | 0.2113 | 14 | 0.2857 | 17 | 0.2353 | 21 | 0.2381 | 18 | 0.3333 | 70 | 0.2714 |
| 2_6547630 | 15 | 0.2667 | 28 | 0.3929 | 24 | 0.2917 | 19 | 0.5789 | 86 | 0.3837 | 16 | 0.4375 | 21 | 0.4762 | 27 | 0.4074 | 20 | 0.7 | 84 | 0.5 |
| 2_6547746 | 15 | 0.2667 | 29 | 0.1724 | 25 | 0.08 | 21 | 0.0476 | 90 | 0.1333 | 17 | 0.1176 | 19 | 0.1053 | 26 | 0.1538 | 21 | 0.381 | 83 | 0.1928 |
| 2_6551481 | 10 | 0.2 | 20 | 0.15 | 9 | 0.4444 | 11 | 0.2727 | 50 | 0.24 | 11 | 0 | 19 | 0.2632 | 17 | 0.2941 | 9 | 0.1111 | 56 | 0.1964 |
| 2_6551567 | 14 | 0 | 23 | 0.0435 | 11 | 0.0909 | 19 | 0.1053 | 67 | 0.0597 | 12 | 0 | 24 | 0.0833 | 22 | 0.1364 | 12 | 0.0833 | 70 | 0.0857 |
| 2_6551804 | 14 | 0.1429 | 20 | 0.1 | 12 | 0.25 | 14 | 0 | 60 | 0.1167 | 10 | 0.2 | 24 | 0.0833 | 24 | 0.125 | 10 | 0.2 | 68 | 0.1324 |
| 2_6877809 | 1 | 1 | NA | NA | NA | NA | 1 | 1 | 2 | 1 | NA | NA | NA | NA | 1 | 1 | NA | NA | 1 | 1 |
| 2_6877872 | 1 | 1 | NA | NA | NA | NA | 1 | 1 | 2 | 1 | NA | NA | NA | NA | 1 | 1 | NA | NA | 1 | 1 |
| 2_9919840 | 1 | 0 | NA | NA | 2 | 0.5 | NA | NA | 3 | 0.3333 | NA | NA | 2 | 1 | 2 | 0.5 | NA | NA | 4 | 0.75 |
| 2_9920142 | 1 | 0 | 1 | 0 | 2 | 0.5 | 1 | 0 | 5 | 0.2 | 2 | 0 | 3 | 0 | 1 | 1 | NA | NA | 6 | 0.1667 |
| 2_9920517 | 1 | 1 | 1 | 1 | 1 | 1 | 1 | 1 | 4 | 1 | 1 | 1 | 1 | 1 | 2 | 1 | 1 | 1 | 5 | 1 |
| 2_9920811 | 1 | 0 | 1 | 1 | NA | NA | NA | NA | 2 | 0.5 | 1 | 0 | NA | NA | 1 | 1 | NA | NA | 2 | 0.5 |
| 2_9920917 | 1 | 0 | 1 | 1 | 1 | 1 | NA | NA | 3 | 0.6667 | 1 | 1 | 1 | 1 | 2 | 0 | NA | NA | 4 | 0.5 |
| 2_9921043 | NA | NA | 2 | 0 | 1 | 1 | NA | NA | 3 | 0.3333 | 1 | 1 | NA | NA | 2 | 0.5 | NA | NA | 3 | 0.6667 |
| 2_9921049 | NA | NA | 1 | 1 | 1 | 1 | NA | NA | 2 | 1 | 1 | 1 | 1 | 0 | 2 | 0.5 | NA | NA | 4 | 0.5 |
| 2_9921077 | NA | NA | 1 | 1 | 1 | 1 | NA | NA | 2 | 1 | 1 | 1 | 1 | 1 | 2 | 1 | NA | NA | 4 | 1 |

| | | | | | | | | | | | | | | | | | | | | |
|---|---|---|---|---|---|---|---|---|---|---|---|---|---|---|---|---|---|---|---|---|
| 2_9921143 | NA | NA | 1 | 1 | 1 | 1 | NA | NA | 2 | 1 | 1 | 0 | NA | NA | 2 | 0.5 | NA | NA | 3 | 0.3333 |
| 2_9956099 | NA | NA | 1 | 1 | 1 | 1 | NA | NA | 2 | 1 | 1 | 1 | 1 | 0 | 2 | 1 | NA | NA | 4 | 0.75 |
| 2_9956579 | NA | NA | 1 | 1 | 1 | 0 | NA | NA | 2 | 0.5 | 1 | 0 | 1 | 0 | 2 | 0.5 | NA | NA | 4 | 0.25 |
| 2_9956770 | NA | NA | 1 | 1 | 1 | 1 | NA | NA | 2 | 1 | 1 | 1 | 1 | 1 | 2 | 0.5 | NA | NA | 4 | 0.75 |
| 2_9957558 | NA | NA | 1 | 1 | NA | NA | NA | NA | 1 | 1 | NA | NA | 1 | 0 | 2 | 0.5 | NA | NA | 3 | 0.3333 |
| 2_1008496 | 2 | 0.5 | 1 | 0 | NA | NA | NA | NA | 3 | 0.3333 | 4 | 0.75 | 1 | 0 | 1 | 1 | NA | NA | 6 | 0.6667 |
| 2_1009673 | 2 | 0.5 | 1 | 1 | 1 | 1 | 1 | 0 | 5 | 0.6 | 2 | 1 | 1 | 1 | NA | NA | 3 | 0.6667 | 6 | 0.8333 |
| 2_1009683 | NA | NA | 1 | 0 | 1 | 1 | 1 | 0 | 3 | 0.3333 | 2 | 0.5 | 1 | 0 | 1 | 0 | 3 | 0 | 7 | 0.1429 |
| 2_1009707 | 2 | 0 | 1 | 1 | 2 | 0 | 1 | 0 | 6 | 0.1667 | 2 | 0 | 1 | 0 | 2 | 0 | 3 | 0.3333 | 8 | 0.125 |
| 2_1009710 | 2 | 0.5 | 1 | 0 | 2 | 0 | 1 | 0 | 6 | 0.1667 | 2 | 0.5 | 1 | 0 | 1 | 0 | 3 | 0 | 7 | 0.1429 |
| 2_1009710 | 2 | 0.5 | 1 | 1 | 2 | 0.5 | 1 | 0 | 6 | 0.5 | 1 | 1 | 1 | 1 | 1 | 1 | 3 | 1 | 6 | 1 |
| 2_1009712 | 2 | 0 | 1 | 0 | 2 | 1 | 1 | 1 | 6 | 0.5 | 2 | 0.5 | 1 | 1 | 2 | 0 | 2 | 1 | 7 | 0.5714 |
| 2_1009712 | 2 | 1 | 1 | 1 | 2 | 0 | 1 | 1 | 6 | 0.6667 | 1 | 0 | 1 | 1 | 2 | 0 | 3 | 0.3333 | 7 | 0.2857 |
| 2_1009713 | 1 | 0 | 1 | 0 | 2 | 0 | 1 | 1 | 5 | 0.2 | 2 | 0.5 | 1 | 0 | 2 | 0 | 3 | 0.3333 | 8 | 0.25 |
| 2_1009714 | 2 | 1 | NA | NA | 2 | 1 | NA | NA | 4 | 1 | 2 | 0.5 | NA | NA | 2 | 0.5 | 3 | 1 | 7 | 0.7143 |
| 2_1009726 | 2 | 1 | 1 | 1 | 2 | 0.5 | 1 | 1 | 6 | 0.8333 | 2 | 0.5 | 1 | 1 | 2 | 1 | 3 | 1 | 8 | 0.875 |
| 2_1009735 | 3 | 0 | 1 | 1 | 2 | 1 | 1 | 1 | 7 | 0.5714 | 2 | 0.5 | 1 | 1 | 2 | 0 | 2 | 0 | 7 | 0.2857 |
| 2_1025939 | 2 | 0 | 5 | 0 | 2 | 0.5 | NA | NA | 9 | 0.1111 | 1 | 0 | NA | NA | 3 | 0.3333 | 1 | 0 | 5 | 0.2 |
| 2_1026092 | 4 | 0 | 5 | 0.4 | 5 | 0.6 | 2 | 0.5 | 16 | 0.375 | 1 | 0 | NA | NA | 3 | 0.3333 | NA | NA | 4 | 0.25 |
| 2_1128859 | 1 | 0 | 1 | 0 | 1 | 0 | 1 | 1 | 4 | 0.25 | 1 | 0 | 1 | 0 | 2 | 0.5 | 1 | 0 | 5 | 0.2 |
| 2_1130491 | 1 | 1 | 2 | 1 | 1 | 0 | NA | NA | 4 | 0.75 | 1 | 0 | 1 | 1 | 1 | 1 | 1 | 1 | 4 | 0.75 |
| 2_1130495 | 1 | 1 | 2 | 1 | 1 | 0 | 1 | 1 | 5 | 0.8 | 1 | 0 | 1 | 0 | 2 | 0 | 1 | 1 | 5 | 0.2 |
| 2_1130498 | 1 | 1 | 2 | 1 | 1 | 1 | 1 | 1 | 5 | 1 | NA | NA | 1 | 1 | 1 | 1 | 1 | 1 | 3 | 1 |
| 2_1130506 | 1 | 1 | 2 | 0.5 | 1 | 1 | 1 | 1 | 5 | 0.8 | NA | NA | 1 | 1 | 2 | 1 | 1 | 1 | 4 | 1 |
| 2_1130508 | 1 | 0 | 1 | 1 | 1 | 0 | NA | NA | 3 | 0.3333 | 1 | 0 | 1 | 1 | 2 | 0.5 | 1 | 1 | 5 | 0.6 |
| 2_1130520 | 1 | 0 | 1 | 1 | 1 | 1 | 1 | 0 | 4 | 0.5 | NA | NA | 2 | 0.5 | 2 | 0.5 | 1 | 1 | 5 | 0.6 |
| 2_1147760 | 3 | 0.3333 | NA | NA | 2 | 0 | 1 | 0 | 6 | 0.1667 | 2 | 0 | 2 | 0 | 2 | 0 | 3 | 0.3333 | 9 | 0.1111 |
| 2_1147764 | 3 | 0.6667 | NA | NA | 2 | 0.5 | 1 | 1 | 6 | 0.6667 | 2 | 0.5 | 2 | 1 | 1 | 0 | 3 | 0.3333 | 8 | 0.5 |
| 2_1147797 | 2 | 0 | 1 | 0 | 3 | 0 | 1 | 1 | 7 | 0.1429 | 2 | 0.5 | 1 | 0 | 2 | 0 | 3 | 0.3333 | 8 | 0.25 |
| 2_1147879 | 2 | 0.5 | 1 | 1 | 3 | 0 | 1 | 0 | 7 | 0.2857 | NA | NA | 2 | 0 | 2 | 0.5 | 2 | 0.5 | 6 | 0.3333 |
| 2_1147909 | 1 | 0 | 1 | 1 | 3 | 0 | 1 | 0 | 6 | 0.1667 | NA | NA | 2 | 0 | 2 | 0 | 2 | 0.5 | 6 | 0.1667 |
| 2_1159032 | 8 | 0.75 | 6 | 0.3333 | 8 | 0.25 | 11 | 0.4545 | 33 | 0.4545 | 7 | 0.2857 | 13 | 0.3846 | 6 | 0.5 | 8 | 0.875 | 34 | 0.5 |
| 2_1159039 | 8 | 0.375 | 6 | 0.3333 | 7 | 0 | 9 | 0.3333 | 30 | 0.2667 | 9 | 0.2222 | 12 | 0.3333 | 6 | 0.5 | 7 | 0.5714 | 34 | 0.3824 |
| 2_1173838 | 1 | 1 | 1 | 1 | NA | NA | 1 | 0 | 3 | 0.6667 | 2 | 1 | NA | NA | NA | NA | NA | NA | 2 | 1 |
| 2_1173861 | NA | NA | 2 | 1 | NA | NA | 1 | 0 | 3 | 0.6667 | 2 | 0.5 | NA | NA | NA | NA | NA | NA | 2 | 0.5 |
| 2_1174749 | 9 | 0.2222 | 7 | 0 | 7 | 0 | 9 | 0.1111 | 32 | 0.0938 | 8 | 0.125 | 10 | 0.1 | 12 | 0 | 6 | 0.1667 | 36 | 0.0833 |
| 2_1174756 | 7 | 0.2857 | 6 | 0.3333 | 6 | 0.1667 | 7 | 0.4286 | 26 | 0.3077 | 8 | 0.375 | 10 | 0.6 | 11 | 0.0909 | 5 | 0.2 | 34 | 0.3235 |
| 2_1174761 | 10 | 0.4 | 7 | 0.2857 | 7 | 0.4286 | 10 | 0.2 | 34 | 0.3235 | 8 | 0.125 | 11 | 0.3636 | 12 | 0.0833 | 5 | 0.4 | 36 | 0.2222 |
| 2_1175408 | 1 | 0 | NA | NA | 1 | 1 | NA | NA | 2 | 0.5 | NA | NA | 1 | 0 | 1 | 1 | NA | NA | 2 | 0.5 |
| 2_1176236 | 7 | 0.1429 | 7 | 0.1429 | 3 | 0 | 6 | 0.1667 | 23 | 0.1304 | 3 | 0 | 6 | 0.3333 | 5 | 0 | 4 | 0 | 18 | 0.1111 |
| 2_1177210 | 5 | 0 | 2 | 0.5 | 2 | 1 | 2 | 0 | 11 | 0.2727 | 2 | 0.5 | 4 | 0.25 | 4 | 0 | 4 | 0.25 | 14 | 0.2143 |
| 2_1177212 | 4 | 0.5 | 4 | 0.5 | 2 | 0.5 | 2 | 0.5 | 12 | 0.5 | 2 | 1 | 3 | 0.3333 | 4 | 1 | 3 | 1 | 12 | 0.8333 |
| 2_1177213 | 4 | 0.25 | 4 | 0.5 | 2 | 0.5 | 2 | 0.5 | 12 | 0.4167 | 2 | 0.5 | 3 | 0 | 3 | 0 | 4 | 0.75 | 12 | 0.3333 |
| 2_1177218 | 3 | 0.3333 | 3 | 0.6667 | 2 | 1 | 2 | 0.5 | 10 | 0.6 | 3 | 0.3333 | 3 | 1 | 3 | 0.6667 | 3 | 0.6667 | 12 | 0.6667 |
| 2_1177242 | 4 | 0 | 3 | 0 | 2 | 0.5 | 2 | 0 | 11 | 0.0909 | 2 | 0.5 | 3 | 0.6667 | 3 | 0.3333 | 3 | 0.3333 | 11 | 0.4545 |
| 2_1179467 | 3 | 0 | 3 | 0.3333 | 3 | 0 | 4 | 0.75 | 13 | 0.3077 | 3 | 0.3333 | 3 | 0.3333 | 3 | 0.3333 | 3 | 0.3333 | 12 | 0.3333 |
| 2_1179476 | 3 | 0 | 2 | 0 | 3 | 0.6667 | 4 | 0 | 12 | 0.1667 | 3 | 0 | 5 | 0.4 | 2 | 0 | 3 | 0.6667 | 13 | 0.3077 |
| 2_1179491 | 3 | 0.3333 | 3 | 1 | 2 | 0.5 | 3 | 0.3333 | 11 | 0.5455 | 3 | 0.6667 | 3 | 0.6667 | NA | NA | 2 | 0.5 | 8 | 0.625 |
| 2_1179493 | 3 | 0.3333 | 3 | 0 | 2 | 0 | 4 | 0.25 | 12 | 0.1667 | 3 | 0 | 5 | 0.4 | 3 | 0.6667 | 2 | 0 | 13 | 0.3077 |
| 2_1179494 | 3 | 1 | 3 | 0.6667 | 3 | 0.6667 | 3 | 0.6667 | 12 | 0.75 | 2 | 0 | 4 | 1 | 3 | 0.6667 | 2 | 0.5 | 11 | 0.6364 |
| 2_1211317 | NA | NA | 2 | 0.5 | NA | NA | 1 | 0 | 3 | 0.3333 | 2 | 1 | NA | NA | NA | NA | 1 | 1 | 3 | 1 |
| 2_1211344 | NA | NA | 2 | 0.5 | NA | NA | 1 | 1 | 3 | 0.6667 | 2 | 1 | NA | NA | NA | NA | 1 | 1 | 3 | 1 |
| 2_1211346 | NA | NA | 2 | 1 | NA | NA | 1 | 1 | 3 | 1 | 2 | 1 | NA | NA | NA | NA | 1 | 1 | 3 | 1 |
| 2_1211356 | NA | NA | 1 | 1 | NA | NA | 1 | 1 | 2 | 1 | 2 | 1 | NA | NA | NA | NA | 1 | 0 | 3 | 0.6667 |
| 2_1211357 | NA | NA | 2 | 1 | NA | NA | 1 | 1 | 3 | 1 | 1 | 1 | NA | NA | NA | NA | 1 | 1 | 2 | 1 |
| 2_1211359 | NA | NA | 1 | 0 | NA | NA | 1 | 1 | 2 | 0.5 | 2 | 1 | NA | NA | NA | NA | NA | NA | 2 | 1 |
| 2_1211360 | NA | NA | 2 | 1 | NA | NA | 1 | 1 | 3 | 1 | 1 | 1 | NA | NA | NA | NA | 1 | 1 | 2 | 1 |
| 2_1211386 | NA | NA | 2 | 0.5 | NA | NA | 1 | 1 | 3 | 0.6667 | 1 | 1 | NA | NA | NA | NA | 1 | 1 | 2 | 1 |
| 2_1235289 | 7 | 0.1429 | 10 | 0.1 | 8 | 0 | 2 | 0 | 27 | 0.0741 | NA | NA | 5 | 0 | 10 | 0.1 | 5 | 0 | 20 | 0.05 |
| 2_1235299 | 8 | 0.25 | 9 | 0.1111 | 10 | 0.1 | 5 | 0 | 32 | 0.125 | NA | NA | 5 | 0.2 | 9 | 0.1111 | 5 | 0.2 | 19 | 0.1579 |
| 2_1242400 | 5 | 0 | 8 | 0.125 | 8 | 0.25 | 6 | 0 | 27 | 0.1111 | NA | NA | 3 | 0 | 9 | 0.1111 | 3 | 0 | 15 | 0.0667 |
| 2_1300670 | 11 | 0.2727 | 18 | 0.3889 | 10 | 0.3 | 7 | 0.5714 | 46 | 0.3696 | 12 | 0.1667 | 2 | 0.5 | 3 | 0.6667 | 9 | 0.2222 | 26 | 0.2692 |
| 2_1300680 | 10 | 0 | 16 | 0.375 | 11 | 0.0909 | 7 | 0.2857 | 44 | 0.2045 | 10 | 0.4 | 2 | 0.5 | 2 | 0.5 | 11 | 0.5455 | 25 | 0.48 |
| 2_1300877 | 7 | 0.2857 | 12 | 0.25 | 4 | 0.25 | 4 | 0.5 | 27 | 0.2963 | 11 | 0.1818 | 1 | 1 | 3 | 0 | 7 | 0.1429 | 22 | 0.1818 |
| 2_1300880 | 7 | 0.1429 | 12 | 0.25 | 6 | 0 | 4 | 0.25 | 29 | 0.1724 | 10 | 0.1 | 1 | 1 | 3 | 0 | 8 | 0 | 22 | 0.0909 |

| | | | | | | | | | | | | | | | | | | | | |
|---|---|---|---|---|---|---|---|---|---|---|---|---|---|---|---|---|---|---|---|---|
| 2_1300891 | 6 | 0.3333 | 12 | 0.1667 | 5 | 0.2 | 4 | 0.5 | 27 | 0.2593 | 9 | 0.2222 | 1 | 1 | 2 | 0 | 8 | 0.25 | 20 | 0.25 |
| 2_1300893 | 6 | 0.3333 | 11 | 0.4545 | 5 | 0.4 | 4 | 0.25 | 26 | 0.3846 | 8 | 0.375 | 1 | 1 | 3 | 0.6667 | 7 | 0.1429 | 19 | 0.3684 |
| 2_1300906 | 5 | 1 | 9 | 1 | 4 | 0.75 | 3 | 1 | 21 | 0.9524 | 8 | 1 | 1 | 1 | 3 | 1 | 6 | 1 | 18 | 1 |
| 2_1300908 | 3 | 1 | 7 | 1 | 4 | 1 | 2 | 1 | 16 | 1 | 8 | 1 | 1 | 1 | NA | NA | 5 | 1 | 14 | 1 |
| 2_1301551 | 2 | 0.5 | 8 | 0.375 | 4 | 0.5 | 2 | 0.5 | 16 | 0.4375 | 4 | 0.25 | 1 | 1 | 1 | 0 | 4 | 0 | 10 | 0.2 |
| 2_1301591 | 2 | 1 | 4 | 1 | 3 | 1 | 1 | 1 | 10 | 1 | 2 | 1 | 1 | 1 | 2 | 0.5 | 5 | 1 | 10 | 0.9 |
| 2_1301613 | 2 | 0.5 | 4 | 0.75 | 3 | 0.3333 | 2 | 0.5 | 11 | 0.5455 | 2 | 0.5 | NA | NA | 2 | 0 | 3 | 0.3333 | 7 | 0.2857 |
| 2_1301621 | 1 | 1 | 5 | 0.2 | 3 | 0.6667 | 2 | 0.5 | 11 | 0.4545 | NA | NA | 1 | 0 | 2 | 0.5 | 3 | 0.3333 | 6 | 0.3333 |
| 2_1301621 | 2 | 0 | 5 | 0 | 3 | 0.3333 | 2 | 0 | 12 | 0.0833 | 2 | 0 | 1 | 0 | 2 | 0 | 4 | 0.25 | 9 | 0.1111 |
| 2_1357949 | 7 | 0.2857 | 9 | 0.1111 | 5 | 0 | 6 | 0.5 | 27 | 0.2222 | 8 | 0.375 | 3 | 0 | 3 | 0.6667 | 6 | 0.3333 | 20 | 0.35 |
| 2_1415117 | 7 | 0.7143 | 10 | 0.5 | 1 | 0 | 7 | 0.4286 | 25 | 0.52 | 7 | 0.5714 | 4 | 0.25 | 6 | 0.6667 | 6 | 0.6667 | 23 | 0.5652 |
| 2_1415119 | 7 | 0.2857 | 8 | 0.125 | 1 | 0 | 8 | 0.125 | 24 | 0.1667 | 5 | 0.4 | 4 | 0 | 5 | 0.2 | 5 | 0.2 | 19 | 0.2105 |
| 2_1415129 | 8 | 1 | 9 | 0.8889 | NA | NA | 8 | 1 | 25 | 0.96 | 7 | 1 | 5 | 1 | 4 | 1 | 5 | 1 | 21 | 1 |
| 2_1415133 | 8 | 1 | 9 | 0.6667 | NA | NA | 8 | 0.875 | 25 | 0.84 | 6 | 1 | 5 | 1 | 6 | 0.6667 | 6 | 1 | 23 | 0.913 |
| 2_1415143 | 7 | 1 | 10 | 1 | 1 | 1 | 7 | 1 | 25 | 1 | 7 | 1 | 4 | 1 | 6 | 1 | 6 | 0.8333 | 23 | 0.9565 |
| 2_1415148 | 9 | 0.1111 | 11 | 0.1818 | NA | NA | 8 | 0.25 | 28 | 0.1786 | 6 | 0.3333 | 7 | 0.1429 | 9 | 0.4444 | 6 | 0.8333 | 28 | 0.4286 |
| 2_1415304 | 3 | 0 | 5 | 0.2 | 2 | 0 | 3 | 0 | 13 | 0.0769 | 4 | 0 | 2 | 0 | 4 | 0.25 | 3 | 0.3333 | 13 | 0.1538 |
| 2_1439413 | 7 | 0 | 2 | 0 | 8 | 0.125 | 3 | 0 | 20 | 0.05 | 3 | 0 | 3 | 0 | 5 | 0.2 | 5 | 0 | 16 | 0.0625 |
| 2_1439434 | 7 | 0 | 2 | 0.5 | 5 | 0.2 | 3 | 0.3333 | 17 | 0.1765 | 5 | 0.2 | 4 | 0.25 | 5 | 0.2 | 6 | 0.1667 | 20 | 0.2 |
| 2_1439436 | 7 | 0.1429 | 2 | 0 | 8 | 0.125 | 3 | 0 | 20 | 0.1 | 4 | 0.25 | 3 | 0 | 4 | 0 | 6 | 0.1667 | 17 | 0.1176 |
| 2_1445679 | 9 | 0.1111 | 2 | 0 | 6 | 0.1667 | 7 | 0 | 24 | 0.0833 | 5 | 0.2 | 2 | 0 | 5 | 0.4 | 6 | 0.3333 | 18 | 0.2778 |
| 2_1445680 | 9 | 0 | 2 | 0.5 | 8 | 0 | 7 | 0.1429 | 26 | 0.0769 | 6 | 0.1667 | 4 | 0 | 6 | 0 | 7 | 0.1429 | 23 | 0.087 |
| 2_1445685 | 10 | 0.1 | 2 | 0 | 8 | 0.25 | 7 | 0.2857 | 27 | 0.1852 | 5 | 0 | 3 | 0 | 6 | 0.1667 | 6 | 0.1667 | 20 | 0.1 |
| 2_1449678 | 1 | 1 | NA | NA | 1 | 0 | NA | NA | 2 | 0.5 | 1 | 1 | NA | NA | NA | NA | NA | NA | 1 | 1 |
| 2_1449703 | 1 | 0 | NA | NA | 1 | 1 | NA | NA | 2 | 0.5 | 1 | 1 | NA | NA | NA | NA | NA | NA | 1 | 1 |
| 2_1449720 | 1 | 1 | NA | NA | NA | NA | NA | NA | 1 | 1 | 1 | 1 | NA | NA | NA | NA | NA | NA | 1 | 1 |
| 2_1449738 | 1 | 1 | NA | NA | NA | NA | NA | NA | 1 | 1 | 1 | 1 | NA | NA | NA | NA | NA | NA | 1 | 1 |
| 2_1449764 | 1 | 1 | NA | NA | 1 | 1 | NA | NA | 2 | 1 | 1 | 1 | NA | NA | NA | NA | NA | NA | 1 | 1 |
| 2_1463176 | 1 | 1 | NA | NA | NA | NA | NA | NA | 1 | 1 | NA | NA | 1 | 1 | NA | NA | NA | NA | 1 | 1 |
| 2_1463188 | 1 | 1 | NA | NA | NA | NA | NA | NA | 1 | 1 | NA | NA | 1 | 1 | NA | NA | NA | NA | 1 | 1 |
| 2_1463197 | 1 | 1 | NA | NA | NA | NA | NA | NA | 1 | 1 | NA | NA | 1 | 1 | NA | NA | NA | NA | 1 | 1 |
| 2_1463224 | 1 | 1 | NA | NA | NA | NA | NA | NA | 1 | 1 | NA | NA | 1 | 1 | NA | NA | NA | NA | 1 | 1 |
| 2_1463235 | 1 | 1 | NA | NA | NA | NA | NA | NA | 1 | 1 | NA | NA | 1 | 1 | NA | NA | NA | NA | 1 | 1 |
| 2_1463241 | 1 | 1 | NA | NA | NA | NA | NA | NA | 1 | 1 | NA | NA | 1 | 1 | NA | NA | NA | NA | 1 | 1 |
| 2_1499053 | 3 | 0 | 7 | 0.2857 | 3 | 0.3333 | 6 | 0.1667 | 19 | 0.2105 | 1 | 0 | 5 | 0.8 | 6 | 0.3333 | 2 | 1 | 14 | 0.5714 |
| 2_1499057 | 3 | 0.3333 | 6 | 0.5 | 3 | 1 | 4 | 0.5 | 16 | 0.5625 | 2 | 0 | 6 | 0.8333 | 8 | 0.5 | 3 | 0.3333 | 19 | 0.5263 |
| 2_1499059 | 4 | 0.25 | 8 | 0.375 | 3 | 0 | 7 | 0.4286 | 22 | 0.3182 | 2 | 0 | 6 | 0 | 7 | 0.4286 | 3 | 0.3333 | 18 | 0.2222 |
| 2_1499063 | 2 | 0 | 2 | 0.5 | 1 | 0 | 3 | 0.3333 | 8 | 0.25 | 2 | 1 | 1 | 0 | 4 | 0.5 | 2 | 0 | 9 | 0.4444 |
| 2_1499064 | 3 | 0.3333 | 6 | 0.3333 | 2 | 0.5 | 6 | 0 | 17 | 0.2353 | 3 | 0.6667 | 6 | 0.1667 | 6 | 0.1667 | 3 | 0 | 18 | 0.2222 |
| 2_1499070 | 4 | 0.5 | 5 | 0.2 | 4 | 0.25 | 5 | 0 | 18 | 0.2222 | 2 | 0 | 5 | 0.4 | 7 | 0.4286 | 3 | 0 | 17 | 0.2941 |
| 2_1499073 | 4 | 1 | 8 | 0.875 | 3 | 1 | 6 | 1 | 21 | 0.9524 | 3 | 1 | 6 | 0.6667 | 8 | 1 | 2 | 1 | 19 | 0.8947 |
| 2_1499075 | 4 | 0.5 | 8 | 0.375 | 3 | 0 | 7 | 0 | 22 | 0.2273 | 3 | 0 | 5 | 0.4 | 7 | 0.1429 | 3 | 0.6667 | 18 | 0.2778 |
| 2_1499081 | 3 | 0.3333 | 9 | 0.1111 | 3 | 0.3333 | 5 | 0 | 20 | 0.15 | 2 | 0 | 4 | 0.25 | 7 | 0.2857 | 3 | 0 | 16 | 0.1875 |
| 2_1499086 | 4 | 0 | 7 | 0.2857 | 2 | 0.5 | 5 | 0.2 | 18 | 0.2222 | 2 | 0 | 2 | 0.5 | 5 | 0 | 1 | 0 | 10 | 0.1 |
| 2_1499107 | 6 | 0 | 8 | 0.375 | 2 | 0.5 | 7 | 0.1429 | 23 | 0.2174 | 4 | 0 | 6 | 0.3333 | 7 | 0.1429 | 4 | 0 | 21 | 0.1429 |
| 2_1499109 | 4 | 0 | 7 | 0 | 3 | 0.3333 | 6 | 0 | 20 | 0.05 | 2 | 0 | 5 | 0.2 | 5 | 0.4 | 4 | 0.25 | 16 | 0.25 |
| 2_1499124 | 5 | 0.6 | 8 | 0.75 | 3 | 0.6667 | 7 | 0.5714 | 23 | 0.6522 | 3 | 0.3333 | 5 | 0.4 | 10 | 0.6 | 5 | 0.4 | 23 | 0.4783 |
| 2_1499156 | 6 | 0.1667 | 12 | 0.25 | 3 | 0 | 10 | 0.3 | 31 | 0.2258 | 5 | 0 | 8 | 0.5 | 9 | 0 | 8 | 0.25 | 30 | 0.2 |
| 2_1499163 | 5 | 0.4 | 15 | 0.0667 | 3 | 0 | 10 | 0 | 33 | 0.0909 | 6 | 0 | 8 | 0.125 | 9 | 0.2222 | 8 | 0.125 | 31 | 0.129 |
| 2_1499191 | 5 | 0.2 | 11 | 0.0909 | 5 | 0 | 7 | 0 | 28 | 0.0714 | 4 | 0 | 9 | 0.3333 | 11 | 0.0909 | 7 | 0.2857 | 31 | 0.1935 |
| 2_1499198 | 5 | 0.4 | 11 | 0.0909 | 4 | 0.5 | 6 | 0.1667 | 26 | 0.2308 | 6 | 0.3333 | 9 | 0.2222 | 10 | 0.4 | 7 | 0.2857 | 32 | 0.3125 |
| 2_1499214 | 5 | 0.8 | 12 | 0.9167 | 6 | 0.5 | 8 | 0.75 | 31 | 0.7742 | 5 | 1 | 11 | 1 | 10 | 0.9 | 5 | 1 | 31 | 0.9677 |
| 2_1499244 | 3 | 0 | 6 | 0.3333 | 3 | 0 | 6 | 0.6667 | 18 | 0.3333 | 1 | 1 | 3 | 0 | 5 | 0.2 | 3 | 0.3333 | 12 | 0.25 |
| 2_1499251 | 5 | 0.4 | 5 | 0.2 | 3 | 0.3333 | 5 | 0.4 | 18 | 0.3333 | 1 | 0 | 3 | 0.6667 | 5 | 0.2 | 5 | 0 | 14 | 0.2143 |
| 2_1499253 | 5 | 0 | 6 | 0.1667 | 2 | 0.5 | 6 | 0.3333 | 19 | 0.2105 | 1 | 1 | 3 | 0.6667 | 5 | 0 | 5 | 0 | 14 | 0.2143 |
| 2_1499259 | 4 | 0.5 | 6 | 0.3333 | 3 | 0.6667 | 6 | 0.6667 | 19 | 0.5263 | 1 | 1 | 3 | 0.3333 | 3 | 0.3333 | 3 | 0.3333 | 10 | 0.4 |
| 2_1499355 | 2 | 0 | 5 | 0.2 | 2 | 0 | 5 | 0.2 | 14 | 0.1429 | 1 | 0 | 2 | 0 | 4 | 0.25 | 2 | 0.5 | 9 | 0.2222 |
| 2_1500072 | 1 | 1 | 1 | 1 | 1 | 1 | 1 | 0 | 4 | 0.75 | NA | NA | NA | NA | 1 | 1 | NA | NA | 1 | 1 |
| 2_1500119 | 2 | 0.5 | 2 | 1 | 2 | 1 | 1 | 1 | 7 | 0.8571 | NA | NA | NA | NA | 1 | 1 | NA | NA | 1 | 1 |
| 2_1508137 | 1 | 1 | NA | NA | NA | NA | NA | NA | 1 | 1 | NA | NA | 1 | 1 | NA | NA | 2 | 0.5 | 3 | 0.6667 |
| 2_1508690 | 2 | 0.5 | 3 | 0 | NA | NA | 2 | 0 | 7 | 0.1429 | 1 | 0 | 5 | 0.2 | 1 | 0 | 4 | 0.5 | 11 | 0.2727 |
| 2_1508714 | 3 | 0.3333 | 1 | 0 | NA | NA | 2 | 0.5 | 6 | 0.3333 | 1 | 0 | 5 | 0.4 | 1 | 1 | 4 | 0.25 | 11 | 0.3636 |
| 2_1636162 | NA | NA | 1 | 1 | 1 | 0 | 1 | 0 | 3 | 0.3333 | 1 | 0 | 3 | 0 | 2 | 1 | 1 | 0 | 7 | 0.2857 |
| 2_1636179 | NA | NA | 2 | 0.5 | NA | NA | 3 | 0.3333 | 5 | 0.4 | 2 | 0.5 | 3 | 0.3333 | 2 | 0 | 1 | 1 | 8 | 0.375 |
| 2_1675612 | 2 | 0.5 | 2 | 1 | 1 | 1 | 5 | 0.4 | 10 | 0.6 | 5 | 0.2 | 3 | 0.6667 | 2 | 0.5 | 1 | 0 | 11 | 0.3636 |

| | | | | | | | | | | | | | | | | | | | | |
|---|---|---|---|---|---|---|---|---|---|---|---|---|---|---|---|---|---|---|---|---|
| 2_16756164 | 2 | 0.5 | 3 | 0.3333 | 3 | 0.3333 | 5 | 0.4 | 13 | 0.3846 | 4 | 0.75 | 2 | 1 | 2 | 1 | 1 | 0 | 9 | 0.7778 |
| 2_16756170 | 1 | 0 | 2 | 0 | 1 | 0 | 4 | 0.5 | 8 | 0.25 | 5 | 0.2 | 3 | 1 | 2 | 0 | 1 | 0 | 11 | 0.3636 |
| 2_16756198 | 1 | 0 | 3 | 0.3333 | 1 | 0 | 2 | 0 | 7 | 0.1429 | 5 | 0.2 | 4 | 0.25 | 2 | 0 | 1 | 0 | 12 | 0.1667 |
| 2_16756268 | 2 | 0.5 | 4 | 0.5 | 2 | 0 | 5 | 0 | 13 | 0.2308 | 5 | 0 | 4 | 0.25 | 2 | 0 | 1 | 1 | 12 | 0.1667 |
| 2_16756285 | 2 | 0.5 | 3 | 0.3333 | 2 | 0 | 4 | 0.5 | 11 | 0.3636 | 4 | 0.25 | 4 | 0.5 | 1 | 0 | NA | NA | 9 | 0.3333 |
| 2_16756299 | 3 | 0 | 4 | 0 | 2 | 0.5 | 5 | 0 | 14 | 0.0714 | 5 | 0.2 | 3 | 0 | 1 | 0 | 1 | 1 | 10 | 0.2 |
| 2_16756305 | 3 | 0.6667 | 4 | 0.5 | 2 | 0.5 | 5 | 0.4 | 14 | 0.5 | 5 | 0.4 | 3 | 0.3333 | 2 | 0 | 1 | 1 | 11 | 0.3636 |
| 2_16756388 | 3 | 0.3333 | 2 | 0.5 | 3 | 0.3333 | 4 | 0.5 | 12 | 0.4167 | 5 | 1 | 2 | 1 | 2 | 0.5 | 1 | 1 | 10 | 0.9 |
| 2_16756672 | 1 | 0 | 1 | 1 | 1 | 0 | 2 | 0.5 | 5 | 0.4 | NA | NA | 1 | 1 | NA | NA | NA | NA | 1 | 1 |
| 2_16756769 | 1 | 0 | 1 | 1 | 1 | 0 | 3 | 0 | 6 | 0.1667 | NA | NA | 2 | 1 | NA | NA | NA | NA | 2 | 1 |
| 2_16756975 | 1 | 1 | NA | NA | 1 | 0 | 2 | 0 | 4 | 0.25 | NA | NA | 2 | 1 | 1 | 0 | NA | NA | 3 | 0.6667 |
| 2_16773846 | NA | NA | NA | NA | 1 | 1 | 1 | 0 | 2 | 0.5 | NA | NA | NA | NA | NA | NA | 1 | 1 | 1 | 1 |
| 2_16774058 | 1 | 1 | NA | NA | 1 | 1 | 1 | 0 | 3 | 0.6667 | NA | NA | NA | NA | NA | NA | 1 | 1 | 1 | 1 |
| 2_16774154 | 1 | 1 | NA | NA | 1 | 0 | 1 | 0 | 3 | 0.3333 | NA | NA | NA | NA | NA | NA | 1 | 1 | 1 | 1 |
| 2_16774199 | 1 | 1 | NA | NA | 1 | 0 | 1 | 0 | 3 | 0.3333 | NA | NA | NA | NA | NA | NA | 1 | 1 | 1 | 1 |
| 2_17065148 | NA | NA | 3 | 0 | NA | NA | 4 | 0.25 | 7 | 0.1429 | 1 | 1 | NA | NA | NA | NA | 1 | 0 | 2 | 0.5 |
| 2_17065398 | NA | NA | 1 | 0 | NA | NA | 4 | 0.5 | 5 | 0.4 | 1 | 1 | NA | NA | NA | NA | NA | NA | 1 | 1 |
| 2_17066504 | NA | NA | 2 | 0 | NA | NA | 3 | 0.3333 | 5 | 0.2 | 1 | 1 | NA | NA | NA | NA | NA | NA | 1 | 1 |
| 2_17174386 | NA | NA | NA | NA | 1 | 1 | NA | NA | 1 | 1 | NA | NA | NA | NA | 1 | 1 | NA | NA | 1 | 1 |
| 2_17174523 | NA | NA | NA | NA | 1 | 1 | NA | NA | 1 | 1 | NA | NA | NA | NA | 1 | 1 | NA | NA | 1 | 1 |
| 2_17348561 | 2 | 0 | 4 | 0 | 6 | 0.1667 | 1 | 0 | 13 | 0.0769 | 1 | 0 | 4 | 0.25 | 1 | 0 | 2 | 0 | 8 | 0.125 |
| 2_17348596 | 4 | 0.25 | 5 | 0.2 | 7 | 0.1429 | 1 | 0 | 17 | 0.1765 | 2 | 0 | 4 | 0.25 | 2 | 0 | 2 | 0.5 | 10 | 0.2 |
| 2_17348784 | 4 | 0.25 | 5 | 0.4 | 7 | 0.2857 | NA | NA | 16 | 0.3125 | 2 | 0.5 | 4 | 0.75 | 2 | 0 | 2 | 0 | 10 | 0.4 |
| 2_17348805 | 4 | 0 | 3 | 0 | 6 | 0.1667 | NA | NA | 13 | 0.0769 | 2 | 0.5 | 4 | 0 | 2 | 0 | 2 | 0.5 | 10 | 0.2 |
| 2_17349128 | 9 | 0.1111 | 9 | 0.3333 | 12 | 0.25 | 3 | 0.3333 | 33 | 0.2424 | 7 | 0 | 7 | 0.2857 | 10 | 0.2 | 5 | 0.2 | 29 | 0.1724 |
| 2_17353347 | 1 | 0 | 4 | 0.25 | 4 | 0 | 3 | 0 | 12 | 0.0833 | 3 | 0.3333 | 4 | 0 | 3 | 0 | NA | NA | 10 | 0.1 |
| 2_18032885 | NA | NA | NA | NA | 3 | 0.6667 | NA | NA | 3 | 0.6667 | NA | NA | 1 | 1 | NA | NA | NA | NA | 1 | 1 |
| 2_18032943 | 2 | 0.5 | NA | NA | 2 | 1 | NA | NA | 4 | 0.75 | NA | NA | 1 | 1 | 1 | 0 | NA | NA | 2 | 0.5 |
| 2_18032998 | 2 | 0 | NA | NA | 3 | 0.3333 | NA | NA | 5 | 0.2 | NA | NA | 2 | 0.5 | 1 | 0 | NA | NA | 3 | 0.3333 |
| 2_18033067 | 2 | 1 | NA | NA | 3 | 1 | NA | NA | 5 | 1 | NA | NA | 2 | 1 | 1 | 1 | NA | NA | 3 | 1 |
| 2_18050173 | NA | NA | 1 | 1 | 1 | 1 | NA | NA | 2 | 1 | NA | NA | NA | NA | NA | NA | 2 | 1 | 2 | 1 |
| 2_18050392 | NA | NA | NA | NA | 1 | 1 | NA | NA | 1 | 1 | NA | NA | NA | NA | NA | NA | 1 | 1 | 1 | 1 |
| 2_18050693 | NA | NA | NA | NA | 1 | 1 | NA | NA | 1 | 1 | NA | NA | NA | NA | NA | NA | 1 | 1 | 1 | 1 |
| 2_18050801 | NA | NA | NA | NA | 1 | 1 | NA | NA | 1 | 1 | NA | NA | NA | NA | 1 | 1 | 1 | 1 | 2 | 1 |
| 2_18050984 | NA | NA | NA | NA | 1 | 1 | NA | NA | 1 | 1 | NA | NA | NA | NA | 1 | 0 | 1 | 1 | 2 | 0.5 |
| 2_18051116 | NA | NA | NA | NA | 1 | 1 | NA | NA | 1 | 1 | NA | NA | NA | NA | 1 | 1 | 1 | 0 | 2 | 0.5 |
| 2_18051464 | NA | NA | NA | NA | 1 | 1 | NA | NA | 1 | 1 | NA | NA | NA | NA | 1 | 1 | 1 | 1 | 2 | 1 |
| 2_18051575 | NA | NA | NA | NA | 1 | 1 | NA | NA | 1 | 1 | NA | NA | NA | NA | 1 | 1 | 1 | 0 | 2 | 0.5 |
| 2_18052418 | NA | NA | NA | NA | 1 | 1 | NA | NA | 1 | 1 | NA | NA | NA | NA | 1 | 1 | 1 | 0 | 2 | 0.5 |
| 2_18167194 | NA | NA | 1 | 1 | 1 | 0 | NA | NA | 2 | 0.5 | NA | NA | NA | NA | NA | NA | 1 | 1 | 1 | 1 |
| 2_18215826 | 1 | 0 | 3 | 0 | 2 | 0 | 7 | 0.1429 | 13 | 0.0769 | 2 | 0.5 | 1 | 0 | 3 | 0 | 6 | 0 | 12 | 0.0833 |
| 2_18215958 | 1 | 0 | 2 | 0.5 | 2 | 0 | 8 | 0.25 | 13 | 0.2308 | 3 | 0.3333 | 1 | 0 | 2 | 0.5 | 6 | 0.1667 | 12 | 0.25 |
| 2_18216064 | 1 | 0 | 2 | 0 | 2 | 0 | 8 | 0.25 | 13 | 0.1538 | 3 | 0.3333 | NA | NA | 3 | 0 | 6 | 0.1667 | 12 | 0.1667 |
| 2_18216397 | 3 | 0 | 3 | 0 | 2 | 0 | 7 | 0.1429 | 15 | 0.0667 | 4 | 0.25 | 1 | 0 | 4 | 0 | 5 | 0 | 14 | 0.0714 |
| 2_18217286 | 1 | 1 | 1 | 0 | NA | NA | NA | NA | 2 | 0.5 | NA | NA | 1 | 1 | NA | NA | NA | NA | 1 | 1 |
| 2_18221999 | NA | NA | 2 | 0.5 | NA | NA | NA | NA | 2 | 0.5 | 1 | 1 | 1 | 0 | NA | NA | 3 | 0 | 5 | 0.2 |
| 2_18222101 | NA | NA | 1 | 1 | NA | NA | NA | NA | 1 | 1 | 1 | 0 | 1 | 0 | NA | NA | 3 | 0.3333 | 5 | 0.2 |
| 2_18224449 | NA | NA | 1 | 1 | NA | NA | NA | NA | 1 | 1 | 1 | 1 | 1 | 1 | NA | NA | 2 | 1 | 4 | 1 |
| 2_18680574 | 2 | 1 | 1 | 0 | NA | NA | NA | NA | 3 | 0.6667 | NA | NA | NA | NA | 1 | 0 | 1 | 1 | 2 | 0.5 |
| 2_18680679 | 19 | 0.4737 | 10 | 0.6 | 6 | 0.1667 | 11 | 0.3636 | 46 | 0.4348 | 6 | 0.5 | 14 | 0.1429 | 11 | 0.3636 | 13 | 0.6154 | 44 | 0.3864 |
| 2_18680921 | 20 | 0.3 | 10 | 0.3 | 8 | 0.125 | 14 | 0.0714 | 52 | 0.2115 | 6 | 0.3333 | 14 | 0.2857 | 14 | 0.2143 | 13 | 0.3077 | 47 | 0.2766 |
| 2_18689037 | 2 | 1 | 2 | 0.5 | 5 | 0.6 | 7 | 0.4286 | 16 | 0.5625 | 4 | 0.5 | 3 | 0.3333 | 3 | 0.3333 | 9 | 0.7778 | 19 | 0.5789 |
| 2_18689066 | 2 | 0 | 2 | 1 | 7 | 0.4286 | 8 | 0.5 | 19 | 0.4737 | 4 | 0.5 | 4 | 0.75 | 5 | 0.4 | 8 | 0.375 | 21 | 0.4762 |
| 2_18689294 | 3 | 0.6667 | 1 | 1 | 6 | 0.6667 | 8 | 0.5 | 18 | 0.6111 | 4 | 1 | 4 | 0.75 | 3 | 1 | 9 | 0.8889 | 20 | 0.9 |
| 2_18689374 | 3 | 0 | 2 | 0.5 | 5 | 0.4 | 4 | 0.5 | 14 | 0.3571 | 2 | 1 | 3 | 0.3333 | 4 | 0.5 | 8 | 0.625 | 17 | 0.5882 |
| 2_18689405 | 4 | 0.25 | 2 | 0.5 | 7 | 0.4286 | 8 | 0.5 | 21 | 0.4286 | 3 | 0.3333 | 4 | 0.5 | 5 | 0.2 | 11 | 0.7273 | 23 | 0.5217 |
| 2_18689431 | 3 | 0.6667 | 3 | 0.3333 | 7 | 0.8571 | 7 | 0.7143 | 20 | 0.7 | 3 | 1 | 5 | 0.6 | 5 | 0.8 | 11 | 0.7273 | 24 | 0.75 |
| 2_18691236 | 3 | 0 | 4 | 0.25 | 4 | 0.5 | 1 | 0 | 12 | 0.25 | 1 | 0 | 4 | 0 | 8 | 0.125 | 6 | 0 | 19 | 0.0526 |
| 2_18691257 | 1 | 1 | 3 | 0.6667 | 6 | 0.6667 | 1 | 1 | 11 | 0.7273 | 2 | 0.5 | 3 | 1 | 9 | 0.8889 | 7 | 0.7143 | 21 | 0.8095 |
| 2_18848538 | 5 | 0.2 | 10 | 0.1 | 7 | 0 | 9 | 0.2222 | 31 | 0.129 | 7 | 0.2857 | 5 | 0.2 | 6 | 0 | 4 | 0 | 22 | 0.1364 |
| 2_18848627 | 7 | 0.1429 | 10 | 0.3 | 5 | 0.2 | 9 | 0.2222 | 31 | 0.2258 | 6 | 0 | 7 | 0.8571 | 6 | 0.5 | 6 | 0.3333 | 25 | 0.44 |
| 2_18848757 | 7 | 0 | 11 | 0.1818 | 8 | 0.375 | 9 | 0.2222 | 35 | 0.2 | 8 | 0 | 6 | 0.3333 | 6 | 0 | 7 | 0 | 27 | 0.0741 |
| 2_18848780 | 7 | 0.1429 | 10 | 0.2 | 7 | 0.2857 | 9 | 0.5556 | 33 | 0.303 | 7 | 0 | 6 | 0.3333 | 6 | 0.3333 | 6 | 0.3333 | 25 | 0.24 |
| 2_18848913 | 3 | 0 | 7 | 0.2857 | 3 | 0 | 5 | 0.2 | 18 | 0.1667 | 7 | 0.1429 | 6 | 0.5 | 4 | 0.25 | 6 | 0.1667 | 23 | 0.2609 |
| 2_18848993 | 8 | 0.625 | 4 | 0.25 | 5 | 0.4 | 5 | 0.8 | 22 | 0.5455 | 4 | 0.5 | 2 | 0.5 | 4 | 1 | 3 | 1 | 13 | 0.7692 |

| | | | | | | | | | | | | | | | | | | | | |
|---|---|---|---|---|---|---|---|---|---|---|---|---|---|---|---|---|---|---|---|---|
| 2_1884905 | 5 | 0.6 | 7 | 0.7143 | 5 | 0 | 6 | 0.5 | 23 | 0.4783 | 5 | 0.6 | 5 | 0 | 6 | 0.5 | 6 | 0.3333 | 22 | 0.3636 |
| 2_1884916 | 6 | 0.6667 | 6 | 0.5 | 3 | 0.3333 | 6 | 0.6667 | 21 | 0.5714 | 6 | 0.3333 | 1 | 1 | 3 | 0.6667 | 4 | 0.75 | 14 | 0.5714 |
| 2_1884921 | 7 | 0.5714 | 9 | 0.7778 | 4 | 1 | 7 | 0.8571 | 27 | 0.7778 | 5 | 0.6 | 5 | 0.8 | 6 | 0.3333 | 5 | 0.8 | 21 | 0.619 |
| 2_1900295 | 2 | 0.5 | 1 | 0 | 1 | 0 | 1 | 1 | 5 | 0.4 | 1 | 0 | 2 | 0 | 3 | 0 | 2 | 0.5 | 8 | 0.125 |
| 2_1935379 | NA | NA | NA | NA | NA | NA | 1 | 1 | 1 | 1 | NA | NA | NA | NA | 1 | 1 | NA | NA | 1 | 1 |
| 2_1935543 | NA | NA | 1 | 1 | 1 | 0 | 2 | 0 | 4 | 0.25 | NA | NA | NA | NA | 1 | 1 | NA | NA | 1 | 1 |
| 2_1935590 | NA | NA | 2 | 1 | 2 | 0.5 | 4 | 0.5 | 8 | 0.625 | 3 | 0.3333 | 2 | 1 | 1 | 0 | NA | NA | 6 | 0.5 |
| 2_1935594 | 1 | 0 | 2 | 1 | 1 | 0 | 4 | 0.75 | 8 | 0.625 | 3 | 0.6667 | 1 | 0 | 2 | 0.5 | NA | NA | 6 | 0.5 |
| 2_1935597 | 1 | 1 | 2 | 0.5 | 1 | 1 | 4 | 0.75 | 8 | 0.75 | 3 | 0.3333 | 1 | 1 | 2 | 0.5 | NA | NA | 6 | 0.5 |
| 2_1935604 | 1 | 0 | 2 | 0 | 3 | 0.3333 | 4 | 0 | 10 | 0.1 | 3 | 0 | 2 | 0.5 | 2 | 0.5 | NA | NA | 7 | 0.2857 |
| 2_1935606 | 1 | 0 | 1 | 1 | 2 | 0.5 | 3 | 0.6667 | 7 | 0.5714 | 3 | 0.6667 | 2 | 1 | 2 | 0.5 | NA | NA | 7 | 0.7143 |
| 2_1935635 | 1 | 1 | 2 | 0 | 2 | 1 | 3 | 0.3333 | 8 | 0.5 | 3 | 0.6667 | 2 | 0.5 | 2 | 0.5 | NA | NA | 7 | 0.5714 |
| 2_1935641 | 2 | 0.5 | 3 | 0.6667 | 3 | 0.6667 | 3 | 1 | 11 | 0.7273 | 3 | 0.3333 | 2 | 1 | 1 | 1 | NA | NA | 6 | 0.6667 |
| 2_1935652 | 1 | 0 | 3 | 0 | 3 | 1 | 3 | 0.3333 | 10 | 0.4 | 3 | 0.3333 | 2 | 0.5 | 2 | 0.5 | NA | NA | 7 | 0.4286 |
| 2_1935667 | 2 | 0.5 | 3 | 0 | 3 | 0.3333 | 4 | 0.25 | 12 | 0.25 | 3 | 0.3333 | 2 | 0.5 | 2 | 0.5 | NA | NA | 7 | 0.4286 |
| 2_1939916 | 3 | 0 | 5 | 0 | 1 | 1 | 3 | 0.6667 | 12 | 0.25 | 5 | 0.4 | NA | NA | 1 | 0 | 4 | 0.75 | 10 | 0.5 |
| 2_1939938 | 3 | 0 | 5 | 0 | 1 | 0 | 4 | 0.25 | 13 | 0.0769 | 7 | 0 | NA | NA | 1 | 0 | 4 | 0.25 | 12 | 0.0833 |
| 2_1939941 | 2 | 0.5 | 4 | 1 | 1 | 1 | 4 | 1 | 11 | 0.9091 | 6 | 1 | NA | NA | 1 | 1 | 4 | 1 | 11 | 1 |
| 2_1939957 | 3 | 0 | 5 | 0.2 | 1 | 0 | 3 | 0.3333 | 12 | 0.1667 | 6 | 0.1667 | NA | NA | 1 | 0 | 3 | 0.6667 | 10 | 0.3 |
| 2_1939959 | 2 | 1 | 3 | 0 | 1 | 0 | 4 | 0.25 | 10 | 0.3 | 5 | 0.2 | NA | NA | 1 | 1 | 4 | 0.5 | 10 | 0.4 |
| 2_1939962 | 3 | 0.3333 | 5 | 0 | 1 | 0 | 4 | 0.5 | 13 | 0.2308 | 7 | 0.2857 | NA | NA | 1 | 0 | 3 | 0.3333 | 11 | 0.2727 |
| 2_1939965 | 3 | 0.3333 | 5 | 0.2 | 2 | 1 | 3 | 0.3333 | 13 | 0.3846 | 6 | 0.3333 | NA | NA | 1 | 0 | 3 | 0.3333 | 10 | 0.3 |
| 2_1966333 | 2 | 0.5 | 2 | 0 | 2 | 0 | 1 | 0 | 7 | 0.1429 | 1 | 0 | 1 | 0 | 1 | 0 | 3 | 0.3333 | 6 | 0.1667 |
| 2_1966369 | 2 | 0.5 | 2 | 1 | 2 | 0.5 | 1 | 0 | 7 | 0.5714 | 2 | 0.5 | 2 | 0 | 2 | 0 | 2 | 0 | 8 | 0.125 |
| 2_1966377 | 2 | 0.5 | 1 | 1 | 2 | 0.5 | 1 | 1 | 6 | 0.6667 | 2 | 0.5 | 2 | 1 | 2 | 0.5 | 3 | 0.6667 | 9 | 0.6667 |
| 2_1966383 | 1 | 1 | 2 | 0 | 2 | 0 | 1 | 0 | 6 | 0.1667 | 1 | 0 | NA | NA | NA | NA | 3 | 0.3333 | 4 | 0.25 |
| 2_1966384 | 2 | 0.5 | 2 | 1 | 2 | 1 | NA | NA | 6 | 0.8333 | 1 | 1 | NA | NA | 1 | 0 | 3 | 0.3333 | 5 | 0.4 |
| 2_1966397 | 1 | 0 | 1 | 1 | 1 | 0 | 1 | 1 | 4 | 0.5 | 1 | 1 | 1 | 0 | 1 | 0 | 2 | 0 | 5 | 0.2 |
| 2_1966408 | 1 | 1 | 2 | 1 | 2 | 1 | 1 | 1 | 6 | 1 | 2 | 1 | 3 | 1 | 2 | 1 | 4 | 1 | 11 | 1 |
| 2_1966412 | 1 | 0 | 4 | 0.25 | 2 | 0.5 | 1 | 1 | 8 | 0.375 | 2 | 0 | 3 | 0 | 1 | 0 | 4 | 0.75 | 10 | 0.3 |
| 2_1966417 | 2 | 1 | 4 | 0.75 | 1 | 1 | 1 | 1 | 8 | 0.875 | 2 | 1 | 2 | 1 | 2 | 1 | 4 | 0.75 | 10 | 0.9 |
| 2_1966421 | 1 | 1 | 3 | 1 | 2 | 0 | NA | NA | 6 | 0.6667 | 2 | 0 | 2 | 1 | 2 | 0.5 | 4 | 0.75 | 10 | 0.6 |
| 2_1966424 | 2 | 1 | 4 | 0.75 | 2 | 0.5 | 1 | 1 | 9 | 0.7778 | 2 | 1 | 3 | 1 | 1 | 1 | 3 | 1 | 9 | 1 |
| 2_1966425 | 1 | 0 | 3 | 0.3333 | 2 | 0 | NA | NA | 6 | 0.1667 | 2 | 0 | NA | NA | 1 | 0 | 3 | 0.3333 | 6 | 0.1667 |
| 2_1966430 | 1 | 1 | 5 | 1 | 2 | 1 | 1 | 1 | 9 | 1 | 2 | 1 | 1 | 1 | 1 | 1 | 3 | 1 | 7 | 1 |
| 2_1966434 | NA | NA | 4 | 1 | 2 | 0.5 | 1 | 1 | 7 | 0.8571 | 2 | 0.5 | 1 | 0 | 1 | 1 | 4 | 0.5 | 8 | 0.5 |
| 2_1966502 | 3 | 0 | 7 | 0.7143 | 4 | 0.25 | 2 | 0.5 | 16 | 0.4375 | 4 | 0.25 | 4 | 0.75 | 4 | 0.5 | 3 | 0.6667 | 15 | 0.5333 |
| 2_2074846 | 1 | 1 | 3 | 0.6667 | 2 | 0.5 | 3 | 1 | 9 | 0.7778 | 2 | 1 | NA | NA | 4 | 0.5 | NA | NA | 6 | 0.6667 |
| 2_2074847 | 1 | 1 | 2 | 0 | 2 | 0.5 | 3 | 0.3333 | 8 | 0.375 | 3 | 0.3333 | 1 | 1 | 4 | 0.75 | NA | NA | 8 | 0.625 |
| 2_2074877 | NA | NA | 3 | 0 | 1 | 0 | 4 | 0.25 | 8 | 0.125 | 3 | 0.3333 | 1 | 1 | 4 | 0.25 | 2 | 0.5 | 10 | 0.4 |
| 2_2074886 | 1 | 0 | 3 | 0 | 2 | 0 | 4 | 0.25 | 10 | 0.1 | 3 | 0.6667 | 1 | 0 | 4 | 0.5 | 2 | 0 | 10 | 0.4 |
| 2_2074887 | 1 | 0 | 4 | 0.5 | 3 | 1 | 4 | 0.25 | 12 | 0.5 | 3 | 0.3333 | 1 | 1 | 4 | 0.25 | 1 | 0 | 9 | 0.3333 |
| 2_2081286 | NA | NA | 1 | 1 | 2 | 1 | 2 | 1 | 5 | 1 | 2 | 1 | NA | NA | NA | NA | 2 | 0.5 | 4 | 0.75 |
| 2_2081355 | 2 | 0 | 7 | 0.1429 | 5 | 0 | 5 | 0.2 | 19 | 0.1053 | 5 | 0 | 3 | 0.3333 | 6 | 0 | 6 | 0 | 20 | 0.05 |
| 2_2081364 | 2 | 0 | 6 | 0 | 5 | 0 | 4 | 0.5 | 17 | 0.1176 | 4 | 0 | 2 | 0 | 4 | 0.25 | 6 | 0.3333 | 16 | 0.1875 |
| 2_2081383 | 2 | 0.5 | 5 | 0.4 | 4 | 0 | 4 | 0.25 | 15 | 0.2667 | 4 | 0 | 3 | 0 | 4 | 0.25 | 6 | 0.5 | 17 | 0.2353 |
| 2_2081393 | 2 | 0.5 | 8 | 0.375 | 3 | 0.3333 | 5 | 0.2 | 18 | 0.3333 | 5 | 0.2 | 3 | 0.3333 | 5 | 0.4 | 7 | 0.2857 | 20 | 0.3 |
| 2_2081439 | 5 | 0 | 11 | 0.0909 | 7 | 0 | 6 | 0.1667 | 29 | 0.069 | 5 | 0 | 3 | 0 | 6 | 0 | 6 | 0.1667 | 20 | 0.05 |
| 2_2081502 | 6 | 0 | 12 | 0.25 | 6 | 0.1667 | 6 | 0 | 30 | 0.1333 | 7 | 0 | 3 | 0.3333 | 7 | 0.1429 | 7 | 0 | 24 | 0.0833 |
| 2_2081512 | 9 | 0 | 12 | 0.0833 | 10 | 0.1 | 7 | 0 | 38 | 0.0526 | 8 | 0.25 | 3 | 0 | 7 | 0.1429 | 8 | 0.5 | 26 | 0.2692 |
| 2_2085282 | 10 | 0.5 | 16 | 0.375 | 11 | 0.7273 | 12 | 0.5833 | 49 | 0.5306 | 11 | 0.6364 | 9 | 0.4444 | 7 | 1 | 13 | 0.6154 | 40 | 0.65 |
| 2_2085286 | 11 | 0.0909 | 16 | 0.125 | 13 | 0.2308 | 12 | 0.1667 | 52 | 0.1538 | 12 | 0.1667 | 10 | 0.3 | 6 | 0.3333 | 13 | 0.3077 | 41 | 0.2683 |
| 2_2085302 | 10 | 0.1 | 14 | 0.0714 | 12 | 0 | 14 | 0.1429 | 50 | 0.08 | 9 | 0 | 9 | 0.1111 | 5 | 0 | 14 | 0.1429 | 37 | 0.0811 |
| 2_2085307 | 13 | 0.7692 | 12 | 0.5 | 11 | 0.6364 | 14 | 0.6429 | 50 | 0.64 | 9 | 0.5556 | 10 | 0.7 | 6 | 0.6667 | 14 | 0.7143 | 39 | 0.6667 |
| 2_2085335 | 13 | 0.0769 | 9 | 0.2222 | 10 | 0.1 | 12 | 0.0833 | 44 | 0.1136 | 7 | 0.2857 | 8 | 0.125 | 4 | 0.25 | 16 | 0.125 | 35 | 0.1714 |
| 2_2085346 | 12 | 0.3333 | 11 | 0.5455 | 11 | 0.2727 | 9 | 0.4444 | 43 | 0.3953 | 5 | 0.8 | 5 | 0.2 | 3 | 0.3333 | 13 | 0.5385 | 26 | 0.5 |
| 2_2085348 | 10 | 0.2 | 10 | 0.1 | 10 | 0.1 | 11 | 0 | 41 | 0.0976 | 4 | 0.25 | 5 | 0.2 | 2 | 0 | 10 | 0.2 | 21 | 0.1905 |
| 2_2085352 | 11 | 0.4545 | 11 | 0.3636 | 12 | 0.5833 | 10 | 0.5 | 44 | 0.4773 | 4 | 0.5 | 8 | 0.375 | 3 | 0 | 9 | 0.1111 | 24 | 0.25 |
| 2_2085353 | 13 | 1 | 12 | 0.9167 | 11 | 0.9091 | 11 | 1 | 47 | 0.9574 | 6 | 0.8333 | 7 | 0.8571 | 3 | 1 | 14 | 0.9286 | 30 | 0.9 |
| 2_2085361 | 10 | 0.1 | 10 | 0.2 | 10 | 0 | 9 | 0 | 39 | 0.0769 | 6 | 0 | 7 | 0.2857 | 4 | 0.25 | 15 | 0.2667 | 32 | 0.2188 |
| 2_2085362 | 7 | 0 | 8 | 0.125 | 5 | 0 | 8 | 0.25 | 28 | 0.1071 | 4 | 0 | 5 | 0 | 1 | 1 | 6 | 0.1667 | 16 | 0.125 |
| 2_2085375 | 9 | 0 | 8 | 0.125 | 12 | 0 | 7 | 0.1429 | 36 | 0.0556 | 5 | 0.2 | 6 | 0.1667 | 3 | 0 | 14 | 0.0714 | 28 | 0.1071 |
| 2_2085384 | 10 | 0.9 | 11 | 0.8182 | 8 | 0.875 | 8 | 0.375 | 37 | 0.7568 | 4 | 0.5 | 6 | 0.6667 | 3 | 0.3333 | 14 | 1 | 27 | 0.7778 |
| 2_2085386 | 8 | 1 | 10 | 0.9 | 10 | 0.9 | 9 | 0.8889 | 37 | 0.9189 | 4 | 1 | 3 | 1 | 2 | 1 | 12 | 0.8333 | 21 | 0.9048 |
| 2_2085390 | 10 | 1 | 9 | 0.7778 | 8 | 1 | 8 | 1 | 35 | 0.9429 | 3 | 1 | 5 | 1 | 1 | 1 | 15 | 0.9333 | 24 | 0.9583 |

| | | | | | | | | | | | | | | | | | | | | |
|---|---|---|---|---|---|---|---|---|---|---|---|---|---|---|---|---|---|---|---|---|
| 2_2085392 | 11 | 0.7273 | 9 | 0.8889 | 9 | 0.3333 | 6 | 0.6667 | 35 | 0.6571 | 2 | 1 | 4 | 0.5 | 2 | 0.5 | 13 | 0.4615 | 21 | 0.5238 |
| 2_2085399 | 10 | 0.1 | 8 | 0 | 7 | 0 | 8 | 0.125 | 33 | 0.0606 | 5 | 0.2 | 4 | 0 | 4 | 0 | 14 | 0.0714 | 27 | 0.0741 |
| 2_2085408 | 9 | 0.3333 | 9 | 0.2222 | 8 | 0.125 | 6 | 0.3333 | 32 | 0.25 | 5 | 0 | 4 | 0 | 2 | 0 | 13 | 0.1538 | 24 | 0.0833 |
| 2_2085409 | 5 | 0.8 | 9 | 0.6667 | 7 | 0.7143 | 2 | 0.5 | 23 | 0.6957 | 1 | 1 | 3 | 0.6667 | 1 | 1 | 7 | 0.1429 | 12 | 0.4167 |
| 2_2085423 | 9 | 0.8889 | 9 | 0.8889 | 9 | 0.5556 | 7 | 0.5714 | 34 | 0.7353 | 3 | 0.6667 | 6 | 0.5 | 2 | 1 | 14 | 0.7857 | 25 | 0.72 |
| 2_2085431 | 8 | 0.75 | 6 | 1 | 7 | 1 | 5 | 1 | 26 | 0.9231 | 4 | 1 | 4 | 1 | 2 | 0.5 | 9 | 0.8889 | 19 | 0.8947 |
| 2_2085460 | 8 | 0.75 | 7 | 0 | 9 | 0.6667 | 6 | 0.6667 | 30 | 0.5333 | 4 | 0.25 | 5 | 0.2 | NA | NA | 11 | 0.9091 | 20 | 0.6 |
| 2_2085477 | 7 | 0.1429 | 7 | 0.1429 | 10 | 0.1 | 6 | 0.5 | 30 | 0.2 | 4 | 0 | 6 | 0.1667 | 3 | 0.3333 | 10 | 0.2 | 23 | 0.1739 |
| 2_2085493 | 5 | 0.8 | 7 | 0.5714 | 9 | 0.4444 | 4 | 1 | 25 | 0.64 | 4 | 0 | 6 | 0.6667 | 1 | 0 | 11 | 0.4545 | 22 | 0.4091 |
| 2_2085506 | 4 | 0.5 | 7 | 0.7143 | 10 | 0.7 | 5 | 0.8 | 26 | 0.6923 | 4 | 0.75 | 6 | 0.6667 | 2 | 0.5 | 8 | 0.625 | 20 | 0.65 |
| 2_2088493 | 1 | 0 | 2 | 0.5 | 1 | 0 | 3 | 0 | 7 | 0.1429 | 2 | 0 | 4 | 0.25 | 1 | 0 | 5 | 0.2 | 12 | 0.1667 |
| 2_2088504 | 1 | 0 | NA | NA | 1 | 0 | 3 | 0.3333 | 5 | 0.2 | 2 | 0.5 | 4 | 0.75 | NA | NA | 5 | 0.2 | 11 | 0.4545 |
| 2_2088508 | 1 | 0 | 1 | 0 | 1 | 0 | 4 | 0.25 | 7 | 0.1429 | 2 | 0.5 | 4 | 0.25 | 1 | 0 | 6 | 0.1667 | 13 | 0.2308 |
| 2_2088513 | 1 | 0 | 2 | 0.5 | 1 | 0 | 3 | 0 | 7 | 0.1429 | 2 | 0 | 4 | 0.25 | 1 | 0 | 6 | 0 | 13 | 0.0769 |
| 2_2088542 | 1 | 1 | 1 | 0 | 1 | 0 | 4 | 0 | 7 | 0.1429 | 2 | 0 | 3 | 0 | 1 | 0 | 5 | 0.6 | 11 | 0.2727 |
| 2_2088551 | 1 | 1 | NA | NA | NA | NA | 4 | 1 | 5 | 1 | 2 | 1 | 2 | 1 | NA | NA | 5 | 0.8 | 9 | 0.8889 |
| 2_2088561 | 1 | 1 | 1 | 1 | 1 | 0 | 3 | 0.6667 | 6 | 0.6667 | 1 | 1 | 2 | 0 | NA | NA | 4 | 0.75 | 7 | 0.5714 |
| 2_2088568 | 1 | 0 | 1 | 0 | 1 | 0 | 3 | 0.3333 | 6 | 0.1667 | NA | NA | 2 | 0.5 | 1 | 0 | 4 | 0 | 7 | 0.1429 |
| 2_2088595 | 1 | 0 | 1 | 1 | NA | NA | 3 | 0 | 5 | 0.2 | 1 | 0 | 3 | 0.6667 | 1 | 0 | 3 | 0 | 8 | 0.25 |
| 2_2088597 | 1 | 1 | 1 | 1 | NA | NA | 3 | 1 | 5 | 1 | 1 | 0 | 1 | 1 | 1 | 0 | 2 | 1 | 5 | 0.6 |
| 2_2088602 | 1 | 1 | 1 | 0 | NA | NA | 3 | 0.6667 | 5 | 0.6 | 1 | 0 | 3 | 0.3333 | 1 | 0 | 3 | 0.3333 | 8 | 0.25 |
| 2_2088603 | NA | NA | 1 | 0 | NA | NA | 3 | 1 | 4 | 0.75 | NA | NA | 3 | 0.6667 | 1 | 0 | 3 | 0.3333 | 7 | 0.4286 |
| 2_2088613 | 1 | 0 | 1 | 1 | NA | NA | 2 | 0.5 | 4 | 0.5 | NA | NA | 3 | 0.3333 | 1 | 1 | 3 | 0.3333 | 7 | 0.4286 |
| 2_2088629 | 1 | 1 | 1 | 0 | NA | NA | NA | NA | 2 | 0.5 | NA | NA | NA | NA | NA | NA | 3 | 0.6667 | 3 | 0.6667 |
| 2_2088634 | 1 | 1 | 1 | 0 | NA | NA | 1 | 1 | 3 | 0.6667 | NA | NA | 3 | 0.6667 | 1 | 0 | 3 | 0.6667 | 7 | 0.5714 |
| 2_2097220 | 1 | 0 | 2 | 1 | NA | NA | 1 | 1 | 4 | 0.75 | NA | NA | 1 | 1 | NA | NA | 2 | 0.5 | 3 | 0.6667 |
| 2_2097225 | 1 | 1 | 2 | 0 | NA | NA | 1 | 1 | 4 | 0.5 | NA | NA | 1 | 0 | NA | NA | 2 | 1 | 3 | 0.6667 |
| 2_2097240 | NA | NA | 1 | 1 | NA | NA | 1 | 1 | 2 | 1 | NA | NA | NA | NA | NA | NA | 2 | 0.5 | 2 | 0.5 |
| 2_2097241 | NA | NA | 1 | 0 | NA | NA | 1 | 1 | 2 | 0.5 | NA | NA | NA | NA | NA | NA | 2 | 0.5 | 2 | 0.5 |
| 2_2097483 | NA | NA | 5 | 1 | 4 | 0.75 | 2 | 0.5 | 11 | 0.8182 | 4 | 1 | 1 | 1 | NA | NA | 2 | 0.5 | 7 | 0.8571 |
| 2_2097492 | NA | NA | 5 | 0.2 | 4 | 0.25 | 2 | 0 | 11 | 0.1818 | 4 | 0.25 | NA | NA | 1 | 0 | 3 | 0 | 8 | 0.125 |
| 2_2121648 | 2 | 0 | 1 | 0 | 1 | 1 | 1 | 0 | 5 | 0.2 | 1 | 0 | 5 | 0.4 | 4 | 0 | 2 | 0 | 12 | 0.1667 |
| 2_2121659 | NA | NA | 1 | 1 | 1 | 0 | 1 | 0 | 3 | 0.3333 | 1 | 0 | 5 | 0 | 2 | 0 | 1 | 1 | 9 | 0.1111 |
| 2_2121679 | 2 | 0.5 | 2 | 1 | 1 | 1 | 1 | 1 | 6 | 0.8333 | 1 | 1 | 6 | 1 | 4 | 1 | 2 | 1 | 13 | 1 |
| 2_2121681 | 2 | 1 | 2 | 1 | 1 | 1 | 1 | 1 | 6 | 1 | NA | NA | 5 | 1 | 4 | 1 | 2 | 1 | 11 | 1 |
| 2_2121693 | 2 | 1 | 2 | 1 | 1 | 1 | 1 | 1 | 6 | 1 | 1 | 1 | 5 | 1 | 4 | 0.75 | 2 | 1 | 12 | 0.9167 |
| 2_2121715 | 2 | 1 | 2 | 1 | 1 | 1 | 2 | 0.5 | 7 | 0.8571 | 1 | 1 | 6 | 1 | 4 | 1 | 2 | 1 | 13 | 1 |
| 2_2121719 | 2 | 1 | 2 | 1 | 1 | 1 | 2 | 1 | 7 | 1 | NA | NA | 5 | 1 | 3 | 0.6667 | 2 | 1 | 10 | 0.9 |
| 2_2121738 | 2 | 0.5 | 2 | 0.5 | 1 | 1 | 2 | 1 | 7 | 0.7143 | 1 | 0 | 6 | 0 | 4 | 0.5 | 2 | 0 | 13 | 0.1538 |
| 2_2121763 | 2 | 0.5 | 2 | 1 | 1 | 1 | 2 | 0 | 7 | 0.5714 | 1 | 0 | 5 | 0.4 | 4 | 0 | 2 | 0.5 | 12 | 0.25 |
| 2_2121777 | 2 | 0 | 4 | 0 | 1 | 0 | 2 | 0.5 | 9 | 0.1111 | 2 | 0 | 5 | 0.6 | 5 | 0.2 | 2 | 0 | 14 | 0.2857 |
| 2_2121786 | 4 | 0.75 | 4 | 0.25 | 1 | 0 | 2 | 0 | 11 | 0.3636 | 1 | 0 | 5 | 0.4 | 5 | 0.8 | 2 | 0 | 13 | 0.4615 |
| 2_2121787 | 4 | 0 | 5 | 0.4 | 1 | 1 | 2 | 0.5 | 12 | 0.3333 | 2 | 0 | 6 | 0.3333 | 4 | 0 | 2 | 0 | 14 | 0.1429 |
| 2_2121788 | 3 | 0.3333 | 4 | 0.5 | NA | NA | 1 | 1 | 8 | 0.5 | 1 | 1 | 2 | 1 | 4 | 1 | 1 | 1 | 8 | 1 |
| 2_2121788 | 4 | 0.75 | 4 | 1 | 2 | 1 | 2 | 0.5 | 12 | 0.8333 | 2 | 1 | 5 | 1 | 4 | 1 | 2 | 1 | 13 | 1 |
| 2_2121799 | 4 | 1 | 4 | 0.75 | 2 | 1 | 2 | 1 | 12 | 0.9167 | 1 | 1 | 6 | 1 | 4 | 1 | 1 | 1 | 12 | 1 |
| 2_2121821 | 4 | 0 | 6 | 0 | 2 | 0.5 | 2 | 0 | 14 | 0.0714 | 2 | 0 | 6 | 0 | 6 | 0.1667 | 2 | 0 | 16 | 0.0625 |
| 2_2121829 | 4 | 1 | 6 | 1 | 2 | 0.5 | 2 | 1 | 14 | 0.9286 | 2 | 0.5 | 5 | 1 | 6 | 0.8333 | 2 | 0.5 | 15 | 0.8 |
| 2_2121845 | 3 | 0.3333 | 6 | 0.6667 | 3 | 0.6667 | 2 | 0 | 14 | 0.5 | 3 | 0.3333 | 5 | 0.2 | 6 | 0.5 | 2 | 1 | 16 | 0.4375 |
| 2_2121852 | 3 | 0 | 6 | 0.1667 | 3 | 0 | 2 | 0 | 14 | 0.0714 | 2 | 0 | 3 | 0 | 4 | 0.25 | 1 | 1 | 10 | 0.2 |
| 2_2121854 | 2 | 1 | 6 | 1 | 3 | 1 | 1 | 1 | 12 | 1 | 2 | 1 | 4 | 1 | 4 | 1 | 2 | 1 | 12 | 1 |
| 2_2121859 | 2 | 1 | 4 | 1 | 2 | 1 | 1 | 1 | 9 | 1 | 2 | 1 | 3 | 1 | 2 | 1 | 2 | 1 | 9 | 1 |
| 2_2121863 | 2 | 0.5 | 3 | 0.3333 | 2 | 0 | 1 | 0 | 8 | 0.25 | 2 | 0 | 3 | 0 | 2 | 0 | 2 | 0.5 | 9 | 0.1111 |
| 2_2121888 | 3 | 0.3333 | 4 | 0.5 | 2 | 0.5 | 1 | 1 | 10 | 0.5 | 1 | 1 | 2 | 1 | 2 | 0.5 | 1 | 1 | 6 | 0.8333 |
| 2_2121890 | 3 | 0.3333 | 4 | 0 | 3 | 0 | 1 | 0 | 11 | 0.0909 | 1 | 0 | 4 | 0.25 | 2 | 0 | 1 | 0 | 8 | 0.125 |
| 2_2121901 | 2 | 0.5 | 3 | 0.6667 | 3 | 0 | 1 | 0 | 9 | 0.3333 | NA | NA | 4 | 0.75 | 2 | 1 | 2 | 0.5 | 8 | 0.75 |
| 2_2123230 | NA | NA | 2 | 0 | NA | NA | 1 | 1 | 3 | 0.3333 | 1 | 1 | NA | NA | NA | NA | NA | NA | 1 | 1 |
| 2_2123231 | NA | NA | 2 | 0.5 | NA | NA | NA | NA | 2 | 0.5 | 1 | 1 | NA | NA | NA | NA | NA | NA | 1 | 1 |
| 2_2123239 | NA | NA | 2 | 1 | NA | NA | 1 | 1 | 3 | 1 | 1 | 1 | 1 | 0 | NA | NA | NA | NA | 2 | 0.5 |
| 2_2123247 | NA | NA | 2 | 0.5 | NA | NA | 1 | 0 | 3 | 0.3333 | NA | NA | 1 | 1 | NA | NA | NA | NA | 1 | 1 |
| 2_2123251 | NA | NA | 2 | 1 | NA | NA | 1 | 1 | 3 | 1 | 1 | 1 | 1 | 0 | NA | NA | NA | NA | 2 | 0.5 |
| 2_2123280 | NA | NA | 2 | 0 | NA | NA | 2 | 1 | 4 | 0.5 | 1 | 0 | 1 | 1 | NA | NA | NA | NA | 2 | 0.5 |
| 2_2137200 | 1 | 1 | NA | NA | NA | NA | NA | NA | 1 | 1 | NA | NA | 1 | 1 | 1 | 1 | 1 | 1 | 3 | 1 |
| 2_2137210 | 1 | 1 | NA | NA | NA | NA | NA | NA | 1 | 1 | NA | NA | 1 | 0 | 1 | 1 | 1 | 1 | 3 | 0.6667 |
| 2_2137294 | 1 | 0 | 1 | 1 | 3 | 0.3333 | NA | NA | 5 | 0.4 | 1 | 0 | 2 | 0 | NA | NA | 2 | 0.5 | 5 | 0.2 |

| | | | | | | | | | | | | | | | | | | | | |
|---|---|---|---|---|---|---|---|---|---|---|---|---|---|---|---|---|---|---|---|---|
| 2_2137301 | 2 | 0 | 2 | 0 | 5 | 0.6 | 1 | 1 | 10 | 0.4 | 1 | 0 | 6 | 0.6667 | NA | NA | 2 | 0.5 | 9 | 0.5556 |
| 2_2137303 | 1 | 0 | 1 | 0 | 5 | 0 | 1 | 1 | 8 | 0.125 | 1 | 0 | 6 | 0.1667 | NA | NA | 2 | 0 | 9 | 0.1111 |
| 2_2137318 | NA | NA | 3 | 0.6667 | 2 | 0 | 3 | 0.3333 | 8 | 0.375 | 1 | 0 | 4 | 0.75 | NA | NA | 2 | 0.5 | 7 | 0.5714 |
| 2_2137333 | 1 | 1 | 3 | 0.6667 | 4 | 0.75 | 2 | 1 | 10 | 0.8 | 2 | 1 | 3 | 0.6667 | NA | NA | 3 | 0 | 8 | 0.5 |
| 2_2137344 | 2 | 1 | 3 | 0.6667 | 5 | 0.6 | 2 | 1 | 12 | 0.75 | 2 | 0.5 | 6 | 0.8333 | NA | NA | 4 | 0.75 | 12 | 0.75 |
| 2_2137350 | 1 | 0 | 1 | 1 | 2 | 0 | 2 | 0 | 6 | 0.1667 | 1 | 0 | 3 | 0.3333 | NA | NA | 2 | 1 | 6 | 0.5 |
| 2_2137368 | 3 | 0 | 3 | 0 | 6 | 0.1667 | 4 | 0.25 | 16 | 0.125 | 2 | 0.5 | 7 | 0.4286 | 1 | 0 | 7 | 0.2857 | 17 | 0.3529 |
| 2_2137372 | 3 | 0 | 3 | 0.6667 | 3 | 0.3333 | 4 | 0.75 | 13 | 0.4615 | 2 | 0 | 6 | 0.5 | 1 | 0 | 5 | 0.2 | 14 | 0.2857 |
| 2_2178799 | 7 | 0.2857 | 5 | 0.6 | 8 | 0.125 | 4 | 0 | 24 | 0.25 | 3 | 0.6667 | 10 | 0.3 | 3 | 0 | 8 | 0.125 | 24 | 0.25 |
| 2_2183105 | 8 | 0.25 | 8 | 0.125 | 7 | 0 | 7 | 0 | 30 | 0.1 | 5 | 0.2 | 7 | 0 | 9 | 0 | 7 | 0.2857 | 28 | 0.1071 |
| 2_2183106 | 8 | 0 | 8 | 0.125 | 9 | 0.1111 | 7 | 0.1429 | 32 | 0.0938 | 5 | 0 | 10 | 0.1 | 9 | 0.1111 | 8 | 0 | 32 | 0.0625 |
| 2_2183119 | 8 | 0.125 | 7 | 0.2857 | 5 | 0.2 | 6 | 0.1667 | 26 | 0.1923 | 4 | 0.5 | 8 | 0.375 | 7 | 0.4286 | 4 | 0.25 | 23 | 0.3913 |
| 2_2183137 | 9 | 0.1111 | 6 | 0.1667 | 9 | 0.3333 | 9 | 0.1111 | 33 | 0.1818 | 7 | 0.1429 | 11 | 0.2727 | 10 | 0.1 | 8 | 0.25 | 36 | 0.1944 |
| 2_2183146 | 11 | 0.1818 | 7 | 0.5714 | 7 | 0.4286 | 9 | 0.2222 | 34 | 0.3235 | 6 | 0.5 | 11 | 0.0909 | 9 | 0.3333 | 7 | 0.2857 | 33 | 0.2727 |
| 2_2183163 | 14 | 0 | 7 | 0.2857 | 8 | 0 | 10 | 0 | 39 | 0.0513 | 7 | 0 | 11 | 0.0909 | 10 | 0.1 | 7 | 0.1429 | 35 | 0.0857 |
| 2_2183166 | 11 | 0.4545 | 8 | 0.375 | 7 | 0.2857 | 7 | 0.1429 | 33 | 0.3333 | 6 | 0.1667 | 11 | 0.5455 | 12 | 0.6667 | 8 | 0.5 | 37 | 0.5135 |
| 2_2183171 | 13 | 0.0769 | 7 | 0 | 8 | 0.25 | 10 | 0.1 | 38 | 0.1053 | 6 | 0 | 13 | 0 | 11 | 0.0909 | 9 | 0.1111 | 39 | 0.0513 |
| 2_2183175 | 14 | 0.9286 | 8 | 1 | 10 | 0.9 | 10 | 0.8 | 42 | 0.9048 | 7 | 0.7143 | 12 | 0.9167 | 12 | 0.9167 | 10 | 0.9 | 41 | 0.878 |
| 2_2183187 | 11 | 1 | 9 | 0.8889 | 10 | 1 | 8 | 1 | 38 | 0.9737 | 5 | 0.8 | 13 | 0.9231 | 12 | 0.9167 | 10 | 1 | 40 | 0.925 |
| 2_2183215 | 12 | 0.4167 | 7 | 0.4286 | 10 | 0.5 | 9 | 0.3333 | 38 | 0.4211 | 8 | 0.375 | 11 | 0.4545 | 10 | 0.6 | 7 | 0.4286 | 36 | 0.4722 |
| 2_2183224 | 13 | 0.7692 | 9 | 0.2222 | 9 | 0.5556 | 10 | 0.6 | 41 | 0.561 | 6 | 0.8333 | 10 | 0.4 | 10 | 0.5 | 10 | 0 | 36 | 0.3889 |
| 2_2183231 | 11 | 0.6364 | 10 | 0.7 | 10 | 0.8 | 10 | 0.9 | 41 | 0.7561 | 10 | 0.9 | 15 | 0.7333 | 11 | 0.4545 | 11 | 0.7273 | 47 | 0.7021 |
| 2_2183235 | 11 | 0.1818 | 10 | 0.2 | 11 | 0.0909 | 10 | 0.3 | 42 | 0.1905 | 9 | 0.2222 | 14 | 0.2857 | 12 | 0.25 | 8 | 0.25 | 43 | 0.2558 |
| 2_2183236 | 17 | 0.3529 | 8 | 0.25 | 11 | 0 | 6 | 0.3333 | 42 | 0.2381 | 8 | 0.375 | 14 | 0.4286 | 11 | 0.3636 | 9 | 0.2222 | 42 | 0.3571 |
| 2_2183242 | 15 | 0.2 | 8 | 0 | 8 | 0 | 9 | 0.1111 | 40 | 0.1 | 10 | 0 | 16 | 0.125 | 13 | 0.1538 | 9 | 0.1111 | 48 | 0.1042 |
| 2_2183244 | 14 | 0.0714 | 9 | 0.1111 | 13 | 0.3077 | 10 | 0.4 | 46 | 0.2174 | 10 | 0.4 | 16 | 0.375 | 15 | 0.2 | 12 | 0.0833 | 53 | 0.2642 |
| 2_2183251 | 15 | 1 | 10 | 1 | 12 | 0.8333 | 11 | 0.9091 | 48 | 0.9375 | 10 | 1 | 15 | 0.9333 | 12 | 0.8333 | 11 | 0.9091 | 48 | 0.9167 |
| 2_2183253 | 13 | 0.5385 | 11 | 0.3636 | 13 | 0.1538 | 6 | 0.8333 | 43 | 0.4186 | 9 | 0.4444 | 13 | 0.4615 | 14 | 0.5 | 9 | 0.3333 | 45 | 0.4444 |
| 2_2183270 | 15 | 0.2667 | 10 | 0.4 | 12 | 0.0833 | 11 | 0.1818 | 48 | 0.2292 | 8 | 0.25 | 15 | 0.3333 | 13 | 0.3077 | 13 | 0.3846 | 49 | 0.3265 |
| 2_2183273 | 17 | 0.8824 | 11 | 0.9091 | 14 | 1 | 11 | 0.9091 | 53 | 0.9245 | 10 | 1 | 15 | 1 | 16 | 0.875 | 12 | 0.9167 | 53 | 0.9434 |
| 2_2183276 | 17 | 1 | 11 | 1 | 14 | 1 | 9 | 1 | 51 | 1 | 12 | 1 | 16 | 1 | 15 | 1 | 14 | 1 | 57 | 1 |
| 2_2183286 | 18 | 0.0556 | 11 | 0.0909 | 14 | 0.0714 | 11 | 0.0909 | 54 | 0.0741 | 12 | 0 | 16 | 0.1875 | 17 | 0.1176 | 13 | 0.2308 | 58 | 0.1379 |
| 2_2183288 | 16 | 1 | 10 | 1 | 13 | 1 | 10 | 1 | 49 | 1 | 11 | 1 | 12 | 0.9167 | 15 | 0.9333 | 14 | 1 | 52 | 0.9615 |
| 2_2183290 | 17 | 0.8235 | 10 | 0.7 | 13 | 0.6154 | 9 | 0.6667 | 49 | 0.7143 | 12 | 0.8333 | 16 | 0.5625 | 16 | 0.5625 | 12 | 0.5 | 56 | 0.6071 |
| 2_2183306 | 16 | 0.1875 | 10 | 0.4 | 11 | 0 | 10 | 0.1 | 47 | 0.1702 | 10 | 0.1 | 12 | 0.25 | 15 | 0.2 | 11 | 0.0909 | 48 | 0.1667 |
| 2_2183335 | 19 | 0.2632 | 12 | 0.1667 | 14 | 0 | 11 | 0.0909 | 56 | 0.1429 | 12 | 0 | 17 | 0.1176 | 16 | 0.0625 | 13 | 0.1538 | 58 | 0.0862 |
| 2_2183346 | 19 | 0.1579 | 12 | 0.25 | 14 | 0.2143 | 13 | 0.0769 | 58 | 0.1724 | 10 | 0.1 | 16 | 0.125 | 14 | 0.0714 | 13 | 0.1538 | 53 | 0.1132 |
| 2_2183419 | 20 | 0.1 | 13 | 0.0769 | 14 | 0.1429 | 11 | 0.2727 | 58 | 0.1379 | 12 | 0 | 15 | 0.1333 | 13 | 0.2308 | 13 | 0 | 53 | 0.0943 |
| 2_2183482 | 32 | 0.25 | 12 | 0.1667 | 20 | 0.2 | 15 | 0.3333 | 79 | 0.2405 | 16 | 0.125 | 20 | 0.2 | 23 | 0.2174 | 17 | 0.1176 | 76 | 0.1711 |
| 2_2183514 | 35 | 0.2857 | 16 | 0.25 | 26 | 0.0769 | 18 | 0.1111 | 95 | 0.1895 | 16 | 0 | 25 | 0.2 | 26 | 0.1154 | 19 | 0.1053 | 86 | 0.1163 |
| 2_2251487 | 22 | 0.1364 | 8 | 0 | 19 | 0.0526 | 19 | 0.1053 | 68 | 0.0882 | 8 | 0.25 | 21 | 0.0476 | 27 | 0.037 | 16 | 0.0625 | 72 | 0.0694 |
| 2_2258031 | 24 | 0.0417 | 15 | 0.1333 | 19 | 0.1579 | 34 | 0.0294 | 92 | 0.0761 | 10 | 0.1 | 33 | 0.1212 | 36 | 0.1111 | 23 | 0 | 102 | 0.0882 |
| 2_2258037 | 24 | 0.1667 | 15 | 0.0667 | 19 | 0.1053 | 37 | 0.2703 | 95 | 0.1789 | 12 | 0.5 | 30 | 0.1333 | 37 | 0.1081 | 23 | 0.2174 | 102 | 0.1863 |
| 2_2258041 | 21 | 0.1905 | 17 | 0.1765 | 23 | 0.087 | 39 | 0.0256 | 100 | 0.1 | 12 | 0.0833 | 33 | 0.0606 | 39 | 0.0513 | 27 | 0.0741 | 111 | 0.0631 |
| 2_2258207 | 30 | 0.2667 | 23 | 0.2174 | 34 | 0.2647 | 60 | 0.15 | 147 | 0.2109 | 22 | 0.3182 | 47 | 0.1915 | 52 | 0.1923 | 39 | 0.2051 | 160 | 0.2125 |
| 2_2258351 | 47 | 0.2553 | 37 | 0.2162 | 45 | 0.1556 | 81 | 0.1975 | 210 | 0.2048 | 24 | 0.2083 | 63 | 0.2698 | 77 | 0.1299 | 56 | 0.1071 | 220 | 0.1727 |
| 2_2258356 | 52 | 0.3846 | 38 | 0.2632 | 53 | 0.3585 | 84 | 0.2857 | 227 | 0.3216 | 24 | 0.375 | 65 | 0.2615 | 77 | 0.2468 | 50 | 0.4 | 216 | 0.3009 |
| 2_2258359 | 59 | 0.0508 | 48 | 0.0208 | 55 | 0.0909 | 94 | 0.117 | 256 | 0.0781 | 27 | 0.037 | 70 | 0.1429 | 88 | 0.0795 | 63 | 0.0952 | 248 | 0.0968 |
| 2_2276524 | 2 | 1 | NA | NA | 1 | 1 | NA | NA | 3 | 1 | 1 | 1 | 1 | 1 | 2 | 0.5 | 1 | 0 | 5 | 0.6 |
| 2_2276533 | 2 | 1 | NA | NA | NA | NA | NA | NA | 2 | 1 | 1 | 1 | NA | NA | 2 | 0 | 1 | 0 | 4 | 0.25 |
| 2_2283178 | 20 | 0.1 | 14 | 0.0714 | 17 | 0.1765 | 19 | 0.1053 | 70 | 0.1143 | 10 | 0.2 | 18 | 0.2222 | 16 | 0.0625 | 23 | 0.1304 | 67 | 0.1493 |
| 2_2283191 | 19 | 0 | 16 | 0.0625 | 17 | 0.1176 | 20 | 0.1 | 72 | 0.0694 | 9 | 0.2222 | 16 | 0.0625 | 15 | 0.0667 | 18 | 0.1667 | 58 | 0.1207 |
| 2_2283193 | 18 | 0.8333 | 15 | 0.6 | 17 | 0.5882 | 18 | 0.7222 | 68 | 0.6912 | 9 | 0.3333 | 15 | 0.6667 | 14 | 0.7143 | 22 | 0.6364 | 60 | 0.6167 |
| 2_2283193 | 20 | 0.2 | 16 | 0 | 18 | 0.1111 | 18 | 0.1111 | 72 | 0.1111 | 8 | 0.125 | 17 | 0.1765 | 17 | 0 | 21 | 0 | 63 | 0.0635 |
| 2_2290025 | 10 | 0 | 4 | 0.25 | 1 | 0 | 1 | 1 | 16 | 0.125 | NA | NA | 4 | 0.25 | 2 | 0 | 5 | 0.2 | 11 | 0.1818 |
| 2_2290030 | 9 | 0.4444 | 3 | 1 | 1 | 1 | 2 | 1 | 15 | 0.6667 | NA | NA | 2 | 0.5 | 2 | 0.5 | 4 | 0.5 | 8 | 0.5 |
| 2_2290039 | 9 | 0.1111 | 4 | 0 | 1 | 1 | 2 | 0.5 | 16 | 0.1875 | 1 | 1 | 4 | 0.25 | 3 | 0.6667 | 5 | 0.2 | 13 | 0.3846 |
| 2_2290097 | 7 | 0.1429 | 4 | 0 | 1 | 0 | 2 | 0 | 14 | 0.0714 | 1 | 0 | 4 | 0.25 | 3 | 0 | 4 | 0 | 12 | 0.0833 |
| 2_2290113 | 6 | 0.8333 | 4 | 0.25 | 1 | 0 | 2 | 0.5 | 13 | 0.5385 | 1 | 1 | 4 | 0.5 | 2 | 0.5 | 4 | 0.25 | 11 | 0.4545 |
| 2_2290142 | 5 | 0 | 3 | 0.3333 | 1 | 0 | 2 | 0 | 11 | 0.0909 | 1 | 0 | 3 | 0 | 2 | 0 | 3 | 0.3333 | 9 | 0.1111 |
| 2_2290159 | 4 | 0 | 2 | 0.5 | 1 | 0 | NA | NA | 7 | 0.1429 | 1 | 1 | 3 | 0.3333 | 2 | 0 | 3 | 0.3333 | 9 | 0.3333 |
| 2_2290168 | 4 | 0 | 2 | 0.5 | 1 | 0 | 1 | 0 | 8 | 0.125 | 1 | 1 | 2 | 0 | 2 | 0 | 2 | 0 | 7 | 0.1429 |
| 2_2290171 | 5 | 0.4 | 2 | 0 | 1 | 0 | 1 | 0 | 9 | 0.2222 | 1 | 0 | 3 | 0.6667 | 2 | 0.5 | 3 | 0 | 9 | 0.3333 |
| 2_2290177 | 5 | 0.6 | 2 | 0.5 | 1 | 0 | 1 | 0 | 9 | 0.4444 | 1 | 1 | 3 | 0.3333 | 1 | 1 | 2 | 0.5 | 7 | 0.5714 |
| 2_2290192 | 2 | 0 | 2 | 1 | 1 | 0 | NA | NA | 5 | 0.4 | 1 | 1 | 3 | 0 | 1 | 0 | 3 | 0.3333 | 8 | 0.25 |

| | | | | | | | | | | | | | | | | | | | | |
|---|---|---|---|---|---|---|---|---|---|---|---|---|---|---|---|---|---|---|---|---|
| 2_22901938 | 3 | 0 | 2 | 0.5 | 1 | 0 | 1 | 0 | 7 | 0.1429 | 1 | 1 | 3 | 0.3333 | 1 | 0 | 3 | 0 | 8 | 0.25 |
| 2_22902143 | 2 | 0.5 | 2 | 0 | 1 | 0 | 1 | 1 | 6 | 0.3333 | 1 | 1 | 3 | 0 | 1 | 1 | 3 | 0.6667 | 8 | 0.5 |
| 2_22959136 | 1 | 0 | 3 | 0.3333 | NA | NA | 1 | 0 | 5 | 0.2 | 1 | 0 | 3 | 0.3333 | 2 | 0.5 | NA | NA | 6 | 0.3333 |
| 2_23002584 | 10 | 0.3 | 5 | 0.2 | 7 | 0.4286 | 6 | 0.3333 | 28 | 0.3214 | 6 | 0 | 9 | 0.1111 | 7 | 0.4286 | 8 | 0.375 | 30 | 0.2333 |
| 2_23002708 | 10 | 0.2 | 4 | 0.75 | 7 | 0.2857 | 6 | 0.1667 | 27 | 0.2963 | 5 | 0.4 | 10 | 0.5 | 7 | 0.4286 | 7 | 0.4286 | 29 | 0.4483 |
| 2_23002767 | 10 | 0.1 | 5 | 0.2 | 7 | 0.1429 | 4 | 0 | 26 | 0.1154 | 5 | 0.4 | 12 | 0.3333 | 8 | 0.375 | 8 | 0.375 | 33 | 0.3636 |
| 2_23002791 | 9 | 0.7778 | 5 | 0.2 | 6 | 0.5 | 6 | 0.3333 | 26 | 0.5 | 5 | 0.4 | 11 | 0.4545 | 10 | 0.4 | 7 | 0.5714 | 33 | 0.4545 |
| 2_23002989 | 10 | 0.1 | 2 | 0 | 5 | 0 | 4 | 0.5 | 21 | 0.1429 | 5 | 0.2 | 10 | 0.2 | 8 | 0.375 | 6 | 0.3333 | 29 | 0.2759 |
| 2_23003037 | 10 | 0.1 | 5 | 0 | 7 | 0 | 6 | 0.1667 | 28 | 0.0714 | 6 | 0 | 13 | 0.0769 | 10 | 0.1 | 8 | 0 | 37 | 0.0541 |
| 2_23003140 | 10 | 0.2 | 5 | 0 | 6 | 0.5 | 6 | 0.3333 | 27 | 0.2593 | 5 | 0.8 | 13 | 0.3077 | 9 | 0.2222 | 8 | 0.25 | 35 | 0.3429 |
| 2_23003400 | 6 | 0.1667 | 5 | 0.2 | 7 | 0.7143 | 5 | 0.4 | 23 | 0.3913 | 4 | 1 | 13 | 0.3846 | 9 | 0.1111 | 7 | 0.2857 | 33 | 0.3636 |
| 2_23086984 | 15 | 0.3333 | 13 | 0.4615 | 11 | 0.1818 | 18 | 0.1667 | 57 | 0.2807 | 7 | 0.1429 | 14 | 0.3571 | 9 | 0.4444 | 11 | 0.5455 | 41 | 0.3902 |
| 2_23087046 | 12 | 0 | 12 | 0.0833 | 11 | 0.0909 | 18 | 0.0556 | 53 | 0.0566 | 6 | 0 | 14 | 0.1429 | 8 | 0.25 | 9 | 0.1111 | 37 | 0.1351 |
| 2_23087262 | 12 | 0.75 | 8 | 0.5 | 8 | 0.625 | 11 | 0.5455 | 39 | 0.6154 | 6 | 0.8333 | 11 | 0.7273 | 4 | 0.75 | 9 | 0.4444 | 30 | 0.6667 |
| 2_23087282 | 13 | 0.0769 | 11 | 0.1818 | 10 | 0.2 | 13 | 0.1538 | 47 | 0.1489 | 7 | 0.1429 | 12 | 0.0833 | 9 | 0.3333 | 9 | 0.2222 | 37 | 0.1892 |
| 2_23289761 | NA | NA | 2 | 1 | 1 | 0 | NA | NA | 3 | 0.6667 | NA | NA | 1 | 1 | NA | NA | NA | NA | 1 | 1 |
| 2_23289876 | NA | NA | 1 | 1 | NA | NA | NA | NA | 1 | 1 | NA | NA | 1 | 1 | NA | NA | NA | NA | 1 | 1 |
| 2_23290197 | NA | NA | 2 | 1 | 1 | 0 | NA | NA | 3 | 0.6667 | NA | NA | 1 | 1 | NA | NA | NA | NA | 1 | 1 |
| 2_23291150 | NA | NA | 2 | 1 | 2 | 0 | NA | NA | 4 | 0.5 | NA | NA | 1 | 0 | NA | NA | 1 | 1 | 2 | 0.5 |
| 2_23291515 | NA | NA | 2 | 0.5 | 2 | 0 | NA | NA | 4 | 0.25 | NA | NA | NA | NA | NA | NA | 1 | 1 | 1 | 1 |
| 2_23384862 | 1 | 1 | NA | NA | NA | NA | NA | NA | 1 | 1 | NA | NA | NA | NA | NA | NA | 1 | 1 | 1 | 1 |
| 2_23397093 | NA | NA | 1 | 0 | 1 | 0 | 2 | 0.5 | 4 | 0.25 | NA | NA | 1 | 0 | 5 | 0 | 2 | 0.5 | 8 | 0.125 |
| 2_23397883 | NA | NA | 1 | 1 | 1 | 0 | 1 | 1 | 3 | 0.6667 | NA | NA | 1 | 0 | 4 | 0 | 2 | 0.5 | 7 | 0.1429 |
| 2_23398410 | NA | NA | 1 | 1 | 1 | 1 | NA | NA | 2 | 1 | NA | NA | 1 | 0 | 2 | 0.5 | 1 | 1 | 4 | 0.5 |
| 2_23401114 | NA | NA | NA | NA | 2 | 0.5 | 1 | 0 | 3 | 0.3333 | NA | NA | NA | NA | 3 | 0.3333 | 2 | 1 | 5 | 0.6 |
| 2_23401154 | NA | NA | NA | NA | 2 | 0.5 | 1 | 1 | 3 | 0.6667 | NA | NA | NA | NA | 3 | 0.3333 | 2 | 1 | 5 | 0.6 |
| 2_23401310 | NA | NA | NA | NA | 1 | 1 | 1 | 1 | 2 | 1 | NA | NA | NA | NA | 2 | 1 | 2 | 1 | 4 | 1 |
| 2_23798692 | 2 | 1 | 1 | 1 | NA | NA | NA | NA | 3 | 1 | NA | NA | 1 | 0 | 1 | 1 | 1 | 1 | 3 | 0.6667 |
| 2_23893807 | 4 | 1 | 5 | 1 | 1 | 1 | 4 | 0.75 | 14 | 0.9286 | 2 | 1 | 3 | 1 | 2 | 1 | 6 | 0.8333 | 13 | 0.9231 |
| 2_23893816 | 3 | 0.3333 | 5 | 0 | 2 | 0.5 | 3 | 0.3333 | 13 | 0.2308 | 2 | 0 | 1 | 1 | 4 | 0.25 | 5 | 0.6 | 12 | 0.4167 |
| 2_24297288 | 1 | 1 | NA | NA | NA | NA | NA | NA | 1 | 1 | NA | NA | 1 | 1 | NA | NA | NA | NA | 1 | 1 |
| 2_24297400 | 1 | 1 | NA | NA | NA | NA | NA | NA | 1 | 1 | NA | NA | 1 | 1 | NA | NA | NA | NA | 1 | 1 |
| 2_24494418 | 5 | 0.6 | 4 | 0.75 | 3 | 0.3333 | 7 | 0.5714 | 19 | 0.5789 | 3 | 0.3333 | 6 | 0.1667 | 5 | 0.6 | 5 | 1 | 19 | 0.5263 |
| 2_24494554 | 4 | 0.5 | 4 | 0 | 4 | 0 | 7 | 0.2857 | 19 | 0.2105 | 3 | 0 | 5 | 0.2 | 5 | 0.2 | 2 | 0 | 15 | 0.1333 |
| 2_24494688 | 3 | 0.3333 | 3 | 0.3333 | 3 | 0.3333 | 6 | 0.1667 | 15 | 0.2667 | 3 | 0.3333 | 4 | 0.25 | 6 | 0.5 | 4 | 0.25 | 17 | 0.3529 |
| 2_24496223 | 3 | 0.3333 | 2 | 0.5 | 3 | 0 | 1 | 0 | 9 | 0.2222 | 1 | 1 | 2 | 0 | 2 | 0 | 2 | 0.5 | 7 | 0.2857 |
| 2_24496239 | 1 | 0 | 2 | 1 | 1 | 0 | 1 | 0 | 5 | 0.4 | 1 | 0 | 1 | 0 | 2 | 0 | 2 | 0.5 | 6 | 0.1667 |
| 2_24496386 | 3 | 0.6667 | 3 | 0.3333 | 3 | 0 | 1 | 0 | 10 | 0.3 | 2 | 0.5 | 2 | 0 | 3 | 0 | 2 | 0.5 | 9 | 0.2222 |
| 2_24496411 | 2 | 0.5 | 3 | 1 | 5 | 0.2 | 3 | 0.3333 | 13 | 0.4615 | 3 | 0.3333 | 3 | 0.3333 | 4 | 0.5 | 1 | 1 | 11 | 0.4545 |
| 2_24496645 | 4 | 1 | 2 | 1 | 3 | 0 | 3 | 1 | 12 | 0.75 | 2 | 0 | 2 | 1 | 5 | 0.2 | 2 | 1 | 11 | 0.4545 |
| 2_24496722 | 4 | 0.5 | 2 | 1 | 4 | 1 | 3 | 0.6667 | 13 | 0.7692 | 3 | 0.6667 | 2 | 1 | 4 | 1 | 3 | 0.6667 | 12 | 0.8333 |
| 2_24496808 | 4 | 0.75 | 1 | 1 | 2 | 1 | 2 | 1 | 9 | 0.8889 | 1 | 0 | 2 | 1 | 1 | 1 | 3 | 0.6667 | 7 | 0.7143 |
| 2_24496900 | 3 | 1 | 2 | 0.5 | 2 | 0.5 | 3 | 0 | 10 | 0.5 | 1 | 0 | 2 | 1 | 4 | 0.75 | 3 | 0.3333 | 10 | 0.6 |
| 2_24681256 | 5 | 0.6 | 4 | 0.75 | 4 | 0.5 | 5 | 0.2 | 18 | 0.5 | 2 | 0.5 | 4 | 0.5 | 4 | 0 | 1 | 1 | 11 | 0.3636 |
| 2_24681264 | 5 | 0.2 | 3 | 0.3333 | 3 | 0 | 5 | 0 | 16 | 0.125 | 2 | 0 | 4 | 0.5 | 4 | 0.25 | 1 | 0 | 11 | 0.2727 |
| 2_24681290 | 5 | 0.4 | 4 | 0.5 | 5 | 0.4 | 5 | 0.4 | 19 | 0.4211 | 2 | 0 | 5 | 0.4 | 4 | 0.25 | 2 | 0 | 13 | 0.2308 |
| 2_24681532 | 4 | 1 | 3 | 1 | 4 | 1 | 4 | 1 | 15 | 1 | 2 | 1 | 4 | 1 | 4 | 1 | 2 | 1 | 12 | 1 |
| 2_24681727 | 2 | 0 | 3 | 0 | 3 | 0 | 5 | 0.2 | 13 | 0.0769 | 1 | 0 | 5 | 0 | 4 | 0.25 | 1 | 0 | 11 | 0.0909 |
| 2_24681748 | 4 | 0 | 3 | 0.3333 | 4 | 0 | 6 | 0.1667 | 17 | 0.1176 | 1 | 1 | 4 | 0 | 4 | 0.25 | 2 | 0.5 | 11 | 0.2727 |
| 2_24681763 | 4 | 1 | 3 | 1 | 4 | 1 | 6 | 0.8333 | 17 | 0.9412 | 2 | 1 | 5 | 1 | 4 | 1 | 2 | 1 | 13 | 1 |
| 2_24681770 | 4 | 1 | 2 | 1 | 4 | 1 | 6 | 1 | 16 | 1 | 2 | 1 | 5 | 1 | 5 | 1 | 2 | 1 | 14 | 1 |
| 2_24681860 | 4 | 1 | 3 | 1 | 4 | 1 | 6 | 1 | 17 | 1 | 2 | 1 | 5 | 0.6 | 3 | 0.6667 | 2 | 1 | 12 | 0.75 |
| 2_24681895 | 3 | 0.3333 | 3 | 0.3333 | 4 | 0.25 | 5 | 0.4 | 15 | 0.3333 | 1 | 0 | 2 | 0 | 3 | 0.6667 | 1 | 0 | 7 | 0.2857 |
| 2_24681917 | 3 | 1 | 3 | 1 | 3 | 1 | 6 | 0.8333 | 15 | 0.9333 | 2 | 1 | 3 | 1 | 5 | 1 | 2 | 1 | 12 | 1 |
| 2_24681940 | 3 | 1 | 3 | 1 | 4 | 1 | 6 | 0.8333 | 16 | 0.9375 | 1 | 1 | 4 | 1 | 5 | 0.8 | 2 | 1 | 12 | 0.9167 |
| 2_24849000 | 3 | 0 | 4 | 0.25 | 3 | 0 | 2 | 0 | 12 | 0.0833 | NA | NA | 1 | 0 | 1 | 0 | 6 | 0.1667 | 8 | 0.125 |
| 2_24849142 | 2 | 1 | 4 | 1 | 3 | 1 | 1 | 1 | 10 | 1 | 1 | 1 | 1 | 1 | NA | NA | 5 | 0.8 | 7 | 0.8571 |
| 2_24849154 | 3 | 0 | 4 | 0.25 | 2 | 0 | 2 | 0 | 11 | 0.0909 | 2 | 0 | 2 | 0 | 2 | 0.5 | 6 | 0 | 12 | 0.0833 |
| 2_24849205 | 2 | 0 | 4 | 0 | 3 | 0 | 2 | 0.5 | 11 | 0.0909 | 2 | 0 | 2 | 0 | 2 | 0 | 6 | 0.1667 | 12 | 0.0833 |
| 2_24849233 | 3 | 0 | 5 | 0.4 | 2 | 0 | 2 | 0 | 12 | 0.1667 | 1 | 0 | 2 | 1 | 2 | 0.5 | 6 | 0.3333 | 11 | 0.4545 |
| 2_24849266 | 4 | 0 | 3 | 0 | 3 | 0.3333 | 2 | 0.5 | 12 | 0.1667 | 1 | 0 | 1 | 0 | 2 | 0 | 6 | 0.6667 | 10 | 0.4 |
| 2_24849287 | 4 | 0.25 | 5 | 0.8 | 3 | 0.6667 | 2 | 0.5 | 14 | 0.5714 | 2 | 0.5 | 2 | 1 | 2 | 0.5 | 6 | 0.6667 | 12 | 0.6667 |
| 2_24849430 | 1 | 1 | 3 | 0.6667 | 2 | 1 | 3 | 0.3333 | 9 | 0.6667 | NA | NA | 1 | 1 | 2 | 0.5 | 4 | 0.5 | 7 | 0.5714 |
| 2_24849459 | NA | NA | 2 | 0.5 | 1 | 1 | 1 | 1 | 4 | 0.75 | NA | NA | NA | NA | 1 | 1 | 4 | 1 | 5 | 1 |
| 2_24849740 | NA | NA | 1 | 0 | 2 | 0 | 4 | 1 | 7 | 0.5714 | NA | NA | 1 | 0 | 1 | 1 | 4 | 0.75 | 6 | 0.6667 |

| | | | | | | | | | | | | | | | | | | | | |
|---|---|---|---|---|---|---|---|---|---|---|---|---|---|---|---|---|---|---|---|---|
| 2_2485115 | 1 | 0 | 1 | 1 | NA | NA | 1 | 1 | 3 | 0.6667 | NA | NA | 2 | 0 | NA | NA | 2 | 0.5 | 4 | 0.25 |
| 2_2485117 | 1 | 0 | 1 | 1 | 1 | 1 | NA | NA | 3 | 0.6667 | NA | NA | 2 | 0.5 | 1 | 0 | 3 | 0.3333 | 6 | 0.3333 |
| 2_2485164 | 1 | 1 | NA | NA | 1 | 0 | 1 | 0 | 3 | 0.3333 | NA | NA | 2 | 0 | 1 | 1 | 3 | 0.6667 | 6 | 0.5 |
| 2_2485176 | 1 | 1 | NA | NA | NA | NA | NA | NA | 1 | 1 | NA | NA | NA | NA | NA | NA | 1 | 1 | 1 | 1 |
| 2_2485194 | 1 | 1 | NA | NA | NA | NA | NA | NA | 1 | 1 | NA | NA | NA | NA | NA | NA | 1 | 1 | 1 | 1 |
| 2_2485255 | 5 | 0 | 5 | 0.2 | 1 | 0 | 6 | 0.1667 | 17 | 0.1176 | 1 | 0 | 5 | 0.2 | 1 | 0 | 1 | 0 | 8 | 0.125 |
| 2_2485270 | 1 | 0 | 2 | 1 | 1 | 1 | 4 | 0 | 8 | 0.375 | NA | NA | 2 | 0.5 | NA | NA | NA | NA | 2 | 0.5 |
| 2_2485314 | 4 | 0.25 | 4 | 0 | 1 | 0 | 4 | 0.5 | 13 | 0.2308 | 1 | 0 | 3 | 0 | 2 | 0.5 | 1 | 0 | 7 | 0.1429 |
| 2_2486154 | 4 | 0 | 5 | 0.2 | 2 | 0 | 8 | 0 | 19 | 0.0526 | 2 | 0 | 6 | 0.1667 | 5 | 0 | 1 | 0 | 14 | 0.0714 |
| 2_2486160 | 4 | 0 | 6 | 0.1667 | 2 | 0 | 8 | 0 | 20 | 0.05 | 1 | 0 | 5 | 0.2 | 4 | 0.25 | 1 | 0 | 11 | 0.1818 |
| 2_2486166 | 4 | 0 | 5 | 0 | 1 | 0 | 9 | 0.1111 | 19 | 0.0526 | 2 | 0 | 5 | 0 | 4 | 0.25 | 1 | 0 | 12 | 0.0833 |
| 2_2486178 | 3 | 0.3333 | 6 | 0.1667 | 2 | 0.5 | 7 | 0.5714 | 18 | 0.3889 | 1 | 0 | 4 | 1 | 5 | 0.6 | 1 | 0 | 11 | 0.6364 |
| 2_2486181 | 4 | 0.5 | 4 | 0.25 | 2 | 1 | 7 | 0.5714 | 17 | 0.5294 | 1 | 0 | 4 | 0.5 | 3 | 1 | 1 | 1 | 9 | 0.6667 |
| 2_2486183 | 6 | 1 | 5 | 1 | 1 | 0 | 9 | 0.8889 | 21 | 0.9048 | 2 | 1 | 5 | 0.8 | 5 | 1 | NA | NA | 12 | 0.9167 |
| 2_2486185 | 6 | 0.6667 | 6 | 0.6667 | 2 | 1 | 9 | 0.4444 | 23 | 0.6087 | 2 | 0.5 | 5 | 1 | 5 | 0.6 | 1 | 0 | 13 | 0.6923 |
| 2_2486189 | 6 | 0.1667 | 6 | 0.3333 | 2 | 0 | 8 | 0.125 | 22 | 0.1818 | 1 | 1 | 5 | 0 | 4 | 0 | NA | NA | 10 | 0.1 |
| 2_2486193 | 6 | 0.1667 | 6 | 0 | 2 | 0 | 9 | 0.1111 | 23 | 0.087 | 2 | 0 | 5 | 0.2 | 4 | 0.25 | 1 | 0 | 12 | 0.1667 |
| 2_2486202 | 6 | 1 | 6 | 1 | 3 | 1 | 9 | 1 | 24 | 1 | 2 | 1 | 5 | 1 | 5 | 1 | 1 | 1 | 13 | 1 |
| 2_2486422 | 18 | 0.2778 | 23 | 0.087 | 14 | 0.1429 | 16 | 0.125 | 71 | 0.1549 | 16 | 0.125 | 21 | 0.0952 | 14 | 0.0714 | 13 | 0.1538 | 64 | 0.1094 |
| 2_2486435 | 18 | 0.9444 | 24 | 0.8333 | 10 | 0.8 | 16 | 0.8125 | 68 | 0.8529 | 16 | 0.8125 | 18 | 1 | 13 | 1 | 14 | 1 | 61 | 0.9508 |
| 2_2486436 | 18 | 0.8889 | 22 | 0.8636 | 10 | 1 | 16 | 0.9375 | 66 | 0.9091 | 14 | 0.9286 | 16 | 1 | 14 | 0.7857 | 11 | 1 | 55 | 0.9273 |
| 2_2486442 | 18 | 0.3333 | 19 | 0.3158 | 10 | 0.1 | 16 | 0.375 | 63 | 0.3016 | 16 | 0.0625 | 17 | 0.4706 | 14 | 0.3571 | 14 | 0.4286 | 61 | 0.3279 |
| 2_2486448 | 15 | 0.8 | 19 | 0.8947 | 8 | 1 | 13 | 0.8462 | 55 | 0.8727 | 11 | 1 | 11 | 1 | 9 | 1 | 11 | 1 | 42 | 1 |
| 2_2486504 | 15 | 0 | 8 | 0 | 5 | 0 | 12 | 0.1667 | 40 | 0.05 | 6 | 0 | 12 | 0.1667 | 5 | 0 | 8 | 0 | 31 | 0.0645 |
| 2_2486532 | NA | NA | NA | NA | NA | NA | 2 | 1 | 2 | 1 | 1 | 1 | 1 | 1 | NA | NA | NA | NA | 2 | 1 |
| 2_2486540 | NA | NA | NA | NA | NA | NA | 3 | 0.3333 | 3 | 0.3333 | 1 | 1 | 1 | 0 | NA | NA | NA | NA | 2 | 0.5 |
| 2_2486547 | NA | NA | NA | NA | NA | NA | 1 | 1 | 1 | 1 | 1 | 1 | 1 | 1 | NA | NA | NA | NA | 2 | 1 |
| 2_2494596 | 7 | 0.4286 | 32 | 0.125 | 24 | 0.0833 | 11 | 0.0909 | 74 | 0.1351 | 24 | 0.125 | 18 | 0.3333 | 17 | 0.2353 | 18 | 0.3333 | 77 | 0.2468 |
| 2_2495176 | 8 | 0.125 | 39 | 0.1795 | 35 | 0.1429 | 16 | 0.3125 | 98 | 0.1837 | 33 | 0.2424 | 34 | 0.2647 | 24 | 0.25 | 20 | 0.25 | 111 | 0.2523 |
| 2_2495183 | 11 | 0.2727 | 47 | 0.1064 | 35 | 0.1714 | 15 | 0.1333 | 108 | 0.1481 | 31 | 0.1935 | 31 | 0.0323 | 29 | 0.069 | 24 | 0.0417 | 115 | 0.087 |
| 2_2495186 | 11 | 0.0909 | 47 | 0.3191 | 36 | 0.1667 | 14 | 0.5714 | 108 | 0.2778 | 32 | 0.375 | 32 | 0.4688 | 27 | 0.4815 | 22 | 0.1364 | 113 | 0.3805 |
| 2_2495190 | 11 | 0.2727 | 50 | 0.24 | 40 | 0.225 | 17 | 0.1765 | 118 | 0.2288 | 38 | 0.2368 | 36 | 0.2222 | 28 | 0.2143 | 25 | 0.28 | 127 | 0.2362 |
| 2_2495267 | 17 | 0.0588 | 58 | 0.1897 | 52 | 0.0385 | 17 | 0.1176 | 144 | 0.1111 | 48 | 0.1458 | 38 | 0.1316 | 31 | 0.129 | 22 | 0.0455 | 139 | 0.1223 |
| 2_2507123 | 4 | 0.25 | 2 | 0 | 1 | 0 | NA | NA | 7 | 0.1429 | 2 | 0 | 5 | 0 | 4 | 0.25 | 3 | 0.3333 | 14 | 0.1429 |
| 2_2507125 | NA | NA | 2 | 1 | 1 | 0 | NA | NA | 3 | 0.6667 | 1 | 0 | 1 | 0 | 4 | 0.25 | 2 | 1 | 8 | 0.375 |
| 2_2507141 | 4 | 0.75 | 2 | 0 | 1 | 1 | NA | NA | 7 | 0.5714 | 2 | 0.5 | 5 | 0 | 5 | 0.4 | 5 | 0.6 | 17 | 0.3529 |
| 2_2507147 | 5 | 1 | 1 | 1 | 1 | 1 | NA | NA | 7 | 1 | 3 | 1 | 5 | 1 | 6 | 1 | 3 | 1 | 17 | 1 |
| 2_2507162 | 3 | 1 | 1 | 1 | 1 | 1 | NA | NA | 5 | 1 | 2 | 0 | 3 | 0.6667 | 8 | 0.875 | 4 | 1 | 17 | 0.7647 |
| 2_2507184 | 3 | 0.6667 | 3 | 1 | 2 | 1 | NA | NA | 8 | 0.875 | 3 | 0.6667 | 5 | 0.8 | 8 | 0.75 | 4 | 1 | 20 | 0.8 |
| 2_2507187 | 5 | 0.2 | 3 | 0 | 2 | 0 | NA | NA | 10 | 0.1 | 3 | 0 | 5 | 0 | 7 | 0.1429 | 5 | 0.2 | 20 | 0.1 |
| 2_2507263 | 5 | 0.2 | 5 | 0.6 | 1 | 1 | NA | NA | 11 | 0.4545 | 3 | 0 | 6 | 0.5 | 6 | 0.6667 | 5 | 0.4 | 20 | 0.45 |
| 2_2507299 | 5 | 0.4 | 5 | 0.6 | NA | NA | 1 | 1 | 11 | 0.5455 | 2 | 0 | 2 | 0.5 | 4 | 0.5 | 6 | 0.6667 | 14 | 0.5 |
| 2_2507305 | 4 | 0 | 4 | 0.5 | 1 | 0 | 1 | 0 | 10 | 0.2 | 4 | 0.25 | 3 | 0.3333 | 7 | 0.1429 | 5 | 0.4 | 19 | 0.2632 |
| 2_2509013 | 10 | 0.2 | 7 | 0.1429 | 6 | 0 | 10 | 0.1 | 33 | 0.1212 | 5 | 0 | 5 | 0 | 2 | 0 | 7 | 0.2857 | 19 | 0.1053 |
| 2_2509031 | 10 | 0.1 | 7 | 0.1429 | 7 | 0.4286 | 9 | 0.3333 | 33 | 0.2424 | 5 | 0.4 | 5 | 0.4 | 3 | 0 | 8 | 0.125 | 21 | 0.2381 |
| 2_2509032 | 10 | 0.2 | 6 | 0.5 | 7 | 0.2857 | 10 | 0.2 | 33 | 0.2727 | 6 | 0.5 | 5 | 0.4 | 3 | 0.3333 | 6 | 0.1667 | 20 | 0.35 |
| 2_2510117 | 3 | 0.3333 | 4 | 0 | 3 | 0 | 5 | 0 | 15 | 0.0667 | 1 | 0 | 3 | 0 | 2 | 0 | 6 | 0.1667 | 12 | 0.0833 |
| 2_2511228 | 50 | 0.82 | 58 | 0.7414 | 50 | 0.72 | 35 | 0.8 | 193 | 0.7668 | 34 | 0.8235 | 35 | 0.8 | 20 | 0.55 | 43 | 0.7907 | 132 | 0.7652 |
| 2_2511259 | 46 | 0.1739 | 46 | 0.1957 | 39 | 0.3333 | 31 | 0.2581 | 162 | 0.2346 | 31 | 0.3548 | 30 | 0.3667 | 18 | 0.4444 | 41 | 0.3171 | 120 | 0.3583 |
| 2_2511260 | 60 | 0.1167 | 55 | 0.0545 | 48 | 0.0625 | 41 | 0.0732 | 204 | 0.0784 | 39 | 0.0256 | 38 | 0.0526 | 21 | 0.0476 | 50 | 0.08 | 148 | 0.0541 |
| 2_2511264 | 45 | 0.0222 | 42 | 0 | 40 | 0.075 | 35 | 0.1429 | 162 | 0.0556 | 30 | 0.0333 | 31 | 0.0968 | 16 | 0.0625 | 42 | 0.0952 | 119 | 0.0756 |
| 2_2511267 | 37 | 0.4054 | 39 | 0.3846 | 30 | 0.4333 | 29 | 0.3793 | 135 | 0.4 | 24 | 0.375 | 26 | 0.2308 | 12 | 0.3333 | 35 | 0.4 | 97 | 0.3402 |
| 2_2511272 | 50 | 0.2 | 43 | 0.1395 | 40 | 0.1 | 39 | 0.2051 | 172 | 0.1628 | 31 | 0.129 | 34 | 0.2647 | 21 | 0.0952 | 42 | 0.2381 | 128 | 0.1953 |
| 2_2511276 | 46 | 0.1522 | 40 | 0.05 | 37 | 0.0541 | 32 | 0.125 | 155 | 0.0968 | 30 | 0.0333 | 33 | 0.1818 | 17 | 0.1176 | 38 | 0.2105 | 118 | 0.1441 |
| 2_2511312 | 45 | 0.2 | 40 | 0.225 | 38 | 0.2105 | 34 | 0.2647 | 157 | 0.2229 | 31 | 0.0968 | 29 | 0.2759 | 16 | 0.1875 | 40 | 0.225 | 116 | 0.1983 |
| 2_2511316 | 28 | 0.1071 | 24 | 0.0833 | 25 | 0.16 | 23 | 0.087 | 100 | 0.11 | 13 | 0 | 17 | 0.1176 | 12 | 0.25 | 23 | 0.3043 | 65 | 0.1846 |
| 2_2511323 | 42 | 0.1905 | 32 | 0.0938 | 29 | 0.1724 | 26 | 0.1154 | 129 | 0.1473 | 27 | 0.1111 | 22 | 0.0909 | 13 | 0.1538 | 30 | 0.2333 | 92 | 0.1522 |
| 2_2511434 | NA | NA | NA | NA | NA | NA | 1 | 1 | 1 | 1 | 1 | 1 | 1 | 0 | NA | NA | NA | NA | 2 | 0.5 |
| 2_2513125 | 2 | 0.5 | NA | NA | NA | NA | NA | NA | 2 | 0.5 | 1 | 1 | NA | NA | NA | NA | NA | NA | 1 | 1 |
| 2_2513134 | 1 | 0 | 1 | 1 | NA | NA | NA | NA | 2 | 0.5 | 1 | 1 | NA | NA | NA | NA | NA | NA | 1 | 1 |
| 2_2513167 | 2 | 1 | NA | NA | NA | NA | NA | NA | 2 | 1 | 1 | 1 | NA | NA | NA | NA | NA | NA | 1 | 1 |
| 2_2513190 | 2 | 0.5 | 1 | 0 | NA | NA | NA | NA | 3 | 0.3333 | 1 | 1 | NA | NA | NA | NA | NA | NA | 1 | 1 |
| 2_2513295 | 28 | 0.1071 | 19 | 0.0526 | 25 | 0.2 | 18 | 0.1667 | 90 | 0.1333 | 12 | 0.25 | 15 | 0 | 14 | 0.0714 | 14 | 0.1429 | 55 | 0.1091 |
| 2_2513301 | 26 | 0.1923 | 19 | 0.0526 | 25 | 0.44 | 17 | 0.0588 | 87 | 0.2069 | 14 | 0.2143 | 12 | 0.25 | 12 | 0.1667 | 15 | 0.2 | 53 | 0.2075 |
| 2_2513311 | 24 | 0.8333 | 21 | 0.9524 | 25 | 0.96 | 17 | 0.8824 | 87 | 0.908 | 14 | 0.7143 | 15 | 0.6667 | 12 | 1 | 11 | 0.8182 | 52 | 0.7885 |

| | | | | | | | | | | | | | | | | | | | | |
|---|---|---|---|---|---|---|---|---|---|---|---|---|---|---|---|---|---|---|---|---|
| 2_2513329 | 33 | 0.3333 | 22 | 0.2273 | 29 | 0.2759 | 22 | 0.3182 | 106 | 0.2925 | 14 | 0.3571 | 19 | 0.3158 | 16 | 0.1875 | 13 | 0.1538 | 62 | 0.2581 |
| 2_2513351 | 35 | 0.2857 | 22 | 0.0455 | 32 | 0.125 | 27 | 0.0741 | 116 | 0.1466 | 19 | 0.1579 | 20 | 0.05 | 18 | 0.1667 | 16 | 0.125 | 73 | 0.1233 |
| 2_2513355 | 33 | 0.2424 | 20 | 0.2 | 28 | 0.25 | 23 | 0.2609 | 104 | 0.2404 | 17 | 0.3529 | 19 | 0.2105 | 16 | 0.25 | 14 | 0.2143 | 66 | 0.2576 |
| 2_2513389 | 32 | 0.9062 | 29 | 0.931 | 29 | 0.8621 | 25 | 0.84 | 115 | 0.887 | 18 | 0.8889 | 24 | 0.9167 | 18 | 0.8889 | 16 | 0.6875 | 76 | 0.8553 |
| 2_2513391 | 27 | 0.3333 | 23 | 0.5217 | 30 | 0.4 | 22 | 0.4545 | 102 | 0.4216 | 20 | 0.3 | 22 | 0.1364 | 15 | 0.4 | 15 | 0.0667 | 72 | 0.2222 |
| 2_2513399 | 27 | 0.8889 | 20 | 1 | 22 | 0.9091 | 20 | 0.95 | 89 | 0.9326 | 14 | 1 | 19 | 0.8421 | 16 | 0.8125 | 11 | 0.8182 | 60 | 0.8667 |
| 2_2513415 | 31 | 0.1613 | 25 | 0.16 | 27 | 0.1481 | 29 | 0.2759 | 112 | 0.1875 | 19 | 0.4737 | 22 | 0.1364 | 17 | 0.2941 | 14 | 0.1429 | 72 | 0.2639 |
| 2_2513425 | 23 | 0.2174 | 24 | 0.2917 | 25 | 0.32 | 33 | 0.2424 | 105 | 0.2667 | 18 | 0.3333 | 23 | 0.2609 | 19 | 0.5789 | 13 | 0.1538 | 73 | 0.3425 |
| 2_2513516 | 3 | 0.3333 | 3 | 0.6667 | 1 | 1 | 2 | 0 | 9 | 0.4444 | 2 | 0.5 | 1 | 1 | 2 | 0.5 | 1 | 1 | 6 | 0.6667 |
| 2_2513540 | 3 | 0 | 2 | 1 | 2 | 0.5 | 3 | 0.3333 | 10 | 0.4 | 2 | 0.5 | 1 | 0 | 2 | 0 | 1 | 1 | 6 | 0.3333 |
| 2_2513547 | 3 | 0.6667 | 3 | 0.6667 | NA | NA | 2 | 1 | 8 | 0.75 | 2 | 1 | NA | NA | 2 | 1 | 1 | 0 | 5 | 0.8 |
| 2_2513556 | 2 | 0.5 | 3 | 0.6667 | 2 | 0 | 2 | 0 | 9 | 0.3333 | 1 | 1 | 1 | 1 | 2 | 0 | 1 | 1 | 5 | 0.6 |
| 2_2513561 | 3 | 0.6667 | 2 | 1 | 1 | 1 | NA | NA | 6 | 0.8333 | 1 | 1 | 1 | 0 | 2 | 1 | NA | NA | 4 | 0.75 |
| 2_2513562 | 2 | 1 | 3 | 1 | 2 | 0.5 | 1 | 0 | 8 | 0.75 | NA | NA | 1 | 0 | 2 | 1 | NA | NA | 3 | 0.6667 |
| 2_2513569 | 3 | 0 | 3 | 0.3333 | 2 | 0.5 | 1 | 0 | 9 | 0.2222 | 1 | 1 | 1 | 0 | 2 | 0.5 | 1 | 1 | 5 | 0.6 |
| 2_2513572 | 3 | 0.3333 | 3 | 0.3333 | 1 | 1 | 1 | 1 | 8 | 0.5 | 1 | 1 | 1 | 1 | 2 | 1 | 1 | 1 | 5 | 1 |
| 2_2513586 | NA | NA | 2 | 1 | 1 | 1 | 1 | 0 | 4 | 0.75 | 1 | 1 | 1 | 1 | 1 | 1 | 1 | 1 | 4 | 1 |
| 2_2513608 | NA | NA | 2 | 1 | NA | NA | 1 | 1 | 3 | 1 | 1 | 1 | 1 | 1 | 2 | 1 | 1 | 1 | 5 | 1 |
| 2_2513611 | 1 | 1 | 3 | 0 | NA | NA | 1 | 0 | 5 | 0.2 | NA | NA | 1 | 0 | 1 | 1 | 1 | 1 | 3 | 0.6667 |
| 2_2513625 | NA | NA | 3 | 0.6667 | 1 | 0 | 1 | 1 | 5 | 0.6 | NA | NA | 1 | 1 | NA | NA | 1 | 1 | 2 | 1 |
| 2_2513655 | 1 | 0 | 2 | 0.5 | NA | NA | 1 | 1 | 4 | 0.5 | 1 | 1 | 1 | 1 | 2 | 0 | 1 | 1 | 5 | 0.6 |
| 2_2513696 | 1 | 1 | 1 | 0 | NA | NA | NA | NA | 2 | 0.5 | NA | NA | 1 | 1 | 1 | 0 | NA | NA | 2 | 0.5 |
| 2_2513741 | 1 | 1 | 5 | 0.6 | 2 | 1 | 2 | 1 | 10 | 0.8 | NA | NA | 2 | 1 | 1 | 1 | 2 | 1 | 5 | 1 |
| 2_2513750 | 1 | 0 | 4 | 0 | 2 | 0.5 | 1 | 0 | 8 | 0.125 | 1 | 1 | 2 | 0 | 1 | 0 | 2 | 0 | 6 | 0.1667 |
| 2_2513773 | 1 | 0 | 5 | 0.2 | 2 | 0 | 1 | 1 | 9 | 0.2222 | 1 | 0 | 2 | 0 | 1 | 0 | 1 | 1 | 5 | 0.2 |
| 2_2514600 | NA | NA | 1 | 0 | 1 | 1 | 2 | 0 | 4 | 0.25 | 1 | 1 | 4 | 0.5 | 1 | 0 | 1 | 1 | 7 | 0.5714 |
| 2_2514614 | NA | NA | 1 | 0 | 2 | 1 | 2 | 0.5 | 5 | 0.6 | NA | NA | 5 | 0.2 | 1 | 0 | 2 | 0 | 8 | 0.125 |
| 2_2514629 | NA | NA | 1 | 0 | 2 | 0.5 | 2 | 0 | 5 | 0.2 | 1 | 0 | 5 | 0.6 | NA | NA | 2 | 1 | 8 | 0.625 |
| 2_2514633 | NA | NA | NA | NA | 1 | 1 | 1 | 1 | 2 | 1 | NA | NA | 1 | 1 | NA | NA | 2 | 1 | 3 | 1 |
| 2_2514635 | 1 | 1 | NA | NA | 1 | 1 | 1 | 1 | 3 | 1 | NA | NA | 1 | 1 | NA | NA | 2 | 1 | 3 | 1 |
| 2_2514946 | NA | NA | 1 | 1 | NA | NA | 2 | 1 | 3 | 1 | 1 | 1 | 2 | 1 | 3 | 0.6667 | 1 | 1 | 7 | 0.8571 |
| 2_2514959 | NA | NA | NA | NA | NA | NA | 1 | 1 | 1 | 1 | 1 | 1 | NA | NA | 2 | 0.5 | NA | NA | 3 | 0.6667 |
| 2_2514982 | NA | NA | 1 | 0 | NA | NA | 2 | 0.5 | 3 | 0.3333 | 1 | 1 | 2 | 0.5 | 3 | 0 | 2 | 1 | 8 | 0.5 |
| 2_2515347 | 1 | 1 | 1 | 1 | 1 | 1 | 1 | 0 | 4 | 0.75 | NA | NA | NA | NA | 1 | 1 | NA | NA | 1 | 1 |
| 2_2515359 | 1 | 1 | 1 | 1 | 1 | 1 | 1 | 0 | 4 | 0.75 | NA | NA | 1 | 1 | 1 | 1 | NA | NA | 2 | 1 |
| 2_2515430 | 3 | 0.3333 | 1 | 1 | 1 | 0 | 2 | 0.5 | 7 | 0.4286 | NA | NA | 1 | 1 | 1 | 0 | NA | NA | 2 | 0.5 |
| 2_2515435 | 3 | 0 | 1 | 1 | 2 | 0 | 2 | 0.5 | 8 | 0.25 | NA | NA | 1 | 0 | 1 | 1 | NA | NA | 2 | 0.5 |
| 2_2515945 | 4 | 0.25 | 1 | 0 | 1 | 0 | 2 | 0 | 8 | 0.125 | 1 | 0 | 3 | 0 | 2 | 0.5 | NA | NA | 6 | 0.1667 |
| 2_2516127 | 11 | 0.1818 | 19 | 0.1579 | 9 | 0.2222 | 8 | 0 | 47 | 0.1489 | 10 | 0.1 | 5 | 0 | 12 | 0.0833 | 10 | 0.1 | 37 | 0.0811 |
| 2_2516133 | 7 | 0.1429 | 16 | 0.4375 | 6 | 0.3333 | 5 | 0.2 | 34 | 0.3235 | 5 | 0.2 | 4 | 0 | 7 | 0.4286 | 10 | 0.4 | 26 | 0.3077 |
| 2_2516149 | 10 | 0.1 | 16 | 0.25 | 7 | 0.1429 | 6 | 0.3333 | 39 | 0.2051 | 9 | 0.3333 | 5 | 0.4 | 11 | 0.1818 | 8 | 0.125 | 33 | 0.2424 |
| 2_2516224 | 3 | 0.3333 | 6 | 0 | 2 | 0 | 2 | 0.5 | 13 | 0.1538 | 1 | 0 | 1 | 0 | 6 | 0.1667 | 2 | 0 | 10 | 0.1 |
| 2_2516321 | 2 | 0.5 | 2 | 1 | 1 | 1 | 2 | 1 | 7 | 0.8571 | 4 | 0.75 | NA | NA | 4 | 0.75 | 5 | 0.8 | 13 | 0.7692 |
| 2_2516331 | 2 | 0.5 | 3 | 0.6667 | 1 | 1 | 3 | 0.3333 | 9 | 0.5556 | 2 | 0.5 | NA | NA | 3 | 0.3333 | 6 | 0.5 | 11 | 0.4545 |
| 2_2516340 | 3 | 0 | 3 | 0.3333 | NA | NA | 2 | 0 | 8 | 0.125 | 3 | 0 | NA | NA | 7 | 0.1429 | 4 | 0 | 14 | 0.0714 |
| 2_2516838 | 3 | 0 | 6 | 0 | 5 | 0.2 | 4 | 0.25 | 18 | 0.1111 | 5 | 0.2 | 3 | 0 | 10 | 0.1 | 7 | 0 | 25 | 0.08 |
| 2_2517458 | 6 | 0.3333 | 5 | 0.2 | 6 | 0 | 7 | 0 | 24 | 0.125 | 5 | 0.2 | 5 | 0 | 12 | 0 | 9 | 0.1111 | 31 | 0.0645 |
| 2_2517535 | 7 | 0 | 4 | 0.25 | 7 | 0 | 7 | 0.2857 | 25 | 0.12 | 5 | 0.4 | 4 | 0.5 | 7 | 0.2857 | 6 | 0.1667 | 22 | 0.3182 |
| 2_2517553 | 8 | 0.75 | 6 | 0.3333 | 7 | 0.4286 | 9 | 0.7778 | 30 | 0.6 | 8 | 0.625 | 9 | 0.5556 | 12 | 0.3333 | 13 | 0.3077 | 42 | 0.4286 |
| 2_2517966 | 2 | 1 | NA | NA | NA | NA | NA | NA | 2 | 1 | 1 | 0 | NA | NA | 1 | 1 | NA | NA | 2 | 0.5 |
| 2_2517993 | 1 | 1 | NA | NA | NA | NA | NA | NA | 1 | 1 | NA | NA | NA | NA | 1 | 1 | NA | NA | 1 | 1 |
| 2_2518000 | 2 | 1 | NA | NA | NA | NA | NA | NA | 2 | 1 | NA | NA | NA | NA | 1 | 1 | NA | NA | 1 | 1 |
| 2_2518084 | 1 | 1 | NA | NA | NA | NA | NA | NA | 1 | 1 | NA | NA | NA | NA | 1 | 1 | NA | NA | 1 | 1 |
| 2_2518126 | 46 | 0.1304 | 38 | 0.1579 | 58 | 0.1034 | 37 | 0.1081 | 179 | 0.1229 | 64 | 0.1562 | 51 | 0.098 | 59 | 0.1525 | 31 | 0.1935 | 205 | 0.1463 |
| 2_2518133 | 44 | 0 | 43 | 0.0233 | 51 | 0.1373 | 33 | 0.0303 | 171 | 0.0526 | 56 | 0.0179 | 49 | 0.102 | 48 | 0.0417 | 33 | 0.0909 | 186 | 0.0591 |
| 2_2518148 | 42 | 0.0952 | 36 | 0.1111 | 44 | 0.1136 | 30 | 0.0667 | 152 | 0.0987 | 47 | 0.0851 | 45 | 0.1333 | 48 | 0.0208 | 26 | 0.0385 | 166 | 0.0723 |
| 2_2518164 | 42 | 0.0952 | 35 | 0.0857 | 37 | 0 | 25 | 0.04 | 139 | 0.0576 | 53 | 0.0943 | 40 | 0.075 | 47 | 0.0851 | 29 | 0.0345 | 169 | 0.0769 |
| 2_2518190 | 38 | 0.2632 | 35 | 0.1714 | 33 | 0.2121 | 25 | 0.08 | 131 | 0.1908 | 50 | 0.14 | 39 | 0.0769 | 47 | 0.2553 | 25 | 0.12 | 161 | 0.1553 |
| 2_2518191 | 43 | 0.093 | 38 | 0 | 43 | 0.1163 | 29 | 0.1379 | 153 | 0.085 | 56 | 0.0893 | 44 | 0.0455 | 46 | 0.0217 | 29 | 0.069 | 175 | 0.0571 |
| 2_2518197 | 35 | 0.7143 | 35 | 0.6857 | 35 | 0.7143 | 25 | 0.6 | 130 | 0.6846 | 46 | 0.7609 | 39 | 0.641 | 37 | 0.7838 | 22 | 0.7727 | 144 | 0.7361 |
| 2_2518202 | 41 | 0.4146 | 35 | 0.4 | 41 | 0.439 | 30 | 0.5 | 147 | 0.4354 | 51 | 0.3725 | 42 | 0.5952 | 45 | 0.4 | 27 | 0.3704 | 165 | 0.4364 |
| 2_2518232 | 19 | 0.0526 | 15 | 0.0667 | 29 | 0.069 | 16 | 0 | 79 | 0.0506 | 34 | 0.0882 | 26 | 0.1154 | 30 | 0 | 16 | 0 | 106 | 0.0566 |
| 2_2518497 | 2 | 0.5 | NA | NA | 2 | 0 | 1 | 0 | 5 | 0.2 | NA | NA | NA | NA | 2 | 0.5 | 1 | 0 | 3 | 0.3333 |
| 2_2518570 | 1 | 0 | 3 | 0.3333 | 1 | 0 | 1 | 0 | 6 | 0.1667 | NA | NA | 1 | 0 | 2 | 0 | 2 | 0.5 | 5 | 0.2 |
| 2_2523955 | 4 | 0 | 4 | 0 | 5 | 0.2 | 3 | 0 | 16 | 0.0625 | 5 | 0 | NA | NA | 10 | 0 | 3 | 0.3333 | 18 | 0.0556 |

| | | | | | | | | | | | | | | | | | | | | |
|---|---|---|---|---|---|---|---|---|---|---|---|---|---|---|---|---|---|---|---|---|
| 2_2524052 | 4 | 1 | 4 | 1 | 4 | 0.5 | 5 | 0.4 | 17 | 0.7059 | 5 | 0.8 | NA | NA | 10 | 0.9 | 3 | 0.6667 | 18 | 0.8333 |
| 2_2524061 | 4 | 0.75 | 3 | 1 | 5 | 0.6 | 4 | 1 | 16 | 0.8125 | 4 | 0.5 | NA | NA | 11 | 0.7273 | 2 | 1 | 17 | 0.7059 |
| 2_2524080 | 4 | 0 | 4 | 0 | 5 | 0 | 5 | 0.2 | 18 | 0.0556 | 7 | 0.1429 | NA | NA | 12 | 0.0833 | 4 | 0 | 23 | 0.087 |
| 2_2524108 | 2 | 0.5 | 5 | 0.4 | 2 | 0 | 8 | 0.375 | 17 | 0.3529 | 3 | 0 | 1 | 0 | 5 | 0 | 4 | 0.25 | 13 | 0.0769 |
| 2_2524156 | 4 | 0 | 4 | 0.25 | 6 | 0 | 5 | 0 | 19 | 0.0526 | 5 | 0 | 1 | 0 | 11 | 0 | 4 | 0.5 | 21 | 0.0952 |
| 2_2524170 | 5 | 0 | 6 | 0.3333 | 6 | 0 | 7 | 0 | 24 | 0.0833 | 9 | 0.2222 | 2 | 0 | 13 | 0 | 6 | 0.1667 | 30 | 0.1 |
| 2_2524235 | 4 | 0.75 | 3 | 0.6667 | 3 | 0.6667 | 3 | 0.6667 | 13 | 0.6923 | 5 | 0.6 | 3 | 0.6667 | 4 | 0.5 | 3 | 0 | 15 | 0.4667 |
| 2_2524453 | 5 | 0 | 2 | 0 | 2 | 0 | 2 | 1 | 11 | 0.1818 | 4 | 0 | 1 | 1 | 2 | 0 | NA | NA | 7 | 0.1429 |
| 2_2524462 | 3 | 0 | 3 | 0.3333 | 2 | 0 | 2 | 0 | 10 | 0.1 | 5 | 0.2 | 1 | 0 | 3 | 0 | 3 | 0 | 12 | 0.0833 |
| 2_2524626 | 2 | 0 | 2 | 0 | 1 | 0 | 1 | 1 | 6 | 0.1667 | 3 | 0.3333 | 1 | 0 | 1 | 0 | 3 | 0 | 8 | 0.125 |
| 2_2525156 | 5 | 0.2 | 5 | 0.2 | 2 | 0.5 | 3 | 0 | 15 | 0.2 | 4 | 0 | 2 | 0 | 2 | 1 | 5 | 0 | 13 | 0.1538 |
| 2_2525166 | 5 | 0 | 5 | 0 | NA | NA | 3 | 0.3333 | 13 | 0.0769 | 2 | 0.5 | 2 | 0 | 2 | 0 | 4 | 0 | 10 | 0.1 |
| 2_2529657 | NA | NA | 1 | 0 | 1 | 0 | 3 | 0.3333 | 5 | 0.2 | NA | NA | NA | NA | 5 | 0.2 | 2 | 0.5 | 7 | 0.2857 |
| 2_2529711 | 1 | 1 | 2 | 0.5 | 3 | 1 | 5 | 0.4 | 11 | 0.6364 | NA | NA | NA | NA | 1 | 0 | 2 | 0.5 | 3 | 0.3333 |
| 2_2529933 | 17 | 0.0588 | 20 | 0.2 | 14 | 0 | 16 | 0.125 | 67 | 0.1045 | 17 | 0.0588 | 17 | 0.1176 | 12 | 0.0833 | 21 | 0.0952 | 67 | 0.0896 |
| 2_2529940 | 43 | 0.1395 | 43 | 0.1628 | 29 | 0 | 43 | 0.093 | 158 | 0.1076 | 36 | 0.1111 | 31 | 0.0968 | 26 | 0.0385 | 43 | 0.1163 | 136 | 0.0956 |
| 2_2533799 | 18 | 0.1667 | 30 | 0.0333 | 20 | 0.2 | 19 | 0.1579 | 87 | 0.1264 | 13 | 0.1538 | 12 | 0.1667 | 22 | 0.2273 | 12 | 0.0833 | 59 | 0.1695 |
| 2_2533817 | 23 | 0.2174 | 34 | 0.2059 | 28 | 0.1071 | 26 | 0.1154 | 111 | 0.1622 | 20 | 0.15 | 14 | 0.0714 | 26 | 0.1154 | 13 | 0.2308 | 73 | 0.137 |
| 2_2533824 | 22 | 0.4091 | 25 | 0.52 | 25 | 0.28 | 19 | 0.3158 | 91 | 0.3846 | 18 | 0.3333 | 11 | 0.1818 | 17 | 0.7059 | 13 | 0.6923 | 59 | 0.4915 |
| 2_2533825 | 21 | 0.2381 | 32 | 0.1562 | 28 | 0.25 | 23 | 0.1304 | 104 | 0.1923 | 18 | 0.2222 | 15 | 0.5333 | 22 | 0.1818 | 13 | 0.1538 | 68 | 0.2647 |
| 2_2534529 | 2 | 0.5 | 3 | 0.6667 | NA | NA | 2 | 1 | 7 | 0.7143 | NA | NA | 1 | 1 | 2 | 1 | 1 | 0 | 4 | 0.75 |
| 2_2534532 | 2 | 0.5 | NA | NA | NA | NA | 1 | 1 | 3 | 0.6667 | NA | NA | 1 | 1 | 1 | 1 | 1 | 1 | 3 | 1 |
| 2_2534538 | 2 | 0.5 | 3 | 0.3333 | NA | NA | 2 | 1 | 7 | 0.5714 | NA | NA | 1 | 1 | 2 | 1 | 1 | 1 | 4 | 1 |
| 2_2534545 | 2 | 1 | 3 | 0.6667 | NA | NA | 2 | 1 | 7 | 0.8571 | NA | NA | 1 | 1 | 1 | 1 | 2 | 0.5 | 4 | 0.75 |
| 2_2534553 | 2 | 0.5 | 3 | 0 | NA | NA | 2 | 0 | 7 | 0.1429 | NA | NA | 1 | 1 | 1 | 0 | 2 | 0.5 | 4 | 0.5 |
| 2_2539356 | 10 | 0 | 10 | 0.1 | 8 | 0.125 | 16 | 0.1875 | 44 | 0.1136 | 5 | 0.2 | 6 | 0.1667 | 9 | 0.2222 | 14 | 0.1429 | 34 | 0.1765 |
| 2_2539362 | 10 | 0.2 | 8 | 0.125 | 10 | 0.6 | 13 | 0.3077 | 41 | 0.3171 | 2 | 0 | 7 | 0.1429 | 7 | 0.1429 | 12 | 0.3333 | 28 | 0.2143 |
| 2_2539372 | 10 | 0.4 | 9 | 0.4444 | 11 | 0.3636 | 16 | 0.4375 | 46 | 0.413 | 4 | 0.5 | 6 | 0.5 | 10 | 0.6 | 13 | 0.5385 | 33 | 0.5455 |
| 2_2539373 | 8 | 0.375 | 10 | 0.5 | 11 | 0.5455 | 14 | 0.7857 | 43 | 0.5814 | 3 | 0.3333 | 6 | 0.5 | 9 | 0.7778 | 13 | 0.6923 | 31 | 0.6452 |
| 2_2539410 | NA | NA | 3 | 0 | 2 | 0.5 | 2 | 0 | 7 | 0.1429 | NA | NA | 1 | 0 | 1 | 0 | 6 | 0.3333 | 8 | 0.25 |
| 2_2539475 | 14 | 0.2143 | 11 | 0.3636 | 12 | 0.0833 | 15 | 0.1333 | 52 | 0.1923 | 5 | 0 | 10 | 0.2 | 12 | 0.3333 | 15 | 0.0667 | 42 | 0.1667 |
| 2_2539483 | 16 | 0.625 | 13 | 0.6923 | 13 | 0.8462 | 14 | 0.3571 | 56 | 0.625 | 4 | 0.5 | 9 | 0.7778 | 12 | 0.5 | 19 | 0.5789 | 44 | 0.5909 |
| 2_2539500 | 15 | 0.0667 | 14 | 0.0714 | 12 | 0.0833 | 17 | 0 | 58 | 0.0517 | 5 | 0.2 | 10 | 0.1 | 10 | 0.1 | 18 | 0 | 43 | 0.0698 |
| 2_2539542 | 13 | 0.6923 | 14 | 0.5714 | 18 | 0.7222 | 19 | 0.7368 | 64 | 0.6875 | 9 | 0.6667 | 12 | 0.5833 | 14 | 0.6429 | 20 | 0.65 | 55 | 0.6364 |
| 2_2539544 | 12 | 0.5 | 14 | 0.6429 | 16 | 0.8125 | 15 | 0.6 | 57 | 0.6491 | 8 | 0.375 | 10 | 0.5 | 13 | 0.6923 | 21 | 0.5714 | 52 | 0.5577 |
| 2_2540906 | 2 | 0.5 | 1 | 0 | 1 | 1 | 1 | 1 | 5 | 0.6 | 1 | 0 | 1 | 1 | NA | NA | 1 | 0 | 3 | 0.3333 |
| 2_2540942 | 2 | 1 | 1 | 1 | 1 | 1 | 1 | 1 | 5 | 1 | 2 | 1 | 1 | 1 | NA | NA | 1 | 1 | 4 | 1 |
| 2_2540943 | 2 | 0.5 | 1 | 1 | NA | NA | 1 | 1 | 4 | 0.75 | 2 | 0.5 | 1 | 0 | NA | NA | 1 | 1 | 4 | 0.5 |
| 2_2545206 | 31 | 0.0645 | 36 | 0.1111 | 24 | 0 | 30 | 0.0333 | 121 | 0.0579 | 29 | 0.069 | 26 | 0.1538 | 35 | 0.1143 | 36 | 0 | 126 | 0.0794 |
| 2_2546359 | 15 | 0.2 | 17 | 0.0588 | 15 | 0.2667 | 16 | 0.0625 | 63 | 0.1429 | 17 | 0.0588 | 16 | 0.1875 | 18 | 0.1667 | 8 | 0.125 | 59 | 0.1356 |
| 2_2546408 | 1 | 0 | 1 | 1 | 2 | 1 | 1 | 0 | 5 | 0.6 | NA | NA | NA | NA | 2 | 0.5 | NA | NA | 2 | 0.5 |
| 2_2546432 | 13 | 0.0769 | 17 | 0.1176 | 13 | 0 | 14 | 0.1429 | 57 | 0.0877 | 16 | 0.125 | 11 | 0 | 17 | 0.1176 | 7 | 0.2857 | 51 | 0.1176 |
| 2_2546503 | 1 | 1 | NA | NA | 1 | 1 | NA | NA | 2 | 1 | NA | NA | NA | NA | 2 | 1 | 1 | 1 | 3 | 1 |
| 2_2546506 | 1 | 1 | 1 | 1 | 2 | 1 | NA | NA | 4 | 1 | NA | NA | 1 | 1 | 2 | 1 | 1 | 1 | 4 | 1 |
| 2_2546520 | 1 | 1 | 1 | 1 | 1 | 1 | NA | NA | 3 | 1 | NA | NA | 1 | 0 | 1 | 1 | 1 | 1 | 3 | 0.6667 |
| 2_2546545 | 7 | 0.5714 | 11 | 0.4545 | 6 | 0.8333 | 12 | 0.75 | 36 | 0.6389 | 8 | 0.375 | 10 | 0.5 | 8 | 0.75 | 9 | 0.5556 | 35 | 0.5429 |
| 2_2546546 | 8 | 0.25 | 11 | 0.2727 | 9 | 0.2222 | 12 | 0.0833 | 40 | 0.2 | 9 | 0 | 8 | 0.125 | 10 | 0.1 | 9 | 0.2222 | 36 | 0.1111 |
| 2_2546550 | 12 | 0.1667 | 8 | 0.25 | 9 | 0.2222 | 8 | 0.375 | 37 | 0.2432 | 7 | 0 | 9 | 0.2222 | 7 | 0.1429 | 9 | 0.2222 | 32 | 0.1562 |
| 2_2546563 | 11 | 0.5455 | 10 | 0.5 | 8 | 0.5 | 11 | 0.4545 | 40 | 0.5 | 8 | 0.375 | 9 | 0.3333 | 8 | 0.625 | 7 | 0.5714 | 32 | 0.4688 |
| 2_2546569 | 9 | 1 | 10 | 1 | 6 | 0.3333 | 10 | 0.9 | 35 | 0.8571 | 6 | 0.5 | 9 | 0.4444 | 6 | 1 | 7 | 0.7143 | 28 | 0.6429 |
| 2_2546569 | 11 | 0.0909 | 7 | 0.4286 | 8 | 0.125 | 10 | 0 | 36 | 0.1389 | 7 | 0 | 10 | 0.2 | 9 | 0.1111 | 8 | 0.125 | 34 | 0.1176 |
| 2_2546580 | 10 | 1 | 7 | 1 | 5 | 0.8 | 11 | 1 | 33 | 0.9697 | 7 | 1 | 8 | 0.875 | 5 | 0.8 | 7 | 0.8571 | 27 | 0.8889 |
| 2_2546601 | 9 | 0.7778 | 8 | 0.875 | 6 | 0.8333 | 10 | 0.9 | 33 | 0.8485 | 6 | 0.6667 | 8 | 0.5 | 6 | 0.6667 | 6 | 0.5 | 26 | 0.5769 |
| 2_2546601 | 5 | 0.4 | 5 | 0.8 | 3 | 0.6667 | 7 | 0.5714 | 20 | 0.6 | 6 | 0.5 | 10 | 0.4 | 6 | 0.5 | 6 | 0 | 28 | 0.3571 |
| 2_2546606 | 12 | 0.4167 | 7 | 0.5714 | 6 | 0.3333 | 12 | 0.5833 | 37 | 0.4865 | 9 | 0.1111 | 10 | 0.2 | 6 | 0.1667 | 8 | 0.25 | 33 | 0.1818 |
| 2_2546656 | 11 | 0.2727 | 8 | 0.25 | 3 | 0 | 10 | 0.2 | 32 | 0.2188 | 8 | 0.25 | 9 | 0.3333 | 2 | 0.5 | 8 | 0.125 | 27 | 0.2593 |
| 2_2546657 | 12 | 0.0833 | 8 | 0.25 | 3 | 0 | 12 | 0.25 | 35 | 0.1714 | 7 | 0 | 10 | 0.2 | 4 | 0.25 | 10 | 0.1 | 31 | 0.129 |
| 2_2546660 | 13 | 0.6923 | 8 | 0.75 | 3 | 0.6667 | 12 | 0.9167 | 36 | 0.7778 | 8 | 0.375 | 9 | 0.8889 | 4 | 1 | 9 | 0.7778 | 30 | 0.7333 |
| 2_2546661 | 13 | 0.3077 | 8 | 0.375 | 3 | 0 | 12 | 0.0833 | 36 | 0.2222 | 8 | 0.125 | 11 | 0.2727 | 4 | 0.25 | 8 | 0.25 | 31 | 0.2258 |
| 2_2546689 | 13 | 0.2308 | 6 | 0 | 2 | 0.5 | 9 | 0.3333 | 30 | 0.2333 | 6 | 0.1667 | 10 | 0.4 | 4 | 0 | 9 | 0.2222 | 29 | 0.2414 |
| 2_2546697 | 11 | 0.9091 | 4 | 1 | 2 | 1 | 6 | 1 | 23 | 0.9565 | 4 | 1 | 8 | 1 | 3 | 1 | 5 | 0.8 | 20 | 0.95 |
| 2_2546698 | 10 | 0.5 | 5 | 0.4 | 2 | 1 | 7 | 0.8571 | 24 | 0.625 | 6 | 0.6667 | 8 | 0.125 | 1 | 0 | 8 | 0.5 | 23 | 0.3913 |
| 2_2546780 | 6 | 1 | 4 | 1 | 3 | 1 | 6 | 0.6667 | 19 | 0.8947 | 6 | 1 | 10 | 1 | 1 | 1 | 6 | 0.8333 | 23 | 0.9565 |
| 2_2546788 | 8 | 0.25 | 4 | 0 | 3 | 0 | 8 | 0.125 | 23 | 0.1304 | 6 | 0.1667 | 7 | 0.1429 | 2 | 0 | 7 | 0.2857 | 22 | 0.1818 |
| 2_2546789 | 6 | 0 | 4 | 0.25 | 2 | 0.5 | 3 | 0 | 15 | 0.1333 | 4 | 0.75 | 7 | 0.2857 | 2 | 0 | 6 | 0 | 19 | 0.2632 |

| | | | | | | | | | | | | | | | | | | | | |
|---|---|---|---|---|---|---|---|---|---|---|---|---|---|---|---|---|---|---|---|---|
| 2_25468069 | 6 | 0.8333 | 4 | 1 | 3 | 1 | 4 | 1 | 17 | 0.9412 | 3 | 1 | 8 | 0.875 | 1 | 1 | 5 | 1 | 17 | 0.9412 |
| 2_25468647 | 5 | 0.4 | 4 | 0 | 2 | 0.5 | 2 | 0.5 | 13 | 0.3077 | 4 | 0 | 5 | 0.2 | 1 | 0 | 2 | 0 | 12 | 0.0833 |
| 2_25468739 | 8 | 0.625 | 3 | 1 | 1 | 0 | 2 | 1 | 14 | 0.7143 | 4 | 0.5 | 8 | 0.875 | 1 | 1 | 2 | 1 | 15 | 0.8 |
| 2_25468777 | 8 | 0.875 | 5 | 1 | 2 | 1 | 2 | 1 | 17 | 0.9412 | 3 | 1 | 9 | 1 | 1 | 1 | 3 | 1 | 16 | 1 |
| 2_25469122 | 8 | 0.375 | 5 | 0.4 | 2 | 0 | 2 | 0.5 | 17 | 0.3529 | 4 | 0.75 | 8 | 0.375 | 1 | 0 | 2 | 1 | 15 | 0.5333 |
| 2_25469159 | 5 | 0.8 | 1 | 1 | 2 | 0.5 | 1 | 1 | 9 | 0.7778 | 1 | 0 | 5 | 0.8 | NA | NA | 1 | 1 | 7 | 0.7143 |
| 2_25469182 | 9 | 0.8889 | 4 | 1 | 2 | 1 | 2 | 1 | 17 | 0.9412 | 4 | 0.75 | 8 | 0.875 | 1 | 1 | 2 | 0.5 | 15 | 0.8 |
| 2_25469344 | 3 | 0.3333 | 5 | 0.2 | 2 | 0 | NA | NA | 10 | 0.2 | 3 | 0.3333 | 8 | 0.375 | NA | NA | 1 | 1 | 12 | 0.4167 |
| 2_25469373 | 6 | 0.1667 | 3 | 0 | 1 | 1 | 2 | 0 | 12 | 0.1667 | 4 | 0.25 | 8 | 0.25 | 1 | 0 | NA | NA | 13 | 0.2308 |
| 2_25469659 | 8 | 0.5 | 3 | 0.3333 | 2 | 0.5 | 2 | 0.5 | 15 | 0.4667 | 4 | 0.25 | 6 | 0.5 | 1 | 0 | 1 | 0 | 12 | 0.3333 |
| 2_25469714 | 9 | 0.1111 | 3 | 0.3333 | 1 | 0 | 2 | 0 | 15 | 0.1333 | 4 | 0 | 6 | 0.1667 | 1 | 0 | 1 | 0 | 12 | 0.0833 |
| 2_25469845 | 7 | 0.1429 | 3 | 0.3333 | 2 | 0 | 2 | 0 | 14 | 0.1429 | 4 | 0.75 | 5 | 0.4 | NA | NA | 1 | 0 | 10 | 0.5 |
| 2_25469860 | 7 | 0 | 4 | 0.5 | 2 | 0 | 2 | 0 | 15 | 0.1333 | 4 | 0.5 | 4 | 0 | NA | NA | 1 | 0 | 9 | 0.2222 |
| 2_25469887 | 8 | 0.125 | 4 | 0.25 | 2 | 0 | 2 | 0 | 16 | 0.125 | 4 | 0 | 5 | 0.4 | NA | NA | 1 | 0 | 10 | 0.2 |
| 2_25469942 | 6 | 0 | 4 | 0 | 2 | 0 | 2 | 0.5 | 14 | 0.0714 | 4 | 0 | 5 | 0.2 | NA | NA | 1 | 0 | 10 | 0.1 |
| 2_25470156 | 3 | 0.3333 | 4 | 0 | 2 | 0 | 2 | 0 | 11 | 0.0909 | 4 | 0 | 4 | 0 | NA | NA | 1 | 1 | 9 | 0.1111 |
| 2_25473318 | 19 | 0.3684 | 12 | 0.3333 | 15 | 0.3333 | 11 | 0.4545 | 57 | 0.3684 | 14 | 0.3571 | 16 | 0.25 | 22 | 0.4091 | 13 | 0.3846 | 65 | 0.3538 |
| 2_25473819 | 16 | 0.6875 | 11 | 0.8182 | 16 | 0.625 | 8 | 0.75 | 51 | 0.7059 | 12 | 0.5833 | 17 | 0.8235 | 23 | 0.5652 | 11 | 0.7273 | 63 | 0.6667 |
| 2_25475692 | 9 | 0.1111 | 7 | 0 | 9 | 0 | 6 | 0.3333 | 31 | 0.0968 | 8 | 0.125 | 7 | 0 | 10 | 0 | 5 | 0.4 | 30 | 0.1 |
| 2_25498176 | 1 | 1 | NA | NA | NA | NA | 1 | 1 | 2 | 1 | NA | NA | NA | NA | 1 | 1 | NA | NA | 1 | 1 |
| 2_25498501 | 1 | 1 | NA | NA | NA | NA | 1 | 1 | 2 | 1 | NA | NA | NA | NA | 1 | 1 | NA | NA | 1 | 1 |
| 2_25498915 | 1 | 1 | NA | NA | NA | NA | 2 | 0 | 3 | 0.3333 | 1 | 1 | 1 | 0 | 1 | 0 | NA | NA | 3 | 0.3333 |
| 2_25498928 | 1 | 1 | NA | NA | NA | NA | 2 | 1 | 3 | 1 | 1 | 1 | 1 | 1 | 1 | 1 | NA | NA | 3 | 1 |
| 2_25510199 | 2 | 1 | 3 | 0 | 3 | 0.3333 | 3 | 0.6667 | 11 | 0.4545 | 2 | 0.5 | 1 | 0 | 5 | 0.2 | 1 | 0 | 9 | 0.2222 |
| 2_25596813 | 2 | 0 | 3 | 0 | 1 | 0 | 2 | 0.5 | 8 | 0.125 | 5 | 0 | 5 | 0.6 | 3 | 0.3333 | 2 | 0.5 | 15 | 0.3333 |
| 2_25596981 | 17 | 0.2353 | 14 | 0.3571 | 9 | 0.1111 | 7 | 0.4286 | 47 | 0.2766 | 9 | 0.4444 | 12 | 0.3333 | 18 | 0.1667 | 12 | 0.5833 | 51 | 0.3529 |
| 2_25597305 | 20 | 1 | 20 | 0.95 | 12 | 0.9167 | 13 | 0.9231 | 65 | 0.9538 | 12 | 1 | 19 | 0.9474 | 25 | 0.84 | 19 | 0.9474 | 75 | 0.92 |
| 2_25597356 | 19 | 0.3158 | 18 | 0.2778 | 10 | 0.2 | 12 | 0.1667 | 59 | 0.2542 | 10 | 0.4 | 12 | 0.0833 | 19 | 0.3684 | 20 | 0.25 | 61 | 0.2787 |
| 2_25597431 | 19 | 0.6842 | 17 | 0.6471 | 13 | 0.2308 | 9 | 0.5556 | 58 | 0.5517 | 11 | 0.5455 | 21 | 0.7619 | 23 | 0.6087 | 22 | 0.5 | 77 | 0.6104 |
| 2_25597474 | 16 | 0.9375 | 16 | 1 | 10 | 1 | 12 | 1 | 54 | 0.9815 | 11 | 1 | 20 | 1 | 23 | 1 | 22 | 0.9545 | 76 | 0.9868 |
| 2_25597496 | 16 | 0 | 13 | 0.0769 | 11 | 0 | 9 | 0.2222 | 49 | 0.0612 | 12 | 0.0833 | 18 | 0 | 19 | 0 | 21 | 0.1429 | 70 | 0.0571 |
| 2_25597640 | 19 | 0.2105 | 21 | 0.1429 | 12 | 0.1667 | 15 | 0.2 | 67 | 0.1791 | 12 | 0.25 | 19 | 0.1053 | 25 | 0.2 | 22 | 0.1818 | 78 | 0.1795 |
| 2_25791889 | 3 | 0 | 4 | 0.25 | 1 | 0 | 1 | 0 | 9 | 0.1111 | 3 | 0 | 2 | 0.5 | 1 | 0 | 1 | 0 | 7 | 0.1429 |
| 2_25801063 | 12 | 0.1667 | 8 | 0.375 | 6 | 0.1667 | 6 | 0.1667 | 32 | 0.2188 | 4 | 0 | 5 | 0.2 | 6 | 0.1667 | 6 | 0 | 21 | 0.0952 |
| 2_25801091 | 13 | 0.0769 | 9 | 0 | 8 | 0.375 | 7 | 0.2857 | 37 | 0.1622 | 4 | 0 | 6 | 0 | 7 | 0.2857 | 9 | 0.1111 | 26 | 0.1154 |
| 2_25806216 | 13 | 0.5385 | 8 | 0.625 | 8 | 1 | 7 | 0.5714 | 36 | 0.6667 | 3 | 0.6667 | 5 | 1 | 8 | 0.625 | 8 | 0.375 | 24 | 0.625 |
| 2_25806237 | 14 | 0.0714 | 8 | 0.25 | 7 | 0.1429 | 7 | 0 | 36 | 0.1111 | 3 | 0.3333 | 5 | 0.2 | 8 | 0.125 | 8 | 0 | 24 | 0.125 |
| 2_25806383 | 8 | 0.125 | 8 | 0.375 | 5 | 0.4 | 4 | 0 | 25 | 0.24 | 1 | 0 | 3 | 0 | 5 | 0.2 | 5 | 0 | 14 | 0.0714 |
| 2_25806426 | 7 | 0.1429 | 9 | 0.3333 | 6 | 0.1667 | 4 | 0.25 | 26 | 0.2308 | 1 | 0 | 6 | 0 | 5 | 0 | 8 | 0.125 | 20 | 0.05 |
| 2_25806540 | 9 | 0.2222 | 8 | 0.375 | 6 | 0.6667 | 4 | 0.25 | 27 | 0.3704 | 1 | 0 | 6 | 0.1667 | 7 | 0.4286 | 7 | 0.2857 | 21 | 0.2857 |
| 2_25806789 | 1 | 1 | 4 | 0 | 3 | 0 | 2 | 0 | 10 | 0.1 | 1 | 0 | 5 | 0 | 4 | 0.5 | 2 | 0 | 12 | 0.1667 |
| 2_25806887 | 4 | 0 | 6 | 0.1667 | 3 | 0.3333 | 3 | 0 | 16 | 0.125 | 1 | 1 | 5 | 0.2 | 3 | 0 | 5 | 0 | 14 | 0.1429 |
| 2_25806894 | 3 | 0.3333 | 4 | 0 | 2 | 0.5 | 3 | 0.3333 | 12 | 0.25 | 1 | 0 | 6 | 0.1667 | 2 | 0 | 3 | 0.3333 | 12 | 0.1667 |
| 2_25806919 | 5 | 0.2 | 6 | 0 | 4 | 0 | 4 | 0 | 19 | 0.0526 | 1 | 0 | 6 | 0.3333 | 2 | 0 | 3 | 0 | 12 | 0.1667 |
| 2_25806935 | 3 | 0.3333 | 6 | 0.1667 | 4 | 0.5 | 4 | 0.25 | 17 | 0.2941 | 1 | 0 | 3 | 0 | 3 | 0.3333 | 5 | 0 | 12 | 0.0833 |
| 2_25807059 | 4 | 0.25 | 5 | 0.4 | 4 | 0.5 | 4 | 0 | 17 | 0.2941 | 1 | 0 | 6 | 0 | 4 | 0.25 | 4 | 0.25 | 15 | 0.1333 |
| 2_25817590 | 5 | 0 | 2 | 0 | 2 | 0.5 | 2 | 0 | 11 | 0.0909 | NA | NA | 2 | 0 | 3 | 0.3333 | 6 | 0.1667 | 11 | 0.1818 |
| 2_25818425 | 6 | 0 | 9 | 0.1111 | 4 | 0.25 | 5 | 0 | 24 | 0.0833 | 5 | 0 | 6 | 0 | 7 | 0.2857 | 11 | 0 | 29 | 0.069 |
| 2_25819250 | 6 | 0.8333 | 9 | 0.8889 | 6 | 1 | 8 | 0.75 | 29 | 0.8621 | 8 | 0.875 | 7 | 1 | 9 | 1 | 10 | 0.7 | 34 | 0.8824 |
| 2_25819496 | 7 | 1 | 10 | 1 | 5 | 1 | 6 | 1 | 28 | 1 | 10 | 1 | 6 | 1 | 8 | 1 | 12 | 1 | 36 | 1 |
| 2_25819521 | 7 | 0.1429 | 10 | 0 | 6 | 0.1667 | 7 | 0 | 30 | 0.0667 | 9 | 0 | 7 | 0 | 10 | 0.1 | 11 | 0.0909 | 37 | 0.0541 |
| 2_25819622 | 6 | 0.3333 | 7 | 0.2857 | 6 | 0 | 6 | 0.1667 | 25 | 0.2 | 7 | 0 | 5 | 0.4 | 7 | 0.2857 | 10 | 0.2 | 29 | 0.2069 |
| 2_25819649 | 8 | 1 | 9 | 1 | 6 | 0.8333 | 7 | 0.8571 | 30 | 0.9333 | 8 | 1 | 7 | 1 | 10 | 1 | 12 | 1 | 37 | 1 |
| 2_25819723 | 6 | 1 | 8 | 1 | 6 | 0.8333 | 5 | 1 | 25 | 0.96 | 9 | 1 | 7 | 1 | 9 | 1 | 10 | 1 | 35 | 1 |
| 2_25819738 | 8 | 0 | 9 | 0 | 6 | 0.1667 | 4 | 0.5 | 27 | 0.1111 | 9 | 0.3333 | 7 | 0 | 9 | 0.2222 | 10 | 0.3 | 35 | 0.2286 |
| 2_25819749 | 4 | 0.75 | 9 | 0.3333 | 4 | 0.75 | 4 | 0.25 | 21 | 0.4762 | 7 | 0.7143 | 6 | 0.6667 | 8 | 0.375 | 9 | 0.5556 | 30 | 0.5667 |
| 2_25819885 | 6 | 0.8333 | 9 | 0.4444 | 3 | 1 | 5 | 0 | 23 | 0.5217 | 7 | 0.7143 | 4 | 0.5 | 9 | 0.6667 | 8 | 0.75 | 28 | 0.6786 |
| 2_25819889 | 6 | 0 | 8 | 0.125 | 3 | 0.3333 | 5 | 0.4 | 22 | 0.1818 | 8 | 0.125 | 7 | 0.1429 | 8 | 0 | 8 | 0.125 | 31 | 0.0968 |
| 2_25819933 | 4 | 0.25 | 7 | 0 | 2 | 0.5 | 4 | 0 | 17 | 0.1176 | 5 | 0 | 6 | 0.1667 | 6 | 0 | 7 | 0.1429 | 24 | 0.0833 |
| 2_25828193 | 4 | 0.75 | 5 | 0.8 | 3 | 1 | 2 | 1 | 14 | 0.8571 | 5 | 0.6 | 6 | 0.8333 | 5 | 1 | 5 | 1 | 21 | 0.8571 |
| 2_25828201 | 6 | 0.1667 | 5 | 0 | 5 | 0 | 2 | 0 | 18 | 0.0556 | 7 | 0 | 6 | 0.3333 | 5 | 0 | 6 | 0.3333 | 24 | 0.1667 |
| 2_25828245 | 5 | 1 | 6 | 0.8333 | 4 | 1 | 2 | 1 | 17 | 0.9412 | 4 | 1 | 6 | 1 | 5 | 1 | 6 | 1 | 21 | 1 |
| 2_25828303 | 6 | 0.1667 | 4 | 0 | 2 | 0 | 2 | 0 | 14 | 0.0714 | 6 | 0.1667 | 5 | 0 | 3 | 0 | 3 | 0 | 17 | 0.0588 |
| 2_25828421 | 4 | 0 | 6 | 0 | 3 | 0.3333 | 2 | 0 | 15 | 0.0667 | 4 | 0.25 | 3 | 0.3333 | 5 | 0.2 | 4 | 0 | 16 | 0.1875 |
| 2_25828438 | 4 | 0.25 | 6 | 0.3333 | 4 | 0.5 | 2 | 0 | 16 | 0.3125 | 5 | 0 | 5 | 0.4 | 4 | 0.25 | 4 | 0.25 | 18 | 0.2222 |

| | | | | | | | | | | | | | | | | | | | | |
|---|---|---|---|---|---|---|---|---|---|---|---|---|---|---|---|---|---|---|---|---|
| 2_2582855 | 5 | 0.2 | 6 | 0.5 | 3 | 0.3333 | 2 | 0 | 16 | 0.3125 | 5 | 0 | 3 | 1 | 4 | 0 | 4 | 0.5 | 16 | 0.3125 |
| 2_2582856 | 2 | 0 | 5 | 0.4 | 3 | 0 | 2 | 0.5 | 12 | 0.25 | 5 | 0 | 4 | 0 | 5 | 0 | 4 | 0.5 | 18 | 0.1111 |
| 2_2582878 | 6 | 0.1667 | 6 | 0 | 4 | 0 | 2 | 0.5 | 18 | 0.1111 | 6 | 0.1667 | 3 | 0.3333 | 4 | 0 | 5 | 0.2 | 18 | 0.1667 |
| 2_2582889 | 5 | 0.6 | 5 | 0.4 | 3 | 0.3333 | 2 | 0 | 15 | 0.4 | 6 | 0.5 | 3 | 0.6667 | 4 | 0.5 | 5 | 1 | 18 | 0.6667 |
| 2_2582893 | 4 | 0.25 | 2 | 0 | 3 | 0.3333 | 1 | 0 | 10 | 0.2 | 5 | 0 | 2 | 0 | 3 | 0.3333 | 1 | 0 | 11 | 0.0909 |
| 2_2582912 | 6 | 0 | 4 | 0 | 4 | 0 | 2 | 0.5 | 16 | 0.0625 | 5 | 0.2 | 2 | 0 | 2 | 0.5 | 5 | 0.4 | 14 | 0.2857 |
| 2_2582951 | 3 | 1 | 3 | 1 | 3 | 1 | 2 | 1 | 11 | 1 | 3 | 1 | 2 | 1 | 1 | 1 | 4 | 1 | 10 | 1 |
| 2_2582954 | 5 | 0 | 3 | 0.3333 | 3 | 0 | 2 | 0 | 13 | 0.0769 | 2 | 0 | 2 | 0 | 1 | 0 | 4 | 0.5 | 9 | 0.2222 |
| 2_2582968 | 5 | 0.2 | 2 | 0 | 3 | 0 | 2 | 0.5 | 12 | 0.1667 | 2 | 0 | 1 | 1 | 1 | 0 | 3 | 0 | 7 | 0.1429 |
| 2_2582977 | 4 | 1 | 3 | 0.3333 | 3 | 0.6667 | 2 | 0.5 | 12 | 0.6667 | 1 | 1 | 1 | 1 | 1 | 1 | 3 | 1 | 6 | 1 |
| 2_2582986 | 4 | 0 | 2 | 1 | 2 | 0 | 2 | 0 | 10 | 0.2 | 2 | 1 | 1 | 1 | NA | NA | 3 | 0.3333 | 6 | 0.6667 |
| 2_2582999 | 4 | 0.25 | 2 | 0 | 3 | 0.3333 | 2 | 0 | 11 | 0.1818 | 2 | 0 | 1 | 0 | NA | NA | 4 | 0.25 | 7 | 0.1429 |
| 2_2583004 | 2 | 0 | NA | NA | 3 | 0.3333 | 2 | 0.5 | 7 | 0.2857 | NA | NA | 1 | 0 | NA | NA | 3 | 0.3333 | 4 | 0.25 |
| 2_2583008 | 3 | 0.3333 | 2 | 0.5 | 2 | 0.5 | 1 | 1 | 8 | 0.5 | 1 | 0 | 1 | 1 | NA | NA | 3 | 0.3333 | 5 | 0.4 |
| 2_2588860 | 9 | 0.2222 | 14 | 0.1429 | 10 | 0.1 | 8 | 0 | 41 | 0.122 | 9 | 0.1111 | 7 | 0.1429 | 2 | 0 | 6 | 0.1667 | 24 | 0.125 |
| 2_2588869 | 9 | 0.1111 | 12 | 0.0833 | 10 | 0 | 7 | 0.2857 | 38 | 0.1053 | 9 | 0.1111 | 9 | 0.1111 | 2 | 0 | 7 | 0 | 27 | 0.0741 |
| 2_2588886 | 9 | 0.8889 | 13 | 0.9231 | 9 | 0.7778 | 7 | 0.8571 | 38 | 0.8684 | 7 | 0.8571 | 7 | 1 | 1 | 1 | 7 | 1 | 22 | 0.9545 |
| 2_2594577 | 9 | 0 | 14 | 0.1429 | 19 | 0.2105 | 14 | 0.2143 | 56 | 0.1607 | 12 | 0 | 11 | 0.0909 | 11 | 0.0909 | 5 | 0.2 | 39 | 0.0769 |
| 2_2594857 | 4 | 0.25 | 6 | 0.1667 | 2 | 0.5 | 3 | 0.3333 | 15 | 0.2667 | 4 | 0.25 | 4 | 0 | 4 | 0.25 | 3 | 0 | 15 | 0.1333 |
| 2_2594866 | 3 | 0 | 5 | 0.2 | 1 | 0 | 3 | 0 | 12 | 0.0833 | 4 | 0.25 | 4 | 0.5 | 3 | 0.6667 | NA | NA | 11 | 0.4545 |
| 2_2594952 | 4 | 0.25 | 3 | 0 | NA | NA | 3 | 0.3333 | 10 | 0.2 | 1 | 1 | 1 | 0 | 3 | 0 | NA | NA | 5 | 0.2 |
| 2_2594987 | 6 | 0.8333 | 4 | 0.75 | 2 | 1 | 2 | 1 | 14 | 0.8571 | 2 | 1 | 4 | 0.5 | 3 | 0.6667 | 1 | 0 | 10 | 0.6 |
| 2_2594994 | 5 | 0.4 | 4 | 0.5 | NA | NA | 3 | 0.3333 | 12 | 0.4167 | 5 | 0 | 3 | 0.3333 | 3 | 0 | 2 | 0 | 13 | 0.0769 |
| 2_2594998 | 6 | 1 | 4 | 1 | 2 | 1 | NA | NA | 12 | 1 | 4 | 1 | 3 | 1 | 3 | 1 | 2 | 1 | 12 | 1 |
| 2_2595021 | 6 | 1 | 5 | 1 | 2 | 1 | 3 | 1 | 16 | 1 | 3 | 1 | 4 | 1 | 4 | 1 | 2 | 1 | 13 | 1 |
| 2_2595030 | 6 | 0 | 5 | 0 | 2 | 0 | 3 | 0.3333 | 16 | 0.0625 | 5 | 0.2 | 4 | 0.25 | 4 | 0.25 | 2 | 0.5 | 15 | 0.2667 |
| 2_2597970 | 2 | 0.5 | 3 | 0 | 1 | 1 | NA | NA | 6 | 0.3333 | 1 | 0 | 4 | 0.5 | 3 | 0.6667 | 1 | 1 | 9 | 0.5556 |
| 2_2597981 | 4 | 0.75 | 4 | 0.5 | 1 | 0 | 1 | 1 | 10 | 0.6 | 2 | 0 | 4 | 0.5 | 3 | 0.6667 | 1 | 0 | 10 | 0.4 |
| 2_2598004 | 3 | 0 | 2 | 0 | 2 | 0.5 | 2 | 0.5 | 9 | 0.2222 | 1 | 0 | 4 | 0.25 | 3 | 0.6667 | 1 | 0 | 9 | 0.3333 |
| 2_2598026 | 4 | 0.25 | 3 | 0 | 1 | 0 | 1 | 0 | 9 | 0.1111 | 2 | 0.5 | 4 | 0.25 | 2 | 0 | 1 | 0 | 9 | 0.2222 |
| 2_2602709 | 9 | 0.1111 | 9 | 0.4444 | 9 | 0.1111 | 5 | 0 | 32 | 0.1875 | 6 | 0.3333 | 6 | 0 | 15 | 0.0667 | 14 | 0.2143 | 41 | 0.1463 |
| 2_2602721 | 7 | 0.2857 | 13 | 0.1538 | 8 | 0.625 | 4 | 0 | 32 | 0.2812 | 5 | 0.6 | 7 | 0.4286 | 13 | 0.3077 | 11 | 0.2727 | 36 | 0.3611 |
| 2_2602733 | 6 | 0 | 12 | 0 | 7 | 0.2857 | 3 | 0 | 28 | 0.0714 | 5 | 0.4 | 6 | 0.1667 | 10 | 0 | 12 | 0 | 33 | 0.0909 |
| 2_2602740 | 8 | 0 | 12 | 0.1667 | 7 | 0 | 2 | 0 | 29 | 0.069 | 4 | 0.25 | 5 | 0 | 8 | 0.25 | 8 | 0 | 25 | 0.12 |
| 2_2602767 | 5 | 0.2 | 13 | 0 | 5 | 0.4 | 2 | 0.5 | 25 | 0.16 | 5 | 0.4 | 4 | 0 | 11 | 0.1818 | 7 | 0.1429 | 27 | 0.1852 |
| 2_2602789 | 6 | 0.3333 | 8 | 0.5 | 5 | 0.4 | 2 | 0.5 | 21 | 0.4286 | 3 | 0.6667 | 3 | 0 | 7 | 0.2857 | 5 | 0.4 | 18 | 0.3333 |
| 2_2602795 | 8 | 0.125 | 9 | 0 | 7 | 0.1429 | 3 | 0 | 27 | 0.0741 | 3 | 0 | 3 | 0 | 11 | 0.1818 | 7 | 0 | 24 | 0.0833 |
| 2_2602805 | 6 | 0.1667 | 7 | 0 | 5 | 0 | 2 | 0 | 20 | 0.05 | 4 | 0.5 | 3 | 0 | 7 | 0.1429 | 6 | 0 | 20 | 0.15 |
| 2_2602823 | 7 | 0 | 4 | 0 | 4 | 0.25 | 1 | 0 | 16 | 0.0625 | 3 | 0.6667 | 2 | 0 | 8 | 0 | 5 | 0 | 18 | 0.1111 |
| 2_2602840 | 1 | 1 | 1 | 1 | 2 | 1 | 1 | 0 | 5 | 0.8 | NA | NA | 1 | 1 | 6 | 0.8333 | 4 | 1 | 11 | 0.9091 |
| 2_2602845 | 1 | 1 | 3 | 0 | 3 | 0.3333 | 1 | 0 | 8 | 0.25 | 1 | 0 | 1 | 0 | 6 | 0.3333 | 6 | 0 | 14 | 0.1429 |
| 2_2602849 | 2 | 1 | 3 | 0.6667 | 3 | 0.6667 | NA | NA | 8 | 0.75 | 1 | 1 | 1 | 1 | 7 | 1 | 6 | 1 | 15 | 1 |
| 2_2602867 | 2 | 0 | 3 | 0 | 2 | 0.5 | NA | NA | 7 | 0.1429 | NA | NA | 1 | 0 | 6 | 0.1667 | 6 | 0 | 13 | 0.0769 |
| 2_2602873 | 2 | 0 | 3 | 0.3333 | 2 | 0 | NA | NA | 7 | 0.1429 | NA | NA | 1 | 0 | 5 | 0.4 | 5 | 0.4 | 11 | 0.3636 |
| 2_2602885 | 2 | 0.5 | 3 | 0 | 2 | 0.5 | NA | NA | 7 | 0.2857 | NA | NA | 1 | 1 | 6 | 0 | 5 | 0.4 | 12 | 0.25 |
| 2_2609941 | 7 | 1 | 3 | 0.6667 | 2 | 0.5 | 4 | 0.75 | 16 | 0.8125 | 4 | 0.75 | 3 | 1 | 8 | 0.625 | 4 | 0.75 | 19 | 0.7368 |
| 2_2609943 | 8 | 0.625 | 3 | 0 | 3 | 0.6667 | 5 | 0.2 | 19 | 0.4211 | 6 | 0.1667 | 4 | 0.5 | 9 | 0.6667 | 7 | 0.4286 | 26 | 0.4615 |
| 2_2609960 | 6 | 0.8333 | 3 | 0.6667 | 4 | 0.75 | 5 | 0.8 | 18 | 0.7778 | 6 | 0.6667 | 4 | 1 | 9 | 0.7778 | 5 | 0.6 | 24 | 0.75 |
| 2_2609964 | 7 | 0.2857 | 1 | 0 | 4 | 0 | 4 | 0.25 | 16 | 0.1875 | 4 | 0 | 2 | 0 | 9 | 0.2222 | 5 | 0.4 | 20 | 0.2 |
| 2_2609985 | 7 | 0.1429 | 2 | 0 | 4 | 0 | 3 | 0 | 16 | 0.0625 | 6 | 0.1667 | 3 | 0.3333 | 9 | 0.1111 | 6 | 0.3333 | 24 | 0.2083 |
| 2_2610015 | 5 | 0.8 | 2 | 1 | 3 | 0.6667 | 2 | 0 | 12 | 0.6667 | 5 | 0.8 | 2 | 0.5 | 8 | 0.625 | 5 | 0.8 | 20 | 0.7 |
| 2_2610016 | 6 | 0.5 | 1 | 0 | 3 | 0.3333 | 1 | 1 | 11 | 0.4545 | 4 | 0.5 | 3 | 1 | 6 | 0.3333 | 5 | 0.4 | 18 | 0.5 |
| 2_2610048 | 7 | 0.1429 | 2 | 0 | 3 | 0.3333 | 2 | 0 | 14 | 0.1429 | 5 | 0 | 3 | 0.3333 | 9 | 0.3333 | 5 | 0.2 | 22 | 0.2273 |
| 2_2610072 | 7 | 0.1429 | 2 | 0 | 3 | 0 | 2 | 0 | 14 | 0.0714 | 4 | 0 | 3 | 0 | 8 | 0.125 | 4 | 0 | 19 | 0.0526 |
| 2_2610078 | 6 | 0.5 | 2 | 0.5 | 2 | 0 | 1 | 0 | 11 | 0.3636 | 3 | 0.3333 | 3 | 0.6667 | 6 | 0.1667 | 3 | 0.3333 | 15 | 0.3333 |
| 2_2610092 | 7 | 0.5714 | 2 | 0 | 3 | 0.6667 | 2 | 0 | 14 | 0.4286 | 3 | 1 | 2 | 0.5 | 8 | 0.625 | 3 | 0.6667 | 16 | 0.6875 |
| 2_2610104 | 5 | 0.8 | 2 | 1 | 3 | 0.6667 | 1 | 0 | 11 | 0.7273 | 1 | 1 | 2 | 0.5 | 8 | 0.625 | 3 | 0.3333 | 14 | 0.5714 |
| 2_2610112 | 3 | 0.6667 | 1 | 1 | 3 | 1 | 2 | 0.5 | 9 | 0.7778 | 1 | 1 | 2 | 0.5 | 7 | 0.8571 | 3 | 0.6667 | 13 | 0.7692 |
| 2_2610113 | 5 | 1 | 1 | 1 | 2 | 0.5 | 2 | 1 | 10 | 0.9 | 1 | 1 | 1 | 1 | 6 | 0.8333 | 1 | 1 | 9 | 0.8889 |
| 2_2610116 | 5 | 0.8 | 1 | 1 | 2 | 1 | 2 | 0.5 | 10 | 0.8 | 2 | 1 | 2 | 1 | 6 | 1 | 3 | 1 | 13 | 1 |
| 2_2610132 | 6 | 0.3333 | 2 | 0 | 1 | 1 | 3 | 0 | 12 | 0.25 | 2 | 0 | 2 | 0 | 6 | 0.1667 | 2 | 0 | 12 | 0.0833 |
| 2_2610138 | 5 | 0 | 1 | 0 | 1 | 1 | 3 | 0.3333 | 10 | 0.2 | 1 | 0 | 1 | 1 | 5 | 0.2 | 2 | 0 | 9 | 0.2222 |
| 2_2611224 | 3 | 0 | 2 | 0 | 1 | 1 | 2 | 0.5 | 8 | 0.25 | 1 | 0 | 1 | 0 | 2 | 1 | 2 | 0 | 6 | 0.3333 |
| 2_2613701 | 1 | 0 | NA | NA | NA | NA | 1 | 1 | 2 | 0.5 | NA | NA | 1 | 0 | NA | NA | 1 | 1 | 2 | 0.5 |
| 2_2613794 | 1 | 1 | NA | NA | NA | NA | 1 | 1 | 2 | 1 | NA | NA | NA | NA | NA | NA | 1 | 1 | 1 | 1 |

| | | | | | | | | | | | | | | | | | | | | |
|---|---|---|---|---|---|---|---|---|---|---|---|---|---|---|---|---|---|---|---|---|
| 2_26236949 | 5 | 0 | 8 | 0.25 | 5 | 0 | 7 | 0 | 25 | 0.08 | 2 | 0 | 5 | 0.2 | 5 | 0.2 | 4 | 0 | 16 | 0.125 |
| 2_26237062 | 5 | 1 | 11 | 0.8182 | 4 | 0.5 | 6 | 0.8333 | 26 | 0.8077 | 2 | 1 | 5 | 1 | 5 | 0.8 | 4 | 0.75 | 16 | 0.875 |
| 2_26237135 | 4 | 0.75 | 10 | 0.6 | 5 | 0.4 | 7 | 0.8571 | 26 | 0.6538 | 2 | 0.5 | 6 | 0.6667 | 6 | 0.6667 | 7 | 0.2857 | 21 | 0.5238 |
| 2_26237424 | 6 | 1 | 13 | 0.7692 | 3 | 0.6667 | 9 | 0.3333 | 31 | 0.6774 | 4 | 1 | 7 | 0.8571 | 6 | 0.6667 | 5 | 0.4 | 22 | 0.7273 |
| 2_26237849 | 9 | 0.1111 | 11 | 0.1818 | 4 | 0 | 8 | 0 | 32 | 0.0938 | 7 | 0.1429 | 8 | 0 | 8 | 0.125 | 4 | 0.25 | 27 | 0.1111 |
| 2_26238238 | 2 | 0 | 6 | 0.3333 | 1 | 0 | 4 | 0.25 | 13 | 0.2308 | 5 | 0.2 | 2 | 1 | 4 | 0.25 | 2 | 0.5 | 13 | 0.3846 |
| 2_26238554 | 2 | 0 | 5 | 0.4 | 1 | 0 | 3 | 0 | 11 | 0.1818 | 6 | 0.3333 | 2 | 0.5 | 5 | 0 | 2 | 0 | 15 | 0.2 |
| 2_26238613 | 1 | 1 | 4 | 0.75 | 1 | 1 | 2 | 0 | 8 | 0.625 | 4 | 0.25 | 2 | 0 | 1 | 0 | 2 | 0 | 9 | 0.1111 |
| 2_26238672 | 2 | 0.5 | 5 | 0.4 | NA | NA | 2 | 0.5 | 9 | 0.4444 | 6 | 0.3333 | 2 | 0 | 5 | 0.6 | 2 | 1 | 15 | 0.4667 |
| 2_26239183 | NA | NA | 3 | 0.6667 | 1 | 0 | 1 | 1 | 5 | 0.6 | 5 | 0.6 | 2 | 0.5 | 5 | 0.8 | 2 | 0.5 | 14 | 0.6429 |
| 2_26239427 | 1 | 1 | 4 | 0 | 3 | 0.3333 | 2 | 0.5 | 10 | 0.3 | 6 | 0.1667 | 2 | 0 | 4 | 0.5 | 2 | 0 | 14 | 0.2143 |
| 2_26239599 | 2 | 0 | 3 | 0.3333 | 3 | 0.3333 | 2 | 0.5 | 10 | 0.3 | 6 | 0.1667 | 1 | 0 | 5 | 0 | 2 | 0 | 14 | 0.0714 |
| 2_26240393 | 11 | 0.0909 | 12 | 0.1667 | 17 | 0.0588 | 13 | 0.1538 | 53 | 0.1132 | 17 | 0 | 16 | 0.0625 | 16 | 0.125 | 14 | 0.0714 | 63 | 0.0635 |
| 2_26240426 | 8 | 0.25 | 10 | 0.4 | 13 | 0.4615 | 11 | 0.1818 | 42 | 0.3333 | 11 | 0.4545 | 14 | 0.2857 | 12 | 0.4167 | 13 | 0.3846 | 50 | 0.38 |
| 2_26240501 | 8 | 0.5 | 12 | 0.0833 | 18 | 0.1667 | 11 | 0.1818 | 49 | 0.2041 | 16 | 0.125 | 12 | 0.0833 | 14 | 0.0714 | 15 | 0.1333 | 57 | 0.1053 |
| 2_26241911 | 6 | 0.3333 | 10 | 0.5 | 11 | 0.4545 | 7 | 0.4286 | 34 | 0.4412 | 11 | 0.3636 | 10 | 0 | 4 | 0.5 | 12 | 0.5 | 37 | 0.3243 |
| 2_26252835 | 2 | 0.5 | 6 | 0.3333 | 3 | 0.6667 | 3 | 1 | 14 | 0.5714 | 1 | 0 | 1 | 0 | 2 | 1 | 2 | 0.5 | 6 | 0.5 |
| 2_26254684 | 2 | 0.5 | 4 | 0.75 | 2 | 0.5 | 3 | 0 | 11 | 0.4545 | NA | NA | NA | NA | NA | NA | 3 | 0.6667 | 3 | 0.6667 |
| 2_26254867 | 2 | 0.5 | 5 | 0.6 | 3 | 0.6667 | 3 | 0 | 13 | 0.4615 | 1 | 1 | 1 | 0 | 2 | 0.5 | 2 | 0 | 6 | 0.3333 |
| 2_26254981 | 2 | 0.5 | 5 | 0.8 | 2 | 0.5 | 3 | 0.6667 | 12 | 0.6667 | 1 | 0 | 1 | 1 | 2 | 0.5 | 3 | 0.6667 | 7 | 0.5714 |
| 2_26254986 | 2 | 0 | 4 | 0 | 2 | 0 | 2 | 0.5 | 10 | 0.1 | 1 | 1 | NA | NA | 2 | 0 | 3 | 0 | 6 | 0.1667 |
| 2_26255140 | 2 | 1 | 5 | 1 | 3 | 1 | 2 | 1 | 12 | 1 | 1 | 1 | 1 | 1 | 2 | 1 | 3 | 1 | 7 | 1 |
| 2_26255193 | 2 | 0.5 | 4 | 1 | 3 | 1 | 3 | 0.3333 | 12 | 0.75 | 1 | 1 | 1 | 1 | 2 | 0.5 | 3 | 0.6667 | 7 | 0.7143 |
| 2_26255202 | 2 | 0.5 | 4 | 0 | 3 | 0 | 3 | 0 | 12 | 0.0833 | 1 | 0 | 1 | 1 | 2 | 0 | 3 | 0.3333 | 7 | 0.2857 |
| 2_26255209 | 2 | 0 | 5 | 0.2 | 3 | 0 | 3 | 0 | 13 | 0.0769 | 1 | 0 | 1 | 0 | 2 | 0 | 3 | 0.3333 | 7 | 0.1429 |
| 2_26255229 | 2 | 0 | 4 | 0 | 1 | 0 | 3 | 0.3333 | 10 | 0.1 | 1 | 1 | 1 | 0 | 1 | 0 | 2 | 0 | 5 | 0.2 |
| 2_26255246 | 2 | 0 | 5 | 0.4 | 3 | 0 | 3 | 0 | 13 | 0.1538 | 1 | 0 | 1 | 1 | 2 | 0.5 | 3 | 0 | 7 | 0.2857 |
| 2_26255280 | 2 | 0 | 5 | 0.6 | 3 | 0.3333 | 3 | 0 | 13 | 0.3077 | 1 | 0 | 1 | 0 | 2 | 0 | 3 | 0.3333 | 7 | 0.1429 |
| 2_26255406 | 1 | 0 | 4 | 0.75 | 2 | 0 | 2 | 0.5 | 9 | 0.4444 | NA | NA | 1 | 1 | 2 | 1 | 3 | 0.6667 | 6 | 0.8333 |
| 2_26255511 | 2 | 0 | 5 | 0.4 | 3 | 0.3333 | 3 | 0 | 13 | 0.2308 | NA | NA | 1 | 0 | 1 | 0 | 3 | 0.3333 | 5 | 0.2 |
| 2_26255597 | 2 | 0.5 | 4 | 0 | 3 | 0.3333 | 3 | 0.3333 | 12 | 0.25 | 1 | 1 | NA | NA | 2 | 1 | 3 | 0.3333 | 6 | 0.6667 |
| 2_26255730 | 1 | 0 | 4 | 0.25 | 2 | 0 | NA | NA | 7 | 0.1429 | 1 | 1 | NA | NA | 2 | 1 | 3 | 0.3333 | 6 | 0.6667 |
| 2_26255906 | 2 | 0.5 | 2 | 0.5 | 2 | 0.5 | 3 | 0.3333 | 9 | 0.4444 | 1 | 1 | 1 | 1 | NA | NA | 1 | 1 | 3 | 1 |
| 2_26255969 | 2 | 0 | 4 | 0.5 | 2 | 1 | 3 | 0.6667 | 11 | 0.5455 | 1 | 0 | 1 | 1 | 1 | 0 | 2 | 0.5 | 5 | 0.4 |
| 2_26259188 | 1 | 1 | 3 | 0.3333 | 1 | 0 | 1 | 0 | 6 | 0.3333 | 1 | 0 | NA | NA | 1 | 0 | 2 | 0.5 | 4 | 0.25 |
| 2_26272975 | 9 | 0.4444 | 8 | 0.625 | 11 | 0.3636 | 6 | 0.6667 | 34 | 0.5 | 10 | 0.6 | 5 | 0.4 | 11 | 0.6364 | 3 | 0.3333 | 29 | 0.5517 |
| 2_26273048 | 8 | 0.5 | 9 | 0.6667 | 11 | 0.5455 | 6 | 0.8333 | 34 | 0.6176 | 9 | 0.6667 | 5 | 0.4 | 12 | 0.75 | 4 | 0.5 | 30 | 0.6333 |
| 2_26273186 | 8 | 0.125 | 9 | 0.3333 | 8 | 0.375 | 6 | 0.3333 | 31 | 0.2903 | 7 | 0.4286 | 5 | 0.2 | 12 | 0.4167 | 4 | 0 | 28 | 0.3214 |
| 2_26273194 | 8 | 0.25 | 9 | 0 | 10 | 0.1 | 4 | 0.5 | 31 | 0.1613 | 8 | 0.125 | 4 | 0.25 | 11 | 0.2727 | 4 | 0.25 | 27 | 0.2222 |
| 2_26273398 | 9 | 0.4444 | 9 | 0 | 10 | 0.3 | 6 | 0.1667 | 34 | 0.2353 | 6 | 0.1667 | 5 | 0 | 9 | 0.2222 | 3 | 0 | 23 | 0.1304 |
| 2_26273418 | 10 | 0.9 | 10 | 1 | 8 | 1 | 6 | 0.6667 | 34 | 0.9118 | 6 | 1 | 4 | 1 | 9 | 1 | 3 | 1 | 22 | 1 |
| 2_26273454 | 9 | 0.3333 | 9 | 0.3333 | 7 | 0 | 6 | 0.1667 | 31 | 0.2258 | 6 | 0.1667 | 4 | 0 | 9 | 0.3333 | 2 | 0.5 | 21 | 0.2381 |
| 2_26273494 | 9 | 0.1111 | 9 | 0.1111 | 10 | 0.2 | 5 | 0.6 | 33 | 0.2121 | 7 | 0.2857 | 4 | 0 | 10 | 0.1 | 3 | 0 | 24 | 0.125 |
| 2_26273617 | 8 | 0.5 | 10 | 0.4 | 9 | 0.1111 | 3 | 0.6667 | 30 | 0.3667 | 6 | 0.6667 | 3 | 0 | 6 | 0.3333 | 2 | 0.5 | 17 | 0.4118 |
| 2_26275352 | 5 | 0 | 1 | 0 | NA | NA | 2 | 0.5 | 8 | 0.125 | 3 | 0 | 2 | 0.5 | 2 | 0 | 1 | 0 | 8 | 0.125 |
| 2_26275376 | 4 | 0.75 | 2 | 0.5 | NA | NA | 1 | 1 | 7 | 0.7143 | 2 | 0.5 | 2 | 1 | 3 | 0.6667 | 1 | 1 | 8 | 0.75 |
| 2_26275519 | 7 | 0.8571 | 6 | 0.5 | 6 | 0.5 | 5 | 0.8 | 24 | 0.6667 | 3 | 0 | 4 | 1 | 5 | 0.4 | 3 | 0 | 15 | 0.4 |
| 2_26275531 | 8 | 0.125 | 7 | 0.1429 | 6 | 0 | 4 | 0 | 25 | 0.08 | 4 | 0 | 3 | 0 | 6 | 0.1667 | 2 | 0 | 15 | 0.0667 |
| 2_26275568 | 8 | 0.25 | 9 | 0.2222 | 6 | 0 | 5 | 0.4 | 28 | 0.2143 | 5 | 0 | 4 | 0.25 | 7 | 0.1429 | 3 | 0 | 19 | 0.1053 |
| 2_26275614 | 5 | 0.4 | 7 | 0.5714 | 5 | 0.6 | 4 | 0.5 | 21 | 0.5238 | 4 | 0.25 | 4 | 0.5 | 7 | 0.2857 | 2 | 0.5 | 17 | 0.3529 |
| 2_26275690 | 4 | 0.25 | 4 | 0.25 | 3 | 0.3333 | 3 | 0 | 14 | 0.2143 | 2 | 0 | 2 | 0 | 3 | 0.3333 | 1 | 1 | 8 | 0.25 |
| 2_26275854 | 2 | 0 | 5 | 0.4 | 2 | 0.5 | 3 | 0.3333 | 12 | 0.3333 | 2 | 0 | 2 | 1 | 2 | 0 | 1 | 0 | 7 | 0.2857 |
| 2_26276094 | 5 | 0.4 | 4 | 0.75 | 1 | 1 | 1 | 0 | 11 | 0.5455 | 1 | 0 | 2 | 1 | 1 | 0 | NA | NA | 4 | 0.5 |
| 2_26276229 | 4 | 0.5 | 4 | 0.5 | 1 | 1 | 2 | 0.5 | 11 | 0.5455 | 1 | 1 | 2 | 0.5 | 1 | 1 | NA | NA | 4 | 0.75 |
| 2_26276240 | 4 | 0.25 | 4 | 0.5 | 1 | 1 | 2 | 0.5 | 11 | 0.4545 | 1 | 0 | 1 | 1 | 2 | 0.5 | NA | NA | 4 | 0.5 |
| 2_26276953 | 4 | 0.25 | 7 | 0.1429 | 6 | 0.1667 | 4 | 0.5 | 21 | 0.2381 | 4 | 0.25 | 4 | 0 | 8 | 0.125 | 2 | 0 | 18 | 0.1111 |
| 2_26276983 | 5 | 0.2 | 8 | 0.125 | 6 | 0.1667 | 5 | 0.4 | 24 | 0.2083 | 5 | 0.2 | 4 | 0.5 | 8 | 0.25 | 2 | 0 | 19 | 0.2632 |
| 2_26277124 | 5 | 0.6 | 8 | 0.5 | 6 | 0.6667 | 4 | 0.75 | 23 | 0.6087 | 5 | 1 | 4 | 0.75 | 7 | 0.1429 | 2 | 0.5 | 18 | 0.5556 |
| 2_26277133 | 3 | 0.3333 | 7 | 0 | 5 | 0.2 | 4 | 0.5 | 19 | 0.2105 | 5 | 0.4 | 3 | 0 | 8 | 0.125 | 1 | 0 | 17 | 0.1765 |
| 2_26277149 | 5 | 0 | 6 | 0.1667 | 6 | 0.3333 | 5 | 0.4 | 22 | 0.2273 | 4 | 0.25 | 4 | 0.5 | 8 | 0.125 | 2 | 0 | 18 | 0.2222 |
| 2_26277300 | 4 | 0.25 | 8 | 0.625 | 5 | 0.8 | 2 | 1 | 19 | 0.6316 | 5 | 0.8 | 4 | 0.5 | 7 | 0.5714 | 1 | 0 | 17 | 0.5882 |
| 2_26277448 | NA | NA | NA | NA | 2 | 1 | 1 | 1 | 3 | 1 | 1 | 0 | 1 | 1 | NA | NA | NA | NA | 2 | 0.5 |
| 2_26279678 | NA | NA | NA | NA | NA | NA | 1 | 1 | 1 | 1 | 1 | 1 | NA | NA | 1 | 1 | 2 | 0.5 | 4 | 0.75 |
| 2_26279874 | NA | NA | NA | NA | NA | NA | 1 | 1 | 1 | 1 | 1 | 1 | NA | NA | NA | NA | 2 | 0.5 | 3 | 0.6667 |
| 2_26280059 | 4 | 0 | 5 | 0 | 3 | 0 | 4 | 0.25 | 16 | 0.0625 | 2 | 0.5 | 1 | 0 | 8 | 0.25 | 9 | 0.1111 | 20 | 0.2 |

| | | | | | | | | | | | | | | | | | | | | |
|---|---|---|---|---|---|---|---|---|---|---|---|---|---|---|---|---|---|---|---|---|
| 2_2628670 | 1 | 1 | NA | NA | 2 | 1 | 1 | 1 | 4 | 1 | 3 | 1 | NA | NA | NA | NA | 1 | 1 | 4 | 1 |
| 2_2628675 | NA | NA | NA | NA | 1 | 1 | 1 | 1 | 2 | 1 | NA | NA | NA | NA | NA | NA | 1 | 1 | 1 | 1 |
| 2_2628685 | 1 | 1 | NA | NA | 1 | 1 | 1 | 0 | 3 | 0.6667 | 3 | 0.3333 | NA | NA | NA | NA | 1 | 0 | 4 | 0.25 |
| 2_2628690 | 4 | 0 | 4 | 0 | 4 | 0.25 | 5 | 0 | 17 | 0.0588 | 4 | 0.5 | 4 | 0.25 | 8 | 0.125 | 11 | 0 | 27 | 0.1481 |
| 2_2628693 | 3 | 0.3333 | 4 | 0 | 3 | 0 | 3 | 0.3333 | 13 | 0.1538 | 4 | 0 | 5 | 0.2 | 8 | 0.5 | 11 | 0.1818 | 28 | 0.25 |
| 2_2628698 | 1 | 1 | NA | NA | 1 | 1 | NA | NA | 2 | 1 | 1 | 1 | NA | NA | NA | NA | 1 | 1 | 2 | 1 |
| 2_2628745 | 1 | 1 | 1 | 1 | 2 | 0 | NA | NA | 4 | 0.5 | 4 | 0.25 | NA | NA | 1 | 1 | 2 | 1 | 7 | 0.5714 |
| 2_2628766 | 1 | 0 | NA | NA | 2 | 0 | 2 | 0.5 | 5 | 0.2 | 3 | 0.6667 | NA | NA | 1 | 1 | 2 | 0 | 6 | 0.5 |
| 2_2628770 | 1 | 1 | 1 | 1 | 1 | 1 | 2 | 1 | 5 | 1 | 3 | 0.6667 | NA | NA | 1 | 1 | 2 | 0.5 | 6 | 0.6667 |
| 2_2628783 | 1 | 0 | 1 | 1 | 2 | 0 | 2 | 0.5 | 6 | 0.3333 | 4 | 0.5 | NA | NA | NA | NA | 2 | 0.5 | 6 | 0.5 |
| 2_2629793 | 1 | 0 | 1 | 1 | 1 | 0 | 1 | 0 | 4 | 0.25 | 2 | 0 | NA | NA | NA | NA | 1 | 1 | 3 | 0.3333 |
| 2_2629799 | 1 | 1 | 1 | 1 | 1 | 1 | 1 | 0 | 4 | 0.75 | 3 | 1 | NA | NA | 1 | 1 | NA | NA | 4 | 1 |
| 2_2629807 | 1 | 1 | NA | NA | 1 | 0 | 1 | 0 | 3 | 0.3333 | 2 | 0.5 | NA | NA | 1 | 1 | NA | NA | 3 | 0.6667 |
| 2_2629811 | 1 | 1 | 1 | 1 | 1 | 0 | 1 | 0 | 4 | 0.5 | 2 | 0 | NA | NA | 1 | 1 | NA | NA | 3 | 0.3333 |
| 2_2629823 | 1 | 1 | 1 | 1 | 1 | 1 | 1 | 1 | 4 | 1 | 3 | 0.3333 | NA | NA | 1 | 0 | NA | NA | 4 | 0.25 |
| 2_2629958 | 7 | 0 | 4 | 0.25 | 4 | 0 | 6 | 0.1667 | 21 | 0.0952 | 4 | 0.5 | 6 | 0.1667 | 5 | 0.2 | 8 | 0 | 23 | 0.1739 |
| 2_2629966 | 5 | 0 | 6 | 0.5 | 3 | 0 | 7 | 0.1429 | 21 | 0.1905 | 6 | 0.3333 | 7 | 0.2857 | 7 | 0 | 8 | 0.125 | 28 | 0.1786 |
| 2_2629971 | 8 | 0.125 | 7 | 0.2857 | 3 | 0 | 6 | 0 | 24 | 0.125 | 4 | 0.25 | 7 | 0 | 6 | 0 | 8 | 0.125 | 25 | 0.08 |
| 2_2629988 | 8 | 0.5 | 7 | 0.7143 | 2 | 0.5 | 4 | 0.5 | 21 | 0.5714 | 5 | 0.8 | 7 | 0.4286 | 4 | 0.75 | 8 | 0.75 | 24 | 0.6667 |
| 2_2629996 | 7 | 0.4286 | 7 | 0.1429 | 1 | 0 | 5 | 0.4 | 20 | 0.3 | 3 | 0 | 7 | 0.2857 | 7 | 0.2857 | 9 | 0.2222 | 26 | 0.2308 |
| 2_2630008 | 7 | 0.1429 | 7 | 0 | 2 | 0 | 4 | 0 | 20 | 0.05 | 5 | 0 | 7 | 0 | 7 | 0.1429 | 8 | 0.125 | 27 | 0.0741 |
| 2_2630014 | 7 | 0.1429 | 5 | 0 | 2 | 0 | 4 | 0 | 18 | 0.0556 | 4 | 0.25 | 6 | 0.1667 | 3 | 0.3333 | 8 | 0 | 21 | 0.1429 |
| 2_2630038 | 6 | 0.1667 | 6 | 0.3333 | 1 | 0 | 3 | 0 | 16 | 0.1875 | 2 | 0.5 | 5 | 0.4 | 4 | 0.5 | 7 | 0.1429 | 18 | 0.3333 |
| 2_2630041 | 7 | 0 | 6 | 0 | 1 | 1 | 3 | 0.3333 | 17 | 0.1176 | 3 | 0.3333 | 6 | 0 | 4 | 0 | 5 | 0 | 18 | 0.0556 |
| 2_2630045 | 5 | 0.6 | 6 | 0 | 1 | 0 | 4 | 0.25 | 16 | 0.25 | 4 | 0.25 | 5 | 0 | 6 | 0.1667 | 9 | 0.4444 | 24 | 0.25 |
| 2_2630045 | 6 | 0.8333 | 5 | 0.4 | 1 | 0 | 4 | 0 | 16 | 0.4375 | 4 | 0.75 | 6 | 0.8333 | 4 | 0.75 | 9 | 0.6667 | 23 | 0.7391 |
| 2_2630054 | 6 | 0.3333 | 7 | 0.2857 | 1 | 1 | 4 | 0 | 18 | 0.2778 | 2 | 1 | 6 | 0.3333 | 4 | 0.25 | 8 | 0.375 | 20 | 0.4 |
| 2_2630065 | 7 | 0 | 7 | 0 | 1 | 1 | 2 | 0 | 17 | 0.0588 | 2 | 0.5 | 4 | 0.25 | 5 | 0 | 8 | 0.25 | 19 | 0.2105 |
| 2_2630079 | 7 | 0.5714 | 7 | 0.2857 | 2 | 0.5 | 4 | 0.75 | 20 | 0.5 | 3 | 0.6667 | 3 | 0.6667 | 5 | 0.8 | 8 | 0.5 | 19 | 0.6316 |
| 2_2634833 | 2 | 0.5 | 2 | 1 | NA | NA | 2 | 0 | 6 | 0.5 | NA | NA | 4 | 0.5 | 1 | 0 | 5 | 1 | 10 | 0.7 |
| 2_2634858 | 2 | 0.5 | 3 | 0.6667 | NA | NA | 3 | 0.3333 | 8 | 0.5 | NA | NA | 3 | 0 | 1 | 0 | 4 | 0.75 | 8 | 0.375 |
| 2_2634861 | 2 | 1 | 2 | 1 | NA | NA | 3 | 1 | 7 | 1 | NA | NA | 2 | 0.5 | 1 | 1 | 4 | 0.75 | 7 | 0.7143 |
| 2_2634867 | 2 | 0 | 2 | 1 | NA | NA | 3 | 1 | 7 | 0.7143 | NA | NA | 3 | 0.6667 | 1 | 1 | 4 | 1 | 8 | 0.875 |
| 2_2634868 | 1 | 1 | 3 | 0.3333 | NA | NA | 3 | 0 | 7 | 0.2857 | NA | NA | 4 | 0 | 1 | 0 | 4 | 0.25 | 9 | 0.1111 |
| 2_2634873 | 2 | 1 | 3 | 1 | NA | NA | 3 | 0.6667 | 8 | 0.875 | NA | NA | 3 | 0.6667 | 1 | 1 | 2 | 1 | 6 | 0.8333 |
| 2_2634893 | 2 | 0.5 | 2 | 0 | NA | NA | 1 | 1 | 5 | 0.4 | NA | NA | 3 | 0.6667 | 1 | 0 | 3 | 0.6667 | 7 | 0.5714 |
| 2_2634901 | 2 | 0 | 2 | 0 | NA | NA | 2 | 0.5 | 6 | 0.1667 | NA | NA | 3 | 0 | 1 | 0 | 2 | 0.5 | 6 | 0.1667 |
| 2_2634972 | 1 | 0 | 1 | 0 | NA | NA | 2 | 0.5 | 4 | 0.25 | NA | NA | 3 | 0 | 1 | 0 | 2 | 0.5 | 6 | 0.1667 |
| 2_2635000 | 1 | 0 | 1 | 0 | NA | NA | 2 | 1 | 4 | 0.5 | NA | NA | 1 | 1 | 1 | 0 | 1 | 0 | 3 | 0.3333 |
| 2_2635056 | NA | NA | 1 | 1 | NA | NA | 2 | 1 | 3 | 1 | NA | NA | 1 | 1 | NA | NA | 2 | 1 | 3 | 1 |
| 2_2635094 | NA | NA | NA | NA | NA | NA | 2 | 0.5 | 2 | 0.5 | NA | NA | 1 | 0 | NA | NA | 1 | 1 | 2 | 0.5 |
| 2_2648258 | 3 | 0.3333 | 5 | 0.2 | 6 | 0.5 | 4 | 0.25 | 18 | 0.3333 | 7 | 0.5714 | 4 | 0.5 | 6 | 0.3333 | 5 | 0.4 | 22 | 0.4545 |
| 2_2652561 | 6 | 0.3333 | 9 | 0.1111 | 7 | 0.4286 | 8 | 0.125 | 30 | 0.2333 | 12 | 0.1667 | 9 | 0.4444 | 9 | 0.2222 | 9 | 0.3333 | 39 | 0.2821 |
| 2_2652574 | 3 | 0.6667 | 11 | 0.4545 | 8 | 0.125 | 8 | 0.625 | 30 | 0.4333 | 15 | 0.3333 | 9 | 0.4444 | 11 | 0.2727 | 8 | 0.5 | 43 | 0.3721 |
| 2_2652586 | 9 | 0.7778 | 8 | 0.875 | 9 | 0.6667 | 9 | 0.7778 | 35 | 0.7714 | 12 | 0.8333 | 9 | 0.7778 | 8 | 0.875 | 8 | 0.75 | 37 | 0.8108 |
| 2_2652595 | 10 | 0.4 | 12 | 0.3333 | 7 | 0.1429 | 7 | 0.4286 | 36 | 0.3333 | 12 | 0.4167 | 9 | 0.3333 | 12 | 0.4167 | 9 | 0.3333 | 42 | 0.381 |
| 2_2652608 | 6 | 0.5 | 8 | 0.375 | 7 | 0.2857 | 6 | 0.5 | 27 | 0.4074 | 13 | 0.4615 | 6 | 0.5 | 11 | 0.4545 | 9 | 0.4444 | 39 | 0.4615 |
| 2_2652611 | 9 | 0 | 12 | 0.1667 | 8 | 0 | 8 | 0 | 37 | 0.0541 | 18 | 0.1667 | 10 | 0.2 | 11 | 0 | 12 | 0 | 51 | 0.098 |
| 2_2652619 | 8 | 1 | 13 | 0.6923 | 11 | 1 | 8 | 0.625 | 40 | 0.825 | 18 | 0.7778 | 7 | 0.7143 | 11 | 0.8182 | 11 | 0.6364 | 47 | 0.7447 |
| 2_2652627 | 7 | 0.8571 | 13 | 0.9231 | 10 | 0.8 | 5 | 0.8 | 35 | 0.8571 | 18 | 0.9444 | 10 | 0.8 | 11 | 0.6364 | 11 | 0.8182 | 50 | 0.82 |
| 2_2652632 | 9 | 0.6667 | 10 | 0.9 | 10 | 0.5 | 8 | 0.75 | 37 | 0.7027 | 15 | 0.5333 | 8 | 0.625 | 10 | 0.8 | 11 | 0.5455 | 44 | 0.6136 |
| 2_2677921 | NA | NA | 1 | 1 | NA | NA | NA | NA | 1 | 1 | NA | NA | NA | NA | 1 | 1 | NA | NA | 1 | 1 |
| 2_2678152 | NA | NA | 1 | 0 | 2 | 0 | 1 | 1 | 4 | 0.25 | 2 | 0.5 | NA | NA | NA | NA | 2 | 1 | 4 | 0.75 |
| 2_2678162 | 1 | 1 | 1 | 1 | 2 | 0 | 1 | 1 | 5 | 0.6 | 3 | 0 | NA | NA | NA | NA | 3 | 0.3333 | 6 | 0.1667 |
| 2_2678167 | 1 | 1 | NA | NA | 2 | 0 | NA | NA | 3 | 0.3333 | 3 | 0.6667 | 1 | 0 | NA | NA | 3 | 0 | 7 | 0.2857 |
| 2_2678196 | 1 | 1 | NA | NA | 2 | 1 | NA | NA | 3 | 1 | 2 | 0.5 | 1 | 1 | NA | NA | 2 | 1 | 5 | 0.8 |
| 2_2678205 | 1 | 1 | NA | NA | 2 | 0.5 | NA | NA | 3 | 0.6667 | 2 | 1 | 1 | 1 | NA | NA | 3 | 0.6667 | 6 | 0.8333 |
| 2_2678209 | 1 | 1 | NA | NA | 2 | 1 | NA | NA | 3 | 1 | 2 | 0 | 1 | 0 | NA | NA | 3 | 1 | 6 | 0.5 |
| 2_2678215 | 1 | 1 | NA | NA | 1 | 1 | NA | NA | 2 | 1 | 2 | 1 | 1 | 1 | NA | NA | 2 | 1 | 5 | 1 |
| 2_2678228 | 1 | 1 | NA | NA | 1 | 0 | NA | NA | 2 | 0.5 | 2 | 0.5 | 1 | 1 | NA | NA | 3 | 0.6667 | 6 | 0.6667 |
| 2_2678253 | 1 | 0 | NA | NA | 1 | 1 | NA | NA | 2 | 0.5 | 2 | 1 | NA | NA | NA | NA | 3 | 1 | 5 | 1 |
| 2_2679570 | 6 | 0.1667 | 4 | 0.25 | 2 | 0 | 5 | 0.4 | 17 | 0.2353 | 4 | 0.5 | 2 | 0 | 5 | 0 | 6 | 0.1667 | 17 | 0.1765 |
| 2_2680351 | 15 | 0.2 | 18 | 0.0556 | 13 | 0.3077 | 16 | 0.375 | 62 | 0.2258 | 12 | 0 | 14 | 0.3571 | 9 | 0.5556 | 20 | 0.2 | 55 | 0.2545 |
| 2_2680987 | 4 | 0 | 2 | 0.5 | 2 | 0 | 2 | 0.5 | 10 | 0.2 | 3 | 0 | 4 | 0 | 8 | 0.125 | 3 | 0 | 18 | 0.0556 |
| 2_2680993 | 4 | 0.75 | 5 | 0.4 | 4 | 0.75 | 2 | 0.5 | 15 | 0.6 | 3 | 0.6667 | 4 | 0.5 | 7 | 0.4286 | 5 | 0.4 | 19 | 0.4737 |

| | | | | | | | | | | | | | | | | | | | | |
|---|---|---|---|---|---|---|---|---|---|---|---|---|---|---|---|---|---|---|---|---|
| 2_26809969 | 4 | 1 | 5 | 0.8 | 5 | 0.6 | 3 | 1 | 17 | 0.8235 | 3 | 0.6667 | 3 | 0.6667 | 7 | 0.4286 | 3 | 0.6667 | 16 | 0.5625 |
| 2_26810041 | 2 | 1 | 4 | 0.25 | 4 | 0.25 | 3 | 0 | 13 | 0.3077 | 4 | 0 | 4 | 0.25 | 7 | 0 | 3 | 0.3333 | 18 | 0.1111 |
| 2_26810612 | 7 | 0.1429 | 7 | 0.2857 | 7 | 0 | 3 | 0 | 24 | 0.125 | 10 | 0.1 | 4 | 0.25 | 4 | 0 | 8 | 0 | 26 | 0.0769 |
| 2_26811785 | 6 | 0.8333 | 7 | 0.4286 | 11 | 0.8182 | 5 | 0.8 | 29 | 0.7241 | 13 | 0.6923 | 8 | 0.875 | 9 | 0.7778 | 11 | 0.8182 | 41 | 0.7805 |
| 2_26811849 | 5 | 0.2 | 4 | 0 | 7 | 0 | 5 | 0.6 | 21 | 0.1905 | 11 | 0.1818 | 7 | 0.4286 | 5 | 0.2 | 7 | 0.7143 | 30 | 0.3667 |
| 2_26812029 | 3 | 0.3333 | 4 | 0 | 11 | 0 | 4 | 0.25 | 22 | 0.0909 | 10 | 0.4 | 6 | 0.1667 | 8 | 0 | 6 | 0 | 30 | 0.1667 |
| 2_26812057 | 4 | 1 | 4 | 0.75 | 5 | 1 | 3 | 1 | 16 | 0.9375 | 10 | 0.6 | 7 | 0.5714 | 4 | 0.75 | 5 | 0.8 | 26 | 0.6538 |
| 2_26812108 | 4 | 0.25 | 6 | 0 | 9 | 0.1111 | 5 | 0 | 24 | 0.0833 | 9 | 0.3333 | 8 | 0 | 8 | 0.25 | 6 | 0.3333 | 31 | 0.2258 |
| 2_26843665 | 5 | 0.6 | 1 | 1 | 5 | 0.6 | 6 | 0.6667 | 17 | 0.6471 | 6 | 0.1667 | 5 | 0.4 | 2 | 0.5 | 8 | 0.5 | 21 | 0.381 |
| 2_26843739 | 6 | 0.5 | 2 | 0.5 | 4 | 0.75 | 8 | 0.625 | 20 | 0.6 | 7 | 0.4286 | 8 | 0.5 | 6 | 0.3333 | 8 | 0.625 | 29 | 0.4828 |
| 2_26843748 | 6 | 0.1667 | 2 | 0.5 | 4 | 0.25 | 7 | 0.5714 | 19 | 0.3684 | 7 | 0.1429 | 7 | 0.1429 | 5 | 0.2 | 7 | 0.2857 | 26 | 0.1923 |
| 2_26843985 | 3 | 1 | 2 | 0.5 | 4 | 0.75 | 6 | 0.8333 | 15 | 0.8 | 5 | 0.8 | 6 | 0.5 | 6 | 0 | 6 | 0.5 | 23 | 0.4348 |
| 2_26844384 | 4 | 0.75 | 1 | 1 | 4 | 0.75 | 5 | 0.6 | 14 | 0.7143 | 7 | 0.5714 | 7 | 0.8571 | 6 | 0.6667 | 6 | 0.6667 | 26 | 0.6923 |
| 2_26844448 | 3 | 0.3333 | 2 | 1 | 3 | 0.3333 | 6 | 0.6667 | 14 | 0.5714 | 7 | 0.4286 | 7 | 0.7143 | 6 | 0.6667 | 6 | 0.3333 | 26 | 0.5385 |
| 2_26844496 | 2 | 0 | NA | NA | 3 | 0 | 4 | 0.5 | 9 | 0.2222 | 4 | 0 | 6 | 0.1667 | 6 | 0 | 6 | 0.3333 | 22 | 0.1364 |
| 2_26852557 | 2 | 0.5 | NA | NA | 3 | 0 | 1 | 0 | 6 | 0.1667 | 4 | 0 | 5 | 0 | 3 | 0 | 1 | 1 | 13 | 0.0769 |
| 2_26852606 | 2 | 0.5 | 1 | 0 | 3 | 0.3333 | 2 | 0 | 8 | 0.25 | 3 | 0 | 5 | 0.4 | 2 | 0 | 1 | 0 | 11 | 0.1818 |
| 2_26854095 | 2 | 0.5 | NA | NA | 2 | 0 | 2 | 0.5 | 6 | 0.3333 | 2 | 0.5 | 5 | 0.4 | 2 | 1 | 1 | 1 | 10 | 0.6 |
| 2_26906064 | NA | NA | 4 | 0.25 | 1 | 0 | 2 | 0 | 7 | 0.1429 | 4 | 0 | 2 | 0 | 3 | 0.3333 | NA | NA | 9 | 0.1111 |
| 2_26906699 | NA | NA | 4 | 0.25 | 2 | 0 | 2 | 0 | 8 | 0.125 | 3 | 0 | 3 | 0 | 2 | 0.5 | NA | NA | 8 | 0.125 |
| 2_26907852 | NA | NA | NA | NA | NA | NA | 1 | 1 | 1 | 1 | NA | NA | 1 | 1 | 1 | 0 | NA | NA | 2 | 0.5 |
| 2_26908472 | 4 | 0.75 | NA | NA | 4 | 0.75 | 2 | 0.5 | 10 | 0.7 | 1 | 0 | 3 | 0 | 4 | 0.75 | 2 | 1 | 10 | 0.5 |
| 2_26908970 | 4 | 0.25 | 1 | 0 | 6 | 0 | 2 | 0 | 13 | 0.0769 | 2 | 0 | 6 | 0.1667 | 5 | 0.2 | 2 | 1 | 15 | 0.2667 |
| 2_26908984 | 4 | 0 | 1 | 0 | 5 | 0.2 | 2 | 0.5 | 12 | 0.1667 | 1 | 1 | 6 | 0.3333 | 3 | 0.6667 | 2 | 0.5 | 12 | 0.5 |
| 2_26909054 | 3 | 0.6667 | 1 | 0 | 4 | 0.25 | 2 | 0.5 | 10 | 0.4 | 1 | 1 | 4 | 0.25 | 5 | 0.6 | 2 | 1 | 12 | 0.5833 |
| 2_26909142 | 4 | 1 | NA | NA | 5 | 1 | 2 | 1 | 11 | 1 | NA | NA | 5 | 1 | 4 | 1 | 1 | 1 | 10 | 1 |
| 2_26909203 | 4 | 0.75 | NA | NA | 4 | 1 | 2 | 1 | 10 | 0.9 | 1 | 1 | 4 | 1 | 6 | 1 | 2 | 1 | 13 | 1 |
| 2_26909255 | 4 | 0 | NA | NA | 5 | 0.2 | 2 | 0 | 11 | 0.0909 | 1 | 0 | 5 | 0 | 5 | 0 | 2 | 0.5 | 13 | 0.0769 |
| 2_26909277 | 3 | 0.3333 | NA | NA | 3 | 0.3333 | 1 | 0 | 7 | 0.2857 | NA | NA | 1 | 0 | 5 | 0.4 | 2 | 0.5 | 8 | 0.375 |
| 2_26910061 | 4 | 0.25 | NA | NA | 8 | 0 | 4 | 0.25 | 16 | 0.125 | 1 | 0 | 6 | 0.1667 | 6 | 0.1667 | 2 | 0.5 | 15 | 0.2 |
| 2_26910081 | 4 | 0.5 | NA | NA | 5 | 0.6 | 2 | 0.5 | 11 | 0.5455 | 1 | 1 | 5 | 0.8 | 3 | 0.3333 | 2 | 1 | 11 | 0.7273 |
| 2_26910134 | 2 | 0 | NA | NA | 8 | 0.375 | 3 | 0 | 13 | 0.2308 | 1 | 0 | 5 | 0.6 | 6 | 0.6667 | 2 | 0.5 | 14 | 0.5714 |
| 2_26910171 | 4 | 0.5 | NA | NA | 8 | 0.375 | 4 | 0.5 | 16 | 0.4375 | 1 | 1 | 6 | 0.5 | 6 | 0.3333 | 2 | 1 | 15 | 0.5333 |
| 2_26910337 | 4 | 0 | NA | NA | 7 | 0.1429 | 3 | 0.3333 | 14 | 0.1429 | 1 | 1 | 6 | 0 | 5 | 0 | 2 | 0 | 14 | 0.0714 |
| 2_26910349 | 3 | 0 | NA | NA | 7 | 0.4286 | 4 | 0.5 | 14 | 0.3571 | 1 | 1 | 6 | 0.5 | 5 | 0.4 | 1 | 1 | 13 | 0.5385 |
| 2_26910432 | 4 | 0 | NA | NA | 7 | 0.4286 | 3 | 0.6667 | 14 | 0.3571 | 1 | 0 | 6 | 0.5 | 5 | 0.2 | 2 | 0.5 | 14 | 0.3571 |
| 2_26910470 | 4 | 1 | NA | NA | 8 | 1 | 4 | 1 | 16 | 1 | 1 | 1 | 5 | 1 | 6 | 1 | 2 | 1 | 14 | 1 |
| 2_26910515 | 4 | 0.75 | NA | NA | 8 | 0.5 | 3 | 1 | 15 | 0.6667 | 1 | 0 | 6 | 0.1667 | 6 | 0.1667 | 2 | 0 | 15 | 0.1333 |
| 2_26910733 | 4 | 0.25 | NA | NA | 6 | 0.1667 | 4 | 0 | 14 | 0.1429 | 1 | 0 | 6 | 0 | 5 | 0.2 | 2 | 0 | 14 | 0.0714 |
| 2_26911485 | NA | NA | 6 | 0.6667 | NA | NA | 1 | 1 | 7 | 0.7143 | 3 | 0.6667 | 1 | 1 | 3 | 1 | 4 | 0.75 | 11 | 0.8182 |
| 2_26911492 | NA | NA | 6 | 0.1667 | NA | NA | 1 | 1 | 7 | 0.2857 | 3 | 0 | 1 | 1 | 4 | 0 | 3 | 0.3333 | 11 | 0.1818 |
| 2_26911532 | 1 | 0 | 6 | 0 | NA | NA | 1 | 1 | 8 | 0.125 | 3 | 0.3333 | NA | NA | 3 | 0 | 4 | 0.5 | 10 | 0.3 |
| 2_26911571 | 1 | 1 | 7 | 0.2857 | NA | NA | 1 | 0 | 9 | 0.3333 | 3 | 0.3333 | 1 | 1 | 4 | 0 | 3 | 0.6667 | 11 | 0.3636 |
| 2_26911614 | 1 | 0 | 7 | 0.2857 | NA | NA | NA | NA | 8 | 0.25 | 3 | 0 | 1 | 0 | 3 | 0.6667 | 3 | 0.6667 | 10 | 0.4 |
| 2_26911674 | 1 | 0 | 7 | 0.1429 | NA | NA | 1 | 0 | 9 | 0.1111 | 3 | 0 | 1 | 0 | 3 | 0 | 3 | 0.3333 | 10 | 0.1 |
| 2_26911801 | NA | NA | 4 | 0.5 | NA | NA | 1 | 1 | 5 | 0.6 | 2 | 0.5 | 1 | 1 | 2 | 0.5 | 3 | 0.6667 | 8 | 0.625 |
| 2_26998696 | NA | NA | NA | NA | 1 | 1 | NA | NA | 1 | 1 | NA | NA | NA | NA | NA | NA | 1 | 1 | 1 | 1 |
| 2_27000443 | 16 | 0.0625 | 33 | 0.0606 | 19 | 0.0526 | 11 | 0.0909 | 79 | 0.0633 | 25 | 0.08 | 7 | 0 | 15 | 0.0667 | 12 | 0.1667 | 59 | 0.0847 |
| 2_27000454 | 16 | 0.5625 | 26 | 0.5769 | 12 | 0.5 | 10 | 0.6 | 64 | 0.5625 | 19 | 0.5263 | 4 | 0.75 | 12 | 0.5833 | 10 | 0.5 | 45 | 0.5556 |
| 2_27001492 | 15 | 0.1333 | 14 | 0.2857 | 11 | 0.3636 | 10 | 0.1 | 50 | 0.22 | 16 | 0.125 | 4 | 0.5 | 10 | 0 | 8 | 0.125 | 38 | 0.1316 |
| 2_27002037 | 14 | 0.0714 | 15 | 0.2667 | 10 | 0.2 | 10 | 0.2 | 49 | 0.1837 | 18 | 0.1111 | 4 | 0.25 | 7 | 0 | 6 | 0.1667 | 35 | 0.1143 |
| 2_27002044 | 12 | 0.1667 | 14 | 0.2143 | 8 | 0.375 | 10 | 0.5 | 44 | 0.2955 | 18 | 0.4444 | 4 | 0.25 | 8 | 0.5 | 7 | 0.5714 | 37 | 0.4595 |
| 2_27002057 | 8 | 0.125 | 15 | 0.0667 | 7 | 0 | 9 | 0.1111 | 39 | 0.0769 | 15 | 0.1333 | 3 | 0 | 8 | 0.125 | 6 | 0 | 32 | 0.0938 |
| 2_27002260 | 7 | 0.2857 | 7 | 0.1429 | 3 | 0.3333 | 4 | 0.5 | 21 | 0.2857 | 6 | 0.1667 | 3 | 0 | 2 | 0.5 | 3 | 0 | 14 | 0.1429 |
| 2_27002288 | 10 | 0.4 | 15 | 0.0667 | 5 | 0 | 8 | 0 | 38 | 0.1316 | 10 | 0.3 | 4 | 0.5 | 4 | 0.25 | 4 | 0.25 | 22 | 0.3182 |
| 2_27002314 | 11 | 0.0909 | 14 | 0.0714 | 8 | 0.125 | 8 | 0 | 41 | 0.0732 | 12 | 0.0833 | 4 | 0.25 | 6 | 0 | 5 | 0.2 | 27 | 0.1111 |
| 2_27002406 | 11 | 0 | 14 | 0.1429 | 8 | 0.5 | 7 | 0.2857 | 40 | 0.2 | 11 | 0.0909 | 2 | 0.5 | 4 | 0 | 5 | 0 | 22 | 0.0909 |
| 2_27002492 | 11 | 0.1818 | 14 | 0.0714 | 6 | 0 | 6 | 0 | 37 | 0.0811 | 14 | 0.1429 | 3 | 0 | 4 | 0.25 | 5 | 0.2 | 26 | 0.1538 |
| 2_27002664 | 9 | 0.4444 | 15 | 0.7333 | 6 | 0.8333 | 3 | 0.3333 | 33 | 0.6364 | 13 | 0.2308 | 3 | 0.3333 | 5 | 0.8 | 3 | 0.6667 | 24 | 0.4167 |
| 2_27078840 | 1 | 1 | 1 | 1 | NA | NA | NA | NA | 2 | 1 | 1 | 0 | 1 | 1 | 2 | 1 | 1 | 0 | 5 | 0.6 |
| 2_27154089 | 1 | 1 | NA | NA | NA | NA | NA | NA | 1 | 1 | 1 | 1 | NA | NA | 2 | 0.5 | NA | NA | 3 | 0.6667 |
| 2_27332768 | 2 | 0 | 2 | 0 | 6 | 0 | 4 | 0.5 | 14 | 0.1429 | 5 | 0.2 | 5 | 0.4 | 5 | 0 | 3 | 0 | 18 | 0.1667 |
| 2_27332845 | NA | NA | NA | NA | 2 | 0.5 | 3 | 0.6667 | 5 | 0.6 | 2 | 1 | 2 | 1 | 3 | 0.6667 | 1 | 0 | 8 | 0.75 |
| 2_27332981 | 2 | 0.5 | 2 | 0.5 | 5 | 0.4 | 4 | 0.5 | 13 | 0.4615 | 5 | 0.8 | 5 | 0.6 | 3 | 1 | 3 | 0 | 16 | 0.625 |
| 2_27333031 | 1 | 0 | 3 | 0 | 4 | 0 | 3 | 0.3333 | 11 | 0.0909 | 4 | 0 | 3 | 1 | 4 | 0 | 2 | 0 | 13 | 0.2308 |

| | | | | | | | | | | | | | | | | | | | | |
|---|---|---|---|---|---|---|---|---|---|---|---|---|---|---|---|---|---|---|---|---|
| 2_2733310 | 2 | 0.5 | 3 | 0.6667 | 4 | 0.5 | 4 | 0.5 | 13 | 0.5385 | 5 | 0.8 | 3 | 0.6667 | 6 | 0.6667 | 3 | 0.3333 | 17 | 0.6471 |
| 2_2733327 | 2 | 0 | 3 | 0 | 5 | 0.2 | 3 | 1 | 13 | 0.3077 | 4 | 0.25 | 4 | 0 | 5 | 0.2 | 3 | 0 | 16 | 0.125 |
| 2_2733328 | 1 | 1 | 2 | 1 | 5 | 1 | 5 | 1 | 13 | 1 | 5 | 1 | 5 | 1 | 5 | 1 | 3 | 1 | 18 | 1 |
| 2_2733358 | 2 | 0.5 | 4 | 0.25 | 5 | 0.8 | 5 | 0 | 16 | 0.375 | 5 | 0 | 8 | 0 | 6 | 0.5 | 3 | 0 | 22 | 0.1364 |
| 2_2733361 | 2 | 1 | 4 | 0.5 | 6 | 1 | 5 | 1 | 17 | 0.8824 | 5 | 1 | 8 | 0.875 | 8 | 0.75 | 2 | 1 | 23 | 0.8696 |
| 2_2733381 | 2 | 1 | 4 | 0.75 | 7 | 0.7143 | 4 | 0.75 | 17 | 0.7647 | 5 | 1 | 8 | 0.75 | 7 | 0.8571 | 3 | 0.6667 | 23 | 0.8261 |
| 2_2733389 | 2 | 1 | 4 | 0 | 6 | 0 | 4 | 0.5 | 16 | 0.25 | 6 | 0.3333 | 6 | 0 | 7 | 0.2857 | 3 | 0 | 22 | 0.1818 |
| 2_2733396 | 2 | 0.5 | 4 | 0.25 | 6 | 0.3333 | 5 | 0.4 | 17 | 0.3529 | 7 | 0.2857 | 8 | 0.25 | 7 | 0.2857 | 3 | 0.3333 | 25 | 0.28 |
| 2_2733611 | 5 | 0 | 5 | 0.2 | 6 | 0.3333 | 8 | 0.375 | 24 | 0.25 | 7 | 0 | 5 | 0.4 | 2 | 0 | 4 | 0 | 18 | 0.1111 |
| 2_2733686 | 6 | 0.1667 | 4 | 0.25 | 5 | 0.4 | 7 | 0.2857 | 22 | 0.2727 | 8 | 0.625 | 4 | 0.25 | 2 | 0.5 | 2 | 0 | 16 | 0.4375 |
| 2_2733698 | 5 | 0.6 | 4 | 0 | 6 | 0.3333 | 7 | 0.1429 | 22 | 0.2727 | 8 | 0.25 | 4 | 0.25 | 2 | 0 | 2 | 0.5 | 16 | 0.25 |
| 2_2733742 | 6 | 0 | 3 | 0 | 4 | 0.25 | 2 | 0 | 15 | 0.0667 | 7 | 0.2857 | 4 | 0.25 | 2 | 0 | 1 | 1 | 14 | 0.2857 |
| 2_2733820 | 3 | 0 | 3 | 0.6667 | 4 | 0.25 | 2 | 0.5 | 12 | 0.3333 | 3 | 0.3333 | 4 | 0.5 | NA | NA | NA | NA | 7 | 0.4286 |
| 2_2733824 | 4 | 0.25 | 4 | 0 | 3 | 0 | 3 | 0.3333 | 14 | 0.1429 | 4 | 0 | 3 | 0.3333 | NA | NA | NA | NA | 7 | 0.1429 |
| 2_2733845 | 3 | 0.6667 | 3 | 0.3333 | 4 | 0.75 | 2 | 0.5 | 12 | 0.5833 | 3 | 1 | 2 | 0.5 | 1 | 0 | NA | NA | 6 | 0.6667 |
| 2_2733846 | 3 | 0 | 2 | 0.5 | 3 | 0.3333 | 1 | 0 | 9 | 0.2222 | 3 | 0.3333 | 4 | 0 | 1 | 1 | NA | NA | 8 | 0.25 |
| 2_2733847 | 3 | 0.6667 | 4 | 1 | 3 | 1 | 2 | 0.5 | 12 | 0.8333 | 4 | 0.75 | 3 | 0.6667 | 1 | 1 | NA | NA | 8 | 0.75 |
| 2_2733847 | 2 | 0 | 4 | 0.25 | 4 | 0.25 | 2 | 0 | 12 | 0.1667 | 4 | 0.25 | 3 | 0.3333 | 1 | 1 | NA | NA | 8 | 0.375 |
| 2_2733848 | 3 | 0 | 4 | 0 | 3 | 0.3333 | 2 | 0.5 | 12 | 0.1667 | 3 | 0.3333 | 4 | 0 | 1 | 0 | NA | NA | 8 | 0.125 |
| 2_2734038 | 1 | 1 | 4 | 0.25 | 3 | 0 | 2 | 0.5 | 10 | 0.3 | 4 | 0.25 | 4 | 0 | NA | NA | NA | NA | 8 | 0.125 |
| 2_2734963 | NA | NA | 4 | 0.25 | 2 | 0 | 2 | 0 | 8 | 0.125 | 1 | 0 | 2 | 0.5 | 1 | 0 | NA | NA | 4 | 0.25 |
| 2_2742545 | 6 | 0.1667 | 5 | 0 | 3 | 0 | 3 | 0 | 17 | 0.0588 | 2 | 0 | 1 | 0 | 2 | 0 | 2 | 0.5 | 7 | 0.1429 |
| 2_2742584 | 9 | 0.2222 | 5 | 0 | 3 | 0 | 3 | 0.3333 | 20 | 0.15 | 3 | 0 | 1 | 1 | 2 | 0 | 4 | 0.25 | 10 | 0.2 |
| 2_2742586 | 9 | 0.2222 | 5 | 0 | 3 | 0.3333 | 2 | 0 | 19 | 0.1579 | 2 | 0 | 1 | 0 | 2 | 0 | 4 | 0.25 | 9 | 0.1111 |
| 2_2764973 | 2 | 1 | 3 | 0.3333 | 2 | 0.5 | 1 | 1 | 8 | 0.625 | 1 | 1 | 1 | 0 | 1 | 0 | 2 | 0.5 | 5 | 0.4 |
| 2_2765024 | 2 | 0 | 2 | 1 | 2 | 1 | 1 | 0 | 7 | 0.5714 | 2 | 0 | 1 | 1 | 2 | 0 | 2 | 0 | 7 | 0.1429 |
| 2_2765048 | 2 | 1 | 2 | 0.5 | 1 | 1 | 2 | 0.5 | 7 | 0.7143 | 2 | 1 | NA | NA | 4 | 1 | 1 | 1 | 7 | 1 |
| 2_2765055 | 2 | 0.5 | 2 | 0 | 2 | 0.5 | 2 | 0.5 | 8 | 0.375 | 2 | 0.5 | 1 | 1 | 4 | 0.5 | 3 | 0 | 10 | 0.4 |
| 2_2765062 | 2 | 1 | 1 | 0 | 2 | 1 | 2 | 0.5 | 7 | 0.7143 | 2 | 1 | NA | NA | 4 | 1 | 1 | 1 | 7 | 1 |
| 2_2765064 | 1 | 1 | 3 | 0.3333 | 2 | 0.5 | 1 | 0 | 7 | 0.4286 | 1 | 0 | 1 | 1 | 3 | 0.6667 | 1 | 1 | 6 | 0.6667 |
| 2_2765068 | 2 | 1 | 3 | 0 | 2 | 0.5 | 2 | 1 | 9 | 0.5556 | 2 | 1 | 1 | 1 | 4 | 0.75 | 3 | 1 | 10 | 0.9 |
| 2_2765069 | 2 | 1 | 3 | 0.6667 | 1 | 1 | 2 | 0.5 | 8 | 0.75 | 2 | 1 | NA | NA | 4 | 0.75 | 2 | 1 | 8 | 0.875 |
| 2_2765072 | 2 | 0.5 | 2 | 0 | 2 | 0.5 | 2 | 0 | 8 | 0.25 | 1 | 1 | 1 | 1 | 4 | 1 | 2 | 0.5 | 8 | 0.875 |
| 2_2765107 | 3 | 0 | 4 | 0.25 | 2 | 0 | 3 | 0.3333 | 12 | 0.1667 | 2 | 0 | 2 | 0 | 4 | 0.25 | 2 | 0 | 10 | 0.1 |
| 2_2765206 | 4 | 0.5 | 4 | 0 | 4 | 0.25 | 4 | 0 | 16 | 0.1875 | 2 | 0 | 3 | 0 | 3 | 0.3333 | 2 | 0 | 10 | 0.1 |
| 2_2765215 | 4 | 0.5 | 3 | 1 | 3 | 0.3333 | 4 | 0.25 | 14 | 0.5 | 2 | 0.5 | 3 | 0.6667 | 3 | 0.3333 | 3 | 0.3333 | 11 | 0.4545 |
| 2_2765216 | 3 | 0.3333 | 3 | 0 | 3 | 0 | 5 | 0 | 14 | 0.0714 | 3 | 0 | 3 | 0.3333 | 3 | 0.3333 | 2 | 0 | 11 | 0.1818 |
| 2_2765221 | 4 | 1 | 3 | 1 | 4 | 1 | 5 | 0.8 | 16 | 0.9375 | 3 | 1 | 3 | 0.6667 | 3 | 1 | 3 | 1 | 12 | 0.9167 |
| 2_2765228 | 4 | 0.75 | 3 | 0.3333 | 2 | 0.5 | 6 | 0.5 | 15 | 0.5333 | 2 | 0.5 | 2 | 0.5 | 3 | 0.3333 | 3 | 0.6667 | 10 | 0.5 |
| 2_2765232 | 4 | 0.75 | 3 | 0.3333 | 4 | 0.75 | 5 | 0.2 | 16 | 0.5 | 3 | 0.3333 | 3 | 0 | 3 | 0.6667 | 3 | 0.6667 | 12 | 0.4167 |
| 2_2765236 | 4 | 1 | 3 | 1 | 4 | 1 | 4 | 0.5 | 15 | 0.8667 | 3 | 1 | 3 | 0.6667 | 2 | 1 | 2 | 1 | 10 | 0.9 |
| 2_2765261 | 3 | 0.3333 | 5 | 0.2 | 4 | 0 | 5 | 0.4 | 17 | 0.2353 | 3 | 0 | 3 | 0.6667 | 3 | 0.6667 | 2 | 0.5 | 11 | 0.4545 |
| 2_2765262 | 4 | 0 | 5 | 0.2 | 4 | 0 | 4 | 0 | 17 | 0.0588 | 3 | 0 | 2 | 0 | 3 | 0.3333 | 2 | 0.5 | 10 | 0.2 |
| 2_2765281 | 3 | 0.3333 | 5 | 0.2 | 2 | 0 | 5 | 0 | 15 | 0.1333 | 3 | 0.6667 | 2 | 0 | 3 | 0 | 2 | 0.5 | 10 | 0.3 |
| 2_2765281 | 4 | 0.5 | 2 | 1 | 2 | 0.5 | 6 | 0.5 | 14 | 0.5714 | 3 | 0.3333 | 1 | 1 | 2 | 0 | NA | NA | 6 | 0.3333 |
| 2_2765284 | 4 | 1 | 4 | 0.75 | 3 | 0.6667 | 5 | 0.6 | 16 | 0.75 | 3 | 1 | 3 | 0.6667 | 2 | 1 | 2 | 1 | 10 | 0.9 |
| 2_2811912 | 17 | 0.1765 | 49 | 0 | 28 | 0.1071 | 13 | 0.1538 | 107 | 0.0748 | 26 | 0.1154 | 16 | 0.0625 | 13 | 0.0769 | 21 | 0.0476 | 76 | 0.0789 |
| 2_2811914 | 17 | 0.1765 | 45 | 0.1111 | 29 | 0.1379 | 13 | 0.3846 | 104 | 0.1635 | 27 | 0 | 14 | 0.2143 | 15 | 0.1333 | 18 | 0.2222 | 74 | 0.1216 |
| 2_2811929 | 19 | 0.8947 | 51 | 0.902 | 28 | 0.8929 | 11 | 1 | 109 | 0.9083 | 28 | 1 | 17 | 0.8824 | 12 | 0.8333 | 20 | 1 | 77 | 0.9481 |
| 2_2811934 | 15 | 0.1333 | 40 | 0.05 | 23 | 0.1739 | 11 | 0.0909 | 89 | 0.1011 | 16 | 0.125 | 12 | 0.0833 | 8 | 0.125 | 16 | 0.125 | 52 | 0.1154 |
| 2_2811959 | 23 | 0.1304 | 56 | 0.1964 | 31 | 0.0968 | 12 | 0 | 122 | 0.1393 | 31 | 0.0968 | 15 | 0 | 14 | 0.1429 | 28 | 0.0714 | 88 | 0.0795 |
| 2_2811961 | 16 | 0.0625 | 47 | 0.0851 | 25 | 0.12 | 11 | 0 | 99 | 0.0808 | 31 | 0.0968 | 16 | 0.1875 | 11 | 0.0909 | 22 | 0.1818 | 80 | 0.1375 |
| 2_2811964 | 20 | 0.9 | 54 | 0.9444 | 27 | 0.8519 | 11 | 1 | 112 | 0.9196 | 33 | 0.9394 | 17 | 0.8824 | 13 | 0.9231 | 24 | 0.9167 | 87 | 0.9195 |
| 2_2811965 | 21 | 0.0952 | 56 | 0.2679 | 29 | 0.3448 | 13 | 0.1538 | 119 | 0.2437 | 33 | 0.2424 | 14 | 0.2857 | 12 | 0 | 22 | 0.1818 | 81 | 0.1975 |
| 2_2811967 | 17 | 0.1765 | 50 | 0.12 | 27 | 0.2593 | 11 | 0.0909 | 105 | 0.1619 | 24 | 0.25 | 16 | 0.1875 | 15 | 0.2 | 22 | 0.1818 | 77 | 0.2078 |
| 2_2811969 | 22 | 0.3636 | 60 | 0.25 | 35 | 0.1429 | 13 | 0.3846 | 130 | 0.2538 | 33 | 0.1212 | 18 | 0.3889 | 12 | 0.0833 | 28 | 0.1071 | 91 | 0.1648 |
| 2_2811978 | 23 | 0.6957 | 57 | 0.6491 | 37 | 0.6757 | 11 | 0.9091 | 128 | 0.6875 | 35 | 0.8 | 19 | 0.6316 | 11 | 0.5455 | 25 | 0.72 | 90 | 0.7111 |
| 2_2811979 | 19 | 0.2105 | 62 | 0.1452 | 33 | 0.2121 | 11 | 0.2727 | 125 | 0.184 | 33 | 0.2727 | 17 | 0.2941 | 12 | 0.3333 | 27 | 0.1852 | 89 | 0.2584 |
| 2_2811982 | 8 | 0.125 | 23 | 0.1304 | 14 | 0.0714 | 5 | 0 | 50 | 0.1 | 14 | 0.0714 | 10 | 0.3 | 8 | 0.125 | 12 | 0.0833 | 44 | 0.1364 |
| 2_2811995 | 22 | 0.7727 | 49 | 0.7755 | 34 | 0.6471 | 8 | 0.625 | 113 | 0.7257 | 30 | 0.8 | 17 | 0.7647 | 13 | 0.3846 | 25 | 0.68 | 85 | 0.6941 |
| 2_2812001 | 24 | 0.2083 | 57 | 0.1404 | 37 | 0.1622 | 9 | 0.1111 | 127 | 0.1575 | 29 | 0.2069 | 20 | 0.15 | 10 | 0.1 | 26 | 0.3846 | 85 | 0.2353 |
| 2_2812003 | 17 | 0.8235 | 49 | 0.7755 | 36 | 0.8611 | 10 | 0.8 | 112 | 0.8125 | 29 | 0.8621 | 18 | 1 | 11 | 0.5455 | 31 | 0.8065 | 89 | 0.8315 |
| 2_2812008 | 27 | 1 | 61 | 1 | 43 | 0.9535 | 17 | 1 | 148 | 0.9865 | 35 | 1 | 20 | 1 | 16 | 1 | 32 | 1 | 103 | 1 |
| 2_2812023 | 28 | 0.2143 | 56 | 0.25 | 37 | 0.1622 | 12 | 0.0833 | 133 | 0.203 | 37 | 0.1622 | 22 | 0.1364 | 11 | 0.2727 | 30 | 0.2 | 100 | 0.18 |
| 2_2812024 | 27 | 0.1481 | 65 | 0.1231 | 40 | 0.175 | 15 | 0.2667 | 147 | 0.1565 | 34 | 0.2059 | 17 | 0.2353 | 15 | 0.2 | 36 | 0.1667 | 102 | 0.1961 |

| | | | | | | | | | | | | | | | | | | | | |
|---|---|---|---|---|---|---|---|---|---|---|---|---|---|---|---|---|---|---|---|---|
| 2_28120263 | 31 | 0.9032 | 64 | 0.8906 | 48 | 0.9583 | 18 | 0.9444 | 161 | 0.9193 | 40 | 0.9 | 21 | 0.9048 | 17 | 0.8824 | 38 | 0.9737 | 116 | 0.9224 |
| 2_28120334 | 30 | 0.2 | 66 | 0.3333 | 45 | 0.2667 | 16 | 0.125 | 157 | 0.2675 | 41 | 0.0244 | 22 | 0.3636 | 14 | 0.4286 | 33 | 0.1818 | 110 | 0.1909 |
| 2_28120547 | 28 | 0.3571 | 65 | 0.3385 | 48 | 0.3125 | 19 | 0.4737 | 160 | 0.35 | 42 | 0.3095 | 19 | 0.4737 | 16 | 0.25 | 36 | 0.3056 | 113 | 0.3274 |
| 2_28120633 | 38 | 0.0263 | 64 | 0.0781 | 49 | 0.0816 | 19 | 0.0526 | 170 | 0.0647 | 43 | 0.093 | 22 | 0.0455 | 15 | 0 | 34 | 0.0882 | 114 | 0.0702 |
| 2_28343238 | NA | NA | NA | NA | 1 | 1 | 1 | 1 | 2 | 1 | NA | NA | 1 | 1 | NA | NA | NA | NA | 1 | 1 |
| 2_28343253 | NA | NA | NA | NA | 1 | 1 | 1 | 1 | 2 | 1 | NA | NA | 1 | 1 | NA | NA | NA | NA | 1 | 1 |
| 2_28343349 | 1 | 1 | NA | NA | NA | NA | 1 | 1 | 2 | 1 | NA | NA | 1 | 1 | NA | NA | NA | NA | 1 | 1 |
| 2_28343372 | 1 | 1 | NA | NA | 1 | 1 | 1 | 0 | 3 | 0.6667 | NA | NA | 1 | 1 | NA | NA | NA | NA | 1 | 1 |
| 2_28343640 | 1 | 1 | NA | NA | 1 | 1 | 1 | 0 | 3 | 0.6667 | NA | NA | 1 | 1 | NA | NA | NA | NA | 1 | 1 |
| 2_28343675 | 1 | 0 | 1 | 1 | 1 | 0 | 1 | 0 | 4 | 0.25 | NA | NA | 1 | 1 | NA | NA | NA | NA | 1 | 1 |
| 2_28343691 | 1 | 1 | 1 | 0 | 1 | 1 | 1 | 0 | 4 | 0.5 | NA | NA | 1 | 1 | NA | NA | NA | NA | 1 | 1 |
| 2_28343715 | 1 | 1 | 1 | 0 | 1 | 0 | 1 | 0 | 4 | 0.25 | NA | NA | 1 | 1 | NA | NA | NA | NA | 1 | 1 |
| 2_28345033 | 2 | 0.5 | 1 | 0 | 1 | 1 | 1 | 0 | 5 | 0.4 | NA | NA | 1 | 1 | 1 | 0 | NA | NA | 2 | 0.5 |
| 2_28345270 | 1 | 0 | 1 | 1 | 1 | 1 | NA | NA | 3 | 0.6667 | 1 | 0 | 1 | 1 | 1 | 1 | NA | NA | 3 | 0.6667 |
| 2_28352304 | NA | NA | NA | NA | 1 | 1 | NA | NA | 1 | 1 | 1 | 1 | NA | NA | NA | NA | 1 | 1 | 2 | 1 |
| 2_28359524 | 1 | 0 | 5 | 0.2 | 1 | 0 | 3 | 0.3333 | 10 | 0.2 | 1 | 1 | 3 | 0.3333 | 2 | 0 | 3 | 0 | 9 | 0.2222 |
| 2_28359604 | 1 | 1 | 4 | 0.75 | 2 | 0.5 | 3 | 0.6667 | 10 | 0.7 | 1 | 1 | 2 | 1 | 3 | 0.3333 | 4 | 0.75 | 10 | 0.7 |
| 2_28359664 | 1 | 0 | 4 | 0.5 | 2 | 0 | 2 | 0 | 9 | 0.2222 | 1 | 0 | 3 | 0.6667 | 2 | 0.5 | 4 | 0.75 | 10 | 0.6 |
| 2_28359707 | 1 | 0 | 5 | 0.2 | 2 | 0.5 | 3 | 0 | 11 | 0.1818 | NA | NA | 3 | 0.3333 | 3 | 0.3333 | 4 | 0 | 10 | 0.2 |
| 2_28359782 | 1 | 1 | 4 | 0.75 | 1 | 0 | 3 | 0.6667 | 9 | 0.6667 | NA | NA | 1 | 1 | 2 | 1 | 4 | 0.75 | 7 | 0.8571 |
| 2_28359806 | 1 | 1 | 5 | 1 | 1 | 1 | 2 | 1 | 9 | 1 | 2 | 1 | 4 | 1 | 2 | 1 | 3 | 1 | 11 | 1 |
| 2_28359898 | NA | NA | 4 | 0 | 2 | 0.5 | NA | NA | 6 | 0.1667 | 2 | 0 | 4 | 0 | 3 | 0.6667 | 5 | 0.4 | 14 | 0.2857 |
| 2_28359985 | 1 | 0 | 4 | 0.75 | 2 | 0 | 3 | 0.6667 | 10 | 0.5 | 2 | 0.5 | 5 | 0.4 | 4 | 0.5 | 5 | 0.6 | 16 | 0.5 |
| 2_28360253 | 2 | 0 | 5 | 0.4 | 3 | 0 | 4 | 0.75 | 14 | 0.3571 | 3 | 0.6667 | 4 | 0.25 | 5 | 0 | 5 | 0.4 | 17 | 0.2941 |
| 2_28360472 | 2 | 0 | 5 | 0.4 | 4 | 0.25 | 5 | 0.2 | 16 | 0.25 | 3 | 0 | 4 | 0.25 | 4 | 0 | 4 | 0.25 | 15 | 0.1333 |
| 2_28360565 | 1 | 0 | 5 | 0.2 | 4 | 0.25 | 6 | 0.1667 | 16 | 0.1875 | 3 | 0 | 4 | 0 | 3 | 0.3333 | 4 | 0 | 14 | 0.0714 |
| 2_28360901 | 1 | 0 | 5 | 0.4 | 3 | 0.3333 | 5 | 0.6 | 14 | 0.4286 | 4 | 0.25 | 3 | 0.3333 | 4 | 0.75 | 6 | 0.8333 | 17 | 0.5882 |
| 2_28434254 | 1 | 0 | 4 | 0 | 5 | 0.2 | 3 | 0 | 13 | 0.0769 | 4 | 0 | 4 | 0 | 7 | 0 | 5 | 0.2 | 20 | 0.05 |
| 2_28434326 | 2 | 0 | 5 | 0 | 5 | 0.4 | 3 | 0 | 15 | 0.1333 | 4 | 0.5 | 6 | 0.3333 | 12 | 0.4167 | 3 | 0.3333 | 25 | 0.4 |
| 2_28442146 | 5 | 0.2 | 5 | 0.2 | 6 | 0.5 | 8 | 0.375 | 24 | 0.3333 | 2 | 1 | 10 | 0.4 | 18 | 0.5 | 4 | 0.25 | 34 | 0.4706 |
| 2_28442252 | 6 | 0.5 | 8 | 0.625 | 7 | 0.8571 | 7 | 0.8571 | 28 | 0.7143 | 5 | 0.8 | 10 | 0.9 | 15 | 0.4667 | 10 | 0.5 | 40 | 0.625 |
| 2_28442313 | 7 | 0.1429 | 9 | 0.3333 | 6 | 0.3333 | 7 | 0.1429 | 29 | 0.2414 | 6 | 0.6667 | 11 | 0.2727 | 17 | 0.3529 | 9 | 0.3333 | 43 | 0.3721 |
| 2_28442328 | 6 | 0.8333 | 7 | 0.4286 | 7 | 0.7143 | 7 | 1 | 27 | 0.7407 | 4 | 1 | 8 | 0.625 | 16 | 0.875 | 6 | 0.8333 | 34 | 0.8235 |
| 2_28442465 | 8 | 0.375 | 9 | 0 | 6 | 0.1667 | 9 | 0.2222 | 32 | 0.1875 | 6 | 0.5 | 12 | 0.1667 | 15 | 0.2667 | 9 | 0.2222 | 42 | 0.2619 |
| 2_28442509 | 5 | 0.8 | 7 | 0.8571 | 7 | 0.7143 | 8 | 0.625 | 27 | 0.7407 | 5 | 1 | 11 | 0.9091 | 18 | 0.8333 | 8 | 0.875 | 42 | 0.881 |
| 2_28442517 | 7 | 0.5714 | 6 | 0.5 | 6 | 0.3333 | 5 | 0.4 | 24 | 0.4583 | 4 | 0.75 | 10 | 0.7 | 14 | 0.3571 | 6 | 0.8333 | 34 | 0.5882 |
| 2_28442552 | 8 | 0.75 | 9 | 0.3333 | 8 | 0.75 | 9 | 0.5556 | 34 | 0.5882 | 6 | 0.8333 | 13 | 0.6154 | 19 | 0.4737 | 9 | 0.8889 | 47 | 0.6383 |
| 2_28442568 | 8 | 1 | 9 | 1 | 7 | 1 | 8 | 0.875 | 32 | 0.9688 | 6 | 0.8333 | 13 | 0.9231 | 18 | 0.8889 | 10 | 0.9 | 47 | 0.8936 |
| 2_28442642 | 8 | 0.125 | 10 | 0.3 | 6 | 0.3333 | 9 | 0.4444 | 33 | 0.303 | 6 | 0.3333 | 13 | 0.4615 | 19 | 0.4737 | 10 | 0.6 | 48 | 0.4792 |
| 2_28442708 | 7 | 1 | 7 | 1 | 4 | 1 | 6 | 1 | 24 | 1 | 5 | 1 | 10 | 0.9 | 15 | 0.9333 | 9 | 1 | 39 | 0.9487 |
| 2_28442859 | 3 | 0.6667 | 7 | 0.4286 | 5 | 0.6 | 8 | 0.625 | 23 | 0.5652 | 4 | 0.75 | 10 | 0.7 | 17 | 0.5294 | 9 | 0.4444 | 40 | 0.575 |
| 2_28442975 | 5 | 0.4 | 11 | 0.1818 | 7 | 0.1429 | 8 | 0.25 | 31 | 0.2258 | 8 | 0.125 | 12 | 0.0833 | 18 | 0.1667 | 10 | 0.2 | 48 | 0.1458 |
| 2_28456390 | 4 | 0 | 1 | 1 | 4 | 0 | 6 | 0.5 | 15 | 0.2667 | 5 | 0 | 4 | 0 | 4 | 0.25 | 6 | 0 | 19 | 0.0526 |
| 2_28456428 | 4 | 1 | 1 | 1 | 4 | 1 | 7 | 0.5714 | 16 | 0.8125 | 4 | 0.75 | 4 | 1 | 4 | 0.75 | 6 | 0.6667 | 18 | 0.7778 |
| 2_28456437 | 4 | 0 | 1 | 0 | 4 | 0.25 | 7 | 0.2857 | 16 | 0.1875 | 5 | 0.2 | 3 | 0 | 4 | 0 | 6 | 0.1667 | 18 | 0.1111 |
| 2_28456441 | 4 | 0.25 | 1 | 1 | 4 | 0.75 | 3 | 0.3333 | 12 | 0.5 | 1 | 0 | 3 | 0.6667 | 3 | 1 | 2 | 0 | 9 | 0.5556 |
| 2_28456539 | 3 | 0 | 1 | 1 | 3 | 0.3333 | 4 | 0.75 | 11 | 0.4545 | 2 | 0 | 4 | 0.5 | 3 | 0.6667 | 4 | 0.25 | 13 | 0.3846 |
| 2_28456661 | 3 | 1 | NA | NA | 2 | 1 | 7 | 0.8571 | 12 | 0.9167 | 5 | 1 | 4 | 0.75 | 3 | 1 | 4 | 1 | 16 | 0.9375 |
| 2_28456777 | 3 | 0.3333 | NA | NA | 4 | 0 | 5 | 0 | 12 | 0.0833 | 5 | 0.2 | 4 | 0 | 3 | 0.3333 | 4 | 0.25 | 16 | 0.1875 |
| 2_28456802 | 3 | 0.6667 | NA | NA | 3 | 0 | 7 | 0.1429 | 13 | 0.2308 | 5 | 0 | 3 | 0.3333 | 2 | 0.5 | 5 | 0 | 15 | 0.1333 |
| 2_28457026 | 3 | 0 | NA | NA | 3 | 0 | 7 | 0.2857 | 13 | 0.1538 | 4 | 0.25 | 2 | 0.5 | 1 | 1 | 5 | 0.6 | 12 | 0.5 |
| 2_28457074 | 4 | 0.5 | NA | NA | 4 | 1 | 6 | 0.6667 | 14 | 0.7143 | 4 | 1 | 2 | 0.5 | 2 | 1 | 5 | 1 | 13 | 0.9231 |
| 2_28457156 | 3 | 0.3333 | NA | NA | 3 | 0 | 6 | 0 | 12 | 0.0833 | 4 | 0 | 3 | 0 | 3 | 0 | 5 | 0.2 | 15 | 0.0667 |
| 2_28457165 | 3 | 0.3333 | NA | NA | 4 | 0 | 5 | 0.2 | 12 | 0.1667 | 4 | 0 | 2 | 0 | 3 | 0.3333 | 4 | 0 | 13 | 0.0769 |
| 2_28566149 | NA | NA | 2 | 0.5 | NA | NA | 2 | 1 | 4 | 0.75 | 2 | 0.5 | 1 | 1 | 1 | 0 | 1 | 0 | 5 | 0.4 |
| 2_28566195 | NA | NA | 2 | 0.5 | NA | NA | 1 | 1 | 3 | 0.6667 | 2 | 0.5 | 1 | 0 | NA | NA | 1 | 0 | 4 | 0.25 |
| 2_28566416 | NA | NA | 1 | 1 | 1 | 0 | 2 | 1 | 4 | 0.75 | 2 | 0 | 1 | 1 | 1 | 1 | 1 | 1 | 5 | 0.6 |
| 2_28576060 | NA | NA | 1 | 0 | 1 | 1 | 3 | 0 | 5 | 0.2 | 2 | 0.5 | NA | NA | 1 | 0 | 2 | 0 | 5 | 0.2 |
| 2_28576708 | NA | NA | 1 | 1 | 1 | 1 | 2 | 0.5 | 4 | 0.75 | 2 | 0.5 | NA | NA | NA | NA | NA | NA | 2 | 0.5 |
| 2_28577001 | NA | NA | 2 | 0.5 | 1 | 1 | 3 | 0.6667 | 6 | 0.6667 | 1 | 1 | 2 | 1 | 1 | 1 | 2 | 1 | 6 | 1 |
| 2_28577045 | NA | NA | 2 | 0.5 | 1 | 0 | 3 | 0 | 6 | 0.1667 | 1 | 0 | 2 | 0.5 | 1 | 1 | 2 | 0.5 | 6 | 0.5 |
| 2_28577050 | NA | NA | 1 | 1 | 1 | 1 | 3 | 1 | 5 | 1 | 2 | 1 | 1 | 1 | NA | NA | 2 | 0.5 | 5 | 0.8 |
| 2_28577061 | NA | NA | 2 | 1 | NA | NA | 3 | 1 | 5 | 1 | 2 | 1 | 2 | 1 | 1 | 1 | 2 | 1 | 7 | 1 |
| 2_28577081 | NA | NA | 1 | 0 | 1 | 1 | 3 | 0.3333 | 5 | 0.4 | 1 | 1 | 2 | 1 | 1 | 1 | 2 | 0.5 | 6 | 0.8333 |
| 2_28577154 | NA | NA | 1 | 0 | 1 | 0 | 3 | 0.6667 | 5 | 0.4 | 2 | 1 | 2 | 1 | 1 | 1 | 1 | 1 | 6 | 1 |

| | | | | | | | | | | | | | | | | | | | | |
|---|---|---|---|---|---|---|---|---|---|---|---|---|---|---|---|---|---|---|---|---|
| 2_2857716 | NA | NA | 2 | 1 | 1 | 1 | 2 | 1 | 5 | 1 | 2 | 1 | 2 | 1 | 1 | 1 | 1 | 1 | 6 | 1 |
| 2_2857730 | NA | NA | 1 | 0 | 1 | 1 | 2 | 0.5 | 4 | 0.5 | 2 | 0 | 2 | 0 | 1 | 1 | 2 | 0.5 | 7 | 0.2857 |
| 2_2857731 | NA | NA | 1 | 1 | 1 | 1 | 3 | 0 | 5 | 0.4 | 2 | 0 | 2 | 0 | 1 | 1 | 2 | 0 | 7 | 0.1429 |
| 2_2857912 | 2 | 0 | 1 | 0 | 2 | 0.5 | 2 | 0 | 7 | 0.1429 | 4 | 0 | 5 | 0.4 | 1 | 0 | 1 | 1 | 11 | 0.2727 |
| 2_2857935 | 2 | 0 | 3 | 0.3333 | 2 | 0.5 | 2 | 0 | 9 | 0.2222 | 4 | 0.5 | 4 | 0.25 | 2 | 0.5 | 2 | 0 | 12 | 0.3333 |
| 2_2858070 | 5 | 0.2 | 7 | 0.5714 | 10 | 0.5 | 5 | 0.2 | 27 | 0.4074 | 4 | 0 | 8 | 0.5 | 7 | 0.1429 | 5 | 0.6 | 24 | 0.3333 |
| 2_2871251 | 4 | 0.75 | 7 | 0.7143 | 2 | 0 | 3 | 0.6667 | 16 | 0.625 | 4 | 0.25 | 4 | 0.25 | 1 | 1 | 4 | 0.5 | 13 | 0.3846 |
| 2_2871359 | 3 | 0 | 3 | 0 | 1 | 0 | 1 | 1 | 8 | 0.125 | 4 | 0.25 | 3 | 0.3333 | 1 | 0 | 2 | 0 | 10 | 0.2 |
| 2_2871465 | 2 | 0.5 | 2 | 0 | NA | NA | 1 | 0 | 5 | 0.2 | 2 | 1 | 1 | 0 | NA | NA | 1 | 1 | 4 | 0.75 |
| 2_2871471 | 1 | 0 | 3 | 0.3333 | NA | NA | 1 | 1 | 5 | 0.4 | 2 | 1 | 1 | 1 | NA | NA | NA | NA | 3 | 1 |
| 2_2871491 | 2 | 1 | 2 | 1 | NA | NA | 2 | 1 | 6 | 1 | 2 | 1 | 1 | 1 | NA | NA | 1 | 1 | 4 | 1 |
| 2_2871538 | 1 | 1 | 3 | 1 | NA | NA | 2 | 1 | 6 | 1 | 1 | 1 | NA | NA | NA | NA | 1 | 1 | 2 | 1 |
| 2_2871565 | 2 | 0.5 | 2 | 0.5 | NA | NA | 2 | 1 | 6 | 0.6667 | 1 | 1 | NA | NA | NA | NA | 2 | 1 | 3 | 1 |
| 2_2871573 | 2 | 0.5 | 1 | 1 | NA | NA | 2 | 1 | 5 | 0.8 | 1 | 1 | NA | NA | NA | NA | 2 | 0 | 3 | 0.3333 |
| 2_2871589 | 2 | 1 | 2 | 0.5 | NA | NA | 2 | 0.5 | 6 | 0.6667 | 1 | 1 | NA | NA | NA | NA | 2 | 1 | 3 | 1 |
| 2_2871593 | 2 | 0 | 2 | 0 | NA | NA | 2 | 0.5 | 6 | 0.1667 | 1 | 1 | NA | NA | NA | NA | 2 | 0.5 | 3 | 0.6667 |
| 2_2871623 | 2 | 0.5 | 1 | 0 | NA | NA | 2 | 1 | 5 | 0.6 | 1 | 1 | NA | NA | NA | NA | 1 | 1 | 2 | 1 |
| 2_2871636 | 2 | 1 | 1 | 0 | NA | NA | 2 | 1 | 5 | 0.8 | 1 | 1 | NA | NA | NA | NA | 1 | 1 | 2 | 1 |
| 2_2871640 | 2 | 1 | 1 | 1 | NA | NA | 2 | 1 | 5 | 1 | 1 | 1 | NA | NA | NA | NA | 1 | 1 | 2 | 1 |
| 2_2871643 | 2 | 0.5 | NA | NA | NA | NA | 2 | 0 | 4 | 0.25 | 1 | 1 | NA | NA | NA | NA | 1 | 0 | 2 | 0.5 |
| 2_2871653 | 2 | 0 | 1 | 1 | NA | NA | 2 | 0 | 5 | 0.2 | NA | NA | NA | NA | NA | NA | 1 | 1 | 1 | 1 |
| 2_2871658 | 1 | 1 | 1 | 0 | NA | NA | 2 | 1 | 4 | 0.75 | NA | NA | NA | NA | NA | NA | 1 | 1 | 1 | 1 |
| 2_2872379 | 2 | 1 | 1 | 1 | NA | NA | 1 | 0 | 4 | 0.75 | 1 | 1 | NA | NA | NA | NA | 1 | 1 | 2 | 1 |
| 2_2872385 | 1 | 1 | 1 | 1 | NA | NA | 1 | 0 | 3 | 0.6667 | 1 | 1 | NA | NA | NA | NA | 1 | 1 | 2 | 1 |
| 2_2872391 | 2 | 0.5 | 1 | 0 | NA | NA | 1 | 0 | 4 | 0.25 | 1 | 0 | NA | NA | NA | NA | 1 | 1 | 2 | 0.5 |
| 2_2872393 | 2 | 1 | 1 | 1 | NA | NA | 1 | 1 | 4 | 1 | 1 | 1 | NA | NA | NA | NA | 1 | 1 | 2 | 1 |
| 2_2872464 | 2 | 1 | NA | NA | NA | NA | 1 | 1 | 3 | 1 | 1 | 1 | NA | NA | NA | NA | 1 | 1 | 2 | 1 |
| 2_2872472 | 2 | 1 | NA | NA | NA | NA | 1 | 1 | 3 | 1 | 1 | 1 | NA | NA | NA | NA | 1 | 1 | 2 | 1 |
| 2_2872507 | 2 | 1 | NA | NA | NA | NA | 1 | 1 | 3 | 1 | 1 | 1 | NA | NA | NA | NA | 1 | 1 | 2 | 1 |
| 2_2872514 | 1 | 1 | NA | NA | NA | NA | 1 | 0 | 2 | 0.5 | 1 | 1 | NA | NA | NA | NA | 1 | 1 | 2 | 1 |
| 2_2872518 | 2 | 1 | NA | NA | NA | NA | NA | NA | 2 | 1 | 1 | 1 | NA | NA | NA | NA | NA | NA | 1 | 1 |
| 2_2879484 | 1 | 0 | 1 | 0 | NA | NA | 1 | 1 | 3 | 0.3333 | 1 | 1 | 1 | 1 | 1 | 0 | 2 | 0.5 | 5 | 0.6 |
| 2_2879492 | 1 | 1 | 1 | 1 | NA | NA | 1 | 1 | 3 | 1 | 1 | 0 | 1 | 1 | 1 | 0 | 2 | 0 | 5 | 0.2 |
| 2_2879494 | 1 | 1 | 1 | 1 | NA | NA | 2 | 0.5 | 4 | 0.75 | 2 | 1 | 1 | 1 | 1 | 1 | 1 | 1 | 5 | 1 |
| 2_2879507 | 1 | 1 | 1 | 1 | NA | NA | 1 | 1 | 3 | 1 | 2 | 1 | 1 | 1 | 2 | 0.5 | 2 | 0.5 | 7 | 0.7143 |
| 2_2895530 | NA | NA | 1 | 1 | NA | NA | NA | NA | 1 | 1 | 1 | 1 | NA | NA | NA | NA | NA | NA | 1 | 1 |
| 2_2895581 | NA | NA | 2 | 1 | NA | NA | NA | NA | 2 | 1 | 1 | 1 | NA | NA | NA | NA | NA | NA | 1 | 1 |
| 2_2897634 | NA | NA | 2 | 0.5 | 1 | 1 | 2 | 1 | 5 | 0.8 | 1 | 1 | 2 | 1 | 1 | 1 | 1 | 1 | 5 | 1 |
| 2_2897674 | NA | NA | 2 | 0 | 1 | 1 | 2 | 0 | 5 | 0.2 | 1 | 0 | 2 | 0.5 | 1 | 0 | 1 | 0 | 5 | 0.2 |
| 2_2897688 | NA | NA | 2 | 0.5 | 1 | 1 | 2 | 0 | 5 | 0.4 | 1 | 0 | 2 | 0 | 1 | 1 | 1 | 1 | 5 | 0.4 |
| 2_2903670 | NA | NA | NA | NA | 1 | 1 | NA | NA | 1 | 1 | 2 | 0.5 | NA | NA | 1 | 0 | NA | NA | 3 | 0.3333 |
| 2_2903703 | NA | NA | NA | NA | 1 | 1 | NA | NA | 1 | 1 | 2 | 1 | NA | NA | 1 | 0 | NA | NA | 3 | 0.6667 |
| 2_2903716 | NA | NA | NA | NA | 1 | 1 | NA | NA | 1 | 1 | 2 | 0 | NA | NA | 1 | 1 | NA | NA | 3 | 0.3333 |
| 2_2903747 | NA | NA | NA | NA | 1 | 1 | NA | NA | 1 | 1 | 2 | 0.5 | NA | NA | 1 | 0 | NA | NA | 3 | 0.3333 |
| 2_2903756 | NA | NA | NA | NA | 1 | 1 | NA | NA | 1 | 1 | 1 | 1 | NA | NA | 1 | 1 | NA | NA | 2 | 1 |
| 2_2903811 | NA | NA | NA | NA | 1 | 1 | NA | NA | 1 | 1 | 1 | 1 | NA | NA | 1 | 1 | NA | NA | 2 | 1 |
| 2_2903844 | NA | NA | NA | NA | 1 | 1 | NA | NA | 1 | 1 | 2 | 0.5 | NA | NA | 3 | 0.3333 | NA | NA | 5 | 0.4 |
| 2_2906947 | 1 | 0 | 2 | 0 | 1 | 1 | 1 | 0 | 5 | 0.2 | 3 | 0 | NA | NA | 6 | 0.1667 | 1 | 0 | 10 | 0.1 |
| 2_2906949 | 1 | 0 | 2 | 0.5 | 1 | 0 | 1 | 0 | 5 | 0.2 | 3 | 0 | NA | NA | 4 | 0.5 | 1 | 0 | 8 | 0.25 |
| 2_2906960 | NA | NA | NA | NA | NA | NA | 2 | 1 | 2 | 1 | 2 | 1 | NA | NA | 1 | 1 | NA | NA | 3 | 1 |
| 2_2906961 | 1 | 1 | 2 | 1 | NA | NA | 2 | 1 | 5 | 1 | 2 | 0.5 | NA | NA | 5 | 0.8 | 1 | 1 | 8 | 0.75 |
| 2_2906961 | 1 | 1 | 2 | 1 | 1 | 1 | 2 | 1 | 6 | 1 | 3 | 1 | NA | NA | 6 | 1 | 1 | 1 | 10 | 1 |
| 2_2906968 | 1 | 0 | 1 | 1 | 1 | 1 | 2 | 1 | 5 | 0.8 | 3 | 1 | NA | NA | 6 | 0.8333 | 1 | 0 | 10 | 0.8 |
| 2_2906976 | 1 | 1 | NA | NA | 1 | 0 | 2 | 0 | 4 | 0.25 | 3 | 0.6667 | NA | NA | 5 | 0.4 | NA | NA | 8 | 0.5 |
| 2_2906985 | 1 | 1 | 2 | 1 | 1 | 1 | 2 | 1 | 6 | 1 | 3 | 1 | NA | NA | 6 | 0.6667 | 1 | 0 | 10 | 0.7 |
| 2_2907005 | 1 | 1 | 1 | 1 | 1 | 1 | 3 | 1 | 6 | 1 | 2 | 0.5 | 2 | 0.5 | 5 | 0.6 | 1 | 1 | 10 | 0.6 |
| 2_2907008 | 1 | 0 | 2 | 0 | 1 | 0 | 3 | 0.3333 | 7 | 0.1429 | 3 | 0.3333 | 2 | 0.5 | 5 | 0 | 1 | 0 | 11 | 0.1818 |
| 2_2907008 | NA | NA | 1 | 1 | 1 | 1 | 3 | 1 | 5 | 1 | 3 | 1 | 1 | 1 | 2 | 1 | NA | NA | 6 | 1 |
| 2_2907012 | 1 | 0 | 2 | 0 | 1 | 1 | 3 | 0.6667 | 7 | 0.4286 | 3 | 0 | 2 | 0.5 | 6 | 0.1667 | 1 | 1 | 12 | 0.25 |
| 2_2907015 | 1 | 1 | 2 | 0.5 | 1 | 0 | 2 | 1 | 6 | 0.6667 | 3 | 1 | 2 | 1 | 5 | 0.6 | 1 | 0 | 11 | 0.7273 |
| 2_2907180 | 6 | 0.3333 | 5 | 0.2 | 4 | 0 | 5 | 0.2 | 20 | 0.2 | 3 | 0 | 4 | 0 | 9 | 0.1111 | 2 | 0 | 18 | 0.0556 |
| 2_2907195 | 5 | 0 | 6 | 0 | 6 | 0.1667 | 4 | 0.25 | 21 | 0.0952 | 5 | 0.2 | 3 | 0 | 9 | 0 | 2 | 0 | 19 | 0.0526 |
| 2_2908413 | 5 | 0.2 | 5 | 0.2 | 9 | 0 | 1 | 0 | 20 | 0.1 | 4 | 0.25 | 1 | 0 | NA | NA | 7 | 0 | 12 | 0.0833 |
| 2_2908416 | 4 | 0 | 4 | 0 | 10 | 0.3 | 1 | 1 | 19 | 0.2105 | 4 | 0.5 | 1 | 0 | NA | NA | 8 | 0.125 | 13 | 0.2308 |
| 2_2908427 | 5 | 0 | 5 | 0 | 8 | 0 | 1 | 1 | 19 | 0.0526 | 2 | 0.5 | 1 | 0 | NA | NA | 6 | 0 | 9 | 0.1111 |

| | | | | | | | | | | | | | | | | | | | | |
|---|---|---|---|---|---|---|---|---|---|---|---|---|---|---|---|---|---|---|---|---|
| 2_2908431 | 6 | 0 | 5 | 0.2 | 9 | 0.1111 | 1 | 0 | 21 | 0.0952 | 2 | 0.5 | 1 | 0 | NA | NA | 6 | 0 | 9 | 0.1111 |
| 2_2908436 | 5 | 0.2 | 2 | 0.5 | 6 | 0.3333 | 1 | 0 | 14 | 0.2857 | 1 | 1 | 1 | 1 | NA | NA | 5 | 0.2 | 7 | 0.4286 |
| 2_2908442 | 5 | 0.4 | 4 | 0 | 9 | 0.1111 | 1 | 0 | 19 | 0.1579 | 1 | 1 | 1 | 0 | NA | NA | 5 | 0 | 7 | 0.1429 |
| 2_2908465 | 4 | 0.25 | 3 | 0.3333 | 8 | 0.25 | 1 | 0 | 16 | 0.25 | 1 | 1 | 1 | 1 | NA | NA | 6 | 0.3333 | 8 | 0.5 |
| 2_2908492 | 4 | 0.5 | 3 | 0.3333 | 3 | 0.6667 | 1 | 1 | 11 | 0.5455 | 1 | 0 | 1 | 1 | NA | NA | 4 | 0.75 | 6 | 0.6667 |
| 2_2911775 | 2 | 1 | 1 | 0 | 6 | 1 | 1 | 0 | 10 | 0.8 | 1 | 0 | 1 | 1 | NA | NA | 4 | 1 | 6 | 0.8333 |
| 2_2911779 | 2 | 1 | 2 | 0.5 | 6 | 0.8333 | 1 | 1 | 11 | 0.8182 | 1 | 1 | 1 | 0 | NA | NA | 5 | 1 | 7 | 0.8571 |
| 2_2911789 | 2 | 0.5 | 1 | 0 | 4 | 0 | 1 | 0 | 8 | 0.125 | 1 | 1 | 1 | 1 | NA | NA | 3 | 0.6667 | 5 | 0.8 |
| 2_2911793 | 1 | 1 | 2 | 1 | 8 | 0.875 | 1 | 0 | 12 | 0.8333 | 1 | 1 | 1 | 0 | NA | NA | 4 | 0.75 | 6 | 0.6667 |
| 2_2911798 | 1 | 1 | 1 | 1 | 6 | 1 | 1 | 1 | 9 | 1 | 1 | 1 | 1 | 1 | NA | NA | 5 | 1 | 7 | 1 |
| 2_2911802 | 2 | 0.5 | 2 | 0.5 | 6 | 1 | 1 | 1 | 11 | 0.8182 | 1 | 1 | 1 | 1 | NA | NA | 3 | 1 | 5 | 1 |
| 2_2911814 | 2 | 1 | 2 | 0.5 | 6 | 0.6667 | 1 | 1 | 11 | 0.7273 | NA | NA | 1 | 1 | NA | NA | 4 | 0.5 | 5 | 0.6 |
| 2_2913800 | 2 | 1 | 2 | 1 | 5 | 1 | 1 | 1 | 10 | 1 | 1 | 1 | 1 | 1 | NA | NA | 4 | 1 | 6 | 1 |
| 2_2913806 | 1 | 1 | NA | NA | 2 | 1 | 1 | 1 | 4 | 1 | NA | NA | 1 | 1 | NA | NA | 2 | 0 | 3 | 0.3333 |
| 2_2913814 | 2 | 0.5 | 2 | 0.5 | 6 | 0.3333 | 1 | 1 | 11 | 0.4545 | 1 | 1 | 1 | 0 | NA | NA | 4 | 0.5 | 6 | 0.5 |
| 2_2913822 | 1 | 0 | 1 | 0 | 6 | 0.1667 | 1 | 0 | 9 | 0.1111 | 1 | 0 | 1 | 1 | NA | NA | 1 | 0 | 3 | 0.3333 |
| 2_2913830 | 1 | 1 | 2 | 1 | 5 | 0.8 | 1 | 1 | 9 | 0.8889 | 1 | 0 | 1 | 0 | NA | NA | 4 | 0.75 | 6 | 0.5 |
| 2_2913845 | 1 | 1 | 2 | 1 | 4 | 1 | NA | NA | 7 | 1 | 1 | 1 | NA | NA | NA | NA | 3 | 0.3333 | 4 | 0.5 |
| 2_2913858 | NA | NA | NA | NA | 3 | 0.6667 | NA | NA | 3 | 0.6667 | 1 | 1 | NA | NA | NA | NA | 1 | 1 | 2 | 1 |
| 2_2941309 | 1 | 0 | NA | NA | 1 | 0 | 1 | 1 | 3 | 0.3333 | NA | NA | NA | NA | NA | NA | 2 | 0.5 | 2 | 0.5 |
| 2_2941347 | 1 | 0 | NA | NA | NA | NA | 1 | 1 | 2 | 0.5 | NA | NA | NA | NA | NA | NA | 2 | 1 | 2 | 1 |
| 2_2941461 | 3 | 0.3333 | 4 | 0.25 | 5 | 0.2 | 4 | 0 | 16 | 0.1875 | 4 | 0.25 | 3 | 0 | 7 | 0 | 5 | 0.2 | 19 | 0.1053 |
| 2_2959244 | 6 | 0.5 | 2 | 1 | 2 | 0.5 | 5 | 0.4 | 15 | 0.5333 | 4 | 0.5 | 4 | 0.25 | 4 | 0.25 | 1 | 0 | 13 | 0.3077 |
| 2_2962801 | 5 | 0.4 | 6 | 0 | 6 | 0.5 | 8 | 0.125 | 25 | 0.24 | 7 | 0 | 5 | 0.2 | 6 | 0 | 4 | 0.25 | 22 | 0.0909 |
| 2_2962812 | 8 | 0.375 | 6 | 0.1667 | 5 | 0.2 | 10 | 0.4 | 29 | 0.3103 | 6 | 0.1667 | 5 | 0.4 | 7 | 0.2857 | 4 | 0.25 | 22 | 0.2727 |
| 2_2962813 | 6 | 0.8333 | 6 | 0.1667 | 7 | 0.7143 | 9 | 0.2222 | 28 | 0.4643 | 6 | 0.5 | 5 | 0.4 | 6 | 0.3333 | 5 | 0.6 | 22 | 0.4545 |
| 2_2962845 | 10 | 0 | 7 | 0 | 6 | 0.3333 | 11 | 0.1818 | 34 | 0.1176 | 8 | 0.25 | 8 | 0 | 7 | 0 | 6 | 0 | 29 | 0.069 |
| 2_2962847 | 8 | 0 | 6 | 0.1667 | 5 | 0.2 | 10 | 0 | 29 | 0.069 | 9 | 0 | 9 | 0.3333 | 6 | 0 | 6 | 0 | 30 | 0.1 |
| 2_2962882 | 11 | 0.4545 | 5 | 1 | 6 | 0.3333 | 10 | 0.6 | 32 | 0.5625 | 8 | 0.5 | 8 | 0.75 | 8 | 0.375 | 8 | 0.625 | 32 | 0.5625 |
| 2_2962884 | 10 | 0.2 | 6 | 0.1667 | 7 | 0 | 8 | 0.25 | 31 | 0.1613 | 10 | 0.1 | 9 | 0 | 9 | 0.2222 | 8 | 0.25 | 36 | 0.1389 |
| 2_2962927 | 10 | 0.3 | 8 | 0.625 | 7 | 0.2857 | 11 | 0.2727 | 36 | 0.3611 | 11 | 0.0909 | 12 | 0.3333 | 6 | 0.1667 | 8 | 0.5 | 37 | 0.2703 |
| 2_2962946 | 9 | 0.3333 | 9 | 0.7778 | 7 | 0.2857 | 11 | 0.4545 | 36 | 0.4722 | 9 | 0.5556 | 11 | 0.4545 | 9 | 0.5556 | 8 | 0.625 | 37 | 0.5405 |
| 2_2963015 | 11 | 0.7273 | 4 | 0.75 | 9 | 0.8889 | 11 | 0.6364 | 35 | 0.7429 | 14 | 0.6429 | 15 | 0.6 | 10 | 0.7 | 9 | 0.6667 | 48 | 0.6458 |
| 2_2966680 | 5 | 0.4 | NA | NA | 6 | 0.5 | 5 | 0.4 | 16 | 0.4375 | 4 | 0.5 | NA | NA | 1 | 1 | 3 | 0.6667 | 8 | 0.625 |
| 2_2966697 | 7 | 0.5714 | 1 | 0 | 4 | 0.25 | 5 | 0.2 | 17 | 0.3529 | 4 | 0.5 | 1 | 0 | 2 | 0.5 | 3 | 0.6667 | 10 | 0.5 |
| 2_2966704 | 5 | 0.2 | 1 | 0 | 4 | 0 | 4 | 0.25 | 14 | 0.1429 | 4 | 0 | 1 | 0 | 2 | 0.5 | 3 | 0 | 10 | 0.1 |
| 2_2966714 | 6 | 0 | 1 | 0 | 3 | 0.3333 | 3 | 0 | 13 | 0.0769 | 1 | 0 | 1 | 0 | 1 | 0 | 3 | 0.3333 | 6 | 0.1667 |
| 2_2966752 | 4 | 0 | 1 | 0 | 4 | 0.25 | 3 | 0 | 12 | 0.0833 | 2 | 0 | 1 | 0 | 2 | 0 | 3 | 0.3333 | 8 | 0.125 |
| 2_2966759 | 5 | 0.6 | NA | NA | 3 | 0.6667 | 2 | 1 | 10 | 0.7 | 2 | 1 | 1 | 1 | 1 | 0 | 2 | 1 | 6 | 0.8333 |
| 2_2966774 | 2 | 1 | 1 | 1 | 2 | 1 | 1 | 1 | 6 | 1 | 3 | 0.3333 | 1 | 1 | 1 | 1 | 1 | 1 | 6 | 0.6667 |
| 2_2966776 | 5 | 1 | 1 | 1 | 2 | 0.5 | 3 | 1 | 11 | 0.9091 | 2 | 1 | 1 | 1 | 1 | 1 | 3 | 1 | 7 | 1 |
| 2_2966781 | 5 | 0.6 | 1 | 1 | 3 | 0 | 3 | 0 | 12 | 0.3333 | 3 | 0 | 1 | 0 | 2 | 0.5 | 3 | 0.6667 | 9 | 0.3333 |
| 2_2966786 | 4 | 0.5 | 1 | 0 | 3 | 0.6667 | 3 | 0.6667 | 11 | 0.5455 | 3 | 0.3333 | 1 | 1 | 1 | 1 | 3 | 0.6667 | 8 | 0.625 |
| 2_2968669 | 1 | 0 | 4 | 0 | 5 | 0.4 | 1 | 0 | 11 | 0.1818 | NA | NA | 6 | 0 | 3 | 0.3333 | 3 | 0.3333 | 12 | 0.1667 |
| 2_2968669 | 1 | 0 | 5 | 0.2 | 4 | 0.25 | 1 | 1 | 11 | 0.2727 | NA | NA | 5 | 0.6 | 3 | 0.3333 | 3 | 0.3333 | 11 | 0.4545 |
| 2_2968686 | 2 | 1 | 5 | 0.6 | NA | NA | 1 | 1 | 8 | 0.75 | NA | NA | 5 | 1 | 2 | 0.5 | 2 | 0 | 9 | 0.6667 |
| 2_2968687 | 1 | 1 | 4 | 0.5 | 4 | 0.5 | 1 | 1 | 10 | 0.6 | NA | NA | 5 | 0.4 | 2 | 0.5 | 3 | 0.3333 | 10 | 0.4 |
| 2_2968694 | 3 | 0.6667 | 6 | 0 | 5 | 0.4 | NA | NA | 14 | 0.2857 | NA | NA | 5 | 0.2 | 3 | 0.3333 | 3 | 0.6667 | 11 | 0.3636 |
| 2_2968707 | NA | NA | 1 | 1 | NA | NA | NA | NA | 1 | 1 | NA | NA | NA | NA | 1 | 1 | NA | NA | 1 | 1 |
| 2_2969576 | 6 | 0.1667 | 4 | 0.25 | 2 | 0 | 7 | 0.1429 | 19 | 0.1579 | 4 | 0.5 | 3 | 0.3333 | 3 | 0.3333 | 4 | 0.5 | 14 | 0.4286 |
| 2_2969577 | 6 | 0.1667 | 3 | 0.6667 | NA | NA | 5 | 0.4 | 14 | 0.3571 | 3 | 0.3333 | 3 | 1 | 2 | 0.5 | 3 | 0.3333 | 11 | 0.5455 |
| 2_2970569 | 11 | 0.0909 | 11 | 0 | 5 | 0 | 11 | 0.0909 | 38 | 0.0526 | 10 | 0.2 | 4 | 0 | 6 | 0 | 5 | 0 | 25 | 0.08 |
| 2_2970589 | 12 | 0.8333 | 11 | 0.5455 | 6 | 0.5 | 11 | 0.7273 | 40 | 0.675 | 11 | 0.8182 | 6 | 0.8333 | 6 | 0.6667 | 9 | 0.6667 | 32 | 0.75 |
| 2_2970611 | 12 | 0.1667 | 12 | 0.0833 | 5 | 0.2 | 9 | 0.1111 | 38 | 0.1316 | 11 | 0.3636 | 6 | 0 | 4 | 0 | 9 | 0 | 30 | 0.1333 |
| 2_2970613 | 12 | 0.3333 | 13 | 0.0769 | 7 | 0.2857 | 12 | 0.4167 | 44 | 0.2727 | 12 | 0.5 | 6 | 0.8333 | 6 | 0.1667 | 10 | 0.1 | 34 | 0.3824 |
| 2_2970623 | 10 | 0.1 | 14 | 0.0714 | 6 | 0.1667 | 9 | 0.1111 | 39 | 0.1026 | 10 | 0.3 | 4 | 1 | 6 | 0.3333 | 8 | 0.375 | 28 | 0.4286 |
| 2_2970677 | 1 | 1 | NA | NA | NA | NA | NA | NA | 1 | 1 | NA | NA | NA | NA | 1 | 1 | NA | NA | 1 | 1 |
| 2_2973356 | 5 | 0.2 | 3 | 0 | 2 | 0 | 3 | 0 | 13 | 0.0769 | 5 | 0.2 | 6 | 0 | 3 | 0 | 5 | 0 | 19 | 0.0526 |
| 2_2973366 | 6 | 0.1667 | 4 | 0 | 2 | 0 | 4 | 0 | 16 | 0.0625 | 6 | 0.3333 | 6 | 0.1667 | 3 | 0.3333 | 3 | 0.3333 | 18 | 0.2778 |
| 2_2973406 | 6 | 0 | 5 | 0.2 | 1 | 0 | 3 | 0 | 15 | 0.0667 | 4 | 0.25 | 6 | 0.3333 | 5 | 0 | 3 | 0.3333 | 18 | 0.2222 |
| 2_2973421 | 4 | 0.25 | 1 | 0 | NA | NA | 2 | 0 | 7 | 0.1429 | NA | NA | 3 | 0.3333 | NA | NA | NA | NA | 3 | 0.3333 |
| 2_2973506 | 3 | 0.6667 | 2 | 0 | 5 | 0.2 | 2 | 0 | 12 | 0.25 | 2 | 0 | 3 | 0.3333 | 1 | 0 | 2 | 1 | 8 | 0.375 |
| 2_2973534 | 1 | 0 | 1 | 0 | 4 | 0.25 | 3 | 0.3333 | 9 | 0.2222 | 2 | 0 | 1 | 0 | 1 | 0 | 2 | 0.5 | 6 | 0.1667 |
| 2_2977248 | NA | NA | NA | NA | 1 | 1 | NA | NA | 1 | 1 | NA | NA | 1 | 1 | 2 | 0.5 | 1 | 0 | 4 | 0.5 |
| 2_2977256 | NA | NA | NA | NA | 1 | 1 | NA | NA | 1 | 1 | NA | NA | NA | NA | 1 | 1 | 1 | 1 | 2 | 1 |

| | | | | | | | | | | | | | | | | | | | | |
|---|---|---|---|---|---|---|---|---|---|---|---|---|---|---|---|---|---|---|---|---|
| 2_2977260 | NA | NA | NA | NA | 1 | 1 | NA | NA | 1 | 1 | NA | NA | NA | NA | 1 | 1 | NA | NA | 1 | 1 |
| 2_2981239 | NA | NA | 2 | 1 | 1 | 1 | 3 | 0.6667 | 6 | 0.8333 | 3 | 0.6667 | 1 | 1 | 1 | 0 | 5 | 0.6 | 10 | 0.6 |
| 2_2981241 | NA | NA | 1 | 1 | 1 | 0 | 3 | 0.6667 | 5 | 0.6 | 2 | 1 | 1 | 1 | 1 | 1 | 5 | 0.2 | 9 | 0.5556 |
| 2_2981246 | NA | NA | 2 | 1 | 1 | 1 | 3 | 0 | 6 | 0.5 | 3 | 0.6667 | 1 | 1 | 1 | 0 | 5 | 0.2 | 10 | 0.4 |
| 2_2981369 | NA | NA | NA | NA | 1 | 1 | NA | NA | 1 | 1 | 1 | 0 | NA | NA | 1 | 1 | 2 | 0.5 | 4 | 0.5 |
| 2_2981396 | NA | NA | 1 | 1 | 1 | 0 | NA | NA | 2 | 0.5 | 1 | 1 | 1 | 1 | NA | NA | 1 | 1 | 3 | 1 |
| 2_2981408 | NA | NA | 1 | 1 | 1 | 0 | NA | NA | 2 | 0.5 | 1 | 0 | 1 | 1 | NA | NA | 2 | 1 | 4 | 0.75 |
| 2_2982093 | 2 | 0.5 | 5 | 0.6 | 3 | 0.6667 | 1 | 0 | 11 | 0.5455 | 2 | 0.5 | 5 | 0.2 | 4 | 0.5 | 3 | 0 | 14 | 0.2857 |
| 2_2982098 | 4 | 0.5 | 3 | 0.6667 | 3 | 0.6667 | 1 | 0 | 11 | 0.5455 | 3 | 1 | 4 | 0.5 | 4 | 1 | 5 | 1 | 16 | 0.875 |
| 2_2982135 | 2 | 0.5 | 4 | 0.75 | 4 | 0.25 | 1 | 0 | 11 | 0.4545 | 3 | 1 | 4 | 0.75 | 4 | 0.5 | 4 | 0.75 | 15 | 0.7333 |
| 2_2982142 | 4 | 0 | 5 | 0.2 | 6 | 0.1667 | 4 | 0 | 19 | 0.1053 | 3 | 0 | 4 | 0 | 4 | 0.25 | 6 | 0.3333 | 17 | 0.1765 |
| 2_2983901 | 3 | 0 | 2 | 0 | 3 | 0 | 4 | 0.25 | 12 | 0.0833 | 4 | 0.25 | 4 | 0 | 4 | 0 | 5 | 0 | 17 | 0.0588 |
| 2_2984857 | 2 | 0.5 | 3 | 0 | 5 | 0.2 | 5 | 0.2 | 15 | 0.2 | 3 | 0.3333 | 5 | 0.2 | 4 | 0 | 9 | 0 | 21 | 0.0952 |
| 2_2992178 | 4 | 0 | 5 | 0.4 | 2 | 1 | 3 | 0.3333 | 14 | 0.3571 | 2 | 0 | 2 | 0.5 | 5 | 0.6 | 4 | 0.75 | 13 | 0.5385 |
| 2_2996201 | 26 | 0.1154 | 24 | 0.375 | 16 | 0.3125 | 30 | 0.2667 | 96 | 0.2604 | 14 | 0.2143 | 10 | 0.3 | 15 | 0.2 | 25 | 0.08 | 64 | 0.1719 |
| 2_2996217 | 24 | 0.3333 | 22 | 0.1818 | 14 | 0.5 | 27 | 0.2222 | 87 | 0.2874 | 13 | 0.3846 | 9 | 0.3333 | 15 | 0.3333 | 26 | 0.0769 | 63 | 0.2381 |
| 2_2996225 | 24 | 0.0833 | 22 | 0.1364 | 19 | 0.0526 | 32 | 0.1875 | 97 | 0.1237 | 16 | 0.0625 | 11 | 0.2727 | 14 | 0.0714 | 26 | 0.1154 | 67 | 0.1194 |
| 2_2996239 | 26 | 0.1923 | 24 | 0.2083 | 18 | 0.0556 | 30 | 0.2333 | 98 | 0.1837 | 18 | 0.1111 | 11 | 0.1818 | 17 | 0.1765 | 26 | 0.1538 | 72 | 0.1528 |
| 2_2998130 | 3 | 0.6667 | NA | NA | NA | NA | 3 | 0.6667 | 6 | 0.6667 | 1 | 0 | 3 | 1 | 1 | 0 | 1 | 1 | 6 | 0.6667 |
| 2_2998738 | 22 | 0.5455 | 18 | 0.5556 | 19 | 0.4737 | 16 | 0.5625 | 75 | 0.5333 | 18 | 0.5 | 19 | 0.2105 | 20 | 0.5 | 12 | 0.3333 | 69 | 0.3913 |
| 2_2998751 | 17 | 0.4118 | 14 | 0.3571 | 20 | 0.3 | 13 | 0.3077 | 64 | 0.3438 | 15 | 0 | 14 | 0.3571 | 15 | 0.2667 | 9 | 0.2222 | 53 | 0.2075 |
| 2_2998780 | 16 | 0.4375 | 8 | 0.25 | 14 | 0.2857 | 9 | 0.5556 | 47 | 0.383 | 12 | 0.4167 | 13 | 0.2308 | 16 | 0.4375 | 10 | 0.4 | 51 | 0.3725 |
| 2_2998869 | 18 | 0.3333 | 11 | 0.1818 | 20 | 0.1 | 13 | 0.0769 | 62 | 0.1774 | 12 | 0.3333 | 9 | 0.4444 | 14 | 0.2143 | 11 | 0.2727 | 46 | 0.3043 |
| 2_3000072 | 6 | 0 | 5 | 0.4 | 11 | 0.1818 | 6 | 0.5 | 28 | 0.25 | 5 | 0.2 | 4 | 0.25 | 8 | 0.125 | 7 | 0.2857 | 24 | 0.2083 |
| 2_3000075 | 5 | 0.2 | 3 | 0.3333 | 11 | 0.3636 | 5 | 1 | 24 | 0.4583 | 2 | 0 | 2 | 0 | 4 | 0.5 | 5 | 0.2 | 13 | 0.2308 |
| 2_3000080 | 7 | 0.7143 | 4 | 0.25 | 9 | 0.5556 | 5 | 1 | 25 | 0.64 | 2 | 1 | 4 | 0.75 | 7 | 0.7143 | 6 | 0.1667 | 19 | 0.5789 |
| 2_3000091 | 4 | 0.5 | 4 | 0.5 | 12 | 0.5 | 3 | 0.3333 | 23 | 0.4783 | 3 | 0.3333 | 3 | 0.6667 | 6 | 0.3333 | 4 | 0.5 | 16 | 0.4375 |
| 2_3000098 | 4 | 0 | 4 | 0 | 10 | 0.2 | 3 | 0 | 21 | 0.0952 | 2 | 0.5 | 4 | 0 | 6 | 0 | 5 | 0 | 17 | 0.0588 |
| 2_3000129 | 4 | 0.5 | 4 | 0 | 9 | 0.1111 | 5 | 0.2 | 22 | 0.1818 | 3 | 0 | 4 | 0.25 | 5 | 0 | 7 | 0 | 19 | 0.0526 |
| 2_3003984 | 1 | 0 | NA | NA | NA | NA | 1 | 1 | 2 | 0.5 | 1 | 1 | NA | NA | 1 | 0 | 4 | 0.5 | 6 | 0.5 |
| 2_3004001 | 1 | 1 | NA | NA | NA | NA | 1 | 1 | 2 | 1 | 1 | 0 | 2 | 0.5 | 1 | 0 | 4 | 0.25 | 8 | 0.25 |
| 2_3004002 | 1 | 0 | NA | NA | NA | NA | 1 | 1 | 2 | 0.5 | 1 | 0 | 2 | 0.5 | 1 | 1 | 3 | 0.3333 | 7 | 0.4286 |
| 2_3004005 | 1 | 1 | NA | NA | NA | NA | 1 | 1 | 2 | 1 | NA | NA | 1 | 1 | 1 | 1 | 2 | 1 | 4 | 1 |
| 2_3004025 | 1 | 0 | NA | NA | NA | NA | 1 | 1 | 2 | 0.5 | 1 | 0 | 2 | 0.5 | 1 | 0 | 3 | 0 | 7 | 0.1429 |
| 2_3004109 | NA | NA | 3 | 0.3333 | 1 | 0 | 2 | 0 | 6 | 0.1667 | 3 | 0 | 4 | 0.25 | 3 | 0.3333 | 5 | 0.2 | 15 | 0.2 |
| 2_3004112 | 1 | 0 | 3 | 0 | 3 | 0.3333 | 4 | 0.25 | 11 | 0.1818 | 3 | 0 | 5 | 0 | 5 | 0 | 6 | 0.6667 | 19 | 0.2105 |
| 2_3004133 | 2 | 0 | 4 | 0.25 | 2 | 0 | 3 | 0 | 11 | 0.0909 | 3 | 0 | 4 | 0 | 6 | 0.1667 | 6 | 0 | 19 | 0.0526 |
| 2_3004162 | 2 | 0 | 1 | 0 | 2 | 0 | 3 | 0.3333 | 8 | 0.125 | 6 | 0.1667 | 5 | 0.4 | 6 | 0.1667 | 6 | 0.1667 | 23 | 0.2174 |
| 2_3004185 | 1 | 0 | 4 | 0 | 2 | 0 | 3 | 0.3333 | 10 | 0.1 | 8 | 0.125 | 5 | 0.2 | 6 | 0 | 7 | 0.1429 | 26 | 0.1154 |
| 2_3004185 | 2 | 0.5 | 4 | 0.75 | 2 | 0 | 4 | 0.25 | 12 | 0.4167 | 8 | 0.5 | 4 | 0.5 | 5 | 0.8 | 8 | 0.375 | 25 | 0.52 |
| 2_3006196 | 5 | 0.2 | 2 | 0 | 5 | 0.6 | 6 | 0.1667 | 18 | 0.2778 | 2 | 0 | 8 | 0.5 | 2 | 0.5 | 2 | 1 | 14 | 0.5 |
| 2_3006355 | 2 | 0 | 7 | 0.4286 | 6 | 0.1667 | 5 | 0.2 | 20 | 0.25 | 7 | 0.2857 | 5 | 0.2 | 5 | 0.2 | 8 | 0.5 | 25 | 0.32 |
| 2_3006357 | 2 | 0.5 | 5 | 0.8 | 7 | 0.7143 | 4 | 0.75 | 18 | 0.7222 | 5 | 0.4 | 4 | 1 | 4 | 0.5 | 9 | 0.5556 | 22 | 0.5909 |
| 2_3006372 | NA | NA | 5 | 1 | 4 | 1 | 5 | 0.8 | 14 | 0.9286 | 6 | 0.8333 | 5 | 1 | 5 | 1 | 8 | 0.875 | 24 | 0.9167 |
| 2_3006387 | 1 | 0 | 4 | 0.25 | 5 | 0 | 4 | 0 | 14 | 0.0714 | 8 | 0.125 | 5 | 0 | 4 | 0 | 6 | 0.1667 | 23 | 0.087 |
| 2_3006399 | 10 | 0.9 | 3 | 1 | 6 | 0.5 | 8 | 1 | 27 | 0.8519 | 4 | 1 | 8 | 1 | 3 | 1 | 4 | 1 | 19 | 1 |
| 2_3006404 | 1 | 0 | 4 | 0.5 | 4 | 0.25 | 3 | 0.6667 | 12 | 0.4167 | 7 | 0.5714 | 2 | 0.5 | 4 | 0.5 | 3 | 0 | 16 | 0.4375 |
| 2_3007533 | 3 | 0 | 2 | 0.5 | 3 | 0.6667 | NA | NA | 8 | 0.375 | 5 | 0.2 | 2 | 0.5 | 4 | 0 | 2 | 0 | 13 | 0.1538 |
| 2_3007537 | 3 | 1 | NA | NA | 4 | 0.25 | 1 | 1 | 8 | 0.625 | 3 | 0.3333 | 2 | 1 | 3 | 0.6667 | 2 | 1 | 10 | 0.7 |
| 2_3007571 | 4 | 0 | 2 | 0 | 3 | 0.3333 | 1 | 0 | 10 | 0.1 | 5 | 0.2 | 2 | 0 | 4 | 0.25 | 2 | 0 | 13 | 0.1538 |
| 2_3007670 | 4 | 0.25 | 2 | 0 | 2 | 0 | 1 | 0 | 9 | 0.1111 | 5 | 0 | 2 | 0 | 4 | 0 | 2 | 0.5 | 13 | 0.0769 |
| 2_3014537 | 2 | 1 | 4 | 0.75 | 1 | 1 | 2 | 1 | 9 | 0.8889 | 3 | 1 | 2 | 0.5 | NA | NA | 1 | 1 | 6 | 0.8333 |
| 2_3014564 | 4 | 0.25 | 3 | 0.6667 | 2 | 0.5 | 1 | 0 | 10 | 0.4 | 3 | 0 | 1 | 0 | 2 | 0.5 | 1 | 0 | 7 | 0.1429 |
| 2_3014577 | 4 | 0 | 4 | 0.25 | 2 | 0 | 2 | 0 | 12 | 0.0833 | 3 | 0 | 1 | 0 | 3 | 0.3333 | 1 | 1 | 8 | 0.25 |
| 2_3014583 | 5 | 0.4 | 5 | 0.2 | 2 | 0 | 2 | 0 | 14 | 0.2143 | 2 | 0.5 | 2 | 0 | 3 | 0 | 1 | 1 | 8 | 0.25 |
| 2_3014588 | 5 | 0.4 | 3 | 0 | NA | NA | 2 | 0 | 10 | 0.2 | 3 | 0.6667 | 2 | 0 | 2 | 0 | 1 | 1 | 8 | 0.375 |
| 2_3014594 | 5 | 0.4 | 4 | 0.5 | 3 | 0 | 2 | 0 | 14 | 0.2857 | 3 | 0.6667 | 1 | 0 | 3 | 0.3333 | 1 | 1 | 8 | 0.5 |
| 2_3014596 | 3 | 1 | 5 | 0.8 | 2 | 0.5 | 2 | 0 | 12 | 0.6667 | 3 | 0.3333 | 2 | 1 | 3 | 0.3333 | 1 | 1 | 9 | 0.5556 |
| 2_3014606 | 4 | 0.25 | 4 | 0.25 | 3 | 0.6667 | 2 | 0.5 | 13 | 0.3846 | 2 | 0.5 | 2 | 1 | 3 | 0.6667 | NA | NA | 7 | 0.7143 |
| 2_3014650 | 5 | 1 | 5 | 0.8 | 2 | 0.5 | 2 | 1 | 14 | 0.8571 | 3 | 1 | 1 | 1 | 3 | 0.6667 | 1 | 1 | 8 | 0.875 |
| 2_3014669 | 3 | 0.3333 | 5 | 0.4 | 3 | 0.3333 | 1 | 0 | 12 | 0.3333 | 3 | 0.3333 | 1 | 0 | 3 | 0 | NA | NA | 7 | 0.1429 |
| 2_3014705 | 5 | 0.4 | 5 | 0 | 2 | 0 | 3 | 0 | 15 | 0.1333 | 3 | 0.6667 | 1 | 1 | 4 | 0.25 | 1 | 1 | 9 | 0.5556 |
| 2_3014725 | 5 | 0.2 | 6 | 0.1667 | 3 | 0 | 3 | 0 | 17 | 0.1176 | 3 | 0.3333 | 2 | 0.5 | 5 | 0 | NA | NA | 10 | 0.2 |
| 2_3015206 | 7 | 1 | 6 | 1 | 3 | 1 | 3 | 1 | 19 | 1 | 3 | 1 | 3 | 1 | 7 | 0.8571 | 1 | 1 | 14 | 0.9286 |
| 2_3015220 | 7 | 0.8571 | 6 | 1 | 3 | 1 | 3 | 1 | 19 | 0.9474 | 3 | 0.6667 | 3 | 1 | 7 | 1 | 1 | 1 | 14 | 0.9286 |

| | | | | | | | | | | | | | | | | | | | | |
|---|---|---|---|---|---|---|---|---|---|---|---|---|---|---|---|---|---|---|---|---|
| 2_30152249 | 7 | 0.8571 | 4 | 0.75 | 2 | 0.5 | 2 | 1 | 15 | 0.8 | 3 | 1 | 3 | 0.6667 | 6 | 1 | 1 | 1 | 13 | 0.9231 |
| 2_30152281 | 6 | 0.5 | 5 | 0.4 | 3 | 0.3333 | 3 | 0.3333 | 17 | 0.4118 | 3 | 0.3333 | 2 | 1 | 7 | 0.5714 | NA | NA | 12 | 0.5833 |
| 2_30152287 | 5 | 1 | 5 | 0.6 | 3 | 1 | 3 | 0.6667 | 16 | 0.8125 | 3 | 0.6667 | 3 | 1 | 6 | 0.6667 | 1 | 1 | 13 | 0.7692 |
| 2_30152296 | 5 | 0.2 | 6 | 0.6667 | 3 | 0.6667 | 3 | 0.3333 | 17 | 0.4706 | 3 | 0.3333 | 2 | 0.5 | 5 | 0 | 1 | 1 | 11 | 0.2727 |
| 2_30152353 | 5 | 1 | 6 | 1 | 3 | 1 | 3 | 1 | 17 | 1 | 2 | 1 | 2 | 1 | 5 | 1 | 1 | 1 | 10 | 1 |
| 2_30152421 | 4 | 0.5 | 5 | 0.2 | 3 | 0.6667 | 4 | 0.5 | 16 | 0.4375 | 2 | 0.5 | 2 | 0 | 5 | 0.6 | NA | NA | 9 | 0.4444 |
| 2_30152587 | 7 | 0 | 6 | 0 | 2 | 0.5 | 4 | 0 | 19 | 0.0526 | 2 | 0 | 2 | 0.5 | 7 | 0 | 1 | 0 | 12 | 0.0833 |
| 2_30152624 | 5 | 0 | 5 | 0.4 | 1 | 0 | 3 | 0 | 14 | 0.1429 | 3 | 0.3333 | 2 | 0 | 6 | 0.1667 | 1 | 0 | 12 | 0.1667 |
| 2_30153605 | 10 | 0.4 | 11 | 0.5455 | 5 | 0.2 | 3 | 0.6667 | 29 | 0.4483 | 3 | 0 | 5 | 0.2 | 7 | 0.5714 | 2 | 0.5 | 17 | 0.3529 |
| 2_30171750 | 1 | 1 | 2 | 0.5 | 1 | 1 | NA | NA | 4 | 0.75 | NA | NA | NA | NA | 1 | 1 | NA | NA | 1 | 1 |
| 2_30174297 | 2 | 0 | 3 | 0.6667 | NA | NA | 4 | 0.5 | 9 | 0.4444 | 1 | 1 | 4 | 0.25 | 1 | 0 | 4 | 0.75 | 10 | 0.5 |
| 2_30174368 | 1 | 0 | 3 | 0.6667 | NA | NA | 4 | 0.5 | 8 | 0.5 | 1 | 0 | 3 | 0.6667 | 1 | 0 | 4 | 0.5 | 9 | 0.4444 |
| 2_30174485 | 1 | 1 | 3 | 1 | NA | NA | 4 | 1 | 8 | 1 | NA | NA | 3 | 1 | NA | NA | 4 | 1 | 7 | 1 |
| 2_30174557 | 2 | 0.5 | 2 | 0 | NA | NA | 3 | 0.6667 | 7 | 0.4286 | NA | NA | 2 | 1 | NA | NA | 2 | 1 | 4 | 1 |
| 2_30174575 | 2 | 0.5 | 3 | 0.3333 | NA | NA | 3 | 0.3333 | 8 | 0.375 | NA | NA | 3 | 0.6667 | NA | NA | 3 | 0 | 6 | 0.3333 |
| 2_30174630 | 2 | 1 | 2 | 1 | NA | NA | 3 | 1 | 7 | 1 | NA | NA | 3 | 1 | NA | NA | 4 | 1 | 7 | 1 |
| 2_30174651 | 2 | 0.5 | 3 | 0.3333 | NA | NA | 3 | 0.3333 | 8 | 0.375 | NA | NA | 3 | 1 | NA | NA | 3 | 0.6667 | 6 | 0.8333 |
| 2_30174767 | 1 | 1 | 3 | 0.6667 | NA | NA | 2 | 1 | 6 | 0.8333 | NA | NA | 2 | 0.5 | NA | NA | 3 | 1 | 5 | 0.8 |
| 2_30174850 | 2 | 1 | 3 | 0.6667 | NA | NA | 3 | 0.6667 | 8 | 0.75 | NA | NA | 3 | 0 | NA | NA | 2 | 0.5 | 5 | 0.2 |
| 2_30174857 | 1 | 1 | 3 | 0.6667 | NA | NA | 2 | 0 | 6 | 0.5 | NA | NA | 2 | 0 | NA | NA | 3 | 0.6667 | 5 | 0.4 |
| 2_30175385 | 2 | 1 | 1 | 1 | NA | NA | 3 | 1 | 6 | 1 | NA | NA | 2 | 1 | NA | NA | 3 | 1 | 5 | 1 |
| 2_30175410 | 2 | 0 | 1 | 0 | NA | NA | 1 | 1 | 4 | 0.25 | NA | NA | 2 | 1 | NA | NA | 2 | 0 | 4 | 0.5 |
| 2_30175749 | 2 | 0.5 | NA | NA | NA | NA | 3 | 0.6667 | 5 | 0.6 | NA | NA | 2 | 0.5 | NA | NA | 3 | 1 | 5 | 0.8 |
| 2_30176045 | 2 | 0.5 | 1 | 0 | NA | NA | 1 | 0 | 4 | 0.25 | NA | NA | 2 | 0.5 | NA | NA | 3 | 0.3333 | 5 | 0.4 |
| 2_30234017 | 3 | 0 | 3 | 0.3333 | 2 | 0.5 | 1 | 0 | 9 | 0.2222 | 2 | 0.5 | 4 | 0 | 1 | 0 | 1 | 1 | 8 | 0.25 |
| 2_30234083 | 3 | 0.3333 | 2 | 0 | 2 | 0 | 1 | 1 | 8 | 0.25 | 2 | 0 | 5 | 0.4 | 1 | 1 | 2 | 0 | 10 | 0.3 |
| 2_30235057 | 19 | 0.1053 | 21 | 0.1905 | 18 | 0.1667 | 13 | 0.1538 | 71 | 0.1549 | 19 | 0.0526 | 21 | 0 | 23 | 0.1739 | 19 | 0.0526 | 82 | 0.0732 |
| 2_30235265 | 23 | 0.5217 | 26 | 0.5 | 22 | 0.4545 | 25 | 0.44 | 96 | 0.4792 | 29 | 0.2759 | 20 | 0.35 | 37 | 0.4865 | 23 | 0.3478 | 109 | 0.3761 |
| 2_30235344 | 23 | 0.2174 | 22 | 0.1364 | 20 | 0.05 | 19 | 0 | 84 | 0.1071 | 23 | 0.0435 | 18 | 0.1111 | 25 | 0 | 19 | 0.1053 | 85 | 0.0588 |
| 2_30271985 | 8 | 0.125 | 2 | 0 | 2 | 0 | NA | NA | 12 | 0.0833 | 4 | 0 | 1 | 0 | 3 | 0 | 6 | 0.3333 | 14 | 0.1429 |
| 2_30274475 | 10 | 0.1 | 5 | 0.2 | 4 | 0 | 7 | 0.1429 | 26 | 0.1154 | 9 | 0.1111 | 4 | 0 | 7 | 0.2857 | 8 | 0 | 28 | 0.1071 |
| 2_30274793 | 12 | 0.1667 | 4 | 0.25 | 3 | 0 | 5 | 0.4 | 24 | 0.2083 | 11 | 0.0909 | 4 | 0 | 10 | 0.1 | 9 | 0.1111 | 34 | 0.0882 |
| 2_30274965 | 10 | 0.1 | 5 | 0.2 | 6 | 0.1667 | 8 | 0.5 | 29 | 0.2414 | 9 | 0.6667 | 4 | 0 | 8 | 0.5 | 9 | 0.2222 | 30 | 0.4 |
| 2_30289190 | 4 | 0.5 | 2 | 0 | 3 | 0.3333 | 2 | 0.5 | 11 | 0.3636 | NA | NA | NA | NA | NA | NA | 2 | 1 | 2 | 1 |
| 2_30289342 | 4 | 0.75 | 1 | 1 | 2 | 1 | 1 | 1 | 8 | 0.875 | NA | NA | NA | NA | NA | NA | 2 | 1 | 2 | 1 |
| 2_30290194 | 5 | 0.2 | 2 | 1 | 3 | 0 | 3 | 0 | 13 | 0.2308 | 3 | 0 | NA | NA | NA | NA | 3 | 0.3333 | 6 | 0.1667 |
| 2_30290239 | 6 | 0.3333 | 2 | 0 | 4 | 0.5 | 3 | 0.3333 | 15 | 0.3333 | 3 | 0 | NA | NA | 1 | 0 | 3 | 0.3333 | 7 | 0.1429 |
| 2_30290809 | NA | NA | 1 | 0 | 1 | 1 | NA | NA | 2 | 0.5 | 1 | 1 | 1 | 0 | NA | NA | NA | NA | 2 | 0.5 |
| 2_30292057 | 9 | 0.3333 | 6 | 0.1667 | 6 | 0.1667 | 1 | 0 | 22 | 0.2273 | 4 | 0.25 | 3 | 0 | 6 | 0.1667 | 2 | 0 | 15 | 0.1333 |
| 2_30292123 | 8 | 0.625 | 7 | 0.5714 | 4 | 0.25 | 1 | 0 | 20 | 0.5 | 4 | 0.5 | 3 | 0.3333 | 6 | 0.5 | 2 | 1 | 15 | 0.5333 |
| 2_30292203 | 8 | 0.75 | 7 | 0.7143 | 3 | 0 | 1 | 0 | 19 | 0.5789 | 3 | 0.6667 | 4 | 0.75 | 5 | 0.4 | 2 | 0.5 | 14 | 0.5714 |
| 2_30292340 | 7 | 0.1429 | 6 | 0.1667 | 4 | 0 | 1 | 0 | 18 | 0.1111 | 3 | 0 | 4 | 0 | 4 | 0.25 | 1 | 0 | 12 | 0.0833 |
| 2_30292379 | 6 | 0.6667 | 6 | 0.3333 | 6 | 0.3333 | 1 | 1 | 19 | 0.4737 | 4 | 0.5 | 3 | 0 | 5 | 0 | 2 | 0 | 14 | 0.1429 |
| 2_30292540 | 7 | 1 | 7 | 1 | 4 | 0.75 | 1 | 0 | 19 | 0.8947 | 4 | 1 | 4 | 1 | 5 | 1 | 2 | 1 | 15 | 1 |
| 2_30292577 | 7 | 0 | 6 | 0 | 6 | 0.1667 | 1 | 0 | 20 | 0.05 | 4 | 0 | 4 | 0 | 5 | 0.4 | 2 | 0 | 15 | 0.1333 |
| 2_30292633 | 5 | 0.6 | 2 | 0 | 1 | 0 | NA | NA | 8 | 0.375 | 1 | 1 | 1 | 1 | 3 | 0.6667 | 2 | 0.5 | 7 | 0.7143 |
| 2_30292789 | 7 | 0.4286 | 7 | 0.4286 | 6 | 0 | 1 | 0 | 21 | 0.2857 | 2 | 0 | 3 | 0 | 4 | 0.25 | 2 | 0 | 11 | 0.0909 |
| 2_30336636 | 16 | 0.3125 | 13 | 0.1538 | 11 | 0.4545 | 14 | 0.0714 | 54 | 0.2407 | 12 | 0.0833 | 15 | 0.3333 | 20 | 0.35 | 11 | 0.0909 | 58 | 0.2414 |
| 2_30336753 | 14 | 0 | 18 | 0.1111 | 15 | 0 | 18 | 0.1111 | 65 | 0.0615 | 14 | 0 | 18 | 0.1667 | 24 | 0.0417 | 17 | 0 | 73 | 0.0548 |
| 2_30337211 | 21 | 0.2381 | 21 | 0.0476 | 19 | 0 | 20 | 0.05 | 81 | 0.0864 | 17 | 0.0588 | 21 | 0.0476 | 25 | 0.2 | 19 | 0.1053 | 82 | 0.1098 |
| 2_30337596 | 20 | 0.15 | 26 | 0.0769 | 19 | 0 | 25 | 0.2 | 90 | 0.1111 | 19 | 0.0526 | 23 | 0.1304 | 30 | 0.1 | 23 | 0.087 | 95 | 0.0947 |
| 2_30361687 | 3 | 0.6667 | 1 | 0 | 4 | 0.5 | 1 | 1 | 9 | 0.5556 | 2 | 1 | 1 | 1 | 3 | 0.6667 | 3 | 1 | 9 | 0.8889 |
| 2_30361912 | 3 | 0 | 1 | 0 | 3 | 0.3333 | 2 | 0 | 9 | 0.1111 | 2 | 0.5 | 1 | 0 | 3 | 0 | 4 | 0.25 | 10 | 0.2 |
| 2_30362628 | NA | NA | 1 | 1 | 1 | 1 | 1 | 1 | 3 | 1 | 1 | 1 | 1 | 0 | 1 | 0 | 2 | 1 | 5 | 0.6 |
| 2_30362748 | 7 | 0.7143 | 6 | 1 | 5 | 1 | 3 | 1 | 21 | 0.9048 | 3 | 1 | 4 | 1 | 6 | 0.8333 | 4 | 0.75 | 17 | 0.8824 |
| 2_30362782 | 7 | 0 | 7 | 0.1429 | 8 | 0.25 | 6 | 0.5 | 28 | 0.2143 | 6 | 0.3333 | 6 | 0.6667 | 10 | 0.3 | 5 | 0.2 | 27 | 0.3704 |
| 2_30362800 | 7 | 0.4286 | 10 | 0.2 | 7 | 0.2857 | 6 | 0.5 | 30 | 0.3333 | 6 | 0 | 6 | 0.1667 | 10 | 0.2 | 7 | 0.2857 | 29 | 0.1724 |
| 2_30362826 | 5 | 1 | 7 | 1 | 5 | 1 | 6 | 1 | 23 | 1 | 5 | 1 | 4 | 0.75 | 8 | 1 | 4 | 1 | 21 | 0.9524 |
| 2_30362900 | 7 | 1 | 7 | 1 | 8 | 0.75 | 5 | 0.8 | 27 | 0.8889 | 5 | 0.6 | 5 | 0.8 | 10 | 0.6 | 6 | 1 | 26 | 0.7308 |
| 2_30362954 | 7 | 0.4286 | 9 | 0 | 7 | 0.1429 | 5 | 0 | 28 | 0.1429 | 5 | 0 | 4 | 0 | 11 | 0.0909 | 6 | 0.1667 | 26 | 0.0769 |
| 2_30362984 | 8 | 0.625 | 9 | 0.4444 | 7 | 0.5714 | 3 | 0.6667 | 27 | 0.5556 | 5 | 0.8 | 4 | 0.5 | 12 | 0.3333 | 3 | 0 | 24 | 0.4167 |
| 2_30362996 | 6 | 1 | 6 | 0.8333 | 3 | 0.6667 | 3 | 1 | 18 | 0.8889 | 3 | 1 | 3 | 1 | 5 | 1 | 4 | 1 | 15 | 1 |
| 2_30363142 | 6 | 0.1667 | 9 | 0.1111 | 6 | 0.1667 | 4 | 0.25 | 25 | 0.16 | 5 | 0.2 | 5 | 0.2 | 9 | 0.1111 | 4 | 0.5 | 23 | 0.2174 |
| 2_30363191 | 6 | 1 | 9 | 0.6667 | 6 | 0.8333 | 4 | 0.75 | 25 | 0.8 | 5 | 0.4 | 5 | 1 | 8 | 0.875 | 4 | 0.75 | 22 | 0.7727 |
| 2_30363236 | 4 | 0.5 | 5 | 0.6 | 6 | 0.5 | 4 | 1 | 19 | 0.6316 | 5 | 0.8 | 2 | 1 | 7 | 0.7143 | 2 | 1 | 16 | 0.8125 |

| | | | | | | | | | | | | | | | | | | | | |
|---|---|---|---|---|---|---|---|---|---|---|---|---|---|---|---|---|---|---|---|---|
| 2_3036324 | 4 | 0.25 | 7 | 0.2857 | 5 | 0.2 | 3 | 0.3333 | 19 | 0.2632 | 5 | 0.2 | 5 | 0.2 | 9 | 0.4444 | 3 | 0.3333 | 22 | 0.3182 |
| 2_3036328 | 6 | 0.8333 | 9 | 0.6667 | 5 | 0.8 | 4 | 1 | 24 | 0.7917 | 4 | 0.5 | 4 | 0.5 | 7 | 0.5714 | 4 | 1 | 19 | 0.6316 |
| 2_3069115 | NA | NA | 1 | 1 | 1 | 0 | 1 | 0 | 3 | 0.3333 | NA | NA | 1 | 1 | NA | NA | 1 | 0 | 2 | 0.5 |
| 2_3069166 | NA | NA | 1 | 1 | 1 | 1 | NA | NA | 2 | 1 | 1 | 1 | NA | NA | 1 | 1 | 1 | 1 | 3 | 1 |
| 2_3071049 | NA | NA | 4 | 0.5 | 2 | 0 | 2 | 0 | 8 | 0.25 | 2 | 0 | 2 | 0 | 2 | 0.5 | 4 | 0.25 | 10 | 0.2 |
| 2_3071066 | 3 | 0 | 5 | 0.2 | 3 | 0 | 2 | 0 | 13 | 0.0769 | 2 | 0 | 1 | 0 | 3 | 0 | 6 | 0.1667 | 12 | 0.0833 |
| 2_3071245 | 3 | 0.6667 | 4 | 0.25 | 3 | 0.6667 | 1 | 0 | 11 | 0.4545 | 1 | 0 | 2 | 1 | 4 | 0.5 | 6 | 0.1667 | 13 | 0.3846 |
| 2_3071249 | 2 | 0 | 5 | 0 | 3 | 0.3333 | 2 | 0 | 12 | 0.0833 | 2 | 1 | 2 | 0.5 | 3 | 0.6667 | 5 | 0.4 | 12 | 0.5833 |
| 2_3071251 | 4 | 0.25 | 5 | 0.4 | 3 | 0 | 2 | 0 | 14 | 0.2143 | 2 | 0 | 2 | 0.5 | 3 | 0 | 5 | 0 | 12 | 0.0833 |
| 2_3071267 | 4 | 0.75 | 6 | 0.6667 | 4 | 0.75 | 2 | 0.5 | 16 | 0.6875 | 2 | 1 | 2 | 0.5 | 3 | 0.6667 | 5 | 0.8 | 12 | 0.75 |
| 2_3071268 | 4 | 1 | 6 | 1 | 4 | 1 | 2 | 1 | 16 | 1 | 2 | 1 | 2 | 1 | 2 | 1 | 5 | 0.8 | 11 | 0.9091 |
| 2_3071283 | 5 | 0.4 | 6 | 0.8333 | 4 | 0.75 | 2 | 1 | 17 | 0.7059 | 3 | 1 | 2 | 0 | 4 | 0.5 | 5 | 0.8 | 14 | 0.6429 |
| 2_3071285 | 4 | 0.75 | 5 | 0.6 | 4 | 0.75 | 2 | 1 | 15 | 0.7333 | 2 | 1 | 2 | 0.5 | 4 | 0.25 | 6 | 0.8333 | 14 | 0.6429 |
| 2_3071286 | 4 | 1 | 5 | 0.8 | 4 | 0.75 | 2 | 1 | 15 | 0.8667 | 3 | 0.6667 | 2 | 0.5 | 4 | 0.75 | 6 | 0.5 | 15 | 0.6 |
| 2_3071288 | 5 | 0.2 | 6 | 0.1667 | 5 | 0.2 | 2 | 0 | 18 | 0.1667 | 3 | 0 | 2 | 0 | 5 | 0 | 6 | 0.1667 | 16 | 0.0625 |
| 2_3071322 | 2 | 0 | 7 | 0.1429 | NA | NA | 8 | 0 | 17 | 0.0588 | 2 | 0 | 5 | 0 | 3 | 0 | 13 | 0.1538 | 23 | 0.087 |
| 2_3071340 | 2 | 0 | 5 | 0.2 | NA | NA | 3 | 0.3333 | 10 | 0.2 | 1 | 0 | 4 | 0.75 | 3 | 0.6667 | 7 | 0.5714 | 15 | 0.6 |
| 2_3071340 | 1 | 1 | 6 | 0.8333 | NA | NA | 4 | 1 | 11 | 0.9091 | 2 | 1 | 3 | 0.3333 | 3 | 0 | 10 | 0.6 | 18 | 0.5 |
| 2_3071380 | 2 | 0.5 | 5 | 0.2 | NA | NA | 5 | 0.6 | 12 | 0.4167 | 1 | 1 | 5 | 0.4 | 3 | 0 | 9 | 0.1111 | 18 | 0.2222 |
| 2_3071385 | 1 | 0 | 4 | 0 | NA | NA | 5 | 0.2 | 10 | 0.1 | 1 | 0 | 3 | 0 | 1 | 0 | 10 | 0.2 | 15 | 0.1333 |
| 2_3071386 | 1 | 0 | 5 | 0.8 | NA | NA | 7 | 0.8571 | 13 | 0.7692 | 1 | 1 | 5 | 0.8 | 4 | 1 | 9 | 0.5556 | 19 | 0.7368 |
| 2_3071393 | 1 | 1 | 5 | 0.6 | NA | NA | 7 | 0.8571 | 13 | 0.7692 | 1 | 1 | 5 | 1 | 4 | 1 | 7 | 0.8571 | 17 | 0.9412 |
| 2_3071408 | 1 | 0 | 4 | 0.75 | NA | NA | 6 | 0.6667 | 11 | 0.6364 | 1 | 0 | 4 | 0.25 | 4 | 0.25 | 6 | 0.8333 | 15 | 0.4667 |
| 2_3071412 | NA | NA | 5 | 0.4 | NA | NA | 7 | 0.8571 | 12 | 0.6667 | 1 | 1 | 4 | 0.5 | 4 | 1 | 11 | 0.6364 | 20 | 0.7 |
| 2_3071413 | 1 | 1 | 5 | 0 | NA | NA | 6 | 0.3333 | 12 | 0.25 | 1 | 0 | 4 | 0.25 | 3 | 0.3333 | 11 | 0.0909 | 19 | 0.1579 |
| 2_3071444 | 1 | 0 | 3 | 0.3333 | NA | NA | 6 | 0 | 10 | 0.1 | 1 | 1 | 3 | 0.3333 | 3 | 0 | 10 | 0.4 | 17 | 0.3529 |
| 2_3077102 | 1 | 1 | NA | NA | NA | NA | NA | NA | 1 | 1 | NA | NA | 1 | 1 | NA | NA | NA | NA | 1 | 1 |
| 2_3077119 | 1 | 1 | NA | NA | NA | NA | NA | NA | 1 | 1 | NA | NA | 1 | 1 | NA | NA | NA | NA | 1 | 1 |
| 2_3078252 | 1 | 1 | 1 | 1 | NA | NA | 1 | 1 | 3 | 1 | 1 | 1 | 1 | 1 | NA | NA | NA | NA | 2 | 1 |
| 2_3078253 | 1 | 0 | 1 | 1 | NA | NA | 1 | 1 | 3 | 0.6667 | 1 | 1 | 1 | 1 | NA | NA | NA | NA | 2 | 1 |
| 2_3084377 | 1 | 1 | 2 | 0 | 2 | 0.5 | 3 | 0.3333 | 8 | 0.375 | NA | NA | 1 | 0 | 1 | 1 | NA | NA | 2 | 0.5 |
| 2_3084383 | 1 | 0 | 2 | 0 | 3 | 0.6667 | 3 | 0.3333 | 9 | 0.3333 | 1 | 1 | 1 | 0 | 2 | 0.5 | 1 | 0 | 5 | 0.4 |
| 2_3084387 | 1 | 1 | 3 | 1 | 3 | 1 | 3 | 1 | 10 | 1 | 1 | 1 | 1 | 1 | 2 | 1 | 2 | 0.5 | 6 | 0.8333 |
| 2_3084517 | 2 | 1 | 1 | 1 | NA | NA | NA | NA | 3 | 1 | 1 | 0 | 1 | 1 | NA | NA | NA | NA | 2 | 0.5 |
| 2_3084531 | NA | NA | 1 | 1 | NA | NA | 1 | 1 | 2 | 1 | NA | NA | 1 | 1 | NA | NA | NA | NA | 1 | 1 |
| 2_3084551 | 1 | 0 | 1 | 1 | NA | NA | 1 | 0 | 3 | 0.3333 | NA | NA | 1 | 1 | NA | NA | NA | NA | 1 | 1 |
| 2_3084617 | NA | NA | 1 | 1 | NA | NA | 1 | 1 | 2 | 1 | NA | NA | 1 | 1 | NA | NA | NA | NA | 1 | 1 |
| 2_3084624 | 1 | 1 | 1 | 1 | NA | NA | 1 | 1 | 3 | 1 | NA | NA | 1 | 1 | NA | NA | NA | NA | 1 | 1 |
| 2_3084635 | NA | NA | NA | NA | NA | NA | 1 | 1 | 1 | 1 | NA | NA | 1 | 1 | NA | NA | NA | NA | 1 | 1 |
| 2_3084659 | 1 | 1 | 1 | 1 | NA | NA | 1 | 0 | 3 | 0.6667 | NA | NA | 1 | 1 | NA | NA | NA | NA | 1 | 1 |
| 2_3084674 | 1 | 1 | 1 | 1 | NA | NA | 1 | 1 | 3 | 1 | NA | NA | 1 | 1 | NA | NA | NA | NA | 1 | 1 |
| 2_3084717 | 2 | 1 | 3 | 1 | 2 | 1 | 4 | 1 | 11 | 1 | NA | NA | 2 | 1 | 3 | 1 | 1 | 1 | 6 | 1 |
| 2_3084721 | 1 | 1 | 3 | 1 | 2 | 1 | 2 | 1 | 8 | 1 | 1 | 1 | 2 | 1 | 3 | 1 | 1 | 1 | 7 | 1 |
| 2_3084727 | 1 | 1 | 2 | 1 | 1 | 1 | 3 | 1 | 7 | 1 | NA | NA | 2 | 0.5 | 2 | 1 | NA | NA | 4 | 0.75 |
| 2_3084749 | 1 | 1 | 2 | 1 | 3 | 1 | 4 | 1 | 10 | 1 | 1 | 1 | 2 | 1 | 3 | 1 | 3 | 1 | 9 | 1 |
| 2_3084755 | 3 | 0.6667 | 4 | 0.75 | 3 | 0.3333 | 2 | 0.5 | 12 | 0.5833 | 1 | 1 | 1 | 0 | 2 | 1 | 3 | 1 | 7 | 0.8571 |
| 2_3084774 | 3 | 0.3333 | 4 | 0.25 | 3 | 0.3333 | 4 | 0 | 14 | 0.2143 | 1 | 0 | 2 | 0 | 3 | 0.3333 | 3 | 0.6667 | 9 | 0.3333 |
| 2_3084779 | 1 | 0 | 4 | 0.25 | 1 | 0 | 4 | 0 | 10 | 0.1 | 1 | 0 | 2 | 0 | 2 | 0 | 1 | 1 | 6 | 0.1667 |
| 2_3084789 | 2 | 1 | 3 | 0.3333 | 2 | 0 | 3 | 0.6667 | 10 | 0.5 | NA | NA | 2 | 0 | 2 | 0 | 1 | 1 | 5 | 0.2 |
| 2_3084861 | 1 | 1 | NA | NA | NA | NA | NA | NA | 1 | 1 | 1 | 0 | NA | NA | 2 | 0.5 | 1 | 0 | 4 | 0.25 |
| 2_3084883 | 1 | 0 | 4 | 0.75 | 2 | 0 | 3 | 0.3333 | 10 | 0.4 | 1 | 1 | 2 | 0.5 | 3 | 0.6667 | 1 | 1 | 7 | 0.7143 |
| 2_3084892 | 2 | 0.5 | 5 | 0.8 | 3 | 0.3333 | 4 | 0.5 | 14 | 0.5714 | 1 | 1 | 2 | 1 | 3 | 0.6667 | 3 | 1 | 9 | 0.8889 |
| 2_3085032 | 1 | 1 | NA | NA | NA | NA | 1 | 0 | 2 | 0.5 | NA | NA | NA | NA | 2 | 0.5 | NA | NA | 2 | 0.5 |
| 2_3085044 | 1 | 1 | NA | NA | NA | NA | 1 | 1 | 2 | 1 | NA | NA | NA | NA | 2 | 1 | 1 | 1 | 3 | 1 |
| 2_3085047 | 1 | 0 | NA | NA | NA | NA | 1 | 1 | 2 | 0.5 | NA | NA | NA | NA | 2 | 1 | 1 | 1 | 3 | 1 |
| 2_3085061 | 1 | 1 | NA | NA | NA | NA | 1 | 1 | 2 | 1 | NA | NA | NA | NA | 2 | 0.5 | 1 | 1 | 3 | 0.6667 |
| 2_3085076 | 4 | 1 | 6 | 1 | 8 | 1 | 5 | 1 | 23 | 1 | 3 | 1 | 11 | 1 | 10 | 1 | 9 | 0.8889 | 33 | 0.9697 |
| 2_3085081 | 3 | 1 | 5 | 0.4 | 7 | 0.5714 | 5 | 0.8 | 20 | 0.65 | 3 | 0.6667 | 11 | 0.8182 | 8 | 0.625 | 7 | 0.7143 | 29 | 0.7241 |
| 2_3085083 | 4 | 0.25 | 5 | 0.2 | 7 | 0.2857 | 5 | 0.6 | 21 | 0.3333 | 2 | 0.5 | 10 | 0.1 | 11 | 0.0909 | 7 | 0.1429 | 30 | 0.1333 |
| 2_3085090 | 2 | 1 | 4 | 1 | 6 | 0.5 | 6 | 0.8333 | 18 | 0.7778 | 3 | 0.6667 | 11 | 1 | 10 | 0.7 | 9 | 0.7778 | 33 | 0.8182 |
| 2_3085096 | 4 | 0.25 | 6 | 0 | 6 | 0 | 6 | 0.3333 | 22 | 0.1364 | 2 | 0 | 10 | 0.2 | 11 | 0.2727 | 8 | 0.125 | 31 | 0.1935 |
| 2_3085097 | 3 | 1 | 6 | 0.8333 | 6 | 0.6667 | 5 | 0.6 | 20 | 0.75 | 3 | 1 | 9 | 0.7778 | 10 | 0.8 | 7 | 0.4286 | 29 | 0.7241 |
| 2_3085099 | 4 | 0.5 | 6 | 0.6667 | 6 | 0.3333 | 6 | 0.6667 | 22 | 0.5455 | 3 | 0.3333 | 11 | 0.3636 | 10 | 0.3 | 8 | 0.5 | 32 | 0.375 |
| 2_3085716 | 4 | 0.25 | 4 | 0.5 | 5 | 0.4 | 4 | 0 | 17 | 0.2941 | 1 | 1 | 8 | 0.5 | 9 | 0.4444 | 9 | 0.6667 | 27 | 0.5556 |
| 2_3085717 | 4 | 1 | 3 | 0.6667 | 5 | 0.4 | 3 | 0.3333 | 15 | 0.6 | 1 | 1 | 6 | 0.3333 | 10 | 0.6 | 8 | 0.875 | 25 | 0.64 |

| | | | | | | | | | | | | | | | | | | | | |
|---|---|---|---|---|---|---|---|---|---|---|---|---|---|---|---|---|---|---|---|---|
| 2_3085733 | 3 | 0.3333 | 4 | 0.5 | 4 | 0.25 | 3 | 0 | 14 | 0.2857 | 1 | 1 | 6 | 0.5 | 9 | 0.6667 | 7 | 0.2857 | 23 | 0.5217 |
| 2_3086266 | 2 | 0.5 | NA | NA | 2 | 0 | 1 | 0 | 5 | 0.2 | NA | NA | NA | NA | 1 | 1 | 1 | 0 | 2 | 0.5 |
| 2_3086301 | 6 | 0.1667 | 12 | 0.1667 | 12 | 0.0833 | 7 | 0.1429 | 37 | 0.1351 | 9 | 0.1111 | 8 | 0.125 | 7 | 0 | 6 | 0.1667 | 30 | 0.1 |
| 2_3090020 | 4 | 0.25 | 3 | 0 | 2 | 0 | 5 | 0.2 | 14 | 0.1429 | 4 | 0 | 2 | 0.5 | NA | NA | 2 | 0 | 8 | 0.125 |
| 2_3090027 | 4 | 0 | 4 | 0 | 2 | 0.5 | 4 | 0 | 14 | 0.0714 | 4 | 0.25 | 3 | 0.3333 | NA | NA | 3 | 0.3333 | 10 | 0.3 |
| 2_3090035 | 4 | 0.5 | 4 | 0 | 1 | 0 | 8 | 0.25 | 17 | 0.2353 | 4 | 0.5 | 3 | 0 | NA | NA | 1 | 0 | 8 | 0.25 |
| 2_3090036 | 3 | 0 | 2 | 0.5 | 2 | 0.5 | 5 | 0 | 12 | 0.1667 | 3 | 0 | 3 | 0.3333 | NA | NA | NA | NA | 6 | 0.1667 |
| 2_3091149 | 8 | 0.125 | 5 | 0 | 2 | 0 | 14 | 0.0714 | 29 | 0.069 | 6 | 0.3333 | 4 | 0.25 | 6 | 0 | 9 | 0.1111 | 25 | 0.16 |
| 2_3091172 | 5 | 0.8 | 6 | 0.5 | 4 | 0.75 | 14 | 0.8571 | 29 | 0.7586 | 3 | 1 | 5 | 1 | 6 | 0.8333 | 8 | 1 | 22 | 0.9545 |
| 2_3091190 | 5 | 0.2 | 7 | 0.1429 | 3 | 0.6667 | 13 | 0.2308 | 28 | 0.25 | 5 | 0.4 | 5 | 0.2 | 5 | 0.2 | 7 | 0.2857 | 22 | 0.2727 |
| 2_3091202 | 4 | 0 | 4 | 0 | 2 | 0 | 8 | 0.125 | 18 | 0.0556 | 3 | 0 | 8 | 0.125 | 7 | 0 | 7 | 0.1429 | 25 | 0.08 |
| 2_3091331 | NA | NA | NA | NA | NA | NA | 2 | 0.5 | 2 | 0.5 | NA | NA | NA | NA | NA | NA | 1 | 1 | 1 | 1 |
| 2_3091698 | 5 | 0.2 | 6 | 0.3333 | 2 | 0 | 6 | 0.6667 | 19 | 0.3684 | 3 | 0.3333 | 6 | 0.3333 | 4 | 0.75 | 2 | 0.5 | 15 | 0.4667 |
| 2_3091699 | 5 | 0 | 6 | 0.3333 | 3 | 0 | 5 | 0.4 | 19 | 0.2105 | 4 | 0 | 4 | 0 | 3 | 0 | 5 | 0.2 | 16 | 0.0625 |
| 2_3091717 | 4 | 0 | 8 | 0.125 | 1 | 0 | 3 | 0.3333 | 16 | 0.125 | 4 | 0.25 | 6 | 0.1667 | 4 | 0 | 5 | 0 | 19 | 0.1053 |
| 2_3091718 | 5 | 0.4 | 8 | 0.125 | 2 | 0.5 | 4 | 0.25 | 19 | 0.2632 | 5 | 0 | 7 | 0.4286 | 3 | 0 | 5 | 0 | 20 | 0.15 |
| 2_3091726 | 3 | 0 | 7 | 0.1429 | 1 | 0 | 4 | 0.25 | 15 | 0.1333 | 4 | 0 | 5 | 0.2 | 3 | 0 | 5 | 0 | 17 | 0.0588 |
| 2_3091741 | 5 | 0.2 | 7 | 0.1429 | 2 | 0 | 3 | 0 | 17 | 0.1176 | 5 | 0 | 6 | 0.1667 | 4 | 0 | 5 | 0 | 20 | 0.05 |
| 2_3091803 | 5 | 0.2 | 6 | 0 | 1 | 1 | 5 | 0.6 | 17 | 0.2941 | 4 | 0.25 | 6 | 0.6667 | 4 | 0 | 4 | 0.5 | 18 | 0.3889 |
| 2_3091805 | 5 | 0.2 | 7 | 0.8571 | 2 | 1 | 5 | 0.8 | 19 | 0.6842 | 5 | 0.8 | 6 | 0.5 | 4 | 0.5 | 5 | 0.6 | 20 | 0.6 |
| 2_3091839 | 3 | 0 | 6 | 0.1667 | 1 | 1 | 5 | 0 | 15 | 0.1333 | 5 | 0 | 7 | 0.2857 | 4 | 0 | 3 | 0 | 19 | 0.1053 |
| 2_3091843 | 5 | 0.2 | 6 | 0 | 1 | 0 | 5 | 0.2 | 17 | 0.1176 | 5 | 0 | 7 | 0.1429 | 4 | 0.25 | 2 | 0 | 18 | 0.1111 |
| 2_3092220 | 1 | 0 | 2 | 0.5 | 1 | 0 | NA | NA | 4 | 0.25 | 1 | 0 | 1 | 0 | 1 | 0 | 2 | 0.5 | 5 | 0.2 |
| 2_3092225 | 1 | 0 | 2 | 1 | NA | NA | NA | NA | 3 | 0.6667 | 1 | 1 | 1 | 1 | 1 | 1 | 2 | 0.5 | 5 | 0.8 |
| 2_3092233 | 1 | 0 | 2 | 0.5 | 1 | 0 | NA | NA | 4 | 0.25 | 1 | 0 | 1 | 1 | 1 | 0 | 2 | 0.5 | 5 | 0.4 |
| 2_3092245 | NA | NA | 2 | 0.5 | 1 | 0 | NA | NA | 3 | 0.3333 | 1 | 0 | 1 | 1 | 1 | 0 | 2 | 0 | 5 | 0.2 |
| 2_3092286 | 1 | 1 | 3 | 0.3333 | 1 | 0 | NA | NA | 5 | 0.4 | 1 | 0 | 1 | 0 | 1 | 1 | 2 | 0.5 | 5 | 0.4 |
| 2_3092291 | 1 | 0 | 2 | 0.5 | 1 | 1 | NA | NA | 4 | 0.5 | 1 | 0 | 1 | 0 | 1 | 1 | 2 | 0.5 | 5 | 0.4 |
| 2_3092303 | 1 | 1 | 2 | 1 | 1 | 0 | NA | NA | 4 | 0.75 | 1 | 1 | 1 | 0 | 1 | 0 | 2 | 0.5 | 5 | 0.4 |
| 2_3092312 | 4 | 0.25 | 7 | 0 | 2 | 0.5 | 5 | 0 | 18 | 0.1111 | 4 | 0 | 6 | 0 | 4 | 0 | 4 | 0.5 | 18 | 0.1111 |
| 2_3104214 | 2 | 0.5 | 3 | 0 | 3 | 0.3333 | NA | NA | 8 | 0.25 | 2 | 0.5 | NA | NA | 1 | 0 | NA | NA | 3 | 0.3333 |
| 2_3110436 | 4 | 0.75 | 2 | 1 | NA | NA | 4 | 0.25 | 10 | 0.6 | 1 | 1 | 1 | 1 | 1 | 0 | 6 | 0.8333 | 9 | 0.7778 |
| 2_3110438 | 4 | 0.25 | 2 | 0 | NA | NA | 4 | 0.25 | 10 | 0.2 | 1 | 0 | 2 | 0.5 | 2 | 0 | 6 | 0.1667 | 11 | 0.1818 |
| 2_3110449 | 3 | 0.6667 | 3 | 0.6667 | NA | NA | 4 | 0.75 | 10 | 0.7 | 1 | 1 | 2 | 0.5 | 2 | 1 | 5 | 1 | 10 | 0.9 |
| 2_3110455 | 4 | 0.25 | 2 | 0 | NA | NA | 4 | 0 | 10 | 0.1 | NA | NA | 1 | 0 | NA | NA | 5 | 0.2 | 6 | 0.1667 |
| 2_3110459 | 4 | 0.75 | 3 | 1 | NA | NA | 4 | 0.5 | 11 | 0.7273 | 1 | 1 | 2 | 1 | 2 | 0.5 | 7 | 0.8571 | 12 | 0.8333 |
| 2_3110503 | 2 | 0 | 1 | 0 | NA | NA | 2 | 0.5 | 5 | 0.2 | 1 | 0 | NA | NA | NA | NA | 4 | 0.25 | 5 | 0.2 |
| 2_3110510 | 3 | 0.3333 | 1 | 0 | NA | NA | 2 | 0 | 6 | 0.1667 | 1 | 0 | 1 | 0 | NA | NA | 5 | 0.2 | 7 | 0.1429 |
| 2_3110521 | 3 | 0.3333 | 1 | 0 | NA | NA | 1 | 0 | 5 | 0.2 | 1 | 1 | 1 | 0 | NA | NA | 5 | 0.2 | 7 | 0.2857 |
| 2_3110535 | 3 | 0 | 1 | 1 | NA | NA | 2 | 0.5 | 6 | 0.3333 | 1 | 1 | 1 | 1 | NA | NA | 6 | 0.3333 | 8 | 0.5 |
| 2_3110547 | 3 | 0.3333 | 1 | 0 | NA | NA | 2 | 0.5 | 6 | 0.3333 | 1 | 1 | 1 | 1 | 1 | 1 | 4 | 0.25 | 7 | 0.5714 |
| 2_3110569 | 3 | 0.6667 | 2 | 0.5 | NA | NA | 1 | 1 | 6 | 0.6667 | 1 | 1 | 1 | 0 | 1 | 1 | 6 | 0.3333 | 9 | 0.4444 |
| 2_3110649 | 5 | 0.2 | 5 | 0.2 | 2 | 0.5 | 4 | 0 | 16 | 0.1875 | 3 | 0 | 2 | 0 | 5 | 0.2 | 5 | 0 | 15 | 0.0667 |
| 2_3118254 | 8 | 0.25 | 11 | 0.4545 | 8 | 0.25 | 8 | 0.25 | 35 | 0.3143 | 7 | 0 | 12 | 0.0833 | 14 | 0.4286 | 12 | 0.25 | 45 | 0.2222 |
| 2_3118262 | 12 | 1 | 14 | 0.9286 | 11 | 1 | 13 | 1 | 50 | 0.98 | 11 | 0.9091 | 16 | 0.9375 | 19 | 1 | 20 | 1 | 66 | 0.9697 |
| 2_3118264 | 8 | 0.25 | 13 | 0.3077 | 11 | 0.3636 | 11 | 0.0909 | 43 | 0.2558 | 12 | 0.25 | 13 | 0.2308 | 16 | 0.1875 | 15 | 0.0667 | 56 | 0.1786 |
| 2_3118267 | 10 | 0.2 | 13 | 0.3077 | 10 | 0.2 | 11 | 0.2727 | 44 | 0.25 | 11 | 0.1818 | 15 | 0.4 | 24 | 0.1667 | 15 | 0.2 | 65 | 0.2308 |
| 2_3118270 | 8 | 0.75 | 10 | 0.5 | 11 | 0.4545 | 10 | 0.7 | 39 | 0.5897 | 9 | 0.7778 | 10 | 0.4 | 21 | 0.6667 | 15 | 0.3333 | 55 | 0.5455 |
| 2_3118281 | 13 | 0.1538 | 13 | 0.3077 | 14 | 0.4286 | 16 | 0.1875 | 56 | 0.2679 | 13 | 0.2308 | 17 | 0.1176 | 26 | 0.1923 | 17 | 0.2941 | 73 | 0.2055 |
| 2_3118285 | 12 | 0.25 | 13 | 0 | 14 | 0.2143 | 16 | 0.1875 | 55 | 0.1636 | 11 | 0 | 17 | 0.0588 | 24 | 0.0833 | 16 | 0.125 | 68 | 0.0735 |
| 2_3118348 | 15 | 0.1333 | 19 | 0.3684 | 16 | 0.0625 | 24 | 0.0417 | 74 | 0.1486 | 19 | 0.1053 | 22 | 0.1818 | 33 | 0.1212 | 22 | 0.0909 | 96 | 0.125 |
| 2_3118355 | 14 | 0.1429 | 17 | 0.5294 | 19 | 0.1579 | 23 | 0.1739 | 73 | 0.2466 | 17 | 0.5294 | 24 | 0.25 | 32 | 0.1875 | 20 | 0.3 | 93 | 0.2903 |
| 2_3118454 | 24 | 0.125 | 30 | 0 | 31 | 0.0645 | 37 | 0.0541 | 122 | 0.0574 | 23 | 0.087 | 31 | 0.0323 | 36 | 0.0556 | 29 | 0.069 | 119 | 0.0588 |
| 2_3118535 | 29 | 0.0345 | 34 | 0.0882 | 39 | 0.1026 | 43 | 0.1163 | 145 | 0.0897 | 31 | 0.129 | 41 | 0.1463 | 46 | 0.1304 | 33 | 0.0606 | 151 | 0.1192 |
| 2_3118802 | NA | NA | NA | NA | 1 | 1 | NA | NA | 1 | 1 | NA | NA | 1 | 1 | NA | NA | NA | NA | 1 | 1 |
| 2_3150706 | 3 | 0 | 4 | 0.25 | 2 | 0.5 | 2 | 0 | 11 | 0.1818 | NA | NA | 3 | 0.3333 | NA | NA | 4 | 0 | 7 | 0.1429 |
| 2_3154476 | 1 | 1 | 1 | 0 | NA | NA | NA | NA | 2 | 0.5 | NA | NA | NA | NA | 1 | 1 | NA | NA | 1 | 1 |
| 2_3154512 | 1 | 1 | 1 | 0 | NA | NA | NA | NA | 2 | 0.5 | NA | NA | NA | NA | 1 | 1 | NA | NA | 1 | 1 |
| 2_3169071 | NA | NA | 2 | 0.5 | 1 | 0 | NA | NA | 3 | 0.3333 | 2 | 1 | 6 | 0.8333 | 3 | 1 | 2 | 1 | 13 | 0.9231 |
| 2_3169106 | NA | NA | 2 | 0.5 | 1 | 1 | NA | NA | 3 | 0.6667 | 2 | 0 | 7 | 0.5714 | 3 | 0.3333 | 1 | 1 | 13 | 0.4615 |
| 2_3169130 | NA | NA | 2 | 0 | 1 | 1 | NA | NA | 3 | 0.3333 | 2 | 0.5 | 6 | 0 | 4 | 0 | 2 | 0.5 | 14 | 0.1429 |
| 2_3169137 | NA | NA | 1 | 0 | 1 | 1 | NA | NA | 2 | 0.5 | 2 | 1 | 7 | 1 | 4 | 1 | 2 | 1 | 15 | 1 |
| 2_3169143 | NA | NA | 3 | 0 | 1 | 1 | NA | NA | 4 | 0.25 | 2 | 0.5 | 6 | 0.1667 | 4 | 0.25 | 2 | 0.5 | 14 | 0.2857 |
| 2_3169161 | NA | NA | 1 | 1 | 1 | 1 | NA | NA | 2 | 1 | 2 | 1 | 2 | 1 | 2 | 1 | 1 | 1 | 7 | 1 |
| 2_3169165 | NA | NA | 2 | 0.5 | 1 | 0 | NA | NA | 3 | 0.3333 | 1 | 0 | 6 | 0.5 | 4 | 0.25 | 1 | 1 | 12 | 0.4167 |

| | | | | | | | | | | | | | | | | | | | | |
|---|---|---|---|---|---|---|---|---|---|---|---|---|---|---|---|---|---|---|---|---|
| 2_3169170 | NA | NA | 3 | 0.3333 | 1 | 0 | NA | NA | 4 | 0.25 | 2 | 0 | 7 | 0.1429 | 5 | 0.6 | 2 | 0 | 16 | 0.25 |
| 2_3169173 | NA | NA | 3 | 1 | 1 | 1 | NA | NA | 4 | 1 | 2 | 1 | 7 | 1 | 4 | 1 | 2 | 1 | 15 | 1 |
| 2_3169197 | NA | NA | 1 | 1 | 1 | 0 | 1 | 1 | 3 | 0.6667 | 2 | 1 | 6 | 0.8333 | 3 | 0.6667 | 2 | 1 | 13 | 0.8462 |
| 2_3169198 | NA | NA | 2 | 0 | 2 | 1 | 1 | 1 | 5 | 0.6 | 2 | 1 | 7 | 1 | 5 | 0.8 | 3 | 1 | 17 | 0.9412 |
| 2_3169202 | NA | NA | 3 | 0.6667 | 1 | 1 | 2 | 0.5 | 6 | 0.6667 | 2 | 1 | 7 | 0.8571 | 3 | 0.3333 | 3 | 1 | 15 | 0.8 |
| 2_3169217 | NA | NA | 3 | 0.3333 | 2 | 0.5 | 2 | 0.5 | 7 | 0.4286 | 2 | 1 | 7 | 0.4286 | 6 | 0.1667 | 3 | 0.3333 | 18 | 0.3889 |
| 2_3169229 | NA | NA | 2 | 1 | 2 | 1 | 2 | 1 | 6 | 1 | 2 | 1 | 8 | 1 | 7 | 1 | 3 | 1 | 20 | 1 |
| 2_3169235 | NA | NA | 3 | 1 | 2 | 0.5 | NA | NA | 5 | 0.8 | 1 | 1 | 4 | 1 | 2 | 1 | 1 | 1 | 8 | 1 |
| 2_3169253 | NA | NA | 1 | 1 | 2 | 1 | 1 | 1 | 4 | 1 | 2 | 0 | 8 | 0.875 | 4 | 0.75 | 3 | 1 | 17 | 0.7647 |
| 2_3169281 | NA | NA | 2 | 0 | 2 | 1 | 1 | 0 | 5 | 0.4 | 1 | 0 | 8 | 0.375 | 8 | 0.375 | 3 | 1 | 20 | 0.45 |
| 2_3169290 | NA | NA | 4 | 0.25 | 1 | 0 | 1 | 1 | 6 | 0.3333 | 1 | 0 | 6 | 0 | 6 | 0.1667 | 2 | 0 | 15 | 0.0667 |
| 2_3169338 | NA | NA | 6 | 0.5 | 2 | 0 | 2 | 0.5 | 10 | 0.4 | 2 | 0 | 11 | 0.1818 | 7 | 0 | 2 | 1 | 22 | 0.1818 |
| 2_3170364 | NA | NA | 1 | 0 | 4 | 0.5 | 2 | 0.5 | 7 | 0.4286 | 1 | 1 | 3 | 1 | 1 | 1 | 1 | 0 | 6 | 0.8333 |
| 2_3170376 | NA | NA | 1 | 1 | 3 | 0.3333 | 2 | 1 | 6 | 0.6667 | 1 | 0 | 3 | 0.3333 | 1 | 0 | 1 | 0 | 6 | 0.1667 |
| 2_3170385 | NA | NA | 1 | 0 | 3 | 0.3333 | 2 | 0.5 | 6 | 0.3333 | 1 | 1 | 3 | 0.6667 | 1 | 1 | 1 | 0 | 6 | 0.6667 |
| 2_3170397 | NA | NA | 1 | 0 | 3 | 0 | 2 | 1 | 6 | 0.3333 | 1 | 1 | 2 | 0.5 | 1 | 0 | 1 | 0 | 5 | 0.4 |
| 2_3170481 | NA | NA | NA | NA | 1 | 0 | 1 | 1 | 2 | 0.5 | NA | NA | 1 | 1 | 1 | 1 | NA | NA | 2 | 1 |
| 2_3170496 | NA | NA | 1 | 0 | 2 | 0 | 2 | 0.5 | 5 | 0.2 | NA | NA | 2 | 0.5 | NA | NA | 1 | 0 | 3 | 0.3333 |
| 2_3170512 | NA | NA | 1 | 1 | 1 | 1 | 2 | 1 | 4 | 1 | 1 | 1 | 1 | 1 | NA | NA | 1 | 1 | 3 | 1 |
| 2_3170522 | NA | NA | NA | NA | 1 | 1 | 1 | 1 | 2 | 1 | 1 | 0 | NA | NA | 1 | 1 | 1 | 1 | 3 | 0.6667 |
| 2_3170525 | NA | NA | 1 | 1 | 2 | 0 | 2 | 0.5 | 5 | 0.4 | 1 | 0 | 1 | 1 | 1 | 1 | 1 | 0 | 4 | 0.5 |
| 2_3170532 | NA | NA | 1 | 1 | 2 | 1 | NA | NA | 3 | 1 | 1 | 1 | 1 | 1 | 1 | 1 | 1 | 1 | 4 | 1 |
| 2_3170545 | NA | NA | 1 | 1 | 1 | 1 | 1 | 1 | 3 | 1 | 1 | 0 | NA | NA | NA | NA | 1 | 1 | 2 | 0.5 |
| 2_3170546 | NA | NA | 1 | 1 | 1 | 0 | 2 | 1 | 4 | 0.75 | 1 | 1 | 1 | 1 | 1 | 1 | 1 | 0 | 4 | 0.75 |
| 2_3170554 | NA | NA | 1 | 1 | 1 | 1 | 2 | 1 | 4 | 1 | 1 | 1 | 1 | 1 | NA | NA | 1 | 1 | 3 | 1 |
| 2_3170556 | NA | NA | 1 | 1 | NA | NA | 2 | 1 | 3 | 1 | 1 | 1 | 1 | 1 | NA | NA | 1 | 1 | 3 | 1 |
| 2_3170560 | NA | NA | 1 | 1 | 1 | 1 | 2 | 1 | 4 | 1 | 1 | 1 | 1 | 1 | NA | NA | 1 | 0 | 3 | 0.6667 |
| 2_3170564 | NA | NA | NA | NA | 1 | 1 | 1 | 1 | 2 | 1 | 1 | 1 | 1 | 1 | NA | NA | 1 | 1 | 3 | 1 |
| 2_3170569 | NA | NA | NA | NA | 1 | 0 | 2 | 0.5 | 3 | 0.3333 | 1 | 0 | 1 | 1 | NA | NA | 1 | 1 | 3 | 0.6667 |
| 2_3170628 | NA | NA | 1 | 1 | 1 | 0 | NA | NA | 2 | 0.5 | 1 | 1 | NA | NA | NA | NA | 1 | 0 | 2 | 0.5 |
| 2_3170665 | 1 | 0 | 1 | 0 | 1 | 0 | 2 | 0.5 | 5 | 0.2 | 1 | 0 | NA | NA | NA | NA | 1 | 1 | 2 | 0.5 |
| 2_3192336 | NA | NA | 2 | 0.5 | 3 | 0.3333 | 1 | 0 | 6 | 0.3333 | NA | NA | 1 | 1 | 3 | 0.6667 | 2 | 0.5 | 6 | 0.6667 |
| 2_3192376 | NA | NA | 2 | 1 | 2 | 1 | 1 | 1 | 5 | 1 | NA | NA | 1 | 1 | 2 | 1 | 2 | 1 | 5 | 1 |
| 2_3192384 | NA | NA | 3 | 0.3333 | 3 | 0 | 2 | 0.5 | 8 | 0.25 | NA | NA | 1 | 1 | 2 | 0 | 2 | 0 | 5 | 0.2 |
| 2_3192412 | NA | NA | 2 | 0 | 2 | 0.5 | 2 | 0.5 | 6 | 0.3333 | NA | NA | 2 | 0 | 4 | 0.75 | 2 | 0 | 8 | 0.375 |
| 2_3192416 | NA | NA | 3 | 0 | 3 | 0.3333 | 2 | 0 | 8 | 0.125 | NA | NA | 2 | 0 | 4 | 0 | 2 | 0.5 | 8 | 0.125 |
| 2_3192427 | NA | NA | 3 | 0.6667 | 2 | 1 | 2 | 0.5 | 7 | 0.7143 | NA | NA | 2 | 1 | 4 | 0.75 | 2 | 1 | 8 | 0.875 |
| 2_3192439 | NA | NA | 3 | 0.3333 | 3 | 0 | 2 | 0.5 | 8 | 0.25 | NA | NA | 2 | 1 | 3 | 0.6667 | 2 | 0.5 | 7 | 0.7143 |
| 2_3192440 | NA | NA | 3 | 1 | 3 | 1 | 2 | 1 | 8 | 1 | NA | NA | 1 | 1 | 4 | 1 | 2 | 1 | 7 | 1 |
| 2_3192462 | NA | NA | 1 | 1 | 2 | 0.5 | 2 | 1 | 5 | 0.8 | NA | NA | 4 | 1 | 4 | 1 | 3 | 0.6667 | 11 | 0.9091 |
| 2_3192463 | NA | NA | 2 | 0.5 | 4 | 0.25 | 3 | 0 | 9 | 0.2222 | NA | NA | 4 | 0 | 2 | 0.5 | 2 | 0 | 8 | 0.125 |
| 2_3192482 | NA | NA | 3 | 1 | 1 | 1 | 2 | 0.5 | 6 | 0.8333 | NA | NA | 4 | 0.75 | 2 | 0.5 | 3 | 1 | 9 | 0.7778 |
| 2_3194222 | NA | NA | 4 | 0.25 | 7 | 0.1429 | 6 | 0.1667 | 17 | 0.1765 | NA | NA | 8 | 0.5 | 4 | 0.25 | 6 | 0.1667 | 18 | 0.3333 |
| 2_3194223 | NA | NA | 6 | 0.1667 | 6 | 0.5 | 4 | 0.5 | 16 | 0.375 | NA | NA | 8 | 0.25 | 6 | 0.5 | 6 | 0.5 | 20 | 0.4 |
| 2_3194230 | NA | NA | 5 | 0.2 | 5 | 0.6 | 6 | 0 | 16 | 0.25 | 1 | 0 | 4 | 0.25 | 5 | 0.6 | 4 | 0.25 | 14 | 0.3571 |
| 2_3194238 | NA | NA | NA | NA | NA | NA | 1 | 1 | 1 | 1 | NA | NA | 1 | 1 | 1 | 0 | 2 | 0.5 | 4 | 0.5 |
| 2_3194241 | NA | NA | NA | NA | 1 | 1 | 1 | 1 | 2 | 1 | NA | NA | NA | NA | 1 | 1 | 2 | 1 | 3 | 1 |
| 2_3194282 | NA | NA | NA | NA | 1 | 1 | 1 | 1 | 2 | 1 | NA | NA | 1 | 1 | 1 | 1 | 2 | 1 | 4 | 1 |
| 2_3194298 | NA | NA | NA | NA | 1 | 1 | 1 | 1 | 2 | 1 | NA | NA | 1 | 1 | NA | NA | 2 | 1 | 3 | 1 |
| 2_3198578 | 2 | 0.5 | NA | NA | 2 | 0.5 | NA | NA | 4 | 0.5 | 1 | 0 | NA | NA | 1 | 0 | 2 | 0.5 | 4 | 0.25 |
| 2_3199970 | 3 | 0.3333 | NA | NA | 4 | 0.5 | 1 | 0 | 8 | 0.375 | 2 | 1 | 1 | 1 | 1 | 1 | 4 | 0.5 | 8 | 0.75 |
| 2_3199972 | 2 | 0.5 | NA | NA | 4 | 1 | 2 | 1 | 8 | 0.875 | 2 | 0.5 | 1 | 0 | 2 | 1 | 4 | 0.75 | 9 | 0.6667 |
| 2_3200006 | 3 | 0.6667 | 1 | 0 | 4 | 1 | 3 | 1 | 11 | 0.8182 | 1 | 1 | 2 | 0.5 | 2 | 1 | 5 | 1 | 10 | 0.9 |
| 2_3200013 | NA | NA | NA | NA | 1 | 1 | 1 | 1 | 2 | 1 | 1 | 1 | NA | NA | NA | NA | 1 | 1 | 2 | 1 |
| 2_3200016 | 2 | 0.5 | NA | NA | 5 | 0 | 2 | 0.5 | 9 | 0.2222 | 2 | 0.5 | 1 | 0 | 2 | 0 | 4 | 0 | 9 | 0.1111 |
| 2_3200056 | 4 | 0.5 | 1 | 1 | 6 | 0.8333 | 2 | 1 | 13 | 0.7692 | 2 | 0.5 | 3 | 1 | 2 | 0.5 | 4 | 0.5 | 11 | 0.6364 |
| 2_3200058 | 4 | 0.5 | 2 | 0 | 4 | 0.25 | 4 | 0.5 | 14 | 0.3571 | 2 | 0 | 3 | 0.3333 | 1 | 0 | 2 | 0.5 | 8 | 0.25 |
| 2_3200075 | 5 | 1 | 1 | 1 | 4 | 1 | 3 | 1 | 13 | 1 | 2 | 1 | 3 | 1 | 2 | 1 | 4 | 1 | 11 | 1 |
| 2_3200080 | 3 | 0.6667 | 2 | 0 | 5 | 0.8 | 3 | 0.6667 | 13 | 0.6154 | 2 | 0.5 | 3 | 0.3333 | NA | NA | 4 | 0.75 | 9 | 0.5556 |
| 2_3210301 | 1 | 1 | 4 | 0.5 | 3 | 0.3333 | 2 | 1 | 10 | 0.6 | 3 | 0.3333 | 1 | 1 | 1 | 1 | 1 | 1 | 6 | 0.6667 |
| 2_3210314 | 1 | 0 | 3 | 0.3333 | 3 | 0.3333 | 2 | 0.5 | 9 | 0.3333 | 3 | 0.3333 | NA | NA | 2 | 0.5 | 1 | 1 | 6 | 0.5 |
| 2_3210332 | NA | NA | 3 | 0 | 4 | 0.25 | 2 | 0.5 | 9 | 0.2222 | 2 | 0.5 | 2 | 0 | 2 | 0.5 | 1 | 0 | 7 | 0.2857 |
| 2_3210339 | 1 | 0 | 4 | 0 | 4 | 0 | 2 | 0.5 | 11 | 0.0909 | 2 | 0.5 | 1 | 0 | 2 | 0 | 1 | 0 | 6 | 0.1667 |
| 2_3210350 | 1 | 0 | 3 | 0 | 2 | 0.5 | 2 | 0 | 8 | 0.125 | 2 | 0 | 1 | 0 | 1 | 1 | 1 | 0 | 5 | 0.2 |
| 2_3210357 | 2 | 0 | 4 | 0.25 | 4 | 0 | 2 | 0 | 12 | 0.0833 | 2 | 0.5 | 2 | 0 | 2 | 1 | 1 | 0 | 7 | 0.4286 |

| | | | | | | | | | | | | | | | | | | | | |
|---|---|---|---|---|---|---|---|---|---|---|---|---|---|---|---|---|---|---|---|---|
| 2_321036 | 2 | 0.5 | 5 | 0.2 | 4 | 0 | 2 | 0 | 13 | 0.1538 | 2 | 0 | 2 | 1 | 2 | 0 | 1 | 0 | 7 | 0.2857 |
| 2_321040 | 2 | 0 | 6 | 0.8333 | 4 | 0.75 | 2 | 1 | 14 | 0.7143 | 2 | 1 | 3 | 0.6667 | 4 | 0.75 | 1 | 0 | 10 | 0.7 |
| 2_321041 | 2 | 1 | 3 | 1 | 3 | 0 | 2 | 0.5 | 10 | 0.6 | 3 | 0.3333 | 2 | 0 | 2 | 0.5 | 1 | 1 | 8 | 0.375 |
| 2_321041 | 3 | 0.3333 | 6 | 0.1667 | 4 | 0.25 | 1 | 0 | 14 | 0.2143 | 3 | 0 | 3 | 0 | 3 | 0.3333 | 1 | 0 | 10 | 0.1 |
| 2_321041 | NA | NA | 5 | 0.2 | 1 | 0 | 1 | 0 | 7 | 0.1429 | 2 | 0 | 3 | 0 | 4 | 0.25 | 1 | 0 | 10 | 0.1 |
| 2_321042 | 3 | 0.3333 | 6 | 0 | 4 | 0 | 3 | 0 | 16 | 0.0625 | 2 | 0 | 3 | 0 | 5 | 0.2 | 1 | 0 | 11 | 0.0909 |
| 2_321042 | 2 | 0 | 6 | 0.1667 | 3 | 0.3333 | 3 | 0 | 14 | 0.1429 | 3 | 0 | 1 | 0 | 3 | 0.3333 | 1 | 0 | 8 | 0.125 |
| 2_321043 | 3 | 0.6667 | 6 | 0.6667 | 4 | 0.75 | 3 | 0.3333 | 16 | 0.625 | 3 | 0.6667 | 3 | 0.3333 | 5 | 0.4 | 1 | 1 | 12 | 0.5 |
| 2_321043 | 3 | 0 | 6 | 0.1667 | 2 | 0 | 3 | 0.3333 | 14 | 0.1429 | 3 | 0 | 2 | 0 | 5 | 0.4 | 1 | 0 | 11 | 0.1818 |
| 2_321043 | 3 | 0 | 6 | 0.1667 | 3 | 0 | 2 | 0 | 14 | 0.0714 | 1 | 0 | 3 | 0.3333 | 4 | 0 | 1 | 0 | 9 | 0.1111 |
| 2_321043 | 3 | 0.3333 | 6 | 0 | 4 | 0.25 | 3 | 0 | 16 | 0.125 | 2 | 0 | 3 | 0.3333 | 5 | 0 | NA | NA | 10 | 0.1 |
| 2_321044 | 3 | 0.3333 | 6 | 0.1667 | 4 | 0 | 3 | 0.3333 | 16 | 0.1875 | 2 | 0 | 3 | 0 | 2 | 0.5 | 1 | 0 | 8 | 0.125 |
| 2_321045 | 3 | 0 | 5 | 0 | 4 | 0.25 | 3 | 0 | 15 | 0.0667 | 2 | 0 | 3 | 0.3333 | 3 | 0.3333 | 1 | 0 | 9 | 0.2222 |
| 2_321046 | 3 | 0.3333 | 6 | 0.3333 | 3 | 0 | 3 | 0 | 15 | 0.2 | 3 | 0 | 4 | 0.25 | 6 | 0 | 1 | 0 | 14 | 0.0714 |
| 2_321048 | 3 | 0 | 5 | 0.4 | 2 | 0 | 4 | 0.25 | 14 | 0.2143 | 3 | 0 | 3 | 0 | 6 | 0.1667 | 2 | 0 | 14 | 0.0714 |
| 2_321095 | 4 | 0 | 7 | 0 | 6 | 0.1667 | 2 | 0 | 19 | 0.0526 | 4 | 0 | 7 | 0.2857 | 10 | 0 | 3 | 0 | 24 | 0.0833 |
| 2_321208 | 7 | 0.4286 | 13 | 0.6154 | 11 | 0.5455 | 11 | 0.3636 | 42 | 0.5 | 3 | 0.3333 | 12 | 0.5833 | 9 | 0.8889 | 8 | 0.5 | 32 | 0.625 |
| 2_321208 | 14 | 0.6429 | 12 | 0.6667 | 10 | 0.8 | 12 | 0.4167 | 48 | 0.625 | 6 | 0.6667 | 12 | 0.8333 | 13 | 0.7692 | 14 | 0.7143 | 45 | 0.7556 |
| 2_321209 | 13 | 0.6154 | 9 | 0.8889 | 8 | 0.75 | 10 | 0.9 | 40 | 0.775 | 6 | 1 | 10 | 0.6 | 17 | 0.7647 | 12 | 0.6667 | 45 | 0.7333 |
| 2_321209 | 15 | 0.4 | 12 | 0.25 | 12 | 0.4167 | 13 | 0.2308 | 52 | 0.3269 | 7 | 0.1429 | 14 | 0.5714 | 14 | 0.2143 | 12 | 0.4167 | 47 | 0.3617 |
| 2_321209 | 15 | 0.8 | 11 | 1 | 12 | 0.9167 | 12 | 0.9167 | 50 | 0.9 | 5 | 1 | 13 | 0.9231 | 14 | 1 | 13 | 0.7692 | 45 | 0.9111 |
| 2_321211 | 18 | 0.3889 | 16 | 0.3125 | 12 | 0.1667 | 17 | 0.1176 | 63 | 0.254 | 9 | 0.4444 | 14 | 0.1429 | 18 | 0.0556 | 17 | 0.2941 | 58 | 0.2069 |
| 2_321213 | 15 | 0.4667 | 15 | 0.4667 | 6 | 0.6667 | 14 | 0.2143 | 50 | 0.42 | 9 | 0.2222 | 15 | 0.4 | 15 | 0.2667 | 12 | 0.1667 | 51 | 0.2745 |
| 2_321214 | 20 | 0.1 | 19 | 0.1579 | 9 | 0.1111 | 18 | 0.1667 | 66 | 0.1364 | 10 | 0.1 | 15 | 0.2 | 20 | 0.15 | 16 | 0.1875 | 61 | 0.1639 |
| 2_321215 | 14 | 0.1429 | 13 | 0 | 9 | 0 | 16 | 0.0625 | 52 | 0.0577 | 9 | 0.3333 | 11 | 0 | 13 | 0 | 12 | 0.0833 | 45 | 0.0889 |
| 2_321217 | 17 | 0.0588 | 21 | 0.0476 | 10 | 0.1 | 19 | 0.0526 | 67 | 0.0597 | 11 | 0.0909 | 20 | 0.1 | 21 | 0.0952 | 19 | 0.1579 | 71 | 0.1127 |
| 2_321217 | 20 | 0 | 24 | 0.0833 | 12 | 0.1667 | 21 | 0.0476 | 77 | 0.0649 | 15 | 0.2 | 23 | 0.0435 | 24 | 0.0833 | 23 | 0.0435 | 85 | 0.0824 |
| 2_321233 | 32 | 0.0938 | 30 | 0.1 | 23 | 0.1739 | 31 | 0.1613 | 116 | 0.1293 | 22 | 0.0455 | 25 | 0.12 | 28 | 0.1071 | 24 | 0.1667 | 99 | 0.1111 |
| 2_321242 | 41 | 0.0976 | 39 | 0.1795 | 25 | 0.2 | 31 | 0.1935 | 136 | 0.1618 | 25 | 0.04 | 34 | 0.0882 | 35 | 0.2286 | 31 | 0.0968 | 125 | 0.12 |
| 2_321445 | 2 | 1 | 2 | 0.5 | 1 | 0 | NA | NA | 5 | 0.6 | 2 | 0.5 | NA | NA | NA | NA | 1 | 0 | 3 | 0.3333 |
| 2_321457 | 3 | 0.3333 | 2 | 0.5 | 1 | 1 | NA | NA | 6 | 0.5 | 2 | 1 | NA | NA | NA | NA | 1 | 0 | 3 | 0.6667 |
| 2_321455 | 9 | 0.2222 | 7 | 0.1429 | 7 | 0 | 8 | 0 | 31 | 0.0968 | 5 | 0.2 | 5 | 0 | 3 | 0 | 6 | 0 | 19 | 0.0526 |
| 2_321455 | 11 | 0.2727 | 6 | 0.5 | 6 | 0.3333 | 7 | 0 | 30 | 0.2667 | 4 | 0.25 | 5 | 0.6 | 3 | 0.3333 | 5 | 0.4 | 17 | 0.4118 |
| 2_321457 | 7 | 0 | 6 | 0.3333 | 5 | 0 | 6 | 0 | 24 | 0.0833 | 4 | 0.5 | 5 | 0 | 3 | 0.6667 | 4 | 0 | 16 | 0.25 |
| 2_321458 | 8 | 0.25 | 5 | 0 | 6 | 0.3333 | 8 | 0.125 | 27 | 0.1852 | 4 | 0.25 | 5 | 0 | 3 | 0 | 5 | 0.2 | 17 | 0.1176 |
| 2_321496 | 7 | 0 | 3 | 0 | 5 | 0.2 | 5 | 0.2 | 20 | 0.1 | 3 | 0.6667 | 2 | 0 | 1 | 1 | 5 | 0.4 | 11 | 0.4545 |
| 2_321966 | 1 | 1 | 5 | 0.8 | 4 | 0.75 | NA | NA | 10 | 0.8 | 3 | 0.6667 | 7 | 0.4286 | 5 | 0.8 | 4 | 0.75 | 19 | 0.6316 |
| 2_321969 | 3 | 0 | 7 | 0.4286 | 6 | 0.3333 | 1 | 0 | 17 | 0.2941 | 3 | 0 | 10 | 0.1 | 7 | 0.1429 | 6 | 0 | 26 | 0.0769 |
| 2_321970 | 3 | 0.3333 | 6 | 0.1667 | 5 | 0 | 1 | 0 | 15 | 0.1333 | 3 | 0 | 9 | 0 | 7 | 0.1429 | 7 | 0.1429 | 26 | 0.0769 |
| 2_321971 | 3 | 0 | 7 | 0.2857 | 5 | 0.2 | 1 | 0 | 16 | 0.1875 | 3 | 0.3333 | 9 | 0 | 6 | 0.1667 | 7 | 0.2857 | 25 | 0.16 |
| 2_321989 | 118 | 0.0763 | 137 | 0.0584 | 87 | 0.0575 | 96 | 0.0312 | 438 | 0.0571 | 84 | 0.0952 | 88 | 0.0795 | 76 | 0.0921 | 76 | 0.0658 | 324 | 0.0833 |
| 2_321989 | 107 | 0.0561 | 117 | 0.0855 | 79 | 0.0633 | 90 | 0.0667 | 393 | 0.0687 | 86 | 0.0814 | 86 | 0.1163 | 67 | 0.0299 | 68 | 0.0441 | 307 | 0.0717 |
| 2_321989 | 119 | 0.1176 | 138 | 0.1087 | 94 | 0.1702 | 99 | 0.1313 | 450 | 0.1289 | 83 | 0.1205 | 90 | 0.1778 | 75 | 0.12 | 76 | 0.1579 | 324 | 0.1451 |
| 2_321993 | 26 | 0.2308 | 36 | 0.1111 | 17 | 0.1765 | 26 | 0.3846 | 105 | 0.219 | 14 | 0.2143 | 22 | 0.2273 | 18 | 0.3333 | 20 | 0.2 | 74 | 0.2432 |
| 2_321994 | 17 | 0.0588 | 25 | 0 | 14 | 0.2143 | 21 | 0.1429 | 77 | 0.0909 | 12 | 0.1667 | 15 | 0.2 | 14 | 0.2143 | 8 | 0.375 | 49 | 0.2245 |
| 2_321995 | 25 | 0.04 | 35 | 0.0857 | 19 | 0.0526 | 24 | 0.0417 | 103 | 0.0583 | 18 | 0 | 24 | 0 | 21 | 0.1905 | 20 | 0.15 | 83 | 0.0843 |
| 2_321997 | NA | NA | 9 | 0 | 4 | 0 | 5 | 0.4 | 18 | 0.1111 | 2 | 0 | 6 | 0 | 9 | 0.3333 | 6 | 0.1667 | 23 | 0.1739 |
| 2_321999 | 4 | 0.25 | 14 | 0.0714 | 4 | 0 | 6 | 0.3333 | 28 | 0.1429 | 2 | 0.5 | 11 | 0.0909 | 8 | 0.25 | 7 | 0.1429 | 28 | 0.1786 |
| 2_322007 | 1 | 0 | 11 | 0.0909 | 3 | 0 | 5 | 0 | 20 | 0.05 | 2 | 0.5 | 10 | 0.2 | 5 | 0 | 6 | 0.1667 | 23 | 0.1739 |
| 2_322001 | 2 | 0 | 10 | 0.1 | 4 | 0.25 | 5 | 0 | 21 | 0.0952 | 2 | 0 | 9 | 0.1111 | 8 | 0 | 7 | 0.1429 | 26 | 0.0769 |
| 2_322001 | 1 | 0 | 10 | 0.1 | 5 | 0.2 | 5 | 0 | 21 | 0.0952 | NA | NA | 6 | 0.5 | 8 | 0.25 | 6 | 0 | 20 | 0.25 |
| 2_322002 | 2 | 0 | 10 | 0.3 | 5 | 0.2 | 5 | 0.2 | 22 | 0.2273 | 2 | 0.5 | 11 | 0.6364 | 8 | 0.375 | 6 | 0.6667 | 27 | 0.5556 |
| 2_322003 | 2 | 0.5 | 6 | 0 | 2 | 0.5 | 4 | 0.5 | 14 | 0.2857 | 2 | 0.5 | 8 | 0.125 | 5 | 0.2 | 6 | 0 | 21 | 0.1429 |
| 2_322672 | 3 | 0.6667 | 1 | 0 | NA | NA | 2 | 0.5 | 6 | 0.5 | 1 | 1 | NA | NA | 1 | 0 | 1 | 1 | 3 | 0.6667 |
| 2_322681 | 5 | 0.2 | 2 | 0 | 4 | 0 | 5 | 0 | 16 | 0.0625 | 5 | 0 | 3 | 0 | 6 | 0 | 5 | 0.4 | 19 | 0.1053 |
| 2_322691 | 24 | 0.0417 | 38 | 0.1579 | 18 | 0 | 15 | 0.2 | 95 | 0.1053 | 31 | 0.1613 | 13 | 0 | 20 | 0 | 25 | 0 | 89 | 0.0562 |
| 2_322692 | 23 | 0.0435 | 39 | 0.1282 | 19 | 0.1579 | 12 | 0 | 93 | 0.0968 | 26 | 0.1154 | 10 | 0 | 15 | 0.1333 | 21 | 0.0476 | 72 | 0.0833 |
| 2_322923 | 5 | 0.4 | 10 | 0.3 | 6 | 0.5 | 7 | 0.2857 | 28 | 0.3571 | 3 | 0.6667 | 15 | 0.0667 | 7 | 0.1429 | 2 | 0 | 27 | 0.1481 |
| 2_322923 | 6 | 0.1667 | 8 | 0 | 7 | 0.2857 | 8 | 0 | 29 | 0.1034 | 3 | 0 | 13 | 0.1538 | 8 | 0.125 | 2 | 0 | 26 | 0.1154 |
| 2_322924 | 7 | 0.8571 | 9 | 0.8889 | 6 | 0.8333 | 6 | 1 | 28 | 0.8929 | 3 | 1 | 10 | 0.9 | 7 | 0.8571 | 2 | 1 | 22 | 0.9091 |
| 2_322925 | 7 | 0.1429 | 10 | 0.4 | 8 | 0.125 | 8 | 0 | 33 | 0.1818 | 4 | 0.25 | 13 | 0.2308 | 6 | 0.1667 | 2 | 0.5 | 25 | 0.24 |
| 2_322926 | 7 | 0.1429 | 10 | 0 | 7 | 0.2857 | 9 | 0.2222 | 33 | 0.1515 | 4 | 0.25 | 14 | 0.1429 | 6 | 0 | 2 | 0 | 26 | 0.1154 |
| 2_322927 | 7 | 0.1429 | 9 | 0.1111 | 8 | 0.125 | 6 | 0.1667 | 30 | 0.1333 | 5 | 0.2 | 13 | 0.3077 | 8 | 0.625 | 2 | 0.5 | 28 | 0.3929 |
| 2_323047 | 7 | 0.1429 | 9 | 0.3333 | 3 | 0 | 5 | 0.4 | 24 | 0.25 | 5 | 0.2 | 16 | 0.125 | 14 | 0.2143 | 8 | 0.5 | 43 | 0.2326 |
| 2_323052 | 5 | 0.8 | 10 | 0.6 | 5 | 0.6 | 5 | 1 | 25 | 0.72 | 4 | 0.5 | 14 | 0.6429 | 15 | 0.4667 | 9 | 0.6667 | 42 | 0.5714 |

| | | | | | | | | | | | | | | | | | | | | |
|---|---|---|---|---|---|---|---|---|---|---|---|---|---|---|---|---|---|---|---|---|
| 2_32305279 | 4 | 0 | 10 | 0.3 | 5 | 0.2 | 5 | 0.2 | 24 | 0.2083 | 4 | 0 | 13 | 0.3846 | 16 | 0.25 | 9 | 0.2222 | 42 | 0.2619 |
| 2_32305429 | 5 | 0.2 | 7 | 0.5714 | 5 | 0 | 7 | 0.5714 | 24 | 0.375 | 2 | 0.5 | 14 | 0.4286 | 17 | 0.2941 | 8 | 0.5 | 41 | 0.3902 |
| 2_32305486 | 5 | 0.4 | 7 | 0.5714 | 5 | 0.2 | 7 | 0.4286 | 24 | 0.4167 | 1 | 1 | 14 | 0.5 | 17 | 0.3529 | 7 | 0.2857 | 39 | 0.4103 |
| 2_32305502 | 5 | 0.2 | 7 | 0.2857 | 5 | 0 | 7 | 0.1429 | 24 | 0.1667 | 2 | 0 | 13 | 0.0769 | 16 | 0.1875 | 7 | 0.2857 | 38 | 0.1579 |
| 2_32305897 | 5 | 0.2 | 5 | 0 | 5 | 0 | 5 | 0.2 | 20 | 0.1 | NA | NA | 13 | 0.2308 | 16 | 0.0625 | 8 | 0.125 | 37 | 0.1351 |
| 2_32306064 | 4 | 0 | 6 | 0.3333 | 5 | 0 | 6 | 0 | 21 | 0.0952 | 1 | 0 | 14 | 0 | 16 | 0.1875 | 6 | 0.3333 | 37 | 0.1351 |
| 2_32306135 | 2 | 0.5 | 4 | 0.75 | 4 | 0.5 | 4 | 1 | 14 | 0.7143 | 1 | 1 | 10 | 0.8 | 13 | 0.6154 | 6 | 0.8333 | 30 | 0.7333 |
| 2_32306275 | 2 | 1 | 3 | 0.6667 | 4 | 0.25 | 2 | 0.5 | 11 | 0.5455 | NA | NA | 7 | 0.4286 | 13 | 0.3846 | 3 | 0.3333 | 23 | 0.3913 |
| 2_32346571 | NA | NA | NA | NA | NA | NA | 1 | 1 | 1 | 1 | NA | NA | 1 | 1 | NA | NA | 1 | 1 | 2 | 1 |
| 2_32425508 | 1 | 0 | 3 | 0.6667 | 2 | 0.5 | 1 | 0 | 7 | 0.4286 | 1 | 1 | 1 | 1 | NA | NA | NA | NA | 2 | 1 |
| 2_32425913 | 1 | 0 | 3 | 0.6667 | 1 | 1 | 2 | 0 | 7 | 0.4286 | 2 | 0.5 | 1 | 1 | 1 | 0 | NA | NA | 4 | 0.5 |
| 2_32426082 | 3 | 0 | 3 | 0.3333 | 2 | 0 | 2 | 0 | 10 | 0.1 | 2 | 0 | 1 | 0 | 2 | 0.5 | 1 | 1 | 6 | 0.3333 |
| 2_32426091 | 3 | 0.3333 | 3 | 0.3333 | 2 | 0 | 2 | 0 | 10 | 0.2 | 2 | 0 | 2 | 0 | 2 | 0 | 1 | 1 | 7 | 0.1429 |
| 2_32457652 | 4 | 0.25 | 3 | 0 | 4 | 0.25 | 2 | 0 | 13 | 0.1538 | 2 | 0 | 3 | 0 | 2 | 0 | 2 | 0.5 | 9 | 0.1111 |
| 2_32457688 | 5 | 0.6 | 4 | 0.5 | 4 | 0.75 | 1 | 1 | 14 | 0.6429 | 3 | 0.6667 | 5 | 0.8 | 2 | 0.5 | 4 | 0.5 | 14 | 0.6429 |
| 2_32457697 | 7 | 0 | 4 | 0.5 | 5 | 0 | 2 | 0 | 18 | 0.1111 | 4 | 0.25 | 6 | 0 | 1 | 0 | 3 | 0.3333 | 14 | 0.1429 |
| 2_32457707 | 6 | 0 | 4 | 0.25 | 5 | 0.4 | 3 | 0 | 18 | 0.1667 | 4 | 0.25 | 5 | 0.4 | 2 | 0 | 4 | 0.25 | 15 | 0.2667 |
| 2_32457725 | 6 | 0.8333 | 4 | 0.5 | 3 | 0.6667 | 3 | 0.3333 | 16 | 0.625 | 3 | 1 | 3 | 0.3333 | 2 | 1 | 3 | 0.3333 | 11 | 0.6364 |
| 2_32458382 | 5 | 0 | 4 | 0 | 5 | 0.2 | 3 | 0 | 17 | 0.0588 | 6 | 0 | 9 | 0.2222 | 3 | 0 | 3 | 0.6667 | 21 | 0.1905 |
| 2_32458743 | 9 | 0 | 3 | 0.3333 | 5 | 0.2 | 3 | 0 | 20 | 0.1 | 8 | 0.125 | 9 | 0.1111 | 3 | 0 | 3 | 0 | 23 | 0.087 |
| 2_32459012 | 13 | 0.0769 | 5 | 0 | 7 | 0.1429 | 5 | 0 | 30 | 0.0667 | 7 | 0 | 12 | 0.25 | 6 | 0 | 4 | 0.25 | 29 | 0.1379 |
| 2_32461975 | NA | NA | 1 | 0 | NA | NA | 1 | 1 | 2 | 0.5 | NA | NA | NA | NA | 1 | 0 | 3 | 0.3333 | 4 | 0.25 |
| 2_32463086 | 8 | 0.375 | 10 | 0.2 | 6 | 0 | 10 | 0.4 | 34 | 0.2647 | 6 | 0.5 | 10 | 0.1 | 11 | 0.3636 | 9 | 0.2222 | 36 | 0.2778 |
| 2_32463143 | 9 | 1 | 9 | 0.5556 | 6 | 0.6667 | 10 | 0.9 | 34 | 0.7941 | 6 | 0.8333 | 10 | 0.6 | 11 | 0.9091 | 8 | 1 | 35 | 0.8286 |
| 2_32463265 | 7 | 0.5714 | 9 | 0.3333 | 7 | 0.4286 | 10 | 0.5 | 33 | 0.4545 | 5 | 0.4 | 8 | 0 | 9 | 0.5556 | 9 | 0.3333 | 31 | 0.3226 |
| 2_32463454 | 10 | 0.7 | 10 | 0.6 | 7 | 0.7143 | 13 | 0.7692 | 40 | 0.7 | 6 | 0.3333 | 12 | 0.5833 | 10 | 0.6 | 9 | 0.6667 | 37 | 0.5676 |
| 2_32463462 | 6 | 0 | 9 | 0.1111 | 6 | 0.1667 | 12 | 0 | 33 | 0.0606 | 2 | 0 | 10 | 0 | 11 | 0.2727 | 7 | 0 | 30 | 0.1 |
| 2_32463484 | 10 | 0.6 | 9 | 0.3333 | 8 | 0.375 | 12 | 0.8333 | 39 | 0.5641 | 5 | 0.4 | 13 | 0.4615 | 9 | 0.6667 | 5 | 0.6 | 32 | 0.5312 |
| 2_32477484 | NA | NA | 1 | 0 | 1 | 1 | 1 | 1 | 3 | 0.6667 | 1 | 0 | 2 | 1 | 1 | 1 | NA | NA | 4 | 0.75 |
| 2_32477505 | NA | NA | 3 | 0.6667 | 2 | 0.5 | 1 | 1 | 6 | 0.6667 | 1 | 1 | 2 | 0 | 1 | 0 | NA | NA | 4 | 0.25 |
| 2_32477527 | NA | NA | 3 | 0.6667 | 2 | 0.5 | 1 | 0 | 6 | 0.5 | 1 | 0 | 2 | 1 | 1 | 0 | NA | NA | 4 | 0.5 |
| 2_32484276 | 1 | 0 | 2 | 0.5 | 4 | 0.75 | 4 | 0.25 | 11 | 0.4545 | 2 | 0.5 | 1 | 0 | 1 | 1 | NA | NA | 4 | 0.5 |
| 2_32487231 | 1 | 1 | 2 | 1 | NA | NA | 3 | 1 | 6 | 1 | 2 | 1 | 1 | 1 | NA | NA | NA | NA | 3 | 1 |
| 2_32487298 | 2 | 1 | 4 | 1 | NA | NA | 4 | 0.75 | 10 | 0.9 | 3 | 1 | 1 | 1 | 1 | 1 | NA | NA | 5 | 1 |
| 2_32487326 | 3 | 0 | 3 | 0 | 4 | 0 | 5 | 0.2 | 15 | 0.0667 | 3 | 0 | 2 | 0 | 1 | 1 | NA | NA | 6 | 0.1667 |
| 2_32487383 | 3 | 0 | 5 | 0.2 | 4 | 0.25 | 5 | 0 | 17 | 0.1176 | 2 | 0.5 | 2 | 0.5 | 1 | 0 | 1 | 0 | 6 | 0.3333 |
| 2_32487485 | 3 | 0.6667 | 4 | 0.5 | 3 | 1 | 4 | 0.75 | 14 | 0.7143 | 4 | 0.75 | 2 | 1 | 2 | 0.5 | 1 | 1 | 9 | 0.7778 |
| 2_32487549 | 2 | 0.5 | 5 | 0.6 | 2 | 0.5 | 4 | 0.5 | 13 | 0.5385 | 4 | 1 | 1 | 1 | 1 | 1 | NA | NA | 6 | 1 |
| 2_32487623 | 2 | 1 | 4 | 1 | 3 | 1 | 7 | 0.8571 | 16 | 0.9375 | 4 | 1 | 2 | 1 | 3 | 1 | 1 | 1 | 10 | 1 |
| 2_32487671 | 3 | 1 | 6 | 1 | 3 | 0.6667 | 6 | 0.6667 | 18 | 0.8333 | 5 | 0.8 | 2 | 1 | 3 | 1 | 1 | 1 | 11 | 0.9091 |
| 2_32487675 | 1 | 0 | 6 | 0.1667 | 4 | 0.25 | 6 | 0 | 17 | 0.1176 | 3 | 0 | 2 | 0 | 3 | 0.3333 | 1 | 0 | 9 | 0.1111 |
| 2_32487714 | 1 | 1 | 3 | 1 | 1 | 1 | 2 | 1 | 7 | 1 | NA | NA | 1 | 1 | NA | NA | NA | NA | 1 | 1 |
| 2_32487852 | 2 | 0.5 | 2 | 1 | NA | NA | 1 | 1 | 5 | 0.8 | NA | NA | NA | NA | 1 | 1 | NA | NA | 1 | 1 |
| 2_32487919 | 2 | 1 | 2 | 1 | 1 | 1 | 1 | 1 | 6 | 1 | NA | NA | NA | NA | 1 | 1 | NA | NA | 1 | 1 |
| 2_32509536 | 4 | 0.5 | 7 | 0.7143 | 5 | 0.6 | 1 | 0 | 17 | 0.5882 | 5 | 0.4 | 3 | 0.3333 | 7 | 0.7143 | 9 | 0.4444 | 24 | 0.5 |
| 2_32509564 | 8 | 0.5 | 10 | 0.3 | 13 | 0.6154 | 3 | 0.3333 | 34 | 0.4706 | 6 | 0.5 | 5 | 0.4 | 7 | 0.7143 | 9 | 0.4444 | 27 | 0.5185 |
| 2_32509948 | 9 | 0.3333 | 14 | 0.2143 | 13 | 0.4615 | 8 | 0.125 | 44 | 0.2955 | 11 | 0.2727 | 8 | 0.125 | 7 | 0.4286 | 12 | 0.1667 | 38 | 0.2368 |
| 2_32510465 | 9 | 0.3333 | 18 | 0.3889 | 14 | 0.5714 | 8 | 0.25 | 49 | 0.4082 | 8 | 0.25 | 5 | 0.4 | 11 | 0.2727 | 14 | 0.2143 | 38 | 0.2632 |
| 2_32510694 | 8 | 0 | 14 | 0 | 13 | 0.2308 | 8 | 0.25 | 43 | 0.1163 | 7 | 0 | 7 | 0.1429 | 10 | 0.1 | 14 | 0.0714 | 38 | 0.0789 |
| 2_32523854 | 14 | 0.9286 | 12 | 0.75 | 16 | 0.9375 | 16 | 0.75 | 58 | 0.8448 | 11 | 0.7273 | 20 | 0.8 | 18 | 0.7778 | 15 | 0.6667 | 64 | 0.75 |
| 2_32571665 | 6 | 0.6667 | 4 | 0.5 | 2 | 1 | 4 | 0.75 | 16 | 0.6875 | 3 | 0 | NA | NA | 1 | 1 | 3 | 1 | 7 | 0.5714 |
| 2_32571766 | 1 | 1 | NA | NA | 1 | 1 | 2 | 0.5 | 4 | 0.75 | NA | NA | NA | NA | NA | NA | 2 | 0.5 | 2 | 0.5 |
| 2_32571903 | 5 | 0.6 | 4 | 0.75 | 2 | 1 | 4 | 0.5 | 15 | 0.6667 | 3 | 0.6667 | NA | NA | 1 | 0 | 3 | 1 | 7 | 0.7143 |
| 2_32571967 | 5 | 1 | 4 | 1 | 2 | 1 | 4 | 0.75 | 15 | 0.9333 | 2 | 0.5 | NA | NA | 1 | 1 | 3 | 0.6667 | 6 | 0.6667 |
| 2_32572123 | 6 | 0.1667 | 3 | 0 | 3 | 0 | 4 | 0 | 16 | 0.0625 | 3 | 0 | NA | NA | 1 | 0 | 3 | 0.3333 | 7 | 0.1429 |
| 2_32572702 | NA | NA | 1 | 0 | 4 | 0.25 | 1 | 1 | 6 | 0.3333 | 1 | 1 | 1 | 0 | NA | NA | 1 | 0 | 3 | 0.3333 |
| 2_32573034 | NA | NA | NA | NA | 4 | 0.25 | NA | NA | 4 | 0.25 | 1 | 1 | 1 | 1 | NA | NA | 1 | 0 | 3 | 0.6667 |
| 2_32573064 | NA | NA | 1 | 0 | 3 | 0.3333 | NA | NA | 4 | 0.25 | 1 | 0 | 1 | 1 | NA | NA | 1 | 0 | 3 | 0.3333 |
| 2_32573098 | NA | NA | 1 | 0 | 4 | 0.5 | NA | NA | 5 | 0.4 | 1 | 1 | 1 | 0 | NA | NA | 1 | 0 | 3 | 0.3333 |
| 2_32597508 | 3 | 0 | 17 | 0.1176 | 9 | 0.3333 | 8 | 0.125 | 37 | 0.1622 | 9 | 0.1111 | 9 | 0.1111 | 11 | 0.1818 | 12 | 0.25 | 41 | 0.1707 |
| 2_32597560 | 1 | 1 | 16 | 0.25 | 6 | 0.1667 | 9 | 0.3333 | 32 | 0.2812 | 8 | 0.25 | 11 | 0.0909 | 10 | 0.1 | 14 | 0.0714 | 43 | 0.1163 |
| 2_32597585 | 2 | 1 | 12 | 0.9167 | 9 | 0.6667 | 8 | 1 | 31 | 0.871 | 6 | 0.6667 | 8 | 0.875 | 10 | 0.9 | 11 | 0.9091 | 35 | 0.8571 |
| 2_32597622 | 2 | 0 | 17 | 0.3529 | 6 | 0.3333 | 5 | 0.4 | 30 | 0.3333 | 7 | 0.1429 | 10 | 0.4 | 10 | 0.1 | 11 | 0.4545 | 38 | 0.2895 |
| 2_32597696 | 1 | 1 | 15 | 0.6667 | 5 | 1 | 7 | 0.7143 | 28 | 0.75 | 6 | 0.8333 | 7 | 0.2857 | 9 | 1 | 11 | 0.8182 | 33 | 0.7576 |
| 2_32597707 | 2 | 0.5 | 12 | 0.5833 | 4 | 0.5 | 5 | 1 | 23 | 0.6522 | 6 | 0.5 | 5 | 0.4 | 7 | 0.8571 | 9 | 0.6667 | 27 | 0.6296 |

| | | | | | | | | | | | | | | | | | | | | |
|---|---|---|---|---|---|---|---|---|---|---|---|---|---|---|---|---|---|---|---|---|
| 2_3259774 | 3 | 0.6667 | 13 | 1 | 6 | 0.6667 | 7 | 0.8571 | 29 | 0.8621 | 9 | 0.6667 | 8 | 0.875 | 9 | 0.5556 | 11 | 0.9091 | 37 | 0.7568 |
| 2_3259786 | 3 | 1 | 16 | 0.9375 | 8 | 1 | 8 | 1 | 35 | 0.9714 | 8 | 1 | 9 | 0.8889 | 7 | 1 | 11 | 0.9091 | 35 | 0.9429 |
| 2_3259787 | 1 | 1 | 17 | 0.1176 | 6 | 0.5 | 7 | 0.1429 | 31 | 0.2258 | 7 | 0.2857 | 10 | 0.2 | 10 | 0.3 | 13 | 0.3077 | 40 | 0.275 |
| 2_3259787 | 2 | 0 | 13 | 0.6154 | 8 | 0.625 | 8 | 0.625 | 31 | 0.5806 | 5 | 0.6 | 9 | 0.1111 | 9 | 0.3333 | 11 | 0.5455 | 34 | 0.3824 |
| 2_3259792 | 2 | 0 | 15 | 0.1333 | 5 | 0.4 | 6 | 0.1667 | 28 | 0.1786 | 3 | 0 | 8 | 0.125 | 9 | 0.1111 | 10 | 0.4 | 30 | 0.2 |
| 2_3259808 | 3 | 0.6667 | 13 | 0.3846 | 8 | 0.375 | 5 | 0.4 | 29 | 0.4138 | 6 | 0.1667 | 10 | 0.3 | 6 | 0.3333 | 13 | 0.3846 | 35 | 0.3143 |
| 2_3259810 | 3 | 0.6667 | 15 | 0.1333 | 6 | 0.3333 | 7 | 0.2857 | 31 | 0.2581 | 7 | 0 | 9 | 0.1111 | 9 | 0.1111 | 11 | 0.3636 | 36 | 0.1667 |
| 2_3259956 | 2 | 0 | 12 | 0.1667 | 7 | 0.2857 | 6 | 0 | 27 | 0.1481 | 4 | 0.25 | 8 | 0.25 | 8 | 0.375 | 11 | 0.1818 | 31 | 0.2581 |
| 2_3262281 | NA | NA | 1 | 1 | NA | NA | NA | NA | 1 | 1 | NA | NA | 1 | 1 | NA | NA | NA | NA | 1 | 1 |
| 2_3263591 | NA | NA | 3 | 0 | 1 | 1 | 1 | 1 | 5 | 0.4 | NA | NA | NA | NA | 1 | 1 | 1 | 1 | 2 | 1 |
| 2_3263937 | NA | NA | 3 | 0.3333 | 2 | 0 | NA | NA | 5 | 0.2 | NA | NA | NA | NA | 1 | 0 | 1 | 1 | 2 | 0.5 |
| 2_3263947 | NA | NA | 2 | 0.5 | 1 | 0 | 2 | 1 | 5 | 0.6 | NA | NA | NA | NA | 1 | 1 | 1 | 1 | 2 | 1 |
| 2_3264306 | 1 | 0 | 3 | 0 | 2 | 0 | 3 | 0.3333 | 9 | 0.1111 | 2 | 0.5 | NA | NA | NA | NA | 3 | 0 | 5 | 0.2 |
| 2_3264464 | 2 | 1 | 4 | 0.5 | 1 | 0 | 3 | 0.6667 | 10 | 0.6 | 2 | 0 | NA | NA | 1 | 1 | 3 | 0.3333 | 6 | 0.3333 |
| 2_3264466 | 2 | 0 | 4 | 0.75 | 2 | 1 | 2 | 0.5 | 10 | 0.6 | 2 | 0.5 | NA | NA | NA | NA | 2 | 1 | 4 | 0.75 |
| 2_3264479 | 1 | 1 | 3 | 1 | 2 | 0 | 3 | 0.3333 | 9 | 0.5556 | 2 | 0.5 | 1 | 0 | 1 | 1 | 3 | 0.3333 | 7 | 0.4286 |
| 2_3264500 | NA | NA | 3 | 0.3333 | NA | NA | 3 | 0.3333 | 6 | 0.3333 | 1 | 0 | NA | NA | NA | NA | 3 | 0.3333 | 4 | 0.25 |
| 2_3265185 | 3 | 0.3333 | 9 | 0.2222 | 8 | 0.375 | 5 | 0 | 25 | 0.24 | 2 | 0 | 3 | 0 | 5 | 0 | 3 | 0.3333 | 13 | 0.0769 |
| 2_3265193 | 3 | 0.3333 | 9 | 0.2222 | 8 | 0.125 | 5 | 0 | 25 | 0.16 | 2 | 0 | 3 | 0.3333 | 5 | 0 | 3 | 0 | 13 | 0.0769 |
| 2_3265205 | 2 | 0.5 | 7 | 0.8571 | 6 | 0.8333 | 5 | 1 | 20 | 0.85 | 2 | 0.5 | 3 | 0.3333 | 4 | 1 | 3 | 1 | 12 | 0.75 |
| 2_3265210 | 3 | 0.3333 | 8 | 0.75 | 8 | 0.5 | 5 | 0.4 | 24 | 0.5417 | 2 | 0.5 | 3 | 0 | 5 | 0.6 | 3 | 1 | 13 | 0.5385 |
| 2_3265216 | 3 | 0.3333 | 9 | 0.4444 | 7 | 0.7143 | 5 | 0.6 | 24 | 0.5417 | 2 | 0 | 4 | 0.75 | 4 | 1 | 3 | 1 | 13 | 0.7692 |
| 2_3267792 | 2 | 0 | 9 | 0.4444 | 2 | 0 | 3 | 0.6667 | 16 | 0.375 | 1 | 0 | 4 | 0 | 4 | 0.25 | 1 | 0 | 10 | 0.1 |
| 2_3267826 | 95 | 0.1789 | 106 | 0.1132 | 83 | 0.1205 | 81 | 0.2346 | 365 | 0.1589 | 73 | 0.1644 | 94 | 0.1702 | 72 | 0.1944 | 79 | 0.1392 | 318 | 0.1667 |
| 2_3267847 | 93 | 0.086 | 107 | 0.0841 | 82 | 0 | 82 | 0.0366 | 364 | 0.0549 | 71 | 0.0423 | 93 | 0.043 | 75 | 0.0667 | 73 | 0.0548 | 312 | 0.0513 |
| 2_3267929 | 78 | 0.0641 | 94 | 0.0745 | 57 | 0.0877 | 65 | 0.0769 | 294 | 0.0748 | 60 | 0.1 | 78 | 0.0769 | 67 | 0.1194 | 58 | 0.1034 | 263 | 0.0989 |
| 2_3281374 | 1 | 1 | NA | NA | NA | NA | 1 | 1 | 2 | 1 | 5 | 0.8 | 2 | 1 | NA | NA | NA | NA | 7 | 0.8571 |
| 2_3281392 | 1 | 0 | 1 | 1 | NA | NA | 1 | 1 | 3 | 0.6667 | 5 | 1 | 2 | 1 | NA | NA | NA | NA | 7 | 1 |
| 2_3281395 | 1 | 1 | 1 | 0 | NA | NA | 1 | 1 | 3 | 0.6667 | 4 | 0.25 | 2 | 0.5 | NA | NA | NA | NA | 6 | 0.3333 |
| 2_3281401 | 1 | 1 | NA | NA | NA | NA | NA | NA | 1 | 1 | 4 | 0.5 | 2 | 1 | NA | NA | NA | NA | 6 | 0.6667 |
| 2_3281406 | 1 | 1 | 1 | 1 | NA | NA | 1 | 1 | 3 | 1 | 5 | 1 | 2 | 1 | NA | NA | NA | NA | 7 | 1 |
| 2_3281514 | 3 | 0 | 2 | 0.5 | 1 | 0 | 1 | 1 | 7 | 0.2857 | 7 | 0.2857 | 2 | 0 | 1 | 0 | 1 | 0 | 11 | 0.1818 |
| 2_3281574 | 4 | 0.5 | 2 | 0.5 | 1 | 0 | 6 | 0.1667 | 13 | 0.3077 | 3 | 0 | NA | NA | 1 | 1 | 1 | 0 | 5 | 0.2 |
| 2_3281650 | 3 | 0.3333 | 3 | 0.6667 | 1 | 0 | 4 | 0.25 | 11 | 0.3636 | 2 | 0 | 1 | 1 | 1 | 0 | 1 | 0 | 5 | 0.2 |
| 2_3281669 | 3 | 0 | 3 | 0.3333 | 1 | 0 | 3 | 0 | 10 | 0.1 | 3 | 0.3333 | NA | NA | 1 | 0 | 1 | 0 | 5 | 0.2 |
| 2_3281673 | 2 | 1 | 3 | 1 | 1 | 1 | 4 | 0.5 | 10 | 0.8 | 3 | 0.6667 | NA | NA | NA | NA | 1 | 1 | 4 | 0.75 |
| 2_3285781 | NA | NA | NA | NA | 1 | 1 | NA | NA | 1 | 1 | NA | NA | NA | NA | 1 | 1 | NA | NA | 1 | 1 |
| 2_3285969 | NA | NA | NA | NA | 1 | 1 | NA | NA | 1 | 1 | NA | NA | NA | NA | 1 | 0 | 1 | 1 | 2 | 0.5 |
| 2_3286004 | NA | NA | NA | NA | 1 | 1 | NA | NA | 1 | 1 | NA | NA | NA | NA | 1 | 1 | 1 | 1 | 2 | 1 |
| 2_3286013 | NA | NA | NA | NA | 1 | 1 | NA | NA | 1 | 1 | NA | NA | NA | NA | NA | NA | 1 | 1 | 1 | 1 |
| 2_3286016 | NA | NA | NA | NA | 1 | 1 | NA | NA | 1 | 1 | NA | NA | NA | NA | NA | NA | 1 | 1 | 1 | 1 |
| 2_3286051 | NA | NA | NA | NA | 1 | 1 | NA | NA | 1 | 1 | NA | NA | NA | NA | 1 | 1 | NA | NA | 1 | 1 |
| 2_3286057 | NA | NA | NA | NA | 1 | 1 | NA | NA | 1 | 1 | NA | NA | NA | NA | NA | NA | 1 | 1 | 1 | 1 |
| 2_3286058 | NA | NA | NA | NA | 1 | 1 | NA | NA | 1 | 1 | NA | NA | NA | NA | 1 | 1 | NA | NA | 1 | 1 |
| 2_3286061 | NA | NA | NA | NA | 1 | 1 | NA | NA | 1 | 1 | NA | NA | NA | NA | 1 | 1 | NA | NA | 1 | 1 |
| 2_3286071 | NA | NA | NA | NA | 1 | 1 | NA | NA | 1 | 1 | NA | NA | NA | NA | NA | NA | 1 | 1 | 1 | 1 |
| 2_3286072 | NA | NA | NA | NA | 1 | 1 | NA | NA | 1 | 1 | NA | NA | NA | NA | 1 | 1 | 1 | 1 | 2 | 1 |
| 2_3286091 | NA | NA | NA | NA | 1 | 1 | NA | NA | 1 | 1 | NA | NA | NA | NA | NA | NA | 1 | 1 | 1 | 1 |
| 2_3286313 | 3 | 0 | 4 | 0.25 | 3 | 0.3333 | 4 | 0 | 14 | 0.1429 | 1 | 1 | 2 | 0.5 | 2 | 0 | NA | NA | 5 | 0.4 |
| 2_3286323 | 3 | 0.3333 | 5 | 0.2 | 2 | 0 | 2 | 0.5 | 12 | 0.25 | 1 | 0 | 2 | 0 | 2 | 1 | NA | NA | 5 | 0.4 |
| 2_3286332 | 3 | 0.3333 | 5 | 0.4 | 3 | 0 | 3 | 0 | 14 | 0.2143 | 1 | 0 | 3 | 0.3333 | 1 | 1 | NA | NA | 5 | 0.4 |
| 2_3286334 | 3 | 0.3333 | 5 | 0.4 | 3 | 0 | 3 | 0.3333 | 14 | 0.2857 | 1 | 1 | 1 | 1 | 1 | 0 | NA | NA | 3 | 0.6667 |
| 2_3286339 | 2 | 0 | 3 | 0.3333 | 2 | 0.5 | 3 | 0 | 10 | 0.2 | 1 | 1 | 3 | 0.3333 | 2 | 0 | NA | NA | 6 | 0.3333 |
| 2_3286343 | 1 | 0 | 5 | 0.2 | 3 | 0 | 4 | 0 | 13 | 0.0769 | 1 | 0 | 3 | 0.3333 | 2 | 0.5 | NA | NA | 6 | 0.3333 |
| 2_3286364 | 2 | 0.5 | 4 | 0.5 | 3 | 0.3333 | 3 | 0.3333 | 12 | 0.4167 | 1 | 0 | 3 | 0.6667 | 2 | 0.5 | NA | NA | 6 | 0.5 |
| 2_3287655 | 3 | 0.3333 | 10 | 0 | 6 | 0.1667 | 3 | 0.6667 | 22 | 0.1818 | 9 | 0.2222 | 4 | 0 | 6 | 0 | 7 | 0.1429 | 26 | 0.1154 |
| 2_3287667 | 3 | 1 | 11 | 0.1818 | 7 | 0.5714 | 3 | 0.3333 | 24 | 0.4167 | 9 | 0.1111 | 3 | 0.3333 | 7 | 0.5714 | 8 | 0.375 | 27 | 0.3333 |
| 2_3302365 | 5 | 0 | 5 | 0.6 | 4 | 0.5 | 6 | 0.3333 | 20 | 0.35 | 3 | 0.3333 | 6 | 0.1667 | 3 | 0.6667 | 5 | 0 | 17 | 0.2353 |
| 2_3302391 | 5 | 0.6 | 3 | 0.6667 | 4 | 0.25 | 5 | 0.8 | 17 | 0.5882 | 4 | 0.75 | 6 | 0.3333 | 4 | 0.75 | 5 | 0.4 | 19 | 0.5263 |
| 2_3302397 | 4 | 0.25 | 2 | 0 | 3 | 0 | 4 | 0.25 | 13 | 0.1538 | 4 | 0 | 3 | 0.3333 | 2 | 0.5 | 5 | 0 | 14 | 0.1429 |
| 2_3302402 | 4 | 0 | 3 | 0.3333 | 4 | 0.25 | 5 | 0.2 | 16 | 0.1875 | 3 | 0 | 5 | 0.2 | 4 | 0.25 | 4 | 0 | 16 | 0.125 |
| 2_3302403 | 4 | 0.25 | 4 | 0.25 | 3 | 0.6667 | 6 | 0 | 17 | 0.2353 | 2 | 0 | 6 | 0 | 3 | 0.3333 | 5 | 0.2 | 16 | 0.125 |
| 2_3302425 | 5 | 0 | 5 | 0.2 | 4 | 0 | 6 | 0.6667 | 20 | 0.25 | 3 | 0.3333 | 6 | 0.1667 | 4 | 0 | 5 | 0 | 18 | 0.1111 |
| 2_3302498 | 4 | 0.5 | 4 | 0 | 2 | 0 | 3 | 0 | 13 | 0.1538 | 2 | 0.5 | 4 | 0 | 4 | 0 | 4 | 0 | 14 | 0.0714 |
| 2_3302519 | 4 | 0 | 4 | 0.25 | 3 | 0 | 5 | 0 | 16 | 0.0625 | 2 | 0 | 4 | 0 | 4 | 0 | 4 | 0.25 | 14 | 0.0714 |

| | | | | | | | | | | | | | | | | | | | | |
|---|---|---|---|---|---|---|---|---|---|---|---|---|---|---|---|---|---|---|---|---|
| 2_3302526 | 4 | 0.5 | 4 | 0.25 | 3 | 0 | 5 | 0 | 16 | 0.1875 | 2 | 0.5 | 5 | 0 | 3 | 0 | 4 | 0.25 | 14 | 0.1429 |
| 2_3302544 | 3 | 0.3333 | 4 | 0.25 | 3 | 0 | 4 | 0.75 | 14 | 0.3571 | 2 | 0 | 5 | 0.2 | 3 | 0.3333 | 3 | 0.6667 | 13 | 0.3077 |
| 2_3302547 | 1 | 1 | 3 | 0 | 3 | 0.3333 | 3 | 0.3333 | 10 | 0.3 | 2 | 0.5 | 3 | 0 | 3 | 0.3333 | 1 | 0 | 9 | 0.2222 |
| 2_3302549 | 4 | 0.25 | 2 | 0 | 2 | 0.5 | 2 | 1 | 10 | 0.4 | 2 | 0.5 | 3 | 0.3333 | 3 | 0 | 2 | 0.5 | 10 | 0.3 |
| 2_3302610 | 4 | 0 | 3 | 0 | 3 | 0.3333 | 1 | 0 | 11 | 0.0909 | 2 | 0.5 | 3 | 0.6667 | 3 | 0 | 2 | 0 | 10 | 0.3 |
| 2_3302616 | 4 | 0.25 | 3 | 0.6667 | 3 | 0.6667 | 1 | 0 | 11 | 0.4545 | 1 | 1 | NA | NA | 2 | 0 | 2 | 1 | 5 | 0.6 |
| 2_3302644 | 4 | 0 | 3 | 0.6667 | 2 | 0.5 | 1 | 0 | 10 | 0.3 | 2 | 0 | 1 | 0 | 1 | 0 | 4 | 0.25 | 8 | 0.125 |
| 2_3302728 | 3 | 0.6667 | 3 | 0.3333 | 1 | 1 | 1 | 0 | 8 | 0.5 | 2 | 1 | 1 | 1 | 1 | 1 | 4 | 0.25 | 8 | 0.625 |
| 2_3302744 | 1 | 0 | 2 | 1 | NA | NA | NA | NA | 3 | 0.6667 | 2 | 0.5 | NA | NA | 2 | 0 | 2 | 0 | 6 | 0.1667 |
| 2_3302745 | 3 | 0.3333 | 3 | 0.3333 | 1 | 0 | 1 | 0 | 8 | 0.25 | 2 | 0 | 2 | 0.5 | 2 | 0.5 | 4 | 0 | 10 | 0.2 |
| 2_3302757 | 3 | 0.6667 | 3 | 0.6667 | 1 | 1 | 1 | 1 | 8 | 0.75 | 1 | 1 | 1 | 1 | 2 | 0.5 | 4 | 1 | 8 | 0.875 |
| 2_3302767 | 3 | 0.3333 | 2 | 0 | 1 | 0 | 1 | 1 | 7 | 0.2857 | 1 | 0 | 2 | 1 | 1 | 0 | 3 | 0.3333 | 7 | 0.4286 |
| 2_3302768 | 4 | 0.5 | 3 | 0 | 1 | 0 | 1 | 0 | 9 | 0.2222 | 2 | 0.5 | 2 | 0 | 2 | 0.5 | 3 | 0 | 9 | 0.2222 |
| 2_3313639 | NA | NA | NA | NA | 1 | 1 | NA | NA | 1 | 1 | 1 | 1 | NA | NA | NA | NA | NA | NA | 1 | 1 |
| 2_3329629 | 4 | 0 | 4 | 0.5 | NA | NA | 2 | 0 | 10 | 0.2 | NA | NA | 3 | 0.3333 | 4 | 0 | 2 | 0 | 9 | 0.1111 |
| 2_3329704 | 3 | 0.6667 | 3 | 0 | NA | NA | 2 | 0 | 8 | 0.25 | NA | NA | 2 | 0.5 | 3 | 0.3333 | 2 | 0 | 7 | 0.2857 |
| 2_3329842 | 3 | 0.3333 | 2 | 1 | NA | NA | 2 | 0.5 | 7 | 0.5714 | NA | NA | 1 | 0 | 1 | 1 | 2 | 0 | 4 | 0.25 |
| 2_3329933 | 2 | 0.5 | 1 | 1 | NA | NA | 1 | 1 | 4 | 0.75 | NA | NA | 1 | 1 | 1 | 1 | 1 | 1 | 3 | 1 |
| 2_3329973 | 1 | 1 | 1 | 0 | NA | NA | 1 | 0 | 3 | 0.3333 | NA | NA | 1 | 0 | 1 | 0 | 1 | 1 | 3 | 0.3333 |
| 2_3330034 | 1 | 1 | 1 | 0 | NA | NA | NA | NA | 2 | 0.5 | NA | NA | 1 | 0 | 1 | 1 | 1 | 1 | 3 | 0.6667 |
| 2_3330090 | 1 | 1 | 1 | 1 | NA | NA | NA | NA | 2 | 1 | NA | NA | 1 | 1 | 1 | 1 | NA | NA | 2 | 1 |
| 2_3330107 | 1 | 1 | NA | NA | NA | NA | NA | NA | 1 | 1 | NA | NA | 1 | 1 | NA | NA | NA | NA | 1 | 1 |
| 2_3330108 | NA | NA | 1 | 1 | NA | NA | NA | NA | 1 | 1 | NA | NA | 1 | 1 | 1 | 1 | 1 | 1 | 3 | 1 |
| 2_3330124 | 1 | 0 | 1 | 1 | NA | NA | NA | NA | 2 | 0.5 | NA | NA | 1 | 0 | 1 | 1 | 1 | 0 | 3 | 0.3333 |
| 2_3330144 | NA | NA | 1 | 1 | NA | NA | NA | NA | 1 | 1 | NA | NA | 1 | 1 | NA | NA | NA | NA | 1 | 1 |
| 2_3330181 | 1 | 1 | 1 | 1 | NA | NA | NA | NA | 2 | 1 | NA | NA | 1 | 0 | 1 | 1 | NA | NA | 2 | 0.5 |
| 2_3330208 | 1 | 1 | NA | NA | NA | NA | NA | NA | 1 | 1 | NA | NA | 1 | 1 | NA | NA | NA | NA | 1 | 1 |
| 2_3334051 | NA | NA | 1 | 0 | 3 | 0 | 1 | 1 | 5 | 0.2 | 9 | 0.4444 | 1 | 1 | 2 | 0 | 6 | 0 | 18 | 0.2778 |
| 2_3334097 | NA | NA | 1 | 0 | 3 | 0.6667 | 2 | 0.5 | 6 | 0.5 | 7 | 0.2857 | NA | NA | 2 | 0 | 5 | 0.4 | 14 | 0.2857 |
| 2_3334209 | NA | NA | NA | NA | 5 | 0.6 | 2 | 0.5 | 7 | 0.5714 | 4 | 0.75 | NA | NA | 1 | 1 | 5 | 0.8 | 10 | 0.8 |
| 2_3334236 | NA | NA | NA | NA | 5 | 0.2 | 1 | 0 | 6 | 0.1667 | 4 | 0 | NA | NA | NA | NA | 4 | 0.25 | 8 | 0.125 |
| 2_3334326 | NA | NA | NA | NA | 4 | 0.75 | 1 | 0 | 5 | 0.6 | 1 | 1 | NA | NA | NA | NA | 3 | 0 | 4 | 0.25 |
| 2_3334343 | NA | NA | NA | NA | 4 | 0.25 | NA | NA | 4 | 0.25 | 1 | 0 | NA | NA | NA | NA | 2 | 0.5 | 3 | 0.3333 |
| 2_3334366 | NA | NA | NA | NA | 2 | 0.5 | NA | NA | 2 | 0.5 | 1 | 0 | NA | NA | 1 | 0 | 3 | 1 | 5 | 0.6 |
| 2_3334377 | NA | NA | NA | NA | 4 | 0.75 | NA | NA | 4 | 0.75 | 1 | 0 | NA | NA | 1 | 0 | 2 | 0.5 | 4 | 0.25 |
| 2_3334379 | NA | NA | NA | NA | 4 | 1 | NA | NA | 4 | 1 | 1 | 0 | NA | NA | NA | NA | 3 | 1 | 4 | 0.75 |
| 2_3334410 | NA | NA | NA | NA | 3 | 0.6667 | NA | NA | 3 | 0.6667 | NA | NA | NA | NA | NA | NA | 1 | 1 | 1 | 1 |
| 2_3334416 | NA | NA | NA | NA | 3 | 1 | NA | NA | 3 | 1 | NA | NA | NA | NA | NA | NA | 2 | 1 | 2 | 1 |
| 2_3334417 | NA | NA | NA | NA | 3 | 0.6667 | NA | NA | 3 | 0.6667 | NA | NA | NA | NA | NA | NA | 2 | 1 | 2 | 1 |
| 2_3334476 | NA | NA | NA | NA | 2 | 0.5 | NA | NA | 2 | 0.5 | NA | NA | NA | NA | NA | NA | 3 | 0.6667 | 3 | 0.6667 |
| 2_3334486 | NA | NA | NA | NA | 1 | 1 | NA | NA | 1 | 1 | NA | NA | NA | NA | NA | NA | 3 | 1 | 3 | 1 |
| 2_3334502 | NA | NA | NA | NA | 1 | 1 | NA | NA | 1 | 1 | NA | NA | NA | NA | NA | NA | 3 | 1 | 3 | 1 |
| 2_3362004 | 6 | 0.1667 | 14 | 0.1429 | 4 | 0 | 10 | 0.2 | 34 | 0.1471 | 5 | 0.2 | 8 | 0 | 13 | 0 | 7 | 0.1429 | 33 | 0.0606 |
| 2_3362008 | 5 | 0.4 | 13 | 0.2308 | 8 | 0.125 | 8 | 0.625 | 34 | 0.3235 | 5 | 1 | 10 | 0.5 | 14 | 0.3571 | 6 | 0.3333 | 35 | 0.4857 |
| 2_3362009 | 5 | 0.2 | 14 | 0.0714 | 7 | 0 | 10 | 0.2 | 36 | 0.1111 | 6 | 0.1667 | 9 | 0.2222 | 13 | 0 | 6 | 0 | 34 | 0.0882 |
| 2_3362011 | 1 | 1 | 5 | 0.8 | 4 | 1 | 2 | 0.5 | 12 | 0.8333 | 2 | 1 | 4 | 1 | 5 | 1 | 2 | 0.5 | 13 | 0.9231 |
| 2_3362017 | 5 | 0.2 | 13 | 0.3846 | 5 | 0.4 | 8 | 0.375 | 31 | 0.3548 | 4 | 0.25 | 9 | 0.2222 | 12 | 0.1667 | 7 | 0.1429 | 32 | 0.1875 |
| 2_3362031 | 6 | 0 | 15 | 0 | 6 | 0 | 9 | 0.3333 | 36 | 0.0833 | 4 | 0.25 | 11 | 0.0909 | 12 | 0.0833 | 4 | 0 | 31 | 0.0968 |
| 2_3362043 | 5 | 0 | 12 | 0.0833 | 6 | 0.1667 | 9 | 0.2222 | 32 | 0.125 | 5 | 0.2 | 12 | 0.0833 | 12 | 0.1667 | 5 | 0 | 34 | 0.1176 |
| 2_3362105 | 5 | 0 | 11 | 0.1818 | 4 | 0.25 | 6 | 0.3333 | 26 | 0.1923 | 4 | 0 | 8 | 0.125 | 12 | 0.0833 | 5 | 0.4 | 29 | 0.1379 |
| 2_3362125 | 3 | 0 | 10 | 0.6 | 3 | 0.3333 | 7 | 0.7143 | 23 | 0.5217 | 3 | 0.6667 | 9 | 0.4444 | 13 | 0.4615 | 5 | 0.6 | 30 | 0.5 |
| 2_3362138 | 2 | 0 | 9 | 0.1111 | 2 | 0 | 6 | 0.3333 | 19 | 0.1579 | 4 | 0.25 | 7 | 0 | 11 | 0.0909 | 4 | 0 | 26 | 0.0769 |
| 2_3362140 | 1 | 0 | 7 | 0.4286 | 2 | 1 | 7 | 0.5714 | 17 | 0.5294 | 4 | 0.5 | 8 | 0.75 | 9 | 0.5556 | 3 | 0 | 24 | 0.5417 |
| 2_3362151 | 2 | 1 | 8 | 1 | 1 | 1 | 5 | 1 | 16 | 1 | 4 | 1 | 8 | 1 | 10 | 1 | 4 | 1 | 26 | 1 |
| 2_3362160 | 2 | 0.5 | 8 | 1 | 2 | 1 | 4 | 0.75 | 16 | 0.875 | 4 | 1 | 5 | 1 | 11 | 1 | 1 | 1 | 21 | 1 |
| 2_3362177 | 2 | 1 | 8 | 0.875 | 2 | 1 | 5 | 1 | 17 | 0.9412 | 3 | 1 | 9 | 1 | 10 | 1 | 2 | 1 | 24 | 1 |
| 2_3362191 | NA | NA | 8 | 1 | 2 | 0.5 | 5 | 0.6 | 15 | 0.8 | 3 | 1 | 7 | 1 | 13 | 0.9231 | 4 | 1 | 27 | 0.963 |
| 2_3362192 | 1 | 0 | 7 | 0.8571 | 1 | 1 | 3 | 0.6667 | 12 | 0.75 | 2 | 1 | 5 | 1 | 8 | 1 | 3 | 1 | 18 | 1 |
| 2_3362197 | 2 | 0.5 | 5 | 0.6 | 2 | 0.5 | 6 | 0.5 | 15 | 0.5333 | 2 | 0.5 | 8 | 0.75 | 13 | 0.5385 | 3 | 0.6667 | 26 | 0.6154 |
| 2_3362223 | 1 | 0 | 9 | 0 | 2 | 0 | 4 | 0.25 | 16 | 0.0625 | 2 | 0 | 6 | 0.1667 | 9 | 0 | 2 | 0 | 19 | 0.0526 |
| 2_3362227 | 2 | 0 | 8 | 0.625 | 2 | 0.5 | 5 | 0.6 | 17 | 0.5294 | 2 | 0 | 7 | 0.7143 | 11 | 0.5455 | 3 | 0.6667 | 23 | 0.5652 |
| 2_3362229 | 1 | 0 | 7 | 0.1429 | 2 | 0 | 5 | 0 | 15 | 0.0667 | 1 | 0 | 5 | 0 | 10 | 0.2 | 2 | 0 | 18 | 0.1111 |
| 2_3362238 | 1 | 0 | 6 | 0 | 2 | 0 | 4 | 0.5 | 13 | 0.1538 | 2 | 0 | 6 | 0.1667 | 11 | 0.5455 | 3 | 0 | 22 | 0.3182 |
| 2_3362245 | 1 | 1 | 5 | 0.4 | 2 | 0.5 | 4 | 0.5 | 12 | 0.5 | 2 | 0.5 | 5 | 0.2 | 9 | 0.4444 | 3 | 0.3333 | 19 | 0.3684 |
| 2_3362248 | 1 | 0 | 4 | 0.25 | 2 | 0 | 5 | 0.2 | 12 | 0.1667 | 2 | 0 | 7 | 0.1429 | 10 | 0.1 | 3 | 0 | 22 | 0.0909 |

| | | | | | | | | | | | | | | | | | | | | |
|---|---|---|---|---|---|---|---|---|---|---|---|---|---|---|---|---|---|---|---|---|
| 2_33622518 | 1 | 1 | 6 | 0.8333 | 2 | 0 | 4 | 1 | 13 | 0.7692 | 2 | 0.5 | 3 | 1 | 9 | 0.6667 | 3 | 0.6667 | 17 | 0.7059 |
| 2_33622639 | 2 | 0.5 | 5 | 0 | 2 | 0.5 | 4 | 0 | 13 | 0.1538 | 1 | 1 | 6 | 0.3333 | 8 | 0.375 | 2 | 0 | 17 | 0.3529 |
| 2_33622823 | 2 | 0 | 4 | 0.25 | 1 | 0 | 4 | 0.25 | 11 | 0.1818 | 2 | 0.5 | 6 | 0.1667 | 11 | 0.3636 | 2 | 0 | 21 | 0.2857 |
| 2_33622870 | 1 | 1 | 5 | 1 | 2 | 1 | 3 | 1 | 11 | 1 | 2 | 1 | 5 | 1 | 10 | 1 | 2 | 1 | 19 | 1 |
| 2_33622897 | 1 | 1 | 4 | 0.75 | 1 | 1 | 4 | 0.5 | 10 | 0.7 | 2 | 1 | 6 | 0.6667 | 11 | 0.8182 | 2 | 0.5 | 21 | 0.7619 |
| 2_33623009 | 2 | 0.5 | 6 | 0.5 | 2 | 0 | 4 | 0 | 14 | 0.2857 | 2 | 0.5 | 7 | 0.1429 | 10 | 0.3 | 2 | 0 | 21 | 0.2381 |
| 2_33623038 | 2 | 1 | 4 | 0.5 | 2 | 0 | 3 | 0.3333 | 11 | 0.4545 | 2 | 0 | 6 | 0.6667 | 10 | 0.7 | 2 | 0.5 | 20 | 0.6 |
| 2_33623087 | NA | NA | 4 | 0.25 | 2 | 0 | 3 | 0 | 9 | 0.1111 | 1 | 0 | 6 | 0.3333 | 9 | 0 | 1 | 0 | 17 | 0.1176 |
| 2_33623155 | NA | NA | 2 | 1 | NA | NA | 2 | 1 | 4 | 1 | 1 | 1 | 1 | 1 | 6 | 0.8333 | NA | NA | 8 | 0.875 |
| 2_33623240 | 1 | 0 | 5 | 0.2 | 1 | 0 | 2 | 1 | 9 | 0.3333 | 1 | 0 | 4 | 0.5 | 9 | 0.6667 | 1 | 1 | 15 | 0.6 |
| 2_33623259 | 2 | 1 | 4 | 1 | 2 | 1 | 3 | 0.6667 | 11 | 0.9091 | 2 | 1 | 5 | 0.6 | 11 | 0.6364 | 2 | 1 | 20 | 0.7 |
| 2_33623278 | 1 | 0 | 3 | 0.3333 | 1 | 0 | 2 | 0.5 | 7 | 0.2857 | 1 | 0 | 4 | 0.5 | 11 | 0.3636 | 1 | 0 | 17 | 0.3529 |
| 2_33623327 | 1 | 0 | 4 | 0.5 | 2 | 0.5 | 2 | 0.5 | 9 | 0.4444 | 1 | 1 | 5 | 0.6 | 8 | 1 | 2 | 1 | 16 | 0.875 |
| 2_33623407 | 1 | 1 | 3 | 0 | 1 | 0 | 3 | 0.3333 | 8 | 0.25 | 2 | 0 | 4 | 0.5 | 7 | 0.1429 | 1 | 0 | 14 | 0.2143 |
| 2_34062076 | 6 | 0 | 4 | 0.75 | 3 | 0 | 7 | 0.1429 | 20 | 0.2 | 1 | 0 | 8 | 0.125 | 6 | 0.1667 | 6 | 0.5 | 21 | 0.2381 |
| 2_34062205 | 4 | 0 | 7 | 0.1429 | 5 | 0 | 7 | 0.7143 | 23 | 0.2609 | 1 | 0 | 10 | 0.2 | 5 | 0 | 5 | 0.2 | 21 | 0.1429 |
| 2_34062361 | 6 | 0.5 | 6 | 0.3333 | 4 | 0.5 | 7 | 0.7143 | 23 | 0.5217 | 1 | 0 | 10 | 0.1 | 6 | 0.5 | 5 | 0.2 | 22 | 0.2273 |
| 2_34062414 | 6 | 0.6667 | 5 | 0.8 | 4 | 0.25 | 5 | 0.8 | 20 | 0.65 | 1 | 0 | 10 | 0.7 | 6 | 0.1667 | 5 | 0.8 | 22 | 0.5455 |
| 2_34062777 | 4 | 0 | 4 | 0.25 | 4 | 0 | 4 | 0.25 | 16 | 0.125 | 1 | 0 | 8 | 0 | 6 | 0.1667 | 3 | 0.3333 | 18 | 0.1111 |
| 2_34261889 | 2 | 0.5 | 4 | 0 | 3 | 0 | 3 | 0 | 12 | 0.0833 | NA | NA | 6 | 0 | 2 | 0 | 3 | 0.3333 | 11 | 0.0909 |
| 2_34261905 | 2 | 0 | 1 | 0 | 2 | 0.5 | 3 | 0 | 8 | 0.125 | NA | NA | 5 | 0 | 1 | 0 | 2 | 0.5 | 8 | 0.125 |
| 2_34322067 | 2 | 1 | 1 | 0 | NA | NA | 4 | 0.25 | 7 | 0.4286 | 3 | 0.3333 | 1 | 0 | 4 | 0.25 | 1 | 1 | 9 | 0.3333 |
| 2_34513704 | 4 | 0.75 | 3 | 1 | 1 | 1 | 12 | 0.6667 | 20 | 0.75 | 5 | 1 | 8 | 0.875 | 8 | 0.75 | 4 | 1 | 25 | 0.88 |
| 2_34513811 | 5 | 0.4 | 3 | 1 | 2 | 0.5 | 12 | 0.3333 | 22 | 0.4545 | 4 | 0.25 | 9 | 0.7778 | 10 | 0.2 | 4 | 0.75 | 27 | 0.4815 |
| 2_34514667 | 5 | 0.8 | 1 | 1 | 3 | 0.6667 | 9 | 0.4444 | 18 | 0.6111 | 2 | 0.5 | 7 | 0.5714 | 5 | 0.6 | 2 | 1 | 16 | 0.625 |
| 2_34567291 | 5 | 0 | 5 | 0.6 | 2 | 0.5 | 2 | 0 | 14 | 0.2857 | 2 | 0 | 5 | 0.2 | 6 | 0 | 8 | 0.25 | 21 | 0.1429 |
| 2_34567334 | 3 | 0.3333 | 4 | 0.5 | 4 | 0.75 | 3 | 0.3333 | 14 | 0.5 | 2 | 0 | 4 | 0.75 | 7 | 0.2857 | 5 | 0.6 | 18 | 0.4444 |
| 2_34567505 | 4 | 0.25 | 6 | 0.3333 | 4 | 0.75 | 3 | 0.3333 | 17 | 0.4118 | 1 | 0 | 6 | 0.5 | 6 | 0.3333 | 10 | 0.5 | 23 | 0.4348 |
| 2_34567524 | 4 | 0 | 6 | 0 | 6 | 0.1667 | 4 | 0 | 20 | 0.05 | 2 | 0 | 7 | 0.1429 | 7 | 0.1429 | 11 | 0.0909 | 27 | 0.1111 |
| 2_34567697 | 2 | 0.5 | 4 | 0.25 | 6 | 0.3333 | 2 | 0 | 14 | 0.2857 | 1 | 0 | 5 | 0 | 6 | 0.1667 | 9 | 0.1111 | 21 | 0.0952 |
| 2_34567809 | 5 | 0.4 | 6 | 0.5 | 4 | 0.5 | 4 | 0.5 | 19 | 0.4737 | 3 | 0.6667 | 7 | 0.5714 | 6 | 0.1667 | 9 | 0.1111 | 25 | 0.32 |
| 2_34567966 | 3 | 0.6667 | 6 | 0.1667 | 4 | 0 | 3 | 0.3333 | 16 | 0.25 | 1 | 1 | 3 | 0.6667 | 4 | 0.25 | 10 | 0.1 | 18 | 0.2778 |
| 2_34567993 | 2 | 0 | 6 | 0 | 3 | 0.3333 | 4 | 0.25 | 15 | 0.1333 | 2 | 0.5 | 6 | 0.1667 | 8 | 0.125 | 10 | 0.1 | 26 | 0.1538 |
| 2_34568021 | 5 | 0.6 | 4 | 0.75 | 3 | 1 | 4 | 0 | 16 | 0.5625 | 3 | 0 | 6 | 0.5 | 7 | 0.2857 | 11 | 0.7273 | 27 | 0.4815 |
| 2_34594379 | 3 | 0 | 3 | 0.3333 | 3 | 0.6667 | 2 | 0 | 11 | 0.2727 | 2 | 0.5 | 6 | 0.1667 | 2 | 0 | 8 | 0.25 | 18 | 0.2222 |
| 2_34618584 | 1 | 1 | 2 | 0 | 2 | 0 | 1 | 0 | 6 | 0.1667 | NA | NA | 4 | 0.25 | 1 | 0 | 5 | 0.2 | 10 | 0.2 |
| 2_34618808 | 1 | 1 | 2 | 1 | 2 | 1 | 1 | 0 | 6 | 0.8333 | NA | NA | 4 | 0.75 | 2 | 1 | 4 | 1 | 10 | 0.9 |
| 2_34618849 | 1 | 1 | 2 | 0.5 | 2 | 1 | 1 | 1 | 6 | 0.8333 | NA | NA | 4 | 0.75 | 2 | 0.5 | 5 | 0.6 | 11 | 0.6364 |
| 2_34618892 | 1 | 0 | 2 | 0 | 2 | 0.5 | 1 | 1 | 6 | 0.3333 | NA | NA | 4 | 0.75 | 1 | 1 | 5 | 0.4 | 10 | 0.6 |
| 2_34619015 | NA | NA | 2 | 0.5 | 1 | 1 | NA | NA | 3 | 0.6667 | NA | NA | 3 | 0.6667 | 1 | 0 | 3 | 0.3333 | 7 | 0.4286 |
| 2_34664254 | 23 | 0.087 | 40 | 0.15 | 27 | 0.1481 | 29 | 0 | 119 | 0.1008 | 27 | 0.037 | 35 | 0.1429 | 29 | 0.069 | 44 | 0.0227 | 135 | 0.0667 |
| 2_34665584 | 29 | 0 | 46 | 0.087 | 32 | 0.125 | 33 | 0.0303 | 140 | 0.0643 | 30 | 0.1 | 43 | 0.186 | 39 | 0.1026 | 46 | 0.0435 | 158 | 0.1076 |
| 2_34665613 | 32 | 0.0625 | 53 | 0.1321 | 33 | 0.1515 | 36 | 0.1111 | 154 | 0.1169 | 32 | 0.0625 | 48 | 0.1667 | 37 | 0.027 | 54 | 0.0926 | 171 | 0.0936 |
| 2_34665694 | 36 | 0.0833 | 54 | 0.0926 | 33 | 0 | 41 | 0.0244 | 164 | 0.0549 | 35 | 0.0571 | 48 | 0.0417 | 42 | 0.0714 | 55 | 0.0364 | 180 | 0.05 |
| 2_34665725 | 32 | 0.0625 | 47 | 0.1064 | 33 | 0.0303 | 38 | 0.1053 | 150 | 0.08 | 30 | 0 | 47 | 0.1064 | 41 | 0.0976 | 50 | 0.08 | 168 | 0.0774 |
| 2_34665746 | 35 | 0.1143 | 52 | 0.0962 | 35 | 0.1429 | 39 | 0.0513 | 161 | 0.0994 | 34 | 0.0588 | 51 | 0.0784 | 41 | 0.0244 | 54 | 0.1111 | 180 | 0.0722 |
| 2_34665839 | 35 | 0.2 | 54 | 0.0741 | 37 | 0.027 | 39 | 0.1282 | 165 | 0.103 | 35 | 0.0286 | 53 | 0.0943 | 44 | 0.1136 | 55 | 0.0909 | 187 | 0.0856 |
| 2_34665881 | 38 | 0.1579 | 53 | 0.0943 | 36 | 0.1111 | 41 | 0.122 | 168 | 0.119 | 36 | 0.0556 | 47 | 0.1702 | 43 | 0.093 | 54 | 0.1111 | 180 | 0.1111 |
| 2_34665915 | 37 | 0.1892 | 54 | 0.1852 | 36 | 0.1111 | 35 | 0.2 | 162 | 0.1728 | 33 | 0.1818 | 47 | 0.1277 | 42 | 0.3095 | 52 | 0.1923 | 174 | 0.2011 |
| 2_34665995 | 43 | 0.9767 | 55 | 1 | 35 | 0.9429 | 37 | 0.973 | 170 | 0.9765 | 36 | 0.9722 | 49 | 0.9592 | 45 | 0.9556 | 52 | 0.9808 | 182 | 0.967 |
| 2_34666004 | 40 | 0.625 | 58 | 0.6207 | 34 | 0.6765 | 40 | 0.825 | 172 | 0.6802 | 37 | 0.6757 | 55 | 0.7818 | 48 | 0.5833 | 52 | 0.7308 | 192 | 0.6979 |
| 2_34666026 | 39 | 0.1795 | 59 | 0.1525 | 33 | 0.1515 | 41 | 0.1463 | 172 | 0.157 | 41 | 0.122 | 52 | 0.2115 | 47 | 0.1702 | 50 | 0.14 | 190 | 0.1632 |
| 2_34666059 | 38 | 0.2105 | 55 | 0.1636 | 31 | 0.0968 | 35 | 0.2 | 159 | 0.1698 | 37 | 0.1892 | 49 | 0.1633 | 39 | 0.2051 | 54 | 0.2037 | 179 | 0.1899 |
| 2_34666166 | 34 | 1 | 47 | 1 | 31 | 0.9677 | 31 | 1 | 143 | 0.993 | 30 | 1 | 48 | 1 | 37 | 1 | 45 | 0.9778 | 160 | 0.9938 |
| 2_34666174 | 40 | 0.95 | 58 | 0.8966 | 36 | 0.9167 | 42 | 0.881 | 176 | 0.9091 | 36 | 0.9722 | 57 | 0.9649 | 47 | 0.8511 | 54 | 0.9259 | 194 | 0.9278 |
| 2_34666200 | 44 | 0.6136 | 58 | 0.7069 | 38 | 0.7105 | 42 | 0.7857 | 182 | 0.7033 | 43 | 0.6977 | 56 | 0.9286 | 48 | 0.7917 | 53 | 0.6981 | 200 | 0.785 |
| 2_34666425 | 36 | 0.1667 | 64 | 0.25 | 38 | 0.1053 | 45 | 0.1778 | 183 | 0.1858 | 47 | 0.1064 | 61 | 0.2131 | 53 | 0.0943 | 56 | 0.1786 | 217 | 0.1521 |
| 2_34667742 | NA | NA | 1 | 1 | NA | NA | NA | NA | 1 | 1 | NA | NA | 2 | 0.5 | NA | NA | NA | NA | 2 | 0.5 |
| 2_34668101 | NA | NA | 1 | 0 | 2 | 1 | NA | NA | 3 | 0.6667 | NA | NA | NA | NA | 2 | 0.5 | NA | NA | 2 | 0.5 |
| 2_34669222 | NA | NA | NA | NA | 1 | 1 | NA | NA | 1 | 1 | 1 | 0 | NA | NA | 1 | 1 | NA | NA | 2 | 0.5 |
| 2_34669253 | 1 | 0 | 3 | 0.3333 | 3 | 0.3333 | 2 | 0.5 | 9 | 0.3333 | 3 | 0 | NA | NA | 5 | 0.4 | 3 | 0.3333 | 11 | 0.2727 |
| 2_34669263 | NA | NA | 2 | 0.5 | 2 | 0 | 2 | 0 | 6 | 0.1667 | 4 | 0.25 | NA | NA | 5 | 0 | 2 | 0 | 11 | 0.0909 |
| 2_34669430 | 2 | 0 | 4 | 0.25 | 3 | 1 | NA | NA | 9 | 0.4444 | 4 | 0.25 | NA | NA | 5 | 0.6 | 4 | 0.25 | 13 | 0.3846 |
| 2_34695638 | 11 | 0 | 7 | 0 | 4 | 0.25 | 5 | 0.2 | 27 | 0.0741 | 5 | 0 | 9 | 0 | 11 | 0 | 7 | 0.2857 | 32 | 0.0625 |
| 2_34695697 | 10 | 1 | 5 | 1 | 6 | 1 | 4 | 1 | 25 | 1 | 5 | 0.8 | 9 | 0.8889 | 8 | 1 | 2 | 1 | 24 | 0.9167 |

| | | | | | | | | | | | | | | | | | | | | |
|---|---|---|---|---|---|---|---|---|---|---|---|---|---|---|---|---|---|---|---|---|
| 2_3469570 | 10 | 0.1 | 7 | 0 | 5 | 0 | 4 | 0.25 | 26 | 0.0769 | 5 | 0.2 | 7 | 0 | 10 | 0 | 5 | 0.2 | 27 | 0.0741 |
| 2_3469590 | 11 | 0.3636 | 7 | 0.1429 | 3 | 0 | 4 | 0 | 25 | 0.2 | 8 | 0.25 | 14 | 0.3571 | 9 | 0.1111 | 8 | 0.125 | 39 | 0.2308 |
| 2_3469591 | 10 | 0.9 | 5 | 1 | 4 | 1 | 5 | 1 | 24 | 0.9583 | 6 | 1 | 14 | 0.8571 | 8 | 1 | 6 | 1 | 34 | 0.9412 |
| 2_3471262 | 4 | 0 | 5 | 0 | 6 | 0.1667 | 7 | 0.2857 | 22 | 0.1364 | 3 | 0 | 11 | 0.2727 | 6 | 0.1667 | 12 | 0.3333 | 32 | 0.25 |
| 2_3471271 | 4 | 0.75 | 5 | 1 | 7 | 0.8571 | 7 | 0.7143 | 23 | 0.8261 | 6 | 0.8333 | 14 | 0.7143 | 7 | 0.7143 | 12 | 1 | 39 | 0.8205 |
| 2_3471276 | 4 | 0.25 | 2 | 0.5 | 4 | 0.5 | 7 | 0.2857 | 17 | 0.3529 | 4 | 0.75 | 12 | 0.6667 | 3 | 0.3333 | 10 | 0.8 | 29 | 0.6897 |
| 2_3471283 | 4 | 1 | 5 | 0.8 | 5 | 0.8 | 7 | 0.8571 | 21 | 0.8571 | 5 | 0.8 | 14 | 0.8571 | 6 | 0.6667 | 8 | 0.75 | 33 | 0.7879 |
| 2_3471289 | 3 | 0.6667 | 5 | 0.8 | 5 | 1 | 7 | 1 | 20 | 0.9 | 6 | 0.6667 | 15 | 0.6667 | 6 | 0.6667 | 11 | 0.8182 | 38 | 0.7105 |
| 2_3471291 | 4 | 0.5 | 6 | 0.1667 | 5 | 0.2 | 7 | 0.2857 | 22 | 0.2727 | 4 | 0.25 | 14 | 0.2143 | 7 | 0.4286 | 10 | 0.1 | 35 | 0.2286 |
| 2_3471297 | 3 | 0.6667 | 2 | 0 | 4 | 0.25 | 6 | 0.1667 | 15 | 0.2667 | 1 | 1 | 11 | 0.4545 | 5 | 0 | 4 | 0 | 21 | 0.2857 |
| 2_3471297 | 3 | 0 | 6 | 0 | 5 | 0 | 6 | 0.1667 | 20 | 0.05 | 5 | 0.2 | 13 | 0.1538 | 6 | 0 | 10 | 0.2 | 34 | 0.1471 |
| 2_3471299 | 3 | 0.6667 | 5 | 1 | 5 | 1 | 4 | 1 | 17 | 0.9412 | 6 | 0.8333 | 12 | 0.9167 | 5 | 0.8 | 7 | 0.8571 | 30 | 0.8667 |
| 2_3471304 | 3 | 1 | 5 | 0.2 | 5 | 0.6 | 4 | 0 | 17 | 0.4118 | 6 | 0.5 | 11 | 0.5455 | 5 | 0.6 | 8 | 0.375 | 30 | 0.5 |
| 2_3474245 | 7 | 0 | 9 | 0.2222 | 5 | 0 | 12 | 0 | 33 | 0.0606 | 9 | 0.2222 | 7 | 0.1429 | 10 | 0 | 9 | 0 | 35 | 0.0857 |
| 2_3474259 | 6 | 0 | 7 | 0.1429 | 5 | 0 | 11 | 0.0909 | 29 | 0.069 | 8 | 0 | 7 | 0.1429 | 9 | 0 | 9 | 0.1111 | 33 | 0.0606 |
| 2_3474267 | 6 | 0.6667 | 9 | 0.3333 | 5 | 0.6 | 10 | 0.5 | 30 | 0.5 | 8 | 0.625 | 7 | 0.8571 | 7 | 0.4286 | 5 | 0.8 | 27 | 0.6667 |
| 2_3474270 | 7 | 0.5714 | 7 | 0.1429 | 5 | 0 | 10 | 0 | 29 | 0.1724 | 8 | 0.125 | 7 | 0.4286 | 7 | 0.1429 | 8 | 0.125 | 30 | 0.2 |
| 2_3474272 | 6 | 1 | 6 | 1 | 4 | 1 | 9 | 1 | 25 | 1 | 8 | 1 | 5 | 0.6 | 7 | 1 | 7 | 1 | 27 | 0.9259 |
| 2_3474277 | 7 | 0.1429 | 8 | 0.375 | 5 | 0 | 11 | 0.1818 | 31 | 0.1935 | 8 | 0.5 | 4 | 0 | 8 | 0.25 | 7 | 0.1429 | 27 | 0.2593 |
| 2_3474281 | 6 | 0.5 | 9 | 0.5556 | 4 | 0.25 | 11 | 0.5455 | 30 | 0.5 | 8 | 0.625 | 7 | 0.5714 | 8 | 0.875 | 9 | 0.5556 | 32 | 0.6562 |
| 2_3507041 | 33 | 0.0606 | 40 | 0.1 | 49 | 0.0204 | 31 | 0.129 | 153 | 0.0719 | 22 | 0.0909 | 33 | 0.0606 | 26 | 0.1154 | 31 | 0.129 | 112 | 0.0982 |
| 2_3507139 | 24 | 0.25 | 33 | 0.2424 | 26 | 0.3462 | 25 | 0.28 | 108 | 0.2778 | 17 | 0.2941 | 24 | 0.3333 | 23 | 0.2174 | 27 | 0.2593 | 91 | 0.2747 |
| 2_3507153 | 22 | 0.1364 | 34 | 0.0882 | 31 | 0.1935 | 27 | 0.1111 | 114 | 0.1316 | 18 | 0 | 21 | 0.0476 | 23 | 0.1304 | 31 | 0.0645 | 93 | 0.0645 |
| 2_3507184 | 30 | 0.0333 | 36 | 0.1667 | 34 | 0.1471 | 25 | 0.16 | 125 | 0.128 | 20 | 0.25 | 21 | 0.2381 | 23 | 0.1739 | 34 | 0.0588 | 98 | 0.1633 |
| 2_3507197 | 30 | 0.1 | 33 | 0.1212 | 32 | 0.0312 | 26 | 0.0769 | 121 | 0.0826 | 18 | 0.1667 | 19 | 0.1053 | 20 | 0.15 | 31 | 0.1935 | 88 | 0.1591 |
| 2_3529037 | 1 | 1 | NA | NA | NA | NA | NA | NA | 1 | 1 | 2 | 1 | 1 | 1 | NA | NA | 1 | 1 | 4 | 1 |
| 2_3529107 | 1 | 1 | NA | NA | NA | NA | NA | NA | 1 | 1 | 1 | 1 | 1 | 0 | NA | NA | NA | NA | 2 | 0.5 |
| 2_3529116 | 1 | 1 | NA | NA | NA | NA | NA | NA | 1 | 1 | 3 | 0.6667 | 1 | 0 | NA | NA | NA | NA | 4 | 0.5 |
| 2_3529129 | 1 | 1 | NA | NA | NA | NA | NA | NA | 1 | 1 | 3 | 1 | 1 | 0 | NA | NA | NA | NA | 4 | 0.75 |
| 2_3529145 | 1 | 1 | NA | NA | NA | NA | NA | NA | 1 | 1 | 3 | 0.6667 | NA | NA | NA | NA | NA | NA | 3 | 0.6667 |
| 2_3529147 | 1 | 1 | NA | NA | NA | NA | NA | NA | 1 | 1 | 3 | 0.6667 | NA | NA | NA | NA | NA | NA | 3 | 0.6667 |
| 2_3544487 | NA | NA | 2 | 1 | 1 | 0 | NA | NA | 3 | 0.6667 | 2 | 0.5 | NA | NA | NA | NA | NA | NA | 2 | 0.5 |
| 2_3560004 | NA | NA | 1 | 0 | 2 | 1 | NA | NA | 3 | 0.6667 | 3 | 0 | 3 | 1 | 2 | 0 | 1 | 1 | 9 | 0.4444 |
| 2_3560991 | 1 | 1 | 3 | 0.3333 | 3 | 0.6667 | 1 | 0 | 8 | 0.5 | NA | NA | 3 | 0.6667 | 3 | 0.6667 | 1 | 0 | 7 | 0.5714 |
| 2_3561003 | 1 | 1 | 3 | 0 | 4 | 0 | 1 | 1 | 9 | 0.2222 | 2 | 0 | 3 | 0.3333 | 2 | 0 | 1 | 0 | 8 | 0.125 |
| 2_3561032 | 1 | 0 | 3 | 0 | 5 | 0.2 | 1 | 1 | 10 | 0.2 | 3 | 0.6667 | 4 | 0.5 | 3 | 0 | 2 | 0.5 | 12 | 0.4167 |
| 2_3561045 | 1 | 0 | 3 | 1 | 3 | 1 | 1 | 1 | 8 | 0.875 | 3 | 0.6667 | 4 | 0.25 | 3 | 0.3333 | 2 | 0.5 | 12 | 0.4167 |
| 2_3561047 | NA | NA | 1 | 1 | 1 | 1 | 1 | 0 | 3 | 0.6667 | 2 | 0.5 | 3 | 1 | NA | NA | 2 | 1 | 7 | 0.8571 |
| 2_3561825 | 2 | 0.5 | 4 | 0.75 | 6 | 0.6667 | 2 | 1 | 14 | 0.7143 | 3 | 1 | 4 | 0.75 | 3 | 0.6667 | 1 | 0 | 11 | 0.7273 |
| 2_3561827 | 2 | 0 | 4 | 0.25 | 5 | 0 | 2 | 0.5 | 13 | 0.1538 | 3 | 0.3333 | 4 | 0 | 3 | 0 | NA | NA | 10 | 0.1 |
| 2_3561829 | 2 | 0 | 4 | 0.25 | 5 | 0 | 3 | 0 | 14 | 0.0714 | 2 | 0 | 4 | 0 | 2 | 0 | 1 | 1 | 9 | 0.1111 |
| 2_3561833 | 1 | 0 | 4 | 0.75 | 5 | 0.4 | 3 | 0.3333 | 13 | 0.4615 | 3 | 1 | 3 | 1 | 2 | 1 | 1 | 0 | 9 | 0.8889 |
| 2_3561835 | 1 | 0 | 4 | 0.75 | 4 | 0 | 1 | 1 | 10 | 0.4 | 1 | 0 | 2 | 0.5 | 2 | 0.5 | 1 | 1 | 6 | 0.5 |
| 2_3561839 | 2 | 0 | 3 | 0.6667 | 7 | 0.1429 | 2 | 0.5 | 14 | 0.2857 | 2 | 0 | 2 | 0 | 3 | 0.3333 | 1 | 0 | 8 | 0.125 |
| 2_3561861 | 1 | 0 | 4 | 0.25 | 6 | 0.1667 | 4 | 0.25 | 15 | 0.2 | 4 | 0 | 4 | 0.25 | 4 | 0 | 3 | 0 | 15 | 0.0667 |
| 2_3561872 | 3 | 0 | 5 | 0 | 7 | 0.2857 | 4 | 0 | 19 | 0.1053 | 4 | 0 | 4 | 0 | 4 | 0.25 | 3 | 0.3333 | 15 | 0.1333 |
| 2_3562006 | 3 | 0 | 10 | 0.2 | 8 | 0.25 | 5 | 0 | 26 | 0.1538 | 5 | 0 | 7 | 0.2857 | 8 | 0.125 | 7 | 0 | 27 | 0.1111 |
| 2_3562008 | 5 | 0 | 10 | 0 | 9 | 0.3333 | 7 | 0.1429 | 31 | 0.129 | 6 | 0 | 9 | 0.3333 | 10 | 0 | 7 | 0.2857 | 32 | 0.1562 |
| 2_3562018 | 5 | 0 | 9 | 0.1111 | 7 | 0 | 6 | 0.1667 | 27 | 0.0741 | 6 | 0.1667 | 8 | 0 | 9 | 0.1111 | 7 | 0 | 30 | 0.0667 |
| 2_3562060 | 7 | 0 | 11 | 0 | 10 | 0.1 | 7 | 0.1429 | 35 | 0.0571 | 11 | 0.1818 | 9 | 0.1111 | 9 | 0.1111 | 7 | 0 | 36 | 0.1111 |
| 2_3562070 | 8 | 0 | 12 | 0 | 12 | 0 | 7 | 0.4286 | 39 | 0.0769 | 10 | 0.1 | 9 | 0.3333 | 9 | 0 | 8 | 0 | 36 | 0.1111 |
| 2_3562074 | 8 | 0 | 10 | 0.2 | 12 | 0.25 | 6 | 0.3333 | 36 | 0.1944 | 11 | 0.0909 | 9 | 0.3333 | 11 | 0.2727 | 7 | 0.2857 | 38 | 0.2368 |
| 2_3563883 | 1 | 0 | 1 | 1 | NA | NA | 1 | 0 | 3 | 0.3333 | NA | NA | NA | NA | 2 | 0.5 | NA | NA | 2 | 0.5 |
| 2_3563937 | 1 | 0 | 1 | 1 | NA | NA | 1 | 0 | 3 | 0.3333 | 1 | 0 | NA | NA | 2 | 0.5 | NA | NA | 3 | 0.3333 |
| 2_3564131 | 6 | 0.3333 | 5 | 0 | 4 | 0.25 | 3 | 0 | 18 | 0.1667 | 5 | 0 | 1 | 0 | 6 | 0.3333 | 2 | 0 | 14 | 0.1429 |
| 2_3564157 | 6 | 0.3333 | 4 | 0.5 | 5 | 0.2 | 4 | 0.25 | 19 | 0.3158 | 3 | 0 | 1 | 0 | 5 | 0.2 | 2 | 0.5 | 11 | 0.1818 |
| 2_3564171 | 5 | 1 | 6 | 1 | 5 | 1 | 4 | 1 | 20 | 1 | 4 | 0.75 | 1 | 0 | 5 | 1 | 2 | 1 | 12 | 0.8333 |
| 2_3564174 | 5 | 0 | 7 | 0.1429 | 6 | 0.1667 | 4 | 0 | 22 | 0.0909 | 3 | 0.3333 | 2 | 0 | 4 | 0 | 2 | 0 | 11 | 0.0909 |
| 2_3564185 | 4 | 1 | 3 | 1 | 2 | 0.5 | 4 | 0.25 | 13 | 0.6923 | 2 | 0 | 1 | 0 | 4 | 0.5 | 1 | 0 | 8 | 0.25 |
| 2_3564189 | 5 | 0.8 | 5 | 0.8 | 1 | 1 | 5 | 0.6 | 16 | 0.75 | 4 | 0.75 | 1 | 0 | 5 | 0.8 | 2 | 1 | 12 | 0.75 |
| 2_3564199 | 5 | 0.4 | 6 | 0.1667 | 5 | 0.2 | 3 | 0 | 19 | 0.2105 | 4 | 0 | 2 | 0 | 4 | 0.75 | 2 | 0.5 | 12 | 0.3333 |
| 2_3564224 | 5 | 0 | 4 | 0.25 | 5 | 0 | 4 | 0 | 18 | 0.0556 | 3 | 0.3333 | 3 | 0 | 4 | 0 | 2 | 0.5 | 12 | 0.1667 |
| 2_3564231 | 3 | 0.3333 | 2 | 0.5 | 3 | 1 | 2 | 0 | 10 | 0.5 | 2 | 0.5 | 2 | 1 | 1 | 1 | 1 | 0 | 6 | 0.6667 |
| 2_3564240 | 5 | 0 | 5 | 0 | 4 | 0.25 | 4 | 0 | 18 | 0.0556 | 3 | 0 | 2 | 0.5 | 5 | 0.2 | 2 | 0 | 12 | 0.1667 |
| 2_3579266 | 4 | 0.75 | 2 | 0.5 | 3 | 0.3333 | 2 | 0.5 | 11 | 0.5455 | 1 | 1 | 2 | 1 | 5 | 0.8 | 1 | 1 | 9 | 0.8889 |

| | | | | | | | | | | | | | | | | | | | | |
|---|---|---|---|---|---|---|---|---|---|---|---|---|---|---|---|---|---|---|---|---|
| 2_35792709 | 4 | 0.25 | 2 | 0.5 | 3 | 0 | 2 | 0.5 | 11 | 0.2727 | 1 | 1 | 2 | 0.5 | 3 | 0.3333 | 1 | 1 | 7 | 0.5714 |
| 2_35792729 | 4 | 0.25 | 1 | 1 | 3 | 0.6667 | 2 | 1 | 10 | 0.6 | 1 | 0 | 1 | 1 | 4 | 0.75 | 1 | 1 | 7 | 0.7143 |
| 2_35792752 | 4 | 1 | 2 | 1 | 3 | 1 | 2 | 0.5 | 11 | 0.9091 | 1 | 1 | 2 | 1 | 4 | 0.75 | 1 | 1 | 8 | 0.875 |
| 2_35792864 | 3 | 0.3333 | 2 | 0.5 | 3 | 0 | NA | NA | 8 | 0.25 | NA | NA | 1 | 0 | 4 | 0.25 | 1 | 0 | 6 | 0.1667 |
| 2_35792897 | 4 | 0.75 | 2 | 1 | 3 | 1 | 2 | 1 | 11 | 0.9091 | 1 | 1 | 1 | 1 | 3 | 1 | 1 | 1 | 6 | 1 |
| 2_35792972 | 3 | 0 | 2 | 0 | 3 | 0 | 2 | 0.5 | 10 | 0.1 | 1 | 0 | 1 | 0 | 5 | 0.2 | 1 | 0 | 8 | 0.125 |
| 2_35793034 | 4 | 0.5 | 2 | 0.5 | 3 | 0.3333 | 2 | 0 | 11 | 0.3636 | 1 | 0 | NA | NA | 5 | 0.2 | 1 | 0 | 7 | 0.1429 |
| 2_35869251 | 3 | 0 | 2 | 0 | 2 | 0.5 | 1 | 0 | 8 | 0.125 | 1 | 0 | 2 | 0.5 | 4 | 0 | 1 | 0 | 8 | 0.125 |
| 2_35869831 | NA | NA | NA | NA | NA | NA | 1 | 1 | 1 | 1 | 1 | 1 | 1 | 1 | NA | NA | NA | NA | 2 | 1 |
| 2_35869962 | NA | NA | NA | NA | NA | NA | 1 | 1 | 1 | 1 | 1 | 1 | NA | NA | NA | NA | NA | NA | 1 | 1 |
| 2_35926415 | 30 | 0.2 | 33 | 0.2424 | 38 | 0.1053 | 40 | 0.225 | 141 | 0.1915 | 31 | 0.1935 | 33 | 0.2424 | 36 | 0.1667 | 41 | 0.1707 | 141 | 0.1915 |
| 2_35926524 | 17 | 0.1176 | 22 | 0.0909 | 33 | 0.0606 | 28 | 0.0714 | 100 | 0.08 | 24 | 0.125 | 22 | 0 | 28 | 0.0357 | 34 | 0.0588 | 108 | 0.0556 |
| 2_35972409 | 2 | 0 | 3 | 1 | 2 | 0.5 | 2 | 0 | 9 | 0.4444 | 3 | 0 | 1 | 0 | 5 | 0.2 | NA | NA | 9 | 0.1111 |
| 2_35972441 | 2 | 0 | 4 | 0.5 | 2 | 0 | 2 | 0.5 | 10 | 0.3 | 3 | 0.3333 | NA | NA | 3 | 0 | NA | NA | 6 | 0.1667 |
| 2_35972516 | 1 | 0 | 3 | 0 | 2 | 0 | 2 | 1 | 8 | 0.25 | 3 | 0 | 1 | 0 | 4 | 0.25 | 1 | 1 | 9 | 0.2222 |
| 2_35972523 | 2 | 1 | 4 | 0.5 | 2 | 0.5 | 2 | 1 | 10 | 0.7 | 3 | 0.3333 | 1 | 0 | 4 | 0 | 1 | 0 | 9 | 0.1111 |
| 2_35972599 | 2 | 0 | 2 | 0.5 | 2 | 0 | 1 | 1 | 7 | 0.2857 | 2 | 0.5 | 1 | 0 | 4 | 0.75 | 1 | 1 | 8 | 0.625 |
| 2_35972752 | 2 | 0 | 4 | 0.25 | 2 | 0 | 2 | 0 | 10 | 0.1 | 2 | 0 | NA | NA | 3 | 0 | 1 | 1 | 6 | 0.1667 |
| 2_35972841 | 2 | 0 | 4 | 0.25 | 3 | 0 | 2 | 0 | 11 | 0.0909 | 3 | 0.3333 | 1 | 1 | 2 | 0 | 1 | 0 | 7 | 0.2857 |
| 2_35973352 | 2 | 1 | 3 | 1 | 2 | 1 | 2 | 1 | 9 | 1 | 2 | 1 | 1 | 1 | 1 | 1 | NA | NA | 4 | 1 |
| 2_35973366 | 2 | 1 | 3 | 1 | 3 | 1 | 2 | 1 | 10 | 1 | 2 | 0.5 | NA | NA | 1 | 1 | NA | NA | 3 | 0.6667 |
| 2_35973487 | 2 | 0.5 | 3 | 0.6667 | 3 | 1 | 1 | 1 | 9 | 0.7778 | 2 | 0 | 1 | 1 | 1 | 1 | NA | NA | 4 | 0.5 |
| 2_35973587 | 2 | 1 | 3 | 0.3333 | 2 | 0.5 | NA | NA | 7 | 0.5714 | 2 | 1 | NA | NA | 1 | 1 | NA | NA | 3 | 1 |
| 2_35973654 | 2 | 0 | 2 | 0.5 | 3 | 0 | 1 | 0 | 8 | 0.125 | 2 | 0 | NA | NA | 1 | 1 | NA | NA | 3 | 0.3333 |
| 2_35992879 | 2 | 0 | 2 | 0.5 | 1 | 0 | NA | NA | 5 | 0.2 | 1 | 0 | 1 | 1 | NA | NA | NA | NA | 2 | 0.5 |
| 2_36006489 | 7 | 0.1429 | 12 | 0.1667 | 10 | 0.1 | 10 | 0.2 | 39 | 0.1538 | 6 | 0 | 12 | 0.0833 | 8 | 0.25 | 11 | 0.0909 | 37 | 0.1081 |
| 2_36006527 | 7 | 0.2857 | 12 | 0.1667 | 11 | 0 | 11 | 0 | 41 | 0.0976 | 5 | 0 | 11 | 0.0909 | 7 | 0 | 12 | 0.1667 | 35 | 0.0857 |
| 2_36006612 | 7 | 0.2857 | 13 | 0.1538 | 10 | 0.4 | 10 | 0.1 | 40 | 0.225 | 5 | 0.4 | 11 | 0.0909 | 5 | 0.2 | 11 | 0.2727 | 32 | 0.2188 |
| 2_36007378 | 7 | 0.1429 | 12 | 0.5 | 10 | 0.2 | 7 | 0 | 36 | 0.25 | 5 | 0 | 8 | 0.125 | 4 | 0.25 | 6 | 0.1667 | 23 | 0.1304 |
| 2_36007603 | 6 | 0.1667 | 7 | 0.1429 | 7 | 0 | 7 | 0.1429 | 27 | 0.1111 | 3 | 0.3333 | 7 | 0 | 3 | 0 | 6 | 0 | 19 | 0.0526 |
| 2_36007615 | 7 | 0.5714 | 11 | 0.2727 | 11 | 0.1818 | 8 | 0.25 | 37 | 0.2973 | 4 | 0.25 | 10 | 0.3 | 5 | 0 | 7 | 0.5714 | 26 | 0.3077 |
| 2_36007663 | 4 | 0.25 | 10 | 0.3 | 9 | 0.1111 | 8 | 0.125 | 31 | 0.1935 | 4 | 0 | 10 | 0 | 5 | 0.2 | 6 | 0.1667 | 25 | 0.08 |
| 2_36007705 | 5 | 0 | 8 | 0.125 | 11 | 0.0909 | 8 | 0 | 32 | 0.0625 | 3 | 0 | 10 | 0 | 5 | 0 | 8 | 0.25 | 26 | 0.0769 |
| 2_36007868 | 6 | 0.8333 | 11 | 1 | 11 | 1 | 8 | 0.875 | 36 | 0.9444 | 4 | 1 | 9 | 1 | 4 | 0.75 | 7 | 0.8571 | 24 | 0.9167 |
| 2_36079394 | NA | NA | 1 | 1 | 3 | 0.6667 | 2 | 1 | 6 | 0.8333 | 4 | 1 | 1 | 1 | 2 | 1 | 3 | 1 | 10 | 1 |
| 2_36079412 | NA | NA | 2 | 0.5 | 5 | 0 | 2 | 0.5 | 9 | 0.2222 | 5 | 0.2 | 1 | 0 | 2 | 0.5 | 2 | 0 | 10 | 0.2 |
| 2_36079418 | NA | NA | 2 | 0 | 4 | 0.25 | 2 | 0 | 8 | 0.125 | 4 | 0.25 | 1 | 0 | 3 | 0.3333 | 3 | 0.6667 | 11 | 0.3636 |
| 2_36079495 | NA | NA | 1 | 1 | 1 | 1 | 2 | 1 | 4 | 1 | 4 | 1 | 1 | 1 | 2 | 1 | 3 | 0.6667 | 10 | 0.9 |
| 2_36079525 | NA | NA | 2 | 0 | 5 | 0.2 | 2 | 0 | 9 | 0.1111 | 5 | 0.2 | 1 | 0 | 2 | 0 | 4 | 0 | 12 | 0.0833 |
| 2_36079557 | NA | NA | 2 | 0 | 4 | 0.5 | 2 | 1 | 8 | 0.5 | 4 | 0.5 | 1 | 0 | 3 | 0 | 4 | 0.25 | 12 | 0.25 |
| 2_36079640 | NA | NA | 1 | 1 | NA | NA | 1 | 0 | 2 | 0.5 | 1 | 1 | NA | NA | 2 | 0 | 2 | 0.5 | 5 | 0.4 |
| 2_36141413 | 2 | 0 | 2 | 1 | 1 | 0 | 2 | 0 | 7 | 0.2857 | 2 | 0.5 | NA | NA | 1 | 0 | 1 | 1 | 4 | 0.5 |
| 2_36141473 | 2 | 1 | 4 | 0.75 | NA | NA | 3 | 1 | 9 | 0.8889 | 2 | 1 | 1 | 1 | 1 | 1 | 1 | 1 | 5 | 1 |
| 2_36142149 | 3 | 0 | 4 | 0.25 | NA | NA | 3 | 0 | 10 | 0.1 | 4 | 0 | 1 | 0 | 1 | 1 | 1 | 0 | 7 | 0.1429 |
| 2_36142181 | 3 | 1 | 4 | 0.5 | 1 | 1 | 3 | 0.3333 | 11 | 0.6364 | 4 | 0.5 | NA | NA | NA | NA | 1 | 1 | 5 | 0.6 |
| 2_37240651 | 2 | 0 | 9 | 0.1111 | 4 | 0 | 3 | 0 | 18 | 0.0556 | 4 | 0.25 | 3 | 0 | 4 | 0.25 | 2 | 0.5 | 13 | 0.2308 |
| 2_37241137 | 2 | 0.5 | 10 | 0.5 | 3 | 0 | 3 | 0 | 18 | 0.3333 | 3 | 0 | 2 | 0 | 5 | 0.2 | 2 | 0 | 12 | 0.0833 |
| 2_37241797 | 2 | 0.5 | 8 | 0 | 4 | 0.25 | 3 | 0 | 17 | 0.1176 | 3 | 0 | 2 | 1 | 5 | 0.2 | 2 | 0.5 | 12 | 0.3333 |
| 2_37241842 | 1 | 1 | 8 | 0.875 | 2 | 1 | 3 | 1 | 14 | 0.9286 | 2 | 1 | 2 | 1 | 5 | 1 | 2 | 1 | 11 | 1 |
| 2_37241889 | 2 | 0.5 | 6 | 0.3333 | 4 | 0.25 | 3 | 0 | 15 | 0.2667 | 2 | 0 | 2 | 0 | 5 | 0.4 | 2 | 0 | 11 | 0.1818 |
| 2_37241961 | 2 | 0.5 | 8 | 0.125 | 4 | 0 | 3 | 0.3333 | 17 | 0.1765 | 2 | 0 | 2 | 0 | 5 | 0.2 | 2 | 0 | 11 | 0.0909 |
| 2_37242166 | 1 | 1 | 7 | 1 | 4 | 1 | 3 | 0.3333 | 15 | 0.8667 | 1 | 1 | 2 | 1 | 4 | 1 | 2 | 0.5 | 9 | 0.8889 |
| 2_37242218 | 1 | 1 | 5 | 1 | 4 | 1 | 3 | 1 | 13 | 1 | 1 | 0 | 2 | 1 | 4 | 0.25 | 2 | 0.5 | 9 | 0.4444 |
| 2_37261867 | 6 | 0 | 2 | 0.5 | 4 | 0 | 6 | 0 | 18 | 0.0556 | 5 | 0 | 8 | 0.25 | 5 | 0.2 | 6 | 0.1667 | 24 | 0.1667 |
| 2_37262240 | 4 | 0 | 2 | 0 | 5 | 0.2 | 6 | 0.1667 | 17 | 0.1176 | 4 | 0 | 7 | 0.4286 | 3 | 0 | 4 | 0 | 18 | 0.1667 |
| 2_37263322 | 6 | 0.3333 | NA | NA | 3 | 0 | 3 | 0 | 12 | 0.1667 | 3 | 0.3333 | 7 | 0.1429 | 5 | 0.2 | 4 | 0 | 19 | 0.1579 |
| 2_37265007 | 2 | 0 | NA | NA | 2 | 0.5 | 2 | 0 | 6 | 0.1667 | 1 | 0 | 3 | 0 | 3 | 0.3333 | 2 | 0 | 9 | 0.1111 |
| 2_37265108 | 2 | 1 | NA | NA | 2 | 1 | 2 | 1 | 6 | 1 | 1 | 1 | 3 | 1 | 3 | 1 | 1 | 1 | 8 | 1 |
| 2_37272949 | 1 | 0 | NA | NA | 1 | 0 | 2 | 0.5 | 4 | 0.25 | 1 | 0 | 2 | 0 | 2 | 0.5 | NA | NA | 5 | 0.2 |
| 2_37273035 | 1 | 0 | NA | NA | 2 | 0 | 2 | 1 | 5 | 0.4 | 1 | 0 | 1 | 1 | 2 | 0.5 | NA | NA | 4 | 0.5 |
| 2_37279679 | 1 | 1 | NA | NA | 2 | 1 | 2 | 0.5 | 5 | 0.8 | 1 | 0 | 1 | 0 | 1 | 1 | NA | NA | 3 | 0.3333 |
| 2_37315243 | NA | NA | 1 | 1 | NA | NA | NA | NA | 1 | 1 | NA | NA | NA | NA | 1 | 1 | NA | NA | 1 | 1 |
| 2_37317899 | 3 | 0.3333 | 1 | 0 | 1 | 1 | NA | NA | 5 | 0.4 | 1 | 1 | 1 | 0 | 1 | 0 | 2 | 1 | 5 | 0.6 |
| 2_37318544 | 2 | 0.5 | 1 | 0 | 1 | 0 | NA | NA | 4 | 0.25 | 1 | 1 | 1 | 0 | 1 | 0 | 1 | 0 | 4 | 0.25 |
| 2_37318643 | 2 | 1 | 1 | 1 | NA | NA | NA | NA | 3 | 1 | 1 | 1 | NA | NA | 2 | 1 | 1 | 1 | 4 | 1 |

| | | | | | | | | | | | | | | | | | | | | |
|---|---|---|---|---|---|---|---|---|---|---|---|---|---|---|---|---|---|---|---|---|
| 2_37318818 | 2 | 0.5 | 1 | 0 | 1 | 1 | NA | NA | 4 | 0.5 | 1 | 0 | NA | NA | 2 | 1 | 2 | 1 | 5 | 0.8 |
| 2_37318846 | 2 | 1 | 1 | 1 | 1 | 1 | NA | NA | 4 | 1 | 1 | 1 | NA | NA | 2 | 1 | 2 | 1 | 5 | 1 |
| 2_37318889 | 1 | 1 | 1 | 1 | 1 | 1 | NA | NA | 3 | 1 | NA | NA | NA | NA | 2 | 1 | 1 | 1 | 3 | 1 |
| 2_37318958 | 1 | 1 | 1 | 1 | 1 | 1 | NA | NA | 3 | 1 | 1 | 1 | NA | NA | 2 | 1 | 1 | 1 | 4 | 1 |
| 2_37318986 | 1 | 1 | 1 | 0 | 1 | 1 | NA | NA | 3 | 0.6667 | 1 | 0 | NA | NA | 1 | 0 | 2 | 0.5 | 4 | 0.25 |
| 2_37319137 | 2 | 0 | NA | NA | 1 | 1 | NA | NA | 3 | 0.3333 | 1 | 1 | NA | NA | 1 | 0 | 1 | 0 | 3 | 0.3333 |
| 2_37319145 | 2 | 0.5 | NA | NA | 1 | 1 | NA | NA | 3 | 0.6667 | 1 | 0 | NA | NA | 1 | 0 | 1 | 1 | 3 | 0.3333 |
| 2_37319356 | 2 | 0.5 | 1 | 1 | 1 | 1 | NA | NA | 4 | 0.75 | 1 | 1 | NA | NA | 1 | 1 | 2 | 1 | 4 | 1 |
| 2_37319451 | 1 | 1 | 1 | 1 | 1 | 1 | NA | NA | 3 | 1 | NA | NA | NA | NA | NA | NA | 2 | 1 | 2 | 1 |
| 2_37319538 | 2 | 0.5 | NA | NA | 1 | 1 | NA | NA | 3 | 0.6667 | 1 | 0 | NA | NA | 1 | 1 | 1 | 1 | 3 | 0.6667 |
| 2_37319614 | 2 | 0.5 | 1 | 0 | 1 | 1 | NA | NA | 4 | 0.5 | 1 | 1 | NA | NA | NA | NA | 2 | 0 | 3 | 0.3333 |
| 2_37325782 | NA | NA | NA | NA | NA | NA | 1 | 1 | 1 | 1 | NA | NA | NA | NA | NA | NA | 2 | 1 | 2 | 1 |
| 2_37325809 | NA | NA | NA | NA | NA | NA | 1 | 1 | 1 | 1 | 1 | 1 | NA | NA | NA | NA | 2 | 1 | 3 | 1 |
| 2_37325974 | NA | NA | NA | NA | NA | NA | 1 | 1 | 1 | 1 | 1 | 1 | NA | NA | NA | NA | 2 | 0.5 | 3 | 0.6667 |
| 2_37326114 | NA | NA | NA | NA | NA | NA | 1 | 1 | 1 | 1 | NA | NA | NA | NA | NA | NA | 1 | 1 | 1 | 1 |
| 2_37326309 | NA | NA | NA | NA | NA | NA | 1 | 1 | 1 | 1 | NA | NA | NA | NA | NA | NA | 2 | 1 | 2 | 1 |
| 2_37326472 | NA | NA | NA | NA | NA | NA | 1 | 1 | 1 | 1 | NA | NA | NA | NA | NA | NA | 2 | 1 | 2 | 1 |
| 2_37326517 | NA | NA | NA | NA | NA | NA | 1 | 1 | 1 | 1 | 1 | 1 | NA | NA | NA | NA | 1 | 1 | 2 | 1 |
| 2_37326705 | NA | NA | NA | NA | NA | NA | 1 | 1 | 1 | 1 | 1 | 1 | NA | NA | NA | NA | 1 | 1 | 2 | 1 |
| 2_37332923 | 1 | 0 | NA | NA | 1 | 1 | NA | NA | 2 | 0.5 | NA | NA | NA | NA | NA | NA | 3 | 0.3333 | 3 | 0.3333 |
| 2_37407227 | NA | NA | 1 | 1 | NA | NA | NA | NA | 1 | 1 | NA | NA | NA | NA | 1 | 1 | NA | NA | 1 | 1 |
| 2_37407525 | NA | NA | 1 | 1 | NA | NA | NA | NA | 1 | 1 | 1 | 1 | NA | NA | 1 | 1 | NA | NA | 2 | 1 |
| 2_37408320 | NA | NA | 1 | 1 | NA | NA | NA | NA | 1 | 1 | 1 | 1 | NA | NA | 1 | 0 | NA | NA | 2 | 0.5 |
| 2_37408397 | NA | NA | 1 | 1 | NA | NA | NA | NA | 1 | 1 | 1 | 1 | NA | NA | 1 | 1 | NA | NA | 2 | 1 |
| 2_37408456 | NA | NA | 1 | 1 | NA | NA | NA | NA | 1 | 1 | 1 | 1 | NA | NA | 1 | 1 | NA | NA | 2 | 1 |
| 2_37408481 | NA | NA | 1 | 1 | NA | NA | NA | NA | 1 | 1 | 1 | 1 | NA | NA | 1 | 1 | NA | NA | 2 | 1 |
| 2_37408620 | NA | NA | 1 | 1 | NA | NA | NA | NA | 1 | 1 | 1 | 1 | NA | NA | 1 | 1 | NA | NA | 2 | 1 |
| 2_37408937 | NA | NA | 1 | 1 | NA | NA | NA | NA | 1 | 1 | 1 | 1 | NA | NA | 1 | 1 | NA | NA | 2 | 1 |
| 2_37409509 | NA | NA | 1 | 1 | NA | NA | NA | NA | 1 | 1 | NA | NA | NA | NA | 1 | 1 | NA | NA | 1 | 1 |
| 2_37433143 | 3 | 0 | 2 | 0.5 | 3 | 0 | 5 | 0 | 13 | 0.0769 | 2 | 0 | 3 | 0 | 6 | 0.1667 | 4 | 0 | 15 | 0.0667 |
| 2_37454778 | 5 | 0.2 | 3 | 0 | 3 | 0 | 5 | 0 | 16 | 0.0625 | 2 | 0 | 5 | 0.2 | 8 | 0 | 5 | 0 | 20 | 0.05 |
| 2_37454864 | 5 | 0 | 3 | 0 | 3 | 0.3333 | 3 | 0 | 14 | 0.0714 | 2 | 0.5 | 6 | 0 | 8 | 0.125 | 5 | 0 | 21 | 0.0952 |
| 2_37454891 | 5 | 0.2 | 4 | 0.25 | 3 | 0 | 5 | 0.4 | 17 | 0.2353 | 2 | 0 | 7 | 0.5714 | 8 | 0 | 5 | 0.2 | 22 | 0.2273 |
| 2_37459974 | 6 | 0.1667 | 20 | 0.3 | 15 | 0.4667 | 10 | 0.4 | 51 | 0.3529 | 8 | 0.5 | 7 | 0 | 6 | 0.3333 | 10 | 0.4 | 31 | 0.3226 |
| 2_37459986 | 8 | 0.875 | 20 | 1 | 14 | 0.9286 | 11 | 1 | 53 | 0.9623 | 8 | 0.875 | 5 | 0.8 | 6 | 1 | 9 | 0.8889 | 28 | 0.8929 |
| 2_37460051 | 9 | 0.1111 | 21 | 0.4762 | 16 | 0.25 | 10 | 0.3 | 56 | 0.3214 | 9 | 0.4444 | 7 | 0.5714 | 6 | 0 | 9 | 0.6667 | 31 | 0.4516 |
| 2_37460099 | 6 | 0.8333 | 21 | 0.8571 | 14 | 0.9286 | 10 | 0.8 | 51 | 0.8627 | 9 | 0.8889 | 7 | 0.7143 | 6 | 0.6667 | 7 | 0.8571 | 29 | 0.7931 |
| 2_37460145 | 8 | 0.875 | 18 | 0.8889 | 13 | 0.9231 | 11 | 1 | 50 | 0.92 | 7 | 1 | 7 | 0.8571 | 6 | 1 | 10 | 0.9 | 30 | 0.9333 |
| 2_37460152 | 6 | 0.5 | 16 | 0.4375 | 14 | 0.4286 | 10 | 0.4 | 46 | 0.4348 | 6 | 0.3333 | 6 | 0.3333 | 6 | 0.5 | 10 | 0.6 | 28 | 0.4643 |
| 2_37460189 | 7 | 0.7143 | 18 | 0.6667 | 13 | 0.7692 | 7 | 1 | 45 | 0.7556 | 7 | 0.8571 | 6 | 0.8333 | 5 | 0.8 | 9 | 0.8889 | 27 | 0.8519 |
| 2_37460793 | 4 | 0 | 10 | 0.4 | 7 | 0.2857 | 9 | 0.5556 | 30 | 0.3667 | 8 | 0.25 | 6 | 0.3333 | 2 | 1 | 8 | 0.75 | 24 | 0.5 |
| 2_37461271 | 2 | 0 | 11 | 0.0909 | 8 | 0.125 | 8 | 0 | 29 | 0.069 | 6 | 0.1667 | 5 | 0 | 1 | 0 | 5 | 0 | 17 | 0.0588 |
| 2_37473918 | 4 | 0.25 | 10 | 0 | 2 | 0 | 4 | 0.25 | 20 | 0.1 | 1 | 0 | 5 | 0 | 3 | 0.3333 | 4 | 0 | 13 | 0.0769 |
| 2_37686644 | NA | NA | NA | NA | 1 | 1 | NA | NA | 1 | 1 | 1 | 1 | NA | NA | NA | NA | NA | NA | 1 | 1 |
| 2_37687720 | NA | NA | 1 | 1 | 1 | 1 | 1 | 1 | 3 | 1 | 1 | 1 | NA | NA | NA | NA | NA | NA | 1 | 1 |
| 2_37687790 | NA | NA | 1 | 1 | 1 | 1 | NA | NA | 2 | 1 | 1 | 1 | NA | NA | NA | NA | NA | NA | 1 | 1 |
| 2_37687985 | NA | NA | 1 | 1 | 1 | 0 | 1 | 1 | 3 | 0.6667 | 1 | 1 | NA | NA | NA | NA | NA | NA | 1 | 1 |
| 2_37688104 | NA | NA | 1 | 1 | 1 | 1 | 1 | 1 | 3 | 1 | 1 | 1 | NA | NA | NA | NA | NA | NA | 1 | 1 |
| 2_37688448 | NA | NA | 1 | 1 | 1 | 0 | 1 | 1 | 3 | 0.6667 | 1 | 1 | NA | NA | NA | NA | NA | NA | 1 | 1 |
| 2_37688532 | NA | NA | 1 | 0 | 1 | 1 | 1 | 1 | 3 | 0.6667 | 1 | 1 | NA | NA | NA | NA | NA | NA | 1 | 1 |
| 2_37689311 | 5 | 0.2 | 8 | 0.375 | 8 | 0.5 | 9 | 0.1111 | 30 | 0.3 | 2 | 0.5 | 5 | 0.2 | 7 | 0.1429 | 1 | 0 | 15 | 0.2 |
| 2_37689329 | 8 | 0.25 | 9 | 0.6667 | 8 | 0.75 | 7 | 0.2857 | 32 | 0.5 | 2 | 0.5 | 5 | 1 | 8 | 0.75 | 1 | 1 | 16 | 0.8125 |
| 2_37689350 | 8 | 0 | 9 | 0.1111 | 9 | 0.1111 | 9 | 0 | 35 | 0.0571 | 3 | 0 | 5 | 0 | 8 | 0.125 | 1 | 0 | 17 | 0.0588 |
| 2_37689712 | 3 | 0 | 4 | 0 | 5 | 0.4 | 6 | 0.3333 | 18 | 0.2222 | 1 | 1 | 4 | 0.75 | 4 | 0.5 | 1 | 1 | 10 | 0.7 |
| 2_37689853 | 4 | 0 | 4 | 0.25 | 8 | 0.375 | 8 | 0.25 | 24 | 0.25 | 2 | 0 | 5 | 0.6 | 6 | 0.5 | 1 | 0 | 14 | 0.4286 |
| 2_37689973 | 5 | 0.4 | 5 | 0.8 | 8 | 0.875 | 8 | 0.625 | 26 | 0.6923 | 1 | 1 | 5 | 1 | 6 | 0.6667 | 1 | 0 | 13 | 0.7692 |
| 2_37690069 | 5 | 0.2 | 5 | 0 | 7 | 0.2857 | 8 | 0.375 | 25 | 0.24 | NA | NA | 5 | 0.2 | 7 | 0.1429 | 1 | 0 | 13 | 0.1538 |
| 2_37690169 | 5 | 0 | 6 | 0.1667 | 9 | 0 | 8 | 0.125 | 28 | 0.0714 | 1 | 0 | 5 | 0.4 | 6 | 0.1667 | 1 | 0 | 13 | 0.2308 |
| 2_37690795 | 5 | 0 | 4 | 0.5 | 5 | 0.4 | 6 | 0.5 | 20 | 0.35 | 1 | 1 | 4 | 0 | 6 | 0.3333 | NA | NA | 11 | 0.2727 |
| 2_37690864 | 4 | 0.5 | 5 | 0 | 5 | 0.2 | 6 | 0 | 20 | 0.15 | NA | NA | 5 | 0.2 | 5 | 0 | 1 | 0 | 11 | 0.0909 |
| 2_37690874 | 5 | 0.2 | 4 | 0.5 | 5 | 1 | 6 | 0.5 | 20 | 0.55 | NA | NA | 4 | 0.5 | 3 | 0.3333 | NA | NA | 7 | 0.4286 |
| 2_37690901 | 2 | 0 | 3 | 0.6667 | 3 | 0 | 4 | 0.25 | 12 | 0.25 | NA | NA | 4 | 0.25 | 4 | 0.25 | NA | NA | 8 | 0.25 |
| 2_37691034 | 4 | 0.25 | 5 | 0 | 4 | 0 | 6 | 0 | 19 | 0.0526 | NA | NA | 4 | 0 | 3 | 0.3333 | NA | NA | 7 | 0.1429 |
| 2_37691260 | 4 | 0.25 | 3 | 0 | 4 | 0 | 5 | 0 | 16 | 0.0625 | 1 | 0 | 5 | 0.2 | 3 | 0 | NA | NA | 9 | 0.1111 |
| 2_38244535 | 2 | 0.5 | 4 | 0.5 | 6 | 0.3333 | 7 | 0.4286 | 19 | 0.4211 | 5 | 0.2 | 1 | 0 | 3 | 0.6667 | 5 | 0.4 | 14 | 0.3571 |

| | | | | | | | | | | | | | | | | | | | | |
|---|---|---|---|---|---|---|---|---|---|---|---|---|---|---|---|---|---|---|---|---|
| 2_38244616 | 2 | 0 | 3 | 0.3333 | 5 | 0.4 | 7 | 0.4286 | 17 | 0.3529 | 5 | 0.4 | 2 | 0.5 | 3 | 0 | 4 | 1 | 14 | 0.5 |
| 2_38259026 | 4 | 1 | 7 | 0.8571 | 6 | 0.8333 | 10 | 0.2 | 27 | 0.6296 | 7 | 0.4286 | 2 | 0 | 5 | 0.6 | 5 | 0.6 | 19 | 0.4737 |
| 2_38259031 | 4 | 0 | 8 | 0.25 | 6 | 0 | 9 | 0.1111 | 27 | 0.1111 | 8 | 0.125 | 2 | 0 | 3 | 0 | 5 | 0 | 18 | 0.0556 |
| 2_38259144 | 3 | 0.3333 | 7 | 0.5714 | 6 | 0.3333 | 7 | 0.7143 | 23 | 0.5217 | 5 | 0.8 | 2 | 0.5 | 3 | 0.6667 | 4 | 0.5 | 14 | 0.6429 |
| 2_38259387 | 3 | 1 | 7 | 0.8571 | 7 | 0.8571 | 11 | 1 | 28 | 0.9286 | 9 | 1 | 1 | 1 | 4 | 1 | 5 | 0.8 | 19 | 0.9474 |
| 2_38259446 | 3 | 1 | 6 | 1 | 5 | 0.8 | 10 | 0.8 | 24 | 0.875 | 6 | 0.6667 | 1 | 1 | 4 | 0.75 | 5 | 0.8 | 16 | 0.75 |
| 2_38475388 | 4 | 0.25 | 4 | 0 | 4 | 0 | 1 | 0 | 13 | 0.0769 | 4 | 0 | 7 | 0.1429 | 2 | 0 | 2 | 0 | 15 | 0.0667 |
| 2_38475487 | 6 | 0.3333 | 4 | 0 | 4 | 0 | 2 | 0 | 16 | 0.125 | 3 | 0 | 7 | 0.1429 | 3 | 0.3333 | 2 | 0 | 15 | 0.1333 |
| 2_38475495 | 6 | 0.8333 | 3 | 1 | 4 | 1 | 2 | 1 | 15 | 0.9333 | 3 | 1 | 6 | 0.8333 | 3 | 0.6667 | 2 | 1 | 14 | 0.8571 |
| 2_38475542 | 6 | 0.3333 | 4 | 0.25 | 4 | 0.25 | 1 | 0 | 15 | 0.2667 | 4 | 0 | 6 | 0.1667 | 3 | 0 | 2 | 0 | 15 | 0.0667 |
| 2_38475563 | 4 | 0.75 | 4 | 1 | 3 | 1 | 1 | 1 | 12 | 0.9167 | 2 | 1 | 2 | 1 | 3 | 1 | 2 | 1 | 9 | 1 |
| 2_38475573 | 5 | 0.2 | 4 | 0 | 3 | 0 | 2 | 0 | 14 | 0.0714 | 3 | 0 | 5 | 0.2 | 3 | 0 | 2 | 0 | 13 | 0.0769 |
| 2_38475692 | 4 | 0 | 4 | 0 | 3 | 0 | 2 | 0.5 | 13 | 0.0769 | 3 | 0 | 6 | 0.1667 | 4 | 0.25 | 2 | 0.5 | 15 | 0.2 |
| 2_38476112 | 5 | 0 | 3 | 0 | 4 | 0.25 | 2 | 0.5 | 14 | 0.1429 | 5 | 0 | 7 | 0.2857 | 4 | 0 | 2 | 0.5 | 18 | 0.1667 |
| 2_38476157 | 6 | 0.1667 | 3 | 0 | 3 | 0.3333 | 2 | 0.5 | 14 | 0.2143 | 4 | 0.25 | 6 | 0 | 4 | 0.5 | 2 | 1 | 16 | 0.3125 |
| 2_38476323 | 6 | 0.1667 | 3 | 0.6667 | 3 | 0.6667 | 3 | 0.3333 | 15 | 0.4 | 4 | 0.75 | 6 | 0.3333 | 2 | 0 | 2 | 1 | 14 | 0.5 |
| 2_38476454 | 7 | 0.5714 | 3 | 0.3333 | 2 | 1 | 3 | 0.6667 | 15 | 0.6 | 5 | 0.8 | 7 | 0.5714 | 3 | 0 | 2 | 0.5 | 17 | 0.5294 |
| 2_38476771 | 3 | 0 | NA | NA | 2 | 0 | 1 | 1 | 6 | 0.1667 | 3 | 0.3333 | 1 | 0 | NA | NA | 1 | 0 | 5 | 0.2 |
| 2_38849898 | 1 | 1 | NA | NA | NA | NA | NA | NA | 1 | 1 | NA | NA | NA | NA | 1 | 1 | NA | NA | 1 | 1 |
| 2_38849931 | 1 | 1 | NA | NA | NA | NA | NA | NA | 1 | 1 | NA | NA | NA | NA | 1 | 1 | NA | NA | 1 | 1 |
| 2_38850102 | 1 | 1 | NA | NA | NA | NA | NA | NA | 1 | 1 | NA | NA | NA | NA | 1 | 1 | NA | NA | 1 | 1 |
| 2_38850138 | 1 | 1 | NA | NA | NA | NA | NA | NA | 1 | 1 | NA | NA | NA | NA | 1 | 1 | NA | NA | 1 | 1 |
| 2_38850288 | 1 | 1 | NA | NA | NA | NA | NA | NA | 1 | 1 | NA | NA | NA | NA | 1 | 1 | NA | NA | 1 | 1 |
| 2_38851515 | 1 | 1 | NA | NA | NA | NA | NA | NA | 1 | 1 | NA | NA | NA | NA | 1 | 1 | NA | NA | 1 | 1 |
| 2_38903814 | 8 | 0 | 4 | 0.25 | 9 | 0.1111 | 7 | 0 | 28 | 0.0714 | 6 | 0 | 4 | 0 | 4 | 0.25 | 5 | 0 | 19 | 0.0526 |
| 2_38905087 | 23 | 0.0435 | 19 | 0 | 16 | 0.125 | 22 | 0.0455 | 80 | 0.05 | 11 | 0.0909 | 12 | 0 | 24 | 0.0833 | 12 | 0 | 59 | 0.0508 |
| 2_38905124 | 21 | 0.5238 | 18 | 0.7778 | 15 | 0.6 | 25 | 0.68 | 79 | 0.6456 | 11 | 0.6364 | 11 | 0.5455 | 22 | 0.6818 | 13 | 0.7692 | 57 | 0.6667 |
| 2_38905189 | 22 | 0.4545 | 20 | 0.45 | 16 | 0.4375 | 21 | 0.381 | 79 | 0.4304 | 10 | 0.5 | 12 | 0.3333 | 24 | 0.3333 | 14 | 0.5 | 60 | 0.4 |
| 2_38905200 | 18 | 0.1667 | 13 | 0.4615 | 17 | 0.2353 | 18 | 0.2222 | 66 | 0.2576 | 9 | 0 | 9 | 0.3333 | 24 | 0.1667 | 13 | 0.2308 | 55 | 0.1818 |
| 2_38905214 | 20 | 0 | 16 | 0.3125 | 12 | 0.0833 | 19 | 0 | 67 | 0.0896 | 11 | 0 | 10 | 0 | 22 | 0.0455 | 12 | 0.1667 | 55 | 0.0545 |
| 2_38905236 | 24 | 0.2083 | 17 | 0.0588 | 17 | 0.2353 | 24 | 0.125 | 82 | 0.1585 | 11 | 0.0909 | 13 | 0.2308 | 25 | 0.24 | 13 | 0.0769 | 62 | 0.1774 |
| 2_38905430 | 21 | 0.1905 | 21 | 0.381 | 16 | 0.125 | 20 | 0.35 | 78 | 0.2692 | 10 | 0 | 12 | 0 | 19 | 0.2632 | 14 | 0.2143 | 55 | 0.1455 |
| 2_38906621 | 5 | 0.4 | 1 | 1 | 2 | 0.5 | 4 | 0.25 | 12 | 0.4167 | 1 | 0 | 1 | 0 | NA | NA | 5 | 0.2 | 7 | 0.1429 |
| 2_38906653 | 6 | 0 | 2 | 0 | 1 | 1 | 4 | 0 | 13 | 0.0769 | 1 | 0 | 1 | 0 | 1 | 0 | 5 | 0.2 | 8 | 0.125 |
| 2_38908059 | 4 | 0 | 2 | 0 | 2 | 0.5 | 3 | 0 | 11 | 0.0909 | 1 | 0 | 1 | 0 | 1 | 0 | 5 | 0.2 | 8 | 0.125 |
| 2_38908256 | 3 | 0 | 2 | 0.5 | 1 | 1 | 3 | 0.3333 | 9 | 0.3333 | 1 | 0 | 1 | 1 | 1 | 0 | 3 | 0.3333 | 6 | 0.3333 |
| 2_38908267 | 5 | 0 | 2 | 0 | 2 | 0.5 | 3 | 0 | 12 | 0.0833 | 1 | 0 | 1 | 0 | 1 | 0 | 4 | 0.5 | 7 | 0.2857 |
| 2_38956649 | 3 | 0.3333 | 3 | 0 | 3 | 0 | 5 | 0 | 14 | 0.0714 | 3 | 0.3333 | 3 | 0 | 3 | 0 | 1 | 0 | 10 | 0.1 |
| 2_38956843 | 2 | 0.5 | 4 | 0 | 4 | 0 | 5 | 0 | 15 | 0.0667 | 3 | 0 | 4 | 0.25 | 5 | 0 | 1 | 0 | 13 | 0.0769 |
| 2_38956857 | 3 | 0.3333 | 4 | 0 | 4 | 0.25 | 5 | 0 | 16 | 0.125 | 3 | 0.6667 | 4 | 0.25 | 5 | 0 | 1 | 0 | 13 | 0.2308 |
| 2_38964704 | 1 | 0 | 1 | 1 | 1 | 0 | 1 | 0 | 4 | 0.25 | 1 | 1 | NA | NA | NA | NA | NA | NA | 1 | 1 |
| 2_39084730 | 3 | 0 | 3 | 0 | 5 | 0.4 | 4 | 0.25 | 15 | 0.2 | 2 | 0.5 | 3 | 0.3333 | 3 | 0.3333 | 3 | 0.3333 | 11 | 0.3636 |
| 2_39084876 | 1 | 0 | 2 | 0.5 | 5 | 0.2 | 3 | 0.3333 | 11 | 0.2727 | 1 | 0 | 2 | 0 | 3 | 0.3333 | 4 | 0.75 | 10 | 0.4 |
| 2_39084925 | 1 | 0 | 3 | 0 | 4 | 0.25 | 3 | 0 | 11 | 0.0909 | 1 | 1 | 1 | 1 | 3 | 0.3333 | 4 | 0 | 9 | 0.3333 |
| 2_39085164 | 9 | 0.2222 | 9 | 0.6667 | 8 | 0.25 | 9 | 0.5556 | 35 | 0.4286 | 3 | 0.6667 | 3 | 0.3333 | 2 | 0.5 | 5 | 0.4 | 13 | 0.4615 |
| 2_39086923 | 10 | 0.5 | 9 | 0.4444 | 10 | 0.4 | 9 | 0.4444 | 38 | 0.4474 | 8 | 0.625 | 11 | 0.4545 | 6 | 0.3333 | 4 | 0.25 | 29 | 0.4483 |
| 2_39087127 | 10 | 0.2 | 11 | 0 | 10 | 0 | 9 | 0 | 40 | 0.05 | 8 | 0.125 | 9 | 0.1111 | 6 | 0 | 5 | 0 | 28 | 0.0714 |
| 2_39087141 | 7 | 0.5714 | 6 | 0.5 | 7 | 0.4286 | 8 | 0 | 28 | 0.3571 | 6 | 0.1667 | 7 | 0.4286 | 4 | 0 | 2 | 0 | 19 | 0.2105 |
| 2_39089736 | 9 | 0.2222 | 10 | 0 | 7 | 0 | 9 | 0.1111 | 35 | 0.0857 | 5 | 0 | 5 | 0 | 4 | 0 | 4 | 0.25 | 18 | 0.0556 |
| 2_39089812 | 9 | 0 | 8 | 0.125 | 7 | 0 | 8 | 0.125 | 32 | 0.0625 | 5 | 0 | 5 | 0.2 | 4 | 0 | 3 | 0.3333 | 17 | 0.1176 |
| 2_40447188 | 7 | 0.7143 | 9 | 0.8889 | 7 | 0.4286 | 10 | 0.5 | 33 | 0.6364 | 8 | 0.625 | 7 | 0.5714 | 15 | 0.7333 | 10 | 0.6 | 40 | 0.65 |
| 2_40447229 | 10 | 0.8 | 9 | 0.4444 | 8 | 0.5 | 12 | 0.5833 | 39 | 0.5897 | 9 | 0.8889 | 12 | 0.6667 | 15 | 0.7333 | 12 | 0.5833 | 48 | 0.7083 |
| 2_40447286 | 10 | 0.3 | 8 | 0.125 | 9 | 0.1111 | 13 | 0.2308 | 40 | 0.2 | 12 | 0.1667 | 13 | 0 | 17 | 0.2353 | 10 | 0.1 | 52 | 0.1346 |
| 2_40486939 | 1 | 0 | 4 | 0.5 | 2 | 0.5 | 1 | 1 | 8 | 0.5 | NA | NA | 4 | 0.5 | 2 | 0.5 | 1 | 1 | 7 | 0.5714 |
| 2_41877245 | 1 | 0 | NA | NA | NA | NA | 1 | 1 | 2 | 0.5 | 3 | 0.6667 | NA | NA | NA | NA | NA | NA | 3 | 0.6667 |
| 2_41877302 | NA | NA | NA | NA | NA | NA | 1 | 1 | 1 | 1 | 1 | 1 | NA | NA | NA | NA | NA | NA | 1 | 1 |
| 2_44277397 | NA | NA | 3 | 0 | 2 | 0.5 | 1 | 0 | 6 | 0.1667 | 1 | 0 | 1 | 0 | 1 | 1 | 3 | 0.3333 | 6 | 0.3333 |
| 2_44454665 | 4 | 0 | 3 | 0.3333 | 4 | 0 | 4 | 0.25 | 15 | 0.1333 | 4 | 0.25 | 2 | 0 | 2 | 0 | 6 | 0 | 14 | 0.0714 |
| 2_44454713 | 5 | 0 | 3 | 0.3333 | 5 | 0 | 6 | 0.3333 | 19 | 0.1579 | 4 | 0.25 | 2 | 0 | 2 | 1 | 6 | 0 | 14 | 0.2143 |
| 2_44454844 | 4 | 1 | 3 | 0 | 6 | 0.1667 | 6 | 0.8333 | 19 | 0.5263 | 3 | 0.6667 | 1 | 0 | 1 | 1 | 4 | 0.5 | 9 | 0.5556 |
| 2_44454918 | 5 | 0.4 | 2 | 0.5 | 3 | 0.3333 | 6 | 0 | 16 | 0.25 | 5 | 0 | 2 | 0 | 1 | 1 | 3 | 0.3333 | 11 | 0.1818 |
| 2_44455039 | 6 | 0.1667 | 4 | 0.25 | 6 | 0 | 7 | 0.1429 | 23 | 0.1304 | 5 | 0.2 | 2 | 0 | 1 | 0 | 5 | 0 | 13 | 0.0769 |
| 2_44455195 | 5 | 0.8 | 4 | 0.75 | 5 | 0.6 | 6 | 0.6667 | 20 | 0.7 | 6 | 0.8333 | 2 | 0 | 2 | 1 | 4 | 0.5 | 14 | 0.6429 |
| 2_44455414 | 6 | 0.3333 | 4 | 0 | 4 | 0.5 | 4 | 0.25 | 18 | 0.2778 | 4 | 0.5 | 2 | 0.5 | 2 | 0.5 | 5 | 0.6 | 13 | 0.5385 |
| 2_44525200 | 4 | 0.25 | 3 | 0.3333 | 4 | 0 | 4 | 0 | 15 | 0.1333 | 3 | 0.3333 | 2 | 0 | 1 | 0 | 5 | 0.2 | 11 | 0.1818 |

| | | | | | | | | | | | | | | | | | | | | |
|---|---|---|---|---|---|---|---|---|---|---|---|---|---|---|---|---|---|---|---|---|
| 2_4487442 | 2 | 0.5 | 7 | 0 | 6 | 0.1667 | 1 | 0 | 16 | 0.125 | 2 | 0 | 4 | 0 | 2 | 0.5 | 8 | 0 | 16 | 0.0625 |
| 2_4487545 | 2 | 0.5 | 7 | 0.2857 | 4 | 0 | 1 | 0 | 14 | 0.2143 | 2 | 0 | 2 | 0 | 2 | 0 | 7 | 0.1429 | 13 | 0.0769 |
| 2_4487726 | 6 | 0.5 | 14 | 0.8571 | 10 | 0.9 | 4 | 0.75 | 34 | 0.7941 | 8 | 0.875 | 4 | 0.5 | 7 | 0.7143 | 8 | 0.875 | 27 | 0.7778 |
| 2_4487743 | 5 | 0.2 | 11 | 0.0909 | 8 | 0.125 | 2 | 0 | 26 | 0.1154 | 5 | 0.2 | 4 | 0 | 5 | 0 | 7 | 0.1429 | 21 | 0.0952 |
| 2_4487846 | 2 | 0.5 | 8 | 0.5 | 5 | 0.2 | 2 | 0 | 17 | 0.3529 | 4 | 0.75 | 5 | 0 | 4 | 0.25 | 5 | 0 | 18 | 0.2222 |
| 2_4487852 | 2 | 0 | 9 | 0 | 7 | 0.1429 | 2 | 0 | 20 | 0.05 | 3 | 0.3333 | 4 | 0 | 1 | 0 | 5 | 0 | 13 | 0.0769 |
| 2_4487858 | 2 | 0 | 10 | 0.4 | 5 | 0.2 | 3 | 0.3333 | 20 | 0.3 | 3 | 0.3333 | 5 | 0.2 | 4 | 0 | 5 | 0.2 | 17 | 0.1765 |
| 2_4487862 | 2 | 0 | 10 | 0 | 6 | 0.1667 | 2 | 0 | 20 | 0.05 | 3 | 0 | 5 | 0.2 | 4 | 0 | 5 | 0 | 17 | 0.0588 |
| 2_4487868 | 1 | 1 | 10 | 1 | 5 | 1 | 3 | 1 | 19 | 1 | 3 | 1 | 5 | 1 | 4 | 1 | 5 | 1 | 17 | 1 |
| 2_4487886 | 2 | 0.5 | 7 | 0.2857 | 5 | 0.8 | 2 | 0.5 | 16 | 0.5 | 2 | 0.5 | 5 | 1 | 5 | 0.8 | 4 | 0.75 | 16 | 0.8125 |
| 2_4487890 | 2 | 0 | 7 | 0.1429 | 5 | 0.4 | 3 | 0.3333 | 17 | 0.2353 | 3 | 0 | 5 | 0.4 | 4 | 0.25 | 4 | 0.25 | 16 | 0.25 |
| 2_4487892 | 1 | 0 | 8 | 0 | 6 | 0 | 3 | 0.3333 | 18 | 0.0556 | 3 | 0 | 4 | 0 | 4 | 0.25 | 6 | 0.1667 | 17 | 0.1176 |
| 2_4487906 | 2 | 0 | 6 | 0.1667 | 6 | 0.8333 | 3 | 0.3333 | 17 | 0.4118 | 3 | 0 | 5 | 0.4 | 4 | 0 | 5 | 0 | 17 | 0.1176 |
| 2_4488614 | 2 | 0 | 6 | 0.1667 | 5 | 0 | 2 | 1 | 15 | 0.2 | 3 | 0.6667 | 4 | 0.5 | 3 | 0 | 5 | 0.2 | 15 | 0.3333 |
| 2_4488629 | 1 | 1 | 4 | 1 | 3 | 1 | 2 | 1 | 10 | 1 | 3 | 0.6667 | 3 | 1 | 2 | 1 | 4 | 1 | 12 | 0.9167 |
| 2_4488633 | 1 | 0 | 4 | 0 | 5 | 0 | 2 | 1 | 12 | 0.1667 | 3 | 0.3333 | 3 | 0.3333 | 2 | 0 | 4 | 0.5 | 12 | 0.3333 |
| 2_4488647 | 1 | 1 | 5 | 0 | 5 | 0.4 | 2 | 0.5 | 13 | 0.3077 | 2 | 0 | 4 | 0.5 | 2 | 0.5 | 3 | 0.6667 | 11 | 0.4545 |
| 2_4488654 | NA | NA | NA | NA | NA | NA | 1 | 1 | 1 | 1 | NA | NA | NA | NA | NA | NA | 1 | 1 | 1 | 1 |
| 2_4488660 | NA | NA | 4 | 0.25 | 5 | 0.6 | 2 | 0.5 | 11 | 0.4545 | 2 | 0 | 2 | 0 | 2 | 0.5 | 3 | 0.3333 | 9 | 0.2222 |
| 2_4488668 | NA | NA | 3 | 0.3333 | 3 | 0 | 2 | 0 | 8 | 0.125 | 2 | 0 | 2 | 0 | 3 | 0 | 4 | 0.25 | 11 | 0.0909 |
| 2_4488671 | 1 | 0 | 3 | 1 | 5 | 0.6 | 2 | 1 | 11 | 0.7273 | 1 | 0 | 3 | 1 | 3 | 0.6667 | 4 | 1 | 11 | 0.8182 |
| 2_4488675 | 1 | 1 | 3 | 1 | 5 | 1 | 2 | 0.5 | 11 | 0.9091 | 2 | 0.5 | 2 | 1 | 3 | 0.6667 | 4 | 1 | 11 | 0.8182 |
| 2_4488690 | 1 | 1 | 2 | 1 | 4 | 0.75 | 2 | 0.5 | 9 | 0.7778 | 2 | 1 | 2 | 1 | 3 | 1 | 3 | 1 | 10 | 1 |
| 2_4488696 | 1 | 0 | 3 | 0 | 4 | 0.25 | 2 | 0 | 10 | 0.1 | 2 | 0 | NA | NA | 3 | 0.3333 | 2 | 0.5 | 7 | 0.2857 |
| 2_4488699 | 1 | 1 | 3 | 1 | 3 | 0.6667 | 2 | 0.5 | 9 | 0.7778 | 2 | 1 | 2 | 1 | 3 | 1 | 3 | 1 | 10 | 1 |
| 2_4488749 | 1 | 1 | 3 | 0 | 3 | 0.3333 | 2 | 0 | 9 | 0.2222 | 2 | 1 | 1 | 0 | 2 | 1 | 2 | 1 | 7 | 0.8571 |
| 2_4488752 | 1 | 0 | 3 | 1 | 1 | 1 | 1 | 1 | 6 | 0.8333 | 2 | 0 | 1 | 0 | 1 | 0 | 2 | 1 | 6 | 0.3333 |
| 2_4488772 | NA | NA | 3 | 0.3333 | 3 | 0.6667 | 2 | 0 | 8 | 0.375 | 2 | 0.5 | 1 | 0 | 2 | 0.5 | 2 | 0 | 7 | 0.2857 |
| 2_4500343 | NA | NA | 2 | 0 | 1 | 1 | NA | NA | 3 | 0.3333 | NA | NA | 1 | 1 | NA | NA | NA | NA | 1 | 1 |
| 2_4870432 | NA | NA | 4 | 0.75 | 2 | 1 | 1 | 0 | 7 | 0.7143 | NA | NA | 1 | 1 | NA | NA | 2 | 0.5 | 3 | 0.6667 |
| 2_4878971 | 2 | 1 | 6 | 1 | 3 | 1 | 2 | 0.5 | 13 | 0.9231 | 3 | 0.3333 | 3 | 1 | 2 | 1 | 5 | 0.6 | 13 | 0.6923 |
| 2_4878973 | 2 | 1 | 6 | 0.6667 | 1 | 1 | NA | NA | 9 | 0.7778 | 3 | 1 | 3 | 1 | 2 | 1 | 4 | 0.5 | 12 | 0.8333 |
| 2_4878977 | 1 | 1 | 5 | 1 | 4 | 1 | 3 | 0.6667 | 13 | 0.9231 | 3 | 1 | 4 | 0.75 | 2 | 1 | 5 | 1 | 14 | 0.9286 |
| 2_4878982 | 2 | 1 | 4 | 1 | 3 | 0.6667 | 2 | 1 | 11 | 0.9091 | 2 | 1 | 3 | 1 | 1 | 0 | 5 | 0.8 | 11 | 0.8182 |
| 2_4878983 | 1 | 1 | 3 | 1 | 4 | 1 | 3 | 1 | 11 | 1 | 3 | 1 | 3 | 1 | 2 | 1 | 6 | 0.8333 | 14 | 0.9286 |
| 2_4878984 | NA | NA | 6 | 0.6667 | 1 | 0 | 1 | 1 | 8 | 0.625 | 2 | 0.5 | 4 | 0.75 | 1 | 0 | 1 | 1 | 8 | 0.625 |
| 2_4878991 | 2 | 0.5 | 6 | 0.3333 | 4 | 0.5 | 3 | 0.3333 | 15 | 0.4 | 3 | 0.3333 | 4 | 1 | 2 | 0.5 | 6 | 0.6667 | 15 | 0.6667 |
| 2_4879002 | 2 | 0 | 7 | 0.1429 | 3 | 0 | 3 | 0.3333 | 15 | 0.1333 | 4 | 0 | 5 | 0.6 | 2 | 0 | 4 | 0 | 15 | 0.2 |
| 2_4879003 | 2 | 0.5 | 6 | 0.6667 | 3 | 0.6667 | 3 | 0 | 14 | 0.5 | 4 | 0.5 | 5 | 1 | 2 | 1 | 6 | 0.6667 | 17 | 0.7647 |
| 2_4879008 | 2 | 1 | 6 | 0.8333 | 4 | 0.25 | 3 | 0.6667 | 15 | 0.6667 | 4 | 0.5 | 5 | 1 | 2 | 1 | 6 | 0.6667 | 17 | 0.7647 |
| 2_4879011 | 2 | 1 | 7 | 0.5714 | 4 | 0.5 | 3 | 0.3333 | 16 | 0.5625 | 3 | 0.6667 | 3 | 0.6667 | 2 | 0 | 4 | 0.5 | 12 | 0.5 |
| 2_4879014 | 2 | 0.5 | 6 | 0.3333 | 4 | 0.25 | 3 | 0.3333 | 15 | 0.3333 | 4 | 0 | 5 | 0.2 | 2 | 0 | 6 | 0 | 17 | 0.0588 |
| 2_4879016 | 1 | 1 | 7 | 0.8571 | 3 | 1 | 2 | 1 | 13 | 0.9231 | 3 | 1 | 5 | 1 | 2 | 1 | 3 | 1 | 13 | 1 |
| 2_4879019 | 2 | 0 | 5 | 0.8 | 1 | 1 | 3 | 0.3333 | 11 | 0.5455 | 4 | 0.25 | 5 | 0.6 | 2 | 0 | 6 | 0.6667 | 17 | 0.4706 |
| 2_4879229 | 6 | 0 | 12 | 0.25 | 4 | 0 | 6 | 0.1667 | 28 | 0.1429 | 2 | 0 | 7 | 0.2857 | 7 | 0.2857 | 8 | 0.25 | 24 | 0.25 |
| 2_4879236 | 6 | 0.1667 | 12 | 0.0833 | 4 | 0.5 | 6 | 0.3333 | 28 | 0.2143 | 2 | 0 | 9 | 0.5556 | 9 | 0.4444 | 8 | 0.25 | 28 | 0.3929 |
| 2_4879252 | 6 | 0.3333 | 12 | 0.25 | 5 | 0.4 | 6 | 0.3333 | 29 | 0.3103 | 1 | 0 | 10 | 0.3 | 7 | 0.4286 | 8 | 0 | 26 | 0.2308 |
| 2_4879262 | 7 | 0 | 11 | 0.4545 | 4 | 0.5 | 6 | 0.1667 | 28 | 0.2857 | NA | NA | 11 | 0.1818 | 8 | 0.25 | 9 | 0.5556 | 28 | 0.3214 |
| 2_4879368 | 1 | 0 | 1 | 0 | 4 | 0 | 1 | 1 | 7 | 0.1429 | 2 | 0.5 | NA | NA | NA | NA | 1 | 1 | 3 | 0.6667 |
| 2_4879382 | 1 | 1 | 2 | 0.5 | 3 | 0.6667 | 1 | 1 | 7 | 0.7143 | 1 | 0 | NA | NA | NA | NA | 1 | 1 | 2 | 0.5 |
| 2_4879528 | 5 | 0.2 | 9 | 0.4444 | 5 | 0 | 6 | 0.5 | 25 | 0.32 | 4 | 0.25 | 6 | 0.5 | 2 | 0 | 7 | 0.7143 | 19 | 0.4737 |
| 2_4879556 | 5 | 0 | 9 | 0.1111 | 4 | 0.25 | 7 | 0 | 25 | 0.08 | 4 | 0.5 | 5 | 0.2 | 3 | 0 | 7 | 0 | 19 | 0.1579 |
| 2_4913595 | NA | NA | 1 | 1 | NA | NA | NA | NA | 1 | 1 | NA | NA | 1 | 1 | 1 | 1 | NA | NA | 2 | 1 |
| 2_4914630 | NA | NA | 1 | 1 | NA | NA | NA | NA | 1 | 1 | 1 | 1 | 1 | 1 | 2 | 0.5 | NA | NA | 4 | 0.75 |
| 2_4914641 | NA | NA | 2 | 1 | NA | NA | NA | NA | 2 | 1 | 1 | 1 | NA | NA | 2 | 1 | NA | NA | 3 | 1 |
| 2_4914651 | NA | NA | 2 | 0.5 | NA | NA | NA | NA | 2 | 0.5 | 1 | 1 | 1 | 1 | 2 | 1 | NA | NA | 4 | 1 |
| 2_4914660 | NA | NA | 1 | 1 | NA | NA | NA | NA | 1 | 1 | 2 | 1 | 1 | 1 | 1 | 1 | NA | NA | 4 | 1 |
| 2_4914667 | NA | NA | 2 | 1 | NA | NA | NA | NA | 2 | 1 | 2 | 1 | 1 | 1 | 1 | 1 | 1 | 0 | 5 | 0.8 |
| 2_4914690 | NA | NA | 1 | 1 | NA | NA | NA | NA | 1 | 1 | 1 | 1 | 1 | 1 | 2 | 0.5 | NA | NA | 4 | 0.75 |
| 2_4914713 | NA | NA | 2 | 1 | NA | NA | NA | NA | 2 | 1 | NA | NA | 1 | 0 | 1 | 1 | NA | NA | 2 | 0.5 |
| 2_4914722 | 1 | 0 | 2 | 1 | NA | NA | NA | NA | 3 | 0.6667 | 2 | 0.5 | 1 | 1 | 2 | 0 | NA | NA | 5 | 0.4 |
| 2_4914733 | 1 | 1 | 2 | 1 | 1 | 1 | NA | NA | 4 | 1 | 1 | 1 | NA | NA | 2 | 1 | 1 | 1 | 4 | 1 |
| 2_4914757 | 1 | 1 | 1 | 0 | 2 | 0.5 | NA | NA | 4 | 0.5 | 1 | 0 | 1 | 0 | 3 | 0.3333 | NA | NA | 5 | 0.2 |
| 2_4914763 | 1 | 1 | 2 | 0.5 | 2 | 1 | NA | NA | 5 | 0.8 | 1 | 1 | 1 | 1 | 3 | 1 | 1 | 1 | 6 | 1 |
| 2_4914775 | 1 | 1 | 2 | 1 | 2 | 1 | NA | NA | 5 | 1 | 1 | 1 | 1 | 1 | 2 | 1 | NA | NA | 4 | 1 |

| | | | | | | | | | | | | | | | | | | | | |
|---|---|---|---|---|---|---|---|---|---|---|---|---|---|---|---|---|---|---|---|---|
| 2_4914780 | 1 | 1 | 2 | 0.5 | 2 | 0 | NA | NA | 5 | 0.4 | 1 | 0 | 1 | 1 | 3 | 0 | 1 | 0 | 6 | 0.1667 |
| 2_4914784 | 2 | 1 | 3 | 1 | 3 | 1 | NA | NA | 8 | 1 | 1 | 1 | 1 | 1 | 3 | 1 | NA | NA | 5 | 1 |
| 2_4914805 | 1 | 0 | 3 | 0 | 3 | 0.3333 | NA | NA | 7 | 0.1429 | 1 | 1 | 1 | 0 | 4 | 0 | 1 | 0 | 7 | 0.1429 |
| 2_4914810 | 1 | 1 | 3 | 0.6667 | 3 | 0.6667 | NA | NA | 7 | 0.7143 | 1 | 1 | NA | NA | 2 | 0.5 | NA | NA | 3 | 0.6667 |
| 2_4914818 | 2 | 1 | 3 | 1 | 3 | 1 | NA | NA | 8 | 1 | 1 | 1 | 1 | 1 | 4 | 0.75 | NA | NA | 6 | 0.8333 |
| 2_4916219 | 2 | 1 | 4 | 0.75 | 1 | 0 | NA | NA | 7 | 0.7143 | 1 | 1 | 1 | 1 | 2 | 0 | 2 | 0 | 6 | 0.3333 |
| 2_4916271 | 1 | 1 | 2 | 1 | NA | NA | NA | NA | 3 | 1 | 1 | 0 | 2 | 0 | NA | NA | 1 | 1 | 4 | 0.25 |
| 2_4916293 | 1 | 1 | 1 | 0 | 1 | 0 | 1 | 0 | 4 | 0.25 | 2 | 0.5 | 1 | 0 | NA | NA | 1 | 0 | 4 | 0.25 |
| 2_4916904 | 2 | 1 | 5 | 0.4 | 3 | 0 | 1 | 0 | 11 | 0.3636 | 2 | 1 | 1 | 0 | 3 | 0.3333 | 1 | 0 | 7 | 0.4286 |
| 2_4916911 | 2 | 0 | 3 | 0 | 1 | 0 | 2 | 0.5 | 8 | 0.125 | 2 | 0.5 | 2 | 0 | 2 | 0 | 1 | 0 | 7 | 0.1429 |
| 2_4916929 | 2 | 0.5 | 5 | 1 | 3 | 1 | 2 | 1 | 12 | 0.9167 | 2 | 0 | 2 | 1 | 5 | 0.8 | 2 | 0.5 | 11 | 0.6364 |
| 2_4916934 | 2 | 1 | 4 | 0.75 | 2 | 1 | 2 | 1 | 10 | 0.9 | 1 | 1 | 2 | 1 | 7 | 0.8571 | 1 | 1 | 11 | 0.9091 |
| 2_4916936 | 2 | 0.5 | 6 | 0.8333 | 3 | 0.6667 | 2 | 0.5 | 13 | 0.6923 | 2 | 1 | 2 | 1 | 5 | 0.8 | 2 | 0.5 | 11 | 0.8182 |
| 2_4916941 | 2 | 0 | 6 | 0 | 3 | 0 | 3 | 0.3333 | 14 | 0.0714 | 3 | 0 | 1 | 0 | 7 | 0.1429 | 2 | 0 | 13 | 0.0769 |
| 2_4916946 | NA | NA | 6 | 0 | 3 | 0 | 2 | 0.5 | 11 | 0.0909 | 3 | 0 | 2 | 0 | 8 | 0.125 | 1 | 0 | 14 | 0.0714 |
| 2_4916952 | 2 | 1 | 6 | 0.8333 | 3 | 0.3333 | 3 | 0.3333 | 14 | 0.6429 | 3 | 0.6667 | 2 | 1 | 6 | 0.6667 | 2 | 0.5 | 13 | 0.6923 |
| 2_4916954 | 2 | 1 | 6 | 0.8333 | 3 | 1 | 3 | 1 | 14 | 0.9286 | 2 | 1 | 2 | 1 | 6 | 0.8333 | 2 | 0.5 | 12 | 0.8333 |
| 2_4916963 | 2 | 0 | 6 | 0.1667 | 2 | 0 | 4 | 0 | 14 | 0.0714 | 3 | 0 | 2 | 0 | 8 | 0.125 | 3 | 0.3333 | 16 | 0.125 |
| 2_4916966 | 2 | 0 | 5 | 0.2 | 3 | 0 | 3 | 0 | 13 | 0.0769 | 1 | 0 | 2 | 0 | 8 | 0.125 | 2 | 0 | 13 | 0.0769 |
| 2_4916971 | 1 | 1 | 3 | 0.6667 | 3 | 0.3333 | 1 | 1 | 8 | 0.625 | 2 | 1 | 2 | 0.5 | 5 | 1 | 2 | 1 | 11 | 0.9091 |
| 2_4920647 | 3 | 1 | 5 | 0.8 | 3 | 1 | 4 | 1 | 15 | 0.9333 | 4 | 1 | 3 | 1 | 9 | 0.8889 | 2 | 1 | 18 | 0.9444 |
| 2_4920694 | 3 | 0 | 8 | 0 | 7 | 0.2857 | 3 | 0 | 21 | 0.0952 | 4 | 0.5 | 4 | 0 | 10 | 0.2 | 3 | 0 | 21 | 0.1905 |
| 2_4942684 | 1 | 0 | 2 | 0.5 | 4 | 0.25 | 3 | 0.3333 | 10 | 0.3 | 1 | 0 | 2 | 0 | 2 | 0 | 5 | 0.4 | 10 | 0.2 |
| 2_4942693 | 1 | 1 | 2 | 1 | 4 | 1 | 2 | 1 | 9 | 1 | 1 | 1 | 4 | 0.5 | 1 | 1 | 7 | 0.8571 | 13 | 0.7692 |
| 2_4942699 | 1 | 1 | 2 | 1 | 3 | 1 | 2 | 1 | 8 | 1 | 1 | 1 | 4 | 0.75 | 2 | 1 | 6 | 0.6667 | 13 | 0.7692 |
| 2_4942707 | 1 | 1 | 2 | 0.5 | 2 | 0.5 | 2 | 1 | 7 | 0.7143 | 1 | 1 | 3 | 0.3333 | 2 | 0.5 | 6 | 0.5 | 12 | 0.5 |
| 2_4942711 | 2 | 0 | 2 | 0.5 | 4 | 0 | 3 | 0 | 11 | 0.0909 | 1 | 0 | 5 | 0.2 | 2 | 0.5 | 7 | 0.1429 | 15 | 0.2 |
| 2_4943852 | 3 | 1 | 3 | 0.6667 | 4 | 0.75 | 3 | 0.6667 | 13 | 0.7692 | 2 | 0.5 | 5 | 1 | 5 | 0.6 | 7 | 0.8571 | 19 | 0.7895 |
| 2_4943858 | 1 | 0 | 3 | 1 | 4 | 0.5 | 3 | 0.3333 | 11 | 0.5455 | 2 | 0 | 5 | 0.2 | 4 | 0.25 | 7 | 0.5714 | 18 | 0.3333 |
| 2_4943859 | NA | NA | NA | NA | 3 | 1 | 1 | 1 | 4 | 1 | 2 | 0.5 | 2 | 1 | 1 | 0 | 5 | 0.8 | 10 | 0.7 |
| 2_4943867 | 4 | 0.75 | 3 | 0.3333 | 4 | 0.75 | 3 | 1 | 14 | 0.7143 | 3 | 0.3333 | 3 | 0.6667 | 4 | 1 | 5 | 0.8 | 15 | 0.7333 |
| 2_4943870 | 4 | 0 | 3 | 0 | 3 | 0.3333 | 3 | 0 | 13 | 0.0769 | 3 | 0.6667 | 4 | 0 | 4 | 0 | 7 | 0 | 18 | 0.1111 |
| 2_4943872 | 3 | 0.3333 | 3 | 0.3333 | 4 | 0 | 3 | 0 | 13 | 0.1538 | 1 | 0 | 3 | 0 | 4 | 0.25 | 4 | 0 | 12 | 0.0833 |
| 2_4943995 | 5 | 0.6 | 3 | 0.3333 | 3 | 0.6667 | 2 | 0.5 | 13 | 0.5385 | 3 | 0.6667 | 4 | 0.25 | 6 | 0.3333 | 6 | 0.3333 | 19 | 0.3684 |
| 2_4943997 | 3 | 1 | 3 | 1 | 4 | 1 | 3 | 0.6667 | 13 | 0.9231 | 3 | 0.6667 | 5 | 1 | 6 | 0.6667 | 7 | 0.8571 | 21 | 0.8095 |
| 2_4943998 | 5 | 0.2 | 3 | 0 | 4 | 0 | 3 | 0 | 15 | 0.0667 | 3 | 0 | 6 | 0 | 5 | 0.2 | 5 | 0 | 19 | 0.0526 |
| 2_4944010 | 5 | 0.2 | 3 | 0 | 5 | 0 | 3 | 0 | 16 | 0.0625 | 3 | 0.3333 | 5 | 0 | 5 | 0 | 7 | 0.1429 | 20 | 0.1 |
| 2_4944011 | 4 | 1 | 3 | 1 | 5 | 1 | 2 | 1 | 14 | 1 | 3 | 1 | 4 | 0.75 | 4 | 1 | 7 | 0.8571 | 18 | 0.8889 |
| 2_4944052 | 4 | 0.25 | 3 | 0.6667 | 6 | 0.6667 | 3 | 0 | 16 | 0.4375 | 2 | 0.5 | 5 | 0.4 | 6 | 0.1667 | 7 | 0.2857 | 20 | 0.3 |
| 2_4966120 | 11 | 0.9091 | 23 | 0.6957 | 10 | 0.8 | 15 | 0.8 | 59 | 0.7797 | 15 | 0.8667 | 15 | 0.7333 | 8 | 0.75 | 14 | 0.7143 | 52 | 0.7692 |
| 2_4966150 | 6 | 1 | 18 | 0.8889 | 11 | 1 | 11 | 0.9091 | 46 | 0.9348 | 8 | 1 | 11 | 0.8182 | 7 | 1 | 13 | 0.8462 | 39 | 0.8974 |
| 2_4966156 | 11 | 0.3636 | 26 | 0.1923 | 9 | 0 | 13 | 0.2308 | 59 | 0.2034 | 16 | 0.0625 | 15 | 0.0667 | 8 | 0.125 | 15 | 0.2 | 54 | 0.1111 |
| 2_4966163 | 10 | 0.8 | 23 | 0.8261 | 8 | 0.25 | 12 | 0.75 | 53 | 0.717 | 11 | 0.9091 | 12 | 0.8333 | 8 | 1 | 13 | 0.6154 | 44 | 0.8182 |
| 2_4966170 | 11 | 0.6364 | 27 | 0.5926 | 10 | 0.3 | 16 | 0.875 | 64 | 0.625 | 15 | 0.5333 | 20 | 0.65 | 7 | 0.7143 | 16 | 0.6875 | 58 | 0.6379 |
| 2_4966175 | 13 | 0.4615 | 25 | 0.16 | 12 | 0 | 16 | 0.4375 | 66 | 0.2576 | 14 | 0.1429 | 13 | 0.1538 | 8 | 0.375 | 15 | 0.2 | 50 | 0.2 |
| 2_4966176 | 13 | 0.3077 | 25 | 0.16 | 11 | 0.0909 | 15 | 0.1333 | 64 | 0.1719 | 12 | 0.25 | 21 | 0.0476 | 7 | 0.1429 | 12 | 0.1667 | 52 | 0.1346 |
| 2_4966179 | 13 | 0.1538 | 28 | 0.1786 | 13 | 0 | 16 | 0.125 | 70 | 0.1286 | 18 | 0.2222 | 19 | 0.1579 | 8 | 0 | 13 | 0.0769 | 58 | 0.1379 |
| 2_4966186 | 14 | 0.5714 | 26 | 0.5 | 13 | 0.4615 | 19 | 0.6316 | 72 | 0.5417 | 17 | 0.6471 | 17 | 0.5294 | 8 | 0.875 | 14 | 0.6429 | 56 | 0.6429 |
| 2_4966189 | 11 | 0.3636 | 22 | 0.3182 | 15 | 0.3333 | 18 | 0.3889 | 66 | 0.3485 | 16 | 0.3125 | 20 | 0.35 | 7 | 0.7143 | 14 | 0.5714 | 57 | 0.4386 |
| 2_4966257 | 20 | 0.05 | 38 | 0.0789 | 19 | 0.1053 | 24 | 0.0417 | 101 | 0.0693 | 19 | 0.0526 | 22 | 0.0909 | 10 | 0.1 | 26 | 0.0385 | 77 | 0.0649 |
| 2_4966417 | 40 | 0.125 | 57 | 0.0351 | 38 | 0.0789 | 42 | 0.1429 | 177 | 0.0904 | 27 | 0.0741 | 37 | 0.027 | 26 | 0.0769 | 33 | 0.1212 | 123 | 0.0732 |
| 2_4966439 | 40 | 0.275 | 60 | 0.2 | 36 | 0.1389 | 50 | 0.2 | 186 | 0.2043 | 35 | 0.2 | 39 | 0.1282 | 38 | 0.1316 | 35 | 0.2571 | 147 | 0.1769 |
| 2_4983766 | NA | NA | NA | NA | 1 | 1 | NA | NA | 1 | 1 | 1 | 1 | 1 | 1 | NA | NA | NA | NA | 2 | 1 |
| 2_4983771 | NA | NA | NA | NA | 1 | 1 | NA | NA | 1 | 1 | NA | NA | 1 | 1 | NA | NA | NA | NA | 1 | 1 |
| 2_4983778 | NA | NA | NA | NA | 1 | 1 | NA | NA | 1 | 1 | 1 | 1 | 1 | 1 | NA | NA | NA | NA | 2 | 1 |
| 2_4983794 | NA | NA | NA | NA | 1 | 1 | NA | NA | 1 | 1 | 1 | 1 | 1 | 1 | NA | NA | NA | NA | 2 | 1 |
| 2_4983796 | NA | NA | NA | NA | 1 | 1 | NA | NA | 1 | 1 | NA | NA | 1 | 1 | NA | NA | NA | NA | 1 | 1 |
| 2_5008081 | 1 | 1 | NA | NA | 1 | 0 | 1 | 0 | 3 | 0.3333 | 2 | 0 | 1 | 0 | 1 | 1 | 3 | 0.6667 | 7 | 0.4286 |
| 2_5017008 | 8 | 0.25 | 8 | 0.125 | 8 | 0.375 | 3 | 0.3333 | 27 | 0.2593 | 4 | 0.5 | 6 | 0.1667 | 6 | 0.6667 | 3 | 0.6667 | 19 | 0.4737 |
| 2_5017015 | 4 | 0 | 7 | 0.5714 | 9 | 0.6667 | 2 | 0 | 22 | 0.4545 | 3 | 0.6667 | 4 | 1 | 8 | 0.875 | 2 | 1 | 17 | 0.8824 |
| 2_5102068 | 1 | 0 | 3 | 0.3333 | 2 | 0 | 1 | 1 | 7 | 0.2857 | 1 | 0 | 1 | 0 | 1 | 0 | 2 | 0.5 | 5 | 0.2 |
| 2_5102106 | 1 | 1 | 4 | 1 | 1 | 1 | 1 | 1 | 7 | 1 | NA | NA | 1 | 1 | 1 | 1 | 1 | 1 | 3 | 1 |
| 2_5102133 | 1 | 0 | 4 | 0.75 | 1 | 0 | NA | NA | 6 | 0.5 | NA | NA | 1 | 1 | 1 | 0 | 1 | 0 | 3 | 0.3333 |
| 2_5102167 | 1 | 0 | 3 | 0.3333 | 1 | 0 | NA | NA | 5 | 0.2 | 1 | 0 | 1 | 0 | 1 | 1 | 1 | 0 | 4 | 0.25 |
| 2_5102179 | 1 | 0 | 4 | 0.25 | 1 | 0 | NA | NA | 6 | 0.1667 | 1 | 0 | 1 | 0 | 1 | 0 | 1 | 1 | 4 | 0.25 |

| | | | | | | | | | | | | | | | | | | | | |
|---|---|---|---|---|---|---|---|---|---|---|---|---|---|---|---|---|---|---|---|---|
| 2_5102181( | 1 | 0 | 4 | 0.75 | 1 | 1 | NA | NA | 6 | 0.6667 | NA | NA | 1 | 0 | 1 | 1 | 1 | 1 | 3 | 0.6667 |
| 2_5102196! | 1 | 0 | 4 | 0.25 | 1 | 0 | NA | NA | 6 | 0.1667 | 1 | 0 | 1 | 0 | 1 | 1 | 1 | 0 | 4 | 0.25 |
| 2_5102201: | 1 | 0 | 2 | 0.5 | 1 | 0 | NA | NA | 4 | 0.25 | 1 | 1 | 1 | 0 | 1 | 0 | 1 | 0 | 4 | 0.25 |
| 2_5102204! | 1 | 0 | 4 | 0.5 | 1 | 1 | NA | NA | 6 | 0.5 | 1 | 0 | 1 | 0 | 1 | 1 | 1 | 0 | 4 | 0.25 |
| 2_5102210' | 1 | 1 | 3 | 0 | 1 | 0 | NA | NA | 5 | 0.2 | 1 | 0 | 1 | 1 | 1 | 0 | 1 | 0 | 4 | 0.25 |
| 2_5102233- | 1 | 0 | 4 | 0.75 | NA | NA | NA | NA | 5 | 0.6 | 1 | 0 | 1 | 1 | NA | NA | NA | NA | 2 | 0.5 |
| 2_5102234( | NA | NA | 4 | 1 | NA | NA | NA | NA | 4 | 1 | 1 | 1 | 1 | 1 | 1 | 1 | 1 | 1 | 4 | 1 |
| 2_5102236( | 1 | 1 | 4 | 0.75 | NA | NA | NA | NA | 5 | 0.8 | 1 | 1 | 1 | 0 | 1 | 1 | 1 | 1 | 4 | 0.75 |
| 2_5102241! | 1 | 0 | 2 | 0.5 | NA | NA | NA | NA | 3 | 0.3333 | 1 | 0 | 1 | 1 | 1 | 0 | 1 | 0 | 4 | 0.25 |
| 2_5102244( | NA | NA | 4 | 0.5 | NA | NA | NA | NA | 4 | 0.5 | 1 | 0 | 1 | 0 | 1 | 1 | 1 | 0 | 4 | 0.25 |
| 2_5102247! | 1 | 0 | 4 | 1 | NA | NA | NA | NA | 5 | 0.8 | 1 | 1 | 1 | 1 | 1 | 1 | 1 | 1 | 4 | 1 |
| 2_5176114! | 9 | 0.2222 | 7 | 0 | 7 | 0.1429 | 4 | 0.25 | 27 | 0.1481 | 7 | 0.2857 | 5 | 0.2 | 3 | 0.6667 | 6 | 0 | 21 | 0.2381 |
| 2_5181451( | 3 | 0 | 1 | 0 | 1 | 0 | 4 | 0.25 | 9 | 0.1111 | 3 | 0.3333 | 1 | 1 | 3 | 0 | 1 | 0 | 8 | 0.25 |
| 2_5181462- | 3 | 0.3333 | 1 | 0 | 1 | 0 | 4 | 0 | 9 | 0.1111 | 2 | 0.5 | 1 | 1 | 1 | 0 | 1 | 0 | 5 | 0.4 |
| 2_5181464! | 3 | 0.3333 | 1 | 1 | 1 | 1 | 4 | 0.75 | 9 | 0.6667 | 3 | 1 | 1 | 1 | 3 | 1 | 1 | 1 | 8 | 1 |
| 2_5181554- | 2 | 0 | 1 | 1 | NA | NA | 4 | 0 | 7 | 0.1429 | 3 | 0 | 1 | 0 | 2 | 0.5 | 1 | 0 | 7 | 0.1429 |
| 2_5181558! | 2 | 1 | 1 | 1 | NA | NA | 4 | 1 | 7 | 1 | 2 | 1 | 1 | 1 | 2 | 1 | NA | NA | 5 | 1 |
| 2_5181564( | 2 | 1 | 1 | 1 | NA | NA | 4 | 0.5 | 7 | 0.7143 | 3 | 0.6667 | 1 | 1 | 3 | 0.3333 | NA | NA | 7 | 0.5714 |
| 2_5181582' | 1 | 0 | 1 | 1 | NA | NA | 4 | 0 | 6 | 0.1667 | 3 | 0.3333 | 1 | 1 | 3 | 0 | 1 | 1 | 8 | 0.375 |
| 2_5183857: | NA | NA | 2 | 0.5 | NA | NA | 1 | 0 | 3 | 0.3333 | 1 | 1 | NA | NA | 1 | 0 | 1 | 1 | 3 | 0.6667 |
| 2_5183870! | NA | NA | 3 | 0.3333 | NA | NA | 1 | 0 | 4 | 0.25 | 2 | 1 | NA | NA | 1 | 0 | 1 | 0 | 4 | 0.5 |
| 2_5194099( | 2 | 0.5 | 2 | 0.5 | 2 | 0 | 2 | 0.5 | 8 | 0.375 | 3 | 0 | 3 | 0.3333 | 1 | 1 | NA | NA | 7 | 0.2857 |
| 2_5194625' | 2 | 0 | 5 | 0.2 | 2 | 0 | 3 | 0.3333 | 12 | 0.1667 | 2 | 0 | 2 | 0.5 | 1 | 0 | NA | NA | 5 | 0.2 |
| 2_5200004! | NA | NA | 1 | 1 | NA | NA | NA | NA | 1 | 1 | NA | NA | 1 | 0 | NA | NA | 2 | 0.5 | 3 | 0.3333 |
| 2_5200208' | NA | NA | 1 | 0 | 2 | 0.5 | NA | NA | 3 | 0.3333 | NA | NA | 1 | 1 | 1 | 1 | 3 | 0.6667 | 5 | 0.8 |
| 2_5200235' | NA | NA | 1 | 1 | 2 | 1 | NA | NA | 3 | 1 | NA | NA | 1 | 1 | 1 | 0 | 3 | 1 | 5 | 0.8 |
| 2_5200241( | NA | NA | 1 | 0 | 2 | 0.5 | NA | NA | 3 | 0.3333 | NA | NA | 1 | 1 | 1 | 1 | 3 | 0.3333 | 5 | 0.6 |
| 2_5200255: | NA | NA | NA | NA | 2 | 0.5 | NA | NA | 2 | 0.5 | NA | NA | 1 | 1 | 1 | 1 | 3 | 0.3333 | 5 | 0.6 |
| 2_5200265( | NA | NA | 1 | 1 | 2 | 1 | NA | NA | 3 | 1 | NA | NA | 1 | 1 | 1 | 1 | 4 | 1 | 6 | 1 |
| 2_5200267' | NA | NA | 1 | 0 | 2 | 0.5 | NA | NA | 3 | 0.3333 | NA | NA | 1 | 0 | NA | NA | 4 | 0.25 | 5 | 0.2 |
| 2_5200268! | NA | NA | 1 | 1 | 2 | 1 | NA | NA | 3 | 1 | NA | NA | 1 | 0 | 1 | 0 | 4 | 1 | 6 | 0.6667 |
| 2_5201034' | 1 | 0 | 2 | 0.5 | 2 | 0 | 1 | 0 | 6 | 0.1667 | NA | NA | 1 | 0 | 1 | 0 | 4 | 0.25 | 6 | 0.1667 |
| 2_5201108! | 1 | 0 | 1 | 0 | 2 | 0.5 | 1 | 1 | 5 | 0.4 | NA | NA | 1 | 1 | 1 | 0 | 4 | 0.75 | 6 | 0.6667 |
| 2_5201129' | 1 | 0 | 1 | 1 | 2 | 1 | 1 | 1 | 5 | 0.8 | NA | NA | 1 | 1 | 1 | 0 | 4 | 0.75 | 6 | 0.6667 |
| 2_5228912! | 9 | 0.1111 | 10 | 0 | 12 | 0.0833 | 13 | 0.2308 | 44 | 0.1136 | 6 | 0 | 10 | 0.2 | 13 | 0.1538 | 11 | 0 | 40 | 0.1 |
| 2_5228933( | 8 | 0.25 | 11 | 0.3636 | 12 | 0 | 11 | 0.0909 | 42 | 0.1667 | 7 | 0 | 12 | 0 | 13 | 0.2308 | 10 | 0.1 | 42 | 0.0952 |
| 2_5229209( | 3 | 0 | 6 | 0 | 7 | 0 | 4 | 0.25 | 20 | 0.05 | 2 | 0 | 4 | 0.5 | 8 | 0.125 | 6 | 0 | 20 | 0.15 |
| 2_5229211: | 5 | 0.8 | 7 | 0.7143 | 7 | 0.8571 | 4 | 1 | 23 | 0.8261 | 4 | 0.25 | 4 | 0.5 | 8 | 0.875 | 5 | 0.8 | 21 | 0.6667 |
| 2_5229212( | 4 | 0.75 | 5 | 1 | 6 | 0.3333 | 6 | 0.6667 | 21 | 0.6667 | 2 | 1 | 3 | 0.6667 | 4 | 0.75 | 5 | 0.8 | 14 | 0.7857 |
| 2_5229216! | 4 | 0.25 | 7 | 0.7143 | 8 | 0.625 | 7 | 0.2857 | 26 | 0.5 | 4 | 0.25 | 4 | 0.25 | 6 | 0.3333 | 6 | 0.3333 | 20 | 0.3 |
| 2_5229219' | 7 | 0.7143 | 3 | 0.6667 | 5 | 0.4 | 5 | 0.6 | 20 | 0.6 | 4 | 0.75 | 2 | 1 | 7 | 0.8571 | 7 | 0.7143 | 20 | 0.8 |
| 2_5229229' | 3 | 0.3333 | 4 | 0.5 | 6 | 0.5 | 3 | 0.3333 | 16 | 0.4375 | 3 | 0.3333 | 3 | 0.6667 | 6 | 0.5 | 6 | 0.6667 | 18 | 0.5556 |
| 2_5229230' | NA | NA | 1 | 1 | 3 | 1 | 4 | 0.75 | 8 | 0.875 | 3 | 0.3333 | 1 | 0 | 4 | 1 | 1 | 1 | 9 | 0.6667 |
| 2_5229234! | 5 | 1 | 5 | 0.8 | 7 | 0.8571 | 6 | 1 | 23 | 0.913 | 4 | 1 | 4 | 1 | 6 | 0.8333 | 6 | 1 | 20 | 0.95 |
| 2_5229237: | 5 | 0.8 | 5 | 0.4 | 5 | 0.8 | 7 | 0.8571 | 22 | 0.7273 | 5 | 0.4 | 4 | 1 | 6 | 0.6667 | 7 | 0.4286 | 22 | 0.5909 |
| 2_5229243: | 5 | 0 | 5 | 0 | 5 | 0.4 | 7 | 0 | 22 | 0.0909 | 4 | 0 | 4 | 0.25 | 8 | 0.125 | 7 | 0.2857 | 23 | 0.1739 |
| 2_5231909: | 16 | 0.0625 | 9 | 0.2222 | 16 | 0 | 19 | 0.1579 | 60 | 0.1 | 8 | 0 | 16 | 0.125 | 21 | 0.0952 | 21 | 0.0952 | 66 | 0.0909 |
| 2_5231976! | 9 | 0 | 7 | 0 | 12 | 0.0833 | 15 | 0.1333 | 43 | 0.0698 | 3 | 0 | 11 | 0.0909 | 7 | 0.1429 | 18 | 0.1111 | 39 | 0.1026 |
| 2_5232409- | 3 | 0 | 5 | 0.4 | 3 | 0 | 6 | 0.1667 | 17 | 0.1765 | 5 | 0.4 | 8 | 0.125 | 4 | 0.25 | 8 | 0.125 | 25 | 0.2 |
| 2_5232457' | 3 | 1 | 4 | 0.5 | 3 | 1 | 2 | 1 | 12 | 0.8333 | 3 | 0.6667 | 6 | 0.6667 | 3 | 0.6667 | 8 | 1 | 20 | 0.8 |
| 2_5232458: | 3 | 0.3333 | 5 | 0.4 | 2 | 1 | 1 | 1 | 11 | 0.5455 | 3 | 0.6667 | 6 | 0.6667 | 3 | 1 | 6 | 0.8333 | 18 | 0.7778 |
| 2_5251044! | NA | NA | NA | NA | NA | NA | 1 | 1 | 1 | 1 | NA | NA | NA | NA | 1 | 1 | NA | NA | 1 | 1 |
| 2_5258462' | NA | NA | 2 | 1 | 2 | 0.5 | 3 | 0.3333 | 7 | 0.5714 | 4 | 0 | 5 | 0 | 4 | 0.5 | 4 | 0.25 | 17 | 0.1765 |
| 2_5258470! | NA | NA | 2 | 0.5 | 3 | 0.3333 | 3 | 0 | 8 | 0.25 | 4 | 0 | 4 | 1 | 4 | 0.25 | 4 | 0 | 16 | 0.3125 |
| 2_5274817' | 2 | 1 | NA | NA | 1 | 0 | 3 | 0.6667 | 6 | 0.6667 | 2 | 0 | 1 | 1 | 2 | 0 | NA | NA | 5 | 0.2 |
| 2_5299747: | 3 | 0 | 1 | 1 | 1 | 0 | 3 | 0.3333 | 8 | 0.25 | 2 | 0 | 1 | 0 | 4 | 0.25 | 1 | 1 | 8 | 0.25 |
| 2_5302196' | 7 | 0.1429 | 2 | 0 | 1 | 0 | 7 | 0 | 17 | 0.0588 | 5 | 0.4 | 3 | 0 | 4 | 0.25 | 2 | 0.5 | 14 | 0.2857 |
| 2_5302199( | 10 | 0.1 | 2 | 0 | 3 | 0.3333 | 8 | 0 | 23 | 0.087 | 3 | 0.3333 | 3 | 0 | 5 | 0.2 | 2 | 0 | 13 | 0.1538 |
| 2_5302237: | 10 | 0.3 | 3 | 0.6667 | 4 | 0 | 8 | 0.125 | 25 | 0.24 | 6 | 0.3333 | 3 | 0 | 4 | 0 | 2 | 0 | 15 | 0.1333 |
| 2_5302257( | 6 | 0.1667 | 2 | 0.5 | 3 | 0.3333 | 6 | 0.3333 | 17 | 0.2941 | 5 | 0 | 5 | 0 | 5 | 0 | 4 | 0.25 | 19 | 0.0526 |
| 2_5302303! | 10 | 0 | 2 | 0.5 | 6 | 0 | 9 | 0.1111 | 27 | 0.0741 | 6 | 0 | 4 | 0.25 | 3 | 0 | 5 | 0 | 18 | 0.0556 |
| 2_5302876! | 1 | 0 | 1 | 1 | 2 | 1 | 1 | 0 | 5 | 0.6 | 3 | 0 | 4 | 0.25 | 4 | 0.5 | 4 | 0.25 | 15 | 0.2667 |
| 2_5302905( | 1 | 1 | 1 | 1 | 1 | 1 | 2 | 1 | 5 | 1 | 2 | 0 | 3 | 0.3333 | 2 | 0 | 3 | 0 | 10 | 0.1 |
| 2_5308179' | 1 | 0 | NA | NA | 1 | 1 | 1 | 1 | 3 | 0.6667 | NA | NA | 2 | 0.5 | NA | NA | 1 | 0 | 3 | 0.3333 |
| 2_5310731! | 2 | 0.5 | 2 | 0.5 | 4 | 0 | 3 | 0.3333 | 11 | 0.2727 | 1 | 1 | 2 | 0 | 5 | 0.4 | 2 | 0.5 | 10 | 0.4 |

| | | | | | | | | | | | | | | | | | | | | |
|---|---|---|---|---|---|---|---|---|---|---|---|---|---|---|---|---|---|---|---|---|
| 2_5310747( | 2 | 0.5 | 1 | 1 | 4 | 0.25 | 3 | 0 | 10 | 0.3 | 1 | 0 | 1 | 0 | 4 | 0 | 3 | 0.3333 | 9 | 0.1111 |
| 2_5310747! | 2 | 0.5 | 2 | 0 | 5 | 0 | 3 | 0 | 12 | 0.0833 | 1 | 0 | 2 | 0 | 5 | 0.2 | 3 | 0.3333 | 11 | 0.1818 |
| 2_5397424 | 5 | 0 | 7 | 0.1429 | 4 | 0.5 | 2 | 0 | 18 | 0.1667 | 6 | 0.1667 | 2 | 0 | 6 | 0 | 2 | 0 | 16 | 0.0625 |
| 2_5397433( | 4 | 0 | 6 | 0.1667 | 5 | 0 | 1 | 0 | 16 | 0.0625 | 6 | 0.1667 | 2 | 0 | 3 | 0 | 1 | 0 | 12 | 0.0833 |
| 2_5397449( | 3 | 0.3333 | 3 | 0 | 4 | 0.25 | 1 | 1 | 11 | 0.2727 | 6 | 0.3333 | 2 | 0 | 2 | 0.5 | 1 | 0 | 11 | 0.2727 |
| 2_5397453: | 4 | 0 | 4 | 0 | 5 | 0.4 | 1 | 1 | 14 | 0.2143 | 5 | 0.4 | 2 | 0 | 2 | 0.5 | 1 | 0 | 10 | 0.3 |
| 2_5397457 | 4 | 1 | 4 | 1 | 3 | 1 | 1 | 1 | 12 | 1 | 6 | 1 | 1 | 1 | 1 | 1 | NA | NA | 8 | 1 |
| 2_5397458 | 4 | 0.25 | 5 | 0 | 4 | 0.25 | 1 | 0 | 14 | 0.1429 | 6 | 0.3333 | 2 | 0 | 2 | 0.5 | 1 | 0 | 11 | 0.2727 |
| 2_5397464: | 3 | 0.3333 | 2 | 1 | 2 | 1 | 1 | 1 | 8 | 0.75 | 4 | 1 | 2 | 0.5 | 2 | 0.5 | NA | NA | 8 | 0.75 |
| 2_5397469: | 4 | 0 | 3 | 0 | 4 | 0.5 | 1 | 0 | 12 | 0.1667 | 5 | 0.2 | 2 | 0 | 1 | 0 | NA | NA | 8 | 0.125 |
| 2_5397473: | 4 | 1 | 3 | 1 | 5 | 1 | 1 | 1 | 13 | 1 | 5 | 1 | 2 | 1 | 1 | 1 | NA | NA | 8 | 1 |
| 2_5397477 | 4 | 0.25 | 4 | 0.5 | 5 | 0.4 | 1 | 0 | 14 | 0.3571 | 5 | 0.6 | 2 | 0 | 2 | 0.5 | NA | NA | 9 | 0.4444 |
| 2_5397479( | 3 | 0.6667 | 3 | 1 | 5 | 1 | 1 | 1 | 12 | 0.9167 | 5 | 1 | 2 | 1 | 2 | 1 | NA | NA | 9 | 1 |
| 2_5397483 | 4 | 0.75 | 3 | 0.3333 | 5 | 0.4 | 1 | 1 | 13 | 0.5385 | 4 | 0.5 | 2 | 0 | 2 | 0.5 | NA | NA | 8 | 0.375 |
| 2_5397485 | 4 | 0.25 | 4 | 0.5 | 3 | 0.3333 | NA | NA | 11 | 0.3636 | 3 | 0.6667 | 2 | 1 | 1 | 0 | NA | NA | 6 | 0.6667 |
| 2_5397490( | 4 | 0.75 | 4 | 0.5 | 3 | 0.3333 | 1 | 1 | 12 | 0.5833 | 5 | 0.8 | 2 | 0.5 | 2 | 1 | NA | NA | 9 | 0.7778 |
| 2_5397503( | 4 | 0.75 | 4 | 0 | 4 | 0.75 | 1 | 0 | 13 | 0.4615 | 4 | 0.25 | 2 | 0.5 | 2 | 0 | NA | NA | 8 | 0.25 |
| 2_5397545 | 2 | 0.5 | 2 | 0.5 | 2 | 0.5 | 1 | 1 | 7 | 0.5714 | 4 | 0.75 | 1 | 1 | 1 | 1 | NA | NA | 6 | 0.8333 |
| 2_5432584( | NA | NA | NA | NA | 1 | 1 | NA | NA | 1 | 1 | NA | NA | NA | NA | 1 | 1 | NA | NA | 1 | 1 |
| 2_5432612 | NA | NA | NA | NA | 1 | 1 | NA | NA | 1 | 1 | NA | NA | NA | NA | 1 | 1 | NA | NA | 1 | 1 |
| 2_5500283 | 5 | 0.2 | 3 | 0 | 2 | 0 | 3 | 0.3333 | 13 | 0.1538 | 1 | 0 | 4 | 0 | 5 | 0 | 2 | 0.5 | 12 | 0.0833 |
| 2_5500287 | 4 | 0 | 4 | 0 | 2 | 0.5 | 3 | 0 | 13 | 0.0769 | 2 | 0 | 6 | 0.3333 | 5 | 0 | 2 | 0 | 15 | 0.1333 |
| 2_5500291 | 4 | 0.25 | 3 | 0 | 2 | 0.5 | 3 | 0.3333 | 12 | 0.25 | 2 | 0 | 5 | 0.6 | 6 | 0 | 2 | 0.5 | 15 | 0.2667 |
| 2_5500296 | 5 | 0 | 4 | 0.25 | 2 | 0 | 3 | 0 | 14 | 0.0714 | 1 | 0 | 6 | 0.1667 | 6 | 0 | 2 | 0 | 15 | 0.0667 |
| 2_5500320 | 3 | 0 | 3 | 0 | 2 | 0.5 | 2 | 0.5 | 10 | 0.2 | 2 | 0 | 2 | 0.5 | 7 | 0.1429 | 1 | 1 | 12 | 0.25 |
| 2_5500410 | 3 | 0.3333 | 2 | 0 | 2 | 0 | 3 | 0 | 10 | 0.1 | 3 | 0 | 7 | 0.1429 | 3 | 0 | 1 | 0 | 14 | 0.0714 |
| 2_5500423 | 1 | 0 | 2 | 0 | 1 | 1 | 2 | 0 | 6 | 0.1667 | 1 | 0 | 5 | 0.2 | 4 | 0 | 1 | 0 | 11 | 0.0909 |
| 2_5500649 | 11 | 0.0909 | 3 | 0.3333 | 6 | 0.1667 | 5 | 0.2 | 25 | 0.16 | 3 | 0.3333 | 8 | 0 | 8 | 0 | 1 | 1 | 20 | 0.1 |
| 2_5500674 | 9 | 0.5556 | 2 | 0.5 | 4 | 0.25 | 3 | 0.3333 | 18 | 0.4444 | 2 | 0.5 | 4 | 0.25 | 6 | 0 | 1 | 0 | 13 | 0.1538 |
| 2_5500686 | 6 | 0.1667 | 2 | 0.5 | 2 | 0 | 5 | 0 | 15 | 0.1333 | 2 | 0.5 | 5 | 0 | 6 | 0.1667 | 1 | 0 | 14 | 0.1429 |
| 2_5500690 | 9 | 0.2222 | 3 | 0.3333 | 4 | 0.25 | 4 | 0 | 20 | 0.2 | 2 | 0.5 | 5 | 0.2 | 6 | 0 | 1 | 0 | 14 | 0.1429 |
| 2_5532704( | 1 | 0 | 1 | 1 | NA | NA | 1 | 1 | 3 | 0.6667 | 1 | 1 | NA | NA | NA | NA | 1 | 1 | 2 | 1 |
| 2_5532730 | NA | NA | 4 | 0.5 | 1 | 0 | 4 | 0.25 | 9 | 0.3333 | 2 | 0.5 | NA | NA | 1 | 0 | 5 | 0.4 | 8 | 0.375 |
| 2_5532742 | NA | NA | 5 | 0.8 | 2 | 0.5 | 5 | 0.6 | 12 | 0.6667 | 2 | 0.5 | NA | NA | 2 | 0 | 4 | 0.75 | 8 | 0.5 |
| 2_5532746( | 1 | 0 | 5 | 0.8 | 2 | 0 | 3 | 0 | 11 | 0.3636 | 2 | 0 | NA | NA | 2 | 0 | 5 | 0.4 | 9 | 0.2222 |
| 2_5532778 | 1 | 0 | 7 | 0.1429 | 3 | 0 | 3 | 0 | 14 | 0.0714 | 3 | 0.3333 | 1 | 0 | 2 | 0 | 4 | 0 | 10 | 0.1 |
| 2_5533688 | 1 | 0 | 5 | 0.2 | 2 | 0.5 | 2 | 0 | 10 | 0.2 | 2 | 1 | NA | NA | 1 | 0 | 4 | 0.25 | 7 | 0.4286 |
| 2_5548504 | 1 | 1 | 10 | 1 | 4 | 1 | 5 | 0.4 | 20 | 0.85 | 4 | 0.75 | 1 | 0 | 4 | 1 | 5 | 0.6 | 14 | 0.7143 |
| 2_5548507( | NA | NA | 10 | 1 | 4 | 1 | 5 | 1 | 19 | 1 | 3 | 1 | 1 | 1 | 4 | 1 | 6 | 1 | 14 | 1 |
| 2_5548508 | 1 | 0 | 10 | 0.5 | 3 | 0.3333 | 5 | 0.6 | 19 | 0.4737 | 4 | 0 | 1 | 1 | 3 | 0.6667 | 6 | 0.8333 | 14 | 0.5714 |
| 2_5548509 | 1 | 0 | 10 | 0 | 5 | 0.4 | 4 | 0.5 | 20 | 0.2 | 3 | 0 | 1 | 0 | 3 | 0.3333 | 5 | 0.2 | 12 | 0.1667 |
| 2_5548513 | 1 | 0 | 11 | 0.8182 | 5 | 0.6 | 5 | 0.6 | 22 | 0.6818 | 3 | 1 | 1 | 1 | 4 | 1 | 5 | 1 | 13 | 1 |
| 2_5548514 | 1 | 0 | 7 | 0.7143 | 2 | 1 | 3 | 0.6667 | 13 | 0.6923 | 3 | 1 | 1 | 1 | 3 | 0.3333 | 4 | 0.5 | 11 | 0.6364 |
| 2_5548521 | 1 | 0 | 10 | 0.7 | 4 | 0.25 | 5 | 0.6 | 20 | 0.55 | 4 | 1 | 1 | 0 | 4 | 0.75 | 5 | 0.6 | 14 | 0.7143 |
| 2_5548525 | 1 | 1 | 9 | 0.7778 | 5 | 0.8 | 3 | 0.6667 | 18 | 0.7778 | 3 | 1 | 1 | 1 | 3 | 1 | 3 | 1 | 10 | 1 |
| 2_5548534 | 1 | 1 | 8 | 0.625 | 4 | 1 | 3 | 0.3333 | 16 | 0.6875 | 4 | 0.75 | 1 | 0 | 3 | 0.6667 | 3 | 0.3333 | 11 | 0.5455 |
| 2_5548537( | 1 | 0 | 11 | 0.1818 | 3 | 0 | 5 | 0 | 20 | 0.1 | 4 | 0 | 1 | 0 | 4 | 0 | 6 | 0.1667 | 15 | 0.0667 |
| 2_5548539( | 1 | 1 | 11 | 1 | 3 | 1 | 4 | 1 | 19 | 1 | 4 | 1 | 1 | 1 | 3 | 1 | 4 | 1 | 12 | 1 |
| 2_5548553 | 1 | 0 | 8 | 0.875 | 3 | 1 | 5 | 0.4 | 17 | 0.7059 | 4 | 0.25 | 1 | 1 | 4 | 1 | 6 | 0.8333 | 15 | 0.7333 |
| 2_5548620 | 1 | 0 | 14 | 0.2143 | 3 | 0.6667 | 3 | 0.3333 | 21 | 0.2857 | 5 | 0.4 | 1 | 0 | 4 | 0.25 | 7 | 0.2857 | 17 | 0.2941 |
| 2_5548649 | 1 | 1 | 18 | 0.1111 | 7 | 0.2857 | 5 | 0.2 | 31 | 0.1935 | 8 | 0.375 | 1 | 0 | 5 | 0.2 | 5 | 0.2 | 19 | 0.2632 |
| 2_5548657 | NA | NA | 15 | 0.4667 | 7 | 0.7143 | 6 | 0.6667 | 28 | 0.5714 | 8 | 0.5 | 1 | 0 | 5 | 0.8 | 6 | 0.5 | 20 | 0.55 |
| 2_5548662 | 1 | 1 | 15 | 0.5333 | 5 | 0.2 | 5 | 0.4 | 26 | 0.4615 | 8 | 0.375 | 1 | 1 | 4 | 0.25 | 8 | 0.375 | 21 | 0.381 |
| 2_5699729 | NA | NA | 3 | 0.3333 | NA | NA | 2 | 0.5 | 5 | 0.4 | 7 | 0.1429 | 3 | 0 | 2 | 0 | 3 | 0 | 15 | 0.0667 |
| 2_5699791 | NA | NA | 2 | 0.5 | 1 | 1 | 2 | 1 | 5 | 0.8 | 5 | 1 | 2 | 0 | 3 | 0.3333 | 2 | 0.5 | 12 | 0.5833 |
| 2_5699799 | NA | NA | 2 | 1 | NA | NA | NA | NA | 2 | 1 | 4 | 1 | 1 | 0 | 1 | 1 | 1 | 1 | 7 | 0.8571 |
| 2_5699814 | NA | NA | 2 | 0.5 | 1 | 1 | 2 | 1 | 5 | 0.8 | 7 | 0.7143 | 2 | 1 | 3 | 0.6667 | 1 | 1 | 13 | 0.7692 |
| 2_5699818 | NA | NA | 1 | 1 | 1 | 0 | 2 | 1 | 4 | 0.75 | 6 | 0.5 | 2 | 1 | 3 | 0.3333 | 2 | 1 | 13 | 0.6154 |
| 2_5699819 | NA | NA | 1 | 1 | NA | NA | 2 | 0.5 | 3 | 0.6667 | 6 | 0.3333 | 1 | 1 | 3 | 1 | 1 | 0 | 11 | 0.5455 |
| 2_5699830 | NA | NA | 2 | 0.5 | NA | NA | 2 | 0 | 4 | 0.25 | 6 | 0.1667 | 2 | 0 | 3 | 0 | 1 | 0 | 12 | 0.0833 |
| 2_5699834 | NA | NA | 2 | 1 | NA | NA | 1 | 1 | 3 | 1 | 5 | 0.8 | 2 | 1 | 3 | 1 | 1 | 1 | 11 | 0.9091 |
| 2_5700219 | NA | NA | NA | NA | NA | NA | 1 | 1 | 1 | 1 | 3 | 0 | 1 | 1 | 2 | 1 | 2 | 0 | 8 | 0.375 |
| 2_5700225 | NA | NA | NA | NA | 1 | 1 | 1 | 0 | 2 | 0.5 | 2 | 0.5 | 1 | 0 | 2 | 1 | 2 | 1 | 7 | 0.7143 |
| 2_5700231 | NA | NA | NA | NA | 1 | 1 | 1 | 0 | 2 | 0.5 | 3 | 1 | 2 | 1 | 2 | 0.5 | 1 | 1 | 8 | 0.875 |
| 2_5725746 | 2 | 0 | 5 | 0.2 | 6 | 0 | 3 | 0 | 16 | 0.0625 | 3 | 0 | 1 | 1 | 2 | 0 | 1 | 0 | 7 | 0.1429 |

| | | | | | | | | | | | | | | | | | | | | |
|---|---|---|---|---|---|---|---|---|---|---|---|---|---|---|---|---|---|---|---|---|
| 2_5725752 | 2 | 0 | 6 | 0.1667 | 5 | 0 | 4 | 0 | 17 | 0.0588 | 4 | 0 | 2 | 0 | 4 | 0.25 | 2 | 0 | 12 | 0.0833 |
| 2_5725755 | 2 | 0 | 6 | 0.1667 | 6 | 0 | 4 | 0 | 18 | 0.0556 | 4 | 0.25 | 2 | 0 | 4 | 0 | 2 | 0 | 12 | 0.0833 |
| 2_5725790 | 2 | 0 | 6 | 0.1667 | 8 | 0.375 | 4 | 0 | 20 | 0.2 | 4 | 0.25 | 2 | 0 | 4 | 0 | 2 | 0 | 12 | 0.0833 |
| 2_5725895 | 3 | 0 | 7 | 0 | 8 | 0.25 | 5 | 0 | 23 | 0.087 | 3 | 0 | 1 | 0 | 4 | 0 | 4 | 0.25 | 12 | 0.0833 |
| 2_5726046 | 4 | 0.25 | 8 | 0 | 9 | 0.2222 | 5 | 0 | 26 | 0.1154 | 4 | 0 | 4 | 0 | 5 | 0 | 6 | 0.1667 | 19 | 0.0526 |
| 2_5726053 | 4 | 0.5 | 7 | 0.2857 | 8 | 0.25 | 2 | 0 | 21 | 0.2857 | 5 | 0.2 | 4 | 0.5 | 6 | 0.3333 | 5 | 0.2 | 20 | 0.3 |
| 2_5726056 | 4 | 0.25 | 7 | 0.4286 | 9 | 0.1111 | 3 | 0.3333 | 23 | 0.2609 | 6 | 0.5 | 5 | 0.2 | 5 | 0.4 | 4 | 0 | 20 | 0.3 |
| 2_5793546 | 5 | 0.2 | 9 | 0.3333 | 6 | 0.1667 | 6 | 0.5 | 26 | 0.3077 | 5 | 0 | 3 | 0 | 5 | 0.2 | 7 | 0.2857 | 20 | 0.15 |
| 2_5793577 | 5 | 0.4 | 8 | 0.25 | 4 | 0.25 | 5 | 0.2 | 22 | 0.2727 | 5 | 0.6 | 2 | 1 | 3 | 0 | 4 | 0.5 | 14 | 0.5 |
| 2_5793581 | 4 | 0.5 | 9 | 0.2222 | 4 | 0.5 | 5 | 0.2 | 22 | 0.3182 | 4 | 0.5 | 3 | 0.6667 | 3 | 0.3333 | 5 | 0.2 | 15 | 0.4 |
| 2_5793588 | 4 | 1 | 9 | 0.4444 | 3 | 0.6667 | 5 | 0.8 | 21 | 0.6667 | 4 | 1 | 2 | 1 | 4 | 0.5 | 3 | 1 | 13 | 0.8462 |
| 2_5793598 | 5 | 0 | 8 | 0 | 4 | 0.25 | 4 | 0.25 | 21 | 0.0952 | 4 | 0 | 2 | 0.5 | 4 | 0.25 | 4 | 0 | 14 | 0.1429 |
| 2_5793600 | 5 | 0 | 9 | 0.2222 | 2 | 0 | 5 | 0.2 | 21 | 0.1429 | 5 | 0.2 | 2 | 0 | 3 | 0 | 3 | 0 | 13 | 0.0769 |
| 2_5793613 | 4 | 0.75 | 9 | 0.5556 | 4 | 0 | 4 | 0.25 | 21 | 0.4286 | 4 | 1 | 3 | 0.3333 | 3 | 1 | 5 | 0.6 | 15 | 0.7333 |
| 2_5793632 | 5 | 0 | 9 | 0.2222 | 4 | 0.25 | 4 | 0.25 | 22 | 0.1818 | 5 | 0.2 | 3 | 0.3333 | 4 | 0.75 | 5 | 0.2 | 17 | 0.3529 |
| 2_5793659 | 5 | 0.6 | 7 | 0.1429 | 5 | 0.2 | 4 | 0 | 21 | 0.2381 | 6 | 0.5 | 4 | 0 | 4 | 0.25 | 6 | 0.5 | 20 | 0.35 |
| 2_5793675 | 5 | 0 | 10 | 0.3 | 4 | 0.5 | 5 | 0.4 | 24 | 0.2917 | 5 | 0 | 4 | 0 | 3 | 0.3333 | 6 | 0.1667 | 18 | 0.1111 |
| 2_5793688 | 5 | 0.2 | 6 | 0.1667 | 5 | 0.2 | 3 | 0.3333 | 19 | 0.2105 | 6 | 0 | 4 | 0.25 | 4 | 0.5 | 6 | 0 | 20 | 0.15 |
| 2_5793754 | 6 | 0.1667 | 10 | 0.6 | 6 | 0.3333 | 7 | 0.2857 | 29 | 0.3793 | 4 | 0.75 | 4 | 0.25 | 4 | 0 | 8 | 0.125 | 20 | 0.25 |
| 2_5793761 | 3 | 0 | 8 | 0.375 | 6 | 0.3333 | 4 | 0.25 | 21 | 0.2857 | 3 | 0.3333 | 4 | 0.5 | 1 | 0 | 6 | 0.3333 | 14 | 0.3571 |
| 2_5793761 | 6 | 0.6667 | 10 | 0.3 | 7 | 0.2857 | 6 | 0.6667 | 29 | 0.4483 | 3 | 0.3333 | 3 | 0.6667 | 3 | 0.6667 | 5 | 0.4 | 14 | 0.5 |
| 2_5793771 | 3 | 0 | 6 | 0.1667 | 5 | 0 | 6 | 0 | 20 | 0.05 | 4 | 0 | 4 | 0 | 4 | 0 | 5 | 0.2 | 17 | 0.0588 |
| 2_5793772 | 4 | 0 | 11 | 0.1818 | 8 | 0.25 | 7 | 0.1429 | 30 | 0.1667 | 4 | 0 | 4 | 0.25 | 4 | 0.25 | 6 | 0 | 18 | 0.1111 |
| 2_5793775 | 2 | 1 | 7 | 0.8571 | 8 | 0.875 | 6 | 0.5 | 23 | 0.7826 | 5 | 1 | 4 | 1 | 4 | 1 | 5 | 1 | 18 | 1 |
| 2_5793776 | 5 | 1 | 10 | 0.8 | 6 | 0.6667 | 3 | 0.3333 | 24 | 0.75 | 4 | 1 | 4 | 1 | 4 | 1 | 3 | 0.6667 | 15 | 0.9333 |
| 2_5793794 | 3 | 1 | 11 | 1 | 6 | 1 | 5 | 1 | 25 | 1 | 3 | 1 | 4 | 1 | 2 | 1 | 5 | 1 | 14 | 1 |
| 2_5793821 | 5 | 0 | 12 | 0.0833 | 4 | 0.5 | 5 | 0.2 | 26 | 0.1538 | 4 | 0 | 4 | 0.25 | 4 | 0 | 7 | 0.4286 | 19 | 0.2105 |
| 2_5794269 | 4 | 0 | 8 | 0 | 4 | 0 | 4 | 0.25 | 20 | 0.05 | 2 | 1 | 2 | 0 | 4 | 0.25 | 5 | 0.2 | 13 | 0.3077 |
| 2_5794280 | 5 | 0 | 12 | 0.25 | 5 | 0.2 | 5 | 0 | 27 | 0.1481 | 2 | 0.5 | 3 | 0 | 4 | 0.25 | 6 | 0.1667 | 15 | 0.2 |
| 2_5815773 | 1 | 0 | 4 | 0.25 | 4 | 0 | 5 | 0 | 14 | 0.0714 | 1 | 0 | 2 | 0 | 6 | 0.1667 | 4 | 0 | 13 | 0.0769 |
| 2_5815793 | 1 | 0 | 3 | 0 | 4 | 0 | 5 | 0.2 | 13 | 0.0769 | 1 | 0 | 1 | 1 | 5 | 0 | 4 | 0.25 | 11 | 0.1818 |
| 2_5815797 | 1 | 0 | 3 | 0.3333 | 4 | 0.5 | 5 | 0.6 | 13 | 0.4615 | 1 | 1 | 2 | 0.5 | 4 | 0.75 | 3 | 1 | 10 | 0.8 |
| 2_5815963 | NA | NA | 1 | 0 | 3 | 0 | 4 | 0.25 | 8 | 0.125 | NA | NA | 1 | 0 | 1 | 0 | 3 | 1 | 5 | 0.6 |
| 2_5816152 | NA | NA | 1 | 0 | 2 | 0.5 | 2 | 1 | 5 | 0.6 | 1 | 1 | 1 | 0 | NA | NA | 1 | 0 | 3 | 0.3333 |
| 2_5816170 | NA | NA | 1 | 0 | 2 | 1 | 2 | 0.5 | 5 | 0.6 | 1 | 0 | 1 | 0 | 1 | 1 | 1 | 1 | 4 | 0.5 |
| 2_5833678 | 3 | 0.3333 | 5 | 0.8 | 2 | 1 | 4 | 1 | 14 | 0.7857 | 4 | 0.75 | 1 | 1 | 3 | 0.6667 | 3 | 0.6667 | 11 | 0.7273 |
| 2_5833693 | 1 | 0 | 4 | 0.25 | 1 | 0 | 3 | 0.6667 | 9 | 0.3333 | 4 | 0.25 | NA | NA | 3 | 0.3333 | 3 | 0.3333 | 10 | 0.3 |
| 2_5833698 | 3 | 1 | 3 | 1 | 1 | 1 | 3 | 1 | 10 | 1 | 4 | 1 | NA | NA | 3 | 1 | 3 | 1 | 10 | 1 |
| 2_5833707 | 3 | 0.6667 | 3 | 1 | NA | NA | 4 | 0.5 | 10 | 0.7 | 4 | 0.25 | 1 | 0 | 3 | 0.6667 | 3 | 0.3333 | 11 | 0.3636 |
| 2_5833723 | 3 | 0 | 2 | 0.5 | 1 | 0 | 3 | 0 | 9 | 0.1111 | 4 | 0.25 | NA | NA | 3 | 0 | 3 | 0.3333 | 10 | 0.2 |
| 2_5833731 | 3 | 1 | 3 | 1 | 1 | 1 | 3 | 0.6667 | 10 | 0.9 | 4 | 0.75 | NA | NA | 2 | 1 | 2 | 1 | 8 | 0.875 |
| 2_5833733 | NA | NA | 3 | 1 | 1 | 1 | 3 | 1 | 7 | 1 | 4 | 1 | NA | NA | 3 | 1 | 3 | 1 | 10 | 1 |
| 2_5833735 | 2 | 0 | 3 | 0.3333 | 1 | 0 | 1 | 0 | 7 | 0.1429 | 3 | 0 | 1 | 0 | 3 | 0 | 3 | 0.3333 | 10 | 0.1 |
| 2_5833735 | 2 | 1 | 2 | 0.5 | 1 | 1 | 2 | 1 | 7 | 0.8571 | 4 | 1 | NA | NA | 3 | 1 | 3 | 1 | 10 | 1 |
| 2_5833746 | 2 | 0.5 | 3 | 0.6667 | 1 | 0 | 1 | 0 | 7 | 0.4286 | 4 | 0.25 | 1 | 0 | 2 | 0 | 2 | 0 | 9 | 0.1111 |
| 2_5833749 | 3 | 0.3333 | 3 | 0 | NA | NA | 2 | 0.5 | 8 | 0.25 | 4 | 1 | 1 | 0 | 3 | 0.6667 | 2 | 0 | 10 | 0.6 |
| 2_5833755 | 2 | 1 | 2 | 1 | NA | NA | 2 | 0.5 | 6 | 0.8333 | 4 | 0.75 | 1 | 1 | 2 | 0.5 | 1 | 1 | 8 | 0.75 |
| 2_5833756 | NA | NA | 1 | 1 | NA | NA | 2 | 0 | 3 | 0.3333 | 3 | 0.3333 | NA | NA | 1 | 0 | NA | NA | 4 | 0.25 |
| 2_5833760 | 3 | 1 | 3 | 1 | 1 | 0 | 2 | 1 | 9 | 0.8889 | 4 | 0.75 | 1 | 1 | 3 | 1 | 2 | 1 | 10 | 0.9 |
| 2_5847347 | NA | NA | 2 | 0.5 | 3 | 0.6667 | 2 | 0 | 7 | 0.4286 | 1 | 0 | 1 | 0 | 1 | 1 | NA | NA | 3 | 0.3333 |
| 2_5847349 | 1 | 1 | 2 | 1 | 3 | 0.6667 | 2 | 1 | 8 | 0.875 | 1 | 1 | 1 | 0 | 1 | 0 | NA | NA | 3 | 0.3333 |
| 2_5847353 | 1 | 1 | NA | NA | 3 | 0.6667 | 2 | 0.5 | 6 | 0.6667 | 2 | 1 | 1 | 1 | 1 | 0 | NA | NA | 4 | 0.75 |
| 2_5847369 | 1 | 1 | 2 | 1 | 2 | 1 | 2 | 1 | 7 | 1 | 2 | 1 | 1 | 1 | 1 | 1 | NA | NA | 4 | 1 |
| 2_5847393 | NA | NA | NA | NA | 1 | 1 | NA | NA | 1 | 1 | NA | NA | 1 | 1 | NA | NA | NA | NA | 1 | 1 |
| 2_5852297 | NA | NA | 1 | 1 | 1 | 1 | 1 | 1 | 3 | 1 | 2 | 1 | NA | NA | NA | NA | NA | NA | 2 | 1 |
| 2_5852302 | NA | NA | 1 | 1 | 1 | 1 | 1 | 1 | 3 | 1 | 1 | 1 | NA | NA | NA | NA | NA | NA | 1 | 1 |
| 2_5852312 | NA | NA | NA | NA | 1 | 0 | 1 | 1 | 2 | 0.5 | 2 | 0.5 | NA | NA | NA | NA | NA | NA | 2 | 0.5 |
| 2_5852319 | NA | NA | 1 | 1 | 1 | 1 | 1 | 1 | 3 | 1 | 2 | 1 | NA | NA | NA | NA | NA | NA | 2 | 1 |
| 2_5852326 | NA | NA | 1 | 1 | 1 | 1 | 1 | 1 | 3 | 1 | 2 | 0.5 | NA | NA | NA | NA | NA | NA | 2 | 0.5 |
| 2_5868017 | 1 | 0 | NA | NA | NA | NA | 3 | 0.6667 | 4 | 0.5 | 1 | 1 | 1 | 1 | NA | NA | NA | NA | 2 | 1 |
| 2_5871349 | 2 | 1 | NA | NA | NA | NA | 1 | 1 | 3 | 1 | NA | NA | NA | NA | 1 | 1 | NA | NA | 1 | 1 |
| 2_5871366 | 2 | 0.5 | 1 | 0 | NA | NA | 1 | 1 | 4 | 0.5 | NA | NA | 1 | 0 | 1 | 1 | NA | NA | 2 | 0.5 |
| 2_5898965 | 1 | 1 | NA | NA | NA | NA | NA | NA | 1 | 1 | 1 | 1 | NA | NA | NA | NA | NA | NA | 1 | 1 |
| 2_5913846 | 7 | 0.2857 | 4 | 0.25 | 2 | 0 | 7 | 0.2857 | 20 | 0.25 | 3 | 0.6667 | 6 | 0.1667 | 3 | 0.3333 | 3 | 0.3333 | 15 | 0.3333 |
| 2_5913850 | 7 | 0.4286 | 5 | 0.6 | 4 | 0.75 | 10 | 0.6 | 26 | 0.5769 | 5 | 0.8 | 6 | 0.3333 | 6 | 0.3333 | 9 | 0.4444 | 26 | 0.4615 |

| | | | | | | | | | | | | | | | | | | | | |
|---|---|---|---|---|---|---|---|---|---|---|---|---|---|---|---|---|---|---|---|---|
| 2_5913865 | 5 | 0.6 | 6 | 0.5 | 4 | 0.5 | 12 | 0.3333 | 27 | 0.4444 | 7 | 0.7143 | 6 | 0.1667 | 7 | 0.5714 | 9 | 0.4444 | 29 | 0.4828 |
| 2_5913871 | 9 | 0.2222 | 8 | 0 | 4 | 0 | 14 | 0.2143 | 35 | 0.1429 | 7 | 0.2857 | 7 | 0.2857 | 8 | 0.125 | 11 | 0.1818 | 33 | 0.2121 |
| 2_5913881 | 9 | 0.6667 | 8 | 0.75 | 6 | 0.5 | 14 | 0.5714 | 37 | 0.6216 | 7 | 0.5714 | 7 | 0.4286 | 7 | 0.7143 | 12 | 0.5833 | 33 | 0.5758 |
| 2_5918460 | 14 | 0.0714 | 21 | 0.2381 | 15 | 0.3333 | 23 | 0.1304 | 73 | 0.1918 | 20 | 0.1 | 12 | 0.1667 | 17 | 0.1765 | 26 | 0.3077 | 75 | 0.2 |
| 2_5918469 | 14 | 0.7857 | 19 | 0.8421 | 15 | 0.8 | 23 | 0.8261 | 71 | 0.8169 | 20 | 0.9 | 10 | 0.9 | 14 | 0.8571 | 24 | 0.7917 | 68 | 0.8529 |
| 2_5918472 | 15 | 0.8 | 20 | 0.8 | 14 | 0.6429 | 20 | 0.8 | 69 | 0.7681 | 19 | 0.6316 | 12 | 0.8333 | 18 | 0.6667 | 24 | 0.75 | 73 | 0.7123 |
| 2_5918475 | 8 | 0 | 14 | 0.2857 | 14 | 0.1429 | 16 | 0.1875 | 52 | 0.1731 | 6 | 0.3333 | 4 | 0.25 | 8 | 0.125 | 14 | 0.0714 | 32 | 0.1562 |
| 2_5918477 | 16 | 0.375 | 21 | 0.1905 | 18 | 0 | 24 | 0.25 | 79 | 0.2025 | 21 | 0.1905 | 9 | 0.1111 | 17 | 0.2353 | 28 | 0.1786 | 75 | 0.1867 |
| 2_5918483 | 14 | 0.0714 | 22 | 0.2273 | 15 | 0.4 | 22 | 0.0909 | 73 | 0.1918 | 20 | 0.25 | 12 | 0.1667 | 13 | 0.3077 | 29 | 0.1379 | 74 | 0.2027 |
| 2_5918498 | 16 | 0.1875 | 21 | 0.5714 | 18 | 0.4444 | 23 | 0.3913 | 78 | 0.4103 | 20 | 0.45 | 13 | 0.3077 | 17 | 0.5294 | 26 | 0.5769 | 76 | 0.4868 |
| 2_5918533 | 20 | 0.35 | 22 | 0.3636 | 21 | 0.2857 | 23 | 0.2609 | 86 | 0.314 | 21 | 0.3333 | 12 | 0.1667 | 18 | 0.3889 | 29 | 0.2759 | 80 | 0.3 |
| 2_5918537 | 19 | 0.3158 | 24 | 0.4167 | 22 | 0.4091 | 24 | 0.4167 | 89 | 0.3933 | 17 | 0.2353 | 12 | 0.5 | 23 | 0.4348 | 32 | 0.4062 | 84 | 0.3929 |
| 2_5918539 | 19 | 0.3158 | 21 | 0.1429 | 20 | 0.25 | 28 | 0.2143 | 88 | 0.2273 | 23 | 0.1739 | 13 | 0 | 20 | 0.25 | 35 | 0.1714 | 91 | 0.1648 |
| 2_5918543 | 18 | 0.0556 | 24 | 0 | 21 | 0.0952 | 29 | 0.069 | 92 | 0.0543 | 24 | 0.125 | 12 | 0.0833 | 24 | 0.0417 | 32 | 0.0312 | 92 | 0.0652 |
| 2_5918548 | 15 | 0.2 | 20 | 0.35 | 20 | 0.35 | 25 | 0.12 | 80 | 0.25 | 25 | 0.24 | 12 | 0.3333 | 20 | 0.15 | 32 | 0.2188 | 89 | 0.2247 |
| 2_5939441 | 1 | 0 | 3 | 0.3333 | NA | NA | NA | NA | 4 | 0.25 | 1 | 0 | 1 | 0 | 1 | 1 | NA | NA | 3 | 0.3333 |
| 2_5963038 | 1 | 0 | 1 | 0 | 1 | 0 | 1 | 1 | 4 | 0.25 | NA | NA | 2 | 0 | 1 | 1 | NA | NA | 3 | 0.3333 |
| 2_5963043 | 1 | 0 | 1 | 0 | NA | NA | 1 | 1 | 3 | 0.3333 | NA | NA | 2 | 0.5 | 1 | 0 | NA | NA | 3 | 0.3333 |
| 2_5967320 | 3 | 0.3333 | 2 | 0 | 2 | 0.5 | 1 | 0 | 8 | 0.25 | NA | NA | 3 | 0 | 2 | 0.5 | 1 | 0 | 6 | 0.1667 |
| 2_5967324 | 2 | 1 | 2 | 1 | 2 | 1 | 1 | 1 | 7 | 1 | NA | NA | 4 | 0.75 | 2 | 1 | 1 | 1 | 7 | 0.8571 |
| 2_5967343 | 3 | 1 | 5 | 1 | 2 | 1 | 1 | 1 | 11 | 1 | NA | NA | 4 | 1 | 2 | 1 | 1 | 1 | 7 | 1 |
| 2_5967353 | 3 | 0.3333 | 4 | 0.25 | 1 | 0 | 1 | 1 | 9 | 0.3333 | NA | NA | 4 | 0.25 | 2 | 0.5 | 1 | 0 | 7 | 0.2857 |
| 2_5967356 | 3 | 0.6667 | 4 | 0 | 2 | 0.5 | 1 | 0 | 10 | 0.3 | NA | NA | 3 | 0.3333 | 2 | 0 | 1 | 0 | 6 | 0.1667 |
| 2_5967366 | 2 | 1 | 5 | 1 | 2 | 1 | NA | NA | 9 | 1 | NA | NA | 3 | 0.6667 | 2 | 1 | 1 | 1 | 6 | 0.8333 |
| 2_5967380 | 2 | 1 | 3 | 1 | 1 | 1 | 1 | 1 | 7 | 1 | NA | NA | 4 | 1 | 2 | 0.5 | NA | NA | 6 | 0.8333 |
| 2_5967381 | 3 | 0.3333 | 5 | 0 | 2 | 0 | 1 | 0 | 11 | 0.0909 | NA | NA | 4 | 0 | 2 | 0.5 | 1 | 0 | 7 | 0.1429 |
| 2_5967397 | 3 | 0.6667 | 5 | 0.4 | 2 | 0.5 | 1 | 1 | 11 | 0.5455 | NA | NA | 4 | 0.25 | 2 | 0.5 | 1 | 1 | 7 | 0.4286 |
| 2_5967398 | 2 | 0.5 | 3 | 0.3333 | 2 | 0.5 | 1 | 0 | 8 | 0.375 | NA | NA | 3 | 0.6667 | 2 | 1 | NA | NA | 5 | 0.8 |
| 2_5967399 | 2 | 1 | 5 | 0.8 | 2 | 1 | 1 | 1 | 10 | 0.9 | NA | NA | 4 | 0.5 | 2 | 0 | NA | NA | 6 | 0.3333 |
| 2_5967403 | 3 | 0 | 3 | 0 | 2 | 0.5 | NA | NA | 8 | 0.125 | NA | NA | 3 | 0 | 1 | 0 | 2 | 0.5 | 6 | 0.1667 |
| 2_5967407 | 2 | 1 | 5 | 1 | 3 | 1 | 1 | 1 | 11 | 1 | NA | NA | 4 | 1 | 2 | 1 | 2 | 1 | 8 | 1 |
| 2_5967410 | 2 | 0.5 | 3 | 0 | 3 | 0 | NA | NA | 8 | 0.125 | NA | NA | 4 | 0.25 | 2 | 0 | 1 | 0 | 7 | 0.1429 |
| 2_5967414 | 1 | 0 | 5 | 0.2 | 3 | 0.3333 | 1 | 0 | 10 | 0.2 | NA | NA | 3 | 0 | 2 | 0.5 | 2 | 1 | 7 | 0.4286 |
| 2_5967418 | 3 | 0.3333 | 2 | 0 | 1 | 1 | 1 | 0 | 7 | 0.2857 | NA | NA | 2 | 1 | 2 | 0.5 | 2 | 0 | 6 | 0.5 |
| 2_5967419 | 2 | 0 | 3 | 0.3333 | 3 | 0 | 1 | 0 | 9 | 0.1111 | NA | NA | 4 | 0 | 2 | 0.5 | 2 | 0 | 8 | 0.125 |
| 2_5967429 | 3 | 0.6667 | 6 | 0.5 | 2 | 0 | 1 | 1 | 12 | 0.5 | NA | NA | 3 | 0 | 1 | 1 | 2 | 0 | 6 | 0.1667 |
| 2_5967433 | 3 | 0.6667 | 3 | 0 | 2 | 0.5 | 1 | 0 | 9 | 0.3333 | NA | NA | 2 | 0 | 2 | 0.5 | 1 | 0 | 5 | 0.2 |
| 2_5967460 | 3 | 0.6667 | 6 | 0 | 3 | 0.6667 | 1 | 0 | 13 | 0.3077 | NA | NA | 4 | 0 | 2 | 0.5 | 3 | 0.3333 | 9 | 0.2222 |
| 2_5967461 | 3 | 0.3333 | 6 | 0.1667 | 3 | 0.3333 | 1 | 0 | 13 | 0.2308 | NA | NA | 3 | 0 | 2 | 0 | 3 | 0.6667 | 8 | 0.25 |
| 2_5967462 | 3 | 0 | 5 | 0 | 3 | 0.3333 | 1 | 0 | 12 | 0.0833 | NA | NA | 3 | 0 | 1 | 0 | 2 | 0.5 | 6 | 0.1667 |
| 2_5967468 | 2 | 0 | 7 | 0.5714 | 3 | 0.6667 | 1 | 0 | 13 | 0.4615 | NA | NA | 3 | 0.3333 | 1 | 0 | 2 | 0.5 | 6 | 0.3333 |
| 2_5967496 | 3 | 0.6667 | 10 | 0.4 | 3 | 0.3333 | 1 | 0 | 17 | 0.4118 | NA | NA | 4 | 0 | 2 | 0 | 3 | 0.6667 | 9 | 0.2222 |
| 2_5967508 | 3 | 0.3333 | 11 | 0.0909 | 3 | 0.3333 | 1 | 0 | 18 | 0.1667 | NA | NA | 4 | 0.25 | 2 | 0 | 3 | 0.3333 | 9 | 0.2222 |
| 2_5967523 | 2 | 0.5 | 5 | 0.2 | 3 | 0.3333 | NA | NA | 10 | 0.3 | NA | NA | 3 | 0 | 2 | 0.5 | 3 | 0.3333 | 8 | 0.25 |
| 2_5967542 | 3 | 0 | 10 | 0.1 | 3 | 0 | 1 | 0 | 17 | 0.0588 | NA | NA | 3 | 0.3333 | 2 | 0 | 2 | 0 | 7 | 0.1429 |
| 2_5967556 | 2 | 0.5 | 10 | 0.5 | 3 | 0.3333 | 1 | 1 | 16 | 0.5 | NA | NA | 3 | 0.3333 | 1 | 1 | 3 | 0.6667 | 7 | 0.5714 |
| 2_5967559 | 3 | 0 | 9 | 0.1111 | 2 | 0 | 1 | 0 | 15 | 0.0667 | NA | NA | 4 | 0 | 1 | 0 | 2 | 0.5 | 7 | 0.1429 |
| 2_5967581 | 3 | 0 | 10 | 0 | 5 | 0 | 1 | 1 | 19 | 0.0526 | 1 | 0 | 4 | 0 | 2 | 0 | 2 | 0.5 | 9 | 0.1111 |
| 2_5968302 | NA | NA | 1 | 1 | NA | NA | 1 | 1 | 2 | 1 | NA | NA | NA | NA | 2 | 1 | 1 | 0 | 3 | 0.6667 |
| 2_5970875 | 1 | 1 | 1 | 0 | NA | NA | 1 | 0 | 3 | 0.3333 | 1 | 0 | NA | NA | 2 | 0.5 | 1 | 0 | 4 | 0.25 |
| 2_5973038 | 1 | 0 | 4 | 0.25 | 3 | 0 | 1 | 1 | 9 | 0.2222 | 1 | 1 | 1 | 0 | NA | NA | 2 | 0.5 | 4 | 0.5 |
| 2_5973042 | 1 | 0 | 4 | 0 | 3 | 0.3333 | 1 | 0 | 9 | 0.1111 | 1 | 1 | NA | NA | NA | NA | 1 | 0 | 2 | 0.5 |
| 2_5973092 | 1 | 0 | 3 | 0 | 4 | 0.25 | NA | NA | 8 | 0.125 | 1 | 1 | 1 | 0 | NA | NA | 2 | 0 | 4 | 0.25 |
| 2_5973095 | 1 | 0 | 2 | 0.5 | 3 | 1 | NA | NA | 6 | 0.6667 | NA | NA | 1 | 0 | NA | NA | 2 | 0.5 | 3 | 0.3333 |
| 2_5973101 | 1 | 0 | 3 | 1 | 4 | 0.75 | NA | NA | 8 | 0.75 | 1 | 1 | 1 | 1 | NA | NA | 2 | 0.5 | 4 | 0.75 |
| 2_5973106 | NA | NA | 4 | 0.25 | 3 | 0.6667 | NA | NA | 7 | 0.4286 | NA | NA | NA | NA | NA | NA | 2 | 0.5 | 2 | 0.5 |
| 2_5973182 | 1 | 0 | 3 | 0.3333 | 4 | 0.5 | NA | NA | 8 | 0.375 | 1 | 0 | 1 | 0 | NA | NA | 2 | 0.5 | 4 | 0.25 |
| 2_6003718 | NA | NA | 1 | 1 | 1 | 0 | NA | NA | 2 | 0.5 | NA | NA | 1 | 1 | 2 | 0.5 | NA | NA | 3 | 0.6667 |
| 2_6003719 | NA | NA | 1 | 1 | 1 | 0 | NA | NA | 2 | 0.5 | NA | NA | 1 | 0 | 2 | 1 | NA | NA | 3 | 0.6667 |
| 2_6003944 | NA | NA | 1 | 1 | NA | NA | NA | NA | 1 | 1 | NA | NA | 1 | 1 | 1 | 1 | NA | NA | 2 | 1 |
| 2_6006438 | NA | NA | 5 | 0.2 | 2 | 0.5 | 2 | 0.5 | 9 | 0.3333 | 1 | 1 | 4 | 0.5 | 3 | 0.3333 | 1 | 1 | 9 | 0.5556 |
| 2_6006448 | NA | NA | 5 | 0.2 | 2 | 0 | 2 | 0.5 | 9 | 0.2222 | 1 | 0 | 4 | 0 | 4 | 0.5 | 1 | 0 | 10 | 0.2 |
| 2_6006452 | NA | NA | 5 | 0.8 | 2 | 0 | 2 | 0.5 | 9 | 0.5556 | 1 | 0 | 4 | 0.5 | 4 | 0.25 | 1 | 1 | 10 | 0.4 |
| 2_6006468 | NA | NA | 5 | 0 | 2 | 0.5 | 2 | 0 | 9 | 0.1111 | 1 | 0 | 4 | 0 | 3 | 0.3333 | 1 | 0 | 9 | 0.1111 |
| 2_6006471 | NA | NA | 4 | 0.25 | 1 | 0 | 2 | 0.5 | 7 | 0.2857 | NA | NA | 3 | 0 | 4 | 0.25 | 1 | 0 | 8 | 0.125 |

| | | | | | | | | | | | | | | | | | | | | |
|---|---|---|---|---|---|---|---|---|---|---|---|---|---|---|---|---|---|---|---|---|
| 2_6006487 | NA | NA | 5 | 0.2 | 2 | 0.5 | 2 | 0.5 | 9 | 0.3333 | 1 | 0 | 4 | 0.75 | 4 | 0.25 | 2 | 0 | 11 | 0.3636 |
| 2_6006515 | NA | NA | 4 | 0.25 | 1 | 1 | 3 | 1 | 8 | 0.625 | NA | NA | 3 | 1 | 4 | 0.75 | 2 | 1 | 9 | 0.8889 |
| 2_6007834 | 5 | 0 | 6 | 0 | 2 | 0.5 | 7 | 0 | 20 | 0.05 | 2 | 0 | 5 | 0.2 | 5 | 0 | 7 | 0.1429 | 19 | 0.1053 |
| 2_6007847 | 5 | 0 | 6 | 0.1667 | 3 | 0 | 5 | 0.2 | 19 | 0.1053 | 4 | 0 | 5 | 0 | 6 | 0.5 | 5 | 0 | 20 | 0.15 |
| 2_6008259 | 4 | 0 | 3 | 0.3333 | 6 | 0.1667 | 3 | 0 | 16 | 0.125 | 5 | 0.2 | 6 | 0 | 4 | 0 | 5 | 0 | 20 | 0.05 |
| 2_6008264 | 6 | 0.5 | 1 | 0 | 4 | 0.5 | 2 | 1 | 13 | 0.5385 | 2 | 0.5 | 6 | 0.5 | 3 | 0.3333 | 4 | 0.75 | 15 | 0.5333 |
| 2_6058175 | 2 | 1 | 4 | 0.5 | NA | NA | 2 | 0.5 | 8 | 0.625 | 1 | 1 | NA | NA | 2 | 0.5 | 2 | 0.5 | 5 | 0.6 |
| 2_6148020 | 7 | 0.5714 | 2 | 0 | 5 | 0.4 | 5 | 0.6 | 19 | 0.4737 | 4 | 0.75 | 2 | 1 | 3 | 0 | 4 | 0.5 | 13 | 0.5385 |
| 2_6148028 | 6 | 1 | 2 | 1 | 5 | 1 | 5 | 0.8 | 18 | 0.9444 | 4 | 1 | 1 | 1 | 3 | 1 | 4 | 1 | 12 | 1 |
| 2_6148028 | 7 | 1 | 2 | 0.5 | 6 | 1 | 4 | 0.75 | 19 | 0.8947 | 3 | 0.3333 | 2 | 0.5 | 3 | 0.6667 | 3 | 1 | 11 | 0.6364 |
| 2_6148038 | 8 | 0.5 | 4 | 0 | 6 | 0.8333 | 4 | 0 | 22 | 0.4091 | 4 | 1 | 3 | 0.3333 | 3 | 0.3333 | 4 | 0.5 | 14 | 0.5714 |
| 2_6148039 | 8 | 0.375 | 4 | 0.25 | 6 | 0.3333 | 5 | 0.2 | 23 | 0.3043 | 5 | 1 | 3 | 0.3333 | 3 | 0.3333 | 4 | 0.75 | 15 | 0.6667 |
| 2_6148041 | 7 | 1 | 3 | 0.6667 | 6 | 0.5 | 5 | 0.6 | 21 | 0.7143 | 5 | 0.4 | 4 | 0.5 | 2 | 0.5 | 3 | 0.3333 | 14 | 0.4286 |
| 2_6148046 | 8 | 0.125 | 4 | 0 | 6 | 0 | 5 | 0.2 | 23 | 0.087 | 5 | 0 | 4 | 0.25 | 3 | 0 | 4 | 0 | 16 | 0.0625 |
| 2_6148048 | 6 | 0.3333 | 5 | 0.4 | 5 | 0 | 5 | 0 | 21 | 0.1905 | 5 | 0.2 | 4 | 0.25 | 4 | 0.25 | 3 | 0 | 16 | 0.1875 |
| 2_6148375 | 7 | 0 | 5 | 0.2 | 5 | 0.2 | 6 | 0.3333 | 23 | 0.1739 | 5 | 0 | 4 | 0.25 | 4 | 0 | 3 | 0 | 16 | 0.0625 |
| 2_6148378 | 5 | 0.6 | 5 | 0.4 | 6 | 0.3333 | 5 | 0.4 | 21 | 0.4286 | 5 | 0 | 4 | 0 | 3 | 0.3333 | 5 | 0.2 | 17 | 0.1176 |
| 2_6148389 | 8 | 0.375 | 5 | 0 | 6 | 0.3333 | 6 | 0.3333 | 25 | 0.28 | 5 | 0.4 | 4 | 0.5 | 4 | 0 | 5 | 0.4 | 18 | 0.3333 |
| 2_6148391 | 6 | 0.1667 | 5 | 0.2 | 6 | 0.1667 | 5 | 0.4 | 22 | 0.2273 | 5 | 0.2 | 3 | 0 | 4 | 0.25 | 4 | 0.25 | 16 | 0.1875 |
| 2_6163500 | NA | NA | 4 | 0.25 | 2 | 0.5 | NA | NA | 6 | 0.3333 | 1 | 1 | NA | NA | NA | NA | 2 | 0.5 | 3 | 0.6667 |
| 2_6163524 | 2 | 0.5 | 5 | 0.4 | 3 | 0.6667 | NA | NA | 10 | 0.5 | 4 | 0 | 1 | 0 | NA | NA | 3 | 1 | 8 | 0.375 |
| 2_6163526 | 2 | 0 | 5 | 0.8 | 4 | 0.5 | NA | NA | 11 | 0.5455 | 4 | 0.25 | 1 | 1 | NA | NA | 2 | 1 | 7 | 0.5714 |
| 2_6164210 | 8 | 0 | 11 | 0.0909 | 6 | 0.1667 | 2 | 0.5 | 27 | 0.1111 | 7 | 0 | 2 | 0 | 2 | 0.5 | 10 | 0.3 | 21 | 0.1905 |
| 2_6164216 | 10 | 0.1 | 13 | 0.3077 | 8 | 0 | 2 | 0.5 | 33 | 0.1818 | 5 | 0.2 | 2 | 0 | 2 | 0.5 | 11 | 0.2727 | 20 | 0.25 |
| 2_6164221 | 9 | 0.8889 | 13 | 0.9231 | 8 | 0.75 | 2 | 1 | 32 | 0.875 | 7 | 1 | 3 | 1 | 2 | 1 | 12 | 1 | 24 | 1 |
| 2_6164225 | 9 | 0.8889 | 12 | 1 | 7 | 1 | 2 | 1 | 30 | 0.9667 | 7 | 1 | 2 | 1 | 2 | 1 | 11 | 0.9091 | 22 | 0.9545 |
| 2_6164255 | 10 | 0.8 | 12 | 0.75 | 7 | 1 | 3 | 0.6667 | 32 | 0.8125 | 7 | 0.7143 | 2 | 1 | 3 | 1 | 9 | 0.8889 | 21 | 0.8571 |
| 2_6164272 | 11 | 0.5455 | 11 | 0.6364 | 8 | 0.75 | 3 | 0.6667 | 33 | 0.6364 | 9 | 0.6667 | 3 | 0.6667 | 2 | 0.5 | 11 | 0.5455 | 25 | 0.6 |
| 2_6164273 | 11 | 0.3636 | 12 | 0.5833 | 6 | 0.6667 | 3 | 0.3333 | 32 | 0.5 | 9 | 0.5556 | 2 | 1 | 2 | 1 | 11 | 0.7273 | 24 | 0.7083 |
| 2_6164274 | 12 | 0.75 | 13 | 0.7692 | 8 | 0.875 | 2 | 1 | 35 | 0.8 | 7 | 0.5714 | 3 | 0.6667 | 3 | 1 | 13 | 0.9231 | 26 | 0.8077 |
| 2_6164293 | 12 | 0.25 | 16 | 0.0625 | 8 | 0 | 3 | 0.3333 | 39 | 0.1282 | 9 | 0.3333 | 3 | 0.3333 | 2 | 0 | 12 | 0.1667 | 26 | 0.2308 |
| 2_6164321 | 13 | 0.1538 | 15 | 0.2 | 8 | 0.125 | 3 | 0 | 39 | 0.1538 | 10 | 0.2 | 4 | 0 | 4 | 0 | 14 | 0.1429 | 32 | 0.125 |
| 2_6164369 | 14 | 0.4286 | 18 | 0.2222 | 9 | 0.5556 | 3 | 0.3333 | 44 | 0.3636 | 14 | 0.3571 | 5 | 0.4 | 4 | 0.25 | 14 | 0.2143 | 37 | 0.2973 |
| 2_6164383 | 15 | 0.2 | 21 | 0.2857 | 12 | 0.3333 | 3 | 0 | 51 | 0.2549 | 14 | 0.2143 | 5 | 0.8 | 5 | 0.2 | 13 | 0.1538 | 37 | 0.2703 |
| 2_6164408 | 16 | 0.0625 | 21 | 0.0476 | 15 | 0.0667 | 2 | 0 | 54 | 0.0556 | 19 | 0.1579 | 6 | 0 | 5 | 0.2 | 12 | 0.0833 | 42 | 0.119 |
| 2_6209905 | 2 | 0.5 | 6 | 0 | 2 | 0.5 | 8 | 0 | 18 | 0.1111 | 1 | 0 | 2 | 0 | 10 | 0.1 | 1 | 1 | 14 | 0.1429 |
| 2_6209923 | 3 | 0.3333 | 8 | 0 | 3 | 0.3333 | 10 | 0.1 | 24 | 0.125 | 4 | 0.25 | 3 | 0.3333 | 12 | 0 | 1 | 1 | 20 | 0.15 |
| 2_6215519 | 12 | 0.1667 | 15 | 0.2 | 5 | 0.2 | 15 | 0.2 | 47 | 0.1915 | 7 | 0.1429 | 7 | 0 | 14 | 0.0714 | 9 | 0.3333 | 37 | 0.1351 |
| 2_6252199 | 1 | 0 | 3 | 0.3333 | 2 | 0 | NA | NA | 6 | 0.1667 | NA | NA | 1 | 0 | 3 | 0.3333 | NA | NA | 4 | 0.25 |
| 2_6252202 | 2 | 0 | 3 | 0.3333 | 2 | 0 | 1 | 0 | 8 | 0.125 | NA | NA | 1 | 0 | 3 | 0.6667 | NA | NA | 4 | 0.5 |
| 2_6252406 | NA | NA | 2 | 0 | 2 | 0 | 2 | 0.5 | 6 | 0.1667 | NA | NA | NA | NA | 3 | 0 | 1 | 1 | 4 | 0.25 |
| 2_6252426 | 4 | 0.25 | 8 | 0.375 | 4 | 0.25 | 1 | 0 | 17 | 0.2941 | 5 | 0.2 | 2 | 1 | 2 | 1 | 2 | 1 | 11 | 0.6364 |
| 2_6252465 | 5 | 0.4 | 7 | 0.4286 | 4 | 0.5 | 1 | 1 | 17 | 0.4706 | 3 | 0.3333 | 2 | 0 | 2 | 0 | 2 | 0.5 | 9 | 0.2222 |
| 2_6252497 | 2 | 0.5 | 4 | 0.25 | 2 | 0.5 | 1 | 0 | 9 | 0.3333 | 2 | 0.5 | 2 | 0.5 | 1 | 0 | 1 | 1 | 6 | 0.5 |
| 2_6252502 | 5 | 0 | 4 | 0 | 3 | 0.3333 | 1 | 0 | 13 | 0.0769 | 3 | 0 | 1 | 0 | 2 | 0.5 | 2 | 0 | 8 | 0.125 |
| 2_6252502 | 5 | 0.2 | 6 | 0.1667 | 3 | 0 | NA | NA | 14 | 0.1429 | 3 | 0 | 2 | 0.5 | 2 | 0 | 3 | 0.6667 | 10 | 0.3 |
| 2_6252516 | 4 | 0 | 6 | 0.1667 | 3 | 0 | 1 | 1 | 14 | 0.1429 | 3 | 0 | 1 | 0 | 2 | 0 | 3 | 0.3333 | 9 | 0.1111 |
| 2_6252550 | NA | NA | 2 | 1 | 2 | 0.5 | 1 | 1 | 5 | 0.8 | 2 | 1 | NA | NA | 1 | 1 | NA | NA | 3 | 1 |
| 2_6261789 | NA | NA | NA | NA | 1 | 1 | NA | NA | 1 | 1 | NA | NA | NA | NA | NA | NA | 1 | 1 | 1 | 1 |
| 2_6261809 | 1 | 1 | NA | NA | 1 | 1 | NA | NA | 2 | 1 | NA | NA | NA | NA | NA | NA | 1 | 1 | 1 | 1 |
| 2_6261812 | 1 | 1 | NA | NA | 1 | 0 | NA | NA | 2 | 0.5 | NA | NA | NA | NA | NA | NA | 1 | 1 | 1 | 1 |
| 2_6380296 | NA | NA | 2 | 1 | NA | NA | NA | NA | 2 | 1 | 1 | 0 | NA | NA | 1 | 1 | NA | NA | 2 | 0.5 |
| 2_6380301 | NA | NA | 2 | 1 | 1 | 1 | NA | NA | 3 | 1 | NA | NA | NA | NA | 1 | 1 | NA | NA | 1 | 1 |
| 2_6380308 | NA | NA | 2 | 1 | 1 | 1 | NA | NA | 3 | 1 | 1 | 0 | NA | NA | 1 | 1 | NA | NA | 2 | 0.5 |
| 2_6474304 | 5 | 0.4 | 7 | 0.1429 | 5 | 0.4 | 6 | 0 | 23 | 0.2174 | 9 | 0 | 5 | 0.2 | 6 | 0 | 3 | 0.3333 | 23 | 0.087 |
| 2_6492837 | 1 | 1 | NA | NA | NA | NA | NA | NA | 1 | 1 | NA | NA | 2 | 0.5 | NA | NA | 3 | 0.3333 | 5 | 0.4 |
| 2_6492852 | 1 | 1 | NA | NA | NA | NA | NA | NA | 1 | 1 | NA | NA | 2 | 1 | NA | NA | 3 | 1 | 5 | 1 |
| 2_6492885 | 1 | 1 | NA | NA | NA | NA | NA | NA | 1 | 1 | NA | NA | 2 | 0.5 | NA | NA | 3 | 1 | 5 | 0.8 |
| 2_6492917 | 1 | 1 | NA | NA | NA | NA | NA | NA | 1 | 1 | NA | NA | 1 | 1 | NA | NA | 2 | 0.5 | 3 | 0.6667 |
| 2_6492957 | 1 | 1 | NA | NA | NA | NA | NA | NA | 1 | 1 | NA | NA | 1 | 1 | NA | NA | 3 | 0.6667 | 4 | 0.75 |
| 2_6528877 | 4 | 0.25 | 1 | 1 | 6 | 0.8333 | 3 | 0.6667 | 14 | 0.6429 | 2 | 0.5 | 6 | 0.6667 | 2 | 1 | 4 | 0.25 | 14 | 0.5714 |
| 2_6528885 | 2 | 1 | 1 | 1 | 4 | 0.5 | 5 | 0.6 | 12 | 0.6667 | 1 | 0 | 2 | 1 | 3 | 0.3333 | 4 | 0.5 | 10 | 0.5 |
| 2_6528930 | 2 | 0.5 | NA | NA | 3 | 0 | 5 | 0 | 10 | 0.1 | 2 | 0.5 | 6 | 0.5 | 1 | 0 | 7 | 0.2857 | 16 | 0.375 |
| 2_6529020 | 1 | 0 | NA | NA | 3 | 0.3333 | 7 | 0.1429 | 11 | 0.1818 | 2 | 0 | 2 | 0 | 2 | 0.5 | 7 | 0.1429 | 13 | 0.1538 |
| 2_6529060 | 1 | 0 | NA | NA | 3 | 0.3333 | 7 | 0.1429 | 11 | 0.1818 | 2 | 0 | 2 | 0 | 2 | 0.5 | 7 | 0 | 13 | 0.0769 |

| | | | | | | | | | | | | | | | | | | | | |
|---|---|---|---|---|---|---|---|---|---|---|---|---|---|---|---|---|---|---|---|---|
| 2_6529062( | 1 | 0 | NA | NA | 3 | 0.3333 | 6 | 0 | 10 | 0.1 | 2 | 0 | 1 | 1 | 2 | 0 | 7 | 0.1429 | 12 | 0.1667 |
| 2_6529066! | 1 | 0 | NA | NA | 3 | 0.3333 | 5 | 0.2 | 9 | 0.2222 | 2 | 0 | 2 | 0 | 2 | 0.5 | 6 | 0.1667 | 12 | 0.1667 |
| 2_6529068: | 1 | 0 | NA | NA | 3 | 0.3333 | 7 | 0.1429 | 11 | 0.1818 | 2 | 0.5 | 2 | 0 | 2 | 0.5 | 6 | 0.3333 | 12 | 0.3333 |
| 2_6529075: | 1 | 1 | NA | NA | 2 | 1 | 4 | 1 | 7 | 1 | 2 | 1 | 1 | 1 | NA | NA | 4 | 1 | 7 | 1 |
| 2_6529078( | 1 | 1 | NA | NA | 3 | 1 | 6 | 1 | 10 | 1 | 2 | 1 | 2 | 1 | 2 | 0.5 | 6 | 1 | 12 | 0.9167 |
| 2_6529093· | 1 | 0 | NA | NA | 3 | 0 | 5 | 0.2 | 9 | 0.1111 | 2 | 0 | 2 | 0 | 2 | 0.5 | 7 | 0 | 13 | 0.0769 |
| 2_6529093! | 1 | 1 | NA | NA | 2 | 1 | 5 | 0.8 | 8 | 0.875 | 2 | 0.5 | 2 | 0 | 2 | 0.5 | 7 | 0.4286 | 13 | 0.3846 |
| 2_6529097 | 1 | 1 | NA | NA | 3 | 1 | 6 | 0.5 | 10 | 0.7 | 2 | 0 | 2 | 0.5 | 2 | 0.5 | 6 | 0.5 | 12 | 0.4167 |
| 2_6529101 | 1 | 1 | NA | NA | 3 | 1 | 5 | 0.8 | 9 | 0.8889 | 2 | 1 | 1 | 1 | 2 | 0.5 | 6 | 1 | 11 | 0.9091 |
| 2_6529104! | 1 | 1 | NA | NA | 1 | 0 | 5 | 0.6 | 7 | 0.5714 | 1 | 1 | 1 | 0 | 2 | 1 | 6 | 0.3333 | 10 | 0.5 |
| 2_6529109! | 1 | 1 | NA | NA | 2 | 1 | 5 | 1 | 8 | 1 | 1 | 1 | NA | NA | 1 | 1 | 5 | 0.8 | 7 | 0.8571 |
| 2_6529110( | 1 | 0 | NA | NA | 3 | 0.3333 | 5 | 0 | 9 | 0.1111 | 1 | 0 | 1 | 0 | 2 | 0 | 7 | 0.1429 | 11 | 0.0909 |
| 2_6529112· | 1 | 1 | NA | NA | 2 | 0.5 | 6 | 0.6667 | 9 | 0.6667 | 2 | 0.5 | 2 | 0.5 | 2 | 1 | 6 | 0.1667 | 12 | 0.4167 |
| 2_6529133 | 1 | 0 | NA | NA | 3 | 1 | 5 | 1 | 9 | 0.8889 | 1 | 1 | 2 | 1 | 2 | 1 | 5 | 1 | 10 | 1 |
| 2_6529137( | NA | NA | NA | NA | 3 | 1 | 4 | 1 | 7 | 1 | 1 | 1 | 1 | 1 | 1 | 1 | 5 | 1 | 8 | 1 |
| 2_6529144! | 1 | 0 | NA | NA | 2 | 0.5 | 6 | 1 | 9 | 0.7778 | 2 | 0.5 | 1 | 0 | 2 | 0.5 | 4 | 0.5 | 9 | 0.4444 |
| 2_6529157 | 1 | 1 | NA | NA | 3 | 0 | 6 | 0.1667 | 10 | 0.2 | 2 | 0.5 | 2 | 0 | 1 | 0 | 5 | 0 | 10 | 0.1 |
| 2_6529159! | 1 | 0 | NA | NA | 2 | 0.5 | 5 | 0 | 8 | 0.125 | 2 | 1 | 1 | 1 | 1 | 0 | 5 | 0 | 9 | 0.3333 |
| 2_6529167( | 1 | 1 | NA | NA | 3 | 1 | 5 | 1 | 9 | 1 | 2 | 1 | 2 | 1 | 2 | 1 | 5 | 1 | 11 | 1 |
| 2_6529170! | 1 | 1 | NA | NA | 3 | 0 | 5 | 0 | 9 | 0.1111 | 2 | 0 | 2 | 0 | 2 | 0 | 5 | 0.2 | 11 | 0.0909 |
| 2_6529171! | 1 | 0 | NA | NA | 3 | 0.6667 | 6 | 0.6667 | 10 | 0.6 | 2 | 1 | 2 | 0.5 | 2 | 0.5 | 5 | 0.4 | 11 | 0.5455 |
| 2_6529175( | 1 | 1 | NA | NA | 3 | 1 | 6 | 1 | 10 | 1 | 2 | 1 | 2 | 1 | 2 | 0.5 | 5 | 1 | 11 | 0.9091 |
| 2_6529180! | NA | NA | NA | NA | 2 | 0 | 5 | 1 | 7 | 0.7143 | 2 | 1 | 2 | 1 | 2 | 1 | 3 | 1 | 9 | 1 |
| 2_6532552( | NA | NA | NA | NA | 2 | 0.5 | 2 | 0 | 4 | 0.25 | 1 | 1 | NA | NA | NA | NA | 1 | 0 | 2 | 0.5 |
| 2_6532778 | NA | NA | NA | NA | 1 | 1 | 1 | 0 | 2 | 0.5 | 1 | 0 | NA | NA | NA | NA | 1 | 1 | 2 | 0.5 |
| 2_6550100: | 1 | 0 | NA | NA | NA | NA | 1 | 1 | 2 | 0.5 | 1 | 1 | 2 | 0 | NA | NA | 1 | 1 | 4 | 0.5 |
| 2_6553216· | 5 | 0.2 | 1 | 0 | NA | NA | 3 | 0.3333 | 9 | 0.2222 | 1 | 0 | 2 | 0 | 1 | 0 | 2 | 1 | 6 | 0.3333 |
| 2_6553221! | 5 | 0 | 1 | 1 | NA | NA | 3 | 0.3333 | 9 | 0.2222 | 1 | 1 | 2 | 0.5 | 1 | 1 | 2 | 1 | 6 | 0.8333 |
| 2_6554212( | 5 | 0.4 | 2 | 0.5 | NA | NA | 5 | 0.6 | 12 | 0.5 | 1 | 0 | 2 | 0.5 | 1 | 0 | 1 | 1 | 5 | 0.4 |
| 2_6554627 | 5 | 0.2 | 2 | 0 | 2 | 0 | 4 | 0 | 13 | 0.0769 | NA | NA | 2 | 0 | 2 | 0 | 3 | 0.3333 | 7 | 0.1429 |
| 2_6554628 | 5 | 0.6 | 1 | 1 | 2 | 0 | 6 | 0.8333 | 14 | 0.6429 | 1 | 0 | 3 | 0.3333 | 2 | 0.5 | 1 | 0 | 7 | 0.2857 |
| 2_6554796 | 5 | 0.6 | 2 | 0.5 | 3 | 0.3333 | 5 | 0.8 | 15 | 0.6 | 1 | 1 | 4 | 0.75 | 2 | 0.5 | 3 | 0.6667 | 10 | 0.7 |
| 2_6554800: | 6 | 0.6667 | 3 | 1 | 3 | 0.3333 | 5 | 1 | 17 | 0.7647 | 1 | 1 | 4 | 0.75 | 2 | 0.5 | 4 | 0.75 | 11 | 0.7273 |
| 2_6554812: | 3 | 0.3333 | 3 | 1 | 2 | 0 | 4 | 0.75 | 12 | 0.5833 | 1 | 1 | 4 | 0.5 | 2 | 1 | 4 | 0.5 | 11 | 0.6364 |
| 2_6554815( | 6 | 0.5 | 3 | 0.6667 | 3 | 0 | 5 | 0.4 | 17 | 0.4118 | 2 | 0 | 4 | 0.25 | 2 | 0 | 3 | 0.3333 | 11 | 0.1818 |
| 2_6554815! | 5 | 0 | 3 | 0 | 3 | 0 | 7 | 0.2857 | 18 | 0.1111 | 2 | 0 | 4 | 0.25 | 2 | 0 | 5 | 0 | 13 | 0.0769 |
| 2_6559423: | 7 | 0 | 8 | 0.25 | 3 | 0 | 8 | 0.125 | 26 | 0.1154 | 4 | 0 | 9 | 0.2222 | 5 | 0 | 6 | 0.3333 | 24 | 0.1667 |
| 2_6559425( | 9 | 0.6667 | 6 | 0.1667 | 3 | 0.6667 | 6 | 0.5 | 24 | 0.5 | 3 | 0.6667 | 9 | 0.4444 | 5 | 0.6 | 6 | 0.8333 | 23 | 0.6087 |
| 2_6559430! | 8 | 1 | 8 | 1 | 2 | 1 | 7 | 1 | 25 | 1 | 4 | 1 | 9 | 1 | 3 | 1 | 7 | 1 | 23 | 1 |
| 2_6559433( | 7 | 0.7143 | 7 | 1 | 2 | 1 | 8 | 0.875 | 24 | 0.875 | 3 | 0.6667 | 7 | 0.7143 | 4 | 0.5 | 6 | 1 | 20 | 0.75 |
| 2_6559435( | 9 | 0.1111 | 9 | 0.2222 | 2 | 0.5 | 9 | 0.3333 | 29 | 0.2414 | 4 | 0.25 | 7 | 0 | 4 | 0 | 7 | 0.2857 | 22 | 0.1364 |
| 2_6559438! | 9 | 1 | 9 | 1 | 3 | 1 | 9 | 1 | 30 | 1 | 3 | 1 | 10 | 0.9 | 5 | 1 | 7 | 1 | 25 | 0.96 |
| 2_6559446 | 9 | 0.3333 | 7 | 0.1429 | 3 | 0.3333 | 8 | 0.125 | 27 | 0.2222 | 3 | 0.3333 | 9 | 0 | 4 | 0 | 5 | 0.2 | 21 | 0.0952 |
| 2_6559448! | 8 | 0.5 | 8 | 0.375 | 3 | 0.3333 | 10 | 0.3 | 29 | 0.3793 | 4 | 0 | 9 | 0.2222 | 4 | 0.5 | 6 | 0.3333 | 23 | 0.2609 |
| 2_6559461· | 10 | 0.4 | 9 | 0.5556 | 3 | 0.6667 | 7 | 0.7143 | 29 | 0.5517 | 5 | 0.6 | 7 | 0.1429 | 5 | 0.4 | 6 | 0.6667 | 23 | 0.4348 |
| 2_6559468: | 7 | 1 | 3 | 1 | 1 | 1 | 8 | 1 | 19 | 1 | 5 | 1 | 8 | 0.875 | 6 | 1 | 7 | 1 | 26 | 0.9615 |
| 2_6559472: | 9 | 1 | 8 | 1 | 2 | 1 | 12 | 1 | 31 | 1 | 4 | 1 | 10 | 0.9 | 6 | 1 | 8 | 1 | 28 | 0.9643 |
| 2_6559495! | 12 | 0.3333 | 10 | 0.3 | 4 | 0.25 | 14 | 0.2857 | 40 | 0.3 | 5 | 0.4 | 10 | 0.3 | 6 | 0 | 9 | 0.5556 | 30 | 0.3333 |
| 2_6559501· | 9 | 0.3333 | 8 | 0.25 | 5 | 0.4 | 12 | 0.3333 | 34 | 0.3235 | 5 | 0.2 | 10 | 0.2 | 5 | 0.6 | 7 | 0.5714 | 27 | 0.3704 |
| 2_6559504: | 12 | 0.25 | 10 | 0.5 | 4 | 0.5 | 13 | 0.4615 | 39 | 0.4103 | 6 | 0.5 | 12 | 0.1667 | 6 | 0.1667 | 9 | 0.8889 | 33 | 0.4242 |
| 2_6559512 | 12 | 0.9167 | 8 | 1 | 6 | 0.8333 | 15 | 0.8667 | 41 | 0.9024 | 6 | 0.8333 | 12 | 1 | 6 | 1 | 8 | 0.875 | 32 | 0.9375 |
| 2_6559513! | 9 | 1 | 9 | 1 | 4 | 0.75 | 15 | 1 | 37 | 0.973 | 6 | 1 | 12 | 1 | 5 | 1 | 9 | 1 | 32 | 1 |
| 2_6559529· | 8 | 0.5 | 8 | 0.375 | 5 | 0.2 | 12 | 0.25 | 33 | 0.3333 | 5 | 0.2 | 9 | 0.1111 | 5 | 0 | 8 | 0.5 | 27 | 0.2222 |
| 2_6559529! | 8 | 1 | 8 | 1 | 5 | 1 | 10 | 0.9 | 31 | 0.9677 | 6 | 1 | 7 | 1 | 6 | 1 | 7 | 0.8571 | 26 | 0.9615 |
| 2_6559534 | 11 | 0.8182 | 10 | 1 | 6 | 1 | 15 | 0.9333 | 42 | 0.9286 | 7 | 1 | 13 | 1 | 7 | 1 | 11 | 1 | 38 | 1 |
| 2_6559536! | 10 | 0.9 | 10 | 1 | 5 | 1 | 14 | 0.7143 | 39 | 0.8718 | 7 | 1 | 12 | 0.8333 | 7 | 1 | 12 | 0.9167 | 38 | 0.9211 |
| 2_6559546: | 13 | 0.0769 | 11 | 0.0909 | 7 | 0.2857 | 14 | 0.0714 | 45 | 0.1111 | 7 | 0.1429 | 12 | 0.1667 | 8 | 0.125 | 12 | 0.0833 | 39 | 0.1282 |
| 2_6559547: | 10 | 0.9 | 11 | 1 | 6 | 1 | 16 | 0.8125 | 43 | 0.907 | 7 | 0.7143 | 12 | 1 | 5 | 1 | 12 | 0.9167 | 36 | 0.9167 |
| 2_6559552: | 11 | 0.4545 | 10 | 0.4 | 6 | 0.3333 | 13 | 0.0769 | 40 | 0.3 | 6 | 0.3333 | 12 | 0.5 | 8 | 0.125 | 11 | 0.1818 | 37 | 0.2973 |
| 2_6559553! | 11 | 0.0909 | 9 | 0 | 6 | 0.1667 | 11 | 0.1818 | 37 | 0.1081 | 6 | 0.1667 | 8 | 0.25 | 6 | 0.5 | 11 | 0.1818 | 31 | 0.2581 |
| 2_6559556: | 10 | 0.1 | 5 | 0.2 | 4 | 0 | 7 | 0.2857 | 26 | 0.1538 | 3 | 0.3333 | 6 | 0 | 3 | 0 | 8 | 0 | 20 | 0.05 |
| 2_6559560: | 13 | 0.4615 | 10 | 0.6 | 7 | 0.4286 | 14 | 0.3571 | 44 | 0.4545 | 8 | 0.375 | 12 | 0.3333 | 9 | 0.5556 | 12 | 0.4167 | 41 | 0.4146 |
| 2_6559697· | 20 | 0.15 | 20 | 0.35 | 14 | 0.1429 | 20 | 0.25 | 74 | 0.2297 | 14 | 0.0714 | 16 | 0.25 | 12 | 0.1667 | 18 | 0.1667 | 60 | 0.1667 |
| 2_6585000 | 1 | 0 | 1 | 1 | 4 | 0.25 | 2 | 0 | 8 | 0.25 | 2 | 0.5 | 4 | 0.25 | 3 | 0 | 6 | 0.1667 | 15 | 0.2 |
| 2_6585240! | 2 | 1 | 1 | 1 | 5 | 1 | 3 | 1 | 11 | 1 | 2 | 1 | 5 | 1 | 5 | 1 | 6 | 0.8333 | 18 | 0.9444 |

| | | | | | | | | | | | | | | | | | | | | |
|---|---|---|---|---|---|---|---|---|---|---|---|---|---|---|---|---|---|---|---|---|
| 2_6585243 | 2 | 1 | NA | NA | 4 | 0.75 | 2 | 1 | 8 | 0.875 | 3 | 0.3333 | 2 | 1 | 5 | 0.6 | 3 | 1 | 13 | 0.6923 |
| 2_6585255 | 2 | 0 | 2 | 0 | 5 | 0 | 3 | 0.6667 | 12 | 0.1667 | 3 | 0 | 5 | 0 | 4 | 0.25 | 5 | 0.2 | 17 | 0.1176 |
| 2_6585260 | 2 | 0 | 2 | 0 | 3 | 0.3333 | 3 | 0 | 10 | 0.1 | 3 | 0.3333 | 6 | 0.1667 | 4 | 0.5 | 5 | 0.4 | 18 | 0.3333 |
| 2_6585261 | 1 | 1 | 2 | 1 | 2 | 1 | 3 | 0.3333 | 8 | 0.75 | 2 | 1 | 6 | 0.8333 | 2 | 0.5 | 4 | 1 | 14 | 0.8571 |
| 2_6585294 | 2 | 1 | 2 | 1 | 4 | 0.75 | 3 | 1 | 11 | 0.9091 | 3 | 1 | 7 | 0.8571 | 2 | 1 | 4 | 1 | 16 | 0.9375 |
| 2_6585304 | 2 | 0.5 | 3 | 0 | 4 | 0.5 | 4 | 0 | 13 | 0.2308 | 4 | 0.5 | 6 | 0.1667 | 6 | 0.1667 | 5 | 0.2 | 21 | 0.2381 |
| 2_6585317 | 2 | 1 | 2 | 0.5 | 2 | 1 | 2 | 0 | 8 | 0.625 | 4 | 0.25 | 5 | 0.4 | 6 | 0.5 | 4 | 0.5 | 19 | 0.4211 |
| 2_6585339 | 2 | 0.5 | 4 | 1 | 6 | 1 | 4 | 1 | 16 | 0.9375 | 5 | 0.8 | 7 | 1 | 5 | 1 | 4 | 0.75 | 21 | 0.9048 |
| 2_6585347 | 3 | 1 | 5 | 1 | 6 | 1 | 4 | 1 | 18 | 1 | 6 | 1 | 7 | 1 | 6 | 1 | 6 | 1 | 25 | 1 |
| 2_6585349 | 2 | 1 | 5 | 0.8 | 4 | 0.75 | 4 | 0.5 | 15 | 0.7333 | 5 | 0.4 | 5 | 0.6 | 5 | 1 | 6 | 1 | 21 | 0.7619 |
| 2_6585357 | 3 | 1 | 5 | 1 | 5 | 1 | 4 | 1 | 17 | 1 | 5 | 1 | 6 | 1 | 5 | 0.8 | 3 | 1 | 19 | 0.9474 |
| 2_6585364 | 3 | 0.6667 | 3 | 0.3333 | 5 | 0.8 | 3 | 0.6667 | 14 | 0.6429 | 3 | 0.6667 | 4 | 0.75 | 5 | 0.6 | 6 | 0.8333 | 18 | 0.7222 |
| 2_6585372 | 4 | 0 | 5 | 0 | 5 | 0.2 | 4 | 0 | 18 | 0.0556 | 5 | 0.2 | 7 | 0.5714 | 5 | 0.4 | 6 | 0.6667 | 23 | 0.4783 |
| 2_6585377 | 3 | 0 | 4 | 0.25 | 7 | 0.1429 | 5 | 0 | 19 | 0.1053 | 6 | 0.3333 | 7 | 0.1429 | 5 | 0.6 | 5 | 0.6 | 23 | 0.3913 |
| 2_6585472 | 3 | 0 | 4 | 0 | 4 | 0.25 | 2 | 0 | 13 | 0.0769 | 1 | 0 | 3 | 0 | 1 | 1 | 2 | 0 | 7 | 0.1429 |
| 2_6585494 | 2 | 1 | 5 | 0.2 | 3 | 0 | 2 | 0 | 12 | 0.25 | NA | NA | 2 | 0.5 | 1 | 0 | 2 | 0 | 5 | 0.2 |
| 2_6585509 | 3 | 0.6667 | 5 | 0.4 | 2 | 0 | 2 | 0 | 12 | 0.3333 | NA | NA | 3 | 0.3333 | 1 | 1 | 2 | 0 | 6 | 0.3333 |
| 2_6585513 | 1 | 1 | 1 | 1 | 1 | 1 | 2 | 1 | 5 | 1 | NA | NA | 1 | 1 | 1 | 1 | 1 | 1 | 3 | 1 |
| 2_6585800 | 5 | 0.2 | 2 | 0 | 2 | 0 | 7 | 0 | 16 | 0.0625 | 1 | 0 | 3 | 0 | 1 | 1 | NA | NA | 5 | 0.2 |
| 2_6585821 | 5 | 0.2 | 3 | 0 | 2 | 0 | 8 | 0 | 18 | 0.0556 | 1 | 0 | 3 | 0.3333 | NA | NA | NA | NA | 4 | 0.25 |
| 2_6585868 | 5 | 0.4 | 3 | 0 | 4 | 0.25 | 7 | 0.1429 | 19 | 0.2105 | NA | NA | 3 | 0.3333 | 1 | 0 | NA | NA | 4 | 0.25 |
| 2_6586103 | 6 | 0 | 5 | 0 | 4 | 0.75 | 11 | 0.1818 | 26 | 0.1923 | 2 | 0 | 5 | 0 | 3 | 0 | 1 | 1 | 11 | 0.0909 |
| 2_6586109 | 5 | 0.2 | 4 | 0.75 | 4 | 0.75 | 12 | 0.25 | 25 | 0.4 | 1 | 1 | 4 | 0.5 | 3 | 0.3333 | 1 | 1 | 9 | 0.5556 |
| 2_6586137 | 7 | 0 | 5 | 0.2 | 4 | 0.25 | 11 | 0.0909 | 27 | 0.1111 | 1 | 0 | 5 | 0.4 | 4 | 0 | 1 | 1 | 11 | 0.2727 |
| 2_6586155 | 7 | 0.2857 | 6 | 0 | 3 | 0 | 13 | 0 | 29 | 0.069 | 2 | 0 | 6 | 0.1667 | 4 | 0.5 | 1 | 0 | 13 | 0.2308 |
| 2_6610121 | 11 | 0.0909 | 13 | 0.1538 | 7 | 0 | 16 | 0.125 | 47 | 0.1064 | 3 | 0 | 6 | 0 | 13 | 0.3077 | 7 | 0 | 29 | 0.1379 |
| 2_6610238 | 8 | 0 | 10 | 0.1 | 6 | 0.1667 | 11 | 0.2727 | 35 | 0.1429 | 4 | 0 | 6 | 0.5 | 8 | 0.125 | 6 | 0 | 24 | 0.1667 |
| 2_6610261 | 3 | 0.3333 | 9 | 0.3333 | 6 | 0.1667 | 8 | 0 | 26 | 0.1923 | 3 | 0.3333 | 4 | 0.25 | 7 | 0.2857 | 5 | 0.4 | 19 | 0.3158 |
| 2_6610274 | 6 | 0.5 | 6 | 0.8333 | 2 | 1 | 9 | 0.4444 | 23 | 0.6087 | 2 | 0.5 | 6 | 0.5 | 4 | 0.75 | 4 | 0 | 16 | 0.4375 |
| 2_6610275 | 8 | 0.125 | 8 | 0.125 | 5 | 0 | 10 | 0 | 31 | 0.0645 | 3 | 0.3333 | 7 | 0.2857 | 6 | 0.1667 | 5 | 0 | 21 | 0.1905 |
| 2_6610284 | 6 | 0.5 | 8 | 0.5 | 5 | 0.8 | 12 | 0.4167 | 31 | 0.5161 | 3 | 0.3333 | 7 | 0.4286 | 5 | 0.8 | 5 | 0.6 | 20 | 0.55 |
| 2_6610289 | 6 | 0 | 7 | 0.1429 | 5 | 0 | 12 | 0.0833 | 30 | 0.0667 | 3 | 0 | 6 | 0.1667 | 6 | 0.1667 | 5 | 0 | 20 | 0.1 |
| 2_6610297 | 5 | 0.4 | 6 | 0.6667 | 5 | 0.6 | 10 | 0.7 | 26 | 0.6154 | 3 | 0.6667 | 6 | 0.8333 | 5 | 0.8 | 5 | 1 | 19 | 0.8421 |
| 2_6610298 | 6 | 0.6667 | 7 | 0.7143 | 5 | 0.2 | 11 | 0.7273 | 29 | 0.6207 | 3 | 0.3333 | 7 | 0.5714 | 5 | 1 | 5 | 0.6 | 20 | 0.65 |
| 2_6610306 | 5 | 0.8 | 7 | 1 | 3 | 1 | 8 | 0.875 | 23 | 0.913 | 2 | 1 | 7 | 0.8571 | 4 | 0.75 | 3 | 1 | 16 | 0.875 |
| 2_6610306 | 6 | 0.3333 | 7 | 0.5714 | 6 | 0.1667 | 11 | 0.3636 | 30 | 0.3667 | 2 | 0 | 5 | 0 | 5 | 0.4 | 4 | 0.25 | 16 | 0.1875 |
| 2_6610309 | 5 | 0.8 | 6 | 0.3333 | 3 | 0.6667 | 9 | 0.5556 | 23 | 0.5652 | 3 | 0.6667 | 6 | 0.3333 | 5 | 0.2 | 5 | 0.6 | 19 | 0.4211 |
| 2_6610313 | 4 | 1 | 8 | 1 | 4 | 1 | 10 | 1 | 26 | 1 | 2 | 1 | 5 | 1 | 4 | 1 | 4 | 1 | 15 | 1 |
| 2_6610316 | 7 | 0.7143 | 8 | 1 | 6 | 1 | 10 | 1 | 31 | 0.9355 | 2 | 1 | 7 | 1 | 5 | 1 | 5 | 0.8 | 19 | 0.9474 |
| 2_6610319 | 5 | 0 | 8 | 0 | 5 | 0.2 | 11 | 0.0909 | 29 | 0.069 | 2 | 0.5 | 7 | 0 | 3 | 0.3333 | 4 | 0.25 | 16 | 0.1875 |
| 2_6610322 | 4 | 0.75 | 6 | 0.3333 | 6 | 0.5 | 10 | 0.4 | 26 | 0.4615 | 1 | 1 | 6 | 0.6667 | 4 | 0.5 | 5 | 0.6 | 16 | 0.625 |
| 2_6610333 | 5 | 0 | 5 | 0.2 | 4 | 0.25 | 9 | 0 | 23 | 0.087 | 2 | 0 | 6 | 0.1667 | 4 | 0 | 4 | 0 | 16 | 0.0625 |
| 2_6610333 | 6 | 0.1667 | 7 | 0.2857 | 4 | 0 | 10 | 0.1 | 27 | 0.1481 | 2 | 0 | 6 | 0.1667 | 4 | 0 | 5 | 0.2 | 17 | 0.1176 |
| 2_6610334 | 4 | 0.5 | 6 | 0.6667 | 4 | 0.75 | 9 | 0.4444 | 23 | 0.5652 | 2 | 1 | 5 | 0.4 | 2 | 0.5 | 4 | 0.25 | 13 | 0.4615 |
| 2_6610335 | 5 | 0.2 | 6 | 1 | 4 | 0.75 | 6 | 0.6667 | 21 | 0.6667 | 2 | 0.5 | 6 | 0.8333 | 4 | 0.5 | 4 | 0.75 | 16 | 0.6875 |
| 2_6610337 | 6 | 0.1667 | 7 | 0.4286 | 4 | 0.75 | 9 | 0.4444 | 26 | 0.4231 | 1 | 1 | 5 | 0.4 | 4 | 0.75 | 4 | 0.25 | 14 | 0.5 |
| 2_6610342 | 4 | 0.25 | 3 | 0.6667 | 5 | 0.4 | 5 | 0.2 | 17 | 0.3529 | 1 | 0 | 3 | 0.6667 | 2 | 0 | 1 | 0 | 7 | 0.2857 |
| 2_6610342 | 5 | 0.6 | 6 | 0.3333 | 2 | 0.5 | 6 | 0.3333 | 19 | 0.4211 | 2 | 0 | 3 | 0 | 2 | 0.5 | 2 | 0 | 9 | 0.1111 |
| 2_6610345 | 4 | 0.75 | 2 | 1 | 3 | 1 | 6 | 0.5 | 15 | 0.7333 | NA | NA | 3 | 0.6667 | 2 | 0.5 | 4 | 0.75 | 9 | 0.6667 |
| 2_6610366 | 5 | 1 | 5 | 1 | 4 | 1 | 10 | 1 | 24 | 1 | 2 | 1 | 4 | 1 | 4 | 1 | 5 | 1 | 15 | 1 |
| 2_6610370 | 1 | 1 | 4 | 1 | 1 | 1 | 4 | 1 | 10 | 1 | 1 | 1 | 2 | 1 | 4 | 1 | 2 | 1 | 9 | 1 |
| 2_6610372 | 3 | 0.3333 | 5 | 0.4 | 3 | 1 | 8 | 0.625 | 19 | 0.5789 | 1 | 1 | 4 | 0.5 | 3 | 0.3333 | 2 | 0 | 10 | 0.4 |
| 2_6610375 | 4 | 0 | 5 | 0.2 | 3 | 0 | 10 | 0.1 | 22 | 0.0909 | 2 | 0 | 4 | 0 | 4 | 0 | 5 | 0.2 | 15 | 0.0667 |
| 2_6610389 | 2 | 0.5 | 6 | 0.3333 | 3 | 0.6667 | 10 | 0.1 | 21 | 0.2857 | 2 | 0 | 3 | 0 | 3 | 0 | 4 | 0.25 | 12 | 0.0833 |
| 2_6610399 | 2 | 1 | 5 | 1 | 4 | 1 | 9 | 0.8889 | 20 | 0.95 | 2 | 1 | 3 | 1 | 4 | 1 | 5 | 1 | 14 | 1 |
| 2_6610401 | 3 | 0 | 5 | 0 | 3 | 0 | 9 | 0.1111 | 20 | 0.05 | 1 | 0 | 3 | 0 | 3 | 0.3333 | 3 | 0 | 10 | 0.1 |
| 2_6610403 | 2 | 0 | 6 | 0.3333 | 4 | 0 | 9 | 0.1111 | 21 | 0.1429 | 2 | 0 | 3 | 0 | 4 | 0.25 | 4 | 0 | 13 | 0.0769 |
| 2_6610404 | 2 | 1 | 5 | 1 | 3 | 0.6667 | 9 | 1 | 19 | 0.9474 | 2 | 1 | 2 | 1 | 4 | 0.75 | 5 | 1 | 13 | 0.9231 |
| 2_6610407 | 3 | 1 | 5 | 0.8 | 2 | 1 | 10 | 0.9 | 20 | 0.9 | 2 | 1 | 3 | 1 | 4 | 1 | 5 | 1 | 14 | 1 |
| 2_6610413 | 3 | 0.3333 | 5 | 0 | 2 | 0.5 | 8 | 0.25 | 18 | 0.2222 | 1 | 0 | 3 | 0 | 4 | 0.25 | 5 | 0 | 13 | 0.0769 |
| 2_6614820 | 3 | 0.6667 | 1 | 1 | 4 | 0.25 | 2 | 0 | 10 | 0.4 | 1 | 0 | 1 | 1 | 2 | 0.5 | 4 | 0 | 8 | 0.25 |
| 2_6614820 | 3 | 0 | 1 | 0 | 3 | 0 | 2 | 0.5 | 9 | 0.1111 | 1 | 0 | NA | NA | 2 | 0 | 4 | 0.25 | 7 | 0.1429 |
| 2_6614823 | 3 | 0.3333 | 1 | 1 | 3 | 0.3333 | 2 | 0.5 | 9 | 0.4444 | NA | NA | 1 | 1 | 2 | 1 | 4 | 1 | 7 | 1 |
| 2_6614835 | 3 | 1 | 1 | 1 | 3 | 1 | 1 | 0 | 8 | 0.875 | NA | NA | 1 | 1 | 2 | 0.5 | 4 | 0.5 | 7 | 0.5714 |
| 2_6614840 | 3 | 1 | 1 | 0 | 3 | 0.6667 | 2 | 0.5 | 9 | 0.6667 | NA | NA | 1 | 0 | 2 | 0.5 | 4 | 0.75 | 7 | 0.5714 |

| | | | | | | | | | | | | | | | | | | | | |
|---|---|---|---|---|---|---|---|---|---|---|---|---|---|---|---|---|---|---|---|---|
| 2_6614852 | 2 | 0.5 | 1 | 0 | 3 | 0 | 1 | 1 | 7 | 0.2857 | NA | NA | 1 | 0 | 2 | 0 | 3 | 0.3333 | 6 | 0.1667 |
| 2_6614989 | 1 | 0 | 1 | 1 | 1 | 0 | 2 | 0 | 5 | 0.2 | NA | NA | NA | NA | 2 | 0.5 | 4 | 0.5 | 6 | 0.5 |
| 2_6615002 | 1 | 1 | 1 | 0 | 1 | 0 | 1 | 0 | 4 | 0.25 | NA | NA | NA | NA | 2 | 0 | 3 | 0.3333 | 5 | 0.2 |
| 2_6615276 | 1 | 0 | 1 | 1 | 2 | 0 | 2 | 0.5 | 6 | 0.3333 | NA | NA | NA | NA | 1 | 1 | 3 | 0.3333 | 4 | 0.5 |
| 2_6650396 | NA | NA | 1 | 1 | NA | NA | NA | NA | 1 | 1 | 1 | 1 | NA | NA | NA | NA | NA | NA | 1 | 1 |
| 2_6739524 | NA | NA | NA | NA | 1 | 1 | NA | NA | 1 | 1 | NA | NA | NA | NA | NA | NA | 1 | 1 | 1 | 1 |
| 2_6739538 | NA | NA | NA | NA | 1 | 1 | NA | NA | 1 | 1 | NA | NA | NA | NA | NA | NA | 1 | 1 | 1 | 1 |
| 2_6794815 | 1 | 0 | NA | NA | 3 | 0.3333 | 3 | 0.6667 | 7 | 0.4286 | 1 | 1 | 1 | 0 | NA | NA | 2 | 0 | 4 | 0.25 |
| 2_6794822 | 1 | 1 | NA | NA | 3 | 1 | 2 | 1 | 6 | 1 | 1 | 1 | 2 | 1 | NA | NA | 2 | 1 | 5 | 1 |
| 2_6794828 | 1 | 0 | NA | NA | 3 | 1 | 2 | 0 | 6 | 0.5 | 1 | 0 | 1 | 1 | 1 | 0 | 2 | 0 | 5 | 0.2 |
| 2_6794832 | NA | NA | NA | NA | 3 | 0.3333 | 2 | 0 | 5 | 0.2 | 1 | 0 | 1 | 1 | 1 | 0 | 1 | 1 | 4 | 0.5 |
| 2_6794842 | 2 | 0 | NA | NA | NA | NA | 3 | 0.3333 | 5 | 0.2 | 1 | 1 | 2 | 0 | 1 | 0 | 2 | 0 | 6 | 0.1667 |
| 2_6794844 | 2 | 1 | NA | NA | 2 | 1 | 1 | 1 | 5 | 1 | 1 | 1 | 1 | 1 | 1 | 1 | 2 | 1 | 5 | 1 |
| 2_6794846 | 2 | 0.5 | NA | NA | 3 | 0.3333 | 3 | 0.3333 | 8 | 0.375 | 1 | 0 | 2 | 1 | 1 | 0 | 2 | 0.5 | 6 | 0.5 |
| 2_6794858 | 2 | 1 | 1 | 0 | 2 | 1 | 3 | 0.3333 | 8 | 0.625 | 1 | 1 | 2 | 0 | 1 | 0 | 2 | 0 | 6 | 0.1667 |
| 2_6794866 | 2 | 0.5 | 1 | 0 | 3 | 0 | 2 | 0 | 8 | 0.125 | 1 | 0 | 2 | 0 | 1 | 0 | 2 | 0.5 | 6 | 0.1667 |
| 2_6794871 | 1 | 1 | 1 | 1 | 2 | 1 | 3 | 1 | 7 | 1 | 1 | 1 | NA | NA | 1 | 0 | 2 | 1 | 4 | 0.75 |
| 2_6794872 | 2 | 1 | 1 | 1 | 2 | 1 | 1 | 0 | 6 | 0.8333 | 1 | 1 | 2 | 1 | 1 | 0 | 2 | 0.5 | 6 | 0.6667 |
| 2_6794882 | 2 | 1 | NA | NA | 1 | 1 | 2 | 0.5 | 5 | 0.8 | 1 | 1 | 2 | 1 | 1 | 1 | 2 | 1 | 6 | 1 |
| 2_6794883 | 2 | 0 | 1 | 0 | 1 | 0 | 2 | 1 | 6 | 0.3333 | 1 | 1 | NA | NA | 1 | 0 | 2 | 0 | 4 | 0.25 |
| 2_6794884 | 2 | 1 | 1 | 1 | 3 | 1 | 2 | 1 | 8 | 1 | 1 | 1 | 2 | 1 | 1 | 1 | 2 | 1 | 6 | 1 |
| 2_6794900 | 2 | 1 | 2 | 1 | 3 | 1 | 3 | 1 | 10 | 1 | NA | NA | 1 | 1 | 1 | 1 | 2 | 1 | 4 | 1 |
| 2_6794903 | 2 | 1 | 2 | 1 | 2 | 1 | 3 | 0.6667 | 9 | 0.8889 | 1 | 1 | 1 | 1 | 1 | 0 | 2 | 1 | 5 | 0.8 |
| 2_6794904 | 2 | 0 | 2 | 0.5 | 4 | 0 | 3 | 0 | 11 | 0.0909 | 1 | 0 | 2 | 0 | 1 | 0 | 2 | 0.5 | 6 | 0.1667 |
| 2_6794909 | 2 | 1 | 2 | 1 | 4 | 1 | 1 | 1 | 9 | 1 | NA | NA | 2 | 1 | 1 | 1 | 2 | 1 | 5 | 1 |
| 2_6794926 | 2 | 1 | 2 | 1 | 4 | 1 | 2 | 1 | 10 | 1 | 1 | 1 | 2 | 1 | 1 | 1 | 2 | 1 | 6 | 1 |
| 2_6794928 | 2 | 0 | 1 | 1 | 4 | 0 | 1 | 0 | 8 | 0.125 | 1 | 0 | 1 | 1 | NA | NA | 2 | 0.5 | 4 | 0.5 |
| 2_6794931 | 2 | 0.5 | 2 | 1 | 3 | 1 | 2 | 1 | 9 | 0.8889 | NA | NA | 1 | 1 | 1 | 1 | 2 | 1 | 4 | 1 |
| 2_6794935 | 2 | 0.5 | 3 | 1 | 4 | 0.75 | 3 | 0.6667 | 12 | 0.75 | 1 | 1 | 1 | 1 | 1 | 0 | 2 | 0.5 | 5 | 0.6 |
| 2_6794951 | 2 | 0 | 3 | 0.3333 | 4 | 0 | 1 | 0 | 10 | 0.1 | 1 | 0 | NA | NA | 1 | 0 | 2 | 0.5 | 4 | 0.25 |
| 2_6795160 | 4 | 0.75 | 4 | 0.75 | 4 | 0.5 | 5 | 0.6 | 17 | 0.6471 | 2 | 1 | 1 | 1 | 3 | 0.3333 | 3 | 1 | 9 | 0.7778 |
| 2_6795161 | 5 | 0.2 | 5 | 0 | 3 | 0 | 4 | 0 | 17 | 0.0588 | 1 | 0 | NA | NA | 1 | 0 | 3 | 0.3333 | 5 | 0.2 |
| 2_6795242 | 4 | 0.75 | 5 | 0.6 | 3 | 0 | 8 | 0.125 | 20 | 0.35 | 1 | 0 | 2 | 0 | 4 | 0.25 | 3 | 0.6667 | 10 | 0.3 |
| 2_6795248 | 5 | 0.8 | 3 | 0.6667 | 5 | 0.4 | 7 | 0.5714 | 20 | 0.6 | 2 | 1 | 2 | 0.5 | 2 | 0.5 | 2 | 1 | 8 | 0.75 |
| 2_6795298 | 7 | 0.1429 | 4 | 0.25 | 8 | 0.5 | 7 | 0.2857 | 26 | 0.3077 | 1 | 1 | 3 | 0.3333 | 1 | 1 | 3 | 0.3333 | 8 | 0.5 |
| 2_6795301 | 6 | 0 | 4 | 0 | 7 | 0.4286 | 8 | 0 | 25 | 0.12 | 2 | 0 | 2 | 0 | 2 | 0 | 4 | 0.5 | 10 | 0.2 |
| 2_6804137 | 4 | 0.25 | 8 | 0.25 | 8 | 0.25 | 4 | 0.25 | 24 | 0.25 | 2 | 0 | 3 | 0 | NA | NA | 7 | 0.1429 | 12 | 0.0833 |
| 2_6804208 | 1 | 0 | 5 | 0.4 | NA | NA | 4 | 0.75 | 10 | 0.5 | 1 | 1 | 2 | 1 | NA | NA | 1 | 0 | 4 | 0.75 |
| 2_6804208 | 1 | 1 | NA | NA | 1 | 1 | NA | NA | 2 | 1 | NA | NA | 1 | 1 | NA | NA | 2 | 1 | 3 | 1 |
| 2_6822500 | 1 | 1 | NA | NA | NA | NA | NA | NA | 1 | 1 | NA | NA | 1 | 1 | NA | NA | NA | NA | 1 | 1 |
| 2_6893904 | NA | NA | 1 | 1 | 1 | 0 | 1 | 0 | 3 | 0.3333 | 1 | 0 | NA | NA | 3 | 0.6667 | NA | NA | 4 | 0.5 |
| 2_6894240 | 2 | 0 | 5 | 0.2 | 1 | 0 | 4 | 0 | 12 | 0.0833 | 1 | 1 | 3 | 0.3333 | 6 | 0 | NA | NA | 10 | 0.2 |
| 2_6894308 | 2 | 0 | 7 | 0.1429 | 2 | 0 | 3 | 0 | 14 | 0.0714 | 3 | 0 | 5 | 0.2 | 6 | 0.3333 | NA | NA | 14 | 0.2143 |
| 2_6894332 | 4 | 0 | 8 | 0.125 | 4 | 0 | 4 | 0.25 | 20 | 0.1 | 3 | 0.3333 | 5 | 0 | 7 | 0 | NA | NA | 15 | 0.0667 |
| 2_6901949 | 2 | 0.5 | 2 | 0 | NA | NA | 1 | 0 | 5 | 0.2 | NA | NA | 2 | 0.5 | 2 | 1 | 1 | 0 | 5 | 0.6 |
| 2_6949752 | 1 | 0 | 2 | 0 | 5 | 0.4 | 2 | 0.5 | 10 | 0.3 | 1 | 0 | 4 | 0.5 | 5 | 0.4 | NA | NA | 10 | 0.4 |
| 2_6949757 | 1 | 0 | 1 | 1 | 4 | 0.75 | 2 | 0.5 | 8 | 0.625 | 1 | 0 | 3 | 0.3333 | 6 | 0.6667 | NA | NA | 10 | 0.5 |
| 2_6949761 | 2 | 0.5 | 2 | 0.5 | 5 | 0.2 | 2 | 0 | 11 | 0.2727 | 1 | 0 | 4 | 0.5 | 4 | 0.5 | NA | NA | 9 | 0.4444 |
| 2_6951724 | 1 | 0 | NA | NA | 3 | 0.6667 | 1 | 0 | 5 | 0.4 | 1 | 0 | 1 | 0 | 2 | 0.5 | 5 | 0.4 | 9 | 0.3333 |
| 2_6951733 | 2 | 0.5 | 1 | 0 | 4 | 0.25 | 2 | 0 | 9 | 0.2222 | 1 | 0 | 2 | 0 | 3 | 0.3333 | 5 | 0.2 | 11 | 0.1818 |
| 2_6951735 | 1 | 1 | NA | NA | 3 | 0.3333 | 1 | 1 | 5 | 0.6 | 2 | 0.5 | NA | NA | 2 | 0 | 5 | 1 | 9 | 0.6667 |
| 2_6951736 | 1 | 0 | NA | NA | 3 | 0.3333 | 2 | 0 | 6 | 0.1667 | NA | NA | 1 | 0 | 2 | 0.5 | 4 | 0 | 7 | 0.1429 |
| 2_6953297 | 2 | 0 | 1 | 1 | 2 | 1 | 2 | 0.5 | 7 | 0.5714 | 1 | 0 | 1 | 0 | 2 | 0.5 | 4 | 0.25 | 8 | 0.25 |
| 2_6953305 | 2 | 0.5 | NA | NA | 1 | 1 | 1 | 0 | 4 | 0.5 | 1 | 0 | 1 | 0 | 2 | 1 | 3 | 0.6667 | 7 | 0.5714 |
| 2_6953305 | 2 | 0.5 | 1 | 1 | 2 | 1 | 2 | 1 | 7 | 0.8571 | 1 | 1 | NA | NA | 2 | 0.5 | 4 | 1 | 7 | 0.8571 |
| 2_6953306 | 2 | 0 | 1 | 0 | 2 | 0.5 | 1 | 0 | 6 | 0.1667 | 1 | 0 | 1 | 0 | 2 | 0 | 4 | 0.25 | 8 | 0.125 |
| 2_6954305 | NA | NA | NA | NA | 2 | 0.5 | 1 | 0 | 3 | 0.3333 | 2 | 0.5 | NA | NA | 2 | 0 | 2 | 0 | 6 | 0.1667 |
| 2_6954309 | NA | NA | NA | NA | 2 | 0.5 | 1 | 0 | 3 | 0.3333 | 1 | 0 | NA | NA | 1 | 0 | 2 | 0.5 | 4 | 0.25 |
| 2_6954330 | NA | NA | NA | NA | 1 | 0 | 1 | 1 | 2 | 0.5 | 1 | 1 | NA | NA | 1 | 1 | 2 | 1 | 4 | 1 |
| 2_6954343 | NA | NA | NA | NA | 1 | 1 | 1 | 1 | 2 | 1 | 1 | 1 | NA | NA | 2 | 1 | 1 | 1 | 4 | 1 |
| 2_6957974 | 1 | 0 | 6 | 0 | 2 | 0 | 5 | 0.4 | 14 | 0.1429 | 2 | 0.5 | 5 | 0.2 | 7 | 0.2857 | 3 | 0.3333 | 17 | 0.2941 |
| 2_6957977 | 1 | 1 | 6 | 0.1667 | 2 | 0 | 5 | 0.2 | 14 | 0.2143 | 2 | 0 | 4 | 0 | 7 | 0.1429 | 3 | 0 | 16 | 0.0625 |
| 2_6957995 | 1 | 0 | 5 | 0 | 2 | 0 | 5 | 0.2 | 13 | 0.0769 | 2 | 0 | 5 | 0.4 | 6 | 0 | 3 | 0 | 16 | 0.125 |
| 2_6958002 | 2 | 0 | 6 | 0 | 2 | 0.5 | 4 | 0 | 14 | 0.0714 | 2 | 1 | 3 | 0 | 6 | 0.1667 | 2 | 0 | 13 | 0.2308 |
| 2_6958005 | 1 | 1 | 6 | 0.1667 | 1 | 1 | 5 | 0 | 13 | 0.2308 | NA | NA | 5 | 0 | 5 | 0.6 | 2 | 1 | 12 | 0.4167 |

| | | | | | | | | | | | | | | | | | | | | |
|---|---|---|---|---|---|---|---|---|---|---|---|---|---|---|---|---|---|---|---|---|
| 2_6958055 | 2 | 0 | 6 | 0.6667 | 2 | 1 | 5 | 0.4 | 15 | 0.5333 | 4 | 0.25 | 5 | 0.2 | 7 | 0.5714 | 3 | 0.6667 | 19 | 0.4211 |
| 2_6958066 | 4 | 0 | 6 | 0.1667 | 1 | 0 | 5 | 0 | 16 | 0.0625 | 4 | 0 | 5 | 0 | 6 | 0 | 2 | 0.5 | 17 | 0.0588 |
| 2_6958077 | 4 | 0.25 | 6 | 0.5 | 5 | 0.4 | 5 | 0.2 | 20 | 0.35 | 4 | 0 | 4 | 0 | 5 | 0.2 | 3 | 0 | 16 | 0.0625 |
| 2_6958087 | 5 | 0.2 | 4 | 0.5 | 3 | 0.6667 | 4 | 0.25 | 16 | 0.375 | 3 | 0.6667 | 3 | 0 | 3 | 0.3333 | 2 | 0 | 11 | 0.2727 |
| 2_6958425 | 7 | 0.1429 | 3 | 0 | 3 | 0 | NA | NA | 13 | 0.0769 | 1 | 0 | 2 | 0 | 6 | 0.1667 | 6 | 0 | 15 | 0.0667 |
| 2_6962597 | 2 | 1 | 6 | 1 | 3 | 1 | 5 | 0.6 | 16 | 0.875 | 2 | 1 | 2 | 1 | 4 | 1 | 6 | 1 | 14 | 1 |
| 2_6962603 | 1 | 1 | 2 | 1 | 2 | 1 | 2 | 1 | 7 | 1 | 2 | 1 | 2 | 1 | 2 | 1 | 2 | 0.5 | 8 | 0.875 |
| 2_6962608 | 3 | 0 | 8 | 0.125 | 3 | 0.6667 | 4 | 0.5 | 18 | 0.2778 | 3 | 0.6667 | 1 | 1 | 5 | 0.2 | 7 | 0.2857 | 16 | 0.375 |
| 2_6962611 | 3 | 0.3333 | 8 | 0.875 | 1 | 1 | 5 | 0.4 | 17 | 0.6471 | 3 | 0.6667 | 2 | 0.5 | 6 | 0.8333 | 6 | 0.8333 | 17 | 0.7647 |
| 2_6962612 | 3 | 0 | 7 | 0.2857 | 2 | 0 | 5 | 0 | 17 | 0.1176 | 2 | 0 | 1 | 1 | 5 | 0.2 | 7 | 0.2857 | 15 | 0.2667 |
| 2_6962616 | 3 | 1 | 7 | 1 | 3 | 1 | 3 | 0.6667 | 16 | 0.9375 | 2 | 0.5 | 2 | 0.5 | 5 | 1 | 5 | 0.8 | 14 | 0.7857 |
| 2_6962640 | 1 | 0 | 8 | 0.375 | 3 | 0 | 5 | 0.2 | 17 | 0.2353 | 2 | 0 | 2 | 0 | 5 | 0 | 7 | 0.4286 | 16 | 0.1875 |
| 2_6962646 | 2 | 0 | 3 | 0 | 2 | 1 | 4 | 1 | 11 | 0.5455 | 4 | 0.25 | 1 | 0 | 5 | 0.4 | 6 | 0.5 | 16 | 0.375 |
| 2_6962718 | 2 | 0.5 | 8 | 0.625 | 1 | 1 | 5 | 0.4 | 16 | 0.5625 | 5 | 1 | 2 | 0.5 | 8 | 0.625 | 8 | 0.625 | 23 | 0.6957 |
| 2_6962743 | 3 | 1 | 8 | 0.875 | 3 | 0.6667 | 5 | 1 | 19 | 0.8947 | 6 | 1 | 2 | 1 | 9 | 1 | 6 | 1 | 23 | 1 |
| 2_6962749 | 2 | 0 | 7 | 0.1429 | 2 | 0 | 2 | 0 | 13 | 0.0769 | 2 | 0.5 | 2 | 0 | 8 | 0 | 7 | 0.2857 | 19 | 0.1579 |
| 2_6965423 | 4 | 0.75 | 2 | 1 | 4 | 0.75 | 6 | 0.8333 | 16 | 0.8125 | 2 | 1 | 5 | 0.8 | 6 | 1 | 10 | 0.9 | 23 | 0.913 |
| 2_6965436 | 4 | 0.5 | 3 | 0.6667 | 4 | 0.75 | 5 | 0.4 | 16 | 0.5625 | 2 | 1 | 5 | 1 | 4 | 0.75 | 9 | 0.4444 | 20 | 0.7 |
| 2_6965453 | 5 | 1 | 4 | 0.75 | 4 | 1 | 7 | 0.7143 | 20 | 0.85 | 3 | 0.6667 | 4 | 0.75 | 5 | 0.4 | 10 | 0.7 | 22 | 0.6364 |
| 2_6965475 | 5 | 1 | 3 | 1 | 5 | 1 | 6 | 0.8333 | 19 | 0.9474 | 2 | 1 | 4 | 0.75 | 5 | 0.6 | 8 | 0.75 | 19 | 0.7368 |
| 2_6965481 | 5 | 0.2 | 4 | 0 | 4 | 0 | 7 | 0 | 20 | 0.05 | 3 | 0 | 5 | 0 | 7 | 0.1429 | 10 | 0.2 | 25 | 0.12 |
| 2_6965508 | 5 | 0.2 | 4 | 0.25 | 5 | 0.2 | 8 | 0.25 | 22 | 0.2273 | 3 | 0 | 5 | 0.4 | 6 | 0.1667 | 11 | 0.1818 | 25 | 0.2 |
| 2_6965510 | 4 | 0 | 4 | 0.25 | 5 | 0.4 | 7 | 0.2857 | 20 | 0.25 | 3 | 0 | 5 | 0 | 5 | 0.2 | 11 | 0.2727 | 24 | 0.1667 |
| 2_6965511 | 5 | 0.6 | 4 | 0 | 4 | 0.25 | 7 | 0.4286 | 20 | 0.35 | 3 | 0.3333 | 6 | 0.1667 | 6 | 0.3333 | 10 | 0 | 25 | 0.16 |
| 2_6965517 | 3 | 0.3333 | 3 | 0.3333 | 5 | 0.2 | 7 | 0 | 18 | 0.1667 | 3 | 0 | 5 | 0.2 | 7 | 0.2857 | 10 | 0.1 | 25 | 0.16 |
| 2_6966402 | 6 | 0.3333 | 4 | 0.25 | 6 | 0.5 | 8 | 0.625 | 24 | 0.4583 | 2 | 1 | 6 | 0.1667 | 7 | 0.1429 | 10 | 0 | 25 | 0.16 |
| 2_6966433 | 5 | 0 | 5 | 0 | 6 | 0.1667 | 7 | 0.1429 | 23 | 0.087 | 3 | 0 | 6 | 0 | 8 | 0.125 | 12 | 0.25 | 29 | 0.1379 |
| 2_6966476 | 7 | 0 | 5 | 0.2 | 6 | 0.1667 | 7 | 0.1429 | 25 | 0.12 | 3 | 0 | 6 | 0.3333 | 9 | 0 | 14 | 0 | 32 | 0.0625 |
| 2_6966511 | 6 | 0.3333 | 5 | 0 | 5 | 0.2 | 7 | 0.2857 | 23 | 0.2174 | 2 | 0 | 7 | 0.1429 | 7 | 0 | 14 | 0.0714 | 30 | 0.0667 |
| 2_6966557 | 9 | 0.1111 | 6 | 0 | 7 | 0 | 10 | 0.2 | 32 | 0.0938 | 3 | 0 | 9 | 0.1111 | 11 | 0.0909 | 14 | 0.1429 | 37 | 0.1081 |
| 2_6966577 | 3 | 0 | 6 | 0.1667 | 6 | 0 | 8 | 0.125 | 23 | 0.087 | 3 | 0 | 7 | 0.1429 | 9 | 0.1111 | 12 | 0 | 31 | 0.0645 |
| 2_6966586 | 6 | 0.1667 | 8 | 0.5 | 10 | 0.3 | 6 | 0.5 | 30 | 0.3667 | 3 | 0.3333 | 5 | 0.4 | 10 | 0.3 | 12 | 0.1667 | 30 | 0.2667 |
| 2_6980369 | 1 | 0 | 3 | 0 | 1 | 0 | 1 | 1 | 6 | 0.1667 | 3 | 0.3333 | 5 | 0.2 | NA | NA | 1 | 0 | 9 | 0.2222 |
| 2_6980370 | 1 | 1 | 4 | 0.75 | 1 | 0 | NA | NA | 6 | 0.6667 | 3 | 0.3333 | 3 | 0.3333 | 2 | 0 | NA | NA | 8 | 0.25 |
| 2_6980372 | 1 | 0 | 4 | 0.25 | 1 | 0 | 2 | 0 | 8 | 0.125 | 4 | 0.5 | 5 | 0.8 | 2 | 0.5 | 1 | 1 | 12 | 0.6667 |
| 2_6980374 | 1 | 0 | 4 | 0 | 2 | 0 | 2 | 0.5 | 9 | 0.1111 | 4 | 0 | 6 | 0 | 2 | 0.5 | 2 | 0 | 14 | 0.0714 |
| 2_6985219 | 6 | 0.3333 | 10 | 0.2 | 4 | 0 | 5 | 0 | 25 | 0.16 | 6 | 0 | 9 | 0.1111 | 11 | 0.1818 | 9 | 0.1111 | 35 | 0.1143 |
| 2_6985300 | 7 | 0.5714 | 7 | 0.7143 | 5 | 0.4 | 5 | 0.6 | 24 | 0.5833 | 6 | 0.5 | 3 | 0.3333 | 8 | 0.375 | 5 | 0.6 | 22 | 0.4545 |
| 2_6985385 | 15 | 0.1333 | 15 | 0.0667 | 11 | 0.0909 | 16 | 0 | 57 | 0.0702 | 12 | 0.0833 | 14 | 0.0714 | 13 | 0 | 10 | 0.1 | 49 | 0.0612 |
| 2_7025735 | NA | NA | 1 | 0 | NA | NA | 1 | 1 | 2 | 0.5 | 1 | 1 | NA | NA | NA | NA | NA | NA | 1 | 1 |
| 2_7025740 | NA | NA | 1 | 1 | NA | NA | 1 | 1 | 2 | 1 | 1 | 1 | NA | NA | NA | NA | 1 | 1 | 2 | 1 |
| 2_7039228 | NA | NA | NA | NA | 1 | 1 | 2 | 1 | 3 | 1 | NA | NA | NA | NA | 3 | 1 | NA | NA | 3 | 1 |
| 2_7052125 | 8 | 0 | 11 | 0.0909 | 18 | 0.0556 | 11 | 0.0909 | 48 | 0.0625 | 10 | 0.1 | 5 | 0.2 | 16 | 0.125 | 12 | 0.25 | 43 | 0.1628 |
| 2_7054331 | 7 | 0 | 7 | 0.1429 | 9 | 0 | 9 | 0.1111 | 32 | 0.0625 | 12 | 0 | 9 | 0.2222 | 10 | 0.1 | 12 | 0 | 43 | 0.0698 |
| 2_7054371 | 6 | 0.5 | 6 | 0.8333 | 8 | 0.875 | 6 | 0.6667 | 26 | 0.7308 | 9 | 0.5556 | 9 | 0.5556 | 9 | 0.5556 | 11 | 0.7273 | 38 | 0.6053 |
| 2_7054381 | 7 | 0.1429 | 6 | 0.6667 | 7 | 0 | 8 | 0 | 28 | 0.1786 | 11 | 0.0909 | 9 | 0.1111 | 9 | 0.1111 | 12 | 0.1667 | 41 | 0.122 |
| 2_7054383 | 7 | 0.1429 | 7 | 0.4286 | 6 | 0.3333 | 7 | 0.2857 | 27 | 0.2963 | 12 | 0.1667 | 7 | 0.2857 | 10 | 0.3 | 11 | 0.2727 | 40 | 0.25 |
| 2_7054384 | 4 | 0.75 | 6 | 1 | 6 | 0.8333 | 7 | 0.8571 | 23 | 0.8696 | 11 | 0.5455 | 9 | 0.7778 | 9 | 0.7778 | 9 | 0.8889 | 38 | 0.7368 |
| 2_7054390 | 6 | 0.6667 | 5 | 0.2 | 4 | 1 | 6 | 1 | 21 | 0.7143 | 11 | 0.4545 | 4 | 1 | 9 | 0.8889 | 9 | 0.6667 | 33 | 0.697 |
| 2_7054398 | 6 | 0 | 5 | 0 | 7 | 0.2857 | 5 | 0.2 | 23 | 0.1304 | 10 | 0 | 9 | 0.1111 | 8 | 0.125 | 12 | 0.0833 | 39 | 0.0769 |
| 2_7054409 | 4 | 1 | 7 | 0.8571 | 7 | 0.8571 | 5 | 1 | 23 | 0.913 | 9 | 1 | 9 | 0.7778 | 10 | 1 | 11 | 0.8182 | 39 | 0.8974 |
| 2_7054416 | 5 | 0.4 | 6 | 0.5 | 6 | 0.1667 | 5 | 0.6 | 22 | 0.4091 | 9 | 0.1111 | 8 | 0.375 | 8 | 0.25 | 9 | 0.3333 | 34 | 0.2647 |
| 2_7054417 | 3 | 1 | 6 | 0.6667 | 6 | 0.8333 | 5 | 1 | 20 | 0.85 | 10 | 1 | 9 | 0.8889 | 9 | 1 | 10 | 1 | 38 | 0.9737 |
| 2_7054421 | 4 | 0 | 5 | 0.4 | 6 | 0.3333 | 4 | 0.5 | 19 | 0.3158 | 9 | 0.4444 | 9 | 0.3333 | 9 | 0.2222 | 10 | 0.6 | 37 | 0.4054 |
| 2_7054423 | 5 | 1 | 7 | 1 | 6 | 1 | 5 | 1 | 23 | 1 | 10 | 1 | 9 | 1 | 8 | 1 | 10 | 0.9 | 37 | 0.973 |
| 2_7054426 | 3 | 0.6667 | 4 | 1 | 7 | 0.8571 | 5 | 1 | 19 | 0.8947 | 10 | 1 | 7 | 1 | 10 | 1 | 7 | 1 | 34 | 1 |
| 2_7054427 | 5 | 0.4 | 8 | 0.5 | 7 | 0.2857 | 5 | 0.4 | 25 | 0.4 | 10 | 0 | 6 | 0.5 | 10 | 0.4 | 11 | 0.3636 | 37 | 0.2973 |
| 2_7054433 | 3 | 1 | 5 | 1 | 6 | 0.6667 | 4 | 0.75 | 18 | 0.8333 | 9 | 0.3333 | 6 | 0.6667 | 9 | 0.6667 | 9 | 1 | 33 | 0.6667 |
| 2_7054437 | 3 | 0 | 5 | 0 | 5 | 0.2 | 5 | 0.2 | 18 | 0.1111 | 10 | 0.1 | 9 | 0 | 10 | 0.1 | 10 | 0.1 | 39 | 0.0769 |
| 2_7073354 | NA | NA | NA | NA | NA | NA | 1 | 1 | 1 | 1 | 1 | 1 | NA | NA | NA | NA | NA | NA | 1 | 1 |
| 2_7073373 | NA | NA | NA | NA | NA | NA | 1 | 1 | 1 | 1 | 1 | 1 | NA | NA | NA | NA | NA | NA | 1 | 1 |
| 2_7079566 | 3 | 0 | 2 | 0.5 | 1 | 1 | 1 | 1 | 7 | 0.4286 | NA | NA | 1 | 1 | 3 | 0 | 2 | 0.5 | 6 | 0.3333 |
| 2_7079571 | 3 | 0.6667 | 2 | 1 | 1 | 1 | 2 | 1 | 8 | 0.875 | NA | NA | 1 | 1 | 3 | 1 | 2 | 0.5 | 6 | 0.8333 |
| 2_7079578 | 2 | 0.5 | 1 | 0 | NA | NA | 1 | 0 | 4 | 0.25 | NA | NA | 1 | 0 | 3 | 0.6667 | 1 | 0 | 5 | 0.4 |
| 2_7079581 | 3 | 0.3333 | 2 | 0 | NA | NA | 2 | 0 | 7 | 0.1429 | NA | NA | 1 | 1 | 3 | 0 | 2 | 0 | 6 | 0.1667 |

| | | | | | | | | | | | | | | | | | | | | |
|---|---|---|---|---|---|---|---|---|---|---|---|---|---|---|---|---|---|---|---|---|
| 2_7079583 | 2 | 0.5 | 1 | 1 | NA | NA | 1 | 1 | 4 | 0.75 | NA | NA | 1 | 1 | 2 | 0.5 | 2 | 0 | 5 | 0.4 |
| 2_7079584 | 3 | 0.6667 | 2 | 0.5 | NA | NA | 2 | 1 | 7 | 0.7143 | NA | NA | 1 | 0 | 3 | 0.6667 | 2 | 0 | 6 | 0.3333 |
| 2_7092029 | 1 | 1 | 1 | 0 | 1 | 0 | NA | NA | 3 | 0.3333 | 1 | 1 | 2 | 0.5 | NA | NA | NA | NA | 3 | 0.6667 |
| 2_7092035 | 1 | 1 | 1 | 1 | 1 | 0 | NA | NA | 3 | 0.6667 | 1 | 1 | 2 | 0 | NA | NA | NA | NA | 3 | 0.3333 |
| 2_7092040 | NA | NA | 1 | 1 | 1 | 0 | NA | NA | 2 | 0.5 | 1 | 0 | 2 | 0.5 | NA | NA | NA | NA | 3 | 0.3333 |
| 2_7092043 | 1 | 1 | 1 | 0 | 1 | 0 | NA | NA | 3 | 0.3333 | 1 | 0 | 2 | 1 | NA | NA | NA | NA | 3 | 0.6667 |
| 2_7092437 | 1 | 1 | NA | NA | 1 | 0 | NA | NA | 2 | 0.5 | 1 | 0 | 2 | 0.5 | NA | NA | NA | NA | 3 | 0.3333 |
| 2_7096917 | 1 | 1 | NA | NA | NA | NA | NA | NA | 1 | 1 | NA | NA | 1 | 1 | NA | NA | NA | NA | 1 | 1 |
| 2_7110515 | 12 | 0.1667 | 21 | 0.1905 | 19 | 0.3158 | 27 | 0.3704 | 79 | 0.2785 | 13 | 0.3077 | 21 | 0.0952 | 16 | 0.375 | 20 | 0.1 | 70 | 0.2 |
| 2_7110521 | 13 | 0.0769 | 23 | 0.087 | 19 | 0.0526 | 30 | 0.1333 | 85 | 0.0941 | 11 | 0.0909 | 22 | 0 | 16 | 0.1875 | 21 | 0.0476 | 70 | 0.0714 |
| 2_7110524 | 11 | 0.4545 | 20 | 0.3 | 19 | 0.5263 | 27 | 0.5185 | 77 | 0.4545 | 8 | 0.625 | 19 | 0.3158 | 15 | 0.6667 | 19 | 0.1579 | 61 | 0.3934 |
| 2_7110527 | 11 | 0.2727 | 23 | 0.2609 | 18 | 0.2778 | 27 | 0.1481 | 79 | 0.2278 | 10 | 0.3 | 20 | 0.15 | 16 | 0.125 | 20 | 0 | 66 | 0.1212 |
| 2_7110549 | 11 | 0.1818 | 21 | 0.0476 | 16 | 0 | 29 | 0.0345 | 77 | 0.0519 | 9 | 0.2222 | 17 | 0.0588 | 11 | 0.0909 | 21 | 0 | 58 | 0.069 |
| 2_7110561 | 8 | 0.25 | 13 | 0.1538 | 12 | 0.1667 | 23 | 0.2609 | 56 | 0.2143 | 9 | 0.4444 | 14 | 0.2857 | 13 | 0.0769 | 19 | 0.0526 | 55 | 0.1818 |
| 2_7135269 | 1 | 1 | NA | NA | NA | NA | NA | NA | 1 | 1 | NA | NA | 1 | 1 | NA | NA | NA | NA | 1 | 1 |
| 2_7135511 | 2 | 0 | 2 | 0.5 | 1 | 0 | NA | NA | 5 | 0.2 | NA | NA | 2 | 0.5 | NA | NA | 1 | 0 | 3 | 0.3333 |
| 2_7135533 | 2 | 0 | 2 | 0 | 1 | 1 | NA | NA | 5 | 0.2 | NA | NA | 2 | 0.5 | NA | NA | 1 | 0 | 3 | 0.3333 |
| 2_7135541 | 2 | 0 | 2 | 1 | 1 | 0 | NA | NA | 5 | 0.4 | NA | NA | 2 | 1 | NA | NA | 1 | 1 | 3 | 1 |
| 2_7136019 | NA | NA | NA | NA | NA | NA | 1 | 1 | 1 | 1 | NA | NA | 1 | 1 | NA | NA | 1 | 0 | 2 | 0.5 |
| 2_7136116 | 1 | 1 | 5 | 0.8 | 2 | 0.5 | 1 | 1 | 9 | 0.7778 | NA | NA | 1 | 1 | 3 | 1 | NA | NA | 4 | 1 |
| 2_7136127 | 1 | 0 | 5 | 0.4 | 2 | 0 | 1 | 1 | 9 | 0.3333 | NA | NA | 1 | 0 | 4 | 0.5 | NA | NA | 5 | 0.4 |
| 2_7136148 | 2 | 0.5 | 2 | 0.5 | 1 | 1 | 1 | 1 | 6 | 0.6667 | 1 | 1 | NA | NA | 2 | 0 | 1 | 0 | 4 | 0.25 |
| 2_7136196 | 1 | 0 | 4 | 0.5 | 1 | 0 | 1 | 0 | 7 | 0.2857 | NA | NA | NA | NA | 3 | 0.6667 | NA | NA | 3 | 0.6667 |
| 2_7136236 | 1 | 0 | 4 | 0.25 | 1 | 0 | 1 | 0 | 7 | 0.1429 | NA | NA | NA | NA | 3 | 0.3333 | NA | NA | 3 | 0.3333 |
| 2_7136248 | 1 | 0 | 3 | 1 | NA | NA | NA | NA | 4 | 0.75 | NA | NA | NA | NA | 2 | 1 | NA | NA | 2 | 1 |
| 2_7136257 | 1 | 1 | 3 | 1 | 1 | 1 | 1 | 1 | 6 | 1 | NA | NA | NA | NA | 2 | 1 | NA | NA | 2 | 1 |
| 2_7136258 | 1 | 0 | 3 | 1 | 1 | 1 | 1 | 0 | 6 | 0.6667 | NA | NA | NA | NA | 2 | 0.5 | NA | NA | 2 | 0.5 |
| 2_7136275 | 2 | 0.5 | 1 | 1 | 1 | 0 | 1 | 0 | 5 | 0.4 | 1 | 1 | NA | NA | NA | NA | 1 | 0 | 2 | 0.5 |
| 2_7136287 | 1 | 1 | 3 | 1 | 1 | 1 | NA | NA | 5 | 1 | NA | NA | NA | NA | 1 | 1 | NA | NA | 1 | 1 |
| 2_7136292 | 1 | 1 | 4 | 0.75 | 1 | 1 | NA | NA | 6 | 0.8333 | NA | NA | NA | NA | 2 | 0.5 | NA | NA | 2 | 0.5 |
| 2_7136302 | 2 | 0.5 | 2 | 1 | 1 | 1 | NA | NA | 5 | 0.8 | NA | NA | 1 | 1 | 2 | 1 | NA | NA | 3 | 1 |
| 2_7136306 | 1 | 1 | 3 | 1 | NA | NA | NA | NA | 4 | 1 | NA | NA | NA | NA | 2 | 1 | NA | NA | 2 | 1 |
| 2_7136307 | 2 | 0 | 4 | 0.5 | 1 | 0 | 2 | 0 | 9 | 0.2222 | 1 | 0 | 1 | 0 | 2 | 0 | 1 | 1 | 5 | 0.2 |
| 2_7136318 | 1 | 1 | 4 | 0 | 1 | 0 | 1 | 1 | 7 | 0.2857 | 1 | 0 | 1 | 1 | 2 | 0.5 | 1 | 0 | 5 | 0.4 |
| 2_7136319 | 2 | 0.5 | 3 | 0.6667 | NA | NA | 1 | 0 | 6 | 0.5 | 1 | 1 | NA | NA | 2 | 0.5 | 1 | 0 | 4 | 0.5 |
| 2_7136319 | 2 | 0 | 3 | 0.6667 | NA | NA | 1 | 1 | 6 | 0.5 | 1 | 1 | NA | NA | 2 | 0 | 1 | 0 | 4 | 0.25 |
| 2_7136321 | 2 | 1 | 5 | 0.6 | 1 | 1 | 1 | 1 | 9 | 0.7778 | 1 | 0 | 1 | 1 | 2 | 1 | 1 | 1 | 5 | 0.8 |
| 2_7136326 | 1 | 1 | 2 | 1 | 1 | 1 | NA | NA | 4 | 1 | NA | NA | NA | NA | 2 | 0.5 | NA | NA | 2 | 0.5 |
| 2_7136331 | 1 | 0 | 4 | 0.25 | 1 | 0 | NA | NA | 6 | 0.1667 | NA | NA | NA | NA | 2 | 0.5 | NA | NA | 2 | 0.5 |
| 2_7136340 | 1 | 1 | 3 | 0.6667 | 1 | 1 | NA | NA | 5 | 0.8 | NA | NA | NA | NA | 2 | 1 | NA | NA | 2 | 1 |
| 2_7136341 | 2 | 0 | 5 | 0.6 | 1 | 0 | 1 | 0 | 9 | 0.3333 | 1 | 1 | 1 | 0 | 2 | 0 | 1 | 1 | 5 | 0.4 |
| 2_7136351 | 2 | 1 | 4 | 1 | 1 | 1 | NA | NA | 7 | 1 | 1 | 1 | 1 | 0 | 2 | 1 | 1 | 1 | 5 | 0.8 |
| 2_7136356 | 2 | 1 | 5 | 1 | 1 | 1 | 2 | 1 | 10 | 1 | 1 | 1 | 1 | 1 | 2 | 1 | 1 | 1 | 5 | 1 |
| 2_7136362 | 1 | 0 | 4 | 0.25 | 1 | 0 | 1 | 1 | 7 | 0.2857 | NA | NA | NA | NA | 2 | 0.5 | 1 | 0 | 3 | 0.3333 |
| 2_7136363 | 2 | 1 | 4 | 1 | 1 | 1 | 1 | 1 | 8 | 1 | 1 | 1 | 1 | 1 | 2 | 1 | 1 | 1 | 5 | 1 |
| 2_7136365 | 2 | 0.5 | 3 | 0.3333 | 1 | 0 | 2 | 0 | 8 | 0.25 | 1 | 0 | 1 | 1 | 1 | 1 | 1 | 1 | 4 | 0.75 |
| 2_7136374 | 2 | 0.5 | 4 | 0.25 | 1 | 1 | 1 | 0 | 8 | 0.375 | 2 | 0.5 | 1 | 0 | 2 | 0.5 | 1 | 0 | 6 | 0.3333 |
| 2_7136380 | 2 | 0.5 | 4 | 0.5 | 2 | 0.5 | 1 | 1 | 9 | 0.5556 | 2 | 0.5 | 1 | 1 | 1 | 1 | 1 | 0 | 5 | 0.6 |
| 2_7136385 | 1 | 0 | 5 | 0.2 | 2 | 1 | 2 | 0 | 10 | 0.3 | 2 | 0.5 | 1 | 0 | 2 | 0.5 | 1 | 0 | 6 | 0.3333 |
| 2_7136393 | 1 | 1 | 5 | 0.2 | 2 | 0 | 2 | 0 | 10 | 0.2 | 2 | 0.5 | 1 | 0 | 2 | 0 | 1 | 0 | 6 | 0.1667 |
| 2_7137410 | 6 | 1 | 5 | 1 | 3 | 1 | 1 | 1 | 15 | 1 | 3 | 0.6667 | 2 | 0.5 | 3 | 1 | 4 | 1 | 12 | 0.8333 |
| 2_7137417 | 2 | 0.5 | NA | NA | 1 | 1 | NA | NA | 3 | 0.6667 | 1 | 0 | 2 | 0 | 2 | 0.5 | 2 | 0.5 | 7 | 0.2857 |
| 2_7137417 | 2 | 0.5 | 3 | 0.3333 | 1 | 0 | 1 | 0 | 7 | 0.2857 | NA | NA | 1 | 0 | NA | NA | 1 | 1 | 2 | 0.5 |
| 2_7137427 | 3 | 0 | 2 | 0.5 | 3 | 0 | 1 | 0 | 9 | 0.1111 | 1 | 0 | 1 | 0 | 3 | 0.3333 | 2 | 0 | 7 | 0.1429 |
| 2_7137458 | 5 | 1 | 4 | 1 | 2 | 1 | 1 | 1 | 12 | 1 | 2 | 1 | 1 | 1 | 3 | 1 | 4 | 1 | 10 | 1 |
| 2_7137483 | 4 | 0.5 | 3 | 1 | 1 | 1 | 1 | 1 | 9 | 0.7778 | 2 | 0.5 | 1 | 1 | 3 | 0.3333 | 3 | 1 | 9 | 0.6667 |
| 2_7137484 | 4 | 0.5 | 5 | 0.4 | 1 | 0 | 1 | 0 | 11 | 0.3636 | 2 | 0.5 | 2 | 1 | 3 | 0.6667 | 4 | 0.25 | 11 | 0.5455 |
| 2_7137485 | 3 | 0.6667 | 4 | 0.5 | 1 | 1 | 1 | 0 | 9 | 0.5556 | 2 | 1 | 2 | 0.5 | 2 | 0 | 4 | 0.5 | 10 | 0.5 |
| 2_7137512 | 4 | 1 | 2 | 1 | 1 | 1 | 1 | 0 | 8 | 0.875 | 2 | 1 | 3 | 1 | 2 | 1 | 4 | 0.75 | 11 | 0.9091 |
| 2_7168638 | NA | NA | NA | NA | 2 | 1 | 1 | 0 | 3 | 0.6667 | 2 | 1 | NA | NA | NA | NA | 1 | 1 | 3 | 1 |
| 2_7168647 | 1 | 1 | 1 | 1 | 2 | 1 | 1 | 1 | 5 | 1 | 3 | 1 | NA | NA | NA | NA | 1 | 1 | 4 | 1 |
| 2_7168649 | 1 | 1 | 1 | 1 | 2 | 1 | 1 | 1 | 5 | 1 | 3 | 1 | NA | NA | NA | NA | 1 | 1 | 4 | 1 |
| 2_7168656 | 1 | 1 | 1 | 1 | 2 | 1 | 1 | 1 | 5 | 1 | 2 | 1 | NA | NA | NA | NA | 1 | 1 | 3 | 1 |
| 2_7168658 | 1 | 1 | 1 | 0 | 2 | 0 | 1 | 1 | 5 | 0.4 | 3 | 0.6667 | NA | NA | NA | NA | NA | NA | 3 | 0.6667 |
| 2_7168676 | 1 | 0 | NA | NA | 1 | 0 | 1 | 1 | 3 | 0.3333 | 3 | 0.3333 | NA | NA | NA | NA | 1 | 0 | 4 | 0.25 |

| | | | | | | | | | | | | | | | | | | | | |
|---|---|---|---|---|---|---|---|---|---|---|---|---|---|---|---|---|---|---|---|---|
| 2_7168687 | 1 | 1 | 1 | 0 | 2 | 1 | 1 | 1 | 5 | 0.8 | 2 | 0.5 | NA | NA | NA | NA | 1 | 1 | 3 | 0.6667 |
| 2_7168693 | 1 | 1 | 1 | 0 | 2 | 1 | 1 | 1 | 5 | 0.8 | 3 | 0.6667 | NA | NA | NA | NA | NA | NA | 3 | 0.6667 |
| 2_7168729 | 2 | 0 | 1 | 0 | 2 | 0.5 | NA | NA | 5 | 0.2 | 4 | 0.5 | NA | NA | NA | NA | 2 | 0 | 6 | 0.3333 |
| 2_7168732 | 1 | 0 | 1 | 0 | 2 | 1 | 1 | 0 | 5 | 0.4 | 4 | 0.75 | NA | NA | NA | NA | 1 | 0 | 5 | 0.6 |
| 2_7168734 | 2 | 1 | NA | NA | 1 | 0 | 1 | 1 | 4 | 0.75 | 3 | 0.6667 | NA | NA | NA | NA | 2 | 0.5 | 5 | 0.6 |
| 2_7168739 | 2 | 0 | 1 | 1 | 2 | 1 | 1 | 0 | 6 | 0.5 | 3 | 0.6667 | NA | NA | NA | NA | 2 | 0 | 5 | 0.4 |
| 2_7168759 | 2 | 1 | 1 | 0 | 2 | 0.5 | NA | NA | 5 | 0.6 | 3 | 1 | NA | NA | NA | NA | 1 | 1 | 4 | 1 |
| 2_7173096 | 2 | 0.5 | 1 | 1 | NA | NA | 1 | 1 | 4 | 0.75 | NA | NA | NA | NA | NA | NA | 2 | 0.5 | 2 | 0.5 |
| 2_7173133 | 1 | 1 | 2 | 0 | NA | NA | 2 | 0.5 | 5 | 0.4 | NA | NA | NA | NA | NA | NA | 2 | 0.5 | 2 | 0.5 |
| 2_7173137 | 3 | 0 | 4 | 0.25 | NA | NA | 2 | 0.5 | 9 | 0.2222 | NA | NA | 1 | 1 | NA | NA | 1 | 0 | 2 | 0.5 |
| 2_7208667 | 19 | 0.7895 | 38 | 0.8947 | 22 | 0.6818 | 24 | 0.7083 | 103 | 0.7864 | 28 | 0.7857 | 13 | 0.7692 | 10 | 0.6 | 25 | 0.92 | 76 | 0.8026 |
| 2_7208672 | 24 | 0.3333 | 45 | 0.1111 | 38 | 0.1579 | 29 | 0.1379 | 136 | 0.1691 | 41 | 0.1951 | 19 | 0.1053 | 17 | 0.0588 | 36 | 0.2778 | 113 | 0.1858 |
| 2_7208679 | 22 | 0.8182 | 43 | 0.8837 | 34 | 0.9412 | 28 | 0.8214 | 127 | 0.874 | 36 | 0.8056 | 17 | 0.9412 | 15 | 0.8 | 33 | 0.9394 | 101 | 0.8713 |
| 2_7208683 | 20 | 0.3 | 46 | 0.4565 | 36 | 0.3333 | 29 | 0.4828 | 131 | 0.4046 | 32 | 0.375 | 19 | 0.4211 | 14 | 0.2857 | 35 | 0.3714 | 100 | 0.37 |
| 2_7208689 | 17 | 0.5882 | 40 | 0.65 | 33 | 0.3939 | 28 | 0.5714 | 118 | 0.5508 | 35 | 0.4571 | 21 | 0.4286 | 16 | 0.375 | 37 | 0.4595 | 109 | 0.4404 |
| 2_7208699 | 21 | 0.0952 | 37 | 0.1892 | 33 | 0.2121 | 25 | 0.2 | 116 | 0.181 | 30 | 0.1667 | 17 | 0.2353 | 12 | 0.3333 | 26 | 0.2308 | 85 | 0.2235 |
| 2_7208705 | 25 | 0.88 | 45 | 0.7778 | 40 | 0.875 | 29 | 0.7931 | 139 | 0.8273 | 37 | 0.8919 | 20 | 0.95 | 16 | 0.875 | 34 | 0.8529 | 107 | 0.8879 |
| 2_7276781 | 4 | 0.25 | 4 | 0 | 5 | 0.2 | 8 | 0.25 | 21 | 0.1905 | 3 | 0 | 7 | 0.2857 | 3 | 0 | 9 | 0.1111 | 22 | 0.1364 |
| 2_7276842 | 4 | 0.25 | 2 | 0.5 | 5 | 0.4 | 9 | 0.2222 | 20 | 0.3 | 4 | 0.25 | 5 | 0.4 | 1 | 1 | 7 | 0.2857 | 17 | 0.3529 |
| 2_7276850 | 3 | 1 | 3 | 0.6667 | 4 | 0.75 | 8 | 1 | 18 | 0.8889 | 4 | 1 | 6 | 0.8333 | 2 | 0.5 | 6 | 1 | 18 | 0.8889 |
| 2_7276855 | 2 | 0.5 | 2 | 0 | 5 | 0 | 7 | 0 | 16 | 0.0625 | 2 | 0 | 6 | 0.1667 | 2 | 0 | 5 | 0 | 15 | 0.0667 |
| 2_7276865 | 3 | 0.3333 | 3 | 0 | 4 | 0 | 6 | 0 | 16 | 0.0625 | 4 | 0 | 5 | 0.2 | 1 | 0 | 7 | 0.1429 | 17 | 0.1176 |
| 2_7280057 | 2 | 0.5 | 1 | 0 | 4 | 0.5 | 4 | 0 | 11 | 0.2727 | 1 | 0 | 5 | 0.6 | NA | NA | 4 | 0.5 | 10 | 0.5 |
| 2_7280065 | 1 | 0 | 2 | 1 | 4 | 0.5 | 2 | 0.5 | 9 | 0.5556 | 1 | 1 | 3 | 0 | 1 | 0 | 5 | 0.4 | 10 | 0.3 |
| 2_7280072 | 2 | 1 | 2 | 0.5 | 4 | 0.5 | 4 | 0.5 | 12 | 0.5833 | 1 | 0 | 3 | 0.3333 | 1 | 0 | 5 | 1 | 10 | 0.6 |
| 2_7280074 | 2 | 0.5 | 1 | 1 | 3 | 0 | 4 | 0.25 | 10 | 0.3 | 1 | 0 | 3 | 0.3333 | 1 | 0 | 5 | 0 | 10 | 0.1 |
| 2_7280076 | 2 | 0.5 | 2 | 1 | 3 | 1 | 4 | 1 | 11 | 0.9091 | 1 | 0 | 2 | 1 | 1 | 1 | 4 | 1 | 8 | 0.875 |
| 2_7280077 | 2 | 0 | 2 | 0 | 3 | 0.3333 | 3 | 0.3333 | 10 | 0.2 | 1 | 0 | 4 | 0.25 | NA | NA | 4 | 0 | 9 | 0.1111 |
| 2_7280088 | 2 | 0 | 1 | 0 | 3 | 0.3333 | 3 | 1 | 9 | 0.4444 | NA | NA | 3 | 0.3333 | 1 | 0 | 4 | 0.75 | 8 | 0.5 |
| 2_7280120 | 2 | 1 | 1 | 1 | 3 | 1 | 4 | 1 | 10 | 1 | NA | NA | 2 | 1 | 1 | 1 | 4 | 0.75 | 7 | 0.8571 |
| 2_7280123 | 2 | 0 | 1 | 1 | 4 | 0 | 4 | 0 | 11 | 0.0909 | 1 | 0 | 3 | 0.3333 | 1 | 0 | 4 | 0 | 9 | 0.1111 |
| 2_7280125 | 2 | 1 | 1 | 1 | 3 | 1 | 4 | 1 | 10 | 1 | 1 | 1 | 3 | 0.6667 | 1 | 1 | 4 | 1 | 9 | 0.8889 |
| 2_7280127 | 2 | 0.5 | NA | NA | 4 | 0.75 | 3 | 0.6667 | 9 | 0.6667 | NA | NA | 2 | 0.5 | 1 | 0 | 4 | 0.5 | 7 | 0.4286 |
| 2_7280130 | 2 | 1 | 1 | 1 | 3 | 1 | 4 | 1 | 10 | 1 | 1 | 1 | 3 | 0.6667 | 1 | 1 | 5 | 1 | 10 | 0.9 |
| 2_7280139 | 2 | 0 | 1 | 0 | 4 | 0 | 4 | 0.25 | 11 | 0.0909 | 1 | 0 | 2 | 0 | 1 | 0 | 5 | 0.2 | 9 | 0.1111 |
| 2_7280144 | 1 | 1 | NA | NA | 2 | 0.5 | 3 | 1 | 6 | 0.8333 | NA | NA | 3 | 0 | 1 | 0 | 4 | 0.75 | 8 | 0.375 |
| 2_7280937 | 1 | 0 | 1 | 0 | 4 | 0 | 4 | 0.25 | 10 | 0.1 | 1 | 0 | 2 | 0 | 1 | 0 | 4 | 0.25 | 8 | 0.125 |
| 2_7281074 | NA | NA | NA | NA | 1 | 1 | NA | NA | 1 | 1 | 1 | 1 | NA | NA | 1 | 1 | 2 | 0.5 | 4 | 0.75 |
| 2_7281094 | NA | NA | NA | NA | 1 | 1 | NA | NA | 1 | 1 | 1 | 1 | NA | NA | 1 | 1 | 2 | 0 | 4 | 0.5 |
| 2_7281118 | NA | NA | NA | NA | 1 | 1 | NA | NA | 1 | 1 | 1 | 1 | NA | NA | 1 | 1 | 2 | 0.5 | 4 | 0.75 |
| 2_7281145 | NA | NA | NA | NA | 1 | 1 | NA | NA | 1 | 1 | 1 | 1 | NA | NA | 1 | 1 | 2 | 0.5 | 4 | 0.75 |
| 2_7281885 | 3 | 0.6667 | 5 | 0.6 | 3 | 0.6667 | 4 | 0.75 | 15 | 0.6667 | 3 | 0.6667 | 4 | 0.25 | 4 | 0.5 | 1 | 0 | 12 | 0.4167 |
| 2_7281893 | 1 | 1 | 5 | 0.2 | 3 | 0.3333 | 4 | 0 | 13 | 0.2308 | 3 | 0.3333 | 4 | 0 | 3 | 0.3333 | NA | NA | 10 | 0.2 |
| 2_7292332 | 4 | 0.5 | 2 | 0.5 | 1 | 1 | 5 | 0.2 | 12 | 0.4167 | 4 | 0.5 | 2 | 0 | 1 | 0 | 2 | 0.5 | 9 | 0.3333 |
| 2_7292386 | 18 | 0.0556 | 22 | 0.2727 | 11 | 0.0909 | 16 | 0.125 | 67 | 0.1493 | 15 | 0.1333 | 23 | 0.1304 | 12 | 0 | 16 | 0.125 | 66 | 0.1061 |
| 2_7309523 | 1 | 1 | 1 | 1 | 1 | 1 | NA | NA | 3 | 1 | NA | NA | 1 | 1 | NA | NA | NA | NA | 1 | 1 |
| 2_7309529 | 1 | 1 | NA | NA | NA | NA | NA | NA | 1 | 1 | NA | NA | 1 | 1 | NA | NA | NA | NA | 1 | 1 |
| 2_7310425 | 3 | 1 | 2 | 1 | 3 | 1 | 5 | 1 | 13 | 1 | 1 | 1 | 1 | 1 | 4 | 1 | 2 | 1 | 8 | 1 |
| 2_7310460 | 4 | 1 | 2 | 1 | 2 | 1 | 6 | 0.8333 | 14 | 0.9286 | 2 | 1 | 2 | 1 | 7 | 0.8571 | 2 | 1 | 13 | 0.9231 |
| 2_7310472 | 4 | 0 | 2 | 0.5 | 3 | 0 | 7 | 0 | 16 | 0.0625 | 2 | 0 | 6 | 0 | 7 | 0.1429 | 2 | 0 | 17 | 0.0588 |
| 2_7310477 | 4 | 0.25 | 2 | 0.5 | 3 | 0 | 8 | 0 | 17 | 0.1176 | 1 | 0 | 5 | 0.2 | 6 | 0.1667 | 3 | 0.3333 | 15 | 0.2 |
| 2_7310493 | 4 | 0.75 | 2 | 0 | 3 | 0.3333 | 7 | 0.2857 | 16 | 0.375 | 2 | 0.5 | 6 | 0.5 | 6 | 0.6667 | 3 | 0.6667 | 17 | 0.5882 |
| 2_7310505 | 3 | 0 | 2 | 0.5 | 1 | 0 | 8 | 0.25 | 14 | 0.2143 | 2 | 0.5 | 4 | 0 | 5 | 0 | 2 | 0 | 13 | 0.0769 |
| 2_7310528 | 3 | 0 | 2 | 0 | 2 | 0 | 6 | 0.1667 | 13 | 0.0769 | 2 | 0.5 | 4 | 0.25 | 5 | 0 | 1 | 0 | 12 | 0.1667 |
| 2_7310540 | 5 | 1 | 2 | 1 | 3 | 0.6667 | 8 | 1 | 18 | 0.9444 | 2 | 1 | 4 | 1 | 9 | 1 | 2 | 1 | 17 | 1 |
| 2_7310541 | 4 | 1 | 2 | 1 | 3 | 1 | 8 | 0.875 | 17 | 0.9412 | 2 | 1 | 5 | 0.8 | 7 | 0.8571 | 3 | 0.6667 | 17 | 0.8235 |
| 2_7310543 | 5 | 0.8 | 2 | 0.5 | 4 | 0 | 8 | 0.125 | 19 | 0.3158 | 2 | 0.5 | 7 | 0.2857 | 9 | 0.2222 | 3 | 0 | 21 | 0.2381 |
| 2_7310545 | 4 | 0.25 | 2 | 0.5 | 3 | 0 | 7 | 0.2857 | 16 | 0.25 | 2 | 0 | 7 | 0.4286 | 9 | 0.3333 | 2 | 0.5 | 20 | 0.35 |
| 2_7310551 | 5 | 1 | 1 | 0 | 4 | 0.75 | 8 | 0.625 | 18 | 0.7222 | 2 | 1 | 8 | 0.875 | 9 | 1 | 3 | 1 | 22 | 0.9545 |
| 2_7310589 | 3 | 1 | 1 | 1 | 1 | 1 | 1 | 1 | 6 | 1 | NA | NA | 5 | 0.6 | 2 | 1 | NA | NA | 7 | 0.7143 |
| 2_7311456 | 4 | 0 | 3 | 0.3333 | 4 | 0 | 2 | 0 | 13 | 0.0769 | 1 | 1 | 3 | 0 | 4 | 0.25 | 4 | 0.5 | 12 | 0.3333 |
| 2_7311463 | 4 | 0.25 | 3 | 0.6667 | 3 | 0.3333 | 4 | 0.5 | 14 | 0.4286 | 1 | 1 | 3 | 1 | 4 | 0.75 | 5 | 0.6 | 13 | 0.7692 |
| 2_7311494 | 4 | 0.25 | 1 | 1 | 4 | 0 | 4 | 0 | 13 | 0.1538 | 1 | 0 | 4 | 0.5 | 3 | 0.3333 | 4 | 0 | 12 | 0.25 |
| 2_7311505 | 3 | 0.6667 | 2 | 0.5 | 4 | 0.25 | 3 | 0.6667 | 12 | 0.5 | NA | NA | 3 | 0.6667 | 2 | 1 | 3 | 0.3333 | 8 | 0.625 |
| 2_7311506 | 3 | 0.3333 | 2 | 0.5 | 4 | 0 | 2 | 0 | 11 | 0.1818 | 1 | 0 | 3 | 0.3333 | 3 | 0 | 4 | 0 | 11 | 0.0909 |

| | | | | | | | | | | | | | | | | | | | | |
|---|---|---|---|---|---|---|---|---|---|---|---|---|---|---|---|---|---|---|---|---|
| 2_7311508 | 3 | 0.3333 | 3 | 0 | 4 | 0 | 3 | 0.3333 | 13 | 0.1538 | 1 | 0 | 4 | 0.5 | 3 | 0.3333 | 4 | 0.25 | 12 | 0.3333 |
| 2_7311509 | 4 | 1 | 3 | 0.6667 | 3 | 0.6667 | 1 | 0 | 11 | 0.7273 | 1 | 1 | 4 | 1 | 3 | 0.3333 | 4 | 0.75 | 12 | 0.75 |
| 2_7314990 | NA | NA | NA | NA | NA | NA | 1 | 1 | 1 | 1 | NA | NA | 1 | 1 | NA | NA | 1 | 0 | 2 | 0.5 |
| 2_7314999 | NA | NA | NA | NA | 1 | 1 | NA | NA | 1 | 1 | NA | NA | 1 | 1 | NA | NA | NA | NA | 1 | 1 |
| 2_7316613 | 1 | 1 | 4 | 0 | 2 | 0 | 3 | 0 | 10 | 0.1 | 1 | 0 | 3 | 0 | NA | NA | 2 | 0.5 | 6 | 0.1667 |
| 2_7317268 | 4 | 0 | 4 | 0 | 8 | 0.25 | 8 | 0.25 | 24 | 0.1667 | 2 | 0.5 | 5 | 0.4 | 10 | 0.4 | 7 | 0.1429 | 24 | 0.3333 |
| 2_7317345 | 3 | 0 | 3 | 0 | 8 | 0.125 | 6 | 0 | 20 | 0.05 | 2 | 0.5 | 6 | 0 | 5 | 0.2 | 5 | 0 | 18 | 0.1111 |
| 2_7317380 | 4 | 0.75 | 2 | 0.5 | 5 | 0.6 | 6 | 0.5 | 17 | 0.5882 | 2 | 0.5 | 4 | 1 | 6 | 0.5 | 7 | 0.5714 | 19 | 0.6316 |
| 2_7317386 | 3 | 0.3333 | 1 | 0 | 5 | 0.2 | 7 | 0.1429 | 16 | 0.1875 | 2 | 0 | 5 | 0 | 6 | 0 | 6 | 0.1667 | 19 | 0.0526 |
| 2_7317399 | 3 | 0.6667 | NA | NA | 6 | 0.5 | 7 | 0.2857 | 16 | 0.4375 | 1 | 1 | 4 | 0.75 | 5 | 1 | 6 | 0.8333 | 16 | 0.875 |
| 2_7317405 | 2 | 0 | 2 | 0 | 6 | 0 | 6 | 0.1667 | 16 | 0.0625 | 2 | 0 | 3 | 0 | 6 | 0 | 6 | 0.1667 | 17 | 0.0588 |
| 2_7320416 | 3 | 0.3333 | 2 | 0 | 6 | 0.1667 | 3 | 0.3333 | 14 | 0.2143 | 1 | 0 | 2 | 0 | 2 | 0 | 5 | 0.2 | 10 | 0.1 |
| 2_7320419 | 2 | 0.5 | 2 | 0 | 4 | 0.75 | 3 | 0.6667 | 11 | 0.5455 | 1 | 1 | 2 | 1 | 2 | 1 | 5 | 0.8 | 10 | 0.9 |
| 2_7320423 | 3 | 0.3333 | 2 | 0.5 | 5 | 0.6 | 3 | 0.6667 | 13 | 0.5385 | 1 | 0 | 1 | 1 | 2 | 0.5 | 5 | 0.8 | 9 | 0.6667 |
| 2_7321222 | 3 | 1 | 1 | 0 | 5 | 0.8 | 2 | 0.5 | 11 | 0.7273 | 1 | 1 | 2 | 1 | 2 | 0.5 | 4 | 0.5 | 9 | 0.6667 |
| 2_7321233 | 2 | 1 | 1 | 1 | 4 | 0.75 | 3 | 1 | 10 | 0.9 | 1 | 1 | 2 | 1 | 1 | 1 | 3 | 0.6667 | 7 | 0.8571 |
| 2_7321241 | 3 | 1 | 1 | 1 | 4 | 1 | 1 | 0 | 9 | 0.8889 | 1 | 0 | 2 | 1 | 2 | 1 | 2 | 1 | 7 | 0.8571 |
| 2_7326248 | NA | NA | NA | NA | 1 | 1 | 1 | 1 | 2 | 1 | NA | NA | 2 | 0.5 | 1 | 1 | 2 | 1 | 5 | 0.8 |
| 2_7344224 | 11 | 0.1818 | 24 | 0.5 | 16 | 0.3125 | 13 | 0.4615 | 64 | 0.3906 | 17 | 0.1176 | 21 | 0.3333 | 19 | 0.2105 | 15 | 0.4667 | 72 | 0.2778 |
| 2_7344230 | 11 | 0.0909 | 24 | 0.0833 | 19 | 0.1053 | 11 | 0.1818 | 65 | 0.1077 | 16 | 0.0625 | 17 | 0.0588 | 15 | 0.0667 | 17 | 0.0588 | 65 | 0.0615 |
| 2_7344323 | 133 | 0.0602 | 156 | 0.1346 | 135 | 0.0963 | 127 | 0.0787 | 551 | 0.0944 | 121 | 0.124 | 124 | 0.0968 | 147 | 0.068 | 124 | 0.121 | 516 | 0.1008 |
| 2_7344341 | 135 | 0.4667 | 149 | 0.4966 | 132 | 0.5379 | 121 | 0.4463 | 537 | 0.4879 | 106 | 0.5094 | 120 | 0.4667 | 137 | 0.4964 | 115 | 0.4957 | 478 | 0.4916 |
| 2_7349012 | 96 | 0.1458 | 105 | 0.1524 | 89 | 0.1798 | 85 | 0.1412 | 375 | 0.1547 | 71 | 0.0845 | 82 | 0.1707 | 90 | 0.1444 | 84 | 0.2024 | 327 | 0.1529 |
| 2_7349054 | 70 | 0.2429 | 71 | 0.0704 | 66 | 0.2576 | 62 | 0.2419 | 269 | 0.2007 | 60 | 0.1833 | 67 | 0.1493 | 73 | 0.2877 | 67 | 0.2388 | 267 | 0.2172 |
| 2_7349055 | 70 | 0.4429 | 75 | 0.3867 | 66 | 0.4091 | 65 | 0.4769 | 276 | 0.4275 | 55 | 0.3273 | 68 | 0.4118 | 75 | 0.4 | 66 | 0.4697 | 264 | 0.4053 |
| 2_7349060 | 63 | 0.6032 | 67 | 0.4925 | 56 | 0.6071 | 49 | 0.6735 | 235 | 0.5872 | 49 | 0.6122 | 49 | 0.4898 | 65 | 0.5077 | 61 | 0.5738 | 224 | 0.5446 |
| 2_7349067 | 52 | 0.25 | 58 | 0.3103 | 47 | 0.4255 | 43 | 0.3721 | 200 | 0.335 | 39 | 0.2308 | 49 | 0.449 | 56 | 0.3929 | 57 | 0.3684 | 201 | 0.3682 |
| 2_7351015 | 11 | 0.4545 | 9 | 0.3333 | 6 | 0.1667 | 3 | 0.3333 | 29 | 0.3448 | 7 | 0.2857 | 4 | 0 | 7 | 0.1429 | 6 | 0.8333 | 24 | 0.3333 |
| 2_7364830 | 15 | 0 | 20 | 0.2 | 16 | 0.4375 | 10 | 0.1 | 61 | 0.1967 | 5 | 0 | 9 | 0.3333 | 10 | 0.1 | 8 | 0.25 | 32 | 0.1875 |
| 2_7364907 | 14 | 0.2857 | 19 | 0.1579 | 16 | 0.1875 | 10 | 0.2 | 59 | 0.2034 | 5 | 0.2 | 8 | 0.125 | 13 | 0.2308 | 6 | 0.1667 | 32 | 0.1875 |
| 2_7364910 | 15 | 0.3333 | 18 | 0.1667 | 15 | 0.2 | 9 | 0.2222 | 57 | 0.2281 | 5 | 0.4 | 7 | 0.5714 | 11 | 0.2727 | 6 | 0.5 | 29 | 0.4138 |
| 2_7364920 | 15 | 0.1333 | 16 | 0.4375 | 13 | 0 | 9 | 0.1111 | 53 | 0.1887 | 5 | 0 | 6 | 0 | 13 | 0.3846 | 6 | 0 | 30 | 0.1667 |
| 2_7364922 | 15 | 0.5333 | 17 | 0.5294 | 14 | 0.4286 | 8 | 0.5 | 54 | 0.5 | 5 | 0.8 | 5 | 0.6 | 12 | 0.4167 | 6 | 0.5 | 28 | 0.5357 |
| 2_7364933 | 13 | 0 | 15 | 0.1333 | 10 | 0.2 | 9 | 0.1111 | 47 | 0.1064 | 5 | 0.2 | 6 | 0 | 11 | 0.0909 | 6 | 0 | 28 | 0.0714 |
| 2_7434547 | 1 | 0 | 4 | 0.5 | 6 | 0 | 2 | 0 | 13 | 0.1538 | 3 | 0.3333 | 1 | 0 | 2 | 0 | 2 | 0.5 | 8 | 0.25 |
| 2_7434556 | 2 | 0 | 5 | 0.2 | 5 | 0 | 2 | 0 | 14 | 0.0714 | 3 | 0.3333 | 1 | 0 | 2 | 0 | 3 | 0 | 9 | 0.1111 |
| 2_7434621 | NA | NA | 4 | 0.5 | 4 | 0 | 1 | 0 | 9 | 0.2222 | 3 | 0.3333 | 1 | 0 | 1 | 0 | 4 | 0 | 9 | 0.1111 |
| 2_7434782 | NA | NA | 3 | 0 | 3 | 0.3333 | 1 | 1 | 7 | 0.2857 | 2 | 0.5 | NA | NA | 1 | 0 | 4 | 0 | 7 | 0.1429 |
| 2_7434839 | 1 | 0 | 2 | 0 | 2 | 0.5 | 1 | 1 | 6 | 0.3333 | 1 | 0 | NA | NA | 1 | 0 | 5 | 0.4 | 7 | 0.2857 |
| 2_7435250 | 3 | 0.6667 | 3 | 0 | 7 | 0.2857 | 3 | 0.3333 | 16 | 0.3125 | 1 | 0 | 3 | 1 | 4 | 0.75 | 2 | 0 | 10 | 0.6 |
| 2_7470671 | 10 | 0.1 | 9 | 0 | 9 | 0.1111 | 7 | 0.1429 | 35 | 0.0857 | 3 | 0 | 7 | 0.1429 | 11 | 0.0909 | 2 | 1 | 23 | 0.1739 |
| 2_7470679 | 11 | 0.1818 | 7 | 0 | 9 | 0 | 9 | 0 | 36 | 0.0556 | 5 | 0 | 7 | 0 | 12 | 0 | 3 | 0.6667 | 27 | 0.0741 |
| 2_7549583 | 15 | 0.6667 | 16 | 0.4375 | 11 | 0.6364 | 16 | 0.75 | 58 | 0.6207 | 7 | 0.8571 | 25 | 0.52 | 15 | 0.4667 | 15 | 0.3333 | 62 | 0.5 |
| 2_7549591 | 17 | 0.5294 | 19 | 0.7895 | 10 | 0.8 | 16 | 0.875 | 62 | 0.7419 | 7 | 0.7143 | 24 | 0.7917 | 14 | 0.5714 | 17 | 0.6471 | 62 | 0.6935 |
| 2_7549699 | 9 | 0.2222 | 12 | 0.0833 | 6 | 0 | 13 | 0.0769 | 40 | 0.1 | 6 | 0.1667 | 14 | 0 | 10 | 0 | 16 | 0.125 | 46 | 0.0652 |
| 2_7549739 | 11 | 0.1818 | 12 | 0.3333 | 9 | 0.1111 | 12 | 0 | 44 | 0.1591 | 9 | 0.1111 | 13 | 0.3077 | 11 | 0.1818 | 16 | 0.125 | 49 | 0.1837 |
| 2_7549756 | 10 | 0.2 | 12 | 0.0833 | 7 | 0.1429 | 10 | 0.1 | 39 | 0.1282 | 5 | 0 | 11 | 0 | 9 | 0 | 18 | 0.2778 | 43 | 0.1163 |
| 2_7549788 | 3 | 0.3333 | 9 | 0.2222 | 4 | 0 | 8 | 0.5 | 24 | 0.2917 | NA | NA | 6 | 0 | 6 | 0.1667 | 12 | 0.5 | 24 | 0.2917 |
| 2_7549793 | 4 | 0.25 | 6 | 0.6667 | 5 | 0.2 | 7 | 0.7143 | 22 | 0.5 | NA | NA | 5 | 0.4 | 5 | 0.4 | 8 | 0.75 | 18 | 0.5556 |
| 2_7549798 | 3 | 0 | 8 | 0.25 | 6 | 0.5 | 6 | 0.1667 | 23 | 0.2609 | NA | NA | 4 | 0.5 | 7 | 0.1429 | 13 | 0.3077 | 24 | 0.2917 |
| 2_7549801 | 5 | 0.4 | 9 | 0.4444 | 7 | 0.4286 | 8 | 0.25 | 29 | 0.3793 | NA | NA | 6 | 0.5 | 6 | 0.1667 | 12 | 0.4167 | 24 | 0.375 |
| 2_7549886 | 7 | 0 | 9 | 0.2222 | 6 | 0 | 12 | 0.0833 | 34 | 0.0882 | 2 | 0 | 5 | 0 | 7 | 0.1429 | 10 | 0.1 | 24 | 0.0833 |
| 2_7549938 | 10 | 0.1 | 13 | 0.3077 | 9 | 0 | 13 | 0 | 45 | 0.1111 | 1 | 0 | 8 | 0.125 | 10 | 0.2 | 15 | 0.1333 | 34 | 0.1471 |
| 2_7550602 | 2 | 0.5 | 6 | 0.3333 | 5 | 0 | 6 | 0.3333 | 19 | 0.2632 | 5 | 0.6 | 3 | 0 | 5 | 0.6 | 6 | 0 | 19 | 0.3158 |
| 2_7550605 | 2 | 0.5 | 7 | 0.2857 | 5 | 0.2 | 4 | 0.25 | 18 | 0.2778 | 3 | 0.3333 | 3 | 0 | 5 | 0.2 | 6 | 0 | 17 | 0.1176 |
| 2_7550616 | 2 | 0 | 6 | 0.1667 | 5 | 0 | 6 | 0.1667 | 19 | 0.1053 | 5 | 0 | 3 | 0 | 4 | 0.5 | 5 | 0 | 17 | 0.1176 |
| 2_7550621 | 2 | 0 | 7 | 0 | 4 | 0.25 | 7 | 0.1429 | 20 | 0.1 | 5 | 0 | 3 | 0.3333 | 4 | 0 | 5 | 0.2 | 17 | 0.1176 |
| 2_7550622 | 2 | 0 | 7 | 0.1429 | 5 | 0 | 7 | 0.1429 | 21 | 0.0952 | 5 | 0 | 4 | 0 | 5 | 0.2 | 5 | 0.2 | 19 | 0.1053 |
| 2_7550630 | 1 | 0 | 4 | 0.25 | 3 | 0 | 4 | 0 | 12 | 0.0833 | 2 | 0.5 | 3 | 0.3333 | 3 | 0 | 3 | 0.3333 | 11 | 0.2727 |
| 2_7550650 | 1 | 1 | 5 | 0 | 3 | 0 | 5 | 0 | 14 | 0.0714 | NA | NA | 2 | 0 | 4 | 0.25 | 2 | 0 | 8 | 0.125 |
| 2_7550654 | 1 | 0 | 7 | 0.1429 | 4 | 0.25 | 7 | 0.2857 | 19 | 0.2105 | 5 | 0.4 | 4 | 0.5 | 3 | 0 | 4 | 0.5 | 16 | 0.375 |
| 2_7550656 | 1 | 0 | 6 | 0.3333 | 4 | 0.25 | 6 | 0 | 17 | 0.1765 | 4 | 0.25 | 4 | 0.5 | 3 | 0 | 2 | 0.5 | 13 | 0.3077 |
| 2_7550658 | 1 | 1 | 6 | 0.5 | 4 | 0.25 | 5 | 0.6 | 16 | 0.5 | 4 | 0.5 | 4 | 0.75 | 4 | 0.75 | 4 | 0.5 | 16 | 0.625 |
| 2_7550668 | 1 | 0 | 5 | 0 | 4 | 0 | 7 | 0.1429 | 17 | 0.0588 | 5 | 0.2 | 4 | 0.5 | 2 | 0 | 4 | 0.25 | 15 | 0.2667 |
| 2_7550679 | 1 | 1 | 7 | 0.5714 | 4 | 0.75 | 6 | 0.3333 | 18 | 0.5556 | 5 | 0.6 | 2 | 0 | 2 | 0.5 | 4 | 0.25 | 13 | 0.3846 |

| | | | | | | | | | | | | | | | | | | | | |
|---|---|---|---|---|---|---|---|---|---|---|---|---|---|---|---|---|---|---|---|---|
| 2_7550700 | 1 | 0 | 5 | 0.4 | 3 | 0.6667 | 7 | 0.1429 | 16 | 0.3125 | 3 | 1 | 4 | 0.5 | 3 | 0.6667 | 2 | 0 | 12 | 0.5833 |
| 2_7550709 | 1 | 0 | 5 | 0.2 | 3 | 0.6667 | 5 | 0.4 | 14 | 0.3571 | 4 | 0.25 | 3 | 0.6667 | 3 | 0 | 3 | 0.3333 | 13 | 0.3077 |
| 2_7550710 | 1 | 1 | 4 | 0.75 | 2 | 0.5 | 6 | 0.6667 | 13 | 0.6923 | 4 | 0.75 | 3 | 1 | 2 | 0.5 | 2 | 0.5 | 11 | 0.7273 |
| 2_7550718 | 1 | 1 | 5 | 1 | 3 | 1 | 7 | 1 | 16 | 1 | 5 | 0.8 | 4 | 1 | 3 | 1 | 4 | 0.75 | 16 | 0.875 |
| 2_7550719 | 1 | 1 | 3 | 0.6667 | 1 | 1 | 3 | 0.3333 | 8 | 0.625 | 3 | 1 | NA | NA | 1 | 1 | 1 | 0 | 5 | 0.8 |
| 2_7550731 | 1 | 0 | 6 | 0.8333 | 3 | 0.6667 | 7 | 0.8571 | 17 | 0.7647 | 5 | 0.6 | 2 | 0 | 3 | 0.6667 | 4 | 0.5 | 14 | 0.5 |
| 2_7550732 | 1 | 0 | 6 | 0 | 3 | 0 | 6 | 0.1667 | 16 | 0.0625 | 4 | 0 | 3 | 0 | 2 | 0 | 4 | 0.25 | 13 | 0.0769 |
| 2_7550745 | 1 | 1 | 6 | 1 | 3 | 1 | 7 | 0.5714 | 17 | 0.8235 | 4 | 0.5 | 2 | 0 | 2 | 1 | 4 | 0.5 | 12 | 0.5 |
| 2_7550968 | 1 | 0 | 5 | 0.2 | 3 | 0.3333 | 6 | 0.3333 | 15 | 0.2667 | 5 | 0.2 | 2 | 0.5 | 2 | 0.5 | 3 | 0 | 12 | 0.25 |
| 2_7553472 | 1 | 0 | 5 | 0.2 | 3 | 0 | 6 | 0.1667 | 15 | 0.1333 | 4 | 0.25 | 1 | 0 | 1 | 0 | 3 | 0 | 9 | 0.1111 |
| 2_7575635 | 3 | 0 | NA | NA | 3 | 0.6667 | 2 | 0 | 8 | 0.25 | 2 | 0.5 | 1 | 0 | 4 | 0 | 4 | 0 | 11 | 0.0909 |
| 2_7575646 | 3 | 0.3333 | NA | NA | 2 | 1 | 2 | 0.5 | 7 | 0.5714 | 2 | 1 | 1 | 0 | 4 | 0.75 | 3 | 0.6667 | 10 | 0.7 |
| 2_7575805 | 6 | 0 | 3 | 0.3333 | 4 | 0 | 3 | 0 | 16 | 0.0625 | 2 | 0 | 2 | 0.5 | 6 | 0 | 6 | 0 | 16 | 0.0625 |
| 2_7575854 | 8 | 0 | 2 | 0 | 6 | 0.1667 | 3 | 0 | 19 | 0.0526 | 2 | 0 | 2 | 0 | 6 | 0.1667 | 7 | 0 | 17 | 0.0588 |
| 2_7575884 | 6 | 0.1667 | 5 | 0 | 6 | 0.1667 | 3 | 0 | 20 | 0.1 | 1 | 0 | 4 | 0 | 9 | 0.1111 | 8 | 0.25 | 22 | 0.1364 |
| 2_7576655 | 1 | 1 | NA | NA | 4 | 0 | 3 | 0.3333 | 8 | 0.25 | 3 | 0.3333 | 2 | 0 | 3 | 0 | 2 | 0 | 10 | 0.1 |
| 2_7576666 | 1 | 1 | 1 | 0 | 4 | 0 | 5 | 0 | 11 | 0.0909 | 1 | 1 | 1 | 0 | 3 | 0 | 2 | 0 | 7 | 0.1429 |
| 2_7576672 | 1 | 0 | 1 | 1 | 4 | 0 | 5 | 0.4 | 11 | 0.2727 | 2 | 0.5 | 1 | 0 | 3 | 0 | 2 | 0 | 8 | 0.125 |
| 2_7576688 | NA | NA | 1 | 1 | 3 | 0.6667 | 3 | 1 | 7 | 0.8571 | 3 | 0.3333 | 1 | 0 | 3 | 0 | 2 | 0 | 9 | 0.1111 |
| 2_7576689 | NA | NA | 1 | 1 | 4 | 1 | 5 | 1 | 10 | 1 | NA | NA | NA | NA | 2 | 1 | 1 | 1 | 3 | 1 |
| 2_7576693 | NA | NA | 1 | 1 | 4 | 1 | 5 | 1 | 10 | 1 | 3 | 1 | 1 | 1 | 3 | 1 | 2 | 1 | 9 | 1 |
| 2_7576696 | NA | NA | 1 | 1 | 2 | 0.5 | 4 | 0.25 | 7 | 0.4286 | 2 | 0.5 | 1 | 0 | 3 | 0.3333 | 1 | 1 | 7 | 0.4286 |
| 2_7576707 | NA | NA | 1 | 1 | 3 | 0.3333 | 5 | 0.6 | 9 | 0.5556 | 2 | 1 | 1 | 1 | 2 | 0.5 | 1 | 0 | 6 | 0.6667 |
| 2_7576710 | NA | NA | 1 | 1 | 1 | 1 | 5 | 1 | 7 | 1 | 1 | 1 | NA | NA | 2 | 1 | 1 | 1 | 4 | 1 |
| 2_7576738 | NA | NA | 1 | 1 | 4 | 0 | 5 | 0.2 | 10 | 0.2 | 2 | 0.5 | 1 | 0 | 1 | 0 | 2 | 0 | 6 | 0.1667 |
| 2_7576744 | NA | NA | 1 | 1 | 3 | 0.6667 | 5 | 0.8 | 9 | 0.7778 | 2 | 1 | NA | NA | 2 | 1 | 2 | 1 | 6 | 1 |
| 2_7576750 | NA | NA | 1 | 1 | 3 | 0.3333 | 5 | 0.4 | 9 | 0.4444 | 1 | 1 | 1 | 0 | 1 | 0 | 1 | 0 | 4 | 0.25 |
| 2_7576759 | NA | NA | 1 | 0 | 4 | 0.5 | 4 | 0.5 | 9 | 0.4444 | 2 | 0.5 | 1 | 0 | 2 | 0.5 | 1 | 0 | 6 | 0.3333 |
| 2_7576767 | NA | NA | 1 | 0 | 3 | 0 | 4 | 0.5 | 8 | 0.25 | 2 | 0 | 1 | 0 | 2 | 0.5 | 1 | 0 | 6 | 0.1667 |
| 2_7576770 | NA | NA | 1 | 1 | 4 | 0.5 | 4 | 0.25 | 9 | 0.4444 | 2 | 1 | 1 | 0 | 2 | 1 | 1 | 1 | 6 | 0.8333 |
| 2_7576779 | NA | NA | 1 | 1 | 4 | 0.75 | 4 | 0.75 | 9 | 0.7778 | 2 | 1 | 1 | 1 | 2 | 1 | 1 | 0 | 6 | 0.8333 |
| 2_7576786 | NA | NA | 1 | 1 | 3 | 0.3333 | 4 | 0.25 | 8 | 0.375 | 1 | 0 | 1 | 0 | 1 | 0 | 1 | 1 | 4 | 0.25 |
| 2_7576825 | NA | NA | 1 | 1 | 4 | 0.25 | 3 | 0.6667 | 8 | 0.5 | 1 | 0 | 1 | 1 | 2 | 0 | 1 | 1 | 5 | 0.4 |
| 2_7576831 | NA | NA | 1 | 1 | 4 | 0.5 | 4 | 0.5 | 9 | 0.5556 | 1 | 0 | 1 | 0 | 2 | 0.5 | 1 | 1 | 5 | 0.4 |
| 2_7581007 | 3 | 0 | 2 | 0.5 | NA | NA | 1 | 0 | 6 | 0.1667 | 2 | 1 | NA | NA | NA | NA | NA | NA | 2 | 1 |
| 2_7581013 | 1 | 1 | 1 | 0 | 1 | 0 | NA | NA | 3 | 0.3333 | 2 | 1 | NA | NA | NA | NA | 1 | 0 | 3 | 0.6667 |
| 2_7581022 | 1 | 1 | 2 | 1 | 1 | 1 | NA | NA | 4 | 1 | 3 | 1 | NA | NA | NA | NA | 1 | 1 | 4 | 1 |
| 2_7581047 | 1 | 0 | 2 | 0.5 | NA | NA | NA | NA | 3 | 0.3333 | 3 | 0.3333 | NA | NA | NA | NA | 1 | 0 | 4 | 0.25 |
| 2_7581071 | 1 | 0 | 2 | 0.5 | 1 | 0 | NA | NA | 4 | 0.25 | 2 | 0 | NA | NA | NA | NA | 1 | 1 | 3 | 0.3333 |
| 2_7581083 | 1 | 1 | 2 | 0.5 | 1 | 0 | 1 | 0 | 5 | 0.4 | 2 | 0.5 | NA | NA | NA | NA | 1 | 1 | 3 | 0.6667 |
| 2_7581090 | 1 | 0 | 2 | 0.5 | NA | NA | NA | NA | 3 | 0.3333 | 2 | 1 | NA | NA | NA | NA | 1 | 1 | 3 | 1 |
| 2_7581112 | 1 | 0 | 1 | 1 | NA | NA | 1 | 1 | 3 | 0.6667 | 2 | 0.5 | NA | NA | NA | NA | 1 | 1 | 3 | 0.6667 |
| 2_7581126 | 1 | 0 | 2 | 0 | NA | NA | 1 | 1 | 4 | 0.25 | 2 | 0.5 | NA | NA | NA | NA | 1 | 0 | 3 | 0.3333 |
| 2_7581134 | 1 | 0 | 2 | 0.5 | NA | NA | 1 | 0 | 4 | 0.25 | 2 | 0.5 | NA | NA | NA | NA | 1 | 0 | 3 | 0.3333 |
| 2_7581141 | 1 | 0 | 1 | 1 | NA | NA | NA | NA | 2 | 0.5 | 2 | 0 | NA | NA | NA | NA | 1 | 1 | 3 | 0.3333 |
| 2_7581158 | 1 | 1 | 2 | 0.5 | NA | NA | NA | NA | 3 | 0.6667 | 2 | 1 | NA | NA | NA | NA | NA | NA | 2 | 1 |
| 2_7582097 | NA | NA | NA | NA | NA | NA | 1 | 1 | 1 | 1 | NA | NA | NA | NA | NA | NA | 1 | 1 | 1 | 1 |
| 2_7582129 | NA | NA | NA | NA | NA | NA | 1 | 1 | 1 | 1 | NA | NA | NA | NA | NA | NA | 1 | 1 | 1 | 1 |
| 2_7621186 | NA | NA | NA | NA | NA | NA | 1 | 1 | 1 | 1 | NA | NA | NA | NA | 1 | 0 | 1 | 1 | 2 | 0.5 |
| 2_7621380 | NA | NA | 5 | 0.2 | 5 | 0 | 3 | 0.3333 | 13 | 0.1538 | 1 | 0 | 1 | 1 | 5 | 0 | 4 | 0.25 | 11 | 0.1818 |
| 2_7621386 | NA | NA | 3 | 0 | 4 | 0.25 | 4 | 0 | 11 | 0.0909 | NA | NA | 1 | 1 | 5 | 0.4 | 4 | 0.25 | 10 | 0.4 |
| 2_7642628 | NA | NA | 1 | 1 | 5 | 1 | 5 | 0.6 | 11 | 0.8182 | 2 | 0.5 | 6 | 0.8333 | NA | NA | 4 | 0.75 | 12 | 0.75 |
| 2_7642632 | NA | NA | 1 | 0 | 5 | 0.4 | 4 | 0.25 | 10 | 0.3 | 2 | 0 | 6 | 0 | NA | NA | 5 | 0.2 | 13 | 0.0769 |
| 2_7642633 | NA | NA | 1 | 0 | 5 | 0.2 | 5 | 0 | 11 | 0.0909 | 2 | 0.5 | 6 | 0.1667 | NA | NA | 5 | 0 | 13 | 0.1538 |
| 2_7642637 | NA | NA | 1 | 1 | 5 | 0.4 | 3 | 0.6667 | 9 | 0.5556 | 1 | 1 | 5 | 0.4 | NA | NA | 2 | 1 | 8 | 0.625 |
| 2_7642664 | NA | NA | 1 | 1 | 3 | 0.6667 | 3 | 0.3333 | 7 | 0.5714 | 2 | 1 | 4 | 0.75 | NA | NA | 3 | 0.6667 | 9 | 0.7778 |
| 2_7642669 | NA | NA | 1 | 1 | 4 | 0 | 3 | 0.6667 | 8 | 0.375 | 2 | 0 | 4 | 0.25 | NA | NA | 3 | 0.3333 | 9 | 0.2222 |
| 2_7642677 | NA | NA | NA | NA | 5 | 0.4 | 5 | 0.2 | 10 | 0.3 | 2 | 1 | 5 | 0.4 | NA | NA | 4 | 0.5 | 11 | 0.5455 |
| 2_7642690 | NA | NA | 1 | 0 | 5 | 0 | 5 | 0.8 | 11 | 0.3636 | 2 | 0.5 | 5 | 0.6 | NA | NA | 3 | 0.6667 | 10 | 0.6 |
| 2_7642694 | NA | NA | 1 | 0 | 6 | 0.1667 | 3 | 0.3333 | 10 | 0.2 | NA | NA | 4 | 0.25 | NA | NA | 2 | 1 | 6 | 0.5 |
| 2_7642888 | NA | NA | 2 | 0.5 | 6 | 0.1667 | 4 | 0.25 | 12 | 0.25 | 1 | 0 | 7 | 0.1429 | NA | NA | 5 | 0.2 | 13 | 0.1538 |
| 2_7642957 | NA | NA | 1 | 1 | 8 | 0.375 | 3 | 0 | 12 | 0.3333 | 2 | 0.5 | 6 | 0.5 | 1 | 0 | 5 | 0.2 | 14 | 0.3571 |
| 2_7643033 | NA | NA | 2 | 0.5 | 4 | 0.25 | 4 | 0 | 10 | 0.2 | 2 | 0.5 | 3 | 0 | 1 | 0 | 2 | 1 | 8 | 0.375 |
| 2_7643034 | NA | NA | 2 | 0 | 3 | 0.3333 | 4 | 0 | 9 | 0.1111 | 2 | 0 | 4 | 0.25 | 1 | 0 | 2 | 0 | 9 | 0.1111 |
| 2_7643036 | NA | NA | 2 | 0 | 4 | 0.25 | 4 | 0 | 10 | 0.1 | 2 | 0 | 4 | 0.25 | 1 | 0 | 2 | 0.5 | 9 | 0.2222 |

| | | | | | | | | | | | | | | | | | | | | |
|---|---|---|---|---|---|---|---|---|---|---|---|---|---|---|---|---|---|---|---|---|
| 2_7643057 | NA | NA | 2 | 0.5 | 4 | 0 | 3 | 0.3333 | 9 | 0.2222 | 2 | 0 | 3 | 0 | 1 | 1 | 3 | 0 | 9 | 0.1111 |
| 2_7646047 | 1 | 0 | 1 | 0 | 3 | 0.3333 | 2 | 0.5 | 7 | 0.2857 | 3 | 0 | 3 | 0 | 3 | 0 | 4 | 0.25 | 13 | 0.0769 |
| 2_7684045 | 1 | 1 | 4 | 1 | 1 | 0 | 3 | 0.6667 | 9 | 0.7778 | 1 | 1 | 2 | 0.5 | 2 | 0.5 | 2 | 0.5 | 7 | 0.5714 |
| 2_7684079 | 1 | 0 | 3 | 0.6667 | 1 | 1 | 3 | 0.3333 | 8 | 0.5 | 1 | 0 | 2 | 0.5 | 3 | 0.6667 | 1 | 0 | 7 | 0.4286 |
| 2_7684093 | 1 | 1 | 2 | 1 | 1 | 1 | 1 | 1 | 5 | 1 | 1 | 0 | 2 | 0 | 3 | 0.6667 | 2 | 1 | 8 | 0.5 |
| 2_7684107 | NA | NA | 2 | 0 | 1 | 1 | 1 | 0 | 4 | 0.25 | NA | NA | 1 | 0 | 2 | 0.5 | 1 | 0 | 4 | 0.25 |
| 2_7684136 | 1 | 0 | 3 | 0.3333 | 1 | 0 | 1 | 0 | 6 | 0.1667 | NA | NA | 2 | 0 | 3 | 0.3333 | 2 | 0 | 7 | 0.1429 |
| 2_7684140 | 1 | 0 | 3 | 0.3333 | 1 | 0 | 1 | 0 | 6 | 0.1667 | 1 | 0 | 2 | 0.5 | 2 | 0 | 1 | 0 | 6 | 0.1667 |
| 2_7684179 | 1 | 0 | 3 | 0 | 1 | 1 | 1 | 0 | 6 | 0.1667 | 1 | 1 | 1 | 1 | 2 | 1 | 1 | 0 | 5 | 0.8 |
| 2_7684181 | 1 | 0 | 3 | 1 | 1 | 0 | 1 | 0 | 6 | 0.5 | 1 | 1 | 1 | 1 | 2 | 1 | 1 | 1 | 5 | 1 |
| 2_7684190 | NA | NA | 2 | 0 | 1 | 0 | 1 | 1 | 4 | 0.25 | 1 | 0 | 1 | 1 | 2 | 0.5 | 1 | 0 | 5 | 0.4 |
| 2_7684204 | 1 | 0 | 3 | 0.3333 | 1 | 0 | 1 | 1 | 6 | 0.3333 | 1 | 0 | 1 | 0 | NA | NA | 1 | 1 | 3 | 0.3333 |
| 2_7684206 | 1 | 0 | 3 | 0.3333 | 1 | 0 | 1 | 1 | 6 | 0.3333 | 1 | 0 | 1 | 0 | 1 | 1 | 1 | 0 | 4 | 0.25 |
| 2_7684478 | 1 | 0 | 3 | 0 | 1 | 0 | 2 | 0.5 | 7 | 0.1429 | 1 | 1 | 1 | 0 | 1 | 0 | 1 | 0 | 4 | 0.25 |
| 2_7684488 | NA | NA | 3 | 0.3333 | 1 | 0 | 1 | 0 | 5 | 0.2 | 1 | 0 | 1 | 1 | 1 | 1 | 1 | 1 | 4 | 0.75 |
| 2_7684521 | NA | NA | 2 | 1 | 1 | 0 | 1 | 0 | 4 | 0.5 | 1 | 1 | 1 | 0 | NA | NA | 1 | 0 | 3 | 0.3333 |
| 2_7684529 | NA | NA | 3 | 0.3333 | 1 | 0 | 1 | 0 | 5 | 0.2 | 1 | 0 | 1 | 1 | NA | NA | 1 | 1 | 3 | 0.6667 |
| 2_7684552 | NA | NA | 3 | 1 | 1 | 0 | 1 | 1 | 5 | 0.8 | 1 | 1 | NA | NA | NA | NA | 1 | 0 | 2 | 0.5 |
| 2_7701727 | 2 | 0.5 | 2 | 0 | 2 | 0 | 2 | 0 | 8 | 0.125 | 1 | 1 | 2 | 0.5 | 4 | 0 | 2 | 0.5 | 9 | 0.3333 |
| 2_7701728 | 3 | 1 | 2 | 0.5 | 2 | 1 | 2 | 1 | 9 | 0.8889 | 1 | 1 | 2 | 1 | 4 | 1 | 2 | 0.5 | 9 | 0.8889 |
| 2_7701753 | 2 | 0 | 2 | 0.5 | 1 | 0 | 3 | 0.3333 | 8 | 0.25 | 1 | 1 | 2 | 0 | 2 | 0.5 | 2 | 1 | 7 | 0.5714 |
| 2_7701754 | 1 | 1 | 1 | 1 | 1 | 0 | 3 | 0.3333 | 6 | 0.5 | 1 | 1 | 2 | 0.5 | 1 | 0 | 2 | 0.5 | 6 | 0.5 |
| 2_7701760 | 3 | 0.3333 | 1 | 0 | 1 | 0 | 3 | 0 | 8 | 0.125 | 1 | 1 | 2 | 0.5 | 3 | 0 | 2 | 0 | 8 | 0.25 |
| 2_7724112 | 10 | 0 | 8 | 0.125 | 9 | 0 | 13 | 0.0769 | 40 | 0.05 | 4 | 0 | 9 | 0.1111 | 13 | 0.0769 | 2 | 0 | 28 | 0.0714 |
| 2_7724193 | 8 | 0.375 | 6 | 0.1667 | 5 | 0.4 | 13 | 0.2308 | 32 | 0.2812 | 1 | 0 | 6 | 0.5 | 7 | 0.4286 | 2 | 1 | 16 | 0.5 |
| 2_7724218 | 7 | 0.5714 | 5 | 0.2 | 5 | 0.4 | 10 | 0.7 | 27 | 0.5185 | 2 | 1 | 7 | 0.5714 | 9 | 0.4444 | 2 | 0.5 | 20 | 0.55 |
| 2_7772627 | 1 | 0 | 3 | 0.6667 | 2 | 1 | 3 | 0.3333 | 9 | 0.5556 | NA | NA | 2 | 0 | 3 | 1 | 1 | 1 | 6 | 0.6667 |
| 2_7772633 | 1 | 0 | 4 | 0.25 | 2 | 0 | 2 | 0 | 9 | 0.1111 | NA | NA | 3 | 0.3333 | 3 | 0 | 3 | 0.3333 | 9 | 0.2222 |
| 2_7772641 | 1 | 0 | 3 | 0.3333 | 2 | 0 | 2 | 0 | 8 | 0.125 | NA | NA | 3 | 0 | 3 | 0.3333 | 3 | 0 | 9 | 0.1111 |
| 2_7772655 | 1 | 1 | 4 | 1 | 2 | 0.5 | 3 | 0.6667 | 10 | 0.8 | NA | NA | 2 | 0 | 3 | 0.6667 | 3 | 0.3333 | 8 | 0.375 |
| 2_7772673 | 1 | 0 | 4 | 0.25 | 2 | 0.5 | 2 | 0 | 9 | 0.2222 | NA | NA | 3 | 0.3333 | 3 | 0 | 3 | 0 | 9 | 0.1111 |
| 2_7772679 | 1 | 1 | 4 | 0.75 | 2 | 1 | 2 | 1 | 9 | 0.8889 | NA | NA | 3 | 1 | 2 | 1 | 3 | 1 | 8 | 1 |
| 2_7772681 | 1 | 0 | 3 | 0 | 2 | 0 | 2 | 0.5 | 8 | 0.125 | NA | NA | 3 | 0 | 2 | 0 | 3 | 0.3333 | 8 | 0.125 |
| 2_7875048 | 10 | 1 | 17 | 0.8824 | 14 | 0.9286 | 5 | 0.8 | 46 | 0.913 | 8 | 0.75 | 13 | 1 | 12 | 1 | 9 | 1 | 42 | 0.9524 |
| 2_7875053 | 13 | 0 | 19 | 0.1053 | 13 | 0.3077 | 6 | 0 | 51 | 0.1176 | 8 | 0.125 | 17 | 0.0588 | 14 | 0.1429 | 10 | 0 | 49 | 0.0816 |
| 2_7928425 | NA | NA | 2 | 1 | NA | NA | 2 | 1 | 4 | 1 | 1 | 1 | NA | NA | 1 | 1 | NA | NA | 2 | 1 |
| 2_7928438 | NA | NA | 4 | 0.5 | NA | NA | 1 | 1 | 5 | 0.6 | 1 | 1 | NA | NA | 1 | 1 | NA | NA | 2 | 1 |
| 2_7928451 | NA | NA | 2 | 0.5 | NA | NA | NA | NA | 2 | 0.5 | 1 | 1 | NA | NA | 1 | 1 | NA | NA | 2 | 1 |
| 2_7928456 | NA | NA | 4 | 0.25 | NA | NA | 1 | 0 | 5 | 0.2 | 1 | 1 | NA | NA | 1 | 1 | NA | NA | 2 | 1 |
| 2_7928459 | NA | NA | 3 | 1 | NA | NA | 1 | 1 | 4 | 1 | NA | NA | NA | NA | 1 | 1 | NA | NA | 1 | 1 |
| 2_7928467 | NA | NA | 3 | 0.3333 | NA | NA | 1 | 1 | 4 | 0.5 | 1 | 1 | NA | NA | 1 | 1 | NA | NA | 2 | 1 |
| 2_7928470 | NA | NA | 1 | 1 | NA | NA | 1 | 0 | 2 | 0.5 | 1 | 1 | NA | NA | 1 | 1 | NA | NA | 2 | 1 |
| 2_7928480 | NA | NA | 3 | 1 | NA | NA | 1 | 1 | 4 | 1 | 1 | 1 | NA | NA | 1 | 1 | NA | NA | 2 | 1 |
| 2_7928493 | NA | NA | 2 | 0.5 | NA | NA | NA | NA | 2 | 0.5 | NA | NA | NA | NA | 1 | 1 | NA | NA | 1 | 1 |
| 2_7928495 | NA | NA | 4 | 0.5 | NA | NA | 1 | 1 | 5 | 0.6 | NA | NA | NA | NA | 1 | 1 | NA | NA | 1 | 1 |
| 2_7928497 | NA | NA | 2 | 1 | NA | NA | NA | NA | 2 | 1 | NA | NA | NA | NA | 1 | 1 | NA | NA | 1 | 1 |
| 2_7947494 | 1 | 1 | 3 | 0 | NA | NA | 1 | 0 | 5 | 0.2 | NA | NA | 1 | 1 | NA | NA | 2 | 0.5 | 3 | 0.6667 |
| 2_7947494 | 1 | 1 | 3 | 0.6667 | 1 | 1 | 2 | 0.5 | 7 | 0.7143 | 1 | 0 | 1 | 0 | NA | NA | 2 | 1 | 4 | 0.5 |
| 2_7947501 | 1 | 1 | 3 | 0.6667 | 1 | 1 | 2 | 0 | 7 | 0.5714 | 1 | 1 | 1 | 1 | NA | NA | 1 | 0 | 3 | 0.6667 |
| 2_7947517 | 1 | 1 | 3 | 0.3333 | NA | NA | 2 | 0.5 | 6 | 0.5 | 1 | 0 | 1 | 1 | NA | NA | 2 | 0.5 | 4 | 0.5 |
| 2_7947528 | 1 | 0 | 3 | 0.3333 | 1 | 0 | 2 | 0 | 7 | 0.1429 | 1 | 1 | 1 | 0 | NA | NA | 2 | 0.5 | 4 | 0.5 |
| 2_7948795 | 1 | 0 | 3 | 0.3333 | 1 | 1 | 2 | 0 | 7 | 0.2857 | 1 | 0 | 1 | 1 | 1 | 1 | 2 | 0.5 | 5 | 0.6 |
| 2_7948798 | 1 | 1 | 1 | 1 | NA | NA | 1 | 0 | 3 | 0.6667 | 1 | 1 | 1 | 0 | 1 | 1 | 1 | 1 | 4 | 0.75 |
| 2_7948799 | 1 | 1 | 2 | 0.5 | NA | NA | 2 | 0 | 5 | 0.4 | 1 | 0 | 1 | 1 | 1 | 0 | 3 | 0.3333 | 6 | 0.3333 |
| 2_7948827 | 1 | 1 | 2 | 0.5 | NA | NA | 2 | 0.5 | 5 | 0.6 | 1 | 1 | 2 | 0.5 | 1 | 0 | 1 | 1 | 5 | 0.6 |
| 2_7948847 | 2 | 0.5 | 3 | 0.3333 | 1 | 0 | NA | NA | 6 | 0.3333 | 1 | 1 | 1 | 0 | 1 | 0 | 2 | 0 | 5 | 0.2 |
| 2_7948848 | 1 | 1 | 3 | 0.3333 | NA | NA | 1 | 1 | 5 | 0.6 | 1 | 0 | 1 | 1 | 2 | 0.5 | 3 | 0.3333 | 7 | 0.4286 |
| 2_7948849 | 2 | 1 | 3 | 0.6667 | 1 | 1 | 2 | 0.5 | 8 | 0.75 | 2 | 1 | 2 | 0 | 2 | 0.5 | 3 | 1 | 9 | 0.6667 |
| 2_7949292 | 2 | 0.5 | 4 | 0 | 3 | 0 | 2 | 0 | 11 | 0.0909 | 3 | 0 | 4 | 0.5 | 2 | 0 | 3 | 0 | 12 | 0.1667 |
| 2_7950221 | 2 | 1 | 4 | 1 | 5 | 0.4 | 4 | 0.5 | 15 | 0.6667 | 4 | 0.5 | 5 | 0.8 | 2 | 1 | 2 | 1 | 13 | 0.7692 |
| 2_7950225 | 2 | 0 | 4 | 0 | 5 | 0.2 | 4 | 0 | 15 | 0.0667 | 4 | 0.25 | 5 | 0 | 2 | 0 | 2 | 0.5 | 13 | 0.1538 |
| 2_7950229 | 2 | 0 | 5 | 0 | 6 | 0 | 4 | 0.25 | 17 | 0.0588 | 4 | 0 | 6 | 0 | 1 | 1 | 3 | 0 | 14 | 0.0714 |
| 2_7950231 | 1 | 0 | 4 | 0 | 5 | 0.2 | 3 | 0.3333 | 13 | 0.1538 | 4 | 0 | 5 | 0 | 2 | 0.5 | 2 | 0 | 13 | 0.0769 |
| 2_7950231 | 2 | 0.5 | 5 | 0 | 5 | 0.2 | 3 | 0 | 15 | 0.1333 | 4 | 0.25 | 6 | 0 | 2 | 0 | 2 | 0.5 | 14 | 0.1429 |
| 2_7966491 | 13 | 0.3846 | 16 | 0.3125 | 12 | 0.5833 | 16 | 0.25 | 57 | 0.3684 | 10 | 0.5 | 12 | 0.3333 | 11 | 0.5455 | 14 | 0.4286 | 47 | 0.4468 |

| | | | | | | | | | | | | | | | | | | | | |
|---|---|---|---|---|---|---|---|---|---|---|---|---|---|---|---|---|---|---|---|---|
| 2_7966501 | 10 | 0.9 | 14 | 0.8571 | 9 | 1 | 12 | 0.9167 | 45 | 0.9111 | 10 | 0.9 | 10 | 0.9 | 12 | 1 | 17 | 0.9412 | 49 | 0.9388 |
| 2_7966511 | 11 | 0.1818 | 11 | 0.0909 | 11 | 0 | 15 | 0.4 | 48 | 0.1875 | 7 | 0.1429 | 13 | 0.2308 | 6 | 0.3333 | 17 | 0.0588 | 43 | 0.1628 |
| 2_7966522 | 9 | 0.2222 | 10 | 0.1 | 10 | 0.1 | 11 | 0.0909 | 40 | 0.125 | 7 | 0 | 11 | 0.0909 | 9 | 0.1111 | 16 | 0.1875 | 43 | 0.1163 |
| 2_7966530 | 11 | 0 | 13 | 0 | 10 | 0.2 | 14 | 0.0714 | 48 | 0.0625 | 6 | 0 | 12 | 0.1667 | 7 | 0 | 16 | 0.0625 | 41 | 0.0732 |
| 2_7966563 | 8 | 0.125 | 11 | 0 | 6 | 0 | 13 | 0.1538 | 38 | 0.0789 | 7 | 0.1429 | 11 | 0.0909 | 4 | 0.25 | 17 | 0.1176 | 39 | 0.1282 |
| 2_7966596 | 7 | 0 | 13 | 0.1538 | 4 | 0.5 | 11 | 0.0909 | 35 | 0.1429 | 5 | 0.4 | 9 | 0 | 5 | 0 | 14 | 0.2143 | 33 | 0.1515 |
| 2_7966608 | 7 | 0.1429 | 15 | 0.0667 | 4 | 0.25 | 10 | 0 | 36 | 0.0833 | 3 | 0 | 6 | 0.1667 | 4 | 0 | 14 | 0.1429 | 27 | 0.1111 |
| 2_7966611 | 9 | 0 | 14 | 0.0714 | 4 | 0.25 | 10 | 0.3 | 37 | 0.1351 | 5 | 0 | 10 | 0 | 5 | 0.2 | 16 | 0.125 | 36 | 0.0833 |
| 2_7966624 | 7 | 0.5714 | 11 | 0.4545 | 2 | 0.5 | 11 | 0.7273 | 31 | 0.5806 | 3 | 1 | 7 | 0.2857 | 3 | 0.3333 | 8 | 0.375 | 21 | 0.4286 |
| 2_7966627 | 8 | 0.125 | 14 | 0.0714 | 4 | 0 | 9 | 0.1111 | 35 | 0.0857 | 5 | 0.4 | 10 | 0 | 4 | 0 | 14 | 0.1429 | 33 | 0.1212 |
| 2_7966678 | 8 | 0.25 | 10 | 0.3 | 1 | 1 | 8 | 0.25 | 27 | 0.2963 | 5 | 0.2 | 7 | 0.8571 | 3 | 0.3333 | 13 | 0.7692 | 28 | 0.6429 |
| 2_7966680 | 6 | 1 | 10 | 1 | 1 | 1 | 9 | 1 | 26 | 1 | 4 | 1 | 6 | 0.8333 | 3 | 1 | 12 | 1 | 25 | 0.96 |
| 2_7966683 | 6 | 0 | 10 | 0.1 | 1 | 0 | 10 | 0.1 | 27 | 0.0741 | 5 | 0 | 7 | 0.1429 | 4 | 0.25 | 13 | 0.1538 | 29 | 0.1379 |
| 2_7966688 | 4 | 0 | 7 | 0.1429 | 1 | 0 | 8 | 0.125 | 20 | 0.1 | 5 | 0 | 6 | 0.3333 | 3 | 0 | 11 | 0 | 25 | 0.08 |
| 2_7966688 | 7 | 0 | 12 | 0.0833 | 1 | 0 | 8 | 0.125 | 28 | 0.0714 | 5 | 0 | 7 | 0 | 4 | 0 | 12 | 0.1667 | 28 | 0.0714 |
| 2_7966690 | 7 | 0.8571 | 12 | 0.3333 | 1 | 1 | 9 | 0.5556 | 29 | 0.5517 | 5 | 0.2 | 7 | 0.7143 | 4 | 0.75 | 11 | 0.6364 | 27 | 0.5926 |
| 2_7966714 | 4 | 0.25 | 9 | 0.1111 | 1 | 0 | 7 | 0.1429 | 21 | 0.1429 | 2 | 0 | 5 | 0.2 | 3 | 0.6667 | 8 | 0.625 | 18 | 0.4444 |
| 2_7966718 | 5 | 1 | 12 | 1 | 1 | 1 | 8 | 1 | 26 | 1 | 3 | 1 | 4 | 1 | 4 | 0.75 | 8 | 1 | 19 | 0.9474 |
| 2_7966723 | 5 | 0.4 | 10 | 0.3 | 1 | 1 | 8 | 0.375 | 24 | 0.375 | 4 | 0 | 5 | 0.6 | 4 | 1 | 11 | 0.8182 | 24 | 0.6667 |
| 2_8014772 | 4 | 0.25 | 3 | 0.6667 | 2 | 0.5 | 4 | 0.75 | 13 | 0.5385 | 4 | 0.5 | 3 | 0.3333 | 4 | 0.25 | 5 | 0.6 | 16 | 0.4375 |
| 2_8018340 | 7 | 0.4286 | 7 | 0.4286 | 8 | 0.625 | 7 | 0.1429 | 29 | 0.4138 | 7 | 0.2857 | 10 | 0.3 | 8 | 0.375 | 10 | 0.4 | 35 | 0.3429 |
| 2_8018353 | 6 | 0.1667 | 8 | 0.625 | 9 | 0 | 5 | 0.2 | 28 | 0.25 | 8 | 0.125 | 10 | 0.4 | 11 | 0.2727 | 11 | 0.4545 | 40 | 0.325 |
| 2_8018355 | 5 | 0.2 | 8 | 0.5 | 7 | 0.2857 | 5 | 0.2 | 25 | 0.32 | 7 | 0.2857 | 8 | 0.375 | 9 | 0.3333 | 10 | 0.4 | 34 | 0.3529 |
| 2_8018356 | 5 | 0.2 | 8 | 0.125 | 7 | 0.2857 | 7 | 0.1429 | 27 | 0.1852 | 7 | 0.2857 | 8 | 0.25 | 12 | 0.25 | 11 | 0.1818 | 38 | 0.2368 |
| 2_8018360 | 6 | 0.1667 | 8 | 0.125 | 9 | 0.3333 | 6 | 0.8333 | 29 | 0.3448 | 7 | 0.1429 | 9 | 0.5556 | 12 | 0.25 | 11 | 0.3636 | 39 | 0.3333 |
| 2_8018361 | 6 | 0.1667 | 8 | 0 | 9 | 0.1111 | 7 | 0 | 30 | 0.0667 | 8 | 0 | 10 | 0.2 | 12 | 0 | 11 | 0.0909 | 41 | 0.0732 |
| 2_8018365 | 6 | 0.1667 | 8 | 0.375 | 7 | 0.4286 | 6 | 0.1667 | 27 | 0.2963 | 8 | 0.25 | 9 | 0.3333 | 11 | 0.2727 | 10 | 0.3 | 38 | 0.2895 |
| 2_8018424 | 3 | 1 | 3 | 1 | 5 | 0.4 | 2 | 1 | 13 | 0.7692 | 4 | 0.5 | 4 | 1 | 5 | 0.8 | 4 | 0.25 | 17 | 0.6471 |
| 2_8024297 | 2 | 0 | 5 | 0.2 | 1 | 0 | 1 | 1 | 9 | 0.2222 | 1 | 0 | 2 | 0.5 | 1 | 0 | NA | NA | 4 | 0.25 |
| 2_8024342 | 1 | 0 | 6 | 0.5 | 1 | 0 | NA | NA | 8 | 0.375 | 1 | 1 | 2 | 1 | 1 | 0 | NA | NA | 4 | 0.75 |
| 2_8024348 | 1 | 0 | 6 | 0.8333 | 1 | 1 | 1 | 1 | 9 | 0.7778 | 1 | 0 | 2 | 0.5 | 1 | 0 | NA | NA | 4 | 0.25 |
| 2_8024364 | 1 | 1 | 5 | 0.8 | 1 | 0 | NA | NA | 7 | 0.7143 | 1 | 1 | 2 | 0 | NA | NA | NA | NA | 3 | 0.3333 |
| 2_8024366 | 1 | 0 | 5 | 0.4 | 1 | 0 | 1 | 1 | 8 | 0.375 | 2 | 0.5 | 2 | 0 | 1 | 0 | NA | NA | 5 | 0.2 |
| 2_8024367 | 1 | 1 | 5 | 0.8 | 1 | 1 | 1 | 1 | 8 | 0.875 | 2 | 0.5 | 2 | 0.5 | 1 | 1 | NA | NA | 5 | 0.6 |
| 2_8024378 | 1 | 0 | 6 | 0.1667 | 1 | 0 | NA | NA | 8 | 0.125 | 2 | 0.5 | 2 | 0.5 | 1 | 1 | NA | NA | 5 | 0.6 |
| 2_8033112 | 1 | 1 | NA | NA | NA | NA | NA | NA | 1 | 1 | 1 | 1 | NA | NA | NA | NA | NA | NA | 1 | 1 |
| 2_8033117 | 45 | 0.4889 | 76 | 0.4737 | 51 | 0.5294 | 54 | 0.6667 | 226 | 0.5354 | 30 | 0.4667 | 47 | 0.5106 | 36 | 0.3889 | 42 | 0.6429 | 155 | 0.5097 |
| 2_8033133 | 51 | 0.1961 | 78 | 0.0385 | 54 | 0.0926 | 69 | 0.0435 | 252 | 0.0833 | 36 | 0.0278 | 52 | 0.1154 | 42 | 0.0952 | 59 | 0.1356 | 189 | 0.1005 |
| 2_8033157 | 45 | 0.4222 | 74 | 0.2703 | 52 | 0.4423 | 55 | 0.2182 | 226 | 0.3274 | 29 | 0.3103 | 40 | 0.275 | 38 | 0.3421 | 51 | 0.2745 | 158 | 0.2975 |
| 2_8033158 | 45 | 0.1778 | 74 | 0.1622 | 51 | 0.1373 | 59 | 0.0678 | 229 | 0.1354 | 29 | 0.069 | 44 | 0.2045 | 41 | 0.0732 | 52 | 0.1538 | 166 | 0.1325 |
| 2_8041174 | NA | NA | 2 | 0 | NA | NA | 1 | 1 | 3 | 0.3333 | 1 | 0 | 1 | 0 | NA | NA | 1 | 1 | 3 | 0.3333 |
| 2_8041203 | 1 | 0 | 4 | 0.25 | 1 | 1 | NA | NA | 6 | 0.3333 | 1 | 0 | 1 | 1 | NA | NA | 1 | 1 | 3 | 0.6667 |
| 2_8043373 | 2 | 1 | 2 | 1 | NA | NA | NA | NA | 4 | 1 | NA | NA | 2 | 1 | NA | NA | 1 | 1 | 3 | 1 |
| 2_8046141 | 1 | 1 | 5 | 1 | 2 | 0.5 | 4 | 1 | 12 | 0.9167 | 5 | 1 | 5 | 0.4 | 2 | 1 | 1 | 1 | 13 | 0.7692 |
| 2_8046144 | 1 | 0 | 5 | 0.4 | 2 | 0 | 4 | 0 | 12 | 0.1667 | 4 | 0.25 | 4 | 0 | 2 | 0 | 1 | 0 | 11 | 0.0909 |
| 2_8046145 | 1 | 0 | 5 | 0.2 | 2 | 0 | 3 | 0 | 11 | 0.0909 | 4 | 0.25 | 5 | 0 | 2 | 0 | 1 | 0 | 12 | 0.0833 |
| 2_8046297 | NA | NA | 2 | 0.5 | 3 | 0.3333 | 1 | 0 | 6 | 0.3333 | 2 | 1 | 3 | 0.3333 | 3 | 0 | 2 | 1 | 10 | 0.5 |
| 2_8046405 | NA | NA | 4 | 0 | 2 | 0.5 | 2 | 0 | 8 | 0.125 | 2 | 0 | 3 | 0 | 5 | 0.2 | 2 | 0 | 12 | 0.0833 |
| 2_8048875 | 6 | 0.8333 | 2 | 0.5 | 1 | 0 | 3 | 1 | 12 | 0.75 | 3 | 0.3333 | 3 | 0.6667 | 2 | 0.5 | 5 | 0.6 | 13 | 0.5385 |
| 2_8188418 | NA | NA | 2 | 0 | 2 | 0 | 4 | 0.25 | 8 | 0.125 | NA | NA | 2 | 0.5 | NA | NA | 2 | 0 | 4 | 0.25 |
| 2_8208672 | NA | NA | 1 | 0 | 2 | 1 | 3 | 0.6667 | 6 | 0.6667 | NA | NA | 2 | 1 | NA | NA | 3 | 0.6667 | 5 | 0.8 |
| 2_8208673 | NA | NA | 1 | 1 | 1 | 0 | 2 | 0 | 4 | 0.25 | NA | NA | 1 | 0 | NA | NA | 2 | 0.5 | 3 | 0.3333 |
| 2_8208678 | NA | NA | 2 | 0 | 1 | 1 | 4 | 0.25 | 7 | 0.2857 | NA | NA | 2 | 0.5 | 1 | 0 | 4 | 0 | 7 | 0.1429 |
| 2_8208720 | NA | NA | 1 | 1 | 2 | 1 | 5 | 0.8 | 8 | 0.875 | NA | NA | 2 | 1 | 1 | 1 | 3 | 1 | 6 | 1 |
| 2_8208741 | NA | NA | 2 | 1 | 2 | 1 | 5 | 0.8 | 9 | 0.8889 | NA | NA | 2 | 1 | 1 | 1 | 4 | 1 | 7 | 1 |
| 2_8208742 | NA | NA | 1 | 0 | 2 | 0.5 | 5 | 0 | 8 | 0.125 | NA | NA | 2 | 0 | 1 | 1 | 3 | 0 | 6 | 0.1667 |
| 2_8208745 | NA | NA | NA | NA | 2 | 0.5 | 4 | 0.5 | 6 | 0.5 | NA | NA | 2 | 0.5 | 1 | 0 | 4 | 0.25 | 7 | 0.2857 |
| 2_8208754 | 1 | 0 | 1 | 1 | 2 | 0 | 5 | 0 | 9 | 0.1111 | NA | NA | 2 | 0 | 1 | 1 | 4 | 0 | 7 | 0.1429 |
| 2_8208762 | 1 | 1 | NA | NA | 1 | 1 | 3 | 1 | 5 | 1 | NA | NA | 2 | 1 | 1 | 1 | 3 | 1 | 6 | 1 |
| 2_8208765 | 1 | 0 | NA | NA | 2 | 0.5 | 5 | 0.2 | 8 | 0.25 | NA | NA | 2 | 0 | 1 | 0 | 4 | 0.75 | 7 | 0.4286 |
| 2_8208769 | NA | NA | 1 | 1 | 2 | 1 | 3 | 0.3333 | 6 | 0.6667 | NA | NA | 2 | 1 | NA | NA | 4 | 0.5 | 6 | 0.6667 |
| 2_8208777 | 1 | 0 | 2 | 0 | 2 | 0.5 | 4 | 0.5 | 9 | 0.3333 | NA | NA | 2 | 0.5 | 1 | 0 | 4 | 0.75 | 7 | 0.5714 |
| 2_8208781 | 1 | 0 | 1 | 0 | 2 | 0 | 4 | 0.25 | 8 | 0.125 | NA | NA | 2 | 0 | 1 | 1 | 4 | 0 | 7 | 0.1429 |
| 2_8208782 | 1 | 0 | 2 | 0 | 1 | 0 | 5 | 0.6 | 9 | 0.3333 | NA | NA | 2 | 0.5 | NA | NA | 4 | 0.25 | 6 | 0.3333 |
| 2_8208818 | 1 | 0 | NA | NA | 2 | 0 | 3 | 0.3333 | 6 | 0.1667 | NA | NA | 2 | 0 | 1 | 0 | 4 | 0.5 | 7 | 0.2857 |

| | | | | | | | | | | | | | | | | | | | | |
|---|---|---|---|---|---|---|---|---|---|---|---|---|---|---|---|---|---|---|---|---|
| 2_8208822 | 1 | 0 | 2 | 0 | 1 | 1 | 4 | 0 | 8 | 0.125 | NA | NA | NA | NA | 1 | 0 | 5 | 0.4 | 6 | 0.3333 |
| 2_8208831 | 1 | 1 | 1 | 0 | 2 | 1 | 5 | 0.6 | 9 | 0.6667 | NA | NA | 1 | 0 | NA | NA | 6 | 0.8333 | 7 | 0.7143 |
| 2_8208833 | NA | NA | 1 | 0 | 1 | 0 | 5 | 0.2 | 7 | 0.1429 | NA | NA | 2 | 0.5 | 1 | 0 | 4 | 0 | 7 | 0.1429 |
| 2_8208844 | 1 | 1 | 2 | 0.5 | 2 | 1 | 5 | 0 | 10 | 0.4 | NA | NA | 1 | 0 | 2 | 1 | 6 | 0.3333 | 9 | 0.4444 |
| 2_8208869 | 1 | 1 | 1 | 1 | 2 | 1 | 6 | 1 | 10 | 1 | NA | NA | 2 | 1 | 2 | 1 | 5 | 1 | 9 | 1 |
| 2_8208871 | 1 | 1 | 2 | 0.5 | 2 | 0 | 6 | 1 | 11 | 0.7273 | NA | NA | 2 | 0 | 2 | 0.5 | 5 | 0.4 | 9 | 0.3333 |
| 2_8208874 | 1 | 0 | 1 | 0 | 2 | 0.5 | 6 | 0.1667 | 10 | 0.2 | 1 | 0 | 2 | 0 | 2 | 0 | 6 | 0.3333 | 11 | 0.1818 |
| 2_8208884 | 1 | 0 | 1 | 0 | 2 | 0.5 | 6 | 0.5 | 10 | 0.4 | 2 | 1 | 2 | 0.5 | 2 | 0 | 4 | 0.5 | 10 | 0.5 |
| 2_8208891 | 1 | 1 | 2 | 0.5 | 2 | 0.5 | 4 | 0.75 | 9 | 0.6667 | 2 | 0 | 2 | 0.5 | 2 | 0.5 | 5 | 0.4 | 11 | 0.3636 |
| 2_8208893 | NA | NA | NA | NA | 1 | 0 | 6 | 0.1667 | 7 | 0.1429 | 2 | 0.5 | 2 | 0 | 1 | 0 | 6 | 0.1667 | 11 | 0.1818 |
| 2_8208896 | 1 | 0 | 2 | 0 | 2 | 0 | 6 | 0.3333 | 11 | 0.1818 | 2 | 0 | 2 | 0 | 2 | 0 | 5 | 0.2 | 11 | 0.0909 |
| 2_8208902 | 1 | 0 | 1 | 1 | 2 | 1 | 2 | 0.5 | 6 | 0.6667 | 1 | 1 | 2 | 1 | 2 | 1 | 5 | 1 | 10 | 1 |
| 2_8208903 | NA | NA | 1 | 0 | 3 | 1 | 5 | 0 | 9 | 0.3333 | 1 | 0 | 2 | 0.5 | 1 | 0 | 4 | 0.75 | 8 | 0.5 |
| 2_8208907 | 1 | 0 | 2 | 0 | 2 | 0.5 | 5 | 0.2 | 10 | 0.2 | 2 | 0 | 2 | 0 | 2 | 0 | 2 | 0.5 | 8 | 0.125 |
| 2_8208932 | 1 | 0 | NA | NA | 1 | 0 | 3 | 0.6667 | 5 | 0.4 | 2 | 1 | 2 | 1 | 2 | 0.5 | 5 | 0 | 11 | 0.4545 |
| 2_8208950 | 1 | 0 | 2 | 0.5 | 3 | 0.6667 | 5 | 0.4 | 11 | 0.4545 | 2 | 0.5 | 2 | 1 | 3 | 0.6667 | 5 | 0.2 | 12 | 0.5 |
| 2_8208956 | 2 | 0 | 3 | 0 | 3 | 0.3333 | 6 | 0.3333 | 14 | 0.2143 | 2 | 0 | 2 | 0 | 3 | 0 | 6 | 0.1667 | 13 | 0.0769 |
| 2_8208959 | NA | NA | 3 | 1 | 4 | 1 | 4 | 1 | 11 | 1 | 1 | 1 | 2 | 1 | 3 | 0.6667 | 5 | 1 | 11 | 0.9091 |
| 2_8208960 | 2 | 1 | 3 | 0.6667 | 3 | 0.3333 | 8 | 0.375 | 16 | 0.5 | 2 | 0.5 | 2 | 0.5 | 3 | 1 | 6 | 0.3333 | 13 | 0.5385 |
| 2_8208979 | 3 | 0.6667 | 4 | 1 | 4 | 1 | 8 | 0.875 | 19 | 0.8947 | 2 | 1 | 2 | 1 | 3 | 1 | 8 | 0.875 | 15 | 0.9333 |
| 2_8208984 | 2 | 1 | 4 | 0.75 | 3 | 0.6667 | 7 | 0.7143 | 16 | 0.75 | 2 | 0.5 | 1 | 1 | 3 | 0.6667 | 7 | 0.7143 | 13 | 0.6923 |
| 2_8208989 | 1 | 0 | 2 | 1 | 2 | 0.5 | 6 | 0.1667 | 11 | 0.3636 | 2 | 0.5 | 2 | 0.5 | 2 | 0 | 6 | 0.8333 | 12 | 0.5833 |
| 2_8209526 | NA | NA | NA | NA | 2 | 0.5 | NA | NA | 2 | 0.5 | NA | NA | 1 | 1 | NA | NA | NA | NA | 1 | 1 |
| 2_8349423 | 8 | 0 | 13 | 0.1538 | 8 | 0.5 | 13 | 0.3077 | 42 | 0.2381 | 9 | 0.2222 | 10 | 0 | 14 | 0.0714 | 10 | 0.1 | 43 | 0.093 |
| 2_8349433 | 8 | 0.125 | 13 | 0.2308 | 10 | 0.1 | 13 | 0.1538 | 44 | 0.1591 | 9 | 0 | 13 | 0 | 17 | 0.1765 | 13 | 0 | 52 | 0.0577 |
| 2_8349439 | 11 | 1 | 17 | 1 | 11 | 1 | 15 | 1 | 54 | 1 | 8 | 1 | 13 | 1 | 15 | 1 | 15 | 0.9333 | 51 | 0.9804 |
| 2_8349440 | 9 | 0.6667 | 14 | 0.7857 | 9 | 0.8889 | 16 | 1 | 48 | 0.8542 | 9 | 0.8889 | 12 | 0.9167 | 14 | 0.7143 | 9 | 0.8889 | 44 | 0.8409 |
| 2_8349469 | 12 | 0.0833 | 17 | 0.0588 | 10 | 0.1 | 18 | 0 | 57 | 0.0526 | 10 | 0.1 | 17 | 0.1176 | 14 | 0 | 17 | 0.0588 | 58 | 0.069 |
| 2_8349475 | 12 | 0.0833 | 19 | 0.2105 | 12 | 0.0833 | 17 | 0.3529 | 60 | 0.2 | 10 | 0.4 | 20 | 0.1 | 15 | 0.2 | 18 | 0.2778 | 63 | 0.2222 |
| 2_8349483 | 11 | 0.3636 | 15 | 0.2667 | 8 | 0.125 | 15 | 0.1333 | 49 | 0.2245 | 11 | 0.1818 | 17 | 0.2353 | 13 | 0.1538 | 13 | 0.1538 | 54 | 0.1852 |
| 2_8349484 | 13 | 0.0769 | 17 | 0.1765 | 8 | 0.125 | 16 | 0.25 | 54 | 0.1667 | 11 | 0.3636 | 17 | 0.1176 | 12 | 0.1667 | 17 | 0.1765 | 57 | 0.193 |
| 2_8349485 | 14 | 0.2143 | 19 | 0.3684 | 12 | 0.4167 | 18 | 0.1111 | 63 | 0.2698 | 12 | 0.25 | 18 | 0.6111 | 16 | 0.5 | 18 | 0.2222 | 64 | 0.4062 |
| 2_8355437 | 1 | 1 | NA | NA | NA | NA | NA | NA | 1 | 1 | 1 | 0 | NA | NA | 1 | 0 | 1 | 1 | 3 | 0.3333 |
| 2_8355450 | 1 | 1 | NA | NA | NA | NA | NA | NA | 1 | 1 | NA | NA | NA | NA | NA | NA | 1 | 1 | 1 | 1 |
| 2_8355457 | 1 | 1 | NA | NA | NA | NA | NA | NA | 1 | 1 | NA | NA | NA | NA | 1 | 0 | 1 | 1 | 2 | 0.5 |
| 2_8355462 | 1 | 1 | NA | NA | NA | NA | NA | NA | 1 | 1 | NA | NA | NA | NA | 1 | 1 | 1 | 1 | 2 | 1 |
| 2_8355464 | 1 | 1 | NA | NA | NA | NA | NA | NA | 1 | 1 | NA | NA | NA | NA | NA | NA | 1 | 1 | 1 | 1 |
| 2_8363375 | 5 | 0 | 4 | 0.5 | 2 | 0 | 5 | 0.2 | 16 | 0.1875 | 3 | 0 | 7 | 0 | 2 | 0.5 | 4 | 0.5 | 16 | 0.1875 |
| 2_8363377 | 6 | 0.8333 | 7 | 1 | 3 | 1 | 5 | 0.8 | 21 | 0.9048 | 5 | 1 | 7 | 0.8571 | 2 | 1 | 7 | 0.8571 | 21 | 0.9048 |
| 2_8363390 | 6 | 0.3333 | 5 | 0.2 | 3 | 0.3333 | 7 | 0.1429 | 21 | 0.2381 | 5 | 0.4 | 8 | 0.125 | 2 | 0 | 7 | 0.2857 | 22 | 0.2273 |
| 2_8363401 | 7 | 0.2857 | 8 | 0.375 | 3 | 0.3333 | 5 | 0.2 | 23 | 0.3043 | 6 | 0.3333 | 8 | 0.375 | 2 | 0 | 8 | 0.5 | 24 | 0.375 |
| 2_8363415 | 7 | 0.1429 | 8 | 0.125 | 3 | 0 | 6 | 0 | 24 | 0.0833 | 6 | 0.3333 | 8 | 0 | 2 | 0 | 7 | 0 | 23 | 0.087 |
| 2_8363417 | 7 | 0.4286 | 9 | 0.4444 | 3 | 0.3333 | 6 | 0.6667 | 25 | 0.48 | 6 | 0.5 | 8 | 0.5 | 2 | 0.5 | 8 | 0.75 | 24 | 0.5833 |
| 2_8363420 | 6 | 0.1667 | 8 | 0.25 | 3 | 0 | 6 | 0.3333 | 23 | 0.2174 | 6 | 0.5 | 8 | 0 | 2 | 0.5 | 8 | 0.125 | 24 | 0.2083 |
| 2_8363424 | 7 | 0.2857 | 9 | 0.7778 | 1 | 0 | 7 | 0.5714 | 24 | 0.5417 | 6 | 0.5 | 7 | 0.2857 | 1 | 0 | 5 | 0 | 19 | 0.2632 |
| 2_8363464 | 7 | 0 | 9 | 0.1111 | 3 | 0.6667 | 6 | 0 | 25 | 0.12 | 8 | 0.25 | 7 | 0.4286 | 4 | 0.5 | 10 | 0.2 | 29 | 0.3103 |
| 2_8363466 | 7 | 0.1429 | 9 | 0 | 3 | 0.3333 | 6 | 0 | 25 | 0.08 | 8 | 0.125 | 8 | 0 | 4 | 0 | 10 | 0.1 | 30 | 0.0667 |
| 2_8363500 | 7 | 0 | 6 | 0.1667 | 2 | 1 | 5 | 0 | 20 | 0.15 | 9 | 0.1111 | 10 | 0.1 | 4 | 0.25 | 7 | 0 | 30 | 0.1 |
| 2_8363662 | 9 | 0 | 11 | 0 | 11 | 0.1818 | 9 | 0.1111 | 40 | 0.075 | 13 | 0.0769 | 10 | 0.1 | 10 | 0 | 13 | 0.0769 | 46 | 0.0652 |
| 2_8363671 | 12 | 0.25 | 12 | 0.25 | 12 | 0.5 | 10 | 0.4 | 46 | 0.3478 | 15 | 0.4667 | 12 | 0.25 | 11 | 0.1818 | 17 | 0.2353 | 55 | 0.2909 |
| 2_8368387 | 9 | 0 | 12 | 0.25 | 9 | 0 | 13 | 0.0769 | 43 | 0.093 | 4 | 0 | 10 | 0 | 8 | 0.125 | 13 | 0.1538 | 35 | 0.0857 |
| 2_8370956 | 11 | 0.3636 | 15 | 0.4667 | 13 | 0.3077 | 16 | 0.25 | 55 | 0.3455 | 10 | 0.4 | 14 | 0.2857 | 15 | 0.5333 | 15 | 0.6 | 54 | 0.463 |
| 2_8370969 | 9 | 0.2222 | 15 | 0.3333 | 13 | 0.4615 | 17 | 0.6471 | 54 | 0.4444 | 12 | 0.4167 | 15 | 0.2667 | 16 | 0.5625 | 14 | 0.4286 | 57 | 0.4211 |
| 2_8370979 | 11 | 0.0909 | 16 | 0.0625 | 14 | 0 | 12 | 0.1667 | 53 | 0.0755 | 8 | 0 | 12 | 0.0833 | 14 | 0.0714 | 14 | 0.0714 | 48 | 0.0625 |
| 2_8370986 | 13 | 0.1538 | 14 | 0 | 13 | 0.2308 | 17 | 0.2353 | 57 | 0.1579 | 10 | 0.3 | 14 | 0.4286 | 13 | 0.6154 | 12 | 0.25 | 49 | 0.4082 |
| 2_8370995 | 13 | 0.0769 | 17 | 0.0588 | 16 | 0.0625 | 18 | 0.2778 | 64 | 0.125 | 13 | 0.1538 | 21 | 0.0476 | 17 | 0.0588 | 14 | 0 | 65 | 0.0615 |
| 2_8371001 | 8 | 0.5 | 14 | 0.6429 | 13 | 0.7692 | 12 | 0.25 | 47 | 0.5532 | 12 | 0.75 | 18 | 0.6111 | 15 | 0.7333 | 13 | 0.7692 | 58 | 0.7069 |
| 2_8371004 | 11 | 0.0909 | 13 | 0.1538 | 11 | 0.3636 | 15 | 0.2667 | 50 | 0.22 | 11 | 0.1818 | 16 | 0.25 | 14 | 0 | 11 | 0 | 52 | 0.1154 |
| 2_8371005 | 10 | 1 | 16 | 0.875 | 13 | 0.9231 | 16 | 0.875 | 55 | 0.9091 | 16 | 1 | 18 | 0.8889 | 16 | 1 | 13 | 0.6154 | 63 | 0.8889 |
| 2_8371006 | 10 | 0.3 | 14 | 0.5 | 13 | 0.1538 | 16 | 0.25 | 53 | 0.3019 | 10 | 0.3 | 18 | 0 | 13 | 0.4615 | 13 | 0.1538 | 54 | 0.2037 |
| 2_8371022 | 12 | 0.9167 | 14 | 0.8571 | 13 | 0.9231 | 10 | 1 | 49 | 0.9184 | 12 | 0.75 | 16 | 0.9375 | 14 | 0.9286 | 8 | 1 | 50 | 0.9 |
| 2_8371026 | 13 | 0 | 17 | 0.0588 | 14 | 0 | 18 | 0.2222 | 62 | 0.0806 | 15 | 0 | 20 | 0.1 | 18 | 0.1111 | 14 | 0.0714 | 67 | 0.0746 |
| 2_8371036 | 14 | 0.2143 | 16 | 0.1875 | 16 | 0.3125 | 17 | 0.2941 | 63 | 0.254 | 14 | 0.3571 | 23 | 0.1739 | 17 | 0.5882 | 17 | 0.2941 | 71 | 0.338 |
| 2_8371037 | 1 | 1 | NA | NA | NA | NA | NA | NA | 1 | 1 | NA | NA | NA | NA | NA | NA | 1 | 1 | 1 | 1 |
| 2_8371038 | 12 | 0.6667 | 17 | 0.8235 | 12 | 0.9167 | 17 | 0.8824 | 58 | 0.8276 | 11 | 0.7273 | 19 | 0.7895 | 16 | 0.8125 | 16 | 0.75 | 62 | 0.7742 |

| | | | | | | | | | | | | | | | | | | | | |
|---|---|---|---|---|---|---|---|---|---|---|---|---|---|---|---|---|---|---|---|---|
| 2_83710428 | 14 | 0.7857 | 16 | 0.9375 | 15 | 0.5333 | 15 | 0.8667 | 60 | 0.7833 | 15 | 0.7333 | 16 | 0.6875 | 15 | 0.8 | 17 | 0.8235 | 63 | 0.7619 |
| 2_83710530 | 14 | 0.1429 | 18 | 0.3333 | 14 | 0.0714 | 17 | 0.2941 | 63 | 0.2222 | 15 | 0.1333 | 23 | 0.087 | 16 | 0.1875 | 21 | 0.1905 | 75 | 0.1467 |
| 2_83710536 | 15 | 0.2 | 19 | 0.0526 | 17 | 0.0588 | 18 | 0.0556 | 69 | 0.087 | 15 | 0.1333 | 23 | 0.0435 | 16 | 0 | 19 | 0.1579 | 73 | 0.0822 |
| 2_83710745 | 16 | 0.5 | 21 | 0.5714 | 16 | 0.375 | 18 | 0.4444 | 71 | 0.4789 | 15 | 0.4 | 22 | 0.3182 | 17 | 0.5294 | 19 | 0.2632 | 73 | 0.3699 |
| 2_83710791 | 16 | 0.0625 | 19 | 0.1053 | 15 | 0.1333 | 18 | 0.1111 | 68 | 0.1029 | 15 | 0.2 | 20 | 0.25 | 16 | 0.125 | 16 | 0 | 67 | 0.1493 |
| 2_83710860 | 16 | 0.9375 | 19 | 1 | 14 | 0.9286 | 16 | 0.9375 | 65 | 0.9538 | 17 | 1 | 22 | 0.8636 | 13 | 1 | 11 | 1 | 63 | 0.9524 |
| 2_83710878 | 18 | 1 | 22 | 1 | 19 | 1 | 19 | 1 | 78 | 1 | 17 | 1 | 26 | 0.9615 | 19 | 0.9474 | 20 | 1 | 82 | 0.9756 |
| 2_83710902 | 16 | 0.8125 | 22 | 0.5 | 17 | 0.4118 | 19 | 0.5263 | 74 | 0.5541 | 17 | 0.6471 | 23 | 0.6957 | 19 | 0.5789 | 19 | 0.5789 | 78 | 0.6282 |
| 2_83711032 | 18 | 0.1111 | 20 | 0.25 | 19 | 0.1053 | 16 | 0.0625 | 73 | 0.137 | 18 | 0.1111 | 25 | 0.08 | 16 | 0.0625 | 15 | 0 | 74 | 0.0676 |
| 2_83711197 | 17 | 0.9412 | 24 | 1 | 20 | 1 | 17 | 0.9412 | 78 | 0.9744 | 17 | 1 | 29 | 1 | 15 | 1 | 18 | 0.9444 | 79 | 0.9873 |
| 2_83711262 | 20 | 0.15 | 26 | 0.1154 | 19 | 0.3158 | 19 | 0.2105 | 84 | 0.1905 | 19 | 0.0526 | 28 | 0.1429 | 20 | 0.25 | 19 | 0.0526 | 86 | 0.1279 |
| 2_83711426 | 19 | 0.1053 | 26 | 0.1923 | 21 | 0.2381 | 18 | 0.0556 | 84 | 0.1548 | 20 | 0.2 | 25 | 0.12 | 19 | 0.2105 | 21 | 0.1429 | 85 | 0.1647 |
| 2_83711608 | 20 | 0.3 | 29 | 0.4483 | 23 | 0.3913 | 22 | 0.1364 | 94 | 0.3298 | 20 | 0.35 | 28 | 0.2857 | 20 | 0.2 | 24 | 0.3333 | 92 | 0.2935 |
| 2_83711661 | 22 | 0.2727 | 24 | 0.5 | 23 | 0.5652 | 23 | 0.3043 | 92 | 0.413 | 19 | 0.6316 | 24 | 0.4167 | 20 | 0.6 | 21 | 0.4762 | 84 | 0.5238 |
| 2_83711915 | 20 | 0.05 | 26 | 0.1154 | 22 | 0.0909 | 23 | 0.1739 | 91 | 0.1099 | 17 | 0 | 27 | 0.1111 | 22 | 0.2273 | 18 | 0.1667 | 84 | 0.131 |
| 2_83712015 | 21 | 0.1429 | 30 | 0.2 | 24 | 0 | 22 | 0 | 97 | 0.0928 | 24 | 0.2083 | 29 | 0.1034 | 22 | 0.0909 | 22 | 0.1364 | 97 | 0.134 |
| 2_83712132 | 23 | 0 | 33 | 0.0606 | 26 | 0.0769 | 24 | 0.0833 | 106 | 0.0566 | 23 | 0.1304 | 32 | 0.0938 | 24 | 0 | 23 | 0.0435 | 102 | 0.0686 |
| 2_83712204 | 24 | 0.0417 | 33 | 0.0909 | 24 | 0.0833 | 24 | 0.0417 | 105 | 0.0667 | 23 | 0.1739 | 34 | 0.0588 | 24 | 0.0833 | 26 | 0.0385 | 107 | 0.0841 |
| 2_83712442 | 25 | 0.44 | 31 | 0.4194 | 26 | 0.4615 | 25 | 0.2 | 107 | 0.3832 | 25 | 0.52 | 37 | 0.3514 | 25 | 0.36 | 25 | 0.56 | 112 | 0.4375 |
| 2_83712509 | 22 | 0.0455 | 27 | 0.1111 | 24 | 0.125 | 20 | 0.15 | 93 | 0.1075 | 24 | 0.1667 | 31 | 0.0968 | 21 | 0.1429 | 21 | 0.0476 | 97 | 0.1134 |
| 2_83713501 | 22 | 0.0455 | 31 | 0.129 | 30 | 0.1 | 23 | 0 | 106 | 0.0755 | 22 | 0.1364 | 30 | 0.0333 | 29 | 0.069 | 24 | 0.2083 | 105 | 0.1048 |
| 2_84163408 | 2 | 1 | NA | NA | NA | NA | NA | NA | 2 | 1 | NA | NA | 2 | 1 | 1 | 1 | NA | NA | 3 | 1 |
| 2_84163443 | 1 | 1 | NA | NA | NA | NA | NA | NA | 1 | 1 | NA | NA | 1 | 1 | 1 | 1 | NA | NA | 2 | 1 |
| 2_84163600 | 2 | 1 | NA | NA | NA | NA | NA | NA | 2 | 1 | NA | NA | 2 | 1 | NA | NA | NA | NA | 2 | 1 |
| 2_84163773 | 1 | 1 | NA | NA | NA | NA | NA | NA | 1 | 1 | NA | NA | 2 | 0.5 | 1 | 1 | NA | NA | 3 | 0.6667 |
| 2_84163876 | 2 | 1 | NA | NA | NA | NA | NA | NA | 2 | 1 | NA | NA | 2 | 1 | 1 | 1 | NA | NA | 3 | 1 |
| 2_84336012 | NA | NA | 1 | 1 | NA | NA | 1 | 1 | 2 | 1 | 1 | 0 | 1 | 1 | 3 | 0.6667 | NA | NA | 5 | 0.6 |
| 2_84432241 | 5 | 0 | 6 | 0.1667 | NA | NA | 4 | 0 | 15 | 0.0667 | 4 | 0.25 | NA | NA | 3 | 0 | 3 | 0 | 10 | 0.1 |
| 2_84432397 | 4 | 0 | 7 | 0.1429 | NA | NA | 3 | 0 | 14 | 0.0714 | 4 | 0.5 | 2 | 0 | NA | NA | 3 | 0 | 9 | 0.2222 |
| 2_84432592 | 4 | 0.5 | 6 | 0.6667 | NA | NA | 4 | 0.25 | 14 | 0.5 | 4 | 0.5 | 2 | 0.5 | 1 | 1 | 2 | 0 | 9 | 0.4444 |
| 2_84432600 | 4 | 0 | 6 | 0.5 | NA | NA | 3 | 0.6667 | 13 | 0.3846 | 5 | 0.6 | 1 | 0 | NA | NA | 3 | 0.6667 | 9 | 0.5556 |
| 2_84432638 | 3 | 0.3333 | 5 | 0.4 | NA | NA | 1 | 0 | 9 | 0.3333 | 2 | 1 | 2 | 0.5 | NA | NA | 3 | 0.6667 | 7 | 0.7143 |
| 2_84432785 | 4 | 0.25 | 6 | 0 | NA | NA | 3 | 0 | 13 | 0.0769 | 5 | 0 | 2 | 0.5 | 1 | 0 | 2 | 0.5 | 10 | 0.2 |
| 2_84432809 | 4 | 0.5 | 5 | 0.4 | NA | NA | 4 | 0 | 13 | 0.3077 | 3 | 0.3333 | 2 | 0 | 1 | 0 | 1 | 0 | 7 | 0.1429 |
| 2_84432822 | 4 | 0.25 | 6 | 0.5 | NA | NA | 4 | 0 | 14 | 0.2857 | 4 | 0.5 | 1 | 1 | 1 | 0 | 2 | 0 | 8 | 0.375 |
| 2_84432829 | 2 | 0.5 | 3 | 0 | NA | NA | 3 | 0.6667 | 8 | 0.375 | 4 | 0 | 1 | 0 | 1 | 1 | 2 | 1 | 8 | 0.375 |
| 2_84432874 | 3 | 1 | 5 | 0.8 | NA | NA | 4 | 1 | 12 | 0.9167 | 4 | 0.75 | 2 | 0 | 1 | 1 | 2 | 1 | 9 | 0.6667 |
| 2_84432900 | 4 | 0.25 | 6 | 0.5 | NA | NA | 4 | 0.25 | 14 | 0.3571 | 2 | 1 | 2 | 0 | 1 | 1 | 1 | 0 | 6 | 0.5 |
| 2_84432934 | 2 | 1 | 3 | 0.6667 | NA | NA | 3 | 1 | 8 | 0.875 | 3 | 1 | 1 | 1 | 1 | 1 | 1 | 1 | 6 | 1 |
| 2_84432973 | 4 | 0.75 | 4 | 0.75 | NA | NA | 4 | 0.75 | 12 | 0.75 | 2 | 0 | 2 | 0 | 1 | 0 | 2 | 0.5 | 7 | 0.1429 |
| 2_84432999 | 3 | 0.3333 | 5 | 0 | NA | NA | 4 | 0.5 | 12 | 0.25 | 2 | 1 | 1 | 0 | 1 | 1 | 2 | 0.5 | 6 | 0.6667 |
| 2_84433142 | 3 | 0.6667 | 5 | 0.8 | NA | NA | 2 | 1 | 10 | 0.8 | 2 | 0 | 1 | 0 | 1 | 1 | 1 | 0 | 5 | 0.2 |
| 2_84433171 | 4 | 1 | 6 | 1 | NA | NA | 4 | 1 | 14 | 1 | 2 | 1 | 2 | 1 | 1 | 1 | 1 | 1 | 6 | 1 |
| 2_84447040 | 1 | 1 | 2 | 0 | NA | NA | 1 | 0 | 4 | 0.25 | NA | NA | 2 | 0 | 1 | 1 | NA | NA | 3 | 0.3333 |
| 2_84501672 | 31 | 0.129 | 34 | 0.1471 | 32 | 0.125 | 26 | 0.1923 | 123 | 0.1463 | 35 | 0.1429 | 33 | 0.0909 | 29 | 0.2069 | 33 | 0.2121 | 130 | 0.1615 |
| 2_84501717 | 28 | 0.1429 | 32 | 0.25 | 32 | 0.125 | 27 | 0.3333 | 119 | 0.2101 | 38 | 0.2105 | 24 | 0.1667 | 25 | 0.2 | 29 | 0.2759 | 116 | 0.2155 |
| 2_84501790 | 38 | 0.3421 | 40 | 0.375 | 40 | 0.35 | 27 | 0.4815 | 145 | 0.3793 | 37 | 0.3514 | 33 | 0.303 | 38 | 0.3158 | 42 | 0.4524 | 150 | 0.36 |
| 2_84502379 | 28 | 0 | 41 | 0.122 | 39 | 0.0513 | 29 | 0.0345 | 137 | 0.0584 | 28 | 0.0714 | 35 | 0.0571 | 37 | 0.0541 | 35 | 0.0286 | 135 | 0.0519 |
| 2_84502690 | 25 | 0.28 | 36 | 0.2778 | 31 | 0.2581 | 28 | 0.1429 | 120 | 0.2417 | 21 | 0.2857 | 32 | 0.25 | 32 | 0.25 | 30 | 0.2667 | 115 | 0.2609 |
| 2_84502740 | 28 | 0.4643 | 38 | 0.5 | 32 | 0.25 | 27 | 0.5185 | 125 | 0.432 | 24 | 0.5417 | 34 | 0.5 | 33 | 0.4545 | 33 | 0.3939 | 124 | 0.4677 |
| 2_84502782 | 27 | 0.0741 | 37 | 0.0541 | 32 | 0.0312 | 27 | 0.1111 | 123 | 0.065 | 24 | 0.0833 | 32 | 0.1562 | 35 | 0.0286 | 34 | 0.1765 | 125 | 0.112 |
| 2_84517283 | 9 | 0 | 5 | 0 | 5 | 0.2 | 8 | 0.125 | 27 | 0.0741 | 5 | 0.2 | 6 | 0.1667 | 6 | 0.3333 | 4 | 0.25 | 21 | 0.2381 |
| 2_84517291 | 5 | 0.2 | 3 | 0 | 3 | 0 | 6 | 0.3333 | 17 | 0.1765 | 2 | 0.5 | 6 | 0.3333 | 3 | 0.3333 | 3 | 0.3333 | 14 | 0.3571 |
| 2_84517338 | 7 | 0 | 6 | 0.1667 | 4 | 0.5 | 9 | 0.2222 | 26 | 0.1923 | 2 | 0 | 5 | 0.2 | 6 | 0 | 3 | 0.3333 | 16 | 0.125 |
| 2_84517437 | 8 | 0 | 6 | 0 | 6 | 0.1667 | 10 | 0.2 | 30 | 0.1 | 5 | 0 | 6 | 0 | 6 | 0 | 4 | 0.5 | 21 | 0.0952 |
| 2_84517541 | 9 | 0.1111 | 6 | 0 | 5 | 0.4 | 11 | 0.1818 | 31 | 0.1613 | 5 | 0.2 | 8 | 0.25 | 6 | 0.1667 | 2 | 0 | 21 | 0.1905 |
| 2_84517700 | 8 | 0.25 | 6 | 0.3333 | 6 | 0.5 | 13 | 0.2308 | 33 | 0.303 | 5 | 0.4 | 9 | 0 | 5 | 0.2 | 3 | 0.3333 | 22 | 0.1818 |
| 2_84518814 | 6 | 0.1667 | 6 | 0 | 4 | 0 | 3 | 0 | 19 | 0.0526 | 1 | 0 | 7 | 0 | 4 | 0 | 8 | 0.125 | 20 | 0.05 |
| 2_84519369 | 2 | 1 | 5 | 0.8 | 4 | 0.75 | 2 | 0 | 13 | 0.6923 | 1 | 0 | 10 | 0.5 | 6 | 0.6667 | 7 | 0.7143 | 24 | 0.5833 |
| 2_84519383 | 3 | 0.3333 | 6 | 0 | 2 | 0 | 3 | 0 | 14 | 0.0714 | 1 | 0 | 7 | 0.1429 | 5 | 0 | 7 | 0 | 20 | 0.05 |
| 2_84519443 | 3 | 0.3333 | 6 | 0.1667 | 3 | 0 | 3 | 0 | 15 | 0.1333 | 1 | 0 | 10 | 0.4 | 7 | 0.1429 | 7 | 0.2857 | 25 | 0.28 |
| 2_84519451 | 3 | 0 | 5 | 0 | 4 | 0.25 | 3 | 0 | 15 | 0.0667 | 1 | 0 | 9 | 0.1111 | 7 | 0.1429 | 5 | 0.2 | 22 | 0.1364 |
| 2_84519525 | 3 | 1 | 4 | 1 | 4 | 1 | 3 | 0.3333 | 14 | 0.8571 | 1 | 0 | 8 | 0.875 | 6 | 1 | 6 | 1 | 21 | 0.9048 |
| 2_84520322 | 2 | 0 | 4 | 0.5 | 3 | 0 | 2 | 0 | 11 | 0.1818 | 1 | 0 | 6 | 0 | 3 | 0 | 6 | 0.1667 | 16 | 0.0625 |
| 2_84520384 | 2 | 0 | 5 | 0.6 | 4 | 0.25 | 3 | 0.6667 | 14 | 0.4286 | 1 | 1 | 5 | 0.8 | 2 | 0 | 6 | 0.6667 | 14 | 0.6429 |

| | | | | | | | | | | | | | | | | | | | | |
|---|---|---|---|---|---|---|---|---|---|---|---|---|---|---|---|---|---|---|---|---|
| 2_84520473 | 2 | 0.5 | 5 | 0.2 | 4 | 0.25 | 3 | 0 | 14 | 0.2143 | 1 | 0 | 7 | 0.1429 | 5 | 0 | 5 | 0.2 | 18 | 0.1111 |
| 2_84520575 | 2 | 0.5 | 3 | 0.3333 | 4 | 0 | 2 | 0 | 11 | 0.1818 | 1 | 0 | 6 | 0.3333 | 5 | 0 | 3 | 0 | 15 | 0.1333 |
| 2_84520595 | 2 | 0 | 5 | 0 | 4 | 0 | 2 | 0.5 | 13 | 0.0769 | 1 | 1 | 6 | 0 | 4 | 0.25 | 3 | 0.6667 | 14 | 0.2857 |
| 2_84520789 | 2 | 0 | 5 | 0.2 | 3 | 0.3333 | 3 | 0.3333 | 13 | 0.2308 | 1 | 0 | 7 | 0 | 5 | 0.2 | 4 | 0 | 17 | 0.0588 |
| 2_84541305 | 2 | 0 | 3 | 0.3333 | 1 | 0 | NA | NA | 6 | 0.1667 | NA | NA | 1 | 0 | 2 | 0.5 | 4 | 0 | 7 | 0.1429 |
| 2_84541347 | 2 | 1 | 2 | 1 | 1 | 1 | NA | NA | 5 | 1 | NA | NA | 1 | 1 | 1 | 1 | 4 | 1 | 6 | 1 |
| 2_84541446 | 2 | 0.5 | 3 | 0.6667 | 1 | 1 | NA | NA | 6 | 0.6667 | NA | NA | NA | NA | 2 | 1 | 4 | 0.5 | 6 | 0.6667 |
| 2_84541602 | 2 | 1 | 3 | 1 | 1 | 1 | NA | NA | 6 | 1 | NA | NA | 1 | 1 | 2 | 1 | 3 | 1 | 6 | 1 |
| 2_84541648 | 2 | 1 | 3 | 0.6667 | 1 | 1 | NA | NA | 6 | 0.8333 | NA | NA | 1 | 1 | 2 | 0.5 | 4 | 0.75 | 7 | 0.7143 |
| 2_84541697 | 2 | 0.5 | 1 | 0 | 1 | 0 | NA | NA | 4 | 0.25 | NA | NA | NA | NA | 1 | 0 | 3 | 0.3333 | 4 | 0.25 |
| 2_84541721 | 2 | 1 | 2 | 1 | 1 | 0 | NA | NA | 5 | 0.8 | NA | NA | 1 | 1 | 2 | 1 | 4 | 0.5 | 7 | 0.7143 |
| 2_84541732 | 2 | 0 | 3 | 0.6667 | 1 | 0 | NA | NA | 6 | 0.3333 | NA | NA | 1 | 0 | 2 | 0.5 | 4 | 0.75 | 7 | 0.5714 |
| 2_84541756 | 2 | 1 | 3 | 0.3333 | 1 | 0 | NA | NA | 6 | 0.5 | NA | NA | 1 | 1 | 1 | 1 | 3 | 0.6667 | 5 | 0.8 |
| 2_84553695 | 1 | 1 | 1 | 0 | NA | NA | 2 | 0 | 4 | 0.25 | 1 | 0 | 1 | 1 | 1 | 0 | 2 | 0.5 | 5 | 0.4 |
| 2_84553754 | NA | NA | 1 | 1 | NA | NA | 1 | 0 | 2 | 0.5 | 1 | 0 | 2 | 1 | NA | NA | NA | NA | 3 | 0.6667 |
| 2_84553796 | 1 | 1 | 1 | 0 | NA | NA | 2 | 0 | 4 | 0.25 | 1 | 0 | 2 | 0 | 1 | 0 | 2 | 0.5 | 6 | 0.1667 |
| 2_84553823 | 1 | 1 | 1 | 1 | NA | NA | 1 | 1 | 3 | 1 | 1 | 1 | 2 | 1 | 1 | 1 | 2 | 1 | 6 | 1 |
| 2_84553933 | 1 | 1 | 1 | 1 | NA | NA | 2 | 1 | 4 | 1 | 1 | 1 | 2 | 1 | 1 | 1 | 1 | 1 | 5 | 1 |
| 2_84554038 | NA | NA | 1 | 1 | NA | NA | 2 | 0.5 | 3 | 0.6667 | 1 | 0 | 2 | 0.5 | 1 | 0 | NA | NA | 4 | 0.25 |
| 2_84554296 | 1 | 1 | 1 | 1 | NA | NA | 2 | 1 | 4 | 1 | 1 | 1 | 2 | 1 | 1 | 1 | 1 | 1 | 5 | 1 |
| 2_84554342 | 1 | 1 | NA | NA | NA | NA | 2 | 1 | 3 | 1 | 1 | 1 | 2 | 1 | 1 | 1 | NA | NA | 4 | 1 |
| 2_84554362 | 1 | 1 | 1 | 0 | NA | NA | 2 | 1 | 4 | 0.75 | 1 | 1 | 2 | 0.5 | 1 | 0 | 1 | 1 | 5 | 0.6 |
| 2_84556219 | 1 | 1 | 1 | 1 | NA | NA | 2 | 1 | 4 | 1 | 1 | 0 | 1 | 1 | 1 | 1 | NA | NA | 3 | 0.6667 |
| 2_84556244 | 1 | 1 | 1 | 1 | NA | NA | 2 | 1 | 4 | 1 | NA | NA | 1 | 1 | 1 | 1 | 1 | 1 | 3 | 1 |
| 2_84565958 | NA | NA | 3 | 1 | NA | NA | NA | NA | 3 | 1 | NA | NA | NA | NA | NA | NA | 1 | 1 | 1 | 1 |
| 2_84566450 | NA | NA | 3 | 0.3333 | NA | NA | NA | NA | 3 | 0.3333 | NA | NA | NA | NA | NA | NA | 1 | 1 | 1 | 1 |
| 2_84566478 | NA | NA | 3 | 1 | NA | NA | NA | NA | 3 | 1 | NA | NA | NA | NA | NA | NA | 1 | 1 | 1 | 1 |
| 2_84566667 | NA | NA | 2 | 1 | NA | NA | NA | NA | 2 | 1 | NA | NA | NA | NA | NA | NA | 1 | 1 | 1 | 1 |
| 2_84566825 | NA | NA | 3 | 0.3333 | NA | NA | NA | NA | 3 | 0.3333 | NA | NA | NA | NA | NA | NA | 2 | 0.5 | 2 | 0.5 |
| 2_84566835 | NA | NA | 2 | 1 | NA | NA | NA | NA | 2 | 1 | NA | NA | NA | NA | NA | NA | 1 | 1 | 1 | 1 |
| 2_84568384 | 1 | 0 | 6 | 0.1667 | 6 | 0.5 | 3 | 0 | 16 | 0.25 | 2 | 0 | 3 | 0.3333 | 4 | 0.5 | 4 | 0.25 | 13 | 0.3077 |
| 2_84595814 | 8 | 0.75 | 2 | 0.5 | 1 | 0 | NA | NA | 11 | 0.6364 | 3 | 1 | NA | NA | 3 | 0.3333 | NA | NA | 6 | 0.6667 |
| 2_84595916 | 5 | 0.2 | 1 | 0 | 2 | 0 | NA | NA | 8 | 0.125 | 2 | 0.5 | NA | NA | 2 | 0 | 1 | 0 | 5 | 0.2 |
| 2_84596077 | 8 | 0.25 | 1 | 0 | 2 | 0 | NA | NA | 11 | 0.1818 | 3 | 0.3333 | NA | NA | 3 | 0 | NA | NA | 6 | 0.1667 |
| 2_84603618 | 3 | 0.6667 | 2 | 0.5 | 2 | 0.5 | NA | NA | 7 | 0.5714 | 3 | 0.3333 | NA | NA | NA | NA | 1 | 0 | 4 | 0.25 |
| 2_84660217 | 7 | 0 | 10 | 0.1 | 7 | 0 | 4 | 0.25 | 28 | 0.0714 | 6 | 0 | 7 | 0.1429 | 6 | 0.1667 | 7 | 0.1429 | 26 | 0.1154 |
| 2_84660241 | 7 | 0.1429 | 9 | 0.4444 | 7 | 0 | 4 | 0.25 | 27 | 0.2222 | 5 | 0 | 6 | 0.1667 | 5 | 0 | 5 | 0.2 | 21 | 0.0952 |
| 2_84660488 | 6 | 0 | 13 | 0.1538 | 6 | 0 | 4 | 0 | 29 | 0.069 | 7 | 0.2857 | 8 | 0 | 10 | 0.2 | 9 | 0.1111 | 34 | 0.1471 |
| 2_84701742 | 7 | 0.1429 | 6 | 0.3333 | 5 | 0 | 1 | 0 | 19 | 0.1579 | 7 | 0 | 3 | 0 | 1 | 0 | 9 | 0.1111 | 20 | 0.05 |
| 2_84701762 | 7 | 0.8571 | 6 | 0.6667 | 4 | 0.5 | NA | NA | 17 | 0.7059 | 9 | 0.6667 | 4 | 0.75 | 1 | 0 | 9 | 0.5556 | 23 | 0.6087 |
| 2_84701838 | 6 | 0.5 | 7 | 0.5714 | 3 | 0.3333 | 1 | 1 | 17 | 0.5294 | 7 | 0.4286 | 4 | 0.5 | 1 | 0 | 8 | 0.5 | 20 | 0.45 |
| 2_84701962 | 2 | 1 | 6 | 1 | 3 | 1 | NA | NA | 11 | 1 | 6 | 1 | 2 | 1 | 1 | 1 | 8 | 1 | 17 | 1 |
| 2_84702013 | 5 | 0.8 | 6 | 1 | 2 | 1 | 1 | 1 | 14 | 0.9286 | 6 | 1 | 4 | 0.5 | NA | NA | 6 | 0.6667 | 16 | 0.75 |
| 2_84702041 | 5 | 0.6 | 3 | 0.3333 | 3 | 0 | 1 | 1 | 12 | 0.4167 | 4 | 0.25 | 4 | 0.5 | NA | NA | 4 | 0 | 12 | 0.25 |
| 2_84702112 | 5 | 0.8 | 6 | 0.8333 | 4 | 0.75 | 1 | 1 | 16 | 0.8125 | 7 | 0.7143 | 3 | 0.3333 | NA | NA | 8 | 0.875 | 18 | 0.7222 |
| 2_84702147 | 5 | 0.4 | 4 | 0.75 | 2 | 0 | 1 | 1 | 12 | 0.5 | 6 | 0.5 | 4 | 0.25 | NA | NA | 6 | 0.3333 | 16 | 0.375 |
| 2_84702290 | 4 | 0.25 | 4 | 0.75 | 1 | 1 | 1 | 1 | 10 | 0.6 | 8 | 0.625 | 3 | 0 | NA | NA | 6 | 0.3333 | 17 | 0.4118 |
| 2_84702369 | 5 | 0.6 | 6 | 0.3333 | 4 | 0 | 1 | 1 | 16 | 0.375 | 3 | 0.6667 | 4 | 0.25 | NA | NA | 5 | 0.2 | 12 | 0.3333 |
| 2_84702489 | 3 | 1 | 5 | 1 | 4 | 0.75 | 1 | 1 | 13 | 0.9231 | 7 | 0.8571 | 4 | 0.75 | NA | NA | 2 | 0.5 | 13 | 0.7692 |
| 2_84702660 | 4 | 0.25 | 5 | 0.4 | 3 | 0.3333 | 1 | 0 | 13 | 0.3077 | 5 | 0.2 | 3 | 0.3333 | NA | NA | 3 | 0 | 11 | 0.1818 |
| 2_84702936 | 4 | 0.25 | 5 | 0.6 | 4 | 0.5 | 1 | 0 | 14 | 0.4286 | 7 | 0.4286 | 3 | 0.6667 | NA | NA | 5 | 0.6 | 15 | 0.5333 |
| 2_84710915 | 4 | 0 | 4 | 0.5 | 2 | 0 | 1 | 0 | 11 | 0.1818 | 4 | 0.25 | 4 | 0.25 | NA | NA | 4 | 0 | 12 | 0.1667 |
| 2_84711053 | 4 | 1 | 3 | 0.6667 | 2 | 1 | 1 | 1 | 10 | 0.9 | 6 | 0.8333 | 4 | 1 | NA | NA | 4 | 1 | 14 | 0.9286 |
| 2_84871118 | 5 | 0.2 | 3 | 0 | 8 | 0.25 | 5 | 0.2 | 21 | 0.1905 | NA | NA | 8 | 0.375 | 5 | 0 | 8 | 0 | 21 | 0.1429 |
| 2_84871827 | 5 | 0.2 | 3 | 0 | 6 | 0 | 5 | 0 | 19 | 0.0526 | NA | NA | 9 | 0.3333 | 6 | 0 | 7 | 0 | 22 | 0.1364 |
| 2_84877129 | 8 | 0.25 | 6 | 0.3333 | 8 | 0 | 5 | 0.4 | 27 | 0.2222 | 2 | 0.5 | 8 | 0.125 | 9 | 0.1111 | 5 | 0.4 | 24 | 0.2083 |
| 2_84877223 | 11 | 0.5455 | 10 | 0.8 | 14 | 0.5714 | 10 | 0.5 | 45 | 0.6 | 3 | 1 | 16 | 0.5625 | 12 | 0.5 | 10 | 0.7 | 41 | 0.6098 |
| 2_84877267 | 13 | 0.2308 | 11 | 0.0909 | 14 | 0.0714 | 12 | 0.0833 | 50 | 0.12 | 3 | 0.3333 | 16 | 0.1875 | 13 | 0.0769 | 10 | 0.2 | 42 | 0.1667 |
| 2_84877300 | 11 | 0.3636 | 10 | 0.3 | 12 | 0.3333 | 10 | 0.2 | 43 | 0.3023 | 2 | 0 | 13 | 0.3846 | 8 | 0.375 | 9 | 0.2222 | 32 | 0.3125 |
| 2_84893537 | 1 | 1 | NA | NA | 2 | 1 | 1 | 0 | 4 | 0.75 | NA | NA | NA | NA | 1 | 1 | NA | NA | 1 | 1 |
| 2_84893559 | 1 | 1 | NA | NA | 2 | 0.5 | 1 | 0 | 4 | 0.5 | NA | NA | NA | NA | 1 | 1 | NA | NA | 1 | 1 |
| 2_84893730 | NA | NA | NA | NA | 3 | 0.6667 | NA | NA | 3 | 0.6667 | NA | NA | NA | NA | 1 | 1 | NA | NA | 1 | 1 |
| 2_84894072 | 1 | 1 | NA | NA | 2 | 1 | NA | NA | 3 | 1 | 2 | 1 | 1 | 0 | 1 | 1 | NA | NA | 4 | 0.75 |
| 2_84894091 | 1 | 0 | NA | NA | 3 | 0.6667 | 1 | 0 | 5 | 0.4 | 2 | 0.5 | 1 | 0 | 1 | 0 | NA | NA | 4 | 0.25 |
| 2_85018816 | 4 | 0.5 | 2 | 0 | 4 | 0.25 | 4 | 0.75 | 14 | 0.4286 | NA | NA | 2 | 1 | 3 | 0.6667 | 2 | 0 | 7 | 0.5714 |

| | | | | | | | | | | | | | | | | | | | | |
|---|---|---|---|---|---|---|---|---|---|---|---|---|---|---|---|---|---|---|---|---|
| 2_8501911 | 2 | 0.5 | 2 | 0 | 3 | 0 | 2 | 0 | 9 | 0.1111 | NA | NA | 2 | 0.5 | 3 | 0 | 2 | 0 | 7 | 0.1429 |
| 2_8501955 | 2 | 0 | 2 | 0.5 | 3 | 0.6667 | 2 | 0.5 | 9 | 0.4444 | NA | NA | 2 | 0 | 3 | 0.3333 | 2 | 0 | 7 | 0.1429 |
| 2_8501963 | 2 | 0 | 2 | 0 | 3 | 0 | 2 | 0.5 | 9 | 0.1111 | NA | NA | 2 | 0 | 4 | 0.25 | 2 | 0 | 8 | 0.125 |
| 2_8501970 | 2 | 0 | 2 | 0 | 3 | 0 | 2 | 0.5 | 9 | 0.1111 | NA | NA | 2 | 0.5 | 4 | 0 | 2 | 0 | 8 | 0.125 |
| 2_8501978 | 2 | 0 | 2 | 0.5 | 3 | 0.3333 | 2 | 0 | 9 | 0.2222 | NA | NA | 1 | 0 | 2 | 0.5 | 2 | 0 | 5 | 0.2 |
| 2_8501987 | 2 | 0 | 2 | 0.5 | 2 | 1 | 2 | 0.5 | 8 | 0.5 | NA | NA | 2 | 0.5 | 3 | 0.6667 | 2 | 0.5 | 7 | 0.5714 |
| 2_8501997 | 2 | 0 | 2 | 0 | 2 | 0.5 | 2 | 0.5 | 8 | 0.25 | NA | NA | 1 | 1 | 4 | 0.25 | 2 | 0 | 7 | 0.2857 |
| 2_8502012 | 2 | 0 | 2 | 0 | 2 | 0 | 2 | 0.5 | 8 | 0.125 | NA | NA | 2 | 0.5 | 3 | 0.3333 | 2 | 0 | 7 | 0.2857 |
| 2_8502030 | 2 | 0 | 2 | 0 | 3 | 0.3333 | 2 | 0.5 | 9 | 0.2222 | NA | NA | 1 | 0 | 2 | 0 | 2 | 0.5 | 5 | 0.2 |
| 2_8502038 | 2 | 0 | 2 | 0 | 3 | 0.6667 | 2 | 0 | 9 | 0.2222 | NA | NA | 1 | 1 | 2 | 0 | 2 | 0 | 5 | 0.2 |
| 2_8502043 | 2 | 0 | 2 | 0 | 3 | 1 | 2 | 0 | 9 | 0.3333 | NA | NA | 1 | 0 | 2 | 0.5 | 2 | 0.5 | 5 | 0.4 |
| 2_8502049 | 2 | 0.5 | 1 | 1 | 2 | 1 | 2 | 1 | 7 | 0.8571 | NA | NA | NA | NA | 1 | 1 | 1 | 0 | 2 | 0.5 |
| 2_8502065 | 1 | 0 | NA | NA | 1 | 1 | 1 | 1 | 3 | 0.6667 | NA | NA | 1 | 1 | NA | NA | 2 | 0.5 | 3 | 0.6667 |
| 2_8502065 | 2 | 0 | 2 | 0.5 | 1 | 1 | 2 | 0 | 7 | 0.2857 | NA | NA | 1 | 0 | 1 | 1 | 2 | 0.5 | 4 | 0.5 |
| 2_8502087 | 2 | 0.5 | 2 | 1 | 1 | 1 | 2 | 1 | 7 | 0.8571 | NA | NA | 1 | 1 | 2 | 1 | 2 | 1 | 5 | 1 |
| 2_8502107 | 2 | 0 | 1 | 1 | 1 | 1 | 2 | 0 | 6 | 0.3333 | NA | NA | 1 | 0 | 1 | 1 | 2 | 0 | 4 | 0.25 |
| 2_8502149 | 1 | 1 | 1 | 1 | 2 | 1 | 2 | 1 | 6 | 1 | NA | NA | NA | NA | 2 | 1 | 2 | 0.5 | 4 | 0.75 |
| 2_8502186 | 1 | 1 | 1 | 0 | 2 | 0 | 2 | 0.5 | 6 | 0.3333 | NA | NA | NA | NA | 1 | 0 | 2 | 0.5 | 3 | 0.3333 |
| 2_8502208 | 1 | 1 | NA | NA | 2 | 1 | 2 | 1 | 5 | 1 | NA | NA | NA | NA | 2 | 0.5 | 2 | 1 | 4 | 0.75 |
| 2_8502220 | 1 | 0 | 1 | 1 | 2 | 0.5 | 2 | 1 | 6 | 0.6667 | NA | NA | NA | NA | 1 | 1 | 2 | 1 | 3 | 1 |
| 2_8502648 | 1 | 1 | 1 | 1 | 2 | 0.5 | 1 | 1 | 5 | 0.8 | NA | NA | NA | NA | 2 | 0.5 | 1 | 1 | 3 | 0.6667 |
| 2_8502648 | 1 | 1 | 1 | 1 | 2 | 1 | 1 | 1 | 5 | 1 | NA | NA | NA | NA | 2 | 1 | 2 | 1 | 4 | 1 |
| 2_8502657 | NA | NA | 1 | 1 | 2 | 1 | 1 | 1 | 4 | 1 | NA | NA | NA | NA | 1 | 1 | 2 | 1 | 3 | 1 |
| 2_8502678 | 1 | 1 | 1 | 1 | 2 | 0.5 | 1 | 1 | 5 | 0.8 | NA | NA | NA | NA | 1 | 0 | 2 | 1 | 3 | 0.6667 |
| 2_8502679 | 1 | 0 | 1 | 0 | 2 | 0.5 | 1 | 0 | 5 | 0.2 | NA | NA | NA | NA | 1 | 1 | 1 | 1 | 2 | 1 |
| 2_8502683 | NA | NA | 1 | 1 | 2 | 1 | 1 | 1 | 4 | 1 | NA | NA | NA | NA | 1 | 1 | 1 | 1 | 2 | 1 |
| 2_8502703 | 1 | 0 | NA | NA | 2 | 0.5 | NA | NA | 3 | 0.3333 | NA | NA | NA | NA | 1 | 1 | 2 | 0.5 | 3 | 0.6667 |
| 2_8502704 | 1 | 1 | 1 | 1 | 1 | 1 | 1 | 1 | 4 | 1 | NA | NA | NA | NA | 1 | 1 | 2 | 0.5 | 3 | 0.6667 |
| 2_8502704 | NA | NA | NA | NA | 1 | 1 | NA | NA | 1 | 1 | NA | NA | NA | NA | 1 | 0 | 2 | 1 | 3 | 0.6667 |
| 2_8502718 | 1 | 0 | 1 | 0 | 1 | 1 | NA | NA | 3 | 0.3333 | NA | NA | NA | NA | 1 | 0 | 2 | 1 | 3 | 0.6667 |
| 2_8502743 | 1 | 1 | NA | NA | 2 | 1 | 1 | 0 | 4 | 0.75 | NA | NA | NA | NA | NA | NA | 2 | 1 | 2 | 1 |
| 2_8502756 | NA | NA | NA | NA | 2 | 1 | 1 | 1 | 3 | 1 | NA | NA | NA | NA | NA | NA | 2 | 1 | 2 | 1 |
| 2_8502770 | NA | NA | NA | NA | 2 | 1 | 1 | 1 | 3 | 1 | NA | NA | NA | NA | NA | NA | 2 | 0.5 | 2 | 0.5 |
| 2_9026341 | 10 | 0.3 | 20 | 0.25 | 10 | 0.4 | 13 | 0.4615 | 53 | 0.3396 | 4 | 0.5 | 7 | 0.8571 | 12 | 0.25 | 7 | 0.5714 | 30 | 0.5 |
| 2_9026353 | 9 | 0.3333 | 19 | 0.4211 | 10 | 0.4 | 13 | 0.3077 | 51 | 0.3725 | 6 | 0.3333 | 7 | 0.5714 | 14 | 0.3571 | 9 | 0.2222 | 36 | 0.3611 |
| 2_9026372 | 8 | 0.875 | 16 | 0.9375 | 9 | 1 | 11 | 0.8182 | 44 | 0.9091 | 4 | 1 | 8 | 1 | 13 | 0.9231 | 9 | 1 | 34 | 0.9706 |
| 2_9026374 | 6 | 0 | 14 | 0.0714 | 9 | 0.3333 | 14 | 0.2143 | 43 | 0.1628 | 5 | 0.6 | 8 | 0.25 | 15 | 0 | 9 | 0 | 37 | 0.1351 |
| 2_9026378 | 9 | 0.6667 | 15 | 0.8 | 9 | 0.5556 | 13 | 0.9231 | 46 | 0.7609 | 3 | 1 | 7 | 0.8571 | 14 | 0.7143 | 6 | 0.6667 | 30 | 0.7667 |
| 2_9026385 | 8 | 0.75 | 12 | 0.4167 | 8 | 0.75 | 12 | 0.5833 | 40 | 0.6 | 4 | 1 | 8 | 0.5 | 11 | 0.8182 | 7 | 0.5714 | 30 | 0.7 |
| 2_9026387 | 7 | 0.8571 | 10 | 0.6 | 9 | 0.8889 | 13 | 0.8462 | 39 | 0.7949 | 4 | 0.75 | 7 | 1 | 14 | 0.9286 | 6 | 1 | 31 | 0.9355 |
| 2_9026389 | 9 | 0.6667 | 13 | 0.3846 | 7 | 0.4286 | 11 | 0.5455 | 40 | 0.5 | 5 | 0.2 | 8 | 0.5 | 13 | 0.4615 | 8 | 0.5 | 34 | 0.4412 |
| 2_9026392 | 9 | 0.1111 | 13 | 0.0769 | 10 | 0.3 | 12 | 0.1667 | 44 | 0.1591 | 5 | 0.4 | 7 | 0.2857 | 13 | 0.2308 | 7 | 0.4286 | 32 | 0.3125 |
| 2_9026394 | 9 | 0 | 11 | 0.1818 | 5 | 0.2 | 12 | 0.1667 | 37 | 0.1351 | 5 | 0 | 5 | 0.2 | 15 | 0.2 | 7 | 0 | 32 | 0.125 |
| 2_9026400 | 8 | 0.125 | 13 | 0 | 5 | 0.2 | 4 | 0 | 30 | 0.0667 | 5 | 0.2 | 6 | 0.1667 | 8 | 0.125 | 4 | 0.25 | 23 | 0.1739 |
| 2_9026400 | 6 | 0.6667 | 13 | 0.7692 | 6 | 0.6667 | 11 | 0.6364 | 36 | 0.6944 | 5 | 0.8 | 5 | 0.8 | 13 | 0.6923 | 8 | 0.75 | 31 | 0.7419 |
| 2_9026412 | 7 | 0.8571 | 13 | 0.6923 | 8 | 0.875 | 8 | 0.5 | 36 | 0.7222 | 4 | 0.75 | 6 | 0.8333 | 13 | 0.7692 | 9 | 0.6667 | 32 | 0.75 |
| 2_9026438 | 8 | 0.125 | 12 | 0.1667 | 10 | 0.1 | 9 | 0.1111 | 39 | 0.1282 | 4 | 0 | 6 | 0.3333 | 12 | 0.1667 | 5 | 0.4 | 27 | 0.2222 |
| 2_9026495 | 8 | 0.375 | 7 | 0.1429 | 7 | 0.1429 | 8 | 0.5 | 30 | 0.3 | 2 | 0 | 4 | 0.25 | 12 | 0.25 | 6 | 0.1667 | 24 | 0.2083 |
| 2_9026496 | 8 | 0.125 | 7 | 0 | 7 | 0.1429 | 7 | 0 | 29 | 0.069 | 2 | 0 | 3 | 0 | 13 | 0.1538 | 6 | 0 | 24 | 0.0833 |
| 2_9026502 | 6 | 0.1667 | 7 | 0.4286 | 7 | 0.1429 | 8 | 0.125 | 28 | 0.2143 | 1 | 0 | 3 | 1 | 10 | 0.6 | 4 | 0.5 | 18 | 0.6111 |
| 2_9026504 | 7 | 0 | 8 | 0.125 | 6 | 0.5 | 7 | 0.2857 | 28 | 0.2143 | 1 | 0 | 4 | 0.5 | 13 | 0.2308 | 6 | 0.1667 | 24 | 0.25 |
| 2_9026508 | 5 | 0.6 | 8 | 0.75 | 6 | 0.8333 | 6 | 0.8333 | 25 | 0.76 | 2 | 1 | 3 | 1 | 11 | 0.6364 | 5 | 1 | 21 | 0.8095 |
| 2_9026513 | 6 | 0.3333 | 5 | 0.4 | 6 | 0.6667 | 7 | 0.5714 | 24 | 0.5 | 2 | 0.5 | 3 | 0.3333 | 11 | 0.7273 | 5 | 0.6 | 21 | 0.619 |
| 2_9026593 | 3 | 0 | 5 | 0.2 | 7 | 0 | 3 | 0.3333 | 18 | 0.1111 | 1 | 0 | 1 | 0 | 9 | 0 | 3 | 0.3333 | 14 | 0.0714 |
| 2_9026609 | 4 | 0 | 5 | 0.2 | 6 | 0 | 1 | 0 | 16 | 0.0625 | 1 | 0 | NA | NA | 8 | 0.25 | 3 | 0 | 12 | 0.1667 |
| 2_9026663 | 2 | 1 | 2 | 0.5 | 3 | 0.3333 | 1 | 1 | 8 | 0.625 | 1 | 0 | NA | NA | 5 | 0.2 | 2 | 0 | 8 | 0.125 |
| 2_9026668 | 2 | 0.5 | 3 | 0 | 4 | 0.25 | 1 | 0 | 10 | 0.2 | 1 | 0 | 1 | 0 | 5 | 0.2 | 2 | 0 | 9 | 0.1111 |
| 2_9026672 | 2 | 0 | 3 | 0 | 3 | 0.3333 | 1 | 1 | 9 | 0.2222 | 1 | 0 | 1 | 1 | 2 | 0 | 3 | 0.3333 | 7 | 0.2857 |
| 2_9026677 | 2 | 1 | 3 | 0.6667 | 5 | 0.4 | 1 | 0 | 11 | 0.5455 | 1 | 1 | 1 | 0 | 6 | 0.8333 | 3 | 0.6667 | 11 | 0.7273 |
| 2_9026685 | 1 | 0 | 3 | 0.3333 | 5 | 0.4 | NA | NA | 9 | 0.3333 | 1 | 1 | 1 | 1 | 6 | 0.6667 | 3 | 0.3333 | 11 | 0.6364 |
| 2_9028864 | 2 | 0 | 1 | 1 | NA | NA | NA | NA | 3 | 0.3333 | NA | NA | 1 | 0 | 2 | 0 | 1 | 1 | 4 | 0.25 |
| 2_9061005 | 7 | 0.2857 | 2 | 1 | 2 | 0.5 | 3 | 0.3333 | 14 | 0.4286 | NA | NA | 2 | 0 | 4 | 0.25 | 4 | 0.25 | 10 | 0.2 |
| 2_9061025 | 8 | 0.25 | 2 | 0 | 2 | 0.5 | 3 | 0 | 15 | 0.2 | 2 | 0.5 | 2 | 0.5 | 4 | 0.25 | 4 | 0.25 | 12 | 0.3333 |
| 2_9071891 | 2 | 0 | 1 | 1 | 1 | 0 | 1 | 0 | 5 | 0.2 | 3 | 0 | 1 | 1 | 1 | 0 | NA | NA | 5 | 0.2 |
| 2_9071989 | 48 | 0.2292 | 62 | 0.0645 | 44 | 0.1818 | 37 | 0.1351 | 191 | 0.1466 | 51 | 0.098 | 59 | 0.0847 | 56 | 0.1786 | 41 | 0.0976 | 207 | 0.1159 |

| | | | | | | | | | | | | | | | | | | | | |
|---|---|---|---|---|---|---|---|---|---|---|---|---|---|---|---|---|---|---|---|---|
| 2_90720348 | 53 | 0.0755 | 66 | 0.0909 | 43 | 0.1163 | 48 | 0.0208 | 210 | 0.0762 | 53 | 0.0755 | 55 | 0.0909 | 57 | 0.0526 | 44 | 0.0455 | 209 | 0.067 |
| 2_90720663 | 65 | 0.0308 | 79 | 0.0886 | 70 | 0.0429 | 62 | 0.0323 | 276 | 0.0507 | 74 | 0.0405 | 72 | 0.0417 | 76 | 0.0526 | 66 | 0.0758 | 288 | 0.0521 |
| 2_90720763 | 75 | 0.16 | 87 | 0.1609 | 74 | 0.1486 | 68 | 0.1618 | 304 | 0.1579 | 78 | 0.1667 | 84 | 0.1548 | 87 | 0.2184 | 73 | 0.1918 | 322 | 0.1832 |
| 2_90720797 | 77 | 0.3117 | 89 | 0.2584 | 68 | 0.3382 | 66 | 0.197 | 300 | 0.2767 | 77 | 0.1948 | 81 | 0.284 | 89 | 0.2584 | 71 | 0.3521 | 318 | 0.2704 |
| 2_90720819 | 68 | 1 | 82 | 1 | 72 | 0.9861 | 65 | 1 | 287 | 0.9965 | 79 | 1 | 77 | 0.987 | 87 | 0.9885 | 77 | 0.987 | 320 | 0.9906 |
| 2_90720875 | 2 | 1 | NA | NA | NA | NA | NA | NA | 2 | 1 | 1 | 0 | 1 | 1 | NA | NA | 3 | 0.6667 | 5 | 0.6 |
| 2_90720940 | 1 | 1 | 1 | 1 | 1 | 1 | 1 | 0 | 4 | 0.75 | NA | NA | NA | NA | NA | NA | 1 | 1 | 1 | 1 |
| 2_90721528 | 1 | 0 | 1 | 0 | 1 | 1 | 1 | 0 | 4 | 0.25 | 1 | 0 | 1 | 1 | NA | NA | NA | NA | 2 | 0.5 |
| 2_90721617 | 2 | 0 | 1 | 1 | 1 | 0 | 1 | 0 | 5 | 0.2 | 1 | 1 | 1 | 0 | NA | NA | NA | NA | 2 | 0.5 |
| 2_90724985 | 23 | 0.1739 | 35 | 0.2286 | 25 | 0.16 | 30 | 0.1667 | 113 | 0.1858 | 21 | 0.3333 | 27 | 0.1481 | 19 | 0.1579 | 28 | 0.25 | 95 | 0.2211 |
| 2_90734678 | NA | NA | NA | NA | NA | NA | 1 | 1 | 1 | 1 | NA | NA | NA | NA | NA | NA | 1 | 1 | 1 | 1 |
| 2_90735080 | NA | NA | NA | NA | NA | NA | 1 | 1 | 1 | 1 | NA | NA | NA | NA | NA | NA | 3 | 0.3333 | 3 | 0.3333 |
| 2_90736091 | 1 | 1 | NA | NA | 1 | 1 | 2 | 1 | 4 | 1 | 2 | 0.5 | NA | NA | 1 | 1 | 4 | 0.75 | 7 | 0.7143 |
| 2_90736146 | 4 | 1 | 2 | 1 | 1 | 0 | 2 | 1 | 9 | 0.8889 | NA | NA | NA | NA | 1 | 0 | 4 | 1 | 5 | 0.8 |
| 2_90736340 | 4 | 1 | 3 | 0.6667 | 1 | 0 | 2 | 1 | 10 | 0.8 | 3 | 0.3333 | 1 | 0 | 3 | 0.3333 | 4 | 0.75 | 11 | 0.4545 |
| 2_90736357 | 4 | 1 | 2 | 1 | NA | NA | 2 | 0.5 | 8 | 0.875 | 3 | 1 | 1 | 1 | 2 | 1 | 4 | 1 | 10 | 1 |
| 2_90736365 | 5 | 0 | 3 | 0 | 1 | 0 | 2 | 0.5 | 11 | 0.0909 | 2 | 0 | 1 | 0 | 2 | 0.5 | 3 | 0 | 8 | 0.125 |
| 2_90736506 | 5 | 0.4 | 3 | 0.3333 | 2 | 0 | 2 | 0 | 12 | 0.25 | 2 | 0.5 | 1 | 1 | 3 | 0.3333 | 4 | 0 | 10 | 0.3 |
| 2_90739679 | 6 | 0.5 | 4 | 0 | 3 | 0.3333 | 3 | 0.3333 | 16 | 0.3125 | 2 | 0 | 1 | 0 | 3 | 0 | 5 | 0.4 | 11 | 0.1818 |
| 2_90739739 | 3 | 0.6667 | 4 | 0.75 | 2 | 0.5 | 3 | 1 | 12 | 0.75 | 3 | 0.6667 | 1 | 0 | 2 | 1 | 4 | 0.75 | 10 | 0.7 |
| 2_90739745 | 6 | 0.3333 | 4 | 0.5 | 3 | 0.3333 | 3 | 0 | 16 | 0.3125 | 3 | 0 | 1 | 0 | 3 | 0 | 5 | 0.2 | 12 | 0.0833 |
| 2_90739836 | 6 | 0.1667 | 3 | 0 | 3 | 0.3333 | 3 | 0.3333 | 15 | 0.2 | 3 | 0.3333 | 1 | 0 | 3 | 0.6667 | 5 | 0.2 | 12 | 0.3333 |
| 2_90739883 | 4 | 0 | 3 | 0.6667 | 3 | 0 | 3 | 0.3333 | 13 | 0.2308 | 2 | 0 | 1 | 0 | 3 | 0.3333 | 4 | 0.25 | 10 | 0.2 |
| 2_90739980 | 5 | 1 | 4 | 1 | 3 | 0.3333 | 3 | 1 | 15 | 0.8667 | 3 | 0.6667 | 1 | 1 | 3 | 0.6667 | 5 | 0.6 | 12 | 0.6667 |
| 2_90740245 | 6 | 0.1667 | 5 | 0 | 5 | 0 | 3 | 0 | 19 | 0.0526 | 3 | 0 | 1 | 0 | 3 | 0.3333 | 5 | 0 | 12 | 0.0833 |
| 2_90740408 | 4 | 0.5 | 4 | 0.75 | 5 | 0.2 | 1 | 1 | 14 | 0.5 | 3 | 0 | 1 | 0 | 2 | 0 | 5 | 0.6 | 11 | 0.2727 |
| 2_90740448 | 6 | 0.1667 | 5 | 0 | 5 | 0.2 | 3 | 0.6667 | 19 | 0.2105 | 3 | 0.3333 | 1 | 0 | 2 | 0.5 | 6 | 0.6667 | 12 | 0.5 |
| 2_90740627 | 6 | 0.8333 | 4 | 0.5 | 6 | 0.5 | 3 | 0.6667 | 19 | 0.6316 | 3 | 0.6667 | 1 | 0 | 2 | 0.5 | 5 | 0.2 | 11 | 0.3636 |
| 2_90740634 | 5 | 0.2 | 4 | 1 | 6 | 0.5 | 2 | 0 | 17 | 0.4706 | 3 | 1 | NA | NA | 2 | 0.5 | 5 | 0.6 | 10 | 0.7 |
| 2_90740727 | 5 | 0.4 | 6 | 0.5 | 6 | 0.1667 | 2 | 0.5 | 19 | 0.3684 | 4 | 0.25 | 1 | 0 | 4 | 0.5 | 6 | 0.3333 | 15 | 0.3333 |
| 2_90740803 | 3 | 0 | 5 | 0.6 | 6 | 0.1667 | 1 | 0 | 15 | 0.2667 | 3 | 0 | 1 | 0 | 4 | 0 | 6 | 0.5 | 14 | 0.2143 |
| 2_90741151 | 4 | 0.25 | 7 | 0 | 6 | 0 | 6 | 0.1667 | 23 | 0.087 | 4 | 0 | 4 | 0 | 4 | 0.25 | 4 | 0 | 16 | 0.0625 |
| 2_90741227 | 5 | 0.4 | 8 | 0.25 | 6 | 0.1667 | 6 | 0 | 25 | 0.2 | 4 | 0.25 | 3 | 0 | 3 | 0.6667 | 4 | 0.5 | 14 | 0.3571 |
| 2_90741397 | 8 | 0.375 | 12 | 0.1667 | 8 | 0.375 | 10 | 0.1 | 38 | 0.2368 | 8 | 0.375 | 5 | 0.4 | 4 | 0.5 | 5 | 0.8 | 22 | 0.5 |
| 2_90741435 | 10 | 0.4 | 14 | 0.3571 | 10 | 0.4 | 12 | 0.3333 | 46 | 0.3696 | 10 | 0.1 | 6 | 0 | 5 | 0.4 | 6 | 0.6667 | 27 | 0.2593 |
| 2_90741473 | 12 | 0.6667 | 11 | 0.5455 | 8 | 0.75 | 10 | 0.6 | 41 | 0.6341 | 10 | 0.4 | 6 | 0.6667 | 4 | 0.75 | 5 | 0.4 | 25 | 0.52 |
| 2_90741490 | 12 | 0.5 | 13 | 0.3846 | 10 | 0.6 | 11 | 0.4545 | 46 | 0.4783 | 9 | 0.3333 | 5 | 0.6 | 5 | 0.6 | 5 | 0.4 | 24 | 0.4583 |
| 2_90741529 | 10 | 0.2 | 9 | 0.1111 | 9 | 0.1111 | 10 | 0.1 | 38 | 0.1316 | 8 | 0.25 | 5 | 0 | 5 | 0.2 | 5 | 0 | 23 | 0.1304 |
| 2_90748285 | 3 | 0.3333 | 2 | 0 | 4 | 0 | 3 | 0.3333 | 12 | 0.1667 | 2 | 0.5 | NA | NA | NA | NA | 1 | 0 | 3 | 0.3333 |
| 2_90748406 | NA | NA | NA | NA | NA | NA | 1 | 1 | 1 | 1 | 1 | 1 | 1 | 1 | 1 | 0 | 1 | 1 | 4 | 0.75 |
| 2_90748899 | NA | NA | NA | NA | 1 | 1 | 1 | 0 | 2 | 0.5 | 1 | 1 | 1 | 1 | 1 | 0 | 1 | 1 | 4 | 0.75 |
| 2_90750017 | NA | NA | NA | NA | 1 | 1 | NA | NA | 1 | 1 | 1 | 1 | 1 | 1 | NA | NA | 1 | 1 | 3 | 1 |
| 2_90750109 | NA | NA | NA | NA | 1 | 0 | 2 | 0.5 | 3 | 0.3333 | 1 | 0 | 1 | 1 | 1 | 0 | 2 | 0 | 5 | 0.2 |
| 2_90750375 | NA | NA | NA | NA | 1 | 0 | 2 | 0.5 | 3 | 0.3333 | NA | NA | 1 | 1 | 1 | 0 | 2 | 0.5 | 4 | 0.5 |
| 2_90750471 | NA | NA | NA | NA | 1 | 1 | 2 | 1 | 3 | 1 | 1 | 0 | 1 | 1 | NA | NA | 1 | 1 | 3 | 0.6667 |
| 2_90750491 | NA | NA | NA | NA | 1 | 0 | 1 | 1 | 2 | 0.5 | 1 | 0 | NA | NA | NA | NA | 1 | 1 | 2 | 0.5 |
| 2_90750685 | NA | NA | NA | NA | 1 | 1 | 2 | 1 | 3 | 1 | 1 | 1 | 1 | 1 | 1 | 1 | 2 | 0.5 | 5 | 0.8 |
| 2_90843726 | 5 | 0.4 | 1 | 1 | 2 | 1 | 4 | 0.5 | 12 | 0.5833 | 1 | 1 | 3 | 0.3333 | 4 | 1 | 3 | 1 | 11 | 0.8182 |
| 2_90843811 | 2 | 1 | 1 | 1 | 1 | 1 | 2 | 1 | 6 | 1 | 1 | 1 | 1 | 1 | 2 | 1 | 3 | 0.6667 | 7 | 0.8571 |
| 2_90843854 | 2 | 0 | 1 | 1 | 2 | 0 | 4 | 0 | 9 | 0.1111 | 1 | 1 | NA | NA | 3 | 0.3333 | 3 | 0 | 7 | 0.2857 |
| 2_90843897 | 2 | 0 | 1 | 0 | 2 | 0 | 4 | 0.25 | 9 | 0.1111 | 1 | 0 | 2 | 0 | 3 | 0 | 3 | 0.3333 | 9 | 0.1111 |
| 2_90845545 | 3 | 0 | 3 | 0 | 5 | 0.2 | 4 | 0 | 15 | 0.0667 | 1 | 0 | 2 | 0 | 5 | 0 | 4 | 0.25 | 12 | 0.0833 |
| 2_90845615 | 5 | 0.2 | 3 | 0 | 6 | 0 | 5 | 0 | 19 | 0.0526 | 1 | 0 | 2 | 0 | 5 | 0 | 4 | 0.25 | 12 | 0.0833 |
| 2_90848854 | 13 | 0 | 10 | 0 | 14 | 0.0714 | 15 | 0.4 | 52 | 0.1346 | 8 | 0 | 11 | 0.0909 | 13 | 0.1538 | 13 | 0.2308 | 45 | 0.1333 |
| 2_90849084 | 9 | 0.5556 | 8 | 0.5 | 13 | 0.4615 | 11 | 0.3636 | 41 | 0.4634 | 5 | 0.6 | 10 | 0.5 | 17 | 0.5882 | 13 | 0.6154 | 45 | 0.5778 |
| 2_90849225 | 16 | 0.0625 | 13 | 0.0769 | 21 | 0.1905 | 22 | 0.1364 | 72 | 0.125 | 7 | 0 | 14 | 0.1429 | 19 | 0.1579 | 14 | 0.1429 | 54 | 0.1296 |
| 2_90849691 | 17 | 0.4118 | 15 | 0.4667 | 21 | 0.3333 | 23 | 0.3913 | 76 | 0.3947 | 7 | 0.5714 | 18 | 0.3333 | 23 | 0.3478 | 12 | 0.5833 | 60 | 0.4167 |
| 2_90892357 | 1 | 0 | 3 | 0 | 4 | 0.5 | 2 | 0 | 10 | 0.2 | 2 | 0 | 4 | 0.25 | 4 | 0 | 1 | 0 | 11 | 0.0909 |
| 2_90892925 | 1 | 0 | 1 | 0 | 4 | 0.25 | 2 | 0.5 | 8 | 0.25 | 1 | 1 | 3 | 0.3333 | 4 | 0 | 1 | 0 | 9 | 0.2222 |
| 2_90893209 | 1 | 1 | 2 | 0.5 | 5 | 0.2 | 3 | 0.6667 | 11 | 0.4545 | 1 | 1 | 4 | 0 | 5 | 0 | 1 | 0 | 11 | 0.0909 |
| 2_90945807 | 1 | 1 | NA | NA | 3 | 0.3333 | NA | NA | 4 | 0.5 | NA | NA | NA | NA | 1 | 1 | 2 | 0.5 | 3 | 0.6667 |
| 2_90945818 | 1 | 1 | NA | NA | 2 | 0.5 | NA | NA | 3 | 0.6667 | NA | NA | NA | NA | 1 | 1 | 2 | 0.5 | 3 | 0.6667 |
| 2_90945908 | 1 | 1 | NA | NA | 3 | 1 | NA | NA | 4 | 1 | NA | NA | NA | NA | 1 | 1 | 2 | 1 | 3 | 1 |
| 2_90967797 | 7 | 0.7143 | 11 | 0.7273 | 14 | 0.9286 | 12 | 0.75 | 44 | 0.7955 | 7 | 0.7143 | 13 | 0.8462 | 11 | 0.9091 | 11 | 0.6364 | 42 | 0.7857 |
| 2_90968171 | 12 | 0.0833 | 17 | 0.1765 | 14 | 0.2857 | 11 | 0.0909 | 54 | 0.1667 | 6 | 0 | 15 | 0.0667 | 14 | 0.1429 | 15 | 0.0667 | 50 | 0.08 |

| | | | | | | | | | | | | | | | | | | | | |
|---|---|---|---|---|---|---|---|---|---|---|---|---|---|---|---|---|---|---|---|---|
| 2_90968190 | 10 | 0.6 | 13 | 0.2308 | 9 | 0.1111 | 8 | 0.125 | 40 | 0.275 | 4 | 0 | 8 | 0.25 | 6 | 0 | 15 | 0.0667 | 33 | 0.0909 |
| 2_90968565 | 9 | 0.6667 | 12 | 0.6667 | 8 | 0.75 | 11 | 0.3636 | 40 | 0.6 | 7 | 0.7143 | 10 | 0.8 | 11 | 0.4545 | 13 | 0.6923 | 41 | 0.6585 |
| 2_91007311 | 2 | 0 | 4 | 0.25 | 2 | 0.5 | 3 | 0 | 11 | 0.1818 | 2 | 0.5 | 1 | 0 | 1 | 0 | 2 | 1 | 6 | 0.5 |
| 2_91008302 | 2 | 0 | 2 | 0 | 2 | 0.5 | 3 | 0 | 9 | 0.1111 | 2 | 0 | 1 | 0 | 1 | 0 | 2 | 0.5 | 6 | 0.1667 |
| 2_91008559 | 2 | 0 | 1 | 0 | 1 | 0 | 3 | 0.3333 | 7 | 0.1429 | 2 | 0 | 1 | 0 | 1 | 1 | 1 | 0 | 5 | 0.2 |
| 2_91014681 | 1 | 0 | NA | NA | 1 | 1 | NA | NA | 2 | 0.5 | NA | NA | NA | NA | 1 | 0 | 1 | 1 | 2 | 0.5 |
| 2_91041170 | 3 | 0.3333 | 6 | 1 | 4 | 0.5 | 2 | 0.5 | 15 | 0.6667 | 1 | 1 | 7 | 0.5714 | 3 | 1 | 4 | 0.75 | 15 | 0.7333 |
| 2_91041285 | 3 | 1 | 4 | 1 | 5 | 1 | 1 | 1 | 13 | 1 | 2 | 1 | 7 | 0.8571 | 4 | 1 | 4 | 1 | 17 | 0.9412 |
| 2_91041333 | 2 | 1 | 5 | 1 | 5 | 1 | 2 | 1 | 14 | 1 | 2 | 1 | 8 | 1 | 4 | 0.75 | 5 | 1 | 19 | 0.9474 |
| 2_91041377 | 3 | 1 | 6 | 1 | 6 | 0.8333 | 1 | 1 | 16 | 0.9375 | 2 | 1 | 8 | 1 | 3 | 1 | 5 | 1 | 18 | 1 |
| 2_91041390 | 3 | 0 | 5 | 0 | 6 | 0.1667 | 1 | 0 | 15 | 0.0667 | 2 | 0.5 | 7 | 0 | 4 | 0 | 4 | 0 | 17 | 0.0588 |
| 2_91041502 | 3 | 1 | 7 | 1 | 5 | 0.8 | 2 | 1 | 17 | 0.9412 | 2 | 1 | 7 | 1 | 5 | 0.8 | 6 | 0.8333 | 20 | 0.9 |
| 2_91041535 | 3 | 1 | 7 | 1 | 4 | 0.5 | 1 | 1 | 15 | 0.8667 | 2 | 1 | 7 | 1 | 5 | 0.6 | 6 | 0.8333 | 20 | 0.85 |
| 2_91041574 | 2 | 1 | 5 | 0.4 | 4 | 0.5 | 2 | 1 | 13 | 0.6154 | 2 | 1 | 4 | 1 | 3 | 0.3333 | 6 | 0.6667 | 15 | 0.7333 |
| 2_91041671 | 3 | 0 | 6 | 0.1667 | 5 | 0 | 2 | 0 | 16 | 0.0625 | 2 | 0 | 7 | 0 | 5 | 0 | 6 | 0.1667 | 20 | 0.05 |
| 2_91041682 | 2 | 1 | 7 | 0.7143 | 5 | 0.4 | 1 | 0 | 15 | 0.6 | 2 | 0 | 6 | 0.3333 | 4 | 0.25 | 6 | 0.3333 | 18 | 0.2778 |
| 2_91041719 | 3 | 0.3333 | 6 | 0.1667 | 6 | 0.3333 | 2 | 0 | 17 | 0.2353 | 2 | 0.5 | 6 | 0.1667 | 5 | 0.4 | 5 | 0 | 18 | 0.2222 |
| 2_91041739 | 2 | 1 | 6 | 0.6667 | 5 | 0.8 | 1 | 1 | 14 | 0.7857 | 2 | 1 | 7 | 0.7143 | 5 | 0.8 | 6 | 0.8333 | 20 | 0.8 |
| 2_91041750 | 3 | 0.3333 | 7 | 0.5714 | 6 | 0.5 | 1 | 0 | 17 | 0.4706 | 2 | 0.5 | 8 | 0.375 | 6 | 0.1667 | 6 | 0.3333 | 22 | 0.3182 |
| 2_91041813 | 2 | 0.5 | 6 | 0.1667 | 5 | 0.4 | 2 | 1 | 15 | 0.4 | 2 | 0 | 7 | 0.4286 | 6 | 0.3333 | 4 | 0.5 | 19 | 0.3684 |
| 2_91042154 | NA | NA | 2 | 0.5 | 2 | 0.5 | 1 | 0 | 5 | 0.4 | 1 | 1 | NA | NA | 1 | 0 | 3 | 0 | 5 | 0.2 |
| 2_91042180 | NA | NA | 1 | 1 | 2 | 0.5 | 3 | 0.6667 | 6 | 0.6667 | 1 | 1 | NA | NA | NA | NA | 3 | 0.3333 | 4 | 0.5 |
| 2_91042439 | 3 | 0.6667 | 5 | 0.2 | 4 | 0 | 2 | 0 | 14 | 0.2143 | 3 | 0.3333 | 7 | 0.2857 | 6 | 0 | 6 | 0.3333 | 22 | 0.2273 |
| 2_91043017 | 4 | 0.25 | 5 | 0 | 5 | 0 | 2 | 0 | 16 | 0.0625 | 5 | 0.2 | 3 | 0 | 4 | 0 | 5 | 0 | 17 | 0.0588 |
| 2_91044160 | 5 | 0 | 5 | 0 | 6 | 0.1667 | 3 | 0 | 19 | 0.0526 | 6 | 0.1667 | 7 | 0 | 4 | 0 | 4 | 0.25 | 21 | 0.0952 |
| 2_91044212 | 5 | 0 | 5 | 0.2 | 7 | 0 | 1 | 0 | 18 | 0.0556 | 6 | 0 | 5 | 0 | 4 | 0.25 | 5 | 0 | 20 | 0.05 |
| 2_91094713 | 3 | 0.3333 | 3 | 0.3333 | 2 | 1 | 1 | 1 | 9 | 0.5556 | 1 | 1 | 3 | 0.3333 | 2 | 0.5 | 3 | 0.3333 | 9 | 0.4444 |
| 2_91106597 | 8 | 0.625 | 10 | 0.1 | 6 | 0.6667 | 9 | 0.3333 | 33 | 0.3939 | 4 | 0 | 6 | 0.1667 | 9 | 0.3333 | 5 | 0.6 | 24 | 0.2917 |
| 2_91106685 | 9 | 0.4444 | 13 | 0.2308 | 7 | 0.2857 | 7 | 0.2857 | 36 | 0.3056 | 4 | 0 | 5 | 0.4 | 7 | 0.2857 | 5 | 0.4 | 21 | 0.2857 |
| 2_91106935 | 8 | 0.625 | 15 | 0.3333 | 8 | 0.25 | 11 | 0.5455 | 42 | 0.4286 | 6 | 0.1667 | 8 | 0.375 | 10 | 0.1 | 6 | 0.5 | 30 | 0.2667 |
| 2_91107001 | 7 | 0.1429 | 11 | 0.1818 | 6 | 0 | 6 | 0.1667 | 30 | 0.1333 | 5 | 0 | 9 | 0.1111 | 9 | 0 | 6 | 0.1667 | 29 | 0.069 |
| 2_91109064 | 9 | 0.2222 | 15 | 0.0667 | 14 | 0.0714 | 14 | 0 | 52 | 0.0769 | 10 | 0 | 12 | 0 | 14 | 0.2143 | 15 | 0.2667 | 51 | 0.1373 |
| 2_91109133 | 8 | 0.25 | 14 | 0.2143 | 12 | 0.4167 | 15 | 0.3333 | 49 | 0.3061 | 10 | 0.3 | 12 | 0.25 | 10 | 0.1 | 15 | 0.2 | 47 | 0.2128 |
| 2_91109148 | 8 | 0.25 | 14 | 0.2143 | 12 | 0.4167 | 12 | 0.1667 | 46 | 0.2609 | 8 | 0 | 10 | 0.3 | 13 | 0.2308 | 14 | 0.1429 | 45 | 0.1778 |
| 2_91112972 | 6 | 0 | 14 | 0.0714 | 13 | 0.0769 | 12 | 0.1667 | 45 | 0.0889 | 7 | 0 | 12 | 0 | 14 | 0.0714 | 15 | 0.1333 | 48 | 0.0625 |
| 2_91342057 | NA | NA | 1 | 1 | 1 | 1 | 2 | 0 | 4 | 0.5 | 2 | 0 | NA | NA | NA | NA | 2 | 1 | 4 | 0.5 |
| 2_91342150 | NA | NA | NA | NA | 1 | 1 | 1 | 0 | 2 | 0.5 | 2 | 1 | NA | NA | NA | NA | 2 | 0.5 | 4 | 0.75 |
| 2_91342250 | NA | NA | 1 | 0 | 1 | 1 | 1 | 0 | 3 | 0.3333 | 2 | 0.5 | NA | NA | NA | NA | 2 | 0 | 4 | 0.25 |
| 2_91342285 | NA | NA | NA | NA | 1 | 1 | 2 | 0 | 3 | 0.3333 | 2 | 0.5 | NA | NA | NA | NA | 2 | 0.5 | 4 | 0.5 |
| 2_91342335 | NA | NA | NA | NA | 1 | 1 | 2 | 1 | 3 | 1 | 2 | 0.5 | NA | NA | NA | NA | 1 | 1 | 3 | 0.6667 |
| 2_91342473 | NA | NA | 2 | 0.5 | 1 | 0 | 2 | 1 | 5 | 0.6 | 2 | 0.5 | NA | NA | NA | NA | 2 | 1 | 4 | 0.75 |
| 2_91342518 | NA | NA | 2 | 0.5 | 1 | 1 | 2 | 0 | 5 | 0.4 | 1 | 1 | NA | NA | NA | NA | 2 | 0.5 | 3 | 0.6667 |
| 2_91342539 | NA | NA | 1 | 0 | 1 | 0 | 2 | 0.5 | 4 | 0.25 | 1 | 1 | NA | NA | NA | NA | 3 | 0.6667 | 4 | 0.75 |
| 2_91342568 | NA | NA | 1 | 1 | 1 | 1 | 2 | 0.5 | 4 | 0.75 | 1 | 1 | NA | NA | NA | NA | 1 | 0 | 2 | 0.5 |
| 2_91342653 | NA | NA | 1 | 1 | NA | NA | 2 | 1 | 3 | 1 | 2 | 1 | NA | NA | NA | NA | 1 | 0 | 3 | 0.6667 |
| 2_91342706 | NA | NA | 2 | 0 | 1 | 0 | 2 | 0.5 | 5 | 0.2 | 1 | 1 | NA | NA | NA | NA | 3 | 0.3333 | 4 | 0.5 |
| 2_91342928 | NA | NA | 2 | 0 | 1 | 1 | 1 | 1 | 4 | 0.5 | 1 | 1 | NA | NA | NA | NA | 2 | 0.5 | 3 | 0.6667 |
| 2_91344166 | 3 | 0.3333 | 4 | 0.75 | 1 | 1 | 4 | 0.25 | 12 | 0.5 | 2 | 0 | NA | NA | NA | NA | 3 | 0.3333 | 5 | 0.2 |
| 2_91450526 | 9 | 0.2222 | 13 | 0.1538 | 23 | 0.0435 | 9 | 0.1111 | 54 | 0.1111 | 10 | 0.1 | 10 | 0 | 12 | 0.1667 | 14 | 0.0714 | 46 | 0.087 |
| 2_91450539 | 11 | 0.8182 | 9 | 0.8889 | 21 | 1 | 9 | 0.8889 | 50 | 0.92 | 10 | 1 | 8 | 0.875 | 12 | 0.9167 | 17 | 0.8824 | 47 | 0.9149 |
| 2_91450658 | 10 | 0.7 | 14 | 0.5714 | 21 | 0.619 | 9 | 0.6667 | 54 | 0.6296 | 10 | 0.9 | 8 | 0.875 | 10 | 0.6 | 13 | 0.6154 | 41 | 0.7317 |
| 2_91450676 | 11 | 0.3636 | 11 | 0.5455 | 22 | 0.4545 | 8 | 0.375 | 52 | 0.4423 | 8 | 0.25 | 9 | 0.4444 | 12 | 0.5 | 17 | 0.2941 | 46 | 0.3696 |
| 2_91450733 | 12 | 0.3333 | 13 | 0.1538 | 23 | 0.0435 | 9 | 0.3333 | 57 | 0.1754 | 10 | 0.3 | 11 | 0.0909 | 14 | 0.2857 | 17 | 0.3529 | 52 | 0.2692 |
| 2_91450737 | 9 | 0.3333 | 11 | 0.2727 | 23 | 0.3478 | 9 | 0.4444 | 52 | 0.3462 | 9 | 0.2222 | 10 | 0.3 | 10 | 0.5 | 13 | 0.2308 | 42 | 0.3095 |
| 2_91450761 | 11 | 0.3636 | 12 | 0.3333 | 19 | 0.4737 | 9 | 0.5556 | 51 | 0.4314 | 10 | 0.5 | 9 | 0.7778 | 14 | 0.4286 | 17 | 0.4118 | 50 | 0.5 |
| 2_91450932 | 10 | 0 | 14 | 0.1429 | 25 | 0.12 | 9 | 0 | 58 | 0.0862 | 12 | 0 | 14 | 0.1429 | 15 | 0.1333 | 19 | 0.2632 | 60 | 0.15 |
| 2_91450980 | 8 | 0.25 | 11 | 0.2727 | 22 | 0.1818 | 10 | 0.1 | 51 | 0.1961 | 13 | 0.3846 | 13 | 0.3077 | 14 | 0.2143 | 18 | 0.3889 | 58 | 0.3276 |
| 2_91501286 | 3 | 0.6667 | 8 | 0.875 | 5 | 1 | 7 | 1 | 23 | 0.913 | 7 | 1 | 8 | 1 | 7 | 1 | 10 | 1 | 32 | 1 |
| 2_91501448 | 4 | 0.5 | 7 | 0.1429 | 5 | 0.2 | 7 | 0.2857 | 23 | 0.2609 | 9 | 0.3333 | 6 | 0.6667 | 8 | 0.125 | 10 | 0.3 | 33 | 0.3333 |
| 2_91502514 | 7 | 0 | 10 | 0.3 | 5 | 0.2 | 5 | 0 | 27 | 0.1481 | 13 | 0.0769 | 6 | 0 | 7 | 0.1429 | 12 | 0.0833 | 38 | 0.0789 |
| 2_91505792 | 6 | 1 | 7 | 0.8571 | 9 | 1 | 6 | 1 | 28 | 0.9643 | 7 | 1 | 11 | 0.8182 | 5 | 0.8 | 7 | 1 | 30 | 0.9 |
| 2_91506009 | 9 | 0 | 10 | 0.3 | 9 | 0.2222 | 8 | 0.125 | 36 | 0.1667 | 12 | 0.3333 | 14 | 0.2857 | 10 | 0.3 | 10 | 0.4 | 46 | 0.3261 |
| 2_91506134 | 10 | 0.1 | 14 | 0.0714 | 9 | 0.2222 | 8 | 0.25 | 41 | 0.1463 | 16 | 0.0625 | 20 | 0.2 | 15 | 0.2 | 12 | 0.0833 | 63 | 0.1429 |
| 2_91506164 | 9 | 0.1111 | 15 | 0.1333 | 11 | 0 | 9 | 0.1111 | 44 | 0.0909 | 15 | 0.0667 | 20 | 0.15 | 12 | 0 | 10 | 0.1 | 57 | 0.0877 |
| 2_91506270 | 4 | 0.25 | 12 | 0.3333 | 9 | 0.5556 | 8 | 0.375 | 33 | 0.3939 | 10 | 0.3 | 15 | 0.2667 | 10 | 0.2 | 8 | 0.5 | 43 | 0.3023 |

| | | | | | | | | | | | | | | | | | | | | |
|---|---|---|---|---|---|---|---|---|---|---|---|---|---|---|---|---|---|---|---|---|
| 2_9150636 | 10 | 0.2 | 14 | 0 | 9 | 0 | 5 | 0.2 | 38 | 0.0789 | 12 | 0.25 | 13 | 0.0769 | 10 | 0.2 | 7 | 0 | 42 | 0.1429 |
| 2_9150648 | 9 | 0.6667 | 15 | 0.5333 | 11 | 0.8182 | 9 | 0.5556 | 44 | 0.6364 | 12 | 0.75 | 18 | 0.6111 | 12 | 0.75 | 9 | 0.7778 | 51 | 0.7059 |
| 2_9150653 | 9 | 0.6667 | 13 | 0.9231 | 11 | 0.5455 | 8 | 0.375 | 41 | 0.6585 | 12 | 0.8333 | 15 | 0.8 | 12 | 0.6667 | 9 | 0.5556 | 48 | 0.7292 |
| 2_9150654 | 8 | 0.875 | 13 | 0.8462 | 9 | 0.8889 | 8 | 0.875 | 38 | 0.8684 | 13 | 0.7692 | 17 | 0.6471 | 14 | 0.8571 | 10 | 0.9 | 54 | 0.7778 |
| 2_9150657 | 9 | 0.4444 | 14 | 0.7143 | 10 | 0.8 | 9 | 0.6667 | 42 | 0.6667 | 11 | 0.9091 | 18 | 1 | 13 | 0.6923 | 8 | 0.875 | 50 | 0.88 |
| 2_9150658 | 7 | 0.7143 | 12 | 0.75 | 8 | 0.625 | 6 | 0.3333 | 33 | 0.6364 | 12 | 0.8333 | 14 | 0.6429 | 9 | 0.7778 | 8 | 0.875 | 43 | 0.7674 |
| 2_9150662 | 9 | 0.1111 | 13 | 0.0769 | 9 | 0.2222 | 9 | 0.1111 | 40 | 0.125 | 11 | 0.4545 | 18 | 0.1111 | 12 | 0.5 | 9 | 0.4444 | 50 | 0.34 |
| 2_9150669 | 10 | 0 | 13 | 0.4615 | 8 | 0.125 | 9 | 0.1111 | 40 | 0.2 | 14 | 0.0714 | 17 | 0.1765 | 12 | 0.5 | 11 | 0.1818 | 54 | 0.2222 |
| 2_9154263 | 1 | 1 | NA | NA | NA | NA | NA | NA | 1 | 1 | NA | NA | 1 | 1 | NA | NA | NA | NA | 1 | 1 |
| 2_9154313 | 1 | 0 | NA | NA | NA | NA | 1 | 1 | 2 | 0.5 | NA | NA | NA | NA | NA | NA | 1 | 1 | 1 | 1 |
| 2_9155097 | 1 | 1 | NA | NA | NA | NA | 1 | 1 | 2 | 1 | NA | NA | 1 | 1 | NA | NA | 1 | 1 | 2 | 1 |
| 2_9155102 | 1 | 1 | NA | NA | NA | NA | 1 | 1 | 2 | 1 | NA | NA | 1 | 1 | NA | NA | 1 | 1 | 2 | 1 |
| 2_9160286 | 1 | 1 | 1 | 1 | 1 | 0 | NA | NA | 3 | 0.6667 | 2 | 0.5 | 4 | 0.75 | 1 | 0 | 1 | 1 | 8 | 0.625 |
| 2_9160300 | 1 | 1 | 1 | 1 | 1 | 1 | NA | NA | 3 | 1 | 2 | 1 | 4 | 0.75 | 1 | 1 | 1 | 1 | 8 | 0.875 |
| 2_9160324 | 2 | 0 | 2 | 0.5 | 1 | 1 | NA | NA | 5 | 0.4 | 2 | 0.5 | 3 | 1 | 1 | 1 | 2 | 0.5 | 8 | 0.75 |
| 2_9160327 | 2 | 0 | 2 | 0 | 1 | 1 | 1 | 1 | 6 | 0.3333 | 3 | 0.3333 | 2 | 0.5 | 1 | 0 | 1 | 0 | 7 | 0.2857 |
| 2_9160328 | 1 | 1 | 1 | 0 | 1 | 1 | 1 | 1 | 4 | 0.75 | NA | NA | 4 | 1 | 1 | 0 | 2 | 0.5 | 7 | 0.7143 |
| 2_9160330 | 1 | 1 | 1 | 0 | 1 | 0 | NA | NA | 3 | 0.3333 | 3 | 1 | 4 | 1 | NA | NA | 1 | 1 | 8 | 1 |
| 2_9160333 | 2 | 0.5 | 2 | 0 | 1 | 0 | 1 | 0 | 6 | 0.1667 | 2 | 0 | 3 | 0.3333 | 1 | 0 | 2 | 0 | 8 | 0.125 |
| 2_9160337 | 2 | 0 | 2 | 0.5 | 1 | 0 | 1 | 0 | 6 | 0.1667 | 3 | 0 | 3 | 0.3333 | 1 | 0 | 2 | 0 | 9 | 0.1111 |
| 2_9160364 | 2 | 0 | 2 | 0.5 | 1 | 0 | 1 | 1 | 6 | 0.3333 | 3 | 0 | 4 | 0.25 | 1 | 0 | 2 | 0.5 | 10 | 0.2 |
| 2_9160388 | NA | NA | 2 | 0.5 | NA | NA | 1 | 1 | 3 | 0.6667 | 3 | 0.6667 | 3 | 1 | 2 | 1 | 2 | 0.5 | 10 | 0.8 |
| 2_9160395 | 1 | 1 | 2 | 1 | 2 | 1 | 1 | 1 | 6 | 1 | 2 | 1 | 4 | 1 | 2 | 1 | 2 | 0.5 | 10 | 0.9 |
| 2_9160402 | 2 | 1 | 2 | 1 | 1 | 1 | 1 | 1 | 6 | 1 | 3 | 0.6667 | 4 | 0.75 | 2 | 1 | 2 | 0.5 | 11 | 0.7273 |
| 2_9174790 | 2 | 0.5 | 8 | 0.5 | 2 | 0.5 | 4 | 1 | 16 | 0.625 | 6 | 0.6667 | 4 | 1 | 8 | 0.5 | 2 | 0 | 20 | 0.6 |
| 2_9174795 | 2 | 0 | 5 | 0.2 | NA | NA | 3 | 0 | 10 | 0.1 | 4 | 0.25 | 3 | 0.6667 | 6 | 0.6667 | 3 | 0 | 16 | 0.4375 |
| 2_9174806 | 3 | 0.3333 | 9 | 0.3333 | 5 | 0.4 | 4 | 0.25 | 21 | 0.3333 | 9 | 0.6667 | 6 | 0.1667 | 10 | 0.4 | 4 | 0.5 | 29 | 0.4483 |
| 2_9174817 | 4 | 0.5 | 10 | 0.2 | 5 | 0.2 | 5 | 0.4 | 24 | 0.2917 | 9 | 0.2222 | 6 | 0.1667 | 11 | 0.1818 | 6 | 0 | 32 | 0.1562 |
| 2_9174823 | 4 | 0.5 | 10 | 0.2 | 5 | 0 | 5 | 0 | 24 | 0.1667 | 9 | 0.1111 | 6 | 0 | 12 | 0.0833 | 5 | 0 | 32 | 0.0625 |
| 2_9174823 | 4 | 0.5 | 9 | 0.4444 | 4 | 0.75 | 3 | 0.6667 | 20 | 0.55 | 9 | 0.5556 | 5 | 0.4 | 12 | 0.6667 | 5 | 1 | 31 | 0.6452 |
| 2_9174827 | 2 | 0.5 | 9 | 0.2222 | 5 | 0.2 | 4 | 0 | 20 | 0.2 | 7 | 0.2857 | 4 | 0.25 | 11 | 0.1818 | 5 | 0.2 | 27 | 0.2222 |
| 2_9174847 | 5 | 0.4 | 9 | 0.6667 | 5 | 0.6 | 6 | 0.3333 | 25 | 0.52 | 7 | 0.7143 | 6 | 0.5 | 13 | 0.3846 | 6 | 0.8333 | 32 | 0.5625 |
| 2_9174854 | 4 | 0.75 | 8 | 0.375 | 5 | 0.4 | 2 | 0.5 | 19 | 0.4737 | 7 | 0.4286 | 4 | 0 | 4 | 0 | 6 | 0.8333 | 21 | 0.381 |
| 2_9174855 | 6 | 0.5 | 8 | 0.375 | 5 | 0.2 | 6 | 0.3333 | 25 | 0.36 | 7 | 0.1429 | 7 | 0.5714 | 11 | 0.7273 | 6 | 0.1667 | 31 | 0.4516 |
| 2_9174858 | 6 | 0 | 8 | 0.125 | 4 | 0.5 | 6 | 0.5 | 24 | 0.25 | 8 | 0.25 | 5 | 0.4 | 11 | 0.5455 | 7 | 0.2857 | 31 | 0.3871 |
| 2_9174871 | 6 | 0.6667 | 9 | 0.6667 | 6 | 0.6667 | 5 | 0.4 | 26 | 0.6154 | 6 | 0.5 | 6 | 0.5 | 12 | 0.4167 | 6 | 0.5 | 30 | 0.4667 |
| 2_9174873 | 4 | 0.25 | 9 | 0.2222 | 5 | 0.4 | 4 | 0.25 | 22 | 0.2727 | 7 | 0.7143 | 5 | 0.6 | 9 | 0.5556 | 6 | 0.5 | 27 | 0.5926 |
| 2_9174900 | 6 | 0 | 11 | 0.1818 | 7 | 0.1429 | 5 | 0.4 | 29 | 0.1724 | 9 | 0.2222 | 9 | 0 | 13 | 0.3077 | 6 | 0 | 37 | 0.1622 |
| 2_9175770 | 4 | 0.5 | 3 | 0.3333 | NA | NA | 9 | 0.4444 | 16 | 0.4375 | NA | NA | 8 | 0.5 | 3 | 1 | 4 | 0.25 | 15 | 0.5333 |
| 2_9175772 | 5 | 0.6 | 3 | 0.3333 | 1 | 0 | 9 | 0.1111 | 18 | 0.2778 | NA | NA | 6 | 0.3333 | 6 | 0.6667 | 3 | 0.3333 | 15 | 0.4667 |
| 2_9175778 | 5 | 0.4 | 2 | 0 | 1 | 0 | 10 | 0.1 | 18 | 0.1667 | NA | NA | 5 | 0 | 3 | 0 | 4 | 0.25 | 12 | 0.0833 |
| 2_9175785 | 5 | 0.6 | 3 | 0.3333 | 1 | 0 | 8 | 0.75 | 17 | 0.5882 | NA | NA | 8 | 0.625 | 6 | 0.6667 | 2 | 1 | 16 | 0.6875 |
| 2_9175792 | 5 | 1 | 3 | 1 | 1 | 1 | 11 | 1 | 20 | 1 | 1 | 1 | 7 | 1 | 6 | 1 | 4 | 1 | 18 | 1 |
| 2_9175795 | 5 | 0.8 | 4 | 0.25 | 2 | 0 | 10 | 0.2 | 21 | 0.3333 | 1 | 0 | 7 | 0.5714 | 6 | 0.1667 | 4 | 0 | 18 | 0.2778 |
| 2_9175812 | 4 | 1 | 3 | 0.6667 | 3 | 0.3333 | 12 | 0.75 | 22 | 0.7273 | 1 | 1 | 8 | 1 | 6 | 1 | 3 | 0.6667 | 18 | 0.9444 |
| 2_9175838 | 5 | 0.2 | 2 | 0 | 2 | 0 | 10 | 0.1 | 19 | 0.1053 | NA | NA | 8 | 0.125 | 6 | 0.1667 | 5 | 0.2 | 19 | 0.1579 |
| 2_9175846 | 7 | 1 | 3 | 0.3333 | 3 | 0.6667 | 11 | 0.4545 | 24 | 0.625 | 1 | 1 | 8 | 0.875 | 6 | 0.5 | 5 | 0.6 | 20 | 0.7 |
| 2_9175858 | 7 | 0 | 4 | 0 | 3 | 0.3333 | 12 | 0.0833 | 26 | 0.0769 | 1 | 0 | 10 | 0.1 | 9 | 0.1111 | 8 | 0 | 28 | 0.0714 |
| 2_9175862 | 7 | 0 | 3 | 0 | 3 | 0.3333 | 11 | 0.0909 | 24 | 0.0833 | 1 | 0 | 6 | 0.1667 | 6 | 0 | 6 | 0 | 19 | 0.0526 |
| 2_9175865 | 5 | 0.2 | 4 | 0.25 | 3 | 0 | 10 | 0.3 | 22 | 0.2273 | 1 | 0 | 11 | 0.2727 | 8 | 0.375 | 7 | 0.1429 | 27 | 0.2593 |
| 2_9175878 | 6 | 0.6667 | 4 | 0.5 | 3 | 0.6667 | 10 | 0.6 | 23 | 0.6087 | NA | NA | 10 | 1 | 8 | 0.625 | 7 | 0.4286 | 25 | 0.72 |
| 2_9175879 | 4 | 1 | 4 | 1 | 2 | 1 | 8 | 1 | 18 | 1 | NA | NA | 10 | 1 | 6 | 0.8333 | 5 | 1 | 21 | 0.9524 |
| 2_9175898 | 5 | 0.6 | 4 | 0.75 | 2 | 0.5 | 10 | 0.4 | 21 | 0.5238 | 2 | 0.5 | 9 | 0.6667 | 5 | 0.8 | 5 | 0.4 | 21 | 0.619 |
| 2_9175901 | 8 | 0.25 | 3 | 0 | 6 | 0.1667 | 12 | 0.1667 | 29 | 0.1724 | 2 | 1 | 10 | 0.1 | 6 | 0.1667 | 5 | 0.2 | 23 | 0.2174 |
| 2_9175915 | 5 | 0.2 | 4 | 0 | 5 | 0 | 12 | 0.1667 | 26 | 0.1154 | 2 | 0 | 8 | 0 | 10 | 0.2 | 7 | 0 | 27 | 0.0741 |
| 2_9175925 | 7 | 0.8571 | 6 | 0.8333 | 4 | 1 | 7 | 1 | 24 | 0.9167 | 2 | 0.5 | 9 | 1 | 6 | 0.8333 | 7 | 0.8571 | 24 | 0.875 |
| 2_9175937 | 7 | 0.4286 | 5 | 0.4 | 6 | 1 | 12 | 0.6667 | 30 | 0.6333 | 1 | 1 | 12 | 0.5833 | 13 | 0.5385 | 8 | 0.625 | 34 | 0.5882 |
| 2_9175953 | 8 | 0.25 | 5 | 0 | 5 | 0.4 | 14 | 0.2857 | 32 | 0.25 | 2 | 0.5 | 16 | 0.125 | 11 | 0.4545 | 9 | 0.2222 | 38 | 0.2632 |
| 2_9175961 | 5 | 0.2 | 3 | 0.3333 | 6 | 0 | 11 | 0.0909 | 25 | 0.12 | 1 | 0 | 9 | 0.2222 | 11 | 0.1818 | 5 | 0 | 26 | 0.1538 |
| 2_9175964 | 9 | 0.2222 | 4 | 0.5 | 5 | 0.6 | 13 | 0.5385 | 31 | 0.4516 | 2 | 0.5 | 12 | 0.1667 | 14 | 0.7143 | 8 | 0.5 | 36 | 0.4722 |
| 2_9175965 | 9 | 0.6667 | 5 | 0.6 | 6 | 0.6667 | 13 | 0.6154 | 33 | 0.6364 | 2 | 1 | 15 | 0.8 | 13 | 0.6923 | 7 | 0.4286 | 37 | 0.7027 |
| 2_9175973 | 7 | 0.1429 | 6 | 0.1667 | 5 | 0 | 11 | 0.2727 | 29 | 0.1724 | 2 | 0.5 | 9 | 0.3333 | 9 | 0 | 6 | 0 | 26 | 0.1538 |
| 2_9176325 | 32 | 0.75 | 38 | 0.5789 | 37 | 0.6216 | 19 | 0.7895 | 126 | 0.6667 | 35 | 0.6 | 29 | 0.6207 | 32 | 0.6562 | 35 | 0.6 | 131 | 0.6183 |
| 2_9176334 | 31 | 0.0645 | 35 | 0.2 | 36 | 0.1111 | 22 | 0.0909 | 124 | 0.121 | 31 | 0.2258 | 29 | 0.1034 | 30 | 0.0333 | 32 | 0.0312 | 122 | 0.0984 |
| 2_9176343 | 31 | 0.5484 | 36 | 0.3889 | 38 | 0.5 | 20 | 0.55 | 125 | 0.488 | 37 | 0.3784 | 28 | 0.5 | 33 | 0.4545 | 32 | 0.4375 | 130 | 0.4385 |

| | | | | | | | | | | | | | | | | | | | | |
|---|---|---|---|---|---|---|---|---|---|---|---|---|---|---|---|---|---|---|---|---|
| 2_9176348 | 25 | 0.24 | 23 | 0.1739 | 28 | 0.1786 | 9 | 0 | 85 | 0.1765 | 26 | 0.1923 | 20 | 0.3 | 27 | 0.2222 | 30 | 0.2333 | 103 | 0.233 |
| 2_9176358 | 26 | 1 | 34 | 0.9706 | 31 | 0.9677 | 19 | 0.9474 | 110 | 0.9727 | 31 | 0.7742 | 25 | 0.96 | 29 | 0.9655 | 31 | 0.8387 | 116 | 0.8793 |
| 2_9176360 | 26 | 0.8846 | 27 | 0.963 | 31 | 0.871 | 18 | 0.8889 | 102 | 0.902 | 29 | 0.7931 | 19 | 0.8421 | 32 | 0.9062 | 28 | 0.7143 | 108 | 0.8148 |
| 2_9176371 | 28 | 0.3929 | 35 | 0.4286 | 27 | 0.5556 | 19 | 0.7895 | 109 | 0.5138 | 30 | 0.6333 | 28 | 0.2857 | 33 | 0.5758 | 34 | 0.4706 | 125 | 0.496 |
| 2_9176372 | 22 | 0.0455 | 33 | 0.2121 | 31 | 0.2258 | 16 | 0.125 | 102 | 0.1667 | 27 | 0.2222 | 26 | 0.1154 | 25 | 0.08 | 33 | 0.1212 | 111 | 0.1351 |
| 2_9176497 | NA | NA | 1 | 1 | 1 | 0 | 1 | 0 | 3 | 0.3333 | NA | NA | 1 | 0 | 2 | 0.5 | 1 | 0 | 4 | 0.25 |
| 2_9176563 | 1 | 0 | 1 | 0 | 6 | 0.1667 | 4 | 0 | 12 | 0.0833 | 5 | 0 | 3 | 0.3333 | 6 | 0 | 3 | 0 | 17 | 0.0588 |
| 2_9176582 | 1 | 0 | 1 | 0 | 8 | 0.25 | 3 | 0.3333 | 13 | 0.2308 | 5 | 0.8 | 3 | 0.3333 | 3 | 0.3333 | 5 | 0.4 | 16 | 0.5 |
| 2_9180609 | 4 | 0.5 | 3 | 0.3333 | 2 | 0.5 | 3 | 0.6667 | 12 | 0.5 | 4 | 0.5 | 3 | 0.3333 | NA | NA | 1 | 1 | 8 | 0.5 |
| 2_9180611 | 4 | 0 | 3 | 0 | 3 | 0.6667 | 4 | 0.75 | 14 | 0.3571 | 2 | 0.5 | 3 | 0 | 1 | 0 | 1 | 1 | 7 | 0.2857 |
| 2_9180615 | 3 | 0.3333 | 4 | 0 | 3 | 0.3333 | 3 | 0.6667 | 13 | 0.3077 | 2 | 0.5 | 3 | 0 | 1 | 0 | 1 | 0 | 7 | 0.1429 |
| 2_9180617 | 3 | 0.6667 | 3 | 0.6667 | 3 | 0.6667 | 3 | 0.6667 | 12 | 0.6667 | 3 | 0.3333 | 2 | 0.5 | 1 | 0 | NA | NA | 6 | 0.3333 |
| 2_9180618 | 4 | 0.75 | 4 | 0.5 | 3 | 0.3333 | 5 | 0.4 | 16 | 0.5 | 4 | 0 | 4 | 0.5 | 1 | 1 | 1 | 0 | 10 | 0.3 |
| 2_9180635 | 4 | 0.25 | 2 | 0.5 | 3 | 0 | 5 | 0.2 | 14 | 0.2143 | 3 | 0.6667 | 2 | 0.5 | 1 | 1 | 1 | 1 | 7 | 0.7143 |
| 2_9180650 | 4 | 0.5 | 3 | 0.3333 | 3 | 0 | 5 | 0 | 15 | 0.2 | 3 | 0 | 3 | 0.3333 | 1 | 0 | 1 | 0 | 8 | 0.125 |
| 2_9188929 | 1 | 0 | 2 | 0.5 | 1 | 1 | 2 | 0.5 | 6 | 0.5 | NA | NA | 1 | 1 | NA | NA | 3 | 0.6667 | 4 | 0.75 |
| 2_9188947 | 1 | 0 | 2 | 0.5 | 2 | 0.5 | 4 | 0 | 9 | 0.2222 | NA | NA | 1 | 1 | NA | NA | 3 | 0.3333 | 4 | 0.5 |
| 2_9188962 | 1 | 1 | 1 | 1 | NA | NA | 2 | 1 | 4 | 1 | NA | NA | 1 | 1 | NA | NA | 3 | 1 | 4 | 1 |
| 2_9188964 | 1 | 0 | 2 | 1 | 1 | 1 | 4 | 1 | 8 | 0.875 | NA | NA | 1 | 1 | NA | NA | 2 | 1 | 3 | 1 |
| 2_9188977 | 1 | 1 | 1 | 1 | 2 | 1 | 1 | 1 | 5 | 1 | NA | NA | NA | NA | NA | NA | 2 | 0.5 | 2 | 0.5 |
| 2_9188987 | 1 | 0 | 3 | 0.6667 | 3 | 0.6667 | 4 | 0.75 | 11 | 0.6364 | NA | NA | 1 | 0 | NA | NA | 2 | 1 | 3 | 0.6667 |
| 2_9189028 | 1 | 0 | 2 | 0.5 | 3 | 0 | 3 | 0.3333 | 9 | 0.2222 | NA | NA | NA | NA | NA | NA | 2 | 0.5 | 2 | 0.5 |
| 2_9189029 | NA | NA | NA | NA | 3 | 1 | 3 | 0.6667 | 6 | 0.8333 | NA | NA | 1 | 1 | NA | NA | 3 | 0.6667 | 4 | 0.75 |
| 2_9189051 | 1 | 0 | 2 | 1 | 1 | 0 | 5 | 0.6 | 9 | 0.5556 | NA | NA | 1 | 1 | NA | NA | 4 | 0.5 | 5 | 0.6 |
| 2_9189070 | 1 | 0 | 3 | 0.3333 | 3 | 0 | 4 | 0.25 | 11 | 0.1818 | NA | NA | 1 | 0 | NA | NA | 4 | 0.25 | 5 | 0.2 |
| 2_9189087 | 1 | 1 | 3 | 1 | 3 | 0.3333 | 3 | 1 | 10 | 0.8 | NA | NA | NA | NA | NA | NA | 3 | 1 | 3 | 1 |
| 2_9189110 | 1 | 1 | 3 | 0.3333 | 4 | 0.25 | 4 | 0.5 | 12 | 0.4167 | NA | NA | 1 | 0 | NA | NA | 3 | 0.6667 | 4 | 0.5 |
| 2_9189125 | 1 | 0 | 3 | 0 | 4 | 0 | 4 | 0.25 | 12 | 0.0833 | NA | NA | 1 | 0 | NA | NA | 4 | 0.25 | 5 | 0.2 |
| 2_9189136 | 1 | 0 | 2 | 1 | 3 | 0.3333 | 5 | 0.6 | 11 | 0.5455 | NA | NA | 1 | 0 | NA | NA | 4 | 0.25 | 5 | 0.2 |
| 2_9189221 | 3 | 0 | 3 | 0.3333 | 5 | 0 | 4 | 0 | 15 | 0.0667 | NA | NA | 2 | 0.5 | NA | NA | 4 | 0 | 6 | 0.1667 |
| 2_9189253 | 1 | 1 | 1 | 1 | NA | NA | NA | NA | 2 | 1 | NA | NA | 2 | 0.5 | NA | NA | 1 | 0 | 3 | 0.3333 |
| 2_9189260 | 3 | 0.6667 | 2 | 1 | NA | NA | NA | NA | 5 | 0.8 | NA | NA | 2 | 1 | NA | NA | 1 | 1 | 3 | 1 |
| 2_9219063 | 3 | 1 | 2 | 1 | 1 | 1 | 4 | 1 | 10 | 1 | 1 | 1 | 1 | 0 | 3 | 1 | 1 | 1 | 6 | 0.8333 |
| 2_9219067 | 4 | 0.5 | 2 | 0 | NA | NA | 4 | 0.5 | 10 | 0.4 | 1 | 0 | 1 | 0 | 3 | 0.6667 | 3 | 0.3333 | 8 | 0.375 |
| 2_9219074 | 3 | 1 | 2 | 1 | NA | NA | 3 | 1 | 8 | 1 | 1 | 1 | 1 | 1 | 3 | 1 | 2 | 1 | 7 | 1 |
| 2_9219076 | 4 | 0.5 | 1 | 1 | NA | NA | 4 | 0.75 | 9 | 0.6667 | 1 | 1 | 1 | 1 | 3 | 1 | 2 | 0.5 | 7 | 0.8571 |
| 2_9219078 | 4 | 0.5 | 1 | 0 | 1 | 0 | 4 | 0.5 | 10 | 0.4 | 1 | 0 | 1 | 1 | 3 | 0.6667 | 2 | 0.5 | 7 | 0.5714 |
| 2_9219095 | 3 | 0.6667 | 1 | 1 | NA | NA | 2 | 1 | 6 | 0.8333 | 1 | 1 | 1 | 1 | 2 | 1 | 3 | 1 | 7 | 1 |
| 2_9219096 | 3 | 1 | 2 | 1 | NA | NA | 4 | 1 | 9 | 1 | NA | NA | 1 | 1 | 2 | 1 | 2 | 1 | 5 | 1 |
| 2_9219101 | 3 | 0.6667 | 2 | 0.5 | NA | NA | 4 | 1 | 9 | 0.7778 | 1 | 1 | 1 | 1 | 3 | 0.3333 | 2 | 0.5 | 7 | 0.5714 |
| 2_9220394 | NA | NA | 1 | 1 | NA | NA | NA | NA | 1 | 1 | 1 | 1 | NA | NA | NA | NA | NA | NA | 1 | 1 |
| 2_9220422 | NA | NA | 1 | 1 | NA | NA | NA | NA | 1 | 1 | 1 | 1 | NA | NA | NA | NA | NA | NA | 1 | 1 |
| 2_9220516 | NA | NA | 1 | 1 | NA | NA | NA | NA | 1 | 1 | 1 | 1 | NA | NA | NA | NA | NA | NA | 1 | 1 |
| 2_9220529 | NA | NA | 1 | 1 | NA | NA | NA | NA | 1 | 1 | 1 | 1 | NA | NA | NA | NA | NA | NA | 1 | 1 |
| 2_9221139 | 2 | 0 | 1 | 1 | 2 | 0 | 5 | 0 | 10 | 0.1 | 1 | 0 | 1 | 1 | 2 | 0.5 | 2 | 1 | 6 | 0.6667 |
| 2_9221425 | 70 | 0.7429 | 78 | 0.8205 | 65 | 0.8308 | 58 | 0.7414 | 271 | 0.786 | 62 | 0.8548 | 56 | 0.6607 | 77 | 0.8312 | 53 | 0.8302 | 248 | 0.7984 |
| 2_9221443 | 57 | 0.1228 | 65 | 0.1231 | 58 | 0.0517 | 54 | 0.037 | 234 | 0.0855 | 55 | 0.0909 | 49 | 0.0408 | 70 | 0.1429 | 44 | 0.1818 | 218 | 0.1147 |
| 2_9221450 | 57 | 0.0526 | 61 | 0.0656 | 49 | 0.0816 | 50 | 0.08 | 217 | 0.0691 | 41 | 0.0488 | 40 | 0.025 | 61 | 0.0656 | 40 | 0.075 | 182 | 0.0549 |
| 2_9221699 | 42 | 0.3333 | 40 | 0.175 | 46 | 0.3696 | 35 | 0.3714 | 163 | 0.3129 | 29 | 0.2069 | 25 | 0.28 | 42 | 0.2143 | 32 | 0.4062 | 128 | 0.2734 |
| 2_9221717 | 33 | 0.1515 | 29 | 0.1724 | 36 | 0.1111 | 28 | 0.2143 | 126 | 0.1587 | 22 | 0.0455 | 14 | 0 | 31 | 0.129 | 26 | 0.1923 | 93 | 0.1075 |
| 2_9221722 | 37 | 0.1351 | 35 | 0.2 | 39 | 0.2051 | 30 | 0.1667 | 141 | 0.1773 | 28 | 0.1786 | 23 | 0 | 40 | 0.125 | 31 | 0.129 | 122 | 0.1148 |
| 2_9221737 | 26 | 0.1923 | 18 | 0.1667 | 27 | 0.1111 | 19 | 0.2105 | 90 | 0.1667 | 16 | 0.25 | 12 | 0.1667 | 21 | 0.0952 | 18 | 0.2222 | 67 | 0.1791 |
| 2_9221973 | NA | NA | NA | NA | 2 | 1 | NA | NA | 2 | 1 | NA | NA | 1 | 1 | 1 | 1 | NA | NA | 2 | 1 |
| 2_9223412 | 27 | 0.1481 | 16 | 0.0625 | 18 | 0.0556 | 22 | 0.2273 | 83 | 0.1325 | 16 | 0.0625 | 17 | 0.0588 | 19 | 0.1053 | 15 | 0.0667 | 67 | 0.0746 |
| 2_9223416 | 24 | 0.6667 | 15 | 0.6667 | 17 | 0.5294 | 25 | 0.72 | 81 | 0.6543 | 17 | 0.8235 | 16 | 0.5 | 20 | 0.6 | 10 | 0.6 | 63 | 0.6349 |
| 2_9223437 | 23 | 0.1304 | 17 | 0.1176 | 16 | 0.0625 | 21 | 0.1429 | 77 | 0.1169 | 16 | 0.0625 | 17 | 0.0588 | 16 | 0.3125 | 11 | 0.0909 | 60 | 0.1333 |
| 2_9223521 | 21 | 0.5238 | 12 | 0.5 | 11 | 0.3636 | 14 | 0.1429 | 58 | 0.3966 | 12 | 0.1667 | 13 | 0.5385 | 9 | 0.2222 | 11 | 0.3636 | 45 | 0.3333 |
| 2_9223522 | 21 | 0.2857 | 13 | 0.2308 | 11 | 0.1818 | 15 | 0.2 | 60 | 0.2333 | 12 | 0.0833 | 14 | 0.4286 | 11 | 0.1818 | 11 | 0.1818 | 48 | 0.2292 |
| 2_9223524 | 20 | 0.25 | 14 | 0 | 12 | 0.1667 | 16 | 0.0625 | 62 | 0.129 | 12 | 0.1667 | 14 | 0.3571 | 9 | 0.1111 | 12 | 0.4167 | 47 | 0.2766 |
| 2_9228426 | 2 | 1 | 4 | 1 | NA | NA | 1 | 1 | 7 | 1 | 3 | 1 | 2 | 1 | 3 | 1 | 5 | 0.8 | 13 | 0.9231 |
| 2_9228461 | 2 | 0 | 4 | 0.25 | NA | NA | 1 | 0 | 7 | 0.1429 | 2 | 0 | 2 | 0.5 | 3 | 0 | 4 | 0 | 11 | 0.0909 |
| 2_9228468 | 1 | 1 | 2 | 0.5 | NA | NA | 1 | 1 | 4 | 0.75 | 1 | 1 | 1 | 0 | 3 | 1 | 4 | 0.75 | 9 | 0.7778 |
| 2_9228473 | 2 | 0.5 | 2 | 0 | 1 | 1 | 1 | 0 | 6 | 0.3333 | 2 | 0 | 2 | 0 | 3 | 0 | 5 | 0.4 | 12 | 0.1667 |
| 2_9228480 | 1 | 1 | 2 | 1 | NA | NA | 1 | 1 | 4 | 1 | 2 | 1 | 1 | 1 | 3 | 1 | 3 | 1 | 9 | 1 |
| 2_9228480 | 2 | 1 | 1 | 1 | NA | NA | 1 | 0 | 4 | 0.75 | 2 | 0 | 2 | 0 | 3 | 0.6667 | 4 | 0 | 11 | 0.1818 |

| | | | | | | | | | | | | | | | | | | | | |
|---|---|---|---|---|---|---|---|---|---|---|---|---|---|---|---|---|---|---|---|---|
| 2_9228921 | NA | NA | 1 | 1 | NA | NA | NA | NA | 1 | 1 | 2 | 1 | 1 | 1 | 1 | 1 | 2 | 1 | 6 | 1 |
| 2_9228949 | 1 | 1 | 1 | 0 | NA | NA | NA | NA | 2 | 0.5 | 1 | 0 | 2 | 0 | 1 | 1 | 3 | 0 | 7 | 0.1429 |
| 2_9228958 | 1 | 0 | 1 | 0 | NA | NA | 1 | 1 | 3 | 0.3333 | 2 | 0.5 | 1 | 1 | 2 | 1 | 3 | 0 | 8 | 0.5 |
| 2_9228968 | 1 | 1 | 1 | 0 | NA | NA | 1 | 0 | 3 | 0.3333 | 1 | 1 | 2 | 0 | 2 | 1 | 2 | 0 | 7 | 0.4286 |
| 2_9229085 | NA | NA | 1 | 1 | NA | NA | NA | NA | 1 | 1 | NA | NA | 1 | 1 | NA | NA | NA | NA | 1 | 1 |
| 2_9229101 | NA | NA | 1 | 1 | NA | NA | NA | NA | 1 | 1 | NA | NA | 1 | 1 | NA | NA | NA | NA | 1 | 1 |
| 2_9244389 | 25 | 0.36 | 33 | 0.4848 | 28 | 0.4286 | 26 | 0.5 | 112 | 0.4464 | 28 | 0.5714 | 23 | 0.3913 | 28 | 0.5357 | 30 | 0.3333 | 109 | 0.4587 |
| 2_9244393 | 22 | 0.3182 | 25 | 0.24 | 27 | 0.3704 | 22 | 0.1818 | 96 | 0.2812 | 26 | 0.1154 | 20 | 0.25 | 25 | 0.32 | 26 | 0.3462 | 97 | 0.2577 |
| 2_9244406 | 31 | 1 | 34 | 1 | 27 | 1 | 26 | 1 | 118 | 1 | 32 | 1 | 23 | 1 | 29 | 1 | 30 | 0.9333 | 114 | 0.9825 |
| 2_9244409 | 31 | 0.1935 | 38 | 0.0789 | 30 | 0.2333 | 27 | 0.1481 | 126 | 0.1587 | 29 | 0.1379 | 23 | 0.1304 | 29 | 0.2759 | 31 | 0.2903 | 112 | 0.2143 |
| 2_9244410 | 30 | 0.0667 | 36 | 0.0833 | 28 | 0.0357 | 31 | 0.0968 | 125 | 0.072 | 31 | 0.0968 | 22 | 0.0455 | 34 | 0.1176 | 32 | 0.125 | 119 | 0.1008 |
| 2_9244414 | 28 | 0.8929 | 35 | 0.8857 | 28 | 0.9643 | 30 | 0.8333 | 121 | 0.8926 | 31 | 0.9355 | 26 | 0.8462 | 27 | 0.8519 | 28 | 0.8571 | 112 | 0.875 |
| 2_9244417 | 33 | 0.3636 | 31 | 0.5484 | 27 | 0.2593 | 28 | 0.3214 | 119 | 0.3782 | 31 | 0.3226 | 26 | 0.4231 | 31 | 0.3871 | 30 | 0.3667 | 118 | 0.3729 |
| 2_9244426 | 35 | 0.0857 | 34 | 0.0294 | 26 | 0 | 32 | 0.0938 | 127 | 0.0551 | 34 | 0.1176 | 30 | 0.0333 | 32 | 0.0312 | 33 | 0.0606 | 129 | 0.062 |
| 2_9244427 | 32 | 0.4062 | 35 | 0.4571 | 26 | 0.6154 | 28 | 0.3929 | 121 | 0.4628 | 32 | 0.4375 | 28 | 0.4286 | 32 | 0.3438 | 34 | 0.3824 | 126 | 0.3968 |
| 2_9244444 | 37 | 0.7838 | 39 | 0.7692 | 33 | 0.7879 | 35 | 0.8 | 144 | 0.7847 | 33 | 0.7879 | 28 | 0.8214 | 35 | 0.7714 | 36 | 0.8333 | 132 | 0.803 |
| 2_9244450 | 29 | 0.9655 | 32 | 0.9688 | 22 | 1 | 31 | 1 | 114 | 0.9825 | 25 | 0.96 | 25 | 0.96 | 30 | 0.9667 | 27 | 1 | 107 | 0.972 |
| 2_9244453 | 21 | 0.5714 | 24 | 0.5417 | 21 | 0.2857 | 20 | 0.4 | 86 | 0.4535 | 19 | 0.4737 | 17 | 0.4118 | 19 | 0.3684 | 22 | 0.4545 | 77 | 0.4286 |
| 2_9244462 | 37 | 0.1351 | 41 | 0.0976 | 30 | 0.0333 | 36 | 0.1389 | 144 | 0.1042 | 34 | 0.1176 | 30 | 0.1667 | 36 | 0.0556 | 35 | 0.1143 | 135 | 0.1111 |
| 2_9244470 | 33 | 0.8788 | 30 | 0.8 | 24 | 0.7917 | 28 | 0.7857 | 115 | 0.8174 | 30 | 0.8333 | 24 | 0.9583 | 23 | 0.8696 | 24 | 0.8333 | 101 | 0.8713 |
| 2_9244474 | 41 | 0.0976 | 36 | 0.2778 | 29 | 0.069 | 38 | 0.2105 | 144 | 0.1667 | 34 | 0.1471 | 31 | 0.129 | 38 | 0.1053 | 33 | 0.1818 | 136 | 0.1397 |
| 2_9244476 | 14 | 0.2143 | 14 | 0.4286 | 10 | 0.4 | 14 | 0.2857 | 52 | 0.3269 | 11 | 0.2727 | 13 | 0.4615 | 16 | 0.1875 | 9 | 0.3333 | 49 | 0.3061 |
| 2_9244481 | 36 | 0.1111 | 34 | 0.1176 | 29 | 0 | 28 | 0.1071 | 127 | 0.0866 | 29 | 0.069 | 26 | 0.0769 | 27 | 0.037 | 31 | 0.0968 | 113 | 0.0708 |
| 2_9244501 | 33 | 0 | 28 | 0.1786 | 25 | 0 | 32 | 0.0312 | 118 | 0.0508 | 28 | 0.0714 | 27 | 0.1111 | 33 | 0.0303 | 31 | 0.0323 | 119 | 0.0588 |
| 2_9244522 | 43 | 0.0465 | 40 | 0.1 | 33 | 0.0606 | 38 | 0.1053 | 154 | 0.0779 | 38 | 0.1316 | 28 | 0.1429 | 36 | 0.1389 | 37 | 0.027 | 139 | 0.1079 |
| 2_9244536 | 43 | 0.2093 | 43 | 0.186 | 38 | 0.2368 | 41 | 0.0732 | 165 | 0.1758 | 38 | 0.2895 | 31 | 0.2903 | 40 | 0.125 | 43 | 0.2326 | 152 | 0.2303 |
| 2_9259219 | NA | NA | NA | NA | NA | NA | 1 | 1 | 1 | 1 | NA | NA | NA | NA | NA | NA | 1 | 1 | 1 | 1 |
| 2_9277611 | NA | NA | 1 | 1 | 1 | 0 | NA | NA | 2 | 0.5 | 1 | 1 | NA | NA | NA | NA | 1 | 0 | 2 | 0.5 |
| 2_9278405 | 6 | 0.8333 | 16 | 0.875 | 8 | 1 | 4 | 1 | 34 | 0.9118 | 7 | 0.8571 | 5 | 1 | NA | NA | 10 | 0.7 | 22 | 0.8182 |
| 2_9278411 | 7 | 0.8571 | 14 | 1 | 7 | 0.8571 | 5 | 1 | 33 | 0.9394 | 5 | 0.8 | 3 | 1 | 2 | 0.5 | 10 | 0.9 | 20 | 0.85 |
| 2_9278420 | 7 | 0.8571 | 20 | 0.9 | 10 | 1 | 4 | 0.75 | 41 | 0.9024 | 9 | 0.5556 | 6 | 0.8333 | 1 | 1 | 13 | 0.9231 | 29 | 0.7931 |
| 2_9278421 | 8 | 1 | 20 | 0.7 | 8 | 0.875 | 3 | 1 | 39 | 0.8205 | 9 | 0.8889 | 5 | 1 | 2 | 0.5 | 16 | 0.75 | 32 | 0.8125 |
| 2_9278423 | 8 | 0.75 | 24 | 0.9167 | 11 | 0.8182 | 5 | 0.8 | 48 | 0.8542 | 9 | 1 | 6 | 1 | 2 | 1 | 16 | 0.875 | 33 | 0.9394 |
| 2_9278441 | 8 | 0.125 | 18 | 0.1111 | 7 | 0.1429 | NA | NA | 33 | 0.1212 | 6 | 0 | 6 | 0 | 2 | 0 | 14 | 0.2857 | 28 | 0.1429 |
| 2_9278445 | 7 | 0 | 22 | 0.3182 | 9 | 0.4444 | 4 | 0.75 | 42 | 0.3333 | 8 | 0.125 | 7 | 0.2857 | 3 | 0.3333 | 17 | 0.1765 | 35 | 0.2 |
| 2_9278468 | 6 | 0.1667 | 21 | 0.0952 | 11 | 0 | 2 | 0 | 40 | 0.075 | 7 | 0 | 6 | 0.1667 | 3 | 0.3333 | 16 | 0.1875 | 32 | 0.1562 |
| 2_9278504 | 8 | 0.25 | 19 | 0.1579 | 11 | 0.2727 | 7 | 0.2857 | 45 | 0.2222 | 12 | 0.1667 | 6 | 0.1667 | 5 | 0 | 18 | 0.1111 | 41 | 0.122 |
| 2_9278580 | 7 | 0 | 22 | 0.0909 | 14 | 0.0714 | 7 | 0 | 50 | 0.06 | 14 | 0.0714 | 8 | 0.25 | 8 | 0.375 | 19 | 0.0526 | 49 | 0.1429 |
| 2_9278586 | 11 | 0.2727 | 25 | 0.08 | 16 | 0.0625 | 10 | 0 | 62 | 0.0968 | 19 | 0.1579 | 8 | 0 | 9 | 0.1111 | 23 | 0.087 | 59 | 0.1017 |
| 2_9278871 | NA | NA | 1 | 1 | NA | NA | NA | NA | 1 | 1 | NA | NA | NA | NA | NA | NA | 1 | 1 | 1 | 1 |
| 2_9278874 | NA | NA | 2 | 0.5 | NA | NA | NA | NA | 2 | 0.5 | NA | NA | NA | NA | NA | NA | 1 | 1 | 1 | 1 |
| 2_9303031 | 8 | 0 | 6 | 0.3333 | 3 | 0.3333 | 4 | 0.5 | 21 | 0.2381 | 5 | 0.4 | 5 | 0 | 3 | 0 | 4 | 0 | 17 | 0.1176 |
| 2_9303158 | 12 | 0 | 8 | 0.125 | 7 | 0.1429 | 8 | 0.125 | 35 | 0.0857 | 6 | 0 | 9 | 0.1111 | 6 | 0.1667 | 11 | 0 | 32 | 0.0625 |
| 2_9303158 | 12 | 0 | 10 | 0 | 7 | 0 | 9 | 0.3333 | 38 | 0.0789 | 6 | 0.1667 | 9 | 0.1111 | 6 | 0 | 11 | 0.0909 | 32 | 0.0938 |
| 2_9303413 | NA | NA | 1 | 1 | NA | NA | NA | NA | 1 | 1 | 1 | 1 | NA | NA | NA | NA | NA | NA | 1 | 1 |
| 2_9303423 | NA | NA | 1 | 1 | NA | NA | NA | NA | 1 | 1 | 1 | 1 | NA | NA | NA | NA | NA | NA | 1 | 1 |
| 2_9303430 | NA | NA | 1 | 1 | NA | NA | NA | NA | 1 | 1 | 1 | 1 | NA | NA | NA | NA | NA | NA | 1 | 1 |
| 2_9303443 | NA | NA | 1 | 1 | NA | NA | NA | NA | 1 | 1 | 1 | 1 | NA | NA | NA | NA | NA | NA | 1 | 1 |
| 2_9303451 | NA | NA | 1 | 1 | NA | NA | NA | NA | 1 | 1 | 1 | 1 | NA | NA | NA | NA | NA | NA | 1 | 1 |
| 2_9324958 | 1 | 1 | 3 | 0 | 2 | 0 | NA | NA | 6 | 0.1667 | 4 | 0 | 4 | 0.75 | NA | NA | NA | NA | 8 | 0.375 |
| 2_9352609 | 3 | 0.3333 | 7 | 0.4286 | 3 | 0.6667 | 4 | 0.75 | 17 | 0.5294 | 4 | 0.5 | 7 | 0.7143 | 3 | 0.6667 | 4 | 0.75 | 18 | 0.6667 |
| 2_9352610 | 1 | 1 | 6 | 1 | 2 | 1 | 3 | 1 | 12 | 1 | 2 | 1 | 5 | 1 | 2 | 1 | 4 | 1 | 13 | 1 |
| 2_9352616 | 2 | 0.5 | 6 | 0.5 | 2 | 1 | 4 | 0.25 | 14 | 0.5 | 2 | 1 | 7 | 0.8571 | 3 | 1 | 2 | 0.5 | 14 | 0.8571 |
| 2_9352631 | 3 | 0.6667 | 7 | 0.2857 | 3 | 1 | 4 | 0.25 | 17 | 0.4706 | 4 | 0.5 | 7 | 0.7143 | 3 | 0.6667 | 5 | 0.6 | 19 | 0.6316 |
| 2_9352635 | 3 | 1 | 4 | 1 | 3 | 1 | 4 | 1 | 14 | 1 | 2 | 1 | 6 | 1 | 3 | 1 | 4 | 1 | 15 | 1 |
| 2_9352651 | 3 | 0.3333 | 6 | 0 | 3 | 0.3333 | 2 | 0.5 | 14 | 0.2143 | 3 | 0 | 6 | 0 | 2 | 0 | 3 | 0.3333 | 14 | 0.0714 |
| 2_9352654 | 3 | 0 | 6 | 0 | 3 | 0.3333 | 3 | 0 | 15 | 0.0667 | 4 | 0 | 7 | 0.1429 | 3 | 0 | 5 | 0.2 | 19 | 0.1053 |
| 2_9353346 | NA | NA | NA | NA | 2 | 1 | NA | NA | 2 | 1 | NA | NA | NA | NA | 1 | 1 | 1 | 1 | 2 | 1 |
| 2_9364388 | 1 | 1 | NA | NA | 2 | 0 | NA | NA | 3 | 0.3333 | 1 | 1 | 2 | 0.5 | 1 | 1 | 2 | 1 | 6 | 0.8333 |
| 2_9366596 | 1 | 1 | NA | NA | NA | NA | NA | NA | 1 | 1 | NA | NA | 2 | 1 | NA | NA | NA | NA | 2 | 1 |
| 2_9378618 | 9 | 0.7778 | 16 | 0.6875 | 7 | 0.5714 | 10 | 0.8 | 42 | 0.7143 | 13 | 0.6154 | 14 | 0.7143 | 14 | 0.7857 | 11 | 0.6364 | 52 | 0.6923 |
| 2_9378631 | 10 | 0.8 | 14 | 0.9286 | 6 | 0.8333 | 8 | 1 | 38 | 0.8947 | 13 | 0.8462 | 14 | 0.9286 | 12 | 0.9167 | 11 | 1 | 50 | 0.92 |
| 2_9378632 | 6 | 0 | 13 | 0.1538 | 4 | 0.25 | 8 | 0.25 | 31 | 0.1613 | 11 | 0.3636 | 11 | 0.4545 | 12 | 0.25 | 11 | 0.3636 | 45 | 0.3556 |
| 2_9378634 | 6 | 0.6667 | 8 | 0.25 | 4 | 0.25 | 3 | 0.3333 | 21 | 0.381 | 7 | 0.2857 | 3 | 0 | 11 | 0.3636 | 7 | 0.7143 | 28 | 0.3929 |
| 2_9378649 | 6 | 0.5 | 9 | 0.8889 | 3 | 0.3333 | 3 | 0.3333 | 21 | 0.619 | 9 | 0.7778 | 7 | 0.8571 | 9 | 0.8889 | 6 | 0.8333 | 31 | 0.8387 |

| | | | | | | | | | | | | | | | | | | | | |
|---|---|---|---|---|---|---|---|---|---|---|---|---|---|---|---|---|---|---|---|---|
| 2_9378659( | 6 | 0 | 14 | 0.1429 | 7 | 0 | 6 | 0 | 33 | 0.0606 | 11 | 0 | 11 | 0.0909 | 13 | 0.0769 | 11 | 0.1818 | 46 | 0.087 |
| 2_9378660{ | 5 | 1 | 11 | 1 | 6 | 1 | 4 | 1 | 26 | 1 | 9 | 1 | 12 | 1 | 9 | 1 | 10 | 1 | 40 | 1 |
| 2_9378671: | 9 | 0.7778 | 13 | 0.7692 | 7 | 1 | 7 | 0.7143 | 36 | 0.8056 | 10 | 0.9 | 13 | 0.6923 | 10 | 0.9 | 11 | 0.7273 | 44 | 0.7955 |
| 2_9378676( | 8 | 0.5 | 14 | 0.3571 | 7 | 0.2857 | 5 | 0 | 34 | 0.3235 | 11 | 0.1818 | 11 | 0.2727 | 12 | 0.5 | 9 | 0.5556 | 43 | 0.3721 |
| 2_9378676! | 7 | 0 | 14 | 0.1429 | 5 | 0 | 6 | 0 | 32 | 0.0625 | 6 | 0.1667 | 10 | 0.1 | 10 | 0 | 8 | 0 | 34 | 0.0588 |
| 2_9381105: | NA | NA | 3 | 0.3333 | 4 | 0.25 | 3 | 0 | 10 | 0.2 | 1 | 0 | 1 | 0 | 2 | 0 | 1 | 1 | 5 | 0.2 |
| 2_9386315! | 18 | 0.0556 | 16 | 0.125 | 13 | 0.0769 | 14 | 0.0714 | 61 | 0.082 | 13 | 0.0769 | 8 | 0 | 12 | 0.0833 | 14 | 0.0714 | 47 | 0.0638 |
| 2_9386335! | 14 | 0.0714 | 10 | 0.1 | 8 | 0 | 11 | 0.1818 | 43 | 0.093 | 10 | 0 | 4 | 0.25 | 11 | 0.0909 | 12 | 0 | 37 | 0.0541 |
| 2_9408234{ | NA | NA | 1 | 1 | 2 | 1 | 1 | 1 | 4 | 1 | NA | NA | 1 | 1 | NA | NA | 1 | 0 | 2 | 0.5 |
| 2_9413130: | 2 | 0.5 | 6 | 0 | 1 | 0 | NA | NA | 9 | 0.1111 | NA | NA | NA | NA | NA | NA | 5 | 0.2 | 5 | 0.2 |
| 2_9413154( | 4 | 0 | 5 | 0.2 | 3 | 0 | NA | NA | 12 | 0.0833 | NA | NA | 1 | 0 | 1 | 0 | 3 | 0.3333 | 5 | 0.2 |
| 2_9413185 | 6 | 0 | 6 | 0 | 4 | 0.25 | NA | NA | 16 | 0.0625 | NA | NA | 1 | 1 | 2 | 0.5 | 4 | 0 | 7 | 0.2857 |
| 2_9424097! | 2 | 1 | NA | NA | NA | NA | NA | NA | 2 | 1 | 1 | 1 | NA | NA | NA | NA | NA | NA | 1 | 1 |
| 2_9424106! | 1 | 1 | NA | NA | 1 | 1 | NA | NA | 2 | 1 | NA | NA | NA | NA | 1 | 1 | 1 | 0 | 2 | 0.5 |
| 2_9424143! | 2 | 0.5 | NA | NA | 1 | 1 | NA | NA | 3 | 0.6667 | 1 | 1 | NA | NA | 1 | 1 | 1 | 0 | 3 | 0.6667 |
| 2_9424192 | 3 | 0.3333 | NA | NA | 1 | 1 | NA | NA | 4 | 0.5 | 1 | 0 | NA | NA | 1 | 1 | 1 | 0 | 3 | 0.3333 |
| 2_9424196 | 3 | 0.3333 | NA | NA | 1 | 0 | NA | NA | 4 | 0.25 | 1 | 0 | NA | NA | 1 | 1 | 1 | 0 | 3 | 0.3333 |
| 2_9424238( | 11 | 0.2727 | 17 | 0.7059 | 7 | 0.5714 | 22 | 0.4545 | 57 | 0.5088 | 6 | 0.5 | 8 | 0.625 | 13 | 0.6154 | 11 | 0.2727 | 38 | 0.5 |
| 2_9424243( | 13 | 0.2308 | 17 | 0.1176 | 6 | 0 | 21 | 0.2381 | 57 | 0.1754 | 6 | 0.3333 | 7 | 0.5714 | 12 | 0.3333 | 15 | 0.1333 | 40 | 0.3 |
| 2_9424253! | 10 | 0 | 14 | 0.2143 | 7 | 0 | 15 | 0.0667 | 46 | 0.087 | 5 | 0 | 5 | 0 | 9 | 0 | 12 | 0.1667 | 31 | 0.0645 |
| 2_9424389 | 10 | 0.1 | 13 | 0.1538 | 6 | 0 | 19 | 0 | 48 | 0.0625 | 3 | 0 | 5 | 0.2 | 10 | 0.2 | 10 | 0.2 | 28 | 0.1786 |
| 2_9424395! | 10 | 0.6 | 12 | 1 | 6 | 0.8333 | 19 | 0.9474 | 47 | 0.8723 | 3 | 1 | 5 | 0.8 | 9 | 0.8889 | 9 | 1 | 26 | 0.9231 |
| 2_9729702( | 1 | 1 | 2 | 1 | 1 | 1 | NA | NA | 4 | 1 | NA | NA | 3 | 0.6667 | 1 | 0 | NA | NA | 4 | 0.5 |
| 2_9729738{ | 2 | 1 | 3 | 1 | 2 | 0.5 | 1 | 1 | 8 | 0.875 | NA | NA | 1 | 1 | 1 | 1 | NA | NA | 2 | 1 |
| 2_9729742: | 3 | 1 | 3 | 0 | 2 | 0 | 1 | 0 | 9 | 0.3333 | NA | NA | 2 | 0.5 | 1 | 0 | NA | NA | 3 | 0.3333 |
| 2_9729751 | 3 | 0.3333 | 2 | 1 | 2 | 0.5 | 1 | 1 | 8 | 0.625 | NA | NA | 3 | 1 | 1 | 1 | NA | NA | 4 | 1 |
| 2_9729817: | 7 | 0.2857 | 3 | 0 | 3 | 0.3333 | 6 | 0 | 19 | 0.1579 | 1 | 0 | 5 | 0.2 | 5 | 0 | 4 | 0 | 15 | 0.0667 |
| 2_9729824{ | 7 | 0.2857 | 4 | 0.75 | 3 | 0.3333 | 6 | 0.3333 | 20 | 0.4 | 1 | 1 | 5 | 0.4 | 5 | 0.4 | 3 | 0.6667 | 14 | 0.5 |
| 2_9729827: | 7 | 0.4286 | 4 | 0.5 | 4 | 0.25 | 7 | 0.1429 | 22 | 0.3182 | 1 | 0 | 3 | 0 | 7 | 0.1429 | 3 | 0 | 14 | 0.0714 |
| 2_9729832! | 6 | 0.8333 | 4 | 0.75 | 3 | 1 | 6 | 0.8333 | 19 | 0.8421 | 1 | 1 | 5 | 1 | 5 | 1 | 5 | 0.6 | 16 | 0.875 |
| 2_9729844( | 6 | 0.5 | 5 | 1 | 4 | 0.75 | 6 | 0.6667 | 21 | 0.7143 | 1 | 1 | 4 | 0.5 | 4 | 0.5 | 6 | 0.8333 | 15 | 0.6667 |
| 2_9729858: | 7 | 0.5714 | 4 | 1 | 4 | 0.5 | 5 | 0.4 | 20 | 0.6 | 1 | 1 | 5 | 0.8 | 6 | 0.6667 | 7 | 0.5714 | 19 | 0.6842 |
| 2_9729860 | 6 | 0 | 4 | 0 | 4 | 0.5 | 4 | 0.25 | 18 | 0.1667 | 1 | 0 | 6 | 0.5 | 6 | 0.6667 | 6 | 0.5 | 19 | 0.5263 |
| 2_9729870( | 7 | 0.1429 | 6 | 0 | 4 | 0.25 | 7 | 0 | 24 | 0.0833 | 1 | 0 | 6 | 0.1667 | 7 | 0 | 6 | 0 | 20 | 0.05 |
| 2_9745965( | 7 | 0.1429 | 6 | 0.1667 | 4 | 0 | 7 | 0 | 24 | 0.0833 | 1 | 1 | 6 | 0 | 9 | 0.1111 | 6 | 0.1667 | 22 | 0.1364 |
| 2_9745970 | 8 | 0.25 | 7 | 0 | 4 | 0 | 9 | 0.1111 | 28 | 0.1071 | 2 | 0 | 7 | 0.1429 | 10 | 0.1 | 8 | 0 | 27 | 0.0741 |
| 2_9745971! | 6 | 0.1667 | 6 | 0 | 2 | 0.5 | 6 | 0 | 20 | 0.1 | 1 | 0 | 8 | 0.25 | 7 | 0.1429 | 8 | 0 | 24 | 0.125 |
| 2_9745976! | 8 | 0.875 | 6 | 1 | 4 | 1 | 7 | 1 | 25 | 0.96 | 2 | 1 | 8 | 1 | 8 | 1 | 8 | 1 | 26 | 1 |
| 2_9745982: | 6 | 0.3333 | 6 | 0.5 | 3 | 0.3333 | 9 | 0.2222 | 24 | 0.3333 | 2 | 1 | 8 | 0.25 | 9 | 0.1111 | 7 | 0.1429 | 26 | 0.2308 |
| 2_9745990{ | 7 | 0.8571 | 7 | 1 | 4 | 1 | 9 | 0.7778 | 27 | 0.8889 | 2 | 1 | 7 | 1 | 9 | 1 | 8 | 0.625 | 26 | 0.8846 |
| 2_9745998: | 8 | 0.5 | 7 | 0.5714 | 4 | 0.25 | 11 | 0.3636 | 30 | 0.4333 | 3 | 0 | 8 | 0.5 | 11 | 0.2727 | 10 | 0.6 | 32 | 0.4062 |
| 2_9746011( | 5 | 1 | 6 | 0.8333 | 4 | 1 | 12 | 1 | 27 | 0.963 | 3 | 1 | 5 | 1 | 10 | 0.9 | 7 | 0.8571 | 25 | 0.92 |
| 2_9746019: | 10 | 0.4 | 6 | 0.3333 | 3 | 0.3333 | 11 | 0.1818 | 30 | 0.3 | 2 | 0.5 | 7 | 0.1429 | 12 | 0.25 | 8 | 0.375 | 29 | 0.2759 |
| 2_9746028( | 10 | 1 | 3 | 1 | 3 | 1 | 6 | 0.8333 | 22 | 0.9545 | 2 | 1 | 6 | 1 | 7 | 1 | 5 | 1 | 20 | 1 |
| 2_9746029: | 8 | 0.375 | 7 | 0 | 3 | 0.3333 | 8 | 0.375 | 26 | 0.2692 | 3 | 0 | 7 | 0.1429 | 9 | 0.2222 | 9 | 0.4444 | 28 | 0.25 |
| 2_9746032{ | 11 | 0.6364 | 7 | 0.4286 | 4 | 0.75 | 11 | 0.8182 | 33 | 0.6667 | 4 | 0.75 | 8 | 0.875 | 12 | 0.5 | 10 | 0.6 | 34 | 0.6471 |
| 2_9746040! | 10 | 0.4 | 6 | 0.6667 | 4 | 0.5 | 12 | 0.4167 | 32 | 0.4688 | 4 | 0.5 | 7 | 0.1429 | 10 | 0.7 | 11 | 0.7273 | 32 | 0.5625 |
| 2_9746046( | 11 | 0.0909 | 6 | 0.1667 | 3 | 0.3333 | 12 | 0 | 32 | 0.0938 | 4 | 0 | 7 | 0.4286 | 11 | 0.1818 | 7 | 0 | 29 | 0.1724 |
| 2_9746054! | 5 | 0.4 | 6 | 0.1667 | 2 | 1 | 7 | 0.4286 | 20 | 0.4 | 4 | 0.25 | 6 | 0.8333 | 8 | 0.875 | 6 | 0.6667 | 24 | 0.7083 |
| 2_9746062 | 9 | 0.3333 | 5 | 0.2 | 2 | 0.5 | 10 | 0.2 | 26 | 0.2692 | 3 | 0.3333 | 7 | 0.2857 | 7 | 0.1429 | 9 | 0.1111 | 26 | 0.1923 |
| 2_9746075( | 10 | 1 | 6 | 1 | 4 | 1 | 13 | 0.8462 | 33 | 0.9394 | 4 | 0.75 | 8 | 1 | 12 | 1 | 11 | 1 | 35 | 0.9714 |
| 2_9746076 | 11 | 0.0909 | 7 | 0.1429 | 2 | 0 | 13 | 0.2308 | 33 | 0.1515 | 4 | 0 | 8 | 0.5 | 12 | 0 | 11 | 0.1818 | 35 | 0.1714 |
| 2_9746078( | 10 | 0.2 | 5 | 0 | 4 | 0 | 11 | 0.3636 | 30 | 0.2 | 5 | 0 | 8 | 0.5 | 11 | 0.3636 | 11 | 0.3636 | 35 | 0.3429 |
| 2_9746088! | 12 | 0.0833 | 8 | 0.25 | 4 | 0.25 | 13 | 0.3077 | 37 | 0.2162 | 4 | 0 | 7 | 0.2857 | 9 | 0.2222 | 11 | 0.0909 | 31 | 0.1613 |
| 2_9746090: | 10 | 0.4 | 7 | 0.4286 | 1 | 0 | 10 | 0.4 | 28 | 0.3929 | 2 | 0 | 8 | 0.125 | 12 | 0.0833 | 10 | 0.3 | 32 | 0.1562 |
| 2_9746106{ | 14 | 1 | 8 | 1 | 6 | 1 | 14 | 1 | 42 | 1 | 5 | 1 | 8 | 1 | 14 | 1 | 11 | 0.9091 | 38 | 0.9737 |
| 2_9746110{ | 10 | 0.6 | 6 | 0.1667 | 6 | 0.6667 | 12 | 0.6667 | 34 | 0.5588 | 4 | 0.25 | 7 | 0.5714 | 13 | 0.5385 | 8 | 0.5 | 32 | 0.5 |
| 2_9746117 | 7 | 0.4286 | 6 | 0.5 | 4 | 0.25 | 9 | 0.4444 | 26 | 0.4231 | 5 | 0.6 | 7 | 0.5714 | 11 | 0.5455 | 10 | 0.4 | 33 | 0.5152 |
| 2_9746124 | 14 | 0.1429 | 8 | 0.25 | 7 | 0.4286 | 13 | 0.3846 | 42 | 0.2857 | 3 | 0 | 8 | 0.5 | 14 | 0.4286 | 10 | 0.4 | 35 | 0.4 |
| 2_9746126 | 11 | 1 | 7 | 1 | 6 | 1 | 8 | 1 | 32 | 1 | 3 | 1 | 6 | 0.8333 | 10 | 1 | 8 | 1 | 27 | 0.963 |
| 2_9746127: | 14 | 0.9286 | 9 | 1 | 8 | 1 | 14 | 0.9286 | 45 | 0.9556 | 5 | 1 | 9 | 0.8889 | 14 | 0.8571 | 12 | 1 | 40 | 0.925 |
| 2_9746128 | 14 | 1 | 10 | 1 | 8 | 1 | 14 | 1 | 46 | 1 | 6 | 1 | 9 | 1 | 14 | 1 | 12 | 1 | 41 | 1 |
| 2_9746130( | 11 | 0.2727 | 10 | 0.2 | 8 | 0.375 | 9 | 0.1111 | 38 | 0.2368 | 5 | 0.2 | 9 | 0.1111 | 13 | 0.2308 | 13 | 0.2308 | 40 | 0.2 |
| 2_9746132: | 14 | 0.0714 | 7 | 0.4286 | 8 | 0 | 12 | 0.1667 | 41 | 0.1463 | 5 | 0.2 | 7 | 0.2857 | 13 | 0.1538 | 12 | 0.0833 | 37 | 0.1622 |
| 2_9746134 | 13 | 1 | 7 | 1 | 8 | 0.875 | 13 | 0.9231 | 41 | 0.9512 | 4 | 1 | 6 | 0.8333 | 10 | 1 | 12 | 1 | 32 | 0.9688 |

| | | | | | | | | | | | | | | | | | | | | |
|---|---|---|---|---|---|---|---|---|---|---|---|---|---|---|---|---|---|---|---|---|
| 2_9746136 | 13 | 0.3077 | 9 | 0.2222 | 8 | 0.375 | 15 | 0.0667 | 45 | 0.2222 | 5 | 0 | 8 | 0.125 | 13 | 0.0769 | 14 | 0.1429 | 40 | 0.1 |
| 2_9746137 | 11 | 0.9091 | 8 | 0.875 | 7 | 1 | 13 | 0.7692 | 39 | 0.8718 | 6 | 0.8333 | 9 | 0.8889 | 13 | 0.8462 | 10 | 0.9 | 38 | 0.8684 |
| 2_9746158 | 13 | 0.3077 | 7 | 0.7143 | 8 | 0.5 | 12 | 0.5833 | 40 | 0.5 | 4 | 0.75 | 5 | 0.4 | 13 | 0.3077 | 8 | 0.5 | 30 | 0.4333 |
| 2_9746185 | 18 | 0.1667 | 7 | 0.2857 | 9 | 0 | 11 | 0.0909 | 45 | 0.1333 | 6 | 0.1667 | 5 | 0.6 | 13 | 0.0769 | 11 | 0.2727 | 35 | 0.2286 |
| 2_1013922 | NA | NA | 2 | 1 | 1 | 1 | 1 | 0 | 4 | 0.75 | 1 | 1 | NA | NA | NA | NA | NA | NA | 1 | 1 |
| 2_1013925 | 9 | 0.1111 | 5 | 0.4 | 9 | 0.1111 | 7 | 0 | 30 | 0.1333 | 7 | 0.1429 | 5 | 0.2 | 12 | 0.1667 | 7 | 0 | 31 | 0.129 |
| 2_1013926 | 10 | 0 | 4 | 0.25 | 9 | 0.1111 | 7 | 0.2857 | 30 | 0.1333 | 4 | 0.25 | 3 | 0 | 10 | 0.3 | 8 | 0.25 | 25 | 0.24 |
| 2_1015270 | 1 | 1 | 1 | 1 | NA | NA | NA | NA | 2 | 1 | NA | NA | 1 | 1 | 1 | 1 | NA | NA | 2 | 1 |
| 2_1015272 | 1 | 1 | 1 | 1 | 1 | 0 | NA | NA | 3 | 0.6667 | NA | NA | 1 | 0 | 1 | 1 | NA | NA | 2 | 0.5 |
| 2_1015272 | 1 | 1 | 1 | 0 | 1 | 0 | NA | NA | 3 | 0.3333 | NA | NA | 1 | 1 | 1 | 1 | NA | NA | 2 | 1 |
| 2_1015272 | 1 | 0 | 1 | 1 | 1 | 0 | NA | NA | 3 | 0.3333 | NA | NA | 1 | 1 | 1 | 1 | NA | NA | 2 | 1 |
| 2_1015273 | 1 | 1 | 1 | 0 | 1 | 0 | NA | NA | 3 | 0.3333 | NA | NA | 1 | 1 | 1 | 1 | NA | NA | 2 | 1 |
| 2_1015274 | 1 | 1 | 1 | 1 | 1 | 1 | NA | NA | 3 | 1 | NA | NA | 1 | 1 | 1 | 1 | NA | NA | 2 | 1 |
| 2_1015275 | 1 | 0 | NA | NA | 1 | 1 | NA | NA | 2 | 0.5 | NA | NA | 1 | 1 | 1 | 0 | NA | NA | 2 | 0.5 |
| 2_1015276 | 1 | 1 | 1 | 1 | 1 | 1 | NA | NA | 3 | 1 | NA | NA | 1 | 1 | 1 | 1 | NA | NA | 2 | 1 |
| 2_1015276 | 1 | 0 | 1 | 1 | 1 | 1 | NA | NA | 3 | 0.6667 | NA | NA | NA | NA | 1 | 1 | NA | NA | 1 | 1 |
| 2_1015284 | 1 | 0 | 1 | 1 | 1 | 0 | 1 | 0 | 4 | 0.25 | NA | NA | 1 | 1 | 1 | 0 | NA | NA | 2 | 0.5 |
| 2_1015308 | 3 | 0 | 1 | 0 | 1 | 0 | 3 | 0.3333 | 8 | 0.125 | NA | NA | 2 | 0.5 | 3 | 0 | NA | NA | 5 | 0.2 |
| 2_1015311 | 2 | 0.5 | 1 | 0 | 1 | 0 | 3 | 0 | 7 | 0.1429 | NA | NA | 3 | 0.3333 | 4 | 0.25 | 1 | 0 | 8 | 0.25 |
| 2_1015311 | 3 | 0 | 1 | 0 | 1 | 0 | 3 | 0.6667 | 8 | 0.25 | NA | NA | 3 | 0.3333 | 5 | 0.4 | 1 | 1 | 9 | 0.4444 |
| 2_1015312 | 2 | 0.5 | 1 | 0 | 1 | 0 | 2 | 0.5 | 6 | 0.3333 | NA | NA | 3 | 0.6667 | 5 | 0.4 | 1 | 0 | 9 | 0.4444 |
| 2_1015316 | 3 | 0.6667 | NA | NA | 1 | 0 | NA | NA | 4 | 0.5 | NA | NA | 2 | 0.5 | 5 | 0.4 | 1 | 1 | 8 | 0.5 |
| 2_1015321 | 1 | 1 | NA | NA | NA | NA | 1 | 0 | 2 | 0.5 | NA | NA | NA | NA | 1 | 1 | 1 | 0 | 2 | 0.5 |
| 2_1015326 | 2 | 1 | NA | NA | NA | NA | 1 | 0 | 3 | 0.6667 | NA | NA | 1 | 1 | 1 | 1 | 1 | 0 | 3 | 0.6667 |
| 2_1015470 | 1 | 1 | 1 | 0 | NA | NA | NA | NA | 2 | 0.5 | 1 | 0 | 1 | 0 | NA | NA | 1 | 1 | 3 | 0.3333 |
| 2_1015471 | 1 | 1 | 1 | 1 | NA | NA | NA | NA | 2 | 1 | 1 | 1 | 1 | 1 | NA | NA | 1 | 1 | 3 | 1 |
| 2_1015474 | 1 | 1 | 1 | 1 | NA | NA | NA | NA | 2 | 1 | 1 | 1 | 1 | 1 | NA | NA | NA | NA | 2 | 1 |
| 2_1015474 | 1 | 1 | 1 | 1 | 1 | 1 | NA | NA | 3 | 1 | 1 | 1 | 1 | 1 | NA | NA | 1 | 1 | 3 | 1 |
| 2_1015474 | 1 | 1 | NA | NA | 1 | 1 | NA | NA | 2 | 1 | 1 | 1 | NA | NA | NA | NA | 1 | 1 | 2 | 1 |
| 2_1015477 | 1 | 0 | NA | NA | 1 | 1 | NA | NA | 2 | 0.5 | 1 | 0 | 1 | 0 | NA | NA | 1 | 1 | 3 | 0.3333 |
| 2_1017314 | 2 | 0 | 3 | 0 | 5 | 0 | 8 | 0.125 | 18 | 0.0556 | 5 | 0.2 | 1 | 0 | 4 | 0 | 5 | 0 | 15 | 0.0667 |
| 2_1017318 | 2 | 0 | 5 | 0.2 | 6 | 0.1667 | 12 | 0.0833 | 25 | 0.12 | 9 | 0 | 2 | 0 | 6 | 0.3333 | 6 | 0 | 23 | 0.087 |
| 2_1017832 | NA | NA | 1 | 1 | NA | NA | 1 | 1 | 2 | 1 | NA | NA | NA | NA | NA | NA | 2 | 1 | 2 | 1 |
| 2_1017832 | NA | NA | 1 | 1 | NA | NA | 1 | 1 | 2 | 1 | NA | NA | NA | NA | NA | NA | 1 | 1 | 1 | 1 |
| 2_1020724 | 2 | 0.5 | NA | NA | NA | NA | NA | NA | 2 | 0.5 | NA | NA | NA | NA | NA | NA | 1 | 1 | 1 | 1 |
| 2_1020727 | 2 | 1 | NA | NA | NA | NA | NA | NA | 2 | 1 | NA | NA | NA | NA | NA | NA | 1 | 1 | 1 | 1 |
| 2_1020729 | 2 | 1 | NA | NA | NA | NA | NA | NA | 2 | 1 | NA | NA | NA | NA | NA | NA | 1 | 1 | 1 | 1 |
| 2_1021341 | 13 | 0.2308 | 9 | 0.1111 | 6 | 0.1667 | 16 | 0.0625 | 44 | 0.1364 | 7 | 0 | 7 | 0 | 20 | 0.1 | 14 | 0.2143 | 48 | 0.1042 |
| 2_1021345 | 11 | 0.0909 | 5 | 0.2 | 6 | 0.1667 | 15 | 0.0667 | 37 | 0.1081 | 8 | 0.25 | 4 | 0.5 | 14 | 0.2143 | 11 | 0.1818 | 37 | 0.2432 |
| 2_1021346 | 10 | 0.1 | 7 | 0.1429 | 8 | 0.25 | 12 | 0.0833 | 37 | 0.1351 | 7 | 0.4286 | 6 | 0.3333 | 13 | 0.1538 | 14 | 0.3571 | 40 | 0.3 |
| 2_1021346 | 9 | 0.3333 | 6 | 0.1667 | 4 | 0.75 | 12 | 0.3333 | 31 | 0.3548 | 7 | 0.4286 | 6 | 0.5 | 16 | 0.4375 | 14 | 0.5714 | 43 | 0.4884 |
| 2_1021878 | 9 | 0.1111 | 5 | 0.2 | 7 | 0 | 13 | 0.2308 | 34 | 0.1471 | 7 | 0.1429 | 4 | 0.25 | 12 | 0 | 13 | 0 | 36 | 0.0556 |
| 2_1022303 | 9 | 0.3333 | 4 | 0 | 7 | 0 | 12 | 0 | 32 | 0.0938 | 6 | 0 | 4 | 0 | 11 | 0.0909 | 9 | 0.3333 | 30 | 0.1333 |
| 2_1022799 | 14 | 0.0714 | 17 | 0.1176 | 12 | 0 | 6 | 0.1667 | 49 | 0.0816 | 14 | 0 | 13 | 0.2308 | 6 | 0 | 12 | 0.1667 | 45 | 0.1111 |
| 2_1022799 | 15 | 0.4667 | 19 | 0.6842 | 15 | 0.6667 | 4 | 0.5 | 53 | 0.6038 | 11 | 0.4545 | 10 | 0.6 | 4 | 0.75 | 14 | 0.6429 | 39 | 0.5897 |
| 2_1022800 | 9 | 0.1111 | 8 | 0.25 | 9 | 0.2222 | 3 | 0 | 29 | 0.1724 | 11 | 0.3636 | 8 | 0.125 | 3 | 0.6667 | 10 | 0.1 | 32 | 0.25 |
| 2_1022800 | 14 | 0.1429 | 17 | 0.1765 | 15 | 0.2667 | 6 | 0 | 52 | 0.1731 | 13 | 0.0769 | 11 | 0.3636 | 7 | 0.2857 | 13 | 0.1538 | 44 | 0.2045 |
| 2_1022800 | 12 | 0.1667 | 15 | 0.3333 | 7 | 0.4286 | 3 | 0 | 37 | 0.2703 | 9 | 0.1111 | 9 | 0.2222 | 3 | 0 | 9 | 0.1111 | 30 | 0.1333 |
| 2_1022805 | 10 | 1 | 14 | 1 | 13 | 0.9231 | 5 | 1 | 42 | 0.9762 | 6 | 1 | 6 | 1 | 5 | 0.8 | 8 | 1 | 25 | 0.96 |
| 2_1022806 | 8 | 1 | 13 | 1 | 10 | 1 | 6 | 1 | 37 | 1 | 6 | 1 | 7 | 1 | 4 | 1 | 10 | 0.8 | 27 | 0.9259 |
| 2_1022806 | 8 | 0.125 | 9 | 0 | 9 | 0.2222 | 4 | 0.25 | 30 | 0.1333 | 6 | 0 | 8 | 0.125 | 4 | 0 | 8 | 0.125 | 26 | 0.0769 |
| 2_1022807 | 7 | 0.1429 | 8 | 0.25 | 9 | 0.2222 | 5 | 0.4 | 29 | 0.2414 | 6 | 0.1667 | 8 | 0.125 | 6 | 0.1667 | 10 | 0 | 30 | 0.1 |
| 2_1022808 | 6 | 1 | 9 | 0.8889 | 11 | 1 | 4 | 1 | 30 | 0.9667 | 6 | 1 | 6 | 1 | 6 | 0.8333 | 8 | 1 | 26 | 0.9615 |
| 2_1022808 | 6 | 0.3333 | 8 | 0.125 | 9 | 0.2222 | 4 | 0.5 | 27 | 0.2593 | 5 | 0.4 | 7 | 0.2857 | 6 | 0.3333 | 9 | 0.2222 | 27 | 0.2963 |
| 2_1022808 | 5 | 1 | 8 | 0.875 | 9 | 1 | 4 | 1 | 26 | 0.9615 | 4 | 1 | 6 | 1 | 5 | 1 | 7 | 1 | 22 | 1 |
| 2_1022809 | 6 | 1 | 6 | 0.8333 | 8 | 1 | 2 | 1 | 22 | 0.9545 | 5 | 0.8 | 5 | 1 | 3 | 1 | 7 | 1 | 20 | 0.95 |
| 2_1022809 | 7 | 0.2857 | 7 | 0.5714 | 10 | 0.4 | 4 | 0.25 | 28 | 0.3929 | 5 | 0 | 6 | 0.1667 | 6 | 0.5 | 10 | 0.1 | 27 | 0.1852 |
| 2_1022809 | 5 | 0.2 | 10 | 0.2 | 12 | 0.3333 | 5 | 0.4 | 32 | 0.2812 | 7 | 0 | 6 | 0 | 7 | 0.4286 | 8 | 0.125 | 28 | 0.1429 |
| 2_1022810 | 6 | 0.6667 | 8 | 1 | 11 | 0.9091 | 4 | 1 | 29 | 0.8966 | 4 | 0.75 | 5 | 1 | 7 | 0.8571 | 9 | 0.8889 | 25 | 0.88 |
| 2_1022810 | 5 | 1 | 6 | 1 | 10 | 0.9 | 4 | 1 | 25 | 0.96 | 5 | 1 | 5 | 0.8 | 4 | 1 | 9 | 1 | 23 | 0.9565 |
| 2_1022812 | 4 | 0.75 | 6 | 0.5 | 6 | 0.8333 | 2 | 0.5 | 18 | 0.6667 | 5 | 0.4 | 5 | 0.6 | 4 | 0.75 | 4 | 1 | 18 | 0.6667 |
| 2_1022816 | 4 | 0.5 | 3 | 0.6667 | 6 | 0.1667 | 3 | 0.6667 | 16 | 0.4375 | 3 | 0 | 4 | 0.25 | 2 | 0.5 | 4 | 0.75 | 13 | 0.3846 |
| 2_1022818 | 4 | 0 | 7 | 0.2857 | 7 | 0.4286 | 3 | 0 | 21 | 0.2381 | 3 | 0.3333 | 4 | 0 | 2 | 0.5 | 5 | 0.2 | 14 | 0.2143 |
| 2_1024724 | 1 | 1 | NA | NA | NA | NA | NA | NA | 1 | 1 | NA | NA | 1 | 1 | NA | NA | 1 | 0 | 2 | 0.5 |
| 2_1024729 | 1 | 1 | NA | NA | 2 | 1 | NA | NA | 3 | 1 | NA | NA | 1 | 1 | 1 | 0 | 1 | 1 | 3 | 0.6667 |

| | | | | | | | | | | | | | | | | | | | | |
|---|---|---|---|---|---|---|---|---|---|---|---|---|---|---|---|---|---|---|---|---|
| 2_1024729 | 1 | 1 | NA | NA | 2 | 1 | NA | NA | 3 | 1 | NA | NA | 1 | 0 | 1 | 0 | 1 | 1 | 3 | 0.3333 |
| 2_1024731( | NA | NA | NA | NA | 2 | 0.5 | NA | NA | 2 | 0.5 | NA | NA | 1 | 0 | 1 | 1 | 1 | 1 | 3 | 0.6667 |
| 2_1024732 | 1 | 0 | NA | NA | 2 | 0.5 | NA | NA | 3 | 0.3333 | 1 | 0 | NA | NA | 1 | 1 | 1 | 0 | 3 | 0.3333 |
| 2_1026080 | 2 | 1 | 3 | 1 | 1 | 1 | NA | NA | 6 | 1 | NA | NA | NA | NA | 2 | 0.5 | 1 | 1 | 3 | 0.6667 |
| 2_1026081 | 2 | 0.5 | 2 | 0 | 1 | 0 | NA | NA | 5 | 0.2 | NA | NA | NA | NA | 2 | 0.5 | 1 | 1 | 3 | 0.6667 |
| 2_1026081 | 1 | 0 | 3 | 0.3333 | 1 | 0 | NA | NA | 5 | 0.2 | 1 | 1 | NA | NA | 2 | 0 | 1 | 1 | 4 | 0.5 |
| 2_1026082 | 2 | 0.5 | 3 | 0.3333 | 1 | 1 | NA | NA | 6 | 0.5 | 1 | 1 | NA | NA | 2 | 0 | 1 | 1 | 4 | 0.5 |
| 2_1026083 | 2 | 0 | 3 | 0 | 1 | 1 | NA | NA | 6 | 0.1667 | 1 | 0 | NA | NA | 3 | 0.3333 | 1 | 1 | 5 | 0.4 |
| 2_1026118 | 16 | 0.1875 | 24 | 0.0833 | 16 | 0 | 15 | 0.0667 | 71 | 0.0845 | 19 | 0 | 12 | 0.0833 | 27 | 0.0741 | 16 | 0.0625 | 74 | 0.0541 |
| 2_1026119 | 10 | 0.2 | 20 | 0.1 | 14 | 0.0714 | 14 | 0.1429 | 58 | 0.1207 | 13 | 0.3077 | 10 | 0.2 | 24 | 0.125 | 14 | 0.2143 | 61 | 0.1967 |
| 2_1026120 | 12 | 0.3333 | 18 | 0.2222 | 9 | 0.3333 | 16 | 0.25 | 55 | 0.2727 | 18 | 0.3889 | 11 | 0.3636 | 24 | 0.375 | 14 | 0.2857 | 67 | 0.3582 |
| 2_1026122 | 11 | 0.6364 | 16 | 0.3125 | 9 | 0.4444 | 15 | 0.2667 | 51 | 0.3922 | 13 | 0.5385 | 10 | 0.7 | 22 | 0.4545 | 9 | 0.5556 | 54 | 0.537 |
| 2_1026122 | 10 | 0.7 | 15 | 0.6 | 8 | 0.75 | 14 | 0.7857 | 47 | 0.7021 | 15 | 0.8 | 13 | 0.6154 | 23 | 0.7826 | 10 | 1 | 61 | 0.7869 |
| 2_1026122 | 13 | 0.0769 | 17 | 0 | 8 | 0.125 | 16 | 0.0625 | 54 | 0.0556 | 18 | 0.0556 | 14 | 0.1429 | 24 | 0.1667 | 10 | 0 | 66 | 0.1061 |
| 2_1026124 | 13 | 0.1538 | 18 | 0 | 10 | 0.1 | 16 | 0.1875 | 57 | 0.1053 | 18 | 0.3333 | 14 | 0.3571 | 23 | 0.3478 | 10 | 0.2 | 65 | 0.3231 |
| 2_1026131 | 17 | 0.2353 | 25 | 0.16 | 8 | 0.25 | 17 | 0.1176 | 67 | 0.1791 | 21 | 0.2381 | 10 | 0.1 | 27 | 0.1111 | 10 | 0.1 | 68 | 0.1471 |
| 2_1026138 | 17 | 0.1176 | 24 | 0.125 | 13 | 0 | 21 | 0 | 75 | 0.0667 | 22 | 0.0909 | 11 | 0.0909 | 30 | 0.0333 | 11 | 0.0909 | 74 | 0.0676 |
| 2_1026177 | 84 | 0.0476 | 97 | 0.1134 | 77 | 0.1039 | 110 | 0.0545 | 368 | 0.0788 | 73 | 0.1507 | 70 | 0.0714 | 99 | 0.0707 | 86 | 0.0581 | 328 | 0.0854 |
| 2_1026188 | 107 | 0.0374 | 126 | 0.0794 | 96 | 0.0938 | 122 | 0.082 | 451 | 0.0732 | 98 | 0.0816 | 89 | 0.0112 | 120 | 0.1083 | 101 | 0.0396 | 408 | 0.0637 |
| 2_1026197 | 149 | 0.3289 | 175 | 0.3429 | 137 | 0.292 | 167 | 0.3772 | 628 | 0.3376 | 126 | 0.3333 | 133 | 0.3759 | 156 | 0.4295 | 131 | 0.4046 | 546 | 0.3883 |
| 2_1026199 | 128 | 0.5625 | 155 | 0.5613 | 123 | 0.6504 | 141 | 0.6454 | 547 | 0.6033 | 112 | 0.5804 | 114 | 0.6842 | 130 | 0.6 | 120 | 0.6917 | 476 | 0.6387 |
| 2_1026199 | 151 | 0.2185 | 190 | 0.0789 | 148 | 0.0676 | 157 | 0.0573 | 646 | 0.1037 | 124 | 0.1694 | 131 | 0.1145 | 155 | 0.1226 | 137 | 0.1314 | 547 | 0.1335 |
| 2_1026199 | 3 | 0.6667 | NA | NA | 2 | 1 | NA | NA | 5 | 0.8 | 2 | 0.5 | NA | NA | NA | NA | 1 | 0 | 3 | 0.3333 |
| 2_1026199 | 153 | 0.915 | 166 | 0.9096 | 145 | 0.9103 | 156 | 0.9487 | 620 | 0.921 | 125 | 0.88 | 130 | 0.9231 | 148 | 0.9189 | 142 | 0.9437 | 545 | 0.9174 |
| 2_1026199 | 155 | 0.8258 | 190 | 0.7211 | 148 | 0.723 | 175 | 0.7543 | 668 | 0.7545 | 136 | 0.7794 | 134 | 0.7388 | 156 | 0.7885 | 144 | 0.7917 | 570 | 0.7754 |
| 2_1026200 | 135 | 0.1185 | 167 | 0.1078 | 141 | 0.0638 | 160 | 0.0625 | 603 | 0.0879 | 124 | 0.0806 | 113 | 0.0708 | 140 | 0.1071 | 122 | 0.082 | 499 | 0.0862 |
| 2_1026201 | 164 | 0.1341 | 198 | 0.1313 | 158 | 0.1013 | 171 | 0.1813 | 691 | 0.1375 | 138 | 0.1812 | 142 | 0.1972 | 169 | 0.1302 | 148 | 0.2162 | 597 | 0.1792 |
| 2_1026201 | 153 | 0.1961 | 178 | 0.1573 | 141 | 0.1702 | 155 | 0.1419 | 627 | 0.1659 | 128 | 0.1172 | 127 | 0.189 | 147 | 0.1973 | 138 | 0.1884 | 540 | 0.1741 |
| 2_1026203 | 156 | 0.2051 | 192 | 0.2188 | 147 | 0.1769 | 168 | 0.1667 | 663 | 0.1931 | 130 | 0.2 | 140 | 0.2071 | 157 | 0.2229 | 140 | 0.2643 | 567 | 0.224 |
| 2_1026203 | 172 | 0.2558 | 203 | 0.266 | 164 | 0.2134 | 180 | 0.2722 | 719 | 0.2531 | 142 | 0.2887 | 150 | 0.3267 | 170 | 0.2529 | 151 | 0.2848 | 613 | 0.2871 |
| 2_1026203 | 142 | 0.8944 | 165 | 0.9273 | 125 | 0.944 | 145 | 0.9241 | 577 | 0.922 | 113 | 0.9115 | 127 | 0.937 | 146 | 0.9041 | 121 | 0.9256 | 507 | 0.9191 |
| 2_1026204 | 167 | 0.0539 | 187 | 0.0642 | 145 | 0.0621 | 162 | 0.0432 | 661 | 0.056 | 122 | 0.0492 | 129 | 0.0775 | 158 | 0.0316 | 139 | 0.0647 | 548 | 0.0547 |
| 2_1026204 | 154 | 0.7013 | 188 | 0.617 | 153 | 0.6797 | 168 | 0.6964 | 663 | 0.6712 | 132 | 0.6894 | 133 | 0.6917 | 155 | 0.7226 | 141 | 0.7305 | 561 | 0.7094 |
| 2_1026205 | 183 | 0.0601 | 216 | 0.0694 | 169 | 0.0769 | 187 | 0.0749 | 755 | 0.0702 | 149 | 0.0671 | 157 | 0.0318 | 174 | 0.1092 | 162 | 0.0988 | 642 | 0.0779 |
| 2_1026206 | 192 | 0.3906 | 225 | 0.4 | 178 | 0.382 | 190 | 0.3684 | 785 | 0.386 | 144 | 0.4097 | 163 | 0.3988 | 180 | 0.4611 | 164 | 0.4512 | 651 | 0.4316 |
| 2_1026207 | 192 | 0.0417 | 222 | 0.0676 | 175 | 0.0571 | 185 | 0.0486 | 774 | 0.0543 | 149 | 0.0537 | 161 | 0.0621 | 183 | 0.0437 | 164 | 0.1159 | 657 | 0.0685 |
| 2_1026208 | 202 | 0.2178 | 234 | 0.2094 | 189 | 0.2063 | 197 | 0.2437 | 822 | 0.219 | 150 | 0.1733 | 162 | 0.2654 | 191 | 0.2827 | 172 | 0.2558 | 675 | 0.2474 |
| 2_1026208 | 192 | 0.3906 | 216 | 0.375 | 174 | 0.3621 | 185 | 0.4054 | 767 | 0.3833 | 144 | 0.2986 | 155 | 0.3935 | 179 | 0.3408 | 163 | 0.3129 | 641 | 0.337 |
| 2_1028522 | 7 | 0.5714 | 9 | 0.4444 | 6 | 0.3333 | 12 | 0.5833 | 34 | 0.5 | 4 | 0.75 | 10 | 0.4 | 8 | 0.375 | 6 | 0.3333 | 28 | 0.4286 |
| 2_1028523 | 5 | 0.2 | 8 | 0.125 | 5 | 0.4 | 13 | 0.3846 | 31 | 0.2903 | 4 | 0.5 | 7 | 0.2857 | 9 | 0.1111 | 6 | 0.1667 | 26 | 0.2308 |
| 2_1028524 | 6 | 0.5 | 6 | 0.3333 | 6 | 0.3333 | 6 | 0.5 | 24 | 0.4167 | 4 | 0.75 | 6 | 0.5 | 6 | 0.3333 | 3 | 0.6667 | 19 | 0.5263 |
| 2_1032848 | 7 | 0.2857 | 7 | 0.2857 | 5 | 0.2 | 7 | 0.2857 | 26 | 0.2692 | 8 | 0.5 | 7 | 0.2857 | 3 | 0.3333 | 3 | 0.3333 | 21 | 0.381 |
| 2_1033967( | NA | NA | 2 | 0 | NA | NA | 1 | 1 | 3 | 0.3333 | NA | NA | 1 | 0 | NA | NA | 1 | 1 | 2 | 0.5 |
| 2_1033971 | NA | NA | 1 | 1 | NA | NA | 1 | 1 | 2 | 1 | NA | NA | 1 | 1 | 1 | 1 | 1 | 1 | 3 | 1 |
| 2_1033972 | NA | NA | 2 | 0.5 | NA | NA | 1 | 0 | 3 | 0.3333 | NA | NA | 1 | 0 | 1 | 0 | 1 | 1 | 3 | 0.3333 |
| 2_1033977 | NA | NA | 1 | 1 | NA | NA | 1 | 1 | 2 | 1 | NA | NA | 1 | 0 | 1 | 1 | 1 | 0 | 3 | 0.3333 |
| 2_1035477 | 1 | 1 | 1 | 1 | 1 | 1 | 2 | 1 | 5 | 1 | 1 | 0 | NA | NA | 3 | 0.6667 | 1 | 1 | 5 | 0.6 |
| 2_1035480 | 1 | 0 | 3 | 1 | 1 | 0 | 2 | 0 | 7 | 0.4286 | 1 | 1 | 2 | 0.5 | 2 | 0 | 1 | 0 | 6 | 0.3333 |
| 2_1035482 | 1 | 0 | 3 | 1 | 1 | 0 | 2 | 0.5 | 7 | 0.5714 | 1 | 0 | 2 | 0.5 | 3 | 0.6667 | NA | NA | 6 | 0.5 |
| 2_1035485 | NA | NA | 3 | 0.3333 | 1 | 0 | 2 | 0.5 | 6 | 0.3333 | 2 | 0 | 1 | 0 | 4 | 0.25 | 1 | 1 | 8 | 0.25 |
| 2_1035516( | NA | NA | NA | NA | 2 | 0 | 3 | 0.3333 | 5 | 0.2 | 2 | 0 | 3 | 0.3333 | 3 | 0.3333 | NA | NA | 8 | 0.25 |
| 2_1035516 | 2 | 1 | 1 | 1 | 4 | 0.5 | 4 | 1 | 11 | 0.8182 | 2 | 1 | 3 | 1 | 6 | 1 | NA | NA | 11 | 1 |
| 2_1035516 | 2 | 0 | 1 | 0 | 4 | 0.5 | 1 | 1 | 8 | 0.375 | 2 | 0 | 4 | 0.25 | 6 | 0.1667 | 1 | 0 | 13 | 0.1538 |
| 2_1035519 | 2 | 0.5 | 1 | 0 | 4 | 0.5 | 3 | 1 | 10 | 0.6 | 2 | 0.5 | 4 | 0.5 | 6 | 0.8333 | NA | NA | 12 | 0.6667 |
| 2_1035520 | 1 | 0 | 1 | 1 | 4 | 0.5 | NA | NA | 6 | 0.5 | 1 | 0 | 3 | 0 | 3 | 0.6667 | NA | NA | 7 | 0.2857 |
| 2_1035520 | 1 | 0 | 1 | 1 | 3 | 0 | 3 | 0.3333 | 8 | 0.25 | 2 | 0.5 | 2 | 0.5 | 3 | 0.3333 | 1 | 0 | 8 | 0.375 |
| 2_1035521( | 2 | 1 | 1 | 1 | 4 | 1 | 3 | 1 | 10 | 1 | 1 | 1 | 4 | 1 | 5 | 0.8 | NA | NA | 10 | 0.9 |
| 2_1035521 | 2 | 0 | 1 | 0 | 4 | 0.5 | 3 | 0 | 10 | 0.2 | 2 | 0 | 4 | 0 | 4 | 0.25 | NA | NA | 10 | 0.1 |
| 2_1035956 | 16 | 0.1875 | 12 | 0.25 | 11 | 0.1818 | 23 | 0.1304 | 62 | 0.1774 | 12 | 0.1667 | 19 | 0.1053 | 13 | 0.0769 | 10 | 0.2 | 54 | 0.1296 |
| 2_1035963 | 14 | 0.1429 | 13 | 0.0769 | 13 | 0.2308 | 21 | 0.1429 | 61 | 0.1475 | 9 | 0 | 19 | 0.1579 | 17 | 0.0588 | 12 | 0.0833 | 57 | 0.0877 |
| 2_1035970 | 13 | 0.3846 | 11 | 0.4545 | 12 | 0.25 | 19 | 0.2632 | 55 | 0.3273 | 8 | 0.5 | 18 | 0.5 | 16 | 0.5 | 12 | 0.25 | 54 | 0.4444 |
| 2_1035970 | 17 | 0.6471 | 12 | 0.5 | 11 | 0.5455 | 20 | 0.55 | 60 | 0.5667 | 8 | 0.625 | 18 | 0.5556 | 18 | 0.8889 | 13 | 0.5385 | 57 | 0.6667 |
| 2_1035986 | 6 | 0 | 6 | 0.3333 | 5 | 0.6 | 7 | 0.2857 | 24 | 0.2917 | 3 | 0.6667 | 8 | 0.375 | 6 | 0.3333 | 8 | 0.625 | 25 | 0.48 |
| 2_1038008 | 1 | 0 | 1 | 0 | 3 | 0.6667 | 4 | 0 | 9 | 0.2222 | 3 | 0.3333 | 3 | 0 | 2 | 0 | 3 | 0 | 11 | 0.0909 |
| 2_1038065 | 3 | 0.3333 | 1 | 1 | 4 | 0.25 | 6 | 0.1667 | 14 | 0.2857 | 4 | 0.25 | 4 | 0.5 | 4 | 0.5 | 4 | 0.25 | 16 | 0.375 |

| | | | | | | | | | | | | | | | | | | | | |
|---|---|---|---|---|---|---|---|---|---|---|---|---|---|---|---|---|---|---|---|---|
| 2_10381155 | 3 | 1 | 2 | 1 | 3 | 1 | 4 | 0.75 | 12 | 0.9167 | 4 | 1 | 3 | 1 | 4 | 1 | 4 | 1 | 15 | 1 |
| 2_10381178 | 3 | 0 | 2 | 0.5 | 4 | 0 | 5 | 0 | 14 | 0.0714 | 4 | 0.25 | 2 | 0 | 5 | 0.2 | 3 | 0 | 14 | 0.1429 |
| 2_10388650 | 6 | 0.1667 | 8 | 0.5 | 7 | 0.1429 | 4 | 0.25 | 25 | 0.28 | 1 | 0 | 2 | 0 | NA | NA | 5 | 0.2 | 8 | 0.125 |
| 2_10388654 | 4 | 0.5 | 9 | 0.8889 | 7 | 0.8571 | 5 | 0.2 | 25 | 0.68 | NA | NA | 2 | 0.5 | 2 | 0 | 6 | 0.8333 | 10 | 0.6 |
| 2_10388778 | 8 | 0.125 | 10 | 0.2 | 9 | 0 | 7 | 0 | 34 | 0.0882 | 1 | 0 | 6 | 0 | 6 | 0.1667 | 7 | 0.1429 | 20 | 0.1 |
| 2_10389059 | 6 | 0.3333 | 5 | 0.4 | 5 | 0 | 6 | 0.1667 | 22 | 0.2273 | 6 | 0.1667 | 6 | 0.1667 | 6 | 0 | 10 | 0 | 28 | 0.0714 |
| 2_10389079 | 6 | 0.3333 | 6 | 0 | 6 | 0.1667 | 8 | 0 | 26 | 0.1154 | 5 | 0.2 | 4 | 0 | 8 | 0.25 | 13 | 0.1538 | 30 | 0.1667 |
| 2_10389080 | 8 | 0.25 | 5 | 0 | 3 | 0 | 7 | 0.2857 | 23 | 0.1739 | 6 | 0.1667 | 6 | 0.3333 | 9 | 0.2222 | 13 | 0.2308 | 34 | 0.2353 |
| 2_10389182 | 2 | 0 | 3 | 0 | 3 | 0.3333 | 4 | 0 | 12 | 0.0833 | 1 | 0 | 7 | 0.2857 | 6 | 0 | 2 | 0 | 16 | 0.125 |
| 2_10389211 | NA | NA | NA | NA | 1 | 1 | NA | NA | 1 | 1 | 1 | 0 | 3 | 0 | 3 | 0.3333 | 1 | 1 | 8 | 0.25 |
| 2_10389310 | NA | NA | NA | NA | 1 | 1 | NA | NA | 1 | 1 | NA | NA | 1 | 0 | 1 | 0 | 1 | 1 | 3 | 0.3333 |
| 2_10394438 | 2 | 0.5 | NA | NA | 2 | 0 | NA | NA | 4 | 0.25 | NA | NA | 2 | 0.5 | 3 | 0.6667 | 2 | 0.5 | 7 | 0.5714 |
| 2_10394447 | 2 | 0.5 | NA | NA | 2 | 0 | NA | NA | 4 | 0.25 | NA | NA | 1 | 0 | 2 | 0 | 4 | 0.25 | 7 | 0.1429 |
| 2_10394547 | 2 | 0 | NA | NA | 2 | 0.5 | 1 | 0 | 5 | 0.2 | 1 | 0 | 2 | 0 | 2 | 0 | 2 | 0.5 | 7 | 0.1429 |
| 2_10397361 | 13 | 0.0769 | 16 | 0.25 | 8 | 0.125 | 9 | 0.2222 | 46 | 0.1739 | 5 | 0.2 | 13 | 0 | 6 | 0.1667 | 7 | 0 | 31 | 0.0645 |
| 2_10397364 | 6 | 0.1667 | 7 | 0.4286 | 3 | 0 | 7 | 0.1429 | 23 | 0.2174 | 4 | 0.75 | 6 | 0 | 1 | 0 | 2 | 0.5 | 13 | 0.3077 |
| 2_10397389 | 10 | 0.2 | 16 | 0.1875 | 6 | 0.1667 | 7 | 0 | 39 | 0.1538 | 3 | 0.3333 | 10 | 0.1 | 5 | 0.6 | 6 | 0.3333 | 24 | 0.2917 |
| 2_10397399 | 10 | 0 | 16 | 0.125 | 6 | 0 | 8 | 0 | 40 | 0.05 | 4 | 0.25 | 10 | 0 | 5 | 0.2 | 6 | 0.1667 | 25 | 0.12 |
| 2_10397399 | 11 | 0.2727 | 16 | 0.25 | 6 | 0.1667 | 8 | 0.125 | 41 | 0.2195 | 3 | 1 | 10 | 0.4 | 4 | 0.75 | 6 | 0.5 | 23 | 0.5652 |
| 2_10397420 | 4 | 0.25 | 13 | 0.2308 | 6 | 0.1667 | 4 | 0.25 | 27 | 0.2222 | 3 | 0.6667 | 9 | 0.3333 | 3 | 0.3333 | 6 | 0.3333 | 21 | 0.381 |
| 2_10397447 | 6 | 0 | 13 | 0.0769 | 6 | 0.1667 | 7 | 0.2857 | 32 | 0.125 | 3 | 0 | 8 | 0.125 | 3 | 0 | 4 | 0.25 | 18 | 0.1111 |
| 2_10397605 | 4 | 0 | 6 | 0 | 1 | 0 | 3 | 0.3333 | 14 | 0.0714 | 1 | 0 | 6 | 0.1667 | 3 | 0 | 2 | 0 | 12 | 0.0833 |
| 2_10397611 | 5 | 0.2 | 5 | 0.2 | 1 | 0 | 4 | 0.25 | 15 | 0.2 | 1 | 0 | 6 | 0 | 3 | 0.3333 | 4 | 0 | 14 | 0.0714 |
| 2_10397637 | 4 | 0.25 | 7 | 0.4286 | 1 | 0 | 4 | 0 | 16 | 0.25 | NA | NA | 6 | 0.1667 | 2 | 0.5 | 4 | 0.25 | 12 | 0.25 |
| 2_10397640 | 3 | 0.3333 | 7 | 0.1429 | 1 | 0 | 4 | 0.25 | 15 | 0.2 | NA | NA | 6 | 0.1667 | 1 | 0 | 3 | 0.6667 | 10 | 0.3 |
| 2_10397645 | 3 | 0 | 6 | 0.3333 | 1 | 0 | 1 | 0 | 11 | 0.1818 | NA | NA | 6 | 0.1667 | NA | NA | 5 | 0 | 11 | 0.0909 |
| 2_10397658 | 4 | 0 | 6 | 0.1667 | 1 | 0 | 2 | 0 | 13 | 0.0769 | NA | NA | 6 | 0 | 2 | 0.5 | 5 | 0 | 13 | 0.0769 |
| 2_10397676 | 4 | 0.75 | 5 | 0.8 | 1 | 0 | 3 | 0 | 13 | 0.5385 | NA | NA | 4 | 0.75 | 2 | 1 | 4 | 0.5 | 10 | 0.7 |
| 2_10397690 | 4 | 0.5 | 6 | 0.6667 | 1 | 0 | 3 | 0.3333 | 14 | 0.5 | NA | NA | 4 | 0.75 | 2 | 1 | 5 | 0.6 | 11 | 0.7273 |
| 2_10397704 | 4 | 0.5 | 7 | 0 | 1 | 0 | 3 | 0 | 15 | 0.1333 | NA | NA | 3 | 0.3333 | 2 | 0.5 | 5 | 0 | 10 | 0.2 |
| 2_10397706 | 5 | 0.2 | 7 | 0.1429 | 1 | 0 | 4 | 0.25 | 17 | 0.1765 | NA | NA | 3 | 0.6667 | 2 | 0.5 | 4 | 0.25 | 9 | 0.4444 |
| 2_10397710 | 5 | 0.2 | 7 | 0.5714 | 1 | 0 | 4 | 0.25 | 17 | 0.3529 | NA | NA | 3 | 0.6667 | 2 | 1 | 5 | 0.8 | 10 | 0.8 |
| 2_10397711 | 5 | 0.2 | 6 | 0.5 | 1 | 0 | 4 | 0.5 | 16 | 0.375 | NA | NA | 3 | 1 | 1 | 1 | 4 | 0.5 | 8 | 0.75 |
| 2_10397718 | 5 | 0 | 7 | 0.1429 | 1 | 0 | 4 | 0 | 17 | 0.0588 | NA | NA | 4 | 0.25 | 2 | 0.5 | 4 | 0 | 10 | 0.2 |
| 2_10397728 | 5 | 0.2 | 5 | 0 | 1 | 0 | 4 | 0 | 15 | 0.0667 | NA | NA | 4 | 0.5 | 2 | 0 | 5 | 0.2 | 11 | 0.2727 |
| 2_10397760 | 5 | 0.2 | 5 | 0.2 | 1 | 0 | 2 | 0 | 13 | 0.1538 | NA | NA | 3 | 0 | 2 | 0 | 5 | 0.2 | 10 | 0.1 |
| 2_10397763 | 5 | 0.2 | 6 | 0.5 | NA | NA | 2 | 0 | 13 | 0.3077 | 1 | 1 | 1 | 0 | 2 | 0 | 3 | 0.3333 | 7 | 0.2857 |
| 2_10397779 | 4 | 0.5 | 5 | 0.6 | 1 | 0 | 2 | 0.5 | 12 | 0.5 | 1 | 1 | 3 | 0.6667 | 2 | 0.5 | 4 | 0.5 | 10 | 0.6 |
| 2_10397787 | 3 | 0.6667 | 2 | 0.5 | 1 | 0 | 2 | 1 | 8 | 0.625 | NA | NA | 1 | 1 | 2 | 0.5 | 2 | 0.5 | 5 | 0.6 |
| 2_10397793 | 5 | 0 | 6 | 0.1667 | 1 | 0 | 3 | 0 | 15 | 0.0667 | 1 | 0 | 3 | 0.6667 | 2 | 0 | 3 | 0 | 9 | 0.2222 |
| 2_10397798 | 5 | 0 | 5 | 0.6 | 1 | 0 | 3 | 0 | 14 | 0.2143 | 1 | 0 | 2 | 0.5 | 2 | 0 | 4 | 0.5 | 9 | 0.3333 |
| 2_10397800 | 3 | 0.6667 | 6 | 0.8333 | 1 | 1 | 2 | 1 | 12 | 0.8333 | 1 | 1 | 1 | 0 | 2 | 1 | 4 | 1 | 8 | 0.875 |
| 2_10397815 | 3 | 1 | 3 | 0.3333 | 1 | 1 | NA | NA | 7 | 0.7143 | 1 | 1 | 2 | 1 | 1 | 0 | 1 | 1 | 5 | 0.8 |
| 2_10397816 | 5 | 0 | 4 | 0.25 | 1 | 0 | 3 | 0 | 13 | 0.0769 | 1 | 0 | 3 | 0.6667 | 1 | 0 | 4 | 0 | 9 | 0.2222 |
| 2_10397824 | 5 | 0.4 | 3 | 0.3333 | 1 | 0 | 3 | 0 | 12 | 0.25 | 1 | 0 | 3 | 0.3333 | 1 | 1 | 3 | 0.6667 | 8 | 0.5 |
| 2_10397826 | 3 | 0 | 3 | 0.3333 | 1 | 0 | 3 | 0.3333 | 10 | 0.2 | 1 | 0 | 3 | 0 | 1 | 0 | 4 | 0.25 | 9 | 0.1111 |
| 2_10397835 | 4 | 1 | 4 | 0.75 | NA | NA | 3 | 1 | 11 | 0.9091 | 1 | 0 | 2 | 1 | 1 | 1 | 3 | 0.6667 | 7 | 0.7143 |
| 2_10397843 | 5 | 0.6 | 3 | 1 | 1 | 1 | 3 | 0.3333 | 12 | 0.6667 | 1 | 0 | 2 | 1 | NA | NA | 3 | 0.6667 | 6 | 0.6667 |
| 2_10397849 | 4 | 0.75 | 2 | 0 | NA | NA | 3 | 0.3333 | 9 | 0.4444 | 1 | 0 | 3 | 0.3333 | 1 | 0 | 4 | 1 | 9 | 0.5556 |
| 2_10397858 | 5 | 0.4 | 3 | 0.3333 | 1 | 0 | 3 | 0.3333 | 12 | 0.3333 | 2 | 0 | 3 | 0.6667 | 1 | 0 | 4 | 0.75 | 10 | 0.5 |
| 2_10397860 | 4 | 0.5 | 2 | 0.5 | 1 | 1 | 3 | 0.6667 | 10 | 0.6 | 2 | 0.5 | 3 | 1 | 1 | 1 | 3 | 1 | 9 | 0.8889 |
| 2_10397865 | 5 | 0 | 3 | 0 | 1 | 0 | 2 | 0.5 | 11 | 0.0909 | 2 | 0 | 3 | 0.3333 | 1 | 0 | 4 | 0 | 10 | 0.1 |
| 2_10397868 | 5 | 1 | 3 | 1 | 1 | 1 | 2 | 1 | 11 | 1 | 1 | 1 | 2 | 1 | 1 | 1 | 4 | 1 | 8 | 1 |
| 2_10397886 | 5 | 1 | 1 | 1 | NA | NA | 3 | 1 | 9 | 1 | 2 | 1 | 3 | 1 | 1 | 1 | 3 | 0.6667 | 9 | 0.8889 |
| 2_10397890 | 5 | 1 | 1 | 1 | 1 | 1 | 3 | 1 | 10 | 1 | 2 | 1 | 3 | 1 | 1 | 1 | 2 | 1 | 8 | 1 |
| 2_10397896 | 5 | 0.4 | 1 | 0 | 1 | 0 | 3 | 0 | 10 | 0.2 | 2 | 0 | 3 | 0.3333 | 1 | 0 | 2 | 0.5 | 8 | 0.25 |
| 2_10397902 | 3 | 0.6667 | 1 | 1 | NA | NA | 2 | 1 | 6 | 0.8333 | 2 | 1 | 3 | 1 | 1 | 1 | 2 | 1 | 8 | 1 |
| 2_10397914 | 5 | 0.8 | 1 | 1 | 1 | 1 | 3 | 0 | 10 | 0.6 | 2 | 0.5 | 3 | 1 | NA | NA | 1 | 1 | 6 | 0.8333 |
| 2_10397923 | 5 | 0.8 | 1 | 0 | 1 | 1 | 3 | 0 | 10 | 0.5 | NA | NA | 3 | 0.3333 | 1 | 0 | 2 | 0 | 6 | 0.1667 |
| 2_10408636 | NA | NA | NA | NA | NA | NA | 1 | 1 | 1 | 1 | NA | NA | 2 | 1 | 2 | 0.5 | 1 | 1 | 5 | 0.8 |
| 2_10408669 | NA | NA | NA | NA | NA | NA | 1 | 1 | 1 | 1 | NA | NA | 2 | 1 | 2 | 1 | 1 | 1 | 5 | 1 |
| 2_10408677 | NA | NA | NA | NA | NA | NA | 1 | 1 | 1 | 1 | NA | NA | 2 | 0.5 | 2 | 0 | 1 | 0 | 5 | 0.2 |
| 2_10408679 | NA | NA | NA | NA | NA | NA | 1 | 1 | 1 | 1 | NA | NA | 2 | 1 | 2 | 0 | 1 | 0 | 5 | 0.4 |
| 2_10425703 | 12 | 0.5833 | 23 | 0.4783 | 15 | 0.4 | 7 | 0.1429 | 57 | 0.4386 | 3 | 0 | 7 | 0.4286 | 9 | 0.1111 | 13 | 0.3077 | 32 | 0.25 |
| 2_10425958 | 4 | 0.25 | 11 | 0.1818 | 9 | 0.1111 | 3 | 0 | 27 | 0.1481 | 1 | 0 | 4 | 0 | 5 | 0.2 | 3 | 0 | 13 | 0.0769 |

| | | | | | | | | | | | | | | | | | | | | |
|---|---|---|---|---|---|---|---|---|---|---|---|---|---|---|---|---|---|---|---|---|
| 2_10426008 | 3 | 0 | 12 | 0.1667 | 9 | 0.1111 | 5 | 0 | 29 | 0.1034 | 4 | 0.25 | 4 | 0 | 5 | 0 | 5 | 0 | 18 | 0.0556 |
| 2_10426019 | 4 | 0 | 8 | 0 | 7 | 0.1429 | 5 | 0.4 | 24 | 0.125 | 3 | 0 | 4 | 0 | 5 | 0.4 | 4 | 0 | 16 | 0.125 |
| 2_10426022 | 4 | 0 | 10 | 0.2 | 9 | 0.1111 | 4 | 0 | 27 | 0.1111 | 4 | 0.25 | 4 | 0 | 5 | 0 | 5 | 0.2 | 18 | 0.1111 |
| 2_10426030 | 3 | 0.3333 | 11 | 0.9091 | 9 | 0.8889 | 4 | 1 | 27 | 0.8519 | 3 | 1 | 4 | 1 | 5 | 0.6 | 5 | 0.6 | 17 | 0.7647 |
| 2_10426050 | 1 | 0 | 11 | 0.3636 | 8 | 0.375 | 4 | 0 | 24 | 0.2917 | 4 | 0.5 | 3 | 0.3333 | 5 | 0.4 | 4 | 0.75 | 16 | 0.5 |
| 2_10426063 | 2 | 0 | 9 | 0 | 7 | 0.2857 | 5 | 0 | 23 | 0.087 | 2 | 0 | 3 | 0.3333 | 5 | 0.2 | 2 | 0 | 12 | 0.1667 |
| 2_10430113 | NA | NA | 4 | 0.5 | 5 | 0.6 | 5 | 1 | 14 | 0.7143 | 1 | 0 | 2 | 1 | 1 | 1 | 2 | 1 | 6 | 0.8333 |
| 2_10430116 | NA | NA | 2 | 1 | 4 | 0.75 | 5 | 1 | 11 | 0.9091 | 1 | 1 | 2 | 1 | NA | NA | 2 | 1 | 5 | 1 |
| 2_10430134 | NA | NA | 4 | 0.25 | 5 | 0 | 5 | 0 | 14 | 0.0714 | 1 | 0 | 2 | 0.5 | 1 | 0 | 2 | 0.5 | 6 | 0.3333 |
| 2_10430134 | NA | NA | 4 | 0 | 5 | 0.2 | 5 | 0.2 | 14 | 0.1429 | 2 | 0 | 2 | 0 | 1 | 0 | 2 | 0.5 | 7 | 0.1429 |
| 2_10430142 | NA | NA | 4 | 0.25 | 1 | 0 | 5 | 0 | 10 | 0.1 | 1 | 1 | 2 | 0 | 1 | 0 | 1 | 0 | 5 | 0.2 |
| 2_10430153 | NA | NA | 4 | 0.75 | 5 | 0.6 | 5 | 1 | 14 | 0.7857 | 1 | 0 | 2 | 1 | 1 | 1 | 2 | 0.5 | 6 | 0.6667 |
| 2_10430167 | NA | NA | 3 | 0 | 5 | 0.4 | 5 | 0.4 | 13 | 0.3077 | 2 | 0 | 1 | 0 | 1 | 1 | 2 | 0 | 6 | 0.1667 |
| 2_10430179 | NA | NA | 3 | 1 | 3 | 1 | 3 | 1 | 9 | 1 | 1 | 1 | 1 | 1 | NA | NA | NA | NA | 2 | 1 |
| 2_10430193 | NA | NA | 4 | 0.25 | 5 | 0.4 | 5 | 0 | 14 | 0.2143 | 1 | 1 | 2 | 0.5 | NA | NA | 1 | 0 | 4 | 0.5 |
| 2_10430197 | NA | NA | 3 | 0.6667 | 5 | 0.4 | 4 | 0.5 | 12 | 0.5 | 2 | 0 | 2 | 1 | 1 | 1 | 1 | 0 | 6 | 0.5 |
| 2_10430204 | NA | NA | 4 | 0.5 | 5 | 0.4 | 5 | 0.2 | 14 | 0.3571 | 2 | 0 | 2 | 0.5 | 1 | 0 | 2 | 0.5 | 7 | 0.2857 |
| 2_10443951 | 3 | 0 | 6 | 0 | 6 | 0 | 6 | 0.3333 | 21 | 0.0952 | 4 | 0 | 5 | 0.2 | 5 | 0 | 2 | 0.5 | 16 | 0.125 |
| 2_10449522 | 2 | 0 | 4 | 0 | 7 | 0 | 5 | 0.2 | 18 | 0.0556 | 2 | 0.5 | 2 | 0 | 4 | 0 | NA | NA | 8 | 0.125 |
| 2_10449529 | 3 | 0 | 3 | 0.6667 | 7 | 0.2857 | 5 | 0.2 | 18 | 0.2778 | 2 | 0 | 2 | 0 | 3 | 0.3333 | NA | NA | 7 | 0.1429 |
| 2_10449531 | 3 | 0.6667 | 3 | 1 | 6 | 0.8333 | 5 | 0.8 | 17 | 0.8235 | 2 | 1 | 3 | 0.6667 | 4 | 1 | NA | NA | 9 | 0.8889 |
| 2_10449534 | 2 | 0 | 4 | 0.25 | 7 | 0.4286 | 4 | 0.25 | 17 | 0.2941 | 2 | 0.5 | 3 | 0 | 4 | 0.25 | NA | NA | 9 | 0.2222 |
| 2_10464749 | NA | NA | NA | NA | 1 | 1 | NA | NA | 1 | 1 | NA | NA | NA | NA | NA | NA | 1 | 1 | 1 | 1 |
| 2_10464750 | NA | NA | NA | NA | 2 | 1 | NA | NA | 2 | 1 | NA | NA | NA | NA | NA | NA | 1 | 1 | 1 | 1 |
| 2_10464762 | NA | NA | NA | NA | 2 | 1 | NA | NA | 2 | 1 | NA | NA | NA | NA | NA | NA | 1 | 1 | 1 | 1 |
| 2_10464790 | NA | NA | NA | NA | 2 | 1 | NA | NA | 2 | 1 | NA | NA | NA | NA | NA | NA | 1 | 1 | 1 | 1 |
| 2_10464799 | NA | NA | NA | NA | 2 | 1 | NA | NA | 2 | 1 | NA | NA | NA | NA | NA | NA | 1 | 1 | 1 | 1 |
| 2_10464804 | NA | NA | NA | NA | 2 | 1 | NA | NA | 2 | 1 | NA | NA | NA | NA | NA | NA | 1 | 1 | 1 | 1 |
| 2_10464830 | NA | NA | NA | NA | 2 | 1 | NA | NA | 2 | 1 | NA | NA | NA | NA | NA | NA | 1 | 1 | 1 | 1 |
| 2_10464844 | NA | NA | NA | NA | 2 | 1 | NA | NA | 2 | 1 | NA | NA | NA | NA | NA | NA | 1 | 1 | 1 | 1 |
| 2_10482151 | 1 | 1 | 1 | 1 | NA | NA | 3 | 0.6667 | 5 | 0.8 | NA | NA | NA | NA | NA | NA | 1 | 1 | 1 | 1 |
| 2_10482216 | 1 | 0 | 1 | 1 | NA | NA | 3 | 0 | 5 | 0.2 | NA | NA | NA | NA | NA | NA | 1 | 1 | 1 | 1 |
| 2_10482230 | 1 | 0 | 1 | 1 | NA | NA | 2 | 0.5 | 4 | 0.5 | NA | NA | NA | NA | NA | NA | 1 | 1 | 1 | 1 |
| 2_10482234 | 1 | 0 | 1 | 1 | NA | NA | 1 | 0 | 3 | 0.3333 | NA | NA | NA | NA | NA | NA | 1 | 1 | 1 | 1 |
| 2_10482238 | 1 | 1 | 1 | 1 | NA | NA | 2 | 0.5 | 4 | 0.75 | NA | NA | NA | NA | NA | NA | 1 | 1 | 1 | 1 |
| 2_10482248 | 1 | 0 | 1 | 1 | NA | NA | 3 | 0.3333 | 5 | 0.4 | NA | NA | NA | NA | NA | NA | 1 | 1 | 1 | 1 |
| 2_10482268 | 1 | 1 | 1 | 1 | NA | NA | 1 | 0 | 3 | 0.6667 | NA | NA | NA | NA | NA | NA | 1 | 1 | 1 | 1 |
| 2_10482273 | 1 | 1 | NA | NA | NA | NA | NA | NA | 1 | 1 | NA | NA | NA | NA | NA | NA | 1 | 1 | 1 | 1 |
| 2_10521758 | 3 | 0 | 4 | 0 | 5 | 0.2 | 7 | 0 | 19 | 0.0526 | 1 | 0 | 3 | 0 | 6 | 0.1667 | 8 | 0 | 18 | 0.0556 |
| 2_10521826 | 3 | 0.3333 | 4 | 0.5 | 4 | 0.25 | 4 | 0 | 15 | 0.2667 | 1 | 0 | 2 | 0.5 | 3 | 0 | 3 | 0.3333 | 9 | 0.2222 |
| 2_10521827 | 3 | 0.6667 | 4 | 1 | 5 | 0.6 | 2 | 0.5 | 14 | 0.7143 | 1 | 0 | 1 | 1 | 3 | 0.6667 | 2 | 1 | 7 | 0.7143 |
| 2_10522936 | 2 | 1 | 4 | 0.25 | 2 | 0 | 2 | 0.5 | 10 | 0.4 | NA | NA | 1 | 0 | 1 | 0 | 2 | 0.5 | 4 | 0.25 |
| 2_10522952 | 1 | 0 | 4 | 0.5 | 3 | 0 | 1 | 1 | 9 | 0.3333 | NA | NA | NA | NA | NA | NA | 1 | 1 | 1 | 1 |
| 2_10552675 | 2 | 0.5 | NA | NA | 4 | 0 | 2 | 0 | 8 | 0.125 | 2 | 1 | 2 | 0.5 | NA | NA | NA | NA | 4 | 0.75 |
| 2_10552676 | 2 | 0 | NA | NA | 4 | 0.75 | 2 | 0 | 8 | 0.375 | 2 | 0.5 | 2 | 0 | NA | NA | NA | NA | 4 | 0.25 |
| 2_10552685 | 2 | 1 | NA | NA | 4 | 0.75 | 2 | 1 | 8 | 0.875 | 2 | 0.5 | 2 | 1 | NA | NA | NA | NA | 4 | 0.75 |
| 2_10552686 | 2 | 0.5 | NA | NA | 4 | 0.75 | 1 | 1 | 7 | 0.7143 | 2 | 1 | 2 | 0 | NA | NA | NA | NA | 4 | 0.5 |
| 2_10552713 | 2 | 0.5 | NA | NA | 4 | 0.5 | 2 | 1 | 8 | 0.625 | 1 | 1 | 1 | 1 | NA | NA | NA | NA | 2 | 1 |
| 2_10553302 | 1 | 1 | 1 | 1 | 2 | 0.5 | NA | NA | 4 | 0.75 | 3 | 0.3333 | 1 | 1 | NA | NA | 1 | 1 | 5 | 0.6 |
| 2_10579715 | 1 | 0 | 2 | 1 | 4 | 0.75 | 4 | 1 | 11 | 0.8182 | NA | NA | 3 | 1 | 2 | 1 | 1 | 1 | 6 | 1 |
| 2_10579727 | 1 | 1 | 2 | 0.5 | 3 | 0.3333 | 3 | 0 | 9 | 0.3333 | NA | NA | 3 | 0.3333 | 3 | 0.6667 | 1 | 0 | 7 | 0.4286 |
| 2_10661402 | 5 | 0.4 | 1 | 0 | 7 | 0.4286 | 3 | 0.3333 | 16 | 0.375 | 2 | 0 | 6 | 0.5 | 9 | 0.4444 | 3 | 0 | 20 | 0.35 |
| 2_10661406 | 4 | 0 | 1 | 0 | 7 | 0.2857 | 5 | 0 | 17 | 0.1176 | 2 | 0.5 | 7 | 0.1429 | 9 | 0.2222 | 3 | 0 | 21 | 0.1905 |
| 2_10661411 | 5 | 0.6 | 1 | 0 | 7 | 0.7143 | 4 | 0.25 | 17 | 0.5294 | 3 | 0.3333 | 6 | 0.6667 | 9 | 0.3333 | 3 | 0.3333 | 21 | 0.4286 |
| 2_10669745 | 2 | 0.5 | NA | NA | 7 | 0.2857 | 6 | 0.1667 | 15 | 0.2667 | 2 | 1 | 7 | 0.2857 | 9 | 0.1111 | 3 | 0.3333 | 21 | 0.2857 |
| 2_10669753 | 2 | 0 | 1 | 0 | 5 | 0.2 | 8 | 0.125 | 16 | 0.125 | 4 | 0.25 | 4 | 0 | 8 | 0 | 2 | 0 | 18 | 0.0556 |
| 2_10669763 | 4 | 0 | 1 | 0 | 9 | 0 | 6 | 0.1667 | 20 | 0.05 | 5 | 0.2 | 9 | 0.1111 | 8 | 0.25 | 3 | 0 | 25 | 0.16 |
| 2_10669770 | 4 | 0.25 | NA | NA | 9 | 0.2222 | 5 | 0 | 18 | 0.1667 | 3 | 0 | 8 | 0.25 | 9 | 0.1111 | 1 | 0 | 21 | 0.1429 |
| 2_10669801 | 4 | 0.25 | 1 | 0 | 10 | 0.3 | 6 | 0.3333 | 21 | 0.2857 | 4 | 0.75 | 7 | 0.4286 | 9 | 0.3333 | 2 | 1 | 22 | 0.5 |
| 2_10669822 | 3 | 0 | 1 | 0 | 9 | 0 | 6 | 0.1667 | 19 | 0.0526 | 3 | 0 | 7 | 0 | 6 | 0.1667 | 2 | 0 | 18 | 0.0556 |
| 2_10669827 | 5 | 0.4 | 1 | 1 | 9 | 0.5556 | 9 | 0.4444 | 24 | 0.5 | 4 | 1 | 6 | 0.6667 | 9 | 0.6667 | 2 | 1 | 21 | 0.7619 |
| 2_10669832 | 4 | 0 | 1 | 1 | 9 | 0.2222 | 6 | 0.3333 | 20 | 0.25 | 5 | 0.2 | 7 | 0.2857 | 7 | 0.1429 | 2 | 0.5 | 21 | 0.2381 |
| 2_10669836 | 5 | 0.2 | 1 | 0 | 11 | 0.0909 | 8 | 0 | 25 | 0.08 | 5 | 0.2 | 8 | 0 | 8 | 0.125 | 2 | 0 | 23 | 0.087 |
| 2_10669840 | 5 | 0 | 1 | 1 | 11 | 0.1818 | 8 | 0 | 25 | 0.12 | 5 | 0.2 | 8 | 0 | 10 | 0.2 | 3 | 0.3333 | 26 | 0.1538 |
| 2_10669842 | 6 | 0 | 1 | 0 | 11 | 0.0909 | 8 | 0.125 | 26 | 0.0769 | 4 | 0 | 8 | 0.125 | 7 | 0.1429 | 4 | 0 | 23 | 0.087 |

| | | | | | | | | | | | | | | | | | | | | |
|---|---|---|---|---|---|---|---|---|---|---|---|---|---|---|---|---|---|---|---|---|
| 2_1067933 | 6 | 0.5 | 12 | 0.5 | 3 | 0.3333 | 6 | 0.6667 | 27 | 0.5185 | 1 | 1 | 4 | 0.75 | 5 | 0.8 | 8 | 0.375 | 18 | 0.6111 |
| 2_1067939 | 8 | 0 | 12 | 0.0833 | 7 | 0 | 6 | 0.1667 | 33 | 0.0606 | 1 | 0 | 3 | 0.3333 | 6 | 0.1667 | 12 | 0 | 22 | 0.0909 |
| 2_1067948 | 6 | 0 | 9 | 0.1111 | 7 | 0.2857 | 6 | 0.1667 | 28 | 0.1429 | 1 | 0 | 3 | 0.3333 | 5 | 0 | 11 | 0 | 20 | 0.05 |
| 2_1067948 | 6 | 1 | 7 | 1 | 5 | 1 | 6 | 1 | 24 | 1 | 1 | 1 | 3 | 1 | 3 | 1 | 11 | 0.8182 | 18 | 0.8889 |
| 2_1067949 | 6 | 0 | 6 | 0.3333 | 4 | 0 | 4 | 0 | 20 | 0.1 | 1 | 0 | 3 | 0.3333 | 5 | 0.2 | 6 | 0 | 15 | 0.1333 |
| 2_1067949 | 5 | 0.4 | 7 | 0.1429 | 6 | 0 | 6 | 0.3333 | 24 | 0.2083 | 1 | 1 | 2 | 0 | 5 | 0.4 | 8 | 0.125 | 16 | 0.25 |
| 2_1067949 | 6 | 0.1667 | 7 | 0.1429 | 6 | 0 | 5 | 0.2 | 24 | 0.125 | 1 | 1 | 3 | 0 | 5 | 0.4 | 10 | 0.2 | 19 | 0.2632 |
| 2_1067950 | 5 | 0 | 6 | 0 | 5 | 0.2 | 6 | 0.1667 | 22 | 0.0909 | 1 | 1 | 3 | 0 | 5 | 0.2 | 10 | 0.2 | 19 | 0.2105 |
| 2_1067951 | 4 | 0.75 | 4 | 0.25 | 6 | 0.8333 | 6 | 0.5 | 20 | 0.6 | 1 | 0 | NA | NA | 2 | 1 | 8 | 0.375 | 11 | 0.4545 |
| 2_1067959 | 4 | 0.25 | 7 | 0.7143 | 5 | 0.6 | 5 | 0.4 | 21 | 0.5238 | 1 | 0 | 2 | 0.5 | 5 | 0.6 | 7 | 0.2857 | 15 | 0.4 |
| 2_1067959 | 5 | 0.2 | 7 | 0.1429 | 5 | 0.2 | 4 | 0 | 21 | 0.1429 | 1 | 0 | 2 | 0.5 | 5 | 0.2 | 8 | 0 | 16 | 0.125 |
| 2_1067995 | 4 | 0.25 | 6 | 0.3333 | 4 | 0 | 3 | 0 | 17 | 0.1765 | 1 | 0 | 1 | 0 | 4 | 0.5 | 6 | 0.3333 | 12 | 0.3333 |
| 2_1067995 | 5 | 0 | 6 | 0.1667 | 4 | 0 | 3 | 0 | 18 | 0.0556 | 1 | 0 | 1 | 1 | 5 | 0 | 5 | 0.2 | 12 | 0.1667 |
| 2_1067996 | 5 | 0.4 | 6 | 0.1667 | 4 | 0 | 3 | 0 | 18 | 0.1667 | 1 | 0 | 1 | 0 | 5 | 0 | 6 | 0.1667 | 13 | 0.0769 |
| 2_1067996 | 5 | 0.2 | 5 | 0.2 | 4 | 0.5 | 3 | 0.3333 | 17 | 0.2941 | 1 | 0 | 1 | 0 | 5 | 0 | 6 | 0.3333 | 13 | 0.1538 |
| 2_1067996 | 5 | 0.6 | 6 | 0.5 | 4 | 0.5 | 2 | 0 | 17 | 0.4706 | 1 | 0 | 1 | 1 | 4 | 0.75 | 6 | 0.5 | 12 | 0.5833 |
| 2_1067997 | 3 | 1 | 4 | 0.75 | 3 | 1 | 2 | 1 | 12 | 0.9167 | NA | NA | NA | NA | 4 | 0.75 | 4 | 1 | 8 | 0.875 |
| 2_1067997 | 3 | 0.3333 | 6 | 0.1667 | 4 | 0 | 3 | 0.3333 | 16 | 0.1875 | 1 | 0 | 1 | 0 | 5 | 0 | 5 | 0.2 | 12 | 0.0833 |
| 2_1067997 | 4 | 0.5 | 6 | 0.3333 | 3 | 0 | 2 | 0 | 15 | 0.2667 | 1 | 0 | 1 | 0 | 5 | 0.4 | 5 | 0 | 12 | 0.1667 |
| 2_1071211 | 1 | 1 | 1 | 1 | NA | NA | 1 | 1 | 3 | 1 | NA | NA | 2 | 0.5 | 1 | 1 | NA | NA | 3 | 0.6667 |
| 2_1071247 | 1 | 1 | 1 | 1 | NA | NA | 2 | 1 | 4 | 1 | NA | NA | 2 | 1 | 1 | 1 | NA | NA | 3 | 1 |
| 2_1071247 | NA | NA | 1 | 0 | NA | NA | 2 | 1 | 3 | 0.6667 | NA | NA | 2 | 0 | 1 | 1 | NA | NA | 3 | 0.3333 |
| 2_1071247 | NA | NA | NA | NA | NA | NA | 2 | 1 | 2 | 1 | NA | NA | 2 | 1 | 1 | 1 | NA | NA | 3 | 1 |
| 2_1071248 | 1 | 1 | 1 | 1 | NA | NA | NA | NA | 2 | 1 | NA | NA | 2 | 0.5 | NA | NA | NA | NA | 2 | 0.5 |
| 2_1071249 | 1 | 0 | 1 | 1 | NA | NA | 2 | 1 | 4 | 0.75 | NA | NA | 3 | 0.3333 | NA | NA | NA | NA | 3 | 0.3333 |
| 2_1071250 | 1 | 0 | 1 | 1 | NA | NA | 1 | 0 | 3 | 0.3333 | NA | NA | 2 | 0.5 | 1 | 0 | NA | NA | 3 | 0.3333 |
| 2_1071251 | 1 | 1 | 1 | 1 | NA | NA | 2 | 1 | 4 | 1 | NA | NA | 4 | 1 | NA | NA | NA | NA | 4 | 1 |
| 2_1071253 | 1 | 0 | 1 | 0 | 1 | 1 | 2 | 0 | 5 | 0.2 | NA | NA | 4 | 0.5 | 1 | 0 | 1 | 1 | 6 | 0.5 |
| 2_1071257 | 1 | 1 | 1 | 0 | 1 | 1 | 2 | 1 | 5 | 0.8 | NA | NA | 3 | 1 | 3 | 0.3333 | 1 | 0 | 7 | 0.5714 |
| 2_1071257 | 1 | 1 | 1 | 1 | 2 | 1 | 2 | 1 | 6 | 1 | NA | NA | 4 | 1 | 3 | 1 | 1 | 1 | 8 | 1 |
| 2_1071258 | 1 | 0 | 1 | 1 | NA | NA | 2 | 0.5 | 4 | 0.5 | NA | NA | 4 | 0.5 | 2 | 0 | 1 | 0 | 7 | 0.2857 |
| 2_1071258 | 1 | 0 | 1 | 1 | 2 | 0 | 2 | 1 | 6 | 0.5 | NA | NA | 5 | 0.2 | 3 | 0.6667 | 1 | 0 | 9 | 0.3333 |
| 2_1071258 | 1 | 1 | 1 | 1 | 2 | 0 | 2 | 1 | 6 | 0.6667 | NA | NA | 5 | 1 | 3 | 0.6667 | 1 | 1 | 9 | 0.8889 |
| 2_1071259 | 1 | 1 | 1 | 0 | 2 | 0 | 2 | 0 | 6 | 0.1667 | NA | NA | 4 | 0.25 | 3 | 0 | NA | NA | 7 | 0.1429 |
| 2_1071259 | NA | NA | 1 | 0 | 1 | 1 | 1 | 0 | 3 | 0.3333 | NA | NA | 3 | 0.6667 | 2 | 0.5 | 1 | 0 | 6 | 0.5 |
| 2_1071262 | 1 | 1 | 1 | 1 | 2 | 1 | 2 | 1 | 6 | 1 | NA | NA | 5 | 1 | 3 | 1 | 1 | 1 | 9 | 1 |
| 2_1071263 | 1 | 0 | 1 | 0 | 2 | 0.5 | 2 | 0 | 6 | 0.1667 | NA | NA | 6 | 0.1667 | 3 | 0 | 1 | 0 | 10 | 0.1 |
| 2_1071263 | 1 | 1 | 1 | 1 | 1 | 1 | 2 | 1 | 5 | 1 | NA | NA | 6 | 1 | 3 | 1 | 1 | 1 | 10 | 1 |
| 2_1071263 | 1 | 1 | 1 | 1 | 1 | 1 | 2 | 1 | 5 | 1 | NA | NA | 6 | 0 | 3 | 0.6667 | 1 | 0 | 10 | 0.2 |
| 2_1071263 | 1 | 1 | 1 | 0 | 1 | 1 | 2 | 1 | 5 | 0.8 | NA | NA | 6 | 0.5 | 3 | 0.3333 | 1 | 0 | 10 | 0.4 |
| 2_1071264 | 1 | 1 | 1 | 1 | 2 | 0.5 | 2 | 0 | 6 | 0.5 | NA | NA | 5 | 0.4 | 1 | 0 | 1 | 1 | 7 | 0.4286 |
| 2_1071264 | 1 | 1 | 1 | 0 | 1 | 1 | 2 | 1 | 5 | 0.8 | NA | NA | 6 | 1 | 2 | 1 | NA | NA | 8 | 1 |
| 2_1071265 | 2 | 0.5 | 1 | 0 | 1 | 0 | 2 | 0 | 6 | 0.1667 | NA | NA | 5 | 0.2 | 3 | 0.3333 | 1 | 0 | 9 | 0.2222 |
| 2_1071266 | 2 | 0.5 | 1 | 1 | 2 | 0 | 2 | 1 | 7 | 0.5714 | NA | NA | 4 | 1 | 4 | 0.5 | 1 | 1 | 9 | 0.7778 |
| 2_1071267 | 2 | 1 | 1 | 1 | 2 | 1 | 2 | 1 | 7 | 1 | NA | NA | 5 | 1 | 4 | 1 | 1 | 1 | 10 | 1 |
| 2_1071267 | 2 | 1 | 2 | 0.5 | 2 | 0.5 | 2 | 0.5 | 8 | 0.625 | NA | NA | 6 | 0.3333 | 3 | 0.3333 | 1 | 1 | 10 | 0.4 |
| 2_1071268 | 2 | 0.5 | 1 | 0 | 1 | 0 | 1 | 0 | 5 | 0.2 | NA | NA | 5 | 0 | 4 | 0.75 | 1 | 0 | 10 | 0.3 |
| 2_1071268 | 2 | 1 | 2 | 1 | 1 | 1 | 2 | 1 | 7 | 1 | NA | NA | 5 | 0.8 | 4 | 0.75 | 1 | 1 | 10 | 0.8 |
| 2_1071269 | 2 | 0.5 | 3 | 0 | 2 | 0.5 | 1 | 0 | 8 | 0.25 | NA | NA | 6 | 0 | 3 | 0.3333 | 1 | 0 | 10 | 0.1 |
| 2_1071269 | 2 | 1 | 3 | 1 | 1 | 1 | 2 | 1 | 8 | 1 | NA | NA | 4 | 0.75 | 3 | 1 | NA | NA | 7 | 0.8571 |
| 2_1071269 | 1 | 1 | 1 | 0 | NA | NA | 1 | 0 | 3 | 0.3333 | NA | NA | 4 | 0.5 | 1 | 0 | NA | NA | 5 | 0.4 |
| 2_1071270 | 2 | 0.5 | 3 | 0.6667 | 2 | 0.5 | 2 | 0 | 9 | 0.4444 | NA | NA | 5 | 0.6 | 4 | 0.25 | NA | NA | 9 | 0.4444 |
| 2_1071272 | 2 | 0.5 | 3 | 0.3333 | 3 | 0.3333 | 2 | 1 | 10 | 0.5 | NA | NA | 5 | 0.8 | 4 | 0.25 | 1 | 0 | 10 | 0.5 |
| 2_1071273 | 2 | 0.5 | 3 | 0 | 2 | 0 | 2 | 0 | 9 | 0.1111 | NA | NA | 6 | 0.1667 | 4 | 0 | 1 | 0 | 11 | 0.0909 |
| 2_1071280 | 1 | 1 | 1 | 1 | 3 | 1 | 1 | 1 | 6 | 1 | 1 | 1 | 6 | 1 | 5 | 1 | 1 | 1 | 13 | 1 |
| 2_1071280 | 1 | 0 | 1 | 1 | 3 | 0.6667 | 1 | 1 | 6 | 0.6667 | 1 | 0 | 5 | 0.2 | 4 | 0.25 | 1 | 0 | 11 | 0.1818 |
| 2_1071281 | NA | NA | 1 | 1 | 2 | 1 | 1 | 0 | 4 | 0.75 | 2 | 0.5 | 6 | 0.6667 | 4 | 1 | 1 | 0 | 13 | 0.6923 |
| 2_1071281 | 1 | 1 | 1 | 1 | 3 | 1 | 1 | 1 | 6 | 1 | 1 | 1 | 5 | 1 | 5 | 1 | 2 | 1 | 13 | 1 |
| 2_1071557 | 1 | 0 | 2 | 0 | 1 | 0 | 6 | 0.1667 | 10 | 0.1 | NA | NA | 4 | 0 | 5 | 0 | 5 | 0.2 | 14 | 0.0714 |
| 2_1071569 | 2 | 0 | 2 | 0.5 | 1 | 0 | 5 | 0.4 | 10 | 0.3 | NA | NA | 4 | 0.25 | 4 | 0.75 | 5 | 0.4 | 13 | 0.4615 |
| 2_1071571 | 2 | 0 | 3 | 0 | 1 | 0 | 10 | 0.2 | 16 | 0.125 | 1 | 0 | 6 | 0 | 6 | 0.1667 | 5 | 0 | 18 | 0.0556 |
| 2_1087808 | NA | NA | 1 | 0 | 3 | 1 | 3 | 1 | 7 | 0.8571 | 2 | 1 | NA | NA | 1 | 0 | 3 | 0 | 6 | 0.3333 |
| 2_1089233 | 1 | 1 | 1 | 0 | 2 | 0.5 | NA | NA | 4 | 0.5 | 1 | 0 | NA | NA | NA | NA | 1 | 1 | 2 | 0.5 |
| 2_1089235 | 2 | 1 | 1 | 1 | 1 | 1 | NA | NA | 4 | 1 | 1 | 1 | NA | NA | NA | NA | 1 | 1 | 2 | 1 |
| 2_1091049 | 1 | 1 | NA | NA | 2 | 0.5 | NA | NA | 3 | 0.6667 | 1 | 1 | NA | NA | NA | NA | 1 | 0 | 2 | 0.5 |

| | | | | | | | | | | | | | | | | | | | | |
|---|---|---|---|---|---|---|---|---|---|---|---|---|---|---|---|---|---|---|---|---|
| 2_10955380 | NA | NA | 4 | 0.25 | 1 | 1 | 1 | 0 | 6 | 0.3333 | 2 | 0 | 2 | 0 | 1 | 0 | 2 | 1 | 7 | 0.2857 |
| 2_10955387 | NA | NA | 4 | 1 | 1 | 1 | NA | NA | 5 | 1 | 3 | 1 | 2 | 1 | 1 | 0 | 2 | 1 | 8 | 0.875 |
| 2_10955401 | NA | NA | 4 | 1 | NA | NA | 1 | 1 | 5 | 1 | 1 | 1 | 2 | 0.5 | 1 | 1 | 2 | 1 | 6 | 0.8333 |
| 2_10955417 | NA | NA | 5 | 0.8 | 1 | 0 | 1 | 1 | 7 | 0.7143 | 2 | 0.5 | 2 | 0.5 | 1 | 0 | 2 | 0.5 | 7 | 0.4286 |
| 2_10955421 | NA | NA | 4 | 1 | 1 | 1 | 1 | 1 | 6 | 1 | 1 | 1 | 2 | 1 | 1 | 1 | NA | NA | 4 | 1 |
| 2_10955422 | NA | NA | 4 | 1 | 1 | 1 | NA | NA | 5 | 1 | 3 | 1 | 1 | 1 | 1 | 1 | NA | NA | 5 | 1 |
| 2_10955435 | NA | NA | 4 | 0.5 | 1 | 1 | 1 | 0 | 6 | 0.5 | 3 | 0.3333 | 2 | 0 | 2 | 0.5 | 2 | 0.5 | 9 | 0.3333 |
| 2_10955436 | NA | NA | 2 | 1 | NA | NA | NA | NA | 2 | 1 | 2 | 1 | NA | NA | 1 | 1 | 2 | 0.5 | 5 | 0.8 |
| 2_10955445 | 1 | 1 | 4 | 0.5 | 1 | 0 | 1 | 0 | 7 | 0.4286 | 3 | 0.3333 | 2 | 0.5 | 2 | 0.5 | 1 | 0 | 8 | 0.375 |
| 2_10955459 | 1 | 1 | 4 | 0.5 | 1 | 0 | 1 | 1 | 7 | 0.5714 | 4 | 0.5 | 2 | 0 | 1 | 1 | 2 | 0.5 | 9 | 0.4444 |
| 2_10955465 | 1 | 0 | 4 | 0.25 | 1 | 0 | 1 | 1 | 7 | 0.2857 | 3 | 0.6667 | 2 | 0 | 2 | 0.5 | 2 | 1 | 9 | 0.5556 |
| 2_10972774 | 7 | 0 | 8 | 0 | 6 | 0.3333 | 6 | 0 | 27 | 0.0741 | 5 | 0.2 | 6 | 0 | 6 | 0 | 3 | 0.3333 | 20 | 0.1 |
| 2_10972775 | 6 | 0.3333 | 8 | 0.375 | 6 | 0.6667 | 5 | 0.4 | 25 | 0.44 | 5 | 0.6 | 6 | 0.6667 | 6 | 0.6667 | 4 | 0.75 | 21 | 0.6667 |
| 2_10972889 | 5 | 0.6 | 10 | 0.1 | 8 | 0.25 | 8 | 0.125 | 31 | 0.2258 | 7 | 0.1429 | 10 | 0.4 | 6 | 0 | 5 | 0 | 28 | 0.1786 |
| 2_10973017 | 7 | 0.2857 | 14 | 0.5 | 9 | 0.4444 | 7 | 0.7143 | 37 | 0.4865 | 6 | 0.3333 | 11 | 0.6364 | 11 | 0.1818 | 10 | 0.4 | 38 | 0.3947 |
| 2_10973019 | 9 | 0.1111 | 11 | 0.1818 | 10 | 0.2 | 6 | 0.3333 | 36 | 0.1944 | 7 | 0 | 12 | 0.0833 | 10 | 0.1 | 8 | 0 | 37 | 0.0541 |
| 2_10973078 | 11 | 0.2727 | 15 | 0.3333 | 10 | 0 | 8 | 0 | 44 | 0.1818 | 8 | 0.125 | 13 | 0.0769 | 13 | 0.1538 | 9 | 0.2222 | 43 | 0.1395 |
| 2_10984176 | 3 | 0 | 1 | 1 | 2 | 1 | NA | NA | 6 | 0.5 | 2 | 0.5 | NA | NA | 1 | 0 | 3 | 0 | 6 | 0.1667 |
| 2_10984211 | 4 | 0.5 | 3 | 0.3333 | 2 | 0.5 | 1 | 0 | 10 | 0.4 | 2 | 0.5 | 1 | 0 | 2 | 0.5 | 3 | 0.6667 | 8 | 0.5 |
| 2_10984214 | 3 | 0.3333 | 3 | 0 | 3 | 0.3333 | 1 | 1 | 10 | 0.3 | 2 | 0.5 | 1 | 1 | 2 | 0.5 | 3 | 0 | 8 | 0.375 |
| 2_10984215 | 3 | 0.3333 | 1 | 1 | 3 | 0.6667 | 1 | 1 | 8 | 0.625 | 1 | 0 | 1 | 0 | 1 | 0 | 3 | 0.3333 | 6 | 0.1667 |
| 2_10984219 | 3 | 0.3333 | 1 | 0 | 2 | 0.5 | NA | NA | 6 | 0.3333 | 1 | 0 | 1 | 0 | 2 | 0.5 | 3 | 0.3333 | 7 | 0.2857 |
| 2_10984223 | 2 | 0 | 3 | 0 | 2 | 1 | 1 | 0 | 8 | 0.25 | 2 | 0 | 1 | 0 | 2 | 0 | 3 | 0.3333 | 8 | 0.125 |
| 2_10984228 | 1 | 1 | 2 | 1 | 2 | 0.5 | 1 | 1 | 6 | 0.8333 | 1 | 1 | 1 | 0 | 1 | 1 | 2 | 1 | 5 | 0.8 |
| 2_10984258 | 2 | 1 | 3 | 1 | 1 | 0 | NA | NA | 6 | 0.8333 | 1 | 1 | NA | NA | 3 | 1 | 2 | 1 | 6 | 1 |
| 2_10984259 | 2 | 1 | 3 | 1 | 3 | 1 | NA | NA | 8 | 1 | 2 | 1 | 1 | 1 | 2 | 1 | 2 | 1 | 7 | 1 |
| 2_10984263 | 2 | 0.5 | 3 | 0.3333 | 2 | 0.5 | NA | NA | 7 | 0.4286 | 2 | 0.5 | 1 | 0 | 3 | 0.3333 | 3 | 0.3333 | 9 | 0.3333 |
| 2_10984266 | 3 | 0 | 2 | 1 | 1 | 1 | 1 | 1 | 7 | 0.5714 | 2 | 1 | NA | NA | 3 | 0.6667 | 3 | 0 | 8 | 0.5 |
| 2_10984273 | 3 | 0.6667 | 3 | 0.6667 | 3 | 0.3333 | 1 | 0 | 10 | 0.5 | 1 | 0 | 1 | 1 | 2 | 1 | 3 | 1 | 7 | 0.8571 |
| 2_10984276 | 3 | 1 | 3 | 1 | 4 | 0.5 | 1 | 1 | 11 | 0.8182 | 2 | 1 | NA | NA | 3 | 1 | 3 | 1 | 8 | 1 |
| 2_10984286 | 3 | 1 | 3 | 1 | 3 | 1 | 1 | 1 | 10 | 1 | 1 | 1 | 1 | 1 | 3 | 0.6667 | 3 | 1 | 8 | 0.875 |
| 2_10984287 | 2 | 0.5 | 3 | 0.6667 | 1 | 0 | 1 | 1 | 7 | 0.5714 | 1 | 0 | NA | NA | 1 | 1 | 2 | 0.5 | 4 | 0.5 |
| 2_10984294 | 3 | 0 | 2 | 0 | 4 | 0.25 | 1 | 0 | 10 | 0.1 | 2 | 0.5 | 1 | 0 | 3 | 0.3333 | 3 | 0.3333 | 9 | 0.3333 |
| 2_10984295 | 2 | 1 | 3 | 1 | 4 | 1 | 1 | 1 | 10 | 1 | 2 | 1 | 1 | 1 | 2 | 1 | 3 | 1 | 8 | 1 |
| 2_10984323 | 2 | 0.5 | 4 | 0.25 | 5 | 0.2 | 1 | 1 | 12 | 0.3333 | 2 | 0.5 | 1 | 1 | 3 | 0.3333 | 2 | 0.5 | 8 | 0.5 |
| 2_10984335 | 1 | 0 | 3 | 0.6667 | 3 | 0.6667 | NA | NA | 7 | 0.5714 | NA | NA | 1 | 0 | 3 | 0.6667 | 3 | 0 | 7 | 0.2857 |
| 2_10984406 | 2 | 0 | 5 | 0 | 4 | 0 | 2 | 0.5 | 13 | 0.0769 | 1 | 0 | 1 | 0 | 3 | 0.3333 | 3 | 0 | 8 | 0.125 |
| 2_10987473 | 1 | 1 | 2 | 0.5 | 2 | 0.5 | 1 | 0 | 6 | 0.5 | 1 | 1 | 1 | 1 | 1 | 1 | 2 | 0 | 5 | 0.6 |
| 2_11014474 | 1 | 1 | NA | NA | NA | NA | NA | NA | 1 | 1 | NA | NA | NA | NA | NA | NA | 1 | 1 | 1 | 1 |
| 2_11019221 | 1 | 1 | NA | NA | 2 | 1 | NA | NA | 3 | 1 | NA | NA | 1 | 1 | NA | NA | 1 | 1 | 2 | 1 |
| 2_11019258 | 1 | 1 | NA | NA | 2 | 1 | 1 | 0 | 4 | 0.75 | NA | NA | NA | NA | NA | NA | 1 | 1 | 1 | 1 |
| 2_11019265 | 1 | 1 | NA | NA | NA | NA | 1 | 1 | 2 | 1 | NA | NA | NA | NA | NA | NA | 1 | 1 | 1 | 1 |
| 2_11019267 | 1 | 1 | NA | NA | 2 | 1 | 1 | 1 | 4 | 1 | NA | NA | NA | NA | NA | NA | 1 | 1 | 1 | 1 |
| 2_11019304 | 1 | 1 | NA | NA | 2 | 1 | 1 | 1 | 4 | 1 | NA | NA | NA | NA | NA | NA | 1 | 1 | 1 | 1 |
| 2_11019310 | NA | NA | NA | NA | 2 | 0.5 | 1 | 1 | 3 | 0.6667 | NA | NA | NA | NA | NA | NA | 1 | 1 | 1 | 1 |
| 2_11048815 | 8 | 0.125 | 6 | 0.3333 | 7 | 0.2857 | 14 | 0 | 35 | 0.1429 | 2 | 0.5 | 6 | 0 | 12 | 0.0833 | 8 | 0 | 28 | 0.0714 |
| 2_11048830 | 8 | 0.625 | 5 | 0.4 | 7 | 0.4286 | 15 | 0.4667 | 35 | 0.4857 | 2 | 0.5 | 3 | 0.3333 | 11 | 0.6364 | 8 | 0.625 | 24 | 0.5833 |
| 2_11048832 | 8 | 0.125 | 6 | 0.1667 | 7 | 0.2857 | 13 | 0.1538 | 34 | 0.1765 | 2 | 0 | 6 | 0 | 11 | 0.0909 | 8 | 0.125 | 27 | 0.0741 |
| 2_11048843 | 7 | 0.2857 | 5 | 0.2 | 7 | 0.2857 | 13 | 0.2308 | 32 | 0.25 | 2 | 0 | 4 | 0.25 | 11 | 0.2727 | 7 | 0.1429 | 24 | 0.2083 |
| 2_11048854 | 7 | 0.2857 | 6 | 0.3333 | 5 | 0.2 | 12 | 0.0833 | 30 | 0.2 | 2 | 0 | 5 | 0 | 6 | 0 | 6 | 0.6667 | 19 | 0.2105 |
| 2_11048854 | 7 | 0.1429 | 6 | 0.1667 | 7 | 0.1429 | 12 | 0.25 | 32 | 0.1875 | 2 | 0 | 4 | 0 | 10 | 0.3 | 8 | 0 | 24 | 0.125 |
| 2_11062538 | 6 | 0 | 3 | 0.3333 | 4 | 0 | 5 | 0 | 18 | 0.0556 | NA | NA | 3 | 0 | 4 | 0 | 7 | 0.1429 | 14 | 0.0714 |
| 2_11207720 | 13 | 0.3077 | 12 | 0.25 | 15 | 0.3333 | 14 | 0.2857 | 54 | 0.2963 | 16 | 0.25 | 17 | 0.2941 | 7 | 0.5714 | 21 | 0.2381 | 61 | 0.2951 |
| 2_11207740 | 13 | 0.1538 | 12 | 0 | 17 | 0.0588 | 11 | 0 | 53 | 0.0566 | 15 | 0.1333 | 18 | 0.0556 | 8 | 0.125 | 19 | 0.0526 | 60 | 0.0833 |
| 2_11207758 | NA | NA | 1 | 1 | NA | NA | 1 | 0 | 2 | 0.5 | NA | NA | 1 | 1 | NA | NA | 1 | 1 | 2 | 1 |
| 2_11209295 | 21 | 0.3333 | 20 | 0.4 | 19 | 0.3684 | 17 | 0.4706 | 77 | 0.3896 | 11 | 0.0909 | 21 | 0.381 | 15 | 0.2 | 17 | 0.4706 | 64 | 0.3125 |
| 2_11209300 | 16 | 0.0625 | 18 | 0.1667 | 11 | 0.0909 | 19 | 0.1579 | 64 | 0.125 | 11 | 0.0909 | 15 | 0.2 | 6 | 0 | 15 | 0.2 | 47 | 0.1489 |
| 2_11209637 | 1 | 0 | 2 | 0.5 | NA | NA | NA | NA | 3 | 0.3333 | 1 | 0 | NA | NA | NA | NA | 3 | 0.3333 | 4 | 0.25 |
| 2_11209675 | 1 | 1 | 2 | 0.5 | NA | NA | NA | NA | 3 | 0.6667 | 1 | 1 | NA | NA | 1 | 0 | 3 | 1 | 5 | 0.8 |
| 2_11209677 | 1 | 1 | 2 | 0.5 | NA | NA | NA | NA | 3 | 0.6667 | 1 | 1 | NA | NA | 1 | 1 | 3 | 1 | 5 | 1 |
| 2_11209679 | 1 | 1 | 1 | 1 | NA | NA | NA | NA | 2 | 1 | 1 | 1 | NA | NA | 1 | 1 | 2 | 1 | 4 | 1 |
| 2_11209686 | 1 | 0 | 2 | 0.5 | NA | NA | NA | NA | 3 | 0.3333 | 1 | 0 | NA | NA | 1 | 1 | 3 | 0.6667 | 5 | 0.6 |
| 2_11209726 | 1 | 1 | 3 | 0 | NA | NA | NA | NA | 4 | 0.25 | 1 | 0 | NA | NA | 1 | 0 | 3 | 0.6667 | 5 | 0.4 |
| 2_11209735 | 1 | 0 | 3 | 1 | NA | NA | 1 | 1 | 5 | 0.8 | 1 | 1 | NA | NA | NA | NA | 3 | 1 | 4 | 1 |
| 2_11218885 | NA | NA | NA | NA | 1 | 1 | NA | NA | 1 | 1 | 2 | 1 | NA | NA | 3 | 1 | NA | NA | 5 | 1 |

| | | | | | | | | | | | | | | | | | | | | |
|---|---|---|---|---|---|---|---|---|---|---|---|---|---|---|---|---|---|---|---|---|
| 2_1121889( | NA | NA | 1 | 1 | 1 | 1 | NA | NA | 2 | 1 | 2 | 0.5 | NA | NA | 2 | 0.5 | 1 | 1 | 5 | 0.6 |
| 2_1121889' | NA | NA | NA | NA | 1 | 1 | NA | NA | 1 | 1 | 2 | 0.5 | NA | NA | 3 | 1 | NA | NA | 5 | 0.8 |
| 2_1121890! | NA | NA | 1 | 1 | 1 | 1 | NA | NA | 2 | 1 | 2 | 1 | NA | NA | 3 | 0.6667 | NA | NA | 5 | 0.8 |
| 2_1121890( | NA | NA | NA | NA | 1 | 1 | NA | NA | 1 | 1 | 2 | 0.5 | NA | NA | 3 | 0.3333 | 1 | 0 | 6 | 0.3333 |
| 2_1121890! | NA | NA | 1 | 1 | 1 | 1 | NA | NA | 2 | 1 | 1 | 0 | NA | NA | 3 | 0.3333 | NA | NA | 4 | 0.25 |
| 2_1121908( | NA | NA | 1 | 0 | 1 | 1 | NA | NA | 2 | 0.5 | 2 | 0.5 | NA | NA | 1 | 1 | 1 | 1 | 4 | 0.75 |
| 2_1121908! | 4 | 0.75 | 8 | 0.625 | 2 | 1 | 8 | 0.625 | 22 | 0.6818 | 4 | 0.5 | 1 | 0 | 6 | 0.5 | 3 | 0.6667 | 14 | 0.5 |
| 2_1121910: | NA | NA | 1 | 0 | 1 | 1 | NA | NA | 2 | 0.5 | 1 | 0 | NA | NA | 1 | 1 | NA | NA | 2 | 0.5 |
| 2_1121911! | 4 | 0 | 9 | 0.1111 | 2 | 0 | 5 | 0 | 20 | 0.05 | 5 | 0 | 2 | 0 | 6 | 0.1667 | 2 | 0 | 15 | 0.0667 |
| 2_1121925· | 4 | 0 | 16 | 0.1875 | 4 | 0 | 9 | 0 | 33 | 0.0909 | 5 | 0 | 4 | 0.25 | 7 | 0 | 6 | 0.1667 | 22 | 0.0909 |
| 2_1121927' | 2 | 0 | 15 | 0.2 | 3 | 0 | 9 | 0.1111 | 29 | 0.1379 | 4 | 0.25 | 4 | 0 | 8 | 0.125 | 5 | 0 | 21 | 0.0952 |
| 2_1121933: | 4 | 0 | 19 | 0.0526 | 6 | 0 | 7 | 0.2857 | 36 | 0.0833 | 9 | 0.1111 | 6 | 0 | 7 | 0.1429 | 6 | 0 | 28 | 0.0714 |
| 2_1121934· | 26 | 0.1923 | 31 | 0.3871 | 25 | 0.16 | 27 | 0.0741 | 109 | 0.211 | 21 | 0.1905 | 26 | 0.2308 | 14 | 0.0714 | 22 | 0.2727 | 83 | 0.2048 |
| 2_1121934! | 23 | 0 | 32 | 0 | 24 | 0.1667 | 28 | 0.1429 | 107 | 0.0748 | 21 | 0.0952 | 22 | 0.0455 | 14 | 0 | 21 | 0.0476 | 78 | 0.0513 |
| 2_1121935( | 23 | 0.1304 | 33 | 0.1515 | 24 | 0.0833 | 29 | 0.2759 | 109 | 0.1651 | 21 | 0.2381 | 26 | 0.1154 | 16 | 0.125 | 21 | 0.1429 | 84 | 0.1548 |
| 2_1121936' | 23 | 0.0435 | 36 | 0.0833 | 27 | 0.037 | 30 | 0.0667 | 116 | 0.0603 | 20 | 0.05 | 25 | 0.04 | 17 | 0.0588 | 23 | 0.087 | 85 | 0.0588 |
| 2_1121936: | 27 | 0 | 34 | 0.0588 | 27 | 0.037 | 31 | 0.0968 | 119 | 0.0504 | 22 | 0.0909 | 26 | 0.2308 | 16 | 0.125 | 23 | 0.1304 | 87 | 0.1494 |
| 2_1122429! | 2 | 1 | 1 | 0 | 5 | 0.2 | 4 | 0.25 | 12 | 0.3333 | NA | NA | 1 | 0 | NA | NA | 2 | 1 | 3 | 0.6667 |
| 2_1122430: | 1 | 1 | 1 | 1 | 4 | 1 | 4 | 1 | 10 | 1 | NA | NA | 2 | 1 | NA | NA | 2 | 1 | 4 | 1 |
| 2_1122430! | 2 | 0.5 | NA | NA | 3 | 0.3333 | 3 | 0.3333 | 8 | 0.375 | NA | NA | NA | NA | NA | NA | 2 | 0.5 | 2 | 0.5 |
| 2_1122431' | 3 | 1 | 1 | 1 | 5 | 1 | 4 | 1 | 13 | 1 | NA | NA | 2 | 1 | NA | NA | 3 | 1 | 5 | 1 |
| 2_1122431: | 3 | 0.6667 | 1 | 1 | 5 | 0.6 | 4 | 0.25 | 13 | 0.5385 | NA | NA | 2 | 0.5 | NA | NA | 3 | 0.6667 | 5 | 0.6 |
| 2_1122432' | 3 | 0.3333 | 1 | 0 | 4 | 0.25 | 4 | 0.25 | 12 | 0.25 | NA | NA | 3 | 1 | NA | NA | 3 | 0 | 6 | 0.5 |
| 2_1122973! | 17 | 0.1765 | 21 | 0.0952 | 15 | 0.0667 | 23 | 0.087 | 76 | 0.1053 | 9 | 0 | 19 | 0.1579 | 17 | 0.0588 | 25 | 0.16 | 70 | 0.1143 |
| 2_1122975( | 17 | 0.1176 | 14 | 0.2143 | 17 | 0.0588 | 22 | 0.1364 | 70 | 0.1286 | 10 | 0.1 | 14 | 0.0714 | 17 | 0.1765 | 20 | 0.15 | 61 | 0.1311 |
| 2_1122975! | 21 | 0.0952 | 17 | 0.0588 | 17 | 0.0588 | 27 | 0.0741 | 82 | 0.0732 | 9 | 0 | 18 | 0.0556 | 20 | 0.1 | 28 | 0.0357 | 75 | 0.0533 |
| 2_1124612' | 2 | 0 | 2 | 0.5 | 1 | 0 | 1 | 1 | 6 | 0.3333 | NA | NA | 2 | 1 | 2 | 0.5 | NA | NA | 4 | 0.75 |
| 2_1124617( | NA | NA | 1 | 1 | NA | NA | NA | NA | 1 | 1 | NA | NA | NA | NA | 2 | 1 | NA | NA | 2 | 1 |
| 2_1124619! | 3 | 0 | 4 | 0.5 | 8 | 0.375 | 4 | 0 | 19 | 0.2632 | NA | NA | 6 | 0 | 4 | 0.25 | 1 | 0 | 11 | 0.0909 |
| 2_1124620! | 4 | 0.25 | 4 | 0.25 | 7 | 0.4286 | 4 | 0.25 | 19 | 0.3158 | NA | NA | 5 | 0.2 | 4 | 0.5 | 2 | 0 | 11 | 0.2727 |
| 2_1124620: | 4 | 0 | 4 | 0 | 8 | 0.25 | 4 | 0.75 | 20 | 0.25 | NA | NA | 5 | 0.2 | 4 | 0.25 | 3 | 0.3333 | 12 | 0.25 |
| 2_1124621: | 4 | 0.75 | 3 | 1 | 8 | 1 | 4 | 0.75 | 19 | 0.8947 | NA | NA | 4 | 0.75 | 4 | 0.75 | 3 | 0.6667 | 11 | 0.7273 |
| 2_1124621! | 2 | 0 | 4 | 0 | 8 | 0.25 | 4 | 0 | 18 | 0.1111 | NA | NA | 5 | 0.2 | 4 | 0 | 3 | 0 | 12 | 0.0833 |
| 2_1124622( | 3 | 0 | 4 | 0.75 | 7 | 0.7143 | 3 | 1 | 17 | 0.6471 | NA | NA | 6 | 0.5 | 4 | 1 | 3 | 0.6667 | 13 | 0.6923 |
| 2_1124622' | 4 | 0.75 | 3 | 0.6667 | 7 | 0.7143 | 4 | 0.25 | 18 | 0.6111 | NA | NA | 5 | 0.6 | 4 | 0.75 | 3 | 1 | 12 | 0.75 |
| 2_1124622· | 4 | 0.25 | 2 | 0 | 6 | 0.1667 | 4 | 0 | 16 | 0.125 | NA | NA | 6 | 0 | 4 | 0.25 | 3 | 0.3333 | 13 | 0.1538 |
| 2_1124622! | 3 | 0.3333 | 4 | 0.25 | 6 | 0.1667 | 3 | 1 | 16 | 0.375 | NA | NA | 4 | 0 | 4 | 0.5 | 3 | 0.6667 | 11 | 0.3636 |
| 2_1124623' | 3 | 1 | 3 | 0.6667 | 7 | 0.4286 | 4 | 0.5 | 17 | 0.5882 | NA | NA | 4 | 0.25 | 4 | 0.25 | 3 | 0.3333 | 11 | 0.2727 |
| 2_1124623( | 3 | 0 | 3 | 0.6667 | 6 | 0.5 | 4 | 0.25 | 16 | 0.375 | NA | NA | 6 | 0.1667 | 3 | 0 | 2 | 0 | 11 | 0.0909 |
| 2_1124625( | 3 | 0 | 4 | 0 | 7 | 0.1429 | 3 | 0 | 17 | 0.0588 | NA | NA | 6 | 0.1667 | 3 | 0 | 3 | 0 | 12 | 0.0833 |
| 2_1124625: | 4 | 0.25 | 3 | 0 | 8 | 0 | 3 | 0 | 18 | 0.0556 | NA | NA | 5 | 0.2 | 2 | 0 | 3 | 0.3333 | 10 | 0.2 |
| 2_1124830' | 2 | 0.5 | 2 | 0.5 | 2 | 0 | 2 | 1 | 8 | 0.5 | NA | NA | 2 | 0.5 | 1 | 1 | 1 | 0 | 4 | 0.5 |
| 2_1124830: | 2 | 1 | 2 | 0.5 | 1 | 0 | 2 | 1 | 7 | 0.7143 | NA | NA | 1 | 0 | 2 | 0.5 | 1 | 1 | 4 | 0.5 |
| 2_1124830! | 1 | 1 | 1 | 1 | 2 | 0.5 | 2 | 1 | 6 | 0.8333 | NA | NA | 1 | 0 | 2 | 0.5 | 1 | 1 | 4 | 0.5 |
| 2_1124832· | 1 | 1 | 2 | 1 | 2 | 0.5 | 2 | 1 | 7 | 0.8571 | NA | NA | 2 | 0.5 | 2 | 0.5 | NA | NA | 4 | 0.5 |
| 2_1124834( | 2 | 0 | 2 | 0.5 | 2 | 0 | 2 | 0.5 | 8 | 0.25 | NA | NA | 2 | 0 | 2 | 0.5 | NA | NA | 4 | 0.25 |
| 2_1124835: | 2 | 0 | 2 | 0.5 | 2 | 1 | 2 | 0.5 | 8 | 0.5 | NA | NA | 2 | 0.5 | 2 | 1 | NA | NA | 4 | 0.75 |
| 2_1124836: | 2 | 0.5 | NA | NA | 2 | 1 | 1 | 0 | 5 | 0.6 | NA | NA | 2 | 0.5 | 2 | 0.5 | NA | NA | 4 | 0.5 |
| 2_1124838' | 2 | 0.5 | 2 | 1 | 2 | 0.5 | 2 | 0.5 | 8 | 0.625 | NA | NA | 1 | 0 | 1 | 1 | NA | NA | 2 | 0.5 |
| 2_1124840: | 2 | 0 | 2 | 0.5 | 1 | 1 | 2 | 0 | 7 | 0.2857 | NA | NA | 1 | 0 | 2 | 1 | NA | NA | 3 | 0.6667 |
| 2_1131619: | NA | NA | 1 | 1 | NA | NA | NA | NA | 1 | 1 | 1 | 1 | NA | NA | NA | NA | NA | NA | 1 | 1 |
| 2_1131620: | NA | NA | 1 | 1 | NA | NA | NA | NA | 1 | 1 | 1 | 1 | NA | NA | NA | NA | NA | NA | 1 | 1 |
| 2_1131620! | NA | NA | 2 | 1 | NA | NA | NA | NA | 2 | 1 | 1 | 1 | NA | NA | NA | NA | NA | NA | 1 | 1 |
| 2_1131621: | NA | NA | 2 | 0.5 | NA | NA | NA | NA | 2 | 0.5 | 1 | 1 | NA | NA | NA | NA | NA | NA | 1 | 1 |
| 2_1131621! | NA | NA | 2 | 1 | NA | NA | NA | NA | 2 | 1 | 1 | 1 | NA | NA | NA | NA | NA | NA | 1 | 1 |
| 2_1131623· | NA | NA | 3 | 1 | NA | NA | NA | NA | 3 | 1 | 1 | 1 | NA | NA | NA | NA | NA | NA | 1 | 1 |
| 2_1131624! | NA | NA | 3 | 1 | NA | NA | NA | NA | 3 | 1 | 1 | 1 | NA | NA | NA | NA | NA | NA | 1 | 1 |
| 2_1131627' | NA | NA | 3 | 0.6667 | NA | NA | NA | NA | 3 | 0.6667 | 1 | 1 | NA | NA | NA | NA | NA | NA | 1 | 1 |
| 2_1131627: | NA | NA | 4 | 1 | NA | NA | NA | NA | 4 | 1 | 1 | 1 | NA | NA | NA | NA | NA | NA | 1 | 1 |
| 2_1131627! | NA | NA | 4 | 0.25 | NA | NA | NA | NA | 4 | 0.25 | 1 | 1 | NA | NA | NA | NA | NA | NA | 1 | 1 |
| 2_1136072( | 26 | 0.1923 | 22 | 0.4091 | 33 | 0.0909 | 31 | 0.1935 | 112 | 0.2054 | 21 | 0.0952 | 38 | 0.2368 | 32 | 0.0625 | 27 | 0.2593 | 118 | 0.1695 |
| 2_1136416: | 1 | 0 | 2 | 0 | 1 | 0 | 3 | 0.3333 | 7 | 0.1429 | 1 | 1 | NA | NA | 2 | 0 | 1 | 0 | 4 | 0.25 |
| 2_1136577! | 26 | 0.0769 | 18 | 0.0556 | 34 | 0.0294 | 26 | 0.0769 | 104 | 0.0577 | 19 | 0.1053 | 38 | 0 | 26 | 0.0385 | 27 | 0.1481 | 110 | 0.0636 |
| 2_1138406! | NA | NA | 1 | 0 | 1 | 1 | 1 | 1 | 3 | 0.6667 | 1 | 1 | NA | NA | NA | NA | NA | NA | 1 | 1 |
| 2_1138412( | NA | NA | 1 | 1 | NA | NA | NA | NA | 1 | 1 | 1 | 1 | NA | NA | NA | NA | NA | NA | 1 | 1 |

| | | | | | | | | | | | | | | | | | | | | |
|---|---|---|---|---|---|---|---|---|---|---|---|---|---|---|---|---|---|---|---|---|
| 2_11384135 | NA | NA | 1 | 1 | NA | NA | NA | NA | 1 | 1 | 1 | 1 | NA | NA | NA | NA | NA | NA | 1 | 1 |
| 2_11384152 | NA | NA | 1 | 1 | NA | NA | NA | NA | 1 | 1 | 1 | 1 | NA | NA | NA | NA | NA | NA | 1 | 1 |
| 2_11393188 | 4 | 0.5 | 7 | 0.2857 | 6 | 0 | 8 | 0.375 | 25 | 0.28 | 6 | 0.1667 | 3 | 1 | 4 | 0.75 | 9 | 0.1111 | 22 | 0.3636 |
| 2_11393189 | 6 | 0.5 | 7 | 0.7143 | 6 | 0.8333 | 8 | 0.625 | 27 | 0.6667 | 7 | 0.8571 | 2 | 1 | 7 | 0.5714 | 7 | 0.8571 | 23 | 0.7826 |
| 2_11393192 | 6 | 0 | 5 | 0.2 | 7 | 0 | 9 | 0.1111 | 27 | 0.0741 | 7 | 0.4286 | 3 | 0 | 7 | 0.4286 | 8 | 0.375 | 25 | 0.36 |
| 2_11393508 | 1 | 0 | NA | NA | 1 | 0 | 1 | 1 | 3 | 0.3333 | NA | NA | 1 | 1 | NA | NA | 3 | 0 | 4 | 0.25 |
| 2_11396120 | 2 | 0 | 3 | 0 | 3 | 0.3333 | 4 | 0.25 | 12 | 0.1667 | 4 | 0.25 | 1 | 0 | 4 | 0 | 3 | 0 | 12 | 0.0833 |
| 2_11402447 | 2 | 0.5 | 4 | 0 | 1 | 0 | 1 | 0 | 8 | 0.125 | 2 | 0 | 5 | 0 | 1 | 0 | 6 | 0.1667 | 14 | 0.0714 |
| 2_11402452 | 2 | 0.5 | 4 | 0.5 | 1 | 0 | 2 | 0 | 9 | 0.3333 | 2 | 0 | 5 | 0 | 1 | 0 | 3 | 0.3333 | 11 | 0.0909 |
| 2_11402577 | NA | NA | 3 | 0.3333 | 1 | 0 | 3 | 0 | 7 | 0.1429 | 1 | 0 | 4 | 0.5 | NA | NA | 5 | 0 | 10 | 0.2 |
| 2_11402580 | 1 | 0 | 4 | 0.25 | 1 | 0 | 3 | 0 | 9 | 0.1111 | 1 | 0 | 5 | 0.2 | NA | NA | 5 | 0 | 11 | 0.0909 |
| 2_11402588 | 1 | 0 | 4 | 0.25 | 1 | 0 | 3 | 0 | 9 | 0.1111 | 1 | 0 | 5 | 0.2 | NA | NA | 3 | 0 | 9 | 0.1111 |
| 2_11402596 | 1 | 0 | 3 | 0.3333 | 1 | 0 | 2 | 0 | 7 | 0.1429 | 1 | 0 | 5 | 0.6 | NA | NA | 5 | 0.2 | 11 | 0.3636 |
| 2_11402604 | 1 | 0 | 3 | 0.3333 | 1 | 0 | 2 | 0 | 7 | 0.1429 | 1 | 0 | 4 | 0.25 | NA | NA | 4 | 0 | 9 | 0.1111 |
| 2_11402609 | 1 | 0 | 3 | 0.3333 | 1 | 0 | 2 | 0 | 7 | 0.1429 | 1 | 0 | 4 | 0.5 | NA | NA | 5 | 0.4 | 10 | 0.4 |
| 2_11402612 | 1 | 0 | 3 | 0.3333 | 1 | 0 | 2 | 0 | 7 | 0.1429 | 1 | 0 | 2 | 0.5 | NA | NA | 4 | 0 | 7 | 0.1429 |
| 2_11402642 | 1 | 1 | 1 | 1 | 1 | 1 | 2 | 1 | 5 | 1 | 1 | 1 | 4 | 1 | NA | NA | 3 | 1 | 8 | 1 |
| 2_11402647 | 1 | 1 | 3 | 0.3333 | NA | NA | 2 | 0 | 6 | 0.3333 | 1 | 0 | 2 | 0 | NA | NA | 3 | 0.3333 | 6 | 0.1667 |
| 2_11402653 | 1 | 1 | 3 | 0.6667 | 1 | 0 | 3 | 0 | 8 | 0.375 | 1 | 1 | 4 | 0.25 | NA | NA | 4 | 0.75 | 9 | 0.5556 |
| 2_11402654 | NA | NA | 2 | 1 | 1 | 1 | 1 | 1 | 4 | 1 | 1 | 0 | 3 | 0.6667 | NA | NA | 4 | 0.75 | 8 | 0.625 |
| 2_11402662 | 1 | 1 | 1 | 1 | 1 | 1 | 2 | 1 | 5 | 1 | 1 | 1 | 4 | 0.75 | NA | NA | 3 | 1 | 8 | 0.875 |
| 2_11402668 | 1 | 0 | 2 | 0.5 | NA | NA | 2 | 0 | 5 | 0.2 | 1 | 1 | 3 | 0.6667 | NA | NA | 2 | 0.5 | 6 | 0.6667 |
| 2_11402671 | 1 | 1 | 2 | 1 | 1 | 1 | 2 | 1 | 6 | 1 | 1 | 1 | 4 | 0.75 | NA | NA | 4 | 1 | 9 | 0.8889 |
| 2_11402684 | 1 | 0 | 1 | 0 | 1 | 0 | 2 | 0.5 | 5 | 0.2 | 1 | 0 | 4 | 0.25 | NA | NA | 2 | 0 | 7 | 0.1429 |
| 2_11402686 | 1 | 1 | 2 | 0 | 1 | 1 | 3 | 0.3333 | 7 | 0.4286 | 1 | 1 | 3 | 0.6667 | NA | NA | 3 | 0.3333 | 7 | 0.5714 |
| 2_11402688 | 1 | 0 | 2 | 0 | 1 | 0 | 3 | 0.3333 | 7 | 0.1429 | 1 | 1 | 3 | 0 | NA | NA | 2 | 0 | 6 | 0.1667 |
| 2_11402690 | 1 | 0 | 2 | 0 | 1 | 1 | 2 | 0.5 | 6 | 0.3333 | 1 | 0 | 2 | 0.5 | NA | NA | 2 | 0 | 5 | 0.2 |
| 2_11402691 | 1 | 0 | 1 | 0 | 1 | 0 | 2 | 0.5 | 5 | 0.2 | 1 | 0 | 3 | 0 | NA | NA | 3 | 1 | 7 | 0.4286 |
| 2_11402692 | NA | NA | 2 | 0.5 | 1 | 0 | 2 | 0.5 | 5 | 0.4 | 1 | 0 | 4 | 0 | NA | NA | 4 | 0.25 | 9 | 0.1111 |
| 2_11402697 | 1 | 0 | 2 | 0.5 | 1 | 0 | 3 | 0.3333 | 7 | 0.2857 | 1 | 1 | 4 | 0.75 | NA | NA | 3 | 1 | 8 | 0.875 |
| 2_11402705 | 1 | 1 | 2 | 1 | 1 | 1 | 2 | 1 | 6 | 1 | 1 | 1 | 4 | 0 | NA | NA | 4 | 0.25 | 9 | 0.2222 |
| 2_11402708 | 1 | 1 | 2 | 0.5 | 1 | 1 | 1 | 1 | 5 | 0.8 | 1 | 0 | 4 | 1 | NA | NA | 4 | 0.75 | 9 | 0.7778 |
| 2_11402712 | 1 | 1 | 2 | 1 | 1 | 1 | 1 | 1 | 5 | 1 | 1 | 0 | 4 | 1 | NA | NA | 4 | 1 | 9 | 0.8889 |
| 2_11402732 | 1 | 1 | 1 | 1 | 1 | 1 | 2 | 1 | 5 | 1 | 1 | 1 | 4 | 1 | NA | NA | 1 | 1 | 6 | 1 |
| 2_11402741 | 1 | 1 | 2 | 0.5 | 1 | 1 | 2 | 1 | 6 | 0.8333 | 1 | 1 | 4 | 0.5 | NA | NA | 3 | 0.3333 | 8 | 0.5 |
| 2_11402786 | 1 | 1 | 2 | 0 | 1 | 0 | 1 | 0 | 5 | 0.2 | NA | NA | 1 | 0 | NA | NA | 3 | 0.3333 | 4 | 0.25 |
| 2_11402786 | NA | NA | 2 | 1 | NA | NA | 1 | 1 | 3 | 1 | 1 | 1 | NA | NA | NA | NA | 1 | 1 | 2 | 1 |
| 2_11434860 | 3 | 0 | 2 | 0 | 2 | 0 | 3 | 0.3333 | 10 | 0.1 | NA | NA | 2 | 0 | 4 | 0 | 5 | 0.2 | 11 | 0.0909 |
| 2_11434863 | 4 | 0.25 | 2 | 0 | 2 | 0 | 3 | 0 | 11 | 0.0909 | NA | NA | 1 | 0 | 5 | 0 | 4 | 0.25 | 10 | 0.1 |
| 2_11560759 | NA | NA | NA | NA | 1 | 0 | 8 | 1 | 9 | 0.8889 | NA | NA | 3 | 0.6667 | 1 | 1 | 1 | 1 | 5 | 0.8 |
| 2_11560760 | NA | NA | NA | NA | 1 | 1 | 6 | 0.5 | 7 | 0.5714 | NA | NA | 3 | 0.6667 | 1 | 1 | NA | NA | 4 | 0.75 |
| 2_11560769 | NA | NA | NA | NA | 1 | 1 | 6 | 0.5 | 7 | 0.5714 | NA | NA | 3 | 0.6667 | 1 | 1 | 1 | 0 | 5 | 0.6 |
| 2_11560830 | NA | NA | NA | NA | 1 | 0 | 6 | 0.3333 | 7 | 0.2857 | NA | NA | 3 | 0.6667 | 1 | 0 | 2 | 0.5 | 6 | 0.5 |
| 2_11569338 | 1 | 0 | 1 | 1 | 1 | 1 | 3 | 0 | 6 | 0.3333 | 2 | 0 | 6 | 0.1667 | 2 | 1 | 2 | 0 | 12 | 0.25 |
| 2_11569362 | 41 | 0.0488 | 38 | 0.0789 | 35 | 0.0857 | 53 | 0.1509 | 167 | 0.0958 | 28 | 0.1786 | 50 | 0.04 | 47 | 0.0851 | 32 | 0.2188 | 157 | 0.1146 |
| 2_11569389 | 33 | 0.2424 | 34 | 0.3824 | 24 | 0.3333 | 45 | 0.2889 | 136 | 0.3088 | 21 | 0.1905 | 43 | 0.3023 | 34 | 0.1471 | 29 | 0.2414 | 127 | 0.2283 |
| 2_11569399 | 36 | 0.5 | 36 | 0.25 | 25 | 0.36 | 43 | 0.3953 | 140 | 0.3786 | 22 | 0.4091 | 42 | 0.4762 | 39 | 0.3846 | 28 | 0.4643 | 131 | 0.4351 |
| 2_11569403 | 41 | 0.2683 | 32 | 0.1562 | 25 | 0.12 | 48 | 0.2083 | 146 | 0.1986 | 25 | 0.2 | 42 | 0.3333 | 41 | 0.122 | 32 | 0.1875 | 140 | 0.2143 |
| 2_11569407 | 30 | 0.7667 | 33 | 0.7576 | 20 | 0.95 | 43 | 0.6977 | 126 | 0.7698 | 23 | 0.7391 | 43 | 0.7442 | 35 | 0.7143 | 30 | 0.9 | 131 | 0.771 |
| 2_11569422 | 37 | 0.7568 | 33 | 0.8182 | 23 | 0.8696 | 49 | 0.8776 | 142 | 0.831 | 22 | 0.8182 | 43 | 0.8605 | 39 | 0.7436 | 28 | 0.9286 | 132 | 0.8333 |
| 2_11569428 | 34 | 0.8824 | 32 | 0.9062 | 25 | 0.92 | 40 | 0.925 | 131 | 0.9084 | 25 | 0.92 | 40 | 0.95 | 35 | 0.9429 | 29 | 0.9655 | 129 | 0.9457 |
| 2_11569432 | 30 | 0.6333 | 28 | 0.7857 | 18 | 0.8889 | 33 | 0.7879 | 109 | 0.7615 | 18 | 0.7222 | 30 | 0.8667 | 29 | 0.8621 | 22 | 0.7727 | 99 | 0.8182 |
| 2_11569443 | 32 | 0.0625 | 30 | 0.1 | 23 | 0.1739 | 40 | 0.075 | 125 | 0.096 | 21 | 0.0476 | 40 | 0.075 | 31 | 0.0968 | 27 | 0.1111 | 119 | 0.084 |
| 2_11569444 | 35 | 0.8 | 32 | 0.9375 | 22 | 1 | 43 | 0.814 | 132 | 0.8712 | 18 | 0.7778 | 37 | 0.8649 | 35 | 0.7143 | 28 | 0.9286 | 118 | 0.822 |
| 2_11569448 | 32 | 0.4688 | 29 | 0.5172 | 25 | 0.48 | 38 | 0.3158 | 124 | 0.4355 | 23 | 0.3478 | 38 | 0.4474 | 32 | 0.5 | 26 | 0.3846 | 119 | 0.4286 |
| 2_11569455 | 35 | 0.0857 | 31 | 0.129 | 26 | 0.2308 | 41 | 0.0488 | 133 | 0.1128 | 23 | 0.2174 | 39 | 0 | 31 | 0.0968 | 26 | 0.1538 | 119 | 0.1008 |
| 2_11569456 | 33 | 0.1212 | 33 | 0.1212 | 25 | 0.04 | 42 | 0.0476 | 133 | 0.0827 | 21 | 0.0476 | 38 | 0.0526 | 33 | 0.0606 | 23 | 0.0435 | 115 | 0.0522 |
| 2_11569463 | 34 | 0.2059 | 29 | 0.069 | 22 | 0.1364 | 41 | 0.0488 | 126 | 0.1111 | 21 | 0.0952 | 39 | 0.0769 | 29 | 0.1724 | 23 | 0.087 | 112 | 0.1071 |
| 2_11569464 | 27 | 0.6667 | 30 | 0.6333 | 20 | 0.7 | 32 | 0.6875 | 109 | 0.6697 | 16 | 0.75 | 34 | 0.7647 | 26 | 0.8462 | 25 | 0.8 | 101 | 0.7921 |
| 2_11569465 | 26 | 0.7308 | 27 | 0.8519 | 21 | 0.7619 | 36 | 0.7222 | 110 | 0.7636 | 18 | 0.8333 | 33 | 0.7273 | 25 | 0.76 | 17 | 0.9412 | 93 | 0.7957 |
| 2_11569466 | 23 | 0.1739 | 21 | 0.1429 | 19 | 0.2105 | 37 | 0.0811 | 100 | 0.14 | 18 | 0.1111 | 30 | 0.1667 | 26 | 0.1154 | 22 | 0.1818 | 96 | 0.1458 |
| 2_11569469 | 32 | 0.2188 | 29 | 0.1724 | 21 | 0.0476 | 39 | 0.1538 | 121 | 0.157 | 20 | 0.1 | 37 | 0.1351 | 29 | 0.1034 | 25 | 0.28 | 111 | 0.1532 |
| 2_11569472 | 31 | 0.9677 | 29 | 0.9655 | 21 | 1 | 41 | 1 | 122 | 0.9836 | 17 | 1 | 38 | 0.9737 | 29 | 0.9655 | 23 | 0.9565 | 107 | 0.972 |
| 2_11569473 | 29 | 0.3448 | 27 | 0.5185 | 22 | 0.5455 | 40 | 0.425 | 118 | 0.4492 | 18 | 0.4444 | 37 | 0.4054 | 28 | 0.5 | 22 | 0.5909 | 105 | 0.4762 |
| 2_11569479 | 28 | 0.1429 | 31 | 0.2258 | 23 | 0.1739 | 40 | 0.225 | 122 | 0.1967 | 21 | 0.1429 | 37 | 0.0811 | 30 | 0.1667 | 25 | 0.28 | 113 | 0.1593 |

| | | | | | | | | | | | | | | | | | | | | |
|---|---|---|---|---|---|---|---|---|---|---|---|---|---|---|---|---|---|---|---|---|
| 2_1156948 | 26 | 0.2308 | 30 | 0.2667 | 20 | 0.35 | 40 | 0.225 | 116 | 0.2586 | 18 | 0.4444 | 35 | 0.1429 | 26 | 0.1538 | 22 | 0.3182 | 101 | 0.2376 |
| 2_1158953 | 2 | 0 | 5 | 0.2 | 4 | 0.5 | 5 | 0 | 16 | 0.1875 | 4 | 0.5 | 3 | 0 | 7 | 0.2857 | 1 | 0 | 15 | 0.2667 |
| 2_1158953 | 2 | 0.5 | 2 | 1 | 4 | 0.5 | 4 | 0.75 | 12 | 0.6667 | 2 | 1 | 2 | 0 | 5 | 0.8 | 1 | 1 | 10 | 0.7 |
| 2_1158954 | 1 | 0 | 3 | 0.3333 | 2 | 0.5 | 2 | 0 | 8 | 0.25 | 2 | 0 | 3 | 0 | 4 | 0.25 | 1 | 0 | 10 | 0.1 |
| 2_1159055 | 1 | 1 | NA | NA | NA | NA | NA | NA | 1 | 1 | 1 | 1 | 2 | 0.5 | 2 | 0 | 1 | 0 | 6 | 0.3333 |
| 2_1167419 | NA | NA | 1 | 1 | NA | NA | 1 | 1 | 2 | 1 | NA | NA | NA | NA | 1 | 1 | NA | NA | 1 | 1 |
| 2_1167449 | NA | NA | NA | NA | NA | NA | 1 | 1 | 1 | 1 | NA | NA | NA | NA | 1 | 1 | NA | NA | 1 | 1 |
| 2_1167450 | NA | NA | NA | NA | NA | NA | 1 | 1 | 1 | 1 | NA | NA | NA | NA | 1 | 1 | NA | NA | 1 | 1 |
| 2_1170078 | 6 | 0 | 16 | 0.1875 | 11 | 0.3636 | 10 | 0.4 | 43 | 0.2558 | 1 | 0 | 14 | 0.3571 | 5 | 0 | 8 | 0.25 | 28 | 0.25 |
| 2_1170079 | 7 | 0.2857 | 13 | 0.5385 | 10 | 0.5 | 11 | 0.5455 | 41 | 0.4878 | 2 | 0 | 16 | 0.4375 | 6 | 0.3333 | 6 | 0.5 | 30 | 0.4 |
| 2_1170079 | 6 | 0.5 | 13 | 0.2308 | 11 | 0.5455 | 12 | 0.5 | 42 | 0.4286 | 2 | 0 | 16 | 0.4375 | 6 | 0.3333 | 8 | 0.375 | 32 | 0.375 |
| 2_1170079 | 5 | 0.4 | 12 | 0.5833 | 9 | 0.5556 | 6 | 0.6667 | 32 | 0.5625 | 1 | 0 | 9 | 0.5556 | 3 | 0.6667 | 7 | 0.4286 | 20 | 0.5 |
| 2_1170079 | 6 | 0.8333 | 12 | 0.6667 | 10 | 0.6 | 11 | 0.7273 | 39 | 0.6923 | 1 | 1 | 11 | 0.6364 | 6 | 0.6667 | 10 | 0.7 | 28 | 0.6786 |
| 2_1170079 | 6 | 0.5 | 12 | 0.6667 | 11 | 0.5455 | 10 | 0.3 | 39 | 0.5128 | 2 | 0.5 | 15 | 0.4 | 7 | 0.5714 | 10 | 0.7 | 34 | 0.5294 |
| 2_1170080 | 6 | 1 | 16 | 1 | 11 | 0.9091 | 12 | 1 | 45 | 0.9778 | 1 | 1 | 15 | 0.9333 | 8 | 1 | 9 | 1 | 33 | 0.9697 |
| 2_1170080 | 7 | 0 | 16 | 0.1875 | 11 | 0.0909 | 12 | 0.4167 | 46 | 0.1957 | 2 | 0 | 18 | 0.1111 | 8 | 0.375 | 10 | 0.2 | 38 | 0.1842 |
| 2_1170082 | 8 | 0.75 | 10 | 0.3 | 11 | 0.8182 | 12 | 0.5833 | 41 | 0.6098 | 1 | 1 | 17 | 0.7647 | 6 | 0.6667 | 10 | 0.8 | 34 | 0.7647 |
| 2_1170082 | 7 | 0.2857 | 14 | 0.2857 | 12 | 0.1667 | 14 | 0.5 | 47 | 0.3191 | 3 | 0.3333 | 18 | 0.1111 | 8 | 0.5 | 12 | 0.4167 | 41 | 0.2927 |
| 2_1170082 | 9 | 0 | 14 | 0 | 12 | 0.0833 | 13 | 0.1538 | 48 | 0.0625 | 4 | 0 | 18 | 0.0556 | 7 | 0.2857 | 11 | 0.0909 | 40 | 0.1 |
| 2_1170085 | 8 | 0.125 | 15 | 0.1333 | 11 | 0.0909 | 14 | 0.1429 | 48 | 0.125 | 5 | 0 | 18 | 0.1667 | 10 | 0.1 | 10 | 0.1 | 43 | 0.1163 |
| 2_1170086 | 8 | 0.25 | 15 | 0.2667 | 11 | 0.4545 | 14 | 0.2143 | 48 | 0.2917 | 4 | 0 | 19 | 0.1579 | 9 | 0.4444 | 9 | 0.1111 | 41 | 0.1951 |
| 2_1170087 | 9 | 0.7778 | 16 | 0.75 | 11 | 0.4545 | 15 | 0.4667 | 51 | 0.6078 | 5 | 0.2 | 16 | 0.625 | 10 | 0.7 | 13 | 0.5385 | 44 | 0.5682 |
| 2_1170087 | 9 | 0.2222 | 17 | 0.1765 | 10 | 0 | 16 | 0.125 | 52 | 0.1346 | 5 | 0.4 | 19 | 0.1053 | 9 | 0.2222 | 14 | 0 | 47 | 0.1277 |
| 2_1170088 | 8 | 0.125 | 15 | 0.1333 | 11 | 0.1818 | 16 | 0.125 | 50 | 0.14 | 3 | 0.3333 | 17 | 0.2353 | 8 | 0.25 | 12 | 0.25 | 40 | 0.25 |
| 2_1170103 | 15 | 0.0667 | 19 | 0.1579 | 12 | 0 | 25 | 0.12 | 71 | 0.0986 | 10 | 0 | 23 | 0.1739 | 16 | 0 | 17 | 0 | 66 | 0.0606 |
| 2_1170108 | 13 | 0.1538 | 12 | 0.25 | 9 | 0.1111 | 23 | 0.2609 | 57 | 0.2105 | 8 | 0.25 | 17 | 0.1176 | 14 | 0.1429 | 8 | 0.125 | 47 | 0.1489 |
| 2_1170114 | 12 | 0 | 12 | 0.0833 | 11 | 0.1818 | 19 | 0 | 54 | 0.0556 | 10 | 0.2 | 20 | 0.15 | 17 | 0 | 15 | 0.1333 | 62 | 0.1129 |
| 2_1171106 | 63 | 0.0635 | 59 | 0.0678 | 51 | 0.098 | 55 | 0.0364 | 228 | 0.0658 | 29 | 0.1034 | 63 | 0.0794 | 78 | 0.0256 | 58 | 0.1034 | 228 | 0.0702 |
| 2_1171106 | 46 | 0.0217 | 40 | 0.125 | 35 | 0.0571 | 39 | 0 | 160 | 0.05 | 24 | 0.0417 | 47 | 0.0638 | 50 | 0.04 | 37 | 0.0541 | 158 | 0.0506 |
| 2_1171106 | 68 | 0.7941 | 56 | 0.8393 | 51 | 0.7255 | 53 | 0.6792 | 228 | 0.7632 | 26 | 0.8846 | 63 | 0.8095 | 76 | 0.8421 | 51 | 0.8824 | 216 | 0.8472 |
| 2_1171112 | 50 | 0.26 | 43 | 0.1163 | 41 | 0.0488 | 43 | 0.1628 | 177 | 0.1525 | 20 | 0.1 | 52 | 0.1923 | 56 | 0.2143 | 47 | 0.2128 | 175 | 0.1943 |
| 2_1171113 | 51 | 0.7255 | 46 | 0.6957 | 44 | 0.4545 | 44 | 0.6591 | 185 | 0.6378 | 22 | 0.6364 | 51 | 0.6667 | 59 | 0.5932 | 49 | 0.6939 | 181 | 0.6464 |
| 2_1171113 | 47 | 0.617 | 41 | 0.6098 | 46 | 0.3261 | 42 | 0.4524 | 176 | 0.5 | 20 | 0.6 | 49 | 0.4286 | 52 | 0.5192 | 49 | 0.6735 | 170 | 0.5471 |
| 2_1171114 | 45 | 0.0222 | 36 | 0.0278 | 39 | 0.0769 | 37 | 0.0811 | 157 | 0.051 | 19 | 0.0526 | 49 | 0.0408 | 49 | 0.0612 | 39 | 0.0769 | 156 | 0.0577 |
| 2_1171115 | 44 | 0.0455 | 43 | 0.1628 | 41 | 0.0244 | 43 | 0.093 | 171 | 0.0819 | 21 | 0.0952 | 54 | 0.1111 | 57 | 0.1228 | 40 | 0.175 | 172 | 0.1279 |
| 2_1171116 | 45 | 0.0444 | 46 | 0.087 | 42 | 0.0476 | 40 | 0.025 | 173 | 0.052 | 21 | 0.0952 | 49 | 0.0408 | 55 | 0.0727 | 47 | 0.0638 | 172 | 0.064 |
| 2_1171126 | 33 | 0.1212 | 32 | 0.125 | 33 | 0 | 32 | 0.0625 | 130 | 0.0769 | 17 | 0 | 39 | 0.0513 | 39 | 0.0256 | 39 | 0.1282 | 134 | 0.0597 |
| 2_1171129 | 24 | 0.5 | 22 | 0.3636 | 27 | 0.3704 | 17 | 0.4706 | 90 | 0.4222 | 15 | 0.7333 | 30 | 0.5333 | 28 | 0.5714 | 27 | 0.4444 | 100 | 0.55 |
| 2_1171129 | 25 | 0.28 | 25 | 0.48 | 28 | 0.4643 | 22 | 0.3182 | 100 | 0.39 | 13 | 0.4615 | 25 | 0.28 | 33 | 0.4242 | 29 | 0.5517 | 100 | 0.43 |
| 2_1171129 | 25 | 0.04 | 23 | 0.087 | 27 | 0.0741 | 19 | 0.0526 | 94 | 0.0638 | 12 | 0 | 28 | 0.0714 | 28 | 0.1071 | 29 | 0.1379 | 97 | 0.0928 |
| 2_1171130 | 31 | 0.6452 | 31 | 0.3548 | 32 | 0.25 | 31 | 0.3226 | 125 | 0.392 | 19 | 0.4737 | 36 | 0.5278 | 38 | 0.3421 | 32 | 0.375 | 125 | 0.424 |
| 2_1171130 | 30 | 0.8333 | 31 | 0.6129 | 27 | 0.5926 | 31 | 0.6129 | 119 | 0.6639 | 18 | 0.7778 | 38 | 0.6053 | 34 | 0.6176 | 33 | 0.697 | 123 | 0.6585 |
| 2_1171153 | 30 | 0.1333 | 29 | 0.2069 | 26 | 0.0769 | 26 | 0.1538 | 111 | 0.1441 | 16 | 0.0625 | 35 | 0.1429 | 32 | 0.125 | 34 | 0.0882 | 117 | 0.1111 |
| 2_1171155 | 27 | 0.8519 | 29 | 0.7586 | 26 | 0.6154 | 25 | 0.64 | 107 | 0.7196 | 13 | 0.9231 | 35 | 0.9143 | 29 | 0.7931 | 28 | 0.8929 | 105 | 0.8762 |
| 2_1171156 | 21 | 0.1429 | 22 | 0.0455 | 23 | 0.087 | 23 | 0.1304 | 89 | 0.1011 | 14 | 0 | 28 | 0.1786 | 23 | 0.0435 | 21 | 0.0476 | 86 | 0.0814 |
| 2_1181079 | 1 | 0 | 1 | 0 | 2 | 0.5 | 2 | 0 | 6 | 0.1667 | 3 | 0.6667 | 3 | 0 | 2 | 0.5 | 1 | 0 | 9 | 0.3333 |
| 2_1181079 | NA | NA | 1 | 0 | 2 | 1 | 2 | 1 | 5 | 0.8 | 3 | 0.6667 | 3 | 0.6667 | 1 | 1 | 1 | 0 | 8 | 0.625 |
| 2_1181099 | NA | NA | NA | NA | 1 | 1 | NA | NA | 1 | 1 | NA | NA | 1 | 1 | NA | NA | NA | NA | 1 | 1 |
| 2_1182525 | 1 | 1 | NA | NA | NA | NA | NA | NA | 1 | 1 | NA | NA | NA | NA | 2 | 1 | NA | NA | 2 | 1 |
| 2_1182527 | 1 | 1 | NA | NA | NA | NA | NA | NA | 1 | 1 | NA | NA | NA | NA | 2 | 0.5 | NA | NA | 2 | 0.5 |
| 2_1183065 | 25 | 0.64 | 26 | 0.2308 | 43 | 0.3023 | 27 | 0.5185 | 121 | 0.405 | 21 | 0.4286 | 23 | 0.4348 | 22 | 0.5 | 23 | 0.7391 | 89 | 0.5281 |
| 2_1183066 | 23 | 0.1739 | 20 | 0.2 | 37 | 0.2703 | 29 | 0.2414 | 109 | 0.2294 | 22 | 0.1818 | 25 | 0.04 | 28 | 0.1429 | 19 | 0.1053 | 94 | 0.117 |
| 2_1184942 | 3 | 0.3333 | 3 | 0.3333 | 3 | 0 | 2 | 0.5 | 11 | 0.2727 | 4 | 0.25 | 3 | 0 | NA | NA | 1 | 0 | 8 | 0.125 |
| 2_1184942 | 5 | 0 | 4 | 0.25 | 5 | 0 | 3 | 0.3333 | 17 | 0.1176 | 5 | 0.6 | 2 | 0.5 | 1 | 1 | 1 | 0 | 9 | 0.5556 |
| 2_1184943 | 5 | 0.2 | 5 | 0.2 | 5 | 0 | 4 | 0 | 19 | 0.1053 | 6 | 0 | 3 | 0 | 1 | 1 | 1 | 0 | 11 | 0.0909 |
| 2_1185204 | 4 | 0 | 5 | 0 | 6 | 0.1667 | 6 | 0.1667 | 21 | 0.0952 | 7 | 0 | 3 | 0.3333 | 1 | 0 | 2 | 0.5 | 13 | 0.1538 |
| 2_1185237 | 5 | 0.6 | 5 | 0.6 | 7 | 0.4286 | 5 | 0.2 | 22 | 0.4545 | 5 | 0.2 | 4 | 0.5 | 1 | 1 | 4 | 0.25 | 14 | 0.3571 |
| 2_1185240 | 4 | 0.5 | 3 | 0 | 5 | 0.2 | 4 | 0.25 | 16 | 0.25 | 4 | 0.25 | 2 | 0 | 2 | 0.5 | 3 | 0.3333 | 11 | 0.2727 |
| 2_1185240 | 8 | 0.25 | 6 | 0 | 8 | 0.25 | 7 | 0 | 29 | 0.1379 | 8 | 0.125 | 5 | 0 | 3 | 0 | 3 | 0.3333 | 19 | 0.1053 |
| 2_1185240 | 6 | 0 | 7 | 0.1429 | 8 | 0.125 | 6 | 0.1667 | 27 | 0.1111 | 7 | 0 | 6 | 0.1667 | 2 | 0 | 3 | 0.6667 | 18 | 0.1667 |
| 2_1185241 | 7 | 0.2857 | 8 | 0.125 | 7 | 0.2857 | 7 | 0 | 29 | 0.1724 | 8 | 0.125 | 4 | 0.25 | 2 | 0 | 3 | 0 | 17 | 0.1176 |
| 2_1185271 | 6 | 1 | 7 | 0.8571 | 10 | 1 | 5 | 1 | 28 | 0.9643 | 7 | 0.8571 | 7 | 1 | 2 | 1 | 7 | 1 | 23 | 0.9565 |
| 2_1185271 | 7 | 0.8571 | 7 | 1 | 9 | 0.7778 | 6 | 0.6667 | 29 | 0.8276 | 6 | 0.6667 | 7 | 0.8571 | 4 | 1 | 7 | 1 | 24 | 0.875 |
| 2_1185272 | 9 | 0.7778 | 9 | 0.8889 | 8 | 0.75 | 6 | 0.5 | 32 | 0.75 | 10 | 0.6 | 9 | 0.7778 | 4 | 0.75 | 8 | 0.75 | 31 | 0.7097 |
| 2_1185272 | 8 | 0.125 | 7 | 0.4286 | 11 | 0.0909 | 6 | 0 | 32 | 0.1562 | 9 | 0.2222 | 9 | 0.2222 | 3 | 0.3333 | 5 | 0 | 26 | 0.1923 |

| | | | | | | | | | | | | | | | | | | | | |
|---|---|---|---|---|---|---|---|---|---|---|---|---|---|---|---|---|---|---|---|---|
| 2_11852727 | 9 | 0.1111 | 9 | 0.1111 | 11 | 0.0909 | 8 | 0.25 | 37 | 0.1351 | 11 | 0.0909 | 11 | 0.0909 | 4 | 0.25 | 7 | 0.1429 | 33 | 0.1212 |
| 2_11852741 | 9 | 0.6667 | 9 | 0.6667 | 11 | 0.1818 | 9 | 0.7778 | 38 | 0.5526 | 11 | 0.6364 | 8 | 0.75 | 3 | 0.3333 | 7 | 0.1429 | 29 | 0.5172 |
| 2_11852745 | 9 | 1 | 9 | 0.8889 | 10 | 1 | 9 | 1 | 37 | 0.973 | 12 | 0.9167 | 10 | 1 | 4 | 1 | 8 | 1 | 34 | 0.9706 |
| 2_11852755 | 9 | 0.1111 | 8 | 0 | 10 | 0 | 9 | 0.1111 | 36 | 0.0556 | 12 | 0.1667 | 8 | 0 | 4 | 0 | 7 | 0.1429 | 31 | 0.0968 |
| 2_11852762 | 9 | 0.8889 | 9 | 0.8889 | 9 | 0.8889 | 9 | 0.7778 | 36 | 0.8611 | 12 | 0.6667 | 10 | 0.9 | 4 | 0.75 | 7 | 1 | 33 | 0.8182 |
| 2_11852791 | 6 | 0.6667 | 8 | 0.125 | 7 | 0 | 9 | 0.1111 | 30 | 0.2 | 8 | 0.375 | 12 | 0.1667 | 1 | 1 | 6 | 0 | 27 | 0.2222 |
| 2_11852811 | 9 | 0 | 7 | 0 | 10 | 0.1 | 10 | 0.2 | 36 | 0.0833 | 12 | 0 | 13 | 0.1538 | 1 | 0 | 8 | 0 | 34 | 0.0588 |
| 2_11852829 | 5 | 0.2 | 6 | 0.1667 | 8 | 0.25 | 9 | 0 | 28 | 0.1429 | 8 | 0.25 | 8 | 0 | 1 | 0 | 6 | 0.1667 | 23 | 0.1304 |
| 2_11852963 | 10 | 0.9 | 5 | 1 | 15 | 0.6 | 14 | 0.8571 | 44 | 0.7955 | 15 | 0.8 | 15 | 0.8 | 16 | 0.5 | 14 | 0.7857 | 60 | 0.7167 |
| 2_11852978 | 17 | 0.8235 | 8 | 0.875 | 12 | 0.8333 | 14 | 1 | 51 | 0.8824 | 17 | 0.8824 | 19 | 0.8421 | 18 | 0.8889 | 14 | 0.7143 | 68 | 0.8382 |
| 2_11852984 | 16 | 0.875 | 5 | 1 | 12 | 1 | 14 | 0.9286 | 47 | 0.9362 | 14 | 0.7857 | 16 | 0.875 | 17 | 0.8824 | 13 | 0.6923 | 60 | 0.8167 |
| 2_11852987 | 14 | 0 | 7 | 0.1429 | 15 | 0 | 14 | 0.1429 | 50 | 0.06 | 11 | 0.1818 | 18 | 0.2222 | 19 | 0.1579 | 14 | 0.2857 | 62 | 0.2097 |
| 2_11853017 | 13 | 0.3846 | 5 | 0.2 | 10 | 0.3 | 13 | 0.3077 | 41 | 0.3171 | 12 | 0.5 | 16 | 0.5625 | 17 | 0.3529 | 9 | 0.5556 | 54 | 0.4815 |
| 2_11853025 | 17 | 0.5882 | 6 | 0.6667 | 13 | 0.6923 | 15 | 0.6667 | 51 | 0.6471 | 12 | 0.75 | 16 | 0.625 | 18 | 0.7778 | 13 | 0.7692 | 59 | 0.7288 |
| 2_11853030 | 14 | 0.1429 | 3 | 0.3333 | 11 | 0.5455 | 6 | 1 | 34 | 0.4412 | 7 | 0.4286 | 12 | 0.4167 | 12 | 0.5 | 9 | 0.4444 | 40 | 0.45 |
| 2_11853037 | 17 | 0.1176 | 6 | 0 | 10 | 0.1 | 14 | 0.0714 | 47 | 0.0851 | 11 | 0.0909 | 17 | 0.2353 | 18 | 0.1111 | 10 | 0.2 | 56 | 0.1607 |
| 2_11853038 | 14 | 0.2143 | 5 | 0.4 | 10 | 0.4 | 13 | 0.2308 | 42 | 0.2857 | 8 | 0.125 | 17 | 0.4118 | 15 | 0.3333 | 11 | 0.0909 | 51 | 0.2745 |
| 2_11853047 | 14 | 0.0714 | 5 | 0 | 8 | 0.375 | 13 | 0.2308 | 40 | 0.175 | 9 | 0.3333 | 13 | 0.1538 | 15 | 0.2667 | 12 | 0.1667 | 49 | 0.2245 |
| 2_11853056 | 14 | 0.2143 | 5 | 0 | 11 | 0.0909 | 15 | 0.2667 | 45 | 0.1778 | 10 | 0.2 | 16 | 0.125 | 13 | 0.0769 | 10 | 0.3 | 49 | 0.1633 |
| 2_11853062 | 11 | 0.3636 | 6 | 0.5 | 7 | 0.1429 | 8 | 0.25 | 32 | 0.3125 | 6 | 0.1667 | 9 | 0.4444 | 12 | 0.4167 | 9 | 0.2222 | 36 | 0.3333 |
| 2_11853076 | 14 | 0.5 | 6 | 0.6667 | 10 | 0.4 | 14 | 0.7143 | 44 | 0.5682 | 9 | 0.6667 | 13 | 0.7692 | 16 | 0.5 | 10 | 0.5 | 48 | 0.6042 |
| 2_11853082 | 12 | 0.8333 | 4 | 1 | 10 | 0.7 | 11 | 0.9091 | 37 | 0.8378 | 6 | 1 | 15 | 0.7333 | 16 | 0.5625 | 9 | 0.8889 | 46 | 0.7391 |
| 2_11853094 | 10 | 0.5 | 4 | 0.25 | 10 | 0.2 | 12 | 0.5 | 36 | 0.3889 | 12 | 0.4167 | 14 | 0.2857 | 16 | 0.1875 | 9 | 0.3333 | 51 | 0.2941 |
| 2_11853097 | 13 | 0.2308 | 4 | 0.5 | 8 | 0 | 13 | 0.1538 | 38 | 0.1842 | 8 | 0.25 | 12 | 0.3333 | 14 | 0.1429 | 10 | 0.1 | 44 | 0.2045 |
| 2_11853156 | 4 | 0.75 | 2 | 0 | 7 | 0.4286 | 8 | 0.75 | 21 | 0.5714 | 5 | 0.8 | 10 | 0.6 | 5 | 0.2 | 5 | 1 | 25 | 0.64 |
| 2_11853161 | 8 | 0.625 | 2 | 1 | 8 | 0.375 | 9 | 0.1111 | 27 | 0.4074 | 5 | 0.4 | 9 | 0.3333 | 9 | 0.2222 | 9 | 0.1111 | 32 | 0.25 |
| 2_11853172 | 7 | 0.1429 | 3 | 0 | 5 | 0.4 | 9 | 0.1111 | 24 | 0.1667 | 10 | 0.1 | 11 | 0.0909 | 7 | 0 | 8 | 0.125 | 36 | 0.0833 |
| 2_11853184 | 7 | 0.2857 | 2 | 0 | 5 | 0.2 | 9 | 0.4444 | 23 | 0.3043 | 9 | 0.2222 | 11 | 0.1818 | 7 | 0 | 9 | 0.1111 | 36 | 0.1389 |
| 2_11853198 | 5 | 0.2 | 1 | 0 | 6 | 0.5 | 6 | 0 | 18 | 0.2222 | 5 | 0.2 | 9 | 0.1111 | 8 | 0.125 | 6 | 0 | 28 | 0.1071 |
| 2_11853204 | 5 | 0.6 | 2 | 1 | 5 | 0.8 | 9 | 0.5556 | 21 | 0.6667 | 9 | 0.6667 | 10 | 0.3 | 6 | 0.6667 | 7 | 0.4286 | 32 | 0.5 |
| 2_11853214 | 6 | 0.5 | 3 | 0.3333 | 6 | 0.3333 | 8 | 0.25 | 23 | 0.3478 | 10 | 0.5 | 10 | 0.2 | 6 | 0.5 | 6 | 0.3333 | 32 | 0.375 |
| 2_11853219 | 6 | 0.6667 | 3 | 0.6667 | 5 | 0.4 | 8 | 0.25 | 22 | 0.4545 | 8 | 0.125 | 8 | 0.25 | 7 | 0.1429 | 6 | 0.1667 | 29 | 0.1724 |
| 2_11853222 | 6 | 0 | 3 | 0.3333 | 4 | 0 | 9 | 0.1111 | 22 | 0.0909 | 7 | 0 | 10 | 0.1 | 6 | 0 | 5 | 0.2 | 28 | 0.0714 |
| 2_11853249 | 4 | 0.25 | 2 | 0 | 4 | 0 | 7 | 0.1429 | 17 | 0.1176 | 7 | 0.2857 | 7 | 0 | 6 | 0 | 3 | 0.3333 | 23 | 0.1304 |
| 2_11853304 | 3 | 0.6667 | 1 | 0 | NA | NA | 6 | 0.3333 | 10 | 0.4 | 6 | 0.1667 | 2 | 0 | 6 | 0 | 3 | 0.3333 | 17 | 0.1176 |
| 2_11853308 | 4 | 0.5 | 2 | 1 | NA | NA | 6 | 0.5 | 12 | 0.5833 | 4 | 0 | 4 | 0.25 | 5 | 0.4 | 4 | 0.25 | 17 | 0.2353 |
| 2_11853309 | 3 | 0 | 2 | 0 | NA | NA | 6 | 0.1667 | 11 | 0.0909 | 7 | 0.2857 | 5 | 0 | 6 | 0 | 4 | 0 | 22 | 0.0909 |
| 2_11853314 | 3 | 0.3333 | 2 | 1 | NA | NA | 5 | 0.6 | 10 | 0.6 | 6 | 0.5 | 5 | 0.2 | 6 | 0.6667 | 2 | 1 | 19 | 0.5263 |
| 2_11853341 | 2 | 1 | 1 | 1 | 1 | 0 | 6 | 1 | 10 | 0.9 | 7 | 0.8571 | 4 | 1 | 5 | 1 | 4 | 1 | 20 | 0.95 |
| 2_11853357 | 3 | 1 | 1 | 0 | NA | NA | 6 | 1 | 10 | 0.9 | 7 | 0.8571 | 4 | 1 | 5 | 0.8 | 3 | 1 | 19 | 0.8947 |
| 2_11853368 | 3 | 0.6667 | 1 | 0 | NA | NA | 6 | 0.1667 | 10 | 0.3 | 7 | 0.1429 | 5 | 0.2 | 4 | 0.5 | 4 | 0 | 20 | 0.2 |
| 2_11853390 | 1 | 1 | 1 | 1 | NA | NA | 2 | 1 | 4 | 1 | 4 | 0.75 | NA | NA | 1 | 0 | 3 | 0 | 8 | 0.375 |
| 2_11853394 | 3 | 0.6667 | 1 | 1 | NA | NA | 6 | 1 | 10 | 0.9 | 5 | 1 | 2 | 0.5 | 2 | 1 | 3 | 1 | 12 | 0.9167 |
| 2_11853405 | 2 | 1 | NA | NA | NA | NA | 7 | 0.7143 | 9 | 0.7778 | 5 | 0.8 | 1 | 1 | 2 | 0.5 | 2 | 1 | 10 | 0.8 |
| 2_11853406 | 2 | 0.5 | NA | NA | NA | NA | 6 | 0 | 8 | 0.125 | 6 | 0 | 2 | 0 | 3 | 0.3333 | 2 | 0.5 | 13 | 0.1538 |
| 2_11853423 | 2 | 0.5 | NA | NA | NA | NA | 4 | 0.25 | 6 | 0.3333 | 2 | 0 | 2 | 0 | 3 | 0.3333 | 1 | 0 | 8 | 0.125 |
| 2_11853604 | NA | NA | 1 | 1 | NA | NA | 1 | 1 | 2 | 1 | 1 | 1 | NA | NA | NA | NA | 1 | 1 | 2 | 1 |
| 2_11853646 | NA | NA | 1 | 1 | NA | NA | NA | NA | 1 | 1 | 1 | 0 | NA | NA | NA | NA | 1 | 1 | 2 | 0.5 |
| 2_11853917 | 1 | 1 | 1 | 0 | NA | NA | NA | NA | 2 | 0.5 | NA | NA | 2 | 0.5 | 1 | 1 | NA | NA | 3 | 0.6667 |
| 2_11853926 | NA | NA | 1 | 1 | NA | NA | NA | NA | 1 | 1 | NA | NA | 2 | 0.5 | 1 | 1 | NA | NA | 3 | 0.6667 |
| 2_11853936 | 1 | 1 | 1 | 1 | NA | NA | NA | NA | 2 | 1 | NA | NA | 2 | 1 | 1 | 1 | NA | NA | 3 | 1 |
| 2_11857665 | 4 | 0 | 1 | 0 | 7 | 0 | 3 | 0.3333 | 15 | 0.0667 | 1 | 0 | 1 | 0 | 4 | 0 | 6 | 0.1667 | 12 | 0.0833 |
| 2_11857679 | 3 | 1 | 1 | 1 | 7 | 0.7143 | 3 | 0.6667 | 14 | 0.7857 | 1 | 1 | 1 | 0 | 3 | 0.3333 | 6 | 0.6667 | 11 | 0.5455 |
| 2_11857685 | 3 | 0.6667 | 2 | 1 | 6 | 1 | 4 | 1 | 15 | 0.9333 | 1 | 1 | NA | NA | 3 | 0.6667 | 7 | 0.8571 | 11 | 0.8182 |
| 2_11857711 | 4 | 1 | 4 | 1 | 7 | 1 | 4 | 1 | 19 | 1 | 3 | 1 | 1 | 1 | 4 | 1 | 5 | 1 | 13 | 1 |
| 2_11857723 | 4 | 0.5 | 4 | 0.5 | 9 | 0.6667 | 6 | 0.3333 | 23 | 0.5217 | 4 | 0.25 | 1 | 0 | 3 | 0.6667 | 6 | 0.6667 | 14 | 0.5 |
| 2_11857725 | 3 | 1 | 4 | 1 | 8 | 0.875 | 6 | 1 | 21 | 0.9524 | 4 | 1 | 1 | 1 | 3 | 1 | 8 | 1 | 16 | 1 |
| 2_11857731 | 1 | 0 | 1 | 1 | 5 | 0 | 3 | 0.6667 | 10 | 0.3 | 3 | 0 | 1 | 0 | 5 | 0.2 | 4 | 0.25 | 13 | 0.1538 |
| 2_11857741 | 4 | 0 | 4 | 0.25 | 7 | 0.1429 | 4 | 0.5 | 19 | 0.2105 | 3 | 0 | 2 | 0.5 | 6 | 0.3333 | 7 | 0.1429 | 18 | 0.2222 |
| 2_11857786 | 5 | 0 | 4 | 0.25 | 8 | 0.25 | 8 | 0.5 | 25 | 0.28 | 3 | 0.3333 | 2 | 0.5 | 7 | 0 | 7 | 0.4286 | 19 | 0.2632 |
| 2_11857880 | 10 | 0.2 | 4 | 0.25 | 10 | 0.5 | 11 | 0.2727 | 35 | 0.3143 | 6 | 0.3333 | 4 | 0.25 | 7 | 0.4286 | 10 | 0.5 | 27 | 0.4074 |
| 2_11857883 | 10 | 0.5 | 6 | 0.1667 | 10 | 0.2 | 10 | 0.3 | 36 | 0.3056 | 6 | 0.1667 | 3 | 0.6667 | 10 | 0.4 | 11 | 0.7273 | 30 | 0.5 |
| 2_11857885 | 8 | 0.25 | 4 | 0.25 | 11 | 0.2727 | 11 | 0.2727 | 34 | 0.2647 | 7 | 0.2857 | 4 | 0.25 | 11 | 0.3636 | 10 | 0.3 | 32 | 0.3125 |
| 2_11857899 | 7 | 1 | 2 | 0.5 | 11 | 0.9091 | 8 | 0.375 | 28 | 0.75 | 4 | 0.75 | 4 | 1 | 10 | 0.6 | 11 | 0.8182 | 29 | 0.7586 |
| 2_11857904 | 11 | 1 | 6 | 1 | 11 | 1 | 12 | 1 | 40 | 1 | 3 | 0.6667 | 4 | 1 | 11 | 0.9091 | 12 | 1 | 30 | 0.9333 |

| | | | | | | | | | | | | | | | | | | | | |
|---|---|---|---|---|---|---|---|---|---|---|---|---|---|---|---|---|---|---|---|---|
| 2_11857911 | 10 | 0.3 | 5 | 0.2 | 12 | 0.1667 | 12 | 0.25 | 39 | 0.2308 | 6 | 0.1667 | 5 | 0.4 | 13 | 0.2308 | 12 | 0.5833 | 36 | 0.3611 |
| 2_11857914 | 12 | 0.9167 | 7 | 0.8571 | 13 | 0.8462 | 11 | 0.8182 | 43 | 0.8605 | 6 | 1 | 6 | 1 | 13 | 0.9231 | 12 | 0.8333 | 37 | 0.9189 |
| 2_11857917 | 12 | 0.9167 | 7 | 0.7143 | 11 | 0.9091 | 12 | 0.5833 | 42 | 0.7857 | 5 | 0.6 | 6 | 0.8333 | 13 | 0.7692 | 13 | 0.9231 | 37 | 0.8108 |
| 2_11857928 | 7 | 0.1429 | 7 | 0.1429 | 9 | 0 | 10 | 0.2 | 33 | 0.1212 | 5 | 0.2 | 5 | 0.2 | 13 | 0.3077 | 11 | 0.0909 | 34 | 0.2059 |
| 2_11858484 | NA | NA | 1 | 1 | 2 | 0.5 | NA | NA | 3 | 0.6667 | 1 | 0 | NA | NA | NA | NA | 3 | 0.3333 | 4 | 0.25 |
| 2_11858546 | NA | NA | 1 | 0 | 2 | 0.5 | NA | NA | 3 | 0.3333 | 1 | 0 | NA | NA | NA | NA | 1 | 1 | 2 | 0.5 |
| 2_11858707 | 1 | 0 | NA | NA | 1 | 1 | NA | NA | 2 | 0.5 | NA | NA | NA | NA | NA | NA | 1 | 1 | 1 | 1 |
| 2_11862039 | 2 | 1 | 3 | 1 | 1 | 1 | 2 | 1 | 8 | 1 | 2 | 1 | 1 | 1 | 4 | 1 | 1 | 1 | 8 | 1 |
| 2_11862040 | 2 | 0.5 | 2 | 0 | 1 | 0 | 2 | 0.5 | 7 | 0.2857 | 3 | 0.6667 | 1 | 0 | 4 | 0.25 | NA | NA | 8 | 0.375 |
| 2_11862046 | 1 | 1 | 2 | 0.5 | 2 | 0.5 | 2 | 0 | 7 | 0.4286 | NA | NA | 1 | 0 | 4 | 0.25 | NA | NA | 5 | 0.2 |
| 2_11862055 | 1 | 1 | 1 | 1 | 1 | 1 | 3 | 1 | 6 | 1 | 2 | 1 | 1 | 1 | 2 | 0.5 | 1 | 1 | 6 | 0.8333 |
| 2_11862057 | 2 | 0.5 | 2 | 0 | 2 | 1 | 2 | 0.5 | 8 | 0.5 | 3 | 0.6667 | 1 | 1 | 1 | 0 | 1 | 0 | 6 | 0.5 |
| 2_11862059 | 3 | 0 | 2 | 0 | 2 | 0.5 | 2 | 0 | 9 | 0.1111 | 3 | 0.3333 | 1 | 0 | 3 | 0.3333 | 1 | 0 | 8 | 0.25 |
| 2_11862147 | 4 | 1 | 6 | 1 | 2 | 1 | 3 | 0.6667 | 15 | 0.9333 | 4 | 1 | 1 | 1 | 6 | 1 | 5 | 1 | 16 | 1 |
| 2_11862149 | 4 | 0.25 | 4 | 0.5 | 2 | 0.5 | 3 | 0.3333 | 13 | 0.3846 | 2 | 0 | NA | NA | 5 | 0.2 | 5 | 0 | 12 | 0.0833 |
| 2_11862153 | 4 | 0.25 | 5 | 0 | 2 | 0.5 | 2 | 0.5 | 13 | 0.2308 | 3 | 0.3333 | 1 | 0 | 7 | 0.1429 | 5 | 0.2 | 16 | 0.1875 |
| 2_11862171 | 4 | 1 | 5 | 0.8 | 2 | 1 | 3 | 1 | 14 | 0.9286 | 5 | 1 | 1 | 1 | 5 | 1 | 5 | 1 | 16 | 1 |
| 2_11862180 | 5 | 0.8 | 6 | 0.6667 | 1 | 0 | 2 | 1 | 14 | 0.7143 | 5 | 0.4 | 2 | 1 | 6 | 0.5 | 5 | 0.2 | 18 | 0.4444 |
| 2_11862222 | 4 | 1 | 6 | 1 | 3 | 1 | 4 | 0.75 | 17 | 0.9412 | 5 | 1 | 4 | 1 | 6 | 0.8333 | 6 | 1 | 21 | 0.9524 |
| 2_11862223 | 3 | 0.3333 | 3 | 0.3333 | 2 | 0 | 2 | 0.5 | 10 | 0.3 | 4 | 0 | 5 | 0.2 | 5 | 0.2 | 3 | 0 | 17 | 0.1176 |
| 2_11862227 | 2 | 0.5 | 7 | 0.8571 | 3 | 0.3333 | 3 | 0.3333 | 15 | 0.6 | 4 | 0.75 | 4 | 0.75 | 3 | 0 | 4 | 0.25 | 15 | 0.4667 |
| 2_11862249 | 2 | 1 | 3 | 1 | 2 | 1 | 1 | 0 | 8 | 0.875 | 6 | 1 | 4 | 1 | 5 | 1 | 6 | 1 | 21 | 1 |
| 2_11862254 | 2 | 0 | 9 | 0.4444 | 3 | 0.3333 | 5 | 0.2 | 19 | 0.3158 | 6 | 0.5 | 6 | 0.6667 | 8 | 0.5 | 8 | 0.5 | 28 | 0.5357 |
| 2_11862259 | 5 | 0.2 | 8 | 0.375 | 3 | 0 | 5 | 0.2 | 21 | 0.2381 | 6 | 0.1667 | 7 | 0.4286 | 8 | 0.375 | 7 | 0.2857 | 28 | 0.3214 |
| 2_11862277 | 5 | 0.2 | 11 | 0.0909 | 3 | 0 | 9 | 0 | 28 | 0.0714 | 7 | 0.4286 | 7 | 0.1429 | 7 | 0.1429 | 8 | 0 | 29 | 0.1724 |
| 2_11862286 | 6 | 0.8333 | 13 | 0.5385 | 2 | 1 | 8 | 0.875 | 29 | 0.7241 | 5 | 1 | 7 | 0.7143 | 10 | 0.8 | 10 | 0.9 | 32 | 0.8438 |
| 2_11862287 | 7 | 0 | 12 | 0.4167 | 3 | 0.6667 | 7 | 0.2857 | 29 | 0.3103 | 7 | 0.2857 | 6 | 0.3333 | 8 | 0.5 | 8 | 0.125 | 29 | 0.3103 |
| 2_11862298 | 6 | 1 | 13 | 1 | 3 | 1 | 10 | 1 | 32 | 1 | 7 | 1 | 7 | 0.7143 | 8 | 1 | 9 | 1 | 31 | 0.9355 |
| 2_11862307 | 8 | 0.125 | 14 | 0.1429 | 3 | 0.3333 | 9 | 0.1111 | 34 | 0.1471 | 8 | 0.125 | 8 | 0.25 | 8 | 0.25 | 10 | 0.1 | 34 | 0.1765 |
| 2_11862340 | 6 | 0.3333 | 20 | 0.3 | 3 | 0.6667 | 12 | 0.4167 | 41 | 0.3659 | 9 | 0 | 8 | 0.375 | 7 | 0.5714 | 11 | 0.4545 | 35 | 0.3429 |
| 2_11862355 | 9 | 1 | 19 | 1 | 4 | 1 | 11 | 1 | 43 | 1 | 10 | 1 | 7 | 0.8571 | 8 | 1 | 10 | 1 | 35 | 0.9714 |
| 2_11862358 | 8 | 0.75 | 20 | 0.7 | 4 | 0 | 11 | 0.7273 | 43 | 0.6512 | 9 | 0.4444 | 9 | 0.8889 | 9 | 0.7778 | 11 | 0.5455 | 38 | 0.6579 |
| 2_11862365 | 8 | 0 | 18 | 0 | 2 | 0 | 11 | 0.1818 | 39 | 0.0513 | 12 | 0.0833 | 10 | 0.1 | 9 | 0.1111 | 11 | 0 | 42 | 0.0714 |
| 2_11862372 | 9 | 0.5556 | 16 | 0.3125 | 4 | 0.5 | 10 | 0.6 | 39 | 0.4615 | 11 | 0.6364 | 9 | 0.4444 | 10 | 0.6 | 12 | 0.5833 | 42 | 0.5714 |
| 2_11862372 | 10 | 0 | 16 | 0.125 | 4 | 0.5 | 12 | 0.1667 | 42 | 0.1429 | 9 | 0.1111 | 10 | 0.1 | 8 | 0 | 10 | 0 | 37 | 0.0541 |
| 2_11862373 | 9 | 0.3333 | 17 | 0.1765 | 3 | 0.3333 | 8 | 0.375 | 37 | 0.2703 | 11 | 0.3636 | 10 | 0.2 | 8 | 0.5 | 12 | 0.3333 | 41 | 0.3415 |
| 2_11862378 | 6 | 1 | 18 | 1 | 4 | 1 | 6 | 0.8333 | 34 | 0.9706 | 11 | 1 | 8 | 1 | 6 | 1 | 11 | 1 | 36 | 1 |
| 2_11862382 | 8 | 0.375 | 17 | 0.2353 | 3 | 0.6667 | 11 | 0.0909 | 39 | 0.2564 | 10 | 0.6 | 7 | 0.1429 | 8 | 0 | 6 | 0.3333 | 31 | 0.2903 |
| 2_11862390 | 10 | 0.2 | 16 | 0.1875 | 6 | 0.1667 | 13 | 0.1538 | 45 | 0.1778 | 9 | 0.5556 | 10 | 0.2 | 10 | 0.4 | 11 | 0.5455 | 40 | 0.425 |
| 2_11862399 | 10 | 0 | 19 | 0.1053 | 6 | 0 | 13 | 0.1538 | 48 | 0.0833 | 11 | 0.0909 | 9 | 0.2222 | 10 | 0.2 | 13 | 0 | 43 | 0.1163 |
| 2_11862406 | 11 | 0.9091 | 19 | 1 | 5 | 0.6 | 13 | 0.9231 | 48 | 0.9167 | 11 | 0.9091 | 11 | 1 | 12 | 1 | 12 | 1 | 46 | 0.9783 |
| 2_11862410 | 11 | 0.1818 | 19 | 0.1579 | 6 | 0.1667 | 12 | 0.4167 | 48 | 0.2292 | 10 | 0.3 | 9 | 0.1111 | 12 | 0.0833 | 13 | 0.3077 | 44 | 0.2045 |
| 2_11862439 | 10 | 0.2 | 18 | 0.0556 | 7 | 0 | 12 | 0 | 47 | 0.0638 | 10 | 0.1 | 9 | 0.2222 | 12 | 0.1667 | 14 | 0.0714 | 45 | 0.1333 |
| 2_11862452 | 11 | 0.2727 | 20 | 0 | 6 | 0.1667 | 12 | 0.1667 | 49 | 0.1224 | 14 | 0.2857 | 10 | 0.2 | 11 | 0.0909 | 14 | 0.2143 | 49 | 0.2041 |
| 2_11862469 | 14 | 0 | 24 | 0.0833 | 6 | 0 | 14 | 0.1429 | 58 | 0.069 | 15 | 0.0667 | 11 | 0 | 12 | 0.0833 | 15 | 0.0667 | 53 | 0.0566 |
| 2_11862485 | 10 | 0 | 24 | 0.0833 | 5 | 0 | 15 | 0.0667 | 54 | 0.0556 | 12 | 0 | 9 | 0.1111 | 12 | 0.0833 | 14 | 0.0714 | 47 | 0.0638 |
| 2_11862487 | 4 | 0 | 9 | 0.3333 | 3 | 0.3333 | 7 | 0 | 23 | 0.1739 | 10 | 0.1 | 5 | 0.4 | 8 | 0.125 | 6 | 0.1667 | 29 | 0.1724 |
| 2_11862505 | 14 | 0.1429 | 24 | 0.375 | 8 | 0.625 | 15 | 0.3333 | 61 | 0.3443 | 14 | 0.5 | 13 | 0.4615 | 14 | 0.5 | 19 | 0.3684 | 60 | 0.45 |
| 2_11862528 | 16 | 0.125 | 28 | 0.1786 | 8 | 0.25 | 18 | 0.1667 | 70 | 0.1714 | 20 | 0.1 | 15 | 0.0667 | 15 | 0.1333 | 21 | 0.0476 | 71 | 0.0845 |
| 2_11862543 | 13 | 0.1538 | 25 | 0.2 | 11 | 0.1818 | 15 | 0.2 | 64 | 0.1875 | 16 | 0.1875 | 16 | 0.3125 | 11 | 0.0909 | 18 | 0.1667 | 61 | 0.1967 |
| 2_11865209 | NA | NA | NA | NA | 1 | 1 | NA | NA | 1 | 1 | 1 | 1 | NA | NA | NA | NA | 1 | 1 | 2 | 1 |
| 2_11871105 | 5 | 0 | 3 | 0 | 5 | 0.2 | 10 | 0.1 | 23 | 0.087 | 3 | 0 | 12 | 0.0833 | 5 | 0.2 | 5 | 0.2 | 25 | 0.12 |
| 2_11871124 | 7 | 0.1429 | 4 | 0 | 7 | 0.4286 | 14 | 0 | 32 | 0.125 | 4 | 0 | 13 | 0.2308 | 5 | 0 | 5 | 0.2 | 27 | 0.1481 |
| 2_11871126 | 7 | 0.2857 | 4 | 0.25 | 8 | 0.625 | 15 | 0.2667 | 34 | 0.3529 | 4 | 0.25 | 9 | 0.3333 | 5 | 0.4 | 5 | 0.6 | 23 | 0.3913 |
| 2_11871150 | 6 | 0 | 4 | 0.25 | 8 | 0 | 14 | 0.1429 | 32 | 0.0938 | 6 | 0.1667 | 14 | 0 | 7 | 0.2857 | 5 | 0.4 | 32 | 0.1562 |
| 2_11871155 | 7 | 0 | 5 | 0.2 | 7 | 0.2857 | 14 | 0.1429 | 33 | 0.1515 | 6 | 0.1667 | 14 | 0.1429 | 7 | 0 | 6 | 0 | 33 | 0.0909 |
| 2_11871270 | 2 | 0 | 7 | 0.1429 | 4 | 0 | 5 | 0 | 18 | 0.0556 | 3 | 0 | 6 | 0.1667 | 1 | 0 | 2 | 1 | 12 | 0.25 |
| 2_11871312 | 1 | 0 | 9 | 0.1111 | 5 | 0 | 5 | 0.2 | 20 | 0.1 | 4 | 0 | 7 | 0.1429 | 1 | 0 | 2 | 0 | 14 | 0.0714 |
| 2_11874654 | NA | NA | 1 | 0 | NA | NA | 2 | 1 | 3 | 0.6667 | NA | NA | 2 | 0.5 | 2 | 0.5 | 1 | 0 | 5 | 0.4 |
| 2_11874692 | NA | NA | 1 | 0 | NA | NA | 2 | 0.5 | 3 | 0.3333 | NA | NA | 1 | 0 | 3 | 0.6667 | 1 | 0 | 5 | 0.4 |
| 2_11874698 | NA | NA | 1 | 1 | NA | NA | 2 | 0.5 | 3 | 0.6667 | NA | NA | 2 | 0.5 | 2 | 1 | 1 | 1 | 5 | 0.8 |
| 2_11874713 | NA | NA | NA | NA | NA | NA | 1 | 1 | 1 | 1 | NA | NA | 1 | 0 | 1 | 1 | NA | NA | 2 | 0.5 |
| 2_11874738 | NA | NA | 1 | 1 | NA | NA | 2 | 1 | 3 | 1 | NA | NA | 2 | 0.5 | 3 | 0.6667 | 1 | 1 | 6 | 0.6667 |
| 2_11875327 | 1 | 0 | 3 | 0.3333 | NA | NA | 4 | 0.25 | 8 | 0.25 | 1 | 0 | 2 | 0 | 5 | 0 | 3 | 0.3333 | 11 | 0.0909 |
| 2_11875334 | 1 | 0 | 3 | 0.3333 | NA | NA | 4 | 0 | 8 | 0.125 | 1 | 0 | 2 | 0 | 5 | 0.4 | 2 | 0 | 10 | 0.2 |

| | | | | | | | | | | | | | | | | | | | | |
|---|---|---|---|---|---|---|---|---|---|---|---|---|---|---|---|---|---|---|---|---|
| 2_1187535 | 2 | 0.5 | 3 | 0 | NA | NA | 5 | 0.4 | 10 | 0.3 | 1 | 0 | 4 | 0 | 5 | 0.2 | 1 | 0 | 11 | 0.0909 |
| 2_1187535 | 2 | 1 | 4 | 1 | NA | NA | 5 | 0.6 | 11 | 0.8182 | 1 | 0 | 2 | 0 | 4 | 1 | 3 | 1 | 10 | 0.7 |
| 2_1187538 | 1 | 0 | 4 | 0.25 | 1 | 0 | 6 | 0.3333 | 12 | 0.25 | 1 | 0 | 4 | 0 | 5 | 0.2 | 5 | 0 | 15 | 0.0667 |
| 2_1187544 | 3 | 0.6667 | 3 | 0 | 4 | 0.25 | 6 | 0.1667 | 16 | 0.25 | 2 | 0 | 4 | 0 | 5 | 0 | 6 | 0.5 | 17 | 0.1765 |
| 2_1187546 | 4 | 0.5 | 5 | 0.2 | 5 | 0.2 | 7 | 0.4286 | 21 | 0.3333 | 3 | 0 | 5 | 0.2 | 4 | 0 | 6 | 0.6667 | 18 | 0.2778 |
| 2_1188484 | 5 | 0.6 | 5 | 0.2 | 4 | 0.5 | 6 | 0.8333 | 20 | 0.55 | 2 | 0 | 2 | 0.5 | 2 | 0.5 | 2 | 1 | 8 | 0.5 |
| 2_1188485 | 5 | 0 | 5 | 0.2 | 4 | 0 | 6 | 0 | 20 | 0.05 | 1 | 0 | 2 | 0 | 2 | 0.5 | 3 | 0 | 8 | 0.125 |
| 2_1188487 | 18 | 0.2222 | 13 | 0.3077 | 12 | 0.1667 | 14 | 0.2857 | 57 | 0.2456 | 3 | 0 | 6 | 0.3333 | 7 | 0.4286 | 5 | 0.4 | 21 | 0.3333 |
| 2_1188488 | 16 | 0.4375 | 13 | 0.3077 | 12 | 0.5833 | 13 | 0.3077 | 54 | 0.4074 | 3 | 0 | 5 | 0.6 | 7 | 0.4286 | 7 | 0.2857 | 22 | 0.3636 |
| 2_1188488 | 16 | 0.3125 | 11 | 0.2727 | 11 | 0.4545 | 13 | 0.3077 | 51 | 0.3333 | 3 | 0.6667 | 6 | 0.3333 | 7 | 0.4286 | 6 | 0.1667 | 22 | 0.3636 |
| 2_1188489 | 14 | 0.1429 | 9 | 0.1111 | 12 | 0 | 11 | 0.0909 | 46 | 0.087 | 3 | 0 | 6 | 0 | 7 | 0.1429 | 3 | 0 | 19 | 0.0526 |
| 2_1188489 | 12 | 0.8333 | 9 | 1 | 9 | 1 | 10 | 1 | 40 | 0.95 | 3 | 1 | 4 | 1 | 7 | 0.8571 | 4 | 0.75 | 18 | 0.8889 |
| 2_1188490 | 14 | 0 | 12 | 0.0833 | 10 | 0.2 | 11 | 0.1818 | 47 | 0.1064 | 1 | 1 | 5 | 0 | 6 | 0 | 3 | 0 | 15 | 0.0667 |
| 2_1188491 | 14 | 0.6429 | 11 | 0.6364 | 10 | 1 | 10 | 0.7 | 45 | 0.7333 | 1 | 0 | 5 | 1 | 7 | 0.7143 | 4 | 1 | 17 | 0.8235 |
| 2_1188492 | 15 | 0.1333 | 12 | 0.25 | 9 | 0.1111 | 10 | 0.1 | 46 | 0.1522 | 2 | 0 | 5 | 0 | 6 | 0.1667 | 3 | 0 | 16 | 0.0625 |
| 2_1188492 | 14 | 0.2857 | 9 | 0.5556 | 10 | 0.2 | 7 | 0.4286 | 40 | 0.35 | 2 | 0 | 2 | 1 | 7 | 0.1429 | 4 | 0.75 | 15 | 0.4 |
| 2_1188493 | 14 | 0.0714 | 11 | 0.0909 | 12 | 0.0833 | 10 | 0 | 47 | 0.0638 | 2 | 0 | 5 | 0 | 7 | 0.1429 | 4 | 0.25 | 18 | 0.1111 |
| 2_1188493 | 9 | 0.2222 | 7 | 0 | 10 | 0 | 7 | 0 | 33 | 0.0606 | 1 | 0 | 3 | 0 | 3 | 0.3333 | 2 | 0 | 9 | 0.1111 |
| 2_1188493 | 10 | 0.7 | 11 | 0.9091 | 9 | 1 | 6 | 0.8333 | 36 | 0.8611 | 2 | 1 | 4 | 1 | 3 | 0.6667 | 3 | 0.6667 | 12 | 0.8333 |
| 2_1188493 | 14 | 1 | 12 | 1 | 11 | 0.9091 | 10 | 1 | 47 | 0.9787 | 2 | 1 | 4 | 1 | 7 | 1 | 3 | 1 | 16 | 1 |
| 2_1188494 | 12 | 0.0833 | 10 | 0.1 | 8 | 0.125 | 9 | 0 | 39 | 0.0769 | 2 | 0 | 4 | 0 | 5 | 0.4 | 4 | 0 | 15 | 0.1333 |
| 2_1188495 | 11 | 0 | 8 | 0.25 | 9 | 0.2222 | 10 | 0.4 | 38 | 0.2105 | 1 | 0 | 3 | 0.6667 | 7 | 0.1429 | 4 | 0.25 | 15 | 0.2667 |
| 2_1188525 | 13 | 0.1538 | 9 | 0.2222 | 11 | 0.1818 | 10 | 0.2 | 43 | 0.186 | 3 | 0.6667 | 2 | 0 | 7 | 0.4286 | 2 | 0 | 14 | 0.3571 |
| 2_1188577 | 6 | 0.1667 | 6 | 0.1667 | 5 | 0 | 4 | 0 | 21 | 0.0952 | 2 | 1 | 2 | 0.5 | 2 | 0 | 2 | 0 | 8 | 0.375 |
| 2_1188577 | 10 | 0.1 | 10 | 0 | 7 | 0.1429 | 8 | 0.125 | 35 | 0.0857 | 2 | 0.5 | 2 | 0 | 4 | 0 | 2 | 0 | 10 | 0.1 |
| 2_1188657 | 3 | 0.3333 | NA | NA | NA | NA | NA | NA | 3 | 0.3333 | NA | NA | NA | NA | 1 | 1 | NA | NA | 1 | 1 |
| 2_1188686 | 3 | 1 | NA | NA | NA | NA | NA | NA | 3 | 1 | NA | NA | 1 | 1 | 1 | 1 | NA | NA | 2 | 1 |
| 2_1188688 | 3 | 1 | NA | NA | NA | NA | NA | NA | 3 | 1 | NA | NA | 1 | 0 | 1 | 1 | NA | NA | 2 | 0.5 |
| 2_1188688 | 2 | 1 | NA | NA | NA | NA | NA | NA | 2 | 1 | NA | NA | 1 | 1 | 1 | 1 | NA | NA | 2 | 1 |
| 2_1188689 | 3 | 1 | NA | NA | NA | NA | NA | NA | 3 | 1 | NA | NA | 1 | 1 | 1 | 1 | NA | NA | 2 | 1 |
| 2_1188690 | 3 | 1 | 1 | 1 | NA | NA | NA | NA | 4 | 1 | NA | NA | 1 | 1 | NA | NA | NA | NA | 1 | 1 |
| 2_1188690 | 3 | 0.6667 | 1 | 0 | NA | NA | NA | NA | 4 | 0.5 | NA | NA | NA | NA | 1 | 1 | NA | NA | 1 | 1 |
| 2_1188691 | 2 | 1 | NA | NA | NA | NA | NA | NA | 2 | 1 | NA | NA | 1 | 1 | 1 | 1 | NA | NA | 2 | 1 |
| 2_1188692 | 3 | 0.6667 | 1 | 0 | NA | NA | NA | NA | 4 | 0.5 | NA | NA | 1 | 1 | 1 | 1 | NA | NA | 2 | 1 |
| 2_1188693 | 3 | 1 | 1 | 1 | NA | NA | NA | NA | 4 | 1 | NA | NA | 1 | 1 | 1 | 1 | NA | NA | 2 | 1 |
| 2_1188698 | 3 | 0.6667 | 1 | 1 | NA | NA | NA | NA | 4 | 0.75 | NA | NA | 1 | 1 | 1 | 0 | NA | NA | 2 | 0.5 |
| 2_1189676 | 5 | 0.6 | 11 | 0.3636 | 9 | 0.3333 | 5 | 0.6 | 30 | 0.4333 | 6 | 0.1667 | 3 | 0.6667 | 6 | 0.3333 | 9 | 0.4444 | 24 | 0.375 |
| 2_1189677 | 5 | 0.6 | 16 | 0.1875 | 13 | 0.3077 | 6 | 0.1667 | 40 | 0.275 | 6 | 0.3333 | 3 | 0.6667 | 6 | 0 | 10 | 0.2 | 25 | 0.24 |
| 2_1189678 | 6 | 0.3333 | 15 | 0.1333 | 13 | 0.3846 | 5 | 0 | 39 | 0.2308 | 7 | 0.1429 | 4 | 0 | 8 | 0.125 | 11 | 0.2727 | 30 | 0.1667 |
| 2_1189680 | 4 | 0 | 13 | 0 | 12 | 0.1667 | 5 | 0.2 | 34 | 0.0882 | 5 | 0.2 | 2 | 0 | 7 | 0.1429 | 9 | 0.2222 | 23 | 0.1739 |
| 2_1189779 | 1 | 0 | 5 | 0 | 5 | 0.2 | 4 | 0 | 15 | 0.0667 | 6 | 0 | 1 | 0 | 6 | 0 | 4 | 0.25 | 17 | 0.0588 |
| 2_1190472 | NA | NA | NA | NA | 3 | 0.3333 | 2 | 0 | 5 | 0.2 | NA | NA | 3 | 0 | 1 | 0 | 1 | 1 | 5 | 0.2 |
| 2_1190474 | 1 | 1 | NA | NA | 4 | 0.25 | 2 | 0.5 | 7 | 0.4286 | NA | NA | 3 | 0.3333 | 1 | 0 | 1 | 1 | 5 | 0.4 |
| 2_1190685 | 1 | 1 | 1 | 1 | NA | NA | NA | NA | 2 | 1 | 1 | 1 | NA | NA | 1 | 0 | NA | NA | 2 | 0.5 |
| 2_1190686 | 1 | 1 | 2 | 0.5 | NA | NA | NA | NA | 3 | 0.6667 | 1 | 0 | NA | NA | 1 | 1 | NA | NA | 2 | 0.5 |
| 2_1190794 | 1 | 1 | 2 | 1 | 3 | 0.6667 | 1 | 0 | 7 | 0.7143 | 1 | 0 | 3 | 0.3333 | 2 | 0.5 | 1 | 1 | 7 | 0.4286 |
| 2_1190794 | 2 | 1 | 2 | 0 | 3 | 0.6667 | 1 | 1 | 8 | 0.625 | NA | NA | 3 | 1 | 2 | 0 | 1 | 1 | 6 | 0.6667 |
| 2_1190795 | 2 | 1 | 2 | 0.5 | 3 | 0.3333 | 1 | 0 | 8 | 0.5 | NA | NA | 3 | 0.3333 | 1 | 0 | 1 | 1 | 5 | 0.4 |
| 2_1190795 | 2 | 0.5 | 2 | 1 | 3 | 0.6667 | 1 | 1 | 8 | 0.75 | 1 | 1 | 3 | 1 | 2 | 0 | 1 | 1 | 7 | 0.7143 |
| 2_1190795 | 2 | 1 | 1 | 0 | 2 | 0.5 | 1 | 1 | 6 | 0.6667 | 1 | 1 | 2 | 0.5 | 1 | 0 | 1 | 0 | 5 | 0.4 |
| 2_1190796 | 2 | 1 | 2 | 1 | 3 | 1 | 1 | 1 | 8 | 1 | 1 | 1 | 3 | 1 | 2 | 0.5 | 1 | 1 | 7 | 0.8571 |
| 2_1190797 | 2 | 1 | 2 | 0.5 | 3 | 1 | 1 | 1 | 8 | 0.875 | 1 | 1 | 3 | 0.6667 | 2 | 0.5 | 1 | 1 | 7 | 0.7143 |
| 2_1191001 | 2 | 0.5 | NA | NA | NA | NA | NA | NA | 2 | 0.5 | 1 | 1 | 1 | 0 | NA | NA | NA | NA | 2 | 0.5 |
| 2_1191003 | 2 | 1 | NA | NA | 1 | 1 | NA | NA | 3 | 1 | 1 | 0 | NA | NA | 1 | 0 | 1 | 1 | 3 | 0.3333 |
| 2_1191004 | NA | NA | NA | NA | 1 | 1 | NA | NA | 1 | 1 | 1 | 1 | 1 | 1 | 1 | 0 | 1 | 1 | 4 | 0.75 |
| 2_1191004 | 1 | 1 | NA | NA | 1 | 1 | NA | NA | 2 | 1 | 1 | 1 | 1 | 1 | 1 | 1 | 1 | 1 | 4 | 1 |
| 2_1191004 | 2 | 0.5 | NA | NA | 1 | 0 | NA | NA | 3 | 0.3333 | 1 | 0 | 1 | 0 | 1 | 0 | 1 | 1 | 4 | 0.25 |
| 2_1191004 | 2 | 1 | NA | NA | NA | NA | NA | NA | 2 | 1 | NA | NA | 1 | 0 | 1 | 1 | 1 | 1 | 3 | 0.6667 |
| 2_1191005 | 2 | 0.5 | NA | NA | NA | NA | NA | NA | 2 | 0.5 | 1 | 1 | 1 | 0 | 1 | 0 | 1 | 1 | 4 | 0.5 |
| 2_1191007 | 2 | 0.5 | NA | NA | NA | NA | NA | NA | 2 | 0.5 | 1 | 0 | 1 | 1 | NA | NA | 1 | 1 | 3 | 0.6667 |
| 2_1191193 | 4 | 0.25 | 1 | 0 | 1 | 0 | 2 | 0 | 8 | 0.125 | 1 | 0 | 3 | 0.3333 | 1 | 0 | 1 | 1 | 6 | 0.3333 |
| 2_1191193 | 1 | 0 | 1 | 0 | 1 | 0 | 2 | 0.5 | 5 | 0.2 | 1 | 1 | 2 | 0 | 1 | 1 | 1 | 0 | 5 | 0.4 |
| 2_1191234 | 4 | 0 | 1 | 0 | 1 | 0 | 2 | 0.5 | 8 | 0.125 | 1 | 0 | 2 | 0 | 1 | 1 | 1 | 0 | 5 | 0.2 |
| 2_1191235 | 3 | 1 | NA | NA | 1 | 1 | 2 | 0.5 | 6 | 0.8333 | 2 | 1 | 2 | 1 | 1 | 1 | NA | NA | 5 | 1 |
| 2_1191238 | 4 | 1 | 1 | 1 | 2 | 1 | 2 | 1 | 9 | 1 | 2 | 1 | 3 | 1 | 1 | 1 | 1 | 1 | 7 | 1 |

| | | | | | | | | | | | | | | | | | | | | |
|---|---|---|---|---|---|---|---|---|---|---|---|---|---|---|---|---|---|---|---|---|
| 2_11912396 | 4 | 1 | 1 | 1 | 3 | 1 | 2 | 1 | 10 | 1 | 2 | 1 | 2 | 1 | NA | NA | 1 | 1 | 5 | 1 |
| 2_11912416 | 4 | 0.25 | 1 | 0 | 1 | 1 | 2 | 0.5 | 8 | 0.375 | 2 | 0.5 | 3 | 0.3333 | NA | NA | 1 | 0 | 6 | 0.3333 |
| 2_11912418 | 4 | 0.75 | 1 | 1 | 3 | 1 | 2 | 1 | 10 | 0.9 | 2 | 1 | 2 | 1 | 1 | 1 | 1 | 1 | 6 | 1 |
| 2_11912428 | 4 | 1 | 1 | 0 | 3 | 0.6667 | 2 | 1 | 10 | 0.8 | 2 | 1 | 3 | 0.6667 | 1 | 1 | 1 | 0 | 7 | 0.7143 |
| 2_11912450 | 1 | 0 | NA | NA | 2 | 0.5 | 2 | 0 | 5 | 0.2 | 1 | 0 | 3 | 0.3333 | 1 | 0 | NA | NA | 5 | 0.2 |
| 2_11912460 | 4 | 0 | 1 | 0 | 3 | 0 | 2 | 0.5 | 10 | 0.1 | 2 | 0 | 3 | 0 | 2 | 0 | 2 | 0.5 | 9 | 0.1111 |
| 2_11912461 | 3 | 0.6667 | 1 | 1 | 1 | 1 | 2 | 1 | 7 | 0.8571 | NA | NA | 2 | 0.5 | 2 | 1 | 1 | 1 | 5 | 0.8 |
| 2_11912462 | 4 | 0.5 | NA | NA | 2 | 0 | 2 | 0.5 | 8 | 0.375 | 1 | 1 | 3 | 0.3333 | 2 | 1 | 1 | 1 | 7 | 0.7143 |
| 2_11912476 | 4 | 0.75 | NA | NA | 3 | 1 | 2 | 1 | 9 | 0.8889 | 2 | 1 | 3 | 1 | 3 | 1 | 2 | 1 | 10 | 1 |
| 2_11912484 | 3 | 0.3333 | 1 | 0 | 2 | 0 | 2 | 0 | 8 | 0.125 | 2 | 0 | 3 | 0 | 2 | 0.5 | 1 | 0 | 8 | 0.125 |
| 2_11912496 | 4 | 0.25 | 1 | 0 | 1 | 0 | 2 | 0 | 8 | 0.125 | 1 | 0 | 1 | 1 | 1 | 1 | 1 | 0 | 4 | 0.5 |
| 2_11912755 | 6 | 0.3333 | 2 | 0 | 4 | 0.75 | 2 | 0.5 | 14 | 0.4286 | 2 | 0 | 3 | 0 | 3 | 0 | 3 | 0.3333 | 11 | 0.0909 |
| 2_11912773 | 6 | 0.6667 | 2 | 0.5 | 4 | 0.75 | 2 | 0.5 | 14 | 0.6429 | 2 | 0 | 3 | 0.6667 | 3 | 0.3333 | 3 | 0.3333 | 11 | 0.3636 |
| 2_11912774 | 5 | 1 | 2 | 1 | 5 | 0.8 | 2 | 1 | 14 | 0.9286 | 2 | 1 | 3 | 1 | 2 | 1 | 3 | 1 | 10 | 1 |
| 2_11912780 | 3 | 1 | 2 | 1 | 4 | 1 | 2 | 1 | 11 | 1 | 2 | 1 | 2 | 1 | 3 | 1 | 2 | 1 | 9 | 1 |
| 2_11912786 | 6 | 0.5 | 2 | 0.5 | 6 | 0.5 | 2 | 0 | 16 | 0.4375 | 1 | 1 | 3 | 0 | 3 | 1 | 4 | 0 | 11 | 0.3636 |
| 2_11912786 | 6 | 0.6667 | 2 | 1 | 4 | 0.75 | 2 | 0.5 | 14 | 0.7143 | 2 | 1 | 3 | 0.6667 | 3 | 0.6667 | 3 | 0.3333 | 11 | 0.6364 |
| 2_11912812 | 5 | 0.8 | 4 | 0.75 | 4 | 0.75 | 4 | 0.5 | 17 | 0.7059 | 1 | 1 | 2 | 0.5 | 5 | 0.6 | 4 | 0.75 | 12 | 0.6667 |
| 2_11912827 | 7 | 0.1429 | 4 | 0.25 | 6 | 0.1667 | 3 | 0 | 20 | 0.15 | 2 | 0.5 | 4 | 0.25 | 5 | 0 | 3 | 0.3333 | 14 | 0.2143 |
| 2_11912838 | 5 | 1 | 5 | 1 | 5 | 1 | 4 | 1 | 19 | 1 | 2 | 1 | 5 | 0.8 | 5 | 0.8 | 4 | 1 | 16 | 0.875 |
| 2_11912843 | 7 | 0.5714 | 2 | 0.5 | 5 | 0.4 | 3 | 0 | 17 | 0.4118 | 1 | 0 | 5 | 0.4 | 5 | 0.4 | 5 | 0.4 | 16 | 0.375 |
| 2_11912848 | 7 | 0.1429 | 4 | 0 | 6 | 0 | 3 | 0.3333 | 20 | 0.1 | 2 | 0 | 4 | 0.25 | 5 | 0 | 5 | 0 | 16 | 0.0625 |
| 2_11912853 | 7 | 0.7143 | 4 | 0.5 | 6 | 0.6667 | 6 | 1 | 23 | 0.7391 | 2 | 0.5 | 5 | 0.2 | 4 | 0.5 | 5 | 0 | 16 | 0.25 |
| 2_11912858 | 6 | 1 | 4 | 1 | 4 | 1 | 4 | 1 | 18 | 1 | 3 | 1 | 7 | 0.8571 | 5 | 1 | 4 | 1 | 19 | 0.9474 |
| 2_11912863 | 4 | 1 | 7 | 1 | 3 | 1 | 4 | 1 | 18 | 1 | 1 | 1 | 6 | 1 | 3 | 1 | 5 | 1 | 15 | 1 |
| 2_11912873 | 2 | 0 | 4 | 0.25 | 3 | 0.3333 | 2 | 0 | 11 | 0.1818 | NA | NA | 4 | 0 | 3 | 0.3333 | 3 | 0 | 10 | 0.1 |
| 2_11918202 | NA | NA | 3 | 0.3333 | 1 | 0 | NA | NA | 4 | 0.25 | 3 | 0.6667 | 2 | 0 | 1 | 1 | 1 | 0 | 7 | 0.4286 |
| 2_11918384 | NA | NA | 3 | 0.3333 | 1 | 0 | NA | NA | 4 | 0.25 | 3 | 0.6667 | 1 | 0 | 1 | 0 | 2 | 0 | 7 | 0.2857 |
| 2_11918395 | NA | NA | 3 | 0.6667 | 1 | 0 | NA | NA | 4 | 0.5 | 1 | 1 | 2 | 0.5 | NA | NA | 1 | 0 | 4 | 0.5 |
| 2_11918411 | NA | NA | 4 | 1 | NA | NA | NA | NA | 4 | 1 | 3 | 1 | 2 | 1 | NA | NA | 1 | 1 | 6 | 1 |
| 2_11918414 | NA | NA | 4 | 1 | 1 | 1 | NA | NA | 5 | 1 | 2 | 1 | 1 | 1 | 1 | 1 | 2 | 1 | 6 | 1 |
| 2_11918424 | NA | NA | 4 | 0.25 | 1 | 1 | NA | NA | 5 | 0.4 | 1 | 1 | 2 | 0.5 | 1 | 0 | 2 | 0 | 6 | 0.3333 |
| 2_11918426 | NA | NA | 2 | 0.5 | NA | NA | NA | NA | 2 | 0.5 | NA | NA | NA | NA | NA | NA | 1 | 1 | 1 | 1 |
| 2_11918442 | NA | NA | 4 | 0.25 | 1 | 1 | NA | NA | 5 | 0.4 | 3 | 0.3333 | 1 | 1 | NA | NA | 2 | 1 | 6 | 0.6667 |
| 2_11918451 | NA | NA | 3 | 0.6667 | 1 | 1 | NA | NA | 4 | 0.75 | 2 | 0 | 2 | 0.5 | 1 | 0 | 2 | 0 | 7 | 0.1429 |
| 2_11918461 | NA | NA | 2 | 0.5 | 1 | 1 | NA | NA | 3 | 0.6667 | 4 | 0.5 | 1 | 1 | 2 | 0.5 | 2 | 0.5 | 9 | 0.5556 |
| 2_11918462 | NA | NA | 4 | 0.5 | NA | NA | NA | NA | 4 | 0.5 | 2 | 1 | 2 | 0 | 1 | 0 | 1 | 0 | 6 | 0.3333 |
| 2_11920399 | 1 | 1 | NA | NA | 3 | 0.3333 | 4 | 0.75 | 8 | 0.625 | 1 | 0 | 3 | 0.3333 | 1 | 0 | 2 | 0 | 7 | 0.1429 |
| 2_11920416 | 1 | 1 | NA | NA | 4 | 0.25 | 5 | 0.4 | 10 | 0.4 | 1 | 0 | 3 | 0 | 2 | 0 | 3 | 0.3333 | 9 | 0.1111 |
| 2_11920456 | NA | NA | NA | NA | 4 | 0.25 | 4 | 0.25 | 8 | 0.25 | 1 | 0 | 3 | 0.3333 | 2 | 0 | 3 | 0 | 9 | 0.1111 |
| 2_11920471 | NA | NA | NA | NA | 2 | 0.5 | 4 | 0.25 | 6 | 0.3333 | 1 | 1 | 2 | 0 | 1 | 0 | 4 | 0 | 8 | 0.125 |
| 2_11920489 | NA | NA | NA | NA | 1 | 1 | 4 | 0.25 | 5 | 0.4 | 1 | 0 | 3 | 0 | 1 | 0 | 4 | 0.5 | 9 | 0.2222 |
| 2_11920493 | NA | NA | NA | NA | 2 | 1 | 3 | 0.6667 | 5 | 0.8 | 1 | 1 | 3 | 0.6667 | 1 | 1 | 4 | 0.75 | 9 | 0.7778 |
| 2_11930855 | 1 | 1 | NA | NA | NA | NA | 1 | 1 | 2 | 1 | 1 | 0 | 1 | 0 | NA | NA | 1 | 1 | 3 | 0.3333 |
| 2_11930858 | 1 | 1 | NA | NA | NA | NA | 1 | 1 | 2 | 1 | 1 | 0 | 1 | 0 | NA | NA | 1 | 1 | 3 | 0.3333 |
| 2_11941568 | 13 | 0.3077 | 19 | 0.3158 | 12 | 0.1667 | 18 | 0.2778 | 62 | 0.2742 | 11 | 0.0909 | 17 | 0.1765 | 13 | 0.3077 | 12 | 0.4167 | 53 | 0.2453 |
| 2_11943008 | 1 | 1 | 3 | 0.3333 | 3 | 0 | 3 | 0.6667 | 10 | 0.4 | 3 | 0.3333 | 2 | 0 | 1 | 1 | 6 | 0.3333 | 12 | 0.3333 |
| 2_11943018 | 2 | 1 | 4 | 1 | 4 | 1 | 4 | 1 | 14 | 1 | 3 | 1 | 3 | 1 | 2 | 1 | 6 | 1 | 14 | 1 |
| 2_11943020 | 2 | 0 | 3 | 0.3333 | 3 | 0.6667 | 3 | 0.3333 | 11 | 0.3636 | 4 | 0.25 | 1 | 0 | 2 | 1 | 7 | 0.7143 | 14 | 0.5714 |
| 2_11943026 | 2 | 0 | 4 | 0.5 | 4 | 0 | 4 | 0 | 14 | 0.1429 | 2 | 0 | 2 | 0.5 | 2 | 0 | 6 | 0 | 12 | 0.0833 |
| 2_11943026 | 2 | 1 | 4 | 1 | 3 | 1 | 4 | 1 | 13 | 1 | 3 | 1 | 3 | 1 | 2 | 1 | 6 | 1 | 14 | 1 |
| 2_11943039 | 2 | 1 | 2 | 1 | 3 | 0.6667 | 2 | 1 | 9 | 0.8889 | 4 | 0.75 | 3 | 1 | 2 | 1 | 6 | 0.8333 | 15 | 0.8667 |
| 2_11943042 | 2 | 0 | 4 | 0.5 | 3 | 0 | 2 | 0 | 11 | 0.1818 | 3 | 0 | 4 | 0 | 2 | 0 | 7 | 0.2857 | 16 | 0.125 |
| 2_11943056 | 3 | 0.6667 | 4 | 0.5 | 4 | 0.25 | 4 | 0.25 | 15 | 0.4 | 4 | 0.5 | 2 | 0 | 1 | 0 | 6 | 0 | 13 | 0.1538 |
| 2_11943071 | 4 | 0.75 | 4 | 1 | 4 | 0.5 | 3 | 0.6667 | 15 | 0.7333 | 2 | 1 | 1 | 1 | 1 | 1 | 7 | 0.4286 | 11 | 0.6364 |
| 2_11943081 | 3 | 0 | 4 | 0.5 | 3 | 0 | 4 | 0.25 | 14 | 0.2143 | 2 | 0.5 | 4 | 0 | 1 | 1 | 6 | 0 | 13 | 0.1538 |
| 2_11943083 | 2 | 0 | 3 | 0.3333 | 5 | 0.2 | 5 | 0 | 15 | 0.1333 | 3 | 0 | 2 | 0.5 | NA | NA | 6 | 0.1667 | 11 | 0.1818 |
| 2_11943088 | 3 | 0 | 4 | 0.25 | 5 | 0.2 | 3 | 0 | 15 | 0.1333 | 2 | 0 | 3 | 0.3333 | 1 | 0 | 6 | 0 | 12 | 0.0833 |
| 2_11943097 | 2 | 0.5 | 5 | 0.2 | 4 | 0 | 4 | 0.75 | 15 | 0.3333 | 4 | 0.25 | 4 | 0.25 | 1 | 1 | 7 | 0 | 16 | 0.1875 |
| 2_11943099 | 3 | 0.3333 | 4 | 0 | 3 | 0 | 5 | 0 | 15 | 0.0667 | 4 | 0.25 | 4 | 0.25 | 1 | 1 | 8 | 0 | 17 | 0.1765 |
| 2_11943103 | 4 | 0.25 | 6 | 0.1667 | 4 | 0.25 | 5 | 0.2 | 19 | 0.2105 | 4 | 0.5 | 3 | 0 | 1 | 0 | 8 | 0 | 16 | 0.125 |
| 2_11943106 | 1 | 1 | 5 | 0 | 5 | 0.2 | 4 | 0.25 | 15 | 0.2 | 4 | 0.25 | 2 | 0 | 1 | 0 | 6 | 0.1667 | 13 | 0.1538 |
| 2_11943128 | 2 | 0 | 7 | 0.2857 | 4 | 0 | 4 | 0 | 17 | 0.1176 | 4 | 0 | 3 | 0.3333 | 1 | 0 | 7 | 0 | 15 | 0.0667 |
| 2_11943157 | 4 | 0 | 8 | 0.125 | 5 | 0 | 5 | 0.2 | 22 | 0.0909 | 3 | 0 | 4 | 0 | 1 | 0 | 8 | 0.125 | 16 | 0.0625 |
| 2_11943756 | 31 | 0.1935 | 59 | 0.1017 | 41 | 0.0732 | 26 | 0.1154 | 157 | 0.1146 | 27 | 0.2222 | 12 | 0.1667 | 24 | 0.0833 | 28 | 0.0714 | 91 | 0.1319 |

| | | | | | | | | | | | | | | | | | | | | |
|---|---|---|---|---|---|---|---|---|---|---|---|---|---|---|---|---|---|---|---|---|
| 2_1194379( | 34 | 0.0882 | 65 | 0.2 | 52 | 0.0962 | 30 | 0.2 | 181 | 0.1492 | 41 | 0.1951 | 19 | 0.2632 | 26 | 0.1538 | 35 | 0.2286 | 121 | 0.2066 |
| 2_1194861: | 7 | 0.2857 | 6 | 0.1667 | 7 | 0.4286 | 5 | 0.2 | 25 | 0.28 | 1 | 0 | 6 | 0.3333 | 3 | 0 | 2 | 0 | 12 | 0.1667 |
| 2_1194861; | 7 | 0.1429 | 5 | 0.4 | 8 | 0.375 | 4 | 0 | 24 | 0.25 | 2 | 0.5 | 6 | 0.1667 | 3 | 0.3333 | 3 | 0 | 14 | 0.2143 |
| 2_1194905: | 5 | 0.2 | 4 | 0.5 | 7 | 0.4286 | 3 | 0 | 19 | 0.3158 | 2 | 0 | 6 | 0.1667 | 2 | 0 | 2 | 0 | 12 | 0.0833 |
| 2_119490( | 7 | 0.7143 | 5 | 0.4 | 8 | 0.25 | 3 | 0.3333 | 23 | 0.4348 | 2 | 0.5 | 4 | 0.25 | 2 | 0 | 3 | 0.6667 | 11 | 0.3636 |
| 2_1194906: | 6 | 0.1667 | 5 | 0.4 | 8 | 0.25 | 3 | 0.6667 | 22 | 0.3182 | 2 | 0.5 | 4 | 0.75 | 2 | 0 | 3 | 0 | 11 | 0.3636 |
| 2_1194906‹ | 6 | 0.3333 | 5 | 0.4 | 8 | 0.5 | 4 | 0.25 | 23 | 0.3913 | 2 | 0.5 | 5 | 0.2 | 2 | 0 | 3 | 1 | 12 | 0.4167 |
| 2_1194907‹ | 5 | 0.6 | 4 | 0.75 | 7 | 0.5714 | 3 | 0.6667 | 19 | 0.6316 | 2 | 0.5 | 5 | 0.8 | 2 | 0.5 | 2 | 1 | 11 | 0.7273 |
| 2_1194908: | 6 | 0.3333 | 5 | 0 | 6 | 0.1667 | 3 | 1 | 20 | 0.3 | 2 | 0.5 | 6 | 0.1667 | 2 | 0 | 3 | 0 | 13 | 0.1538 |
| 2_1194908' | 5 | 0.6 | 5 | 0.4 | 7 | 0.2857 | 3 | 0.3333 | 20 | 0.4 | 2 | 0 | 6 | 0.5 | 2 | 0 | 3 | 0.6667 | 13 | 0.3846 |
| 2_1194908‹ | 2 | 0 | 5 | 0.4 | 6 | 0.6667 | 3 | 0.3333 | 16 | 0.4375 | 2 | 1 | 5 | 0.4 | 2 | 1 | 3 | 0.3333 | 12 | 0.5833 |
| 2_1194909: | 6 | 0.3333 | 4 | 0.25 | 7 | 0.1429 | 3 | 0.3333 | 20 | 0.25 | 2 | 0.5 | 6 | 0.1667 | 2 | 1 | 3 | 1 | 13 | 0.5385 |
| 2_1194909 | 5 | 0.4 | 4 | 0.25 | 7 | 0 | 2 | 0.5 | 18 | 0.2222 | 2 | 0 | 5 | 0.2 | NA | NA | 1 | 0 | 8 | 0.125 |
| 2_1194910: | 6 | 0.6667 | 4 | 1 | 5 | 0.4 | 3 | 1 | 18 | 0.7222 | 1 | 1 | 4 | 0.75 | 1 | 1 | 3 | 0.6667 | 9 | 0.7778 |
| 2_1195634' | 7 | 0.2857 | 8 | 0.5 | 2 | 0.5 | 8 | 0.375 | 25 | 0.4 | 1 | 1 | 4 | 0 | 3 | 0.6667 | 8 | 0.375 | 16 | 0.375 |
| 2_1195634‹ | 6 | 0.8333 | 8 | 0.875 | 4 | 0.5 | 11 | 1 | 29 | 0.8621 | 2 | 1 | 4 | 1 | 4 | 1 | 11 | 0.8182 | 21 | 0.9048 |
| 2_1195635' | 7 | 0.1429 | 8 | 0.125 | 3 | 0 | 11 | 0.0909 | 29 | 0.1034 | 2 | 0 | 5 | 0.2 | 3 | 0 | 9 | 0 | 19 | 0.0526 |
| 2_1195635: | 7 | 0.1429 | 7 | 0.1429 | 1 | 0 | 11 | 0.3636 | 26 | 0.2308 | 2 | 0.5 | 5 | 0.2 | 4 | 0.25 | 11 | 0.1818 | 22 | 0.2273 |
| 2_1195637' | 6 | 0 | 8 | 0.375 | NA | NA | 8 | 0.375 | 22 | 0.2727 | 2 | 0 | 5 | 0.2 | 3 | 0.3333 | 9 | 0.3333 | 19 | 0.2632 |
| 2_1195638( | 4 | 0.25 | 8 | 0.25 | 3 | 0 | 10 | 0.6 | 25 | 0.36 | 4 | 0.25 | 4 | 0.25 | 4 | 0.5 | 9 | 0.5556 | 21 | 0.4286 |
| 2_1195643‹ | 4 | 0 | 7 | 0.1429 | 2 | 0 | 6 | 0 | 19 | 0.0526 | 1 | 0 | 3 | 0 | 5 | 0 | 10 | 0.1 | 19 | 0.0526 |
| 2_1195649' | 7 | 0 | 4 | 0.25 | 3 | 0 | 8 | 0.125 | 22 | 0.0909 | 1 | 0 | 3 | 0 | 7 | 0.2857 | 9 | 0.2222 | 20 | 0.2 |
| 2_1195650 | 7 | 0.1429 | 4 | 0.5 | 5 | 0 | 7 | 0.1429 | 23 | 0.1739 | NA | NA | 2 | 0 | 6 | 0.3333 | 8 | 0.375 | 16 | 0.3125 |
| 2_1195651( | 7 | 0.2857 | 3 | 0.6667 | 5 | 0.4 | 8 | 0.75 | 23 | 0.5217 | 1 | 0 | 3 | 0.6667 | 7 | 0.1429 | 8 | 0.75 | 19 | 0.4737 |
| 2_1195651: | 8 | 0.375 | 3 | 0.3333 | 4 | 0.5 | 8 | 0.375 | 23 | 0.3913 | 1 | 0 | 1 | 0 | 7 | 0.4286 | 6 | 0.5 | 15 | 0.4 |
| 2_1195652( | 8 | 0.875 | 4 | 0.5 | 5 | 0.6 | 6 | 0.8333 | 23 | 0.7391 | 1 | 1 | 3 | 0.6667 | 8 | 0.625 | 10 | 0.8 | 22 | 0.7273 |
| 2_1195652: | 6 | 0.5 | 3 | 0 | 3 | 0.3333 | 7 | 0.1429 | 19 | 0.2632 | 2 | 0.5 | 2 | 0.5 | 5 | 0.4 | 9 | 0.2222 | 18 | 0.3333 |
| 2_1195655' | 4 | 0 | 2 | 0 | 2 | 1 | 4 | 0.25 | 12 | 0.25 | 2 | 0 | 3 | 0.6667 | 4 | 0.25 | 3 | 0.6667 | 12 | 0.4167 |
| 2_1195656 | 5 | 0 | 3 | 0.3333 | 3 | 0 | 6 | 0 | 17 | 0.0588 | 2 | 0 | 2 | 0 | 4 | 0.25 | 3 | 0 | 11 | 0.0909 |
| 2_1195732( | 12 | 0.0833 | 14 | 0.0714 | 13 | 0.0769 | 17 | 0.1176 | 56 | 0.0893 | 7 | 0.1429 | 26 | 0.1538 | 18 | 0.1111 | 6 | 0.1667 | 57 | 0.1404 |
| 2_1195813( | 21 | 0.7143 | 24 | 0.5 | 17 | 0.4706 | 20 | 0.35 | 82 | 0.5122 | 13 | 0.5385 | 30 | 0.6 | 22 | 0.5455 | 12 | 0.8333 | 77 | 0.6104 |
| 2_1195814: | 20 | 0.8 | 30 | 0.9333 | 25 | 0.72 | 27 | 0.7778 | 102 | 0.8137 | 17 | 0.7647 | 41 | 0.9024 | 33 | 0.7273 | 19 | 0.7895 | 110 | 0.8091 |
| 2_1195815 | 20 | 0.1 | 26 | 0.1154 | 22 | 0.0909 | 24 | 0.0833 | 92 | 0.0978 | 19 | 0.1053 | 41 | 0.1463 | 31 | 0.0645 | 17 | 0.1176 | 108 | 0.1111 |
| 2_1195816' | 20 | 0.25 | 29 | 0.1034 | 30 | 0.0333 | 29 | 0.0345 | 108 | 0.0926 | 15 | 0.1333 | 46 | 0.087 | 35 | 0.0571 | 20 | 0.05 | 116 | 0.0776 |
| 2_1195821: | 2 | 1 | 4 | 0.5 | NA | NA | NA | NA | 6 | 0.6667 | 2 | 0.5 | NA | NA | NA | NA | 1 | 0 | 3 | 0.3333 |
| 2_1195946‹ | 3 | 0.6667 | 1 | 1 | 2 | 0.5 | 2 | 0 | 8 | 0.5 | 1 | 1 | 5 | 0.4 | 3 | 0.3333 | 3 | 0.6667 | 12 | 0.5 |
| 2_1195947: | 1 | 1 | 2 | 0.5 | 2 | 0 | 1 | 1 | 6 | 0.5 | 1 | 0 | 3 | 0.6667 | 2 | 0 | 3 | 0.3333 | 9 | 0.3333 |
| 2_1195981: | 2 | 0.5 | 1 | 0 | 1 | 0 | 3 | 0 | 7 | 0.1429 | 1 | 0 | 3 | 0 | 1 | 0 | 2 | 0.5 | 7 | 0.1429 |
| 2_1195981' | 2 | 1 | 1 | 1 | 1 | 1 | 3 | 1 | 7 | 1 | 1 | 1 | 2 | 1 | 1 | 1 | 2 | 1 | 6 | 1 |
| 2_1195982 | 1 | 1 | 1 | 1 | 1 | 1 | 3 | 1 | 6 | 1 | 1 | 1 | 3 | 1 | 1 | 1 | 2 | 1 | 7 | 1 |
| 2_1195983 | 1 | 0 | 1 | 1 | 1 | 0 | 1 | 0 | 4 | 0.25 | 1 | 0 | 3 | 0.3333 | 1 | 0 | 2 | 0.5 | 7 | 0.2857 |
| 2_1195984( | 1 | 1 | 1 | 0 | 1 | 0 | 2 | 0 | 5 | 0.2 | NA | NA | 3 | 0.3333 | 1 | 0 | NA | NA | 4 | 0.25 |
| 2_1195986‹ | NA | NA | NA | NA | 1 | 1 | 2 | 0.5 | 3 | 0.6667 | NA | NA | 3 | 0.6667 | NA | NA | 2 | 1 | 5 | 0.8 |
| 2_1196433: | 9 | 0.4444 | 5 | 0.4 | 6 | 0.1667 | 15 | 0.3333 | 35 | 0.3429 | 8 | 0.25 | 5 | 0.2 | 15 | 0.7333 | 6 | 0.1667 | 34 | 0.4412 |
| 2_1196433‹ | 12 | 0.1667 | 4 | 0 | 7 | 0.2857 | 15 | 0.1333 | 38 | 0.1579 | 9 | 0.1111 | 7 | 0.1429 | 19 | 0.3158 | 8 | 0 | 43 | 0.186 |
| 2_1196471 | 14 | 0.5 | 12 | 0 | 8 | 0.25 | 19 | 0.0526 | 53 | 0.1887 | 16 | 0.0625 | 5 | 0.4 | 20 | 0 | 12 | 0.25 | 53 | 0.1132 |
| 2_1196472( | 15 | 0.2 | 6 | 0.1667 | 8 | 0.25 | 19 | 0.1579 | 48 | 0.1875 | 16 | 0.1875 | 6 | 0.3333 | 20 | 0.35 | 12 | 0.25 | 54 | 0.2778 |
| 2_1196472‹ | 16 | 0.25 | 12 | 0 | 9 | 0.1111 | 25 | 0.04 | 62 | 0.0968 | 14 | 0.0714 | 6 | 0 | 22 | 0.0455 | 13 | 0.0769 | 55 | 0.0545 |
| 2_1196473' | 17 | 0.7647 | 12 | 0.5833 | 11 | 0.7273 | 23 | 0.8261 | 63 | 0.746 | 14 | 0.8571 | 9 | 0.8889 | 20 | 0.9 | 15 | 0.8 | 58 | 0.8621 |
| 2_1196474( | 16 | 0.0625 | 15 | 0.2 | 12 | 0.25 | 24 | 0.125 | 67 | 0.1493 | 15 | 0.0667 | 8 | 0.25 | 21 | 0.2381 | 13 | 0.0769 | 57 | 0.1579 |
| 2_1196476 | 19 | 0.6316 | 17 | 0.5882 | 13 | 0.3846 | 25 | 0.64 | 74 | 0.5811 | 16 | 0.4375 | 10 | 0.6 | 22 | 0.5909 | 16 | 0.625 | 64 | 0.5625 |
| 2_1196479' | 19 | 0.1053 | 15 | 0.0667 | 15 | 0.2 | 27 | 0.1852 | 76 | 0.1447 | 17 | 0.2941 | 10 | 0.1 | 28 | 0.1071 | 15 | 0 | 70 | 0.1286 |
| 2_1196481: | 17 | 0.1765 | 14 | 0.0714 | 12 | 0.1667 | 26 | 0.1154 | 69 | 0.1304 | 13 | 0 | 10 | 0.2 | 26 | 0.2308 | 14 | 0.2143 | 63 | 0.1746 |
| 2_1196481 | 23 | 0.1304 | 16 | 0.125 | 17 | 0.0588 | 30 | 0.2333 | 86 | 0.1512 | 16 | 0.1875 | 11 | 0.2727 | 27 | 0.2222 | 18 | 0.4444 | 72 | 0.2778 |
| 2_1196483‹ | 22 | 0.5 | 20 | 0.35 | 19 | 0.3158 | 29 | 0.5172 | 90 | 0.4333 | 21 | 0.3333 | 16 | 0.4375 | 29 | 0.4828 | 14 | 0.5 | 80 | 0.4375 |
| 2_1197719( | 1 | 1 | 1 | 1 | NA | NA | NA | NA | 2 | 1 | NA | NA | 1 | 1 | NA | NA | 1 | 1 | 2 | 1 |
| 2_1197775( | 2 | 1 | 1 | 1 | NA | NA | NA | NA | 3 | 1 | NA | NA | 1 | 1 | 1 | 1 | 1 | 1 | 3 | 1 |
| 2_1197779 | 1 | 1 | 1 | 1 | NA | NA | NA | NA | 2 | 1 | NA | NA | 1 | 1 | 1 | 1 | 1 | 1 | 3 | 1 |
| 2_1197780: | 1 | 1 | NA | NA | NA | NA | NA | NA | 1 | 1 | NA | NA | 1 | 1 | 1 | 0 | 1 | 1 | 3 | 0.6667 |
| 2_1197781 | 1 | 1 | 1 | 0 | 1 | 0 | NA | NA | 3 | 0.3333 | NA | NA | 1 | 0 | 1 | 1 | 1 | 0 | 3 | 0.3333 |
| 2_1197782: | 2 | 0.5 | 1 | 0 | 1 | 0 | NA | NA | 4 | 0.25 | NA | NA | 1 | 0 | 1 | 1 | 1 | 0 | 3 | 0.3333 |
| 2_1197782‹ | 1 | 1 | 1 | 0 | NA | NA | NA | NA | 2 | 0.5 | NA | NA | 1 | 1 | 1 | 1 | NA | NA | 2 | 1 |
| 2_1197783 | 2 | 0.5 | 1 | 0 | NA | NA | NA | NA | 3 | 0.3333 | NA | NA | NA | NA | 1 | 1 | 1 | 1 | 2 | 1 |
| 2_1197784: | 2 | 0.5 | 1 | 0 | 1 | 0 | NA | NA | 4 | 0.25 | NA | NA | 1 | 0 | 1 | 0 | 1 | 1 | 3 | 0.3333 |
| 2_1197784 | 2 | 1 | 1 | 0 | 1 | 1 | NA | NA | 4 | 0.75 | NA | NA | 1 | 1 | NA | NA | NA | NA | 1 | 1 |

| | | | | | | | | | | | | | | | | | | | | |
|---|---|---|---|---|---|---|---|---|---|---|---|---|---|---|---|---|---|---|---|---|
| 2_1197784 | 2 | 1 | 1 | 1 | 1 | 0 | NA | NA | 4 | 0.75 | NA | NA | 1 | 1 | 2 | 1 | 1 | 1 | 4 | 1 |
| 2_1197786 | 3 | 0.3333 | 1 | 0 | NA | NA | NA | NA | 4 | 0.25 | NA | NA | NA | NA | 1 | 1 | 1 | 1 | 2 | 1 |
| 2_1197787 | 2 | 0.5 | 2 | 0.5 | 1 | 0 | NA | NA | 5 | 0.4 | NA | NA | 1 | 0 | 2 | 0.5 | 1 | 0 | 4 | 0.25 |
| 2_1197859 | NA | NA | NA | NA | 1 | 1 | NA | NA | 1 | 1 | NA | NA | 1 | 1 | NA | NA | NA | NA | 1 | 1 |
| 2_1197943 | 2 | 1 | 2 | 0.5 | 2 | 0.5 | 1 | 1 | 7 | 0.7143 | 1 | 1 | NA | NA | 3 | 1 | NA | NA | 4 | 1 |
| 2_1197945 | 4 | 0.5 | 2 | 0.5 | 2 | 1 | 1 | 0 | 9 | 0.5556 | 1 | 1 | 1 | 0 | 3 | 0.6667 | 2 | 1 | 7 | 0.7143 |
| 2_1197946 | 4 | 1 | 2 | 0 | 2 | 1 | 2 | 1 | 10 | 0.8 | 1 | 1 | 1 | 1 | 3 | 0.6667 | 2 | 1 | 7 | 0.8571 |
| 2_1197946 | 4 | 0.75 | 2 | 0.5 | 2 | 0.5 | 1 | 1 | 9 | 0.6667 | 1 | 1 | 1 | 1 | 2 | 1 | 1 | 1 | 5 | 1 |
| 2_1197946 | 3 | 0 | 2 | 0 | 3 | 0 | 2 | 0.5 | 10 | 0.1 | 1 | 1 | 1 | 0 | 3 | 0.3333 | 2 | 0 | 7 | 0.2857 |
| 2_1197947 | 4 | 1 | 2 | 1 | 3 | 1 | 2 | 1 | 11 | 1 | 1 | 1 | 1 | 1 | 3 | 1 | 2 | 1 | 7 | 1 |
| 2_1197947 | 3 | 1 | 2 | 0.5 | 2 | 1 | 2 | 1 | 9 | 0.8889 | 1 | 1 | NA | NA | 1 | 1 | 1 | 1 | 3 | 1 |
| 2_1197947 | 3 | 1 | 2 | 1 | 3 | 1 | 2 | 1 | 10 | 1 | 1 | 1 | NA | NA | 3 | 0.6667 | 2 | 1 | 6 | 0.8333 |
| 2_1197948 | 3 | 0.6667 | 2 | 0 | 3 | 0 | 1 | 0 | 9 | 0.2222 | 1 | 1 | 1 | 0 | 3 | 0 | 1 | 0 | 6 | 0.1667 |
| 2_1197948 | 4 | 0 | 1 | 0 | 2 | 0 | 1 | 1 | 8 | 0.125 | NA | NA | NA | NA | 2 | 1 | 1 | 0 | 3 | 0.6667 |
| 2_1197948 | 4 | 1 | 2 | 1 | 2 | 1 | 2 | 1 | 10 | 1 | 1 | 1 | 1 | 1 | 1 | 1 | 1 | 1 | 4 | 1 |
| 2_1197949 | 4 | 0.75 | 1 | 0 | 2 | 1 | 2 | 1 | 9 | 0.7778 | 1 | 1 | NA | NA | 1 | 1 | 1 | 1 | 3 | 1 |
| 2_1197950 | 3 | 0 | 2 | 1 | 2 | 0.5 | 2 | 0 | 9 | 0.3333 | 1 | 0 | NA | NA | 3 | 1 | 2 | 0.5 | 6 | 0.6667 |
| 2_1197950 | 4 | 0.5 | 2 | 0 | 2 | 0.5 | 2 | 0 | 10 | 0.3 | 1 | 1 | 1 | 0 | 3 | 0.3333 | 2 | 0.5 | 7 | 0.4286 |
| 2_1197951 | 3 | 0.6667 | 2 | 0 | 2 | 0.5 | 1 | 1 | 8 | 0.5 | 1 | 0 | 2 | 0.5 | 2 | 0.5 | 2 | 0.5 | 7 | 0.4286 |
| 2_1197953 | 4 | 1 | 3 | 1 | 3 | 1 | 2 | 1 | 12 | 1 | 1 | 1 | 2 | 1 | 4 | 1 | 2 | 1 | 9 | 1 |
| 2_1197955 | 5 | 0.4 | 3 | 0.3333 | 1 | 1 | 2 | 0.5 | 11 | 0.4545 | 1 | 1 | 2 | 0.5 | 4 | 0 | 2 | 0 | 9 | 0.2222 |
| 2_1197955 | 5 | 0.2 | 3 | 0 | 1 | 0 | 2 | 0 | 11 | 0.0909 | 1 | 0 | 2 | 0 | 4 | 0 | 2 | 0.5 | 9 | 0.1111 |
| 2_1197958 | 5 | 0.2 | 3 | 0 | 3 | 0 | 1 | 0 | 12 | 0.0833 | 1 | 0 | 2 | 0 | 4 | 0.25 | 2 | 0 | 9 | 0.1111 |
| 2_1197966 | 4 | 0 | 3 | 0 | 3 | 0.3333 | 2 | 0 | 12 | 0.0833 | 1 | 0 | 3 | 0.3333 | 5 | 0 | 2 | 0 | 11 | 0.0909 |
| 2_1197968 | 5 | 0.2 | 6 | 0 | 3 | 0.3333 | 2 | 0 | 16 | 0.125 | 1 | 0 | 3 | 0 | 4 | 0.25 | 2 | 0.5 | 10 | 0.2 |
| 2_1197975 | 6 | 0 | 5 | 0 | 4 | 0 | 4 | 0.25 | 19 | 0.0526 | 1 | 1 | 4 | 0 | 4 | 0 | 1 | 0 | 10 | 0.1 |
| 2_1198542 | NA | NA | 1 | 1 | NA | NA | 1 | 0 | 2 | 0.5 | 1 | 0 | 1 | 0 | 2 | 0 | 2 | 0.5 | 6 | 0.1667 |
| 2_1198558 | NA | NA | 1 | 1 | NA | NA | 1 | 1 | 2 | 1 | 1 | 0 | NA | NA | 3 | 1 | 1 | 1 | 5 | 0.8 |
| 2_1198559 | NA | NA | 2 | 0.5 | NA | NA | 2 | 0.5 | 4 | 0.5 | 1 | 1 | 1 | 0 | 3 | 0.3333 | 2 | 1 | 7 | 0.5714 |
| 2_1198559 | NA | NA | 1 | 0 | NA | NA | 2 | 0.5 | 3 | 0.3333 | 1 | 1 | 1 | 0 | 3 | 0.3333 | 2 | 0.5 | 7 | 0.4286 |
| 2_1198560 | NA | NA | NA | NA | NA | NA | 2 | 1 | 2 | 1 | 1 | 1 | NA | NA | 4 | 1 | 2 | 1 | 7 | 1 |
| 2_1198561 | NA | NA | 1 | 0 | NA | NA | 2 | 0.5 | 3 | 0.3333 | 1 | 1 | 2 | 0.5 | 3 | 0.6667 | 1 | 0 | 7 | 0.5714 |
| 2_1198565 | NA | NA | 1 | 0 | 2 | 0 | 2 | 0.5 | 5 | 0.2 | 2 | 0 | 3 | 0.3333 | 4 | 0 | 2 | 0 | 11 | 0.0909 |
| 2_1198566 | NA | NA | 1 | 0 | 2 | 0 | 3 | 0.3333 | 6 | 0.1667 | 2 | 0 | 3 | 0 | 4 | 0.25 | 1 | 0 | 10 | 0.1 |
| 2_1198570 | NA | NA | NA | NA | 1 | 0 | 1 | 1 | 2 | 0.5 | 3 | 0 | 5 | 0 | 5 | 0 | 2 | 0.5 | 15 | 0.0667 |
| 2_1198720 | 1 | 1 | 1 | 1 | NA | NA | NA | NA | 2 | 1 | 1 | 1 | NA | NA | 1 | 1 | 1 | 1 | 3 | 1 |
| 2_1198721 | 1 | 1 | 2 | 0 | NA | NA | NA | NA | 3 | 0.3333 | 1 | 0 | NA | NA | NA | NA | 4 | 0.25 | 5 | 0.2 |
| 2_1198721 | 1 | 0 | 2 | 0.5 | NA | NA | NA | NA | 3 | 0.3333 | 1 | 0 | NA | NA | 1 | 1 | 3 | 0 | 5 | 0.2 |
| 2_1198721 | NA | NA | 1 | 1 | NA | NA | NA | NA | 1 | 1 | 1 | 1 | NA | NA | NA | NA | 2 | 0.5 | 3 | 0.6667 |
| 2_1198722 | 1 | 1 | 2 | 0.5 | NA | NA | NA | NA | 3 | 0.6667 | 1 | 1 | NA | NA | 1 | 1 | 2 | 1 | 4 | 1 |
| 2_1198724 | 1 | 1 | 1 | 1 | NA | NA | NA | NA | 2 | 1 | 1 | 1 | NA | NA | 1 | 1 | NA | NA | 2 | 1 |
| 2_1198724 | 1 | 1 | 2 | 1 | NA | NA | NA | NA | 3 | 1 | 1 | 1 | NA | NA | 1 | 1 | 1 | 1 | 3 | 1 |
| 2_1199209 | 2 | 0 | 3 | 0.3333 | 1 | 0 | 1 | 0 | 7 | 0.1429 | NA | NA | 1 | 1 | NA | NA | 1 | 0 | 2 | 0.5 |
| 2_1199212 | 3 | 0.3333 | 3 | 0.6667 | 1 | 1 | 1 | 1 | 8 | 0.625 | NA | NA | 2 | 0 | NA | NA | 1 | 1 | 3 | 0.3333 |
| 2_1199212 | 2 | 1 | 3 | 0.6667 | 1 | 0 | 1 | 0 | 7 | 0.5714 | NA | NA | 1 | 1 | NA | NA | 1 | 1 | 2 | 1 |
| 2_1199213 | 2 | 0.5 | 3 | 1 | 1 | 0 | 1 | 0 | 7 | 0.5714 | NA | NA | 2 | 0.5 | NA | NA | 1 | 1 | 3 | 0.6667 |
| 2_1199216 | 2 | 1 | 1 | 1 | 1 | 1 | NA | NA | 4 | 1 | NA | NA | 2 | 1 | NA | NA | 1 | 1 | 3 | 1 |
| 2_1199218 | 2 | 1 | 1 | 1 | 1 | 1 | NA | NA | 4 | 1 | NA | NA | 2 | 1 | NA | NA | 1 | 1 | 3 | 1 |
| 2_1199220 | 2 | 0.5 | 1 | 0 | 1 | 1 | NA | NA | 4 | 0.5 | NA | NA | NA | NA | NA | NA | 1 | 1 | 1 | 1 |
| 2_1199973 | 1 | 1 | NA | NA | NA | NA | NA | NA | 1 | 1 | NA | NA | NA | NA | NA | NA | 1 | 1 | 1 | 1 |
| 2_1199976 | 1 | 1 | NA | NA | NA | NA | NA | NA | 1 | 1 | NA | NA | NA | NA | NA | NA | 1 | 1 | 1 | 1 |
| 2_1201471 | 9 | 0 | 5 | 0 | 6 | 0 | 14 | 0.1429 | 34 | 0.0588 | 10 | 0.1 | 8 | 0.125 | 9 | 0.1111 | 5 | 0 | 32 | 0.0938 |
| 2_1201478 | 8 | 0 | 6 | 0 | 5 | 0 | 11 | 0.1818 | 30 | 0.0667 | 8 | 0.125 | 7 | 0.7143 | 11 | 0.1818 | 4 | 0 | 30 | 0.2667 |
| 2_1201478 | 9 | 0.6667 | 4 | 0.5 | 4 | 0.5 | 10 | 0.9 | 27 | 0.7037 | 8 | 0.875 | 7 | 1 | 7 | 0.8571 | 4 | 0.5 | 26 | 0.8462 |
| 2_1201481 | 9 | 0.6667 | 4 | 0.75 | 5 | 0.4 | 12 | 1 | 30 | 0.7667 | 4 | 0.75 | 7 | 0.8571 | 10 | 1 | 5 | 0.8 | 26 | 0.8846 |
| 2_1201482 | 7 | 0.1429 | 4 | 0.25 | 1 | 1 | 9 | 0.6667 | 21 | 0.4286 | 5 | 0 | 4 | 0.25 | 9 | 0.2222 | 5 | 0.2 | 23 | 0.1739 |
| 2_1201482 | 9 | 0.6667 | 5 | 0.4 | 5 | 0.6 | 12 | 0.4167 | 31 | 0.5161 | 5 | 0.4 | 6 | 0.1667 | 10 | 0.4 | 5 | 1 | 26 | 0.4615 |
| 2_1201483 | 7 | 0.5714 | 5 | 0.8 | 4 | 0.5 | 13 | 0.2308 | 29 | 0.4483 | 6 | 0.5 | 7 | 0.4286 | 9 | 0.7778 | 3 | 0.6667 | 25 | 0.6 |
| 2_1201483 | 7 | 1 | 5 | 0.8 | 4 | 1 | 10 | 1 | 26 | 0.9615 | 3 | 1 | 5 | 1 | 10 | 0.9 | 2 | 1 | 20 | 0.95 |
| 2_1201483 | 5 | 0.4 | 4 | 0.25 | 3 | 0.3333 | 9 | 0.6667 | 21 | 0.4762 | 3 | 0.3333 | 4 | 0.25 | 7 | 0.5714 | 2 | 0.5 | 16 | 0.4375 |
| 2_1201483 | 8 | 0.125 | 3 | 0.3333 | 4 | 0.25 | 12 | 0 | 27 | 0.1111 | 6 | 0.1667 | 7 | 0.2857 | 6 | 0 | 4 | 0 | 23 | 0.1304 |
| 2_1201484 | 9 | 0.2222 | 4 | 0 | 3 | 0 | 8 | 0.125 | 24 | 0.125 | 4 | 0 | 2 | 0 | 9 | 0.1111 | 3 | 0.3333 | 18 | 0.1111 |
| 2_1201866 | 3 | 1 | 4 | 0.75 | 4 | 1 | 1 | 0 | 12 | 0.8333 | 6 | 0.8333 | 2 | 0.5 | 4 | 0.75 | 4 | 1 | 16 | 0.8125 |
| 2_1201867 | 3 | 0.3333 | 5 | 0.2 | 4 | 0.5 | 1 | 1 | 13 | 0.3846 | 4 | 0.75 | 2 | 0 | 3 | 1 | 5 | 0.4 | 14 | 0.5714 |
| 2_1201869 | 3 | 0 | 3 | 0.3333 | 4 | 0.25 | NA | NA | 10 | 0.2 | 4 | 0 | 2 | 0 | 2 | 0.5 | 4 | 0 | 12 | 0.0833 |

| | | | | | | | | | | | | | | | | | | | | |
|---|---|---|---|---|---|---|---|---|---|---|---|---|---|---|---|---|---|---|---|---|
| 2_12033742 | 12 | 0.0833 | 23 | 0.1304 | 12 | 0 | 24 | 0.0417 | 71 | 0.0704 | 8 | 0 | 13 | 0 | 15 | 0 | 10 | 0.3 | 46 | 0.0652 |
| 2_12033777 | 9 | 0 | 19 | 0.0526 | 10 | 0.1 | 22 | 0.0455 | 60 | 0.05 | 9 | 0 | 12 | 0.0833 | 15 | 0.0667 | 11 | 0.1818 | 47 | 0.0851 |
| 2_12033951 | 7 | 0 | 13 | 0 | 7 | 0.1429 | 11 | 0.2727 | 38 | 0.1053 | 6 | 0.1667 | 5 | 0.4 | 14 | 0.0714 | 6 | 0 | 31 | 0.129 |
| 2_12033997 | 5 | 0.2 | 12 | 0.0833 | 5 | 0 | 9 | 0.1111 | 31 | 0.0968 | 6 | 0.1667 | 5 | 0 | 12 | 0.1667 | 6 | 0.3333 | 29 | 0.1724 |
| 2_12034036 | 3 | 0.3333 | 13 | 0.1538 | 6 | 0.1667 | 9 | 0.1111 | 31 | 0.1613 | 7 | 0.1429 | 9 | 0.1111 | 10 | 0.2 | 7 | 0.1429 | 33 | 0.1515 |
| 2_12035458 | 1 | 1 | 7 | 0.2857 | 4 | 0 | 3 | 1 | 15 | 0.4 | 7 | 0.2857 | 6 | 0 | 9 | 0.2222 | 5 | 0 | 27 | 0.1481 |
| 2_12035460 | 1 | 0 | 9 | 0.1111 | 4 | 0 | 5 | 0 | 19 | 0.0526 | 7 | 0 | 6 | 0.1667 | 9 | 0 | 6 | 0.1667 | 28 | 0.0714 |
| 2_12035464 | NA | NA | 9 | 0.1111 | 4 | 0 | 4 | 0 | 17 | 0.0588 | 5 | 0.4 | 6 | 0 | 10 | 0.1 | 5 | 0.2 | 26 | 0.1538 |
| 2_12035470 | NA | NA | 6 | 0 | 3 | 0.3333 | 4 | 0 | 13 | 0.0769 | 5 | 0.2 | 1 | 0 | 5 | 0.2 | 4 | 0 | 15 | 0.1333 |
| 2_12035491 | NA | NA | 8 | 0.125 | 4 | 0.25 | 3 | 0 | 15 | 0.1333 | 4 | 0.25 | 4 | 0 | 8 | 0 | 4 | 0 | 20 | 0.05 |
| 2_12035503 | NA | NA | 9 | 0 | 2 | 0.5 | 2 | 0 | 13 | 0.0769 | 4 | 0 | 4 | 0.25 | 9 | 0.1111 | 3 | 0 | 20 | 0.1 |
| 2_12036240 | NA | NA | 3 | 0 | 3 | 0 | 2 | 1 | 8 | 0.25 | 4 | 0 | 5 | 0 | 9 | 0.1111 | 1 | 0 | 19 | 0.0526 |
| 2_12036253 | NA | NA | 5 | 0.8 | 3 | 0.6667 | 4 | 0.75 | 12 | 0.75 | 5 | 1 | 2 | 1 | 9 | 0.8889 | 2 | 1 | 18 | 0.9444 |
| 2_12036266 | NA | NA | 5 | 0.2 | 3 | 0 | 5 | 0 | 13 | 0.0769 | 5 | 0 | 2 | 0.5 | 6 | 0 | 2 | 0 | 15 | 0.0667 |
| 2_12036427 | NA | NA | 3 | 0.3333 | 1 | 0 | 4 | 0.25 | 8 | 0.25 | 2 | 0 | NA | NA | 4 | 0.5 | 1 | 0 | 7 | 0.2857 |
| 2_12036434 | NA | NA | 2 | 0 | 1 | 1 | 4 | 0.5 | 7 | 0.4286 | 2 | 1 | 2 | 1 | 6 | 0.6667 | 1 | 1 | 11 | 0.8182 |
| 2_12036436 | NA | NA | 3 | 0.6667 | 1 | 1 | 3 | 1 | 7 | 0.8571 | 3 | 1 | 2 | 0.5 | 5 | 1 | NA | NA | 10 | 0.9 |
| 2_12036452 | NA | NA | 2 | 0.5 | 1 | 0 | 4 | 0 | 7 | 0.1429 | 2 | 0 | NA | NA | 7 | 0.1429 | 1 | 0 | 10 | 0.1 |
| 2_12036469 | NA | NA | 2 | 0.5 | 1 | 0 | 4 | 0.5 | 7 | 0.4286 | 3 | 0.3333 | 2 | 0 | 5 | 0 | 1 | 0 | 11 | 0.0909 |
| 2_12036484 | NA | NA | 2 | 0 | 1 | 0 | 4 | 0.25 | 7 | 0.1429 | 3 | 0 | 2 | 0 | 6 | 0.1667 | 1 | 0 | 12 | 0.0833 |
| 2_12036486 | NA | NA | 2 | 1 | 1 | 0 | 4 | 0.25 | 7 | 0.4286 | 2 | 0.5 | 2 | 0.5 | 7 | 0.7143 | NA | NA | 11 | 0.6364 |
| 2_12036494 | NA | NA | 3 | 0.3333 | 1 | 0 | 2 | 0.5 | 6 | 0.3333 | NA | NA | 2 | 0.5 | 5 | 0.2 | NA | NA | 7 | 0.2857 |
| 2_12036500 | NA | NA | 3 | 0 | 1 | 0 | 4 | 0.25 | 8 | 0.125 | 2 | 0 | 2 | 0 | 6 | 0.1667 | NA | NA | 10 | 0.1 |
| 2_12036525 | NA | NA | 3 | 0.3333 | 1 | 1 | 4 | 0.75 | 8 | 0.625 | 2 | 1 | 2 | 1 | 6 | 0.6667 | NA | NA | 10 | 0.8 |
| 2_12036530 | NA | NA | 3 | 0 | 1 | 0 | 3 | 0.3333 | 7 | 0.1429 | 1 | 0 | 2 | 0 | 4 | 0.25 | NA | NA | 7 | 0.1429 |
| 2_12036537 | NA | NA | 2 | 0 | NA | NA | 3 | 0.3333 | 5 | 0.2 | 2 | 0 | 2 | 0 | 5 | 0.2 | NA | NA | 9 | 0.1111 |
| 2_12036584 | NA | NA | 3 | 0 | 1 | 0 | 3 | 0.3333 | 7 | 0.1429 | 1 | 0 | 2 | 0 | 2 | 0.5 | NA | NA | 5 | 0.2 |
| 2_12036594 | NA | NA | 3 | 0 | 1 | 0 | 4 | 0.25 | 8 | 0.125 | 1 | 1 | 2 | 0 | 1 | 0 | NA | NA | 4 | 0.25 |
| 2_12036601 | NA | NA | 2 | 1 | 1 | 1 | 4 | 0.5 | 7 | 0.7143 | 1 | 0 | 1 | 1 | NA | NA | NA | NA | 2 | 0.5 |
| 2_12042978 | 3 | 0 | 4 | 0 | 3 | 0.3333 | 3 | 0 | 13 | 0.0769 | 2 | 0.5 | NA | NA | NA | NA | 1 | 0 | 3 | 0.3333 |
| 2_12042985 | 2 | 0 | 5 | 0 | 2 | 0 | 3 | 0.3333 | 12 | 0.0833 | 2 | 0.5 | NA | NA | 1 | 0 | 1 | 0 | 4 | 0.25 |
| 2_12043539 | 4 | 0.25 | 7 | 0.4286 | 4 | 0 | 1 | 1 | 16 | 0.3125 | 1 | 0 | 1 | 0 | 4 | 0.75 | 3 | 0 | 9 | 0.3333 |
| 2_12043557 | 4 | 0.25 | 6 | 0.1667 | 3 | 0 | 1 | 1 | 14 | 0.2143 | NA | NA | 1 | 0 | 2 | 0 | 3 | 1 | 6 | 0.5 |
| 2_12043559 | 3 | 0 | 9 | 0.3333 | 4 | 0.25 | 1 | 0 | 17 | 0.2353 | 1 | 0 | 1 | 1 | 3 | 0.3333 | 3 | 0 | 8 | 0.25 |
| 2_12043565 | 4 | 0 | 9 | 0.2222 | 4 | 0 | NA | NA | 17 | 0.1176 | 1 | 0 | 1 | 0 | 4 | 0.25 | 3 | 0 | 9 | 0.1111 |
| 2_12044959 | 2 | 1 | 1 | 1 | 1 | 1 | 1 | 0 | 5 | 0.8 | 1 | 1 | 2 | 0.5 | 1 | 0 | 5 | 0.8 | 9 | 0.6667 |
| 2_12045046 | 3 | 0.6667 | 1 | 0 | 3 | 0.6667 | 1 | 0 | 8 | 0.5 | NA | NA | 2 | 0 | 1 | 0 | 4 | 0.25 | 7 | 0.1429 |
| 2_12054519 | NA | NA | 1 | 1 | NA | NA | NA | NA | 1 | 1 | NA | NA | 1 | 1 | 1 | 1 | 2 | 1 | 4 | 1 |
| 2_12054541 | NA | NA | 1 | 1 | NA | NA | NA | NA | 1 | 1 | NA | NA | 1 | 1 | 1 | 1 | 2 | 1 | 4 | 1 |
| 2_12054543 | NA | NA | 1 | 1 | NA | NA | NA | NA | 1 | 1 | NA | NA | 1 | 0 | NA | NA | 2 | 1 | 3 | 0.6667 |
| 2_12054557 | NA | NA | 1 | 1 | NA | NA | NA | NA | 1 | 1 | NA | NA | 1 | 1 | 1 | 1 | 2 | 1 | 4 | 1 |
| 2_12054565 | NA | NA | 1 | 1 | NA | NA | NA | NA | 1 | 1 | NA | NA | NA | NA | NA | NA | 2 | 0.5 | 2 | 0.5 |
| 2_12054725 | NA | NA | 1 | 1 | NA | NA | NA | NA | 1 | 1 | NA | NA | 1 | 0 | NA | NA | 1 | 1 | 2 | 0.5 |
| 2_12054879 | 1 | 1 | 2 | 0.5 | 2 | 0.5 | 1 | 0 | 6 | 0.5 | 1 | 1 | 2 | 0.5 | 2 | 0.5 | 4 | 0 | 9 | 0.3333 |
| 2_12054883 | 1 | 1 | 3 | 0.3333 | 2 | 0 | 1 | 1 | 7 | 0.4286 | NA | NA | 3 | 0.3333 | 2 | 1 | 3 | 0.3333 | 8 | 0.5 |
| 2_12054890 | 1 | 1 | 2 | 1 | 1 | 1 | 1 | 1 | 5 | 1 | 1 | 1 | 3 | 0.6667 | 2 | 0 | 4 | 1 | 10 | 0.7 |
| 2_12054893 | 1 | 1 | 2 | 1 | 2 | 1 | 1 | 1 | 6 | 1 | 1 | 1 | 3 | 1 | 2 | 1 | 4 | 1 | 10 | 1 |
| 2_12054903 | 1 | 0 | 1 | 1 | NA | NA | 1 | 0 | 3 | 0.3333 | 1 | 0 | 2 | 0 | 1 | 0 | 4 | 0.25 | 8 | 0.125 |
| 2_12054904 | 1 | 1 | 3 | 0.3333 | 2 | 0.5 | 1 | 1 | 7 | 0.5714 | 1 | 1 | 3 | 1 | 2 | 0.5 | 4 | 0.5 | 10 | 0.7 |
| 2_12054930 | NA | NA | 2 | 0.5 | 2 | 1 | 1 | 1 | 5 | 0.8 | 1 | 1 | 1 | 1 | 2 | 0 | 3 | 0.6667 | 7 | 0.5714 |
| 2_12054931 | 1 | 0 | 2 | 0.5 | 2 | 0 | NA | NA | 5 | 0.2 | 1 | 1 | 2 | 0 | 2 | 0 | 4 | 0 | 9 | 0.1111 |
| 2_12054932 | NA | NA | 2 | 0.5 | 2 | 0.5 | 1 | 1 | 5 | 0.6 | 1 | 0 | 2 | 0.5 | 2 | 1 | 4 | 0.75 | 9 | 0.6667 |
| 2_12056564 | 2 | 0.5 | 7 | 0.1429 | 2 | 0 | 2 | 0 | 13 | 0.1538 | 2 | 0 | 5 | 0 | 1 | 0 | 2 | 0.5 | 10 | 0.1 |
| 2_12056758 | 1 | 1 | 4 | 1 | 4 | 0.25 | 2 | 0 | 11 | 0.5455 | 1 | 1 | 2 | 0.5 | 1 | 0 | 1 | 0 | 5 | 0.4 |
| 2_12056860 | 1 | 1 | 4 | 0.5 | 1 | 0 | 2 | 1 | 8 | 0.625 | 1 | 1 | 1 | 1 | 1 | 0 | NA | NA | 3 | 0.6667 |
| 2_12056873 | 1 | 0 | 3 | 0 | 3 | 0 | 3 | 0.3333 | 10 | 0.1 | 1 | 0 | NA | NA | 1 | 1 | NA | NA | 2 | 0.5 |
| 2_12056876 | 2 | 1 | 3 | 0 | 2 | 0 | 2 | 0 | 9 | 0.2222 | 1 | 0 | 1 | 0 | 1 | 1 | NA | NA | 3 | 0.3333 |
| 2_12056886 | 2 | 1 | 3 | 1 | 3 | 0.6667 | 1 | 1 | 9 | 0.8889 | 1 | 1 | NA | NA | 1 | 1 | NA | NA | 2 | 1 |
| 2_12056887 | 2 | 0.5 | 3 | 0.3333 | 3 | 0 | 3 | 0 | 11 | 0.1818 | 1 | 1 | 1 | 0 | NA | NA | NA | NA | 2 | 0.5 |
| 2_12056979 | 1 | 0 | 4 | 0 | NA | NA | 3 | 0.3333 | 8 | 0.125 | 1 | 0 | 1 | 1 | NA | NA | NA | NA | 2 | 0.5 |
| 2_12057022 | NA | NA | 3 | 0.6667 | NA | NA | 4 | 0 | 7 | 0.2857 | 1 | 1 | 1 | 0 | NA | NA | NA | NA | 2 | 0.5 |
| 2_12057063 | NA | NA | 4 | 0.75 | NA | NA | 3 | 0 | 7 | 0.4286 | 1 | 1 | NA | NA | NA | NA | NA | NA | 1 | 1 |
| 2_12057597 | NA | NA | 3 | 0.3333 | 1 | 0 | 2 | 0 | 6 | 0.1667 | 1 | 0 | NA | NA | 1 | 1 | NA | NA | 2 | 0.5 |
| 2_12057628 | NA | NA | 4 | 0.75 | NA | NA | 2 | 1 | 6 | 0.8333 | 1 | 1 | NA | NA | 1 | 0 | NA | NA | 2 | 0.5 |
| 2_12057649 | NA | NA | 4 | 0.25 | NA | NA | 2 | 0 | 6 | 0.1667 | 1 | 1 | NA | NA | 1 | 1 | NA | NA | 2 | 1 |

| | | | | | | | | | | | | | | | | | | | | |
|---|---|---|---|---|---|---|---|---|---|---|---|---|---|---|---|---|---|---|---|---|
| 2_12057657 | NA | NA | 4 | 0.75 | NA | NA | 1 | 1 | 5 | 0.8 | NA | NA | NA | NA | 1 | 1 | NA | NA | 1 | 1 |
| 2_12057910 | NA | NA | 3 | 1 | NA | NA | 2 | 1 | 5 | 1 | NA | NA | NA | NA | 1 | 1 | NA | NA | 1 | 1 |
| 2_12057919 | NA | NA | 4 | 0.5 | NA | NA | 2 | 0 | 6 | 0.3333 | NA | NA | NA | NA | 1 | 1 | NA | NA | 1 | 1 |
| 2_12069277 | 2 | 0 | 5 | 0.2 | 1 | 0 | 3 | 0 | 11 | 0.0909 | 3 | 0.3333 | 6 | 0.1667 | 4 | 0.25 | 1 | 1 | 14 | 0.2857 |
| 2_12069306 | 3 | 0.3333 | 8 | 0.5 | 2 | 1 | 5 | 0.2 | 18 | 0.4444 | 4 | 0.5 | 4 | 0.25 | 5 | 0.6 | 1 | 1 | 14 | 0.5 |
| 2_12069308 | 3 | 0.6667 | 8 | 0.75 | 2 | 1 | 3 | 0.6667 | 16 | 0.75 | 3 | 1 | 3 | 0.3333 | 4 | 1 | NA | NA | 10 | 0.8 |
| 2_12069317 | 3 | 0 | 9 | 0.1111 | 2 | 0 | 6 | 0.1667 | 20 | 0.1 | 4 | 0.25 | 6 | 0 | 4 | 0 | NA | NA | 14 | 0.0714 |
| 2_12083748 | 3 | 0 | 2 | 0.5 | 9 | 0.2222 | 5 | 0.4 | 19 | 0.2632 | 2 | 0.5 | 2 | 0 | 4 | 0.25 | 1 | 0 | 9 | 0.2222 |
| 2_12083751 | 3 | 1 | 2 | 1 | 10 | 0.7 | 6 | 0.5 | 21 | 0.7143 | 2 | 1 | 2 | 1 | 3 | 0.3333 | 1 | 1 | 8 | 0.75 |
| 2_12083752 | 3 | 0 | 3 | 0.3333 | 11 | 0.2727 | 6 | 0.5 | 23 | 0.3043 | 3 | 1 | 3 | 0.3333 | 5 | 0.2 | 1 | 0 | 12 | 0.4167 |
| 2_12083756 | 1 | 1 | 3 | 1 | 9 | 0.8889 | 5 | 0.8 | 18 | 0.8889 | 3 | 1 | 2 | 0.5 | 3 | 0.6667 | 1 | 1 | 9 | 0.7778 |
| 2_12083757 | 3 | 0.6667 | 2 | 1 | 9 | 0.1111 | 3 | 0 | 17 | 0.2941 | 2 | 0 | 3 | 0.3333 | 5 | 0.4 | 1 | 0 | 11 | 0.2727 |
| 2_12083758 | 3 | 0.3333 | 2 | 0.5 | 7 | 0.4286 | 5 | 0.6 | 17 | 0.4706 | 3 | 0.6667 | 3 | 0.3333 | 3 | 1 | 1 | 0 | 10 | 0.6 |
| 2_12083775 | 3 | 1 | 3 | 1 | 11 | 0.9091 | 5 | 0.8 | 22 | 0.9091 | 3 | 0.6667 | 3 | 0.3333 | 5 | 1 | 1 | 1 | 12 | 0.75 |
| 2_12083780 | 2 | 1 | 3 | 1 | 10 | 1 | 6 | 1 | 21 | 1 | 3 | 1 | 3 | 1 | 4 | 1 | 1 | 1 | 11 | 1 |
| 2_12083794 | 3 | 0 | 3 | 0.3333 | 7 | 0.2857 | 3 | 0 | 16 | 0.1875 | 3 | 0 | 2 | 0 | 5 | 0.4 | 1 | 0 | 11 | 0.1818 |
| 2_12083800 | 3 | 1 | 3 | 1 | 10 | 0.6 | 6 | 0.6667 | 22 | 0.7273 | 3 | 0.6667 | 2 | 1 | 4 | 1 | 1 | 0 | 10 | 0.8 |
| 2_12083802 | 3 | 0.6667 | 2 | 1 | 9 | 0.4444 | 5 | 0.2 | 19 | 0.4737 | 3 | 0.3333 | 3 | 0.3333 | 5 | 0.6 | 1 | 1 | 12 | 0.5 |
| 2_12083802 | 3 | 0 | 3 | 0.3333 | 11 | 0.1818 | 3 | 0 | 20 | 0.15 | 3 | 0.6667 | 3 | 0 | 4 | 0.25 | 1 | 0 | 11 | 0.2727 |
| 2_12083812 | 2 | 1 | NA | NA | 5 | 0.4 | 2 | 0.5 | 9 | 0.5556 | 2 | 1 | NA | NA | 3 | 0.6667 | NA | NA | 5 | 0.8 |
| 2_12083827 | 1 | 0 | 2 | 0.5 | 10 | 0.1 | 5 | 0.2 | 18 | 0.1667 | 3 | 0 | NA | NA | 4 | 0.75 | NA | NA | 7 | 0.4286 |
| 2_12084328 | 6 | 0.1667 | 9 | 0 | 10 | 0.1 | 13 | 0.1538 | 38 | 0.1053 | 6 | 0.1667 | 10 | 0 | 10 | 0.1 | 12 | 0.0833 | 38 | 0.0789 |
| 2_12096901 | 1 | 0 | 4 | 1 | NA | NA | 1 | 1 | 6 | 0.8333 | NA | NA | NA | NA | 1 | 1 | 1 | 1 | 2 | 1 |
| 2_12096909 | 1 | 0 | 4 | 0.5 | NA | NA | 2 | 0.5 | 7 | 0.4286 | NA | NA | NA | NA | 1 | 1 | 2 | 1 | 3 | 1 |
| 2_12096922 | 1 | 0 | 3 | 0.3333 | NA | NA | 2 | 0 | 6 | 0.1667 | NA | NA | NA | NA | 1 | 0 | 2 | 0.5 | 3 | 0.3333 |
| 2_12096923 | 2 | 0.5 | 3 | 1 | NA | NA | 1 | 1 | 6 | 0.8333 | NA | NA | NA | NA | NA | NA | 1 | 1 | 1 | 1 |
| 2_12096941 | 1 | 1 | 3 | 1 | NA | NA | 2 | 1 | 6 | 1 | NA | NA | NA | NA | 1 | 1 | 2 | 1 | 3 | 1 |
| 2_12096953 | 1 | 1 | 2 | 1 | NA | NA | 2 | 1 | 5 | 1 | NA | NA | NA | NA | 1 | 1 | 1 | 1 | 2 | 1 |
| 2_12096955 | 1 | 1 | 3 | 1 | NA | NA | 2 | 1 | 6 | 1 | NA | NA | NA | NA | 1 | 1 | 1 | 1 | 2 | 1 |
| 2_12096985 | 1 | 1 | 1 | 1 | NA | NA | 2 | 1 | 4 | 1 | NA | NA | NA | NA | 1 | 1 | NA | NA | 1 | 1 |
| 2_12097013 | 1 | 0 | 2 | 0.5 | NA | NA | 2 | 0.5 | 5 | 0.4 | NA | NA | NA | NA | 1 | 1 | 1 | 0 | 2 | 0.5 |
| 2_12097029 | 1 | 1 | 1 | 1 | NA | NA | 2 | 0.5 | 4 | 0.75 | NA | NA | NA | NA | 1 | 0 | 1 | 1 | 2 | 0.5 |
| 2_12097062 | 1 | 0 | 1 | 1 | NA | NA | NA | NA | 2 | 0.5 | NA | NA | NA | NA | 1 | 0 | 1 | 1 | 2 | 0.5 |
| 2_12098596 | NA | NA | 1 | 0 | 3 | 1 | 3 | 0.6667 | 7 | 0.7143 | NA | NA | NA | NA | 4 | 1 | 1 | 1 | 5 | 1 |
| 2_12098597 | NA | NA | 1 | 1 | 6 | 0.8333 | 3 | 1 | 10 | 0.9 | 1 | 1 | 1 | 1 | 4 | 1 | 1 | 1 | 7 | 1 |
| 2_12098600 | NA | NA | 2 | 0 | 6 | 0.3333 | 3 | 0.6667 | 11 | 0.3636 | 1 | 0 | 1 | 0 | 4 | 0.75 | 1 | 0 | 7 | 0.4286 |
| 2_12098601 | NA | NA | 1 | 1 | 5 | 0 | 3 | 0.3333 | 9 | 0.2222 | 1 | 0 | 1 | 0 | 4 | 0.25 | 1 | 0 | 7 | 0.1429 |
| 2_12098623 | NA | NA | 2 | 0.5 | 6 | 0.3333 | 3 | 0 | 11 | 0.2727 | 1 | 0 | NA | NA | 3 | 0.3333 | 1 | 1 | 5 | 0.4 |
| 2_12098901 | 1 | 0 | 1 | 1 | 1 | 0 | NA | NA | 3 | 0.3333 | NA | NA | NA | NA | 1 | 1 | NA | NA | 1 | 1 |
| 2_12099113 | 1 | 0 | NA | NA | 1 | 1 | NA | NA | 2 | 0.5 | NA | NA | NA | NA | 1 | 1 | NA | NA | 1 | 1 |
| 2_12099120 | 1 | 1 | NA | NA | 1 | 1 | NA | NA | 2 | 1 | NA | NA | NA | NA | 1 | 1 | NA | NA | 1 | 1 |
| 2_12099177 | NA | NA | NA | NA | 1 | 1 | NA | NA | 1 | 1 | NA | NA | NA | NA | 1 | 1 | NA | NA | 1 | 1 |
| 2_12099664 | 1 | 0 | NA | NA | 1 | 1 | 1 | 1 | 3 | 0.6667 | NA | NA | NA | NA | 1 | 1 | NA | NA | 1 | 1 |
| 2_12099692 | NA | NA | NA | NA | 1 | 1 | 1 | 0 | 2 | 0.5 | NA | NA | NA | NA | 1 | 1 | NA | NA | 1 | 1 |
| 2_12099698 | NA | NA | NA | NA | 1 | 1 | 1 | 1 | 2 | 1 | NA | NA | NA | NA | 1 | 1 | NA | NA | 1 | 1 |
| 2_12100040 | NA | NA | NA | NA | 1 | 1 | 1 | 1 | 2 | 1 | NA | NA | NA | NA | 1 | 1 | NA | NA | 1 | 1 |
| 2_12100048 | NA | NA | NA | NA | 1 | 1 | NA | NA | 1 | 1 | NA | NA | NA | NA | 1 | 1 | NA | NA | 1 | 1 |
| 2_12100114 | NA | NA | NA | NA | 1 | 1 | 1 | 1 | 2 | 1 | NA | NA | NA | NA | 1 | 1 | NA | NA | 1 | 1 |
| 2_12100137 | NA | NA | NA | NA | 1 | 1 | 1 | 0 | 2 | 0.5 | NA | NA | NA | NA | 1 | 1 | NA | NA | 1 | 1 |
| 2_12101500 | 1 | 1 | NA | NA | NA | NA | NA | NA | 1 | 1 | NA | NA | 1 | 1 | NA | NA | NA | NA | 1 | 1 |
| 2_12101518 | 1 | 1 | NA | NA | NA | NA | NA | NA | 1 | 1 | NA | NA | 1 | 1 | NA | NA | NA | NA | 1 | 1 |
| 2_12102671 | 1 | 0 | 2 | 0.5 | 3 | 0 | 4 | 0 | 10 | 0.1 | NA | NA | 3 | 0 | 3 | 0.3333 | 2 | 0 | 8 | 0.125 |
| 2_12102731 | 1 | 0 | 2 | 1 | 2 | 1 | 4 | 0 | 9 | 0.4444 | 1 | 0 | 4 | 0 | 4 | 0.25 | 2 | 1 | 11 | 0.2727 |
| 2_12102738 | 1 | 0 | 1 | 0 | 3 | 0.3333 | 2 | 0 | 7 | 0.1429 | 1 | 0 | 4 | 0.25 | 4 | 0 | 1 | 0 | 10 | 0.1 |
| 2_12102743 | 1 | 0 | 2 | 0 | 3 | 0.6667 | 4 | 0 | 10 | 0.2 | NA | NA | 3 | 0 | 5 | 0 | 2 | 0.5 | 10 | 0.1 |
| 2_12102747 | 1 | 0 | 2 | 1 | 2 | 1 | 3 | 0.3333 | 8 | 0.625 | 1 | 1 | 4 | 0.5 | 4 | 0.25 | 2 | 0.5 | 11 | 0.4545 |
| 2_12102751 | 1 | 0 | 2 | 0.5 | 4 | 0 | 3 | 0 | 10 | 0.1 | 1 | 0 | 3 | 0.3333 | 3 | 0 | 3 | 0 | 10 | 0.1 |
| 2_12102764 | 1 | 0 | 2 | 0.5 | 4 | 0.25 | 2 | 0 | 9 | 0.2222 | 1 | 0 | 4 | 0.5 | 4 | 0.5 | 2 | 0 | 11 | 0.3636 |
| 2_12102877 | 7 | 0.1429 | 8 | 0 | 7 | 0 | 5 | 0.2 | 27 | 0.0741 | 5 | 0.2 | 4 | 0 | 7 | 0.1429 | 3 | 0 | 19 | 0.1053 |
| 2_12102908 | 9 | 0.1111 | 6 | 0 | 5 | 0 | 5 | 0.2 | 25 | 0.08 | 5 | 0 | 3 | 0.3333 | 7 | 0.1429 | 3 | 0 | 18 | 0.1111 |
| 2_12103822 | 5 | 0 | 6 | 0.1667 | 5 | 0.4 | 5 | 0 | 21 | 0.1429 | 2 | 0 | 4 | 0 | 3 | 0.3333 | 3 | 0 | 12 | 0.0833 |
| 2_12103825 | 7 | 0.2857 | 7 | 0.2857 | 5 | 0 | 6 | 0.3333 | 25 | 0.24 | 2 | 0 | 4 | 0.5 | 4 | 0.25 | 3 | 0.3333 | 13 | 0.3077 |
| 2_12103918 | 1 | 0 | 1 | 1 | NA | NA | 2 | 0 | 4 | 0.25 | NA | NA | NA | NA | NA | NA | 1 | 1 | 1 | 1 |
| 2_12103961 | 4 | 0.25 | 4 | 0 | 2 | 0 | 1 | 0 | 11 | 0.0909 | 2 | 0 | 2 | 0 | 3 | 0 | 3 | 0.3333 | 10 | 0.1 |
| 2_12103977 | 4 | 0.25 | 4 | 0.25 | 3 | 0 | 3 | 0.6667 | 14 | 0.2857 | 2 | 0.5 | 2 | 0.5 | 3 | 0 | 1 | 0 | 8 | 0.25 |

| | | | | | | | | | | | | | | | | | | | | |
|---|---|---|---|---|---|---|---|---|---|---|---|---|---|---|---|---|---|---|---|---|
| 2_12104624 | 2 | 0.5 | 3 | 0.3333 | 2 | 0.5 | 3 | 0 | 10 | 0.3 | 1 | 1 | 1 | 1 | 3 | 0 | 2 | 1 | 7 | 0.5714 |
| 2_12104629 | 3 | 0.3333 | 3 | 0 | 3 | 0.3333 | 3 | 0.3333 | 12 | 0.25 | 1 | 1 | NA | NA | 3 | 0.3333 | 2 | 0 | 6 | 0.3333 |
| 2_12104631 | 2 | 0 | 3 | 0.3333 | 2 | 1 | 3 | 0.3333 | 10 | 0.4 | 1 | 1 | 1 | 0 | 3 | 0 | 2 | 0 | 7 | 0.1429 |
| 2_12104633 | 3 | 0 | 2 | 0 | 2 | 0 | 3 | 0.3333 | 10 | 0.1 | NA | NA | 1 | 0 | 3 | 0.3333 | 2 | 0 | 6 | 0.1667 |
| 2_12104637 | 2 | 1 | 3 | 0 | 2 | 0 | 2 | 0.5 | 9 | 0.3333 | NA | NA | 1 | 1 | 2 | 0 | 2 | 1 | 5 | 0.6 |
| 2_12104639 | 3 | 0 | 2 | 0 | 2 | 0.5 | 3 | 0 | 10 | 0.1 | 1 | 0 | 1 | 0 | 3 | 0.3333 | 2 | 0 | 7 | 0.1429 |
| 2_12104641 | 3 | 0.6667 | 3 | 1 | 2 | 0.5 | 3 | 1 | 11 | 0.8182 | NA | NA | 1 | 1 | 3 | 0.6667 | 1 | 1 | 5 | 0.8 |
| 2_12104652 | 3 | 0.3333 | 2 | 0.5 | 1 | 1 | 3 | 0.6667 | 9 | 0.5556 | 1 | 1 | 1 | 1 | 3 | 1 | NA | NA | 5 | 1 |
| 2_12106616 | 2 | 0.5 | 2 | 0 | 1 | 1 | 1 | 0 | 6 | 0.3333 | NA | NA | 1 | 1 | 2 | 0 | 1 | 1 | 4 | 0.5 |
| 2_12106652 | 2 | 0 | 2 | 0 | 2 | 0.5 | 1 | 1 | 7 | 0.2857 | NA | NA | 1 | 0 | 3 | 0.3333 | 1 | 0 | 5 | 0.2 |
| 2_12106680 | 1 | 1 | 1 | 0 | 2 | 0 | 1 | 1 | 5 | 0.4 | NA | NA | 1 | 0 | 3 | 1 | 1 | 0 | 5 | 0.6 |
| 2_12106695 | 2 | 0.5 | 2 | 0 | 1 | 0 | 1 | 1 | 6 | 0.3333 | NA | NA | NA | NA | 2 | 0.5 | NA | NA | 2 | 0.5 |
| 2_12106707 | 1 | 0 | 2 | 0 | 1 | 1 | 1 | 0 | 5 | 0.2 | NA | NA | 1 | 0 | 2 | 0.5 | NA | NA | 3 | 0.3333 |
| 2_12106711 | 2 | 0.5 | 2 | 0.5 | 1 | 1 | NA | NA | 5 | 0.6 | NA | NA | 1 | 1 | 2 | 0.5 | 1 | 0 | 4 | 0.5 |
| 2_12112985 | 2 | 0.5 | 1 | 0 | NA | NA | NA | NA | 3 | 0.3333 | 2 | 0.5 | 2 | 0 | 2 | 0 | 1 | 1 | 7 | 0.2857 |
| 2_12113004 | 2 | 1 | 1 | 1 | NA | NA | NA | NA | 3 | 1 | 2 | 1 | 1 | 1 | 2 | 1 | 1 | 1 | 6 | 1 |
| 2_12113013 | 2 | 0.5 | 1 | 0 | NA | NA | NA | NA | 3 | 0.3333 | 1 | 0 | 2 | 1 | 1 | 1 | 1 | 1 | 5 | 0.8 |
| 2_12113014 | 3 | 0.3333 | 1 | 1 | NA | NA | NA | NA | 4 | 0.5 | 2 | 0 | 2 | 0.5 | 2 | 0 | 1 | 1 | 7 | 0.2857 |
| 2_12113049 | 2 | 0.5 | 1 | 1 | NA | NA | NA | NA | 3 | 0.6667 | 2 | 1 | 2 | 0.5 | 2 | 0 | 1 | 1 | 7 | 0.5714 |
| 2_12113052 | 3 | 0.6667 | 2 | 1 | NA | NA | NA | NA | 5 | 0.8 | NA | NA | 2 | 1 | 2 | 1 | 1 | 1 | 5 | 1 |
| 2_12113075 | 3 | 0.3333 | 2 | 0.5 | NA | NA | 1 | 0 | 6 | 0.3333 | 2 | 0 | 2 | 0.5 | 2 | 0 | 1 | 1 | 7 | 0.2857 |
| 2_12113079 | 3 | 0.3333 | 1 | 0 | NA | NA | NA | NA | 4 | 0.25 | 2 | 0.5 | 2 | 0 | 2 | 0.5 | 1 | 0 | 7 | 0.2857 |
| 2_12113140 | 4 | 0.25 | 2 | 0 | 1 | 0 | 1 | 0 | 8 | 0.125 | 1 | 0 | 2 | 0.5 | 3 | 0 | 2 | 0 | 8 | 0.125 |
| 2_12113148 | 4 | 0.25 | 2 | 0 | 1 | 0 | 1 | 0 | 8 | 0.125 | 2 | 0.5 | 2 | 0 | 1 | 0 | 1 | 1 | 6 | 0.3333 |
| 2_12113150 | 4 | 0.75 | 1 | 1 | 1 | 0 | 1 | 1 | 7 | 0.7143 | 2 | 1 | 2 | 1 | 3 | 0.6667 | 2 | 1 | 9 | 0.8889 |
| 2_12113155 | 3 | 0.3333 | 2 | 0 | 1 | 0 | NA | NA | 6 | 0.1667 | 2 | 0 | 1 | 0 | 3 | 0.6667 | 2 | 0.5 | 8 | 0.375 |
| 2_12113155 | 4 | 0 | 1 | 1 | 1 | 1 | 1 | 0 | 7 | 0.2857 | 3 | 0.3333 | 2 | 1 | 2 | 0.5 | 2 | 0.5 | 9 | 0.5556 |
| 2_12113200 | 4 | 0.25 | 1 | 0 | 3 | 0.3333 | 1 | 0 | 9 | 0.2222 | 3 | 0 | 2 | 0.5 | 2 | 0 | 2 | 0 | 9 | 0.1111 |
| 2_12113213 | 5 | 0.6 | 2 | 0.5 | 2 | 0.5 | 1 | 0 | 10 | 0.5 | 3 | 0.6667 | 4 | 0.5 | 2 | 1 | 3 | 1 | 12 | 0.75 |
| 2_12113229 | 5 | 0.2 | 1 | 0 | 3 | 0 | 1 | 0 | 10 | 0.1 | 3 | 0.6667 | 4 | 0 | 4 | 0 | 2 | 0 | 13 | 0.1538 |
| 2_12113231 | 5 | 1 | 1 | 1 | 3 | 1 | 1 | 0 | 10 | 0.9 | 4 | 1 | 4 | 1 | 3 | 0.6667 | 2 | 1 | 13 | 0.9231 |
| 2_12113232 | 4 | 0.5 | 2 | 0 | 3 | 0.3333 | 1 | 0 | 10 | 0.3 | 4 | 0.25 | 4 | 0.5 | 2 | 0.5 | 2 | 0.5 | 12 | 0.4167 |
| 2_12113255 | 6 | 0.5 | 2 | 1 | 2 | 1 | 1 | 1 | 11 | 0.7273 | 3 | 0.6667 | 4 | 1 | 3 | 1 | 2 | 0.5 | 12 | 0.8333 |
| 2_12113274 | 7 | 0.1429 | 2 | 0 | 4 | 0.5 | 3 | 0.3333 | 16 | 0.25 | 6 | 0.1667 | 6 | 0.3333 | 3 | 0.3333 | 3 | 0 | 18 | 0.2222 |
| 2_12113278 | 5 | 0.2 | 1 | 0 | 3 | 0.3333 | 2 | 0.5 | 11 | 0.2727 | 2 | 0 | 2 | 0.5 | 1 | 0 | 1 | 0 | 6 | 0.1667 |
| 2_12113302 | 8 | 0 | 1 | 1 | 5 | 0 | 3 | 0 | 17 | 0.0588 | 6 | 0.1667 | 5 | 0.2 | 4 | 0 | 2 | 0 | 17 | 0.1176 |
| 2_12113309 | 8 | 0.5 | 1 | 1 | 5 | 0.8 | 3 | 0 | 17 | 0.5294 | 6 | 0.6667 | 6 | 0.3333 | 3 | 0.3333 | 4 | 0.5 | 19 | 0.4737 |
| 2_12113312 | 8 | 0 | 2 | 0 | 5 | 0.4 | 2 | 0.5 | 17 | 0.1765 | 6 | 0.6667 | 5 | 0.4 | 4 | 0.5 | 4 | 0.5 | 19 | 0.5263 |
| 2_12113315 | 6 | 0.1667 | NA | NA | 5 | 0 | 3 | 0 | 14 | 0.0714 | 5 | 0 | 4 | 0 | 3 | 0 | 4 | 0.25 | 16 | 0.0625 |
| 2_12113318 | 6 | 0.5 | 2 | 1 | 5 | 1 | 2 | 1 | 15 | 0.8 | 7 | 1 | 3 | 0.6667 | 4 | 0.75 | 2 | 1 | 16 | 0.875 |
| 2_12113323 | 7 | 0 | 2 | 0 | 4 | 0.5 | 3 | 0.3333 | 16 | 0.1875 | 6 | 0.3333 | 6 | 0.3333 | 2 | 1 | 3 | 0.3333 | 17 | 0.4118 |
| 2_12113334 | 8 | 0.25 | 1 | 0 | 4 | 0 | 3 | 0.3333 | 16 | 0.1875 | 7 | 0.1429 | 4 | 0 | 4 | 0.25 | 4 | 0.25 | 19 | 0.1579 |
| 2_12113341 | 7 | 0.1429 | 2 | 0.5 | 4 | 0.25 | 1 | 0 | 14 | 0.2143 | 6 | 0.5 | 4 | 0.25 | 4 | 0.25 | 3 | 0.6667 | 17 | 0.4118 |
| 2_12113379 | 8 | 0.625 | 3 | 1 | 4 | 0.5 | 2 | 0.5 | 17 | 0.6471 | 7 | 0.5714 | 4 | 0.75 | 3 | 0 | 4 | 0.25 | 18 | 0.4444 |
| 2_12113386 | 4 | 0 | 3 | 0 | 4 | 0.25 | 4 | 0 | 15 | 0.0667 | 7 | 0 | 5 | 0 | 3 | 0 | 3 | 0.3333 | 18 | 0.0556 |
| 2_12113396 | 10 | 0 | 2 | 0 | 6 | 0.1667 | 4 | 0.25 | 22 | 0.0909 | 8 | 0.125 | 7 | 0.1429 | 3 | 0 | 4 | 0.25 | 22 | 0.1364 |
| 2_12113400 | 10 | 0.1 | 3 | 0.3333 | 6 | 0 | 4 | 0.25 | 23 | 0.1304 | 9 | 0.1111 | 7 | 0.1429 | 4 | 0 | 4 | 0.25 | 24 | 0.125 |
| 2_12113409 | 11 | 0.0909 | 4 | 0.5 | 5 | 0 | 4 | 0.25 | 24 | 0.1667 | 9 | 0.3333 | 7 | 0.4286 | 4 | 0.5 | 4 | 0.5 | 24 | 0.4167 |
| 2_12113415 | 9 | 0.4444 | 3 | 1 | 4 | 0.25 | 2 | 0.5 | 18 | 0.5 | 6 | 0.6667 | 6 | 0.3333 | 3 | 0 | 3 | 0 | 18 | 0.3333 |
| 2_12113419 | 11 | 0.2727 | 4 | 0 | 6 | 0.3333 | 4 | 0.5 | 25 | 0.28 | 9 | 0.2222 | 7 | 0.1429 | 3 | 0.3333 | 4 | 0 | 23 | 0.1739 |
| 2_12113447 | 12 | 0.25 | 3 | 0.3333 | 6 | 0.5 | 3 | 0.3333 | 24 | 0.3333 | 10 | 0.4 | 8 | 0.125 | 4 | 0.25 | 4 | 0.5 | 26 | 0.3077 |
| 2_12113453 | 10 | 0.2 | 2 | 0 | 4 | 0 | 4 | 0.5 | 20 | 0.2 | 7 | 0.2857 | 5 | 0.2 | 4 | 0.25 | 4 | 0.25 | 20 | 0.25 |
| 2_12113487 | 15 | 0.2667 | 6 | 0 | 7 | 0 | 7 | 0.2857 | 35 | 0.1714 | 12 | 0.1667 | 9 | 0.2222 | 7 | 0.1429 | 6 | 0.1667 | 34 | 0.1765 |
| 2_12113487 | 16 | 0.25 | 7 | 0.2857 | 7 | 0 | 6 | 0.5 | 36 | 0.25 | 11 | 0.2727 | 9 | 0.4444 | 7 | 0.4286 | 6 | 0.3333 | 33 | 0.3636 |
| 2_12113495 | 15 | 0.3333 | 6 | 0.1667 | 6 | 0.1667 | 8 | 0.5 | 35 | 0.3143 | 13 | 0.4615 | 8 | 0.125 | 7 | 0.1429 | 6 | 0.3333 | 34 | 0.2941 |
| 2_12113502 | 13 | 0.2308 | 5 | 0.2 | 4 | 0.25 | 7 | 0.2857 | 29 | 0.2414 | 10 | 0.1 | 10 | 0.2 | 5 | 0.4 | 5 | 0.4 | 30 | 0.2333 |
| 2_12113516 | 16 | 0.375 | 6 | 0.1667 | 5 | 0.4 | 7 | 0.4286 | 34 | 0.3529 | 13 | 0.2308 | 9 | 0.1111 | 6 | 0.3333 | 7 | 0.5714 | 35 | 0.2857 |
| 2_12113517 | 15 | 0.4 | 7 | 0.2857 | 7 | 0.2857 | 6 | 0.1667 | 35 | 0.3143 | 9 | 0.1111 | 8 | 0.25 | 4 | 0.75 | 5 | 0 | 26 | 0.2308 |
| 2_12113517 | 15 | 0.2667 | 4 | 0.25 | 6 | 0.3333 | 8 | 0.75 | 33 | 0.3939 | 11 | 0.5455 | 8 | 0.625 | 3 | 0 | 6 | 0.5 | 28 | 0.5 |
| 2_12113523 | 12 | 0.25 | 6 | 0.5 | 5 | 0.4 | 4 | 0.75 | 27 | 0.4074 | 13 | 0.3846 | 10 | 0.5 | 6 | 0.5 | 7 | 0.4286 | 36 | 0.4444 |
| 2_12113525 | 15 | 0.1333 | 7 | 0 | 7 | 0.1429 | 8 | 0.25 | 37 | 0.1351 | 12 | 0.25 | 10 | 0.3 | 9 | 0 | 7 | 0.1429 | 38 | 0.1842 |
| 2_12113527 | 15 | 0.1333 | 7 | 0.2857 | 7 | 0 | 9 | 0.1111 | 38 | 0.1316 | 16 | 0.125 | 7 | 0.2857 | 9 | 0.1111 | 7 | 0.2857 | 39 | 0.1795 |
| 2_12113530 | 13 | 0.1538 | 5 | 0 | 7 | 0.2857 | 8 | 0.25 | 33 | 0.1818 | 12 | 0.0833 | 8 | 0.125 | 8 | 0.5 | 5 | 0.4 | 33 | 0.2424 |
| 2_12113535 | 14 | 0.4286 | 5 | 0.8 | 7 | 0.4286 | 8 | 0.75 | 34 | 0.5588 | 15 | 0.4667 | 9 | 0.6667 | 10 | 0.5 | 6 | 0.8333 | 40 | 0.575 |
| 2_12113546 | 14 | 0 | 3 | 0 | 7 | 0.2857 | 7 | 0.2857 | 31 | 0.129 | 14 | 0.2143 | 5 | 0.2 | 7 | 0.1429 | 6 | 0.1667 | 32 | 0.1875 |

| | | | | | | | | | | | | | | | | | | | | |
|---|---|---|---|---|---|---|---|---|---|---|---|---|---|---|---|---|---|---|---|---|
| 2_12113553 | 19 | 0.8947 | 7 | 0.7143 | 7 | 0.5714 | 8 | 1 | 41 | 0.8293 | 12 | 0.9167 | 12 | 0.8333 | 10 | 0.8 | 6 | 0.6667 | 40 | 0.825 |
| 2_12113565 | 17 | 0.1765 | 9 | 0.1111 | 10 | 0.1 | 11 | 0.3636 | 47 | 0.1915 | 17 | 0.2353 | 14 | 0.1429 | 12 | 0.1667 | 8 | 0.375 | 51 | 0.2157 |
| 2_12113572 | 15 | 0.2 | 4 | 0.5 | 8 | 0.25 | 11 | 0.1818 | 38 | 0.2368 | 12 | 0.0833 | 9 | 0.1111 | 11 | 0.1818 | 3 | 0 | 35 | 0.1143 |
| 2_12113593 | 19 | 0.2105 | 13 | 0.0769 | 12 | 0.1667 | 13 | 0.3077 | 57 | 0.193 | 18 | 0.1667 | 13 | 0.1538 | 12 | 0 | 7 | 0.2857 | 50 | 0.14 |
| 2_12113595 | 18 | 0.0556 | 11 | 0.1818 | 10 | 0.1 | 12 | 0.1667 | 51 | 0.1176 | 17 | 0.1765 | 11 | 0.0909 | 11 | 0.1818 | 8 | 0 | 47 | 0.1277 |
| 2_12113614 | 14 | 0.2857 | 10 | 0.6 | 8 | 0.375 | 10 | 0.5 | 42 | 0.4286 | 13 | 0.3846 | 10 | 0.6 | 11 | 0.3636 | 8 | 0.375 | 42 | 0.4286 |
| 2_12113614 | 15 | 0.1333 | 7 | 0 | 6 | 0 | 9 | 0 | 37 | 0.0541 | 10 | 0 | 8 | 0.25 | 10 | 0.1 | 7 | 0.1429 | 35 | 0.1143 |
| 2_12113618 | 24 | 0.0833 | 13 | 0.1538 | 14 | 0.0714 | 15 | 0.2 | 66 | 0.1212 | 21 | 0.0952 | 16 | 0.3125 | 16 | 0.125 | 11 | 0.0909 | 64 | 0.1562 |
| 2_12113624 | 20 | 0.15 | 15 | 0 | 13 | 0 | 12 | 0.0833 | 60 | 0.0667 | 19 | 0.1053 | 14 | 0.0714 | 13 | 0.1538 | 13 | 0.0769 | 59 | 0.1017 |
| 2_12113627 | 19 | 0.6316 | 13 | 0.6154 | 14 | 0.7143 | 11 | 0.7273 | 57 | 0.6667 | 16 | 0.5 | 12 | 0.5 | 14 | 0.4286 | 9 | 0.3333 | 51 | 0.451 |
| 2_12113628 | 23 | 0.1739 | 12 | 0.3333 | 15 | 0.2667 | 18 | 0.1667 | 68 | 0.2206 | 22 | 0.0909 | 14 | 0.2143 | 18 | 0.1111 | 13 | 0.1538 | 67 | 0.1343 |
| 2_12113630 | 21 | 0.6667 | 16 | 0.625 | 14 | 0.6429 | 15 | 0.6 | 66 | 0.6364 | 21 | 0.2857 | 17 | 0.6471 | 15 | 0.5333 | 12 | 0.8333 | 65 | 0.5385 |
| 2_12113641 | 12 | 0.1667 | 15 | 0.1333 | 11 | 0 | 12 | 0.0833 | 50 | 0.1 | 17 | 0.0588 | 13 | 0.1538 | 11 | 0.0909 | 8 | 0.25 | 49 | 0.1224 |
| 2_12113644 | 25 | 0.36 | 18 | 0.2778 | 16 | 0.0625 | 16 | 0.125 | 75 | 0.2267 | 20 | 0.15 | 17 | 0.4706 | 18 | 0.1111 | 16 | 0.25 | 71 | 0.2394 |
| 2_12113667 | 24 | 0.0833 | 17 | 0.0588 | 18 | 0.1111 | 17 | 0.1765 | 76 | 0.1053 | 20 | 0.15 | 13 | 0 | 16 | 0.0625 | 10 | 0 | 59 | 0.0678 |
| 2_12113672 | 25 | 0.32 | 20 | 0.4 | 17 | 0.1765 | 18 | 0.3333 | 80 | 0.3125 | 22 | 0.1364 | 20 | 0.35 | 15 | 0.2667 | 17 | 0.6471 | 74 | 0.3378 |
| 2_12113676 | 27 | 0.3704 | 24 | 0.5417 | 16 | 0.375 | 19 | 0.5263 | 86 | 0.4535 | 23 | 0.3913 | 21 | 0.619 | 18 | 0.6111 | 15 | 0.4667 | 77 | 0.5195 |
| 2_12113685 | 26 | 0.1538 | 23 | 0.087 | 18 | 0.1667 | 20 | 0.1 | 87 | 0.1264 | 22 | 0.2727 | 19 | 0.0526 | 18 | 0.1667 | 15 | 0.0667 | 74 | 0.1486 |
| 2_12113712 | 24 | 0.2917 | 26 | 0.3846 | 22 | 0.1818 | 21 | 0.2857 | 93 | 0.2903 | 27 | 0.2222 | 24 | 0.3333 | 22 | 0.2727 | 19 | 0.2105 | 92 | 0.2609 |
| 2_12113718 | 29 | 0.1379 | 23 | 0.0435 | 21 | 0.2381 | 21 | 0.1429 | 94 | 0.1383 | 24 | 0.125 | 19 | 0.0526 | 15 | 0.2 | 18 | 0.1667 | 76 | 0.1316 |
| 2_12113722 | 30 | 0.0667 | 26 | 0.1538 | 23 | 0.1739 | 25 | 0.12 | 104 | 0.125 | 29 | 0.1034 | 24 | 0.25 | 22 | 0.1364 | 21 | 0.0952 | 96 | 0.1458 |
| 2_12113738 | 33 | 0.4545 | 24 | 0.3333 | 22 | 0.2273 | 20 | 0.35 | 99 | 0.3535 | 30 | 0.3333 | 23 | 0.2609 | 20 | 0.3 | 18 | 0.3333 | 91 | 0.3077 |
| 2_12113742 | 33 | 0.0909 | 32 | 0.125 | 23 | 0.2609 | 21 | 0.0476 | 109 | 0.1284 | 32 | 0.0625 | 27 | 0.0741 | 27 | 0.037 | 19 | 0.1579 | 105 | 0.0762 |
| 2_12113744 | 26 | 0.6923 | 26 | 0.7308 | 20 | 0.65 | 17 | 0.7647 | 89 | 0.7079 | 31 | 0.7419 | 26 | 0.7692 | 23 | 0.8261 | 19 | 0.6316 | 99 | 0.7475 |
| 2_12113745 | 21 | 0.8571 | 23 | 0.7391 | 15 | 0.9333 | 12 | 0.75 | 71 | 0.8169 | 21 | 0.7619 | 16 | 0.9375 | 19 | 0.9474 | 17 | 0.8235 | 73 | 0.863 |
| 2_12113747 | 26 | 0.4231 | 26 | 0.4615 | 14 | 0.3571 | 19 | 0.3684 | 85 | 0.4118 | 29 | 0.3448 | 24 | 0.5417 | 22 | 0.3636 | 19 | 0.6316 | 94 | 0.4574 |
| 2_12113752 | 30 | 0.2667 | 31 | 0.2903 | 26 | 0.1538 | 23 | 0.2174 | 110 | 0.2364 | 31 | 0.2258 | 25 | 0.24 | 23 | 0.1304 | 16 | 0.0625 | 95 | 0.1789 |
| 2_12113760 | 30 | 0.8667 | 32 | 0.9688 | 26 | 0.9231 | 30 | 0.9333 | 118 | 0.9237 | 32 | 0.9062 | 30 | 0.8667 | 25 | 0.92 | 24 | 1 | 111 | 0.9189 |
| 2_12113808 | 28 | 0.2143 | 37 | 0.1622 | 37 | 0.1081 | 30 | 0.3333 | 132 | 0.197 | 38 | 0.1842 | 31 | 0.1935 | 31 | 0.1935 | 24 | 0.1667 | 124 | 0.1855 |
| 2_12113868 | 35 | 0.8 | 47 | 0.7447 | 39 | 0.7949 | 44 | 0.6818 | 165 | 0.7515 | 49 | 0.7959 | 35 | 0.8857 | 47 | 0.6809 | 44 | 0.8864 | 175 | 0.8057 |
| 2_12113954 | 60 | 0.2833 | 59 | 0.1017 | 55 | 0.1273 | 49 | 0.1429 | 223 | 0.1659 | 52 | 0.1154 | 46 | 0.2391 | 52 | 0.1538 | 49 | 0.1633 | 199 | 0.1658 |
| 2_12113971 | 84 | 0.2381 | 72 | 0.2639 | 70 | 0.2143 | 64 | 0.125 | 290 | 0.2138 | 74 | 0.2432 | 58 | 0.1552 | 65 | 0.2769 | 61 | 0.2131 | 258 | 0.2248 |
| 2_12114097 | NA | NA | 4 | 0.25 | 6 | 0.1667 | NA | NA | 10 | 0.2 | 2 | 0 | 4 | 0 | 1 | 0 | 5 | 0.4 | 12 | 0.1667 |
| 2_12114143 | 6 | 0.5 | 6 | 0.5 | 7 | 0.1429 | 6 | 0.3333 | 25 | 0.36 | 6 | 0.1667 | 3 | 0 | 7 | 0.1429 | 4 | 0.25 | 20 | 0.15 |
| 2_12114144 | 6 | 0.1667 | 9 | 0.1111 | 6 | 0 | 6 | 0 | 27 | 0.0741 | 6 | 0.3333 | 4 | 0.25 | 5 | 0 | 4 | 0 | 19 | 0.1579 |
| 2_12114150 | 6 | 0.1667 | 9 | 0.3333 | 6 | 0 | 5 | 0.4 | 26 | 0.2308 | 6 | 0.6667 | 4 | 0.25 | 7 | 0.2857 | 5 | 0.4 | 22 | 0.4091 |
| 2_12114160 | 4 | 0.5 | 7 | 0.1429 | 8 | 0.25 | 6 | 0.3333 | 25 | 0.28 | 6 | 0.1667 | 4 | 0.25 | 6 | 0.3333 | 4 | 0.25 | 20 | 0.25 |
| 2_12114172 | 4 | 0.5 | 7 | 0.4286 | 7 | 0.5714 | 6 | 0.5 | 24 | 0.5 | 7 | 0.7143 | 4 | 1 | 6 | 0.8333 | 5 | 0.8 | 22 | 0.8182 |
| 2_12114176 | 3 | 0 | 5 | 0 | 5 | 0 | 4 | 0.25 | 17 | 0.0588 | 7 | 0.1429 | 4 | 0.25 | 6 | 0.1667 | 4 | 0 | 21 | 0.1429 |
| 2_12114179 | 3 | 0 | 7 | 0.2857 | 7 | 0 | 6 | 0.5 | 23 | 0.2174 | 6 | 0 | 3 | 0 | 7 | 0.1429 | 5 | 0.2 | 21 | 0.0952 |
| 2_12114180 | 5 | 0.8 | 5 | 0.8 | 7 | 1 | 6 | 0.6667 | 23 | 0.8261 | 6 | 0.8333 | 4 | 0.5 | 7 | 0.8571 | 5 | 1 | 22 | 0.8182 |
| 2_12114191 | 3 | 1 | 5 | 0.8 | 6 | 0.8333 | 5 | 1 | 19 | 0.8947 | 6 | 0.8333 | 4 | 0.5 | 7 | 0.8571 | 5 | 1 | 22 | 0.8182 |
| 2_12114198 | 4 | 0.25 | 5 | 0 | 4 | 0 | 4 | 0 | 17 | 0.0588 | 3 | 0.3333 | 3 | 0 | 7 | 0 | 4 | 0.25 | 17 | 0.1176 |
| 2_12114205 | 3 | 1 | 6 | 1 | 7 | 0.4286 | 4 | 0.75 | 20 | 0.75 | 6 | 1 | 3 | 1 | 6 | 1 | 3 | 1 | 18 | 1 |
| 2_12114211 | 5 | 0 | 5 | 0.2 | 7 | 0.1429 | 3 | 0 | 20 | 0.1 | 4 | 0.25 | 4 | 0.5 | 3 | 0 | 4 | 0.5 | 15 | 0.3333 |
| 2_12114222 | 4 | 0.25 | 5 | 0 | 5 | 0.4 | 2 | 0 | 16 | 0.1875 | 3 | 0.6667 | 3 | 0.6667 | 5 | 0.2 | 4 | 0 | 15 | 0.3333 |
| 2_12114224 | 3 | 0 | 5 | 0.6 | 5 | 0 | 4 | 0.25 | 17 | 0.2353 | 6 | 0.1667 | 4 | 0.5 | 4 | 0 | 5 | 0.2 | 19 | 0.2105 |
| 2_12114225 | 5 | 0.2 | 6 | 0 | 6 | 0 | 4 | 0.5 | 21 | 0.1429 | 6 | 0.1667 | 4 | 0 | 5 | 0 | 4 | 0 | 19 | 0.0526 |
| 2_12114226 | 2 | 0.5 | 6 | 0.3333 | 7 | 0.1429 | 4 | 0.25 | 19 | 0.2632 | 6 | 0.1667 | 3 | 0 | 6 | 0.3333 | 5 | 0.2 | 20 | 0.2 |
| 2_12114257 | 3 | 1 | 4 | 0.75 | 6 | 0.8333 | 4 | 0.5 | 17 | 0.7647 | 4 | 0.75 | 4 | 0.75 | 6 | 0.5 | 5 | 0.8 | 19 | 0.6842 |
| 2_12114277 | 4 | 0.25 | 2 | 0 | 4 | 0.25 | 4 | 0.25 | 14 | 0.2143 | 2 | 0 | 3 | 0.3333 | 6 | 0.3333 | 4 | 0.5 | 15 | 0.3333 |
| 2_12114279 | 2 | 0 | 3 | 0 | 5 | 0 | 4 | 0.25 | 14 | 0.0714 | 4 | 0.25 | 4 | 0.25 | 6 | 0.1667 | 3 | 0.3333 | 17 | 0.2353 |
| 2_12114285 | 3 | 0.6667 | 4 | 0.25 | 5 | 0.4 | 4 | 0.25 | 16 | 0.375 | 3 | 0.3333 | 2 | 0 | 4 | 0.5 | 4 | 0.75 | 13 | 0.4615 |
| 2_12114289 | 4 | 0.25 | 4 | 0 | 6 | 0 | 4 | 0 | 18 | 0.0556 | 3 | 0 | 4 | 0 | 5 | 0.2 | 4 | 0.5 | 16 | 0.1875 |
| 2_12117772 | NA | NA | 1 | 1 | 2 | 0.5 | NA | NA | 3 | 0.6667 | 1 | 1 | NA | NA | NA | NA | 4 | 0.25 | 5 | 0.4 |
| 2_12118937 | 18 | 0.0556 | 17 | 0 | 13 | 0.2308 | 26 | 0.0385 | 74 | 0.0676 | 13 | 0 | 15 | 0 | 16 | 0.125 | 15 | 0.0667 | 59 | 0.0508 |
| 2_12119413 | 40 | 0.175 | 43 | 0.186 | 33 | 0.1515 | 42 | 0.0714 | 158 | 0.1456 | 38 | 0.0789 | 25 | 0.28 | 35 | 0.2 | 29 | 0.2759 | 127 | 0.1969 |
| 2_12119417 | 37 | 0.027 | 38 | 0.1316 | 31 | 0.0645 | 42 | 0.0238 | 148 | 0.0608 | 35 | 0 | 26 | 0.1154 | 32 | 0.0625 | 28 | 0.1429 | 121 | 0.0744 |
| 2_12119420 | 32 | 0.1562 | 35 | 0.1429 | 28 | 0.1071 | 32 | 0.1562 | 127 | 0.1417 | 39 | 0.2051 | 25 | 0.12 | 34 | 0.2059 | 26 | 0.1538 | 124 | 0.1774 |
| 2_12126821 | 35 | 0.9143 | 21 | 0.8095 | 19 | 0.8947 | 19 | 0.9474 | 94 | 0.8936 | 28 | 0.9286 | 38 | 0.8684 | 26 | 0.9231 | 25 | 1 | 117 | 0.9231 |
| 2_12126824 | 37 | 0.3784 | 25 | 0.24 | 23 | 0.4348 | 21 | 0.3333 | 106 | 0.3491 | 30 | 0.2 | 35 | 0.1714 | 24 | 0.2083 | 27 | 0.4815 | 116 | 0.2586 |
| 2_12126828 | 36 | 0.2222 | 23 | 0.2609 | 20 | 0.3 | 21 | 0.2381 | 100 | 0.25 | 31 | 0.1935 | 38 | 0.1053 | 25 | 0.16 | 25 | 0.28 | 119 | 0.1765 |
| 2_12126832 | 37 | 0.0811 | 25 | 0.12 | 21 | 0.1429 | 26 | 0.3846 | 109 | 0.1743 | 32 | 0.1875 | 40 | 0.175 | 26 | 0.1923 | 27 | 0.4444 | 125 | 0.24 |
| 2_12126980 | 1 | 0 | 1 | 0 | 1 | 1 | NA | NA | 3 | 0.3333 | 3 | 0 | 1 | 0 | 2 | 0.5 | 2 | 0.5 | 8 | 0.25 |
| 2_12126985 | 1 | 1 | 1 | 1 | 1 | 0 | NA | NA | 3 | 0.6667 | 4 | 0.5 | 1 | 1 | 2 | 0.5 | 2 | 1 | 9 | 0.6667 |

| | | | | | | | | | | | | | | | | | | | | |
|---|---|---|---|---|---|---|---|---|---|---|---|---|---|---|---|---|---|---|---|---|
| 2_12126991 | 1 | 1 | 1 | 1 | 1 | 1 | NA | NA | 3 | 1 | 3 | 1 | 1 | 1 | 2 | 1 | 1 | 0 | 7 | 0.8571 |
| 2_12126995 | 1 | 1 | 1 | 0 | 1 | 0 | NA | NA | 3 | 0.3333 | 3 | 0 | 1 | 1 | 2 | 0 | 2 | 0.5 | 8 | 0.25 |
| 2_12126997 | 1 | 1 | 1 | 1 | 1 | 1 | NA | NA | 3 | 1 | 3 | 0.6667 | NA | NA | 2 | 1 | 2 | 1 | 7 | 0.8571 |
| 2_12127011 | NA | NA | NA | NA | 1 | 1 | NA | NA | 1 | 1 | 1 | 1 | NA | NA | 1 | 1 | NA | NA | 2 | 1 |
| 2_12127013 | NA | NA | NA | NA | 1 | 1 | NA | NA | 1 | 1 | 1 | 1 | NA | NA | 1 | 1 | NA | NA | 2 | 1 |
| 2_12128155 | 3 | 0 | 3 | 0 | 2 | 0 | 4 | 0.25 | 12 | 0.0833 | 2 | 0.5 | 3 | 0 | 2 | 0 | 4 | 0 | 11 | 0.0909 |
| 2_12128174 | 3 | 0 | 5 | 0 | 2 | 0 | 4 | 0.5 | 14 | 0.1429 | 4 | 0.25 | 6 | 0 | 2 | 0 | 4 | 0 | 16 | 0.0625 |
| 2_12128175 | 3 | 0 | 5 | 0 | 2 | 0 | 5 | 0.2 | 15 | 0.0667 | 3 | 0.3333 | 6 | 0 | 2 | 0 | 4 | 0.75 | 15 | 0.2667 |
| 2_12128875 | 13 | 0.2308 | 10 | 0.4 | 15 | 0.4667 | 17 | 0.0588 | 55 | 0.2727 | 18 | 0.2222 | 12 | 0.3333 | 14 | 0.2143 | 16 | 0.375 | 60 | 0.2833 |
| 2_12129167 | 9 | 0.1111 | 13 | 0.0769 | 11 | 0.1818 | 14 | 0.2143 | 47 | 0.1489 | 8 | 0 | 11 | 0.0909 | 11 | 0.2727 | 8 | 0.125 | 38 | 0.1316 |
| 2_12129168 | 7 | 0.1429 | 13 | 0.1538 | 10 | 0.2 | 14 | 0.1429 | 44 | 0.1591 | 8 | 0.25 | 9 | 0.3333 | 10 | 0.2 | 6 | 0.1667 | 33 | 0.2424 |
| 2_12129230 | 4 | 0.25 | 11 | 0.2727 | 10 | 0.4 | 13 | 0.2308 | 38 | 0.2895 | 6 | 0.3333 | 7 | 0.2857 | 10 | 0.3 | 9 | 0.3333 | 32 | 0.3125 |
| 2_12129234 | 6 | 0.6667 | 13 | 0.7692 | 9 | 0.8889 | 11 | 0.6364 | 39 | 0.7436 | 6 | 0.8333 | 7 | 0.7143 | 9 | 0.5556 | 7 | 0.7143 | 29 | 0.6897 |
| 2_12129235 | 7 | 0 | 11 | 0 | 10 | 0.1 | 13 | 0.1538 | 41 | 0.0732 | 6 | 0 | 9 | 0.2222 | 7 | 0.1429 | 8 | 0.125 | 30 | 0.1333 |
| 2_12129246 | 7 | 1 | 10 | 1 | 12 | 1 | 14 | 1 | 43 | 1 | 4 | 1 | 8 | 1 | 9 | 0.8889 | 9 | 1 | 30 | 0.9667 |
| 2_12129251 | 8 | 0.25 | 12 | 0.4167 | 11 | 0.1818 | 13 | 0.3077 | 44 | 0.2955 | 4 | 0.25 | 9 | 0.4444 | 10 | 0.4 | 9 | 0.3333 | 32 | 0.375 |
| 2_12129257 | 6 | 1 | 9 | 0.7778 | 9 | 0.8889 | 11 | 0.9091 | 35 | 0.8857 | 2 | 1 | 7 | 1 | 7 | 0.7143 | 7 | 1 | 23 | 0.913 |
| 2_12129271 | 5 | 0 | 8 | 0.125 | 8 | 0.125 | 9 | 0 | 30 | 0.0667 | NA | NA | 5 | 0 | 6 | 0.3333 | 4 | 0.5 | 15 | 0.2667 |
| 2_12130064 | 4 | 0.25 | 7 | 0.1429 | 10 | 0.3 | 8 | 0 | 29 | 0.1724 | 2 | 0 | 4 | 0.5 | 5 | 0.4 | 8 | 0.25 | 19 | 0.3158 |
| 2_12131203 | NA | NA | NA | NA | 1 | 1 | NA | NA | 1 | 1 | NA | NA | 2 | 1 | NA | NA | NA | NA | 2 | 1 |
| 2_12132485 | NA | NA | 4 | 0.5 | 4 | 0.25 | 7 | 0.2857 | 15 | 0.3333 | NA | NA | 8 | 0.375 | 5 | 0.4 | 5 | 0.6 | 18 | 0.4444 |
| 2_12132489 | 2 | 1 | 6 | 1 | 3 | 0.6667 | 4 | 1 | 15 | 0.9333 | 1 | 1 | 8 | 1 | 6 | 0.3333 | 4 | 1 | 19 | 0.7895 |
| 2_12132489 | 2 | 0.5 | 6 | 0.1667 | 4 | 0.25 | 6 | 0.1667 | 18 | 0.2222 | 1 | 0 | 9 | 0.2222 | 6 | 0 | 5 | 0.2 | 21 | 0.1429 |
| 2_12132500 | 2 | 1 | 6 | 1 | 4 | 1 | 7 | 1 | 19 | 1 | NA | NA | 9 | 1 | 5 | 1 | 4 | 1 | 18 | 1 |
| 2_12132505 | 3 | 0.6667 | 6 | 0.5 | 4 | 0.25 | 6 | 0.6667 | 19 | 0.5263 | NA | NA | 8 | 0.75 | 6 | 0.1667 | 5 | 0.2 | 19 | 0.4211 |
| 2_12132512 | 3 | 1 | 3 | 1 | 3 | 0.6667 | 4 | 0.75 | 13 | 0.8462 | NA | NA | 4 | 1 | 5 | 1 | 3 | 1 | 12 | 1 |
| 2_12132522 | 1 | 0 | 5 | 0.6 | 4 | 0 | 6 | 0 | 16 | 0.1875 | NA | NA | 8 | 0.125 | 6 | 0 | 4 | 0 | 18 | 0.0556 |
| 2_12132525 | 2 | 0.5 | 4 | 0.25 | 2 | 1 | 1 | 0 | 9 | 0.4444 | NA | NA | 3 | 0.6667 | 1 | 1 | 3 | 0 | 7 | 0.4286 |
| 2_12133319 | 2 | 0.5 | 4 | 0.25 | 3 | 0 | 3 | 0.6667 | 12 | 0.3333 | NA | NA | 9 | 0.3333 | 4 | 0 | 3 | 0.3333 | 16 | 0.25 |
| 2_12137622 | 1 | 1 | 2 | 0.5 | 1 | 1 | 2 | 0 | 6 | 0.5 | 1 | 0 | 2 | 1 | NA | NA | 3 | 0.3333 | 6 | 0.5 |
| 2_12137646 | 1 | 0 | 2 | 0.5 | 1 | 0 | 2 | 0 | 6 | 0.1667 | 1 | 0 | 2 | 0.5 | NA | NA | 4 | 0.25 | 7 | 0.2857 |
| 2_12137672 | 1 | 1 | 2 | 0 | 1 | 1 | 2 | 0.5 | 6 | 0.5 | 1 | 0 | 2 | 0.5 | NA | NA | 4 | 0 | 7 | 0.1429 |
| 2_12137676 | 1 | 1 | 2 | 0 | 1 | 1 | 2 | 0.5 | 6 | 0.5 | 1 | 0 | 2 | 0.5 | NA | NA | 4 | 0.5 | 7 | 0.4286 |
| 2_12137685 | 1 | 1 | 2 | 0 | NA | NA | 2 | 0.5 | 5 | 0.4 | 1 | 0 | 2 | 0 | NA | NA | 4 | 0.5 | 7 | 0.2857 |
| 2_12137712 | 1 | 1 | 2 | 1 | 1 | 1 | 1 | 0 | 5 | 0.8 | 1 | 0 | 2 | 0.5 | NA | NA | 3 | 0.3333 | 6 | 0.3333 |
| 2_12137776 | 1 | 1 | 2 | 0 | 1 | 0 | 1 | 0 | 5 | 0.2 | 1 | 0 | 1 | 0 | NA | NA | 2 | 1 | 4 | 0.5 |
| 2_12137873 | 1 | 0 | 2 | 0 | 2 | 0.5 | NA | NA | 5 | 0.2 | 1 | 0 | 2 | 0.5 | 1 | 1 | 1 | 0 | 5 | 0.4 |
| 2_12137885 | 2 | 0.5 | 3 | 0 | 2 | 0 | NA | NA | 7 | 0.1429 | 3 | 0 | 6 | 0.1667 | 4 | 0 | 3 | 0 | 16 | 0.0625 |
| 2_12137894 | 3 | 0.3333 | 3 | 1 | 2 | 1 | 1 | 1 | 9 | 0.7778 | 3 | 1 | 5 | 0.8 | 4 | 0.25 | 3 | 0.3333 | 15 | 0.6 |
| 2_12137902 | 3 | 0 | 1 | 1 | 2 | 0 | 1 | 0 | 7 | 0.1429 | 2 | 0.5 | 3 | 0 | 3 | 0 | 1 | 0 | 9 | 0.1111 |
| 2_12137906 | 3 | 0 | 2 | 0.5 | 3 | 0 | 1 | 0 | 9 | 0.1111 | 3 | 0 | 6 | 0.1667 | 4 | 0 | 2 | 0 | 15 | 0.0667 |
| 2_12137910 | 3 | 0.3333 | 2 | 0 | 3 | 0 | 1 | 0 | 9 | 0.1111 | 3 | 0 | 6 | 0.6667 | 3 | 0.3333 | 2 | 0 | 14 | 0.3571 |
| 2_12137912 | 3 | 0.3333 | 2 | 0 | 2 | 0.5 | 1 | 0 | 8 | 0.25 | 2 | 0 | 6 | 0.3333 | 3 | 0 | 2 | 0 | 13 | 0.1538 |
| 2_12137922 | 3 | 0.3333 | 1 | 0 | 1 | 1 | NA | NA | 5 | 0.4 | 1 | 1 | 2 | 0.5 | 1 | 0 | 2 | 0.5 | 6 | 0.5 |
| 2_12137931 | 3 | 1 | 1 | 1 | 2 | 1 | NA | NA | 6 | 1 | 1 | 1 | 2 | 1 | 1 | 1 | 2 | 1 | 6 | 1 |
| 2_12137932 | 3 | 0.6667 | 1 | 0 | NA | NA | NA | NA | 4 | 0.5 | 2 | 0.5 | 3 | 0.3333 | 1 | 0 | 1 | 1 | 7 | 0.4286 |
| 2_12137937 | 4 | 0.25 | 2 | 0 | 2 | 0.5 | 1 | 0 | 9 | 0.2222 | 3 | 0.3333 | 4 | 0.5 | 2 | 0 | 2 | 0 | 11 | 0.2727 |
| 2_12137956 | 4 | 0 | 2 | 0 | 2 | 0.5 | 1 | 0 | 9 | 0.1111 | 3 | 0.3333 | 6 | 0.5 | 3 | 0 | 1 | 0 | 13 | 0.3077 |
| 2_12169789 | 5 | 0.4 | 6 | 0.5 | 4 | 0 | 5 | 0.2 | 20 | 0.3 | 2 | 0 | 4 | 0 | 2 | 0.5 | 5 | 0.2 | 13 | 0.1538 |
| 2_12169790 | 6 | 0 | 5 | 0 | 3 | 0.3333 | 5 | 0 | 19 | 0.0526 | 2 | 0 | 3 | 0 | 2 | 0 | 5 | 0.2 | 12 | 0.0833 |
| 2_12176409 | 10 | 0.2 | 11 | 0.0909 | 5 | 0 | 9 | 0 | 35 | 0.0857 | 3 | 0 | 4 | 0.25 | 6 | 0 | 6 | 0 | 19 | 0.0526 |
| 2_12176418 | 8 | 0.375 | 11 | 0.4545 | 6 | 0.3333 | 9 | 0.1111 | 34 | 0.3235 | 3 | 0.6667 | 5 | 0.4 | 4 | 0.25 | 8 | 0.125 | 20 | 0.3 |
| 2_12176481 | 12 | 0.25 | 12 | 0 | 8 | 0.125 | 8 | 0.125 | 40 | 0.125 | 3 | 0.6667 | 5 | 0.2 | 5 | 0.2 | 8 | 0.5 | 21 | 0.381 |
| 2_12184111 | NA | NA | 1 | 1 | 3 | 0.6667 | 1 | 0 | 5 | 0.6 | NA | NA | 1 | 1 | 3 | 0.3333 | 3 | 0.6667 | 7 | 0.5714 |
| 2_12184116 | NA | NA | 1 | 1 | 3 | 1 | 1 | 1 | 5 | 1 | NA | NA | 1 | 1 | 2 | 1 | 2 | 0.5 | 5 | 0.8 |
| 2_12184127 | NA | NA | 1 | 1 | 3 | 1 | 1 | 0 | 5 | 0.8 | NA | NA | 1 | 1 | 2 | 0.5 | 3 | 0.6667 | 6 | 0.6667 |
| 2_12184144 | NA | NA | 1 | 1 | 2 | 0.5 | 1 | 1 | 4 | 0.75 | NA | NA | 1 | 0 | 3 | 0.3333 | 3 | 0.3333 | 7 | 0.2857 |
| 2_12184243 | 1 | 0 | 1 | 1 | 3 | 0.3333 | 1 | 0 | 6 | 0.3333 | NA | NA | 3 | 0 | 3 | 0.3333 | 3 | 0 | 9 | 0.1111 |
| 2_12184427 | 2 | 0 | 2 | 0.5 | 1 | 0 | NA | NA | 5 | 0.2 | NA | NA | 2 | 0.5 | 2 | 0 | 2 | 0 | 6 | 0.1667 |
| 2_12184438 | 1 | 1 | 2 | 0 | 3 | 0.6667 | NA | NA | 6 | 0.5 | NA | NA | 2 | 0 | 3 | 0.3333 | 3 | 0 | 8 | 0.125 |
| 2_12184460 | 1 | 0 | 2 | 1 | 2 | 1 | NA | NA | 5 | 0.8 | NA | NA | 2 | 0 | 3 | 0.6667 | 3 | 0.6667 | 8 | 0.5 |
| 2_12184565 | 3 | 0.6667 | 4 | 0.25 | 4 | 0 | 1 | 0 | 12 | 0.25 | 1 | 0 | 2 | 0 | 4 | 0 | 3 | 0.3333 | 10 | 0.1 |
| 2_12188257 | 7 | 0 | 9 | 0.2222 | 5 | 0 | 2 | 0.5 | 23 | 0.1304 | 4 | 0.25 | 7 | 0 | 2 | 0 | 5 | 0 | 18 | 0.0556 |
| 2_12188274 | 7 | 0 | 9 | 0.2222 | 4 | 0.5 | 1 | 0 | 21 | 0.1905 | 5 | 0 | 7 | 0.1429 | 2 | 0 | 5 | 0.2 | 19 | 0.1053 |
| 2_12188295 | 4 | 0.25 | 10 | 0 | 5 | 0 | 2 | 0.5 | 21 | 0.0952 | 5 | 0.4 | 4 | 0.25 | 2 | 0.5 | 5 | 0 | 16 | 0.25 |

| | | | | | | | | | | | | | | | | | | | | |
|---|---|---|---|---|---|---|---|---|---|---|---|---|---|---|---|---|---|---|---|---|
| 2_12188320 | 7 | 0.7143 | 13 | 0.4615 | 4 | 0.5 | 3 | 0.3333 | 27 | 0.5185 | 7 | 0.7143 | 8 | 0.75 | 4 | 0.5 | 5 | 0.2 | 24 | 0.5833 |
| 2_12188326 | 7 | 0.4286 | 12 | 0.5 | 5 | 1 | 3 | 0.3333 | 27 | 0.5556 | 9 | 0.3333 | 10 | 0.5 | 4 | 0.25 | 6 | 0.1667 | 29 | 0.3448 |
| 2_12188369 | 4 | 0.25 | 7 | 0 | 3 | 0 | 1 | 0 | 15 | 0.0667 | 1 | 0 | 6 | 0 | 2 | 0 | 4 | 0.25 | 13 | 0.0769 |
| 2_12188440 | 3 | 0 | 1 | 0 | 3 | 0 | 1 | 0 | 8 | 0.125 | 2 | 0 | 2 | 0.5 | NA | NA | 3 | 0 | 7 | 0.1429 |
| 2_12188440 | 3 | 0 | 1 | 1 | 3 | 0 | 1 | 0 | 8 | 0.125 | 2 | 0 | 2 | 0.5 | NA | NA | 3 | 0 | 7 | 0.1429 |
| 2_12188440 | 3 | 0 | 1 | 0 | 3 | 0 | 1 | 0 | 8 | 0.125 | 2 | 0 | 2 | 0.5 | NA | NA | 3 | 0 | 7 | 0.1429 |
| 2_12188440 | 3 | 0 | 1 | 1 | 3 | 0 | 1 | 0 | 8 | 0.125 | 2 | 0 | 2 | 0.5 | NA | NA | 3 | 0 | 7 | 0.1429 |
| 2_12188653 | 1 | 1 | 3 | 0 | NA | NA | NA | NA | 4 | 0.25 | 2 | 0 | 1 | 1 | 2 | 0 | 1 | 0 | 6 | 0.1667 |
| 2_12188697 | 1 | 1 | 2 | 0 | NA | NA | NA | NA | 3 | 0.3333 | 2 | 0 | 1 | 0 | 2 | 0 | 1 | 1 | 6 | 0.1667 |
| 2_12189565 | 2 | 0.5 | NA | NA | 1 | 0 | 1 | 0 | 4 | 0.25 | 2 | 0 | 1 | 1 | NA | NA | NA | NA | 3 | 0.3333 |
| 2_12189571 | 2 | 0 | NA | NA | 2 | 1 | NA | NA | 4 | 0.5 | 1 | 0 | 2 | 0 | NA | NA | 1 | 1 | 4 | 0.25 |
| 2_12189573 | NA | NA | 1 | 1 | NA | NA | NA | NA | 1 | 1 | NA | NA | NA | NA | 1 | 1 | NA | NA | 1 | 1 |
| 2_12189576 | NA | NA | NA | NA | 1 | 1 | 1 | 1 | 2 | 1 | 2 | 0 | 2 | 1 | NA | NA | 1 | 1 | 5 | 0.6 |
| 2_12189603 | 2 | 1 | NA | NA | NA | NA | NA | NA | 2 | 1 | NA | NA | 1 | 1 | NA | NA | 2 | 0.5 | 3 | 0.6667 |
| 2_12189607 | 2 | 0 | NA | NA | 1 | 1 | 1 | 0 | 4 | 0.25 | 2 | 0.5 | 2 | 0 | NA | NA | 2 | 0.5 | 6 | 0.3333 |
| 2_12189621 | NA | NA | 1 | 1 | NA | NA | NA | NA | 1 | 1 | NA | NA | NA | NA | 1 | 1 | NA | NA | 1 | 1 |
| 2_12190042 | 2 | 0.5 | NA | NA | 1 | 0 | NA | NA | 3 | 0.3333 | 1 | 0 | NA | NA | NA | NA | 1 | 1 | 2 | 0.5 |
| 2_12198152 | 26 | 0.5385 | 45 | 0.3333 | 19 | 0.2105 | 16 | 0.3125 | 106 | 0.3585 | 35 | 0.3429 | 19 | 0.2105 | 17 | 0.2353 | 32 | 0.1562 | 103 | 0.2427 |
| 2_12198156 | 27 | 0.1481 | 41 | 0.1951 | 19 | 0.0526 | 15 | 0.0667 | 102 | 0.1373 | 34 | 0.0882 | 19 | 0.2105 | 16 | 0.0625 | 32 | 0.0938 | 101 | 0.1089 |
| 2_12198594 | NA | NA | NA | NA | 1 | 1 | NA | NA | 1 | 1 | 1 | 0 | NA | NA | 1 | 1 | NA | NA | 2 | 0.5 |
| 2_12198600 | NA | NA | NA | NA | 1 | 1 | NA | NA | 1 | 1 | 1 | 0 | NA | NA | 1 | 1 | NA | NA | 2 | 0.5 |
| 2_12209476 | 2 | 0 | 4 | 0.25 | 2 | 0 | 1 | 0 | 9 | 0.1111 | 4 | 0.25 | 2 | 0 | 6 | 0.1667 | 4 | 0.25 | 16 | 0.1875 |
| 2_12209574 | 1 | 0 | 2 | 0.5 | 2 | 0 | 2 | 0 | 7 | 0.1429 | 6 | 0 | 1 | 1 | 5 | 0 | 6 | 0 | 18 | 0.0556 |
| 2_12217958 | 1 | 1 | NA | NA | 2 | 0 | NA | NA | 3 | 0.3333 | NA | NA | 3 | 0.3333 | 2 | 0.5 | 2 | 0 | 7 | 0.2857 |
| 2_12217965 | 3 | 1 | 4 | 1 | 4 | 1 | 5 | 0.8 | 16 | 0.9375 | 5 | 1 | 4 | 0.75 | 7 | 1 | 8 | 1 | 24 | 0.9583 |
| 2_12217975 | 2 | 0.5 | 2 | 0.5 | 5 | 0.8 | 6 | 1 | 15 | 0.8 | 3 | 0.6667 | 5 | 1 | 7 | 1 | 7 | 0.8571 | 22 | 0.9091 |
| 2_12217980 | 2 | 0 | 3 | 0.6667 | 4 | 0 | 5 | 0.2 | 14 | 0.2143 | 3 | 0 | 4 | 0 | 6 | 0.1667 | 6 | 0.1667 | 19 | 0.1053 |
| 2_12217980 | 3 | 0 | 2 | 0.5 | 4 | 0 | 4 | 0.25 | 13 | 0.1538 | 4 | 0.25 | 4 | 0.25 | 7 | 0.1429 | 7 | 0 | 22 | 0.1364 |
| 2_12217984 | 3 | 1 | 3 | 1 | 4 | 1 | 4 | 1 | 14 | 1 | 5 | 0.6 | 5 | 1 | 6 | 1 | 6 | 1 | 22 | 0.9091 |
| 2_12242589 | 7 | 0.2857 | 18 | 0.7222 | 7 | 0.4286 | 9 | 0.4444 | 41 | 0.5366 | 7 | 0.7143 | 9 | 0.8889 | 9 | 0.7778 | 14 | 0.7857 | 39 | 0.7949 |
| 2_12247467 | NA | NA | 1 | 1 | 1 | 1 | 2 | 0.5 | 4 | 0.75 | NA | NA | 2 | 0 | 1 | 1 | NA | NA | 3 | 0.3333 |
| 2_12252439 | 5 | 0 | 3 | 0 | 2 | 0.5 | 6 | 0 | 16 | 0.0625 | 2 | 0 | NA | NA | 1 | 1 | 5 | 0.2 | 8 | 0.25 |
| 2_12252588 | 3 | 0.3333 | 3 | 0 | 3 | 0.6667 | 5 | 0 | 14 | 0.2143 | 2 | 0 | 4 | 0 | 1 | 1 | 5 | 0.2 | 12 | 0.1667 |
| 2_12252658 | 3 | 0.3333 | 3 | 0.3333 | 2 | 0.5 | 6 | 0.5 | 14 | 0.4286 | 1 | 0 | 3 | 0 | NA | NA | 6 | 0.1667 | 10 | 0.1 |
| 2_12252673 | 2 | 1 | 3 | 0.3333 | 1 | 0 | 6 | 0.5 | 12 | 0.5 | 1 | 1 | 3 | 0.3333 | 1 | 1 | 5 | 0.4 | 10 | 0.5 |
| 2_12252681 | 2 | 0 | 1 | 0 | NA | NA | 6 | 0.3333 | 9 | 0.2222 | 1 | 0 | 4 | 0.75 | 1 | 1 | 4 | 0.5 | 10 | 0.6 |
| 2_12252684 | 2 | 0 | 3 | 0.3333 | 1 | 0 | 6 | 0.1667 | 12 | 0.1667 | 1 | 0 | 3 | 0.3333 | 1 | 0 | 6 | 0.5 | 11 | 0.3636 |
| 2_12252692 | 2 | 0.5 | 1 | 1 | 1 | 1 | 4 | 1 | 8 | 0.875 | 1 | 1 | 4 | 1 | 1 | 1 | 3 | 1 | 9 | 1 |
| 2_12252693 | 2 | 0 | 3 | 0 | 1 | 0 | 6 | 0.1667 | 12 | 0.0833 | 1 | 0 | 4 | 0.25 | 1 | 0 | 4 | 0 | 10 | 0.1 |
| 2_12252695 | 2 | 0.5 | 3 | 0.6667 | 1 | 1 | 5 | 0.6 | 11 | 0.6364 | 1 | 1 | 4 | 0.75 | 1 | 1 | 6 | 0 | 12 | 0.4167 |
| 2_12252696 | 2 | 1 | 2 | 1 | 1 | 1 | 3 | 0.6667 | 8 | 0.875 | 1 | 1 | 4 | 0.5 | 1 | 1 | 5 | 0.8 | 11 | 0.7273 |
| 2_12252704 | 2 | 1 | 1 | 1 | 1 | 1 | 6 | 1 | 10 | 1 | 1 | 1 | NA | NA | 1 | 1 | 5 | 1 | 7 | 1 |
| 2_12252713 | 2 | 1 | 2 | 0 | 1 | 0 | 5 | 1 | 10 | 0.7 | 1 | 1 | 4 | 0.5 | 1 | 1 | 5 | 0.4 | 11 | 0.5455 |
| 2_12252719 | 2 | 0.5 | NA | NA | 1 | 0 | 5 | 0.4 | 8 | 0.375 | 1 | 0 | 2 | 0.5 | NA | NA | 4 | 0.25 | 7 | 0.2857 |
| 2_12254406 | 1 | 1 | 1 | 0 | 1 | 0 | 5 | 0 | 8 | 0.125 | 1 | 0 | 4 | 0 | NA | NA | 4 | 0.25 | 9 | 0.1111 |
| 2_12254410 | 2 | 1 | 1 | 0 | 1 | 1 | 4 | 1 | 8 | 0.875 | 1 | 1 | 4 | 1 | NA | NA | 4 | 0.75 | 9 | 0.8889 |
| 2_12254418 | 1 | 0 | 1 | 0 | 1 | 0 | 4 | 0.25 | 7 | 0.1429 | 1 | 1 | 3 | 0 | NA | NA | 4 | 0 | 8 | 0.125 |
| 2_12258873 | 3 | 0.3333 | 3 | 0 | 3 | 0 | 9 | 0.1111 | 18 | 0.1111 | 3 | 0 | 1 | 0 | 4 | 0.25 | 5 | 0.2 | 13 | 0.1538 |
| 2_12258898 | 3 | 0.3333 | 4 | 0 | 4 | 0.25 | 11 | 0.0909 | 22 | 0.1364 | 2 | 0 | NA | NA | 2 | 0 | 5 | 0.4 | 9 | 0.2222 |
| 2_12265133 | 2 | 0.5 | 1 | 1 | 1 | 1 | 1 | 0 | 5 | 0.6 | NA | NA | NA | NA | 4 | 0.75 | 1 | 0 | 5 | 0.6 |
| 2_12450619 | 1 | 1 | NA | NA | 1 | 0 | 4 | 0.25 | 6 | 0.3333 | 3 | 0.3333 | 2 | 0.5 | 4 | 0.5 | 1 | 0 | 10 | 0.4 |
| 2_12450624 | 1 | 0 | 1 | 0 | NA | NA | 4 | 0.75 | 6 | 0.5 | 3 | 0.6667 | 3 | 0.6667 | 2 | 0.5 | 1 | 1 | 9 | 0.6667 |
| 2_12450641 | 1 | 0 | 1 | 0 | 1 | 0 | 3 | 1 | 6 | 0.5 | 3 | 0 | 3 | 0 | 3 | 0.3333 | 1 | 0 | 10 | 0.1 |
| 2_12450667 | 2 | 0.5 | 2 | 0.5 | 3 | 0.3333 | 6 | 0.1667 | 13 | 0.3077 | 3 | 0 | 2 | 0.5 | 4 | 0.25 | 1 | 0 | 10 | 0.2 |
| 2_12450672 | 1 | 0 | 2 | 0 | 3 | 0 | 4 | 0.25 | 10 | 0.1 | 3 | 0.3333 | 2 | 0 | 3 | 0 | 1 | 0 | 9 | 0.1111 |
| 2_12450673 | 2 | 0.5 | 2 | 0 | 1 | 0 | 4 | 0.5 | 9 | 0.3333 | 2 | 0 | 2 | 0.5 | 3 | 0.6667 | 1 | 1 | 8 | 0.5 |
| 2_12450674 | 2 | 0 | 2 | 0.5 | 3 | 0 | 5 | 0 | 12 | 0.0833 | 4 | 0 | 3 | 0.3333 | 4 | 0.5 | 1 | 0 | 12 | 0.25 |
| 2_12450687 | 2 | 1 | 2 | 1 | 4 | 0.75 | 6 | 1 | 14 | 0.9286 | 3 | 1 | 2 | 0.5 | 2 | 1 | 2 | 1 | 9 | 0.8889 |
| 2_12450688 | 2 | 1 | 2 | 1 | 3 | 1 | 5 | 1 | 12 | 1 | 3 | 1 | 3 | 0.6667 | 4 | 1 | 4 | 1 | 14 | 0.9286 |
| 2_12450694 | 2 | 1 | 1 | 1 | 1 | 1 | 4 | 1 | 8 | 1 | 4 | 0.75 | 3 | 1 | 2 | 1 | 2 | 1 | 11 | 0.9091 |
| 2_12450695 | 1 | 0 | 1 | 0 | 3 | 0.3333 | 3 | 0 | 8 | 0.125 | 3 | 0.3333 | 2 | 0.5 | 2 | 0 | 2 | 0 | 9 | 0.2222 |
| 2_12450698 | 2 | 0.5 | 2 | 0 | 3 | 0.3333 | 4 | 0 | 11 | 0.1818 | 4 | 0.25 | 3 | 0 | 4 | 1 | 3 | 0 | 14 | 0.3571 |
| 2_12450699 | 2 | 1 | 2 | 1 | 3 | 0.6667 | 6 | 0.8333 | 13 | 0.8462 | 4 | 0.5 | 2 | 1 | 4 | 0.75 | 1 | 1 | 11 | 0.7273 |
| 2_12450703 | 2 | 1 | 2 | 1 | NA | NA | 4 | 1 | 8 | 1 | 4 | 1 | 3 | 1 | 4 | 0.75 | 1 | 1 | 12 | 0.9167 |
| 2_12450705 | 2 | 0 | 2 | 0 | 4 | 0 | 5 | 0.2 | 13 | 0.0769 | 4 | 0.25 | 3 | 0 | 2 | 0 | 3 | 0.3333 | 12 | 0.1667 |

| | | | | | | | | | | | | | | | | | | | | |
|---|---|---|---|---|---|---|---|---|---|---|---|---|---|---|---|---|---|---|---|---|
| 2_12450713 | 2 | 0.5 | 2 | 1 | 3 | 1 | 5 | 1 | 12 | 0.9167 | 4 | 0.25 | 3 | 1 | 3 | 1 | 2 | 1 | 12 | 0.75 |
| 2_12450714 | 2 | 0.5 | 2 | 1 | 4 | 1 | 5 | 1 | 13 | 0.9231 | 4 | 0.75 | 3 | 1 | 3 | 0.6667 | 2 | 1 | 12 | 0.8333 |
| 2_12450720 | 1 | 0 | 2 | 0 | 5 | 0 | 6 | 0.1667 | 14 | 0.0714 | 4 | 0.25 | 2 | 0 | 4 | 0 | 1 | 0 | 11 | 0.0909 |
| 2_12450733 | 2 | 0 | 1 | 0 | 3 | 0 | 4 | 0.25 | 10 | 0.1 | 3 | 0 | 1 | 0 | 3 | 0.3333 | 3 | 0.3333 | 10 | 0.2 |
| 2_12450734 | 2 | 1 | 2 | 1 | 5 | 1 | 6 | 1 | 15 | 1 | 3 | 0.6667 | 3 | 0.6667 | 3 | 1 | 2 | 1 | 11 | 0.8182 |
| 2_12450737 | 2 | 1 | 2 | 0.5 | 3 | 1 | 4 | 1 | 11 | 0.9091 | 1 | 1 | 2 | 1 | 2 | 0.5 | NA | NA | 5 | 0.8 |
| 2_12450753 | 2 | 1 | 2 | 1 | 5 | 0.8 | 5 | 0.4 | 14 | 0.7143 | 3 | 1 | 3 | 0.6667 | 4 | 1 | 1 | 0 | 11 | 0.8182 |
| 2_12450756 | 3 | 0.6667 | 2 | 0 | 5 | 0.6 | 5 | 0.6 | 15 | 0.5333 | 4 | 0.75 | 2 | 0.5 | 4 | 0.75 | 2 | 1 | 12 | 0.75 |
| 2_12450760 | 1 | 1 | 2 | 1 | 4 | 1 | 5 | 0.4 | 12 | 0.75 | 2 | 1 | 2 | 0.5 | 3 | 0.6667 | 2 | 1 | 9 | 0.7778 |
| 2_12450781 | 3 | 0 | 2 | 0 | 6 | 0.1667 | 5 | 0 | 16 | 0.0625 | 3 | 0 | 4 | 0 | 3 | 0.3333 | 3 | 0 | 13 | 0.0769 |
| 2_12450806 | 3 | 0.6667 | 2 | 0 | 5 | 0.6 | 6 | 0.1667 | 16 | 0.375 | 5 | 0 | 4 | 0 | 4 | 0.75 | 3 | 0 | 16 | 0.1875 |
| 2_12450809 | 3 | 0.3333 | 2 | 0 | 4 | 0 | 6 | 0 | 15 | 0.0667 | 5 | 0 | 3 | 0.3333 | 4 | 0.25 | 3 | 0 | 15 | 0.1333 |
| 2_12450817 | 3 | 0 | 2 | 0 | 5 | 0.2 | 6 | 0 | 16 | 0.0625 | 5 | 0 | 5 | 0 | 5 | 0.2 | 3 | 0.3333 | 18 | 0.1111 |
| 2_12450853 | 3 | 0 | 3 | 0 | 6 | 0.1667 | 6 | 0 | 18 | 0.0556 | 3 | 0 | 4 | 0 | 6 | 0 | 3 | 0.3333 | 16 | 0.0625 |
| 2_12450869 | 3 | 0.3333 | 3 | 0.3333 | 8 | 0.25 | 7 | 0.4286 | 21 | 0.3333 | 5 | 0 | 5 | 0.2 | 6 | 0.5 | 4 | 0.25 | 20 | 0.25 |
| 2_12450905 | 2 | 0.5 | 3 | 0.3333 | 6 | 0.1667 | 6 | 0 | 17 | 0.1765 | 6 | 0 | 5 | 0.4 | 5 | 0.2 | 4 | 0.25 | 20 | 0.2 |
| 2_12450915 | 4 | 0.25 | 1 | 0 | 6 | 0.1667 | 4 | 0.25 | 15 | 0.2 | 6 | 0.3333 | 6 | 0.1667 | 6 | 0.3333 | 5 | 0.4 | 23 | 0.3043 |
| 2_12492309 | 1 | 0 | 4 | 0.25 | NA | NA | 2 | 0 | 7 | 0.1429 | 1 | 0 | 2 | 0 | NA | NA | 2 | 0.5 | 5 | 0.2 |
| 2_1249232 | NA | NA | 3 | 0.3333 | NA | NA | 1 | 0 | 4 | 0.25 | NA | NA | 2 | 0 | NA | NA | 1 | 1 | 3 | 0.3333 |
| 2_12492326 | 1 | 0 | 3 | 0.6667 | 1 | 0 | 3 | 0 | 8 | 0.25 | 1 | 1 | 3 | 0 | NA | NA | 2 | 0.5 | 6 | 0.3333 |
| 2_12492336 | 1 | 0 | 4 | 0 | 1 | 0 | 2 | 0.5 | 8 | 0.125 | NA | NA | 3 | 0.3333 | NA | NA | 2 | 0.5 | 5 | 0.4 |
| 2_12492340 | NA | NA | 4 | 0.25 | 1 | 0 | 2 | 0 | 7 | 0.1429 | 1 | 1 | 3 | 0.3333 | NA | NA | 1 | 0 | 5 | 0.4 |
| 2_1249271 | NA | NA | 1 | 0 | 1 | 1 | 2 | 0 | 4 | 0.25 | NA | NA | 1 | 0 | NA | NA | 1 | 1 | 2 | 0.5 |
| 2_12492857 | NA | NA | 1 | 1 | 1 | 0 | 4 | 0.5 | 6 | 0.5 | NA | NA | 1 | 0 | NA | NA | 1 | 1 | 2 | 0.5 |
| 2_12492903 | NA | NA | NA | NA | 1 | 0 | 3 | 0.6667 | 4 | 0.5 | NA | NA | 1 | 0 | NA | NA | 1 | 1 | 2 | 0.5 |
| 2_12493007 | 7 | 0.5714 | 12 | 0.25 | 9 | 0.2222 | 11 | 0.0909 | 39 | 0.2564 | 7 | 0.4286 | 14 | 0.0714 | 10 | 0.5 | 11 | 0.0909 | 42 | 0.2381 |
| 2_12493053 | 8 | 0.25 | 10 | 0.5 | 9 | 0.1111 | 11 | 0.2727 | 38 | 0.2895 | 7 | 0.5714 | 15 | 0.2 | 9 | 0.2222 | 8 | 0.25 | 39 | 0.2821 |
| 2_12493069 | 6 | 0.8333 | 4 | 0.75 | 9 | 0.7778 | 10 | 0.6 | 29 | 0.7241 | 5 | 1 | 13 | 0.5385 | 6 | 0.6667 | 5 | 0.4 | 29 | 0.6207 |
| 2_12493075 | 7 | 0 | 4 | 0 | 7 | 0.1429 | 9 | 0.1111 | 27 | 0.0741 | 6 | 0 | 10 | 0.4 | 7 | 0.7143 | 5 | 0.4 | 28 | 0.3929 |
| 2_12493077 | 8 | 0.125 | 7 | 0.2857 | 9 | 0.2222 | 10 | 0 | 34 | 0.1471 | 6 | 0 | 15 | 0.2 | 8 | 0 | 7 | 0 | 36 | 0.0833 |
| 2_12493080 | 7 | 0.2857 | 7 | 0.2857 | 9 | 0.2222 | 8 | 0.375 | 31 | 0.2903 | 5 | 0 | 12 | 0.1667 | 7 | 0.1429 | 5 | 0 | 29 | 0.1034 |
| 2_12493090 | 8 | 0.375 | 6 | 0.1667 | 8 | 0.375 | 7 | 0.4286 | 29 | 0.3448 | 5 | 0.6 | 14 | 0.3571 | 6 | 0.3333 | 5 | 0.2 | 30 | 0.3667 |
| 2_12496634 | 2 | 0.5 | NA | NA | 2 | 0 | 2 | 0 | 6 | 0.1667 | 1 | 0 | 1 | 1 | NA | NA | 1 | 0 | 3 | 0.3333 |
| 2_12497370 | NA | NA | NA | NA | NA | NA | 2 | 0.5 | 2 | 0.5 | NA | NA | NA | NA | NA | NA | 1 | 1 | 1 | 1 |
| 2_12497378 | NA | NA | NA | NA | NA | NA | 2 | 0.5 | 2 | 0.5 | NA | NA | NA | NA | NA | NA | 1 | 1 | 1 | 1 |
| 2_12497384 | NA | NA | NA | NA | NA | NA | 1 | 1 | 1 | 1 | NA | NA | NA | NA | NA | NA | 1 | 1 | 1 | 1 |
| 2_12498950 | 7 | 1 | NA | NA | 2 | 1 | 1 | 1 | 10 | 1 | 4 | 1 | 5 | 1 | 5 | 1 | 4 | 1 | 18 | 1 |
| 2_12498960 | 7 | 0.1429 | NA | NA | 3 | 0.3333 | 2 | 0.5 | 12 | 0.25 | 5 | 0.2 | 5 | 0.4 | 4 | 0.5 | 7 | 0.7143 | 21 | 0.4762 |
| 2_12509933 | 10 | 0.1 | 5 | 0.2 | 4 | 0.25 | 7 | 0.1429 | 26 | 0.1538 | 5 | 0.2 | 5 | 0 | 2 | 0.5 | 4 | 0 | 16 | 0.125 |
| 2_12509951 | 10 | 0 | 6 | 0.1667 | 6 | 0 | 8 | 0.125 | 30 | 0.0667 | 5 | 0.2 | 7 | 0 | 2 | 0 | 4 | 0 | 18 | 0.0556 |
| 2_12509989 | 4 | 0 | 1 | 0 | 3 | 0 | 3 | 0.3333 | 11 | 0.0909 | NA | NA | 1 | 0 | 1 | 0 | 1 | 1 | 3 | 0.3333 |
| 2_12510024 | 3 | 0 | 1 | 0 | 3 | 0.3333 | 4 | 0 | 11 | 0.0909 | NA | NA | 2 | 0 | 2 | 0.5 | 1 | 0 | 5 | 0.2 |
| 2_12514317 | 1 | 1 | NA | NA | NA | NA | NA | NA | 1 | 1 | 1 | 1 | NA | NA | 1 | 1 | NA | NA | 2 | 1 |
| 2_12514325 | 1 | 1 | NA | NA | NA | NA | NA | NA | 1 | 1 | 1 | 1 | NA | NA | 1 | 1 | NA | NA | 2 | 1 |
| 2_12514355 | 1 | 1 | NA | NA | NA | NA | NA | NA | 1 | 1 | 1 | 1 | NA | NA | NA | NA | NA | NA | 1 | 1 |
| 2_12514359 | 1 | 1 | NA | NA | NA | NA | NA | NA | 1 | 1 | 1 | 1 | NA | NA | NA | NA | NA | NA | 1 | 1 |
| 2_12514364 | 1 | 1 | NA | NA | NA | NA | NA | NA | 1 | 1 | 1 | 1 | NA | NA | NA | NA | NA | NA | 1 | 1 |
| 2_12551530 | 27 | 0.0741 | 34 | 0.0882 | 27 | 0.0741 | 30 | 0.0333 | 118 | 0.0678 | 20 | 0.25 | 31 | 0.1613 | 29 | 0.069 | 26 | 0 | 106 | 0.1132 |
| 2_12551533 | 27 | 0.2593 | 31 | 0.1935 | 26 | 0.0769 | 30 | 0.0333 | 114 | 0.1404 | 20 | 0.1 | 29 | 0.2414 | 27 | 0.037 | 24 | 0.25 | 100 | 0.16 |
| 2_12551561 | 25 | 0.12 | 26 | 0.1923 | 21 | 0.1429 | 25 | 0.28 | 97 | 0.1856 | 15 | 0.1333 | 28 | 0.1786 | 22 | 0.2273 | 22 | 0.0455 | 87 | 0.1494 |
| 2_12551568 | 19 | 1 | 25 | 0.96 | 18 | 1 | 18 | 1 | 80 | 0.9875 | 14 | 1 | 24 | 0.9583 | 19 | 1 | 20 | 0.9 | 77 | 0.961 |
| 2_12551572 | 26 | 0.9615 | 34 | 1 | 26 | 1 | 31 | 0.9677 | 117 | 0.9829 | 22 | 1 | 30 | 1 | 29 | 1 | 21 | 1 | 102 | 1 |
| 2_12551573 | 28 | 0.2143 | 36 | 0.1944 | 26 | 0.1923 | 30 | 0.0667 | 120 | 0.1667 | 24 | 0 | 34 | 0.2059 | 30 | 0.1 | 28 | 0.1429 | 116 | 0.1207 |
| 2_12551580 | 34 | 0.2353 | 37 | 0.1892 | 29 | 0.3103 | 34 | 0.2941 | 134 | 0.2537 | 24 | 0.4583 | 33 | 0.4545 | 29 | 0.2069 | 29 | 0.3793 | 115 | 0.3739 |
| 2_12551584 | 31 | 0.0968 | 35 | 0.0857 | 24 | 0.0833 | 29 | 0.1724 | 119 | 0.1092 | 22 | 0.0909 | 31 | 0.0968 | 27 | 0.1852 | 24 | 0.1667 | 104 | 0.1346 |
| 2_12551585 | 32 | 1 | 25 | 1 | 23 | 1 | 30 | 1 | 110 | 1 | 22 | 1 | 32 | 0.9688 | 24 | 1 | 26 | 0.9231 | 104 | 0.9712 |
| 2_12551594 | 30 | 0.4667 | 34 | 0.4412 | 27 | 0.4815 | 32 | 0.4688 | 123 | 0.4634 | 24 | 0.5417 | 32 | 0.5625 | 29 | 0.5172 | 27 | 0.6296 | 112 | 0.5625 |
| 2_12551600 | 24 | 0.9583 | 28 | 1 | 20 | 1 | 28 | 0.9286 | 100 | 0.97 | 22 | 1 | 31 | 0.9677 | 23 | 1 | 25 | 0.96 | 101 | 0.9802 |
| 2_12551603 | 24 | 0.9583 | 32 | 0.8438 | 21 | 1 | 18 | 0.8333 | 95 | 0.9053 | 21 | 1 | 23 | 0.7391 | 22 | 0.9091 | 24 | 0.8333 | 90 | 0.8667 |
| 2_12551608 | 32 | 0.0625 | 35 | 0.0286 | 27 | 0.1111 | 28 | 0.0714 | 122 | 0.0656 | 26 | 0.0385 | 37 | 0.0811 | 32 | 0.0625 | 29 | 0.1379 | 124 | 0.0806 |
| 2_12551621 | 35 | 0.0286 | 38 | 0.0526 | 29 | 0.1379 | 33 | 0.1212 | 135 | 0.0815 | 28 | 0.0714 | 36 | 0.0833 | 35 | 0.1714 | 32 | 0.0312 | 131 | 0.0916 |
| 2_12551636 | 25 | 0.12 | 32 | 0.2188 | 17 | 0.1176 | 22 | 0.1364 | 96 | 0.1562 | 17 | 0.2353 | 27 | 0.1481 | 26 | 0.2308 | 16 | 0.1875 | 86 | 0.1977 |
| 2_12551643 | 35 | 0.0857 | 42 | 0.0476 | 29 | 0.0345 | 34 | 0.0294 | 140 | 0.05 | 28 | 0.0357 | 36 | 0.1111 | 32 | 0.0312 | 28 | 0.1071 | 124 | 0.0726 |
| 2_12551646 | 34 | 0.4118 | 40 | 0.3 | 29 | 0.2759 | 35 | 0.4571 | 138 | 0.3623 | 27 | 0.3333 | 36 | 0.2778 | 31 | 0.3548 | 31 | 0.3548 | 125 | 0.328 |
| 2_12558011 | 6 | 0.1667 | 9 | 0.2222 | 9 | 0.1111 | 14 | 0.2143 | 38 | 0.1842 | 8 | 0 | 7 | 0 | 7 | 0.1429 | 14 | 0.2143 | 36 | 0.1111 |

| | | | | | | | | | | | | | | | | | | | | |
|---|---|---|---|---|---|---|---|---|---|---|---|---|---|---|---|---|---|---|---|---|
| 2_12558038 | 7 | 0.2857 | 9 | 0 | 7 | 0 | 14 | 0.1429 | 37 | 0.1081 | 6 | 0.1667 | 5 | 0 | 8 | 0.125 | 14 | 0 | 33 | 0.0606 |
| 2_12558080 | 5 | 0.2 | 6 | 0.3333 | 4 | 0.25 | 9 | 0.2222 | 24 | 0.25 | 4 | 0.5 | 4 | 0.5 | 8 | 0.5 | 11 | 0.3636 | 27 | 0.4444 |
| 2_12558088 | 5 | 0.2 | 7 | 0.2857 | 5 | 0.6 | 9 | 0.4444 | 26 | 0.3846 | 5 | 0.4 | 4 | 0.75 | 5 | 0.6 | 10 | 0.2 | 24 | 0.4167 |
| 2_12558120 | 5 | 0.2 | 6 | 0 | 6 | 0 | 10 | 0.1 | 27 | 0.0741 | 5 | 0.2 | 3 | 0 | 4 | 0 | 6 | 0 | 18 | 0.0556 |
| 2_12558135 | 3 | 0.3333 | 7 | 0.1429 | 5 | 0.2 | 9 | 0.6667 | 24 | 0.375 | 5 | 0.2 | 4 | 0.25 | 7 | 0.1429 | 8 | 0.5 | 24 | 0.2917 |
| 2_12558141 | 4 | 0 | 7 | 0 | 5 | 0.2 | 7 | 0.4286 | 23 | 0.1739 | 5 | 0.2 | 3 | 0 | 5 | 0.4 | 5 | 0.2 | 18 | 0.2222 |
| 2_12558153 | 4 | 0.25 | 8 | 0.25 | 5 | 0.4 | 8 | 0.125 | 25 | 0.24 | 5 | 0.8 | 3 | 0 | 6 | 0.3333 | 7 | 0 | 21 | 0.2857 |
| 2_12558164 | 3 | 1 | 7 | 1 | 6 | 1 | 6 | 1 | 22 | 1 | 4 | 1 | 4 | 1 | 7 | 1 | 8 | 1 | 23 | 1 |
| 2_12558173 | 3 | 0.3333 | 5 | 0.6 | 4 | 0.75 | 6 | 0.8333 | 18 | 0.6667 | 3 | 0.6667 | 3 | 0.6667 | 6 | 0.5 | 5 | 0.6 | 17 | 0.5882 |
| 2_12558173 | 4 | 0 | 8 | 0.125 | 6 | 0 | 7 | 0.2857 | 25 | 0.12 | 5 | 0 | 3 | 0 | 7 | 0.2857 | 7 | 0.1429 | 22 | 0.1364 |
| 2_12558175 | 3 | 1 | 7 | 1 | 6 | 1 | 7 | 1 | 23 | 1 | 3 | 1 | 3 | 1 | 7 | 1 | 7 | 1 | 20 | 1 |
| 2_12558184 | 4 | 1 | 6 | 1 | 6 | 0.8333 | 6 | 0.8333 | 22 | 0.9091 | 5 | 1 | 3 | 1 | 6 | 1 | 8 | 0.875 | 22 | 0.9545 |
| 2_12558190 | 3 | 0.3333 | 4 | 0.25 | 5 | 0 | 3 | 0 | 15 | 0.1333 | 5 | 0.4 | 2 | 0 | 5 | 0.2 | 7 | 0 | 19 | 0.1579 |
| 2_12558191 | 4 | 0 | 4 | 0.5 | 6 | 0.1667 | 4 | 0 | 18 | 0.1667 | 5 | 0 | 3 | 0.6667 | 5 | 0 | 6 | 0 | 19 | 0.1053 |
| 2_12558193 | 4 | 0 | 6 | 0.1667 | 6 | 0.1667 | 6 | 0.5 | 22 | 0.2273 | 5 | 0.2 | 4 | 0.25 | 7 | 0.1429 | 8 | 0 | 24 | 0.125 |
| 2_12558194 | 3 | 1 | 4 | 0.75 | 3 | 0.6667 | 5 | 0.8 | 15 | 0.8 | 4 | 1 | 2 | 1 | 7 | 0.8571 | 7 | 0.5714 | 20 | 0.8 |
| 2_12558198 | 4 | 1 | 6 | 1 | 6 | 1 | 6 | 1 | 22 | 1 | 4 | 0.5 | 3 | 1 | 6 | 1 | 6 | 1 | 19 | 0.8947 |
| 2_12558202 | 4 | 0 | 2 | 0.5 | 1 | 0 | 3 | 0 | 10 | 0.1 | 3 | 0 | 1 | 0 | 4 | 0.25 | 6 | 0 | 14 | 0.0714 |
| 2_12558203 | 3 | 1 | 5 | 0.8 | 4 | 0.75 | 3 | 0.6667 | 15 | 0.8 | 4 | 1 | 3 | 1 | 6 | 1 | 7 | 0.7143 | 20 | 0.9 |
| 2_12558207 | 3 | 1 | 6 | 1 | 6 | 1 | 6 | 1 | 21 | 1 | 5 | 1 | 3 | 0.6667 | 7 | 0.8571 | 8 | 1 | 23 | 0.913 |
| 2_12558218 | 3 | 0.3333 | 5 | 0 | 6 | 0 | 4 | 0.25 | 18 | 0.1111 | 4 | 0 | 2 | 0.5 | 4 | 0.25 | 6 | 0 | 16 | 0.125 |
| 2_12558226 | 3 | 1 | 5 | 1 | 4 | 0.75 | 5 | 1 | 17 | 0.9412 | 3 | 0.3333 | 4 | 1 | 5 | 0.8 | 8 | 1 | 20 | 0.85 |
| 2_12558227 | 3 | 0 | 5 | 0 | 5 | 0 | 5 | 0.2 | 18 | 0.0556 | 4 | 0.25 | 3 | 0 | 6 | 0.3333 | 8 | 0 | 21 | 0.1429 |
| 2_12558227 | 2 | 0.5 | 4 | 0.5 | 4 | 0.25 | 5 | 0 | 15 | 0.2667 | 2 | 0 | 3 | 0.3333 | 5 | 0.4 | 7 | 0.4286 | 17 | 0.3529 |
| 2_12558234 | 1 | 0 | 6 | 0.5 | 4 | 0.5 | 4 | 0.25 | 15 | 0.4 | 2 | 0 | 4 | 0 | 5 | 0.2 | 7 | 0.1429 | 18 | 0.1111 |
| 2_12558238 | 3 | 1 | 6 | 0.8333 | 6 | 0.5 | 3 | 0 | 18 | 0.6111 | 2 | 1 | 4 | 0.5 | 6 | 0.8333 | 8 | 0.5 | 20 | 0.65 |
| 2_12558239 | 4 | 1 | 6 | 1 | 6 | 1 | 5 | 0.8 | 21 | 0.9524 | 3 | 0.6667 | 4 | 1 | 6 | 1 | 8 | 1 | 21 | 0.9524 |
| 2_12558254 | 3 | 0.6667 | 6 | 0.6667 | 5 | 0.8 | 3 | 1 | 17 | 0.7647 | 3 | 0.6667 | 1 | 0 | 3 | 0.3333 | 6 | 0.5 | 13 | 0.4615 |
| 2_12558258 | 4 | 0 | 6 | 0.1667 | 6 | 0.1667 | 5 | 0 | 21 | 0.0952 | 3 | 0 | 3 | 0 | 6 | 0.1667 | 6 | 0 | 18 | 0.0556 |
| 2_12558273 | 3 | 0.6667 | 5 | 0.2 | 4 | 0.25 | 5 | 0.6 | 17 | 0.4118 | 3 | 0 | 3 | 0 | 6 | 0.1667 | 7 | 0.4286 | 19 | 0.2105 |
| 2_12567260 | 5 | 0.2 | 8 | 0.125 | 7 | 0.2857 | 8 | 0.125 | 28 | 0.1786 | 2 | 0.5 | 10 | 0.4 | 8 | 0 | 9 | 0.4444 | 29 | 0.3103 |
| 2_12567326 | 2 | 0 | 7 | 0 | 6 | 0 | 6 | 0.3333 | 21 | 0.0952 | NA | NA | 9 | 0.1111 | 8 | 0.125 | 8 | 0 | 25 | 0.08 |
| 2_12567370 | 6 | 0.5 | 15 | 0.0667 | 13 | 0.1538 | 11 | 0 | 45 | 0.1333 | 5 | 0.4 | 6 | 0.1667 | 10 | 0.3 | 11 | 0.2727 | 32 | 0.2812 |
| 2_12567372 | 9 | 0 | 18 | 0 | 15 | 0.2667 | 10 | 0 | 52 | 0.0769 | 7 | 0.1429 | 12 | 0 | 13 | 0.0769 | 13 | 0.0769 | 45 | 0.0667 |
| 2_12567380 | 7 | 0.2857 | 11 | 0.3636 | 11 | 0.0909 | 10 | 0.1 | 39 | 0.2051 | 6 | 0.1667 | 11 | 0.3636 | 8 | 0.375 | 10 | 0.3 | 35 | 0.3143 |
| 2_12567380 | 2 | 1 | 2 | 0 | 2 | 0 | 1 | 0 | 7 | 0.2857 | NA | NA | 1 | 0 | 2 | 0.5 | 1 | 0 | 4 | 0.25 |
| 2_12567388 | 12 | 0.0833 | 15 | 0.1333 | 16 | 0 | 13 | 0.0769 | 56 | 0.0714 | 7 | 0 | 13 | 0.1538 | 13 | 0.2308 | 12 | 0.25 | 45 | 0.1778 |
| 2_12567404 | 10 | 0.2 | 14 | 0.2143 | 14 | 0.2857 | 13 | 0.0769 | 51 | 0.1961 | 6 | 0 | 11 | 0.1818 | 12 | 0 | 9 | 0.2222 | 38 | 0.1053 |
| 2_12567405 | 10 | 0.7 | 15 | 0.8667 | 13 | 0.9231 | 12 | 0.8333 | 50 | 0.84 | 6 | 0.8333 | 11 | 0.9091 | 11 | 1 | 11 | 1 | 39 | 0.9487 |
| 2_12567409 | 11 | 0.7273 | 13 | 0.9231 | 15 | 0.8 | 12 | 0.6667 | 51 | 0.7843 | 7 | 0.8571 | 12 | 0.6667 | 10 | 0.8 | 11 | 1 | 40 | 0.825 |
| 2_12567412 | 11 | 0.0909 | 13 | 0.3077 | 15 | 0.0667 | 11 | 0.1818 | 50 | 0.16 | 5 | 0.2 | 13 | 0.2308 | 9 | 0.2222 | 10 | 0.3 | 37 | 0.2432 |
| 2_12567416 | 11 | 1 | 15 | 0.5333 | 13 | 0.6923 | 13 | 0.8462 | 52 | 0.75 | 6 | 0.1667 | 12 | 0.5833 | 10 | 0.6 | 11 | 0.9091 | 39 | 0.6154 |
| 2_12567421 | 9 | 0 | 14 | 0.1429 | 13 | 0 | 12 | 0.1667 | 48 | 0.0833 | 6 | 0 | 12 | 0 | 9 | 0.2222 | 10 | 0 | 37 | 0.0541 |
| 2_12582520 | 5 | 0.4 | 6 | 0.5 | 8 | 0 | 5 | 0.2 | 24 | 0.25 | 1 | 0 | 10 | 0.3 | 3 | 0.3333 | 5 | 0.2 | 19 | 0.2632 |
| 2_12600679 | 2 | 1 | 1 | 1 | 3 | 1 | 1 | 1 | 7 | 1 | 3 | 0.6667 | 4 | 0.75 | 2 | 1 | NA | NA | 9 | 0.7778 |
| 2_12600697 | 2 | 0.5 | 1 | 0 | 3 | 0 | 1 | 1 | 7 | 0.2857 | 2 | 0.5 | 3 | 0 | 2 | 0 | NA | NA | 7 | 0.1429 |
| 2_12600707 | 2 | 0 | 1 | 1 | 3 | 0 | 1 | 0 | 7 | 0.1429 | 3 | 0.3333 | 5 | 0.2 | 2 | 0 | NA | NA | 10 | 0.2 |
| 2_12642970 | 1 | 0 | 3 | 0.3333 | 1 | 0 | 1 | 0 | 6 | 0.1667 | 1 | 1 | 2 | 0.5 | 1 | 0 | 2 | 0.5 | 6 | 0.5 |
| 2_12642980 | 1 | 0 | 3 | 0.3333 | NA | NA | 1 | 0 | 5 | 0.2 | 1 | 0 | 1 | 0 | 2 | 0 | 2 | 0.5 | 6 | 0.1667 |
| 2_12642984 | 1 | 0 | 3 | 1 | 1 | 0 | 1 | 0 | 6 | 0.5 | 1 | 0 | 2 | 0.5 | 2 | 0 | 2 | 0 | 7 | 0.1429 |
| 2_12647103 | 3 | 0.3333 | 2 | 0 | 2 | 0.5 | 3 | 0.6667 | 10 | 0.4 | 2 | 0.5 | 2 | 0.5 | 5 | 0 | 2 | 0 | 11 | 0.1818 |
| 2_12647121 | 3 | 1 | 2 | 1 | 2 | 0.5 | 3 | 0.6667 | 10 | 0.8 | 2 | 0.5 | 1 | 1 | 4 | 1 | 2 | 1 | 9 | 0.8889 |
| 2_12647133 | 3 | 0.3333 | 2 | 1 | 2 | 0.5 | 4 | 0.5 | 11 | 0.5455 | 2 | 1 | 2 | 0.5 | 6 | 0.3333 | 2 | 1 | 12 | 0.5833 |
| 2_12647171 | 4 | 0.25 | 2 | 0.5 | 2 | 0 | 3 | 0.6667 | 11 | 0.3636 | NA | NA | 2 | 1 | 6 | 0.3333 | 2 | 0.5 | 10 | 0.5 |
| 2_12647175 | 3 | 1 | 2 | 1 | 2 | 1 | 4 | 0.5 | 11 | 0.8182 | 1 | 1 | 2 | 1 | 3 | 1 | 2 | 1 | 8 | 1 |
| 2_12647198 | 3 | 0.3333 | 1 | 0 | 2 | 0 | 4 | 0.25 | 10 | 0.2 | 1 | 1 | 2 | 0 | 4 | 0 | 1 | 1 | 8 | 0.25 |
| 2_12647202 | 3 | 1 | 1 | 1 | 1 | 1 | 4 | 0.75 | 9 | 0.8889 | 1 | 1 | 2 | 1 | 6 | 1 | NA | NA | 9 | 1 |
| 2_12647207 | 3 | 0 | 1 | 1 | 2 | 1 | 4 | 0.5 | 10 | 0.5 | 1 | 0 | 2 | 0.5 | 6 | 0.5 | 1 | 1 | 10 | 0.5 |
| 2_12647209 | 1 | 0 | 1 | 1 | 2 | 1 | 3 | 0.6667 | 7 | 0.7143 | 1 | 1 | 2 | 1 | 4 | 1 | 2 | 1 | 9 | 1 |
| 2_12655113 | 23 | 0.6087 | 20 | 0.75 | 9 | 0.7778 | 12 | 0.8333 | 64 | 0.7188 | 23 | 0.5217 | 14 | 0.5 | 15 | 0.6 | 17 | 0.5294 | 69 | 0.5362 |
| 2_12655121 | 24 | 0.375 | 15 | 0.3333 | 8 | 0.375 | 11 | 0.4545 | 58 | 0.3793 | 17 | 0.1176 | 13 | 0.3846 | 8 | 0.375 | 15 | 0.2667 | 53 | 0.2642 |
| 2_12656001 | 1 | 1 | 1 | 0 | 1 | 0 | NA | NA | 3 | 0.3333 | 1 | 0 | 3 | 0 | 2 | 0.5 | 1 | 0 | 7 | 0.1429 |
| 2_12657990 | 3 | 0 | 4 | 0.5 | 1 | 0 | 5 | 0 | 13 | 0.1538 | 1 | 0 | 4 | 0 | 4 | 0.25 | 4 | 0 | 13 | 0.0769 |
| 2_12658425 | 3 | 0.6667 | 6 | 0.5 | 1 | 1 | 6 | 0.6667 | 16 | 0.625 | NA | NA | 4 | 0.25 | 6 | 0.3333 | 5 | 0.6 | 15 | 0.4 |
| 2_12658425 | 3 | 0 | 6 | 0.3333 | 1 | 0 | 7 | 0.1429 | 17 | 0.1765 | 1 | 0 | 5 | 0.2 | 6 | 0 | 5 | 0.2 | 17 | 0.1176 |

| | | | | | | | | | | | | | | | | | | | | |
|---|---|---|---|---|---|---|---|---|---|---|---|---|---|---|---|---|---|---|---|---|
| 2_1265843 | 3 | 0 | 6 | 0 | NA | NA | 7 | 0.1429 | 16 | 0.0625 | 2 | 0 | 5 | 0 | 6 | 0.1667 | 5 | 0.2 | 18 | 0.1111 |
| 2_1265844 | 4 | 0 | 6 | 0.1667 | NA | NA | 7 | 0.1429 | 17 | 0.1176 | 1 | 0 | 5 | 0 | 6 | 0.1667 | 5 | 0 | 17 | 0.0588 |
| 2_1265844 | 4 | 0 | 6 | 0.3333 | 1 | 0 | 7 | 0.1429 | 18 | 0.1667 | 2 | 0 | 6 | 0 | 6 | 0 | 5 | 0.2 | 19 | 0.0526 |
| 2_1265929 | 5 | 0 | 4 | 0.75 | NA | NA | 6 | 0.3333 | 15 | 0.3333 | 2 | 0 | 6 | 0 | 6 | 0.1667 | 7 | 0.1429 | 21 | 0.0952 |
| 2_1265930 | 5 | 0.4 | 4 | 0.25 | 1 | 0 | 7 | 0.2857 | 17 | 0.2941 | 3 | 0 | 4 | 0 | 6 | 0 | 6 | 0.1667 | 19 | 0.0526 |
| 2_1265942 | 6 | 0.1667 | 7 | 0 | 1 | 0 | 8 | 0.25 | 22 | 0.1364 | 3 | 0.3333 | 6 | 0.1667 | 8 | 0.125 | 6 | 0.3333 | 23 | 0.2174 |
| 2_1266006 | 2 | 0 | 5 | 0.2 | NA | NA | 9 | 0.1111 | 16 | 0.125 | 4 | 0 | 5 | 0.2 | 8 | 0.125 | 6 | 0 | 23 | 0.087 |
| 2_1266341 | 1 | 1 | 5 | 0.6 | 6 | 0.3333 | 2 | 0.5 | 14 | 0.5 | NA | NA | 2 | 0 | 7 | 0.2857 | 8 | 0.125 | 17 | 0.1765 |
| 2_1266344 | 1 | 1 | 5 | 0.2 | 6 | 0.1667 | 1 | 0 | 13 | 0.2308 | NA | NA | 3 | 0 | 6 | 0.3333 | 8 | 0 | 17 | 0.1176 |
| 2_1266347 | 1 | 0 | 3 | 0 | 3 | 0.3333 | NA | NA | 7 | 0.1429 | NA | NA | 2 | 0.5 | 3 | 0.3333 | 5 | 0 | 10 | 0.2 |
| 2_1266549 | NA | NA | 2 | 0 | 2 | 0.5 | NA | NA | 4 | 0.25 | NA | NA | 2 | 0 | 4 | 0.25 | 3 | 0 | 9 | 0.1111 |
| 2_1267343 | 3 | 0 | 4 | 0 | 3 | 0.3333 | 5 | 0 | 15 | 0.0667 | 3 | 0.3333 | 2 | 0 | 4 | 0 | 2 | 0 | 11 | 0.0909 |
| 2_1267354 | NA | NA | 4 | 0.25 | 1 | 0 | 5 | 0.2 | 10 | 0.2 | 2 | 0 | 3 | 0.3333 | 4 | 0.5 | 4 | 0 | 13 | 0.2308 |
| 2_1267359 | 2 | 0 | 3 | 0 | 2 | 0.5 | 4 | 0 | 11 | 0.0909 | 3 | 0 | 3 | 0 | 3 | 0 | 5 | 0.4 | 14 | 0.1429 |
| 2_1267361 | 2 | 0 | 4 | 0.75 | 2 | 0 | 5 | 0.2 | 13 | 0.3077 | 2 | 0.5 | 3 | 0.6667 | 4 | 0 | 1 | 0 | 10 | 0.3 |
| 2_1269052 | NA | NA | 1 | 1 | 1 | 1 | NA | NA | 2 | 1 | NA | NA | NA | NA | NA | NA | 2 | 0.5 | 2 | 0.5 |
| 2_1269088 | NA | NA | NA | NA | 1 | 1 | NA | NA | 1 | 1 | NA | NA | NA | NA | NA | NA | 2 | 0.5 | 2 | 0.5 |
| 2_1269387 | 5 | 0.4 | 2 | 0.5 | 6 | 0 | 1 | 0 | 14 | 0.2143 | 2 | 1 | 3 | 0 | 3 | 1 | 6 | 0.8333 | 14 | 0.7143 |
| 2_1269822 | 1 | 1 | NA | NA | NA | NA | NA | NA | 1 | 1 | 1 | 1 | NA | NA | NA | NA | NA | NA | 1 | 1 |
| 2_1269824 | 1 | 1 | NA | NA | NA | NA | NA | NA | 1 | 1 | 1 | 1 | NA | NA | NA | NA | NA | NA | 1 | 1 |
| 2_1269828 | 1 | 1 | NA | NA | NA | NA | NA | NA | 1 | 1 | 1 | 1 | NA | NA | NA | NA | NA | NA | 1 | 1 |
| 2_1270721 | 1 | 0 | 4 | 0 | 7 | 0 | 3 | 0.3333 | 15 | 0.0667 | 5 | 0 | 2 | 0.5 | 4 | 0 | 1 | 0 | 12 | 0.0833 |
| 2_1270743 | 3 | 0 | 5 | 0 | 10 | 0.1 | 9 | 0.1111 | 27 | 0.0741 | 5 | 0.2 | 4 | 0 | 6 | 0 | 1 | 0 | 16 | 0.0625 |
| 2_1270800 | 6 | 0 | 9 | 0.1111 | 20 | 0.05 | 11 | 0.0909 | 46 | 0.0652 | 13 | 0.1538 | 6 | 0 | 11 | 0.1818 | 6 | 0 | 36 | 0.1111 |
| 2_1270821 | 8 | 0.125 | 11 | 0.2727 | 13 | 0.3077 | 7 | 0.2857 | 39 | 0.2564 | 10 | 0.3 | 3 | 0.6667 | 7 | 0.2857 | 6 | 0.3333 | 26 | 0.3462 |
| 2_1270822 | 7 | 0 | 15 | 0.0667 | 22 | 0.1818 | 12 | 0.3333 | 56 | 0.1607 | 14 | 0.2143 | 6 | 0.1667 | 14 | 0.1429 | 6 | 0.1667 | 40 | 0.175 |
| 2_1270830 | 1 | 0 | 2 | 0.5 | 2 | 0 | 3 | 0.6667 | 8 | 0.375 | NA | NA | 4 | 0.25 | 2 | 0 | 1 | 0 | 7 | 0.1429 |
| 2_1270831 | 1 | 0 | 1 | 0 | 2 | 0.5 | 2 | 0 | 6 | 0.1667 | NA | NA | 4 | 0 | 2 | 0.5 | 1 | 1 | 7 | 0.2857 |
| 2_1270832 | 1 | 0 | 2 | 0 | 2 | 0.5 | 2 | 0 | 7 | 0.1429 | NA | NA | 4 | 0.25 | 2 | 0 | 1 | 0 | 7 | 0.1429 |
| 2_1270846 | 8 | 0.875 | 11 | 1 | 8 | 0.875 | 4 | 1 | 31 | 0.9355 | 6 | 1 | 10 | 0.9 | 6 | 1 | 10 | 1 | 32 | 0.9688 |
| 2_1270846 | 9 | 0.4444 | 7 | 0.4286 | 7 | 0.2857 | 4 | 0 | 27 | 0.3333 | 6 | 0.5 | 8 | 0.25 | 6 | 0.3333 | 11 | 0.4545 | 31 | 0.3871 |
| 2_1270847 | 10 | 0.2 | 8 | 0.125 | 8 | 0.25 | 2 | 0 | 28 | 0.1786 | 4 | 0 | 6 | 0.3333 | 5 | 0.4 | 8 | 0.25 | 23 | 0.2609 |
| 2_1270900 | NA | NA | 1 | 0 | 1 | 1 | NA | NA | 2 | 0.5 | NA | NA | NA | NA | 1 | 1 | NA | NA | 1 | 1 |
| 2_1270993 | 4 | 0.25 | 1 | 0 | 3 | 0.3333 | 3 | 0 | 11 | 0.1818 | 2 | 0 | 2 | 0 | 3 | 0.3333 | 1 | 0 | 8 | 0.125 |
| 2_1270993 | 4 | 1 | 1 | 0 | 3 | 0.6667 | 3 | 0.3333 | 11 | 0.6364 | 2 | 1 | 2 | 0 | 3 | 0.6667 | 2 | 0.5 | 9 | 0.5556 |
| 2_1270994 | 5 | 0.6 | 1 | 0 | 1 | 0 | 3 | 0.6667 | 10 | 0.5 | 1 | 0 | 2 | 1 | 3 | 1 | 1 | 1 | 7 | 0.8571 |
| 2_1270994 | 3 | 1 | 1 | 0 | 2 | 1 | 3 | 0.6667 | 9 | 0.7778 | 2 | 0.5 | 1 | 0 | 2 | 0 | 2 | 1 | 7 | 0.4286 |
| 2_1270995 | 4 | 1 | 1 | 0 | 2 | 0.5 | 3 | 0 | 10 | 0.5 | 2 | 0.5 | 2 | 1 | 3 | 0.3333 | 2 | 0 | 9 | 0.4444 |
| 2_1270998 | 4 | 0.5 | 1 | 1 | 2 | 0.5 | 3 | 1 | 10 | 0.7 | 1 | 1 | 2 | 0.5 | 3 | 0.6667 | 1 | 1 | 7 | 0.7143 |
| 2_1270998 | 4 | 0 | 1 | 1 | 2 | 0 | 3 | 0 | 10 | 0.1 | 1 | 0 | 3 | 0.3333 | 3 | 0 | 1 | 0 | 8 | 0.125 |
| 2_1271000 | 5 | 0 | 1 | 0 | 2 | 0.5 | 2 | 0 | 10 | 0.1 | 2 | 0 | 4 | 0.25 | 3 | 0 | 2 | 0 | 11 | 0.0909 |
| 2_1271001 | 4 | 0.25 | 1 | 0 | 1 | 0 | 2 | 0.5 | 8 | 0.25 | 2 | 0.5 | 3 | 0 | 3 | 0.3333 | 1 | 0 | 9 | 0.2222 |
| 2_1271003 | 6 | 0.1667 | 1 | 0 | 2 | 0 | 2 | 0 | 11 | 0.0909 | 1 | 0 | 2 | 0 | 3 | 0.3333 | 1 | 0 | 7 | 0.1429 |
| 2_1271007 | 8 | 0 | 5 | 0 | 4 | 0 | 2 | 0.5 | 19 | 0.0526 | 3 | 0 | 5 | 0.2 | 2 | 0 | 4 | 0 | 14 | 0.0714 |
| 2_1271007 | 9 | 0.8889 | 3 | 0.6667 | 4 | 0.75 | 3 | 0.6667 | 19 | 0.7895 | 3 | 0.3333 | 4 | 0.5 | 4 | 0.5 | 3 | 1 | 14 | 0.5714 |
| 2_1271007 | 7 | 0.8571 | 5 | 1 | 4 | 1 | 3 | 1 | 19 | 0.9474 | 2 | 1 | 5 | 0.8 | 2 | 0.5 | 2 | 1 | 11 | 0.8182 |
| 2_1271008 | 7 | 1 | 5 | 0.6 | 5 | 0.4 | 3 | 1 | 20 | 0.75 | 3 | 0.6667 | 5 | 0.8 | 4 | 1 | 3 | 0.6667 | 15 | 0.8 |
| 2_1271011 | 7 | 1 | 6 | 1 | 5 | 1 | 3 | 1 | 21 | 1 | 3 | 1 | 9 | 1 | 4 | 1 | 5 | 1 | 21 | 1 |
| 2_1271011 | 8 | 0.125 | 5 | 0.6 | 4 | 0.75 | 5 | 0.8 | 22 | 0.5 | 3 | 0.3333 | 7 | 0.4286 | 5 | 1 | 4 | 0.5 | 19 | 0.5789 |
| 2_1271012 | 8 | 0.625 | 6 | 0 | 5 | 0.8 | 5 | 0 | 24 | 0.375 | 2 | 0 | 8 | 0.5 | 5 | 0.6 | 5 | 0.4 | 20 | 0.45 |
| 2_1271012 | 9 | 0.7778 | 5 | 0.2 | 4 | 0.5 | 4 | 0.75 | 22 | 0.5909 | 3 | 0.6667 | 6 | 0.5 | 5 | 0.4 | 6 | 0.1667 | 20 | 0.4 |
| 2_1271014 | 9 | 1 | 4 | 1 | 4 | 1 | 5 | 1 | 22 | 1 | 3 | 1 | 9 | 1 | 4 | 1 | 5 | 1 | 21 | 1 |
| 2_1271015 | 8 | 0.25 | 3 | 0.3333 | 3 | 0.3333 | 5 | 0.6 | 19 | 0.3684 | 3 | 0.6667 | 6 | 0.1667 | 3 | 0.6667 | 4 | 0 | 16 | 0.3125 |
| 2_1271017 | 8 | 0.125 | 4 | 0 | 3 | 0.3333 | 3 | 0.3333 | 18 | 0.1667 | 2 | 0 | 7 | 0.2857 | 3 | 0.3333 | 5 | 0.2 | 17 | 0.2353 |
| 2_1271018 | 9 | 0.2222 | 6 | 0.5 | 4 | 0.5 | 6 | 0.3333 | 25 | 0.36 | 3 | 0.3333 | 8 | 0.375 | 4 | 0.25 | 5 | 0.2 | 20 | 0.3 |
| 2_1271019 | 9 | 0.4444 | 5 | 0.2 | 5 | 0.4 | 7 | 0.2857 | 26 | 0.3462 | 7 | 0.5714 | 8 | 0.25 | 1 | 0 | 7 | 0.2857 | 23 | 0.3478 |
| 2_1271019 | 9 | 0.6667 | 6 | 1 | 5 | 0.6 | 6 | 0.8333 | 26 | 0.7692 | 6 | 1 | 9 | 0.6667 | 4 | 1 | 7 | 1 | 26 | 0.8846 |
| 2_1271020 | 9 | 0.6667 | 6 | 0.5 | 5 | 0.4 | 5 | 0.8 | 25 | 0.6 | 4 | 0.5 | 7 | 0.5714 | 2 | 0 | 4 | 0.5 | 17 | 0.4706 |
| 2_1271024 | 7 | 0.4286 | 9 | 0.4444 | 8 | 0.375 | 7 | 0.1429 | 31 | 0.3548 | 9 | 0.3333 | 7 | 0.1429 | 7 | 0 | 7 | 0 | 30 | 0.1333 |
| 2_1271026 | 10 | 0.2 | 10 | 0.2 | 9 | 0.1111 | 8 | 0 | 37 | 0.1351 | 12 | 0 | 8 | 0.125 | 9 | 0.1111 | 7 | 0.1429 | 36 | 0.0833 |
| 2_1271028 | 1 | 1 | 1 | 1 | NA | NA | NA | NA | 2 | 1 | 1 | 1 | NA | NA | 3 | 0.3333 | 1 | 1 | 5 | 0.6 |
| 2_1271124 | 1 | 0 | 3 | 0.3333 | 1 | 0 | 3 | 0.3333 | 8 | 0.25 | 1 | 0 | 3 | 0.3333 | 1 | 0 | 2 | 0.5 | 7 | 0.2857 |
| 2_1271390 | 6 | 0.3333 | 8 | 0 | 6 | 0 | 8 | 0.25 | 28 | 0.1429 | 8 | 0.125 | 8 | 0.25 | 9 | 0.1111 | 9 | 0.3333 | 34 | 0.2059 |
| 2_1271392 | 5 | 0 | 9 | 0.5556 | 6 | 0.1667 | 6 | 0.1667 | 26 | 0.2692 | 9 | 0.2222 | 10 | 0.1 | 8 | 0.375 | 10 | 0.4 | 37 | 0.2703 |
| 2_1271392 | 4 | 0.25 | 10 | 0.2 | 7 | 0 | 9 | 0.2222 | 30 | 0.1667 | 10 | 0.1 | 10 | 0 | 9 | 0 | 9 | 0.2222 | 38 | 0.0789 |

| | | | | | | | | | | | | | | | | | | | | |
|---|---|---|---|---|---|---|---|---|---|---|---|---|---|---|---|---|---|---|---|---|
| 2_12713928 | 4 | 0.75 | 9 | 0.6667 | 6 | 0.8333 | 8 | 0.375 | 27 | 0.6296 | 8 | 0.5 | 9 | 0.5556 | 8 | 0.5 | 10 | 0.8 | 35 | 0.6 |
| 2_12713931 | 5 | 0.2 | 9 | 0.2222 | 6 | 0.5 | 9 | 0.3333 | 29 | 0.3103 | 8 | 0.125 | 10 | 0.2 | 9 | 0.4444 | 7 | 0.4286 | 34 | 0.2941 |
| 2_12713934 | 4 | 1 | 9 | 0.8889 | 6 | 0.6667 | 9 | 0.8889 | 28 | 0.8571 | 9 | 0.8889 | 9 | 0.8889 | 7 | 0.7143 | 7 | 1 | 32 | 0.875 |
| 2_12713949 | 5 | 0.2 | 10 | 0.1 | 7 | 0 | 9 | 0.1111 | 31 | 0.0968 | 9 | 0.1111 | 10 | 0 | 7 | 0 | 10 | 0.1 | 36 | 0.0556 |
| 2_12713953 | 6 | 0.1667 | 9 | 0.4444 | 4 | 0 | 8 | 0.125 | 27 | 0.2222 | 10 | 0.4 | 10 | 0.2 | 8 | 0.125 | 10 | 0.2 | 38 | 0.2368 |
| 2_12713970 | 7 | 0 | 12 | 0.0833 | 6 | 0.1667 | 6 | 0 | 31 | 0.0645 | 9 | 0.1111 | 9 | 0.1111 | 8 | 0 | 9 | 0 | 35 | 0.0571 |
| 2_12720698 | 1 | 1 | NA | NA | NA | NA | NA | NA | 1 | 1 | 1 | 1 | 1 | 1 | NA | NA | NA | NA | 2 | 1 |
| 2_12720708 | 1 | 1 | NA | NA | NA | NA | NA | NA | 1 | 1 | 1 | 1 | 1 | 1 | NA | NA | NA | NA | 2 | 1 |
| 2_12727824 | 2 | 0 | 4 | 0 | 4 | 0.25 | 3 | 0 | 13 | 0.0769 | 2 | 0.5 | 4 | 0.25 | 4 | 0.25 | 3 | 0.3333 | 13 | 0.3077 |
| 2_12728260 | NA | NA | NA | NA | NA | NA | 1 | 1 | 1 | 1 | NA | NA | NA | NA | 1 | 1 | NA | NA | 1 | 1 |
| 2_12729854 | 21 | 0.5714 | 25 | 0.32 | 16 | 0.5 | 17 | 0.3529 | 79 | 0.4304 | 11 | 0.7273 | 11 | 0.2727 | 13 | 0.1538 | 20 | 0.35 | 55 | 0.3636 |
| 2_12729854 | 18 | 0.0556 | 23 | 0.0435 | 16 | 0.1875 | 18 | 0.0556 | 75 | 0.08 | 9 | 0.2222 | 12 | 0.0833 | 15 | 0.0667 | 20 | 0.2 | 56 | 0.1429 |
| 2_12729867 | 20 | 0.1 | 23 | 0.0435 | 13 | 0.0769 | 15 | 0.0667 | 71 | 0.0704 | 11 | 0.1818 | 13 | 0.0769 | 12 | 0 | 17 | 0 | 53 | 0.0566 |
| 2_12729882 | 19 | 0.0526 | 22 | 0.0909 | 11 | 0.0909 | 15 | 0.0667 | 67 | 0.0746 | 8 | 0.25 | 12 | 0.1667 | 12 | 0 | 22 | 0.0455 | 54 | 0.0926 |
| 2_12729897 | 20 | 0 | 20 | 0.2 | 11 | 0.0909 | 14 | 0.0714 | 65 | 0.0923 | 7 | 0.1429 | 12 | 0.0833 | 9 | 0.2222 | 16 | 0.125 | 44 | 0.1364 |
| 2_12729906 | 19 | 0.1579 | 22 | 0.0909 | 14 | 0.0714 | 14 | 0.0714 | 69 | 0.1014 | 10 | 0.1 | 10 | 0.1 | 11 | 0.2727 | 20 | 0.2 | 51 | 0.1765 |
| 2_12729933 | 18 | 0.3889 | 22 | 0.2727 | 14 | 0.3571 | 13 | 0.3077 | 67 | 0.3284 | 8 | 0.5 | 12 | 0.25 | 11 | 0.1818 | 18 | 0.4444 | 49 | 0.3469 |
| 2_12729946 | 10 | 0.5 | 13 | 0.6923 | 8 | 0.375 | 7 | 0.1429 | 38 | 0.4737 | 8 | 0.875 | 6 | 0.5 | 6 | 0.5 | 12 | 0.5 | 32 | 0.5938 |
| 2_12729965 | 10 | 0.5 | 10 | 0.5 | 7 | 0.2857 | 8 | 0.25 | 35 | 0.4 | 7 | 0.2857 | 10 | 0.3 | 8 | 0.5 | 7 | 0.2857 | 32 | 0.3438 |
| 2_12729983 | 17 | 0.2941 | 18 | 0.1111 | 9 | 0.3333 | 10 | 0.3 | 54 | 0.2407 | 7 | 0.2857 | 11 | 0.2727 | 8 | 0.125 | 12 | 0.0833 | 38 | 0.1842 |
| 2_12729992 | 15 | 0.1333 | 20 | 0.05 | 9 | 0.1111 | 8 | 0.25 | 52 | 0.1154 | 5 | 0 | 8 | 0.125 | 8 | 0 | 15 | 0.2 | 36 | 0.1111 |
| 2_12730025 | 11 | 0.1818 | 16 | 0.25 | 4 | 0 | 7 | 0.1429 | 38 | 0.1842 | 7 | 0 | 9 | 0.3333 | 3 | 0 | 10 | 0.4 | 29 | 0.2414 |
| 2_12741870 | 1 | 1 | NA | NA | NA | NA | 1 | 0 | 2 | 0.5 | NA | NA | NA | NA | 1 | 1 | NA | NA | 1 | 1 |
| 2_12741887 | 1 | 1 | NA | NA | NA | NA | 1 | 1 | 2 | 1 | NA | NA | NA | NA | 2 | 1 | NA | NA | 2 | 1 |
| 2_12741920 | NA | NA | NA | NA | NA | NA | 1 | 1 | 1 | 1 | NA | NA | NA | NA | 1 | 1 | NA | NA | 1 | 1 |
| 2_12741936 | NA | NA | NA | NA | NA | NA | 1 | 1 | 1 | 1 | NA | NA | NA | NA | 1 | 1 | NA | NA | 1 | 1 |
| 2_12741954 | NA | NA | NA | NA | NA | NA | 1 | 1 | 1 | 1 | NA | NA | NA | NA | 1 | 1 | NA | NA | 1 | 1 |
| 2_12744570 | 7 | 0.5714 | 6 | 0.6667 | 7 | 0.7143 | 6 | 0.6667 | 26 | 0.6538 | 5 | 0.6 | 6 | 0.1667 | 11 | 0.7273 | 5 | 0.8 | 27 | 0.5926 |
| 2_12744572 | 6 | 0.1667 | 7 | 0.2857 | 5 | 0.6 | 7 | 0.1429 | 25 | 0.28 | 5 | 0.8 | 7 | 0.4286 | 8 | 0.75 | 5 | 0.2 | 25 | 0.56 |
| 2_12744582 | 8 | 0.125 | 8 | 0 | 8 | 0.25 | 8 | 0.375 | 32 | 0.1875 | 5 | 0 | 7 | 0.1429 | 9 | 0.4444 | 6 | 0.3333 | 27 | 0.2593 |
| 2_12747553 | 35 | 0.4857 | 36 | 0.4722 | 32 | 0.5625 | 38 | 0.6053 | 141 | 0.5319 | 26 | 0.6154 | 21 | 0.5238 | 30 | 0.7 | 37 | 0.6757 | 114 | 0.6404 |
| 2_12747580 | 35 | 0.2571 | 33 | 0.2424 | 34 | 0.3235 | 36 | 0.2778 | 138 | 0.2754 | 24 | 0.25 | 24 | 0.125 | 29 | 0.2759 | 31 | 0.1935 | 108 | 0.213 |
| 2_12747597 | 34 | 0.0882 | 37 | 0.2162 | 35 | 0.1714 | 36 | 0.1389 | 142 | 0.1549 | 23 | 0.087 | 22 | 0.1818 | 29 | 0.1379 | 30 | 0.1 | 104 | 0.125 |
| 2_12747600 | 23 | 0.2609 | 26 | 0.3077 | 23 | 0.4348 | 28 | 0.2857 | 100 | 0.32 | 19 | 0.3684 | 12 | 0.25 | 22 | 0.0909 | 28 | 0.2857 | 81 | 0.2469 |
| 2_12747621 | 27 | 0.4074 | 26 | 0.1154 | 20 | 0.35 | 34 | 0.1765 | 107 | 0.2523 | 18 | 0.1111 | 20 | 0.25 | 20 | 0.45 | 23 | 0.2609 | 81 | 0.2716 |
| 2_12747626 | 30 | 0.5 | 26 | 0.5 | 25 | 0.32 | 32 | 0.5312 | 113 | 0.469 | 20 | 0.55 | 20 | 0.5 | 17 | 0.3529 | 24 | 0.4167 | 81 | 0.4568 |
| 2_12747631 | 32 | 0.1562 | 37 | 0.0811 | 30 | 0.1667 | 36 | 0.0833 | 135 | 0.1185 | 23 | 0.087 | 22 | 0.1364 | 25 | 0.28 | 30 | 0.1333 | 100 | 0.16 |
| 2_12747635 | 29 | 0.1379 | 32 | 0.1562 | 26 | 0.0769 | 32 | 0.125 | 119 | 0.1261 | 16 | 0.125 | 21 | 0.1429 | 20 | 0.05 | 26 | 0.2308 | 83 | 0.1446 |
| 2_12747660 | 21 | 0.1905 | 27 | 0.3704 | 25 | 0.24 | 29 | 0.2414 | 102 | 0.2647 | 18 | 0.2778 | 19 | 0.3158 | 16 | 0.1875 | 24 | 0.125 | 77 | 0.2208 |
| 2_12747662 | 24 | 0.7917 | 30 | 0.8 | 26 | 0.8846 | 32 | 0.9375 | 112 | 0.8571 | 17 | 0.8235 | 10 | 0.8 | 20 | 0.75 | 22 | 0.8182 | 69 | 0.7971 |
| 2_12747664 | 26 | 0.1923 | 28 | 0.2143 | 26 | 0.1923 | 30 | 0.1 | 110 | 0.1727 | 19 | 0.1053 | 16 | 0.0625 | 18 | 0.1111 | 22 | 0.2727 | 75 | 0.1467 |
| 2_12747682 | 29 | 0.5517 | 29 | 0.6552 | 23 | 0.7826 | 31 | 0.6452 | 112 | 0.6518 | 21 | 0.4762 | 18 | 0.6111 | 17 | 0.7059 | 23 | 0.3913 | 79 | 0.5316 |
| 2_12758275 | 2 | 0.5 | 3 | 1 | 2 | 1 | 3 | 0.6667 | 10 | 0.8 | NA | NA | 6 | 0.5 | 5 | 0.4 | 3 | 1 | 14 | 0.5714 |
| 2_12758276 | 2 | 0 | 4 | 0 | 2 | 0 | 3 | 0.3333 | 11 | 0.0909 | NA | NA | 7 | 0 | 4 | 0 | 3 | 0.3333 | 14 | 0.0714 |
| 2_12758280 | 1 | 0 | 3 | 0 | 1 | 0 | 4 | 0.25 | 9 | 0.1111 | NA | NA | 7 | 0.1429 | 5 | 0.2 | 2 | 0 | 14 | 0.1429 |
| 2_12758284 | 1 | 1 | 3 | 1 | NA | NA | 2 | 1 | 6 | 1 | NA | NA | 6 | 0.3333 | 3 | 0.6667 | 2 | 1 | 11 | 0.5455 |
| 2_12758287 | 2 | 0 | 3 | 0.3333 | 1 | 1 | 4 | 0 | 10 | 0.2 | NA | NA | 7 | 0 | 5 | 0.2 | 3 | 0.3333 | 15 | 0.1333 |
| 2_12758287 | 2 | 0 | 3 | 0.3333 | 2 | 0 | 3 | 0.3333 | 10 | 0.2 | NA | NA | 7 | 0 | 4 | 0.25 | 3 | 0.3333 | 14 | 0.1429 |
| 2_12763334 | 6 | 0.3333 | 9 | 0.1111 | 7 | 0.1429 | 10 | 0.1 | 32 | 0.1562 | 6 | 0 | 9 | 0.2222 | 5 | 0 | 7 | 0 | 27 | 0.0741 |
| 2_12763349 | 6 | 0.1667 | 9 | 0 | 7 | 0 | 11 | 0.1818 | 33 | 0.0909 | 4 | 0 | 10 | 0.2 | 4 | 0.25 | 8 | 0 | 26 | 0.1154 |
| 2_12763352 | 5 | 0.6 | 7 | 0.1429 | 7 | 0.5714 | 11 | 0.5455 | 30 | 0.4667 | 5 | 0.6 | 11 | 0.5455 | 4 | 0 | 7 | 0.5714 | 27 | 0.4815 |
| 2_12763357 | 6 | 1 | 8 | 0.875 | 6 | 0.8333 | 11 | 0.9091 | 31 | 0.9032 | 5 | 0.6 | 9 | 1 | 6 | 0.6667 | 8 | 0.75 | 28 | 0.7857 |
| 2_12800131 | NA | NA | NA | NA | 1 | 1 | NA | NA | 1 | 1 | 1 | 1 | NA | NA | NA | NA | NA | NA | 1 | 1 |
| 2_12800352 | 1 | 1 | NA | NA | 1 | 0 | NA | NA | 2 | 0.5 | 2 | 1 | NA | NA | NA | NA | NA | NA | 2 | 1 |
| 2_12800385 | 1 | 1 | NA | NA | 1 | 1 | NA | NA | 2 | 1 | 2 | 1 | NA | NA | NA | NA | NA | NA | 2 | 1 |
| 2_12845269 | 8 | 0 | 5 | 0.4 | 3 | 0 | 11 | 0 | 27 | 0.0741 | 3 | 0 | 2 | 1 | 1 | 0 | 3 | 0.3333 | 9 | 0.3333 |
| 2_12845283 | 6 | 0.1667 | 6 | 0.1667 | 3 | 0.3333 | 9 | 0.3333 | 24 | 0.25 | 2 | 1 | 2 | 0.5 | 1 | 0 | 3 | 0 | 8 | 0.375 |
| 2_12845290 | 6 | 0.1667 | 6 | 0.3333 | 2 | 0 | 9 | 0.2222 | 23 | 0.2174 | 2 | 0 | NA | NA | 1 | 0 | 4 | 0.25 | 7 | 0.1429 |
| 2_12845293 | 6 | 0.8333 | 5 | 1 | 3 | 0.6667 | 7 | 0.8571 | 21 | 0.8571 | 2 | 1 | 2 | 1 | 1 | 1 | 2 | 1 | 7 | 1 |
| 2_12845303 | 8 | 1 | 5 | 1 | 2 | 1 | 7 | 1 | 22 | 1 | 3 | 1 | 1 | 1 | 1 | 1 | 4 | 1 | 9 | 1 |
| 2_12845308 | 5 | 0 | 4 | 0.25 | 2 | 0 | 7 | 0.2857 | 18 | 0.1667 | 2 | 1 | 2 | 0 | 1 | 0 | 2 | 0 | 7 | 0.2857 |
| 2_12845317 | 5 | 0.2 | 6 | 0.1667 | 2 | 0 | 7 | 0.4286 | 20 | 0.25 | 1 | 0 | 2 | 1 | 1 | 0 | 3 | 0.3333 | 7 | 0.4286 |
| 2_12845318 | 7 | 0.2857 | 6 | 0 | 2 | 0 | 10 | 0.2 | 25 | 0.16 | 3 | 0 | 2 | 0 | NA | NA | 4 | 0.25 | 9 | 0.1111 |
| 2_12845322 | 5 | 0.2 | 5 | 0 | 2 | 0 | 11 | 0.0909 | 23 | 0.087 | 3 | 0.3333 | 1 | 0 | 1 | 0 | 4 | 0.25 | 9 | 0.2222 |
| 2_12845328 | 7 | 0.2857 | 3 | 0 | 2 | 0 | 10 | 0 | 22 | 0.0909 | 3 | 0.3333 | 1 | 0 | NA | NA | 4 | 0.25 | 8 | 0.25 |

| | | | | | | | | | | | | | | | | | | | | |
|---|---|---|---|---|---|---|---|---|---|---|---|---|---|---|---|---|---|---|---|---|
| 2_1284534 | 5 | 0.2 | 6 | 1 | 2 | 0.5 | 8 | 0.375 | 21 | 0.5238 | 3 | 0.3333 | 1 | 0 | 1 | 1 | 4 | 0.25 | 9 | 0.3333 |
| 2_1284535 | 7 | 0.7143 | 5 | 1 | 2 | 1 | 9 | 1 | 23 | 0.913 | 3 | 1 | 1 | 1 | 1 | 1 | 4 | 0.75 | 9 | 0.8889 |
| 2_1284535 | 5 | 0.8 | 4 | 0.75 | 2 | 1 | 11 | 0.8182 | 22 | 0.8182 | 3 | 1 | 1 | 1 | 1 | 1 | 4 | 0.5 | 9 | 0.7778 |
| 2_1284536 | 5 | 1 | 4 | 1 | 2 | 1 | 8 | 0.75 | 19 | 0.8947 | 3 | 1 | 1 | 1 | 1 | 1 | 4 | 1 | 9 | 1 |
| 2_1284538 | 6 | 0.5 | 4 | 0 | 2 | 1 | 10 | 0.3 | 22 | 0.3636 | 3 | 0 | 1 | 0 | 1 | 0 | 4 | 0.25 | 9 | 0.1111 |
| 2_1284538 | 7 | 0.5714 | 3 | 1 | 2 | 1 | 8 | 1 | 20 | 0.85 | 2 | 1 | 1 | 1 | 1 | 1 | 1 | 1 | 5 | 1 |
| 2_1284539 | 7 | 0.4286 | 3 | 0 | 2 | 0 | 9 | 0.4444 | 21 | 0.3333 | 3 | 0.6667 | 1 | 0 | NA | NA | 3 | 0.6667 | 7 | 0.5714 |
| 2_1284539 | 5 | 0.6 | 4 | 0.5 | 1 | 1 | 8 | 0.125 | 18 | 0.3889 | 3 | 0 | 1 | 1 | NA | NA | 2 | 0 | 6 | 0.1667 |
| 2_1284540 | 5 | 0.4 | 3 | 0.6667 | 2 | 0.5 | 8 | 0.5 | 18 | 0.5 | 1 | 0 | 1 | 0 | 1 | 0 | 4 | 0.5 | 7 | 0.2857 |
| 2_1284541 | 6 | 0.3333 | 2 | 0 | 2 | 0.5 | 8 | 0.125 | 18 | 0.2222 | 2 | 0 | 1 | 0 | 1 | 0 | 4 | 0.5 | 8 | 0.25 |
| 2_1284541 | 5 | 0.6 | 2 | 0 | 1 | 1 | 7 | 0.5714 | 15 | 0.5333 | 2 | 0 | 1 | 0 | 1 | 0 | 2 | 0.5 | 6 | 0.1667 |
| 2_1284543 | 6 | 1 | 2 | 1 | 1 | 1 | 8 | 0.875 | 17 | 0.9412 | NA | NA | 1 | 1 | 1 | 1 | 3 | 1 | 5 | 1 |
| 2_1284545 | 6 | 0.8333 | 2 | 1 | NA | NA | 9 | 0.7778 | 17 | 0.8235 | 2 | 1 | 1 | 1 | NA | NA | 3 | 1 | 6 | 1 |
| 2_1284545 | 6 | 0.6667 | 3 | 1 | 1 | 0 | 6 | 0.3333 | 16 | 0.5625 | 2 | 1 | 1 | 0 | 1 | 1 | 3 | 0.3333 | 7 | 0.5714 |
| 2_1284546 | 5 | 1 | 2 | 1 | 1 | 1 | 6 | 0.6667 | 14 | 0.8571 | 2 | 1 | 1 | 1 | 1 | 1 | 3 | 0 | 7 | 0.5714 |
| 2_1284546 | 6 | 0 | 2 | 0.5 | 2 | 0 | 8 | 0.25 | 18 | 0.1667 | 4 | 0.5 | 2 | 0 | 4 | 0.25 | 3 | 0 | 13 | 0.2308 |
| 2_1284547 | 6 | 0.1667 | 4 | 0.5 | 3 | 0.3333 | 7 | 0.1429 | 20 | 0.25 | 3 | 0 | 1 | 0 | 4 | 0.25 | 2 | 0 | 10 | 0.1 |
| 2_1284547 | 5 | 0.8 | 4 | 1 | 1 | 1 | 8 | 0.75 | 18 | 0.8333 | 5 | 1 | 1 | 1 | 2 | 0.5 | 2 | 1 | 10 | 0.9 |
| 2_1284549 | 7 | 0.1429 | 3 | 0 | 3 | 0.3333 | 7 | 0.2857 | 20 | 0.2 | 5 | 0.4 | 1 | 0 | 3 | 0 | 2 | 0 | 11 | 0.1818 |
| 2_1285713 | 1 | 1 | 4 | 0.5 | 2 | 0 | 2 | 0 | 9 | 0.3333 | 2 | 0.5 | 6 | 0.1667 | 2 | 0.5 | 2 | 0 | 12 | 0.25 |
| 2_1285713 | 2 | 0 | 5 | 0.2 | 2 | 0.5 | 3 | 0.3333 | 12 | 0.25 | 2 | 0 | 6 | 0.1667 | 1 | 1 | 2 | 0 | 11 | 0.1818 |
| 2_1286150 | NA | NA | NA | NA | 1 | 1 | NA | NA | 1 | 1 | NA | NA | 1 | 1 | 1 | 1 | NA | NA | 2 | 1 |
| 2_1286150 | NA | NA | NA | NA | 1 | 1 | NA | NA | 1 | 1 | NA | NA | 1 | 0 | 1 | 1 | NA | NA | 2 | 0.5 |
| 2_1286152 | NA | NA | NA | NA | 1 | 1 | NA | NA | 1 | 1 | NA | NA | 1 | 1 | 1 | 1 | NA | NA | 2 | 1 |
| 2_1286152 | NA | NA | NA | NA | 1 | 1 | NA | NA | 1 | 1 | NA | NA | 1 | 1 | 1 | 1 | NA | NA | 2 | 1 |
| 2_1286154 | NA | NA | NA | NA | 1 | 1 | NA | NA | 1 | 1 | NA | NA | 2 | 1 | 1 | 1 | NA | NA | 3 | 1 |
| 2_1286155 | NA | NA | NA | NA | 1 | 1 | NA | NA | 1 | 1 | NA | NA | 2 | 1 | 1 | 1 | NA | NA | 3 | 1 |
| 2_1286157 | NA | NA | NA | NA | 1 | 1 | NA | NA | 1 | 1 | NA | NA | 2 | 0.5 | 1 | 1 | NA | NA | 3 | 0.6667 |
| 2_1286432 | 4 | 0 | 11 | 0.4545 | 5 | 0.2 | 5 | 0 | 25 | 0.24 | 4 | 0 | 9 | 0.4444 | 4 | 0 | 6 | 0.1667 | 23 | 0.2174 |
| 2_1286432 | 4 | 0.5 | 10 | 0.9 | 5 | 0.8 | 4 | 0.75 | 23 | 0.7826 | 2 | 0 | 7 | 0.5714 | 3 | 0.6667 | 2 | 0.5 | 14 | 0.5 |
| 2_1286432 | 3 | 0 | 11 | 0.2727 | 5 | 0 | 6 | 0.3333 | 25 | 0.2 | 4 | 0.25 | 9 | 0.3333 | 4 | 0.25 | 5 | 0.2 | 22 | 0.2727 |
| 2_1286434 | 3 | 0.6667 | 9 | 1 | 4 | 1 | 7 | 0.4286 | 23 | 0.7826 | 2 | 0.5 | 7 | 0.8571 | 4 | 0.75 | 4 | 1 | 17 | 0.8235 |
| 2_1286434 | 3 | 0.6667 | 11 | 0.5455 | 3 | 0.3333 | 6 | 0.3333 | 23 | 0.4783 | 3 | 0.3333 | 6 | 0.5 | 4 | 0.75 | 3 | 1 | 16 | 0.625 |
| 2_1286435 | 5 | 0.8 | 11 | 0.9091 | 4 | 1 | 4 | 0.75 | 24 | 0.875 | 4 | 0.75 | 8 | 0.75 | 3 | 1 | 2 | 1 | 17 | 0.8235 |
| 2_1286435 | 6 | 0 | 15 | 0.3333 | 6 | 0.3333 | 4 | 0.25 | 31 | 0.2581 | 4 | 0.5 | 11 | 0.2727 | 4 | 0.75 | 7 | 0.4286 | 26 | 0.4231 |
| 2_1286436 | 6 | 0.3333 | 12 | 0.1667 | 5 | 0.2 | 7 | 0 | 30 | 0.1667 | 5 | 0.2 | 6 | 0 | 3 | 0 | 7 | 0.1429 | 21 | 0.0952 |
| 2_1286441 | 7 | 0 | 14 | 0.2143 | 6 | 0.1667 | 8 | 0.25 | 35 | 0.1714 | 4 | 0 | 16 | 0.125 | 4 | 0 | 7 | 0.1429 | 31 | 0.0968 |
| 2_1286443 | 4 | 0 | 11 | 0.0909 | 4 | 1 | 7 | 0.5714 | 26 | 0.3462 | 3 | 0.6667 | 13 | 0.3846 | 5 | 0.6 | 6 | 0.6667 | 27 | 0.5185 |
| 2_1286933 | 2 | 0 | 6 | 0.1667 | 5 | 0.6 | 6 | 0.1667 | 19 | 0.2632 | 3 | 0.3333 | 1 | 1 | 4 | 0.25 | 2 | 0.5 | 10 | 0.4 |
| 2_1286934 | 5 | 0.4 | 5 | 0.6 | 3 | 0 | 5 | 0.8 | 18 | 0.5 | 3 | 1 | 1 | 0 | 4 | 0.5 | 1 | 1 | 9 | 0.6667 |
| 2_1286934 | 5 | 0.4 | 6 | 0.3333 | 5 | 0.2 | 7 | 0.4286 | 23 | 0.3478 | 3 | 0.6667 | 1 | 0 | 4 | 0 | 3 | 0.6667 | 11 | 0.3636 |
| 2_1286935 | 5 | 0.8 | 5 | 0.6 | 5 | 0.4 | 6 | 0.6667 | 21 | 0.619 | 3 | 1 | 1 | 1 | 4 | 0.75 | 3 | 1 | 11 | 0.9091 |
| 2_1286936 | 5 | 0.2 | 6 | 0.3333 | 5 | 0 | 6 | 0.3333 | 22 | 0.2273 | 3 | 1 | 1 | 0 | 4 | 0 | 3 | 1 | 11 | 0.5455 |
| 2_1286938 | 7 | 0.4286 | 6 | 0.6667 | 4 | 0.25 | 8 | 0 | 25 | 0.32 | 3 | 0.6667 | 1 | 0 | 5 | 0.2 | 3 | 0 | 12 | 0.25 |
| 2_1286938 | 6 | 0.8333 | 7 | 0.7143 | 4 | 0.25 | 8 | 0.625 | 25 | 0.64 | 3 | 1 | 1 | 1 | 5 | 0.6 | 3 | 1 | 12 | 0.8333 |
| 2_1286940 | 4 | 0.75 | 6 | 0.5 | 4 | 0.5 | 8 | 0.5 | 22 | 0.5455 | 3 | 0.3333 | 1 | 1 | 4 | 0.5 | 3 | 1 | 11 | 0.6364 |
| 2_1286940 | 7 | 0.1429 | 8 | 0.375 | 4 | 0 | 7 | 0.2857 | 26 | 0.2308 | 3 | 0 | 1 | 0 | 5 | 0.2 | 3 | 0 | 12 | 0.0833 |
| 2_1286941 | 8 | 0.25 | 8 | 0.25 | 6 | 0.3333 | 9 | 0.4444 | 31 | 0.3226 | 3 | 0.6667 | 1 | 1 | 5 | 0.2 | 4 | 0.5 | 13 | 0.4615 |
| 2_1286943 | 10 | 0 | 8 | 0.125 | 9 | 0.1111 | 7 | 0.2857 | 34 | 0.1176 | 1 | 0 | 1 | 0 | 5 | 0.2 | 2 | 0 | 9 | 0.1111 |
| 2_1286944 | 9 | 0.3333 | 7 | 0.2857 | 10 | 0.2 | 7 | 0.1429 | 33 | 0.2424 | 2 | 0 | 1 | 0 | 6 | 0.3333 | 3 | 0 | 12 | 0.1667 |
| 2_1286950 | 3 | 0 | 2 | 0.5 | 2 | 0.5 | 2 | 0 | 9 | 0.2222 | NA | NA | NA | NA | 2 | 0.5 | NA | NA | 2 | 0.5 |
| 2_1288075 | NA | NA | 1 | 1 | NA | NA | NA | NA | 1 | 1 | NA | NA | NA | NA | 1 | 1 | NA | NA | 1 | 1 |
| 2_1288075 | NA | NA | 1 | 1 | NA | NA | NA | NA | 1 | 1 | NA | NA | NA | NA | 1 | 1 | NA | NA | 1 | 1 |
| 2_1288083 | 1 | 0 | 1 | 1 | NA | NA | 2 | 0 | 4 | 0.25 | 1 | 1 | NA | NA | 2 | 0 | 2 | 0 | 5 | 0.2 |
| 2_1288086 | NA | NA | 1 | 0 | NA | NA | 1 | 1 | 2 | 0.5 | NA | NA | NA | NA | 2 | 0.5 | 3 | 0 | 5 | 0.2 |
| 2_1288087 | 2 | 1 | 1 | 1 | NA | NA | 1 | 0 | 4 | 0.75 | 1 | 1 | NA | NA | 2 | 0.5 | 3 | 0.3333 | 6 | 0.5 |
| 2_1288088 | 2 | 0.5 | 1 | 1 | NA | NA | 1 | 0 | 4 | 0.5 | 1 | 1 | NA | NA | 2 | 1 | 2 | 0.5 | 5 | 0.8 |
| 2_1288089 | 2 | 1 | NA | NA | NA | NA | 2 | 1 | 4 | 1 | 1 | 1 | NA | NA | 2 | 1 | 2 | 0.5 | 5 | 0.8 |
| 2_1288089 | 2 | 0.5 | 1 | 1 | NA | NA | 1 | 0 | 4 | 0.5 | 1 | 1 | NA | NA | 2 | 1 | 3 | 0.6667 | 6 | 0.8333 |
| 2_1288092 | 1 | 1 | 1 | 1 | NA | NA | 1 | 1 | 3 | 1 | NA | NA | NA | NA | NA | NA | 2 | 0.5 | 2 | 0.5 |
| 2_1288093 | 1 | 1 | 1 | 1 | NA | NA | NA | NA | 2 | 1 | NA | NA | NA | NA | NA | NA | 2 | 1 | 2 | 1 |
| 2_1288095 | NA | NA | 1 | 1 | NA | NA | 2 | 0.5 | 3 | 0.6667 | NA | NA | 3 | 0.6667 | 1 | 0 | NA | NA | 4 | 0.5 |
| 2_1288095 | 1 | 1 | 1 | 0 | NA | NA | NA | NA | 2 | 0.5 | 1 | 1 | NA | NA | 2 | 0.5 | 1 | 0 | 4 | 0.5 |
| 2_1288095 | 1 | 1 | 1 | 0 | NA | NA | 1 | 0 | 3 | 0.3333 | 1 | 0 | NA | NA | 2 | 0 | 2 | 0.5 | 5 | 0.2 |
| 2_1288097 | 1 | 1 | 1 | 0 | NA | NA | NA | NA | 2 | 0.5 | 1 | 1 | NA | NA | 1 | 0 | 2 | 0 | 4 | 0.25 |

| | | | | | | | | | | | | | | | | | | | | |
|---|---|---|---|---|---|---|---|---|---|---|---|---|---|---|---|---|---|---|---|---|
| 2_1288565 | 1 | 1 | 2 | 0 | 1 | 1 | 2 | 0 | 6 | 0.3333 | NA | NA | 2 | 1 | NA | NA | 1 | 1 | 3 | 1 |
| 2_1288581 | NA | NA | 1 | 1 | 1 | 1 | 1 | 1 | 3 | 1 | NA | NA | 2 | 0.5 | NA | NA | 1 | 1 | 3 | 0.6667 |
| 2_1288581 | 1 | 1 | 2 | 0.5 | 1 | 0 | 1 | 0 | 5 | 0.4 | NA | NA | 2 | 1 | NA | NA | 1 | 0 | 3 | 0.6667 |
| 2_1288582 | NA | NA | 2 | 1 | 1 | 1 | 3 | 0.6667 | 6 | 0.8333 | NA | NA | 2 | 1 | NA | NA | 1 | 1 | 3 | 1 |
| 2_1288582 | 1 | 1 | 2 | 1 | 1 | 1 | 2 | 1 | 6 | 1 | NA | NA | 2 | 1 | NA | NA | 1 | 0 | 3 | 0.6667 |
| 2_1288582 | 1 | 0 | 2 | 0.5 | 1 | 1 | 3 | 0.3333 | 7 | 0.4286 | NA | NA | 2 | 0.5 | NA | NA | 1 | 0 | 3 | 0.3333 |
| 2_1288583 | 1 | 0 | 2 | 0 | NA | NA | 3 | 0.6667 | 6 | 0.3333 | NA | NA | 2 | 0.5 | NA | NA | 1 | 0 | 3 | 0.3333 |
| 2_1288584 | NA | NA | 1 | 0 | 1 | 1 | 2 | 0.5 | 4 | 0.5 | NA | NA | 2 | 0.5 | NA | NA | 1 | 0 | 3 | 0.3333 |
| 2_1288585 | 1 | 1 | 2 | 1 | 1 | 1 | 3 | 0.3333 | 7 | 0.7143 | NA | NA | 2 | 1 | NA | NA | 1 | 1 | 3 | 1 |
| 2_1288585 | NA | NA | 2 | 1 | 1 | 1 | 3 | 1 | 6 | 1 | 1 | 1 | 2 | 1 | NA | NA | 1 | 1 | 4 | 1 |
| 2_1288586 | 1 | 0 | 2 | 1 | 1 | 1 | 3 | 0.3333 | 7 | 0.5714 | 1 | 1 | 2 | 0 | NA | NA | 1 | 0 | 4 | 0.25 |
| 2_1288587 | 1 | 1 | 2 | 0 | 1 | 0 | 1 | 1 | 5 | 0.4 | NA | NA | 3 | 0.3333 | NA | NA | 1 | 0 | 4 | 0.25 |
| 2_1288588 | 1 | 1 | 2 | 1 | 1 | 1 | 2 | 0 | 6 | 0.6667 | 1 | 1 | 3 | 0.6667 | NA | NA | 1 | 1 | 5 | 0.8 |
| 2_1288588 | 1 | 1 | 2 | 1 | 1 | 1 | 1 | 1 | 5 | 1 | 1 | 1 | 3 | 1 | NA | NA | 1 | 1 | 5 | 1 |
| 2_1288589 | 1 | 0 | 1 | 1 | 1 | 0 | 1 | 1 | 4 | 0.5 | NA | NA | 3 | 1 | NA | NA | 1 | 1 | 4 | 1 |
| 2_1288590 | 1 | 1 | 2 | 0 | NA | NA | 3 | 1 | 6 | 0.6667 | 2 | 0.5 | 3 | 1 | NA | NA | 1 | 0 | 6 | 0.6667 |
| 2_1288590 | 1 | 1 | 2 | 0 | 1 | 0 | 2 | 0 | 6 | 0.1667 | 2 | 1 | 3 | 0.6667 | NA | NA | 1 | 0 | 6 | 0.6667 |
| 2_1288592 | 1 | 0 | 1 | 1 | 1 | 0 | 1 | 1 | 4 | 0.5 | 1 | 0 | 3 | 0.6667 | NA | NA | 1 | 1 | 5 | 0.6 |
| 2_1288592 | 1 | 1 | 1 | 0 | 1 | 0 | 2 | 0 | 5 | 0.2 | NA | NA | 3 | 0.6667 | NA | NA | 1 | 0 | 4 | 0.5 |
| 2_1288592 | 2 | 0 | 2 | 0 | 1 | 0 | 3 | 1 | 8 | 0.375 | 2 | 0.5 | 3 | 0.3333 | NA | NA | 1 | 1 | 6 | 0.5 |
| 2_1288592 | 2 | 0.5 | 1 | 0 | 1 | 0 | 3 | 0 | 7 | 0.1429 | 1 | 1 | 3 | 0.3333 | NA | NA | 1 | 0 | 5 | 0.4 |
| 2_1288594 | 2 | 0.5 | 2 | 0 | 2 | 0.5 | 4 | 0.25 | 10 | 0.3 | 2 | 0.5 | 3 | 0.3333 | NA | NA | 1 | 0 | 6 | 0.3333 |
| 2_1288596 | 2 | 1 | 2 | 0 | 1 | 0 | 3 | 1 | 8 | 0.625 | 1 | 1 | 2 | 1 | NA | NA | 1 | 0 | 4 | 0.75 |
| 2_1288596 | 2 | 1 | 2 | 0.5 | 2 | 0.5 | 4 | 0.75 | 10 | 0.7 | 2 | 1 | 2 | 1 | NA | NA | 1 | 1 | 5 | 1 |
| 2_1288597 | 2 | 0.5 | 2 | 0.5 | 3 | 1 | 3 | 1 | 10 | 0.8 | 2 | 0.5 | 3 | 0.6667 | NA | NA | 1 | 0 | 6 | 0.5 |
| 2_1288602 | 5 | 0 | 2 | 0.5 | 4 | 0.25 | 3 | 0 | 14 | 0.1429 | 2 | 0.5 | 3 | 0 | NA | NA | 1 | 0 | 6 | 0.1667 |
| 2_1288604 | 4 | 0.25 | 3 | 0 | 4 | 0 | 2 | 0.5 | 13 | 0.1538 | 1 | 1 | 3 | 0 | NA | NA | 1 | 0 | 5 | 0.2 |
| 2_1289240 | 4 | 0.25 | 3 | 0.3333 | 2 | 0 | 2 | 0 | 11 | 0.1818 | 3 | 0.3333 | 4 | 0 | 3 | 0.3333 | 7 | 0.5714 | 17 | 0.3529 |
| 2_1289348 | 2 | 0.5 | 2 | 0 | 1 | 0 | 2 | 0 | 7 | 0.1429 | 2 | 0 | 5 | 0.2 | 1 | 0 | 4 | 0 | 12 | 0.0833 |
| 2_1289349 | 4 | 0 | 6 | 0.1667 | 2 | 0 | 2 | 0 | 14 | 0.0714 | 3 | 0 | 5 | 0.2 | 5 | 0.2 | 7 | 0.1429 | 20 | 0.15 |
| 2_1289350 | 4 | 0.25 | 6 | 0.3333 | 2 | 0 | 2 | 0 | 14 | 0.2143 | 3 | 0.3333 | 5 | 0.4 | 7 | 0.2857 | 7 | 0.2857 | 22 | 0.3182 |
| 2_1289350 | 5 | 0.8 | 6 | 1 | 3 | 1 | 1 | 1 | 15 | 0.9333 | 3 | 1 | 6 | 1 | 6 | 1 | 8 | 1 | 23 | 1 |
| 2_1289350 | 4 | 0.5 | 5 | 0.4 | 3 | 0.6667 | 1 | 1 | 13 | 0.5385 | 4 | 0.75 | 5 | 0.6 | 6 | 0.5 | 6 | 0.6667 | 21 | 0.619 |
| 2_1289351 | 5 | 1 | 7 | 0.7143 | 2 | 1 | 2 | 1 | 16 | 0.875 | 4 | 0.75 | 6 | 0.8333 | 7 | 1 | 9 | 1 | 26 | 0.9231 |
| 2_1289352 | 3 | 0.3333 | 7 | 0.1429 | 3 | 0 | 1 | 0 | 14 | 0.1429 | 3 | 0 | 5 | 0.2 | 6 | 0.5 | 8 | 0.25 | 22 | 0.2727 |
| 2_1289353 | 5 | 0.8 | 5 | 1 | 2 | 1 | 2 | 1 | 14 | 0.9286 | 2 | 0.5 | 5 | 1 | 5 | 1 | 7 | 1 | 19 | 0.9474 |
| 2_1289353 | 4 | 1 | 1 | 1 | NA | NA | 1 | 1 | 6 | 1 | 1 | 1 | NA | NA | 3 | 0.6667 | 4 | 0.75 | 8 | 0.75 |
| 2_1289353 | 5 | 0.8 | 6 | 1 | 2 | 1 | 1 | 1 | 14 | 0.9286 | 3 | 1 | 4 | 1 | 6 | 1 | 10 | 0.9 | 23 | 0.9565 |
| 2_1289354 | 1 | 1 | 5 | 1 | 1 | 1 | 2 | 1 | 9 | 1 | 3 | 0.6667 | 5 | 0.8 | 5 | 0.8 | 8 | 0.875 | 21 | 0.8095 |
| 2_1289354 | 4 | 0.5 | 8 | 0.625 | 4 | 0.5 | 2 | 0.5 | 18 | 0.5556 | 4 | 0.75 | 5 | 0.8 | 4 | 0.75 | 4 | 0.5 | 17 | 0.7059 |
| 2_1289357 | NA | NA | 1 | 1 | NA | NA | NA | NA | 1 | 1 | NA | NA | NA | NA | NA | NA | 1 | 1 | 1 | 1 |
| 2_1289357 | NA | NA | 1 | 1 | NA | NA | NA | NA | 1 | 1 | NA | NA | NA | NA | NA | NA | 1 | 1 | 1 | 1 |
| 2_1289360 | NA | NA | 1 | 1 | 1 | 0 | 1 | 0 | 3 | 0.3333 | NA | NA | NA | NA | NA | NA | 1 | 1 | 1 | 1 |
| 2_1289361 | NA | NA | 1 | 1 | 1 | 1 | 1 | 1 | 3 | 1 | NA | NA | NA | NA | NA | NA | 1 | 1 | 1 | 1 |
| 2_1289378 | 4 | 0 | 6 | 0 | 3 | 0.3333 | 4 | 0.25 | 17 | 0.1176 | 5 | 0 | 6 | 0.1667 | 7 | 0.1429 | 3 | 0.3333 | 21 | 0.1429 |
| 2_1289673 | NA | NA | 1 | 1 | NA | NA | NA | NA | 1 | 1 | 2 | 0.5 | 1 | 0 | 2 | 1 | 1 | 1 | 6 | 0.6667 |
| 2_1289673 | NA | NA | 1 | 1 | NA | NA | 1 | 1 | 2 | 1 | 2 | 0.5 | 1 | 1 | 2 | 1 | 2 | 1 | 7 | 0.8571 |
| 2_1289674 | NA | NA | 1 | 1 | NA | NA | 1 | 1 | 2 | 1 | 2 | 0.5 | 1 | 0 | 2 | 0.5 | 1 | 0 | 6 | 0.3333 |
| 2_1289675 | NA | NA | 1 | 1 | NA | NA | 1 | 1 | 2 | 1 | 2 | 1 | 1 | 1 | 2 | 1 | 2 | 1 | 7 | 1 |
| 2_1289676 | NA | NA | 1 | 1 | NA | NA | 1 | 0 | 2 | 0.5 | 2 | 0.5 | 1 | 0 | 2 | 0.5 | 2 | 0.5 | 7 | 0.4286 |
| 2_1289680 | NA | NA | 1 | 0 | NA | NA | 1 | 1 | 2 | 0.5 | 1 | 1 | 1 | 0 | 2 | 0.5 | 2 | 0 | 6 | 0.3333 |
| 2_1289752 | NA | NA | 2 | 0 | 3 | 0.3333 | 3 | 0.3333 | 8 | 0.25 | 1 | 1 | 1 | 0 | 4 | 0.25 | 1 | 0 | 7 | 0.2857 |
| 2_1289753 | NA | NA | 2 | 0.5 | 3 | 0 | 3 | 0.6667 | 8 | 0.375 | 1 | 1 | 2 | 1 | 4 | 0.5 | 1 | 0 | 8 | 0.625 |
| 2_1289753 | NA | NA | 2 | 1 | 2 | 0.5 | 2 | 1 | 6 | 0.8333 | 1 | 1 | NA | NA | 3 | 1 | 1 | 1 | 5 | 1 |
| 2_1289758 | NA | NA | 2 | 1 | 2 | 1 | 2 | 1 | 6 | 1 | 3 | 0.6667 | 4 | 0.75 | 5 | 0.6 | NA | NA | 12 | 0.6667 |
| 2_1289758 | NA | NA | 2 | 0 | NA | NA | 2 | 1 | 4 | 0.5 | 3 | 0.3333 | 4 | 0 | 4 | 0 | NA | NA | 11 | 0.0909 |
| 2_1290392 | NA | NA | 3 | 0.3333 | 1 | 1 | NA | NA | 4 | 0.5 | NA | NA | 1 | 1 | 2 | 0.5 | 1 | 0 | 4 | 0.5 |
| 2_1290399 | NA | NA | 3 | 1 | NA | NA | 1 | 1 | 4 | 1 | NA | NA | NA | NA | 2 | 1 | 2 | 1 | 4 | 1 |
| 2_1290400 | NA | NA | 1 | 1 | NA | NA | 1 | 0 | 2 | 0.5 | NA | NA | NA | NA | 2 | 0.5 | 2 | 0 | 4 | 0.25 |
| 2_1290403 | NA | NA | 2 | 1 | 1 | 1 | 1 | 1 | 4 | 1 | NA | NA | NA | NA | 1 | 1 | 1 | 1 | 2 | 1 |
| 2_1290403 | NA | NA | 3 | 1 | 1 | 1 | 1 | 1 | 5 | 1 | NA | NA | NA | NA | 2 | 0.5 | 2 | 1 | 4 | 0.75 |
| 2_1290404 | NA | NA | 3 | 1 | 1 | 1 | 1 | 1 | 5 | 1 | NA | NA | NA | NA | 2 | 0.5 | 2 | 0.5 | 4 | 0.5 |
| 2_1290404 | NA | NA | 2 | 0.5 | 1 | 1 | 1 | 1 | 4 | 0.75 | NA | NA | NA | NA | 2 | 0.5 | 1 | 1 | 3 | 0.6667 |
| 2_1290405 | NA | NA | 3 | 0.6667 | 1 | 1 | 1 | 0 | 5 | 0.6 | NA | NA | NA | NA | 1 | 1 | 2 | 0.5 | 3 | 0.6667 |
| 2_1290406 | NA | NA | 3 | 0.3333 | 1 | 0 | 1 | 0 | 5 | 0.2 | NA | NA | NA | NA | 2 | 0.5 | 2 | 0 | 4 | 0.25 |

| | | | | | | | | | | | | | | | | | | | | |
|---|---|---|---|---|---|---|---|---|---|---|---|---|---|---|---|---|---|---|---|---|
| 2_1290408( | NA | NA | 2 | 1 | 1 | 1 | 1 | 1 | 4 | 1 | NA | NA | NA | NA | 1 | 1 | 2 | 1 | 3 | 1 |
| 2_1290409 | NA | NA | 2 | 0.5 | 1 | 0 | 1 | 0 | 4 | 0.25 | NA | NA | NA | NA | 1 | 0 | 2 | 0.5 | 3 | 0.3333 |
| 2_1290418( | 5 | 0.2 | 3 | 0 | 5 | 0.2 | 5 | 0 | 18 | 0.1111 | 3 | 0.3333 | 5 | 0.2 | 4 | 0.25 | 3 | 0 | 15 | 0.2 |
| 2_1290462( | 3 | 0 | NA | NA | 3 | 0.3333 | 1 | 0 | 7 | 0.1429 | NA | NA | 2 | 0.5 | 2 | 0.5 | 1 | 0 | 5 | 0.4 |
| 2_1290462( | 3 | 0.3333 | NA | NA | 3 | 0 | 1 | 0 | 7 | 0.1429 | NA | NA | 2 | 0 | 2 | 0.5 | 1 | 0 | 5 | 0.2 |
| 2_1290463 | 3 | 0.3333 | NA | NA | 1 | 0 | 1 | 0 | 5 | 0.2 | NA | NA | 1 | 1 | 1 | 1 | 1 | 0 | 3 | 0.6667 |
| 2_1290465 | 3 | 0.3333 | NA | NA | 3 | 0.3333 | NA | NA | 6 | 0.3333 | NA | NA | 1 | 1 | 2 | 1 | 1 | 1 | 4 | 1 |
| 2_1290466 | 3 | 1 | NA | NA | 2 | 0.5 | 1 | 1 | 6 | 0.8333 | NA | NA | 1 | 1 | 2 | 1 | 1 | 1 | 4 | 1 |
| 2_1290467 | 3 | 1 | NA | NA | 3 | 0.3333 | 1 | 1 | 7 | 0.7143 | NA | NA | 2 | 1 | 2 | 1 | 1 | 1 | 5 | 1 |
| 2_1290468( | 2 | 0.5 | NA | NA | 3 | 0.3333 | 1 | 1 | 6 | 0.5 | NA | NA | 1 | 1 | 2 | 1 | 1 | 1 | 4 | 1 |
| 2_1290468 | 2 | 1 | NA | NA | 3 | 0.6667 | NA | NA | 5 | 0.8 | NA | NA | 1 | 1 | 1 | 1 | 1 | 1 | 3 | 1 |
| 2_1290499 | 6 | 0.1667 | 6 | 0 | 13 | 0.0769 | 8 | 0 | 33 | 0.0606 | 4 | 0 | 9 | 0 | 3 | 0 | 9 | 0.2222 | 25 | 0.08 |
| 2_1290500 | 6 | 0.1667 | 6 | 0.1667 | 13 | 0.2308 | 8 | 0.125 | 33 | 0.1818 | 4 | 0 | 10 | 0.2 | 3 | 0 | 9 | 0.3333 | 26 | 0.1923 |
| 2_1290500 | 6 | 0 | 6 | 0 | 15 | 0.3333 | 8 | 0.125 | 35 | 0.1714 | 4 | 0.25 | 9 | 0.1111 | 3 | 0 | 9 | 0.1111 | 25 | 0.12 |
| 2_1290502 | 6 | 0.5 | 6 | 0.8333 | 16 | 0.625 | 6 | 0.6667 | 34 | 0.6471 | 3 | 0.6667 | 8 | 0.5 | 3 | 0.6667 | 7 | 0.5714 | 21 | 0.5714 |
| 2_1290502 | 4 | 0.25 | 6 | 0.8333 | 12 | 0.5833 | 8 | 0.75 | 30 | 0.6333 | 2 | 0 | 8 | 0.25 | 2 | 1 | 8 | 0.5 | 20 | 0.4 |
| 2_1290527 | 6 | 0.1667 | 7 | 0.2857 | 16 | 0.125 | 12 | 0 | 41 | 0.122 | 6 | 0 | 7 | 0 | 4 | 0.25 | 13 | 0.0769 | 30 | 0.0667 |
| 2_1290533 | 12 | 0.1667 | 7 | 0.1429 | 16 | 0.3125 | 13 | 0.3077 | 48 | 0.25 | 5 | 0 | 9 | 0.1111 | 7 | 0 | 12 | 0.1667 | 33 | 0.0909 |
| 2_1290534 | 12 | 0.0833 | 8 | 0 | 22 | 0.1364 | 13 | 0.3846 | 55 | 0.1636 | 5 | 0 | 8 | 0.5 | 8 | 0.25 | 15 | 0.3333 | 36 | 0.3056 |
| 2_1294346 | 1 | 1 | NA | NA | NA | NA | NA | NA | 1 | 1 | 1 | 0 | NA | NA | 2 | 0.5 | NA | NA | 3 | 0.3333 |
| 2_1294348( | 1 | 1 | 1 | 0 | NA | NA | 1 | 0 | 3 | 0.3333 | 1 | 0 | 1 | 0 | 2 | 1 | NA | NA | 4 | 0.5 |
| 2_1294509( | 9 | 0.4444 | 14 | 0.5 | 13 | 0.1538 | 15 | 0.2 | 51 | 0.3137 | 14 | 0.2143 | 10 | 0.1 | 15 | 0.1333 | 14 | 0.2857 | 53 | 0.1887 |
| 2_1294509 | 12 | 0.0833 | 18 | 0.2222 | 13 | 0.3846 | 19 | 0.2105 | 62 | 0.2258 | 16 | 0.375 | 11 | 0.3636 | 17 | 0.1176 | 16 | 0.0625 | 60 | 0.2167 |
| 2_1294510 | 12 | 0.1667 | 16 | 0.1875 | 10 | 0.1 | 17 | 0.0588 | 55 | 0.1273 | 14 | 0.0714 | 6 | 0 | 16 | 0.1875 | 16 | 0.0625 | 52 | 0.0962 |
| 2_1294510 | 11 | 0.9091 | 17 | 0.7647 | 16 | 0.875 | 17 | 0.8824 | 61 | 0.8525 | 14 | 1 | 11 | 0.8182 | 17 | 0.8235 | 17 | 0.8824 | 59 | 0.8814 |
| 2_1294575( | 6 | 0 | 15 | 0.1333 | 14 | 0.0714 | 15 | 0 | 50 | 0.06 | 13 | 0.1538 | 8 | 0.125 | 16 | 0.0625 | 15 | 0.1333 | 52 | 0.1154 |
| 2_1294583( | 14 | 0 | 19 | 0.0526 | 19 | 0.1053 | 24 | 0.0417 | 76 | 0.0526 | 23 | 0.0435 | 14 | 0.0714 | 22 | 0.0909 | 20 | 0 | 79 | 0.0506 |
| 2_1294583 | 12 | 0.6667 | 19 | 0.4737 | 18 | 0.4444 | 21 | 0.5714 | 70 | 0.5286 | 22 | 0.5909 | 12 | 0.75 | 21 | 0.4286 | 20 | 0.5 | 75 | 0.5467 |
| 2_1294583 | 12 | 0.25 | 17 | 0.1765 | 18 | 0.1667 | 20 | 0.25 | 67 | 0.209 | 21 | 0.2381 | 11 | 0.4545 | 21 | 0.381 | 18 | 0.3333 | 71 | 0.338 |
| 2_1294617( | NA | NA | 1 | 1 | NA | NA | NA | NA | 1 | 1 | NA | NA | NA | NA | 1 | 0 | 2 | 0.5 | 3 | 0.3333 |
| 2_1294713 | 2 | 0.5 | NA | NA | NA | NA | NA | NA | 2 | 0.5 | 2 | 0.5 | NA | NA | NA | NA | NA | NA | 2 | 0.5 |
| 2_1294719 | 23 | 0.1304 | 26 | 0.0769 | 26 | 0.3462 | 29 | 0.2069 | 104 | 0.1923 | 32 | 0.1875 | 12 | 0 | 23 | 0.087 | 31 | 0.1613 | 98 | 0.1327 |
| 2_1294720 | 20 | 0.7 | 24 | 0.625 | 21 | 0.7143 | 26 | 0.5385 | 91 | 0.6374 | 28 | 0.6071 | 13 | 0.6923 | 22 | 0.5909 | 25 | 0.64 | 88 | 0.625 |
| 2_1294721 | 21 | 0.7619 | 23 | 0.7391 | 23 | 0.8261 | 29 | 0.7586 | 96 | 0.7708 | 31 | 0.7742 | 12 | 0.9167 | 21 | 0.8571 | 27 | 0.7778 | 91 | 0.8132 |
| 2_1294722 | 22 | 0.9545 | 26 | 1 | 27 | 1 | 31 | 1 | 106 | 0.9906 | 30 | 0.9333 | 11 | 1 | 27 | 1 | 29 | 1 | 97 | 0.9794 |
| 2_1294723 | 24 | 0.125 | 28 | 0.0357 | 30 | 0.1 | 35 | 0.0286 | 117 | 0.0684 | 36 | 0.1111 | 17 | 0.1765 | 29 | 0 | 34 | 0.1176 | 116 | 0.0948 |
| 2_1294723 | 19 | 0.9474 | 18 | 0.6667 | 24 | 0.875 | 29 | 0.931 | 90 | 0.8667 | 26 | 0.9615 | 14 | 0.9286 | 22 | 0.8636 | 26 | 0.9231 | 88 | 0.9205 |
| 2_1294725( | 25 | 0.8 | 27 | 0.7037 | 32 | 0.8438 | 36 | 0.8611 | 120 | 0.8083 | 37 | 0.8649 | 17 | 0.6471 | 22 | 0.8182 | 31 | 0.7097 | 107 | 0.7757 |
| 2_1294725( | 22 | 0.6818 | 26 | 0.6538 | 31 | 0.6774 | 34 | 0.6176 | 113 | 0.6549 | 34 | 0.7353 | 16 | 0.6875 | 28 | 0.6786 | 27 | 0.5556 | 105 | 0.6667 |
| 2_1294725 | 23 | 0.4783 | 26 | 0.3077 | 31 | 0.4194 | 30 | 0.4333 | 110 | 0.4091 | 39 | 0.4103 | 13 | 0.3846 | 24 | 0.3333 | 27 | 0.3333 | 103 | 0.3689 |
| 2_1294726 | 27 | 0.4815 | 33 | 0.5455 | 33 | 0.4545 | 34 | 0.6471 | 127 | 0.5354 | 40 | 0.45 | 15 | 0.4 | 28 | 0.3571 | 34 | 0.6765 | 117 | 0.4872 |
| 2_1294726 | 27 | 1 | 29 | 0.9655 | 36 | 0.9722 | 37 | 0.9459 | 129 | 0.969 | 41 | 1 | 16 | 0.9375 | 29 | 1 | 35 | 0.9429 | 121 | 0.9752 |
| 2_1294727 | 27 | 0.3333 | 27 | 0.2593 | 31 | 0.0323 | 35 | 0.1429 | 120 | 0.1833 | 37 | 0.1622 | 14 | 0 | 25 | 0.2 | 29 | 0.1724 | 105 | 0.1524 |
| 2_1294727( | 25 | 1 | 26 | 1 | 29 | 0.9655 | 34 | 1 | 114 | 0.9912 | 36 | 1 | 15 | 1 | 23 | 0.9565 | 22 | 0.9545 | 96 | 0.9792 |
| 2_1294730 | 30 | 0.1 | 37 | 0.027 | 35 | 0.0857 | 37 | 0.0541 | 139 | 0.0647 | 47 | 0.1064 | 18 | 0.1111 | 34 | 0 | 36 | 0.0278 | 135 | 0.0593 |
| 2_1294731 | 18 | 0.2222 | 32 | 0.0938 | 29 | 0.069 | 32 | 0.0938 | 111 | 0.1081 | 32 | 0.0625 | 12 | 0.1667 | 25 | 0.16 | 26 | 0.0385 | 95 | 0.0947 |
| 2_1294732 | 24 | 0.7917 | 34 | 0.8529 | 33 | 0.7879 | 32 | 0.5938 | 123 | 0.7561 | 37 | 0.7297 | 16 | 0.75 | 30 | 0.8 | 28 | 0.8214 | 111 | 0.7748 |
| 2_1294732 | 5 | 1 | NA | NA | 2 | 1 | 3 | 0.6667 | 10 | 0.9 | 5 | 1 | NA | NA | 4 | 1 | 6 | 0.8333 | 15 | 0.9333 |
| 2_1294733 | 22 | 0.0909 | 32 | 0.0625 | 34 | 0.0588 | 31 | 0.0968 | 119 | 0.0756 | 40 | 0.025 | 17 | 0.1176 | 30 | 0.1 | 30 | 0.1333 | 117 | 0.0855 |
| 2_1294733 | 29 | 0.1034 | 41 | 0.0488 | 41 | 0.0488 | 41 | 0.0244 | 152 | 0.0526 | 47 | 0.0213 | 18 | 0.1111 | 34 | 0.0882 | 39 | 0.0769 | 138 | 0.0652 |
| 2_1294734 | 24 | 0.0833 | 29 | 0.1379 | 32 | 0.1562 | 32 | 0.0625 | 117 | 0.1111 | 32 | 0.1875 | 16 | 0.125 | 27 | 0.0741 | 31 | 0.1613 | 106 | 0.1415 |
| 2_1294734 | 31 | 0.0323 | 41 | 0.1707 | 40 | 0.25 | 38 | 0.0789 | 150 | 0.14 | 49 | 0.1633 | 16 | 0.25 | 35 | 0.1429 | 37 | 0 | 137 | 0.1241 |
| 2_1294737 | 38 | 0.1842 | 40 | 0.125 | 48 | 0.0833 | 41 | 0.1463 | 167 | 0.1317 | 48 | 0.1875 | 18 | 0.1111 | 30 | 0.2 | 41 | 0.1951 | 137 | 0.1825 |
| 2_1295285 | 35 | 0.0571 | 7 | 0 | 27 | 0.0741 | 44 | 0.0682 | 113 | 0.0619 | 6 | 0 | 42 | 0.2143 | 48 | 0.1667 | 22 | 0.0909 | 118 | 0.161 |
| 2_1295285 | 42 | 0.119 | 8 | 0 | 32 | 0.0625 | 55 | 0.0545 | 137 | 0.073 | 8 | 0 | 54 | 0.1111 | 55 | 0.0727 | 31 | 0.129 | 148 | 0.0946 |
| 2_1295287( | 38 | 0.5526 | 7 | 0.2857 | 29 | 0.5172 | 51 | 0.5686 | 125 | 0.536 | 8 | 0.625 | 52 | 0.5385 | 52 | 0.5962 | 24 | 0.5833 | 136 | 0.5735 |
| 2_1295287 | 33 | 0.2727 | 7 | 0.2857 | 24 | 0.375 | 44 | 0.2727 | 108 | 0.2963 | 7 | 0.1429 | 43 | 0.186 | 41 | 0.3415 | 21 | 0.2857 | 112 | 0.2589 |
| 2_1295287 | 39 | 0.2308 | 8 | 0 | 26 | 0.2692 | 51 | 0.1765 | 124 | 0.2016 | 6 | 0.3333 | 55 | 0.1636 | 53 | 0.2075 | 26 | 0.1538 | 140 | 0.1857 |
| 2_1295289 | 35 | 0.9143 | 7 | 0.8571 | 26 | 0.9231 | 48 | 0.9167 | 116 | 0.9138 | 5 | 1 | 50 | 0.86 | 48 | 0.9583 | 21 | 0.9048 | 124 | 0.9113 |
| 2_1295290 | 30 | 0.4667 | 6 | 0.6667 | 21 | 0.5714 | 37 | 0.4054 | 94 | 0.4787 | 5 | 0.8 | 41 | 0.6341 | 42 | 0.7143 | 21 | 0.6667 | 109 | 0.6789 |
| 2_1295291 | 31 | 0.0968 | 6 | 0 | 25 | 0.08 | 49 | 0.0408 | 111 | 0.0631 | 4 | 0 | 50 | 0.04 | 52 | 0.0192 | 26 | 0.1538 | 132 | 0.053 |
| 2_1295293 | 27 | 0.3333 | 5 | 0.4 | 24 | 0.2083 | 43 | 0.1395 | 99 | 0.2222 | 4 | 0.5 | 47 | 0.234 | 47 | 0.4043 | 21 | 0.2381 | 119 | 0.3109 |
| 2_1295295 | 28 | 0.5714 | 7 | 0.7143 | 22 | 0.7727 | 40 | 0.625 | 97 | 0.6495 | 3 | 1 | 43 | 0.7907 | 46 | 0.7826 | 21 | 0.8571 | 113 | 0.8053 |
| 2_1295296 | 33 | 0.2727 | 6 | 0 | 22 | 0.0909 | 42 | 0.119 | 103 | 0.1553 | 3 | 0.6667 | 47 | 0.2128 | 50 | 0.18 | 24 | 0.25 | 124 | 0.2177 |
| 2_1295296 | 29 | 0.2759 | 7 | 0.1429 | 20 | 0.4 | 40 | 0.2 | 96 | 0.2604 | 3 | 0.6667 | 42 | 0.2857 | 45 | 0.2889 | 23 | 0.1739 | 113 | 0.2743 |

| | | | | | | | | | | | | | | | | | | | | |
|---|---|---|---|---|---|---|---|---|---|---|---|---|---|---|---|---|---|---|---|---|
| 2_1295297( | 22 | 0 | 3 | 0.3333 | 14 | 0.0714 | 33 | 0.1515 | 72 | 0.0972 | 2 | 0 | 32 | 0.0625 | 36 | 0.1111 | 21 | 0.2381 | 91 | 0.1209 |
| 2_1295300: | 32 | 0.4688 | 6 | 0.1667 | 17 | 0.4118 | 36 | 0.4167 | 91 | 0.4176 | 4 | 0.25 | 38 | 0.3158 | 45 | 0.4667 | 19 | 0.4211 | 106 | 0.3962 |
| 2_1295300 | 32 | 1 | 6 | 0.8333 | 21 | 1 | 41 | 0.9756 | 100 | 0.98 | 4 | 0.75 | 39 | 0.9744 | 45 | 0.9333 | 21 | 0.9524 | 109 | 0.945 |
| 2_1295301 | 26 | 0.0769 | 5 | 0 | 18 | 0.1111 | 23 | 0.0435 | 72 | 0.0694 | 3 | 0 | 29 | 0.069 | 37 | 0.0811 | 14 | 0.0714 | 83 | 0.0723 |
| 2_1295301 | 19 | 1 | 6 | 1 | 15 | 1 | 25 | 1 | 65 | 1 | 2 | 1 | 27 | 0.963 | 30 | 1 | 12 | 1 | 71 | 0.9859 |
| 2_1295301 | NA | NA | NA | NA | NA | NA | 1 | 1 | 1 | 1 | NA | NA | NA | NA | 1 | 1 | NA | NA | 1 | 1 |
| 2_1295303( | 29 | 0.1379 | 6 | 0 | 20 | 0.05 | 32 | 0.125 | 87 | 0.1034 | 3 | 0.3333 | 30 | 0 | 43 | 0.1163 | 18 | 0.0556 | 94 | 0.0745 |
| 2_1296416 | NA | NA | 1 | 0 | NA | NA | 2 | 0.5 | 3 | 0.3333 | NA | NA | NA | NA | 1 | 1 | NA | NA | 1 | 1 |
| 2_1296419 | NA | NA | NA | NA | 1 | 1 | 1 | 1 | 2 | 1 | NA | NA | 1 | 1 | NA | NA | NA | NA | 1 | 1 |
| 2_1296422 | NA | NA | 1 | 1 | 1 | 1 | 2 | 1 | 4 | 1 | NA | NA | 1 | 1 | NA | NA | NA | NA | 1 | 1 |
| 2_1296428 | NA | NA | NA | NA | 1 | 1 | 1 | 1 | 2 | 1 | NA | NA | 1 | 1 | NA | NA | NA | NA | 1 | 1 |
| 2_1296429( | NA | NA | NA | NA | 1 | 0 | 1 | 1 | 2 | 0.5 | NA | NA | 1 | 1 | NA | NA | NA | NA | 1 | 1 |
| 2_1296432( | NA | NA | NA | NA | 1 | 1 | NA | NA | 1 | 1 | NA | NA | 1 | 1 | NA | NA | NA | NA | 1 | 1 |
| 2_1296434( | NA | NA | 1 | 1 | NA | NA | NA | NA | 1 | 1 | NA | NA | NA | NA | 2 | 0.5 | 1 | 0 | 3 | 0.3333 |
| 2_1296525 | NA | NA | 1 | 1 | NA | NA | NA | NA | 1 | 1 | NA | NA | 2 | 0.5 | 2 | 0.5 | 1 | 1 | 5 | 0.6 |
| 2_1296526 | NA | NA | 1 | 1 | NA | NA | NA | NA | 1 | 1 | NA | NA | 1 | 1 | 1 | 1 | NA | NA | 2 | 1 |
| 2_1296528( | NA | NA | 1 | 1 | NA | NA | NA | NA | 1 | 1 | NA | NA | 1 | 1 | 2 | 1 | 1 | 1 | 4 | 1 |
| 2_1296528 | NA | NA | 1 | 1 | NA | NA | NA | NA | 1 | 1 | NA | NA | 2 | 1 | 2 | 1 | 1 | 1 | 5 | 1 |
| 2_1296528 | NA | NA | 1 | 1 | NA | NA | NA | NA | 1 | 1 | NA | NA | 1 | 1 | 2 | 1 | 1 | 0 | 4 | 0.75 |
| 2_1296529 | NA | NA | 1 | 1 | NA | NA | NA | NA | 1 | 1 | NA | NA | 1 | 1 | 2 | 1 | 1 | 1 | 4 | 1 |
| 2_1296530( | NA | NA | 1 | 1 | NA | NA | NA | NA | 1 | 1 | NA | NA | 1 | 1 | 2 | 1 | 1 | 1 | 4 | 1 |
| 2_1296530 | NA | NA | 1 | 1 | NA | NA | NA | NA | 1 | 1 | NA | NA | NA | NA | 2 | 0.5 | 1 | 1 | 3 | 0.6667 |
| 2_1296699( | NA | NA | NA | NA | NA | NA | 1 | 1 | 1 | 1 | NA | NA | 1 | 1 | 2 | 0.5 | NA | NA | 3 | 0.6667 |
| 2_1296703( | NA | NA | 1 | 0 | 1 | 1 | NA | NA | 2 | 0.5 | NA | NA | 2 | 0.5 | 2 | 0 | NA | NA | 4 | 0.25 |
| 2_1301194 | 5 | 0.2 | 9 | 0 | 12 | 0.25 | 6 | 0.3333 | 32 | 0.1875 | 8 | 0 | 10 | 0.1 | 6 | 0 | 9 | 0.1111 | 33 | 0.0606 |
| 2_1301211 | 6 | 0.5 | 12 | 0.5833 | 13 | 0.6923 | 7 | 0.5714 | 38 | 0.6053 | 11 | 0.6364 | 14 | 0.9286 | 9 | 0.7778 | 14 | 0.6429 | 48 | 0.75 |
| 2_1301214 | 35 | 0.0857 | 40 | 0.075 | 45 | 0.0222 | 44 | 0.0455 | 164 | 0.0549 | 27 | 0.1111 | 38 | 0.0789 | 39 | 0.2308 | 27 | 0.1111 | 131 | 0.1374 |
| 2_1301214 | 38 | 0.1053 | 41 | 0.0976 | 47 | 0.1064 | 44 | 0.0682 | 170 | 0.0941 | 28 | 0.0357 | 37 | 0.0541 | 42 | 0.0476 | 26 | 0.1154 | 133 | 0.0602 |
| 2_1301215 | 34 | 0.0882 | 38 | 0.1053 | 48 | 0.0625 | 40 | 0.1 | 160 | 0.0875 | 27 | 0 | 37 | 0.1892 | 40 | 0.05 | 25 | 0.08 | 129 | 0.0853 |
| 2_1302070 | 1 | 1 | 1 | 0 | 1 | 1 | NA | NA | 3 | 0.6667 | NA | NA | NA | NA | NA | NA | 1 | 1 | 1 | 1 |
| 2_1302117 | 1 | 0 | NA | NA | 1 | 1 | NA | NA | 2 | 0.5 | NA | NA | 1 | 0 | 1 | 1 | 2 | 0.5 | 4 | 0.5 |
| 2_1302119 | 1 | 0 | 1 | 0 | 1 | 1 | NA | NA | 3 | 0.3333 | NA | NA | 1 | 1 | 1 | 1 | 1 | 1 | 3 | 1 |
| 2_1302119( | NA | NA | 1 | 0 | 1 | 1 | 1 | 0 | 3 | 0.3333 | NA | NA | NA | NA | 1 | 0 | 2 | 0.5 | 3 | 0.3333 |
| 2_1302327 | 1 | 0 | 1 | 1 | NA | NA | NA | NA | 2 | 0.5 | NA | NA | 1 | 0 | 2 | 1 | NA | NA | 3 | 0.6667 |
| 2_1302328 | 1 | 0 | 2 | 1 | NA | NA | NA | NA | 3 | 0.6667 | NA | NA | 1 | 0 | 1 | 1 | NA | NA | 2 | 0.5 |
| 2_1302601 | 1 | 0 | 2 | 0 | NA | NA | 2 | 0.5 | 5 | 0.2 | 4 | 0.25 | 1 | 0 | NA | NA | 1 | 0 | 6 | 0.1667 |
| 2_1302606 | 2 | 0.5 | 1 | 0 | NA | NA | 2 | 0 | 5 | 0.2 | 3 | 0 | 1 | 0 | NA | NA | 1 | 1 | 5 | 0.2 |
| 2_1302610 | 3 | 0.3333 | 2 | 0.5 | NA | NA | 2 | 0 | 7 | 0.2857 | 5 | 0.2 | 2 | 0 | 1 | 1 | 1 | 0 | 9 | 0.2222 |
| 2_1302610 | 3 | 1 | 2 | 0 | NA | NA | 2 | 0 | 7 | 0.4286 | 5 | 0.4 | 1 | 0 | 1 | 0 | 1 | 0 | 8 | 0.25 |
| 2_1302612 | 3 | 0.3333 | 1 | 0 | NA | NA | 2 | 0 | 6 | 0.1667 | 5 | 0.6 | 2 | 0.5 | 1 | 0 | NA | NA | 8 | 0.5 |
| 2_1302640 | 1 | 1 | NA | NA | NA | NA | NA | NA | 1 | 1 | 1 | 1 | NA | NA | NA | NA | NA | NA | 1 | 1 |
| 2_1302641 | 1 | 1 | NA | NA | NA | NA | NA | NA | 1 | 1 | 1 | 1 | NA | NA | NA | NA | NA | NA | 1 | 1 |
| 2_1302641 | 1 | 1 | 1 | 0 | NA | NA | NA | NA | 2 | 0.5 | 2 | 0 | 2 | 0.5 | NA | NA | 2 | 0 | 6 | 0.1667 |
| 2_1302642 | 1 | 1 | 1 | 1 | NA | NA | NA | NA | 2 | 1 | 4 | 0.25 | 2 | 0.5 | 1 | 1 | 2 | 0 | 9 | 0.3333 |
| 2_1302642 | NA | NA | 1 | 1 | NA | NA | NA | NA | 1 | 1 | 5 | 0.2 | 2 | 0.5 | 1 | 0 | 2 | 0 | 10 | 0.2 |
| 2_1302644 | 1 | 1 | 1 | 1 | NA | NA | NA | NA | 2 | 1 | 4 | 0 | 2 | 0.5 | 1 | 0 | 2 | 0 | 9 | 0.1111 |
| 2_1302644( | 1 | 1 | NA | NA | NA | NA | NA | NA | 1 | 1 | 4 | 0 | 2 | 0.5 | 1 | 0 | 2 | 0 | 9 | 0.1111 |
| 2_1302645 | 1 | 1 | 1 | 1 | NA | NA | NA | NA | 2 | 1 | 4 | 0.5 | 1 | 1 | 1 | 0 | 2 | 0 | 8 | 0.375 |
| 2_1302648( | NA | NA | 1 | 1 | NA | NA | NA | NA | 1 | 1 | NA | NA | 1 | 1 | NA | NA | 1 | 1 | 2 | 1 |
| 2_1302653 | NA | NA | 1 | 1 | NA | NA | NA | NA | 1 | 1 | 1 | 0 | 1 | 1 | 1 | 0 | 1 | 1 | 4 | 0.5 |
| 2_1302922 | 5 | 0.2 | 6 | 0.1667 | 4 | 0 | 4 | 0 | 19 | 0.1053 | 2 | 0 | 6 | 0.1667 | 5 | 0 | 3 | 0 | 16 | 0.0625 |
| 2_1303460 | 10 | 0 | 11 | 0.1818 | 9 | 0.1111 | 13 | 0.0769 | 43 | 0.093 | 4 | 0 | 10 | 0.1 | 14 | 0.0714 | 9 | 0 | 37 | 0.0541 |
| 2_1303460 | 9 | 0 | 12 | 0 | 8 | 0.125 | 11 | 0.0909 | 40 | 0.05 | 4 | 0.25 | 9 | 0.1111 | 15 | 0 | 7 | 0 | 35 | 0.0571 |
| 2_1303956 | 4 | 0 | 3 | 0.3333 | 4 | 0 | 4 | 0 | 15 | 0.0667 | 4 | 0.25 | 3 | 0.3333 | 3 | 0 | 4 | 0 | 14 | 0.1429 |
| 2_1303998 | 2 | 0.5 | NA | NA | 1 | 0 | NA | NA | 3 | 0.3333 | 1 | 0 | 1 | 0 | 1 | 1 | NA | NA | 3 | 0.3333 |
| 2_1304061 | 3 | 0 | 1 | 0 | 2 | 0 | 7 | 0.1429 | 13 | 0.0769 | 2 | 0 | 1 | 1 | 2 | 0 | 2 | 0 | 7 | 0.1429 |
| 2_1304069 | 14 | 0.5 | 10 | 0.2 | 14 | 0.4286 | 13 | 0.0769 | 51 | 0.3137 | 11 | 0 | 14 | 0.2857 | 15 | 0.3333 | 16 | 0.4375 | 56 | 0.2857 |
| 2_1304069 | 11 | 0.8182 | 8 | 0.625 | 15 | 0.9333 | 12 | 0.9167 | 46 | 0.8478 | 11 | 1 | 16 | 0.875 | 16 | 0.9375 | 14 | 0.9286 | 57 | 0.9298 |
| 2_1304071 | 15 | 0.1333 | 10 | 0.1 | 13 | 0.0769 | 12 | 0.0833 | 50 | 0.1 | 12 | 0 | 16 | 0.1875 | 14 | 0.1429 | 12 | 0.0833 | 54 | 0.1111 |
| 2_1304071 | 13 | 0.3846 | 10 | 0.3 | 11 | 0.2727 | 9 | 0.3333 | 43 | 0.3256 | 9 | 0.2222 | 14 | 0.2143 | 15 | 0.1333 | 12 | 0.1667 | 50 | 0.18 |
| 2_1304072 | 14 | 0.2143 | 8 | 0.375 | 12 | 0.5 | 11 | 0.2727 | 45 | 0.3333 | 11 | 0.2727 | 13 | 0.3077 | 13 | 0.4615 | 12 | 0.3333 | 49 | 0.3469 |
| 2_1304072 | 10 | 0.5 | 9 | 0.5556 | 12 | 0.5833 | 12 | 0.6667 | 43 | 0.5814 | 11 | 0.8182 | 11 | 0.4545 | 13 | 0.3077 | 12 | 0.6667 | 47 | 0.5532 |
| 2_1304332( | NA | NA | 2 | 0 | 2 | 0.5 | NA | NA | 4 | 0.25 | NA | NA | 2 | 0.5 | 4 | 0 | 1 | 0 | 7 | 0.1429 |
| 2_1304333 | NA | NA | 2 | 1 | 2 | 1 | 1 | 1 | 5 | 1 | 1 | 1 | 2 | 0.5 | 3 | 1 | NA | NA | 6 | 0.8333 |
| 2_1304333 | NA | NA | 2 | 1 | 1 | 1 | 1 | 1 | 4 | 1 | NA | NA | 2 | 1 | 2 | 1 | NA | NA | 4 | 1 |

| | | | | | | | | | | | | | | | | | | | | |
|---|---|---|---|---|---|---|---|---|---|---|---|---|---|---|---|---|---|---|---|---|
| 2_1304336 | NA | NA | 2 | 0.5 | 2 | 1 | 1 | 1 | 5 | 0.8 | 1 | 0 | 2 | 1 | 2 | 0.5 | 1 | 1 | 6 | 0.6667 |
| 2_1304336 | NA | NA | 2 | 0.5 | 2 | 0.5 | 1 | 0 | 5 | 0.4 | 1 | 0 | 2 | 0.5 | 3 | 0 | 1 | 0 | 7 | 0.1429 |
| 2_1304416 | NA | NA | NA | NA | 1 | 0 | 1 | 1 | 2 | 0.5 | NA | NA | 2 | 1 | 1 | 0 | 1 | 0 | 4 | 0.5 |
| 2_1304419 | NA | NA | NA | NA | 1 | 0 | 1 | 1 | 2 | 0.5 | NA | NA | 2 | 0.5 | 1 | 1 | 1 | 0 | 4 | 0.5 |
| 2_1304420 | NA | NA | NA | NA | 1 | 1 | 1 | 1 | 2 | 1 | NA | NA | 1 | 0 | NA | NA | 1 | 1 | 2 | 0.5 |
| 2_1304422 | NA | NA | NA | NA | 1 | 1 | 1 | 1 | 2 | 1 | NA | NA | 1 | 1 | 1 | 0 | NA | NA | 2 | 0.5 |
| 2_1304423 | NA | NA | NA | NA | 1 | 1 | 1 | 1 | 2 | 1 | NA | NA | 1 | 1 | 1 | 1 | 1 | 0 | 3 | 0.6667 |
| 2_1304559 | 2 | 1 | 2 | 0.5 | 1 | 1 | 2 | 1 | 7 | 0.8571 | NA | NA | 4 | 0.5 | 1 | 1 | 2 | 1 | 7 | 0.7143 |
| 2_1304560 | 1 | 1 | 2 | 0.5 | 1 | 1 | 2 | 1 | 6 | 0.8333 | NA | NA | 2 | 1 | 1 | 1 | 2 | 1 | 5 | 1 |
| 2_1304561 | NA | NA | 1 | 0 | 1 | 1 | 2 | 1 | 4 | 0.75 | NA | NA | 4 | 0.5 | NA | NA | 2 | 0 | 6 | 0.3333 |
| 2_1304562 | 2 | 0.5 | 2 | 0.5 | NA | NA | 2 | 0.5 | 6 | 0.5 | NA | NA | 3 | 0.6667 | 1 | 1 | 1 | 1 | 5 | 0.8 |
| 2_1304564 | 2 | 0.5 | 1 | 1 | 1 | 0 | 1 | 1 | 5 | 0.6 | NA | NA | 2 | 1 | 1 | 1 | 2 | 1 | 5 | 1 |
| 2_1304564 | 2 | 1 | 2 | 1 | 1 | 1 | 2 | 1 | 7 | 1 | NA | NA | 3 | 1 | NA | NA | 2 | 1 | 5 | 1 |
| 2_1304565 | 2 | 1 | 2 | 0.5 | 1 | 1 | 2 | 1 | 7 | 0.8571 | NA | NA | 3 | 1 | 1 | 1 | 2 | 1 | 6 | 1 |
| 2_1304567 | 1 | 1 | 1 | 0 | 2 | 0 | 1 | 0 | 5 | 0.2 | NA | NA | 2 | 0 | NA | NA | 1 | 1 | 3 | 0.3333 |
| 2_1304567 | 2 | 1 | 1 | 0 | 2 | 0.5 | 1 | 1 | 6 | 0.6667 | NA | NA | 3 | 1 | 1 | 1 | 2 | 1 | 6 | 1 |
| 2_1304569 | 1 | 1 | 1 | 1 | 1 | 1 | 1 | 1 | 4 | 1 | NA | NA | 3 | 1 | 1 | 1 | 2 | 0.5 | 6 | 0.8333 |
| 2_1304569 | 2 | 1 | NA | NA | 1 | 1 | 1 | 0 | 4 | 0.75 | NA | NA | 1 | 1 | 1 | 1 | 2 | 1 | 4 | 1 |
| 2_1304571 | 2 | 1 | 1 | 1 | 1 | 1 | 1 | 1 | 5 | 1 | NA | NA | 3 | 1 | 1 | 1 | 2 | 1 | 6 | 1 |
| 2_1304576 | 1 | 0 | NA | NA | 2 | 0.5 | NA | NA | 3 | 0.3333 | NA | NA | 3 | 0.3333 | 1 | 0 | 1 | 1 | 5 | 0.4 |
| 2_1304579 | 1 | 1 | 1 | 0 | 2 | 0 | 1 | 0 | 5 | 0.2 | NA | NA | 3 | 0 | 1 | 1 | 1 | 0 | 5 | 0.2 |
| 2_1304580 | 2 | 0.5 | 1 | 0 | 1 | 1 | NA | NA | 4 | 0.5 | NA | NA | 1 | 1 | 1 | 1 | 1 | 1 | 3 | 1 |
| 2_1304580 | 2 | 0.5 | 1 | 0 | 1 | 1 | 1 | 0 | 5 | 0.4 | NA | NA | 2 | 0.5 | 1 | 1 | 1 | 0 | 4 | 0.5 |
| 2_1304581 | 1 | 0 | 1 | 0 | 2 | 0.5 | 1 | 0 | 5 | 0.2 | NA | NA | 2 | 0 | 1 | 0 | 1 | 1 | 4 | 0.25 |
| 2_1304583 | 2 | 0 | 1 | 1 | 1 | 1 | 1 | 0 | 5 | 0.4 | NA | NA | 2 | 0 | 1 | 1 | 1 | 0 | 4 | 0.25 |
| 2_1304584 | 2 | 0.5 | 1 | 0 | 1 | 0 | 1 | 1 | 5 | 0.4 | NA | NA | 2 | 0.5 | 1 | 0 | 1 | 0 | 4 | 0.25 |
| 2_1304585 | 2 | 0 | 1 | 1 | 1 | 0 | 1 | 1 | 5 | 0.4 | NA | NA | 1 | 1 | NA | NA | NA | NA | 1 | 1 |
| 2_1304748 | 32 | 0.0938 | 17 | 0.1765 | 22 | 0.1364 | 44 | 0.1364 | 115 | 0.1304 | 13 | 0.2308 | 41 | 0.0732 | 53 | 0.0755 | 23 | 0.1304 | 130 | 0.1 |
| 2_1304750 | 25 | 0.56 | 13 | 0.5385 | 14 | 0.4286 | 36 | 0.4444 | 88 | 0.4886 | 10 | 0.6 | 31 | 0.4516 | 40 | 0.55 | 14 | 0.6429 | 95 | 0.5368 |
| 2_1304755 | 20 | 0 | 11 | 0.0909 | 14 | 0.0714 | 30 | 0.1 | 75 | 0.0667 | 10 | 0 | 28 | 0.1071 | 35 | 0.0286 | 19 | 0.1579 | 92 | 0.0761 |
| 2_1304760 | 18 | 0.6111 | 9 | 0.3333 | 13 | 0.4615 | 26 | 0.6538 | 66 | 0.5606 | 9 | 0.3333 | 25 | 0.52 | 32 | 0.375 | 18 | 0.6667 | 84 | 0.4762 |
| 2_1304765 | 19 | 0.0526 | 8 | 0.25 | 10 | 0 | 23 | 0.0435 | 60 | 0.0667 | 7 | 0.1429 | 20 | 0.1 | 29 | 0.0345 | 18 | 0 | 74 | 0.0541 |
| 2_1304767 | 16 | 0.1875 | 6 | 0.1667 | 9 | 0.1111 | 18 | 0.2222 | 49 | 0.1837 | 6 | 0 | 17 | 0.1765 | 29 | 0.1379 | 15 | 0 | 67 | 0.1045 |
| 2_1304770 | 11 | 0.1818 | 5 | 0 | 6 | 0 | 11 | 0.1818 | 33 | 0.1212 | 4 | 0.25 | 11 | 0 | 19 | 0.1053 | 7 | 0 | 41 | 0.0732 |
| 2_1304771 | 8 | 1 | 5 | 0.8 | 7 | 1 | 11 | 0.8182 | 31 | 0.9032 | 3 | 0.3333 | 10 | 1 | 16 | 1 | 8 | 0.875 | 37 | 0.9189 |
| 2_1304772 | 8 | 1 | 4 | 1 | 7 | 0.8571 | 13 | 1 | 32 | 0.9688 | 4 | 1 | 11 | 0.9091 | 16 | 0.75 | 9 | 1 | 40 | 0.875 |
| 2_1304773 | 10 | 0.9 | 5 | 1 | 7 | 1 | 14 | 1 | 36 | 0.9722 | 4 | 0.5 | 10 | 0.9 | 15 | 0.7333 | 7 | 0.7143 | 36 | 0.75 |
| 2_1304781 | 2 | 0.5 | 2 | 0 | 5 | 0 | 3 | 0 | 12 | 0.0833 | 1 | 0 | 2 | 0 | 3 | 0.3333 | 2 | 0 | 8 | 0.125 |
| 2_1304781 | 5 | 0.4 | 4 | 0.25 | 7 | 0.2857 | 5 | 0 | 21 | 0.2381 | NA | NA | 3 | 0 | 8 | 0.25 | 6 | 0.3333 | 17 | 0.2353 |
| 2_1304783 | 5 | 0.4 | 2 | 0 | 6 | 0 | 5 | 0.2 | 18 | 0.1667 | 1 | 0 | 4 | 0 | 8 | 0.375 | 5 | 0.2 | 18 | 0.2222 |
| 2_1304783 | 4 | 0 | 3 | 0.3333 | 4 | 0.25 | 4 | 0.5 | 15 | 0.2667 | 1 | 0 | 4 | 0 | 8 | 0.25 | 5 | 0.2 | 18 | 0.1667 |
| 2_1304784 | 5 | 0.4 | 3 | 0 | 4 | 0 | 3 | 0.3333 | 15 | 0.2 | 1 | 0 | 4 | 0.25 | 7 | 0.1429 | 5 | 0.4 | 17 | 0.2353 |
| 2_1304784 | 5 | 0.6 | 3 | 0.6667 | 5 | 0 | 6 | 0.5 | 19 | 0.4211 | NA | NA | 4 | 0.75 | 8 | 0.625 | 5 | 0.2 | 17 | 0.5294 |
| 2_1304784 | 5 | 0.2 | 3 | 0.3333 | 5 | 0.2 | 5 | 0.2 | 18 | 0.2222 | 1 | 0 | 4 | 0.25 | 9 | 0 | 4 | 0 | 18 | 0.0556 |
| 2_1304784 | 3 | 0.3333 | 3 | 0 | 5 | 0 | 6 | 0.3333 | 17 | 0.1765 | 1 | 0 | 2 | 0 | 7 | 0 | 4 | 0.25 | 14 | 0.0714 |
| 2_1304792 | 3 | 0.3333 | 3 | 0 | 5 | 0.4 | 3 | 0 | 14 | 0.2143 | 1 | 0 | 3 | 0.3333 | 7 | 0 | 3 | 0.3333 | 14 | 0.1429 |
| 2_1305055 | NA | NA | NA | NA | NA | NA | 1 | 1 | 1 | 1 | NA | NA | 1 | 1 | NA | NA | NA | NA | 1 | 1 |
| 2_1305502 | 4 | 0 | 2 | 0 | 2 | 0 | 7 | 0.4286 | 15 | 0.2 | 1 | 0 | 1 | 0 | 5 | 0.2 | 2 | 0 | 9 | 0.1111 |
| 2_1305507 | 4 | 0 | 2 | 0 | 2 | 0 | 8 | 0.25 | 16 | 0.125 | 1 | 0 | 1 | 0 | 3 | 0.3333 | 1 | 0 | 6 | 0.1667 |
| 2_1305508 | 4 | 0.25 | 2 | 0 | 1 | 0 | 7 | 0.1429 | 14 | 0.1429 | 1 | 1 | NA | NA | 3 | 0.3333 | 1 | 1 | 5 | 0.6 |
| 2_1305508 | 4 | 0.25 | 2 | 0 | 1 | 1 | 7 | 0.2857 | 14 | 0.2857 | 1 | 1 | 1 | 0 | 2 | 0 | 1 | 0 | 5 | 0.2 |
| 2_1305509 | 3 | 0 | 1 | 0 | 2 | 0.5 | 5 | 0 | 11 | 0.0909 | 1 | 1 | 1 | 0 | 3 | 1 | 1 | 0 | 6 | 0.6667 |
| 2_1305511 | 4 | 0 | 1 | 0 | 1 | 0 | 6 | 0.1667 | 12 | 0.0833 | 1 | 1 | 1 | 0 | 3 | 0 | 1 | 0 | 6 | 0.1667 |
| 2_1306453 | NA | NA | 2 | 0.5 | 1 | 0 | 1 | 1 | 4 | 0.5 | 1 | 0 | 1 | 1 | NA | NA | NA | NA | 2 | 0.5 |
| 2_1307150 | 1 | 0 | 2 | 0.5 | NA | NA | 1 | 0 | 4 | 0.25 | NA | NA | 1 | 1 | NA | NA | NA | NA | 1 | 1 |
| 2_1307162 | 1 | 1 | NA | NA | NA | NA | NA | NA | 1 | 1 | NA | NA | 1 | 1 | NA | NA | NA | NA | 1 | 1 |
| 2_1307163 | 1 | 1 | NA | NA | NA | NA | NA | NA | 1 | 1 | NA | NA | 1 | 1 | NA | NA | NA | NA | 1 | 1 |
| 2_1307168 | 1 | 1 | NA | NA | NA | NA | NA | NA | 1 | 1 | NA | NA | 1 | 1 | NA | NA | NA | NA | 1 | 1 |
| 2_1308677 | NA | NA | 2 | 0 | 4 | 0.25 | 4 | 0 | 10 | 0.1 | 2 | 0.5 | 3 | 0.3333 | 2 | 0 | 3 | 0 | 10 | 0.2 |
| 2_1308679 | 1 | 1 | 4 | 0.25 | 3 | 0 | 3 | 0.6667 | 11 | 0.3636 | 4 | 0.75 | 6 | 0.3333 | 3 | 0.3333 | 1 | 0 | 14 | 0.4286 |
| 2_1308679 | 1 | 0 | 4 | 0.5 | 4 | 0.5 | 4 | 0.5 | 13 | 0.4615 | 4 | 0.75 | 6 | 0.3333 | 3 | 0.6667 | 3 | 0.3333 | 16 | 0.5 |
| 2_1308682 | 1 | 1 | 3 | 1 | 3 | 1 | 4 | 1 | 11 | 1 | 2 | 1 | 5 | 1 | 2 | 1 | 3 | 1 | 12 | 1 |
| 2_1308684 | 2 | 0.5 | 5 | 0 | 5 | 0 | 4 | 0.25 | 16 | 0.125 | 4 | 0 | 6 | 0 | 3 | 0.3333 | 5 | 0.2 | 18 | 0.1111 |
| 2_1308684 | 1 | 0 | 5 | 0.8 | 5 | 0.8 | 4 | 0.75 | 15 | 0.7333 | 5 | 0.4 | 6 | 0.1667 | 4 | 0.5 | 5 | 0 | 20 | 0.25 |
| 2_1308684 | 2 | 0 | 5 | 0 | 6 | 0.1667 | 4 | 0.5 | 17 | 0.1765 | 5 | 0.2 | 7 | 0.1429 | 4 | 0 | 5 | 0 | 21 | 0.0952 |

| | | | | | | | | | | | | | | | | | | | | |
|---|---|---|---|---|---|---|---|---|---|---|---|---|---|---|---|---|---|---|---|---|
| 2_1308690 | 3 | 0 | 5 | 0 | 6 | 0.1667 | 7 | 0.2857 | 21 | 0.1429 | 3 | 0 | 6 | 0.1667 | 7 | 0.2857 | 5 | 0 | 21 | 0.1429 |
| 2_1308710 | 8 | 0.625 | 19 | 0.6842 | 6 | 0.3333 | 14 | 0.5714 | 47 | 0.5957 | 6 | 1 | 9 | 0.5556 | 13 | 0.5385 | 6 | 0.8333 | 34 | 0.6765 |
| 2_1308710 | 10 | 0.3 | 21 | 0.2381 | 9 | 0.1111 | 13 | 0.3077 | 53 | 0.2453 | 5 | 0 | 11 | 0.0909 | 13 | 0.3846 | 5 | 0 | 34 | 0.1765 |
| 2_1308711 | 12 | 0 | 23 | 0.0435 | 9 | 0.1111 | 14 | 0.0714 | 58 | 0.0517 | 8 | 0.25 | 10 | 0 | 12 | 0 | 6 | 0 | 36 | 0.0556 |
| 2_1308712 | 11 | 0.1818 | 22 | 0.2727 | 13 | 0.2308 | 16 | 0.25 | 62 | 0.2419 | 8 | 0.375 | 11 | 0.1818 | 13 | 0.1538 | 6 | 0 | 38 | 0.1842 |
| 2_1308715 | 12 | 0.1667 | 18 | 0.0556 | 11 | 0 | 17 | 0 | 58 | 0.0517 | 8 | 0.125 | 9 | 0.2222 | 10 | 0.1 | 6 | 0 | 33 | 0.1212 |
| 2_1308715 | 12 | 0.8333 | 24 | 0.9167 | 13 | 0.9231 | 16 | 0.875 | 65 | 0.8923 | 7 | 0.8571 | 12 | 0.8333 | 13 | 1 | 6 | 0.8333 | 38 | 0.8947 |
| 2_1308717 | 14 | 0.0714 | 22 | 0.1818 | 16 | 0 | 18 | 0.1667 | 70 | 0.1143 | 9 | 0.3333 | 13 | 0.1538 | 15 | 0.2 | 9 | 0.1111 | 46 | 0.1957 |
| 2_1308718 | 14 | 0.1429 | 24 | 0.2083 | 14 | 0.2143 | 16 | 0 | 68 | 0.1471 | 7 | 0.2857 | 11 | 0.0909 | 11 | 0.2727 | 9 | 0.3333 | 38 | 0.2368 |
| 2_1308718 | 14 | 0.9286 | 21 | 0.9048 | 15 | 0.8 | 19 | 0.8947 | 69 | 0.8841 | 9 | 1 | 12 | 0.9167 | 14 | 0.8571 | 8 | 0.75 | 43 | 0.8837 |
| 2_1308718 | 15 | 0.3333 | 26 | 0.0385 | 16 | 0.1875 | 17 | 0 | 74 | 0.1216 | 10 | 0.1 | 13 | 0.1538 | 12 | 0.1667 | 8 | 0.125 | 43 | 0.1395 |
| 2_1308719 | 14 | 0.1429 | 25 | 0.64 | 17 | 0.3529 | 15 | 0.4 | 71 | 0.4225 | 8 | 0.375 | 13 | 0.4615 | 14 | 0.3571 | 6 | 0 | 41 | 0.3415 |
| 2_1308720 | 14 | 0.1429 | 24 | 0.125 | 17 | 0.0588 | 18 | 0 | 73 | 0.0822 | 10 | 0.4 | 13 | 0 | 15 | 0.1333 | 10 | 0.3 | 48 | 0.1875 |
| 2_1308816 | 16 | 0.125 | 9 | 0.1111 | 10 | 0 | 11 | 0.1818 | 46 | 0.1087 | 13 | 0.1538 | 8 | 0.125 | 10 | 0 | 8 | 0 | 39 | 0.0769 |
| 2_1308816 | 14 | 0.1429 | 8 | 0.125 | 8 | 0 | 11 | 0 | 41 | 0.0732 | 13 | 0.1538 | 6 | 0 | 9 | 0 | 5 | 0 | 33 | 0.0606 |
| 2_1308817 | 11 | 0 | 8 | 0.125 | 10 | 0.2 | 5 | 0 | 34 | 0.0882 | 9 | 0 | 6 | 0 | 8 | 0.25 | 6 | 0 | 29 | 0.069 |
| 2_1308818 | 16 | 0.3125 | 11 | 0.1818 | 9 | 0.4444 | 10 | 0.2 | 46 | 0.2826 | 9 | 0.6667 | 7 | 0.1429 | 9 | 0.3333 | 7 | 0.5714 | 32 | 0.4375 |
| 2_1308818 | 17 | 0.1176 | 11 | 0.0909 | 9 | 0 | 12 | 0.1667 | 49 | 0.102 | 12 | 0.0833 | 8 | 0.25 | 10 | 0.3 | 8 | 0.125 | 38 | 0.1842 |
| 2_1308819 | 17 | 0.1176 | 9 | 0 | 10 | 0.2 | 12 | 0.0833 | 48 | 0.1042 | 11 | 0.0909 | 8 | 0.375 | 8 | 0.125 | 6 | 0.1667 | 33 | 0.1818 |
| 2_1308828 | 6 | 1 | 1 | 1 | NA | NA | 1 | 1 | 8 | 1 | 2 | 1 | 3 | 0.6667 | 2 | 1 | 1 | 0 | 8 | 0.75 |
| 2_1309871 | NA | NA | 1 | 0 | 2 | 0.5 | NA | NA | 3 | 0.3333 | NA | NA | NA | NA | 1 | 1 | 1 | 0 | 2 | 0.5 |
| 2_1309872 | NA | NA | 2 | 1 | 2 | 1 | NA | NA | 4 | 1 | NA | NA | NA | NA | 1 | 1 | 1 | 1 | 2 | 1 |
| 2_1309873 | 1 | 1 | 2 | 0 | 1 | 0 | NA | NA | 4 | 0.25 | NA | NA | NA | NA | 1 | 1 | NA | NA | 1 | 1 |
| 2_1309873 | 1 | 0 | 2 | 1 | 2 | 0.5 | NA | NA | 5 | 0.6 | NA | NA | NA | NA | 1 | 0 | 1 | 1 | 2 | 0.5 |
| 2_1309874 | 1 | 0 | 1 | 1 | 2 | 1 | NA | NA | 4 | 0.75 | NA | NA | 2 | 1 | 1 | 0 | 1 | 1 | 4 | 0.75 |
| 2_1309876 | 1 | 0 | 3 | 0 | 2 | 0.5 | NA | NA | 6 | 0.1667 | NA | NA | 2 | 0 | 2 | 0.5 | 1 | 0 | 5 | 0.2 |
| 2_1309876 | 1 | 1 | 3 | 0 | 2 | 0 | NA | NA | 6 | 0.1667 | NA | NA | 2 | 0.5 | 2 | 0 | 1 | 0 | 5 | 0.2 |
| 2_1309876 | 1 | 0 | 3 | 0.6667 | 2 | 0 | NA | NA | 6 | 0.3333 | NA | NA | 2 | 0.5 | 2 | 0 | 1 | 0 | 5 | 0.2 |
| 2_1309877 | 1 | 0 | 3 | 0.3333 | 2 | 0 | NA | NA | 6 | 0.1667 | NA | NA | 1 | 0 | 2 | 0.5 | NA | NA | 3 | 0.3333 |
| 2_1310134 | NA | NA | 4 | 0.5 | NA | NA | 1 | 1 | 5 | 0.6 | 1 | 1 | NA | NA | NA | NA | 1 | 0 | 2 | 0.5 |
| 2_1310198 | 10 | 0.1 | 10 | 0.1 | 17 | 0.1176 | 7 | 0 | 44 | 0.0909 | 7 | 0 | 8 | 0.125 | 11 | 0.0909 | 5 | 0 | 31 | 0.0645 |
| 2_1310199 | 14 | 1 | 11 | 0.9091 | 11 | 1 | 12 | 1 | 48 | 0.9792 | 8 | 1 | 10 | 1 | 10 | 0.9 | 6 | 1 | 34 | 0.9706 |
| 2_1310199 | 17 | 0.9412 | 17 | 0.7647 | 19 | 0.8947 | 11 | 0.8182 | 64 | 0.8594 | 11 | 0.7273 | 12 | 0.75 | 12 | 1 | 7 | 0.8571 | 42 | 0.8333 |
| 2_1310200 | 17 | 0.7059 | 16 | 0.875 | 22 | 0.8636 | 14 | 0.8571 | 69 | 0.8261 | 12 | 0.8333 | 11 | 0.8182 | 15 | 0.8667 | 6 | 0.8333 | 44 | 0.8409 |
| 2_1310200 | 15 | 0.2 | 12 | 0.1667 | 16 | 0.0625 | 9 | 0 | 52 | 0.1154 | 8 | 0.125 | 10 | 0.2 | 12 | 0 | 5 | 0 | 35 | 0.0857 |
| 2_1310200 | 12 | 0.4167 | 11 | 0.0909 | 17 | 0.0588 | 6 | 0.3333 | 46 | 0.1957 | 11 | 0.1818 | 7 | 0.4286 | 13 | 0.3846 | 1 | 0 | 32 | 0.3125 |
| 2_1310202 | 13 | 0.2308 | 14 | 0.0714 | 16 | 0.0625 | 9 | 0.2222 | 52 | 0.1346 | 10 | 0.3 | 7 | 0.2857 | 8 | 0.125 | 4 | 0 | 29 | 0.2069 |
| 2_1310202 | 12 | 0.6667 | 14 | 0.5 | 17 | 0.7059 | 10 | 0.9 | 53 | 0.6792 | 10 | 0.9 | 7 | 0.7143 | 9 | 0.7778 | 3 | 1 | 29 | 0.8276 |
| 2_1310203 | 8 | 0 | 12 | 0 | 15 | 0.2 | 9 | 0.1111 | 44 | 0.0909 | 7 | 0 | 7 | 0 | 7 | 0.1429 | 2 | 0.5 | 23 | 0.087 |
| 2_1310206 | 5 | 0.6 | 2 | 1 | 4 | 1 | 5 | 1 | 16 | 0.875 | 1 | 1 | 1 | 1 | 7 | 1 | 1 | 1 | 10 | 1 |
| 2_1310208 | 4 | 1 | 2 | 1 | 4 | 1 | 3 | 0.6667 | 13 | 0.9231 | 1 | 1 | NA | NA | 3 | 1 | 1 | 1 | 5 | 1 |
| 2_1310209 | 4 | 1 | 2 | 1 | 3 | 1 | 2 | 1 | 11 | 1 | 1 | 0 | 1 | 1 | 4 | 0.75 | 1 | 1 | 7 | 0.7143 |
| 2_1310211 | 4 | 0.75 | NA | NA | NA | NA | 3 | 1 | 7 | 0.8571 | 1 | 0 | 1 | 1 | 4 | 0.75 | 1 | 1 | 7 | 0.7143 |
| 2_1310392 | 1 | 0 | 2 | 0.5 | NA | NA | 1 | 0 | 4 | 0.25 | 1 | 0 | 1 | 0 | 1 | 0 | 2 | 0.5 | 5 | 0.2 |
| 2_1310545 | 6 | 0.5 | 5 | 0.4 | 1 | 0 | 3 | 0 | 15 | 0.3333 | 2 | 0.5 | 1 | 0 | 4 | 0 | NA | NA | 7 | 0.1429 |
| 2_1310545 | 6 | 1 | 5 | 0.8 | 1 | 1 | 3 | 1 | 15 | 0.9333 | 1 | 1 | 1 | 1 | 5 | 0.6 | 1 | 0 | 8 | 0.625 |
| 2_1310545 | 6 | 0.3333 | 4 | 0.75 | 1 | 1 | 5 | 0.4 | 16 | 0.5 | 1 | 0 | NA | NA | 5 | 0.8 | 1 | 1 | 7 | 0.7143 |
| 2_1310545 | 5 | 0 | 3 | 0 | 1 | 1 | 5 | 0 | 14 | 0.0714 | 2 | 0.5 | 1 | 0 | 4 | 0 | 1 | 1 | 8 | 0.25 |
| 2_1310546 | 5 | 0.2 | 4 | 0 | 1 | 0 | 5 | 0.2 | 15 | 0.1333 | 1 | 1 | 1 | 0 | 4 | 0 | 2 | 0 | 8 | 0.125 |
| 2_1310548 | 5 | 0.2 | 5 | 0 | 2 | 0 | 4 | 0 | 16 | 0.0625 | 2 | 0 | 1 | 0 | 6 | 0 | 3 | 0.3333 | 12 | 0.0833 |
| 2_1310549 | 7 | 1 | 5 | 1 | 2 | 1 | 6 | 1 | 20 | 1 | 2 | 1 | 1 | 1 | 6 | 0.8333 | 3 | 1 | 12 | 0.9167 |
| 2_1310549 | 7 | 0.8571 | 5 | 1 | 2 | 0.5 | 4 | 1 | 18 | 0.8889 | 2 | 0.5 | 1 | 1 | 6 | 0.6667 | 3 | 0.6667 | 12 | 0.6667 |
| 2_1310550 | 6 | 0.1667 | 4 | 0 | 2 | 0 | 6 | 0 | 18 | 0.0556 | 3 | 0.3333 | 1 | 0 | 4 | 0 | 2 | 0 | 10 | 0.1 |
| 2_1310550 | 8 | 0.125 | 5 | 0.2 | 2 | 0 | 7 | 0.1429 | 22 | 0.1364 | 3 | 0 | 1 | 0 | 6 | 0.5 | 3 | 0 | 13 | 0.2308 |
| 2_1310551 | 7 | 0.4286 | 4 | 0.25 | 1 | 0 | 5 | 0.2 | 17 | 0.2941 | 4 | 0.75 | 1 | 0 | 6 | 0.1667 | 3 | 0.3333 | 14 | 0.3571 |
| 2_1310554 | 4 | 0 | 3 | 0.3333 | 4 | 0 | 5 | 0 | 16 | 0.0625 | 4 | 0.25 | 1 | 0 | 3 | 0.3333 | 2 | 1 | 10 | 0.4 |
| 2_1310871 | NA | NA | 1 | 0 | 2 | 1 | 1 | 0 | 4 | 0.5 | NA | NA | NA | NA | NA | NA | 2 | 0.5 | 2 | 0.5 |
| 2_1310873 | NA | NA | 1 | 0 | 2 | 0.5 | NA | NA | 3 | 0.3333 | NA | NA | NA | NA | NA | NA | 2 | 0.5 | 2 | 0.5 |
| 2_1310883 | 1 | 1 | 1 | 0 | 2 | 0 | 1 | 0 | 5 | 0.2 | NA | NA | NA | NA | NA | NA | 2 | 0.5 | 2 | 0.5 |
| 2_1310884 | 1 | 1 | NA | NA | 2 | 0.5 | 1 | 1 | 4 | 0.75 | NA | NA | NA | NA | NA | NA | 2 | 0.5 | 2 | 0.5 |
| 2_1310888 | 1 | 0 | 1 | 0 | 2 | 0.5 | NA | NA | 4 | 0.25 | NA | NA | NA | NA | NA | NA | 2 | 0.5 | 2 | 0.5 |
| 2_1311160 | 3 | 0.3333 | 3 | 1 | 7 | 0.2857 | 2 | 0.5 | 15 | 0.4667 | 2 | 1 | 6 | 0.3333 | 7 | 0.7143 | 1 | 0 | 16 | 0.5625 |
| 2_1311160 | 3 | 0.3333 | 4 | 0.25 | 6 | 0.1667 | 2 | 0 | 15 | 0.2 | 4 | 0.5 | 7 | 0 | 6 | 0 | 1 | 0 | 18 | 0.1111 |
| 2_1311161 | 3 | 0 | 3 | 0.3333 | 6 | 0 | 1 | 0 | 13 | 0.0769 | 2 | 0.5 | 5 | 0.6 | 4 | 0.5 | 1 | 0 | 12 | 0.5 |
| 2_1311382 | 3 | 0 | 5 | 0.2 | 10 | 0.2 | 3 | 0 | 21 | 0.1429 | 3 | 0 | 6 | 0 | 6 | 0.3333 | 2 | 0 | 17 | 0.1176 |

| | | | | | | | | | | | | | | | | | | | | |
|---|---|---|---|---|---|---|---|---|---|---|---|---|---|---|---|---|---|---|---|---|
| 2_1311383 | 3 | 1 | 5 | 0.8 | 10 | 0.7 | 3 | 1 | 21 | 0.8095 | 3 | 0.6667 | 7 | 0.5714 | 6 | 0.8333 | 2 | 1 | 18 | 0.7222 |
| 2_1311384 | 2 | 0 | 4 | 0.75 | 8 | 0.75 | 3 | 0.3333 | 17 | 0.5882 | 6 | 0.5 | 6 | 0.5 | 6 | 0.1667 | 2 | 0 | 20 | 0.35 |
| 2_1311385 | 2 | 0 | 2 | 0.5 | 6 | 0.5 | 2 | 0.5 | 12 | 0.4167 | 3 | 0.6667 | 5 | 0.6 | 3 | 1 | NA | NA | 11 | 0.7273 |
| 2_1311385 | 3 | 0.3333 | 5 | 0 | 5 | 0.4 | 2 | 0 | 15 | 0.2 | 5 | 0.2 | 5 | 0.6 | 6 | 0.3333 | NA | NA | 16 | 0.375 |
| 2_1311388 | 2 | 0 | 1 | 1 | 4 | 1 | 1 | 0 | 8 | 0.625 | 1 | 1 | 4 | 0.75 | 4 | 1 | NA | NA | 9 | 0.8889 |
| 2_1311389 | 3 | 0 | 1 | 1 | 3 | 0.6667 | NA | NA | 7 | 0.4286 | 2 | 0 | 5 | 0.4 | 4 | 0.25 | NA | NA | 11 | 0.2727 |
| 2_1311394 | 3 | 0.3333 | 1 | 1 | 2 | 0 | 1 | 0 | 7 | 0.2857 | 2 | 0 | 5 | 0.2 | 4 | 0 | NA | NA | 11 | 0.0909 |
| 2_1311408 | 1 | 0 | 1 | 0 | 4 | 0.5 | 6 | 0.5 | 12 | 0.4167 | 3 | 0.3333 | 3 | 0.3333 | 2 | 0.5 | 7 | 0.2857 | 15 | 0.3333 |
| 2_1311409 | 6 | 0 | 2 | 0 | 4 | 0.25 | 4 | 0.25 | 16 | 0.125 | 2 | 0.5 | 3 | 0 | 3 | 0 | 2 | 0 | 10 | 0.1 |
| 2_1311422 | NA | NA | 1 | 1 | 2 | 0 | 6 | 0 | 9 | 0.1111 | 3 | 0.3333 | 2 | 0 | 2 | 0 | 4 | 0.25 | 11 | 0.1818 |
| 2_1311422 | 1 | 0 | 1 | 0 | 1 | 0 | 6 | 0.3333 | 9 | 0.2222 | 3 | 0.3333 | 2 | 0 | 2 | 0 | 4 | 0 | 11 | 0.0909 |
| 2_1311428 | 1 | 1 | 2 | 0 | 2 | 0 | 6 | 0.1667 | 11 | 0.1818 | 2 | 0 | 2 | 0 | 2 | 0.5 | 5 | 0.2 | 11 | 0.1818 |
| 2_1311435 | NA | NA | 1 | 0 | 3 | 0 | 5 | 0.2 | 9 | 0.1111 | 1 | 0 | 3 | 0.3333 | 3 | 0 | 3 | 0 | 10 | 0.1 |
| 2_1311441 | NA | NA | NA | NA | 2 | 0.5 | 2 | 1 | 4 | 0.75 | 1 | 1 | 3 | 0.6667 | 3 | 0.3333 | 3 | 0.6667 | 10 | 0.6 |
| 2_1311441 | NA | NA | NA | NA | 2 | 0 | 2 | 0.5 | 4 | 0.25 | 1 | 1 | 2 | 0 | 3 | 0.3333 | 3 | 0 | 9 | 0.2222 |
| 2_1311442 | NA | NA | NA | NA | 2 | 1 | 2 | 1 | 4 | 1 | 1 | 1 | 2 | 1 | 2 | 1 | 3 | 1 | 8 | 1 |
| 2_1311444 | NA | NA | NA | NA | 2 | 0 | 1 | 1 | 3 | 0.3333 | 1 | 0 | 2 | 1 | 3 | 0 | 3 | 0 | 9 | 0.2222 |
| 2_1311447 | NA | NA | NA | NA | 2 | 0 | 2 | 0.5 | 4 | 0.25 | NA | NA | 3 | 0.3333 | NA | NA | 2 | 0.5 | 5 | 0.4 |
| 2_1311448 | NA | NA | NA | NA | 1 | 1 | 1 | 1 | 2 | 1 | NA | NA | 2 | 1 | 2 | 1 | 1 | 0 | 5 | 0.8 |
| 2_1311448 | NA | NA | NA | NA | 1 | 1 | 2 | 1 | 3 | 1 | 1 | 1 | 3 | 1 | 2 | 1 | 2 | 1 | 8 | 1 |
| 2_1311449 | NA | NA | NA | NA | 1 | 1 | 1 | 1 | 2 | 1 | 1 | 1 | 2 | 0.5 | 3 | 1 | 2 | 0.5 | 8 | 0.75 |
| 2_1311450 | NA | NA | NA | NA | 1 | 1 | 1 | 1 | 2 | 1 | 1 | 1 | 2 | 1 | 3 | 1 | 2 | 1 | 8 | 1 |
| 2_1311451 | NA | NA | NA | NA | 1 | 1 | 1 | 0 | 2 | 0.5 | 1 | 0 | 2 | 0 | 3 | 0.6667 | 2 | 0.5 | 8 | 0.375 |
| 2_1311452 | NA | NA | NA | NA | 1 | 1 | NA | NA | 1 | 1 | 1 | 1 | 2 | 0.5 | 2 | 1 | 2 | 0.5 | 7 | 0.7143 |
| 2_1311453 | NA | NA | NA | NA | 1 | 1 | 1 | 0 | 2 | 0.5 | 1 | 1 | 2 | 1 | 2 | 0 | 2 | 0.5 | 7 | 0.5714 |
| 2_1311453 | NA | NA | NA | NA | 1 | 1 | 1 | 1 | 2 | 1 | 1 | 1 | 1 | 0 | 2 | 1 | 2 | 1 | 6 | 0.8333 |
| 2_1313668 | 8 | 0 | 8 | 0.25 | 5 | 0.2 | 10 | 0.3 | 31 | 0.1935 | 3 | 0 | 5 | 0.2 | 9 | 0.1111 | 8 | 0.25 | 25 | 0.16 |
| 2_1313669 | 7 | 0.1429 | 9 | 0.1111 | 7 | 0 | 8 | 0 | 31 | 0.0645 | 2 | 0 | 7 | 0.1429 | 9 | 0.1111 | 7 | 0 | 25 | 0.08 |
| 2_1313874 | NA | NA | 2 | 1 | 1 | 1 | NA | NA | 3 | 1 | NA | NA | NA | NA | NA | NA | 1 | 1 | 1 | 1 |
| 2_1313875 | NA | NA | 2 | 1 | 1 | 1 | NA | NA | 3 | 1 | NA | NA | NA | NA | NA | NA | 1 | 1 | 1 | 1 |
| 2_1313876 | NA | NA | 2 | 1 | 1 | 1 | NA | NA | 3 | 1 | NA | NA | NA | NA | NA | NA | 1 | 1 | 1 | 1 |
| 2_1313877 | NA | NA | 2 | 1 | 1 | 1 | NA | NA | 3 | 1 | NA | NA | NA | NA | NA | NA | 1 | 1 | 1 | 1 |
| 2_1313878 | NA | NA | 3 | 1 | 1 | 1 | NA | NA | 4 | 1 | NA | NA | NA | NA | NA | NA | 1 | 1 | 1 | 1 |
| 2_1317783 | 31 | 0.4839 | 56 | 0.4286 | 17 | 0.2941 | 42 | 0.381 | 146 | 0.411 | 35 | 0.2286 | 26 | 0.4231 | 28 | 0.3929 | 35 | 0.5714 | 124 | 0.4032 |
| 2_1317783 | 32 | 0.9688 | 49 | 1 | 19 | 1 | 41 | 0.9268 | 141 | 0.9716 | 33 | 0.9697 | 29 | 0.9655 | 28 | 0.9643 | 35 | 0.9714 | 125 | 0.968 |
| 2_1317785 | 38 | 0.3947 | 53 | 0.3962 | 18 | 0.1667 | 40 | 0.375 | 149 | 0.3624 | 33 | 0.3939 | 31 | 0.2903 | 33 | 0.3636 | 39 | 0.4103 | 136 | 0.3676 |
| 2_1317785 | 40 | 0.55 | 50 | 0.46 | 20 | 0.4 | 44 | 0.3182 | 154 | 0.4351 | 41 | 0.5366 | 29 | 0.3793 | 29 | 0.2759 | 40 | 0.425 | 139 | 0.4173 |
| 2_1317786 | 46 | 0.1522 | 59 | 0.0678 | 22 | 0 | 43 | 0.0233 | 170 | 0.0706 | 40 | 0.05 | 37 | 0.1351 | 32 | 0.0938 | 41 | 0.122 | 150 | 0.1 |
| 2_1317786 | 43 | 0.0698 | 60 | 0.0833 | 27 | 0.1481 | 41 | 0.1463 | 171 | 0.1053 | 39 | 0.0513 | 27 | 0.0741 | 36 | 0.1944 | 44 | 0.2045 | 146 | 0.137 |
| 2_1317787 | 61 | 0.3279 | 77 | 0.3766 | 35 | 0.3143 | 68 | 0.3088 | 241 | 0.3361 | 60 | 0.3167 | 54 | 0.2778 | 62 | 0.4355 | 57 | 0.2982 | 233 | 0.3348 |
| 2_1317789 | 62 | 0.129 | 79 | 0.0506 | 33 | 0.1212 | 68 | 0.1471 | 242 | 0.1074 | 58 | 0.1897 | 57 | 0.0877 | 60 | 0.1333 | 52 | 0.1731 | 227 | 0.1454 |
| 2_1317793 | 72 | 0.0833 | 76 | 0.0789 | 35 | 0.0857 | 69 | 0.1159 | 252 | 0.0913 | 73 | 0.0959 | 63 | 0.1111 | 63 | 0.0794 | 61 | 0.1475 | 260 | 0.1077 |
| 2_1317798 | 94 | 0.1064 | 104 | 0.0673 | 51 | 0.0588 | 85 | 0.1176 | 334 | 0.0898 | 93 | 0.172 | 78 | 0.1538 | 85 | 0.1176 | 86 | 0.1279 | 342 | 0.1433 |
| 2_1317798 | 103 | 0.0291 | 101 | 0.0891 | 58 | 0.069 | 104 | 0.0577 | 366 | 0.0601 | 96 | 0.1042 | 86 | 0.0465 | 92 | 0.0652 | 95 | 0.0842 | 369 | 0.0759 |
| 2_1318348 | 6 | 0.1667 | 3 | 0 | 8 | 0.125 | 2 | 0.5 | 19 | 0.1579 | 8 | 0 | 3 | 0 | 4 | 0.25 | 2 | 0.5 | 17 | 0.1176 |
| 2_1318349 | 6 | 0.1667 | 2 | 0.5 | 7 | 0.4286 | 2 | 0 | 17 | 0.2941 | 7 | 0 | 2 | 0.5 | 4 | 0.75 | 2 | 0 | 15 | 0.2667 |
| 2_1318351 | 2 | 0 | 1 | 1 | 3 | 0.6667 | 2 | 0.5 | 8 | 0.5 | 5 | 0.6 | 2 | 1 | 1 | 0 | 3 | 0.3333 | 11 | 0.5455 |
| 2_1318351 | 5 | 0.6 | 3 | 0 | 6 | 0.3333 | 2 | 0.5 | 16 | 0.375 | 7 | 0.1429 | 2 | 0 | 4 | 0.75 | 4 | 0 | 17 | 0.2353 |
| 2_1318351 | 7 | 0.1429 | 2 | 0 | 3 | 0 | 2 | 0 | 14 | 0.0714 | 6 | 0.1667 | 3 | 0 | 4 | 0 | 4 | 0 | 17 | 0.0588 |
| 2_1318354 | 5 | 0.6 | 2 | 0.5 | 4 | 0 | 2 | 0.5 | 13 | 0.3846 | 5 | 0.4 | 3 | 0 | 3 | 0.3333 | 4 | 0 | 15 | 0.2 |
| 2_1318356 | 3 | 0.6667 | 2 | 0.5 | 3 | 0 | 2 | 0 | 10 | 0.3 | 1 | 1 | 1 | 0 | 2 | 0.5 | 1 | 1 | 5 | 0.6 |
| 2_1318357 | 4 | 0.25 | 3 | 0 | 3 | 0 | 3 | 0 | 13 | 0.0769 | 6 | 0.3333 | 2 | 0 | 2 | 0 | 3 | 0.3333 | 13 | 0.2308 |
| 2_1318357 | 5 | 0 | 4 | 0 | 5 | 0.2 | 3 | 0 | 17 | 0.0588 | 5 | 0.2 | 2 | 0 | 2 | 0 | 4 | 0 | 13 | 0.0769 |
| 2_1318358 | 5 | 0 | 2 | 0 | 3 | 0.3333 | 3 | 0 | 13 | 0.0769 | 4 | 0.25 | 3 | 0 | 2 | 0 | 3 | 0 | 12 | 0.0833 |
| 2_1318362 | 3 | 0.6667 | 3 | 0 | 4 | 0 | 3 | 0 | 13 | 0.1538 | 5 | 0.6 | 1 | 1 | 1 | 1 | 2 | 0 | 9 | 0.5556 |
| 2_1318363 | 2 | 0 | 3 | 0.3333 | 3 | 0.3333 | 2 | 0 | 10 | 0.2 | 5 | 0.8 | 1 | 0 | 1 | 0 | 2 | 0 | 9 | 0.4444 |
| 2_1318363 | 2 | 0.5 | 3 | 0.6667 | 3 | 0.3333 | 3 | 0.3333 | 11 | 0.4545 | 4 | 0.75 | 1 | 1 | 1 | 1 | 3 | 0.6667 | 9 | 0.7778 |
| 2_1318366 | 2 | 0 | 2 | 0 | 1 | 1 | 4 | 0.5 | 9 | 0.3333 | 4 | 0.5 | NA | NA | NA | NA | 3 | 0.3333 | 7 | 0.4286 |
| 2_1318366 | 2 | 0.5 | 4 | 0 | 2 | 1 | 4 | 0.25 | 12 | 0.3333 | 3 | 0.6667 | 1 | 1 | 1 | 0 | 3 | 0.3333 | 8 | 0.5 |
| 2_1318367 | NA | NA | 2 | 1 | 1 | 0 | 2 | 0.5 | 5 | 0.6 | 2 | 0.5 | 1 | 1 | 1 | 0 | 2 | 1 | 6 | 0.6667 |
| 2_1318368 | 2 | 0.5 | 4 | 0.5 | 3 | 0 | 3 | 0.6667 | 12 | 0.4167 | 4 | 0.5 | 1 | 1 | NA | NA | 4 | 0.75 | 9 | 0.6667 |
| 2_1318369 | 2 | 1 | 3 | 1 | 3 | 1 | 4 | 1 | 12 | 1 | 4 | 1 | 1 | 1 | 1 | 1 | 3 | 1 | 9 | 1 |
| 2_1318370 | NA | NA | 3 | 0.6667 | 2 | 0.5 | 3 | 0.3333 | 8 | 0.5 | 5 | 0.6 | 1 | 0 | 1 | 0 | 3 | 0.3333 | 10 | 0.4 |
| 2_1318371 | 2 | 1 | 1 | 0 | 1 | 0 | 2 | 0 | 6 | 0.3333 | 2 | 0.5 | 1 | 1 | 1 | 1 | 3 | 0.3333 | 7 | 0.5714 |
| 2_1318378 | 1 | 1 | 2 | 1 | NA | NA | 4 | 1 | 7 | 1 | 1 | 1 | NA | NA | 1 | 1 | 2 | 0.5 | 4 | 0.75 |

| | | | | | | | | | | | | | | | | | | | | |
|---|---|---|---|---|---|---|---|---|---|---|---|---|---|---|---|---|---|---|---|---|
| 2_13183789 | 1 | 1 | 4 | 0 | NA | NA | 2 | 0 | 7 | 0.1429 | 1 | 0 | NA | NA | 1 | 0 | 2 | 0.5 | 4 | 0.25 |
| 2_13183808 | 1 | 1 | 4 | 1 | NA | NA | 4 | 1 | 9 | 1 | NA | NA | NA | NA | 1 | 1 | 3 | 1 | 4 | 1 |
| 2_13183809 | 1 | 1 | 3 | 1 | NA | NA | 4 | 1 | 8 | 1 | 1 | 1 | NA | NA | 1 | 1 | 4 | 1 | 6 | 1 |
| 2_13183812 | 1 | 0 | 3 | 1 | 1 | 1 | 3 | 0.3333 | 8 | 0.625 | 1 | 1 | NA | NA | NA | NA | 4 | 0.5 | 5 | 0.6 |
| 2_13183814 | 1 | 0 | 4 | 0 | NA | NA | 4 | 0.25 | 9 | 0.1111 | 1 | 1 | NA | NA | 1 | 0 | 3 | 0 | 5 | 0.2 |
| 2_13183816 | 1 | 1 | 4 | 0.75 | 1 | 0 | 4 | 0.25 | 10 | 0.5 | 1 | 1 | NA | NA | 1 | 0 | 4 | 0.25 | 6 | 0.3333 |
| 2_13189725 | 4 | 0.25 | 4 | 0 | 2 | 0 | 1 | 1 | 11 | 0.1818 | 7 | 0.1429 | 1 | 0 | 1 | 0 | NA | NA | 9 | 0.1111 |
| 2_13189835 | 4 | 0 | 6 | 0 | 2 | 0.5 | 1 | 0 | 13 | 0.0769 | 6 | 0.1667 | NA | NA | NA | NA | NA | NA | 6 | 0.1667 |
| 2_13189871 | 2 | 1 | 2 | 1 | 2 | 1 | 1 | 0 | 7 | 0.8571 | 4 | 0.75 | NA | NA | NA | NA | NA | NA | 4 | 0.75 |
| 2_13208158 | 7 | 0.1429 | 7 | 0.1429 | 4 | 0 | 4 | 0.25 | 22 | 0.1364 | 4 | 0 | 2 | 0 | 5 | 0.2 | 9 | 0.1111 | 20 | 0.1 |
| 2_13208223 | 5 | 0.4 | 5 | 0.8 | 6 | 0.6667 | 4 | 0 | 20 | 0.5 | 2 | 0.5 | 4 | 0.25 | 3 | 0.6667 | 7 | 1 | 16 | 0.6875 |
| 2_13208234 | 1 | 0 | 3 | 1 | 5 | 1 | 2 | 0.5 | 11 | 0.8182 | 2 | 0.5 | 3 | 0.3333 | 3 | 0.3333 | 5 | 0.8 | 13 | 0.5385 |
| 2_13208241 | 4 | 0.5 | 4 | 0 | 7 | 0.2857 | 4 | 0.25 | 19 | 0.2632 | 3 | 0 | 4 | 0.25 | 2 | 0 | 7 | 0.1429 | 16 | 0.125 |
| 2_13209483 | 14 | 0.2143 | 13 | 0 | 6 | 0.1667 | 9 | 0 | 42 | 0.0952 | 10 | 0.2 | 9 | 0 | 6 | 0 | 7 | 0.1429 | 32 | 0.0938 |
| 2_13213804 | 1 | 1 | NA | NA | NA | NA | NA | NA | 1 | 1 | 1 | 0 | 1 | 1 | NA | NA | NA | NA | 2 | 0.5 |
| 2_13214727 | 4 | 1 | 6 | 0.6667 | 6 | 0.6667 | 4 | 0.5 | 20 | 0.7 | 8 | 0.5 | 9 | 0.5556 | 6 | 0.8333 | 6 | 0.5 | 29 | 0.5862 |
| 2_13214826 | 6 | 0 | 9 | 0 | 7 | 0 | 7 | 0.2857 | 29 | 0.069 | 8 | 0.125 | 11 | 0 | 8 | 0.125 | 8 | 0.125 | 35 | 0.0857 |
| 2_13215354 | 30 | 0.1333 | 39 | 0.0769 | 34 | 0.0588 | 30 | 0.0333 | 133 | 0.0752 | 38 | 0.0526 | 33 | 0 | 34 | 0.0588 | 28 | 0.1786 | 133 | 0.0677 |
| 2_13215369 | 22 | 0.4091 | 30 | 0.2667 | 30 | 0.2333 | 26 | 0.3077 | 108 | 0.2963 | 31 | 0.4516 | 27 | 0.2963 | 28 | 0.3214 | 24 | 0.4167 | 110 | 0.3727 |
| 2_13215373 | 24 | 0.1667 | 33 | 0.1818 | 32 | 0.0938 | 29 | 0.2069 | 118 | 0.161 | 30 | 0.0333 | 31 | 0.0968 | 30 | 0.0667 | 26 | 0.1538 | 117 | 0.0855 |
| 2_13221384 | 1 | 1 | 1 | 0 | NA | NA | NA | NA | 2 | 0.5 | 1 | 1 | NA | NA | NA | NA | NA | NA | 1 | 1 |
| 2_13221401 | NA | NA | 1 | 1 | NA | NA | NA | NA | 1 | 1 | 1 | 1 | NA | NA | NA | NA | NA | NA | 1 | 1 |
| 2_13237245 | 8 | 0.875 | 10 | 0.9 | 9 | 0.7778 | 7 | 0.7143 | 34 | 0.8235 | 3 | 1 | 6 | 1 | 8 | 0.875 | 7 | 0.7143 | 24 | 0.875 |
| 2_13237257 | 6 | 0.6667 | 10 | 0.8 | 10 | 0.6 | 10 | 0.5 | 36 | 0.6389 | 3 | 1 | 6 | 0.8333 | 8 | 0.875 | 7 | 0.8571 | 24 | 0.875 |
| 2_13259217 | 1 | 0 | 4 | 0.5 | NA | NA | 3 | 0.6667 | 8 | 0.5 | 1 | 0 | 1 | 1 | 3 | 0.3333 | 3 | 0 | 8 | 0.25 |
| 2_13259223 | 1 | 0 | 3 | 0.3333 | 1 | 0 | 3 | 0 | 8 | 0.125 | 1 | 1 | 1 | 1 | 3 | 0 | 2 | 0.5 | 7 | 0.4286 |
| 2_13259233 | 2 | 0 | 4 | 0.25 | 3 | 0 | 3 | 0.6667 | 12 | 0.25 | 1 | 0 | 1 | 0 | 3 | 0.3333 | 2 | 0.5 | 7 | 0.2857 |
| 2_13259245 | 1 | 0 | 4 | 0.75 | 3 | 0.6667 | 2 | 1 | 10 | 0.7 | 1 | 1 | 1 | 0 | 4 | 1 | 3 | 0.6667 | 9 | 0.7778 |
| 2_13259251 | 1 | 1 | 4 | 1 | 3 | 0.6667 | 3 | 1 | 11 | 0.9091 | 1 | 1 | 1 | 1 | 3 | 1 | 2 | 1 | 7 | 1 |
| 2_13259255 | 2 | 1 | 4 | 0.75 | 2 | 0.5 | 3 | 0.6667 | 11 | 0.7273 | 1 | 1 | 1 | 0 | 3 | 0.6667 | 4 | 0.75 | 9 | 0.6667 |
| 2_13259256 | 2 | 0.5 | 4 | 1 | 2 | 1 | 2 | 1 | 10 | 0.9 | 1 | 0 | 1 | 1 | 3 | 1 | 3 | 1 | 8 | 0.875 |
| 2_13259287 | 2 | 0 | 3 | 0.3333 | 3 | 0.3333 | 3 | 0.3333 | 11 | 0.2727 | 1 | 1 | NA | NA | 3 | 0.6667 | 3 | 0.3333 | 7 | 0.5714 |
| 2_13263438 | 15 | 0.0667 | 10 | 0 | 7 | 0 | 8 | 0.125 | 40 | 0.05 | 9 | 0.1111 | 12 | 0.0833 | 6 | 0 | 7 | 0.1429 | 34 | 0.0882 |
| 2_13263442 | 14 | 0.1429 | 4 | 0 | 5 | 0 | 5 | 0 | 28 | 0.0714 | 8 | 0 | 10 | 0.1 | 5 | 0.2 | 5 | 0 | 28 | 0.0714 |
| 2_13263513 | 21 | 0 | 15 | 0.0667 | 16 | 0.0625 | 9 | 0.4444 | 61 | 0.0984 | 15 | 0.2 | 18 | 0.1111 | 12 | 0 | 12 | 0.0833 | 57 | 0.1053 |
| 2_13263523 | 26 | 0.2692 | 25 | 0.08 | 19 | 0.1053 | 13 | 0.2308 | 83 | 0.1687 | 14 | 0.1429 | 22 | 0.1364 | 15 | 0.2 | 15 | 0.1333 | 66 | 0.1515 |
| 2_13263548 | 25 | 0.08 | 23 | 0.087 | 20 | 0.25 | 12 | 0.0833 | 80 | 0.125 | 14 | 0.2143 | 19 | 0.1579 | 11 | 0.0909 | 16 | 0.125 | 60 | 0.15 |
| 2_13263556 | 28 | 0.1071 | 28 | 0.1071 | 24 | 0.2083 | 16 | 0.25 | 96 | 0.1562 | 18 | 0 | 26 | 0.1538 | 16 | 0.0625 | 18 | 0 | 78 | 0.0641 |
| 2_13263565 | 34 | 0.3824 | 29 | 0.4483 | 28 | 0.2857 | 13 | 0.3077 | 104 | 0.3654 | 20 | 0.55 | 23 | 0.2174 | 17 | 0.2941 | 22 | 0.5909 | 82 | 0.4146 |
| 2_13263576 | 30 | 0.3 | 33 | 0.1818 | 30 | 0.4 | 14 | 0.5 | 107 | 0.3178 | 18 | 0.4444 | 24 | 0.4167 | 17 | 0.2941 | 21 | 0.1905 | 80 | 0.3375 |
| 2_13263678 | 37 | 0.1081 | 32 | 0.0938 | 30 | 0.1667 | 16 | 0.3125 | 115 | 0.1478 | 19 | 0.2632 | 25 | 0.2 | 20 | 0.15 | 24 | 0.2083 | 88 | 0.2045 |
| 2_13264631 | 3 | 0 | 5 | 0.2 | 7 | 0.1429 | 2 | 0 | 17 | 0.1176 | 3 | 0.3333 | 6 | 0 | 3 | 0 | 3 | 0.6667 | 15 | 0.2 |
| 2_13264655 | 3 | 0 | 5 | 0.2 | 6 | 0.1667 | 2 | 0.5 | 16 | 0.1875 | 3 | 0 | 5 | 0.2 | 2 | 0 | 3 | 0 | 13 | 0.0769 |
| 2_13264661 | 3 | 0.3333 | 5 | 0.2 | 7 | 0.1429 | 2 | 0 | 17 | 0.1765 | 2 | 0 | 6 | 0.3333 | 3 | 0 | 3 | 0.3333 | 14 | 0.2143 |
| 2_13264716 | 2 | 0.5 | 2 | 1 | 6 | 0.8333 | 2 | 0.5 | 12 | 0.75 | NA | NA | 3 | 1 | 3 | 1 | 1 | 1 | 7 | 1 |
| 2_13265079 | 2 | 1 | 1 | 1 | 7 | 0.7143 | 2 | 1 | 12 | 0.8333 | NA | NA | 3 | 1 | 1 | 1 | 2 | 0.5 | 6 | 0.8333 |
| 2_13270781 | 3 | 0.3333 | 7 | 0 | 7 | 0.1429 | 3 | 0 | 20 | 0.1 | 2 | 0 | 3 | 0.3333 | 4 | 0 | 5 | 0 | 14 | 0.0714 |
| 2_13270791 | 3 | 0.6667 | 7 | 0.4286 | 7 | 0.4286 | 3 | 0.3333 | 20 | 0.45 | 2 | 0.5 | 3 | 0.6667 | 6 | 0.6667 | 4 | 0.5 | 15 | 0.6 |
| 2_13270805 | 2 | 0.5 | 7 | 0 | 8 | 0.125 | 3 | 0 | 20 | 0.1 | NA | NA | 4 | 0 | 7 | 0 | 5 | 0.2 | 16 | 0.0625 |
| 2_13339454 | NA | NA | 1 | 1 | 2 | 0.5 | NA | NA | 3 | 0.6667 | 1 | 1 | NA | NA | 1 | 1 | NA | NA | 2 | 1 |
| 2_13339460 | NA | NA | 1 | 1 | 2 | 0.5 | NA | NA | 3 | 0.6667 | 1 | 1 | NA | NA | 1 | 0 | NA | NA | 2 | 0.5 |
| 2_13339525 | NA | NA | 1 | 1 | 2 | 0 | NA | NA | 3 | 0.3333 | 1 | 1 | NA | NA | 1 | 0 | NA | NA | 2 | 0.5 |
| 2_13340301 | NA | NA | 1 | 0 | 2 | 1 | 2 | 0 | 5 | 0.4 | 1 | 0 | NA | NA | 2 | 1 | NA | NA | 3 | 0.6667 |
| 2_13340302 | NA | NA | 1 | 0 | 3 | 1 | 2 | 1 | 6 | 0.8333 | NA | NA | NA | NA | 2 | 1 | NA | NA | 2 | 1 |
| 2_13340309 | NA | NA | 1 | 0 | 3 | 0.6667 | 1 | 0 | 5 | 0.4 | NA | NA | NA | NA | 2 | 0.5 | NA | NA | 2 | 0.5 |
| 2_13340310 | NA | NA | 1 | 1 | 2 | 1 | 1 | 1 | 4 | 1 | 1 | 1 | NA | NA | 1 | 1 | NA | NA | 2 | 1 |
| 2_13340317 | NA | NA | 1 | 1 | 3 | 1 | 2 | 1 | 6 | 1 | 1 | 1 | NA | NA | 2 | 1 | NA | NA | 3 | 1 |
| 2_13340320 | NA | NA | 1 | 0 | 3 | 0.3333 | 1 | 0 | 5 | 0.2 | 1 | 1 | NA | NA | 2 | 0.5 | NA | NA | 3 | 0.6667 |
| 2_13340324 | NA | NA | NA | NA | 1 | 1 | 2 | 1 | 3 | 1 | 1 | 1 | NA | NA | 1 | 1 | NA | NA | 2 | 1 |
| 2_13340324 | NA | NA | 1 | 1 | 3 | 1 | 2 | 0.5 | 6 | 0.8333 | 1 | 1 | NA | NA | 2 | 1 | NA | NA | 3 | 1 |
| 2_13340326 | NA | NA | 1 | 0 | 3 | 0.6667 | 2 | 0.5 | 6 | 0.5 | 1 | 1 | NA | NA | 2 | 1 | NA | NA | 3 | 1 |
| 2_13340333 | NA | NA | 1 | 0 | 3 | 0.3333 | 2 | 0.5 | 6 | 0.3333 | 1 | 1 | NA | NA | 2 | 0.5 | NA | NA | 3 | 0.6667 |
| 2_13340335 | NA | NA | NA | NA | 3 | 1 | 1 | 1 | 4 | 1 | NA | NA | NA | NA | 2 | 1 | NA | NA | 2 | 1 |
| 2_13340337 | NA | NA | 1 | 1 | 2 | 1 | 1 | 1 | 4 | 1 | 1 | 1 | NA | NA | 1 | 1 | NA | NA | 2 | 1 |
| 2_13340349 | NA | NA | 1 | 1 | 2 | 0.5 | 2 | 0.5 | 5 | 0.6 | 1 | 1 | NA | NA | NA | NA | NA | NA | 1 | 1 |

| | | | | | | | | | | | | | | | | | | | | |
|---|---|---|---|---|---|---|---|---|---|---|---|---|---|---|---|---|---|---|---|---|
| 2_13340348 | NA | NA | NA | NA | 2 | 0 | 1 | 1 | 3 | 0.3333 | 1 | 0 | NA | NA | 2 | 0.5 | NA | NA | 3 | 0.3333 |
| 2_13340349 | NA | NA | 1 | 1 | 3 | 1 | 1 | 1 | 5 | 1 | 1 | 1 | NA | NA | 2 | 1 | NA | NA | 3 | 1 |
| 2_13340360 | NA | NA | NA | NA | 3 | 0.3333 | 2 | 1 | 5 | 0.6 | 1 | 0 | NA | NA | 2 | 0.5 | NA | NA | 3 | 0.3333 |
| 2_13340439 | NA | NA | 1 | 1 | 4 | 0 | 1 | 0 | 6 | 0.1667 | 1 | 0 | NA | NA | 3 | 0.3333 | NA | NA | 4 | 0.25 |
| 2_13443624 | 2 | 0 | 2 | 0 | 1 | 1 | NA | NA | 5 | 0.2 | 2 | 0 | 2 | 0.5 | 1 | 0 | 2 | 0 | 7 | 0.1429 |
| 2_13443636 | 1 | 1 | 2 | 0 | NA | NA | NA | NA | 3 | 0.3333 | 2 | 0.5 | 1 | 0 | 1 | 0 | 2 | 1 | 6 | 0.5 |
| 2_13443689 | 32 | 0.0312 | 36 | 0.0556 | 36 | 0.0556 | 30 | 0.1333 | 134 | 0.0672 | 23 | 0 | 36 | 0.1111 | 29 | 0.069 | 32 | 0 | 120 | 0.05 |
| 2_13443694 | 26 | 0.1923 | 29 | 0.1379 | 27 | 0.1111 | 21 | 0.2381 | 103 | 0.165 | 17 | 0.2941 | 24 | 0.2083 | 22 | 0.0455 | 24 | 0.1667 | 87 | 0.1724 |
| 2_13443706 | 28 | 0.0357 | 34 | 0.0294 | 31 | 0.0968 | 27 | 0.037 | 120 | 0.05 | 20 | 0.1 | 33 | 0.1515 | 24 | 0.0417 | 27 | 0.037 | 104 | 0.0865 |
| 2_13444017 | 9 | 0.2222 | 13 | 0.2308 | 10 | 0 | 17 | 0.1765 | 49 | 0.1633 | 7 | 0.2857 | 16 | 0.1875 | 11 | 0.0909 | 12 | 0.0833 | 46 | 0.1522 |
| 2_13444043 | 6 | 0 | 11 | 0.0909 | 9 | 0.2222 | 18 | 0.0556 | 44 | 0.0909 | 6 | 0 | 15 | 0.0667 | 12 | 0.25 | 10 | 0 | 43 | 0.093 |
| 2_13444065 | 6 | 0.5 | 10 | 0.5 | 8 | 0.625 | 18 | 0.6667 | 42 | 0.5952 | 5 | 0.6 | 13 | 0.5385 | 10 | 0.7 | 12 | 0.6667 | 40 | 0.625 |
| 2_13444069 | 5 | 0.4 | 7 | 0 | 9 | 0.2222 | 16 | 0.1875 | 37 | 0.1892 | 4 | 0.25 | 12 | 0.25 | 9 | 0.5556 | 9 | 0.4444 | 34 | 0.3824 |
| 2_13464638 | NA | NA | NA | NA | NA | NA | 1 | 1 | 1 | 1 | NA | NA | NA | NA | 1 | 1 | NA | NA | 1 | 1 |
| 2_13510247 | 1 | 0 | 3 | 0 | 4 | 0 | 4 | 0.25 | 12 | 0.0833 | 1 | 0 | 3 | 0 | 5 | 0 | 11 | 0.0909 | 20 | 0.05 |
| 2_13531407 | 17 | 0.4118 | 21 | 0.0476 | 13 | 0.1538 | 10 | 0.2 | 61 | 0.1967 | 6 | 0.8333 | 18 | 0.1111 | 16 | 0.0625 | 28 | 0.0714 | 68 | 0.1471 |
| 2_13531413 | 13 | 0 | 13 | 0.1538 | 9 | 0 | 10 | 0.1 | 45 | 0.0667 | 3 | 0.3333 | 15 | 0.0667 | 15 | 0.0667 | 22 | 0.1818 | 55 | 0.1273 |
| 2_13531416 | 16 | 0.0625 | 20 | 0.05 | 15 | 0.0667 | 12 | 0.0833 | 63 | 0.0635 | 6 | 0 | 19 | 0.1053 | 17 | 0 | 27 | 0.0741 | 69 | 0.058 |
| 2_13531422 | 18 | 0.1111 | 20 | 0.15 | 15 | 0.1333 | 12 | 0.0833 | 65 | 0.1231 | 7 | 0.1429 | 18 | 0.1111 | 18 | 0.2222 | 31 | 0.1613 | 74 | 0.1622 |
| 2_13531425 | 18 | 1 | 21 | 1 | 14 | 0.9286 | 12 | 1 | 65 | 0.9846 | 6 | 1 | 19 | 1 | 19 | 1 | 31 | 1 | 75 | 1 |
| 2_13531425 | 17 | 0.2353 | 21 | 0.3333 | 15 | 0.1333 | 8 | 0.125 | 61 | 0.2295 | 5 | 0.4 | 17 | 0.1765 | 17 | 0.1176 | 30 | 0.2667 | 69 | 0.2174 |
| 2_13531432 | 12 | 0.1667 | 18 | 0.2778 | 14 | 0.2143 | 10 | 0.3 | 54 | 0.2407 | 6 | 0 | 20 | 0.3 | 15 | 0.2667 | 29 | 0.2414 | 70 | 0.2429 |
| 2_13531438 | 15 | 0.3333 | 21 | 0.1905 | 14 | 0.2857 | 10 | 0.4 | 60 | 0.2833 | 8 | 0.25 | 20 | 0.35 | 19 | 0.2632 | 30 | 0.1333 | 77 | 0.2338 |
| 2_13531440 | 16 | 0.25 | 20 | 0.1 | 16 | 0.1875 | 13 | 0.0769 | 65 | 0.1538 | 9 | 0.1111 | 18 | 0.2222 | 16 | 0.125 | 23 | 0.087 | 66 | 0.1364 |
| 2_13531449 | 15 | 0.8667 | 21 | 0.9524 | 16 | 0.9375 | 11 | 1 | 63 | 0.9365 | 8 | 1 | 20 | 1 | 20 | 0.9 | 26 | 0.9231 | 74 | 0.9459 |
| 2_13531461 | 15 | 0.6 | 19 | 0.5789 | 11 | 0.6364 | 10 | 0.8 | 55 | 0.6364 | 8 | 0.625 | 19 | 0.6842 | 14 | 0.7143 | 26 | 0.4615 | 67 | 0.597 |
| 2_13531479 | 17 | 0.1765 | 22 | 0.4091 | 19 | 0.1579 | 13 | 0.3077 | 71 | 0.2676 | 9 | 0.1111 | 20 | 0.15 | 18 | 0.2778 | 31 | 0.2258 | 78 | 0.2051 |
| 2_13531622 | 11 | 0.6364 | 22 | 0.3636 | 15 | 0.5333 | 15 | 0.3333 | 63 | 0.4444 | 6 | 0.3333 | 10 | 0.5 | 9 | 0.7778 | 21 | 0.4286 | 46 | 0.5 |
| 2_13531629 | 10 | 0.7 | 22 | 0.5455 | 16 | 0.5 | 14 | 0.4286 | 62 | 0.5323 | 7 | 0.1429 | 10 | 0.5 | 10 | 0.6 | 26 | 0.5 | 53 | 0.4717 |
| 2_13531632 | 10 | 0.2 | 23 | 0 | 14 | 0 | 14 | 0.1429 | 61 | 0.0656 | 7 | 0.1429 | 10 | 0 | 11 | 0 | 21 | 0.1429 | 49 | 0.0816 |
| 2_13531634 | 9 | 0.1111 | 24 | 0.125 | 17 | 0.2941 | 12 | 0.1667 | 62 | 0.1774 | 4 | 0.25 | 8 | 0.125 | 8 | 0.375 | 21 | 0 | 41 | 0.122 |
| 2_13531637 | 11 | 0.8182 | 24 | 0.5417 | 14 | 0.6429 | 15 | 0.6 | 64 | 0.625 | 5 | 0.4 | 13 | 0.6154 | 10 | 0.6 | 24 | 0.5833 | 52 | 0.5769 |
| 2_13585507 | 1 | 0 | 2 | 0 | 3 | 0.3333 | 4 | 0 | 10 | 0.1 | 4 | 0.25 | 4 | 0 | 1 | 0 | 2 | 0 | 11 | 0.0909 |
| 2_13585509 | 1 | 1 | 1 | 1 | 3 | 0.6667 | 3 | 1 | 8 | 0.875 | 3 | 0.6667 | 4 | 1 | 1 | 1 | 2 | 1 | 10 | 0.9 |
| 2_13590002 | 1 | 0 | 1 | 0 | 2 | 0 | 5 | 0.2 | 9 | 0.1111 | 2 | 0 | 4 | 0.25 | 4 | 0 | 2 | 0 | 12 | 0.0833 |
| 2_13591105 | 13 | 0.2308 | 9 | 0.2222 | 13 | 0.0769 | 20 | 0.1 | 55 | 0.1455 | 10 | 0.4 | 10 | 0.2 | 16 | 0.125 | 14 | 0.2857 | 50 | 0.24 |
| 2_13591108 | 21 | 0.3333 | 11 | 0.3636 | 13 | 0.0769 | 23 | 0.3043 | 68 | 0.2794 | 9 | 0.3333 | 12 | 0.0833 | 17 | 0.1176 | 17 | 0.1176 | 55 | 0.1455 |
| 2_13591140 | 18 | 0.0556 | 9 | 0.2222 | 11 | 0.0909 | 21 | 0.0476 | 59 | 0.0847 | 7 | 0.1429 | 15 | 0.0667 | 21 | 0.0952 | 16 | 0.125 | 59 | 0.1017 |
| 2_13591168 | 21 | 0.2857 | 12 | 0.3333 | 14 | 0.5714 | 26 | 0.4615 | 73 | 0.411 | 11 | 0.3636 | 15 | 0.4 | 26 | 0.4615 | 17 | 0.4118 | 69 | 0.4203 |
| 2_13592320 | 3 | 0.6667 | 6 | 0.8333 | 1 | 0 | 2 | 0 | 12 | 0.5833 | 1 | 1 | 1 | 1 | 5 | 0.6 | 3 | 0.6667 | 10 | 0.7 |
| 2_13592330 | 4 | 0.25 | 5 | 0.2 | 1 | 1 | 1 | 1 | 11 | 0.3636 | 1 | 0 | 1 | 1 | 4 | 0.25 | 4 | 0.5 | 10 | 0.4 |
| 2_13592364 | 3 | 0.6667 | 5 | 0 | 1 | 0 | 3 | 0 | 12 | 0.1667 | 1 | 1 | 2 | 0 | 4 | 0 | 4 | 0 | 11 | 0.0909 |
| 2_13592406 | 3 | 0 | 5 | 0.4 | NA | NA | 2 | 0 | 10 | 0.2 | 2 | 1 | 1 | 0 | 3 | 0.3333 | 3 | 0 | 9 | 0.3333 |
| 2_13592416 | 3 | 0 | 5 | 0.2 | NA | NA | 2 | 0.5 | 10 | 0.2 | 2 | 1 | NA | NA | 2 | 0 | 2 | 0.5 | 6 | 0.5 |
| 2_13592441 | 3 | 0 | 3 | 0.3333 | NA | NA | 2 | 0.5 | 8 | 0.25 | 2 | 0 | 1 | 0 | 3 | 0 | 4 | 0.25 | 10 | 0.1 |
| 2_13592485 | 3 | 0 | 3 | 0.3333 | NA | NA | 2 | 0.5 | 8 | 0.25 | 1 | 1 | 1 | 0 | 4 | 0 | 1 | 0 | 7 | 0.1429 |
| 2_13592486 | 3 | 1 | 3 | 1 | NA | NA | 2 | 1 | 8 | 1 | 2 | 1 | 1 | 1 | 4 | 1 | 4 | 1 | 11 | 1 |
| 2_13592502 | 2 | 0 | 2 | 0 | NA | NA | 2 | 0.5 | 6 | 0.1667 | 2 | 0.5 | NA | NA | 2 | 0 | 4 | 0 | 8 | 0.125 |
| 2_13592505 | 2 | 0 | 3 | 0 | NA | NA | 1 | 1 | 6 | 0.1667 | 2 | 0.5 | 1 | 1 | 4 | 0.25 | 4 | 0.25 | 11 | 0.3636 |
| 2_13592508 | 3 | 0.3333 | 2 | 0 | NA | NA | 1 | 0 | 6 | 0.1667 | 2 | 0.5 | 1 | 0 | 4 | 0 | 4 | 0 | 11 | 0.0909 |
| 2_13592510 | 1 | 0 | 3 | 0.6667 | NA | NA | 2 | 0.5 | 6 | 0.5 | 2 | 0.5 | 1 | 0 | 4 | 1 | 4 | 0.75 | 11 | 0.7273 |
| 2_13592515 | 3 | 0.3333 | 3 | 0.3333 | NA | NA | 2 | 0.5 | 8 | 0.375 | 2 | 1 | NA | NA | 3 | 0.6667 | 2 | 0 | 7 | 0.5714 |
| 2_13592521 | 3 | 0.3333 | 3 | 0.6667 | NA | NA | 2 | 0.5 | 8 | 0.5 | 1 | 1 | 1 | 1 | 4 | 0.25 | 4 | 0.5 | 10 | 0.5 |
| 2_13592522 | 2 | 0.5 | 3 | 0.3333 | NA | NA | 2 | 0.5 | 7 | 0.4286 | 1 | 1 | 1 | 0 | 3 | 0.3333 | 4 | 0 | 9 | 0.2222 |
| 2_13594284 | 1 | 1 | 1 | 1 | NA | NA | 1 | 0 | 3 | 0.6667 | 1 | 1 | NA | NA | 1 | 1 | NA | NA | 2 | 1 |
| 2_13595815 | 2 | 0 | 2 | 1 | NA | NA | 2 | 0.5 | 6 | 0.5 | 1 | 1 | 1 | 0 | 4 | 0.5 | 2 | 0.5 | 8 | 0.5 |
| 2_13595829 | 2 | 0 | 2 | 0.5 | NA | NA | 2 | 0.5 | 6 | 0.3333 | 1 | 0 | 1 | 1 | 4 | 0 | 2 | 0 | 8 | 0.125 |
| 2_13595836 | 2 | 0 | 2 | 0.5 | NA | NA | 2 | 0 | 6 | 0.1667 | NA | NA | 1 | 1 | 4 | 0.25 | 2 | 0.5 | 7 | 0.4286 |
| 2_13595837 | 1 | 0 | 2 | 0.5 | NA | NA | 2 | 0.5 | 5 | 0.4 | 1 | 0 | 1 | 0 | 3 | 0 | 2 | 0.5 | 7 | 0.1429 |
| 2_13595840 | 2 | 1 | 2 | 1 | NA | NA | 1 | 1 | 5 | 1 | 1 | 1 | 1 | 1 | 4 | 0.75 | 2 | 1 | 8 | 0.875 |
| 2_13595844 | 2 | 0 | 2 | 0 | NA | NA | 2 | 0.5 | 6 | 0.1667 | 1 | 0 | 1 | 0 | 3 | 0.3333 | 2 | 0.5 | 7 | 0.2857 |
| 2_13595847 | 2 | 1 | 2 | 1 | NA | NA | 2 | 1 | 6 | 1 | 1 | 1 | 1 | 1 | 2 | 0.5 | 2 | 1 | 6 | 0.8333 |
| 2_13595866 | 2 | 1 | 2 | 1 | NA | NA | 1 | 1 | 5 | 1 | 1 | 1 | 1 | 1 | 2 | 1 | 2 | 0.5 | 6 | 0.8333 |
| 2_13661446 | 3 | 0 | 4 | 0.25 | 1 | 0 | 3 | 0 | 11 | 0.0909 | 1 | 0 | 4 | 0 | 2 | 0 | 5 | 0.2 | 12 | 0.0833 |
| 2_13662366 | 238 | 0.1008 | 399 | 0.1103 | 283 | 0.1025 | 251 | 0.0916 | 1171 | 0.1025 | 206 | 0.1165 | 210 | 0.0952 | 254 | 0.0866 | 250 | 0.124 | 920 | 0.1054 |

| | | | | | | | | | | | | | | | | | | | | |
|---|---|---|---|---|---|---|---|---|---|---|---|---|---|---|---|---|---|---|---|---|
| 2_1366237 | 242 | 0.124 | 401 | 0.1272 | 284 | 0.1373 | 248 | 0.121 | 1175 | 0.1277 | 209 | 0.134 | 211 | 0.1659 | 244 | 0.1393 | 245 | 0.151 | 909 | 0.1474 |
| 2_1366238 | 248 | 0.0806 | 410 | 0.0561 | 290 | 0.0897 | 255 | 0.0745 | 1203 | 0.0732 | 221 | 0.0814 | 219 | 0.0594 | 259 | 0.0618 | 240 | 0.0625 | 939 | 0.066 |
| 2_1367162 | 1 | 1 | 1 | 1 | 5 | 1 | 2 | 1 | 9 | 1 | 3 | 1 | 4 | 1 | 4 | 1 | 3 | 1 | 14 | 1 |
| 2_1367162 | 1 | 0 | 1 | 1 | 5 | 0 | 2 | 0 | 9 | 0.1111 | 3 | 0.3333 | 3 | 0.6667 | 4 | 0 | 3 | 0.6667 | 13 | 0.3846 |
| 2_1367164 | NA | NA | 1 | 0 | 3 | 0.3333 | 1 | 0 | 5 | 0.2 | 2 | 0 | 3 | 0.3333 | 3 | 0.6667 | 2 | 0.5 | 10 | 0.4 |
| 2_1367164 | 1 | 0 | 1 | 1 | 4 | 0.75 | 2 | 0.5 | 8 | 0.625 | 3 | 0.6667 | 3 | 0.6667 | 4 | 1 | 3 | 1 | 13 | 0.8462 |
| 2_1367165 | 1 | 0 | 1 | 0 | 4 | 0.25 | 2 | 0 | 8 | 0.125 | 3 | 0 | 4 | 0.25 | 4 | 0.5 | 3 | 0 | 14 | 0.2143 |
| 2_1367225 | 1 | 1 | 1 | 1 | 2 | 0 | 2 | 1 | 6 | 0.6667 | 2 | 0 | 3 | 1 | 4 | 0.5 | 3 | 0.6667 | 12 | 0.5833 |
| 2_1367228 | 3 | 0.6667 | 1 | 1 | 3 | 0.3333 | 3 | 0.6667 | 10 | 0.6 | 2 | 0.5 | 4 | 1 | 4 | 0.25 | 4 | 0.5 | 14 | 0.5714 |
| 2_1367228 | 3 | 0.3333 | 1 | 0 | 3 | 0.6667 | 2 | 0 | 9 | 0.3333 | 1 | 0 | 4 | 0.5 | 4 | 0.5 | 4 | 0.75 | 13 | 0.5385 |
| 2_1367231 | 3 | 0 | 1 | 0 | 2 | 0.5 | 3 | 0.6667 | 9 | 0.3333 | 2 | 0 | 4 | 0.5 | 4 | 0 | 5 | 0.2 | 15 | 0.2 |
| 2_1367231 | 6 | 0.3333 | 3 | 0 | 5 | 0 | 2 | 0.5 | 16 | 0.1875 | 7 | 0.1429 | 2 | 0.5 | 7 | 0 | 6 | 0.1667 | 22 | 0.1364 |
| 2_1367232 | 7 | 0.2857 | 6 | 0.3333 | 7 | 0.2857 | 5 | 0.2 | 25 | 0.28 | 6 | 0.1667 | 5 | 0.8 | 9 | 0.2222 | 6 | 0.6667 | 26 | 0.4231 |
| 2_1367233 | 12 | 0.9167 | 9 | 0.7778 | 11 | 0.7273 | 8 | 1 | 40 | 0.85 | 8 | 0.875 | 11 | 0.6364 | 16 | 0.875 | 10 | 0.8 | 45 | 0.8 |
| 2_1367234 | 12 | 0 | 9 | 0.1111 | 15 | 0.1333 | 7 | 0 | 43 | 0.0698 | 9 | 0.1111 | 12 | 0 | 15 | 0.0667 | 11 | 0.0909 | 47 | 0.0638 |
| 2_1367234 | 12 | 0.5833 | 7 | 0.5714 | 14 | 0.3571 | 8 | 0.25 | 41 | 0.439 | 7 | 0.4286 | 13 | 0.3077 | 12 | 0.4167 | 10 | 0.4 | 42 | 0.381 |
| 2_1369136 | 4 | 0.25 | NA | NA | 1 | 0 | 1 | 0 | 6 | 0.1667 | NA | NA | NA | NA | NA | NA | 1 | 1 | 1 | 1 |
| 2_1369235 | 1 | 0 | 1 | 1 | NA | NA | 2 | 0 | 4 | 0.25 | NA | NA | 2 | 0.5 | 1 | 1 | 1 | 0 | 4 | 0.5 |
| 2_1369249 | 1 | 1 | 1 | 1 | NA | NA | 1 | 1 | 3 | 1 | NA | NA | 1 | 1 | NA | NA | 1 | 1 | 2 | 1 |
| 2_1369251 | 1 | 0 | 1 | 1 | NA | NA | 1 | 1 | 3 | 0.6667 | NA | NA | 1 | 1 | NA | NA | 1 | 1 | 2 | 1 |
| 2_1369251 | 1 | 1 | 1 | 0 | NA | NA | NA | NA | 2 | 0.5 | NA | NA | 1 | 1 | NA | NA | 1 | 1 | 2 | 1 |
| 2_1381208 | 4 | 0.5 | 6 | 0.1667 | 10 | 0.2 | 6 | 0.1667 | 26 | 0.2308 | 3 | 0 | 6 | 0.6667 | 6 | 0.3333 | 7 | 0.5714 | 22 | 0.4545 |
| 2_1381208 | 3 | 0 | 7 | 0 | 9 | 0.3333 | 5 | 0.2 | 24 | 0.1667 | 3 | 0 | 2 | 0.5 | 5 | 0.4 | 5 | 0.2 | 15 | 0.2667 |
| 2_1381208 | 5 | 0.2 | 8 | 0.25 | 10 | 0.2 | 5 | 0 | 28 | 0.1786 | 3 | 0.6667 | 7 | 0.2857 | 7 | 0.2857 | 8 | 0.125 | 25 | 0.28 |
| 2_1381254 | 7 | 0 | 10 | 0.1 | 11 | 0.2727 | 8 | 0.125 | 36 | 0.1389 | 5 | 0.4 | 11 | 0.1818 | 10 | 0.2 | 9 | 0.1111 | 35 | 0.2 |
| 2_1381255 | 7 | 0.2857 | 13 | 0.6154 | 12 | 0.0833 | 9 | 0.3333 | 41 | 0.3415 | 7 | 0.5714 | 12 | 0.5833 | 13 | 0.3077 | 12 | 0.25 | 44 | 0.4091 |
| 2_1381256 | 8 | 0.125 | 12 | 0.1667 | 11 | 0.4545 | 9 | 0.3333 | 40 | 0.275 | 7 | 0.1429 | 11 | 0.2727 | 11 | 0.1818 | 11 | 0.2727 | 40 | 0.225 |
| 2_1381256 | 7 | 0.2857 | 15 | 0.4 | 10 | 0.4 | 9 | 0.3333 | 41 | 0.3659 | 6 | 0.6667 | 13 | 0.5385 | 13 | 0.6154 | 13 | 0.6923 | 45 | 0.6222 |
| 2_1381257 | 9 | 0.1111 | 12 | 0.25 | 11 | 0.2727 | 9 | 0.1111 | 41 | 0.1951 | 6 | 0.3333 | 11 | 0.3636 | 12 | 0.1667 | 12 | 0.1667 | 41 | 0.2439 |
| 2_1381258 | 8 | 0.625 | 13 | 0.6923 | 13 | 0.4615 | 8 | 0.875 | 42 | 0.6429 | 7 | 0.7143 | 9 | 0.8889 | 9 | 0.5556 | 12 | 0.6667 | 37 | 0.7027 |
| 2_1381258 | 8 | 1 | 15 | 0.9333 | 10 | 1 | 8 | 1 | 41 | 0.9756 | 7 | 1 | 12 | 0.9167 | 11 | 0.9091 | 12 | 0.9167 | 42 | 0.9286 |
| 2_1381258 | 10 | 0.3 | 15 | 0.2667 | 10 | 0.5 | 9 | 0.3333 | 44 | 0.3409 | 6 | 0.3333 | 12 | 0.5833 | 14 | 0.4286 | 13 | 0.5385 | 45 | 0.4889 |
| 2_1381258 | 8 | 1 | 13 | 0.6923 | 13 | 0.4615 | 7 | 0.2857 | 41 | 0.6098 | 7 | 0.7143 | 12 | 0.75 | 13 | 0.4615 | 11 | 0.6364 | 43 | 0.6279 |
| 2_1381261 | 6 | 1 | 9 | 1 | 6 | 1 | 5 | 1 | 26 | 1 | 5 | 1 | 5 | 1 | 8 | 1 | 6 | 1 | 24 | 1 |
| 2_1381261 | 11 | 0.7273 | 14 | 0.7143 | 14 | 1 | 8 | 1 | 47 | 0.8511 | 6 | 0.6667 | 12 | 0.9167 | 14 | 0.8571 | 13 | 0.9231 | 45 | 0.8667 |
| 2_1381262 | 9 | 0.5556 | 18 | 0.3333 | 14 | 0.5714 | 10 | 0.8 | 51 | 0.5294 | 6 | 0.3333 | 15 | 0.7333 | 13 | 0.6154 | 17 | 0.7059 | 51 | 0.6471 |
| 2_1381262 | 9 | 0.8889 | 13 | 0.9231 | 9 | 1 | 10 | 0.9 | 41 | 0.9268 | 6 | 0.6667 | 13 | 1 | 12 | 1 | 13 | 1 | 44 | 0.9545 |
| 2_1381263 | 10 | 0.9 | 17 | 0.8824 | 15 | 0.8667 | 11 | 0.9091 | 53 | 0.8868 | 5 | 1 | 13 | 0.9231 | 17 | 0.8235 | 16 | 0.875 | 51 | 0.8824 |
| 2_1381263 | 8 | 0.125 | 17 | 0.0588 | 12 | 0.1667 | 12 | 0.1667 | 49 | 0.1224 | 7 | 0.4286 | 15 | 0.3333 | 12 | 0.3333 | 17 | 0.1176 | 51 | 0.2745 |
| 2_1381263 | 9 | 0.8889 | 15 | 0.6667 | 13 | 0.9231 | 9 | 0.8889 | 46 | 0.8261 | 5 | 1 | 14 | 0.7143 | 14 | 0.6429 | 17 | 0.7647 | 50 | 0.74 |
| 2_1381264 | 7 | 0.7143 | 11 | 0.8182 | 4 | 0.75 | 7 | 0.7143 | 29 | 0.7586 | 3 | 1 | 8 | 1 | 12 | 1 | 7 | 1 | 30 | 1 |
| 2_1381265 | 10 | 0 | 17 | 0.2353 | 14 | 0.2857 | 8 | 0 | 49 | 0.1633 | 6 | 0.1667 | 12 | 0.3333 | 15 | 0.0667 | 14 | 0.2143 | 47 | 0.1915 |
| 2_1381275 | 11 | 0.9091 | 13 | 0.6154 | 11 | 0.4545 | 12 | 0.9167 | 47 | 0.7234 | 5 | 0.4 | 11 | 0.5455 | 17 | 0.6471 | 14 | 0.5 | 47 | 0.5532 |
| 2_1381276 | 15 | 0.0667 | 20 | 0.05 | 20 | 0.15 | 15 | 0.1333 | 70 | 0.1 | 8 | 0 | 12 | 0.0833 | 15 | 0.0667 | 14 | 0.0714 | 49 | 0.0612 |
| 2_1381281 | 13 | 0.0769 | 18 | 0.1667 | 16 | 0 | 13 | 0 | 60 | 0.0667 | 7 | 0 | 19 | 0.1053 | 18 | 0 | 16 | 0.25 | 60 | 0.1 |
| 2_1395156 | NA | NA | 2 | 1 | NA | NA | 1 | 0 | 3 | 0.6667 | NA | NA | 1 | 0 | 1 | 1 | 1 | 0 | 3 | 0.3333 |
| 2_1395159 | NA | NA | 2 | 0.5 | 1 | 0 | NA | NA | 3 | 0.3333 | NA | NA | 1 | 1 | 2 | 0 | 2 | 0 | 5 | 0.2 |
| 2_1395161 | NA | NA | 2 | 0.5 | 1 | 0 | 1 | 1 | 4 | 0.5 | NA | NA | 1 | 0 | 2 | 0.5 | 2 | 0.5 | 5 | 0.4 |
| 2_1395163 | NA | NA | 2 | 1 | 1 | 1 | 1 | 0 | 4 | 0.75 | NA | NA | 1 | 1 | 2 | 0 | 2 | 0 | 5 | 0.2 |
| 2_1395178 | NA | NA | 1 | 1 | NA | NA | NA | NA | 1 | 1 | NA | NA | NA | NA | 2 | 0.5 | 1 | 0 | 3 | 0.3333 |
| 2_1395182 | NA | NA | 3 | 0.3333 | 1 | 0 | NA | NA | 4 | 0.25 | 3 | 0 | 1 | 1 | 6 | 0 | 1 | 0 | 11 | 0.0909 |
| 2_1395182 | 1 | 0 | 2 | 0 | 2 | 1 | NA | NA | 5 | 0.4 | 3 | 0.6667 | 3 | 0.6667 | 5 | 0.6 | 1 | 1 | 12 | 0.6667 |
| 2_1395190 | 3 | 1 | 2 | 1 | 3 | 0.6667 | 1 | 1 | 9 | 0.8889 | NA | NA | 5 | 0.8 | 6 | 0.6667 | 3 | 1 | 14 | 0.7857 |
| 2_1395191 | 2 | 1 | 1 | 1 | 3 | 0.3333 | NA | NA | 6 | 0.6667 | 1 | 1 | 4 | 1 | 4 | 1 | 1 | 1 | 10 | 1 |
| 2_1395193 | 1 | 1 | NA | NA | 2 | 1 | NA | NA | 3 | 1 | NA | NA | 3 | 0.6667 | 3 | 1 | 2 | 1 | 8 | 0.875 |
| 2_1395193 | 1 | 1 | NA | NA | 2 | 1 | 1 | 0 | 4 | 0.75 | NA | NA | 1 | 0 | 2 | 1 | 1 | 1 | 4 | 0.75 |
| 2_1395195 | 1 | 0 | NA | NA | 2 | 0.5 | 1 | 0 | 4 | 0.25 | 1 | 0 | 3 | 0.3333 | 5 | 0 | 3 | 0 | 12 | 0.0833 |
| 2_1395198 | 1 | 1 | NA | NA | 3 | 1 | NA | NA | 4 | 1 | NA | NA | 2 | 1 | 2 | 0.5 | 1 | 1 | 5 | 0.8 |
| 2_1395199 | 1 | 1 | NA | NA | 3 | 0.6667 | 1 | 1 | 5 | 0.8 | NA | NA | 1 | 0 | 5 | 0.6 | 2 | 1 | 8 | 0.625 |
| 2_1395202 | 1 | 1 | NA | NA | 1 | 0 | NA | NA | 2 | 0.5 | 1 | 1 | 1 | 1 | 1 | 1 | 1 | 1 | 4 | 1 |
| 2_1399625 | NA | NA | 2 | 0 | 5 | 0.4 | 3 | 0.3333 | 10 | 0.3 | 3 | 0.3333 | 4 | 0.25 | 2 | 0.5 | 3 | 0.3333 | 12 | 0.3333 |
| 2_1399625 | NA | NA | 2 | 0 | 4 | 0.5 | 2 | 0.5 | 8 | 0.375 | 3 | 0 | 2 | 0.5 | 2 | 0 | 1 | 0 | 8 | 0.125 |
| 2_1399626 | NA | NA | 2 | 0 | 5 | 0.2 | 3 | 0 | 10 | 0.1 | 3 | 0.6667 | 5 | 0.2 | 3 | 0 | 3 | 0 | 14 | 0.2143 |
| 2_1400097 | NA | NA | NA | NA | 2 | 0 | 3 | 0.3333 | 5 | 0.2 | 1 | 0 | 2 | 0.5 | 1 | 0 | 1 | 1 | 5 | 0.4 |
| 2_1400099 | NA | NA | 1 | 0 | 2 | 0.5 | 1 | 1 | 4 | 0.5 | 1 | 0 | 2 | 0 | 1 | 1 | 1 | 1 | 5 | 0.4 |

| | | | | | | | | | | | | | | | | | | | | |
|---|---|---|---|---|---|---|---|---|---|---|---|---|---|---|---|---|---|---|---|---|
| 2_1400101( | NA | NA | 1 | 0 | 2 | 0 | 3 | 0.3333 | 6 | 0.1667 | 1 | 1 | 2 | 1 | 1 | 1 | 1 | 1 | 5 | 1 |
| 2_1400101ε | NA | NA | 1 | 1 | 2 | 1 | 3 | 0.6667 | 6 | 0.8333 | 1 | 0 | 1 | 1 | 1 | 1 | 1 | 1 | 4 | 0.75 |
| 2_1400103 | NA | NA | 1 | 1 | 1 | 0 | 3 | 0.6667 | 5 | 0.6 | 1 | 0 | 2 | 0 | 1 | 1 | 1 | 0 | 5 | 0.2 |
| 2_1402618 | 2 | 1 | 2 | 0 | 2 | 0 | 3 | 0.6667 | 9 | 0.4444 | 1 | 0 | 3 | 0.3333 | 4 | 0.25 | 2 | 0 | 10 | 0.2 |
| 2_1402619 | 2 | 1 | 1 | 0 | 2 | 0 | 2 | 0.5 | 7 | 0.4286 | 1 | 0 | 2 | 0.5 | 5 | 0.6 | 2 | 0 | 10 | 0.4 |
| 2_1402620 | 3 | 1 | 2 | 0.5 | 2 | 0.5 | 3 | 0.6667 | 10 | 0.7 | 1 | 0 | 3 | 0.6667 | 5 | 0.6 | 2 | 0.5 | 11 | 0.5455 |
| 2_1402620 | 3 | 1 | 1 | 1 | 2 | 1 | 3 | 1 | 9 | 1 | 1 | 0 | 1 | 1 | 5 | 0.8 | 1 | 0 | 8 | 0.625 |
| 2_1402621 | 3 | 0.6667 | 2 | 0.5 | 1 | 0 | 4 | 0.75 | 10 | 0.6 | 1 | 1 | 3 | 0.6667 | 3 | 0.6667 | 2 | 1 | 9 | 0.7778 |
| 2_1402621 | 3 | 1 | 2 | 0 | 2 | 0 | 3 | 0.3333 | 10 | 0.4 | 1 | 0 | 3 | 0 | 4 | 0.25 | 2 | 0.5 | 10 | 0.2 |
| 2_1402622 | 3 | 0 | 2 | 0.5 | 3 | 0 | 4 | 0.25 | 12 | 0.1667 | 1 | 0 | 2 | 0 | 5 | 0.2 | 2 | 0 | 10 | 0.1 |
| 2_1405002 | 5 | 0.6 | 4 | 0.75 | 5 | 0.6 | 7 | 0.4286 | 21 | 0.5714 | 2 | 0 | 5 | 0.4 | 6 | 0.3333 | 3 | 0.3333 | 16 | 0.3125 |
| 2_1405010 | 4 | 0.25 | 5 | 0.2 | 5 | 0 | 8 | 0.125 | 22 | 0.1364 | 2 | 0 | 7 | 0 | 7 | 0.2857 | 5 | 0 | 21 | 0.0952 |
| 2_1405013 | 3 | 1 | 5 | 0.8 | 5 | 0.6 | 9 | 0.5556 | 22 | 0.6818 | 2 | 1 | 8 | 0.75 | 7 | 0.7143 | 5 | 0.8 | 22 | 0.7727 |
| 2_1405013 | 4 | 0.25 | 7 | 0.4286 | 5 | 0 | 10 | 0.2 | 26 | 0.2308 | 2 | 0 | 8 | 0.125 | 7 | 0.1429 | 4 | 0.5 | 21 | 0.1905 |
| 2_1405013 | 4 | 0 | 6 | 0 | 5 | 0 | 10 | 0.2 | 25 | 0.08 | 2 | 1 | 8 | 0 | 7 | 0 | 3 | 0 | 20 | 0.1 |
| 2_1405015 | 3 | 0.6667 | 5 | 1 | 4 | 1 | 9 | 1 | 21 | 0.9524 | 2 | 1 | 8 | 0.875 | 7 | 1 | 2 | 1 | 19 | 0.9474 |
| 2_1405015 | 4 | 1 | 6 | 1 | 4 | 1 | 8 | 1 | 22 | 1 | 2 | 1 | 9 | 1 | 6 | 1 | 2 | 1 | 19 | 1 |
| 2_1405016 | 4 | 1 | 6 | 0.8333 | 4 | 1 | 7 | 0.8571 | 21 | 0.9048 | 1 | 1 | 7 | 0.8571 | 6 | 1 | 3 | 1 | 17 | 0.9412 |
| 2_1405016 | 4 | 1 | 3 | 1 | 3 | 1 | 5 | 1 | 15 | 1 | 1 | 1 | 4 | 1 | 3 | 1 | 2 | 1 | 10 | 1 |
| 2_1405016 | 2 | 0 | 5 | 0.6 | 2 | 0.5 | 5 | 0.2 | 14 | 0.3571 | 1 | 0 | 9 | 0.3333 | 4 | 0.5 | 1 | 0 | 15 | 0.3333 |
| 2_1405016 | 4 | 0 | 6 | 0.3333 | 4 | 0.25 | 8 | 0.5 | 22 | 0.3182 | 2 | 0.5 | 9 | 0.1111 | 5 | 0 | 3 | 0 | 19 | 0.1053 |
| 2_1405017 | 4 | 1 | 6 | 0.6667 | 3 | 1 | 8 | 1 | 21 | 0.9048 | 2 | 0.5 | 8 | 1 | 5 | 0.8 | 1 | 1 | 16 | 0.875 |
| 2_1405017 | 4 | 0.25 | 6 | 0 | 4 | 0 | 7 | 0.1429 | 21 | 0.0952 | NA | NA | 8 | 0.125 | 5 | 0 | 3 | 0.3333 | 16 | 0.125 |
| 2_1405018 | 2 | 1 | 4 | 0.75 | 5 | 0.8 | 5 | 0.6 | 16 | 0.75 | 1 | 0 | 9 | 0.6667 | 1 | 0 | 1 | 1 | 12 | 0.5833 |
| 2_1405018 | 4 | 0.5 | 6 | 0 | 5 | 0.2 | 8 | 0.25 | 23 | 0.2174 | NA | NA | 8 | 0.25 | 5 | 0.4 | 2 | 0.5 | 15 | 0.3333 |
| 2_1405019 | 3 | 0 | 6 | 0.5 | 5 | 0 | 5 | 0 | 19 | 0.1579 | 1 | 0 | 9 | 0.5556 | 4 | 0.25 | 3 | 0.6667 | 17 | 0.4706 |
| 2_1405019 | 4 | 1 | 5 | 1 | 2 | 1 | 7 | 1 | 18 | 1 | 2 | 1 | 5 | 1 | 4 | 1 | 1 | 1 | 12 | 1 |
| 2_1405020 | 4 | 1 | 6 | 0.6667 | 2 | 1 | 5 | 1 | 17 | 0.8824 | NA | NA | 6 | 0.8333 | 4 | 1 | 3 | 0.6667 | 13 | 0.8462 |
| 2_1405021 | 4 | 0 | 5 | 0.4 | 4 | 0.25 | 7 | 0.2857 | 20 | 0.25 | 2 | 0.5 | 9 | 0.4444 | 5 | 0.4 | 3 | 1 | 19 | 0.5263 |
| 2_1405021 | 4 | 0.25 | 5 | 0 | 4 | 0.25 | 7 | 0 | 20 | 0.1 | 2 | 0 | 8 | 0.125 | 4 | 0.25 | 3 | 0 | 17 | 0.1176 |
| 2_1405021 | 4 | 0.25 | 5 | 0.2 | 4 | 0.25 | 7 | 0 | 20 | 0.15 | 2 | 0 | 8 | 0.25 | 5 | 0.2 | 2 | 0 | 17 | 0.1765 |
| 2_1405022 | 4 | 0 | 4 | 0 | 4 | 0 | 7 | 0.1429 | 19 | 0.0526 | 2 | 0 | 9 | 0.3333 | 4 | 0 | 3 | 0.3333 | 18 | 0.2222 |
| 2_1405022 | 4 | 0.25 | 5 | 0.2 | 2 | 0.5 | 6 | 0.3333 | 17 | 0.2941 | 1 | 0 | 8 | 0.25 | 4 | 0.25 | 3 | 0 | 16 | 0.1875 |
| 2_1405023 | 4 | 0.5 | 4 | 0.5 | 2 | 0.5 | 7 | 0.5714 | 17 | 0.5294 | 2 | 0.5 | 8 | 0.625 | 3 | 0.6667 | 3 | 0 | 16 | 0.5 |
| 2_1405023 | 4 | 0.25 | 5 | 0.6 | 4 | 0.25 | 7 | 0.2857 | 20 | 0.35 | 2 | 0 | 8 | 0.75 | 4 | 0.25 | 3 | 1 | 17 | 0.5882 |
| 2_1405023 | 4 | 0 | 5 | 0 | 4 | 0 | 6 | 0.1667 | 19 | 0.0526 | 2 | 0.5 | 8 | 0 | 4 | 0 | 3 | 0 | 17 | 0.0588 |
| 2_1405024 | 3 | 0 | 5 | 0.2 | 3 | 0.3333 | 7 | 0 | 18 | 0.1111 | 2 | 0 | 7 | 0.1429 | 4 | 0.25 | 3 | 0.3333 | 16 | 0.1875 |
| 2_1405024 | 3 | 0.3333 | 3 | 1 | 2 | 1 | 7 | 0.8571 | 15 | 0.8 | 1 | 0 | 6 | 1 | 4 | 1 | 3 | 1 | 14 | 0.9286 |
| 2_1405025 | 3 | 0 | 3 | 0.6667 | NA | NA | 5 | 0.6 | 11 | 0.4545 | NA | NA | 5 | 1 | 2 | 1 | 1 | 1 | 8 | 1 |
| 2_1405025 | 2 | 1 | 5 | 1 | 2 | 1 | 7 | 1 | 16 | 1 | 1 | 1 | 7 | 1 | 3 | 1 | 2 | 1 | 13 | 1 |
| 2_1405025 | 3 | 0.6667 | 5 | 0.4 | 2 | 0 | 7 | 0.2857 | 17 | 0.3529 | 1 | 0 | 7 | 0.5714 | 4 | 0 | 3 | 0.3333 | 15 | 0.3333 |
| 2_1405026 | 3 | 0 | 4 | 0.5 | 2 | 0.5 | 7 | 0.2857 | 16 | 0.3125 | 1 | 0 | 6 | 0.3333 | 3 | 0 | 3 | 1 | 13 | 0.3846 |
| 2_1405027 | 3 | 1 | 4 | 1 | 2 | 1 | 6 | 1 | 15 | 1 | 1 | 1 | 6 | 1 | 3 | 1 | 3 | 1 | 13 | 1 |
| 2_1405028 | 4 | 0 | 5 | 0.2 | 2 | 0 | 6 | 0.1667 | 17 | 0.1176 | 1 | 0 | 7 | 0 | 3 | 0 | 3 | 0.3333 | 14 | 0.0714 |
| 2_1405029 | 3 | 0.3333 | 3 | 0 | 1 | 0 | 2 | 0 | 9 | 0.1111 | NA | NA | 7 | 0.2857 | 2 | 0.5 | 3 | 0 | 12 | 0.25 |
| 2_1405030 | 4 | 0 | 5 | 0.2 | 2 | 0 | 6 | 0.1667 | 17 | 0.1176 | 1 | 1 | 7 | 0.2857 | 3 | 0.3333 | 3 | 0.6667 | 14 | 0.4286 |
| 2_1405030 | 4 | 0 | 4 | 0 | 2 | 0.5 | 6 | 0 | 16 | 0.0625 | 1 | 0 | 7 | 0.1429 | 3 | 0 | 2 | 0 | 13 | 0.0769 |
| 2_1405031 | 3 | 1 | 4 | 1 | 2 | 1 | 6 | 0.8333 | 15 | 0.9333 | NA | NA | 6 | 1 | 3 | 1 | 3 | 1 | 12 | 1 |
| 2_1405031 | 3 | 0 | 3 | 0 | 2 | 0.5 | 5 | 0.2 | 13 | 0.1538 | 1 | 0 | 5 | 0.2 | 3 | 0.3333 | 3 | 0.3333 | 12 | 0.25 |
| 2_1405031 | 4 | 0.25 | 4 | 0.25 | 2 | 0 | 6 | 0 | 16 | 0.125 | 1 | 0 | 7 | 0.1429 | 3 | 0 | 2 | 0 | 13 | 0.0769 |
| 2_1405032 | 4 | 1 | 4 | 1 | 2 | 1 | 5 | 1 | 15 | 1 | 1 | 1 | 7 | 1 | 3 | 1 | 3 | 1 | 14 | 1 |
| 2_1405033 | 4 | 1 | 3 | 0.6667 | 2 | 0 | 6 | 0.6667 | 15 | 0.6667 | 1 | 0 | 6 | 0.8333 | 3 | 1 | 3 | 0.6667 | 13 | 0.7692 |
| 2_1405034 | 3 | 0 | 3 | 0 | 2 | 0.5 | 6 | 0 | 14 | 0.0714 | 1 | 0 | 7 | 0.2857 | 3 | 0 | 3 | 0 | 14 | 0.1429 |
| 2_1405035 | 4 | 1 | 3 | 0.3333 | NA | NA | 6 | 0.5 | 13 | 0.6154 | 1 | 1 | 7 | 0.5714 | 3 | 0.6667 | 3 | 1 | 14 | 0.7143 |
| 2_1405052 | 4 | 0 | 2 | 0 | 1 | 1 | 5 | 0.4 | 12 | 0.25 | 1 | 0 | 6 | 0.3333 | 1 | 0 | 2 | 0 | 10 | 0.2 |
| 2_1422340 | 4 | 0.25 | 1 | 0 | 2 | 0 | 5 | 0 | 12 | 0.0833 | 2 | 0 | 2 | 0 | 6 | 0.1667 | 6 | 0 | 16 | 0.0625 |
| 2_1422342 | 4 | 0.25 | 2 | 0 | 2 | 0 | 6 | 0 | 14 | 0.0714 | 2 | 0 | 3 | 0 | 6 | 0 | 6 | 0.1667 | 17 | 0.0588 |
| 2_1424613 | NA | NA | NA | NA | 3 | 1 | 1 | 1 | 4 | 1 | NA | NA | 1 | 1 | NA | NA | NA | NA | 1 | 1 |
| 2_1424613 | NA | NA | NA | NA | 3 | 1 | 1 | 1 | 4 | 1 | NA | NA | 1 | 1 | NA | NA | NA | NA | 1 | 1 |
| 2_1424613 | NA | NA | NA | NA | 3 | 1 | 1 | 1 | 4 | 1 | NA | NA | 1 | 1 | NA | NA | NA | NA | 1 | 1 |
| 2_1425540 | NA | NA | NA | NA | 1 | 1 | NA | NA | 1 | 1 | NA | NA | 1 | 1 | NA | NA | NA | NA | 1 | 1 |
| 2_1425543 | NA | NA | NA | NA | 1 | 1 | NA | NA | 1 | 1 | NA | NA | 1 | 1 | NA | NA | NA | NA | 1 | 1 |
| 2_1429134 | 9 | 1 | 10 | 0.7 | 8 | 0.75 | 9 | 0.7778 | 36 | 0.8056 | 15 | 0.8 | 6 | 1 | 7 | 1 | 6 | 0.6667 | 34 | 0.8529 |
| 2_1436351 | 28 | 0.0357 | 17 | 0.1176 | 22 | 0.0455 | 31 | 0.2581 | 98 | 0.1224 | 9 | 0.1111 | 37 | 0.2703 | 27 | 0.1111 | 34 | 0.2059 | 107 | 0.1963 |
| 2_1436430 | 34 | 0.2941 | 20 | 0.2 | 28 | 0.25 | 35 | 0.2857 | 117 | 0.265 | 10 | 0.2 | 43 | 0.2791 | 29 | 0.2069 | 33 | 0.1818 | 115 | 0.2261 |

| | | | | | | | | | | | | | | | | | | | | |
|---|---|---|---|---|---|---|---|---|---|---|---|---|---|---|---|---|---|---|---|---|
| 2_14365115 | NA | NA | NA | NA | NA | NA | 1 | 1 | 1 | 1 | NA | NA | 1 | 1 | NA | NA | NA | NA | 1 | 1 |
| 2_14365136 | NA | NA | NA | NA | NA | NA | 1 | 1 | 1 | 1 | NA | NA | 1 | 1 | NA | NA | NA | NA | 1 | 1 |
| 2_14365137 | NA | NA | NA | NA | NA | NA | 1 | 1 | 1 | 1 | NA | NA | 1 | 1 | NA | NA | NA | NA | 1 | 1 |
| 2_14365551 | 32 | 0.7812 | 17 | 0.5882 | 28 | 0.6429 | 36 | 0.6944 | 113 | 0.6903 | 10 | 0.6 | 38 | 0.6316 | 26 | 0.7308 | 32 | 0.875 | 106 | 0.7264 |
| 2_14365551 | 39 | 0.8462 | 22 | 0.8182 | 27 | 0.8148 | 38 | 0.8421 | 126 | 0.8333 | 13 | 0.5385 | 45 | 0.8222 | 27 | 0.8889 | 37 | 0.8108 | 122 | 0.8033 |
| 2_14365557 | 40 | 0.975 | 22 | 0.9545 | 31 | 0.9677 | 39 | 0.9744 | 132 | 0.9697 | 15 | 0.9333 | 46 | 0.913 | 28 | 0.9643 | 41 | 0.9756 | 130 | 0.9462 |
| 2_14365572 | 36 | 0.4444 | 19 | 0.4211 | 31 | 0.3548 | 33 | 0.3939 | 119 | 0.4034 | 11 | 0.1818 | 38 | 0.3158 | 26 | 0.3462 | 36 | 0.4167 | 111 | 0.3423 |
| 2_14365596 | 12 | 0.4167 | 4 | 0 | 9 | 0.1111 | 17 | 0.1176 | 42 | 0.1905 | 4 | 0.25 | 16 | 0.5 | 11 | 0.3636 | 22 | 0.4545 | 53 | 0.434 |
| 2_14365610 | 13 | 0.2308 | 5 | 0 | 12 | 0.1667 | 17 | 0.2353 | 47 | 0.1915 | 3 | 0.3333 | 14 | 0.2143 | 8 | 0.125 | 21 | 0.3333 | 46 | 0.2609 |
| 2_14365614 | 12 | 0.25 | 6 | 0 | 11 | 0.0909 | 21 | 0.1429 | 50 | 0.14 | 5 | 0 | 20 | 0.05 | 14 | 0.2143 | 22 | 0.0455 | 61 | 0.082 |
| 2_14365627 | 14 | 0.0714 | 7 | 0 | 9 | 0.2222 | 15 | 0 | 45 | 0.0667 | 4 | 0 | 19 | 0.0526 | 10 | 0 | 20 | 0.1 | 53 | 0.0566 |
| 2_14365694 | 25 | 0.04 | 9 | 0.1111 | 13 | 0.0769 | 24 | 0.125 | 71 | 0.0845 | 9 | 0 | 25 | 0.12 | 15 | 0 | 23 | 0.0435 | 72 | 0.0556 |
| 2_14365750 | 17 | 0.3529 | 6 | 0.1667 | 11 | 0.2727 | 18 | 0.2222 | 52 | 0.2692 | 10 | 0.4 | 18 | 0.2222 | 10 | 0.3 | 19 | 0.4211 | 57 | 0.3333 |
| 2_14365806 | 26 | 0.1923 | 8 | 0.125 | 16 | 0.125 | 21 | 0.1429 | 71 | 0.1549 | 8 | 0.125 | 24 | 0.2083 | 16 | 0.0625 | 20 | 0.2 | 68 | 0.1618 |
| 2_14375733 | 2 | 0 | 1 | 1 | NA | NA | NA | NA | 3 | 0.3333 | 1 | 0 | NA | NA | 1 | 1 | NA | NA | 2 | 0.5 |
| 2_14378419 | 26 | 0.1923 | 29 | 0.0345 | 11 | 0 | 21 | 0.0476 | 87 | 0.0805 | 17 | 0.0588 | 19 | 0.0526 | 15 | 0.1333 | 21 | 0.0476 | 72 | 0.0694 |
| 2_14378511 | 33 | 0.0606 | 38 | 0.1316 | 20 | 0.1 | 24 | 0.0417 | 115 | 0.087 | 25 | 0.2 | 24 | 0.125 | 19 | 0.1579 | 26 | 0.1154 | 94 | 0.1489 |
| 2_14378935 | 7 | 0 | 2 | 0.5 | 1 | 0 | 1 | 0 | 11 | 0.0909 | NA | NA | 4 | 0 | NA | NA | 2 | 0.5 | 6 | 0.1667 |
| 2_14378937 | 8 | 0.25 | 3 | 0.3333 | 1 | 0 | 1 | 1 | 13 | 0.3077 | NA | NA | 3 | 0.6667 | 1 | 1 | 2 | 0.5 | 6 | 0.6667 |
| 2_14378953 | 6 | 0.5 | 3 | 1 | NA | NA | 1 | 1 | 10 | 0.7 | NA | NA | 3 | 0.6667 | NA | NA | 2 | 1 | 5 | 0.8 |
| 2_14378956 | 8 | 0.25 | 2 | 0 | NA | NA | 1 | 0 | 11 | 0.1818 | NA | NA | 2 | 0 | NA | NA | 2 | 1 | 4 | 0.5 |
| 2_14378963 | 4 | 0 | 2 | 0.5 | NA | NA | 1 | 0 | 7 | 0.1429 | NA | NA | 2 | 0 | NA | NA | 2 | 0.5 | 4 | 0.25 |
| 2_14378965 | 8 | 0.375 | 2 | 0 | NA | NA | 1 | 0 | 11 | 0.2727 | NA | NA | 4 | 0.25 | NA | NA | 2 | 0.5 | 6 | 0.3333 |
| 2_14409217 | 4 | 0 | 7 | 0.2857 | 8 | 0 | 4 | 0 | 23 | 0.087 | 4 | 0.25 | 5 | 0.2 | 5 | 0 | 5 | 0 | 19 | 0.1053 |
| 2_14409226 | 4 | 0 | 7 | 0.1429 | 8 | 0.125 | 3 | 0 | 22 | 0.0909 | 4 | 0 | 4 | 0 | 4 | 0 | 6 | 0.1667 | 18 | 0.0556 |
| 2_14409271 | 14 | 0.2857 | 10 | 0.1 | 11 | 0.1818 | 7 | 0.1429 | 42 | 0.1905 | 8 | 0 | 10 | 0.2 | 4 | 0 | 12 | 0.25 | 34 | 0.1471 |
| 2_14409285 | 16 | 0.125 | 10 | 0.1 | 11 | 0 | 7 | 0 | 44 | 0.0682 | 8 | 0.125 | 9 | 0.1111 | 5 | 0 | 13 | 0.0769 | 35 | 0.0857 |
| 2_14409292 | 8 | 0.375 | 8 | 0.25 | 11 | 0.0909 | 3 | 0 | 30 | 0.2 | 6 | 0.3333 | 5 | 0 | 3 | 0 | 9 | 0 | 23 | 0.087 |
| 2_14411401 | 2 | 0 | 2 | 0.5 | NA | NA | 1 | 0 | 5 | 0.2 | 2 | 0 | 2 | 0.5 | 1 | 1 | 1 | 1 | 6 | 0.5 |
| 2_14411413 | 2 | 0.5 | 3 | 0.3333 | NA | NA | 1 | 1 | 6 | 0.5 | 2 | 1 | 2 | 0 | 1 | 1 | 1 | 1 | 6 | 0.6667 |
| 2_14412153 | 3 | 0 | 3 | 0.3333 | NA | NA | 1 | 0 | 7 | 0.1429 | 1 | 0 | 2 | 0 | 1 | 0 | 1 | 1 | 5 | 0.2 |
| 2_14412165 | 3 | 0.6667 | 2 | 1 | NA | NA | 1 | 1 | 6 | 0.8333 | 1 | 1 | 2 | 0.5 | NA | NA | 1 | 1 | 4 | 0.75 |
| 2_14412174 | 3 | 0.3333 | 3 | 0 | NA | NA | 1 | 1 | 7 | 0.2857 | 2 | 0 | 2 | 0 | 1 | 1 | 1 | 0 | 6 | 0.1667 |
| 2_14412178 | 2 | 1 | 3 | 1 | NA | NA | NA | NA | 5 | 1 | 2 | 1 | 1 | 1 | 1 | 1 | 1 | 1 | 5 | 1 |
| 2_14412181 | 2 | 0.5 | 3 | 1 | NA | NA | 1 | 0 | 6 | 0.6667 | 2 | 0 | 2 | 0.5 | 1 | 1 | 1 | 1 | 6 | 0.5 |
| 2_14412204 | 3 | 0.3333 | 1 | 0 | NA | NA | 1 | 0 | 5 | 0.2 | 2 | 0 | 1 | 0 | 1 | 0 | 2 | 1 | 6 | 0.3333 |
| 2_14412221 | 2 | 1 | 3 | 1 | NA | NA | 2 | 1 | 7 | 1 | 2 | 1 | 2 | 1 | 1 | 1 | 2 | 1 | 7 | 1 |
| 2_14412224 | 3 | 0.6667 | 2 | 0 | 1 | 0 | 2 | 0.5 | 8 | 0.375 | 2 | 0.5 | 2 | 0 | 1 | 0 | 2 | 0.5 | 7 | 0.2857 |
| 2_14412225 | 3 | 0.3333 | 3 | 0.3333 | 1 | 0 | 2 | 0 | 9 | 0.2222 | 2 | 0.5 | 2 | 0 | 1 | 0 | 2 | 0 | 7 | 0.1429 |
| 2_14421529 | NA | NA | 3 | 0.3333 | NA | NA | NA | NA | 3 | 0.3333 | NA | NA | NA | NA | 1 | 1 | NA | NA | 1 | 1 |
| 2_14421538 | 2 | 1 | 3 | 0.6667 | NA | NA | NA | NA | 5 | 0.8 | NA | NA | NA | NA | 1 | 1 | NA | NA | 1 | 1 |
| 2_14421562 | 2 | 0 | 3 | 1 | NA | NA | NA | NA | 5 | 0.6 | NA | NA | 1 | 0 | 1 | 1 | NA | NA | 2 | 0.5 |
| 2_14423526 | 2 | 1 | 2 | 0.5 | 1 | 0 | NA | NA | 5 | 0.6 | NA | NA | 2 | 0.5 | 1 | 0 | 1 | 0 | 4 | 0.25 |
| 2_14423598 | 1 | 1 | 2 | 1 | 1 | 1 | 1 | 0 | 5 | 0.8 | 1 | 1 | 1 | 1 | 1 | 0 | 2 | 0.5 | 5 | 0.6 |
| 2_14423610 | 2 | 0.5 | 1 | 0 | 2 | 0.5 | 1 | 1 | 6 | 0.5 | 1 | 0 | 2 | 0.5 | 2 | 0 | 3 | 0 | 8 | 0.125 |
| 2_14423615 | 2 | 0.5 | 2 | 0.5 | 1 | 1 | 1 | 0 | 6 | 0.5 | 1 | 1 | 1 | 1 | 1 | 0 | 3 | 0.6667 | 6 | 0.6667 |
| 2_14423627 | 2 | 0.5 | 1 | 1 | 2 | 1 | NA | NA | 5 | 0.8 | 1 | 0 | 2 | 0 | 2 | 0.5 | 3 | 0.6667 | 8 | 0.375 |
| 2_14424893 | 5 | 1 | 4 | 1 | 3 | 1 | 1 | 1 | 13 | 1 | 1 | 1 | 2 | 1 | 3 | 1 | 3 | 1 | 9 | 1 |
| 2_14424894 | 5 | 0.2 | 4 | 0 | 3 | 0 | 2 | 0 | 14 | 0.0714 | 2 | 0 | 3 | 0 | 3 | 0.3333 | 3 | 0 | 11 | 0.0909 |
| 2_14424895 | 5 | 0.8 | 4 | 1 | 3 | 0.6667 | 2 | 1 | 14 | 0.8571 | 2 | 1 | 3 | 1 | 3 | 1 | 4 | 1 | 12 | 1 |
| 2_14424916 | 5 | 0 | 5 | 0.2 | 3 | 0 | 3 | 0 | 16 | 0.0625 | 2 | 0 | 2 | 0.5 | 2 | 0 | 3 | 0.3333 | 9 | 0.2222 |
| 2_14424917 | 5 | 1 | 3 | 1 | 3 | 1 | 1 | 1 | 12 | 1 | 2 | 1 | 2 | 1 | 3 | 1 | 4 | 1 | 11 | 1 |
| 2_14424921 | 5 | 1 | 3 | 1 | 2 | 0.5 | 2 | 1 | 12 | 0.9167 | 2 | 1 | 3 | 1 | 2 | 1 | 3 | 1 | 10 | 1 |
| 2_14424930 | 7 | 1 | 5 | 1 | 2 | 0.5 | 3 | 1 | 17 | 0.9412 | 3 | 1 | 3 | 1 | 5 | 1 | 4 | 1 | 15 | 1 |
| 2_14424930 | 7 | 0.1429 | 5 | 0 | 3 | 0.3333 | 3 | 0 | 18 | 0.1111 | 3 | 0 | 3 | 0.3333 | 5 | 0.2 | 4 | 0 | 15 | 0.1333 |
| 2_14424936 | 6 | 0.6667 | 6 | 0.5 | 3 | 0.3333 | 3 | 0.3333 | 18 | 0.5 | 1 | 1 | 3 | 0.3333 | 4 | 0.75 | 4 | 0.25 | 12 | 0.5 |
| 2_14424939 | 5 | 1 | 5 | 0.8 | 3 | 0.6667 | 3 | 1 | 16 | 0.875 | 2 | 1 | 2 | 1 | 5 | 1 | 4 | 1 | 13 | 1 |
| 2_14424949 | 7 | 1 | 5 | 1 | 2 | 1 | 3 | 1 | 17 | 1 | 2 | 1 | 3 | 1 | 4 | 1 | 4 | 1 | 13 | 1 |
| 2_14431323 | 20 | 0.15 | 10 | 0.2 | 10 | 0.3 | 20 | 0.4 | 60 | 0.2667 | 10 | 0.4 | 20 | 0.15 | 24 | 0.375 | 13 | 0.0769 | 67 | 0.2537 |
| 2_14431537 | 9 | 0.5556 | 4 | 0.5 | 3 | 0.3333 | 11 | 0.3636 | 27 | 0.4444 | 2 | 0 | 9 | 0.3333 | 6 | 0.1667 | 9 | 0.4444 | 26 | 0.3077 |
| 2_14431549 | 7 | 0.8571 | 3 | 0.6667 | 3 | 1 | 3 | 0.6667 | 16 | 0.8125 | 2 | 1 | 3 | 1 | 5 | 0.4 | 6 | 0.3333 | 16 | 0.5625 |
| 2_14431675 | 3 | 0 | 1 | 0 | 2 | 0.5 | 3 | 0 | 9 | 0.1111 | 1 | 1 | 6 | 0.1667 | 2 | 0 | 3 | 0 | 12 | 0.1667 |
| 2_14431681 | 1 | 0 | 1 | 0 | NA | NA | 2 | 0.5 | 4 | 0.25 | 1 | 0 | 3 | 0.3333 | NA | NA | 1 | 0 | 5 | 0.2 |
| 2_14437919 | 1 | 1 | 1 | 0 | 2 | 0 | 1 | 0 | 5 | 0.2 | NA | NA | 2 | 0 | NA | NA | 1 | 1 | 3 | 0.3333 |
| 2_14438064 | 1 | 1 | 1 | 0 | 1 | 1 | 1 | 1 | 4 | 0.75 | NA | NA | 2 | 0.5 | NA | NA | 1 | 1 | 3 | 0.6667 |

| | | | | | | | | | | | | | | | | | | | | |
|---|---|---|---|---|---|---|---|---|---|---|---|---|---|---|---|---|---|---|---|---|
| 2_14438064 | 1 | 0 | 1 | 1 | 2 | 0.5 | 1 | 0 | 5 | 0.4 | NA | NA | 2 | 0.5 | NA | NA | 1 | 0 | 3 | 0.3333 |
| 2_14438065 | NA | NA | 1 | 1 | 1 | 0 | 1 | 0 | 3 | 0.3333 | NA | NA | 1 | 0 | NA | NA | 1 | 1 | 2 | 0.5 |
| 2_14438066 | NA | NA | 1 | 0 | 1 | 1 | 1 | 0 | 3 | 0.3333 | NA | NA | 2 | 0.5 | NA | NA | 1 | 0 | 3 | 0.3333 |
| 2_14438072 | 1 | 1 | NA | NA | 2 | 1 | 1 | 1 | 4 | 1 | NA | NA | 2 | 1 | NA | NA | NA | NA | 2 | 1 |
| 2_14438100 | 2 | 0 | 1 | 0 | 2 | 0.5 | 1 | 0 | 6 | 0.1667 | NA | NA | 1 | 0 | NA | NA | 2 | 0.5 | 3 | 0.3333 |
| 2_14438100 | 1 | 0 | 1 | 1 | NA | NA | 1 | 1 | 3 | 0.6667 | NA | NA | 2 | 0.5 | NA | NA | 1 | 0 | 3 | 0.3333 |
| 2_14438109 | NA | NA | 1 | 1 | 1 | 1 | 1 | 1 | 3 | 1 | NA | NA | 1 | 1 | NA | NA | 1 | 1 | 2 | 1 |
| 2_14438189 | 2 | 0.5 | 2 | 0.5 | 2 | 0 | 1 | 0 | 7 | 0.2857 | NA | NA | 3 | 0.3333 | 1 | 0 | 3 | 0.3333 | 7 | 0.2857 |
| 2_14438323 | 1 | 0 | 2 | 0.5 | 1 | 0 | NA | NA | 4 | 0.25 | NA | NA | 1 | 0 | 2 | 0.5 | 2 | 0 | 5 | 0.2 |
| 2_14438574 | 6 | 0.1667 | 6 | 0.1667 | 7 | 0.7143 | 3 | 0.6667 | 22 | 0.4091 | 3 | 0.6667 | 5 | 0.4 | 7 | 0.4286 | 6 | 0.3333 | 21 | 0.4286 |
| 2_14443715 | 5 | 1 | 3 | 1 | 10 | 0.8 | 9 | 0.8889 | 27 | 0.8889 | 6 | 0.8333 | 9 | 0.7778 | 6 | 0.6667 | 5 | 0.8 | 26 | 0.7692 |
| 2_14443720 | 6 | 0.5 | 5 | 0.8 | 13 | 0.8462 | 12 | 0.6667 | 36 | 0.7222 | 8 | 0.5 | 11 | 0.7273 | 10 | 0.7 | 9 | 0.4444 | 38 | 0.6053 |
| 2_14461020 | 14 | 0.4286 | 14 | 0.4286 | 9 | 0.4444 | 11 | 0.8182 | 48 | 0.5208 | 10 | 0.2 | 11 | 0.4545 | 12 | 0.5 | 11 | 0.6364 | 44 | 0.4545 |
| 2_14461027 | 13 | 0.2308 | 12 | 0.1667 | 10 | 0.2 | 12 | 0.25 | 47 | 0.2128 | 10 | 0.3 | 10 | 0.2 | 11 | 0.0909 | 12 | 0.0833 | 43 | 0.1628 |
| 2_14544852 | 6 | 0.1667 | 9 | 0 | 9 | 0.1111 | 4 | 0 | 28 | 0.0714 | 6 | 0 | 1 | 0 | 9 | 0.2222 | 6 | 0 | 22 | 0.0909 |
| 2_14548225 | 4 | 0.5 | 9 | 0.5556 | 3 | 0.3333 | 1 | 1 | 17 | 0.5294 | 4 | 0.5 | 2 | 0.5 | 3 | 0.3333 | 2 | 0.5 | 11 | 0.4545 |
| 2_14548227 | 5 | 1 | 13 | 1 | 5 | 1 | 1 | 1 | 24 | 1 | 5 | 1 | 3 | 1 | 7 | 1 | 7 | 1 | 22 | 1 |
| 2_14548257 | 4 | 0.5 | 10 | 0.2 | 7 | 0.4286 | 2 | 0.5 | 23 | 0.3478 | 6 | 0.1667 | 7 | 0.2857 | 6 | 0.3333 | 8 | 0.5 | 27 | 0.3333 |
| 2_14548258 | 9 | 0.7778 | 13 | 0.8462 | 5 | 1 | 3 | 1 | 30 | 0.8667 | 9 | 1 | 5 | 0.6 | 8 | 0.875 | 9 | 0.8889 | 31 | 0.871 |
| 2_14548259 | 7 | 0 | 12 | 0.0833 | 9 | 0.2222 | 4 | 0 | 32 | 0.0938 | 10 | 0.1 | 8 | 0 | 10 | 0 | 8 | 0.375 | 36 | 0.1111 |
| 2_14548269 | 11 | 0.8182 | 15 | 0.7333 | 8 | 1 | 4 | 1 | 38 | 0.8421 | 8 | 1 | 6 | 0.8333 | 7 | 0.8571 | 9 | 0.7778 | 30 | 0.8667 |
| 2_14548320 | 7 | 0.2857 | 16 | 0.25 | 12 | 0.3333 | 4 | 0.5 | 39 | 0.3077 | 13 | 0.4615 | 10 | 0.5 | 8 | 0.375 | 9 | 0.4444 | 40 | 0.45 |
| 2_14548323 | 10 | 0.3 | 13 | 0.3077 | 13 | 0.2308 | 5 | 0 | 41 | 0.2439 | 15 | 0.2667 | 8 | 0.625 | 7 | 0.1429 | 11 | 0.2727 | 41 | 0.3171 |
| 2_14548332 | 9 | 0.6667 | 15 | 0.8 | 15 | 0.5333 | 4 | 0.5 | 43 | 0.6512 | 11 | 0.6364 | 7 | 0.5714 | 8 | 0.375 | 10 | 0.6 | 36 | 0.5556 |
| 2_14548334 | 11 | 0.5455 | 20 | 0.45 | 17 | 0.6471 | 6 | 0.5 | 54 | 0.537 | 15 | 0.5333 | 9 | 0.4444 | 10 | 0.7 | 11 | 0.5455 | 45 | 0.5556 |
| 2_14548343 | 11 | 0.8182 | 16 | 0.9375 | 16 | 0.8125 | 6 | 1 | 49 | 0.8776 | 13 | 0.6923 | 10 | 1 | 11 | 0.8182 | 12 | 0.75 | 46 | 0.8043 |
| 2_14548355 | 11 | 0.8182 | 19 | 0.7368 | 15 | 0.7333 | 7 | 0.8571 | 52 | 0.7692 | 18 | 0.8333 | 8 | 1 | 14 | 0.5714 | 13 | 0.8462 | 53 | 0.7925 |
| 2_14548359 | 10 | 0.7 | 20 | 0.65 | 10 | 0.6 | 7 | 0.4286 | 47 | 0.617 | 15 | 0.5333 | 10 | 0.6 | 10 | 0.6 | 10 | 0.6 | 45 | 0.5778 |
| 2_14548362 | 11 | 0.8182 | 16 | 0.8125 | 16 | 0.625 | 8 | 1 | 51 | 0.7843 | 13 | 0.7692 | 10 | 0.9 | 12 | 0.6667 | 12 | 0.6667 | 47 | 0.7447 |
| 2_14548374 | 13 | 0.4615 | 20 | 0.45 | 19 | 0.2632 | 8 | 0.375 | 60 | 0.3833 | 14 | 0.2857 | 11 | 0.4545 | 15 | 0.5333 | 11 | 0.5455 | 51 | 0.451 |
| 2_14548388 | 8 | 0.375 | 19 | 0.3684 | 15 | 0.4667 | 5 | 0.6 | 47 | 0.4255 | 17 | 0.4706 | 11 | 0.3636 | 13 | 0.6154 | 15 | 0.4 | 56 | 0.4643 |
| 2_14548394 | 12 | 0 | 21 | 0.0952 | 16 | 0.0625 | 6 | 0 | 55 | 0.0545 | 19 | 0.1579 | 11 | 0.0909 | 13 | 0.0769 | 14 | 0 | 57 | 0.0877 |
| 2_14570272 | NA | NA | 2 | 0.5 | 1 | 0 | NA | NA | 3 | 0.3333 | 2 | 0.5 | 1 | 0 | 2 | 0.5 | NA | NA | 5 | 0.4 |
| 2_14570290 | 1 | 0 | 5 | 0.4 | 2 | 0.5 | 2 | 0.5 | 10 | 0.4 | 3 | 0.3333 | NA | NA | 2 | 0.5 | NA | NA | 5 | 0.4 |
| 2_14570293 | 1 | 0 | 3 | 0.3333 | 2 | 1 | 2 | 0 | 8 | 0.375 | 3 | 0.6667 | 1 | 0 | 2 | 1 | NA | NA | 6 | 0.6667 |
| 2_14570302 | 1 | 0 | 3 | 0 | 2 | 0.5 | 2 | 1 | 8 | 0.375 | 3 | 0.3333 | 1 | 0 | 2 | 0.5 | NA | NA | 6 | 0.3333 |
| 2_14572764 | 1 | 1 | 4 | 0.5 | 4 | 0.75 | 2 | 0.5 | 11 | 0.6364 | 4 | 0.75 | 1 | 0 | 2 | 0 | 1 | 1 | 8 | 0.5 |
| 2_14572768 | 1 | 1 | 3 | 0 | 3 | 0 | 2 | 0 | 9 | 0.1111 | 5 | 0 | 1 | 1 | 2 | 0.5 | 1 | 0 | 9 | 0.2222 |
| 2_14572770 | 1 | 1 | 4 | 1 | 3 | 1 | 1 | 1 | 9 | 1 | 3 | 1 | 1 | 1 | 2 | 1 | 1 | 1 | 7 | 1 |
| 2_14572784 | 1 | 1 | 3 | 1 | 1 | 1 | 2 | 1 | 7 | 1 | 5 | 1 | 1 | 0 | 1 | 1 | 1 | 1 | 8 | 0.875 |
| 2_14572785 | 2 | 1 | 2 | 1 | 4 | 1 | 1 | 1 | 9 | 1 | 4 | 1 | 1 | 1 | 2 | 1 | NA | NA | 7 | 1 |
| 2_14572796 | 3 | 0.6667 | 2 | 0 | 4 | 0.25 | 2 | 0 | 11 | 0.2727 | 5 | 0 | 1 | 0 | 1 | 1 | 1 | 0 | 8 | 0.125 |
| 2_14572805 | 3 | 0.3333 | 4 | 0 | 4 | 0 | 2 | 0 | 13 | 0.0769 | 5 | 0.2 | NA | NA | 2 | 0.5 | 1 | 1 | 8 | 0.375 |
| 2_14572822 | 1 | 0 | 4 | 0.25 | 3 | 0.3333 | 2 | 0.5 | 10 | 0.3 | 4 | 0.25 | 1 | 0 | 2 | 0 | 1 | 0 | 8 | 0.125 |
| 2_14572857 | 3 | 0.3333 | 4 | 0 | 5 | 0.2 | 2 | 0.5 | 14 | 0.2143 | 4 | 0.5 | 1 | 0 | 2 | 0 | 2 | 0 | 9 | 0.2222 |
| 2_14572898 | 3 | 0 | 3 | 0 | 4 | 0.25 | 2 | 0.5 | 12 | 0.1667 | 6 | 0 | 1 | 1 | 3 | 0 | 2 | 0.5 | 12 | 0.1667 |
| 2_14572926 | 3 | 0.3333 | 5 | 0.2 | 5 | 0.2 | 4 | 0 | 17 | 0.1765 | 5 | 0.2 | 1 | 0 | 3 | 0 | 2 | 1 | 11 | 0.2727 |
| 2_14572938 | 5 | 0.2 | 7 | 0 | 5 | 0.2 | 5 | 0 | 22 | 0.0909 | 6 | 0.1667 | 1 | 0 | 3 | 0.3333 | 2 | 0 | 12 | 0.1667 |
| 2_14575779 | 13 | 0 | 22 | 0.0455 | 15 | 0.0667 | 8 | 0.125 | 58 | 0.0517 | 10 | 0.1 | 11 | 0.0909 | 12 | 0.0833 | 13 | 0.0769 | 46 | 0.087 |
| 2_14575798 | 15 | 0.2667 | 22 | 0.1818 | 16 | 0.1875 | 10 | 0.2 | 63 | 0.2063 | 9 | 0.1111 | 13 | 0.0769 | 13 | 0.2308 | 11 | 0.3636 | 46 | 0.1957 |
| 2_14576037 | 2 | 0 | 2 | 0 | 4 | 0.25 | 2 | 1 | 10 | 0.3 | NA | NA | 1 | 1 | 2 | 0.5 | 2 | 0.5 | 5 | 0.6 |
| 2_14606227 | 1 | 0 | 1 | 1 | NA | NA | NA | NA | 2 | 0.5 | NA | NA | 1 | 0 | 1 | 1 | 1 | 0 | 3 | 0.3333 |
| 2_14606259 | 1 | 0 | 1 | 1 | NA | NA | NA | NA | 2 | 0.5 | NA | NA | 1 | 0 | 1 | 1 | 1 | 1 | 3 | 0.6667 |
| 2_14606263 | 1 | 0 | 1 | 1 | NA | NA | NA | NA | 2 | 0.5 | NA | NA | 1 | 0 | NA | NA | 1 | 1 | 2 | 0.5 |
| 2_14606307 | 1 | 0 | 1 | 1 | NA | NA | NA | NA | 2 | 0.5 | NA | NA | 1 | 1 | 1 | 1 | 1 | 0 | 3 | 0.6667 |
| 2_14606314 | NA | NA | 1 | 1 | NA | NA | NA | NA | 1 | 1 | NA | NA | 1 | 1 | 1 | 1 | 1 | 1 | 3 | 1 |
| 2_14606322 | 1 | 1 | NA | NA | NA | NA | NA | NA | 1 | 1 | NA | NA | 1 | 1 | 1 | 1 | 1 | 1 | 3 | 1 |
| 2_14606387 | 1 | 1 | NA | NA | NA | NA | NA | NA | 1 | 1 | NA | NA | NA | NA | NA | NA | 1 | 1 | 1 | 1 |
| 2_14606407 | 1 | 1 | NA | NA | NA | NA | NA | NA | 1 | 1 | NA | NA | NA | NA | NA | NA | 1 | 1 | 1 | 1 |
| 2_14606422 | 1 | 1 | NA | NA | NA | NA | NA | NA | 1 | 1 | NA | NA | NA | NA | NA | NA | 1 | 1 | 1 | 1 |
| 2_14606449 | 1 | 1 | NA | NA | NA | NA | NA | NA | 1 | 1 | NA | NA | NA | NA | NA | NA | 1 | 1 | 1 | 1 |
| 2_14606457 | 1 | 1 | NA | NA | NA | NA | NA | NA | 1 | 1 | NA | NA | NA | NA | NA | NA | 1 | 1 | 1 | 1 |
| 2_14606525 | 1 | 0 | NA | NA | 2 | 1 | NA | NA | 3 | 0.6667 | NA | NA | 3 | 0 | 1 | 0 | 6 | 0.5 | 10 | 0.3 |
| 2_14606543 | 1 | 1 | NA | NA | 4 | 0.75 | NA | NA | 5 | 0.8 | NA | NA | 3 | 0.6667 | 1 | 1 | 7 | 1 | 11 | 0.9091 |
| 2_14606570 | 1 | 1 | 1 | 0 | 6 | 0.5 | NA | NA | 8 | 0.5 | NA | NA | 3 | 0.6667 | 2 | 0.5 | 8 | 0.625 | 13 | 0.6154 |

| | | | | | | | | | | | | | | | | | | | | |
|---|---|---|---|---|---|---|---|---|---|---|---|---|---|---|---|---|---|---|---|---|
| 2_1460657 | 1 | 1 | NA | NA | 5 | 1 | NA | NA | 6 | 1 | NA | NA | 3 | 1 | 1 | 1 | 8 | 1 | 12 | 1 |
| 2_1460659 | 1 | 1 | 1 | 0 | 5 | 0.4 | NA | NA | 7 | 0.4286 | NA | NA | 4 | 1 | 2 | 0.5 | 8 | 1 | 14 | 0.9286 |
| 2_1460659 | 1 | 1 | 1 | 0 | 5 | 0.6 | NA | NA | 7 | 0.5714 | NA | NA | 2 | 1 | 1 | 1 | 8 | 0.625 | 11 | 0.7273 |
| 2_1460662 | NA | NA | 1 | 0 | 5 | 0.6 | NA | NA | 6 | 0.5 | NA | NA | 4 | 0.25 | 2 | 0.5 | 8 | 0.25 | 14 | 0.2857 |
| 2_1460664 | 1 | 1 | 1 | 0 | 5 | 0 | 1 | 0 | 8 | 0.125 | NA | NA | 4 | 0.5 | 3 | 0 | 7 | 0 | 14 | 0.1429 |
| 2_1460837 | NA | NA | 3 | 0.3333 | 1 | 0 | NA | NA | 4 | 0.25 | 2 | 0.5 | NA | NA | 2 | 0.5 | NA | NA | 4 | 0.5 |
| 2_1460855 | NA | NA | 3 | 0.6667 | 1 | 0 | NA | NA | 4 | 0.5 | 1 | 1 | 1 | 0 | NA | NA | NA | NA | 2 | 0.5 |
| 2_1460858 | NA | NA | 3 | 0.3333 | NA | NA | NA | NA | 3 | 0.3333 | 1 | 0 | 1 | 1 | 1 | 0 | 1 | 1 | 4 | 0.5 |
| 2_1460864 | NA | NA | 1 | 1 | NA | NA | NA | NA | 1 | 1 | NA | NA | 1 | 1 | 1 | 0 | 1 | 1 | 3 | 0.6667 |
| 2_1461481 | NA | NA | 1 | 1 | NA | NA | 1 | 0 | 2 | 0.5 | NA | NA | NA | NA | NA | NA | 1 | 1 | 1 | 1 |
| 2_1461482 | NA | NA | 1 | 1 | NA | NA | 1 | 1 | 2 | 1 | NA | NA | NA | NA | NA | NA | 1 | 1 | 1 | 1 |
| 2_1461484 | NA | NA | 2 | 0.5 | NA | NA | 1 | 0 | 3 | 0.3333 | NA | NA | NA | NA | NA | NA | 1 | 1 | 1 | 1 |
| 2_1461842 | NA | NA | 1 | 1 | NA | NA | NA | NA | 1 | 1 | 1 | 1 | NA | NA | NA | NA | NA | NA | 1 | 1 |
| 2_1461844 | NA | NA | 1 | 1 | NA | NA | NA | NA | 1 | 1 | 1 | 1 | NA | NA | NA | NA | NA | NA | 1 | 1 |
| 2_1467310 | NA | NA | 3 | 1 | 1 | 0 | 2 | 1 | 6 | 0.8333 | 1 | 1 | 1 | 1 | 2 | 0 | 2 | 0.5 | 6 | 0.5 |
| 2_1467311 | NA | NA | 3 | 0 | 1 | 0 | 2 | 0.5 | 6 | 0.1667 | 1 | 1 | 1 | 0 | 3 | 0 | 2 | 0.5 | 7 | 0.2857 |
| 2_1467311 | NA | NA | 3 | 0.3333 | 1 | 0 | 1 | 0 | 5 | 0.2 | 1 | 0 | 1 | 0 | 3 | 0.3333 | 2 | 0 | 7 | 0.1429 |
| 2_1467331 | NA | NA | 3 | 0 | 2 | 0.5 | 3 | 0 | 8 | 0.125 | 1 | 0 | 1 | 1 | 3 | 0.3333 | 2 | 0.5 | 7 | 0.4286 |
| 2_1469197 | 5 | 0.2 | 8 | 0.125 | 2 | 0 | 10 | 0.6 | 25 | 0.32 | 5 | 0.6 | 5 | 0.4 | 9 | 0.4444 | 11 | 0.2727 | 30 | 0.4 |
| 2_1471326 | NA | NA | NA | NA | 1 | 1 | NA | NA | 1 | 1 | 1 | 1 | NA | NA | NA | NA | NA | NA | 1 | 1 |
| 2_1471326 | NA | NA | NA | NA | 1 | 1 | NA | NA | 1 | 1 | 1 | 1 | NA | NA | NA | NA | NA | NA | 1 | 1 |
| 2_1471332 | NA | NA | NA | NA | 1 | 1 | NA | NA | 1 | 1 | 1 | 1 | NA | NA | NA | NA | NA | NA | 1 | 1 |
| 2_1482384 | 1 | 1 | NA | NA | 1 | 1 | 2 | 0.5 | 4 | 0.75 | NA | NA | 1 | 1 | 1 | 1 | NA | NA | 2 | 1 |
| 2_1482386 | 1 | 1 | NA | NA | 1 | 0 | 2 | 0 | 4 | 0.25 | NA | NA | 1 | 0 | 1 | 0 | 1 | 1 | 3 | 0.3333 |
| 2_1482386 | 1 | 1 | NA | NA | 2 | 1 | 3 | 1 | 6 | 1 | NA | NA | 1 | 1 | 1 | 1 | 1 | 1 | 3 | 1 |
| 2_1482387 | 1 | 1 | NA | NA | NA | NA | 3 | 1 | 4 | 1 | NA | NA | 1 | 1 | 1 | 1 | 1 | 1 | 3 | 1 |
| 2_1482387 | 1 | 1 | NA | NA | 1 | 1 | 2 | 1 | 4 | 1 | NA | NA | 1 | 1 | 1 | 1 | 1 | 1 | 3 | 1 |
| 2_1482387 | 1 | 1 | NA | NA | 2 | 1 | 2 | 1 | 5 | 1 | NA | NA | 1 | 1 | 1 | 1 | NA | NA | 2 | 1 |
| 2_1482388 | 1 | 1 | NA | NA | 2 | 0 | 2 | 0.5 | 5 | 0.4 | NA | NA | 1 | 0 | 1 | 0 | 1 | 1 | 3 | 0.3333 |
| 2_1482388 | NA | NA | NA | NA | NA | NA | 2 | 1 | 2 | 1 | NA | NA | 1 | 1 | 1 | 1 | 1 | 1 | 3 | 1 |
| 2_1482391 | 1 | 1 | NA | NA | 1 | 0 | 2 | 0 | 4 | 0.25 | NA | NA | 1 | 0 | 1 | 1 | 2 | 1 | 4 | 0.75 |
| 2_1482401 | 1 | 0 | 1 | 0 | 1 | 1 | 2 | 0 | 5 | 0.2 | NA | NA | 1 | 0 | 1 | 0 | 2 | 0.5 | 4 | 0.25 |
| 2_1482413 | 3 | 0.3333 | 1 | 0 | 1 | 0 | 2 | 0 | 7 | 0.1429 | NA | NA | 2 | 0.5 | 2 | 1 | 1 | 0 | 5 | 0.6 |
| 2_1482423 | 2 | 1 | 3 | 1 | 1 | 0 | 4 | 0.5 | 10 | 0.7 | 1 | 1 | 3 | 0.6667 | 1 | 1 | 4 | 0.5 | 9 | 0.6667 |
| 2_1482424 | 3 | 0 | 2 | 0.5 | 1 | 0 | 4 | 0 | 10 | 0.1 | 2 | 0 | 3 | 0 | 3 | 0 | 5 | 0.2 | 13 | 0.0769 |
| 2_1482469 | 3 | 0.3333 | 3 | 0.3333 | NA | NA | NA | NA | 6 | 0.3333 | 1 | 0 | NA | NA | 1 | 0 | 4 | 0.5 | 6 | 0.3333 |
| 2_1482469 | 3 | 0.3333 | 4 | 0 | NA | NA | 1 | 0 | 8 | 0.125 | NA | NA | NA | NA | 1 | 0 | 4 | 0.25 | 5 | 0.2 |
| 2_1482471 | 2 | 0.5 | 4 | 0 | NA | NA | 1 | 0 | 7 | 0.1429 | 1 | 1 | NA | NA | 1 | 0 | 4 | 0.25 | 6 | 0.3333 |
| 2_1482479 | 2 | 0.5 | 3 | 1 | NA | NA | NA | NA | 5 | 0.8 | 1 | 0 | NA | NA | 1 | 0 | 4 | 0.75 | 6 | 0.5 |
| 2_1482479 | 2 | 0 | 3 | 0.3333 | NA | NA | NA | NA | 5 | 0.2 | 1 | 0 | NA | NA | NA | NA | 3 | 0.6667 | 4 | 0.5 |
| 2_1482481 | 2 | 0 | 3 | 1 | NA | NA | NA | NA | 5 | 0.6 | NA | NA | NA | NA | 1 | 1 | 4 | 0.25 | 5 | 0.4 |
| 2_1482481 | 2 | 1 | 3 | 1 | NA | NA | NA | NA | 5 | 1 | NA | NA | NA | NA | 1 | 1 | 3 | 1 | 4 | 1 |
| 2_1482484 | 2 | 1 | 1 | 1 | NA | NA | NA | NA | 3 | 1 | NA | NA | NA | NA | 1 | 1 | 4 | 1 | 5 | 1 |
| 2_1482485 | 1 | 1 | 2 | 1 | NA | NA | NA | NA | 3 | 1 | NA | NA | NA | NA | 1 | 1 | 3 | 1 | 4 | 1 |
| 2_1482486 | 1 | 0 | 2 | 1 | NA | NA | NA | NA | 3 | 0.6667 | NA | NA | NA | NA | 1 | 0 | 3 | 0.6667 | 4 | 0.5 |
| 2_1482490 | 1 | 1 | 2 | 1 | NA | NA | NA | NA | 3 | 1 | NA | NA | NA | NA | 1 | 1 | 2 | 0.5 | 3 | 0.6667 |
| 2_1482490 | 1 | 1 | 2 | 1 | NA | NA | NA | NA | 3 | 1 | NA | NA | NA | NA | 1 | 1 | 3 | 0.3333 | 4 | 0.5 |
| 2_1482491 | 1 | 0 | 2 | 1 | NA | NA | NA | NA | 3 | 0.6667 | NA | NA | NA | NA | 1 | 1 | 3 | 0.6667 | 4 | 0.75 |
| 2_1482492 | NA | NA | 2 | 1 | NA | NA | NA | NA | 2 | 1 | NA | NA | NA | NA | NA | NA | 2 | 1 | 2 | 1 |
| 2_1482493 | 1 | 0 | 2 | 0.5 | NA | NA | NA | NA | 3 | 0.3333 | NA | NA | NA | NA | 1 | 0 | 2 | 0.5 | 3 | 0.3333 |
| 2_1482496 | 1 | 1 | 2 | 1 | NA | NA | NA | NA | 3 | 1 | NA | NA | NA | NA | 1 | 0 | 2 | 1 | 3 | 0.6667 |
| 2_1482564 | NA | NA | 1 | 0 | NA | NA | 1 | 1 | 2 | 0.5 | NA | NA | NA | NA | 1 | 1 | 1 | 1 | 2 | 1 |
| 2_1482566 | NA | NA | 1 | 0 | NA | NA | 1 | 1 | 2 | 0.5 | NA | NA | NA | NA | 1 | 0 | 1 | 1 | 2 | 0.5 |
| 2_1482795 | NA | NA | NA | NA | 1 | 1 | 2 | 0.5 | 3 | 0.6667 | NA | NA | NA | NA | NA | NA | 1 | 1 | 1 | 1 |
| 2_1485174 | 2 | 0.5 | 2 | 0 | 3 | 1 | 1 | 0 | 8 | 0.5 | 2 | 0.5 | 1 | 0 | 3 | 0.6667 | 3 | 0.6667 | 9 | 0.5556 |
| 2_1485174 | 2 | 0.5 | 2 | 0 | 3 | 1 | 1 | 0 | 8 | 0.5 | 3 | 1 | 2 | 0.5 | 1 | 0 | 3 | 1 | 9 | 0.7778 |
| 2_1485175 | 1 | 0 | NA | NA | 3 | 0.3333 | NA | NA | 4 | 0.25 | 3 | 0.6667 | 2 | 0 | 2 | 0 | 1 | 0 | 8 | 0.25 |
| 2_1485175 | 2 | 0.5 | 2 | 0.5 | 4 | 0.75 | 1 | 1 | 9 | 0.6667 | 3 | 0.6667 | 1 | 1 | 2 | 0.5 | 3 | 0.6667 | 9 | 0.6667 |
| 2_1485176 | 2 | 0.5 | 1 | 0 | 2 | 0 | 1 | 0 | 6 | 0.1667 | 2 | 0 | 1 | 0 | 2 | 0 | 3 | 0.3333 | 8 | 0.125 |
| 2_1485176 | 2 | 0.5 | 1 | 1 | 4 | 1 | 1 | 0 | 8 | 0.75 | 3 | 1 | 1 | 1 | 2 | 1 | 3 | 1 | 9 | 1 |
| 2_1485176 | 2 | 0.5 | 2 | 0.5 | 4 | 1 | 1 | 1 | 9 | 0.7778 | 4 | 0.75 | 2 | 0.5 | 3 | 0.3333 | 3 | 0.6667 | 12 | 0.5833 |
| 2_1485177 | 2 | 0 | 2 | 0 | 4 | 0.25 | 1 | 1 | 9 | 0.2222 | 3 | 0.6667 | 1 | 0 | 3 | 0 | 3 | 0.3333 | 10 | 0.3 |
| 2_1485178 | 2 | 1 | 2 | 1 | 4 | 1 | 1 | 1 | 9 | 1 | 4 | 1 | 2 | 1 | 2 | 1 | 3 | 1 | 11 | 1 |
| 2_1485179 | 1 | 0 | 1 | 1 | 4 | 0.5 | 1 | 0 | 7 | 0.4286 | 4 | 0.75 | 1 | 1 | 3 | 0.3333 | 2 | 0.5 | 10 | 0.6 |
| 2_1485257 | 3 | 0.6667 | 1 | 1 | 3 | 1 | 1 | 1 | 8 | 0.875 | 1 | 1 | 3 | 0.6667 | 1 | 1 | 3 | 1 | 8 | 0.875 |

| | | | | | | | | | | | | | | | | | | | | |
|---|---|---|---|---|---|---|---|---|---|---|---|---|---|---|---|---|---|---|---|---|
| 2_14852578 | 3 | 0.6667 | 1 | 1 | 3 | 0.3333 | 1 | 0 | 8 | 0.5 | 1 | 0 | 3 | 0.6667 | 1 | 0 | 2 | 0.5 | 7 | 0.4286 |
| 2_14852588 | 3 | 1 | 1 | 1 | 2 | 1 | NA | NA | 6 | 1 | 1 | 1 | 3 | 1 | 1 | 1 | 2 | 0.5 | 7 | 0.8571 |
| 2_14852609 | 3 | 0.6667 | 1 | 1 | 3 | 0.6667 | 1 | 1 | 8 | 0.75 | 2 | 1 | 2 | 1 | 1 | 1 | 3 | 0.6667 | 8 | 0.875 |
| 2_14852614 | 3 | 0.3333 | 2 | 0 | 3 | 0.3333 | 1 | 0 | 9 | 0.2222 | 2 | 0.5 | 3 | 0 | 1 | 0 | 3 | 0 | 9 | 0.1111 |
| 2_14852634 | 3 | 0 | 3 | 0.3333 | 4 | 0 | 1 | 1 | 11 | 0.1818 | 1 | 1 | 3 | 0 | 1 | 0 | 3 | 0.6667 | 8 | 0.375 |
| 2_14852639 | 3 | 0 | 3 | 0.3333 | 3 | 0 | 1 | 0 | 10 | 0.1 | 1 | 1 | 4 | 0.75 | 1 | 0 | 4 | 0.25 | 10 | 0.5 |
| 2_14852657 | 4 | 0.5 | 3 | 0.3333 | 4 | 0.25 | 1 | 1 | 12 | 0.4167 | 2 | 1 | 4 | 0.5 | 1 | 0 | 4 | 0.75 | 11 | 0.6364 |
| 2_14852678 | 4 | 1 | 2 | 1 | 4 | 0.75 | 1 | 1 | 11 | 0.9091 | 3 | 1 | 4 | 0.5 | 1 | 1 | 2 | 0.5 | 10 | 0.7 |
| 2_14852699 | 1 | 1 | NA | NA | NA | NA | NA | NA | 1 | 1 | 1 | 1 | NA | NA | NA | NA | NA | NA | 1 | 1 |
| 2_14852705 | 1 | 1 | 1 | 1 | NA | NA | NA | NA | 2 | 1 | 1 | 1 | NA | NA | NA | NA | NA | NA | 1 | 1 |
| 2_14853211 | 6 | 0.3333 | 3 | 0 | 5 | 0 | 1 | 0 | 15 | 0.1333 | 2 | 0 | 4 | 0.25 | 2 | 0 | 3 | 0 | 11 | 0.0909 |
| 2_14853274 | 4 | 0 | 2 | 0.5 | 5 | 0 | 1 | 0 | 12 | 0.0833 | 2 | 0.5 | 4 | 0 | 2 | 0 | 4 | 0.5 | 12 | 0.25 |
| 2_14853297 | 5 | 0.4 | 3 | 0.3333 | 5 | 0.4 | 2 | 0.5 | 15 | 0.4 | 2 | 1 | 3 | 0.6667 | 2 | 0.5 | 4 | 0.5 | 11 | 0.6364 |
| 2_14853306 | 7 | 0 | 3 | 0.3333 | 5 | 0 | 2 | 0 | 17 | 0.0588 | 2 | 0.5 | 3 | 0 | 2 | 0 | 4 | 0 | 11 | 0.0909 |
| 2_14853308 | 8 | 0.125 | 3 | 0.3333 | 5 | 0.4 | 2 | 0 | 18 | 0.2222 | 1 | 0 | 4 | 0 | 2 | 0 | 4 | 0.25 | 11 | 0.0909 |
| 2_14853312 | 8 | 0.125 | 3 | 0.3333 | 2 | 0 | 2 | 0.5 | 15 | 0.2 | NA | NA | 4 | 0.25 | 2 | 0 | 2 | 1 | 8 | 0.375 |
| 2_14853407 | 6 | 0.1667 | 3 | 0 | 7 | 0.1429 | 5 | 0.2 | 21 | 0.1429 | 3 | 0.3333 | 4 | 0.5 | 4 | 0 | 3 | 0 | 14 | 0.2143 |
| 2_14853449 | 7 | 0 | 2 | 0 | 7 | 0 | 4 | 0.25 | 20 | 0.05 | 3 | 0 | 3 | 0 | 5 | 0.2 | 4 | 0 | 15 | 0.0667 |
| 2_14853468 | 8 | 0.25 | 4 | 0.25 | 5 | 0 | 4 | 0 | 21 | 0.1429 | 2 | 0 | 4 | 0 | 4 | 0 | 6 | 0.1667 | 16 | 0.0625 |
| 2_14853667 | 27 | 0 | 52 | 0.1538 | 42 | 0.0714 | 35 | 0.1429 | 156 | 0.1026 | 33 | 0.1818 | 29 | 0.069 | 35 | 0.0571 | 25 | 0.24 | 122 | 0.1311 |
| 2_14853818 | 21 | 0 | 40 | 0.1 | 40 | 0.05 | 35 | 0.0571 | 136 | 0.0588 | 29 | 0.1034 | 23 | 0.2174 | 32 | 0.1875 | 24 | 0.0417 | 108 | 0.1389 |
| 2_14853861 | 17 | 0 | 33 | 0.0909 | 36 | 0.0278 | 34 | 0.0882 | 120 | 0.0583 | 25 | 0.04 | 19 | 0.2105 | 30 | 0.0667 | 16 | 0.0625 | 90 | 0.0889 |
| 2_14871365 | 3 | 0 | 3 | 0.3333 | 1 | 0 | 3 | 0 | 10 | 0.1 | 1 | 0 | NA | NA | 3 | 0 | 5 | 0.2 | 9 | 0.1111 |
| 2_14871385 | 624 | 0.2356 | 479 | 0.2338 | 529 | 0.2382 | 497 | 0.2233 | 2129 | 0.233 | 614 | 0.1987 | 492 | 0.2215 | 486 | 0.1996 | 453 | 0.2693 | 2045 | 0.22 |
| 2_14967217 | 2 | 0.5 | NA | NA | NA | NA | NA | NA | 2 | 0.5 | NA | NA | NA | NA | 1 | 1 | NA | NA | 1 | 1 |
| 2_14967239 | 1 | 1 | NA | NA | NA | NA | NA | NA | 1 | 1 | NA | NA | NA | NA | 1 | 1 | NA | NA | 1 | 1 |
| 2_14967263 | 1 | 1 | 1 | 0 | 2 | 0 | 1 | 0 | 5 | 0.2 | NA | NA | NA | NA | 1 | 1 | NA | NA | 1 | 1 |
| 2_14974150 | 11 | 0.6364 | 6 | 0.1667 | 9 | 0.2222 | 20 | 0.25 | 46 | 0.3261 | 3 | 0.3333 | 8 | 0.125 | 12 | 0.1667 | 7 | 0.4286 | 30 | 0.2333 |
| 2_14974163 | 14 | 0.6429 | 6 | 0.1667 | 11 | 0.1818 | 24 | 0.625 | 55 | 0.4909 | 3 | 0.6667 | 12 | 0.5833 | 15 | 0.5333 | 8 | 0.5 | 38 | 0.5526 |
| 2_14974174 | 12 | 0.5 | 6 | 0.3333 | 11 | 0.2727 | 21 | 0.3333 | 50 | 0.36 | 4 | 0.5 | 10 | 0.4 | 15 | 0.3333 | 6 | 0.3333 | 35 | 0.3714 |
| 2_14974176 | 12 | 0.0833 | 6 | 0 | 11 | 0 | 23 | 0.087 | 52 | 0.0577 | 4 | 0.25 | 12 | 0.25 | 15 | 0.1333 | 6 | 0.1667 | 37 | 0.1892 |
| 2_14974178 | 12 | 0.0833 | 5 | 0 | 10 | 0.1 | 22 | 0.0455 | 49 | 0.0612 | 4 | 0 | 11 | 0.1818 | 17 | 0.1765 | 8 | 0.125 | 40 | 0.15 |
| 2_14984519 | 12 | 0.9167 | 6 | 0.8333 | 8 | 1 | 23 | 1 | 49 | 0.9592 | 4 | 1 | 8 | 0.875 | 12 | 0.9167 | 5 | 1 | 29 | 0.931 |
| 2_14984531 | 15 | 0.6667 | 7 | 0.5714 | 13 | 0.4615 | 25 | 0.6 | 60 | 0.5833 | 5 | 0.8 | 11 | 0.7273 | 16 | 0.5 | 7 | 0.4286 | 39 | 0.5897 |
| 2_14984550 | 15 | 0.1333 | 9 | 0.2222 | 13 | 0.2308 | 26 | 0.0385 | 63 | 0.127 | 5 | 0.2 | 11 | 0 | 16 | 0.125 | 7 | 0.2857 | 39 | 0.1282 |
| 2_14984556 | 13 | 0.8462 | 9 | 0.8889 | 13 | 1 | 25 | 0.88 | 60 | 0.9 | 5 | 1 | 13 | 0.9231 | 15 | 0.9333 | 8 | 1 | 41 | 0.9512 |
| 2_14984559 | 6 | 0.8333 | 4 | 0.75 | 7 | 1 | 13 | 0.8462 | 30 | 0.8667 | 1 | 1 | 5 | 1 | 9 | 1 | 2 | 0.5 | 17 | 0.9412 |
| 2_14984564 | 14 | 0.9286 | 10 | 1 | 14 | 1 | 25 | 1 | 63 | 0.9841 | 4 | 1 | 14 | 1 | 16 | 1 | 8 | 1 | 42 | 1 |
| 2_14984572 | 14 | 0.3571 | 9 | 0.1111 | 13 | 0.3846 | 23 | 0.087 | 59 | 0.2203 | 3 | 0 | 12 | 0.25 | 14 | 0.1429 | 7 | 0 | 36 | 0.1389 |
| 2_14984595 | 6 | 0.3333 | 11 | 0.2727 | 11 | 0.4545 | 17 | 0.2353 | 45 | 0.3111 | 3 | 0.3333 | 13 | 0.3077 | 16 | 0.125 | 6 | 0.3333 | 38 | 0.2368 |
| 2_14984596 | 11 | 0.2727 | 6 | 0 | 10 | 0.2 | 16 | 0.0625 | 43 | 0.1395 | 5 | 0.4 | 7 | 0.2857 | 11 | 0.2727 | 5 | 0.2 | 28 | 0.2857 |
| 2_14984600 | 16 | 0.125 | 11 | 0 | 15 | 0.1333 | 30 | 0.1333 | 72 | 0.1111 | 5 | 0.2 | 17 | 0.0588 | 19 | 0.2632 | 8 | 0.125 | 49 | 0.1633 |
| 2_14984603 | 14 | 1 | 9 | 1 | 13 | 1 | 27 | 1 | 63 | 1 | 5 | 1 | 12 | 1 | 16 | 1 | 4 | 1 | 37 | 1 |
| 2_14984604 | 16 | 0.6875 | 8 | 0.5 | 14 | 0.5714 | 25 | 0.48 | 63 | 0.5556 | 4 | 0.75 | 13 | 0.6923 | 13 | 0.4615 | 7 | 0.5714 | 37 | 0.5946 |
| 2_14984630 | 2 | 1 | 1 | 1 | 1 | 1 | NA | NA | 4 | 1 | 1 | 1 | NA | NA | NA | NA | 1 | 1 | 2 | 1 |
| 2_14984636 | 2 | 1 | 1 | 1 | NA | NA | NA | NA | 3 | 1 | NA | NA | NA | NA | NA | NA | 1 | 1 | 1 | 1 |
| 2_14995655 | 1 | 0 | 1 | 1 | 2 | 0 | 3 | 0 | 7 | 0.1429 | NA | NA | 2 | 0.5 | 1 | 0 | 4 | 0.5 | 7 | 0.4286 |
| 2_14995823 | 1 | 0 | NA | NA | 2 | 0.5 | NA | NA | 3 | 0.3333 | NA | NA | 2 | 0.5 | NA | NA | 2 | 0 | 4 | 0.25 |
| 2_14995922 | 1 | 1 | NA | NA | 2 | 0.5 | 1 | 1 | 4 | 0.75 | NA | NA | 2 | 0.5 | NA | NA | 4 | 1 | 6 | 0.8333 |
| 2_14995943 | 1 | 1 | NA | NA | 2 | 0 | NA | NA | 3 | 0.3333 | NA | NA | 1 | 0 | NA | NA | 2 | 0.5 | 3 | 0.3333 |
| 2_14995950 | 1 | 1 | NA | NA | 2 | 1 | 1 | 0 | 4 | 0.75 | NA | NA | 2 | 0 | NA | NA | 4 | 0.25 | 6 | 0.1667 |
| 2_14995969 | 1 | 1 | NA | NA | 2 | 0 | 1 | 1 | 4 | 0.5 | NA | NA | 2 | 0.5 | NA | NA | 3 | 0.3333 | 5 | 0.4 |
| 2_14995972 | 1 | 0 | NA | NA | 2 | 0.5 | 1 | 0 | 4 | 0.25 | NA | NA | 2 | 0.5 | NA | NA | 4 | 0 | 6 | 0.1667 |
| 2_14995974 | NA | NA | NA | NA | 2 | 0.5 | 1 | 1 | 3 | 0.6667 | NA | NA | 1 | 1 | NA | NA | 1 | 1 | 2 | 1 |
| 2_14995978 | 1 | 1 | NA | NA | 2 | 1 | 1 | 0 | 4 | 0.75 | NA | NA | 2 | 0.5 | NA | NA | 4 | 0.75 | 6 | 0.6667 |
| 2_14995984 | 1 | 0 | NA | NA | 2 | 0 | 1 | 1 | 4 | 0.25 | NA | NA | 2 | 0 | NA | NA | 3 | 0.3333 | 5 | 0.2 |
| 2_14995994 | 1 | 0 | NA | NA | 2 | 0.5 | 1 | 0 | 4 | 0.25 | NA | NA | 2 | 1 | NA | NA | 3 | 0 | 5 | 0.4 |
| 2_14995998 | 1 | 0 | NA | NA | 2 | 1 | 1 | 1 | 4 | 0.75 | NA | NA | 2 | 1 | NA | NA | 3 | 0.3333 | 5 | 0.6 |
| 2_14996006 | 1 | 1 | NA | NA | 2 | 1 | 1 | 1 | 4 | 1 | NA | NA | 2 | 1 | NA | NA | 3 | 0.6667 | 5 | 0.8 |
| 2_15017389 | 1 | 0 | NA | NA | NA | NA | 1 | 1 | 2 | 0.5 | NA | NA | 1 | 1 | 1 | 1 | NA | NA | 2 | 1 |
| 2_15017392 | 1 | 0 | NA | NA | NA | NA | 1 | 1 | 2 | 0.5 | NA | NA | NA | NA | 1 | 1 | NA | NA | 1 | 1 |
| 2_15017397 | 1 | 0 | NA | NA | NA | NA | 1 | 1 | 2 | 0.5 | NA | NA | 1 | 0 | 1 | 1 | NA | NA | 2 | 0.5 |
| 2_15024445 | 1 | 1 | NA | NA | NA | NA | NA | NA | 1 | 1 | NA | NA | NA | NA | NA | NA | 1 | 1 | 1 | 1 |
| 2_15024582 | 1 | 0 | 1 | 1 | NA | NA | 2 | 0 | 4 | 0.25 | NA | NA | 2 | 0 | 1 | 0 | 3 | 0.3333 | 6 | 0.1667 |
| 2_15024659 | NA | NA | 1 | 0 | NA | NA | 2 | 0.5 | 3 | 0.3333 | NA | NA | 2 | 0.5 | NA | NA | 3 | 0.6667 | 5 | 0.6 |

| | | | | | | | | | | | | | | | | | | | | |
|---|---|---|---|---|---|---|---|---|---|---|---|---|---|---|---|---|---|---|---|---|
| 2_1502466 | 1 | 0 | 1 | 0 | NA | NA | 2 | 1 | 4 | 0.5 | NA | NA | 2 | 0.5 | NA | NA | 3 | 0.6667 | 5 | 0.6 |
| 2_1502466 | 1 | 1 | 1 | 1 | NA | NA | 2 | 1 | 4 | 1 | NA | NA | 2 | 0.5 | NA | NA | 3 | 0.6667 | 5 | 0.6 |
| 2_1502467 | 1 | 1 | NA | NA | NA | NA | 2 | 1 | 3 | 1 | NA | NA | NA | NA | NA | NA | 2 | 0.5 | 2 | 0.5 |
| 2_1502468 | 1 | 0 | 1 | 1 | NA | NA | 2 | 0 | 4 | 0.25 | NA | NA | 2 | 0 | NA | NA | 2 | 0.5 | 4 | 0.25 |
| 2_1502470 | 1 | 1 | 1 | 1 | NA | NA | 1 | 1 | 3 | 1 | NA | NA | 2 | 1 | NA | NA | 1 | 1 | 3 | 1 |
| 2_1502471 | 1 | 0 | 1 | 1 | NA | NA | 2 | 0 | 4 | 0.25 | NA | NA | 2 | 0 | NA | NA | 2 | 0.5 | 4 | 0.25 |
| 2_1502472 | 1 | 0 | NA | NA | NA | NA | 1 | 1 | 2 | 0.5 | NA | NA | 1 | 0 | NA | NA | 2 | 0.5 | 3 | 0.3333 |
| 2_1502473 | 1 | 0 | 1 | 1 | NA | NA | 2 | 1 | 4 | 0.75 | NA | NA | 1 | 1 | NA | NA | 2 | 1 | 3 | 1 |
| 2_1502473 | 1 | 0 | 1 | 1 | NA | NA | 1 | 0 | 3 | 0.3333 | NA | NA | 1 | 1 | NA | NA | 1 | 1 | 2 | 1 |
| 2_1502475 | NA | NA | 1 | 0 | NA | NA | 2 | 1 | 3 | 0.6667 | NA | NA | 2 | 0 | NA | NA | 2 | 0.5 | 4 | 0.25 |
| 2_1502475 | NA | NA | 1 | 1 | NA | NA | 2 | 1 | 3 | 1 | NA | NA | 1 | 1 | NA | NA | NA | NA | 1 | 1 |
| 2_1502496 | NA | NA | 1 | 1 | NA | NA | NA | NA | 1 | 1 | NA | NA | 2 | 0.5 | NA | NA | 1 | 1 | 3 | 0.6667 |
| 2_1502499 | NA | NA | 1 | 1 | NA | NA | NA | NA | 1 | 1 | NA | NA | 2 | 1 | NA | NA | 1 | 1 | 3 | 1 |
| 2_1502503 | NA | NA | 1 | 1 | NA | NA | NA | NA | 1 | 1 | NA | NA | 2 | 1 | NA | NA | 1 | 1 | 3 | 1 |
| 2_1502513 | NA | NA | 1 | 1 | NA | NA | NA | NA | 1 | 1 | NA | NA | 2 | 1 | NA | NA | 1 | 1 | 3 | 1 |
| 2_1502515 | NA | NA | 1 | 1 | NA | NA | NA | NA | 1 | 1 | NA | NA | 1 | 1 | NA | NA | 1 | 1 | 2 | 1 |
| 2_1504253 | 6 | 0.1667 | 4 | 0.25 | 1 | 0 | 13 | 0.5385 | 24 | 0.375 | 3 | 1 | 10 | 0.4 | 7 | 0.5714 | 5 | 0.4 | 25 | 0.52 |
| 2_1504257 | 8 | 0.625 | 1 | 0 | 1 | 1 | 8 | 0.375 | 18 | 0.5 | 2 | 0.5 | 7 | 0.7143 | 5 | 1 | 4 | 0 | 18 | 0.6111 |
| 2_1504258 | 7 | 0.4286 | 4 | 0.5 | 1 | 1 | 11 | 0.3636 | 23 | 0.4348 | 3 | 1 | 7 | 0.4286 | 5 | 0.8 | 3 | 0.6667 | 18 | 0.6667 |
| 2_1504260 | 9 | 0.5556 | 3 | 0 | NA | NA | 12 | 0.25 | 24 | 0.3333 | 2 | 0 | 8 | 0.25 | 6 | 0.3333 | 4 | 0.5 | 20 | 0.3 |
| 2_1504604 | 7 | 0.4286 | 2 | 1 | 1 | 1 | 4 | 0.75 | 14 | 0.6429 | 3 | 0.6667 | 2 | 1 | NA | NA | 2 | 0.5 | 7 | 0.7143 |
| 2_1504605 | 8 | 0.375 | 2 | 1 | 1 | 0 | 4 | 0.5 | 15 | 0.4667 | 3 | 0.3333 | 3 | 0 | NA | NA | 2 | 0.5 | 8 | 0.25 |
| 2_1504605 | 8 | 0.125 | 2 | 0 | 1 | 1 | 4 | 0.5 | 15 | 0.2667 | 3 | 0 | 3 | 0.6667 | NA | NA | 1 | 1 | 7 | 0.4286 |
| 2_1504605 | 8 | 0.5 | 2 | 0.5 | 1 | 1 | 3 | 1 | 14 | 0.6429 | 2 | 0.5 | 3 | 0.6667 | NA | NA | 1 | 1 | 6 | 0.6667 |
| 2_1504608 | 5 | 0.8 | 1 | 1 | NA | NA | 3 | 1 | 9 | 0.8889 | 2 | 0.5 | 3 | 0.6667 | NA | NA | 1 | 1 | 6 | 0.6667 |
| 2_1504608 | 5 | 0.2 | 1 | 1 | NA | NA | 4 | 0.25 | 10 | 0.3 | 3 | 0.6667 | 3 | 0 | NA | NA | NA | NA | 6 | 0.3333 |
| 2_1504610 | 4 | 0 | 1 | 1 | NA | NA | 4 | 0 | 9 | 0.1111 | 2 | 0 | 3 | 0.3333 | NA | NA | NA | NA | 5 | 0.2 |
| 2_1504612 | 5 | 0.4 | 1 | 0 | NA | NA | 2 | 0.5 | 8 | 0.375 | 2 | 0 | 3 | 0.3333 | NA | NA | NA | NA | 5 | 0.2 |
| 2_1504614 | 5 | 0 | 1 | 1 | NA | NA | 3 | 0 | 9 | 0.1111 | 2 | 0 | 3 | 0.3333 | NA | NA | NA | NA | 5 | 0.2 |
| 2_1505950 | 8 | 0.75 | 2 | 1 | 3 | 0 | 4 | 0.75 | 17 | 0.6471 | 5 | 1 | 2 | 1 | 1 | 0 | 4 | 0.25 | 12 | 0.6667 |
| 2_1505951 | 8 | 0.125 | 2 | 0 | 2 | 0 | 6 | 0 | 18 | 0.0556 | 5 | 0.2 | 3 | 0 | 1 | 0 | 3 | 0 | 12 | 0.0833 |
| 2_1506126 | 10 | 0.1 | 5 | 0 | 5 | 0 | 8 | 0.25 | 28 | 0.1071 | 8 | 0.125 | 5 | 0.2 | 1 | 0 | 9 | 0.1111 | 23 | 0.1304 |
| 2_1506127 | 11 | 0.2727 | 6 | 0 | 4 | 0 | 9 | 0.1111 | 30 | 0.1333 | 9 | 0.3333 | 5 | 0.4 | NA | NA | 8 | 0.375 | 22 | 0.3636 |
| 2_1506127 | 10 | 0.6 | 5 | 0.6 | 3 | 0 | 8 | 0.5 | 26 | 0.5 | 8 | 0.625 | 5 | 0.6 | 1 | 1 | 9 | 0.5556 | 23 | 0.6087 |
| 2_1506131 | 9 | 0.1111 | 7 | 0.1429 | 4 | 0 | 10 | 0.4 | 30 | 0.2 | 8 | 0.125 | 5 | 0.2 | 1 | 1 | 7 | 0 | 21 | 0.1429 |
| 2_1506132 | 9 | 0.5556 | 3 | 0 | 3 | 0 | 7 | 0 | 22 | 0.2273 | 5 | 0.2 | 3 | 0 | 1 | 0 | 5 | 0 | 14 | 0.0714 |
| 2_1506132 | 10 | 0.5 | 7 | 0.4286 | 4 | 0.5 | 8 | 0.5 | 29 | 0.4828 | 11 | 0.5455 | 5 | 0.2 | 1 | 0 | 9 | 0.2222 | 26 | 0.3462 |
| 2_1506133 | 9 | 0.2222 | 6 | 0 | 5 | 0 | 14 | 0 | 34 | 0.0588 | 10 | 0.2 | 5 | 0 | 2 | 0 | 7 | 0 | 24 | 0.0833 |
| 2_1506724 | 13 | 0.8462 | 15 | 0.8667 | 22 | 0.6364 | 10 | 0.7 | 60 | 0.75 | 19 | 0.7895 | 17 | 0.6471 | 8 | 0.625 | 17 | 0.7647 | 61 | 0.7213 |
| 2_1506724 | 13 | 0.3077 | 17 | 0 | 20 | 0.1 | 14 | 0.2143 | 64 | 0.1406 | 18 | 0.0556 | 18 | 0.1667 | 6 | 0.1667 | 18 | 0.2222 | 60 | 0.15 |
| 2_1506725 | 11 | 0.7273 | 16 | 0.8125 | 19 | 0.4211 | 10 | 0.5 | 56 | 0.6071 | 18 | 0.5 | 19 | 0.6316 | 8 | 0.5 | 17 | 0.5294 | 62 | 0.5484 |
| 2_1506725 | 14 | 0.2857 | 19 | 0.0526 | 21 | 0.0476 | 13 | 0.0769 | 67 | 0.1045 | 19 | 0.2632 | 20 | 0.1 | 8 | 0 | 18 | 0.2222 | 65 | 0.1692 |
| 2_1506726 | 11 | 0.3636 | 15 | 0.4667 | 17 | 0.5294 | 13 | 0.3077 | 56 | 0.4286 | 15 | 0.4667 | 19 | 0.5789 | 6 | 0.5 | 16 | 0.4375 | 56 | 0.5 |
| 2_1506729 | 14 | 0.1429 | 20 | 0 | 23 | 0.1304 | 14 | 0 | 71 | 0.0704 | 17 | 0.0588 | 19 | 0.1579 | 8 | 0 | 16 | 0.0625 | 60 | 0.0833 |
| 2_1506730 | 16 | 0.125 | 22 | 0.0909 | 23 | 0.0435 | 15 | 0 | 76 | 0.0658 | 20 | 0.1 | 20 | 0.05 | 8 | 0.375 | 18 | 0 | 66 | 0.0909 |
| 2_1506730 | 17 | 0.4118 | 20 | 0.35 | 22 | 0.1818 | 15 | 0.2 | 74 | 0.2838 | 18 | 0.3889 | 20 | 0.4 | 8 | 0.375 | 18 | 0.2222 | 64 | 0.3438 |
| 2_1506731 | 16 | 0.125 | 18 | 0.1667 | 22 | 0.1818 | 13 | 0 | 69 | 0.1304 | 19 | 0.1579 | 19 | 0.0526 | 8 | 0.125 | 16 | 0.3125 | 62 | 0.1613 |
| 2_1506731 | 12 | 0.0833 | 19 | 0.1053 | 19 | 0.1579 | 12 | 0 | 62 | 0.0968 | 14 | 0.2143 | 15 | 0.0667 | 6 | 0.1667 | 15 | 0.1333 | 50 | 0.14 |
| 2_1506734 | 17 | 0.1765 | 20 | 0.3 | 22 | 0.1818 | 15 | 0.2 | 74 | 0.2162 | 22 | 0.0909 | 19 | 0.3158 | 7 | 0.4286 | 21 | 0.2381 | 69 | 0.2319 |
| 2_1506746 | 47 | 0.0426 | 60 | 0.15 | 43 | 0.2093 | 49 | 0.102 | 199 | 0.1256 | 48 | 0.2083 | 49 | 0.1429 | 53 | 0.0943 | 47 | 0.1489 | 197 | 0.1472 |
| 2_1506747 | 35 | 0 | 47 | 0.0851 | 36 | 0.1111 | 33 | 0.0303 | 151 | 0.0596 | 41 | 0.0732 | 45 | 0.0444 | 48 | 0.0625 | 39 | 0.0256 | 173 | 0.052 |
| 2_1506748 | 45 | 0.3333 | 54 | 0.2963 | 39 | 0.2308 | 43 | 0.2558 | 181 | 0.2818 | 43 | 0.3023 | 49 | 0.1837 | 50 | 0.34 | 44 | 0.4318 | 186 | 0.3118 |
| 2_1506748 | 37 | 0.4595 | 48 | 0.625 | 38 | 0.5263 | 35 | 0.4571 | 158 | 0.5253 | 43 | 0.5116 | 42 | 0.3333 | 51 | 0.5098 | 44 | 0.4545 | 180 | 0.4556 |
| 2_1506772 | NA | NA | NA | NA | NA | NA | 1 | 1 | 1 | 1 | 1 | 1 | NA | NA | NA | NA | NA | NA | 1 | 1 |
| 2_1506777 | NA | NA | NA | NA | 1 | 1 | 1 | 0 | 2 | 0.5 | 1 | 1 | 1 | 0 | NA | NA | 4 | 0.25 | 6 | 0.3333 |
| 2_1506781 | NA | NA | NA | NA | 1 | 0 | 1 | 1 | 2 | 0.5 | 1 | 1 | 1 | 0 | NA | NA | 3 | 0.6667 | 5 | 0.6 |
| 2_1507730 | NA | NA | 1 | 1 | NA | NA | NA | NA | 1 | 1 | 1 | 1 | NA | NA | NA | NA | NA | NA | 1 | 1 |
| 2_1507770 | NA | NA | 2 | 0.5 | NA | NA | 1 | 1 | 3 | 0.6667 | NA | NA | NA | NA | NA | NA | 1 | 1 | 1 | 1 |
| 2_1508718 | 3 | 0 | 3 | 0.3333 | 2 | 0 | 2 | 0.5 | 10 | 0.2 | 7 | 0 | 5 | 0.2 | 5 | 0 | 3 | 0 | 20 | 0.05 |
| 2_1508718 | 5 | 0.8 | 3 | 0.6667 | 2 | 1 | 3 | 1 | 13 | 0.8462 | 7 | 1 | 6 | 0.8333 | 6 | 0.5 | 3 | 0.6667 | 22 | 0.7727 |
| 2_1508719 | 5 | 0.2 | 3 | 0 | 1 | 0 | 2 | 0 | 11 | 0.0909 | 6 | 0 | 6 | 0.3333 | 5 | 0 | 2 | 0 | 19 | 0.1053 |
| 2_1508721 | 5 | 0 | 3 | 0.3333 | 2 | 0.5 | 2 | 0 | 12 | 0.1667 | 6 | 0.5 | 3 | 0.3333 | 4 | 0 | 2 | 0.5 | 15 | 0.3333 |
| 2_1508721 | 5 | 0.2 | 3 | 0.3333 | 2 | 0.5 | 3 | 0 | 13 | 0.2308 | 6 | 0.1667 | 5 | 0.2 | 5 | 0.2 | 1 | 1 | 17 | 0.2353 |
| 2_1508723 | 4 | 1 | 2 | 1 | 2 | 1 | 2 | 1 | 10 | 1 | 5 | 1 | 5 | 1 | 4 | 1 | 2 | 1 | 16 | 1 |
| 2_1508723 | 5 | 0 | 3 | 0 | 2 | 0 | 2 | 0.5 | 12 | 0.0833 | 5 | 0 | 6 | 0 | 4 | 0.25 | 3 | 0 | 18 | 0.0556 |

| | | | | | | | | | | | | | | | | | | | | |
|---|---|---|---|---|---|---|---|---|---|---|---|---|---|---|---|---|---|---|---|---|
| 2_1508723 | 5 | 1 | 3 | 1 | 2 | 1 | 1 | 1 | 11 | 1 | 5 | 1 | 6 | 1 | 4 | 1 | 3 | 1 | 18 | 1 |
| 2_1508724 | 3 | 0.6667 | 1 | 0 | 1 | 0 | 2 | 1 | 7 | 0.5714 | 5 | 0.4 | 6 | 0.5 | 3 | 0.3333 | 3 | 0.6667 | 17 | 0.4706 |
| 2_1508725 | 3 | 0.6667 | 2 | 1 | NA | NA | 1 | 1 | 6 | 0.8333 | 3 | 1 | 3 | 1 | 2 | 1 | 2 | 1 | 10 | 1 |
| 2_1508725 | 4 | 0.25 | 2 | 0.5 | NA | NA | 1 | 0 | 7 | 0.2857 | 4 | 0 | 5 | 0.2 | 4 | 0 | 3 | 0 | 16 | 0.0625 |
| 2_1508725 | 2 | 0.5 | 3 | 0.6667 | NA | NA | 1 | 1 | 6 | 0.6667 | 5 | 1 | 4 | 0.75 | 4 | 0.25 | 3 | 0.6667 | 16 | 0.6875 |
| 2_1508726 | 5 | 0.2 | 3 | 0 | NA | NA | 2 | 0 | 10 | 0.1 | 5 | 0 | 6 | 0.1667 | 4 | 0 | 3 | 0 | 18 | 0.0556 |
| 2_1508727 | 5 | 1 | 3 | 1 | NA | NA | 1 | 1 | 9 | 1 | 5 | 1 | 5 | 1 | 3 | 0.6667 | 3 | 1 | 16 | 0.9375 |
| 2_1508727 | 3 | 0.3333 | 3 | 0 | NA | NA | 1 | 0 | 7 | 0.1429 | 3 | 0 | 5 | 0.4 | 2 | 0 | 3 | 0 | 13 | 0.1538 |
| 2_1513363 | 10 | 0 | 11 | 0.0909 | 8 | 0.125 | 17 | 0.1176 | 46 | 0.087 | 10 | 0 | 8 | 0.25 | 11 | 0.0909 | 9 | 0.1111 | 38 | 0.1053 |
| 2_1513530 | 19 | 0.2105 | 14 | 0.2143 | 14 | 0.2143 | 23 | 0.2174 | 70 | 0.2143 | 15 | 0.0667 | 12 | 0.1667 | 17 | 0.1176 | 17 | 0.0588 | 61 | 0.0984 |
| 2_1513531 | 11 | 0.0909 | 14 | 0.4286 | 6 | 0.6667 | 20 | 0.5 | 51 | 0.4118 | 13 | 0.3846 | 9 | 0.4444 | 12 | 0.4167 | 15 | 0.5333 | 49 | 0.449 |
| 2_1513531 | 13 | 0.4615 | 14 | 0.7143 | 10 | 0.4 | 17 | 0.7059 | 54 | 0.5926 | 11 | 0.3636 | 12 | 0.8333 | 14 | 0.5 | 13 | 0.6923 | 50 | 0.6 |
| 2_1514031 | 122 | 0.1311 | 97 | 0.1959 | 77 | 0.1558 | 99 | 0.1919 | 395 | 0.1671 | 70 | 0.1429 | 131 | 0.1985 | 115 | 0.1739 | 112 | 0.1518 | 428 | 0.1706 |
| 2_1514032 | 155 | 0.1677 | 97 | 0.2371 | 101 | 0.2178 | 129 | 0.2481 | 482 | 0.2137 | 86 | 0.1977 | 161 | 0.236 | 139 | 0.2446 | 137 | 0.3066 | 523 | 0.2505 |
| 2_1514326 | 24 | 0.0417 | 37 | 0.1081 | 18 | 0.2222 | 29 | 0.069 | 108 | 0.1019 | 25 | 0.24 | 22 | 0.2727 | 30 | 0.2333 | 18 | 0 | 95 | 0.2 |
| 2_1514326 | 19 | 0.2105 | 37 | 0.0811 | 17 | 0.2941 | 28 | 0.1786 | 101 | 0.1683 | 24 | 0.125 | 21 | 0.0952 | 27 | 0.2222 | 22 | 0.0909 | 94 | 0.1383 |
| 2_1514330 | 16 | 0.125 | 30 | 0.0333 | 15 | 0.1333 | 20 | 0 | 81 | 0.0617 | 21 | 0.1429 | 18 | 0.1111 | 13 | 0 | 17 | 0.0588 | 69 | 0.087 |
| 2_1514334 | 18 | 0.0556 | 33 | 0.1515 | 17 | 0.0588 | 24 | 0.1667 | 92 | 0.1196 | 21 | 0.1429 | 17 | 0.0588 | 20 | 0.1 | 20 | 0.3 | 78 | 0.1538 |
| 2_1514335 | 21 | 0.0476 | 37 | 0.2432 | 20 | 0.2 | 30 | 0.3 | 108 | 0.213 | 26 | 0.4231 | 21 | 0.1429 | 22 | 0.4091 | 21 | 0.2381 | 90 | 0.3111 |
| 2_1514335 | 17 | 0.2941 | 27 | 0.3333 | 14 | 0.2857 | 23 | 0.3478 | 81 | 0.321 | 21 | 0.3333 | 19 | 0.2105 | 17 | 0.4118 | 15 | 0.2667 | 72 | 0.3056 |
| 2_1514336 | 22 | 0.5 | 38 | 0.5526 | 20 | 0.45 | 29 | 0.6552 | 109 | 0.5505 | 24 | 0.5417 | 25 | 0.56 | 23 | 0.6087 | 26 | 0.5 | 98 | 0.551 |
| 2_1514337 | 22 | 0.0455 | 42 | 0.119 | 25 | 0.08 | 37 | 0.1081 | 126 | 0.0952 | 27 | 0.1481 | 28 | 0.1429 | 27 | 0.1481 | 28 | 0.0714 | 110 | 0.1273 |
| 2_1514337 | 21 | 0.0952 | 39 | 0.1282 | 23 | 0.1304 | 36 | 0.2222 | 119 | 0.1513 | 26 | 0.1154 | 26 | 0.0769 | 27 | 0.2222 | 25 | 0.16 | 104 | 0.1442 |
| 2_1514337 | 16 | 0.0625 | 34 | 0.1471 | 19 | 0.0526 | 29 | 0.2414 | 98 | 0.1429 | 21 | 0.1429 | 22 | 0.1818 | 19 | 0.2105 | 21 | 0.2381 | 83 | 0.1928 |
| 2_1514337 | 21 | 0.0476 | 40 | 0.1 | 24 | 0.0833 | 39 | 0.0256 | 124 | 0.0645 | 27 | 0.037 | 27 | 0.1111 | 23 | 0.1304 | 25 | 0.16 | 102 | 0.1078 |
| 2_1514338 | 18 | 0.2778 | 37 | 0.4054 | 24 | 0.4167 | 36 | 0.4444 | 115 | 0.4 | 26 | 0.4615 | 26 | 0.3462 | 22 | 0.4545 | 24 | 0.5417 | 98 | 0.449 |
| 2_1514340 | 19 | 0.0526 | 29 | 0.1724 | 14 | 0.1429 | 36 | 0.25 | 98 | 0.1735 | 18 | 0.2778 | 21 | 0.1429 | 22 | 0.1364 | 20 | 0.05 | 81 | 0.1481 |
| 2_1514344 | 20 | 0.65 | 32 | 0.75 | 15 | 0.8 | 31 | 0.7419 | 98 | 0.7347 | 19 | 0.6842 | 20 | 0.6 | 22 | 0.8182 | 21 | 0.5714 | 82 | 0.6707 |
| 2_1514348 | 16 | 0.0625 | 26 | 0.1154 | 14 | 0.0714 | 24 | 0 | 80 | 0.0625 | 9 | 0.1111 | 18 | 0.2222 | 17 | 0.0588 | 22 | 0.2273 | 66 | 0.1667 |
| 2_1514348 | 18 | 0.1667 | 31 | 0.3226 | 16 | 0.3125 | 24 | 0.1667 | 89 | 0.2472 | 14 | 0.4286 | 18 | 0.2222 | 19 | 0.2632 | 20 | 0.1 | 71 | 0.2394 |
| 2_1514349 | 18 | 0.3333 | 25 | 0.24 | 12 | 0.0833 | 23 | 0.3478 | 78 | 0.2692 | 14 | 0.0714 | 21 | 0.2381 | 18 | 0.1667 | 18 | 0.1111 | 71 | 0.1549 |
| 2_1514350 | 13 | 0.0769 | 29 | 0.1379 | 15 | 0 | 24 | 0.125 | 81 | 0.0988 | 14 | 0.0714 | 21 | 0.0952 | 15 | 0.1333 | 19 | 0.0526 | 69 | 0.087 |
| 2_1514351 | 15 | 0.1333 | 23 | 0.3043 | 13 | 0.3077 | 22 | 0.1818 | 73 | 0.2329 | 14 | 0.0714 | 16 | 0.375 | 18 | 0.1111 | 18 | 0.2778 | 66 | 0.2121 |
| 2_1514355 | 13 | 0.3077 | 21 | 0.0952 | 11 | 0.2727 | 19 | 0.2632 | 64 | 0.2188 | 13 | 0.1538 | 15 | 0.0667 | 12 | 0.1667 | 16 | 0.4375 | 56 | 0.2143 |
| 2_1514356 | 10 | 0.9 | 18 | 1 | 7 | 1 | 14 | 0.9286 | 49 | 0.9592 | 12 | 1 | 16 | 0.8125 | 14 | 0.9286 | 12 | 0.9167 | 54 | 0.9074 |
| 2_1514357 | 13 | 0 | 21 | 0.2857 | 8 | 0.25 | 16 | 0.3125 | 58 | 0.2241 | 10 | 0.2 | 16 | 0.3125 | 12 | 0.0833 | 15 | 0.2 | 53 | 0.2075 |
| 2_1514357 | 13 | 0.6923 | 21 | 0.619 | 10 | 0.7 | 18 | 0.7222 | 62 | 0.6774 | 14 | 0.7143 | 18 | 0.6111 | 15 | 0.5333 | 15 | 0.8667 | 62 | 0.6774 |
| 2_1514357 | 12 | 0.5833 | 21 | 0.5714 | 8 | 0.875 | 18 | 0.5556 | 59 | 0.6102 | 8 | 0.625 | 14 | 0.5 | 13 | 0.4615 | 14 | 0.5714 | 49 | 0.5306 |
| 2_1514358 | 14 | 0.1429 | 17 | 0.1176 | 6 | 0 | 16 | 0 | 53 | 0.0755 | 11 | 0.2727 | 16 | 0 | 12 | 0 | 11 | 0.0909 | 50 | 0.08 |
| 2_1514358 | 10 | 0.7 | 16 | 0.9375 | 10 | 0.9 | 14 | 0.9286 | 50 | 0.88 | 12 | 0.9167 | 16 | 0.9375 | 10 | 0.8 | 13 | 0.8462 | 51 | 0.8824 |
| 2_1514358 | 10 | 0.2 | 15 | 0.1333 | 9 | 0.2222 | 17 | 0.1765 | 51 | 0.1765 | 11 | 0.0909 | 13 | 0.0769 | 12 | 0.0833 | 13 | 0.0769 | 49 | 0.0816 |
| 2_1514360 | 11 | 0.9091 | 15 | 1 | 9 | 1 | 12 | 1 | 47 | 0.9787 | 10 | 1 | 15 | 1 | 13 | 0.9231 | 10 | 1 | 48 | 0.9792 |
| 2_1514362 | 11 | 0.0909 | 17 | 0 | 6 | 0.1667 | 16 | 0.125 | 50 | 0.08 | 11 | 0.2727 | 18 | 0.0556 | 10 | 0.1 | 12 | 0.0833 | 51 | 0.1176 |
| 2_1514363 | 11 | 0.4545 | 16 | 0.6875 | 8 | 0.5 | 15 | 0.7333 | 50 | 0.62 | 11 | 0.2727 | 17 | 0.4706 | 10 | 0.6 | 13 | 0.6154 | 51 | 0.4902 |
| 2_1514363 | 11 | 0.3636 | 17 | 0.4118 | 7 | 0.2857 | 15 | 0.2 | 50 | 0.32 | 11 | 0.4545 | 19 | 0.3684 | 10 | 0.5 | 12 | 0.25 | 52 | 0.3846 |
| 2_1514364 | 11 | 0.0909 | 16 | 0.25 | 7 | 0.1429 | 13 | 0.0769 | 47 | 0.1489 | 9 | 0.1111 | 19 | 0.2105 | 9 | 0.1111 | 11 | 0 | 48 | 0.125 |
| 2_1514391 | NA | NA | 1 | 0 | 1 | 1 | 1 | 0 | 3 | 0.3333 | NA | NA | 1 | 1 | 1 | 0 | NA | NA | 2 | 0.5 |
| 2_1514432 | 1 | 1 | 1 | 0 | 1 | 1 | 2 | 0.5 | 5 | 0.6 | NA | NA | 3 | 0.3333 | 1 | 0 | 2 | 0.5 | 6 | 0.3333 |
| 2_1514432 | 2 | 1 | 2 | 1 | 1 | 1 | 2 | 1 | 7 | 1 | NA | NA | 4 | 1 | 1 | 1 | 2 | 1 | 7 | 1 |
| 2_1514432 | 2 | 0 | 2 | 0.5 | NA | NA | 2 | 1 | 6 | 0.5 | NA | NA | 3 | 1 | 1 | 1 | 2 | 1 | 6 | 1 |
| 2_1514433 | 2 | 0 | 2 | 0 | 1 | 1 | 1 | 1 | 6 | 0.3333 | NA | NA | 3 | 0.3333 | NA | NA | 2 | 0 | 5 | 0.2 |
| 2_1514433 | 1 | 0 | 1 | 0 | 1 | 1 | 1 | 0 | 4 | 0.25 | NA | NA | 4 | 0.25 | 1 | 0 | 2 | 0 | 7 | 0.1429 |
| 2_1514434 | 1 | 0 | 2 | 0.5 | NA | NA | 1 | 0 | 4 | 0.25 | NA | NA | 2 | 0.5 | 1 | 1 | 2 | 0 | 5 | 0.4 |
| 2_1515484 | NA | NA | 2 | 0.5 | 2 | 0.5 | NA | NA | 4 | 0.5 | NA | NA | NA | NA | 2 | 1 | NA | NA | 2 | 1 |
| 2_1515487 | NA | NA | 2 | 1 | 3 | 1 | NA | NA | 5 | 1 | NA | NA | NA | NA | 2 | 1 | NA | NA | 2 | 1 |
| 2_1515488 | NA | NA | 1 | 0 | 3 | 0.6667 | NA | NA | 4 | 0.5 | NA | NA | NA | NA | 2 | 0.5 | NA | NA | 2 | 0.5 |
| 2_1515489 | NA | NA | 2 | 0.5 | 3 | 0.3333 | NA | NA | 5 | 0.4 | NA | NA | NA | NA | 2 | 0.5 | NA | NA | 2 | 0.5 |
| 2_1515589 | 1 | 0 | 1 | 0 | 5 | 0.2 | 1 | 0 | 8 | 0.125 | 1 | 1 | 2 | 1 | 1 | 1 | 1 | 0 | 5 | 0.8 |
| 2_1515593 | 1 | 1 | NA | NA | 4 | 0.5 | 1 | 1 | 6 | 0.6667 | NA | NA | 3 | 0.3333 | 1 | 1 | NA | NA | 4 | 0.5 |
| 2_1515593 | 1 | 0 | NA | NA | 6 | 0 | 1 | 1 | 8 | 0.125 | NA | NA | 3 | 0 | 1 | 0 | 1 | 1 | 5 | 0.2 |
| 2_1515599 | 1 | 1 | NA | NA | 5 | 0.2 | 1 | 0 | 7 | 0.2857 | NA | NA | 3 | 0.3333 | 1 | 0 | 2 | 0 | 6 | 0.1667 |
| 2_1515599 | 1 | 0 | NA | NA | 6 | 0.3333 | NA | NA | 7 | 0.2857 | NA | NA | 1 | 1 | 1 | 0 | 1 | 0 | 3 | 0.3333 |
| 2_1515601 | 1 | 1 | NA | NA | 5 | 1 | 1 | 1 | 7 | 1 | NA | NA | 3 | 1 | NA | NA | NA | NA | 3 | 1 |
| 2_1515601 | 1 | 1 | NA | NA | 4 | 0.75 | 1 | 1 | 6 | 0.8333 | NA | NA | 2 | 0.5 | 1 | 0 | 1 | 0 | 4 | 0.25 |
| 2_1515602 | 6 | 0.1667 | 2 | 0.5 | 12 | 0.1667 | 3 | 0 | 23 | 0.1739 | 5 | 0 | 9 | 0.4444 | 6 | 0 | 3 | 0.3333 | 23 | 0.2174 |

| | | | | | | | | | | | | | | | | | | | | |
|---|---|---|---|---|---|---|---|---|---|---|---|---|---|---|---|---|---|---|---|---|
| 2_15156024 | 5 | 0.4 | 2 | 0 | 13 | 0.1538 | 3 | 0.6667 | 23 | 0.2609 | 4 | 0.5 | 7 | 0.4286 | 6 | 0.1667 | 2 | 0 | 19 | 0.3158 |
| 2_15181138 | 1 | 0 | 1 | 1 | 2 | 1 | NA | NA | 4 | 0.75 | 1 | 1 | 1 | 0 | NA | NA | NA | NA | 2 | 0.5 |
| 2_15181139 | 1 | 0 | 1 | 1 | 2 | 0 | NA | NA | 4 | 0.25 | 2 | 0 | 1 | 1 | NA | NA | 1 | 0 | 4 | 0.25 |
| 2_15181151 | 1 | 1 | 1 | 1 | 2 | 0 | NA | NA | 4 | 0.5 | 1 | 1 | 1 | 0 | NA | NA | 1 | 0 | 3 | 0.3333 |
| 2_15181171 | 1 | 1 | 1 | 1 | 2 | 1 | NA | NA | 4 | 1 | 2 | 1 | 1 | 0 | NA | NA | 1 | 1 | 4 | 0.75 |
| 2_15181174 | 1 | 0 | 1 | 0 | 2 | 0.5 | NA | NA | 4 | 0.25 | 1 | 1 | 1 | 0 | NA | NA | 1 | 1 | 3 | 0.6667 |
| 2_15181177 | 1 | 1 | 1 | 1 | 2 | 1 | NA | NA | 4 | 1 | 1 | 1 | 1 | 1 | NA | NA | 1 | 1 | 3 | 1 |
| 2_15181183 | 1 | 1 | 1 | 0 | 2 | 1 | NA | NA | 4 | 0.75 | 1 | 1 | 1 | 1 | NA | NA | 1 | 1 | 3 | 1 |
| 2_15181186 | 1 | 1 | 1 | 1 | 2 | 1 | NA | NA | 4 | 1 | 1 | 1 | 1 | 1 | NA | NA | 1 | 1 | 3 | 1 |
| 2_15181204 | NA | NA | 1 | 0 | 2 | 1 | NA | NA | 3 | 0.6667 | 1 | 1 | 1 | 0 | NA | NA | 1 | 1 | 3 | 0.6667 |
| 2_15181207 | 1 | 0 | NA | NA | 1 | 1 | NA | NA | 2 | 0.5 | 1 | 1 | 1 | 1 | NA | NA | 1 | 0 | 3 | 0.6667 |
| 2_15181219 | 1 | 1 | 1 | 1 | 2 | 1 | NA | NA | 4 | 1 | 1 | 1 | NA | NA | NA | NA | 1 | 1 | 2 | 1 |
| 2_15192304 | NA | NA | NA | NA | NA | NA | 1 | 1 | 1 | 1 | 1 | 1 | NA | NA | NA | NA | 1 | 0 | 2 | 0.5 |
| 2_15192366 | 2 | 0 | 1 | 0 | NA | NA | 1 | 1 | 4 | 0.25 | 1 | 1 | 3 | 0 | 1 | 0 | 1 | 0 | 6 | 0.1667 |
| 2_15193540 | 5 | 0.8 | 1 | 1 | 1 | 0 | 3 | 1 | 10 | 0.8 | 3 | 1 | 5 | 0.6 | 2 | 1 | 4 | 1 | 14 | 0.8571 |
| 2_15193589 | 6 | 1 | 4 | 1 | 3 | 0.6667 | 9 | 0.7778 | 22 | 0.8636 | 5 | 1 | 10 | 0.9 | 11 | 0.9091 | 8 | 1 | 34 | 0.9412 |
| 2_15193598 | 6 | 0.1667 | 4 | 0 | 3 | 0.3333 | 8 | 0.25 | 21 | 0.1905 | 5 | 0 | 10 | 0.1 | 10 | 0.1 | 9 | 0.2222 | 34 | 0.1176 |
| 2_15193659 | 3 | 0.6667 | 6 | 0.1667 | 3 | 0 | 7 | 0 | 19 | 0.1579 | 6 | 0 | 9 | 0.3333 | 14 | 0 | 7 | 0.1429 | 36 | 0.1111 |
| 2_15193662 | 4 | 0.75 | 4 | 0 | 2 | 0 | 7 | 0.4286 | 17 | 0.3529 | 5 | 0.6 | 9 | 0.6667 | 13 | 0.2308 | 6 | 0.3333 | 33 | 0.4242 |
| 2_15193667 | 4 | 0.75 | 3 | 0 | 3 | 1 | 5 | 0.8 | 15 | 0.6667 | 5 | 0.2 | 8 | 0.5 | 12 | 0.6667 | 5 | 0.8 | 30 | 0.5667 |
| 2_15193677 | 3 | 1 | 6 | 0.1667 | 2 | 1 | 4 | 0.75 | 15 | 0.6 | 4 | 0 | 7 | 0.5714 | 9 | 0.5556 | 7 | 0.4286 | 27 | 0.4444 |
| 2_15193710 | 5 | 0 | 6 | 0 | 3 | 0 | 6 | 0.1667 | 20 | 0.05 | 8 | 0.125 | 12 | 0.0833 | 15 | 0.0667 | 8 | 0 | 43 | 0.0698 |
| 2_15193720 | 3 | 0.3333 | 5 | 0.4 | 3 | 0 | 5 | 0.6 | 16 | 0.375 | 8 | 0.25 | 10 | 0.5 | 9 | 0.5556 | 6 | 0.1667 | 33 | 0.3939 |
| 2_15193737 | 5 | 0.8 | 6 | 0.6667 | 3 | 0.6667 | 4 | 0.5 | 18 | 0.6667 | 8 | 0.875 | 12 | 0.5833 | 14 | 0.5 | 7 | 0.5714 | 41 | 0.6098 |
| 2_15193770 | 4 | 0.25 | 6 | 0.3333 | 3 | 0.3333 | 8 | 0.125 | 21 | 0.2381 | 9 | 0.1111 | 9 | 0.3333 | 18 | 0.3889 | 8 | 0.25 | 44 | 0.2955 |
| 2_15195103 | 6 | 0.3333 | 6 | 0.3333 | 7 | 0 | 6 | 0 | 25 | 0.16 | 10 | 0.1 | 11 | 0.1818 | 14 | 0.0714 | 8 | 0.125 | 43 | 0.1163 |
| 2_15195110 | 6 | 1 | 4 | 0.75 | 7 | 0.8571 | 7 | 0.8571 | 24 | 0.875 | 11 | 0.8182 | 11 | 0.9091 | 15 | 0.9333 | 7 | 0.8571 | 44 | 0.8864 |
| 2_15195117 | 6 | 0.1667 | 7 | 0.4286 | 5 | 0 | 5 | 0 | 23 | 0.1739 | 11 | 0.1818 | 11 | 0.3636 | 13 | 0.0769 | 9 | 0.2222 | 44 | 0.2045 |
| 2_15195137 | 9 | 1 | 5 | 1 | 5 | 1 | 9 | 1 | 28 | 1 | 13 | 0.9231 | 11 | 1 | 15 | 0.9333 | 10 | 0.9 | 49 | 0.9388 |
| 2_15195137 | 9 | 0 | 6 | 0.1667 | 7 | 0 | 10 | 0.2 | 32 | 0.0938 | 13 | 0.0769 | 12 | 0 | 15 | 0.0667 | 11 | 0.0909 | 51 | 0.0588 |
| 2_15195138 | 9 | 0.8889 | 7 | 1 | 7 | 1 | 10 | 1 | 33 | 0.9697 | 14 | 1 | 12 | 1 | 15 | 1 | 11 | 1 | 52 | 1 |
| 2_15195142 | 7 | 1 | 7 | 1 | 5 | 1 | 9 | 0.8889 | 28 | 0.9643 | 10 | 1 | 12 | 1 | 15 | 1 | 10 | 1 | 47 | 1 |
| 2_15195150 | 4 | 0.75 | 3 | 1 | 7 | 1 | 6 | 1 | 20 | 0.95 | 8 | 0.875 | 7 | 0.8571 | 10 | 1 | 6 | 0.8333 | 31 | 0.9032 |
| 2_15195153 | 10 | 0.5 | 7 | 0.2857 | 5 | 0.4 | 8 | 0.625 | 30 | 0.4667 | 13 | 0.6923 | 12 | 0.5833 | 15 | 0.3333 | 12 | 0.5833 | 52 | 0.5385 |
| 2_15195164 | 7 | 0.2857 | 8 | 0.125 | 5 | 0 | 10 | 0.2 | 30 | 0.1667 | 14 | 0.2143 | 12 | 0.0833 | 13 | 0.2308 | 12 | 0.25 | 51 | 0.1961 |
| 2_15195167 | 8 | 0.625 | 9 | 0.5556 | 4 | 0.5 | 9 | 0.4444 | 30 | 0.5333 | 14 | 0.4286 | 10 | 0.8 | 13 | 0.4615 | 9 | 0.2222 | 46 | 0.4783 |
| 2_15195172 | 7 | 0.8571 | 8 | 1 | 7 | 0.8571 | 9 | 1 | 31 | 0.9355 | 11 | 1 | 12 | 0.9167 | 13 | 0.8462 | 12 | 1 | 48 | 0.9375 |
| 2_15195181 | 6 | 0 | 7 | 0 | 7 | 0 | 11 | 0.1818 | 31 | 0.0645 | 10 | 0.1 | 12 | 0.1667 | 14 | 0 | 12 | 0 | 48 | 0.0625 |
| 2_15195183 | 6 | 0.1667 | 5 | 0.2 | 4 | 0 | 9 | 0.3333 | 24 | 0.2083 | 9 | 0.4444 | 9 | 0.3333 | 15 | 0.2 | 10 | 0.5 | 43 | 0.3488 |
| 2_15195185 | 6 | 0.6667 | 5 | 0.8 | 6 | 0 | 5 | 0.6 | 22 | 0.5 | 8 | 0.625 | 7 | 0.8571 | 8 | 0.625 | 7 | 0.5714 | 30 | 0.6667 |
| 2_15195223 | 8 | 0 | 9 | 0.1111 | 9 | 0.1111 | 14 | 0 | 40 | 0.05 | 11 | 0.1818 | 9 | 0 | 18 | 0.0556 | 11 | 0 | 49 | 0.0612 |
| 2_15195261 | 9 | 0.2222 | 6 | 0.1667 | 7 | 0.1429 | 9 | 0.4444 | 31 | 0.2581 | 8 | 0.625 | 6 | 0.1667 | 13 | 0.1538 | 12 | 0.1667 | 39 | 0.2564 |
| 2_15195264 | 7 | 0.2857 | 4 | 0.25 | 7 | 0.1429 | 11 | 0.3636 | 29 | 0.2759 | 8 | 0.5 | 8 | 0.625 | 13 | 0.2308 | 10 | 0.2 | 39 | 0.359 |
| 2_15195268 | 10 | 0.1 | 7 | 0.1429 | 7 | 0 | 11 | 0.0909 | 35 | 0.0857 | 9 | 0.2222 | 11 | 0.1818 | 15 | 0.1333 | 11 | 0.0909 | 46 | 0.1522 |
| 2_15195296 | 11 | 0 | 9 | 0.2222 | 8 | 0 | 13 | 0.0769 | 41 | 0.0732 | 11 | 0.2727 | 10 | 0.1 | 16 | 0.0625 | 13 | 0.0769 | 50 | 0.12 |
| 2_15195309 | 12 | 0 | 9 | 0 | 8 | 0.125 | 15 | 0.2 | 44 | 0.0909 | 8 | 0.25 | 10 | 0.2 | 18 | 0.1667 | 10 | 0 | 46 | 0.1522 |
| 2_15195310 | 12 | 0.1667 | 11 | 0.2727 | 6 | 0 | 13 | 0.3077 | 42 | 0.2143 | 10 | 0.1 | 11 | 0.3636 | 19 | 0.1053 | 11 | 0.1818 | 51 | 0.1765 |
| 2_15195314 | 11 | 0.5455 | 10 | 0.4 | 9 | 0.7778 | 15 | 0.6667 | 45 | 0.6 | 12 | 0.4167 | 13 | 0.6154 | 19 | 0.4737 | 12 | 0.5833 | 56 | 0.5179 |
| 2_15195647 | NA | NA | NA | NA | NA | NA | 3 | 0.3333 | 3 | 0.3333 | NA | NA | NA | NA | 1 | 1 | NA | NA | 1 | 1 |
| 2_15195788 | NA | NA | NA | NA | NA | NA | 2 | 0.5 | 2 | 0.5 | NA | NA | NA | NA | 1 | 1 | NA | NA | 1 | 1 |
| 2_15202718 | NA | NA | NA | NA | NA | NA | 2 | 0.5 | 2 | 0.5 | 1 | 0 | 3 | 0.6667 | 1 | 1 | 2 | 0 | 7 | 0.4286 |
| 2_15212109 | 6 | 0 | 9 | 0.3333 | 6 | 0 | 4 | 0.25 | 25 | 0.16 | 4 | 0.25 | 7 | 0 | 6 | 0.1667 | 6 | 0 | 23 | 0.087 |
| 2_15212122 | 7 | 1 | 6 | 0.8333 | 3 | 0.6667 | 5 | 0.8 | 21 | 0.8571 | 1 | 1 | 2 | 1 | 6 | 0.8333 | 4 | 1 | 13 | 0.9231 |
| 2_15212128 | 9 | 0.6667 | 11 | 0.6364 | 7 | 0.7143 | 6 | 0.5 | 33 | 0.6364 | 4 | 0.5 | 7 | 0.8571 | 8 | 0.75 | 7 | 0.5714 | 26 | 0.6923 |
| 2_15212134 | 12 | 0.0833 | 11 | 0.0909 | 4 | 0.5 | 6 | 0 | 33 | 0.1212 | 4 | 0 | 6 | 0 | 6 | 0 | 8 | 0.25 | 24 | 0.0833 |
| 2_15212155 | 11 | 0.1818 | 10 | 0 | 7 | 0 | 6 | 0.1667 | 34 | 0.0882 | 5 | 0 | 6 | 0 | 6 | 0.1667 | 9 | 0.1111 | 26 | 0.0769 |
| 2_15212319 | 13 | 0.2308 | 12 | 0 | 9 | 0.1111 | 6 | 0.1667 | 40 | 0.125 | 11 | 0.0909 | 7 | 0 | 9 | 0 | 5 | 0.4 | 32 | 0.0938 |
| 2_15212331 | 12 | 0.0833 | 14 | 0.2143 | 8 | 0.125 | 7 | 0.1429 | 41 | 0.1463 | 11 | 0.3636 | 8 | 0.25 | 9 | 0 | 4 | 0 | 32 | 0.1875 |
| 2_15212931 | 2 | 0.5 | NA | NA | 1 | 0 | 1 | 0 | 4 | 0.25 | 1 | 1 | 3 | 0.3333 | 1 | 0 | NA | NA | 5 | 0.4 |
| 2_15215365 | 2 | 0.5 | 11 | 0.5455 | 2 | 1 | 7 | 0.1429 | 22 | 0.4545 | 6 | 0.1667 | 7 | 0.1429 | 8 | 0.625 | 6 | 0.6667 | 27 | 0.4074 |
| 2_15215371 | 1 | 1 | 9 | 0.6667 | 4 | 0.5 | 6 | 0.3333 | 20 | 0.55 | 6 | 0.3333 | 7 | 0.7143 | 8 | 0.75 | 5 | 0.4 | 26 | 0.5769 |
| 2_15215376 | 2 | 0.5 | 12 | 0.3333 | 4 | 0.75 | 8 | 0.25 | 26 | 0.3846 | 6 | 0.1667 | 6 | 0 | 6 | 0 | 4 | 0.5 | 22 | 0.1364 |
| 2_15215378 | 2 | 0.5 | 8 | 0.625 | 3 | 0.3333 | 5 | 0.6 | 18 | 0.5556 | 5 | 0.2 | 5 | 0.6 | 8 | 0.625 | 5 | 0.2 | 23 | 0.4348 |
| 2_15215406 | 3 | 0.3333 | 14 | 0.2143 | 3 | 0.6667 | 8 | 0.375 | 28 | 0.3214 | 6 | 0.1667 | 8 | 0.25 | 9 | 0.4444 | 6 | 0.3333 | 29 | 0.3103 |
| 2_15215444 | 3 | 0 | 13 | 0 | 5 | 0.6 | 12 | 0 | 33 | 0.0909 | 7 | 0.2857 | 9 | 0.1111 | 9 | 0.2222 | 6 | 0 | 31 | 0.1613 |

| | | | | | | | | | | | | | | | | | | | | |
|---|---|---|---|---|---|---|---|---|---|---|---|---|---|---|---|---|---|---|---|---|
| 2_1521548 | 5 | 0.2 | 15 | 0.1333 | 7 | 0.2857 | 13 | 0.1538 | 40 | 0.175 | 8 | 0.5 | 10 | 0.2 | 11 | 0.1818 | 5 | 0 | 34 | 0.2353 |
| 2_1521555 | 9 | 0.2222 | 21 | 0.0952 | 10 | 0 | 16 | 0.0625 | 56 | 0.0893 | 9 | 0 | 10 | 0.1 | 13 | 0 | 7 | 0.1429 | 39 | 0.0513 |
| 2_1521556 | 7 | 0 | 17 | 0.1176 | 7 | 0.4286 | 12 | 0.1667 | 43 | 0.1628 | 9 | 0.1111 | 9 | 0 | 15 | 0.3333 | 6 | 0.1667 | 39 | 0.1795 |
| 2_1521557 | 9 | 0.8889 | 20 | 0.7 | 6 | 0.1667 | 14 | 0.5714 | 49 | 0.6327 | 7 | 0.5714 | 9 | 0.1111 | 14 | 0.2143 | 7 | 0.2857 | 37 | 0.2703 |
| 2_1521584 | 18 | 0.1667 | 13 | 0.2308 | 11 | 0.1818 | 20 | 0.4 | 62 | 0.2581 | 9 | 0.1111 | 27 | 0.1111 | 29 | 0.2759 | 21 | 0.1905 | 86 | 0.186 |
| 2_1521586 | 15 | 0.3333 | 9 | 0.2222 | 9 | 0.5556 | 12 | 0.1667 | 45 | 0.3111 | 5 | 0.2 | 21 | 0.0476 | 19 | 0.2632 | 18 | 0.0556 | 63 | 0.127 |
| 2_1521633 | 1 | 1 | 1 | 0 | NA | NA | 1 | 0 | 3 | 0.3333 | NA | NA | 1 | 0 | 2 | 0.5 | 1 | 0 | 4 | 0.25 |
| 2_1521660 | 4 | 0.25 | NA | NA | NA | NA | 2 | 0 | 6 | 0.1667 | NA | NA | 2 | 0 | 1 | 1 | 2 | 0 | 5 | 0.2 |
| 2_1521661 | 2 | 0.5 | NA | NA | NA | NA | 2 | 0 | 4 | 0.25 | 1 | 1 | 1 | 0 | 1 | 0 | 1 | 0 | 4 | 0.25 |
| 2_1521741 | 10 | 0.2 | 5 | 0 | 4 | 0 | 8 | 0.375 | 27 | 0.1852 | 6 | 0.1667 | 5 | 0 | 5 | 0 | 10 | 0.3 | 26 | 0.1538 |
| 2_1521794 | 1 | 1 | 1 | 1 | NA | NA | NA | NA | 2 | 1 | NA | NA | NA | NA | 1 | 1 | 1 | 1 | 2 | 1 |
| 2_1521794 | 1 | 1 | 1 | 1 | NA | NA | NA | NA | 2 | 1 | NA | NA | NA | NA | 2 | 0 | 1 | 1 | 3 | 0.3333 |
| 2_1521797 | NA | NA | 1 | 1 | NA | NA | NA | NA | 1 | 1 | NA | NA | NA | NA | 1 | 0 | 1 | 1 | 2 | 0.5 |
| 2_1521798 | 1 | 1 | 1 | 1 | NA | NA | NA | NA | 2 | 1 | NA | NA | NA | NA | 1 | 1 | 1 | 1 | 2 | 1 |
| 2_1521800 | 1 | 1 | 1 | 1 | NA | NA | NA | NA | 2 | 1 | NA | NA | NA | NA | 1 | 1 | 1 | 0 | 2 | 0.5 |
| 2_1521803 | 1 | 0 | 1 | 1 | NA | NA | NA | NA | 2 | 0.5 | NA | NA | NA | NA | NA | NA | 1 | 1 | 1 | 1 |
| 2_1522109 | 38 | 0.4211 | 19 | 0.3158 | 19 | 0.6316 | 40 | 0.3 | 116 | 0.3966 | 15 | 0.1333 | 43 | 0.3721 | 42 | 0.5 | 27 | 0.4074 | 127 | 0.3937 |
| 2_1522112 | 32 | 0.1875 | 21 | 0.0952 | 18 | 0.1667 | 37 | 0.1892 | 108 | 0.1667 | 16 | 0.0625 | 33 | 0.1212 | 43 | 0.2093 | 23 | 0.2609 | 115 | 0.1739 |
| 2_1522112 | 33 | 0.5152 | 15 | 0.6 | 17 | 0.7647 | 39 | 0.641 | 104 | 0.6154 | 14 | 0.5 | 34 | 0.6765 | 35 | 0.7429 | 22 | 0.6364 | 105 | 0.6667 |
| 2_1522113 | 36 | 0.1667 | 19 | 0.1053 | 18 | 0.2222 | 38 | 0.0789 | 111 | 0.1351 | 16 | 0.0625 | 38 | 0.1316 | 37 | 0.2162 | 25 | 0.16 | 116 | 0.1552 |
| 2_1522113 | 34 | 0.1176 | 19 | 0.1579 | 16 | 0.1875 | 40 | 0.075 | 109 | 0.1193 | 12 | 0.25 | 31 | 0.2903 | 36 | 0.25 | 22 | 0.0455 | 101 | 0.2178 |
| 2_1522114 | 37 | 0.7838 | 20 | 0.95 | 16 | 0.625 | 36 | 0.8611 | 109 | 0.8165 | 14 | 0.7143 | 36 | 0.8611 | 40 | 0.875 | 21 | 0.9048 | 111 | 0.8559 |
| 2_1522115 | 32 | 0.0625 | 20 | 0.05 | 18 | 0 | 39 | 0.0769 | 109 | 0.055 | 13 | 0.1538 | 36 | 0.0278 | 39 | 0.1282 | 20 | 0 | 108 | 0.0741 |
| 2_1522115 | 32 | 0.1562 | 18 | 0.3333 | 12 | 0.0833 | 28 | 0.1429 | 90 | 0.1778 | 10 | 0.1 | 26 | 0.2308 | 31 | 0.0968 | 16 | 0.1875 | 83 | 0.1566 |
| 2_1522115 | 36 | 0.2222 | 19 | 0.1579 | 15 | 0.3333 | 39 | 0.1282 | 109 | 0.1927 | 13 | 0.2308 | 36 | 0.1389 | 37 | 0.1351 | 20 | 0.1 | 106 | 0.1415 |
| 2_1522368 | NA | NA | 2 | 1 | 3 | 1 | 4 | 0.75 | 9 | 0.8889 | 3 | 1 | 3 | 0.3333 | 5 | 0.4 | 3 | 1 | 14 | 0.6429 |
| 2_1522368 | NA | NA | 1 | 0 | 2 | 0.5 | 4 | 0.25 | 7 | 0.2857 | 3 | 0.6667 | 3 | 0.6667 | 4 | 0.25 | 3 | 1 | 13 | 0.6154 |
| 2_1522369 | 2 | 1 | 3 | 0.6667 | 3 | 0.6667 | 3 | 0.3333 | 11 | 0.6364 | 3 | 0.6667 | 2 | 0.5 | 5 | 0.4 | 3 | 0.3333 | 13 | 0.4615 |
| 2_1522375 | 2 | 0.5 | 3 | 0.3333 | 3 | 0 | 3 | 0 | 11 | 0.1818 | 2 | 0 | 3 | 0.3333 | 1 | 1 | 3 | 0 | 9 | 0.2222 |
| 2_1522387 | 2 | 1 | 1 | 1 | 3 | 1 | 2 | 1 | 8 | 1 | 2 | 1 | NA | NA | 1 | 1 | 2 | 0.5 | 5 | 0.8 |
| 2_1522387 | 1 | 1 | 1 | 1 | 3 | 1 | 1 | 1 | 6 | 1 | 2 | 1 | NA | NA | 1 | 1 | 1 | 1 | 4 | 1 |
| 2_1522387 | 2 | 1 | 1 | 1 | 1 | 0 | 2 | 0.5 | 6 | 0.6667 | 2 | 1 | NA | NA | 1 | 1 | 2 | 1 | 5 | 1 |
| 2_1522388 | 2 | 1 | 1 | 1 | NA | NA | 2 | 0.5 | 5 | 0.8 | 1 | 1 | NA | NA | 1 | 0 | 2 | 0.5 | 4 | 0.5 |
| 2_1522388 | 2 | 1 | NA | NA | 1 | 1 | 2 | 1 | 5 | 1 | 2 | 1 | NA | NA | 1 | 1 | 2 | 1 | 5 | 1 |
| 2_1522391 | 2 | 1 | 1 | 1 | 1 | 0 | 1 | 0 | 5 | 0.6 | 2 | 1 | NA | NA | 1 | 1 | 2 | 0.5 | 5 | 0.8 |
| 2_1522393 | 2 | 0.5 | NA | NA | 1 | 0 | NA | NA | 3 | 0.3333 | 2 | 0.5 | NA | NA | 1 | 0 | 2 | 0.5 | 5 | 0.4 |
| 2_1522393 | 2 | 0.5 | 1 | 0 | 1 | 0 | NA | NA | 4 | 0.25 | 2 | 0 | NA | NA | 1 | 0 | 2 | 1 | 5 | 0.4 |
| 2_1522394 | 2 | 1 | 1 | 0 | 1 | 0 | NA | NA | 4 | 0.5 | 1 | 0 | NA | NA | 1 | 0 | 2 | 0.5 | 4 | 0.25 |
| 2_1522549 | NA | NA | 4 | 1 | 1 | 1 | 1 | 0 | 6 | 0.8333 | 2 | 1 | 1 | 1 | 1 | 1 | NA | NA | 4 | 1 |
| 2_1522550 | NA | NA | 3 | 0.6667 | 1 | 0 | 1 | 0 | 5 | 0.4 | 2 | 1 | 1 | 1 | 2 | 0 | NA | NA | 5 | 0.6 |
| 2_1522552 | NA | NA | 3 | 0.3333 | 1 | 0 | NA | NA | 4 | 0.25 | 1 | 1 | 1 | 1 | NA | NA | NA | NA | 2 | 1 |
| 2_1522554 | NA | NA | 3 | 0.3333 | 1 | 0 | NA | NA | 4 | 0.25 | 2 | 1 | 1 | 1 | 2 | 0 | NA | NA | 5 | 0.6 |
| 2_1522555 | NA | NA | 4 | 0.25 | 1 | 0 | 1 | 0 | 6 | 0.1667 | 2 | 0.5 | 1 | 0 | 2 | 0.5 | NA | NA | 5 | 0.4 |
| 2_1522556 | NA | NA | 4 | 1 | NA | NA | 1 | 1 | 5 | 1 | 2 | 1 | 1 | 1 | 2 | 1 | NA | NA | 5 | 1 |
| 2_1522556 | NA | NA | 3 | 0.3333 | 1 | 0 | 1 | 1 | 5 | 0.4 | 1 | 0 | 1 | 1 | 2 | 0.5 | NA | NA | 4 | 0.5 |
| 2_1522558 | NA | NA | 3 | 1 | 1 | 1 | NA | NA | 4 | 1 | 2 | 0.5 | 1 | 1 | 2 | 1 | NA | NA | 5 | 0.8 |
| 2_1522559 | NA | NA | 1 | 1 | NA | NA | 1 | 1 | 2 | 1 | 2 | 1 | 1 | 1 | 2 | 1 | NA | NA | 5 | 1 |
| 2_1522559 | NA | NA | NA | NA | NA | NA | 1 | 1 | 1 | 1 | 2 | 1 | 1 | 1 | 2 | 1 | NA | NA | 5 | 1 |
| 2_1522561 | NA | NA | 2 | 0.5 | NA | NA | 1 | 1 | 3 | 0.6667 | 2 | 0.5 | 1 | 1 | 2 | 0.5 | NA | NA | 5 | 0.6 |
| 2_1522561 | NA | NA | 2 | 0.5 | NA | NA | 1 | 0 | 3 | 0.3333 | 2 | 0 | 1 | 0 | 1 | 1 | NA | NA | 4 | 0.25 |
| 2_1524949 | NA | NA | NA | NA | NA | NA | 1 | 1 | 1 | 1 | NA | NA | 1 | 1 | NA | NA | NA | NA | 1 | 1 |
| 2_1524950 | NA | NA | NA | NA | NA | NA | 1 | 1 | 1 | 1 | NA | NA | 1 | 1 | NA | NA | NA | NA | 1 | 1 |
| 2_1524951 | NA | NA | NA | NA | NA | NA | 1 | 1 | 1 | 1 | NA | NA | 1 | 1 | NA | NA | NA | NA | 1 | 1 |
| 2_1525292 | NA | NA | 1 | 0 | 4 | 0.75 | 1 | 1 | 6 | 0.6667 | 2 | 0 | 5 | 0.6 | 4 | 0.5 | 1 | 1 | 12 | 0.5 |
| 2_1525294 | NA | NA | 2 | 0.5 | 3 | 1 | 1 | 0 | 6 | 0.6667 | 2 | 0.5 | 7 | 0.8571 | 5 | 0.8 | 2 | 1 | 16 | 0.8125 |
| 2_1525294 | NA | NA | 2 | 0.5 | 3 | 0.3333 | 1 | 1 | 6 | 0.5 | 1 | 1 | 6 | 0.3333 | 4 | 0.5 | 2 | 0.5 | 13 | 0.4615 |
| 2_1525466 | 2 | 0 | 8 | 0.375 | 4 | 0.25 | 8 | 0.25 | 22 | 0.2727 | 6 | 0.1667 | 11 | 0.2727 | 3 | 0 | 5 | 0.2 | 25 | 0.2 |
| 2_1525469 | 4 | 0.75 | 6 | 0.6667 | 4 | 1 | 7 | 1 | 21 | 0.8571 | 8 | 0.875 | 12 | 0.6667 | 4 | 0.75 | 4 | 1 | 28 | 0.7857 |
| 2_1525492 | 2 | 0.5 | NA | NA | NA | NA | NA | NA | 2 | 0.5 | NA | NA | 1 | 1 | 1 | 1 | NA | NA | 2 | 1 |
| 2_1525493 | 2 | 0.5 | NA | NA | NA | NA | NA | NA | 2 | 0.5 | NA | NA | 2 | 0.5 | 1 | 0 | NA | NA | 3 | 0.3333 |
| 2_1525493 | 1 | 1 | NA | NA | NA | NA | 1 | 1 | 2 | 1 | NA | NA | 2 | 0.5 | 1 | 1 | NA | NA | 3 | 0.6667 |
| 2_1525493 | 2 | 1 | NA | NA | NA | NA | NA | NA | 2 | 1 | NA | NA | 2 | 1 | 1 | 1 | NA | NA | 3 | 1 |
| 2_1525495 | 1 | 1 | NA | NA | NA | NA | NA | NA | 1 | 1 | NA | NA | 1 | 1 | 1 | 1 | NA | NA | 2 | 1 |
| 2_1525498 | 2 | 1 | NA | NA | NA | NA | NA | NA | 2 | 1 | NA | NA | 2 | 0 | 1 | 1 | NA | NA | 3 | 0.3333 |
| 2_1525789 | 4 | 0.25 | 6 | 0.3333 | 4 | 0.5 | 5 | 0 | 19 | 0.2632 | 4 | 0.25 | 6 | 0 | 4 | 0 | 3 | 0 | 17 | 0.0588 |

| | | | | | | | | | | | | | | | | | | | | |
|---|---|---|---|---|---|---|---|---|---|---|---|---|---|---|---|---|---|---|---|---|
| 2_1525790 | 7 | 0.1429 | 4 | 0 | 3 | 0 | 5 | 0.2 | 19 | 0.1053 | 6 | 0 | 6 | 0.1667 | 3 | 0.3333 | 3 | 0 | 18 | 0.1111 |
| 2_1525790 | 7 | 0.4286 | 6 | 0.8333 | 3 | 0.3333 | 5 | 0.8 | 21 | 0.619 | 5 | 0.4 | 7 | 0.7143 | 4 | 0.5 | 3 | 0.6667 | 19 | 0.5789 |
| 2_1525791 | 6 | 1 | 4 | 1 | 3 | 1 | 5 | 1 | 18 | 1 | 5 | 1 | 7 | 0.8571 | 4 | 1 | 3 | 1 | 19 | 0.9474 |
| 2_1525791 | 8 | 0 | 5 | 0.2 | 4 | 0.5 | 5 | 0.6 | 22 | 0.2727 | 5 | 0.2 | 7 | 0.2857 | 4 | 0 | 3 | 0 | 19 | 0.1579 |
| 2_1526227 | 14 | 0.4286 | 20 | 0.35 | 15 | 0.5333 | 23 | 0.5217 | 72 | 0.4583 | 22 | 0.5909 | 13 | 0.3846 | 23 | 0.3913 | 17 | 0.5882 | 75 | 0.4933 |
| 2_1526231 | 13 | 0.0769 | 17 | 0.1176 | 13 | 0.3077 | 18 | 0.2222 | 61 | 0.1803 | 14 | 0.2857 | 13 | 0.0769 | 16 | 0.1875 | 17 | 0.1765 | 60 | 0.1833 |
| 2_1526232 | 10 | 0.1 | 14 | 0 | 15 | 0 | 20 | 0.1 | 59 | 0.0508 | 16 | 0.125 | 12 | 0.1667 | 18 | 0 | 15 | 0 | 61 | 0.0656 |
| 2_1526233 | 9 | 0 | 14 | 0.0714 | 9 | 0 | 18 | 0.1111 | 50 | 0.06 | 12 | 0.1667 | 11 | 0.0909 | 18 | 0.0556 | 12 | 0 | 53 | 0.0755 |
| 2_1526237 | 9 | 0.1111 | 12 | 0.1667 | 9 | 0 | 16 | 0.0625 | 46 | 0.087 | 12 | 0.1667 | 8 | 0.125 | 17 | 0.1176 | 14 | 0.0714 | 51 | 0.1176 |
| 2_1526241 | 8 | 0 | 10 | 0 | 10 | 0.2 | 11 | 0 | 39 | 0.0513 | 8 | 0 | 8 | 0 | 13 | 0.0769 | 15 | 0.2 | 44 | 0.0909 |
| 2_1526715 | 10 | 0 | 7 | 0 | 11 | 0.1818 | 13 | 0.1538 | 41 | 0.0976 | 7 | 0.1429 | 7 | 0 | 12 | 0.1667 | 10 | 0.2 | 36 | 0.1389 |
| 2_1526715 | 9 | 0.8889 | 7 | 1 | 6 | 1 | 11 | 0.8182 | 33 | 0.9091 | 8 | 1 | 2 | 1 | 12 | 0.9167 | 10 | 1 | 32 | 0.9688 |
| 2_1526715 | 8 | 0.75 | 8 | 0.25 | 10 | 0.5 | 15 | 0.6 | 41 | 0.5366 | 9 | 0.4444 | 8 | 0.875 | 12 | 0.5 | 12 | 0.8333 | 41 | 0.6585 |
| 2_1526716 | 7 | 0.7143 | 5 | 0.8 | 6 | 0.6667 | 11 | 0.6364 | 29 | 0.6897 | 8 | 0.5 | 6 | 0.3333 | 9 | 0.3333 | 8 | 0.5 | 31 | 0.4194 |
| 2_1526717 | 7 | 0.1429 | 7 | 0 | 8 | 0.25 | 10 | 0 | 32 | 0.0938 | 7 | 0.2857 | 6 | 0 | 10 | 0.2 | 7 | 0 | 30 | 0.1333 |
| 2_1526718 | 6 | 0.6667 | 7 | 0.8571 | 7 | 0.8571 | 13 | 0.6923 | 33 | 0.7576 | 8 | 0.75 | 5 | 0.6 | 9 | 0.4444 | 8 | 0.875 | 30 | 0.6667 |
| 2_1526719 | 5 | 1 | 8 | 1 | 9 | 0.8889 | 12 | 0.75 | 34 | 0.8824 | 6 | 0.8333 | 7 | 1 | 9 | 0.6667 | 8 | 0.875 | 30 | 0.8333 |
| 2_1526719 | 7 | 0.7143 | 8 | 0.875 | 7 | 0.5714 | 11 | 0.6364 | 33 | 0.697 | 4 | 0.75 | 7 | 0.7143 | 10 | 0.6 | 9 | 0.7778 | 30 | 0.7 |
| 2_1527831 | NA | NA | NA | NA | 1 | 1 | NA | NA | 1 | 1 | NA | NA | 1 | 1 | 1 | 1 | NA | NA | 2 | 1 |
| 2_1527846 | NA | NA | NA | NA | 1 | 1 | NA | NA | 1 | 1 | NA | NA | NA | NA | NA | NA | 1 | 1 | 1 | 1 |
| 2_1527848 | 1 | 0 | 1 | 0 | 3 | 1 | NA | NA | 5 | 0.6 | 1 | 1 | NA | NA | 1 | 0 | 1 | 1 | 3 | 0.6667 |
| 2_1527848 | 1 | 1 | 1 | 1 | 1 | 1 | 1 | 1 | 4 | 1 | 1 | 1 | NA | NA | 1 | 1 | 1 | 1 | 3 | 1 |
| 2_1527851 | 1 | 0 | 1 | 0 | 3 | 1 | 1 | 1 | 6 | 0.6667 | 1 | 1 | NA | NA | NA | NA | 1 | 0 | 2 | 0.5 |
| 2_1528511 | 11 | 0.0909 | 6 | 0.3333 | 3 | 0 | 12 | 0.0833 | 32 | 0.125 | 12 | 0.25 | 11 | 0 | 8 | 0.125 | 8 | 0 | 39 | 0.1026 |
| 2_1528513 | 8 | 0 | 7 | 0.1429 | 3 | 0.3333 | 11 | 0.1818 | 29 | 0.1379 | 14 | 0.2143 | 10 | 0.4 | 9 | 0.1111 | 8 | 0 | 41 | 0.1951 |
| 2_1528514 | 6 | 0.3333 | 8 | 0.375 | 3 | 0 | 11 | 0.0909 | 28 | 0.2143 | 14 | 0.2143 | 6 | 0 | 8 | 0.25 | 7 | 0.1429 | 35 | 0.1714 |
| 2_1528515 | 8 | 0.25 | 8 | 0.375 | 3 | 0 | 11 | 0.1818 | 30 | 0.2333 | 15 | 0.0667 | 9 | 0.1111 | 9 | 0.1111 | 8 | 0 | 41 | 0.0732 |
| 2_1529016 | 1 | 1 | NA | NA | 2 | 0.5 | NA | NA | 3 | 0.6667 | 1 | 0 | 2 | 0.5 | 1 | 1 | 1 | 1 | 5 | 0.6 |
| 2_1529932 | NA | NA | NA | NA | 1 | 1 | 1 | 0 | 2 | 0.5 | 1 | 0 | NA | NA | NA | NA | 1 | 1 | 2 | 0.5 |
| 2_1530505 | 2 | 0.5 | 11 | 0 | 4 | 0 | 2 | 0 | 19 | 0.0526 | 4 | 0 | 7 | 0.1429 | 5 | 0 | 3 | 0.3333 | 19 | 0.1053 |
| 2_1530506 | 1 | 0 | 9 | 0.2222 | 8 | 0.125 | 3 | 0.6667 | 21 | 0.2381 | 2 | 0 | 6 | 0.5 | 6 | 0.5 | 4 | 0.25 | 18 | 0.3889 |
| 2_1530506 | 2 | 0.5 | 12 | 0.6667 | 5 | 0.8 | 2 | 1 | 21 | 0.7143 | 4 | 0.25 | 4 | 0.75 | 4 | 0.5 | 5 | 0.4 | 17 | 0.4706 |
| 2_1530507 | 2 | 1 | 11 | 0.9091 | 7 | 1 | 4 | 0.75 | 24 | 0.9167 | 4 | 0.75 | 7 | 1 | 6 | 1 | 4 | 0.75 | 21 | 0.9048 |
| 2_1530507 | 2 | 1 | 11 | 0.2727 | 8 | 0.875 | 4 | 0.75 | 25 | 0.6 | 4 | 0.25 | 6 | 0.3333 | 5 | 0.8 | 4 | 0.75 | 19 | 0.5263 |
| 2_1530511 | 1 | 0 | 8 | 0 | 6 | 0.1667 | 2 | 0.5 | 17 | 0.1176 | 4 | 0 | 6 | 0 | 7 | 0.2857 | 5 | 0 | 22 | 0.0909 |
| 2_1530512 | 2 | 0.5 | 9 | 0.2222 | 7 | 0.4286 | 3 | 0.6667 | 21 | 0.381 | 5 | 0.2 | 7 | 0.4286 | 6 | 0.3333 | 3 | 0.6667 | 21 | 0.381 |
| 2_1530513 | 1 | 1 | 13 | 0.6154 | 8 | 0.625 | 3 | 0.6667 | 25 | 0.64 | 6 | 0.8333 | 7 | 0.5714 | 6 | 0.8333 | 4 | 0.5 | 23 | 0.6957 |
| 2_1530513 | 2 | 0 | 13 | 0.1538 | 6 | 0 | 3 | 0.3333 | 24 | 0.125 | 5 | 0 | 8 | 0.125 | 5 | 0.4 | 6 | 0.3333 | 24 | 0.2083 |
| 2_1530513 | 2 | 0 | 17 | 0.0588 | 8 | 0.125 | 3 | 0 | 30 | 0.0667 | 6 | 0 | 7 | 0 | 8 | 0 | 6 | 0.3333 | 27 | 0.0741 |
| 2_1530513 | 1 | 1 | 15 | 0.8667 | 7 | 1 | 3 | 0.6667 | 26 | 0.8846 | 6 | 1 | 7 | 0.8571 | 7 | 1 | 5 | 1 | 25 | 0.96 |
| 2_1530514 | 2 | 0.5 | 16 | 0.125 | 6 | 0.5 | 4 | 0 | 28 | 0.2143 | 4 | 0 | 7 | 0.4286 | 7 | 0.2857 | 7 | 0.1429 | 25 | 0.24 |
| 2_1530521 | 2 | 0 | 21 | 0.0952 | 8 | 0.125 | 6 | 0.3333 | 37 | 0.1351 | 5 | 0.2 | 10 | 0 | 9 | 0.1111 | 9 | 0.1111 | 33 | 0.0909 |
| 2_1530644 | 3 | 0 | 5 | 0.8 | 3 | 0.3333 | 5 | 0.6 | 16 | 0.5 | 4 | 0.75 | 6 | 0.6667 | 3 | 1 | 5 | 0.4 | 18 | 0.6667 |
| 2_1530645 | 3 | 0.3333 | 3 | 0.6667 | 3 | 0.3333 | 3 | 1 | 12 | 0.5833 | 4 | 0.25 | 5 | 0 | 4 | 0.25 | 5 | 0.4 | 18 | 0.2222 |
| 2_1530646 | 2 | 0.5 | 3 | 0 | 3 | 0 | 3 | 0 | 11 | 0.0909 | 3 | 0 | 6 | 0 | 5 | 0.2 | 6 | 0 | 20 | 0.05 |
| 2_1530646 | 3 | 1 | 3 | 1 | 3 | 0.3333 | 4 | 0.75 | 13 | 0.7692 | 3 | 0.6667 | 3 | 1 | 5 | 1 | 4 | 0.75 | 15 | 0.8667 |
| 2_1530646 | 2 | 0.5 | 2 | 0 | 2 | 0 | 4 | 0.25 | 10 | 0.2 | 3 | 0.3333 | 4 | 0 | 4 | 0 | 5 | 0.2 | 16 | 0.125 |
| 2_1530647 | 2 | 0.5 | 3 | 0.3333 | 3 | 0 | 3 | 0.3333 | 11 | 0.2727 | 4 | 0.25 | 5 | 0.4 | 5 | 0.2 | 5 | 0.2 | 19 | 0.2632 |
| 2_1530648 | 3 | 0.3333 | 3 | 0.3333 | 3 | 0 | 5 | 0.2 | 14 | 0.2143 | 1 | 0 | 5 | 0.2 | 4 | 0 | 6 | 0.1667 | 16 | 0.125 |
| 2_1530650 | 3 | 0 | 2 | 0.5 | 3 | 0 | 4 | 0.5 | 12 | 0.25 | 2 | 0.5 | 2 | 0 | 5 | 0 | 5 | 0.2 | 14 | 0.1429 |
| 2_1530651 | 2 | 1 | 2 | 1 | 3 | 0 | 4 | 1 | 11 | 0.7273 | NA | NA | 3 | 0.6667 | 2 | 1 | 5 | 1 | 10 | 0.9 |
| 2_1530651 | 3 | 0 | 3 | 1 | 3 | 0 | 4 | 0 | 13 | 0.2308 | 3 | 0 | 4 | 0.25 | 4 | 0.25 | 5 | 0.2 | 16 | 0.1875 |
| 2_1530651 | 3 | 0.3333 | 2 | 0.5 | 3 | 0 | 5 | 0.8 | 13 | 0.4615 | 3 | 0.3333 | 3 | 0.3333 | 3 | 0.6667 | 5 | 0.4 | 14 | 0.4286 |
| 2_1530652 | 1 | 0 | 3 | 0.3333 | 2 | 0 | 4 | 1 | 10 | 0.5 | 3 | 0 | 4 | 0 | 3 | 0 | 3 | 0.3333 | 13 | 0.0769 |
| 2_1530653 | 2 | 0 | 3 | 0.3333 | 3 | 0 | 5 | 0 | 13 | 0.0769 | 3 | 0.3333 | 3 | 0.3333 | 4 | 0 | 4 | 0 | 14 | 0.1429 |
| 2_1530653 | 2 | 0 | 3 | 0 | 2 | 0 | 4 | 0.25 | 11 | 0.0909 | 3 | 0 | 2 | 0 | 4 | 0.25 | 3 | 0.3333 | 12 | 0.1667 |
| 2_1530653 | 2 | 0 | 2 | 0 | 3 | 0.3333 | 5 | 0.4 | 12 | 0.25 | 3 | 0.3333 | 3 | 0.3333 | 4 | 0.5 | 4 | 0.5 | 14 | 0.4286 |
| 2_1530656 | 2 | 0 | 3 | 0 | 3 | 0.3333 | 5 | 0.4 | 13 | 0.2308 | 3 | 0.3333 | 2 | 0 | 4 | 0.75 | 4 | 0 | 13 | 0.3077 |
| 2_1530657 | 1 | 0 | 3 | 0.3333 | 2 | 0 | 5 | 0.2 | 11 | 0.1818 | 3 | 0.3333 | 3 | 0 | 3 | 0 | 3 | 0.3333 | 12 | 0.1667 |
| 2_1530657 | 2 | 1 | 1 | 1 | 2 | 1 | 5 | 1 | 10 | 1 | 3 | 1 | 1 | 1 | 3 | 1 | 4 | 0.75 | 11 | 0.9091 |
| 2_1530658 | 2 | 0.5 | 3 | 0.3333 | 2 | 0 | 4 | 0.25 | 11 | 0.2727 | 3 | 0.6667 | 3 | 0 | 3 | 0.6667 | 3 | 0.3333 | 12 | 0.4167 |
| 2_1530658 | 2 | 0 | 3 | 0.3333 | 2 | 1 | 2 | 0 | 9 | 0.3333 | 3 | 0 | 2 | 0 | 3 | 0.3333 | 3 | 0 | 11 | 0.0909 |
| 2_1530658 | 2 | 0.5 | 3 | 1 | 2 | 1 | 4 | 0.75 | 11 | 0.8182 | 2 | 1 | 1 | 1 | 2 | 1 | 4 | 1 | 9 | 1 |
| 2_1530659 | 2 | 0.5 | 3 | 0.6667 | 1 | 1 | 3 | 1 | 9 | 0.7778 | 3 | 0 | 3 | 1 | 3 | 0 | 4 | 0.5 | 13 | 0.3846 |
| 2_1530659 | 1 | 1 | 3 | 0.6667 | 2 | 1 | 3 | 1 | 9 | 0.8889 | 2 | 0.5 | 1 | 1 | 2 | 1 | 4 | 0.5 | 9 | 0.6667 |

| | | | | | | | | | | | | | | | | | | | | |
|---|---|---|---|---|---|---|---|---|---|---|---|---|---|---|---|---|---|---|---|---|
| 2_1530661( | 1 | 1 | 3 | 1 | 2 | 1 | 2 | 1 | 8 | 1 | 1 | 1 | 3 | 1 | 3 | 0.6667 | 4 | 1 | 11 | 0.9091 |
| 2_1530661: | 1 | 1 | 2 | 0 | 1 | 0 | 2 | 0.5 | 6 | 0.3333 | 2 | 1 | 3 | 1 | 3 | 0.3333 | 4 | 0.5 | 12 | 0.6667 |
| 2_1530662( | 1 | 0 | 2 | 1 | 2 | 1 | 2 | 1 | 7 | 0.8571 | 2 | 1 | 2 | 1 | 2 | 1 | 4 | 1 | 10 | 1 |
| 2_1530668( | NA | NA | 1 | 1 | 1 | 0 | 2 | 0.5 | 4 | 0.5 | 2 | 1 | 1 | 1 | 2 | 0.5 | 2 | 0.5 | 7 | 0.7143 |
| 2_1530669! | NA | NA | NA | NA | 1 | 0 | 2 | 0.5 | 3 | 0.3333 | 1 | 0 | 2 | 0 | 1 | 0 | 2 | 0.5 | 6 | 0.1667 |
| 2_1530670( | NA | NA | 1 | 0 | 1 | 1 | 2 | 0 | 4 | 0.25 | 1 | 0 | 2 | 0.5 | 1 | 0 | 1 | 0 | 5 | 0.2 |
| 2_1530701! | 1 | 0 | 4 | 0 | 1 | 1 | 1 | 0 | 7 | 0.1429 | 1 | 1 | 3 | 0 | 1 | 0 | 1 | 1 | 6 | 0.3333 |
| 2_1530709! | 1 | 0 | 5 | 0.4 | NA | NA | 1 | 0 | 7 | 0.2857 | 1 | 1 | 1 | 0 | 1 | 0 | 1 | 0 | 4 | 0.25 |
| 2_1530712( | NA | NA | 3 | 0.3333 | NA | NA | 1 | 0 | 4 | 0.25 | NA | NA | 1 | 1 | 1 | 1 | 1 | 0 | 3 | 0.6667 |
| 2_1530713' | NA | NA | 5 | 0.4 | NA | NA | 1 | 1 | 6 | 0.5 | 1 | 0 | 1 | 0 | 1 | 1 | 1 | 0 | 4 | 0.25 |
| 2_1530717: | NA | NA | 5 | 0.2 | NA | NA | NA | NA | 5 | 0.2 | 1 | 1 | 1 | 0 | 1 | 0 | 1 | 0 | 4 | 0.25 |
| 2_1530719‹ | NA | NA | 5 | 0.2 | NA | NA | 1 | 0 | 6 | 0.1667 | NA | NA | 1 | 0 | 1 | 1 | 1 | 0 | 3 | 0.3333 |
| 2_1530721' | NA | NA | 5 | 1 | NA | NA | 1 | 1 | 6 | 1 | 1 | 1 | 1 | 1 | 1 | 1 | 1 | 1 | 4 | 1 |
| 2_1530722! | NA | NA | 3 | 1 | NA | NA | 1 | 1 | 4 | 1 | 1 | 1 | 1 | 1 | 1 | 1 | 1 | 1 | 4 | 1 |
| 2_1530722( | NA | NA | 3 | 0.6667 | NA | NA | 1 | 0 | 4 | 0.5 | 1 | 1 | 1 | 1 | 1 | 1 | 1 | 1 | 4 | 1 |
| 2_1530723 | NA | NA | 4 | 0.75 | NA | NA | 1 | 1 | 5 | 0.8 | 1 | 1 | 1 | 0 | 1 | 1 | 1 | 1 | 4 | 0.75 |
| 2_1530725 | NA | NA | 1 | 1 | NA | NA | 1 | 1 | 2 | 1 | 1 | 1 | 1 | 1 | 1 | 1 | 1 | 1 | 4 | 1 |
| 2_1530726( | NA | NA | 4 | 1 | NA | NA | 1 | 1 | 5 | 1 | 1 | 1 | 1 | 1 | 1 | 1 | NA | NA | 3 | 1 |
| 2_1530726( | NA | NA | 3 | 0.3333 | NA | NA | 1 | 0 | 4 | 0.25 | 1 | 1 | 1 | 1 | 1 | 1 | NA | NA | 3 | 1 |
| 2_1530726' | NA | NA | 3 | 0.3333 | NA | NA | 1 | 1 | 4 | 0.5 | 1 | 1 | NA | NA | 1 | 1 | NA | NA | 2 | 1 |
| 2_1530727 | NA | NA | 3 | 0.3333 | NA | NA | NA | NA | 3 | 0.3333 | 1 | 1 | 1 | 0 | 1 | 0 | 1 | 1 | 4 | 0.5 |
| 2_1530729! | NA | NA | 4 | 0.25 | NA | NA | 1 | 1 | 5 | 0.4 | 1 | 1 | 1 | 0 | 1 | 0 | 1 | 0 | 4 | 0.25 |
| 2_1530731! | NA | NA | 2 | 1 | NA | NA | 1 | 1 | 3 | 1 | 1 | 1 | 1 | 1 | 1 | 1 | NA | NA | 3 | 1 |
| 2_1530732( | NA | NA | 2 | 0.5 | NA | NA | 1 | 1 | 3 | 0.6667 | 1 | 1 | 1 | 1 | 1 | 0 | NA | NA | 3 | 0.6667 |
| 2_1530734! | NA | NA | 2 | 1 | NA | NA | NA | NA | 2 | 1 | 1 | 1 | 1 | 1 | 1 | 1 | NA | NA | 3 | 1 |
| 2_1530736( | NA | NA | 2 | 1 | NA | NA | NA | NA | 2 | 1 | 1 | 1 | NA | NA | 1 | 1 | NA | NA | 2 | 1 |
| 2_1530738‹ | NA | NA | 2 | 1 | NA | NA | NA | NA | 2 | 1 | NA | NA | NA | NA | 1 | 1 | NA | NA | 1 | 1 |
| 2_1530741( | NA | NA | 1 | 1 | NA | NA | NA | NA | 1 | 1 | 1 | 1 | NA | NA | 1 | 1 | NA | NA | 2 | 1 |
| 2_1530742‹ | NA | NA | 1 | 1 | NA | NA | NA | NA | 1 | 1 | 1 | 1 | NA | NA | 1 | 1 | NA | NA | 2 | 1 |
| 2_1531114( | NA | NA | 1 | 0 | NA | NA | 4 | 0.75 | 5 | 0.6 | 2 | 1 | 2 | 1 | 1 | 0 | NA | NA | 5 | 0.8 |
| 2_1531115 | NA | NA | 1 | 1 | 1 | 1 | 4 | 0.75 | 6 | 0.8333 | 2 | 1 | 2 | 1 | 1 | 0 | 1 | 1 | 6 | 0.8333 |
| 2_1531116( | NA | NA | 1 | 0 | 1 | 0 | 2 | 0.5 | 4 | 0.25 | 1 | 0 | 2 | 1 | 1 | 0 | 1 | 1 | 5 | 0.6 |
| 2_1531116( | NA | NA | 1 | 0 | 1 | 0 | 3 | 0.3333 | 5 | 0.2 | 2 | 0.5 | 2 | 0 | 1 | 0 | 1 | 0 | 6 | 0.1667 |
| 2_1531118( | NA | NA | 1 | 1 | 1 | 1 | 4 | 1 | 6 | 1 | 1 | 1 | 1 | 1 | 1 | 1 | 1 | 1 | 4 | 1 |
| 2_1531119: | NA | NA | 1 | 0 | 1 | 0 | 4 | 0.25 | 6 | 0.1667 | 1 | 0 | 3 | 0.3333 | 1 | 0 | 1 | 1 | 6 | 0.3333 |
| 2_1531119: | NA | NA | NA | NA | 1 | 1 | 2 | 1 | 3 | 1 | 1 | 0 | 3 | 0.6667 | 1 | 1 | 1 | 1 | 6 | 0.6667 |
| 2_1531119‹ | NA | NA | 1 | 0 | 1 | 1 | 3 | 0.6667 | 5 | 0.6 | 1 | 0 | 3 | 0.6667 | NA | NA | 1 | 1 | 5 | 0.6 |
| 2_1531119 | NA | NA | NA | NA | 1 | 1 | 4 | 1 | 5 | 1 | NA | NA | 1 | 1 | 1 | 0 | 1 | 1 | 3 | 0.6667 |
| 2_1531120: | NA | NA | 1 | 1 | 1 | 1 | 4 | 1 | 6 | 1 | NA | NA | 3 | 0.6667 | 1 | 0 | NA | NA | 4 | 0.5 |
| 2_1531120: | NA | NA | 1 | 0 | NA | NA | 2 | 0.5 | 3 | 0.3333 | 1 | 1 | 2 | 1 | 1 | 1 | 1 | 1 | 5 | 1 |
| 2_1531120( | NA | NA | 1 | 1 | 1 | 1 | 3 | 0.6667 | 5 | 0.8 | 1 | 1 | 2 | 1 | 1 | 1 | NA | NA | 4 | 1 |
| 2_1531120! | NA | NA | NA | NA | 1 | 0 | 2 | 0.5 | 3 | 0.3333 | 1 | 1 | 3 | 0 | 1 | 0 | 1 | 0 | 6 | 0.1667 |
| 2_1531584( | NA | NA | NA | NA | 1 | 1 | NA | NA | 1 | 1 | NA | NA | 1 | 1 | NA | NA | NA | NA | 1 | 1 |
| 2_1531590( | NA | NA | NA | NA | 1 | 1 | NA | NA | 1 | 1 | NA | NA | 1 | 1 | 1 | 1 | NA | NA | 2 | 1 |
| 2_1531594( | 2 | 0 | 1 | 0 | 1 | 1 | NA | NA | 4 | 0.25 | NA | NA | 1 | 0 | 2 | 0.5 | NA | NA | 3 | 0.3333 |
| 2_1531721‹ | 2 | 0 | 2 | 0.5 | 1 | 0 | 1 | 0 | 6 | 0.1667 | NA | NA | 2 | 0.5 | 2 | 0 | NA | NA | 4 | 0.25 |
| 2_1531722: | 2 | 0 | 2 | 0 | 1 | 1 | NA | NA | 5 | 0.2 | NA | NA | NA | NA | 2 | 1 | NA | NA | 2 | 1 |
| 2_1531724( | 2 | 1 | 2 | 1 | 1 | 1 | 2 | 1 | 7 | 1 | NA | NA | 2 | 1 | 3 | 0.6667 | 1 | 1 | 6 | 0.8333 |
| 2_1531726! | 2 | 1 | 2 | 0.5 | 1 | 0 | 2 | 0.5 | 7 | 0.5714 | 1 | 1 | 4 | 0.5 | 3 | 0.3333 | 2 | 1 | 10 | 0.6 |
| 2_1531727' | 2 | 1 | 2 | 1 | 2 | 1 | 2 | 1 | 8 | 1 | 1 | 1 | 3 | 0 | 3 | 0 | 1 | 0 | 8 | 0.125 |
| 2_1531735 | 3 | 0 | 3 | 0.3333 | 4 | 0.25 | 3 | 0 | 13 | 0.1538 | 2 | 0.5 | 4 | 0.25 | 4 | 0.25 | 4 | 0 | 14 | 0.2143 |
| 2_1531737( | NA | NA | NA | NA | 3 | 0.3333 | 2 | 0 | 5 | 0.2 | 3 | 0.3333 | 2 | 0.5 | 2 | 0.5 | 1 | 0 | 8 | 0.375 |
| 2_1531739! | 4 | 0 | 3 | 0.3333 | 4 | 0 | 3 | 0 | 14 | 0.0714 | 2 | 0 | 4 | 0 | 2 | 0 | 4 | 0.25 | 12 | 0.0833 |
| 2_1531741' | 3 | 0.3333 | 3 | 0 | 5 | 0.4 | 3 | 0 | 14 | 0.2143 | 3 | 0.6667 | 5 | 0.6 | 4 | 0.5 | 6 | 0.3333 | 18 | 0.5 |
| 2_1531742' | 4 | 0.5 | 3 | 0.3333 | 7 | 0.5714 | 3 | 0.3333 | 17 | 0.4706 | 3 | 0.6667 | 4 | 0.75 | 4 | 0.25 | 5 | 0.2 | 16 | 0.4375 |
| 2_1531744( | 4 | 0.25 | 3 | 0.3333 | 4 | 1 | 2 | 0.5 | 13 | 0.5385 | 2 | 0.5 | 4 | 0.75 | 4 | 1 | 6 | 0.5 | 16 | 0.6875 |
| 2_1531745! | 5 | 1 | 2 | 1 | 5 | 1 | 3 | 1 | 15 | 1 | 2 | 1 | 4 | 1 | 4 | 1 | 6 | 1 | 16 | 1 |
| 2_1531746 | 3 | 1 | 1 | 1 | 6 | 0.5 | 1 | 1 | 11 | 0.7273 | 2 | 1 | 4 | 0.75 | 4 | 1 | 6 | 0.6667 | 16 | 0.8125 |
| 2_1531746( | 6 | 0.6667 | 4 | 0.5 | 7 | 0.5714 | 4 | 0.25 | 21 | 0.5238 | 2 | 0.5 | 4 | 0 | 4 | 0.75 | 6 | 0.3333 | 16 | 0.375 |
| 2_1531748: | 7 | 0.8571 | 5 | 0.8 | 6 | 0.5 | 6 | 1 | 24 | 0.7917 | 3 | 0.6667 | 3 | 0.6667 | 4 | 0.75 | 6 | 0.6667 | 16 | 0.6875 |
| 2_1531749: | 8 | 0.125 | 4 | 0.5 | 5 | 0.4 | 4 | 0 | 21 | 0.2381 | 2 | 0 | 4 | 0.5 | 5 | 0.2 | 6 | 0.5 | 17 | 0.3529 |
| 2_1531751! | 6 | 1 | 4 | 1 | 6 | 0.8333 | 5 | 1 | 21 | 0.9524 | 3 | 0.6667 | 2 | 1 | 7 | 1 | 6 | 1 | 18 | 0.9444 |
| 2_1531838( | 3 | 0 | NA | NA | 1 | 1 | NA | NA | 4 | 0.25 | 1 | 1 | 1 | 0 | 2 | 0.5 | 1 | 0 | 5 | 0.4 |
| 2_1531838' | 3 | 0.6667 | NA | NA | 1 | 0 | 1 | 1 | 5 | 0.6 | 1 | 0 | 2 | 0 | 2 | 0.5 | 1 | 1 | 6 | 0.3333 |
| 2_1531839 | 3 | 0 | NA | NA | 1 | 0 | 1 | 1 | 5 | 0.2 | 1 | 0 | 2 | 0 | 2 | 0 | 1 | 1 | 6 | 0.1667 |

| | | | | | | | | | | | | | | | | | | | | |
|---|---|---|---|---|---|---|---|---|---|---|---|---|---|---|---|---|---|---|---|---|
| 2_1531841 | 1 | 1 | NA | NA | 1 | 1 | NA | NA | 2 | 1 | 1 | 1 | 1 | 1 | 2 | 1 | 1 | 1 | 5 | 1 |
| 2_1531843 | 3 | 1 | NA | NA | 1 | 0 | 1 | 1 | 5 | 0.8 | 1 | 1 | 2 | 1 | 2 | 1 | NA | NA | 5 | 1 |
| 2_1531844 | 2 | 1 | NA | NA | 1 | 1 | 1 | 1 | 4 | 1 | 1 | 1 | 2 | 1 | 2 | 1 | 1 | 1 | 6 | 1 |
| 2_1531845 | 3 | 0.3333 | NA | NA | 1 | 0 | NA | NA | 4 | 0.25 | 1 | 0 | 2 | 0.5 | 2 | 0.5 | 1 | 0 | 6 | 0.3333 |
| 2_1531846 | 3 | 0.6667 | NA | NA | 1 | 1 | 1 | 1 | 5 | 0.8 | 1 | 1 | 2 | 1 | 2 | 1 | 1 | 1 | 6 | 1 |
| 2_1531846 | 3 | 0 | NA | NA | 1 | 1 | 1 | 0 | 5 | 0.2 | 1 | 1 | 2 | 0.5 | 2 | 0.5 | 1 | 1 | 6 | 0.6667 |
| 2_1531846 | 3 | 0.6667 | NA | NA | 1 | 0 | 1 | 0 | 5 | 0.4 | 1 | 0 | 2 | 0.5 | 2 | 0 | 1 | 1 | 6 | 0.3333 |
| 2_1531847 | 3 | 0.3333 | NA | NA | 1 | 1 | 1 | 1 | 5 | 0.6 | 1 | 1 | 2 | 0.5 | 2 | 1 | 1 | 1 | 6 | 0.8333 |
| 2_1532394 | NA | NA | NA | NA | 1 | 1 | 2 | 1 | 3 | 1 | 1 | 1 | 1 | 1 | 1 | 1 | NA | NA | 3 | 1 |
| 2_1532396 | NA | NA | 1 | 1 | 1 | 0 | 2 | 0.5 | 4 | 0.5 | 2 | 0.5 | 2 | 0.5 | 1 | 1 | NA | NA | 5 | 0.6 |
| 2_1532412 | NA | NA | 1 | 1 | 1 | 1 | 2 | 1 | 4 | 1 | 1 | 1 | 2 | 1 | 1 | 1 | 1 | 1 | 5 | 1 |
| 2_1532412 | NA | NA | 1 | 0 | 1 | 1 | 2 | 1 | 4 | 0.75 | 1 | 0 | 2 | 0.5 | 1 | 1 | 1 | 0 | 5 | 0.4 |
| 2_1532412 | NA | NA | NA | NA | 1 | 0 | 2 | 1 | 3 | 0.6667 | 1 | 0 | 2 | 0.5 | NA | NA | 1 | 1 | 4 | 0.5 |
| 2_1532414 | NA | NA | 1 | 0 | 1 | 1 | 3 | 0.6667 | 5 | 0.6 | 1 | 1 | 2 | 0 | 1 | 1 | 1 | 1 | 5 | 0.6 |
| 2_1532417 | 1 | 1 | 1 | 1 | 1 | 1 | 2 | 1 | 5 | 1 | NA | NA | 2 | 1 | NA | NA | 1 | 1 | 3 | 1 |
| 2_1532418 | 1 | 1 | 1 | 1 | 2 | 0.5 | 2 | 1 | 6 | 0.8333 | 3 | 0.6667 | 2 | 0 | 1 | 0 | 2 | 0.5 | 8 | 0.375 |
| 2_1532421 | 1 | 0 | 1 | 0 | 2 | 0.5 | 5 | 0 | 9 | 0.1111 | 2 | 0 | 3 | 0.3333 | 1 | 0 | 1 | 1 | 7 | 0.2857 |
| 2_1532422 | 1 | 1 | 1 | 1 | 2 | 0.5 | 4 | 1 | 8 | 0.875 | 2 | 1 | 3 | 1 | NA | NA | 1 | 1 | 6 | 1 |
| 2_1532422 | 1 | 0 | 2 | 0.5 | 1 | 1 | 5 | 0.2 | 9 | 0.3333 | NA | NA | 2 | 0.5 | 1 | 0 | 1 | 0 | 4 | 0.25 |
| 2_1532423 | 1 | 0 | 2 | 0 | 2 | 0 | 5 | 0.6 | 10 | 0.3 | 2 | 0.5 | 4 | 0 | 1 | 1 | 2 | 0.5 | 9 | 0.3333 |
| 2_1532423 | 1 | 1 | 2 | 1 | 2 | 0.5 | 4 | 1 | 9 | 0.8889 | 2 | 0.5 | 3 | 1 | 1 | 0 | 1 | 1 | 7 | 0.7143 |
| 2_1532424 | 1 | 0 | 2 | 0 | 3 | 0.3333 | 5 | 0.2 | 11 | 0.1818 | 2 | 0.5 | 4 | 0.5 | 1 | 0 | 2 | 0.5 | 9 | 0.4444 |
| 2_1532424 | 1 | 0 | 2 | 0.5 | 4 | 0.25 | 5 | 0.4 | 12 | 0.3333 | 3 | 0.3333 | 4 | 0.25 | 1 | 0 | 1 | 0 | 9 | 0.2222 |
| 2_1532424 | 1 | 0 | 2 | 0.5 | 4 | 0 | 4 | 0.5 | 11 | 0.2727 | 3 | 0.3333 | 4 | 0.5 | 1 | 0 | 1 | 0 | 9 | 0.3333 |
| 2_1532425 | 1 | 0 | 2 | 0.5 | 4 | 0.5 | 5 | 0 | 12 | 0.25 | 3 | 0.3333 | 3 | 0.6667 | 1 | 0 | 2 | 0.5 | 9 | 0.4444 |
| 2_1532425 | 1 | 0 | 2 | 1 | 4 | 1 | 3 | 1 | 10 | 0.9 | 1 | 0 | 4 | 1 | 1 | 1 | 1 | 1 | 7 | 0.8571 |
| 2_1532426 | 1 | 1 | 2 | 0 | 4 | 0 | 6 | 0.3333 | 13 | 0.2308 | 3 | 0.3333 | 4 | 0.25 | 1 | 0 | 1 | 0 | 9 | 0.2222 |
| 2_1532430 | 1 | 1 | 2 | 0 | 4 | 0.5 | 5 | 0.6 | 12 | 0.5 | 2 | 1 | 4 | 0.25 | 1 | 0 | 1 | 1 | 8 | 0.5 |
| 2_1532430 | 1 | 1 | NA | NA | 2 | 0.5 | 3 | 0 | 6 | 0.3333 | 1 | 1 | 2 | 0.5 | 1 | 1 | NA | NA | 4 | 0.75 |
| 2_1532432 | 1 | 0 | 2 | 0.5 | 4 | 0.5 | 6 | 0 | 13 | 0.2308 | 2 | 0 | 3 | 0 | 1 | 1 | NA | NA | 6 | 0.1667 |
| 2_1532432 | 1 | 0 | 2 | 0.5 | 4 | 0.25 | 6 | 0.6667 | 13 | 0.4615 | 3 | 0.3333 | 4 | 0.5 | 1 | 0 | 1 | 0 | 9 | 0.3333 |
| 2_1532432 | 1 | 0 | 1 | 1 | 3 | 0 | 6 | 0.3333 | 11 | 0.2727 | 1 | 1 | 4 | 0.75 | NA | NA | 1 | 0 | 6 | 0.6667 |
| 2_1532432 | 1 | 1 | 2 | 0 | 3 | 1 | 6 | 0 | 12 | 0.3333 | 2 | 1 | 2 | 0.5 | 1 | 0 | 1 | 0 | 6 | 0.5 |
| 2_1532433 | NA | NA | NA | NA | 4 | 0.5 | 4 | 1 | 8 | 0.75 | 2 | 0.5 | 2 | 1 | 1 | 1 | NA | NA | 5 | 0.8 |
| 2_1532434 | 1 | 0 | 2 | 0 | 4 | 0 | 6 | 0.1667 | 13 | 0.0769 | 1 | 0 | 3 | 0 | 1 | 0 | 1 | 1 | 6 | 0.1667 |
| 2_1532434 | 2 | 0.5 | 2 | 0 | 4 | 0 | 5 | 0.2 | 13 | 0.1538 | 2 | 0 | 4 | 0.25 | 1 | 0 | 1 | 0 | 8 | 0.125 |
| 2_1532434 | 2 | 0.5 | 2 | 0.5 | 4 | 0.5 | 6 | 0.8333 | 14 | 0.6429 | 2 | 0.5 | 3 | 1 | 1 | 0 | NA | NA | 6 | 0.6667 |
| 2_1532435 | 2 | 0.5 | 2 | 0.5 | 4 | 0.5 | 6 | 0.1667 | 14 | 0.3571 | 2 | 0 | 3 | 0.3333 | 1 | 0 | 1 | 0 | 7 | 0.1429 |
| 2_1532435 | 2 | 0.5 | 2 | 1 | 4 | 1 | 6 | 0.8333 | 14 | 0.8571 | 3 | 1 | 3 | 1 | 1 | 1 | 1 | 0 | 8 | 0.875 |
| 2_1532437 | 2 | 0 | 2 | 0 | 4 | 0.5 | 6 | 0.3333 | 14 | 0.2857 | 3 | 0.3333 | 4 | 0.25 | 2 | 0.5 | 1 | 0 | 10 | 0.3 |
| 2_1532438 | 1 | 0 | 2 | 0 | 4 | 0.25 | 3 | 0.3333 | 10 | 0.2 | 3 | 0 | 4 | 0 | 2 | 0 | 2 | 0.5 | 11 | 0.0909 |
| 2_1532439 | 2 | 0.5 | 1 | 0 | 4 | 0.5 | 6 | 0.3333 | 13 | 0.3846 | 2 | 0.5 | 4 | 0.5 | 2 | 1 | NA | NA | 8 | 0.625 |
| 2_1532439 | 2 | 1 | 2 | 1 | 3 | 0.6667 | 6 | 0.8333 | 13 | 0.8462 | 4 | 1 | 4 | 0.75 | 1 | 1 | 1 | 1 | 10 | 0.9 |
| 2_1532440 | NA | NA | 2 | 0 | 3 | 0.3333 | 4 | 0.25 | 9 | 0.2222 | 3 | 0.6667 | 2 | 0.5 | 2 | 0.5 | 1 | 1 | 8 | 0.625 |
| 2_1532442 | NA | NA | 2 | 1 | 2 | 0.5 | 3 | 1 | 7 | 0.8571 | 4 | 1 | 2 | 1 | 2 | 1 | 1 | 1 | 9 | 1 |
| 2_1532443 | NA | NA | 3 | 0 | 3 | 0 | 4 | 0.5 | 10 | 0.2 | 2 | 0 | 4 | 0 | 2 | 0 | 1 | 1 | 9 | 0.1111 |
| 2_1532443 | NA | NA | 3 | 0 | 3 | 0.3333 | 5 | 0 | 11 | 0.0909 | 3 | 0.3333 | 3 | 0 | 2 | 0.5 | 1 | 1 | 9 | 0.3333 |
| 2_1532445 | 1 | 1 | 3 | 0.6667 | 2 | 0.5 | 4 | 0.25 | 10 | 0.5 | 3 | 1 | 1 | 1 | 1 | 1 | 2 | 0 | 7 | 0.7143 |
| 2_1532446 | 1 | 1 | 2 | 1 | 3 | 0.6667 | 4 | 1 | 10 | 0.9 | 3 | 1 | 1 | 1 | 1 | 1 | 1 | 1 | 6 | 1 |
| 2_1532446 | 1 | 1 | 3 | 0 | 3 | 0 | 5 | 0 | 12 | 0.0833 | 3 | 0 | 4 | 0.25 | 3 | 0 | 1 | 1 | 11 | 0.1818 |
| 2_1532448 | 2 | 0 | 3 | 0 | 2 | 0 | 4 | 0.25 | 11 | 0.0909 | 5 | 0.2 | 4 | 0.5 | 4 | 0.25 | 3 | 0.3333 | 16 | 0.3125 |
| 2_1532449 | 2 | 0 | 3 | 0 | 3 | 0 | 4 | 0.25 | 12 | 0.0833 | 5 | 0 | 4 | 0 | 4 | 0.25 | 1 | 0 | 14 | 0.0714 |
| 2_1532450 | 3 | 0.3333 | NA | NA | 2 | 0 | 4 | 0 | 9 | 0.1111 | 4 | 0.25 | 4 | 0 | 4 | 0.75 | 2 | 0.5 | 14 | 0.3571 |
| 2_1532454 | 2 | 0 | NA | NA | 3 | 0.3333 | 4 | 0 | 9 | 0.1111 | 4 | 0 | 4 | 0 | 4 | 0 | 1 | 1 | 13 | 0.0769 |
| 2_1532507 | 3 | 0.6667 | 6 | 0.1667 | 8 | 0.125 | 4 | 0 | 21 | 0.1905 | 1 | 0 | 6 | 0.1667 | 4 | 0.25 | 5 | 0.4 | 16 | 0.25 |
| 2_1532507 | 1 | 0 | 5 | 0.8 | 6 | 0.3333 | 4 | 0.25 | 16 | 0.4375 | 1 | 0 | 5 | 0 | 3 | 0.3333 | 4 | 0.5 | 13 | 0.2308 |
| 2_1532531 | 2 | 0.5 | 2 | 0 | 2 | 0 | 3 | 0.6667 | 9 | 0.3333 | 2 | 0.5 | 2 | 0.5 | 3 | 0.3333 | 4 | 0.5 | 11 | 0.4545 |
| 2_1532533 | 3 | 0.6667 | 3 | 0.6667 | 1 | 1 | 2 | 0.5 | 9 | 0.6667 | 1 | 0 | 1 | 0 | 1 | 1 | 4 | 1 | 7 | 0.7143 |
| 2_1532533 | 3 | 1 | 3 | 1 | 1 | 1 | 2 | 1 | 9 | 1 | 1 | 1 | 1 | 1 | 1 | 1 | 4 | 1 | 7 | 1 |
| 2_1532534 | 3 | 1 | 3 | 0 | 1 | 0 | 2 | 0 | 9 | 0.3333 | 1 | 1 | 1 | 0 | 1 | 0 | 4 | 0.5 | 7 | 0.4286 |
| 2_1532535 | 1 | 1 | NA | NA | 1 | 1 | 2 | 0 | 4 | 0.5 | NA | NA | 1 | 1 | 2 | 0 | 3 | 0.3333 | 6 | 0.3333 |
| 2_1532535 | 1 | 1 | NA | NA | 1 | 1 | 2 | 1 | 4 | 1 | 1 | 1 | 1 | 0 | 1 | 1 | 3 | 0.6667 | 6 | 0.6667 |
| 2_1534608 | 10 | 0.1 | 9 | 0.3333 | 8 | 0.125 | 8 | 0.125 | 35 | 0.1714 | 6 | 0.6667 | 4 | 0.75 | 5 | 0.8 | 7 | 0.5714 | 22 | 0.6818 |
| 2_1534610 | 10 | 0.1 | 8 | 0 | 6 | 0.3333 | 7 | 0.1429 | 31 | 0.129 | 3 | 0 | 5 | 0.2 | 5 | 0 | 3 | 0 | 16 | 0.0625 |
| 2_1534611 | 10 | 0.9 | 10 | 0.7 | 7 | 1 | 9 | 1 | 36 | 0.8889 | 6 | 0.8333 | 5 | 1 | 6 | 1 | 7 | 0.8571 | 24 | 0.9167 |

| | | | | | | | | | | | | | | | | | | | | |
|---|---|---|---|---|---|---|---|---|---|---|---|---|---|---|---|---|---|---|---|---|
| 2_1534611 | 9 | 0.3333 | 11 | 0.4545 | 8 | 0 | 7 | 0.2857 | 35 | 0.2857 | 7 | 0 | 5 | 0 | 6 | 0.1667 | 6 | 0.3333 | 24 | 0.125 |
| 2_1534612 | 9 | 0.1111 | 10 | 0.1 | 7 | 0 | 6 | 0 | 32 | 0.0625 | 6 | 0.1667 | 2 | 0.5 | 6 | 0 | 7 | 0.1429 | 21 | 0.1429 |
| 2_1535998 | 12 | 0 | 7 | 0.1429 | 6 | 0 | 10 | 0.1 | 35 | 0.0571 | 6 | 0 | 7 | 0 | 15 | 0.1333 | 9 | 0 | 37 | 0.0541 |
| 2_1536089 | 13 | 0.3077 | 8 | 0.25 | 7 | 0 | 9 | 0.1111 | 37 | 0.1892 | 6 | 0.1667 | 8 | 0.25 | 16 | 0.125 | 8 | 0.25 | 38 | 0.1842 |
| 2_1536091 | 10 | 0.4 | 9 | 0.2222 | 6 | 0.3333 | 6 | 0.5 | 31 | 0.3548 | 7 | 0.7143 | 7 | 0.7143 | 12 | 0.5833 | 9 | 0.4444 | 35 | 0.6 |
| 2_1536091 | 17 | 0.8235 | 10 | 0.8 | 7 | 0.7143 | 11 | 0.5455 | 45 | 0.7333 | 8 | 0.875 | 8 | 1 | 17 | 0.8824 | 9 | 0.7778 | 42 | 0.881 |
| 2_1536093 | 8 | 0.125 | 4 | 0 | 5 | 0.2 | 4 | 0 | 21 | 0.0952 | 4 | 0 | 8 | 0 | 9 | 0.1111 | 9 | 0.1111 | 30 | 0.0667 |
| 2_1536098 | 5 | 0 | 3 | 0 | 3 | 1 | 5 | 0 | 16 | 0.1875 | 4 | 0.25 | 5 | 0 | 6 | 0.1667 | 3 | 0.3333 | 18 | 0.1667 |
| 2_1536100 | 4 | 0 | 2 | 0 | 5 | 0 | 6 | 0.1667 | 17 | 0.0588 | 6 | 0 | 5 | 0.2 | 6 | 0.1667 | 7 | 0.1429 | 24 | 0.125 |
| 2_1536149 | 3 | 0 | 3 | 0.3333 | 1 | 0 | 1 | 0 | 8 | 0.125 | 2 | 0 | 2 | 0 | 4 | 0 | 2 | 1 | 10 | 0.2 |
| 2_1541774 | NA | NA | 2 | 0.5 | NA | NA | NA | NA | 2 | 0.5 | 2 | 0 | 1 | 0 | 1 | 0 | 2 | 0.5 | 6 | 0.1667 |
| 2_1542184 | 9 | 0.1111 | 9 | 0.1111 | 6 | 0 | 4 | 0.25 | 28 | 0.1071 | 5 | 0 | 3 | 0.3333 | 2 | 0.5 | 5 | 0 | 15 | 0.1333 |
| 2_1542185 | 7 | 0.2857 | 7 | 0.4286 | 6 | 0.3333 | 4 | 0.25 | 24 | 0.3333 | 3 | 0.3333 | 2 | 0 | 1 | 0 | 3 | 0.3333 | 9 | 0.2222 |
| 2_1542185 | 7 | 0.2857 | 8 | 0.25 | 6 | 0.5 | 4 | 0.5 | 25 | 0.36 | 4 | 0.25 | 4 | 0.5 | 2 | 0.5 | 4 | 0.25 | 14 | 0.3571 |
| 2_1542186 | 8 | 0 | 7 | 0.1429 | 5 | 0.4 | 4 | 0 | 24 | 0.125 | 5 | 0 | 4 | 0.25 | 3 | 0 | 3 | 0 | 15 | 0.0667 |
| 2_1542186 | 6 | 0 | 5 | 0.2 | 4 | 0.5 | 3 | 0 | 18 | 0.1667 | 5 | 0 | 4 | 0.25 | 1 | 0 | 3 | 0 | 13 | 0.0769 |
| 2_1542189 | 2 | 0 | 3 | 0.6667 | 4 | 0.5 | 2 | 0 | 11 | 0.3636 | 3 | 0 | 2 | 0 | 1 | 0 | 2 | 0.5 | 8 | 0.125 |
| 2_1543768 | NA | NA | NA | NA | 3 | 1 | NA | NA | 3 | 1 | 2 | 0.5 | 2 | 1 | 4 | 1 | NA | NA | 8 | 0.875 |
| 2_1543770 | 1 | 0 | NA | NA | 3 | 0.3333 | 1 | 0 | 5 | 0.2 | 2 | 0.5 | 3 | 0 | 4 | 0.75 | NA | NA | 9 | 0.4444 |
| 2_1543770 | 1 | 0 | NA | NA | 2 | 0.5 | 1 | 0 | 4 | 0.25 | 2 | 0 | 3 | 0 | 5 | 0.2 | NA | NA | 10 | 0.1 |
| 2_1543771 | 1 | 1 | NA | NA | 2 | 0.5 | 1 | 1 | 4 | 0.75 | 1 | 1 | 3 | 1 | 3 | 1 | NA | NA | 7 | 1 |
| 2_1543771 | 1 | 1 | NA | NA | 2 | 1 | 1 | 1 | 4 | 1 | NA | NA | 1 | 0 | 4 | 0.75 | NA | NA | 5 | 0.6 |
| 2_1543772 | 1 | 1 | NA | NA | 2 | 1 | 1 | 1 | 4 | 1 | 2 | 1 | 4 | 1 | 5 | 1 | NA | NA | 11 | 1 |
| 2_1543774 | 1 | 1 | NA | NA | 3 | 1 | 1 | 1 | 5 | 1 | 1 | 1 | 4 | 0.75 | 5 | 1 | NA | NA | 10 | 0.9 |
| 2_1543775 | 1 | 1 | 1 | 1 | 4 | 0.75 | 1 | 0 | 7 | 0.7143 | 2 | 0.5 | 4 | 0.75 | 6 | 0.6667 | NA | NA | 12 | 0.6667 |
| 2_1543775 | 1 | 1 | NA | NA | 3 | 1 | 1 | 1 | 5 | 1 | 2 | 1 | 4 | 1 | 6 | 0.8333 | NA | NA | 12 | 0.9167 |
| 2_1543775 | 1 | 0 | 1 | 1 | 4 | 0.5 | 1 | 1 | 7 | 0.5714 | NA | NA | 4 | 0.5 | 5 | 0.4 | NA | NA | 9 | 0.4444 |
| 2_1543777 | 1 | 1 | 1 | 1 | 1 | 1 | 1 | 0 | 4 | 0.75 | 1 | 1 | 4 | 1 | 5 | 1 | NA | NA | 10 | 1 |
| 2_1543779 | 1 | 0 | 1 | 1 | 3 | 0.6667 | 1 | 0 | 6 | 0.5 | 2 | 0 | 3 | 0.3333 | 6 | 0.6667 | NA | NA | 11 | 0.4545 |
| 2_1543779 | 1 | 1 | 1 | 1 | 3 | 1 | 1 | 1 | 6 | 1 | 2 | 0.5 | 4 | 1 | 5 | 1 | NA | NA | 11 | 0.9091 |
| 2_1543781 | 1 | 0 | 1 | 1 | 2 | 0.5 | 1 | 0 | 5 | 0.4 | 2 | 0 | 4 | 0.75 | 6 | 0.5 | NA | NA | 12 | 0.5 |
| 2_1543782 | 1 | 0 | 1 | 1 | 2 | 0 | 1 | 0 | 5 | 0.2 | 2 | 0 | 4 | 0 | 3 | 0.3333 | NA | NA | 9 | 0.1111 |
| 2_1543783 | 1 | 0 | 1 | 0 | 3 | 0.3333 | 1 | 1 | 6 | 0.3333 | 2 | 0 | 4 | 0.25 | 5 | 0 | NA | NA | 11 | 0.0909 |
| 2_1543797 | 5 | 0 | 2 | 0 | 6 | 0.1667 | 1 | 0 | 14 | 0.0714 | 2 | 0 | 4 | 0.25 | 8 | 0 | 1 | 0 | 15 | 0.0667 |
| 2_1543803 | 5 | 0.4 | 2 | 0.5 | 7 | 0.1429 | 1 | 0 | 15 | 0.2667 | 3 | 0 | 5 | 0.4 | 8 | 0.25 | 4 | 0 | 20 | 0.2 |
| 2_1543804 | 5 | 0.2 | 2 | 0 | 4 | 0.25 | 1 | 0 | 12 | 0.1667 | 2 | 0 | 5 | 0.4 | 6 | 0.1667 | 1 | 0 | 14 | 0.2143 |
| 2_1543804 | 5 | 0.4 | 3 | 0.3333 | 4 | 0.25 | NA | NA | 12 | 0.3333 | 3 | 0.6667 | 4 | 0.75 | 4 | 0.25 | 4 | 0.25 | 15 | 0.4667 |
| 2_1543804 | 4 | 0.25 | 3 | 0.3333 | 4 | 0 | 1 | 0 | 12 | 0.1667 | 3 | 0.3333 | 5 | 0.8 | 8 | 0.125 | 3 | 0.3333 | 19 | 0.3684 |
| 2_1543805 | 4 | 0 | 3 | 0 | 7 | 0.1429 | 1 | 0 | 15 | 0.0667 | 3 | 0 | 5 | 0.2 | 7 | 0 | 3 | 0 | 18 | 0.0556 |
| 2_1543805 | 4 | 0.5 | 3 | 0.3333 | 8 | 0.5 | 1 | 0 | 16 | 0.4375 | 3 | 0.6667 | 5 | 0.8 | 7 | 0.2857 | 3 | 0.6667 | 18 | 0.5556 |
| 2_1543805 | 5 | 0.2 | 2 | 0 | 6 | 0.1667 | 1 | 0 | 14 | 0.1429 | 2 | 0 | 5 | 0.2 | 7 | 0 | 3 | 0.3333 | 17 | 0.1176 |
| 2_1543805 | 5 | 0.4 | 3 | 0 | 8 | 0.25 | 1 | 0 | 17 | 0.2353 | 3 | 0 | 5 | 0.2 | 6 | 0.3333 | 2 | 0 | 16 | 0.1875 |
| 2_1543806 | 3 | 0.3333 | 3 | 0.3333 | 6 | 0 | 1 | 0 | 13 | 0.1538 | 3 | 0 | 5 | 0 | 7 | 0.1429 | 4 | 0 | 19 | 0.0526 |
| 2_1543807 | 5 | 0 | 3 | 0.3333 | 6 | 0 | 1 | 0 | 15 | 0.0667 | 3 | 0.3333 | 7 | 0 | 7 | 0 | 3 | 0 | 20 | 0.05 |
| 2_1543808 | 5 | 0.6 | 2 | 0.5 | 6 | 0.3333 | 1 | 1 | 14 | 0.5 | 3 | 1 | 5 | 0.8 | 7 | 0.2857 | 3 | 0.6667 | 18 | 0.6111 |
| 2_1543808 | 5 | 0.4 | 3 | 0 | 8 | 0 | 2 | 0 | 18 | 0.1111 | 3 | 0.3333 | 4 | 0.25 | 6 | 0 | 4 | 0.25 | 17 | 0.1765 |
| 2_1543811 | 5 | 0.4 | 3 | 0.3333 | 8 | 0.25 | 2 | 0.5 | 18 | 0.3333 | 4 | 0 | 7 | 0 | 9 | 0.2222 | 4 | 0.5 | 24 | 0.1667 |
| 2_1543866 | 7 | 0 | 13 | 0.2308 | 11 | 0.0909 | 4 | 0 | 35 | 0.1143 | 12 | 0.1667 | 4 | 0.25 | 4 | 0.25 | 5 | 0.2 | 25 | 0.2 |
| 2_1543869 | 6 | 0.3333 | 10 | 0.2 | 9 | 0.2222 | 4 | 0.25 | 29 | 0.2414 | 11 | 0.0909 | 4 | 0.5 | 3 | 0 | 5 | 0 | 23 | 0.1304 |
| 2_1544026 | 1 | 1 | 2 | 0.5 | NA | NA | NA | NA | 3 | 0.6667 | NA | NA | 2 | 1 | 1 | 1 | NA | NA | 3 | 1 |
| 2_1544026 | 1 | 1 | 2 | 1 | 2 | 1 | NA | NA | 5 | 1 | NA | NA | NA | NA | 1 | 1 | 1 | 1 | 2 | 1 |
| 2_1544026 | 1 | 0 | 2 | 0.5 | 2 | 0 | NA | NA | 5 | 0.2 | NA | NA | 2 | 0 | NA | NA | 1 | 1 | 3 | 0.3333 |
| 2_1544279 | 1 | 1 | NA | NA | NA | NA | NA | NA | 1 | 1 | 2 | 0.5 | 1 | 1 | 2 | 1 | 2 | 1 | 7 | 0.8571 |
| 2_1544293 | 2 | 0 | 1 | 1 | 2 | 0.5 | NA | NA | 5 | 0.4 | 2 | 0 | NA | NA | 2 | 0.5 | 4 | 0.25 | 8 | 0.25 |
| 2_1544297 | 3 | 0 | 3 | 0 | 2 | 0.5 | NA | NA | 8 | 0.125 | 2 | 0 | NA | NA | 1 | 0 | 4 | 0.25 | 7 | 0.1429 |
| 2_1544297 | 3 | 0.3333 | 3 | 0.3333 | 1 | 0 | NA | NA | 7 | 0.2857 | 2 | 0.5 | NA | NA | 1 | 0 | 4 | 0.5 | 7 | 0.4286 |
| 2_1544297 | 3 | 0.3333 | 3 | 0.3333 | 2 | 0 | NA | NA | 8 | 0.25 | 2 | 0.5 | NA | NA | 1 | 0 | 3 | 0 | 6 | 0.1667 |
| 2_1544298 | 3 | 0.3333 | 2 | 0 | 1 | 0 | NA | NA | 6 | 0.1667 | 2 | 0 | NA | NA | 1 | 0 | 4 | 0.25 | 7 | 0.1429 |
| 2_1544373 | NA | NA | NA | NA | 3 | 0.6667 | 1 | 1 | 4 | 0.75 | NA | NA | NA | NA | 1 | 1 | 1 | 0 | 2 | 0.5 |
| 2_1544878 | NA | NA | 1 | 1 | 2 | 0 | 1 | 1 | 4 | 0.5 | NA | NA | NA | NA | 2 | 0.5 | 1 | 0 | 3 | 0.3333 |
| 2_1544886 | NA | NA | 1 | 1 | 3 | 0 | 1 | 0 | 5 | 0.2 | NA | NA | 1 | 1 | 1 | 0 | 3 | 0 | 5 | 0.2 |
| 2_1545359 | 15 | 0.4667 | 14 | 0.7143 | 18 | 0.5 | 24 | 0.4583 | 71 | 0.5211 | 18 | 0.4444 | 26 | 0.5 | 21 | 0.619 | 13 | 0.6154 | 78 | 0.5385 |
| 2_1545361 | 26 | 0.0385 | 29 | 0.069 | 26 | 0.0769 | 35 | 0.1143 | 116 | 0.0776 | 28 | 0.1071 | 37 | 0.1351 | 31 | 0 | 23 | 0.2174 | 119 | 0.1092 |
| 2_1545362 | 20 | 0.05 | 30 | 0.2333 | 22 | 0 | 27 | 0.0741 | 99 | 0.101 | 19 | 0.1579 | 24 | 0.1667 | 27 | 0.037 | 20 | 0.1 | 90 | 0.1111 |
| 2_1547090 | 13 | 0.3846 | 13 | 0.0769 | 12 | 0.0833 | 13 | 0.1538 | 51 | 0.1765 | 8 | 0.125 | 15 | 0.1333 | 14 | 0.2143 | 14 | 0.1429 | 51 | 0.1569 |

| | | | | | | | | | | | | | | | | | | | | |
|---|---|---|---|---|---|---|---|---|---|---|---|---|---|---|---|---|---|---|---|---|
| 2_1547136 | 1 | 0 | NA | NA | 2 | 0.5 | 3 | 0 | 6 | 0.1667 | 1 | 0 | 3 | 0 | 1 | 1 | 3 | 0 | 8 | 0.125 |
| 2_1547143 | 5 | 0.2 | 17 | 0.0588 | 13 | 0.0769 | 12 | 0.1667 | 47 | 0.1064 | 7 | 0.1429 | 12 | 0.0833 | 15 | 0 | 12 | 0.1667 | 46 | 0.087 |
| 2_1547145 | 4 | 0.5 | 15 | 0.4667 | 10 | 0.6 | 10 | 0.4 | 39 | 0.4872 | 6 | 0.1667 | 11 | 0.6364 | 16 | 0.4375 | 12 | 0.3333 | 45 | 0.4222 |
| 2_1549022 | 9 | 0.7778 | 5 | 0.4 | 6 | 0.8333 | 9 | 0.5556 | 29 | 0.6552 | 3 | 1 | 4 | 0.75 | 12 | 0.75 | 4 | 0.75 | 23 | 0.7826 |
| 2_1550664 | 5 | 0.4 | 4 | 0.25 | 5 | 0.2 | 8 | 0.5 | 22 | 0.3636 | 2 | 0.5 | 5 | 0.2 | 5 | 0.4 | 2 | 0.5 | 14 | 0.3571 |
| 2_1550664 | 6 | 0.6667 | 3 | 1 | 4 | 1 | 8 | 0.875 | 21 | 0.8571 | 1 | 0 | 4 | 1 | 5 | 0.4 | 1 | 1 | 11 | 0.6364 |
| 2_1550669 | 6 | 0 | 6 | 0 | 6 | 0.1667 | 9 | 0.3333 | 27 | 0.1481 | 2 | 0 | 6 | 0.1667 | 5 | 0 | 4 | 0 | 17 | 0.0588 |
| 2_1551199 | 34 | 0.0294 | 62 | 0.0323 | 53 | 0.0566 | 48 | 0.0833 | 197 | 0.0508 | 44 | 0.0682 | 28 | 0.1071 | 49 | 0.102 | 47 | 0.0426 | 168 | 0.0774 |
| 2_1551202 | 2 | 0 | 8 | 0.25 | 4 | 0.25 | 5 | 0.4 | 19 | 0.2632 | 3 | 0.6667 | 4 | 0.25 | 1 | 0 | 2 | 0.5 | 10 | 0.4 |
| 2_1551202 | 1 | 0 | 9 | 0 | 3 | 0.3333 | 5 | 0.2 | 18 | 0.1111 | 3 | 0.3333 | 3 | 0.3333 | 1 | 0 | 1 | 1 | 8 | 0.375 |
| 2_1551203 | 1 | 1 | 6 | 0.8333 | 4 | 1 | 5 | 0.8 | 16 | 0.875 | 1 | 1 | 4 | 0.75 | 1 | 1 | 2 | 1 | 8 | 0.875 |
| 2_1551204 | NA | NA | 6 | 0.8333 | 3 | 0.3333 | 3 | 1 | 12 | 0.75 | 1 | 1 | 3 | 0.6667 | 1 | 1 | 1 | 1 | 6 | 0.8333 |
| 2_1552309 | 16 | 0.125 | 25 | 0.16 | 19 | 0.0526 | 17 | 0 | 77 | 0.0909 | 18 | 0 | 14 | 0 | 10 | 0 | 9 | 0.3333 | 51 | 0.0588 |
| 2_1552314 | 18 | 0 | 25 | 0.08 | 23 | 0.3043 | 16 | 0.125 | 82 | 0.1341 | 20 | 0.1 | 18 | 0.1667 | 12 | 0.0833 | 9 | 0.2222 | 59 | 0.1356 |
| 2_1552314 | 17 | 0.3529 | 23 | 0.3043 | 22 | 0.3636 | 20 | 0.3 | 82 | 0.3293 | 17 | 0.2353 | 15 | 0.3333 | 11 | 0.1818 | 8 | 0.125 | 51 | 0.2353 |
| 2_1552315 | 18 | 0.2778 | 27 | 0.1852 | 23 | 0.087 | 22 | 0.0909 | 90 | 0.1556 | 22 | 0 | 17 | 0.1765 | 12 | 0.0833 | 12 | 0.25 | 63 | 0.1111 |
| 2_1552316 | 18 | 0 | 26 | 0 | 21 | 0.1905 | 19 | 0.0526 | 84 | 0.0595 | 21 | 0.1429 | 15 | 0 | 12 | 0 | 10 | 0.2 | 58 | 0.0862 |
| 2_1552316 | 18 | 0 | 26 | 0.1154 | 23 | 0.087 | 22 | 0 | 89 | 0.0562 | 23 | 0.0435 | 18 | 0.1111 | 11 | 0 | 12 | 0.1667 | 64 | 0.0781 |
| 2_1552316 | 16 | 0.25 | 26 | 0.1538 | 23 | 0.2174 | 19 | 0.1053 | 84 | 0.1786 | 18 | 0.2222 | 13 | 0 | 11 | 0.0909 | 9 | 0 | 51 | 0.098 |
| 2_1552475 | 5 | 0.4 | 5 | 0.2 | 4 | 0 | 10 | 0.3 | 24 | 0.25 | 5 | 0.2 | 7 | 0 | 4 | 0.5 | 4 | 0.5 | 20 | 0.25 |
| 2_1552476 | 4 | 0.25 | 4 | 0 | 4 | 0.25 | 8 | 0.25 | 20 | 0.2 | 4 | 0 | 6 | 0 | 2 | 0 | 4 | 0.25 | 16 | 0.0625 |
| 2_1552476 | 4 | 0.5 | 4 | 0.75 | 4 | 1 | 6 | 0.8333 | 18 | 0.7778 | 5 | 1 | 6 | 1 | 2 | 0.5 | 2 | 1 | 15 | 0.9333 |
| 2_1552476 | 4 | 0 | 4 | 0 | 5 | 0 | 8 | 0.25 | 21 | 0.0952 | 3 | 0 | 6 | 0.1667 | 4 | 0 | 2 | 0.5 | 15 | 0.1333 |
| 2_1552477 | 2 | 0.5 | 4 | 1 | 4 | 0.75 | 8 | 1 | 18 | 0.8889 | 6 | 0.8333 | 5 | 1 | 3 | 1 | 1 | 1 | 15 | 0.9333 |
| 2_1552477 | 4 | 0.25 | 1 | 0 | 5 | 0.2 | 9 | 0 | 19 | 0.1053 | 6 | 0.1667 | 4 | 0.25 | 3 | 0 | 2 | 0 | 15 | 0.1333 |
| 2_1552478 | 1 | 1 | 3 | 1 | 5 | 1 | 7 | 1 | 16 | 1 | 6 | 0.6667 | 6 | 1 | 4 | 1 | 2 | 0.5 | 18 | 0.8333 |
| 2_1552479 | 2 | 0.5 | 3 | 0.6667 | 4 | 0.5 | 5 | 0.4 | 14 | 0.5 | 3 | 0.3333 | 4 | 0.25 | 3 | 0 | 2 | 0 | 12 | 0.1667 |
| 2_1552481 | 2 | 0.5 | 2 | 0 | 4 | 0.25 | 8 | 0 | 16 | 0.125 | 4 | 0.25 | 6 | 0 | 2 | 0.5 | 1 | 0 | 13 | 0.1538 |
| 2_1552481 | 1 | 0 | 2 | 0 | 3 | 0.3333 | 8 | 0.125 | 14 | 0.1429 | 3 | 0 | 3 | 0.3333 | 3 | 0 | 1 | 0 | 10 | 0.1 |
| 2_1552481 | 1 | 1 | 4 | 0.75 | 4 | 0.75 | 7 | 0.8571 | 16 | 0.8125 | 6 | 0.8333 | 6 | 1 | 4 | 1 | 1 | 1 | 17 | 0.9412 |
| 2_1552483 | 1 | 1 | 3 | 1 | 3 | 0.6667 | 6 | 1 | 13 | 0.9231 | 6 | 0.8333 | 6 | 0.8333 | 4 | 1 | 2 | 0.5 | 18 | 0.8333 |
| 2_1552484 | 2 | 1 | 2 | 1 | 2 | 0.5 | 8 | 0.375 | 14 | 0.5714 | 3 | 1 | 4 | 0.75 | 2 | 0.5 | 2 | 1 | 11 | 0.8182 |
| 2_1552486 | 2 | 0 | 4 | 0 | 3 | 0 | 9 | 0.2222 | 18 | 0.1111 | 5 | 0 | 4 | 0.5 | 3 | 0.3333 | 1 | 0 | 13 | 0.2308 |
| 2_1552487 | 2 | 0 | 3 | 0.3333 | 3 | 0.3333 | 7 | 0.7143 | 15 | 0.4667 | 4 | 0.25 | 4 | 0.75 | 4 | 0.25 | 2 | 0.5 | 14 | 0.4286 |
| 2_1552487 | 2 | 0 | 4 | 0 | 3 | 0.3333 | 8 | 0 | 17 | 0.0588 | 5 | 0 | 4 | 0.25 | 4 | 0.25 | 2 | 0 | 15 | 0.1333 |
| 2_1552488 | 2 | 0 | 2 | 0 | 3 | 0 | 7 | 0.1429 | 14 | 0.0714 | 3 | 0.3333 | 2 | 0 | 4 | 0 | 2 | 0 | 11 | 0.0909 |
| 2_1552490 | 2 | 0 | 4 | 0.5 | 2 | 0 | 8 | 0.125 | 16 | 0.1875 | 3 | 0 | 2 | 0.5 | 2 | 0.5 | 1 | 0 | 8 | 0.25 |
| 2_1552490 | 2 | 0 | 4 | 0.25 | 3 | 0.3333 | 7 | 0.2857 | 16 | 0.25 | 3 | 0.3333 | 4 | 0 | 4 | 0 | 2 | 0.5 | 13 | 0.1538 |
| 2_1552490 | 2 | 1 | 4 | 0.5 | 3 | 0.3333 | 9 | 0.2222 | 18 | 0.3889 | 3 | 0.6667 | 2 | 1 | 4 | 0.75 | 2 | 0.5 | 11 | 0.7273 |
| 2_1552495 | 1 | 0 | 3 | 0.3333 | 2 | 1 | 8 | 0.625 | 14 | 0.5714 | 4 | 0.5 | 3 | 1 | 2 | 0.5 | 1 | 1 | 10 | 0.7 |
| 2_1552495 | 2 | 0.5 | 4 | 0.25 | 3 | 0.3333 | 8 | 0.875 | 17 | 0.5882 | 4 | 0.5 | 2 | 0.5 | 4 | 0.75 | 1 | 1 | 11 | 0.6364 |
| 2_1552497 | 1 | 0 | 3 | 0.3333 | 2 | 1 | 7 | 0.2857 | 13 | 0.3846 | 4 | 0.5 | 2 | 0 | 3 | 0.3333 | 1 | 0 | 10 | 0.3 |
| 2_1552500 | 1 | 0 | 2 | 0 | 3 | 0 | 5 | 0.2 | 11 | 0.0909 | 3 | 0.3333 | 3 | 0 | 3 | 0 | 1 | 0 | 10 | 0.1 |
| 2_1552500 | 1 | 0 | 2 | 0.5 | 3 | 0.6667 | 6 | 0.8333 | 12 | 0.6667 | 4 | 0.25 | 3 | 0.3333 | 4 | 0.5 | 1 | 1 | 12 | 0.4167 |
| 2_1552568 | 2 | 0.5 | 1 | 0 | 2 | 1 | 6 | 0.3333 | 11 | 0.4545 | 2 | 0 | 2 | 0.5 | 1 | 0 | NA | NA | 5 | 0.2 |
| 2_1552571 | 2 | 0 | 2 | 0 | 4 | 0 | 5 | 0.2 | 13 | 0.0769 | 2 | 0 | 3 | 0.3333 | 2 | 0 | 1 | 0 | 8 | 0.125 |
| 2_1552572 | 2 | 0.5 | 2 | 0.5 | 4 | 1 | 5 | 0.6 | 13 | 0.6923 | 2 | 0 | 3 | 0.3333 | 2 | 0.5 | 1 | 0 | 8 | 0.25 |
| 2_1552572 | 2 | 0 | 2 | 1 | 3 | 0.6667 | 4 | 0.5 | 11 | 0.5455 | 2 | 1 | 2 | 0.5 | 2 | 1 | 1 | 1 | 7 | 0.8571 |
| 2_1552572 | 2 | 1 | 2 | 0 | 2 | 0.5 | 3 | 0.3333 | 9 | 0.4444 | 2 | 0.5 | 3 | 0.6667 | 2 | 0.5 | NA | NA | 7 | 0.5714 |
| 2_1552572 | 2 | 0 | 2 | 0 | 3 | 0.3333 | 5 | 0 | 12 | 0.0833 | 2 | 0.5 | 2 | 0 | 2 | 0 | 1 | 0 | 7 | 0.1429 |
| 2_1552573 | 1 | 0 | 1 | 0 | 3 | 0.3333 | 5 | 0 | 10 | 0.1 | 2 | 0.5 | 1 | 0 | 1 | 0 | 1 | 0 | 5 | 0.2 |
| 2_1552573 | 1 | 0 | 2 | 0 | 4 | 0.25 | 5 | 0 | 12 | 0.0833 | 1 | 0 | 2 | 0 | 2 | 0.5 | 1 | 0 | 6 | 0.1667 |
| 2_1552575 | 2 | 1 | 2 | 1 | 3 | 1 | 5 | 1 | 12 | 1 | 1 | 0 | 2 | 0.5 | 1 | 1 | NA | NA | 4 | 0.5 |
| 2_1552575 | 2 | 0 | 1 | 0 | 4 | 0.25 | 4 | 0.25 | 11 | 0.1818 | 1 | 0 | 2 | 0.5 | 2 | 0 | 1 | 0 | 6 | 0.1667 |
| 2_1552576 | 2 | 0.5 | 2 | 0 | 4 | 1 | 3 | 1 | 11 | 0.7273 | 1 | 1 | 1 | 1 | 2 | 1 | 1 | 1 | 5 | 1 |
| 2_1552632 | 1 | 1 | 1 | 1 | 1 | 1 | 3 | 1 | 6 | 1 | 1 | 1 | 1 | 1 | 1 | 1 | 1 | 1 | 4 | 1 |
| 2_1552633 | 1 | 0 | 2 | 0 | 2 | 0 | 4 | 0.5 | 9 | 0.2222 | 1 | 1 | 1 | 0 | 1 | 0 | 1 | 0 | 4 | 0.25 |
| 2_1552633 | 1 | 1 | 2 | 0 | 3 | 0 | 4 | 0 | 10 | 0.1 | 1 | 0 | 1 | 0 | 1 | 1 | 1 | 0 | 4 | 0.25 |
| 2_1553566 | 5 | 0.2 | 4 | 0 | 7 | 0 | 3 | 0 | 19 | 0.0526 | 4 | 0 | 2 | 0 | 1 | 1 | 2 | 0 | 9 | 0.1111 |
| 2_1554766 | 10 | 0.7 | 22 | 0.7273 | 14 | 0.6429 | 18 | 0.7222 | 64 | 0.7031 | 19 | 0.6842 | 16 | 0.625 | 19 | 0.7895 | 14 | 0.6429 | 68 | 0.6912 |
| 2_1554769 | 9 | 0.2222 | 19 | 0.4737 | 15 | 0.4 | 16 | 0.4375 | 59 | 0.4068 | 17 | 0.3529 | 16 | 0.3125 | 19 | 0.4211 | 14 | 0.2143 | 66 | 0.3333 |
| 2_1554769 | 10 | 0.4 | 16 | 0.3125 | 16 | 0.3125 | 18 | 0.2778 | 60 | 0.3167 | 15 | 0.3333 | 16 | 0.375 | 18 | 0.2778 | 13 | 0.3077 | 62 | 0.3226 |
| 2_1554771 | 9 | 0.1111 | 20 | 0 | 13 | 0 | 16 | 0.1875 | 58 | 0.069 | 17 | 0.1176 | 16 | 0.0625 | 15 | 0.0667 | 14 | 0.0714 | 62 | 0.0806 |
| 2_1555442 | 3 | 0.6667 | 3 | 1 | 1 | 1 | 4 | 1 | 11 | 0.9091 | 1 | 1 | 2 | 1 | 4 | 0.75 | 2 | 1 | 9 | 0.8889 |
| 2_1555443 | 2 | 1 | 2 | 0.5 | 1 | 1 | 3 | 0.6667 | 8 | 0.75 | NA | NA | 2 | 1 | 1 | 0 | 1 | 0 | 4 | 0.5 |

| | | | | | | | | | | | | | | | | | | | | |
|---|---|---|---|---|---|---|---|---|---|---|---|---|---|---|---|---|---|---|---|---|
| 2_1555444 | 3 | 0.3333 | 3 | 0.6667 | 1 | 0 | 3 | 0 | 10 | 0.3 | 1 | 0 | 2 | 0 | 3 | 0.3333 | 1 | 0 | 7 | 0.1429 |
| 2_1556525 | NA | NA | 1 | 1 | NA | NA | NA | NA | 1 | 1 | NA | NA | 1 | 1 | NA | NA | NA | NA | 1 | 1 |
| 2_1556559 | 2 | 0 | 5 | 0.2 | 1 | 0 | 3 | 0 | 11 | 0.0909 | 2 | 0 | NA | NA | 6 | 0.1667 | 1 | 0 | 9 | 0.1111 |
| 2_1556576 | 1 | 1 | 4 | 0.75 | 1 | 0 | 2 | 0 | 8 | 0.5 | 2 | 0 | NA | NA | 4 | 0.25 | 1 | 0 | 7 | 0.1429 |
| 2_1556576 | 2 | 0.5 | 5 | 0.2 | 1 | 0 | 2 | 0 | 10 | 0.2 | 2 | 0.5 | NA | NA | 5 | 0.4 | 1 | 0 | 8 | 0.375 |
| 2_1556577 | 2 | 0 | 5 | 0 | 1 | 0 | 2 | 0.5 | 10 | 0.1 | 2 | 0 | NA | NA | 5 | 0.2 | 1 | 0 | 8 | 0.125 |
| 2_1556577 | 2 | 1 | 4 | 0.5 | 1 | 1 | 1 | 1 | 8 | 0.75 | 1 | 1 | NA | NA | 4 | 0.75 | 1 | 1 | 6 | 0.8333 |
| 2_1556583 | 2 | 0 | 5 | 0.2 | NA | NA | 1 | 0 | 8 | 0.125 | 2 | 0.5 | NA | NA | 7 | 0 | 1 | 0 | 10 | 0.1 |
| 2_1556585 | 3 | 0 | 6 | 0.1667 | 1 | 0 | 2 | 0 | 12 | 0.0833 | 2 | 0 | NA | NA | 7 | 0.1429 | 1 | 0 | 10 | 0.1 |
| 2_1556587 | 2 | 0.5 | 6 | 0 | 2 | 1 | 3 | 0.3333 | 13 | 0.3077 | 2 | 0.5 | NA | NA | 7 | 0.2857 | 1 | 0 | 10 | 0.3 |
| 2_1556594 | 3 | 0 | 5 | 0.2 | 4 | 0.75 | 3 | 0.3333 | 15 | 0.3333 | 2 | 0.5 | NA | NA | 6 | 0.3333 | 1 | 0 | 9 | 0.3333 |
| 2_1556596 | 11 | 0.2727 | 7 | 0.1429 | 14 | 0.0714 | 16 | 0.0625 | 48 | 0.125 | 14 | 0 | 15 | 0.0667 | 16 | 0.125 | 4 | 0 | 49 | 0.0612 |
| 2_1556598 | 12 | 0.25 | 7 | 0 | 14 | 0.3571 | 13 | 0 | 46 | 0.1739 | 9 | 0.3333 | 14 | 0 | 16 | 0.1875 | 4 | 0.25 | 43 | 0.1628 |
| 2_1556598 | 11 | 0.1818 | 6 | 0.1667 | 12 | 0 | 15 | 0.0667 | 44 | 0.0909 | 15 | 0.1333 | 13 | 0 | 15 | 0.2 | 5 | 0.2 | 48 | 0.125 |
| 2_1556598 | 5 | 0.4 | 5 | 0 | 1 | 0 | 2 | 0 | 13 | 0.1538 | 2 | 0 | 1 | 0 | 5 | 0.2 | 2 | 0 | 10 | 0.1 |
| 2_1556598 | 9 | 0.5556 | 5 | 0.6 | 11 | 0.0909 | 13 | 0.5385 | 38 | 0.4211 | 13 | 0.3077 | 15 | 0.4 | 16 | 0.4375 | 4 | 0.25 | 48 | 0.375 |
| 2_1556598 | 11 | 0.3636 | 5 | 0.4 | 15 | 0.2 | 15 | 0.2667 | 46 | 0.2826 | 14 | 0.0714 | 10 | 0.2 | 14 | 0.2143 | 5 | 0.6 | 43 | 0.2093 |
| 2_1556598 | 6 | 0.3333 | 7 | 0.2857 | 12 | 0.0833 | 15 | 0.3333 | 40 | 0.25 | 10 | 0.5 | 8 | 0.125 | 12 | 0.4167 | 4 | 0.25 | 34 | 0.3529 |
| 2_1556599 | 8 | 0.75 | 4 | 0 | 11 | 0.0909 | 13 | 0.0769 | 36 | 0.2222 | 13 | 0.2308 | 12 | 0.1667 | 14 | 0.5714 | 3 | 0 | 42 | 0.3095 |
| 2_1556601 | 9 | 0.5556 | 6 | 0.5 | 12 | 0.6667 | 13 | 0.6154 | 40 | 0.6 | 11 | 0.5455 | 12 | 0.6667 | 15 | 0.4 | 5 | 1 | 43 | 0.5814 |
| 2_1556601 | 8 | 0.25 | 7 | 0.2857 | 9 | 0.1111 | 15 | 0.3333 | 39 | 0.2564 | 11 | 0.0909 | 12 | 0.1667 | 15 | 0.2667 | 4 | 0 | 42 | 0.1667 |
| 2_1556604 | 10 | 0.4 | 3 | 0.3333 | 13 | 0.4615 | 13 | 0.6154 | 39 | 0.4872 | 12 | 0.3333 | 12 | 0.3333 | 15 | 0.4 | 7 | 0.7143 | 46 | 0.413 |
| 2_1556605 | 10 | 1 | 1 | 1 | 9 | 0.6667 | 8 | 1 | 28 | 0.8929 | 9 | 0.7778 | 9 | 0.6667 | 12 | 0.5833 | 5 | 1 | 35 | 0.7143 |
| 2_1556606 | 9 | 0.4444 | 2 | 0.5 | 12 | 0.1667 | 15 | 0.3333 | 38 | 0.3158 | 12 | 0.3333 | 11 | 0.1818 | 14 | 0.0714 | 7 | 0.5714 | 44 | 0.25 |
| 2_1556607 | 10 | 0.4 | 1 | 0 | 10 | 0.7 | 8 | 0.375 | 29 | 0.4828 | 12 | 0.4167 | 11 | 0.4545 | 14 | 0.3571 | 7 | 0.5714 | 44 | 0.4318 |
| 2_1556617 | 5 | 0.2 | 7 | 0 | 8 | 0.125 | 8 | 0 | 28 | 0.0714 | 6 | 0 | 9 | 0.1111 | 8 | 0 | 7 | 0.2857 | 30 | 0.1 |
| 2_1556619 | 3 | 0 | 7 | 0.1429 | 11 | 0 | 11 | 0.3636 | 32 | 0.1562 | 7 | 0.1429 | 12 | 0.0833 | 9 | 0.2222 | 7 | 0.2857 | 35 | 0.1714 |
| 2_1556682 | 4 | 0 | 8 | 0.125 | 6 | 0.3333 | 6 | 0 | 24 | 0.125 | 5 | 0 | 12 | 0.0833 | 8 | 0.125 | 7 | 0 | 32 | 0.0625 |
| 2_1556685 | 3 | 0 | 3 | 0.3333 | 3 | 0 | 3 | 0.3333 | 12 | 0.1667 | 3 | 0 | 8 | 0.125 | 5 | 0.2 | 3 | 0 | 19 | 0.1053 |
| 2_1556685 | 3 | 0 | 4 | 0.25 | 3 | 0 | 2 | 0 | 12 | 0.0833 | 2 | 0 | 7 | 0 | 4 | 0 | 3 | 0.3333 | 16 | 0.0625 |
| 2_1556888 | 6 | 0.6667 | 6 | 0 | 9 | 0.3333 | 6 | 0.1667 | 27 | 0.2963 | 1 | 0 | 5 | 0.4 | 5 | 0.8 | 7 | 0.5714 | 18 | 0.5556 |
| 2_1556891 | 4 | 0.5 | 6 | 0 | 9 | 0 | 6 | 0.1667 | 25 | 0.12 | 2 | 0.5 | 5 | 0 | 6 | 0.1667 | 7 | 0 | 20 | 0.1 |
| 2_1556892 | 5 | 0.4 | 6 | 0 | 9 | 0.1111 | 4 | 0 | 24 | 0.125 | 2 | 0 | 4 | 0.5 | 6 | 0.6667 | 7 | 0.1429 | 19 | 0.3684 |
| 2_1556900 | 6 | 0.1667 | 7 | 0 | 7 | 0.1429 | 5 | 0.2 | 25 | 0.12 | 4 | 0.25 | 7 | 0.2857 | 5 | 0.4 | 5 | 0 | 21 | 0.2381 |
| 2_1556902 | 3 | 0.6667 | 3 | 0 | 6 | 0.5 | 2 | 0.5 | 14 | 0.4286 | NA | NA | 4 | 0.25 | 4 | 0.5 | 4 | 0.75 | 12 | 0.5 |
| 2_1556902 | 5 | 0.2 | 7 | 0.4286 | 8 | 0.25 | 5 | 0.2 | 25 | 0.28 | 4 | 0 | 7 | 0.2857 | 5 | 0.2 | 6 | 0.5 | 22 | 0.2727 |
| 2_1556902 | 4 | 0.5 | 6 | 0.1667 | 7 | 0.2857 | 5 | 0.6 | 22 | 0.3636 | 3 | 0.6667 | 6 | 0.5 | 4 | 0.25 | 6 | 0.3333 | 19 | 0.4211 |
| 2_1556902 | 6 | 0 | 6 | 0.3333 | 8 | 0.125 | 5 | 0.2 | 25 | 0.16 | 5 | 0 | 7 | 0.2857 | 5 | 0.2 | 7 | 0.2857 | 24 | 0.2083 |
| 2_1556902 | 6 | 0.3333 | 6 | 0.1667 | 7 | 0.5714 | 2 | 0 | 21 | 0.3333 | 4 | 0 | 7 | 0 | 4 | 0.5 | 6 | 0.1667 | 21 | 0.1429 |
| 2_1556903 | 6 | 0.6667 | 6 | 0.6667 | 8 | 0.5 | 4 | 0.75 | 24 | 0.625 | 5 | 0.8 | 7 | 0.7143 | 5 | 1 | 7 | 0.5714 | 24 | 0.75 |
| 2_1558327 | NA | NA | 1 | 1 | NA | NA | NA | NA | 1 | 1 | 1 | 0 | NA | NA | 1 | 0 | 1 | 1 | 3 | 0.3333 |
| 2_1558335 | NA | NA | 1 | 1 | NA | NA | NA | NA | 1 | 1 | 1 | 1 | NA | NA | 2 | 0.5 | NA | NA | 3 | 0.6667 |
| 2_1558337 | NA | NA | 1 | 1 | 1 | 0 | NA | NA | 2 | 0.5 | 1 | 1 | NA | NA | 3 | 0 | 1 | 1 | 5 | 0.4 |
| 2_1558342 | 1 | 0 | 1 | 1 | 1 | 0 | 1 | 0 | 4 | 0.25 | NA | NA | NA | NA | 3 | 0.3333 | 1 | 1 | 4 | 0.5 |
| 2_1558343 | 1 | 0 | 2 | 0 | 1 | 1 | 1 | 0 | 5 | 0.2 | 1 | 0 | NA | NA | 3 | 0.3333 | 1 | 0 | 5 | 0.2 |
| 2_1558345 | 1 | 0 | 3 | 0.3333 | 1 | 1 | 1 | 0 | 6 | 0.3333 | 1 | 0 | NA | NA | 3 | 1 | 1 | 0 | 5 | 0.6 |
| 2_1558403 | NA | NA | 3 | 0.6667 | 4 | 0.5 | 1 | 1 | 8 | 0.625 | 1 | 1 | NA | NA | 3 | 1 | 1 | 0 | 5 | 0.8 |
| 2_1558403 | 1 | 0 | 3 | 0 | 3 | 0.3333 | 1 | 0 | 8 | 0.125 | 1 | 1 | NA | NA | 3 | 0.6667 | NA | NA | 4 | 0.75 |
| 2_1558403 | 1 | 0 | 2 | 0 | 3 | 0.3333 | 1 | 0 | 7 | 0.1429 | 1 | 0 | NA | NA | 3 | 0.3333 | 1 | 0 | 5 | 0.2 |
| 2_1558405 | 1 | 1 | 3 | 0.3333 | 4 | 0.75 | 1 | 0 | 9 | 0.5556 | 1 | 0 | NA | NA | 3 | 0.3333 | 1 | 0 | 5 | 0.2 |
| 2_1558581 | 1 | 1 | NA | NA | NA | NA | NA | NA | 1 | 1 | NA | NA | NA | NA | 1 | 1 | NA | NA | 1 | 1 |
| 2_1559099 | NA | NA | NA | NA | NA | NA | 1 | 1 | 1 | 1 | NA | NA | NA | NA | 1 | 1 | NA | NA | 1 | 1 |
| 2_1559152 | 4 | 0 | 2 | 0 | 4 | 0 | 5 | 0.2 | 15 | 0.0667 | NA | NA | 1 | 0 | 8 | 0.125 | 4 | 0.5 | 13 | 0.2308 |
| 2_1559153 | 3 | 0.3333 | 2 | 0 | 4 | 0 | 6 | 0 | 15 | 0.0667 | NA | NA | 1 | 0 | 8 | 0.125 | 5 | 0 | 14 | 0.0714 |
| 2_1559377 | 1 | 1 | 1 | 0 | 2 | 0 | NA | NA | 4 | 0.25 | 3 | 0.3333 | 4 | 0.25 | 3 | 0.6667 | 3 | 0.3333 | 13 | 0.3846 |
| 2_1559377 | 2 | 0.5 | 1 | 0 | 2 | 0 | 1 | 1 | 6 | 0.3333 | 2 | 0.5 | 4 | 0.25 | 4 | 0 | 2 | 1 | 12 | 0.3333 |
| 2_1559379 | 2 | 0.5 | 1 | 1 | 1 | 0 | 1 | 0 | 5 | 0.4 | 2 | 1 | 3 | 0 | 4 | 0 | 1 | 0 | 10 | 0.2 |
| 2_1559384 | 2 | 0 | 1 | 1 | 1 | 0 | NA | NA | 4 | 0.25 | 1 | 1 | 3 | 1 | 4 | 1 | 2 | 1 | 10 | 1 |
| 2_1559384 | 2 | 0.5 | 1 | 0 | 1 | 1 | 1 | 0 | 5 | 0.4 | 2 | 0.5 | 1 | 0 | 4 | 0 | 2 | 0 | 9 | 0.1111 |
| 2_1559385 | NA | NA | 1 | 1 | 1 | 0 | NA | NA | 2 | 0.5 | 1 | 0 | 3 | 0.3333 | 2 | 0 | 2 | 0 | 8 | 0.125 |
| 2_1559386 | NA | NA | 1 | 1 | 1 | 1 | NA | NA | 2 | 1 | 2 | 1 | 2 | 0 | 2 | 0.5 | 1 | 0 | 7 | 0.4286 |
| 2_1559386 | 2 | 0.5 | 1 | 1 | 1 | 1 | NA | NA | 4 | 0.75 | 1 | 1 | 3 | 0.3333 | 3 | 0.3333 | 1 | 1 | 8 | 0.5 |
| 2_1559387 | 2 | 0.5 | 1 | 0 | NA | NA | 1 | 1 | 4 | 0.5 | 1 | 0 | 4 | 1 | 4 | 0.75 | 2 | 0.5 | 11 | 0.7273 |
| 2_1559387 | 1 | 1 | 1 | 0 | 1 | 0 | NA | NA | 3 | 0.3333 | NA | NA | 2 | 0 | 4 | 0.25 | 1 | 0 | 7 | 0.1429 |
| 2_1559388 | 2 | 0.5 | 1 | 1 | NA | NA | NA | NA | 3 | 0.6667 | 1 | 0 | 2 | 0.5 | 4 | 0.25 | NA | NA | 7 | 0.2857 |

| | | | | | | | | | | | | | | | | | | | | |
|---|---|---|---|---|---|---|---|---|---|---|---|---|---|---|---|---|---|---|---|---|
| 2_15593887 | 2 | 1 | 1 | 1 | NA | NA | NA | NA | 3 | 1 | 1 | 1 | 3 | 1 | 4 | 1 | NA | NA | 8 | 1 |
| 2_15593913 | 1 | 1 | 1 | 1 | NA | NA | NA | NA | 2 | 1 | NA | NA | 4 | 1 | 3 | 1 | 1 | 1 | 8 | 1 |
| 2_15593946 | 1 | 1 | 1 | 0 | NA | NA | NA | NA | 2 | 0.5 | NA | NA | 2 | 1 | 4 | 0.75 | NA | NA | 6 | 0.8333 |
| 2_15593982 | 1 | 1 | 1 | 0 | NA | NA | NA | NA | 2 | 0.5 | NA | NA | 1 | 1 | 2 | 1 | 1 | 1 | 4 | 1 |
| 2_15593991 | 1 | 0 | 1 | 1 | NA | NA | NA | NA | 2 | 0.5 | NA | NA | 2 | 0.5 | 3 | 0 | NA | NA | 5 | 0.2 |
| 2_15593999 | 1 | 0 | 1 | 1 | NA | NA | NA | NA | 2 | 0.5 | NA | NA | 2 | 1 | 2 | 0 | 1 | 1 | 5 | 0.6 |
| 2_15594001 | 1 | 1 | 1 | 1 | NA | NA | NA | NA | 2 | 1 | NA | NA | 2 | 1 | NA | NA | NA | NA | 2 | 1 |
| 2_15594021 | 1 | 1 | NA | NA | NA | NA | NA | NA | 1 | 1 | NA | NA | 1 | 0 | 5 | 0.8 | 1 | 1 | 7 | 0.7143 |
| 2_15596574 | 10 | 0.1 | 7 | 0.2857 | 6 | 0 | 10 | 0.2 | 33 | 0.1515 | 2 | 0 | 8 | 0.125 | 9 | 0.1111 | 7 | 0 | 26 | 0.0769 |
| 2_15596577 | 10 | 0.6 | 8 | 0.125 | 5 | 0.4 | 9 | 0.4444 | 32 | 0.4062 | 3 | 0 | 7 | 0.8571 | 9 | 0.1111 | 7 | 0.5714 | 26 | 0.4231 |
| 2_15596579 | 10 | 0.2 | 8 | 0 | 6 | 0.3333 | 8 | 0.375 | 32 | 0.2188 | 3 | 0 | 8 | 0 | 7 | 0.2857 | 7 | 0.1429 | 25 | 0.12 |
| 2_15596580 | 7 | 0.5714 | 6 | 1 | 2 | 0.5 | 8 | 0.75 | 23 | 0.7391 | 1 | 0 | 4 | 0.5 | 6 | 0.6667 | 5 | 1 | 16 | 0.6875 |
| 2_15596584 | 9 | 0.4444 | 7 | 0.5714 | 5 | 0.2 | 8 | 0.625 | 29 | 0.4828 | 3 | 0.6667 | 8 | 0.875 | 9 | 0.6667 | 7 | 0.8571 | 27 | 0.7778 |
| 2_15596587 | 9 | 0.1111 | 8 | 0.5 | 6 | 0.5 | 9 | 0.4444 | 32 | 0.375 | 3 | 0 | 7 | 0.2857 | 8 | 0.375 | 5 | 0.2 | 23 | 0.2609 |
| 2_15596589 | 10 | 0.3 | 4 | 0.25 | 5 | 0.4 | 8 | 0.25 | 27 | 0.2963 | 2 | 0 | 5 | 0 | 6 | 0.3333 | 5 | 0.2 | 18 | 0.1667 |
| 2_15596606 | 9 | 0 | 6 | 0.3333 | 6 | 0.5 | 9 | 0.3333 | 30 | 0.2667 | 3 | 0.3333 | 7 | 0.2857 | 8 | 0 | 6 | 0.3333 | 24 | 0.2083 |
| 2_15596616 | 9 | 0.1111 | 5 | 0.2 | 5 | 0 | 10 | 0.2 | 29 | 0.1379 | 3 | 0.3333 | 8 | 0.25 | 9 | 0.1111 | 6 | 0.1667 | 26 | 0.1923 |
| 2_15598403 | NA | NA | 2 | 0.5 | NA | NA | NA | NA | 2 | 0.5 | 1 | 1 | NA | NA | NA | NA | NA | NA | 1 | 1 |
| 2_15601583 | 3 | 0 | 3 | 0 | 3 | 0 | 3 | 0.3333 | 12 | 0.0833 | 2 | 0 | 1 | 0 | 3 | 0.3333 | 5 | 0 | 11 | 0.0909 |
| 2_15601965 | NA | NA | NA | NA | 3 | 0.3333 | 2 | 0.5 | 5 | 0.4 | NA | NA | 1 | 0 | 1 | 1 | 2 | 0 | 4 | 0.25 |
| 2_15601990 | NA | NA | NA | NA | 3 | 0.3333 | NA | NA | 3 | 0.3333 | NA | NA | 1 | 0 | NA | NA | 2 | 0.5 | 3 | 0.3333 |
| 2_15612981 | 2 | 0.5 | 4 | 0.5 | 4 | 0.25 | NA | NA | 10 | 0.4 | 3 | 0.6667 | 1 | 1 | 3 | 0.6667 | 3 | 0.6667 | 10 | 0.7 |
| 2_15614483 | 1 | 0 | 2 | 0.5 | 2 | 0.5 | NA | NA | 5 | 0.4 | 1 | 1 | NA | NA | 1 | 1 | 1 | 0 | 3 | 0.6667 |
| 2_15614923 | 1 | 0 | 5 | 0.2 | 3 | 0.3333 | NA | NA | 9 | 0.2222 | NA | NA | 1 | 0 | 2 | 1 | 3 | 0 | 6 | 0.3333 |
| 2_15614930 | 2 | 0 | 7 | 0.1429 | 3 | 0 | 2 | 1 | 14 | 0.2143 | 4 | 0.25 | 1 | 0 | 4 | 0.5 | 4 | 0 | 13 | 0.2308 |
| 2_15614937 | 2 | 0.5 | 7 | 0.2857 | 3 | 0.3333 | 2 | 0.5 | 14 | 0.3571 | 6 | 0.6667 | 1 | 0 | 4 | 0.25 | 5 | 0.6 | 16 | 0.5 |
| 2_15614943 | 2 | 0.5 | 7 | 0.5714 | 4 | 0.25 | 2 | 0.5 | 15 | 0.4667 | 4 | 0.75 | 1 | 0 | 5 | 0.6 | 3 | 0.6667 | 13 | 0.6154 |
| 2_15614945 | 2 | 0 | 7 | 0.1429 | 4 | 0 | 3 | 0 | 16 | 0.0625 | 6 | 0.1667 | 1 | 0 | 5 | 0.2 | 4 | 0 | 16 | 0.125 |
| 2_15614946 | 1 | 0 | 5 | 1 | 1 | 0 | 3 | 0.6667 | 10 | 0.7 | 3 | 0.3333 | 1 | 0 | 4 | 0.5 | 4 | 0.5 | 12 | 0.4167 |
| 2_15614946 | 2 | 0 | 5 | 0.2 | 3 | 0 | 2 | 0 | 12 | 0.0833 | 5 | 0.2 | 1 | 0 | 4 | 0 | 5 | 0 | 15 | 0.0667 |
| 2_15614949 | 2 | 1 | 8 | 1 | 4 | 1 | 3 | 0.6667 | 17 | 0.9412 | 5 | 0.8 | 1 | 0 | 6 | 0.6667 | 5 | 0.8 | 17 | 0.7059 |
| 2_15614950 | 2 | 1 | 6 | 0.6667 | 5 | 0.4 | 3 | 0.6667 | 16 | 0.625 | 4 | 0.5 | 1 | 0 | 4 | 1 | 5 | 0.8 | 14 | 0.7143 |
| 2_15614951 | 1 | 1 | 7 | 0.7143 | 5 | 0.2 | 3 | 1 | 16 | 0.625 | 5 | 0.8 | 2 | 0.5 | 6 | 0.5 | 4 | 0.75 | 17 | 0.6471 |
| 2_15614953 | 2 | 1 | 8 | 0.875 | 5 | 0.2 | 2 | 0.5 | 17 | 0.6471 | 6 | 0.5 | 2 | 0 | 5 | 1 | 5 | 0.8 | 18 | 0.6667 |
| 2_15614965 | 2 | 0 | 9 | 0.2222 | 5 | 0 | 3 | 0 | 19 | 0.1053 | 6 | 0.3333 | 2 | 0 | 6 | 0 | 4 | 0 | 18 | 0.1111 |
| 2_15614968 | 2 | 0.5 | 9 | 0.5556 | 4 | 0.25 | 3 | 0.6667 | 18 | 0.5 | 6 | 0.1667 | 1 | 0 | 5 | 0.4 | 5 | 0.4 | 17 | 0.2941 |
| 2_15614969 | 2 | 1 | 7 | 0 | 2 | 0.5 | 3 | 0.3333 | 14 | 0.2857 | 3 | 0.3333 | 1 | 1 | 4 | 0 | 3 | 0 | 11 | 0.1818 |
| 2_15614973 | 2 | 1 | 8 | 0.875 | 2 | 1 | 3 | 1 | 15 | 0.9333 | 4 | 1 | NA | NA | 4 | 1 | 5 | 1 | 13 | 1 |
| 2_15614984 | 2 | 1 | 7 | 1 | 5 | 1 | 3 | 1 | 17 | 1 | 5 | 1 | 1 | 1 | 6 | 1 | 4 | 0.75 | 16 | 0.9375 |
| 2_15614990 | 2 | 0 | 8 | 0.125 | 4 | 0.5 | 3 | 0 | 17 | 0.1765 | 4 | 0.25 | 1 | 1 | 5 | 0.4 | 5 | 0.4 | 15 | 0.4 |
| 2_15614995 | 2 | 1 | 9 | 0.7778 | 4 | 0.75 | 3 | 0.6667 | 18 | 0.7778 | 4 | 0.5 | 1 | 1 | 6 | 0.5 | 4 | 0.75 | 15 | 0.6 |
| 2_15615001 | 2 | 1 | 6 | 0.5 | 4 | 0.5 | 2 | 1 | 14 | 0.6429 | 7 | 0.5714 | 1 | 1 | 5 | 0.6 | 5 | 0.2 | 18 | 0.5 |
| 2_15615010 | 2 | 0.5 | 9 | 1 | 6 | 1 | 3 | 0.6667 | 20 | 0.9 | 7 | 0.8571 | 1 | 1 | 7 | 0.8571 | 5 | 1 | 20 | 0.9 |
| 2_15615018 | 2 | 1 | 6 | 0.8333 | 4 | 0.75 | 1 | 1 | 13 | 0.8462 | 4 | 1 | NA | NA | 5 | 0.8 | 5 | 1 | 14 | 0.9286 |
| 2_15615052 | 3 | 0.3333 | 10 | 0.1 | 6 | 0 | 3 | 0.3333 | 22 | 0.1364 | 8 | 0.25 | 2 | 0 | 7 | 0.1429 | 5 | 0 | 22 | 0.1364 |
| 2_15615141 | 5 | 0 | 9 | 0.2222 | 7 | 0.1429 | 5 | 0.2 | 26 | 0.1538 | 8 | 0.125 | 2 | 0 | 6 | 0.1667 | 5 | 0 | 21 | 0.0952 |
| 2_15615381 | 23 | 0.6087 | 22 | 0.7273 | 19 | 0.6842 | 17 | 0.8235 | 81 | 0.7037 | 17 | 0.5882 | 20 | 0.85 | 22 | 0.8182 | 19 | 0.5263 | 78 | 0.7051 |
| 2_15615396 | 21 | 0.0476 | 22 | 0.0455 | 18 | 0.1111 | 16 | 0 | 77 | 0.0519 | 13 | 0 | 20 | 0.05 | 20 | 0.1 | 17 | 0.0588 | 70 | 0.0571 |
| 2_15615430 | 23 | 0.087 | 20 | 0.05 | 15 | 0.1333 | 14 | 0.2857 | 72 | 0.125 | 14 | 0.0714 | 18 | 0.1667 | 20 | 0.2 | 14 | 0.0714 | 66 | 0.1364 |
| 2_15620053 | NA | NA | NA | NA | NA | NA | 1 | 1 | 1 | 1 | 2 | 1 | NA | NA | NA | NA | NA | NA | 2 | 1 |
| 2_15620074 | NA | NA | NA | NA | NA | NA | 2 | 0.5 | 2 | 0.5 | 2 | 0.5 | NA | NA | NA | NA | NA | NA | 2 | 0.5 |
| 2_15623225 | 75 | 0.16 | 91 | 0.1978 | 66 | 0.2121 | 90 | 0.0778 | 322 | 0.1584 | 44 | 0.0909 | 79 | 0.2152 | 73 | 0.1233 | 64 | 0.1562 | 260 | 0.1538 |
| 2_15623254 | 75 | 0.1467 | 88 | 0.25 | 59 | 0.1186 | 85 | 0.1529 | 307 | 0.1726 | 39 | 0.2308 | 77 | 0.0909 | 60 | 0.15 | 57 | 0.0877 | 233 | 0.1288 |
| 2_15623305 | 64 | 0.1562 | 70 | 0.2 | 50 | 0.14 | 75 | 0.16 | 259 | 0.166 | 30 | 0.1333 | 66 | 0.1818 | 47 | 0.1277 | 55 | 0.2182 | 198 | 0.1717 |
| 2_15623312 | 69 | 0.1449 | 71 | 0.0986 | 45 | 0.2 | 77 | 0.1948 | 262 | 0.1565 | 34 | 0.1765 | 66 | 0.1061 | 47 | 0.1277 | 52 | 0.1731 | 199 | 0.1407 |
| 2_15623338 | 55 | 0.1455 | 55 | 0.1273 | 35 | 0.3429 | 59 | 0.0847 | 204 | 0.1569 | 27 | 0.1111 | 58 | 0.0345 | 30 | 0.1333 | 39 | 0.1282 | 154 | 0.0909 |
| 2_15623403 | 42 | 0.2381 | 41 | 0.3171 | 28 | 0.2857 | 50 | 0.26 | 161 | 0.2733 | 21 | 0.2381 | 36 | 0.2778 | 28 | 0.3214 | 34 | 0.2353 | 119 | 0.2689 |
| 2_15623409 | 38 | 0.7632 | 39 | 0.9231 | 24 | 0.9583 | 47 | 0.8085 | 148 | 0.8514 | 21 | 0.9524 | 37 | 0.8919 | 28 | 0.9286 | 32 | 0.9062 | 118 | 0.9153 |
| 2_15623417 | 24 | 0.5833 | 21 | 0.381 | 16 | 0.625 | 23 | 0.6522 | 84 | 0.5595 | 15 | 0.8 | 18 | 0.6667 | 19 | 0.4211 | 18 | 0.5556 | 70 | 0.6 |
| 2_15623419 | 36 | 0.9722 | 37 | 1 | 26 | 0.9615 | 40 | 0.975 | 139 | 0.9784 | 19 | 1 | 28 | 0.9643 | 25 | 1 | 29 | 1 | 101 | 0.9901 |
| 2_15623441 | 31 | 0.0323 | 32 | 0.125 | 20 | 0.05 | 38 | 0.1316 | 121 | 0.0909 | 17 | 0.2353 | 27 | 0.037 | 25 | 0.08 | 23 | 0.1739 | 92 | 0.1196 |
| 2_15623442 | 38 | 0.1316 | 37 | 0.0541 | 24 | 0 | 43 | 0.0233 | 142 | 0.0563 | 19 | 0.0526 | 33 | 0.0606 | 26 | 0.0769 | 26 | 0.1154 | 104 | 0.0769 |
| 2_15623447 | 25 | 0.8 | 33 | 0.6667 | 21 | 0.6667 | 34 | 0.4706 | 113 | 0.6372 | 16 | 0.5625 | 28 | 0.75 | 17 | 0.7059 | 25 | 0.6 | 86 | 0.6628 |
| 2_15636341 | NA | NA | NA | NA | NA | NA | 1 | 1 | 1 | 1 | NA | NA | 1 | 1 | NA | NA | 1 | 1 | 2 | 1 |
| 2_15636387 | NA | NA | NA | NA | NA | NA | 2 | 1 | 2 | 1 | NA | NA | 1 | 0 | NA | NA | 1 | 1 | 2 | 0.5 |

| | | | | | | | | | | | | | | | | | | | | |
|---|---|---|---|---|---|---|---|---|---|---|---|---|---|---|---|---|---|---|---|---|
| 2_15636413 | NA | NA | NA | NA | NA | NA | 2 | 1 | 2 | 1 | NA | NA | NA | NA | 1 | 1 | 1 | 1 | 2 | 1 |
| 2_15636488 | NA | NA | NA | NA | 1 | 0 | 2 | 1 | 3 | 0.6667 | NA | NA | 1 | 1 | 2 | 0.5 | NA | NA | 3 | 0.6667 |
| 2_15636504 | NA | NA | 1 | 0 | 1 | 0 | 3 | 0.3333 | 5 | 0.2 | NA | NA | 1 | 1 | 2 | 0.5 | 1 | 0 | 4 | 0.5 |
| 2_15636542 | NA | NA | NA | NA | NA | NA | 2 | 1 | 2 | 1 | NA | NA | 1 | 1 | 1 | 1 | NA | NA | 2 | 1 |
| 2_15636561 | NA | NA | 1 | 0 | 1 | 0 | 3 | 0.3333 | 5 | 0.2 | NA | NA | 1 | 0 | 2 | 0.5 | 1 | 1 | 4 | 0.5 |
| 2_15636699 | 16 | 0.3125 | 20 | 0.3 | 15 | 0.2667 | 17 | 0.4118 | 68 | 0.3235 | 15 | 0.4667 | 13 | 0.3077 | 16 | 0.25 | 21 | 0.4762 | 65 | 0.3846 |
| 2_15636707 | 15 | 0.2667 | 23 | 0.3913 | 14 | 0.4286 | 19 | 0.4211 | 71 | 0.3803 | 17 | 0.4706 | 11 | 0.1818 | 19 | 0.4737 | 18 | 0.5556 | 65 | 0.4462 |
| 2_15638186 | 21 | 0.9524 | 28 | 1 | 21 | 1 | 34 | 1 | 104 | 0.9904 | 19 | 0.9474 | 21 | 0.8095 | 25 | 1 | 31 | 1 | 96 | 0.9479 |
| 2_15638201 | 24 | 0.2083 | 24 | 0.2083 | 18 | 0.1111 | 33 | 0.2121 | 99 | 0.1919 | 23 | 0.1739 | 21 | 0.2381 | 24 | 0.125 | 29 | 0.3103 | 97 | 0.2165 |
| 2_15638205 | 19 | 0.2105 | 24 | 0.125 | 19 | 0.0526 | 26 | 0.1923 | 88 | 0.1477 | 18 | 0.1667 | 17 | 0.1176 | 26 | 0.0769 | 28 | 0.2857 | 89 | 0.1685 |
| 2_15638209 | 24 | 0.1667 | 27 | 0.1481 | 19 | 0.2105 | 34 | 0.1765 | 104 | 0.1731 | 23 | 0.087 | 22 | 0.0455 | 29 | 0.1034 | 31 | 0.1935 | 105 | 0.1143 |
| 2_15638211 | 26 | 0 | 30 | 0.0667 | 23 | 0.087 | 40 | 0.125 | 119 | 0.0756 | 23 | 0.1304 | 22 | 0 | 30 | 0.0667 | 33 | 0.0909 | 108 | 0.0741 |
| 2_15638212 | 24 | 1 | 29 | 0.6897 | 23 | 0.6522 | 36 | 0.6944 | 112 | 0.75 | 26 | 0.7308 | 23 | 0.7391 | 24 | 0.5833 | 32 | 0.7812 | 105 | 0.7143 |
| 2_15638215 | 26 | 0.4231 | 30 | 0.4667 | 24 | 0.4583 | 37 | 0.5946 | 117 | 0.4957 | 26 | 0.3462 | 26 | 0.4231 | 32 | 0.375 | 37 | 0.4054 | 121 | 0.3884 |
| 2_15638226 | 22 | 0.1364 | 31 | 0.129 | 25 | 0.08 | 41 | 0.0488 | 119 | 0.0924 | 26 | 0.1538 | 26 | 0.0769 | 29 | 0.0345 | 35 | 0.0857 | 116 | 0.0862 |
| 2_15638240 | 25 | 0.32 | 32 | 0.5312 | 22 | 0.5 | 38 | 0.4474 | 117 | 0.453 | 26 | 0.4615 | 19 | 0.4737 | 33 | 0.5455 | 35 | 0.4571 | 113 | 0.4867 |
| 2_15638256 | 18 | 0.2778 | 26 | 0.2692 | 21 | 0.3333 | 31 | 0.2258 | 96 | 0.2708 | 20 | 0.25 | 22 | 0.3182 | 27 | 0.5185 | 30 | 0.2333 | 99 | 0.3333 |
| 2_15638258 | 29 | 0.1034 | 37 | 0.1351 | 25 | 0.28 | 44 | 0.1591 | 135 | 0.163 | 25 | 0.04 | 28 | 0.1786 | 35 | 0.0571 | 39 | 0.1026 | 127 | 0.0945 |
| 2_15638262 | 28 | 0.3214 | 34 | 0.3235 | 26 | 0.1538 | 37 | 0.2703 | 125 | 0.272 | 28 | 0.2143 | 24 | 0.4167 | 32 | 0.125 | 39 | 0.2564 | 123 | 0.2439 |
| 2_15638267 | 24 | 0.8333 | 34 | 0.8235 | 19 | 0.8421 | 38 | 0.8684 | 115 | 0.8435 | 29 | 0.8276 | 28 | 0.6071 | 35 | 0.8857 | 35 | 0.6286 | 127 | 0.7402 |
| 2_15638273 | 24 | 0.2917 | 29 | 0.3448 | 20 | 0.35 | 39 | 0.3077 | 112 | 0.3214 | 27 | 0.2222 | 24 | 0.25 | 35 | 0.2571 | 37 | 0.3243 | 123 | 0.2683 |
| 2_15638278 | 11 | 0.1818 | 22 | 0.2727 | 15 | 0.1333 | 22 | 0.2273 | 70 | 0.2143 | 10 | 0.1 | 15 | 0 | 16 | 0.4375 | 22 | 0.3182 | 63 | 0.2381 |
| 2_15638283 | 5 | 0 | 12 | 0.1667 | 11 | 0.1818 | 10 | 0.1 | 38 | 0.1316 | 7 | 0.2857 | 9 | 0.1111 | 10 | 0.1 | 14 | 0.3571 | 40 | 0.225 |
| 2_15638284 | 10 | 1 | 20 | 1 | 15 | 1 | 18 | 1 | 63 | 1 | 8 | 1 | 12 | 0.9167 | 14 | 1 | 18 | 0.9444 | 52 | 0.9615 |
| 2_15638438 | 20 | 0.25 | 22 | 0.0909 | 18 | 0.1667 | 23 | 0.2174 | 83 | 0.1807 | 11 | 0.4545 | 16 | 0.1875 | 20 | 0.15 | 23 | 0.1739 | 70 | 0.2143 |
| 2_15638491 | 26 | 0.0385 | 39 | 0.1795 | 29 | 0.0345 | 28 | 0.0357 | 122 | 0.082 | 20 | 0.1 | 25 | 0.08 | 25 | 0.04 | 29 | 0.1034 | 99 | 0.0808 |
| 2_15639268 | 3 | 0.3333 | 6 | 0.1667 | 3 | 0.3333 | 6 | 0.1667 | 18 | 0.2222 | 2 | 0 | 3 | 0 | 2 | 0 | 6 | 0.1667 | 13 | 0.0769 |
| 2_15639270 | 2 | 0.5 | 8 | 0.75 | 4 | 0.5 | 7 | 0.4286 | 21 | 0.5714 | 2 | 1 | 3 | 0.3333 | 5 | 0.6 | 4 | 0.25 | 14 | 0.5 |
| 2_15639271 | 2 | 0.5 | 8 | 0.125 | 4 | 0.25 | 6 | 0.3333 | 20 | 0.25 | 2 | 0.5 | 4 | 0.25 | 4 | 0 | 7 | 0.5714 | 17 | 0.3529 |
| 2_15639280 | 3 | 0.6667 | 6 | 0.3333 | 3 | 0.6667 | 6 | 0.5 | 18 | 0.5 | 2 | 0 | 6 | 0.1667 | 4 | 0.5 | 6 | 0.3333 | 18 | 0.2778 |
| 2_15639298 | 3 | 0.3333 | 7 | 0.5714 | 6 | 0.8333 | 5 | 0.4 | 21 | 0.5714 | 3 | 0.3333 | 4 | 0.5 | 5 | 0.4 | 7 | 0.5714 | 19 | 0.4737 |
| 2_15639312 | 3 | 1 | 3 | 1 | 6 | 0.8333 | 3 | 0.6667 | 15 | 0.8667 | 1 | 1 | 3 | 1 | 5 | 0.8 | 5 | 1 | 14 | 0.9286 |
| 2_15639322 | 4 | 0.75 | 9 | 0.2222 | 7 | 0.1429 | 8 | 0.25 | 28 | 0.2857 | 3 | 0 | 5 | 0.8 | 2 | 0 | 6 | 0.5 | 16 | 0.4375 |
| 2_15639330 | 4 | 0.75 | 8 | 0.625 | 7 | 0.8571 | 6 | 0.8333 | 25 | 0.76 | 3 | 1 | 4 | 1 | 5 | 0.6 | 7 | 1 | 19 | 0.8947 |
| 2_15639332 | 3 | 0 | 8 | 0 | 6 | 0.3333 | 7 | 0 | 24 | 0.0833 | 3 | 0 | 5 | 0.2 | 5 | 0 | 7 | 0 | 20 | 0.05 |
| 2_15639793 | 4 | 0.25 | 4 | 0 | 5 | 0 | 5 | 0.2 | 18 | 0.1111 | 2 | 0 | 4 | 0.5 | 3 | 0 | 8 | 0.25 | 17 | 0.2353 |
| 2_15640888 | 5 | 0.4 | 7 | 0 | 4 | 0 | 7 | 0 | 23 | 0.087 | 4 | 0.25 | 4 | 0.25 | 4 | 0 | 7 | 0 | 19 | 0.1053 |
| 2_15640890 | 6 | 0.8333 | 8 | 0.875 | 6 | 0.8333 | 7 | 0.5714 | 27 | 0.7778 | 4 | 1 | 7 | 1 | 4 | 0.75 | 10 | 0.6 | 25 | 0.8 |
| 2_15640902 | 5 | 0 | 8 | 0.125 | 6 | 0.3333 | 6 | 0.1667 | 25 | 0.16 | 4 | 0.25 | 5 | 0.4 | 5 | 0.4 | 10 | 0.1 | 24 | 0.25 |
| 2_15640914 | 4 | 1 | 8 | 1 | 6 | 1 | 6 | 0.8333 | 24 | 0.9583 | 3 | 1 | 7 | 0.8571 | 5 | 1 | 7 | 1 | 22 | 0.9545 |
| 2_15640921 | 7 | 0.1429 | 12 | 0 | 8 | 0.5 | 8 | 0.125 | 35 | 0.1714 | 4 | 0.25 | 6 | 0 | 6 | 0.1667 | 11 | 0.2727 | 27 | 0.1852 |
| 2_15640923 | 6 | 1 | 9 | 1 | 7 | 0.8571 | 7 | 0.8571 | 29 | 0.931 | 3 | 1 | 6 | 0.8333 | 3 | 1 | 7 | 1 | 19 | 0.9474 |
| 2_15640925 | 5 | 0.8 | 10 | 0.8 | 8 | 1 | 7 | 0.7143 | 30 | 0.8333 | 3 | 0.6667 | 5 | 1 | 5 | 1 | 11 | 0.7273 | 24 | 0.8333 |
| 2_15640941 | 7 | 0.8571 | 10 | 0.7 | 7 | 1 | 4 | 0.5 | 28 | 0.7857 | 4 | 0.5 | 4 | 0.5 | 6 | 0.8333 | 9 | 0.5556 | 23 | 0.6087 |
| 2_15640956 | 8 | 0.625 | 11 | 0.7273 | 7 | 0.8571 | 8 | 0.875 | 34 | 0.7647 | 5 | 1 | 7 | 0.8571 | 9 | 0.7778 | 9 | 0.7778 | 30 | 0.8333 |
| 2_15640967 | 7 | 0 | 10 | 0.1 | 8 | 0.25 | 7 | 0 | 32 | 0.0938 | 4 | 0.25 | 7 | 0.1429 | 8 | 0 | 10 | 0.2 | 29 | 0.1379 |
| 2_15640975 | 7 | 0.2857 | 10 | 0.2 | 7 | 0.2857 | 7 | 0.2857 | 31 | 0.2581 | 7 | 0 | 5 | 0 | 6 | 0.3333 | 11 | 0.3636 | 29 | 0.2069 |
| 2_15640991 | 8 | 0.125 | 11 | 0.1818 | 8 | 0 | 8 | 0.25 | 35 | 0.1429 | 7 | 0 | 8 | 0 | 9 | 0 | 12 | 0.25 | 36 | 0.0833 |
| 2_15641000 | 7 | 0.7143 | 10 | 0.9 | 5 | 1 | 9 | 0.7778 | 31 | 0.8387 | 6 | 0.6667 | 8 | 0.875 | 5 | 1 | 10 | 0.8 | 29 | 0.8276 |
| 2_15641007 | 3 | 1 | 6 | 0.8333 | 4 | 0.5 | 7 | 0.4286 | 20 | 0.65 | 2 | 0 | 6 | 0.8333 | 6 | 0.6667 | 6 | 0.1667 | 20 | 0.5 |
| 2_15645136 | 1 | 1 | 1 | 1 | 1 | 1 | NA | NA | 3 | 1 | NA | NA | NA | NA | NA | NA | 1 | 1 | 1 | 1 |
| 2_15654267 | 2 | 0 | 11 | 0.8182 | 5 | 0.6 | 8 | 0.625 | 26 | 0.6538 | 7 | 0.7143 | 5 | 0.6 | 12 | 0.6667 | 1 | 0 | 25 | 0.64 |
| 2_15654291 | 3 | 0.3333 | 12 | 0.4167 | 5 | 0.6 | 9 | 0.3333 | 29 | 0.4138 | 6 | 0.5 | 5 | 0.2 | 13 | 0.3077 | NA | NA | 24 | 0.3333 |
| 2_15654328 | 4 | 0.25 | 16 | 0.25 | 6 | 0.1667 | 8 | 0.375 | 34 | 0.2647 | 11 | 0.3636 | 7 | 0.1429 | 12 | 0.5 | 2 | 0.5 | 32 | 0.375 |
| 2_15654337 | 3 | 1 | 13 | 0.6154 | 4 | 0.75 | 9 | 0.6667 | 29 | 0.6897 | 10 | 0.3 | 7 | 0.5714 | 10 | 0.4 | 2 | 0.5 | 29 | 0.4138 |
| 2_15654338 | 4 | 0 | 16 | 0 | 7 | 0.1429 | 10 | 0.1 | 37 | 0.0541 | 11 | 0.2727 | 6 | 0.1667 | 10 | 0.1 | 2 | 0 | 29 | 0.1724 |
| 2_15658787 | 13 | 0.2308 | 22 | 0 | 8 | 0.125 | 20 | 0.05 | 63 | 0.0794 | 14 | 0.1429 | 18 | 0.2222 | 16 | 0.125 | 8 | 0 | 56 | 0.1429 |
| 2_15658794 | 14 | 0.1429 | 21 | 0.0476 | 7 | 0.1429 | 24 | 0.1667 | 66 | 0.1212 | 14 | 0.2143 | 21 | 0.0952 | 19 | 0.1053 | 8 | 0 | 62 | 0.1129 |
| 2_15658804 | 15 | 0.4 | 26 | 0.4231 | 8 | 0.25 | 26 | 0.3846 | 75 | 0.3867 | 16 | 0.3125 | 26 | 0.3462 | 23 | 0.3478 | 11 | 0.7273 | 76 | 0.3947 |
| 2_15658809 | 13 | 0.0769 | 23 | 0.1739 | 10 | 0 | 26 | 0.0385 | 72 | 0.0833 | 14 | 0.0714 | 23 | 0.1304 | 22 | 0 | 8 | 0.125 | 67 | 0.0746 |
| 2_15659096 | 16 | 0 | 28 | 0.1429 | 14 | 0.1429 | 30 | 0.1667 | 88 | 0.125 | 18 | 0.2222 | 30 | 0.2333 | 28 | 0.1071 | 13 | 0.0769 | 89 | 0.1685 |
| 2_15659121 | 14 | 0 | 26 | 0.1538 | 13 | 0.1538 | 26 | 0.1538 | 79 | 0.1266 | 18 | 0.0556 | 30 | 0.1333 | 26 | 0.0769 | 12 | 0.0833 | 86 | 0.093 |
| 2_15660469 | 14 | 0.0714 | 26 | 0.0769 | 12 | 0.0833 | 30 | 0.0667 | 82 | 0.0732 | 21 | 0.0952 | 27 | 0.037 | 29 | 0.1034 | 14 | 0 | 91 | 0.0659 |
| 2_15660472 | 18 | 0.1667 | 31 | 0.2581 | 14 | 0.3571 | 29 | 0.1724 | 92 | 0.2283 | 23 | 0.3913 | 32 | 0.2812 | 29 | 0.2069 | 16 | 0.25 | 100 | 0.28 |
| 2_15660478 | 16 | 0.1875 | 33 | 0.1818 | 17 | 0.1765 | 31 | 0.1613 | 97 | 0.1753 | 23 | 0.1739 | 34 | 0.2059 | 30 | 0.1667 | 17 | 0.2353 | 104 | 0.1923 |

| | | | | | | | | | | | | | | | | | | | | |
|---|---|---|---|---|---|---|---|---|---|---|---|---|---|---|---|---|---|---|---|---|
| 2_1566048( | NA | NA | 1 | 1 | NA | NA | NA | NA | 1 | 1 | NA | NA | 1 | 1 | NA | NA | NA | NA | 1 | 1 |
| 2_1566048( | 14 | 0.2143 | 32 | 0.4062 | 11 | 0.4545 | 32 | 0.4375 | 89 | 0.3933 | 20 | 0.55 | 30 | 0.4667 | 27 | 0.4815 | 15 | 0.5333 | 92 | 0.5 |
| 2_1566048: | 14 | 0.0714 | 25 | 0.04 | 11 | 0 | 23 | 0.087 | 73 | 0.0548 | 21 | 0.4286 | 20 | 0.2 | 25 | 0.24 | 11 | 0.3636 | 77 | 0.2987 |
| 2_1566049: | 13 | 0.5385 | 26 | 0.6538 | 13 | 0.3846 | 24 | 0.5417 | 76 | 0.5526 | 22 | 0.4091 | 23 | 0.6087 | 28 | 0.3571 | 12 | 0.5 | 85 | 0.4588 |
| 2_1566049· | 11 | 0.2727 | 19 | 0.1053 | 7 | 0.1429 | 18 | 0.2222 | 55 | 0.1818 | 15 | 0.5333 | 14 | 0.2143 | 18 | 0.0556 | 12 | 0.1667 | 59 | 0.2373 |
| 2_1566049· | 1 | 0 | 5 | 0.2 | NA | NA | 1 | 0 | 7 | 0.1429 | 4 | 0 | 1 | 1 | 2 | 0.5 | NA | NA | 7 | 0.2857 |
| 2_1566050( | 15 | 0.6 | 28 | 0.4286 | 12 | 0.3333 | 24 | 0.4583 | 79 | 0.4557 | 23 | 0.3478 | 22 | 0.5455 | 28 | 0.4286 | 14 | 0.4286 | 87 | 0.4368 |
| 2_1566050( | 13 | 0.2308 | 23 | 0.1739 | 11 | 0.0909 | 18 | 0.2222 | 65 | 0.1846 | 21 | 0.1905 | 22 | 0.1818 | 28 | 0.2857 | 12 | 0 | 83 | 0.1928 |
| 2_1566050! | 7 | 0.1429 | 12 | 0.0833 | 8 | 0.125 | 17 | 0.2941 | 44 | 0.1818 | 13 | 0.0769 | 14 | 0 | 18 | 0.1111 | 9 | 0.1111 | 54 | 0.0741 |
| 2_1566050: | 12 | 0.0833 | 28 | 0.1071 | 12 | 0.0833 | 26 | 0.1154 | 78 | 0.1026 | 23 | 0.2174 | 22 | 0.0909 | 28 | 0.0714 | 15 | 0.1333 | 88 | 0.125 |
| 2_1566051! | 15 | 0 | 24 | 0.1667 | 13 | 0.1538 | 26 | 0.1154 | 78 | 0.1154 | 23 | 0.2174 | 25 | 0.16 | 29 | 0.2069 | 14 | 0.2143 | 91 | 0.1978 |
| 2_1566052! | 7 | 0 | 13 | 0 | 11 | 0.1818 | 20 | 0.1 | 51 | 0.0784 | 11 | 0.0909 | 17 | 0.1765 | 18 | 0.1111 | 9 | 0 | 55 | 0.1091 |
| 2_1566053: | 15 | 0.0667 | 28 | 0.1429 | 13 | 0.0769 | 26 | 0.2308 | 82 | 0.1463 | 23 | 0.1304 | 23 | 0.1304 | 28 | 0.1071 | 15 | 0.2 | 89 | 0.1348 |
| 2_1566055: | 15 | 0.1333 | 29 | 0.1034 | 13 | 0.0769 | 24 | 0.125 | 81 | 0.1111 | 19 | 0.1579 | 24 | 0.0833 | 30 | 0.1333 | 15 | 0.1333 | 88 | 0.125 |
| 2_1566231! | 7 | 0.2857 | 3 | 0.3333 | 3 | 0.3333 | 7 | 0.7143 | 20 | 0.45 | 3 | 0 | NA | NA | 3 | 0.3333 | 4 | 0.25 | 10 | 0.2 |
| 2_1566234' | 8 | 0.625 | 3 | 0.3333 | 4 | 0.25 | 5 | 0.6 | 20 | 0.5 | 3 | 0.3333 | 1 | 1 | 2 | 0 | 4 | 0.25 | 10 | 0.3 |
| 2_1566858! | NA | NA | NA | NA | 1 | 1 | NA | NA | 1 | 1 | NA | NA | 1 | 1 | NA | NA | NA | NA | 1 | 1 |
| 2_1566955! | 1 | 0 | 1 | 0 | 3 | 0.3333 | NA | NA | 5 | 0.2 | 1 | 0 | 1 | 1 | NA | NA | 1 | 0 | 3 | 0.3333 |
| 2_1566956( | 1 | 1 | 1 | 1 | 1 | 1 | NA | NA | 3 | 1 | 1 | 0 | 1 | 1 | NA | NA | NA | NA | 2 | 0.5 |
| 2_1566956; | NA | NA | 1 | 1 | 2 | 1 | NA | NA | 3 | 1 | 1 | 0 | 1 | 1 | NA | NA | 1 | 1 | 3 | 0.6667 |
| 2_1566958( | 1 | 1 | 1 | 1 | 3 | 1 | NA | NA | 5 | 1 | 1 | 1 | 1 | 1 | NA | NA | 1 | 1 | 3 | 1 |
| 2_1567051: | 5 | 0.2 | 11 | 0 | 10 | 0 | 12 | 0.0833 | 38 | 0.0526 | 5 | 0.2 | 12 | 0 | 14 | 0.0714 | 7 | 0.1429 | 38 | 0.0789 |
| 2_1567055: | NA | NA | NA | NA | 1 | 1 | NA | NA | 1 | 1 | NA | NA | NA | NA | 1 | 1 | NA | NA | 1 | 1 |
| 2_1567150· | 8 | 0.625 | 17 | 0.1765 | 14 | 0.5714 | 17 | 0.2941 | 56 | 0.375 | 11 | 0.1818 | 17 | 0.2941 | 17 | 0.2941 | 11 | 0.1818 | 56 | 0.25 |
| 2_1567151: | 11 | 0.4545 | 18 | 0.4444 | 16 | 0.375 | 17 | 0.3529 | 62 | 0.4032 | 10 | 0.2 | 18 | 0.1667 | 15 | 0.4 | 10 | 0.3 | 53 | 0.2642 |
| 2_1567151· | 11 | 0.6364 | 16 | 0.625 | 16 | 0.75 | 15 | 0.5333 | 58 | 0.6379 | 10 | 0.8 | 13 | 0.6154 | 18 | 0.5556 | 10 | 0.6 | 51 | 0.6275 |
| 2_1567151( | 12 | 0.8333 | 18 | 0.7222 | 12 | 0.5 | 17 | 0.6471 | 59 | 0.678 | 11 | 0.2727 | 18 | 0.7778 | 20 | 0.5 | 11 | 0.7273 | 60 | 0.5833 |
| 2_1567151: | 10 | 0.2 | 16 | 0.25 | 14 | 0.1429 | 13 | 0.3077 | 53 | 0.2264 | 8 | 0.125 | 10 | 0.5 | 14 | 0.2143 | 11 | 0.4545 | 43 | 0.3256 |
| 2_1567154! | 9 | 0.2222 | 18 | 0.2222 | 15 | 0.2 | 16 | 0.125 | 58 | 0.1897 | 10 | 0 | 17 | 0.1176 | 18 | 0.0556 | 11 | 0.1818 | 56 | 0.0893 |
| 2_1567694: | 67 | 0.3433 | 93 | 0.3118 | 76 | 0.2763 | 65 | 0.2308 | 301 | 0.2924 | 74 | 0.2297 | 48 | 0.2292 | 49 | 0.1224 | 69 | 0.3333 | 240 | 0.2375 |
| 2_1567698: | 55 | 0.0909 | 78 | 0.0256 | 60 | 0.0667 | 46 | 0.1522 | 239 | 0.0753 | 56 | 0.0893 | 37 | 0.027 | 41 | 0.0244 | 55 | 0.0909 | 189 | 0.0635 |
| 2_1567702: | 55 | 0.2 | 84 | 0.1667 | 62 | 0.2581 | 43 | 0.1163 | 244 | 0.1885 | 59 | 0.2712 | 43 | 0.1395 | 35 | 0.1429 | 51 | 0.2157 | 188 | 0.2021 |
| 2_1567705! | 37 | 0.3784 | 62 | 0.3065 | 46 | 0.3043 | 36 | 0.3056 | 181 | 0.3204 | 38 | 0.3684 | 29 | 0.3793 | 27 | 0.3704 | 46 | 0.2174 | 140 | 0.3214 |
| 2_1567706( | 44 | 0.0909 | 75 | 0.12 | 52 | 0.0769 | 41 | 0.0732 | 212 | 0.0943 | 42 | 0.0952 | 35 | 0.0857 | 28 | 0.1429 | 53 | 0.0755 | 158 | 0.0949 |
| 2_1567706! | 44 | 0.2955 | 69 | 0.1594 | 45 | 0.2889 | 34 | 0.1471 | 192 | 0.2188 | 42 | 0.0952 | 34 | 0.1765 | 30 | 0.1 | 43 | 0.3023 | 149 | 0.1745 |
| 2_1567706: | 44 | 0.0455 | 70 | 0.1 | 49 | 0.1429 | 36 | 0.0556 | 199 | 0.0905 | 46 | 0.1087 | 35 | 0.1143 | 31 | 0.0968 | 52 | 0.1154 | 164 | 0.1098 |
| 2_1568335· | NA | NA | NA | NA | NA | NA | 1 | 1 | 1 | 1 | NA | NA | 1 | 1 | NA | NA | NA | NA | 1 | 1 |
| 2_1568378( | 1 | 1 | NA | NA | NA | NA | NA | NA | 1 | 1 | 1 | 1 | NA | NA | NA | NA | 1 | 0 | 2 | 0.5 |
| 2_1568527· | NA | NA | 2 | 0.5 | 3 | 0 | NA | NA | 5 | 0.2 | 6 | 0.1667 | 5 | 0 | 5 | 0 | 2 | 0 | 18 | 0.0556 |
| 2_1568529' | NA | NA | 4 | 0 | 3 | 0.3333 | NA | NA | 7 | 0.1429 | 4 | 0.25 | 6 | 0 | 6 | 0.1667 | 2 | 0 | 18 | 0.1111 |
| 2_1568578: | 1 | 0 | 2 | 0.5 | NA | NA | 1 | 1 | 4 | 0.5 | 1 | 0 | 1 | 1 | 2 | 0.5 | NA | NA | 4 | 0.5 |
| 2_1568584· | NA | NA | 2 | 0 | NA | NA | 2 | 0.5 | 4 | 0.25 | 1 | 0 | 1 | 0 | 2 | 0.5 | NA | NA | 4 | 0.25 |
| 2_1568585: | 1 | 1 | 1 | 1 | NA | NA | 2 | 1 | 4 | 1 | NA | NA | 1 | 1 | 3 | 1 | NA | NA | 4 | 1 |
| 2_1568585: | 1 | 0 | 2 | 0.5 | NA | NA | 2 | 0 | 5 | 0.2 | NA | NA | 1 | 1 | 2 | 0 | NA | NA | 3 | 0.3333 |
| 2_1568588' | 1 | 1 | 2 | 1 | NA | NA | 1 | 1 | 4 | 1 | NA | NA | 1 | 1 | 2 | 1 | NA | NA | 3 | 1 |
| 2_1568588: | 1 | 0 | 2 | 0.5 | NA | NA | NA | NA | 3 | 0.3333 | NA | NA | NA | NA | 2 | 1 | NA | NA | 2 | 1 |
| 2_1568589! | 1 | 0 | 2 | 0.5 | NA | NA | 1 | 1 | 4 | 0.5 | NA | NA | 1 | 0 | 2 | 1 | NA | NA | 3 | 0.6667 |
| 2_1568599' | 1 | 1 | NA | NA | NA | NA | 1 | 0 | 2 | 0.5 | NA | NA | 1 | 1 | 1 | 0 | NA | NA | 2 | 0.5 |
| 2_1568600· | 1 | 1 | NA | NA | NA | NA | 1 | 1 | 2 | 1 | NA | NA | 1 | 1 | 1 | 1 | NA | NA | 2 | 1 |
| 2_1568601' | 1 | 1 | NA | NA | NA | NA | 1 | 1 | 2 | 1 | NA | NA | 1 | 0 | 1 | 1 | NA | NA | 2 | 0.5 |
| 2_1568820( | 1 | 1 | NA | NA | NA | NA | NA | NA | 1 | 1 | 1 | 1 | NA | NA | NA | NA | NA | NA | 1 | 1 |
| 2_1569399' | 1 | 1 | 5 | 0.2 | 1 | 0 | 2 | 0 | 9 | 0.2222 | 3 | 0 | 2 | 0.5 | 1 | 0 | NA | NA | 6 | 0.1667 |
| 2_1569401' | 1 | 1 | 4 | 1 | 1 | 1 | 2 | 1 | 8 | 1 | 4 | 1 | 2 | 1 | 2 | 0 | NA | NA | 8 | 0.75 |
| 2_1569401' | 1 | 0 | 6 | 0.1667 | 1 | 1 | 3 | 0 | 11 | 0.1818 | 4 | 0 | 3 | 0 | 2 | 0.5 | NA | NA | 9 | 0.1111 |
| 2_1569402! | 1 | 0 | 5 | 0.4 | 1 | 1 | 3 | 0 | 10 | 0.3 | 3 | 0.3333 | 3 | 0 | 1 | 0 | NA | NA | 7 | 0.1429 |
| 2_1569406( | 1 | 1 | 5 | 0 | NA | NA | 3 | 0 | 9 | 0.1111 | 3 | 0.3333 | NA | NA | 1 | 0 | NA | NA | 4 | 0.25 |
| 2_1569406; | 1 | 1 | 5 | 0.2 | NA | NA | 2 | 0 | 8 | 0.25 | 3 | 0.3333 | 3 | 0.3333 | 1 | 0 | NA | NA | 7 | 0.2857 |
| 2_1569407: | 1 | 1 | 4 | 0.25 | NA | NA | 3 | 0.3333 | 8 | 0.375 | 2 | 0 | 3 | 0.3333 | 2 | 0 | NA | NA | 7 | 0.1429 |
| 2_1569410( | 1 | 1 | 4 | 0 | NA | NA | 3 | 0.6667 | 8 | 0.375 | 3 | 0.6667 | 3 | 0.6667 | 2 | 0.5 | NA | NA | 8 | 0.625 |
| 2_1569412: | 1 | 0 | 4 | 0 | NA | NA | 2 | 0.5 | 7 | 0.1429 | 3 | 0.3333 | 3 | 0 | 1 | 0 | NA | NA | 7 | 0.1429 |
| 2_1571668' | NA | NA | NA | NA | 1 | 1 | NA | NA | 1 | 1 | 1 | 0 | NA | NA | 1 | 1 | 1 | 1 | 3 | 0.6667 |
| 2_1571677( | 18 | 0.1111 | 24 | 0.0417 | 14 | 0.0714 | 14 | 0.3571 | 70 | 0.1286 | 18 | 0.1111 | 16 | 0.0625 | 24 | 0.2083 | 19 | 0.0526 | 77 | 0.1169 |
| 2_1572379: | 7 | 0.1429 | 7 | 0.2857 | 7 | 0.1429 | 9 | 0.2222 | 30 | 0.2 | 4 | 0.25 | 6 | 0.1667 | 8 | 0.25 | 3 | 0 | 21 | 0.1905 |
| 2_1572380: | 11 | 0.9091 | 12 | 1 | 8 | 1 | 8 | 1 | 39 | 0.9744 | 6 | 1 | 9 | 0.8889 | 11 | 0.9091 | 4 | 0.75 | 30 | 0.9 |
| 2_1572380: | 3 | 0 | 5 | 0.4 | 5 | 0 | 6 | 0.3333 | 19 | 0.2105 | 5 | 0.4 | 5 | 0.2 | 9 | 0.3333 | 2 | 0.5 | 21 | 0.3333 |

| | | | | | | | | | | | | | | | | | | | | |
|---|---|---|---|---|---|---|---|---|---|---|---|---|---|---|---|---|---|---|---|---|
| 2_1572383 | 13 | 0.2308 | 13 | 0.1538 | 10 | 0.4 | 9 | 0.3333 | 45 | 0.2667 | 8 | 0.25 | 6 | 0.3333 | 13 | 0.0769 | 5 | 0 | 32 | 0.1562 |
| 2_1572384 | 4 | 0 | 12 | 0.0833 | 8 | 0.125 | 6 | 0.1667 | 30 | 0.1 | 5 | 0 | 8 | 0.25 | 11 | 0.0909 | 6 | 0 | 30 | 0.1 |
| 2_1572385 | 13 | 0.6154 | 10 | 0.5 | 9 | 0.2222 | 10 | 0.6 | 42 | 0.5 | 8 | 0.375 | 9 | 0.6667 | 13 | 0.4615 | 7 | 0.1429 | 37 | 0.4324 |
| 2_1572390 | 2 | 0.5 | NA | NA | NA | NA | NA | NA | 2 | 0.5 | NA | NA | 1 | 1 | NA | NA | NA | NA | 1 | 1 |
| 2_1573118 | 2 | 0.5 | 4 | 1 | 1 | 0 | 6 | 1 | 13 | 0.8462 | 8 | 0.875 | 7 | 1 | 1 | 1 | 2 | 1 | 18 | 0.9444 |
| 2_1573118 | NA | NA | 1 | 1 | NA | NA | 4 | 0.75 | 5 | 0.8 | 5 | 0.8 | 4 | 1 | NA | NA | 3 | 0.6667 | 12 | 0.8333 |
| 2_1573120 | 2 | 0.5 | 3 | 1 | NA | NA | 7 | 1 | 12 | 0.9167 | 7 | 0.7143 | 3 | 0.6667 | 1 | 1 | 3 | 1 | 14 | 0.7857 |
| 2_1573120 | 2 | 0.5 | 3 | 1 | 1 | 1 | 7 | 0.7143 | 13 | 0.7692 | 8 | 0.75 | 6 | 0.5 | 1 | 1 | 2 | 1 | 17 | 0.7059 |
| 2_1573121 | 3 | 0.6667 | 2 | 1 | NA | NA | 8 | 0.875 | 13 | 0.8462 | 4 | 0.75 | 5 | 0.6 | 1 | 1 | 3 | 0.3333 | 13 | 0.6154 |
| 2_1573123 | 3 | 0 | 5 | 0.4 | 2 | 0 | 6 | 0.1667 | 16 | 0.1875 | 7 | 0.2857 | 5 | 0.2 | 1 | 1 | 2 | 0.5 | 15 | 0.3333 |
| 2_1573124 | 3 | 0 | 6 | 0.1667 | 3 | 0 | 9 | 0.3333 | 21 | 0.1905 | 11 | 0.5455 | 8 | 0.25 | 1 | 1 | 5 | 0.6 | 25 | 0.48 |
| 2_1573125 | 4 | 0 | 5 | 0.6 | 2 | 0 | 8 | 0.375 | 19 | 0.3158 | 8 | 0.5 | 5 | 0 | 2 | 0.5 | 4 | 0.5 | 19 | 0.3684 |
| 2_1573135 | 6 | 0 | 14 | 0.1429 | 9 | 0.1111 | 15 | 0 | 44 | 0.0682 | 17 | 0 | 11 | 0.1818 | 9 | 0 | 8 | 0.375 | 45 | 0.1111 |
| 2_1573861 | 1 | 1 | NA | NA | NA | NA | NA | NA | 1 | 1 | NA | NA | 1 | 1 | 3 | 1 | 2 | 1 | 6 | 1 |
| 2_1573861 | 1 | 0 | NA | NA | NA | NA | 1 | 1 | 2 | 0.5 | NA | NA | NA | NA | 1 | 1 | 2 | 1 | 3 | 1 |
| 2_1573863 | 1 | 0 | NA | NA | NA | NA | 2 | 0.5 | 3 | 0.3333 | NA | NA | 1 | 1 | 3 | 0.3333 | 2 | 0.5 | 6 | 0.5 |
| 2_1573983 | 6 | 0.6667 | 9 | 0.2222 | 13 | 0.5385 | 12 | 0.3333 | 40 | 0.425 | 4 | 0 | 17 | 0.3529 | 19 | 0.3158 | 13 | 0.4615 | 53 | 0.3396 |
| 2_1573983 | 8 | 0.5 | 15 | 0.2 | 15 | 0.1333 | 13 | 0.0769 | 51 | 0.1961 | 9 | 0.1111 | 20 | 0.1 | 18 | 0.1111 | 13 | 0.2308 | 60 | 0.1333 |
| 2_1573984 | 6 | 0.5 | 14 | 0.3571 | 14 | 0.5 | 13 | 0.3077 | 47 | 0.4043 | 8 | 0.5 | 20 | 0.6 | 18 | 0.1667 | 11 | 0.0909 | 57 | 0.3509 |
| 2_1573985 | 7 | 1 | 15 | 1 | 12 | 0.9167 | 12 | 0.9167 | 46 | 0.9565 | 9 | 1 | 20 | 0.95 | 20 | 0.9 | 15 | 1 | 64 | 0.9531 |
| 2_1573985 | 8 | 0.625 | 15 | 0.7333 | 15 | 0.6667 | 14 | 0.7143 | 52 | 0.6923 | 10 | 0.4 | 19 | 0.6842 | 21 | 0.619 | 16 | 0.625 | 66 | 0.6061 |
| 2_1573986 | 5 | 0.8 | 15 | 0.8 | 14 | 0.8571 | 13 | 0.6154 | 47 | 0.766 | 9 | 0.8889 | 18 | 0.7778 | 19 | 0.6842 | 13 | 1 | 59 | 0.8136 |
| 2_1573987 | 7 | 0.2857 | 12 | 0.3333 | 13 | 0.6923 | 13 | 0.3077 | 45 | 0.4222 | 10 | 0.4 | 19 | 0.5263 | 21 | 0.2381 | 13 | 0.5385 | 63 | 0.4127 |
| 2_1573997 | 4 | 0 | 9 | 0 | 9 | 0 | 11 | 0.1818 | 33 | 0.0606 | 7 | 0 | 16 | 0.125 | 15 | 0.0667 | 12 | 0 | 50 | 0.06 |
| 2_1574039 | 65 | 0.7231 | 53 | 0.6792 | 45 | 0.6667 | 58 | 0.7759 | 221 | 0.7149 | 42 | 0.8333 | 63 | 0.6984 | 60 | 0.6833 | 43 | 0.7907 | 208 | 0.7404 |
| 2_1576310 | 6 | 0.3333 | 7 | 0.2857 | 3 | 0.3333 | 4 | 0 | 20 | 0.25 | 3 | 0 | 4 | 0 | 8 | 0 | 3 | 0.3333 | 18 | 0.0556 |
| 2_1577331 | 3 | 0.3333 | 1 | 1 | 2 | 0 | 4 | 0 | 10 | 0.2 | 2 | 0 | 4 | 0.5 | 3 | 0 | 1 | 0 | 10 | 0.2 |
| 2_1577335 | 2 | 0.5 | NA | NA | 1 | 1 | 2 | 0 | 5 | 0.4 | 2 | 1 | 2 | 0.5 | 2 | 0 | 1 | 0 | 7 | 0.4286 |
| 2_1577864 | 26 | 0.0769 | 28 | 0.0714 | 26 | 0.0769 | 53 | 0.0189 | 133 | 0.0526 | 16 | 0.0625 | 33 | 0.0606 | 39 | 0.1026 | 24 | 0 | 112 | 0.0625 |
| 2_1577865 | 27 | 0.037 | 27 | 0.1852 | 22 | 0.0455 | 57 | 0.1579 | 133 | 0.1203 | 16 | 0.0625 | 33 | 0.1212 | 39 | 0.1795 | 19 | 0.1053 | 107 | 0.1308 |
| 2_1578240 | 1 | 1 | 3 | 0.6667 | 4 | 0.75 | 4 | 1 | 12 | 0.8333 | 1 | 1 | 2 | 1 | 3 | 0.6667 | 1 | 1 | 7 | 0.8571 |
| 2_1578241 | 1 | 0 | 3 | 0.3333 | 4 | 0 | 4 | 0 | 12 | 0.0833 | 1 | 0 | 3 | 0.3333 | 4 | 0.25 | 1 | 1 | 9 | 0.3333 |
| 2_1578490 | NA | NA | 1 | 0 | 3 | 1 | 2 | 0.5 | 6 | 0.6667 | 1 | 0 | 1 | 1 | NA | NA | NA | NA | 2 | 0.5 |
| 2_1578491 | NA | NA | NA | NA | 2 | 0.5 | 2 | 0.5 | 4 | 0.5 | 2 | 0 | 1 | 1 | 1 | 1 | NA | NA | 4 | 0.5 |
| 2_1578493 | NA | NA | NA | NA | 2 | 1 | 2 | 1 | 4 | 1 | 2 | 0.5 | 1 | 1 | 1 | 1 | NA | NA | 4 | 0.75 |
| 2_1578493 | NA | NA | NA | NA | 2 | 1 | 1 | 0 | 3 | 0.6667 | 1 | 1 | 1 | 0 | NA | NA | NA | NA | 2 | 0.5 |
| 2_1578497 | NA | NA | NA | NA | 2 | 0.5 | 2 | 1 | 4 | 0.75 | NA | NA | 1 | 1 | NA | NA | NA | NA | 1 | 1 |
| 2_1578500 | NA | NA | NA | NA | 2 | 0.5 | 1 | 1 | 3 | 0.6667 | NA | NA | 1 | 1 | NA | NA | NA | NA | 1 | 1 |
| 2_1578503 | NA | NA | NA | NA | 2 | 0.5 | 1 | 1 | 3 | 0.6667 | NA | NA | 1 | 1 | NA | NA | NA | NA | 1 | 1 |
| 2_1578508 | NA | NA | NA | NA | 1 | 0 | 1 | 1 | 2 | 0.5 | NA | NA | 1 | 1 | NA | NA | NA | NA | 1 | 1 |
| 2_1578508 | NA | NA | NA | NA | 2 | 0.5 | 1 | 0 | 3 | 0.3333 | NA | NA | 1 | 1 | NA | NA | NA | NA | 1 | 1 |
| 2_1578508 | NA | NA | NA | NA | 2 | 1 | 1 | 1 | 3 | 1 | NA | NA | 1 | 1 | NA | NA | NA | NA | 1 | 1 |
| 2_1578509 | NA | NA | NA | NA | 2 | 1 | 1 | 0 | 3 | 0.6667 | NA | NA | 1 | 1 | NA | NA | NA | NA | 1 | 1 |
| 2_1579168 | 2 | 0 | 5 | 0.4 | 3 | 0 | 3 | 0 | 13 | 0.1538 | NA | NA | 4 | 0.25 | 2 | 0 | 1 | 0 | 7 | 0.1429 |
| 2_1579169 | 2 | 0 | 4 | 0.5 | 3 | 0 | 3 | 0.3333 | 12 | 0.25 | 1 | 1 | 4 | 0.25 | 2 | 0 | 1 | 0 | 8 | 0.25 |
| 2_1579170 | 2 | 0.5 | 5 | 0.4 | 2 | 0 | 3 | 0.6667 | 12 | 0.4167 | 1 | 1 | 4 | 0.75 | 1 | 1 | 1 | 0 | 7 | 0.7143 |
| 2_1579171 | 2 | 1 | 4 | 0.25 | 3 | 0 | 3 | 0.3333 | 12 | 0.3333 | NA | NA | 3 | 0 | 1 | 0 | 1 | 1 | 5 | 0.2 |
| 2_1579171 | 2 | 0 | 5 | 0.2 | 3 | 0.3333 | 3 | 0 | 13 | 0.1538 | 2 | 0 | 3 | 0.3333 | 1 | 0 | 1 | 0 | 7 | 0.1429 |
| 2_1579171 | 2 | 1 | 5 | 1 | 2 | 1 | 1 | 1 | 10 | 1 | 1 | 1 | 4 | 1 | 2 | 1 | 1 | 1 | 8 | 1 |
| 2_1579172 | 2 | 0 | 4 | 0 | 3 | 0 | 3 | 0.3333 | 12 | 0.0833 | 1 | 0 | 4 | 0.5 | 2 | 0 | 1 | 0 | 8 | 0.25 |
| 2_1579172 | 2 | 1 | 6 | 0.5 | 3 | 0 | 3 | 0.6667 | 14 | 0.5 | 1 | 0 | 3 | 0.3333 | 2 | 1 | 1 | 1 | 7 | 0.5714 |
| 2_1579174 | 4 | 0.5 | 5 | 1 | 3 | 0.3333 | 3 | 0.3333 | 15 | 0.6 | 2 | 0 | 3 | 0 | 2 | 1 | 1 | 1 | 8 | 0.375 |
| 2_1579177 | 4 | 0.25 | 5 | 0 | 2 | 0 | 4 | 0 | 15 | 0.0667 | 3 | 0 | 5 | 0 | 2 | 0 | 1 | 1 | 11 | 0.0909 |
| 2_1579178 | 4 | 0.25 | 5 | 0 | 4 | 0.5 | 5 | 0.4 | 18 | 0.2778 | 4 | 0 | 6 | 0.5 | 2 | 1 | 1 | 1 | 13 | 0.4615 |
| 2_1579179 | 4 | 0 | 5 | 0 | 4 | 0.25 | 5 | 0.4 | 18 | 0.1667 | 4 | 0 | 5 | 0.2 | 2 | 0 | 1 | 0 | 12 | 0.0833 |
| 2_1579183 | 6 | 0 | 5 | 0 | 4 | 0.25 | 4 | 0 | 19 | 0.0526 | 5 | 0 | 6 | 0.1667 | 2 | 0.5 | 1 | 0 | 14 | 0.1429 |
| 2_1579186 | 6 | 0.3333 | 5 | 0.2 | 4 | 0.5 | 5 | 0.2 | 20 | 0.3 | 6 | 0.1667 | 6 | 0.5 | 3 | 0.6667 | 1 | 0 | 16 | 0.375 |
| 2_1579196 | NA | NA | NA | NA | 1 | 1 | 3 | 1 | 4 | 1 | 1 | 1 | NA | NA | NA | NA | NA | NA | 1 | 1 |
| 2_1579591 | 3 | 0.6667 | 6 | 0.8333 | 3 | 1 | NA | NA | 12 | 0.8333 | 1 | 0 | 2 | 1 | 3 | 0.3333 | NA | NA | 6 | 0.5 |
| 2_1579592 | 2 | 0 | 6 | 0.1667 | 3 | 0.6667 | NA | NA | 11 | 0.2727 | 1 | 1 | 2 | 0 | 2 | 0 | NA | NA | 5 | 0.2 |
| 2_1579592 | 3 | 0.3333 | 6 | 0.1667 | 2 | 0 | NA | NA | 11 | 0.1818 | 1 | 1 | 2 | 0 | 2 | 0.5 | NA | NA | 5 | 0.4 |
| 2_1579594 | 2 | 1 | NA | NA | 2 | 0.5 | NA | NA | 4 | 0.75 | 1 | 1 | 2 | 0.5 | 2 | 0.5 | NA | NA | 5 | 0.6 |
| 2_1579594 | 3 | 1 | 4 | 1 | 2 | 1 | NA | NA | 9 | 1 | 1 | 1 | 1 | 1 | 3 | 1 | NA | NA | 5 | 1 |
| 2_1579595 | 3 | 0.3333 | 6 | 0.5 | 3 | 0.3333 | NA | NA | 12 | 0.4167 | 1 | 0 | 2 | 0.5 | 3 | 0 | NA | NA | 6 | 0.1667 |
| 2_1579597 | 2 | 0.5 | 5 | 0.4 | 1 | 0 | NA | NA | 8 | 0.375 | NA | NA | 1 | 0 | 3 | 0.3333 | NA | NA | 4 | 0.25 |

| | | | | | | | | | | | | | | | | | | | | |
|---|---|---|---|---|---|---|---|---|---|---|---|---|---|---|---|---|---|---|---|---|
| 2_1579597 | 3 | 0.3333 | 6 | 0.1667 | 2 | 0 | NA | NA | 11 | 0.1818 | NA | NA | 2 | 0 | 3 | 0.3333 | NA | NA | 5 | 0.2 |
| 2_1581977 | 1 | 1 | NA | NA | NA | NA | 1 | 1 | 2 | 1 | NA | NA | NA | NA | 1 | 0 | 2 | 1 | 3 | 0.6667 |
| 2_1582266 | 18 | 0 | 42 | 0.0476 | 23 | 0.087 | 40 | 0.1 | 123 | 0.065 | 13 | 0.2308 | 29 | 0.1034 | 36 | 0.1111 | 33 | 0.1818 | 111 | 0.1441 |
| 2_1582273 | 39 | 0.0769 | 66 | 0.1212 | 38 | 0.2632 | 61 | 0.0656 | 204 | 0.1225 | 19 | 0.0526 | 36 | 0.0278 | 52 | 0.0769 | 44 | 0.1364 | 151 | 0.0795 |
| 2_1582273 | 38 | 0.1579 | 59 | 0.2542 | 34 | 0.2353 | 57 | 0.2281 | 188 | 0.2234 | 19 | 0.1579 | 37 | 0.1892 | 53 | 0.2075 | 40 | 0.3 | 149 | 0.2215 |
| 2_1582273 | 39 | 0.2051 | 65 | 0.1846 | 38 | 0.2632 | 61 | 0.1803 | 203 | 0.202 | 20 | 0 | 39 | 0.2308 | 55 | 0.1091 | 47 | 0.1915 | 161 | 0.1491 |
| 2_1582274 | 35 | 0.4 | 59 | 0.5254 | 32 | 0.5625 | 56 | 0.6964 | 182 | 0.5604 | 17 | 0.7059 | 34 | 0.6765 | 48 | 0.5208 | 39 | 0.5385 | 138 | 0.587 |
| 2_1582275 | 38 | 0.0526 | 66 | 0.0909 | 40 | 0.1 | 61 | 0.0984 | 205 | 0.0878 | 19 | 0.0526 | 39 | 0.1282 | 53 | 0.1698 | 48 | 0.1458 | 159 | 0.1384 |
| 2_1582276 | 38 | 0 | 60 | 0.0667 | 35 | 0.0571 | 61 | 0.082 | 194 | 0.0567 | 21 | 0 | 37 | 0.0541 | 53 | 0.0566 | 44 | 0.1136 | 155 | 0.0645 |
| 2_1582280 | 26 | 0.2308 | 56 | 0.2321 | 30 | 0.2333 | 50 | 0.1 | 162 | 0.1914 | 13 | 0.3077 | 35 | 0.3429 | 43 | 0.2093 | 38 | 0.2632 | 129 | 0.2713 |
| 2_1583087 | 1 | 0 | 4 | 0 | 2 | 0.5 | 7 | 0.1429 | 14 | 0.1429 | 3 | 0 | 1 | 0 | 4 | 0.25 | 1 | 0 | 9 | 0.1111 |
| 2_1583374 | 2 | 0 | 6 | 0.3333 | 2 | 0.5 | 8 | 0.125 | 18 | 0.2222 | 3 | 0.3333 | 3 | 0.3333 | 5 | 0.8 | 2 | 0 | 13 | 0.4615 |
| 2_1583374 | 2 | 0 | 6 | 0.3333 | 3 | 0.6667 | 7 | 0.7143 | 18 | 0.5 | 3 | 0.3333 | 3 | 0.3333 | 6 | 0.3333 | 1 | 0 | 13 | 0.3077 |
| 2_1583375 | 2 | 1 | 4 | 1 | 3 | 1 | 7 | 1 | 16 | 1 | 3 | 1 | 3 | 1 | 5 | 1 | 2 | 0.5 | 13 | 0.9231 |
| 2_1583375 | 2 | 0.5 | 7 | 0.8571 | 3 | 0.6667 | 5 | 0.8 | 17 | 0.7647 | 4 | 1 | 3 | 0.6667 | 5 | 1 | 1 | 1 | 13 | 0.9231 |
| 2_1583375 | NA | NA | 6 | 0.3333 | 2 | 1 | 5 | 0.8 | 13 | 0.6154 | 2 | 0.5 | 2 | 1 | 3 | 1 | 2 | 0 | 9 | 0.6667 |
| 2_1583376 | 2 | 0.5 | 4 | 0.75 | 2 | 0.5 | 6 | 0.3333 | 14 | 0.5 | 3 | 0.6667 | 2 | 1 | 6 | 0.5 | 2 | 0.5 | 13 | 0.6154 |
| 2_1583377 | 3 | 0.3333 | 7 | 0.1429 | 4 | 0.75 | 7 | 0.4286 | 21 | 0.381 | 4 | 0.5 | 5 | 0.8 | 5 | 0.4 | 3 | 0.6667 | 17 | 0.5882 |
| 2_1583377 | 2 | 0.5 | 6 | 0.6667 | 1 | 1 | 4 | 1 | 13 | 0.7692 | 3 | 0.6667 | 5 | 1 | 6 | 1 | 3 | 0.6667 | 17 | 0.8824 |
| 2_1583378 | 2 | 1 | 4 | 0.75 | 4 | 0.75 | 5 | 1 | 15 | 0.8667 | 2 | 0.5 | 5 | 0.8 | 7 | 0.8571 | 2 | 0.5 | 16 | 0.75 |
| 2_1583378 | 4 | 0.25 | 8 | 0.25 | 2 | 0.5 | 7 | 0.1429 | 21 | 0.2381 | 4 | 0.25 | 5 | 0 | 8 | 0.125 | 5 | 0.2 | 22 | 0.1364 |
| 2_1583379 | 5 | 0.8 | 7 | 0.5714 | 4 | 1 | 5 | 0.8 | 21 | 0.7619 | 4 | 0.5 | 5 | 0.8 | 6 | 0.6667 | 6 | 0.5 | 21 | 0.619 |
| 2_1583386 | 6 | 0.3333 | 6 | 0 | 7 | 0.2857 | 11 | 0 | 30 | 0.1333 | 4 | 0.25 | 8 | 0.125 | 7 | 0.2857 | 3 | 0.6667 | 22 | 0.2727 |
| 2_1583393 | 5 | 0.4 | 6 | 0.1667 | 6 | 0.3333 | 9 | 0 | 26 | 0.1923 | 4 | 0.5 | 8 | 0.125 | 9 | 0.4444 | 5 | 0.4 | 26 | 0.3462 |
| 2_1583398 | 7 | 0.8571 | 5 | 0.6 | 9 | 0.6667 | 11 | 0.8182 | 32 | 0.75 | 4 | 0.75 | 8 | 0.875 | 11 | 0.6364 | 7 | 0.7143 | 30 | 0.7333 |
| 2_1583398 | 6 | 0.6667 | 7 | 0.7143 | 10 | 0.3 | 9 | 0.4444 | 32 | 0.5 | 4 | 0.25 | 8 | 0.25 | 11 | 0.3636 | 4 | 0.75 | 27 | 0.3704 |
| 2_1583402 | 4 | 0 | 6 | 0 | 8 | 0.25 | 9 | 0 | 27 | 0.0741 | 2 | 0.5 | 8 | 0.125 | 11 | 0.0909 | 3 | 0.3333 | 24 | 0.1667 |
| 2_1584375 | NA | NA | NA | NA | 1 | 1 | NA | NA | 1 | 1 | 1 | 1 | 3 | 0.6667 | NA | NA | NA | NA | 4 | 0.75 |
| 2_1584375 | NA | NA | NA | NA | 1 | 1 | NA | NA | 1 | 1 | 1 | 1 | 3 | 0.3333 | 1 | 1 | NA | NA | 5 | 0.6 |
| 2_1584376 | NA | NA | NA | NA | 1 | 1 | NA | NA | 1 | 1 | 1 | 1 | 2 | 1 | 1 | 1 | NA | NA | 4 | 1 |
| 2_1584379 | NA | NA | NA | NA | 2 | 0.5 | NA | NA | 2 | 0.5 | NA | NA | 1 | 1 | 1 | 1 | NA | NA | 2 | 1 |
| 2_1584392 | NA | NA | 2 | 0.5 | 3 | 0.3333 | 1 | 0 | 6 | 0.3333 | 1 | 1 | NA | NA | 1 | 0 | NA | NA | 2 | 0.5 |
| 2_1584438 | NA | NA | NA | NA | NA | NA | 1 | 1 | 1 | 1 | NA | NA | 1 | 1 | NA | NA | NA | NA | 1 | 1 |
| 2_1584529 | 28 | 0.1429 | 17 | 0.1176 | 11 | 0.1818 | 23 | 0.087 | 79 | 0.1266 | 12 | 0 | 27 | 0.1852 | 23 | 0.0435 | 15 | 0.2 | 77 | 0.1169 |
| 2_1584533 | 34 | 0.2647 | 17 | 0.2353 | 14 | 0.2143 | 27 | 0.2963 | 92 | 0.2609 | 17 | 0.4118 | 31 | 0.2581 | 28 | 0.4643 | 18 | 0.3333 | 94 | 0.3617 |
| 2_1584534 | 32 | 0.0312 | 20 | 0.15 | 16 | 0.25 | 25 | 0.12 | 93 | 0.1183 | 14 | 0 | 35 | 0.2286 | 28 | 0.1429 | 17 | 0.0588 | 94 | 0.1383 |
| 2_1584558 | 35 | 0.9714 | 25 | 1 | 17 | 0.9412 | 32 | 0.9688 | 109 | 0.9725 | 16 | 0.9375 | 42 | 0.9762 | 35 | 1 | 19 | 0.9474 | 112 | 0.9732 |
| 2_1584563 | 32 | 0.875 | 25 | 0.8 | 16 | 1 | 24 | 0.625 | 97 | 0.8144 | 17 | 0.6471 | 38 | 0.6842 | 30 | 0.7333 | 15 | 0.7333 | 100 | 0.7 |
| 2_1584563 | 32 | 0.5312 | 30 | 0.5333 | 22 | 0.4091 | 32 | 0.2812 | 116 | 0.4397 | 17 | 0.5294 | 42 | 0.381 | 29 | 0.4483 | 17 | 0.5294 | 105 | 0.4476 |
| 2_1584569 | 43 | 0.093 | 32 | 0.0938 | 27 | 0.037 | 38 | 0.1053 | 140 | 0.0857 | 23 | 0.1304 | 50 | 0.16 | 40 | 0.175 | 22 | 0.0455 | 135 | 0.1407 |
| 2_1584569 | 22 | 1 | 16 | 1 | 18 | 1 | 23 | 1 | 79 | 1 | 15 | 0.9333 | 32 | 1 | 29 | 1 | 15 | 1 | 91 | 0.989 |
| 2_1584572 | 43 | 0.1395 | 36 | 0.0833 | 30 | 0.1667 | 34 | 0.1765 | 143 | 0.1399 | 33 | 0.0909 | 51 | 0.1373 | 50 | 0.2 | 24 | 0.125 | 158 | 0.1456 |
| 2_1584572 | 43 | 0.2791 | 37 | 0.2973 | 30 | 0.3667 | 35 | 0.2286 | 145 | 0.2897 | 26 | 0.4231 | 50 | 0.3 | 47 | 0.3191 | 23 | 0.4783 | 146 | 0.3562 |
| 2_1584573 | 46 | 1 | 37 | 1 | 34 | 1 | 38 | 0.9474 | 155 | 0.9871 | 32 | 1 | 51 | 0.9804 | 44 | 0.9545 | 23 | 1 | 150 | 0.98 |
| 2_1584575 | 51 | 0.3333 | 41 | 0.2195 | 39 | 0.1538 | 44 | 0.1818 | 175 | 0.2286 | 34 | 0.2353 | 60 | 0.25 | 55 | 0.2 | 32 | 0.375 | 181 | 0.2541 |
| 2_1584575 | 48 | 0.0208 | 41 | 0.0976 | 38 | 0.0526 | 40 | 0.05 | 167 | 0.0539 | 31 | 0.0645 | 59 | 0.0847 | 54 | 0.0556 | 30 | 0.0667 | 174 | 0.069 |
| 2_1584575 | 37 | 0.6757 | 35 | 0.8286 | 31 | 0.871 | 37 | 0.8108 | 140 | 0.7929 | 30 | 0.7667 | 46 | 0.6957 | 42 | 0.6429 | 25 | 0.72 | 143 | 0.6993 |
| 2_1584806 | 2 | 1 | 1 | 1 | 2 | 0.5 | 1 | 1 | 6 | 0.8333 | NA | NA | NA | NA | 1 | 1 | 1 | 1 | 2 | 1 |
| 2_1584807 | 2 | 0.5 | NA | NA | 1 | 0 | 1 | 0 | 4 | 0.25 | NA | NA | NA | NA | 1 | 1 | NA | NA | 1 | 1 |
| 2_1584808 | 2 | 0.5 | 1 | 1 | 2 | 0.5 | 1 | 1 | 6 | 0.6667 | NA | NA | NA | NA | 2 | 1 | 1 | 1 | 3 | 1 |
| 2_1584808 | 1 | 1 | 1 | 1 | 2 | 1 | 1 | 1 | 5 | 1 | NA | NA | NA | NA | NA | NA | 1 | 1 | 1 | 1 |
| 2_1584809 | 2 | 1 | 1 | 0 | 2 | 1 | NA | NA | 5 | 0.8 | NA | NA | NA | NA | NA | NA | 1 | 1 | 1 | 1 |
| 2_1584809 | 2 | 0.5 | NA | NA | 3 | 1 | 1 | 1 | 6 | 0.8333 | NA | NA | NA | NA | 1 | 1 | NA | NA | 1 | 1 |
| 2_1584810 | 1 | 1 | 1 | 1 | 1 | 1 | 1 | 1 | 4 | 1 | NA | NA | NA | NA | 1 | 1 | NA | NA | 1 | 1 |
| 2_1584810 | NA | NA | 1 | 0 | 1 | 0 | 1 | 1 | 3 | 0.3333 | NA | NA | NA | NA | 1 | 0 | 1 | 1 | 2 | 0.5 |
| 2_1584810 | 1 | 1 | 1 | 1 | 1 | 1 | 1 | 0 | 4 | 0.75 | NA | NA | NA | NA | 2 | 0.5 | 1 | 1 | 3 | 0.6667 |
| 2_1586033 | 3 | 0.3333 | 2 | 0.5 | 2 | 0.5 | 4 | 0.5 | 11 | 0.4545 | 4 | 0.75 | 5 | 0.8 | 7 | 0.2857 | 2 | 0.5 | 18 | 0.5556 |
| 2_1586034 | 3 | 1 | 5 | 1 | 2 | 1 | 6 | 0.8333 | 16 | 0.9375 | 4 | 0.75 | 7 | 0.7143 | 7 | 1 | 4 | 0.75 | 22 | 0.8182 |
| 2_1586036 | 3 | 0.6667 | 4 | 0.5 | 1 | 0 | 6 | 0.6667 | 14 | 0.5714 | 4 | 0.75 | 4 | 0.75 | 7 | 0.7143 | 3 | 0.6667 | 18 | 0.7222 |
| 2_1586036 | 4 | 0.75 | 5 | 1 | 2 | 0.5 | 9 | 0.7778 | 20 | 0.8 | 5 | 1 | 8 | 1 | 7 | 0.8571 | 4 | 1 | 24 | 0.9583 |
| 2_1586037 | 4 | 1 | 4 | 1 | 2 | 1 | 9 | 1 | 19 | 1 | 4 | 1 | 7 | 1 | 6 | 1 | 5 | 1 | 22 | 1 |
| 2_1586042 | 6 | 0.3333 | 6 | 0.8333 | 2 | 0.5 | 8 | 0.375 | 22 | 0.5 | 2 | 1 | 8 | 0.625 | 8 | 0.625 | 7 | 0.4286 | 25 | 0.6 |
| 2_1586042 | 4 | 0.25 | 2 | 0.5 | 1 | 1 | 7 | 0.1429 | 14 | 0.2857 | 4 | 0 | 9 | 0.2222 | 8 | 0.25 | 7 | 0.2857 | 28 | 0.2143 |
| 2_1586043 | 4 | 0 | 5 | 0.4 | 4 | 0 | 10 | 0.1 | 23 | 0.1304 | 5 | 0.2 | 10 | 0.3 | 9 | 0.2222 | 7 | 0.1429 | 31 | 0.2258 |
| 2_1586054 | 12 | 0.1667 | 8 | 0.125 | 6 | 0.1667 | 15 | 0.0667 | 41 | 0.122 | 5 | 0 | 14 | 0.1429 | 17 | 0.0588 | 10 | 0.1 | 46 | 0.087 |

| | | | | | | | | | | | | | | | | | | | | |
|---|---|---|---|---|---|---|---|---|---|---|---|---|---|---|---|---|---|---|---|---|
| 2_1586055 | 13 | 0 | 8 | 0.125 | 7 | 0.1429 | 16 | 0.0625 | 44 | 0.0682 | 5 | 0.2 | 16 | 0.125 | 17 | 0 | 12 | 0.0833 | 50 | 0.08 |
| 2_1586056 | 12 | 0.0833 | 8 | 0.25 | 7 | 0.1429 | 17 | 0.0588 | 44 | 0.1136 | 5 | 0.2 | 15 | 0.3333 | 18 | 0.2778 | 13 | 0.3846 | 51 | 0.3137 |
| 2_1586057 | 9 | 0.1111 | 6 | 0 | 9 | 0.1111 | 19 | 0.0526 | 43 | 0.0698 | 4 | 0 | 14 | 0.0714 | 21 | 0.0476 | 15 | 0.1333 | 54 | 0.0741 |
| 2_1586060 | 10 | 0.3 | 9 | 0.1111 | 11 | 0.1818 | 19 | 0.0526 | 49 | 0.1429 | 4 | 0.25 | 20 | 0.1 | 23 | 0.2609 | 15 | 0.0667 | 62 | 0.1613 |
| 2_1586062 | 11 | 0.5455 | 8 | 0.625 | 10 | 0.1 | 17 | 0.2941 | 46 | 0.3696 | 4 | 0.25 | 15 | 0.5333 | 22 | 0.3636 | 12 | 0.3333 | 53 | 0.3962 |
| 2_1586068 | 14 | 0.0714 | 17 | 0.1765 | 16 | 0.1875 | 21 | 0.0952 | 68 | 0.1324 | 6 | 0 | 22 | 0.2273 | 26 | 0.1923 | 9 | 0 | 63 | 0.1587 |
| 2_1586076 | 18 | 0.7222 | 19 | 0.4737 | 24 | 0.7083 | 27 | 0.4815 | 88 | 0.5909 | 11 | 0.5455 | 26 | 0.4615 | 33 | 0.6364 | 18 | 0.3889 | 88 | 0.5227 |
| 2_1586076 | 17 | 0.4118 | 16 | 0.4375 | 23 | 0.3913 | 25 | 0.24 | 81 | 0.358 | 11 | 0.2727 | 25 | 0.16 | 32 | 0.4375 | 19 | 0.2105 | 87 | 0.2874 |
| 2_1586079 | 19 | 0.1053 | 21 | 0.0952 | 27 | 0.1852 | 27 | 0.037 | 94 | 0.1064 | 13 | 0 | 31 | 0.0968 | 40 | 0.1 | 22 | 0.0909 | 106 | 0.0849 |
| 2_1586273 | NA | NA | NA | NA | 1 | 0 | 1 | 1 | 2 | 0.5 | NA | NA | NA | NA | NA | NA | 2 | 1 | 2 | 1 |
| 2_1586275 | NA | NA | NA | NA | NA | NA | 1 | 1 | 1 | 1 | NA | NA | NA | NA | NA | NA | 1 | 1 | 1 | 1 |
| 2_1586276 | NA | NA | NA | NA | 1 | 1 | NA | NA | 1 | 1 | NA | NA | NA | NA | NA | NA | 1 | 1 | 1 | 1 |
| 2_1586278 | NA | NA | NA | NA | 1 | 1 | 1 | 1 | 2 | 1 | NA | NA | NA | NA | 1 | 1 | 2 | 0.5 | 3 | 0.6667 |
| 2_1586278 | NA | NA | NA | NA | 1 | 1 | 1 | 1 | 2 | 1 | NA | NA | NA | NA | NA | NA | 2 | 0.5 | 2 | 0.5 |
| 2_1586280 | NA | NA | NA | NA | 1 | 1 | 1 | 1 | 2 | 1 | NA | NA | NA | NA | 1 | 1 | 2 | 1 | 3 | 1 |
| 2_1586280 | NA | NA | NA | NA | 1 | 1 | NA | NA | 1 | 1 | NA | NA | NA | NA | NA | NA | 2 | 1 | 2 | 1 |
| 2_1586996 | 6 | 0.5 | 2 | 0.5 | NA | NA | NA | NA | 8 | 0.5 | 2 | 0 | 2 | 0.5 | 2 | 0.5 | NA | NA | 6 | 0.3333 |
| 2_1586997 | 6 | 0 | 3 | 0.3333 | NA | NA | NA | NA | 9 | 0.1111 | 2 | 0.5 | 2 | 0 | 2 | 0.5 | 1 | 1 | 7 | 0.4286 |
| 2_1586999 | 6 | 0.1667 | 3 | 0 | 1 | 0 | NA | NA | 10 | 0.1 | 1 | 0 | 2 | 0.5 | 2 | 0 | 1 | 0 | 6 | 0.1667 |
| 2_1602060 | 4 | 1 | NA | NA | 1 | 1 | 1 | 1 | 6 | 1 | NA | NA | 2 | 1 | 3 | 0.3333 | NA | NA | 5 | 0.6 |
| 2_1602070 | 3 | 1 | 1 | 1 | 1 | 1 | 2 | 1 | 7 | 1 | NA | NA | 1 | 1 | 3 | 0.6667 | NA | NA | 4 | 0.75 |
| 2_1602072 | 3 | 0.3333 | 1 | 1 | 1 | 0 | 1 | 0 | 6 | 0.3333 | NA | NA | 1 | 1 | 2 | 0 | NA | NA | 3 | 0.3333 |
| 2_1602072 | 3 | 1 | 1 | 1 | 1 | 1 | 2 | 1 | 7 | 1 | NA | NA | 1 | 1 | 2 | 0 | NA | NA | 3 | 0.3333 |
| 2_1602072 | 2 | 1 | 1 | 1 | 1 | 1 | 1 | 1 | 5 | 1 | NA | NA | 1 | 1 | 2 | 0.5 | NA | NA | 3 | 0.6667 |
| 2_1602072 | 3 | 1 | 1 | 1 | 1 | 1 | NA | NA | 5 | 1 | NA | NA | 1 | 1 | 1 | 1 | NA | NA | 2 | 1 |
| 2_1602074 | 3 | 1 | 1 | 1 | NA | NA | NA | NA | 4 | 1 | NA | NA | NA | NA | 2 | 0.5 | NA | NA | 2 | 0.5 |
| 2_1602074 | 3 | 0.3333 | 1 | 1 | 1 | 0 | NA | NA | 5 | 0.4 | NA | NA | 1 | 1 | 2 | 0 | NA | NA | 3 | 0.3333 |
| 2_1602075 | 3 | 1 | NA | NA | 1 | 1 | NA | NA | 4 | 1 | NA | NA | 1 | 1 | 2 | 1 | NA | NA | 3 | 1 |
| 2_1602075 | 2 | 0.5 | NA | NA | 1 | 1 | 1 | 0 | 4 | 0.5 | NA | NA | 1 | 0 | 2 | 0.5 | NA | NA | 3 | 0.3333 |
| 2_1605634 | 4 | 0.25 | 8 | 0.25 | 16 | 0.3125 | 11 | 0.1818 | 39 | 0.2564 | 2 | 0 | 13 | 0.3077 | 22 | 0.1364 | 8 | 0 | 45 | 0.1556 |
| 2_1605635 | 3 | 0 | 3 | 0 | 12 | 0.0833 | 12 | 0.0833 | 30 | 0.0667 | 1 | 0 | 11 | 0.1818 | 16 | 0.25 | 5 | 0.6 | 33 | 0.2727 |
| 2_1605635 | 4 | 0 | 8 | 0 | 11 | 0.1818 | 11 | 0.0909 | 34 | 0.0882 | 2 | 0 | 12 | 0.25 | 22 | 0.0455 | 9 | 0.1111 | 45 | 0.1111 |
| 2_1605897 | NA | NA | 1 | 1 | 1 | 0 | NA | NA | 2 | 0.5 | NA | NA | NA | NA | 1 | 1 | 2 | 0 | 3 | 0.3333 |
| 2_1606035 | 2 | 0 | 2 | 0 | 3 | 0 | 3 | 0.3333 | 10 | 0.1 | 4 | 0 | 5 | 0.4 | 3 | 0.3333 | 4 | 0 | 16 | 0.1875 |
| 2_1606038 | NA | NA | NA | NA | NA | NA | 1 | 1 | 1 | 1 | 1 | 1 | 1 | 1 | NA | NA | NA | NA | 2 | 1 |
| 2_1606039 | 1 | 1 | NA | NA | NA | NA | 1 | 1 | 2 | 1 | 2 | 0.5 | 1 | 1 | NA | NA | NA | NA | 3 | 0.6667 |
| 2_1606040 | 1 | 0 | 2 | 0 | 3 | 1 | 2 | 1 | 8 | 0.625 | 3 | 0.6667 | 4 | 0.75 | 3 | 1 | 4 | 1 | 14 | 0.8571 |
| 2_1606041 | 4 | 1 | 2 | 1 | 3 | 1 | 1 | 1 | 10 | 1 | 2 | 1 | 5 | 0.8 | 3 | 0.6667 | 5 | 1 | 15 | 0.8667 |
| 2_1606041 | 4 | 0 | 3 | 0.3333 | 3 | 0 | 3 | 0 | 13 | 0.0769 | 6 | 0 | 6 | 0 | 3 | 0.3333 | 5 | 0 | 20 | 0.05 |
| 2_1606042 | 3 | 1 | 3 | 0.6667 | 2 | 1 | 2 | 1 | 10 | 0.9 | 4 | 0.75 | 6 | 1 | 3 | 1 | 4 | 1 | 17 | 0.9412 |
| 2_1606045 | 3 | 0.3333 | 2 | 0.5 | 3 | 0.3333 | 3 | 0.6667 | 11 | 0.4545 | 5 | 0.4 | 6 | 1 | 3 | 1 | 4 | 1 | 18 | 0.8333 |
| 2_1606114 | NA | NA | 2 | 0 | 4 | 0.5 | 1 | 0 | 7 | 0.2857 | 2 | 0.5 | 1 | 0 | 2 | 0.5 | NA | NA | 5 | 0.4 |
| 2_1606114 | NA | NA | 2 | 0 | 4 | 0.25 | 1 | 0 | 7 | 0.1429 | 2 | 0.5 | 1 | 0 | 2 | 0 | 1 | 0 | 6 | 0.1667 |
| 2_1606125 | NA | NA | NA | NA | 4 | 0.25 | NA | NA | 4 | 0.25 | 1 | 1 | NA | NA | 1 | 0 | 2 | 0 | 4 | 0.25 |
| 2_1606127 | NA | NA | NA | NA | 3 | 0.3333 | NA | NA | 3 | 0.3333 | 2 | 0.5 | 1 | 0 | 1 | 0 | 2 | 0.5 | 6 | 0.3333 |
| 2_1606128 | NA | NA | NA | NA | 4 | 1 | NA | NA | 4 | 1 | 2 | 1 | 1 | 1 | 1 | 0 | 2 | 0.5 | 6 | 0.6667 |
| 2_1606129 | NA | NA | NA | NA | 3 | 0.3333 | NA | NA | 3 | 0.3333 | 1 | 1 | 1 | 1 | 1 | 0 | NA | NA | 3 | 0.6667 |
| 2_1606129 | NA | NA | NA | NA | 1 | 1 | NA | NA | 1 | 1 | 1 | 0 | 1 | 0 | 1 | 1 | 1 | 0 | 4 | 0.25 |
| 2_1606129 | NA | NA | NA | NA | 4 | 0.5 | NA | NA | 4 | 0.5 | 2 | 1 | NA | NA | 1 | 1 | 1 | 1 | 4 | 1 |
| 2_1606130 | NA | NA | NA | NA | 4 | 1 | NA | NA | 4 | 1 | 2 | 1 | 1 | 1 | 1 | 1 | NA | NA | 4 | 1 |
| 2_1606131 | NA | NA | NA | NA | 3 | 1 | NA | NA | 3 | 1 | 2 | 1 | 1 | 1 | 1 | 0 | 1 | 1 | 5 | 0.8 |
| 2_1606131 | NA | NA | NA | NA | 4 | 0.75 | NA | NA | 4 | 0.75 | 2 | 0.5 | 1 | 1 | 1 | 1 | 1 | 1 | 5 | 0.8 |
| 2_1606131 | NA | NA | NA | NA | 2 | 1 | NA | NA | 2 | 1 | 2 | 1 | 1 | 1 | 1 | 1 | 1 | 1 | 5 | 1 |
| 2_1606132 | NA | NA | NA | NA | 5 | 0.8 | NA | NA | 5 | 0.8 | 2 | 1 | 2 | 1 | 1 | 1 | 1 | 1 | 6 | 1 |
| 2_1606133 | NA | NA | NA | NA | 4 | 0.75 | NA | NA | 4 | 0.75 | 2 | 1 | 2 | 0.5 | 1 | 1 | 1 | 1 | 6 | 0.8333 |
| 2_1606213 | NA | NA | NA | NA | 5 | 0.6 | NA | NA | 5 | 0.6 | 2 | 0.5 | 2 | 0 | 1 | 0 | 1 | 0 | 6 | 0.1667 |
| 2_1606216 | NA | NA | NA | NA | 6 | 0.6667 | NA | NA | 6 | 0.6667 | 1 | 1 | 2 | 0.5 | 1 | 0 | 1 | 0 | 5 | 0.4 |
| 2_1606217 | NA | NA | NA | NA | 4 | 0.25 | NA | NA | 4 | 0.25 | 1 | 1 | 2 | 0.5 | 1 | 0 | NA | NA | 4 | 0.5 |
| 2_1606218 | NA | NA | NA | NA | 5 | 1 | NA | NA | 5 | 1 | 1 | 1 | 2 | 0.5 | 1 | 1 | 1 | 1 | 5 | 0.8 |
| 2_1606224 | NA | NA | NA | NA | 2 | 1 | NA | NA | 2 | 1 | 1 | 1 | 2 | 1 | 1 | 1 | 1 | 1 | 5 | 1 |
| 2_1606229 | NA | NA | NA | NA | 4 | 0.5 | NA | NA | 4 | 0.5 | 1 | 1 | 1 | 0 | NA | NA | 1 | 1 | 3 | 0.6667 |
| 2_1606231 | NA | NA | NA | NA | 3 | 0.6667 | NA | NA | 3 | 0.6667 | 1 | 1 | 2 | 0.5 | NA | NA | 1 | 1 | 4 | 0.75 |
| 2_1606232 | NA | NA | NA | NA | 4 | 1 | NA | NA | 4 | 1 | NA | NA | 2 | 1 | NA | NA | NA | NA | 2 | 1 |
| 2_1606237 | NA | NA | NA | NA | 3 | 1 | NA | NA | 3 | 1 | NA | NA | 1 | 1 | NA | NA | NA | NA | 1 | 1 |
| 2_1606238 | NA | NA | NA | NA | 3 | 0.3333 | NA | NA | 3 | 0.3333 | NA | NA | 1 | 1 | NA | NA | NA | NA | 1 | 1 |

| | | | | | | | | | | | | | | | | | | | | |
|---|---|---|---|---|---|---|---|---|---|---|---|---|---|---|---|---|---|---|---|---|
| 2_1606238 | NA | NA | NA | NA | 3 | 1 | NA | NA | 3 | 1 | NA | NA | 1 | 1 | NA | NA | NA | NA | 1 | 1 |
| 2_1607476 | NA | NA | 1 | 1 | NA | NA | NA | NA | 1 | 1 | NA | NA | NA | NA | 1 | 1 | NA | NA | 1 | 1 |
| 2_1607890 | 6 | 0 | 6 | 0.3333 | 2 | 0 | 2 | 0 | 16 | 0.125 | 2 | 0 | 7 | 0 | 5 | 0.2 | 5 | 0 | 19 | 0.0526 |
| 2_1607896 | 6 | 0.3333 | 8 | 0.25 | 5 | 0.4 | 5 | 0.6 | 24 | 0.375 | 3 | 0.6667 | 9 | 0.2222 | 9 | 0.1111 | 7 | 0.1429 | 28 | 0.2143 |
| 2_1607900 | 4 | 0 | 4 | 0.25 | 5 | 0 | 5 | 0 | 18 | 0.0556 | 3 | 0 | 6 | 0.3333 | 6 | 0 | 4 | 0 | 19 | 0.1053 |
| 2_1607910 | 5 | 0.8 | 5 | 0.8 | 5 | 0.4 | 4 | 0.75 | 19 | 0.6842 | 1 | 0 | 8 | 0.625 | 5 | 0.8 | 5 | 0.8 | 19 | 0.6842 |
| 2_1607912 | 5 | 1 | 6 | 1 | 5 | 1 | 4 | 1 | 20 | 1 | 2 | 1 | 7 | 1 | 5 | 1 | 5 | 0.8 | 19 | 0.9474 |
| 2_1607912 | 4 | 0.75 | 6 | 0.8333 | 4 | 0.5 | 4 | 0.75 | 18 | 0.7222 | 2 | 1 | 7 | 1 | 5 | 0.6 | 7 | 0.7143 | 21 | 0.8095 |
| 2_1607912 | 4 | 0 | 5 | 0.2 | 3 | 0 | 3 | 0.3333 | 15 | 0.1333 | 2 | 0 | 7 | 0.1429 | 3 | 0 | 6 | 0.1667 | 18 | 0.1111 |
| 2_1607913 | 4 | 1 | 6 | 0.8333 | 2 | 1 | 3 | 1 | 15 | 0.9333 | 1 | 1 | 8 | 1 | 5 | 1 | 5 | 1 | 19 | 1 |
| 2_1607913 | 2 | 0 | 4 | 0 | 2 | 0 | 3 | 0.3333 | 11 | 0.0909 | NA | NA | 5 | 0 | 5 | 0.4 | 5 | 0 | 15 | 0.1333 |
| 2_1607914 | 4 | 0.5 | 5 | 0.6 | 3 | 1 | 2 | 1 | 14 | 0.7143 | 2 | 1 | 7 | 0.4286 | 6 | 0.1667 | 7 | 0.5714 | 22 | 0.4545 |
| 2_1607915 | 3 | 1 | 6 | 1 | 3 | 1 | 3 | 1 | 15 | 1 | 2 | 1 | 8 | 1 | 6 | 1 | 7 | 1 | 23 | 1 |
| 2_1607915 | 2 | 0.5 | 4 | 0.25 | 4 | 0.5 | 2 | 1 | 12 | 0.5 | 2 | 1 | 5 | 0.4 | 4 | 0.5 | 4 | 0.5 | 15 | 0.5333 |
| 2_1607915 | 4 | 0 | 5 | 0.2 | 4 | 0.25 | 2 | 0.5 | 15 | 0.2 | 1 | 1 | 8 | 0.375 | 6 | 0.3333 | 6 | 0.3333 | 21 | 0.381 |
| 2_1607916 | 4 | 0 | 6 | 0.1667 | 4 | 0 | 2 | 0 | 16 | 0.0625 | NA | NA | 5 | 0.2 | 5 | 0 | 5 | 0 | 15 | 0.0667 |
| 2_1607916 | 3 | 0.3333 | 5 | 0.8 | 4 | 0.5 | 2 | 1 | 14 | 0.6429 | 2 | 0.5 | 8 | 0.875 | 4 | 0.75 | 5 | 0.8 | 19 | 0.7895 |
| 2_1607917 | 3 | 0.6667 | 6 | 0.8333 | 3 | 0.6667 | 2 | 1 | 14 | 0.7857 | 2 | 1 | 8 | 1 | 6 | 0.5 | 5 | 0.8 | 21 | 0.8095 |
| 2_1607918 | 3 | 0.3333 | 3 | 0.6667 | 3 | 0.6667 | 1 | 0 | 10 | 0.5 | 2 | 1 | 7 | 0.8571 | 5 | 0.4 | 5 | 0.8 | 19 | 0.7368 |
| 2_1607919 | 2 | 1 | 5 | 1 | 4 | 1 | 2 | 1 | 13 | 1 | 2 | 1 | 7 | 1 | 5 | 0.8 | 5 | 1 | 19 | 0.9474 |
| 2_1607919 | 3 | 0.6667 | 5 | 0.6 | 4 | 1 | 1 | 1 | 13 | 0.7692 | 2 | 1 | 5 | 0.8 | 5 | 0.2 | 3 | 0.6667 | 15 | 0.6 |
| 2_1607919 | 3 | 1 | 5 | 1 | 3 | 1 | 2 | 1 | 13 | 1 | 2 | 1 | 7 | 1 | 4 | 1 | 4 | 1 | 17 | 1 |
| 2_1607920 | 2 | 0.5 | 4 | 0.75 | 1 | 1 | 2 | 1 | 9 | 0.7778 | 2 | 0 | 5 | 0.6 | 4 | 0.25 | 3 | 0.6667 | 14 | 0.4286 |
| 2_1608020 | 1 | 1 | 2 | 0.5 | NA | NA | NA | NA | 3 | 0.6667 | 2 | 1 | 6 | 0.8333 | 3 | 0.3333 | 4 | 1 | 15 | 0.8 |
| 2_1608072 | 2 | 0.5 | 1 | 0 | 1 | 0 | NA | NA | 4 | 0.25 | 1 | 1 | 3 | 0.6667 | 3 | 0.3333 | 3 | 0 | 10 | 0.4 |
| 2_1608073 | 2 | 0 | 1 | 1 | 1 | 1 | NA | NA | 4 | 0.5 | 1 | 0 | 3 | 0.6667 | 2 | 0.5 | 2 | 0.5 | 8 | 0.5 |
| 2_1608076 | 1 | 0 | NA | NA | 1 | 1 | NA | NA | 2 | 0.5 | 1 | 1 | 3 | 0.3333 | 2 | 0 | 2 | 0 | 8 | 0.25 |
| 2_1608077 | 1 | 1 | NA | NA | 1 | 0 | NA | NA | 2 | 0.5 | NA | NA | 2 | 0 | 1 | 0 | 2 | 0.5 | 5 | 0.2 |
| 2_1608080 | 1 | 0 | NA | NA | 1 | 1 | NA | NA | 2 | 0.5 | 1 | 0 | 3 | 0.3333 | 2 | 1 | 1 | 1 | 7 | 0.5714 |
| 2_1608081 | 1 | 1 | NA | NA | 1 | 0 | NA | NA | 2 | 0.5 | 1 | 0 | 2 | 0 | 2 | 0.5 | 2 | 0 | 7 | 0.1429 |
| 2_1608082 | 1 | 1 | NA | NA | 1 | 1 | NA | NA | 2 | 1 | 1 | 0 | 3 | 0.3333 | 2 | 1 | 2 | 0.5 | 8 | 0.5 |
| 2_1608085 | 1 | 1 | NA | NA | NA | NA | NA | NA | 1 | 1 | 1 | 1 | 1 | 0 | 1 | 0 | 2 | 0.5 | 5 | 0.4 |
| 2_1608561 | 1 | 1 | NA | NA | NA | NA | NA | NA | 1 | 1 | NA | NA | NA | NA | 2 | 0 | 1 | 1 | 3 | 0.3333 |
| 2_1608946 | 1 | 0 | 1 | 1 | NA | NA | NA | NA | 2 | 0.5 | NA | NA | 1 | 1 | NA | NA | NA | NA | 1 | 1 |
| 2_1613711 | 1 | 1 | 1 | 1 | NA | NA | NA | NA | 2 | 1 | 1 | 1 | NA | NA | 1 | 1 | NA | NA | 2 | 1 |
| 2_1613713 | 1 | 1 | NA | NA | NA | NA | NA | NA | 1 | 1 | 1 | 1 | NA | NA | NA | NA | NA | NA | 1 | 1 |
| 2_1625152 | NA | NA | NA | NA | NA | NA | 1 | 1 | 1 | 1 | NA | NA | NA | NA | NA | NA | 1 | 1 | 1 | 1 |
| 2_1625154 | NA | NA | NA | NA | NA | NA | 1 | 1 | 1 | 1 | NA | NA | NA | NA | NA | NA | 1 | 1 | 1 | 1 |
| 2_1625157 | 2 | 0.5 | NA | NA | NA | NA | 1 | 0 | 3 | 0.3333 | NA | NA | 1 | 1 | 1 | 1 | 1 | 1 | 3 | 1 |
| 2_1625157 | 1 | 1 | NA | NA | NA | NA | 1 | 1 | 2 | 1 | NA | NA | 1 | 1 | 2 | 1 | 1 | 1 | 4 | 1 |
| 2_1625166 | 4 | 0.75 | NA | NA | NA | NA | 1 | 0 | 5 | 0.6 | NA | NA | 1 | 0 | 2 | 0.5 | 2 | 0.5 | 5 | 0.4 |
| 2_1625168 | 2 | 1 | NA | NA | NA | NA | 2 | 1 | 4 | 1 | NA | NA | 1 | 1 | 2 | 1 | 2 | 1 | 5 | 1 |
| 2_1625168 | 4 | 1 | NA | NA | NA | NA | 1 | 0 | 5 | 0.8 | NA | NA | 1 | 0 | 1 | 1 | 2 | 1 | 4 | 0.75 |
| 2_1625170 | 4 | 0.25 | NA | NA | NA | NA | 1 | 0 | 5 | 0.2 | NA | NA | 1 | 0 | 1 | 1 | 2 | 0.5 | 4 | 0.5 |
| 2_1625170 | 3 | 0.3333 | NA | NA | NA | NA | 2 | 0 | 5 | 0.2 | NA | NA | 1 | 0 | 2 | 0.5 | 1 | 0 | 4 | 0.25 |
| 2_1627767 | 2 | 1 | 13 | 0.8462 | 4 | 0.5 | 9 | 0.4444 | 28 | 0.6786 | 4 | 0.5 | 8 | 0.5 | 8 | 0.25 | 8 | 0.75 | 28 | 0.5 |
| 2_1627773 | 2 | 0 | 7 | 0.1429 | 4 | 0 | 2 | 0.5 | 15 | 0.1333 | NA | NA | 1 | 0 | NA | NA | 3 | 0.6667 | 4 | 0.5 |
| 2_1627786 | 6 | 0.1667 | 7 | 0.1429 | 7 | 0.2857 | 6 | 0.1667 | 26 | 0.1923 | 2 | 0 | 4 | 0 | 7 | 0.1429 | 7 | 0.1429 | 20 | 0.1 |
| 2_1628187 | NA | NA | NA | NA | NA | NA | 1 | 1 | 1 | 1 | 1 | 1 | NA | NA | NA | NA | 2 | 1 | 3 | 1 |
| 2_1628872 | 1 | 0 | 5 | 0.4 | NA | NA | 3 | 0.3333 | 9 | 0.3333 | 4 | 0.75 | NA | NA | 1 | 1 | NA | NA | 5 | 0.8 |
| 2_1628873 | 1 | 0 | 6 | 0.1667 | 1 | 0 | 3 | 0 | 11 | 0.0909 | 8 | 0 | NA | NA | 2 | 0.5 | NA | NA | 10 | 0.1 |
| 2_1628873 | 1 | 1 | 6 | 0.3333 | 1 | 0 | 3 | 0.6667 | 11 | 0.4545 | 7 | 1 | NA | NA | 2 | 0.5 | NA | NA | 9 | 0.8889 |
| 2_1628874 | NA | NA | 4 | 0.25 | 1 | 0 | 2 | 0 | 7 | 0.1429 | 7 | 0.1429 | NA | NA | 1 | 0 | NA | NA | 8 | 0.125 |
| 2_1628876 | 2 | 0.5 | 6 | 0.1667 | 1 | 0 | 1 | 0 | 10 | 0.2 | 7 | 0.1429 | NA | NA | NA | NA | NA | NA | 7 | 0.1429 |
| 2_1628876 | 1 | 0 | 5 | 0.4 | NA | NA | 3 | 0.3333 | 9 | 0.3333 | 4 | 0.75 | NA | NA | 1 | 0 | NA | NA | 5 | 0.6 |
| 2_1628878 | 2 | 0.5 | 8 | 0.625 | NA | NA | 1 | 0 | 11 | 0.5455 | 3 | 0.6667 | NA | NA | 1 | 1 | NA | NA | 4 | 0.75 |
| 2_1632479 | 2 | 1 | 6 | 0.3333 | 4 | 0.25 | 2 | 0 | 14 | 0.3571 | 4 | 0 | NA | NA | 4 | 0.25 | NA | NA | 8 | 0.125 |
| 2_1632479 | 2 | 0.5 | 6 | 0.5 | 4 | 0.75 | 3 | 0.3333 | 15 | 0.5333 | 5 | 0.2 | NA | NA | 4 | 0.5 | NA | NA | 9 | 0.3333 |
| 2_1632479 | 1 | 0 | 3 | 1 | 3 | 0.3333 | 2 | 0 | 9 | 0.4444 | 5 | 0.4 | NA | NA | 3 | 0.6667 | NA | NA | 8 | 0.5 |
| 2_1632479 | 2 | 1 | 6 | 0.1667 | 4 | 0.25 | 3 | 0.3333 | 15 | 0.3333 | 3 | 0 | NA | NA | 4 | 0.25 | NA | NA | 7 | 0.1429 |
| 2_1632480 | 2 | 1 | 5 | 0.2 | 4 | 0 | 3 | 0.3333 | 14 | 0.2857 | 4 | 0.5 | NA | NA | 4 | 0.5 | NA | NA | 8 | 0.5 |
| 2_1632480 | 2 | 0 | 5 | 0 | 2 | 0 | 3 | 0.3333 | 12 | 0.0833 | 3 | 0 | NA | NA | 4 | 0.5 | NA | NA | 7 | 0.2857 |
| 2_1632480 | 2 | 0.5 | 6 | 1 | 3 | 0.6667 | 3 | 1 | 14 | 0.8571 | 5 | 0.8 | NA | NA | 3 | 0.3333 | NA | NA | 8 | 0.625 |
| 2_1632480 | 2 | 0 | 6 | 0.3333 | 4 | 0 | 3 | 0 | 15 | 0.1333 | 4 | 0 | NA | NA | 3 | 0.3333 | NA | NA | 7 | 0.1429 |
| 2_1632482 | NA | NA | 5 | 1 | 3 | 0.6667 | 3 | 0.6667 | 11 | 0.8182 | 3 | 0.3333 | NA | NA | 1 | 1 | NA | NA | 4 | 0.5 |

| | | | | | | | | | | | | | | | | | | | | |
|---|---|---|---|---|---|---|---|---|---|---|---|---|---|---|---|---|---|---|---|---|
| 2_1632484( | 2 | 1 | 5 | 0.6 | 4 | 0.25 | 3 | 0.6667 | 14 | 0.5714 | 3 | 0.6667 | NA | NA | 4 | 1 | NA | NA | 7 | 0.8571 |
| 2_1632484 | 1 | 0 | 5 | 0.8 | 2 | 1 | 1 | 0 | 9 | 0.6667 | 3 | 0.3333 | NA | NA | 3 | 0.6667 | NA | NA | 6 | 0.5 |
| 2_1632484 | 1 | 0 | 4 | 0.25 | 4 | 0.5 | 3 | 0.3333 | 12 | 0.3333 | 3 | 1 | NA | NA | 3 | 0.3333 | NA | NA | 6 | 0.6667 |
| 2_1632486 | 2 | 0.5 | 4 | 1 | 2 | 1 | 1 | 1 | 9 | 0.8889 | 4 | 0.75 | NA | NA | 1 | 0 | NA | NA | 5 | 0.6 |
| 2_1632955 | 9 | 0.2222 | 12 | 0.0833 | 15 | 0.4 | 3 | 0.3333 | 39 | 0.2564 | 9 | 0.1111 | 7 | 0.1429 | 8 | 0 | 10 | 0.2 | 34 | 0.1176 |
| 2_1632958 | 4 | 0.75 | 5 | 0.2 | 6 | 0.8333 | 4 | 0.25 | 19 | 0.5263 | 5 | 0.2 | 4 | 0.5 | 5 | 0.8 | 4 | 0.5 | 18 | 0.5 |
| 2_1632959 | 6 | 1 | 3 | 0.6667 | 7 | 0.5714 | 3 | 1 | 19 | 0.7895 | 3 | 1 | 5 | 0.4 | 2 | 1 | 3 | 1 | 13 | 0.7692 |
| 2_1632960 | 5 | 0 | 4 | 0 | 7 | 0.2857 | 4 | 0 | 20 | 0.1 | 5 | 0.2 | 4 | 0.25 | 4 | 0 | 4 | 0 | 17 | 0.1176 |
| 2_1632960 | 6 | 0.3333 | 5 | 0.2 | 7 | 0.1429 | 4 | 0 | 22 | 0.1818 | 5 | 0 | 5 | 0 | 3 | 0.3333 | 4 | 0.5 | 17 | 0.1765 |
| 2_1632961 | 4 | 0.75 | 5 | 0.6 | 8 | 0.25 | 3 | 0 | 20 | 0.4 | 5 | 0.4 | 4 | 0.5 | 4 | 0.5 | 4 | 0.25 | 17 | 0.4118 |
| 2_1632961 | 6 | 0.6667 | 5 | 0.4 | 10 | 0.3 | 3 | 0.3333 | 24 | 0.4167 | 6 | 0.8333 | 5 | 0.2 | 4 | 0.5 | 4 | 0.25 | 19 | 0.4737 |
| 2_1632962( | 3 | 0.6667 | 1 | 0 | 2 | 0 | 1 | 1 | 7 | 0.4286 | 2 | 0.5 | 3 | 0.3333 | 1 | 0 | NA | NA | 6 | 0.3333 |
| 2_1632962( | 6 | 0.1667 | 3 | 0 | 8 | 0.125 | 3 | 0 | 20 | 0.1 | 6 | 0 | 4 | 0.25 | 4 | 0.25 | 4 | 0 | 18 | 0.1111 |
| 2_1632963 | 7 | 0.2857 | 4 | 0 | 8 | 0.125 | 4 | 0.5 | 23 | 0.2174 | 5 | 0.6 | 5 | 0.4 | 2 | 0 | 3 | 0 | 15 | 0.3333 |
| 2_1632963 | 4 | 0.75 | 6 | 0.8333 | 9 | 0.7778 | 3 | 1 | 22 | 0.8182 | 4 | 0.75 | 5 | 0.8 | 2 | 1 | 3 | 1 | 14 | 0.8571 |
| 2_1633085( | 3 | 0 | 4 | 0 | 5 | 0.4 | 4 | 0 | 16 | 0.125 | 4 | 0.25 | 2 | 0 | 6 | 0 | 1 | 0 | 13 | 0.0769 |
| 2_1633091 | 3 | 0.3333 | 4 | 0.5 | 5 | 0.6 | 2 | 0 | 14 | 0.4286 | 2 | 0.5 | NA | NA | 4 | 0 | NA | NA | 6 | 0.1667 |
| 2_1633101 | 1 | 0 | 2 | 0 | 4 | 0 | 3 | 0.3333 | 10 | 0.1 | 2 | 0.5 | 1 | 0 | 5 | 0.4 | 1 | 0 | 9 | 0.3333 |
| 2_1633106 | 1 | 1 | 2 | 0.5 | 3 | 0.3333 | 2 | 0.5 | 8 | 0.5 | 1 | 1 | 1 | 0 | 3 | 0.3333 | NA | NA | 5 | 0.4 |
| 2_1633107 | 2 | 0.5 | 3 | 0 | 3 | 0 | 2 | 0 | 10 | 0.1 | 1 | 0 | 1 | 0 | 3 | 0.3333 | 2 | 0 | 7 | 0.1429 |
| 2_1633108 | 2 | 1 | 2 | 0 | 2 | 0 | 2 | 0.5 | 8 | 0.375 | 1 | 1 | NA | NA | 2 | 0 | 2 | 0 | 5 | 0.2 |
| 2_1633109 | 2 | 1 | 1 | 1 | 3 | 0.6667 | 2 | 0.5 | 8 | 0.75 | 1 | 1 | 1 | 1 | 4 | 0.75 | 2 | 1 | 8 | 0.875 |
| 2_1633112( | 1 | 0 | 1 | 0 | 2 | 0 | 2 | 0.5 | 6 | 0.1667 | 1 | 1 | NA | NA | 4 | 0.25 | 2 | 0 | 7 | 0.2857 |
| 2_1633113 | 1 | 1 | 1 | 1 | 3 | 0.3333 | 1 | 1 | 6 | 0.6667 | 1 | 1 | 1 | 1 | 3 | 1 | 2 | 1 | 7 | 1 |
| 2_1633114( | 1 | 0 | 1 | 0 | 3 | 0.3333 | 1 | 1 | 6 | 0.3333 | 1 | 1 | 1 | 1 | 3 | 0.3333 | 1 | 1 | 6 | 0.6667 |
| 2_1633114 | NA | NA | 1 | 1 | 3 | 0.6667 | NA | NA | 4 | 0.75 | 1 | 1 | 1 | 0 | 3 | 0.6667 | 2 | 1 | 7 | 0.7143 |
| 2_1633114 | 1 | 1 | 1 | 0 | 3 | 0.3333 | NA | NA | 5 | 0.4 | 1 | 1 | 1 | 0 | 4 | 0.75 | 2 | 0.5 | 8 | 0.625 |
| 2_1633114 | 1 | 1 | 1 | 1 | 2 | 1 | NA | NA | 4 | 1 | 1 | 1 | 1 | 1 | 4 | 1 | 2 | 0.5 | 8 | 0.875 |
| 2_1633115 | NA | NA | NA | NA | 3 | 0.6667 | NA | NA | 3 | 0.6667 | 1 | 1 | 1 | 1 | 2 | 1 | 1 | 1 | 5 | 1 |
| 2_1633116 | 1 | 1 | 1 | 1 | 3 | 0 | NA | NA | 5 | 0.4 | 1 | 1 | 1 | 1 | 4 | 0.75 | 2 | 1 | 8 | 0.875 |
| 2_1633117 | 1 | 0 | 1 | 0 | 3 | 0.3333 | NA | NA | 5 | 0.2 | 1 | 1 | NA | NA | 3 | 0.3333 | 1 | 1 | 5 | 0.6 |
| 2_1633118 | 1 | 0 | 1 | 0 | 3 | 0.3333 | NA | NA | 5 | 0.2 | 1 | 0 | NA | NA | 4 | 0 | 2 | 0.5 | 7 | 0.1429 |
| 2_1633120 | 1 | 0 | 1 | 0 | 3 | 0.6667 | NA | NA | 5 | 0.4 | NA | NA | NA | NA | 3 | 0 | 2 | 1 | 5 | 0.4 |
| 2_1634491 | NA | NA | 4 | 0 | 3 | 0.3333 | 3 | 0 | 10 | 0.1 | 1 | 0 | 1 | 0 | 2 | 0 | 4 | 0.25 | 8 | 0.125 |
| 2_1634491( | NA | NA | 4 | 0.5 | 2 | 0 | 4 | 0 | 10 | 0.2 | 1 | 0 | 1 | 0 | 2 | 0 | 4 | 0.75 | 8 | 0.375 |
| 2_1634492 | 1 | 0 | 4 | 0.25 | 3 | 0 | 4 | 0 | 12 | 0.0833 | 1 | 1 | 2 | 0 | 3 | 0 | 4 | 0 | 10 | 0.1 |
| 2_1634573 | 2 | 0.5 | 5 | 0.6 | 3 | 1 | 3 | 1 | 13 | 0.7692 | 1 | 0 | 3 | 0.6667 | 1 | 1 | 5 | 0.8 | 10 | 0.7 |
| 2_1634575 | 2 | 0 | 4 | 0.25 | 3 | 0 | 6 | 0.3333 | 15 | 0.2 | 1 | 0 | 3 | 0 | 3 | 0.3333 | 7 | 0.2857 | 14 | 0.2143 |
| 2_1634576 | 2 | 1 | 5 | 1 | 3 | 1 | 7 | 0.5714 | 17 | 0.8235 | 1 | 1 | 3 | 1 | 3 | 1 | 7 | 0.8571 | 14 | 0.9286 |
| 2_1634577 | 2 | 0.5 | 5 | 0 | 3 | 0.3333 | 3 | 0.3333 | 13 | 0.2308 | 1 | 0 | 2 | 0 | 2 | 0.5 | 5 | 0.4 | 10 | 0.3 |
| 2_1634577 | 1 | 1 | 4 | 1 | 3 | 1 | 5 | 1 | 13 | 1 | NA | NA | 3 | 0.6667 | NA | NA | 5 | 0.8 | 8 | 0.75 |
| 2_1634577 | 2 | 0 | 5 | 0.4 | 3 | 0 | 7 | 0 | 17 | 0.1176 | 1 | 1 | 4 | 0 | 4 | 0 | 9 | 0.1111 | 18 | 0.1111 |
| 2_1634578 | 1 | 1 | 2 | 1 | 1 | 1 | 4 | 1 | 8 | 1 | 1 | 1 | 3 | 1 | 4 | 1 | 4 | 0.5 | 12 | 0.8333 |
| 2_1634578 | 2 | 0.5 | 5 | 0 | 3 | 0 | 7 | 0 | 17 | 0.0588 | 1 | 0 | 5 | 0 | 4 | 0 | 9 | 0.3333 | 19 | 0.1579 |
| 2_1634578( | 2 | 1 | 5 | 1 | 3 | 1 | 7 | 0.8571 | 17 | 0.9412 | 1 | 0 | 5 | 1 | 4 | 1 | 6 | 0.8333 | 16 | 0.875 |
| 2_1634578 | NA | NA | 3 | 0 | 2 | 0.5 | 5 | 0 | 10 | 0.1 | 1 | 0 | 4 | 0.25 | 4 | 0 | 6 | 0.1667 | 15 | 0.1333 |
| 2_1634579 | 2 | 1 | 5 | 1 | 3 | 1 | 6 | 1 | 16 | 1 | 1 | 1 | 5 | 1 | 4 | 1 | 6 | 0.6667 | 16 | 0.875 |
| 2_1634580 | 3 | 0 | 5 | 0.6 | 3 | 0 | 6 | 0 | 17 | 0.1765 | 1 | 0 | 5 | 0.4 | 2 | 0.5 | 7 | 0.1429 | 15 | 0.2667 |
| 2_1634581 | 3 | 0.3333 | 5 | 0.4 | 2 | 0 | 7 | 0.7143 | 17 | 0.4706 | 1 | 1 | 5 | 0.2 | 4 | 0.5 | 8 | 0.125 | 18 | 0.2778 |
| 2_1634582( | 4 | 1 | 5 | 0.8 | 3 | 1 | 6 | 1 | 18 | 0.9444 | 1 | 1 | 4 | 1 | 5 | 0.8 | 9 | 0.8889 | 19 | 0.8947 |
| 2_1634582 | 4 | 0 | 5 | 0.2 | 3 | 0 | 5 | 0.4 | 17 | 0.1765 | 1 | 0 | 4 | 0.25 | 5 | 0.2 | 7 | 0.2857 | 17 | 0.2353 |
| 2_1634582 | 3 | 0.6667 | 5 | 0 | 3 | 0 | 6 | 0 | 17 | 0.1176 | 1 | 0 | 5 | 0.2 | 4 | 0.25 | 8 | 0 | 18 | 0.1111 |
| 2_1634583 | 5 | 0.2 | 5 | 0.2 | 3 | 0 | 8 | 0 | 21 | 0.0952 | 1 | 0 | 5 | 0 | 5 | 0 | 8 | 0.125 | 19 | 0.0526 |
| 2_1634585( | 4 | 0 | 5 | 0.4 | 4 | 0 | 7 | 0 | 20 | 0.1 | 1 | 0 | 6 | 0 | 4 | 0.25 | 10 | 0.1 | 21 | 0.0952 |
| 2_1634587 | 4 | 0.75 | 7 | 0.8571 | 4 | 0.5 | 4 | 1 | 19 | 0.7895 | 2 | 1 | 7 | 0.4286 | 4 | 1 | 9 | 0.5556 | 22 | 0.6364 |
| 2_1634588 | 4 | 0 | 8 | 0.625 | 5 | 0.4 | 6 | 0.3333 | 23 | 0.3913 | 1 | 0 | 7 | 0.5714 | 1 | 0 | 8 | 0.25 | 17 | 0.3529 |
| 2_1634590 | 4 | 0.5 | 7 | 0.2857 | 5 | 0.4 | 6 | 0.6667 | 22 | 0.4545 | 2 | 0.5 | 5 | 0.2 | 4 | 0.5 | 10 | 0.4 | 21 | 0.381 |
| 2_1634594 | 6 | 0.3333 | 4 | 0.5 | 6 | 0.5 | 7 | 0.5714 | 23 | 0.4783 | 2 | 1 | 10 | 0.6 | 4 | 0.25 | 6 | 0 | 22 | 0.4091 |
| 2_1634596 | 6 | 0.3333 | 9 | 0.3333 | 6 | 0.3333 | 7 | 0.5714 | 28 | 0.3929 | 2 | 0.5 | 10 | 0.3 | 6 | 0 | 9 | 0.2222 | 27 | 0.2222 |
| 2_1634604 | 9 | 0 | 11 | 0.0909 | 7 | 0 | 9 | 0.3333 | 36 | 0.1111 | 2 | 0 | 12 | 0.1667 | 8 | 0.125 | 8 | 0.125 | 30 | 0.1333 |
| 2_1634607( | 10 | 0 | 11 | 0.1818 | 9 | 0.1111 | 10 | 0.2 | 40 | 0.125 | 5 | 0 | 12 | 0.0833 | 10 | 0 | 11 | 0.0909 | 38 | 0.0526 |
| 2_1634672 | 54 | 0.1111 | 49 | 0.0816 | 52 | 0.0962 | 31 | 0.0968 | 186 | 0.0968 | 47 | 0.1489 | 27 | 0.1111 | 41 | 0.0732 | 45 | 0.1556 | 160 | 0.125 |
| 2_1634673 | 65 | 0.1077 | 53 | 0.0755 | 63 | 0.0952 | 37 | 0.0541 | 218 | 0.0872 | 52 | 0.0962 | 32 | 0.0938 | 42 | 0.1667 | 52 | 0.1154 | 178 | 0.118 |
| 2_1634673 | 57 | 0.2456 | 48 | 0.125 | 52 | 0.2115 | 32 | 0.1875 | 189 | 0.1958 | 44 | 0.1818 | 32 | 0.1562 | 40 | 0.125 | 43 | 0.1628 | 159 | 0.1572 |
| 2_1634674( | 38 | 0.7895 | 29 | 0.5862 | 39 | 0.3846 | 24 | 0.5833 | 130 | 0.5846 | 33 | 0.6364 | 30 | 0.6333 | 31 | 0.5161 | 40 | 0.625 | 134 | 0.6045 |

| | | | | | | | | | | | | | | | | | | | | |
|---|---|---|---|---|---|---|---|---|---|---|---|---|---|---|---|---|---|---|---|---|
| 2_16356750 | 18 | 0 | 21 | 0.1905 | 17 | 0.1765 | 18 | 0.1667 | 74 | 0.1351 | 19 | 0.1053 | 18 | 0.1667 | 27 | 0.1852 | 20 | 0.2 | 84 | 0.1667 |
| 2_16356752 | 19 | 0.2105 | 23 | 0.3478 | 18 | 0.5 | 17 | 0.2353 | 77 | 0.3247 | 22 | 0.2727 | 21 | 0.3333 | 28 | 0.4643 | 19 | 0.1579 | 90 | 0.3222 |
| 2_16356772 | 23 | 0.1304 | 23 | 0.1304 | 21 | 0.2381 | 17 | 0.1765 | 84 | 0.1667 | 23 | 0.087 | 19 | 0.4737 | 27 | 0.2222 | 20 | 0.15 | 89 | 0.2247 |
| 2_16356798 | 17 | 0.5294 | 18 | 0.4444 | 22 | 0.4545 | 12 | 0.4167 | 69 | 0.4638 | 24 | 0.4167 | 17 | 0.3529 | 22 | 0.5 | 17 | 0.4706 | 80 | 0.4375 |
| 2_16356803 | 23 | 0.3043 | 23 | 0.3043 | 19 | 0.2105 | 17 | 0.4118 | 82 | 0.3049 | 21 | 0.0952 | 24 | 0.4583 | 30 | 0.2 | 21 | 0.4762 | 96 | 0.3021 |
| 2_16370502 | 2 | 0 | 1 | 0 | 2 | 0.5 | 2 | 0 | 7 | 0.1429 | 2 | 0 | 1 | 0 | 3 | 0.3333 | 1 | 1 | 7 | 0.2857 |
| 2_16370504 | 2 | 1 | 1 | 1 | 2 | 1 | 1 | 1 | 6 | 1 | 2 | 1 | 1 | 1 | 1 | 0 | 1 | 0 | 5 | 0.6 |
| 2_16370507 | 2 | 1 | 1 | 1 | 2 | 1 | 2 | 1 | 7 | 1 | 2 | 1 | 1 | 1 | 1 | 1 | 1 | 0 | 5 | 0.8 |
| 2_16370508 | 1 | 1 | NA | NA | 2 | 1 | 2 | 1 | 5 | 1 | 1 | 1 | NA | NA | 1 | 1 | 1 | 1 | 3 | 1 |
| 2_16370517 | 1 | 0 | 1 | 0 | 2 | 0.5 | 2 | 0 | 6 | 0.1667 | 2 | 0 | 1 | 0 | 2 | 0.5 | 1 | 0 | 6 | 0.1667 |
| 2_16370555 | 1 | 1 | 1 | 1 | 1 | 1 | 2 | 1 | 5 | 1 | 2 | 1 | NA | NA | 1 | 1 | 1 | 1 | 4 | 1 |
| 2_16370560 | 2 | 0 | 1 | 0 | 1 | 0 | 2 | 0.5 | 6 | 0.1667 | 2 | 0 | 1 | 1 | 2 | 0.5 | 1 | 0 | 6 | 0.3333 |
| 2_16370573 | 1 | 1 | 1 | 1 | NA | NA | 2 | 1 | 4 | 1 | 1 | 1 | 1 | 1 | 2 | 1 | 1 | 1 | 5 | 1 |
| 2_16370596 | 1 | 0 | 1 | 0 | 1 | 0 | 2 | 1 | 5 | 0.4 | 1 | 1 | 1 | 1 | 2 | 0 | 1 | 0 | 5 | 0.4 |
| 2_16385362 | 38 | 0.0789 | 41 | 0.0488 | 28 | 0.1071 | 42 | 0.0476 | 149 | 0.0671 | 20 | 0 | 46 | 0.1304 | 37 | 0.0541 | 38 | 0.1316 | 141 | 0.0922 |
| 2_16385363 | 28 | 0.0357 | 29 | 0.1379 | 22 | 0.1364 | 32 | 0.0312 | 111 | 0.0811 | 17 | 0.0588 | 34 | 0.1176 | 25 | 0.08 | 28 | 0.1429 | 104 | 0.1058 |
| 2_16385873 | 46 | 0.3043 | 51 | 0.3529 | 39 | 0.2564 | 44 | 0.3864 | 180 | 0.3278 | 30 | 0.3667 | 52 | 0.3654 | 43 | 0.4884 | 47 | 0.383 | 172 | 0.4012 |
| 2_16385907 | 41 | 0.0732 | 49 | 0.0204 | 42 | 0.0952 | 43 | 0.0698 | 175 | 0.0629 | 23 | 0.1304 | 46 | 0.087 | 42 | 0.0238 | 36 | 0.0556 | 147 | 0.068 |
| 2_16389612 | 10 | 0.2 | 11 | 0.1818 | 7 | 0.4286 | 7 | 0 | 35 | 0.2 | 16 | 0.0625 | 12 | 0 | 9 | 0.1111 | 12 | 0.0833 | 49 | 0.0612 |
| 2_16389632 | 10 | 0.4 | 6 | 0.1667 | 8 | 0.25 | 6 | 0.1667 | 30 | 0.2667 | 11 | 0.1818 | 7 | 0.1429 | 8 | 0.125 | 10 | 0.1 | 36 | 0.1389 |
| 2_16399379 | 1 | 1 | 1 | 0 | NA | NA | 1 | 1 | 3 | 0.6667 | NA | NA | 1 | 1 | NA | NA | 2 | 0.5 | 3 | 0.6667 |
| 2_16399384 | NA | NA | 1 | 1 | NA | NA | 1 | 1 | 2 | 1 | NA | NA | 1 | 1 | NA | NA | 1 | 0 | 2 | 0.5 |
| 2_16399407 | 1 | 1 | 1 | 0 | NA | NA | 1 | 0 | 3 | 0.3333 | NA | NA | 1 | 0 | 1 | 1 | 2 | 0.5 | 4 | 0.5 |
| 2_16399440 | 1 | 1 | 1 | 1 | NA | NA | 1 | 1 | 3 | 1 | NA | NA | 1 | 0 | 1 | 0 | 3 | 0.3333 | 5 | 0.2 |
| 2_16399455 | 1 | 1 | 1 | 0 | NA | NA | 1 | 0 | 3 | 0.3333 | NA | NA | 1 | 0 | 1 | 1 | 3 | 0.6667 | 5 | 0.6 |
| 2_16399457 | 1 | 1 | 1 | 0 | NA | NA | 2 | 0.5 | 4 | 0.5 | NA | NA | 1 | 1 | 1 | 0 | 3 | 0.3333 | 5 | 0.4 |
| 2_16399628 | 1 | 1 | 2 | 0 | NA | NA | 2 | 0 | 5 | 0.2 | NA | NA | 2 | 0.5 | 1 | 1 | 5 | 0.2 | 8 | 0.375 |
| 2_16399675 | 1 | 0 | 2 | 0.5 | 1 | 0 | 3 | 0 | 7 | 0.1429 | NA | NA | 6 | 0 | 2 | 0 | 6 | 0.3333 | 14 | 0.1429 |
| 2_16399686 | 1 | 1 | 3 | 0 | 1 | 0 | 3 | 0.3333 | 8 | 0.25 | NA | NA | 5 | 0.2 | 2 | 0 | 5 | 0 | 12 | 0.0833 |
| 2_16400590 | 1 | 1 | NA | NA | 1 | 1 | 1 | 1 | 3 | 1 | 2 | 1 | 1 | 1 | NA | NA | 2 | 1 | 5 | 1 |
| 2_16400654 | NA | NA | NA | NA | 1 | 1 | 1 | 1 | 2 | 1 | 1 | 0 | 1 | 1 | NA | NA | 2 | 0.5 | 4 | 0.5 |
| 2_16400659 | NA | NA | NA | NA | 1 | 1 | 1 | 1 | 2 | 1 | NA | NA | 1 | 1 | NA | NA | 1 | 1 | 2 | 1 |
| 2_16400675 | NA | NA | NA | NA | 1 | 1 | 1 | 1 | 2 | 1 | 1 | 1 | 1 | 1 | NA | NA | 1 | 1 | 3 | 1 |
| 2_16400983 | NA | NA | NA | NA | 1 | 1 | 1 | 0 | 2 | 0.5 | 1 | 0 | 1 | 0 | NA | NA | 1 | 1 | 3 | 0.3333 |
| 2_16423140 | NA | NA | 3 | 0.3333 | 1 | 0 | NA | NA | 4 | 0.25 | 2 | 0.5 | NA | NA | NA | NA | 1 | 0 | 3 | 0.3333 |
| 2_16426797 | 5 | 1 | 8 | 0.875 | 7 | 0.5714 | 7 | 0.5714 | 27 | 0.7407 | 13 | 0.6923 | 9 | 0.6667 | 9 | 0.6667 | 7 | 1 | 38 | 0.7368 |
| 2_16430653 | 15 | 0.7333 | 6 | 1 | 11 | 0.8182 | 8 | 0.75 | 40 | 0.8 | 9 | 0.8889 | 15 | 0.8667 | 8 | 0.875 | 5 | 0.8 | 37 | 0.8649 |
| 2_16430653 | NA | NA | NA | NA | 1 | 1 | NA | NA | 1 | 1 | 1 | 1 | NA | NA | NA | NA | NA | NA | 1 | 1 |
| 2_16430674 | 13 | 0.2308 | 1 | 0 | 4 | 0 | 5 | 0.2 | 23 | 0.1739 | 4 | 0.5 | 9 | 0.1111 | 7 | 0.2857 | 2 | 0.5 | 22 | 0.2727 |
| 2_16430710 | 4 | 0 | 2 | 0 | 4 | 0.25 | NA | NA | 10 | 0.1 | 3 | 0.3333 | 4 | 0 | 2 | 0.5 | 1 | 0 | 10 | 0.2 |
| 2_16430712 | 3 | 0.6667 | 2 | 1 | 2 | 0.5 | NA | NA | 7 | 0.7143 | 2 | 0.5 | 3 | 0.6667 | 1 | 1 | 1 | 1 | 7 | 0.7143 |
| 2_16430716 | 5 | 0.2 | 2 | 0 | 4 | 0 | NA | NA | 11 | 0.0909 | 2 | 0 | 4 | 0.25 | 2 | 0 | 1 | 0 | 9 | 0.1111 |
| 2_16431571 | NA | NA | NA | NA | NA | NA | 1 | 1 | 1 | 1 | NA | NA | NA | NA | 1 | 1 | 1 | 1 | 2 | 1 |
| 2_16432751 | 2 | 0.5 | 1 | 0 | 1 | 0 | NA | NA | 4 | 0.25 | 2 | 0 | NA | NA | 1 | 0 | 1 | 1 | 4 | 0.25 |
| 2_16432804 | 2 | 1 | 1 | 0 | 1 | 0 | NA | NA | 4 | 0.5 | NA | NA | 1 | 1 | 1 | 0 | 2 | 0.5 | 4 | 0.5 |
| 2_16432806 | 2 | 1 | 1 | 1 | NA | NA | NA | NA | 3 | 1 | 2 | 1 | 1 | 1 | 1 | 0 | 1 | 1 | 5 | 0.8 |
| 2_16433224 | 31 | 0.129 | 28 | 0.0357 | 27 | 0.0741 | 26 | 0.1154 | 112 | 0.0893 | 26 | 0.0385 | 29 | 0.1379 | 29 | 0 | 33 | 0.0909 | 117 | 0.0684 |
| 2_16433227 | 23 | 0.7391 | 27 | 0.7407 | 23 | 0.6957 | 28 | 0.6429 | 101 | 0.703 | 27 | 0.5556 | 26 | 0.8077 | 24 | 0.7083 | 27 | 0.5556 | 104 | 0.6538 |
| 2_16433233 | 39 | 0.0513 | 31 | 0.129 | 31 | 0.0968 | 31 | 0.0968 | 132 | 0.0909 | 27 | 0.1111 | 34 | 0.0294 | 33 | 0.0909 | 33 | 0.0303 | 127 | 0.063 |
| 2_16433247 | 32 | 0.375 | 30 | 0.1333 | 27 | 0.1852 | 27 | 0.2963 | 116 | 0.25 | 28 | 0.1786 | 28 | 0.3214 | 28 | 0.25 | 32 | 0.3125 | 116 | 0.2672 |
| 2_16433248 | 36 | 0.2778 | 29 | 0.2759 | 31 | 0.2581 | 28 | 0.25 | 124 | 0.2661 | 25 | 0.16 | 33 | 0.1515 | 29 | 0.3448 | 27 | 0.3333 | 114 | 0.2456 |
| 2_16433317 | NA | NA | 4 | 0.5 | 4 | 0 | 1 | 0 | 9 | 0.2222 | 1 | 0 | 3 | 0 | 2 | 0.5 | 2 | 0 | 8 | 0.125 |
| 2_16433402 | 1 | 0 | 5 | 0.2 | 4 | 0 | 1 | 0 | 11 | 0.0909 | 1 | 0 | 3 | 0.3333 | 3 | 0.3333 | 4 | 0 | 11 | 0.1818 |
| 2_16433456 | 1 | 0 | 7 | 0.4286 | 2 | 0.5 | 2 | 0 | 12 | 0.3333 | 4 | 0.25 | 4 | 0 | 6 | 0.3333 | 3 | 0.3333 | 17 | 0.2353 |
| 2_16433512 | 3 | 0.3333 | 9 | 0.2222 | 4 | 0 | 5 | 0 | 21 | 0.1429 | 5 | 0 | 4 | 0.5 | 6 | 0 | 5 | 0.2 | 20 | 0.15 |
| 2_16434445 | 10 | 0.2 | 5 | 0 | 2 | 0 | 3 | 0 | 20 | 0.1 | 4 | 0.25 | 7 | 0.2857 | 3 | 0 | 4 | 0.25 | 18 | 0.2222 |
| 2_16434927 | 15 | 0.1333 | 8 | 0.125 | 8 | 0.375 | 9 | 0.2222 | 40 | 0.2 | 10 | 0 | 10 | 0.1 | 12 | 0.1667 | 5 | 0.6 | 37 | 0.1622 |
| 2_16434998 | 16 | 0.0625 | 11 | 0 | 9 | 0 | 8 | 0.25 | 44 | 0.0682 | 13 | 0 | 11 | 0 | 17 | 0.1176 | 8 | 0.25 | 49 | 0.0816 |
| 2_16435006 | 17 | 0.1176 | 9 | 0.1111 | 8 | 0.25 | 9 | 0.2222 | 43 | 0.1628 | 14 | 0.0714 | 9 | 0 | 17 | 0.2353 | 7 | 0.4286 | 47 | 0.1702 |
| 2_16435009 | 10 | 0.6 | 7 | 0.5714 | 4 | 0.5 | 6 | 0.6667 | 27 | 0.5926 | 8 | 0.625 | 9 | 0.6667 | 9 | 0.7778 | 5 | 1 | 31 | 0.7419 |
| 2_16460965 | 7 | 0 | 6 | 0 | 1 | 0 | 3 | 0.3333 | 17 | 0.0588 | 6 | 0.1667 | 4 | 0 | 2 | 0 | 5 | 0 | 17 | 0.0588 |
| 2_16461001 | 5 | 0.6 | 6 | 0.3333 | NA | NA | 4 | 1 | 15 | 0.6 | 5 | 0.4 | 3 | 0.6667 | NA | NA | 5 | 0.4 | 13 | 0.4615 |
| 2_16461005 | 7 | 0.1429 | 6 | 0 | NA | NA | 4 | 0.25 | 17 | 0.1176 | 3 | 0.3333 | 2 | 0.5 | 2 | 0.5 | 4 | 0 | 11 | 0.2727 |
| 2_16461007 | 5 | 0 | 6 | 0 | NA | NA | 5 | 0.2 | 16 | 0.0625 | 4 | 0 | 3 | 0.3333 | 2 | 0 | 4 | 0.25 | 13 | 0.1538 |
| 2_16462864 | 2 | 0 | 1 | 0 | 2 | 0.5 | 4 | 0 | 9 | 0.1111 | 5 | 0.2 | 3 | 0 | 1 | 0 | 3 | 0 | 12 | 0.0833 |

| | | | | | | | | | | | | | | | | | | | | |
|---|---|---|---|---|---|---|---|---|---|---|---|---|---|---|---|---|---|---|---|---|
| 2_1646337( | 2 | 1 | 2 | 0.5 | 2 | 1 | 5 | 0.8 | 11 | 0.8182 | 4 | 0.75 | 2 | 1 | 4 | 1 | 2 | 0.5 | 12 | 0.8333 |
| 2_1646337! | 2 | 0.5 | 2 | 0.5 | 2 | 1 | 6 | 0.5 | 12 | 0.5833 | 5 | 0.4 | 3 | 0.6667 | 3 | 0.3333 | 3 | 1 | 14 | 0.5714 |
| 2_1646338( | 3 | 0.3333 | 2 | 0 | 1 | 0 | 5 | 0 | 11 | 0.0909 | 5 | 0 | 3 | 0 | 3 | 0.3333 | 3 | 0 | 14 | 0.0714 |
| 2_1646338: | NA | NA | 1 | 0 | 2 | 1 | 6 | 1 | 9 | 0.8889 | 3 | 0.6667 | 3 | 1 | 4 | 0.75 | 1 | 1 | 11 | 0.8182 |
| 2_1646338! | 4 | 0.25 | 3 | 0.3333 | 3 | 0.3333 | 5 | 0.2 | 15 | 0.2667 | 7 | 0 | 3 | 0.3333 | 3 | 0.3333 | 3 | 0 | 16 | 0.125 |
| 2_1646338! | 3 | 1 | 3 | 0.6667 | 3 | 1 | 5 | 0.8 | 14 | 0.8571 | 6 | 1 | 3 | 1 | 4 | 0.75 | 3 | 0.6667 | 16 | 0.875 |
| 2_1646340: | 4 | 0 | 3 | 0 | 3 | 0.3333 | 6 | 0 | 16 | 0.0625 | 6 | 0 | 4 | 0.25 | 3 | 0 | 3 | 0 | 16 | 0.0625 |
| 2_1646340: | 5 | 0.8 | 2 | 0.5 | 3 | 0.6667 | 5 | 0.6 | 15 | 0.6667 | 6 | 0.6667 | 4 | 0.25 | 3 | 0.6667 | 2 | 0 | 15 | 0.4667 |
| 2_1646341: | 5 | 0 | 3 | 0 | 3 | 0 | 6 | 0.1667 | 17 | 0.0588 | 6 | 0.1667 | 4 | 0.5 | 4 | 0.25 | 3 | 0 | 17 | 0.2353 |
| 2_1646342( | 4 | 0.75 | 2 | 1 | 3 | 0 | 6 | 0.1667 | 15 | 0.4 | 8 | 0.5 | 4 | 0.5 | 4 | 0 | 3 | 0.3333 | 19 | 0.3684 |
| 2_1646343( | 4 | 1 | 4 | 1 | 3 | 1 | 5 | 1 | 16 | 1 | 8 | 1 | 4 | 1 | 2 | 1 | 3 | 1 | 17 | 1 |
| 2_1646343( | 5 | 0.2 | 3 | 0 | 3 | 0 | 5 | 0.2 | 16 | 0.125 | 6 | 0.1667 | 5 | 0.2 | 3 | 0 | 2 | 0 | 16 | 0.125 |
| 2_1646343' | 5 | 0.8 | 3 | 0.3333 | 3 | 0.6667 | 6 | 0.6667 | 17 | 0.6471 | 8 | 0.5 | 4 | 0.25 | 3 | 0 | 2 | 0 | 17 | 0.2941 |
| 2_1646344: | 5 | 0.2 | 4 | 0.5 | 2 | 0.5 | 6 | 0.3333 | 17 | 0.3529 | 6 | 0.3333 | 5 | 0.6 | 4 | 0 | 3 | 0.3333 | 18 | 0.3333 |
| 2_1646344: | 6 | 0.3333 | 4 | 0.5 | 3 | 0.6667 | 6 | 0.1667 | 19 | 0.3684 | 8 | 0.375 | 5 | 0 | 4 | 0.25 | 3 | 0 | 20 | 0.2 |
| 2_1646344! | 6 | 0.8333 | 5 | 0.2 | 3 | 0.3333 | 7 | 0.7143 | 21 | 0.5714 | 8 | 0.375 | 4 | 1 | 4 | 0.5 | 3 | 0.6667 | 19 | 0.5789 |
| 2_1646346: | 4 | 0.25 | 4 | 0.25 | 3 | 0 | 4 | 0 | 15 | 0.1333 | 8 | 0 | 3 | 0.3333 | 2 | 0 | 1 | 0 | 14 | 0.0714 |
| 2_1646347· | 1 | 0 | 1 | 1 | 1 | 0 | 1 | 0 | 4 | 0.25 | 1 | 0 | 2 | 0.5 | 2 | 0 | 1 | 0 | 6 | 0.1667 |
| 2_1646347' | 4 | 0.5 | 5 | 0.4 | 4 | 0.5 | 2 | 0.5 | 15 | 0.4667 | 6 | 0.8333 | 1 | 0 | 2 | 1 | 1 | 1 | 10 | 0.8 |
| 2_1646352' | 1 | 0 | NA | NA | 1 | 0 | 1 | 1 | 3 | 0.3333 | 1 | 0 | 3 | 0.3333 | 3 | 0 | 1 | 0 | 8 | 0.125 |
| 2_1646391! | 2 | 1 | 1 | 1 | NA | NA | NA | NA | 3 | 1 | NA | NA | 1 | 1 | NA | NA | NA | NA | 1 | 1 |
| 2_1646617! | NA | NA | 2 | 1 | NA | NA | NA | NA | 2 | 1 | NA | NA | NA | NA | NA | NA | 1 | 1 | 1 | 1 |
| 2_1646620! | 1 | 1 | 2 | 1 | NA | NA | NA | NA | 3 | 1 | NA | NA | NA | NA | NA | NA | 1 | 1 | 1 | 1 |
| 2_1646621' | 1 | 1 | 2 | 0 | NA | NA | NA | NA | 3 | 0.3333 | NA | NA | NA | NA | NA | NA | 1 | 1 | 1 | 1 |
| 2_1646626: | 1 | 1 | 3 | 1 | NA | NA | NA | NA | 4 | 1 | NA | NA | NA | NA | NA | NA | 1 | 1 | 1 | 1 |
| 2_1646626! | 1 | 1 | 1 | 0 | NA | NA | NA | NA | 2 | 0.5 | NA | NA | NA | NA | NA | NA | 1 | 1 | 1 | 1 |
| 2_1646627! | 1 | 1 | 3 | 0.3333 | NA | NA | NA | NA | 4 | 0.5 | NA | NA | NA | NA | NA | NA | 1 | 1 | 1 | 1 |
| 2_1646627! | 1 | 1 | 3 | 1 | NA | NA | NA | NA | 4 | 1 | NA | NA | NA | NA | NA | NA | 1 | 1 | 1 | 1 |
| 2_1646633! | NA | NA | 3 | 0.3333 | 1 | 1 | NA | NA | 4 | 0.5 | NA | NA | NA | NA | NA | NA | 1 | 1 | 1 | 1 |
| 2_1646634( | 1 | 1 | 2 | 1 | 1 | 1 | 1 | 0 | 5 | 0.8 | NA | NA | NA | NA | NA | NA | 1 | 1 | 1 | 1 |
| 2_1646636! | 1 | 1 | 3 | 1 | 2 | 0.5 | NA | NA | 6 | 0.8333 | 1 | 1 | NA | NA | NA | NA | 1 | 1 | 2 | 1 |
| 2_1646636' | 1 | 0 | 2 | 0.5 | 1 | 0 | 1 | 0 | 5 | 0.2 | 1 | 1 | NA | NA | NA | NA | 1 | 1 | 2 | 1 |
| 2_1646637· | 1 | 0 | 3 | 0.3333 | 2 | 0 | 1 | 0 | 7 | 0.1429 | 1 | 0 | NA | NA | NA | NA | 1 | 1 | 2 | 0.5 |
| 2_1646638! | 1 | 0 | 3 | 0.3333 | 2 | 0 | 1 | 0 | 7 | 0.1429 | 1 | 1 | NA | NA | NA | NA | 1 | 0 | 2 | 0.5 |
| 2_1646639· | 1 | 0 | 3 | 0 | 2 | 0.5 | 1 | 0 | 7 | 0.1429 | 1 | 0 | NA | NA | NA | NA | 1 | 1 | 2 | 0.5 |
| 2_1646667! | 1 | 1 | 5 | 1 | 1 | 0 | 1 | 1 | 8 | 0.875 | 1 | 1 | 2 | 0 | 1 | 1 | 1 | 1 | 5 | 0.6 |
| 2_1646669( | NA | NA | 4 | 1 | 1 | 1 | 1 | 1 | 6 | 1 | 1 | 1 | 2 | 1 | 2 | 1 | 1 | 0 | 6 | 0.8333 |
| 2_1646669! | 1 | 0 | 5 | 0.6 | 1 | 1 | 1 | 0 | 8 | 0.5 | 1 | 0 | 2 | 1 | 2 | 0.5 | 2 | 1 | 7 | 0.7143 |
| 2_1646675: | 2 | 0.5 | 4 | 0.25 | 1 | 0 | 1 | 1 | 8 | 0.375 | 3 | 0 | 2 | 0.5 | 3 | 0.6667 | 3 | 0.3333 | 11 | 0.3636 |
| 2_1646675: | 1 | 1 | 5 | 0 | 2 | 0 | 1 | 1 | 9 | 0.2222 | 3 | 0.3333 | 2 | 0.5 | 3 | 0.3333 | 4 | 0.25 | 12 | 0.3333 |
| 2_1646675' | 2 | 1 | 4 | 1 | 2 | 1 | 1 | 1 | 9 | 1 | 2 | 1 | 2 | 1 | 2 | 1 | 2 | 1 | 8 | 1 |
| 2_1646677( | 2 | 0 | 7 | 0 | 2 | 0 | 1 | 1 | 12 | 0.0833 | 3 | 0 | 1 | 0 | 3 | 0 | 5 | 0.2 | 12 | 0.0833 |
| 2_1646679' | 3 | 0.6667 | 6 | 0.3333 | 2 | 0.5 | 1 | 1 | 12 | 0.5 | 2 | 1 | 2 | 0.5 | 2 | 1 | 5 | 1 | 11 | 0.9091 |
| 2_1646679: | 2 | 1 | 2 | 0 | NA | NA | 1 | 1 | 5 | 0.6 | 2 | 0 | 1 | 0 | 3 | 0.3333 | 2 | 0.5 | 8 | 0.25 |
| 2_1646679! | 3 | 0.6667 | 7 | 0.5714 | 2 | 0 | 1 | 0 | 13 | 0.4615 | 3 | 0 | 1 | 0 | 4 | 0.25 | 3 | 0.3333 | 11 | 0.1818 |
| 2_1646681· | 1 | 1 | 4 | 1 | 2 | 1 | NA | NA | 7 | 1 | 3 | 1 | 2 | 1 | 4 | 1 | 2 | 1 | 11 | 1 |
| 2_1646682· | 2 | 0 | 4 | 0.75 | 1 | 1 | NA | NA | 7 | 0.5714 | 1 | 1 | 2 | 1 | 2 | 1 | 2 | 0.5 | 7 | 0.8571 |
| 2_1646682' | 3 | 0.6667 | 8 | 0.375 | 3 | 0.3333 | 1 | 0 | 15 | 0.4 | 3 | 0.3333 | 2 | 0.5 | 5 | 0.8 | 5 | 0.6 | 15 | 0.6 |
| 2_1646687' | 3 | 0.6667 | 8 | 0.625 | 4 | 0.75 | 1 | 0 | 16 | 0.625 | 5 | 0.8 | 2 | 1 | 5 | 1 | 4 | 0.75 | 16 | 0.875 |
| 2_1646749( | 3 | 0.3333 | 1 | 1 | NA | NA | 2 | 0 | 6 | 0.3333 | 1 | 0 | 2 | 1 | 5 | 0.4 | 1 | 0 | 9 | 0.4444 |
| 2_1646761( | 10 | 0.1 | 7 | 0.1429 | 10 | 0 | 6 | 0 | 33 | 0.0606 | 6 | 0.3333 | 9 | 0 | 7 | 0 | 8 | 0 | 30 | 0.0667 |
| 2_1646813: | 22 | 0.0909 | 18 | 0.0556 | 14 | 0 | 12 | 0.0833 | 66 | 0.0606 | 13 | 0 | 17 | 0 | 16 | 0.125 | 13 | 0.0769 | 59 | 0.0508 |
| 2_1646813! | 13 | 0.1538 | 6 | 0.1667 | 6 | 0 | 5 | 0 | 30 | 0.1 | 7 | 0.1429 | 5 | 0.2 | 8 | 0.25 | 8 | 0.25 | 28 | 0.2143 |
| 2_1647212' | NA | NA | NA | NA | 1 | 1 | NA | NA | 1 | 1 | NA | NA | NA | NA | NA | NA | 1 | 1 | 1 | 1 |
| 2_1647248! | 5 | 0.2 | 1 | 0 | 5 | 0 | 4 | 0 | 15 | 0.0667 | 4 | 0.25 | 3 | 0 | 2 | 0.5 | 3 | 0 | 12 | 0.1667 |
| 2_1647329! | 1 | 1 | 1 | 0 | 2 | 0 | 2 | 0.5 | 6 | 0.3333 | 3 | 0.6667 | 3 | 0.6667 | 3 | 1 | 1 | 1 | 10 | 0.8 |
| 2_1647330! | 1 | 1 | 1 | 1 | 2 | 1 | 2 | 0.5 | 6 | 0.8333 | 3 | 0.3333 | 3 | 0.3333 | 4 | 0.5 | 1 | 0 | 11 | 0.3636 |
| 2_1647331: | 1 | 1 | 1 | 1 | 3 | 0 | 3 | 0 | 8 | 0.25 | 2 | 0.5 | 4 | 0.5 | 4 | 0.75 | NA | NA | 10 | 0.6 |
| 2_1647331· | 1 | 0 | 1 | 0 | 3 | 0 | 3 | 0.3333 | 8 | 0.125 | 3 | 0.6667 | 3 | 0.6667 | 5 | 0 | 1 | 0 | 12 | 0.3333 |
| 2_1647338' | 1 | 1 | 1 | 0 | NA | NA | NA | NA | 2 | 0.5 | NA | NA | 2 | 0 | 5 | 0.4 | 1 | 0 | 8 | 0.25 |
| 2_1647339' | 7 | 0.4286 | 4 | 0 | 2 | 1 | 3 | 0.6667 | 16 | 0.4375 | 2 | 1 | 1 | 1 | 1 | 0 | NA | NA | 4 | 0.75 |
| 2_1647339' | 1 | 1 | 1 | 0 | NA | NA | NA | NA | 2 | 0.5 | NA | NA | 2 | 0 | 6 | 0.1667 | 1 | 1 | 9 | 0.2222 |
| 2_1647339! | 4 | 0.25 | 6 | 0.3333 | 3 | 0 | 1 | 0 | 14 | 0.2143 | 2 | 0.5 | 2 | 0.5 | 2 | 0 | NA | NA | 6 | 0.3333 |
| 2_1647343( | 1 | 0 | 1 | 1 | NA | NA | NA | NA | 2 | 0.5 | NA | NA | 1 | 0 | 6 | 0.1667 | 1 | 0 | 8 | 0.125 |
| 2_1647344( | 1 | 1 | 2 | 0 | NA | NA | NA | NA | 3 | 0.3333 | NA | NA | 1 | 1 | 6 | 0.5 | 1 | 0 | 8 | 0.5 |

| | | | | | | | | | | | | | | | | | | | | |
|---|---|---|---|---|---|---|---|---|---|---|---|---|---|---|---|---|---|---|---|---|
| 2_16473452 | 1 | 1 | 1 | 1 | NA | NA | NA | NA | 2 | 1 | NA | NA | 1 | 0 | 5 | 0.6 | 1 | 1 | 7 | 0.5714 |
| 2_16473462 | 1 | 0 | 2 | 0.5 | NA | NA | NA | NA | 3 | 0.3333 | NA | NA | NA | NA | 6 | 0.1667 | 1 | 1 | 7 | 0.2857 |
| 2_16473477 | 1 | 1 | 2 | 0.5 | NA | NA | NA | NA | 3 | 0.6667 | NA | NA | NA | NA | 5 | 0.6 | 1 | 1 | 6 | 0.6667 |
| 2_16473525 | 1 | 1 | 2 | 0.5 | 1 | 0 | NA | NA | 4 | 0.5 | NA | NA | 1 | 0 | 4 | 0.5 | 1 | 1 | 6 | 0.5 |
| 2_16473584 | 1 | 0 | 3 | 0 | 1 | 1 | NA | NA | 5 | 0.2 | NA | NA | 3 | 0 | 6 | 0.5 | 2 | 0.5 | 11 | 0.3636 |
| 2_16475363 | NA | NA | 1 | 0 | NA | NA | 3 | 0.3333 | 4 | 0.25 | NA | NA | NA | NA | 1 | 1 | NA | NA | 1 | 1 |
| 2_16483940 | 19 | 0 | 21 | 0.1429 | 19 | 0 | 29 | 0.069 | 88 | 0.0568 | 18 | 0.2222 | 26 | 0.0385 | 26 | 0 | 23 | 0.1304 | 93 | 0.086 |
| 2_16483948 | 25 | 0.12 | 22 | 0.0455 | 21 | 0.0476 | 30 | 0.2 | 98 | 0.1122 | 23 | 0.1739 | 31 | 0.0968 | 32 | 0.1562 | 27 | 0.0741 | 113 | 0.1239 |
| 2_16483972 | 20 | 0.3 | 21 | 0.2381 | 24 | 0.2083 | 29 | 0.2759 | 94 | 0.2553 | 21 | 0.381 | 27 | 0.5185 | 32 | 0.25 | 24 | 0.375 | 104 | 0.375 |
| 2_16483983 | 23 | 0.8261 | 22 | 0.8636 | 24 | 0.7083 | 31 | 0.7097 | 100 | 0.77 | 23 | 0.8696 | 31 | 0.7097 | 34 | 0.7353 | 30 | 0.8667 | 118 | 0.7881 |
| 2_16484000 | 28 | 0.1071 | 25 | 0.12 | 31 | 0.3871 | 35 | 0.2571 | 119 | 0.2269 | 28 | 0.1786 | 32 | 0.0312 | 38 | 0.1316 | 31 | 0.0645 | 129 | 0.1008 |
| 2_16484009 | 25 | 0.2 | 24 | 0.0833 | 29 | 0.1379 | 33 | 0.0606 | 111 | 0.1171 | 24 | 0.2083 | 29 | 0.1724 | 34 | 0.1471 | 29 | 0.069 | 116 | 0.1466 |
| 2_16484020 | 23 | 0.8261 | 25 | 0.84 | 28 | 0.75 | 25 | 0.6 | 101 | 0.7525 | 24 | 0.7917 | 33 | 0.6364 | 30 | 0.9333 | 28 | 0.6071 | 115 | 0.7391 |
| 2_16484025 | 22 | 0.4091 | 21 | 0.381 | 27 | 0.3333 | 32 | 0.375 | 102 | 0.3725 | 29 | 0.4138 | 38 | 0.4737 | 39 | 0.4615 | 32 | 0.4062 | 138 | 0.442 |
| 2_16484051 | 26 | 0.0385 | 26 | 0.1538 | 30 | 0.2333 | 34 | 0.1765 | 116 | 0.1552 | 25 | 0.16 | 36 | 0.25 | 37 | 0.2703 | 32 | 0.125 | 130 | 0.2077 |
| 2_16484083 | 36 | 0.2222 | 28 | 0.0357 | 40 | 0.125 | 44 | 0.1818 | 148 | 0.1486 | 31 | 0.0645 | 42 | 0.119 | 44 | 0.1136 | 39 | 0.1282 | 156 | 0.109 |
| 2_16484093 | 34 | 0.2647 | 34 | 0.2059 | 39 | 0.2564 | 47 | 0.234 | 154 | 0.2403 | 37 | 0.2162 | 44 | 0.25 | 51 | 0.2353 | 39 | 0.2821 | 171 | 0.2456 |
| 2_16484220 | NA | NA | NA | NA | 1 | 1 | 1 | 0 | 2 | 0.5 | NA | NA | NA | NA | 2 | 0.5 | NA | NA | 2 | 0.5 |
| 2_16484265 | 7 | 0.2857 | 7 | 0.2857 | 9 | 0.5556 | 6 | 0.1667 | 29 | 0.3448 | 8 | 0.25 | 9 | 0.3333 | 6 | 0.1667 | 5 | 0 | 28 | 0.2143 |
| 2_16484271 | 7 | 0.2857 | 6 | 0.1667 | 7 | 0.1429 | 6 | 0.3333 | 26 | 0.2308 | 7 | 0.2857 | 7 | 0.4286 | 5 | 0.4 | 4 | 0 | 23 | 0.3043 |
| 2_16484286 | 6 | 0.3333 | 6 | 0.6667 | 7 | 0.4286 | 6 | 0.6667 | 25 | 0.52 | 8 | 0.625 | 8 | 0.625 | 6 | 0.5 | 3 | 0.3333 | 25 | 0.56 |
| 2_16484287 | 3 | 0.6667 | 4 | 1 | 7 | 0.8571 | 2 | 1 | 16 | 0.875 | 6 | 0.5 | 8 | 0.875 | 6 | 0.6667 | 2 | 1 | 22 | 0.7273 |
| 2_16484297 | 7 | 0.2857 | 5 | 0.2 | 7 | 0.4286 | 5 | 0.6 | 24 | 0.375 | 7 | 0.2857 | 7 | 0.1429 | 6 | 0.3333 | 5 | 0.2 | 25 | 0.24 |
| 2_16484298 | 8 | 0.75 | 7 | 1 | 6 | 0.8333 | 6 | 1 | 27 | 0.8889 | 7 | 1 | 8 | 0.75 | 7 | 0.8571 | 3 | 0.6667 | 25 | 0.84 |
| 2_16484301 | 7 | 0.7143 | 5 | 1 | 7 | 1 | 6 | 1 | 25 | 0.92 | 7 | 0.7143 | 8 | 0.875 | 6 | 0.8333 | 3 | 1 | 24 | 0.8333 |
| 2_16484303 | 7 | 0.7143 | 7 | 1 | 7 | 0.7143 | 6 | 1 | 27 | 0.8519 | 7 | 0.7143 | 9 | 0.7778 | 5 | 0.6 | 4 | 0.5 | 25 | 0.68 |
| 2_16484304 | 8 | 0 | 7 | 0 | 8 | 0.25 | 5 | 0.2 | 28 | 0.1071 | 7 | 0 | 9 | 0.2222 | 6 | 0.1667 | 4 | 0.5 | 26 | 0.1923 |
| 2_16484307 | 8 | 0 | 7 | 0.1429 | 8 | 0.25 | 7 | 0.1429 | 30 | 0.1333 | 8 | 0.125 | 8 | 0.25 | 7 | 0 | 2 | 0 | 25 | 0.12 |
| 2_16484318 | 6 | 0.1667 | 6 | 0 | 4 | 0 | 4 | 0.25 | 20 | 0.1 | 7 | 0.1429 | 6 | 0.1667 | 4 | 0 | 2 | 0 | 19 | 0.1053 |
| 2_16484321 | 8 | 0 | 7 | 0 | 8 | 0.125 | 5 | 0.2 | 28 | 0.0714 | 7 | 0.1429 | 9 | 0 | 7 | 0 | 3 | 0.3333 | 26 | 0.0769 |
| 2_16484332 | 6 | 0.3333 | 5 | 0.8 | 6 | 0.6667 | 4 | 1 | 21 | 0.6667 | 6 | 0.3333 | 8 | 0.375 | 7 | 0.8571 | 3 | 0.6667 | 24 | 0.5417 |
| 2_16484333 | 6 | 0.6667 | 6 | 0.8333 | 8 | 0.625 | 5 | 0.8 | 25 | 0.72 | 6 | 0.3333 | 5 | 0.2 | 6 | 0.6667 | 3 | 0.6667 | 20 | 0.45 |
| 2_16484356 | 5 | 0.4 | 6 | 0 | 8 | 0 | 6 | 0.3333 | 25 | 0.16 | 8 | 0.125 | 8 | 0.25 | 6 | 0 | 2 | 0 | 24 | 0.125 |
| 2_16484360 | 5 | 0.6 | 7 | 0.5714 | 9 | 0.6667 | 5 | 0.6 | 26 | 0.6154 | 8 | 0.5 | 8 | 0.375 | 6 | 0.6667 | 2 | 0.5 | 24 | 0.5 |
| 2_16484363 | 5 | 0.4 | 7 | 0.1429 | 7 | 0.2857 | 5 | 0.6 | 24 | 0.3333 | 5 | 0.2 | 6 | 0.3333 | 4 | 0.5 | 1 | 0 | 16 | 0.3125 |
| 2_16484364 | 4 | 0.75 | 5 | 0.6 | 5 | 0.8 | 6 | 0.8333 | 20 | 0.75 | 5 | 0.4 | 4 | 1 | 4 | 0.75 | 1 | 1 | 14 | 0.7143 |
| 2_16484375 | 4 | 0 | 6 | 0.6667 | 5 | 0.2 | 5 | 0.2 | 20 | 0.3 | 8 | 0.375 | 5 | 0.2 | 2 | 0.5 | 1 | 0 | 16 | 0.3125 |
| 2_16484377 | 6 | 0.6667 | 6 | 1 | 7 | 0.5714 | 5 | 0.6 | 24 | 0.7083 | 6 | 0.8333 | 5 | 0.8 | 3 | 1 | NA | NA | 14 | 0.8571 |
| 2_16484378 | 6 | 0 | 7 | 0.2857 | 6 | 0 | 5 | 0 | 24 | 0.0833 | 6 | 0 | 5 | 0.2 | 2 | 0.5 | 1 | 0 | 14 | 0.1429 |
| 2_16484381 | 6 | 0 | 7 | 0 | 7 | 0.1429 | 5 | 0.2 | 25 | 0.08 | 8 | 0.125 | 6 | 0 | 4 | 0.5 | 1 | 0 | 19 | 0.1579 |
| 2_16484481 | 5 | 0 | 4 | 0.5 | 4 | 0 | 2 | 0 | 15 | 0.1333 | 6 | 0 | 5 | 0 | 1 | 0 | 1 | 1 | 13 | 0.0769 |
| 2_16485476 | 4 | 0 | 4 | 0.25 | 3 | 0 | 1 | 0 | 12 | 0.0833 | 4 | 0 | 4 | 0 | 2 | 0.5 | 1 | 0 | 11 | 0.0909 |
| 2_16485483 | 4 | 0.75 | 3 | 0.3333 | 3 | 1 | 1 | 1 | 11 | 0.7273 | 4 | 0.5 | 3 | 0 | 2 | 1 | 1 | 1 | 10 | 0.5 |
| 2_16485485 | 4 | 0.5 | 4 | 0.5 | 3 | 0.6667 | 1 | 0 | 12 | 0.5 | 4 | 0.25 | 4 | 0.25 | 2 | 0.5 | 1 | 0 | 11 | 0.2727 |
| 2_16485491 | 4 | 0 | 4 | 0.25 | 2 | 0 | 1 | 0 | 11 | 0.0909 | 3 | 0 | 4 | 0.25 | 1 | 0 | 1 | 0 | 9 | 0.1111 |
| 2_16495387 | 5 | 1 | 6 | 1 | 6 | 0.8333 | 4 | 1 | 21 | 0.9524 | 1 | 1 | 6 | 0.8333 | 6 | 0.8333 | 2 | 0.5 | 15 | 0.8 |
| 2_16495390 | 5 | 0.4 | 7 | 0.2857 | 6 | 0.5 | 5 | 0 | 23 | 0.3043 | 2 | 0.5 | 8 | 0.25 | 5 | 0 | 4 | 0 | 19 | 0.1579 |
| 2_16495398 | 3 | 0.6667 | 7 | 0.4286 | 6 | 0.3333 | 6 | 0.3333 | 22 | 0.4091 | NA | NA | 7 | 0.5714 | 6 | 0.8333 | 3 | 0.3333 | 16 | 0.625 |
| 2_16495417 | 5 | 0.6 | 5 | 0.8 | 5 | 0.6 | 5 | 0.8 | 20 | 0.7 | 1 | 1 | 8 | 0.75 | 6 | 0.8333 | 5 | 0.8 | 20 | 0.8 |
| 2_16495428 | 5 | 0.2 | 5 | 0.4 | 5 | 0.2 | 4 | 0.25 | 19 | 0.2632 | 1 | 1 | 8 | 0.125 | 6 | 0.1667 | 5 | 0.4 | 20 | 0.25 |
| 2_16495431 | 4 | 1 | 6 | 1 | 5 | 0.8 | 4 | 0.75 | 19 | 0.8947 | 1 | 1 | 6 | 0.6667 | 3 | 0.6667 | 3 | 0.6667 | 13 | 0.6923 |
| 2_16495433 | 5 | 1 | 4 | 1 | 4 | 1 | 5 | 1 | 18 | 1 | 1 | 1 | 7 | 0.8571 | 5 | 0.8 | 3 | 1 | 16 | 0.875 |
| 2_16495449 | 4 | 0 | 6 | 0.1667 | 3 | 0.3333 | 3 | 0 | 16 | 0.125 | 1 | 1 | 8 | 0.125 | 4 | 0 | 4 | 0 | 17 | 0.1176 |
| 2_16495453 | 5 | 0.4 | 5 | 0 | 4 | 0.25 | 5 | 0.6 | 19 | 0.3158 | 1 | 0 | 7 | 0.2857 | 5 | 0.6 | 3 | 0.3333 | 16 | 0.375 |
| 2_16502324 | 1 | 0 | 2 | 0 | 1 | 0 | 3 | 0.3333 | 7 | 0.1429 | NA | NA | 2 | 0.5 | 3 | 0 | 3 | 0 | 8 | 0.125 |
| 2_16502329 | 1 | 0 | 2 | 0.5 | 1 | 0 | 3 | 0.3333 | 7 | 0.2857 | NA | NA | 3 | 0 | 3 | 0 | 1 | 1 | 7 | 0.1429 |
| 2_16511850 | 7 | 0.4286 | 6 | 0.5 | 5 | 0.6 | 5 | 0.6 | 23 | 0.5217 | 7 | 0.4286 | 5 | 0.4 | 6 | 0.6667 | 1 | 1 | 19 | 0.5263 |
| 2_16511871 | 10 | 0.8 | 10 | 0.7 | 5 | 0.6 | 3 | 0.6667 | 28 | 0.7143 | 9 | 0.8889 | 8 | 0.875 | 7 | 0.8571 | 2 | 1 | 26 | 0.8846 |
| 2_16511877 | 10 | 0.9 | 10 | 1 | 6 | 1 | 5 | 1 | 31 | 0.9677 | 11 | 1 | 8 | 1 | 9 | 1 | 3 | 0.6667 | 31 | 0.9677 |
| 2_16511887 | 11 | 0.3636 | 9 | 0.5556 | 6 | 0.3333 | 3 | 0.3333 | 29 | 0.4138 | 12 | 0.5 | 8 | 0.75 | 8 | 0.75 | 3 | 0.3333 | 31 | 0.6129 |
| 2_16511893 | 10 | 1 | 9 | 1 | 6 | 1 | 3 | 1 | 28 | 1 | 11 | 0.8182 | 8 | 0.875 | 8 | 0.875 | 3 | 1 | 30 | 0.8667 |
| 2_16512544 | 4 | 0 | 5 | 0.4 | 2 | 0.5 | 1 | 0 | 12 | 0.25 | 4 | 0 | 6 | 0 | 5 | 0.2 | 1 | 0 | 16 | 0.0625 |
| 2_16513163 | 9 | 0.3333 | 13 | 0.2308 | 10 | 0.3 | 13 | 0.3077 | 45 | 0.2889 | 6 | 0 | 7 | 0.2857 | 14 | 0.2857 | 12 | 0.25 | 39 | 0.2308 |
| 2_16513189 | 13 | 0.9231 | 20 | 0.8 | 13 | 0.7692 | 15 | 0.8667 | 61 | 0.8361 | 11 | 0.9091 | 11 | 0.4545 | 17 | 0.6471 | 9 | 0.8889 | 48 | 0.7083 |
| 2_16513194 | 16 | 0.25 | 18 | 0.3333 | 14 | 0.2857 | 12 | 0.1667 | 60 | 0.2667 | 9 | 0.3333 | 8 | 0.375 | 15 | 0.2 | 12 | 0.3333 | 44 | 0.2955 |

| | | | | | | | | | | | | | | | | | | | | |
|---|---|---|---|---|---|---|---|---|---|---|---|---|---|---|---|---|---|---|---|---|
| 2_16513203 | 16 | 0.5 | 21 | 0.619 | 17 | 0.7059 | 19 | 0.3158 | 73 | 0.5342 | 12 | 0.4167 | 11 | 0.5455 | 21 | 0.4762 | 17 | 0.5294 | 61 | 0.4918 |
| 2_16513208 | 14 | 0.7143 | 16 | 0.5625 | 12 | 0.5833 | 9 | 0.3333 | 51 | 0.5686 | 6 | 0.8333 | 8 | 0.625 | 15 | 0.6667 | 10 | 0.5 | 39 | 0.641 |
| 2_16513218 | 16 | 0.0625 | 23 | 0.1304 | 18 | 0.2222 | 20 | 0.1 | 77 | 0.1299 | 11 | 0.1818 | 10 | 0.2 | 20 | 0.1 | 17 | 0.2353 | 58 | 0.1724 |
| 2_16513222 | 12 | 0.8333 | 19 | 0.8947 | 15 | 1 | 16 | 0.9375 | 62 | 0.9194 | 10 | 1 | 10 | 0.7 | 19 | 0.6842 | 13 | 1 | 52 | 0.8269 |
| 2_16513225 | 16 | 0.8125 | 20 | 0.65 | 15 | 0.4 | 17 | 0.2941 | 68 | 0.5441 | 7 | 0.5714 | 9 | 0.3333 | 20 | 0.5 | 14 | 0.6429 | 50 | 0.52 |
| 2_16513227 | 9 | 1 | 17 | 0.8824 | 16 | 0.875 | 15 | 0.8667 | 57 | 0.8947 | 8 | 0.875 | 5 | 1 | 18 | 0.7222 | 13 | 0.9231 | 44 | 0.8409 |
| 2_16513235 | 12 | 0.25 | 16 | 0.1875 | 14 | 0.0714 | 16 | 0.0625 | 58 | 0.1379 | 8 | 0.375 | 8 | 0.25 | 19 | 0.2105 | 15 | 0.3333 | 50 | 0.28 |
| 2_16513236 | 13 | 0.6923 | 18 | 0.4444 | 12 | 0.5 | 17 | 0.4706 | 60 | 0.5167 | 9 | 0.3333 | 6 | 0.3333 | 19 | 0.2632 | 13 | 0.4615 | 47 | 0.3404 |
| 2_16513246 | 13 | 0.2308 | 18 | 0.3333 | 14 | 0.3571 | 15 | 0.2 | 60 | 0.2833 | 9 | 0.2222 | 9 | 0.4444 | 17 | 0.1176 | 12 | 0.25 | 47 | 0.234 |
| 2_16514015 | 11 | 0 | 11 | 0.0909 | 8 | 0 | 11 | 0.1818 | 41 | 0.0732 | 8 | 0 | 5 | 0 | 15 | 0 | 14 | 0.2143 | 42 | 0.0714 |
| 2_16521793 | 1 | 1 | NA | NA | 2 | 0 | NA | NA | 3 | 0.3333 | NA | NA | 1 | 1 | NA | NA | NA | NA | 1 | 1 |
| 2_16521941 | NA | NA | 2 | 0.5 | 2 | 0.5 | 2 | 0 | 6 | 0.3333 | 1 | 0 | 1 | 1 | NA | NA | 2 | 1 | 4 | 0.75 |
| 2_16521952 | 1 | 1 | 2 | 0.5 | 4 | 0.75 | 2 | 0 | 9 | 0.5556 | 1 | 1 | 1 | 1 | NA | NA | 1 | 1 | 3 | 1 |
| 2_16521977 | 1 | 0 | 2 | 0.5 | 2 | 0.5 | 2 | 0.5 | 7 | 0.4286 | 1 | 1 | 1 | 1 | NA | NA | 2 | 1 | 4 | 1 |
| 2_16521986 | 1 | 0 | 2 | 0.5 | 4 | 0.25 | 1 | 0 | 8 | 0.25 | 1 | 0 | 1 | 0 | NA | NA | 2 | 0.5 | 4 | 0.25 |
| 2_16521993 | 1 | 0 | 1 | 1 | 3 | 0.6667 | 1 | 0 | 6 | 0.5 | 1 | 0 | 1 | 1 | NA | NA | 1 | 1 | 3 | 0.6667 |
| 2_16521995 | 1 | 0 | 2 | 0.5 | 4 | 0.75 | 1 | 0 | 8 | 0.5 | 1 | 1 | NA | NA | NA | NA | 1 | 1 | 2 | 1 |
| 2_16521997 | 1 | 0 | 2 | 0.5 | 3 | 0.3333 | NA | NA | 6 | 0.3333 | 1 | 1 | NA | NA | NA | NA | 2 | 1 | 3 | 1 |
| 2_16521998 | 1 | 0 | 2 | 0 | 4 | 0.25 | 1 | 0 | 8 | 0.125 | 1 | 1 | 1 | 0 | NA | NA | 2 | 0 | 4 | 0.25 |
| 2_16522141 | NA | NA | NA | NA | 2 | 0.5 | NA | NA | 2 | 0.5 | 1 | 1 | 1 | 0 | NA | NA | 2 | 0 | 4 | 0.25 |
| 2_16522163 | NA | NA | 2 | 1 | 2 | 0.5 | NA | NA | 4 | 0.75 | 1 | 1 | 1 | 1 | NA | NA | 1 | 1 | 3 | 1 |
| 2_16522178 | NA | NA | 2 | 0.5 | 2 | 0 | NA | NA | 4 | 0.25 | 1 | 0 | 1 | 1 | NA | NA | 2 | 1 | 4 | 0.75 |
| 2_16522182 | NA | NA | 2 | 1 | 2 | 1 | NA | NA | 4 | 1 | 1 | 1 | 1 | 1 | NA | NA | 2 | 0.5 | 4 | 0.75 |
| 2_16522184 | NA | NA | 1 | 1 | 2 | 0.5 | NA | NA | 3 | 0.6667 | 1 | 1 | 1 | 1 | NA | NA | 2 | 1 | 4 | 1 |
| 2_16522191 | NA | NA | 2 | 1 | 2 | 1 | NA | NA | 4 | 1 | 1 | 1 | 1 | 1 | NA | NA | 2 | 1 | 4 | 1 |
| 2_16522211 | NA | NA | 2 | 0.5 | 2 | 0 | NA | NA | 4 | 0.25 | 1 | 0 | NA | NA | NA | NA | 1 | 1 | 2 | 0.5 |
| 2_16522211 | NA | NA | 2 | 0 | 1 | 1 | NA | NA | 3 | 0.3333 | 1 | 0 | 1 | 1 | NA | NA | 1 | 0 | 3 | 0.3333 |
| 2_16522226 | NA | NA | 2 | 1 | 2 | 1 | NA | NA | 4 | 1 | 1 | 1 | 1 | 1 | NA | NA | 2 | 1 | 4 | 1 |
| 2_16522361 | NA | NA | 1 | 1 | NA | NA | NA | NA | 1 | 1 | 1 | 0 | 1 | 1 | NA | NA | NA | NA | 2 | 0.5 |
| 2_16533218 | 3 | 0 | 5 | 0 | 5 | 0 | 4 | 0.25 | 17 | 0.0588 | 2 | 0 | 1 | 0 | 5 | 0.2 | 3 | 0 | 11 | 0.0909 |
| 2_16533241 | 6 | 0.3333 | 3 | 0 | 5 | 0.2 | 1 | 0 | 15 | 0.2 | 2 | 0 | 1 | 0 | 4 | 0 | 3 | 0.3333 | 10 | 0.1 |
| 2_16533244 | 4 | 0.75 | 4 | 1 | 3 | 1 | 3 | 1 | 14 | 0.9286 | 2 | 0.5 | NA | NA | 3 | 1 | 2 | 1 | 7 | 0.8571 |
| 2_16533257 | 4 | 0.25 | 4 | 0.75 | 5 | 0.6 | 3 | 0.6667 | 16 | 0.5625 | 2 | 0.5 | NA | NA | 3 | 0.6667 | 3 | 0.6667 | 8 | 0.625 |
| 2_16533264 | 5 | 0.8 | 3 | 1 | 5 | 1 | 3 | 0.6667 | 16 | 0.875 | 1 | 1 | 1 | 1 | 3 | 0.6667 | 3 | 0.6667 | 8 | 0.75 |
| 2_16533271 | 5 | 0.8 | 3 | 0.6667 | 5 | 0.6 | 2 | 1 | 15 | 0.7333 | 2 | 0 | 1 | 1 | 4 | 0.75 | 1 | 1 | 8 | 0.625 |
| 2_16533272 | 5 | 0.8 | 4 | 0.5 | 4 | 0.75 | 3 | 0.3333 | 16 | 0.625 | 2 | 0.5 | 1 | 1 | 3 | 0.3333 | 3 | 0.6667 | 9 | 0.5556 |
| 2_16533274 | 6 | 0 | 4 | 0.25 | 6 | 0 | 3 | 0 | 19 | 0.0526 | 2 | 0 | 1 | 1 | 3 | 0.3333 | 3 | 0 | 9 | 0.2222 |
| 2_16533278 | 6 | 0 | 4 | 0.25 | 5 | 0.2 | 3 | 0 | 18 | 0.1111 | 2 | 0 | 1 | 0 | 3 | 0.3333 | 2 | 0.5 | 8 | 0.25 |
| 2_16533293 | 5 | 0.6 | 3 | 1 | 5 | 0.8 | 2 | 1 | 15 | 0.8 | 2 | 0.5 | 1 | 1 | 3 | 0.6667 | 3 | 1 | 9 | 0.7778 |
| 2_16533310 | 6 | 0.5 | 3 | 1 | 3 | 1 | 2 | 1 | 14 | 0.7857 | 1 | 0 | 1 | 1 | 1 | 1 | 2 | 0.5 | 5 | 0.6 |
| 2_16533313 | 6 | 1 | 3 | 0.6667 | 5 | 0.6 | 3 | 1 | 17 | 0.8235 | 1 | 1 | 1 | 1 | 3 | 1 | 3 | 1 | 8 | 1 |
| 2_16533334 | 4 | 0.5 | 2 | 0.5 | 5 | 0.8 | 2 | 0.5 | 13 | 0.6154 | 1 | 1 | NA | NA | 2 | 1 | 2 | 0.5 | 5 | 0.8 |
| 2_16533343 | 4 | 0.25 | 3 | 0 | 2 | 1 | 1 | 0 | 10 | 0.3 | 1 | 1 | NA | NA | 2 | 0.5 | 2 | 0.5 | 5 | 0.6 |
| 2_16533349 | 4 | 0.5 | 2 | 0 | 5 | 0.2 | 2 | 0 | 13 | 0.2308 | 1 | 1 | NA | NA | 2 | 1 | 2 | 0.5 | 5 | 0.8 |
| 2_16533351 | 5 | 0.4 | 1 | 0 | 5 | 0 | 3 | 0.6667 | 14 | 0.2857 | 1 | 1 | NA | NA | 3 | 0.6667 | 3 | 0 | 7 | 0.4286 |
| 2_16533352 | 3 | 0.6667 | 2 | 1 | 4 | 0.75 | 3 | 0.6667 | 12 | 0.75 | 1 | 1 | NA | NA | 2 | 1 | 2 | 1 | 5 | 1 |
| 2_16533355 | 5 | 0.4 | 2 | 0 | 3 | 0.6667 | 3 | 0.6667 | 13 | 0.4615 | 1 | 0 | NA | NA | 2 | 1 | 3 | 0.6667 | 6 | 0.6667 |
| 2_16533356 | 4 | 0 | 2 | 0.5 | 4 | 0 | 3 | 0.3333 | 13 | 0.1538 | 1 | 0 | NA | NA | 3 | 0 | 3 | 0.3333 | 7 | 0.1429 |
| 2_16533360 | 3 | 0.6667 | 2 | 1 | 4 | 0.75 | 3 | 0 | 12 | 0.5833 | 1 | 1 | NA | NA | 3 | 0.3333 | 3 | 0.3333 | 7 | 0.4286 |
| 2_16561319 | NA | NA | 3 | 0.6667 | NA | NA | NA | NA | 3 | 0.6667 | NA | NA | 1 | 1 | NA | NA | 1 | 1 | 2 | 1 |
| 2_16561326 | NA | NA | 3 | 1 | NA | NA | NA | NA | 3 | 1 | NA | NA | 1 | 1 | NA | NA | 1 | 1 | 2 | 1 |
| 2_16562639 | 6 | 0.3333 | 7 | 0 | 10 | 0.2 | 13 | 0.1538 | 36 | 0.1667 | 8 | 0 | 8 | 0.375 | 8 | 0.25 | 8 | 0.25 | 32 | 0.2188 |
| 2_16562650 | 7 | 0.1429 | 7 | 0 | 12 | 0.1667 | 15 | 0.3333 | 41 | 0.1951 | 8 | 0.125 | 10 | 0.3 | 9 | 0.1111 | 10 | 0.1 | 37 | 0.1622 |
| 2_16562664 | 5 | 1 | 5 | 1 | 11 | 0.8182 | 14 | 0.8571 | 35 | 0.8857 | 5 | 0.8 | 10 | 0.9 | 7 | 0.8571 | 7 | 0.8571 | 29 | 0.8621 |
| 2_16562690 | 6 | 0.8333 | 7 | 0.4286 | 13 | 0.8462 | 15 | 0.6 | 41 | 0.6829 | 9 | 0.8889 | 10 | 0.7 | 7 | 0.8571 | 10 | 0.9 | 36 | 0.8333 |
| 2_16562736 | 3 | 1 | 7 | 0.8571 | 11 | 1 | 11 | 0.9091 | 32 | 0.9375 | 7 | 0.8571 | 8 | 0.875 | 4 | 1 | 7 | 0.8571 | 26 | 0.8846 |
| 2_16562737 | 3 | 0.3333 | 7 | 0.4286 | 10 | 0.2 | 13 | 0.0769 | 33 | 0.2121 | 7 | 0.2857 | 8 | 0.125 | 3 | 0.6667 | 8 | 0.5 | 26 | 0.3462 |
| 2_16562743 | 5 | 0.2 | 6 | 0.5 | 10 | 0.2 | 11 | 0.2727 | 32 | 0.2812 | 6 | 0.3333 | 7 | 0.2857 | 4 | 0 | 6 | 0.5 | 23 | 0.3043 |
| 2_16562751 | 5 | 1 | 6 | 1 | 11 | 0.9091 | 12 | 0.9167 | 34 | 0.9412 | 6 | 0.8333 | 7 | 0.7143 | 5 | 0.8 | 7 | 0.8571 | 25 | 0.8 |
| 2_16564963 | 1 | 0 | 2 | 0 | 6 | 0 | 9 | 0.1111 | 18 | 0.0556 | 4 | 0.25 | 3 | 0 | 4 | 0 | 6 | 0.3333 | 17 | 0.1765 |
| 2_16565434 | 2 | 0.5 | 2 | 0.5 | 4 | 1 | 6 | 0.6667 | 14 | 0.7143 | 4 | 0 | 2 | 0 | 3 | 0.3333 | 6 | 0.1667 | 15 | 0.1333 |
| 2_16565435 | 2 | 0 | 2 | 0.5 | 4 | 0.25 | 8 | 0.375 | 16 | 0.3125 | 2 | 0 | 3 | 0 | 4 | 0.75 | 6 | 0.5 | 15 | 0.4 |
| 2_16565437 | 2 | 0.5 | 2 | 1 | 5 | 0.8 | 8 | 0.75 | 17 | 0.7647 | 4 | 0.5 | 2 | 1 | 4 | 0.75 | 6 | 1 | 16 | 0.8125 |
| 2_16565443 | 2 | 0 | 2 | 0.5 | 4 | 0.75 | 7 | 0.4286 | 15 | 0.4667 | 4 | 0.75 | 3 | 0.6667 | 4 | 0.25 | 6 | 0.3333 | 17 | 0.4706 |
| 2_16565447 | 2 | 0.5 | 2 | 0.5 | 5 | 1 | 6 | 0.6667 | 15 | 0.7333 | 3 | 0.6667 | 3 | 0.6667 | 4 | 1 | 6 | 1 | 16 | 0.875 |

| | | | | | | | | | | | | | | | | | | | | |
|---|---|---|---|---|---|---|---|---|---|---|---|---|---|---|---|---|---|---|---|---|
| 2_16565461 | 2 | 0 | 1 | 0 | 3 | 0.3333 | 5 | 0.2 | 11 | 0.1818 | 3 | 0.3333 | 3 | 0.3333 | 4 | 0 | 4 | 0 | 14 | 0.1429 |
| 2_16565463 | 2 | 0.5 | 2 | 0 | 5 | 0.2 | 7 | 0 | 16 | 0.125 | 4 | 0 | 3 | 0 | 4 | 0.25 | 5 | 0.2 | 16 | 0.125 |
| 2_16566959 | 2 | 0 | 1 | 0 | 1 | 1 | 1 | 0 | 5 | 0.2 | 1 | 0 | 2 | 0.5 | NA | NA | 2 | 0 | 5 | 0.2 |
| 2_16567014 | 3 | 0.3333 | 3 | 1 | 5 | 0.2 | 7 | 0.5714 | 18 | 0.5 | 3 | 0.6667 | 4 | 0.75 | 1 | 0 | 5 | 0.4 | 13 | 0.5385 |
| 2_16567023 | 3 | 1 | 1 | 1 | 4 | 0.75 | 6 | 0.8333 | 14 | 0.8571 | 2 | 0.5 | 3 | 1 | 1 | 0 | 2 | 0.5 | 8 | 0.625 |
| 2_16567049 | 2 | 0.5 | 3 | 0 | 3 | 0 | 8 | 0.125 | 16 | 0.125 | 2 | 0 | 4 | 0.25 | 1 | 1 | 4 | 0 | 11 | 0.1818 |
| 2_16567054 | 2 | 0.5 | 3 | 0 | 4 | 0.75 | 8 | 0.5 | 17 | 0.4706 | 3 | 0.3333 | 3 | 0 | 1 | 0 | 3 | 1 | 10 | 0.4 |
| 2_16568796 | 2 | 0 | 2 | 0.5 | 3 | 0.3333 | 5 | 0.2 | 12 | 0.25 | 1 | 0 | 3 | 0.3333 | 1 | 0 | 4 | 0.25 | 9 | 0.2222 |
| 2_16568798 | 3 | 0.3333 | 2 | 0 | 3 | 0.3333 | 6 | 0.1667 | 14 | 0.2143 | 2 | 0.5 | 3 | 0 | 1 | 0 | 4 | 0 | 10 | 0.1 |
| 2_16589688 | 1 | 1 | 2 | 1 | NA | NA | 2 | 1 | 5 | 1 | NA | NA | 1 | 1 | 1 | 1 | 1 | 1 | 3 | 1 |
| 2_16589712 | 1 | 1 | 2 | 1 | NA | NA | 2 | 1 | 5 | 1 | NA | NA | 1 | 0 | 1 | 1 | 2 | 0.5 | 4 | 0.5 |
| 2_16589718 | 1 | 1 | 2 | 1 | NA | NA | 2 | 1 | 5 | 1 | NA | NA | NA | NA | 1 | 1 | 1 | 1 | 2 | 1 |
| 2_16589747 | 2 | 0.5 | 2 | 0 | NA | NA | 2 | 0 | 6 | 0.1667 | NA | NA | 1 | 0 | 1 | 1 | 3 | 0 | 5 | 0.2 |
| 2_16590781 | 2 | 0 | 1 | 0 | NA | NA | 2 | 1 | 5 | 0.4 | 1 | 1 | 1 | 0 | 2 | 1 | 2 | 0.5 | 6 | 0.6667 |
| 2_16591207 | 2 | 0 | 2 | 0.5 | NA | NA | 3 | 0.6667 | 7 | 0.4286 | 1 | 0 | 1 | 0 | 2 | 0 | 4 | 0.5 | 8 | 0.25 |
| 2_16591208 | 2 | 0.5 | 3 | 0 | NA | NA | 1 | 0 | 6 | 0.1667 | 1 | 0 | 1 | 0 | 2 | 0.5 | 4 | 0 | 8 | 0.125 |
| 2_16591274 | 2 | 0.5 | 3 | 0 | NA | NA | 4 | 0.25 | 9 | 0.2222 | 2 | 0 | 2 | 0 | 4 | 0.25 | 4 | 0 | 12 | 0.0833 |
| 2_16591275 | 2 | 0 | 3 | 0.3333 | NA | NA | 4 | 0 | 9 | 0.1111 | 2 | 0 | 2 | 0.5 | 2 | 0 | 4 | 0 | 10 | 0.1 |
| 2_16591284 | 2 | 0 | 3 | 0.3333 | NA | NA | 3 | 0 | 8 | 0.125 | 1 | 1 | 2 | 0 | 3 | 0.3333 | 4 | 0 | 10 | 0.2 |
| 2_16591290 | 1 | 1 | 2 | 0 | NA | NA | 4 | 0.25 | 7 | 0.2857 | 2 | 0.5 | 2 | 0.5 | 1 | 0 | 3 | 0.6667 | 8 | 0.5 |
| 2_16591498 | 2 | 0.5 | 3 | 0.6667 | 1 | 0 | 7 | 0.8571 | 13 | 0.6923 | 1 | 1 | 4 | 0.25 | 5 | 0.4 | 2 | 0 | 12 | 0.3333 |
| 2_16591508 | 3 | 0 | 4 | 0.5 | 1 | 1 | 5 | 0 | 13 | 0.2308 | 2 | 0 | 4 | 0.25 | 4 | 0 | 2 | 0 | 12 | 0.0833 |
| 2_16591600 | 9 | 0.4444 | 12 | 0.5 | 11 | 0.6364 | 7 | 0.4286 | 39 | 0.5128 | 9 | 0.3333 | 4 | 0.5 | 7 | 0.2857 | 11 | 0.3636 | 31 | 0.3548 |
| 2_16591604 | 10 | 0.4 | 13 | 0.1538 | 13 | 0.3077 | 7 | 0.5714 | 43 | 0.3256 | 11 | 0.4545 | 4 | 0.25 | 6 | 0.1667 | 12 | 0.25 | 33 | 0.303 |
| 2_16591629 | 6 | 0.6667 | 10 | 0.8 | 7 | 0.8571 | 5 | 0.8 | 28 | 0.7857 | 8 | 1 | 3 | 1 | 6 | 0.6667 | 11 | 0.8182 | 28 | 0.8571 |
| 2_16591640 | 6 | 0.8333 | 10 | 1 | 8 | 0.875 | 7 | 0.7143 | 31 | 0.871 | 8 | 1 | 5 | 0.8 | 4 | 1 | 11 | 0.9091 | 28 | 0.9286 |
| 2_16591642 | 5 | 0.8 | 12 | 0.9167 | 7 | 0.7143 | 8 | 0.625 | 32 | 0.7812 | 8 | 0.5 | 6 | 0.5 | 6 | 0.6667 | 12 | 0.8333 | 32 | 0.6562 |
| 2_16591643 | 6 | 0.5 | 11 | 0.4545 | 8 | 0.25 | 8 | 0.25 | 33 | 0.3636 | 7 | 0.2857 | 6 | 0.5 | 4 | 0.25 | 12 | 0.1667 | 29 | 0.2759 |
| 2_16591654 | 6 | 0.6667 | 8 | 0.625 | 7 | 1 | 7 | 0.7143 | 28 | 0.75 | 7 | 0.8571 | 6 | 0.6667 | 4 | 0.75 | 11 | 0.7273 | 28 | 0.75 |
| 2_16591655 | 5 | 0 | 9 | 0.1111 | 7 | 0.2857 | 4 | 0.25 | 25 | 0.16 | 8 | 0.25 | 4 | 0 | 6 | 0 | 12 | 0 | 30 | 0.0667 |
| 2_16591657 | 7 | 0.2857 | 9 | 0.2222 | 8 | 0.25 | 6 | 0.1667 | 30 | 0.2333 | 6 | 0.5 | 4 | 0.25 | 5 | 0.4 | 12 | 0.25 | 27 | 0.3333 |
| 2_16591661 | 7 | 0.7143 | 10 | 0.5 | 8 | 0.125 | 7 | 0.5714 | 32 | 0.4688 | 6 | 0.1667 | 6 | 0.5 | 6 | 0.5 | 11 | 0.4545 | 29 | 0.4138 |
| 2_16599713 | NA | NA | NA | NA | 1 | 1 | NA | NA | 1 | 1 | NA | NA | NA | NA | NA | NA | 1 | 1 | 1 | 1 |
| 2_16640860 | 10 | 0 | 13 | 0.1538 | 7 | 0 | 9 | 0 | 39 | 0.0513 | 15 | 0.0667 | 13 | 0.0769 | 11 | 0.1818 | 9 | 0 | 48 | 0.0833 |
| 2_16640877 | 11 | 0.4545 | 10 | 0.2 | 6 | 0.5 | 9 | 0.2222 | 36 | 0.3333 | 16 | 0.3125 | 11 | 0.2727 | 13 | 0.5385 | 8 | 0.125 | 48 | 0.3333 |
| 2_16640889 | 13 | 0.1538 | 12 | 0.1667 | 7 | 0.1429 | 8 | 0.125 | 40 | 0.15 | 18 | 0.3333 | 11 | 0.2727 | 14 | 0.1429 | 7 | 0.1429 | 50 | 0.24 |
| 2_16640895 | 12 | 0.0833 | 15 | 0.1333 | 8 | 0 | 9 | 0.2222 | 44 | 0.1136 | 18 | 0.1111 | 16 | 0.0625 | 13 | 0 | 9 | 0 | 56 | 0.0536 |
| 2_16640919 | 9 | 0.5556 | 9 | 0.7778 | 7 | 0.5714 | 6 | 0.8333 | 31 | 0.6774 | 8 | 0.625 | 8 | 0.875 | 7 | 0.5714 | 6 | 1 | 29 | 0.7586 |
| 2_16640921 | 8 | 0.75 | 13 | 0.9231 | 7 | 0.8571 | 8 | 0.875 | 36 | 0.8611 | 14 | 0.9286 | 14 | 0.9286 | 8 | 1 | 9 | 1 | 45 | 0.9556 |
| 2_16640924 | 6 | 0.3333 | 10 | 0.2 | 8 | 0.25 | 2 | 0.5 | 26 | 0.2692 | 15 | 0.2667 | 9 | 0.3333 | 7 | 0.8571 | 6 | 0.8333 | 37 | 0.4865 |
| 2_16640924 | 7 | 0.2857 | 11 | 0.3636 | 7 | 0.1429 | 7 | 0 | 32 | 0.2188 | 15 | 0 | 15 | 0.1333 | 10 | 0.2 | 10 | 0.1 | 50 | 0.1 |
| 2_16640926 | 8 | 0.25 | 9 | 0.4444 | 6 | 0.3333 | 5 | 0.2 | 28 | 0.3214 | 14 | 0.5 | 14 | 0.3571 | 9 | 0.2222 | 7 | 0.5714 | 44 | 0.4091 |
| 2_16640928 | 11 | 0.6364 | 11 | 0.3636 | 8 | 0.125 | 7 | 0.4286 | 37 | 0.4054 | 17 | 0.8235 | 13 | 0.3846 | 10 | 0.6 | 8 | 0.625 | 48 | 0.625 |
| 2_16640931 | 9 | 0.7778 | 11 | 0.8182 | 6 | 0.8333 | 7 | 0.8571 | 33 | 0.8182 | 15 | 0.8 | 16 | 1 | 10 | 0.9 | 7 | 0.7143 | 48 | 0.875 |
| 2_16640935 | 5 | 0 | 9 | 0.1111 | 6 | 0.3333 | 5 | 0 | 25 | 0.12 | 12 | 0.4167 | 12 | 0.1667 | 8 | 0 | 9 | 0.3333 | 41 | 0.2439 |
| 2_16640955 | 9 | 0.6667 | 9 | 0.5556 | 7 | 1 | 4 | 0.25 | 29 | 0.6552 | 14 | 0.6429 | 11 | 0.6364 | 8 | 0.875 | 6 | 0.8333 | 39 | 0.7179 |
| 2_16640958 | 7 | 0.1429 | 11 | 0.1818 | 7 | 0.2857 | 6 | 0.1667 | 31 | 0.1935 | 11 | 0.4545 | 10 | 0 | 8 | 0.375 | 9 | 0.2222 | 38 | 0.2632 |
| 2_16640967 | 10 | 0 | 10 | 0.1 | 6 | 0 | 6 | 0.1667 | 32 | 0.0625 | 16 | 0.25 | 14 | 0.0714 | 10 | 0.1 | 9 | 0.1111 | 49 | 0.1429 |
| 2_16663759 | 1 | 1 | 2 | 0.5 | NA | NA | 2 | 0.5 | 5 | 0.6 | NA | NA | NA | NA | 1 | 1 | NA | NA | 1 | 1 |
| 2_16663779 | 1 | 1 | 2 | 1 | NA | NA | 3 | 1 | 6 | 1 | NA | NA | NA | NA | 1 | 1 | NA | NA | 1 | 1 |
| 2_16663783 | 1 | 1 | 2 | 0.5 | NA | NA | 3 | 1 | 6 | 0.8333 | NA | NA | NA | NA | 1 | 1 | NA | NA | 1 | 1 |
| 2_16663794 | 1 | 1 | 2 | 1 | NA | NA | 3 | 0.3333 | 6 | 0.6667 | NA | NA | NA | NA | 1 | 1 | NA | NA | 1 | 1 |
| 2_16663803 | 1 | 1 | 2 | 1 | NA | NA | 3 | 1 | 6 | 1 | NA | NA | NA | NA | 1 | 1 | NA | NA | 1 | 1 |
| 2_16664128 | NA | NA | NA | NA | 1 | 1 | NA | NA | 1 | 1 | NA | NA | NA | NA | 1 | 1 | NA | NA | 1 | 1 |
| 2_16673680 | 4 | 0.75 | 2 | 1 | 3 | 1 | 1 | 1 | 10 | 0.9 | 2 | 1 | 2 | 1 | 4 | 0.5 | 1 | 0 | 9 | 0.6667 |
| 2_16673682 | 4 | 0 | 4 | 0 | 4 | 0.25 | 1 | 0 | 13 | 0.0769 | 1 | 0 | 2 | 0 | 3 | 0.3333 | 1 | 0 | 7 | 0.1429 |
| 2_16673688 | NA | NA | 3 | 0.3333 | 5 | 0.4 | 1 | 0 | 9 | 0.3333 | 1 | 1 | 1 | 0 | 2 | 0 | 1 | 0 | 5 | 0.2 |
| 2_16673693 | 5 | 0 | 4 | 0.25 | 5 | 0 | NA | NA | 14 | 0.0714 | 3 | 0 | 2 | 0 | 4 | 0.25 | 1 | 0 | 10 | 0.1 |
| 2_16673697 | 3 | 1 | 3 | 0.3333 | 4 | 1 | 1 | 1 | 11 | 0.8182 | 1 | 1 | 2 | 1 | 4 | 0.5 | NA | NA | 7 | 0.7143 |
| 2_16673712 | 3 | 0.3333 | 2 | 0 | 3 | 0 | 1 | 0 | 9 | 0.1111 | 1 | 0 | 2 | 0 | 1 | 1 | NA | NA | 4 | 0.25 |
| 2_16673765 | 3 | 0 | 4 | 0.25 | 3 | 0 | 1 | 1 | 11 | 0.1818 | 1 | 0 | 3 | 0.3333 | 2 | 0.5 | NA | NA | 6 | 0.3333 |
| 2_16673767 | 2 | 0.5 | 3 | 0 | 2 | 0 | 1 | 1 | 8 | 0.25 | 1 | 0 | 2 | 0 | 2 | 1 | NA | NA | 5 | 0.4 |
| 2_16673793 | 4 | 0 | 3 | 0 | 3 | 0.3333 | 1 | 1 | 11 | 0.1818 | 1 | 0 | 3 | 0.3333 | 2 | 0 | NA | NA | 6 | 0.1667 |
| 2_16673795 | 4 | 0 | 4 | 0.5 | 3 | 0 | 1 | 0 | 12 | 0.1667 | 1 | 0 | 3 | 0 | 2 | 0.5 | NA | NA | 6 | 0.1667 |
| 2_16673834 | 4 | 0 | 6 | 0.1667 | 4 | 0 | 2 | 0 | 16 | 0.0625 | 2 | 0 | 2 | 0 | 3 | 0.3333 | NA | NA | 7 | 0.1429 |

| | | | | | | | | | | | | | | | | | | | | |
|---|---|---|---|---|---|---|---|---|---|---|---|---|---|---|---|---|---|---|---|---|
| 2_16678302 | NA | NA | 1 | 1 | 1 | 1 | NA | NA | 2 | 1 | NA | NA | NA | NA | 1 | 1 | NA | NA | 1 | 1 |
| 2_16678324 | NA | NA | NA | NA | 1 | 1 | NA | NA | 1 | 1 | NA | NA | NA | NA | 1 | 1 | NA | NA | 1 | 1 |
| 2_16678337 | 5 | 0.2 | 3 | 0.3333 | 3 | 0 | 4 | 0 | 15 | 0.1333 | 4 | 0 | 5 | 0.2 | 3 | 0 | 4 | 0 | 16 | 0.0625 |
| 2_16678776 | 5 | 0.4 | 5 | 1 | 4 | 0.5 | 7 | 0.5714 | 21 | 0.619 | 5 | 0.4 | 6 | 0.6667 | 6 | 0.5 | 6 | 0.6667 | 23 | 0.5652 |
| 2_16678781 | 4 | 0.75 | 2 | 0.5 | 3 | 0 | 6 | 0.1667 | 15 | 0.3333 | 5 | 0 | 4 | 0 | 2 | 0.5 | 5 | 0.2 | 16 | 0.125 |
| 2_16678782 | 5 | 0.2 | 4 | 0 | 3 | 0.6667 | 7 | 0.1429 | 19 | 0.2105 | 4 | 0.25 | 4 | 0.25 | 5 | 0.6 | 6 | 0.1667 | 19 | 0.3158 |
| 2_16678784 | 5 | 0.4 | 4 | 0.75 | 2 | 0.5 | 8 | 0.125 | 19 | 0.3684 | 5 | 0.2 | 6 | 0.3333 | 6 | 0.3333 | 5 | 0.2 | 22 | 0.2727 |
| 2_16678789 | 6 | 0.1667 | 5 | 0 | 4 | 0 | 7 | 0.1429 | 22 | 0.0909 | 5 | 0 | 6 | 0.1667 | 5 | 0 | 5 | 0.4 | 21 | 0.1429 |
| 2_16678796 | 5 | 0.4 | 4 | 0.5 | 3 | 0 | 3 | 0 | 15 | 0.2667 | 4 | 0.25 | 5 | 0.2 | 4 | 0 | 5 | 0.2 | 18 | 0.1667 |
| 2_16678948 | 1 | 0 | NA | NA | NA | NA | 3 | 0.3333 | 4 | 0.25 | 1 | 0 | 1 | 0 | 1 | 0 | 2 | 0.5 | 5 | 0.2 |
| 2_16678959 | 2 | 1 | NA | NA | 1 | 0 | 2 | 0.5 | 5 | 0.6 | 1 | 0 | 1 | 0 | NA | NA | 2 | 0.5 | 4 | 0.25 |
| 2_16679009 | 7 | 0.1429 | 3 | 0 | 3 | 0 | 6 | 0 | 19 | 0.0526 | 1 | 0 | 7 | 0 | 2 | 0 | 8 | 0.125 | 18 | 0.0556 |
| 2_16679060 | 9 | 0.2222 | 5 | 0.2 | 9 | 0.1111 | 8 | 0.375 | 31 | 0.2258 | 2 | 0 | 7 | 0.4286 | 7 | 0.4286 | 12 | 0.1667 | 28 | 0.2857 |
| 2_16679061 | 10 | 0.7 | 4 | 0.25 | 7 | 0 | 6 | 0 | 27 | 0.2963 | 3 | 0.3333 | 4 | 0.25 | 5 | 0.2 | 10 | 0.2 | 22 | 0.2273 |
| 2_16679087 | 10 | 0.4 | 4 | 0.25 | 8 | 0.625 | 5 | 0.8 | 27 | 0.5185 | 2 | 0 | 3 | 0.3333 | 7 | 0.4286 | 8 | 0.5 | 20 | 0.4 |
| 2_16679087 | 6 | 0.5 | 4 | 0.75 | 5 | 0.4 | 5 | 0.4 | 20 | 0.5 | NA | NA | 2 | 0.5 | 7 | 0.1429 | 5 | 0.2 | 14 | 0.2143 |
| 2_16679095 | 8 | 0.875 | 3 | 1 | 8 | 0.875 | 6 | 0.8333 | 25 | 0.88 | 2 | 1 | 4 | 1 | 6 | 0.8333 | 9 | 1 | 21 | 0.9524 |
| 2_16679112 | 10 | 0.3 | 4 | 0 | 8 | 0.125 | 8 | 0.25 | 30 | 0.2 | 1 | 0 | 5 | 0.2 | 5 | 0.2 | 10 | 0.1 | 21 | 0.1429 |
| 2_16679120 | 8 | 0.875 | 3 | 1 | 7 | 0.5714 | 8 | 0.875 | 26 | 0.8077 | 1 | 1 | 5 | 1 | 7 | 0.8571 | 8 | 0.875 | 21 | 0.9048 |
| 2_16687830 | 13 | 0.1538 | 3 | 0 | 11 | 0 | 11 | 0 | 38 | 0.0526 | 10 | 0 | 3 | 0 | 9 | 0.1111 | 10 | 0.1 | 32 | 0.0625 |
| 2_16687841 | 11 | 1 | 1 | 1 | 9 | 0.6667 | 6 | 1 | 27 | 0.8889 | 8 | 0.75 | NA | NA | 5 | 1 | 2 | 1 | 15 | 0.8667 |
| 2_16687867 | 11 | 1 | 3 | 0.6667 | 10 | 1 | 10 | 1 | 34 | 0.9706 | 5 | 1 | 3 | 1 | 7 | 1 | 7 | 1 | 22 | 1 |
| 2_16687898 | 10 | 0.1 | 3 | 0 | 9 | 0.3333 | 10 | 0.5 | 32 | 0.2812 | 8 | 0.125 | 3 | 1 | 6 | 0.3333 | 9 | 0 | 26 | 0.2308 |
| 2_16687902 | 12 | 0.25 | 2 | 0 | 8 | 0.25 | 8 | 0.125 | 30 | 0.2 | 8 | 0.25 | 1 | 0 | 5 | 0.2 | 8 | 0.25 | 22 | 0.2273 |
| 2_16687913 | 11 | 0.0909 | 2 | 0 | 11 | 0.0909 | 8 | 0.25 | 32 | 0.125 | 9 | 0 | 2 | 0.5 | 7 | 0.4286 | 8 | 0 | 26 | 0.1538 |
| 2_16687914 | 12 | 1 | 2 | 1 | 11 | 1 | 9 | 1 | 34 | 1 | 7 | 1 | 2 | 0 | 7 | 1 | 8 | 1 | 24 | 0.9167 |
| 2_16687920 | 12 | 0.5833 | 2 | 1 | 11 | 0.5455 | 9 | 0.6667 | 34 | 0.6176 | 9 | 1 | 2 | 0.5 | 8 | 0.75 | 8 | 0.625 | 27 | 0.7778 |
| 2_16687930 | 10 | 0.2 | 2 | 0.5 | 10 | 0.3 | 9 | 0 | 31 | 0.1935 | 9 | 0.2222 | 3 | 0.6667 | 7 | 0.4286 | 8 | 0 | 27 | 0.2593 |
| 2_16687959 | 9 | 0 | 2 | 0.5 | 8 | 0 | 7 | 0.1429 | 26 | 0.0769 | 5 | 0 | 3 | 0 | 5 | 0.2 | 8 | 0.125 | 21 | 0.0952 |
| 2_16687982 | 6 | 0.6667 | 2 | 0.5 | 8 | 0.125 | 8 | 0.25 | 24 | 0.3333 | 6 | 0.1667 | 2 | 0 | 5 | 0.2 | 7 | 0.2857 | 20 | 0.2 |
| 2_16687992 | 8 | 0.125 | 2 | 0 | 8 | 0.125 | 9 | 0.1111 | 27 | 0.1111 | 7 | 0.1429 | 2 | 0 | 5 | 0.2 | 5 | 0 | 19 | 0.1053 |
| 2_16687993 | 5 | 0 | 2 | 0 | 7 | 0.2857 | 9 | 0.1111 | 23 | 0.1304 | 8 | 0.375 | 2 | 0.5 | 2 | 0.5 | 6 | 0 | 18 | 0.2778 |
| 2_16688009 | 6 | 0 | 2 | 0.5 | 7 | 0.1429 | 9 | 0.1111 | 24 | 0.125 | 7 | 0.2857 | 2 | 0.5 | 5 | 0.4 | 7 | 0.1429 | 21 | 0.2857 |
| 2_16688018 | 7 | 0.7143 | 1 | 1 | 7 | 0.8571 | 6 | 0.6667 | 21 | 0.7619 | 6 | 1 | 1 | 1 | 4 | 0.5 | 4 | 1 | 15 | 0.8667 |
| 2_16688032 | 5 | 0.8 | NA | NA | 6 | 0 | 8 | 0 | 19 | 0.2105 | 5 | 0.2 | 1 | 0 | 4 | 0 | 5 | 0.4 | 15 | 0.2 |
| 2_16688046 | 5 | 0.2 | 1 | 1 | 3 | 0.3333 | 5 | 0.2 | 14 | 0.2857 | 6 | 0.5 | 1 | 0 | 5 | 0.2 | 4 | 0 | 16 | 0.25 |
| 2_16688058 | 7 | 0.1429 | 1 | 0 | 6 | 0.3333 | 8 | 0 | 22 | 0.1364 | 8 | 0.125 | 1 | 0 | 5 | 0 | 5 | 0 | 19 | 0.0526 |
| 2_16688063 | 6 | 0 | 1 | 0 | 6 | 0.1667 | 5 | 0 | 18 | 0.0556 | 7 | 0.1429 | 1 | 0 | 5 | 0 | 6 | 0 | 19 | 0.0526 |
| 2_16688069 | 7 | 0.2857 | 1 | 0 | 5 | 0 | 5 | 0.2 | 18 | 0.1667 | 6 | 0.1667 | 1 | 0 | 4 | 0 | 5 | 0 | 16 | 0.0625 |
| 2_16688084 | 5 | 0.2 | NA | NA | 3 | 1 | 5 | 0.4 | 13 | 0.4615 | 6 | 0.5 | 1 | 1 | 3 | 0.3333 | 3 | 0.6667 | 13 | 0.5385 |
| 2_16688088 | 7 | 0.1429 | 1 | 0 | 6 | 0.1667 | 5 | 0.2 | 19 | 0.1579 | 5 | 0.6 | 1 | 0 | 4 | 0.5 | 3 | 0.3333 | 13 | 0.4615 |
| 2_16689721 | 3 | 0.3333 | NA | NA | 3 | 0.3333 | 2 | 0.5 | 8 | 0.375 | 2 | 0.5 | NA | NA | 1 | 0 | 1 | 0 | 4 | 0.25 |
| 2_16689727 | 2 | 0 | NA | NA | 1 | 0 | 2 | 1 | 5 | 0.4 | 2 | 0.5 | NA | NA | 1 | 0 | NA | NA | 3 | 0.3333 |
| 2_16689734 | 4 | 1 | NA | NA | 3 | 1 | 3 | 1 | 10 | 1 | 2 | 0.5 | NA | NA | 1 | 0 | NA | NA | 3 | 0.3333 |
| 2_16689744 | 4 | 1 | NA | NA | 3 | 0.6667 | 2 | 1 | 9 | 0.8889 | 2 | 1 | NA | NA | 1 | 1 | 1 | 1 | 4 | 1 |
| 2_16689756 | 4 | 0.25 | NA | NA | 3 | 0 | 3 | 0.3333 | 10 | 0.2 | 2 | 0.5 | NA | NA | 1 | 1 | 1 | 1 | 4 | 0.75 |
| 2_16689842 | 3 | 1 | NA | NA | 1 | 1 | 3 | 0.6667 | 7 | 0.8571 | 2 | 1 | NA | NA | 1 | 0 | 1 | 1 | 4 | 0.75 |
| 2_16689851 | 4 | 1 | NA | NA | 1 | 0 | 3 | 0.3333 | 8 | 0.625 | 2 | 1 | NA | NA | NA | NA | 1 | 1 | 3 | 1 |
| 2_16693802 | 18 | 0.1667 | 16 | 0.25 | 14 | 0.2143 | 11 | 0.1818 | 59 | 0.2034 | 9 | 0.5556 | 18 | 0.4444 | 14 | 0.2857 | 11 | 0.1818 | 52 | 0.3654 |
| 2_16693967 | 9 | 0.1111 | 8 | 0.125 | 5 | 0 | 2 | 0.5 | 24 | 0.125 | 3 | 0 | 7 | 0.1429 | 7 | 0.2857 | 7 | 0 | 24 | 0.125 |
| 2_16694011 | 6 | 0.3333 | 2 | 0 | 3 | 0.3333 | 1 | 0 | 12 | 0.25 | 3 | 0.3333 | 5 | 0.2 | 4 | 0.25 | 4 | 0 | 16 | 0.1875 |
| 2_16694045 | 5 | 0 | 3 | 0.3333 | 3 | 0 | 1 | 0 | 12 | 0.0833 | NA | NA | 3 | 0.3333 | 4 | 0 | 3 | 0 | 10 | 0.1 |
| 2_16694087 | 7 | 0.4286 | 9 | 0.1111 | 4 | 0 | 8 | 0.125 | 28 | 0.1786 | 4 | 0.25 | 7 | 0.1429 | 7 | 0.1429 | 3 | 0 | 21 | 0.1429 |
| 2_16694099 | 7 | 0.2857 | 9 | 0.2222 | 4 | 0 | 8 | 0.125 | 28 | 0.1786 | 4 | 0.25 | 5 | 0.2 | 6 | 0.1667 | 4 | 0.25 | 19 | 0.2105 |
| 2_16694105 | 7 | 1 | 9 | 1 | 5 | 1 | 8 | 0.875 | 29 | 0.9655 | 3 | 1 | 5 | 0.8 | 7 | 1 | 3 | 0.6667 | 18 | 0.8889 |
| 2_16694111 | 6 | 0.5 | 7 | 0.8571 | 3 | 0.3333 | 8 | 0.875 | 24 | 0.7083 | 4 | 0.75 | 5 | 0.2 | 7 | 1 | 4 | 0.5 | 20 | 0.65 |
| 2_16694116 | 5 | 1 | 6 | 0.8333 | 3 | 0 | 7 | 0.8571 | 21 | 0.7619 | 3 | 0.6667 | 3 | 0.6667 | 5 | 0.4 | 4 | 0.5 | 15 | 0.5333 |
| 2_16694119 | 6 | 0 | 6 | 0.1667 | 4 | 0.5 | 6 | 0.3333 | 22 | 0.2273 | 2 | 0.5 | 3 | 0.3333 | 6 | 0.6667 | 4 | 0 | 15 | 0.4 |
| 2_16694120 | 6 | 0.1667 | 5 | 0.2 | 3 | 0.3333 | 6 | 0.6667 | 20 | 0.35 | 4 | 0.25 | 5 | 0 | 6 | 0.1667 | 2 | 0.5 | 17 | 0.1765 |
| 2_16694124 | 5 | 0 | 6 | 0.1667 | 3 | 0.3333 | 8 | 0 | 22 | 0.0909 | 4 | 0 | 4 | 0 | 6 | 0.3333 | 3 | 0.3333 | 17 | 0.1765 |
| 2_16694131 | 6 | 0.6667 | 7 | 0.5714 | 4 | 1 | 7 | 0.5714 | 24 | 0.6667 | 3 | 0.3333 | 4 | 0.25 | 5 | 0.8 | 4 | 0.5 | 16 | 0.5 |
| 2_16694136 | 7 | 0.2857 | 8 | 0.25 | 4 | 0 | 6 | 0.3333 | 25 | 0.24 | 4 | 0 | 2 | 0 | 5 | 0 | 3 | 0.3333 | 14 | 0.0714 |
| 2_16694136 | 5 | 0.2 | 7 | 0.1429 | 3 | 0.3333 | 8 | 0.125 | 23 | 0.1739 | 4 | 0.25 | 4 | 0.25 | 3 | 0 | 3 | 0 | 14 | 0.1429 |
| 2_16694139 | 7 | 0.1429 | 8 | 0.125 | 4 | 0 | 7 | 0 | 26 | 0.0769 | 4 | 0.25 | 5 | 0.4 | 5 | 0 | 4 | 0 | 18 | 0.1667 |
| 2_16694148 | 7 | 0.1429 | 9 | 0.4444 | 4 | 1 | 7 | 0.5714 | 27 | 0.4815 | 3 | 0.3333 | 5 | 0.2 | 6 | 0.6667 | 4 | 0.5 | 18 | 0.4444 |

| | | | | | | | | | | | | | | | | | | | | |
|---|---|---|---|---|---|---|---|---|---|---|---|---|---|---|---|---|---|---|---|---|
| 2_16694164 | 6 | 1 | 5 | 1 | 4 | 1 | 7 | 1 | 22 | 1 | 2 | 1 | 5 | 1 | 3 | 1 | 4 | 1 | 14 | 1 |
| 2_16694170 | 6 | 0.1667 | 7 | 0.2857 | 2 | 0 | 5 | 0 | 20 | 0.15 | 2 | 0.5 | 5 | 0.4 | 4 | 0 | 4 | 0 | 15 | 0.2 |
| 2_16694187 | 6 | 0 | 5 | 0.4 | 3 | 0 | 3 | 0.3333 | 17 | 0.1765 | 3 | 0 | 5 | 0.2 | 2 | 1 | 3 | 0.6667 | 13 | 0.3846 |
| 2_16694218 | 6 | 0 | 3 | 0 | 2 | 0.5 | 3 | 0 | 14 | 0.0714 | 3 | 0 | 3 | 0 | 2 | 0 | 3 | 0.3333 | 11 | 0.0909 |
| 2_16694229 | 5 | 0 | 2 | 0.5 | 1 | 1 | 3 | 0.3333 | 11 | 0.2727 | 2 | 0.5 | 2 | 0 | 2 | 0 | 1 | 0 | 7 | 0.1429 |
| 2_16694230 | 5 | 0.2 | 2 | 0 | 2 | 0 | 3 | 0 | 12 | 0.0833 | 2 | 0 | 2 | 0 | 2 | 0 | 2 | 0.5 | 8 | 0.125 |
| 2_16694236 | 6 | 0.1667 | 3 | 0 | 1 | 1 | 3 | 0.3333 | 13 | 0.2308 | 3 | 0.3333 | 2 | 0 | 1 | 0 | 2 | 0.5 | 8 | 0.25 |
| 2_16694280 | 3 | 0 | 2 | 0 | NA | NA | 2 | 0.5 | 7 | 0.1429 | 1 | 0 | 1 | 0 | 2 | 0.5 | NA | NA | 4 | 0.25 |
| 2_16694300 | 3 | 0 | 1 | 0 | 2 | 0.5 | 2 | 0.5 | 8 | 0.25 | 1 | 1 | 2 | 0 | 2 | 0.5 | NA | NA | 5 | 0.4 |
| 2_16694301 | 4 | 0 | 6 | 0.5 | 3 | 0.6667 | 5 | 0.2 | 18 | 0.3333 | 1 | 0 | 2 | 0.5 | 2 | 0 | NA | NA | 5 | 0.2 |
| 2_16694305 | 5 | 0.4 | 2 | 1 | 1 | 1 | 4 | 0.75 | 12 | 0.6667 | 2 | 0.5 | 2 | 0.5 | 1 | 1 | NA | NA | 5 | 0.6 |
| 2_16694312 | 4 | 0.5 | 6 | 0.6667 | 2 | 0.5 | 4 | 1 | 16 | 0.6875 | 2 | 0.5 | 2 | 1 | 2 | 0.5 | NA | NA | 6 | 0.6667 |
| 2_16694322 | 4 | 0 | 2 | 0 | NA | NA | 3 | 0.3333 | 9 | 0.1111 | 1 | 0 | 1 | 0 | 2 | 0.5 | NA | NA | 4 | 0.25 |
| 2_16694325 | 2 | 0.5 | 5 | 0.4 | 2 | 1 | 3 | 0 | 12 | 0.4167 | NA | NA | 2 | 0.5 | 2 | 0.5 | NA | NA | 4 | 0.5 |
| 2_16694347 | 4 | 0.75 | 4 | 0.75 | 2 | 0.5 | 4 | 0.25 | 14 | 0.5714 | 2 | 0 | 2 | 0 | 1 | 1 | NA | NA | 5 | 0.2 |
| 2_16694358 | 4 | 0 | 6 | 0.5 | 1 | 0 | 3 | 0 | 14 | 0.2143 | 1 | 1 | 2 | 0 | 2 | 0 | NA | NA | 5 | 0.2 |
| 2_16694366 | 4 | 0 | 5 | 0 | 1 | 0 | 3 | 0.3333 | 13 | 0.0769 | 2 | 0.5 | 2 | 0 | 2 | 0 | NA | NA | 6 | 0.1667 |
| 2_16694409 | 3 | 0.3333 | 3 | 0.3333 | 1 | 1 | 2 | 0.5 | 9 | 0.4444 | 2 | 0 | 1 | 0 | 2 | 0.5 | NA | NA | 5 | 0.2 |
| 2_16694526 | 2 | 0 | 4 | 1 | 1 | 1 | 2 | 1 | 9 | 0.7778 | 2 | 1 | NA | NA | 1 | 1 | NA | NA | 3 | 1 |
| 2_16694545 | 3 | 0.6667 | 2 | 1 | 2 | 1 | 2 | 1 | 9 | 0.8889 | 1 | 0 | NA | NA | 2 | 1 | 1 | 1 | 4 | 0.75 |
| 2_16694554 | 3 | 1 | 2 | 1 | 3 | 1 | 1 | 1 | 9 | 1 | 1 | 1 | 2 | 0.5 | 3 | 1 | 2 | 0.5 | 8 | 0.75 |
| 2_16694555 | 3 | 1 | 2 | 1 | 3 | 1 | 1 | 1 | 9 | 1 | 1 | 1 | 2 | 1 | 3 | 1 | 2 | 1 | 8 | 1 |
| 2_16694560 | 3 | 1 | 1 | 1 | 3 | 1 | 2 | 1 | 9 | 1 | 1 | 1 | 2 | 1 | 3 | 0.3333 | 1 | 1 | 7 | 0.7143 |
| 2_16694571 | 3 | 0.3333 | 2 | 0 | 3 | 0 | 1 | 0 | 9 | 0.1111 | 1 | 1 | 2 | 0.5 | 3 | 0.3333 | 1 | 0 | 7 | 0.4286 |
| 2_16694586 | 2 | 1 | 2 | 1 | 3 | 1 | 1 | 1 | 8 | 1 | NA | NA | 2 | 0.5 | 2 | 1 | 2 | 1 | 6 | 0.8333 |
| 2_16694607 | 2 | 0.5 | 2 | 1 | 3 | 0.6667 | 1 | 1 | 8 | 0.75 | 1 | 0 | 1 | 1 | 3 | 0.6667 | 2 | 0.5 | 7 | 0.5714 |
| 2_16694609 | 2 | 0.5 | 2 | 0 | 3 | 0.6667 | 1 | 0 | 8 | 0.375 | NA | NA | 1 | 1 | 3 | 0.6667 | 1 | 1 | 5 | 0.8 |
| 2_16694617 | 2 | 1 | 2 | 0.5 | 3 | 0.6667 | 1 | 1 | 8 | 0.75 | 1 | 1 | 2 | 1 | 3 | 1 | 1 | 1 | 7 | 1 |
| 2_16694619 | 2 | 0 | 2 | 0.5 | 3 | 0 | 1 | 0 | 8 | 0.125 | NA | NA | 2 | 0 | 3 | 0.3333 | 2 | 0 | 7 | 0.1429 |
| 2_16694634 | NA | NA | 2 | 1 | 1 | 0 | 1 | 0 | 4 | 0.5 | 1 | 0 | 2 | 0.5 | 3 | 0.3333 | 1 | 0 | 7 | 0.2857 |
| 2_16694641 | 2 | 1 | NA | NA | 1 | 1 | 1 | 1 | 4 | 1 | 1 | 1 | 1 | 1 | NA | NA | 2 | 1 | 4 | 1 |
| 2_16694653 | 2 | 1 | 2 | 1 | 1 | 1 | 1 | 1 | 6 | 1 | NA | NA | 2 | 0.5 | 3 | 0.6667 | 2 | 0.5 | 7 | 0.5714 |
| 2_16694658 | 2 | 0 | 2 | 0.5 | NA | NA | 1 | 0 | 5 | 0.2 | 1 | 0 | 2 | 0 | 3 | 0.3333 | 2 | 0.5 | 8 | 0.25 |
| 2_16694672 | 1 | 1 | 1 | 1 | NA | NA | 1 | 1 | 3 | 1 | NA | NA | 1 | 1 | 3 | 1 | 2 | 1 | 6 | 1 |
| 2_16694705 | 2 | 1 | 1 | 1 | NA | NA | 1 | 0 | 4 | 0.75 | NA | NA | NA | NA | 3 | 0.6667 | NA | NA | 3 | 0.6667 |
| 2_16694768 | 2 | 1 | NA | NA | NA | NA | 1 | 1 | 3 | 1 | NA | NA | NA | NA | 2 | 1 | NA | NA | 2 | 1 |
| 2_16694805 | 1 | 0 | NA | NA | NA | NA | 1 | 1 | 2 | 0.5 | NA | NA | NA | NA | 3 | 0.6667 | NA | NA | 3 | 0.6667 |
| 2_16703018 | 1 | 1 | NA | NA | NA | NA | 1 | 1 | 2 | 1 | NA | NA | NA | NA | 1 | 1 | NA | NA | 1 | 1 |
| 2_16703375 | 1 | 0 | 1 | 1 | 1 | 0 | 2 | 0.5 | 5 | 0.4 | 2 | 0.5 | NA | NA | 1 | 0 | 1 | 1 | 4 | 0.5 |
| 2_16703394 | 1 | 0 | 1 | 0 | 1 | 1 | 1 | 0 | 4 | 0.25 | 2 | 0.5 | NA | NA | 1 | 0 | 1 | 0 | 4 | 0.25 |
| 2_16703395 | 1 | 1 | 1 | 1 | 1 | 0 | 2 | 1 | 5 | 0.8 | 1 | 0 | NA | NA | 1 | 1 | 1 | 0 | 3 | 0.3333 |
| 2_16703397 | 1 | 0 | 1 | 0 | NA | NA | 1 | 1 | 3 | 0.3333 | 1 | 1 | NA | NA | 1 | 0 | 1 | 0 | 3 | 0.3333 |
| 2_16703402 | 1 | 0 | 1 | 1 | 1 | 0 | 2 | 0 | 5 | 0.2 | 2 | 0.5 | NA | NA | 1 | 0 | 1 | 0 | 4 | 0.25 |
| 2_16703405 | 1 | 0 | 1 | 1 | 1 | 0 | 1 | 1 | 4 | 0.5 | 2 | 1 | NA | NA | 1 | 1 | 1 | 0 | 4 | 0.75 |
| 2_16703876 | 1 | 1 | 1 | 0 | NA | NA | NA | NA | 2 | 0.5 | 1 | 1 | NA | NA | NA | NA | NA | NA | 1 | 1 |
| 2_16704984 | 1 | 1 | NA | NA | NA | NA | NA | NA | 1 | 1 | NA | NA | NA | NA | NA | NA | 1 | 1 | 1 | 1 |
| 2_16705005 | 1 | 1 | NA | NA | NA | NA | NA | NA | 1 | 1 | NA | NA | NA | NA | NA | NA | 1 | 1 | 1 | 1 |
| 2_16705025 | 1 | 1 | NA | NA | NA | NA | NA | NA | 1 | 1 | NA | NA | NA | NA | NA | NA | 1 | 1 | 1 | 1 |
| 2_16705089 | 1 | 1 | NA | NA | NA | NA | NA | NA | 1 | 1 | NA | NA | NA | NA | NA | NA | 1 | 1 | 1 | 1 |
| 2_16714068 | 5 | 0.2 | 4 | 0.25 | 1 | 0 | 7 | 0 | 17 | 0.1176 | 3 | 0.3333 | 4 | 0.25 | 3 | 0 | 5 | 0 | 15 | 0.1333 |
| 2_16714162 | 6 | 0 | 3 | 0 | 1 | 1 | 7 | 0 | 17 | 0.0588 | 3 | 0 | 2 | 0.5 | 2 | 0.5 | 5 | 0 | 12 | 0.1667 |
| 2_16714234 | 5 | 0.4 | 1 | 0 | NA | NA | 3 | 0 | 9 | 0.2222 | NA | NA | 1 | 1 | 2 | 1 | 2 | 0.5 | 5 | 0.8 |
| 2_16714308 | 1 | 0 | 1 | 0 | NA | NA | 2 | 0.5 | 4 | 0.25 | NA | NA | 1 | 0 | 2 | 1 | 2 | 0 | 5 | 0.4 |
| 2_16714313 | 1 | 0 | 1 | 0 | NA | NA | 2 | 1 | 4 | 0.5 | NA | NA | 1 | 0 | 2 | 0 | 3 | 0.6667 | 6 | 0.3333 |
| 2_16714315 | 1 | 1 | 1 | 1 | NA | NA | 1 | 0 | 3 | 0.6667 | NA | NA | 1 | 1 | 2 | 1 | 3 | 0.6667 | 6 | 0.8333 |
| 2_16714317 | 1 | 1 | 1 | 0 | NA | NA | 1 | 0 | 3 | 0.3333 | NA | NA | 1 | 1 | 2 | 1 | 3 | 0 | 6 | 0.5 |
| 2_16714326 | 1 | 1 | 1 | 0 | NA | NA | 2 | 0 | 4 | 0.25 | NA | NA | 1 | 1 | 2 | 0.5 | 3 | 0.6667 | 6 | 0.6667 |
| 2_16731647 | 4 | 1 | NA | NA | 1 | 1 | 2 | 1 | 7 | 1 | 4 | 1 | NA | NA | NA | NA | 2 | 1 | 6 | 1 |
| 2_16731689 | 4 | 0.75 | NA | NA | 1 | 1 | 1 | 1 | 6 | 0.8333 | 3 | 0.6667 | NA | NA | 1 | 1 | 2 | 1 | 6 | 0.8333 |
| 2_16731717 | 4 | 0 | NA | NA | 1 | 1 | 1 | 0 | 6 | 0.1667 | 2 | 0 | NA | NA | 1 | 0 | 2 | 0.5 | 5 | 0.2 |
| 2_16731732 | 4 | 0.25 | NA | NA | 1 | 1 | 1 | 0 | 6 | 0.3333 | 3 | 0 | NA | NA | 1 | 1 | 2 | 0.5 | 6 | 0.3333 |
| 2_16731764 | 4 | 0 | NA | NA | NA | NA | 2 | 0.5 | 6 | 0.1667 | 3 | 0 | NA | NA | NA | NA | 3 | 0.3333 | 6 | 0.1667 |
| 2_16732348 | 4 | 0.5 | 7 | 0.1429 | 9 | 0.1111 | 7 | 0.4286 | 27 | 0.2593 | 7 | 0 | 4 | 0 | 8 | 0.125 | 5 | 0.2 | 24 | 0.0833 |
| 2_16732355 | 2 | 0.5 | 7 | 0 | 10 | 0.4 | 6 | 0.3333 | 25 | 0.28 | 7 | 0.2857 | 3 | 0 | 9 | 0.3333 | 3 | 0.3333 | 22 | 0.2727 |
| 2_16732368 | 3 | 0.3333 | 6 | 0 | 9 | 0.1111 | 7 | 0 | 25 | 0.08 | 8 | 0 | 4 | 0 | 8 | 0.25 | 4 | 0.25 | 24 | 0.125 |

| | | | | | | | | | | | | | | | | | | | | |
|---|---|---|---|---|---|---|---|---|---|---|---|---|---|---|---|---|---|---|---|---|
| 2_16732438 | 3 | 0.3333 | 7 | 0.2857 | 7 | 0.5714 | 5 | 0.6 | 22 | 0.4545 | 7 | 0.2857 | 4 | 0 | 6 | 0.5 | 3 | 0.6667 | 20 | 0.35 |
| 2_16732462 | 1 | 1 | 8 | 1 | 7 | 1 | 4 | 1 | 20 | 1 | 8 | 1 | 4 | 1 | 4 | 1 | 4 | 1 | 20 | 1 |
| 2_16732463 | 2 | 1 | 7 | 1 | 6 | 1 | 3 | 1 | 18 | 1 | 8 | 1 | 4 | 1 | 3 | 1 | 4 | 1 | 19 | 1 |
| 2_16732474 | 2 | 0.5 | 8 | 0.75 | 5 | 0.2 | 5 | 0.8 | 20 | 0.6 | 8 | 0.625 | 4 | 0.5 | 4 | 0 | 3 | 0.6667 | 19 | 0.4737 |
| 2_16732492 | 2 | 1 | 8 | 1 | 5 | 1 | 5 | 1 | 20 | 1 | 9 | 1 | 4 | 1 | 2 | 1 | 4 | 1 | 19 | 1 |
| 2_16732496 | 2 | 1 | 7 | 1 | 5 | 1 | 5 | 1 | 19 | 1 | 8 | 1 | 4 | 1 | 3 | 1 | 4 | 1 | 19 | 1 |
| 2_16732507 | 3 | 1 | 7 | 1 | 5 | 0.8 | 5 | 0.8 | 20 | 0.9 | 7 | 0.8571 | 4 | 0.75 | 3 | 1 | 4 | 1 | 18 | 0.8889 |
| 2_16732517 | 3 | 1 | 7 | 1 | 5 | 1 | 4 | 1 | 19 | 1 | 7 | 1 | 4 | 1 | 3 | 1 | 5 | 1 | 19 | 1 |
| 2_16732522 | 3 | 0 | 8 | 0.125 | 4 | 0 | 5 | 0 | 20 | 0.05 | 7 | 0.1429 | 2 | 0 | 3 | 0.3333 | 3 | 0.3333 | 15 | 0.2 |
| 2_16732531 | 2 | 0 | 7 | 0.5714 | 5 | 0.4 | 4 | 0.25 | 18 | 0.3889 | 6 | 0.1667 | 3 | 0 | 3 | 0.6667 | 2 | 0.5 | 14 | 0.2857 |
| 2_16732552 | 1 | 1 | 6 | 0.8333 | 3 | 1 | 3 | 0.6667 | 13 | 0.8462 | 6 | 0.8333 | 3 | 1 | 2 | 1 | 2 | 0 | 13 | 0.7692 |
| 2_16732564 | 1 | 0 | 8 | 0 | 5 | 0.4 | 3 | 0.3333 | 17 | 0.1765 | 6 | 0.1667 | 1 | 0 | 2 | 0 | NA | NA | 9 | 0.1111 |
| 2_16732568 | 2 | 0 | 4 | 0 | 5 | 0.2 | 4 | 0 | 15 | 0.0667 | 3 | 0.3333 | 2 | 0.5 | 1 | 0 | 2 | 0.5 | 8 | 0.375 |
| 2_16732581 | 3 | 0 | 6 | 0.1667 | 5 | 0 | 4 | 0 | 18 | 0.0556 | 6 | 0.1667 | 3 | 0 | 3 | 0 | 1 | 0 | 13 | 0.0769 |
| 2_16732582 | 3 | 0.6667 | 6 | 0.3333 | 5 | 0.8 | 4 | 0.75 | 18 | 0.6111 | 6 | 1 | 3 | 0.6667 | 3 | 1 | 2 | 0.5 | 14 | 0.8571 |
| 2_16732597 | 2 | 0 | 4 | 0.5 | 5 | 0.4 | 3 | 0.3333 | 14 | 0.3571 | 6 | 0 | 2 | 0.5 | 3 | 0.3333 | 2 | 0.5 | 13 | 0.2308 |
| 2_16732611 | 2 | 0.5 | 6 | 0.8333 | 5 | 0.8 | 3 | 1 | 16 | 0.8125 | 4 | 1 | 2 | 1 | 3 | 0.6667 | 1 | 1 | 10 | 0.9 |
| 2_16732628 | 2 | 0.5 | 5 | 0.2 | 3 | 1 | 3 | 0.3333 | 13 | 0.4615 | 3 | 0.6667 | 2 | 0.5 | 3 | 0.6667 | 2 | 0.5 | 10 | 0.6 |
| 2_16732638 | 2 | 0 | 4 | 0.25 | 4 | 0.25 | 3 | 0 | 13 | 0.1538 | 4 | 0.5 | 2 | 0.5 | 3 | 0.3333 | 2 | 0 | 11 | 0.3636 |
| 2_16732640 | 2 | 1 | 6 | 1 | 4 | 1 | 3 | 1 | 15 | 1 | 4 | 1 | 2 | 1 | 3 | 1 | 2 | 1 | 11 | 1 |
| 2_16732644 | 2 | 0.5 | 6 | 0.5 | 4 | 0.5 | 2 | 0.5 | 14 | 0.5 | 3 | 0.6667 | 2 | 0 | 2 | 0 | 2 | 0.5 | 9 | 0.3333 |
| 2_16732676 | NA | NA | NA | NA | 1 | 1 | NA | NA | 1 | 1 | 1 | 1 | NA | NA | NA | NA | NA | NA | 1 | 1 |
| 2_16732724 | 2 | 0.5 | 5 | 0.8 | 3 | 1 | 3 | 0 | 13 | 0.6154 | 4 | 1 | 2 | 0.5 | 3 | 0.6667 | 2 | 0.5 | 11 | 0.7273 |
| 2_16735671 | 8 | 0.125 | 9 | 0 | 7 | 0.1429 | 5 | 0 | 29 | 0.069 | 5 | 0 | 8 | 0.125 | 6 | 0.1667 | 5 | 0.2 | 24 | 0.125 |
| 2_16735703 | 8 | 0.375 | 8 | 0.375 | 9 | 0.6667 | 6 | 0.5 | 31 | 0.4839 | 6 | 0.1667 | 10 | 0.3 | 8 | 0.75 | 6 | 0.1667 | 30 | 0.3667 |
| 2_16735767 | 10 | 0 | 9 | 0.1111 | 9 | 0.1111 | 7 | 0 | 35 | 0.0571 | 6 | 0 | 9 | 0.1111 | 10 | 0.1 | 5 | 0 | 30 | 0.0667 |
| 2_16735774 | 10 | 0.3 | 10 | 0.2 | 9 | 0.3333 | 10 | 0.2 | 39 | 0.2564 | 9 | 0.2222 | 11 | 0.1818 | 10 | 0.4 | 8 | 0.25 | 38 | 0.2632 |
| 2_16735783 | 9 | 0.1111 | 11 | 0 | 9 | 0 | 11 | 0.0909 | 40 | 0.05 | 8 | 0.125 | 11 | 0 | 10 | 0.2 | 10 | 0.3 | 39 | 0.1538 |
| 2_16735797 | 11 | 0 | 12 | 0.1667 | 9 | 0 | 11 | 0.1818 | 43 | 0.093 | 8 | 0 | 12 | 0.0833 | 9 | 0.3333 | 10 | 0 | 39 | 0.1026 |
| 2_16744982 | 39 | 0.0769 | 39 | 0.0769 | 38 | 0.1316 | 46 | 0.1087 | 162 | 0.0988 | 26 | 0.1154 | 40 | 0.025 | 26 | 0 | 29 | 0.1034 | 121 | 0.0579 |
| 2_16744991 | 38 | 0.1316 | 39 | 0.0256 | 39 | 0.0513 | 44 | 0.0909 | 160 | 0.075 | 25 | 0.04 | 42 | 0.0714 | 25 | 0.12 | 27 | 0.1111 | 119 | 0.084 |
| 2_16744995 | 38 | 0.5526 | 39 | 0.5128 | 39 | 0.5897 | 42 | 0.4762 | 158 | 0.5316 | 29 | 0.4828 | 41 | 0.6341 | 26 | 0.5769 | 29 | 0.4138 | 125 | 0.536 |
| 2_16745016 | 29 | 0.069 | 36 | 0.0278 | 31 | 0.0968 | 35 | 0.0857 | 131 | 0.0687 | 26 | 0.0385 | 35 | 0.1143 | 20 | 0.05 | 26 | 0.0769 | 107 | 0.0748 |
| 2_16745028 | 30 | 0.1 | 29 | 0.069 | 32 | 0.25 | 30 | 0.0333 | 121 | 0.1157 | 26 | 0.1538 | 29 | 0.1034 | 23 | 0.1739 | 27 | 0.2593 | 105 | 0.1714 |
| 2_16745059 | 34 | 0.8529 | 30 | 0.9 | 35 | 1 | 40 | 0.875 | 139 | 0.9065 | 26 | 0.8846 | 41 | 0.9512 | 20 | 0.9 | 27 | 0.963 | 114 | 0.9298 |
| 2_16745072 | 38 | 0.0789 | 34 | 0.2059 | 32 | 0.2812 | 37 | 0.2703 | 141 | 0.2057 | 27 | 0.1481 | 35 | 0.1714 | 25 | 0.28 | 25 | 0.28 | 112 | 0.2143 |
| 2_16745072 | 23 | 0.0435 | 19 | 0.2105 | 19 | 0.0526 | 25 | 0.2 | 86 | 0.1279 | 20 | 0.25 | 22 | 0.2727 | 17 | 0.2353 | 19 | 0.2105 | 78 | 0.2436 |
| 2_16745079 | 27 | 0.7778 | 32 | 0.75 | 28 | 0.8929 | 40 | 0.725 | 127 | 0.7795 | 21 | 0.8571 | 29 | 0.7241 | 15 | 0.8 | 25 | 0.84 | 90 | 0.8 |
| 2_16745087 | 32 | 0.25 | 25 | 0.4 | 27 | 0.1852 | 37 | 0.3243 | 121 | 0.2893 | 21 | 0.1905 | 32 | 0.3438 | 17 | 0.3529 | 22 | 0.5 | 92 | 0.3478 |
| 2_16745089 | 33 | 0.5758 | 28 | 0.3214 | 30 | 0.4667 | 36 | 0.5556 | 127 | 0.4882 | 24 | 0.3333 | 37 | 0.6216 | 20 | 0.4 | 23 | 0.5217 | 104 | 0.4904 |
| 2_16745093 | 29 | 0.069 | 29 | 0.069 | 24 | 0 | 33 | 0.0909 | 115 | 0.0609 | 23 | 0.1304 | 30 | 0.0667 | 18 | 0 | 26 | 0.0385 | 97 | 0.0619 |
| 2_16745103 | 32 | 0.875 | 28 | 0.7143 | 29 | 0.7931 | 37 | 0.8649 | 126 | 0.8175 | 23 | 0.6087 | 34 | 0.6471 | 24 | 0.75 | 26 | 0.6154 | 107 | 0.6542 |
| 2_16745977 | 29 | 0.1034 | 29 | 0.069 | 26 | 0.0385 | 35 | 0.0857 | 119 | 0.0756 | 22 | 0.1364 | 37 | 0.027 | 18 | 0.0556 | 23 | 0 | 100 | 0.05 |
| 2_16748459 | 3 | 0.3333 | 4 | 0 | 2 | 0 | NA | NA | 9 | 0.1111 | 4 | 0.25 | 3 | 0.3333 | 3 | 0.3333 | 2 | 0 | 12 | 0.25 |
| 2_16748460 | 3 | 0.3333 | 5 | 0 | 1 | 0 | NA | NA | 9 | 0.1111 | 4 | 0 | 3 | 0 | 4 | 0.25 | 2 | 0 | 13 | 0.0769 |
| 2_16748561 | 12 | 0.0833 | 11 | 0 | 4 | 0 | 8 | 0.125 | 35 | 0.0571 | 8 | 0.125 | 3 | 0 | 5 | 0 | 2 | 0 | 18 | 0.0556 |
| 2_16748574 | 10 | 0 | 12 | 0.1667 | 6 | 0 | 6 | 0.1667 | 34 | 0.0882 | 8 | 0.125 | 6 | 0.1667 | 5 | 0 | 3 | 0 | 22 | 0.0909 |
| 2_16748584 | 9 | 0.1111 | 12 | 0.0833 | 6 | 0.1667 | 8 | 0.25 | 35 | 0.1429 | 7 | 0 | 8 | 0.375 | 5 | 0.2 | 2 | 0 | 22 | 0.1818 |
| 2_16748594 | 8 | 0.125 | 9 | 0.2222 | 6 | 0.1667 | 5 | 0.2 | 28 | 0.1786 | 3 | 0.3333 | 4 | 0 | 3 | 0 | 2 | 0 | 12 | 0.0833 |
| 2_16748597 | 9 | 0.8889 | 11 | 0.6364 | 5 | 0.6 | 6 | 0.8333 | 31 | 0.7419 | 6 | 0.6667 | 6 | 1 | 5 | 0.6 | 3 | 1 | 20 | 0.8 |
| 2_16753136 | 4 | 0.25 | 2 | 1 | 1 | 1 | 3 | 0.6667 | 10 | 0.6 | 1 | 1 | 6 | 0.3333 | 3 | 1 | 4 | 0.75 | 14 | 0.6429 |
| 2_16753178 | 6 | 0.8333 | 2 | 1 | 2 | 0 | 3 | 0.3333 | 13 | 0.6154 | 3 | 0.6667 | 6 | 0.8333 | 3 | 0.6667 | 2 | 0.5 | 14 | 0.7143 |
| 2_16753192 | 2 | 1 | 1 | 1 | NA | NA | NA | NA | 3 | 1 | 2 | 0.5 | 6 | 0.8333 | 2 | 0.5 | 3 | 0.3333 | 13 | 0.6154 |
| 2_16753196 | 5 | 0.4 | 1 | 1 | 2 | 0 | 4 | 0 | 12 | 0.25 | 3 | 0 | 5 | 0.2 | 4 | 0 | 2 | 0 | 14 | 0.0714 |
| 2_16753202 | 8 | 1 | 3 | 1 | 2 | 1 | 4 | 1 | 17 | 1 | 3 | 1 | 7 | 1 | 4 | 0.75 | 5 | 1 | 19 | 0.9474 |
| 2_16753206 | 6 | 0 | 4 | 0.5 | 2 | 0.5 | 4 | 0.25 | 16 | 0.25 | 3 | 0.3333 | 7 | 0.1429 | 3 | 0.6667 | 5 | 0.4 | 18 | 0.3333 |
| 2_16753207 | 9 | 1 | 3 | 1 | 2 | 1 | 4 | 1 | 18 | 1 | 2 | 1 | 7 | 1 | 5 | 0.8 | 5 | 1 | 19 | 0.9474 |
| 2_16753230 | 8 | 0 | 2 | 0 | 3 | 0 | 3 | 0.3333 | 16 | 0.0625 | 4 | 0 | 6 | 0.6667 | 5 | 0.2 | 4 | 0.25 | 19 | 0.3158 |
| 2_16753231 | 7 | 0.1429 | 1 | 0 | 2 | 0.5 | 2 | 0.5 | 12 | 0.25 | 4 | 0 | 5 | 0.4 | 3 | 0 | 1 | 0 | 13 | 0.1538 |
| 2_16781856 | 2 | 1 | 2 | 0.5 | 2 | 1 | 4 | 1 | 10 | 0.9 | 7 | 0.5714 | 2 | 1 | 4 | 1 | NA | NA | 13 | 0.7692 |
| 2_16781865 | 3 | 0.6667 | 3 | 1 | 1 | 1 | 4 | 0.75 | 11 | 0.8182 | 6 | 0.6667 | 3 | 1 | 6 | 0.6667 | NA | NA | 15 | 0.7333 |
| 2_16781872 | 3 | 0.3333 | NA | NA | 1 | 0 | 4 | 0.25 | 8 | 0.25 | 3 | 0.3333 | 1 | 1 | 4 | 0.5 | NA | NA | 8 | 0.5 |
| 2_16781901 | 4 | 0 | 1 | 0 | NA | NA | 4 | 0.25 | 9 | 0.1111 | 5 | 0.2 | 2 | 0 | 3 | 0 | NA | NA | 10 | 0.1 |
| 2_16781936 | 4 | 0 | 1 | 1 | NA | NA | 6 | 0.5 | 11 | 0.3636 | 6 | 0.3333 | 4 | 0.5 | 4 | 0.5 | 1 | 0 | 15 | 0.4 |
| 2_16781941 | 5 | 0.4 | 1 | 0 | NA | NA | 6 | 0.1667 | 12 | 0.25 | 6 | 0.1667 | 4 | 0 | 3 | 0.3333 | 1 | 1 | 14 | 0.2143 |

| | | | | | | | | | | | | | | | | | | | | |
|---|---|---|---|---|---|---|---|---|---|---|---|---|---|---|---|---|---|---|---|---|
| 2_1678194 | 3 | 0.3333 | 1 | 0 | NA | NA | 3 | 0 | 7 | 0.1429 | 4 | 0 | 2 | 0 | 3 | 0.3333 | NA | NA | 9 | 0.1111 |
| 2_1678194 | 4 | 0.75 | 1 | 0 | NA | NA | 4 | 0.25 | 9 | 0.4444 | 5 | 0.6 | 3 | 1 | 3 | 0.3333 | 1 | 0 | 12 | 0.5833 |
| 2_1678194 | 5 | 0.8 | NA | NA | NA | NA | 5 | 1 | 10 | 0.9 | 6 | 0.5 | 3 | 1 | 4 | 0.5 | NA | NA | 13 | 0.6154 |
| 2_1678195 | 2 | 1 | NA | NA | NA | NA | 5 | 1 | 7 | 1 | 2 | 1 | 2 | 1 | 4 | 1 | 1 | 1 | 9 | 1 |
| 2_1678200 | 4 | 0.5 | 2 | 0 | 1 | 0 | 6 | 0.1667 | 13 | 0.2308 | 6 | 0 | 3 | 0.3333 | 6 | 0.1667 | 1 | 1 | 16 | 0.1875 |
| 2_1678200 | 6 | 0 | 2 | 0 | 1 | 1 | 7 | 0.4286 | 16 | 0.25 | 5 | 0.2 | 3 | 0.3333 | 7 | 0.1429 | 1 | 0 | 16 | 0.1875 |
| 2_1678205 | 6 | 0.1667 | 2 | 0 | 1 | 0 | 3 | 0 | 12 | 0.0833 | 6 | 0.5 | 3 | 0.3333 | 7 | 0 | 1 | 0 | 17 | 0.2353 |
| 2_1678205 | 5 | 0.2 | NA | NA | 1 | 0 | 7 | 0 | 13 | 0.0769 | 5 | 0.2 | 3 | 0.3333 | 3 | 0 | NA | NA | 11 | 0.1818 |
| 2_1678211 | 6 | 0.1667 | 3 | 0.3333 | 2 | 0.5 | 6 | 0.3333 | 17 | 0.2941 | 7 | 0 | 3 | 0.3333 | 6 | 0.1667 | 1 | 0 | 17 | 0.1176 |
| 2_1679293 | 1 | 1 | 1 | 0 | 1 | 0 | 1 | 0 | 4 | 0.25 | 1 | 0 | NA | NA | 2 | 0.5 | NA | NA | 3 | 0.3333 |
| 2_1679407 | 1 | 1 | 1 | 1 | NA | NA | 2 | 0.5 | 4 | 0.75 | 1 | 1 | NA | NA | 2 | 1 | NA | NA | 3 | 1 |
| 2_1679407 | 1 | 1 | NA | NA | 1 | 0 | 2 | 1 | 4 | 0.75 | 1 | 1 | NA | NA | 1 | 1 | NA | NA | 2 | 1 |
| 2_1679410 | 1 | 1 | 1 | 1 | 1 | 0 | 2 | 1 | 5 | 0.8 | 1 | 1 | NA | NA | 1 | 1 | NA | NA | 2 | 1 |
| 2_1679410 | 1 | 1 | 1 | 0 | 1 | 1 | NA | NA | 3 | 0.6667 | 1 | 1 | NA | NA | NA | NA | NA | NA | 1 | 1 |
| 2_1679410 | 1 | 1 | 1 | 0 | 1 | 1 | 2 | 0 | 5 | 0.4 | 1 | 1 | NA | NA | 2 | 1 | NA | NA | 3 | 1 |
| 2_1679410 | 1 | 1 | 1 | 1 | 1 | 1 | 2 | 0 | 5 | 0.6 | 1 | 1 | NA | NA | 1 | 0 | NA | NA | 2 | 0.5 |
| 2_1679410 | 1 | 1 | 1 | 0 | 1 | 0 | 2 | 1 | 5 | 0.6 | 1 | 1 | NA | NA | 2 | 0.5 | NA | NA | 3 | 0.6667 |
| 2_1679411 | 1 | 1 | 1 | 1 | 1 | 1 | 2 | 1 | 5 | 1 | 1 | 1 | NA | NA | 2 | 1 | 1 | 1 | 4 | 1 |
| 2_1679411 | NA | NA | 1 | 1 | 1 | 1 | 2 | 0.5 | 4 | 0.75 | 1 | 1 | NA | NA | 2 | 0.5 | 1 | 1 | 4 | 0.75 |
| 2_1679411 | 1 | 1 | 1 | 0 | NA | NA | 1 | 1 | 3 | 0.6667 | NA | NA | NA | NA | 1 | 0 | 1 | 1 | 2 | 0.5 |
| 2_1679412 | 1 | 1 | 1 | 1 | 1 | 1 | 2 | 1 | 5 | 1 | NA | NA | NA | NA | 1 | 1 | 1 | 1 | 2 | 1 |
| 2_1679413 | 1 | 0 | 1 | 0 | 1 | 1 | 2 | 0 | 5 | 0.2 | 1 | 1 | NA | NA | 1 | 1 | NA | NA | 2 | 1 |
| 2_1679414 | 1 | 1 | 1 | 0 | 1 | 0 | 2 | 0.5 | 5 | 0.4 | 1 | 0 | NA | NA | 2 | 0.5 | NA | NA | 3 | 0.3333 |
| 2_1679421 | 1 | 0 | 1 | 0 | 1 | 0 | 2 | 1 | 5 | 0.4 | 1 | 1 | NA | NA | 2 | 0.5 | 1 | 1 | 4 | 0.75 |
| 2_1679422 | 1 | 0 | 1 | 1 | 1 | 0 | 2 | 0.5 | 5 | 0.4 | 1 | 0 | NA | NA | 1 | 1 | 1 | 1 | 3 | 0.6667 |
| 2_1679431 | 1 | 0 | 1 | 0 | NA | NA | 2 | 0.5 | 4 | 0.25 | NA | NA | NA | NA | 2 | 1 | 1 | 0 | 3 | 0.6667 |
| 2_1680232 | 12 | 0.5 | 15 | 0.5333 | 13 | 0.6923 | 8 | 0.5 | 48 | 0.5625 | 13 | 0.4615 | 13 | 0.2308 | 15 | 0.4 | 13 | 0.4615 | 54 | 0.3889 |
| 2_1680233 | 10 | 0.3 | 12 | 0.25 | 12 | 0.4167 | 9 | 0.4444 | 43 | 0.3488 | 11 | 0.1818 | 12 | 0.1667 | 12 | 0.0833 | 11 | 0.5455 | 46 | 0.2391 |
| 2_1680233 | 14 | 0.4286 | 16 | 0.4375 | 14 | 0.3571 | 10 | 0.7 | 54 | 0.463 | 15 | 0.4667 | 14 | 0.3571 | 13 | 0.2308 | 15 | 0.3333 | 57 | 0.3509 |
| 2_1680235 | 11 | 0.4545 | 15 | 0.4667 | 13 | 0.6923 | 10 | 0.4 | 49 | 0.5102 | 13 | 0.6923 | 10 | 0.4 | 15 | 0.6667 | 15 | 0.6667 | 53 | 0.6226 |
| 2_1680235 | 12 | 0 | 16 | 0.25 | 13 | 0.0769 | 11 | 0 | 52 | 0.0962 | 14 | 0 | 11 | 0.1818 | 14 | 0.0714 | 14 | 0.0714 | 53 | 0.0755 |
| 2_1680237 | 12 | 0.25 | 13 | 0.3846 | 10 | 0.5 | 7 | 0.5714 | 42 | 0.4048 | 13 | 0.3077 | 9 | 0.4444 | 14 | 0.2143 | 12 | 0.1667 | 48 | 0.2708 |
| 2_1680237 | 10 | 0.8 | 14 | 0.5714 | 9 | 0.7778 | 8 | 0.875 | 41 | 0.7317 | 10 | 0.6 | 10 | 0.8 | 13 | 0.8462 | 14 | 0.5714 | 47 | 0.7021 |
| 2_1680237 | 11 | 0.4545 | 13 | 0.3077 | 9 | 0.3333 | 8 | 0.125 | 41 | 0.3171 | 8 | 0.375 | 9 | 0.5556 | 12 | 0.25 | 13 | 0.0769 | 42 | 0.2857 |
| 2_1680238 | 8 | 1 | 13 | 1 | 10 | 1 | 9 | 0.8889 | 40 | 0.975 | 11 | 1 | 8 | 1 | 11 | 1 | 12 | 1 | 42 | 1 |
| 2_1680238 | 10 | 0.3 | 12 | 0.1667 | 6 | 0 | 5 | 0.2 | 33 | 0.1818 | 10 | 0 | 9 | 0.2222 | 13 | 0 | 12 | 0.0833 | 44 | 0.0682 |
| 2_1680238 | 11 | 0.2727 | 14 | 0.2857 | 8 | 0.5 | 9 | 0.3333 | 42 | 0.3333 | 11 | 0.1818 | 9 | 0.4444 | 14 | 0.5714 | 11 | 0.3636 | 45 | 0.4 |
| 2_1680239 | 7 | 1 | 13 | 0.8462 | 11 | 1 | 7 | 1 | 38 | 0.9474 | 11 | 1 | 9 | 1 | 13 | 1 | 13 | 1 | 46 | 1 |
| 2_1680239 | 9 | 0 | 14 | 0.0714 | 9 | 0.1111 | 8 | 0.125 | 40 | 0.075 | 8 | 0 | 6 | 0.1667 | 11 | 0.0909 | 10 | 0.1 | 35 | 0.0857 |
| 2_1680240 | 7 | 0.5714 | 14 | 0.4286 | 5 | 0.6 | 5 | 0.4 | 31 | 0.4839 | 10 | 0.4 | 8 | 0.75 | 10 | 0.7 | 9 | 0.2222 | 37 | 0.5135 |
| 2_1680240 | 9 | 0.8889 | 9 | 0.5556 | 5 | 0.8 | 8 | 0.875 | 31 | 0.7742 | 10 | 0.8 | 7 | 0.4286 | 11 | 0.6364 | 10 | 0.8 | 38 | 0.6842 |
| 2_1680241 | 11 | 0.4545 | 13 | 0.3077 | 9 | 0.7778 | 8 | 0.5 | 41 | 0.4878 | 10 | 0.1 | 8 | 0.625 | 11 | 0.3636 | 12 | 0.4167 | 41 | 0.3659 |
| 2_1680242 | 10 | 0 | 13 | 0.0769 | 6 | 0.3333 | 7 | 0.1429 | 36 | 0.1111 | 11 | 0.3636 | 8 | 0.375 | 10 | 0 | 11 | 0.0909 | 40 | 0.2 |
| 2_1680243 | 9 | 1 | 14 | 0.8571 | 7 | 1 | 6 | 1 | 36 | 0.9444 | 12 | 1 | 6 | 0.8333 | 10 | 1 | 12 | 1 | 40 | 0.975 |
| 2_1680244 | 9 | 0.1111 | 11 | 0 | 6 | 0 | 3 | 0.3333 | 29 | 0.069 | 10 | 0.1 | 7 | 0 | 11 | 0.1818 | 9 | 0.1111 | 37 | 0.1081 |
| 2_1680244 | 6 | 1 | 10 | 0.9 | 4 | 1 | 6 | 0.8333 | 26 | 0.9231 | 9 | 0.7778 | 8 | 0.75 | 9 | 1 | 9 | 1 | 35 | 0.8857 |
| 2_1680244 | 7 | 0.1429 | 11 | 0.2727 | 5 | 0.2 | 7 | 0 | 30 | 0.1667 | 11 | 0.2727 | 7 | 0.4286 | 11 | 0.0909 | 9 | 0.3333 | 38 | 0.2632 |
| 2_1680246 | 5 | 0 | 9 | 0.4444 | 5 | 0 | 4 | 0.25 | 23 | 0.2174 | 11 | 0.3636 | 7 | 0.2857 | 9 | 0.4444 | 8 | 0.5 | 35 | 0.4 |
| 2_1680247 | 6 | 0.5 | 11 | 0.7273 | 5 | 1 | 3 | 1 | 25 | 0.76 | 10 | 0.7 | 8 | 0.875 | 10 | 1 | 10 | 0.8 | 38 | 0.8421 |
| 2_1680247 | 8 | 0.5 | 12 | 0.5 | 3 | 0.3333 | 4 | 0.5 | 27 | 0.4815 | 10 | 0.5 | 7 | 0.4286 | 10 | 0.2 | 8 | 0.25 | 35 | 0.3429 |
| 2_1680249 | 7 | 0 | 10 | 0.2 | 5 | 0.4 | 5 | 0 | 27 | 0.1481 | 11 | 0.0909 | 6 | 0.3333 | 9 | 0.2222 | 8 | 0.25 | 34 | 0.2059 |
| 2_1680253 | 4 | 0 | 7 | 0 | 1 | 0 | 6 | 0.1667 | 18 | 0.0556 | 9 | 0.2222 | 6 | 0 | 6 | 0.1667 | 9 | 0.1111 | 30 | 0.1333 |
| 2_1680253 | 6 | 0.3333 | 7 | 0.2857 | 2 | 0.5 | 5 | 0.2 | 20 | 0.3 | 6 | 0.3333 | 4 | 0.5 | 4 | 0.5 | 6 | 0.5 | 20 | 0.45 |
| 2_1680253 | 4 | 0.5 | 8 | 0.125 | 3 | 0.3333 | 5 | 0.2 | 20 | 0.25 | 10 | 0.1 | 5 | 0.2 | 6 | 0 | 9 | 0.1111 | 30 | 0.1 |
| 2_1680253 | 5 | 0.8 | 8 | 0.75 | 3 | 1 | 3 | 1 | 19 | 0.8421 | 9 | 0.7778 | 5 | 1 | 6 | 1 | 9 | 0.6667 | 29 | 0.8276 |
| 2_1680256 | 5 | 0 | 8 | 0.5 | 3 | 0.6667 | 4 | 0 | 20 | 0.3 | 7 | 0 | 5 | 0.2 | 5 | 0 | 8 | 0.125 | 25 | 0.08 |
| 2_1680261 | 4 | 0.75 | 8 | 0.625 | 2 | 0.5 | 3 | 0.6667 | 17 | 0.6471 | 4 | 0.25 | 4 | 0.25 | 3 | 0.3333 | 7 | 0.2857 | 18 | 0.2778 |
| 2_1680261 | 2 | 0.5 | 7 | 0 | 1 | 1 | 2 | 0 | 12 | 0.1667 | 4 | 0.25 | 2 | 0 | 5 | 0.2 | 6 | 0.1667 | 17 | 0.1765 |
| 2_1680262 | 3 | 1 | 7 | 0.8571 | 1 | 1 | 5 | 0.8 | 16 | 0.875 | 4 | 0.5 | 4 | 0.75 | 4 | 0.75 | 6 | 0.5 | 18 | 0.6111 |
| 2_1680264 | 3 | 1 | 6 | 0.6667 | 1 | 1 | 5 | 0.6 | 15 | 0.7333 | 3 | 1 | 3 | 0.6667 | 3 | 1 | 5 | 1 | 14 | 0.9286 |
| 2_1680264 | 3 | 0.3333 | 5 | 0 | 1 | 0 | 5 | 0 | 14 | 0.0714 | 3 | 0 | 4 | 0.25 | 3 | 0 | 5 | 0 | 15 | 0.0667 |
| 2_1680265 | 3 | 1 | 3 | 1 | 1 | 1 | 4 | 1 | 11 | 1 | 2 | 1 | 4 | 0.75 | 4 | 1 | 5 | 0.8 | 15 | 0.8667 |
| 2_1680265 | 3 | 0.3333 | 5 | 0.4 | 1 | 1 | 4 | 0.5 | 13 | 0.4615 | 3 | 0.6667 | 4 | 0.5 | 3 | 0.3333 | 6 | 0.5 | 16 | 0.5 |
| 2_1680269 | 3 | 0.6667 | 3 | 1 | 1 | 1 | 3 | 1 | 10 | 0.9 | 2 | 1 | 3 | 1 | 1 | 1 | 5 | 1 | 11 | 1 |
| 2_1680269 | 2 | 0.5 | 4 | 0.5 | 1 | 0 | 4 | 0.25 | 11 | 0.3636 | 2 | 1 | 4 | 0.25 | 3 | 0 | 4 | 0.25 | 13 | 0.3077 |

| | | | | | | | | | | | | | | | | | | | | |
|---|---|---|---|---|---|---|---|---|---|---|---|---|---|---|---|---|---|---|---|---|
| 2_1680295 | 3 | 0 | 4 | 0 | NA | NA | 3 | 0.3333 | 10 | 0.1 | NA | NA | 3 | 0 | 3 | 0 | 4 | 0.25 | 10 | 0.1 |
| 2_1680524 | 13 | 0.3846 | 13 | 0.1538 | 6 | 0.1667 | 9 | 0.1111 | 41 | 0.2195 | 3 | 0 | 5 | 0.6 | 10 | 0.4 | 9 | 0.4444 | 27 | 0.4074 |
| 2_1680730 | 2 | 0 | 1 | 1 | 1 | 0 | 1 | 0 | 5 | 0.2 | 3 | 0 | 2 | 0.5 | 4 | 0 | 1 | 0 | 10 | 0.1 |
| 2_1680734 | 2 | 0.5 | 1 | 0 | 1 | 1 | 3 | 0.3333 | 7 | 0.4286 | 3 | 0.3333 | 2 | 0.5 | 4 | 0.5 | 1 | 1 | 10 | 0.5 |
| 2_1680736 | 2 | 1 | 1 | 1 | NA | NA | 3 | 0.3333 | 6 | 0.6667 | 3 | 0.3333 | 4 | 0.5 | 4 | 0.5 | 1 | 0 | 12 | 0.4167 |
| 2_1680737 | 2 | 1 | 1 | 0 | 1 | 0 | 3 | 0 | 7 | 0.2857 | 1 | 1 | 3 | 0.6667 | 5 | 0.6 | 1 | 0 | 10 | 0.6 |
| 2_1680737 | 2 | 1 | 1 | 1 | 1 | 0 | 4 | 0.5 | 8 | 0.625 | 2 | 0.5 | 2 | 0.5 | 4 | 0.5 | 1 | 1 | 9 | 0.5556 |
| 2_1680738 | 1 | 1 | 1 | 1 | NA | NA | 4 | 0.75 | 6 | 0.8333 | 2 | 1 | 3 | 1 | 4 | 1 | 1 | 1 | 10 | 1 |
| 2_1680738 | 1 | 1 | NA | NA | 1 | 0 | 4 | 0.25 | 6 | 0.3333 | 2 | 0 | 4 | 0.25 | 3 | 0 | NA | NA | 9 | 0.1111 |
| 2_1680738 | 2 | 1 | 1 | 1 | 1 | 1 | 4 | 1 | 8 | 1 | 3 | 1 | 4 | 1 | 4 | 1 | NA | NA | 11 | 1 |
| 2_1680738 | 2 | 1 | 1 | 1 | 1 | 1 | 4 | 0.75 | 8 | 0.875 | 2 | 1 | 4 | 1 | 5 | 1 | 1 | 1 | 12 | 1 |
| 2_1680739 | 2 | 1 | 1 | 1 | 1 | 1 | 4 | 1 | 8 | 1 | 3 | 1 | 3 | 1 | 5 | 1 | 1 | 1 | 12 | 1 |
| 2_1680739 | 2 | 1 | NA | NA | 1 | 1 | 4 | 0.25 | 7 | 0.5714 | 3 | 0.6667 | 4 | 1 | 5 | 1 | 1 | 1 | 13 | 0.9231 |
| 2_1680741 | 2 | 1 | 1 | 1 | 2 | 0.5 | 3 | 1 | 8 | 0.875 | 2 | 0.5 | 5 | 0.2 | 5 | 0.8 | 1 | 1 | 13 | 0.5385 |
| 2_1680741 | 1 | 0 | 1 | 1 | 2 | 0.5 | 3 | 0 | 7 | 0.2857 | 3 | 0.3333 | 4 | 0.25 | 6 | 0.1667 | 1 | 0 | 14 | 0.2143 |
| 2_1680743 | NA | NA | 1 | 1 | 1 | 1 | 2 | 0 | 4 | 0.5 | 1 | 1 | 3 | 0.3333 | 5 | 0 | 1 | 0 | 10 | 0.2 |
| 2_1680744 | 1 | 0 | 1 | 1 | 1 | 1 | 1 | 1 | 4 | 0.75 | 1 | 1 | 1 | 1 | 4 | 1 | 1 | 1 | 7 | 1 |
| 2_1681021 | 6 | 0.1667 | 5 | 0.2 | 4 | 0.25 | 2 | 0.5 | 17 | 0.2353 | 5 | 0.2 | 4 | 0 | 4 | 0 | 8 | 0.125 | 21 | 0.0952 |
| 2_1681022 | 5 | 0 | 6 | 0.3333 | 4 | 0 | 2 | 0 | 17 | 0.1176 | 5 | 0 | 4 | 0 | 4 | 0 | 6 | 0.1667 | 19 | 0.0526 |
| 2_1681022 | 4 | 1 | 5 | 1 | 1 | 1 | 2 | 1 | 12 | 1 | 4 | 1 | 3 | 1 | 5 | 0.8 | 5 | 1 | 17 | 0.9412 |
| 2_1681022 | 3 | 0.6667 | 3 | 0.6667 | 4 | 0.25 | 1 | 0 | 11 | 0.4545 | 5 | 0.2 | 1 | 0 | 4 | 0.25 | 6 | 0.3333 | 16 | 0.25 |
| 2_1681023 | 5 | 0.6 | 4 | 0.25 | 4 | 0 | 2 | 0.5 | 15 | 0.3333 | 4 | 0.5 | 3 | 0 | 3 | 0 | 7 | 0.2857 | 17 | 0.2353 |
| 2_1681023 | 5 | 0 | 6 | 0.3333 | 4 | 0.25 | 2 | 0.5 | 17 | 0.2353 | 3 | 0 | 3 | 0.3333 | 5 | 0 | 7 | 0 | 18 | 0.0556 |
| 2_1681024 | 5 | 0.8 | 6 | 0.8333 | 4 | 1 | 2 | 1 | 17 | 0.8824 | 4 | 1 | 4 | 0.75 | 5 | 1 | 7 | 0.8571 | 20 | 0.9 |
| 2_1681025 | 5 | 0.2 | 5 | 0.2 | 4 | 0 | 2 | 0 | 16 | 0.125 | 4 | 0 | 4 | 0 | 4 | 0.25 | 6 | 0 | 18 | 0.0556 |
| 2_1681026 | 5 | 0.2 | 6 | 0.1667 | 4 | 0 | 2 | 0.5 | 17 | 0.1765 | 4 | 0 | 3 | 0.3333 | 4 | 0 | 5 | 0 | 16 | 0.0625 |
| 2_1681036 | 3 | 0.3333 | 2 | 0 | 2 | 0.5 | 2 | 0 | 9 | 0.2222 | 4 | 0.25 | 3 | 0.3333 | 4 | 0 | 4 | 0.25 | 15 | 0.2 |
| 2_1681037 | 2 | 0.5 | 2 | 0 | 2 | 0 | 2 | 0 | 8 | 0.125 | 2 | 0 | 2 | 0.5 | 4 | 0.25 | 4 | 0 | 12 | 0.1667 |
| 2_1681038 | 2 | 0.5 | 2 | 0.5 | 1 | 0 | 2 | 0 | 7 | 0.2857 | 4 | 0.5 | 2 | 0.5 | 4 | 0.5 | 2 | 0.5 | 12 | 0.5 |
| 2_1681038 | 3 | 0.6667 | 1 | 1 | 1 | 1 | 2 | 0 | 7 | 0.5714 | 3 | 0.6667 | 3 | 0.6667 | 4 | 0.5 | 4 | 0.75 | 14 | 0.6429 |
| 2_1681038 | 2 | 1 | 1 | 0 | 2 | 1 | 2 | 1 | 7 | 0.8571 | 2 | 1 | 3 | 1 | 4 | 0.75 | 2 | 1 | 11 | 0.9091 |
| 2_1681039 | 3 | 1 | 2 | 1 | 2 | 1 | 2 | 1 | 9 | 1 | 3 | 1 | 3 | 1 | 4 | 1 | 4 | 1 | 14 | 1 |
| 2_1681039 | 2 | 0.5 | 2 | 0 | 1 | 0 | 1 | 1 | 6 | 0.3333 | 2 | 1 | 3 | 0.6667 | 4 | 0.75 | 3 | 1 | 12 | 0.8333 |
| 2_1681039 | 3 | 0 | NA | NA | 2 | 0.5 | 2 | 0 | 7 | 0.1429 | 3 | 0.3333 | 2 | 0.5 | 4 | 0.5 | 3 | 0 | 12 | 0.3333 |
| 2_1681041 | 3 | 0.6667 | 1 | 1 | 1 | 1 | 2 | 1 | 7 | 0.8571 | 3 | 1 | 3 | 1 | 2 | 1 | 4 | 1 | 12 | 1 |
| 2_1681042 | 2 | 1 | 1 | 1 | 1 | 1 | 2 | 1 | 6 | 1 | 3 | 1 | 2 | 1 | 4 | 1 | 3 | 1 | 12 | 1 |
| 2_1681043 | 1 | 1 | 1 | 0 | 2 | 0.5 | 1 | 0 | 5 | 0.4 | 3 | 1 | NA | NA | 4 | 0.75 | 3 | 0.6667 | 10 | 0.8 |
| 2_1681043 | 3 | 0 | 1 | 1 | 2 | 0 | 2 | 0.5 | 8 | 0.25 | 3 | 0.3333 | 3 | 0.3333 | 4 | 0.25 | 3 | 0 | 13 | 0.2308 |
| 2_1681043 | 3 | 0.3333 | 1 | 0 | 2 | 0 | 1 | 0 | 7 | 0.1429 | 3 | 0.3333 | 3 | 0 | 4 | 0.5 | 3 | 0.3333 | 13 | 0.3077 |
| 2_1681043 | 3 | 1 | 1 | 1 | 1 | 1 | 2 | 1 | 7 | 1 | 2 | 1 | 3 | 1 | 4 | 0.5 | 4 | 1 | 13 | 0.8462 |
| 2_1681047 | 2 | 0 | 1 | 0 | 1 | 1 | 2 | 0.5 | 6 | 0.3333 | 2 | 0.5 | 3 | 0.6667 | 4 | 0.5 | 4 | 0.25 | 13 | 0.4615 |
| 2_1681048 | 2 | 0 | 1 | 0 | 1 | 1 | 2 | 0.5 | 6 | 0.3333 | 2 | 1 | 3 | 0.3333 | 4 | 0.75 | 4 | 1 | 13 | 0.7692 |
| 2_1681242 | NA | NA | NA | NA | NA | NA | 1 | 1 | 1 | 1 | NA | NA | NA | NA | NA | NA | 1 | 1 | 1 | 1 |
| 2_1681243 | NA | NA | NA | NA | NA | NA | 1 | 1 | 1 | 1 | NA | NA | NA | NA | NA | NA | 1 | 1 | 1 | 1 |
| 2_1681243 | NA | NA | NA | NA | NA | NA | 2 | 0.5 | 2 | 0.5 | NA | NA | NA | NA | NA | NA | 1 | 1 | 1 | 1 |
| 2_1681249 | NA | NA | NA | NA | NA | NA | 2 | 0.5 | 2 | 0.5 | NA | NA | NA | NA | NA | NA | 1 | 1 | 1 | 1 |
| 2_1683186 | 3 | 1 | 1 | 1 | 3 | 1 | NA | NA | 7 | 1 | NA | NA | 2 | 1 | 1 | 1 | 1 | 1 | 4 | 1 |
| 2_1683187 | 3 | 0.6667 | 1 | 1 | 2 | 1 | NA | NA | 6 | 0.8333 | NA | NA | 2 | 1 | 1 | 1 | NA | NA | 3 | 1 |
| 2_1683187 | 2 | 0.5 | 1 | 0 | 3 | 0.3333 | NA | NA | 6 | 0.3333 | NA | NA | 2 | 0 | 1 | 1 | NA | NA | 3 | 0.3333 |
| 2_1683188 | 3 | 0 | 1 | 0 | 3 | 0.3333 | NA | NA | 7 | 0.1429 | NA | NA | 2 | 0.5 | NA | NA | NA | NA | 2 | 0.5 |
| 2_1683189 | 2 | 1 | NA | NA | 2 | 1 | NA | NA | 4 | 1 | NA | NA | 2 | 1 | NA | NA | NA | NA | 2 | 1 |
| 2_1683189 | 2 | 0.5 | NA | NA | 2 | 0 | NA | NA | 4 | 0.25 | NA | NA | 1 | 1 | 1 | 0 | NA | NA | 2 | 0.5 |
| 2_1683190 | 1 | 1 | NA | NA | 3 | 1 | NA | NA | 4 | 1 | NA | NA | 2 | 1 | 1 | 1 | NA | NA | 3 | 1 |
| 2_1683190 | 1 | 0 | NA | NA | 3 | 0.3333 | NA | NA | 4 | 0.25 | NA | NA | 1 | 1 | 1 | 1 | NA | NA | 2 | 1 |
| 2_1683194 | 2 | 0 | NA | NA | 3 | 0.3333 | NA | NA | 5 | 0.2 | NA | NA | 2 | 0.5 | 1 | 1 | NA | NA | 3 | 0.6667 |
| 2_1683194 | 2 | 0.5 | NA | NA | 3 | 1 | NA | NA | 5 | 0.8 | NA | NA | 2 | 1 | 1 | 1 | NA | NA | 3 | 1 |
| 2_1683195 | 2 | 0.5 | NA | NA | 3 | 1 | NA | NA | 5 | 0.8 | NA | NA | 2 | 1 | 1 | 1 | NA | NA | 3 | 1 |
| 2_1683196 | 1 | 1 | NA | NA | 2 | 0.5 | NA | NA | 3 | 0.6667 | NA | NA | 1 | 1 | NA | NA | NA | NA | 1 | 1 |
| 2_1683206 | 2 | 0.5 | NA | NA | 2 | 0 | NA | NA | 4 | 0.25 | NA | NA | 2 | 0.5 | 1 | 0 | NA | NA | 3 | 0.3333 |
| 2_1683208 | 2 | 0 | NA | NA | 2 | 0.5 | NA | NA | 4 | 0.25 | NA | NA | 1 | 1 | 1 | 1 | NA | NA | 2 | 1 |
| 2_1683208 | 1 | 1 | NA | NA | 1 | 1 | NA | NA | 2 | 1 | NA | NA | 1 | 1 | 1 | 1 | NA | NA | 2 | 1 |
| 2_1683208 | 2 | 0.5 | NA | NA | 2 | 0 | NA | NA | 4 | 0.25 | NA | NA | 1 | 0 | 1 | 1 | NA | NA | 2 | 0.5 |
| 2_1683208 | 2 | 1 | NA | NA | 2 | 1 | NA | NA | 4 | 1 | NA | NA | 1 | 1 | 1 | 1 | NA | NA | 2 | 1 |
| 2_1683210 | 2 | 0 | NA | NA | 2 | 1 | NA | NA | 4 | 0.5 | NA | NA | 1 | 1 | 1 | 1 | NA | NA | 2 | 1 |
| 2_1683210 | 2 | 1 | NA | NA | 1 | 1 | NA | NA | 3 | 1 | NA | NA | 1 | 1 | 1 | 1 | NA | NA | 2 | 1 |

| | | | | | | | | | | | | | | | | | | | | |
|---|---|---|---|---|---|---|---|---|---|---|---|---|---|---|---|---|---|---|---|---|
| 2_1683213 | 1 | 1 | NA | NA | 1 | 1 | NA | NA | 2 | 1 | NA | NA | 1 | 1 | 1 | 1 | NA | NA | 2 | 1 |
| 2_1683213 | 2 | 0.5 | NA | NA | 1 | 1 | NA | NA | 3 | 0.6667 | NA | NA | 1 | 1 | NA | NA | NA | NA | 1 | 1 |
| 2_1683214 | 1 | 0 | NA | NA | 1 | 1 | NA | NA | 2 | 0.5 | NA | NA | 1 | 1 | NA | NA | NA | NA | 1 | 1 |
| 2_1683214 | 2 | 1 | NA | NA | 1 | 1 | NA | NA | 3 | 1 | NA | NA | 1 | 1 | NA | NA | NA | NA | 1 | 1 |
| 2_1683217 | 2 | 0 | NA | NA | 1 | 1 | NA | NA | 3 | 0.3333 | NA | NA | 1 | 1 | NA | NA | NA | NA | 1 | 1 |
| 2_1683218 | 2 | 0 | NA | NA | 1 | 1 | NA | NA | 3 | 0.3333 | NA | NA | 1 | 1 | NA | NA | NA | NA | 1 | 1 |
| 2_1683604 | NA | NA | 1 | 1 | 1 | 0 | NA | NA | 2 | 0.5 | 1 | 1 | NA | NA | NA | NA | NA | NA | 1 | 1 |
| 2_1683606 | NA | NA | 1 | 1 | 1 | 1 | NA | NA | 2 | 1 | 1 | 1 | NA | NA | NA | NA | NA | NA | 1 | 1 |
| 2_1683608 | NA | NA | 1 | 1 | NA | NA | NA | NA | 1 | 1 | 1 | 1 | NA | NA | NA | NA | NA | NA | 1 | 1 |
| 2_1683610 | 1 | 1 | 2 | 1 | 1 | 1 | 2 | 1 | 6 | 1 | 3 | 1 | 1 | 1 | 1 | 1 | NA | NA | 5 | 1 |
| 2_1683613 | 1 | 0 | 2 | 0.5 | NA | NA | 3 | 0 | 6 | 0.1667 | 3 | 0.3333 | NA | NA | 2 | 0.5 | NA | NA | 5 | 0.4 |
| 2_1684764 | 43 | 0.7209 | 47 | 0.7021 | 38 | 0.6053 | 51 | 0.8039 | 179 | 0.7151 | 48 | 0.7917 | 48 | 0.7708 | 52 | 0.6923 | 38 | 0.8158 | 186 | 0.7634 |
| 2_1684765 | 43 | 0.0698 | 36 | 0.0556 | 32 | 0.0625 | 46 | 0.0217 | 157 | 0.051 | 42 | 0.0714 | 40 | 0.025 | 43 | 0.1163 | 35 | 0.0571 | 160 | 0.0688 |
| 2_1684765 | 43 | 0.3721 | 44 | 0.5 | 35 | 0.5714 | 49 | 0.3469 | 171 | 0.4386 | 45 | 0.4222 | 43 | 0.3721 | 46 | 0.5435 | 37 | 0.5135 | 171 | 0.462 |
| 2_1685836 | 5 | 0 | 4 | 0.25 | 4 | 0 | 3 | 0 | 16 | 0.0625 | 4 | 0 | 4 | 0 | 8 | 0.125 | 4 | 0 | 20 | 0.05 |
| 2_1687354 | NA | NA | 1 | 1 | NA | NA | NA | NA | 1 | 1 | NA | NA | 2 | 1 | NA | NA | NA | NA | 2 | 1 |
| 2_1687354 | NA | NA | 1 | 1 | NA | NA | NA | NA | 1 | 1 | NA | NA | 1 | 1 | NA | NA | NA | NA | 1 | 1 |
| 2_1687355 | NA | NA | 1 | 1 | NA | NA | NA | NA | 1 | 1 | NA | NA | 2 | 0.5 | NA | NA | NA | NA | 2 | 0.5 |
| 2_1687356 | NA | NA | 1 | 1 | NA | NA | NA | NA | 1 | 1 | NA | NA | 1 | 1 | NA | NA | NA | NA | 1 | 1 |
| 2_1687358 | NA | NA | 1 | 1 | NA | NA | NA | NA | 1 | 1 | NA | NA | 1 | 1 | NA | NA | NA | NA | 1 | 1 |
| 2_1687358 | NA | NA | 1 | 1 | NA | NA | NA | NA | 1 | 1 | NA | NA | 1 | 1 | NA | NA | NA | NA | 1 | 1 |
| 2_1687358 | NA | NA | 1 | 1 | NA | NA | NA | NA | 1 | 1 | NA | NA | 1 | 1 | NA | NA | NA | NA | 1 | 1 |
| 2_1687360 | NA | NA | 1 | 1 | NA | NA | NA | NA | 1 | 1 | NA | NA | 1 | 1 | NA | NA | NA | NA | 1 | 1 |
| 2_1687673 | 1 | 1 | 1 | 0 | 3 | 1 | 1 | 1 | 6 | 0.8333 | 1 | 1 | 2 | 0.5 | 2 | 1 | 2 | 1 | 7 | 0.8571 |
| 2_1687675 | NA | NA | NA | NA | 4 | 0.5 | 1 | 1 | 5 | 0.6 | 1 | 0 | 3 | 0.6667 | 1 | 1 | 2 | 0 | 7 | 0.4286 |
| 2_1687675 | 1 | 0 | 1 | 0 | 4 | 0.25 | 1 | 0 | 7 | 0.1429 | 1 | 0 | 4 | 0 | 2 | 0 | 2 | 0.5 | 9 | 0.1111 |
| 2_1687676 | NA | NA | NA | NA | 4 | 0.25 | NA | NA | 4 | 0.25 | 1 | 0 | 3 | 0.3333 | 1 | 0 | 1 | 1 | 6 | 0.3333 |
| 2_1687676 | 1 | 1 | NA | NA | 3 | 0.3333 | NA | NA | 4 | 0.5 | 1 | 0 | 3 | 1 | 2 | 0 | 1 | 0 | 7 | 0.4286 |
| 2_1687677 | 1 | 1 | 1 | 0 | 2 | 1 | NA | NA | 4 | 0.75 | 1 | 1 | 4 | 0 | 2 | 0 | 2 | 0 | 9 | 0.1111 |
| 2_1687677 | 1 | 1 | 1 | 1 | 2 | 1 | NA | NA | 4 | 1 | NA | NA | 3 | 1 | 2 | 1 | 2 | 1 | 7 | 1 |
| 2_1687677 | 1 | 1 | 1 | 1 | 3 | 1 | NA | NA | 5 | 1 | 1 | 1 | 4 | 0.25 | 2 | 0.5 | 2 | 1 | 9 | 0.5556 |
| 2_1687678 | NA | NA | 1 | 1 | 3 | 0.6667 | NA | NA | 4 | 0.75 | 1 | 0 | 3 | 0.6667 | 1 | 0 | 2 | 1 | 7 | 0.5714 |
| 2_1687678 | 1 | 0 | 1 | 0 | 2 | 0.5 | NA | NA | 4 | 0.25 | 1 | 1 | 4 | 0.25 | 2 | 0 | 2 | 0 | 9 | 0.2222 |
| 2_1687682 | NA | NA | 1 | 1 | 3 | 0 | NA | NA | 4 | 0.25 | 1 | 0 | 4 | 0.5 | 1 | 0 | 2 | 0.5 | 8 | 0.375 |
| 2_1687683 | 1 | 1 | NA | NA | 3 | 0.3333 | NA | NA | 4 | 0.5 | 1 | 1 | 4 | 0.5 | 1 | 0 | 2 | 0.5 | 8 | 0.5 |
| 2_1687687 | 1 | 1 | NA | NA | 3 | 1 | NA | NA | 4 | 1 | 1 | 1 | 3 | 1 | 1 | 1 | 1 | 1 | 6 | 1 |
| 2_1697280 | NA | NA | 1 | 1 | NA | NA | 1 | 0 | 2 | 0.5 | 1 | 1 | 1 | 1 | 2 | 1 | NA | NA | 4 | 1 |
| 2_1697281 | NA | NA | 1 | 1 | NA | NA | 1 | 0 | 2 | 0.5 | 1 | 0 | 2 | 0 | 3 | 0.3333 | NA | NA | 6 | 0.1667 |
| 2_1697281 | NA | NA | 1 | 1 | NA | NA | 1 | 1 | 2 | 1 | 1 | 1 | 2 | 1 | 2 | 1 | NA | NA | 5 | 1 |
| 2_1697281 | NA | NA | NA | NA | NA | NA | 1 | 1 | 1 | 1 | 1 | 1 | 2 | 1 | 2 | 1 | NA | NA | 5 | 1 |
| 2_1697282 | NA | NA | 1 | 0 | NA | NA | 1 | 1 | 2 | 0.5 | 1 | 0 | 1 | 1 | 2 | 1 | NA | NA | 4 | 0.75 |
| 2_1697283 | NA | NA | 1 | 1 | NA | NA | 1 | 0 | 2 | 0.5 | 1 | 1 | 2 | 1 | 2 | 1 | NA | NA | 5 | 1 |
| 2_1697283 | NA | NA | 1 | 1 | NA | NA | 1 | 1 | 2 | 1 | 1 | 1 | 2 | 1 | 3 | 1 | NA | NA | 6 | 1 |
| 2_1697287 | NA | NA | 1 | 1 | NA | NA | 1 | 0 | 2 | 0.5 | 1 | 0 | 2 | 0.5 | 3 | 1 | NA | NA | 6 | 0.6667 |
| 2_1697288 | NA | NA | 1 | 1 | NA | NA | 1 | 0 | 2 | 0.5 | 1 | 1 | 1 | 0 | 3 | 0.6667 | NA | NA | 5 | 0.6 |
| 2_1697294 | NA | NA | 1 | 0 | NA | NA | 1 | 1 | 2 | 0.5 | 1 | 1 | 2 | 0 | 2 | 0 | 1 | 0 | 6 | 0.1667 |
| 2_1697300 | 1 | 1 | 1 | 0 | NA | NA | 1 | 0 | 3 | 0.3333 | 1 | 1 | 2 | 0.5 | 2 | 0.5 | 1 | 0 | 6 | 0.5 |
| 2_1699146 | NA | NA | 1 | 1 | NA | NA | NA | NA | 1 | 1 | NA | NA | 1 | 1 | NA | NA | NA | NA | 1 | 1 |
| 2_1701833 | NA | NA | NA | NA | NA | NA | 1 | 1 | 1 | 1 | NA | NA | 1 | 1 | 1 | 1 | NA | NA | 2 | 1 |
| 2_1701836 | NA | NA | NA | NA | NA | NA | 1 | 1 | 1 | 1 | NA | NA | 1 | 1 | 1 | 0 | NA | NA | 2 | 0.5 |
| 2_1701890 | 2 | 0 | 1 | 0 | 4 | 0.25 | NA | NA | 7 | 0.1429 | 3 | 0.6667 | 2 | 1 | 2 | 0 | 3 | 0.3333 | 10 | 0.5 |
| 2_1701891 | 19 | 0 | 28 | 0.1429 | 21 | 0.0476 | 9 | 0.2222 | 77 | 0.0909 | 29 | 0.1034 | 14 | 0.1429 | 11 | 0.1818 | 15 | 0 | 69 | 0.1014 |
| 2_1701895 | 15 | 0.0667 | 21 | 0.1429 | 17 | 0.2941 | 9 | 0.1111 | 62 | 0.1613 | 26 | 0.1923 | 10 | 0.3 | 8 | 0.125 | 8 | 0.25 | 52 | 0.2115 |
| 2_1701900 | 16 | 0.125 | 22 | 0 | 13 | 0.0769 | 8 | 0 | 59 | 0.0508 | 24 | 0.2083 | 11 | 0.0909 | 8 | 0.125 | 8 | 0 | 51 | 0.1373 |
| 2_1704215 | 1 | 1 | NA | NA | NA | NA | NA | NA | 1 | 1 | 1 | 1 | 1 | 1 | NA | NA | NA | NA | 2 | 1 |
| 2_1704219 | 1 | 1 | NA | NA | NA | NA | NA | NA | 1 | 1 | 1 | 1 | 1 | 1 | NA | NA | NA | NA | 2 | 1 |
| 2_1707215 | 2 | 0.5 | 2 | 0 | 2 | 0 | 1 | 0 | 7 | 0.1429 | 1 | 1 | 3 | 0.3333 | 3 | 0 | 3 | 0 | 10 | 0.2 |
| 2_1707215 | 2 | 0.5 | 3 | 0.6667 | 1 | 1 | 2 | 0.5 | 8 | 0.625 | 3 | 1 | 2 | 1 | 5 | 0.8 | 3 | 1 | 13 | 0.9231 |
| 2_1707216 | 2 | 0 | 1 | 0 | 3 | 0.3333 | 1 | 1 | 7 | 0.2857 | 2 | 0 | 2 | 0 | 5 | 0.4 | 2 | 0 | 11 | 0.1818 |
| 2_1718788 | NA | NA | 2 | 0.5 | NA | NA | 1 | 0 | 3 | 0.3333 | NA | NA | 2 | 0.5 | 2 | 0 | 1 | 0 | 5 | 0.2 |
| 2_1718790 | 1 | 1 | 3 | 1 | NA | NA | 1 | 1 | 5 | 1 | NA | NA | 2 | 1 | 2 | 1 | 1 | 1 | 5 | 1 |
| 2_1718790 | 1 | 1 | 3 | 1 | NA | NA | 1 | 1 | 5 | 1 | NA | NA | 2 | 1 | 1 | 1 | 1 | 1 | 4 | 1 |
| 2_1718791 | 1 | 1 | 3 | 1 | NA | NA | 1 | 1 | 5 | 1 | NA | NA | 1 | 1 | 1 | 1 | 1 | 1 | 3 | 1 |
| 2_1718844 | NA | NA | NA | NA | NA | NA | 1 | 1 | 1 | 1 | NA | NA | 1 | 1 | 1 | 1 | NA | NA | 2 | 1 |
| 2_1718845 | NA | NA | NA | NA | NA | NA | 1 | 1 | 1 | 1 | NA | NA | 1 | 1 | 1 | 0 | NA | NA | 2 | 0.5 |

| | | | | | | | | | | | | | | | | | | | | |
|---|---|---|---|---|---|---|---|---|---|---|---|---|---|---|---|---|---|---|---|---|
| 2_1718845 | NA | NA | NA | NA | NA | NA | 1 | 1 | 1 | 1 | NA | NA | 2 | 1 | NA | NA | NA | NA | 2 | 1 |
| 2_1718847 | NA | NA | NA | NA | NA | NA | 1 | 1 | 1 | 1 | NA | NA | 2 | 1 | 1 | 1 | NA | NA | 3 | 1 |
| 2_1721946 | 3 | 1 | 2 | 1 | 1 | 1 | 1 | 1 | 7 | 1 | 1 | 1 | 1 | 1 | 4 | 1 | 2 | 1 | 8 | 1 |
| 2_1721950 | 3 | 0.3333 | 4 | 0.5 | 1 | 0 | 1 | 0 | 9 | 0.3333 | 2 | 0.5 | 1 | 0 | 4 | 0.5 | 4 | 0.25 | 11 | 0.3636 |
| 2_1721951 | 2 | 0 | 3 | 1 | 1 | 0 | 1 | 0 | 7 | 0.4286 | 3 | 1 | 1 | 0 | 3 | 0.3333 | 2 | 0 | 9 | 0.4444 |
| 2_1721954 | 3 | 0 | 5 | 0.2 | 1 | 0 | 2 | 0 | 11 | 0.0909 | 1 | 0 | NA | NA | 3 | 0 | 4 | 0.25 | 8 | 0.125 |
| 2_1721956 | 2 | 0.5 | 5 | 0 | 1 | 0 | 3 | 0 | 11 | 0.0909 | 2 | 0 | 1 | 0 | 2 | 0 | 5 | 0.4 | 10 | 0.2 |
| 2_1721961 | 2 | 0.5 | 3 | 0.3333 | NA | NA | 3 | 0.3333 | 8 | 0.375 | NA | NA | NA | NA | 3 | 0.6667 | 1 | 0 | 4 | 0.5 |
| 2_1721963 | 2 | 0.5 | 4 | 0 | 1 | 0 | 1 | 0 | 8 | 0.125 | NA | NA | NA | NA | 1 | 1 | 2 | 1 | 3 | 1 |
| 2_1721969 | 2 | 0.5 | 4 | 0.25 | 2 | 0 | 3 | 0 | 11 | 0.1818 | NA | NA | NA | NA | 4 | 0.5 | 2 | 0 | 6 | 0.3333 |
| 2_1722177 | 3 | 1 | 3 | 0 | 1 | 0 | 2 | 0 | 9 | 0.3333 | 2 | 0 | 1 | 0 | 2 | 0.5 | 3 | 0.3333 | 8 | 0.25 |
| 2_1722179 | 4 | 0.25 | 4 | 0 | 1 | 0 | 3 | 0 | 12 | 0.0833 | 2 | 0.5 | 1 | 0 | 3 | 0 | 3 | 0 | 9 | 0.1111 |
| 2_1722180 | 4 | 0.5 | 3 | 0 | NA | NA | 3 | 0.3333 | 10 | 0.3 | 1 | 0 | 1 | 0 | 3 | 0.3333 | 2 | 1 | 7 | 0.4286 |
| 2_1722198 | 3 | 0.3333 | 4 | 0.75 | 1 | 1 | 4 | 0.75 | 12 | 0.6667 | 3 | 0.6667 | 1 | 1 | 3 | 0.3333 | 3 | 1 | 10 | 0.7 |
| 2_1722223 | 4 | 0.75 | 3 | 0.6667 | 2 | 1 | 3 | 0.6667 | 12 | 0.75 | 3 | 0.6667 | 1 | 0 | 2 | 1 | 2 | 0 | 8 | 0.5 |
| 2_1722223 | 3 | 0.6667 | 2 | 0 | 1 | 0 | 4 | 0 | 10 | 0.2 | 2 | 0 | 1 | 0 | 1 | 0 | 3 | 0.3333 | 7 | 0.1429 |
| 2_1722223 | 4 | 0.75 | 5 | 0 | 1 | 1 | 3 | 0.6667 | 13 | 0.4615 | 3 | 1 | 1 | 0 | 2 | 0 | 4 | 0.5 | 10 | 0.5 |
| 2_1722225 | 4 | 1 | 3 | 1 | 1 | 0 | 4 | 1 | 12 | 0.9167 | 1 | 1 | 1 | 1 | 3 | 1 | 3 | 1 | 8 | 1 |
| 2_1722225 | 1 | 1 | 2 | 0.5 | 1 | 1 | 1 | 1 | 5 | 0.8 | 2 | 0.5 | NA | NA | 1 | 1 | 1 | 1 | 4 | 0.75 |
| 2_1722225 | 4 | 1 | 4 | 1 | 2 | 1 | 4 | 1 | 14 | 1 | 4 | 1 | 1 | 1 | 4 | 0.75 | 4 | 0.75 | 13 | 0.8462 |
| 2_1722227 | 3 | 1 | 4 | 1 | 3 | 1 | 5 | 1 | 15 | 1 | 5 | 1 | 3 | 1 | 4 | 0.75 | 4 | 1 | 16 | 0.9375 |
| 2_1722238 | 1 | 0 | NA | NA | 1 | 1 | NA | NA | 2 | 0.5 | 2 | 0 | NA | NA | 3 | 0.3333 | 1 | 0 | 6 | 0.1667 |
| 2_1723109 | 3 | 0.3333 | 2 | 0.5 | 6 | 0 | 4 | 0 | 15 | 0.1333 | 4 | 0 | 8 | 0.125 | 4 | 0 | 1 | 0 | 17 | 0.0588 |
| 2_1723109 | 2 | 0.5 | 2 | 0.5 | 5 | 0.4 | 6 | 0.6667 | 15 | 0.5333 | 5 | 0.2 | 8 | 0.5 | 2 | 0 | NA | NA | 15 | 0.3333 |
| 2_1723109 | 3 | 0.3333 | 2 | 0.5 | 6 | 0.1667 | 6 | 0 | 17 | 0.1765 | 5 | 0 | 9 | 0 | 4 | 0.5 | 1 | 0 | 19 | 0.1053 |
| 2_1723110 | 3 | 0.6667 | 2 | 1 | 6 | 0.1667 | 5 | 0.4 | 16 | 0.4375 | 5 | 0.2 | 8 | 0.125 | 4 | 0.5 | 1 | 0 | 18 | 0.2222 |
| 2_1723112 | 3 | 0.6667 | 2 | 1 | 3 | 0.3333 | 5 | 0.4 | 13 | 0.5385 | 5 | 0.2 | 7 | 0.2857 | 4 | 0.5 | 2 | 0.5 | 18 | 0.3333 |
| 2_1723117 | 3 | 0.6667 | 2 | 0 | 5 | 0.2 | 6 | 0.3333 | 16 | 0.3125 | 7 | 0.5714 | 8 | 0.125 | 5 | 0.6 | 3 | 0 | 23 | 0.3478 |
| 2_1727117 | 3 | 0.6667 | 2 | 0.5 | NA | NA | 1 | 1 | 6 | 0.6667 | 2 | 0.5 | NA | NA | 3 | 0.6667 | 4 | 0.25 | 9 | 0.4444 |
| 2_1727118 | 3 | 0.3333 | 2 | 0 | NA | NA | 1 | 0 | 6 | 0.1667 | 2 | 0 | NA | NA | 2 | 0 | 3 | 0.3333 | 7 | 0.1429 |
| 2_1727815 | 1 | 0 | 2 | 0.5 | NA | NA | 1 | 0 | 4 | 0.25 | NA | NA | NA | NA | 3 | 0.3333 | 2 | 1 | 5 | 0.6 |
| 2_1727816 | 2 | 0 | 2 | 0.5 | NA | NA | 1 | 0 | 5 | 0.2 | NA | NA | NA | NA | 3 | 0.3333 | 2 | 0 | 5 | 0.2 |
| 2_1727820 | 1 | 1 | 2 | 0.5 | NA | NA | 1 | 0 | 4 | 0.5 | NA | NA | NA | NA | 2 | 0 | 2 | 0.5 | 4 | 0.25 |
| 2_1727821 | 1 | 1 | NA | NA | NA | NA | 1 | 1 | 2 | 1 | NA | NA | NA | NA | 1 | 1 | 1 | 1 | 2 | 1 |
| 2_1728569 | 1 | 1 | NA | NA | NA | NA | NA | NA | 1 | 1 | NA | NA | NA | NA | 2 | 0.5 | NA | NA | 2 | 0.5 |
| 2_1728571 | 6 | 1 | 3 | 1 | 2 | 1 | 4 | 1 | 15 | 1 | 2 | 1 | 3 | 1 | 4 | 1 | 6 | 1 | 15 | 1 |
| 2_1728572 | 6 | 0.8333 | 3 | 1 | 3 | 0.6667 | 4 | 0.5 | 16 | 0.75 | 1 | 1 | 6 | 0.8333 | 3 | 0.6667 | 8 | 0.75 | 18 | 0.7778 |
| 2_1728573 | 7 | 0.1429 | 3 | 0 | 3 | 0 | 4 | 0 | 17 | 0.0588 | 2 | 0 | 7 | 0 | 3 | 0.3333 | 8 | 0 | 20 | 0.05 |
| 2_1728757 | NA | NA | 1 | 1 | 1 | 1 | 2 | 1 | 4 | 1 | 1 | 1 | NA | NA | 1 | 0 | 1 | 1 | 3 | 0.6667 |
| 2_1728758 | NA | NA | 1 | 1 | NA | NA | 2 | 1 | 3 | 1 | NA | NA | NA | NA | 1 | 1 | 1 | 1 | 2 | 1 |
| 2_1728759 | NA | NA | 1 | 0 | NA | NA | 2 | 1 | 3 | 0.6667 | 1 | 1 | NA | NA | 1 | 0 | 1 | 1 | 3 | 0.6667 |
| 2_1730665 | 8 | 0.625 | 8 | 0.75 | 3 | 0.3333 | 5 | 0.6 | 24 | 0.625 | 6 | 0.8333 | 7 | 0.5714 | 7 | 0.2857 | 3 | 0.6667 | 23 | 0.5652 |
| 2_1730671 | 7 | 0 | 5 | 0 | 2 | 0 | 5 | 0.2 | 19 | 0.0526 | 4 | 0 | 5 | 0.2 | 6 | 0 | 3 | 0 | 18 | 0.0556 |
| 2_1730681 | 1 | 0 | 3 | 0 | 1 | 1 | 4 | 0 | 9 | 0.1111 | 1 | 0 | 1 | 1 | 5 | 0.4 | 3 | 0 | 10 | 0.3 |
| 2_1730686 | 12 | 0.1667 | 7 | 0 | 5 | 0.2 | 12 | 0.0833 | 36 | 0.1111 | 11 | 0 | 12 | 0 | 16 | 0.125 | 9 | 0.1111 | 48 | 0.0625 |
| 2_1730687 | 6 | 0.3333 | 7 | 0.1429 | 3 | 0.3333 | 13 | 0.1538 | 29 | 0.2069 | 7 | 0 | 8 | 0.125 | 13 | 0.0769 | 6 | 0.3333 | 34 | 0.1176 |
| 2_1730691 | 8 | 0 | 7 | 0 | 6 | 0.1667 | 13 | 0.0769 | 34 | 0.0588 | 8 | 0.125 | 11 | 0.0909 | 13 | 0.0769 | 8 | 0.125 | 40 | 0.1 |
| 2_1730692 | 7 | 0 | 9 | 0 | 4 | 0 | 10 | 0.3 | 30 | 0.1 | 10 | 0 | 11 | 0.0909 | 15 | 0.2 | 8 | 0.125 | 44 | 0.1136 |
| 2_1730695 | 4 | 0.25 | 6 | 0 | 3 | 0.3333 | 5 | 0 | 18 | 0.1111 | 7 | 0.4286 | 6 | 0.1667 | 8 | 0.125 | 4 | 0.25 | 25 | 0.24 |
| 2_1730695 | 9 | 0 | 9 | 0.1111 | 2 | 0 | 11 | 0.0909 | 31 | 0.0645 | 7 | 0.4286 | 12 | 0 | 11 | 0 | 8 | 0 | 38 | 0.0789 |
| 2_1730696 | 6 | 0.1667 | 8 | 0.25 | 2 | 0 | 6 | 0.1667 | 22 | 0.1818 | 6 | 0 | 13 | 0.1538 | 8 | 0 | 5 | 0 | 32 | 0.0625 |
| 2_1730696 | 5 | 0.2 | 6 | 0 | 2 | 0 | 8 | 0.375 | 21 | 0.1905 | 8 | 0.25 | 11 | 0.0909 | 9 | 0 | 5 | 0.2 | 33 | 0.1212 |
| 2_1730698 | 5 | 0 | 2 | 0 | 2 | 0 | 5 | 0.2 | 14 | 0.0714 | 5 | 0 | 12 | 0.0833 | 6 | 0.1667 | 2 | 0 | 25 | 0.08 |
| 2_1730698 | 3 | 1 | 1 | 1 | 1 | 1 | 4 | 0.75 | 9 | 0.8889 | 3 | 1 | 5 | 1 | 5 | 1 | 3 | 0.3333 | 16 | 0.875 |
| 2_1730698 | 7 | 0.1429 | 7 | 0.2857 | 2 | 0 | 7 | 0.1429 | 23 | 0.1739 | 8 | 0.375 | 13 | 0.3846 | 9 | 0.5556 | 9 | 0.4444 | 39 | 0.4359 |
| 2_1730698 | 8 | 0.5 | 7 | 0.2857 | 3 | 0 | 6 | 0 | 24 | 0.25 | 6 | 0.5 | 10 | 0.3 | 8 | 0.125 | 8 | 0.375 | 32 | 0.3125 |
| 2_1730699 | 6 | 0.3333 | 7 | 0.4286 | 3 | 0 | 6 | 0.5 | 22 | 0.3636 | 8 | 0.375 | 14 | 0.4286 | 7 | 0.5714 | 8 | 0.625 | 37 | 0.4865 |
| 2_1734253 | NA | NA | 1 | 1 | NA | NA | NA | NA | 1 | 1 | NA | NA | NA | NA | 1 | 1 | NA | NA | 1 | 1 |
| 2_1734274 | 1 | 0 | 1 | 0 | 1 | 1 | NA | NA | 3 | 0.3333 | NA | NA | 1 | 1 | 1 | 1 | NA | NA | 2 | 1 |
| 2_1734274 | 1 | 0 | 1 | 1 | NA | NA | NA | NA | 2 | 0.5 | NA | NA | NA | NA | 1 | 1 | NA | NA | 1 | 1 |
| 2_1734274 | 1 | 0 | 1 | 1 | 1 | 1 | NA | NA | 3 | 0.6667 | NA | NA | 1 | 1 | 1 | 0 | NA | NA | 2 | 0.5 |
| 2_1735430 | 5 | 1 | 4 | 1 | 4 | 1 | 4 | 1 | 17 | 1 | 4 | 1 | 3 | 0.6667 | 3 | 1 | 2 | 1 | 12 | 0.9167 |
| 2_1735430 | 5 | 0.4 | 4 | 0.75 | 5 | 0.8 | 5 | 0.4 | 19 | 0.5789 | 3 | 0.3333 | 3 | 0.6667 | 3 | 0.6667 | 3 | 0.6667 | 12 | 0.5833 |
| 2_1735430 | 5 | 1 | 3 | 0.6667 | 5 | 1 | 5 | 1 | 18 | 0.9444 | 4 | 1 | 2 | 1 | 3 | 1 | 4 | 1 | 13 | 1 |
| 2_1735431 | 5 | 0.2 | 3 | 0 | 4 | 0.5 | 5 | 0.4 | 17 | 0.2941 | 4 | 0.5 | 3 | 0.3333 | 2 | 0 | 3 | 0 | 12 | 0.25 |

| | | | | | | | | | | | | | | | | | | | | |
|---|---|---|---|---|---|---|---|---|---|---|---|---|---|---|---|---|---|---|---|---|
| 2_1735449( | 2 | 1 | NA | NA | 3 | 0.3333 | NA | NA | 5 | 0.6 | 2 | 1 | 1 | 0 | 1 | 1 | NA | NA | 4 | 0.75 |
| 2_1735485 | 2 | 0.5 | 1 | 1 | 5 | 0 | 5 | 0.2 | 13 | 0.2308 | 5 | 0.2 | 1 | 0 | 3 | 0.3333 | 1 | 0 | 10 | 0.2 |
| 2_1736206 | 10 | 0 | 13 | 0.1538 | 8 | 0 | 12 | 0.25 | 43 | 0.1163 | 4 | 0 | 5 | 0.4 | 9 | 0.1111 | 7 | 0 | 25 | 0.12 |
| 2_1736235 | NA | NA | 4 | 0.25 | 1 | 0 | 4 | 0.5 | 9 | 0.3333 | 1 | 1 | 3 | 0 | 3 | 0.6667 | 4 | 0.5 | 11 | 0.4545 |
| 2_1736237 | NA | NA | 4 | 0.5 | 1 | 0 | 4 | 0 | 9 | 0.2222 | 1 | 0 | 3 | 0.3333 | 2 | 0 | 4 | 0 | 10 | 0.1 |
| 2_1736243( | 1 | 0 | 3 | 0.3333 | 1 | 0 | 4 | 0.25 | 9 | 0.2222 | 1 | 0 | 5 | 0.4 | 2 | 0.5 | 4 | 0.25 | 12 | 0.3333 |
| 2_1736246 | 1 | 0 | 5 | 0.4 | 1 | 0 | 5 | 0.2 | 12 | 0.25 | 1 | 1 | 7 | 0.1429 | 3 | 0.3333 | 4 | 0.25 | 15 | 0.2667 |
| 2_1736247( | 1 | 0 | 5 | 0.6 | 1 | 0 | 4 | 0.25 | 11 | 0.3636 | 1 | 0 | 7 | 0 | 3 | 0 | 4 | 0.25 | 15 | 0.0667 |
| 2_1736257 | 4 | 0 | 4 | 0 | 2 | 0 | 5 | 0.2 | 15 | 0.0667 | 2 | 0 | 7 | 0.1429 | 3 | 0 | 4 | 0 | 16 | 0.0625 |
| 2_1736258( | 1 | 1 | 3 | 0 | 3 | 0.3333 | 4 | 0 | 11 | 0.1818 | 1 | 0 | 3 | 0 | 3 | 0 | 4 | 0.25 | 11 | 0.0909 |
| 2_1736258 | 4 | 0.5 | 4 | 0.5 | NA | NA | 5 | 0.2 | 13 | 0.3846 | 1 | 0 | 6 | 0.5 | 3 | 0.3333 | 2 | 0.5 | 12 | 0.4167 |
| 2_1739388( | 5 | 1 | 2 | 1 | 5 | 1 | 7 | 1 | 19 | 1 | 1 | 1 | 3 | 1 | 5 | 1 | 2 | 1 | 11 | 1 |
| 2_1739389 | 5 | 0 | 3 | 0.3333 | 4 | 0 | 7 | 0 | 19 | 0.0526 | 1 | 0 | 2 | 0.5 | 5 | 0.2 | 2 | 0 | 10 | 0.2 |
| 2_1739389 | 5 | 0.4 | 1 | 0 | 5 | 0.2 | 4 | 0.25 | 15 | 0.2667 | NA | NA | 2 | 0 | 5 | 0.4 | NA | NA | 7 | 0.2857 |
| 2_1739390 | 5 | 1 | 3 | 1 | 5 | 0.8 | 7 | 0.8571 | 20 | 0.9 | 1 | 1 | 3 | 0.6667 | 5 | 1 | 2 | 0.5 | 11 | 0.8182 |
| 2_1739390 | 4 | 0.5 | 3 | 0.6667 | 5 | 1 | 5 | 0.8 | 17 | 0.7647 | 1 | 1 | 2 | 0.5 | 5 | 0.8 | 2 | 0.5 | 10 | 0.7 |
| 2_1739396( | 5 | 0.4 | 3 | 0.3333 | 6 | 0.3333 | 8 | 0.25 | 22 | 0.3182 | NA | NA | 5 | 0.2 | 7 | 0.4286 | 3 | 0 | 15 | 0.2667 |
| 2_1739402( | 5 | 0.8 | 2 | 0.5 | 7 | 0.1429 | 4 | 0 | 18 | 0.3333 | 2 | 0 | 5 | 0.4 | 6 | 0.5 | 4 | 0 | 17 | 0.2941 |
| 2_1739403 | 7 | 0 | 5 | 0.4 | 6 | 0.1667 | 9 | 0.2222 | 27 | 0.1852 | 1 | 0 | 3 | 0.3333 | 8 | 0.125 | 5 | 0 | 17 | 0.1176 |
| 2_1739407( | 7 | 0.2857 | 3 | 0 | 5 | 0.2 | 7 | 0.1429 | 22 | 0.1818 | 2 | 0 | 6 | 0.1667 | 8 | 0.125 | 5 | 0.2 | 21 | 0.1429 |
| 2_1739409 | 10 | 0.6 | 5 | 0.2 | 9 | 0.6667 | 13 | 0.4615 | 37 | 0.5135 | 3 | 1 | 4 | 0.5 | 10 | 0.6 | 5 | 0.4 | 22 | 0.5909 |
| 2_1739410 | 10 | 0.3 | 5 | 0.2 | 9 | 0.5556 | 9 | 0.3333 | 33 | 0.3636 | 4 | 0.25 | 4 | 0.25 | 9 | 0.4444 | 6 | 0.3333 | 23 | 0.3478 |
| 2_1739410 | 10 | 0.1 | 5 | 0 | 8 | 0 | 12 | 0.0833 | 35 | 0.0571 | 4 | 0 | 6 | 0 | 7 | 0.1429 | 5 | 0.2 | 22 | 0.0909 |
| 2_1739411 | 8 | 0.375 | 3 | 0.3333 | 8 | 0.375 | 8 | 0.5 | 27 | 0.4074 | 3 | 0 | 3 | 0.3333 | 9 | 0.4444 | 4 | 0.5 | 19 | 0.3684 |
| 2_1739640 | 3 | 0.3333 | NA | NA | 1 | 1 | 3 | 0.6667 | 7 | 0.5714 | 2 | 0.5 | NA | NA | 2 | 0.5 | 2 | 0.5 | 6 | 0.5 |
| 2_1741708 | 2 | 1 | 3 | 1 | NA | NA | NA | NA | 5 | 1 | NA | NA | 3 | 1 | 2 | 1 | 1 | 1 | 6 | 1 |
| 2_1741713 | 1 | 0 | 3 | 0.3333 | NA | NA | 1 | 0 | 5 | 0.2 | NA | NA | 3 | 0.3333 | 3 | 0 | 1 | 0 | 7 | 0.1429 |
| 2_1741716 | 7 | 0 | 5 | 0.2 | 3 | 0 | 2 | 0 | 17 | 0.0588 | 3 | 0 | 5 | 0.2 | 3 | 0 | 4 | 0 | 15 | 0.0667 |
| 2_1742689 | 8 | 0.625 | 1 | 1 | 3 | 1 | 4 | 1 | 16 | 0.8125 | 5 | 0.6 | 2 | 0 | 4 | 0.75 | 4 | 0.75 | 15 | 0.6 |
| 2_1742690 | 9 | 0.2222 | 2 | 0 | 4 | 0.5 | 4 | 0.25 | 19 | 0.2632 | 5 | 0.2 | 3 | 0 | 4 | 0.25 | 5 | 0.2 | 17 | 0.1765 |
| 2_1742691 | 9 | 0.3333 | 1 | 0 | 4 | 0.75 | 3 | 0.3333 | 17 | 0.4118 | 5 | 0 | 3 | 0 | 4 | 0.25 | 5 | 0 | 17 | 0.0588 |
| 2_1742693 | 6 | 0.1667 | 15 | 0.2667 | 6 | 0.3333 | 9 | 0.2222 | 36 | 0.25 | 14 | 0.1429 | 11 | 0.1818 | 5 | 0.4 | 5 | 0 | 35 | 0.1714 |
| 2_1742693 | 4 | 0.5 | 13 | 0.5385 | 6 | 0.5 | 10 | 0.1 | 33 | 0.3939 | 12 | 0.25 | 8 | 0.25 | 4 | 0 | 5 | 0.4 | 29 | 0.2414 |
| 2_1742694 | 6 | 0.1667 | 9 | 0.3333 | 7 | 0.1429 | 10 | 0.1 | 32 | 0.1875 | 14 | 0.2143 | 9 | 0.1111 | 4 | 0.75 | 6 | 0 | 33 | 0.2121 |
| 2_1742695 | 4 | 0.75 | 11 | 0.8182 | 6 | 0.8333 | 9 | 0.6667 | 30 | 0.7667 | 11 | 0.6364 | 9 | 0.7778 | 5 | 0.4 | 5 | 1 | 30 | 0.7 |
| 2_1742695 | 6 | 0 | 15 | 0.1333 | 7 | 0 | 10 | 0.1 | 38 | 0.0789 | 13 | 0 | 10 | 0.2 | 5 | 0.2 | 5 | 0.2 | 33 | 0.1212 |
| 2_1742695 | 5 | 0.2 | 9 | 0.3333 | 7 | 0.4286 | 6 | 0.5 | 27 | 0.3704 | 12 | 0.25 | 11 | 0.0909 | 4 | 0.25 | 5 | 0 | 32 | 0.1562 |
| 2_1742695 | 5 | 0.6 | 11 | 0.1818 | 4 | 0.5 | 9 | 0.4444 | 29 | 0.3793 | 13 | 0.1538 | 11 | 0.3636 | 5 | 0.4 | 5 | 0.4 | 34 | 0.2941 |
| 2_1742696 | 4 | 0 | 10 | 0.2 | 3 | 0 | 8 | 0.375 | 25 | 0.2 | 13 | 0 | 8 | 0 | 3 | 0 | 4 | 0.5 | 28 | 0.0714 |
| 2_1742696 | 5 | 0.2 | 12 | 0.3333 | 7 | 0.5714 | 7 | 0.1429 | 31 | 0.3226 | 12 | 0.4167 | 6 | 0.1667 | NA | NA | 4 | 0.5 | 22 | 0.3636 |
| 2_1742696 | 5 | 0 | 12 | 0.0833 | 6 | 0.3333 | 10 | 0.1 | 33 | 0.1212 | 14 | 0.2143 | 10 | 0.2 | 4 | 0.25 | 6 | 0 | 34 | 0.1765 |
| 2_1743071 | 9 | 0 | 14 | 0.1429 | 9 | 0 | 10 | 0.1 | 42 | 0.0714 | 9 | 0.1111 | 9 | 0 | 12 | 0.0833 | 11 | 0.0909 | 41 | 0.0732 |
| 2_1743071 | 9 | 0.5556 | 12 | 0.3333 | 10 | 0.4 | 10 | 0.5 | 41 | 0.439 | 8 | 0.375 | 9 | 0.2222 | 9 | 0.3333 | 10 | 0.2 | 36 | 0.2778 |
| 2_1743072 | 9 | 0 | 13 | 0 | 7 | 0.1429 | 8 | 0.375 | 37 | 0.1081 | 9 | 0.3333 | 9 | 0.1111 | 10 | 0 | 9 | 0.1111 | 37 | 0.1351 |
| 2_1743072 | 7 | 0 | 10 | 0.1 | 8 | 0.125 | 9 | 0 | 34 | 0.0588 | 8 | 0.125 | 7 | 0.1429 | 9 | 0 | 8 | 0 | 32 | 0.0625 |
| 2_1743074 | 7 | 0 | 11 | 0.0909 | 7 | 0.1429 | 6 | 0 | 31 | 0.0645 | 9 | 0 | 6 | 0.1667 | 10 | 0.1 | 10 | 0.1 | 35 | 0.0857 |
| 2_1743074 | 8 | 0 | 12 | 0.25 | 8 | 0.125 | 8 | 0 | 36 | 0.1111 | 6 | 0 | 8 | 0.25 | 9 | 0 | 10 | 0.1 | 33 | 0.0909 |
| 2_1743075 | 9 | 0.1111 | 11 | 0.0909 | 7 | 0.1429 | 7 | 0 | 34 | 0.0882 | 6 | 0 | 9 | 0 | 9 | 0.2222 | 9 | 0.2222 | 33 | 0.1212 |
| 2_1743075 | 9 | 0.4444 | 10 | 0.3 | 8 | 0.5 | 8 | 0.25 | 35 | 0.3714 | 6 | 0 | 9 | 0.6667 | 9 | 0.4444 | 8 | 0.625 | 32 | 0.4688 |
| 2_1744872 | NA | NA | NA | NA | NA | NA | 1 | 1 | 1 | 1 | NA | NA | NA | NA | 1 | 1 | NA | NA | 1 | 1 |
| 2_1744878 | NA | NA | NA | NA | 1 | 0 | 1 | 1 | 2 | 0.5 | NA | NA | NA | NA | 1 | 1 | NA | NA | 1 | 1 |
| 2_1744878 | 1 | 0 | NA | NA | 1 | 1 | 1 | 1 | 3 | 0.6667 | NA | NA | NA | NA | 1 | 1 | NA | NA | 1 | 1 |
| 2_1744879 | 1 | 1 | NA | NA | 1 | 1 | 1 | 0 | 3 | 0.6667 | NA | NA | NA | NA | 1 | 1 | NA | NA | 1 | 1 |
| 2_1744881 | 1 | 1 | NA | NA | 1 | 1 | 1 | 1 | 3 | 1 | NA | NA | 1 | 1 | 1 | 1 | NA | NA | 2 | 1 |
| 2_1744883 | NA | NA | NA | NA | 1 | 0 | 1 | 1 | 2 | 0.5 | NA | NA | 1 | 1 | 1 | 0 | NA | NA | 2 | 0.5 |
| 2_1744885 | 1 | 0 | NA | NA | 1 | 1 | 1 | 0 | 3 | 0.3333 | NA | NA | 1 | 1 | 1 | 1 | NA | NA | 2 | 1 |
| 2_1744887 | 1 | 1 | NA | NA | NA | NA | NA | NA | 1 | 1 | NA | NA | 1 | 1 | 1 | 0 | NA | NA | 2 | 0.5 |
| 2_1744888 | 1 | 1 | NA | NA | 1 | 1 | 1 | 1 | 3 | 1 | NA | NA | 1 | 0 | 1 | 1 | NA | NA | 2 | 0.5 |
| 2_1745469 | 1 | 1 | NA | NA | 1 | 0 | 1 | 0 | 3 | 0.3333 | NA | NA | 1 | 0 | 1 | 1 | NA | NA | 2 | 0.5 |
| 2_1745469 | 1 | 0 | NA | NA | 1 | 0 | 1 | 1 | 3 | 0.3333 | NA | NA | 1 | 1 | 1 | 1 | NA | NA | 2 | 1 |
| 2_1745472 | 1 | 1 | NA | NA | 1 | 1 | 1 | 0 | 3 | 0.6667 | NA | NA | 1 | 1 | 1 | 0 | NA | NA | 2 | 0.5 |
| 2_1745473 | 1 | 1 | NA | NA | 1 | 0 | 1 | 0 | 3 | 0.3333 | NA | NA | 1 | 1 | 1 | 0 | NA | NA | 2 | 0.5 |
| 2_1745475 | 1 | 1 | NA | NA | 1 | 1 | 1 | 1 | 3 | 1 | NA | NA | 1 | 1 | 1 | 1 | NA | NA | 2 | 1 |
| 2_1745475 | 1 | 1 | NA | NA | 1 | 1 | 1 | 0 | 3 | 0.6667 | NA | NA | 1 | 1 | 1 | 1 | NA | NA | 2 | 1 |
| 2_1745476 | 1 | 1 | 1 | 0 | 1 | 1 | 1 | 1 | 4 | 0.75 | NA | NA | 1 | 1 | NA | NA | NA | NA | 1 | 1 |

| | | | | | | | | | | | | | | | | | | | | |
|---|---|---|---|---|---|---|---|---|---|---|---|---|---|---|---|---|---|---|---|---|
| 2_17454778 | 1 | 1 | 1 | 1 | 1 | 1 | 1 | 1 | 4 | 1 | NA | NA | NA | NA | 1 | 1 | 1 | 1 | 2 | 1 |
| 2_17454781 | 1 | 1 | 1 | 1 | 1 | 0 | 1 | 0 | 4 | 0.5 | NA | NA | 1 | 0 | 1 | 1 | 1 | 1 | 3 | 0.6667 |
| 2_17454791 | 1 | 1 | 1 | 1 | 1 | 1 | 1 | 1 | 4 | 1 | NA | NA | 1 | 1 | 1 | 1 | 1 | 1 | 3 | 1 |
| 2_17454803 | 1 | 1 | NA | NA | 1 | 1 | 1 | 1 | 3 | 1 | NA | NA | 1 | 1 | 1 | 1 | NA | NA | 2 | 1 |
| 2_17454809 | 1 | 1 | NA | NA | NA | NA | 1 | 0 | 2 | 0.5 | NA | NA | 1 | 0 | 1 | 0 | 1 | 1 | 3 | 0.3333 |
| 2_17454812 | 1 | 1 | 1 | 1 | NA | NA | 1 | 1 | 3 | 1 | NA | NA | 1 | 1 | 1 | 1 | 1 | 1 | 3 | 1 |
| 2_17454824 | 1 | 1 | 1 | 0 | NA | NA | NA | NA | 2 | 0.5 | NA | NA | NA | NA | NA | NA | 1 | 1 | 1 | 1 |
| 2_17454830 | NA | NA | NA | NA | 1 | 1 | 1 | 1 | 2 | 1 | NA | NA | 1 | 1 | 1 | 1 | 1 | 1 | 3 | 1 |
| 2_17454833 | 1 | 1 | NA | NA | 1 | 1 | NA | NA | 2 | 1 | NA | NA | 1 | 1 | 1 | 1 | NA | NA | 2 | 1 |
| 2_17454852 | 1 | 1 | 1 | 0 | NA | NA | NA | NA | 2 | 0.5 | NA | NA | 1 | 0 | 1 | 1 | NA | NA | 2 | 0.5 |
| 2_17454905 | 1 | 1 | 1 | 0 | NA | NA | NA | NA | 2 | 0.5 | NA | NA | 1 | 0 | 1 | 1 | NA | NA | 2 | 0.5 |
| 2_17455198 | 2 | 0 | 4 | 0 | 3 | 0.3333 | 2 | 0.5 | 11 | 0.1818 | NA | NA | NA | NA | 1 | 0 | 1 | 1 | 2 | 0.5 |
| 2_17455239 | 1 | 0 | 4 | 0.25 | 3 | 0.3333 | 2 | 0 | 10 | 0.2 | 3 | 0.3333 | NA | NA | 1 | 0 | 1 | 0 | 5 | 0.2 |
| 2_17462410 | 1 | 1 | NA | NA | NA | NA | NA | NA | 1 | 1 | 1 | 1 | NA | NA | NA | NA | NA | NA | 1 | 1 |
| 2_17462428 | 1 | 1 | NA | NA | NA | NA | NA | NA | 1 | 1 | 1 | 1 | NA | NA | NA | NA | NA | NA | 1 | 1 |
| 2_17477767 | NA | NA | 1 | 1 | NA | NA | NA | NA | 1 | 1 | NA | NA | 1 | 0 | NA | NA | 1 | 1 | 2 | 0.5 |
| 2_17496915 | 2 | 0 | NA | NA | 1 | 1 | NA | NA | 3 | 0.3333 | NA | NA | NA | NA | NA | NA | 1 | 1 | 1 | 1 |
| 2_17496943 | 2 | 0 | NA | NA | 1 | 1 | NA | NA | 3 | 0.3333 | 1 | 0 | NA | NA | NA | NA | 1 | 1 | 2 | 0.5 |
| 2_17496961 | 2 | 0 | NA | NA | 1 | 1 | NA | NA | 3 | 0.3333 | 1 | 1 | NA | NA | NA | NA | 1 | 0 | 2 | 0.5 |
| 2_17496967 | 2 | 0.5 | NA | NA | 1 | 0 | NA | NA | 3 | 0.3333 | 1 | 0 | NA | NA | NA | NA | 1 | 1 | 2 | 0.5 |
| 2_17496987 | 1 | 0 | NA | NA | 1 | 1 | NA | NA | 2 | 0.5 | 1 | 1 | NA | NA | NA | NA | 1 | 1 | 2 | 1 |
| 2_17497002 | 2 | 0.5 | NA | NA | NA | NA | NA | NA | 2 | 0.5 | 1 | 1 | NA | NA | NA | NA | 1 | 1 | 2 | 1 |
| 2_17501329 | NA | NA | 1 | 1 | NA | NA | NA | NA | 1 | 1 | NA | NA | 1 | 1 | NA | NA | NA | NA | 1 | 1 |
| 2_17503529 | 1 | 1 | 3 | 0 | 1 | 0 | 2 | 0.5 | 7 | 0.2857 | NA | NA | NA | NA | 1 | 1 | NA | NA | 1 | 1 |
| 2_17503921 | 2 | 0 | 1 | 1 | NA | NA | 2 | 0.5 | 5 | 0.4 | 1 | 0 | NA | NA | 2 | 0.5 | NA | NA | 3 | 0.3333 |
| 2_17513118 | 2 | 0 | NA | NA | NA | NA | 5 | 0.2 | 7 | 0.1429 | 1 | 1 | NA | NA | NA | NA | NA | NA | 1 | 1 |
| 2_17513134 | 2 | 0 | NA | NA | NA | NA | 5 | 0.6 | 7 | 0.4286 | 1 | 1 | NA | NA | NA | NA | NA | NA | 1 | 1 |
| 2_17513144 | 1 | 0 | NA | NA | NA | NA | 3 | 0.3333 | 4 | 0.25 | 1 | 1 | NA | NA | NA | NA | NA | NA | 1 | 1 |
| 2_17513149 | 2 | 1 | NA | NA | NA | NA | 5 | 0.8 | 7 | 0.8571 | 1 | 1 | NA | NA | NA | NA | 1 | 1 | 2 | 1 |
| 2_17513177 | 2 | 0 | NA | NA | NA | NA | 5 | 0.2 | 7 | 0.1429 | 1 | 1 | NA | NA | NA | NA | 1 | 1 | 2 | 1 |
| 2_17513202 | 2 | 0.5 | NA | NA | NA | NA | 5 | 1 | 7 | 0.8571 | 1 | 1 | NA | NA | NA | NA | 1 | 0 | 2 | 0.5 |
| 2_17513202 | 2 | 0.5 | NA | NA | NA | NA | 5 | 0.8 | 7 | 0.7143 | 1 | 1 | NA | NA | NA | NA | 1 | 1 | 2 | 1 |
| 2_17513205 | 2 | 0 | NA | NA | NA | NA | 5 | 1 | 7 | 0.7143 | 1 | 1 | NA | NA | NA | NA | 1 | 1 | 2 | 1 |
| 2_17513206 | 2 | 1 | NA | NA | NA | NA | 5 | 1 | 7 | 1 | 1 | 1 | NA | NA | NA | NA | 1 | 1 | 2 | 1 |
| 2_17513234 | 2 | 1 | NA | NA | NA | NA | 5 | 0.4 | 7 | 0.5714 | 1 | 1 | NA | NA | NA | NA | 2 | 0.5 | 3 | 0.6667 |
| 2_17516827 | NA | NA | 2 | 0 | 1 | 1 | 1 | 0 | 4 | 0.25 | 2 | 0.5 | 1 | 0 | 1 | 0 | NA | NA | 4 | 0.25 |
| 2_17516887 | 1 | 1 | 2 | 1 | 1 | 0 | 1 | 0 | 5 | 0.6 | 3 | 0 | 1 | 1 | NA | NA | NA | NA | 4 | 0.25 |
| 2_17522991 | NA | NA | 1 | 1 | NA | NA | NA | NA | 1 | 1 | 1 | 0 | NA | NA | 2 | 1 | 1 | 1 | 4 | 0.75 |
| 2_17526583 | 1 | 0 | 5 | 0.2 | NA | NA | NA | NA | 6 | 0.1667 | NA | NA | 4 | 0.25 | 3 | 0.3333 | 1 | 0 | 8 | 0.25 |
| 2_17526643 | 1 | 0 | 7 | 0.2857 | NA | NA | NA | NA | 8 | 0.25 | NA | NA | 3 | 0 | 4 | 0.5 | 1 | 1 | 8 | 0.375 |
| 2_17532431 | 1 | 1 | 1 | 1 | NA | NA | NA | NA | 2 | 1 | NA | NA | 1 | 1 | NA | NA | NA | NA | 1 | 1 |
| 2_17532432 | 2 | 1 | 1 | 1 | NA | NA | NA | NA | 3 | 1 | NA | NA | NA | NA | 1 | 1 | NA | NA | 1 | 1 |
| 2_17532432 | 2 | 1 | 1 | 0 | NA | NA | NA | NA | 3 | 0.6667 | NA | NA | 1 | 1 | 1 | 1 | NA | NA | 2 | 1 |
| 2_17532434 | 2 | 0.5 | 1 | 0 | NA | NA | NA | NA | 3 | 0.3333 | NA | NA | 1 | 0 | 1 | 1 | NA | NA | 2 | 0.5 |
| 2_17532436 | 2 | 1 | NA | NA | NA | NA | NA | NA | 2 | 1 | NA | NA | 1 | 1 | 1 | 0 | NA | NA | 2 | 0.5 |
| 2_17532445 | 2 | 1 | 1 | 1 | NA | NA | NA | NA | 3 | 1 | NA | NA | NA | NA | 1 | 1 | NA | NA | 1 | 1 |
| 2_17532458 | 2 | 1 | 1 | 0 | NA | NA | NA | NA | 3 | 0.6667 | NA | NA | 1 | 0 | 1 | 1 | NA | NA | 2 | 0.5 |
| 2_17532488 | 1 | 0 | 1 | 1 | 1 | 1 | NA | NA | 3 | 0.6667 | NA | NA | 1 | 1 | NA | NA | NA | NA | 1 | 1 |
| 2_17532515 | 2 | 1 | 1 | 1 | 1 | 1 | NA | NA | 4 | 1 | NA | NA | 1 | 0 | 1 | 1 | NA | NA | 2 | 0.5 |
| 2_17532521 | 2 | 0.5 | 1 | 0 | 1 | 1 | NA | NA | 4 | 0.5 | NA | NA | 1 | 0 | 1 | 1 | NA | NA | 2 | 0.5 |
| 2_17532523 | 2 | 1 | 1 | 0 | 1 | 1 | NA | NA | 4 | 0.75 | NA | NA | 1 | 0 | 1 | 1 | NA | NA | 2 | 0.5 |
| 2_17532524 | 2 | 1 | 1 | 0 | 1 | 1 | NA | NA | 4 | 0.75 | NA | NA | 1 | 1 | 1 | 1 | NA | NA | 2 | 1 |
| 2_17532548 | 2 | 1 | 1 | 0 | 2 | 1 | NA | NA | 5 | 0.8 | NA | NA | 1 | 1 | 1 | 1 | NA | NA | 2 | 1 |
| 2_17532566 | 2 | 1 | 1 | 1 | 2 | 0 | NA | NA | 5 | 0.6 | NA | NA | 1 | 1 | 1 | 1 | NA | NA | 2 | 1 |
| 2_17532655 | 1 | 1 | 1 | 0 | 2 | 1 | 1 | 0 | 5 | 0.6 | NA | NA | 1 | 0 | 1 | 1 | NA | NA | 2 | 0.5 |
| 2_17532669 | 3 | 1 | 1 | 1 | 2 | 1 | 1 | 1 | 7 | 1 | NA | NA | 1 | 1 | 1 | 1 | NA | NA | 2 | 1 |
| 2_17532698 | 3 | 0 | 1 | 0 | 2 | 0.5 | 1 | 0 | 7 | 0.1429 | NA | NA | 1 | 1 | 2 | 0 | NA | NA | 3 | 0.3333 |
| 2_17532722 | 3 | 0.6667 | 1 | 1 | 2 | 0.5 | 1 | 0 | 7 | 0.5714 | NA | NA | 1 | 1 | 2 | 1 | NA | NA | 3 | 1 |
| 2_17532723 | 2 | 1 | 1 | 1 | 2 | 1 | 1 | 1 | 6 | 1 | NA | NA | 2 | 1 | 1 | 1 | NA | NA | 3 | 1 |
| 2_17532725 | 2 | 1 | 1 | 1 | 2 | 0.5 | 1 | 1 | 6 | 0.8333 | NA | NA | 1 | 1 | 1 | 1 | NA | NA | 2 | 1 |
| 2_17532726 | 3 | 1 | 1 | 1 | 2 | 1 | 1 | 1 | 7 | 1 | NA | NA | 2 | 1 | 1 | 1 | NA | NA | 3 | 1 |
| 2_17532739 | 2 | 0.5 | 1 | 0 | 1 | 0 | 1 | 0 | 5 | 0.2 | NA | NA | 2 | 0 | 2 | 0.5 | NA | NA | 4 | 0.25 |
| 2_17532754 | 2 | 1 | 1 | 0 | 2 | 0.5 | NA | NA | 5 | 0.6 | NA | NA | 1 | 1 | 2 | 0 | NA | NA | 3 | 0.3333 |
| 2_17532760 | 3 | 0.3333 | 1 | 0 | 1 | 1 | 1 | 0 | 6 | 0.3333 | NA | NA | 2 | 0.5 | 3 | 0.3333 | NA | NA | 5 | 0.4 |
| 2_17544331 | 1 | 0 | 1 | 0 | 1 | 1 | 1 | 0 | 4 | 0.25 | 1 | 0 | 2 | 0 | 5 | 0.6 | 2 | 0.5 | 10 | 0.4 |

| | | | | | | | | | | | | | | | | | | | | |
|---|---|---|---|---|---|---|---|---|---|---|---|---|---|---|---|---|---|---|---|---|
| 2_1754436 | 2 | 0 | 1 | 0 | 1 | 0 | 1 | 1 | 5 | 0.2 | 1 | 0 | 2 | 0 | 3 | 0 | 3 | 0.3333 | 9 | 0.1111 |
| 2_1754439 | 1 | 1 | 1 | 0 | 1 | 0 | 2 | 0 | 5 | 0.2 | NA | NA | 1 | 0 | 4 | 0 | 2 | 0.5 | 7 | 0.1429 |
| 2_1754798 | NA | NA | NA | NA | NA | NA | 1 | 1 | 1 | 1 | NA | NA | NA | NA | 2 | 0.5 | 1 | 1 | 3 | 0.6667 |
| 2_1755408 | 2 | 0.5 | 2 | 0.5 | NA | NA | NA | NA | 4 | 0.5 | 1 | 0 | 1 | 0 | 3 | 0.3333 | 1 | 0 | 6 | 0.1667 |
| 2_1755414 | NA | NA | 2 | 0.5 | NA | NA | NA | NA | 2 | 0.5 | NA | NA | NA | NA | 3 | 0.3333 | 1 | 0 | 4 | 0.25 |
| 2_1755774 | 1 | 0 | 1 | 1 | NA | NA | 2 | 0 | 4 | 0.25 | NA | NA | 1 | 0 | NA | NA | 1 | 1 | 2 | 0.5 |
| 2_1755776 | 1 | 1 | 1 | 1 | NA | NA | 2 | 1 | 4 | 1 | NA | NA | 1 | 1 | NA | NA | NA | NA | 1 | 1 |
| 2_1755776 | NA | NA | 1 | 1 | NA | NA | 2 | 1 | 3 | 1 | NA | NA | 1 | 1 | NA | NA | 1 | 1 | 2 | 1 |
| 2_1755777 | 1 | 1 | 1 | 1 | NA | NA | 2 | 1 | 4 | 1 | NA | NA | 1 | 1 | NA | NA | 1 | 1 | 2 | 1 |
| 2_1755777 | 1 | 0 | 1 | 1 | NA | NA | 2 | 1 | 4 | 0.75 | NA | NA | 1 | 1 | NA | NA | NA | NA | 1 | 1 |
| 2_1755777 | 1 | 0 | 1 | 1 | NA | NA | 2 | 0 | 4 | 0.25 | NA | NA | 1 | 0 | NA | NA | 1 | 1 | 2 | 0.5 |
| 2_1755779 | 1 | 0 | 1 | 1 | NA | NA | 1 | 0 | 3 | 0.3333 | NA | NA | 1 | 0 | NA | NA | 1 | 1 | 2 | 0.5 |
| 2_1755782 | 1 | 1 | 1 | 1 | NA | NA | NA | NA | 2 | 1 | NA | NA | NA | NA | NA | NA | 1 | 1 | 1 | 1 |
| 2_1755782 | 1 | 1 | 1 | 1 | NA | NA | 1 | 1 | 3 | 1 | NA | NA | NA | NA | NA | NA | 1 | 1 | 1 | 1 |
| 2_1755784 | NA | NA | 1 | 1 | NA | NA | 1 | 0 | 2 | 0.5 | NA | NA | NA | NA | NA | NA | 1 | 1 | 1 | 1 |
| 2_1755792 | 1 | 1 | NA | NA | NA | NA | 1 | 1 | 2 | 1 | NA | NA | NA | NA | NA | NA | 1 | 1 | 1 | 1 |
| 2_1755794 | 1 | 1 | NA | NA | NA | NA | 1 | 0 | 2 | 0.5 | NA | NA | NA | NA | NA | NA | 1 | 1 | 1 | 1 |
| 2_1755794 | 1 | 1 | NA | NA | NA | NA | 1 | 1 | 2 | 1 | NA | NA | NA | NA | NA | NA | 1 | 1 | 1 | 1 |
| 2_1755797 | 1 | 1 | NA | NA | NA | NA | 1 | 1 | 2 | 1 | NA | NA | NA | NA | NA | NA | 1 | 1 | 1 | 1 |
| 2_1755797 | 1 | 1 | NA | NA | NA | NA | 1 | 1 | 2 | 1 | NA | NA | NA | NA | NA | NA | 1 | 1 | 1 | 1 |
| 2_1755797 | 1 | 1 | NA | NA | NA | NA | 1 | 1 | 2 | 1 | NA | NA | NA | NA | NA | NA | 1 | 1 | 1 | 1 |
| 2_1755798 | 1 | 0 | NA | NA | NA | NA | 1 | 1 | 2 | 0.5 | NA | NA | NA | NA | NA | NA | 1 | 1 | 1 | 1 |
| 2_1755800 | 1 | 1 | NA | NA | NA | NA | 1 | 1 | 2 | 1 | NA | NA | NA | NA | NA | NA | 1 | 1 | 1 | 1 |
| 2_1755803 | 1 | 0 | NA | NA | NA | NA | 1 | 1 | 2 | 0.5 | NA | NA | NA | NA | NA | NA | 1 | 1 | 1 | 1 |
| 2_1755804 | NA | NA | NA | NA | NA | NA | 1 | 1 | 1 | 1 | NA | NA | NA | NA | NA | NA | 1 | 1 | 1 | 1 |
| 2_1755805 | 1 | 0 | NA | NA | NA | NA | 1 | 1 | 2 | 0.5 | NA | NA | NA | NA | NA | NA | 1 | 1 | 1 | 1 |
| 2_1755806 | 1 | 1 | NA | NA | NA | NA | 1 | 1 | 2 | 1 | NA | NA | NA | NA | NA | NA | 1 | 1 | 1 | 1 |
| 2_1755806 | 1 | 1 | NA | NA | NA | NA | 1 | 1 | 2 | 1 | NA | NA | NA | NA | NA | NA | 1 | 1 | 1 | 1 |
| 2_1755806 | 1 | 1 | NA | NA | NA | NA | NA | NA | 1 | 1 | NA | NA | NA | NA | NA | NA | 1 | 1 | 1 | 1 |
| 2_1755835 | NA | NA | NA | NA | 1 | 0 | 1 | 1 | 2 | 0.5 | NA | NA | NA | NA | NA | NA | 2 | 0.5 | 2 | 0.5 |
| 2_1756705 | NA | NA | 1 | 0 | 1 | 1 | 4 | 0.5 | 6 | 0.5 | 2 | 0.5 | NA | NA | NA | NA | NA | NA | 2 | 0.5 |
| 2_1757072 | 1 | 1 | NA | NA | NA | NA | 1 | 1 | 2 | 1 | NA | NA | 1 | 0 | 1 | 1 | NA | NA | 2 | 0.5 |
| 2_1757075 | NA | NA | NA | NA | NA | NA | 1 | 1 | 1 | 1 | NA | NA | 1 | 1 | 1 | 1 | NA | NA | 2 | 1 |
| 2_1757075 | NA | NA | NA | NA | NA | NA | 1 | 1 | 1 | 1 | NA | NA | 1 | 1 | 1 | 1 | NA | NA | 2 | 1 |
| 2_1757075 | NA | NA | NA | NA | NA | NA | 1 | 1 | 1 | 1 | NA | NA | 1 | 1 | 1 | 1 | NA | NA | 2 | 1 |
| 2_1757075 | NA | NA | NA | NA | NA | NA | 1 | 1 | 1 | 1 | NA | NA | NA | NA | 1 | 1 | NA | NA | 1 | 1 |
| 2_1757075 | NA | NA | NA | NA | NA | NA | 1 | 1 | 1 | 1 | NA | NA | 1 | 1 | 1 | 0 | NA | NA | 2 | 0.5 |
| 2_1757080 | NA | NA | NA | NA | NA | NA | 1 | 1 | 1 | 1 | NA | NA | 1 | 0 | 1 | 1 | NA | NA | 2 | 0.5 |
| 2_1757081 | NA | NA | NA | NA | NA | NA | 1 | 1 | 1 | 1 | NA | NA | 1 | 1 | NA | NA | NA | NA | 1 | 1 |
| 2_1757082 | NA | NA | NA | NA | NA | NA | 1 | 1 | 1 | 1 | NA | NA | 1 | 1 | 1 | 1 | NA | NA | 2 | 1 |
| 2_1757682 | NA | NA | 3 | 0.3333 | 2 | 0.5 | 3 | 0.3333 | 8 | 0.375 | NA | NA | NA | NA | 1 | 1 | NA | NA | 1 | 1 |
| 2_1757777 | 1 | 0 | 1 | 1 | NA | NA | NA | NA | 2 | 0.5 | NA | NA | NA | NA | 1 | 1 | NA | NA | 1 | 1 |
| 2_1758812 | NA | NA | 1 | 1 | NA | NA | NA | NA | 1 | 1 | NA | NA | 1 | 1 | 1 | 0 | NA | NA | 2 | 0.5 |
| 2_1758813 | NA | NA | 1 | 1 | NA | NA | NA | NA | 1 | 1 | NA | NA | 1 | 1 | 1 | 0 | NA | NA | 2 | 0.5 |
| 2_1758815 | NA | NA | 1 | 1 | NA | NA | NA | NA | 1 | 1 | NA | NA | 1 | 1 | 1 | 0 | NA | NA | 2 | 0.5 |
| 2_1758816 | NA | NA | 1 | 1 | NA | NA | NA | NA | 1 | 1 | NA | NA | 1 | 1 | 1 | 1 | NA | NA | 2 | 1 |
| 2_1758819 | NA | NA | 1 | 1 | NA | NA | NA | NA | 1 | 1 | NA | NA | 1 | 1 | 1 | 0 | 1 | 0 | 3 | 0.3333 |
| 2_1759993 | 2 | 0.5 | 2 | 1 | NA | NA | NA | NA | 4 | 0.75 | NA | NA | NA | NA | NA | NA | 1 | 1 | 1 | 1 |
| 2_1759993 | 2 | 0 | 2 | 0 | 1 | 1 | NA | NA | 5 | 0.2 | NA | NA | NA | NA | NA | NA | 1 | 1 | 1 | 1 |
| 2_1759994 | 1 | 1 | 1 | 1 | 1 | 1 | NA | NA | 3 | 1 | NA | NA | NA | NA | NA | NA | 1 | 1 | 1 | 1 |
| 2_1759996 | 2 | 0.5 | 2 | 1 | 1 | 1 | NA | NA | 5 | 0.8 | NA | NA | NA | NA | NA | NA | 1 | 1 | 1 | 1 |
| 2_1759996 | 2 | 0 | 1 | 1 | 1 | 0 | NA | NA | 4 | 0.25 | NA | NA | NA | NA | NA | NA | 1 | 1 | 1 | 1 |
| 2_1760010 | 3 | 0.6667 | 2 | 1 | 1 | 1 | NA | NA | 6 | 0.8333 | NA | NA | NA | NA | NA | NA | 1 | 1 | 1 | 1 |
| 2_1760020 | 3 | 0.3333 | 2 | 0 | 1 | 0 | NA | NA | 6 | 0.1667 | NA | NA | NA | NA | NA | NA | 1 | 1 | 1 | 1 |
| 2_1760029 | 3 | 0 | 2 | 0.5 | 2 | 0 | 1 | 0 | 8 | 0.125 | 1 | 0 | NA | NA | NA | NA | 1 | 1 | 2 | 0.5 |
| 2_1760030 | 3 | 0 | 2 | 1 | 2 | 0 | 1 | 0 | 8 | 0.25 | 1 | 0 | NA | NA | NA | NA | 1 | 1 | 2 | 0.5 |
| 2_1760030 | 1 | 0 | 2 | 1 | 1 | 0 | 1 | 0 | 5 | 0.4 | 1 | 1 | NA | NA | NA | NA | 1 | 0 | 2 | 0.5 |
| 2_1760032 | 3 | 0.3333 | 2 | 1 | 2 | 0.5 | NA | NA | 7 | 0.5714 | 1 | 1 | NA | NA | NA | NA | 1 | 1 | 2 | 1 |
| 2_1760033 | 3 | 1 | 2 | 1 | 2 | 1 | 1 | 1 | 8 | 1 | 1 | 1 | NA | NA | NA | NA | 1 | 1 | 2 | 1 |
| 2_1760033 | 3 | 0 | 2 | 0.5 | 2 | 0 | 1 | 0 | 8 | 0.125 | 1 | 1 | NA | NA | NA | NA | 1 | 0 | 2 | 0.5 |
| 2_1760034 | 3 | 0.3333 | 2 | 0 | 2 | 0.5 | 1 | 0 | 8 | 0.25 | 1 | 0 | NA | NA | 1 | 1 | 1 | 0 | 3 | 0.3333 |
| 2_1761241 | NA | NA | NA | NA | NA | NA | 1 | 1 | 1 | 1 | NA | NA | NA | NA | 2 | 1 | NA | NA | 2 | 1 |
| 2_1761244 | NA | NA | NA | NA | NA | NA | 2 | 0.5 | 2 | 0.5 | NA | NA | NA | NA | 2 | 0.5 | NA | NA | 2 | 0.5 |
| 2_1761276 | NA | NA | 2 | 1 | NA | NA | 1 | 1 | 3 | 1 | NA | NA | NA | NA | NA | NA | 1 | 1 | 1 | 1 |
| 2_1761276 | NA | NA | 2 | 1 | NA | NA | 1 | 1 | 3 | 1 | NA | NA | NA | NA | NA | NA | 1 | 1 | 1 | 1 |

| | | | | | | | | | | | | | | | | | | | | |
|---|---|---|---|---|---|---|---|---|---|---|---|---|---|---|---|---|---|---|---|---|
| 2_1761276 | NA | NA | 2 | 1 | NA | NA | NA | NA | 2 | 1 | NA | NA | NA | NA | NA | NA | 1 | 1 | 1 | 1 |
| 2_1761276 | NA | NA | 2 | 0 | NA | NA | 1 | 1 | 3 | 0.3333 | NA | NA | NA | NA | NA | NA | 1 | 1 | 1 | 1 |
| 2_1761276 | NA | NA | 2 | 1 | NA | NA | 1 | 0 | 3 | 0.6667 | NA | NA | NA | NA | NA | NA | 1 | 1 | 1 | 1 |
| 2_1761277 | NA | NA | 2 | 1 | NA | NA | 1 | 1 | 3 | 1 | NA | NA | NA | NA | NA | NA | 1 | 1 | 1 | 1 |
| 2_1761279 | NA | NA | 2 | 1 | NA | NA | 1 | 0 | 3 | 0.6667 | NA | NA | NA | NA | NA | NA | 1 | 1 | 1 | 1 |
| 2_1761284 | NA | NA | 2 | 1 | NA | NA | 1 | 1 | 3 | 1 | NA | NA | NA | NA | NA | NA | 1 | 1 | 1 | 1 |
| 2_1761284 | NA | NA | 1 | 1 | NA | NA | 1 | 1 | 2 | 1 | NA | NA | NA | NA | NA | NA | 1 | 1 | 1 | 1 |
| 2_1761285 | NA | NA | 2 | 1 | NA | NA | 1 | 1 | 3 | 1 | NA | NA | NA | NA | NA | NA | 1 | 1 | 1 | 1 |
| 2_1761285 | NA | NA | 1 | 1 | NA | NA | NA | NA | 1 | 1 | NA | NA | NA | NA | NA | NA | 1 | 1 | 1 | 1 |
| 2_1761285 | NA | NA | 2 | 1 | NA | NA | 1 | 1 | 3 | 1 | NA | NA | NA | NA | NA | NA | 1 | 1 | 1 | 1 |
| 2_1761288 | NA | NA | 2 | 1 | NA | NA | 1 | 1 | 3 | 1 | NA | NA | NA | NA | NA | NA | 1 | 1 | 1 | 1 |
| 2_1761289 | NA | NA | 2 | 0.5 | NA | NA | 1 | 1 | 3 | 0.6667 | NA | NA | NA | NA | NA | NA | 1 | 1 | 1 | 1 |
| 2_1761299 | NA | NA | 2 | 1 | NA | NA | 1 | 0 | 3 | 0.6667 | NA | NA | NA | NA | 1 | 1 | NA | NA | 1 | 1 |
| 2_1761300 | NA | NA | 1 | 1 | NA | NA | 1 | 1 | 2 | 1 | NA | NA | NA | NA | NA | NA | 1 | 1 | 1 | 1 |
| 2_1761302 | NA | NA | 2 | 0.5 | NA | NA | 1 | 0 | 3 | 0.3333 | NA | NA | NA | NA | 1 | 1 | 1 | 0 | 2 | 0.5 |
| 2_1761305 | NA | NA | 2 | 1 | NA | NA | 1 | 1 | 3 | 1 | NA | NA | NA | NA | NA | NA | 1 | 1 | 1 | 1 |
| 2_1761305 | NA | NA | 2 | 1 | NA | NA | NA | NA | 2 | 1 | NA | NA | NA | NA | 1 | 1 | 1 | 1 | 2 | 1 |
| 2_1761305 | NA | NA | 1 | 1 | NA | NA | 1 | 0 | 2 | 0.5 | NA | NA | NA | NA | 1 | 1 | 1 | 1 | 2 | 1 |
| 2_1761305 | NA | NA | 2 | 1 | NA | NA | 1 | 1 | 3 | 1 | NA | NA | NA | NA | 1 | 1 | 1 | 1 | 2 | 1 |
| 2_1761307 | NA | NA | 2 | 0.5 | NA | NA | 1 | 0 | 3 | 0.3333 | NA | NA | NA | NA | 1 | 1 | 1 | 1 | 2 | 1 |
| 2_1761308 | 1 | 0 | 2 | 0.5 | NA | NA | 1 | 0 | 4 | 0.25 | NA | NA | NA | NA | NA | NA | 1 | 1 | 1 | 1 |
| 2_1761636 | NA | NA | NA | NA | NA | NA | 1 | 1 | 1 | 1 | NA | NA | 3 | 0 | NA | NA | 1 | 1 | 4 | 0.25 |
| 2_1761674 | 1 | 0 | 1 | 1 | 1 | 1 | 2 | 1 | 5 | 0.8 | 1 | 1 | 1 | 1 | 2 | 0.5 | NA | NA | 4 | 0.75 |
| 2_1763163 | NA | NA | 1 | 1 | 3 | 0 | 1 | 1 | 5 | 0.4 | 1 | 0 | 1 | 0 | NA | NA | 1 | 1 | 3 | 0.3333 |
| 2_1763169 | NA | NA | 3 | 0 | 2 | 1 | 1 | 0 | 6 | 0.3333 | 1 | 0 | 1 | 0 | NA | NA | 2 | 0.5 | 4 | 0.25 |
| 2_1763748 | NA | NA | 1 | 1 | 1 | 1 | NA | NA | 2 | 1 | 1 | 0 | NA | NA | 2 | 0.5 | NA | NA | 3 | 0.3333 |
| 2_1763748 | NA | NA | 1 | 0 | 1 | 1 | NA | NA | 2 | 0.5 | 1 | 1 | NA | NA | 2 | 1 | NA | NA | 3 | 1 |
| 2_1763748 | NA | NA | 1 | 1 | 1 | 1 | NA | NA | 2 | 1 | 1 | 1 | NA | NA | 2 | 1 | NA | NA | 3 | 1 |
| 2_1763749 | NA | NA | 1 | 1 | 1 | 1 | NA | NA | 2 | 1 | 1 | 1 | NA | NA | 1 | 1 | NA | NA | 2 | 1 |
| 2_1763751 | NA | NA | 1 | 1 | 1 | 1 | NA | NA | 2 | 1 | NA | NA | NA | NA | 2 | 1 | NA | NA | 2 | 1 |
| 2_1763756 | NA | NA | 1 | 1 | 1 | 1 | NA | NA | 2 | 1 | 1 | 1 | NA | NA | 3 | 1 | NA | NA | 4 | 1 |
| 2_1763756 | NA | NA | 1 | 1 | 1 | 1 | NA | NA | 2 | 1 | NA | NA | NA | NA | 3 | 0.6667 | NA | NA | 3 | 0.6667 |
| 2_1763756 | 1 | 1 | 1 | 0 | 1 | 1 | NA | NA | 3 | 0.6667 | 1 | 1 | NA | NA | 3 | 0.6667 | NA | NA | 4 | 0.75 |
| 2_1763757 | 1 | 1 | 1 | 1 | 1 | 0 | NA | NA | 3 | 0.6667 | 1 | 0 | NA | NA | 2 | 0.5 | 1 | 0 | 4 | 0.25 |
| 2_1763757 | 1 | 1 | 1 | 1 | 1 | 1 | NA | NA | 3 | 1 | 1 | 0 | NA | NA | 2 | 0.5 | NA | NA | 3 | 0.3333 |
| 2_1763759 | NA | NA | 1 | 0 | 1 | 1 | NA | NA | 2 | 0.5 | 1 | 0 | NA | NA | 2 | 0.5 | 1 | 0 | 4 | 0.25 |
| 2_1763760 | 1 | 1 | 1 | 1 | 1 | 1 | NA | NA | 3 | 1 | 1 | 1 | 1 | 1 | 3 | 1 | NA | NA | 5 | 1 |
| 2_1763761 | 1 | 1 | 1 | 1 | 1 | 1 | NA | NA | 3 | 1 | 1 | 0 | NA | NA | 2 | 1 | 1 | 0 | 4 | 0.5 |
| 2_1763762 | 1 | 1 | 2 | 0.5 | 1 | 0 | NA | NA | 4 | 0.5 | 1 | 0 | NA | NA | 3 | 1 | 1 | 0 | 5 | 0.6 |
| 2_1763762 | 1 | 1 | 1 | 1 | 1 | 0 | NA | NA | 3 | 0.6667 | 1 | 0 | NA | NA | 3 | 1 | 1 | 0 | 5 | 0.6 |
| 2_1763770 | 1 | 1 | 2 | 0.5 | 2 | 1 | NA | NA | 5 | 0.8 | 1 | 0 | NA | NA | 3 | 0.6667 | 1 | 1 | 5 | 0.6 |
| 2_1763771 | 1 | 1 | 3 | 1 | 2 | 1 | NA | NA | 6 | 1 | NA | NA | NA | NA | 3 | 1 | NA | NA | 3 | 1 |
| 2_1763772 | 1 | 1 | 3 | 1 | 2 | 1 | NA | NA | 6 | 1 | 1 | 1 | 1 | 1 | 3 | 1 | 1 | 1 | 6 | 1 |
| 2_1763775 | 1 | 0 | 4 | 0.25 | 2 | 0 | NA | NA | 7 | 0.1429 | 1 | 0 | NA | NA | 3 | 0 | 1 | 1 | 5 | 0.2 |
| 2_1763777 | 1 | 1 | 4 | 0.75 | 2 | 1 | NA | NA | 7 | 0.8571 | 1 | 1 | 1 | 1 | 3 | 0.6667 | NA | NA | 5 | 0.8 |
| 2_1763777 | 1 | 1 | 4 | 1 | 3 | 1 | NA | NA | 8 | 1 | 1 | 1 | NA | NA | 2 | 1 | 1 | 1 | 4 | 1 |
| 2_1763777 | 1 | 1 | 4 | 0.75 | 2 | 0.5 | NA | NA | 7 | 0.7143 | NA | NA | 1 | 1 | 3 | 0.6667 | 1 | 1 | 5 | 0.8 |
| 2_1763778 | 1 | 1 | 3 | 1 | 3 | 1 | NA | NA | 7 | 1 | NA | NA | NA | NA | 3 | 1 | NA | NA | 3 | 1 |
| 2_1763779 | 2 | 0 | 4 | 0.25 | 3 | 0 | NA | NA | 9 | 0.1111 | 1 | 0 | NA | NA | 3 | 0.3333 | NA | NA | 4 | 0.25 |
| 2_1763780 | 2 | 1 | 2 | 1 | 2 | 0 | NA | NA | 6 | 0.6667 | 1 | 0 | 2 | 0.5 | 4 | 0 | NA | NA | 7 | 0.1429 |
| 2_1763781 | 1 | 0 | 4 | 0 | 3 | 0.3333 | NA | NA | 8 | 0.125 | NA | NA | 2 | 0.5 | 3 | 0.3333 | NA | NA | 5 | 0.4 |
| 2_1763913 | NA | NA | NA | NA | 2 | 1 | NA | NA | 2 | 1 | NA | NA | NA | NA | 2 | 1 | NA | NA | 2 | 1 |
| 2_1763914 | NA | NA | NA | NA | 2 | 1 | NA | NA | 2 | 1 | NA | NA | NA | NA | 2 | 0.5 | NA | NA | 2 | 0.5 |
| 2_1763915 | NA | NA | NA | NA | 2 | 0.5 | NA | NA | 2 | 0.5 | NA | NA | NA | NA | 2 | 0.5 | NA | NA | 2 | 0.5 |
| 2_1763915 | NA | NA | NA | NA | 2 | 1 | NA | NA | 2 | 1 | NA | NA | NA | NA | 2 | 1 | NA | NA | 2 | 1 |
| 2_1763918 | NA | NA | NA | NA | 2 | 1 | NA | NA | 2 | 1 | NA | NA | NA | NA | 1 | 1 | NA | NA | 1 | 1 |
| 2_1763919 | NA | NA | NA | NA | 1 | 1 | NA | NA | 1 | 1 | NA | NA | NA | NA | 2 | 0.5 | NA | NA | 2 | 0.5 |
| 2_1763921 | NA | NA | NA | NA | 2 | 1 | NA | NA | 2 | 1 | NA | NA | NA | NA | 2 | 1 | NA | NA | 2 | 1 |
| 2_1763923 | NA | NA | NA | NA | 2 | 0.5 | NA | NA | 2 | 0.5 | NA | NA | NA | NA | 2 | 0.5 | NA | NA | 2 | 0.5 |
| 2_1763923 | NA | NA | NA | NA | 2 | 1 | NA | NA | 2 | 1 | NA | NA | NA | NA | 2 | 1 | NA | NA | 2 | 1 |
| 2_1763924 | NA | NA | NA | NA | 2 | 0.5 | NA | NA | 2 | 0.5 | NA | NA | NA | NA | 2 | 1 | NA | NA | 2 | 1 |
| 2_1763924 | NA | NA | NA | NA | 2 | 1 | NA | NA | 2 | 1 | NA | NA | NA | NA | 2 | 1 | NA | NA | 2 | 1 |
| 2_1763926 | NA | NA | NA | NA | 1 | 1 | NA | NA | 1 | 1 | NA | NA | NA | NA | 2 | 0.5 | NA | NA | 2 | 0.5 |
| 2_1763927 | NA | NA | NA | NA | 2 | 1 | NA | NA | 2 | 1 | NA | NA | NA | NA | 2 | 1 | NA | NA | 2 | 1 |
| 2_1763927 | NA | NA | NA | NA | 1 | 1 | NA | NA | 1 | 1 | NA | NA | NA | NA | 1 | 1 | NA | NA | 1 | 1 |

| | | | | | | | | | | | | | | | | | | | | |
|---|---|---|---|---|---|---|---|---|---|---|---|---|---|---|---|---|---|---|---|---|
| 2_1764112( | 4 | 0.25 | 6 | 0.5 | 4 | 1 | 2 | 0.5 | 16 | 0.5625 | NA | NA | 5 | 0 | 5 | 1 | 4 | 1 | 14 | 0.6429 |
| 2_1764113: | 4 | 0 | 6 | 0 | 5 | 0 | 2 | 0.5 | 17 | 0.0588 | NA | NA | 7 | 0 | 5 | 0 | 3 | 0.3333 | 15 | 0.0667 |
| 2_1764118( | 2 | 0 | 6 | 0 | 3 | 0 | 3 | 0.3333 | 14 | 0.0714 | NA | NA | 6 | 0.1667 | 4 | 0 | 3 | 0 | 13 | 0.0769 |
| 2_1764118! | 3 | 0 | 4 | 0 | 3 | 0 | 2 | 0.5 | 12 | 0.0833 | NA | NA | 5 | 0.2 | 4 | 0 | 3 | 0 | 12 | 0.0833 |
| 2_1764118( | 2 | 0.5 | 4 | 0.25 | 3 | 0 | 3 | 0.3333 | 12 | 0.25 | NA | NA | 6 | 0.5 | 3 | 0.6667 | 3 | 0.3333 | 12 | 0.5 |
| 2_1764124: | NA | NA | 1 | 0 | 3 | 0.6667 | NA | NA | 4 | 0.5 | NA | NA | 2 | 0.5 | NA | NA | 3 | 0.3333 | 5 | 0.4 |
| 2_1764124( | NA | NA | 1 | 0 | 3 | 0.6667 | NA | NA | 4 | 0.5 | 1 | 1 | 2 | 0 | NA | NA | 2 | 0.5 | 5 | 0.4 |
| 2_1764125( | NA | NA | 1 | 0 | 3 | 0.3333 | NA | NA | 4 | 0.25 | 1 | 0 | 2 | 0 | NA | NA | 3 | 0.3333 | 6 | 0.1667 |
| 2_1764134( | NA | NA | 2 | 0 | 2 | 0.5 | 1 | 0 | 5 | 0.2 | 1 | 1 | 3 | 0.3333 | NA | NA | 1 | 0 | 5 | 0.4 |
| 2_1764145! | NA | NA | 5 | 0 | 3 | 0.3333 | 3 | 0 | 11 | 0.0909 | 2 | 0 | 3 | 0 | NA | NA | 4 | 0.25 | 9 | 0.1111 |
| 2_1764150: | NA | NA | 2 | 0 | 1 | 0 | 2 | 1 | 5 | 0.4 | 1 | 0 | 2 | 0 | NA | NA | 4 | 0.25 | 7 | 0.1429 |
| 2_1764150( | NA | NA | 2 | 0 | 1 | 0 | 3 | 0.6667 | 6 | 0.3333 | 1 | 0 | 1 | 0 | NA | NA | 4 | 0.5 | 6 | 0.3333 |
| 2_1764996 | 4 | 0.5 | 3 | 0 | 4 | 0.25 | 4 | 0 | 15 | 0.2 | 1 | 0 | 2 | 0 | 5 | 0.6 | 3 | 0.3333 | 11 | 0.3636 |
| 2_1764997: | 4 | 0.25 | 3 | 0 | 4 | 0 | 4 | 0 | 15 | 0.0667 | 1 | 0 | 2 | 0.5 | 6 | 0 | 3 | 0 | 12 | 0.0833 |
| 2_1765845 | NA | NA | NA | NA | 5 | 0.4 | 1 | 1 | 6 | 0.5 | 3 | 0.6667 | NA | NA | 2 | 0.5 | 2 | 0.5 | 7 | 0.5714 |
| 2_1765846: | NA | NA | NA | NA | 3 | 1 | 1 | 1 | 4 | 1 | 3 | 1 | NA | NA | 2 | 1 | 2 | 1 | 7 | 1 |
| 2_1765846( | NA | NA | NA | NA | 3 | 0.6667 | 1 | 1 | 4 | 0.75 | 3 | 1 | NA | NA | 2 | 0.5 | 1 | 1 | 6 | 0.8333 |
| 2_1765849 | NA | NA | NA | NA | 4 | 0.25 | 1 | 0 | 5 | 0.2 | 3 | 0 | NA | NA | 2 | 0 | 2 | 0.5 | 7 | 0.1429 |
| 2_1765849( | NA | NA | NA | NA | 4 | 0.5 | 1 | 0 | 5 | 0.4 | 3 | 0.6667 | NA | NA | 2 | 0 | 2 | 0 | 7 | 0.2857 |
| 2_1765850: | NA | NA | NA | NA | 4 | 0.25 | 1 | 0 | 5 | 0.2 | 3 | 0.3333 | NA | NA | 1 | 0 | 2 | 0.5 | 6 | 0.3333 |
| 2_1765858! | NA | NA | NA | NA | 3 | 0.3333 | 1 | 0 | 4 | 0.25 | 3 | 0.3333 | NA | NA | 1 | 0 | 2 | 0.5 | 6 | 0.3333 |
| 2_1765859( | NA | NA | NA | NA | 2 | 0.5 | 1 | 0 | 3 | 0.3333 | 3 | 0 | NA | NA | 1 | 0 | 1 | 1 | 5 | 0.2 |
| 2_1765864( | NA | NA | NA | NA | 3 | 0.3333 | 1 | 0 | 4 | 0.25 | 3 | 0 | NA | NA | 1 | 0 | 2 | 0.5 | 6 | 0.1667 |
| 2_1765864! | NA | NA | NA | NA | 3 | 0.3333 | 1 | 0 | 4 | 0.25 | 2 | 0.5 | NA | NA | 1 | 0 | 2 | 0.5 | 5 | 0.4 |
| 2_1765867 | NA | NA | NA | NA | 3 | 0.3333 | 1 | 0 | 4 | 0.25 | 2 | 0 | NA | NA | 1 | 0 | 2 | 0.5 | 5 | 0.2 |
| 2_1765868! | NA | NA | NA | NA | 3 | 0.3333 | 1 | 1 | 4 | 0.5 | 2 | 1 | NA | NA | 1 | 0 | 2 | 0.5 | 5 | 0.6 |
| 2_1765883: | NA | NA | NA | NA | 2 | 0.5 | 1 | 0 | 3 | 0.3333 | 1 | 1 | NA | NA | 1 | 1 | 2 | 1 | 4 | 1 |
| 2_1765884( | NA | NA | NA | NA | 1 | 0 | 1 | 1 | 2 | 0.5 | 1 | 1 | NA | NA | 1 | 0 | 2 | 0.5 | 4 | 0.5 |
| 2_1765888 | NA | NA | NA | NA | 1 | 1 | 1 | 0 | 2 | 0.5 | 1 | 0 | NA | NA | 1 | 0 | 2 | 0.5 | 4 | 0.25 |
| 2_1766064( | NA | NA | NA | NA | 1 | 1 | NA | NA | 1 | 1 | NA | NA | NA | NA | NA | NA | 1 | 1 | 1 | 1 |
| 2_1766069: | NA | NA | NA | NA | 1 | 1 | NA | NA | 1 | 1 | NA | NA | NA | NA | NA | NA | 1 | 1 | 1 | 1 |
| 2_1766073: | NA | NA | NA | NA | 1 | 1 | NA | NA | 1 | 1 | NA | NA | NA | NA | NA | NA | 1 | 1 | 1 | 1 |
| 2_1766073: | NA | NA | NA | NA | 1 | 1 | NA | NA | 1 | 1 | NA | NA | NA | NA | NA | NA | 1 | 1 | 1 | 1 |
| 2_1767059: | NA | NA | 1 | 1 | NA | NA | 1 | 1 | 2 | 1 | 1 | 0 | 1 | 1 | 2 | 1 | NA | NA | 4 | 0.75 |
| 2_1767059! | NA | NA | 1 | 1 | NA | NA | 1 | 1 | 2 | 1 | 1 | 1 | 1 | 1 | 2 | 1 | NA | NA | 4 | 1 |
| 2_1767060( | NA | NA | 1 | 1 | NA | NA | NA | NA | 1 | 1 | 1 | 1 | 1 | 1 | 2 | 1 | NA | NA | 4 | 1 |
| 2_1767060 | NA | NA | NA | NA | NA | NA | 1 | 1 | 1 | 1 | 1 | 1 | 1 | 1 | 2 | 1 | NA | NA | 4 | 1 |
| 2_1767062! | NA | NA | NA | NA | NA | NA | 1 | 1 | 1 | 1 | 1 | 1 | 1 | 1 | 2 | 0 | NA | NA | 4 | 0.5 |
| 2_1767062( | NA | NA | 1 | 1 | NA | NA | 1 | 1 | 2 | 1 | 1 | 1 | 1 | 1 | 2 | 1 | NA | NA | 4 | 1 |
| 2_1767064 | NA | NA | 1 | 1 | NA | NA | 1 | 1 | 2 | 1 | 1 | 1 | 1 | 1 | 2 | 1 | NA | NA | 4 | 1 |
| 2_1767065( | NA | NA | 1 | 1 | NA | NA | 1 | 0 | 2 | 0.5 | 1 | 0 | 1 | 1 | 2 | 0 | NA | NA | 4 | 0.25 |
| 2_1767067! | NA | NA | 1 | 1 | NA | NA | 1 | 1 | 2 | 1 | NA | NA | 1 | 1 | 2 | 1 | NA | NA | 3 | 1 |
| 2_1767067! | NA | NA | NA | NA | NA | NA | 1 | 1 | 1 | 1 | 1 | 1 | 1 | 1 | 2 | 1 | NA | NA | 4 | 1 |
| 2_1767068! | NA | NA | NA | NA | NA | NA | 1 | 1 | 1 | 1 | 1 | 1 | 1 | 1 | 2 | 0.5 | NA | NA | 4 | 0.75 |
| 2_1767071 | NA | NA | 1 | 0 | NA | NA | 1 | 1 | 2 | 0.5 | NA | NA | 1 | 1 | 2 | 0.5 | NA | NA | 3 | 0.6667 |
| 2_1768754( | 1 | 1 | NA | NA | NA | NA | 1 | 0 | 2 | 0.5 | 2 | 0.5 | NA | NA | NA | NA | NA | NA | 2 | 0.5 |
| 2_1768755! | 1 | 1 | NA | NA | NA | NA | 1 | 1 | 2 | 1 | 1 | 1 | NA | NA | NA | NA | NA | NA | 1 | 1 |
| 2_1768825( | NA | NA | 1 | 0 | 3 | 0.3333 | NA | NA | 4 | 0.25 | NA | NA | NA | NA | 2 | 0.5 | NA | NA | 2 | 0.5 |
| 2_1768829: | NA | NA | NA | NA | 3 | 0.3333 | NA | NA | 3 | 0.3333 | NA | NA | 1 | 0 | 2 | 0.5 | NA | NA | 3 | 0.3333 |
| 2_1768833: | NA | NA | 1 | 0 | 2 | 0.5 | NA | NA | 3 | 0.3333 | NA | NA | NA | NA | 2 | 0.5 | NA | NA | 2 | 0.5 |
| 2_1768868( | 3 | 0.3333 | NA | NA | 2 | 0 | 1 | 0 | 6 | 0.1667 | 2 | 1 | 2 | 0.5 | NA | NA | 1 | 1 | 5 | 0.8 |
| 2_1768870: | 3 | 0 | 1 | 0 | 4 | 0.25 | 3 | 0.3333 | 11 | 0.1818 | 3 | 0 | 4 | 0.25 | 2 | 0 | 1 | 0 | 10 | 0.1 |
| 2_1768870( | 3 | 0.3333 | 1 | 0 | 3 | 0.3333 | 3 | 0 | 10 | 0.2 | 3 | 0 | 4 | 0 | 2 | 0.5 | 1 | 0 | 10 | 0.1 |
| 2_1768871( | 3 | 1 | 1 | 0 | 2 | 0.5 | 3 | 1 | 9 | 0.7778 | 3 | 1 | 3 | 0.6667 | 1 | 1 | 1 | 0 | 8 | 0.75 |
| 2_1768871: | 3 | 0 | 1 | 0 | 4 | 0.25 | 3 | 0 | 11 | 0.0909 | 3 | 0 | 3 | 0 | 1 | 1 | 1 | 0 | 8 | 0.125 |
| 2_1770061! | 1 | 1 | NA | NA | NA | NA | NA | NA | 1 | 1 | NA | NA | NA | NA | NA | NA | 1 | 1 | 1 | 1 |
| 2_1771658! | 1 | 0 | 1 | 1 | NA | NA | 2 | 0.5 | 4 | 0.5 | NA | NA | 1 | 1 | 1 | 0 | NA | NA | 2 | 0.5 |
| 2_1771663: | 1 | 1 | NA | NA | NA | NA | NA | NA | 1 | 1 | NA | NA | 1 | 1 | NA | NA | NA | NA | 1 | 1 |
| 2_1771697( | NA | NA | 1 | 1 | 1 | 1 | 1 | 1 | 3 | 1 | 1 | 1 | NA | NA | 2 | 1 | 2 | 0.5 | 5 | 0.8 |
| 2_1771697: | NA | NA | NA | NA | 1 | 1 | 1 | 1 | 2 | 1 | NA | NA | NA | NA | 2 | 1 | 1 | 1 | 3 | 1 |
| 2_1771697! | 1 | 1 | NA | NA | NA | NA | 1 | 1 | 2 | 1 | 1 | 1 | NA | NA | 2 | 1 | 2 | 0.5 | 5 | 0.8 |
| 2_1771697! | 1 | 1 | NA | NA | 1 | 0 | 1 | 1 | 3 | 0.6667 | 1 | 1 | NA | NA | 2 | 1 | 2 | 0.5 | 5 | 0.8 |
| 2_1771698 | NA | NA | NA | NA | NA | NA | 1 | 1 | 1 | 1 | 1 | 1 | NA | NA | 2 | 1 | 2 | 1 | 5 | 1 |
| 2_1771699 | 1 | 1 | NA | NA | 2 | 1 | 1 | 1 | 4 | 1 | 1 | 1 | NA | NA | 1 | 1 | 2 | 1 | 4 | 1 |
| 2_1771700: | 1 | 1 | NA | NA | 1 | 1 | 1 | 1 | 3 | 1 | 1 | 1 | NA | NA | 2 | 1 | 2 | 1 | 5 | 1 |

| | | | | | | | | | | | | | | | | | | | | |
|---|---|---|---|---|---|---|---|---|---|---|---|---|---|---|---|---|---|---|---|---|
| 2_17717018 | 1 | 1 | 1 | 1 | 2 | 1 | 1 | 1 | 5 | 1 | 1 | 1 | NA | NA | 2 | 1 | 2 | 1 | 5 | 1 |
| 2_17717033 | 1 | 1 | 1 | 1 | 2 | 0.5 | 1 | 1 | 5 | 0.8 | 1 | 1 | NA | NA | 2 | 1 | 3 | 0.6667 | 6 | 0.8333 |
| 2_17717042 | 1 | 0 | 1 | 0 | 1 | 1 | 1 | 0 | 4 | 0.25 | 1 | 0 | NA | NA | 2 | 0.5 | 3 | 0.3333 | 6 | 0.3333 |
| 2_17717051 | 1 | 1 | 1 | 0 | 2 | 1 | 1 | 1 | 5 | 0.8 | 1 | 1 | NA | NA | 2 | 0 | 2 | 0.5 | 5 | 0.4 |
| 2_17717054 | 1 | 1 | 1 | 1 | 1 | 1 | 1 | 1 | 4 | 1 | 1 | 0 | NA | NA | 2 | 1 | 3 | 1 | 6 | 0.8333 |
| 2_17717061 | 1 | 1 | 1 | 0 | 1 | 1 | 2 | 1 | 5 | 0.8 | 1 | 1 | NA | NA | 2 | 1 | 3 | 1 | 6 | 1 |
| 2_17717067 | 1 | 0 | 2 | 1 | 1 | 1 | 1 | 0 | 5 | 0.6 | NA | NA | NA | NA | 1 | 1 | 2 | 0 | 3 | 0.3333 |
| 2_17717068 | 1 | 1 | NA | NA | 1 | 1 | 2 | 0.5 | 4 | 0.75 | 1 | 1 | NA | NA | NA | NA | 1 | 0 | 2 | 0.5 |
| 2_17717069 | 1 | 1 | 2 | 1 | 2 | 1 | 1 | 1 | 6 | 1 | 1 | 1 | 1 | 1 | 1 | 1 | 3 | 1 | 6 | 1 |
| 2_17717093 | NA | NA | 2 | 0.5 | 3 | 0.6667 | 2 | 1 | 7 | 0.7143 | 1 | 1 | 1 | 0 | 2 | 1 | 2 | 1 | 6 | 0.8333 |
| 2_17720038 | NA | NA | NA | NA | 3 | 1 | NA | NA | 3 | 1 | NA | NA | 2 | 1 | 1 | 1 | 1 | 0 | 4 | 0.75 |
| 2_17720045 | NA | NA | NA | NA | 3 | 0.3333 | NA | NA | 3 | 0.3333 | NA | NA | 2 | 0.5 | NA | NA | 1 | 0 | 3 | 0.3333 |
| 2_17720078 | NA | NA | NA | NA | 3 | 0.3333 | NA | NA | 3 | 0.3333 | NA | NA | 2 | 0.5 | 1 | 1 | 1 | 0 | 4 | 0.5 |
| 2_17720109 | NA | NA | 1 | 0 | 3 | 0.6667 | NA | NA | 4 | 0.5 | NA | NA | 2 | 0.5 | 1 | 1 | 1 | 0 | 4 | 0.5 |
| 2_17720128 | NA | NA | 1 | 0 | 3 | 0.3333 | NA | NA | 4 | 0.25 | NA | NA | 2 | 0 | 1 | 1 | 1 | 0 | 4 | 0.25 |
| 2_17720274 | NA | NA | 1 | 1 | 1 | 1 | NA | NA | 2 | 1 | NA | NA | 1 | 1 | 1 | 1 | NA | NA | 2 | 1 |
| 2_17720275 | NA | NA | 1 | 1 | 1 | 1 | NA | NA | 2 | 1 | NA | NA | NA | NA | 1 | 1 | NA | NA | 1 | 1 |
| 2_17720278 | NA | NA | 1 | 1 | 1 | 1 | NA | NA | 2 | 1 | NA | NA | 1 | 1 | 1 | 1 | NA | NA | 2 | 1 |
| 2_17720300 | NA | NA | 1 | 1 | 1 | 1 | NA | NA | 2 | 1 | NA | NA | 1 | 1 | 1 | 1 | NA | NA | 2 | 1 |
| 2_17720316 | NA | NA | 1 | 1 | 1 | 1 | NA | NA | 2 | 1 | NA | NA | 2 | 1 | NA | NA | NA | NA | 2 | 1 |
| 2_17720317 | NA | NA | 1 | 0 | 1 | 1 | NA | NA | 2 | 0.5 | NA | NA | 1 | 1 | 1 | 0 | NA | NA | 2 | 0.5 |
| 2_17720342 | NA | NA | 1 | 0 | 2 | 0.5 | NA | NA | 3 | 0.3333 | NA | NA | 2 | 0.5 | 1 | 0 | NA | NA | 3 | 0.3333 |
| 2_17720344 | NA | NA | 1 | 0 | 1 | 1 | NA | NA | 2 | 0.5 | NA | NA | 2 | 0 | 1 | 1 | NA | NA | 3 | 0.3333 |
| 2_17720349 | NA | NA | 1 | 1 | 2 | 1 | NA | NA | 3 | 1 | NA | NA | 2 | 1 | NA | NA | NA | NA | 2 | 1 |
| 2_17720352 | NA | NA | 1 | 0 | 1 | 1 | NA | NA | 2 | 0.5 | NA | NA | 2 | 0.5 | 1 | 1 | NA | NA | 3 | 0.6667 |
| 2_17720359 | NA | NA | 1 | 1 | 1 | 1 | NA | NA | 2 | 1 | NA | NA | 2 | 1 | 1 | 0 | NA | NA | 3 | 0.6667 |
| 2_17741369 | 1 | 1 | NA | NA | NA | NA | 1 | 0 | 2 | 0.5 | NA | NA | 1 | 1 | NA | NA | NA | NA | 1 | 1 |
| 2_17741370 | 1 | 1 | NA | NA | NA | NA | 1 | 0 | 2 | 0.5 | NA | NA | 1 | 1 | NA | NA | NA | NA | 1 | 1 |
| 2_17741395 | 1 | 1 | NA | NA | NA | NA | 1 | 0 | 2 | 0.5 | NA | NA | 1 | 1 | NA | NA | NA | NA | 1 | 1 |
| 2_17741412 | 1 | 1 | NA | NA | NA | NA | 1 | 0 | 2 | 0.5 | NA | NA | 1 | 1 | NA | NA | NA | NA | 1 | 1 |
| 2_17741444 | 1 | 1 | NA | NA | NA | NA | 1 | 0 | 2 | 0.5 | NA | NA | 1 | 1 | NA | NA | NA | NA | 1 | 1 |
| 2_17741447 | 1 | 1 | NA | NA | NA | NA | 1 | 0 | 2 | 0.5 | NA | NA | 1 | 1 | NA | NA | NA | NA | 1 | 1 |
| 2_17741454 | 1 | 1 | NA | NA | NA | NA | 1 | 0 | 2 | 0.5 | NA | NA | 1 | 1 | NA | NA | NA | NA | 1 | 1 |
| 2_17741466 | 1 | 1 | NA | NA | NA | NA | 1 | 0 | 2 | 0.5 | NA | NA | 1 | 1 | NA | NA | NA | NA | 1 | 1 |
| 2_17747298 | NA | NA | 5 | 0.2 | 1 | 0 | 3 | 0 | 9 | 0.1111 | 1 | 0 | 2 | 0.5 | 2 | 0 | 4 | 0 | 9 | 0.1111 |
| 2_17747370 | NA | NA | 3 | 0 | 1 | 0 | 2 | 0.5 | 6 | 0.1667 | 1 | 1 | 1 | 0 | 2 | 0 | 2 | 0 | 6 | 0.1667 |
| 2_17747474 | NA | NA | 1 | 0 | 1 | 0 | 2 | 0.5 | 4 | 0.25 | NA | NA | 1 | 1 | 2 | 0 | 2 | 0.5 | 5 | 0.4 |
| 2_17747532 | 3 | 0 | 4 | 0.25 | 5 | 0.4 | 5 | 0.2 | 17 | 0.2353 | 4 | 0 | 4 | 0 | 9 | 0.3333 | 8 | 0.375 | 25 | 0.24 |
| 2_17754689 | NA | NA | 1 | 1 | 1 | 0 | NA | NA | 2 | 0.5 | NA | NA | NA | NA | 1 | 0 | 2 | 0.5 | 3 | 0.3333 |
| 2_17755125 | NA | NA | NA | NA | NA | NA | 1 | 1 | 1 | 1 | 2 | 1 | 1 | 1 | 4 | 0.5 | 3 | 0.6667 | 10 | 0.7 |
| 2_17758505 | 1 | 0 | 3 | 0.6667 | 1 | 0 | 4 | 0.25 | 9 | 0.3333 | NA | NA | 2 | 0 | 1 | 0 | 2 | 0.5 | 5 | 0.2 |
| 2_17758511 | 1 | 0 | 3 | 0.3333 | 1 | 0 | 4 | 0.25 | 9 | 0.2222 | NA | NA | 1 | 0 | NA | NA | 2 | 0.5 | 3 | 0.3333 |
| 2_17758515 | 1 | 0 | 3 | 0.6667 | 1 | 0 | 3 | 0.3333 | 8 | 0.375 | NA | NA | NA | NA | NA | NA | 2 | 0.5 | 2 | 0.5 |
| 2_17758515 | 1 | 0 | 3 | 0.3333 | 1 | 0 | 4 | 0.25 | 9 | 0.2222 | NA | NA | 1 | 0 | NA | NA | 2 | 0.5 | 3 | 0.3333 |
| 2_17758537 | 1 | 0 | 2 | 0 | 1 | 0 | 4 | 0.25 | 8 | 0.125 | NA | NA | 1 | 0 | NA | NA | 2 | 0.5 | 3 | 0.3333 |
| 2_17758549 | 1 | 0 | 3 | 0.6667 | 1 | 0 | 3 | 0.3333 | 8 | 0.375 | NA | NA | 1 | 0 | NA | NA | 2 | 0.5 | 3 | 0.3333 |
| 2_17758564 | 1 | 0 | 3 | 0.6667 | 1 | 1 | 2 | 0.5 | 7 | 0.5714 | NA | NA | 1 | 1 | NA | NA | 2 | 0.5 | 3 | 0.6667 |
| 2_17758589 | 1 | 0 | 2 | 0.5 | 2 | 0.5 | 4 | 0.5 | 9 | 0.4444 | NA | NA | NA | NA | NA | NA | 2 | 0.5 | 2 | 0.5 |
| 2_17758590 | 1 | 0 | 2 | 1 | 2 | 0 | 4 | 0.5 | 9 | 0.4444 | NA | NA | 1 | 1 | NA | NA | 2 | 0 | 3 | 0.3333 |
| 2_17758744 | 1 | 0 | 1 | 1 | 2 | 0 | 2 | 0 | 6 | 0.1667 | NA | NA | 2 | 0.5 | 1 | 1 | 2 | 0.5 | 5 | 0.6 |
| 2_17758860 | 1 | 0 | 1 | 1 | 1 | 0 | 1 | 0 | 4 | 0.25 | NA | NA | 2 | 0.5 | NA | NA | 1 | 0 | 3 | 0.3333 |
| 2_17758866 | 1 | 0 | 1 | 1 | 1 | 1 | 2 | 1 | 5 | 0.8 | NA | NA | 2 | 0.5 | NA | NA | 1 | 0 | 3 | 0.3333 |
| 2_17758868 | 1 | 0 | 2 | 0.5 | 1 | 0 | 2 | 0 | 6 | 0.1667 | NA | NA | 2 | 0.5 | NA | NA | 1 | 0 | 3 | 0.3333 |
| 2_17758923 | 3 | 0 | 1 | 1 | 6 | 0.3333 | 2 | 0 | 12 | 0.25 | 1 | 0 | 2 | 0 | 1 | 0 | 3 | 0.3333 | 7 | 0.1429 |
| 2_17758953 | 3 | 0.3333 | 2 | 0 | 3 | 0.3333 | 4 | 0.25 | 12 | 0.25 | 1 | 0 | 4 | 1 | 1 | 0 | 3 | 0.6667 | 9 | 0.6667 |
| 2_17767806 | NA | NA | 1 | 1 | 2 | 1 | 1 | 0 | 4 | 0.75 | NA | NA | NA | NA | 1 | 1 | NA | NA | 1 | 1 |
| 2_17767814 | NA | NA | 1 | 1 | 1 | 0 | 1 | 1 | 3 | 0.6667 | NA | NA | NA | NA | 1 | 1 | NA | NA | 1 | 1 |
| 2_17767815 | NA | NA | 1 | 1 | 2 | 0.5 | 1 | 1 | 4 | 0.75 | NA | NA | NA | NA | 1 | 1 | NA | NA | 1 | 1 |
| 2_17767838 | NA | NA | 1 | 1 | 2 | 1 | 1 | 1 | 4 | 1 | NA | NA | NA | NA | 1 | 1 | NA | NA | 1 | 1 |
| 2_17767945 | NA | NA | 1 | 1 | 1 | 1 | 1 | 1 | 3 | 1 | NA | NA | NA | NA | 1 | 1 | NA | NA | 1 | 1 |
| 2_17767959 | NA | NA | 1 | 1 | 2 | 1 | 1 | 1 | 4 | 1 | NA | NA | NA | NA | 2 | 0 | 1 | 1 | 3 | 0.3333 |
| 2_17767993 | NA | NA | 1 | 1 | 3 | 0 | 1 | 0 | 5 | 0.2 | NA | NA | NA | NA | 2 | 0.5 | 1 | 0 | 3 | 0.3333 |
| 2_17768013 | NA | NA | 1 | 1 | 3 | 1 | 1 | 1 | 5 | 1 | NA | NA | NA | NA | 1 | 1 | 1 | 1 | 2 | 1 |
| 2_17768013 | NA | NA | 1 | 1 | 3 | 0.6667 | 1 | 1 | 5 | 0.8 | NA | NA | NA | NA | 1 | 1 | 1 | 1 | 2 | 1 |
| 2_17768016 | NA | NA | 1 | 1 | 1 | 1 | 1 | 1 | 3 | 1 | NA | NA | NA | NA | 2 | 0.5 | NA | NA | 2 | 0.5 |

| | | | | | | | | | | | | | | | | | | | | |
|---|---|---|---|---|---|---|---|---|---|---|---|---|---|---|---|---|---|---|---|---|
| 2_1776801 | NA | NA | 1 | 1 | 2 | 0.5 | 1 | 0 | 4 | 0.5 | NA | NA | NA | NA | 2 | 1 | 1 | 1 | 3 | 1 |
| 2_1776804 | NA | NA | 2 | 0.5 | 2 | 0 | 1 | 0 | 5 | 0.2 | NA | NA | 1 | 1 | 2 | 0 | 1 | 1 | 4 | 0.5 |
| 2_1776805 | NA | NA | 2 | 1 | 2 | 0.5 | 1 | 0 | 5 | 0.6 | NA | NA | 1 | 1 | NA | NA | NA | NA | 1 | 1 |
| 2_1776982 | NA | NA | NA | NA | 1 | 1 | NA | NA | 1 | 1 | NA | NA | NA | NA | NA | NA | 1 | 1 | 1 | 1 |
| 2_1776983 | NA | NA | NA | NA | 1 | 1 | NA | NA | 1 | 1 | NA | NA | NA | NA | NA | NA | 1 | 1 | 1 | 1 |
| 2_1776991 | NA | NA | NA | NA | 1 | 1 | NA | NA | 1 | 1 | NA | NA | NA | NA | NA | NA | 1 | 1 | 1 | 1 |
| 2_1776999 | NA | NA | NA | NA | 1 | 1 | NA | NA | 1 | 1 | NA | NA | NA | NA | NA | NA | 1 | 1 | 1 | 1 |
| 2_1776999 | NA | NA | NA | NA | 1 | 1 | NA | NA | 1 | 1 | NA | NA | NA | NA | NA | NA | 1 | 1 | 1 | 1 |
| 2_1777001 | NA | NA | NA | NA | 1 | 1 | NA | NA | 1 | 1 | NA | NA | NA | NA | NA | NA | 1 | 1 | 1 | 1 |
| 2_1777106 | 2 | 0.5 | 3 | 0.3333 | 1 | 0 | NA | NA | 6 | 0.3333 | 1 | 0 | 4 | 0.25 | 2 | 0 | 2 | 0.5 | 9 | 0.2222 |
| 2_1777108 | 2 | 0 | 4 | 0.25 | 2 | 0 | NA | NA | 8 | 0.125 | 1 | 0 | 4 | 0.5 | 2 | 0 | 3 | 0 | 10 | 0.2 |
| 2_1777111 | 2 | 0 | 3 | 0 | 2 | 0 | 1 | 1 | 8 | 0.125 | 2 | 0 | 4 | 0 | 2 | 0.5 | 3 | 0 | 11 | 0.0909 |
| 2_1777112 | 2 | 0 | 3 | 0.3333 | 2 | 0 | 1 | 1 | 8 | 0.25 | 3 | 0.3333 | 2 | 0.5 | 3 | 0.3333 | 3 | 1 | 11 | 0.5455 |
| 2_1777838 | 10 | 0 | 8 | 0.25 | 4 | 0.25 | 4 | 0 | 26 | 0.1154 | 7 | 0 | 9 | 0.2222 | 5 | 0 | 5 | 0.2 | 26 | 0.1154 |
| 2_1782015 | 2 | 0 | 1 | 1 | NA | NA | 2 | 0.5 | 5 | 0.4 | NA | NA | 1 | 0 | 1 | 1 | NA | NA | 2 | 0.5 |
| 2_1782020 | 2 | 0.5 | 1 | 1 | NA | NA | 2 | 0 | 5 | 0.4 | NA | NA | 1 | 1 | 1 | 0 | NA | NA | 2 | 0.5 |
| 2_1782027 | 1 | 1 | 1 | 1 | NA | NA | 2 | 0 | 4 | 0.5 | NA | NA | 1 | 1 | 1 | 1 | NA | NA | 2 | 1 |
| 2_1779696 | 9 | 0.5556 | 11 | 0.1818 | 3 | 1 | 5 | 0.4 | 28 | 0.4286 | 7 | 0.4286 | 4 | 0.25 | 5 | 0.4 | 5 | 0.4 | 21 | 0.381 |
| 2_1779697 | 8 | 0.25 | 12 | 0.5833 | 3 | 0.3333 | 7 | 0.2857 | 30 | 0.4 | 9 | 0.8889 | 4 | 0.25 | 6 | 0.3333 | 5 | 0.2 | 24 | 0.5 |
| 2_1779697 | 8 | 0.5 | 7 | 0.4286 | 1 | 0 | 7 | 0.5714 | 23 | 0.4783 | 5 | 0.6 | 4 | 0.25 | 4 | 0.5 | 3 | 0.3333 | 16 | 0.4375 |
| 2_1779698 | 5 | 0.4 | 7 | 0.1429 | 1 | 1 | 5 | 0.8 | 18 | 0.4444 | 6 | 0.1667 | 4 | 0.5 | 4 | 0.5 | 2 | 0 | 16 | 0.3125 |
| 2_1779698 | 7 | 0.2857 | 7 | 0.1429 | 3 | 0.3333 | 4 | 0 | 21 | 0.1905 | 9 | 0.2222 | 3 | 0 | 4 | 0 | 4 | 0 | 20 | 0.1 |
| 2_1779773 | 18 | 0.0556 | 19 | 0.2105 | 11 | 0.0909 | 14 | 0.0714 | 62 | 0.1129 | 22 | 0.1818 | 18 | 0.2778 | 14 | 0.1429 | 19 | 0.1579 | 73 | 0.1918 |
| 2_1779773 | 31 | 0.3871 | 28 | 0.3571 | 20 | 0.4 | 26 | 0.3462 | 105 | 0.3714 | 24 | 0.5 | 29 | 0.3103 | 24 | 0.5 | 23 | 0.4348 | 100 | 0.43 |
| 2_1779775 | 28 | 0.3571 | 25 | 0.52 | 15 | 0.3333 | 29 | 0.3793 | 97 | 0.4021 | 20 | 0.45 | 29 | 0.4828 | 20 | 0.25 | 22 | 0.1818 | 91 | 0.3516 |
| 2_1779775 | 26 | 0.6923 | 23 | 0.4348 | 11 | 0.4545 | 27 | 0.3704 | 87 | 0.4943 | 18 | 0.3333 | 20 | 0.6 | 16 | 0.625 | 21 | 0.7143 | 75 | 0.5733 |
| 2_1779775 | 32 | 0.0312 | 32 | 0.0938 | 19 | 0.0526 | 30 | 0.0667 | 113 | 0.0619 | 23 | 0.1304 | 31 | 0.2258 | 25 | 0.12 | 20 | 0.15 | 99 | 0.1616 |
| 2_1779776 | 35 | 0.6286 | 32 | 0.7812 | 22 | 0.7273 | 33 | 0.6667 | 122 | 0.6967 | 26 | 0.8077 | 31 | 0.7419 | 26 | 0.7692 | 23 | 0.5217 | 106 | 0.717 |
| 2_1779776 | 33 | 0.3333 | 33 | 0.2727 | 23 | 0.4348 | 35 | 0.3143 | 124 | 0.3306 | 25 | 0.32 | 31 | 0.2903 | 28 | 0.3571 | 26 | 0.2308 | 110 | 0.3 |
| 2_1779779 | 6 | 0.1667 | 4 | 0.25 | 3 | 0.3333 | 6 | 0.1667 | 19 | 0.2105 | 5 | 0.2 | 5 | 0 | 4 | 0.25 | 9 | 0.2222 | 23 | 0.1739 |
| 2_1779780 | 5 | 0 | 5 | 0.2 | 3 | 0.3333 | 5 | 0 | 18 | 0.1111 | 4 | 0 | 7 | 0.1429 | 6 | 0.1667 | 9 | 0.1111 | 26 | 0.1154 |
| 2_1780534 | NA | NA | 3 | 0.3333 | 1 | 0 | 3 | 0.3333 | 7 | 0.2857 | 3 | 0 | 1 | 0 | 1 | 1 | 3 | 0 | 8 | 0.125 |
| 2_1780535 | NA | NA | 3 | 0.6667 | 1 | 1 | 1 | 1 | 5 | 0.8 | 2 | 0 | NA | NA | 1 | 1 | 2 | 0.5 | 5 | 0.4 |
| 2_1780536 | NA | NA | 3 | 1 | 1 | 1 | 2 | 1 | 6 | 1 | 2 | 1 | 1 | 1 | 1 | 1 | 2 | 1 | 6 | 1 |
| 2_1780536 | NA | NA | 2 | 0.5 | 1 | 0 | 2 | 0 | 5 | 0.2 | 1 | 0 | 1 | 1 | 1 | 0 | 2 | 1 | 5 | 0.6 |
| 2_1780537 | NA | NA | 2 | 0.5 | 1 | 0 | 1 | 0 | 4 | 0.25 | 3 | 0 | 1 | 0 | 1 | 0 | 3 | 0.3333 | 8 | 0.125 |
| 2_1780537 | NA | NA | 3 | 0 | 1 | 0 | 2 | 0.5 | 6 | 0.1667 | 2 | 0 | 1 | 0 | 1 | 1 | 3 | 0.6667 | 7 | 0.4286 |
| 2_1780538 | NA | NA | 3 | 0.6667 | 1 | 1 | 1 | 0 | 5 | 0.6 | 3 | 0.3333 | 1 | 0 | 1 | 1 | 3 | 0.3333 | 8 | 0.375 |
| 2_1780538 | NA | NA | 3 | 1 | 1 | 0 | 1 | 0 | 5 | 0.6 | 3 | 0.3333 | 1 | 0 | 1 | 0 | 3 | 0.3333 | 8 | 0.25 |
| 2_1780538 | NA | NA | 2 | 1 | 2 | 0 | 1 | 0 | 5 | 0.4 | 2 | 0 | 1 | 0 | 1 | 1 | 3 | 0.6667 | 7 | 0.4286 |
| 2_1780544 | NA | NA | 2 | 0 | 2 | 0 | 1 | 1 | 5 | 0.2 | 3 | 0.3333 | 1 | 1 | 1 | 1 | 3 | 0.3333 | 8 | 0.5 |
| 2_1780544 | NA | NA | 2 | 0.5 | 2 | 0 | 2 | 0.5 | 6 | 0.3333 | 3 | 0.3333 | 1 | 1 | 1 | 1 | 4 | 0.5 | 9 | 0.5556 |
| 2_1780545 | NA | NA | 2 | 0 | 2 | 0 | 2 | 0.5 | 6 | 0.1667 | 3 | 0 | 1 | 0 | 1 | 1 | 4 | 0.25 | 9 | 0.2222 |
| 2_1780546 | NA | NA | 2 | 0 | 2 | 0.5 | 2 | 0 | 6 | 0.1667 | 1 | 0 | 1 | 0 | 1 | 0 | 2 | 0.5 | 5 | 0.2 |
| 2_1780551 | NA | NA | 2 | 1 | 2 | 0.5 | 1 | 1 | 5 | 0.8 | 2 | 0 | NA | NA | 1 | 1 | 3 | 0 | 6 | 0.1667 |
| 2_1780552 | NA | NA | 2 | 1 | 2 | 1 | 1 | 1 | 5 | 1 | NA | NA | NA | NA | 1 | 1 | 4 | 1 | 5 | 1 |
| 2_1780553 | NA | NA | 2 | 0.5 | 2 | 0.5 | NA | NA | 4 | 0.5 | 1 | 0 | NA | NA | 1 | 0 | 3 | 0.6667 | 5 | 0.4 |
| 2_1780553 | NA | NA | 2 | 1 | NA | NA | 1 | 0 | 3 | 0.6667 | 1 | 0 | NA | NA | 1 | 1 | 3 | 0.6667 | 5 | 0.6 |
| 2_1780623 | 1 | 1 | 1 | 0 | NA | NA | NA | NA | 2 | 0.5 | 1 | 0 | 1 | 1 | 1 | 1 | NA | NA | 3 | 0.6667 |
| 2_1780623 | 1 | 1 | 1 | 1 | NA | NA | NA | NA | 2 | 1 | 1 | 1 | 1 | 0 | 1 | 1 | NA | NA | 3 | 0.6667 |
| 2_1780624 | NA | NA | 1 | 1 | NA | NA | NA | NA | 1 | 1 | 1 | 1 | 1 | 0 | 1 | 0 | NA | NA | 3 | 0.3333 |
| 2_1780625 | 1 | 1 | 1 | 1 | NA | NA | NA | NA | 2 | 1 | 1 | 1 | NA | NA | 1 | 1 | NA | NA | 2 | 1 |
| 2_1780626 | 1 | 1 | 1 | 1 | NA | NA | NA | NA | 2 | 1 | 1 | 0 | 1 | 0 | 1 | 1 | NA | NA | 3 | 0.3333 |
| 2_1780628 | 1 | 1 | 1 | 1 | NA | NA | NA | NA | 2 | 1 | 1 | 1 | 1 | 1 | 1 | 1 | NA | NA | 3 | 1 |
| 2_1780628 | 1 | 1 | 1 | 0 | NA | NA | NA | NA | 2 | 0.5 | 1 | 0 | 1 | 1 | NA | NA | NA | NA | 2 | 0.5 |
| 2_1780629 | 1 | 1 | NA | NA | NA | NA | NA | NA | 1 | 1 | 1 | 0 | 1 | 1 | 1 | 0 | NA | NA | 3 | 0.3333 |
| 2_1780630 | NA | NA | 1 | 1 | NA | NA | NA | NA | 1 | 1 | 1 | 0 | 2 | 0.5 | 1 | 1 | NA | NA | 4 | 0.5 |
| 2_1780630 | 1 | 0 | 1 | 1 | NA | NA | NA | NA | 2 | 0.5 | 1 | 0 | 1 | 0 | 1 | 1 | NA | NA | 3 | 0.3333 |
| 2_1780630 | 1 | 0 | 1 | 1 | NA | NA | NA | NA | 2 | 0.5 | 1 | 1 | 2 | 1 | 1 | 1 | NA | NA | 4 | 1 |
| 2_1780631 | NA | NA | 1 | 1 | NA | NA | NA | NA | 1 | 1 | 1 | 1 | 1 | 0 | 1 | 1 | NA | NA | 3 | 0.6667 |
| 2_1780632 | NA | NA | 1 | 1 | NA | NA | NA | NA | 1 | 1 | 1 | 1 | 1 | 0 | NA | NA | NA | NA | 2 | 0.5 |
| 2_1780634 | 1 | 0 | 1 | 1 | NA | NA | NA | NA | 2 | 0.5 | 1 | 0 | 2 | 0.5 | 1 | 1 | NA | NA | 4 | 0.5 |
| 2_1780639 | 1 | 1 | 1 | 1 | NA | NA | NA | NA | 2 | 1 | 1 | 0 | 1 | 0 | 1 | 1 | 1 | 1 | 4 | 0.5 |
| 2_1780642 | 3 | 0 | 2 | 1 | NA | NA | 2 | 0 | 7 | 0.2857 | 1 | 0 | 2 | 0.5 | 1 | 0 | 2 | 0 | 6 | 0.1667 |
| 2_1780649 | 5 | 0 | 8 | 0.125 | 4 | 0.25 | 3 | 0 | 20 | 0.1 | 3 | 0 | 5 | 0.2 | 5 | 0.2 | 5 | 0.2 | 18 | 0.1667 |

| | | | | | | | | | | | | | | | | | | | | |
|---|---|---|---|---|---|---|---|---|---|---|---|---|---|---|---|---|---|---|---|---|
| 2_17806498 | 5 | 0 | 7 | 0.1429 | 4 | 0 | 3 | 0 | 19 | 0.0526 | 3 | 0 | 4 | 0.25 | 4 | 0.25 | 5 | 0.2 | 16 | 0.1875 |
| 2_17806617 | 2 | 1 | 4 | 1 | 1 | 0 | 1 | 0 | 8 | 0.75 | 2 | 1 | 3 | 0.6667 | 3 | 1 | 5 | 0.6 | 13 | 0.7692 |
| 2_17864151 | NA | NA | NA | NA | NA | NA | 1 | 1 | 1 | 1 | NA | NA | 1 | 1 | NA | NA | NA | NA | 1 | 1 |
| 2_17864155 | NA | NA | NA | NA | NA | NA | 1 | 1 | 1 | 1 | NA | NA | 1 | 1 | NA | NA | NA | NA | 1 | 1 |
| 2_17864229 | NA | NA | NA | NA | NA | NA | 1 | 1 | 1 | 1 | NA | NA | 1 | 0 | 1 | 1 | NA | NA | 2 | 0.5 |
| 2_17864245 | NA | NA | NA | NA | NA | NA | 1 | 1 | 1 | 1 | NA | NA | NA | NA | 1 | 1 | NA | NA | 1 | 1 |
| 2_17953765 | NA | NA | 2 | 0.5 | 3 | 0.6667 | NA | NA | 5 | 0.6 | 1 | 1 | 1 | 1 | 1 | 1 | NA | NA | 3 | 1 |
| 2_17953768 | NA | NA | 1 | 1 | 2 | 1 | NA | NA | 3 | 1 | 1 | 1 | 1 | 0 | 1 | 0 | 1 | 1 | 4 | 0.5 |
| 2_17953770 | NA | NA | 2 | 0.5 | 3 | 0.6667 | NA | NA | 5 | 0.6 | 1 | 1 | 1 | 1 | 1 | 0 | 1 | 0 | 4 | 0.5 |
| 2_17953775 | NA | NA | 2 | 1 | 3 | 1 | NA | NA | 5 | 1 | 1 | 0 | 1 | 1 | NA | NA | NA | NA | 2 | 0.5 |
| 2_17953784 | NA | NA | NA | NA | 3 | 1 | NA | NA | 3 | 1 | 1 | 1 | NA | NA | NA | NA | NA | NA | 1 | 1 |
| 2_17953803 | NA | NA | 2 | 0.5 | 3 | 0.6667 | NA | NA | 5 | 0.6 | 1 | 1 | 1 | 1 | NA | NA | 1 | 0 | 3 | 0.6667 |
| 2_17953839 | NA | NA | 2 | 0.5 | 1 | 0 | NA | NA | 3 | 0.3333 | NA | NA | 1 | 0 | NA | NA | 2 | 1 | 3 | 0.6667 |
| 2_17953857 | NA | NA | 1 | 0 | 2 | 0.5 | NA | NA | 3 | 0.3333 | 1 | 1 | 1 | 1 | 1 | 1 | 2 | 0.5 | 5 | 0.8 |
| 2_17953876 | NA | NA | 2 | 0.5 | 3 | 0.3333 | NA | NA | 5 | 0.4 | NA | NA | 1 | 0 | 1 | 0 | 2 | 0.5 | 4 | 0.25 |
| 2_17953878 | NA | NA | 2 | 0.5 | 2 | 0.5 | NA | NA | 4 | 0.5 | 1 | 1 | 1 | 1 | 1 | 1 | 1 | 1 | 4 | 1 |
| 2_17953882 | NA | NA | 2 | 1 | 3 | 0.6667 | NA | NA | 5 | 0.8 | 1 | 1 | 1 | 1 | NA | NA | 1 | 1 | 3 | 1 |
| 2_17953884 | NA | NA | 2 | 0.5 | 1 | 1 | NA | NA | 3 | 0.6667 | 1 | 1 | 1 | 0 | 1 | 0 | 2 | 0.5 | 5 | 0.4 |
| 2_17953890 | NA | NA | 2 | 0.5 | 3 | 0.6667 | NA | NA | 5 | 0.6 | 1 | 0 | 1 | 0 | 1 | 0 | 2 | 0.5 | 5 | 0.2 |
| 2_17953947 | 1 | 1 | 2 | 0 | 2 | 0 | NA | NA | 5 | 0.2 | 2 | 0 | 1 | 0 | 2 | 0 | 2 | 0.5 | 7 | 0.1429 |
| 2_17954079 | 3 | 0.3333 | 3 | 0.6667 | 4 | 0.75 | 1 | 1 | 11 | 0.6364 | 2 | 0.5 | 1 | 1 | 2 | 0.5 | 3 | 0.6667 | 8 | 0.625 |
| 2_17955412 | 4 | 0.25 | 3 | 0.3333 | 4 | 0 | 2 | 0.5 | 13 | 0.2308 | 1 | 1 | 4 | 0.25 | 3 | 0.6667 | 5 | 0.6 | 13 | 0.5385 |
| 2_17955416 | 5 | 0 | 2 | 0 | 3 | 0.3333 | 3 | 0 | 13 | 0.0769 | 2 | 0.5 | 4 | 0.25 | 3 | 0 | 6 | 0 | 15 | 0.1333 |
| 2_17955437 | 2 | 1 | 4 | 0.25 | 2 | 0 | 3 | 0.6667 | 11 | 0.4545 | 1 | 1 | 3 | 0.6667 | 2 | 0.5 | 6 | 0.3333 | 12 | 0.5 |
| 2_17955449 | 4 | 0.75 | 3 | 0.3333 | 4 | 0.75 | 3 | 0.6667 | 14 | 0.6429 | NA | NA | 2 | 0.5 | 1 | 0 | 5 | 0.4 | 8 | 0.375 |
| 2_17955453 | 4 | 0.25 | 4 | 0 | 4 | 0.25 | 4 | 0.25 | 16 | 0.1875 | 1 | 0 | 4 | 0.25 | 2 | 0 | 6 | 0.1667 | 13 | 0.1538 |
| 2_17955458 | 4 | 0.25 | 4 | 0 | 4 | 0 | 3 | 0 | 15 | 0.0667 | 1 | 0 | 4 | 0.25 | 2 | 0 | 6 | 0.1667 | 13 | 0.1538 |
| 2_17955479 | 3 | 0.6667 | 3 | 0.6667 | 4 | 0.5 | 2 | 0.5 | 12 | 0.5833 | 1 | 1 | 4 | 0.25 | 1 | 1 | 6 | 0.3333 | 12 | 0.4167 |
| 2_17955480 | 3 | 0.6667 | 4 | 0 | 3 | 0.3333 | 3 | 0.3333 | 13 | 0.3077 | 1 | 0 | 4 | 0 | 2 | 0.5 | 6 | 0.3333 | 13 | 0.2308 |
| 2_17955495 | 2 | 1 | 2 | 1 | 3 | 1 | 3 | 1 | 10 | 1 | NA | NA | 4 | 0.75 | 2 | 0.5 | 4 | 0.25 | 10 | 0.5 |
| 2_17955501 | 4 | 0.5 | 3 | 0.6667 | 3 | 0 | 3 | 0 | 13 | 0.3077 | NA | NA | 3 | 0 | 2 | 0.5 | 6 | 0 | 11 | 0.0909 |
| 2_17955530 | 4 | 0.25 | 3 | 0.3333 | 3 | 0.3333 | 2 | 0 | 12 | 0.25 | NA | NA | 3 | 0.6667 | 2 | 0.5 | 5 | 0.2 | 10 | 0.4 |
| 2_17955640 | 1 | 0 | NA | NA | NA | NA | 1 | 1 | 2 | 0.5 | NA | NA | NA | NA | 1 | 1 | NA | NA | 1 | 1 |
| 2_17967563 | 9 | 0 | 4 | 0 | 1 | 0 | 3 | 0.3333 | 17 | 0.0588 | 5 | 0 | 3 | 0 | 3 | 0 | 3 | 0.3333 | 14 | 0.0714 |
| 2_17967621 | 10 | 0.8 | 4 | 0.75 | 3 | 1 | 3 | 1 | 20 | 0.85 | 5 | 0.4 | 2 | 0.5 | 2 | 1 | 6 | 0.8333 | 15 | 0.6667 |
| 2_17967622 | 10 | 0.4 | 5 | 0.2 | 3 | 0.3333 | 4 | 0.5 | 22 | 0.3636 | 4 | 0.25 | 2 | 1 | 3 | 0 | 6 | 0.3333 | 15 | 0.3333 |
| 2_17967641 | 8 | 0.125 | 3 | 0.3333 | 3 | 0 | 4 | 0 | 18 | 0.1111 | 4 | 0 | 3 | 0 | 3 | 0 | 6 | 0.1667 | 16 | 0.0625 |
| 2_17967724 | 10 | 0 | 16 | 0.125 | 10 | 0.1 | 8 | 0.5 | 44 | 0.1591 | 9 | 0 | 9 | 0.2222 | 13 | 0.1538 | 11 | 0.1818 | 42 | 0.1429 |
| 2_17968355 | NA | NA | NA | NA | 1 | 0 | 3 | 0.3333 | 4 | 0.25 | NA | NA | NA | NA | NA | NA | 1 | 1 | 1 | 1 |
| 2_17968360 | NA | NA | NA | NA | 1 | 0 | 3 | 0.3333 | 4 | 0.25 | NA | NA | 1 | 0 | NA | NA | 1 | 1 | 2 | 0.5 |
| 2_17968397 | 1 | 0 | NA | NA | 2 | 0 | 3 | 0.3333 | 6 | 0.1667 | NA | NA | 2 | 0.5 | NA | NA | 1 | 0 | 3 | 0.3333 |
| 2_17970919 | 3 | 0 | 1 | 1 | 3 | 0.3333 | 3 | 0 | 10 | 0.2 | 2 | 0.5 | NA | NA | 3 | 0 | NA | NA | 5 | 0.2 |
| 2_17970931 | 3 | 0 | 2 | 0 | 3 | 0.3333 | 3 | 0 | 11 | 0.0909 | 2 | 0 | NA | NA | 6 | 0.1667 | 2 | 0 | 10 | 0.1 |
| 2_17970951 | 3 | 0 | 2 | 0 | 3 | 0.3333 | 3 | 0 | 11 | 0.0909 | 2 | 0 | NA | NA | 6 | 0.1667 | 2 | 0 | 10 | 0.1 |
| 2_17970955 | 3 | 1 | 2 | 1 | 2 | 0.5 | 2 | 1 | 9 | 0.8889 | NA | NA | NA | NA | 5 | 0.4 | 2 | 0.5 | 7 | 0.4286 |
| 2_17971074 | 4 | 0.25 | 7 | 0.4286 | 5 | 0.2 | 2 | 0.5 | 18 | 0.3333 | 3 | 0.3333 | 2 | 0.5 | 6 | 0.1667 | 5 | 0.2 | 16 | 0.25 |
| 2_17971077 | 4 | 0.25 | 6 | 0.1667 | 4 | 0 | 3 | 0.3333 | 17 | 0.1765 | 1 | 0 | 2 | 0 | 5 | 0.2 | 2 | 0 | 10 | 0.1 |
| 2_17971094 | 4 | 0.25 | 8 | 0.125 | 5 | 0 | 3 | 0 | 20 | 0.1 | 3 | 0 | 3 | 0 | 6 | 0 | 6 | 0.1667 | 18 | 0.0556 |
| 2_17971159 | 3 | 0 | 10 | 0 | 5 | 0.2 | 2 | 0 | 20 | 0.05 | 3 | 0.3333 | 3 | 0 | 6 | 0 | 6 | 0.1667 | 18 | 0.1111 |
| 2_17971172 | 3 | 0.6667 | 9 | 0.4444 | 4 | 1 | 3 | 1 | 19 | 0.6842 | 3 | 1 | 3 | 0.6667 | 4 | 0.75 | 7 | 0.8571 | 17 | 0.8235 |
| 2_17971178 | 1 | 0 | 9 | 0.1111 | 4 | 0 | 2 | 0.5 | 16 | 0.125 | 4 | 0 | 3 | 0.3333 | 6 | 0 | 6 | 0.1667 | 19 | 0.1053 |
| 2_17972730 | 1 | 1 | 2 | 0.5 | 1 | 1 | 2 | 0.5 | 6 | 0.6667 | NA | NA | 2 | 1 | 2 | 0.5 | 4 | 1 | 8 | 0.875 |
| 2_17972732 | 1 | 1 | 2 | 1 | 1 | 0 | 1 | 1 | 5 | 0.8 | NA | NA | 2 | 1 | 1 | 1 | 2 | 0.5 | 5 | 0.8 |
| 2_17972743 | 1 | 1 | 2 | 0 | 1 | 1 | NA | NA | 4 | 0.5 | 1 | 1 | 1 | 0 | 2 | 0.5 | 4 | 0.5 | 8 | 0.5 |
| 2_17972746 | 1 | 0 | 2 | 0 | 1 | 1 | 2 | 0 | 6 | 0.1667 | 1 | 0 | 2 | 0.5 | 2 | 0 | 5 | 0.2 | 10 | 0.2 |
| 2_17974142 | 22 | 0.5455 | 13 | 0.6154 | 11 | 0.9091 | 19 | 0.5263 | 65 | 0.6154 | 12 | 0.75 | 18 | 0.7778 | 26 | 0.6923 | 23 | 0.7826 | 79 | 0.7468 |
| 2_17974163 | 31 | 0.871 | 20 | 0.85 | 17 | 0.8235 | 24 | 0.7917 | 92 | 0.837 | 17 | 0.8235 | 27 | 0.7778 | 30 | 0.7667 | 23 | 0.7826 | 97 | 0.7835 |
| 2_17974163 | 1 | 0 | 1 | 0 | NA | NA | 2 | 1 | 4 | 0.5 | 1 | 1 | 2 | 0.5 | NA | NA | 2 | 1 | 5 | 0.8 |
| 2_17974167 | 34 | 0.1765 | 21 | 0.1429 | 17 | 0.1765 | 22 | 0.0909 | 94 | 0.1489 | 16 | 0.125 | 23 | 0.0435 | 32 | 0.1562 | 24 | 0.2917 | 95 | 0.1579 |
| 2_17974174 | 34 | 0.1176 | 22 | 0.0909 | 15 | 0.1333 | 19 | 0.1053 | 90 | 0.1111 | 14 | 0.0714 | 23 | 0.0435 | 33 | 0.0909 | 23 | 0.1739 | 93 | 0.0968 |
| 2_17974206 | 27 | 0.8889 | 16 | 0.9375 | 14 | 0.8571 | 19 | 0.8947 | 76 | 0.8947 | 12 | 0.9167 | 22 | 0.8636 | 27 | 0.8519 | 22 | 0.9545 | 83 | 0.8916 |
| 2_17974211 | 29 | 0.7586 | 18 | 0.6667 | 17 | 0.7647 | 19 | 0.6842 | 83 | 0.7229 | 14 | 0.7857 | 26 | 0.4231 | 24 | 0.625 | 24 | 0.7083 | 88 | 0.6136 |
| 2_17974256 | 22 | 0.2727 | 13 | 0.4615 | 15 | 0.2667 | 14 | 0.2857 | 64 | 0.3125 | 7 | 0.2857 | 18 | 0.2222 | 21 | 0.5238 | 19 | 0.6316 | 65 | 0.4462 |
| 2_17974277 | 18 | 0.8889 | 14 | 0.8571 | 13 | 0.8462 | 16 | 0.9375 | 61 | 0.8852 | 11 | 0.6364 | 23 | 0.913 | 22 | 0.7727 | 19 | 0.6842 | 75 | 0.7733 |
| 2_17974290 | 19 | 0 | 18 | 0.1667 | 14 | 0.0714 | 16 | 0 | 67 | 0.0597 | 11 | 0 | 22 | 0 | 21 | 0.1429 | 19 | 0.0526 | 73 | 0.0548 |

| | | | | | | | | | | | | | | | | | | | | |
|---|---|---|---|---|---|---|---|---|---|---|---|---|---|---|---|---|---|---|---|---|
| 2_17974295 | 15 | 0.0667 | 14 | 0.0714 | 14 | 0.0714 | 11 | 0.0909 | 54 | 0.0741 | 10 | 0 | 19 | 0.1053 | 16 | 0.125 | 14 | 0.0714 | 59 | 0.0847 |
| 2_17974584 | 16 | 0 | 10 | 0 | 7 | 0.1429 | 11 | 0.2727 | 44 | 0.0909 | 7 | 0 | 15 | 0 | 7 | 0.2857 | 8 | 0 | 37 | 0.0541 |
| 2_17974597 | 1 | 1 | NA | NA | NA | NA | NA | NA | 1 | 1 | 1 | 1 | NA | NA | NA | NA | NA | NA | 1 | 1 |
| 2_17974608 | 11 | 0.0909 | 7 | 0.1429 | 4 | 0.5 | 5 | 0.2 | 27 | 0.1852 | 7 | 0.2857 | 10 | 0.2 | 8 | 0.25 | 6 | 0.3333 | 31 | 0.2581 |
| 2_17974611 | 15 | 0.0667 | 10 | 0.1 | 6 | 0 | 7 | 0.1429 | 38 | 0.0789 | 7 | 0 | 18 | 0.1111 | 10 | 0 | 10 | 0.2 | 45 | 0.0889 |
| 2_17974612 | 14 | 0.2143 | 10 | 0.2 | 6 | 0.1667 | 7 | 0.1429 | 37 | 0.1892 | 7 | 0 | 15 | 0.3333 | 9 | 0 | 9 | 0.4444 | 40 | 0.225 |
| 2_17974638 | 1 | 1 | NA | NA | NA | NA | 1 | 0 | 2 | 0.5 | 1 | 1 | NA | NA | NA | NA | NA | NA | 1 | 1 |
| 2_17980336 | 5 | 0.4 | 4 | 0.25 | 4 | 0.25 | 11 | 0.1818 | 24 | 0.25 | 3 | 0.3333 | 7 | 0.1429 | 8 | 0.5 | 7 | 0.2857 | 25 | 0.32 |
| 2_17980358 | 6 | 0.1667 | 4 | 0.75 | 6 | 0.3333 | 11 | 0.6364 | 27 | 0.4815 | 3 | 0.6667 | 6 | 0.3333 | 8 | 0.875 | 3 | 1 | 20 | 0.7 |
| 2_17980371 | 7 | 0.1429 | 6 | 0.8333 | 6 | 0.1667 | 9 | 0.2222 | 28 | 0.3214 | 2 | 0.5 | 6 | 0 | 9 | 0.5556 | 11 | 0.6364 | 28 | 0.4643 |
| 2_17980380 | 9 | 0.2222 | 6 | 0 | 6 | 0.1667 | 11 | 0 | 32 | 0.0938 | 3 | 0.3333 | 6 | 0.1667 | 11 | 0 | 11 | 0.0909 | 31 | 0.0968 |
| 2_17980385 | 8 | 0.5 | 6 | 0.1667 | 6 | 0.3333 | 11 | 0.4545 | 31 | 0.3871 | 2 | 0.5 | 7 | 0.1429 | 7 | 0.4286 | 10 | 0.2 | 26 | 0.2692 |
| 2_17980397 | 8 | 0.125 | 6 | 0 | 4 | 0.25 | 13 | 0 | 31 | 0.0645 | 1 | 0 | 6 | 0.1667 | 12 | 0.0833 | 5 | 0 | 24 | 0.0833 |
| 2_17980398 | 9 | 0 | 6 | 0 | 5 | 0 | 12 | 0.25 | 32 | 0.0938 | 4 | 0 | 6 | 0 | 12 | 0.25 | 9 | 0.3333 | 31 | 0.1935 |
| 2_17980406 | 6 | 0.1667 | 6 | 0 | 4 | 0 | 12 | 0.1667 | 28 | 0.1071 | 3 | 0 | 5 | 0 | 8 | 0.25 | 9 | 0 | 25 | 0.08 |
| 2_17980429 | 7 | 0.8571 | 4 | 0.5 | 5 | 0.4 | 12 | 0.6667 | 28 | 0.6429 | 2 | 1 | 6 | 0.8333 | 11 | 1 | 7 | 0.7143 | 26 | 0.8846 |
| 2_17990048 | 2 | 0.5 | 4 | 0 | 3 | 0.3333 | 2 | 1 | 11 | 0.3636 | 2 | 0.5 | 1 | 1 | 4 | 0.25 | 1 | 1 | 8 | 0.5 |
| 2_17990052 | 2 | 1 | 6 | 0.5 | 3 | 0.6667 | 2 | 0.5 | 13 | 0.6154 | 3 | 0.3333 | 1 | 1 | 4 | 0.5 | NA | NA | 8 | 0.5 |
| 2_17990069 | 3 | 0 | 5 | 0.2 | 2 | 0 | 2 | 0.5 | 12 | 0.1667 | 3 | 0 | 1 | 0 | 5 | 0.2 | 1 | 0 | 10 | 0.1 |
| 2_17990071 | 2 | 1 | 4 | 0.25 | 3 | 0.3333 | 2 | 1 | 11 | 0.5455 | 2 | 0 | 1 | 0 | 4 | 0.25 | 1 | 0 | 8 | 0.125 |
| 2_17990098 | 2 | 1 | 5 | 0.4 | 2 | 0 | 1 | 1 | 10 | 0.5 | 2 | 0.5 | 1 | 1 | 2 | 0.5 | 1 | 1 | 6 | 0.6667 |
| 2_17990107 | 1 | 1 | 4 | 0.25 | 3 | 0 | 2 | 0.5 | 10 | 0.3 | 1 | 0 | 1 | 0 | 4 | 0.25 | 2 | 0 | 8 | 0.125 |
| 2_17990114 | 1 | 1 | 4 | 0.75 | 3 | 0.6667 | 2 | 0.5 | 10 | 0.7 | 1 | 1 | 1 | 1 | 5 | 0.8 | 2 | 1 | 9 | 0.8889 |
| 2_17990116 | NA | NA | 5 | 0 | 2 | 0 | 2 | 0.5 | 9 | 0.1111 | 1 | 0 | 1 | 0 | 4 | 0 | 2 | 1 | 8 | 0.25 |
| 2_17990117 | 1 | 1 | 4 | 0.25 | 1 | 0 | 1 | 0 | 7 | 0.2857 | 1 | 1 | 1 | 0 | 2 | 0.5 | 2 | 0.5 | 6 | 0.5 |
| 2_17990123 | 1 | 0 | 2 | 0.5 | 2 | 0.5 | 2 | 0.5 | 7 | 0.4286 | 1 | 0 | 1 | 1 | 3 | 0.3333 | 1 | 1 | 6 | 0.5 |
| 2_17990124 | 1 | 1 | 3 | 1 | 3 | 0.6667 | 2 | 1 | 9 | 0.8889 | 1 | 1 | 1 | 1 | 3 | 1 | 2 | 1 | 7 | 1 |
| 2_17990125 | 1 | 0 | 2 | 1 | 2 | 1 | 2 | 0 | 7 | 0.5714 | 1 | 0 | NA | NA | 5 | 0.4 | 1 | 0 | 7 | 0.2857 |
| 2_17990149 | 1 | 1 | 3 | 0.6667 | 3 | 1 | 2 | 1 | 9 | 0.8889 | 1 | 1 | 1 | 1 | 5 | 1 | 2 | 0.5 | 9 | 0.8889 |
| 2_17990151 | 1 | 1 | 3 | 0.3333 | 3 | 0.3333 | 1 | 0 | 8 | 0.375 | 1 | 0 | 1 | 1 | 4 | 0.25 | 2 | 0.5 | 8 | 0.375 |
| 2_17990162 | 1 | 1 | 1 | 0 | 2 | 0.5 | 2 | 0.5 | 6 | 0.5 | NA | NA | 1 | 1 | 1 | 1 | 2 | 1 | 4 | 1 |
| 2_17990174 | 1 | 0 | 2 | 0 | 3 | 0.3333 | 2 | 0.5 | 8 | 0.25 | 1 | 0 | 1 | 0 | 5 | 0.4 | 1 | 0 | 8 | 0.25 |
| 2_17990177 | 1 | 1 | 3 | 0.6667 | 3 | 1 | 2 | 0.5 | 9 | 0.7778 | 1 | 1 | 1 | 1 | 4 | 0.75 | 2 | 1 | 8 | 0.875 |
| 2_17990190 | 1 | 0 | 3 | 0.3333 | 3 | 0.3333 | 1 | 0 | 8 | 0.25 | 1 | 0 | 1 | 0 | 4 | 0.75 | 1 | 0 | 7 | 0.4286 |
| 2_17990214 | 1 | 0 | 3 | 0.3333 | 3 | 0.3333 | NA | NA | 7 | 0.2857 | 1 | 0 | 1 | 1 | 4 | 0 | 1 | 0 | 7 | 0.1429 |
| 2_17990311 | NA | NA | NA | NA | 2 | 0.5 | NA | NA | 2 | 0.5 | 1 | 0 | NA | NA | 2 | 0.5 | 1 | 0 | 4 | 0.25 |
| 2_18018626 | NA | NA | 1 | 0 | 1 | 1 | NA | NA | 2 | 0.5 | 1 | 1 | NA | NA | NA | NA | NA | NA | 1 | 1 |
| 2_18018634 | NA | NA | 1 | 1 | 1 | 1 | 1 | 0 | 3 | 0.6667 | 1 | 1 | NA | NA | NA | NA | NA | NA | 1 | 1 |
| 2_18018637 | NA | NA | 1 | 1 | 1 | 1 | 1 | 1 | 3 | 1 | 1 | 1 | NA | NA | NA | NA | NA | NA | 1 | 1 |
| 2_18022623 | 1 | 1 | 2 | 0.5 | 2 | 0 | 2 | 0 | 7 | 0.2857 | 4 | 0 | 3 | 0 | 2 | 0 | 2 | 0.5 | 11 | 0.0909 |
| 2_18022742 | 4 | 0.25 | 2 | 0 | 2 | 0 | 2 | 0 | 10 | 0.1 | 4 | 0 | 3 | 0.3333 | 3 | 0 | 2 | 0 | 12 | 0.0833 |
| 2_18022743 | 4 | 0.25 | 1 | 0 | 2 | 0 | 2 | 0 | 9 | 0.1111 | 4 | 0 | 3 | 0.3333 | 2 | 0 | 2 | 0 | 11 | 0.0909 |
| 2_18022764 | 4 | 0.5 | 2 | 1 | 3 | 0.3333 | 2 | 1 | 11 | 0.6364 | 3 | 0.6667 | 2 | 0.5 | 3 | 1 | 2 | 0.5 | 10 | 0.7 |
| 2_18022773 | 4 | 1 | 2 | 1 | 3 | 1 | 2 | 1 | 11 | 1 | 3 | 1 | 3 | 0.6667 | 3 | 1 | 2 | 0.5 | 11 | 0.8182 |
| 2_18022802 | 2 | 1 | 2 | 0 | 4 | 0.25 | 2 | 0 | 10 | 0.3 | 3 | 0 | 2 | 0 | 4 | 0 | 1 | 1 | 10 | 0.1 |
| 2_18023615 | 4 | 0.25 | 3 | 1 | 2 | 0 | 3 | 0.6667 | 12 | 0.5 | 3 | 1 | 3 | 0.3333 | 8 | 0.5 | 3 | 1 | 17 | 0.6471 |
| 2_18023630 | 4 | 0.25 | 3 | 0.3333 | 2 | 0 | 3 | 0.6667 | 12 | 0.3333 | 4 | 0.25 | 4 | 0.25 | 5 | 0.2 | 3 | 0.3333 | 16 | 0.25 |
| 2_18023639 | 5 | 0.2 | 5 | 0.4 | 2 | 0 | 3 | 0 | 15 | 0.2 | 6 | 0.5 | 4 | 0.25 | 8 | 0.25 | 3 | 0.3333 | 21 | 0.3333 |
| 2_18023640 | 5 | 0.2 | 4 | 0 | 2 | 0 | 2 | 0 | 13 | 0.0769 | 5 | 0.2 | 4 | 0 | 7 | 0.1429 | 2 | 0 | 18 | 0.1111 |
| 2_18023701 | 7 | 0.1429 | 7 | 0 | 2 | 0 | 4 | 0 | 20 | 0.05 | 6 | 0.1667 | 6 | 0.1667 | 9 | 0 | 3 | 0 | 24 | 0.0833 |
| 2_18023744 | 5 | 0.2 | 4 | 0 | 2 | 0 | 3 | 0 | 14 | 0.0714 | 4 | 0.25 | 6 | 0 | 9 | 0.1111 | 4 | 0.25 | 23 | 0.1304 |
| 2_18026322 | 1 | 0 | 2 | 0.5 | 1 | 0 | 1 | 1 | 5 | 0.4 | 2 | 0 | 1 | 0 | 1 | 0 | 2 | 1 | 6 | 0.3333 |
| 2_18026331 | 2 | 1 | 2 | 0.5 | 1 | 1 | NA | NA | 5 | 0.8 | 1 | 0 | NA | NA | 1 | 0 | 2 | 1 | 4 | 0.5 |
| 2_18030005 | 7 | 0.1429 | 5 | 0.4 | 4 | 0.5 | 7 | 0.1429 | 23 | 0.2609 | 1 | 0 | 3 | 0 | 3 | 0.6667 | 3 | 0.3333 | 10 | 0.3 |
| 2_18030020 | 9 | 0.3333 | 4 | 0 | 3 | 1 | 8 | 0.625 | 24 | 0.4583 | 1 | 0 | 1 | 0 | 5 | 0.4 | 5 | 0.4 | 12 | 0.3333 |
| 2_18030037 | 8 | 0 | 4 | 0.25 | 2 | 0.5 | 8 | 0.125 | 22 | 0.1364 | 1 | 1 | 2 | 0 | 5 | 0.6 | 6 | 0.3333 | 14 | 0.4286 |
| 2_18030038 | 9 | 0.2222 | 4 | 0 | 3 | 0.6667 | 7 | 0.2857 | 23 | 0.2609 | 1 | 1 | NA | NA | 4 | 0.25 | 6 | 0.5 | 11 | 0.4545 |
| 2_18030043 | 9 | 0 | 3 | 0 | 3 | 0.3333 | 7 | 0.2857 | 22 | 0.1364 | 1 | 0 | 2 | 0 | 4 | 0.25 | 5 | 0.2 | 12 | 0.1667 |
| 2_18030056 | 8 | 0.125 | 3 | 0 | 2 | 1 | 9 | 0.1111 | 22 | 0.1818 | 1 | 1 | 2 | 0 | 5 | 0 | 6 | 0.1667 | 14 | 0.1429 |
| 2_18030075 | 7 | 0 | 2 | 0 | 2 | 0 | 9 | 0.1111 | 20 | 0.05 | 1 | 0 | 1 | 0 | 5 | 0.4 | 5 | 0.2 | 12 | 0.25 |
| 2_18030095 | 7 | 0.5714 | 2 | 1 | 2 | 1 | 8 | 0.5 | 19 | 0.6316 | 1 | 1 | 2 | 0 | 2 | 1 | 5 | 0.8 | 10 | 0.7 |
| 2_18030097 | 6 | 0.6667 | 1 | 1 | 2 | 1 | 7 | 0.5714 | 16 | 0.6875 | 1 | 1 | 2 | 0.5 | 1 | 0 | 5 | 1 | 9 | 0.7778 |
| 2_18030104 | 8 | 0 | 2 | 0.5 | 2 | 0.5 | 8 | 0.25 | 20 | 0.2 | 1 | 1 | 2 | 0 | 3 | 0.3333 | 5 | 0.2 | 11 | 0.2727 |
| 2_18030108 | 8 | 0.25 | 2 | 0.5 | 1 | 0 | 8 | 0.25 | 19 | 0.2632 | 1 | 0 | 2 | 0.5 | 3 | 0.3333 | 4 | 0.5 | 10 | 0.4 |
| 2_18030108 | 8 | 0.25 | 2 | 0 | 2 | 0.5 | 5 | 0.4 | 17 | 0.2941 | 1 | 0 | 2 | 0 | 1 | 0 | 4 | 0.75 | 8 | 0.375 |

| | | | | | | | | | | | | | | | | | | | | |
|---|---|---|---|---|---|---|---|---|---|---|---|---|---|---|---|---|---|---|---|---|
| 2_1803011( | 8 | 0.375 | 1 | 0 | 2 | 0.5 | 7 | 0.2857 | 18 | 0.3333 | NA | NA | 2 | 1 | 2 | 0 | 4 | 0 | 8 | 0.25 |
| 2_1803011( | 7 | 1 | 2 | 1 | 2 | 1 | 7 | 0.4286 | 18 | 0.7778 | NA | NA | 2 | 0.5 | 3 | 0.6667 | 3 | 0.6667 | 8 | 0.625 |
| 2_1803011ł | 8 | 0 | 1 | 0 | 2 | 0 | 7 | 0.4286 | 18 | 0.1667 | NA | NA | 2 | 0.5 | 3 | 0 | 4 | 0.25 | 9 | 0.2222 |
| 2_1803012' | 7 | 0.4286 | 2 | 0.5 | 1 | 1 | 7 | 0.4286 | 17 | 0.4706 | NA | NA | 2 | 0 | 3 | 0 | 3 | 0.6667 | 8 | 0.25 |
| 2_1803012; | 7 | 0.4286 | 1 | 0 | 2 | 0.5 | 6 | 0.3333 | 16 | 0.375 | NA | NA | 2 | 0.5 | 3 | 1 | 3 | 0.6667 | 8 | 0.75 |
| 2_1803012‹ | 7 | 0.4286 | 2 | 0.5 | 1 | 0 | 6 | 0.1667 | 16 | 0.3125 | NA | NA | 2 | 0 | 2 | 0 | 4 | 0.75 | 8 | 0.375 |
| 2_1803012( | 7 | 0.2857 | 2 | 1 | 2 | 1 | 8 | 0.25 | 19 | 0.4211 | NA | NA | 2 | 0.5 | 3 | 0.6667 | 3 | 0 | 8 | 0.375 |
| 2_1803012ł | 7 | 0.7143 | 2 | 0 | 2 | 0.5 | 8 | 0.5 | 19 | 0.5263 | NA | NA | 1 | 0 | 2 | 0 | 4 | 0.5 | 7 | 0.2857 |
| 2_1803013; | 6 | 0.3333 | NA | NA | 2 | 0 | 6 | 0 | 14 | 0.1429 | NA | NA | 1 | 0 | 3 | 0.3333 | 1 | 0 | 5 | 0.2 |
| 2_1803013! | 6 | 0 | 2 | 0 | 2 | 0 | 6 | 0.3333 | 16 | 0.125 | NA | NA | 2 | 0 | 3 | 0 | 3 | 0.3333 | 8 | 0.125 |
| 2_1803013! | 4 | 0 | 2 | 1 | 2 | 0.5 | 7 | 0.2857 | 15 | 0.3333 | NA | NA | 2 | 0 | 2 | 1 | 3 | 0.6667 | 7 | 0.5714 |
| 2_1803014‹ | 4 | 0.5 | 1 | 1 | 2 | 0.5 | 7 | 0.7143 | 14 | 0.6429 | NA | NA | 2 | 0 | 3 | 0.6667 | 2 | 0 | 7 | 0.2857 |
| 2_1803014( | 1 | 1 | 2 | 1 | 2 | 1 | 7 | 1 | 12 | 1 | NA | NA | 2 | 1 | 2 | 1 | 2 | 1 | 6 | 1 |
| 2_1803014ł | 3 | 0.6667 | 1 | 1 | 2 | 1 | 6 | 0.6667 | 12 | 0.75 | NA | NA | 2 | 0 | 2 | 1 | 2 | 1 | 6 | 0.6667 |
| 2_1803016( | 3 | 0 | 1 | 1 | 2 | 0 | 7 | 0.2857 | 13 | 0.2308 | NA | NA | 1 | 0 | 2 | 0.5 | 2 | 1 | 5 | 0.6 |
| 2_1803017! | 3 | 1 | 1 | 1 | 2 | 1 | 6 | 1 | 12 | 1 | NA | NA | 2 | 1 | 1 | 1 | 1 | 1 | 4 | 1 |
| 2_1803018! | 2 | 0.5 | 2 | 0.5 | 2 | 1 | 3 | 1 | 9 | 0.7778 | NA | NA | 1 | 0 | 1 | 1 | 1 | 1 | 3 | 0.6667 |
| 2_1803019! | 3 | 0.3333 | 1 | 1 | 1 | 1 | 3 | 1 | 8 | 0.75 | NA | NA | 1 | 1 | NA | NA | NA | NA | 1 | 1 |
| 2_1803021! | 3 | 0.6667 | 2 | 1 | 1 | 1 | 5 | 1 | 11 | 0.9091 | NA | NA | 2 | 0.5 | 1 | 1 | 1 | 1 | 4 | 0.75 |
| 2_1803021 | 1 | 1 | 1 | 1 | 1 | 1 | 1 | 1 | 4 | 1 | NA | NA | 2 | 0.5 | 1 | 1 | 1 | 1 | 4 | 0.75 |
| 2_1803022! | 2 | 0.5 | 1 | 1 | 1 | 1 | 3 | 1 | 7 | 0.8571 | NA | NA | 2 | 0.5 | 1 | 1 | 1 | 1 | 4 | 0.75 |
| 2_1803023ł | 2 | 1 | 1 | 1 | 1 | 1 | 3 | 1 | 7 | 1 | NA | NA | NA | NA | 1 | 1 | NA | NA | 1 | 1 |
| 2_1803023! | 1 | 1 | 2 | 1 | 1 | 1 | 3 | 0.6667 | 7 | 0.8571 | NA | NA | 2 | 1 | 1 | 1 | 1 | 1 | 4 | 1 |
| 2_1803024! | 2 | 1 | 2 | 0.5 | 1 | 1 | 3 | 0.3333 | 8 | 0.625 | NA | NA | 2 | 1 | 1 | 0 | 1 | 1 | 4 | 0.75 |
| 2_1803025' | 2 | 1 | 1 | 0 | 1 | 1 | 3 | 0.3333 | 7 | 0.5714 | NA | NA | 2 | 0.5 | 1 | 0 | 1 | 0 | 4 | 0.25 |
| 2_1803025‹ | 1 | 1 | 1 | 1 | NA | NA | 1 | 1 | 3 | 1 | NA | NA | 1 | 1 | NA | NA | 1 | 0 | 2 | 0.5 |
| 2_1803027( | 1 | 1 | 1 | 1 | 1 | 1 | 3 | 1 | 6 | 1 | NA | NA | 2 | 1 | 1 | 1 | 1 | 1 | 4 | 1 |
| 2_1803029 | 1 | 0 | 1 | 1 | 1 | 1 | 3 | 1 | 6 | 0.8333 | NA | NA | 1 | 0 | 1 | 1 | 1 | 1 | 3 | 0.6667 |
| 2_1803031; | 1 | 1 | 1 | 1 | 1 | 1 | 2 | 1 | 5 | 1 | NA | NA | 1 | 1 | 1 | 1 | 1 | 1 | 3 | 1 |
| 2_1803032‹ | 2 | 0 | 1 | 1 | 1 | 0 | 3 | 0.3333 | 7 | 0.2857 | NA | NA | 1 | 1 | 1 | 1 | 1 | 1 | 3 | 1 |
| 2_1803032( | 1 | 0 | NA | NA | 1 | 0 | 2 | 0.5 | 4 | 0.25 | NA | NA | 1 | 1 | 1 | 0 | 1 | 0 | 3 | 0.3333 |
| 2_1803035! | 2 | 0 | 1 | 0 | NA | NA | 3 | 0.3333 | 6 | 0.1667 | NA | NA | 1 | 0 | 1 | 1 | 1 | 0 | 3 | 0.3333 |
| 2_1803038( | 1 | 0 | 1 | 1 | NA | NA | 2 | 0 | 4 | 0.25 | NA | NA | 1 | 0 | 1 | 1 | 1 | 1 | 3 | 0.6667 |
| 2_1803041( | NA | NA | 1 | 0 | NA | NA | 2 | 0.5 | 3 | 0.3333 | NA | NA | 1 | 1 | NA | NA | NA | NA | 1 | 1 |
| 2_1803232' | NA | NA | 2 | 1 | 1 | 1 | NA | NA | 3 | 1 | 1 | 1 | NA | NA | 1 | 1 | NA | NA | 2 | 1 |
| 2_1803234' | NA | NA | 1 | 1 | 1 | 1 | NA | NA | 2 | 1 | 1 | 1 | NA | NA | 1 | 0 | NA | NA | 2 | 0.5 |
| 2_1803235; | NA | NA | 1 | 1 | 1 | 1 | NA | NA | 2 | 1 | 1 | 0 | NA | NA | 1 | 1 | NA | NA | 2 | 0.5 |
| 2_1803294; | 1 | 1 | 1 | 1 | 1 | 1 | NA | NA | 3 | 1 | NA | NA | 1 | 1 | NA | NA | NA | NA | 1 | 1 |
| 2_1803597( | 8 | 0.125 | 16 | 0.3125 | 11 | 0.0909 | 13 | 0.0769 | 48 | 0.1667 | 2 | 0.5 | 9 | 0 | 5 | 0.2 | 10 | 0.1 | 26 | 0.1154 |
| 2_1803598' | 7 | 0.4286 | 16 | 0.5625 | 9 | 0.6667 | 11 | 0.5455 | 43 | 0.5581 | 3 | 1 | 10 | 0.4 | 6 | 0.6667 | 9 | 0.3333 | 28 | 0.5 |
| 2_1803598‹ | 8 | 0.25 | 16 | 0.6875 | 11 | 0.7273 | 12 | 0.6667 | 47 | 0.617 | 3 | 0.3333 | 10 | 0.5 | 5 | 0.4 | 8 | 0.5 | 26 | 0.4615 |
| 2_1803598( | 7 | 0.4286 | 13 | 0.2308 | 11 | 0.1818 | 14 | 0.0714 | 45 | 0.2 | 3 | 0.3333 | 10 | 0 | 5 | 0.4 | 9 | 0.1111 | 27 | 0.1481 |
| 2_1803598! | 7 | 0 | 12 | 0.25 | 11 | 0.1818 | 12 | 0.3333 | 42 | 0.2143 | 4 | 0 | 10 | 0.1 | 6 | 0.1667 | 9 | 0.2222 | 29 | 0.1379 |
| 2_1803601; | 8 | 0 | 16 | 0.25 | 12 | 0.1667 | 11 | 0.0909 | 47 | 0.1489 | 4 | 0.75 | 10 | 0 | 8 | 0 | 10 | 0.2 | 32 | 0.1562 |
| 2_1803601; | 5 | 0.4 | 15 | 0.9333 | 11 | 0.7273 | 12 | 0.8333 | 43 | 0.7907 | 3 | 0.6667 | 10 | 0.7 | 8 | 0.75 | 8 | 0.625 | 29 | 0.6897 |
| 2_1803601! | 8 | 0.5 | 15 | 0.6 | 12 | 0.4167 | 10 | 0.3 | 45 | 0.4667 | 4 | 0.5 | 9 | 0.3333 | 8 | 0.625 | 10 | 0.4 | 31 | 0.4516 |
| 2_1803624; | 1 | 0 | 7 | 0.1429 | 5 | 0 | 3 | 0 | 16 | 0.0625 | 2 | 0 | 1 | 0 | 5 | 0 | 3 | 0.3333 | 11 | 0.0909 |
| 2_1803627ł | 1 | 0 | 7 | 0 | 6 | 0.1667 | 3 | 0 | 17 | 0.0588 | 3 | 0 | 1 | 0 | 6 | 0.1667 | 3 | 0 | 13 | 0.0769 |
| 2_1803630 | 1 | 0 | 4 | 0.25 | 3 | 0 | 5 | 0 | 13 | 0.0769 | 3 | 0 | 1 | 0 | 6 | 0.1667 | 3 | 0 | 13 | 0.0769 |
| 2_1803630! | 1 | 0 | 6 | 0.3333 | 6 | 0.3333 | 4 | 0 | 17 | 0.2353 | 2 | 0 | 1 | 1 | 5 | 0.2 | 3 | 0.6667 | 11 | 0.3636 |
| 2_1805353( | NA | NA | 1 | 1 | 1 | 1 | NA | NA | 2 | 1 | 1 | 1 | NA | NA | 1 | 1 | NA | NA | 2 | 1 |
| 2_1805360' | NA | NA | 4 | 0.25 | 1 | 0 | NA | NA | 5 | 0.2 | 1 | 1 | 1 | 0 | 2 | 0.5 | 4 | 0.5 | 8 | 0.5 |
| 2_1805360; | NA | NA | 2 | 0.5 | 1 | 0 | NA | NA | 3 | 0.3333 | 1 | 0 | NA | NA | 1 | 0 | 4 | 0.25 | 6 | 0.1667 |
| 2_1805360ł | NA | NA | 3 | 0.6667 | 2 | 0.5 | NA | NA | 5 | 0.6 | 1 | 1 | NA | NA | 2 | 0.5 | 3 | 0.6667 | 6 | 0.6667 |
| 2_1805368‹ | NA | NA | 4 | 0.25 | 2 | 0 | NA | NA | 6 | 0.1667 | 1 | 0 | 3 | 0.6667 | 2 | 0.5 | 5 | 0.2 | 11 | 0.3636 |
| 2_1805463; | 8 | 0.125 | 9 | 0.1111 | 2 | 0.5 | 6 | 0 | 25 | 0.12 | 14 | 0.2857 | 8 | 0 | 5 | 0 | 8 | 0.125 | 35 | 0.1429 |
| 2_1805463; | 7 | 0 | 9 | 0.1111 | 2 | 0 | 7 | 0.1429 | 25 | 0.08 | 11 | 0.1818 | 10 | 0.1 | 6 | 0 | 10 | 0 | 37 | 0.0811 |
| 2_1805474( | 4 | 0.5 | 4 | 0 | 1 | 0 | 5 | 0.6 | 14 | 0.3571 | 9 | 0.5556 | 7 | 0.1429 | 6 | 0.5 | 7 | 0.4286 | 29 | 0.4138 |
| 2_1805526; | 4 | 0 | 3 | 0.6667 | 1 | 0 | 5 | 0 | 13 | 0.1538 | 7 | 0 | 5 | 0.2 | 3 | 0 | 8 | 0.25 | 23 | 0.1304 |
| 2_1805526! | 3 | 1 | 4 | 0.75 | 1 | 1 | 5 | 0.6 | 13 | 0.7692 | 9 | 0.8889 | 6 | 0.6667 | 5 | 0.6 | 7 | 0.7143 | 27 | 0.7407 |
| 2_1805527; | 2 | 1 | 4 | 1 | NA | NA | 5 | 1 | 11 | 1 | 8 | 0.875 | 5 | 1 | 5 | 0.8 | 7 | 0.8571 | 25 | 0.88 |
| 2_1805527( | 1 | 1 | 3 | 0.6667 | 1 | 1 | 4 | 0.5 | 9 | 0.6667 | 7 | 0.8571 | 5 | 1 | 5 | 1 | 7 | 0.5714 | 24 | 0.8333 |
| 2_1805528( | 3 | 1 | 4 | 1 | 1 | 1 | 3 | 1 | 11 | 1 | 6 | 1 | 2 | 1 | 4 | 1 | 7 | 1 | 19 | 1 |
| 2_1805530; | 2 | 0.5 | 4 | 1 | 1 | 1 | 6 | 0.6667 | 13 | 0.7692 | 7 | 0.4286 | 3 | 0.6667 | 1 | 1 | 7 | 0.8571 | 18 | 0.6667 |
| 2_1805530ł | 1 | 1 | 3 | 0.3333 | NA | NA | 4 | 0 | 8 | 0.25 | 4 | 0.5 | 5 | 0.2 | 3 | 0.3333 | 5 | 0 | 17 | 0.2353 |

| | | | | | | | | | | | | | | | | | | | | |
|---|---|---|---|---|---|---|---|---|---|---|---|---|---|---|---|---|---|---|---|---|
| 2_1805530 | 3 | 0.3333 | 1 | 0 | 1 | 1 | 4 | 0.25 | 9 | 0.3333 | 4 | 0 | 4 | 0.25 | 3 | 0.3333 | 5 | 0.4 | 16 | 0.25 |
| 2_1805533 | 2 | 1 | 1 | 1 | 1 | 1 | 3 | 1 | 7 | 1 | 7 | 1 | 2 | 1 | 1 | 1 | 6 | 0.8333 | 16 | 0.9375 |
| 2_1805534 | 3 | 0.6667 | 2 | 1 | NA | NA | 4 | 0.5 | 9 | 0.6667 | 7 | 0.5714 | 5 | 0.4 | 2 | 1 | 5 | 0.6 | 19 | 0.5789 |
| 2_1805534 | 2 | 0 | 2 | 0.5 | 1 | 0 | 3 | 0 | 8 | 0.125 | 5 | 0.2 | 4 | 0 | 2 | 0 | 4 | 0 | 15 | 0.0667 |
| 2_1805535 | 1 | 1 | NA | NA | NA | NA | 4 | 0 | 5 | 0.2 | 4 | 0.5 | 4 | 0 | 2 | 0 | 4 | 0 | 14 | 0.1429 |
| 2_1805536 | 3 | 0.3333 | 2 | 0.5 | 1 | 0 | 4 | 0 | 10 | 0.2 | 7 | 0.2857 | 4 | 0.25 | 2 | 0 | 6 | 0 | 19 | 0.1579 |
| 2_1805537 | 3 | 0.6667 | 2 | 0 | 1 | 0 | 5 | 0 | 11 | 0.1818 | 5 | 0.2 | 5 | 0.4 | 2 | 1 | 6 | 0.3333 | 18 | 0.3889 |
| 2_1805537 | 3 | 0.3333 | 2 | 0.5 | 1 | 0 | 2 | 0 | 8 | 0.25 | 4 | 0.5 | 4 | 0 | 1 | 0 | 3 | 0.3333 | 12 | 0.25 |
| 2_1805538 | 3 | 1 | 2 | 1 | 1 | 0 | 4 | 1 | 10 | 0.9 | 4 | 0.5 | 3 | 1 | 2 | 1 | 5 | 1 | 14 | 0.8571 |
| 2_1805835 | 1 | 1 | NA | NA | NA | NA | NA | NA | 1 | 1 | NA | NA | NA | NA | 1 | 1 | NA | NA | 1 | 1 |
| 2_1805839 | 1 | 1 | NA | NA | NA | NA | NA | NA | 1 | 1 | NA | NA | NA | NA | 1 | 1 | NA | NA | 1 | 1 |
| 2_1805840 | 1 | 1 | NA | NA | NA | NA | NA | NA | 1 | 1 | NA | NA | NA | NA | 1 | 1 | NA | NA | 1 | 1 |
| 2_1805841 | 1 | 1 | NA | NA | NA | NA | NA | NA | 1 | 1 | NA | NA | NA | NA | 1 | 1 | NA | NA | 1 | 1 |
| 2_1806145 | 4 | 0.25 | 6 | 0 | 1 | 0 | 2 | 0 | 13 | 0.0769 | 3 | 0 | 4 | 0 | 2 | 0 | 3 | 0.3333 | 12 | 0.0833 |
| 2_1806230 | 6 | 0.5 | 5 | 0.4 | 2 | 1 | 5 | 0.6 | 18 | 0.5556 | 9 | 0.7778 | 5 | 1 | 2 | 0.5 | 3 | 0.6667 | 19 | 0.7895 |
| 2_1806230 | 3 | 1 | 5 | 0 | 2 | 0 | 4 | 0.25 | 14 | 0.2857 | 5 | 0 | 3 | 0.3333 | 2 | 0.5 | 3 | 0 | 13 | 0.1538 |
| 2_1806231 | 6 | 0.1667 | 7 | 0.1429 | 3 | 0.6667 | 9 | 0.2222 | 25 | 0.24 | 10 | 0.6 | 7 | 0.1429 | 2 | 0 | 5 | 0 | 24 | 0.2917 |
| 2_1806231 | 6 | 0.8333 | 9 | 0.7778 | 3 | 1 | 10 | 1 | 28 | 0.8929 | 10 | 1 | 4 | 0.75 | 3 | 1 | 5 | 0.8 | 22 | 0.9091 |
| 2_1806231 | 6 | 0 | 11 | 0.3636 | 3 | 0 | 9 | 0.1111 | 29 | 0.1724 | 8 | 0.5 | 6 | 0.3333 | 5 | 0 | 6 | 0.1667 | 25 | 0.28 |
| 2_1806232 | 6 | 0.1667 | 6 | 0.5 | 2 | 0 | 8 | 0.125 | 22 | 0.2273 | 7 | 0.1429 | 4 | 0 | 4 | 0 | 3 | 0.3333 | 18 | 0.1111 |
| 2_1806232 | 8 | 1 | 11 | 0.6364 | 2 | 1 | 12 | 1 | 33 | 0.8788 | 9 | 0.7778 | 8 | 0.5 | 6 | 1 | 6 | 0.6667 | 29 | 0.7241 |
| 2_1806233 | 9 | 0.3333 | 9 | 0 | 4 | 0 | 10 | 0.1 | 32 | 0.125 | 10 | 0.1 | 7 | 0.4286 | 6 | 0.3333 | 6 | 0 | 29 | 0.2069 |
| 2_1806233 | 8 | 1 | 10 | 1 | 5 | 0.8 | 5 | 1 | 28 | 0.9643 | 8 | 1 | 6 | 0.8333 | 2 | 1 | 4 | 1 | 20 | 0.95 |
| 2_1806233 | 9 | 0 | 11 | 0.0909 | 4 | 0.25 | 12 | 0.25 | 36 | 0.1389 | 10 | 0.1 | 7 | 0 | 5 | 0.2 | 6 | 0 | 28 | 0.0714 |
| 2_1806234 | 9 | 0.7778 | 11 | 0.7273 | 3 | 1 | 11 | 0.8182 | 34 | 0.7941 | 10 | 1 | 7 | 0.8571 | 6 | 0.8333 | 5 | 1 | 28 | 0.9286 |
| 2_1806234 | 9 | 0 | 12 | 0.1667 | 6 | 0.3333 | 12 | 0.1667 | 39 | 0.1538 | 11 | 0.0909 | 8 | 0.25 | 7 | 0.1429 | 7 | 0.2857 | 33 | 0.1818 |
| 2_1806234 | 7 | 0 | 12 | 0.25 | 5 | 0 | 12 | 0.1667 | 36 | 0.1389 | 8 | 0.375 | 5 | 0.2 | 7 | 0.1429 | 6 | 0 | 26 | 0.1923 |
| 2_1806236 | 9 | 0.3333 | 11 | 0.0909 | 4 | 0 | 10 | 0.2 | 34 | 0.1765 | 4 | 0.25 | 5 | 0.2 | 7 | 0.1429 | 6 | 0.1667 | 22 | 0.1818 |
| 2_1806236 | 7 | 1 | 11 | 0.7273 | 5 | 1 | 10 | 0.8 | 33 | 0.8485 | 10 | 0.6 | 6 | 1 | 7 | 1 | 6 | 0.8333 | 29 | 0.8276 |
| 2_1806236 | 7 | 0.1429 | 10 | 0.2 | 6 | 0 | 12 | 0.1667 | 35 | 0.1429 | 8 | 0.5 | 6 | 0.1667 | 5 | 0 | 8 | 0.125 | 27 | 0.2222 |
| 2_1806237 | 6 | 0.8333 | 10 | 0.7 | 3 | 0.3333 | 9 | 0.7778 | 28 | 0.7143 | 9 | 0.6667 | 7 | 0.5714 | 5 | 0.8 | 8 | 0.875 | 29 | 0.7241 |
| 2_1806237 | 8 | 0.75 | 10 | 0.8 | 4 | 0.5 | 11 | 1 | 33 | 0.8182 | 11 | 0.9091 | 5 | 1 | 7 | 0.8571 | 6 | 1 | 29 | 0.931 |
| 2_1806237 | 9 | 0.8889 | 8 | 0.875 | 6 | 1 | 11 | 0.9091 | 34 | 0.9118 | 9 | 0.8889 | 6 | 1 | 6 | 0.8333 | 8 | 1 | 29 | 0.931 |
| 2_1806237 | 8 | 1 | 10 | 0.8 | 5 | 1 | 10 | 1 | 33 | 0.9394 | 8 | 1 | 7 | 0.8571 | 7 | 1 | 6 | 1 | 28 | 0.9643 |
| 2_1806239 | 7 | 0.2857 | 6 | 0.1667 | 5 | 0.4 | 14 | 0.3571 | 32 | 0.3125 | 9 | 0.3333 | 5 | 0.2 | 7 | 0.4286 | 8 | 0.5 | 29 | 0.3793 |
| 2_1806240 | 9 | 0.4444 | 10 | 0.3 | 7 | 0.4286 | 12 | 0.0833 | 38 | 0.2895 | 14 | 0.3571 | 7 | 0.1429 | 4 | 0 | 8 | 0.125 | 33 | 0.2121 |
| 2_1806241 | 7 | 0.4286 | 8 | 0.125 | 5 | 0.2 | 12 | 0.3333 | 32 | 0.2812 | 10 | 0.8 | 6 | 0.6667 | 6 | 0.1667 | 6 | 0.6667 | 28 | 0.6071 |
| 2_1806289 | NA | NA | 1 | 0 | 1 | 1 | 1 | 0 | 3 | 0.3333 | 1 | 0 | 1 | 0 | NA | NA | 1 | 1 | 3 | 0.3333 |
| 2_1806603 | NA | NA | NA | NA | 1 | 1 | 1 | 0 | 2 | 0.5 | 1 | 1 | NA | NA | 1 | 0 | 1 | 1 | 3 | 0.6667 |
| 2_1806605 | NA | NA | NA | NA | 1 | 0 | 1 | 1 | 2 | 0.5 | NA | NA | NA | NA | 1 | 1 | 1 | 0 | 2 | 0.5 |
| 2_1806605 | NA | NA | NA | NA | 1 | 1 | 1 | 0 | 2 | 0.5 | NA | NA | NA | NA | 1 | 0 | 1 | 1 | 2 | 0.5 |
| 2_1806643 | NA | NA | 2 | 0.5 | NA | NA | 2 | 0.5 | 4 | 0.5 | NA | NA | NA | NA | 1 | 0 | 2 | 1 | 3 | 0.6667 |
| 2_1806646 | 3 | 0.6667 | 2 | 1 | 2 | 1 | 1 | 0 | 8 | 0.75 | 4 | 1 | NA | NA | 3 | 0.3333 | 1 | 0 | 8 | 0.625 |
| 2_1806646 | 5 | 0.2 | 2 | 0 | 2 | 0 | 2 | 0 | 11 | 0.0909 | 4 | 0 | NA | NA | 4 | 0 | 2 | 0.5 | 10 | 0.1 |
| 2_1806648 | 1 | 1 | 2 | 0 | 2 | 0 | 2 | 0 | 7 | 0.1429 | 3 | 0.3333 | NA | NA | 3 | 0 | 2 | 0.5 | 8 | 0.25 |
| 2_1806648 | 4 | 1 | 2 | 1 | 1 | 1 | 3 | 1 | 10 | 1 | 4 | 1 | NA | NA | 3 | 1 | 2 | 1 | 9 | 1 |
| 2_1806648 | 2 | 0.5 | 3 | 0.6667 | 1 | 0 | 1 | 0 | 7 | 0.4286 | 2 | 0 | NA | NA | 2 | 0.5 | 1 | 0 | 5 | 0.2 |
| 2_1806648 | 4 | 0.75 | 3 | 0.6667 | 2 | 1 | 3 | 1 | 12 | 0.8333 | 3 | 1 | NA | NA | 3 | 1 | 2 | 1 | 8 | 1 |
| 2_1806649 | 4 | 1 | 2 | 0.5 | 2 | 1 | 3 | 1 | 11 | 0.9091 | 3 | 0.6667 | NA | NA | 3 | 1 | 2 | 1 | 8 | 0.875 |
| 2_1806650 | 2 | 1 | 2 | 1 | 1 | 1 | 1 | 1 | 6 | 1 | 2 | 1 | NA | NA | 2 | 0.5 | 2 | 0 | 6 | 0.5 |
| 2_1806650 | 4 | 0.5 | 3 | 0.6667 | 2 | 0 | 3 | 0.3333 | 12 | 0.4167 | 1 | 1 | NA | NA | 3 | 0.6667 | 3 | 0.3333 | 7 | 0.5714 |
| 2_1806650 | 4 | 0.5 | 1 | 1 | 2 | 1 | 3 | 0.6667 | 10 | 0.7 | 2 | 0.5 | NA | NA | 2 | 1 | 3 | 0.3333 | 7 | 0.5714 |
| 2_1806651 | 4 | 0.5 | 3 | 0.3333 | 1 | 0 | 4 | 0.5 | 12 | 0.4167 | 2 | 0 | NA | NA | 3 | 0.6667 | 2 | 0.5 | 7 | 0.4286 |
| 2_1806652 | 4 | 0.25 | 3 | 0 | 2 | 1 | 2 | 0 | 11 | 0.2727 | 2 | 0.5 | NA | NA | 3 | 0.6667 | 2 | 0 | 7 | 0.4286 |
| 2_1806652 | 3 | 0.6667 | 3 | 0.6667 | 2 | 1 | 2 | 0.5 | 10 | 0.7 | 2 | 0.5 | NA | NA | 3 | 0.6667 | 2 | 0 | 7 | 0.4286 |
| 2_1806654 | 4 | 0.5 | 3 | 0 | 1 | 1 | 2 | 0 | 10 | 0.3 | 2 | 0.5 | NA | NA | 3 | 0.3333 | 1 | 1 | 6 | 0.5 |
| 2_1806656 | 2 | 1 | 3 | 0.3333 | 1 | 1 | 2 | 0 | 8 | 0.5 | 2 | 0.5 | NA | NA | 3 | 0.3333 | 2 | 0.5 | 7 | 0.4286 |
| 2_1806656 | 2 | 0.5 | 3 | 1 | 1 | 1 | 3 | 1 | 9 | 0.8889 | 2 | 1 | NA | NA | 3 | 1 | 2 | 0.5 | 7 | 0.8571 |
| 2_1806657 | 2 | 0.5 | 2 | 1 | NA | NA | 2 | 1 | 6 | 0.8333 | 3 | 0.6667 | NA | NA | 3 | 0.3333 | 2 | 0 | 8 | 0.375 |
| 2_1806659 | 3 | 0.6667 | 3 | 0.6667 | NA | NA | 2 | 0.5 | 8 | 0.625 | 3 | 1 | NA | NA | 3 | 0.6667 | 2 | 1 | 8 | 0.875 |
| 2_1806663 | 3 | 0.3333 | 3 | 0.3333 | NA | NA | 2 | 0 | 8 | 0.25 | 3 | 0 | NA | NA | 3 | 0.3333 | 2 | 0 | 8 | 0.125 |
| 2_1806663 | 2 | 0.5 | 3 | 0.6667 | NA | NA | 2 | 0 | 7 | 0.4286 | 1 | 1 | NA | NA | 3 | 0 | 1 | 0 | 5 | 0.2 |
| 2_1806663 | 3 | 1 | 3 | 1 | NA | NA | 2 | 1 | 8 | 1 | 2 | 1 | NA | NA | 3 | 1 | 3 | 0.6667 | 8 | 0.875 |
| 2_1806665 | 1 | 1 | 3 | 1 | NA | NA | 2 | 1 | 6 | 1 | 3 | 1 | NA | NA | 3 | 1 | 3 | 0.6667 | 9 | 0.8889 |
| 2_1806705 | 3 | 0.6667 | 2 | 0.5 | NA | NA | 1 | 1 | 6 | 0.6667 | 1 | 1 | NA | NA | 1 | 0 | 2 | 0.5 | 4 | 0.5 |

| | | | | | | | | | | | | | | | | | | | | |
|---|---|---|---|---|---|---|---|---|---|---|---|---|---|---|---|---|---|---|---|---|
| 2_18067065 | 3 | 0.3333 | 2 | 0.5 | NA | NA | 1 | 0 | 6 | 0.3333 | 1 | 1 | NA | NA | 1 | 0 | 1 | 1 | 3 | 0.6667 |
| 2_18067070 | 3 | 1 | 1 | 1 | NA | NA | NA | NA | 4 | 1 | 1 | 1 | NA | NA | 1 | 1 | 1 | 1 | 3 | 1 |
| 2_18067114 | 1 | 0 | 1 | 1 | NA | NA | NA | NA | 2 | 0.5 | 1 | 1 | NA | NA | NA | NA | 1 | 1 | 2 | 1 |
| 2_18067122 | 1 | 1 | NA | NA | NA | NA | NA | NA | 1 | 1 | 1 | 1 | NA | NA | NA | NA | 1 | 1 | 2 | 1 |
| 2_18071749 | 15 | 0.8667 | 19 | 0.8421 | 26 | 0.8846 | 24 | 0.875 | 84 | 0.869 | 16 | 0.875 | 19 | 0.6316 | 26 | 0.8077 | 13 | 0.6154 | 74 | 0.7432 |
| 2_18071757 | 23 | 0.8696 | 24 | 0.9583 | 31 | 0.9032 | 32 | 0.9062 | 110 | 0.9091 | 17 | 0.8824 | 24 | 0.875 | 37 | 0.8919 | 35 | 0.8857 | 113 | 0.885 |
| 2_18071759 | 19 | 0.7895 | 18 | 0.9444 | 22 | 0.9545 | 23 | 1 | 82 | 0.9268 | 12 | 0.9167 | 15 | 0.8667 | 22 | 0.9091 | 25 | 0.96 | 74 | 0.9189 |
| 2_18071787 | 13 | 0.8462 | 16 | 0.75 | 14 | 0.8571 | 15 | 0.7333 | 58 | 0.7931 | 10 | 0.6 | 12 | 0.5833 | 20 | 0.5 | 14 | 0.5714 | 56 | 0.5536 |
| 2_18071800 | 14 | 0.3571 | 17 | 0.4706 | 13 | 0.5385 | 23 | 0.5652 | 67 | 0.4925 | 11 | 0.4545 | 16 | 0.375 | 22 | 0.3182 | 14 | 0.5714 | 63 | 0.4127 |
| 2_18071801 | 13 | 0.6154 | 20 | 0.45 | 19 | 0.4211 | 29 | 0.7241 | 81 | 0.5679 | 12 | 0.5 | 19 | 0.5263 | 29 | 0.4828 | 19 | 0.3158 | 79 | 0.4557 |
| 2_18071809 | 18 | 0.0556 | 26 | 0.2692 | 20 | 0.15 | 30 | 0.1333 | 94 | 0.1596 | 14 | 0.2143 | 18 | 0.1111 | 35 | 0.1714 | 24 | 0.0833 | 91 | 0.1429 |
| 2_18071814 | 16 | 0.4375 | 22 | 0.7273 | 17 | 0.6471 | 23 | 0.5652 | 78 | 0.6026 | 11 | 0.5455 | 15 | 0.8 | 25 | 0.6 | 21 | 0.5238 | 72 | 0.6111 |
| 2_18071815 | 18 | 0.1111 | 22 | 0.1364 | 17 | 0.2353 | 27 | 0.1481 | 84 | 0.1548 | 14 | 0.4286 | 17 | 0.2353 | 33 | 0.0909 | 23 | 0.1304 | 87 | 0.1839 |
| 2_18071827 | 18 | 0.2778 | 23 | 0.087 | 18 | 0.1667 | 29 | 0.1379 | 88 | 0.1591 | 9 | 0.2222 | 15 | 0.4 | 28 | 0.1786 | 18 | 0.1667 | 70 | 0.2286 |
| 2_18071906 | 14 | 0 | 25 | 0.04 | 15 | 0.0667 | 23 | 0.1304 | 77 | 0.0649 | 9 | 0.1111 | 11 | 0 | 22 | 0.1364 | 17 | 0.0588 | 59 | 0.0847 |
| 2_18071931 | 11 | 0.1818 | 15 | 0 | 15 | 0.2 | 19 | 0.1053 | 60 | 0.1167 | 10 | 0.3 | 12 | 0.25 | 18 | 0.0556 | 15 | 0.2 | 55 | 0.1818 |
| 2_18071974 | 10 | 0.2 | 14 | 0.0714 | 12 | 0.1667 | 17 | 0.1176 | 53 | 0.1321 | 8 | 0 | 10 | 0.1 | 11 | 0 | 8 | 0.125 | 37 | 0.0541 |
| 2_18072056 | 9 | 0.2222 | 14 | 0.1429 | 10 | 0 | 18 | 0.0556 | 51 | 0.098 | 6 | 0 | 8 | 0 | 14 | 0.0714 | 8 | 0.25 | 36 | 0.0833 |
| 2_18072234 | 6 | 0.1667 | 9 | 0 | 11 | 0.0909 | 13 | 0 | 39 | 0.0513 | 5 | 0 | 7 | 0 | 14 | 0.1429 | 4 | 0 | 30 | 0.0667 |
| 2_18072252 | 4 | 0.25 | 10 | 0.2 | 9 | 0.2222 | 11 | 0 | 34 | 0.1471 | 3 | 0.3333 | 7 | 0.1429 | 13 | 0.0769 | 5 | 0.2 | 28 | 0.1429 |
| 2_18072276 | 3 | 0 | 9 | 0.3333 | 9 | 0 | 12 | 0.0833 | 33 | 0.1212 | 5 | 0.2 | 7 | 0.1429 | 16 | 0.0625 | 4 | 0 | 32 | 0.0938 |
| 2_18072279 | 3 | 0 | 5 | 0.2 | 9 | 0.2222 | 10 | 0.1 | 27 | 0.1481 | 5 | 0.4 | 6 | 0.1667 | 10 | 0.2 | 5 | 0.2 | 26 | 0.2308 |
| 2_18072294 | 3 | 0.6667 | 9 | 0.5556 | 9 | 0.2222 | 10 | 0.3 | 31 | 0.3871 | 5 | 0.2 | 8 | 0.5 | 13 | 0.1538 | 5 | 0 | 31 | 0.2258 |
| 2_18072302 | 1 | 0 | 9 | 0.1111 | 9 | 0.2222 | 9 | 0.3333 | 28 | 0.2143 | 5 | 0.2 | 7 | 0.1429 | 11 | 0.2727 | 5 | 0.2 | 28 | 0.2143 |
| 2_18072308 | 2 | 1 | 7 | 0.7143 | 8 | 0.5 | 5 | 1 | 22 | 0.7273 | 2 | 0.5 | 5 | 0.8 | 7 | 0.7143 | 2 | 0.5 | 16 | 0.6875 |
| 2_18072335 | 2 | 0.5 | 8 | 0.125 | 9 | 0.4444 | 5 | 0.6 | 24 | 0.375 | 4 | 0.25 | 6 | 0.3333 | 12 | 0.6667 | 2 | 0.5 | 24 | 0.5 |
| 2_18072343 | 3 | 1 | 5 | 1 | 10 | 0.7 | 3 | 1 | 21 | 0.8571 | 3 | 0 | 2 | 1 | 8 | 0.75 | 3 | 0.6667 | 16 | 0.625 |
| 2_18072349 | 3 | 0.6667 | 8 | 1 | 8 | 0.5 | 5 | 0.8 | 24 | 0.75 | 4 | 0.5 | 4 | 0.75 | 10 | 0.6 | 2 | 1 | 20 | 0.65 |
| 2_18078326 | 1 | 0 | 1 | 1 | 3 | 0.3333 | 5 | 0 | 10 | 0.2 | NA | NA | 1 | 1 | 2 | 0 | NA | NA | 3 | 0.3333 |
| 2_18078404 | 12 | 0.0833 | 9 | 0.2222 | 13 | 0.0769 | 12 | 0 | 46 | 0.087 | 8 | 0.125 | 8 | 0.125 | 7 | 0.1429 | 6 | 0.1667 | 29 | 0.1379 |
| 2_18078642 | 7 | 0.1429 | 2 | 0 | 4 | 0.5 | 5 | 0 | 18 | 0.1667 | 1 | 0 | 2 | 0 | 5 | 0.2 | NA | NA | 8 | 0.125 |
| 2_18078814 | 8 | 0.125 | 12 | 0.4167 | 6 | 0.1667 | 10 | 0.1 | 36 | 0.2222 | 3 | 0 | 4 | 0.5 | 8 | 0.375 | 4 | 0.75 | 19 | 0.4211 |
| 2_18078820 | 9 | 0 | 10 | 0.2 | 6 | 0 | 11 | 0 | 36 | 0.0556 | 4 | 0 | 2 | 0.5 | 5 | 0.2 | 3 | 0 | 14 | 0.1429 |
| 2_18078829 | 8 | 0 | 11 | 0.1818 | 4 | 0.25 | 8 | 0 | 31 | 0.0968 | 2 | 0 | 4 | 0 | 7 | 0.1429 | 4 | 0 | 17 | 0.0588 |
| 2_18078838 | 6 | 0.1667 | 11 | 0.0909 | 4 | 0.5 | 8 | 0 | 29 | 0.1379 | 3 | 0 | 3 | 0.6667 | 7 | 0 | 2 | 0 | 15 | 0.1333 |
| 2_18078853 | 6 | 0 | 12 | 0 | 3 | 0.6667 | 9 | 0.1111 | 30 | 0.1 | 2 | 0.5 | 3 | 0.3333 | 7 | 0 | 4 | 0.25 | 16 | 0.1875 |
| 2_18078854 | 6 | 0 | 12 | 0 | 2 | 0.5 | 9 | 0.1111 | 29 | 0.069 | 2 | 0 | 3 | 0.3333 | 8 | 0 | 4 | 0.25 | 17 | 0.1176 |
| 2_18078856 | 2 | 0 | 10 | 0.6 | 3 | 1 | 7 | 0.2857 | 22 | 0.5 | 1 | 0 | 2 | 0.5 | 6 | 0.1667 | 4 | 1 | 13 | 0.4615 |
| 2_18078872 | 6 | 0.3333 | 9 | 0.3333 | 1 | 1 | 9 | 0.2222 | 25 | 0.32 | 1 | 0 | 1 | 0 | 8 | 0.5 | 3 | 1 | 13 | 0.5385 |
| 2_18078892 | 10 | 0.4 | 14 | 0.2857 | 3 | 0.3333 | 13 | 0.1538 | 40 | 0.275 | 3 | 0.6667 | 4 | 0 | 11 | 0.2727 | 5 | 0.2 | 23 | 0.2609 |
| 2_18078919 | 9 | 0.3333 | 15 | 0.3333 | 3 | 0.3333 | 10 | 0 | 37 | 0.2432 | 5 | 0.2 | 5 | 0.4 | 5 | 0 | 2 | 0 | 17 | 0.1765 |
| 2_18078922 | 8 | 0.625 | 14 | 0.3571 | 4 | 0.5 | 11 | 0.4545 | 37 | 0.4595 | 3 | 0.3333 | 3 | 0 | 8 | 0.25 | 2 | 0.5 | 16 | 0.25 |
| 2_18078923 | 6 | 0.3333 | 10 | 0.4 | 5 | 0.2 | 8 | 0.125 | 29 | 0.2759 | 2 | 1 | 2 | 0.5 | 5 | 0.2 | 1 | 1 | 10 | 0.5 |
| 2_18078923 | 8 | 0 | 17 | 0.1765 | 4 | 0 | 12 | 0.1667 | 41 | 0.122 | 5 | 0.2 | 4 | 0.25 | 9 | 0 | 4 | 0 | 22 | 0.0909 |
| 2_18078938 | 6 | 0.3333 | 9 | 0.2222 | 5 | 0 | 8 | 0 | 28 | 0.1429 | 5 | 0 | 3 | 0 | 10 | 0.3 | 3 | 0 | 21 | 0.1429 |
| 2_18078949 | 150 | 0.2667 | 170 | 0.2765 | 154 | 0.2468 | 151 | 0.2914 | 625 | 0.2704 | 180 | 0.2667 | 172 | 0.2209 | 136 | 0.2868 | 144 | 0.1944 | 632 | 0.2421 |
| 2_18079473 | 135 | 0.2148 | 132 | 0.2045 | 127 | 0.126 | 138 | 0.1884 | 532 | 0.1842 | 119 | 0.1933 | 125 | 0.184 | 115 | 0.1739 | 111 | 0.1802 | 470 | 0.183 |
| 2_18079486 | 119 | 0.1345 | 114 | 0.1404 | 127 | 0.1181 | 128 | 0.0703 | 488 | 0.1148 | 101 | 0.0594 | 119 | 0.084 | 107 | 0.0374 | 103 | 0.0777 | 430 | 0.0651 |
| 2_18083008 | 19 | 0.1053 | 15 | 0.0667 | 12 | 0.25 | 20 | 0.1 | 66 | 0.1212 | 17 | 0.1176 | 14 | 0.2143 | 17 | 0.1765 | 20 | 0.15 | 68 | 0.1618 |
| 2_18083023 | 22 | 0.4091 | 12 | 0.5 | 14 | 0.2857 | 15 | 0.4667 | 63 | 0.4127 | 15 | 0.3333 | 11 | 0.4545 | 14 | 0.2857 | 16 | 0.4375 | 56 | 0.375 |
| 2_18083059 | 1 | 0 | NA | NA | 1 | 0 | 3 | 0.3333 | 5 | 0.2 | 1 | 0 | 5 | 0.2 | 1 | 0 | 3 | 0 | 10 | 0.1 |
| 2_18083190 | 2 | 0.5 | NA | NA | 2 | 0.5 | 3 | 0.3333 | 7 | 0.4286 | 1 | 0 | 5 | 0.2 | 2 | 0 | 2 | 0 | 10 | 0.1 |
| 2_18086971 | 1 | 1 | 1 | 1 | NA | NA | 2 | 0.5 | 4 | 0.75 | NA | NA | 1 | 1 | NA | NA | NA | NA | 1 | 1 |
| 2_18087027 | 1 | 1 | 1 | 1 | NA | NA | NA | NA | 2 | 1 | NA | NA | 1 | 1 | NA | NA | NA | NA | 1 | 1 |
| 2_18089494 | NA | NA | 1 | 1 | 2 | 0 | 1 | 0 | 4 | 0.25 | 1 | 0 | 1 | 0 | NA | NA | 1 | 1 | 3 | 0.3333 |
| 2_18089515 | NA | NA | 1 | 0 | 1 | 1 | 1 | 1 | 3 | 0.6667 | NA | NA | 1 | 0 | NA | NA | 2 | 0.5 | 3 | 0.3333 |
| 2_18089536 | NA | NA | NA | NA | 1 | 1 | 1 | 1 | 2 | 1 | NA | NA | 1 | 1 | NA | NA | 1 | 1 | 2 | 1 |
| 2_18089559 | NA | NA | 1 | 1 | 2 | 1 | 1 | 0 | 4 | 0.75 | NA | NA | 1 | 0 | NA | NA | 2 | 0.5 | 3 | 0.3333 |
| 2_18089563 | NA | NA | 1 | 1 | 2 | 1 | 1 | 1 | 4 | 1 | 1 | 1 | 1 | 1 | NA | NA | 2 | 1 | 4 | 1 |
| 2_18089566 | NA | NA | 1 | 1 | 2 | 0 | 1 | 1 | 4 | 0.5 | 1 | 1 | 1 | 0 | NA | NA | 2 | 0.5 | 4 | 0.5 |
| 2_18089571 | NA | NA | 1 | 1 | 2 | 0.5 | 1 | 1 | 4 | 0.75 | 1 | 1 | 1 | 1 | NA | NA | 2 | 1 | 4 | 1 |
| 2_18089582 | NA | NA | 1 | 1 | 2 | 1 | 1 | 1 | 4 | 1 | 1 | 1 | 1 | 1 | NA | NA | 2 | 1 | 4 | 1 |
| 2_18089606 | NA | NA | 1 | 1 | 2 | 0.5 | 1 | 1 | 4 | 0.75 | NA | NA | 1 | 1 | NA | NA | 2 | 0.5 | 3 | 0.6667 |
| 2_18094841 | 87 | 0.1149 | 92 | 0.1304 | 89 | 0.1124 | 79 | 0.0886 | 347 | 0.1124 | 73 | 0.1233 | 76 | 0.1447 | 79 | 0.0886 | 79 | 0.1013 | 307 | 0.114 |
| 2_18099801 | NA | NA | NA | NA | 1 | 1 | NA | NA | 1 | 1 | NA | NA | 3 | 0.3333 | 2 | 0 | NA | NA | 5 | 0.2 |

| | | | | | | | | | | | | | | | | | | | | |
|---|---|---|---|---|---|---|---|---|---|---|---|---|---|---|---|---|---|---|---|---|
| 2_18100046 | 3 | 0.3333 | 4 | 0.5 | 2 | 0 | 2 | 0 | 11 | 0.2727 | 1 | 0 | 4 | 0 | 5 | 0.2 | 3 | 0.3333 | 13 | 0.1538 |
| 2_18100108 | 3 | 0 | 5 | 0 | 3 | 0.3333 | 3 | 0.3333 | 14 | 0.1429 | NA | NA | 5 | 0 | 6 | 0.3333 | 3 | 0 | 14 | 0.1429 |
| 2_18100119 | 4 | 0.5 | 5 | 0.6 | 4 | 0 | 8 | 0.5 | 21 | 0.4286 | 1 | 1 | 5 | 0.6 | 6 | 0.3333 | 7 | 0.4286 | 19 | 0.4737 |
| 2_18100124 | 6 | 0.5 | 7 | 0.4286 | 4 | 0.5 | 9 | 0.5556 | 26 | 0.5 | 1 | 0 | 5 | 0.4 | 7 | 0.4286 | 4 | 1 | 17 | 0.5294 |
| 2_18100748 | 1 | 0 | 2 | 0.5 | 2 | 0.5 | 4 | 0.5 | 9 | 0.4444 | 2 | 0 | 3 | 0.3333 | NA | NA | 1 | 0 | 6 | 0.1667 |
| 2_18100756 | 2 | 0.5 | 2 | 0 | 2 | 0 | 3 | 0 | 9 | 0.1111 | 2 | 0 | 3 | 0.6667 | NA | NA | 1 | 0 | 6 | 0.3333 |
| 2_18100784 | 2 | 0.5 | 2 | 0.5 | 2 | 0.5 | 4 | 0.75 | 10 | 0.6 | 2 | 0 | 3 | 1 | 1 | 1 | 1 | 1 | 7 | 0.7143 |
| 2_18102176 | 2 | 0 | 5 | 0.2 | 3 | 0.6667 | 4 | 0.25 | 14 | 0.2857 | 3 | 0 | 4 | 0.5 | 3 | 0.3333 | 1 | 0 | 11 | 0.2727 |
| 2_18102177 | 3 | 0.6667 | 4 | 0 | 2 | 0 | 3 | 0 | 12 | 0.1667 | 3 | 1 | 3 | 0.6667 | 1 | 0 | 1 | 1 | 8 | 0.75 |
| 2_18102180 | 3 | 0 | 5 | 0.2 | 2 | 0 | 4 | 0 | 14 | 0.0714 | 2 | 0 | 3 | 0 | 4 | 0.25 | NA | NA | 9 | 0.1111 |
| 2_18102234 | 3 | 0.3333 | 5 | 0.4 | 3 | 0.3333 | 4 | 1 | 15 | 0.5333 | 3 | 0 | 4 | 0.75 | 6 | 0.6667 | 1 | 0 | 14 | 0.5 |
| 2_18102235 | 2 | 0 | 5 | 0 | 2 | 0.5 | 4 | 0.25 | 13 | 0.1538 | 3 | 0.3333 | 3 | 0 | 6 | 0 | 1 | 0 | 13 | 0.0769 |
| 2_18102236 | 3 | 0.3333 | 6 | 0.5 | 3 | 1 | 4 | 0.25 | 16 | 0.5 | 3 | 0.6667 | 4 | 0.25 | 6 | 0.5 | 1 | 0 | 14 | 0.4286 |
| 2_18102246 | 2 | 0.5 | 3 | 0.6667 | 2 | 1 | 3 | 0.3333 | 10 | 0.6 | 3 | 0.6667 | 2 | 0.5 | 4 | 1 | 1 | 1 | 10 | 0.8 |
| 2_18102248 | 3 | 1 | 5 | 0.6 | 3 | 0.6667 | 4 | 0.75 | 15 | 0.7333 | 2 | 0.5 | 4 | 0.5 | 5 | 0.8 | 1 | 1 | 12 | 0.6667 |
| 2_18102250 | 3 | 1 | 6 | 0.8333 | 3 | 1 | 4 | 0.75 | 16 | 0.875 | 1 | 1 | 4 | 0.75 | 5 | 1 | 1 | 1 | 11 | 0.9091 |
| 2_18102255 | 2 | 1 | 5 | 0.8 | 3 | 1 | 3 | 1 | 13 | 0.9231 | 2 | 1 | 4 | 1 | 4 | 1 | 1 | 1 | 11 | 1 |
| 2_18102259 | 2 | 1 | 4 | 0 | 2 | 0 | 3 | 0.3333 | 11 | 0.2727 | 3 | 0.6667 | 2 | 0.5 | 3 | 0.3333 | NA | NA | 8 | 0.5 |
| 2_18102262 | 2 | 0 | 4 | 0 | 2 | 1 | 3 | 1 | 11 | 0.4545 | 3 | 0.6667 | 2 | 0.5 | 3 | 0.6667 | NA | NA | 8 | 0.625 |
| 2_18102272 | 3 | 0 | 3 | 0 | 2 | 0 | 2 | 0.5 | 10 | 0.1 | 3 | 0.3333 | 1 | 0 | 4 | 0 | NA | NA | 8 | 0.125 |
| 2_18102276 | 3 | 0.3333 | 4 | 0 | 2 | 0.5 | 3 | 0.3333 | 12 | 0.25 | 3 | 0 | 2 | 0.5 | 4 | 0.25 | NA | NA | 9 | 0.2222 |
| 2_18102297 | 3 | 0 | 3 | 0.6667 | 2 | 0.5 | 2 | 1 | 10 | 0.5 | 2 | 0 | 3 | 0 | 5 | 0.6 | NA | NA | 10 | 0.3 |
| 2_18102304 | 3 | 0.3333 | 4 | 0.75 | 3 | 0.6667 | 3 | 0.6667 | 13 | 0.6154 | 2 | 0.5 | 3 | 0.6667 | 5 | 0.8 | NA | NA | 10 | 0.7 |
| 2_18102765 | 1 | 1 | NA | NA | NA | NA | NA | NA | 1 | 1 | NA | NA | 3 | 1 | NA | NA | 1 | 1 | 4 | 1 |
| 2_18102784 | 1 | 1 | NA | NA | NA | NA | 1 | 1 | 2 | 1 | NA | NA | 3 | 1 | NA | NA | 1 | 1 | 4 | 1 |
| 2_18102898 | 1 | 1 | NA | NA | NA | NA | 2 | 1 | 3 | 1 | NA | NA | 4 | 0.75 | NA | NA | 2 | 1 | 6 | 0.8333 |
| 2_18102906 | NA | NA | NA | NA | NA | NA | 1 | 1 | 1 | 1 | NA | NA | 3 | 0.6667 | NA | NA | 2 | 0.5 | 5 | 0.6 |
| 2_18106572 | NA | NA | NA | NA | NA | NA | 1 | 1 | 1 | 1 | NA | NA | NA | NA | NA | NA | 1 | 1 | 1 | 1 |
| 2_18106592 | NA | NA | NA | NA | NA | NA | 1 | 1 | 1 | 1 | NA | NA | NA | NA | NA | NA | 1 | 1 | 1 | 1 |
| 2_18115705 | 10 | 0.4 | 13 | 0.0769 | 9 | 0.1111 | 12 | 0.1667 | 44 | 0.1818 | 7 | 0 | 9 | 0.2222 | 9 | 0.2222 | 7 | 0.4286 | 32 | 0.2188 |
| 2_18115708 | 10 | 0.4 | 10 | 0.3 | 7 | 0.2857 | 10 | 0.3 | 37 | 0.3243 | 7 | 0.1429 | 8 | 0.125 | 7 | 0.4286 | 7 | 0 | 29 | 0.1724 |
| 2_18115713 | 10 | 0.6 | 13 | 0.6154 | 6 | 0.3333 | 9 | 0.5556 | 38 | 0.5526 | 9 | 0.4444 | 6 | 0.6667 | 7 | 0.8571 | 7 | 0.5714 | 29 | 0.6207 |
| 2_18115721 | 5 | 1 | 6 | 0.8333 | 4 | 0.5 | 5 | 0.6 | 20 | 0.75 | 3 | 0.3333 | 5 | 0.2 | 5 | 0.6 | 2 | 0 | 15 | 0.3333 |
| 2_18115732 | 4 | 0 | 7 | 0.1429 | 5 | 0 | 5 | 0.2 | 21 | 0.0952 | 2 | 0 | 6 | 0.1667 | 5 | 0 | 2 | 0 | 15 | 0.0667 |
| 2_18115740 | 5 | 0.4 | 7 | 0 | 6 | 0 | 5 | 0 | 23 | 0.087 | 3 | 0 | 8 | 0.125 | 6 | 0 | 3 | 0 | 20 | 0.05 |
| 2_18115822 | 4 | 0 | 4 | 0.25 | 1 | 1 | 1 | 0 | 10 | 0.2 | 2 | 0 | 3 | 0 | 2 | 1 | 1 | 0 | 8 | 0.25 |
| 2_18119123 | 9 | 0.8889 | 8 | 1 | 7 | 1 | 8 | 1 | 32 | 0.9688 | 6 | 1 | 4 | 1 | 8 | 1 | 9 | 0.8889 | 27 | 0.963 |
| 2_18119188 | 16 | 0.625 | 10 | 0.4 | 11 | 0.3636 | 9 | 0.7778 | 46 | 0.5435 | 10 | 0.6 | 6 | 0.6667 | 12 | 0.75 | 13 | 0.4615 | 41 | 0.6098 |
| 2_18119205 | 17 | 0.4706 | 11 | 0.4545 | 8 | 0.375 | 9 | 0.5556 | 45 | 0.4667 | 12 | 0.5 | 8 | 0.375 | 10 | 0.5 | 13 | 0.3846 | 43 | 0.4419 |
| 2_18119399 | 24 | 0.0417 | 21 | 0.0952 | 19 | 0.1579 | 17 | 0.0588 | 81 | 0.0864 | 23 | 0.087 | 20 | 0.05 | 27 | 0.037 | 18 | 0.1111 | 88 | 0.0682 |
| 2_18119439 | 29 | 0.1034 | 20 | 0 | 21 | 0.0952 | 17 | 0.1176 | 87 | 0.0805 | 24 | 0.0833 | 22 | 0.0455 | 24 | 0.0417 | 21 | 0.1429 | 91 | 0.0769 |
| 2_18119514 | 6 | 0.1667 | 6 | 0.1667 | 3 | 0 | 6 | 0.3333 | 21 | 0.1905 | 4 | 0.5 | 2 | 0 | 12 | 0 | 6 | 0 | 24 | 0.0833 |
| 2_18119561 | 2 | 0 | 5 | 0 | 3 | 0.3333 | 4 | 0 | 14 | 0.0714 | 1 | 0 | 4 | 0.25 | 7 | 0.1429 | 5 | 0.2 | 17 | 0.1765 |
| 2_18119653 | 9 | 0.2222 | 12 | 0.0833 | 5 | 0 | 8 | 0 | 34 | 0.0882 | 2 | 1 | 16 | 0.0625 | 8 | 0.125 | 8 | 0 | 34 | 0.1176 |
| 2_18119658 | 6 | 0.1667 | 9 | 0.1111 | 5 | 0.2 | 8 | 0 | 28 | 0.1071 | 2 | 0 | 13 | 0.0769 | 12 | 0.0833 | 6 | 0.3333 | 33 | 0.1212 |
| 2_18119668 | 9 | 0.1111 | 14 | 0 | 6 | 0.1667 | 9 | 0 | 38 | 0.0526 | 2 | 0 | 16 | 0.25 | 14 | 0.1429 | 8 | 0.125 | 40 | 0.175 |
| 2_18119676 | 8 | 0.125 | 14 | 0.2143 | 7 | 0.2857 | 8 | 0 | 37 | 0.1622 | 4 | 0.25 | 15 | 0.2 | 15 | 0.1333 | 9 | 0.3333 | 43 | 0.2093 |
| 2_18119683 | 11 | 0.2727 | 18 | 0 | 6 | 0 | 10 | 0.1 | 45 | 0.0889 | 3 | 0 | 18 | 0.0556 | 13 | 0.3846 | 9 | 0.1111 | 43 | 0.1628 |
| 2_18121154 | 1 | 0 | 2 | 0.5 | 1 | 0 | NA | NA | 4 | 0.25 | 2 | 0 | 3 | 0 | 3 | 0.3333 | 2 | 0 | 10 | 0.1 |
| 2_18122416 | 2 | 0.5 | 6 | 0.6667 | 1 | 1 | 4 | 0.5 | 13 | 0.6154 | NA | NA | 2 | 0.5 | 2 | 0 | 4 | 0.75 | 8 | 0.5 |
| 2_18122416 | 1 | 1 | NA | NA | NA | NA | NA | NA | 1 | 1 | NA | NA | NA | NA | NA | NA | 1 | 1 | 1 | 1 |
| 2_18122474 | 3 | 0 | 5 | 0.2 | 1 | 0 | 4 | 0 | 13 | 0.0769 | NA | NA | 1 | 0 | 3 | 0 | 6 | 0.1667 | 10 | 0.1 |
| 2_18122502 | 3 | 0 | 4 | 0.75 | 1 | 0 | 4 | 0.5 | 12 | 0.4167 | NA | NA | 1 | 1 | 2 | 0.5 | 5 | 0.6 | 8 | 0.625 |
| 2_18122539 | 2 | 0 | 5 | 0.4 | 1 | 0 | 4 | 0 | 12 | 0.1667 | NA | NA | 1 | 1 | 3 | 0.3333 | 4 | 0 | 8 | 0.25 |
| 2_18122576 | NA | NA | 4 | 0.25 | 1 | 0 | 4 | 0 | 9 | 0.1111 | NA | NA | 1 | 1 | 3 | 0 | 4 | 0.25 | 8 | 0.25 |
| 2_18122597 | NA | NA | 5 | 0.4 | 1 | 0 | 4 | 0 | 10 | 0.2 | NA | NA | 1 | 1 | 3 | 0 | 4 | 0 | 8 | 0.125 |
| 2_18122615 | NA | NA | 3 | 0.3333 | 1 | 0 | 3 | 0 | 7 | 0.1429 | NA | NA | NA | NA | 3 | 0 | 3 | 0.3333 | 6 | 0.1667 |
| 2_18122622 | NA | NA | 5 | 1 | 1 | 1 | 3 | 1 | 9 | 1 | NA | NA | 1 | 1 | 2 | 1 | 3 | 1 | 6 | 1 |
| 2_18122626 | NA | NA | 5 | 0.8 | 1 | 1 | 3 | 0.3333 | 9 | 0.6667 | NA | NA | NA | NA | 3 | 0.3333 | 3 | 0.3333 | 6 | 0.3333 |
| 2_18122636 | NA | NA | 3 | 1 | 1 | 1 | 3 | 1 | 7 | 1 | NA | NA | 1 | 1 | 3 | 0.6667 | 3 | 1 | 7 | 0.8571 |
| 2_18122667 | NA | NA | 4 | 0.75 | 1 | 1 | 2 | 0.5 | 7 | 0.7143 | NA | NA | 1 | 1 | 3 | 0.3333 | 2 | 0 | 6 | 0.3333 |
| 2_18122681 | NA | NA | 4 | 0.25 | 1 | 1 | 2 | 0 | 7 | 0.2857 | NA | NA | NA | NA | 2 | 0.5 | 3 | 0.3333 | 5 | 0.4 |
| 2_18123732 | 2 | 1 | 7 | 0.1429 | 5 | 0.4 | 1 | 0 | 15 | 0.3333 | NA | NA | 2 | 0 | 1 | 0 | 3 | 0.3333 | 6 | 0.1667 |
| 2_18123734 | 1 | 1 | 5 | 0.2 | 7 | 0.4286 | 2 | 0.5 | 15 | 0.4 | 1 | 1 | 2 | 0.5 | 2 | 0.5 | 3 | 0.3333 | 8 | 0.5 |
| 2_18123768 | 2 | 0 | 6 | 0 | 5 | 0.2 | 2 | 1 | 15 | 0.2 | 1 | 1 | 1 | 0 | 2 | 0 | 4 | 0 | 8 | 0.125 |

| | | | | | | | | | | | | | | | | | | | | |
|---|---|---|---|---|---|---|---|---|---|---|---|---|---|---|---|---|---|---|---|---|
| 2_18123807 | 2 | 0 | 3 | 0.3333 | 3 | 0 | 2 | 0.5 | 10 | 0.2 | 1 | 0 | 2 | 0 | NA | NA | 2 | 0.5 | 5 | 0.2 |
| 2_18123815 | 2 | 0 | 5 | 0 | 6 | 0.1667 | 2 | 0.5 | 15 | 0.1333 | 1 | 0 | 2 | 0 | 2 | 0 | 3 | 0.3333 | 8 | 0.125 |
| 2_18123823 | 1 | 0 | 5 | 0.2 | 5 | 0 | 2 | 0 | 13 | 0.0769 | 1 | 0 | 2 | 0 | 2 | 0 | 3 | 0.6667 | 8 | 0.25 |
| 2_18129739 | 4 | 0.25 | 4 | 0.75 | 8 | 0.625 | 15 | 0.5333 | 31 | 0.5484 | 6 | 0.8333 | 6 | 0.6667 | 5 | 0.6 | 2 | 1 | 19 | 0.7368 |
| 2_18130100 | NA | NA | NA | NA | NA | NA | 1 | 1 | 1 | 1 | NA | NA | NA | NA | 1 | 1 | NA | NA | 1 | 1 |
| 2_18131171 | 2 | 0 | 2 | 0 | 3 | 0.3333 | 1 | 1 | 8 | 0.25 | 4 | 0.25 | 2 | 0.5 | 1 | 1 | 3 | 0.3333 | 10 | 0.4 |
| 2_18131214 | 3 | 1 | 2 | 0.5 | 2 | 1 | 2 | 0.5 | 9 | 0.7778 | 2 | 0 | 3 | 1 | 1 | 1 | 3 | 0.6667 | 9 | 0.6667 |
| 2_18131218 | 3 | 0.3333 | 2 | 1 | 2 | 0 | 2 | 1 | 9 | 0.5556 | 2 | 0.5 | 3 | 0.3333 | 1 | 0 | 2 | 0 | 8 | 0.25 |
| 2_18131233 | 2 | 0.5 | 2 | 1 | 2 | 0.5 | 1 | 1 | 7 | 0.7143 | 2 | 0.5 | 3 | 0.6667 | 1 | 0 | 2 | 0 | 8 | 0.375 |
| 2_18131238 | 3 | 0.3333 | 2 | 1 | 2 | 0 | 2 | 0 | 9 | 0.3333 | 2 | 0 | 3 | 0.3333 | 2 | 0.5 | 2 | 0.5 | 9 | 0.3333 |
| 2_18131243 | 3 | 1 | 1 | 1 | 3 | 1 | 2 | 1 | 9 | 1 | 2 | 1 | 3 | 1 | 1 | 1 | 1 | 1 | 7 | 1 |
| 2_18132240 | 1 | 1 | NA | NA | 2 | 0 | NA | NA | 3 | 0.3333 | NA | NA | 3 | 0.6667 | NA | NA | NA | NA | 3 | 0.6667 |
| 2_18132871 | 2 | 0 | 4 | 0.25 | 7 | 0 | 2 | 0 | 15 | 0.0667 | 3 | 0 | 2 | 0 | 3 | 0 | 5 | 0.2 | 13 | 0.0769 |
| 2_18132895 | NA | NA | 1 | 1 | 2 | 0.5 | NA | NA | 3 | 0.6667 | NA | NA | NA | NA | 1 | 0 | 1 | 1 | 2 | 0.5 |
| 2_18133037 | 1 | 0 | 4 | 0.25 | 4 | 0 | 3 | 0 | 12 | 0.0833 | 4 | 0 | 1 | 0 | 6 | 0.1667 | 4 | 0.25 | 15 | 0.1333 |
| 2_18133042 | NA | NA | 3 | 0 | 3 | 0 | 3 | 0.3333 | 9 | 0.1111 | 4 | 0.25 | 1 | 0 | 6 | 0.6667 | 4 | 0.25 | 15 | 0.4 |
| 2_18133058 | 1 | 0 | 3 | 0.3333 | 3 | 0.3333 | 3 | 0 | 10 | 0.2 | 4 | 0.25 | 1 | 0 | 4 | 0.5 | 2 | 0 | 11 | 0.2727 |
| 2_18133071 | 1 | 0 | 3 | 0 | 2 | 0 | 2 | 0.5 | 8 | 0.125 | 4 | 0.25 | NA | NA | 5 | 0.2 | 2 | 0 | 11 | 0.1818 |
| 2_18133107 | 1 | 0 | 2 | 0 | 1 | 0 | 3 | 0.3333 | 7 | 0.1429 | 3 | 0 | NA | NA | 2 | 0.5 | 2 | 0 | 7 | 0.1429 |
| 2_18135765 | 3 | 0.3333 | 2 | 0 | 5 | 0 | 3 | 0 | 13 | 0.0769 | 4 | 0.25 | 2 | 0 | 3 | 0 | 5 | 0 | 14 | 0.0714 |
| 2_18136036 | 2 | 0 | 2 | 0 | 5 | 0 | 6 | 0.3333 | 15 | 0.1333 | 4 | 0.5 | 2 | 0 | 4 | 0.5 | 6 | 0.1667 | 16 | 0.3125 |
| 2_18136042 | 3 | 0 | 3 | 0 | 6 | 0.3333 | 6 | 0.5 | 18 | 0.2778 | 4 | 0 | 1 | 0 | 3 | 0.3333 | 8 | 0.375 | 16 | 0.25 |
| 2_18136064 | 4 | 0 | 3 | 0.3333 | 6 | 0.1667 | 5 | 0.6 | 18 | 0.2778 | 5 | 0.2 | 2 | 0 | 3 | 0.3333 | 7 | 0.1429 | 17 | 0.1765 |
| 2_18136078 | 4 | 0.25 | 3 | 0 | 5 | 0.2 | 6 | 0.3333 | 18 | 0.2222 | 5 | 0 | 2 | 0.5 | 3 | 0.3333 | 8 | 0.125 | 18 | 0.1667 |
| 2_18136084 | 4 | 1 | 3 | 0.6667 | 6 | 0.6667 | 6 | 0.8333 | 19 | 0.7895 | 5 | 0.8 | 2 | 1 | 2 | 1 | 8 | 0.875 | 17 | 0.8824 |
| 2_18136087 | 4 | 1 | 2 | 1 | 5 | 1 | 5 | 1 | 16 | 1 | 5 | 0.8 | 1 | 1 | 3 | 1 | 6 | 1 | 15 | 0.9333 |
| 2_18136090 | 4 | 0.75 | 3 | 0.6667 | 6 | 0.6667 | 5 | 0.4 | 18 | 0.6111 | 4 | 0.5 | 2 | 1 | 3 | 0.6667 | 7 | 0.5714 | 16 | 0.625 |
| 2_18136099 | 4 | 0.25 | 2 | 0.5 | 5 | 0.4 | 5 | 1 | 16 | 0.5625 | 5 | 0.8 | 2 | 1 | 2 | 1 | 6 | 1 | 15 | 0.9333 |
| 2_18136110 | 4 | 0.25 | 3 | 0.6667 | 6 | 0.1667 | 5 | 0.6 | 18 | 0.3889 | 5 | 0 | 3 | 0.6667 | 3 | 0.6667 | 7 | 0.1429 | 18 | 0.2778 |
| 2_18136111 | 4 | 1 | 1 | 1 | 5 | 0.2 | 5 | 0.6 | 15 | 0.6 | 4 | 0.75 | 3 | 0.6667 | 3 | 0.6667 | 5 | 0.8 | 15 | 0.7333 |
| 2_18136114 | 4 | 1 | 3 | 0 | 5 | 0.6 | 6 | 0.5 | 18 | 0.5556 | 4 | 0.75 | 3 | 0.6667 | 2 | 0.5 | 6 | 0.8333 | 15 | 0.7333 |
| 2_18136122 | 2 | 0.5 | 2 | 0 | 5 | 0.8 | 5 | 0.6 | 14 | 0.5714 | 4 | 1 | 3 | 0.6667 | 2 | 1 | 6 | 0.8333 | 15 | 0.8667 |
| 2_18136129 | 3 | 0 | 3 | 0 | 5 | 0 | 6 | 0.1667 | 17 | 0.0588 | 4 | 0 | 4 | 0 | 4 | 0 | 8 | 0.125 | 20 | 0.05 |
| 2_18136131 | 4 | 1 | 3 | 1 | 5 | 0.6 | 4 | 0.75 | 16 | 0.8125 | 5 | 1 | 4 | 1 | 5 | 0.8 | 7 | 0.8571 | 21 | 0.9048 |
| 2_18136139 | 4 | 1 | 1 | 1 | 4 | 1 | 5 | 0.8 | 14 | 0.9286 | 2 | 1 | 4 | 1 | 4 | 1 | 6 | 1 | 16 | 1 |
| 2_18136142 | 2 | 1 | 3 | 0.6667 | 4 | 0.75 | 3 | 1 | 12 | 0.8333 | 4 | 0.75 | 2 | 1 | 2 | 1 | 5 | 1 | 13 | 0.9231 |
| 2_18136146 | 4 | 1 | 2 | 1 | 4 | 0.75 | 5 | 1 | 15 | 0.9333 | 3 | 1 | 4 | 1 | 3 | 1 | 8 | 0.625 | 18 | 0.8333 |
| 2_18136159 | 4 | 1 | 1 | 0 | 6 | 0.8333 | 5 | 0.8 | 16 | 0.8125 | 4 | 1 | 4 | 0.75 | 4 | 0.75 | 4 | 1 | 16 | 0.875 |
| 2_18136166 | 3 | 0.3333 | 3 | 0.3333 | 7 | 0.2857 | 5 | 0 | 18 | 0.2222 | 5 | 0.2 | 1 | 0 | 5 | 0 | 9 | 0 | 20 | 0.05 |
| 2_18136172 | 4 | 1 | 3 | 1 | 8 | 0.875 | 5 | 1 | 20 | 0.95 | 5 | 1 | 2 | 0.5 | 6 | 1 | 9 | 0.8889 | 22 | 0.9091 |
| 2_18136177 | 4 | 0.5 | 3 | 1 | 5 | 0.4 | 5 | 1 | 17 | 0.7059 | 3 | 0.6667 | 3 | 1 | 3 | 0.3333 | 8 | 0.875 | 17 | 0.7647 |
| 2_18136181 | 2 | 0.5 | 1 | 0 | 6 | 0 | 4 | 0.5 | 13 | 0.2308 | 5 | 0.4 | 2 | 0.5 | 2 | 0 | 5 | 0.2 | 14 | 0.2857 |
| 2_18136184 | 4 | 0.25 | 3 | 0.3333 | 7 | 0 | 6 | 0 | 20 | 0.1 | 5 | 0.4 | 4 | 0.25 | 6 | 0 | 7 | 0 | 22 | 0.1364 |
| 2_18136189 | 4 | 0.75 | 4 | 1 | 5 | 1 | 4 | 1 | 17 | 0.9412 | 4 | 1 | 3 | 1 | 5 | 0.8 | 6 | 1 | 18 | 0.9444 |
| 2_18136191 | 2 | 1 | 4 | 1 | 7 | 1 | 5 | 1 | 18 | 1 | 4 | 1 | 4 | 1 | 5 | 1 | 8 | 1 | 21 | 1 |
| 2_18136193 | 4 | 0.25 | 3 | 0.6667 | 6 | 0.3333 | 4 | 0.75 | 17 | 0.4706 | 3 | 0.3333 | 3 | 0.6667 | 4 | 0.5 | 9 | 0.4444 | 19 | 0.4737 |
| 2_18136206 | 4 | 0.75 | 3 | 1 | 7 | 0.5714 | 6 | 0.5 | 20 | 0.65 | 4 | 0.25 | 3 | 0.6667 | 5 | 0.2 | 7 | 0.5714 | 19 | 0.4211 |
| 2_18136212 | 4 | 0.5 | 4 | 0 | 7 | 0 | 5 | 0.2 | 20 | 0.15 | 4 | 0 | 3 | 0.3333 | 4 | 0 | 10 | 0.1 | 21 | 0.0952 |
| 2_18136217 | 3 | 1 | 3 | 0.6667 | 7 | 0.7143 | 4 | 1 | 17 | 0.8235 | 5 | 0.2 | 4 | 0.75 | 5 | 0.6 | 7 | 0.4286 | 21 | 0.4762 |
| 2_18136221 | 3 | 0 | 4 | 0.25 | 8 | 0 | 3 | 0 | 18 | 0.0556 | 4 | 0 | 4 | 0.25 | 5 | 0 | 11 | 0.0909 | 24 | 0.0833 |
| 2_18136224 | 4 | 0.75 | 4 | 0.5 | 8 | 0.625 | 6 | 0.6667 | 22 | 0.6364 | 4 | 0.5 | 3 | 0.6667 | 6 | 0.8333 | 9 | 0.7778 | 22 | 0.7273 |
| 2_18136233 | 4 | 1 | 4 | 0.75 | 9 | 1 | 5 | 1 | 22 | 0.9545 | 5 | 1 | 4 | 1 | 6 | 1 | 8 | 1 | 23 | 1 |
| 2_18136235 | 3 | 1 | 3 | 1 | 8 | 0.875 | 6 | 1 | 20 | 0.95 | 4 | 1 | 4 | 1 | 7 | 1 | 9 | 1 | 24 | 1 |
| 2_18136248 | 4 | 1 | 2 | 1 | 9 | 1 | 5 | 0.6 | 20 | 0.9 | 4 | 0.75 | 4 | 1 | 7 | 1 | 8 | 1 | 23 | 0.9565 |
| 2_18136258 | 3 | 0 | 4 | 0 | 6 | 0.3333 | 5 | 0 | 18 | 0.1111 | 7 | 0.5714 | 2 | 0.5 | 7 | 0.5714 | 10 | 0.3 | 26 | 0.4615 |
| 2_18146788 | NA | NA | NA | NA | NA | NA | 1 | 1 | 1 | 1 | NA | NA | 1 | 0 | 1 | 1 | NA | NA | 2 | 0.5 |
| 2_18146804 | NA | NA | NA | NA | NA | NA | 2 | 1 | 2 | 1 | NA | NA | 1 | 1 | 1 | 1 | NA | NA | 2 | 1 |
| 2_18146943 | NA | NA | NA | NA | 1 | 1 | 4 | 0.5 | 5 | 0.6 | NA | NA | 2 | 0.5 | 2 | 0.5 | 2 | 1 | 6 | 0.6667 |
| 2_18146950 | NA | NA | NA | NA | 1 | 0 | 4 | 0.25 | 5 | 0.2 | 1 | 1 | 1 | 0 | 2 | 0 | 2 | 0 | 6 | 0.1667 |
| 2_18148427 | 6 | 0.5 | 3 | 0.6667 | 3 | 0.3333 | 5 | 0.6 | 17 | 0.5294 | 1 | 1 | 3 | 0.3333 | 2 | 0.5 | 3 | 1 | 9 | 0.6667 |
| 2_18148434 | 6 | 0.1667 | 3 | 0.3333 | 3 | 0.3333 | 5 | 0.6 | 17 | 0.3529 | 1 | 0 | 3 | 0.3333 | 2 | 0.5 | 3 | 0.3333 | 9 | 0.3333 |
| 2_18149675 | 7 | 0.1429 | 3 | 0.3333 | 5 | 0.2 | 8 | 0.25 | 23 | 0.2174 | 2 | 0.5 | 4 | 0.25 | 6 | 0.5 | 4 | 0.75 | 16 | 0.5 |
| 2_18149683 | 8 | 0.375 | 2 | 0.5 | 5 | 0 | 9 | 0 | 24 | 0.1667 | 2 | 0.5 | 3 | 0 | 5 | 0 | 3 | 0.6667 | 13 | 0.2308 |
| 2_18149688 | 9 | 0 | 3 | 0 | 6 | 0.3333 | 9 | 0 | 27 | 0.0741 | 2 | 0 | 4 | 0 | 6 | 0.1667 | 4 | 0 | 16 | 0.0625 |
| 2_18149689 | 9 | 0.7778 | 3 | 1 | 6 | 1 | 8 | 0.75 | 26 | 0.8462 | 2 | 1 | 3 | 1 | 6 | 0.6667 | 4 | 1 | 15 | 0.8667 |

| | | | | | | | | | | | | | | | | | | | | |
|---|---|---|---|---|---|---|---|---|---|---|---|---|---|---|---|---|---|---|---|---|
| 2_1814969 | 8 | 0.5 | 3 | 0.3333 | 6 | 0.5 | 9 | 0.6667 | 26 | 0.5385 | 2 | 0.5 | 4 | 0.5 | 5 | 1 | 4 | 1 | 15 | 0.8 |
| 2_1814970 | 7 | 0.8571 | 3 | 0.6667 | 5 | 0.2 | 7 | 0.7143 | 22 | 0.6364 | NA | NA | 3 | 1 | 3 | 0.6667 | 3 | 0.3333 | 9 | 0.6667 |
| 2_1814971 | 10 | 0.3 | 3 | 0 | 6 | 0.1667 | 9 | 0.1111 | 28 | 0.1786 | 2 | 0 | 4 | 0 | 6 | 0.3333 | 4 | 0.25 | 16 | 0.1875 |
| 2_1814971 | 10 | 0.1 | 3 | 0.3333 | 6 | 0.1667 | 9 | 0.2222 | 28 | 0.1786 | 2 | 0.5 | 4 | 0.25 | 5 | 0.4 | 4 | 0.25 | 15 | 0.3333 |
| 2_1814972 | 9 | 0.3333 | 4 | 0.25 | 6 | 0 | 5 | 0.2 | 24 | 0.2083 | 2 | 0.5 | 4 | 0 | 6 | 0.1667 | 4 | 0.5 | 16 | 0.25 |
| 2_1814973 | 8 | 0.625 | 2 | 0.5 | 6 | 0 | 10 | 0.5 | 26 | 0.4231 | 2 | 0 | 6 | 0.3333 | 9 | 0 | 4 | 0.5 | 21 | 0.1905 |
| 2_1814973 | 10 | 0.1 | 6 | 0 | 7 | 0 | 10 | 0.3 | 33 | 0.1212 | 2 | 0 | 6 | 0.1667 | 7 | 0 | 4 | 0 | 19 | 0.0526 |
| 2_1814976 | 9 | 0.5556 | 4 | 0.25 | 4 | 0 | 10 | 0.5 | 27 | 0.4074 | 4 | 0.25 | 5 | 0.2 | 10 | 0.6 | 5 | 0.6 | 24 | 0.4583 |
| 2_1814978 | 11 | 0.3636 | 6 | 0 | 6 | 0.3333 | 12 | 0.25 | 35 | 0.2571 | 4 | 0 | 7 | 0.4286 | 10 | 0.1 | 4 | 0 | 25 | 0.16 |
| 2_1814980 | 14 | 0.0714 | 6 | 0.3333 | 7 | 0 | 14 | 0.2143 | 41 | 0.1463 | 5 | 0.2 | 7 | 0.1429 | 14 | 0.0714 | 5 | 0 | 31 | 0.0968 |
| 2_1814986 | 11 | 0 | 7 | 0 | 7 | 0.1429 | 13 | 0.1538 | 38 | 0.0789 | 3 | 0 | 5 | 0 | 14 | 0.2143 | 4 | 0.25 | 26 | 0.1538 |
| 2_1814987 | 6 | 0.5 | 5 | 0.8 | 6 | 0.5 | 10 | 0.4 | 27 | 0.5185 | 4 | 1 | 9 | 0.8889 | 12 | 0.5833 | 5 | 0.8 | 30 | 0.7667 |
| 2_1815035 | 2 | 1 | 2 | 0.5 | 1 | 1 | 1 | 1 | 6 | 0.8333 | NA | NA | NA | NA | 1 | 1 | 1 | 1 | 2 | 1 |
| 2_1815035 | 2 | 0.5 | 1 | 1 | 1 | 1 | 1 | 0 | 5 | 0.6 | NA | NA | NA | NA | 1 | 1 | 1 | 0 | 2 | 0.5 |
| 2_1815035 | 2 | 0 | 2 | 1 | NA | NA | 1 | 0 | 5 | 0.4 | NA | NA | NA | NA | 1 | 1 | 1 | 0 | 2 | 0.5 |
| 2_1815035 | 1 | 1 | 2 | 1 | 1 | 1 | 1 | 1 | 5 | 1 | NA | NA | NA | NA | 1 | 1 | 1 | 1 | 2 | 1 |
| 2_1815035 | 2 | 1 | 2 | 1 | NA | NA | 1 | 1 | 5 | 1 | NA | NA | NA | NA | 1 | 1 | 1 | 1 | 2 | 1 |
| 2_1815036 | 2 | 1 | 1 | 1 | 1 | 1 | 1 | 1 | 5 | 1 | NA | NA | NA | NA | 1 | 1 | 1 | 1 | 2 | 1 |
| 2_1815036 | 1 | 1 | 2 | 1 | 1 | 1 | 1 | 1 | 5 | 1 | NA | NA | NA | NA | 1 | 1 | 1 | 1 | 2 | 1 |
| 2_1815039 | 2 | 1 | 2 | 1 | NA | NA | 1 | 1 | 5 | 1 | NA | NA | NA | NA | 1 | 1 | 1 | 1 | 2 | 1 |
| 2_1815040 | 1 | 1 | 1 | 1 | NA | NA | 1 | 1 | 3 | 1 | NA | NA | NA | NA | 1 | 1 | 1 | 1 | 2 | 1 |
| 2_1815043 | NA | NA | 2 | 1 | NA | NA | 1 | 1 | 3 | 1 | NA | NA | NA | NA | 1 | 1 | 1 | 1 | 2 | 1 |
| 2_1815043 | 1 | 0 | 1 | 1 | NA | NA | NA | NA | 2 | 0.5 | NA | NA | NA | NA | 1 | 1 | 1 | 0 | 2 | 0.5 |
| 2_1815044 | NA | NA | 2 | 1 | NA | NA | 1 | 1 | 3 | 1 | NA | NA | NA | NA | 1 | 1 | 1 | 1 | 2 | 1 |
| 2_1815044 | 1 | 1 | NA | NA | NA | NA | 1 | 1 | 2 | 1 | NA | NA | NA | NA | NA | NA | 1 | 1 | 1 | 1 |
| 2_1815044 | 1 | 1 | NA | NA | NA | NA | 1 | 1 | 2 | 1 | NA | NA | NA | NA | 1 | 1 | NA | NA | 1 | 1 |
| 2_1815049 | 1 | 1 | 1 | 1 | NA | NA | 1 | 1 | 3 | 1 | NA | NA | NA | NA | 1 | 1 | 1 | 1 | 2 | 1 |
| 2_1815049 | 1 | 0 | 1 | 1 | NA | NA | 1 | 0 | 3 | 0.3333 | NA | NA | NA | NA | 1 | 1 | NA | NA | 1 | 1 |
| 2_1815056 | 1 | 1 | 1 | 1 | NA | NA | 1 | 1 | 3 | 1 | NA | NA | NA | NA | 1 | 1 | NA | NA | 1 | 1 |
| 2_1815057 | NA | NA | 1 | 1 | NA | NA | 1 | 0 | 2 | 0.5 | NA | NA | NA | NA | 1 | 1 | NA | NA | 1 | 1 |
| 2_1815058 | NA | NA | 1 | 1 | NA | NA | 1 | 1 | 2 | 1 | NA | NA | NA | NA | 1 | 1 | NA | NA | 1 | 1 |
| 2_1815101 | 2 | 0 | 6 | 0.1667 | 4 | 0 | 4 | 0.25 | 16 | 0.125 | 4 | 0 | 6 | 0 | 4 | 0.25 | 2 | 0 | 16 | 0.0625 |
| 2_1815101 | 2 | 0.5 | 5 | 0.2 | 4 | 0 | 6 | 0.3333 | 17 | 0.2353 | 4 | 0 | 5 | 0.4 | 3 | 0.6667 | 4 | 0.5 | 16 | 0.375 |
| 2_1815108 | 3 | 0 | 4 | 0.5 | 6 | 0.1667 | 4 | 0 | 17 | 0.1765 | 2 | 0 | 4 | 0.75 | 3 | 0.3333 | 2 | 0 | 11 | 0.3636 |
| 2_1815109 | 5 | 0.2 | 5 | 0 | 6 | 0 | 5 | 0.2 | 21 | 0.0952 | 2 | 0 | 4 | 0.5 | 4 | 0 | 1 | 1 | 11 | 0.2727 |
| 3_4878908 | NA | NA | NA | NA | NA | NA | 1 | 1 | 1 | 1 | NA | NA | 1 | 0 | 1 | 1 | NA | NA | 2 | 0.5 |
| 3_4879021 | NA | NA | NA | NA | NA | NA | 1 | 1 | 1 | 1 | NA | NA | 2 | 1 | NA | NA | NA | NA | 2 | 1 |
| 3_5479410 | NA | NA | 1 | 1 | 1 | 0 | 2 | 0.5 | 4 | 0.5 | NA | NA | NA | NA | NA | NA | 1 | 1 | 1 | 1 |
| 3_5479976 | NA | NA | 1 | 0 | 1 | 0 | 2 | 0.5 | 4 | 0.25 | NA | NA | NA | NA | NA | NA | 1 | 1 | 1 | 1 |
| 3_5480363 | NA | NA | 1 | 1 | 1 | 0 | 2 | 0.5 | 4 | 0.5 | NA | NA | 1 | 1 | NA | NA | 1 | 0 | 2 | 0.5 |
| 3_5480403 | NA | NA | 1 | 0 | 1 | 0 | 1 | 1 | 3 | 0.3333 | NA | NA | 1 | 0 | NA | NA | 1 | 1 | 2 | 0.5 |
| 3_5480452 | NA | NA | 1 | 0 | 1 | 1 | 2 | 1 | 4 | 0.75 | NA | NA | NA | NA | NA | NA | 1 | 1 | 1 | 1 |
| 3_5480493 | NA | NA | 1 | 1 | 1 | 1 | 2 | 1 | 4 | 1 | NA | NA | 1 | 1 | NA | NA | 1 | 1 | 2 | 1 |
| 3_5626226 | 2 | 0 | 3 | 0 | 6 | 0 | 6 | 0.1667 | 17 | 0.0588 | 1 | 0 | 7 | 0 | 5 | 0 | 2 | 0.5 | 15 | 0.0667 |
| 3_5626350 | 2 | 0 | 3 | 0 | 5 | 0.4 | 6 | 0.3333 | 16 | 0.25 | 1 | 0 | 7 | 0 | 5 | 0.4 | 1 | 0 | 14 | 0.1429 |
| 3_5626376 | 3 | 0.6667 | 3 | 1 | 5 | 1 | 6 | 1 | 17 | 0.9412 | NA | NA | 7 | 0.7143 | 5 | 0.8 | 1 | 1 | 13 | 0.7692 |
| 3_5626499 | 2 | 0.5 | 2 | 1 | 6 | 0.6667 | 6 | 0.5 | 16 | 0.625 | 1 | 0 | 7 | 0.7143 | 3 | 0.6667 | 1 | 1 | 12 | 0.6667 |
| 3_5626507 | 3 | 0 | 2 | 0 | 5 | 0.2 | 5 | 0 | 15 | 0.0667 | 1 | 0 | 6 | 0 | 3 | 0.3333 | 2 | 1 | 12 | 0.25 |
| 3_5626667 | 2 | 1 | 3 | 0.6667 | 6 | 1 | 5 | 0.6 | 16 | 0.8125 | 1 | 1 | 6 | 0.8333 | 1 | 1 | 1 | 1 | 9 | 0.8889 |
| 3_5626736 | 2 | 0 | 3 | 0.3333 | 5 | 0.2 | 5 | 0.2 | 15 | 0.2 | NA | NA | 6 | 0 | 2 | 0 | 1 | 1 | 9 | 0.1111 |
| 3_7507180 | 10 | 0 | 14 | 0.0714 | 13 | 0.3077 | 22 | 0.1364 | 59 | 0.1356 | 10 | 0.1 | 22 | 0.0455 | 14 | 0.1429 | 13 | 0.1538 | 59 | 0.1017 |
| 3_7507270 | 13 | 0.3846 | 13 | 0.3077 | 12 | 0.3333 | 22 | 0.5 | 60 | 0.4 | 9 | 0.4444 | 19 | 0.4211 | 11 | 0.3636 | 12 | 0.1667 | 51 | 0.3529 |
| 3_7507559 | 13 | 0 | 16 | 0.125 | 11 | 0.6364 | 25 | 0.16 | 65 | 0.2 | 10 | 0.3 | 22 | 0 | 13 | 0.1538 | 15 | 0.2667 | 60 | 0.15 |
| 3_7509062 | 21 | 0.0476 | 19 | 0.0526 | 18 | 0 | 34 | 0.0882 | 92 | 0.0543 | 14 | 0.2857 | 30 | 0.0333 | 20 | 0.1 | 19 | 0.1053 | 83 | 0.1084 |
| 3_7509516 | 25 | 0.16 | 25 | 0.12 | 19 | 0.1579 | 33 | 0.0303 | 102 | 0.1078 | 13 | 0.0769 | 35 | 0.1429 | 24 | 0.0417 | 22 | 0.0909 | 94 | 0.0957 |
| 3_7616512 | 4 | 0.5 | 4 | 0.5 | 3 | 0 | 8 | 0.375 | 19 | 0.3684 | 3 | 0.3333 | 8 | 0.25 | 5 | 0.6 | 10 | 0.6 | 26 | 0.4615 |
| 3_7616543 | 5 | 0.4 | 6 | 0.1667 | 2 | 0.5 | 7 | 0 | 20 | 0.2 | 3 | 0.3333 | 9 | 0.1111 | 6 | 0.5 | 10 | 0.4 | 28 | 0.3214 |
| 3_7618306 | 7 | 0 | 8 | 0 | 6 | 0.3333 | 13 | 0.0769 | 34 | 0.0882 | 4 | 0 | 10 | 0 | 11 | 0.1818 | 9 | 0.2222 | 34 | 0.1176 |
| 3_8625373 | 32 | 0.0625 | 47 | 0.1277 | 35 | 0.0857 | 24 | 0.0833 | 138 | 0.0942 | 40 | 0.125 | 28 | 0.1071 | 10 | 0.1 | 24 | 0.125 | 102 | 0.1176 |
| 3_8625492 | 33 | 0.2727 | 53 | 0.283 | 39 | 0.2051 | 30 | 0.1667 | 155 | 0.2387 | 40 | 0.175 | 28 | 0.1429 | 10 | 0.1 | 27 | 0.1481 | 105 | 0.1524 |
| 3_8626315 | 29 | 0.1034 | 46 | 0.1304 | 25 | 0.16 | 21 | 0.0952 | 121 | 0.124 | 31 | 0.129 | 21 | 0.2381 | 10 | 0.2 | 15 | 0.1333 | 77 | 0.1688 |
| 3_8626368 | 29 | 0.5862 | 39 | 0.5641 | 26 | 0.4615 | 17 | 0.5294 | 111 | 0.5405 | 29 | 0.4138 | 19 | 0.6316 | 10 | 0.5 | 18 | 0.4444 | 76 | 0.4868 |
| 3_8729097 | 5 | 0 | 11 | 0.0909 | 3 | 0 | 7 | 0.1429 | 26 | 0.0769 | 4 | 0 | 8 | 0.125 | 7 | 0.2857 | 8 | 0.25 | 27 | 0.1852 |
| 3_8729424 | 6 | 0 | 10 | 0.1 | 4 | 0 | 5 | 0.2 | 25 | 0.08 | 3 | 0 | 4 | 0.5 | 5 | 0.4 | 7 | 0.2857 | 19 | 0.3158 |
| 3_8729538 | 3 | 0.3333 | 7 | 0 | 2 | 0 | 4 | 0.25 | 16 | 0.125 | 2 | 0.5 | 5 | 0.4 | 2 | 0.5 | 4 | 0 | 13 | 0.3077 |

| | | | | | | | | | | | | | | | | | | | | |
|---|---|---|---|---|---|---|---|---|---|---|---|---|---|---|---|---|---|---|---|---|
| 3_8729552 | 6 | 1 | 6 | 1 | 4 | 1 | 5 | 1 | 21 | 1 | 1 | 1 | 6 | 1 | 4 | 1 | 6 | 1 | 17 | 1 |
| 3_8729576 | 6 | 0.6667 | 9 | 0.3333 | 3 | 0.6667 | 5 | 0.2 | 23 | 0.4348 | 2 | 0 | 6 | 0.6667 | 4 | 0.5 | 6 | 0.8333 | 18 | 0.6111 |
| 3_8729586 | 6 | 1 | 9 | 0.8889 | 3 | 0.6667 | 4 | 1 | 22 | 0.9091 | 2 | 1 | 6 | 1 | 4 | 1 | 6 | 0.6667 | 18 | 0.8889 |
| 3_8729709 | 4 | 0 | 9 | 0.3333 | 2 | 0 | 4 | 0.75 | 19 | 0.3158 | 1 | 1 | 4 | 0.5 | 5 | 0.2 | 6 | 0.3333 | 16 | 0.375 |
| 3_8729742 | 6 | 0.5 | 7 | 0.5714 | 3 | 0 | 5 | 0.6 | 21 | 0.4762 | 2 | 0 | 5 | 0.2 | 5 | 0.8 | 6 | 0.6667 | 18 | 0.5 |
| 3_8729901 | 5 | 0.4 | 8 | 0.25 | 3 | 0 | 4 | 0.75 | 20 | 0.35 | 2 | 0 | 4 | 0.5 | 4 | 0.5 | 6 | 0.5 | 16 | 0.4375 |
| 3_8729928 | 4 | 0.75 | 6 | 1 | 2 | 0.5 | 5 | 1 | 17 | 0.8824 | 2 | 1 | 3 | 0.6667 | 4 | 0.75 | 6 | 0.8333 | 15 | 0.8 |
| 3_8730027 | 4 | 0.5 | 6 | 0 | 3 | 0.6667 | 5 | 0.8 | 18 | 0.4444 | 1 | 1 | 4 | 0.5 | 4 | 0.5 | 6 | 0.5 | 15 | 0.5333 |
| 3_8730094 | 5 | 0.6 | 4 | 0.25 | 3 | 0 | 4 | 0.75 | 16 | 0.4375 | 2 | 0 | 5 | 0.4 | 4 | 0.75 | 6 | 0.3333 | 17 | 0.4118 |
| 3_8867302 | 1 | 1 | 5 | 0.2 | 3 | 0 | 9 | 0.1111 | 18 | 0.1667 | 2 | 0.5 | 10 | 0.6 | 8 | 0.25 | 6 | 0 | 26 | 0.3462 |
| 3_8867340 | 2 | 0.5 | 4 | 0.25 | 4 | 0 | 8 | 0.125 | 18 | 0.1667 | 3 | 0.3333 | 10 | 0.1 | 9 | 0.3333 | 6 | 0 | 28 | 0.1786 |
| 3_8994555 | 5 | 0.2 | 2 | 0 | 6 | 0 | 3 | 0 | 16 | 0.0625 | 1 | 0 | 2 | 0 | 5 | 0.2 | 2 | 0 | 10 | 0.1 |
| 3_8995032 | 10 | 0.2 | 6 | 0.1667 | 4 | 0.75 | 4 | 0 | 24 | 0.25 | 4 | 0.5 | 5 | 0 | 9 | 0.2222 | 5 | 0.4 | 23 | 0.2609 |
| 3_9492340 | 3 | 1 | 6 | 0 | 7 | 0.2857 | 3 | 0.3333 | 19 | 0.3158 | 5 | 0 | 7 | 0.2857 | 1 | 1 | 4 | 0.5 | 17 | 0.2941 |
| 3_9492689 | 2 | 0.5 | 4 | 1 | 4 | 0.25 | 2 | 0.5 | 12 | 0.5833 | 2 | 0 | 7 | 0.5714 | 1 | 0 | 1 | 0 | 11 | 0.3636 |
| 3_9492872 | 2 | 0 | 4 | 0 | 6 | 0.1667 | 2 | 0 | 14 | 0.0714 | 2 | 0.5 | 7 | 0 | 2 | 0 | 4 | 0 | 15 | 0.0667 |
| 3_9495179 | NA | NA | 2 | 0.5 | 2 | 0 | NA | NA | 4 | 0.25 | NA | NA | 1 | 1 | 1 | 0 | 1 | 0 | 3 | 0.3333 |
| 3_9495558 | NA | NA | 1 | 1 | 2 | 1 | NA | NA | 3 | 1 | NA | NA | 1 | 0 | 1 | 0 | 1 | 1 | 3 | 0.3333 |
| 3_9755663 | NA | NA | 2 | 0.5 | NA | NA | 1 | 1 | 3 | 0.6667 | 1 | 1 | NA | NA | NA | NA | NA | NA | 1 | 1 |
| 3_9755719 | 1 | 1 | 2 | 1 | NA | NA | 1 | 1 | 4 | 1 | 1 | 1 | NA | NA | NA | NA | NA | NA | 1 | 1 |
| 3_9755757 | 1 | 1 | 2 | 1 | NA | NA | 1 | 1 | 4 | 1 | 1 | 1 | NA | NA | NA | NA | NA | NA | 1 | 1 |
| 3_9757096 | 1 | 0 | 2 | 0 | NA | NA | 1 | 1 | 4 | 0.25 | 1 | 1 | NA | NA | NA | NA | NA | NA | 1 | 1 |
| 3_9757337 | 1 | 1 | 2 | 1 | NA | NA | 1 | 1 | 4 | 1 | 1 | 1 | NA | NA | NA | NA | NA | NA | 1 | 1 |
| 3_9757381 | 1 | 1 | 1 | 1 | NA | NA | 1 | 1 | 3 | 1 | 1 | 1 | NA | NA | NA | NA | NA | NA | 1 | 1 |
| 3_9757690 | 1 | 1 | 2 | 1 | NA | NA | 1 | 1 | 4 | 1 | 1 | 1 | NA | NA | NA | NA | NA | NA | 1 | 1 |
| 3_9758264 | 1 | 1 | 2 | 1 | NA | NA | 1 | 1 | 4 | 1 | 1 | 1 | NA | NA | NA | NA | NA | NA | 1 | 1 |
| 3_9758336 | 1 | 0 | 2 | 0 | NA | NA | 1 | 1 | 4 | 0.25 | 1 | 1 | NA | NA | NA | NA | NA | NA | 1 | 1 |
| 3_9758393 | 1 | 1 | 2 | 1 | NA | NA | 1 | 1 | 4 | 1 | 1 | 1 | NA | NA | NA | NA | NA | NA | 1 | 1 |
| 3_10081328 | 29 | 0.1034 | 28 | 0.1786 | 25 | 0.12 | 24 | 0.2083 | 106 | 0.1509 | 19 | 0.0526 | 33 | 0.2424 | 26 | 0.1538 | 37 | 0.0811 | 115 | 0.1391 |
| 3_10379604 | 12 | 0.25 | 14 | 0.2143 | 8 | 0 | 6 | 0.3333 | 40 | 0.2 | 12 | 0.4167 | 8 | 0.125 | 7 | 0.1429 | 9 | 0.1111 | 36 | 0.2222 |
| 3_10380161 | 14 | 0.2857 | 17 | 0.0588 | 9 | 0 | 12 | 0.1667 | 52 | 0.1346 | 10 | 0.1 | 15 | 0.2667 | 10 | 0.2 | 10 | 0.1 | 45 | 0.1778 |
| 3_10380166 | 10 | 0.2 | 17 | 0.1765 | 8 | 0.125 | 10 | 0 | 45 | 0.1333 | 11 | 0.1818 | 16 | 0.125 | 9 | 0.1111 | 9 | 0 | 45 | 0.1111 |
| 3_10405379 | 3 | 0.3333 | 1 | 0 | 2 | 0 | 4 | 0 | 10 | 0.1 | 2 | 0.5 | 3 | 0 | 1 | 0 | 3 | 0.3333 | 9 | 0.2222 |
| 3_10405811 | 1 | 0 | 2 | 0.5 | 3 | 0.3333 | 4 | 0.5 | 10 | 0.4 | 1 | 0 | 4 | 0.75 | 1 | 0 | 4 | 0.25 | 10 | 0.4 |
| 3_10405831 | 3 | 0 | 2 | 0.5 | NA | NA | 4 | 0 | 9 | 0.1111 | 1 | 0 | 4 | 0 | 1 | 0 | 4 | 0.25 | 10 | 0.1 |
| 3_10416343 | NA | NA | 1 | 1 | 2 | 0.5 | 1 | 0 | 4 | 0.5 | NA | NA | 1 | 1 | NA | NA | 3 | 0 | 4 | 0.25 |
| 3_10484125 | 5 | 0.2 | 7 | 0.7143 | 5 | 0.4 | 10 | 0.9 | 27 | 0.6296 | 6 | 0.3333 | 4 | 0.75 | 3 | 1 | 8 | 0.875 | 21 | 0.7143 |
| 3_10503008 | 3 | 0.6667 | 5 | 0.8 | 3 | 0.6667 | 5 | 0.8 | 16 | 0.75 | 5 | 0.2 | 4 | 0.75 | 3 | 0 | 6 | 0.6667 | 18 | 0.4444 |
| 3_10503023 | 3 | 0.6667 | 5 | 0.8 | 2 | 1 | 5 | 1 | 15 | 0.8667 | 5 | 0.6 | 2 | 1 | 3 | 0.3333 | 7 | 0.8571 | 17 | 0.7059 |
| 3_10503061 | 2 | 1 | 5 | 1 | 1 | 1 | 4 | 1 | 12 | 1 | 4 | 0.75 | 4 | 0.75 | 3 | 0.6667 | 8 | 0.75 | 19 | 0.7368 |
| 3_10503109 | 3 | 1 | 5 | 0.8 | 2 | 1 | 3 | 0.6667 | 13 | 0.8462 | 4 | 0.5 | 4 | 0.75 | 1 | 1 | 5 | 0.8 | 14 | 0.7143 |
| 3_10503284 | 2 | 1 | 3 | 1 | 1 | 0 | 3 | 1 | 9 | 0.8889 | 4 | 1 | 4 | 1 | 3 | 0.6667 | 8 | 0.875 | 19 | 0.8947 |
| 3_10503315 | 3 | 0 | 4 | 0.25 | 1 | 0 | 4 | 0.25 | 12 | 0.1667 | 5 | 0.2 | 4 | 0.25 | 3 | 1 | 6 | 0 | 18 | 0.2778 |
| 3_14246198 | 10 | 0.4 | 13 | 0.4615 | 19 | 0.2105 | 20 | 0.45 | 62 | 0.371 | 10 | 0.1 | 12 | 0.3333 | 14 | 0.5 | 7 | 0.2857 | 43 | 0.3256 |
| 3_14246245 | 9 | 0.5556 | 15 | 0.8667 | 20 | 0.55 | 19 | 0.8947 | 63 | 0.7302 | 11 | 0.9091 | 12 | 0.75 | 10 | 0.5 | 10 | 0.8 | 43 | 0.7442 |
| 3_14246685 | 13 | 0.4615 | 19 | 0.8421 | 22 | 0.5455 | 21 | 0.5714 | 75 | 0.6133 | 12 | 0.9167 | 16 | 0.75 | 13 | 0.6923 | 11 | 0.5455 | 52 | 0.7308 |
| 3_14246768 | 13 | 0.0769 | 19 | 0.1579 | 21 | 0.0952 | 22 | 0.0909 | 75 | 0.1067 | 12 | 0.0833 | 14 | 0 | 14 | 0.1429 | 12 | 0 | 52 | 0.0577 |
| 3_14246783 | 11 | 0.4545 | 17 | 0.2941 | 23 | 0.087 | 19 | 0.1053 | 70 | 0.2 | 11 | 0.1818 | 14 | 0.1429 | 11 | 0 | 11 | 0.2727 | 47 | 0.1489 |
| 3_14246795 | 11 | 0.1818 | 20 | 0.25 | 23 | 0.3043 | 21 | 0.3333 | 75 | 0.28 | 14 | 0.3571 | 16 | 0.125 | 15 | 0.4667 | 12 | 0.25 | 57 | 0.2982 |
| 3_14246814 | 11 | 0.1818 | 19 | 0.1053 | 21 | 0.0476 | 19 | 0.1053 | 70 | 0.1 | 15 | 0.1333 | 14 | 0.1429 | 11 | 0.2727 | 12 | 0.1667 | 52 | 0.1731 |
| 3_14630946 | NA | NA | NA | NA | 4 | 0.25 | 1 | 1 | 5 | 0.4 | 1 | 1 | NA | NA | NA | NA | 2 | 0.5 | 3 | 0.6667 |
| 3_14670796 | NA | NA | 3 | 0.3333 | 3 | 0.6667 | 3 | 0 | 9 | 0.3333 | 1 | 0 | 4 | 0 | 1 | 0 | 3 | 0.6667 | 9 | 0.2222 |
| 3_14671604 | 1 | 1 | 1 | 0 | 1 | 0 | 1 | 0 | 4 | 0.25 | 1 | 0 | 2 | 0 | NA | NA | 2 | 0.5 | 5 | 0.2 |
| 3_16258127 | 3 | 1 | 1 | 1 | 2 | 0.5 | 3 | 1 | 9 | 0.8889 | NA | NA | 4 | 1 | 4 | 1 | 3 | 0.6667 | 11 | 0.9091 |
| 3_16258228 | 3 | 0 | 3 | 0 | 2 | 0.5 | 3 | 0 | 11 | 0.0909 | NA | NA | 3 | 0 | 5 | 0 | 6 | 0.1667 | 14 | 0.0714 |
| 3_16258283 | 2 | 0.5 | 3 | 1 | 3 | 0.6667 | 4 | 1 | 12 | 0.8333 | NA | NA | 4 | 1 | 5 | 0.6 | 5 | 0.6 | 14 | 0.7143 |
| 3_16258361 | 1 | 1 | 2 | 0.5 | NA | NA | 2 | 1 | 5 | 0.8 | NA | NA | 2 | 1 | 3 | 1 | NA | NA | 5 | 1 |
| 3_16258428 | 2 | 0 | 3 | 0 | 3 | 0 | 4 | 0.5 | 12 | 0.1667 | NA | NA | 5 | 0.4 | 5 | 0.2 | 7 | 0.1429 | 17 | 0.2353 |
| 3_16258433 | 3 | 1 | 3 | 1 | 3 | 1 | 4 | 1 | 13 | 1 | NA | NA | 4 | 1 | 6 | 0.8333 | 5 | 1 | 15 | 0.9333 |
| 3_16258473 | 3 | 0 | 3 | 0.3333 | 2 | 0.5 | 4 | 0 | 12 | 0.1667 | NA | NA | 4 | 0.25 | 5 | 0 | 6 | 0.5 | 15 | 0.2667 |
| 3_16258506 | 3 | 0 | 4 | 0.25 | 4 | 0 | 4 | 0 | 15 | 0.0667 | NA | NA | 5 | 0 | 5 | 0.2 | 7 | 0.1429 | 17 | 0.1176 |
| 3_16258607 | 1 | 0 | 3 | 0.3333 | 3 | 0 | 3 | 0 | 10 | 0.1 | NA | NA | 3 | 0 | 6 | 0 | 5 | 0.2 | 14 | 0.0714 |
| 3_16258686 | 2 | 1 | 4 | 0.75 | 2 | 1 | 4 | 0.25 | 12 | 0.6667 | NA | NA | 3 | 1 | 6 | 0.6667 | 6 | 0.1667 | 15 | 0.5333 |
| 3_16258853 | 3 | 0.3333 | 4 | 0 | 3 | 0 | 4 | 0.25 | 14 | 0.1429 | NA | NA | 5 | 0 | 6 | 0.1667 | 7 | 0 | 18 | 0.0556 |
| 3_16258954 | 3 | 0 | 4 | 0 | 5 | 0.2 | 4 | 0 | 16 | 0.0625 | NA | NA | 5 | 0.2 | 7 | 0 | 5 | 0 | 17 | 0.0588 |

| | | | | | | | | | | | | | | | | | | | | |
|---|---|---|---|---|---|---|---|---|---|---|---|---|---|---|---|---|---|---|---|---|
| 3_1625897 | 2 | 0.5 | 2 | 0.5 | 3 | 0.3333 | 4 | 0.75 | 11 | 0.5455 | NA | NA | 4 | 0.75 | 6 | 0.5 | 5 | 0.6 | 15 | 0.6 |
| 3_1625904 | 2 | 0 | 4 | 0 | 5 | 0 | 4 | 0.25 | 15 | 0.0667 | NA | NA | 5 | 0 | 9 | 0.1111 | 7 | 0.1429 | 21 | 0.0952 |
| 3_1625905 | 3 | 0 | 3 | 0.3333 | 5 | 0.2 | 4 | 0.25 | 15 | 0.2 | NA | NA | 5 | 0.2 | 8 | 0.125 | 6 | 0.1667 | 19 | 0.1579 |
| 3_1625907 | 3 | 1 | 4 | 1 | 4 | 1 | 4 | 1 | 15 | 1 | NA | NA | 3 | 0.6667 | 8 | 1 | 7 | 0.7143 | 18 | 0.8333 |
| 3_1625926 | 3 | 1 | 3 | 1 | 6 | 1 | 5 | 1 | 17 | 1 | NA | NA | 4 | 1 | 10 | 0.9 | 6 | 0.8333 | 20 | 0.9 |
| 3_1625930 | 3 | 0.3333 | 4 | 0.25 | 6 | 0.5 | 5 | 0.2 | 18 | 0.3333 | NA | NA | 4 | 0.75 | 10 | 0.7 | 5 | 0.6 | 19 | 0.6842 |
| 3_1625942 | 3 | 0.3333 | 3 | 0 | 6 | 0 | 4 | 0 | 16 | 0.0625 | NA | NA | 5 | 0 | 10 | 0.1 | 7 | 0.1429 | 22 | 0.0909 |
| 3_1625945 | 2 | 1 | 4 | 1 | 6 | 1 | 4 | 1 | 16 | 1 | NA | NA | 5 | 0.8 | 9 | 0.5556 | 7 | 0.7143 | 21 | 0.6667 |
| 3_1625953 | 3 | 0.3333 | 3 | 0.3333 | 6 | 0.1667 | 5 | 0 | 17 | 0.1765 | NA | NA | 6 | 0.3333 | 9 | 0.2222 | 7 | 0.4286 | 22 | 0.3182 |
| 3_1626819 | 4 | 0.75 | 3 | 0.6667 | 5 | 0.6 | 3 | 1 | 15 | 0.7333 | NA | NA | 4 | 1 | 7 | 1 | 5 | 0.8 | 16 | 0.9375 |
| 3_1626825 | 3 | 0 | 4 | 0 | 8 | 0.125 | 5 | 0.4 | 20 | 0.15 | NA | NA | 7 | 0.1429 | 10 | 0.2 | 6 | 0.1667 | 23 | 0.1739 |
| 3_1626826 | 4 | 0.75 | 4 | 0.25 | 8 | 0.375 | 6 | 0.1667 | 22 | 0.3636 | NA | NA | 5 | 0.6 | 9 | 0.5556 | 7 | 0.4286 | 21 | 0.5238 |
| 3_1626834 | 4 | 1 | 4 | 1 | 5 | 1 | 6 | 1 | 19 | 1 | NA | NA | 5 | 1 | 9 | 1 | 5 | 1 | 19 | 1 |
| 3_1626843 | 4 | 0.5 | 4 | 0.5 | 9 | 0.4444 | 6 | 0.8333 | 23 | 0.5652 | NA | NA | 6 | 0.8333 | 10 | 0.5 | 7 | 0.5714 | 23 | 0.6087 |
| 3_1626846 | 4 | 0.25 | 4 | 0 | 9 | 0.2222 | 8 | 0 | 25 | 0.12 | NA | NA | 7 | 0.1429 | 7 | 0 | 7 | 0.1429 | 21 | 0.0952 |
| 3_1626873 | 3 | 0 | 4 | 0 | 8 | 0.125 | 7 | 0.1429 | 22 | 0.0909 | 2 | 0 | 5 | 0 | 8 | 0 | 5 | 0.2 | 20 | 0.05 |
| 3_1783895 | NA | NA | NA | NA | NA | NA | 1 | 1 | 1 | 1 | NA | NA | NA | NA | 1 | 1 | NA | NA | 1 | 1 |
| 3_1783925 | NA | NA | NA | NA | NA | NA | 1 | 1 | 1 | 1 | NA | NA | NA | NA | 1 | 1 | NA | NA | 1 | 1 |
| 3_1783927 | NA | NA | NA | NA | NA | NA | 1 | 1 | 1 | 1 | NA | NA | NA | NA | 1 | 1 | NA | NA | 1 | 1 |
| 3_1783928 | NA | NA | NA | NA | NA | NA | 1 | 1 | 1 | 1 | NA | NA | NA | NA | 1 | 1 | NA | NA | 1 | 1 |
| 3_1783945 | NA | NA | NA | NA | NA | NA | 1 | 1 | 1 | 1 | NA | NA | NA | NA | 1 | 1 | NA | NA | 1 | 1 |
| 3_1783957 | NA | NA | NA | NA | NA | NA | 1 | 1 | 1 | 1 | NA | NA | NA | NA | 1 | 1 | NA | NA | 1 | 1 |
| 3_1783961 | NA | NA | NA | NA | NA | NA | 1 | 1 | 1 | 1 | NA | NA | NA | NA | 1 | 1 | NA | NA | 1 | 1 |
| 3_1783965 | NA | NA | NA | NA | NA | NA | 1 | 1 | 1 | 1 | NA | NA | NA | NA | 1 | 1 | NA | NA | 1 | 1 |
| 3_1783971 | NA | NA | NA | NA | NA | NA | 1 | 1 | 1 | 1 | NA | NA | NA | NA | 1 | 1 | NA | NA | 1 | 1 |
| 3_1784899 | NA | NA | NA | NA | NA | NA | 3 | 0.3333 | 3 | 0.3333 | 3 | 0 | 1 | 0 | 3 | 0.3333 | 1 | 0 | 8 | 0.125 |
| 3_1784906 | NA | NA | 1 | 0 | NA | NA | 2 | 0.5 | 3 | 0.3333 | 3 | 0 | 2 | 0.5 | 3 | 0.3333 | 2 | 0 | 10 | 0.2 |
| 3_1784916 | NA | NA | 1 | 1 | NA | NA | 3 | 1 | 4 | 1 | 3 | 0.3333 | 2 | 0.5 | 3 | 0.6667 | 2 | 1 | 10 | 0.6 |
| 3_1785021 | 1 | 0 | 7 | 0.1429 | 2 | 0 | 2 | 0 | 12 | 0.0833 | 1 | 0 | NA | NA | 3 | 0 | 2 | 0.5 | 6 | 0.1667 |
| 3_1785168 | 1 | 0 | 7 | 0 | 2 | 0.5 | 3 | 0 | 13 | 0.0769 | 1 | 0 | NA | NA | 4 | 0.25 | 3 | 0 | 8 | 0.125 |
| 3_1785365 | 3 | 0.6667 | 9 | 0.6667 | 3 | 0.6667 | 7 | 0.4286 | 22 | 0.5909 | 1 | 1 | 1 | 0 | 9 | 0.1111 | 3 | 0 | 14 | 0.1429 |
| 3_1785366 | 2 | 0 | 8 | 0.125 | 2 | 0 | 10 | 0.1 | 22 | 0.0909 | 3 | 0 | 2 | 0.5 | 9 | 0.2222 | 3 | 0 | 17 | 0.1765 |
| 3_1810661 | 1 | 0 | 3 | 0.3333 | 3 | 0 | NA | NA | 7 | 0.1429 | 2 | 0.5 | 1 | 0 | 1 | 0 | 1 | 0 | 5 | 0.2 |
| 3_1810688 | 1 | 1 | 3 | 1 | 3 | 1 | NA | NA | 7 | 1 | 2 | 0.5 | 1 | 1 | 1 | 1 | NA | NA | 4 | 0.75 |
| 3_1810694 | 1 | 1 | 2 | 1 | 3 | 1 | NA | NA | 6 | 1 | 2 | 1 | 1 | 1 | NA | NA | NA | NA | 3 | 1 |
| 3_1810702 | 1 | 0 | 3 | 0 | 2 | 0.5 | NA | NA | 6 | 0.1667 | 2 | 0.5 | 1 | 0 | 1 | 0 | NA | NA | 4 | 0.25 |
| 3_1810728 | 1 | 1 | 2 | 1 | 2 | 1 | NA | NA | 5 | 1 | NA | NA | 1 | 1 | 1 | 1 | NA | NA | 2 | 1 |
| 3_1810816 | 1 | 1 | 1 | 1 | 1 | 1 | NA | NA | 3 | 1 | NA | NA | 1 | 1 | 1 | 1 | NA | NA | 2 | 1 |
| 3_1810851 | 1 | 1 | 1 | 1 | 2 | 1 | NA | NA | 4 | 1 | NA | NA | 1 | 1 | 1 | 1 | NA | NA | 2 | 1 |
| 3_1810860 | 1 | 0 | 2 | 0 | 2 | 0.5 | NA | NA | 5 | 0.2 | NA | NA | NA | NA | 1 | 1 | NA | NA | 1 | 1 |
| 3_1810908 | NA | NA | 1 | 1 | 1 | 1 | NA | NA | 2 | 1 | 1 | 1 | 1 | 0 | 1 | 0 | NA | NA | 3 | 0.3333 |
| 3_1810923 | NA | NA | 1 | 0 | 2 | 1 | NA | NA | 3 | 0.6667 | NA | NA | 1 | 1 | 1 | 0 | NA | NA | 2 | 0.5 |
| 3_1810974 | NA | NA | 1 | 1 | 1 | 1 | NA | NA | 2 | 1 | 1 | 1 | NA | NA | NA | NA | 1 | 1 | 2 | 1 |
| 3_1811012 | NA | NA | 2 | 0 | 2 | 0 | 1 | 1 | 5 | 0.2 | 2 | 0.5 | NA | NA | NA | NA | 2 | 0.5 | 4 | 0.5 |
| 3_1811015 | NA | NA | 3 | 0.6667 | 2 | 1 | 1 | 1 | 6 | 0.8333 | 2 | 0.5 | NA | NA | NA | NA | 2 | 1 | 4 | 0.75 |
| 3_1811017 | NA | NA | 2 | 0.5 | 2 | 1 | 1 | 1 | 5 | 0.8 | 1 | 1 | NA | NA | NA | NA | 2 | 0.5 | 3 | 0.6667 |
| 3_1811017 | NA | NA | 3 | 1 | 2 | 1 | 1 | 1 | 6 | 1 | 2 | 1 | NA | NA | NA | NA | 2 | 1 | 4 | 1 |
| 3_1811047 | NA | NA | 5 | 0.4 | 2 | 0.5 | 1 | 1 | 8 | 0.5 | 2 | 0.5 | 1 | 1 | NA | NA | 3 | 0.6667 | 6 | 0.6667 |
| 3_1811055 | NA | NA | 4 | 0.75 | 2 | 1 | 1 | 0 | 7 | 0.7143 | 1 | 0 | 1 | 1 | NA | NA | 2 | 1 | 4 | 0.75 |
| 3_1811058 | NA | NA | 4 | 0.5 | 1 | 0 | 1 | 0 | 6 | 0.3333 | 2 | 0.5 | 1 | 0 | NA | NA | 3 | 0 | 6 | 0.1667 |
| 3_1811060 | NA | NA | 5 | 0.8 | 3 | 0.6667 | 1 | 1 | 9 | 0.7778 | 1 | 1 | 1 | 1 | NA | NA | 2 | 0.5 | 4 | 0.75 |
| 3_1811128 | NA | NA | 7 | 0.4286 | 3 | 0.3333 | 1 | 0 | 11 | 0.3636 | 1 | 1 | NA | NA | NA | NA | 3 | 0 | 4 | 0.25 |
| 3_1812682 | 1 | 1 | 1 | 0 | 2 | 1 | 1 | 1 | 5 | 0.8 | 1 | 1 | 3 | 1 | NA | NA | 1 | 1 | 5 | 1 |
| 3_1815144 | 2 | 1 | NA | NA | 1 | 0 | 1 | 1 | 4 | 0.75 | NA | NA | 2 | 0 | NA | NA | 1 | 1 | 3 | 0.3333 |
| 3_1918638 | 10 | 0.1 | 18 | 0.2222 | 16 | 0.0625 | 19 | 0.1053 | 63 | 0.127 | 5 | 0.2 | 9 | 0.1111 | 12 | 0.0833 | 16 | 0.1875 | 42 | 0.1429 |
| 3_1918685 | 10 | 0.1 | 16 | 0.1875 | 16 | 0.125 | 15 | 0.1333 | 57 | 0.1404 | 4 | 0 | 8 | 0.125 | 11 | 0.0909 | 13 | 0.0769 | 36 | 0.0833 |
| 3_1918804 | 5 | 1 | 12 | 0.9167 | 11 | 0.9091 | 12 | 0.8333 | 40 | 0.9 | 1 | 1 | 5 | 0.4 | 6 | 1 | 10 | 0.8 | 22 | 0.7727 |
| 3_1918807 | 8 | 0.25 | 13 | 0.2308 | 13 | 0.1538 | 11 | 0.2727 | 45 | 0.2222 | 2 | 0 | 6 | 0 | 6 | 0.3333 | 12 | 0.25 | 26 | 0.1923 |
| 3_1918812 | 5 | 1 | 9 | 1 | 14 | 0.8571 | 10 | 0.9 | 38 | 0.9211 | 2 | 1 | 4 | 1 | 5 | 1 | 13 | 0.9231 | 24 | 0.9583 |
| 3_1918816 | 7 | 1 | 11 | 1 | 13 | 0.9231 | 14 | 0.9286 | 45 | 0.9556 | 3 | 1 | 6 | 1 | 7 | 1 | 14 | 1 | 30 | 1 |
| 3_1918819 | 7 | 0.2857 | 13 | 0 | 14 | 0.1429 | 13 | 0.3077 | 47 | 0.1702 | 3 | 0.3333 | 7 | 0.1429 | 6 | 0.3333 | 13 | 0 | 29 | 0.1379 |
| 3_1918822 | 6 | 0 | 11 | 0 | 14 | 0.2857 | 13 | 0.3846 | 44 | 0.2045 | 3 | 0.3333 | 6 | 0.5 | 7 | 0.1429 | 14 | 0.0714 | 30 | 0.2 |
| 3_1927327 | 2 | 0.5 | NA | NA | 1 | 0 | NA | NA | 3 | 0.3333 | NA | NA | 1 | 1 | 2 | 0.5 | 2 | 0.5 | 5 | 0.6 |
| 3_1927780 | 3 | 0.3333 | 1 | 0 | NA | NA | 1 | 0 | 5 | 0.2 | 2 | 0.5 | 1 | 0 | 1 | 0 | 1 | 0 | 5 | 0.2 |
| 3_1927811 | 3 | 0.3333 | 1 | 0 | NA | NA | 1 | 0 | 5 | 0.2 | 3 | 0 | 1 | 0 | NA | NA | 1 | 1 | 5 | 0.2 |

| | | | | | | | | | | | | | | | | | | | | |
|---|---|---|---|---|---|---|---|---|---|---|---|---|---|---|---|---|---|---|---|---|
| 3_19280304 | 1 | 0 | 3 | 0 | 1 | 0 | 2 | 0.5 | 7 | 0.1429 | 2 | 1 | 3 | 0.3333 | 1 | 0 | 2 | 0 | 8 | 0.375 |
| 3_19280491 | 1 | 0 | 2 | 0.5 | 1 | 0 | 2 | 1 | 6 | 0.5 | 2 | 0.5 | 2 | 1 | 1 | 0 | 2 | 1 | 7 | 0.7143 |
| 3_19280534 | 1 | 0 | 2 | 0.5 | NA | NA | 2 | 0 | 5 | 0.2 | 2 | 0 | 2 | 0 | 1 | 0 | 2 | 0.5 | 7 | 0.1429 |
| 3_19281027 | 2 | 1 | NA | NA | 1 | 0 | 2 | 0 | 5 | 0.4 | 1 | 0 | 3 | 0 | 1 | 1 | NA | NA | 5 | 0.2 |
| 3_19281087 | 2 | 1 | NA | NA | 1 | 0 | 2 | 0.5 | 5 | 0.6 | 2 | 1 | 2 | 1 | 1 | 1 | NA | NA | 5 | 1 |
| 3_19281164 | 2 | 0.5 | 1 | 1 | NA | NA | 2 | 1 | 5 | 0.8 | 2 | 0.5 | NA | NA | 1 | 0 | NA | NA | 3 | 0.3333 |
| 3_19281253 | 2 | 0 | 1 | 0 | 1 | 0 | 2 | 0.5 | 6 | 0.1667 | 1 | 1 | 3 | 0.3333 | 1 | 1 | NA | NA | 5 | 0.6 |
| 3_19281323 | 2 | 1 | NA | NA | 1 | 1 | 2 | 1 | 5 | 1 | 1 | 1 | 3 | 1 | 1 | 1 | NA | NA | 5 | 1 |
| 3_19281477 | 2 | 1 | NA | NA | 1 | 1 | 1 | 1 | 4 | 1 | NA | NA | 1 | 1 | 1 | 1 | NA | NA | 2 | 1 |
| 3_19281634 | 2 | 1 | 1 | 1 | 1 | 0 | 1 | 1 | 5 | 0.8 | 1 | 1 | 2 | 0.5 | 1 | 0 | NA | NA | 4 | 0.5 |
| 3_19281722 | 2 | 1 | 1 | 0 | 1 | 1 | 1 | 1 | 5 | 0.8 | 1 | 1 | 1 | 0 | 1 | 1 | NA | NA | 3 | 0.6667 |
| 3_19281732 | NA | NA | 1 | 1 | 1 | 0 | 1 | 1 | 3 | 0.6667 | 1 | 1 | 2 | 1 | 1 | 0 | NA | NA | 4 | 0.75 |
| 3_19281784 | 2 | 0.5 | NA | NA | 1 | 0 | 1 | 0 | 4 | 0.25 | 1 | 1 | 3 | 1 | 1 | 1 | NA | NA | 5 | 1 |
| 3_19281823 | 1 | 0 | NA | NA | 1 | 1 | 1 | 1 | 3 | 0.6667 | 1 | 1 | 1 | 1 | 1 | 0 | NA | NA | 3 | 0.6667 |
| 3_19281837 | 1 | 1 | 1 | 1 | 1 | 1 | 1 | 1 | 4 | 1 | NA | NA | 2 | 1 | 1 | 1 | NA | NA | 3 | 1 |
| 3_19281890 | 1 | 1 | 1 | 1 | 1 | 1 | NA | NA | 3 | 1 | 1 | 1 | 2 | 0.5 | 1 | 0 | NA | NA | 4 | 0.5 |
| 3_19281923 | 1 | 1 | 1 | 1 | NA | NA | NA | NA | 2 | 1 | 1 | 0 | 1 | 1 | 1 | 0 | NA | NA | 3 | 0.3333 |
| 3_19748101 | 1 | 1 | 3 | 0.3333 | 3 | 1 | NA | NA | 7 | 0.7143 | NA | NA | 2 | 1 | NA | NA | NA | NA | 2 | 1 |
| 3_19748339 | NA | NA | 3 | 0 | 3 | 0.6667 | NA | NA | 6 | 0.3333 | NA | NA | 2 | 0.5 | NA | NA | NA | NA | 2 | 0.5 |
| 3_19949528 | 1 | 1 | 1 | 1 | NA | NA | 2 | 1 | 4 | 1 | NA | NA | 3 | 1 | 1 | 1 | 1 | 1 | 5 | 1 |
| 3_19949854 | 1 | 1 | NA | NA | 1 | 1 | 2 | 1 | 4 | 1 | NA | NA | 1 | 1 | 1 | 1 | 1 | 1 | 3 | 1 |
| 3_19950056 | 2 | 0.5 | NA | NA | 1 | 1 | 2 | 1 | 5 | 0.8 | NA | NA | 3 | 1 | 1 | 1 | NA | NA | 4 | 1 |
| 3_19950273 | NA | NA | NA | NA | 1 | 1 | 2 | 0 | 3 | 0.3333 | NA | NA | 2 | 0.5 | 1 | 0 | 1 | 0 | 4 | 0.25 |
| 3_19950284 | 2 | 1 | NA | NA | 1 | 1 | 2 | 1 | 5 | 1 | NA | NA | 3 | 0.6667 | 1 | 1 | 1 | 0 | 5 | 0.6 |
| 3_19950315 | 2 | 1 | 1 | 1 | 1 | 1 | 2 | 1 | 6 | 1 | NA | NA | 3 | 1 | 1 | 1 | 1 | 1 | 5 | 1 |
| 3_19950338 | 2 | 1 | 1 | 1 | 1 | 1 | 1 | 1 | 5 | 1 | NA | NA | 2 | 1 | 1 | 1 | NA | NA | 3 | 1 |
| 3_19950350 | 2 | 1 | 1 | 1 | 1 | 1 | 2 | 1 | 6 | 1 | NA | NA | 3 | 1 | 1 | 1 | 1 | 1 | 5 | 1 |
| 3_19950371 | 2 | 0.5 | NA | NA | 1 | 0 | 1 | 1 | 4 | 0.5 | NA | NA | 2 | 0.5 | 1 | 1 | 1 | 1 | 4 | 0.75 |
| 3_19950402 | 2 | 0 | 1 | 0 | 1 | 0 | 2 | 0.5 | 6 | 0.1667 | NA | NA | 3 | 0.3333 | 1 | 1 | 1 | 0 | 5 | 0.4 |
| 3_19950438 | 2 | 0.5 | 1 | 1 | NA | NA | 1 | 0 | 4 | 0.5 | NA | NA | 1 | 1 | 1 | 1 | 1 | 1 | 3 | 1 |
| 3_19950933 | NA | NA | 1 | 0 | 1 | 1 | 2 | 1 | 4 | 0.75 | NA | NA | 1 | 0 | 1 | 1 | NA | NA | 2 | 0.5 |
| 3_19950987 | 1 | 1 | NA | NA | 1 | 1 | 1 | 1 | 3 | 1 | NA | NA | 3 | 1 | 1 | 1 | NA | NA | 4 | 1 |
| 3_19951339 | 2 | 0 | 1 | 1 | 1 | 0 | 1 | 0 | 5 | 0.2 | NA | NA | 1 | 1 | NA | NA | NA | NA | 1 | 1 |
| 3_19952979 | 1 | 1 | 2 | 0 | 1 | 0 | 2 | 0 | 6 | 0.1667 | NA | NA | 2 | 0 | 2 | 0.5 | NA | NA | 4 | 0.25 |
| 3_20046899 | 1 | 0 | 2 | 0.5 | NA | NA | 1 | 0 | 4 | 0.25 | 2 | 0.5 | NA | NA | NA | NA | 1 | 0 | 3 | 0.3333 |
| 3_20050455 | 1 | 1 | 1 | 1 | 1 | 1 | 1 | 1 | 4 | 1 | 1 | 1 | 2 | 0 | 4 | 0.5 | NA | NA | 7 | 0.4286 |
| 3_20050512 | 1 | 0 | 1 | 0 | 1 | 0 | 1 | 1 | 4 | 0.25 | 3 | 0.3333 | 2 | 0 | 4 | 0 | NA | NA | 9 | 0.1111 |
| 3_20050730 | 1 | 0 | 1 | 0 | 1 | 1 | 1 | 0 | 4 | 0.25 | 3 | 0.3333 | 2 | 0 | 4 | 0 | NA | NA | 9 | 0.1111 |
| 3_20062893 | 3 | 0 | NA | NA | NA | NA | 2 | 0.5 | 5 | 0.2 | NA | NA | NA | NA | 1 | 1 | 1 | 0 | 2 | 0.5 |
| 3_20170749 | NA | NA | 3 | 0.3333 | 4 | 0 | 1 | 0 | 8 | 0.125 | 3 | 0.3333 | 1 | 0 | 2 | 0 | 2 | 0 | 8 | 0.125 |
| 3_20170871 | NA | NA | 3 | 0 | 2 | 0.5 | 1 | 0 | 6 | 0.1667 | 3 | 0 | 2 | 0.5 | 2 | 0 | 2 | 0 | 9 | 0.1111 |
| 3_20176477 | 1 | 0 | 7 | 0.1429 | 2 | 0.5 | 6 | 0.3333 | 16 | 0.25 | 9 | 0.3333 | 7 | 0.1429 | 4 | 0 | 3 | 0.6667 | 23 | 0.2609 |
| 3_20176678 | 1 | 0 | 6 | 0 | 2 | 0.5 | 4 | 0 | 13 | 0.0769 | 10 | 0.1 | 5 | 0 | 3 | 0 | 3 | 0.3333 | 21 | 0.0952 |
| 3_20176832 | 1 | 0 | 4 | 0.75 | 2 | 0.5 | 5 | 0.2 | 12 | 0.4167 | 7 | 0.1429 | 7 | 0.2857 | 3 | 0.3333 | 3 | 0.6667 | 20 | 0.3 |
| 3_22128715 | NA | NA | 1 | 0 | 2 | 1 | NA | NA | 3 | 0.6667 | 2 | 0.5 | NA | NA | 1 | 0 | NA | NA | 3 | 0.3333 |
| 3_22128755 | NA | NA | 1 | 1 | 2 | 1 | NA | NA | 3 | 1 | 1 | 1 | NA | NA | 1 | 0 | NA | NA | 2 | 0.5 |
| 3_22128906 | NA | NA | 1 | 1 | 2 | 0.5 | NA | NA | 3 | 0.6667 | 2 | 1 | NA | NA | NA | NA | NA | NA | 2 | 1 |
| 3_22129417 | NA | NA | 1 | 1 | 1 | 1 | NA | NA | 2 | 1 | 1 | 1 | NA | NA | NA | NA | NA | NA | 1 | 1 |
| 3_22264614 | NA | NA | 6 | 0.5 | NA | NA | 1 | 0 | 7 | 0.4286 | 1 | 0 | 2 | 0 | 1 | 1 | 2 | 0.5 | 6 | 0.3333 |
| 3_22264644 | 1 | 0 | 6 | 0.3333 | NA | NA | 2 | 0 | 9 | 0.2222 | 1 | 0 | 2 | 0.5 | 1 | 1 | 2 | 0 | 6 | 0.3333 |
| 3_22264682 | 1 | 0 | 4 | 0 | NA | NA | 2 | 0.5 | 7 | 0.1429 | 1 | 1 | 2 | 0 | 1 | 0 | 2 | 0 | 6 | 0.1667 |
| 3_22264741 | 1 | 1 | 6 | 0.5 | 1 | 0 | 2 | 0 | 10 | 0.4 | 1 | 1 | 2 | 0.5 | 1 | 0 | 2 | 0 | 6 | 0.3333 |
| 3_22264921 | 1 | 0 | 6 | 0.3333 | 1 | 1 | 2 | 0 | 10 | 0.3 | 1 | 1 | 2 | 0 | NA | NA | 2 | 0 | 5 | 0.2 |
| 3_22264951 | 2 | 0.5 | 6 | 0.1667 | 1 | 0 | 2 | 0 | 11 | 0.1818 | 1 | 0 | 2 | 0 | 1 | 0 | 2 | 0.5 | 6 | 0.1667 |
| 3_22265013 | 2 | 1 | 5 | 0.8 | 1 | 1 | 2 | 0.5 | 10 | 0.8 | 1 | 1 | 2 | 0.5 | 1 | 1 | 2 | 0.5 | 6 | 0.6667 |
| 3_22267093 | 4 | 0.25 | 9 | 0.1111 | 2 | 0 | 2 | 0 | 17 | 0.1176 | 3 | 0 | 3 | 0 | 4 | 0.25 | 2 | 0 | 12 | 0.0833 |
| 3_22267421 | 6 | 0.1667 | 10 | 0.1 | 2 | 0 | 2 | 0 | 20 | 0.1 | 3 | 0 | 3 | 0.3333 | 4 | 0 | 1 | 0 | 11 | 0.0909 |
| 3_22269173 | 9 | 0.1111 | 11 | 0 | 5 | 0 | 4 | 0.5 | 29 | 0.1034 | 4 | 0 | 6 | 0.5 | 6 | 0.1667 | 3 | 0 | 19 | 0.2105 |
| 3_25476340 | NA | NA | 1 | 0 | 1 | 0 | 2 | 1 | 4 | 0.5 | NA | NA | 1 | 1 | 1 | 0 | 1 | 0 | 3 | 0.3333 |
| 3_25476382 | NA | NA | 2 | 0.5 | NA | NA | 1 | 0 | 3 | 0.3333 | NA | NA | 1 | 0 | 1 | 0 | 3 | 0.3333 | 5 | 0.2 |
| 3_25486233 | 7 | 1 | 4 | 1 | 8 | 1 | 3 | 1 | 22 | 1 | 1 | 1 | 6 | 1 | 3 | 0.6667 | 3 | 1 | 13 | 0.9231 |
| 3_25486368 | 7 | 0.5714 | 4 | 0.75 | 3 | 0.3333 | 3 | 0.3333 | 17 | 0.5294 | 2 | 0 | 6 | 0.3333 | 4 | 0.25 | 3 | 0 | 15 | 0.2 |
| 3_25487174 | 3 | 0.6667 | 4 | 0.75 | 1 | 1 | 2 | 0.5 | 10 | 0.7 | 1 | 0 | 4 | 0 | 5 | 1 | 3 | 0.3333 | 13 | 0.4615 |
| 3_25487235 | 2 | 0 | 3 | 0.6667 | 2 | 1 | 3 | 0.3333 | 10 | 0.5 | 1 | 0 | 4 | 0 | 5 | 0.6 | 3 | 0.6667 | 13 | 0.3846 |
| 3_25487308 | 2 | 0.5 | 4 | 0.75 | 2 | 1 | 3 | 0.3333 | 11 | 0.6364 | 1 | 1 | 3 | 0.3333 | 3 | 1 | 3 | 0.6667 | 10 | 0.7 |

| | | | | | | | | | | | | | | | | | | | | |
|---|---|---|---|---|---|---|---|---|---|---|---|---|---|---|---|---|---|---|---|---|
| 3_2548733 | 3 | 0 | 4 | 0.25 | 2 | 0 | 3 | 0 | 12 | 0.0833 | 1 | 0 | 2 | 0 | 3 | 0.3333 | 3 | 0.3333 | 9 | 0.2222 |
| 3_2548739 | 3 | 1 | 4 | 1 | 2 | 1 | 3 | 0.3333 | 12 | 0.8333 | 1 | 1 | 2 | 0 | 3 | 1 | 3 | 1 | 9 | 0.7778 |
| 3_2548743 | 2 | 0 | 3 | 0.3333 | 2 | 0 | 2 | 0 | 9 | 0.1111 | 1 | 0 | 1 | 0 | 3 | 0 | 3 | 0.3333 | 8 | 0.125 |
| 3_2548744 | 3 | 0.3333 | 2 | 0 | 1 | 0 | 3 | 0 | 9 | 0.1111 | 1 | 0 | 1 | 0 | 3 | 0.3333 | 3 | 0 | 8 | 0.125 |
| 3_2548744 | 2 | 0 | 4 | 0.5 | 2 | 0 | 3 | 0 | 11 | 0.1818 | 1 | 0 | 2 | 0 | 4 | 0.25 | 3 | 0 | 10 | 0.1 |
| 3_2548749 | 3 | 0.6667 | 4 | 0.5 | 2 | 0.5 | 3 | 0 | 12 | 0.4167 | 1 | 0 | 2 | 0 | 2 | 0.5 | 3 | 0.3333 | 8 | 0.25 |
| 3_2548766 | 3 | 1 | 3 | 1 | 2 | 1 | 3 | 1 | 11 | 1 | 1 | 1 | 1 | 1 | 3 | 1 | 3 | 1 | 8 | 1 |
| 3_2548771 | 2 | 1 | 2 | 1 | 1 | 1 | 3 | 1 | 8 | 1 | 1 | 1 | 2 | 0 | 4 | 1 | 3 | 0.6667 | 10 | 0.7 |
| 3_2548778 | 2 | 0.5 | 2 | 1 | 2 | 1 | 3 | 0 | 9 | 0.5556 | 1 | 1 | 2 | 0.5 | 4 | 0.5 | 3 | 1 | 10 | 0.7 |
| 3_2548778 | 2 | 0.5 | 3 | 0 | 2 | 0 | 3 | 0 | 10 | 0.1 | 1 | 1 | 2 | 0 | 4 | 0.25 | 3 | 0 | 10 | 0.2 |
| 3_2548786 | 2 | 1 | 1 | 1 | 2 | 0.5 | 3 | 1 | 8 | 0.875 | 1 | 1 | 2 | 0.5 | 4 | 1 | 2 | 0.5 | 9 | 0.7778 |
| 3_2548794 | 1 | 0 | 3 | 0.3333 | 2 | 0 | 3 | 0 | 9 | 0.1111 | 1 | 0 | 2 | 0 | 4 | 0.5 | 2 | 0 | 9 | 0.2222 |
| 3_2548805 | 1 | 1 | 1 | 0 | 2 | 0.5 | 1 | 0 | 5 | 0.4 | 1 | 1 | 1 | 0 | 4 | 0.5 | 2 | 0.5 | 8 | 0.5 |
| 3_2548826 | 1 | 1 | 3 | 1 | 1 | 1 | 1 | 1 | 6 | 1 | 1 | 1 | 2 | 1 | 4 | 1 | 1 | 1 | 8 | 1 |
| 3_2548836 | 2 | 0 | 3 | 0 | 1 | 0 | 2 | 0.5 | 8 | 0.125 | 1 | 0 | 2 | 0 | 4 | 0.25 | 2 | 0.5 | 9 | 0.2222 |
| 3_2548837 | 2 | 0.5 | 3 | 1 | 1 | 1 | 2 | 1 | 8 | 0.875 | 1 | 0 | 1 | 1 | 4 | 0.75 | 2 | 0.5 | 8 | 0.625 |
| 3_2548842 | 1 | 1 | 3 | 1 | 1 | 1 | 2 | 1 | 7 | 1 | 1 | 1 | 2 | 1 | 4 | 0.75 | 2 | 1 | 9 | 0.8889 |
| 3_2548854 | 2 | 1 | 3 | 0.6667 | 1 | 0 | 2 | 1 | 8 | 0.75 | 1 | 1 | 1 | 0 | 4 | 0.75 | 2 | 1 | 8 | 0.75 |
| 3_2549023 | 2 | 0.5 | 2 | 1 | 1 | 1 | 2 | 1 | 7 | 0.8571 | NA | NA | 2 | 1 | 4 | 1 | 1 | 1 | 7 | 1 |
| 3_2549028 | 1 | 1 | 2 | 1 | 1 | 1 | 2 | 0 | 6 | 0.6667 | NA | NA | 2 | 1 | 4 | 1 | 1 | 1 | 7 | 1 |
| 3_2549034 | 1 | 1 | NA | NA | 1 | 1 | 1 | 1 | 3 | 1 | NA | NA | 2 | 1 | 3 | 1 | 1 | 1 | 6 | 1 |
| 3_2549050 | 2 | 0.5 | 1 | 0 | 1 | 0 | 1 | 0 | 5 | 0.2 | NA | NA | 1 | 0 | 4 | 0.25 | 1 | 1 | 6 | 0.3333 |
| 3_2549056 | 2 | 0 | 3 | 0.3333 | 1 | 0 | 1 | 0 | 7 | 0.1429 | NA | NA | 1 | 0 | 3 | 0.3333 | 1 | 0 | 5 | 0.2 |
| 3_2549059 | 2 | 0.5 | 3 | 0.6667 | 1 | 1 | 1 | 1 | 7 | 0.7143 | NA | NA | 1 | 1 | 3 | 1 | 1 | 1 | 5 | 1 |
| 3_2549064 | 2 | 0 | 3 | 0.6667 | 1 | 0 | 1 | 1 | 7 | 0.4286 | NA | NA | 1 | 0 | 4 | 0.5 | 1 | 0 | 6 | 0.3333 |
| 3_2549080 | NA | NA | 2 | 0.5 | 1 | 0 | NA | NA | 3 | 0.3333 | NA | NA | NA | NA | 3 | 0 | 1 | 1 | 4 | 0.25 |
| 3_2638574 | NA | NA | NA | NA | NA | NA | 1 | 1 | 1 | 1 | NA | NA | 1 | 1 | NA | NA | 1 | 1 | 2 | 1 |
| 3_2638601 | NA | NA | NA | NA | NA | NA | 1 | 1 | 1 | 1 | NA | NA | 1 | 0 | NA | NA | 1 | 1 | 2 | 0.5 |
| 3_2638621 | NA | NA | NA | NA | NA | NA | 1 | 1 | 1 | 1 | NA | NA | 1 | 1 | NA | NA | 1 | 1 | 2 | 1 |
| 3_2729542 | 7 | 0.2857 | 5 | 0.6 | 3 | 0.6667 | 8 | 0.375 | 23 | 0.4348 | 3 | 0.6667 | 7 | 0.5714 | 9 | 0.6667 | 8 | 0.75 | 27 | 0.6667 |
| 3_2729545 | 7 | 0.1429 | 3 | 0 | 2 | 0.5 | 6 | 0.5 | 18 | 0.2778 | 3 | 0.3333 | 5 | 0.2 | 8 | 0.375 | 6 | 0.1667 | 22 | 0.2727 |
| 3_2729550 | 6 | 0.3333 | 6 | 0.5 | 3 | 0.6667 | 7 | 0.2857 | 22 | 0.4091 | 3 | 0 | 5 | 0.6 | 9 | 0.6667 | 5 | 0.4 | 22 | 0.5 |
| 3_2729554 | 4 | 1 | 5 | 0.6 | 3 | 1 | 6 | 0.8333 | 18 | 0.8333 | 5 | 0.6 | 4 | 1 | 6 | 0.6667 | 6 | 0.8333 | 21 | 0.7619 |
| 3_2729560 | 6 | 0.5 | 6 | 0.3333 | 3 | 0.6667 | 7 | 0.4286 | 22 | 0.4545 | 5 | 0.6 | 6 | 0.6667 | 11 | 0.4545 | 7 | 0.7143 | 29 | 0.5862 |
| 3_2729563 | 5 | 0.2 | 5 | 0.6 | 3 | 0 | 6 | 0 | 19 | 0.2105 | 4 | 0.5 | 5 | 0.4 | 9 | 0.1111 | 6 | 0.1667 | 24 | 0.25 |
| 3_2729585 | 7 | 0.5714 | 6 | 0.6667 | 3 | 0.6667 | 7 | 0.4286 | 23 | 0.5652 | 4 | 0 | 5 | 0.2 | 7 | 0.4286 | 6 | 0.3333 | 22 | 0.2727 |
| 3_2729592 | 7 | 0.8571 | 6 | 0.8333 | 3 | 1 | 9 | 0.8889 | 25 | 0.88 | 5 | 0.8 | 7 | 0.8571 | 11 | 1 | 9 | 0.8889 | 32 | 0.9062 |
| 3_2729602 | 7 | 0.2857 | 5 | 0 | 4 | 0.25 | 9 | 0.4444 | 25 | 0.28 | 6 | 0 | 5 | 0.2 | 9 | 0.1111 | 7 | 0.4286 | 27 | 0.1852 |
| 3_2729605 | 5 | 0 | 3 | 0 | 1 | 1 | 4 | 0 | 13 | 0.0769 | 4 | 0 | 4 | 0.75 | 7 | 0 | 2 | 0 | 17 | 0.1765 |
| 3_2729605 | 7 | 0.4286 | 5 | 0.8 | 3 | 0.6667 | 7 | 0.5714 | 22 | 0.5909 | 6 | 0.8333 | 5 | 1 | 9 | 0.6667 | 6 | 0.8333 | 26 | 0.8077 |
| 3_2729612 | 10 | 0.7 | 9 | 0.7778 | 4 | 0.75 | 9 | 0.5556 | 32 | 0.6875 | 5 | 0.8 | 7 | 0.4286 | 11 | 0.4545 | 9 | 1 | 32 | 0.6562 |
| 3_2729614 | 10 | 1 | 7 | 1 | 3 | 1 | 9 | 1 | 29 | 1 | 6 | 1 | 7 | 1 | 9 | 1 | 7 | 1 | 29 | 1 |
| 3_2729615 | 7 | 0 | 8 | 0.125 | 3 | 0.3333 | 8 | 0.375 | 26 | 0.1923 | 5 | 0 | 6 | 0.1667 | 10 | 0.3 | 5 | 0.2 | 26 | 0.1923 |
| 3_2729617 | 8 | 0.375 | 7 | 0.2857 | 4 | 0.25 | 7 | 0.2857 | 26 | 0.3077 | 4 | 0.25 | 7 | 0.5714 | 12 | 0.3333 | 8 | 0.375 | 31 | 0.3871 |
| 3_2729617 | 9 | 0.6667 | 8 | 0.75 | 3 | 1 | 7 | 0.7143 | 27 | 0.7407 | 5 | 1 | 6 | 0.8333 | 11 | 0.7273 | 8 | 0.75 | 30 | 0.8 |
| 3_2729619 | 9 | 1 | 8 | 1 | 4 | 1 | 7 | 0.8571 | 28 | 0.9643 | 6 | 1 | 6 | 0.8333 | 10 | 0.8 | 7 | 1 | 29 | 0.8966 |
| 3_2729634 | 10 | 0.3 | 7 | 0.1429 | 3 | 0 | 6 | 0.1667 | 26 | 0.1923 | 6 | 0 | 7 | 0 | 10 | 0.1 | 8 | 0.125 | 31 | 0.0645 |
| 3_2729642 | 12 | 1 | 8 | 1 | 3 | 1 | 10 | 1 | 33 | 1 | 7 | 1 | 5 | 1 | 10 | 1 | 8 | 1 | 30 | 1 |
| 3_2729644 | 10 | 1 | 9 | 1 | 3 | 1 | 8 | 1 | 30 | 1 | 7 | 1 | 4 | 1 | 8 | 1 | 7 | 1 | 26 | 1 |
| 3_2729693 | 13 | 0.2308 | 9 | 0 | 8 | 0.875 | 8 | 0.375 | 38 | 0.3421 | 8 | 0.125 | 7 | 0.2857 | 12 | 0.25 | 9 | 0.3333 | 36 | 0.25 |
| 3_2729849 | 18 | 0.0556 | 10 | 0 | 15 | 0.0667 | 10 | 0.1 | 53 | 0.0566 | 13 | 0.1538 | 15 | 0 | 16 | 0.0625 | 14 | 0.0714 | 58 | 0.069 |
| 3_2729861 | 15 | 0.4667 | 12 | 0.3333 | 11 | 0.2727 | 7 | 0.4286 | 45 | 0.3778 | 12 | 0.25 | 9 | 0.3333 | 15 | 0.2667 | 11 | 0.1818 | 47 | 0.2553 |
| 3_2729883 | 17 | 0.6471 | 15 | 0.8667 | 15 | 0.8 | 10 | 0.7 | 57 | 0.7544 | 13 | 0.8462 | 15 | 0.8667 | 17 | 0.7647 | 12 | 0.9167 | 57 | 0.8421 |
| 3_2729889 | 15 | 0.7333 | 16 | 0.875 | 15 | 0.8667 | 12 | 0.9167 | 58 | 0.8448 | 13 | 1 | 17 | 0.8235 | 16 | 0.8125 | 12 | 0.9167 | 58 | 0.8793 |
| 3_2729890 | 19 | 0.6842 | 15 | 0.6 | 13 | 0.6154 | 11 | 0.9091 | 58 | 0.6897 | 13 | 0.7692 | 18 | 0.7778 | 17 | 0.7647 | 16 | 0.9375 | 64 | 0.8125 |
| 3_2747052 | 1 | 0 | 6 | 0.1667 | 4 | 0 | 6 | 0.1667 | 17 | 0.1176 | 3 | 0.3333 | 1 | 0 | 4 | 0 | 5 | 0 | 13 | 0.0769 |
| 3_2747205 | NA | NA | 2 | 0 | 1 | 1 | 5 | 0.2 | 8 | 0.25 | 1 | 0 | 3 | 0 | 1 | 0 | 4 | 0.5 | 9 | 0.2222 |
| 3_2747218 | NA | NA | 4 | 0 | 1 | 0 | 5 | 0.2 | 10 | 0.1 | 1 | 0 | 3 | 0 | 2 | 0.5 | 3 | 0 | 9 | 0.1111 |
| 3_2747252 | NA | NA | 3 | 0 | 1 | 1 | 4 | 0 | 8 | 0.125 | 2 | 0 | 3 | 0 | 2 | 0.5 | 4 | 0 | 11 | 0.0909 |
| 3_2747352 | NA | NA | 1 | 1 | NA | NA | 2 | 1 | 3 | 1 | 1 | 1 | 2 | 0.5 | 1 | 1 | NA | NA | 4 | 0.75 |
| 3_2747357 | NA | NA | 2 | 1 | NA | NA | 4 | 0.25 | 6 | 0.5 | 1 | 0 | 1 | 0 | 1 | 1 | 2 | 0 | 5 | 0.2 |
| 3_2747365 | NA | NA | 2 | 1 | NA | NA | 3 | 0.3333 | 5 | 0.6 | 1 | 0 | 2 | 0.5 | NA | NA | 3 | 0.6667 | 6 | 0.5 |
| 3_2750559 | NA | NA | 1 | 0 | NA | NA | 3 | 0.3333 | 4 | 0.25 | 1 | 0 | 1 | 0 | 1 | 0 | 1 | 1 | 4 | 0.25 |
| 3_2818571 | 2 | 0.5 | NA | NA | NA | NA | 1 | 1 | 3 | 0.6667 | NA | NA | NA | NA | 1 | 0 | 1 | 1 | 2 | 0.5 |
| 3_2818588 | 3 | 1 | NA | NA | NA | NA | 1 | 1 | 4 | 1 | NA | NA | NA | NA | NA | NA | 1 | 1 | 1 | 1 |

| | | | | | | | | | | | | | | | | | | | | |
|---|---|---|---|---|---|---|---|---|---|---|---|---|---|---|---|---|---|---|---|---|
| 3_2818595 | 3 | 1 | NA | NA | NA | NA | 1 | 1 | 4 | 1 | NA | NA | NA | NA | 1 | 1 | NA | NA | 1 | 1 |
| 3_2818600 | 1 | 1 | NA | NA | NA | NA | 1 | 1 | 2 | 1 | NA | NA | NA | NA | 1 | 1 | 1 | 1 | 2 | 1 |
| 3_2818604 | 3 | 0.6667 | NA | NA | NA | NA | 1 | 1 | 4 | 0.75 | NA | NA | NA | NA | 1 | 1 | 1 | 1 | 2 | 1 |
| 3_2818605 | 3 | 0.3333 | NA | NA | NA | NA | 1 | 1 | 4 | 0.5 | NA | NA | NA | NA | 1 | 1 | NA | NA | 1 | 1 |
| 3_2818619 | 4 | 0.75 | NA | NA | NA | NA | 1 | 0 | 5 | 0.6 | NA | NA | NA | NA | 1 | 0 | 1 | 1 | 2 | 0.5 |
| 3_2818667 | 4 | 0.5 | NA | NA | NA | NA | 1 | 0 | 5 | 0.4 | NA | NA | 1 | 1 | 1 | 0 | NA | NA | 2 | 0.5 |
| 3_2818690 | 5 | 0.4 | NA | NA | NA | NA | 1 | 1 | 6 | 0.5 | NA | NA | 1 | 1 | 1 | 1 | 1 | 1 | 3 | 1 |
| 3_2818709 | 5 | 0.2 | NA | NA | NA | NA | 1 | 1 | 6 | 0.3333 | NA | NA | 1 | 1 | 1 | 0 | 1 | 1 | 3 | 0.6667 |
| 3_2872010 | 2 | 0 | 1 | 1 | 1 | 0 | NA | NA | 4 | 0.25 | 1 | 0 | 1 | 0 | 2 | 0.5 | 3 | 0 | 7 | 0.1429 |
| 3_2872030 | 7 | 0 | 3 | 0.6667 | 4 | 0.25 | 5 | 0.2 | 19 | 0.2105 | 2 | 0 | 5 | 0.2 | 13 | 0.1538 | 6 | 0 | 26 | 0.1154 |
| 3_2872212 | 6 | 1 | 1 | 1 | 3 | 1 | 3 | 0.6667 | 13 | 0.9231 | 2 | 1 | 4 | 1 | 9 | 1 | 5 | 0.8 | 20 | 0.95 |
| 3_2872218 | 6 | 0 | 2 | 0.5 | 2 | 0.5 | 3 | 0.3333 | 13 | 0.2308 | 4 | 0.25 | 3 | 0.3333 | 7 | 0.7143 | 4 | 1 | 18 | 0.6111 |
| 3_2872218 | 7 | 0.2857 | 2 | 0 | 4 | 0.25 | 5 | 0 | 18 | 0.1667 | 3 | 0 | 4 | 0 | 11 | 0.0909 | 5 | 0.2 | 23 | 0.087 |
| 3_2872223 | 8 | 0.125 | 2 | 0 | 3 | 0 | 5 | 0 | 18 | 0.0556 | 3 | 0.3333 | 5 | 0 | 12 | 0.1667 | 6 | 0.3333 | 26 | 0.1923 |
| 3_2872237 | 9 | 0.4444 | 2 | 0 | 4 | 0.25 | 7 | 0.2857 | 22 | 0.3182 | 3 | 0.6667 | 4 | 0.75 | 13 | 0.3077 | 5 | 0.6 | 25 | 0.48 |
| 3_2872238 | 10 | 0.9 | 2 | 0.5 | 4 | 1 | 6 | 0.8333 | 22 | 0.8636 | 4 | 1 | 3 | 0.3333 | 13 | 0.7692 | 5 | 0.6 | 25 | 0.72 |
| 3_2872241 | 11 | 0.2727 | 3 | 0 | 4 | 0.75 | 8 | 0.375 | 26 | 0.3462 | 4 | 0.25 | 5 | 0.6 | 13 | 0.1538 | 6 | 0.3333 | 28 | 0.2857 |
| 3_2872248 | 10 | 0.3 | 2 | 0.5 | 3 | 0.3333 | 6 | 0.3333 | 21 | 0.3333 | 3 | 0 | 4 | 0.25 | 10 | 0.1 | 6 | 0.3333 | 23 | 0.1739 |
| 3_2872265 | 7 | 0 | 3 | 0 | 4 | 0.75 | 7 | 0.4286 | 21 | 0.2857 | 4 | 0.25 | 4 | 0 | 15 | 0.2 | 6 | 0.3333 | 29 | 0.2069 |
| 3_2872319 | 6 | 0.1667 | 2 | 0 | 3 | 0 | 7 | 0.2857 | 18 | 0.1667 | 4 | 0 | 6 | 0.5 | 11 | 0.3636 | 7 | 0.4286 | 28 | 0.3571 |
| 3_2872328 | 7 | 0 | 2 | 0 | 3 | 0 | 8 | 0.375 | 20 | 0.15 | 5 | 0 | 6 | 0 | 9 | 0.1111 | 5 | 0.2 | 25 | 0.08 |
| 3_2872334 | 6 | 0.3333 | 2 | 0 | 2 | 0 | 6 | 0.1667 | 16 | 0.1875 | 5 | 0 | 4 | 0.25 | 10 | 0.1 | 5 | 0.2 | 24 | 0.125 |
| 3_2872335 | 7 | 0 | 2 | 0 | 1 | 0 | 6 | 0.1667 | 16 | 0.0625 | 5 | 0 | 4 | 0 | 9 | 0.1111 | 5 | 0.2 | 23 | 0.087 |
| 3_2872344 | 8 | 0 | 2 | 0 | 1 | 0 | 5 | 0.4 | 16 | 0.125 | 5 | 0.2 | 3 | 0 | 9 | 0.1111 | 6 | 0.1667 | 23 | 0.1304 |
| 3_2872405 | 6 | 0.1667 | 2 | 0 | 1 | 0 | 4 | 0 | 13 | 0.0769 | 4 | 0 | 4 | 0 | 10 | 0.1 | 4 | 0.25 | 22 | 0.0909 |
| 3_2872452 | 11 | 0.3636 | 4 | 0.25 | 1 | 1 | 7 | 0.1429 | 23 | 0.3043 | 7 | 0.1429 | 11 | 0 | 14 | 0.2143 | 8 | 0 | 40 | 0.1 |
| 3_2872623 | 5 | 0 | 4 | 0.25 | 3 | 0 | 4 | 0 | 16 | 0.0625 | 4 | 0.25 | 5 | 0.2 | 9 | 0.1111 | 7 | 0 | 25 | 0.12 |
| 3_2872624 | 3 | 0 | 4 | 0.25 | 3 | 0 | 4 | 0 | 14 | 0.0714 | 4 | 0 | 4 | 0.25 | 8 | 0 | 7 | 0.1429 | 23 | 0.087 |
| 3_2872636 | 6 | 0.1667 | 5 | 0 | 3 | 0 | 5 | 0 | 19 | 0.0526 | 4 | 0.25 | 5 | 0.2 | 9 | 0 | 7 | 0 | 25 | 0.08 |
| 3_2872637 | 6 | 0.3333 | 5 | 0.2 | 3 | 0.6667 | 5 | 0.6 | 19 | 0.4211 | 3 | 0 | 4 | 0.5 | 6 | 0.5 | 6 | 0.5 | 19 | 0.4211 |
| 3_2872655 | 4 | 0.25 | 4 | 0 | 2 | 0.5 | 4 | 0.5 | 14 | 0.2857 | 4 | 0.25 | 5 | 0.4 | 8 | 0.375 | 7 | 0.2857 | 24 | 0.3333 |
| 3_2872661 | 2 | 0 | 3 | 0.3333 | 1 | 1 | 4 | 0.25 | 10 | 0.3 | 2 | 0.5 | 3 | 0.6667 | 3 | 0.3333 | 3 | 0 | 11 | 0.3636 |
| 3_2872666 | 3 | 0 | 4 | 0.25 | 2 | 0 | 4 | 0 | 13 | 0.0769 | 4 | 0.25 | 3 | 0 | 6 | 0 | 6 | 0.1667 | 19 | 0.1053 |
| 3_2872715 | 4 | 0.25 | 1 | 0 | 2 | 0 | 1 | 0 | 8 | 0.125 | 4 | 0 | 3 | 0 | 6 | 0 | 6 | 0.1667 | 19 | 0.0526 |
| 3_2885059 | 2 | 0 | 5 | 0 | 1 | 1 | 3 | 0 | 11 | 0.0909 | 5 | 0 | 3 | 0 | 5 | 0 | 7 | 0.1429 | 20 | 0.05 |
| 3_2885102 | 6 | 0.1667 | 6 | 0 | 1 | 0 | 3 | 0 | 16 | 0.0625 | 4 | 0 | 2 | 0 | 5 | 0.2 | 6 | 0 | 17 | 0.0588 |
| 3_2885261 | 6 | 0.1667 | 11 | 0.0909 | 2 | 0 | 3 | 0.3333 | 22 | 0.1364 | 10 | 0.2 | 3 | 0 | 7 | 0 | 8 | 0 | 28 | 0.0714 |
| 3_2886153 | 41 | 0.3659 | 48 | 0.3958 | 36 | 0.4167 | 44 | 0.3636 | 169 | 0.3846 | 32 | 0.4062 | 49 | 0.3878 | 47 | 0.4468 | 34 | 0.5294 | 162 | 0.4383 |
| 3_3065181 | NA | NA | NA | NA | 1 | 1 | NA | NA | 1 | 1 | 1 | 1 | NA | NA | NA | NA | NA | NA | 1 | 1 |
| 3_3065197 | 1 | 1 | 1 | 1 | NA | NA | NA | NA | 2 | 1 | 1 | 1 | 1 | 0 | NA | NA | NA | NA | 2 | 0.5 |
| 3_3065201 | 1 | 0 | 1 | 1 | NA | NA | NA | NA | 2 | 0.5 | NA | NA | 1 | 1 | NA | NA | NA | NA | 1 | 1 |
| 3_3066140 | 1 | 1 | 2 | 1 | 1 | 1 | 1 | 1 | 5 | 1 | NA | NA | 1 | 1 | 1 | 1 | NA | NA | 2 | 1 |
| 3_3066162 | 1 | 1 | 2 | 1 | 1 | 0 | 1 | 0 | 5 | 0.6 | NA | NA | 1 | 1 | 1 | 0 | NA | NA | 2 | 0.5 |
| 3_3066167 | 1 | 0 | 2 | 0.5 | 1 | 1 | 1 | 0 | 5 | 0.4 | NA | NA | 1 | 1 | 1 | 0 | NA | NA | 2 | 0.5 |
| 3_3066173 | 1 | 0 | 2 | 0.5 | 1 | 0 | 1 | 1 | 5 | 0.4 | NA | NA | 1 | 1 | 1 | 1 | NA | NA | 2 | 1 |
| 3_3067096 | 2 | 0.5 | 2 | 0.5 | 1 | 1 | 1 | 1 | 6 | 0.6667 | NA | NA | 3 | 0.3333 | NA | NA | NA | NA | 3 | 0.3333 |
| 3_3067097 | 1 | 1 | 4 | 1 | 1 | 1 | 1 | 1 | 7 | 1 | NA | NA | 3 | 0.6667 | 1 | 1 | NA | NA | 4 | 0.75 |
| 3_3087272 | 8 | 0.125 | 11 | 0.4545 | 16 | 0.3125 | 16 | 0.25 | 51 | 0.2941 | 5 | 0 | 16 | 0.0625 | 16 | 0.3125 | 14 | 0.2857 | 51 | 0.1961 |
| 3_3087274 | 10 | 0.3 | 11 | 0.2727 | 20 | 0.3 | 19 | 0.4737 | 60 | 0.35 | 5 | 0.2 | 15 | 0.5333 | 19 | 0.2632 | 20 | 0.1 | 59 | 0.2712 |
| 3_3087278 | 11 | 0.5455 | 11 | 0.9091 | 21 | 0.7619 | 18 | 0.8333 | 61 | 0.7705 | 4 | 1 | 20 | 0.75 | 18 | 0.7778 | 21 | 0.4762 | 63 | 0.6825 |
| 3_3087279 | 9 | 0.5556 | 11 | 0.2727 | 21 | 0.5238 | 16 | 0.5625 | 57 | 0.4912 | 5 | 0.2 | 17 | 0.5294 | 17 | 0.2941 | 17 | 0.5294 | 56 | 0.4286 |
| 3_3087285 | 11 | 0.1818 | 14 | 0 | 22 | 0.1364 | 20 | 0.05 | 67 | 0.0896 | 5 | 0.2 | 19 | 0.1579 | 21 | 0.0952 | 20 | 0.1 | 65 | 0.1231 |
| 3_3087291 | 11 | 0.1818 | 12 | 0.25 | 23 | 0.3478 | 19 | 0.2632 | 65 | 0.2769 | 3 | 0.6667 | 19 | 0.3158 | 22 | 0.4545 | 21 | 0.381 | 65 | 0.4 |
| 3_3087294 | 7 | 0.8571 | 9 | 0.8889 | 17 | 0.8824 | 15 | 1 | 48 | 0.9167 | 5 | 0.8 | 13 | 0.9231 | 17 | 0.9412 | 19 | 1 | 54 | 0.9444 |
| 3_3087295 | 14 | 0 | 13 | 0 | 22 | 0.0909 | 22 | 0.1364 | 71 | 0.0704 | 4 | 0 | 18 | 0.1111 | 20 | 0.05 | 17 | 0.0588 | 59 | 0.0678 |
| 3_3087304 | 13 | 0.1538 | 14 | 0.3571 | 24 | 0.1667 | 20 | 0.3 | 71 | 0.2394 | 5 | 0 | 17 | 0.2941 | 24 | 0.1667 | 19 | 0.1579 | 65 | 0.1846 |
| 3_3087312 | 14 | 0.4286 | 12 | 0.5 | 22 | 0.4091 | 20 | 0.4 | 68 | 0.4265 | 4 | 1 | 18 | 0.4444 | 21 | 0.5238 | 20 | 0.25 | 63 | 0.4444 |
| 3_3087313 | 12 | 0.8333 | 14 | 0.8571 | 22 | 0.8182 | 20 | 0.9 | 68 | 0.8529 | 6 | 1 | 16 | 0.75 | 18 | 0.8333 | 20 | 0.85 | 60 | 0.8333 |
| 3_3087425 | 1 | 0 | 5 | 0.6 | 5 | 0.8 | 4 | 0.75 | 15 | 0.6667 | NA | NA | 5 | 0.4 | 7 | 1 | 4 | 1 | 16 | 0.8125 |
| 3_3087495 | 3 | 0 | 5 | 0.4 | 4 | 0 | 4 | 0.25 | 16 | 0.1875 | 2 | 0 | 4 | 0.5 | 7 | 0.1429 | 4 | 0.25 | 17 | 0.2353 |
| 3_3087522 | 4 | 0.25 | 7 | 0.1429 | 6 | 0 | 4 | 0 | 21 | 0.0952 | 3 | 0 | 6 | 0 | 6 | 0.1667 | 5 | 0 | 20 | 0.05 |
| 3_3095381 | 5 | 0.2 | 10 | 0.1 | 5 | 0.2 | 6 | 0.1667 | 26 | 0.1538 | 4 | 0 | 6 | 0.1667 | 3 | 0.6667 | 1 | 0 | 14 | 0.2143 |
| 3_3095507 | 5 | 0.2 | 3 | 0 | 1 | 0 | 2 | 0.5 | 11 | 0.1818 | 1 | 0 | 3 | 0.3333 | 2 | 0 | NA | NA | 6 | 0.1667 |
| 3_3095544 | 3 | 0.3333 | 5 | 0 | 1 | 0 | 3 | 0 | 12 | 0.0833 | 1 | 0 | 4 | 0.25 | 2 | 0 | NA | NA | 7 | 0.1429 |
| 3_3095549 | 5 | 0 | 7 | 0.1429 | 1 | 0 | 4 | 0.25 | 17 | 0.1176 | 1 | 0 | 3 | 0.6667 | 1 | 0 | NA | NA | 5 | 0.4 |

| | | | | | | | | | | | | | | | | | | | | |
|---|---|---|---|---|---|---|---|---|---|---|---|---|---|---|---|---|---|---|---|---|
| 3_3095551 | 5 | 0.4 | 6 | 0.1667 | 1 | 0 | 4 | 0.25 | 16 | 0.25 | 1 | 0 | 3 | 0.3333 | 3 | 0 | NA | NA | 7 | 0.1429 |
| 3_3095566 | 2 | 0 | 5 | 0.2 | 1 | 0 | 4 | 0 | 12 | 0.0833 | NA | NA | 3 | 0.3333 | 3 | 0 | NA | NA | 6 | 0.1667 |
| 3_3095619 | 3 | 0 | 3 | 0 | NA | NA | 4 | 0.5 | 10 | 0.2 | NA | NA | 3 | 0.6667 | 3 | 0.3333 | NA | NA | 6 | 0.5 |
| 3_3096125 | 2 | 0.5 | NA | NA | NA | NA | NA | NA | 2 | 0.5 | NA | NA | NA | NA | 1 | 1 | NA | NA | 1 | 1 |
| 3_3110455 | 2 | 1 | NA | NA | 3 | 1 | 2 | 1 | 7 | 1 | 2 | 1 | 1 | 1 | 3 | 0.6667 | 3 | 0.6667 | 9 | 0.7778 |
| 3_3110562 | 3 | 0.3333 | 6 | 0.5 | 2 | 0 | 3 | 0.3333 | 14 | 0.3571 | 4 | 0.5 | 4 | 0.25 | 4 | 0.5 | 3 | 0 | 15 | 0.3333 |
| 3_3110594 | 5 | 0 | 6 | 0.1667 | 3 | 0.6667 | 2 | 0 | 16 | 0.1875 | 4 | 0.25 | 4 | 0.25 | 4 | 0 | 4 | 0 | 16 | 0.125 |
| 3_3110603 | 5 | 0 | 7 | 0 | 3 | 0 | 4 | 0.25 | 19 | 0.0526 | 4 | 0 | 4 | 0.25 | 2 | 0 | 4 | 0 | 14 | 0.0714 |
| 3_3110613 | 6 | 0 | 7 | 0 | 3 | 0 | 4 | 0.25 | 20 | 0.05 | 4 | 0 | 4 | 0.25 | 4 | 0 | 4 | 0 | 16 | 0.0625 |
| 3_3110617 | 6 | 0.5 | 4 | 0.25 | 3 | 0.6667 | 4 | 0.25 | 17 | 0.4118 | 5 | 0 | 4 | 0.5 | 3 | 0.3333 | 5 | 0.4 | 17 | 0.2941 |
| 3_3115089 | NA | NA | 1 | 1 | 2 | 0 | NA | NA | 3 | 0.3333 | 1 | 1 | NA | NA | NA | NA | 1 | 0 | 2 | 0.5 |
| 3_3115110 | 1 | 0 | 2 | 0 | 2 | 0.5 | 1 | 0 | 6 | 0.1667 | 1 | 0 | NA | NA | NA | NA | 1 | 1 | 2 | 0.5 |
| 3_3115115 | 1 | 0 | 2 | 0.5 | 3 | 0.6667 | 1 | 0 | 7 | 0.4286 | 1 | 0 | NA | NA | 1 | 0 | 1 | 1 | 3 | 0.3333 |
| 3_3115116 | 2 | 0.5 | 2 | 1 | 4 | 1 | NA | NA | 8 | 0.875 | 1 | 1 | NA | NA | 1 | 1 | 1 | 1 | 3 | 1 |
| 3_3115145 | 2 | 0 | 2 | 0 | 4 | 0.5 | 2 | 0 | 10 | 0.2 | 2 | 0.5 | NA | NA | 1 | 0 | 1 | 0 | 4 | 0.25 |
| 3_3115170 | 2 | 0 | 1 | 1 | 4 | 0.75 | 2 | 1 | 9 | 0.6667 | 2 | 0.5 | NA | NA | 2 | 0.5 | 1 | 0 | 5 | 0.4 |
| 3_3115173 | 2 | 0 | 2 | 0.5 | 3 | 0 | 2 | 0 | 9 | 0.1111 | 2 | 0.5 | NA | NA | 2 | 0 | 1 | 0 | 5 | 0.2 |
| 3_3115177 | 2 | 0 | 2 | 0 | 4 | 0.25 | 2 | 0.5 | 10 | 0.2 | 2 | 0 | NA | NA | 2 | 0.5 | 1 | 0 | 5 | 0.2 |
| 3_3115190 | 2 | 1 | 1 | 0 | 4 | 0.5 | 1 | 1 | 8 | 0.625 | 1 | 1 | NA | NA | 1 | 1 | NA | NA | 2 | 1 |
| 3_3115196 | 2 | 1 | 2 | 1 | 4 | 1 | 1 | 1 | 9 | 1 | 2 | 1 | NA | NA | 2 | 1 | 1 | 0 | 5 | 0.8 |
| 3_3115215 | 2 | 0.5 | 2 | 1 | 4 | 1 | 2 | 1 | 10 | 0.9 | 1 | 1 | 1 | 1 | 2 | 1 | 1 | 1 | 5 | 1 |
| 3_3115218 | 2 | 1 | 2 | 1 | 2 | 1 | 2 | 1 | 8 | 1 | 2 | 1 | 1 | 1 | 2 | 1 | 1 | 0 | 6 | 0.8333 |
| 3_3115226 | 2 | 0.5 | 2 | 1 | 2 | 0.5 | 2 | 0 | 8 | 0.5 | 2 | 0.5 | 1 | 0 | 2 | 0 | 1 | 1 | 6 | 0.3333 |
| 3_3115241 | 3 | 0.6667 | 2 | 0.5 | 4 | 0 | 2 | 1 | 11 | 0.4545 | 2 | 1 | 1 | 1 | 2 | 1 | 1 | 1 | 6 | 1 |
| 3_3115246 | 2 | 0 | 2 | 0 | 2 | 1 | NA | NA | 6 | 0.3333 | 2 | 0 | 1 | 0 | 3 | 0.3333 | 1 | 0 | 7 | 0.1429 |
| 3_3115248 | 2 | 0 | 2 | 0 | 3 | 0.3333 | 2 | 0 | 9 | 0.1111 | 2 | 0 | 1 | 0 | 3 | 0.3333 | 1 | 0 | 7 | 0.1429 |
| 3_3117165 | 4 | 0.75 | 2 | 0.5 | 5 | 0.4 | 2 | 1 | 13 | 0.6154 | 3 | 1 | NA | NA | 2 | 0.5 | 1 | 1 | 6 | 0.8333 |
| 3_3117261 | 5 | 0.2 | 2 | 0 | 5 | 0 | 3 | 0.3333 | 15 | 0.1333 | 3 | 0.3333 | 1 | 0 | 3 | 0.3333 | 2 | 0.5 | 9 | 0.3333 |
| 3_3117266 | 6 | 0.1667 | 2 | 0 | 4 | 0.25 | 3 | 0.3333 | 15 | 0.2 | 3 | 0 | 1 | 0 | 3 | 0 | 1 | 1 | 8 | 0.125 |
| 3_3117275 | 5 | 0.8 | 1 | 1 | 5 | 0.6 | 3 | 0.6667 | 14 | 0.7143 | 3 | 1 | 1 | 0 | 3 | 1 | 2 | 0.5 | 9 | 0.7778 |
| 3_3117316 | 5 | 0.4 | 2 | 0.5 | 3 | 0 | 3 | 0.3333 | 13 | 0.3077 | 4 | 0.5 | 1 | 0 | 3 | 0 | 2 | 0.5 | 10 | 0.3 |
| 3_3117341 | 6 | 0.1667 | 2 | 0 | 5 | 0 | 4 | 0 | 17 | 0.0588 | 4 | 0.25 | 1 | 0 | 2 | 0 | 3 | 0.3333 | 10 | 0.2 |
| 3_3117361 | 4 | 0.25 | 3 | 0.6667 | 3 | 0 | 2 | 0.5 | 12 | 0.3333 | 4 | 0.75 | 1 | 0 | 3 | 0.6667 | 3 | 0.3333 | 11 | 0.5455 |
| 3_3117366 | 4 | 0.25 | 3 | 0.3333 | 5 | 0.4 | 2 | 0 | 14 | 0.2857 | 4 | 0 | 1 | 0 | 3 | 0.3333 | 4 | 0 | 12 | 0.0833 |
| 3_3117450 | 6 | 0.3333 | 3 | 0.6667 | 5 | 0.2 | 7 | 0 | 21 | 0.2381 | 4 | 0.5 | 2 | 0 | 7 | 0.4286 | 8 | 0.25 | 21 | 0.3333 |
| 3_3117501 | 8 | 0.25 | 3 | 0 | 5 | 0.4 | 8 | 0.375 | 24 | 0.2917 | 4 | 0.25 | 2 | 0 | 9 | 0.1111 | 8 | 0.125 | 23 | 0.1304 |
| 3_3117551 | 6 | 0 | 5 | 0.4 | 6 | 0.5 | 7 | 0.1429 | 24 | 0.25 | 5 | 0.4 | 3 | 0 | 9 | 0.2222 | 8 | 0.125 | 25 | 0.2 |
| 3_3117567 | 8 | 0 | 5 | 0 | 7 | 0.2857 | 8 | 0 | 28 | 0.0714 | 5 | 0 | 3 | 0 | 9 | 0.1111 | 8 | 0.125 | 25 | 0.08 |
| 3_3117579 | 6 | 0 | 6 | 0 | 5 | 0 | 7 | 0.2857 | 24 | 0.0833 | 4 | 0.25 | 4 | 0 | 9 | 0 | 7 | 0.1429 | 24 | 0.0833 |
| 3_3117589 | 8 | 0.375 | 5 | 0.6 | 7 | 0.4286 | 8 | 0 | 28 | 0.3214 | 5 | 0.2 | 5 | 0 | 10 | 0.4 | 6 | 0.3333 | 26 | 0.2692 |
| 3_3120443 | 19 | 0.0526 | 24 | 0.1667 | 20 | 0 | 21 | 0.1429 | 84 | 0.0952 | 20 | 0.3 | 8 | 0.25 | 12 | 0.1667 | 6 | 0.1667 | 46 | 0.2391 |
| 3_3121778 | 21 | 0.1429 | 32 | 0.1562 | 24 | 0.125 | 28 | 0.1429 | 105 | 0.1429 | 29 | 0.2069 | 14 | 0.1429 | 17 | 0.1176 | 8 | 0.125 | 68 | 0.1618 |
| 3_3121779 | 23 | 0.1739 | 31 | 0.1613 | 27 | 0.1111 | 26 | 0.1538 | 107 | 0.1495 | 29 | 0.2069 | 15 | 0 | 19 | 0.1053 | 6 | 0 | 69 | 0.1159 |
| 3_3121787 | 17 | 0.0588 | 33 | 0.1515 | 27 | 0.0741 | 27 | 0.037 | 104 | 0.0865 | 33 | 0.0909 | 17 | 0.0588 | 19 | 0.1053 | 8 | 0 | 77 | 0.0779 |
| 3_3125759 | 10 | 0.6 | 6 | 0.5 | 5 | 0.8 | 13 | 0.7692 | 34 | 0.6765 | 9 | 0.3333 | 8 | 0.625 | 6 | 0.3333 | 9 | 0.7778 | 32 | 0.5312 |
| 3_3126274 | NA | NA | NA | NA | NA | NA | 2 | 0.5 | 2 | 0.5 | NA | NA | 3 | 0.6667 | 1 | 1 | NA | NA | 4 | 0.75 |
| 3_3126298 | NA | NA | NA | NA | NA | NA | 2 | 0.5 | 2 | 0.5 | NA | NA | 2 | 1 | 1 | 0 | NA | NA | 3 | 0.6667 |
| 3_3126338 | NA | NA | NA | NA | NA | NA | 2 | 1 | 2 | 1 | NA | NA | 3 | 1 | 1 | 1 | NA | NA | 4 | 1 |
| 3_3127774 | NA | NA | NA | NA | NA | NA | 2 | 0.5 | 2 | 0.5 | NA | NA | 3 | 0.3333 | 1 | 1 | NA | NA | 4 | 0.5 |
| 3_3127799 | NA | NA | NA | NA | NA | NA | 2 | 1 | 2 | 1 | NA | NA | 3 | 0.6667 | 1 | 0 | NA | NA | 4 | 0.5 |
| 3_3225329 | NA | NA | NA | NA | NA | NA | 1 | 1 | 1 | 1 | 1 | 1 | NA | NA | NA | NA | NA | NA | 1 | 1 |
| 3_3225332 | NA | NA | NA | NA | NA | NA | 1 | 1 | 1 | 1 | 1 | 1 | NA | NA | NA | NA | NA | NA | 1 | 1 |
| 3_3238918 | 13 | 0.0769 | 14 | 0.0714 | 10 | 0 | 3 | 0 | 40 | 0.05 | 13 | 0.1538 | 9 | 0 | 11 | 0 | 5 | 0 | 38 | 0.0526 |
| 3_3239028 | 8 | 0 | 10 | 0 | 9 | 0.1111 | 3 | 0.3333 | 30 | 0.0667 | 12 | 0.1667 | 8 | 0 | 7 | 0.1429 | 4 | 0 | 31 | 0.0968 |
| 3_3239097 | 6 | 0 | 8 | 0.125 | 5 | 0.2 | 2 | 0 | 21 | 0.0952 | 11 | 0.3636 | 3 | 0 | 7 | 0.4286 | 2 | 0 | 23 | 0.3043 |
| 3_3239119 | 6 | 0.1667 | 7 | 0.2857 | 7 | 0 | 3 | 0 | 23 | 0.1304 | 11 | 0.2727 | 3 | 0 | 6 | 0.1667 | 4 | 0 | 24 | 0.1667 |
| 3_3239129 | 6 | 0.1667 | 8 | 0.125 | 7 | 0 | 3 | 0 | 24 | 0.0833 | 11 | 0.3636 | 5 | 0.4 | 7 | 0.1429 | 3 | 0.3333 | 26 | 0.3077 |
| 3_3239445 | 5 | 0 | 6 | 0.1667 | 5 | 0.4 | 5 | 0.4 | 21 | 0.2381 | 8 | 0.375 | 2 | 0.5 | 7 | 0.1429 | 2 | 0 | 19 | 0.2632 |
| 3_3239490 | 4 | 0.75 | 2 | 1 | 4 | 0.75 | 3 | 0.6667 | 13 | 0.7692 | 7 | 0.8571 | 4 | 0.5 | 5 | 0.8 | 1 | 1 | 17 | 0.7647 |
| 3_3239501 | 3 | 0 | 4 | 0.25 | 4 | 0 | 4 | 0 | 15 | 0.0667 | 7 | 0.2857 | 1 | 0 | 4 | 0 | 1 | 0 | 13 | 0.1538 |
| 3_3239503 | 5 | 0.2 | 6 | 0.5 | 4 | 0 | 2 | 0.5 | 17 | 0.2941 | 7 | 0.2857 | 3 | 0 | 6 | 0 | 1 | 0 | 17 | 0.1176 |
| 3_3239528 | 5 | 0.2 | 5 | 1 | 3 | 0.3333 | 5 | 0.4 | 18 | 0.5 | 6 | 0.5 | 3 | 0.3333 | 6 | 0.5 | 1 | 0 | 16 | 0.4375 |
| 3_3239565 | 2 | 0 | 3 | 0.3333 | 4 | 0 | 3 | 0 | 12 | 0.0833 | 4 | 0 | 3 | 0 | 5 | 0 | 1 | 1 | 13 | 0.0769 |
| 3_3239578 | 3 | 0 | 3 | 0 | 4 | 0 | 3 | 0.3333 | 13 | 0.0769 | 5 | 0 | 3 | 0 | 3 | 0.3333 | NA | NA | 11 | 0.0909 |
| 3_3241522 | 1 | 1 | 4 | 0 | 2 | 0.5 | 1 | 0 | 8 | 0.25 | 4 | 0.25 | 2 | 0.5 | 2 | 0 | 1 | 0 | 9 | 0.2222 |

| | | | | | | | | | | | | | | | | | | | | |
|---|---|---|---|---|---|---|---|---|---|---|---|---|---|---|---|---|---|---|---|---|
| 3_32419914 | 1 | 0 | NA | NA | 1 | 1 | 1 | 1 | 3 | 0.6667 | NA | NA | 1 | 1 | NA | NA | NA | NA | 1 | 1 |
| 3_32420122 | NA | NA | NA | NA | 2 | 1 | 1 | 1 | 3 | 1 | NA | NA | 1 | 1 | NA | NA | 1 | 1 | 2 | 1 |
| 3_32420161 | 2 | 0.5 | NA | NA | 2 | 1 | 1 | 0 | 5 | 0.6 | NA | NA | 2 | 0 | 1 | 1 | 1 | 1 | 4 | 0.5 |
| 3_32420293 | 2 | 0 | NA | NA | 2 | 0.5 | 1 | 0 | 5 | 0.2 | NA | NA | 2 | 0.5 | NA | NA | 1 | 0 | 3 | 0.3333 |
| 3_32420428 | 2 | 1 | NA | NA | 2 | 1 | 1 | 1 | 5 | 1 | NA | NA | 2 | 1 | 1 | 1 | 1 | 1 | 4 | 1 |
| 3_32420460 | 2 | 0.5 | NA | NA | 3 | 0.3333 | 1 | 1 | 6 | 0.5 | NA | NA | 2 | 0 | 1 | 1 | NA | NA | 3 | 0.3333 |
| 3_32420511 | 1 | 1 | NA | NA | 3 | 1 | 1 | 1 | 5 | 1 | NA | NA | 2 | 1 | NA | NA | 1 | 1 | 3 | 1 |
| 3_32420669 | 1 | 1 | NA | NA | 2 | 1 | 1 | 1 | 4 | 1 | NA | NA | 2 | 1 | NA | NA | 1 | 1 | 3 | 1 |
| 3_32420697 | 2 | 1 | NA | NA | 3 | 0.6667 | NA | NA | 5 | 0.8 | NA | NA | 2 | 0.5 | 1 | 0 | 1 | 0 | 4 | 0.25 |
| 3_32421180 | 2 | 1 | NA | NA | 3 | 0.6667 | 1 | 1 | 6 | 0.8333 | NA | NA | 3 | 0.3333 | 1 | 1 | 1 | 1 | 5 | 0.6 |
| 3_32490451 | 1 | 1 | NA | NA | 2 | 0 | 1 | 0 | 4 | 0.25 | 1 | 0 | 2 | 1 | NA | NA | 2 | 0 | 5 | 0.4 |
| 3_32490498 | 1 | 1 | 1 | 0 | 1 | 1 | 1 | 1 | 4 | 0.75 | NA | NA | 1 | 0 | NA | NA | 2 | 1 | 3 | 0.6667 |
| 3_32516809 | NA | NA | 5 | 0.4 | 4 | 0 | 2 | 0 | 11 | 0.1818 | 1 | 0 | 3 | 0.6667 | 2 | 0.5 | 3 | 0 | 9 | 0.3333 |
| 3_32516826 | 1 | 1 | 5 | 0 | 4 | 0.5 | 2 | 0.5 | 12 | 0.3333 | 2 | 0 | 3 | 0.3333 | 2 | 0.5 | 4 | 0.25 | 11 | 0.2727 |
| 3_32516846 | NA | NA | 6 | 1 | 5 | 1 | 3 | 1 | 14 | 1 | 1 | 1 | 3 | 1 | 2 | 0.5 | 4 | 1 | 10 | 0.9 |
| 3_32516859 | 1 | 1 | 5 | 0.6 | 3 | 0.3333 | 3 | 0.6667 | 12 | 0.5833 | 3 | 0.6667 | 2 | 1 | 2 | 0.5 | 4 | 0.75 | 11 | 0.7273 |
| 3_32516864 | 1 | 0 | 5 | 0 | 5 | 0.2 | 1 | 0 | 12 | 0.0833 | 2 | 0 | 2 | 0.5 | 1 | 0 | 4 | 0 | 9 | 0.1111 |
| 3_32516976 | NA | NA | 6 | 1 | 2 | 1 | 2 | 1 | 10 | 1 | 2 | 1 | 3 | 1 | 2 | 1 | 2 | 1 | 9 | 1 |
| 3_32516994 | 1 | 1 | 6 | 0.8333 | 5 | 0.6 | 2 | 0.5 | 14 | 0.7143 | 3 | 1 | 3 | 0.6667 | 1 | 1 | 4 | 0.75 | 11 | 0.8182 |
| 3_32517088 | 1 | 1 | 5 | 0.6 | 3 | 1 | 3 | 0.6667 | 12 | 0.75 | 3 | 0.6667 | 3 | 1 | 2 | 1 | 4 | 0.75 | 12 | 0.8333 |
| 3_32517100 | 1 | 0 | 5 | 0.2 | 2 | 0 | 2 | 0 | 10 | 0.1 | 3 | 0.6667 | 3 | 0 | 2 | 0.5 | 4 | 0 | 12 | 0.25 |
| 3_32517364 | 2 | 0 | 3 | 0 | 4 | 0 | 3 | 0.3333 | 12 | 0.0833 | 2 | 0 | 3 | 0 | 2 | 0 | 3 | 0.3333 | 10 | 0.1 |
| 3_32517535 | 2 | 0.5 | 5 | 0.6 | 4 | 0.75 | 3 | 0 | 14 | 0.5 | 3 | 0.3333 | 3 | 0.6667 | 3 | 0.3333 | 4 | 0.5 | 13 | 0.4615 |
| 3_32517572 | 2 | 1 | 4 | 1 | 4 | 0.5 | 4 | 1 | 14 | 0.8571 | 3 | 0.6667 | 3 | 0.6667 | 3 | 0.6667 | 4 | 1 | 13 | 0.7692 |
| 3_32517667 | 3 | 0 | 3 | 0.3333 | 2 | 0 | 2 | 0.5 | 10 | 0.2 | 1 | 0 | 1 | 1 | 1 | 0 | 4 | 0.75 | 7 | 0.5714 |
| 3_32518269 | NA | NA | 4 | 0 | 4 | 0.25 | 3 | 0.3333 | 11 | 0.1818 | 1 | 0 | 1 | 0 | 1 | 0 | 2 | 0.5 | 5 | 0.2 |
| 3_32518288 | 1 | 0 | 4 | 0 | 4 | 0 | 3 | 0.3333 | 12 | 0.0833 | 1 | 0 | NA | NA | 1 | 1 | 2 | 0 | 4 | 0.25 |
| 3_32518628 | 1 | 0 | 3 | 0 | 4 | 0 | 3 | 0.6667 | 11 | 0.1818 | NA | NA | 1 | 1 | NA | NA | 2 | 0 | 3 | 0.3333 |
| 3_32519183 | 1 | 1 | 3 | 0.6667 | 3 | 0.6667 | 3 | 0.3333 | 10 | 0.6 | NA | NA | 1 | 1 | 1 | 0 | 2 | 0 | 4 | 0.25 |
| 3_32573825 | 5 | 0.2 | 8 | 0.25 | 2 | 0.5 | 6 | 0.1667 | 21 | 0.2381 | 2 | 0 | 4 | 0 | 4 | 0 | 4 | 0.25 | 14 | 0.0714 |
| 3_32573846 | 6 | 0.1667 | 5 | 0.8 | 1 | 1 | 5 | 0.2 | 17 | 0.4118 | 2 | 0 | 4 | 0.25 | 3 | 0 | 3 | 0.3333 | 12 | 0.1667 |
| 3_32573898 | 7 | 0.1429 | 8 | 0 | 2 | 0 | 5 | 0.4 | 22 | 0.1364 | 2 | 0 | 4 | 0.25 | 4 | 0.25 | 3 | 0.3333 | 13 | 0.2308 |
| 3_32573938 | 5 | 1 | 6 | 0.6667 | NA | NA | 4 | 0.75 | 15 | 0.8 | 2 | 0.5 | 4 | 0.5 | 4 | 1 | 4 | 0.5 | 14 | 0.6429 |
| 3_32573997 | 6 | 1 | 4 | 1 | 2 | 1 | 3 | 1 | 15 | 1 | 1 | 0 | 3 | 1 | 4 | 0.75 | 3 | 1 | 11 | 0.8182 |
| 3_32574068 | 7 | 0.4286 | 8 | 0.5 | 2 | 0 | 5 | 0.2 | 22 | 0.3636 | 2 | 0.5 | 4 | 0.5 | 3 | 0 | 4 | 0 | 13 | 0.2308 |
| 3_32574181 | 7 | 1 | 6 | 0.8333 | 2 | 1 | 6 | 0.6667 | 21 | 0.8571 | 2 | 1 | 2 | 1 | 4 | 0.75 | 2 | 1 | 10 | 0.9 |
| 3_32574359 | 6 | 0.5 | 7 | 0.4286 | 2 | 0.5 | 4 | 0.25 | 19 | 0.4211 | 1 | 0 | 5 | 0.4 | 4 | 0 | 5 | 0.8 | 15 | 0.4 |
| 3_32574368 | 7 | 1 | 5 | 1 | 2 | 1 | 6 | 0.8333 | 20 | 0.95 | 1 | 1 | 5 | 1 | 4 | 0.75 | 4 | 1 | 14 | 0.9286 |
| 3_32574405 | 4 | 0.5 | 4 | 0.75 | 1 | 1 | 4 | 0.25 | 13 | 0.5385 | 2 | 1 | 4 | 0.5 | 3 | 0.3333 | 5 | 0.2 | 14 | 0.4286 |
| 3_32574422 | 7 | 0 | 7 | 0.1429 | 2 | 0 | 5 | 0.2 | 21 | 0.0952 | 2 | 0 | 5 | 0.2 | 3 | 0 | 4 | 0 | 14 | 0.0714 |
| 3_32574528 | 6 | 0.5 | 8 | 0.5 | 2 | 0.5 | 5 | 0 | 21 | 0.381 | 2 | 0.5 | 5 | 0.6 | 3 | 0.3333 | 6 | 0.3333 | 16 | 0.4375 |
| 3_32574549 | 7 | 0.7143 | 7 | 0.7143 | 2 | 0 | 6 | 0.6667 | 22 | 0.6364 | 2 | 0 | 4 | 0.5 | 4 | 0.5 | 5 | 0.2 | 15 | 0.3333 |
| 3_32574596 | 6 | 1 | 7 | 1 | 1 | 1 | 6 | 0.8333 | 20 | 0.95 | 2 | 1 | 5 | 0.8 | 3 | 1 | 4 | 0.75 | 14 | 0.8571 |
| 3_32574699 | 7 | 0.7143 | 7 | 0.4286 | 2 | 1 | 6 | 0.8333 | 22 | 0.6818 | 2 | 1 | 5 | 0.6 | 3 | 1 | 7 | 0.8571 | 17 | 0.8235 |
| 3_32574746 | 7 | 0.2857 | 7 | 0.1429 | 4 | 0.75 | 5 | 0 | 23 | 0.2609 | 2 | 0 | 5 | 0.4 | 4 | 0.25 | 7 | 0.2857 | 18 | 0.2778 |
| 3_32574797 | 6 | 1 | 8 | 1 | 3 | 1 | 5 | 0.8 | 22 | 0.9545 | 2 | 1 | 5 | 0.8 | 3 | 1 | 6 | 1 | 16 | 0.9375 |
| 3_32616269 | NA | NA | 1 | 0 | 2 | 0.5 | NA | NA | 3 | 0.3333 | NA | NA | NA | NA | NA | NA | 1 | 1 | 1 | 1 |
| 3_32631318 | 5 | 0.2 | 12 | 0.0833 | 7 | 0.2857 | 4 | 0 | 28 | 0.1429 | 7 | 0.1429 | 6 | 0.1667 | 7 | 0.1429 | 6 | 0.1667 | 26 | 0.1538 |
| 3_32631371 | 3 | 0.3333 | 12 | 0.0833 | 6 | 0.1667 | 4 | 0 | 25 | 0.12 | 7 | 0.1429 | 7 | 0.2857 | 5 | 0 | 4 | 0 | 23 | 0.1304 |
| 3_32631479 | 5 | 0 | 9 | 0.2222 | 7 | 0.4286 | 3 | 0.3333 | 24 | 0.25 | 6 | 0.1667 | 5 | 0.2 | 7 | 0.4286 | 6 | 0.1667 | 24 | 0.25 |
| 3_32633025 | NA | NA | 1 | 1 | NA | NA | NA | NA | 1 | 1 | NA | NA | NA | NA | NA | NA | 1 | 1 | 1 | 1 |
| 3_32633580 | NA | NA | 1 | 0 | NA | NA | 1 | 1 | 2 | 0.5 | 1 | 1 | NA | NA | NA | NA | NA | NA | 1 | 1 |
| 3_32637465 | 1 | 0 | 1 | 0 | 1 | 1 | 2 | 0 | 5 | 0.2 | NA | NA | 2 | 0.5 | NA | NA | NA | NA | 2 | 0.5 |
| 3_32637805 | 2 | 0 | 2 | 0.5 | 1 | 0 | 4 | 0 | 9 | 0.1111 | 2 | 0.5 | 5 | 0.2 | NA | NA | NA | NA | 7 | 0.2857 |
| 3_32638482 | 5 | 0.2 | 4 | 0.25 | 5 | 0.4 | 8 | 0 | 22 | 0.1818 | 4 | 0 | 6 | 0.1667 | 5 | 0 | 5 | 0.4 | 20 | 0.15 |
| 3_32638784 | 5 | 0.2 | 3 | 0.6667 | 6 | 0.5 | 8 | 0.25 | 22 | 0.3636 | 3 | 0.3333 | 4 | 0.25 | 5 | 0.2 | 5 | 0.2 | 17 | 0.2353 |
| 3_32638903 | 2 | 0 | 2 | 0 | 5 | 0.2 | 6 | 0.3333 | 15 | 0.2 | 2 | 0 | 4 | 0 | 6 | 0 | 5 | 0.8 | 17 | 0.2353 |
| 3_32638914 | 4 | 0 | 3 | 0.6667 | 5 | 0.2 | 8 | 0.625 | 20 | 0.4 | 3 | 1 | 5 | 0.2 | 6 | 0 | 5 | 0 | 19 | 0.2105 |
| 3_32639101 | 4 | 0 | 3 | 0.3333 | 5 | 0.6 | 8 | 0.25 | 20 | 0.3 | 3 | 0 | 5 | 0 | 5 | 0.4 | 3 | 0 | 16 | 0.125 |
| 3_32639145 | 4 | 0.25 | 3 | 0.3333 | 4 | 0 | 6 | 0.1667 | 17 | 0.1765 | 3 | 0.3333 | 5 | 0.2 | 5 | 0 | 2 | 0 | 15 | 0.1333 |
| 3_32639294 | 4 | 0.75 | 2 | 0.5 | 3 | 1 | 5 | 1 | 14 | 0.8571 | 3 | 1 | 5 | 1 | 6 | 0.8333 | 3 | 0.6667 | 17 | 0.8824 |
| 3_32639331 | 4 | 0 | 2 | 0 | 3 | 0.3333 | 5 | 0.2 | 14 | 0.1429 | 3 | 0.6667 | 5 | 0 | 6 | 0.1667 | 3 | 0 | 17 | 0.1765 |
| 3_32780224 | NA | NA | 3 | 0.3333 | NA | NA | NA | NA | 3 | 0.3333 | NA | NA | 1 | 0 | 2 | 0.5 | 2 | 0 | 5 | 0.2 |
| 3_32780308 | NA | NA | 3 | 0.6667 | 1 | 1 | NA | NA | 4 | 0.75 | NA | NA | NA | NA | 2 | 0 | 2 | 0.5 | 4 | 0.25 |
| 3_32780515 | NA | NA | 3 | 0.3333 | 1 | 0 | NA | NA | 4 | 0.25 | NA | NA | 1 | 1 | 2 | 0 | 2 | 0.5 | 5 | 0.4 |

| | | | | | | | | | | | | | | | | | | | | |
|---|---|---|---|---|---|---|---|---|---|---|---|---|---|---|---|---|---|---|---|---|
| 3_3280532 | 39 | 0.0769 | 37 | 0.027 | 28 | 0.1429 | 33 | 0.0909 | 137 | 0.0803 | 28 | 0.0357 | 28 | 0.1071 | 48 | 0.1042 | 35 | 0.1143 | 139 | 0.0935 |
| 3_3280551 | 10 | 0.1 | 7 | 0 | 4 | 0 | 9 | 0.1111 | 30 | 0.0667 | 10 | 0.1 | 6 | 0 | 11 | 0.1818 | 14 | 0 | 41 | 0.0732 |
| 3_3280562 | 10 | 0.1 | 7 | 0.2857 | 4 | 0.25 | 7 | 0.1429 | 28 | 0.1786 | 12 | 0.25 | 5 | 0.2 | 10 | 0.1 | 12 | 0.1667 | 39 | 0.1795 |
| 3_3298749 | NA | NA | NA | NA | 1 | 1 | NA | NA | 1 | 1 | 2 | 1 | NA | NA | NA | NA | 1 | 0 | 3 | 0.6667 |
| 3_3298750 | NA | NA | NA | NA | 1 | 1 | NA | NA | 1 | 1 | 2 | 1 | NA | NA | NA | NA | 1 | 1 | 3 | 1 |
| 3_3298775 | NA | NA | NA | NA | 2 | 1 | NA | NA | 2 | 1 | 2 | 1 | NA | NA | NA | NA | 1 | 1 | 3 | 1 |
| 3_3299832 | 1 | 1 | 6 | 0 | 8 | 0.25 | 1 | 0 | 16 | 0.1875 | 3 | 0.3333 | NA | NA | 1 | 0 | 6 | 0 | 10 | 0.1 |
| 3_3300433 | 6 | 0 | 9 | 0.1111 | 5 | 0.4 | 4 | 0 | 24 | 0.125 | 6 | 0 | 4 | 0 | 6 | 0.3333 | 2 | 0.5 | 18 | 0.1667 |
| 3_3300440 | 6 | 0 | 7 | 0.2857 | 5 | 0.4 | 5 | 0 | 23 | 0.1739 | 5 | 0.2 | 6 | 0.3333 | 5 | 0.6 | 2 | 1 | 18 | 0.4444 |
| 3_3300443 | 7 | 0.1429 | 7 | 0.1429 | 4 | 0.5 | 4 | 0.25 | 22 | 0.2273 | 5 | 0 | 5 | 0.6 | 4 | 0.25 | 2 | 0.5 | 16 | 0.3125 |
| 3_3300446 | 8 | 0.375 | 8 | 0.625 | 5 | 0.4 | 5 | 0.4 | 26 | 0.4615 | 5 | 0.6 | 6 | 0.3333 | 6 | 0.5 | 2 | 0.5 | 19 | 0.4737 |
| 3_3300448 | 7 | 0.1429 | 6 | 0 | 5 | 0.6 | 4 | 0.5 | 22 | 0.2727 | 6 | 0.6667 | 5 | 0.2 | 6 | 0.5 | 2 | 0.5 | 19 | 0.4737 |
| 3_3300457 | 8 | 0.125 | 7 | 0 | 5 | 0.2 | 4 | 0 | 24 | 0.0833 | 5 | 0 | 6 | 0.1667 | 5 | 0.2 | 2 | 1 | 18 | 0.2222 |
| 3_3300458 | 5 | 0.6 | 4 | 0.5 | 4 | 0.5 | 4 | 0.75 | 17 | 0.5882 | 6 | 0.1667 | 5 | 0.2 | 3 | 0.3333 | 2 | 0.5 | 16 | 0.25 |
| 3_3300474 | 6 | 0.5 | 6 | 0.3333 | 3 | 0 | 4 | 0.5 | 19 | 0.3684 | 3 | 0.3333 | 5 | 0.2 | 2 | 0 | NA | NA | 10 | 0.2 |
| 3_3300475 | 7 | 0.5714 | 8 | 0.625 | 4 | 0.25 | 3 | 0.3333 | 22 | 0.5 | 6 | 0.5 | 6 | 0.6667 | 5 | 0.6 | 2 | 0.5 | 19 | 0.5789 |
| 3_3300489 | 7 | 0.2857 | 10 | 0.1 | 4 | 0.5 | 4 | 0.5 | 25 | 0.28 | 4 | 0 | 5 | 0.2 | 5 | 0.2 | 2 | 0.5 | 16 | 0.1875 |
| 3_3386247 | 4 | 0 | NA | NA | 3 | 0.3333 | 1 | 1 | 8 | 0.25 | NA | NA | 4 | 0.25 | 1 | 0 | NA | NA | 5 | 0.2 |
| 3_3386270 | 4 | 0 | NA | NA | 4 | 0.25 | 1 | 1 | 9 | 0.2222 | NA | NA | 5 | 0.2 | 2 | 1 | NA | NA | 7 | 0.4286 |
| 3_3386328 | 3 | 0.6667 | 1 | 1 | 3 | 1 | 3 | 1 | 10 | 0.9 | NA | NA | 3 | 1 | 2 | 0.5 | NA | NA | 5 | 0.8 |
| 3_3386333 | 5 | 0.6 | 1 | 0 | 5 | 0.6 | 4 | 0.5 | 15 | 0.5333 | NA | NA | 6 | 0.6667 | 2 | 1 | NA | NA | 8 | 0.75 |
| 3_3386337 | 5 | 0.8 | 1 | 1 | 4 | 1 | 4 | 1 | 14 | 0.9286 | NA | NA | 6 | 0.6667 | 2 | 0.5 | NA | NA | 8 | 0.625 |
| 3_3386344 | 5 | 0.6 | 1 | 1 | 6 | 0.6667 | 3 | 1 | 15 | 0.7333 | NA | NA | 5 | 1 | 2 | 0 | NA | NA | 7 | 0.7143 |
| 3_3386347 | 4 | 0 | 1 | 0 | 5 | 0.2 | 2 | 0 | 12 | 0.0833 | NA | NA | 5 | 0.4 | 2 | 0 | NA | NA | 7 | 0.2857 |
| 3_3386357 | 4 | 0.75 | 1 | 1 | 4 | 1 | 4 | 1 | 13 | 0.9231 | NA | NA | 5 | 1 | 2 | 1 | NA | NA | 7 | 1 |
| 3_3386391 | 4 | 0 | 1 | 1 | 6 | 0.6667 | 2 | 0.5 | 13 | 0.4615 | NA | NA | 6 | 0.6667 | 2 | 0 | 2 | 0 | 10 | 0.4 |
| 3_3386466 | 1 | 0 | 2 | 0.5 | 4 | 0 | 1 | 0 | 8 | 0.125 | NA | NA | 1 | 1 | 1 | 0 | 2 | 0 | 4 | 0.25 |
| 3_3386489 | 2 | 0.5 | 2 | 1 | 4 | 0.5 | 2 | 1 | 10 | 0.7 | NA | NA | 1 | 1 | 1 | 1 | 2 | 0.5 | 4 | 0.75 |
| 3_3387555 | 1 | 1 | NA | NA | NA | NA | NA | NA | 1 | 1 | NA | NA | 1 | 1 | NA | NA | 1 | 0 | 2 | 0.5 |
| 3_3387558 | 1 | 1 | NA | NA | NA | NA | NA | NA | 1 | 1 | NA | NA | 1 | 1 | NA | NA | NA | NA | 1 | 1 |
| 3_3387574 | 1 | 1 | NA | NA | NA | NA | NA | NA | 1 | 1 | NA | NA | 1 | 1 | NA | NA | NA | NA | 1 | 1 |
| 3_3387588 | 1 | 1 | NA | NA | NA | NA | NA | NA | 1 | 1 | NA | NA | 1 | 1 | NA | NA | NA | NA | 1 | 1 |
| 3_3387598 | 1 | 1 | NA | NA | NA | NA | NA | NA | 1 | 1 | NA | NA | 1 | 1 | NA | NA | NA | NA | 1 | 1 |
| 3_3387601 | 1 | 1 | NA | NA | NA | NA | NA | NA | 1 | 1 | NA | NA | 1 | 1 | NA | NA | NA | NA | 1 | 1 |
| 3_3387606 | 1 | 1 | NA | NA | NA | NA | NA | NA | 1 | 1 | NA | NA | 1 | 1 | NA | NA | NA | NA | 1 | 1 |
| 3_3387671 | 1 | 1 | NA | NA | NA | NA | NA | NA | 1 | 1 | NA | NA | 1 | 1 | NA | NA | NA | NA | 1 | 1 |
| 3_3387734 | 1 | 1 | NA | NA | NA | NA | NA | NA | 1 | 1 | NA | NA | 1 | 1 | NA | NA | NA | NA | 1 | 1 |
| 3_3411198 | NA | NA | NA | NA | 1 | 1 | NA | NA | 1 | 1 | NA | NA | 2 | 0.5 | NA | NA | NA | NA | 2 | 0.5 |
| 3_3411842 | 5 | 0 | 7 | 0.1429 | 5 | 0.2 | 7 | 0.2857 | 24 | 0.1667 | 4 | 0.25 | 3 | 0 | 1 | 0 | 4 | 0 | 12 | 0.0833 |
| 3_3412317 | 6 | 0.3333 | 8 | 0.125 | 3 | 0 | 8 | 0.125 | 25 | 0.16 | 2 | 0 | 4 | 0 | 1 | 0 | 7 | 0.2857 | 14 | 0.1429 |
| 3_3412380 | 3 | 0.6667 | 9 | 0.1111 | 4 | 0 | 6 | 0.1667 | 22 | 0.1818 | 1 | 0 | 6 | 0 | 3 | 0 | 5 | 0.2 | 15 | 0.0667 |
| 3_3413218 | NA | NA | 1 | 1 | 2 | 1 | NA | NA | 3 | 1 | 1 | 1 | NA | NA | NA | NA | NA | NA | 1 | 1 |
| 3_3469258 | NA | NA | 3 | 0 | 2 | 0 | 2 | 0.5 | 7 | 0.1429 | 1 | 0 | 2 | 1 | 2 | 0 | 1 | 1 | 6 | 0.5 |
| 3_3470481 | 3 | 0.3333 | 4 | 0 | 9 | 0.2222 | NA | NA | 16 | 0.1875 | 3 | 0 | 2 | 0.5 | 3 | 0 | 4 | 0.5 | 12 | 0.25 |
| 3_3470498 | 3 | 0.6667 | 5 | 0.4 | 7 | 0.4286 | 2 | 0.5 | 17 | 0.4706 | 2 | 0 | 1 | 1 | 3 | 1 | 3 | 0.3333 | 9 | 0.5556 |
| 3_3470503 | 3 | 0 | 7 | 0.2857 | 8 | 0 | 2 | 0 | 20 | 0.1 | 3 | 0 | 2 | 0 | 3 | 0 | 3 | 0.3333 | 11 | 0.0909 |
| 3_3470504 | 3 | 1 | 7 | 1 | 7 | 1 | 1 | 1 | 18 | 1 | 3 | 1 | 2 | 1 | 3 | 1 | 4 | 1 | 12 | 1 |
| 3_3470521 | 4 | 0.25 | 5 | 0.2 | 8 | 0.125 | 2 | 0 | 19 | 0.1579 | 3 | 0.3333 | NA | NA | 2 | 0 | 4 | 0 | 9 | 0.1111 |
| 3_3470537 | 4 | 0.25 | 7 | 0.4286 | 9 | 0.5556 | 2 | 0 | 22 | 0.4091 | 4 | 0.5 | 1 | 1 | 3 | 0 | 4 | 0.25 | 12 | 0.3333 |
| 3_3470538 | 3 | 0.6667 | 6 | 0.6667 | 8 | 0.75 | 2 | 0.5 | 19 | 0.6842 | 4 | 0.75 | 1 | 1 | 3 | 1 | 5 | 0.8 | 13 | 0.8462 |
| 3_3470544 | 3 | 1 | 5 | 1 | 8 | 1 | 3 | 1 | 19 | 1 | 4 | 1 | 2 | 1 | 3 | 1 | 3 | 1 | 12 | 1 |
| 3_3470553 | 4 | 1 | 7 | 1 | 6 | 1 | 2 | 1 | 19 | 1 | 3 | 1 | 2 | 1 | 3 | 1 | 4 | 1 | 12 | 1 |
| 3_3470553 | 2 | 1 | 5 | 1 | 7 | 0.8571 | 2 | 0.5 | 16 | 0.875 | 4 | 1 | 3 | 1 | 3 | 1 | 4 | 1 | 14 | 1 |
| 3_3470572 | 5 | 0.2 | 5 | 1 | 11 | 0.2727 | 2 | 0 | 23 | 0.3913 | 5 | 0.4 | 3 | 0.3333 | 5 | 0 | 5 | 0.4 | 18 | 0.2778 |
| 3_3470574 | 5 | 1 | 7 | 0.8571 | 11 | 1 | 3 | 1 | 26 | 0.9615 | 5 | 1 | 2 | 1 | 5 | 1 | 4 | 1 | 16 | 1 |
| 3_3470575 | 5 | 0.6 | 5 | 0.4 | 9 | 0.4444 | 3 | 0.3333 | 22 | 0.4545 | 4 | 0 | 3 | 0.6667 | 5 | 0.6 | 4 | 1 | 16 | 0.5625 |
| 3_3470581 | 4 | 0 | 5 | 0 | 10 | 0.2 | 3 | 0 | 22 | 0.0909 | 5 | 0.4 | 2 | 0 | 5 | 0 | 5 | 0.2 | 17 | 0.1765 |
| 3_3470613 | 4 | 0 | 6 | 0.1667 | 10 | 0.1 | 2 | 0 | 22 | 0.0909 | 5 | 0.6 | 2 | 0 | 6 | 0.1667 | 3 | 0.3333 | 16 | 0.3125 |
| 3_3470651 | 4 | 0.5 | 8 | 0.125 | 11 | 0.0909 | 3 | 0.3333 | 26 | 0.1923 | 7 | 0 | 4 | 0 | 6 | 0.3333 | 5 | 0.2 | 22 | 0.1364 |
| 3_3470660 | 3 | 0.3333 | 8 | 0.625 | 9 | 0.1111 | 3 | 0.3333 | 23 | 0.3478 | 7 | 0.2857 | 2 | 0.5 | 2 | 0 | 4 | 0.25 | 15 | 0.2667 |
| 3_3473000 | 2 | 0 | 3 | 0 | 2 | 0.5 | 2 | 0.5 | 9 | 0.2222 | 1 | 0 | 1 | 1 | 2 | 0.5 | 1 | 0 | 5 | 0.4 |
| 3_3473039 | 3 | 0.3333 | 3 | 0.3333 | 2 | 0.5 | 1 | 0 | 9 | 0.3333 | 1 | 0 | 2 | 0 | 2 | 0.5 | 1 | 1 | 6 | 0.3333 |
| 3_3473165 | 2 | 0.5 | 4 | 0 | 1 | 0 | 2 | 0.5 | 9 | 0.2222 | 1 | 0 | 2 | 0 | 3 | 0 | 1 | 1 | 7 | 0.1429 |
| 3_3473174 | 2 | 0 | 3 | 0 | 2 | 0 | 2 | 0.5 | 9 | 0.1111 | 1 | 0 | 2 | 0 | 3 | 0 | 1 | 1 | 7 | 0.1429 |
| 3_3587417 | NA | NA | 6 | 0.1667 | 2 | 0.5 | NA | NA | 8 | 0.25 | 2 | 0 | 1 | 0 | 4 | 0.5 | 4 | 0 | 11 | 0.1818 |

| | | | | | | | | | | | | | | | | | | | | |
|---|---|---|---|---|---|---|---|---|---|---|---|---|---|---|---|---|---|---|---|---|
| 3_35909319 | 1 | 0 | 7 | 0.2857 | 3 | 0.3333 | 2 | 0.5 | 13 | 0.3077 | 4 | 0.75 | 4 | 0.5 | 6 | 0 | 7 | 0.1429 | 21 | 0.2857 |
| 3_35909340 | 2 | 0 | 7 | 0 | 2 | 0.5 | 2 | 0 | 13 | 0.0769 | 3 | 0 | 3 | 0.3333 | 5 | 0 | 7 | 0.1429 | 18 | 0.1111 |
| 3_35909348 | 2 | 0.5 | 9 | 0 | 3 | 0 | 2 | 0.5 | 16 | 0.125 | 5 | 0.6 | 4 | 0.25 | 6 | 0.3333 | 7 | 0.2857 | 22 | 0.3636 |
| 3_35909741 | 2 | 0 | 8 | 0.125 | 3 | 0 | 2 | 0 | 15 | 0.0667 | 7 | 0 | 2 | 0.5 | 6 | 0 | 5 | 0 | 20 | 0.05 |
| 3_35944195 | NA | NA | 1 | 1 | NA | NA | NA | NA | 1 | 1 | NA | NA | NA | NA | 1 | 1 | NA | NA | 1 | 1 |
| 3_35946697 | NA | NA | NA | NA | NA | NA | 1 | 1 | 1 | 1 | 1 | 0 | NA | NA | 1 | 1 | NA | NA | 2 | 0.5 |
| 3_35947753 | 3 | 0 | 5 | 0.2 | 1 | 0 | 5 | 0 | 14 | 0.0714 | 2 | 0 | 4 | 0 | 5 | 0.2 | 4 | 0 | 15 | 0.0667 |
| 3_35947861 | 4 | 0 | 5 | 0 | 5 | 0 | 6 | 0.1667 | 20 | 0.05 | 3 | 0.3333 | 8 | 0.125 | 6 | 0 | 5 | 0 | 22 | 0.0909 |
| 3_35948010 | 3 | 0 | 5 | 0.2 | 7 | 0 | 7 | 0.1429 | 22 | 0.0909 | 2 | 0 | 7 | 0 | 5 | 0 | 6 | 0.3333 | 20 | 0.1 |
| 3_35948350 | 5 | 0.2 | 4 | 0 | 5 | 0.2 | 7 | 0 | 21 | 0.0952 | 1 | 0 | 9 | 0.3333 | 3 | 0 | 3 | 0.3333 | 16 | 0.25 |
| 3_35948456 | 5 | 0 | 5 | 0.2 | 7 | 0.1429 | 7 | 0.4286 | 24 | 0.2083 | 2 | 0.5 | 9 | 0.2222 | 4 | 0 | 4 | 0.5 | 19 | 0.2632 |
| 3_35948552 | 6 | 0 | 5 | 0.2 | 4 | 0 | 8 | 0.25 | 23 | 0.1304 | 2 | 0 | 9 | 0.1111 | 4 | 0 | 5 | 0 | 20 | 0.05 |
| 3_35948672 | 5 | 0.2 | 5 | 0 | 6 | 0 | 7 | 0.2857 | 23 | 0.1304 | 2 | 0 | 8 | 0.375 | 4 | 0.25 | 4 | 0.25 | 18 | 0.2778 |
| 3_35949047 | 4 | 0 | 4 | 0.25 | 5 | 0.4 | 4 | 0 | 17 | 0.1765 | 2 | 0 | 8 | 0.375 | 3 | 0.3333 | 5 | 0.2 | 18 | 0.2778 |
| 3_35949145 | 2 | 1 | 2 | 0 | 5 | 0.6 | 5 | 0.4 | 14 | 0.5 | 1 | 1 | 7 | 0.4286 | 2 | 1 | 4 | 0.25 | 14 | 0.5 |
| 3_35949327 | 2 | 1 | 4 | 1 | 4 | 1 | 7 | 1 | 17 | 1 | 1 | 1 | 8 | 1 | 3 | 1 | 5 | 1 | 17 | 1 |
| 3_35949362 | 2 | 0 | 2 | 0.5 | 2 | 0.5 | 7 | 0.4286 | 13 | 0.3846 | 2 | 0 | 5 | 0.8 | 2 | 1 | 2 | 0.5 | 11 | 0.6364 |
| 3_35949363 | NA | NA | 1 | 1 | 1 | 0 | NA | NA | 2 | 0.5 | NA | NA | 1 | 1 | 2 | 1 | 1 | 0 | 4 | 0.75 |
| 3_35949383 | 1 | 1 | 4 | 0.25 | 4 | 0.5 | 6 | 0.5 | 15 | 0.4667 | 2 | 0 | 7 | 0.5714 | 4 | 0.5 | 6 | 0.5 | 19 | 0.4737 |
| 3_35949416 | 3 | 0 | 5 | 0 | 4 | 0.25 | 6 | 0.1667 | 18 | 0.1111 | 3 | 0 | 6 | 0 | 5 | 0 | 6 | 0.3333 | 20 | 0.1 |
| 3_35949515 | 1 | 1 | 4 | 1 | 5 | 0.8 | 6 | 1 | 16 | 0.9375 | 2 | 1 | 7 | 0.8571 | 5 | 0.8 | 2 | 1 | 16 | 0.875 |
| 3_35949623 | 3 | 1 | 5 | 1 | 4 | 1 | 5 | 1 | 17 | 1 | 2 | 1 | 8 | 0.625 | 4 | 1 | 5 | 0.8 | 19 | 0.7895 |
| 3_35949644 | 2 | 0.5 | 2 | 0.5 | 4 | 0.75 | 4 | 0.25 | 12 | 0.5 | 2 | 0 | 7 | 0.2857 | NA | NA | 3 | 1 | 12 | 0.4167 |
| 3_35949649 | 3 | 0.3333 | 3 | 0.3333 | 2 | 0 | 6 | 0.3333 | 14 | 0.2857 | 3 | 0 | 7 | 0.1429 | 4 | 1 | 1 | 0 | 15 | 0.3333 |
| 3_35949662 | 3 | 0.3333 | 4 | 0.25 | 4 | 0.5 | 5 | 0 | 16 | 0.25 | 3 | 0.3333 | 7 | 0 | 3 | 0.3333 | 5 | 0.2 | 18 | 0.1667 |
| 3_35949688 | 3 | 1 | 3 | 1 | 3 | 1 | 5 | 1 | 14 | 1 | 3 | 0.6667 | 6 | 1 | 3 | 1 | 2 | 1 | 14 | 0.9286 |
| 3_35949758 | 2 | 0 | 1 | 0 | 3 | 0.3333 | 5 | 0.6 | 11 | 0.3636 | 3 | 0.3333 | 4 | 0.25 | 3 | 0 | 3 | 0.6667 | 13 | 0.3077 |
| 3_35949789 | 3 | 0.3333 | 4 | 0.25 | 4 | 0.25 | 7 | 0.7143 | 18 | 0.4444 | 3 | 0 | 6 | 0.3333 | 3 | 0.3333 | 5 | 0.8 | 17 | 0.4118 |
| 3_35949824 | 3 | 0 | 4 | 0 | 4 | 0 | 7 | 0.1429 | 18 | 0.0556 | 3 | 0 | 7 | 0 | 4 | 0.25 | 4 | 0 | 18 | 0.0556 |
| 3_35975193 | 3 | 0 | 4 | 0.25 | 4 | 0 | 5 | 0 | 16 | 0.0625 | 1 | 0 | 5 | 0.4 | 2 | 0.5 | 3 | 0 | 11 | 0.2727 |
| 3_35975239 | 3 | 0 | 4 | 0 | 4 | 0.5 | 4 | 0 | 15 | 0.1333 | 2 | 0 | 6 | 0.1667 | 3 | 0.3333 | 3 | 0 | 14 | 0.1429 |
| 3_36010215 | NA | NA | 2 | 0.5 | 1 | 0 | NA | NA | 3 | 0.3333 | NA | NA | 1 | 0 | 2 | 0.5 | NA | NA | 3 | 0.3333 |
| 3_36010267 | NA | NA | 2 | 0.5 | 1 | 0 | NA | NA | 3 | 0.3333 | NA | NA | 1 | 0 | 2 | 1 | NA | NA | 3 | 0.6667 |
| 3_36010346 | NA | NA | 2 | 0.5 | 1 | 0 | NA | NA | 3 | 0.3333 | NA | NA | 1 | 0 | 2 | 0.5 | NA | NA | 3 | 0.3333 |
| 3_36010594 | NA | NA | 2 | 0.5 | 1 | 1 | NA | NA | 3 | 0.6667 | NA | NA | 1 | 0 | 2 | 1 | NA | NA | 3 | 0.6667 |
| 3_36010875 | NA | NA | 2 | 0.5 | NA | NA | NA | NA | 2 | 0.5 | NA | NA | 1 | 0 | 2 | 0.5 | NA | NA | 3 | 0.3333 |
| 3_36013980 | NA | NA | 4 | 0.25 | NA | NA | NA | NA | 4 | 0.25 | NA | NA | NA | NA | 1 | 0 | 1 | 1 | 2 | 0.5 |
| 3_36014149 | 2 | 0 | 5 | 0.6 | 2 | 1 | 3 | 0.3333 | 12 | 0.5 | NA | NA | 4 | 0.75 | 3 | 0.3333 | NA | NA | 7 | 0.5714 |
| 3_36130891 | 4 | 0 | 6 | 0.1667 | 3 | 0 | 3 | 0 | 16 | 0.0625 | 3 | 0 | 1 | 0 | 2 | 0 | 4 | 0.25 | 10 | 0.1 |
| 3_36142710 | 4 | 0 | 8 | 0 | 2 | 0.5 | 4 | 0 | 18 | 0.0556 | 3 | 0 | 2 | 0.5 | 2 | 0.5 | 4 | 0 | 11 | 0.1818 |
| 3_36145153 | 4 | 0.75 | 8 | 1 | 4 | 1 | 4 | 1 | 20 | 0.95 | 6 | 0.8333 | 3 | 1 | 2 | 1 | 4 | 0.5 | 15 | 0.8 |
| 3_36145180 | 4 | 0.75 | 9 | 0.7778 | 4 | 0.25 | 4 | 0.5 | 21 | 0.619 | 6 | 0.3333 | 2 | 0 | 2 | 0.5 | 3 | 0.6667 | 13 | 0.3846 |
| 3_36145182 | NA | NA | NA | NA | NA | NA | 1 | 1 | 1 | 1 | NA | NA | 1 | 1 | NA | NA | NA | NA | 1 | 1 |
| 3_36145358 | 4 | 0.5 | 10 | 0.6 | 4 | 0.5 | 5 | 0.8 | 23 | 0.6087 | 7 | 0.8571 | 2 | 0.5 | 1 | 1 | 3 | 0.6667 | 13 | 0.7692 |
| 3_36145395 | 3 | 0 | 8 | 0.375 | 4 | 0 | 5 | 0.2 | 20 | 0.2 | 7 | 0.1429 | 2 | 0 | 2 | 0.5 | 4 | 0 | 15 | 0.1333 |
| 3_36145439 | 4 | 0 | 9 | 0.1111 | 4 | 0 | 5 | 0.2 | 22 | 0.0909 | 8 | 0 | 3 | 0 | 2 | 0.5 | 3 | 0 | 16 | 0.0625 |
| 3_36145474 | 4 | 0.5 | 8 | 0.875 | 4 | 0.5 | 6 | 1 | 22 | 0.7727 | 8 | 0.625 | 2 | 1 | 2 | 1 | 3 | 1 | 15 | 0.8 |
| 3_36145535 | 4 | 0.5 | 7 | 0.1429 | 3 | 0.3333 | 6 | 0.8333 | 20 | 0.45 | 6 | 0 | 2 | 0 | 2 | 0.5 | 4 | 0.25 | 14 | 0.1429 |
| 3_36218331 | 1 | 1 | NA | NA | NA | NA | NA | NA | 1 | 1 | 1 | 1 | NA | NA | NA | NA | 1 | 1 | 2 | 1 |
| 3_36218659 | 1 | 1 | NA | NA | NA | NA | NA | NA | 1 | 1 | 1 | 0 | 1 | 1 | NA | NA | 1 | 1 | 3 | 0.6667 |
| 3_36218677 | 1 | 1 | NA | NA | NA | NA | NA | NA | 1 | 1 | 1 | 0 | 1 | 1 | NA | NA | 1 | 1 | 3 | 0.6667 |
| 3_36218703 | 1 | 1 | NA | NA | NA | NA | NA | NA | 1 | 1 | NA | NA | 1 | 0 | NA | NA | 1 | 1 | 2 | 0.5 |
| 3_36218927 | 1 | 1 | NA | NA | NA | NA | NA | NA | 1 | 1 | 1 | 1 | NA | NA | NA | NA | 1 | 1 | 2 | 1 |
| 3_36219049 | 1 | 1 | NA | NA | NA | NA | NA | NA | 1 | 1 | 1 | 1 | 1 | 0 | NA | NA | 1 | 1 | 3 | 0.6667 |
| 3_36219261 | 1 | 1 | NA | NA | 1 | 1 | NA | NA | 2 | 1 | 1 | 1 | NA | NA | NA | NA | NA | NA | 1 | 1 |
| 3_36219513 | 1 | 1 | NA | NA | 1 | 0 | NA | NA | 2 | 0.5 | 1 | 1 | 1 | 1 | NA | NA | 1 | 0 | 3 | 0.6667 |
| 3_36219597 | 1 | 1 | NA | NA | 1 | 1 | NA | NA | 2 | 1 | 1 | 1 | 1 | 1 | NA | NA | 1 | 1 | 3 | 1 |
| 3_36219681 | 1 | 1 | NA | NA | 1 | 1 | NA | NA | 2 | 1 | 1 | 1 | 1 | 1 | NA | NA | 1 | 1 | 3 | 1 |
| 3_36219849 | NA | NA | NA | NA | 1 | 1 | NA | NA | 1 | 1 | 1 | 1 | 1 | 1 | NA | NA | 1 | 1 | 3 | 1 |
| 3_36220017 | 1 | 1 | NA | NA | 1 | 0 | NA | NA | 2 | 0.5 | 1 | 1 | 1 | 1 | NA | NA | 1 | 1 | 3 | 1 |
| 3_36220041 | 1 | 1 | NA | NA | 1 | 0 | NA | NA | 2 | 0.5 | 1 | 0 | 1 | 0 | NA | NA | 1 | 1 | 3 | 0.3333 |
| 3_36220164 | 1 | 1 | NA | NA | 1 | 0 | NA | NA | 2 | 0.5 | NA | NA | 1 | 1 | NA | NA | 1 | 1 | 2 | 1 |
| 3_36220185 | 1 | 1 | NA | NA | 1 | 1 | NA | NA | 2 | 1 | 1 | 1 | 1 | 1 | NA | NA | 1 | 1 | 3 | 1 |
| 3_36220596 | 1 | 1 | NA | NA | 1 | 1 | NA | NA | 2 | 1 | 1 | 1 | 1 | 0 | NA | NA | 1 | 1 | 3 | 0.6667 |
| 3_36221092 | 1 | 1 | NA | NA | 1 | 0 | NA | NA | 2 | 0.5 | NA | NA | 1 | 0 | NA | NA | 1 | 1 | 2 | 0.5 |

| | | | | | | | | | | | | | | | | | | | | |
|---|---|---|---|---|---|---|---|---|---|---|---|---|---|---|---|---|---|---|---|---|
| 3_3622115 | 1 | 1 | NA | NA | 1 | 0 | NA | NA | 2 | 0.5 | NA | NA | 1 | 1 | NA | NA | 1 | 1 | 2 | 1 |
| 3_3622137 | 1 | 1 | NA | NA | 3 | 0.3333 | NA | NA | 4 | 0.5 | NA | NA | 1 | 1 | NA | NA | 1 | 1 | 2 | 1 |
| 3_3622225 | 2 | 0 | NA | NA | 2 | 0.5 | NA | NA | 4 | 0.25 | NA | NA | 1 | 1 | NA | NA | 1 | 1 | 2 | 1 |
| 3_3622412 | 1 | 1 | NA | NA | 2 | 0 | NA | NA | 3 | 0.3333 | 1 | 0 | 4 | 0 | 1 | 1 | 2 | 0.5 | 8 | 0.25 |
| 3_3622416 | 1 | 1 | NA | NA | 2 | 0.5 | NA | NA | 3 | 0.6667 | 1 | 1 | 2 | 0 | 1 | 1 | 2 | 1 | 6 | 0.6667 |
| 3_3622422 | 1 | 1 | NA | NA | 3 | 0.3333 | NA | NA | 4 | 0.5 | 1 | 0 | 4 | 0.25 | NA | NA | 2 | 0.5 | 7 | 0.2857 |
| 3_3622423 | 1 | 0 | 1 | 0 | 3 | 0.3333 | NA | NA | 5 | 0.2 | 1 | 0 | 3 | 0.6667 | 1 | 1 | 2 | 0.5 | 7 | 0.5714 |
| 3_3657047 | 2 | 1 | 1 | 1 | 2 | 0.5 | 1 | 1 | 6 | 0.8333 | 6 | 0.8333 | 1 | 1 | NA | NA | 2 | 1 | 9 | 0.8889 |
| 3_3657049 | 2 | 1 | 4 | 1 | 2 | 1 | 2 | 1 | 10 | 1 | 5 | 1 | 1 | 1 | NA | NA | 4 | 1 | 10 | 1 |
| 3_3657064 | 2 | 1 | 5 | 0.8 | 2 | 0.5 | 2 | 1 | 11 | 0.8182 | 6 | 0.3333 | 1 | 0 | NA | NA | 3 | 0.6667 | 10 | 0.4 |
| 3_3657078 | NA | NA | 4 | 0.75 | 2 | 1 | 1 | 1 | 7 | 0.8571 | 4 | 1 | 1 | 1 | NA | NA | 3 | 1 | 8 | 1 |
| 3_3657089 | NA | NA | 3 | 0.3333 | 2 | 0 | 1 | 0 | 6 | 0.1667 | 5 | 0.2 | 1 | 1 | NA | NA | 3 | 0.6667 | 9 | 0.4444 |
| 3_3657091 | 1 | 1 | 4 | 1 | 2 | 1 | 2 | 1 | 9 | 1 | 4 | 1 | 1 | 1 | NA | NA | 3 | 1 | 8 | 1 |
| 3_3657099 | 1 | 1 | 2 | 0.5 | 2 | 0.5 | 2 | 0 | 7 | 0.4286 | 3 | 0.3333 | NA | NA | NA | NA | 1 | 0 | 4 | 0.25 |
| 3_3657106 | 1 | 1 | 5 | 0.4 | 2 | 0 | 2 | 1 | 10 | 0.5 | 4 | 0.5 | 1 | 1 | NA | NA | 4 | 0.5 | 9 | 0.5556 |
| 3_3657108 | 1 | 0 | 3 | 1 | 2 | 0.5 | 2 | 1 | 8 | 0.75 | 4 | 0.5 | 1 | 1 | NA | NA | 2 | 0.5 | 7 | 0.5714 |
| 3_3657116 | 1 | 0 | 3 | 0.6667 | 2 | 0.5 | 2 | 1 | 8 | 0.625 | 3 | 0 | 1 | 1 | NA | NA | 2 | 0 | 6 | 0.1667 |
| 3_3662761 | 3 | 0.3333 | 4 | 0 | 3 | 0 | 4 | 0 | 14 | 0.0714 | NA | NA | 4 | 0.25 | 4 | 0.5 | 5 | 0.4 | 13 | 0.3846 |
| 3_3667485 | 1 | 1 | 1 | 0 | NA | NA | 2 | 0.5 | 4 | 0.5 | NA | NA | 2 | 0.5 | 2 | 0 | NA | NA | 4 | 0.25 |
| 3_3667532 | 1 | 0 | 1 | 1 | NA | NA | 1 | 0 | 3 | 0.3333 | NA | NA | 3 | 0.3333 | 2 | 0 | NA | NA | 5 | 0.2 |
| 3_3667544 | 1 | 0 | 1 | 0 | NA | NA | 2 | 0.5 | 4 | 0.25 | NA | NA | 2 | 0 | 2 | 0.5 | NA | NA | 4 | 0.25 |
| 3_3672536 | 1 | 1 | NA | NA | NA | NA | 1 | 0 | 2 | 0.5 | NA | NA | 1 | 0 | 1 | 1 | NA | NA | 2 | 0.5 |
| 3_3672557 | NA | NA | NA | NA | NA | NA | 1 | 1 | 1 | 1 | NA | NA | 1 | 1 | NA | NA | NA | NA | 1 | 1 |
| 3_3709096 | 2 | 0 | 7 | 0.1429 | 6 | 0 | 3 | 0 | 18 | 0.0556 | 1 | 0 | 6 | 0.1667 | 8 | 0 | 9 | 0.1111 | 24 | 0.0833 |
| 3_3710648 | 4 | 0 | 7 | 0.1429 | 6 | 0.3333 | 4 | 0.75 | 21 | 0.2857 | 2 | 1 | 7 | 0.2857 | 7 | 0.4286 | 7 | 0.2857 | 23 | 0.3913 |
| 3_3710655 | 2 | 0.5 | 7 | 0.2857 | 10 | 0.4 | 5 | 0.4 | 24 | 0.375 | 3 | 0.3333 | 11 | 0.3636 | 8 | 0.125 | 8 | 0.5 | 30 | 0.3333 |
| 3_3710659 | 4 | 0.25 | 10 | 0.1 | 12 | 0.1667 | 5 | 0.2 | 31 | 0.1613 | 3 | 0.3333 | 14 | 0.1429 | 11 | 0.2727 | 10 | 0.2 | 38 | 0.2105 |
| 3_3710663 | 3 | 0 | 8 | 0.125 | 8 | 0.125 | 6 | 0.3333 | 25 | 0.16 | 2 | 0 | 12 | 0.25 | 9 | 0.2222 | 10 | 0.1 | 33 | 0.1818 |
| 3_3710667 | 4 | 0 | 10 | 0.1 | 9 | 0.1111 | 7 | 0.1429 | 30 | 0.1 | 3 | 0.3333 | 14 | 0.0714 | 11 | 0.0909 | 9 | 0 | 37 | 0.0811 |
| 3_3710671 | 5 | 0.6 | 10 | 0.3 | 11 | 0.1818 | 7 | 0.1429 | 33 | 0.2727 | 3 | 0 | 15 | 0.2 | 10 | 0.1 | 9 | 0.3333 | 37 | 0.1892 |
| 3_3736296 | 4 | 0.25 | 1 | 0 | 1 | 0 | NA | NA | 6 | 0.1667 | 2 | 0.5 | 3 | 0 | 3 | 0 | 2 | 0 | 10 | 0.1 |
| 3_3736302 | 4 | 0.25 | 1 | 0 | 1 | 0 | NA | NA | 6 | 0.1667 | 1 | 0 | 3 | 0 | 4 | 0.25 | NA | NA | 8 | 0.125 |
| 3_3737528 | NA | NA | NA | NA | 1 | 1 | NA | NA | 1 | 1 | NA | NA | NA | NA | NA | NA | 1 | 1 | 1 | 1 |
| 3_3745925 | NA | NA | NA | NA | 2 | 1 | 1 | 0 | 3 | 0.6667 | 1 | 1 | NA | NA | 1 | 1 | NA | NA | 2 | 1 |
| 3_3745930 | NA | NA | NA | NA | 2 | 1 | 1 | 1 | 3 | 1 | 1 | 0 | NA | NA | 1 | 1 | NA | NA | 2 | 0.5 |
| 3_3745937 | NA | NA | NA | NA | 2 | 1 | NA | NA | 2 | 1 | 1 | 1 | NA | NA | NA | NA | NA | NA | 1 | 1 |
| 3_3745942 | NA | NA | NA | NA | 2 | 0.5 | 1 | 0 | 3 | 0.3333 | 1 | 1 | NA | NA | 1 | 1 | NA | NA | 2 | 1 |
| 3_3748593 | 1 | 0 | 2 | 0 | NA | NA | 1 | 1 | 4 | 0.25 | NA | NA | 1 | 0 | 2 | 0.5 | NA | NA | 3 | 0.3333 |
| 3_3748605 | 1 | 1 | 1 | 1 | NA | NA | NA | NA | 2 | 1 | NA | NA | NA | NA | 1 | 1 | NA | NA | 1 | 1 |
| 3_3748611 | 1 | 1 | 2 | 0.5 | NA | NA | 1 | 1 | 4 | 0.75 | NA | NA | 1 | 0 | 2 | 1 | NA | NA | 3 | 0.6667 |
| 3_3748631 | 2 | 0.5 | 1 | 0 | NA | NA | NA | NA | 3 | 0.3333 | NA | NA | 1 | 0 | 1 | 1 | NA | NA | 2 | 0.5 |
| 3_3769684 | 1 | 1 | 2 | 0 | 1 | 1 | NA | NA | 4 | 0.5 | NA | NA | 2 | 1 | 4 | 0.5 | 1 | 0 | 7 | 0.5714 |
| 3_3769685 | 1 | 1 | 2 | 0.5 | 1 | 0 | NA | NA | 4 | 0.5 | NA | NA | 2 | 0.5 | 4 | 0.75 | 1 | 1 | 7 | 0.7143 |
| 3_3769698 | 1 | 1 | 1 | 1 | 1 | 1 | NA | NA | 3 | 1 | NA | NA | 2 | 1 | 4 | 1 | 1 | 1 | 7 | 1 |
| 3_3769728 | 1 | 1 | 1 | 1 | 1 | 1 | NA | NA | 3 | 1 | NA | NA | NA | NA | 3 | 0.6667 | 1 | 1 | 4 | 0.75 |
| 3_3769739 | 1 | 1 | 2 | 1 | 1 | 1 | NA | NA | 4 | 1 | NA | NA | 1 | 1 | 4 | 1 | 1 | 1 | 6 | 1 |
| 3_3769760 | 1 | 1 | 2 | 0.5 | 1 | 1 | NA | NA | 4 | 0.75 | NA | NA | 2 | 0.5 | 4 | 0.75 | NA | NA | 6 | 0.6667 |
| 3_3769763 | 1 | 0 | 2 | 0 | 1 | 1 | NA | NA | 4 | 0.25 | NA | NA | 2 | 0.5 | 4 | 0.25 | 1 | 0 | 7 | 0.2857 |
| 3_3769769 | 1 | 1 | 1 | 1 | 1 | 1 | NA | NA | 3 | 1 | NA | NA | 1 | 1 | 3 | 1 | 1 | 1 | 5 | 1 |
| 3_3769770 | NA | NA | 2 | 0.5 | NA | NA | NA | NA | 2 | 0.5 | NA | NA | 2 | 0.5 | 2 | 1 | NA | NA | 4 | 0.75 |
| 3_3769774 | 1 | 1 | 2 | 1 | 1 | 1 | NA | NA | 4 | 1 | NA | NA | 2 | 0.5 | 4 | 0.5 | 1 | 1 | 7 | 0.5714 |
| 3_3769782 | 1 | 1 | 2 | 1 | 1 | 1 | NA | NA | 4 | 1 | NA | NA | 2 | 1 | 4 | 1 | 1 | 1 | 7 | 1 |
| 3_3769782 | 1 | 1 | 2 | 1 | 1 | 1 | NA | NA | 4 | 1 | NA | NA | 1 | 0 | 3 | 1 | 1 | 0 | 5 | 0.6 |
| 3_3769798 | 1 | 1 | 2 | 1 | 1 | 1 | NA | NA | 4 | 1 | NA | NA | 2 | 1 | 4 | 1 | 2 | 1 | 8 | 1 |
| 3_3769831 | 1 | 0 | 2 | 0.5 | 1 | 0 | NA | NA | 4 | 0.25 | NA | NA | 3 | 0.3333 | 4 | 0.25 | 2 | 1 | 9 | 0.4444 |
| 3_3769861 | 1 | 1 | 1 | 1 | 1 | 0 | NA | NA | 3 | 0.6667 | NA | NA | 3 | 0.3333 | 3 | 0 | 2 | 0.5 | 8 | 0.25 |
| 3_3850405 | 5 | 0 | 3 | 0 | 4 | 0 | 6 | 0.1667 | 18 | 0.0556 | 3 | 0 | 6 | 0 | 3 | 0.3333 | 4 | 0.25 | 16 | 0.125 |
| 3_3850494 | 4 | 0 | 4 | 0.75 | 1 | 0 | 2 | 0.5 | 11 | 0.3636 | 2 | 0 | 4 | 0 | 1 | 1 | 3 | 0.3333 | 10 | 0.2 |
| 3_3850505 | 4 | 0.25 | 4 | 0.25 | 1 | 0 | 4 | 0 | 13 | 0.1538 | 2 | 0 | 4 | 0 | 1 | 1 | 3 | 0 | 10 | 0.1 |
| 3_3850506 | 4 | 0 | 3 | 0.3333 | 1 | 0 | 5 | 0 | 13 | 0.0769 | 2 | 0 | 4 | 0.25 | 1 | 0 | 2 | 0 | 9 | 0.1111 |
| 3_3850555 | 4 | 0 | 4 | 0.25 | 1 | 0 | 2 | 0 | 11 | 0.0909 | 2 | 0 | 3 | 0.3333 | NA | NA | 1 | 0 | 6 | 0.1667 |
| 3_3850586 | 4 | 1 | 4 | 1 | 1 | 1 | 3 | 0.6667 | 12 | 0.9167 | 2 | 1 | 2 | 1 | NA | NA | 3 | 0.6667 | 7 | 0.8571 |
| 3_3850590 | 4 | 0 | 4 | 0.5 | 1 | 1 | 1 | 1 | 10 | 0.4 | 2 | 1 | 2 | 0 | NA | NA | 2 | 1 | 6 | 0.6667 |
| 3_3850591 | 3 | 0.6667 | 3 | 1 | 2 | 1 | 2 | 0.5 | 10 | 0.8 | 2 | 1 | 1 | 1 | NA | NA | 3 | 1 | 6 | 1 |
| 3_3850594 | 4 | 0 | 3 | 0.6667 | NA | NA | 3 | 0.3333 | 10 | 0.3 | 2 | 0.5 | 1 | 1 | NA | NA | 2 | 0.5 | 5 | 0.6 |

| | | | | | | | | | | | | | | | | | | | | |
|---|---|---|---|---|---|---|---|---|---|---|---|---|---|---|---|---|---|---|---|---|
| 3_38506069 | 2 | 0 | 2 | 0.5 | 1 | 0 | 2 | 0 | 7 | 0.1429 | 2 | 0.5 | 1 | 0 | NA | NA | 1 | 1 | 4 | 0.5 |
| 3_38506154 | 2 | 0 | 4 | 0.5 | 1 | 0 | 2 | 0 | 9 | 0.2222 | 2 | 0.5 | 1 | 0 | NA | NA | NA | NA | 3 | 0.3333 |
| 3_38506534 | 3 | 0 | 5 | 0.4 | 2 | 0 | 2 | 0 | 12 | 0.1667 | 2 | 0 | 2 | 0.5 | 1 | 1 | 2 | 0 | 7 | 0.2857 |
| 3_38506564 | 3 | 1 | 5 | 0.6 | 2 | 0.5 | 3 | 0.3333 | 13 | 0.6154 | 1 | 1 | 2 | 0.5 | 1 | 0 | 2 | 1 | 6 | 0.6667 |
| 3_38508392 | 3 | 0 | 2 | 0.5 | 2 | 0 | 3 | 0 | 10 | 0.1 | 2 | 0 | 1 | 0 | NA | NA | 2 | 0.5 | 5 | 0.2 |
| 3_38508553 | 2 | 1 | 4 | 0.5 | 2 | 0 | 3 | 0.3333 | 11 | 0.4545 | 1 | 1 | 1 | 1 | 1 | 1 | 1 | 1 | 4 | 1 |
| 3_38508573 | 2 | 1 | 4 | 1 | 1 | 1 | 3 | 1 | 10 | 1 | 1 | 0 | 1 | 0 | 1 | 0 | 2 | 0.5 | 5 | 0.2 |
| 3_38509536 | 1 | 1 | 3 | 0.3333 | 1 | 1 | 3 | 0.3333 | 8 | 0.5 | 1 | 1 | 1 | 1 | NA | NA | 1 | 0 | 3 | 0.6667 |
| 3_38509658 | 2 | 0.5 | 4 | 0.25 | 2 | 0 | 3 | 0 | 11 | 0.1818 | 1 | 0 | 1 | 1 | NA | NA | 1 | 1 | 3 | 0.6667 |
| 3_38509697 | 2 | 0.5 | 4 | 0.25 | 2 | 1 | 3 | 0.6667 | 11 | 0.5455 | 1 | 0 | 1 | 0 | NA | NA | 1 | 1 | 3 | 0.3333 |
| 3_38509796 | 2 | 0.5 | 5 | 1 | 2 | 0.5 | 3 | 0.6667 | 12 | 0.75 | 1 | 0 | 1 | 1 | NA | NA | 1 | 0 | 3 | 0.3333 |
| 3_38509847 | 2 | 0.5 | 4 | 0.75 | 1 | 0 | 3 | 0.6667 | 10 | 0.6 | NA | NA | 1 | 1 | NA | NA | 1 | 0 | 2 | 0.5 |
| 3_38509973 | 1 | 0 | 2 | 1 | 2 | 1 | NA | NA | 5 | 0.8 | NA | NA | 1 | 1 | NA | NA | NA | NA | 1 | 1 |
| 3_38510456 | 2 | 0 | 4 | 0.25 | 1 | 1 | 3 | 0 | 10 | 0.2 | NA | NA | 1 | 1 | NA | NA | NA | NA | 1 | 1 |
| 3_38510564 | 2 | 1 | 5 | 1 | 1 | 1 | 1 | 1 | 9 | 1 | NA | NA | 1 | 1 | NA | NA | NA | NA | 1 | 1 |
| 3_38510838 | NA | NA | 3 | 1 | 1 | 1 | 1 | 0 | 5 | 0.8 | NA | NA | 1 | 1 | NA | NA | NA | NA | 1 | 1 |
| 3_38510889 | 1 | 1 | 5 | 1 | 1 | 1 | 2 | 1 | 9 | 1 | NA | NA | 1 | 1 | NA | NA | NA | NA | 1 | 1 |
| 3_38511326 | 1 | 0 | 4 | 0.5 | 1 | 0 | 2 | 0 | 8 | 0.25 | NA | NA | 1 | 1 | NA | NA | NA | NA | 1 | 1 |
| 3_38511354 | 1 | 1 | 4 | 0.75 | NA | NA | 2 | 1 | 7 | 0.8571 | NA | NA | 1 | 1 | NA | NA | NA | NA | 1 | 1 |
| 3_38511378 | 1 | 0 | 4 | 0.25 | 1 | 0 | 2 | 0 | 8 | 0.125 | NA | NA | 1 | 1 | NA | NA | NA | NA | 1 | 1 |
| 3_38933838 | NA | NA | NA | NA | NA | NA | 1 | 1 | 1 | 1 | NA | NA | 1 | 1 | NA | NA | NA | NA | 1 | 1 |
| 3_38934104 | NA | NA | NA | NA | NA | NA | 1 | 1 | 1 | 1 | 1 | 0 | 1 | 1 | NA | NA | NA | NA | 2 | 0.5 |
| 3_39064211 | NA | NA | NA | NA | 1 | 1 | NA | NA | 1 | 1 | NA | NA | NA | NA | NA | NA | 1 | 1 | 1 | 1 |
| 3_39064409 | NA | NA | 1 | 0 | 1 | 1 | NA | NA | 2 | 0.5 | NA | NA | NA | NA | NA | NA | 1 | 1 | 1 | 1 |
| 3_39064837 | NA | NA | 1 | 0 | 1 | 1 | NA | NA | 2 | 0.5 | NA | NA | NA | NA | 1 | 0 | 1 | 1 | 2 | 0.5 |
| 3_39064849 | NA | NA | NA | NA | 1 | 1 | NA | NA | 1 | 1 | NA | NA | NA | NA | 1 | 1 | NA | NA | 1 | 1 |
| 3_39064979 | NA | NA | 1 | 1 | 1 | 1 | NA | NA | 2 | 1 | NA | NA | NA | NA | NA | NA | 1 | 1 | 1 | 1 |
| 3_39065076 | NA | NA | 1 | 0 | 1 | 1 | NA | NA | 2 | 0.5 | NA | NA | NA | NA | NA | NA | 1 | 1 | 1 | 1 |
| 3_39065090 | NA | NA | 1 | 1 | 1 | 1 | NA | NA | 2 | 1 | NA | NA | NA | NA | 1 | 1 | 1 | 1 | 2 | 1 |
| 3_39065126 | NA | NA | 1 | 0 | 1 | 1 | NA | NA | 2 | 0.5 | NA | NA | NA | NA | 1 | 1 | 1 | 1 | 2 | 1 |
| 3_39065288 | NA | NA | 1 | 0 | 1 | 1 | NA | NA | 2 | 0.5 | NA | NA | NA | NA | 1 | 1 | 1 | 1 | 2 | 1 |
| 3_39065404 | NA | NA | NA | NA | 1 | 1 | NA | NA | 1 | 1 | NA | NA | NA | NA | 1 | 0 | 1 | 1 | 2 | 0.5 |
| 3_39065713 | NA | NA | 1 | 1 | 1 | 1 | NA | NA | 2 | 1 | NA | NA | NA | NA | NA | NA | 1 | 1 | 1 | 1 |
| 3_40647113 | NA | NA | 1 | 1 | NA | NA | NA | NA | 1 | 1 | NA | NA | NA | NA | 1 | 0 | 1 | 1 | 2 | 0.5 |
| 3_40655662 | NA | NA | 1 | 0 | 1 | 1 | 1 | 1 | 3 | 0.6667 | NA | NA | NA | NA | 1 | 1 | 1 | 1 | 2 | 1 |
| 3_40655787 | NA | NA | 1 | 1 | 1 | 1 | 1 | 1 | 3 | 1 | NA | NA | NA | NA | 1 | 0 | 1 | 1 | 2 | 0.5 |
| 3_40655921 | NA | NA | 1 | 1 | 1 | 1 | 1 | 0 | 3 | 0.6667 | NA | NA | NA | NA | NA | NA | 1 | 1 | 1 | 1 |
| 3_40721284 | 3 | 0 | 2 | 0 | 5 | 0 | 9 | 0.1111 | 19 | 0.0526 | 4 | 0 | 6 | 0 | 5 | 0 | 5 | 0.2 | 20 | 0.05 |
| 3_40743711 | 5 | 0.4 | 6 | 0 | 7 | 0 | 13 | 0 | 31 | 0.0645 | 9 | 0.1111 | 6 | 0 | 4 | 0 | 6 | 0.3333 | 25 | 0.12 |
| 3_40743731 | 5 | 0.4 | 8 | 0.125 | 8 | 0.125 | 13 | 0.0769 | 34 | 0.1471 | 9 | 0.1111 | 8 | 0 | 6 | 0.1667 | 8 | 0.125 | 31 | 0.0968 |
| 3_40743743 | 5 | 0 | 8 | 0 | 9 | 0.3333 | 11 | 0.0909 | 33 | 0.1212 | 8 | 0 | 6 | 0.1667 | 7 | 0 | 8 | 0.125 | 29 | 0.069 |
| 3_40743820 | 5 | 0.8 | 8 | 0.75 | 8 | 0.875 | 13 | 0.5385 | 34 | 0.7059 | 9 | 0.5556 | 8 | 0.5 | 7 | 1 | 9 | 0.7778 | 33 | 0.697 |
| 3_40744063 | 5 | 0.6 | 11 | 0.1818 | 8 | 0.25 | 13 | 0.2308 | 37 | 0.2703 | 9 | 0 | 7 | 0.5714 | 8 | 0.25 | 8 | 0.375 | 32 | 0.2812 |
| 3_40744374 | 5 | 0.2 | 7 | 0 | 9 | 0.2222 | 14 | 0.1429 | 35 | 0.1429 | 9 | 0 | 4 | 0 | 8 | 0.25 | 7 | 0.2857 | 28 | 0.1429 |
| 3_40744452 | 4 | 0.5 | 8 | 0.25 | 10 | 0.3 | 16 | 0.25 | 38 | 0.2895 | 5 | 0.2 | 8 | 0.25 | 9 | 0.2222 | 8 | 0.375 | 30 | 0.2667 |
| 3_40744466 | 5 | 0 | 10 | 0.1 | 9 | 0.1111 | 16 | 0.0625 | 40 | 0.075 | 7 | 0 | 6 | 0 | 8 | 0.125 | 9 | 0.1111 | 30 | 0.0667 |
| 3_40744570 | 6 | 0 | 11 | 0.1818 | 10 | 0.1 | 15 | 0.0667 | 42 | 0.0952 | 9 | 0 | 9 | 0 | 9 | 0.2222 | 9 | 0.1111 | 36 | 0.0833 |
| 3_40750127 | 9 | 0.3333 | 7 | 0.1429 | 8 | 0.25 | 6 | 0.1667 | 30 | 0.2333 | 1 | 1 | 4 | 0 | 13 | 0.5385 | 3 | 0.3333 | 21 | 0.4286 |
| 3_40760100 | NA | NA | NA | NA | NA | NA | 1 | 1 | 1 | 1 | NA | NA | 1 | 1 | 1 | 1 | 1 | 1 | 3 | 1 |
| 3_40772869 | 7 | 0.1429 | 1 | 0 | 5 | 0 | 2 | 0 | 15 | 0.0667 | 1 | 0 | 2 | 0.5 | 3 | 0.3333 | 5 | 0 | 11 | 0.1818 |
| 3_40773038 | 8 | 0.625 | 1 | 0 | 5 | 0.4 | 2 | 0.5 | 16 | 0.5 | 1 | 1 | 2 | 0.5 | 3 | 0.3333 | 5 | 0.2 | 11 | 0.3636 |
| 3_40773047 | 8 | 0.125 | 1 | 0 | 5 | 0.2 | 3 | 0 | 17 | 0.1176 | 1 | 0 | 2 | 0.5 | 3 | 0 | 3 | 0 | 9 | 0.1111 |
| 3_40773360 | 8 | 0.25 | NA | NA | 6 | 0.5 | 3 | 0.3333 | 17 | 0.3529 | 1 | 1 | 2 | 0 | 3 | 0 | 4 | 0 | 10 | 0.1 |
| 3_40773569 | 4 | 0 | NA | NA | 6 | 0.1667 | 3 | 0 | 13 | 0.0769 | 1 | 1 | 2 | 0.5 | 3 | 0 | 4 | 0.25 | 10 | 0.3 |
| 3_40773705 | 6 | 0 | NA | NA | 4 | 0.25 | 1 | 0 | 11 | 0.0909 | 1 | 0 | 1 | 0 | 2 | 0.5 | 3 | 0 | 7 | 0.1429 |
| 3_40773723 | 5 | 1 | NA | NA | 5 | 0.6 | 3 | 0.6667 | 13 | 0.7692 | 1 | 1 | 1 | 1 | 2 | 1 | 3 | 0.3333 | 7 | 0.7143 |
| 3_40773803 | 7 | 0.5714 | NA | NA | 5 | 0.4 | 3 | 0.3333 | 15 | 0.4667 | 1 | 0 | 2 | 0.5 | 2 | 0.5 | 3 | 0 | 8 | 0.25 |
| 3_40773839 | 6 | 0.3333 | NA | NA | 4 | 0.25 | 3 | 0 | 13 | 0.2308 | NA | NA | 1 | 0 | 2 | 0.5 | 3 | 0.3333 | 6 | 0.3333 |
| 3_40773872 | 6 | 0 | NA | NA | 4 | 0.25 | 2 | 0 | 12 | 0.0833 | 1 | 0 | 1 | 0 | 1 | 0 | 3 | 0.3333 | 6 | 0.1667 |
| 3_40773991 | 6 | 0.8333 | 1 | 1 | 5 | 0.8 | 3 | 0.6667 | 15 | 0.8 | 1 | 1 | 2 | 1 | 1 | 1 | 3 | 0.6667 | 7 | 0.8571 |
| 3_40802217 | NA | NA | 1 | 1 | NA | NA | NA | NA | 1 | 1 | NA | NA | NA | NA | NA | NA | 1 | 1 | 1 | 1 |
| 3_40802346 | NA | NA | 1 | 1 | NA | NA | NA | NA | 1 | 1 | 1 | 1 | NA | NA | 1 | 0 | 1 | 0 | 3 | 0.3333 |
| 3_40889969 | 1 | 1 | NA | NA | NA | NA | 2 | 0.5 | 3 | 0.6667 | NA | NA | 2 | 0 | 3 | 0.3333 | 1 | 1 | 6 | 0.3333 |
| 3_40904791 | NA | NA | NA | NA | 2 | 0.5 | NA | NA | 2 | 0.5 | NA | NA | NA | NA | 1 | 0 | 1 | 1 | 2 | 0.5 |
| 3_40931877 | NA | NA | 3 | 1 | 2 | 1 | 1 | 0 | 6 | 0.8333 | NA | NA | NA | NA | 1 | 1 | 2 | 0.5 | 3 | 0.6667 |

| | | | | | | | | | | | | | | | | | | | | |
|---|---|---|---|---|---|---|---|---|---|---|---|---|---|---|---|---|---|---|---|---|
| 3_4093238 | NA | NA | 1 | 1 | NA | NA | NA | NA | 1 | 1 | 1 | 1 | NA | NA | NA | NA | NA | NA | 1 | 1 |
| 3_4102113 | 2 | 0 | 4 | 0.25 | 8 | 0.25 | NA | NA | 14 | 0.2143 | NA | NA | 2 | 0 | 3 | 0 | 3 | 0.3333 | 8 | 0.125 |
| 3_4102135 | 2 | 0 | 5 | 0.2 | 7 | 0.1429 | 1 | 0 | 15 | 0.1333 | 1 | 0 | 3 | 0 | 3 | 0.3333 | 1 | 0 | 8 | 0.125 |
| 3_4102154 | 1 | 0 | 4 | 0.25 | 6 | 0.6667 | 1 | 0 | 12 | 0.4167 | 1 | 0 | 2 | 0 | 2 | 1 | 2 | 0.5 | 7 | 0.4286 |
| 3_4102282 | 12 | 0.3333 | 12 | 0.4167 | 11 | 0.5455 | 9 | 0.3333 | 44 | 0.4091 | 10 | 0.3 | 10 | 0.4 | 12 | 0.1667 | 6 | 0.3333 | 38 | 0.2895 |
| 3_4102302 | 10 | 0.9 | 10 | 0.9 | 10 | 0.7 | 7 | 1 | 37 | 0.8649 | 6 | 0.6667 | 10 | 0.7 | 12 | 0.9167 | 5 | 1 | 33 | 0.8182 |
| 3_4102306 | 9 | 0.6667 | 7 | 0.8571 | 5 | 0.6 | 3 | 1 | 24 | 0.75 | 8 | 1 | 8 | 1 | 11 | 0.6364 | 4 | 0.75 | 31 | 0.8387 |
| 3_4102306 | 10 | 0.9 | 5 | 0.8 | 7 | 0.8571 | 6 | 0.8333 | 28 | 0.8571 | 6 | 1 | 7 | 0.8571 | 9 | 0.8889 | 3 | 1 | 25 | 0.92 |
| 3_4103714 | 6 | 0 | 7 | 0 | 7 | 0.1429 | 2 | 0.5 | 22 | 0.0909 | 6 | 0 | 5 | 0 | 4 | 0.5 | 3 | 0.3333 | 18 | 0.1667 |
| 3_4103719 | 6 | 0 | 4 | 0.25 | 5 | 0 | 2 | 0 | 17 | 0.0588 | 5 | 0 | 5 | 0 | 7 | 0.1429 | 3 | 0 | 20 | 0.05 |
| 3_4155934 | 4 | 0.5 | 1 | 1 | 2 | 0.5 | 4 | 1 | 11 | 0.7273 | NA | NA | 3 | 0.3333 | 3 | 0.3333 | NA | NA | 6 | 0.3333 |
| 3_4155941 | 3 | 0.6667 | 2 | 0 | 2 | 0 | 5 | 1 | 12 | 0.5833 | NA | NA | 4 | 0.5 | 2 | 0.5 | NA | NA | 6 | 0.5 |
| 3_4155945 | 2 | 0.5 | NA | NA | 1 | 0 | 3 | 0.6667 | 6 | 0.5 | NA | NA | 1 | 1 | 1 | 1 | NA | NA | 2 | 1 |
| 3_4155960 | 3 | 0.3333 | 1 | 0 | 1 | 0 | 3 | 0.3333 | 8 | 0.25 | NA | NA | 1 | 0 | 1 | 0 | 1 | 1 | 3 | 0.3333 |
| 3_4155999 | 2 | 0 | 1 | 1 | 2 | 0 | 4 | 0.5 | 9 | 0.3333 | 2 | 0 | 1 | 1 | 1 | 1 | 1 | 0 | 5 | 0.4 |
| 3_4156036 | 3 | 0.3333 | 1 | 0 | 1 | 0 | 3 | 0 | 8 | 0.125 | 2 | 1 | 1 | 0 | NA | NA | 1 | 0 | 4 | 0.5 |
| 3_4156764 | 2 | 1 | 2 | 1 | 1 | 1 | 1 | 1 | 6 | 1 | NA | NA | 3 | 0.6667 | 2 | 1 | NA | NA | 5 | 0.8 |
| 3_4156774 | 1 | 1 | 1 | 1 | 1 | 1 | NA | NA | 3 | 1 | NA | NA | 2 | 1 | 2 | 1 | NA | NA | 4 | 1 |
| 3_4156783 | 2 | 1 | 2 | 1 | 1 | 1 | 1 | 1 | 6 | 1 | NA | NA | 3 | 1 | 2 | 1 | NA | NA | 5 | 1 |
| 3_4156791 | 2 | 0.5 | 2 | 1 | 1 | 1 | 2 | 1 | 7 | 0.8571 | NA | NA | 3 | 1 | 1 | 0 | 1 | 0 | 5 | 0.6 |
| 3_4156795 | 2 | 0 | 2 | 0.5 | 1 | 0 | 2 | 0.5 | 7 | 0.2857 | NA | NA | 3 | 0.3333 | 2 | 0 | NA | NA | 5 | 0.2 |
| 3_4156796 | 2 | 1 | 2 | 1 | 1 | 1 | 1 | 1 | 6 | 1 | NA | NA | 3 | 0.6667 | 1 | 1 | 1 | 1 | 5 | 0.8 |
| 3_4156800 | 1 | 0 | 2 | 1 | 1 | 1 | 2 | 0.5 | 6 | 0.6667 | NA | NA | 3 | 0 | 1 | 1 | 1 | 0 | 5 | 0.2 |
| 3_4156802 | 2 | 1 | 2 | 1 | 1 | 1 | 1 | 0 | 6 | 0.8333 | NA | NA | 3 | 0.3333 | 2 | 1 | 1 | 0 | 6 | 0.5 |
| 3_4156823 | 3 | 1 | NA | NA | 1 | 1 | 2 | 0.5 | 6 | 0.8333 | NA | NA | 3 | 0.3333 | 2 | 1 | NA | NA | 5 | 0.6 |
| 3_4156829 | 2 | 0.5 | 2 | 0 | NA | NA | NA | NA | 4 | 0.25 | NA | NA | 3 | 0.3333 | 1 | 1 | 1 | 0 | 5 | 0.4 |
| 3_4156834 | 2 | 1 | 2 | 1 | NA | NA | 1 | 1 | 5 | 1 | NA | NA | 2 | 0.5 | NA | NA | NA | NA | 2 | 0.5 |
| 3_4156840 | 3 | 1 | 2 | 0.5 | 1 | 1 | 2 | 0.5 | 8 | 0.75 | NA | NA | 3 | 0.3333 | 1 | 0 | 1 | 1 | 5 | 0.4 |
| 3_4156853 | 2 | 0 | 2 | 0.5 | NA | NA | 2 | 0.5 | 6 | 0.3333 | NA | NA | 3 | 0.3333 | 2 | 0.5 | 1 | 1 | 6 | 0.5 |
| 3_4156864 | 3 | 0.6667 | 2 | 0 | 1 | 0 | 2 | 0 | 8 | 0.25 | NA | NA | 2 | 0.5 | 1 | 1 | 1 | 0 | 4 | 0.5 |
| 3_4156865 | 3 | 0.3333 | 2 | 0 | 1 | 0 | 2 | 0 | 8 | 0.125 | NA | NA | 2 | 0.5 | 1 | 0 | 1 | 0 | 4 | 0.25 |
| 3_4156867 | 2 | 0.5 | 1 | 0 | 1 | 0 | 2 | 0 | 6 | 0.1667 | NA | NA | 2 | 0 | 1 | 1 | 1 | 0 | 4 | 0.25 |
| 3_4156877 | 3 | 0.6667 | 1 | 0 | NA | NA | 2 | 0.5 | 6 | 0.5 | NA | NA | 1 | 0 | 1 | 1 | NA | NA | 2 | 0.5 |
| 3_4156910 | 2 | 1 | NA | NA | 1 | 0 | 1 | 0 | 4 | 0.5 | NA | NA | 1 | 0 | 3 | 0.3333 | 1 | 0 | 5 | 0.2 |
| 3_4156953 | 3 | 0.6667 | 1 | 0 | 1 | 0 | 3 | 0 | 8 | 0.25 | 1 | 0 | 2 | 0.5 | 3 | 0.3333 | 1 | 0 | 7 | 0.2857 |
| 3_4157027 | 3 | 0.3333 | 2 | 0 | 2 | 0.5 | 3 | 0 | 10 | 0.2 | 2 | 0.5 | 3 | 0 | 2 | 0.5 | 1 | 0 | 8 | 0.25 |
| 3_4157098 | 1 | 0 | 3 | 0 | 1 | 0 | 3 | 0.3333 | 8 | 0.125 | 2 | 0.5 | 4 | 0 | 4 | 0.5 | NA | NA | 10 | 0.3 |
| 3_4171067 | NA | NA | NA | NA | 1 | 0 | 2 | 0.5 | 3 | 0.3333 | NA | NA | NA | NA | NA | NA | 1 | 1 | 1 | 1 |
| 3_4522915 | 1 | 1 | 1 | 1 | NA | NA | NA | NA | 2 | 1 | NA | NA | 3 | 0.6667 | 1 | 1 | NA | NA | 4 | 0.75 |
| 3_4522919 | 1 | 1 | 1 | 1 | NA | NA | NA | NA | 2 | 1 | NA | NA | 1 | 1 | 1 | 1 | NA | NA | 2 | 1 |
| 3_4522924 | 1 | 1 | 1 | 1 | NA | NA | NA | NA | 2 | 1 | NA | NA | 3 | 0.3333 | 1 | 0 | NA | NA | 4 | 0.25 |
| 3_4522934 | 1 | 1 | 1 | 1 | NA | NA | NA | NA | 2 | 1 | NA | NA | 3 | 1 | 1 | 1 | NA | NA | 4 | 1 |
| 3_4533289 | 3 | 0.3333 | 2 | 0 | 3 | 0.3333 | 2 | 0.5 | 10 | 0.3 | 2 | 0.5 | 2 | 0.5 | 3 | 0 | 3 | 0.6667 | 10 | 0.4 |
| 3_4533331 | 3 | 1 | 2 | 1 | 3 | 0.6667 | 4 | 0.75 | 12 | 0.8333 | 4 | 0.75 | 1 | 1 | 4 | 1 | 2 | 1 | 11 | 0.9091 |
| 3_4533339 | 4 | 0.25 | 1 | 0 | 3 | 0.3333 | 5 | 0.6 | 13 | 0.3846 | 3 | 0.3333 | 3 | 0.3333 | 6 | 0.3333 | 3 | 0 | 15 | 0.2667 |
| 3_4533345 | 3 | 0 | 2 | 0.5 | 3 | 0 | 5 | 0.2 | 13 | 0.1538 | 4 | 0.25 | 3 | 0 | 4 | 0.25 | 4 | 0 | 15 | 0.1333 |
| 3_4533381 | 3 | 1 | 4 | 1 | 2 | 1 | 3 | 0.6667 | 12 | 0.9167 | 3 | 1 | 3 | 1 | 8 | 1 | 4 | 1 | 18 | 1 |
| 3_4533382 | 3 | 0 | 3 | 0 | 3 | 0.3333 | 5 | 0 | 14 | 0.0714 | 1 | 1 | 2 | 0 | 7 | 0 | 1 | 0 | 11 | 0.0909 |
| 3_4533386 | 5 | 0.2 | 4 | 0.25 | 3 | 0 | 6 | 0.1667 | 18 | 0.1667 | 4 | 0 | 4 | 0.5 | 6 | 0.3333 | 4 | 0.5 | 18 | 0.3333 |
| 3_4533388 | 5 | 0.8 | 3 | 1 | 2 | 1 | 5 | 1 | 15 | 0.9333 | 2 | 1 | 4 | 0.5 | 5 | 0.8 | 4 | 1 | 15 | 0.8 |
| 3_4533414 | 5 | 0.2 | 4 | 0.25 | 4 | 0 | 8 | 0.125 | 21 | 0.1429 | 4 | 0.25 | 4 | 0.25 | 8 | 0.125 | 5 | 0 | 21 | 0.1429 |
| 3_4533416 | 4 | 0 | 4 | 0 | 4 | 0 | 8 | 0.125 | 20 | 0.05 | 3 | 0.3333 | 2 | 0 | 9 | 0.1111 | 4 | 0 | 18 | 0.1111 |
| 3_4533423 | 3 | 0.3333 | 3 | 0 | 4 | 0.5 | 5 | 0.2 | 15 | 0.2667 | 3 | 0 | 1 | 0 | 7 | 0.1429 | 3 | 0.3333 | 14 | 0.1429 |
| 3_4533425 | 3 | 0.3333 | 3 | 0.3333 | 4 | 0.25 | 7 | 0.5714 | 17 | 0.4118 | 4 | 0.5 | 4 | 0.75 | 9 | 0.2222 | 5 | 0.2 | 22 | 0.3636 |
| 3_4533429 | 3 | 0.3333 | 4 | 0 | 4 | 0 | 8 | 0 | 19 | 0.0526 | 3 | 0.6667 | 2 | 0 | 7 | 0 | 3 | 0 | 15 | 0.1333 |
| 3_4533456 | NA | NA | NA | NA | 1 | 1 | NA | NA | 1 | 1 | NA | NA | 1 | 1 | NA | NA | 1 | 1 | 2 | 1 |
| 3_4533459 | 6 | 1 | 6 | 0.8333 | 6 | 0.6667 | 8 | 0.75 | 26 | 0.8077 | 5 | 0.8 | 6 | 1 | 10 | 0.8 | 7 | 0.5714 | 28 | 0.7857 |
| 3_4533463 | 4 | 0.75 | 5 | 0.6 | 6 | 0.6667 | 9 | 0.5556 | 24 | 0.625 | 6 | 0.8333 | 8 | 0.625 | 12 | 0.5833 | 8 | 0.75 | 34 | 0.6765 |
| 3_4533467 | NA | NA | NA | NA | 1 | 1 | NA | NA | 1 | 1 | NA | NA | 1 | 1 | NA | NA | 1 | 1 | 2 | 1 |
| 3_4533468 | 4 | 0.75 | 5 | 0.8 | 5 | 0.8 | 7 | 0.4286 | 21 | 0.6667 | 4 | 0.5 | 7 | 0.7143 | 10 | 0.7 | 7 | 0.8571 | 28 | 0.7143 |
| 3_4533484 | 7 | 0.5714 | 7 | 0.4286 | 7 | 0.5714 | 8 | 0.625 | 29 | 0.5517 | 6 | 0.3333 | 7 | 0.4286 | 12 | 0.75 | 7 | 0.2857 | 32 | 0.5 |
| 3_4533495 | 5 | 0.6 | 3 | 0.6667 | 7 | 0.1429 | 6 | 0.1667 | 21 | 0.3333 | 6 | 0.1667 | 9 | 0.2222 | 17 | 0.2353 | 6 | 0 | 38 | 0.1842 |
| 3_4533496 | 6 | 0.1667 | 5 | 0.4 | 7 | 0.4286 | 11 | 0.0909 | 29 | 0.2414 | 7 | 0.7143 | 12 | 0.25 | 16 | 0.4375 | 6 | 0.6667 | 41 | 0.4634 |
| 3_4533506 | 7 | 0 | 8 | 0.125 | 7 | 0.1429 | 11 | 0 | 33 | 0.0606 | 6 | 0.3333 | 11 | 0 | 16 | 0 | 6 | 0 | 39 | 0.0513 |
| 3_4533515 | 7 | 0 | 9 | 0 | 6 | 0.1667 | 11 | 0.0909 | 33 | 0.0606 | 7 | 0.1429 | 11 | 0.1818 | 15 | 0.0667 | 9 | 0.2222 | 42 | 0.1429 |

| | | | | | | | | | | | | | | | | | | | | |
|---|---|---|---|---|---|---|---|---|---|---|---|---|---|---|---|---|---|---|---|---|
| 3_45335300 | 8 | 0.625 | 8 | 0.75 | 8 | 0.75 | 10 | 0.5 | 34 | 0.6471 | 6 | 0.6667 | 8 | 0.625 | 15 | 0.8 | 10 | 0.6 | 39 | 0.6923 |
| 3_45335437 | 8 | 1 | 9 | 0.7778 | 7 | 1 | 9 | 0.7778 | 33 | 0.8788 | 8 | 0.625 | 11 | 0.7273 | 15 | 0.8 | 10 | 0.9 | 44 | 0.7727 |
| 3_45335567 | 8 | 0.5 | 9 | 0.6667 | 7 | 0.5714 | 10 | 0.5 | 34 | 0.5588 | 5 | 0.6 | 8 | 0.5 | 8 | 0.625 | 9 | 0.4444 | 30 | 0.5333 |
| 3_45335582 | 7 | 0.8571 | 7 | 1 | 3 | 0.6667 | 8 | 0.75 | 25 | 0.84 | 6 | 0.8333 | 9 | 0.8889 | 14 | 0.8571 | 9 | 0.7778 | 38 | 0.8421 |
| 3_45335759 | 5 | 0.6 | 8 | 0.75 | 6 | 0.8333 | 6 | 0.5 | 25 | 0.68 | 4 | 0.75 | 8 | 0.75 | 10 | 0.3 | 7 | 0.8571 | 29 | 0.6207 |
| 3_45335825 | 9 | 0.6667 | 10 | 0.7 | 6 | 1 | 11 | 0.7273 | 36 | 0.75 | 8 | 0.875 | 10 | 1 | 14 | 0.7857 | 11 | 0.7273 | 43 | 0.8372 |
| 3_45335922 | 10 | 0.4 | 11 | 0.4545 | 7 | 0.2857 | 12 | 0.25 | 40 | 0.35 | 11 | 0.2727 | 10 | 0.3 | 12 | 0.25 | 14 | 0.2143 | 47 | 0.2553 |
| 3_45336030 | 11 | 0.0909 | 12 | 0.1667 | 8 | 0.125 | 8 | 0.125 | 39 | 0.1282 | 10 | 0.2 | 14 | 0.2143 | 15 | 0.2 | 12 | 0.1667 | 51 | 0.1961 |
| 3_45336140 | 7 | 0.8571 | 9 | 1 | 3 | 1 | 5 | 1 | 24 | 0.9583 | 4 | 0.75 | 8 | 0.625 | 9 | 1 | 11 | 0.8182 | 32 | 0.8125 |
| 3_45336943 | 3 | 1 | NA | NA | 1 | 1 | NA | NA | 4 | 1 | NA | NA | 2 | 0.5 | NA | NA | NA | NA | 2 | 0.5 |
| 3_45337035 | 3 | 0.6667 | NA | NA | 3 | 0.3333 | NA | NA | 6 | 0.5 | NA | NA | 3 | 0.3333 | NA | NA | NA | NA | 3 | 0.3333 |
| 3_45337273 | 3 | 1 | 1 | 1 | 1 | 1 | 1 | 0 | 6 | 0.8333 | 2 | 0.5 | 3 | 0.3333 | NA | NA | NA | NA | 5 | 0.4 |
| 3_45347142 | 10 | 1 | 16 | 0.9375 | 7 | 1 | 10 | 1 | 43 | 0.9767 | 12 | 0.9167 | 19 | 1 | 16 | 1 | 20 | 1 | 67 | 0.9851 |
| 3_45374012 | 7 | 1 | 11 | 0.7273 | 3 | 1 | 7 | 0.8571 | 28 | 0.8571 | 7 | 1 | 13 | 0.9231 | 13 | 0.8462 | 15 | 0.7333 | 48 | 0.8542 |
| 3_45374058 | 12 | 0.9167 | 12 | 1 | 3 | 1 | 7 | 1 | 34 | 0.9706 | 8 | 0.75 | 12 | 0.9167 | 12 | 1 | 15 | 0.9333 | 47 | 0.9149 |
| 3_45374067 | 9 | 0.1111 | 12 | 0.1667 | 2 | 0.5 | 2 | 0 | 25 | 0.16 | 7 | 0.1429 | 12 | 0.25 | 11 | 0.3636 | 8 | 0 | 38 | 0.2105 |
| 3_45374246 | 12 | 0.5833 | 12 | 0.5 | 2 | 0.5 | 6 | 0.3333 | 32 | 0.5 | 9 | 0.3333 | 14 | 0.5 | 13 | 0.5385 | 16 | 0.5625 | 52 | 0.5 |
| 3_45374383 | 9 | 0.5556 | 15 | 0.6 | 4 | 0.5 | 8 | 0.375 | 36 | 0.5278 | 7 | 0.4286 | 14 | 0 | 14 | 0.2143 | 14 | 0.4286 | 49 | 0.2449 |
| 3_45374682 | 7 | 0 | 13 | 0.2308 | 2 | 0.5 | 9 | 0.3333 | 31 | 0.2258 | 8 | 0.25 | 13 | 0.0769 | 15 | 0.1333 | 15 | 0.2667 | 51 | 0.1765 |
| 3_45376216 | 6 | 0.3333 | 7 | 0 | 2 | 0.5 | 5 | 0 | 20 | 0.15 | 5 | 0.2 | 12 | 0 | 12 | 0.0833 | 9 | 0 | 38 | 0.0526 |
| 3_45377531 | 7 | 0.1429 | 7 | 0 | 6 | 0.1667 | 6 | 0 | 26 | 0.0769 | 6 | 0.1667 | 10 | 0.1 | 17 | 0.2353 | 9 | 0 | 42 | 0.1429 |
| 3_45377559 | 9 | 0.1111 | 8 | 0.125 | 6 | 0 | 7 | 0 | 30 | 0.0667 | 5 | 0.2 | 11 | 0 | 19 | 0.1053 | 10 | 0 | 45 | 0.0667 |
| 3_45377895 | 8 | 0 | 7 | 0 | 6 | 0.3333 | 7 | 0.1429 | 28 | 0.1071 | 9 | 0.1111 | 10 | 0.1 | 16 | 0.1875 | 10 | 0.3 | 45 | 0.1778 |
| 3_45387065 | 2 | 1 | 1 | 1 | 1 | 1 | 4 | 0.5 | 8 | 0.75 | 1 | 1 | 4 | 1 | 2 | 0.5 | 2 | 0.5 | 9 | 0.7778 |
| 3_45387112 | 2 | 0.5 | 1 | 0 | 1 | 1 | 4 | 0.75 | 8 | 0.625 | 2 | 1 | 3 | 0.6667 | 1 | 1 | 2 | 1 | 8 | 0.875 |
| 3_45391877 | 1 | 1 | NA | NA | NA | NA | NA | NA | 1 | 1 | 1 | 1 | NA | NA | 1 | 0 | NA | NA | 2 | 0.5 |
| 3_45403100 | 1 | 1 | NA | NA | NA | NA | NA | NA | 1 | 1 | NA | NA | 1 | 1 | NA | NA | NA | NA | 1 | 1 |
| 3_49698983 | 1 | 1 | NA | NA | NA | NA | NA | NA | 1 | 1 | NA | NA | NA | NA | 1 | 1 | NA | NA | 1 | 1 |
| 3_49699083 | 1 | 1 | NA | NA | NA | NA | NA | NA | 1 | 1 | NA | NA | 1 | 0 | 1 | 0 | 1 | 1 | 3 | 0.3333 |
| 3_49699122 | 1 | 1 | NA | NA | NA | NA | NA | NA | 1 | 1 | NA | NA | 1 | 1 | 1 | 1 | NA | NA | 2 | 1 |
| 3_50319462 | 4 | 0.25 | 5 | 0 | 4 | 0 | 1 | 0 | 14 | 0.0714 | 3 | 0 | 6 | 0 | 2 | 0 | 3 | 0.3333 | 14 | 0.0714 |
| 3_50319669 | 2 | 0 | 6 | 0.1667 | 5 | 0.2 | 2 | 0 | 15 | 0.1333 | 3 | 0.3333 | 6 | 0.3333 | 3 | 0.6667 | 3 | 0 | 15 | 0.3333 |
| 3_50319922 | 2 | 0.5 | 6 | 0 | 3 | 0 | 2 | 0 | 13 | 0.0769 | 3 | 0 | 5 | 0.2 | 2 | 0.5 | 2 | 0.5 | 12 | 0.25 |
| 3_50326663 | NA | NA | 1 | 0 | NA | NA | 1 | 1 | 2 | 0.5 | NA | NA | 2 | 0.5 | NA | NA | 1 | 1 | 3 | 0.6667 |
| 3_51155362 | 13 | 0.0769 | 5 | 0 | 6 | 0.1667 | 8 | 0 | 32 | 0.0625 | 2 | 0.5 | 5 | 0 | 3 | 0 | 9 | 0.1111 | 19 | 0.1053 |
| 3_51157247 | 13 | 0.5385 | 6 | 0.5 | 7 | 0.5714 | 8 | 0.5 | 34 | 0.5294 | 2 | 0 | 6 | 0.5 | 3 | 0.3333 | 10 | 0.5 | 21 | 0.4286 |
| 3_51157284 | 13 | 0.7692 | 5 | 0.8 | 7 | 0.8571 | 6 | 0.6667 | 31 | 0.7742 | 1 | 1 | 6 | 0.6667 | 3 | 0.6667 | 10 | 0.8 | 20 | 0.75 |
| 3_51157341 | 10 | 0.7 | 6 | 0.5 | 7 | 0.2857 | 6 | 0.1667 | 29 | 0.4483 | 2 | 0.5 | 6 | 0.1667 | 2 | 0.5 | 10 | 0.3 | 20 | 0.3 |
| 3_51157432 | 12 | 0.0833 | 7 | 0.1429 | 9 | 0.3333 | 8 | 0.25 | 36 | 0.1944 | 2 | 0 | 7 | 0 | 5 | 0 | 12 | 0.25 | 26 | 0.1154 |
| 3_51157441 | 13 | 0.0769 | 7 | 0.1429 | 8 | 0 | 7 | 0.1429 | 35 | 0.0857 | 1 | 0 | 5 | 0.4 | 5 | 0 | 12 | 0.1667 | 23 | 0.1739 |
| 3_51157465 | 12 | 0.1667 | 7 | 0.2857 | 9 | 0.1111 | 8 | 0.125 | 36 | 0.1667 | 3 | 0 | 7 | 0.1429 | 5 | 0.6 | 12 | 0.0833 | 27 | 0.1852 |
| 3_51157565 | 14 | 0.1429 | 7 | 0 | 9 | 0 | 8 | 0 | 38 | 0.0526 | 3 | 0 | 7 | 0.1429 | 5 | 0 | 12 | 0.1667 | 27 | 0.1111 |
| 3_51157628 | 13 | 0 | 6 | 0.6667 | 8 | 0.5 | 8 | 0.5 | 35 | 0.3429 | 3 | 0.6667 | 7 | 0.2857 | 5 | 0.6 | 12 | 0.1667 | 27 | 0.3333 |
| 3_51157662 | 11 | 0.2727 | 7 | 0.1429 | 10 | 0.1 | 7 | 0 | 35 | 0.1429 | 3 | 0.3333 | 7 | 0.2857 | 5 | 0.2 | 11 | 0 | 26 | 0.1538 |
| 3_51157722 | 14 | 0.8571 | 6 | 1 | 8 | 1 | 8 | 0.875 | 36 | 0.9167 | 4 | 1 | 7 | 1 | 4 | 1 | 12 | 1 | 27 | 1 |
| 3_51157759 | 8 | 0.75 | 7 | 0.1429 | 8 | 0.25 | 6 | 0.5 | 29 | 0.4138 | 2 | 0 | 4 | 0.25 | 5 | 0.2 | 10 | 0.3 | 21 | 0.2381 |
| 3_51157787 | 12 | 1 | 6 | 1 | 8 | 0.875 | 9 | 1 | 35 | 0.9714 | 2 | 1 | 6 | 1 | 4 | 1 | 10 | 1 | 22 | 1 |
| 3_51157851 | 9 | 0.5556 | 5 | 0.4 | 7 | 0.2857 | 4 | 0.75 | 25 | 0.48 | 3 | 0.6667 | 4 | 0 | 4 | 0.25 | 8 | 0.375 | 19 | 0.3158 |
| 3_51157895 | 7 | 0 | 7 | 0.4286 | 7 | 0.4286 | 6 | 0.5 | 27 | 0.3333 | 1 | 0 | 6 | 0.6667 | 3 | 1 | 7 | 0.1429 | 17 | 0.4706 |
| 3_51157916 | 11 | 0.7273 | 8 | 0.75 | 10 | 0.4 | 6 | 0.5 | 35 | 0.6 | 3 | 1 | 5 | 1 | 4 | 0.5 | 10 | 0.7 | 22 | 0.7727 |
| 3_51157987 | 10 | 0.7 | 6 | 0.6667 | 9 | 0.6667 | 6 | 0.3333 | 31 | 0.6129 | 2 | 0.5 | 3 | 0.6667 | 5 | 0.6 | 10 | 0.5 | 20 | 0.55 |
| 3_51157992 | 12 | 0.5 | 8 | 0.625 | 7 | 0.5714 | 8 | 0.5 | 35 | 0.5429 | 2 | 0.5 | 5 | 0.4 | 4 | 0.25 | 11 | 0.4545 | 22 | 0.4091 |
| 3_51158020 | 14 | 0.9286 | 9 | 0.7778 | 9 | 0.8889 | 10 | 0.9 | 42 | 0.881 | 4 | 1 | 7 | 0.7143 | 6 | 1 | 12 | 0.75 | 29 | 0.8276 |
| 3_51158072 | 10 | 0.8 | 7 | 0.7143 | 9 | 0.5556 | 6 | 0.8333 | 32 | 0.7188 | 1 | 1 | 7 | 0.7143 | 6 | 0.8333 | 12 | 0.6667 | 26 | 0.7308 |
| 3_51158085 | 13 | 0.1538 | 8 | 0.25 | 9 | 0 | 9 | 0 | 39 | 0.1026 | 1 | 0 | 6 | 0.1667 | 8 | 0 | 12 | 0.0833 | 27 | 0.0741 |
| 3_51158395 | 14 | 0.6429 | 10 | 0.7 | 8 | 0.75 | 5 | 0.6 | 37 | 0.6757 | 3 | 0.3333 | 8 | 0.875 | 8 | 0.75 | 9 | 0.4444 | 28 | 0.6429 |
| 3_51158482 | 14 | 0.0714 | 8 | 0.5 | 7 | 0.4286 | 6 | 0.1667 | 35 | 0.2571 | 3 | 0 | 7 | 0.1429 | 7 | 0.4286 | 9 | 0.3333 | 26 | 0.2692 |
| 3_51158725 | 16 | 0.0625 | 9 | 0 | 8 | 0.125 | 8 | 0.25 | 41 | 0.0976 | 3 | 0 | 6 | 0.5 | 7 | 0 | 12 | 0.1667 | 28 | 0.1786 |
| 3_51158736 | 15 | 0.4667 | 10 | 0.3 | 10 | 0.6 | 8 | 0.375 | 43 | 0.4419 | 4 | 0.25 | 6 | 0.6667 | 4 | 0.5 | 9 | 0.3333 | 23 | 0.4348 |
| 3_51158780 | 16 | 0 | 11 | 0.0909 | 12 | 0.0833 | 9 | 0.2222 | 48 | 0.0833 | 4 | 0.25 | 7 | 0.1429 | 7 | 0 | 12 | 0.0833 | 30 | 0.1 |
| 3_51158850 | 16 | 0.1875 | 11 | 0 | 13 | 0.1538 | 10 | 0.1 | 50 | 0.12 | 4 | 0 | 6 | 0.1667 | 7 | 0.2857 | 12 | 0 | 29 | 0.1034 |
| 3_51158874 | 14 | 0.5714 | 10 | 0.8 | 11 | 0.5455 | 8 | 0.375 | 43 | 0.5814 | 4 | 0.25 | 7 | 0.4286 | 6 | 0.8333 | 12 | 0.5 | 29 | 0.5172 |
| 3_51159034 | 15 | 0.6 | 4 | 0.25 | 12 | 0.3333 | 6 | 0.1667 | 37 | 0.4054 | 2 | 0 | 7 | 0.2857 | 6 | 0.3333 | 10 | 0.4 | 25 | 0.32 |
| 3_51162882 | 2 | 0 | 6 | 0.5 | 2 | 0.5 | 4 | 0.5 | 14 | 0.4286 | 3 | 0.3333 | 1 | 0 | 5 | 0.2 | 6 | 0.1667 | 15 | 0.2 |
| 3_51163305 | 1 | 0 | 2 | 0.5 | 2 | 0.5 | 5 | 0.8 | 10 | 0.6 | 4 | 1 | 1 | 0 | 2 | 0 | 6 | 0.5 | 13 | 0.5385 |

| | | | | | | | | | | | | | | | | | | | | |
|---|---|---|---|---|---|---|---|---|---|---|---|---|---|---|---|---|---|---|---|---|
| 3_51163335 | 2 | 0.5 | 4 | 0.25 | 2 | 1 | 3 | 0.6667 | 11 | 0.5455 | 3 | 0.6667 | 1 | 1 | 3 | 0.6667 | 6 | 0.5 | 13 | 0.6154 |
| 3_51163490 | 2 | 1 | 3 | 0.6667 | 2 | 1 | 4 | 0.75 | 11 | 0.8182 | 3 | 0.6667 | NA | NA | 2 | 0.5 | 4 | 0.5 | 9 | 0.5556 |
| 3_51163533 | 2 | 0.5 | 4 | 0.75 | 1 | 0 | 5 | 0.6 | 12 | 0.5833 | 2 | 0.5 | 1 | 0 | 2 | 0.5 | 3 | 0.3333 | 8 | 0.375 |
| 3_51163560 | 2 | 0 | 4 | 0.25 | 2 | 0 | 5 | 0.2 | 13 | 0.1538 | 3 | 0 | 1 | 0 | 2 | 0.5 | 4 | 0.25 | 10 | 0.2 |
| 3_51163586 | 2 | 0 | 3 | 0 | 1 | 1 | 4 | 0.75 | 10 | 0.4 | 3 | 0.6667 | NA | NA | 2 | 0.5 | 3 | 0 | 8 | 0.375 |
| 3_51163593 | 2 | 0 | 4 | 0 | 2 | 0 | 5 | 0.4 | 13 | 0.1538 | 3 | 0 | 1 | 0 | 1 | 0 | 3 | 0.3333 | 8 | 0.125 |
| 3_51163612 | 2 | 1 | 3 | 0 | 2 | 1 | 4 | 0.75 | 11 | 0.6364 | 2 | 1 | 1 | 1 | 2 | 1 | 3 | 0.3333 | 8 | 0.75 |
| 3_51163640 | 2 | 0 | 4 | 0.5 | 2 | 0 | 5 | 0 | 13 | 0.1538 | 3 | 0 | 1 | 0 | 2 | 0 | 2 | 0.5 | 8 | 0.125 |
| 3_51163753 | 2 | 1 | 3 | 0.3333 | 1 | 1 | 3 | 0.3333 | 9 | 0.5556 | 1 | 0 | 1 | 1 | 2 | 1 | 1 | 1 | 5 | 0.8 |
| 3_51163820 | 1 | 1 | 4 | 1 | 1 | 1 | 2 | 1 | 8 | 1 | 1 | 1 | 1 | 1 | 2 | 0.5 | 3 | 0.6667 | 7 | 0.7143 |
| 3_51163874 | 1 | 1 | 3 | 1 | 1 | 0 | 1 | 1 | 6 | 0.8333 | 2 | 1 | 1 | 1 | 1 | 1 | 2 | 0.5 | 6 | 0.8333 |
| 3_51163929 | 2 | 0 | 4 | 0.25 | 2 | 0 | 4 | 0 | 12 | 0.0833 | 3 | 0.3333 | 1 | 1 | 1 | 0 | 3 | 0 | 8 | 0.25 |
| 3_51163997 | 2 | 0.5 | 3 | 1 | 1 | 0 | 4 | 0.75 | 10 | 0.7 | 3 | 0.6667 | 1 | 0 | 1 | 0 | 3 | 0.3333 | 8 | 0.375 |
| 3_51164031 | 2 | 1 | 3 | 1 | 1 | 1 | 2 | 1 | 8 | 1 | 2 | 1 | 1 | 1 | 1 | 1 | 3 | 1 | 7 | 1 |
| 3_51164104 | 1 | 1 | 2 | 0.5 | 2 | 0.5 | 2 | 0 | 7 | 0.4286 | 2 | 1 | 1 | 0 | NA | NA | 3 | 0.3333 | 6 | 0.5 |
| 3_51164405 | 2 | 0 | 3 | 0 | 2 | 0.5 | 3 | 0 | 10 | 0.1 | 2 | 0 | 1 | 0 | NA | NA | 2 | 0.5 | 5 | 0.2 |
| 3_51165131 | 2 | 0.5 | 2 | 0 | 2 | 0.5 | 2 | 0 | 8 | 0.25 | 2 | 0 | 1 | 0 | NA | NA | 2 | 0.5 | 5 | 0.2 |
| 3_51166674 | 1 | 1 | 2 | 0 | 2 | 0 | 1 | 0 | 6 | 0.1667 | 2 | 0 | 1 | 0 | NA | NA | 1 | 1 | 4 | 0.25 |
| 3_51166708 | 2 | 0.5 | 2 | 0.5 | 2 | 0 | 1 | 0 | 7 | 0.2857 | 2 | 1 | 1 | 0 | NA | NA | NA | NA | 3 | 0.6667 |
| 3_51310168 | NA | NA | NA | NA | 1 | 1 | NA | NA | 1 | 1 | NA | NA | NA | NA | 1 | 0 | 1 | 1 | 2 | 0.5 |
| 3_51312865 | 1 | 1 | 1 | 1 | NA | NA | NA | NA | 2 | 1 | 1 | 1 | 2 | 1 | 2 | 0.5 | NA | NA | 5 | 0.8 |
| 3_51315602 | 39 | 0.0769 | 39 | 0.0769 | 36 | 0.0833 | 45 | 0.0889 | 159 | 0.0818 | 29 | 0.1379 | 28 | 0.1786 | 28 | 0.1071 | 39 | 0.0256 | 124 | 0.1048 |
| 3_51315901 | NA | NA | NA | NA | 1 | 1 | NA | NA | 1 | 1 | 1 | 1 | NA | NA | 1 | 0 | NA | NA | 2 | 0.5 |
| 3_51380067 | 6 | 0.3333 | 5 | 0 | 12 | 0.25 | 11 | 0.1818 | 34 | 0.2059 | 4 | 0.25 | NA | NA | 6 | 0 | 4 | 0.25 | 14 | 0.1429 |
| 3_51380203 | 6 | 0.3333 | 4 | 0 | 12 | 0.25 | 12 | 0.25 | 34 | 0.2353 | 4 | 0.5 | NA | NA | 5 | 0.2 | 4 | 0 | 13 | 0.2308 |
| 3_51380248 | 6 | 0.1667 | 3 | 0.3333 | 11 | 0.1818 | 11 | 0.1818 | 31 | 0.1935 | 3 | 0.6667 | NA | NA | 5 | 0 | 4 | 0 | 12 | 0.1667 |
| 3_51380255 | 5 | 0.2 | 4 | 0.25 | 9 | 0.1111 | 12 | 0.3333 | 30 | 0.2333 | 4 | 0 | NA | NA | 6 | 0.3333 | 4 | 0.5 | 14 | 0.2857 |
| 3_51380781 | 7 | 0.5714 | 5 | 0 | 11 | 0.1818 | 13 | 0.2308 | 36 | 0.25 | 6 | 0.1667 | NA | NA | 7 | 0.1429 | 6 | 0.3333 | 19 | 0.2105 |
| 3_51380787 | 5 | 0.4 | 5 | 0.2 | 12 | 0.1667 | 14 | 0.1429 | 36 | 0.1944 | 6 | 0.5 | NA | NA | 4 | 0.25 | 6 | 0 | 16 | 0.25 |
| 3_51380948 | 8 | 0.125 | 3 | 0 | 11 | 0.1818 | 12 | 0 | 34 | 0.0882 | 5 | 0 | 1 | 1 | 5 | 0.2 | 5 | 0 | 16 | 0.125 |
| 3_51380999 | 8 | 0.25 | 6 | 0.1667 | 11 | 0.3636 | 13 | 0.0769 | 38 | 0.2105 | 6 | 0 | 1 | 0 | 5 | 0.2 | 7 | 0.1429 | 19 | 0.1053 |
| 3_51381116 | 8 | 0 | 5 | 0 | 11 | 0.1818 | 12 | 0.0833 | 36 | 0.0833 | 5 | 0.2 | 1 | 0 | 6 | 0 | 6 | 0.1667 | 18 | 0.1111 |
| 3_51381137 | 8 | 0.125 | 5 | 0.2 | 11 | 0.2727 | 14 | 0.1429 | 38 | 0.1842 | 6 | 0 | 1 | 0 | 6 | 0.1667 | 6 | 0 | 19 | 0.0526 |
| 3_51381149 | 9 | 0.2222 | 5 | 0.4 | 12 | 0.5833 | 14 | 0 | 40 | 0.275 | 6 | 0.3333 | 1 | 1 | 7 | 0.7143 | 7 | 0.1429 | 21 | 0.4286 |
| 3_51381221 | 8 | 0.25 | 5 | 0.4 | 12 | 0.4167 | 11 | 0.1818 | 36 | 0.3056 | 6 | 0.1667 | 1 | 0 | 5 | 0.2 | 6 | 0.3333 | 18 | 0.2222 |
| 3_51382390 | 3 | 0.3333 | 2 | 0 | 5 | 0.2 | 8 | 0 | 18 | 0.1111 | 5 | 0.2 | 1 | 0 | 4 | 0 | 2 | 0 | 12 | 0.0833 |
| 3_51382458 | 3 | 0 | 3 | 0.3333 | 3 | 0 | 7 | 0.1429 | 16 | 0.125 | 5 | 0.4 | 2 | 0.5 | 4 | 0 | 1 | 1 | 12 | 0.3333 |
| 3_51382657 | 3 | 0 | 3 | 0 | 3 | 0.6667 | 7 | 0.4286 | 16 | 0.3125 | 5 | 0.2 | 2 | 1 | 3 | 0 | 1 | 1 | 11 | 0.3636 |
| 3_51400979 | 5 | 1 | 2 | 1 | 1 | 1 | 1 | 1 | 9 | 1 | 2 | 1 | 4 | 1 | 4 | 1 | NA | NA | 10 | 1 |
| 3_51401009 | 6 | 1 | 2 | 1 | 1 | 1 | 1 | 1 | 10 | 1 | 2 | 1 | 3 | 1 | 4 | 1 | NA | NA | 9 | 1 |
| 3_51401030 | 6 | 0.3333 | 2 | 0 | 1 | 0 | NA | NA | 9 | 0.2222 | 2 | 0.5 | 4 | 0 | 4 | 0 | NA | NA | 10 | 0.1 |
| 3_51401056 | 6 | 0.1667 | 2 | 0.5 | 1 | 0 | 1 | 1 | 10 | 0.3 | 2 | 0.5 | 4 | 0 | 4 | 0 | NA | NA | 10 | 0.1 |
| 3_51401069 | 6 | 0 | 2 | 1 | 1 | 0 | 1 | 0 | 10 | 0.2 | 2 | 0.5 | 4 | 0.25 | 4 | 0.5 | NA | NA | 10 | 0.4 |
| 3_51401084 | 4 | 0.25 | 2 | 0.5 | 1 | 0 | 1 | 0 | 8 | 0.25 | 1 | 0 | 4 | 0.25 | 2 | 0 | NA | NA | 7 | 0.1429 |
| 3_51401107 | 4 | 0.25 | 2 | 0 | 1 | 0 | NA | NA | 7 | 0.1429 | 2 | 1 | 4 | 0.25 | 4 | 0.25 | NA | NA | 10 | 0.4 |
| 3_51401188 | 6 | 0 | 2 | 0.5 | 1 | 0 | 1 | 0 | 10 | 0.1 | 2 | 0 | 4 | 0.5 | 5 | 0.2 | NA | NA | 11 | 0.2727 |
| 3_51401203 | 6 | 0.6667 | 2 | 0 | 1 | 1 | 1 | 0 | 10 | 0.5 | 2 | 0.5 | 4 | 0.5 | 5 | 0.6 | NA | NA | 11 | 0.5455 |
| 3_51401245 | 5 | 0 | 2 | 1 | 1 | 0 | 1 | 0 | 9 | 0.2222 | 2 | 0.5 | 4 | 0.75 | 3 | 0.3333 | NA | NA | 9 | 0.5556 |
| 3_51401275 | 5 | 0.4 | 2 | 0 | 1 | 0 | 1 | 0 | 9 | 0.2222 | 2 | 0.5 | 3 | 0.3333 | 4 | 0.25 | NA | NA | 9 | 0.3333 |
| 3_51401306 | 6 | 1 | NA | NA | NA | NA | 1 | 1 | 7 | 1 | 2 | 1 | 3 | 1 | 4 | 1 | NA | NA | 9 | 1 |
| 3_51401387 | 6 | 0 | 3 | 0 | 1 | 0 | 2 | 0.5 | 12 | 0.0833 | 2 | 0 | 4 | 0 | 4 | 0.25 | NA | NA | 10 | 0.1 |
| 3_51401433 | 5 | 0.2 | 3 | 0 | 1 | 0 | 2 | 0 | 11 | 0.0909 | 2 | 0 | 3 | 0 | 5 | 0.2 | NA | NA | 10 | 0.1 |
| 3_51401493 | 6 | 0.8333 | 3 | 0.3333 | 1 | 1 | 2 | 0.5 | 12 | 0.6667 | 2 | 1 | 4 | 0.75 | 5 | 1 | NA | NA | 11 | 0.9091 |
| 3_51401632 | 6 | 0.3333 | 2 | 0.5 | 1 | 0 | 2 | 0 | 11 | 0.2727 | 2 | 1 | 4 | 0.25 | 6 | 0.6667 | NA | NA | 12 | 0.5833 |
| 3_51402238 | 6 | 0 | 2 | 0 | 1 | 1 | 1 | 0 | 10 | 0.1 | 3 | 0 | 2 | 0 | 7 | 0.4286 | 1 | 0 | 13 | 0.2308 |
| 3_51402537 | 7 | 0 | 3 | 0.3333 | 1 | 0 | 2 | 0 | 13 | 0.0769 | 3 | 0.3333 | 4 | 0 | 6 | 0.3333 | 1 | 0 | 14 | 0.2143 |
| 3_51402619 | 5 | 0.6 | 3 | 0.6667 | 1 | 0 | 2 | 0.5 | 11 | 0.5455 | 3 | 0.6667 | 2 | 0 | 4 | 0.75 | 1 | 1 | 10 | 0.6 |
| 3_51402771 | 4 | 0 | 4 | 0.75 | 1 | 0 | 2 | 0.5 | 11 | 0.3636 | 3 | 0.3333 | 3 | 0 | 6 | 0.8333 | NA | NA | 12 | 0.5 |
| 3_51402787 | 6 | 0.3333 | 4 | 0 | 2 | 0 | 2 | 0 | 14 | 0.1429 | 2 | 0 | 3 | 0 | 7 | 0.1429 | 2 | 0 | 14 | 0.0714 |
| 3_51402974 | 6 | 0.1667 | 4 | 0 | 2 | 0 | 2 | 0 | 14 | 0.0714 | 3 | 0 | 3 | 0 | 6 | 0 | 2 | 0.5 | 14 | 0.0714 |
| 3_51403027 | 7 | 0.1429 | 4 | 0.25 | 2 | 1 | 2 | 0 | 15 | 0.2667 | 3 | 0 | 3 | 0.3333 | 6 | 0 | 2 | 1 | 14 | 0.2143 |
| 3_51423338 | 13 | 0 | 14 | 0.1429 | 12 | 0.0833 | 12 | 0.1667 | 51 | 0.098 | 4 | 0.25 | 10 | 0.1 | 11 | 0.0909 | 7 | 0 | 32 | 0.0938 |
| 3_51425328 | 5 | 0 | 8 | 0.25 | 5 | 0 | 9 | 0 | 27 | 0.0741 | 4 | 0 | 5 | 0 | 6 | 0.1667 | 3 | 0 | 18 | 0.0556 |
| 3_51425578 | 5 | 0.2 | 3 | 0 | 4 | 0 | 8 | 0 | 20 | 0.05 | 4 | 0 | 4 | 0.25 | 5 | 0 | 3 | 0 | 16 | 0.0625 |
| 3_51426284 | 5 | 0 | 4 | 0 | 3 | 0.3333 | 4 | 0 | 16 | 0.0625 | 3 | 0 | 5 | 0 | 6 | 0.1667 | 3 | 0 | 17 | 0.0588 |

| | | | | | | | | | | | | | | | | | | | | |
|---|---|---|---|---|---|---|---|---|---|---|---|---|---|---|---|---|---|---|---|---|
| 3_51426878 | 4 | 0 | 5 | 0.2 | 3 | 0 | 6 | 0 | 18 | 0.0556 | 3 | 0 | 4 | 0 | 5 | 0.2 | 3 | 0 | 15 | 0.0667 |
| 3_51426905 | 3 | 0.3333 | 3 | 0 | 1 | 0 | 4 | 0 | 11 | 0.0909 | 1 | 0 | 1 | 0 | 4 | 0.5 | 2 | 0 | 8 | 0.25 |
| 3_51428446 | 2 | 0.5 | 2 | 0 | 2 | 0 | 5 | 0.4 | 11 | 0.2727 | 1 | 0 | 4 | 0 | 3 | 0 | 1 | 1 | 9 | 0.1111 |
| 3_51428516 | 1 | 0 | 2 | 0 | 2 | 0.5 | 3 | 0.3333 | 8 | 0.25 | 1 | 0 | 4 | 0.25 | 3 | 0 | 2 | 0 | 10 | 0.1 |
| 3_51428644 | 1 | 1 | 2 | 0.5 | 3 | 1 | 5 | 0.8 | 11 | 0.8182 | 1 | 1 | 3 | 1 | 4 | 0.75 | 2 | 0.5 | 10 | 0.8 |
| 3_51428654 | 1 | 0 | 2 | 0 | 3 | 0 | 5 | 0.2 | 11 | 0.0909 | 1 | 0 | 2 | 0 | 3 | 0 | 2 | 0.5 | 8 | 0.125 |
| 3_51428690 | 1 | 0 | 2 | 0.5 | 3 | 0.3333 | 5 | 0.2 | 11 | 0.2727 | 1 | 0 | 3 | 0.3333 | 4 | 0.25 | 2 | 0.5 | 10 | 0.3 |
| 3_51428703 | 1 | 0 | 2 | 0.5 | 3 | 0 | 5 | 0 | 11 | 0.0909 | 1 | 0 | 3 | 0.3333 | 4 | 0 | 2 | 0 | 10 | 0.1 |
| 3_51468149 | NA | NA | 1 | 1 | NA | NA | NA | NA | 1 | 1 | NA | NA | NA | NA | 2 | 0.5 | NA | NA | 2 | 0.5 |
| 3_51595447 | NA | NA | 2 | 1 | 1 | 1 | NA | NA | 3 | 1 | NA | NA | 1 | 1 | NA | NA | NA | NA | 1 | 1 |
| 3_51595657 | 1 | 1 | 3 | 0.6667 | 2 | 1 | 1 | 1 | 7 | 0.8571 | NA | NA | 1 | 1 | NA | NA | NA | NA | 1 | 1 |
| 3_51595926 | 1 | 1 | 2 | 1 | 1 | 1 | 1 | 1 | 5 | 1 | NA | NA | 1 | 1 | NA | NA | NA | NA | 1 | 1 |
| 3_51596039 | NA | NA | 3 | 1 | 1 | 1 | 1 | 1 | 5 | 1 | NA | NA | 1 | 1 | NA | NA | NA | NA | 1 | 1 |
| 3_51596211 | 1 | 1 | 3 | 1 | 2 | 0.5 | 1 | 1 | 7 | 0.8571 | NA | NA | 1 | 1 | NA | NA | NA | NA | 1 | 1 |
| 3_51596747 | 1 | 1 | 3 | 1 | 1 | 1 | 1 | 0 | 6 | 0.8333 | NA | NA | 1 | 1 | NA | NA | NA | NA | 1 | 1 |
| 3_52176260 | 1 | 1 | NA | NA | NA | NA | 1 | 1 | 2 | 1 | NA | NA | 1 | 1 | 2 | 1 | 3 | 0.6667 | 6 | 0.8333 |
| 3_52176461 | NA | NA | NA | NA | 2 | 0.5 | 2 | 0 | 4 | 0.25 | NA | NA | 3 | 0 | 3 | 0.6667 | 3 | 0.6667 | 9 | 0.4444 |
| 3_52176530 | 1 | 1 | NA | NA | 1 | 0 | 2 | 0 | 4 | 0.25 | NA | NA | 4 | 0.75 | 2 | 0 | 3 | 0.6667 | 9 | 0.5556 |
| 3_52252520 | 1 | 0 | 3 | 0.3333 | 1 | 0 | 3 | 0 | 8 | 0.125 | 1 | 0 | 5 | 0.2 | 3 | 0.6667 | 3 | 0.3333 | 12 | 0.3333 |
| 3_52252602 | 1 | 0 | 1 | 0 | 3 | 0.6667 | 3 | 0.3333 | 8 | 0.375 | 1 | 1 | 6 | 0.6667 | 3 | 0.6667 | 2 | 1 | 12 | 0.75 |
| 3_52252746 | 2 | 0 | 3 | 0.3333 | 1 | 1 | 2 | 1 | 8 | 0.5 | 1 | 1 | 5 | 0.8 | 2 | 0.5 | 2 | 1 | 10 | 0.8 |
| 3_52253150 | 2 | 0 | 3 | 0.3333 | 4 | 0 | 2 | 0 | 11 | 0.0909 | 2 | 0 | 8 | 0.125 | 3 | 0.3333 | 5 | 0 | 18 | 0.1111 |
| 3_52253160 | 2 | 0.5 | 3 | 0 | 3 | 0.3333 | 3 | 0.6667 | 11 | 0.3636 | 2 | 1 | 7 | 0.5714 | 2 | 1 | 5 | 0.2 | 16 | 0.5625 |
| 3_52253196 | 2 | 0 | 2 | 0 | 3 | 0.6667 | 3 | 0 | 10 | 0.2 | 2 | 0 | 7 | 0.4286 | 2 | 0 | 4 | 0 | 15 | 0.2 |
| 3_52253209 | 2 | 1 | 2 | 1 | 4 | 1 | 3 | 1 | 11 | 1 | 2 | 1 | 9 | 1 | 2 | 1 | 5 | 1 | 18 | 1 |
| 3_52253228 | 2 | 0.5 | 3 | 0 | 2 | 0.5 | 3 | 1 | 10 | 0.5 | 2 | 0 | 6 | 0.3333 | 3 | 0.6667 | 5 | 0.6 | 16 | 0.4375 |
| 3_52253235 | 2 | 0.5 | 3 | 0.6667 | 4 | 0.5 | 1 | 0 | 10 | 0.5 | NA | NA | 8 | 0.625 | 3 | 0.3333 | 3 | 0.6667 | 14 | 0.5714 |
| 3_52253253 | 1 | 1 | 1 | 1 | 1 | 1 | 3 | 0.6667 | 6 | 0.8333 | 2 | 1 | 4 | 0.5 | 3 | 0.3333 | 4 | 0.75 | 13 | 0.6154 |
| 3_52253382 | 2 | 0 | 3 | 0 | 3 | 0.3333 | 2 | 0.5 | 10 | 0.2 | 2 | 0 | 7 | 0 | 2 | 0 | 4 | 0.25 | 15 | 0.0667 |
| 3_52253407 | 1 | 0 | 3 | 1 | 2 | 1 | 3 | 0.6667 | 9 | 0.7778 | 2 | 1 | 6 | 1 | 3 | 1 | 4 | 1 | 15 | 1 |
| 3_52253487 | 2 | 0 | 3 | 0.3333 | 3 | 0.6667 | 3 | 0 | 11 | 0.2727 | 2 | 0.5 | 9 | 0.3333 | 3 | 0.3333 | 5 | 0 | 19 | 0.2632 |
| 3_52253631 | 2 | 1 | 3 | 0.3333 | 2 | 1 | 3 | 1 | 10 | 0.8 | 1 | 1 | 11 | 0.6364 | 3 | 0.6667 | 4 | 0.75 | 19 | 0.6842 |
| 3_52253661 | 2 | 1 | 1 | 1 | 3 | 1 | 3 | 1 | 9 | 1 | NA | NA | 9 | 1 | 3 | 1 | 4 | 1 | 16 | 1 |
| 3_52254456 | 2 | 1 | 3 | 0 | 4 | 0.25 | 3 | 0.3333 | 12 | 0.3333 | 2 | 0 | 11 | 0.1818 | 3 | 0 | 4 | 0 | 20 | 0.1 |
| 3_52254555 | 2 | 1 | 4 | 0.25 | 4 | 0.5 | 2 | 0.5 | 12 | 0.5 | 2 | 0 | 12 | 0.9167 | 3 | 1 | 5 | 1 | 22 | 0.8636 |
| 3_52254633 | 2 | 1 | 5 | 0.8 | 5 | 1 | 3 | 1 | 15 | 0.9333 | 2 | 1 | 14 | 1 | 3 | 1 | 4 | 1 | 23 | 1 |
| 3_52254698 | 2 | 1 | 5 | 1 | 5 | 1 | 3 | 1 | 15 | 1 | 2 | 1 | 13 | 1 | 4 | 1 | 6 | 1 | 25 | 1 |
| 3_52254843 | 2 | 1 | 4 | 0.25 | 5 | 0.2 | 3 | 0 | 14 | 0.2857 | 2 | 0.5 | 14 | 0.5714 | 4 | 0.5 | 5 | 0.4 | 25 | 0.52 |
| 3_52254889 | 2 | 1 | 2 | 0.5 | 2 | 0 | 4 | 0.5 | 10 | 0.5 | 2 | 0.5 | 9 | 0.4444 | 4 | 0.5 | 6 | 0.6667 | 21 | 0.5238 |
| 3_52254993 | 1 | 1 | 4 | 1 | 4 | 1 | 4 | 1 | 13 | 1 | 2 | 1 | 13 | 1 | 3 | 1 | 6 | 1 | 24 | 1 |
| 3_52255002 | 2 | 0.5 | 6 | 0.5 | 3 | 0.6667 | 4 | 0.75 | 15 | 0.6 | 1 | 1 | 11 | 0.6364 | 3 | 0.6667 | 5 | 0.8 | 20 | 0.7 |
| 3_52255131 | 2 | 0.5 | 6 | 0.1667 | 5 | 0.2 | 3 | 0 | 16 | 0.1875 | 2 | 0.5 | 13 | 0 | 6 | 0.1667 | 5 | 0 | 26 | 0.0769 |
| 3_52255152 | 2 | 0 | 5 | 0 | 5 | 0.6 | 4 | 0.25 | 16 | 0.25 | 2 | 0 | 11 | 0.4545 | 6 | 0.1667 | 6 | 0.1667 | 25 | 0.28 |
| 3_52255656 | 3 | 0 | 6 | 0.1667 | 4 | 0.5 | 3 | 0.3333 | 16 | 0.25 | 4 | 0.25 | 14 | 0 | 6 | 0 | 5 | 0.2 | 29 | 0.069 |
| 3_52255947 | 3 | 0.3333 | 6 | 0.1667 | 6 | 0.5 | 4 | 0.5 | 19 | 0.3684 | 3 | 0.3333 | 15 | 0.2 | 6 | 0.1667 | 6 | 0.5 | 30 | 0.2667 |
| 3_52256406 | 7 | 0.1429 | 7 | 0.1429 | 7 | 0 | 3 | 0.6667 | 24 | 0.1667 | 3 | 0.3333 | 18 | 0.1111 | 7 | 0.1429 | 6 | 0 | 34 | 0.1176 |
| 3_52256845 | 8 | 0 | 8 | 0.25 | 8 | 0 | 6 | 0.5 | 30 | 0.1667 | 5 | 0.4 | 18 | 0.0556 | 10 | 0.1 | 6 | 0.1667 | 39 | 0.1282 |
| 3_52257386 | 4 | 0.5 | 6 | 0.5 | 7 | 0.1429 | 5 | 0.2 | 22 | 0.3182 | 5 | 0.8 | 17 | 0.2941 | 8 | 0.375 | 5 | 0.4 | 35 | 0.4 |
| 3_52901570 | NA | NA | NA | NA | NA | NA | 1 | 1 | 1 | 1 | NA | NA | NA | NA | 1 | 1 | 1 | 1 | 2 | 1 |
| 3_52901654 | NA | NA | 1 | 1 | NA | NA | 1 | 1 | 2 | 1 | NA | NA | NA | NA | NA | NA | 1 | 1 | 1 | 1 |
| 3_52901770 | NA | NA | NA | NA | NA | NA | 1 | 1 | 1 | 1 | NA | NA | NA | NA | 1 | 1 | NA | NA | 1 | 1 |
| 3_52903433 | 6 | 0 | 5 | 0 | 1 | 1 | 6 | 0.1667 | 18 | 0.1111 | 3 | 0 | 4 | 0 | 5 | 0 | 2 | 0.5 | 14 | 0.0714 |
| 3_52950661 | 2 | 0 | 4 | 0 | 3 | 0 | 2 | 0.5 | 11 | 0.0909 | 1 | 0 | NA | NA | 3 | 0 | 3 | 0.3333 | 7 | 0.1429 |
| 3_52950702 | 2 | 1 | 3 | 1 | 3 | 0.3333 | 1 | 0 | 9 | 0.6667 | 1 | 0 | NA | NA | 4 | 0.75 | 2 | 0.5 | 7 | 0.5714 |
| 3_52950743 | 2 | 0.5 | 4 | 0.25 | 3 | 0.6667 | 3 | 0.3333 | 12 | 0.4167 | 1 | 0 | NA | NA | 5 | 0.2 | 2 | 0 | 8 | 0.125 |
| 3_52951060 | 3 | 0.6667 | 5 | 0 | 3 | 0 | 3 | 0 | 14 | 0.1429 | 1 | 0 | NA | NA | 5 | 0.2 | 2 | 0 | 8 | 0.125 |
| 3_52951073 | 3 | 0.3333 | 5 | 0 | 3 | 0 | 3 | 0 | 14 | 0.0714 | NA | NA | NA | NA | 5 | 0 | 3 | 0.3333 | 8 | 0.125 |
| 3_53168169 | 1 | 0 | 6 | 0 | 2 | 0 | 4 | 0.5 | 13 | 0.1538 | 2 | 0 | NA | NA | 6 | 0.1667 | 2 | 0 | 10 | 0.1 |
| 3_53386163 | NA | NA | 2 | 0 | 1 | 1 | 1 | 0 | 4 | 0.25 | NA | NA | 3 | 0 | 1 | 1 | NA | NA | 4 | 0.25 |
| 3_53386993 | NA | NA | 3 | 0 | 2 | 0 | 2 | 0.5 | 7 | 0.1429 | 2 | 0 | 3 | 0.3333 | 1 | 0 | NA | NA | 6 | 0.1667 |
| 3_53388100 | NA | NA | 3 | 0.3333 | 2 | 0 | 1 | 0 | 6 | 0.1667 | 2 | 0 | 5 | 0.2 | 1 | 1 | NA | NA | 8 | 0.25 |
| 3_53761269 | 5 | 0.2 | 2 | 0 | 1 | 0 | 1 | 0 | 9 | 0.1111 | 1 | 0 | NA | NA | 1 | 0 | 1 | 1 | 3 | 0.3333 |
| 3_53764881 | 2 | 0 | 9 | 0.1111 | 6 | 0.1667 | 5 | 0 | 22 | 0.0909 | 6 | 0 | 11 | 0.0909 | 2 | 0 | 4 | 0.25 | 23 | 0.087 |
| 3_53765093 | 5 | 1 | 9 | 0.8889 | 7 | 1 | 5 | 1 | 26 | 0.9615 | 6 | 1 | 9 | 1 | 5 | 1 | 7 | 1 | 27 | 1 |
| 3_53765137 | 5 | 0 | 9 | 0 | 8 | 0.25 | 5 | 0 | 27 | 0.0741 | 5 | 0 | 10 | 0 | 5 | 0 | 6 | 0.3333 | 26 | 0.0769 |

| | | | | | | | | | | | | | | | | | | | | |
|---|---|---|---|---|---|---|---|---|---|---|---|---|---|---|---|---|---|---|---|---|
| 3_5376536 | 5 | 0.2 | 11 | 0 | 9 | 0 | 5 | 0.2 | 30 | 0.0667 | 5 | 0.2 | 11 | 0.0909 | 8 | 0 | 10 | 0 | 34 | 0.0588 |
| 3_5376538 | 6 | 0.3333 | 6 | 0 | 5 | 0 | 5 | 0.2 | 22 | 0.1364 | 5 | 0.4 | 10 | 0.1 | 7 | 0.1429 | 8 | 0.125 | 30 | 0.1667 |
| 3_5410223 | 1 | 1 | NA | NA | NA | NA | NA | NA | 1 | 1 | NA | NA | 1 | 1 | NA | NA | 1 | 1 | 2 | 1 |
| 3_5410236 | 1 | 1 | NA | NA | NA | NA | NA | NA | 1 | 1 | NA | NA | NA | NA | 1 | 1 | 1 | 0 | 2 | 0.5 |
| 3_5412999 | 2 | 0.5 | NA | NA | NA | NA | NA | NA | 2 | 0.5 | NA | NA | 1 | 1 | 1 | 1 | 2 | 1 | 4 | 1 |
| 3_5422508 | 1 | 1 | 1 | 1 | NA | NA | NA | NA | 2 | 1 | 3 | 1 | 1 | 0 | 2 | 1 | 1 | 1 | 7 | 0.8571 |
| 3_5422517 | 1 | 1 | 1 | 1 | NA | NA | NA | NA | 2 | 1 | 2 | 1 | 1 | 1 | 1 | 1 | NA | NA | 4 | 1 |
| 3_5422529 | 3 | 0.6667 | 2 | 1 | NA | NA | NA | NA | 5 | 0.8 | 2 | 1 | NA | NA | 1 | 1 | 2 | 1 | 5 | 1 |
| 3_5422535 | 2 | 1 | 2 | 1 | NA | NA | NA | NA | 4 | 1 | 2 | 1 | 1 | 1 | NA | NA | 2 | 1 | 5 | 1 |
| 3_5422539 | 3 | 1 | 2 | 1 | NA | NA | NA | NA | 5 | 1 | 3 | 1 | 1 | 1 | 1 | 1 | NA | NA | 5 | 1 |
| 3_5422550 | 4 | 1 | 3 | 0.6667 | NA | NA | NA | NA | 7 | 0.8571 | 3 | 1 | 1 | 0 | 2 | 1 | 2 | 1 | 8 | 0.875 |
| 3_5422551 | 4 | 1 | 3 | 1 | NA | NA | NA | NA | 7 | 1 | 2 | 1 | 1 | 1 | 2 | 1 | 2 | 1 | 7 | 1 |
| 3_5422553 | 4 | 0 | 4 | 1 | NA | NA | NA | NA | 8 | 0.5 | 2 | 0.5 | 1 | 0 | 1 | 1 | 2 | 0.5 | 6 | 0.5 |
| 3_5422557 | 4 | 0.25 | 4 | 0 | NA | NA | NA | NA | 8 | 0.125 | 3 | 0 | 1 | 0 | 2 | 0.5 | 2 | 0 | 8 | 0.125 |
| 3_5439006 | 3 | 0.6667 | 2 | 0.5 | 3 | 1 | 4 | 0.25 | 12 | 0.5833 | 1 | 0 | NA | NA | 6 | 0.1667 | 4 | 0.25 | 11 | 0.1818 |
| 3_5439014 | 4 | 0.5 | 3 | 0.6667 | 4 | 0.75 | 3 | 1 | 14 | 0.7143 | 1 | 1 | 2 | 0.5 | 6 | 0.3333 | 4 | 0.25 | 13 | 0.3846 |
| 3_5462490 | NA | NA | 1 | 0 | 1 | 0 | 3 | 0.3333 | 5 | 0.2 | 2 | 0 | 1 | 1 | NA | NA | 2 | 0 | 5 | 0.2 |
| 3_5463609 | 1 | 1 | 6 | 0 | 3 | 0.3333 | 8 | 0.25 | 18 | 0.2222 | 4 | 0 | 4 | 0.25 | 3 | 0 | 3 | 0 | 14 | 0.0714 |
| 3_5464175 | 4 | 0 | 2 | 0 | 1 | 0 | 5 | 0.2 | 12 | 0.0833 | NA | NA | 3 | 0 | 3 | 0 | 3 | 0.6667 | 9 | 0.2222 |
| 3_5464199 | NA | NA | 1 | 1 | 1 | 1 | NA | NA | 2 | 1 | NA | NA | 1 | 1 | NA | NA | 2 | 0 | 3 | 0.3333 |
| 3_5464223 | NA | NA | NA | NA | 1 | 1 | NA | NA | 1 | 1 | NA | NA | 1 | 0 | NA | NA | 2 | 1 | 3 | 0.6667 |
| 3_5464230 | NA | NA | NA | NA | 1 | 1 | NA | NA | 1 | 1 | NA | NA | 1 | 0 | NA | NA | 1 | 1 | 2 | 0.5 |
| 3_5465678 | 6 | 0.6667 | 4 | 0.25 | 2 | 1 | 1 | 1 | 13 | 0.6154 | 4 | 0.25 | 2 | 0.5 | 1 | 0 | 3 | 0.3333 | 10 | 0.3 |
| 3_5465701 | 1 | 1 | 1 | 0 | 3 | 0.6667 | 1 | 1 | 6 | 0.6667 | 4 | 0.25 | 1 | 0 | 1 | 0 | 4 | 0.75 | 10 | 0.4 |
| 3_5465715 | 1 | 1 | 1 | 0 | 3 | 0.3333 | 1 | 0 | 6 | 0.3333 | 4 | 0 | 1 | 1 | 1 | 0 | 4 | 0.25 | 10 | 0.2 |
| 3_5470485 | 2 | 0 | 1 | 1 | NA | NA | NA | NA | 3 | 0.3333 | NA | NA | 1 | 1 | NA | NA | NA | NA | 1 | 1 |
| 3_5470495 | 1 | 0 | NA | NA | 1 | 1 | NA | NA | 2 | 0.5 | 1 | 1 | 1 | 1 | 1 | 1 | 1 | 1 | 4 | 1 |
| 3_5470517 | 2 | 0 | 1 | 1 | 1 | 1 | NA | NA | 4 | 0.5 | 1 | 0 | 1 | 1 | 1 | 0 | 1 | 1 | 4 | 0.5 |
| 3_5470617 | 2 | 0 | 1 | 0 | 1 | 1 | NA | NA | 4 | 0.25 | 1 | 0 | 1 | 0 | 2 | 0.5 | 1 | 0 | 5 | 0.2 |
| 3_5471116 | NA | NA | 2 | 0 | 2 | 0 | 5 | 0.2 | 9 | 0.1111 | 2 | 0 | 4 | 0 | 2 | 0.5 | 1 | 0 | 9 | 0.1111 |
| 3_5471144 | 1 | 1 | 1 | 1 | 2 | 1 | 6 | 1 | 10 | 1 | 1 | 0 | 4 | 1 | 2 | 0.5 | 1 | 0 | 8 | 0.625 |
| 3_5471167 | 3 | 0.3333 | 7 | 0 | 4 | 0 | 6 | 0 | 20 | 0.05 | 3 | 0 | 9 | 0.1111 | 3 | 0 | 3 | 0 | 18 | 0.0556 |
| 3_5471177 | 2 | 0.5 | 7 | 0 | 1 | 0 | 6 | 0.3333 | 16 | 0.1875 | 3 | 0 | 6 | 0.1667 | 2 | 0.5 | 3 | 0.3333 | 14 | 0.2143 |
| 3_5471177 | 1 | 0 | NA | NA | 2 | 0 | 1 | 1 | 4 | 0.25 | NA | NA | 1 | 0 | 1 | 1 | NA | NA | 2 | 0.5 |
| 3_5471178 | 3 | 0.3333 | 10 | 0 | 4 | 0.25 | 8 | 0.5 | 25 | 0.24 | 3 | 0.3333 | 10 | 0.2 | 3 | 0 | 4 | 0.5 | 20 | 0.25 |
| 3_5471180 | 3 | 1 | 8 | 0.5 | 4 | 0.75 | 7 | 0.7143 | 22 | 0.6818 | 2 | 0.5 | 6 | 0.5 | 2 | 1 | 4 | 0.75 | 14 | 0.6429 |
| 3_5471182 | 3 | 0.3333 | 8 | 0.25 | 4 | 0.5 | 8 | 0.25 | 23 | 0.3043 | 3 | 0.3333 | 8 | 0.375 | 2 | 0 | 3 | 0.6667 | 16 | 0.375 |
| 3_5471278 | 3 | 0.3333 | 10 | 0 | 4 | 0.25 | 7 | 0.1429 | 24 | 0.125 | 3 | 0.3333 | 9 | 0 | 3 | 0 | 4 | 0 | 19 | 0.0526 |
| 3_5480453 | NA | NA | 3 | 0.6667 | 1 | 0 | 2 | 0 | 6 | 0.3333 | 1 | 0 | NA | NA | 1 | 0 | 2 | 0.5 | 4 | 0.25 |
| 3_5480457 | NA | NA | 2 | 0.5 | 1 | 0 | 2 | 0 | 5 | 0.2 | 1 | 0 | NA | NA | 2 | 0.5 | 3 | 0.3333 | 6 | 0.3333 |
| 3_5480461 | NA | NA | 1 | 0 | 1 | 1 | 1 | 1 | 3 | 0.6667 | NA | NA | NA | NA | NA | NA | 3 | 1 | 3 | 1 |
| 3_5480469 | NA | NA | NA | NA | 1 | 0 | 1 | 1 | 2 | 0.5 | 1 | 1 | NA | NA | 2 | 0 | 3 | 0 | 6 | 0.1667 |
| 3_5480472 | NA | NA | 2 | 1 | 1 | 1 | 2 | 1 | 5 | 1 | NA | NA | NA | NA | 2 | 1 | 2 | 1 | 4 | 1 |
| 3_5480481 | NA | NA | 3 | 0.6667 | 1 | 0 | 2 | 1 | 6 | 0.6667 | 1 | 1 | NA | NA | 1 | 1 | 1 | 0 | 3 | 0.6667 |
| 3_5480490 | NA | NA | 2 | 0.5 | 1 | 0 | 1 | 0 | 4 | 0.25 | NA | NA | NA | NA | 1 | 0 | 3 | 0.6667 | 4 | 0.5 |
| 3_5480505 | NA | NA | 2 | 1 | 1 | 1 | 1 | 1 | 4 | 1 | 1 | 1 | NA | NA | NA | NA | 1 | 1 | 2 | 1 |
| 3_5480531 | NA | NA | 2 | 1 | 1 | 1 | 1 | 0 | 4 | 0.75 | 1 | 0 | NA | NA | 1 | 1 | 3 | 0 | 5 | 0.2 |
| 3_5480548 | NA | NA | 2 | 0.5 | 1 | 0 | NA | NA | 3 | 0.3333 | 1 | 1 | NA | NA | NA | NA | 2 | 0.5 | 3 | 0.6667 |
| 3_5480577 | NA | NA | 3 | 0.6667 | 1 | 0 | 1 | 1 | 5 | 0.6 | 1 | 0 | NA | NA | 1 | 0 | 2 | 0.5 | 4 | 0.25 |
| 3_5480591 | NA | NA | 2 | 1 | 1 | 1 | 1 | 0 | 4 | 0.75 | 1 | 1 | NA | NA | 1 | 0 | 2 | 0 | 4 | 0.25 |
| 3_5480615 | NA | NA | NA | NA | 1 | 1 | 1 | 1 | 2 | 1 | 1 | 1 | NA | NA | 1 | 1 | 2 | 1 | 4 | 1 |
| 3_5480634 | NA | NA | NA | NA | NA | NA | 1 | 1 | 1 | 1 | 1 | 1 | NA | NA | NA | NA | 1 | 1 | 2 | 1 |
| 3_5480640 | NA | NA | NA | NA | NA | NA | 1 | 1 | 1 | 1 | 1 | 1 | NA | NA | NA | NA | 1 | 1 | 2 | 1 |
| 3_5480642 | NA | NA | NA | NA | NA | NA | 1 | 1 | 1 | 1 | 1 | 1 | NA | NA | NA | NA | 1 | 1 | 2 | 1 |
| 3_5502449 | 2 | 0 | 2 | 0.5 | 1 | 1 | 4 | 1 | 9 | 0.6667 | 3 | 0.6667 | 3 | 0.6667 | 2 | 1 | 3 | 0.6667 | 11 | 0.7273 |
| 3_5502453 | 2 | 0.5 | 2 | 0 | 2 | 0.5 | 4 | 0.5 | 10 | 0.4 | 2 | 0.5 | 4 | 0.25 | 2 | 1 | 3 | 0.3333 | 11 | 0.4545 |
| 3_5502455 | 1 | 0 | 1 | 1 | NA | NA | 4 | 0.25 | 6 | 0.3333 | 2 | 0 | 4 | 0.5 | 1 | 0 | 3 | 1 | 10 | 0.5 |
| 3_5502504 | 2 | 0.5 | 2 | 0.5 | 2 | 0 | 6 | 0.6667 | 12 | 0.5 | 4 | 0.75 | 3 | 0.3333 | 2 | 0.5 | 3 | 0.3333 | 12 | 0.5 |
| 3_5504296 | 1 | 0 | 4 | 0.5 | 2 | 0 | 9 | 0.3333 | 16 | 0.3125 | 4 | 0.5 | 6 | 0.5 | 2 | 0 | 4 | 0.5 | 16 | 0.4375 |
| 3_5504297 | 1 | 1 | 4 | 0 | 2 | 0 | 8 | 0.125 | 15 | 0.1333 | 4 | 0.25 | 6 | 0.1667 | 2 | 0.5 | 4 | 0.25 | 16 | 0.25 |
| 3_5504309 | 3 | 1 | 4 | 0.75 | 2 | 1 | 8 | 0.75 | 17 | 0.8235 | 4 | 1 | 6 | 0.6667 | 1 | 1 | 5 | 1 | 16 | 0.875 |
| 3_5504321 | 3 | 1 | 3 | 0.6667 | 2 | 1 | 8 | 1 | 16 | 0.9375 | 3 | 1 | 4 | 1 | 1 | 1 | 5 | 1 | 13 | 1 |
| 3_5504348 | 3 | 1 | 4 | 0.75 | 1 | 0 | 7 | 1 | 15 | 0.8667 | 4 | 1 | 5 | 1 | 3 | 1 | 3 | 1 | 15 | 1 |
| 3_5504356 | 3 | 0.3333 | 4 | 0.5 | 2 | 0 | 6 | 0.8333 | 15 | 0.5333 | 2 | 1 | 6 | 0.1667 | 3 | 0.6667 | 4 | 0.75 | 15 | 0.5333 |
| 3_5504360 | 3 | 0 | 5 | 0.8 | 2 | 0.5 | 6 | 0.3333 | 16 | 0.4375 | 3 | 0.6667 | 5 | 0.2 | 3 | 0.6667 | 3 | 0 | 14 | 0.3571 |

| | | | | | | | | | | | | | | | | | | | | |
|---|---|---|---|---|---|---|---|---|---|---|---|---|---|---|---|---|---|---|---|---|
| 3_5504375 | 4 | 0.25 | 5 | 0.4 | 1 | 0 | 7 | 0.2857 | 17 | 0.2941 | 3 | 0 | 6 | 0 | 2 | 0.5 | 4 | 0 | 15 | 0.0667 |
| 3_5504399 | 4 | 0.5 | 5 | 0 | 2 | 0 | 7 | 0 | 18 | 0.1111 | 4 | 0 | 6 | 0.1667 | 3 | 0 | 5 | 0 | 18 | 0.0556 |
| 3_5504400 | 4 | 0 | 5 | 0 | 2 | 0.5 | 7 | 0 | 18 | 0.0556 | 3 | 0.3333 | 5 | 0 | 2 | 0 | 5 | 0.2 | 15 | 0.1333 |
| 3_5504410 | 3 | 0.3333 | 5 | 0.4 | 2 | 1 | 7 | 0.2857 | 17 | 0.4118 | 4 | 0.25 | 4 | 0.25 | 2 | 0 | 5 | 0.2 | 15 | 0.2 |
| 3_5504419 | 3 | 0.3333 | 4 | 0.75 | 2 | 0.5 | 4 | 0.75 | 13 | 0.6154 | 4 | 0.75 | 4 | 0.75 | 3 | 0.3333 | 5 | 1 | 16 | 0.75 |
| 3_5504427 | 4 | 0 | 5 | 0 | 2 | 0 | 7 | 0.1429 | 18 | 0.0556 | 5 | 0.2 | 5 | 0 | 3 | 0.3333 | 5 | 0.6 | 18 | 0.2778 |
| 3_5504449 | 4 | 0.25 | 7 | 0 | 2 | 0 | 7 | 0.1429 | 20 | 0.1 | 5 | 0 | 6 | 0.1667 | 5 | 0 | 5 | 0.4 | 21 | 0.1429 |
| 3_5504453 | 4 | 0.25 | 7 | 0.1429 | 2 | 0 | 5 | 0.2 | 18 | 0.1667 | 4 | 0 | 6 | 0 | 3 | 0.6667 | 5 | 0.2 | 18 | 0.1667 |
| 3_5515484 | 4 | 0.5 | 3 | 0 | 3 | 0 | 2 | 0 | 12 | 0.1667 | 4 | 0 | 4 | 0 | 2 | 0.5 | 4 | 0 | 14 | 0.0714 |
| 3_5515485 | 2 | 0 | 1 | 1 | 3 | 0 | NA | NA | 6 | 0.1667 | 3 | 1 | 3 | 0.6667 | NA | NA | 2 | 1 | 8 | 0.875 |
| 3_5537546 | NA | NA | NA | NA | 4 | 0 | 4 | 0.25 | 8 | 0.125 | 2 | 0.5 | 2 | 0 | NA | NA | 3 | 0 | 7 | 0.1429 |
| 3_5537571 | NA | NA | 2 | 0 | 4 | 0.5 | 4 | 0.25 | 10 | 0.3 | 2 | 0.5 | 2 | 0.5 | NA | NA | 3 | 0.3333 | 7 | 0.4286 |
| 3_5537575 | NA | NA | 3 | 0.3333 | 4 | 0 | 3 | 1 | 10 | 0.4 | 2 | 0.5 | 2 | 1 | NA | NA | 2 | 1 | 6 | 0.8333 |
| 3_5537577 | NA | NA | 3 | 0.6667 | 4 | 0 | 4 | 0.25 | 11 | 0.2727 | 2 | 0.5 | 2 | 0 | NA | NA | 3 | 0.3333 | 7 | 0.2857 |
| 3_5537836 | 3 | 0.3333 | 8 | 0.625 | 6 | 0.3333 | 4 | 0.5 | 21 | 0.4762 | 4 | 0.75 | 2 | 1 | 5 | 0.4 | 4 | 0.75 | 15 | 0.6667 |
| 3_5542960 | NA | NA | NA | NA | 3 | 0.6667 | 1 | 0 | 4 | 0.5 | 2 | 1 | 5 | 0.6 | 3 | 0.3333 | 1 | 1 | 11 | 0.6364 |
| 3_5543221 | NA | NA | 1 | 1 | NA | NA | NA | NA | 1 | 1 | NA | NA | 1 | 1 | 1 | 0 | 1 | 0 | 3 | 0.3333 |
| 3_5543223 | NA | NA | 1 | 1 | NA | NA | NA | NA | 1 | 1 | NA | NA | 1 | 1 | 1 | 0 | 1 | 0 | 3 | 0.3333 |
| 3_5544112 | 35 | 0.4857 | 57 | 0.4211 | 49 | 0.5306 | 42 | 0.4524 | 183 | 0.4699 | 28 | 0.5 | 32 | 0.4688 | 45 | 0.4889 | 37 | 0.5676 | 142 | 0.507 |
| 3_5544119 | 36 | 0.4167 | 53 | 0.4717 | 48 | 0.5 | 42 | 0.5 | 179 | 0.4749 | 29 | 0.5172 | 30 | 0.4667 | 43 | 0.2558 | 36 | 0.4167 | 138 | 0.3986 |
| 3_5544431 | 10 | 0.2 | 22 | 0.1364 | 11 | 0.0909 | 8 | 0.375 | 51 | 0.1765 | 12 | 0.3333 | 16 | 0.25 | 11 | 0.3636 | 15 | 0.0667 | 54 | 0.2407 |
| 3_5544489 | 16 | 0.0625 | 21 | 0.0476 | 12 | 0.0833 | 9 | 0.1111 | 58 | 0.069 | 14 | 0.0714 | 17 | 0 | 16 | 0.0625 | 16 | 0.1875 | 63 | 0.0794 |
| 3_5544490 | 16 | 0.125 | 20 | 0.05 | 13 | 0 | 8 | 0 | 57 | 0.0526 | 13 | 0.2308 | 18 | 0.0556 | 11 | 0 | 15 | 0.0667 | 57 | 0.0877 |
| 3_5544497 | 12 | 0.25 | 15 | 0.2 | 9 | 0 | 8 | 0 | 44 | 0.1364 | 9 | 0.1111 | 15 | 0.2 | 10 | 0.3 | 13 | 0.0769 | 47 | 0.1702 |
| 3_5544521 | 16 | 0.3125 | 27 | 0.2963 | 15 | 0.2 | 10 | 0.2 | 68 | 0.2647 | 15 | 0.0667 | 21 | 0.0952 | 17 | 0.2941 | 21 | 0.2381 | 74 | 0.1757 |
| 3_5544529 | 14 | 0.2143 | 24 | 0.25 | 17 | 0.3529 | 11 | 0.3636 | 66 | 0.2879 | 10 | 0.3 | 18 | 0.2222 | 16 | 0.1875 | 19 | 0.1579 | 63 | 0.2063 |
| 3_5544530 | 16 | 0.125 | 25 | 0.04 | 19 | 0.1053 | 9 | 0 | 69 | 0.0725 | 16 | 0.0625 | 24 | 0.0833 | 15 | 0.1333 | 21 | 0.0476 | 76 | 0.0789 |
| 3_5553344 | 7 | 0 | 13 | 0.0769 | 8 | 0.125 | 12 | 0.0833 | 40 | 0.075 | 5 | 0 | 9 | 0.1111 | 12 | 0.0833 | 7 | 0 | 33 | 0.0606 |
| 3_5553397 | 8 | 0.25 | 11 | 0 | 7 | 0 | 16 | 0.0625 | 42 | 0.0714 | 6 | 0.1667 | 11 | 0.0909 | 17 | 0 | 11 | 0.0909 | 45 | 0.0667 |
| 3_5553403 | 9 | 0.8889 | 10 | 0.9 | 8 | 0.75 | 17 | 1 | 44 | 0.9091 | 6 | 1 | 11 | 0.9091 | 18 | 0.7222 | 10 | 0.8 | 45 | 0.8222 |
| 3_5553415 | 8 | 0.75 | 12 | 0.8333 | 7 | 0.7143 | 16 | 0.8125 | 43 | 0.7907 | 4 | 1 | 9 | 0.8889 | 15 | 0.8667 | 7 | 0.8571 | 35 | 0.8857 |
| 3_5553441 | 5 | 0.2 | 13 | 0 | 7 | 0 | 18 | 0.1667 | 43 | 0.093 | 6 | 0.6667 | 11 | 0 | 14 | 0.1429 | 10 | 0.1 | 41 | 0.1707 |
| 3_5553596 | 7 | 0.1429 | 8 | 0 | 6 | 0 | 15 | 0.0667 | 36 | 0.0556 | 5 | 0.2 | 8 | 0 | 14 | 0.0714 | 10 | 0.2 | 37 | 0.1081 |
| 3_5583806 | NA | NA | NA | NA | 1 | 1 | NA | NA | 1 | 1 | 3 | 0.3333 | NA | NA | NA | NA | NA | NA | 3 | 0.3333 |
| 3_5587559 | 2 | 0.5 | 3 | 1 | 3 | 0.6667 | 2 | 1 | 10 | 0.8 | 2 | 0.5 | 5 | 0.6 | 3 | 1 | 1 | 0 | 11 | 0.6364 |
| 3_5591210 | 1 | 1 | 2 | 0.5 | 1 | 0 | 3 | 0 | 7 | 0.2857 | NA | NA | 3 | 0.3333 | 2 | 0 | NA | NA | 5 | 0.2 |
| 3_5591219 | 1 | 0 | 3 | 0 | 2 | 0.5 | 4 | 0 | 10 | 0.1 | NA | NA | 5 | 0.4 | 2 | 0 | NA | NA | 7 | 0.2857 |
| 3_5591224 | 1 | 0 | 3 | 0.6667 | 2 | 0 | 3 | 0 | 9 | 0.2222 | NA | NA | 4 | 0.25 | 2 | 0.5 | NA | NA | 6 | 0.3333 |
| 3_5591257 | 1 | 0 | 3 | 0.6667 | 1 | 0 | 5 | 0.6 | 10 | 0.5 | NA | NA | 4 | 0.5 | 1 | 0 | 1 | 1 | 6 | 0.5 |
| 3_5591259 | 1 | 1 | 3 | 1 | 2 | 1 | 5 | 0.6 | 11 | 0.8182 | NA | NA | 4 | 0.75 | 2 | 0.5 | NA | NA | 6 | 0.6667 |
| 3_5591273 | 1 | 0 | 2 | 0.5 | 2 | 0.5 | 3 | 1 | 8 | 0.625 | NA | NA | 3 | 0.3333 | 2 | 0.5 | NA | NA | 5 | 0.4 |
| 3_5599851 | 1 | 0 | 2 | 0.5 | 2 | 0 | NA | NA | 5 | 0.2 | NA | NA | NA | NA | 2 | 0.5 | NA | NA | 2 | 0.5 |
| 3_5719496 | 3 | 1 | 1 | 1 | 4 | 0.5 | NA | NA | 8 | 0.75 | NA | NA | 1 | 1 | 2 | 1 | 4 | 0.75 | 7 | 0.8571 |
| 3_5719510 | 5 | 0.2 | 1 | 0 | 3 | 0.3333 | NA | NA | 9 | 0.2222 | NA | NA | 1 | 0 | 2 | 0 | 4 | 0.25 | 7 | 0.1429 |
| 3_5755377 | 1 | 1 | NA | NA | NA | NA | NA | NA | 1 | 1 | 1 | 1 | 1 | 1 | NA | NA | 1 | 0 | 3 | 0.6667 |
| 3_5755385 | 6 | 1 | 5 | 0.8 | 3 | 1 | 4 | 0.5 | 18 | 0.8333 | 4 | 1 | 9 | 0.8889 | 3 | 1 | 3 | 1 | 19 | 0.9474 |
| 3_5755386 | 5 | 1 | 4 | 0.75 | 3 | 1 | 4 | 0.5 | 16 | 0.8125 | 4 | 1 | 7 | 0.8571 | 3 | 1 | 2 | 1 | 16 | 0.9375 |
| 3_5755407 | 4 | 0.25 | 5 | 0 | 2 | 0 | 5 | 0.2 | 16 | 0.125 | 3 | 0 | 9 | 0.2222 | 3 | 0.3333 | 3 | 0.3333 | 18 | 0.2222 |
| 3_5755758 | 6 | 0 | 2 | 0 | 2 | 0.5 | 4 | 0 | 14 | 0.0714 | 2 | 0 | 6 | 0 | 3 | 0.3333 | 2 | 0.5 | 13 | 0.1538 |
| 3_5774064 | 1 | 0 | 1 | 0 | 1 | 0 | 3 | 0.3333 | 6 | 0.1667 | 1 | 1 | 2 | 0 | 1 | 0 | NA | NA | 4 | 0.25 |
| 3_5774095 | 13 | 0.0769 | 18 | 0.1111 | 19 | 0.1053 | 10 | 0.1 | 60 | 0.1 | 8 | 0.25 | 11 | 0.2727 | 13 | 0.1538 | 10 | 0.2 | 42 | 0.2143 |
| 3_5774099 | 12 | 0.1667 | 17 | 0.3529 | 19 | 0.2105 | 9 | 0.1111 | 57 | 0.2281 | 7 | 0.1429 | 8 | 0.375 | 12 | 0.4167 | 10 | 0.2 | 37 | 0.2973 |
| 3_5774100 | 11 | 0.9091 | 17 | 0.7647 | 19 | 0.6842 | 10 | 0.7 | 57 | 0.7544 | 8 | 0.875 | 10 | 0.9 | 13 | 0.9231 | 12 | 0.8333 | 43 | 0.8837 |
| 3_5774101 | 7 | 0.1429 | 11 | 0 | 16 | 0.125 | 8 | 0 | 42 | 0.0714 | 4 | 0 | 6 | 0.1667 | 11 | 0.1818 | 8 | 0 | 29 | 0.1034 |
| 3_5774127 | 11 | 0.5455 | 14 | 0.3571 | 16 | 0.5 | 11 | 0.1818 | 52 | 0.4038 | 8 | 0.375 | 9 | 0.6667 | 13 | 0.4615 | 9 | 0.5556 | 39 | 0.5128 |
| 3_5774152 | 17 | 0.0588 | 20 | 0.1 | 22 | 0.0909 | 11 | 0 | 70 | 0.0714 | 11 | 0.0909 | 12 | 0.1667 | 13 | 0 | 11 | 0.0909 | 47 | 0.0851 |
| 3_5774953 | 63 | 0.1111 | 97 | 0.0103 | 83 | 0.0964 | 80 | 0.0375 | 323 | 0.0588 | 69 | 0.058 | 74 | 0.0676 | 86 | 0.0349 | 88 | 0.0795 | 317 | 0.0599 |
| 3_5774958 | 53 | 0.2264 | 85 | 0.2824 | 72 | 0.25 | 71 | 0.2535 | 281 | 0.2562 | 65 | 0.2615 | 68 | 0.2794 | 83 | 0.2651 | 74 | 0.2973 | 290 | 0.2759 |
| 3_5774964 | 63 | 0.1111 | 92 | 0.0652 | 79 | 0.0886 | 74 | 0.1351 | 308 | 0.0974 | 72 | 0.0972 | 74 | 0.0946 | 86 | 0.093 | 85 | 0.1176 | 317 | 0.1009 |
| 3_5774983 | 55 | 0.2364 | 92 | 0.2174 | 82 | 0.3049 | 73 | 0.2055 | 302 | 0.2417 | 65 | 0.1538 | 68 | 0.1765 | 85 | 0.2235 | 80 | 0.2125 | 298 | 0.1946 |
| 3_5774994 | 53 | 0.1132 | 73 | 0.0959 | 68 | 0.0441 | 62 | 0.1129 | 256 | 0.0898 | 53 | 0.1132 | 57 | 0.0702 | 76 | 0.0658 | 73 | 0.0822 | 259 | 0.0811 |
| 3_5775040 | 41 | 0.122 | 63 | 0.1587 | 54 | 0.0926 | 46 | 0.2826 | 204 | 0.1618 | 46 | 0.1304 | 41 | 0.0488 | 63 | 0.0952 | 55 | 0.1273 | 205 | 0.1024 |
| 3_5775065 | 43 | 0.4186 | 70 | 0.5714 | 59 | 0.5085 | 48 | 0.625 | 220 | 0.5364 | 56 | 0.5893 | 49 | 0.5918 | 67 | 0.597 | 69 | 0.5797 | 241 | 0.5892 |
| 3_5793849 | NA | NA | 1 | 1 | NA | NA | NA | NA | 1 | 1 | NA | NA | 1 | 1 | NA | NA | NA | NA | 1 | 1 |
| 3_5793858 | NA | NA | 1 | 1 | NA | NA | NA | NA | 1 | 1 | NA | NA | 1 | 1 | NA | NA | NA | NA | 1 | 1 |

| | | | | | | | | | | | | | | | | | | | | |
|---|---|---|---|---|---|---|---|---|---|---|---|---|---|---|---|---|---|---|---|---|
| 3_5793899 | NA | NA | 1 | 1 | NA | NA | NA | NA | 1 | 1 | NA | NA | 1 | 1 | NA | NA | NA | NA | 1 | 1 |
| 3_5793943 | NA | NA | 1 | 1 | NA | NA | NA | NA | 1 | 1 | NA | NA | 1 | 1 | NA | NA | NA | NA | 1 | 1 |
| 3_5801020 | 14 | 0 | 19 | 0.0526 | 15 | 0.2 | 14 | 0.2143 | 62 | 0.1129 | 15 | 0.2667 | 13 | 0 | 20 | 0.15 | 13 | 0.0769 | 61 | 0.1311 |
| 3_5801072 | 5 | 0 | 5 | 0 | 5 | 0.2 | 1 | 0 | 16 | 0.0625 | 4 | 0 | 4 | 0 | 6 | 0 | 4 | 0.25 | 18 | 0.0556 |
| 3_5832282 | 1 | 0 | 1 | 1 | NA | NA | 2 | 0.5 | 4 | 0.5 | NA | NA | NA | NA | 2 | 0.5 | NA | NA | 2 | 0.5 |
| 3_5832302 | 1 | 1 | 1 | 1 | NA | NA | 2 | 1 | 4 | 1 | NA | NA | NA | NA | 2 | 1 | 1 | 1 | 3 | 1 |
| 3_5832357 | NA | NA | NA | NA | NA | NA | 1 | 1 | 1 | 1 | NA | NA | NA | NA | 2 | 0.5 | 1 | 1 | 3 | 0.6667 |
| 3_5832370 | NA | NA | 1 | 1 | NA | NA | 1 | 1 | 2 | 1 | 1 | 0 | NA | NA | 2 | 0.5 | 1 | 0 | 4 | 0.25 |
| 3_5832378 | NA | NA | 1 | 1 | NA | NA | 2 | 1 | 3 | 1 | 1 | 1 | NA | NA | 1 | 1 | 1 | 1 | 3 | 1 |
| 3_5832379 | 1 | 1 | 1 | 1 | NA | NA | 2 | 0.5 | 4 | 0.75 | NA | NA | NA | NA | 2 | 1 | 1 | 1 | 3 | 1 |
| 3_5832482 | 1 | 0 | 1 | 1 | NA | NA | 5 | 0.6 | 7 | 0.5714 | 1 | 1 | NA | NA | 2 | 0.5 | 1 | 1 | 4 | 0.75 |
| 3_5832497 | 1 | 1 | 1 | 1 | NA | NA | 5 | 0.6 | 7 | 0.7143 | 2 | 1 | NA | NA | 2 | 1 | 1 | 1 | 5 | 1 |
| 3_5836751 | 1 | 1 | 2 | 0.5 | 2 | 0 | 5 | 0.4 | 10 | 0.4 | 2 | 0.5 | NA | NA | 3 | 1 | NA | NA | 5 | 0.8 |
| 3_5836755 | 1 | 1 | 2 | 0 | 3 | 0 | 4 | 0 | 10 | 0.1 | 2 | 0.5 | NA | NA | 2 | 0 | 1 | 0 | 5 | 0.2 |
| 3_5836771 | 1 | 0 | 1 | 1 | 1 | 1 | 4 | 0.5 | 7 | 0.5714 | 2 | 1 | NA | NA | 3 | 1 | 1 | 1 | 6 | 1 |
| 3_5836781 | 1 | 0 | 3 | 0 | 3 | 0.6667 | 5 | 0 | 12 | 0.1667 | 2 | 0 | NA | NA | 3 | 0.3333 | 1 | 1 | 6 | 0.3333 |
| 3_5842992 | 8 | 0.125 | 9 | 0 | 4 | 0.25 | 11 | 0.0909 | 32 | 0.0938 | 5 | 0 | 10 | 0.1 | 7 | 0 | 7 | 0.2857 | 29 | 0.1034 |
| 3_5843113 | 6 | 0.1667 | 9 | 0.3333 | 7 | 0.2857 | 9 | 0.3333 | 31 | 0.2903 | 5 | 0.6 | 7 | 0.1429 | 6 | 0.3333 | 6 | 0.3333 | 24 | 0.3333 |
| 3_5843120 | 5 | 0.2 | 8 | 0.625 | 6 | 0.3333 | 8 | 0.75 | 27 | 0.5185 | 5 | 0.6 | 6 | 0.3333 | 5 | 0.2 | 5 | 0.4 | 21 | 0.381 |
| 3_5843124 | 6 | 0.3333 | 11 | 0.2727 | 6 | 0.5 | 10 | 0.4 | 33 | 0.3636 | 5 | 0.2 | 6 | 0.1667 | 6 | 0.1667 | 5 | 0.4 | 22 | 0.2273 |
| 3_5843313 | 4 | 0 | 7 | 0.2857 | 6 | 0 | 4 | 0.25 | 21 | 0.1429 | 6 | 0.1667 | 5 | 0.2 | 3 | 0 | 4 | 0.25 | 18 | 0.1667 |
| 3_5844500 | 3 | 0.3333 | 3 | 0 | 3 | 0 | 9 | 0 | 18 | 0.0556 | 1 | 0 | 5 | 0 | 5 | 0 | 6 | 0.1667 | 17 | 0.0588 |
| 3_5845593 | 7 | 0.5714 | 6 | 0.3333 | 3 | 0.3333 | 11 | 0.3636 | 27 | 0.4074 | 1 | 0 | 5 | 0.6 | 5 | 1 | 8 | 0.375 | 19 | 0.5789 |
| 3_5845594 | 9 | 0.8889 | 6 | 0.8333 | 5 | 0.8 | 10 | 0.8 | 30 | 0.8333 | 1 | 1 | 6 | 0.8333 | 5 | 0.8 | 7 | 1 | 19 | 0.8947 |
| 3_5845598 | 8 | 0.75 | 6 | 1 | 5 | 1 | 8 | 0.75 | 27 | 0.8519 | 1 | 1 | 6 | 0.8333 | 6 | 0.8333 | 9 | 0.5556 | 22 | 0.7273 |
| 3_5845601 | 8 | 0.375 | 6 | 0.3333 | 4 | 0.5 | 10 | 0.4 | 28 | 0.3929 | 1 | 0 | 6 | 0.6667 | 6 | 0 | 10 | 0.2 | 23 | 0.2609 |
| 3_5845604 | 9 | 0 | 5 | 0.2 | 5 | 0.4 | 10 | 0 | 29 | 0.1034 | 1 | 0 | 6 | 0.1667 | 6 | 0.1667 | 9 | 0.1111 | 22 | 0.1364 |
| 3_5849824 | 3 | 0 | 5 | 0.2 | 6 | 0.1667 | 5 | 0.2 | 19 | 0.1579 | 8 | 0.25 | 9 | 0.3333 | 3 | 0 | 3 | 0 | 23 | 0.2174 |
| 3_5849932 | 16 | 0.1875 | 22 | 0.2727 | 23 | 0.2174 | 21 | 0.1429 | 82 | 0.2073 | 20 | 0.3 | 24 | 0.125 | 28 | 0.2143 | 23 | 0.2609 | 95 | 0.2211 |
| 3_5849981 | 24 | 0.0833 | 31 | 0.1935 | 30 | 0.2 | 25 | 0 | 110 | 0.1273 | 22 | 0.0909 | 27 | 0.1111 | 28 | 0.0714 | 26 | 0.1538 | 103 | 0.1068 |
| 3_5858254 | 1 | 0 | 3 | 0 | 5 | 0.2 | 2 | 0 | 11 | 0.0909 | 1 | 1 | 3 | 0.6667 | 2 | 0.5 | 3 | 0.3333 | 9 | 0.5556 |
| 3_5858256 | 2 | 0.5 | 3 | 0.3333 | 5 | 0 | 3 | 0 | 13 | 0.1538 | 1 | 1 | 3 | 0 | 2 | 0.5 | 3 | 0.3333 | 9 | 0.3333 |
| 3_5858270 | 2 | 0.5 | 2 | 0 | 4 | 0.25 | 2 | 0 | 10 | 0.2 | 1 | 1 | 2 | 0 | 2 | 0.5 | 2 | 0.5 | 7 | 0.4286 |
| 3_5858297 | 1 | 1 | 2 | 1 | 3 | 0.3333 | 2 | 0 | 8 | 0.5 | 1 | 1 | 3 | 0.6667 | NA | NA | 2 | 1 | 6 | 0.8333 |
| 3_5858312 | NA | NA | 1 | 1 | 3 | 0.3333 | 2 | 0 | 6 | 0.3333 | 1 | 0 | 3 | 0.3333 | 1 | 0 | 1 | 0 | 6 | 0.1667 |
| 3_5858334 | NA | NA | 2 | 0.5 | 3 | 0 | 3 | 0 | 8 | 0.125 | 1 | 1 | 4 | 0 | 1 | 0 | 1 | 0 | 7 | 0.1429 |
| 3_5858339 | NA | NA | 2 | 1 | 4 | 1 | 3 | 1 | 9 | 1 | 1 | 1 | 1 | 1 | 1 | 1 | 1 | 1 | 4 | 1 |
| 3_5858346 | NA | NA | 2 | 1 | 4 | 0.75 | 3 | 0 | 9 | 0.5556 | 1 | 1 | 4 | 0.5 | NA | NA | NA | NA | 5 | 0.6 |
| 3_5858349 | NA | NA | 2 | 1 | 3 | 1 | 2 | 0 | 7 | 0.7143 | 1 | 1 | 4 | 0.5 | 1 | 1 | 1 | 0 | 7 | 0.5714 |
| 3_5858350 | NA | NA | 2 | 0.5 | 3 | 1 | 3 | 1 | 8 | 0.875 | NA | NA | 3 | 1 | NA | NA | NA | NA | 3 | 1 |
| 3_5858363 | NA | NA | 2 | 1 | 4 | 1 | 3 | 1 | 9 | 1 | 1 | 1 | 4 | 1 | 1 | 1 | 1 | 1 | 7 | 1 |
| 3_5858365 | NA | NA | 2 | 0.5 | 3 | 1 | 2 | 0 | 7 | 0.5714 | 1 | 1 | 3 | 1 | 1 | 1 | 1 | 0 | 6 | 0.8333 |
| 3_5858367 | NA | NA | 2 | 1 | 4 | 1 | 2 | 0 | 8 | 0.75 | 1 | 1 | 4 | 0.75 | 1 | 1 | 1 | 1 | 7 | 0.8571 |
| 3_5902305 | 2 | 1 | NA | NA | 1 | 1 | NA | NA | 3 | 1 | 2 | 1 | NA | NA | NA | NA | NA | NA | 2 | 1 |
| 3_5902324 | 2 | 0.5 | NA | NA | NA | NA | NA | NA | 2 | 0.5 | 2 | 1 | NA | NA | NA | NA | NA | NA | 2 | 1 |
| 3_5902325 | 2 | 0.5 | NA | NA | 1 | 1 | NA | NA | 3 | 0.6667 | 2 | 1 | NA | NA | NA | NA | NA | NA | 2 | 1 |
| 3_5911689 | 1 | 1 | NA | NA | 1 | 1 | 1 | 1 | 3 | 1 | 1 | 1 | 1 | 1 | NA | NA | 3 | 1 | 5 | 1 |
| 3_5911697 | 1 | 1 | NA | NA | NA | NA | NA | NA | 1 | 1 | NA | NA | 1 | 0 | NA | NA | 3 | 0.3333 | 4 | 0.25 |
| 3_5911697 | 1 | 0 | NA | NA | 1 | 1 | 1 | 1 | 3 | 0.6667 | NA | NA | 1 | 0 | NA | NA | 2 | 0.5 | 3 | 0.3333 |
| 3_5911709 | 1 | 0 | NA | NA | 1 | 0 | 1 | 1 | 3 | 0.3333 | NA | NA | 1 | 1 | NA | NA | NA | NA | 1 | 1 |
| 3_5911756 | 1 | 1 | NA | NA | 1 | 1 | 1 | 1 | 3 | 1 | NA | NA | NA | NA | NA | NA | 2 | 0.5 | 2 | 0.5 |
| 3_5911757 | 1 | 0 | NA | NA | 1 | 1 | 1 | 1 | 3 | 0.6667 | 1 | 0 | NA | NA | NA | NA | 2 | 1 | 3 | 0.6667 |
| 3_5911771 | 1 | 0 | NA | NA | 1 | 0 | 1 | 1 | 3 | 0.3333 | NA | NA | NA | NA | NA | NA | 3 | 0.3333 | 3 | 0.3333 |
| 3_5912410 | 8 | 0.25 | 7 | 0.1429 | 5 | 0.4 | 8 | 0.125 | 28 | 0.2143 | 2 | 0 | 6 | 0.1667 | 6 | 0 | 3 | 0 | 17 | 0.0588 |
| 3_5912442 | 2 | 1 | 7 | 0.2857 | 3 | 0.6667 | 4 | 0.5 | 16 | 0.5 | 1 | 1 | 4 | 0.75 | 3 | 1 | 2 | 1 | 10 | 0.9 |
| 3_5912455 | 5 | 0 | 5 | 0 | 3 | 0 | 4 | 0.25 | 17 | 0.0588 | 1 | 0 | 6 | 0 | 7 | 0.1429 | 3 | 0 | 17 | 0.0588 |
| 3_5912459 | 5 | 0 | 6 | 0.1667 | 5 | 0.6 | 7 | 0.2857 | 23 | 0.2609 | 3 | 0 | 5 | 0.6 | 8 | 0.125 | 4 | 0.5 | 20 | 0.3 |
| 3_5912462 | 2 | 1 | 4 | 0.25 | 4 | 0.5 | 2 | 1 | 12 | 0.5833 | 2 | 0.5 | 4 | 0.75 | 4 | 0.75 | 3 | 0.6667 | 13 | 0.6923 |
| 3_5912466 | 4 | 0.5 | 6 | 0.3333 | 5 | 0.2 | 7 | 0.2857 | 22 | 0.3182 | 3 | 0.3333 | 6 | 0.1667 | 7 | 0.1429 | 4 | 0 | 20 | 0.15 |
| 3_5912468 | 5 | 0.4 | 6 | 0.5 | 5 | 0.2 | 5 | 0 | 21 | 0.2857 | 3 | 0.3333 | 6 | 0.1667 | 8 | 0.5 | 4 | 0 | 21 | 0.2857 |
| 3_5912485 | 5 | 0.4 | 5 | 0 | 4 | 0.5 | 6 | 0.3333 | 20 | 0.3 | 3 | 0.3333 | 6 | 0.3333 | 7 | 0.2857 | 4 | 0.5 | 20 | 0.35 |
| 3_5912498 | 4 | 0.25 | 6 | 0 | 3 | 0.3333 | 4 | 0 | 17 | 0.1176 | 2 | 0 | 6 | 0.1667 | 7 | 0.1429 | 3 | 0 | 18 | 0.1111 |
| 3_5912501 | 4 | 1 | 5 | 1 | 4 | 1 | 5 | 1 | 18 | 1 | 2 | 1 | 5 | 1 | 5 | 1 | 2 | 0.5 | 14 | 0.9286 |
| 3_5912529 | 4 | 0.25 | 6 | 0 | 4 | 0.25 | 4 | 0 | 18 | 0.1111 | 3 | 0 | 5 | 0.2 | 6 | 0.1667 | 3 | 0 | 17 | 0.1176 |
| 3_5912537 | 4 | 0.5 | 7 | 0.2857 | 4 | 0.5 | 5 | 0.4 | 20 | 0.4 | 3 | 0.3333 | 4 | 0 | 7 | 0.1429 | 4 | 0 | 18 | 0.1111 |

| | | | | | | | | | | | | | | | | | | | | |
|---|---|---|---|---|---|---|---|---|---|---|---|---|---|---|---|---|---|---|---|---|
| 3_5912539 | 5 | 0.4 | 6 | 0.1667 | 4 | 0.25 | 6 | 0 | 21 | 0.1905 | 3 | 0 | 4 | 0 | 7 | 0.1429 | 4 | 0.25 | 18 | 0.1111 |
| 3_5912543 | 3 | 0.6667 | 6 | 0.3333 | 3 | 0.6667 | 6 | 0.1667 | 18 | 0.3889 | 2 | 0.5 | 4 | 0 | 6 | 0 | 3 | 0.3333 | 15 | 0.1333 |
| 3_5912546 | 4 | 0.25 | 6 | 0.3333 | 2 | 0 | 3 | 0.3333 | 15 | 0.2667 | 1 | 0 | 3 | 0.3333 | 7 | 0.1429 | 3 | 0.6667 | 14 | 0.2857 |
| 3_5912552 | 5 | 0.2 | 4 | 0 | 4 | 0.25 | 7 | 0.2857 | 20 | 0.2 | 2 | 0 | 3 | 0 | 6 | 0 | 4 | 0.25 | 15 | 0.0667 |
| 3_5912559 | 5 | 0.2 | 6 | 0.1667 | 4 | 0 | 7 | 0.1429 | 22 | 0.1364 | 3 | 0.3333 | 4 | 0 | 7 | 0.1429 | 4 | 0 | 18 | 0.1111 |
| 3_5922241 | NA | NA | 1 | 0 | NA | NA | 1 | 1 | 2 | 0.5 | NA | NA | 1 | 0 | 2 | 0.5 | 1 | 0 | 4 | 0.25 |
| 3_5922251 | NA | NA | 1 | 1 | NA | NA | 2 | 0 | 3 | 0.3333 | NA | NA | 3 | 0.3333 | 3 | 0 | 1 | 1 | 7 | 0.2857 |
| 3_5922255 | NA | NA | 1 | 1 | NA | NA | NA | NA | 1 | 1 | NA | NA | 1 | 0 | 3 | 0.3333 | NA | NA | 4 | 0.25 |
| 3_5922266 | NA | NA | 1 | 1 | NA | NA | 2 | 0.5 | 3 | 0.6667 | NA | NA | 3 | 0.3333 | 2 | 0.5 | 1 | 0 | 6 | 0.3333 |
| 3_5922268 | NA | NA | 2 | 1 | NA | NA | 2 | 0.5 | 4 | 0.75 | NA | NA | 2 | 1 | 3 | 1 | 1 | 1 | 6 | 1 |
| 3_5922278 | NA | NA | 2 | 1 | NA | NA | 2 | 1 | 4 | 1 | NA | NA | 3 | 1 | 2 | 1 | 1 | 1 | 6 | 1 |
| 3_5922303 | NA | NA | 1 | 0 | NA | NA | 2 | 0.5 | 3 | 0.3333 | NA | NA | 3 | 1 | 3 | 0 | 1 | 0 | 7 | 0.4286 |
| 3_5922369 | NA | NA | 3 | 0 | NA | NA | 4 | 0.25 | 7 | 0.1429 | 1 | 0 | 3 | 0 | 4 | 0.25 | NA | NA | 8 | 0.125 |
| 3_5922376 | NA | NA | 3 | 0.3333 | NA | NA | 4 | 0.25 | 7 | 0.2857 | 1 | 0 | 3 | 0.6667 | 5 | 0.8 | NA | NA | 9 | 0.6667 |
| 3_5922489 | NA | NA | 4 | 0 | 1 | 0 | 4 | 0.25 | 9 | 0.1111 | 1 | 1 | 3 | 0 | 2 | 0 | NA | NA | 6 | 0.1667 |
| 3_5922512 | NA | NA | 5 | 0.2 | 1 | 0 | 4 | 0 | 10 | 0.1 | 2 | 0.5 | 3 | 0 | 3 | 0 | 1 | 0 | 9 | 0.1111 |
| 3_5922520 | NA | NA | 5 | 0.2 | 1 | 0 | 5 | 0.2 | 11 | 0.1818 | 3 | 0 | 4 | 0.25 | 6 | 0 | 2 | 0 | 15 | 0.0667 |
| 3_6040204 | NA | NA | NA | NA | 3 | 0.6667 | NA | NA | 3 | 0.6667 | NA | NA | NA | NA | 1 | 1 | NA | NA | 1 | 1 |
| 3_6043646 | 2 | 0 | 4 | 0.5 | 4 | 0 | 3 | 0.6667 | 13 | 0.3077 | NA | NA | 1 | 0 | 5 | 0.4 | 1 | 1 | 7 | 0.4286 |
| 3_6043648 | 2 | 0 | 4 | 0 | 4 | 0.25 | 3 | 0 | 13 | 0.0769 | NA | NA | 1 | 1 | 5 | 0 | 1 | 0 | 7 | 0.1429 |
| 3_6043655 | 2 | 0 | 4 | 0 | 4 | 0 | 3 | 0.3333 | 13 | 0.0769 | NA | NA | 1 | 0 | 5 | 0.2 | 1 | 0 | 7 | 0.1429 |
| 3_6043684 | 2 | 0.5 | 2 | 1 | 4 | 1 | 3 | 1 | 11 | 0.9091 | NA | NA | 2 | 1 | 5 | 0.8 | 1 | 1 | 8 | 0.875 |
| 3_6043708 | 2 | 0.5 | 4 | 0 | 4 | 0.5 | 3 | 0.3333 | 13 | 0.3077 | NA | NA | 2 | 0.5 | 3 | 0 | NA | NA | 5 | 0.2 |
| 3_6043712 | 1 | 0 | 4 | 0.25 | 4 | 0 | 5 | 0.8 | 14 | 0.3571 | NA | NA | 2 | 0 | 5 | 0.4 | 2 | 0.5 | 9 | 0.3333 |
| 3_6053382 | 5 | 0.2 | 5 | 0 | 6 | 0 | 9 | 0.2222 | 25 | 0.12 | 2 | 0.5 | 9 | 0.4444 | 7 | 0.4286 | 3 | 0.6667 | 21 | 0.4762 |
| 3_6053412 | 5 | 0.2 | 3 | 0 | 6 | 0.1667 | 9 | 0 | 23 | 0.087 | 1 | 0 | 8 | 0.375 | 11 | 0 | 3 | 0.3333 | 23 | 0.1739 |
| 3_6053412 | 3 | 0.6667 | 4 | 0.25 | 6 | 0.3333 | 10 | 0.5 | 23 | 0.4348 | 1 | 0 | 9 | 0.1111 | 6 | 0.5 | 3 | 0.3333 | 19 | 0.2632 |
| 3_6053444 | 6 | 0.3333 | 5 | 0 | 6 | 0.1667 | 9 | 0.2222 | 26 | 0.1923 | 1 | 0 | 8 | 0 | 13 | 0.1538 | 3 | 0.3333 | 25 | 0.12 |
| 3_6053533 | 7 | 0 | 8 | 0 | 10 | 0.1 | 10 | 0.3 | 35 | 0.1143 | 4 | 0.25 | 16 | 0.1875 | 15 | 0 | 7 | 0 | 42 | 0.0952 |
| 3_6053607 | 8 | 0.25 | 15 | 0.0667 | 12 | 0.1667 | 10 | 0.3 | 45 | 0.1778 | 7 | 0 | 18 | 0.3333 | 16 | 0.3125 | 6 | 0.3333 | 47 | 0.2766 |
| 3_6053631 | 8 | 0.25 | 13 | 0.1538 | 12 | 0.4167 | 10 | 0.6 | 43 | 0.3488 | 5 | 0.2 | 18 | 0.4444 | 14 | 0.5714 | 6 | 0.3333 | 43 | 0.4419 |
| 3_6053639 | 9 | 0.4444 | 16 | 0.5 | 15 | 0.3333 | 11 | 0.6364 | 51 | 0.4706 | 3 | 0.3333 | 16 | 0.1875 | 19 | 0.3158 | 8 | 0.25 | 46 | 0.2609 |
| 3_6053645 | 9 | 0.3333 | 14 | 0.1429 | 12 | 0.1667 | 11 | 0.2727 | 46 | 0.2174 | 6 | 0.1667 | 15 | 0.3333 | 20 | 0.5 | 8 | 0.125 | 49 | 0.3469 |
| 3_6053660 | 8 | 0.375 | 16 | 0.3125 | 17 | 0.4118 | 14 | 0.6429 | 55 | 0.4364 | 7 | 0.5714 | 17 | 0.5882 | 22 | 0.4091 | 9 | 0.2222 | 55 | 0.4545 |
| 3_6053662 | 11 | 0.6364 | 17 | 0.4118 | 17 | 0.4118 | 12 | 0.75 | 57 | 0.5263 | 6 | 0.8333 | 17 | 0.5882 | 23 | 0.6522 | 8 | 0.375 | 54 | 0.6111 |
| 3_6053669 | 10 | 0.5 | 16 | 0.3125 | 17 | 0.1765 | 12 | 0.4167 | 55 | 0.3273 | 5 | 0.4 | 18 | 0.5 | 19 | 0.3684 | 8 | 0.25 | 50 | 0.4 |
| 3_6053690 | 10 | 0.1 | 14 | 0.2143 | 13 | 0.0769 | 16 | 0.0625 | 53 | 0.1132 | 8 | 0.25 | 18 | 0.2222 | 19 | 0 | 10 | 0 | 55 | 0.1091 |
| 3_6053698 | 13 | 0.1538 | 17 | 0.1176 | 19 | 0.1579 | 17 | 0 | 66 | 0.1061 | 8 | 0.25 | 21 | 0.0952 | 25 | 0.04 | 10 | 0.2 | 64 | 0.1094 |
| 3_6078499 | 1 | 1 | NA | NA | NA | NA | NA | NA | 1 | 1 | NA | NA | 1 | 1 | 1 | 1 | NA | NA | 2 | 1 |
| 3_6078523 | 1 | 1 | NA | NA | NA | NA | NA | NA | 1 | 1 | NA | NA | NA | NA | 1 | 1 | NA | NA | 1 | 1 |
| 3_6078586 | 4 | 0 | 3 | 0.3333 | NA | NA | 1 | 0 | 8 | 0.125 | NA | NA | 3 | 0.3333 | 6 | 0.1667 | NA | NA | 9 | 0.2222 |
| 3_6078599 | 3 | 0 | 3 | 0 | 2 | 0 | 4 | 0.25 | 12 | 0.0833 | 1 | 0 | 5 | 0 | 7 | 0.1429 | NA | NA | 13 | 0.0769 |
| 3_6078604 | 5 | 0 | 4 | 0 | 3 | 0 | 3 | 0.3333 | 15 | 0.0667 | 1 | 0 | 3 | 0.3333 | 8 | 0.125 | NA | NA | 12 | 0.1667 |
| 3_6078607 | 6 | 0.5 | 5 | 0.4 | 4 | 0.75 | 4 | 0.75 | 19 | 0.5789 | NA | NA | 5 | 0.2 | 9 | 0.3333 | NA | NA | 14 | 0.2857 |
| 3_6091049 | 1 | 1 | NA | NA | 1 | 0 | NA | NA | 2 | 0.5 | 1 | 1 | 1 | 1 | 1 | 0 | NA | NA | 3 | 0.6667 |
| 3_6091144 | 1 | 1 | NA | NA | 1 | 1 | NA | NA | 2 | 1 | 1 | 1 | 1 | 1 | 1 | 1 | NA | NA | 3 | 1 |
| 3_6091145 | 1 | 1 | NA | NA | 1 | 1 | NA | NA | 2 | 1 | 1 | 0 | 1 | 1 | 1 | 1 | NA | NA | 3 | 0.6667 |
| 3_6091160 | 1 | 1 | NA | NA | 1 | 0 | NA | NA | 2 | 0.5 | 1 | 0 | 1 | 1 | 1 | 0 | NA | NA | 3 | 0.3333 |
| 3_6091161 | 1 | 0 | NA | NA | 1 | 1 | NA | NA | 2 | 0.5 | 1 | 0 | 1 | 1 | 1 | 0 | NA | NA | 3 | 0.3333 |
| 3_6091186 | 1 | 1 | NA | NA | 1 | 1 | NA | NA | 2 | 1 | 1 | 1 | 1 | 1 | 1 | 1 | NA | NA | 3 | 1 |
| 3_6091193 | 1 | 1 | NA | NA | 1 | 1 | NA | NA | 2 | 1 | 1 | 1 | 1 | 1 | 1 | 1 | NA | NA | 3 | 1 |
| 3_6091279 | NA | NA | NA | NA | 1 | 1 | NA | NA | 1 | 1 | NA | NA | NA | NA | NA | NA | 1 | 1 | 1 | 1 |
| 3_6091429 | NA | NA | NA | NA | 2 | 1 | 1 | 0 | 3 | 0.6667 | NA | NA | 1 | 1 | NA | NA | 1 | 0 | 2 | 0.5 |
| 3_6091443 | NA | NA | NA | NA | 2 | 1 | 1 | 1 | 3 | 1 | NA | NA | 2 | 1 | 1 | 1 | NA | NA | 3 | 1 |
| 3_6091445 | 1 | 1 | NA | NA | 1 | 0 | 1 | 0 | 3 | 0.3333 | NA | NA | 2 | 1 | 1 | 0 | 1 | 1 | 4 | 0.75 |
| 3_6091446 | NA | NA | NA | NA | 2 | 1 | 1 | 1 | 3 | 1 | NA | NA | 1 | 1 | 1 | 1 | 1 | 1 | 3 | 1 |
| 3_6091458 | NA | NA | NA | NA | 3 | 1 | 1 | 0 | 4 | 0.75 | NA | NA | 1 | 1 | 1 | 1 | NA | NA | 2 | 1 |
| 3_6091461 | 2 | 1 | NA | NA | 2 | 1 | 1 | 1 | 5 | 1 | NA | NA | 1 | 1 | 1 | 1 | 1 | 1 | 3 | 1 |
| 3_6091473 | 2 | 0.5 | NA | NA | 3 | 0.6667 | 1 | 0 | 6 | 0.5 | NA | NA | 2 | 0 | 1 | 0 | 1 | 1 | 4 | 0.25 |
| 3_6091479 | NA | NA | NA | NA | 3 | 1 | 1 | 1 | 4 | 1 | NA | NA | 2 | 1 | 1 | 1 | 1 | 1 | 4 | 1 |
| 3_6091482 | 2 | 1 | NA | NA | 3 | 1 | 1 | 1 | 6 | 1 | NA | NA | 2 | 1 | 1 | 1 | 1 | 1 | 4 | 1 |
| 3_6091488 | 1 | 0 | NA | NA | 3 | 0.3333 | 1 | 1 | 5 | 0.4 | NA | NA | 2 | 0 | 1 | 0 | 1 | 1 | 4 | 0.25 |
| 3_6091491 | 2 | 1 | NA | NA | 3 | 1 | 1 | 1 | 6 | 1 | NA | NA | 3 | 0.6667 | 1 | 1 | 1 | 1 | 5 | 0.8 |
| 3_6091497 | 2 | 1 | NA | NA | 3 | 1 | 1 | 1 | 6 | 1 | NA | NA | 1 | 0 | 1 | 1 | NA | NA | 2 | 0.5 |
| 3_6091524 | 2 | 0.5 | NA | NA | 3 | 0.6667 | 1 | 0 | 6 | 0.5 | NA | NA | 2 | 0.5 | 1 | 0 | 1 | 0 | 4 | 0.25 |

| | | | | | | | | | | | | | | | | | | | | |
|---|---|---|---|---|---|---|---|---|---|---|---|---|---|---|---|---|---|---|---|---|
| 3_6127259( | 6 | 0.3333 | NA | NA | NA | NA | 4 | 0.75 | 10 | 0.5 | 1 | 1 | 6 | 0.8333 | 1 | 1 | 3 | 0.3333 | 11 | 0.7273 |
| 3_6127261 | 4 | 0 | NA | NA | NA | NA | 4 | 0.25 | 8 | 0.125 | 2 | 0 | 6 | 0.1667 | 1 | 0 | 2 | 0.5 | 11 | 0.1818 |
| 3_6127268 | 6 | 0 | NA | NA | NA | NA | 5 | 0.6 | 11 | 0.2727 | 2 | 0 | 7 | 1 | 2 | 0.5 | 3 | 0.3333 | 14 | 0.6429 |
| 3_6127275 | 5 | 0.4 | NA | NA | NA | NA | 4 | 0.25 | 9 | 0.3333 | 2 | 0 | 6 | 0.6667 | 1 | 0 | 4 | 0.5 | 13 | 0.4615 |
| 3_6127283 | 4 | 1 | NA | NA | NA | NA | 4 | 1 | 8 | 1 | 2 | 1 | 6 | 0.8333 | 2 | 1 | 5 | 1 | 15 | 0.9333 |
| 3_6127291( | 3 | 0 | NA | NA | NA | NA | 5 | 0.2 | 8 | 0.125 | 1 | 0 | 7 | 0.1429 | 2 | 0 | 4 | 0 | 14 | 0.0714 |
| 3_6127297 | 4 | 1 | NA | NA | NA | NA | 5 | 1 | 9 | 1 | 2 | 1 | 7 | 1 | 2 | 1 | 5 | 1 | 16 | 1 |
| 3_6127316 | 6 | 0.3333 | NA | NA | NA | NA | 5 | 0.2 | 11 | 0.2727 | 2 | 0 | 6 | 0.3333 | 1 | 0 | 5 | 0.6 | 14 | 0.3571 |
| 3_6127317( | 7 | 0.1429 | NA | NA | NA | NA | 6 | 0.1667 | 13 | 0.1538 | 2 | 0 | 7 | 0.5714 | 2 | 0.5 | 5 | 0.2 | 16 | 0.375 |
| 3_6127323 | 8 | 0.25 | NA | NA | NA | NA | 6 | 0.5 | 14 | 0.3571 | 4 | 0.25 | 8 | 0.375 | 2 | 1 | 4 | 0.25 | 18 | 0.3889 |
| 3_6127334 | 8 | 0.125 | 1 | 0 | NA | NA | 7 | 0 | 16 | 0.0625 | 4 | 0 | 8 | 0.125 | 2 | 0.5 | 5 | 0 | 19 | 0.1053 |
| 3_6127336 | 8 | 0.25 | 1 | 1 | NA | NA | 7 | 0.1429 | 16 | 0.25 | 4 | 0.25 | 6 | 0 | 2 | 0 | 5 | 0.2 | 17 | 0.1176 |
| 3_6127352 | 11 | 0.6364 | NA | NA | NA | NA | 7 | 0.2857 | 18 | 0.5 | 5 | 0.2 | 8 | 0.5 | 3 | 0.6667 | 8 | 0.5 | 24 | 0.4583 |
| 3_6127356 | 10 | 0.6 | 1 | 1 | NA | NA | 8 | 0.625 | 19 | 0.6316 | 5 | 0.6 | 9 | 0.8889 | 3 | 1 | 8 | 1 | 25 | 0.88 |
| 3_6127360 | 10 | 0.9 | NA | NA | NA | NA | 7 | 1 | 17 | 0.9412 | 6 | 1 | 8 | 1 | 3 | 1 | 9 | 1 | 26 | 1 |
| 3_6127368 | 8 | 0.625 | NA | NA | 1 | 1 | 4 | 1 | 13 | 0.7692 | 4 | 1 | 8 | 0.875 | 1 | 1 | 8 | 1 | 21 | 0.9524 |
| 3_6127369 | 11 | 1 | 1 | 1 | 1 | 1 | 10 | 1 | 23 | 1 | 5 | 1 | 8 | 1 | 4 | 1 | 9 | 1 | 26 | 1 |
| 3_6127399 | 14 | 0.1429 | 2 | 0 | 2 | 0.5 | 12 | 0.3333 | 30 | 0.2333 | 3 | 0 | 10 | 0.9 | 6 | 0.5 | 13 | 0.3077 | 32 | 0.5 |
| 3_6127416 | 21 | 0.0952 | 2 | 0.5 | 5 | 0.2 | 13 | 0.2308 | 41 | 0.1707 | 5 | 0 | 16 | 0.0625 | 7 | 0.2857 | 18 | 0.2222 | 46 | 0.1522 |
| 3_6127442 | 17 | 0 | 4 | 0 | 6 | 0.1667 | 17 | 0.1765 | 44 | 0.0909 | 5 | 0 | 11 | 0.1818 | 8 | 0.125 | 14 | 0.5 | 38 | 0.2632 |
| 3_6127473 | 9 | 0.8889 | 1 | 0 | 2 | 0.5 | 9 | 0.4444 | 21 | 0.619 | 1 | 1 | 7 | 0.5714 | 4 | 0.5 | 4 | 0.25 | 16 | 0.5 |
| 3_6127570( | 14 | 0.3571 | 7 | 0.1429 | 4 | 0.25 | 7 | 0.2857 | 32 | 0.2812 | 2 | 0.5 | 8 | 0.375 | 5 | 0 | 7 | 0.2857 | 22 | 0.2727 |
| 3_6127780 | 1 | 0 | 1 | 1 | 2 | 0 | 1 | 0 | 5 | 0.2 | 2 | 0 | NA | NA | NA | NA | 2 | 0.5 | 4 | 0.25 |
| 3_6224591 | NA | NA | NA | NA | NA | NA | 2 | 1 | 2 | 1 | 1 | 1 | NA | NA | 1 | 1 | 1 | 1 | 3 | 1 |
| 3_6224593 | NA | NA | NA | NA | NA | NA | 2 | 1 | 2 | 1 | 1 | 0 | NA | NA | 1 | 1 | 1 | 1 | 3 | 0.6667 |
| 3_6224597 | NA | NA | NA | NA | NA | NA | 2 | 1 | 2 | 1 | 1 | 1 | NA | NA | 1 | 1 | NA | NA | 2 | 1 |
| 3_6224601 | NA | NA | NA | NA | NA | NA | 2 | 0.5 | 2 | 0.5 | 1 | 1 | NA | NA | 1 | 0 | NA | NA | 2 | 0.5 |
| 3_6224639 | NA | NA | NA | NA | NA | NA | 2 | 0.5 | 2 | 0.5 | 1 | 1 | NA | NA | 1 | 1 | NA | NA | 2 | 1 |
| 3_6224670 | NA | NA | NA | NA | NA | NA | 2 | 0.5 | 2 | 0.5 | 1 | 1 | NA | NA | NA | NA | NA | NA | 1 | 1 |
| 3_6224677 | NA | NA | NA | NA | NA | NA | 1 | 1 | 1 | 1 | 1 | 1 | NA | NA | 1 | 1 | NA | NA | 2 | 1 |
| 3_6224683 | NA | NA | NA | NA | NA | NA | 1 | 1 | 1 | 1 | 3 | 0 | 1 | 1 | NA | NA | 3 | 0 | 7 | 0.1429 |
| 3_6224735 | NA | NA | NA | NA | 1 | 1 | 1 | 0 | 2 | 0.5 | 3 | 0 | 2 | 0 | NA | NA | 3 | 0.6667 | 8 | 0.25 |
| 3_6224738 | NA | NA | NA | NA | 1 | 1 | NA | NA | 1 | 1 | 2 | 1 | 1 | 1 | NA | NA | 2 | 0.5 | 5 | 0.8 |
| 3_6224747( | NA | NA | NA | NA | 1 | 0 | 1 | 1 | 2 | 0.5 | 3 | 0.3333 | 1 | 0 | NA | NA | 2 | 0 | 6 | 0.1667 |
| 3_6224749 | NA | NA | NA | NA | 1 | 1 | 1 | 0 | 2 | 0.5 | 2 | 1 | 2 | 0.5 | NA | NA | 3 | 0.6667 | 7 | 0.7143 |
| 3_6224751 | NA | NA | NA | NA | 1 | 1 | 2 | 0.5 | 3 | 0.6667 | 2 | 1 | 2 | 0.5 | NA | NA | 2 | 1 | 6 | 0.8333 |
| 3_6224754 | NA | NA | NA | NA | NA | NA | 1 | 1 | 1 | 1 | 1 | 1 | NA | NA | NA | NA | NA | NA | 1 | 1 |
| 3_6224755 | NA | NA | NA | NA | 1 | 0 | 2 | 0.5 | 3 | 0.3333 | 1 | 1 | 1 | 0 | NA | NA | 2 | 0 | 4 | 0.25 |
| 3_6224769 | NA | NA | NA | NA | 1 | 0 | 2 | 0.5 | 3 | 0.3333 | 2 | 0.5 | 2 | 0.5 | 1 | 0 | 3 | 0.6667 | 8 | 0.5 |
| 3_6224772 | NA | NA | NA | NA | NA | NA | 1 | 1 | 1 | 1 | 1 | 1 | NA | NA | NA | NA | NA | NA | 1 | 1 |
| 3_6224776 | NA | NA | NA | NA | 1 | 0 | 2 | 0.5 | 3 | 0.3333 | 3 | 0.3333 | 2 | 0 | NA | NA | 3 | 0.3333 | 8 | 0.25 |
| 3_6224781 | NA | NA | NA | NA | NA | NA | 1 | 1 | 1 | 1 | 2 | 0 | 2 | 0.5 | 1 | 0 | 2 | 0 | 7 | 0.1429 |
| 3_6236774 | 1 | 1 | 3 | 1 | 2 | 1 | 3 | 1 | 9 | 1 | 4 | 1 | 2 | 1 | 3 | 1 | 6 | 1 | 15 | 1 |
| 3_6236779 | 1 | 1 | 3 | 1 | 2 | 1 | 3 | 0.3333 | 9 | 0.7778 | 4 | 1 | 3 | 0 | 4 | 1 | 6 | 0.8333 | 17 | 0.7647 |
| 3_6236784( | 1 | 1 | 3 | 0 | 2 | 0 | 3 | 0 | 9 | 0.1111 | 4 | 0 | 3 | 0 | 4 | 0 | 6 | 0.5 | 17 | 0.1765 |
| 3_6236867 | 1 | 0 | 3 | 0 | 2 | 1 | 4 | 0 | 10 | 0.2 | 5 | 0.2 | 3 | 0 | 4 | 0 | 6 | 0 | 18 | 0.0556 |
| 3_6236875 | 2 | 0 | 3 | 0 | 2 | 0 | 4 | 0.25 | 11 | 0.0909 | 4 | 0.5 | 2 | 0 | 4 | 0 | 6 | 0 | 16 | 0.125 |
| 3_6237628 | 1 | 0 | 2 | 0 | 2 | 0.5 | 8 | 0.125 | 13 | 0.1538 | 2 | 0 | 2 | 0 | 2 | 0.5 | 5 | 0.2 | 11 | 0.1818 |
| 3_6237779 | 6 | 0.3333 | 9 | 0.1111 | 9 | 0.1111 | 14 | 0.1429 | 38 | 0.1579 | 6 | 0 | 8 | 0.125 | 10 | 0.4 | 9 | 0.1111 | 33 | 0.1818 |
| 3_6237802( | 5 | 0 | 9 | 0 | 7 | 0.2857 | 11 | 0.4545 | 32 | 0.2188 | 4 | 0 | 5 | 0.2 | 8 | 0 | 6 | 0.1667 | 23 | 0.087 |
| 3_6237805 | 6 | 0 | 8 | 0.125 | 7 | 0.5714 | 12 | 0.3333 | 33 | 0.2727 | 6 | 0 | 3 | 0 | 9 | 0.2222 | 8 | 0.125 | 26 | 0.1154 |
| 3_6237808 | 6 | 1 | 8 | 0.75 | 7 | 1 | 13 | 0.7692 | 34 | 0.8529 | 6 | 0.5 | 5 | 1 | 8 | 0.75 | 8 | 0.875 | 27 | 0.7778 |
| 3_6237816 | 6 | 0.1667 | 8 | 0.125 | 7 | 0.1429 | 12 | 0.25 | 33 | 0.1818 | 3 | 0 | 5 | 0.2 | 9 | 0.2222 | 6 | 0 | 23 | 0.1304 |
| 3_6243722 | 7 | 0.5714 | 1 | 1 | 4 | 0.25 | 10 | 0.6 | 22 | 0.5455 | 2 | 0.5 | 5 | 0.6 | 4 | 0.75 | 2 | 0.5 | 13 | 0.6154 |
| 3_6243745 | 8 | 0.25 | 1 | 1 | 3 | 0.6667 | 14 | 0.6429 | 26 | 0.5385 | 2 | 0 | 5 | 0.2 | 4 | 0.5 | 1 | 0 | 12 | 0.25 |
| 3_6243774 | 5 | 0.2 | 1 | 0 | 2 | 0 | 11 | 0 | 19 | 0.0526 | 2 | 0 | 3 | 0 | 3 | 0.3333 | 1 | 0 | 9 | 0.1111 |
| 3_6243788 | 5 | 0.6 | 1 | 1 | 3 | 0.6667 | 13 | 0.4615 | 22 | 0.5455 | 2 | 1 | 2 | 1 | 3 | 0.6667 | NA | NA | 7 | 0.8571 |
| 3_6243790 | 6 | 0.8333 | 1 | 1 | 3 | 0.6667 | 10 | 0.8 | 20 | 0.8 | 2 | 0 | 1 | 1 | 5 | 0.8 | NA | NA | 8 | 0.625 |
| 3_6243794 | 6 | 0.3333 | 1 | 1 | 2 | 0.5 | 10 | 0.6 | 19 | 0.5263 | 2 | 0.5 | 2 | 0.5 | 5 | 0.2 | NA | NA | 9 | 0.3333 |
| 3_6243805 | 5 | 0.6 | NA | NA | 1 | 0 | 8 | 0 | 14 | 0.2143 | 1 | 0 | 2 | 0.5 | 5 | 0.2 | NA | NA | 8 | 0.25 |
| 3_6243814 | 6 | 0.1667 | 1 | 0 | 3 | 0 | 11 | 0.3636 | 21 | 0.2381 | 2 | 0.5 | 2 | 0 | 5 | 0.8 | NA | NA | 9 | 0.5556 |
| 3_6243825 | 6 | 0.8333 | 1 | 1 | 3 | 1 | 11 | 0.5455 | 21 | 0.7143 | 2 | 0 | 2 | 0.5 | 5 | 0.8 | NA | NA | 9 | 0.5556 |
| 3_6243831 | 6 | 1 | 1 | 1 | 3 | 1 | 11 | 0.5455 | 21 | 0.7619 | 2 | 1 | 2 | 0.5 | 4 | 0.75 | NA | NA | 8 | 0.75 |
| 3_6243834 | 3 | 0.3333 | 1 | 0 | NA | NA | 7 | 0.7143 | 11 | 0.5455 | 1 | 1 | 2 | 1 | 4 | 0.75 | NA | NA | 7 | 0.8571 |
| 3_6243848 | 5 | 0.6 | NA | NA | 3 | 0.3333 | 11 | 0.3636 | 19 | 0.4211 | 2 | 0.5 | 2 | 0 | 5 | 0.8 | NA | NA | 9 | 0.5556 |

| | | | | | | | | | | | | | | | | | | | | |
|---|---|---|---|---|---|---|---|---|---|---|---|---|---|---|---|---|---|---|---|---|
| 3_6250227 | 4 | 0 | NA | NA | 3 | 0 | 9 | 0.2222 | 16 | 0.125 | 2 | 0 | 1 | 0 | 5 | 0.4 | NA | NA | 8 | 0.25 |
| 3_6250235 | 3 | 0.6667 | NA | NA | 2 | 1 | 6 | 0.1667 | 11 | 0.4545 | 1 | 0 | 1 | 1 | 3 | 1 | NA | NA | 5 | 0.8 |
| 3_6250242 | 4 | 0.25 | NA | NA | 3 | 0 | 9 | 0.1111 | 16 | 0.125 | 2 | 0 | 1 | 0 | 5 | 0.2 | NA | NA | 8 | 0.125 |
| 3_6250246 | 4 | 0.25 | NA | NA | 3 | 0 | 8 | 0.375 | 15 | 0.2667 | 2 | 0 | 1 | 1 | 3 | 0.3333 | NA | NA | 6 | 0.3333 |
| 3_6250252 | 4 | 0.25 | NA | NA | 2 | 0 | 9 | 0 | 15 | 0.0667 | 2 | 0 | 1 | 0 | 4 | 0.25 | NA | NA | 7 | 0.1429 |
| 3_6250262 | 3 | 1 | NA | NA | 2 | 1 | 8 | 1 | 13 | 1 | 2 | 1 | NA | NA | 3 | 1 | NA | NA | 5 | 1 |
| 3_6251111 | 2 | 0 | NA | NA | 2 | 1 | 9 | 0.4444 | 13 | 0.4615 | 2 | 0.5 | 1 | 0 | 3 | 0.6667 | NA | NA | 6 | 0.5 |
| 3_6251132 | 3 | 0.3333 | NA | NA | 2 | 0 | 8 | 0.125 | 13 | 0.1538 | 2 | 0 | 1 | 1 | 3 | 0 | NA | NA | 6 | 0.1667 |
| 3_6251153 | 2 | 0.5 | NA | NA | 2 | 1 | 4 | 1 | 8 | 0.875 | 2 | 0.5 | 1 | 0 | 3 | 1 | NA | NA | 6 | 0.6667 |
| 3_6251155 | 2 | 0.5 | NA | NA | 2 | 0 | 7 | 0.4286 | 11 | 0.3636 | 2 | 0 | 1 | 1 | 2 | 0.5 | NA | NA | 5 | 0.4 |
| 3_6251159 | 1 | 0 | NA | NA | 1 | 0 | 4 | 0.25 | 6 | 0.1667 | 2 | 0.5 | 1 | 1 | 2 | 0.5 | NA | NA | 5 | 0.6 |
| 3_6251180 | 1 | 1 | NA | NA | 1 | 1 | 7 | 1 | 9 | 1 | 1 | 1 | 1 | 1 | 3 | 1 | NA | NA | 5 | 1 |
| 3_6251186 | 2 | 1 | NA | NA | 1 | 1 | 6 | 0.8333 | 9 | 0.8889 | 2 | 0.5 | 1 | 1 | 3 | 0.6667 | NA | NA | 6 | 0.6667 |
| 3_6251189 | NA | NA | NA | NA | 1 | 0 | 7 | 0.2857 | 8 | 0.25 | 1 | 0 | 1 | 0 | 3 | 0.6667 | NA | NA | 5 | 0.4 |
| 3_6251192 | 2 | 0.5 | NA | NA | 1 | 1 | 7 | 0.4286 | 10 | 0.5 | 2 | 0 | NA | NA | 3 | 0.6667 | NA | NA | 5 | 0.4 |
| 3_6251196 | 2 | 0.5 | NA | NA | 1 | 1 | 7 | 0.8571 | 10 | 0.8 | 1 | 0 | 1 | 1 | 3 | 0.6667 | NA | NA | 5 | 0.6 |
| 3_6251203 | 2 | 0.5 | NA | NA | 1 | 1 | 5 | 0.2 | 8 | 0.375 | 1 | 0 | 1 | 0 | 3 | 0.3333 | NA | NA | 5 | 0.2 |
| 3_6279674 | NA | NA | NA | NA | 2 | 1 | 2 | 1 | 4 | 1 | NA | NA | NA | NA | 1 | 1 | NA | NA | 1 | 1 |
| 3_6328766 | 2 | 0.5 | 1 | 0 | 4 | 0 | 2 | 1 | 9 | 0.3333 | 2 | 0.5 | 2 | 0.5 | 5 | 0.6 | 1 | 1 | 10 | 0.6 |
| 3_6328781 | 2 | 0 | 1 | 0 | 4 | 0.25 | 2 | 0.5 | 9 | 0.2222 | 2 | 1 | 3 | 0.3333 | 5 | 0.2 | 1 | 1 | 11 | 0.4545 |
| 3_6328808 | 2 | 0 | NA | NA | 4 | 0 | 2 | 0.5 | 8 | 0.125 | 2 | 0 | 2 | 0 | 5 | 0.2 | 1 | 0 | 10 | 0.1 |
| 3_6328821 | 3 | 0 | 1 | 1 | 3 | 0.3333 | 3 | 0 | 10 | 0.2 | 2 | 0.5 | 1 | 0 | 5 | 0.4 | 1 | 0 | 9 | 0.3333 |
| 3_6328824 | 3 | 0 | 1 | 1 | 5 | 0 | 2 | 0 | 11 | 0.0909 | 2 | 0 | 2 | 0 | 5 | 0.2 | NA | NA | 9 | 0.1111 |
| 3_6328859 | 3 | 0.3333 | 1 | 0 | 5 | 0 | 4 | 0.25 | 13 | 0.1538 | 2 | 0 | 1 | 0 | 5 | 0.2 | 2 | 0 | 10 | 0.1 |
| 3_6328864 | 4 | 0.5 | 1 | 0 | 5 | 0 | 3 | 0 | 13 | 0.1538 | 2 | 0.5 | 2 | 0 | 4 | 0.75 | 1 | 1 | 9 | 0.5556 |
| 3_6328894 | 4 | 0.5 | 1 | 0 | 5 | 0.2 | 3 | 0.3333 | 13 | 0.3077 | 2 | 1 | 2 | 0.5 | 6 | 0.6667 | 1 | 0 | 11 | 0.6364 |
| 3_6328949 | 5 | 0.4 | 1 | 0 | 3 | 0 | 3 | 0 | 12 | 0.1667 | 2 | 0.5 | 1 | 0 | 5 | 0.4 | 2 | 0 | 10 | 0.3 |
| 3_6328950 | 5 | 0 | 1 | 0 | 4 | 0.25 | 3 | 0 | 13 | 0.0769 | 3 | 0.3333 | 1 | 1 | 5 | 0.4 | 2 | 0.5 | 11 | 0.4545 |
| 3_6328956 | 5 | 0 | 1 | 0 | 4 | 0.25 | 2 | 0 | 12 | 0.0833 | 3 | 1 | 1 | 0 | 3 | 0.3333 | 2 | 0 | 9 | 0.4444 |
| 3_6382271 | 2 | 0.5 | 1 | 1 | 1 | 1 | 2 | 1 | 6 | 0.8333 | 1 | 1 | 2 | 1 | 1 | 1 | 1 | 1 | 5 | 1 |
| 3_6382280 | 3 | 0 | 1 | 1 | 1 | 0 | 2 | 0 | 7 | 0.1429 | 1 | 0 | 1 | 0 | 1 | 1 | 1 | 0 | 4 | 0.25 |
| 3_6382283 | 3 | 1 | 1 | 0 | 1 | 0 | 2 | 0.5 | 7 | 0.5714 | 1 | 1 | 2 | 0 | 1 | 0 | 1 | 0 | 5 | 0.2 |
| 3_6382287 | 3 | 0.6667 | 1 | 1 | 1 | 1 | 2 | 0.5 | 7 | 0.7143 | NA | NA | 2 | 0.5 | 1 | 1 | 1 | 1 | 4 | 0.75 |
| 3_6382292 | 3 | 1 | 1 | 1 | 1 | 1 | 1 | 1 | 6 | 1 | NA | NA | 2 | 1 | 1 | 1 | 1 | 0 | 4 | 0.75 |
| 3_6382293 | 3 | 0.3333 | 1 | 0 | 1 | 0 | 2 | 0.5 | 7 | 0.2857 | NA | NA | 2 | 0.5 | 1 | 1 | 1 | 1 | 4 | 0.75 |
| 3_6382295 | 3 | 1 | 1 | 1 | 1 | 0 | 1 | 0 | 6 | 0.6667 | NA | NA | 2 | 0 | 1 | 0 | 1 | 1 | 4 | 0.25 |
| 3_6382307 | 3 | 0.6667 | NA | NA | NA | NA | 2 | 0.5 | 5 | 0.6 | NA | NA | 2 | 0 | 1 | 0 | 1 | 1 | 4 | 0.25 |
| 3_6385065 | 5 | 0.6 | 3 | 0.6667 | 3 | 0.3333 | 3 | 0.3333 | 14 | 0.5 | 2 | 1 | 2 | 0.5 | 3 | 1 | 3 | 0.6667 | 10 | 0.8 |
| 3_6385070 | 7 | 0.1429 | 4 | 0 | 2 | 0.5 | 5 | 0 | 18 | 0.1111 | 1 | 0 | 4 | 0 | 6 | 0.1667 | 4 | 0 | 15 | 0.0667 |
| 3_6387089 | 5 | 0.4 | 5 | 0.8 | 2 | 0.5 | 4 | 1 | 16 | 0.6875 | NA | NA | 3 | 0 | 3 | 0.3333 | 2 | 0.5 | 8 | 0.25 |
| 3_6387100 | 4 | 0.25 | 5 | 0.6 | 1 | 0 | 4 | 0.5 | 14 | 0.4286 | NA | NA | 3 | 0.3333 | 3 | 0.3333 | 2 | 0.5 | 8 | 0.375 |
| 3_6387103 | 4 | 0.75 | 4 | 0.5 | 1 | 1 | 3 | 0.3333 | 12 | 0.5833 | NA | NA | 3 | 0 | 3 | 0.6667 | 2 | 0.5 | 8 | 0.375 |
| 3_6387106 | 5 | 1 | 5 | 1 | 1 | 1 | 4 | 1 | 15 | 1 | NA | NA | 3 | 0.6667 | 3 | 1 | 2 | 1 | 8 | 0.875 |
| 3_6387112 | 5 | 0.6 | 5 | 0.4 | 1 | 0 | 4 | 0.25 | 15 | 0.4 | NA | NA | 3 | 0 | 2 | 1 | 2 | 1 | 7 | 0.5714 |
| 3_6387120 | 3 | 1 | 4 | 1 | NA | NA | 3 | 0.6667 | 10 | 0.9 | NA | NA | 2 | 1 | 2 | 1 | 1 | 1 | 5 | 1 |
| 3_6387139 | 4 | 1 | 2 | 1 | 1 | 0 | 1 | 1 | 8 | 0.875 | NA | NA | 2 | 1 | 1 | 1 | 1 | 1 | 4 | 1 |
| 3_6387143 | 4 | 0.5 | 4 | 0.5 | 1 | 1 | 4 | 0.25 | 13 | 0.4615 | NA | NA | 3 | 0.6667 | 2 | 0.5 | 2 | 0.5 | 7 | 0.5714 |
| 3_6387149 | 3 | 1 | 3 | 1 | 1 | 1 | 4 | 0.5 | 11 | 0.8182 | NA | NA | 2 | 1 | 2 | 1 | 2 | 0.5 | 6 | 0.8333 |
| 3_6387158 | 4 | 0.25 | 4 | 0.5 | 1 | 0 | 4 | 0.25 | 13 | 0.3077 | NA | NA | 1 | 0 | 2 | 1 | 2 | 1 | 5 | 0.8 |
| 3_6387166 | 4 | 0.25 | 4 | 0.75 | 1 | 1 | 1 | 1 | 10 | 0.6 | NA | NA | 1 | 1 | 1 | 0 | 2 | 0.5 | 4 | 0.5 |
| 3_6387167 | 5 | 0.2 | 4 | 0.25 | 1 | 0 | 4 | 0.5 | 14 | 0.2857 | NA | NA | 3 | 0 | 2 | 0.5 | 1 | 0 | 6 | 0.1667 |
| 3_6392417 | NA | NA | 3 | 0.3333 | NA | NA | 5 | 0.8 | 8 | 0.625 | NA | NA | 2 | 0.5 | 3 | 0.3333 | 2 | 0 | 7 | 0.2857 |
| 3_6392496 | NA | NA | 3 | 0.3333 | 1 | 0 | 4 | 0 | 8 | 0.125 | 1 | 0 | 2 | 0 | 4 | 0 | 4 | 0.25 | 11 | 0.0909 |
| 3_6392618 | 3 | 0 | 4 | 0 | 1 | 0 | 4 | 0.25 | 12 | 0.0833 | 2 | 0.5 | 3 | 0 | 4 | 0 | 3 | 0 | 12 | 0.0833 |
| 3_6392688 | 4 | 0.25 | 4 | 0 | 3 | 0 | 5 | 0.2 | 16 | 0.125 | 5 | 0.2 | 4 | 0 | 4 | 0 | 4 | 0 | 17 | 0.0588 |
| 3_6392709 | 4 | 0.25 | 4 | 0.25 | 4 | 0 | 4 | 0.25 | 16 | 0.1875 | 5 | 0.4 | 3 | 0.3333 | 5 | 0.2 | 5 | 0.2 | 18 | 0.2778 |
| 3_6392713 | 3 | 0.6667 | 3 | 0.3333 | 4 | 0 | 5 | 0 | 15 | 0.2 | 3 | 0 | 2 | 0 | 6 | 0.1667 | 4 | 0.25 | 15 | 0.1333 |
| 3_6392717 | 4 | 0 | 4 | 0.25 | 4 | 0 | 2 | 0 | 14 | 0.0714 | 5 | 0 | 4 | 0.25 | 3 | 0 | 5 | 0 | 17 | 0.0588 |
| 3_6392727 | 6 | 0.1667 | 4 | 0.25 | 4 | 0 | 5 | 0 | 19 | 0.1053 | 5 | 0.2 | 4 | 0.25 | 6 | 0.3333 | 4 | 0 | 19 | 0.2105 |
| 3_6392734 | 5 | 0 | 4 | 0.25 | 3 | 0 | 5 | 0 | 17 | 0.0588 | 3 | 0 | 3 | 0 | 6 | 0.1667 | 3 | 0 | 15 | 0.0667 |
| 3_6392818 | 9 | 0.1111 | 8 | 0.125 | 8 | 0 | 5 | 0 | 30 | 0.0667 | 5 | 0.2 | 5 | 0.2 | 8 | 0 | 7 | 0.1429 | 25 | 0.12 |
| 3_6392829 | 7 | 0.2857 | 8 | 0.375 | 7 | 0.2857 | 4 | 0 | 26 | 0.2692 | 5 | 0.2 | 4 | 0.25 | 6 | 0.1667 | 6 | 0.1667 | 21 | 0.1905 |
| 3_6392865 | 10 | 0 | 9 | 0.1111 | 10 | 0.1 | 6 | 0 | 35 | 0.0571 | 5 | 0.2 | 7 | 0 | 11 | 0 | 8 | 0.125 | 31 | 0.0645 |
| 3_6501533 | NA | NA | 1 | 1 | NA | NA | 1 | 1 | 2 | 1 | NA | NA | 1 | 1 | 1 | 1 | NA | NA | 2 | 1 |
| 3_6501540 | NA | NA | 1 | 1 | NA | NA | 2 | 1 | 3 | 1 | NA | NA | 1 | 1 | 1 | 1 | NA | NA | 2 | 1 |

| | | | | | | | | | | | | | | | | | | | | |
|---|---|---|---|---|---|---|---|---|---|---|---|---|---|---|---|---|---|---|---|---|
| 3_6501547 | NA | NA | 1 | 1 | NA | NA | 1 | 1 | 2 | 1 | NA | NA | 1 | 1 | 1 | 1 | NA | NA | 2 | 1 |
| 3_6501557 | NA | NA | 1 | 1 | NA | NA | 2 | 1 | 3 | 1 | NA | NA | 1 | 1 | NA | NA | NA | NA | 1 | 1 |
| 3_6501559 | NA | NA | 1 | 0 | NA | NA | 2 | 1 | 3 | 0.6667 | NA | NA | 1 | 1 | 1 | 0 | NA | NA | 2 | 0.5 |
| 3_6501626 | NA | NA | 1 | 1 | NA | NA | 1 | 1 | 2 | 1 | NA | NA | NA | NA | 1 | 1 | NA | NA | 1 | 1 |
| 3_6501640 | NA | NA | 1 | 1 | NA | NA | 1 | 1 | 2 | 1 | NA | NA | 1 | 1 | 1 | 1 | NA | NA | 2 | 1 |
| 3_6501648 | NA | NA | 6 | 0.1667 | 4 | 0 | 4 | 0 | 14 | 0.0714 | 2 | 0 | 8 | 0.125 | 1 | 0 | 4 | 0 | 15 | 0.0667 |
| 3_6501664 | NA | NA | 1 | 1 | NA | NA | NA | NA | 1 | 1 | NA | NA | NA | NA | 1 | 1 | NA | NA | 1 | 1 |
| 3_6501680 | 1 | 1 | 11 | 0.5455 | 9 | 0.4444 | 12 | 0.6667 | 33 | 0.5758 | 7 | 0.4286 | 15 | 0.6667 | 8 | 0.375 | 9 | 0.6667 | 39 | 0.5641 |
| 3_6501700 | 4 | 0.5 | 11 | 0.3636 | 11 | 0.4545 | 14 | 0.6429 | 40 | 0.5 | 9 | 0.4444 | 18 | 0.6667 | 9 | 0.4444 | 9 | 0.3333 | 45 | 0.5111 |
| 3_6501706 | 4 | 0.75 | 13 | 0.2308 | 11 | 0.3636 | 13 | 0.4615 | 41 | 0.3902 | 8 | 0.25 | 15 | 0.2667 | 9 | 0.2222 | 12 | 0.5 | 44 | 0.3182 |
| 3_6501710 | 6 | 0.5 | 17 | 0.3529 | 14 | 0.2143 | 15 | 0.3333 | 52 | 0.3269 | 9 | 0.1111 | 17 | 0.4706 | 13 | 0.3077 | 13 | 0.0769 | 52 | 0.2692 |
| 3_6501717 | 6 | 0.1667 | 18 | 0.1667 | 15 | 0.2667 | 17 | 0.2353 | 56 | 0.2143 | 9 | 0.1111 | 20 | 0.2 | 15 | 0.2 | 15 | 0.0667 | 59 | 0.1525 |
| 3_6528397 | 12 | 0.75 | 19 | 0.2632 | 20 | 0.3 | 23 | 0.3913 | 74 | 0.3919 | 12 | 0.5833 | 31 | 0.3548 | 23 | 0.3913 | 25 | 0.28 | 91 | 0.3736 |
| 3_6528421 | 8 | 0.125 | 11 | 0.2727 | 13 | 0.2308 | 16 | 0.3125 | 48 | 0.25 | 7 | 0.5714 | 18 | 0.2778 | 21 | 0.381 | 14 | 0.2857 | 60 | 0.35 |
| 3_6528847 | 23 | 0.0435 | 14 | 0.2143 | 18 | 0.0556 | 16 | 0 | 71 | 0.0704 | 12 | 0.0833 | 14 | 0 | 22 | 0.0455 | 14 | 0.1429 | 62 | 0.0645 |
| 3_6528907 | 18 | 0.0556 | 14 | 0.0714 | 11 | 0.0909 | 12 | 0.0833 | 55 | 0.0727 | 9 | 0.1111 | 10 | 0.1 | 19 | 0.0526 | 13 | 0 | 51 | 0.0588 |
| 3_6543517 | 1 | 1 | NA | NA | NA | NA | NA | NA | 1 | 1 | NA | NA | NA | NA | 1 | 1 | NA | NA | 1 | 1 |
| 3_6543871 | NA | NA | NA | NA | 1 | 0 | 2 | 0.5 | 3 | 0.3333 | 2 | 0.5 | 2 | 0.5 | 2 | 0 | 2 | 0 | 8 | 0.25 |
| 3_6543882 | 1 | 1 | NA | NA | 1 | 0 | 1 | 0 | 3 | 0.3333 | 1 | 0 | 2 | 0 | 2 | 0.5 | 1 | 1 | 6 | 0.3333 |
| 3_6543883 | 1 | 1 | NA | NA | 1 | 1 | 1 | 1 | 3 | 1 | 2 | 0 | 3 | 0.6667 | 1 | 1 | 2 | 1 | 8 | 0.625 |
| 3_6543891 | 1 | 0 | NA | NA | 1 | 0 | 2 | 0.5 | 4 | 0.25 | 2 | 0.5 | 3 | 0.3333 | 2 | 1 | 2 | 0.5 | 9 | 0.5556 |
| 3_6543894 | 1 | 0 | NA | NA | 1 | 0 | 2 | 0.5 | 4 | 0.25 | 2 | 0 | 2 | 0 | 2 | 0 | 2 | 0.5 | 8 | 0.125 |
| 3_6543903 | 1 | 0 | NA | NA | 1 | 1 | 2 | 0 | 4 | 0.25 | 1 | 0 | 3 | 0.3333 | 2 | 0.5 | 2 | 1 | 8 | 0.5 |
| 3_6543913 | 1 | 0 | NA | NA | 1 | 1 | 1 | 1 | 3 | 0.6667 | 2 | 0.5 | 2 | 1 | 2 | 1 | 2 | 0.5 | 8 | 0.75 |
| 3_6543915 | 1 | 1 | NA | NA | 1 | 1 | 2 | 1 | 4 | 1 | 2 | 1 | 3 | 0.6667 | 2 | 0.5 | 2 | 1 | 9 | 0.7778 |
| 3_6543926 | 1 | 1 | NA | NA | 1 | 0 | 2 | 0 | 4 | 0.25 | 2 | 0 | 3 | 0 | 2 | 0 | 2 | 0.5 | 9 | 0.1111 |
| 3_6543936 | 1 | 1 | NA | NA | NA | NA | 1 | 1 | 2 | 1 | 2 | 1 | 3 | 1 | 2 | 1 | NA | NA | 7 | 1 |
| 3_6543942 | NA | NA | NA | NA | 2 | 0.5 | 2 | 1 | 4 | 0.75 | 2 | 1 | 3 | 1 | 2 | 1 | 2 | 1 | 9 | 1 |
| 3_6543950 | 1 | 1 | NA | NA | 2 | 1 | 2 | 0.5 | 5 | 0.8 | 2 | 0.5 | 3 | 1 | 1 | 1 | 1 | 1 | 7 | 0.8571 |
| 3_6546008 | 1 | 1 | NA | NA | 1 | 0 | 3 | 0.6667 | 5 | 0.6 | 2 | 0.5 | 3 | 0.6667 | 2 | 0 | 2 | 0 | 9 | 0.3333 |
| 3_6546015 | 2 | 0 | NA | NA | 2 | 1 | 1 | 0 | 5 | 0.4 | 2 | 0 | 4 | 0.25 | 2 | 0.5 | NA | NA | 8 | 0.25 |
| 3_6546055 | NA | NA | NA | NA | 1 | 1 | 2 | 0.5 | 3 | 0.6667 | 2 | 0 | 4 | 0.25 | 1 | 0 | 1 | 0 | 8 | 0.125 |
| 3_6546059 | 1 | 1 | NA | NA | 1 | 1 | 2 | 1 | 4 | 1 | 2 | 1 | 4 | 1 | 2 | 1 | 2 | 1 | 10 | 1 |
| 3_6546069 | 1 | 1 | NA | NA | 2 | 0.5 | 4 | 0 | 7 | 0.2857 | 1 | 0 | 4 | 0.25 | 2 | 0.5 | 2 | 0.5 | 9 | 0.3333 |
| 3_6546073 | 1 | 0 | NA | NA | 2 | 0.5 | 3 | 0 | 6 | 0.1667 | 2 | 0 | 2 | 0.5 | 2 | 0.5 | 2 | 1 | 8 | 0.5 |
| 3_6546086 | 2 | 0.5 | NA | NA | 2 | 1 | 3 | 0.3333 | 7 | 0.5714 | 2 | 0.5 | 4 | 0.5 | 1 | 0 | 2 | 0.5 | 9 | 0.4444 |
| 3_6546087 | 1 | 0 | NA | NA | 1 | 1 | 3 | 0.3333 | 5 | 0.4 | 2 | 0.5 | 4 | 0.25 | 2 | 0.5 | 2 | 0.5 | 10 | 0.4 |
| 3_6546101 | 1 | 0 | NA | NA | 2 | 0.5 | 4 | 0.5 | 7 | 0.4286 | 2 | 0.5 | 3 | 0 | 2 | 0 | 2 | 0 | 9 | 0.1111 |
| 3_6546142 | 2 | 0.5 | 1 | 0 | 2 | 0 | 4 | 0 | 9 | 0.1111 | 1 | 0 | 4 | 0.25 | 2 | 0 | 2 | 0 | 9 | 0.1111 |
| 3_6546153 | 1 | 0 | 1 | 0 | 2 | 0 | 4 | 0.25 | 8 | 0.125 | 2 | 0 | 4 | 0.25 | 2 | 0.5 | 2 | 0.5 | 10 | 0.3 |
| 3_6546188 | 2 | 0 | 1 | 0 | 2 | 0 | 2 | 0.5 | 7 | 0.1429 | 2 | 0.5 | 4 | 0.25 | 2 | 0.5 | 2 | 0 | 10 | 0.3 |
| 3_6546197 | 3 | 0.3333 | 1 | 0 | 2 | 0.5 | 2 | 0 | 8 | 0.25 | 2 | 0 | 4 | 0.25 | 2 | 0.5 | 2 | 0 | 10 | 0.2 |
| 3_6585385 | 2 | 1 | 5 | 0 | 2 | 0.5 | 6 | 0.3333 | 15 | 0.3333 | 4 | 0 | 8 | 0.125 | 6 | 0.1667 | 7 | 0 | 25 | 0.08 |
| 3_6585387 | 2 | 0 | 6 | 0.8333 | 2 | 0.5 | 6 | 0.3333 | 16 | 0.5 | 4 | 0.75 | 9 | 0.6667 | 5 | 0.2 | 5 | 0.4 | 23 | 0.5217 |
| 3_6585393 | 1 | 0 | 2 | 0 | 2 | 0.5 | 2 | 1 | 7 | 0.4286 | NA | NA | 1 | 0 | 1 | 1 | 1 | 1 | 3 | 0.6667 |
| 3_6585393 | NA | NA | NA | NA | NA | NA | 2 | 0.5 | 2 | 0.5 | 3 | 0 | 2 | 0 | 1 | 0 | 2 | 0.5 | 8 | 0.125 |
| 3_6585395 | 2 | 1 | 5 | 1 | 2 | 1 | 7 | 1 | 16 | 1 | 4 | 1 | 9 | 1 | 6 | 1 | 7 | 1 | 26 | 1 |
| 3_6585402 | 2 | 0.5 | 6 | 0.3333 | 2 | 1 | 5 | 0.2 | 15 | 0.4 | 4 | 0.25 | 6 | 0 | 5 | 0 | 6 | 0.3333 | 21 | 0.1429 |
| 3_6585404 | 2 | 0 | 6 | 0.1667 | 2 | 0 | 7 | 0.5714 | 17 | 0.2941 | 3 | 0.6667 | 9 | 0.3333 | 5 | 0.6 | 7 | 0.1429 | 24 | 0.375 |
| 3_6585411 | 2 | 0.5 | 4 | 0 | 3 | 0.3333 | 7 | 0.2857 | 16 | 0.25 | 4 | 0.25 | 9 | 0.1111 | 5 | 0 | 7 | 0 | 25 | 0.08 |
| 3_6585412 | 1 | 1 | 4 | 0.5 | 2 | 0.5 | 7 | 0.2857 | 14 | 0.4286 | 3 | 0 | 9 | 0.1111 | 5 | 0.4 | 4 | 0.5 | 21 | 0.2381 |
| 3_6585414 | 2 | 0 | 2 | 0.5 | 2 | 0 | 7 | 0 | 13 | 0.0769 | 4 | 0 | 7 | 0 | 4 | 0 | 5 | 0.2 | 20 | 0.05 |
| 3_6585421 | 2 | 0 | 3 | 0.6667 | 2 | 0 | 6 | 0.5 | 13 | 0.3846 | 3 | 0.3333 | 7 | 0.5714 | 3 | 0.3333 | 5 | 0 | 18 | 0.3333 |
| 3_6586585 | NA | NA | NA | NA | 1 | 1 | NA | NA | 1 | 1 | 1 | 1 | 1 | 1 | NA | NA | NA | NA | 2 | 1 |
| 3_6586594 | NA | NA | NA | NA | 1 | 1 | NA | NA | 1 | 1 | 1 | 1 | 1 | 1 | NA | NA | NA | NA | 2 | 1 |
| 3_6586828 | NA | NA | NA | NA | 1 | 1 | NA | NA | 1 | 1 | NA | NA | 1 | 1 | NA | NA | NA | NA | 1 | 1 |
| 3_6586930 | 1 | 1 | NA | NA | 2 | 1 | NA | NA | 3 | 1 | NA | NA | 1 | 1 | NA | NA | NA | NA | 1 | 1 |
| 3_6726365 | 3 | 0.3333 | 7 | 0.1429 | 6 | 0.6667 | 5 | 0.2 | 21 | 0.3333 | 4 | 0.75 | 3 | 1 | 2 | 0 | 3 | 0.3333 | 12 | 0.5833 |
| 3_6726391 | 3 | 0 | 8 | 0 | 4 | 0.25 | 4 | 0 | 19 | 0.0526 | 3 | 0 | 1 | 0 | NA | NA | 4 | 0.25 | 8 | 0.125 |
| 3_6726416 | 2 | 0 | 6 | 0.1667 | 4 | 0 | 3 | 0.3333 | 15 | 0.1333 | 3 | 0 | 1 | 0 | 1 | 0 | 2 | 1 | 7 | 0.2857 |
| 3_6726507 | 2 | 0 | 8 | 0.125 | 3 | 0 | 4 | 0.25 | 17 | 0.1176 | 4 | 0 | 2 | 0.5 | 2 | 0 | 4 | 0 | 12 | 0.0833 |
| 3_6726512 | 3 | 0 | 7 | 0.1429 | 3 | 0 | 4 | 0 | 17 | 0.0588 | 4 | 0 | 2 | 0 | 1 | 0 | 4 | 0.25 | 11 | 0.0909 |
| 3_6738217 | 6 | 0.3333 | 5 | 0.6 | 6 | 0.1667 | 9 | 0.5556 | 26 | 0.4231 | 2 | 0 | 3 | 0.6667 | 1 | 1 | 3 | 1 | 9 | 0.6667 |
| 3_6738218 | 5 | 0.2 | 5 | 0.6 | 5 | 0.4 | 8 | 0.125 | 23 | 0.3043 | 2 | 0.5 | 3 | 0.3333 | 3 | 0 | 2 | 0 | 10 | 0.2 |
| 3_6738220 | 7 | 0.2857 | 5 | 0.6 | 6 | 0.3333 | 9 | 0.4444 | 27 | 0.4074 | 2 | 1 | 4 | 0.5 | 4 | 1 | 3 | 0.3333 | 13 | 0.6923 |

| | | | | | | | | | | | | | | | | | | | | |
|---|---|---|---|---|---|---|---|---|---|---|---|---|---|---|---|---|---|---|---|---|
| 3_6751017 | 2 | 1 | 5 | 0.4 | 7 | 0.7143 | 2 | 0.5 | 16 | 0.625 | 2 | 0.5 | NA | NA | 2 | 0 | 3 | 0.6667 | 7 | 0.4286 |
| 3_6751025 | 4 | 0 | 4 | 0.25 | 8 | 0 | 3 | 0 | 19 | 0.0526 | 3 | 0.3333 | NA | NA | 4 | 0 | 3 | 0.3333 | 10 | 0.2 |
| 3_6852501 | 29 | 0.0345 | 23 | 0.1304 | 13 | 0 | 13 | 0.0769 | 78 | 0.0641 | 13 | 0.0769 | 16 | 0 | 16 | 0.1875 | 21 | 0.0476 | 66 | 0.0758 |
| 3_6853343 | 38 | 0.2895 | 34 | 0.2059 | 16 | 0.3125 | 22 | 0.1364 | 110 | 0.2364 | 18 | 0.1667 | 23 | 0.3043 | 19 | 0 | 26 | 0.1923 | 86 | 0.1744 |
| 3_6853350 | 37 | 0.1081 | 37 | 0.0541 | 17 | 0.0588 | 21 | 0.0476 | 112 | 0.0714 | 17 | 0 | 23 | 0.087 | 17 | 0.0588 | 23 | 0.1304 | 80 | 0.075 |
| 3_6877725 | NA | NA | NA | NA | NA | NA | 1 | 1 | 1 | 1 | 1 | 1 | NA | NA | 2 | 0 | 1 | 0 | 4 | 0.25 |
| 3_6877750 | NA | NA | NA | NA | NA | NA | 1 | 1 | 1 | 1 | 1 | 1 | 2 | 0.5 | 2 | 1 | 1 | 1 | 6 | 0.8333 |
| 3_6877774 | NA | NA | NA | NA | 1 | 1 | 1 | 1 | 2 | 1 | 2 | 1 | 2 | 1 | 2 | 1 | 1 | 1 | 7 | 1 |
| 3_6877787 | NA | NA | 2 | 0.5 | 1 | 1 | NA | NA | 3 | 0.6667 | 1 | 1 | 2 | 1 | 2 | 1 | NA | NA | 5 | 1 |
| 3_6877802 | NA | NA | 3 | 0.6667 | 1 | 0 | 1 | 1 | 5 | 0.6 | 2 | 1 | 2 | 1 | 2 | 1 | 1 | 1 | 7 | 1 |
| 3_6877821 | NA | NA | 3 | 0.6667 | 1 | 0 | 1 | 0 | 5 | 0.4 | 1 | 1 | 1 | 0 | 1 | 0 | 1 | 1 | 4 | 0.5 |
| 3_6877823 | NA | NA | 3 | 1 | 1 | 1 | 1 | 1 | 5 | 1 | 2 | 1 | 2 | 1 | 1 | 1 | 1 | 1 | 6 | 1 |
| 3_6877831 | NA | NA | 3 | 0.3333 | 1 | 0 | 1 | 0 | 5 | 0.2 | 2 | 0.5 | 2 | 1 | 2 | 1 | 1 | 1 | 7 | 0.8571 |
| 3_6877833 | NA | NA | 2 | 0 | 1 | 0 | 1 | 1 | 4 | 0.25 | 2 | 1 | 1 | 0 | 1 | 0 | 1 | 0 | 5 | 0.4 |
| 3_6877846 | NA | NA | 2 | 1 | 1 | 1 | 2 | 0.5 | 5 | 0.8 | 2 | 1 | 2 | 1 | 2 | 1 | 1 | 1 | 7 | 1 |
| 3_6877848 | NA | NA | 2 | 0.5 | 1 | 0 | 2 | 0 | 5 | 0.2 | 2 | 0.5 | 1 | 0 | 1 | 0 | 1 | 1 | 5 | 0.4 |
| 3_6877879 | NA | NA | 2 | 0.5 | 1 | 0 | 2 | 1 | 5 | 0.6 | 2 | 0 | 2 | 0 | 1 | 1 | NA | NA | 5 | 0.2 |
| 3_6877893 | NA | NA | 2 | 1 | 2 | 1 | 3 | 1 | 7 | 1 | 1 | 1 | 2 | 1 | 2 | 0.5 | 1 | 1 | 6 | 0.8333 |
| 3_6877904 | NA | NA | 1 | 0 | 2 | 0.5 | 3 | 0.6667 | 6 | 0.5 | 1 | 1 | 1 | 1 | NA | NA | NA | NA | 2 | 1 |
| 3_6877937 | 1 | 0 | 3 | 0 | 3 | 0 | 3 | 0.3333 | 10 | 0.1 | 2 | 0 | 2 | 0 | 2 | 0 | 1 | 1 | 7 | 0.1429 |
| 3_6880001 | 1 | 0 | 4 | 0.75 | 3 | 0.3333 | 1 | 0 | 9 | 0.4444 | 3 | 0.3333 | 3 | 0 | 2 | 0 | 2 | 0.5 | 10 | 0.2 |
| 3_6880137 | 1 | 0 | 1 | 1 | 1 | 0 | 2 | 0.5 | 5 | 0.4 | 1 | 0 | NA | NA | 1 | 1 | 2 | 0.5 | 4 | 0.5 |
| 3_6894185 | NA | NA | 5 | 0.6 | 1 | 0 | 1 | 0 | 7 | 0.4286 | NA | NA | NA | NA | 1 | 1 | 1 | 0 | 2 | 0.5 |
| 3_6894349 | 1 | 1 | 5 | 0.8 | 1 | 1 | NA | NA | 7 | 0.8571 | NA | NA | NA | NA | 1 | 1 | NA | NA | 1 | 1 |
| 3_6894436 | NA | NA | 6 | 1 | 2 | 1 | 1 | 1 | 9 | 1 | NA | NA | NA | NA | 1 | 1 | NA | NA | 1 | 1 |
| 3_6894486 | NA | NA | 5 | 0.8 | 1 | 1 | 1 | 1 | 7 | 0.8571 | NA | NA | NA | NA | 1 | 1 | NA | NA | 1 | 1 |
| 3_6894511 | NA | NA | 5 | 0.8 | 2 | 1 | 1 | 1 | 8 | 0.875 | NA | NA | NA | NA | 1 | 1 | NA | NA | 1 | 1 |
| 3_6894521 | NA | NA | 2 | 0.5 | 2 | 1 | NA | NA | 4 | 0.75 | NA | NA | NA | NA | 1 | 1 | NA | NA | 1 | 1 |
| 3_6894524 | NA | NA | 4 | 1 | 1 | 1 | 1 | 1 | 6 | 1 | NA | NA | NA | NA | 1 | 1 | NA | NA | 1 | 1 |
| 3_6898047 | 17 | 0.0588 | 14 | 0.0714 | 9 | 0 | 9 | 0.1111 | 49 | 0.0612 | 4 | 0 | 9 | 0 | 16 | 0.0625 | 16 | 0.1875 | 45 | 0.0889 |
| 3_6898077 | 11 | 0.0909 | 7 | 0.2857 | 5 | 0 | 6 | 0.1667 | 29 | 0.1379 | 3 | 0 | 10 | 0.1 | 14 | 0.1429 | 14 | 0.0714 | 41 | 0.0976 |
| 3_6898138 | 11 | 0.0909 | 7 | 0.1429 | 5 | 0 | 8 | 0.25 | 31 | 0.129 | 3 | 0 | 6 | 0.3333 | 12 | 0.0833 | 11 | 0 | 32 | 0.0938 |
| 3_6922328 | 1 | 1 | NA | NA | NA | NA | NA | NA | 1 | 1 | 1 | 1 | 1 | 1 | NA | NA | 2 | 1 | 4 | 1 |
| 3_6922357 | 1 | 1 | NA | NA | NA | NA | NA | NA | 1 | 1 | 1 | 1 | 1 | 1 | NA | NA | 2 | 1 | 4 | 1 |
| 3_6922369 | 1 | 1 | NA | NA | NA | NA | NA | NA | 1 | 1 | 1 | 1 | 1 | 0 | NA | NA | 2 | 1 | 4 | 0.75 |
| 3_6922453 | 1 | 1 | 1 | 0 | NA | NA | 1 | 0 | 3 | 0.3333 | 1 | 0 | 2 | 0.5 | NA | NA | 2 | 0 | 5 | 0.2 |
| 3_6922491 | 1 | 1 | 1 | 1 | NA | NA | 1 | 1 | 3 | 1 | NA | NA | 2 | 0.5 | NA | NA | 2 | 0 | 4 | 0.25 |
| 3_6946029 | 1 | 0 | 1 | 0 | NA | NA | 1 | 1 | 3 | 0.3333 | NA | NA | 3 | 0.3333 | NA | NA | 2 | 0.5 | 5 | 0.4 |
| 3_6946054 | 1 | 1 | 2 | 1 | NA | NA | 1 | 1 | 4 | 1 | NA | NA | 4 | 1 | NA | NA | 2 | 1 | 6 | 1 |
| 3_6946084 | 2 | 0 | 2 | 0.5 | NA | NA | 1 | 0 | 5 | 0.2 | NA | NA | 4 | 0 | 1 | 0 | 2 | 0.5 | 7 | 0.1429 |
| 3_6946104 | NA | NA | 1 | 1 | NA | NA | NA | NA | 1 | 1 | 1 | 1 | 2 | 1 | NA | NA | 1 | 1 | 4 | 1 |
| 3_6946150 | 2 | 0 | 1 | 0 | 1 | 1 | 1 | 0 | 5 | 0.2 | 1 | 0 | 4 | 0 | 1 | 0 | 2 | 0.5 | 8 | 0.125 |
| 3_6946156 | 1 | 1 | 2 | 0 | NA | NA | 1 | 0 | 4 | 0.25 | 1 | 0 | 4 | 0.5 | 2 | 0 | 2 | 0 | 9 | 0.2222 |
| 3_6946160 | 1 | 1 | 2 | 0 | 1 | 0 | 1 | 0 | 5 | 0.2 | 1 | 1 | 5 | 0.2 | 2 | 0 | 2 | 0 | 10 | 0.2 |
| 3_6946164 | 2 | 1 | 2 | 1 | 1 | 1 | 1 | 1 | 6 | 1 | 2 | 1 | 5 | 1 | 2 | 1 | 2 | 0.5 | 11 | 0.9091 |
| 3_6946168 | 1 | 0 | 2 | 0 | 1 | 1 | NA | NA | 4 | 0.25 | 1 | 0 | 2 | 0 | 1 | 0 | 2 | 1 | 6 | 0.3333 |
| 3_6946169 | 1 | 1 | 2 | 1 | 2 | 1 | 1 | 0 | 6 | 0.8333 | 1 | 1 | 4 | 1 | 2 | 0.5 | 4 | 0.75 | 11 | 0.8182 |
| 3_6946184 | 2 | 0.5 | 2 | 0.5 | 1 | 0 | 1 | 0 | 6 | 0.3333 | 2 | 0.5 | 5 | 0.8 | 2 | 0.5 | 4 | 1 | 13 | 0.7692 |
| 3_6946196 | 2 | 0 | 2 | 0 | 2 | 0.5 | 2 | 0 | 8 | 0.125 | 2 | 0 | 5 | 0.6 | NA | NA | 5 | 0.6 | 12 | 0.5 |
| 3_6946205 | 1 | 1 | 3 | 0.6667 | 1 | 1 | 2 | 0 | 7 | 0.5714 | 2 | 0.5 | 2 | 0 | 1 | 1 | 5 | 0.6 | 10 | 0.5 |
| 3_6946206 | 1 | 1 | 3 | 1 | 2 | 1 | 2 | 1 | 8 | 1 | 2 | 1 | 3 | 1 | 1 | 1 | 2 | 1 | 8 | 1 |
| 3_6946207 | 1 | 1 | 2 | 1 | 2 | 1 | 2 | 1 | 7 | 1 | 2 | 1 | 5 | 1 | 1 | 1 | 5 | 1 | 13 | 1 |
| 3_6946207 | 2 | 0 | 3 | 0.3333 | 2 | 0.5 | 2 | 0 | 9 | 0.2222 | 2 | 0.5 | 3 | 0.6667 | 1 | 1 | 4 | 0.25 | 10 | 0.5 |
| 3_6946209 | 2 | 0.5 | 3 | 0.6667 | 2 | 1 | 2 | 0 | 9 | 0.5556 | 2 | 1 | 5 | 0.6 | 1 | 0 | 4 | 0.75 | 12 | 0.6667 |
| 3_6946306 | 4 | 0 | 2 | 0 | 4 | 0.25 | 3 | 0 | 13 | 0.0769 | 1 | 0 | 4 | 0.25 | 3 | 0 | 4 | 0.25 | 12 | 0.1667 |
| 3_6946330 | 4 | 0 | 5 | 0 | 4 | 0.25 | 3 | 0 | 16 | 0.0625 | 2 | 0.5 | 5 | 0.4 | 3 | 0 | 4 | 0 | 14 | 0.2143 |
| 3_6946345 | 3 | 0.3333 | 4 | 0.25 | 1 | 1 | 2 | 0 | 10 | 0.3 | 1 | 1 | 3 | 0.3333 | 3 | 0 | 4 | 0.5 | 11 | 0.3636 |
| 3_6946354 | 5 | 0.4 | 4 | 1 | 2 | 1 | 1 | 1 | 12 | 0.75 | 1 | 1 | 4 | 0.25 | 3 | 0.6667 | 4 | 0.75 | 12 | 0.5833 |
| 3_6946394 | 4 | 0.5 | 6 | 0.1667 | 4 | 0.25 | 4 | 0 | 18 | 0.2222 | 2 | 0 | 4 | 0 | 3 | 0.3333 | 5 | 0 | 14 | 0.0714 |
| 3_6946419 | 3 | 0.3333 | 2 | 0 | 2 | 0.5 | 4 | 0 | 11 | 0.1818 | 1 | 0 | 5 | 0.2 | 2 | 0 | 4 | 0 | 12 | 0.0833 |
| 3_6946652 | 9 | 0 | 13 | 0.0769 | 9 | 0.1111 | 2 | 0 | 33 | 0.0606 | 3 | 0 | 6 | 0.1667 | 5 | 0.2 | 7 | 0 | 21 | 0.0952 |
| 3_6946701 | 9 | 0 | 16 | 0 | 10 | 0.1 | 4 | 0.25 | 39 | 0.0513 | 5 | 0.2 | 8 | 0.125 | 6 | 0 | 10 | 0 | 29 | 0.069 |
| 3_6946761 | 9 | 0.3333 | 14 | 0 | 11 | 0.2727 | 4 | 0.25 | 38 | 0.1842 | 5 | 0.2 | 4 | 0.5 | 7 | 0.1429 | 11 | 0 | 27 | 0.1481 |
| 3_6946776 | 16 | 0.125 | 19 | 0.1579 | 11 | 0.0909 | 8 | 0.125 | 54 | 0.1296 | 7 | 0.1429 | 9 | 0.1111 | 8 | 0 | 12 | 0 | 36 | 0.0556 |
| 3_6946779 | 12 | 0.1667 | 14 | 0.1429 | 11 | 0.2727 | 6 | 0.1667 | 43 | 0.186 | 8 | 0 | 7 | 0.1429 | 8 | 0.125 | 11 | 0 | 34 | 0.0588 |

| | | | | | | | | | | | | | | | | | | | | |
|---|---|---|---|---|---|---|---|---|---|---|---|---|---|---|---|---|---|---|---|---|
| 3_69482228 | 4 | 0 | 7 | 0.4286 | 2 | 0 | 3 | 0.3333 | 16 | 0.25 | 2 | 0 | 3 | 0 | 3 | 0.6667 | 8 | 0.25 | 16 | 0.25 |
| 3_69482397 | 3 | 0 | 7 | 0 | 2 | 0.5 | 3 | 0 | 15 | 0.0667 | 1 | 0 | 2 | 0.5 | 3 | 0 | 6 | 0 | 12 | 0.0833 |
| 3_69482421 | 3 | 1 | 8 | 1 | 2 | 1 | 3 | 1 | 16 | 1 | 2 | 1 | 3 | 1 | 2 | 1 | 5 | 1 | 12 | 1 |
| 3_69482799 | 3 | 0.3333 | 3 | 0 | NA | NA | 1 | 0 | 7 | 0.1429 | 2 | 0 | 3 | 0.3333 | 3 | 0 | 7 | 0.1429 | 15 | 0.1333 |
| 3_69482820 | 2 | 0.5 | 5 | 0.2 | 1 | 0 | 3 | 0 | 11 | 0.1818 | 2 | 0.5 | 3 | 0.3333 | 3 | 0.3333 | 6 | 0.5 | 14 | 0.4286 |
| 3_69483045 | 1 | 1 | 3 | 1 | NA | NA | 1 | 1 | 5 | 1 | 2 | 1 | 1 | 1 | 1 | 1 | 3 | 0.6667 | 7 | 0.8571 |
| 3_69483075 | 2 | 0 | 5 | 0.6 | 1 | 1 | 1 | 0 | 9 | 0.4444 | 2 | 0 | 3 | 0.6667 | 3 | 0.6667 | 7 | 0.5714 | 15 | 0.5333 |
| 3_69483080 | 2 | 0 | 4 | 0.25 | 1 | 0 | 3 | 0 | 10 | 0.1 | 2 | 0 | 2 | 0 | 2 | 0 | 7 | 0.4286 | 13 | 0.2308 |
| 3_72954861 | 6 | 0.1667 | 6 | 0 | 4 | 0 | 7 | 0.4286 | 23 | 0.1739 | 7 | 0 | 7 | 0.1429 | 9 | 0.2222 | 8 | 0 | 31 | 0.0968 |
| 3_72954892 | 6 | 0 | 10 | 0 | 4 | 0.25 | 7 | 0.1429 | 27 | 0.0741 | 5 | 0.2 | 7 | 0.1429 | 7 | 0.2857 | 10 | 0.5 | 29 | 0.3103 |
| 3_72955314 | 5 | 0 | 4 | 0.25 | 3 | 0 | 5 | 0 | 17 | 0.0588 | 5 | 0.2 | 2 | 0.5 | 6 | 0 | 4 | 0 | 17 | 0.1176 |
| 3_72955531 | 2 | 1 | 4 | 0.5 | 2 | 1 | 5 | 0.4 | 13 | 0.6154 | 4 | 0.25 | 3 | 1 | 2 | 0.5 | 8 | 0.625 | 17 | 0.5882 |
| 3_72955537 | 7 | 0.8571 | 5 | 0.6 | 2 | 1 | 6 | 0.8333 | 20 | 0.8 | 6 | 0.8333 | 3 | 1 | 4 | 0.75 | 8 | 0.875 | 21 | 0.8571 |
| 3_72955544 | 8 | 0 | 6 | 0.5 | 2 | 0 | 6 | 0 | 22 | 0.1364 | 5 | 0 | 3 | 0.3333 | 3 | 0 | 6 | 0.1667 | 17 | 0.1176 |
| 3_72955584 | 6 | 1 | 5 | 1 | 4 | 0.75 | 7 | 0.8571 | 22 | 0.9091 | 5 | 0.6 | 5 | 1 | 3 | 1 | 9 | 0.7778 | 22 | 0.8182 |
| 3_72955691 | 4 | 1 | 4 | 1 | 2 | 1 | 5 | 1 | 15 | 1 | 2 | 1 | 1 | 1 | 5 | 1 | 5 | 1 | 13 | 1 |
| 3_72955696 | 7 | 1 | 4 | 0.75 | 4 | 1 | 5 | 1 | 20 | 0.95 | 2 | 1 | 4 | 1 | 5 | 0.8 | 7 | 1 | 18 | 0.9444 |
| 3_72955726 | 4 | 0 | 5 | 0.2 | 3 | 0 | 6 | 0.1667 | 18 | 0.1111 | 2 | 0 | 2 | 0 | 4 | 0 | 6 | 0.3333 | 14 | 0.1429 |
| 3_72955878 | 4 | 0 | 5 | 0 | 4 | 0 | 5 | 0.2 | 18 | 0.0556 | 5 | 0 | 4 | 0 | 3 | 0 | 8 | 0.125 | 20 | 0.05 |
| 3_72955928 | 5 | 0 | 6 | 0 | 3 | 0.3333 | 9 | 0.3333 | 23 | 0.1739 | 3 | 0 | 3 | 0.6667 | 5 | 0 | 9 | 0.3333 | 20 | 0.25 |
| 3_72955962 | 5 | 0 | 7 | 0.1429 | 4 | 0 | 8 | 0.125 | 24 | 0.0833 | 6 | 0 | 4 | 0 | 6 | 0 | 9 | 0.2222 | 25 | 0.08 |
| 3_72955991 | 4 | 1 | 6 | 1 | 4 | 1 | 7 | 0.8571 | 21 | 0.9524 | 5 | 0.8 | 4 | 1 | 5 | 1 | 7 | 0.5714 | 21 | 0.8095 |
| 3_72956026 | 5 | 1 | 6 | 1 | 2 | 1 | 6 | 1 | 19 | 1 | 3 | 1 | 3 | 1 | 5 | 1 | 6 | 1 | 17 | 1 |
| 3_72956060 | 4 | 1 | 7 | 1 | 4 | 1 | 8 | 1 | 23 | 1 | 4 | 1 | 3 | 1 | 6 | 1 | 8 | 1 | 21 | 1 |
| 3_72956105 | 5 | 0 | 7 | 0.4286 | 3 | 0 | 9 | 0.2222 | 24 | 0.2083 | 4 | 0.25 | 3 | 0 | 6 | 0 | 9 | 0.1111 | 22 | 0.0909 |
| 3_72956195 | 5 | 0.4 | 5 | 0.8 | 2 | 0.5 | 6 | 0.5 | 18 | 0.5556 | 4 | 0.5 | 4 | 0.25 | 6 | 0.3333 | 4 | 0.5 | 18 | 0.3889 |
| 3_72956205 | 6 | 1 | 6 | 1 | 3 | 1 | 8 | 1 | 23 | 1 | 5 | 1 | 4 | 1 | 6 | 0.8333 | 5 | 1 | 20 | 0.95 |
| 3_72956267 | 4 | 0.5 | 2 | 0.5 | 1 | 0 | 8 | 0.625 | 15 | 0.5333 | 5 | 0.2 | 3 | 0 | 7 | 0 | 1 | 1 | 16 | 0.125 |
| 3_72956340 | 5 | 0.8 | 4 | 1 | 2 | 0.5 | 7 | 1 | 18 | 0.8889 | 5 | 1 | 3 | 0.6667 | 7 | 0.4286 | 4 | 1 | 19 | 0.7368 |
| 3_72956456 | 5 | 0.6 | 3 | 0.6667 | 1 | 1 | 8 | 0.625 | 17 | 0.6471 | 4 | 0.25 | 4 | 0.5 | 4 | 0.75 | 4 | 0.75 | 16 | 0.5625 |
| 3_72956526 | 4 | 0.5 | 2 | 0 | 1 | 1 | 4 | 0.25 | 11 | 0.3636 | 1 | 1 | 2 | 0.5 | 5 | 0.8 | 3 | 0 | 11 | 0.5455 |
| 3_72956636 | 5 | 1 | 2 | 1 | 2 | 1 | 4 | 1 | 13 | 1 | 4 | 0.75 | 3 | 0.6667 | 5 | 1 | 3 | 1 | 15 | 0.8667 |
| 3_72956678 | 5 | 0.4 | 2 | 0.5 | 2 | 0.5 | 6 | 0.8333 | 15 | 0.6 | 3 | 0.6667 | 4 | 0.75 | 5 | 0.4 | 3 | 0.6667 | 15 | 0.6 |
| 3_72956863 | 4 | 0 | 2 | 0 | 1 | 0 | 6 | 0.1667 | 13 | 0.0769 | 6 | 0.1667 | 4 | 0.25 | 5 | 0.4 | 3 | 0.6667 | 18 | 0.3333 |
| 3_72956905 | 3 | 0.6667 | 2 | 0.5 | 1 | 1 | 5 | 0.6 | 11 | 0.6364 | 4 | 1 | 3 | 0.6667 | 3 | 0.3333 | 3 | 0.6667 | 13 | 0.6923 |
| 3_72956981 | 4 | 0.25 | 1 | 0 | 2 | 0 | 5 | 0 | 12 | 0.0833 | 3 | 0 | 4 | 0.5 | 4 | 0 | 3 | 0.6667 | 14 | 0.2857 |
| 3_72957074 | 4 | 1 | 2 | 1 | 1 | 1 | 6 | 1 | 13 | 1 | 5 | 0.8 | 3 | 1 | 5 | 0.8 | 2 | 1 | 15 | 0.8667 |
| 3_72957158 | 5 | 0.6 | 2 | 1 | 1 | 1 | 5 | 0.4 | 13 | 0.6154 | 4 | 0.75 | 3 | 1 | 5 | 0.4 | 2 | 1 | 14 | 0.7143 |
| 3_72957216 | 5 | 1 | 2 | 1 | 2 | 1 | 6 | 1 | 15 | 1 | 4 | 1 | 4 | 1 | 6 | 1 | 2 | 1 | 16 | 1 |
| 3_72957227 | 4 | 0 | 1 | 0 | 2 | 0 | 6 | 0.5 | 13 | 0.2308 | 2 | 0.5 | 3 | 0 | 6 | 0.1667 | 1 | 0 | 12 | 0.1667 |
| 3_72957562 | 4 | 0 | 2 | 0.5 | 1 | 1 | 5 | 0.4 | 12 | 0.3333 | 2 | 0.5 | 3 | 0 | 3 | 0.3333 | 1 | 0 | 9 | 0.2222 |
| 3_72957603 | 4 | 0.5 | 1 | 1 | 1 | 1 | 5 | 0.6 | 11 | 0.6364 | 2 | 0.5 | 3 | 0.6667 | 5 | 0.6 | 1 | 1 | 11 | 0.6364 |
| 3_72957614 | 4 | 0 | 1 | 0 | 1 | 1 | 6 | 0.6667 | 12 | 0.4167 | 2 | 0.5 | 3 | 0 | 2 | 0 | 1 | 0 | 8 | 0.125 |
| 3_72957675 | 4 | 0.75 | 1 | 1 | 2 | 1 | 5 | 1 | 12 | 0.9167 | 2 | 1 | 3 | 0.6667 | 4 | 0.75 | 1 | 1 | 10 | 0.8 |
| 3_72957911 | 3 | 0.3333 | NA | NA | 2 | 0 | 5 | 0.4 | 10 | 0.3 | 2 | 0 | NA | NA | 4 | 0.25 | 1 | 0 | 7 | 0.1429 |
| 3_72957931 | 3 | 0.3333 | NA | NA | 1 | 1 | 5 | 0.2 | 9 | 0.3333 | 2 | 0.5 | 3 | 0.6667 | 4 | 0 | NA | NA | 9 | 0.3333 |
| 3_72957936 | 3 | 0.3333 | NA | NA | 2 | 0.5 | 6 | 0.3333 | 11 | 0.3636 | 2 | 0.5 | 3 | 0.6667 | 4 | 0 | NA | NA | 9 | 0.3333 |
| 3_72958064 | 4 | 1 | NA | NA | 2 | 1 | 6 | 0.8333 | 12 | 0.9167 | 1 | 1 | 3 | 1 | 4 | 0.75 | 1 | 1 | 9 | 0.8889 |
| 3_72958163 | 2 | 0.5 | NA | NA | 2 | 0.5 | 6 | 0.3333 | 10 | 0.4 | 1 | 0 | 1 | 1 | 3 | 0 | NA | NA | 5 | 0.2 |
| 3_72958256 | 4 | 0.25 | NA | NA | 2 | 0.5 | 5 | 0.2 | 11 | 0.2727 | 2 | 0 | 3 | 0 | 4 | 0.25 | 1 | 0 | 10 | 0.1 |
| 3_72958381 | 2 | 1 | NA | NA | 2 | 1 | 5 | 1 | 9 | 1 | 1 | 1 | 3 | 1 | 3 | 1 | 1 | 1 | 8 | 1 |
| 3_72958416 | 4 | 0.25 | NA | NA | 1 | 0 | 5 | 0.2 | 10 | 0.2 | 1 | 0 | 3 | 0.3333 | 4 | 0 | 1 | 0 | 9 | 0.1111 |
| 3_72958424 | 3 | 1 | NA | NA | 2 | 1 | 2 | 1 | 7 | 1 | 1 | 1 | 3 | 1 | 4 | 1 | 1 | 1 | 9 | 1 |
| 3_72958439 | 4 | 0.25 | NA | NA | 1 | 0 | 3 | 0 | 8 | 0.125 | 2 | 0 | 1 | 0 | 4 | 0.25 | NA | NA | 7 | 0.1429 |
| 3_72958495 | 4 | 0 | NA | NA | 2 | 1 | 4 | 0 | 10 | 0.2 | 1 | 0 | 3 | 0.3333 | 4 | 0.25 | 1 | 0 | 9 | 0.2222 |
| 3_72958525 | 3 | 0.3333 | NA | NA | 2 | 0.5 | 5 | 0.6 | 10 | 0.5 | 1 | 0 | 3 | 1 | 3 | 0.6667 | 1 | 0 | 8 | 0.625 |
| 3_72958534 | 4 | 0 | NA | NA | 2 | 0.5 | 5 | 0 | 11 | 0.0909 | 2 | 0 | 3 | 0.3333 | 4 | 0 | 1 | 0 | 10 | 0.1 |
| 3_72958661 | 3 | 1 | NA | NA | 2 | 0.5 | 4 | 1 | 9 | 0.8889 | 1 | 1 | 3 | 1 | 4 | 0.5 | 1 | 1 | 9 | 0.7778 |
| 3_72958686 | 4 | 1 | NA | NA | 2 | 0.5 | 4 | 0.75 | 10 | 0.8 | 1 | 1 | 3 | 0.6667 | 4 | 1 | 1 | 1 | 9 | 0.8889 |
| 3_73543191 | 1 | 0 | 1 | 1 | 1 | 1 | 1 | 1 | 4 | 0.75 | NA | NA | NA | NA | NA | NA | 1 | 1 | 1 | 1 |
| 3_73547308 | NA | NA | 2 | 1 | NA | NA | NA | NA | 2 | 1 | NA | NA | NA | NA | NA | NA | 1 | 1 | 1 | 1 |
| 3_73608214 | NA | NA | 2 | 1 | NA | NA | NA | NA | 2 | 1 | NA | NA | 1 | 1 | NA | NA | 1 | 1 | 2 | 1 |
| 3_73608319 | NA | NA | 2 | 1 | NA | NA | NA | NA | 2 | 1 | NA | NA | 1 | 1 | NA | NA | NA | NA | 1 | 1 |
| 3_73608359 | NA | NA | 2 | 1 | NA | NA | NA | NA | 2 | 1 | NA | NA | 1 | 1 | NA | NA | 1 | 1 | 2 | 1 |
| 3_73608974 | NA | NA | 1 | 1 | NA | NA | NA | NA | 1 | 1 | NA | NA | 1 | 1 | NA | NA | 1 | 1 | 2 | 1 |

| | | | | | | | | | | | | | | | | | | | | |
|---|---|---|---|---|---|---|---|---|---|---|---|---|---|---|---|---|---|---|---|---|
| 3_73609262 | NA | NA | 2 | 0.5 | NA | NA | NA | NA | 2 | 0.5 | NA | NA | 1 | 1 | NA | NA | 1 | 1 | 2 | 1 |
| 3_75424752 | 12 | 0 | 11 | 0.1818 | 10 | 0.1 | 5 | 0.4 | 38 | 0.1316 | 10 | 0.1 | 18 | 0.1667 | 17 | 0.1176 | 11 | 0 | 56 | 0.1071 |
| 3_75523905 | 13 | 0 | 39 | 0.0513 | 21 | 0.0952 | 10 | 0.1 | 83 | 0.0602 | 16 | 0 | 10 | 0 | 6 | 0.3333 | 23 | 0.0435 | 55 | 0.0545 |
| 3_75548052 | 19 | 0.1053 | 43 | 0.093 | 24 | 0.1667 | 11 | 0.0909 | 97 | 0.1134 | 19 | 0.1053 | 9 | 0.1111 | 12 | 0 | 31 | 0.2258 | 71 | 0.1408 |
| 3_75549426 | 26 | 0.0385 | 56 | 0.1071 | 39 | 0.0513 | 19 | 0.1053 | 140 | 0.0786 | 30 | 0.0667 | 12 | 0 | 18 | 0.1111 | 39 | 0.0769 | 99 | 0.0707 |
| 3_75784629 | NA | NA | 4 | 0 | NA | NA | 1 | 1 | 5 | 0.2 | 4 | 0.25 | 3 | 0.3333 | 2 | 1 | NA | NA | 9 | 0.4444 |
| 3_75785320 | NA | NA | 4 | 0 | NA | NA | 1 | 1 | 5 | 0.2 | 4 | 0.75 | 1 | 1 | 1 | 1 | NA | NA | 6 | 0.8333 |
| 3_75785333 | NA | NA | 4 | 1 | NA | NA | 1 | 1 | 5 | 1 | 3 | 0 | 1 | 1 | NA | NA | NA | NA | 4 | 0.25 |
| 3_7661487 | 2 | 1 | NA | NA | NA | NA | 4 | 1 | 6 | 1 | 2 | 0.5 | NA | NA | 4 | 0.75 | 2 | 1 | 8 | 0.75 |
| 3_7661497 | 2 | 0.5 | 1 | 1 | 1 | 1 | 3 | 1 | 7 | 0.8571 | 1 | 1 | NA | NA | 3 | 0 | 2 | 0 | 6 | 0.1667 |
| 3_7661498 | 1 | 1 | 1 | 1 | 1 | 1 | 4 | 1 | 7 | 1 | 2 | 0.5 | NA | NA | 3 | 0.6667 | 1 | 0 | 6 | 0.5 |
| 3_7661499 | NA | NA | 1 | 0 | NA | NA | 2 | 1 | 3 | 0.6667 | 2 | 1 | NA | NA | 3 | 1 | 2 | 1 | 7 | 1 |
| 3_7661504 | 2 | 1 | 1 | 1 | 1 | 1 | 5 | 0.8 | 9 | 0.8889 | 3 | 1 | NA | NA | 4 | 1 | 1 | 1 | 8 | 1 |
| 3_7661536 | 1 | 1 | 1 | 1 | 1 | 1 | 5 | 0.8 | 8 | 0.875 | 3 | 1 | NA | NA | 3 | 1 | 3 | 1 | 9 | 1 |
| 3_7661542 | 1 | 1 | 1 | 1 | 1 | 0 | 5 | 0.6 | 8 | 0.625 | 3 | 0.6667 | NA | NA | 4 | 0.25 | 3 | 0.3333 | 10 | 0.4 |
| 3_7661598 | 2 | 0 | 1 | 0 | 3 | 0 | 4 | 0.5 | 10 | 0.2 | 3 | 0.3333 | 1 | 0 | 3 | 0 | 2 | 0 | 9 | 0.1111 |
| 3_7661616 | 3 | 0 | 1 | 0 | 3 | 0.3333 | 5 | 0.2 | 12 | 0.1667 | 3 | 0.3333 | 1 | 0 | 4 | 0 | 2 | 0 | 10 | 0.1 |
| 3_7661673 | 2 | 0 | 1 | 0 | 6 | 0 | 5 | 0.2 | 14 | 0.0714 | 3 | 0 | 1 | 1 | 5 | 0 | 4 | 0.25 | 13 | 0.1538 |
| 3_7897008 | 23 | 0.2609 | 14 | 0.4286 | 19 | 0.3684 | 21 | 0.381 | 77 | 0.3506 | 10 | 0 | 25 | 0.32 | 16 | 0.375 | 9 | 0.4444 | 60 | 0.3 |
| 3_7897017 | 25 | 0.24 | 15 | 0.4667 | 19 | 0.3684 | 23 | 0.3478 | 82 | 0.3415 | 11 | 0.4545 | 26 | 0.3077 | 19 | 0.3684 | 13 | 0.5385 | 69 | 0.3913 |
| 3_7897047 | 22 | 0.1364 | 11 | 0.1818 | 19 | 0.1053 | 20 | 0.1 | 72 | 0.125 | 9 | 0.1111 | 24 | 0.0417 | 17 | 0.1176 | 12 | 0.0833 | 62 | 0.0806 |
| 3_7897052 | 22 | 0.1364 | 11 | 0.2727 | 18 | 0.4444 | 18 | 0.1111 | 69 | 0.2319 | 10 | 0.1 | 26 | 0.1923 | 16 | 0.125 | 13 | 0.1538 | 65 | 0.1538 |
| 3_7897138 | 15 | 0.9333 | 11 | 1 | 14 | 1 | 12 | 0.8333 | 52 | 0.9423 | 7 | 1 | 16 | 1 | 11 | 0.9091 | 9 | 1 | 43 | 0.9767 |
| 3_7897142 | 6 | 0.6667 | 6 | 0.6667 | 10 | 0.7 | 4 | 0.5 | 26 | 0.6538 | 2 | 0.5 | 7 | 0.8571 | 8 | 0.25 | 3 | 0.3333 | 20 | 0.5 |
| 3_7897163 | 14 | 1 | 8 | 1 | 13 | 0.9231 | 11 | 1 | 46 | 0.9783 | 4 | 1 | 11 | 0.9091 | 12 | 0.8333 | 10 | 0.8 | 37 | 0.8649 |
| 3_7897174 | 14 | 0.6429 | 7 | 0.4286 | 15 | 0.6 | 13 | 0.3077 | 49 | 0.5102 | 6 | 0.5 | 13 | 0.3846 | 13 | 0.7692 | 11 | 0.4545 | 43 | 0.5349 |
| 3_7897643 | 6 | 0.3333 | 7 | 0.5714 | 9 | 0.4444 | 8 | 0.625 | 30 | 0.5 | 4 | 0 | 7 | 0.2857 | 11 | 0.3636 | 8 | 0.625 | 30 | 0.3667 |
| 3_7897647 | 12 | 0.9167 | 7 | 1 | 10 | 1 | 10 | 1 | 39 | 0.9744 | 5 | 1 | 8 | 1 | 12 | 1 | 6 | 0.8333 | 31 | 0.9677 |
| 3_7897652 | 7 | 0.5714 | 7 | 0.5714 | 8 | 0.375 | 6 | 0.8333 | 28 | 0.5714 | 1 | 0 | 6 | 0.6667 | 10 | 0.6 | 6 | 0.5 | 23 | 0.5652 |
| 3_7897661 | 10 | 0.3 | 5 | 0.4 | 9 | 0.2222 | 8 | 0.25 | 32 | 0.2812 | 4 | 0 | 7 | 0.7143 | 9 | 0.1111 | 4 | 0.5 | 24 | 0.3333 |
| 3_7899329 | 6 | 1 | 4 | 0.75 | 7 | 0.4286 | 6 | 0.6667 | 23 | 0.6957 | 3 | 0 | 5 | 0 | 4 | 0.25 | 4 | 0.5 | 16 | 0.1875 |
| 3_7899330 | 8 | 0.125 | 4 | 0.25 | 7 | 0.1429 | 6 | 0.1667 | 25 | 0.16 | 3 | 0.3333 | 4 | 0.5 | 5 | 0.4 | 4 | 0.25 | 16 | 0.375 |
| 3_7919395 | NA | NA | 2 | 0.5 | 1 | 1 | NA | NA | 3 | 0.6667 | NA | NA | NA | NA | 1 | 1 | NA | NA | 1 | 1 |
| 3_7919510 | NA | NA | 1 | 1 | NA | NA | NA | NA | 1 | 1 | 1 | 0 | NA | NA | NA | NA | 1 | 1 | 2 | 0.5 |
| 3_7919524 | NA | NA | 1 | 0 | 1 | 1 | NA | NA | 2 | 0.5 | NA | NA | NA | NA | NA | NA | 1 | 1 | 1 | 1 |
| 3_7936369 | 3 | 0 | 3 | 0 | 4 | 0.25 | 8 | 0.5 | 18 | 0.2778 | 1 | 0 | 5 | 0.2 | NA | NA | 6 | 0 | 12 | 0.0833 |
| 3_7936387 | 3 | 0.3333 | 3 | 0 | 4 | 0 | 8 | 0 | 18 | 0.0556 | 1 | 0 | 5 | 0.2 | NA | NA | 5 | 0 | 11 | 0.0909 |
| 3_7936430 | 3 | 0.3333 | 2 | 0.5 | 4 | 0.75 | 7 | 0.8571 | 16 | 0.6875 | 1 | 0 | 5 | 0.8 | NA | NA | 4 | 0.5 | 10 | 0.6 |
| 3_7936455 | 2 | 0.5 | 4 | 0.25 | 2 | 0.5 | 7 | 0.4286 | 15 | 0.4 | 1 | 1 | 4 | 0.5 | NA | NA | 4 | 0.75 | 9 | 0.6667 |
| 3_7936490 | 3 | 1 | 3 | 1 | 2 | 1 | 4 | 1 | 12 | 1 | 1 | 1 | 4 | 1 | NA | NA | 1 | 1 | 6 | 1 |
| 3_7936515 | 3 | 0.3333 | 4 | 0.75 | 2 | 0.5 | 7 | 0.2857 | 16 | 0.4375 | 1 | 1 | 4 | 0.75 | NA | NA | 3 | 0 | 8 | 0.5 |
| 3_7936520 | 2 | 0 | 3 | 0.6667 | 2 | 0.5 | 6 | 0.1667 | 13 | 0.3077 | 1 | 1 | 4 | 0.5 | NA | NA | 3 | 0.3333 | 8 | 0.5 |
| 3_7936529 | 3 | 1 | 2 | 1 | 2 | 1 | 6 | 0.8333 | 13 | 0.9231 | 1 | 1 | 3 | 0.6667 | NA | NA | 3 | 0.6667 | 7 | 0.7143 |
| 3_7936577 | 3 | 0.3333 | 4 | 0 | 2 | 0 | 4 | 0.5 | 13 | 0.2308 | 1 | 1 | 2 | 0.5 | NA | NA | 2 | 0 | 5 | 0.4 |
| 3_7936595 | 3 | 0.6667 | 2 | 1 | 2 | 0 | 2 | 1 | 9 | 0.6667 | NA | NA | 2 | 1 | NA | NA | 2 | 0 | 4 | 0.5 |
| 3_7936612 | 2 | 1 | 1 | 1 | 2 | 1 | 6 | 0.8333 | 11 | 0.9091 | NA | NA | 1 | 1 | NA | NA | 2 | 0.5 | 3 | 0.6667 |
| 3_7936626 | 1 | 0 | 2 | 0.5 | 2 | 1 | 6 | 0.6667 | 11 | 0.6364 | NA | NA | 2 | 0.5 | NA | NA | 2 | 0 | 4 | 0.25 |
| 3_7936630 | 2 | 1 | 1 | 0 | 1 | 0 | 2 | 1 | 6 | 0.6667 | NA | NA | 1 | 0 | NA | NA | 1 | 1 | 2 | 0.5 |
| 3_7936636 | 1 | 1 | 2 | 1 | 2 | 0.5 | 6 | 1 | 11 | 0.9091 | NA | NA | 2 | 1 | NA | NA | 2 | 0.5 | 4 | 0.75 |
| 3_7936647 | 1 | 0 | 2 | 1 | 2 | 0.5 | 1 | 0 | 6 | 0.5 | NA | NA | 1 | 1 | NA | NA | 2 | 0.5 | 3 | 0.6667 |
| 3_7936654 | 2 | 0.5 | NA | NA | 2 | 0.5 | 4 | 1 | 8 | 0.75 | NA | NA | 2 | 1 | NA | NA | 1 | 1 | 3 | 1 |
| 3_7936663 | 1 | 1 | 1 | 1 | 2 | 0.5 | 3 | 1 | 7 | 0.8571 | NA | NA | 1 | 1 | NA | NA | NA | NA | 1 | 1 |
| 3_7936666 | 2 | 1 | 2 | 1 | 2 | 1 | 5 | 0.8 | 11 | 0.9091 | NA | NA | 2 | 1 | NA | NA | 1 | 0 | 3 | 0.6667 |
| 3_7936678 | NA | NA | NA | NA | NA | NA | 1 | 1 | 1 | 1 | NA | NA | 1 | 0 | NA | NA | 1 | 1 | 2 | 0.5 |
| 3_7938790 | NA | NA | 1 | 1 | 1 | 0 | 3 | 0 | 5 | 0.2 | NA | NA | 2 | 0 | NA | NA | 1 | 1 | 3 | 0.3333 |
| 3_7938811 | NA | NA | 1 | 1 | 1 | 1 | 3 | 0.3333 | 5 | 0.6 | NA | NA | 2 | 0 | NA | NA | 1 | 1 | 3 | 0.3333 |
| 3_7938815 | NA | NA | 1 | 1 | 1 | 0 | 3 | 0.3333 | 5 | 0.4 | NA | NA | 1 | 0 | NA | NA | 1 | 1 | 2 | 0.5 |
| 3_7938822 | NA | NA | NA | NA | 1 | 0 | 2 | 0.5 | 3 | 0.3333 | NA | NA | 1 | 0 | NA | NA | 1 | 1 | 2 | 0.5 |
| 3_7938841 | NA | NA | 1 | 1 | 1 | 1 | 1 | 1 | 3 | 1 | NA | NA | NA | NA | NA | NA | 1 | 1 | 1 | 1 |
| 3_7938846 | NA | NA | 1 | 0 | 1 | 0 | 3 | 0.3333 | 5 | 0.2 | NA | NA | 1 | 1 | NA | NA | 1 | 0 | 2 | 0.5 |
| 3_7951082 | 13 | 0.0769 | 15 | 0.0667 | 11 | 0.0909 | 11 | 0.2727 | 50 | 0.12 | 9 | 0.2222 | 4 | 0 | 10 | 0 | 14 | 0.0714 | 37 | 0.0811 |
| 3_7951125 | 3 | 0 | 7 | 0 | 2 | 0.5 | 2 | 0.5 | 14 | 0.1429 | 2 | 0 | 3 | 0 | 4 | 0.25 | 2 | 0 | 11 | 0.0909 |
| 3_7951140 | 3 | 0.6667 | 8 | 0.125 | 2 | 0.5 | 2 | 0.5 | 15 | 0.3333 | 3 | 0.3333 | 3 | 0.3333 | 5 | 0.4 | 3 | 1 | 14 | 0.5 |
| 3_7951236 | 2 | 0 | 7 | 0.1429 | 2 | 0 | 1 | 1 | 12 | 0.1667 | 3 | 0.3333 | 3 | 0 | 4 | 0 | 3 | 0 | 13 | 0.0769 |
| 3_7972004 | 1 | 1 | 1 | 0 | 1 | 1 | NA | NA | 3 | 0.6667 | 1 | 0 | 4 | 0.75 | 4 | 0 | NA | NA | 9 | 0.3333 |

| | | | | | | | | | | | | | | | | | | | | |
|---|---|---|---|---|---|---|---|---|---|---|---|---|---|---|---|---|---|---|---|---|
| 3_7972025 | 1 | 0 | 1 | 1 | 1 | 0 | NA | NA | 3 | 0.3333 | 1 | 0 | 4 | 0.25 | 4 | 0 | 1 | 0 | 10 | 0.1 |
| 3_7972066 | 1 | 0 | NA | NA | 1 | 1 | NA | NA | 2 | 0.5 | NA | NA | 4 | 0.25 | 3 | 0 | NA | NA | 7 | 0.1429 |
| 3_7972099 | 1 | 0 | NA | NA | 1 | 1 | NA | NA | 2 | 0.5 | NA | NA | 3 | 0.3333 | 4 | 0.5 | 1 | 0 | 8 | 0.375 |
| 3_7972113 | 1 | 1 | NA | NA | 1 | 0 | NA | NA | 2 | 0.5 | 1 | 1 | 4 | 0.75 | 4 | 0.75 | 1 | 0 | 10 | 0.7 |
| 3_8059062 | 1 | 0 | 3 | 0.6667 | NA | NA | 1 | 1 | 5 | 0.6 | 1 | 0 | 1 | 1 | NA | NA | NA | NA | 2 | 0.5 |
| 3_8059065 | 1 | 0 | 3 | 0.3333 | NA | NA | 1 | 0 | 5 | 0.2 | NA | NA | 1 | 1 | NA | NA | NA | NA | 1 | 1 |
| 3_8059311 | 3 | 0.3333 | 4 | 0 | 2 | 0 | 3 | 0.3333 | 12 | 0.1667 | 3 | 0.3333 | 1 | 0 | 3 | 0 | 2 | 0 | 9 | 0.1111 |
| 3_8059360 | 37 | 0.1351 | 51 | 0.1373 | 49 | 0.1837 | 55 | 0.2545 | 192 | 0.1823 | 28 | 0.25 | 42 | 0.1667 | 47 | 0.1702 | 50 | 0.26 | 167 | 0.2096 |
| 3_8059363 | 34 | 0.1471 | 53 | 0.0566 | 43 | 0.1628 | 52 | 0.1154 | 182 | 0.1154 | 29 | 0.2069 | 38 | 0.0263 | 43 | 0.1395 | 50 | 0.16 | 160 | 0.1312 |
| 3_8059531 | 23 | 0.6522 | 33 | 0.5455 | 31 | 0.6452 | 41 | 0.5854 | 128 | 0.6016 | 17 | 0.7647 | 30 | 0.8 | 30 | 0.6333 | 32 | 0.7188 | 109 | 0.7248 |
| 3_8059539 | 29 | 0.1034 | 44 | 0.1591 | 39 | 0.1282 | 51 | 0.2157 | 163 | 0.1595 | 21 | 0.0476 | 36 | 0 | 39 | 0.1026 | 44 | 0.0682 | 140 | 0.0571 |
| 3_8059544 | 21 | 0.4762 | 28 | 0.5 | 27 | 0.4815 | 39 | 0.5385 | 115 | 0.5043 | 9 | 0.2222 | 25 | 0.4 | 27 | 0.4815 | 33 | 0.697 | 94 | 0.5106 |
| 3_8059544 | 26 | 0.1923 | 43 | 0.093 | 36 | 0.1667 | 52 | 0.1154 | 157 | 0.1338 | 19 | 0.2632 | 36 | 0.1667 | 36 | 0.1667 | 45 | 0.3333 | 136 | 0.2353 |
| 3_8059548 | 10 | 0.3 | 22 | 0.2273 | 22 | 0.7727 | 21 | 0.4762 | 75 | 0.4667 | 9 | 0.6667 | 15 | 0.6 | 9 | 0.4444 | 14 | 0.7143 | 47 | 0.617 |
| 3_8059548 | 7 | 0.5714 | 5 | 0.2 | 2 | 0.5 | 7 | 0.4286 | 21 | 0.4286 | 2 | 0 | 6 | 0.3333 | 5 | 0.6 | 11 | 0.4545 | 24 | 0.4167 |
| 3_8059548 | 2 | 0.5 | 8 | 0.625 | 4 | 0 | 6 | 0.6667 | 20 | 0.5 | 4 | 0.75 | 7 | 0.5714 | 8 | 0.5 | 4 | 0.75 | 23 | 0.6087 |
| 3_8059555 | 24 | 0.7083 | 37 | 0.5946 | 34 | 0.7353 | 48 | 0.6042 | 143 | 0.6503 | 16 | 0.875 | 35 | 0.5714 | 38 | 0.7105 | 43 | 0.6744 | 132 | 0.6818 |
| 3_8059560 | 26 | 0.2308 | 41 | 0.2195 | 37 | 0.1351 | 51 | 0.1176 | 155 | 0.1677 | 18 | 0.1111 | 38 | 0.2632 | 38 | 0.0789 | 43 | 0.2093 | 137 | 0.1752 |
| 3_8059562 | 19 | 0.2632 | 31 | 0.3548 | 27 | 0.2222 | 31 | 0.129 | 108 | 0.2407 | 17 | 0 | 22 | 0.2727 | 30 | 0.1333 | 28 | 0.2143 | 97 | 0.1649 |
| 3_8059565 | 23 | 0.5652 | 38 | 0.6053 | 34 | 0.7353 | 49 | 0.6327 | 144 | 0.6389 | 17 | 0.7059 | 37 | 0.5405 | 36 | 0.8333 | 41 | 0.7561 | 131 | 0.7099 |
| 3_8059574 | 22 | 0 | 38 | 0.0789 | 33 | 0.0909 | 45 | 0.0889 | 138 | 0.0725 | 17 | 0 | 33 | 0.0909 | 30 | 0.1 | 40 | 0.075 | 120 | 0.075 |
| 3_8059584 | 24 | 0.25 | 37 | 0.2162 | 33 | 0.2727 | 45 | 0.2222 | 139 | 0.2374 | 20 | 0.2 | 35 | 0.2 | 40 | 0.25 | 40 | 0.425 | 135 | 0.2815 |
| 3_8059587 | 20 | 0.7 | 36 | 0.6111 | 29 | 0.6207 | 43 | 0.6744 | 128 | 0.6484 | 18 | 0.7222 | 29 | 0.5862 | 37 | 0.6757 | 40 | 0.825 | 124 | 0.7097 |
| 3_8059620 | NA | NA | 4 | 0 | 3 | 0.3333 | 5 | 0 | 12 | 0.0833 | 1 | 0 | 6 | 0.1667 | 2 | 0.5 | 4 | 0 | 13 | 0.1538 |
| 3_8059627 | 1 | 0 | 9 | 0 | 5 | 0 | 5 | 0.2 | 20 | 0.05 | 2 | 0 | 8 | 0.125 | 5 | 0 | 5 | 0 | 20 | 0.05 |
| 3_8059638 | 1 | 0 | 7 | 0.1429 | 5 | 0 | 3 | 0 | 16 | 0.0625 | 2 | 0 | 8 | 0.125 | 5 | 0 | 5 | 0 | 20 | 0.05 |
| 3_8059952 | 1 | 0 | 4 | 0 | 3 | 0 | 4 | 0.25 | 12 | 0.0833 | NA | NA | NA | NA | 1 | 0 | 2 | 0.5 | 3 | 0.3333 |
| 3_8059976 | 17 | 0.0588 | 22 | 0.0455 | 22 | 0.0455 | 27 | 0.1481 | 88 | 0.0795 | 14 | 0.1429 | 20 | 0.05 | 23 | 0 | 26 | 0.1154 | 83 | 0.0723 |
| 3_8059980 | 13 | 0.6154 | 17 | 0.4118 | 20 | 0.5 | 27 | 0.5185 | 77 | 0.5065 | 12 | 0.5 | 21 | 0.381 | 17 | 0.6471 | 21 | 0.3333 | 71 | 0.4507 |
| 3_8064841 | 7 | 0 | 13 | 0.1538 | 10 | 0.1 | 14 | 0.1429 | 44 | 0.1136 | 6 | 0 | 13 | 0.1538 | 10 | 0.2 | 14 | 0.2143 | 43 | 0.1628 |
| 3_8070964 | 7 | 0.1429 | 10 | 0.1 | 8 | 0 | 11 | 0.0909 | 36 | 0.0833 | 5 | 0 | 11 | 0 | 10 | 0 | 12 | 0.3333 | 38 | 0.1053 |
| 3_8070989 | 6 | 0.3333 | 8 | 0 | 8 | 0.125 | 9 | 0 | 31 | 0.0968 | 5 | 0 | 8 | 0 | 7 | 0.1429 | 10 | 0.1 | 30 | 0.0667 |
| 3_8070990 | 6 | 0.5 | 9 | 0.2222 | 8 | 0 | 10 | 0.1 | 33 | 0.1818 | 4 | 0 | 6 | 0 | 8 | 0.375 | 10 | 0.1 | 28 | 0.1429 |
| 3_8070993 | 7 | 0.1429 | 9 | 0 | 7 | 0.1429 | 9 | 0 | 32 | 0.0625 | 4 | 0 | 7 | 0.1429 | 9 | 0 | 7 | 0.2857 | 27 | 0.1111 |
| 3_8075219 | 19 | 0.0526 | 21 | 0.0952 | 11 | 0.0909 | 30 | 0.0333 | 81 | 0.0617 | 21 | 0.0952 | 16 | 0.0625 | 23 | 0.1739 | 21 | 0.0476 | 81 | 0.0988 |
| 3_8075242 | 17 | 0.8235 | 16 | 0.8125 | 7 | 1 | 30 | 0.9 | 70 | 0.8714 | 19 | 1 | 13 | 0.6154 | 21 | 0.8571 | 18 | 0.8333 | 71 | 0.8451 |
| 3_8075245 | 18 | 0.4444 | 16 | 0.3125 | 8 | 0.625 | 28 | 0.5 | 70 | 0.4571 | 18 | 0.3889 | 13 | 0.3846 | 22 | 0.4091 | 18 | 0.3333 | 71 | 0.3803 |
| 3_8075250 | 19 | 0.1579 | 19 | 0.1579 | 10 | 0.4 | 30 | 0.0333 | 78 | 0.141 | 19 | 0.1053 | 15 | 0.2 | 21 | 0.0952 | 18 | 0.2222 | 73 | 0.1507 |
| 3_8111985 | 2 | 1 | 1 | 1 | NA | NA | NA | NA | 3 | 1 | 4 | 0.75 | 1 | 1 | 2 | 1 | NA | NA | 7 | 0.8571 |
| 3_8112006 | 2 | 1 | 1 | 1 | NA | NA | NA | NA | 3 | 1 | 4 | 1 | 1 | 0 | 1 | 1 | NA | NA | 6 | 0.8333 |
| 3_8112019 | 2 | 0.5 | 1 | 1 | NA | NA | NA | NA | 3 | 0.6667 | 4 | 0.5 | 1 | 0 | 2 | 1 | NA | NA | 7 | 0.5714 |
| 3_8112020 | 1 | 1 | 1 | 0 | NA | NA | NA | NA | 2 | 0.5 | 4 | 0.25 | 1 | 0 | 2 | 0.5 | NA | NA | 7 | 0.2857 |
| 3_8112022 | 2 | 0.5 | 1 | 0 | NA | NA | NA | NA | 3 | 0.3333 | 3 | 0.6667 | 1 | 1 | NA | NA | NA | NA | 4 | 0.75 |
| 3_8112030 | 2 | 1 | 1 | 1 | NA | NA | NA | NA | 3 | 1 | 3 | 0.6667 | 1 | 1 | 1 | 1 | NA | NA | 5 | 0.8 |
| 3_8112035 | 2 | 0.5 | NA | NA | NA | NA | NA | NA | 2 | 0.5 | 2 | 1 | 1 | 0 | 2 | 1 | NA | NA | 5 | 0.8 |
| 3_8112044 | 2 | 1 | 1 | 1 | NA | NA | NA | NA | 3 | 1 | 5 | 1 | NA | NA | 1 | 1 | NA | NA | 6 | 1 |
| 3_8182554 | 2 | 0.5 | NA | NA | 1 | 0 | NA | NA | 3 | 0.3333 | NA | NA | 2 | 1 | 3 | 0.3333 | NA | NA | 5 | 0.6 |
| 3_8182556 | 2 | 0.5 | NA | NA | 1 | 1 | NA | NA | 3 | 0.6667 | NA | NA | 2 | 1 | 3 | 0.6667 | NA | NA | 5 | 0.8 |
| 3_8182561 | 2 | 0.5 | NA | NA | 1 | 0 | NA | NA | 3 | 0.3333 | NA | NA | 1 | 0 | 2 | 0.5 | NA | NA | 3 | 0.3333 |
| 3_8182589 | 2 | 0.5 | NA | NA | 1 | 0 | NA | NA | 3 | 0.3333 | NA | NA | 2 | 0.5 | 1 | 1 | NA | NA | 3 | 0.6667 |
| 3_8182597 | 2 | 0.5 | NA | NA | 1 | 1 | NA | NA | 3 | 0.6667 | NA | NA | 2 | 1 | 3 | 0.6667 | NA | NA | 5 | 0.8 |
| 3_8186173 | 4 | 0.5 | 5 | 0.4 | NA | NA | 3 | 0.3333 | 12 | 0.4167 | 4 | 0.5 | 5 | 0.6 | 1 | 1 | 3 | 0.6667 | 13 | 0.6154 |
| 3_8186175 | 2 | 0.5 | 6 | 0.3333 | NA | NA | 5 | 0.2 | 13 | 0.3077 | 3 | 0.3333 | 6 | 0.6667 | 1 | 1 | 3 | 0.3333 | 13 | 0.5385 |
| 3_8186175 | 2 | 0.5 | 5 | 0.8 | NA | NA | 6 | 0.8333 | 13 | 0.7692 | 3 | 0.3333 | 3 | 0.6667 | 1 | 1 | 2 | 1 | 9 | 0.6667 |
| 3_8186179 | 2 | 0 | 3 | 0 | NA | NA | 4 | 0.25 | 9 | 0.1111 | 4 | 0 | 7 | 0.1429 | 1 | 0 | 2 | 0 | 14 | 0.0714 |
| 3_8186180 | 4 | 0.75 | 6 | 1 | NA | NA | 7 | 0.7143 | 17 | 0.8235 | 4 | 0.75 | 9 | 0.7778 | 1 | 1 | 3 | 1 | 17 | 0.8235 |
| 3_8186188 | 4 | 0.5 | 6 | 0.8333 | NA | NA | 5 | 0.4 | 15 | 0.6 | 3 | 0.6667 | 8 | 0.75 | 1 | 0 | 3 | 1 | 15 | 0.7333 |
| 3_8186198 | 3 | 0.6667 | 6 | 0.6667 | NA | NA | 8 | 0.5 | 17 | 0.5882 | 4 | 0.5 | 8 | 0.375 | 1 | 0 | 3 | 0.6667 | 16 | 0.4375 |
| 3_8186202 | 4 | 1 | 5 | 0.4 | NA | NA | 8 | 0.75 | 17 | 0.7059 | 2 | 1 | 8 | 0.875 | 1 | 1 | 3 | 1 | 14 | 0.9286 |
| 3_8186212 | 4 | 0.5 | 5 | 0.6 | NA | NA | 8 | 0.75 | 17 | 0.6471 | 4 | 0.5 | 9 | 1 | 1 | 1 | 3 | 1 | 17 | 0.8824 |
| 3_8186213 | 4 | 0.25 | 6 | 0 | NA | NA | 7 | 0.1429 | 17 | 0.1176 | 3 | 0 | 8 | 0 | 1 | 0 | 3 | 0.3333 | 15 | 0.0667 |
| 3_8186219 | 4 | 0.5 | 6 | 0.3333 | NA | NA | 5 | 0.4 | 15 | 0.4 | 3 | 0.3333 | 8 | 0.5 | 1 | 0 | 2 | 0.5 | 14 | 0.4286 |
| 3_8186222 | 4 | 0.25 | 6 | 0.1667 | NA | NA | 6 | 0.1667 | 16 | 0.1875 | 2 | 0 | 9 | 0.1111 | 1 | 0 | 3 | 0 | 15 | 0.0667 |
| 3_8186272 | 5 | 0 | 5 | 0 | 1 | 0 | 8 | 0.125 | 19 | 0.0526 | 4 | 0 | 4 | 0 | 1 | 1 | 3 | 0 | 12 | 0.0833 |
| 3_8193956 | 39 | 0.3077 | 26 | 0.5769 | 44 | 0.2727 | 34 | 0.4706 | 143 | 0.3846 | 25 | 0.44 | 36 | 0.4167 | 37 | 0.2703 | 34 | 0.3824 | 132 | 0.3712 |

| | | | | | | | | | | | | | | | | | | | | |
|---|---|---|---|---|---|---|---|---|---|---|---|---|---|---|---|---|---|---|---|---|
| 3_8193971 | 34 | 0 | 17 | 0.1765 | 39 | 0.0256 | 33 | 0.0909 | 123 | 0.0569 | 26 | 0.0769 | 30 | 0.1333 | 36 | 0.1389 | 33 | 0.0909 | 125 | 0.112 |
| 3_8193994 | 33 | 0.0909 | 16 | 0.0625 | 37 | 0.1351 | 22 | 0.0909 | 108 | 0.1019 | 23 | 0.1304 | 29 | 0 | 29 | 0.1034 | 27 | 0.1111 | 108 | 0.0833 |
| 3_8199985 | 12 | 0.0833 | 14 | 0 | 15 | 0.1333 | 25 | 0.08 | 66 | 0.0758 | 11 | 0 | 36 | 0.1111 | 28 | 0 | 17 | 0.0588 | 92 | 0.0543 |
| 3_8200066 | 13 | 0.0769 | 8 | 0.375 | 12 | 0.25 | 27 | 0.1111 | 60 | 0.1667 | 8 | 0.125 | 31 | 0 | 24 | 0.0417 | 14 | 0.2143 | 77 | 0.0649 |
| 3_8200119 | 7 | 0.1429 | 7 | 0 | 9 | 0 | 26 | 0.0769 | 49 | 0.0612 | 9 | 0.1111 | 30 | 0.0333 | 25 | 0.04 | 12 | 0.25 | 76 | 0.0789 |
| 3_8200176 | 5 | 0.2 | 4 | 0 | 7 | 0.1429 | 14 | 0.0714 | 30 | 0.1 | 6 | 0.1667 | 16 | 0.1875 | 12 | 0.1667 | 9 | 0 | 43 | 0.1395 |
| 3_8200178 | 11 | 0.3636 | 6 | 0.5 | 8 | 0.375 | 28 | 0.5 | 53 | 0.4528 | 9 | 0.6667 | 20 | 0.45 | 15 | 0.4667 | 12 | 0.6667 | 56 | 0.5357 |
| 3_8200208 | 10 | 0.7 | 5 | 1 | 8 | 0.625 | 26 | 0.5 | 49 | 0.6122 | 8 | 0.875 | 21 | 0.8095 | 13 | 0.6923 | 8 | 0.75 | 50 | 0.78 |
| 3_8200213 | 7 | 0.1429 | 6 | 0 | 8 | 0.125 | 27 | 0.1111 | 48 | 0.1042 | 9 | 0 | 21 | 0.0952 | 15 | 0.1333 | 11 | 0 | 56 | 0.0714 |
| 3_8200219 | 7 | 0.4286 | 6 | 0.3333 | 8 | 0.625 | 28 | 0.3214 | 49 | 0.3878 | 9 | 0.5556 | 22 | 0.4091 | 14 | 0.2143 | 10 | 0.3 | 55 | 0.3636 |
| 3_8201323 | 7 | 0.5714 | 5 | 0.8 | 7 | 0.7143 | 21 | 0.7619 | 40 | 0.725 | 7 | 0.7143 | 15 | 0.6667 | 11 | 0.6364 | 11 | 0.9091 | 44 | 0.7273 |
| 3_8201329 | 5 | 0.2 | 3 | 0 | 7 | 0.5714 | 19 | 0.5263 | 34 | 0.4412 | 5 | 0.2 | 12 | 0.4167 | 6 | 0.1667 | 9 | 0.6667 | 32 | 0.4062 |
| 3_8201347 | 7 | 0.2857 | 5 | 0 | 6 | 0.8333 | 18 | 0.2778 | 36 | 0.3333 | 4 | 0.5 | 14 | 0.3571 | 8 | 0.375 | 8 | 0.625 | 34 | 0.4412 |
| 3_8201595 | 5 | 0.2 | 5 | 0 | 6 | 0.1667 | 17 | 0 | 33 | 0.0606 | 2 | 0 | 12 | 0.0833 | 8 | 0.125 | 11 | 0 | 33 | 0.0606 |
| 3_8201598 | 6 | 0 | 3 | 0 | 7 | 0.2857 | 17 | 0.0588 | 33 | 0.0909 | 4 | 0 | 10 | 0.1 | 6 | 0.5 | 11 | 0.3636 | 31 | 0.2581 |
| 3_8201604 | 7 | 0.4286 | 5 | 0.6 | 7 | 0.4286 | 16 | 0.5 | 35 | 0.4857 | 4 | 0.25 | 12 | 0.75 | 8 | 0.75 | 11 | 0.3636 | 35 | 0.5714 |
| 3_8201610 | 3 | 0.6667 | 4 | 0.5 | 3 | 0 | 14 | 0.4286 | 24 | 0.4167 | 6 | 0.3333 | 10 | 0.1 | 6 | 0 | 8 | 0.375 | 30 | 0.2 |
| 3_8201613 | 5 | 0.6 | 4 | 0.75 | 5 | 0.4 | 18 | 0.2778 | 32 | 0.4062 | 4 | 0 | 12 | 0.4167 | 7 | 0.1429 | 10 | 0.5 | 33 | 0.3333 |
| 3_8201625 | 5 | 1 | 5 | 1 | 6 | 1 | 17 | 0.9412 | 33 | 0.9697 | 6 | 0.8333 | 14 | 0.8571 | 10 | 0.8 | 10 | 1 | 40 | 0.875 |
| 3_8201639 | 6 | 0.8333 | 2 | 1 | 5 | 1 | 17 | 1 | 30 | 0.9667 | 5 | 1 | 10 | 1 | 9 | 0.8889 | 10 | 1 | 34 | 0.9706 |
| 3_8201640 | 6 | 0 | 2 | 0.5 | 6 | 0.1667 | 18 | 0.1111 | 32 | 0.125 | 5 | 0 | 13 | 0.2308 | 10 | 0.1 | 10 | 0 | 38 | 0.1053 |
| 3_8201650 | 5 | 1 | 2 | 1 | 4 | 1 | 17 | 0.8824 | 28 | 0.9286 | 3 | 1 | 11 | 1 | 9 | 1 | 10 | 1 | 33 | 1 |
| 3_8227070 | 3 | 1 | 1 | 0 | 3 | 0 | 5 | 0.2 | 12 | 0.3333 | 3 | 0.3333 | 2 | 0 | 1 | 0 | 3 | 0.3333 | 9 | 0.2222 |
| 3_8227075 | 1 | 0 | 1 | 0 | 3 | 0 | 3 | 0.6667 | 8 | 0.25 | 2 | 0.5 | 2 | 0 | 1 | 1 | 2 | 0.5 | 7 | 0.4286 |
| 3_8227860 | 2 | 0.5 | 3 | 0 | 2 | 0 | 4 | 0 | 11 | 0.0909 | 3 | 0 | 3 | 0 | 2 | 0.5 | 3 | 0 | 11 | 0.0909 |
| 3_8228430 | 4 | 0.25 | 3 | 0 | 4 | 0 | 6 | 0 | 17 | 0.0588 | 3 | 0.3333 | 2 | 1 | 2 | 0 | 1 | 0 | 8 | 0.375 |
| 3_8229853 | 4 | 0.25 | 1 | 1 | 4 | 0 | 9 | 0 | 18 | 0.1111 | 3 | 0 | 2 | 0 | 2 | 0 | 6 | 0.1667 | 13 | 0.0769 |
| 3_8229864 | 4 | 0.5 | 2 | 1 | 3 | 0.3333 | 9 | 0.6667 | 18 | 0.6111 | 3 | 0.3333 | 3 | 0.6667 | 3 | 0.3333 | 7 | 0.5714 | 16 | 0.5 |
| 3_8229871 | 5 | 0.4 | 3 | 0 | 5 | 0.2 | 7 | 0.7143 | 20 | 0.4 | 3 | 0 | 3 | 0.3333 | 3 | 0 | 6 | 0.3333 | 15 | 0.2 |
| 3_8229874 | 4 | 1 | 3 | 0.6667 | 4 | 0.75 | 8 | 0.875 | 19 | 0.8421 | 2 | 1 | 3 | 0.6667 | 3 | 0.6667 | 6 | 1 | 14 | 0.8571 |
| 3_8229878 | 1 | 0 | 1 | 0 | 2 | 1 | 4 | 0.5 | 8 | 0.5 | NA | NA | 1 | 0 | 1 | 1 | 2 | 0.5 | 4 | 0.5 |
| 3_8229895 | 5 | 0.2 | 3 | 0 | 5 | 0.4 | 9 | 0.1111 | 22 | 0.1818 | 4 | 0.5 | 3 | 0 | 3 | 0 | 7 | 0 | 17 | 0.1176 |
| 3_8229905 | 4 | 0 | 3 | 0 | 4 | 0 | 8 | 0.125 | 19 | 0.0526 | 3 | 0.3333 | 2 | 0 | 2 | 0 | 7 | 0.2857 | 14 | 0.2143 |
| 3_8229907 | 4 | 0.75 | 3 | 0.3333 | 5 | 0.6 | 7 | 0.4286 | 19 | 0.5263 | 4 | 0.25 | 3 | 0.6667 | 1 | 1 | 7 | 0.4286 | 15 | 0.4667 |
| 3_8229934 | 4 | 0 | 3 | 0.3333 | 4 | 0 | 8 | 0 | 19 | 0.0526 | 4 | 0 | 2 | 0 | 2 | 0 | 7 | 0.2857 | 15 | 0.1333 |
| 3_8229954 | 4 | 0.5 | 2 | 0.5 | 3 | 0 | 10 | 0.2 | 19 | 0.2632 | 2 | 0 | 2 | 0 | 4 | 0.25 | 7 | 0.2857 | 15 | 0.2 |
| 3_8229960 | 5 | 0.4 | 5 | 0.2 | 6 | 0.3333 | 10 | 0.4 | 26 | 0.3462 | 3 | 0.3333 | 3 | 0 | 2 | 0 | 7 | 0.7143 | 15 | 0.4 |
| 3_8229968 | 1 | 0 | 4 | 1 | 4 | 0.5 | 9 | 0.5556 | 18 | 0.6111 | NA | NA | 2 | 0 | 4 | 0.5 | 3 | 1 | 9 | 0.5556 |
| 3_8229968 | 5 | 0 | 2 | 0 | 3 | 0 | 9 | 0.1111 | 19 | 0.0526 | 4 | 0.25 | 3 | 0 | 5 | 0.4 | 7 | 0.2857 | 19 | 0.2632 |
| 3_8229974 | 5 | 0.4 | 4 | 0.25 | 5 | 0 | 10 | 0.1 | 24 | 0.1667 | 5 | 0.2 | 3 | 0 | 5 | 0 | 7 | 0.1429 | 20 | 0.1 |
| 3_8229978 | 4 | 1 | 5 | 1 | 6 | 1 | 11 | 1 | 26 | 1 | 6 | 1 | 3 | 1 | 5 | 1 | 8 | 1 | 22 | 1 |
| 3_8229984 | 5 | 0.4 | 5 | 0.4 | 5 | 0.6 | 11 | 0.5455 | 26 | 0.5 | 6 | 1 | 3 | 0.3333 | 3 | 0 | 8 | 0.75 | 20 | 0.65 |
| 3_8230003 | 4 | 1 | 3 | 1 | 4 | 1 | 8 | 1 | 19 | 1 | 6 | 1 | 3 | 1 | 4 | 1 | 6 | 1 | 19 | 1 |
| 3_8230004 | 4 | 0.25 | 5 | 0.4 | 6 | 0.1667 | 9 | 0.3333 | 24 | 0.2917 | 5 | 0.2 | 3 | 0.3333 | 4 | 0.25 | 6 | 0.8333 | 18 | 0.4444 |
| 3_8230010 | 5 | 1 | 4 | 1 | 4 | 1 | 11 | 1 | 24 | 1 | 6 | 0.8333 | 3 | 1 | 5 | 1 | 9 | 1 | 23 | 0.9565 |
| 3_8230022 | 4 | 0.75 | 3 | 0.3333 | 4 | 0.75 | 9 | 0.5556 | 20 | 0.6 | 4 | 0.75 | 3 | 0.6667 | 5 | 0.8 | 9 | 0.8889 | 21 | 0.8095 |
| 3_8230029 | 4 | 0.25 | 2 | 0.5 | 6 | 0.5 | 10 | 0.2 | 22 | 0.3182 | 5 | 0 | 3 | 0 | 4 | 0.75 | 10 | 0.3 | 22 | 0.2727 |
| 3_8230039 | 4 | 1 | 4 | 1 | 6 | 1 | 11 | 0.8182 | 25 | 0.92 | 5 | 1 | 3 | 0.6667 | 5 | 0.6 | 9 | 0.8889 | 22 | 0.8182 |
| 3_8230039 | 5 | 1 | 5 | 0.6 | 7 | 1 | 8 | 1 | 25 | 0.92 | 6 | 1 | 4 | 1 | 5 | 0.8 | 8 | 1 | 23 | 0.9565 |
| 3_8230040 | 4 | 0 | 4 | 0 | 6 | 0.5 | 10 | 0 | 24 | 0.125 | 6 | 0.5 | 4 | 0.25 | 4 | 0 | 10 | 0.3 | 24 | 0.2917 |
| 3_8230045 | 5 | 0.8 | 5 | 1 | 7 | 0.5714 | 11 | 1 | 28 | 0.8571 | 5 | 0.8 | 4 | 0.75 | 5 | 0.8 | 8 | 0.875 | 22 | 0.8182 |
| 3_8230046 | 5 | 0.6 | 5 | 0.4 | 3 | 0.3333 | 7 | 0.1429 | 20 | 0.35 | 6 | 0.6667 | 3 | 1 | 3 | 0.3333 | 9 | 0.4444 | 21 | 0.5714 |
| 3_8230050 | 4 | 0.25 | 4 | 0 | 6 | 0.6667 | 8 | 0.375 | 22 | 0.3636 | 6 | 0.1667 | 4 | 0 | 5 | 0.6 | 8 | 0.5 | 23 | 0.3478 |
| 3_8230053 | 4 | 0.25 | 4 | 0 | 4 | 0.25 | 8 | 0.375 | 20 | 0.25 | 5 | 0 | 2 | 0.5 | 3 | 0 | 6 | 0.3333 | 16 | 0.1875 |
| 3_8230081 | 3 | 1 | 5 | 0.6 | 6 | 1 | 9 | 1 | 23 | 0.913 | 2 | 1 | 2 | 1 | 3 | 0.6667 | 9 | 0.8889 | 16 | 0.875 |
| 3_8230091 | 5 | 0 | 7 | 0.1429 | 10 | 0.3 | 12 | 0.3333 | 34 | 0.2353 | 7 | 0.4286 | 5 | 0 | 4 | 0 | 9 | 0.4444 | 25 | 0.28 |
| 3_8230098 | 3 | 0.3333 | 5 | 0 | 6 | 0.3333 | 10 | 0.1 | 24 | 0.1667 | 6 | 0.1667 | 5 | 0.2 | 6 | 0 | 9 | 0.4444 | 26 | 0.2308 |
| 3_8230128 | 5 | 0.4 | 6 | 0.1667 | 9 | 0.1111 | 13 | 0 | 33 | 0.1212 | 7 | 0.2857 | 6 | 0.1667 | 7 | 0.2857 | 9 | 0.1111 | 29 | 0.2069 |
| 3_8230129 | 4 | 0.25 | 6 | 0.3333 | 10 | 0.1 | 13 | 0 | 33 | 0.1212 | 8 | 0.125 | 6 | 0.1667 | 7 | 0 | 10 | 0 | 31 | 0.0645 |
| 3_8244467 | NA | NA | NA | NA | NA | NA | 1 | 1 | 1 | 1 | 1 | 1 | NA | NA | NA | NA | NA | NA | 1 | 1 |
| 3_8244596 | NA | NA | 2 | 1 | NA | NA | 2 | 1 | 4 | 1 | 1 | 1 | NA | NA | 2 | 1 | NA | NA | 3 | 1 |
| 3_8244598 | NA | NA | 3 | 0.6667 | NA | NA | 2 | 1 | 5 | 0.8 | 1 | 1 | NA | NA | 2 | 1 | NA | NA | 3 | 1 |
| 3_8244599 | NA | NA | 2 | 0.5 | NA | NA | 2 | 0.5 | 4 | 0.5 | 1 | 1 | NA | NA | 2 | 1 | NA | NA | 3 | 1 |
| 3_8244615 | NA | NA | 2 | 0.5 | NA | NA | 2 | 0 | 4 | 0.25 | 1 | 0 | NA | NA | 2 | 0.5 | NA | NA | 3 | 0.3333 |
| 3_8244728 | NA | NA | 2 | 1 | NA | NA | 2 | 1 | 4 | 1 | 1 | 1 | NA | NA | 1 | 1 | NA | NA | 2 | 1 |

| | | | | | | | | | | | | | | | | | | | | |
|---|---|---|---|---|---|---|---|---|---|---|---|---|---|---|---|---|---|---|---|---|
| 3_8244752 | NA | NA | 2 | 1 | NA | NA | 2 | 1 | 3 | 1 | 1 | 1 | NA | NA | 1 | 1 | NA | NA | 2 | 1 |
| 3_8244762 | NA | NA | 2 | 1 | NA | NA | 2 | 1 | 4 | 1 | 1 | 1 | NA | NA | NA | NA | NA | NA | 1 | 1 |
| 3_8244765 | NA | NA | 2 | 1 | NA | NA | 2 | 1 | 4 | 1 | 1 | 1 | NA | NA | 1 | 1 | NA | NA | 2 | 1 |
| 3_8244767 | NA | NA | 2 | 0.5 | NA | NA | 2 | 0 | 4 | 0.25 | NA | NA | NA | NA | 1 | 1 | NA | NA | 1 | 1 |
| 3_8244768 | NA | NA | 2 | 0 | NA | NA | 2 | 1 | 4 | 0.5 | 1 | 1 | NA | NA | 1 | 0 | NA | NA | 2 | 0.5 |
| 3_8244786 | NA | NA | 2 | 0.5 | NA | NA | 2 | 0 | 4 | 0.25 | 1 | 1 | NA | NA | 1 | 0 | NA | NA | 2 | 0.5 |
| 3_8244818 | NA | NA | NA | NA | NA | NA | 1 | 1 | 1 | 1 | 1 | 1 | NA | NA | 1 | 1 | NA | NA | 2 | 1 |
| 3_8244835 | NA | NA | 1 | 1 | NA | NA | 1 | 1 | 2 | 1 | NA | NA | NA | NA | 1 | 1 | NA | NA | 1 | 1 |
| 3_8303264 | 1 | 1 | NA | NA | NA | NA | NA | NA | 1 | 1 | NA | NA | 1 | 1 | NA | NA | NA | NA | 1 | 1 |
| 3_8303285 | 1 | 1 | NA | NA | NA | NA | NA | NA | 1 | 1 | NA | NA | 1 | 1 | NA | NA | NA | NA | 1 | 1 |
| 3_8303383 | 1 | 1 | NA | NA | NA | NA | NA | NA | 1 | 1 | NA | NA | 1 | 1 | NA | NA | NA | NA | 1 | 1 |
| 3_8303407 | NA | NA | NA | NA | 1 | 1 | NA | NA | 1 | 1 | NA | NA | 1 | 1 | NA | NA | NA | NA | 1 | 1 |
| 3_8303514 | NA | NA | NA | NA | 1 | 1 | NA | NA | 1 | 1 | NA | NA | 1 | 1 | NA | NA | NA | NA | 1 | 1 |
| 3_8303586 | NA | NA | NA | NA | NA | NA | 1 | 1 | 1 | 1 | NA | NA | NA | NA | NA | NA | 1 | 1 | 1 | 1 |
| 3_8303597 | 1 | 1 | NA | NA | NA | NA | NA | NA | 1 | 1 | NA | NA | NA | NA | NA | NA | 1 | 1 | 1 | 1 |
| 3_8303670 | 1 | 1 | NA | NA | NA | NA | 1 | 1 | 2 | 1 | NA | NA | NA | NA | NA | NA | 1 | 1 | 1 | 1 |
| 3_8303673 | 1 | 1 | NA | NA | NA | NA | NA | NA | 1 | 1 | NA | NA | NA | NA | NA | NA | 1 | 1 | 1 | 1 |
| 3_8303680 | 1 | 1 | NA | NA | NA | NA | NA | NA | 1 | 1 | NA | NA | NA | NA | NA | NA | 1 | 1 | 1 | 1 |
| 3_8303710 | 1 | 1 | NA | NA | NA | NA | NA | NA | 1 | 1 | NA | NA | NA | NA | NA | NA | 1 | 1 | 1 | 1 |
| 3_8303723 | 1 | 1 | NA | NA | NA | NA | NA | NA | 1 | 1 | NA | NA | NA | NA | NA | NA | 1 | 1 | 1 | 1 |
| 3_8303732 | 1 | 1 | NA | NA | NA | NA | NA | NA | 1 | 1 | NA | NA | NA | NA | NA | NA | 1 | 1 | 1 | 1 |
| 3_8303762 | 1 | 1 | NA | NA | NA | NA | NA | NA | 1 | 1 | NA | NA | NA | NA | NA | NA | 1 | 1 | 1 | 1 |
| 3_8303779 | 1 | 1 | NA | NA | NA | NA | NA | NA | 1 | 1 | NA | NA | NA | NA | NA | NA | 1 | 1 | 1 | 1 |
| 3_8303786 | 1 | 1 | NA | NA | NA | NA | NA | NA | 1 | 1 | NA | NA | NA | NA | NA | NA | 1 | 1 | 1 | 1 |
| 3_8303799 | 1 | 1 | NA | NA | NA | NA | NA | NA | 1 | 1 | NA | NA | NA | NA | NA | NA | 1 | 1 | 1 | 1 |
| 3_8303802 | 1 | 1 | NA | NA | NA | NA | NA | NA | 1 | 1 | NA | NA | NA | NA | NA | NA | 1 | 1 | 1 | 1 |
| 3_8303806 | 1 | 1 | NA | NA | NA | NA | NA | NA | 1 | 1 | NA | NA | NA | NA | NA | NA | 1 | 1 | 1 | 1 |
| 3_8303812 | 1 | 1 | NA | NA | NA | NA | NA | NA | 1 | 1 | NA | NA | NA | NA | NA | NA | 1 | 1 | 1 | 1 |
| 3_8326162 | 1 | 1 | NA | NA | NA | NA | 1 | 1 | 2 | 1 | NA | NA | NA | NA | 2 | 1 | NA | NA | 2 | 1 |
| 3_8326180 | 1 | 1 | NA | NA | NA | NA | 1 | 0 | 2 | 0.5 | NA | NA | NA | NA | 2 | 1 | NA | NA | 2 | 1 |
| 3_8326182 | 1 | 1 | NA | NA | NA | NA | 1 | 0 | 2 | 0.5 | NA | NA | NA | NA | 2 | 1 | NA | NA | 2 | 1 |
| 3_8326216 | 1 | 1 | NA | NA | NA | NA | 1 | 1 | 2 | 1 | NA | NA | NA | NA | 3 | 1 | NA | NA | 3 | 1 |
| 3_8326242 | 1 | 1 | NA | NA | NA | NA | 2 | 0.5 | 3 | 0.6667 | NA | NA | NA | NA | 3 | 0.3333 | NA | NA | 3 | 0.3333 |
| 3_8326261 | 1 | 1 | NA | NA | NA | NA | NA | NA | 1 | 1 | NA | NA | NA | NA | 3 | 0.6667 | NA | NA | 3 | 0.6667 |
| 3_8326289 | 1 | 1 | NA | NA | NA | NA | NA | NA | 1 | 1 | NA | NA | NA | NA | 3 | 0.6667 | NA | NA | 3 | 0.6667 |
| 3_8326291 | 1 | 1 | NA | NA | NA | NA | 1 | 0 | 2 | 0.5 | NA | NA | NA | NA | 3 | 1 | NA | NA | 3 | 1 |
| 3_8326302 | 1 | 1 | NA | NA | NA | NA | 1 | 1 | 2 | 1 | NA | NA | NA | NA | 3 | 1 | NA | NA | 3 | 1 |
| 3_8326308 | 1 | 1 | NA | NA | NA | NA | 1 | 1 | 2 | 1 | NA | NA | NA | NA | 3 | 1 | NA | NA | 3 | 1 |
| 3_8326342 | 1 | 1 | NA | NA | NA | NA | 1 | 1 | 2 | 1 | NA | NA | NA | NA | 3 | 0.3333 | NA | NA | 3 | 0.3333 |
| 3_8326347 | 1 | 1 | NA | NA | NA | NA | 1 | 1 | 2 | 1 | NA | NA | NA | NA | 3 | 1 | NA | NA | 3 | 1 |
| 3_8326366 | 1 | 0 | NA | NA | NA | NA | 1 | 1 | 2 | 0.5 | NA | NA | NA | NA | 1 | 1 | NA | NA | 1 | 1 |
| 3_8326367 | 1 | 0 | NA | NA | 1 | 1 | 1 | 1 | 3 | 0.6667 | NA | NA | NA | NA | 2 | 1 | NA | NA | 2 | 1 |
| 3_8326382 | 1 | 1 | NA | NA | 1 | 1 | 1 | 1 | 3 | 1 | NA | NA | NA | NA | 2 | 1 | NA | NA | 2 | 1 |
| 3_8326387 | NA | NA | NA | NA | 1 | 1 | 1 | 1 | 2 | 1 | NA | NA | NA | NA | 3 | 1 | NA | NA | 3 | 1 |
| 3_8326391 | 1 | 1 | NA | NA | 1 | 0 | NA | NA | 2 | 0.5 | NA | NA | NA | NA | 2 | 0.5 | NA | NA | 2 | 0.5 |
| 3_8326401 | 1 | 1 | NA | NA | 1 | 0 | 1 | 0 | 3 | 0.3333 | NA | NA | NA | NA | 3 | 0.3333 | NA | NA | 3 | 0.3333 |
| 3_8374392 | NA | NA | NA | NA | 1 | 1 | NA | NA | 1 | 1 | 1 | 1 | NA | NA | NA | NA | NA | NA | 1 | 1 |
| 3_8374408 | NA | NA | NA | NA | 1 | 1 | NA | NA | 1 | 1 | 1 | 1 | NA | NA | NA | NA | NA | NA | 1 | 1 |
| 3_8374554 | NA | NA | NA | NA | 1 | 1 | NA | NA | 1 | 1 | 1 | 1 | NA | NA | NA | NA | NA | NA | 1 | 1 |
| 3_8374560 | NA | NA | NA | NA | 1 | 1 | NA | NA | 1 | 1 | 1 | 1 | NA | NA | NA | NA | NA | NA | 1 | 1 |
| 3_8380564 | 1 | 1 | 3 | 0 | NA | NA | 1 | 0 | 5 | 0.2 | NA | NA | 5 | 0.2 | 2 | 1 | NA | NA | 7 | 0.4286 |
| 3_8380567 | 1 | 1 | 3 | 0.6667 | NA | NA | 1 | 0 | 5 | 0.6 | 1 | 1 | 6 | 0.5 | 2 | 0 | NA | NA | 9 | 0.4444 |
| 3_8380571 | 1 | 0 | 3 | 1 | NA | NA | NA | NA | 4 | 0.75 | NA | NA | 3 | 1 | 2 | 1 | NA | NA | 5 | 1 |
| 3_8382960 | 1 | 1 | 1 | 0 | NA | NA | NA | NA | 2 | 0.5 | NA | NA | 3 | 0 | 1 | 1 | NA | NA | 4 | 0.25 |
| 3_8382964 | 1 | 0 | 1 | 1 | NA | NA | NA | NA | 2 | 0.5 | NA | NA | 4 | 0.25 | 1 | 1 | NA | NA | 5 | 0.4 |
| 3_8406837 | 20 | 0.05 | 39 | 0.1538 | 20 | 0.1 | 27 | 0.2593 | 106 | 0.1509 | 13 | 0.2308 | 15 | 0.1333 | 18 | 0.1111 | 13 | 0 | 59 | 0.1186 |
| 3_8406874 | 13 | 0.4615 | 36 | 0.1111 | 19 | 0.1053 | 21 | 0.1429 | 89 | 0.1685 | 16 | 0.125 | 13 | 0.3846 | 16 | 0.125 | 9 | 0.1111 | 54 | 0.1852 |
| 3_8406890 | 13 | 0.3077 | 35 | 0.1429 | 22 | 0.1364 | 23 | 0.1304 | 93 | 0.1613 | 11 | 0 | 12 | 0.0833 | 13 | 0.0769 | 9 | 0.3333 | 45 | 0.1111 |
| 3_8406959 | 12 | 0.1667 | 22 | 0.0909 | 13 | 0.2308 | 20 | 0.2 | 67 | 0.1642 | 7 | 0.4286 | 10 | 0.3 | 13 | 0.2308 | 8 | 0.125 | 38 | 0.2632 |
| 3_8406971 | 12 | 0 | 26 | 0.1538 | 15 | 0 | 20 | 0.2 | 73 | 0.1096 | 9 | 0 | 11 | 0 | 12 | 0.1667 | 8 | 0 | 40 | 0.05 |
| 3_8407014 | 12 | 0.25 | 24 | 0.3333 | 14 | 0.2857 | 18 | 0.3333 | 68 | 0.3088 | 8 | 0.375 | 9 | 0.2222 | 10 | 0.4 | 8 | 0.25 | 35 | 0.3143 |
| 3_8407024 | 12 | 0.25 | 27 | 0.037 | 13 | 0.0769 | 18 | 0 | 70 | 0.0714 | 8 | 0.25 | 8 | 0.125 | 9 | 0.1111 | 7 | 0 | 32 | 0.125 |
| 3_8407089 | 6 | 0.1667 | 19 | 0.1053 | 7 | 0 | 10 | 0.5 | 42 | 0.1905 | 5 | 0.2 | 6 | 0.1667 | 7 | 0 | 4 | 0 | 22 | 0.0909 |
| 3_8407123 | 5 | 0 | 20 | 0 | 7 | 0.1429 | 11 | 0.2727 | 43 | 0.093 | 7 | 0.1429 | 8 | 0.125 | 7 | 0 | 3 | 0 | 25 | 0.08 |
| 3_8407151 | 5 | 0 | 18 | 0.3333 | 8 | 0.125 | 9 | 0.3333 | 40 | 0.25 | 4 | 0.5 | 8 | 0.5 | 6 | 0.3333 | 4 | 0 | 22 | 0.3636 |

| | | | | | | | | | | | | | | | | | | | | |
|---|---|---|---|---|---|---|---|---|---|---|---|---|---|---|---|---|---|---|---|---|
| 3_8407167 | 5 | 1 | 17 | 1 | 7 | 1 | 9 | 0.8889 | 38 | 0.9737 | 4 | 1 | 7 | 1 | 5 | 1 | 4 | 1 | 20 | 1 |
| 3_8407198 | 5 | 1 | 18 | 0.7222 | 6 | 0.6667 | 8 | 0.75 | 37 | 0.7568 | 4 | 1 | 5 | 1 | 8 | 0.875 | 4 | 1 | 21 | 0.9524 |
| 3_8407203 | 5 | 0 | 13 | 0.0769 | 5 | 0.2 | 7 | 0.1429 | 30 | 0.1 | 4 | 0 | 6 | 0.1667 | 8 | 0.125 | 2 | 0 | 20 | 0.1 |
| 3_8407208 | 4 | 0.5 | 17 | 0.2941 | 6 | 0.8333 | 6 | 0.1667 | 33 | 0.3939 | 3 | 0.6667 | 5 | 0.8 | 7 | 0.4286 | 3 | 0 | 18 | 0.5 |
| 3_8407214 | 5 | 0.4 | 17 | 0.1176 | 6 | 0.1667 | 10 | 0.2 | 38 | 0.1842 | 4 | 0.25 | 6 | 0.1667 | 8 | 0.25 | 3 | 0 | 21 | 0.1905 |
| 3_8407216 | 5 | 1 | 14 | 0.7857 | 5 | 1 | 7 | 0.8571 | 31 | 0.871 | 3 | 1 | 5 | 0.8 | 6 | 1 | 2 | 1 | 16 | 0.9375 |
| 3_8407218 | 4 | 0.75 | 12 | 0.4167 | 6 | 0.8333 | 9 | 0.6667 | 31 | 0.6129 | 4 | 0.5 | 5 | 0 | 5 | 0.6 | 3 | 0.6667 | 17 | 0.4118 |
| 3_8407225 | 5 | 1 | 14 | 0.9286 | 3 | 1 | 6 | 1 | 28 | 0.9643 | 3 | 1 | 5 | 1 | 5 | 1 | 2 | 1 | 15 | 1 |
| 3_8407229 | 4 | 0 | 15 | 0.2667 | 5 | 0 | 8 | 0.375 | 32 | 0.2188 | 3 | 0.6667 | 5 | 0.6 | 7 | 0.4286 | 3 | 0 | 18 | 0.4444 |
| 3_8407232 | 5 | 1 | 15 | 0.9333 | 6 | 1 | 6 | 1 | 32 | 0.9688 | 3 | 1 | 5 | 1 | 4 | 1 | 3 | 1 | 15 | 1 |
| 3_8407234 | 4 | 0.5 | 16 | 0.125 | 6 | 0.3333 | 7 | 0.2857 | 33 | 0.2424 | 4 | 0.25 | 4 | 0 | 7 | 0.2857 | 2 | 0.5 | 17 | 0.2353 |
| 3_8407236 | 5 | 0.4 | 12 | 0.4167 | 5 | 0.4 | 5 | 0.4 | 27 | 0.4074 | 4 | 0.25 | 4 | 0.25 | 8 | 0.75 | 3 | 0 | 19 | 0.4211 |
| 3_8407239 | 5 | 1 | 14 | 1 | 3 | 1 | 8 | 0.75 | 30 | 0.9333 | 4 | 0.5 | 3 | 0.6667 | 6 | 0.6667 | 3 | 1 | 16 | 0.6875 |
| 3_8407244 | 5 | 0 | 15 | 0.2 | 6 | 0.1667 | 8 | 0.125 | 34 | 0.1471 | 3 | 0.3333 | 5 | 0 | 7 | 0.1429 | 3 | 0.3333 | 18 | 0.1667 |
| 3_8409805 | 3 | 0 | 9 | 0.2222 | 3 | 0.6667 | 1 | 1 | 16 | 0.3125 | 3 | 0.3333 | 3 | 1 | 5 | 0 | 2 | 1 | 13 | 0.4615 |
| 3_8409810 | 4 | 0 | 9 | 0 | 3 | 0.3333 | 3 | 0.3333 | 19 | 0.1053 | 3 | 0.6667 | 3 | 0.3333 | 6 | 0 | 2 | 0 | 14 | 0.2143 |
| 3_8409819 | 2 | 0.5 | 8 | 0.5 | 4 | 0.5 | 2 | 0 | 16 | 0.4375 | 3 | 0 | 4 | 0.75 | 4 | 0.75 | 1 | 1 | 12 | 0.5833 |
| 3_8409830 | 3 | 0.6667 | 8 | 0.625 | 4 | 0.5 | 2 | 0.5 | 17 | 0.5882 | 3 | 0.3333 | 2 | 0 | 5 | 0.6 | 2 | 0.5 | 12 | 0.4167 |
| 3_8409832 | 2 | 0.5 | 8 | 0.5 | 3 | 1 | 3 | 0.3333 | 16 | 0.5625 | 3 | 0.6667 | 2 | 1 | 5 | 0.6 | 2 | 1 | 12 | 0.75 |
| 3_8409838 | 3 | 0.6667 | 9 | 0.2222 | 5 | 0.4 | 2 | 0 | 19 | 0.3158 | 3 | 0.3333 | 3 | 0.3333 | 5 | 0.8 | 2 | 0.5 | 13 | 0.5385 |
| 3_8409840 | 2 | 0.5 | 7 | 0.1429 | 3 | 0.6667 | 3 | 0.6667 | 15 | 0.4 | 3 | 0 | 2 | 0 | 5 | 0.2 | 2 | 0.5 | 12 | 0.1667 |
| 3_8411555 | 2 | 0 | 6 | 0 | 5 | 0.2 | 1 | 0 | 14 | 0.0714 | 2 | 0 | 2 | 0 | 5 | 0.2 | 1 | 0 | 10 | 0.1 |
| 3_8416687 | 2 | 0.5 | 1 | 1 | 2 | 1 | 1 | 0 | 6 | 0.6667 | NA | NA | 1 | 0 | 2 | 0.5 | 1 | 1 | 4 | 0.5 |
| 3_8435138 | NA | NA | 1 | 1 | NA | NA | NA | NA | 1 | 1 | 1 | 1 | 1 | 1 | NA | NA | NA | NA | 2 | 1 |
| 3_8435145 | NA | NA | 1 | 1 | 1 | 0 | NA | NA | 2 | 0.5 | 1 | 1 | 1 | 1 | NA | NA | NA | NA | 2 | 1 |
| 3_8435150 | NA | NA | 1 | 1 | 1 | 1 | NA | NA | 2 | 1 | 1 | 1 | 1 | 1 | NA | NA | NA | NA | 2 | 1 |
| 3_8435172 | NA | NA | 1 | 1 | NA | NA | NA | NA | 1 | 1 | 1 | 1 | 1 | 1 | NA | NA | NA | NA | 2 | 1 |
| 3_8435196 | NA | NA | 1 | 1 | NA | NA | NA | NA | 1 | 1 | NA | NA | 1 | 1 | NA | NA | NA | NA | 1 | 1 |
| 3_8435201 | NA | NA | 1 | 1 | NA | NA | NA | NA | 1 | 1 | NA | NA | 1 | 1 | NA | NA | NA | NA | 1 | 1 |
| 3_8435238 | NA | NA | 1 | 1 | NA | NA | NA | NA | 1 | 1 | 1 | 0 | 1 | 1 | NA | NA | NA | NA | 2 | 0.5 |
| 3_8435290 | NA | NA | 1 | 1 | NA | NA | NA | NA | 1 | 1 | 1 | 1 | 1 | 1 | NA | NA | NA | NA | 2 | 1 |
| 3_8440450 | 4 | 0.25 | NA | NA | 3 | 0 | 5 | 0 | 12 | 0.0833 | 3 | 0 | 7 | 0.1429 | 3 | 0 | 1 | 0 | 14 | 0.0714 |
| 3_8440499 | 3 | 0.6667 | NA | NA | 2 | 0 | 5 | 0.4 | 10 | 0.4 | 2 | 0 | 4 | 0.5 | 2 | 0.5 | 1 | 1 | 9 | 0.4444 |
| 3_8440511 | 3 | 0.6667 | NA | NA | 3 | 0.6667 | 4 | 0 | 10 | 0.4 | 2 | 0.5 | 5 | 0.2 | 3 | 1 | 1 | 0 | 11 | 0.4545 |
| 3_8471913 | NA | NA | NA | NA | 2 | 0.5 | 2 | 0.5 | 4 | 0.5 | NA | NA | NA | NA | 2 | 1 | NA | NA | 2 | 1 |
| 3_8472009 | NA | NA | NA | NA | 1 | 0 | 2 | 0.5 | 3 | 0.3333 | NA | NA | NA | NA | 2 | 0.5 | NA | NA | 2 | 0.5 |
| 3_8472021 | NA | NA | NA | NA | 2 | 0 | 2 | 0.5 | 4 | 0.25 | NA | NA | NA | NA | 1 | 1 | NA | NA | 1 | 1 |
| 3_8472046 | NA | NA | NA | NA | 2 | 0 | 2 | 0.5 | 4 | 0.25 | NA | NA | NA | NA | 2 | 0.5 | NA | NA | 2 | 0.5 |
| 3_8472061 | NA | NA | NA | NA | 1 | 0 | 2 | 0.5 | 3 | 0.3333 | NA | NA | NA | NA | 2 | 0.5 | NA | NA | 2 | 0.5 |
| 3_8472106 | NA | NA | NA | NA | 1 | 1 | 2 | 0.5 | 3 | 0.6667 | NA | NA | NA | NA | 2 | 0.5 | NA | NA | 2 | 0.5 |
| 3_8472124 | NA | NA | NA | NA | 1 | 1 | 2 | 0.5 | 3 | 0.6667 | NA | NA | NA | NA | 1 | 1 | NA | NA | 1 | 1 |
| 3_8472231 | NA | NA | NA | NA | 1 | 1 | NA | NA | 1 | 1 | NA | NA | 1 | 1 | NA | NA | 1 | 0 | 2 | 0.5 |
| 3_8481091 | 2 | 0 | NA | NA | 2 | 0.5 | 3 | 0.6667 | 7 | 0.4286 | NA | NA | 1 | 1 | NA | NA | 2 | 0.5 | 3 | 0.6667 |
| 3_8481096 | 3 | 0.3333 | NA | NA | 1 | 1 | 3 | 0.3333 | 7 | 0.4286 | NA | NA | 1 | 1 | NA | NA | 2 | 0.5 | 3 | 0.6667 |
| 3_8481105 | 3 | 0.6667 | NA | NA | 1 | 0 | 3 | 0.6667 | 7 | 0.5714 | NA | NA | 1 | 1 | NA | NA | 2 | 0 | 3 | 0.3333 |
| 3_8481110 | 3 | 0.3333 | NA | NA | 2 | 0 | 3 | 0.3333 | 8 | 0.25 | NA | NA | 1 | 1 | NA | NA | 2 | 0.5 | 3 | 0.6667 |
| 3_8481114 | 3 | 0.3333 | NA | NA | 1 | 0 | 3 | 0.6667 | 7 | 0.4286 | NA | NA | 1 | 1 | NA | NA | 2 | 1 | 3 | 1 |
| 3_8481124 | 3 | 0.3333 | NA | NA | 2 | 0 | 4 | 0.25 | 9 | 0.2222 | NA | NA | 1 | 0 | NA | NA | 2 | 0.5 | 3 | 0.3333 |
| 3_8481129 | 3 | 0 | NA | NA | 2 | 0.5 | 1 | 0 | 6 | 0.1667 | NA | NA | 1 | 0 | 1 | 0 | 2 | 0.5 | 4 | 0.25 |
| 3_8481131 | 2 | 0.5 | NA | NA | 2 | 0.5 | 4 | 0.75 | 8 | 0.625 | NA | NA | 1 | 1 | 1 | 0 | 2 | 1 | 4 | 0.75 |
| 3_8481133 | 2 | 0 | NA | NA | 3 | 0 | 3 | 0.3333 | 8 | 0.125 | NA | NA | NA | NA | 1 | 0 | 2 | 0.5 | 3 | 0.3333 |
| 3_8485984 | 2 | 0 | 4 | 0 | 3 | 0.3333 | 4 | 0.5 | 13 | 0.2308 | 4 | 0.25 | 2 | 0 | 3 | 0 | 7 | 0 | 16 | 0.0625 |
| 3_8488466 | 5 | 0.2 | 9 | 0.4444 | 8 | 0.375 | 10 | 0.5 | 32 | 0.4062 | 7 | 0.1429 | 4 | 0.25 | 12 | 0.1667 | 18 | 0.2778 | 41 | 0.2195 |
| 3_8488467 | 5 | 1 | 9 | 0.7778 | 7 | 0.8571 | 10 | 1 | 31 | 0.9032 | 5 | 1 | 4 | 0.75 | 9 | 1 | 18 | 0.8333 | 36 | 0.8889 |
| 3_8488479 | 6 | 0.8333 | 9 | 0.6667 | 7 | 0.8571 | 11 | 0.7273 | 33 | 0.7576 | 7 | 0.8571 | 5 | 0.8 | 13 | 0.9231 | 15 | 0.7333 | 40 | 0.825 |
| 3_8488482 | 6 | 1 | 9 | 1 | 9 | 1 | 11 | 1 | 35 | 1 | 7 | 1 | 6 | 1 | 11 | 1 | 17 | 1 | 41 | 1 |
| 3_8488483 | 7 | 0.8571 | 7 | 1 | 7 | 0.8571 | 8 | 0.875 | 29 | 0.8966 | 5 | 1 | 6 | 0.8333 | 12 | 0.8333 | 16 | 0.875 | 39 | 0.8718 |
| 3_8488488 | 3 | 0 | 6 | 0.5 | 6 | 0.5 | 11 | 0.4545 | 26 | 0.4231 | 8 | 0.625 | 6 | 0.3333 | 12 | 0.6667 | 19 | 0.5789 | 45 | 0.5778 |
| 3_8488491 | 5 | 0.2 | 8 | 0 | 7 | 0.1429 | 10 | 0 | 30 | 0.0667 | 5 | 0.2 | 3 | 0 | 9 | 0 | 15 | 0.0667 | 32 | 0.0625 |
| 3_8488492 | 5 | 0 | 8 | 0.125 | 7 | 0.1429 | 8 | 0 | 28 | 0.0714 | 8 | 0.25 | 3 | 0 | 10 | 0 | 18 | 0.1667 | 39 | 0.1282 |
| 3_8488495 | 4 | 0.25 | 8 | 0.125 | 4 | 0 | 7 | 0.4286 | 23 | 0.2174 | 2 | 0 | 2 | 0.5 | 7 | 0.1429 | 11 | 0.0909 | 22 | 0.1364 |
| 3_8488502 | 7 | 0.1429 | 9 | 0.3333 | 10 | 0.3 | 11 | 0.1818 | 37 | 0.2432 | 8 | 0.25 | 7 | 0.4286 | 14 | 0.4286 | 18 | 0.1667 | 47 | 0.2979 |
| 3_8488503 | 6 | 0.5 | 7 | 0.8571 | 6 | 0.5 | 7 | 0.5714 | 26 | 0.6154 | 5 | 0.4 | 5 | 0.4 | 12 | 0.5 | 13 | 0.5385 | 35 | 0.4857 |
| 3_8488507 | 8 | 0 | 9 | 0.1111 | 7 | 0.2857 | 13 | 0.0769 | 37 | 0.1081 | 7 | 0 | 7 | 0.2857 | 13 | 0.1538 | 18 | 0.1111 | 45 | 0.1333 |
| 3_8488508 | 4 | 0 | 5 | 0 | 6 | 0.1667 | 9 | 0.1111 | 24 | 0.0833 | 5 | 0 | 5 | 0 | 12 | 0 | 14 | 0.1429 | 36 | 0.0556 |

| | | | | | | | | | | | | | | | | | | | | |
|---|---|---|---|---|---|---|---|---|---|---|---|---|---|---|---|---|---|---|---|---|
| 3_8488513 | 7 | 0.8571 | 9 | 0.4444 | 5 | 0.8 | 13 | 0.8462 | 34 | 0.7353 | 9 | 1 | 6 | 0.8333 | 12 | 0.5833 | 18 | 0.6667 | 45 | 0.7333 |
| 3_8488574 | 13 | 0.0769 | 13 | 0.2308 | 15 | 0 | 18 | 0.0556 | 59 | 0.0847 | 10 | 0.4 | 9 | 0 | 17 | 0 | 20 | 0.05 | 56 | 0.0893 |
| 3_8488578 | 10 | 0.3 | 12 | 0.5 | 12 | 0.25 | 11 | 0.6364 | 45 | 0.4222 | 7 | 0.7143 | 9 | 0.6667 | 14 | 0.5 | 17 | 0.6471 | 47 | 0.617 |
| 3_8488614 | 9 | 0.3333 | 12 | 0.3333 | 9 | 0.3333 | 15 | 0.6 | 45 | 0.4222 | 12 | 0.4167 | 9 | 0.5556 | 13 | 0.5385 | 15 | 0.6 | 49 | 0.5306 |
| 3_8488626 | 11 | 0.5455 | 15 | 0.7333 | 14 | 0.8571 | 12 | 0.5833 | 52 | 0.6923 | 11 | 0.8182 | 9 | 0.7778 | 12 | 0.5 | 14 | 1 | 46 | 0.7826 |
| 3_8556119 | 3 | 1 | 2 | 0.5 | 2 | 1 | 2 | 1 | 9 | 0.8889 | 2 | 0.5 | NA | NA | 1 | 1 | NA | NA | 3 | 0.6667 |
| 3_8556122 | 3 | 1 | 2 | 1 | 2 | 1 | 2 | 1 | 9 | 1 | 3 | 0.6667 | NA | NA | 1 | 1 | NA | NA | 4 | 0.75 |
| 3_8613256 | 1 | 0 | 1 | 0 | NA | NA | 1 | 1 | 3 | 0.3333 | NA | NA | NA | NA | NA | NA | 1 | 1 | 1 | 1 |
| 3_8625563 | 1 | 0 | 1 | 1 | NA | NA | NA | NA | 2 | 0.5 | NA | NA | 1 | 1 | NA | NA | 1 | 0 | 2 | 0.5 |
| 3_8625598 | 1 | 1 | NA | NA | NA | NA | NA | NA | 1 | 1 | NA | NA | 1 | 0 | NA | NA | 2 | 0.5 | 3 | 0.3333 |
| 3_8625853 | 1 | 0 | 1 | 0 | NA | NA | 1 | 1 | 3 | 0.3333 | NA | NA | 1 | 0 | NA | NA | 2 | 0.5 | 3 | 0.3333 |
| 3_8626712 | 1 | 0 | 1 | 1 | NA | NA | NA | NA | 2 | 0.5 | NA | NA | 1 | 0 | NA | NA | 2 | 1 | 3 | 0.6667 |
| 3_8626735 | 1 | 0 | 1 | 1 | NA | NA | 1 | 0 | 3 | 0.3333 | NA | NA | 1 | 0 | NA | NA | 3 | 0.3333 | 4 | 0.25 |
| 3_8657915 | 13 | 0.1538 | 12 | 0.3333 | 9 | 0.4444 | 8 | 0.125 | 42 | 0.2619 | 10 | 0.2 | 10 | 0.2 | 9 | 0.3333 | 12 | 0.25 | 41 | 0.2439 |
| 3_8668892 | 2 | 0 | 3 | 0 | 3 | 0.3333 | 1 | 0 | 9 | 0.1111 | 1 | 0 | 3 | 0 | 1 | 0 | 3 | 0.6667 | 8 | 0.25 |
| 3_8668901 | 2 | 0.5 | 3 | 0 | 3 | 0.3333 | 2 | 0 | 10 | 0.2 | 1 | 0 | 4 | 0 | 1 | 0 | 2 | 0.5 | 8 | 0.125 |
| 3_8669387 | 12 | 0.0833 | 9 | 0.2222 | 4 | 0 | 10 | 0.3 | 35 | 0.1714 | 12 | 0 | 12 | 0.0833 | 13 | 0.0769 | 10 | 0.1 | 47 | 0.0638 |
| 3_8669397 | 13 | 0.9231 | 7 | 0.8571 | 3 | 1 | 11 | 0.8182 | 34 | 0.8824 | 15 | 0.6667 | 12 | 0.8333 | 12 | 0.9167 | 11 | 0.4545 | 50 | 0.72 |
| 3_8669399 | 15 | 0.5333 | 8 | 0.375 | 4 | 0 | 12 | 0.4167 | 39 | 0.4103 | 14 | 0.2857 | 11 | 0.7273 | 13 | 0.5385 | 12 | 0.5833 | 50 | 0.52 |
| 3_8669401 | 14 | 0 | 8 | 0 | 5 | 0.4 | 11 | 0.0909 | 38 | 0.0789 | 14 | 0 | 14 | 0.0714 | 12 | 0.3333 | 11 | 0 | 51 | 0.098 |
| 3_8669409 | 12 | 0.9167 | 9 | 0.7778 | 5 | 0.4 | 12 | 1 | 38 | 0.8421 | 12 | 0.6667 | 14 | 0.9286 | 13 | 0.6154 | 11 | 0.5455 | 50 | 0.7 |
| 3_8669411 | 10 | 1 | 6 | 1 | 4 | 1 | 10 | 0.8 | 30 | 0.9333 | 12 | 1 | 12 | 0.8333 | 14 | 0.9286 | 11 | 1 | 49 | 0.9388 |
| 3_8669421 | 14 | 0.5 | 8 | 0.625 | 4 | 0.75 | 7 | 0.5714 | 33 | 0.5758 | 11 | 0.3636 | 9 | 0.3333 | 14 | 0.5 | 10 | 0.3 | 44 | 0.3864 |
| 3_8669425 | 13 | 0.7692 | 7 | 0.7143 | 5 | 1 | 8 | 1 | 33 | 0.8485 | 12 | 1 | 11 | 0.9091 | 16 | 0.9375 | 12 | 0.8333 | 51 | 0.9216 |
| 3_8669426 | 11 | 0 | 10 | 0.1 | 4 | 0 | 8 | 0.125 | 33 | 0.0606 | 12 | 0.1667 | 13 | 0 | 16 | 0.125 | 12 | 0 | 53 | 0.0755 |
| 3_8669437 | 8 | 0.75 | 6 | 0.8333 | 3 | 0.6667 | 4 | 1 | 21 | 0.8095 | 7 | 1 | 9 | 1 | 8 | 1 | 6 | 0.6667 | 30 | 0.9333 |
| 3_8669437 | 6 | 0.6667 | 6 | 0.3333 | 3 | 0.6667 | 4 | 0.5 | 19 | 0.5263 | 6 | 0.3333 | 5 | 0.4 | 6 | 0.3333 | 8 | 0.5 | 25 | 0.4 |
| 3_8669441 | 8 | 0 | 8 | 0.25 | 4 | 0 | 6 | 0.3333 | 26 | 0.1538 | 11 | 0.3636 | 12 | 0.4167 | 11 | 0.1818 | 6 | 0 | 40 | 0.275 |
| 3_8669452 | 10 | 0.5 | 5 | 0.4 | 4 | 0 | 6 | 0.5 | 25 | 0.4 | 9 | 0 | 9 | 0.5556 | 15 | 0.4667 | 9 | 0.4444 | 42 | 0.381 |
| 3_8681329 | 6 | 0.8333 | 2 | 0 | 3 | 0.3333 | 4 | 0.75 | 15 | 0.6 | 7 | 0.7143 | 9 | 0.6667 | 6 | 0.5 | 3 | 1 | 25 | 0.68 |
| 3_8681335 | 5 | 0.6 | 3 | 0.6667 | 4 | 0.75 | 4 | 0.25 | 16 | 0.5625 | 6 | 0.8333 | 6 | 0.3333 | 8 | 0.625 | 2 | 0 | 22 | 0.5455 |
| 3_8681337 | 5 | 0.4 | 3 | 0.3333 | 4 | 0 | 3 | 0.3333 | 15 | 0.2667 | 6 | 0.5 | 8 | 0.5 | 7 | 0.4286 | 3 | 0 | 24 | 0.4167 |
| 3_8681345 | 4 | 0 | 2 | 0.5 | 3 | 0.3333 | 4 | 0.25 | 13 | 0.2308 | 4 | 0 | 6 | 0.6667 | 5 | 0.6 | 2 | 0 | 17 | 0.4118 |
| 3_8682729 | 2 | 0.5 | 3 | 0.6667 | 2 | 0 | 3 | 0.3333 | 10 | 0.4 | 4 | 0 | 6 | 0.5 | 5 | 0.6 | 2 | 0.5 | 17 | 0.4118 |
| 3_8682743 | 4 | 0.5 | 1 | 1 | 3 | 0 | 3 | 0 | 11 | 0.2727 | 3 | 0 | 7 | 0.7143 | 5 | 0.4 | 1 | 0 | 16 | 0.4375 |
| 3_8682744 | 3 | 0.6667 | 1 | 1 | 2 | 0 | 3 | 0 | 9 | 0.3333 | 3 | 0.3333 | 7 | 0.5714 | 5 | 0 | 2 | 0.5 | 17 | 0.3529 |
| 3_8682768 | 3 | 0.3333 | 2 | 0.5 | 1 | 1 | 2 | 0 | 8 | 0.375 | 3 | 0 | 3 | 0.3333 | 1 | 1 | NA | NA | 7 | 0.2857 |
| 3_8698663 | NA | NA | NA | NA | 1 | 1 | NA | NA | 1 | 1 | NA | NA | NA | NA | 3 | 0.3333 | NA | NA | 3 | 0.3333 |
| 3_8699049 | NA | NA | NA | NA | 1 | 1 | NA | NA | 1 | 1 | NA | NA | NA | NA | 1 | 1 | NA | NA | 1 | 1 |
| 3_8699066 | NA | NA | NA | NA | 1 | 1 | NA | NA | 1 | 1 | NA | NA | NA | NA | 1 | 1 | NA | NA | 1 | 1 |
| 3_8711471 | NA | NA | NA | NA | 1 | 1 | 1 | 0 | 2 | 0.5 | NA | NA | 1 | 1 | NA | NA | NA | NA | 1 | 1 |
| 3_8711477 | NA | NA | NA | NA | 1 | 1 | 1 | 1 | 2 | 1 | NA | NA | 1 | 1 | NA | NA | NA | NA | 1 | 1 |
| 3_8715856 | 1 | 0 | 2 | 0.5 | 4 | 0.25 | 2 | 1 | 9 | 0.4444 | 3 | 1 | 6 | 0.8333 | 1 | 1 | 4 | 1 | 14 | 0.9286 |
| 3_8715995 | 1 | 0 | 2 | 0 | 3 | 0.3333 | 2 | 0 | 8 | 0.125 | 1 | 1 | 4 | 0 | 1 | 1 | 2 | 0 | 8 | 0.25 |
| 3_8716684 | NA | NA | 3 | 0 | 4 | 0.25 | 2 | 0.5 | 9 | 0.2222 | 2 | 0 | 3 | 0.3333 | 1 | 1 | 3 | 0.3333 | 9 | 0.3333 |
| 3_8744303 | NA | NA | NA | NA | NA | NA | 1 | 1 | 1 | 1 | NA | NA | 2 | 0.5 | NA | NA | 1 | 1 | 3 | 0.6667 |
| 3_8744336 | NA | NA | NA | NA | NA | NA | 1 | 1 | 1 | 1 | NA | NA | 4 | 0.25 | NA | NA | 1 | 1 | 5 | 0.4 |
| 3_8744364 | NA | NA | 1 | 1 | NA | NA | 1 | 0 | 2 | 0.5 | NA | NA | 3 | 0.3333 | NA | NA | 2 | 1 | 5 | 0.6 |
| 3_8744365 | NA | NA | 1 | 1 | NA | NA | NA | NA | 1 | 1 | 1 | 0 | 3 | 0.3333 | NA | NA | 2 | 1 | 6 | 0.5 |
| 3_8744376 | NA | NA | 1 | 0 | NA | NA | 1 | 1 | 2 | 0.5 | NA | NA | 3 | 1 | NA | NA | 2 | 1 | 5 | 1 |
| 3_8744391 | NA | NA | 1 | 1 | NA | NA | 1 | 0 | 2 | 0.5 | 1 | 1 | 4 | 0.5 | NA | NA | 2 | 0 | 7 | 0.4286 |
| 3_8744407 | NA | NA | 1 | 1 | 1 | 1 | 2 | 0.5 | 4 | 0.75 | 1 | 1 | 4 | 0.5 | NA | NA | 2 | 1 | 7 | 0.7143 |
| 3_8744413 | NA | NA | 2 | 0 | 1 | 0 | 1 | 1 | 4 | 0.25 | 1 | 0 | 2 | 0.5 | 1 | 0 | 2 | 0 | 6 | 0.1667 |
| 3_8744417 | NA | NA | 2 | 1 | 1 | 1 | 1 | 0 | 4 | 0.75 | NA | NA | 3 | 0.3333 | NA | NA | 2 | 1 | 5 | 0.6 |
| 3_8744594 | 1 | 1 | 5 | 0.2 | 1 | 0 | 2 | 0.5 | 9 | 0.3333 | 2 | 0.5 | 5 | 0 | 2 | 0 | 3 | 0.3333 | 12 | 0.1667 |
| 3_8744614 | 1 | 0 | 5 | 0 | 1 | 1 | 2 | 0 | 9 | 0.1111 | 3 | 0 | 5 | 0 | 2 | 0.5 | 4 | 0 | 14 | 0.0714 |
| 3_8744630 | 1 | 1 | 4 | 1 | 1 | 1 | 2 | 1 | 8 | 1 | 2 | 0.5 | 5 | 0.2 | 5 | 0 | 5 | 0.6 | 17 | 0.2941 |
| 3_8744645 | 1 | 0 | 6 | 0.1667 | 2 | 0 | 2 | 0 | 11 | 0.0909 | 3 | 0 | 5 | 0 | 3 | 0 | 5 | 0.2 | 16 | 0.0625 |
| 3_8744705 | 1 | 0 | 6 | 0.1667 | 2 | 0.5 | 2 | 0 | 11 | 0.1818 | 3 | 0.3333 | 6 | 0 | 5 | 0.2 | 5 | 0 | 19 | 0.1053 |
| 3_8752508 | 1 | 1 | 1 | 0 | NA | NA | 1 | 1 | 3 | 0.6667 | NA | NA | 2 | 1 | NA | NA | NA | NA | 2 | 1 |
| 3_8752512 | 2 | 0 | 1 | 1 | NA | NA | 1 | 0 | 4 | 0.25 | NA | NA | 2 | 0.5 | 1 | 1 | NA | NA | 3 | 0.6667 |
| 3_8752569 | 1 | 1 | 1 | 0 | NA | NA | 1 | 1 | 3 | 0.6667 | NA | NA | 3 | 0.3333 | 1 | 1 | NA | NA | 4 | 0.5 |
| 3_8752600 | 2 | 1 | 1 | 0 | NA | NA | 1 | 0 | 4 | 0.5 | NA | NA | 3 | 0.3333 | 1 | 1 | NA | NA | 4 | 0.5 |
| 3_8764073 | 2 | 0 | 3 | 0.3333 | 2 | 0.5 | 2 | 0.5 | 9 | 0.3333 | 1 | 0 | 4 | 0.5 | 5 | 0.8 | 1 | 0 | 11 | 0.5455 |
| 3_8764506 | 2 | 0 | 8 | 0.125 | 1 | 0 | 3 | 0 | 14 | 0.0714 | 4 | 0 | 9 | 0.1111 | 8 | 0.125 | 2 | 0 | 23 | 0.087 |

| | | | | | | | | | | | | | | | | | | | | |
|---|---|---|---|---|---|---|---|---|---|---|---|---|---|---|---|---|---|---|---|---|
| 3_87691015 | 1 | 1 | 1 | 0 | NA | NA | NA | NA | 2 | 0.5 | NA | NA | 2 | 0.5 | 3 | 1 | NA | NA | 5 | 0.8 |
| 3_87766293 | 6 | 0.1667 | 12 | 0.25 | 12 | 0 | 7 | 0.1429 | 37 | 0.1351 | 9 | 0.1111 | 7 | 0.2857 | 9 | 0.1111 | 8 | 0.25 | 33 | 0.1818 |
| 3_87766370 | 6 | 0.5 | 10 | 0.6 | 10 | 0.4 | 9 | 0.3333 | 35 | 0.4571 | 10 | 0.8 | 8 | 0.625 | 9 | 0.4444 | 7 | 0.7143 | 34 | 0.6471 |
| 3_87766436 | 6 | 0 | 12 | 0.3333 | 13 | 0.1538 | 9 | 0.2222 | 40 | 0.2 | 10 | 0.4 | 9 | 0.1111 | 8 | 0 | 8 | 0.5 | 35 | 0.2571 |
| 3_87766462 | 6 | 0.6667 | 9 | 0.4444 | 12 | 0.6667 | 8 | 0.5 | 35 | 0.5714 | 11 | 0.6364 | 8 | 0.625 | 9 | 0.5556 | 7 | 0.5714 | 35 | 0.6 |
| 3_87766477 | 6 | 0.1667 | 12 | 0 | 9 | 0.2222 | 11 | 0.2727 | 38 | 0.1579 | 9 | 0 | 8 | 0.375 | 9 | 0 | 8 | 0 | 34 | 0.0882 |
| 3_87766557 | 5 | 0.4 | 12 | 0.5 | 12 | 0.25 | 11 | 0.3636 | 40 | 0.375 | 10 | 0.1 | 8 | 0.25 | 9 | 0.2222 | 7 | 0.2857 | 34 | 0.2059 |
| 3_87777149 | 4 | 0.5 | 9 | 0.6667 | 8 | 0.875 | 8 | 0.75 | 29 | 0.7241 | 7 | 0.7143 | 5 | 1 | 4 | 0.25 | 4 | 1 | 20 | 0.75 |
| 3_87777390 | 3 | 0.6667 | 5 | 0.8 | 7 | 0.4286 | 10 | 0.6 | 25 | 0.6 | 6 | 0.6667 | 7 | 0.7143 | 5 | 0.8 | 5 | 0.6 | 23 | 0.6957 |
| 3_87777402 | 3 | 0.3333 | 5 | 0.6 | 7 | 0.2857 | 11 | 0.5455 | 26 | 0.4615 | 6 | 0.1667 | 4 | 0.5 | 4 | 0.25 | 3 | 0 | 17 | 0.2353 |
| 3_87792711 | 2 | 0 | 3 | 0.6667 | 3 | 0.3333 | 5 | 0.2 | 13 | 0.3077 | 7 | 0.1429 | 3 | 0 | 3 | 0 | 2 | 0 | 15 | 0.0667 |
| 3_87822669 | 17 | 0.1176 | 18 | 0.1111 | 17 | 0.1765 | 13 | 0.0769 | 65 | 0.1231 | 12 | 0 | 16 | 0.1875 | 17 | 0.0588 | 16 | 0.25 | 61 | 0.1311 |
| 3_87823181 | 15 | 0.1333 | 14 | 0.0714 | 21 | 0.1905 | 9 | 0.3333 | 59 | 0.1695 | 15 | 0 | 18 | 0.2222 | 16 | 0.125 | 17 | 0.1176 | 66 | 0.1212 |
| 3_87826867 | 3 | 0 | 2 | 0 | 3 | 0 | 3 | 0.3333 | 11 | 0.0909 | 3 | 0.3333 | 4 | 0 | 7 | 0 | 1 | 0 | 15 | 0.0667 |
| 3_87827062 | 8 | 0 | 3 | 0 | 3 | 0.3333 | 3 | 0 | 17 | 0.0588 | 5 | 0.2 | 4 | 0.25 | 6 | 0.3333 | NA | NA | 15 | 0.2667 |
| 3_87827177 | NA | NA | NA | NA | NA | NA | 2 | 0.5 | 2 | 0.5 | NA | NA | NA | NA | 1 | 1 | NA | NA | 1 | 1 |
| 3_87830102 | NA | NA | NA | NA | 2 | 0.5 | NA | NA | 2 | 0.5 | NA | NA | 2 | 0.5 | 3 | 1 | NA | NA | 5 | 0.8 |
| 3_87830295 | NA | NA | 1 | 1 | 3 | 1 | NA | NA | 4 | 1 | NA | NA | 2 | 1 | 3 | 1 | NA | NA | 5 | 1 |
| 3_87830355 | NA | NA | 1 | 1 | 3 | 0.6667 | NA | NA | 4 | 0.75 | NA | NA | 2 | 1 | 3 | 0.3333 | NA | NA | 5 | 0.6 |
| 3_87830558 | 15 | 0.0667 | 6 | 0.3333 | 7 | 0 | 8 | 0 | 36 | 0.0833 | 8 | 0.125 | 8 | 0 | 11 | 0.1818 | 5 | 0 | 32 | 0.0938 |
| 3_87830640 | NA | NA | 1 | 0 | 3 | 1 | NA | NA | 4 | 0.75 | NA | NA | 1 | 1 | NA | NA | 1 | 0 | 2 | 0.5 |
| 3_87830731 | 13 | 0.2308 | 7 | 0.4286 | 5 | 0.4 | 10 | 0.5 | 35 | 0.3714 | 8 | 0.375 | 6 | 0 | 17 | 0.2353 | 3 | 0 | 34 | 0.2059 |
| 3_87830749 | NA | NA | 1 | 0 | 4 | 0.25 | NA | NA | 5 | 0.2 | NA | NA | 2 | 0.5 | 1 | 0 | 1 | 0 | 4 | 0.25 |
| 3_87830864 | NA | NA | 2 | 0.5 | 4 | 0.25 | NA | NA | 6 | 0.3333 | NA | NA | 2 | 0.5 | NA | NA | NA | NA | 2 | 0.5 |
| 3_87831930 | NA | NA | 4 | 0.25 | 2 | 0.5 | NA | NA | 6 | 0.3333 | NA | NA | 1 | 0 | 2 | 0.5 | 2 | 0.5 | 5 | 0.4 |
| 3_87832268 | NA | NA | 6 | 0.5 | 5 | 0.6 | 1 | 0 | 12 | 0.5 | NA | NA | 2 | 0.5 | 2 | 0 | 3 | 0.6667 | 7 | 0.4286 |
| 3_87832404 | NA | NA | 6 | 0.5 | 5 | 0.8 | 1 | 1 | 12 | 0.6667 | NA | NA | 2 | 1 | 2 | 0.5 | 3 | 1 | 7 | 0.8571 |
| 3_87832448 | NA | NA | 4 | 0.5 | 3 | 0 | 1 | 1 | 8 | 0.375 | NA | NA | 1 | 0 | NA | NA | 3 | 0.6667 | 4 | 0.5 |
| 3_87832500 | NA | NA | 5 | 0 | 3 | 0.6667 | 1 | 0 | 9 | 0.2222 | NA | NA | 1 | 0 | 2 | 0 | 3 | 0.3333 | 6 | 0.1667 |
| 3_87838702 | 3 | 0.3333 | NA | NA | 1 | 1 | NA | NA | 4 | 0.5 | NA | NA | 1 | 0 | 1 | 1 | NA | NA | 2 | 0.5 |
| 3_87838816 | 2 | 1 | NA | NA | 1 | 1 | 1 | 1 | 4 | 1 | 1 | 0 | 1 | 1 | 2 | 1 | NA | NA | 4 | 0.75 |
| 3_87838895 | 3 | 0.3333 | NA | NA | 1 | 0 | 1 | 0 | 5 | 0.2 | NA | NA | 1 | 0 | 2 | 0.5 | NA | NA | 3 | 0.3333 |
| 3_87839050 | 3 | 0.6667 | NA | NA | 1 | 0 | 1 | 1 | 5 | 0.6 | 1 | 1 | 1 | 1 | 2 | 1 | NA | NA | 4 | 1 |
| 3_87839239 | 3 | 0.3333 | NA | NA | 1 | 0 | 1 | 0 | 5 | 0.2 | 1 | 0 | 1 | 1 | 2 | 0.5 | NA | NA | 4 | 0.5 |
| 3_87839383 | 2 | 1 | NA | NA | 1 | 1 | NA | NA | 3 | 1 | NA | NA | 1 | 1 | 1 | 1 | NA | NA | 2 | 1 |
| 3_87839405 | 3 | 0.6667 | NA | NA | 1 | 0 | 1 | 0 | 5 | 0.4 | 1 | 1 | 1 | 1 | 2 | 0.5 | NA | NA | 4 | 0.75 |
| 3_87894927 | 28 | 0.1071 | 27 | 0.1481 | 19 | 0.1579 | 30 | 0.0333 | 104 | 0.1058 | 26 | 0.1923 | 33 | 0.0606 | 22 | 0.0455 | 26 | 0.0385 | 107 | 0.0841 |
| 3_87895071 | 26 | 0.4231 | 29 | 0.4138 | 19 | 0.3158 | 25 | 0.32 | 99 | 0.3737 | 26 | 0.1538 | 30 | 0.1667 | 19 | 0.2632 | 19 | 0.3684 | 94 | 0.2234 |
| 3_87895086 | 25 | 0.36 | 26 | 0.6154 | 17 | 0.4118 | 23 | 0.5217 | 91 | 0.4835 | 24 | 0.5 | 29 | 0.2414 | 21 | 0.3333 | 24 | 0.5 | 98 | 0.3878 |
| 3_87895968 | NA | NA | 1 | 0 | NA | NA | 2 | 0.5 | 3 | 0.3333 | NA | NA | 5 | 0.2 | 4 | 0.25 | 2 | 0 | 11 | 0.1818 |
| 3_87900452 | 28 | 0.1429 | 33 | 0.0909 | 19 | 0.0526 | 21 | 0.1905 | 101 | 0.1188 | 16 | 0 | 25 | 0.08 | 14 | 0.2143 | 16 | 0.0625 | 71 | 0.0845 |
| 3_87900671 | 30 | 0.6 | 28 | 0.5714 | 23 | 0.6087 | 25 | 0.56 | 106 | 0.5849 | 19 | 0.4211 | 28 | 0.5357 | 16 | 0.5 | 15 | 0.5333 | 78 | 0.5 |
| 3_87900684 | 24 | 0.2917 | 23 | 0.2174 | 20 | 0.4 | 23 | 0.3913 | 90 | 0.3222 | 19 | 0.1579 | 31 | 0.2258 | 17 | 0.4118 | 15 | 0.3333 | 82 | 0.2683 |
| 3_87900829 | 29 | 0.4483 | 29 | 0.4138 | 23 | 0.6087 | 25 | 0.52 | 106 | 0.4906 | 15 | 0.5333 | 30 | 0.5 | 17 | 0.6471 | 17 | 0.5882 | 79 | 0.557 |
| 3_87900865 | 27 | 0.1481 | 28 | 0.1786 | 20 | 0.15 | 25 | 0.24 | 100 | 0.18 | 15 | 0 | 25 | 0.16 | 18 | 0.1111 | 15 | 0.0667 | 73 | 0.0959 |
| 3_87901588 | 27 | 0.1111 | 24 | 0.0833 | 19 | 0.0526 | 23 | 0.0435 | 93 | 0.0753 | 14 | 0.0714 | 25 | 0 | 17 | 0.0588 | 11 | 0.1818 | 67 | 0.0597 |
| 3_87902884 | 20 | 0.1 | 19 | 0.0526 | 16 | 0 | 18 | 0.0556 | 73 | 0.0548 | 9 | 0.1111 | 20 | 0.05 | 10 | 0.1 | 10 | 0 | 49 | 0.0612 |
| 3_87904520 | 19 | 0.1579 | 16 | 0.125 | 19 | 0 | 18 | 0.3333 | 72 | 0.1528 | 12 | 0.3333 | 18 | 0.2222 | 13 | 0.2308 | 6 | 0 | 49 | 0.2245 |
| 3_87908100 | NA | NA | NA | NA | NA | NA | 1 | 1 | 1 | 1 | 1 | 1 | NA | NA | NA | NA | NA | NA | 1 | 1 |
| 3_87928027 | NA | NA | 3 | 0.3333 | 3 | 0 | 1 | 0 | 7 | 0.1429 | NA | NA | 2 | 0.5 | NA | NA | 2 | 0 | 4 | 0.25 |
| 3_87928748 | NA | NA | 2 | 0 | 3 | 0.3333 | NA | NA | 5 | 0.2 | NA | NA | 1 | 1 | NA | NA | 3 | 0 | 4 | 0.25 |
| 3_87929209 | 1 | 1 | 4 | 0.25 | 2 | 0 | NA | NA | 7 | 0.2857 | 1 | 0 | 2 | 0.5 | NA | NA | 2 | 0 | 5 | 0.2 |
| 3_87929216 | 1 | 1 | 6 | 0.6667 | 1 | 0 | NA | NA | 8 | 0.625 | 1 | 0 | 2 | 1 | NA | NA | 3 | 0.6667 | 6 | 0.6667 |
| 3_87929321 | 1 | 1 | 7 | 0.4286 | 3 | 0.3333 | NA | NA | 11 | 0.4545 | 1 | 0 | 2 | 0 | NA | NA | 2 | 0.5 | 5 | 0.2 |
| 3_87929616 | 1 | 0 | 7 | 0.5714 | 3 | 0 | NA | NA | 11 | 0.3636 | 2 | 0.5 | NA | NA | NA | NA | 3 | 0.3333 | 5 | 0.4 |
| 3_87929688 | 1 | 0 | 10 | 0.1 | 3 | 0 | NA | NA | 14 | 0.0714 | 2 | 0 | 1 | 0 | 2 | 0 | 4 | 0.25 | 9 | 0.1111 |
| 3_87929731 | 1 | 0 | 9 | 0.3333 | 6 | 0.1667 | NA | NA | 16 | 0.25 | 4 | 0 | 1 | 0 | 4 | 0.25 | 4 | 0.25 | 13 | 0.1538 |
| 3_87929814 | 4 | 0 | 12 | 0.1667 | 6 | 0 | NA | NA | 22 | 0.0909 | 5 | 0 | 1 | 0 | 5 | 0.2 | 5 | 0 | 16 | 0.0625 |
| 3_87929818 | 3 | 0.3333 | 11 | 0.3636 | 5 | 0.2 | NA | NA | 19 | 0.3158 | 4 | 0.25 | 1 | 0 | 5 | 0.2 | 4 | 0.25 | 14 | 0.2143 |
| 3_87929872 | 4 | 0.25 | 12 | 0.0833 | 6 | 0 | NA | NA | 22 | 0.0909 | 5 | 0 | 2 | 0 | 5 | 0.2 | 6 | 0.1667 | 18 | 0.1111 |
| 3_87930692 | 3 | 0 | 10 | 0.2 | 5 | 0 | NA | NA | 18 | 0.1111 | 5 | 0.2 | 1 | 0 | 4 | 0 | 3 | 0.3333 | 13 | 0.1538 |
| 3_87931233 | 3 | 0 | 7 | 0.1429 | 3 | 0 | NA | NA | 13 | 0.0769 | 4 | 0 | 1 | 1 | 2 | 0 | 3 | 0 | 10 | 0.1 |
| 3_87931247 | 3 | 0.3333 | 10 | 0.6 | 5 | 0.6 | NA | NA | 18 | 0.5556 | 5 | 0.6 | NA | NA | 4 | 0.75 | 4 | 0.75 | 13 | 0.6923 |
| 3_87939843 | 2 | 0 | 6 | 0.5 | 5 | 0.4 | 2 | 0 | 15 | 0.3333 | 4 | 0.5 | 3 | 0.3333 | 2 | 0 | 4 | 0 | 13 | 0.2308 |
| 3_87939943 | 2 | 1 | 7 | 0.5714 | 4 | 1 | 2 | 0.5 | 15 | 0.7333 | 2 | 0.5 | 3 | 1 | 2 | 1 | 3 | 1 | 10 | 0.9 |

| | | | | | | | | | | | | | | | | | | | | |
|---|---|---|---|---|---|---|---|---|---|---|---|---|---|---|---|---|---|---|---|---|
| 3_87939987 | 3 | 0.3333 | 7 | 0.8571 | 5 | 0.4 | 4 | 1 | 19 | 0.6842 | 3 | 0.6667 | 2 | 0.5 | 1 | 1 | 3 | 1 | 9 | 0.7778 |
| 3_87940058 | 2 | 1 | 5 | 0.4 | 5 | 1 | 4 | 0.5 | 16 | 0.6875 | 2 | 0.5 | 1 | 0 | 4 | 0.75 | 5 | 0.4 | 12 | 0.5 |
| 3_87940160 | 1 | 0 | 6 | 0.3333 | 6 | 0 | 4 | 0.25 | 17 | 0.1765 | 3 | 0 | 3 | 0 | 2 | 0.5 | 6 | 0 | 14 | 0.0714 |
| 3_87940212 | NA | NA | 4 | 0.5 | 5 | 0 | 2 | 0 | 11 | 0.1818 | 2 | 1 | 3 | 0.3333 | 2 | 0.5 | 3 | 0.3333 | 10 | 0.5 |
| 3_87940244 | 3 | 0.6667 | 6 | 0.5 | 6 | 0.8333 | 3 | 1 | 18 | 0.7222 | 4 | 0.75 | 3 | 0.3333 | 3 | 0.6667 | 6 | 0.1667 | 16 | 0.4375 |
| 3_87940351 | 4 | 0.75 | 7 | 0.8571 | 5 | 0.6 | 5 | 0.8 | 21 | 0.7619 | 3 | 1 | 3 | 1 | 4 | 0.5 | 5 | 1 | 15 | 0.8667 |
| 3_87940375 | 3 | 1 | 7 | 0.8571 | 4 | 1 | 4 | 1 | 18 | 0.9444 | 3 | 1 | 2 | 1 | 4 | 1 | 4 | 1 | 13 | 1 |
| 3_87962795 | 3 | 0.3333 | 2 | 0.5 | NA | NA | 1 | 0 | 6 | 0.3333 | 1 | 1 | 3 | 0.3333 | 2 | 0 | NA | NA | 6 | 0.3333 |
| 3_87962897 | 2 | 0.5 | 2 | 0 | 2 | 1 | 2 | 0.5 | 8 | 0.5 | 1 | 1 | 2 | 0.5 | 2 | 1 | NA | NA | 5 | 0.8 |
| 3_87962929 | 3 | 0.3333 | 1 | 0 | 1 | 1 | 2 | 0.5 | 7 | 0.4286 | 1 | 1 | 1 | 1 | 2 | 1 | NA | NA | 4 | 1 |
| 3_87963008 | 3 | 0.6667 | 1 | 1 | 1 | 0 | 2 | 0.5 | 7 | 0.5714 | 1 | 0 | 4 | 0.5 | NA | NA | NA | NA | 5 | 0.4 |
| 3_87963152 | 4 | 0.25 | 2 | 0.5 | 2 | 0.5 | 2 | 0.5 | 10 | 0.4 | 1 | 1 | 4 | 0.5 | 2 | 0.5 | NA | NA | 7 | 0.5714 |
| 3_87963169 | 4 | 0.75 | 2 | 0 | 1 | 0 | 2 | 0 | 9 | 0.3333 | 1 | 0 | 3 | 0.3333 | 2 | 0 | NA | NA | 6 | 0.1667 |
| 3_87963289 | 4 | 0 | 1 | 1 | NA | NA | 2 | 0 | 7 | 0.1429 | NA | NA | 4 | 0.25 | 1 | 1 | NA | NA | 5 | 0.4 |
| 3_87964229 | NA | NA | NA | NA | 1 | 0 | 1 | 1 | 2 | 0.5 | NA | NA | 1 | 0 | NA | NA | 1 | 1 | 2 | 0.5 |
| 3_87964330 | 7 | 0 | 8 | 0.125 | 9 | 0 | 19 | 0.1579 | 43 | 0.093 | 8 | 0.125 | 9 | 0.1111 | 9 | 0 | 13 | 0.0769 | 39 | 0.0769 |
| 3_87965062 | 7 | 0.2857 | 8 | 0 | 9 | 0.3333 | 14 | 0.1429 | 38 | 0.1842 | 7 | 0.1429 | 9 | 0.1111 | 8 | 0.125 | 11 | 0.1818 | 35 | 0.1429 |
| 3_88050282 | NA | NA | 1 | 1 | 1 | 0 | NA | NA | 2 | 0.5 | NA | NA | 1 | 0 | NA | NA | 1 | 1 | 2 | 0.5 |
| 3_88063017 | 8 | 0 | 5 | 0.2 | 3 | 0 | 6 | 0.3333 | 22 | 0.1364 | 7 | 0 | 11 | 0.1818 | 8 | 0 | 9 | 0.1111 | 35 | 0.0857 |
| 3_88075535 | 11 | 0.0909 | 10 | 0.1 | 5 | 0 | 9 | 0 | 35 | 0.0571 | 11 | 0.1818 | 16 | 0.0625 | 16 | 0.0625 | 12 | 0 | 55 | 0.0727 |
| 3_88075563 | 9 | 0.5556 | 9 | 0.4444 | 5 | 0.4 | 12 | 0.25 | 35 | 0.4 | 10 | 0.3 | 18 | 0.3333 | 16 | 0.25 | 10 | 0.5 | 54 | 0.3333 |
| 3_88075939 | 11 | 0.0909 | 11 | 0.1818 | 3 | 0.6667 | 10 | 0.2 | 35 | 0.2 | 10 | 0.2 | 17 | 0.0588 | 16 | 0.0625 | 13 | 0.0769 | 56 | 0.0893 |
| 3_88076390 | 12 | 0.25 | 12 | 0.25 | 5 | 0.4 | 14 | 0.2143 | 43 | 0.2558 | 9 | 0.3333 | 16 | 0.1875 | 18 | 0.1111 | 16 | 0.1875 | 59 | 0.1864 |
| 3_88076667 | 14 | 0.0714 | 13 | 0 | 5 | 0.2 | 19 | 0.1053 | 51 | 0.0784 | 12 | 0 | 21 | 0.0952 | 20 | 0.1 | 18 | 0 | 71 | 0.0563 |
| 3_88079294 | NA | NA | 1 | 1 | NA | NA | NA | NA | 1 | 1 | 1 | 1 | 1 | 1 | 2 | 1 | NA | NA | 4 | 1 |
| 3_88107853 | NA | NA | 1 | 1 | 1 | 0 | 1 | 0 | 3 | 0.3333 | 1 | 1 | 2 | 0.5 | NA | NA | NA | NA | 3 | 0.6667 |
| 3_88107906 | 1 | 0 | 2 | 0 | 3 | 0 | 1 | 1 | 7 | 0.1429 | 1 | 0 | 3 | 0.3333 | 1 | 1 | NA | NA | 5 | 0.4 |
| 3_88108229 | 3 | 0.3333 | 4 | 0 | 8 | 0.25 | 10 | 0.2 | 25 | 0.2 | 7 | 0.1429 | 3 | 0 | 4 | 0.25 | 2 | 0.5 | 16 | 0.1875 |
| 3_88108360 | 3 | 0 | 3 | 0.6667 | 5 | 0.4 | 5 | 0.8 | 16 | 0.5 | 7 | 0.1429 | 1 | 0 | 3 | 0.6667 | 1 | 1 | 12 | 0.3333 |
| 3_88108426 | 5 | 0.4 | 5 | 0.4 | 10 | 0.5 | 8 | 0.25 | 28 | 0.3929 | 9 | 0.3333 | 5 | 0.4 | 3 | 0.3333 | 2 | 0 | 19 | 0.3158 |
| 3_88110559 | 6 | 0.6667 | 5 | 0 | 8 | 0.125 | 9 | 0.5556 | 28 | 0.3571 | 10 | 0.3 | 5 | 0.4 | 5 | 0.4 | 1 | 0 | 21 | 0.3333 |
| 3_88110624 | 3 | 0 | 6 | 0 | 12 | 0.1667 | 10 | 0.2 | 31 | 0.129 | 11 | 0 | 6 | 0.3333 | 4 | 0 | 2 | 0.5 | 23 | 0.1304 |
| 3_88228933 | 18 | 0.2778 | 18 | 0.2778 | 8 | 0.375 | 24 | 0.25 | 68 | 0.2794 | 13 | 0.2308 | 30 | 0.1333 | 21 | 0.0952 | 14 | 0.5 | 78 | 0.2051 |
| 3_88235671 | 7 | 0.2857 | 11 | 0.4545 | 11 | 0.3636 | 7 | 0.4286 | 36 | 0.3889 | 4 | 0.25 | 4 | 0 | 10 | 0.2 | 3 | 0 | 21 | 0.1429 |
| 3_88235710 | 6 | 0.8333 | 10 | 0.8 | 8 | 0.75 | 6 | 0.6667 | 30 | 0.7667 | 4 | 0.75 | 6 | 0.8333 | 11 | 0.7273 | 4 | 0.5 | 25 | 0.72 |
| 3_88235889 | 9 | 0.4444 | 9 | 0.7778 | 9 | 0.5556 | 6 | 0.6667 | 33 | 0.6061 | 6 | 0.5 | 8 | 0.5 | 12 | 0.5833 | 4 | 0.75 | 30 | 0.5667 |
| 3_88235920 | 6 | 0.1667 | 8 | 0.25 | 5 | 0.2 | 6 | 0.1667 | 25 | 0.2 | 6 | 0.3333 | 6 | 0 | 11 | 0.1818 | 4 | 0 | 27 | 0.1481 |
| 3_88243426 | NA | NA | 1 | 1 | 1 | 0 | NA | NA | 2 | 0.5 | NA | NA | NA | NA | NA | NA | 1 | 1 | 1 | 1 |
| 3_88258451 | 2 | 0 | 4 | 0 | 5 | 0.2 | 1 | 0 | 12 | 0.0833 | 2 | 0.5 | 5 | 0.2 | 1 | 0 | 2 | 0 | 10 | 0.2 |
| 3_88258475 | 2 | 0 | 4 | 0.25 | 4 | 0 | 1 | 0 | 11 | 0.0909 | 2 | 0 | 5 | 0 | 1 | 0 | 2 | 0.5 | 10 | 0.1 |
| 3_88258571 | 2 | 0 | 3 | 0 | 5 | 0.6 | 1 | 0 | 11 | 0.2727 | 2 | 1 | 4 | 0.25 | 1 | 0 | 2 | 0 | 9 | 0.3333 |
| 3_88258871 | 3 | 0.3333 | 4 | 0 | 4 | 0.25 | 1 | 0 | 12 | 0.1667 | 2 | 0 | 4 | 0.25 | 1 | 1 | 4 | 0 | 11 | 0.1818 |
| 3_88268513 | 7 | 0 | 5 | 0 | 9 | 0.3333 | 5 | 0 | 26 | 0.1154 | 7 | 0.1429 | 15 | 0.1333 | 6 | 0 | 7 | 0.1429 | 35 | 0.1143 |
| 3_88268749 | 7 | 0.2857 | 8 | 0.25 | 8 | 0.375 | 4 | 0.5 | 27 | 0.3333 | 7 | 0.4286 | 17 | 0.4118 | 5 | 0 | 7 | 0.2857 | 36 | 0.3333 |
| 3_88275249 | NA | NA | NA | NA | 3 | 0.6667 | NA | NA | 3 | 0.6667 | 1 | 1 | 2 | 0 | 1 | 0 | NA | NA | 4 | 0.25 |
| 3_88275598 | 1 | 0 | NA | NA | 2 | 1 | NA | NA | 3 | 0.6667 | NA | NA | 1 | 0 | 1 | 1 | NA | NA | 2 | 0.5 |
| 3_88301684 | 2 | 0.5 | 4 | 0.25 | NA | NA | 1 | 1 | 7 | 0.4286 | 3 | 0.3333 | 1 | 0 | NA | NA | NA | NA | 4 | 0.25 |
| 3_88317887 | 14 | 0.2143 | 16 | 0.25 | 16 | 0.125 | 13 | 0.2308 | 59 | 0.2034 | 8 | 0.25 | 6 | 0.5 | 10 | 0.1 | 6 | 0 | 30 | 0.2 |
| 3_88318127 | 17 | 0.1176 | 13 | 0 | 16 | 0.1875 | 14 | 0 | 60 | 0.0833 | 8 | 0.125 | 9 | 0 | 10 | 0.2 | 7 | 0.1429 | 34 | 0.1176 |
| 3_88319045 | 12 | 0.25 | 10 | 0.1 | 12 | 0.0833 | 9 | 0.2222 | 43 | 0.1628 | 4 | 0.25 | 8 | 0.125 | 5 | 0.2 | 6 | 0.1667 | 23 | 0.1739 |
| 3_88319059 | 13 | 0.1538 | 11 | 0.0909 | 11 | 0.0909 | 10 | 0.1 | 45 | 0.1111 | 5 | 0.2 | 7 | 0.1429 | 5 | 0.6 | 7 | 0 | 24 | 0.2083 |
| 3_88319239 | 12 | 0.1667 | 10 | 0.1 | 10 | 0.2 | 10 | 0.1 | 42 | 0.1429 | 4 | 0 | 7 | 0.1429 | 6 | 0.1667 | 5 | 0 | 22 | 0.0909 |
| 3_88319593 | 12 | 0.1667 | 10 | 0 | 9 | 0.1111 | 11 | 0 | 42 | 0.0714 | 4 | 0 | 6 | 0 | 7 | 0.1429 | 3 | 0 | 20 | 0.05 |
| 3_88319758 | 10 | 0 | 9 | 0.1111 | 9 | 0.1111 | 9 | 0 | 37 | 0.0541 | 4 | 0.25 | 8 | 0 | 8 | 0.125 | 4 | 0 | 24 | 0.0833 |
| 3_88319869 | 8 | 1 | 7 | 1 | 8 | 1 | 8 | 0.75 | 31 | 0.9355 | 4 | 1 | 6 | 0.3333 | 8 | 0.875 | 4 | 1 | 22 | 0.7727 |
| 3_88319876 | 11 | 0.1818 | 8 | 0.25 | 9 | 0.2222 | 8 | 0.125 | 36 | 0.1944 | 5 | 0 | 7 | 0 | 8 | 0.25 | 4 | 0.25 | 24 | 0.125 |
| 3_88320029 | 8 | 0.875 | 6 | 0.8333 | 10 | 0.9 | 7 | 1 | 31 | 0.9032 | 5 | 0.8 | 5 | 1 | 6 | 1 | 4 | 0.75 | 20 | 0.9 |
| 3_88327930 | 6 | 0.8333 | 6 | 0.8333 | 7 | 0.7143 | 9 | 0.7778 | 28 | 0.7857 | 4 | 1 | 6 | 0.5 | 5 | 0.6 | 4 | 0.75 | 19 | 0.6842 |
| 3_88327937 | 6 | 0 | 6 | 0.1667 | 7 | 0.4286 | 8 | 0 | 27 | 0.1481 | 5 | 0 | 5 | 0 | 5 | 0 | 4 | 0.25 | 19 | 0.0526 |
| 3_88327962 | 5 | 0.8 | 5 | 1 | 5 | 1 | 7 | 1 | 22 | 0.9545 | 5 | 0.4 | 5 | 0.6 | 3 | 0.6667 | 4 | 1 | 17 | 0.6471 |
| 3_88327972 | 6 | 0.1667 | 4 | 0.75 | 6 | 0.6667 | 6 | 0.6667 | 22 | 0.5455 | 4 | 0.25 | 5 | 0 | 4 | 0.5 | 3 | 0 | 16 | 0.1875 |
| 3_88328110 | 6 | 0 | 6 | 0 | 6 | 0.3333 | 5 | 0 | 23 | 0.087 | 3 | 0 | 3 | 0.3333 | 2 | 0 | 1 | 0 | 9 | 0.1111 |
| 3_88328125 | 7 | 0 | 6 | 0.1667 | 5 | 0.4 | 7 | 0 | 25 | 0.12 | 3 | 0.3333 | 4 | 0 | 3 | 0.3333 | 4 | 0 | 14 | 0.1429 |
| 3_88328264 | 6 | 0.5 | 5 | 0.4 | 6 | 0.3333 | 7 | 0.2857 | 24 | 0.375 | 3 | 0 | 6 | 0.3333 | 4 | 0 | 4 | 0.5 | 17 | 0.2353 |
| 3_88328351 | 6 | 0.1667 | 5 | 0.2 | 5 | 0.2 | 7 | 0 | 23 | 0.1304 | 3 | 0.3333 | 5 | 0.2 | 4 | 0 | 3 | 0 | 15 | 0.1333 |

| | | | | | | | | | | | | | | | | | | | | |
|---|---|---|---|---|---|---|---|---|---|---|---|---|---|---|---|---|---|---|---|---|
| 3_8833206 | 6 | 0.5 | 3 | 0.3333 | 1 | 1 | 4 | 0.25 | 14 | 0.4286 | NA | NA | 1 | 0 | 2 | 0.5 | 4 | 0.25 | 7 | 0.2857 |
| 3_8833210 | 6 | 0.5 | 3 | 0.6667 | 2 | 0.5 | 5 | 0.4 | 16 | 0.5 | 1 | 1 | 3 | 0.3333 | 2 | 1 | 3 | 0.6667 | 9 | 0.6667 |
| 3_8833236 | 6 | 0.3333 | 5 | 0 | 4 | 0.25 | 4 | 0 | 19 | 0.1579 | 2 | 0 | 1 | 0 | 2 | 0 | 3 | 0.3333 | 8 | 0.125 |
| 3_8834330 | 4 | 0.25 | 7 | 0 | 2 | 0 | 5 | 0 | 18 | 0.0556 | 1 | 0 | 3 | 0 | 3 | 0.3333 | 3 | 0 | 10 | 0.1 |
| 3_8834344 | 6 | 0.6667 | 4 | 1 | 3 | 0.6667 | 5 | 0.2 | 18 | 0.6111 | 1 | 1 | 4 | 0.5 | 2 | 0.5 | 4 | 0.5 | 11 | 0.5455 |
| 3_8834362 | 6 | 0.1667 | 5 | 0 | 2 | 0 | 4 | 0 | 17 | 0.0588 | NA | NA | 3 | 0 | 3 | 0 | 3 | 0.3333 | 9 | 0.1111 |
| 3_8834366 | 6 | 0.1667 | 5 | 0 | 2 | 0 | 4 | 0 | 17 | 0.0588 | 1 | 0 | 4 | 0.25 | 3 | 0 | 4 | 0 | 12 | 0.0833 |
| 3_8834373 | 5 | 0 | 6 | 0.3333 | 3 | 0 | 3 | 0 | 17 | 0.1176 | 1 | 0 | 3 | 0 | 3 | 0.3333 | 2 | 0.5 | 9 | 0.2222 |
| 3_8834382 | 6 | 1 | 4 | 0.75 | 2 | 0.5 | 3 | 1 | 15 | 0.8667 | 1 | 1 | 3 | 0.3333 | 3 | 1 | 3 | 0.6667 | 10 | 0.7 |
| 3_8834396 | 5 | 0.6 | 4 | 1 | 2 | 0 | 3 | 0.6667 | 14 | 0.6429 | 1 | 0 | 2 | 0.5 | 3 | 0.3333 | 3 | 0.6667 | 9 | 0.4444 |
| 3_8834402 | 5 | 0.4 | 6 | 0.1667 | 2 | 0 | 4 | 0 | 17 | 0.1765 | 1 | 1 | 2 | 0 | 3 | 0.6667 | 4 | 0.25 | 10 | 0.4 |
| 3_8834403 | 5 | 0.6 | 5 | 0.4 | 2 | 0 | 3 | 0 | 15 | 0.3333 | 1 | 0 | 2 | 0.5 | 3 | 0 | 4 | 0.5 | 10 | 0.3 |
| 3_8834430 | 5 | 0 | 3 | 0 | 1 | 0 | 3 | 0.3333 | 12 | 0.0833 | NA | NA | 2 | 0 | 2 | 0 | 2 | 0.5 | 6 | 0.1667 |
| 3_8845963 | NA | NA | 1 | 1 | 4 | 0.25 | 1 | 0 | 6 | 0.3333 | NA | NA | 2 | 0.5 | NA | NA | 1 | 0 | 3 | 0.3333 |
| 3_8845966 | 9 | 0.3333 | 6 | 0.6667 | 12 | 0.6667 | 5 | 0.4 | 32 | 0.5312 | 6 | 0.5 | 8 | 0.5 | 9 | 0.2222 | 7 | 0.8571 | 30 | 0.5 |
| 3_8845977 | 8 | 0 | 6 | 0.1667 | 10 | 0.2 | 6 | 0 | 30 | 0.1 | 7 | 0.2857 | 9 | 0.2222 | 9 | 0 | 6 | 0 | 31 | 0.129 |
| 3_8845987 | NA | NA | 1 | 0 | 1 | 1 | 2 | 0.5 | 4 | 0.5 | NA | NA | 1 | 0 | NA | NA | 1 | 1 | 2 | 0.5 |
| 3_8845994 | NA | NA | 1 | 1 | 1 | 1 | 2 | 1 | 4 | 1 | NA | NA | 1 | 1 | NA | NA | 1 | 1 | 2 | 1 |
| 3_8846398 | 5 | 0.2 | 3 | 0.3333 | 4 | 0 | 2 | 0 | 14 | 0.1429 | 4 | 0 | 3 | 0 | 6 | 0.1667 | 5 | 0 | 18 | 0.0556 |
| 3_8847549 | 7 | 0.5714 | 10 | 0.3 | 6 | 0.3333 | 9 | 0.1111 | 32 | 0.3125 | 13 | 0.3846 | 4 | 0 | 11 | 0.3636 | 5 | 0.6 | 33 | 0.3636 |
| 3_8847551 | 7 | 0.8571 | 9 | 0.8889 | 8 | 0.875 | 9 | 0.8889 | 33 | 0.8788 | 14 | 0.7143 | 5 | 1 | 11 | 0.8182 | 5 | 1 | 35 | 0.8286 |
| 3_8847564 | 6 | 0 | 7 | 0 | 7 | 0.5714 | 7 | 0.4286 | 27 | 0.2593 | 11 | 0.2727 | 4 | 0.25 | 11 | 0.1818 | 4 | 0 | 30 | 0.2 |
| 3_8847566 | 8 | 0.125 | 11 | 0 | 10 | 0 | 9 | 0.1111 | 38 | 0.0526 | 15 | 0 | 6 | 0 | 12 | 0.1667 | 6 | 0 | 39 | 0.0513 |
| 3_8847584 | 9 | 0.6667 | 11 | 0.5455 | 7 | 0.5714 | 8 | 0.75 | 35 | 0.6286 | 17 | 0.7059 | 6 | 1 | 11 | 0.8182 | 6 | 0.8333 | 40 | 0.8 |
| 3_8847599 | 8 | 0.625 | 10 | 0.2 | 10 | 0.4 | 11 | 0.3636 | 39 | 0.3846 | 17 | 0.3529 | 8 | 0.5 | 10 | 0.5 | 7 | 0.4286 | 42 | 0.4286 |
| 3_8847611 | 10 | 0.5 | 13 | 0.3846 | 6 | 0.3333 | 6 | 0.5 | 35 | 0.4286 | 16 | 0.6875 | 7 | 0.7143 | 9 | 0.6667 | 5 | 0.6 | 37 | 0.6757 |
| 3_8847615 | 9 | 0.5556 | 12 | 0.25 | 12 | 0.4167 | 10 | 0.4 | 43 | 0.3953 | 15 | 0.1333 | 8 | 0.375 | 12 | 0.5833 | 6 | 0.5 | 41 | 0.3659 |
| 3_8847621 | 11 | 0.3636 | 14 | 0.2143 | 13 | 0.2308 | 12 | 0.1667 | 50 | 0.24 | 18 | 0.0556 | 9 | 0.1111 | 12 | 0.25 | 9 | 0.1111 | 48 | 0.125 |
| 3_8847627 | 11 | 0.9091 | 10 | 1 | 9 | 1 | 9 | 1 | 39 | 0.9744 | 20 | 1 | 9 | 1 | 13 | 0.9231 | 8 | 0.875 | 50 | 0.96 |
| 3_8847628 | 9 | 1 | 12 | 0.5 | 10 | 0.9 | 11 | 0.8182 | 42 | 0.7857 | 10 | 0.8 | 7 | 0.8571 | 8 | 0.625 | 8 | 0.875 | 33 | 0.7879 |
| 3_8847631 | 11 | 0.3636 | 15 | 0.3333 | 11 | 0.0909 | 11 | 0.0909 | 48 | 0.2292 | 20 | 0.05 | 10 | 0.5 | 11 | 0.3636 | 7 | 0.2857 | 48 | 0.25 |
| 3_8847635 | 11 | 0.0909 | 12 | 0 | 13 | 0.3077 | 8 | 0.125 | 44 | 0.1364 | 18 | 0.2778 | 8 | 0.125 | 12 | 0.3333 | 8 | 0.125 | 46 | 0.2391 |
| 3_8847638 | 11 | 0.5455 | 14 | 0.1429 | 13 | 0.2308 | 10 | 0.3 | 48 | 0.2917 | 18 | 0.3889 | 10 | 0.3 | 8 | 0.125 | 8 | 0.25 | 44 | 0.2955 |
| 3_8847680 | 12 | 0.0833 | 15 | 0.2 | 15 | 0.0667 | 12 | 0.0833 | 54 | 0.1111 | 18 | 0.1667 | 7 | 0.1429 | 17 | 0.0588 | 10 | 0.2 | 52 | 0.1346 |
| 3_8849541 | 15 | 0.0667 | 7 | 0.1429 | 9 | 0 | 11 | 0.0909 | 42 | 0.0714 | 5 | 0 | 11 | 0.0909 | 5 | 0.2 | 8 | 0.125 | 29 | 0.1034 |
| 3_8849556 | 9 | 0.1111 | 5 | 0 | 8 | 0.125 | 9 | 0 | 31 | 0.0645 | 4 | 0 | 5 | 0 | 6 | 0.1667 | 9 | 0.2222 | 24 | 0.125 |
| 3_8855421 | 7 | 0.1429 | 12 | 0.1667 | 7 | 0.1429 | 5 | 0.2 | 31 | 0.1613 | 7 | 0.1429 | 10 | 0.3 | 9 | 0.1111 | 5 | 0 | 31 | 0.1613 |
| 3_8855424 | 5 | 1 | 10 | 0.8 | 7 | 0.7143 | 3 | 1 | 25 | 0.84 | 6 | 1 | 9 | 0.7778 | 6 | 1 | 5 | 1 | 26 | 0.9231 |
| 3_8855429 | 7 | 0.4286 | 12 | 0.4167 | 10 | 0.6 | 7 | 0.2857 | 36 | 0.4444 | 9 | 0.6667 | 10 | 0.3 | 8 | 0.125 | 2 | 0.5 | 29 | 0.3793 |
| 3_8855430 | 6 | 0.1667 | 13 | 0.0769 | 8 | 0.375 | 8 | 0.25 | 35 | 0.2 | 6 | 0.3333 | 13 | 0.0769 | 7 | 0 | 4 | 0 | 30 | 0.1 |
| 3_8855523 | 11 | 0.6364 | 19 | 0.3684 | 17 | 0.2941 | 10 | 0.4 | 57 | 0.4035 | 15 | 0.4 | 15 | 0.3333 | 10 | 0.2 | 7 | 0.2857 | 47 | 0.3191 |
| 3_8861914 | 3 | 0.6667 | 1 | 1 | 4 | 1 | 4 | 0.75 | 12 | 0.8333 | 4 | 0.5 | 2 | 0.5 | 8 | 0.5 | 1 | 1 | 15 | 0.5333 |
| 3_8861916 | 5 | 0.8 | 4 | 0.75 | 4 | 1 | 3 | 0.3333 | 16 | 0.75 | 5 | 0.8 | 3 | 0.3333 | 8 | 0.875 | 2 | 1 | 18 | 0.7778 |
| 3_8861920 | 4 | 0.75 | 4 | 1 | 4 | 1 | 2 | 1 | 14 | 0.9286 | 4 | 0.5 | 3 | 0.3333 | 8 | 0.75 | 2 | 0.5 | 17 | 0.5882 |
| 3_8861921 | 5 | 0.4 | 4 | 0.25 | 4 | 0.25 | 4 | 0 | 17 | 0.2353 | 5 | 0.2 | 3 | 0.3333 | 9 | 0.3333 | 2 | 0 | 19 | 0.2632 |
| 3_8861940 | 5 | 0.4 | 3 | 0.3333 | 4 | 0.25 | 4 | 0 | 16 | 0.25 | 5 | 0 | 4 | 0 | 8 | 0.125 | 3 | 0 | 20 | 0.05 |
| 3_8861986 | 4 | 0.5 | 4 | 0.5 | 4 | 0.5 | 4 | 0.25 | 16 | 0.4375 | 6 | 0.1667 | 4 | 0.25 | 6 | 0 | 4 | 0.25 | 20 | 0.15 |
| 3_8861991 | 6 | 0.1667 | 5 | 0.2 | 5 | 0.4 | 5 | 0.4 | 21 | 0.2857 | 4 | 0 | 5 | 0.4 | 10 | 0.3 | 5 | 0 | 24 | 0.2083 |
| 3_8861991 | 6 | 0.3333 | 5 | 0.4 | 6 | 0.3333 | 5 | 0.2 | 22 | 0.3182 | 5 | 0.4 | 5 | 0 | 11 | 0.6364 | 5 | 0.2 | 26 | 0.3846 |
| 3_8861997 | 5 | 1 | 2 | 1 | 6 | 0.8333 | 5 | 1 | 18 | 0.9444 | 5 | 0.8 | 4 | 0.75 | 9 | 0.8889 | 4 | 1 | 22 | 0.8636 |
| 3_8863663 | 4 | 0.5 | 6 | 0.3333 | 4 | 0.75 | 2 | 0 | 16 | 0.4375 | 2 | 0.5 | 7 | 0.4286 | 7 | 0.2857 | 6 | 0.6667 | 22 | 0.4545 |
| 3_8863667 | 3 | 0.3333 | 5 | 0.8 | 4 | 0.75 | 3 | 0.6667 | 15 | 0.6667 | 2 | 1 | 9 | 0.7778 | 6 | 1 | 5 | 1 | 22 | 0.9091 |
| 3_8863670 | 3 | 1 | 4 | 1 | 4 | 1 | 3 | 1 | 14 | 1 | 1 | 1 | 8 | 0.875 | 7 | 1 | 6 | 0.8333 | 22 | 0.9091 |
| 3_8863679 | 4 | 0.75 | 8 | 1 | 4 | 1 | 2 | 1 | 18 | 0.9444 | 3 | 1 | 8 | 1 | 8 | 0.75 | 6 | 1 | 25 | 0.92 |
| 3_8863682 | 4 | 0.75 | 6 | 1 | 4 | 1 | 3 | 1 | 17 | 0.9412 | 3 | 1 | 7 | 0.8571 | 8 | 1 | 5 | 0.8 | 23 | 0.913 |
| 3_8863685 | 4 | 0 | 7 | 0.4286 | 4 | 0.5 | 4 | 0 | 19 | 0.2632 | 3 | 0 | 9 | 0.4444 | 7 | 0.4286 | 6 | 0.5 | 25 | 0.4 |
| 3_8863686 | 3 | 1 | 5 | 0.8 | 2 | 1 | 2 | 1 | 12 | 0.9167 | 3 | 1 | 5 | 1 | 2 | 1 | 3 | 1 | 13 | 1 |
| 3_8863707 | NA | NA | 2 | 0.5 | 1 | 0 | 1 | 0 | 4 | 0.25 | NA | NA | 2 | 0.5 | 2 | 0 | NA | NA | 4 | 0.25 |
| 3_8863718 | NA | NA | 2 | 0 | NA | NA | 1 | 1 | 3 | 0.3333 | NA | NA | 2 | 0.5 | 2 | 1 | NA | NA | 4 | 0.75 |
| 3_8863733 | NA | NA | 2 | 0 | 1 | 0 | 1 | 1 | 4 | 0.25 | NA | NA | 3 | 0 | 2 | 0.5 | NA | NA | 5 | 0.2 |
| 3_8863817 | NA | NA | 2 | 0 | 1 | 0 | 2 | 0.5 | 5 | 0.2 | 1 | 0 | NA | NA | 2 | 0.5 | NA | NA | 3 | 0.3333 |
| 3_8863821 | NA | NA | 2 | 0 | 1 | 0 | 2 | 0.5 | 5 | 0.2 | 1 | 1 | NA | NA | 3 | 0.3333 | NA | NA | 4 | 0.5 |
| 3_8865212 | 48 | 0.1458 | 52 | 0.1923 | 63 | 0.1429 | 48 | 0.1667 | 211 | 0.1611 | 48 | 0.1042 | 58 | 0.1034 | 55 | 0.0909 | 43 | 0.1628 | 204 | 0.1127 |
| 3_8865236 | 51 | 0.1961 | 58 | 0.1724 | 73 | 0.1918 | 44 | 0.1364 | 226 | 0.177 | 55 | 0.2909 | 55 | 0.2 | 55 | 0.2182 | 48 | 0.1667 | 213 | 0.2207 |
| 3_8865405 | 34 | 0.1176 | 32 | 0.125 | 47 | 0.0638 | 32 | 0.1562 | 145 | 0.1103 | 38 | 0.0789 | 31 | 0.1613 | 31 | 0.1613 | 34 | 0.1765 | 134 | 0.1418 |

| | | | | | | | | | | | | | | | | | | | | |
|---|---|---|---|---|---|---|---|---|---|---|---|---|---|---|---|---|---|---|---|---|
| 3_8865500 | 24 | 0.2083 | 21 | 0.2381 | 24 | 0.1667 | 27 | 0.0741 | 96 | 0.1667 | 27 | 0.2222 | 18 | 0.1667 | 20 | 0.1 | 24 | 0.1667 | 89 | 0.1685 |
| 3_8865515 | 24 | 0.2917 | 18 | 0.2778 | 29 | 0.1724 | 22 | 0.4545 | 93 | 0.2903 | 24 | 0.375 | 16 | 0.4375 | 18 | 0.3333 | 25 | 0.32 | 83 | 0.3614 |
| 3_8866917 | 27 | 0.2593 | 23 | 0.087 | 26 | 0.2308 | 23 | 0.2609 | 99 | 0.2121 | 24 | 0.2083 | 19 | 0.1579 | 19 | 0.1579 | 18 | 0.2222 | 80 | 0.1875 |
| 3_8866939 | 21 | 0.381 | 14 | 0.7143 | 22 | 0.6818 | 18 | 0.6111 | 75 | 0.5867 | 21 | 0.5714 | 15 | 0.6 | 14 | 0.5714 | 17 | 0.5882 | 67 | 0.5821 |
| 3_8866950 | 26 | 0.8077 | 13 | 0.7692 | 21 | 0.7143 | 20 | 0.9 | 80 | 0.8 | 21 | 0.9048 | 17 | 0.6471 | 17 | 0.7647 | 17 | 0.8235 | 72 | 0.7917 |
| 3_8866951 | 20 | 1 | 16 | 0.8125 | 15 | 0.8 | 21 | 1 | 72 | 0.9167 | 23 | 0.8696 | 19 | 1 | 15 | 1 | 17 | 1 | 74 | 0.9595 |
| 3_8866953 | 25 | 0.88 | 14 | 0.8571 | 21 | 1 | 18 | 1 | 78 | 0.9359 | 24 | 0.9167 | 18 | 0.8333 | 16 | 0.8125 | 16 | 0.9375 | 74 | 0.8784 |
| 3_8866961 | 24 | 0.0417 | 14 | 0 | 18 | 0.1111 | 19 | 0.0526 | 75 | 0.0533 | 23 | 0.0435 | 18 | 0.1667 | 17 | 0.1176 | 15 | 0.1333 | 73 | 0.1096 |
| 3_8866968 | 26 | 0.1538 | 15 | 0.3333 | 19 | 0.2105 | 20 | 0.25 | 80 | 0.225 | 24 | 0.3333 | 17 | 0.2941 | 17 | 0.2941 | 16 | 0.1875 | 74 | 0.2838 |
| 3_8866979 | 20 | 0.55 | 12 | 0.3333 | 13 | 0.6154 | 16 | 0.625 | 61 | 0.541 | 23 | 0.6957 | 13 | 0.5385 | 13 | 0.6154 | 12 | 0.5833 | 61 | 0.623 |
| 3_8868447 | NA | NA | 1 | 1 | NA | NA | 1 | 1 | 2 | 1 | 1 | 1 | NA | NA | 2 | 0.5 | NA | NA | 3 | 0.6667 |
| 3_8868457 | NA | NA | 1 | 0 | NA | NA | 1 | 1 | 2 | 0.5 | NA | NA | NA | NA | 2 | 1 | NA | NA | 2 | 1 |
| 3_8870829 | NA | NA | 1 | 1 | NA | NA | NA | NA | 1 | 1 | NA | NA | NA | NA | 1 | 1 | NA | NA | 1 | 1 |
| 3_8871276 | 2 | 0.5 | 2 | 0.5 | 2 | 0 | 1 | 0 | 7 | 0.2857 | 1 | 1 | 1 | 0 | 2 | 0.5 | 2 | 0.5 | 6 | 0.5 |
| 3_8871449 | 3 | 0.3333 | 4 | 0.5 | 2 | 0.5 | 2 | 0.5 | 11 | 0.4545 | 2 | 0.5 | 3 | 0 | 3 | 0 | 3 | 0.3333 | 11 | 0.1818 |
| 3_8871477 | 2 | 0 | 3 | 0.3333 | 2 | 0 | 2 | 0 | 9 | 0.1111 | 2 | 0.5 | 3 | 0 | 4 | 0.5 | 3 | 0 | 12 | 0.25 |
| 3_8871483 | 2 | 0.5 | 3 | 0 | 2 | 0.5 | 2 | 0.5 | 9 | 0.3333 | 2 | 0.5 | 3 | 0.3333 | 4 | 0.5 | 3 | 0.6667 | 12 | 0.5 |
| 3_8871507 | 4 | 0.25 | 3 | 0 | 6 | 0 | 3 | 0 | 16 | 0.0625 | 4 | 0.25 | 3 | 0 | 4 | 0.25 | 6 | 0 | 17 | 0.1176 |
| 3_8871513 | 3 | 0.3333 | 4 | 0.25 | 6 | 0.3333 | 2 | 0 | 15 | 0.2667 | 4 | 0.75 | 1 | 0 | 4 | 0.25 | 6 | 0.5 | 15 | 0.4667 |
| 3_8871519 | 3 | 0 | 3 | 0.3333 | 5 | 0 | 2 | 0 | 13 | 0.0769 | 4 | 0.25 | 3 | 0.6667 | 4 | 0.5 | 4 | 0 | 15 | 0.3333 |
| 3_8871530 | 3 | 0.3333 | 4 | 0.25 | 7 | 0.1429 | 3 | 0 | 17 | 0.1765 | 4 | 0.25 | 2 | 0 | 5 | 0.2 | 5 | 0.4 | 16 | 0.25 |
| 3_8871543 | 4 | 0.25 | 3 | 0.3333 | 5 | 0 | 2 | 0.5 | 14 | 0.2143 | 4 | 0 | 3 | 0 | 5 | 0.2 | 5 | 0.2 | 17 | 0.1176 |
| 3_8871549 | 3 | 0.3333 | 5 | 0.8 | 3 | 0.6667 | 3 | 0.3333 | 14 | 0.5714 | 3 | 0.3333 | 3 | 0.6667 | 6 | 0.6667 | 5 | 0.6 | 17 | 0.5882 |
| 3_8871550 | 4 | 0 | 3 | 0.3333 | 4 | 0.25 | 3 | 0.6667 | 14 | 0.2857 | 3 | 0.3333 | 1 | 0 | 4 | 0.75 | 3 | 0 | 11 | 0.3636 |
| 3_8871570 | NA | NA | 1 | 1 | 1 | 0 | NA | NA | 2 | 0.5 | NA | NA | NA | NA | 1 | 0 | 1 | 1 | 2 | 0.5 |
| 3_8871796 | NA | NA | NA | NA | NA | NA | 3 | 0.6667 | 3 | 0.6667 | NA | NA | NA | NA | NA | NA | 1 | 1 | 1 | 1 |
| 3_8871803 | NA | NA | NA | NA | NA | NA | 2 | 0.5 | 2 | 0.5 | NA | NA | NA | NA | NA | NA | 1 | 1 | 1 | 1 |
| 3_8871816 | NA | NA | NA | NA | NA | NA | 3 | 0.3333 | 3 | 0.3333 | NA | NA | NA | NA | NA | NA | 1 | 1 | 1 | 1 |
| 3_8871819 | NA | NA | NA | NA | NA | NA | 3 | 1 | 3 | 1 | NA | NA | NA | NA | NA | NA | 1 | 1 | 1 | 1 |
| 3_8872330 | NA | NA | 1 | 1 | NA | NA | NA | NA | 1 | 1 | 2 | 1 | NA | NA | 1 | 0 | NA | NA | 3 | 0.6667 |
| 3_8872332 | NA | NA | 1 | 1 | NA | NA | NA | NA | 1 | 1 | 2 | 1 | NA | NA | 1 | 0 | NA | NA | 3 | 0.6667 |
| 3_8872358 | NA | NA | 1 | 1 | NA | NA | NA | NA | 1 | 1 | 1 | 1 | NA | NA | NA | NA | NA | NA | 1 | 1 |
| 3_8872364 | NA | NA | 1 | 1 | NA | NA | NA | NA | 1 | 1 | 1 | 1 | NA | NA | NA | NA | NA | NA | 1 | 1 |
| 3_8872407 | NA | NA | 1 | 1 | NA | NA | NA | NA | 1 | 1 | 1 | 1 | NA | NA | NA | NA | NA | NA | 1 | 1 |
| 3_8872462 | NA | NA | NA | NA | NA | NA | 2 | 0.5 | 2 | 0.5 | NA | NA | NA | NA | NA | NA | 1 | 1 | 1 | 1 |
| 3_8872785 | NA | NA | NA | NA | NA | NA | 3 | 0.6667 | 3 | 0.6667 | NA | NA | NA | NA | NA | NA | 2 | 0.5 | 2 | 0.5 |
| 3_8872857 | 2 | 1 | NA | NA | NA | NA | 3 | 1 | 5 | 1 | 1 | 1 | 1 | 1 | NA | NA | 2 | 0.5 | 4 | 0.75 |
| 3_8872866 | 3 | 0.6667 | NA | NA | NA | NA | 3 | 0.6667 | 6 | 0.6667 | 1 | 0 | NA | NA | NA | NA | 1 | 1 | 2 | 0.5 |
| 3_8872870 | 2 | 0 | NA | NA | NA | NA | 3 | 0.3333 | 5 | 0.2 | 1 | 0 | 1 | 0 | NA | NA | 2 | 0.5 | 4 | 0.25 |
| 3_8872886 | 3 | 0.6667 | NA | NA | NA | NA | 3 | 1 | 6 | 0.8333 | NA | NA | 1 | 1 | NA | NA | 2 | 1 | 3 | 1 |
| 3_8872893 | 3 | 0 | NA | NA | 1 | 1 | 3 | 0.6667 | 7 | 0.4286 | NA | NA | 1 | 1 | NA | NA | 2 | 0.5 | 3 | 0.6667 |
| 3_8872902 | 3 | 1 | NA | NA | 1 | 0 | 3 | 0.6667 | 7 | 0.7143 | 1 | 1 | 1 | 1 | NA | NA | 2 | 0 | 4 | 0.5 |
| 3_8872935 | NA | NA | 1 | 1 | NA | NA | 1 | 0 | 2 | 0.5 | NA | NA | 1 | 0 | 1 | 0 | 1 | 1 | 3 | 0.3333 |
| 3_8873000 | NA | NA | NA | NA | NA | NA | 1 | 1 | 1 | 1 | NA | NA | 1 | 1 | 1 | 0 | 1 | 0 | 3 | 0.3333 |
| 3_8873089 | 1 | 0 | 3 | 0.6667 | 1 | 1 | 2 | 1 | 7 | 0.7143 | NA | NA | 2 | 0.5 | NA | NA | 2 | 0 | 4 | 0.25 |
| 3_8873090 | 1 | 0 | 3 | 1 | 1 | 1 | NA | NA | 5 | 0.8 | NA | NA | 1 | 1 | NA | NA | 1 | 1 | 2 | 1 |
| 3_8873151 | NA | NA | 1 | 1 | 1 | 1 | NA | NA | 2 | 1 | NA | NA | 2 | 0.5 | NA | NA | 2 | 0.5 | 4 | 0.5 |
| 3_8873161 | 1 | 1 | 2 | 1 | 1 | 1 | NA | NA | 4 | 1 | NA | NA | 2 | 1 | NA | NA | 1 | 1 | 3 | 1 |
| 3_8873169 | 1 | 0 | 3 | 0.6667 | 1 | 0 | 1 | 1 | 6 | 0.5 | NA | NA | 2 | 0.5 | NA | NA | 2 | 1 | 4 | 0.75 |
| 3_8873186 | 1 | 1 | 2 | 0 | 1 | 1 | NA | NA | 4 | 0.5 | NA | NA | 1 | 1 | NA | NA | 1 | 1 | 2 | 1 |
| 3_8873202 | 1 | 1 | 3 | 1 | 1 | 1 | 1 | 1 | 6 | 1 | NA | NA | 1 | 1 | NA | NA | 1 | 1 | 2 | 1 |
| 3_8873248 | 1 | 1 | 1 | 1 | 1 | 1 | 1 | 1 | 4 | 1 | 1 | 1 | 3 | 0.6667 | 1 | 1 | 2 | 1 | 7 | 0.8571 |
| 3_8873250 | 1 | 1 | 1 | 1 | 1 | 1 | 1 | 1 | 4 | 1 | 1 | 0 | 2 | 0 | 2 | 0.5 | 2 | 0 | 7 | 0.1429 |
| 3_8873282 | 3 | 0.3333 | NA | NA | 1 | 0 | 3 | 0 | 7 | 0.1429 | 1 | 0 | 1 | 0 | 2 | 0.5 | 1 | 0 | 5 | 0.2 |
| 3_8873284 | 2 | 1 | NA | NA | 1 | 0 | 3 | 1 | 6 | 0.8333 | 1 | 1 | 1 | 1 | 1 | 0 | 1 | 1 | 4 | 0.75 |
| 3_8873289 | 1 | 1 | NA | NA | 1 | 1 | 3 | 0.3333 | 5 | 0.6 | 1 | 0 | 1 | 0 | 2 | 0.5 | 1 | 0 | 5 | 0.2 |
| 3_8873291 | 2 | 0.5 | NA | NA | 1 | 1 | 2 | 1 | 5 | 0.8 | NA | NA | 1 | 1 | 1 | 1 | 1 | 1 | 3 | 1 |
| 3_8873309 | 3 | 0.6667 | NA | NA | 1 | 1 | 3 | 0.3333 | 7 | 0.5714 | 1 | 1 | 1 | 0 | 2 | 0.5 | 1 | 0 | 5 | 0.4 |
| 3_8880334 | 2 | 1 | 2 | 0.5 | 2 | 0 | NA | NA | 6 | 0.5 | NA | NA | 1 | 1 | 1 | 0 | NA | NA | 2 | 0.5 |
| 3_8880343 | 2 | 0.5 | 2 | 0.5 | 3 | 0.3333 | NA | NA | 7 | 0.4286 | NA | NA | 1 | 1 | NA | NA | NA | NA | 1 | 1 |
| 3_8880363 | 1 | 1 | 3 | 0.3333 | 3 | 0.6667 | NA | NA | 7 | 0.5714 | NA | NA | 1 | 1 | 1 | 1 | NA | NA | 2 | 1 |
| 3_8880370 | 2 | 0.5 | 4 | 0.5 | 3 | 0.6667 | NA | NA | 9 | 0.5556 | NA | NA | 1 | 0 | 1 | 1 | NA | NA | 2 | 0.5 |
| 3_8880379 | 2 | 0 | 3 | 0.6667 | 4 | 0.25 | NA | NA | 9 | 0.3333 | NA | NA | 1 | 0 | 1 | 1 | NA | NA | 2 | 0.5 |
| 3_8880383 | 1 | 0 | 3 | 0 | 4 | 0.25 | NA | NA | 8 | 0.125 | NA | NA | 1 | 1 | NA | NA | NA | NA | 1 | 1 |
| 3_8881538 | 2 | 0.5 | 4 | 0 | 4 | 0 | 1 | 0 | 11 | 0.0909 | 1 | 1 | 4 | 0 | 1 | 0 | 2 | 0 | 8 | 0.125 |

| | | | | | | | | | | | | | | | | | | | | |
|---|---|---|---|---|---|---|---|---|---|---|---|---|---|---|---|---|---|---|---|---|
| 3_8881552 | 2 | 0 | 4 | 0.25 | 5 | 0 | 2 | 0.5 | 13 | 0.1538 | 1 | 0 | 4 | 0.5 | 1 | 0 | 2 | 0 | 8 | 0.25 |
| 3_8881552 | 2 | 0.5 | 4 | 1 | 5 | 0.4 | 2 | 0 | 13 | 0.5385 | 2 | 0.5 | 4 | 1 | 1 | 0 | 2 | 0.5 | 9 | 0.6667 |
| 3_8881556 | 3 | 0.6667 | 3 | 1 | 4 | 0.75 | 2 | 1 | 12 | 0.8333 | 2 | 1 | 4 | 0.5 | 1 | 1 | 2 | 1 | 9 | 0.7778 |
| 3_8881557 | 3 | 0 | 3 | 0.3333 | 5 | 0.2 | 2 | 0.5 | 13 | 0.2308 | 2 | 0.5 | 3 | 0 | 1 | 0 | 2 | 0 | 8 | 0.125 |
| 3_8881562 | 2 | 0.5 | 4 | 0 | 5 | 0.2 | 2 | 0 | 13 | 0.1538 | 2 | 0.5 | 4 | 0 | 1 | 0 | 2 | 0 | 9 | 0.1111 |
| 3_8881562 | 2 | 0.5 | 5 | 0.2 | 5 | 0 | 2 | 0.5 | 14 | 0.2143 | 1 | 0 | 4 | 0.25 | 1 | 0 | 2 | 0 | 8 | 0.125 |
| 3_8881670 | 3 | 0 | 3 | 0 | 6 | 0.1667 | 3 | 0 | 15 | 0.0667 | 3 | 0 | 4 | 0 | 5 | 0.2 | 3 | 0 | 15 | 0.0667 |
| 3_8881676 | 3 | 1 | 4 | 0.5 | 5 | 0.6 | 3 | 0.3333 | 15 | 0.6 | 1 | 0 | 3 | 0.3333 | 4 | 0.25 | 3 | 0.6667 | 11 | 0.3636 |
| 3_8881682 | 3 | 0 | 5 | 0.2 | 7 | 0 | 3 | 0 | 18 | 0.0556 | 3 | 0 | 4 | 0 | 5 | 0.2 | 3 | 0 | 15 | 0.0667 |
| 3_8881700 | 2 | 0.5 | 5 | 1 | 5 | 0.4 | 3 | 0.6667 | 15 | 0.6667 | 3 | 0.6667 | 4 | 0.75 | 4 | 0.75 | 3 | 0.6667 | 14 | 0.7143 |
| 3_8881700 | 1 | 1 | NA | NA | 4 | 0.25 | NA | NA | 5 | 0.4 | 7 | 0.5714 | 1 | 0 | 1 | 1 | 3 | 0.3333 | 12 | 0.5 |
| 3_8881701 | 3 | 0.6667 | 6 | 1 | 5 | 0.8 | 3 | 1 | 17 | 0.8824 | 4 | 1 | 3 | 0.6667 | 5 | 0.8 | 3 | 1 | 15 | 0.8667 |
| 3_8881732 | 1 | 0 | 5 | 0.4 | 5 | 0.4 | 4 | 1 | 15 | 0.5333 | 3 | 0.6667 | 4 | 0.5 | 4 | 0.25 | NA | NA | 11 | 0.4545 |
| 3_8881733 | 2 | 0 | 6 | 0.3333 | 6 | 0.3333 | 2 | 0.5 | 16 | 0.3125 | 2 | 0.5 | 4 | 0.75 | 4 | 0.25 | 2 | 0 | 12 | 0.4167 |
| 3_8881737 | 3 | 0.3333 | 2 | 0 | 5 | 0.2 | 2 | 0.5 | 12 | 0.25 | 4 | 0 | 5 | 0.6 | 4 | 0 | 2 | 0 | 15 | 0.2 |
| 3_8881739 | 1 | 0 | 3 | 0 | 4 | 0.25 | 2 | 0.5 | 10 | 0.2 | 2 | 0 | 2 | 0.5 | 4 | 0.25 | 3 | 0 | 11 | 0.1818 |
| 3_8882642 | 1 | 1 | 1 | 0 | NA | NA | NA | NA | 2 | 0.5 | NA | NA | NA | NA | NA | NA | 1 | 1 | 1 | 1 |
| 3_8882644 | 1 | 1 | 1 | 0 | NA | NA | NA | NA | 2 | 0.5 | NA | NA | NA | NA | NA | NA | 1 | 1 | 1 | 1 |
| 3_8882899 | 1 | 1 | 1 | 1 | 1 | 1 | NA | NA | 3 | 1 | NA | NA | NA | NA | 1 | 0 | 1 | 1 | 2 | 0.5 |
| 3_8883062 | NA | NA | NA | NA | 1 | 1 | NA | NA | 1 | 1 | NA | NA | NA | NA | 1 | 1 | NA | NA | 1 | 1 |
| 3_8883076 | 1 | 0 | NA | NA | 2 | 1 | NA | NA | 3 | 0.6667 | 1 | 0 | NA | NA | 1 | 1 | 2 | 0 | 4 | 0.25 |
| 3_8883085 | 1 | 1 | NA | NA | 2 | 0 | NA | NA | 3 | 0.3333 | 1 | 0 | NA | NA | 2 | 0.5 | 1 | 1 | 4 | 0.5 |
| 3_8883089 | 1 | 1 | NA | NA | 2 | 0.5 | NA | NA | 3 | 0.6667 | 1 | 1 | NA | NA | 2 | 0.5 | 1 | 1 | 4 | 0.75 |
| 3_8887322 | NA | NA | 1 | 1 | NA | NA | NA | NA | 1 | 1 | NA | NA | NA | NA | NA | NA | 1 | 1 | 1 | 1 |
| 3_8887345 | NA | NA | NA | NA | NA | NA | 1 | 1 | 1 | 1 | NA | NA | NA | NA | NA | NA | 1 | 1 | 1 | 1 |
| 3_8887352 | NA | NA | 1 | 1 | NA | NA | 1 | 1 | 2 | 1 | NA | NA | NA | NA | NA | NA | 1 | 1 | 1 | 1 |
| 3_8888887 | NA | NA | 1 | 1 | NA | NA | 1 | 1 | 2 | 1 | NA | NA | NA | NA | NA | NA | 1 | 1 | 1 | 1 |
| 3_8888892 | NA | NA | 1 | 0 | NA | NA | 1 | 1 | 2 | 0.5 | NA | NA | NA | NA | NA | NA | 1 | 1 | 1 | 1 |
| 3_8888902 | NA | NA | 1 | 1 | NA | NA | 1 | 0 | 2 | 0.5 | NA | NA | NA | NA | NA | NA | 1 | 1 | 1 | 1 |
| 3_8888907 | NA | NA | 1 | 1 | NA | NA | 1 | 0 | 2 | 0.5 | NA | NA | NA | NA | NA | NA | 2 | 0.5 | 2 | 0.5 |
| 3_8888911 | NA | NA | 1 | 0 | NA | NA | 1 | 1 | 2 | 0.5 | NA | NA | NA | NA | NA | NA | 1 | 1 | 1 | 1 |
| 3_8888921 | NA | NA | 1 | 1 | NA | NA | NA | NA | 1 | 1 | NA | NA | NA | NA | NA | NA | 3 | 0.6667 | 3 | 0.6667 |
| 3_8888932 | NA | NA | 1 | 1 | NA | NA | 2 | 1 | 3 | 1 | NA | NA | NA | NA | NA | NA | 2 | 1 | 2 | 1 |
| 3_8889001 | NA | NA | 1 | 0 | NA | NA | 2 | 1 | 3 | 0.6667 | NA | NA | NA | NA | NA | NA | 3 | 0.3333 | 3 | 0.3333 |
| 3_8889009 | NA | NA | 2 | 0.5 | NA | NA | 2 | 0 | 4 | 0.25 | NA | NA | NA | NA | NA | NA | 3 | 0.3333 | 3 | 0.3333 |
| 3_8889031 | NA | NA | 2 | 1 | NA | NA | 2 | 1 | 4 | 1 | NA | NA | NA | NA | NA | NA | 3 | 0.3333 | 3 | 0.3333 |
| 3_8889065 | NA | NA | 2 | 1 | NA | NA | 1 | 1 | 3 | 1 | NA | NA | NA | NA | NA | NA | 3 | 1 | 3 | 1 |
| 3_8889104 | NA | NA | 1 | 1 | 1 | 1 | 1 | 1 | 3 | 1 | NA | NA | 1 | 1 | NA | NA | 2 | 1 | 3 | 1 |
| 3_8889145 | 1 | 1 | 2 | 1 | 1 | 0 | 2 | 0 | 6 | 0.5 | NA | NA | 1 | 0 | NA | NA | 3 | 0.6667 | 4 | 0.5 |
| 3_8889155 | 1 | 0 | 1 | 1 | 1 | 0 | 2 | 0 | 5 | 0.2 | NA | NA | 1 | 1 | NA | NA | 3 | 0.6667 | 4 | 0.75 |
| 3_8889275 | 1 | 1 | 2 | 0.5 | 2 | 1 | 3 | 1 | 8 | 0.875 | 4 | 0.5 | 1 | 1 | 1 | 0 | 4 | 0.75 | 10 | 0.6 |
| 3_8889284 | 1 | 1 | NA | NA | 2 | 1 | 4 | 1 | 7 | 1 | 3 | 1 | NA | NA | 1 | 1 | 3 | 1 | 7 | 1 |
| 3_8889291 | 1 | 1 | 1 | 1 | 1 | 0 | 4 | 0.25 | 7 | 0.4286 | 3 | 1 | 1 | 1 | 1 | 1 | 3 | 0.6667 | 8 | 0.875 |
| 3_8889292 | NA | NA | NA | NA | NA | NA | 2 | 1 | 2 | 1 | 3 | 0.3333 | 1 | 1 | 1 | 1 | 4 | 0.75 | 9 | 0.6667 |
| 3_8889299 | 1 | 1 | NA | NA | 1 | 0 | 2 | 0.5 | 4 | 0.5 | 3 | 0.6667 | 1 | 0 | 1 | 1 | 3 | 0.6667 | 8 | 0.625 |
| 3_8891468 | 1 | 1 | 2 | 1 | 1 | 1 | 5 | 1 | 9 | 1 | 4 | 0.75 | 1 | 1 | 1 | 0 | 5 | 1 | 11 | 0.8182 |
| 3_8891482 | 1 | 0 | 3 | 0.6667 | 1 | 0 | 5 | 0 | 10 | 0.2 | 4 | 0.25 | 1 | 0 | 1 | 0 | 4 | 0 | 10 | 0.1 |
| 3_8891483 | 1 | 1 | 3 | 1 | 1 | 1 | 5 | 0.8 | 10 | 0.9 | 4 | 1 | 1 | 1 | 1 | 1 | 4 | 0.75 | 10 | 0.9 |
| 3_8891486 | 1 | 0 | 3 | 0.3333 | NA | NA | 4 | 0.25 | 8 | 0.25 | 4 | 0.25 | 1 | 1 | 1 | 0 | 5 | 0.6 | 11 | 0.4545 |
| 3_8891488 | 1 | 0 | 2 | 1 | 2 | 1 | 5 | 0.2 | 10 | 0.5 | 3 | 0.6667 | 1 | 0 | 1 | 0 | 5 | 0.4 | 10 | 0.4 |
| 3_8898344 | 3 | 0 | 7 | 0.2857 | 2 | 0.5 | 9 | 0.4444 | 21 | 0.3333 | 3 | 0 | 1 | 0 | 1 | 0 | 4 | 0.25 | 9 | 0.1111 |
| 3_8898438 | 3 | 0.3333 | 9 | 0.7778 | 3 | 1 | 9 | 0.5556 | 24 | 0.6667 | 5 | 0.6 | 2 | 0 | 3 | 0.3333 | 8 | 0.625 | 18 | 0.5 |
| 3_8898447 | 3 | 0.6667 | 8 | 1 | 3 | 0.6667 | 9 | 0.8889 | 23 | 0.8696 | 4 | 1 | 1 | 1 | 4 | 1 | 7 | 0.8571 | 16 | 0.9375 |
| 3_8899149 | 16 | 0.375 | 24 | 0.25 | 19 | 0.3684 | 10 | 0.6 | 69 | 0.3623 | 10 | 0.2 | 15 | 0.2 | 11 | 0.8182 | 17 | 0.4118 | 53 | 0.3962 |
| 3_8899167 | 11 | 0.3636 | 17 | 0.4118 | 16 | 0.375 | 10 | 0.4 | 54 | 0.3889 | 7 | 0.4286 | 13 | 0.0769 | 7 | 0.4286 | 11 | 0.1818 | 38 | 0.2368 |
| 3_8899529 | 9 | 0.2222 | 18 | 0.3889 | 16 | 0.1875 | 7 | 0.1429 | 50 | 0.26 | 6 | 0.6667 | 10 | 0.4 | 7 | 0.1429 | 13 | 0.3846 | 36 | 0.3889 |
| 3_8899531 | 9 | 0.6667 | 17 | 0.3529 | 12 | 0.3333 | 5 | 0.2 | 43 | 0.3953 | 5 | 0.2 | 11 | 0.3636 | 7 | 0.2857 | 11 | 0.5455 | 34 | 0.3824 |
| 3_8899536 | 10 | 0.3 | 17 | 0.4706 | 15 | 0.0667 | 8 | 0 | 50 | 0.24 | 6 | 0.1667 | 12 | 0.1667 | 4 | 0.5 | 14 | 0.2857 | 36 | 0.25 |
| 3_8899545 | 4 | 0 | 9 | 0.1111 | 10 | 0.1 | 5 | 0 | 28 | 0.0714 | 2 | 0 | 7 | 0 | 5 | 0 | 8 | 0.25 | 22 | 0.0909 |
| 3_8899863 | 5 | 0 | 5 | 0 | 7 | 0 | 3 | 0.3333 | 20 | 0.05 | 2 | 0 | 4 | 0 | 3 | 0 | 5 | 0.4 | 14 | 0.1429 |
| 3_8899868 | 6 | 0.1667 | 6 | 0.3333 | 9 | 0.2222 | 4 | 0.25 | 25 | 0.24 | 3 | 0 | 3 | 0.3333 | 3 | 0.3333 | 6 | 0.3333 | 15 | 0.2667 |
| 3_8899887 | 8 | 0.875 | 5 | 1 | 8 | 0.75 | 1 | 0 | 22 | 0.8182 | 2 | 1 | 6 | 0.6667 | 2 | 1 | 7 | 1 | 17 | 0.8824 |
| 3_8899892 | 7 | 0.4286 | 4 | 0.25 | 6 | 0.8333 | 2 | 1 | 19 | 0.5789 | 2 | 1 | 5 | 1 | 2 | 1 | 4 | 0.5 | 13 | 0.8462 |
| 3_8899892 | NA | NA | 1 | 1 | NA | NA | NA | NA | 1 | 1 | NA | NA | NA | NA | NA | NA | 1 | 1 | 1 | 1 |
| 3_8899895 | 5 | 0.4 | 4 | 0 | 7 | 0 | 2 | 0 | 18 | 0.1111 | 2 | 0.5 | 4 | 0.75 | 3 | 0.6667 | 4 | 0.25 | 13 | 0.5385 |

| | | | | | | | | | | | | | | | | | | | | |
|---|---|---|---|---|---|---|---|---|---|---|---|---|---|---|---|---|---|---|---|---|
| 3_8899900 | 8 | 0.25 | 6 | 0.3333 | 5 | 0.6 | 3 | 1 | 22 | 0.4545 | 3 | 0.6667 | 5 | 0.8 | 3 | 0.3333 | 5 | 0.6 | 16 | 0.625 |
| 3_8899903 | 6 | 0.8333 | 6 | 0.8333 | 7 | 0.7143 | 3 | 0.6667 | 22 | 0.7727 | 3 | 0.6667 | 5 | 0.6 | 3 | 1 | 6 | 0.6667 | 17 | 0.7059 |
| 3_8899903 | 5 | 0.6 | 5 | 0.6 | 5 | 0.8 | 2 | 0.5 | 17 | 0.6471 | NA | NA | 4 | 0.75 | 2 | 1 | 4 | 0.75 | 10 | 0.8 |
| 3_8899906 | 5 | 0 | 5 | 0 | 5 | 0 | 3 | 0.3333 | 18 | 0.0556 | 2 | 0 | 4 | 0 | 3 | 0.3333 | 5 | 0 | 14 | 0.0714 |
| 3_8899918 | 7 | 0 | 6 | 0 | 5 | 0.2 | 3 | 0.3333 | 21 | 0.0952 | 3 | 0 | 5 | 0.4 | 4 | 0 | 6 | 0 | 18 | 0.1111 |
| 3_8899924 | 3 | 0.3333 | 5 | 0.6 | 4 | 0 | 2 | 0.5 | 14 | 0.3571 | NA | NA | 4 | 0.5 | 4 | 0.25 | 5 | 0.2 | 13 | 0.3077 |
| 3_8899925 | 7 | 0.5714 | 6 | 0.5 | 6 | 0.5 | 2 | 1 | 21 | 0.5714 | 2 | 1 | 4 | 1 | 4 | 1 | 6 | 0.8333 | 16 | 0.9375 |
| 3_8899931 | 6 | 0.3333 | 6 | 0 | 5 | 0.4 | 3 | 0.6667 | 20 | 0.3 | 3 | 0 | 4 | 0 | 3 | 0 | 5 | 0.2 | 15 | 0.0667 |
| 3_8899952 | 5 | 0 | 5 | 0.2 | 4 | 0.25 | 3 | 0 | 17 | 0.1176 | 3 | 0 | 4 | 0 | 3 | 0 | 5 | 0.4 | 15 | 0.1333 |
| 3_8899968 | 6 | 0 | 5 | 0.2 | 4 | 0.5 | 2 | 0 | 17 | 0.1765 | 2 | 0 | 3 | 0.6667 | 2 | 0.5 | 4 | 1 | 11 | 0.6364 |
| 3_8900098 | 15 | 0.0667 | 17 | 0.1176 | 8 | 0 | 11 | 0 | 51 | 0.0588 | 16 | 0 | 5 | 0.2 | 13 | 0.1538 | 9 | 0.2222 | 43 | 0.1163 |
| 3_8905498 | 1 | 0 | NA | NA | 1 | 1 | NA | NA | 2 | 0.5 | 1 | 1 | NA | NA | 1 | 1 | NA | NA | 2 | 1 |
| 3_8908383 | 2 | 0.5 | NA | NA | 1 | 0 | 5 | 0.6 | 8 | 0.5 | 2 | 0.5 | 5 | 0.6 | 5 | 0.2 | 2 | 1 | 14 | 0.5 |
| 3_8908386 | 3 | 0.6667 | NA | NA | NA | NA | 5 | 0.8 | 8 | 0.75 | NA | NA | 4 | 0.5 | 5 | 0.6 | 2 | 1 | 11 | 0.6364 |
| 3_8908394 | 3 | 0.6667 | NA | NA | 1 | 0 | 6 | 0.5 | 10 | 0.5 | 2 | 0.5 | 6 | 0.8333 | 4 | 1 | 2 | 1 | 14 | 0.8571 |
| 3_8909437 | NA | NA | NA | NA | 2 | 0.5 | 1 | 0 | 3 | 0.3333 | 2 | 0.5 | 3 | 0 | 2 | 0.5 | NA | NA | 7 | 0.2857 |
| 3_8909453 | 3 | 0 | 1 | 0 | 1 | 0 | 4 | 0.25 | 9 | 0.1111 | 4 | 0.5 | 7 | 0 | 2 | 0.5 | 2 | 0 | 15 | 0.2 |
| 3_8909470 | 5 | 0 | 2 | 0 | 2 | 0.5 | 6 | 0 | 15 | 0.0667 | 2 | 0 | 10 | 0.1 | 5 | 0.2 | 1 | 0 | 18 | 0.1111 |
| 3_8909603 | 7 | 0.4286 | 4 | 0.5 | 2 | 0 | 7 | 0.4286 | 20 | 0.4 | 3 | 0 | 10 | 0.4 | 9 | 0.3333 | 4 | 0.25 | 26 | 0.3077 |
| 3_8909644 | 6 | 0.3333 | 6 | 0 | 4 | 0 | 10 | 0.2 | 26 | 0.1538 | 7 | 0.2857 | 11 | 0.0909 | 7 | 0.1429 | 3 | 0 | 28 | 0.1429 |
| 3_8911516 | 6 | 0 | 3 | 0 | 9 | 0.1111 | 1 | 0 | 19 | 0.0526 | 5 | 0 | 4 | 0 | 3 | 0.3333 | 5 | 0 | 17 | 0.0588 |
| 3_8911593 | 5 | 0.2 | 2 | 0 | 6 | 0.1667 | 1 | 0 | 14 | 0.1429 | 6 | 0.1667 | 3 | 0 | 3 | 0 | 5 | 0.2 | 17 | 0.1176 |
| 3_8911596 | 5 | 0.4 | 2 | 0.5 | 7 | 0.5714 | NA | NA | 14 | 0.5 | 6 | 0.6667 | 4 | 0.5 | 3 | 0.6667 | 5 | 0.6 | 18 | 0.6111 |
| 3_8911600 | 1 | 1 | NA | NA | 3 | 0.3333 | NA | NA | 4 | 0.5 | NA | NA | 1 | 0 | 1 | 0 | 1 | 1 | 3 | 0.3333 |
| 3_8911603 | NA | NA | NA | NA | 3 | 0.3333 | NA | NA | 3 | 0.3333 | NA | NA | 1 | 1 | 2 | 0 | 1 | 0 | 4 | 0.25 |
| 3_8911639 | 1 | 1 | 1 | 0 | 1 | 0 | 2 | 0 | 5 | 0.2 | 4 | 0.25 | 2 | 0 | 4 | 0.5 | 5 | 0.4 | 15 | 0.3333 |
| 3_8911640 | 8 | 0 | 7 | 0.1429 | 6 | 0.3333 | 3 | 0.3333 | 24 | 0.1667 | 9 | 0.2222 | 5 | 0 | 10 | 0.1 | 5 | 0.2 | 29 | 0.1379 |
| 3_8911649 | 8 | 0.375 | 8 | 0 | 8 | 0.25 | 5 | 0 | 29 | 0.1724 | 9 | 0 | 5 | 0 | 8 | 0.5 | 8 | 0.25 | 30 | 0.2 |
| 3_8911649 | 8 | 0.5 | 10 | 0.8 | 8 | 0.625 | 5 | 0.4 | 31 | 0.6129 | 9 | 0.4444 | 4 | 0.75 | 8 | 0.5 | 8 | 0.625 | 29 | 0.5517 |
| 3_8911651 | 9 | 0.5556 | 8 | 0.5 | 7 | 0.8571 | 5 | 0.2 | 29 | 0.5517 | 11 | 0.5455 | 5 | 0.8 | 9 | 0.6667 | 9 | 0.8889 | 34 | 0.7059 |
| 3_8911718 | NA | NA | NA | NA | 1 | 1 | NA | NA | 1 | 1 | NA | NA | 1 | 1 | 1 | 0 | NA | NA | 2 | 0.5 |
| 3_8911994 | 1 | 1 | NA | NA | 2 | 0.5 | NA | NA | 3 | 0.6667 | 1 | 1 | 1 | 0 | NA | NA | NA | NA | 2 | 0.5 |
| 3_8914186 | 12 | 0 | 7 | 0.1429 | 14 | 0 | 17 | 0.1765 | 50 | 0.08 | 5 | 0 | 12 | 0 | 13 | 0.1538 | 9 | 0.1111 | 39 | 0.0769 |
| 3_8914204 | 10 | 0.5 | 11 | 0.7273 | 16 | 0.5625 | 15 | 0.2667 | 52 | 0.5 | 6 | 0.1667 | 12 | 0.5 | 14 | 0.4286 | 5 | 0.4 | 37 | 0.4054 |
| 3_8914212 | 14 | 0.1429 | 9 | 0 | 17 | 0.1765 | 15 | 0.0667 | 55 | 0.1091 | 5 | 0.2 | 11 | 0 | 11 | 0.2727 | 7 | 0 | 34 | 0.1176 |
| 3_8914221 | 9 | 1 | 8 | 1 | 11 | 1 | 12 | 1 | 40 | 1 | 4 | 1 | 9 | 0.7778 | 10 | 1 | 7 | 1 | 30 | 0.9333 |
| 3_8914221 | 8 | 0.125 | 6 | 0.1667 | 12 | 0.0833 | 12 | 0.0833 | 38 | 0.1053 | 4 | 0 | 8 | 0 | 10 | 0.1 | 7 | 0.2857 | 29 | 0.1034 |
| 3_8914233 | 10 | 0.9 | 8 | 0.75 | 12 | 1 | 7 | 1 | 37 | 0.9189 | 4 | 1 | 7 | 1 | 10 | 1 | 6 | 0.8333 | 27 | 0.963 |
| 3_8914239 | 7 | 0.8571 | 6 | 0.6667 | 9 | 0.8889 | 12 | 0.5833 | 34 | 0.7353 | 3 | 0.6667 | 8 | 0.875 | 8 | 0.75 | 6 | 0.8333 | 25 | 0.8 |
| 3_8916677 | 2 | 0.5 | NA | NA | 2 | 0 | 6 | 0.1667 | 10 | 0.2 | NA | NA | 2 | 0.5 | NA | NA | 3 | 0 | 5 | 0.2 |
| 3_8917565 | 7 | 0.4286 | 7 | 0.2857 | 7 | 0.7143 | 8 | 0.625 | 29 | 0.5172 | 5 | 0.8 | 4 | 0.25 | 6 | 0.6667 | 10 | 0.5 | 25 | 0.56 |
| 3_8917568 | 6 | 0.3333 | 7 | 0.4286 | 6 | 0.1667 | 4 | 0 | 23 | 0.2609 | 5 | 0.4 | 4 | 0.5 | 6 | 0.5 | 10 | 0.4 | 25 | 0.44 |
| 3_8917570 | 5 | 0.4 | 7 | 0.5714 | 7 | 0 | 6 | 0.1667 | 25 | 0.28 | 5 | 0.2 | 3 | 0 | 4 | 0.75 | 9 | 0.3333 | 21 | 0.3333 |
| 3_8917923 | 2 | 0.5 | NA | NA | NA | NA | NA | NA | 2 | 0.5 | 2 | 1 | NA | NA | NA | NA | NA | NA | 2 | 1 |
| 3_8917942 | 2 | 1 | NA | NA | 1 | 1 | NA | NA | 3 | 1 | 1 | 1 | NA | NA | NA | NA | NA | NA | 1 | 1 |
| 3_8917947 | 2 | 1 | NA | NA | 1 | 1 | NA | NA | 3 | 1 | 2 | 1 | NA | NA | NA | NA | NA | NA | 2 | 1 |
| 3_8917976 | 2 | 1 | NA | NA | 1 | 1 | NA | NA | 3 | 1 | 1 | 1 | NA | NA | NA | NA | NA | NA | 1 | 1 |
| 3_8922248 | 5 | 0 | 9 | 0.1111 | 4 | 0.25 | 16 | 0 | 34 | 0.0588 | 2 | 0.5 | 5 | 0 | NA | NA | 1 | 0 | 8 | 0.125 |
| 3_8924699 | 13 | 0.4615 | 13 | 0.3077 | 7 | 0.7143 | 3 | 0 | 36 | 0.4167 | 7 | 0.4286 | 8 | 0.25 | 4 | 0 | 4 | 0 | 23 | 0.2174 |
| 3_8924700 | 13 | 0 | 14 | 0 | 8 | 0.375 | 5 | 0 | 40 | 0.075 | 11 | 0.0909 | 10 | 0.1 | 5 | 0 | 4 | 0 | 30 | 0.0667 |
| 3_8925967 | 1 | 1 | NA | NA | NA | NA | NA | NA | 1 | 1 | NA | NA | 1 | 1 | NA | NA | 1 | 1 | 2 | 1 |
| 3_8925976 | 1 | 1 | NA | NA | NA | NA | NA | NA | 1 | 1 | NA | NA | 1 | 1 | NA | NA | NA | NA | 1 | 1 |
| 3_8925985 | 1 | 1 | NA | NA | NA | NA | NA | NA | 1 | 1 | NA | NA | 1 | 1 | NA | NA | 1 | 1 | 2 | 1 |
| 3_8925994 | 1 | 1 | NA | NA | NA | NA | NA | NA | 1 | 1 | NA | NA | 1 | 1 | NA | NA | 1 | 0 | 2 | 0.5 |
| 3_8926024 | 1 | 1 | NA | NA | NA | NA | NA | NA | 1 | 1 | NA | NA | 1 | 1 | NA | NA | 1 | 0 | 2 | 0.5 |
| 3_8930544 | 2 | 0 | 2 | 0.5 | 2 | 1 | NA | NA | 6 | 0.5 | 2 | 0 | 1 | 1 | NA | NA | NA | NA | 3 | 0.3333 |
| 3_8930928 | 1 | 0 | NA | NA | 2 | 0.5 | 2 | 0 | 5 | 0.2 | 2 | 1 | 2 | 0.5 | 1 | 1 | 2 | 1 | 7 | 0.8571 |
| 3_8930935 | 1 | 1 | 1 | 1 | 3 | 1 | 4 | 1 | 9 | 1 | 2 | 1 | 2 | 1 | 1 | 1 | 5 | 0.8 | 10 | 0.9 |
| 3_8930943 | 1 | 0 | 1 | 0 | 3 | 0.3333 | 4 | 0.25 | 9 | 0.2222 | 2 | 0 | 2 | 0 | 2 | 0.5 | 4 | 0 | 10 | 0.1 |
| 3_8930944 | 1 | 1 | 1 | 0 | 2 | 0 | 4 | 0 | 8 | 0.125 | 2 | 0.5 | 2 | 0.5 | 2 | 0 | 4 | 0.25 | 10 | 0.3 |
| 3_8930956 | 1 | 0 | 1 | 1 | 2 | 0.5 | 3 | 0.3333 | 7 | 0.4286 | 2 | 0.5 | 2 | 0 | 1 | 0 | 4 | 0.25 | 9 | 0.2222 |
| 3_8930977 | 1 | 0 | 1 | 1 | 1 | 1 | 4 | 0.25 | 7 | 0.4286 | 2 | 0.5 | 2 | 1 | 1 | 0 | 4 | 0.75 | 9 | 0.6667 |
| 3_8931032 | 1 | 0 | NA | NA | 3 | 0.6667 | 1 | 0 | 5 | 0.4 | 1 | 0 | 2 | 1 | NA | NA | 3 | 0.3333 | 6 | 0.5 |
| 3_8931045 | 3 | 0.3333 | 5 | 0 | 5 | 0.2 | 4 | 0 | 17 | 0.1176 | 3 | 0.3333 | 4 | 0 | 3 | 0 | 5 | 0 | 15 | 0.0667 |
| 3_8931297 | 4 | 0 | 6 | 0 | 5 | 0 | 5 | 0.2 | 20 | 0.05 | 4 | 0 | 7 | 0.1429 | 2 | 0 | 6 | 0 | 19 | 0.0526 |

| | | | | | | | | | | | | | | | | | | | | |
|---|---|---|---|---|---|---|---|---|---|---|---|---|---|---|---|---|---|---|---|---|
| 3_89315819 | 3 | 0 | 5 | 0.2 | 3 | 0.3333 | 6 | 0.3333 | 17 | 0.2353 | 3 | 0.3333 | 4 | 0.25 | 3 | 0.3333 | 5 | 0.4 | 15 | 0.3333 |
| 3_89315863 | 3 | 0.3333 | 4 | 0 | 1 | 0 | 6 | 0.1667 | 14 | 0.1429 | 3 | 0 | 4 | 0 | 3 | 0 | 6 | 0.1667 | 16 | 0.0625 |
| 3_89315902 | 3 | 0.3333 | 4 | 0 | 2 | 0 | 5 | 0.4 | 14 | 0.2143 | 4 | 0.5 | 5 | 0 | 3 | 0 | 5 | 0.2 | 17 | 0.1765 |
| 3_89316045 | 2 | 1 | 5 | 0.8 | 1 | 1 | 6 | 0.8333 | 14 | 0.8571 | 3 | 1 | 3 | 0.6667 | 3 | 1 | 6 | 1 | 15 | 0.9333 |
| 3_89316137 | 1 | 0 | 4 | 0 | 2 | 0.5 | 4 | 0 | 11 | 0.0909 | 4 | 0.25 | 4 | 0 | 3 | 0 | 4 | 0 | 15 | 0.0667 |
| 3_89316215 | 1 | 0 | 2 | 0 | 2 | 0.5 | 5 | 0.6 | 10 | 0.4 | NA | NA | 1 | 0 | 2 | 0.5 | 3 | 0.3333 | 6 | 0.3333 |
| 3_89318554 | 2 | 0.5 | 2 | 0 | 1 | 0 | 3 | 0 | 8 | 0.125 | 1 | 0 | 1 | 0 | NA | NA | 3 | 0.3333 | 5 | 0.2 |
| 3_89329459 | 1 | 0 | NA | NA | 1 | 1 | NA | NA | 2 | 0.5 | 1 | 1 | NA | NA | NA | NA | NA | NA | 1 | 1 |
| 3_89335682 | 2 | 0 | 2 | 0.5 | 2 | 0 | NA | NA | 6 | 0.1667 | 1 | 1 | NA | NA | 1 | 0 | NA | NA | 2 | 0.5 |
| 3_89335716 | 2 | 0.5 | 2 | 0.5 | 3 | 0.6667 | NA | NA | 7 | 0.5714 | 1 | 1 | NA | NA | 1 | 1 | 1 | 1 | 3 | 1 |
| 3_89337144 | 3 | 0 | 3 | 0.3333 | 3 | 0 | 1 | 0 | 10 | 0.1 | 3 | 0.3333 | 1 | 0 | 2 | 0 | 2 | 0 | 8 | 0.125 |
| 3_89340085 | 1 | 1 | 6 | 0.6667 | 2 | 0.5 | 5 | 1 | 14 | 0.7857 | 3 | 1 | 3 | 1 | 7 | 0.7143 | 5 | 1 | 18 | 0.8889 |
| 3_89340200 | 1 | 0 | 6 | 0.1667 | 2 | 0.5 | 4 | 0 | 13 | 0.1538 | 3 | 0.3333 | 3 | 0 | 5 | 0 | 6 | 0 | 17 | 0.0588 |
| 3_89340287 | 1 | 1 | 5 | 0.2 | 4 | 0 | 5 | 0 | 15 | 0.1333 | 3 | 0 | 3 | 0 | 8 | 0 | 5 | 0.4 | 19 | 0.1053 |
| 3_89340299 | 1 | 1 | 6 | 0.3333 | 4 | 0 | 5 | 0 | 16 | 0.1875 | 4 | 0 | 3 | 0.6667 | 7 | 0.1429 | 6 | 0 | 20 | 0.15 |
| 3_89340821 | 3 | 1 | 6 | 0.8333 | 5 | 0.6 | 2 | 1 | 16 | 0.8125 | 2 | 1 | 4 | 1 | 7 | 0.7143 | 6 | 0.8333 | 19 | 0.8421 |
| 3_89340932 | 3 | 0 | 9 | 0.2222 | 6 | 0 | 7 | 0 | 25 | 0.08 | 4 | 0.25 | 4 | 0 | 6 | 0.1667 | 8 | 0.25 | 22 | 0.1818 |
| 3_89340958 | 3 | 1 | 6 | 1 | 4 | 1 | 4 | 1 | 17 | 1 | 3 | 1 | 4 | 0.75 | 6 | 1 | 7 | 1 | 20 | 0.95 |
| 3_89340968 | 2 | 0 | 9 | 0.1111 | 4 | 0.5 | 5 | 0.2 | 20 | 0.2 | 3 | 0 | 5 | 0 | 6 | 0.1667 | 5 | 0.2 | 19 | 0.1053 |
| 3_89341035 | 3 | 1 | 9 | 0.6667 | 6 | 0.3333 | 6 | 0.5 | 24 | 0.5833 | 2 | 0 | 3 | 0.6667 | 8 | 0.5 | 6 | 0.6667 | 19 | 0.5263 |
| 3_89341599 | 2 | 0.5 | 9 | 0.3333 | 5 | 0.4 | 8 | 0.25 | 24 | 0.3333 | 4 | 0.25 | 4 | 0.5 | 8 | 0.5 | 7 | 0.5714 | 23 | 0.4783 |
| 3_89341638 | 3 | 1 | 11 | 0.2727 | 4 | 0.25 | 6 | 0.5 | 24 | 0.4167 | 4 | 0.25 | 4 | 0.25 | 11 | 0.4545 | 9 | 0.6667 | 28 | 0.4643 |
| 3_89342135 | 3 | 0.3333 | 10 | 0 | 7 | 0.4286 | 7 | 0.2857 | 27 | 0.2222 | 4 | 0.5 | 5 | 0 | 12 | 0.0833 | 5 | 0.2 | 26 | 0.1538 |
| 3_89342179 | 3 | 0.3333 | 11 | 0 | 8 | 0 | 8 | 0.125 | 30 | 0.0667 | 4 | 0 | 5 | 0 | 11 | 0.0909 | 9 | 0.1111 | 29 | 0.069 |
| 3_89354714 | 1 | 0 | NA | NA | 2 | 1 | NA | NA | 3 | 0.6667 | 3 | 0.3333 | NA | NA | 1 | 0 | NA | NA | 4 | 0.25 |
| 3_89354775 | 1 | 1 | NA | NA | 3 | 0.3333 | 1 | 0 | 5 | 0.4 | 2 | 0.5 | NA | NA | 1 | 0 | NA | NA | 3 | 0.3333 |
| 3_89356138 | 4 | 0 | 1 | 0 | 3 | 0.3333 | 1 | 0 | 9 | 0.1111 | 4 | 0.5 | 2 | 0 | 5 | 0.6 | 1 | 0 | 12 | 0.4167 |
| 3_89356314 | 3 | 1 | 2 | 1 | 1 | 1 | 1 | 1 | 7 | 1 | 2 | 0.5 | 2 | 1 | 5 | 0.8 | 1 | 1 | 10 | 0.8 |
| 3_89356327 | 4 | 0.25 | 3 | 0 | 4 | 0 | 1 | 0 | 12 | 0.0833 | 3 | 0.3333 | 2 | 0.5 | 5 | 0 | 1 | 0 | 11 | 0.1818 |
| 3_89356397 | 4 | 0.75 | 4 | 0.5 | 3 | 1 | 1 | 1 | 12 | 0.75 | 4 | 1 | 2 | 1 | 6 | 0.8333 | 1 | 1 | 13 | 0.9231 |
| 3_89356492 | 5 | 0 | 3 | 0.3333 | 4 | 0.25 | 1 | 0 | 13 | 0.1538 | 3 | 0.3333 | 3 | 0.3333 | 4 | 0.25 | 1 | 0 | 11 | 0.2727 |
| 3_89356504 | 3 | 0 | 3 | 0 | 3 | 0.3333 | 1 | 0 | 10 | 0.1 | 4 | 0 | 3 | 0 | 5 | 0.2 | 1 | 0 | 13 | 0.0769 |
| 3_89375985 | 6 | 0.3333 | 3 | 1 | 5 | 0.8 | 8 | 0.875 | 22 | 0.7273 | 6 | 0.3333 | 5 | 0.8 | 2 | 1 | 6 | 0.6667 | 19 | 0.6316 |
| 3_89376148 | 5 | 0 | 4 | 0.25 | 6 | 0 | 5 | 0.2 | 20 | 0.1 | 4 | 0.25 | 5 | 0 | 2 | 0.5 | 4 | 0 | 15 | 0.1333 |
| 3_89575217 | NA | NA | 1 | 1 | NA | NA | NA | NA | 1 | 1 | 1 | 1 | NA | NA | NA | NA | NA | NA | 1 | 1 |
| 3_89575906 | NA | NA | 1 | 0 | 2 | 0.5 | NA | NA | 3 | 0.3333 | 2 | 0.5 | NA | NA | NA | NA | NA | NA | 2 | 0.5 |
| 3_89575981 | NA | NA | 2 | 0 | 2 | 0.5 | NA | NA | 4 | 0.25 | 2 | 1 | NA | NA | NA | NA | NA | NA | 2 | 1 |
| 3_89578960 | 2 | 0 | 4 | 0.25 | 3 | 0.3333 | 2 | 0.5 | 11 | 0.2727 | 1 | 1 | 1 | 0 | 1 | 0 | 1 | 1 | 4 | 0.5 |
| 3_89579232 | 2 | 1 | 4 | 1 | 3 | 1 | 2 | 1 | 11 | 1 | 1 | 1 | 1 | 0 | NA | NA | 1 | 1 | 3 | 0.6667 |
| 3_89579243 | 2 | 0.5 | 4 | 0.75 | 3 | 1 | 2 | 0.5 | 11 | 0.7273 | 1 | 0 | 1 | 0 | 1 | 0 | 2 | 0.5 | 5 | 0.2 |
| 3_89579323 | 2 | 1 | 4 | 0.25 | 2 | 0.5 | 2 | 1 | 10 | 0.6 | 1 | 1 | 1 | 0 | 1 | 0 | 2 | 0.5 | 5 | 0.4 |
| 3_89579374 | 2 | 1 | 3 | 0.6667 | 3 | 1 | 1 | 1 | 9 | 0.8889 | 1 | 0 | NA | NA | 1 | 0 | 2 | 1 | 4 | 0.5 |
| 3_89579380 | 1 | 0 | 3 | 0.6667 | 2 | 0.5 | 1 | 1 | 7 | 0.5714 | 1 | 1 | 1 | 1 | 1 | 0 | 3 | 0.6667 | 6 | 0.6667 |
| 3_89579409 | 1 | 0 | 3 | 0.3333 | 2 | 0.5 | 1 | 0 | 7 | 0.2857 | 1 | 0 | 1 | 1 | 1 | 0 | 1 | 1 | 4 | 0.5 |
| 3_89579452 | 2 | 0 | 4 | 0.75 | 2 | 0.5 | 2 | 0.5 | 10 | 0.5 | 1 | 1 | 1 | 0 | 1 | 0 | 3 | 0 | 6 | 0.1667 |
| 3_89580843 | 3 | 0 | 5 | 0.2 | 3 | 0 | 4 | 0 | 15 | 0.0667 | 1 | 1 | 2 | 0.5 | 4 | 0 | 4 | 0 | 11 | 0.1818 |
| 3_89581044 | 3 | 0 | 5 | 0.2 | 3 | 0 | 4 | 0 | 15 | 0.0667 | 1 | 0 | 2 | 0.5 | 4 | 0 | 4 | 0 | 11 | 0.0909 |
| 3_89642345 | NA | NA | 4 | 1 | 1 | 0 | 1 | 1 | 6 | 0.8333 | NA | NA | 3 | 0.6667 | 1 | 0 | 1 | 0 | 5 | 0.4 |
| 3_89642418 | NA | NA | 3 | 0.3333 | 1 | 1 | NA | NA | 4 | 0.5 | NA | NA | 4 | 1 | 1 | 0 | NA | NA | 5 | 0.8 |
| 3_89642638 | 1 | 0 | 3 | 0.6667 | 1 | 1 | 2 | 0.5 | 7 | 0.5714 | 1 | 1 | 5 | 0.8 | 1 | 1 | 1 | 1 | 8 | 0.875 |
| 3_89642739 | 1 | 1 | 4 | 0.75 | NA | NA | 2 | 1 | 7 | 0.8571 | 1 | 1 | 4 | 0.75 | 2 | 1 | 1 | 1 | 8 | 0.875 |
| 3_89642809 | NA | NA | 4 | 1 | 1 | 1 | 2 | 1 | 7 | 1 | 1 | 1 | 4 | 0.75 | 2 | 1 | NA | NA | 7 | 0.8571 |
| 3_89642846 | NA | NA | 4 | 1 | 1 | 1 | 2 | 0.5 | 7 | 0.8571 | 1 | 1 | 5 | 0.6 | 2 | 0 | 1 | 1 | 9 | 0.5556 |
| 3_89642893 | 1 | 0 | 3 | 0 | 1 | 0 | 2 | 0.5 | 7 | 0.1429 | 1 | 0 | 4 | 0.25 | 2 | 0 | NA | NA | 7 | 0.1429 |
| 3_89642941 | NA | NA | 2 | 1 | NA | NA | 3 | 0.6667 | 5 | 0.8 | NA | NA | 3 | 1 | 1 | 1 | 2 | 0.5 | 6 | 0.8333 |
| 3_89642986 | 1 | 0 | 4 | 0.75 | 1 | 0 | 3 | 0.3333 | 9 | 0.4444 | 1 | 0 | 5 | 0 | 1 | 0 | 2 | 0.5 | 9 | 0.1111 |
| 3_89643049 | 1 | 0 | 4 | 0.5 | 1 | 0 | 3 | 0 | 9 | 0.2222 | 1 | 0 | 4 | 0.25 | 2 | 0.5 | 2 | 1 | 9 | 0.4444 |
| 3_89643263 | 2 | 0 | 1 | 1 | 1 | 1 | 1 | 1 | 5 | 0.6 | NA | NA | 5 | 0.2 | 1 | 1 | 2 | 1 | 8 | 0.5 |
| 3_89643272 | 1 | 0 | 3 | 1 | 1 | 1 | 2 | 1 | 7 | 0.8571 | 1 | 1 | 5 | 1 | 3 | 0.6667 | 3 | 1 | 12 | 0.9167 |
| 3_89658364 | 8 | 0.25 | 11 | 0.0909 | 5 | 0.4 | 7 | 0 | 31 | 0.1613 | 1 | 0 | 9 | 0.2222 | 9 | 0 | 9 | 0.4444 | 28 | 0.2143 |
| 3_89658370 | 8 | 0.125 | 11 | 0.1818 | 4 | 0.25 | 7 | 0.4286 | 30 | 0.2333 | 1 | 0 | 9 | 0.1111 | 9 | 0.1111 | 10 | 0.1 | 29 | 0.1034 |
| 3_89658382 | 7 | 0.1429 | 9 | 0.2222 | 4 | 0 | 7 | 0.1429 | 27 | 0.1481 | 1 | 0 | 9 | 0.2222 | 7 | 0.2857 | 9 | 0.3333 | 26 | 0.2692 |
| 3_89658451 | 8 | 0.875 | 11 | 0.9091 | 4 | 0.5 | 6 | 1 | 29 | 0.8621 | 1 | 0 | 10 | 0.7 | 9 | 0.6667 | 10 | 0.8 | 30 | 0.7 |
| 3_89658478 | 7 | 0.4286 | 6 | 1 | 5 | 0.2 | 5 | 0.6 | 23 | 0.5652 | NA | NA | 6 | 0.5 | 7 | 0.7143 | 8 | 0.375 | 21 | 0.5238 |
| 3_89658615 | 5 | 0.6 | 8 | 0.25 | 3 | 0 | 6 | 0.5 | 22 | 0.3636 | 1 | 0 | 9 | 0.2222 | 6 | 0.5 | 6 | 0.6667 | 22 | 0.4091 |

| | | | | | | | | | | | | | | | | | | | | |
|---|---|---|---|---|---|---|---|---|---|---|---|---|---|---|---|---|---|---|---|---|
| 3_8965912 | 2 | 0 | 7 | 0.4286 | 5 | 0.2 | 8 | 0.125 | 22 | 0.2273 | NA | NA | 8 | 0.25 | 7 | 0.1429 | 7 | 0.1429 | 22 | 0.1818 |
| 3_8965950 | 4 | 0.5 | 12 | 0.4167 | 7 | 0.2857 | 7 | 0.1429 | 30 | 0.3333 | 1 | 0 | 12 | 0.1667 | 11 | 0.4545 | 14 | 0.1429 | 38 | 0.2368 |
| 3_8965951 | 6 | 0.1667 | 13 | 0.3846 | 7 | 0 | 8 | 0.25 | 34 | 0.2353 | 2 | 0 | 12 | 0.0833 | 11 | 0.4545 | 15 | 0.4 | 40 | 0.3 |
| 3_8965972 | 4 | 0.5 | 14 | 0.5714 | 6 | 0.3333 | 5 | 0.4 | 29 | 0.4828 | 4 | 0.25 | 13 | 0.5385 | 13 | 0.6923 | 11 | 0.5455 | 41 | 0.561 |
| 3_8965981 | 5 | 0 | 16 | 0.125 | 7 | 0 | 8 | 0.25 | 36 | 0.1111 | 1 | 0 | 12 | 0.1667 | 14 | 0.0714 | 14 | 0.0714 | 41 | 0.0976 |
| 3_8966000 | 3 | 0.6667 | 11 | 0.0909 | 8 | 0.125 | 6 | 0.3333 | 28 | 0.2143 | 5 | 0 | 9 | 0.1111 | 12 | 0.4167 | 12 | 0 | 38 | 0.1579 |
| 3_8966002 | 6 | 0 | 17 | 0.0588 | 8 | 0.125 | 8 | 0 | 39 | 0.0513 | 6 | 0 | 13 | 0 | 14 | 0.1429 | 11 | 0.0909 | 44 | 0.0682 |
| 3_8966022 | 7 | 0.8571 | 16 | 0.75 | 8 | 0.75 | 6 | 0.8333 | 37 | 0.7838 | 5 | 1 | 15 | 0.7333 | 17 | 0.7059 | 14 | 0.7143 | 51 | 0.7451 |
| 3_8966036 | 7 | 0.8571 | 21 | 0.7143 | 9 | 0.6667 | 7 | 0.8571 | 44 | 0.75 | 8 | 0.625 | 13 | 0.5385 | 17 | 0.8824 | 15 | 0.8667 | 53 | 0.7547 |
| 3_8966060 | 12 | 0.25 | 22 | 0.2727 | 12 | 0 | 12 | 0 | 58 | 0.1552 | 9 | 0 | 14 | 0.0714 | 14 | 0.0714 | 14 | 0.2857 | 51 | 0.1176 |
| 3_8966075 | 3 | 0 | 2 | 0 | 4 | 0.25 | 3 | 0.3333 | 12 | 0.1667 | NA | NA | 2 | 0.5 | 2 | 0.5 | 2 | 0 | 6 | 0.3333 |
| 3_8966112 | 4 | 0.5 | 4 | 0 | 2 | 0.5 | 4 | 0 | 14 | 0.2143 | 4 | 0 | NA | NA | 2 | 0 | 3 | 0.3333 | 9 | 0.1111 |
| 3_8966120 | 3 | 0.3333 | 4 | 0.25 | 2 | 0.5 | 4 | 0.5 | 13 | 0.3846 | 4 | 0.5 | NA | NA | 4 | 0.25 | 3 | 0.3333 | 11 | 0.3636 |
| 3_8966140 | 4 | 0.75 | 3 | 0.3333 | 2 | 0.5 | 5 | 0.6 | 14 | 0.5714 | 4 | 0.25 | NA | NA | 3 | 0.6667 | 3 | 0.6667 | 10 | 0.5 |
| 3_8966146 | 2 | 0.5 | 4 | 0.5 | 1 | 1 | 2 | 0.5 | 9 | 0.5556 | 3 | 0.3333 | NA | NA | 2 | 0.5 | 3 | 0.6667 | 8 | 0.5 |
| 3_8966148 | 3 | 0 | 4 | 0.25 | 2 | 0.5 | 4 | 0.25 | 13 | 0.2308 | 4 | 0.25 | NA | NA | 3 | 0.3333 | 3 | 0.6667 | 10 | 0.4 |
| 3_8966149 | 4 | 0 | 4 | 0 | 2 | 1 | 5 | 0 | 15 | 0.1333 | 4 | 0.25 | NA | NA | 2 | 0.5 | 3 | 0.3333 | 9 | 0.3333 |
| 3_8966185 | 4 | 0 | 3 | 0.3333 | NA | NA | 4 | 0 | 11 | 0.0909 | 4 | 0.25 | NA | NA | 3 | 0 | 2 | 0 | 9 | 0.1111 |
| 3_8966209 | 5 | 0 | 6 | 0.1667 | 3 | 0 | 7 | 0.1429 | 21 | 0.0952 | 6 | 0.3333 | 2 | 0.5 | 7 | 0.4286 | 1 | 0 | 16 | 0.375 |
| 3_8966237 | 6 | 0.1667 | 5 | 0.2 | 3 | 0.3333 | 6 | 0.5 | 20 | 0.3 | 7 | 0.2857 | 2 | 0 | 6 | 0.3333 | 1 | 0 | 16 | 0.25 |
| 3_8966264 | 4 | 0.5 | 6 | 0.5 | 1 | 1 | 5 | 1 | 16 | 0.6875 | 6 | 0.3333 | 2 | 0 | 6 | 0.6667 | 1 | 0 | 15 | 0.4 |
| 3_8966266 | 4 | 0.25 | 6 | 0.3333 | 1 | 0 | 4 | 0.25 | 15 | 0.2667 | 7 | 0.4286 | 2 | 0.5 | 5 | 1 | 1 | 1 | 15 | 0.6667 |
| 3_8966266 | 5 | 0 | 5 | 0.4 | 1 | 0 | 4 | 0 | 15 | 0.1333 | 5 | 0.4 | 2 | 0 | 4 | 0 | NA | NA | 11 | 0.1818 |
| 3_8966284 | 5 | 0.2 | 6 | 0.3333 | 1 | 0 | 4 | 0 | 16 | 0.1875 | 6 | 0.3333 | 2 | 1 | 4 | 0.25 | 1 | 0 | 13 | 0.3846 |
| 3_8966303 | 4 | 0.75 | 4 | 0.5 | 1 | 1 | 4 | 1 | 13 | 0.7692 | 5 | 0.8 | 1 | 1 | 5 | 0.8 | 1 | 0 | 12 | 0.75 |
| 3_8966316 | 4 | 0.25 | 5 | 0 | 1 | 0 | 3 | 0.3333 | 13 | 0.1538 | 6 | 0.1667 | 2 | 0 | 4 | 0 | NA | NA | 12 | 0.0833 |
| 3_8966330 | 4 | 0 | 4 | 0 | 1 | 1 | 2 | 0 | 11 | 0.0909 | 4 | 0.25 | 2 | 0 | 4 | 0 | 1 | 0 | 11 | 0.0909 |
| 3_8966333 | 5 | 0.2 | 4 | 0.25 | 1 | 0 | 1 | 0 | 11 | 0.1818 | 3 | 0 | 2 | 0 | 5 | 0.2 | NA | NA | 10 | 0.1 |
| 3_8966359 | 3 | 0.6667 | 1 | 0 | 1 | 0 | 3 | 0.6667 | 8 | 0.5 | 4 | 0.5 | 2 | 1 | 3 | 0.6667 | 1 | 1 | 10 | 0.7 |
| 3_8966363 | 4 | 0.25 | 2 | 0 | 1 | 1 | 2 | 0.5 | 9 | 0.3333 | 4 | 0.5 | 2 | 0.5 | 4 | 0.75 | 1 | 1 | 11 | 0.6364 |
| 3_8966364 | 4 | 1 | 2 | 0.5 | 1 | 0 | 1 | 0 | 8 | 0.625 | 4 | 0.75 | 2 | 0.5 | 5 | 1 | 1 | 0 | 12 | 0.75 |
| 3_8966373 | 4 | 0.5 | 1 | 0 | 1 | 0 | 2 | 0 | 8 | 0.25 | 4 | 0 | 2 | 0 | 4 | 0.75 | 1 | 0 | 11 | 0.2727 |
| 3_8966427 | 3 | 0 | 1 | 0 | 1 | 1 | 2 | 0 | 7 | 0.1429 | 4 | 0 | 2 | 0 | 3 | 0.3333 | 1 | 0 | 10 | 0.1 |
| 3_8966446 | 4 | 0.5 | NA | NA | 1 | 0 | 2 | 0 | 7 | 0.2857 | 2 | 0 | 1 | 1 | 2 | 0.5 | NA | NA | 5 | 0.4 |
| 3_8966449 | 3 | 0.6667 | 1 | 1 | 1 | 1 | 2 | 0.5 | 7 | 0.7143 | 3 | 1 | 1 | 1 | 1 | 1 | NA | NA | 5 | 1 |
| 3_8966801 | 3 | 0.3333 | NA | NA | 1 | 0 | 1 | 0 | 5 | 0.2 | 3 | 0.3333 | 2 | 0 | 3 | 0.3333 | NA | NA | 8 | 0.25 |
| 3_8966851 | 1 | 1 | 1 | 0 | 1 | 1 | 1 | 1 | 4 | 0.75 | 3 | 0.3333 | 2 | 1 | 2 | 0.5 | NA | NA | 7 | 0.5714 |
| 3_8966853 | 1 | 1 | 1 | 1 | 1 | 1 | 1 | 1 | 4 | 1 | 3 | 1 | 2 | 1 | 2 | 1 | NA | NA | 7 | 1 |
| 3_8966870 | 1 | 0 | 1 | 1 | NA | NA | NA | NA | 2 | 0.5 | 3 | 0.3333 | 2 | 0 | 2 | 0 | NA | NA | 7 | 0.1429 |
| 3_8967516 | NA | NA | 1 | 1 | 2 | 0.5 | NA | NA | 3 | 0.6667 | NA | NA | NA | NA | NA | NA | 1 | 1 | 1 | 1 |
| 3_8967581 | NA | NA | 2 | 0.5 | 2 | 0 | NA | NA | 4 | 0.25 | NA | NA | NA | NA | 1 | 1 | 1 | 0 | 2 | 0.5 |
| 3_8967600 | NA | NA | 2 | 0 | 2 | 0.5 | NA | NA | 4 | 0.25 | NA | NA | NA | NA | 1 | 1 | 1 | 0 | 2 | 0.5 |
| 3_8967865 | NA | NA | 1 | 1 | 1 | 0 | 2 | 0.5 | 4 | 0.5 | NA | NA | NA | NA | 1 | 0 | 1 | 1 | 2 | 0.5 |
| 3_8967880 | NA | NA | 1 | 0 | 1 | 0 | 2 | 0.5 | 4 | 0.25 | NA | NA | NA | NA | 1 | 1 | 1 | 0 | 2 | 0.5 |
| 3_8968973 | 12 | 0.4167 | 12 | 0.5 | 12 | 0.75 | 4 | 0.5 | 40 | 0.55 | 10 | 0.7 | 12 | 0.5 | 11 | 0.5455 | 11 | 0.4545 | 44 | 0.5455 |
| 3_8968978 | 11 | 0.8182 | 10 | 1 | 10 | 0.7 | 1 | 1 | 32 | 0.8438 | 10 | 1 | 9 | 0.7778 | 6 | 0.8333 | 11 | 0.9091 | 36 | 0.8889 |
| 3_8968980 | 11 | 0.1818 | 13 | 0.2308 | 12 | 0.0833 | 5 | 0 | 41 | 0.1463 | 11 | 0.2727 | 13 | 0.1538 | 11 | 0.1818 | 10 | 0 | 45 | 0.1556 |
| 3_8968986 | 9 | 0.3333 | 11 | 0.4545 | 14 | 0.6429 | 3 | 0.6667 | 37 | 0.5135 | 11 | 0.4545 | 12 | 0.3333 | 5 | 0.2 | 10 | 0.5 | 38 | 0.3947 |
| 3_8969029 | 2 | 0 | 2 | 0 | 10 | 0.1 | 1 | 0 | 15 | 0.0667 | 3 | 0 | 4 | 0.25 | 5 | 0 | 2 | 0.5 | 14 | 0.1429 |
| 3_8969037 | 2 | 0.5 | 2 | 0 | 9 | 0.4444 | 1 | 0 | 14 | 0.3571 | 3 | 0.3333 | 2 | 0 | 6 | 0.3333 | 4 | 0 | 15 | 0.2 |
| 3_8969040 | 2 | 1 | 2 | 0.5 | 8 | 0.625 | 1 | 1 | 13 | 0.6923 | 3 | 0.6667 | 2 | 1 | 1 | 0 | 3 | 1 | 9 | 0.7778 |
| 3_8969055 | 4 | 1 | 2 | 0.5 | 9 | 1 | NA | NA | 15 | 0.9333 | 3 | 1 | 3 | 1 | 3 | 0.6667 | 4 | 0.5 | 13 | 0.7692 |
| 3_8969056 | 1 | 1 | 2 | 0.5 | 9 | 0.7778 | 1 | 1 | 13 | 0.7692 | 2 | 1 | 1 | 1 | 4 | 0.75 | 3 | 1 | 10 | 0.9 |
| 3_8969064 | 3 | 0 | 3 | 0 | 9 | 0.1111 | 2 | 0 | 17 | 0.0588 | 3 | 0.3333 | 3 | 0 | 6 | 0.1667 | 3 | 0.3333 | 15 | 0.2 |
| 3_8969066 | 3 | 0 | 3 | 0 | 8 | 0.375 | 1 | 0 | 15 | 0.2 | 2 | 0 | 3 | 0.3333 | 6 | 0.1667 | 4 | 0.5 | 15 | 0.2667 |
| 3_8969071 | 3 | 0.3333 | 3 | 0.3333 | 8 | 0.25 | 2 | 0 | 16 | 0.25 | 3 | 0 | 3 | 0 | 5 | 0.4 | 3 | 0.3333 | 14 | 0.2143 |
| 3_8969072 | 3 | 0.3333 | 4 | 0.25 | 10 | 0.5 | 2 | 0 | 19 | 0.3684 | 2 | 0.5 | 4 | 0.5 | 7 | 0.2857 | 3 | 0 | 16 | 0.3125 |
| 3_8969079 | 3 | 0 | 4 | 0 | 10 | 0.1 | 2 | 0 | 19 | 0.0526 | 3 | 0 | 3 | 0 | 7 | 0 | 4 | 0.25 | 17 | 0.0588 |
| 3_8969080 | 3 | 0 | 4 | 0.25 | 10 | 0.1 | 2 | 0.5 | 19 | 0.1579 | 3 | 0 | 3 | 0 | 7 | 0.1429 | 4 | 0 | 17 | 0.0588 |
| 3_8969082 | 4 | 0.25 | 4 | 0.5 | 7 | 0.2857 | 2 | 0.5 | 17 | 0.3529 | 3 | 0 | 2 | 0 | 5 | 0 | 2 | 0.5 | 12 | 0.0833 |
| 3_8969087 | 3 | 0.3333 | 4 | 0.5 | 7 | 0.2857 | 2 | 0.5 | 16 | 0.375 | 3 | 0.3333 | 4 | 0 | 4 | 0.25 | 3 | 0 | 14 | 0.1429 |
| 3_8969106 | 3 | 0.6667 | 4 | 0 | 10 | 0 | 1 | 0 | 18 | 0.1111 | 3 | 0 | 4 | 0.25 | 7 | 0.2857 | 4 | 0 | 18 | 0.1667 |
| 3_8969110 | 2 | 0.5 | 4 | 1 | 8 | 0.875 | 2 | 1 | 16 | 0.875 | 2 | 0.5 | 3 | 0.6667 | 6 | 0.5 | 3 | 1 | 14 | 0.6429 |
| 3_8969114 | 1 | 1 | 3 | 0.3333 | 9 | 0.4444 | 1 | 1 | 14 | 0.5 | 2 | 0.5 | 4 | 0.5 | 5 | 0.4 | 3 | 0.3333 | 14 | 0.4286 |
| 3_8977505 | 1 | 0 | 1 | 1 | NA | NA | NA | NA | 2 | 0.5 | NA | NA | NA | NA | 1 | 1 | NA | NA | 1 | 1 |

| | | | | | | | | | | | | | | | | | | | | |
|---|---|---|---|---|---|---|---|---|---|---|---|---|---|---|---|---|---|---|---|---|
| 3_8984769 | 6 | 0 | 4 | 0 | 3 | 0 | 6 | 0.1667 | 19 | 0.0526 | 1 | 0 | 3 | 0 | 5 | 0.2 | 1 | 0 | 10 | 0.1 |
| 3_8984782 | 7 | 0 | 6 | 0.3333 | 5 | 0 | 9 | 0.1111 | 27 | 0.1111 | 1 | 0 | 3 | 0 | 6 | 0.1667 | 4 | 0.25 | 14 | 0.1429 |
| 3_8984808 | 4 | 0.75 | 3 | 0.3333 | 5 | 0 | 9 | 0.5556 | 21 | 0.4286 | 2 | 0.5 | 4 | 0.25 | 9 | 0 | 4 | 0.5 | 19 | 0.2105 |
| 3_8984818 | 6 | 0.3333 | 6 | 0.3333 | 5 | 0.4 | 8 | 0.25 | 25 | 0.32 | 2 | 0 | 4 | 0 | 8 | 0.375 | 4 | 0.75 | 18 | 0.3333 |
| 3_8984823 | 6 | 0.8333 | 4 | 1 | 4 | 1 | 5 | 1 | 19 | 0.9474 | 2 | 0.5 | 2 | 1 | 6 | 1 | 4 | 1 | 14 | 0.9286 |
| 3_8984826 | 4 | 0.25 | 4 | 0.25 | 3 | 0.6667 | 7 | 0.2857 | 18 | 0.3333 | 1 | 0 | 3 | 0.3333 | 7 | 0.2857 | 4 | 0.25 | 15 | 0.2667 |
| 3_8984830 | 5 | 0.4 | 5 | 0.8 | 5 | 0.8 | 8 | 0.75 | 23 | 0.6957 | 2 | 1 | 3 | 0 | 8 | 0.625 | 4 | 0.25 | 17 | 0.4706 |
| 3_8984831 | 6 | 0.3333 | 4 | 0 | 5 | 0 | 8 | 0 | 23 | 0.087 | 2 | 0 | 3 | 0.3333 | 9 | 0 | 5 | 0 | 19 | 0.0526 |
| 3_8984835 | 6 | 0.6667 | 4 | 0.75 | 4 | 1 | 8 | 0.75 | 22 | 0.7727 | 2 | 1 | 3 | 1 | 5 | 1 | 3 | 0.6667 | 13 | 0.9231 |
| 3_8984840 | 5 | 0 | 3 | 0.3333 | 3 | 0.3333 | 8 | 0 | 19 | 0.1053 | 2 | 0.5 | 3 | 0 | 7 | 0 | 4 | 0.25 | 16 | 0.125 |
| 3_8984842 | 6 | 0.3333 | 1 | 0 | 4 | 0.25 | 8 | 0.25 | 19 | 0.2632 | 2 | 0.5 | 3 | 0.3333 | 9 | 0.4444 | 3 | 0.3333 | 17 | 0.4118 |
| 3_8984844 | 5 | 0.2 | 3 | 0 | 5 | 0 | 8 | 0.25 | 21 | 0.1429 | 2 | 0 | 3 | 0 | 9 | 0 | 4 | 0.25 | 18 | 0.0556 |
| 3_8984850 | 4 | 0.75 | 1 | 1 | 4 | 0.25 | 7 | 0.5714 | 16 | 0.5625 | 2 | 0.5 | 2 | 0.5 | 9 | 0.6667 | 3 | 1 | 16 | 0.6875 |
| 3_8984859 | 5 | 0 | 2 | 0 | 4 | 0 | 8 | 0.375 | 19 | 0.1579 | 2 | 1 | 3 | 0.3333 | 9 | 0 | 3 | 0.6667 | 17 | 0.2941 |
| 3_8984876 | NA | NA | NA | NA | NA | NA | 1 | 1 | 1 | 1 | NA | NA | 1 | 0 | 2 | 0.5 | 1 | 1 | 4 | 0.5 |
| 3_8984887 | NA | NA | NA | NA | NA | NA | 1 | 1 | 1 | 1 | NA | NA | 1 | 1 | 1 | 1 | NA | NA | 2 | 1 |
| 3_8985357 | 2 | 0 | 2 | 0 | 1 | 0 | 5 | 0.2 | 10 | 0.1 | 1 | 0 | 3 | 0.3333 | 5 | 0 | 3 | 0 | 12 | 0.0833 |
| 3_8990484 | 8 | 0.375 | 9 | 0.2222 | 6 | 0.1667 | 10 | 0.1 | 33 | 0.2121 | 10 | 0.1 | 7 | 0.2857 | 7 | 0.4286 | 11 | 0.1818 | 35 | 0.2286 |
| 3_8990493 | NA | NA | 2 | 0 | 2 | 0.5 | NA | NA | 4 | 0.25 | NA | NA | NA | NA | 2 | 0.5 | 1 | 1 | 3 | 0.6667 |
| 3_8991371 | 3 | 0 | 3 | 0 | 1 | 0 | 1 | 1 | 8 | 0.125 | 2 | 0 | 6 | 0.1667 | 6 | 0 | 2 | 0 | 16 | 0.0625 |
| 3_8996873 | 1 | 0 | 2 | 0 | 1 | 0 | 2 | 0.5 | 6 | 0.1667 | NA | NA | 1 | 0 | 2 | 0.5 | 3 | 0 | 6 | 0.1667 |
| 3_8996910 | 7 | 0.2857 | 21 | 0.1905 | 11 | 0.1818 | 10 | 0.6 | 49 | 0.2857 | 10 | 0.4 | 10 | 0.2 | 7 | 0.4286 | 11 | 0.4545 | 38 | 0.3684 |
| 3_8996922 | 6 | 0.6667 | 11 | 0.6364 | 10 | 0.2 | 6 | 0.6667 | 33 | 0.5152 | 6 | 0.8333 | 5 | 1 | 7 | 0.7143 | 8 | 0.625 | 26 | 0.7692 |
| 3_8997291 | 1 | 1 | 3 | 0 | NA | NA | 1 | 0 | 5 | 0.2 | NA | NA | 3 | 0.6667 | 1 | 1 | 2 | 1 | 6 | 0.8333 |
| 3_8997300 | 1 | 0 | 4 | 0 | NA | NA | 2 | 0.5 | 7 | 0.1429 | NA | NA | 3 | 0.6667 | 1 | 0 | 2 | 0.5 | 6 | 0.5 |
| 3_8997301 | 1 | 1 | 3 | 0.3333 | NA | NA | 2 | 0 | 6 | 0.3333 | NA | NA | 2 | 0 | 1 | 1 | 1 | 1 | 4 | 0.5 |
| 3_8997305 | 1 | 0 | 3 | 0 | NA | NA | 1 | 1 | 5 | 0.2 | NA | NA | 3 | 0.6667 | NA | NA | 1 | 1 | 4 | 0.75 |
| 3_8997543 | NA | NA | 1 | 1 | NA | NA | NA | NA | 1 | 1 | NA | NA | 1 | 1 | NA | NA | NA | NA | 1 | 1 |
| 3_9000575 | NA | NA | 4 | 0.25 | 2 | 0.5 | 7 | 0.7143 | 13 | 0.5385 | 2 | 0.5 | 5 | 0.4 | 4 | 0.25 | 1 | 0 | 12 | 0.3333 |
| 3_9000588 | NA | NA | 5 | 0 | 1 | 0 | 6 | 0.1667 | 12 | 0.0833 | 3 | 0 | 4 | 0.25 | 3 | 0 | 2 | 0 | 12 | 0.0833 |
| 3_9000599 | NA | NA | 5 | 0 | 3 | 0 | 10 | 0.1 | 18 | 0.0556 | 3 | 0.3333 | 6 | 0 | 5 | 0 | 2 | 0 | 16 | 0.0625 |
| 3_9000603 | NA | NA | 5 | 0.2 | 3 | 0 | 8 | 0.125 | 16 | 0.125 | 3 | 0.3333 | 6 | 0 | 5 | 0.2 | 2 | 0 | 16 | 0.125 |
| 3_9000700 | 12 | 0.1667 | 18 | 0 | 15 | 0 | 18 | 0.3333 | 63 | 0.127 | 10 | 0.1 | 12 | 0.0833 | 20 | 0.05 | 17 | 0.2353 | 59 | 0.1186 |
| 3_9000708 | 12 | 0.25 | 16 | 0.1875 | 14 | 0.0714 | 14 | 0 | 56 | 0.125 | 10 | 0 | 13 | 0.2308 | 20 | 0.15 | 15 | 0.1333 | 58 | 0.1379 |
| 3_9000743 | 9 | 0.3333 | 18 | 0.0556 | 10 | 0.1 | 13 | 0.0769 | 50 | 0.12 | 10 | 0.1 | 13 | 0.2308 | 17 | 0.2941 | 10 | 0.2 | 50 | 0.22 |
| 3_9013294 | NA | NA | 1 | 1 | 3 | 0.3333 | 2 | 1 | 6 | 0.6667 | NA | NA | NA | NA | 1 | 1 | NA | NA | 1 | 1 |
| 3_9013297 | NA | NA | 1 | 1 | 3 | 0.6667 | 2 | 1 | 6 | 0.8333 | NA | NA | NA | NA | 1 | 1 | NA | NA | 1 | 1 |
| 3_9013313 | NA | NA | 1 | 0 | 2 | 1 | NA | NA | 3 | 0.6667 | NA | NA | NA | NA | 1 | 1 | NA | NA | 1 | 1 |
| 3_9013919 | 5 | 0.2 | 7 | 0.2857 | 5 | 0.2 | 6 | 0.3333 | 23 | 0.2609 | 4 | 0.5 | 4 | 0.75 | 3 | 0 | 4 | 0.25 | 15 | 0.4 |
| 3_9014280 | 4 | 0.5 | 7 | 0.2857 | 8 | 0.25 | 8 | 0 | 27 | 0.2222 | 3 | 0 | 4 | 0 | 7 | 0.2857 | 7 | 0 | 21 | 0.0952 |
| 3_9014288 | 6 | 1 | 9 | 1 | 12 | 0.9167 | 8 | 0.75 | 35 | 0.9143 | 4 | 0.75 | 6 | 0.6667 | 9 | 0.7778 | 6 | 0.8333 | 25 | 0.76 |
| 3_9014295 | 6 | 0.3333 | 9 | 0.4444 | 13 | 0.3846 | 8 | 0.25 | 36 | 0.3611 | 4 | 0.25 | 7 | 0 | 9 | 0.1111 | 8 | 0.25 | 28 | 0.1429 |
| 3_9014300 | 6 | 0 | 8 | 0.25 | 11 | 0.2727 | 6 | 0.3333 | 31 | 0.2258 | 3 | 0.6667 | 9 | 0.2222 | 8 | 0.25 | 6 | 0.1667 | 26 | 0.2692 |
| 3_9014301 | 7 | 0.1429 | 9 | 0.2222 | 12 | 0.4167 | 8 | 0.25 | 36 | 0.2778 | 4 | 0.5 | 9 | 0 | 10 | 0.1 | 7 | 0.5714 | 30 | 0.2333 |
| 3_9015916 | 2 | 0.5 | 6 | 0.1667 | 6 | 0.3333 | 2 | 0.5 | 16 | 0.3125 | 1 | 0 | 3 | 0.3333 | 2 | 0 | 1 | 0 | 7 | 0.1429 |
| 3_9015955 | 1 | 0 | 8 | 0.125 | 3 | 0 | 2 | 0 | 14 | 0.0714 | 2 | 0 | 3 | 0.3333 | NA | NA | 2 | 0 | 7 | 0.1429 |
| 3_9015980 | 3 | 0.6667 | 8 | 0.25 | 4 | 0.5 | 3 | 0 | 18 | 0.3333 | 4 | 0.5 | 4 | 0.75 | 3 | 0.3333 | 2 | 0.5 | 13 | 0.5385 |
| 3_9016013 | 3 | 0.3333 | 10 | 0 | 6 | 0.1667 | 4 | 0 | 23 | 0.087 | 4 | 0.25 | 3 | 0.3333 | 3 | 0 | 3 | 0 | 13 | 0.1538 |
| 3_9016066 | 5 | 0 | 9 | 0.2222 | 7 | 0 | 4 | 0 | 25 | 0.08 | 4 | 0.25 | 4 | 0.25 | 2 | 0 | 7 | 0 | 17 | 0.1176 |
| 3_9017092 | 1 | 1 | 2 | 1 | 5 | 0.4 | 2 | 0 | 10 | 0.5 | 2 | 0 | 6 | 0.1667 | 4 | 0.75 | 1 | 0 | 13 | 0.3077 |
| 3_9017097 | 2 | 0 | 2 | 0.5 | 5 | 0.8 | 1 | 0 | 10 | 0.5 | 2 | 1 | 6 | 0.1667 | 4 | 0.25 | NA | NA | 12 | 0.3333 |
| 3_9017108 | 2 | 0.5 | 3 | 0 | 5 | 0.2 | 1 | 1 | 11 | 0.2727 | 2 | 0.5 | 6 | 0.5 | 4 | 0.5 | 1 | 0 | 13 | 0.4615 |
| 3_9017113 | 1 | 0 | 3 | 0.3333 | 4 | 0 | 1 | 0 | 9 | 0.1111 | 2 | 0 | 5 | 0.2 | 3 | 0 | NA | NA | 10 | 0.1 |
| 3_9018737 | 3 | 0.3333 | 1 | 0 | 1 | 0 | 3 | 0 | 8 | 0.125 | 3 | 1 | 4 | 0.5 | 5 | 0.2 | 1 | 0 | 13 | 0.4615 |
| 3_9018782 | 3 | 0.3333 | NA | NA | 2 | 0.5 | 2 | 0 | 7 | 0.2857 | 2 | 0 | 4 | 0 | 6 | 0.1667 | 1 | 1 | 13 | 0.1538 |
| 3_9025920 | NA | NA | 1 | 0 | 2 | 0.5 | NA | NA | 3 | 0.3333 | NA | NA | 1 | 0 | 1 | 1 | NA | NA | 2 | 0.5 |
| 3_9026545 | NA | NA | 3 | 0.3333 | 4 | 0.25 | NA | NA | 7 | 0.2857 | 1 | 1 | 2 | 0 | 1 | 0 | 1 | 0 | 5 | 0.2 |
| 3_9026880 | 4 | 0.5 | 7 | 0.2857 | 8 | 0.5 | 7 | 0.2857 | 26 | 0.3846 | 2 | 0 | 6 | 0.3333 | 6 | 0.3333 | 6 | 0.6667 | 20 | 0.4 |
| 3_9026945 | NA | NA | 4 | 0.25 | 4 | 0 | 6 | 0.1667 | 14 | 0.1429 | 3 | 0.3333 | 4 | 0 | 3 | 0 | 5 | 0 | 15 | 0.0667 |
| 3_9027054 | 7 | 0.1429 | 9 | 0.3333 | 6 | 0.1667 | 7 | 0.1429 | 29 | 0.2069 | 6 | 0.1667 | 7 | 0.2857 | 7 | 0.1429 | 9 | 0.1111 | 29 | 0.1724 |
| 3_9029881 | 7 | 0.4286 | 8 | 0.75 | 7 | 0.7143 | 4 | 0 | 26 | 0.5385 | 5 | 0.2 | 5 | 0.6 | 9 | 0.6667 | 7 | 0.4286 | 26 | 0.5 |
| 3_9029892 | 9 | 0 | 10 | 0.2 | 6 | 0 | 4 | 0 | 29 | 0.069 | 6 | 0 | 6 | 0 | 9 | 0.1111 | 7 | 0.4286 | 28 | 0.1429 |
| 3_9029912 | 6 | 0.8333 | 9 | 0.4444 | 5 | 0.6 | 2 | 0 | 22 | 0.5455 | 5 | 0.8 | 4 | 0.75 | 9 | 0.3333 | 6 | 0.8333 | 24 | 0.625 |
| 3_9029933 | 8 | 0.125 | 10 | 0.3 | 5 | 0.2 | 3 | 0 | 26 | 0.1923 | 5 | 0.8 | 6 | 0 | 8 | 0.5 | 8 | 0.125 | 27 | 0.3333 |
| 3_9030010 | 9 | 0.2222 | 7 | 0 | 3 | 0 | 3 | 0 | 22 | 0.0909 | 5 | 0.4 | 6 | 0 | 7 | 0 | 7 | 0 | 25 | 0.08 |

| | | | | | | | | | | | | | | | | | | | | |
|---|---|---|---|---|---|---|---|---|---|---|---|---|---|---|---|---|---|---|---|---|
| 3_90340308 | 2 | 1 | NA | NA | NA | NA | NA | NA | 2 | 1 | NA | NA | NA | NA | 2 | 0.5 | 2 | 0.5 | 4 | 0.5 |
| 3_90340630 | 1 | 0 | NA | NA | 1 | 1 | NA | NA | 2 | 0.5 | 1 | 0 | NA | NA | 2 | 0 | 2 | 1 | 5 | 0.4 |
| 3_90394915 | 11 | 0.0909 | 14 | 0 | 9 | 0 | 23 | 0.087 | 57 | 0.0526 | 4 | 0 | 12 | 0 | 9 | 0.2222 | 7 | 0 | 32 | 0.0625 |
| 3_90395035 | 10 | 0.4 | 13 | 0.3846 | 9 | 0.6667 | 20 | 0.4 | 52 | 0.4423 | 4 | 0.75 | 13 | 0.3846 | 8 | 0.375 | 9 | 0.3333 | 34 | 0.4118 |
| 3_90395099 | 10 | 0.4 | 11 | 0.3636 | 8 | 0.625 | 21 | 0.381 | 50 | 0.42 | 5 | 0.8 | 12 | 0.25 | 9 | 0.3333 | 9 | 0.5556 | 35 | 0.4286 |
| 3_90395525 | 5 | 0 | 1 | 1 | NA | NA | 7 | 0.1429 | 13 | 0.1538 | 1 | 0 | 8 | 0.25 | 3 | 0 | 2 | 0 | 14 | 0.1429 |
| 3_90395790 | 7 | 0 | 3 | 0 | 7 | 0.1429 | 5 | 0.2 | 22 | 0.0909 | 10 | 0 | 3 | 0.3333 | 7 | 0.2857 | 7 | 0.2857 | 27 | 0.1852 |
| 3_90396643 | 5 | 0.4 | 6 | 0.6667 | 8 | 0.75 | 7 | 0.8571 | 26 | 0.6923 | 9 | 0.8889 | 3 | 1 | 6 | 1 | 5 | 1 | 23 | 0.9565 |
| 3_90396710 | 4 | 0.25 | 6 | 0.1667 | 7 | 0.1429 | 7 | 0.2857 | 24 | 0.2083 | 10 | 0.3 | 3 | 0.3333 | 8 | 0.25 | 5 | 0.2 | 26 | 0.2692 |
| 3_90406436 | 29 | 0.3448 | 15 | 0.4667 | 27 | 0.2963 | 27 | 0.3704 | 98 | 0.3571 | 11 | 0.3636 | 23 | 0.4348 | 23 | 0.4783 | 19 | 0.2632 | 76 | 0.3947 |
| 3_90406440 | 24 | 0.1667 | 21 | 0 | 25 | 0 | 25 | 0.04 | 95 | 0.0526 | 9 | 0 | 23 | 0.0435 | 24 | 0.125 | 19 | 0.0526 | 75 | 0.0667 |
| 3_90407082 | 32 | 1 | 17 | 0.9412 | 29 | 0.9655 | 23 | 0.913 | 101 | 0.9604 | 8 | 1 | 30 | 1 | 21 | 1 | 21 | 0.9048 | 80 | 0.975 |
| 3_90407103 | 27 | 0.3333 | 15 | 0.2 | 23 | 0.5217 | 22 | 0.5455 | 87 | 0.4138 | 7 | 0.4286 | 28 | 0.5357 | 18 | 0.5 | 17 | 0.5294 | 70 | 0.5143 |
| 3_90407178 | 31 | 0.8387 | 18 | 0.9444 | 29 | 0.7241 | 22 | 0.8182 | 100 | 0.82 | 10 | 0.5 | 28 | 0.9286 | 21 | 0.8571 | 20 | 0.7 | 79 | 0.7975 |
| 3_90407232 | 31 | 0.129 | 18 | 0.1667 | 28 | 0.1786 | 23 | 0.087 | 100 | 0.14 | 10 | 0.1 | 26 | 0.0769 | 22 | 0.0909 | 20 | 0.05 | 78 | 0.0769 |
| 3_90407269 | 26 | 0.3462 | 18 | 0.6111 | 26 | 0.5 | 19 | 0.4211 | 89 | 0.4607 | 10 | 0.3 | 26 | 0.5 | 17 | 0.4118 | 18 | 0.4444 | 71 | 0.4366 |
| 3_90414796 | 3 | 0.3333 | 5 | 0 | 5 | 0 | 3 | 0 | 16 | 0.0625 | 1 | 0 | 6 | 0.1667 | 2 | 0 | 2 | 0 | 11 | 0.0909 |
| 3_90418601 | 21 | 0.2381 | 21 | 0.0952 | 28 | 0.2143 | 19 | 0.1053 | 89 | 0.1685 | 29 | 0.069 | 14 | 0.3571 | 19 | 0.1579 | 20 | 0.15 | 82 | 0.1585 |
| 3_90431709 | 14 | 0 | 4 | 0.25 | 6 | 0.1667 | 9 | 0.1111 | 33 | 0.0909 | 14 | 0 | 9 | 0.1111 | 9 | 0.2222 | 8 | 0.125 | 40 | 0.1 |
| 3_90449964 | 13 | 0.2308 | 26 | 0.1154 | 13 | 0.3077 | 9 | 0.1111 | 61 | 0.1803 | 12 | 0.0833 | 15 | 0.1333 | 17 | 0.0588 | 9 | 0.1111 | 53 | 0.0943 |
| 3_91987631 | NA | NA | 5 | 0.4 | 2 | 0 | 4 | 0 | 11 | 0.1818 | 3 | 0.3333 | 2 | 0 | 5 | 0.4 | 2 | 0 | 12 | 0.25 |
| 3_91987759 | NA | NA | 5 | 0.2 | 2 | 0 | 3 | 1 | 10 | 0.4 | 3 | 0 | 2 | 0.5 | 2 | 0.5 | 2 | 0.5 | 9 | 0.3333 |
| 3_91989281 | 1 | 0 | 3 | 0.3333 | NA | NA | NA | NA | 4 | 0.25 | NA | NA | NA | NA | 1 | 0 | 1 | 1 | 2 | 0.5 |
| 3_91989429 | NA | NA | 2 | 1 | NA | NA | NA | NA | 2 | 1 | NA | NA | NA | NA | 1 | 1 | 1 | 1 | 2 | 1 |
| 3_91989442 | NA | NA | 1 | 1 | NA | NA | NA | NA | 1 | 1 | NA | NA | NA | NA | NA | NA | 1 | 1 | 1 | 1 |
| 3_91989691 | NA | NA | 2 | 1 | NA | NA | NA | NA | 2 | 1 | NA | NA | NA | NA | NA | NA | 1 | 1 | 1 | 1 |
| 3_91990297 | NA | NA | 1 | 1 | NA | NA | NA | NA | 1 | 1 | NA | NA | NA | NA | NA | NA | 1 | 1 | 1 | 1 |
| 3_91990453 | NA | NA | 3 | 1 | NA | NA | NA | NA | 3 | 1 | NA | NA | NA | NA | NA | NA | 1 | 1 | 1 | 1 |
| 3_91990493 | NA | NA | 3 | 1 | NA | NA | NA | NA | 3 | 1 | NA | NA | NA | NA | NA | NA | 1 | 1 | 1 | 1 |
| 3_91990599 | NA | NA | 2 | 0.5 | NA | NA | NA | NA | 2 | 0.5 | NA | NA | NA | NA | NA | NA | 1 | 1 | 1 | 1 |
| 3_91990777 | NA | NA | 3 | 1 | NA | NA | NA | NA | 3 | 1 | NA | NA | NA | NA | NA | NA | 1 | 1 | 1 | 1 |
| 3_91991190 | NA | NA | 3 | 0.3333 | NA | NA | NA | NA | 3 | 0.3333 | NA | NA | NA | NA | NA | NA | 1 | 1 | 1 | 1 |
| 3_91991227 | NA | NA | 2 | 1 | NA | NA | NA | NA | 2 | 1 | NA | NA | NA | NA | NA | NA | 1 | 1 | 1 | 1 |
| 3_91991833 | NA | NA | 2 | 0.5 | NA | NA | NA | NA | 2 | 0.5 | NA | NA | NA | NA | NA | NA | 1 | 1 | 1 | 1 |
| 3_91992083 | NA | NA | 3 | 0.3333 | NA | NA | NA | NA | 3 | 0.3333 | NA | NA | NA | NA | NA | NA | 2 | 0.5 | 2 | 0.5 |
| 3_93353797 | NA | NA | 1 | 1 | NA | NA | NA | NA | 1 | 1 | NA | NA | NA | NA | NA | NA | 1 | 1 | 1 | 1 |
| 3_93353894 | NA | NA | 1 | 1 | NA | NA | NA | NA | 1 | 1 | NA | NA | NA | NA | NA | NA | 1 | 1 | 1 | 1 |
| 3_93356351 | NA | NA | 1 | 1 | 2 | 0.5 | NA | NA | 3 | 0.6667 | NA | NA | 1 | 1 | NA | NA | 2 | 0 | 3 | 0.3333 |
| 3_93471806 | 2 | 0 | 17 | 0.1176 | 5 | 0 | 1 | 0 | 25 | 0.08 | 7 | 0 | 1 | 0 | 5 | 0 | 5 | 0.2 | 18 | 0.0556 |
| 3_94238892 | 1 | 0 | 1 | 0 | 5 | 0.2 | 3 | 0 | 10 | 0.1 | 3 | 0 | 2 | 0 | 3 | 0.3333 | 4 | 0 | 12 | 0.0833 |
| 3_94239041 | 2 | 1 | 1 | 0 | 4 | 0.75 | 3 | 1 | 10 | 0.8 | 4 | 0.75 | 2 | 1 | 3 | 1 | 5 | 0.8 | 14 | 0.8571 |
| 3_94239119 | 2 | 1 | NA | NA | 5 | 0.6 | 4 | 0.5 | 11 | 0.6364 | 4 | 0.75 | 2 | 1 | 3 | 0.6667 | 5 | 0.8 | 14 | 0.7857 |
| 3_94239148 | 1 | 1 | 1 | 0 | 5 | 1 | 3 | 1 | 10 | 0.9 | 3 | 1 | 2 | 1 | 2 | 0.5 | 4 | 1 | 11 | 0.9091 |
| 3_94239171 | 2 | 0.5 | 1 | 0 | 5 | 0.2 | 4 | 0.5 | 12 | 0.3333 | 3 | 0 | 2 | 0 | 2 | 0.5 | 5 | 0 | 12 | 0.0833 |
| 3_94239187 | 2 | 0.5 | 1 | 1 | 5 | 0.6 | 4 | 0.75 | 12 | 0.6667 | 4 | 0.75 | 2 | 1 | 3 | 0.3333 | 4 | 1 | 13 | 0.7692 |
| 3_94239219 | 2 | 1 | 1 | 0 | 5 | 1 | 3 | 0.6667 | 11 | 0.8182 | 4 | 1 | 2 | 1 | 3 | 1 | 5 | 1 | 14 | 1 |
| 3_94239230 | 1 | 0 | 1 | 1 | 4 | 0 | 4 | 0.25 | 10 | 0.2 | 4 | 0.5 | 2 | 0.5 | 3 | 0.6667 | 4 | 0.25 | 13 | 0.4615 |
| 3_94239236 | 2 | 0.5 | 1 | 1 | 5 | 0.4 | 3 | 0 | 11 | 0.3636 | 4 | 0 | 2 | 0.5 | 3 | 0 | 4 | 0.25 | 13 | 0.1538 |
| 3_94239530 | 1 | 1 | 1 | 1 | 3 | 0.6667 | 2 | 0 | 7 | 0.5714 | 3 | 0.6667 | NA | NA | 2 | 0 | 2 | 0.5 | 7 | 0.4286 |
| 3_94239543 | 1 | 1 | 1 | 0 | 4 | 0 | 3 | 0 | 9 | 0.1111 | 4 | 0 | NA | NA | 3 | 0.3333 | 3 | 0.3333 | 10 | 0.2 |
| 3_94239651 | 2 | 0.5 | 1 | 0 | 4 | 0 | 2 | 0 | 9 | 0.1111 | 4 | 0.5 | NA | NA | 3 | 0 | 3 | 0.6667 | 10 | 0.4 |
| 3_94333271 | 3 | 0.3333 | 1 | 1 | NA | NA | 3 | 0.3333 | 7 | 0.4286 | 4 | 0.75 | NA | NA | 1 | 1 | 5 | 0.8 | 10 | 0.8 |
| 3_94333331 | 2 | 0.5 | 1 | 0 | NA | NA | 3 | 0 | 6 | 0.1667 | 5 | 0.2 | NA | NA | 1 | 1 | 4 | 0.75 | 10 | 0.5 |
| 3_94338474 | 1 | 1 | NA | NA | 1 | 0 | 3 | 0.6667 | 5 | 0.6 | 2 | 0.5 | 2 | 0.5 | NA | NA | 4 | 0 | 8 | 0.25 |
| 3_94338501 | 1 | 1 | NA | NA | 1 | 0 | 3 | 0.6667 | 5 | 0.6 | 2 | 0.5 | 2 | 0.5 | NA | NA | 4 | 0.5 | 8 | 0.5 |
| 3_94342053 | 3 | 1 | 1 | 1 | 2 | 1 | 2 | 1 | 8 | 1 | NA | NA | 1 | 1 | NA | NA | 2 | 1 | 3 | 1 |
| 3_94342155 | 2 | 0 | 2 | 0 | 1 | 0 | 2 | 0.5 | 7 | 0.1429 | NA | NA | 1 | 0 | NA | NA | 1 | 1 | 2 | 0.5 |
| 3_94355302 | 2 | 0 | 4 | 0 | 8 | 0 | 3 | 0.3333 | 17 | 0.0588 | NA | NA | 7 | 0.1429 | 7 | 0 | 5 | 0 | 19 | 0.0526 |
| 3_94355310 | 4 | 0.75 | 4 | 0.75 | 7 | 0.7143 | 1 | 0 | 16 | 0.6875 | 2 | 0.5 | 7 | 1 | 7 | 0.8571 | 4 | 1 | 20 | 0.9 |
| 3_94355319 | 6 | 0.1667 | 2 | 0 | 9 | 0.1111 | 3 | 0 | 20 | 0.1 | 2 | 0.5 | 7 | 0 | 8 | 0.25 | 4 | 0.25 | 21 | 0.1905 |
| 3_94355513 | 5 | 0 | 5 | 0 | 9 | 0.2222 | 3 | 0.3333 | 22 | 0.1364 | 2 | 0 | 7 | 0 | 3 | 0.3333 | 5 | 0 | 17 | 0.0588 |
| 3_94355536 | 6 | 0.5 | 6 | 0.5 | 9 | 0.1111 | 3 | 0 | 24 | 0.2917 | 2 | 0 | 7 | 0.1429 | 6 | 0.3333 | 4 | 0.25 | 19 | 0.2105 |
| 3_94355693 | 6 | 0.1667 | 6 | 0 | 6 | 0 | 2 | 0 | 20 | 0.05 | 1 | 1 | 6 | 0 | 5 | 0.2 | 5 | 0.2 | 17 | 0.1765 |
| 3_94355834 | 5 | 0.6 | 3 | 0 | 9 | 0.2222 | 1 | 1 | 18 | 0.3333 | 2 | 0.5 | 6 | 0.3333 | 7 | 0.1429 | 5 | 0.4 | 20 | 0.3 |
| 3_94395621 | 2 | 0.5 | 2 | 0 | 4 | 0.25 | 3 | 0 | 11 | 0.1818 | 2 | 0 | 2 | 0 | 3 | 0 | 2 | 0.5 | 9 | 0.1111 |

| | | | | | | | | | | | | | | | | | | | | |
|---|---|---|---|---|---|---|---|---|---|---|---|---|---|---|---|---|---|---|---|---|
| 3_94395702 | 2 | 1 | 2 | 1 | 4 | 0.75 | 3 | 0.6667 | 11 | 0.8182 | 1 | 1 | 2 | 0.5 | 2 | 0.5 | 2 | 1 | 7 | 0.7143 |
| 3_94395745 | 1 | 0 | 2 | 1 | 4 | 0 | 2 | 0.5 | 9 | 0.3333 | 1 | 0 | 1 | 1 | 3 | 0.3333 | 2 | 0.5 | 7 | 0.4286 |
| 3_94395885 | 7 | 0.1429 | 10 | 0.2 | 12 | 0.25 | 12 | 0.25 | 41 | 0.2195 | 6 | 0.3333 | 13 | 0.1538 | 12 | 0.25 | 7 | 0.4286 | 38 | 0.2632 |
| 3_94396001 | 1 | 1 | NA | NA | 4 | 0 | 1 | 0 | 6 | 0.1667 | 2 | 0 | 1 | 0 | 2 | 1 | 2 | 1 | 7 | 0.5714 |
| 3_94396012 | NA | NA | 1 | 1 | 4 | 1 | 2 | 1 | 7 | 1 | NA | NA | 1 | 1 | 2 | 1 | 2 | 1 | 5 | 1 |
| 3_94396026 | 1 | 0 | 1 | 0 | 1 | 0 | 2 | 0.5 | 5 | 0.2 | 2 | 1 | NA | NA | 2 | 0.5 | 1 | 1 | 5 | 0.8 |
| 3_94396035 | 1 | 1 | 1 | 1 | 4 | 0.5 | 1 | 1 | 7 | 0.7143 | 2 | 0.5 | 1 | 1 | 1 | 1 | 2 | 1 | 6 | 0.8333 |
| 3_94396076 | 1 | 1 | 1 | 1 | 4 | 0.75 | 1 | 1 | 7 | 0.8571 | 1 | 0 | 1 | 1 | 2 | 1 | 1 | 1 | 5 | 0.8 |
| 3_94396496 | 1 | 0 | 1 | 0 | 2 | 0.5 | 1 | 0 | 5 | 0.2 | 3 | 0.6667 | 1 | 1 | 2 | 0 | 2 | 1 | 8 | 0.625 |
| 3_94396594 | 2 | 0.5 | 1 | 0 | 2 | 0.5 | 2 | 0.5 | 7 | 0.4286 | 3 | 1 | 1 | 0 | 2 | 0 | 2 | 0.5 | 8 | 0.5 |
| 3_94398534 | 9 | 0.6667 | 7 | 0.7143 | 10 | 0.7 | 11 | 0.5455 | 37 | 0.6486 | 5 | 0.4 | 11 | 0.6364 | 9 | 0.4444 | 4 | 0.75 | 29 | 0.5517 |
| 3_94398814 | 13 | 0.1538 | 6 | 0.5 | 12 | 0.3333 | 13 | 0.1538 | 44 | 0.25 | 5 | 0.4 | 15 | 0.1333 | 15 | 0.2667 | 7 | 0.1429 | 42 | 0.2143 |
| 3_94408356 | 1 | 0 | 4 | 0 | 5 | 0 | 1 | 1 | 11 | 0.0909 | 6 | 0 | 9 | 0.2222 | 8 | 0.125 | 4 | 0.25 | 27 | 0.1481 |
| 3_94408864 | NA | NA | 4 | 0.25 | 4 | 0 | NA | NA | 8 | 0.125 | 2 | 0 | 1 | 0 | 2 | 0.5 | 1 | 0 | 6 | 0.1667 |
| 3_94409211 | NA | NA | 4 | 0.25 | 3 | 0.3333 | 1 | 0 | 8 | 0.25 | 4 | 0.25 | 3 | 0 | 3 | 0 | 2 | 0 | 12 | 0.0833 |
| 3_94409357 | NA | NA | 4 | 0.25 | 3 | 0.3333 | 1 | 0 | 8 | 0.25 | 4 | 0.5 | 2 | 0 | 3 | 0 | 2 | 0.5 | 11 | 0.2727 |
| 3_94469166 | NA | NA | 2 | 0 | 2 | 1 | NA | NA | 4 | 0.5 | 3 | 0 | 3 | 0.3333 | NA | NA | 4 | 0 | 10 | 0.1 |
| 3_94786210 | 2 | 1 | 3 | 0.6667 | 10 | 0.7 | 6 | 1 | 21 | 0.8095 | 2 | 1 | 5 | 1 | 10 | 0.8 | 2 | 1 | 19 | 0.8947 |
| 3_94786371 | 2 | 0 | 3 | 0.3333 | 7 | 0 | 5 | 0 | 17 | 0.0588 | 2 | 0 | 4 | 0.25 | 7 | 0 | 4 | 0.25 | 17 | 0.1176 |
| 3_94786383 | 1 | 1 | 1 | 0 | 5 | 0.6 | 5 | 0 | 12 | 0.3333 | 1 | 0 | 4 | 0.25 | 6 | 0.3333 | 1 | 1 | 12 | 0.3333 |
| 3_94786451 | 1 | 1 | 2 | 0.5 | 7 | 0.7143 | 5 | 0.6 | 15 | 0.6667 | 1 | 0 | 5 | 0.4 | 9 | 0.6667 | 3 | 0.3333 | 18 | 0.5 |
| 3_94786491 | 2 | 1 | 3 | 0.6667 | 7 | 0.1429 | 4 | 0 | 16 | 0.3125 | NA | NA | 5 | 0 | 8 | 0.5 | 3 | 0.3333 | 16 | 0.3125 |
| 3_94786546 | 2 | 0 | 3 | 0 | 10 | 0.1 | 5 | 0.2 | 20 | 0.1 | 3 | 0.3333 | 5 | 0 | 10 | 0.1 | 4 | 0 | 22 | 0.0909 |
| 3_94786707 | 3 | 1 | 3 | 1 | 9 | 0.8889 | 6 | 1 | 21 | 0.9524 | 2 | 1 | 4 | 1 | 11 | 1 | 3 | 0.6667 | 20 | 0.95 |
| 3_94786748 | 4 | 1 | 3 | 0.6667 | 10 | 0.8 | 5 | 0.6 | 22 | 0.7727 | 3 | 1 | 4 | 1 | 9 | 0.7778 | 4 | 0.75 | 20 | 0.85 |
| 3_94786799 | 1 | 1 | 2 | 0 | 5 | 0 | 6 | 0.3333 | 14 | 0.2143 | 3 | 0 | 3 | 0 | 7 | 0.1429 | 3 | 0.6667 | 16 | 0.1875 |
| 3_94786846 | 3 | 0.3333 | 3 | 0 | 9 | 0.3333 | 6 | 0 | 21 | 0.1905 | 2 | 0.5 | 5 | 0.2 | 7 | 0.2857 | 3 | 0 | 17 | 0.2353 |
| 3_94786935 | 4 | 0 | 3 | 0.6667 | 8 | 0.25 | 5 | 0 | 20 | 0.2 | 2 | 0.5 | 5 | 0 | 8 | 0.125 | 3 | 0 | 18 | 0.1111 |
| 3_94786964 | 3 | 1 | 3 | 1 | 8 | 0.625 | 5 | 0.6 | 19 | 0.7368 | 2 | 0.5 | 7 | 0.7143 | 9 | 0.2222 | 4 | 1 | 22 | 0.5455 |
| 3_94786988 | 4 | 0.75 | 3 | 0.3333 | 9 | 0.6667 | 5 | 0.6 | 21 | 0.619 | 2 | 0.5 | 5 | 0.8 | 11 | 0.3636 | 4 | 0.75 | 22 | 0.5455 |
| 3_94787087 | 4 | 1 | 3 | 1 | 9 | 1 | 5 | 1 | 21 | 1 | 3 | 1 | 7 | 1 | 12 | 1 | 4 | 1 | 26 | 1 |
| 3_94787212 | 3 | 0.6667 | 1 | 1 | 7 | 1 | 4 | 1 | 15 | 0.9333 | 2 | 1 | 6 | 1 | 10 | 0.8 | 2 | 1 | 20 | 0.9 |
| 3_94787222 | 3 | 0.6667 | 1 | 1 | 6 | 0.8333 | 6 | 1 | 16 | 0.875 | 3 | 1 | 8 | 1 | 9 | 0.7778 | 4 | 1 | 24 | 0.9167 |
| 3_94787247 | 2 | 0 | 2 | 0 | 9 | 0.4444 | 5 | 0 | 18 | 0.2222 | 2 | 0 | 4 | 0 | 9 | 0.2222 | 2 | 0 | 17 | 0.1176 |
| 3_94787342 | 3 | 0.6667 | 2 | 0.5 | 9 | 0.5556 | 5 | 0.8 | 19 | 0.6316 | 2 | 0.5 | 6 | 0.6667 | 9 | 0.6667 | 3 | 0.3333 | 20 | 0.6 |
| 3_94787542 | 4 | 0.75 | 4 | 0 | 8 | 0.75 | 4 | 0.75 | 20 | 0.6 | 2 | 1 | 4 | 0.5 | 12 | 0.5833 | 4 | 0.5 | 22 | 0.5909 |
| 3_94787772 | 5 | 0.6 | 5 | 0.6 | 9 | 0.8889 | 4 | 1 | 23 | 0.7826 | 4 | 0.75 | 5 | 0.8 | 12 | 0.8333 | 4 | 0.5 | 25 | 0.76 |
| 3_94787950 | 5 | 0.4 | 4 | 0.25 | 9 | 0.6667 | 6 | 0.8333 | 24 | 0.5833 | 2 | 0.5 | 7 | 0.8571 | 13 | 0.6154 | 4 | 0.75 | 26 | 0.6923 |
| 3_94788077 | 6 | 0.1667 | 4 | 0 | 7 | 0.2857 | 3 | 0 | 20 | 0.15 | 3 | 0 | 8 | 0.125 | 11 | 0.1818 | 3 | 0.3333 | 25 | 0.16 |
| 3_94788117 | 4 | 0 | 4 | 0.25 | 9 | 0 | 6 | 0.1667 | 23 | 0.087 | 5 | 0 | 8 | 0.25 | 14 | 0.2143 | 4 | 0 | 31 | 0.1613 |
| 3_94788205 | 1 | 1 | NA | NA | 3 | 0 | 2 | 0 | 6 | 0.1667 | 1 | 0 | 3 | 0.6667 | 3 | 0.3333 | 2 | 0 | 9 | 0.3333 |
| 3_94788432 | 2 | 0 | NA | NA | 2 | 0 | 4 | 0.25 | 8 | 0.125 | 1 | 0 | 2 | 0 | 2 | 0.5 | 1 | 1 | 6 | 0.3333 |
| 3_94788555 | 3 | 0 | NA | NA | 5 | 1 | 4 | 0.75 | 12 | 0.6667 | 1 | 0 | 2 | 1 | 4 | 0 | 3 | 0.6667 | 10 | 0.4 |
| 3_94788634 | 2 | 0 | NA | NA | 6 | 0.1667 | 4 | 0 | 12 | 0.0833 | 2 | 0 | 2 | 0 | 4 | 0.25 | 2 | 0 | 10 | 0.1 |
| 3_94788638 | 3 | 0.3333 | NA | NA | 5 | 0.2 | 3 | 0.3333 | 11 | 0.2727 | 1 | 0 | 2 | 0.5 | 3 | 0 | 1 | 0 | 7 | 0.1429 |
| 3_94788730 | 3 | 0 | NA | NA | 5 | 0.2 | 4 | 0.25 | 12 | 0.1667 | 2 | 0 | 2 | 0 | 4 | 0 | 3 | 0.6667 | 11 | 0.1818 |
| 3_94789015 | 4 | 0 | NA | NA | 6 | 0.3333 | 5 | 0.2 | 15 | 0.2 | 3 | 0.3333 | 2 | 0.5 | 4 | 0.25 | 2 | 0.5 | 11 | 0.3636 |
| 3_94789053 | 3 | 0.6667 | NA | NA | 5 | 1 | 5 | 0.4 | 13 | 0.6923 | 3 | 0 | 2 | 0 | 4 | 0.75 | 2 | 1 | 11 | 0.4545 |
| 3_94789178 | 4 | 0.75 | NA | NA | 6 | 0.8333 | 5 | 1 | 15 | 0.8667 | 2 | 0 | 2 | 1 | 3 | 1 | 4 | 1 | 11 | 0.8182 |
| 3_94789229 | 6 | 0.8333 | NA | NA | 7 | 0.5714 | 4 | 0.5 | 17 | 0.6471 | 2 | 1 | 2 | 1 | 4 | 0.5 | 4 | 0.75 | 12 | 0.75 |
| 3_94789290 | 4 | 0.75 | NA | NA | 7 | 1 | 3 | 1 | 14 | 0.9286 | 3 | 0.3333 | 2 | 0.5 | 5 | 0.8 | 4 | 0.75 | 14 | 0.6429 |
| 3_94789334 | 5 | 1 | NA | NA | 7 | 1 | 5 | 0.8 | 17 | 0.9412 | 3 | 0.6667 | 2 | 1 | 5 | 1 | 4 | 1 | 14 | 0.9286 |
| 3_94789433 | 5 | 0.8 | NA | NA | 7 | 1 | 6 | 1 | 18 | 0.9444 | 2 | 1 | 1 | 1 | 3 | 0.6667 | 3 | 1 | 9 | 0.8889 |
| 3_94789545 | 5 | 0.8 | NA | NA | 6 | 0.8333 | 6 | 0.8333 | 17 | 0.8235 | 3 | 0.6667 | 2 | 1 | 5 | 1 | 4 | 0.75 | 14 | 0.8571 |
| 3_94851794 | 3 | 0.6667 | NA | NA | NA | NA | 2 | 0 | 5 | 0.4 | NA | NA | 5 | 0.2 | 4 | 0.5 | 3 | 0.6667 | 12 | 0.4167 |
| 3_94866125 | NA | NA | 1 | 1 | 2 | 1 | 1 | 1 | 4 | 1 | NA | NA | 1 | 1 | NA | NA | 1 | 1 | 2 | 1 |
| 3_94866303 | NA | NA | 1 | 1 | 1 | 0 | 2 | 0.5 | 4 | 0.5 | 1 | 0 | 2 | 1 | NA | NA | NA | NA | 3 | 0.6667 |
| 3_94866382 | NA | NA | 1 | 1 | 2 | 1 | 1 | 1 | 4 | 1 | 1 | 0 | 1 | 1 | 1 | 0 | 1 | 1 | 4 | 0.5 |
| 3_94867534 | NA | NA | 2 | 0 | 2 | 0 | 2 | 0.5 | 6 | 0.1667 | 1 | 0 | 4 | 0.25 | 2 | 0 | 1 | 0 | 8 | 0.125 |
| 3_94867785 | NA | NA | 2 | 0 | 2 | 0 | 3 | 0.3333 | 7 | 0.1429 | 1 | 0 | 5 | 0.6 | 2 | 0 | 1 | 1 | 9 | 0.4444 |
| 3_94867926 | NA | NA | 2 | 0 | 1 | 0 | 4 | 0.25 | 7 | 0.1429 | 1 | 0 | 5 | 0 | 2 | 0 | 1 | 1 | 9 | 0.1111 |
| 3_94868620 | NA | NA | 2 | 0 | 3 | 0 | 4 | 0.25 | 9 | 0.1111 | 1 | 0 | 4 | 0 | 1 | 0 | 1 | 1 | 7 | 0.1429 |
| 3_94882138 | 1 | 1 | NA | NA | NA | NA | NA | NA | 1 | 1 | NA | NA | NA | NA | 1 | 1 | NA | NA | 1 | 1 |
| 3_94891438 | 3 | 1 | 2 | 0.5 | NA | NA | 2 | 1 | 7 | 0.8571 | 1 | 1 | 1 | 1 | 2 | 1 | 1 | 1 | 5 | 1 |
| 3_94891572 | 2 | 0.5 | NA | NA | NA | NA | 2 | 1 | 4 | 0.75 | 1 | 1 | 1 | 1 | 2 | 1 | 1 | 1 | 5 | 1 |

| | | | | | | | | | | | | | | | | | | | | |
|---|---|---|---|---|---|---|---|---|---|---|---|---|---|---|---|---|---|---|---|---|
| 3_9489173 | 2 | 1 | 2 | 0.5 | NA | NA | 2 | 0.5 | 6 | 0.6667 | 1 | 1 | 1 | 1 | 4 | 0.5 | 1 | 0 | 7 | 0.5714 |
| 3_9489174 | 2 | 0.5 | 2 | 0 | NA | NA | 2 | 0 | 6 | 0.1667 | 1 | 0 | 1 | 0 | 4 | 0.25 | 1 | 0 | 7 | 0.1429 |
| 3_9489183 | 3 | 0.6667 | 2 | 1 | NA | NA | 2 | 1 | 7 | 0.8571 | 1 | 1 | 1 | 1 | 3 | 1 | 1 | 1 | 6 | 1 |
| 3_9489202 | 2 | 0.5 | 3 | 0 | NA | NA | 4 | 0.25 | 9 | 0.2222 | 2 | 0.5 | 3 | 0.6667 | 3 | 0 | 1 | 0 | 9 | 0.3333 |
| 3_9489213 | 3 | 0.6667 | 3 | 1 | NA | NA | 4 | 1 | 10 | 0.9 | 2 | 0.5 | 2 | 0.5 | 3 | 0.6667 | 1 | 1 | 8 | 0.625 |
| 3_9491174 | NA | NA | NA | NA | NA | NA | 1 | 1 | 1 | 1 | NA | NA | NA | NA | NA | NA | 1 | 1 | 1 | 1 |
| 3_9491203 | 4 | 0 | 6 | 0 | 1 | 1 | 5 | 0.4 | 16 | 0.1875 | 2 | 0 | 5 | 0.2 | 5 | 0 | 3 | 0 | 15 | 0.0667 |
| 3_9491325 | 6 | 0.1667 | 7 | 0.1429 | 1 | 0 | 5 | 0 | 19 | 0.1053 | 4 | 0 | 3 | 0 | 5 | 0 | 3 | 0.3333 | 15 | 0.0667 |
| 3_9491328 | 6 | 0.6667 | 7 | 0.5714 | 1 | 1 | 4 | 0.75 | 18 | 0.6667 | 4 | 0.75 | 5 | 0.8 | 6 | 1 | 3 | 0.3333 | 18 | 0.7778 |
| 3_9491330 | 6 | 0.5 | 7 | 0.5714 | 1 | 1 | 5 | 0.2 | 19 | 0.4737 | 4 | 0.25 | 4 | 0.75 | 5 | 0.2 | 2 | 0 | 15 | 0.3333 |
| 3_9491344 | 7 | 0.4286 | 8 | 0.125 | 4 | 0 | 3 | 0 | 22 | 0.1818 | 4 | 0 | 5 | 0.2 | 6 | 0.5 | 3 | 0 | 18 | 0.2222 |
| 3_9491350 | 7 | 0.8571 | 8 | 0.75 | 3 | 0 | 5 | 0.6 | 23 | 0.6522 | 5 | 0.8 | 5 | 0.8 | 6 | 1 | 3 | 0.3333 | 19 | 0.7895 |
| 3_9491353 | 6 | 0.1667 | 5 | 0 | 4 | 0.5 | 5 | 0 | 20 | 0.15 | 4 | 0.25 | 5 | 0 | 5 | 0 | 3 | 0 | 17 | 0.0588 |
| 3_9491354 | 7 | 0.4286 | 8 | 0.75 | 4 | 0.5 | 5 | 0.4 | 24 | 0.5417 | 5 | 0.6 | 5 | 0.4 | 7 | 0.5714 | 3 | 0 | 20 | 0.45 |
| 3_9491365 | 7 | 0.4286 | 7 | 0.4286 | 3 | 0.3333 | 6 | 0.1667 | 23 | 0.3478 | 5 | 0.6 | 5 | 0.4 | 7 | 0.2857 | 2 | 1 | 19 | 0.4737 |
| 3_9491378 | 3 | 0.3333 | 7 | 0.2857 | 1 | 1 | 2 | 0 | 13 | 0.3077 | 4 | 0.25 | 2 | 0 | 2 | 0.5 | 2 | 0.5 | 10 | 0.3 |
| 3_9491448 | 2 | 0.5 | 4 | 0.25 | 1 | 0 | 3 | 0.3333 | 10 | 0.3 | 2 | 0 | 2 | 1 | 5 | 0.4 | 2 | 0 | 11 | 0.3636 |
| 3_9491789 | 1 | 1 | 1 | 1 | NA | NA | NA | NA | 2 | 1 | 1 | 1 | 1 | 1 | 1 | 1 | 1 | 0 | 4 | 0.75 |
| 3_9491801 | 1 | 0 | 2 | 1 | NA | NA | 1 | 1 | 4 | 0.75 | 1 | 0 | 1 | 1 | 1 | 1 | 1 | 1 | 4 | 0.75 |
| 3_9491822 | 1 | 1 | 2 | 0 | NA | NA | 1 | 0 | 4 | 0.25 | 1 | 0 | 1 | 1 | 1 | 0 | 1 | 0 | 4 | 0.25 |
| 3_9491840 | 1 | 1 | 1 | 1 | NA | NA | 1 | 1 | 3 | 1 | NA | NA | 1 | 0 | 1 | 1 | 1 | 1 | 3 | 0.6667 |
| 3_9491873 | NA | NA | 1 | 1 | NA | NA | NA | NA | 1 | 1 | NA | NA | 1 | 1 | 1 | 1 | NA | NA | 2 | 1 |
| 3_9491883 | 1 | 0 | 1 | 1 | NA | NA | NA | NA | 2 | 0.5 | 1 | 0 | 1 | 1 | 1 | 1 | 1 | 0 | 4 | 0.5 |
| 3_9491916 | 1 | 1 | 1 | 1 | NA | NA | NA | NA | 2 | 1 | 1 | 0 | 1 | 1 | NA | NA | NA | NA | 2 | 0.5 |
| 3_9491928 | 1 | 1 | 1 | 1 | NA | NA | NA | NA | 2 | 1 | 1 | 1 | 1 | 1 | NA | NA | 1 | 1 | 3 | 1 |
| 3_9491941 | 1 | 1 | 1 | 1 | NA | NA | NA | NA | 2 | 1 | 1 | 1 | 1 | 1 | NA | NA | 1 | 1 | 3 | 1 |
| 3_9491944 | 1 | 1 | 1 | 0 | NA | NA | NA | NA | 2 | 0.5 | 1 | 1 | 1 | 1 | NA | NA | 1 | 1 | 3 | 1 |
| 3_9493953 | 1 | 1 | NA | NA | NA | NA | 1 | 0 | 2 | 0.5 | 1 | 0 | 1 | 0 | 1 | 1 | NA | NA | 3 | 0.3333 |
| 3_9494009 | 21 | 0.4762 | 26 | 0.5385 | 15 | 0.2 | 19 | 0.5789 | 81 | 0.4691 | 17 | 0.2941 | 22 | 0.4545 | 15 | 0.2667 | 15 | 0.4667 | 69 | 0.3768 |
| 3_9496698 | 9 | 0.1111 | 11 | 0.0909 | 2 | 0 | 1 | 0 | 23 | 0.087 | 2 | 0.5 | 7 | 0 | 6 | 0 | 1 | 0 | 16 | 0.0625 |
| 3_9496702 | 6 | 0.3333 | 9 | 0.3333 | 2 | 1 | NA | NA | 17 | 0.4118 | 1 | 0 | 6 | 0.5 | 3 | 0 | 1 | 0 | 11 | 0.2727 |
| 3_9496703 | 8 | 0.625 | 9 | 0.7778 | 1 | 0 | 2 | 1 | 20 | 0.7 | 2 | 0.5 | 7 | 0.7143 | 6 | 0.6667 | 1 | 1 | 16 | 0.6875 |
| 3_9496705 | 6 | 0.6667 | 9 | 1 | 2 | 0.5 | 2 | 1 | 19 | 0.8421 | 2 | 0.5 | 5 | 0.8 | 5 | 1 | 1 | 1 | 13 | 0.8462 |
| 3_9496709 | 8 | 0.875 | 9 | 1 | 1 | 1 | 2 | 1 | 20 | 0.95 | 2 | 1 | 7 | 1 | 5 | 1 | 1 | 1 | 15 | 1 |
| 3_9496721 | 7 | 0.8571 | 9 | 0.6667 | 2 | 0.5 | 1 | 1 | 19 | 0.7368 | 2 | 0.5 | 5 | 0.8 | 4 | 1 | NA | NA | 11 | 0.8182 |
| 3_9496722 | 5 | 0.4 | 6 | 0.1667 | 1 | 0 | 1 | 0 | 13 | 0.2308 | 2 | 0 | 5 | 0 | 3 | 0.3333 | NA | NA | 10 | 0.1 |
| 3_9502990 | 3 | 0 | 7 | 0.2857 | 7 | 0.1429 | 10 | 0.2 | 27 | 0.1852 | 4 | 0.25 | 6 | 0.1667 | 4 | 0.25 | 7 | 0.2857 | 21 | 0.2381 |
| 3_9502998 | 2 | 0 | 7 | 0.1429 | 6 | 0.5 | 8 | 0.25 | 23 | 0.2609 | 3 | 0.3333 | 6 | 0.1667 | 5 | 0.6 | 5 | 0 | 19 | 0.2632 |
| 3_9503009 | 4 | 0.25 | 7 | 0 | 7 | 0.1429 | 10 | 0.1 | 28 | 0.1071 | 5 | 0 | 6 | 0.5 | 5 | 0 | 6 | 0.1667 | 22 | 0.1818 |
| 3_9503669 | NA | NA | 1 | 1 | NA | NA | NA | NA | 1 | 1 | NA | NA | NA | NA | NA | NA | 1 | 1 | 1 | 1 |
| 3_9503709 | 4 | 0.25 | 8 | 0.25 | 5 | 0.4 | 6 | 0.1667 | 23 | 0.2609 | 3 | 0 | 6 | 0.5 | 5 | 0.2 | 4 | 0.25 | 18 | 0.2778 |
| 3_9504162 | NA | NA | NA | NA | NA | NA | 1 | 1 | 1 | 1 | 1 | 1 | 1 | 1 | 2 | 1 | NA | NA | 4 | 1 |
| 3_9504193 | NA | NA | NA | NA | NA | NA | 3 | 0.3333 | 3 | 0.3333 | 3 | 0.6667 | 1 | 0 | 5 | 0.6 | 1 | 1 | 10 | 0.6 |
| 3_9504503 | NA | NA | NA | NA | NA | NA | 4 | 1 | 4 | 1 | 4 | 0.5 | 3 | 0.6667 | 8 | 0.875 | 1 | 1 | 16 | 0.75 |
| 3_9504598 | 1 | 1 | NA | NA | NA | NA | 2 | 1 | 3 | 1 | 2 | 0 | 3 | 0.3333 | 6 | 0.5 | 1 | 0 | 12 | 0.3333 |
| 3_9504637 | 3 | 0 | NA | NA | NA | NA | 5 | 0.2 | 8 | 0.125 | 5 | 0 | 4 | 0.25 | 7 | 0.2857 | 2 | 0.5 | 18 | 0.2222 |
| 3_9504691 | 1 | 1 | NA | NA | NA | NA | 1 | 1 | 2 | 1 | NA | NA | NA | NA | 2 | 1 | NA | NA | 2 | 1 |
| 3_9504718 | 1 | 1 | NA | NA | NA | NA | NA | NA | 1 | 1 | NA | NA | NA | NA | 1 | 1 | NA | NA | 1 | 1 |
| 3_9504728 | 1 | 1 | NA | NA | NA | NA | NA | NA | 1 | 1 | NA | NA | NA | NA | 2 | 1 | NA | NA | 2 | 1 |
| 3_9504753 | 1 | 1 | NA | NA | NA | NA | NA | NA | 1 | 1 | NA | NA | NA | NA | 1 | 1 | NA | NA | 1 | 1 |
| 3_9505228 | NA | NA | 1 | 1 | NA | NA | NA | NA | 1 | 1 | NA | NA | NA | NA | NA | NA | 1 | 1 | 1 | 1 |
| 3_9505252 | NA | NA | 1 | 1 | NA | NA | NA | NA | 1 | 1 | NA | NA | NA | NA | NA | NA | 1 | 1 | 1 | 1 |
| 3_9507997 | 1 | 0 | 1 | 0 | NA | NA | 4 | 0.25 | 6 | 0.1667 | 3 | 0 | 1 | 0 | 1 | 1 | 1 | 1 | 6 | 0.3333 |
| 3_9508011 | 1 | 1 | NA | NA | NA | NA | 4 | 1 | 5 | 1 | 2 | 0.5 | 1 | 1 | 1 | 0 | 1 | 1 | 5 | 0.6 |
| 3_9508016 | 1 | 0 | NA | NA | NA | NA | 4 | 0.25 | 5 | 0.2 | 3 | 0 | 1 | 1 | 1 | 0 | NA | NA | 5 | 0.2 |
| 3_9508023 | 1 | 1 | NA | NA | NA | NA | 3 | 1 | 4 | 1 | 1 | 1 | 1 | 1 | 1 | 1 | 1 | 1 | 4 | 1 |
| 3_9508065 | 1 | 1 | NA | NA | 1 | 1 | 3 | 1 | 5 | 1 | 3 | 1 | 1 | 1 | NA | NA | 2 | 1 | 6 | 1 |
| 3_9508066 | 1 | 1 | NA | NA | 1 | 1 | 2 | 1 | 4 | 1 | 1 | 1 | 1 | 1 | 1 | 1 | 1 | 1 | 4 | 1 |
| 3_9508070 | 1 | 1 | NA | NA | 2 | 1 | 4 | 0.75 | 7 | 0.8571 | 3 | 0.6667 | 1 | 1 | NA | NA | 2 | 1 | 6 | 0.8333 |
| 3_9508073 | 1 | 1 | NA | NA | 1 | 1 | 2 | 0.5 | 4 | 0.75 | 3 | 0.6667 | 1 | 1 | 1 | 1 | 2 | 0.5 | 7 | 0.7143 |
| 3_9508076 | 1 | 0 | 1 | 0 | 1 | 0 | 6 | 0.5 | 9 | 0.3333 | 3 | 1 | 1 | 1 | 1 | 1 | 2 | 0 | 7 | 0.7143 |
| 3_9508082 | 1 | 0 | NA | NA | NA | NA | 4 | 0.5 | 5 | 0.4 | 3 | 1 | 1 | 1 | NA | NA | 2 | 0.5 | 6 | 0.8333 |
| 3_9508083 | 1 | 1 | NA | NA | NA | NA | 4 | 0.5 | 5 | 0.6 | 2 | 1 | NA | NA | NA | NA | 2 | 1 | 4 | 1 |
| 3_9508097 | 1 | 0 | NA | NA | 1 | 1 | 3 | 0.6667 | 5 | 0.6 | 2 | 0 | 1 | 1 | NA | NA | 1 | 0 | 4 | 0.25 |
| 3_9508115 | 1 | 1 | NA | NA | NA | NA | 4 | 0.75 | 5 | 0.8 | 2 | 0.5 | 1 | 0 | NA | NA | 2 | 0.5 | 5 | 0.4 |

| | | | | | | | | | | | | | | | | | | | | |
|---|---|---|---|---|---|---|---|---|---|---|---|---|---|---|---|---|---|---|---|---|
| 3_95124972 | NA | NA | NA | NA | NA | NA | 1 | 1 | 1 | 1 | NA | NA | NA | NA | NA | NA | 1 | 1 | 1 | 1 |
| 3_95125246 | NA | NA | NA | NA | NA | NA | 1 | 1 | 1 | 1 | NA | NA | NA | NA | NA | NA | 2 | 0.5 | 2 | 0.5 |
| 3_95125359 | NA | NA | NA | NA | NA | NA | 1 | 1 | 1 | 1 | NA | NA | NA | NA | NA | NA | 2 | 1 | 2 | 1 |
| 3_95125461 | NA | NA | NA | NA | NA | NA | 1 | 1 | 1 | 1 | NA | NA | NA | NA | NA | NA | 2 | 1 | 2 | 1 |
| 3_95125537 | NA | NA | NA | NA | NA | NA | 1 | 1 | 1 | 1 | NA | NA | NA | NA | NA | NA | 2 | 1 | 2 | 1 |
| 3_95127506 | 39 | 0.9487 | 39 | 0.9744 | 34 | 0.9706 | 39 | 1 | 151 | 0.9735 | 29 | 1 | 44 | 0.9545 | 34 | 1 | 54 | 0.963 | 161 | 0.9752 |
| 3_95127706 | 33 | 0.2121 | 36 | 0.1944 | 33 | 0.1818 | 36 | 0.1944 | 138 | 0.1957 | 31 | 0.1935 | 41 | 0.2195 | 26 | 0.1923 | 46 | 0.2609 | 144 | 0.2222 |
| 3_95135752 | 3 | 0.3333 | 1 | 0 | 4 | 0 | 7 | 0 | 15 | 0.0667 | 3 | 0.3333 | 2 | 0 | 3 | 0 | 1 | 0 | 9 | 0.1111 |
| 3_95135759 | 5 | 0 | 1 | 1 | 6 | 0.5 | 7 | 0 | 19 | 0.2105 | 3 | 0 | 2 | 0 | 2 | 0.5 | 2 | 0.5 | 9 | 0.2222 |
| 3_95135789 | 5 | 0 | 1 | 0 | 5 | 0.2 | 7 | 0.1429 | 18 | 0.1111 | 3 | 0 | 2 | 0 | 3 | 0 | 1 | 1 | 9 | 0.1111 |
| 3_95139170 | 6 | 0.1667 | 3 | 0 | 5 | 0.2 | 4 | 0.25 | 18 | 0.1667 | 2 | 0 | 1 | 0 | NA | NA | 5 | 0.2 | 8 | 0.125 |
| 3_95139240 | 6 | 0 | 2 | 0 | 3 | 0.6667 | 3 | 0.3333 | 14 | 0.2143 | 2 | 0.5 | 1 | 0 | NA | NA | 4 | 0 | 7 | 0.1429 |
| 3_95139276 | 5 | 1 | 2 | 1 | 6 | 0.8333 | 4 | 0.5 | 17 | 0.8235 | 3 | 0.6667 | 1 | 1 | NA | NA | 4 | 0.75 | 8 | 0.75 |
| 3_95141758 | 6 | 0.8333 | 2 | 1 | 6 | 0.8333 | 3 | 0.6667 | 17 | 0.8235 | 8 | 0.875 | 1 | 1 | 1 | 1 | 7 | 1 | 17 | 0.9412 |
| 3_95141770 | 8 | 0.5 | 3 | 0.6667 | 8 | 0.5 | 3 | 0.6667 | 22 | 0.5455 | 8 | 0.375 | 1 | 0 | 1 | 0 | 7 | 0.4286 | 17 | 0.3529 |
| 3_95142536 | 7 | 0.1429 | 4 | 0 | 4 | 0 | 3 | 0 | 18 | 0.0556 | 8 | 0.25 | 3 | 0 | 2 | 0 | 9 | 0.1111 | 22 | 0.1364 |
| 3_95143509 | 10 | 0.2 | 8 | 0 | 7 | 0.1429 | 6 | 0 | 31 | 0.0968 | 11 | 0 | 3 | 0 | 2 | 0.5 | 10 | 0.1 | 26 | 0.0769 |
| 3_95161968 | 1 | 0 | 2 | 0.5 | 4 | 0.75 | 5 | 0.4 | 12 | 0.5 | 5 | 0.8 | 1 | 1 | 5 | 0.2 | 3 | 0.3333 | 14 | 0.5 |
| 3_95162101 | 3 | 0 | 3 | 0 | 4 | 0.25 | 5 | 0 | 15 | 0.0667 | 5 | 0 | 1 | 0 | 5 | 0.2 | 4 | 0 | 15 | 0.0667 |
| 3_95162140 | 2 | 0 | 3 | 0.3333 | 4 | 0.5 | 5 | 0.2 | 14 | 0.2857 | 5 | 0.2 | 1 | 0 | 5 | 0 | 2 | 0.5 | 13 | 0.1538 |
| 3_95162181 | 2 | 0.5 | 2 | 0 | 4 | 0 | 3 | 0 | 11 | 0.0909 | 5 | 0 | 1 | 0 | 4 | 0.25 | 3 | 0 | 13 | 0.0769 |
| 3_95162307 | 3 | 1 | 2 | 0.5 | 4 | 0.5 | 4 | 0.75 | 13 | 0.6923 | 5 | 0.6 | 1 | 1 | 5 | 1 | 3 | 0.6667 | 14 | 0.7857 |
| 3_95162431 | 2 | 0.5 | 1 | 1 | 3 | 0 | 4 | 0.5 | 10 | 0.4 | 5 | 0.4 | 1 | 1 | 3 | 0.3333 | 1 | 1 | 10 | 0.5 |
| 3_95162506 | 1 | 1 | 1 | 0 | 2 | 0.5 | 4 | 0.5 | 8 | 0.5 | 1 | 0 | 1 | 0 | 4 | 0.25 | 1 | 0 | 7 | 0.1429 |
| 3_95162512 | 3 | 1 | 2 | 1 | 3 | 0.6667 | 5 | 1 | 13 | 0.9231 | 5 | 1 | 1 | 1 | 5 | 1 | 2 | 1 | 13 | 1 |
| 3_95162518 | 2 | 1 | 2 | 0.5 | 1 | 0 | 4 | 0.25 | 9 | 0.4444 | 5 | 0.8 | 1 | 1 | 5 | 0.2 | 2 | 0 | 13 | 0.4615 |
| 3_95195125 | 2 | 0 | 2 | 1 | 1 | 1 | NA | NA | 5 | 0.6 | 1 | 1 | NA | NA | 2 | 0.5 | 3 | 0.6667 | 6 | 0.6667 |
| 3_95195159 | 2 | 0.5 | 2 | 0 | 1 | 0 | NA | NA | 5 | 0.2 | 1 | 1 | NA | NA | 2 | 0.5 | 3 | 0 | 6 | 0.3333 |
| 3_95195226 | 2 | 0 | 2 | 0.5 | 1 | 0 | NA | NA | 5 | 0.2 | NA | NA | NA | NA | 1 | 0 | 1 | 1 | 2 | 0.5 |
| 3_95195247 | 2 | 0 | 3 | 0.6667 | 1 | 0 | NA | NA | 6 | 0.3333 | 1 | 1 | NA | NA | 2 | 0.5 | 3 | 0.6667 | 6 | 0.6667 |
| 3_95195274 | 2 | 1 | 2 | 0 | 1 | 0 | NA | NA | 5 | 0.4 | NA | NA | NA | NA | 1 | 1 | 3 | 0.6667 | 4 | 0.75 |
| 3_95195313 | 1 | 1 | 3 | 0.6667 | 1 | 1 | NA | NA | 5 | 0.8 | 1 | 1 | NA | NA | 2 | 1 | 3 | 0.6667 | 6 | 0.8333 |
| 3_95230096 | 9 | 0.1111 | 8 | 0.375 | 9 | 0 | 6 | 0.1667 | 32 | 0.1562 | 11 | 0.2727 | 7 | 0.1429 | 6 | 0.1667 | 7 | 0.4286 | 31 | 0.2581 |
| 3_95231239 | 3 | 0.3333 | 4 | 0.75 | 6 | 0.1667 | 1 | 0 | 14 | 0.3571 | 2 | 0 | 3 | 0.3333 | 5 | 0.2 | 5 | 0.6 | 15 | 0.3333 |
| 3_95231311 | 2 | 0 | 3 | 0.3333 | 5 | 0 | 1 | 1 | 11 | 0.1818 | NA | NA | 3 | 0.3333 | 4 | 0.5 | 5 | 0.4 | 12 | 0.4167 |
| 3_95231319 | 3 | 0.3333 | 5 | 0 | 6 | 0.1667 | 1 | 0 | 15 | 0.1333 | 1 | 0 | 4 | 0.5 | 5 | 0 | 5 | 0.2 | 15 | 0.2 |
| 3_95231333 | 3 | 0 | 4 | 0 | 6 | 0.1667 | 1 | 0 | 14 | 0.0714 | 1 | 0 | 2 | 0 | 5 | 0 | 4 | 0.25 | 12 | 0.0833 |
| 3_95231352 | 3 | 0 | 4 | 0.25 | 5 | 0 | 1 | 1 | 13 | 0.1538 | 1 | 0 | 3 | 0.3333 | 4 | 0.25 | 5 | 0.4 | 13 | 0.3077 |
| 3_95234658 | 2 | 0.5 | 4 | 0 | 5 | 0 | 1 | 0 | 12 | 0.0833 | 1 | 1 | 2 | 0 | 2 | 0 | 4 | 0.25 | 9 | 0.2222 |
| 3_95234722 | 2 | 0 | 4 | 0.5 | 5 | 0.2 | 1 | 0 | 12 | 0.25 | 1 | 0 | 5 | 0.4 | 2 | 0 | 4 | 0 | 12 | 0.1667 |
| 3_95234811 | 2 | 1 | 3 | 1 | 5 | 0.8 | 1 | 0 | 11 | 0.8182 | 1 | 0 | 4 | 1 | 3 | 0.6667 | 4 | 0.75 | 12 | 0.75 |
| 3_95234857 | 2 | 0.5 | 2 | 1 | 3 | 0.3333 | 1 | 0 | 8 | 0.5 | 1 | 1 | 2 | 1 | 2 | 0.5 | 4 | 0.5 | 9 | 0.6667 |
| 3_95234904 | 1 | 1 | 4 | 0.25 | 5 | 0.6 | 1 | 1 | 11 | 0.5455 | 1 | 1 | 5 | 0.4 | 2 | 0.5 | 4 | 0.25 | 12 | 0.4167 |
| 3_95235054 | 2 | 0 | 4 | 0 | 4 | 0.25 | 1 | 0 | 11 | 0.0909 | 1 | 0 | 4 | 0 | 3 | 0 | 3 | 0.3333 | 11 | 0.0909 |
| 3_95245681 | 1 | 0 | 2 | 0.5 | 3 | 0 | NA | NA | 6 | 0.1667 | 1 | 0 | 4 | 0.5 | 1 | 1 | 1 | 1 | 7 | 0.5714 |
| 3_95245828 | 1 | 1 | 2 | 1 | 4 | 0.75 | NA | NA | 7 | 0.8571 | NA | NA | 4 | 0.75 | 1 | 1 | 1 | 1 | 6 | 0.8333 |
| 3_95245918 | 1 | 0 | 2 | 1 | 3 | 1 | NA | NA | 6 | 0.8333 | 1 | 1 | 4 | 0.5 | 1 | 1 | 1 | 1 | 7 | 0.7143 |
| 3_95245928 | 1 | 1 | 2 | 0.5 | 3 | 1 | NA | NA | 6 | 0.8333 | NA | NA | 4 | 0.5 | 1 | 1 | 1 | 1 | 6 | 0.6667 |
| 3_95246050 | 1 | 0 | 2 | 0 | 3 | 0.3333 | NA | NA | 6 | 0.1667 | NA | NA | 4 | 0.25 | 1 | 0 | 1 | 0 | 6 | 0.1667 |
| 3_95246094 | 1 | 1 | NA | NA | 1 | 0 | NA | NA | 2 | 0.5 | 1 | 0 | 1 | 1 | NA | NA | NA | NA | 2 | 0.5 |
| 3_95403481 | 2 | 0.5 | 1 | 0 | 2 | 0 | 3 | 0.3333 | 8 | 0.25 | 1 | 1 | 2 | 0 | 1 | 0 | NA | NA | 4 | 0.25 |
| 3_95403623 | NA | NA | 1 | 0 | 1 | 0 | 3 | 0.3333 | 5 | 0.2 | 1 | 0 | 3 | 0 | 1 | 0 | 1 | 1 | 6 | 0.1667 |
| 3_95416433 | 1 | 0 | NA | NA | 1 | 1 | NA | NA | 2 | 0.5 | NA | NA | NA | NA | 1 | 1 | NA | NA | 1 | 1 |
| 3_95416507 | 1 | 0 | NA | NA | 1 | 1 | NA | NA | 2 | 0.5 | NA | NA | NA | NA | 1 | 1 | NA | NA | 1 | 1 |
| 3_95436819 | 16 | 0.125 | 13 | 0.2308 | 10 | 0 | 6 | 0 | 45 | 0.1111 | 12 | 0.1667 | 12 | 0 | 4 | 0 | 10 | 0.1 | 38 | 0.0789 |
| 3_95452750 | 19 | 0 | 18 | 0.1111 | 8 | 0.125 | 13 | 0.0769 | 58 | 0.069 | 20 | 0.1 | 17 | 0.1176 | 9 | 0 | 14 | 0.1429 | 60 | 0.1 |
| 3_95524565 | 2 | 0 | NA | NA | 4 | 0 | 2 | 0.5 | 8 | 0.125 | NA | NA | 2 | 0.5 | 3 | 0 | 4 | 0.25 | 9 | 0.2222 |
| 3_95524580 | 2 | 1 | NA | NA | 4 | 1 | 2 | 0.5 | 8 | 0.875 | NA | NA | 2 | 1 | 3 | 0.6667 | 4 | 0.75 | 9 | 0.7778 |
| 3_95524588 | 2 | 0 | 1 | 1 | 4 | 0.75 | 1 | 0 | 8 | 0.5 | NA | NA | 2 | 0 | 3 | 0.3333 | 3 | 0 | 8 | 0.125 |
| 3_95524684 | 2 | 0.5 | 1 | 1 | 2 | 0 | 2 | 0 | 7 | 0.2857 | NA | NA | NA | NA | 2 | 0.5 | 4 | 0.25 | 6 | 0.3333 |
| 3_95524691 | 2 | 1 | 1 | 1 | 4 | 1 | 2 | 0.5 | 9 | 0.8889 | NA | NA | 2 | 1 | 3 | 0.6667 | 4 | 0.75 | 9 | 0.7778 |
| 3_95524701 | 1 | 1 | 1 | 1 | 4 | 1 | 1 | 1 | 7 | 1 | NA | NA | NA | NA | 1 | 1 | 2 | 1 | 3 | 1 |
| 3_95524706 | 2 | 0.5 | 1 | 1 | 4 | 0 | 2 | 0 | 9 | 0.2222 | NA | NA | 2 | 1 | 3 | 0.6667 | 2 | 0.5 | 7 | 0.7143 |
| 3_95524723 | 2 | 0 | 1 | 0 | 4 | 0 | 2 | 0.5 | 9 | 0.1111 | NA | NA | 2 | 0 | 3 | 0.6667 | 4 | 0.25 | 9 | 0.3333 |
| 3_95524823 | 2 | 0.5 | 1 | 1 | 4 | 1 | 2 | 1 | 9 | 0.8889 | NA | NA | 2 | 1 | 3 | 0.6667 | 4 | 0.5 | 9 | 0.6667 |

| | | | | | | | | | | | | | | | | | | | | |
|---|---|---|---|---|---|---|---|---|---|---|---|---|---|---|---|---|---|---|---|---|
| 3_9552483 | 2 | 0 | 1 | 1 | 4 | 1 | 2 | 1 | 9 | 0.7778 | NA | NA | 2 | 1 | 3 | 1 | 4 | 0.5 | 9 | 0.7778 |
| 3_9552489 | 2 | 0.5 | 1 | 1 | 4 | 0 | 2 | 1 | 9 | 0.4444 | NA | NA | 1 | 0 | 2 | 0 | 3 | 0.3333 | 6 | 0.1667 |
| 3_9552495 | 2 | 0.5 | 1 | 1 | 4 | 1 | 2 | 1 | 9 | 0.8889 | NA | NA | 2 | 1 | 3 | 1 | 4 | 0.75 | 9 | 0.8889 |
| 3_9552512 | 2 | 0 | 1 | 0 | 3 | 0.3333 | 1 | 0 | 7 | 0.1429 | NA | NA | 1 | 0 | 1 | 1 | 3 | 0.3333 | 5 | 0.4 |
| 3_9552520 | 2 | 0 | 2 | 0 | 4 | 0 | 2 | 0.5 | 10 | 0.1 | NA | NA | 2 | 0 | 4 | 0.25 | 4 | 0 | 10 | 0.1 |
| 3_9552525 | 2 | 0 | 2 | 0 | 4 | 0.25 | 2 | 0.5 | 10 | 0.2 | NA | NA | 2 | 0 | 4 | 0.25 | 4 | 0 | 10 | 0.1 |
| 3_9552561 | 2 | 0 | 1 | 0 | 4 | 0.5 | 2 | 1 | 9 | 0.4444 | NA | NA | 2 | 0 | 4 | 0.25 | 3 | 0 | 9 | 0.1111 |
| 3_9552610 | 2 | 0 | 2 | 0 | 4 | 0.25 | 2 | 0 | 10 | 0.1 | NA | NA | 2 | 0.5 | 3 | 1 | 4 | 0.5 | 9 | 0.6667 |
| 3_9553605 | 4 | 0.75 | 10 | 0.7 | 8 | 0.5 | 3 | 1 | 25 | 0.68 | 3 | 0.6667 | 5 | 0.8 | 7 | 0.8571 | 8 | 0.625 | 23 | 0.7391 |
| 3_9553894 | 31 | 0.1613 | 34 | 0.0882 | 25 | 0.08 | 32 | 0.125 | 122 | 0.1148 | 21 | 0.3333 | 41 | 0.0732 | 33 | 0.0909 | 20 | 0.15 | 115 | 0.1391 |
| 3_9554869 | 1 | 1 | NA | NA | NA | NA | 1 | 0 | 2 | 0.5 | 1 | 1 | 2 | 0 | 1 | 0 | NA | NA | 4 | 0.25 |
| 3_9554871 | 1 | 1 | NA | NA | NA | NA | NA | NA | 1 | 1 | 1 | 1 | 3 | 0.6667 | 1 | 0 | NA | NA | 5 | 0.6 |
| 3_9556648 | 2 | 0 | 6 | 0 | 1 | 1 | 4 | 0.5 | 13 | 0.2308 | 1 | 0 | 1 | 1 | 4 | 0.25 | 2 | 0 | 8 | 0.25 |
| 3_9556669 | 2 | 0.5 | 9 | 0.6667 | 1 | 0 | 4 | 0.5 | 16 | 0.5625 | 4 | 1 | 4 | 0.25 | 5 | 0.6 | 3 | 0.6667 | 16 | 0.625 |
| 3_9556709 | 3 | 0 | 9 | 0.2222 | 3 | 0 | 3 | 0.3333 | 18 | 0.1667 | 2 | 0.5 | 4 | 0 | 5 | 0.8 | 3 | 0.6667 | 14 | 0.5 |
| 3_9556716 | 4 | 0.5 | 10 | 0.5 | 3 | 0.3333 | 4 | 0.25 | 21 | 0.4286 | 4 | 0.25 | 6 | 0 | 5 | 0.4 | 4 | 0.75 | 19 | 0.3158 |
| 3_9556717 | 4 | 0.75 | 9 | 0.7778 | 2 | 1 | 4 | 0.75 | 19 | 0.7895 | 4 | 0.75 | 4 | 0.75 | 4 | 1 | 4 | 0.75 | 16 | 0.8125 |
| 3_9556729 | 4 | 0.25 | 9 | 0.3333 | 1 | 0 | 4 | 0 | 18 | 0.2222 | 3 | 0.3333 | 5 | 0 | 4 | 0.25 | 4 | 0.25 | 16 | 0.1875 |
| 3_9556804 | 4 | 0.25 | 7 | 0.7143 | 2 | 0 | 3 | 0 | 16 | 0.375 | 5 | 0 | 4 | 0 | 4 | 0.5 | 4 | 0.25 | 17 | 0.1765 |
| 3_9556815 | 4 | 0.75 | 11 | 1 | 3 | 1 | 3 | 1 | 21 | 0.9524 | 4 | 1 | 4 | 1 | 5 | 1 | 5 | 1 | 18 | 1 |
| 3_9556821 | 4 | 0.75 | 13 | 0.6923 | 2 | 1 | 3 | 1 | 22 | 0.7727 | 4 | 1 | 3 | 0.6667 | 3 | 0.6667 | 4 | 0.75 | 14 | 0.7857 |
| 3_9556840 | 5 | 0.4 | 10 | 0.6 | 3 | 1 | 3 | 1 | 21 | 0.6667 | 4 | 1 | 3 | 0.6667 | 4 | 1 | 2 | 1 | 13 | 0.9231 |
| 3_9556868 | 3 | 0.3333 | 12 | 0.3333 | 5 | 0.2 | 6 | 0 | 26 | 0.2308 | 5 | 0 | 7 | 0.1429 | 6 | 0.5 | 6 | 0.1667 | 24 | 0.2083 |
| 3_9556902 | 5 | 0 | 12 | 0.1667 | 7 | 0.2857 | 5 | 0.4 | 29 | 0.2069 | 5 | 0.4 | 5 | 0 | 5 | 0.2 | 6 | 0.1667 | 21 | 0.1905 |
| 3_9564332 | 2 | 0.5 | 1 | 0 | NA | NA | 3 | 0.3333 | 6 | 0.3333 | 1 | 0 | 1 | 0 | NA | NA | 3 | 0.6667 | 5 | 0.4 |
| 3_9564732 | 2 | 0.5 | 5 | 0 | 6 | 0.1667 | 6 | 0.1667 | 19 | 0.1579 | 1 | 0 | 3 | 0 | 4 | 0.25 | 4 | 0 | 12 | 0.0833 |
| 3_9564738 | 2 | 0.5 | 4 | 0.25 | 6 | 0.3333 | 6 | 0.1667 | 18 | 0.2778 | 1 | 1 | 2 | 0.5 | 4 | 0.25 | 3 | 0.3333 | 10 | 0.4 |
| 3_9564745 | 2 | 0.5 | 6 | 0 | 6 | 0.1667 | 6 | 0.1667 | 20 | 0.15 | 2 | 0.5 | 3 | 0 | 3 | 0 | 4 | 0.25 | 12 | 0.1667 |
| 3_9566738 | NA | NA | 1 | 1 | NA | NA | 1 | 1 | 2 | 1 | 1 | 0 | 1 | 1 | 2 | 0.5 | 1 | 1 | 5 | 0.6 |
| 3_9566805 | NA | NA | 4 | 0 | 1 | 0 | 6 | 0.1667 | 11 | 0.0909 | 1 | 0 | 2 | 0 | 4 | 0.25 | NA | NA | 7 | 0.1429 |
| 3_9566846 | 1 | 1 | 7 | 0.1429 | 1 | 0 | 7 | 0.4286 | 16 | 0.3125 | NA | NA | 3 | 0.6667 | 6 | 0.1667 | 3 | 0 | 12 | 0.25 |
| 3_9566864 | 1 | 1 | 3 | 0.6667 | 1 | 1 | 5 | 0.8 | 10 | 0.8 | NA | NA | 4 | 1 | 5 | 0.4 | 2 | 1 | 11 | 0.7273 |
| 3_9566932 | NA | NA | 2 | 0 | NA | NA | 5 | 0.4 | 7 | 0.2857 | NA | NA | NA | NA | 2 | 0.5 | NA | NA | 2 | 0.5 |
| 3_9567032 | 1 | 0 | 2 | 0.5 | NA | NA | 3 | 0 | 6 | 0.1667 | NA | NA | 1 | 1 | NA | NA | NA | NA | 1 | 1 |
| 3_9567096 | 3 | 0 | 1 | 0 | 2 | 0 | 4 | 0.25 | 10 | 0.1 | 1 | 1 | NA | NA | 1 | 1 | NA | NA | 2 | 1 |
| 3_9567106 | 3 | 0.3333 | 2 | 0.5 | 1 | 0 | 4 | 0.75 | 10 | 0.5 | 1 | 1 | 2 | 1 | 1 | 1 | NA | NA | 4 | 1 |
| 3_9567108 | 3 | 0 | 2 | 0.5 | 2 | 0.5 | 3 | 0.3333 | 10 | 0.3 | 1 | 0 | 2 | 1 | 1 | 0 | NA | NA | 4 | 0.5 |
| 3_9567111 | 3 | 0 | 2 | 0 | 2 | 0 | 3 | 0.3333 | 10 | 0.1 | NA | NA | 2 | 0.5 | 1 | 1 | NA | NA | 3 | 0.6667 |
| 3_9567134 | 3 | 0.6667 | 1 | 0 | 1 | 0 | 3 | 0.6667 | 8 | 0.5 | 1 | 1 | 2 | 1 | 1 | 1 | NA | NA | 4 | 1 |
| 3_9567137 | 3 | 0.6667 | 2 | 1 | 1 | 0 | 3 | 0.6667 | 9 | 0.6667 | 1 | 1 | 2 | 1 | 1 | 1 | NA | NA | 4 | 1 |
| 3_9567149 | 3 | 0.6667 | 1 | 1 | 2 | 0.5 | 3 | 0.6667 | 9 | 0.6667 | NA | NA | 2 | 1 | 1 | 1 | NA | NA | 3 | 1 |
| 3_9567151 | 2 | 0.5 | NA | NA | 2 | 0 | 3 | 0.3333 | 7 | 0.2857 | NA | NA | 2 | 1 | 1 | 1 | NA | NA | 3 | 1 |
| 3_9567162 | 2 | 0 | 1 | 1 | 2 | 0.5 | 3 | 0.3333 | 8 | 0.375 | 1 | 0 | 1 | 1 | NA | NA | NA | NA | 2 | 0.5 |
| 3_9567174 | 3 | 0.3333 | 2 | 0 | 2 | 0 | 2 | 0 | 9 | 0.1111 | NA | NA | 2 | 1 | 1 | 0 | NA | NA | 3 | 0.6667 |
| 3_9567193 | 2 | 0.5 | 1 | 1 | 2 | 0.5 | 3 | 0.3333 | 8 | 0.5 | 1 | 1 | 2 | 1 | 1 | 0 | NA | NA | 4 | 0.75 |
| 3_9567199 | 2 | 0.5 | 1 | 1 | 1 | 1 | 4 | 1 | 8 | 0.875 | 1 | 1 | 1 | 1 | 1 | 1 | NA | NA | 3 | 1 |
| 3_9567203 | 2 | 1 | 1 | 1 | 2 | 1 | 2 | 1 | 7 | 1 | 1 | 1 | 2 | 1 | 1 | 1 | NA | NA | 4 | 1 |
| 3_9567213 | 3 | 0.6667 | 2 | 0.5 | 2 | 1 | 2 | 1 | 9 | 0.7778 | 1 | 1 | 1 | 1 | 1 | 1 | NA | NA | 3 | 1 |
| 3_9567215 | 3 | 0.6667 | 2 | 0.5 | 1 | 0 | 3 | 0.6667 | 9 | 0.5556 | 1 | 1 | 1 | 0 | NA | NA | NA | NA | 2 | 0.5 |
| 3_9567245 | 2 | 0.5 | 1 | 1 | NA | NA | 1 | 1 | 4 | 0.75 | 1 | 1 | 1 | 1 | 1 | 0 | NA | NA | 3 | 0.6667 |
| 3_9567252 | 1 | 0 | 1 | 0 | 1 | 1 | 2 | 0 | 5 | 0.2 | 1 | 0 | 1 | 1 | NA | NA | NA | NA | 2 | 0.5 |
| 3_9567257 | 2 | 1 | 1 | 1 | 1 | 1 | 2 | 0.5 | 6 | 0.8333 | 1 | 0 | 1 | 1 | 1 | 1 | NA | NA | 3 | 0.6667 |
| 3_9567264 | 1 | 1 | 2 | 1 | NA | NA | 2 | 1 | 5 | 1 | NA | NA | 1 | 1 | 1 | 1 | NA | NA | 2 | 1 |
| 3_9567265 | 2 | 1 | 1 | 1 | 1 | 1 | 2 | 1 | 6 | 1 | 1 | 1 | NA | NA | 1 | 1 | NA | NA | 2 | 1 |
| 3_9567285 | 2 | 0 | 2 | 0 | 1 | 1 | 1 | 0 | 6 | 0.1667 | 1 | 1 | 1 | 0 | 1 | 1 | NA | NA | 3 | 0.6667 |
| 3_9567290 | 2 | 1 | 2 | 1 | 1 | 1 | 2 | 1 | 7 | 1 | 1 | 1 | 1 | 1 | 1 | 1 | NA | NA | 3 | 1 |
| 3_9567295 | 2 | 0 | 1 | 1 | 1 | 1 | 3 | 0.3333 | 7 | 0.4286 | 1 | 0 | 1 | 0 | 1 | 1 | NA | NA | 3 | 0.3333 |
| 3_9567315 | 2 | 0.5 | 2 | 0 | 1 | 0 | 2 | 1 | 7 | 0.4286 | 1 | 0 | 1 | 1 | 1 | 0 | NA | NA | 3 | 0.3333 |
| 3_9573800 | 1 | 1 | 4 | 0 | 2 | 0.5 | 2 | 1 | 9 | 0.4444 | 4 | 0.25 | 6 | 0.1667 | 2 | 0 | 3 | 0.3333 | 15 | 0.2 |
| 3_9573803 | 2 | 1 | 3 | 0 | 2 | 0 | 1 | 0 | 8 | 0.25 | 4 | 0.25 | 4 | 0.5 | 1 | 0 | 3 | 0.3333 | 12 | 0.3333 |
| 3_9576998 | 12 | 0.0833 | 12 | 0 | 14 | 0.0714 | 14 | 0.0714 | 52 | 0.0577 | 6 | 0 | 12 | 0.0833 | 16 | 0 | 18 | 0.1111 | 52 | 0.0577 |
| 3_9577009 | 20 | 0.1 | 24 | 0.0417 | 20 | 0.15 | 19 | 0.1053 | 83 | 0.0964 | 18 | 0.1111 | 23 | 0.1304 | 22 | 0.1364 | 18 | 0.0556 | 81 | 0.1111 |
| 3_9578582 | 2 | 0.5 | 3 | 0 | 4 | 0 | 3 | 0 | 12 | 0.0833 | 4 | 0.25 | 3 | 0 | 1 | 0 | 2 | 0 | 10 | 0.1 |
| 3_9578600 | 5 | 0.2 | 4 | 0.75 | 4 | 0.25 | 4 | 0 | 17 | 0.2941 | 5 | 0.4 | 3 | 0.6667 | 1 | 0 | 6 | 0.3333 | 15 | 0.4 |
| 3_9578606 | 5 | 0.2 | 2 | 0 | 5 | 0.8 | 4 | 0 | 16 | 0.3125 | 3 | 0.3333 | 3 | 0 | 2 | 0 | 6 | 0.3333 | 14 | 0.2143 |

| | | | | | | | | | | | | | | | | | | | | |
|---|---|---|---|---|---|---|---|---|---|---|---|---|---|---|---|---|---|---|---|---|
| 3_9578632 | 5 | 0 | 2 | 0 | 3 | 0 | 3 | 0.3333 | 13 | 0.0769 | 4 | 0 | 3 | 0 | 2 | 0 | 5 | 0.2 | 14 | 0.0714 |
| 3_9578638 | 5 | 1 | 2 | 1 | 4 | 0.5 | 3 | 0 | 14 | 0.6429 | 3 | 0.3333 | 3 | 1 | 2 | 1 | 6 | 0.8333 | 14 | 0.7857 |
| 3_9578693 | NA | NA | NA | NA | NA | NA | 3 | 0.3333 | 3 | 0.3333 | 2 | 0 | 2 | 1 | NA | NA | 1 | 1 | 5 | 0.6 |
| 3_9578875 | 3 | 0 | 2 | 0.5 | 3 | 0.3333 | 4 | 0.25 | 12 | 0.25 | 2 | 0 | 2 | 0.5 | 2 | 0 | 3 | 0.6667 | 9 | 0.3333 |
| 3_9579146 | NA | NA | 1 | 1 | NA | NA | 1 | 1 | 2 | 1 | NA | NA | 1 | 1 | NA | NA | NA | NA | 1 | 1 |
| 3_9580141 | 6 | 0 | 2 | 0 | 6 | 0.1667 | 5 | 0 | 19 | 0.0526 | 3 | 0 | 4 | 0 | NA | NA | 5 | 0.2 | 12 | 0.0833 |
| 3_9580367 | 2 | 1 | 2 | 0.5 | 10 | 0.3 | 3 | 0 | 17 | 0.3529 | 4 | 0.25 | 1 | 1 | 1 | 0 | 8 | 0.375 | 14 | 0.3571 |
| 3_9580369 | 2 | 0 | 2 | 0 | 7 | 0.2857 | 2 | 0 | 13 | 0.1538 | 4 | 0.5 | NA | NA | 2 | 0 | 7 | 0.1429 | 13 | 0.2308 |
| 3_9580388 | 2 | 0.5 | 2 | 0 | 10 | 0.3 | 3 | 0.6667 | 17 | 0.3529 | 4 | 0.5 | 1 | 1 | 2 | 0.5 | 9 | 0.1111 | 16 | 0.3125 |
| 3_9580402 | 4 | 0 | 4 | 0.5 | 7 | 0.2857 | 9 | 0.3333 | 24 | 0.2917 | 3 | 0.3333 | 3 | 0.3333 | 3 | 0 | 8 | 0 | 17 | 0.1176 |
| 3_9580406 | 4 | 0.5 | 5 | 0.4 | 11 | 0.1818 | 11 | 0.3636 | 31 | 0.3226 | 5 | 0 | 5 | 0.4 | 4 | 0.5 | 7 | 0.2857 | 21 | 0.2857 |
| 3_9580425 | 7 | 0.1429 | 5 | 0 | 11 | 0 | 8 | 0.125 | 31 | 0.0645 | 6 | 0 | 5 | 0 | 4 | 0.25 | 10 | 0.1 | 25 | 0.08 |
| 3_9580522 | 3 | 0 | 4 | 0 | 2 | 0 | 4 | 0.25 | 13 | 0.0769 | NA | NA | 1 | 0 | 1 | 1 | 1 | 0 | 3 | 0.3333 |
| 3_9589620 | 4 | 0.5 | 4 | 1 | 4 | 1 | NA | NA | 12 | 0.8333 | 6 | 0.8333 | 3 | 0.6667 | 4 | 1 | 5 | 1 | 18 | 0.8889 |
| 3_9589628 | 4 | 0 | 6 | 0 | 3 | 0.3333 | NA | NA | 13 | 0.0769 | 6 | 0 | 3 | 0.3333 | 4 | 0 | 4 | 0 | 17 | 0.0588 |
| 3_9589629 | 3 | 1 | 6 | 0.5 | 4 | 1 | NA | NA | 13 | 0.7692 | 6 | 0.6667 | 2 | 0 | 2 | 1 | 4 | 1 | 14 | 0.7143 |
| 3_9589631 | 4 | 0.75 | 5 | 0.6 | 4 | 0.75 | NA | NA | 13 | 0.6923 | 8 | 0.5 | 3 | 0.3333 | 3 | 1 | 5 | 0.2 | 19 | 0.4737 |
| 3_9589632 | 4 | 0.25 | 6 | 0.3333 | 4 | 0.75 | NA | NA | 14 | 0.4286 | 8 | 0.125 | 3 | 0.3333 | 4 | 0.25 | 5 | 0.6 | 20 | 0.3 |
| 3_9589645 | 4 | 0.5 | 6 | 0.3333 | 4 | 0 | NA | NA | 14 | 0.2857 | 7 | 0 | 3 | 0 | 4 | 0.5 | 4 | 0 | 18 | 0.1111 |
| 3_9589648 | 3 | 1 | 6 | 0.6667 | 4 | 0.75 | NA | NA | 13 | 0.7692 | 6 | 0.6667 | 3 | 1 | 4 | 0.75 | 4 | 0.5 | 17 | 0.7059 |
| 3_9589651 | 3 | 0.3333 | 6 | 0.1667 | 3 | 0.3333 | NA | NA | 12 | 0.25 | 7 | 0.1429 | 3 | 0 | 4 | 0 | 4 | 0 | 18 | 0.0556 |
| 3_9589654 | 4 | 0.25 | 6 | 0.1667 | 4 | 0.25 | NA | NA | 14 | 0.2143 | 7 | 0.2857 | 3 | 0 | 4 | 0.25 | 4 | 0 | 18 | 0.1667 |
| 3_9589663 | 4 | 1 | 6 | 0.8333 | 3 | 1 | NA | NA | 13 | 0.9231 | 7 | 0.5714 | 3 | 0.6667 | 4 | 1 | 3 | 1 | 17 | 0.7647 |
| 3_9589679 | 3 | 0.3333 | 6 | 0.3333 | 3 | 0.6667 | NA | NA | 12 | 0.4167 | 6 | 0.1667 | 3 | 0 | 4 | 0.75 | 3 | 0.3333 | 16 | 0.3125 |
| 3_9589683 | 3 | 0 | 6 | 0.1667 | 2 | 0 | NA | NA | 11 | 0.0909 | 4 | 0.25 | 2 | 0 | 4 | 0.5 | 2 | 0 | 12 | 0.25 |
| 3_9590722 | 4 | 0.25 | 7 | 0.4286 | 10 | 0.4 | 4 | 0.5 | 25 | 0.4 | 3 | 0 | 10 | 0.2 | 3 | 0.3333 | 8 | 0.375 | 24 | 0.25 |
| 3_9590724 | 3 | 0.6667 | 6 | 0 | 8 | 0.375 | 3 | 0 | 20 | 0.25 | 1 | 0 | 7 | 0.1429 | 3 | 0.3333 | 4 | 0 | 15 | 0.1333 |
| 3_9590727 | 4 | 0.25 | 6 | 0 | 8 | 0.125 | 4 | 0 | 22 | 0.0909 | 3 | 0 | 10 | 0.2 | 2 | 0 | 7 | 0 | 22 | 0.0909 |
| 3_9594874 | 1 | 1 | 5 | 0 | 7 | 0 | 2 | 0 | 15 | 0.0667 | 1 | 0 | 9 | 0.1111 | 2 | 0 | 8 | 0 | 20 | 0.05 |
| 3_9606278 | 1 | 1 | 1 | 1 | 1 | 0 | NA | NA | 3 | 0.6667 | NA | NA | 1 | 1 | NA | NA | NA | NA | 1 | 1 |
| 3_9606299 | 1 | 0 | 1 | 1 | 1 | 1 | NA | NA | 3 | 0.6667 | NA | NA | 1 | 1 | NA | NA | NA | NA | 1 | 1 |
| 3_9606313 | 1 | 1 | 1 | 1 | 1 | 1 | NA | NA | 3 | 1 | NA | NA | 1 | 1 | NA | NA | NA | NA | 1 | 1 |
| 3_9606323 | 1 | 1 | 1 | 1 | NA | NA | 1 | 1 | 3 | 1 | NA | NA | 1 | 1 | NA | NA | NA | NA | 1 | 1 |
| 3_9606329 | 1 | 1 | 1 | 0 | 1 | 0 | 1 | 0 | 4 | 0.25 | NA | NA | 1 | 1 | NA | NA | NA | NA | 1 | 1 |
| 3_9606831 | 4 | 0.25 | NA | NA | 5 | 0 | 1 | 0 | 10 | 0.1 | NA | NA | 1 | 0 | 1 | 0 | 1 | 1 | 3 | 0.3333 |
| 3_9608091 | 13 | 0.0769 | 9 | 0.3333 | 8 | 0.125 | 10 | 0 | 40 | 0.125 | 6 | 0.1667 | 12 | 0.0833 | 9 | 0.1111 | 9 | 0 | 36 | 0.0833 |
| 3_9608101 | 12 | 0.5833 | 14 | 0.2857 | 11 | 0.0909 | 12 | 0 | 49 | 0.2449 | 7 | 0.1429 | 11 | 0.0909 | 8 | 0.25 | 11 | 0.0909 | 37 | 0.1351 |
| 3_9608109 | 13 | 0.0769 | 14 | 0.1429 | 12 | 0.1667 | 12 | 0.0833 | 51 | 0.1176 | 8 | 0.125 | 13 | 0.1538 | 9 | 0 | 10 | 0.1 | 40 | 0.1 |
| 3_9608123 | 11 | 1 | 11 | 1 | 12 | 1 | 11 | 0.9091 | 45 | 0.9778 | 8 | 0.625 | 13 | 0.9231 | 7 | 0.8571 | 10 | 0.9 | 38 | 0.8421 |
| 3_9608127 | 13 | 0.3077 | 14 | 0.3571 | 11 | 0.6364 | 11 | 0.3636 | 49 | 0.4082 | 8 | 0.375 | 13 | 0.4615 | 11 | 0.4545 | 10 | 0.2 | 42 | 0.381 |
| 3_9609219 | 15 | 0.1333 | 7 | 0 | 6 | 0.3333 | 4 | 0.5 | 32 | 0.1875 | 9 | 0.2222 | 6 | 0 | 4 | 0 | 8 | 0.125 | 27 | 0.1111 |
| 3_9609235 | 13 | 0.4615 | 7 | 0.1429 | 5 | 0.4 | 4 | 0.75 | 29 | 0.4138 | 9 | 0.2222 | 8 | 0.25 | 1 | 0 | 10 | 0.3 | 28 | 0.25 |
| 3_9609280 | 16 | 0.75 | 17 | 0.6471 | 9 | 0.4444 | 9 | 0.8889 | 51 | 0.6863 | 11 | 0.8182 | 12 | 0.8333 | 5 | 0.8 | 16 | 0.75 | 44 | 0.7955 |
| 3_9610163 | 27 | 0.0741 | 40 | 0.2 | 31 | 0.129 | 24 | 0.1667 | 122 | 0.1475 | 32 | 0.1875 | 37 | 0.1081 | 26 | 0.1538 | 31 | 0.1935 | 126 | 0.1587 |
| 3_9610259 | 45 | 0.0667 | 56 | 0.0357 | 59 | 0.1017 | 41 | 0.0976 | 201 | 0.0746 | 46 | 0.1304 | 64 | 0.0938 | 46 | 0.087 | 42 | 0.0714 | 198 | 0.096 |
| 3_9610262 | 48 | 0.2292 | 57 | 0.2632 | 57 | 0.3333 | 40 | 0.15 | 202 | 0.2525 | 45 | 0.2889 | 62 | 0.1935 | 47 | 0.2979 | 39 | 0.2308 | 193 | 0.2487 |
| 3_9610418 | 6 | 0.6667 | 4 | 0.5 | 6 | 0.5 | 4 | 1 | 20 | 0.65 | 4 | 0.75 | 2 | 0.5 | 7 | 0.7143 | 5 | 0.4 | 18 | 0.6111 |
| 3_9610419 | 6 | 0 | 3 | 0 | 6 | 0.1667 | 3 | 0 | 18 | 0.0556 | 5 | 0.4 | 1 | 0 | 6 | 0 | 3 | 0.3333 | 15 | 0.2 |
| 3_9610423 | 5 | 1 | 3 | 1 | 5 | 1 | 3 | 1 | 16 | 1 | 5 | 0.6 | 2 | 1 | 5 | 1 | 4 | 1 | 16 | 0.875 |
| 3_9610429 | 6 | 0.1667 | 2 | 0 | 6 | 0.1667 | 4 | 0 | 18 | 0.1111 | 7 | 0 | 2 | 0 | 7 | 0.1429 | 4 | 0.25 | 20 | 0.1 |
| 3_9610450 | 6 | 0 | 2 | 0.5 | 5 | 0 | 5 | 0.2 | 18 | 0.1111 | 6 | 0 | 2 | 0 | 7 | 0.1429 | 5 | 0 | 20 | 0.05 |
| 3_9610462 | 1 | 0 | 1 | 1 | NA | NA | 1 | 0 | 3 | 0.3333 | 1 | 1 | 1 | 0 | 2 | 0.5 | 1 | 1 | 5 | 0.6 |
| 3_9612845 | NA | NA | 1 | 1 | NA | NA | NA | NA | 1 | 1 | NA | NA | 3 | 0.6667 | NA | NA | NA | NA | 3 | 0.6667 |
| 3_9612867 | NA | NA | 1 | 1 | NA | NA | NA | NA | 1 | 1 | NA | NA | 2 | 1 | NA | NA | NA | NA | 2 | 1 |
| 3_9612879 | NA | NA | 1 | 1 | NA | NA | NA | NA | 1 | 1 | NA | NA | 3 | 1 | NA | NA | NA | NA | 3 | 1 |
| 3_9613024 | NA | NA | 1 | 1 | NA | NA | NA | NA | 1 | 1 | NA | NA | 2 | 1 | NA | NA | NA | NA | 2 | 1 |
| 3_9613030 | NA | NA | 2 | 1 | NA | NA | NA | NA | 2 | 1 | NA | NA | 3 | 1 | NA | NA | NA | NA | 3 | 1 |
| 3_9613037 | NA | NA | 2 | 1 | NA | NA | NA | NA | 2 | 1 | NA | NA | 3 | 1 | NA | NA | NA | NA | 3 | 1 |
| 3_9613039 | NA | NA | 2 | 1 | NA | NA | NA | NA | 2 | 1 | NA | NA | 3 | 1 | NA | NA | NA | NA | 3 | 1 |
| 3_9613043 | NA | NA | 2 | 0.5 | NA | NA | NA | NA | 2 | 0.5 | NA | NA | 3 | 0.6667 | NA | NA | NA | NA | 3 | 0.6667 |
| 3_9613057 | NA | NA | 2 | 0.5 | NA | NA | NA | NA | 2 | 0.5 | NA | NA | 3 | 0.6667 | NA | NA | NA | NA | 3 | 0.6667 |
| 3_9613063 | NA | NA | 2 | 1 | NA | NA | NA | NA | 2 | 1 | NA | NA | 3 | 0.6667 | NA | NA | NA | NA | 3 | 0.6667 |
| 3_9614588 | 1 | 1 | NA | NA | 1 | 1 | NA | NA | 2 | 1 | NA | NA | NA | NA | NA | NA | 1 | 1 | 1 | 1 |
| 3_9614602 | 1 | 1 | NA | NA | 1 | 0 | NA | NA | 2 | 0.5 | NA | NA | NA | NA | NA | NA | 1 | 1 | 1 | 1 |
| 3_9614660 | NA | NA | 1 | 0 | 1 | 1 | NA | NA | 2 | 0.5 | NA | NA | NA | NA | NA | NA | 1 | 1 | 1 | 1 |

| | | | | | | | | | | | | | | | | | | | | |
|---|---|---|---|---|---|---|---|---|---|---|---|---|---|---|---|---|---|---|---|---|
| 3_96146812 | NA | NA | 1 | 1 | 1 | 1 | NA | NA | 2 | 1 | NA | NA | NA | NA | NA | NA | 1 | 1 | 1 | 1 |
| 3_96153647 | NA | NA | 2 | 1 | NA | NA | NA | NA | 2 | 1 | NA | NA | 1 | 0 | NA | NA | 1 | 1 | 2 | 0.5 |
| 3_96153653 | 1 | 0 | 1 | 1 | NA | NA | NA | NA | 2 | 0.5 | NA | NA | 1 | 0 | NA | NA | 1 | 1 | 2 | 0.5 |
| 3_96153755 | 1 | 0 | 2 | 0.5 | NA | NA | NA | NA | 3 | 0.3333 | NA | NA | 1 | 0 | NA | NA | 1 | 1 | 2 | 0.5 |
| 3_96153788 | 1 | 0 | 2 | 0.5 | NA | NA | NA | NA | 3 | 0.3333 | NA | NA | 1 | 1 | NA | NA | 1 | 0 | 2 | 0.5 |
| 3_96153800 | NA | NA | 2 | 1 | NA | NA | NA | NA | 2 | 1 | NA | NA | 1 | 1 | NA | NA | 1 | 1 | 2 | 1 |
| 3_96154220 | 1 | 0 | 2 | 0.5 | 1 | 1 | NA | NA | 4 | 0.5 | NA | NA | 1 | 0 | NA | NA | 1 | 1 | 2 | 0.5 |
| 3_96154434 | 1 | 1 | 2 | 1 | 1 | 1 | NA | NA | 4 | 1 | NA | NA | 1 | 1 | NA | NA | 1 | 1 | 2 | 1 |
| 3_96154446 | 2 | 0 | 2 | 0.5 | 1 | 1 | NA | NA | 5 | 0.4 | NA | NA | 1 | 1 | NA | NA | 1 | 1 | 2 | 1 |
| 3_96154521 | 2 | 1 | 2 | 1 | 1 | 1 | NA | NA | 5 | 1 | NA | NA | 1 | 0 | NA | NA | 1 | 1 | 2 | 0.5 |
| 3_96155695 | 1 | 1 | 3 | 0.6667 | NA | NA | NA | NA | 4 | 0.75 | 1 | 0 | 3 | 0.3333 | NA | NA | 1 | 0 | 5 | 0.2 |
| 3_96191049 | NA | NA | 2 | 0.5 | NA | NA | NA | NA | 2 | 0.5 | 1 | 1 | 1 | 0 | 1 | 1 | NA | NA | 3 | 0.6667 |
| 3_96191273 | NA | NA | 2 | 0.5 | NA | NA | NA | NA | 2 | 0.5 | 1 | 1 | 1 | 0 | 1 | 1 | NA | NA | 3 | 0.6667 |
| 3_96191316 | NA | NA | 2 | 0.5 | NA | NA | NA | NA | 2 | 0.5 | 1 | 0 | 1 | 0 | 1 | 1 | NA | NA | 3 | 0.3333 |
| 3_96191388 | NA | NA | 1 | 1 | NA | NA | NA | NA | 1 | 1 | NA | NA | 1 | 0 | 1 | 1 | NA | NA | 2 | 0.5 |
| 3_96191505 | NA | NA | 2 | 1 | NA | NA | NA | NA | 2 | 1 | 1 | 1 | 1 | 0 | 1 | 1 | NA | NA | 3 | 0.6667 |
| 3_96191538 | NA | NA | 1 | 1 | NA | NA | 1 | 1 | 2 | 1 | 1 | 1 | 1 | 1 | 1 | 1 | NA | NA | 3 | 1 |
| 3_96191576 | NA | NA | 3 | 0.6667 | NA | NA | 1 | 1 | 4 | 0.75 | 2 | 0.5 | 1 | 1 | 1 | 0 | NA | NA | 4 | 0.5 |
| 3_96191584 | NA | NA | 3 | 0.6667 | NA | NA | 1 | 1 | 4 | 0.75 | 2 | 1 | 1 | 1 | 1 | 1 | NA | NA | 4 | 1 |
| 3_96467066 | 5 | 0.2 | 5 | 0 | 2 | 0 | 3 | 0.3333 | 15 | 0.1333 | 3 | 0 | 3 | 0.3333 | 5 | 0.2 | 8 | 0.25 | 19 | 0.2105 |
| 3_96467902 | 4 | 0.25 | 3 | 0.3333 | 1 | 1 | 1 | 0 | 9 | 0.3333 | 5 | 0.2 | 2 | 0.5 | 5 | 0.4 | 5 | 0.2 | 17 | 0.2941 |
| 3_96467919 | 5 | 0.4 | 4 | 0.75 | 4 | 0.75 | 3 | 0.3333 | 16 | 0.5625 | 5 | 0.6 | 3 | 1 | 4 | 0.75 | 8 | 0.875 | 20 | 0.8 |
| 3_96468065 | 5 | 0.2 | 5 | 0.2 | 3 | 0 | 3 | 0.3333 | 16 | 0.1875 | 4 | 0 | 2 | 0 | 4 | 0.25 | 8 | 0.625 | 18 | 0.3333 |
| 3_96468085 | 5 | 0.4 | 5 | 0 | 4 | 0 | 4 | 0 | 18 | 0.1111 | 5 | 0 | 3 | 0.3333 | 5 | 0.2 | 10 | 0.1 | 23 | 0.1304 |
| 3_96468119 | 5 | 0.4 | 4 | 0.25 | 3 | 0 | 4 | 0.25 | 16 | 0.25 | 5 | 0.2 | 3 | 0.6667 | 5 | 0.2 | 10 | 0.2 | 23 | 0.2609 |
| 3_96468133 | 5 | 0.4 | 3 | 0 | 3 | 0.3333 | 2 | 0 | 13 | 0.2308 | 4 | 0 | 2 | 0.5 | 4 | 0.25 | 6 | 0.5 | 16 | 0.3125 |
| 3_96468194 | 5 | 1 | 5 | 0.8 | 4 | 1 | 4 | 0.75 | 18 | 0.8889 | 4 | 0.5 | 4 | 1 | 5 | 0.8 | 10 | 0.7 | 23 | 0.7391 |
| 3_96468244 | 7 | 0.1429 | 4 | 0.25 | 4 | 0.75 | 4 | 0.25 | 19 | 0.3158 | 4 | 0.5 | 4 | 0.25 | 5 | 0 | 8 | 0 | 21 | 0.1429 |
| 3_96468344 | 5 | 0.2 | 5 | 0.4 | 4 | 0.25 | 3 | 0 | 17 | 0.2353 | 3 | 0.3333 | NA | NA | 2 | 0 | 7 | 0 | 12 | 0.0833 |
| 3_96468402 | 7 | 0.1429 | 4 | 0 | 4 | 0 | 3 | 0 | 18 | 0.0556 | 4 | 0 | 3 | 0.3333 | 4 | 0 | 10 | 0.1 | 21 | 0.0952 |
| 3_96468636 | 9 | 0.2222 | 4 | 0 | 3 | 0 | 3 | 0 | 19 | 0.1053 | 5 | 0.2 | 3 | 0.3333 | 4 | 0 | 9 | 0 | 21 | 0.0952 |
| 3_96468781 | 7 | 0.2857 | 5 | 0 | 4 | 0 | 3 | 0 | 19 | 0.1053 | 5 | 0 | 5 | 0.2 | 3 | 0.3333 | 7 | 0 | 20 | 0.1 |
| 3_96469006 | 7 | 0.4286 | 4 | 0.25 | 4 | 0.25 | 6 | 0.1667 | 21 | 0.2857 | 5 | 0.4 | 3 | 0.3333 | 4 | 0.5 | 9 | 0.1111 | 21 | 0.2857 |
| 3_96485736 | 2 | 0.5 | 5 | 0.2 | 3 | 0 | 2 | 0 | 12 | 0.1667 | 1 | 0 | 4 | 0.5 | NA | NA | 4 | 0.5 | 9 | 0.4444 |
| 3_96485782 | 2 | 0 | 5 | 0.6 | 4 | 0.75 | 3 | 0.6667 | 14 | 0.5714 | 2 | 0 | 3 | 0.3333 | NA | NA | 3 | 1 | 8 | 0.5 |
| 3_96485802 | 2 | 0 | 2 | 0.5 | 5 | 1 | 2 | 0.5 | 11 | 0.6364 | 1 | 1 | 4 | 0.75 | NA | NA | 3 | 0.6667 | 8 | 0.75 |
| 3_96485864 | 3 | 0.3333 | 4 | 0.25 | 3 | 0.6667 | 2 | 1 | 12 | 0.5 | 2 | 0.5 | 2 | 1 | NA | NA | 3 | 0.6667 | 7 | 0.7143 |
| 3_96489165 | NA | NA | 1 | 0 | 1 | 1 | 1 | 0 | 3 | 0.3333 | NA | NA | 1 | 1 | NA | NA | NA | NA | 1 | 1 |
| 3_96504839 | 3 | 0.3333 | 1 | 1 | 15 | 0.3333 | 8 | 0.125 | 27 | 0.2963 | 1 | 0 | 5 | 0.8 | 3 | 0 | 6 | 0.5 | 15 | 0.4667 |
| 3_96504915 | 4 | 0.25 | 1 | 1 | 11 | 0.5455 | 5 | 0.4 | 21 | 0.4762 | 2 | 0 | 4 | 0.5 | 3 | 0.3333 | 5 | 0.6 | 14 | 0.4286 |
| 3_96505018 | 4 | 0 | 1 | 0 | 14 | 0.0714 | 8 | 0.25 | 27 | 0.1111 | 3 | 0 | 6 | 0.3333 | 3 | 0.3333 | 8 | 0 | 20 | 0.15 |
| 3_96505156 | 3 | 0.3333 | 1 | 0 | 10 | 0.3 | 8 | 0.125 | 22 | 0.2273 | 2 | 0.5 | 5 | 0.4 | 1 | 0 | 4 | 0 | 12 | 0.25 |
| 3_96505306 | 5 | 0.6 | 1 | 1 | 15 | 0.6 | 9 | 0.2222 | 30 | 0.5 | 4 | 0.25 | 6 | 0.3333 | 4 | 0.5 | 10 | 0.3 | 24 | 0.3333 |
| 3_96505368 | 2 | 0 | 1 | 0 | 8 | 0.25 | 7 | 0.2857 | 18 | 0.2222 | 1 | 0 | 4 | 0.25 | 2 | 0.5 | 7 | 0 | 14 | 0.1429 |
| 3_96505429 | 4 | 1 | NA | NA | 16 | 1 | 10 | 1 | 30 | 1 | 4 | 1 | 6 | 1 | 4 | 1 | 10 | 1 | 24 | 1 |
| 3_96505465 | 5 | 0.2 | 1 | 0 | 13 | 0.1538 | 7 | 0 | 26 | 0.1154 | 3 | 0 | 5 | 0.2 | 3 | 0 | 9 | 0.1111 | 20 | 0.1 |
| 3_96505790 | 4 | 0.5 | NA | NA | 12 | 1 | 6 | 1 | 22 | 0.9091 | 3 | 0.3333 | 2 | 1 | 4 | 0.75 | 8 | 0.75 | 17 | 0.7059 |
| 3_96505829 | 4 | 1 | 2 | 1 | 13 | 1 | 9 | 1 | 28 | 1 | 4 | 1 | 5 | 1 | 5 | 1 | 9 | 1 | 23 | 1 |
| 3_96505843 | 5 | 0 | 3 | 0 | 14 | 0.0714 | 11 | 0.0909 | 33 | 0.0606 | 4 | 0 | 6 | 0.1667 | 5 | 0.2 | 10 | 0 | 25 | 0.08 |
| 3_96506025 | 5 | 0.8 | 4 | 0 | 16 | 0.3125 | 11 | 0.1818 | 36 | 0.3056 | 4 | 0.25 | 5 | 0.6 | 6 | 0.1667 | 11 | 0.1818 | 26 | 0.2692 |
| 3_96506038 | 3 | 0 | 2 | 0 | 13 | 0.2308 | 8 | 0.25 | 26 | 0.1923 | 3 | 0.3333 | 3 | 0.3333 | 6 | 0 | 10 | 0.5 | 22 | 0.3182 |
| 3_96506184 | 4 | 0 | 4 | 0 | 14 | 0.0714 | 11 | 0.0909 | 33 | 0.0606 | 3 | 0 | 5 | 0.4 | 6 | 0.1667 | 11 | 0.0909 | 25 | 0.16 |
| 3_96506220 | 4 | 0.75 | 3 | 0.6667 | 12 | 0.6667 | 8 | 1 | 27 | 0.7778 | 4 | 0.75 | 5 | 0.6 | 4 | 0.75 | 9 | 0.6667 | 22 | 0.6818 |
| 3_96506356 | 5 | 0.4 | 5 | 0.2 | 17 | 0.2353 | 10 | 0 | 37 | 0.1892 | 4 | 0.5 | 7 | 0.4286 | 5 | 0.4 | 12 | 0.1667 | 28 | 0.3214 |
| 3_96531044 | 2 | 0.5 | 2 | 0 | 2 | 0 | 2 | 0.5 | 8 | 0.25 | 1 | 0 | 2 | 0.5 | 1 | 0 | NA | NA | 4 | 0.25 |
| 3_96539128 | 7 | 0 | 9 | 0.3333 | 10 | 0.1 | 11 | 0 | 37 | 0.1081 | 13 | 0.1538 | 9 | 0 | 5 | 0 | 5 | 0 | 32 | 0.0625 |
| 3_96539331 | 2 | 0.5 | 2 | 0 | 6 | 0 | 6 | 0 | 16 | 0.0625 | 4 | 0 | 3 | 0.3333 | 2 | 0 | 1 | 0 | 10 | 0.1 |
| 3_96550964 | 3 | 0 | 8 | 0.5 | 5 | 0.4 | 2 | 0.5 | 18 | 0.3889 | 8 | 0.5 | 14 | 0.4286 | 3 | 0.6667 | 6 | 0.5 | 31 | 0.4839 |
| 3_96551024 | 4 | 0.25 | 8 | 0.5 | 5 | 0.6 | 2 | 1 | 19 | 0.5263 | 4 | 0.75 | 13 | 0.3846 | 4 | 0.75 | 5 | 0.8 | 26 | 0.5769 |
| 3_96551032 | 3 | 0 | 8 | 0.375 | 5 | 0.2 | 2 | 0 | 18 | 0.2222 | 9 | 0 | 12 | 0.25 | 2 | 0 | 4 | 0 | 27 | 0.1111 |
| 3_96551048 | 4 | 0.75 | 8 | 0 | 5 | 0.2 | 3 | 0 | 20 | 0.2 | 7 | 0.2857 | 9 | 0.1111 | 3 | 0 | 6 | 0.3333 | 25 | 0.2 |
| 3_96551160 | 5 | 0 | 8 | 0.125 | 4 | 0.25 | 3 | 0 | 20 | 0.1 | 8 | 0 | 14 | 0.0714 | 4 | 0.5 | 6 | 0 | 32 | 0.0938 |
| 3_96551184 | 2 | 0.5 | 6 | 0.8333 | 4 | 1 | 2 | 1 | 14 | 0.8571 | 4 | 0.5 | 12 | 0.8333 | 2 | 0.5 | 5 | 0.8 | 23 | 0.7391 |
| 3_96551296 | 5 | 0.2 | 7 | 0 | 5 | 0.2 | 3 | 0.3333 | 20 | 0.15 | 8 | 0.25 | 12 | 0 | 4 | 0 | 6 | 0 | 30 | 0.0667 |
| 3_96551394 | 4 | 0 | 4 | 0.25 | 4 | 0 | 2 | 0 | 14 | 0.0714 | 7 | 0 | 12 | 0.1667 | 2 | 0 | 4 | 0 | 25 | 0.08 |

| | | | | | | | | | | | | | | | | | | | | |
|---|---|---|---|---|---|---|---|---|---|---|---|---|---|---|---|---|---|---|---|---|
| 3_9659136 | NA | NA | NA | NA | 1 | 0 | 1 | 1 | 2 | 0.5 | NA | NA | 1 | 1 | 3 | 1 | NA | NA | 4 | 1 |
| 3_9659153 | NA | NA | NA | NA | 2 | 1 | 2 | 0.5 | 4 | 0.75 | NA | NA | 1 | 1 | 2 | 1 | NA | NA | 3 | 1 |
| 3_9659156 | NA | NA | NA | NA | 2 | 0.5 | 2 | 1 | 4 | 0.75 | NA | NA | 1 | 1 | 3 | 1 | NA | NA | 4 | 1 |
| 3_9659181 | NA | NA | NA | NA | 2 | 1 | 2 | 1 | 4 | 1 | NA | NA | 1 | 1 | 2 | 1 | NA | NA | 3 | 1 |
| 3_9659230 | 1 | 1 | NA | NA | 2 | 0.5 | 2 | 0 | 5 | 0.4 | NA | NA | 1 | 1 | 4 | 0 | NA | NA | 5 | 0.2 |
| 3_9659231 | 1 | 1 | NA | NA | 1 | 1 | 2 | 0.5 | 4 | 0.75 | NA | NA | 1 | 0 | 3 | 0.6667 | NA | NA | 4 | 0.5 |
| 3_9661242 | 2 | 1 | 1 | 0 | 1 | 1 | 1 | 1 | 5 | 0.8 | 1 | 0 | 3 | 1 | 2 | 1 | 1 | 0 | 7 | 0.7143 |
| 3_9661285 | 3 | 0.3333 | 2 | 0.5 | 1 | 1 | 3 | 0 | 9 | 0.3333 | 1 | 0 | 3 | 0.6667 | 3 | 0.3333 | 1 | 0 | 8 | 0.375 |
| 3_9661305 | NA | NA | 1 | 1 | NA | NA | NA | NA | 1 | 1 | NA | NA | 1 | 1 | NA | NA | 2 | 0 | 3 | 0.3333 |
| 3_9663100 | 1 | 1 | 2 | 0.5 | 1 | 0 | 1 | 1 | 5 | 0.6 | 1 | 1 | 1 | 0 | 1 | 1 | 2 | 0.5 | 5 | 0.6 |
| 3_9663112 | 1 | 0 | 1 | 1 | 2 | 0 | 3 | 0.3333 | 7 | 0.2857 | 1 | 0 | 2 | 0.5 | 1 | 0 | 3 | 0.3333 | 7 | 0.2857 |
| 3_9663470 | 1 | 1 | NA | NA | 2 | 0 | 2 | 0 | 5 | 0.2 | NA | NA | 2 | 0 | 1 | 0 | 3 | 0.6667 | 6 | 0.3333 |
| 3_9669723 | 4 | 0 | 6 | 0 | 15 | 0.0667 | 7 | 0.2857 | 32 | 0.0938 | 2 | 0 | 8 | 0 | 5 | 0.2 | 8 | 0.125 | 23 | 0.087 |
| 3_9669731 | 3 | 0.6667 | 6 | 1 | 16 | 0.75 | 7 | 0.8571 | 32 | 0.8125 | 3 | 1 | 8 | 0.5 | 6 | 0.6667 | 9 | 0.7778 | 26 | 0.6923 |
| 3_9677565 | 3 | 0.6667 | 1 | 1 | 3 | 0.3333 | 3 | 0.6667 | 10 | 0.6 | 1 | 1 | NA | NA | 3 | 0.6667 | 1 | 1 | 5 | 0.8 |
| 3_9677586 | 3 | 0.6667 | NA | NA | 3 | 0.6667 | 2 | 1 | 8 | 0.75 | NA | NA | NA | NA | 3 | 0.3333 | 1 | 1 | 4 | 0.5 |
| 3_9677592 | 3 | 0.6667 | 1 | 0 | 2 | 1 | 2 | 1 | 8 | 0.75 | NA | NA | NA | NA | 3 | 1 | 1 | 1 | 4 | 1 |
| 3_9677595 | 3 | 1 | 1 | 1 | 2 | 1 | 2 | 1 | 8 | 1 | NA | NA | NA | NA | 3 | 1 | 1 | 1 | 4 | 1 |
| 3_9677613 | 1 | 1 | 1 | 1 | 1 | 0 | 1 | 1 | 4 | 0.75 | NA | NA | NA | NA | 2 | 1 | NA | NA | 2 | 1 |
| 3_9677617 | 2 | 0.5 | 1 | 0 | 1 | 0 | 2 | 1 | 6 | 0.5 | NA | NA | NA | NA | 2 | 0 | 1 | 1 | 3 | 0.3333 |
| 3_9677625 | 2 | 0.5 | 1 | 0 | 1 | 0 | 1 | 0 | 5 | 0.2 | NA | NA | NA | NA | 2 | 0.5 | 1 | 0 | 3 | 0.3333 |
| 3_9677651 | 1 | 1 | 1 | 1 | 1 | 1 | 2 | 1 | 5 | 1 | NA | NA | NA | NA | 1 | 1 | 1 | 1 | 2 | 1 |
| 3_9677681 | NA | NA | 1 | 0 | 1 | 0 | 2 | 1 | 4 | 0.5 | NA | NA | NA | NA | NA | NA | 1 | 1 | 1 | 1 |
| 3_9677757 | 1 | 1 | NA | NA | 1 | 1 | 1 | 1 | 3 | 1 | NA | NA | NA | NA | NA | NA | 1 | 1 | 1 | 1 |
| 3_9677765 | 1 | 0 | NA | NA | 1 | 1 | 2 | 0 | 4 | 0.25 | NA | NA | NA | NA | NA | NA | 1 | 1 | 1 | 1 |
| 3_9677775 | 1 | 1 | NA | NA | 1 | 1 | 2 | 1 | 4 | 1 | NA | NA | NA | NA | NA | NA | 1 | 1 | 1 | 1 |
| 3_9677790 | 1 | 1 | NA | NA | 1 | 1 | 1 | 1 | 3 | 1 | NA | NA | NA | NA | NA | NA | 1 | 1 | 1 | 1 |
| 3_9677867 | 3 | 0 | NA | NA | NA | NA | 4 | 0.25 | 7 | 0.1429 | 1 | 0 | 3 | 0 | 5 | 0.4 | 4 | 0.25 | 13 | 0.2308 |
| 3_9677869 | 3 | 0.6667 | NA | NA | NA | NA | 4 | 1 | 7 | 0.8571 | NA | NA | 3 | 0.6667 | 4 | 0.25 | 4 | 0.25 | 11 | 0.3636 |
| 3_9677870 | 2 | 1 | NA | NA | NA | NA | 3 | 1 | 5 | 1 | NA | NA | 3 | 0.3333 | 2 | 0 | 2 | 1 | 7 | 0.4286 |
| 3_9706642 | NA | NA | 2 | 0 | NA | NA | 2 | 0.5 | 4 | 0.25 | 1 | 0 | NA | NA | 3 | 0.6667 | NA | NA | 4 | 0.5 |
| 3_9711164 | NA | NA | 3 | 1 | NA | NA | 1 | 1 | 4 | 1 | 2 | 1 | 3 | 1 | NA | NA | 1 | 1 | 6 | 1 |
| 3_9711183 | NA | NA | 2 | 0.5 | NA | NA | 1 | 1 | 3 | 0.6667 | 1 | 1 | 1 | 0 | NA | NA | NA | NA | 2 | 0.5 |
| 3_9711194 | NA | NA | 2 | 1 | NA | NA | 1 | 1 | 3 | 1 | 1 | 1 | 2 | 0.5 | NA | NA | 1 | 0 | 4 | 0.5 |
| 3_9711206 | NA | NA | 3 | 1 | NA | NA | 2 | 1 | 5 | 1 | 2 | 1 | 3 | 1 | 1 | 1 | 1 | 1 | 7 | 1 |
| 3_9711220 | NA | NA | 3 | 1 | NA | NA | 2 | 0.5 | 5 | 0.8 | 2 | 1 | 3 | 1 | 1 | 1 | 1 | 1 | 7 | 1 |
| 3_9711227 | NA | NA | 2 | 1 | NA | NA | NA | NA | 2 | 1 | 2 | 0 | 3 | 0.6667 | 1 | 0 | 1 | 0 | 7 | 0.2857 |
| 3_9711229 | NA | NA | 3 | 1 | NA | NA | 1 | 1 | 4 | 1 | 3 | 1 | 3 | 0.6667 | 1 | 0 | 1 | 0 | 8 | 0.625 |
| 3_9711247 | NA | NA | 3 | 0.6667 | NA | NA | 1 | 0 | 4 | 0.5 | 2 | 0.5 | 2 | 1 | 1 | 0 | 1 | 1 | 6 | 0.6667 |
| 3_9711250 | NA | NA | 3 | 0.3333 | NA | NA | 1 | 0 | 4 | 0.25 | 1 | 1 | 2 | 0 | 1 | 0 | NA | NA | 4 | 0.25 |
| 3_9711255 | NA | NA | 3 | 1 | NA | NA | NA | NA | 3 | 1 | 2 | 1 | 1 | 1 | 1 | 1 | 1 | 1 | 5 | 1 |
| 3_9711256 | NA | NA | 3 | 1 | NA | NA | 1 | 1 | 4 | 1 | 2 | 0.5 | 2 | 0.5 | 1 | 1 | 1 | 1 | 6 | 0.6667 |
| 3_9711271 | NA | NA | 3 | 0.3333 | NA | NA | 1 | 1 | 4 | 0.5 | 2 | 1 | 1 | 0 | 1 | 1 | 1 | 0 | 5 | 0.6 |
| 3_9711293 | NA | NA | 3 | 0 | NA | NA | 1 | 1 | 4 | 0.25 | 1 | 0 | 1 | 0 | 1 | 1 | 1 | 1 | 4 | 0.5 |
| 3_9711316 | NA | NA | 3 | 0 | NA | NA | 1 | 1 | 4 | 0.25 | 2 | 0.5 | 1 | 0 | 1 | 0 | 1 | 1 | 5 | 0.4 |
| 3_9711603 | NA | NA | 2 | 0.5 | NA | NA | NA | NA | 2 | 0.5 | 1 | 0 | 1 | 1 | NA | NA | NA | NA | 2 | 0.5 |
| 3_9711626 | NA | NA | 2 | 1 | NA | NA | NA | NA | 2 | 1 | 1 | 1 | NA | NA | NA | NA | NA | NA | 1 | 1 |
| 3_9711730 | NA | NA | 1 | 1 | NA | NA | NA | NA | 1 | 1 | 1 | 1 | 1 | 0 | NA | NA | NA | NA | 2 | 0.5 |
| 3_9711750 | NA | NA | 1 | 1 | NA | NA | NA | NA | 1 | 1 | 1 | 1 | NA | NA | NA | NA | NA | NA | 1 | 1 |
| 3_9711759 | NA | NA | 1 | 1 | NA | NA | NA | NA | 1 | 1 | 1 | 1 | 1 | 1 | NA | NA | NA | NA | 2 | 1 |
| 3_9755960 | NA | NA | 1 | 1 | 1 | 1 | NA | NA | 2 | 1 | 1 | 1 | 1 | 1 | NA | NA | NA | NA | 2 | 1 |
| 3_9757760 | 1 | 1 | 2 | 1 | 1 | 0 | 4 | 0.75 | 8 | 0.75 | NA | NA | 1 | 0 | 3 | 0.3333 | 1 | 0 | 5 | 0.2 |
| 3_9757880 | 2 | 0.5 | 5 | 0 | NA | NA | 6 | 0 | 13 | 0.0769 | 1 | 0 | 1 | 0 | 4 | 0.25 | 1 | 1 | 7 | 0.2857 |
| 3_9758029 | 9 | 0.4444 | 5 | 0.6 | 1 | 0 | 8 | 0.5 | 23 | 0.4783 | 3 | 0.6667 | 2 | 0 | 5 | 0.2 | 1 | 1 | 11 | 0.3636 |
| 3_9758032 | 6 | 0.3333 | 5 | 0.2 | 1 | 0 | 7 | 0.1429 | 19 | 0.2105 | 2 | 0 | 3 | 0.3333 | 4 | 0 | 1 | 0 | 10 | 0.1 |
| 3_9758034 | 8 | 0.625 | 5 | 0.8 | 1 | 0 | 8 | 0.75 | 22 | 0.6818 | 2 | 0.5 | 3 | 0.3333 | 5 | 0.4 | 3 | 0.6667 | 13 | 0.4615 |
| 3_9758048 | 8 | 0.25 | 5 | 0.6 | 2 | 0 | 8 | 0.125 | 23 | 0.2609 | 3 | 0.6667 | 3 | 0 | 5 | 0 | 2 | 0.5 | 13 | 0.2308 |
| 3_9758060 | 9 | 0.2222 | 6 | 0.1667 | 2 | 0 | 8 | 0.375 | 25 | 0.24 | 3 | 0.3333 | 3 | 0 | 5 | 0.2 | 2 | 0.5 | 13 | 0.2308 |
| 3_9758095 | 7 | 0 | 5 | 0.4 | 3 | 0 | 7 | 0.1429 | 22 | 0.1364 | 4 | 0.5 | 4 | 0 | 4 | 0 | 2 | 1 | 14 | 0.2857 |
| 3_9758099 | 9 | 0.5556 | 6 | 0.5 | 2 | 0 | 6 | 0.3333 | 23 | 0.4348 | 2 | 0.5 | 4 | 0.25 | 5 | 0 | 2 | 0 | 13 | 0.1538 |
| 3_9758100 | 9 | 0.1111 | 7 | 0.2857 | 2 | 0 | 8 | 0.625 | 26 | 0.3077 | 4 | 0.5 | 2 | 0.5 | 5 | 0 | 3 | 0.3333 | 14 | 0.2857 |
| 3_9758102 | 8 | 0.375 | 5 | 0.2 | 2 | 0 | 5 | 0.4 | 20 | 0.3 | 2 | 0.5 | 3 | 0.3333 | 6 | 0.1667 | 3 | 0 | 14 | 0.2143 |
| 3_9758104 | 8 | 0.125 | 6 | 0.6667 | 2 | 0 | 7 | 0.5714 | 23 | 0.3913 | 2 | 0.5 | 4 | 0.5 | 6 | 0.1667 | 2 | 0.5 | 14 | 0.3571 |
| 3_9758105 | 10 | 0.1 | 6 | 0.3333 | 2 | 0 | 7 | 0.1429 | 25 | 0.16 | 4 | 0 | 4 | 0 | 6 | 0.1667 | 2 | 0 | 16 | 0.0625 |
| 3_9759720 | 14 | 0.1429 | 14 | 0.0714 | 11 | 0.0909 | 10 | 0.1 | 49 | 0.102 | 11 | 0.2727 | 10 | 0.1 | 9 | 0 | 8 | 0.125 | 38 | 0.1316 |

| | | | | | | | | | | | | | | | | | | | | |
|---|---|---|---|---|---|---|---|---|---|---|---|---|---|---|---|---|---|---|---|---|
| 3_97597312 | 10 | 0 | 10 | 0.2 | 7 | 0 | 8 | 0.125 | 35 | 0.0857 | 6 | 0.1667 | 11 | 0.2727 | 9 | 0.3333 | 8 | 0 | 34 | 0.2059 |
| 3_97597460 | 12 | 0.0833 | 12 | 0.0833 | 8 | 0 | 9 | 0.1111 | 41 | 0.0732 | 7 | 0.2857 | 10 | 0.1 | 5 | 0 | 6 | 0 | 28 | 0.1071 |
| 3_97597627 | 13 | 0.0769 | 11 | 0.2727 | 7 | 0.1429 | 8 | 0.375 | 39 | 0.2051 | 9 | 0.3333 | 7 | 0.1429 | 7 | 0.1429 | 7 | 0.2857 | 30 | 0.2333 |
| 3_97597660 | 13 | 0.7692 | 14 | 0.7143 | 7 | 0.7143 | 6 | 1 | 40 | 0.775 | 9 | 1 | 11 | 0.6364 | 7 | 0.7143 | 8 | 0.625 | 35 | 0.7429 |
| 3_97597755 | 10 | 0.3 | 13 | 0.5385 | 6 | 0.5 | 6 | 0.6667 | 35 | 0.4857 | 8 | 0.375 | 10 | 0.6 | 6 | 0.5 | 7 | 0.4286 | 31 | 0.4839 |
| 3_97597839 | 9 | 0 | 12 | 0.0833 | 6 | 0.1667 | 7 | 0 | 34 | 0.0588 | 7 | 0.1429 | 8 | 0.125 | 3 | 0.3333 | 9 | 0 | 27 | 0.1111 |
| 3_97597872 | 8 | 1 | 11 | 0.6364 | 4 | 0.75 | 7 | 0.8571 | 30 | 0.8 | 7 | 1 | 9 | 0.8889 | 5 | 0.6 | 8 | 0.875 | 29 | 0.8621 |
| 3_97597882 | 5 | 0.2 | 9 | 0.2222 | 4 | 0 | 3 | 0.3333 | 21 | 0.1905 | 5 | 0.2 | 6 | 0.1667 | 3 | 0 | 3 | 0.3333 | 17 | 0.1765 |
| 3_97597924 | 10 | 0.7 | 14 | 0.6429 | 5 | 0.6 | 5 | 0.6 | 34 | 0.6471 | 6 | 0.8333 | 10 | 0.7 | 6 | 0.6667 | 6 | 1 | 28 | 0.7857 |
| 3_97597961 | 12 | 0.0833 | 14 | 0.2857 | 6 | 0.5 | 7 | 0.4286 | 39 | 0.2821 | 7 | 0.5714 | 8 | 0.25 | 5 | 0.6 | 5 | 0.2 | 25 | 0.4 |
| 3_97597982 | 11 | 0.2727 | 11 | 0.2727 | 6 | 0 | 6 | 0 | 34 | 0.1765 | 7 | 0.1429 | 9 | 0 | 6 | 0.1667 | 8 | 0.25 | 30 | 0.1333 |
| 3_97598024 | 10 | 0.4 | 12 | 0.5833 | 7 | 0.2857 | 7 | 0.7143 | 36 | 0.5 | 5 | 0.2 | 10 | 0.5 | 4 | 0 | 8 | 0.25 | 27 | 0.2963 |
| 3_97622477 | 1 | 1 | NA | NA | NA | NA | 1 | 0 | 2 | 0.5 | 3 | 1 | 1 | 1 | 2 | 0.5 | 2 | 0.5 | 8 | 0.75 |
| 3_97622552 | 2 | 0.5 | 2 | 0 | NA | NA | 2 | 0.5 | 6 | 0.3333 | 2 | 1 | 1 | 0 | 1 | 1 | 3 | 1 | 7 | 0.8571 |
| 3_97622608 | 3 | 0.6667 | 2 | 1 | NA | NA | 2 | 0 | 7 | 0.5714 | 2 | 0 | 1 | 0 | NA | NA | 1 | 1 | 4 | 0.25 |
| 3_97626731 | 2 | 1 | 1 | 0 | NA | NA | 2 | 1 | 5 | 0.8 | 1 | 0 | NA | NA | 1 | 1 | 1 | 1 | 3 | 0.6667 |
| 3_97636609 | 4 | 0 | 7 | 0.4286 | 1 | 1 | 3 | 0.3333 | 15 | 0.3333 | 2 | 0.5 | 2 | 0.5 | 1 | 0 | NA | NA | 5 | 0.4 |
| 3_97636761 | 5 | 0.6 | 6 | 0.5 | 1 | 1 | 4 | 0.5 | 16 | 0.5625 | 1 | 1 | 2 | 0.5 | 1 | 1 | NA | NA | 4 | 0.75 |
| 3_97636824 | 5 | 0.4 | 5 | 0.4 | NA | NA | 3 | 0 | 13 | 0.3077 | 1 | 0 | 2 | 0.5 | 1 | 0 | NA | NA | 4 | 0.25 |
| 3_97636870 | 5 | 1 | 7 | 1 | NA | NA | 4 | 1 | 16 | 1 | 1 | 1 | 1 | 1 | 1 | 1 | NA | NA | 3 | 1 |
| 3_97636992 | 4 | 1 | 5 | 0.8 | NA | NA | 2 | 1 | 11 | 0.9091 | 1 | 1 | 1 | 1 | 1 | 1 | NA | NA | 3 | 1 |
| 3_97637023 | 4 | 1 | 5 | 0.6 | NA | NA | 4 | 0.75 | 13 | 0.7692 | 1 | 0 | 2 | 0.5 | 1 | 0 | NA | NA | 4 | 0.25 |
| 3_97637107 | 4 | 0.5 | 4 | 0.25 | NA | NA | 3 | 0.6667 | 11 | 0.4545 | 1 | 1 | 2 | 0.5 | 1 | 0 | NA | NA | 4 | 0.5 |
| 3_97637243 | 4 | 0.5 | 6 | 0 | NA | NA | 4 | 0 | 14 | 0.1429 | 1 | 0 | 1 | 1 | 1 | 0 | NA | NA | 3 | 0.3333 |
| 3_97637267 | 4 | 0.25 | 4 | 0.25 | NA | NA | 4 | 0.5 | 12 | 0.3333 | 1 | 1 | 1 | 1 | 1 | 0 | NA | NA | 3 | 0.6667 |
| 3_97637339 | 4 | 0.25 | 4 | 0.25 | NA | NA | 1 | 0 | 9 | 0.2222 | 1 | 1 | 1 | 0 | 1 | 0 | NA | NA | 3 | 0.3333 |
| 3_97637438 | 3 | 1 | 5 | 1 | NA | NA | 2 | 1 | 10 | 1 | 1 | 1 | 1 | 1 | 1 | 1 | NA | NA | 3 | 1 |
| 3_97637443 | 1 | 1 | 4 | 0.5 | NA | NA | 2 | 1 | 7 | 0.7143 | 1 | 1 | 1 | 1 | 1 | 0 | NA | NA | 3 | 0.6667 |
| 3_97637543 | 3 | 0.6667 | 5 | 1 | NA | NA | 3 | 1 | 11 | 0.9091 | 1 | 1 | 1 | 1 | 1 | 1 | NA | NA | 3 | 1 |
| 3_97637590 | 3 | 0.3333 | 5 | 0.6 | NA | NA | 3 | 0.6667 | 11 | 0.5455 | 1 | 0 | 1 | 1 | 1 | 1 | NA | NA | 3 | 0.6667 |
| 3_97637742 | 2 | 0.5 | 6 | 1 | NA | NA | 2 | 1 | 10 | 0.9 | 1 | 1 | 1 | 1 | 1 | 1 | NA | NA | 3 | 1 |
| 3_97646873 | 5 | 0 | 7 | 0 | 5 | 0.2 | 4 | 0.25 | 21 | 0.0952 | 3 | 0 | 6 | 0 | 6 | 0.1667 | 4 | 0 | 19 | 0.0526 |
| 3_97648422 | 4 | 0 | 7 | 0 | 2 | 0 | 3 | 0.3333 | 16 | 0.0625 | 3 | 0 | 4 | 0.25 | 4 | 0 | NA | NA | 11 | 0.0909 |
| 3_97648479 | 2 | 0 | 6 | 0.1667 | 1 | 0 | 1 | 1 | 10 | 0.2 | 3 | 0.6667 | 3 | 0 | 4 | 0 | NA | NA | 10 | 0.2 |
| 3_97648519 | 3 | 0.6667 | 6 | 0.5 | 2 | 1 | 2 | 1 | 13 | 0.6923 | 3 | 0.6667 | 4 | 1 | 4 | 0.75 | NA | NA | 11 | 0.8182 |
| 3_97648626 | 3 | 1 | 4 | 0.75 | 2 | 1 | 3 | 0.6667 | 12 | 0.8333 | 3 | 0.6667 | 3 | 0.6667 | 4 | 0.5 | NA | NA | 10 | 0.6 |
| 3_97674054 | 1 | 0 | 1 | 1 | NA | NA | 1 | 0 | 3 | 0.3333 | 1 | 1 | NA | NA | NA | NA | NA | NA | 1 | 1 |
| 3_97674100 | 2 | 0.5 | 3 | 0.3333 | NA | NA | 1 | 0 | 6 | 0.3333 | 1 | 1 | NA | NA | NA | NA | NA | NA | 1 | 1 |
| 3_97674114 | 2 | 1 | 3 | 1 | NA | NA | 1 | 1 | 6 | 1 | 1 | 1 | NA | NA | NA | NA | NA | NA | 1 | 1 |
| 3_97674206 | 2 | 0 | 3 | 0.6667 | NA | NA | 1 | 1 | 6 | 0.5 | 1 | 1 | NA | NA | NA | NA | NA | NA | 1 | 1 |
| 3_97674256 | 2 | 1 | 2 | 1 | NA | NA | 1 | 1 | 5 | 1 | 1 | 1 | NA | NA | NA | NA | NA | NA | 1 | 1 |
| 3_97828623 | 6 | 0 | 13 | 0 | 6 | 0 | 13 | 0.1538 | 38 | 0.0526 | 14 | 0.1429 | 8 | 0.125 | 9 | 0 | 14 | 0 | 45 | 0.0667 |
| 3_97828648 | 6 | 0.1667 | 12 | 0.0833 | 4 | 0 | 12 | 0.0833 | 34 | 0.0882 | 10 | 0.2 | 7 | 0.1429 | 8 | 0.125 | 13 | 0.1538 | 38 | 0.1579 |
| 3_97828686 | 7 | 0 | 12 | 0 | 5 | 0.4 | 12 | 0.1667 | 36 | 0.1111 | 12 | 0.0833 | 7 | 0.2857 | 10 | 0.2 | 14 | 0.2857 | 43 | 0.2093 |
| 3_97828702 | 9 | 0.3333 | 13 | 0.3077 | 6 | 0.3333 | 14 | 0.2143 | 42 | 0.2857 | 14 | 0.1429 | 8 | 0.375 | 10 | 0 | 16 | 0.25 | 48 | 0.1875 |
| 3_97828717 | 9 | 0 | 14 | 0.1429 | 6 | 0.1667 | 14 | 0.3571 | 43 | 0.186 | 14 | 0.0714 | 7 | 0.1429 | 8 | 0 | 16 | 0.0625 | 45 | 0.0667 |
| 3_97828785 | 6 | 0.1667 | 9 | 0.2222 | 5 | 0 | 12 | 0.5 | 32 | 0.2812 | 9 | 0.5556 | 7 | 0.2857 | 7 | 0.2857 | 13 | 0.1538 | 36 | 0.3056 |
| 3_97828789 | 8 | 0.25 | 14 | 0.2143 | 6 | 0.5 | 13 | 0.3846 | 41 | 0.3171 | 11 | 0.1818 | 7 | 0.1429 | 9 | 0.1111 | 15 | 0.2 | 42 | 0.1667 |
| 3_97828880 | 9 | 0.3333 | 12 | 0.25 | 7 | 0.2857 | 14 | 0.1429 | 42 | 0.2381 | 14 | 0.0714 | 8 | 0.25 | 9 | 0 | 15 | 0.1333 | 46 | 0.1087 |
| 3_97829029 | 6 | 0 | 12 | 0.0833 | 6 | 0 | 14 | 0.0714 | 38 | 0.0526 | 14 | 0.1429 | 8 | 0.25 | 10 | 0.1 | 14 | 0.0714 | 46 | 0.1304 |
| 3_97829943 | 7 | 0.1429 | 10 | 0.1 | 3 | 0 | 7 | 0 | 27 | 0.0741 | 6 | 0.1667 | 2 | 0 | 5 | 0.2 | 5 | 0 | 18 | 0.1111 |
| 3_97830183 | 7 | 0.2857 | 10 | 0.4 | 3 | 0 | 9 | 0.1111 | 29 | 0.2414 | 3 | 0.6667 | 2 | 0.5 | 5 | 0.2 | 5 | 0.6 | 15 | 0.4667 |
| 3_98053780 | 1 | 1 | 3 | 1 | 2 | 1 | NA | NA | 6 | 1 | NA | NA | 1 | 1 | NA | NA | NA | NA | 1 | 1 |
| 3_98054260 | 1 | 0 | 3 | 0.6667 | 2 | 0.5 | NA | NA | 6 | 0.5 | NA | NA | 1 | 1 | 1 | 1 | NA | NA | 2 | 1 |
| 3_98055032 | 2 | 0.5 | 4 | 0 | 2 | 0 | 1 | 0 | 9 | 0.1111 | 1 | 1 | 1 | 0 | 1 | 0 | NA | NA | 3 | 0.3333 |
| 3_98055086 | 1 | 1 | 4 | 1 | 2 | 0.5 | 1 | 1 | 8 | 0.875 | 2 | 1 | 1 | 0 | 1 | 1 | NA | NA | 4 | 0.75 |
| 3_98055145 | 2 | 0.5 | 3 | 1 | 2 | 1 | 1 | 0 | 8 | 0.75 | 2 | 1 | 1 | 1 | 1 | 1 | NA | NA | 4 | 1 |
| 3_98055288 | 1 | 0 | 3 | 0.3333 | 2 | 0.5 | 1 | 0 | 7 | 0.2857 | 2 | 0 | 1 | 1 | 1 | 0 | NA | NA | 4 | 0.25 |
| 3_98055467 | 2 | 1 | 4 | 1 | 2 | 1 | 1 | 1 | 9 | 1 | 2 | 1 | 1 | 1 | 1 | 1 | NA | NA | 4 | 1 |
| 3_98055486 | 2 | 0.5 | 3 | 0.6667 | 1 | 0 | NA | NA | 6 | 0.5 | 2 | 0.5 | 1 | 1 | 1 | 0 | NA | NA | 4 | 0.5 |
| 3_98055594 | 2 | 1 | 3 | 1 | NA | NA | 1 | 1 | 6 | 1 | 2 | 1 | NA | NA | 1 | 1 | NA | NA | 3 | 1 |
| 3_98055728 | 2 | 0.5 | 3 | 1 | 2 | 1 | 1 | 0 | 8 | 0.75 | 2 | 0.5 | 1 | 1 | 1 | 1 | NA | NA | 4 | 0.75 |
| 3_98055760 | 2 | 1 | 4 | 1 | 2 | 1 | 1 | 1 | 9 | 1 | 1 | 1 | 1 | 1 | 1 | 1 | NA | NA | 3 | 1 |
| 3_98055866 | 1 | 1 | 3 | 1 | 2 | 0.5 | NA | NA | 6 | 0.8333 | 2 | 1 | NA | NA | 1 | 0 | NA | NA | 3 | 0.6667 |
| 3_98055899 | 2 | 1 | 2 | 0.5 | 2 | 0.5 | 1 | 1 | 7 | 0.7143 | 2 | 0.5 | 1 | 0 | 1 | 1 | NA | NA | 4 | 0.5 |

| | | | | | | | | | | | | | | | | | | | | |
|---|---|---|---|---|---|---|---|---|---|---|---|---|---|---|---|---|---|---|---|---|
| 3_9805591 | 2 | 0.5 | 4 | 0 | 1 | 1 | 1 | 0 | 8 | 0.25 | 2 | 0.5 | 1 | 1 | 1 | 0 | NA | NA | 4 | 0.5 |
| 3_9805597 | 2 | 1 | 4 | 1 | 2 | 1 | 1 | 1 | 9 | 1 | 2 | 1 | 1 | 1 | 1 | 1 | NA | NA | 4 | 1 |
| 3_9805604 | 2 | 1 | 2 | 1 | 2 | 1 | 1 | 0 | 7 | 0.8571 | 1 | 1 | 1 | 1 | NA | NA | NA | NA | 2 | 1 |
| 3_9805620 | 2 | 0.5 | 4 | 0.75 | 2 | 1 | NA | NA | 8 | 0.75 | 2 | 1 | 1 | 1 | 1 | 1 | NA | NA | 4 | 1 |
| 3_9805621 | 2 | 1 | 2 | 1 | 2 | 0 | 1 | 1 | 7 | 0.7143 | 2 | 0.5 | 1 | 1 | 1 | 0 | NA | NA | 4 | 0.5 |
| 3_9805634 | 2 | 0 | 4 | 0.25 | 2 | 0.5 | 1 | 0 | 9 | 0.2222 | 2 | 0.5 | 1 | 0 | 1 | 0 | NA | NA | 4 | 0.25 |
| 3_9805668 | 2 | 1 | 4 | 1 | 2 | 0 | 1 | 1 | 9 | 0.7778 | 2 | 0.5 | 1 | 0 | 1 | 1 | NA | NA | 4 | 0.5 |
| 3_9805670 | 2 | 0.5 | 3 | 1 | 2 | 1 | 1 | 1 | 8 | 0.875 | 2 | 1 | 1 | 1 | 1 | 1 | NA | NA | 4 | 1 |
| 3_9805724 | 3 | 0.3333 | 4 | 1 | 2 | 1 | 1 | 0 | 10 | 0.7 | 2 | 1 | 1 | 0 | 1 | 1 | NA | NA | 4 | 0.75 |
| 3_9805737 | 4 | 0.75 | 5 | 0.6 | 2 | 0.5 | 1 | 1 | 12 | 0.6667 | 2 | 1 | NA | NA | 1 | 0 | NA | NA | 3 | 0.6667 |
| 3_9805740 | 4 | 1 | 2 | 1 | 2 | 1 | 1 | 1 | 9 | 1 | 2 | 1 | 1 | 0 | 1 | 1 | NA | NA | 4 | 0.75 |
| 3_9819830 | 5 | 0.4 | 3 | 0.3333 | 1 | 1 | 1 | 0 | 10 | 0.4 | NA | NA | 1 | 0 | 2 | 0.5 | 1 | 0 | 4 | 0.25 |
| 3_9819840 | 4 | 0.25 | 4 | 0.25 | 2 | 0 | 1 | 0 | 11 | 0.1818 | 1 | 0 | 2 | 0.5 | 2 | 0.5 | 1 | 0 | 6 | 0.3333 |
| 3_9819853 | 7 | 0.1429 | 4 | 0.5 | 4 | 0.75 | 1 | 0 | 16 | 0.375 | 2 | 0.5 | 2 | 0.5 | 1 | 0 | 1 | 0 | 6 | 0.3333 |
| 3_9821643 | 9 | 0.3333 | 5 | 0.2 | 4 | 1 | 2 | 0.5 | 20 | 0.45 | 2 | 0 | 3 | 0.3333 | 2 | 1 | NA | NA | 7 | 0.4286 |
| 3_9821654 | 6 | 0 | 5 | 0.2 | 3 | 0 | 1 | 0 | 15 | 0.0667 | 2 | 0 | 2 | 0 | 2 | 0.5 | 1 | 0 | 7 | 0.1429 |
| 3_9821666 | 6 | 0.5 | 7 | 0.8571 | 4 | 0.75 | 2 | 1 | 19 | 0.7368 | 2 | 0.5 | 2 | 0.5 | 2 | 1 | 2 | 0 | 8 | 0.5 |
| 3_9821680 | 1 | 0 | 3 | 0.6667 | 2 | 1 | 2 | 0.5 | 8 | 0.625 | 1 | 1 | NA | NA | NA | NA | NA | NA | 1 | 1 |
| 3_9821693 | 1 | 0 | 3 | 0.3333 | 2 | 0 | 2 | 0 | 8 | 0.125 | 1 | 1 | NA | NA | NA | NA | NA | NA | 1 | 1 |
| 3_9821706 | 1 | 0 | 2 | 0 | 2 | 0.5 | 2 | 0 | 7 | 0.1429 | 1 | 1 | NA | NA | NA | NA | NA | NA | 1 | 1 |
| 3_9821758 | 1 | 0 | 3 | 0.6667 | 2 | 0 | 1 | 0 | 7 | 0.2857 | 1 | 1 | 1 | 1 | NA | NA | NA | NA | 2 | 1 |
| 3_9821781 | 2 | 0 | 3 | 0.3333 | 3 | 0 | 1 | 1 | 9 | 0.2222 | 1 | 1 | 1 | 0 | NA | NA | NA | NA | 2 | 0.5 |
| 3_9831790 | NA | NA | NA | NA | 1 | 1 | NA | NA | 1 | 1 | NA | NA | NA | NA | 1 | 1 | NA | NA | 1 | 1 |
| 3_9831840 | NA | NA | NA | NA | 1 | 1 | NA | NA | 1 | 1 | NA | NA | NA | NA | 1 | 1 | 1 | 1 | 2 | 1 |
| 3_9831842 | NA | NA | NA | NA | 1 | 1 | NA | NA | 1 | 1 | NA | NA | NA | NA | 1 | 1 | NA | NA | 1 | 1 |
| 3_9831854 | NA | NA | NA | NA | 1 | 1 | NA | NA | 1 | 1 | NA | NA | NA | NA | 1 | 0 | 1 | 1 | 2 | 0.5 |
| 3_9833456 | NA | NA | NA | NA | NA | NA | 4 | 0.75 | 4 | 0.75 | NA | NA | NA | NA | 1 | 1 | 1 | 1 | 2 | 1 |
| 3_9833479 | NA | NA | NA | NA | NA | NA | 4 | 1 | 4 | 1 | 1 | 1 | NA | NA | 1 | 1 | 1 | 1 | 3 | 1 |
| 3_9833480 | NA | NA | NA | NA | NA | NA | 3 | 1 | 3 | 1 | 1 | 1 | NA | NA | 1 | 1 | 1 | 1 | 3 | 1 |
| 3_9833485 | NA | NA | NA | NA | NA | NA | 4 | 1 | 4 | 1 | 1 | 1 | NA | NA | 1 | 1 | 1 | 1 | 3 | 1 |
| 3_9833512 | NA | NA | 1 | 1 | NA | NA | 4 | 0.5 | 5 | 0.6 | NA | NA | NA | NA | 1 | 1 | 2 | 0 | 3 | 0.3333 |
| 3_9833549 | NA | NA | 1 | 1 | NA | NA | 4 | 1 | 5 | 1 | 1 | 1 | NA | NA | 1 | 1 | 2 | 1 | 4 | 1 |
| 3_9833555 | NA | NA | NA | NA | NA | NA | 3 | 0.6667 | 3 | 0.6667 | 1 | 1 | NA | NA | 1 | 0 | 1 | 1 | 3 | 0.6667 |
| 3_9833602 | NA | NA | 1 | 1 | NA | NA | 4 | 1 | 5 | 1 | 1 | 1 | NA | NA | 1 | 1 | 1 | 1 | 3 | 1 |
| 3_9833613 | NA | NA | 1 | 0 | 1 | 0 | 3 | 0.3333 | 5 | 0.2 | 1 | 1 | NA | NA | NA | NA | 1 | 0 | 2 | 0.5 |
| 3_9833625 | 1 | 0 | 1 | 1 | 1 | 1 | 2 | 0.5 | 5 | 0.6 | 1 | 1 | NA | NA | 1 | 0 | 2 | 1 | 4 | 0.75 |
| 3_9833626 | 1 | 1 | 1 | 0 | 1 | 0 | 3 | 0.3333 | 6 | 0.3333 | 1 | 1 | NA | NA | 1 | 0 | 2 | 0 | 4 | 0.25 |
| 3_9833666 | 1 | 0 | 1 | 0 | 1 | 1 | 4 | 0 | 7 | 0.1429 | 1 | 0 | NA | NA | 1 | 0 | 2 | 0.5 | 4 | 0.25 |
| 3_9833711 | 1 | 0 | 1 | 0 | 1 | 0 | 3 | 0.3333 | 6 | 0.1667 | NA | NA | NA | NA | 2 | 0.5 | NA | NA | 2 | 0.5 |
| 3_9833730 | 1 | 0 | 1 | 1 | 1 | 0 | 4 | 0.5 | 7 | 0.4286 | 2 | 0.5 | NA | NA | 2 | 0 | 2 | 0 | 6 | 0.1667 |
| 3_9833757 | 1 | 0 | 1 | 0 | 1 | 0 | 3 | 0.3333 | 6 | 0.1667 | 2 | 0 | NA | NA | 2 | 0.5 | 2 | 0 | 6 | 0.1667 |
| 3_9833770 | 1 | 0 | 1 | 0 | 1 | 0 | 3 | 0.3333 | 6 | 0.1667 | 1 | 0 | NA | NA | 2 | 0.5 | 2 | 0 | 5 | 0.2 |
| 3_9833776 | 1 | 0 | 1 | 0 | 1 | 1 | 3 | 0 | 6 | 0.1667 | 2 | 0 | NA | NA | 1 | 0 | 2 | 0.5 | 5 | 0.2 |
| 3_9833848 | 1 | 1 | 1 | 1 | 1 | 1 | 2 | 1 | 5 | 1 | 2 | 1 | NA | NA | 2 | 1 | 1 | 1 | 5 | 1 |
| 3_9833856 | 1 | 0 | 2 | 0 | 1 | 1 | 3 | 0 | 7 | 0.1429 | 2 | 0.5 | NA | NA | 1 | 1 | 2 | 0 | 5 | 0.4 |
| 3_9833885 | 1 | 0 | 2 | 0.5 | 1 | 0 | 3 | 0 | 7 | 0.1429 | 2 | 0.5 | NA | NA | 1 | 0 | 3 | 0 | 6 | 0.1667 |
| 3_9833888 | 1 | 0 | 1 | 0 | 1 | 1 | 2 | 0.5 | 5 | 0.4 | 2 | 0.5 | NA | NA | 1 | 0 | 3 | 0.3333 | 6 | 0.3333 |
| 3_9833895 | 1 | 0 | 2 | 0 | 1 | 0 | 3 | 0.3333 | 7 | 0.1429 | 2 | 0 | NA | NA | 2 | 0 | 3 | 0.3333 | 7 | 0.1429 |
| 3_9833919 | 1 | 0 | 2 | 0.5 | 1 | 1 | 1 | 0 | 5 | 0.4 | 2 | 0 | NA | NA | NA | NA | 2 | 0.5 | 4 | 0.25 |
| 3_9872955 | 2 | 1 | 3 | 1 | 3 | 1 | 2 | 1 | 10 | 1 | 3 | 1 | 1 | 0 | 2 | 0.5 | 2 | 1 | 8 | 0.75 |
| 3_9872981 | 7 | 0.2857 | 8 | 0.375 | 8 | 0.25 | 7 | 0.1429 | 30 | 0.2667 | 7 | 0.1429 | 5 | 0 | 10 | 0.1 | 6 | 0.3333 | 28 | 0.1429 |
| 3_1000456 | 4 | 0.5 | 3 | 0 | NA | NA | 1 | 1 | 8 | 0.375 | 1 | 1 | 1 | 0 | 4 | 0.25 | 4 | 0.25 | 10 | 0.3 |
| 3_1000457 | 4 | 0.5 | 3 | 0 | NA | NA | 1 | 1 | 8 | 0.375 | NA | NA | 1 | 0 | 4 | 0.5 | 4 | 0 | 9 | 0.2222 |
| 3_1000459 | 1 | 0 | 2 | 0.5 | NA | NA | 1 | 1 | 4 | 0.5 | NA | NA | NA | NA | 2 | 1 | 3 | 1 | 5 | 1 |
| 3_1000460 | 2 | 1 | 3 | 0.6667 | NA | NA | 1 | 1 | 6 | 0.8333 | NA | NA | NA | NA | 3 | 1 | 5 | 0.4 | 8 | 0.625 |
| 3_1001132 | 4 | 0.25 | 4 | 0.25 | 5 | 0.4 | 5 | 0 | 18 | 0.2222 | 1 | 1 | 7 | 0.1429 | 5 | 0.2 | 4 | 0.25 | 17 | 0.2353 |
| 3_1001133 | 4 | 0 | 4 | 0.25 | 5 | 0 | 5 | 0 | 18 | 0.0556 | 2 | 0.5 | 6 | 0 | 5 | 0 | 4 | 0.25 | 17 | 0.1176 |
| 3_1001133 | 5 | 0.8 | 4 | 0.5 | 4 | 0.25 | 5 | 0.8 | 18 | 0.6111 | 1 | 0 | 7 | 0.5714 | 5 | 0.2 | 4 | 1 | 17 | 0.5294 |
| 3_1001133 | 4 | 0.25 | 4 | 0.5 | 5 | 0.2 | 4 | 0 | 17 | 0.2353 | 2 | 0.5 | 7 | 0 | 5 | 0 | 3 | 0 | 17 | 0.0588 |
| 3_1001134 | 4 | 0.5 | 3 | 0 | 4 | 0.5 | 5 | 0.4 | 16 | 0.375 | 2 | 0.5 | 6 | 0.3333 | 5 | 0.2 | 4 | 0.75 | 17 | 0.4118 |
| 3_1001134 | 3 | 0.6667 | 1 | 0 | 3 | 0 | 5 | 0.6 | 12 | 0.4167 | 1 | 1 | 4 | 0.5 | 3 | 0.3333 | 3 | 0.6667 | 11 | 0.5455 |
| 3_1001134 | 4 | 1 | 4 | 0.5 | 6 | 0.5 | 5 | 0.6 | 19 | 0.6316 | 2 | 1 | 5 | 1 | 5 | 0.6 | 3 | 1 | 15 | 0.8667 |
| 3_1001138 | 5 | 0.8 | 5 | 0.2 | 9 | 0.4444 | 6 | 0.6667 | 25 | 0.52 | 3 | 0 | 7 | 0.2857 | 5 | 0.4 | 3 | 0.3333 | 18 | 0.2778 |
| 3_1001139 | 3 | 0 | 5 | 0 | 7 | 0.2857 | 5 | 0.4 | 20 | 0.2 | 2 | 0 | 6 | 0 | 2 | 0 | 3 | 0.3333 | 13 | 0.0769 |
| 3_1001174 | NA | NA | 1 | 1 | 2 | 0 | 1 | 0 | 4 | 0.25 | 1 | 0 | NA | NA | NA | NA | 3 | 0.3333 | 4 | 0.25 |

| | | | | | | | | | | | | | | | | | | | | |
|---|---|---|---|---|---|---|---|---|---|---|---|---|---|---|---|---|---|---|---|---|
| 3_1003790 | 1 | 1 | NA | NA | NA | NA | NA | NA | 1 | 1 | NA | NA | 1 | 1 | 2 | 1 | NA | NA | 3 | 1 |
| 3_1003790 | 1 | 1 | NA | NA | NA | NA | NA | NA | 1 | 1 | NA | NA | 1 | 1 | 2 | 1 | NA | NA | 2 | 1 |
| 3_1003793 | 1 | 1 | NA | NA | NA | NA | NA | NA | 1 | 1 | NA | NA | 1 | 0 | 1 | 1 | NA | NA | 2 | 0.5 |
| 3_1003793 | 1 | 1 | NA | NA | NA | NA | NA | NA | 1 | 1 | NA | NA | 1 | 0 | 1 | 1 | NA | NA | 2 | 0.5 |
| 3_1003795 | 1 | 1 | NA | NA | NA | NA | NA | NA | 1 | 1 | NA | NA | 1 | 1 | 1 | 1 | NA | NA | 2 | 1 |
| 3_1004701 | 5 | 0.4 | 7 | 0.4286 | 3 | 0 | 3 | 0.3333 | 18 | 0.3333 | 2 | 0.5 | 6 | 0 | 5 | 0.4 | 3 | 0.3333 | 16 | 0.25 |
| 3_1004704 | 4 | 0 | 5 | 0 | 4 | 0 | 2 | 0.5 | 15 | 0.0667 | 2 | 0.5 | 5 | 0.2 | 3 | 0 | 2 | 0 | 12 | 0.1667 |
| 3_1004709 | 3 | 0 | 6 | 0.1667 | 3 | 0.3333 | 1 | 0 | 13 | 0.1538 | 2 | 0.5 | 4 | 0 | 4 | 0 | 3 | 0 | 13 | 0.0769 |
| 3_1004720 | 5 | 0.2 | 5 | 0 | 3 | 0 | 1 | 0 | 14 | 0.0714 | 1 | 0 | 4 | 0 | 3 | 0 | 2 | 0.5 | 10 | 0.1 |
| 3_1004728 | 5 | 0.2 | 2 | 0 | 5 | 0.4 | 2 | 0 | 14 | 0.2143 | NA | NA | 5 | 0.2 | 4 | 0 | 6 | 0.3333 | 15 | 0.2 |
| 3_1004733 | 5 | 0.2 | 3 | 0 | 5 | 0 | 2 | 0 | 15 | 0.0667 | NA | NA | 3 | 0 | 2 | 0 | 6 | 0.1667 | 11 | 0.0909 |
| 3_1004737 | 5 | 0.2 | 4 | 0 | 7 | 0 | 2 | 0.5 | 18 | 0.1111 | NA | NA | 4 | 0 | 3 | 0 | 5 | 0.4 | 12 | 0.1667 |
| 3_1004742 | 4 | 0 | 4 | 0.25 | 5 | 0 | 3 | 0 | 16 | 0.0625 | NA | NA | 4 | 0 | 2 | 0 | 6 | 0.1667 | 12 | 0.0833 |
| 3_1004743 | 4 | 0.5 | 4 | 0.5 | 5 | 0.8 | 2 | 0 | 15 | 0.5333 | NA | NA | 4 | 0 | 2 | 1 | 5 | 0.4 | 11 | 0.3636 |
| 3_1004744 | 4 | 0 | 4 | 0 | 5 | 0 | 3 | 0.3333 | 16 | 0.0625 | NA | NA | 4 | 0 | 2 | 0 | 5 | 0.2 | 11 | 0.0909 |
| 3_1004744 | 4 | 0 | 4 | 0 | 4 | 0.5 | 3 | 0 | 15 | 0.1333 | NA | NA | 4 | 0.5 | 2 | 0 | 6 | 0 | 12 | 0.1667 |
| 3_1004745 | 3 | 0.3333 | 3 | 0 | 5 | 0.2 | 2 | 0.5 | 13 | 0.2308 | NA | NA | 4 | 0 | 2 | 0.5 | 5 | 0 | 11 | 0.0909 |
| 3_1004746 | 3 | 0.3333 | 4 | 0 | 4 | 0.5 | 1 | 1 | 12 | 0.3333 | NA | NA | 3 | 0 | 2 | 0 | 4 | 0.5 | 9 | 0.2222 |
| 3_1004747 | 3 | 0.6667 | 4 | 0.25 | 4 | 1 | 2 | 1 | 13 | 0.6923 | NA | NA | 4 | 0 | 2 | 0.5 | 5 | 0.8 | 11 | 0.4545 |
| 3_1004747 | 3 | 0.6667 | 4 | 0.5 | 4 | 0.75 | 2 | 0 | 13 | 0.5385 | NA | NA | 4 | 0.5 | 2 | 0.5 | 5 | 0.4 | 11 | 0.4545 |
| 3_1004747 | 2 | 0.5 | 4 | 0.5 | 4 | 0.5 | 2 | 0.5 | 12 | 0.5 | NA | NA | 4 | 0.5 | 2 | 0.5 | 4 | 0.75 | 10 | 0.6 |
| 3_1004748 | 2 | 1 | 4 | 1 | 4 | 1 | 2 | 0.5 | 12 | 0.9167 | NA | NA | 3 | 1 | 2 | 1 | 3 | 1 | 8 | 1 |
| 3_1004748 | 2 | 0.5 | 3 | 1 | 4 | 1 | 1 | 1 | 10 | 0.9 | NA | NA | 3 | 1 | 2 | 1 | 2 | 1 | 7 | 1 |
| 3_1004748 | 2 | 1 | 4 | 0.75 | 3 | 0.6667 | 2 | 1 | 11 | 0.8182 | NA | NA | 3 | 0.3333 | 2 | 1 | 3 | 0.3333 | 8 | 0.5 |
| 3_1004749 | 2 | 0.5 | 4 | 0.5 | 4 | 1 | 2 | 0.5 | 12 | 0.6667 | NA | NA | 2 | 0 | 2 | 1 | 3 | 1 | 7 | 0.7143 |
| 3_1005918 | 1 | 0 | 1 | 0 | 2 | 0 | 2 | 1 | 6 | 0.3333 | NA | NA | 2 | 0 | 1 | 1 | 1 | 0 | 4 | 0.25 |
| 3_1005920 | 1 | 0 | 1 | 0 | 1 | 0 | 2 | 1 | 5 | 0.4 | NA | NA | 2 | 0.5 | NA | NA | NA | NA | 2 | 0.5 |
| 3_1005920 | 1 | 1 | 1 | 1 | 1 | 1 | 1 | 1 | 4 | 1 | NA | NA | 2 | 1 | NA | NA | NA | NA | 2 | 1 |
| 3_1005922 | 1 | 0 | NA | NA | 1 | 1 | 1 | 1 | 3 | 0.6667 | NA | NA | 1 | 1 | NA | NA | 1 | 1 | 2 | 1 |
| 3_1005943 | NA | NA | NA | NA | 1 | 1 | 1 | 1 | 2 | 1 | 1 | 1 | NA | NA | NA | NA | 1 | 0 | 2 | 0.5 |
| 3_1005943 | NA | NA | NA | NA | NA | NA | 1 | 1 | 1 | 1 | 1 | 1 | NA | NA | NA | NA | 1 | 0 | 2 | 0.5 |
| 3_1005944 | NA | NA | NA | NA | 1 | 0 | 1 | 1 | 2 | 0.5 | 1 | 0 | NA | NA | NA | NA | 1 | 1 | 2 | 0.5 |
| 3_1008327 | NA | NA | 1 | 1 | NA | NA | NA | NA | 1 | 1 | NA | NA | NA | NA | 1 | 1 | NA | NA | 1 | 1 |
| 3_1008327 | NA | NA | 1 | 1 | NA | NA | NA | NA | 1 | 1 | NA | NA | NA | NA | 1 | 1 | NA | NA | 1 | 1 |
| 3_1009479 | NA | NA | 2 | 0 | 1 | 1 | 5 | 0 | 8 | 0.125 | 1 | 0 | 3 | 0.3333 | 2 | 0.5 | NA | NA | 6 | 0.3333 |
| 3_1009491 | NA | NA | 1 | 0 | 2 | 0 | 6 | 0.1667 | 9 | 0.1111 | 1 | 0 | 3 | 0.3333 | 2 | 0 | NA | NA | 6 | 0.1667 |
| 3_1009496 | NA | NA | 1 | 1 | 2 | 1 | 4 | 0.25 | 7 | 0.5714 | 1 | 0 | 2 | 0.5 | 2 | 1 | NA | NA | 5 | 0.6 |
| 3_1009496 | NA | NA | 1 | 1 | 2 | 0.5 | 3 | 0.3333 | 6 | 0.5 | 1 | 0 | 3 | 0.3333 | 2 | 0.5 | NA | NA | 6 | 0.3333 |
| 3_1009505 | NA | NA | NA | NA | 1 | 0 | 1 | 1 | 2 | 0.5 | 1 | 1 | NA | NA | NA | NA | NA | NA | 1 | 1 |
| 3_1009506 | NA | NA | NA | NA | 1 | 1 | 1 | 1 | 2 | 1 | 2 | 1 | NA | NA | NA | NA | NA | NA | 2 | 1 |
| 3_1009506 | NA | NA | NA | NA | 1 | 1 | NA | NA | 1 | 1 | 2 | 1 | NA | NA | NA | NA | NA | NA | 2 | 1 |
| 3_1009528 | NA | NA | NA | NA | 1 | 1 | 1 | 0 | 2 | 0.5 | 2 | 1 | NA | NA | NA | NA | NA | NA | 2 | 1 |
| 3_1009681 | NA | NA | NA | NA | NA | NA | 1 | 1 | 1 | 1 | 1 | 1 | NA | NA | NA | NA | NA | NA | 1 | 1 |
| 3_1013698 | 1 | 1 | NA | NA | 4 | 1 | NA | NA | 5 | 1 | NA | NA | NA | NA | 1 | 1 | NA | NA | 1 | 1 |
| 3_1013699 | 2 | 1 | NA | NA | 3 | 1 | NA | NA | 5 | 1 | NA | NA | NA | NA | 1 | 1 | NA | NA | 1 | 1 |
| 3_1013700 | 2 | 1 | NA | NA | 4 | 0.5 | NA | NA | 6 | 0.6667 | NA | NA | 1 | 1 | 1 | 1 | NA | NA | 2 | 1 |
| 3_1013701 | 2 | 1 | 1 | 0 | 4 | 0.25 | NA | NA | 7 | 0.4286 | NA | NA | 1 | 1 | 1 | 1 | NA | NA | 2 | 1 |
| 3_1017668 | NA | NA | NA | NA | NA | NA | 1 | 1 | 1 | 1 | 1 | 1 | NA | NA | 1 | 1 | NA | NA | 2 | 1 |
| 3_1017669 | NA | NA | NA | NA | NA | NA | 1 | 1 | 1 | 1 | 1 | 1 | NA | NA | 1 | 1 | 1 | 0 | 3 | 0.6667 |
| 3_1017669 | NA | NA | NA | NA | NA | NA | 1 | 1 | 1 | 1 | NA | NA | NA | NA | 1 | 1 | 1 | 1 | 2 | 1 |
| 3_1017670 | NA | NA | NA | NA | NA | NA | 1 | 1 | 1 | 1 | 1 | 1 | NA | NA | NA | NA | NA | NA | 1 | 1 |
| 3_1017675 | NA | NA | NA | NA | NA | NA | 1 | 1 | 1 | 1 | 1 | 1 | NA | NA | NA | NA | NA | NA | 1 | 1 |
| 3_1017675 | NA | NA | NA | NA | NA | NA | 1 | 1 | 1 | 1 | 1 | 1 | NA | NA | NA | NA | NA | NA | 1 | 1 |
| 3_1020527 | NA | NA | 1 | 1 | NA | NA | NA | NA | 1 | 1 | NA | NA | 1 | 1 | NA | NA | NA | NA | 1 | 1 |
| 3_1020538 | NA | NA | 1 | 1 | NA | NA | 1 | 1 | 2 | 1 | NA | NA | 1 | 1 | NA | NA | NA | NA | 1 | 1 |
| 3_1021864 | NA | NA | 1 | 1 | NA | NA | NA | NA | 1 | 1 | 1 | 1 | NA | NA | NA | NA | NA | NA | 1 | 1 |
| 3_1024276 | NA | NA | 1 | 1 | NA | NA | NA | NA | 1 | 1 | 1 | 0 | NA | NA | 1 | 1 | 1 | 1 | 3 | 0.6667 |
| 3_1024280 | NA | NA | 1 | 1 | NA | NA | NA | NA | 1 | 1 | 1 | 1 | NA | NA | 1 | 1 | 1 | 1 | 3 | 1 |
| 3_1024282 | NA | NA | 1 | 1 | NA | NA | NA | NA | 1 | 1 | 1 | 1 | NA | NA | 1 | 0 | 2 | 1 | 4 | 0.75 |
| 3_1029093 | 1 | 1 | 2 | 0.5 | NA | NA | NA | NA | 3 | 0.6667 | NA | NA | NA | NA | 3 | 0.6667 | 2 | 1 | 5 | 0.8 |
| 3_1029103 | 5 | 0.8 | 4 | 0.25 | 1 | 1 | NA | NA | 10 | 0.6 | NA | NA | NA | NA | 4 | 0.25 | 3 | 0 | 7 | 0.1429 |
| 3_1029653 | 19 | 0.3158 | 32 | 0.0938 | 19 | 0.1579 | 31 | 0.129 | 101 | 0.1584 | 22 | 0.0909 | 19 | 0.2105 | 24 | 0.25 | 20 | 0.05 | 85 | 0.1529 |
| 3_1029717 | 3 | 0.3333 | 3 | 0.3333 | 2 | 0 | 2 | 0 | 10 | 0.2 | 7 | 0 | 6 | 0.3333 | 3 | 0.3333 | 5 | 0 | 21 | 0.1429 |
| 3_1029717 | 3 | 0 | 3 | 0.3333 | 3 | 0 | 2 | 0 | 11 | 0.0909 | 7 | 0.1429 | 6 | 0.1667 | 4 | 0 | 5 | 0.2 | 22 | 0.1364 |
| 3_1029719 | 2 | 0.5 | 1 | 0 | 3 | 0.3333 | 1 | 0 | 7 | 0.2857 | 6 | 0.3333 | 5 | 0.2 | 3 | 0 | 4 | 0 | 18 | 0.1667 |

| | | | | | | | | | | | | | | | | | | | | |
|---|---|---|---|---|---|---|---|---|---|---|---|---|---|---|---|---|---|---|---|---|
| 3_10297218 | 3 | 0 | 3 | 0.3333 | 2 | 0 | 2 | 0 | 10 | 0.1 | 7 | 0.1429 | 6 | 0 | 3 | 0 | 4 | 0.25 | 20 | 0.1 |
| 3_10297230 | 3 | 0.3333 | 3 | 0 | 1 | 0 | 2 | 0 | 9 | 0.1111 | 7 | 0 | 6 | 0.1667 | 4 | 0 | 5 | 0.2 | 22 | 0.0909 |
| 3_10297454 | 6 | 0.1667 | 4 | 0 | 4 | 0 | 6 | 0 | 20 | 0.05 | 9 | 0.1111 | 7 | 0 | 4 | 0 | 8 | 0.125 | 28 | 0.0714 |
| 3_10308580 | 16 | 0.0625 | 16 | 0 | 18 | 0.1667 | 14 | 0 | 64 | 0.0625 | 13 | 0 | 11 | 0.0909 | 16 | 0.125 | 19 | 0.0526 | 59 | 0.0678 |
| 3_10308582 | 16 | 0.0625 | 15 | 0 | 18 | 0.1111 | 15 | 0.0667 | 64 | 0.0625 | 13 | 0.0769 | 11 | 0.0909 | 17 | 0.0588 | 17 | 0 | 58 | 0.0517 |
| 3_10308583 | 16 | 0.25 | 14 | 0.1429 | 17 | 0.3529 | 12 | 0.25 | 59 | 0.2542 | 13 | 0.3846 | 10 | 0.3 | 16 | 0.1875 | 16 | 0.3125 | 55 | 0.2909 |
| 3_10308588 | 15 | 0.4 | 14 | 0.6429 | 18 | 0.4444 | 12 | 0.4167 | 59 | 0.4746 | 11 | 0.5455 | 9 | 0.5556 | 14 | 0.7143 | 18 | 0.5556 | 52 | 0.5962 |
| 3_10325175 | NA | NA | 3 | 1 | 3 | 1 | 3 | 1 | 9 | 1 | 1 | 1 | NA | NA | 2 | 1 | 1 | 1 | 4 | 1 |
| 3_10326115 | 1 | 0 | 3 | 0.3333 | 3 | 0 | 4 | 0 | 11 | 0.0909 | 1 | 0 | 1 | 0 | 4 | 0.25 | 1 | 0 | 7 | 0.1429 |
| 3_10326120 | 1 | 0 | 3 | 0.3333 | 4 | 0 | 4 | 0.5 | 12 | 0.25 | 1 | 0 | 2 | 0 | 3 | 0.3333 | 1 | 1 | 7 | 0.2857 |
| 3_10326123 | 1 | 0 | 3 | 0.6667 | 3 | 0.6667 | 3 | 0.6667 | 10 | 0.6 | 1 | 1 | 2 | 0.5 | 3 | 1 | NA | NA | 6 | 0.8333 |
| 3_10326123 | 1 | 1 | 2 | 1 | 3 | 1 | 4 | 0.75 | 10 | 0.9 | 1 | 1 | 2 | 1 | 3 | 1 | 1 | 1 | 7 | 1 |
| 3_10326138 | NA | NA | 2 | 0 | 3 | 0 | 4 | 0.5 | 9 | 0.2222 | 1 | 0 | 2 | 0.5 | 3 | 0 | 1 | 0 | 7 | 0.1429 |
| 3_10326141 | 1 | 1 | 3 | 1 | 4 | 0.5 | 3 | 1 | 11 | 0.8182 | 1 | 0 | 2 | 0.5 | 4 | 0.75 | 1 | 1 | 8 | 0.625 |
| 3_10326143 | 1 | 0 | 2 | 0.5 | 4 | 0.75 | 4 | 0 | 11 | 0.3636 | 1 | 0 | 2 | 0 | 2 | 0 | 1 | 1 | 6 | 0.1667 |
| 3_10326175 | 1 | 0 | 3 | 0.3333 | 5 | 0 | 5 | 0 | 14 | 0.0714 | 1 | 0 | 2 | 0 | 4 | 0.25 | 1 | 0 | 8 | 0.125 |
| 3_10326196 | NA | NA | 3 | 0 | 7 | 0.2857 | 5 | 0.2 | 15 | 0.2 | 2 | 0 | 3 | 0 | 5 | 0.2 | 1 | 0 | 11 | 0.0909 |
| 3_10326240 | 1 | 0 | 2 | 0.5 | 6 | 0.1667 | 5 | 0.2 | 14 | 0.2143 | 3 | 0.3333 | 3 | 0 | 7 | 0.2857 | 1 | 1 | 14 | 0.2857 |
| 3_10326241 | 2 | 1 | 3 | 0.6667 | 7 | 1 | 5 | 1 | 17 | 0.9412 | 3 | 1 | 2 | 1 | 7 | 1 | 2 | 1 | 14 | 1 |
| 3_10326242 | 2 | 0.5 | 3 | 0.3333 | 7 | 0.1429 | 5 | 0.4 | 17 | 0.2941 | 3 | 0 | 4 | 0.25 | 6 | 0.3333 | 2 | 0.5 | 15 | 0.2667 |
| 3_10326244 | 2 | 0 | 3 | 0.6667 | 7 | 0.4286 | 5 | 0.4 | 17 | 0.4118 | 3 | 0 | 5 | 0.4 | 8 | 0.625 | 2 | 0.5 | 18 | 0.4444 |
| 3_10326249 | 2 | 0 | 4 | 0.25 | 5 | 0 | 3 | 0 | 14 | 0.0714 | 2 | 0 | 3 | 0 | 6 | 0.1667 | 1 | 0 | 12 | 0.0833 |
| 3_10326260 | 2 | 0 | 3 | 0.3333 | 6 | 0.3333 | 4 | 0.5 | 15 | 0.3333 | 3 | 0.6667 | 5 | 0.6 | 6 | 0.8333 | 2 | 0.5 | 16 | 0.6875 |
| 3_10326266 | 2 | 0.5 | 4 | 0.5 | 7 | 0.5714 | 5 | 0.6 | 18 | 0.5556 | 3 | 0.6667 | 5 | 0.6 | 7 | 0.8571 | 2 | 1 | 17 | 0.7647 |
| 3_10326270 | 2 | 0.5 | 4 | 0.25 | 7 | 0.1429 | 3 | 0 | 16 | 0.1875 | 3 | 0.3333 | 5 | 0 | 7 | 0 | 2 | 0 | 17 | 0.0588 |
| 3_10326277 | 3 | 0 | 4 | 0 | 6 | 0.1667 | 5 | 0.2 | 18 | 0.1111 | 3 | 0.3333 | 5 | 0.4 | 7 | 0.1429 | 3 | 0.3333 | 18 | 0.2778 |
| 3_10326520 | 2 | 0 | 2 | 0.5 | 5 | 0 | 3 | 0.3333 | 12 | 0.1667 | NA | NA | 3 | 0.3333 | 7 | 0 | 1 | 0 | 11 | 0.0909 |
| 3_10326567 | 2 | 0 | 3 | 0 | 6 | 0.1667 | 3 | 0 | 14 | 0.0714 | NA | NA | 2 | 0 | 6 | 0.1667 | 2 | 0 | 10 | 0.1 |
| 3_10349280 | 2 | 0 | 1 | 1 | 4 | 0.5 | 8 | 0.125 | 15 | 0.2667 | 1 | 0 | 8 | 0.125 | 6 | 0.1667 | 2 | 0 | 17 | 0.1176 |
| 3_10349460 | 2 | 1 | 1 | 1 | 4 | 0.75 | 6 | 0.5 | 13 | 0.6923 | NA | NA | 6 | 0.6667 | 9 | 0.5556 | 3 | 0.6667 | 18 | 0.6111 |
| 3_10349475 | 3 | 0.3333 | 2 | 0.5 | 5 | 0.4 | 10 | 0.1 | 20 | 0.25 | 2 | 0.5 | 9 | 0.2222 | 9 | 0.1111 | 3 | 1 | 23 | 0.3043 |
| 3_10349481 | 3 | 1 | 2 | 0.5 | 5 | 1 | 8 | 1 | 18 | 0.9444 | 2 | 0.5 | 9 | 0.8889 | 9 | 1 | 3 | 1 | 23 | 0.913 |
| 3_10349492 | 4 | 0.75 | 2 | 0 | 3 | 0.6667 | 10 | 0.4 | 19 | 0.4737 | NA | NA | 9 | 0.5556 | 7 | 0.2857 | 4 | 0.75 | 20 | 0.5 |
| 3_10349499 | 1 | 1 | 1 | 0 | 4 | 0.5 | 7 | 0.4286 | 13 | 0.4615 | 1 | 1 | 7 | 0.4286 | 8 | 0.375 | 3 | 0.3333 | 19 | 0.4211 |
| 3_10349504 | 3 | 0.3333 | 2 | 1 | 4 | 0.5 | 8 | 0.75 | 17 | 0.6471 | 2 | 0.5 | 7 | 0.7143 | 8 | 0.875 | 4 | 0.75 | 21 | 0.7619 |
| 3_10353297 | 4 | 0 | 3 | 0 | 6 | 0 | 12 | 0.1667 | 25 | 0.08 | 2 | 0.5 | 10 | 0.3 | 11 | 0.0909 | 6 | 0.1667 | 29 | 0.2069 |
| 3_10353985 | 4 | 1 | 5 | 0.8 | 7 | 0.4286 | 14 | 0.7143 | 30 | 0.7 | 3 | 0.3333 | 10 | 0.6 | 12 | 0.6667 | 6 | 0.6667 | 31 | 0.6129 |
| 3_10353996 | 2 | 0.5 | 2 | 0 | 4 | 0 | 10 | 0.1 | 18 | 0.1111 | 1 | 0 | 7 | 0.2857 | 4 | 0.25 | 2 | 0.5 | 14 | 0.2857 |
| 3_10353999 | 3 | 0.3333 | 4 | 0.5 | 6 | 0.3333 | 15 | 0.2 | 28 | 0.2857 | 3 | 0.6667 | 10 | 0.1 | 8 | 0.25 | 5 | 0.2 | 26 | 0.2308 |
| 3_10354009 | 4 | 0.5 | 5 | 0 | 7 | 0.1429 | 14 | 0.2143 | 30 | 0.2 | 3 | 0 | 11 | 0.1818 | 13 | 0.3077 | 6 | 0.3333 | 33 | 0.2424 |
| 3_10354019 | 3 | 1 | 2 | 1 | 3 | 0.6667 | 11 | 1 | 19 | 0.9474 | 1 | 1 | 7 | 0.8571 | 7 | 0.8571 | 3 | 1 | 18 | 0.8889 |
| 3_10354029 | 5 | 1 | 3 | 0.6667 | 7 | 0.8571 | 16 | 0.875 | 31 | 0.871 | 3 | 0.6667 | 8 | 1 | 15 | 0.9333 | 6 | 1 | 32 | 0.9375 |
| 3_10354042 | 5 | 1 | 5 | 1 | 8 | 0.875 | 16 | 0.9375 | 34 | 0.9412 | 3 | 1 | 10 | 0.9 | 14 | 0.9286 | 6 | 1 | 33 | 0.9394 |
| 3_10354047 | 4 | 0.5 | 5 | 0.2 | 6 | 0 | 13 | 0.1538 | 28 | 0.1786 | 3 | 0.6667 | 4 | 0.25 | 11 | 0.5455 | 5 | 0.4 | 23 | 0.4783 |
| 3_10354048 | 1 | 0 | 5 | 0.4 | 8 | 0 | 11 | 0.3636 | 25 | 0.24 | 1 | 0 | 5 | 0 | 11 | 0.0909 | 7 | 0.2857 | 24 | 0.125 |
| 3_10354049 | 4 | 0 | 5 | 0.6 | 9 | 0.3333 | 14 | 0.4286 | 32 | 0.375 | 3 | 0.3333 | 10 | 0.4 | 13 | 0.3846 | 7 | 0.5714 | 33 | 0.4242 |
| 3_10354084 | 6 | 0 | 6 | 0 | 9 | 0 | 18 | 0.1111 | 39 | 0.0513 | 3 | 0.3333 | 12 | 0 | 15 | 0 | 7 | 0.1429 | 37 | 0.0541 |
| 3_10354086 | 6 | 0.8333 | 6 | 1 | 9 | 0.5556 | 17 | 0.6471 | 38 | 0.7105 | 3 | 0.6667 | 12 | 0.75 | 14 | 0.6429 | 7 | 1 | 36 | 0.75 |
| 3_10354086 | 5 | 0 | 6 | 0.5 | 7 | 0.1429 | 14 | 0.0714 | 32 | 0.1562 | 3 | 0.3333 | 11 | 0.1818 | 14 | 0.2143 | 4 | 0 | 32 | 0.1875 |
| 3_10354102 | 6 | 0.1667 | 6 | 0.3333 | 9 | 0.1111 | 18 | 0.1667 | 39 | 0.1795 | 3 | 0 | 10 | 0 | 14 | 0.1429 | 7 | 0.4286 | 34 | 0.1471 |
| 3_10354213 | 3 | 0.3333 | 7 | 0.5714 | 10 | 0.5 | 15 | 0.5333 | 35 | 0.5143 | 5 | 0.4 | 12 | 0.6667 | 12 | 0.5 | 11 | 0.1818 | 40 | 0.45 |
| 3_10354217 | 6 | 0.1667 | 9 | 0.1111 | 14 | 0.0714 | 24 | 0.0833 | 53 | 0.0943 | 6 | 0.1667 | 15 | 0.0667 | 19 | 0.1579 | 11 | 0 | 51 | 0.098 |
| 3_10354219 | 4 | 0 | 7 | 0.5714 | 12 | 0.0833 | 23 | 0.3478 | 46 | 0.2826 | 5 | 0.4 | 14 | 0.1429 | 15 | 0.2 | 12 | 0.25 | 46 | 0.2174 |
| 3_10364282 | 4 | 0.25 | 4 | 0.25 | 7 | 0 | NA | NA | 15 | 0.1333 | NA | NA | 2 | 0 | 4 | 0.25 | 1 | 0 | 7 | 0.1429 |
| 3_10364297 | 3 | 0.6667 | 4 | 0.5 | 8 | 0.25 | NA | NA | 15 | 0.4 | NA | NA | 4 | 0 | 2 | 0 | 1 | 1 | 7 | 0.1429 |
| 3_10364303 | 3 | 0 | 5 | 0.8 | 6 | 0.5 | NA | NA | 14 | 0.5 | NA | NA | 3 | 0.6667 | 3 | 1 | 1 | 1 | 7 | 0.8571 |
| 3_10364339 | 3 | 1 | 1 | 1 | 8 | 0.625 | NA | NA | 12 | 0.75 | NA | NA | 2 | 0.5 | 3 | 0.3333 | 1 | 1 | 6 | 0.5 |
| 3_10364346 | 3 | 0 | 2 | 0.5 | 6 | 0.1667 | NA | NA | 11 | 0.1818 | NA | NA | 2 | 0.5 | 3 | 0.3333 | 1 | 0 | 6 | 0.3333 |
| 3_10364346 | 3 | 1 | 2 | 0 | 6 | 0.5 | NA | NA | 11 | 0.5455 | NA | NA | 2 | 0.5 | 2 | 0 | 1 | 1 | 5 | 0.4 |
| 3_10364363 | 3 | 0.6667 | 1 | 0 | 4 | 0.5 | NA | NA | 8 | 0.5 | NA | NA | 2 | 0.5 | 3 | 0.6667 | NA | NA | 5 | 0.6 |
| 3_10364369 | 3 | 1 | 3 | 0.6667 | 7 | 0.7143 | NA | NA | 13 | 0.7692 | NA | NA | 1 | 1 | 3 | 1 | 1 | 0 | 5 | 0.8 |
| 3_10364381 | 3 | 0.3333 | 3 | 0 | 6 | 0 | NA | NA | 12 | 0.0833 | NA | NA | 2 | 0 | 3 | 0 | 1 | 1 | 6 | 0.1667 |
| 3_10364449 | 2 | 0.5 | 3 | 0.6667 | 5 | 0 | NA | NA | 10 | 0.3 | NA | NA | 2 | 0 | 2 | 0.5 | 1 | 0 | 5 | 0.2 |
| 3_10364993 | 14 | 0.1429 | 6 | 0.1667 | 6 | 0.1667 | 5 | 0.2 | 31 | 0.1613 | 8 | 0.75 | 5 | 0.2 | 7 | 1 | 4 | 0.5 | 24 | 0.6667 |
| 3_10365040 | 12 | 0.1667 | 7 | 0 | 6 | 0 | 4 | 0 | 29 | 0.069 | 7 | 0 | 6 | 0.1667 | 11 | 0.0909 | 5 | 0 | 29 | 0.069 |

| | | | | | | | | | | | | | | | | | | | | |
|---|---|---|---|---|---|---|---|---|---|---|---|---|---|---|---|---|---|---|---|---|
| 3_10365058 | 11 | 0.4545 | 7 | 0.7143 | 5 | 0.4 | 4 | 0.5 | 27 | 0.5185 | 5 | 0.8 | 6 | 0.8333 | 11 | 0.7273 | 3 | 0.6667 | 25 | 0.76 |
| 3_10365080 | 11 | 0.0909 | 6 | 0.3333 | 5 | 0.4 | 4 | 0.25 | 26 | 0.2308 | 5 | 0.4 | 6 | 0.1667 | 10 | 0.3 | 5 | 0 | 26 | 0.2308 |
| 3_10365081 | 6 | 0 | 6 | 0.1667 | 2 | 0.5 | 2 | 1 | 16 | 0.25 | 1 | 0 | 3 | 0 | 5 | 0.4 | 1 | 0 | 10 | 0.2 |
| 3_10365087 | 12 | 0.3333 | 5 | 0.4 | 4 | 0.25 | 4 | 0.25 | 25 | 0.32 | 5 | 0.4 | 6 | 0.3333 | 9 | 0.2222 | 5 | 0.2 | 25 | 0.28 |
| 3_10365093 | 12 | 0.25 | 6 | 0.3333 | 5 | 0.2 | 3 | 0.3333 | 26 | 0.2692 | 5 | 0.4 | 6 | 0.5 | 9 | 0.3333 | 5 | 0.6 | 25 | 0.44 |
| 3_10365103 | 9 | 0.2222 | 5 | 0.4 | 5 | 0 | 2 | 0.5 | 21 | 0.2381 | 4 | 0.25 | 5 | 0.2 | 7 | 0 | 5 | 0.2 | 21 | 0.1429 |
| 3_10365109 | 10 | 0.1 | 5 | 0.2 | 5 | 0 | 4 | 0 | 24 | 0.0833 | 2 | 0 | 5 | 0.2 | 7 | 0 | 5 | 0.6 | 19 | 0.2105 |
| 3_10365111 | 8 | 0.5 | 5 | 0.6 | 4 | 0 | 4 | 0.25 | 21 | 0.381 | 4 | 0.5 | 4 | 0.5 | 8 | 0.25 | 5 | 0.4 | 21 | 0.381 |
| 3_10368460 | 3 | 0 | 3 | 0.6667 | 1 | 1 | 3 | 0.6667 | 10 | 0.5 | 1 | 1 | NA | NA | 1 | 1 | 2 | 0 | 4 | 0.5 |
| 3_10368479 | 3 | 1 | 2 | 1 | 1 | 1 | 3 | 0.6667 | 9 | 0.8889 | 1 | 1 | NA | NA | 1 | 1 | 2 | 0.5 | 4 | 0.75 |
| 3_10368486 | 3 | 0 | 2 | 0 | 1 | 1 | 3 | 0.3333 | 9 | 0.2222 | 1 | 1 | NA | NA | 1 | 1 | 2 | 0 | 4 | 0.5 |
| 3_10368492 | 3 | 1 | 2 | 1 | 1 | 0 | 3 | 0.3333 | 9 | 0.6667 | 1 | 1 | NA | NA | 1 | 1 | 2 | 1 | 4 | 1 |
| 3_10368500 | 2 | 1 | 2 | 0.5 | 1 | 0 | 3 | 0.3333 | 8 | 0.5 | 1 | 1 | NA | NA | NA | NA | 1 | 0 | 2 | 0.5 |
| 3_10368501 | 2 | 0.5 | 2 | 1 | 1 | 0 | 3 | 1 | 8 | 0.75 | 1 | 1 | NA | NA | NA | NA | 1 | 1 | 2 | 1 |
| 3_10368554 | 3 | 0 | 3 | 0 | NA | NA | 3 | 0.3333 | 9 | 0.1111 | 1 | 0 | NA | NA | NA | NA | 2 | 0.5 | 3 | 0.3333 |
| 3_10370902 | 1 | 0 | NA | NA | 1 | 0 | 1 | 1 | 3 | 0.3333 | 2 | 0.5 | NA | NA | 1 | 0 | NA | NA | 3 | 0.3333 |
| 3_10371024 | 1 | 0 | NA | NA | 1 | 1 | 3 | 0 | 5 | 0.2 | 3 | 0.6667 | NA | NA | 2 | 0 | NA | NA | 5 | 0.4 |
| 3_10371028 | NA | NA | NA | NA | 1 | 0 | 3 | 0.6667 | 4 | 0.5 | 3 | 0.6667 | NA | NA | 2 | 0 | NA | NA | 5 | 0.4 |
| 3_10371049 | 1 | 1 | NA | NA | 1 | 1 | 2 | 1 | 4 | 1 | 2 | 1 | NA | NA | 1 | 1 | NA | NA | 3 | 1 |
| 3_10371057 | 1 | 0 | NA | NA | 1 | 0 | 4 | 0.25 | 6 | 0.1667 | 2 | 0.5 | NA | NA | 1 | 0 | NA | NA | 3 | 0.3333 |
| 3_10371063 | 1 | 1 | NA | NA | 1 | 1 | 1 | 1 | 3 | 1 | 2 | 0.5 | NA | NA | 2 | 0.5 | NA | NA | 4 | 0.5 |
| 3_10371080 | 1 | 1 | NA | NA | 1 | 1 | 4 | 1 | 6 | 1 | 2 | 1 | NA | NA | 3 | 1 | NA | NA | 5 | 1 |
| 3_10371086 | 1 | 0 | NA | NA | 1 | 1 | 3 | 1 | 5 | 0.8 | 2 | 1 | NA | NA | 2 | 1 | NA | NA | 4 | 1 |
| 3_10371098 | 1 | 1 | NA | NA | 1 | 1 | 4 | 1 | 6 | 1 | 2 | 1 | NA | NA | 2 | 1 | NA | NA | 4 | 1 |
| 3_10371121 | 1 | 0 | NA | NA | 1 | 1 | 2 | 0 | 4 | 0.25 | 2 | 0.5 | NA | NA | 2 | 0 | NA | NA | 4 | 0.25 |
| 3_10371131 | 1 | 0 | NA | NA | 1 | 1 | 4 | 0.5 | 6 | 0.5 | 2 | 0.5 | NA | NA | 1 | 0 | NA | NA | 3 | 0.3333 |
| 3_10371137 | 1 | 1 | NA | NA | 1 | 0 | 2 | 0 | 4 | 0.25 | 1 | 0 | NA | NA | 2 | 0.5 | NA | NA | 3 | 0.3333 |
| 3_10371636 | NA | NA | NA | NA | NA | NA | 3 | 0.6667 | 3 | 0.6667 | 2 | 0.5 | NA | NA | 2 | 0.5 | NA | NA | 4 | 0.5 |
| 3_10372221 | 6 | 0.1667 | NA | NA | 2 | 0 | NA | NA | 8 | 0.125 | NA | NA | 3 | 1 | 2 | 0.5 | 1 | 1 | 6 | 0.8333 |
| 3_10372224 | 5 | 0.6 | NA | NA | 2 | 0.5 | NA | NA | 7 | 0.5714 | NA | NA | 3 | 0.6667 | 3 | 0.6667 | NA | NA | 6 | 0.6667 |
| 3_10372229 | 6 | 0.5 | NA | NA | 1 | 0 | NA | NA | 7 | 0.4286 | NA | NA | 3 | 0 | 2 | 0 | 1 | 1 | 6 | 0.1667 |
| 3_10372233 | 6 | 0.3333 | NA | NA | 2 | 0.5 | NA | NA | 8 | 0.375 | NA | NA | 3 | 0.6667 | 1 | 1 | NA | NA | 4 | 0.75 |
| 3_10372764 | 8 | 0.125 | NA | NA | 2 | 0 | 1 | 0 | 11 | 0.0909 | NA | NA | 3 | 0 | 2 | 0 | 2 | 0.5 | 7 | 0.1429 |
| 3_10372811 | 4 | 0.25 | NA | NA | 1 | 0 | NA | NA | 5 | 0.2 | NA | NA | 1 | 0 | 1 | 0 | 2 | 0.5 | 4 | 0.25 |
| 3_10372885 | 5 | 0.2 | NA | NA | 2 | 1 | 1 | 1 | 8 | 0.5 | NA | NA | 2 | 0.5 | 3 | 1 | 2 | 0 | 7 | 0.5714 |
| 3_10372896 | 8 | 0.125 | NA | NA | 1 | 0 | 1 | 1 | 10 | 0.2 | NA | NA | 3 | 0 | 2 | 0 | 2 | 0.5 | 7 | 0.1429 |
| 3_10372900 | 8 | 0.875 | NA | NA | 2 | 1 | 1 | 1 | 11 | 0.9091 | NA | NA | 3 | 1 | 3 | 1 | 2 | 1 | 8 | 1 |
| 3_10372904 | 9 | 0.2222 | NA | NA | 2 | 0.5 | 1 | 0 | 12 | 0.25 | NA | NA | 2 | 0 | 3 | 0.3333 | 1 | 0 | 6 | 0.1667 |
| 3_10372906 | 9 | 0 | NA | NA | 2 | 0.5 | 1 | 0 | 12 | 0.0833 | NA | NA | 3 | 0.3333 | 1 | 0 | 2 | 0 | 6 | 0.1667 |
| 3_10383585 | 2 | 1 | 1 | 1 | 1 | 1 | 2 | 1 | 6 | 1 | NA | NA | NA | NA | NA | NA | 2 | 1 | 2 | 1 |
| 3_10383596 | 1 | 1 | 1 | 0 | 1 | 0 | 2 | 0 | 5 | 0.2 | NA | NA | NA | NA | NA | NA | 1 | 1 | 1 | 1 |
| 3_10383610 | 1 | 0 | 1 | 1 | 1 | 1 | NA | NA | 3 | 0.6667 | NA | NA | NA | NA | NA | NA | 1 | 1 | 1 | 1 |
| 3_10383623 | 2 | 0.5 | 2 | 0 | 1 | 0 | 1 | 0 | 6 | 0.1667 | NA | NA | NA | NA | NA | NA | 2 | 0.5 | 2 | 0.5 |
| 3_10383624 | 2 | 0.5 | 2 | 1 | 1 | 1 | 2 | 1 | 7 | 0.8571 | NA | NA | NA | NA | NA | NA | 1 | 1 | 1 | 1 |
| 3_10386960 | 3 | 0 | 1 | 0 | NA | NA | 1 | 1 | 5 | 0.2 | NA | NA | NA | NA | 1 | 1 | 1 | 1 | 2 | 1 |
| 3_10386972 | 2 | 0.5 | 2 | 0.5 | NA | NA | 2 | 1 | 6 | 0.6667 | NA | NA | NA | NA | 1 | 1 | 2 | 0 | 3 | 0.3333 |
| 3_10386974 | 1 | 1 | 1 | 0 | NA | NA | 2 | 1 | 4 | 0.75 | NA | NA | NA | NA | 1 | 1 | 1 | 1 | 2 | 1 |
| 3_10386981 | 3 | 0.3333 | 1 | 0 | 1 | 1 | 2 | 0 | 7 | 0.2857 | NA | NA | NA | NA | 1 | 1 | 1 | 0 | 2 | 0.5 |
| 3_10386987 | 3 | 1 | 1 | 1 | 1 | 1 | 2 | 1 | 7 | 1 | NA | NA | NA | NA | 1 | 1 | 2 | 1 | 3 | 1 |
| 3_10387039 | 1 | 0 | 3 | 0.3333 | 1 | 0 | 1 | 1 | 6 | 0.3333 | NA | NA | 1 | 1 | 2 | 0 | 2 | 1 | 5 | 0.6 |
| 3_10387071 | 2 | 0 | 3 | 0.3333 | 1 | 0 | 1 | 0 | 7 | 0.1429 | NA | NA | 1 | 1 | 2 | 0.5 | 2 | 0.5 | 5 | 0.6 |
| 3_10387078 | 2 | 0 | 3 | 0 | 1 | 1 | 1 | 0 | 7 | 0.1429 | NA | NA | 1 | 1 | 2 | 0 | 2 | 0 | 5 | 0.2 |
| 3_10387197 | 3 | 0 | 6 | 0.3333 | 3 | 0.3333 | 2 | 0.5 | 14 | 0.2857 | NA | NA | 3 | 1 | 2 | 0.5 | 1 | 1 | 6 | 0.8333 |
| 3_10387225 | 3 | 0.6667 | 6 | 0.8333 | 3 | 1 | 2 | 0.5 | 14 | 0.7857 | NA | NA | 4 | 0.5 | 2 | 0 | NA | NA | 6 | 0.3333 |
| 3_10387229 | 3 | 0 | 9 | 0.1111 | 2 | 0 | 3 | 0.3333 | 17 | 0.1176 | NA | NA | 5 | 0.4 | 2 | 0 | 1 | 0 | 8 | 0.25 |
| 3_10387336 | 2 | 0 | 9 | 0.1111 | 3 | 0.3333 | 4 | 0 | 18 | 0.1111 | NA | NA | 6 | 0.1667 | 5 | 0.2 | 1 | 0 | 12 | 0.1667 |
| 3_10387368 | 2 | 0.5 | 8 | 0.125 | 4 | 0.5 | 2 | 0 | 16 | 0.25 | NA | NA | 6 | 0.1667 | 6 | 0.1667 | 3 | 0.3333 | 15 | 0.2 |
| 3_10391284 | 2 | 0 | 7 | 0 | 6 | 0 | 2 | 0.5 | 17 | 0.0588 | 5 | 0.2 | 6 | 0 | 4 | 0 | 5 | 0 | 20 | 0.05 |
| 3_10391434 | 2 | 0.5 | 6 | 0.3333 | 6 | 0.3333 | 1 | 0 | 15 | 0.3333 | 3 | 0 | 6 | 0.5 | 4 | 0.25 | 7 | 0.4286 | 20 | 0.35 |
| 3_10391437 | 2 | 0 | 7 | 0 | 7 | 0.1429 | 1 | 0 | 17 | 0.0588 | 4 | 0 | 5 | 0.2 | 4 | 0 | 6 | 0 | 19 | 0.0526 |
| 3_10391456 | 3 | 0.3333 | 8 | 0.375 | 7 | 0.5714 | 2 | 0 | 20 | 0.4 | 6 | 0.3333 | 8 | 0.625 | 5 | 0.4 | 7 | 0.4286 | 26 | 0.4615 |
| 3_10391457 | 2 | 0.5 | 8 | 0.625 | 7 | 0.7143 | 2 | 0.5 | 19 | 0.6316 | 5 | 0.6 | 5 | 0.8 | 5 | 0.8 | 7 | 0.4286 | 22 | 0.6364 |
| 3_10391458 | 3 | 0.6667 | 9 | 0.8889 | 8 | 0.75 | 2 | 0.5 | 22 | 0.7727 | 4 | 0.5 | 6 | 1 | 5 | 0.8 | 7 | 1 | 22 | 0.8636 |
| 3_10391474 | 3 | 0.6667 | 10 | 0.3 | 9 | 0.5556 | 1 | 0 | 23 | 0.4348 | 5 | 0.6 | 7 | 0.5714 | 5 | 0.6 | 8 | 0.5 | 25 | 0.56 |
| 3_10391476 | 3 | 0.3333 | 11 | 0.1818 | 9 | 0.1111 | 1 | 0 | 24 | 0.1667 | 6 | 0.1667 | 7 | 0 | 6 | 0.1667 | 8 | 0.125 | 27 | 0.1111 |

| | | | | | | | | | | | | | | | | | | | | |
|---|---|---|---|---|---|---|---|---|---|---|---|---|---|---|---|---|---|---|---|---|
| 3_10391479 | 2 | 0.5 | 8 | 0.5 | 8 | 0.5 | 1 | 0 | 19 | 0.4737 | 5 | 0.4 | 8 | 0.5 | 5 | 0.4 | 7 | 0.5714 | 25 | 0.48 |
| 3_10392097 | 4 | 0.25 | 4 | 0.5 | 6 | 0.6667 | 8 | 0.375 | 22 | 0.4545 | 1 | 0 | 4 | 0 | 14 | 0.6429 | 4 | 0.25 | 23 | 0.4348 |
| 3_10392127 | 3 | 0.3333 | 4 | 0.5 | 5 | 0.2 | 8 | 0.125 | 20 | 0.25 | 2 | 0 | 4 | 0 | 13 | 0.0769 | 6 | 0.1667 | 25 | 0.08 |
| 3_10392169 | 1 | 0 | 5 | 0 | 2 | 0 | 7 | 0.1429 | 15 | 0.0667 | 2 | 0 | 3 | 0 | 9 | 0.1111 | 3 | 0 | 17 | 0.0588 |
| 3_10392235 | 1 | 1 | 4 | 0.25 | 2 | 0 | 6 | 0.5 | 13 | 0.3846 | 2 | 1 | 3 | 0.6667 | 9 | 0.5556 | 6 | 0.8333 | 20 | 0.7 |
| 3_10392242 | NA | NA | 3 | 1 | 1 | 1 | 5 | 1 | 9 | 1 | 1 | 1 | 1 | 1 | 5 | 0.8 | 5 | 1 | 12 | 0.9167 |
| 3_10392261 | 1 | 1 | 2 | 1 | 2 | 1 | 5 | 0.4 | 10 | 0.7 | 1 | 1 | 3 | 0.3333 | 9 | 0.7778 | 4 | 0.75 | 17 | 0.7059 |
| 3_10392275 | 1 | 1 | 2 | 1 | 2 | 0 | 4 | 0.25 | 9 | 0.4444 | 1 | 0 | 2 | 0.5 | 9 | 0.5556 | 5 | 0.6 | 17 | 0.5294 |
| 3_10392289 | 1 | 1 | 3 | 1 | 1 | 0 | 4 | 0.75 | 9 | 0.7778 | 1 | 1 | 1 | 0 | 8 | 0.625 | 5 | 1 | 15 | 0.7333 |
| 3_10392295 | 1 | 1 | 3 | 0.3333 | 2 | 0 | 6 | 0 | 12 | 0.1667 | 1 | 0 | 1 | 0 | 8 | 0.375 | 5 | 0.4 | 15 | 0.3333 |
| 3_10392301 | 1 | 0 | 3 | 0.3333 | 2 | 0 | 5 | 0.4 | 11 | 0.2727 | NA | NA | 1 | 0 | 10 | 0.7 | 4 | 0.5 | 15 | 0.6 |
| 3_10392314 | 1 | 0 | 2 | 0 | 2 | 0 | 5 | 0.2 | 10 | 0.1 | 1 | 0 | 1 | 0 | 9 | 0.2222 | 5 | 0.2 | 16 | 0.1875 |
| 3_10392321 | 1 | 1 | 3 | 0.6667 | 2 | 0 | 6 | 0.8333 | 12 | 0.6667 | 1 | 1 | 1 | 1 | 8 | 0.875 | 5 | 0.8 | 15 | 0.8667 |
| 3_10392326 | 1 | 0 | 3 | 0 | 2 | 0.5 | 5 | 0.2 | 11 | 0.1818 | 1 | 0 | 1 | 0 | 8 | 0.5 | 5 | 0.4 | 15 | 0.4 |
| 3_10395828 | 1 | 0 | 3 | 0.3333 | 1 | 0 | 4 | 0.5 | 9 | 0.3333 | NA | NA | NA | NA | 4 | 0 | 2 | 0.5 | 6 | 0.1667 |
| 3_10395834 | 1 | 1 | 2 | 1 | NA | NA | 3 | 1 | 6 | 1 | 1 | 1 | NA | NA | 7 | 0.8571 | 2 | 0.5 | 10 | 0.8 |
| 3_10395844 | NA | NA | 2 | 0 | 1 | 0 | 3 | 0.3333 | 6 | 0.1667 | 1 | 1 | NA | NA | 7 | 0.5714 | 2 | 0 | 10 | 0.5 |
| 3_10395850 | 1 | 0 | 3 | 0.3333 | NA | NA | 2 | 0.5 | 6 | 0.3333 | NA | NA | NA | NA | 5 | 0.2 | 2 | 0.5 | 7 | 0.2857 |
| 3_10395852 | NA | NA | 3 | 1 | 1 | 1 | 3 | 1 | 7 | 1 | 1 | 1 | NA | NA | 7 | 0.7143 | 2 | 0.5 | 10 | 0.7 |
| 3_10395855 | 1 | 1 | 2 | 0.5 | NA | NA | 3 | 0.6667 | 6 | 0.6667 | 1 | 1 | NA | NA | 6 | 0.8333 | 2 | 0 | 9 | 0.6667 |
| 3_10436274 | 1 | 1 | 2 | 1 | 1 | 0 | NA | NA | 4 | 0.75 | 2 | 1 | 1 | 0 | 2 | 0 | 2 | 0 | 7 | 0.2857 |
| 3_10436284 | 1 | 0 | 2 | 0.5 | 1 | 0 | NA | NA | 4 | 0.25 | 1 | 1 | 1 | 0 | 2 | 0.5 | 2 | 0 | 6 | 0.3333 |
| 3_10436316 | 1 | 1 | 2 | 1 | 1 | 1 | NA | NA | 4 | 1 | 2 | 1 | 1 | 0 | 2 | 0 | 2 | 1 | 7 | 0.5714 |
| 3_10436330 | NA | NA | 1 | 1 | 1 | 0 | NA | NA | 2 | 0.5 | 1 | 1 | 1 | 0 | 2 | 0.5 | 2 | 0.5 | 6 | 0.5 |
| 3_10436339 | 1 | 1 | 2 | 0.5 | 1 | 0 | NA | NA | 4 | 0.5 | 2 | 1 | 1 | 1 | 2 | 0.5 | 2 | 1 | 7 | 0.8571 |
| 3_10436422 | 1 | 0 | NA | NA | 1 | 1 | NA | NA | 2 | 0.5 | 2 | 0 | NA | NA | 2 | 0 | 2 | 0.5 | 6 | 0.1667 |
| 3_10436485 | 1 | 0 | 1 | 0 | 1 | 1 | NA | NA | 3 | 0.3333 | 1 | 0 | NA | NA | 2 | 0.5 | 2 | 0 | 5 | 0.2 |
| 3_10436505 | 1 | 1 | 3 | 1 | 1 | 1 | NA | NA | 5 | 1 | 2 | 1 | NA | NA | 2 | 1 | 2 | 1 | 6 | 1 |
| 3_10436511 | 1 | 1 | 3 | 1 | 1 | 1 | NA | NA | 5 | 1 | 2 | 1 | NA | NA | 2 | 1 | 2 | 1 | 6 | 1 |
| 3_10436513 | 1 | 1 | 3 | 0 | 1 | 0 | NA | NA | 5 | 0.2 | 2 | 0.5 | NA | NA | 2 | 0.5 | 2 | 0.5 | 6 | 0.5 |
| 3_10436524 | 1 | 0 | 2 | 0.5 | 1 | 0 | NA | NA | 4 | 0.25 | 2 | 0 | NA | NA | 1 | 1 | 1 | 1 | 4 | 0.5 |
| 3_10437137 | 1 | 1 | 1 | 1 | 1 | 1 | NA | NA | 3 | 1 | 2 | 1 | NA | NA | 2 | 1 | 1 | 1 | 5 | 1 |
| 3_10437138 | 1 | 1 | 2 | 1 | 1 | 0 | NA | NA | 4 | 0.75 | 2 | 1 | NA | NA | 1 | 1 | 1 | 0 | 4 | 0.75 |
| 3_10437151 | 1 | 0 | 1 | 1 | 1 | 0 | NA | NA | 3 | 0.3333 | 2 | 0.5 | NA | NA | 2 | 0 | 1 | 0 | 5 | 0.2 |
| 3_10456014 | 3 | 0.3333 | 1 | 0 | 2 | 0 | 2 | 0 | 8 | 0.125 | 2 | 0.5 | 3 | 0 | 1 | 1 | 3 | 0.3333 | 9 | 0.3333 |
| 3_10456021 | 2 | 0.5 | 2 | 1 | 2 | 0.5 | 2 | 1 | 8 | 0.75 | 2 | 0.5 | 3 | 0.3333 | 1 | 1 | 3 | 1 | 9 | 0.6667 |
| 3_10456043 | 3 | 1 | 3 | 1 | 2 | 1 | 2 | 1 | 10 | 1 | 3 | 1 | 2 | 1 | 1 | 1 | 3 | 1 | 9 | 1 |
| 3_10456050 | 2 | 1 | 3 | 1 | 2 | 1 | 1 | 1 | 8 | 1 | 3 | 0.6667 | 3 | 1 | 1 | 1 | 3 | 1 | 10 | 0.9 |
| 3_10456062 | 4 | 0.75 | 2 | 1 | 1 | 1 | 2 | 1 | 9 | 0.8889 | 3 | 0.6667 | 3 | 1 | 1 | 1 | 3 | 1 | 10 | 0.9 |
| 3_10456080 | 4 | 0.25 | 3 | 0.3333 | 2 | 0.5 | 2 | 0.5 | 11 | 0.3636 | 3 | 0.3333 | 3 | 0 | 1 | 0 | 3 | 0.3333 | 10 | 0.2 |
| 3_10456081 | 4 | 0.75 | 2 | 1 | 1 | 1 | 2 | 0.5 | 9 | 0.7778 | 3 | 0 | 2 | 1 | 1 | 1 | 2 | 0.5 | 8 | 0.5 |
| 3_10456288 | 2 | 1 | 2 | 0 | 1 | 1 | 3 | 0 | 8 | 0.375 | 3 | 0 | 1 | 1 | 1 | 1 | 2 | 0.5 | 7 | 0.4286 |
| 3_10456301 | 4 | 0 | 3 | 0.3333 | 2 | 0.5 | 2 | 0.5 | 11 | 0.2727 | 2 | 0 | 3 | 0 | 1 | 0 | 2 | 0.5 | 8 | 0.125 |
| 3_10456313 | 3 | 1 | 3 | 1 | 2 | 1 | 2 | 1 | 10 | 1 | 3 | 1 | 3 | 1 | 1 | 1 | 3 | 1 | 10 | 1 |
| 3_10473168 | 1 | 0 | 9 | 0.2222 | 4 | 0 | 7 | 0.1429 | 21 | 0.1429 | 2 | 0.5 | 2 | 0 | 2 | 0 | 3 | 0.3333 | 9 | 0.2222 |
| 3_10477217 | NA | NA | 1 | 1 | 1 | 0 | NA | NA | 2 | 0.5 | NA | NA | NA | NA | 2 | 1 | NA | NA | 2 | 1 |
| 3_10483381 | 2 | 0.5 | 1 | 1 | 1 | 0 | 2 | 1 | 6 | 0.6667 | 4 | 0.75 | 6 | 0.6667 | 5 | 0.8 | 2 | 0.5 | 17 | 0.7059 |
| 3_10483390 | 2 | 0.5 | 2 | 0.5 | 1 | 1 | 2 | 0.5 | 7 | 0.5714 | 4 | 0.75 | 6 | 1 | 5 | 0.4 | 2 | 0.5 | 17 | 0.7059 |
| 3_10483402 | 2 | 0 | 2 | 0.5 | 2 | 0.5 | 1 | 0 | 7 | 0.2857 | 4 | 1 | 5 | 0.6 | 5 | 0.4 | 2 | 0.5 | 16 | 0.625 |
| 3_10483405 | 1 | 1 | 2 | 1 | 2 | 0 | 1 | 1 | 6 | 0.6667 | 3 | 0.3333 | 4 | 0.5 | 5 | 0.6 | 2 | 0.5 | 14 | 0.5 |
| 3_10483690 | NA | NA | NA | NA | NA | NA | 2 | 0.5 | 2 | 0.5 | 1 | 0 | 1 | 1 | NA | NA | NA | NA | 2 | 0.5 |
| 3_10485647 | NA | NA | 3 | 0.3333 | NA | NA | 1 | 0 | 4 | 0.25 | NA | NA | 1 | 0 | 2 | 0.5 | NA | NA | 3 | 0.3333 |
| 3_10490973 | 4 | 0.25 | 2 | 0 | 1 | 0 | 1 | 1 | 8 | 0.25 | 1 | 0 | 2 | 0 | 2 | 0 | 3 | 0.3333 | 8 | 0.125 |
| 3_10491088 | 4 | 0.25 | 2 | 0.5 | 1 | 0 | NA | NA | 7 | 0.2857 | 2 | 0 | 1 | 0 | 2 | 0.5 | 3 | 0.3333 | 8 | 0.25 |
| 3_10491114 | 2 | 1 | 1 | 1 | 1 | 1 | NA | NA | 4 | 1 | 2 | 1 | 1 | 1 | NA | NA | 3 | 1 | 6 | 1 |
| 3_10491118 | 1 | 1 | NA | NA | 1 | 1 | NA | NA | 2 | 1 | 2 | 1 | 1 | 1 | 1 | 1 | 3 | 0.3333 | 7 | 0.7143 |
| 3_10491127 | 2 | 0.5 | 1 | 1 | 1 | 1 | NA | NA | 4 | 0.75 | 1 | 1 | 1 | 1 | 1 | 0 | 3 | 0.3333 | 6 | 0.5 |
| 3_10491132 | 2 | 1 | 1 | 1 | 1 | 1 | NA | NA | 4 | 1 | 1 | 0 | NA | NA | NA | NA | 2 | 1 | 3 | 0.6667 |
| 3_10491136 | 1 | 1 | 1 | 1 | 1 | 1 | NA | NA | 3 | 1 | 1 | 1 | 1 | 1 | 1 | 1 | 2 | 1 | 5 | 1 |
| 3_10491141 | 1 | 1 | 1 | 0 | NA | NA | NA | NA | 2 | 0.5 | NA | NA | 2 | 0.5 | 1 | 0 | 3 | 0.3333 | 6 | 0.3333 |
| 3_10491144 | 1 | 1 | 1 | 1 | 1 | 1 | NA | NA | 3 | 1 | 1 | 1 | 2 | 1 | 1 | 1 | 3 | 1 | 7 | 1 |
| 3_10491151 | 1 | 0 | 1 | 1 | 1 | 0 | NA | NA | 3 | 0.3333 | 1 | 1 | 2 | 1 | 1 | 0 | 3 | 0.6667 | 7 | 0.7143 |
| 3_10491158 | 2 | 1 | 1 | 1 | 1 | 1 | NA | NA | 4 | 1 | NA | NA | 1 | 1 | 1 | 1 | 1 | 1 | 3 | 1 |
| 3_10491168 | 2 | 0.5 | 1 | 1 | 1 | 1 | NA | NA | 4 | 0.75 | 1 | 0 | 1 | 1 | 1 | 1 | 1 | 1 | 4 | 0.75 |
| 3_10491175 | 2 | 0.5 | 1 | 0 | 1 | 0 | NA | NA | 4 | 0.25 | 1 | 0 | 1 | 1 | 1 | 0 | 2 | 0 | 5 | 0.2 |

| | | | | | | | | | | | | | | | | | | | | |
|---|---|---|---|---|---|---|---|---|---|---|---|---|---|---|---|---|---|---|---|---|
| 3_10491178 | 1 | 1 | 1 | 0 | 1 | 0 | NA | NA | 3 | 0.3333 | 1 | 0 | 1 | 0 | 1 | 0 | 2 | 0.5 | 5 | 0.2 |
| 3_10491192 | 1 | 0 | 1 | 1 | 1 | 1 | NA | NA | 3 | 0.6667 | NA | NA | 1 | 1 | NA | NA | 2 | 1 | 3 | 1 |
| 3_10491193 | 2 | 1 | 1 | 0 | 1 | 1 | NA | NA | 4 | 0.75 | 1 | 1 | 1 | 1 | NA | NA | 1 | 1 | 3 | 1 |
| 3_10491208 | 1 | 0 | 1 | 1 | 1 | 1 | NA | NA | 3 | 0.6667 | 1 | 1 | 1 | 1 | NA | NA | 1 | 1 | 3 | 1 |
| 3_10491229 | 2 | 0.5 | 1 | 1 | 1 | 1 | NA | NA | 4 | 0.75 | 1 | 1 | 1 | 1 | NA | NA | NA | NA | 2 | 1 |
| 3_10491242 | 2 | 1 | 1 | 1 | NA | NA | NA | NA | 3 | 1 | 1 | 1 | 1 | 1 | NA | NA | 1 | 0 | 3 | 0.6667 |
| 3_10491269 | 2 | 0.5 | 1 | 0 | 1 | 0 | NA | NA | 4 | 0.25 | 1 | 1 | 1 | 0 | NA | NA | 1 | 1 | 3 | 0.6667 |
| 3_10491269 | 2 | 0.5 | 1 | 0 | 1 | 1 | NA | NA | 4 | 0.5 | 1 | 1 | NA | NA | NA | NA | NA | NA | 1 | 1 |
| 3_10536630 | 1 | 0 | 1 | 1 | NA | NA | 1 | 1 | 3 | 0.6667 | 1 | 0 | NA | NA | NA | NA | 2 | 0.5 | 3 | 0.3333 |
| 3_10536648 | NA | NA | 1 | 1 | NA | NA | NA | NA | 1 | 1 | 1 | 1 | NA | NA | NA | NA | NA | NA | 1 | 1 |
| 3_10536658 | NA | NA | NA | NA | NA | NA | 1 | 1 | 1 | 1 | 1 | 1 | NA | NA | NA | NA | 2 | 1 | 3 | 1 |
| 3_10536669 | NA | NA | 1 | 1 | NA | NA | NA | NA | 1 | 1 | 1 | 1 | NA | NA | NA | NA | 2 | 1 | 3 | 1 |
| 3_10536682 | 1 | 1 | 1 | 1 | NA | NA | 1 | 1 | 3 | 1 | 1 | 1 | NA | NA | NA | NA | 1 | 1 | 2 | 1 |
| 3_10557635 | 1 | 1 | 2 | 0.5 | NA | NA | 2 | 0 | 5 | 0.4 | 1 | 1 | 1 | 1 | NA | NA | 2 | 0.5 | 4 | 0.75 |
| 3_10557647 | 1 | 1 | 1 | 1 | NA | NA | 2 | 1 | 4 | 1 | 1 | 1 | 1 | 1 | NA | NA | 2 | 1 | 4 | 1 |
| 3_10557653 | NA | NA | 1 | 1 | NA | NA | 1 | 1 | 2 | 1 | NA | NA | 1 | 1 | NA | NA | 2 | 1 | 3 | 1 |
| 3_10557659 | 1 | 1 | 1 | 0 | NA | NA | 1 | 0 | 3 | 0.3333 | 1 | 0 | NA | NA | NA | NA | 2 | 0.5 | 3 | 0.3333 |
| 3_10557665 | 1 | 0 | 2 | 1 | NA | NA | 2 | 0.5 | 5 | 0.6 | 1 | 1 | 1 | 1 | NA | NA | 2 | 1 | 4 | 1 |
| 3_10557671 | 1 | 1 | 2 | 1 | NA | NA | NA | NA | 3 | 1 | NA | NA | NA | NA | NA | NA | 3 | 1 | 3 | 1 |
| 3_10557672 | 1 | 1 | 2 | 0.5 | NA | NA | 2 | 0.5 | 5 | 0.6 | 1 | 0 | 1 | 0 | NA | NA | 3 | 0.3333 | 5 | 0.2 |
| 3_10557704 | 1 | 1 | NA | NA | 1 | 1 | 2 | 0.5 | 4 | 0.75 | NA | NA | 1 | 1 | NA | NA | 3 | 0.6667 | 4 | 0.75 |
| 3_10557712 | 1 | 1 | 1 | 0 | 1 | 1 | 2 | 1 | 5 | 0.8 | 1 | 1 | 1 | 0 | NA | NA | 3 | 0 | 5 | 0.2 |
| 3_10557727 | NA | NA | 1 | 1 | 1 | 1 | 2 | 1 | 4 | 1 | 1 | 1 | 1 | 0 | NA | NA | 4 | 1 | 6 | 0.8333 |
| 3_10557733 | 1 | 1 | 2 | 1 | 1 | 1 | 3 | 1 | 7 | 1 | 1 | 1 | 1 | 1 | 1 | 1 | 3 | 1 | 6 | 1 |
| 3_10557736 | 1 | 1 | NA | NA | 1 | 1 | 2 | 1 | 4 | 1 | 1 | 1 | 1 | 1 | 1 | 1 | 3 | 1 | 6 | 1 |
| 3_10557736 | NA | NA | NA | NA | NA | NA | 3 | 0.3333 | 3 | 0.3333 | 2 | 0.5 | 1 | 0 | 1 | 0 | 3 | 0.3333 | 7 | 0.2857 |
| 3_10557746 | NA | NA | 1 | 1 | 1 | 1 | 2 | 0 | 4 | 0.5 | 1 | 0 | 1 | 0 | 1 | 0 | 3 | 0.3333 | 6 | 0.1667 |
| 3_10557876 | 3 | 0 | 2 | 0 | 2 | 0.5 | 3 | 0 | 10 | 0.1 | 3 | 0 | 2 | 0 | 1 | 0 | 5 | 0.2 | 11 | 0.0909 |
| 3_10557879 | 1 | 0 | 1 | 1 | 2 | 0.5 | 5 | 0.8 | 9 | 0.6667 | 3 | 0 | NA | NA | 1 | 0 | 4 | 0.25 | 8 | 0.125 |
| 3_10557885 | 3 | 0.6667 | 1 | 0 | 2 | 1 | 3 | 0.6667 | 9 | 0.6667 | 2 | 0.5 | 2 | 0.5 | 1 | 1 | 5 | 0.4 | 10 | 0.5 |
| 3_10557888 | 3 | 0 | 2 | 0 | 2 | 0 | 5 | 0.4 | 12 | 0.1667 | 3 | 0.3333 | 2 | 0 | 1 | 0 | 3 | 0 | 9 | 0.1111 |
| 3_10557889 | 3 | 0.3333 | 2 | 0.5 | 2 | 0.5 | 5 | 0.6 | 12 | 0.5 | 3 | 0.6667 | 2 | 0 | 1 | 1 | 4 | 0.25 | 10 | 0.4 |
| 3_10557896 | 3 | 0.3333 | NA | NA | 2 | 1 | 3 | 0.3333 | 8 | 0.5 | 3 | 0.3333 | 2 | 0.5 | 1 | 1 | 4 | 0.25 | 10 | 0.4 |
| 3_10557920 | 3 | 0.3333 | 2 | 0 | 2 | 0.5 | 4 | 0.5 | 11 | 0.3636 | 2 | 0 | 2 | 0 | 1 | 1 | 5 | 0.4 | 10 | 0.3 |
| 3_10557931 | 3 | 0 | 1 | 0 | 2 | 0.5 | 3 | 0 | 9 | 0.1111 | 3 | 0.6667 | 2 | 0 | 1 | 0 | 4 | 0.25 | 10 | 0.3 |
| 3_10557938 | 3 | 0.3333 | 2 | 1 | 1 | 1 | 6 | 1 | 12 | 0.8333 | 3 | 1 | 2 | 1 | 1 | 1 | 3 | 1 | 9 | 1 |
| 3_10557947 | 3 | 0 | 2 | 0 | 3 | 0 | 6 | 0.1667 | 14 | 0.0714 | 3 | 0.6667 | 2 | 0 | 1 | 0 | 5 | 0 | 11 | 0.1818 |
| 3_10557948 | 3 | 0.3333 | 3 | 0 | 3 | 1 | 6 | 0.6667 | 15 | 0.5333 | 3 | 0.6667 | 2 | 1 | 1 | 0 | 4 | 0.75 | 10 | 0.7 |
| 3_10557965 | 3 | 0.6667 | 3 | 0 | 2 | 0 | 5 | 0.4 | 13 | 0.3077 | 2 | 0 | 2 | 0 | 1 | 0 | 5 | 0.2 | 10 | 0.1 |
| 3_10557968 | 2 | 1 | 2 | 0.5 | 3 | 0.3333 | 6 | 0.6667 | 13 | 0.6154 | 3 | 0.6667 | 2 | 0.5 | 1 | 1 | 4 | 0.25 | 10 | 0.5 |
| 3_10557982 | 3 | 0 | 3 | 0 | 3 | 0 | 6 | 0.6667 | 15 | 0.2667 | 2 | 1 | 2 | 0 | 1 | 0 | 3 | 0.3333 | 8 | 0.375 |
| 3_10557982 | 3 | 1 | 3 | 0 | 3 | 0.3333 | 8 | 0.375 | 17 | 0.4118 | 3 | 0.6667 | 1 | 0 | 1 | 0 | 3 | 0 | 8 | 0.25 |
| 3_10558002 | 2 | 0.5 | 3 | 0 | 3 | 0.3333 | 7 | 0 | 15 | 0.1333 | 4 | 0 | 2 | 0.5 | 1 | 0 | 5 | 0 | 12 | 0.0833 |
| 3_10558015 | 2 | 0 | 2 | 0 | 3 | 0 | 8 | 0.25 | 15 | 0.1333 | 3 | 0 | 2 | 0 | 1 | 0 | 5 | 0.2 | 11 | 0.0909 |
| 3_10558075 | 3 | 0 | 5 | 0 | 3 | 0.3333 | 9 | 0.2222 | 20 | 0.15 | 3 | 0 | 2 | 0 | 2 | 0.5 | 5 | 0 | 12 | 0.0833 |
| 3_10558085 | 3 | 0 | 5 | 0.2 | 4 | 0 | 11 | 0.0909 | 23 | 0.087 | 4 | 0.25 | 2 | 0 | 3 | 0 | 7 | 0.1429 | 16 | 0.125 |
| 3_10558093 | 4 | 0.5 | 6 | 0 | 4 | 0 | 11 | 0.2727 | 25 | 0.2 | 5 | 0 | 2 | 0.5 | 4 | 0 | 6 | 0 | 17 | 0.0588 |
| 3_10558094 | 4 | 0.25 | 6 | 0.3333 | 4 | 0 | 10 | 0.3 | 24 | 0.25 | 5 | 0.6 | 2 | 0 | 4 | 0.25 | 5 | 0.8 | 16 | 0.5 |
| 3_10558095 | 3 | 0.6667 | 5 | 1 | 3 | 0.6667 | 10 | 0.7 | 21 | 0.7619 | 4 | 1 | 2 | 1 | 4 | 1 | 4 | 1 | 14 | 1 |
| 3_10558100 | 4 | 0 | 4 | 0.25 | 3 | 0 | 10 | 0.1 | 21 | 0.0952 | 5 | 0.2 | 2 | 0 | 3 | 0 | 6 | 0 | 16 | 0.0625 |
| 3_10558564 | 5 | 0.4 | 1 | 0 | 5 | 0 | 2 | 0.5 | 13 | 0.2308 | 1 | 0 | 1 | 0 | 1 | 1 | 2 | 0 | 5 | 0.2 |
| 3_10558571 | 4 | 1 | 1 | 1 | 5 | 0 | 2 | 0.5 | 12 | 0.5 | 1 | 1 | NA | NA | 1 | 0 | 2 | 0 | 4 | 0.25 |
| 3_10558574 | 5 | 0.6 | 1 | 1 | 5 | 0.6 | 1 | 1 | 12 | 0.6667 | 1 | 1 | NA | NA | 1 | 1 | 2 | 1 | 4 | 1 |
| 3_10558576 | 5 | 1 | 1 | 1 | 4 | 1 | 2 | 1 | 12 | 1 | 1 | 1 | NA | NA | 1 | 1 | 2 | 1 | 4 | 1 |
| 3_10558577 | 5 | 0 | 1 | 0 | 5 | 0.2 | 2 | 0 | 13 | 0.0769 | 1 | 0 | NA | NA | NA | NA | 2 | 0.5 | 3 | 0.3333 |
| 3_10558584 | 2 | 1 | 1 | 1 | 4 | 1 | 1 | 1 | 8 | 1 | 1 | 1 | NA | NA | NA | NA | 2 | 1 | 3 | 1 |
| 3_10558584 | 4 | 0.25 | NA | NA | 4 | 0.5 | 1 | 0 | 9 | 0.3333 | 1 | 1 | NA | NA | 1 | 0 | 2 | 0 | 4 | 0.25 |
| 3_10558593 | 4 | 0.5 | 1 | 1 | 3 | 0.3333 | 1 | 0 | 9 | 0.4444 | 1 | 0 | NA | NA | 1 | 0 | 2 | 0.5 | 4 | 0.25 |
| 3_10558607 | 3 | 0 | NA | NA | 2 | 0.5 | 1 | 1 | 6 | 0.3333 | 1 | 1 | NA | NA | NA | NA | 2 | 0 | 3 | 0.3333 |
| 3_10558635 | 4 | 0.75 | NA | NA | 2 | 1 | 1 | 1 | 7 | 0.8571 | 1 | 1 | NA | NA | 1 | 1 | 2 | 0.5 | 4 | 0.75 |
| 3_10558638 | 4 | 0.25 | NA | NA | 2 | 0 | 1 | 0 | 7 | 0.1429 | 1 | 1 | NA | NA | NA | NA | 2 | 0 | 3 | 0.3333 |
| 3_10558656 | 3 | 0.6667 | NA | NA | 2 | 0.5 | 1 | 0 | 6 | 0.5 | 1 | 0 | NA | NA | NA | NA | 2 | 1 | 3 | 0.6667 |
| 3_10558657 | 3 | 0.3333 | NA | NA | 1 | 0 | 1 | 1 | 5 | 0.4 | 1 | 1 | NA | NA | NA | NA | 2 | 1 | 3 | 1 |
| 3_10562384 | 2 | 0.5 | 1 | 0 | NA | NA | 3 | 0 | 6 | 0.1667 | 2 | 0 | NA | NA | 2 | 0 | 3 | 0.3333 | 7 | 0.1429 |
| 3_10562390 | 3 | 0.3333 | 1 | 0 | 1 | 1 | 4 | 0.75 | 9 | 0.5556 | 2 | 1 | 3 | 0.6667 | 4 | 0.25 | 4 | 0.25 | 13 | 0.4615 |

| | | | | | | | | | | | | | | | | | | | | |
|---|---|---|---|---|---|---|---|---|---|---|---|---|---|---|---|---|---|---|---|---|
| 3_1056240 | 2 | 1 | 1 | 1 | 1 | 1 | 4 | 0.75 | 8 | 0.875 | 1 | 1 | 2 | 1 | 3 | 1 | 3 | 1 | 9 | 1 |
| 3_1056240 | 2 | 0.5 | 2 | 0 | 2 | 0 | 4 | 0 | 10 | 0.1 | 2 | 0 | 3 | 0 | 4 | 0 | 4 | 0.5 | 13 | 0.1538 |
| 3_1056240 | 3 | 0.3333 | 2 | 0 | 2 | 0.5 | 3 | 1 | 10 | 0.5 | 2 | 0.5 | 3 | 0 | 4 | 0.25 | 4 | 0.25 | 13 | 0.2308 |
| 3_1056241 | 3 | 0.6667 | 1 | 1 | 2 | 1 | 3 | 1 | 9 | 0.8889 | 1 | 1 | 4 | 0.75 | 4 | 0.75 | 4 | 1 | 13 | 0.8462 |
| 3_1056241 | 3 | 0.3333 | 1 | 0 | 1 | 0 | 4 | 0 | 9 | 0.1111 | 2 | 0 | 4 | 0.25 | 3 | 0 | 3 | 0 | 12 | 0.0833 |
| 3_1056241 | 3 | 1 | 2 | 1 | 1 | 1 | 3 | 0.6667 | 9 | 0.8889 | 2 | 1 | 1 | 1 | 4 | 1 | 3 | 1 | 10 | 1 |
| 3_1056241 | 4 | 0.5 | 3 | 1 | 2 | 1 | 4 | 0.5 | 13 | 0.6923 | 2 | 0.5 | 5 | 1 | 4 | 0.25 | 6 | 0.6667 | 17 | 0.6471 |
| 3_1056242 | 5 | 0.4 | 2 | 0 | 3 | 0 | 5 | 0 | 15 | 0.1333 | 3 | 0 | 6 | 0 | 4 | 0.25 | 7 | 0.1429 | 20 | 0.1 |
| 3_1056243 | 5 | 0 | 4 | 0 | 2 | 0.5 | 8 | 0 | 19 | 0.0526 | 4 | 0 | 6 | 0.1667 | 6 | 0 | 10 | 0.1 | 26 | 0.0769 |
| 3_1056245 | 4 | 0 | 4 | 0 | 4 | 0.25 | 7 | 0 | 19 | 0.0526 | 2 | 0 | 3 | 0 | 5 | 0 | 9 | 0.1111 | 19 | 0.0526 |
| 3_1056245 | 5 | 0 | 4 | 0.25 | 4 | 0 | 5 | 0.2 | 18 | 0.1111 | 4 | 0 | 4 | 0 | 6 | 0.1667 | 11 | 0.1818 | 25 | 0.12 |
| 3_1056245 | 3 | 0.3333 | 4 | 0.5 | 4 | 0.75 | 8 | 0.25 | 19 | 0.4211 | 2 | 0.5 | 4 | 0.25 | 5 | 0.6 | 8 | 0.5 | 19 | 0.4737 |
| 3_1056253 | 6 | 0.1667 | 5 | 0 | 7 | 0.1429 | 8 | 0 | 26 | 0.0769 | 5 | 0 | 6 | 0.3333 | 7 | 0.1429 | 11 | 0.0909 | 29 | 0.1379 |
| 3_1056258 | 7 | 0 | 8 | 0.125 | 7 | 0 | 7 | 0.1429 | 29 | 0.069 | 12 | 0.0833 | 6 | 0.1667 | 7 | 0 | 11 | 0.0909 | 36 | 0.0833 |
| 3_1056262 | 8 | 0.25 | 10 | 0.3 | 7 | 0.4286 | 14 | 0.3571 | 39 | 0.3333 | 11 | 0.1818 | 8 | 0.25 | 8 | 0.375 | 8 | 0 | 35 | 0.2 |
| 3_1056262 | 10 | 0.3 | 14 | 0.5 | 8 | 0.5 | 18 | 0.4444 | 50 | 0.44 | 10 | 0.1 | 9 | 0.2222 | 11 | 0.2727 | 12 | 0.4167 | 42 | 0.2619 |
| 3_1056263 | 10 | 0.4 | 11 | 0 | 7 | 0.4286 | 13 | 0.4615 | 41 | 0.3171 | 10 | 0.2 | 7 | 0.5714 | 12 | 0.3333 | 12 | 0.25 | 41 | 0.3171 |
| 3_1056263 | 10 | 1 | 13 | 1 | 7 | 1 | 15 | 1 | 45 | 1 | 10 | 1 | 7 | 1 | 8 | 1 | 8 | 1 | 33 | 1 |
| 3_1056264 | 11 | 0.9091 | 13 | 0.9231 | 6 | 0.8333 | 16 | 0.8125 | 46 | 0.8696 | 10 | 0.8 | 9 | 0.8889 | 11 | 0.7273 | 13 | 0.9231 | 43 | 0.8372 |
| 3_1056264 | 11 | 0.2727 | 13 | 0.3846 | 8 | 0.375 | 18 | 0.2222 | 50 | 0.3 | 10 | 0.3 | 9 | 0.1111 | 12 | 0.3333 | 13 | 0.0769 | 44 | 0.2045 |
| 3_1056265 | 9 | 0.2222 | 12 | 0.3333 | 6 | 0.3333 | 18 | 0.3333 | 45 | 0.3111 | 10 | 0.3 | 6 | 0.3333 | 10 | 0.4 | 10 | 0.1 | 36 | 0.2778 |
| 3_1056265 | 3 | 0.6667 | 5 | 0.2 | 3 | 0.6667 | 9 | 0.7778 | 20 | 0.6 | 4 | 1 | 3 | 0 | 5 | 0.4 | 3 | 0 | 15 | 0.4 |
| 3_1056266 | 4 | 0.25 | 1 | 1 | 3 | 1 | 7 | 0.4286 | 15 | 0.5333 | 4 | 0 | 2 | 1 | 5 | 0.2 | 4 | 0.25 | 15 | 0.2667 |
| 3_1056269 | 4 | 0 | 3 | 0.3333 | 2 | 0 | 9 | 0.1111 | 18 | 0.1111 | 1 | 0 | 1 | 0 | 6 | 0.1667 | 3 | 0 | 11 | 0.0909 |
| 3_1056278 | NA | NA | 2 | 1 | NA | NA | 1 | 0 | 3 | 0.6667 | NA | NA | NA | NA | 1 | 1 | 1 | 0 | 2 | 0.5 |
| 3_1056281 | NA | NA | 1 | 0 | NA | NA | 1 | 1 | 2 | 0.5 | 2 | 0 | 1 | 0 | 1 | 1 | 1 | 0 | 5 | 0.2 |
| 3_1058502 | 58 | 0.1034 | 68 | 0.0882 | 62 | 0.0645 | 69 | 0.087 | 257 | 0.0856 | 56 | 0.1429 | 60 | 0.0667 | 77 | 0.0909 | 62 | 0.0806 | 255 | 0.0941 |
| 3_1058745 | 2 | 0 | 9 | 0.2222 | 6 | 0.5 | 8 | 0.25 | 25 | 0.28 | 5 | 0.2 | 9 | 0.2222 | 2 | 0.5 | 7 | 0.4286 | 23 | 0.3043 |
| 3_1058746 | 2 | 0.5 | 7 | 0.2857 | 5 | 0.2 | 8 | 0.25 | 22 | 0.2727 | 6 | 0.1667 | 10 | 0.3 | 2 | 0.5 | 6 | 0.1667 | 24 | 0.25 |
| 3_1058748 | 3 | 0 | 9 | 0.1111 | 6 | 0.1667 | 9 | 0.2222 | 27 | 0.1481 | 7 | 0 | 9 | 0.2222 | 2 | 0.5 | 7 | 0 | 25 | 0.12 |
| 3_1058752 | 3 | 0 | 9 | 0.1111 | 6 | 0 | 10 | 0.2 | 28 | 0.1071 | 7 | 0 | 10 | 0.3 | 4 | 0 | 8 | 0.125 | 29 | 0.1379 |
| 3_1058754 | 3 | 0.3333 | 9 | 0.6667 | 6 | 0.5 | 9 | 0.5556 | 27 | 0.5556 | 7 | 0.2857 | 8 | 0.375 | 1 | 0 | 9 | 0.2222 | 25 | 0.28 |
| 3_1058754 | 3 | 0 | 8 | 0.25 | 6 | 0.6667 | 12 | 0.4167 | 29 | 0.3793 | 8 | 0.25 | 10 | 0.2 | 4 | 0.25 | 10 | 0.3 | 32 | 0.25 |
| 3_1058756 | 1 | 1 | 1 | 0 | 4 | 0.5 | 6 | 0 | 12 | 0.25 | 4 | 0.25 | 4 | 0.25 | 2 | 0.5 | 4 | 0 | 14 | 0.2143 |
| 3_1058766 | 1 | 0 | 1 | 1 | 5 | 0.4 | 6 | 0.1667 | 13 | 0.3077 | 6 | 0.1667 | 3 | 0.3333 | 4 | 0.5 | 8 | 0.125 | 21 | 0.2381 |
| 3_1058766 | 1 | 0 | 1 | 0 | 6 | 0.3333 | 7 | 0.4286 | 15 | 0.3333 | 6 | 0.3333 | 4 | 0.75 | 4 | 0.75 | 8 | 0.25 | 22 | 0.4545 |
| 3_1058936 | NA | NA | NA | NA | 1 | 1 | NA | NA | 1 | 1 | NA | NA | 2 | 0.5 | 1 | 0 | NA | NA | 3 | 0.3333 |
| 3_1058942 | NA | NA | NA | NA | 1 | 1 | NA | NA | 1 | 1 | NA | NA | 2 | 0.5 | 1 | 0 | 1 | 0 | 4 | 0.25 |
| 3_1058943 | NA | NA | NA | NA | 1 | 1 | NA | NA | 1 | 1 | NA | NA | 2 | 0.5 | 1 | 0 | 1 | 0 | 4 | 0.25 |
| 3_1064108 | 10 | 0.6 | 7 | 1 | 5 | 0.4 | 9 | 0.5556 | 31 | 0.6452 | 6 | 0.6667 | 5 | 0.6 | 11 | 0.7273 | 5 | 0.6 | 27 | 0.6667 |
| 3_1064465 | 8 | 0.375 | 4 | 0 | 2 | 0.5 | 8 | 0.75 | 22 | 0.4545 | 4 | 0.75 | 4 | 0.75 | 3 | 0 | 4 | 0.5 | 15 | 0.5333 |
| 3_1064809 | 7 | 0 | 1 | 0 | 3 | 0.3333 | 5 | 0 | 16 | 0.0625 | 3 | 0 | 3 | 0 | 8 | 0.125 | 3 | 0 | 17 | 0.0588 |
| 3_1064809 | 7 | 0 | 2 | 0 | 2 | 0 | 5 | 0.2 | 16 | 0.0625 | 3 | 0 | 4 | 0 | 6 | 0.1667 | 3 | 0 | 16 | 0.0625 |
| 3_1064810 | 5 | 1 | 2 | 1 | 3 | 0.6667 | 4 | 0.5 | 14 | 0.7857 | 2 | 0.5 | 4 | 1 | 9 | 0.7778 | 4 | 1 | 19 | 0.8421 |
| 3_1064811 | 7 | 0 | 2 | 0.5 | 4 | 0.25 | 5 | 0.2 | 18 | 0.1667 | 3 | 0 | 5 | 0.4 | 9 | 0.2222 | 4 | 0 | 21 | 0.1905 |
| 3_1064812 | 7 | 0.1429 | 1 | 0 | 3 | 0 | 4 | 0 | 15 | 0.0667 | 3 | 0 | 4 | 0 | 9 | 0.2222 | 4 | 0 | 20 | 0.1 |
| 3_1064813 | 3 | 0 | NA | NA | 1 | 0 | 3 | 0.3333 | 7 | 0.1429 | 3 | 0.3333 | 5 | 0.6 | 5 | 0.2 | 2 | 0.5 | 15 | 0.4 |
| 3_1066403 | NA | NA | 2 | 0.5 | 1 | 1 | 1 | 1 | 4 | 0.75 | NA | NA | NA | NA | 1 | 1 | NA | NA | 1 | 1 |
| 3_1066428 | 2 | 0.5 | 3 | 1 | 2 | 0.5 | 2 | 1 | 9 | 0.7778 | 1 | 1 | NA | NA | 3 | 0 | 3 | 1 | 7 | 0.5714 |
| 3_1066428 | 2 | 0 | 2 | 0 | 3 | 0.6667 | 2 | 0 | 9 | 0.2222 | 1 | 1 | NA | NA | 2 | 0 | 4 | 0.25 | 7 | 0.2857 |
| 3_1066429 | 2 | 0.5 | 3 | 0 | 3 | 1 | 2 | 0.5 | 10 | 0.5 | 1 | 1 | NA | NA | 3 | 0.3333 | 3 | 0.3333 | 7 | 0.4286 |
| 3_1066430 | 2 | 1 | 3 | 0.3333 | 1 | 1 | 1 | 1 | 7 | 0.7143 | 1 | 1 | NA | NA | 2 | 1 | 1 | 1 | 4 | 1 |
| 3_1066431 | 2 | 0 | 3 | 0 | 3 | 0.3333 | 2 | 0 | 10 | 0.1 | NA | NA | NA | NA | 4 | 0.5 | 3 | 0 | 7 | 0.2857 |
| 3_1066431 | 3 | 0.6667 | 3 | 1 | 3 | 1 | 2 | 1 | 11 | 0.9091 | 1 | 1 | NA | NA | 4 | 0.75 | 5 | 1 | 10 | 0.9 |
| 3_1066433 | 3 | 0 | 3 | 0 | 3 | 0.3333 | 2 | 1 | 11 | 0.2727 | 1 | 0 | NA | NA | 4 | 0.5 | 4 | 0.75 | 9 | 0.5556 |
| 3_1066435 | 3 | 0.6667 | 3 | 1 | 5 | 0.6 | 2 | 1 | 13 | 0.7692 | NA | NA | NA | NA | 3 | 0.6667 | 4 | 0.5 | 7 | 0.5714 |
| 3_1066435 | 3 | 0 | 3 | 0.6667 | 5 | 0 | 2 | 0 | 13 | 0.1538 | 1 | 1 | NA | NA | 3 | 0.3333 | 5 | 0.4 | 9 | 0.4444 |
| 3_1066679 | 1 | 0 | 2 | 0.5 | NA | NA | 2 | 1 | 5 | 0.6 | 1 | 1 | 1 | 0 | 2 | 0.5 | 1 | 1 | 5 | 0.6 |
| 3_1069443 | NA | NA | 1 | 1 | 1 | 1 | NA | NA | 2 | 1 | NA | NA | 1 | 1 | NA | NA | 2 | 1 | 3 | 1 |
| 3_1069444 | NA | NA | 1 | 1 | 1 | 0 | NA | NA | 2 | 0.5 | NA | NA | 1 | 1 | NA | NA | 2 | 1 | 3 | 1 |
| 3_1069445 | NA | NA | NA | NA | 1 | 1 | NA | NA | 1 | 1 | NA | NA | 1 | 1 | NA | NA | 2 | 1 | 3 | 1 |
| 3_1069445 | NA | NA | 1 | 1 | 1 | 1 | NA | NA | 2 | 1 | NA | NA | 1 | 1 | NA | NA | 2 | 1 | 3 | 1 |
| 3_1069447 | NA | NA | 1 | 1 | 1 | 1 | NA | NA | 2 | 1 | NA | NA | 1 | 1 | NA | NA | 2 | 1 | 3 | 1 |
| 3_1069449 | NA | NA | 1 | 0 | 1 | 1 | NA | NA | 2 | 0.5 | NA | NA | NA | NA | NA | NA | 2 | 0.5 | 2 | 0.5 |
| 3_1069452 | NA | NA | 1 | 1 | NA | NA | NA | NA | 1 | 1 | NA | NA | 1 | 1 | NA | NA | NA | NA | 1 | 1 |

| | | | | | | | | | | | | | | | | | | | | |
|---|---|---|---|---|---|---|---|---|---|---|---|---|---|---|---|---|---|---|---|---|
| 3_1069453( | NA | NA | 1 | 1 | 1 | 0 | NA | NA | 2 | 0.5 | NA | NA | NA | NA | NA | NA | 2 | 0.5 | 2 | 0.5 |
| 3_1069456‹ | NA | NA | 1 | 0 | 1 | 1 | NA | NA | 2 | 0.5 | NA | NA | 1 | 0 | NA | NA | 2 | 0.5 | 3 | 0.3333 |
| 3_1069457: | NA | NA | 1 | 1 | 1 | 1 | NA | NA | 2 | 1 | NA | NA | 1 | 1 | NA | NA | 2 | 1 | 3 | 1 |
| 3_1069457‹ | NA | NA | 1 | 1 | 1 | 1 | NA | NA | 2 | 1 | NA | NA | 1 | 1 | NA | NA | 1 | 1 | 2 | 1 |
| 3_1069458 | NA | NA | 1 | 1 | 1 | 1 | NA | NA | 2 | 1 | NA | NA | 1 | 1 | NA | NA | 2 | 1 | 3 | 1 |
| 3_1069461( | NA | NA | 2 | 1 | 2 | 1 | NA | NA | 4 | 1 | NA | NA | 1 | 1 | NA | NA | 2 | 0.5 | 3 | 0.6667 |
| 3_1069463: | NA | NA | 2 | 0.5 | NA | NA | NA | NA | 2 | 0.5 | NA | NA | 1 | 1 | NA | NA | 2 | 0 | 3 | 0.3333 |
| 3_1069463( | NA | NA | 2 | 1 | 2 | 1 | NA | NA | 4 | 1 | NA | NA | 1 | 1 | NA | NA | 2 | 1 | 3 | 1 |
| 3_1069464: | NA | NA | 2 | 1 | NA | NA | NA | NA | 2 | 1 | NA | NA | 1 | 1 | NA | NA | 2 | 0.5 | 3 | 0.6667 |
| 3_1069467 | NA | NA | 2 | 1 | 2 | 1 | NA | NA | 4 | 1 | NA | NA | 1 | 1 | 1 | 0 | 2 | 1 | 4 | 0.75 |
| 3_1069563: | NA | NA | 1 | 1 | 2 | 1 | NA | NA | 3 | 1 | NA | NA | 2 | 0.5 | NA | NA | 2 | 0 | 4 | 0.25 |
| 3_1069563( | NA | NA | 2 | 0 | 2 | 0.5 | NA | NA | 4 | 0.25 | NA | NA | 2 | 0 | 1 | 0 | 2 | 1 | 5 | 0.4 |
| 3_1069590: | 1 | 0 | 3 | 0 | 2 | 0.5 | 3 | 0.3333 | 9 | 0.2222 | 1 | 0 | 1 | 1 | 1 | 0 | 1 | 0 | 4 | 0.25 |
| 3_1069601 | 3 | 0 | 2 | 0 | 3 | 0.3333 | 3 | 0 | 11 | 0.0909 | 3 | 0 | 2 | 0 | 1 | 0 | 2 | 0.5 | 8 | 0.125 |
| 3_1069605 | 4 | 0 | 1 | 0 | 1 | 0 | 3 | 0.6667 | 9 | 0.2222 | 3 | 0.3333 | 1 | 0 | 4 | 0 | 2 | 0 | 10 | 0.1 |
| 3_1070115 | 1 | 0 | NA | NA | NA | NA | 1 | 1 | 2 | 0.5 | 2 | 0.5 | NA | NA | NA | NA | NA | NA | 2 | 0.5 |
| 3_1070118 | 1 | 1 | NA | NA | NA | NA | 1 | 1 | 2 | 1 | 2 | 1 | NA | NA | NA | NA | NA | NA | 2 | 1 |
| 3_1070120 | 1 | 1 | NA | NA | NA | NA | 1 | 1 | 2 | 1 | 1 | 1 | NA | NA | NA | NA | NA | NA | 1 | 1 |
| 3_1070120 | 1 | 1 | NA | NA | NA | NA | 1 | 1 | 2 | 1 | 1 | 1 | NA | NA | NA | NA | NA | NA | 1 | 1 |
| 3_1070121 | 1 | 1 | NA | NA | NA | NA | 1 | 1 | 2 | 1 | 2 | 0.5 | NA | NA | NA | NA | NA | NA | 2 | 0.5 |
| 3_1070128: | 1 | 0 | NA | NA | NA | NA | 1 | 1 | 2 | 0.5 | 1 | 1 | 1 | 0 | 1 | 1 | NA | NA | 3 | 0.6667 |
| 3_1070129 | 1 | 0 | NA | NA | NA | NA | 1 | 1 | 2 | 0.5 | 2 | 0.5 | 1 | 0 | 1 | 1 | NA | NA | 4 | 0.5 |
| 3_1070210 | 12 | 0.25 | 9 | 0.3333 | 17 | 0.1176 | 8 | 0.125 | 46 | 0.1957 | 12 | 0.25 | 9 | 0.5556 | 5 | 0.2 | 10 | 0 | 36 | 0.25 |
| 3_1070210 | 13 | 0.2308 | 11 | 0.2727 | 16 | 0.0625 | 10 | 0.2 | 50 | 0.18 | 13 | 0 | 11 | 0.4545 | 9 | 0.1111 | 10 | 0.1 | 43 | 0.1628 |
| 3_1070216 | 17 | 0.0588 | 16 | 0 | 19 | 0.0526 | 12 | 0.1667 | 64 | 0.0625 | 14 | 0 | 10 | 0.1 | 10 | 0 | 15 | 0.1333 | 49 | 0.0612 |
| 3_1070218 | 17 | 0.2353 | 17 | 0.2941 | 22 | 0.1364 | 10 | 0.3 | 66 | 0.2273 | 14 | 0.0714 | 11 | 0.2727 | 7 | 0.2857 | 15 | 0.3333 | 47 | 0.234 |
| 3_1070218 | 14 | 0 | 19 | 0.1053 | 22 | 0.0455 | 10 | 0.1 | 65 | 0.0615 | 14 | 0 | 9 | 0.1111 | 10 | 0.1 | 16 | 0.125 | 49 | 0.0816 |
| 3_1070219 | 16 | 0.1875 | 18 | 0.2778 | 18 | 0.2778 | 7 | 0.1429 | 59 | 0.2373 | 12 | 0.4167 | 10 | 0.4 | 10 | 0.4 | 15 | 0.2667 | 47 | 0.3617 |
| 3_1070219 | 17 | 0.5294 | 18 | 0.6111 | 20 | 0.6 | 9 | 0.4444 | 64 | 0.5625 | 13 | 0.4615 | 9 | 0.7778 | 9 | 0.6667 | 14 | 0.4286 | 45 | 0.5556 |
| 3_1070219 | 15 | 0.0667 | 14 | 0.3571 | 22 | 0.2273 | 11 | 0.1818 | 62 | 0.2097 | 12 | 0.0833 | 10 | 0 | 9 | 0.1111 | 14 | 0.1429 | 45 | 0.0889 |
| 3_1070220 | 17 | 0.3529 | 21 | 0.381 | 21 | 0.5714 | 14 | 0.2857 | 73 | 0.411 | 14 | 0.2857 | 11 | 0.3636 | 11 | 0.2727 | 17 | 0.4706 | 53 | 0.3585 |
| 3_1070221 | 13 | 0.3077 | 15 | 0.3333 | 18 | 0.2778 | 11 | 0.1818 | 57 | 0.2807 | 8 | 0.25 | 8 | 0 | 9 | 0.4444 | 11 | 0.3636 | 36 | 0.2778 |
| 3_1070221 | 18 | 0.1111 | 22 | 0.1818 | 23 | 0.087 | 13 | 0.2308 | 76 | 0.1447 | 15 | 0.0667 | 11 | 0.0909 | 12 | 0 | 17 | 0.1765 | 55 | 0.0909 |
| 3_1072331: | NA | NA | 1 | 1 | NA | NA | NA | NA | 1 | 1 | NA | NA | NA | NA | 2 | 1 | NA | NA | 2 | 1 |
| 3_1072333 | 1 | 1 | 3 | 0 | 4 | 0 | NA | NA | 8 | 0.125 | 2 | 0.5 | 2 | 0.5 | 3 | 0 | 3 | 0 | 10 | 0.2 |
| 3_1072334 | 2 | 0 | 3 | 0.6667 | 4 | 0.25 | 1 | 0 | 10 | 0.3 | 4 | 0.25 | 2 | 0 | 5 | 0.2 | 3 | 0 | 14 | 0.1429 |
| 3_1072335 | 1 | 1 | 2 | 1 | 4 | 1 | 1 | 1 | 8 | 1 | 4 | 1 | 2 | 1 | 3 | 1 | 3 | 1 | 12 | 1 |
| 3_1072337: | 1 | 1 | 1 | 0 | 4 | 0.5 | 1 | 1 | 7 | 0.5714 | 3 | 0.6667 | 2 | 1 | 2 | 0.5 | 2 | 1 | 9 | 0.7778 |
| 3_1072362 | 1 | 1 | 1 | 1 | 1 | 0 | 1 | 1 | 4 | 0.75 | 2 | 0.5 | 2 | 0 | NA | NA | 2 | 0.5 | 6 | 0.3333 |
| 3_1072376 | 1 | 0 | 1 | 1 | NA | NA | NA | NA | 2 | 0.5 | 2 | 0.5 | 1 | 0 | 1 | 0 | 2 | 0 | 6 | 0.1667 |
| 3_1072376 | 1 | 1 | 2 | 1 | NA | NA | 1 | 1 | 4 | 1 | 2 | 0.5 | 1 | 1 | 1 | 1 | 2 | 1 | 6 | 0.8333 |
| 3_1072377: | 1 | 1 | 1 | 1 | NA | NA | 1 | 1 | 3 | 1 | 1 | 1 | 1 | 1 | 1 | 1 | 2 | 1 | 5 | 1 |
| 3_1072377 | 1 | 1 | 2 | 1 | NA | NA | NA | NA | 3 | 1 | 1 | 1 | 1 | 1 | 1 | 1 | 2 | 1 | 5 | 1 |
| 3_1072378 | 1 | 1 | 2 | 0.5 | NA | NA | 1 | 0 | 4 | 0.5 | 1 | 1 | 1 | 1 | 1 | 1 | 2 | 0.5 | 5 | 0.8 |
| 3_1072380( | NA | NA | 2 | 0.5 | NA | NA | NA | NA | 2 | 0.5 | 2 | 0.5 | 1 | 1 | 1 | 0 | 2 | 0.5 | 6 | 0.5 |
| 3_1072380‹ | 1 | 1 | 1 | 1 | NA | NA | 1 | 0 | 3 | 0.6667 | NA | NA | 1 | 0 | 1 | 0 | 2 | 0.5 | 4 | 0.25 |
| 3_1072380 | 1 | 1 | 2 | 1 | NA | NA | NA | NA | 3 | 1 | NA | NA | NA | NA | 1 | 1 | 1 | 1 | 2 | 1 |
| 3_1072382: | 1 | 0 | 1 | 1 | NA | NA | 1 | 0 | 3 | 0.3333 | 1 | 0 | 1 | 0 | 1 | 0 | 2 | 0.5 | 5 | 0.2 |
| 3_1072385( | 1 | 1 | 2 | 1 | NA | NA | 1 | 0 | 4 | 0.75 | 2 | 0 | 1 | 1 | 1 | 1 | 2 | 0.5 | 6 | 0.5 |
| 3_1072385: | 1 | 1 | 2 | 0.5 | NA | NA | 1 | 0 | 4 | 0.5 | 1 | 1 | 1 | 0 | NA | NA | 2 | 0.5 | 4 | 0.5 |
| 3_1072386( | 1 | 1 | 2 | 0.5 | NA | NA | 1 | 0 | 4 | 0.5 | 2 | 0 | 1 | 0 | NA | NA | 2 | 0.5 | 5 | 0.2 |
| 3_1072386 | 1 | 1 | 2 | 0 | NA | NA | 1 | 0 | 4 | 0.25 | 2 | 0 | 1 | 0 | NA | NA | 2 | 0.5 | 5 | 0.2 |
| 3_1072387( | 1 | 0 | 1 | 1 | NA | NA | 1 | 0 | 3 | 0.3333 | 1 | 1 | 1 | 0 | NA | NA | 1 | 0 | 3 | 0.3333 |
| 3_1072387: | 1 | 1 | 2 | 0.5 | NA | NA | NA | NA | 3 | 0.6667 | 1 | 1 | 1 | 1 | NA | NA | 1 | 1 | 3 | 1 |
| 3_1072387 | 1 | 1 | 2 | 0.5 | NA | NA | 1 | 0 | 4 | 0.5 | 2 | 1 | 1 | 1 | NA | NA | 2 | 0.5 | 5 | 0.8 |
| 3_1072388 | 1 | 0 | 2 | 0.5 | NA | NA | 1 | 0 | 4 | 0.25 | 2 | 1 | 1 | 0 | NA | NA | 2 | 0 | 5 | 0.4 |
| 3_1072389: | 1 | 1 | 1 | 0 | NA | NA | 1 | 0 | 3 | 0.3333 | NA | NA | 1 | 0 | NA | NA | 2 | 0.5 | 3 | 0.3333 |
| 3_1072390 | 1 | 1 | 1 | 0 | NA | NA | 1 | 0 | 3 | 0.3333 | 1 | 1 | 1 | 1 | NA | NA | 2 | 0.5 | 4 | 0.75 |
| 3_1072392 | 1 | 1 | 1 | 1 | NA | NA | 1 | 1 | 3 | 1 | 2 | 1 | 1 | 0 | NA | NA | 2 | 0.5 | 5 | 0.6 |
| 3_1072394( | 1 | 1 | NA | NA | NA | NA | 1 | 0 | 2 | 0.5 | 2 | 0.5 | 1 | 0 | NA | NA | 2 | 0 | 5 | 0.2 |
| 3_1072394( | NA | NA | 1 | 1 | NA | NA | 1 | 1 | 2 | 1 | 1 | 1 | 1 | 1 | NA | NA | 2 | 1 | 4 | 1 |
| 3_1073464( | 8 | 0.125 | 3 | 0 | 3 | 0 | 2 | 0 | 16 | 0.0625 | 8 | 0.125 | 2 | 0 | 3 | 0.3333 | 2 | 0 | 15 | 0.1333 |
| 3_1073465‹ | 8 | 0.25 | 5 | 0.6 | 3 | 0.3333 | 3 | 0.6667 | 19 | 0.4211 | 7 | 0.7143 | 1 | 0 | 3 | 1 | 2 | 1 | 13 | 0.7692 |
| 3_1073465 | 8 | 1 | 5 | 0.8 | 1 | 1 | 1 | 1 | 15 | 0.9333 | 6 | 1 | 2 | 0.5 | 3 | 1 | 2 | 0.5 | 13 | 0.8462 |
| 3_1073466: | 8 | 0 | 6 | 0.1667 | 3 | 0 | 3 | 0 | 20 | 0.05 | 7 | 0 | 2 | 0.5 | 3 | 0 | 2 | 0 | 14 | 0.0714 |

| | | | | | | | | | | | | | | | | | | | | |
|---|---|---|---|---|---|---|---|---|---|---|---|---|---|---|---|---|---|---|---|---|
| 3_1073466 | 7 | 0.4286 | 5 | 0.2 | 2 | 0 | 2 | 0 | 16 | 0.25 | 7 | 0.1429 | 2 | 0 | 3 | 0 | 2 | 0.5 | 14 | 0.1429 |
| 3_1073528 | 8 | 0 | 4 | 0 | 2 | 0.5 | 3 | 0 | 17 | 0.0588 | 5 | 0 | 1 | 0 | 1 | 1 | NA | NA | 7 | 0.1429 |
| 3_1073550 | 5 | 0.6 | 4 | 0.75 | NA | NA | 2 | 1 | 11 | 0.7273 | 5 | 0.8 | 1 | 1 | 1 | 0 | 1 | 1 | 8 | 0.75 |
| 3_1073551 | 5 | 0.8 | 4 | 1 | 2 | 1 | 3 | 1 | 14 | 0.9286 | 5 | 1 | 1 | 1 | 1 | 1 | NA | NA | 7 | 1 |
| 3_1073551 | 4 | 0.25 | 5 | 0 | 1 | 0 | 3 | 0.3333 | 13 | 0.1538 | 4 | 0.25 | 1 | 0 | 1 | 0 | 1 | 0 | 7 | 0.1429 |
| 3_1073555 | 4 | 0.75 | 5 | 0.8 | 1 | 1 | 2 | 0.5 | 12 | 0.75 | 5 | 0.8 | NA | NA | 1 | 1 | 1 | 1 | 7 | 0.8571 |
| 3_1073555 | 3 | 1 | 5 | 1 | 1 | 1 | 2 | 1 | 11 | 1 | 5 | 1 | NA | NA | 1 | 1 | 1 | 1 | 7 | 1 |
| 3_1073657 | 1 | 0 | 4 | 0.5 | NA | NA | NA | NA | 5 | 0.4 | 4 | 0.75 | NA | NA | 1 | 0 | 1 | 0 | 6 | 0.5 |
| 3_1073666 | 2 | 1 | 3 | 1 | NA | NA | NA | NA | 5 | 1 | 3 | 1 | NA | NA | NA | NA | NA | NA | 3 | 1 |
| 3_1073666 | 1 | 1 | 3 | 0.6667 | NA | NA | NA | NA | 4 | 0.75 | 3 | 0.6667 | NA | NA | NA | NA | NA | NA | 3 | 0.6667 |
| 3_1073667 | 1 | 1 | 2 | 0.5 | NA | NA | NA | NA | 3 | 0.6667 | 2 | 1 | NA | NA | NA | NA | NA | NA | 2 | 1 |
| 3_1073671 | NA | NA | 2 | 0.5 | NA | NA | NA | NA | 2 | 0.5 | 2 | 0.5 | NA | NA | NA | NA | NA | NA | 2 | 0.5 |
| 3_1073751 | 26 | 0.4231 | 32 | 0.3125 | 31 | 0.2903 | 24 | 0.2917 | 113 | 0.3274 | 31 | 0.3548 | 19 | 0.2632 | 26 | 0.2308 | 23 | 0.5217 | 99 | 0.3434 |
| 3_1073764 | 18 | 0.0556 | 25 | 0.12 | 21 | 0.1905 | 19 | 0.1053 | 83 | 0.1205 | 15 | 0.2667 | 17 | 0.1765 | 14 | 0.4286 | 15 | 0.0667 | 61 | 0.2295 |
| 3_1073771 | 15 | 0.1333 | 18 | 0.2778 | 15 | 0.1333 | 13 | 0.0769 | 61 | 0.1639 | 12 | 0.5833 | 19 | 0.2632 | 13 | 0.1538 | 10 | 0.6 | 54 | 0.3704 |
| 3_1073780 | 10 | 0.4 | 16 | 0.625 | 6 | 0.3333 | 9 | 0.3333 | 41 | 0.4634 | 10 | 0.6 | 9 | 0.4444 | 8 | 0.625 | 10 | 0.5 | 37 | 0.5405 |
| 3_1073782 | 10 | 0.9 | 10 | 0.8 | 5 | 0.2 | 9 | 0.8889 | 34 | 0.7647 | 10 | 0.5 | 9 | 0.8889 | 8 | 0.875 | 6 | 0.6667 | 33 | 0.7273 |
| 3_1073782 | 10 | 0.3 | 15 | 0.4667 | 5 | 0.2 | 9 | 0.7778 | 39 | 0.4615 | 9 | 0.5556 | 10 | 0.4 | 10 | 0.2 | 7 | 0.2857 | 36 | 0.3611 |
| 3_1073782 | 6 | 0.8333 | 9 | 0.8889 | 5 | 1 | 8 | 1 | 28 | 0.9286 | 7 | 0.8571 | 7 | 1 | 8 | 0.875 | 8 | 0.875 | 30 | 0.9 |
| 3_1073782 | 11 | 0.0909 | 13 | 0.0769 | 6 | 0 | 7 | 0.2857 | 37 | 0.1081 | 9 | 0 | 9 | 0.3333 | 8 | 0.125 | 7 | 0.2857 | 33 | 0.1818 |
| 3_1073783 | 11 | 0.2727 | 13 | 0 | 5 | 0.2 | 8 | 0.125 | 37 | 0.1351 | 7 | 0 | 10 | 0 | 9 | 0.2222 | 8 | 0.25 | 34 | 0.1176 |
| 3_1073783 | 7 | 0.4286 | 10 | 0.1 | 4 | 0 | 2 | 0 | 23 | 0.1739 | 4 | 0 | 5 | 0 | 6 | 0 | 5 | 0.2 | 20 | 0.05 |
| 3_1073783 | 10 | 0.1 | 15 | 0.4 | 6 | 0 | 8 | 0.25 | 39 | 0.2308 | 9 | 0 | 9 | 0.3333 | 10 | 0.1 | 6 | 0.1667 | 34 | 0.1471 |
| 3_1073786 | 6 | 0.6667 | 11 | 0.9091 | 4 | 0.75 | 6 | 0.6667 | 27 | 0.7778 | 6 | 0.8333 | 6 | 0.6667 | 7 | 0.7143 | 7 | 1 | 26 | 0.8077 |
| 3_1073841 | 4 | 0 | 11 | 0.0909 | 3 | 0.3333 | 2 | 0 | 20 | 0.1 | 5 | 0.2 | 4 | 0 | 6 | 0.1667 | 6 | 0.1667 | 21 | 0.1429 |
| 3_1073846 | 2 | 0 | 10 | 0 | 2 | 0 | 3 | 0.3333 | 17 | 0.0588 | 6 | 0 | 4 | 0 | 5 | 0 | 5 | 0.2 | 20 | 0.05 |
| 3_1074078 | 1 | 1 | 6 | 0.8333 | 2 | 0 | 2 | 0 | 11 | 0.5455 | 4 | 0 | NA | NA | 3 | 0.3333 | 2 | 1 | 9 | 0.3333 |
| 3_1075199 | 4 | 0 | 8 | 0 | 8 | 0.25 | 1 | 1 | 21 | 0.1429 | 5 | 0.4 | 3 | 0.3333 | 2 | 0 | 5 | 0 | 15 | 0.2 |
| 3_1075200 | 5 | 0.6 | 7 | 0.7143 | 11 | 0.9091 | 3 | 0.6667 | 26 | 0.7692 | 5 | 0.8 | 3 | 0.3333 | 3 | 0.6667 | 4 | 0.75 | 15 | 0.6667 |
| 3_1075200 | 4 | 0.75 | 8 | 1 | 12 | 0.9167 | 2 | 1 | 26 | 0.9231 | 4 | 1 | 3 | 1 | 3 | 1 | 6 | 1 | 16 | 1 |
| 3_1075201 | 4 | 0.25 | 8 | 0 | 11 | 0.4545 | 2 | 0 | 25 | 0.24 | 3 | 0.3333 | 3 | 0 | 3 | 0 | 5 | 0 | 14 | 0.0714 |
| 3_1075201 | 5 | 0.8 | 7 | 0.5714 | 11 | 0.7273 | 1 | 1 | 24 | 0.7083 | 4 | 1 | 2 | 0.5 | 3 | 1 | 6 | 0.8333 | 15 | 0.8667 |
| 3_1075202 | 4 | 0.75 | 7 | 0.5714 | 8 | 0.75 | 3 | 0.3333 | 22 | 0.6364 | 4 | 0.75 | 1 | 0 | 3 | 1 | 6 | 0.3333 | 14 | 0.5714 |
| 3_1075204 | 4 | 1 | 7 | 1 | 10 | 0.9 | 2 | 0.5 | 23 | 0.913 | 4 | 1 | 3 | 1 | 3 | 1 | 5 | 1 | 15 | 1 |
| 3_1075205 | 4 | 0 | 7 | 0.4286 | 11 | 0.1818 | 3 | 0.3333 | 25 | 0.24 | 2 | 0.5 | 3 | 0 | 3 | 1 | 6 | 0.3333 | 14 | 0.4286 |
| 3_1075206 | 3 | 1 | 4 | 1 | 5 | 1 | 3 | 1 | 15 | 1 | NA | NA | NA | NA | 3 | 1 | 2 | 1 | 5 | 1 |
| 3_1075206 | 4 | 0.5 | 4 | 0.75 | 11 | 0.7273 | 3 | 0.3333 | 22 | 0.6364 | 4 | 0.75 | 3 | 0.6667 | 3 | 0.6667 | 6 | 0.6667 | 16 | 0.6875 |
| 3_1075206 | 4 | 0.75 | 5 | 0.8 | 11 | 0.6364 | 1 | 1 | 21 | 0.7143 | 4 | 0.5 | 3 | 0.3333 | 3 | 0.6667 | 5 | 1 | 15 | 0.6667 |
| 3_1075207 | 3 | 0.3333 | 5 | 0.2 | 7 | 0 | 2 | 0 | 17 | 0.1176 | 2 | 0 | 2 | 0 | 1 | 0 | 5 | 0.2 | 10 | 0.1 |
| 3_1075705 | 40 | 0.075 | 54 | 0.0556 | 46 | 0.1957 | 46 | 0.087 | 186 | 0.1022 | 36 | 0.1111 | 36 | 0.0556 | 37 | 0.1351 | 31 | 0.0968 | 140 | 0.1 |
| 3_1075705 | 38 | 0.1579 | 49 | 0.2245 | 46 | 0.1522 | 39 | 0.2308 | 172 | 0.1919 | 32 | 0.2812 | 35 | 0.2 | 32 | 0.1562 | 27 | 0.1481 | 126 | 0.1984 |
| 3_1075712 | 34 | 0.0588 | 44 | 0.1364 | 39 | 0.1026 | 40 | 0.05 | 157 | 0.0892 | 32 | 0.0312 | 32 | 0 | 31 | 0.0323 | 25 | 0.16 | 120 | 0.05 |
| 3_1076487 | 2 | 0.5 | 3 | 0.6667 | 2 | 0.5 | 3 | 0 | 10 | 0.4 | 3 | 0.3333 | NA | NA | 4 | 0.5 | 2 | 0.5 | 9 | 0.4444 |
| 3_1076487 | 2 | 0 | 2 | 0.5 | 1 | 1 | 2 | 0 | 7 | 0.2857 | 2 | 0.5 | NA | NA | 3 | 0.3333 | 2 | 0 | 7 | 0.2857 |
| 3_1076488 | 2 | 0 | 3 | 0.6667 | 2 | 0.5 | 2 | 0 | 9 | 0.3333 | 2 | 0 | NA | NA | 4 | 0.75 | 2 | 0 | 8 | 0.375 |
| 3_1076493 | 2 | 0.5 | 3 | 0 | 2 | 0 | 2 | 0.5 | 9 | 0.2222 | 3 | 0 | NA | NA | 2 | 0.5 | 1 | 0 | 6 | 0.1667 |
| 3_1076493 | 2 | 0.5 | 2 | 0.5 | 2 | 0 | 2 | 0.5 | 8 | 0.375 | 2 | 0 | NA | NA | 2 | 1 | 2 | 0 | 6 | 0.3333 |
| 3_1076495 | 1 | 1 | NA | NA | 1 | 0 | NA | NA | 2 | 0.5 | 1 | 1 | NA | NA | 3 | 1 | 2 | 0.5 | 6 | 0.8333 |
| 3_1076496 | NA | NA | 3 | 0.3333 | 2 | 0 | 1 | 0 | 6 | 0.1667 | 3 | 0.3333 | NA | NA | 3 | 0.3333 | 1 | 0 | 7 | 0.2857 |
| 3_1076498 | 2 | 1 | 2 | 1 | 2 | 0.5 | NA | NA | 6 | 0.8333 | 3 | 0.6667 | NA | NA | 3 | 0.3333 | 1 | 0 | 7 | 0.4286 |
| 3_1076498 | 2 | 0.5 | 2 | 0 | NA | NA | 1 | 0 | 5 | 0.2 | 3 | 0.3333 | NA | NA | 2 | 0 | 1 | 0 | 6 | 0.1667 |
| 3_1076499 | 2 | 0.5 | 3 | 0.6667 | 2 | 0.5 | 2 | 1 | 9 | 0.6667 | 3 | 1 | NA | NA | 3 | 0.6667 | 1 | 0 | 7 | 0.7143 |
| 3_1076500 | 2 | 0 | 3 | 0.3333 | 2 | 0 | 1 | 1 | 8 | 0.25 | 3 | 0.3333 | NA | NA | 3 | 0.3333 | 1 | 0 | 7 | 0.2857 |
| 3_1076500 | 2 | 0 | 3 | 0.3333 | 2 | 1 | 1 | 0 | 8 | 0.375 | 1 | 0 | NA | NA | 2 | 0 | 1 | 1 | 4 | 0.25 |
| 3_1076501 | 2 | 1 | 3 | 0.6667 | 2 | 1 | 2 | 0.5 | 9 | 0.7778 | 2 | 1 | NA | NA | 3 | 1 | 1 | 1 | 6 | 1 |
| 3_1076501 | 2 | 1 | 3 | 1 | 2 | 0.5 | 1 | 1 | 8 | 0.875 | 3 | 1 | NA | NA | 3 | 1 | 1 | 1 | 7 | 1 |
| 3_1076502 | 1 | 1 | 1 | 1 | 2 | 1 | NA | NA | 4 | 1 | 3 | 0.6667 | NA | NA | 2 | 0.5 | 1 | 1 | 6 | 0.6667 |
| 3_1076503 | 1 | 0 | 3 | 0.6667 | 2 | 0 | 1 | 0 | 7 | 0.2857 | 3 | 0.6667 | NA | NA | 2 | 0.5 | NA | NA | 5 | 0.6 |
| 3_1076504 | 1 | 0 | 3 | 0.3333 | 1 | 1 | 1 | 0 | 6 | 0.3333 | 3 | 0 | NA | NA | 3 | 0.3333 | NA | NA | 6 | 0.1667 |
| 3_1076556 | NA | NA | 3 | 0.6667 | 2 | 0 | NA | NA | 5 | 0.4 | 2 | 0 | NA | NA | 2 | 0.5 | NA | NA | 4 | 0.25 |
| 3_1076558 | NA | NA | 3 | 0 | 2 | 0.5 | NA | NA | 5 | 0.2 | 1 | 0 | NA | NA | 2 | 0.5 | NA | NA | 3 | 0.3333 |
| 3_1076559 | NA | NA | 3 | 0.3333 | 2 | 0 | NA | NA | 5 | 0.2 | 1 | 1 | NA | NA | 2 | 0 | NA | NA | 3 | 0.3333 |
| 3_1076560 | NA | NA | 3 | 1 | 2 | 1 | NA | NA | 5 | 1 | 1 | 1 | NA | NA | 1 | 1 | NA | NA | 2 | 1 |
| 3_1076562 | NA | NA | 3 | 0.6667 | 2 | 0.5 | NA | NA | 5 | 0.6 | 1 | 0 | NA | NA | 2 | 1 | NA | NA | 3 | 0.6667 |
| 3_1076562 | NA | NA | 3 | 0.3333 | NA | NA | NA | NA | 3 | 0.3333 | NA | NA | NA | NA | 1 | 1 | NA | NA | 1 | 1 |

| | | | | | | | | | | | | | | | | | | | | |
|---|---|---|---|---|---|---|---|---|---|---|---|---|---|---|---|---|---|---|---|---|
| 3_1076563 | NA | NA | NA | NA | 1 | 1 | NA | NA | 1 | 1 | 1 | 0 | NA | NA | 2 | 0.5 | NA | NA | 3 | 0.3333 |
| 3_1077978 | 8 | 0.375 | 8 | 0.375 | 4 | 0.5 | 6 | 0.3333 | 26 | 0.3846 | 1 | 1 | 2 | 0 | 3 | 0 | 5 | 0.2 | 11 | 0.1818 |
| 3_1077983 | 3 | 0 | 9 | 0.2222 | 4 | 0.75 | 5 | 0.2 | 21 | 0.2857 | 3 | 0 | 2 | 0.5 | 3 | 0 | 4 | 0.5 | 12 | 0.25 |
| 3_1077987 | 3 | 0.3333 | 8 | 0.125 | 4 | 0 | 6 | 0.3333 | 21 | 0.1905 | 2 | 0.5 | 2 | 0 | 2 | 0 | 2 | 0 | 8 | 0.125 |
| 3_1077987 | 3 | 0 | 8 | 0 | 4 | 0 | 5 | 0.2 | 20 | 0.05 | 1 | 0 | 1 | 0 | 2 | 0.5 | 3 | 0 | 7 | 0.1429 |
| 3_1077988 | 5 | 0 | 8 | 0.125 | 4 | 0.25 | 3 | 0 | 20 | 0.1 | 3 | 0.3333 | 1 | 0 | 3 | 0 | 2 | 0 | 9 | 0.1111 |
| 3_1077996 | 4 | 0 | 6 | 0.1667 | 5 | 0.6 | 6 | 0 | 21 | 0.1905 | 3 | 0 | NA | NA | 2 | 0.5 | 4 | 0.25 | 9 | 0.2222 |
| 3_1077999 | 3 | 1 | 6 | 0.6667 | 4 | 0.75 | 6 | 0.5 | 19 | 0.6842 | 3 | 0.6667 | NA | NA | 2 | 1 | 4 | 0.75 | 9 | 0.7778 |
| 3_1078000 | 3 | 0.3333 | 6 | 0 | 5 | 0 | 5 | 0 | 19 | 0.0526 | 3 | 0.3333 | NA | NA | 2 | 0 | 4 | 0.25 | 9 | 0.2222 |
| 3_1078001 | 2 | 0.5 | 6 | 0 | 5 | 0.6 | 6 | 0.1667 | 19 | 0.2632 | 1 | 0 | NA | NA | 2 | 0 | 3 | 0.3333 | 6 | 0.1667 |
| 3_1078005 | 2 | 0 | 5 | 0.2 | 4 | 0.5 | 6 | 0 | 17 | 0.1765 | 2 | 0 | NA | NA | 3 | 0 | 4 | 0.25 | 9 | 0.1111 |
| 3_1078005 | 2 | 0.5 | 6 | 0.6667 | 4 | 0.25 | 5 | 0.4 | 17 | 0.4706 | 1 | 1 | NA | NA | 2 | 1 | 4 | 0.5 | 7 | 0.7143 |
| 3_1078005 | 2 | 1 | 5 | 1 | 3 | 0.6667 | 6 | 1 | 16 | 0.9375 | 2 | 1 | NA | NA | 3 | 0.3333 | 4 | 1 | 9 | 0.7778 |
| 3_1078006 | 2 | 1 | 6 | 0.3333 | 3 | 0.6667 | 5 | 0.4 | 16 | 0.5 | 2 | 0.5 | NA | NA | 2 | 0.5 | 5 | 0.8 | 9 | 0.6667 |
| 3_1078006 | 1 | 0 | 4 | 0.5 | 3 | 0.3333 | 4 | 0.75 | 12 | 0.5 | 2 | 0 | NA | NA | 4 | 0.5 | 3 | 0.6667 | 9 | 0.4444 |
| 3_1078006 | 2 | 0.5 | 6 | 0.6667 | 4 | 0.75 | 6 | 0.6667 | 18 | 0.6667 | 2 | 1 | NA | NA | 4 | 0.5 | 4 | 0.25 | 10 | 0.5 |
| 3_1078007 | 2 | 0.5 | 3 | 0.6667 | 4 | 0.5 | 6 | 0.6667 | 15 | 0.6 | NA | NA | NA | NA | 3 | 0 | 5 | 1 | 8 | 0.625 |
| 3_1078007 | 2 | 0 | 5 | 0.4 | 4 | 0.5 | 5 | 0.2 | 16 | 0.3125 | 2 | 0 | NA | NA | 4 | 0.25 | 5 | 0.2 | 11 | 0.1818 |
| 3_1078008 | 2 | 0 | 2 | 0 | 4 | 0.25 | 3 | 0 | 11 | 0.0909 | 2 | 0 | NA | NA | 2 | 0.5 | 4 | 0 | 8 | 0.125 |
| 3_1078342 | 2 | 0 | 1 | 0 | 2 | 0.5 | 1 | 0 | 6 | 0.1667 | 1 | 0 | 5 | 0.4 | 3 | 0 | 1 | 0 | 10 | 0.2 |
| 3_1079197 | 70 | 0.0714 | 47 | 0.0851 | 46 | 0.0435 | 52 | 0.0385 | 215 | 0.0605 | 49 | 0.0612 | 42 | 0.0476 | 46 | 0.0435 | 35 | 0.0571 | 172 | 0.0523 |
| 3_1079197 | 66 | 0.2273 | 46 | 0.087 | 44 | 0.1136 | 48 | 0.1458 | 204 | 0.152 | 46 | 0.2609 | 37 | 0.1081 | 44 | 0.0909 | 32 | 0.0625 | 159 | 0.1384 |
| 3_1079482 | 1 | 1 | NA | NA | 3 | 0 | NA | NA | 4 | 0.25 | 2 | 0 | 1 | 0 | 1 | 0 | 3 | 0.3333 | 7 | 0.1429 |
| 3_1079815 | 5 | 0.8 | 7 | 0.2857 | 5 | 0.4 | 6 | 0.3333 | 23 | 0.4348 | 5 | 0.8 | 7 | 0.7143 | 10 | 0.2 | 3 | 0 | 25 | 0.44 |
| 3_1079817 | 5 | 0.2 | 7 | 0.4286 | 6 | 0.1667 | 6 | 0.5 | 24 | 0.3333 | 6 | 0.5 | 8 | 0.625 | 12 | 0.5833 | 3 | 0.3333 | 29 | 0.5517 |
| 3_1079821 | 4 | 0.25 | 6 | 0.1667 | 4 | 0 | 6 | 0.3333 | 20 | 0.2 | 5 | 0.2 | 8 | 0.125 | 8 | 0.5 | 2 | 0 | 23 | 0.2609 |
| 3_1080151 | 4 | 0 | 2 | 0 | 3 | 0.3333 | 6 | 0.1667 | 15 | 0.1333 | 2 | 0 | 3 | 0.3333 | 6 | 0.1667 | 6 | 0 | 17 | 0.1176 |
| 3_1080155 | 3 | 0 | 3 | 0 | 1 | 0 | 4 | 0.25 | 11 | 0.0909 | 2 | 0 | 3 | 0 | 5 | 0.2 | 4 | 0 | 14 | 0.0714 |
| 3_1080158 | 3 | 0.3333 | 3 | 0.3333 | 1 | 1 | 4 | 0 | 11 | 0.2727 | 3 | 0 | 3 | 0 | 5 | 0.6 | 5 | 0 | 16 | 0.1875 |
| 3_1080160 | 3 | 0.6667 | 3 | 0.6667 | 2 | 0.5 | 3 | 0.3333 | 11 | 0.5455 | 2 | 0 | 2 | 1 | 5 | 1 | 6 | 0.6667 | 15 | 0.7333 |
| 3_1080161 | 4 | 0.25 | 3 | 0.3333 | 1 | 0 | 5 | 0 | 13 | 0.1538 | 2 | 0 | 1 | 0 | 4 | 0.5 | 5 | 0.4 | 12 | 0.3333 |
| 3_1080162 | 4 | 0.75 | 2 | 0.5 | 1 | 0 | 3 | 0.6667 | 10 | 0.6 | 2 | 0.5 | 1 | 1 | 4 | 0.75 | 4 | 1 | 11 | 0.8182 |
| 3_1080164 | 4 | 0.75 | 2 | 1 | 1 | 0 | 4 | 1 | 11 | 0.8182 | 1 | 1 | 1 | 1 | 4 | 1 | 5 | 1 | 11 | 1 |
| 3_1080164 | 4 | 0.5 | 1 | 0 | 1 | 0 | 2 | 0 | 8 | 0.25 | 2 | 0 | 1 | 0 | 3 | 0 | 5 | 0.4 | 11 | 0.1818 |
| 3_1080164 | 3 | 0 | 2 | 0.5 | 2 | 0.5 | 4 | 0.25 | 11 | 0.2727 | 2 | 0.5 | 1 | 0 | 5 | 0.6 | 4 | 0 | 12 | 0.3333 |
| 3_1080165 | 2 | 0.5 | 2 | 0 | 2 | 0 | 3 | 0 | 9 | 0.1111 | 2 | 0 | 1 | 0 | 4 | 0 | 5 | 0.2 | 12 | 0.0833 |
| 3_1080563 | 4 | 0 | 2 | 0 | 2 | 0 | 2 | 0.5 | 10 | 0.1 | NA | NA | 2 | 0.5 | NA | NA | 1 | 1 | 3 | 0.6667 |
| 3_1080571 | 3 | 1 | 1 | 1 | NA | NA | 1 | 0 | 5 | 0.8 | NA | NA | 3 | 1 | NA | NA | 1 | 1 | 4 | 1 |
| 3_1080571 | 4 | 1 | 1 | 1 | NA | NA | 1 | 1 | 6 | 1 | NA | NA | 3 | 1 | NA | NA | 1 | 1 | 4 | 1 |
| 3_1080571 | 1 | 1 | 1 | 1 | NA | NA | 1 | 1 | 3 | 1 | NA | NA | 3 | 1 | NA | NA | 1 | 0 | 4 | 0.75 |
| 3_1080572 | 4 | 1 | 1 | 1 | NA | NA | 1 | 1 | 6 | 1 | NA | NA | 3 | 1 | NA | NA | 1 | 1 | 4 | 1 |
| 3_1080572 | 3 | 1 | 1 | 1 | NA | NA | 1 | 1 | 5 | 1 | NA | NA | 3 | 1 | NA | NA | 1 | 1 | 4 | 1 |
| 3_1080572 | 3 | 1 | 1 | 1 | NA | NA | 1 | 1 | 5 | 1 | NA | NA | 3 | 1 | NA | NA | 1 | 1 | 4 | 1 |
| 3_1080573 | 3 | 1 | 1 | 1 | NA | NA | 1 | 1 | 5 | 1 | NA | NA | 3 | 0.6667 | NA | NA | 1 | 1 | 4 | 0.75 |
| 3_1080574 | 3 | 1 | NA | NA | NA | NA | NA | NA | 3 | 1 | NA | NA | 2 | 1 | NA | NA | NA | NA | 2 | 1 |
| 3_1080574 | 1 | 1 | 1 | 1 | NA | NA | NA | NA | 2 | 1 | NA | NA | 3 | 1 | NA | NA | NA | NA | 3 | 1 |
| 3_1080575 | 3 | 0.3333 | 1 | 0 | NA | NA | NA | NA | 4 | 0.25 | NA | NA | 2 | 0.5 | NA | NA | NA | NA | 2 | 0.5 |
| 3_1080577 | 3 | 0.6667 | 1 | 1 | NA | NA | NA | NA | 4 | 0.75 | NA | NA | 3 | 0.6667 | NA | NA | NA | NA | 3 | 0.6667 |
| 3_1080577 | 2 | 0.5 | 1 | 1 | NA | NA | NA | NA | 3 | 0.6667 | NA | NA | 3 | 0.6667 | NA | NA | NA | NA | 3 | 0.6667 |
| 3_1080588 | 2 | 1 | NA | NA | NA | NA | NA | NA | 2 | 1 | NA | NA | 1 | 1 | NA | NA | NA | NA | 1 | 1 |
| 3_1080589 | 1 | 1 | NA | NA | NA | NA | NA | NA | 1 | 1 | NA | NA | 1 | 1 | NA | NA | NA | NA | 1 | 1 |
| 3_1080590 | 2 | 0.5 | NA | NA | NA | NA | NA | NA | 2 | 0.5 | NA | NA | 1 | 1 | NA | NA | NA | NA | 1 | 1 |
| 3_1080944 | 1 | 0 | 2 | 0.5 | 1 | 0 | NA | NA | 4 | 0.25 | 2 | 1 | 1 | 0 | NA | NA | 3 | 0.6667 | 6 | 0.6667 |
| 3_1080947 | 2 | 0 | 1 | 0 | 1 | 1 | NA | NA | 4 | 0.25 | 2 | 1 | NA | NA | NA | NA | 2 | 0.5 | 4 | 0.75 |
| 3_1080948 | 2 | 0.5 | 1 | 0 | 1 | 0 | NA | NA | 4 | 0.25 | 2 | 0.5 | 1 | 1 | NA | NA | 2 | 0 | 5 | 0.4 |
| 3_1080949 | 2 | 0 | 2 | 1 | 1 | 1 | 1 | 1 | 6 | 0.6667 | 2 | 1 | NA | NA | NA | NA | 3 | 1 | 5 | 1 |
| 3_1080949 | 3 | 0.6667 | 2 | 0.5 | 1 | 1 | 1 | 1 | 7 | 0.7143 | 2 | 0.5 | 1 | 1 | NA | NA | 3 | 0.6667 | 6 | 0.6667 |
| 3_1080949 | 2 | 0.5 | 2 | 0.5 | 1 | 1 | 1 | 0 | 6 | 0.5 | 1 | 0 | 1 | 0 | NA | NA | 2 | 0.5 | 4 | 0.25 |
| 3_1080950 | 1 | 1 | NA | NA | NA | NA | NA | NA | 1 | 1 | 2 | 1 | NA | NA | NA | NA | 1 | 1 | 3 | 1 |
| 3_1080951 | 3 | 0 | 2 | 0.5 | 1 | 0 | 1 | 0 | 7 | 0.1429 | 3 | 0.3333 | 1 | 0 | NA | NA | 3 | 0 | 7 | 0.1429 |
| 3_1080952 | 4 | 1 | 2 | 1 | 1 | 1 | 1 | 1 | 8 | 1 | 3 | 1 | 1 | 1 | NA | NA | 3 | 1 | 7 | 1 |
| 3_1080952 | 4 | 0.25 | 2 | 1 | 1 | 1 | NA | NA | 7 | 0.5714 | 2 | 0.5 | NA | NA | NA | NA | 3 | 1 | 5 | 0.8 |
| 3_1080954 | 1 | 1 | 1 | 1 | 1 | 1 | 2 | 1 | 5 | 1 | 1 | 1 | NA | NA | NA | NA | 2 | 1 | 3 | 1 |
| 3_1080955 | 4 | 1 | 2 | 1 | 1 | 1 | 2 | 1 | 9 | 1 | 3 | 1 | 1 | 1 | NA | NA | 3 | 0.3333 | 7 | 0.7143 |
| 3_1080955 | 4 | 0.75 | 2 | 0.5 | 1 | 1 | 3 | 0.3333 | 10 | 0.6 | 2 | 1 | 1 | 1 | NA | NA | 3 | 0.3333 | 6 | 0.6667 |

| | | | | | | | | | | | | | | | | | | | | |
|---|---|---|---|---|---|---|---|---|---|---|---|---|---|---|---|---|---|---|---|---|
| 3_1080955 | 4 | 1 | 2 | 1 | 1 | 1 | 2 | 1 | 9 | 1 | 2 | 1 | 1 | 1 | NA | NA | 3 | 1 | 6 | 1 |
| 3_1080957 | 5 | 1 | 3 | 0.3333 | 1 | 1 | 2 | 1 | 11 | 0.8182 | 3 | 0.6667 | 2 | 0.5 | NA | NA | 2 | 0.5 | 7 | 0.5714 |
| 3_1080958 | 4 | 0.25 | 2 | 0 | 2 | 0 | 3 | 0 | 11 | 0.0909 | 3 | 0 | 1 | 0 | NA | NA | 2 | 0.5 | 6 | 0.1667 |
| 3_1080958 | 3 | 0.3333 | 3 | 0.6667 | 2 | 0.5 | 3 | 0.3333 | 11 | 0.4545 | 4 | 0 | 1 | 1 | NA | NA | 3 | 0.3333 | 8 | 0.25 |
| 3_1080958 | 5 | 0.4 | 2 | 0 | 2 | 0.5 | 2 | 0.5 | 11 | 0.3636 | 2 | 0 | NA | NA | NA | NA | 2 | 1 | 4 | 0.5 |
| 3_1080962 | 5 | 0 | 4 | 0.25 | 2 | 0 | 3 | 0 | 14 | 0.0714 | 3 | 0.3333 | 2 | 0 | NA | NA | 3 | 0 | 8 | 0.125 |
| 3_1080964 | 3 | 0.3333 | 1 | 0 | 1 | 1 | 2 | 0 | 7 | 0.2857 | 2 | 0 | 1 | 0 | 1 | 0 | 1 | 1 | 5 | 0.2 |
| 3_1080980 | 4 | 0 | 8 | 0.125 | 3 | 0 | 5 | 0 | 20 | 0.05 | 6 | 0.1667 | 4 | 0 | 2 | 0 | 3 | 0.3333 | 15 | 0.1333 |
| 3_1080984 | 5 | 0.4 | 9 | 0.1111 | 2 | 0 | 3 | 0.6667 | 19 | 0.2632 | 6 | 0.1667 | 6 | 0 | 4 | 0 | 3 | 0 | 19 | 0.0526 |
| 3_1080985 | 5 | 0 | 9 | 0 | 5 | 0 | 7 | 0.2857 | 26 | 0.0769 | 7 | 0 | 5 | 0.2 | 4 | 0 | 4 | 0.25 | 20 | 0.1 |
| 3_1080989 | 3 | 0 | 10 | 0.2 | 11 | 0.0909 | 9 | 0 | 33 | 0.0909 | 9 | 0 | 7 | 0.1429 | 3 | 0 | 5 | 0.2 | 24 | 0.0833 |
| 3_1081057 | NA | NA | 2 | 0.5 | 1 | 0 | NA | NA | 3 | 0.3333 | 2 | 0.5 | 2 | 0 | NA | NA | 2 | 0 | 6 | 0.1667 |
| 3_1081060 | NA | NA | 2 | 0.5 | 1 | 1 | NA | NA | 3 | 0.6667 | 2 | 0 | 2 | 0.5 | NA | NA | 3 | 0 | 7 | 0.1429 |
| 3_1081065 | NA | NA | 1 | 1 | 1 | 0 | NA | NA | 2 | 0.5 | 1 | 0 | 1 | 1 | NA | NA | 2 | 0 | 4 | 0.25 |
| 3_1081066 | NA | NA | 1 | 1 | 1 | 1 | NA | NA | 2 | 1 | 1 | 1 | 2 | 1 | NA | NA | 3 | 1 | 6 | 1 |
| 3_1081067 | NA | NA | 1 | 1 | 1 | 1 | NA | NA | 2 | 1 | 1 | 1 | 2 | 0.5 | NA | NA | 3 | 0.6667 | 6 | 0.6667 |
| 3_1081108 | 1 | 1 | 1 | 1 | 3 | 1 | 1 | 1 | 6 | 1 | NA | NA | 2 | 1 | 2 | 1 | 1 | 1 | 5 | 1 |
| 3_1081108 | NA | NA | 1 | 0 | 1 | 1 | 1 | 1 | 3 | 0.6667 | NA | NA | 1 | 1 | 1 | 1 | 1 | 1 | 3 | 1 |
| 3_1081108 | 1 | 0 | 1 | 0 | 3 | 0.3333 | 1 | 0 | 6 | 0.1667 | NA | NA | 2 | 0 | 2 | 0.5 | 2 | 0 | 6 | 0.1667 |
| 3_1081109 | 1 | 0 | NA | NA | 3 | 0 | 1 | 1 | 5 | 0.2 | NA | NA | 2 | 0.5 | 2 | 0.5 | 2 | 0 | 6 | 0.3333 |
| 3_1081871 | 12 | 0.1667 | 19 | 0.2105 | 17 | 0 | 12 | 0.0833 | 60 | 0.1167 | 7 | 0.1429 | 18 | 0.1667 | 19 | 0.1579 | 13 | 0.3077 | 57 | 0.193 |
| 3_1082646 | 11 | 0.3636 | 12 | 0.4167 | 11 | 0.1818 | 10 | 0 | 44 | 0.25 | 8 | 0.375 | 9 | 0.2222 | 12 | 0.25 | 9 | 0.1111 | 38 | 0.2368 |
| 3_1082648 | 11 | 0.2727 | 14 | 0.3571 | 10 | 0.2 | 10 | 0.2 | 45 | 0.2667 | 7 | 0.2857 | 10 | 0.5 | 11 | 0.5455 | 9 | 0.4444 | 37 | 0.4595 |
| 3_1082648 | 10 | 0.2 | 13 | 0.3077 | 7 | 0.1429 | 11 | 0.0909 | 41 | 0.1951 | 5 | 0.4 | 8 | 0 | 9 | 0.3333 | 7 | 0 | 29 | 0.1724 |
| 3_1082648 | 9 | 0.3333 | 13 | 0.5385 | 8 | 0.625 | 9 | 0.4444 | 39 | 0.4872 | 7 | 0.7143 | 6 | 0.3333 | 10 | 0.3 | 8 | 0.625 | 31 | 0.4839 |
| 3_1082650 | 9 | 0.1111 | 16 | 0.0625 | 9 | 0 | 11 | 0.1818 | 45 | 0.0889 | 7 | 0.1429 | 9 | 0.1111 | 13 | 0.1538 | 11 | 0 | 40 | 0.1 |
| 3_1082650 | 10 | 0.5 | 16 | 0.1875 | 10 | 0.2 | 9 | 0.1111 | 45 | 0.2444 | 9 | 0.2222 | 9 | 0.2222 | 12 | 0.1667 | 9 | 0 | 39 | 0.1538 |
| 3_1082651 | 11 | 0.0909 | 17 | 0.1765 | 9 | 0.1111 | 14 | 0.3571 | 51 | 0.1961 | 10 | 0.3 | 10 | 0.3 | 13 | 0.1538 | 9 | 0.2222 | 42 | 0.2381 |
| 3_1082654 | 11 | 0.0909 | 18 | 0.0556 | 9 | 0.2222 | 10 | 0 | 48 | 0.0833 | 10 | 0.1 | 7 | 0.1429 | 15 | 0.2 | 5 | 0.2 | 37 | 0.1622 |
| 3_1082655 | 13 | 0.2308 | 17 | 0.5294 | 12 | 0.3333 | 14 | 0.3571 | 56 | 0.375 | 10 | 0.1 | 10 | 0.5 | 16 | 0.1875 | 9 | 0.3333 | 45 | 0.2667 |
| 3_1082681 | 30 | 0.3667 | 42 | 0.2619 | 32 | 0.4062 | 27 | 0.2593 | 131 | 0.3206 | 29 | 0.3448 | 28 | 0.1429 | 32 | 0.25 | 29 | 0.3448 | 118 | 0.2712 |
| 3_1082688 | 15 | 0.0667 | 28 | 0.1786 | 17 | 0.0588 | 26 | 0.0385 | 86 | 0.093 | 29 | 0 | 24 | 0.2083 | 20 | 0.05 | 17 | 0 | 90 | 0.0667 |
| 3_1082696 | NA | NA | 1 | 1 | NA | NA | NA | NA | 1 | 1 | 1 | 1 | 1 | 0 | NA | NA | 1 | 1 | 3 | 0.6667 |
| 3_1082991 | 6 | 0.6667 | 10 | 0.8 | 6 | 0.6667 | 10 | 0.6 | 32 | 0.6875 | 7 | 0.8571 | 10 | 1 | 11 | 0.7273 | 10 | 1 | 38 | 0.8947 |
| 3_1082992 | 7 | 0.4286 | 12 | 0.5833 | 7 | 0.2857 | 14 | 0.5 | 40 | 0.475 | 8 | 0.375 | 11 | 0.5455 | 12 | 0.5 | 9 | 0.4444 | 40 | 0.475 |
| 3_1082992 | 6 | 0.3333 | 9 | 0.2222 | 6 | 0.3333 | 13 | 0 | 34 | 0.1765 | 6 | 0.3333 | 10 | 0.2 | 11 | 0.3636 | 11 | 0.1818 | 38 | 0.2632 |
| 3_1082992 | 6 | 0 | 12 | 0 | 7 | 0.2857 | 9 | 0.1111 | 34 | 0.0882 | 8 | 0.125 | 9 | 0.1111 | 11 | 0 | 11 | 0 | 39 | 0.0513 |
| 3_1082992 | 6 | 1 | 11 | 0.8182 | 8 | 0.75 | 13 | 0.8462 | 38 | 0.8421 | 8 | 0.75 | 10 | 0.8 | 12 | 0.8333 | 12 | 0.8333 | 42 | 0.8095 |
| 3_1082993 | 5 | 0.4 | 11 | 0.2727 | 8 | 0.125 | 9 | 0.6667 | 33 | 0.3636 | 8 | 0.5 | 10 | 0.5 | 10 | 0.7 | 7 | 0.4286 | 35 | 0.5429 |
| 3_1082993 | 7 | 0.5714 | 13 | 0.3846 | 8 | 0.75 | 13 | 0.7692 | 41 | 0.6098 | 8 | 0.625 | 11 | 0.8182 | 12 | 0.4167 | 11 | 0.5455 | 42 | 0.5952 |
| 3_1082994 | 7 | 0.7143 | 8 | 0.75 | 5 | 0.8 | 8 | 1 | 28 | 0.8214 | 5 | 0.8 | 7 | 0.8571 | 11 | 1 | 8 | 1 | 31 | 0.9355 |
| 3_1082994 | 7 | 0.4286 | 11 | 0.7273 | 5 | 0.4 | 12 | 0.6667 | 35 | 0.6 | 7 | 0.4286 | 11 | 0.9091 | 10 | 0.8 | 10 | 0.3 | 38 | 0.6316 |
| 3_1082994 | 5 | 1 | 12 | 0.9167 | 6 | 0.8333 | 12 | 0.75 | 35 | 0.8571 | 7 | 0.8571 | 7 | 0.5714 | 12 | 0.6667 | 9 | 0.5556 | 35 | 0.6571 |
| 3_1082995 | 6 | 1 | 13 | 1 | 8 | 1 | 12 | 1 | 39 | 1 | 6 | 0.8333 | 11 | 1 | 9 | 1 | 9 | 1 | 35 | 0.9714 |
| 3_1082995 | 5 | 0 | 11 | 0.0909 | 7 | 0.2857 | 10 | 0.2 | 33 | 0.1515 | 6 | 0 | 9 | 0.2222 | 11 | 0.0909 | 7 | 0.1429 | 33 | 0.1212 |
| 3_1082995 | 5 | 0 | 10 | 0.4 | 6 | 0.3333 | 12 | 0.25 | 33 | 0.2727 | 6 | 0.5 | 9 | 0.2222 | 11 | 0.2727 | 9 | 0.3333 | 35 | 0.3143 |
| 3_1082996 | 6 | 0.8333 | 11 | 0.8182 | 8 | 0.875 | 10 | 0.9 | 35 | 0.8571 | 7 | 1 | 10 | 0.9 | 11 | 1 | 8 | 0.875 | 36 | 0.9444 |
| 3_1082996 | 6 | 0.1667 | 12 | 0.0833 | 7 | 0.1429 | 12 | 0.1667 | 37 | 0.1351 | 6 | 0.1667 | 7 | 0.2857 | 9 | 0.1111 | 6 | 0 | 28 | 0.1429 |
| 3_1082997 | 6 | 1 | 10 | 0.6 | 8 | 0.75 | 7 | 1 | 31 | 0.8065 | 6 | 0.6667 | 10 | 0.6 | 6 | 0.6667 | 6 | 0.8333 | 28 | 0.6786 |
| 3_1083141 | 1 | 1 | 1 | 1 | 1 | 1 | 1 | 1 | 4 | 1 | NA | NA | NA | NA | 1 | 1 | 3 | 0.6667 | 4 | 0.75 |
| 3_1083141 | 1 | 1 | 2 | 1 | 1 | 0 | 1 | 1 | 5 | 0.8 | NA | NA | NA | NA | NA | NA | 3 | 0.3333 | 3 | 0.3333 |
| 3_1083145 | 1 | 1 | 2 | 1 | 1 | 1 | 1 | 1 | 5 | 1 | NA | NA | NA | NA | 1 | 1 | 2 | 1 | 3 | 1 |
| 3_1083196 | 1 | 0 | 2 | 1 | NA | NA | 1 | 1 | 4 | 0.75 | NA | NA | NA | NA | 1 | 0 | 2 | 0.5 | 3 | 0.3333 |
| 3_1083198 | 1 | 1 | 2 | 0.5 | 1 | 1 | 1 | 1 | 5 | 0.8 | NA | NA | NA | NA | 1 | 1 | 3 | 1 | 4 | 1 |
| 3_1083198 | 1 | 1 | 2 | 1 | 1 | 1 | NA | NA | 4 | 1 | NA | NA | NA | NA | 1 | 1 | 3 | 1 | 4 | 1 |
| 3_1083199 | 1 | 1 | NA | NA | 1 | 1 | 1 | 0 | 3 | 0.6667 | NA | NA | NA | NA | 1 | 1 | 3 | 1 | 4 | 1 |
| 3_1083200 | 1 | 0 | NA | NA | 1 | 1 | 1 | 1 | 3 | 0.6667 | NA | NA | NA | NA | 1 | 1 | 3 | 0.3333 | 4 | 0.5 |
| 3_1083202 | 1 | 1 | NA | NA | 1 | 1 | 1 | 1 | 3 | 1 | NA | NA | NA | NA | 1 | 0 | 3 | 0.6667 | 4 | 0.5 |
| 3_1083204 | 1 | 1 | NA | NA | NA | NA | 1 | 1 | 2 | 1 | NA | NA | NA | NA | 1 | 1 | 3 | 0.6667 | 4 | 0.75 |
| 3_1083204 | 1 | 1 | NA | NA | NA | NA | 1 | 1 | 2 | 1 | NA | NA | NA | NA | NA | NA | 1 | 1 | 1 | 1 |
| 3_1083207 | NA | NA | NA | NA | NA | NA | 1 | 1 | 1 | 1 | NA | NA | NA | NA | NA | NA | 3 | 0.3333 | 3 | 0.3333 |
| 3_1083208 | NA | NA | NA | NA | NA | NA | 1 | 1 | 1 | 1 | NA | NA | NA | NA | NA | NA | 3 | 1 | 3 | 1 |
| 3_1083338 | 4 | 0.5 | 3 | 0.6667 | 7 | 0.1429 | 3 | 0 | 17 | 0.2941 | 4 | 0 | 5 | 0.2 | 9 | 0.2222 | 6 | 0 | 24 | 0.125 |
| 3_1083339 | 26 | 0.0385 | 25 | 0.24 | 34 | 0.2059 | 32 | 0.2188 | 117 | 0.1795 | 26 | 0.1538 | 36 | 0.0833 | 29 | 0.069 | 25 | 0.08 | 116 | 0.0948 |
| 3_1083340 | 25 | 0.44 | 22 | 0.2273 | 28 | 0.0714 | 25 | 0.16 | 100 | 0.22 | 19 | 0.1579 | 31 | 0.1613 | 21 | 0.0952 | 21 | 0.2381 | 92 | 0.163 |

| | | | | | | | | | | | | | | | | | | | | |
|---|---|---|---|---|---|---|---|---|---|---|---|---|---|---|---|---|---|---|---|---|
| 3_1083340 | 24 | 0.2083 | 24 | 0.25 | 30 | 0.2 | 30 | 0.2667 | 108 | 0.2315 | 22 | 0.4091 | 33 | 0.2727 | 22 | 0.1818 | 24 | 0.2917 | 101 | 0.2871 |
| 3_1083340 | 26 | 0.3462 | 28 | 0.1429 | 30 | 0.2 | 27 | 0.2222 | 111 | 0.2252 | 24 | 0.125 | 31 | 0.3226 | 27 | 0.1852 | 21 | 0.1429 | 103 | 0.2039 |
| 3_1083341 | 25 | 0.36 | 23 | 0.4348 | 26 | 0.3846 | 31 | 0.4194 | 105 | 0.4 | 22 | 0.3636 | 26 | 0.4231 | 23 | 0.3913 | 22 | 0.4091 | 93 | 0.3978 |
| 3_1083527 | 1 | 1 | NA | NA | NA | NA | NA | NA | 1 | 1 | NA | NA | 1 | 0 | 1 | 1 | 1 | 0 | 3 | 0.3333 |
| 3_1083631 | 5 | 0.6 | 3 | 0 | 6 | 0.5 | 9 | 0.4444 | 23 | 0.4348 | 3 | 0 | 4 | 0.25 | 2 | 0.5 | 6 | 0.3333 | 15 | 0.2667 |
| 3_1083634 | 4 | 0.5 | 2 | 0 | 6 | 0.3333 | 8 | 0.25 | 20 | 0.3 | 2 | 0 | 4 | 0.25 | 1 | 0 | 4 | 0 | 11 | 0.0909 |
| 3_1083638 | 5 | 0.8 | NA | NA | 2 | 1 | 7 | 0.7143 | 14 | 0.7857 | 2 | 0.5 | 3 | 0.6667 | NA | NA | 5 | 0.8 | 10 | 0.7 |
| 3_1083638 | 5 | 0.6 | NA | NA | 5 | 0.4 | 5 | 0.2 | 15 | 0.4 | 2 | 0 | 4 | 0.25 | 1 | 0 | 4 | 0 | 11 | 0.0909 |
| 3_1083644 | 4 | 0 | NA | NA | 4 | 0.25 | 6 | 0.1667 | 14 | 0.1429 | 3 | 0 | 2 | 0 | 1 | 0 | 5 | 0.2 | 11 | 0.0909 |
| 3_1083652 | 3 | 0.3333 | NA | NA | 2 | 0 | 5 | 0 | 10 | 0.1 | 1 | 0 | 1 | 0 | 1 | 1 | 3 | 0.3333 | 6 | 0.3333 |
| 3_1083654 | 1 | 1 | NA | NA | 3 | 0.3333 | 5 | 0.2 | 9 | 0.3333 | NA | NA | 1 | 1 | NA | NA | 3 | 0 | 4 | 0.25 |
| 3_1083655 | 4 | 0 | 2 | 0 | 6 | 0 | 5 | 0.4 | 17 | 0.1176 | 4 | 0 | 3 | 0.3333 | 2 | 0 | 5 | 0.4 | 14 | 0.2143 |
| 3_1083657 | 2 | 0.5 | 1 | 1 | 4 | 0.25 | 3 | 0.6667 | 10 | 0.5 | 3 | 1 | 1 | 1 | 2 | 0.5 | 2 | 0.5 | 8 | 0.75 |
| 3_1083657 | 4 | 0.5 | 2 | 0 | 6 | 0 | 2 | 0 | 14 | 0.1429 | 3 | 0.3333 | 3 | 0 | 1 | 0 | 3 | 0.3333 | 10 | 0.2 |
| 3_1084646 | 2 | 0 | 7 | 0.2857 | 10 | 0.2 | 10 | 0.4 | 29 | 0.2759 | 6 | 0 | 5 | 0 | 5 | 0.4 | 6 | 0.1667 | 22 | 0.1364 |
| 3_1084647 | 1 | 0 | 7 | 0.1429 | 10 | 0.1 | 10 | 0.1 | 28 | 0.1071 | 4 | 0 | 5 | 0.2 | 3 | 0 | 6 | 0 | 18 | 0.0556 |
| 3_1084654 | 2 | 0.5 | 3 | 0.3333 | 5 | 0.2 | 8 | 0.125 | 18 | 0.2222 | 4 | 0.25 | 3 | 0 | 4 | 0.25 | 4 | 0.5 | 15 | 0.2667 |
| 3_1084655 | 2 | 0.5 | 3 | 0.3333 | 6 | 0 | 10 | 0.2 | 21 | 0.1905 | 5 | 0.2 | 3 | 0.3333 | 3 | 0.3333 | 4 | 0.25 | 15 | 0.2667 |
| 3_1084655 | 2 | 1 | 3 | 0.3333 | 5 | 0.4 | 9 | 0.6667 | 19 | 0.5789 | 5 | 0.2 | 3 | 0.6667 | 4 | 0.5 | 4 | 0.5 | 16 | 0.4375 |
| 3_1084656 | 1 | 1 | 3 | 0 | 5 | 0 | 7 | 0 | 16 | 0.0625 | 4 | 0.25 | 4 | 0 | 4 | 0.25 | 3 | 0 | 15 | 0.1333 |
| 3_1084656 | 1 | 0 | 2 | 1 | 6 | 0.1667 | 5 | 0 | 14 | 0.2143 | 4 | 0 | 4 | 0 | 3 | 0 | 3 | 0.3333 | 14 | 0.0714 |
| 3_1084657 | NA | NA | 3 | 0.3333 | 4 | 0.25 | 8 | 0.25 | 15 | 0.2667 | 4 | 0.25 | 4 | 0.25 | 4 | 0 | 3 | 0 | 15 | 0.1333 |
| 3_1084657 | NA | NA | 3 | 0.6667 | 4 | 0.5 | 5 | 0 | 12 | 0.3333 | 3 | 0.3333 | 2 | 0 | 3 | 0 | 3 | 0.3333 | 11 | 0.1818 |
| 3_1084660 | 1 | 1 | 3 | 1 | 4 | 1 | 5 | 1 | 13 | 1 | 1 | 1 | 2 | 1 | 2 | 1 | 2 | 0.5 | 7 | 0.8571 |
| 3_1084661 | 1 | 1 | 3 | 0.6667 | 6 | 0.5 | 5 | 0.4 | 15 | 0.5333 | 2 | 0 | 3 | 0.3333 | 4 | 0.5 | 3 | 0.6667 | 12 | 0.4167 |
| 3_1084661 | 1 | 0 | 4 | 0.25 | 6 | 0.1667 | 6 | 0.5 | 17 | 0.2941 | 2 | 0 | 1 | 0 | 4 | 0.25 | 3 | 0.3333 | 10 | 0.2 |
| 3_1084884 | 7 | 0.7143 | 4 | 1 | 5 | 0.8 | 2 | 1 | 18 | 0.8333 | 2 | 1 | 2 | 1 | 4 | 0.25 | 5 | 0.4 | 13 | 0.5385 |
| 3_1084884 | 6 | 0 | 5 | 0.4 | 5 | 0 | 2 | 0 | 18 | 0.1111 | 2 | 0 | 2 | 0.5 | 4 | 0.25 | 6 | 0 | 14 | 0.1429 |
| 3_1084886 | 6 | 0 | 5 | 0.2 | 6 | 0.1667 | 3 | 0 | 20 | 0.1 | 4 | 0 | 4 | 0.25 | 3 | 0 | 5 | 0 | 16 | 0.0625 |
| 3_1084888 | 5 | 0 | 3 | 0.3333 | 6 | 0 | 1 | 0 | 15 | 0.0667 | 3 | 0 | 4 | 0 | 5 | 0.2 | 6 | 0.1667 | 18 | 0.1111 |
| 3_1085259 | 4 | 0.75 | 8 | 0.75 | 5 | 0.8 | 5 | 0.4 | 22 | 0.6818 | 3 | 1 | 5 | 0.8 | 6 | 0.5 | 2 | 1 | 16 | 0.75 |
| 3_1085260 | 3 | 0.3333 | 3 | 0.3333 | 3 | 0.3333 | 6 | 0 | 15 | 0.2 | 3 | 0 | 4 | 0.25 | 4 | 0 | 1 | 0 | 12 | 0.0833 |
| 3_1085262 | 4 | 0.5 | 7 | 0.7143 | 7 | 0.5714 | 5 | 0.2 | 23 | 0.5217 | 2 | 1 | 2 | 0.5 | 4 | 0.5 | NA | NA | 8 | 0.625 |
| 3_1085262 | 3 | 0.3333 | 7 | 0.1429 | 6 | 0.1667 | 5 | 0.2 | 21 | 0.1905 | 3 | 0 | 5 | 0 | 6 | 0.1667 | 2 | 0.5 | 16 | 0.125 |
| 3_1085262 | 4 | 0.25 | 5 | 0.4 | 7 | 0.2857 | 5 | 0.4 | 21 | 0.3333 | 3 | 0.6667 | 4 | 0.5 | 6 | 0 | 1 | 0 | 14 | 0.2857 |
| 3_1085263 | 4 | 0.5 | 9 | 0.5556 | 7 | 0.7143 | 5 | 0.6 | 25 | 0.6 | 3 | 0.6667 | 5 | 0.8 | 7 | 0.2857 | 3 | 0.6667 | 18 | 0.5556 |
| 3_1085263 | 1 | 1 | 6 | 0.8333 | 7 | 0.4286 | 4 | 0.75 | 18 | 0.6667 | 2 | 1 | 3 | 0.6667 | 6 | 0.5 | 1 | 0 | 12 | 0.5833 |
| 3_1085264 | 3 | 0.6667 | 9 | 0.5556 | 4 | 0.75 | 6 | 0.5 | 22 | 0.5909 | 2 | 0 | 5 | 1 | 7 | 0 | 3 | 0 | 17 | 0.2941 |
| 3_1085264 | 4 | 0.25 | 10 | 0.2 | 5 | 0.2 | 5 | 0.2 | 24 | 0.2083 | 3 | 0.3333 | 5 | 0.2 | 6 | 0.1667 | 2 | 0 | 16 | 0.1875 |
| 3_1085518 | 10 | 0.2 | 12 | 0.0833 | 10 | 0.2 | 7 | 0.4286 | 39 | 0.2051 | 7 | 0.5714 | 9 | 0.4444 | 12 | 0.25 | 7 | 0.1429 | 35 | 0.3429 |
| 3_1085518 | 6 | 0.3333 | 11 | 0.4545 | 10 | 0.5 | 5 | 0.8 | 32 | 0.5 | 7 | 0.4286 | 8 | 0.25 | 7 | 0.5714 | 5 | 0.8 | 27 | 0.4815 |
| 3_1085518 | 9 | 0.8889 | 9 | 0.7778 | 9 | 0.8889 | 7 | 1 | 34 | 0.8824 | 4 | 1 | 10 | 1 | 8 | 0.875 | 5 | 1 | 27 | 0.963 |
| 3_1085519 | 10 | 0.2 | 12 | 0.0833 | 10 | 0 | 10 | 0.2 | 42 | 0.119 | 10 | 0 | 9 | 0 | 12 | 0.25 | 9 | 0.2222 | 40 | 0.125 |
| 3_1085519 | 8 | 0.625 | 9 | 0.7778 | 10 | 0.9 | 9 | 0.6667 | 36 | 0.75 | 9 | 0.7778 | 9 | 0.6667 | 11 | 0.8182 | 6 | 0.6667 | 35 | 0.7429 |
| 3_1085519 | 11 | 0.2727 | 11 | 0.5455 | 11 | 0.4545 | 9 | 0.3333 | 42 | 0.4048 | 11 | 0.0909 | 11 | 0.2727 | 13 | 0.3077 | 8 | 0.25 | 43 | 0.2326 |
| 3_1085520 | 7 | 0.2857 | 7 | 0.1429 | 6 | 0.3333 | 1 | 0 | 21 | 0.2381 | 6 | 0.3333 | 3 | 0 | 7 | 0 | 2 | 0.5 | 18 | 0.1667 |
| 3_1085523 | 13 | 0.0769 | 13 | 0.0769 | 13 | 0.0769 | 11 | 0.0909 | 50 | 0.08 | 12 | 0 | 11 | 0.2727 | 14 | 0.0714 | 9 | 0.1111 | 46 | 0.1087 |
| 3_1085526 | 13 | 0.1538 | 14 | 0.0714 | 14 | 0.0714 | 14 | 0.2143 | 55 | 0.1273 | 11 | 0.1818 | 10 | 0.1 | 12 | 0 | 9 | 0.3333 | 42 | 0.1429 |
| 3_1085820 | 1 | 0 | 4 | 0.5 | 8 | 0 | 2 | 0 | 15 | 0.1333 | 4 | 0.25 | 1 | 0 | 3 | 0 | 3 | 0 | 11 | 0.0909 |
| 3_1085820 | NA | NA | 5 | 0 | 9 | 0.1111 | 2 | 0 | 16 | 0.0625 | 4 | 0 | 3 | 0 | 2 | 0.5 | 3 | 0 | 12 | 0.0833 |
| 3_1085841 | 2 | 0 | 5 | 0 | 7 | 0 | 3 | 0.3333 | 17 | 0.0588 | 6 | 0 | 2 | 0.5 | 1 | 0 | 2 | 0 | 11 | 0.0909 |
| 3_1085842 | 1 | 1 | 5 | 0.6 | 9 | 0.5556 | 3 | 1 | 18 | 0.6667 | 6 | 0.6667 | 2 | 1 | 3 | 0.6667 | 3 | 1 | 14 | 0.7857 |
| 3_1085843 | NA | NA | NA | NA | 1 | 1 | NA | NA | 1 | 1 | 2 | 0.5 | NA | NA | NA | NA | NA | NA | 2 | 0.5 |
| 3_1085860 | 2 | 0 | 5 | 0 | 8 | 0.125 | 2 | 0 | 17 | 0.0588 | 5 | 0.2 | 3 | 0 | 3 | 0 | 4 | 0 | 15 | 0.0667 |
| 3_1085860 | 2 | 0 | 5 | 0 | 8 | 0.125 | 2 | 0 | 17 | 0.0588 | 1 | 0 | 3 | 0 | 3 | 0 | 4 | 0 | 15 | 0.0667 |
| 3_1085860 | 1 | 0 | 5 | 0 | 7 | 0.1429 | 2 | 0 | 15 | 0.0667 | 6 | 0.1667 | 3 | 0 | 3 | 0 | 4 | 0 | 16 | 0.0625 |
| 3_1085867 | 2 | 0.5 | 4 | 0.25 | 1 | 0 | 2 | 0.5 | 9 | 0.3333 | 4 | 0.25 | 3 | 0.3333 | 2 | 0.5 | 1 | 1 | 10 | 0.4 |
| 3_1085868 | 2 | 1 | 3 | 1 | 1 | 1 | 2 | 1 | 8 | 1 | 4 | 1 | 3 | 1 | 1 | 1 | 1 | 1 | 9 | 1 |
| 3_1085869 | 3 | 0.6667 | 4 | 1 | 1 | 1 | 1 | 1 | 9 | 0.8889 | 4 | 1 | 2 | 1 | NA | NA | 1 | 1 | 7 | 1 |
| 3_1085869 | 3 | 0.6667 | 5 | 0.8 | NA | NA | 1 | 1 | 9 | 0.7778 | 4 | 0.75 | 1 | 1 | NA | NA | 1 | 1 | 6 | 0.8333 |
| 3_1085869 | 3 | 0.6667 | 4 | 0.25 | 1 | 1 | 2 | 1 | 10 | 0.6 | 4 | 0.5 | 3 | 0 | 1 | 0 | 1 | 0 | 9 | 0.2222 |
| 3_1085870 | 2 | 0.5 | 5 | 0.4 | 1 | 1 | 1 | 0 | 9 | 0.4444 | 5 | 0.2 | 2 | 0 | 1 | 1 | 1 | 0 | 9 | 0.2222 |
| 3_1086187 | NA | NA | 2 | 0.5 | 1 | 0 | 1 | 0 | 4 | 0.25 | 2 | 0.5 | 1 | 0 | NA | NA | 1 | 0 | 4 | 0.25 |
| 3_1086187 | NA | NA | 1 | 1 | 1 | 0 | NA | NA | 2 | 0.5 | 1 | 1 | NA | NA | NA | NA | 1 | 0 | 2 | 0.5 |
| 3_1086188 | NA | NA | 1 | 0 | 1 | 0 | 1 | 1 | 3 | 0.3333 | 2 | 0.5 | 1 | 0 | NA | NA | 1 | 0 | 4 | 0.25 |

| | | | | | | | | | | | | | | | | | | | | |
|---|---|---|---|---|---|---|---|---|---|---|---|---|---|---|---|---|---|---|---|---|
| 3_1088106( | 7 | 0.1429 | 4 | 0.5 | 7 | 0 | 4 | 0 | 22 | 0.1364 | 5 | 0.2 | 4 | 0.25 | 4 | 0 | 1 | 1 | 14 | 0.2143 |
| 3_1088108: | 7 | 0.5714 | 5 | 1 | 7 | 0.8571 | 4 | 0.5 | 23 | 0.7391 | 5 | 0.6 | 4 | 0.5 | 4 | 0.75 | 2 | 1 | 15 | 0.6667 |
| 3_1088112‹ | 7 | 0.7143 | 10 | 0.9 | 7 | 0.8571 | 5 | 1 | 29 | 0.8621 | 4 | 1 | 4 | 1 | 7 | 0.8571 | 4 | 1 | 19 | 0.9474 |
| 3_1088113: | 8 | 0.375 | 11 | 0.4545 | 5 | 0.6 | 7 | 0.4286 | 31 | 0.4516 | 3 | 0 | 6 | 0.8333 | 7 | 0.8571 | 6 | 0.3333 | 22 | 0.5909 |
| 3_1088114: | 8 | 0.625 | 11 | 0.1818 | 7 | 0.2857 | 8 | 0.75 | 34 | 0.4412 | 3 | 0.6667 | 6 | 0.5 | 6 | 0.3333 | 7 | 0.4286 | 22 | 0.4545 |
| 3_1088115 | 8 | 0.125 | 9 | 0.5556 | 7 | 0.4286 | 7 | 0.5714 | 31 | 0.4194 | 3 | 0 | 6 | 0.1667 | 6 | 0.1667 | 6 | 0.1667 | 21 | 0.1429 |
| 3_1088116: | 5 | 0.2 | 9 | 0.3333 | 5 | 0.2 | 8 | 0.375 | 27 | 0.2963 | 3 | 0.6667 | 5 | 0.2 | 4 | 0.25 | 2 | 1 | 14 | 0.4286 |
| 3_1088117 | 3 | 0.3333 | 9 | 0.3333 | 6 | 0.5 | 5 | 0.4 | 23 | 0.3913 | 3 | 0.6667 | 5 | 0.2 | 5 | 0.4 | 3 | 0.6667 | 16 | 0.4375 |
| 3_1088118( | 5 | 0 | 8 | 0.25 | 5 | 0 | 7 | 0 | 25 | 0.08 | 3 | 0 | 5 | 0 | 2 | 0 | 3 | 0.6667 | 13 | 0.1538 |
| 3_1088678 | NA | NA | NA | NA | NA | NA | 1 | 1 | 1 | 1 | NA | NA | 1 | 1 | NA | NA | NA | NA | 1 | 1 |
| 3_1088785( | 19 | 0.0526 | 8 | 0 | 12 | 0 | 24 | 0.125 | 63 | 0.0635 | 7 | 0 | 12 | 0.0833 | 17 | 0 | 24 | 0.0833 | 60 | 0.05 |
| 3_1088790( | 12 | 0 | 8 | 0.125 | 7 | 0.1429 | 18 | 0.0556 | 45 | 0.0667 | 2 | 0 | 6 | 0.1667 | 14 | 0.0714 | 18 | 0.2778 | 40 | 0.175 |
| 3_1088816 | 1 | 0 | 2 | 0.5 | 5 | 0 | 11 | 0.0909 | 19 | 0.1053 | 2 | 0 | 3 | 0 | 1 | 0 | 11 | 0.0909 | 17 | 0.0588 |
| 3_1088823 | 1 | 0 | 3 | 0.3333 | 5 | 0.8 | 10 | 0.5 | 19 | 0.5263 | 2 | 1 | 3 | 0.3333 | 2 | 0.5 | 12 | 0.75 | 19 | 0.6842 |
| 3_1088824 | 2 | 0.5 | 2 | 1 | 4 | 0.75 | 7 | 0.5714 | 15 | 0.6667 | 2 | 1 | 2 | 1 | 2 | 0.5 | 9 | 0.5556 | 15 | 0.6667 |
| 3_1088825 | 2 | 0 | 1 | 0 | 4 | 0 | 7 | 0.2857 | 14 | 0.1429 | 2 | 0 | 3 | 0.3333 | 2 | 0.5 | 11 | 0.0909 | 18 | 0.1667 |
| 3_1096885 | 2 | 0 | 1 | 0 | NA | NA | 4 | 0.25 | 7 | 0.1429 | 1 | 1 | 1 | 0 | NA | NA | 1 | 0 | 3 | 0.3333 |
| 3_1096886 | 2 | 1 | 1 | 1 | NA | NA | 4 | 1 | 7 | 1 | 1 | 1 | 1 | 1 | NA | NA | 1 | 1 | 3 | 1 |
| 3_1096887 | 2 | 0 | 1 | 0 | NA | NA | 3 | 0.3333 | 6 | 0.1667 | 1 | 0 | 1 | 0 | NA | NA | 1 | 1 | 3 | 0.3333 |
| 3_1096887 | 2 | 1 | 1 | 1 | NA | NA | 3 | 1 | 6 | 1 | 1 | 1 | 1 | 1 | NA | NA | 1 | 1 | 3 | 1 |
| 3_1096888 | 1 | 0 | NA | NA | NA | NA | 3 | 0.3333 | 4 | 0.25 | 1 | 1 | 1 | 0 | NA | NA | 1 | 1 | 3 | 0.6667 |
| 3_1096888 | 2 | 1 | 1 | 1 | NA | NA | 3 | 1 | 6 | 1 | 1 | 1 | 1 | 1 | NA | NA | 1 | 1 | 3 | 1 |
| 3_1096890 | 2 | 1 | 2 | 1 | NA | NA | 2 | 1 | 6 | 1 | 1 | 1 | 1 | 1 | NA | NA | 1 | 1 | 3 | 1 |
| 3_1096891 | 1 | 0 | 2 | 0 | NA | NA | 2 | 0.5 | 5 | 0.2 | 1 | 1 | NA | NA | NA | NA | NA | NA | 1 | 1 |
| 3_1096896 | 1 | 1 | 2 | 0.5 | NA | NA | 2 | 1 | 5 | 0.8 | 1 | 1 | 1 | 0 | NA | NA | NA | NA | 2 | 0.5 |
| 3_1096898 | 1 | 1 | 2 | 1 | NA | NA | 1 | 1 | 4 | 1 | 1 | 1 | 1 | 1 | NA | NA | NA | NA | 2 | 1 |
| 3_1096898 | 1 | 0 | 2 | 0.5 | NA | NA | 2 | 0 | 5 | 0.2 | 1 | 0 | 1 | 1 | NA | NA | NA | NA | 2 | 0.5 |
| 3_1096900 | NA | NA | 2 | 0 | NA | NA | 2 | 1 | 4 | 0.5 | 1 | 1 | 1 | 0 | NA | NA | NA | NA | 2 | 0.5 |
| 3_1098422 | NA | NA | 1 | 0 | NA | NA | 2 | 0.5 | 3 | 0.3333 | 1 | 1 | NA | NA | NA | NA | NA | NA | 1 | 1 |
| 3_1101569 | 1 | 1 | 1 | 1 | 1 | 1 | NA | NA | 3 | 1 | NA | NA | 2 | 1 | 1 | 1 | NA | NA | 3 | 1 |
| 3_1101570 | 2 | 1 | 1 | 1 | 1 | 1 | NA | NA | 4 | 1 | NA | NA | 2 | 1 | 1 | 1 | NA | NA | 3 | 1 |
| 3_1101570 | 2 | 0 | 1 | 1 | 1 | 1 | 1 | 0 | 5 | 0.4 | NA | NA | 1 | 1 | 1 | 0 | 1 | 1 | 3 | 0.6667 |
| 3_1101571 | 2 | 1 | 2 | 1 | NA | NA | 1 | 1 | 5 | 1 | NA | NA | 2 | 1 | 1 | 1 | 1 | 0 | 4 | 0.75 |
| 3_1101572 | 1 | 1 | 2 | 1 | 1 | 1 | 1 | 1 | 5 | 1 | NA | NA | 1 | 1 | 1 | 1 | 1 | 0 | 3 | 0.6667 |
| 3_1101573 | 1 | 1 | 2 | 0.5 | 1 | 1 | 1 | 1 | 5 | 0.8 | NA | NA | 1 | 0 | 1 | 1 | 1 | 0 | 3 | 0.3333 |
| 3_1101577 | 1 | 0 | 1 | 1 | NA | NA | NA | NA | 2 | 0.5 | NA | NA | NA | NA | NA | NA | 1 | 1 | 1 | 1 |
| 3_1101577 | 1 | 1 | 2 | 0.5 | NA | NA | 1 | 1 | 4 | 0.75 | NA | NA | NA | NA | 1 | 1 | 1 | 1 | 2 | 1 |
| 3_1133992 | 2 | 0 | 1 | 0 | 1 | 0 | 4 | 0.25 | 8 | 0.125 | NA | NA | 2 | 0 | 3 | 0.6667 | 5 | 0 | 10 | 0.2 |
| 3_1146979 | NA | NA | 5 | 0.2 | NA | NA | 3 | 0.3333 | 8 | 0.25 | NA | NA | 2 | 0.5 | 4 | 0 | 3 | 0 | 9 | 0.1111 |
| 3_1149158 | 1 | 1 | 11 | 0.7273 | 3 | 0.6667 | 6 | 0.6667 | 21 | 0.7143 | NA | NA | 6 | 0.6667 | 8 | 0.75 | 6 | 0.5 | 20 | 0.65 |
| 3_1149158 | 1 | 0 | 11 | 0.5455 | 3 | 0.3333 | 6 | 0.3333 | 21 | 0.4286 | 1 | 1 | 5 | 0.4 | 6 | 0.1667 | 5 | 0.4 | 17 | 0.3529 |
| 3_1149159 | 1 | 0 | 11 | 0.9091 | 3 | 1 | 6 | 1 | 21 | 0.9048 | 1 | 1 | 6 | 1 | 8 | 0.875 | 5 | 1 | 20 | 0.95 |
| 3_1149159 | 1 | 0 | 10 | 0.4 | 3 | 1 | 5 | 0.6 | 19 | 0.5263 | NA | NA | 6 | 0.5 | 8 | 0.625 | 6 | 0.5 | 20 | 0.55 |
| 3_1149160 | 1 | 1 | 10 | 0.7 | 3 | 0.3333 | 5 | 0.2 | 19 | 0.5263 | 1 | 0 | 5 | 1 | 8 | 0.375 | 7 | 0.1429 | 21 | 0.4286 |
| 3_1149161( | NA | NA | 11 | 0.6364 | 3 | 1 | 5 | 1 | 19 | 0.7895 | 1 | 1 | 3 | 0.6667 | 4 | 0.5 | 5 | 0.8 | 13 | 0.6923 |
| 3_1149161 | 1 | 0 | 12 | 0.75 | 2 | 0.5 | 5 | 0.2 | 20 | 0.55 | 1 | 0 | 6 | 0.1667 | 7 | 0.5714 | 6 | 0 | 20 | 0.25 |
| 3_1149162 | 2 | 1 | 11 | 1 | 3 | 1 | 8 | 1 | 24 | 1 | 1 | 1 | 7 | 1 | 8 | 0.875 | 6 | 1 | 22 | 0.9545 |
| 3_1149163 | 2 | 0.5 | 12 | 0.25 | 2 | 0.5 | 6 | 0.1667 | 22 | 0.2727 | 1 | 1 | 7 | 0.4286 | 7 | 0.4286 | 7 | 0.2857 | 22 | 0.4091 |
| 3_1149163 | 2 | 0.5 | 13 | 0.9231 | 3 | 1 | 7 | 1 | 25 | 0.92 | 1 | 1 | 7 | 0.8571 | 8 | 0.875 | 7 | 0.8571 | 23 | 0.8696 |
| 3_1149164( | 1 | 1 | 12 | 0.0833 | 2 | 0.5 | 7 | 0.1429 | 22 | 0.1818 | 1 | 0 | 6 | 0 | 8 | 0.25 | 7 | 0 | 22 | 0.0909 |
| 3_1149164 | 2 | 0 | 7 | 0.1429 | 3 | 0 | 6 | 0 | 18 | 0.0556 | NA | NA | 4 | 0.25 | 3 | 0 | 4 | 0 | 11 | 0.0909 |
| 3_1149164 | 2 | 0.5 | 10 | 0.7 | 1 | 1 | 5 | 0.4 | 18 | 0.6111 | 1 | 1 | 5 | 0.6 | 8 | 0.75 | 5 | 0.6 | 19 | 0.6842 |
| 3_1149165 | 2 | 0 | 11 | 0.3636 | 2 | 0.5 | 7 | 0.1429 | 22 | 0.2727 | 1 | 0 | 7 | 0.1429 | 10 | 0.1 | 7 | 0.2857 | 25 | 0.16 |
| 3_1155044 | 30 | 0.0667 | 28 | 0.1786 | 21 | 0.0952 | 23 | 0.2174 | 102 | 0.1373 | 22 | 0.0455 | 20 | 0.15 | 18 | 0.0556 | 23 | 0.1304 | 83 | 0.0964 |
| 3_1155044 | 25 | 0.2 | 22 | 0.1818 | 18 | 0.0556 | 20 | 0.15 | 85 | 0.1529 | 23 | 0.2609 | 18 | 0.2222 | 17 | 0.1765 | 18 | 0.0556 | 76 | 0.1842 |
| 3_1155051( | 24 | 0.25 | 23 | 0.6957 | 18 | 0.5556 | 23 | 0.3913 | 88 | 0.4659 | 23 | 0.4783 | 16 | 0.75 | 16 | 0.5625 | 18 | 0.5556 | 73 | 0.5753 |
| 3_1155052 | 21 | 0.2381 | 21 | 0.3333 | 13 | 0.3846 | 22 | 0.1818 | 77 | 0.2727 | 15 | 0.1333 | 12 | 0.6667 | 12 | 0.3333 | 14 | 0.7857 | 53 | 0.4717 |
| 3_1155054( | 24 | 0.6667 | 25 | 0.64 | 20 | 0.85 | 22 | 0.7273 | 91 | 0.7143 | 23 | 0.7826 | 18 | 0.4444 | 15 | 0.9333 | 19 | 0.6316 | 75 | 0.6933 |
| 3_1155054 | 27 | 0.8148 | 27 | 0.8519 | 22 | 0.9545 | 19 | 0.7895 | 95 | 0.8526 | 23 | 0.8696 | 20 | 0.8 | 16 | 0.75 | 22 | 0.5909 | 81 | 0.7531 |
| 3_1155054 | 26 | 0.1923 | 26 | 0.1538 | 22 | 0.1818 | 22 | 0.1364 | 96 | 0.1667 | 23 | 0.087 | 20 | 0.25 | 15 | 0.2 | 21 | 0.0952 | 79 | 0.1519 |
| 3_1155055 | 25 | 0.04 | 25 | 0.16 | 20 | 0.15 | 21 | 0.1905 | 91 | 0.1319 | 24 | 0.2083 | 19 | 0.2105 | 16 | 0.0625 | 20 | 0.15 | 79 | 0.1646 |
| 3_1155055 | 26 | 0.6923 | 25 | 0.52 | 20 | 0.7 | 24 | 0.6667 | 95 | 0.6421 | 24 | 0.5417 | 20 | 0.7 | 13 | 0.6923 | 21 | 0.619 | 78 | 0.6282 |
| 3_1155056 | 22 | 0.6818 | 25 | 0.36 | 20 | 0.75 | 23 | 0.5217 | 90 | 0.5667 | 22 | 0.5455 | 18 | 0.5556 | 15 | 0.6667 | 21 | 0.5238 | 76 | 0.5658 |
| 3_1155057 | 23 | 0 | 27 | 0.037 | 22 | 0.0455 | 22 | 0.1364 | 94 | 0.0532 | 20 | 0.1 | 20 | 0.1 | 17 | 0.0588 | 16 | 0.125 | 73 | 0.0959 |
| 3_1155058 | 13 | 0.2308 | 17 | 0.2353 | 18 | 0.1111 | 13 | 0.1538 | 61 | 0.1803 | 15 | 0.2 | 14 | 0 | 14 | 0.2143 | 16 | 0.0625 | 59 | 0.1186 |

| | | | | | | | | | | | | | | | | | | | | |
|---|---|---|---|---|---|---|---|---|---|---|---|---|---|---|---|---|---|---|---|---|
| 3_1155059 | 25 | 0.24 | 24 | 0.4583 | 20 | 0.2 | 22 | 0.4091 | 91 | 0.3297 | 18 | 0.2222 | 19 | 0.4211 | 13 | 0.0769 | 15 | 0.2 | 65 | 0.2462 |
| 3_1155060 | 23 | 1 | 23 | 0.913 | 16 | 0.9375 | 16 | 1 | 78 | 0.9615 | 18 | 0.9444 | 16 | 0.9375 | 10 | 1 | 15 | 0.8667 | 59 | 0.9322 |
| 3_1155063 | 17 | 1 | 20 | 1 | 17 | 0.9412 | 17 | 0.9412 | 71 | 0.9718 | 16 | 1 | 13 | 1 | 11 | 1 | 14 | 0.9286 | 54 | 0.9815 |
| 3_1155064 | 19 | 0.3158 | 19 | 0.2632 | 17 | 0.4118 | 18 | 0.3333 | 73 | 0.3288 | 18 | 0.5 | 13 | 0.3846 | 13 | 0.2308 | 13 | 0.1538 | 57 | 0.3333 |
| 3_1155064 | 17 | 0.1765 | 17 | 0.0588 | 9 | 0 | 18 | 0.1111 | 61 | 0.0984 | 13 | 0 | 10 | 0 | 11 | 0.2727 | 9 | 0.2222 | 43 | 0.1163 |
| 3_1155064 | 20 | 0.65 | 21 | 0.5714 | 17 | 0.5294 | 18 | 0.6111 | 76 | 0.5921 | 15 | 0.5333 | 14 | 0.5 | 13 | 0.6154 | 15 | 0.6 | 57 | 0.5614 |
| 3_1155064 | 17 | 0.1176 | 15 | 0.2 | 14 | 0 | 15 | 0.2667 | 61 | 0.1475 | 15 | 0.0667 | 10 | 0 | 13 | 0.2308 | 12 | 0.0833 | 50 | 0.1 |
| 3_1155065 | 19 | 0.8421 | 20 | 0.8 | 17 | 0.7059 | 17 | 0.7647 | 73 | 0.7808 | 13 | 0.8462 | 12 | 0.75 | 11 | 0.5455 | 12 | 0.75 | 48 | 0.7292 |
| 3_1155065 | 15 | 0.7333 | 18 | 0.7778 | 13 | 0.8462 | 15 | 0.7333 | 61 | 0.7705 | 13 | 0.6154 | 10 | 0.9 | 11 | 0.7273 | 10 | 0.7 | 44 | 0.7273 |
| 3_1155066 | 19 | 0.7368 | 23 | 0.6087 | 19 | 0.5263 | 16 | 0.8125 | 77 | 0.6623 | 16 | 0.4375 | 11 | 0.5455 | 10 | 0.6 | 13 | 0.6154 | 50 | 0.54 |
| 3_1155084 | 14 | 0.1429 | 15 | 0.1333 | 11 | 0.0909 | 11 | 0.2727 | 51 | 0.1569 | 11 | 0.3636 | 8 | 0.125 | 9 | 0.1111 | 7 | 0 | 35 | 0.1714 |
| 3_1156752 | 2 | 0.5 | NA | NA | 2 | 0 | 1 | 0 | 5 | 0.2 | 1 | 0 | 3 | 0 | 2 | 0 | 4 | 0.5 | 10 | 0.2 |
| 3_1156815 | 2 | 0.5 | NA | NA | 1 | 0 | 1 | 0 | 4 | 0.25 | 2 | 0 | 4 | 0 | 3 | 0.3333 | 3 | 0.3333 | 12 | 0.1667 |
| 3_1156815 | 1 | 1 | NA | NA | 1 | 0 | NA | NA | 2 | 0.5 | 1 | 0 | 2 | 1 | 2 | 0.5 | 3 | 1 | 8 | 0.75 |
| 3_1157223 | 2 | 1 | NA | NA | NA | NA | NA | NA | 2 | 1 | 2 | 1 | 1 | 1 | NA | NA | 1 | 1 | 4 | 1 |
| 3_1157223 | 2 | 1 | NA | NA | NA | NA | 1 | 1 | 3 | 1 | 1 | 1 | 4 | 1 | NA | NA | 1 | 1 | 6 | 1 |
| 3_1157226 | 2 | 1 | NA | NA | 1 | 1 | 2 | 1 | 5 | 1 | 3 | 0.6667 | 4 | 1 | NA | NA | 1 | 1 | 8 | 0.875 |
| 3_1157226 | 1 | 1 | NA | NA | NA | NA | 1 | 1 | 2 | 1 | 2 | 1 | 2 | 1 | NA | NA | 1 | 1 | 5 | 1 |
| 3_1157226 | 1 | 1 | NA | NA | 1 | 0 | 1 | 1 | 3 | 0.6667 | 3 | 0.6667 | 3 | 0.3333 | NA | NA | 1 | 0 | 7 | 0.4286 |
| 3_1157227 | 1 | 0 | NA | NA | 1 | 1 | 1 | 0 | 3 | 0.3333 | 3 | 0 | 2 | 0.5 | NA | NA | 1 | 0 | 6 | 0.1667 |
| 3_1157329 | 2 | 0.5 | 3 | 0.3333 | 1 | 1 | NA | NA | 6 | 0.5 | NA | NA | 1 | 0 | NA | NA | 3 | 0.3333 | 4 | 0.25 |
| 3_1157331 | NA | NA | 4 | 0.5 | 1 | 0 | NA | NA | 5 | 0.4 | NA | NA | 2 | 1 | NA | NA | 4 | 0.5 | 6 | 0.6667 |
| 3_1157332 | NA | NA | 2 | 0.5 | 1 | 0 | NA | NA | 3 | 0.3333 | NA | NA | 2 | 0.5 | NA | NA | 2 | 0 | 4 | 0.25 |
| 3_1157334 | NA | NA | 3 | 0.3333 | 1 | 0 | NA | NA | 4 | 0.25 | NA | NA | 2 | 0 | NA | NA | 4 | 0.25 | 6 | 0.1667 |
| 3_1157339 | NA | NA | 1 | 1 | 1 | 0 | NA | NA | 2 | 0.5 | NA | NA | 2 | 0.5 | NA | NA | 2 | 0.5 | 4 | 0.5 |
| 3_1157339 | NA | NA | 1 | 1 | 1 | 1 | NA | NA | 2 | 1 | NA | NA | 2 | 0.5 | NA | NA | 3 | 0.6667 | 5 | 0.6 |
| 3_1157363 | 1 | 0 | 1 | 1 | NA | NA | 2 | 0 | 4 | 0.25 | NA | NA | NA | NA | NA | NA | 2 | 1 | 2 | 1 |
| 3_1158013 | 2 | 0.5 | 1 | 1 | 4 | 0 | 4 | 0.5 | 11 | 0.3636 | 1 | 0 | NA | NA | 4 | 0.75 | 2 | 0.5 | 7 | 0.5714 |
| 3_1158178 | 6 | 0.3333 | 8 | 0.125 | 8 | 0 | 5 | 0.2 | 27 | 0.1481 | 1 | 1 | 7 | 0 | 5 | 0.2 | 4 | 0 | 17 | 0.1176 |
| 3_1158178 | 10 | 0.1 | 10 | 0.1 | 11 | 0.1818 | 9 | 0.2222 | 40 | 0.15 | 2 | 0.5 | 9 | 0.1111 | 10 | 0.1 | 8 | 0.375 | 29 | 0.2069 |
| 3_1158178 | 10 | 0.8 | 9 | 0.8889 | 8 | 1 | 6 | 1 | 33 | 0.9091 | 1 | 1 | 12 | 0.9167 | 9 | 1 | 6 | 0.8333 | 28 | 0.9286 |
| 3_1158179 | 8 | 0.375 | 8 | 0.375 | 9 | 0.4444 | 5 | 0.4 | 30 | 0.4 | 2 | 0 | 9 | 0.3333 | 8 | 0.25 | 7 | 0.4286 | 26 | 0.3077 |
| 3_1158208 | 9 | 0.4444 | 10 | 0.5 | 11 | 0.1818 | 8 | 0.5 | 38 | 0.3947 | 1 | 1 | 12 | 0.3333 | 11 | 0.3636 | 7 | 0.2857 | 31 | 0.3548 |
| 3_1158208 | 11 | 0.8182 | 12 | 0.8333 | 14 | 0.5714 | 8 | 0.75 | 45 | 0.7333 | 2 | 1 | 11 | 0.9091 | 12 | 0.5 | 6 | 0.8333 | 31 | 0.7419 |
| 3_1158211 | 10 | 0.2 | 10 | 0.1 | 13 | 0.0769 | 9 | 0.2222 | 42 | 0.1429 | 3 | 0.3333 | 15 | 0.4667 | 9 | 0.2222 | 8 | 0 | 35 | 0.2857 |
| 3_1158211 | 11 | 0.6364 | 11 | 0.3636 | 14 | 0.4286 | 10 | 0.6 | 46 | 0.5 | 4 | 0.25 | 15 | 0.4 | 12 | 0.5 | 7 | 0.5714 | 38 | 0.4474 |
| 3_1158212 | 11 | 1 | 12 | 1 | 12 | 1 | 9 | 1 | 44 | 1 | 4 | 1 | 16 | 1 | 12 | 1 | 7 | 1 | 39 | 1 |
| 3_1158212 | 12 | 0.25 | 13 | 0.0769 | 12 | 0 | 9 | 0 | 46 | 0.087 | 5 | 0 | 15 | 0.0667 | 16 | 0.125 | 8 | 0 | 44 | 0.0682 |
| 3_1158212 | 9 | 0.2222 | 14 | 0.1429 | 14 | 0.2857 | 11 | 0 | 48 | 0.1667 | 5 | 0 | 16 | 0.1875 | 15 | 0 | 10 | 0 | 46 | 0.0652 |
| 3_1158213 | 11 | 0 | 12 | 0 | 13 | 0.0769 | 11 | 0.1818 | 47 | 0.0638 | 5 | 0.2 | 15 | 0.1333 | 14 | 0 | 8 | 0 | 42 | 0.0714 |
| 3_1158213 | 10 | 1 | 15 | 1 | 12 | 1 | 10 | 1 | 47 | 1 | 3 | 1 | 15 | 1 | 13 | 1 | 6 | 1 | 37 | 1 |
| 3_1158214 | 10 | 0.3 | 11 | 0.3636 | 15 | 0.2 | 7 | 0.5714 | 43 | 0.3256 | 4 | 0.75 | 16 | 0.5 | 14 | 0.5 | 10 | 0.4 | 44 | 0.5 |
| 3_1158214 | 5 | 0.8 | 11 | 0.9091 | 9 | 0.8889 | 10 | 0.8 | 35 | 0.8571 | 5 | 1 | 13 | 0.9231 | 10 | 0.8 | 4 | 0.75 | 32 | 0.875 |
| 3_1158214 | 12 | 1 | 14 | 1 | 16 | 1 | 12 | 1 | 54 | 1 | 4 | 1 | 18 | 0.9444 | 14 | 1 | 6 | 1 | 42 | 0.9762 |
| 3_1158215 | 12 | 0.9167 | 14 | 1 | 15 | 1 | 12 | 1 | 53 | 0.9811 | 5 | 1 | 13 | 1 | 15 | 0.9333 | 9 | 1 | 42 | 0.9762 |
| 3_1158216 | 12 | 0.75 | 13 | 1 | 13 | 0.9231 | 8 | 1 | 46 | 0.913 | 4 | 1 | 16 | 0.9375 | 13 | 0.9231 | 8 | 0.875 | 41 | 0.9268 |
| 3_1158216 | 9 | 1 | 10 | 1 | 11 | 1 | 7 | 0.8571 | 37 | 0.973 | 4 | 1 | 11 | 0.8182 | 13 | 0.9231 | 5 | 0.8 | 33 | 0.8788 |
| 3_1158216 | 7 | 0 | 9 | 0.4444 | 8 | 0.125 | 9 | 0.4444 | 33 | 0.2727 | 4 | 0.25 | 11 | 0.2727 | 10 | 0.2 | 8 | 0.25 | 33 | 0.2424 |
| 3_1158217 | 12 | 0.0833 | 16 | 0.25 | 15 | 0.4 | 11 | 0.5455 | 54 | 0.3148 | 5 | 0.4 | 17 | 0.2353 | 15 | 0.2667 | 10 | 0.4 | 47 | 0.2979 |
| 3_1158217 | 10 | 0.3 | 14 | 0.1429 | 16 | 0.3125 | 11 | 0.2727 | 51 | 0.2549 | 4 | 0.25 | 16 | 0.1875 | 16 | 0.1875 | 9 | 0 | 45 | 0.1556 |
| 3_1158221 | 4 | 1 | 8 | 1 | 7 | 0.5714 | 9 | 1 | 28 | 0.8929 | 1 | 1 | 9 | 0.6667 | 8 | 1 | 5 | 0.8 | 23 | 0.8261 |
| 3_1158256 | NA | NA | 1 | 1 | NA | NA | NA | NA | 1 | 1 | NA | NA | NA | NA | NA | NA | 1 | 1 | 1 | 1 |
| 3_1158257 | 1 | 1 | 1 | 1 | NA | NA | NA | NA | 2 | 1 | NA | NA | NA | NA | NA | NA | 1 | 1 | 1 | 1 |
| 3_1159041 | 5 | 0 | 8 | 0.25 | 6 | 0.6667 | 4 | 0.5 | 23 | 0.3478 | 4 | 0.5 | 2 | 0 | 5 | 0.2 | 2 | 0 | 13 | 0.2308 |
| 3_1159042 | 4 | 0 | 8 | 0.25 | 6 | 0.1667 | 5 | 0.4 | 23 | 0.2174 | 4 | 0 | 2 | 0.5 | 5 | 0 | 1 | 0 | 12 | 0.0833 |
| 3_1159043 | 6 | 0.1667 | 8 | 0.25 | 6 | 0.6667 | 5 | 1 | 25 | 0.48 | 5 | 0.4 | 2 | 0.5 | 4 | 0.5 | 2 | 1 | 13 | 0.5385 |
| 3_1159044 | 6 | 0 | 8 | 0.125 | 5 | 0.2 | 4 | 0 | 23 | 0.087 | 3 | 0.3333 | 2 | 0 | 5 | 0 | 2 | 0 | 12 | 0.0833 |
| 3_1159044 | 5 | 0.4 | 7 | 0.2857 | 4 | 0.5 | 5 | 0.6 | 21 | 0.4286 | 4 | 0.5 | NA | NA | 3 | 0.3333 | 1 | 0 | 8 | 0.375 |
| 3_1159044 | 6 | 1 | 7 | 0.8571 | 5 | 1 | 5 | 0.8 | 23 | 0.913 | 4 | 1 | 2 | 0.5 | 5 | 0.8 | 1 | 1 | 12 | 0.8333 |
| 3_1159046 | 5 | 0 | 6 | 0.1667 | 5 | 0 | 4 | 0 | 20 | 0.05 | 4 | 0 | 2 | 0.5 | 5 | 0 | 2 | 0 | 13 | 0.0769 |
| 3_1159046 | 5 | 0 | 4 | 0.25 | 5 | 0 | 5 | 0.4 | 19 | 0.1579 | 4 | 0.25 | 2 | 0 | 5 | 0.2 | 2 | 0 | 13 | 0.1538 |
| 3_1159096 | 3 | 0 | 6 | 0 | 4 | 0.25 | 3 | 0 | 16 | 0.0625 | 5 | 0 | 1 | 1 | 2 | 0 | 2 | 0 | 10 | 0.1 |
| 3_1160435 | 110 | 0.0818 | 39 | 0.0769 | 82 | 0.0976 | 123 | 0.0569 | 354 | 0.0763 | 38 | 0.0263 | 127 | 0.0945 | 131 | 0.0534 | 102 | 0.1078 | 398 | 0.0779 |
| 3_1160435 | 102 | 0.0294 | 35 | 0.1143 | 70 | 0.0429 | 114 | 0.0614 | 321 | 0.053 | 32 | 0.0312 | 109 | 0.0826 | 122 | 0.0902 | 97 | 0.1237 | 360 | 0.0917 |
| 3_1160435 | 93 | 0.0323 | 36 | 0.1667 | 71 | 0.0704 | 104 | 0.0865 | 304 | 0.0757 | 30 | 0 | 101 | 0.0891 | 108 | 0.1296 | 91 | 0.0549 | 330 | 0.0848 |

| | | | | | | | | | | | | | | | | | | | | |
|---|---|---|---|---|---|---|---|---|---|---|---|---|---|---|---|---|---|---|---|---|
| 3_11604377 | 97 | 0.5876 | 36 | 0.6389 | 75 | 0.7333 | 99 | 0.6465 | 307 | 0.6482 | 29 | 0.6207 | 119 | 0.563 | 114 | 0.6404 | 97 | 0.6804 | 359 | 0.624 |
| 3_11604399 | 81 | 0.1111 | 26 | 0.0769 | 59 | 0.1017 | 96 | 0.0729 | 262 | 0.0916 | 27 | 0.0741 | 101 | 0.0792 | 100 | 0.07 | 85 | 0.0706 | 313 | 0.0735 |
| 3_11604411 | 83 | 0.2048 | 31 | 0.0968 | 63 | 0.2222 | 94 | 0.2553 | 271 | 0.214 | 23 | 0.2174 | 97 | 0.1753 | 100 | 0.15 | 83 | 0.2048 | 303 | 0.1782 |
| 3_11604457 | 62 | 0.2903 | 24 | 0.25 | 51 | 0.1961 | 75 | 0.16 | 212 | 0.217 | 25 | 0.28 | 88 | 0.1136 | 79 | 0.1899 | 70 | 0.2714 | 262 | 0.1947 |
| 3_11604474 | 71 | 0.7606 | 31 | 0.7742 | 54 | 0.8148 | 94 | 0.8085 | 250 | 0.792 | 24 | 0.75 | 94 | 0.8617 | 92 | 0.8261 | 83 | 0.9036 | 293 | 0.8532 |
| 3_11604488 | 68 | 0.0294 | 26 | 0.1154 | 53 | 0.0189 | 90 | 0.0667 | 237 | 0.0506 | 24 | 0 | 95 | 0.0737 | 91 | 0.0659 | 79 | 0.1013 | 289 | 0.0727 |
| 3_11604490 | 68 | 0.2353 | 24 | 0.2083 | 53 | 0.1887 | 88 | 0.2159 | 233 | 0.2146 | 25 | 0.16 | 91 | 0.2967 | 85 | 0.2235 | 73 | 0.2877 | 274 | 0.2591 |
| 3_11604491 | 66 | 1 | 26 | 1 | 54 | 1 | 87 | 1 | 233 | 1 | 22 | 1 | 93 | 0.9892 | 87 | 1 | 77 | 0.987 | 279 | 0.9928 |
| 3_11604492 | 64 | 0.75 | 22 | 0.7273 | 48 | 0.7708 | 78 | 0.6795 | 212 | 0.7264 | 22 | 0.7273 | 82 | 0.8537 | 80 | 0.775 | 67 | 0.791 | 251 | 0.8008 |
| 3_11604493 | 67 | 0.1343 | 23 | 0.1304 | 52 | 0.1154 | 84 | 0.1667 | 226 | 0.1416 | 24 | 0.1667 | 89 | 0.1348 | 85 | 0.1882 | 77 | 0.2078 | 275 | 0.1745 |
| 3_11604495 | 64 | 0.4844 | 26 | 0.3846 | 52 | 0.3462 | 86 | 0.4186 | 228 | 0.4167 | 23 | 0.4348 | 89 | 0.4494 | 82 | 0.4512 | 77 | 0.5974 | 271 | 0.4908 |
| 3_11604502 | 63 | 0.381 | 21 | 0.2381 | 40 | 0.325 | 67 | 0.403 | 191 | 0.3613 | 19 | 0.1579 | 77 | 0.2857 | 62 | 0.4032 | 64 | 0.3438 | 222 | 0.3243 |
| 3_11604506 | 55 | 0.9636 | 23 | 1 | 41 | 0.9756 | 70 | 0.9429 | 189 | 0.963 | 23 | 0.9565 | 75 | 0.9467 | 65 | 0.9692 | 66 | 0.9848 | 229 | 0.9651 |
| 3_11604590 | 53 | 0.3962 | 19 | 0.2105 | 30 | 0.4 | 58 | 0.3103 | 160 | 0.3438 | 15 | 0.2 | 56 | 0.4107 | 58 | 0.431 | 53 | 0.283 | 182 | 0.3626 |
| 3_11604599 | 43 | 1 | 21 | 0.9048 | 29 | 1 | 56 | 1 | 149 | 0.9866 | 12 | 0.9167 | 65 | 0.9692 | 68 | 0.9559 | 51 | 0.9804 | 196 | 0.9643 |
| 3_11604677 | 24 | 0.0417 | 11 | 0.0909 | 25 | 0.04 | 33 | 0.0606 | 93 | 0.0538 | 10 | 0.1 | 45 | 0.1111 | 31 | 0.0968 | 30 | 0.1667 | 116 | 0.1207 |
| 3_11606799 | NA | NA | 2 | 0 | NA | NA | 1 | 1 | 3 | 0.3333 | NA | NA | 1 | 1 | 1 | 1 | 1 | 0 | 3 | 0.6667 |
| 3_11606818 | NA | NA | 2 | 0.5 | NA | NA | 1 | 1 | 3 | 0.6667 | NA | NA | 1 | 1 | 1 | 1 | 1 | 1 | 3 | 1 |
| 3_11628270 | 12 | 0.25 | 12 | 0 | 10 | 0.1 | 15 | 0.0667 | 49 | 0.102 | 15 | 0 | 6 | 0 | 7 | 0.2857 | 9 | 0 | 37 | 0.0541 |
| 3_11628279 | 11 | 0.5455 | 13 | 0.3077 | 9 | 0.4444 | 14 | 0.4286 | 47 | 0.4255 | 15 | 0.5333 | 7 | 0.5714 | 8 | 0.5 | 8 | 0.625 | 38 | 0.5526 |
| 3_11628285 | 12 | 0 | 13 | 0.1538 | 9 | 0 | 16 | 0.0625 | 50 | 0.06 | 14 | 0.1429 | 8 | 0.25 | 8 | 0 | 7 | 0.1429 | 37 | 0.1351 |
| 3_11634186 | 5 | 0.4 | 3 | 0.6667 | 1 | 1 | 2 | 0 | 11 | 0.4545 | 3 | 0.3333 | 2 | 0.5 | 4 | 0.25 | 5 | 0.2 | 14 | 0.2857 |
| 3_11634187 | 5 | 0.6 | 3 | 1 | 1 | 1 | 2 | 1 | 11 | 0.8182 | 3 | 1 | 1 | 1 | 4 | 0.75 | 3 | 0.6667 | 11 | 0.8182 |
| 3_11634226 | 5 | 0 | 3 | 0 | 2 | 1 | 3 | 0.3333 | 13 | 0.2308 | 4 | 0.25 | 4 | 0.25 | 3 | 0 | 5 | 0 | 16 | 0.125 |
| 3_11637514 | 2 | 0.5 | 1 | 1 | NA | NA | 2 | 0 | 5 | 0.4 | NA | NA | NA | NA | 2 | 1 | 1 | 1 | 3 | 1 |
| 3_11637519 | 2 | 0.5 | 1 | 0 | NA | NA | 2 | 0 | 5 | 0.2 | NA | NA | NA | NA | 2 | 0.5 | NA | NA | 2 | 0.5 |
| 3_11637632 | 2 | 1 | 1 | 1 | NA | NA | 1 | 0 | 4 | 0.75 | NA | NA | NA | NA | 1 | 1 | 1 | 1 | 2 | 1 |
| 3_11637636 | 2 | 0.5 | 1 | 0 | NA | NA | 1 | 0 | 4 | 0.25 | NA | NA | NA | NA | 1 | 1 | 1 | 0 | 2 | 0.5 |
| 3_11637636 | 2 | 0.5 | 1 | 1 | NA | NA | 1 | 0 | 4 | 0.5 | NA | NA | NA | NA | 1 | 1 | 1 | 0 | 2 | 0.5 |
| 3_11642360 | NA | NA | NA | NA | NA | NA | 1 | 1 | 1 | 1 | NA | NA | 1 | 1 | NA | NA | NA | NA | 1 | 1 |
| 3_11642512 | 11 | 0.0909 | 9 | 0.1111 | 12 | 0.25 | 7 | 0.2857 | 39 | 0.1795 | 9 | 0.3333 | 11 | 0 | 9 | 0.1111 | 12 | 0.1667 | 41 | 0.1463 |
| 3_11642557 | 11 | 0.2727 | 9 | 0.4444 | 9 | 0.6667 | 4 | 0.75 | 33 | 0.4848 | 8 | 0.125 | 10 | 0.2 | 9 | 0.2222 | 10 | 0 | 37 | 0.1351 |
| 3_11642569 | 10 | 0.1 | 8 | 0.125 | 8 | 0 | 5 | 0.4 | 31 | 0.129 | 7 | 0 | 9 | 0 | 9 | 0 | 7 | 0.2857 | 32 | 0.0625 |
| 3_11642576 | 9 | 1 | 7 | 0.8571 | 10 | 0.8 | 5 | 0.8 | 31 | 0.871 | 5 | 1 | 8 | 0.75 | 8 | 1 | 8 | 1 | 29 | 0.931 |
| 3_11642579 | 8 | 0 | 8 | 0.25 | 9 | 0.4444 | 5 | 0.2 | 30 | 0.2333 | 4 | 0.5 | 9 | 0.1111 | 8 | 0.125 | 8 | 0.125 | 29 | 0.1724 |
| 3_11642582 | 9 | 0.8889 | 7 | 0.8571 | 8 | 0.875 | 5 | 1 | 29 | 0.8966 | 7 | 1 | 9 | 0.8889 | 8 | 1 | 9 | 1 | 33 | 0.9697 |
| 3_11642590 | 9 | 0.2222 | 8 | 0.25 | 9 | 0 | 5 | 0 | 31 | 0.129 | 7 | 0 | 9 | 0.1111 | 9 | 0 | 10 | 0.1 | 35 | 0.0571 |
| 3_11642593 | 9 | 0 | 6 | 0.1667 | 8 | 0.125 | 5 | 0.2 | 28 | 0.1071 | 6 | 0.1667 | 9 | 0 | 9 | 0.2222 | 9 | 0.1111 | 33 | 0.1212 |
| 3_11642595 | 9 | 0.1111 | 7 | 0.1429 | 5 | 0 | 5 | 0.4 | 26 | 0.1538 | 5 | 0 | 10 | 0.1 | 9 | 0.1111 | 9 | 0.1111 | 33 | 0.0909 |
| 3_11646524 | 3 | 0 | 2 | 0.5 | 3 | 0 | 3 | 0 | 11 | 0.0909 | NA | NA | 2 | 0.5 | NA | NA | 1 | 0 | 3 | 0.3333 |
| 3_11646531 | 3 | 0 | 2 | 0 | 3 | 0.3333 | 2 | 0.5 | 10 | 0.2 | NA | NA | 2 | 1 | NA | NA | 1 | 0 | 3 | 0.6667 |
| 3_11646702 | 2 | 1 | NA | NA | 1 | 0 | 3 | 0 | 6 | 0.3333 | 1 | 0 | 1 | 1 | NA | NA | 3 | 0 | 5 | 0.2 |
| 3_11646720 | 2 | 0.5 | NA | NA | 1 | 0 | 3 | 1 | 6 | 0.6667 | 1 | 0 | NA | NA | NA | NA | 3 | 0.3333 | 4 | 0.25 |
| 3_11648818 | 2 | 0 | NA | NA | 1 | 1 | 1 | 0 | 4 | 0.25 | NA | NA | NA | NA | NA | NA | 2 | 0.5 | 2 | 0.5 |
| 3_11653746 | 2 | 0 | NA | NA | 1 | 0 | 2 | 0.5 | 5 | 0.2 | 2 | 0 | 1 | 1 | NA | NA | 4 | 0 | 7 | 0.1429 |
| 3_11657443 | 2 | 0.5 | NA | NA | 1 | 0 | NA | NA | 3 | 0.3333 | 1 | 1 | NA | NA | NA | NA | 2 | 0 | 3 | 0.3333 |
| 3_11671201 | 6 | 0.1667 | 2 | 0 | 1 | 0 | 3 | 0.3333 | 12 | 0.1667 | NA | NA | 2 | 0 | 2 | 1 | 2 | 1 | 6 | 0.6667 |
| 3_11671205 | 6 | 0.1667 | 2 | 0 | 1 | 1 | 2 | 0 | 11 | 0.1818 | 1 | 0 | 1 | 1 | 1 | 0 | NA | NA | 3 | 0.3333 |
| 3_11671690 | 5 | 0 | 2 | 0.5 | 1 | 1 | 2 | 0.5 | 10 | 0.3 | 1 | 1 | 2 | 0 | 1 | 0 | 1 | 0 | 5 | 0.2 |
| 3_11671712 | 5 | 0.8 | 1 | 1 | 1 | 1 | 2 | 0.5 | 9 | 0.7778 | 1 | 1 | 1 | 1 | 1 | 0 | 1 | 1 | 4 | 0.75 |
| 3_11671739 | 3 | 0 | 1 | 1 | 1 | 0 | 2 | 0 | 7 | 0.1429 | 1 | 0 | 1 | 1 | 1 | 1 | 1 | 1 | 4 | 0.75 |
| 3_11671761 | 3 | 1 | 1 | 1 | 1 | 1 | 2 | 1 | 7 | 1 | 1 | 1 | 2 | 1 | 1 | 1 | 1 | 1 | 5 | 1 |
| 3_11671775 | 3 | 0.3333 | 1 | 1 | 1 | 0 | 2 | 0.5 | 7 | 0.4286 | 1 | 1 | 2 | 0 | 1 | 0 | NA | NA | 4 | 0.25 |
| 3_11671786 | 3 | 0.3333 | 1 | 1 | 1 | 1 | 2 | 0 | 7 | 0.4286 | 1 | 0 | 2 | 0.5 | 1 | 0 | 1 | 0 | 5 | 0.2 |
| 3_11711345 | 28 | 0.2143 | 33 | 0.1818 | 19 | 0.4737 | 22 | 0.3182 | 102 | 0.2745 | 16 | 0.4375 | 25 | 0.32 | 23 | 0.3043 | 26 | 0.4615 | 90 | 0.3778 |
| 3_11711359 | 31 | 0.3548 | 40 | 0.275 | 22 | 0.1364 | 25 | 0.16 | 118 | 0.2458 | 19 | 0.3684 | 29 | 0.3448 | 27 | 0.2963 | 29 | 0.3793 | 104 | 0.3462 |
| 3_11711364 | 31 | 0.0968 | 39 | 0.1282 | 22 | 0.0455 | 23 | 0 | 115 | 0.0783 | 20 | 0.1 | 30 | 0.1667 | 27 | 0.1111 | 31 | 0.2581 | 108 | 0.1667 |
| 3_11711434 | 23 | 0 | 23 | 0.087 | 18 | 0.0556 | 16 | 0.0625 | 80 | 0.05 | 16 | 0.0625 | 23 | 0.087 | 20 | 0.1 | 20 | 0.1 | 79 | 0.0886 |
| 3_11711472 | 16 | 0.625 | 22 | 0.4091 | 14 | 0.4286 | 16 | 0.3125 | 68 | 0.4412 | 12 | 0.5833 | 16 | 0.75 | 17 | 0.4706 | 20 | 0.5 | 65 | 0.5692 |
| 3_11711490 | 23 | 0.2609 | 25 | 0.08 | 18 | 0.1667 | 17 | 0.1765 | 83 | 0.1687 | 16 | 0.0625 | 22 | 0.0909 | 18 | 0.3889 | 25 | 0.24 | 81 | 0.1975 |
| 3_11711492 | 23 | 0.8696 | 26 | 0.7308 | 19 | 0.7895 | 16 | 0.75 | 84 | 0.7857 | 15 | 0.7333 | 23 | 0.8696 | 19 | 0.7895 | 24 | 0.8333 | 81 | 0.8148 |
| 3_11711499 | 15 | 0.4667 | 22 | 0.2727 | 9 | 0.4444 | 14 | 0.4286 | 60 | 0.3833 | 14 | 0.2143 | 21 | 0.3333 | 13 | 0.1538 | 22 | 0.5455 | 70 | 0.3429 |
| 3_11711500 | 20 | 0.15 | 19 | 0.4211 | 12 | 0.4167 | 18 | 0.3333 | 69 | 0.3188 | 11 | 0.2727 | 20 | 0.45 | 10 | 0.3 | 19 | 0.4737 | 60 | 0.4 |
| 3_11711504 | 15 | 0.4 | 20 | 0.1 | 10 | 0.5 | 17 | 0.3529 | 62 | 0.3065 | 13 | 0.2308 | 19 | 0.4737 | 14 | 0.4286 | 21 | 0.3333 | 67 | 0.3731 |
| 3_11711506 | 17 | 0.2941 | 24 | 0.3333 | 13 | 0.5385 | 14 | 0.5 | 68 | 0.3971 | 14 | 0.3571 | 20 | 0.35 | 14 | 0.2857 | 19 | 0.4737 | 67 | 0.3731 |

| | | | | | | | | | | | | | | | | | | | | |
|---|---|---|---|---|---|---|---|---|---|---|---|---|---|---|---|---|---|---|---|---|
| 3_1171151 | 11 | 0.0909 | 21 | 0.1429 | 13 | 0 | 14 | 0.1429 | 59 | 0.1017 | 11 | 0.3636 | 20 | 0.25 | 16 | 0.1875 | 15 | 0.2667 | 62 | 0.2581 |
| 3_1171152 | 17 | 0.0588 | 21 | 0.1429 | 15 | 0.2667 | 17 | 0.1176 | 70 | 0.1429 | 14 | 0.0714 | 19 | 0.2632 | 16 | 0.1875 | 19 | 0.2632 | 68 | 0.2059 |
| 3_1171152 | 13 | 0.8462 | 16 | 0.4375 | 10 | 0.6 | 10 | 0.6 | 49 | 0.6122 | 9 | 0.6667 | 16 | 0.5 | 14 | 0.5714 | 14 | 0.7857 | 53 | 0.6226 |
| 3_1171152 | 14 | 0.5714 | 14 | 0.5714 | 13 | 0.4615 | 13 | 0.5385 | 54 | 0.537 | 12 | 0.5833 | 15 | 0.6 | 15 | 0.5333 | 17 | 0.7647 | 59 | 0.6271 |
| 3_1171152 | 20 | 0.05 | 24 | 0.0833 | 15 | 0.0667 | 14 | 0 | 73 | 0.0548 | 15 | 0 | 21 | 0.0952 | 18 | 0 | 21 | 0.0952 | 75 | 0.0533 |
| 3_1171153 | 17 | 1 | 17 | 1 | 11 | 0.8182 | 11 | 1 | 56 | 0.9643 | 13 | 0.9231 | 17 | 0.9412 | 13 | 0.9231 | 18 | 1 | 61 | 0.9508 |
| 3_1171154 | 16 | 0.3125 | 17 | 0.0588 | 12 | 0.1667 | 15 | 0.0667 | 60 | 0.15 | 14 | 0.3571 | 19 | 0.0526 | 17 | 0.1176 | 19 | 0.2105 | 69 | 0.1739 |
| 3_1171155 | 7 | 0.4286 | 10 | 0.3 | 7 | 0.7143 | 7 | 0.2857 | 31 | 0.4194 | 9 | 0.3333 | 12 | 0.3333 | 11 | 0.3636 | 15 | 0.2667 | 47 | 0.3191 |
| 3_1171158 | 12 | 0.75 | 18 | 0.8333 | 12 | 0.8333 | 12 | 0.75 | 54 | 0.7963 | 10 | 0.9 | 18 | 0.6667 | 13 | 0.5385 | 13 | 0.7692 | 54 | 0.7037 |
| 3_1171158 | 14 | 0.8571 | 16 | 0.8125 | 13 | 1 | 13 | 0.9231 | 56 | 0.8929 | 12 | 0.8333 | 17 | 0.7647 | 16 | 0.9375 | 13 | 0.9231 | 58 | 0.8621 |
| 3_1171160 | 11 | 0.6364 | 14 | 0.3571 | 13 | 0.4615 | 10 | 0.3 | 48 | 0.4375 | 10 | 0.4 | 15 | 0.4667 | 11 | 0.4545 | 13 | 0.6154 | 49 | 0.4898 |
| 3_1171164 | 9 | 1 | 14 | 0.8571 | 10 | 1 | 10 | 0.8 | 43 | 0.907 | 9 | 1 | 15 | 0.9333 | 10 | 1 | 14 | 1 | 48 | 0.9792 |
| 3_1171168 | 8 | 0.25 | 12 | 0 | 12 | 0 | 6 | 0.1667 | 38 | 0.0789 | 6 | 0.1667 | 10 | 0.1 | 10 | 0.1 | 10 | 0.1 | 36 | 0.1111 |
| 3_1171539 | 2 | 1 | 1 | 1 | 2 | 0 | 2 | 0 | 7 | 0.4286 | 1 | 1 | 2 | 0 | 4 | 0.25 | 1 | 1 | 8 | 0.375 |
| 3_1173687 | 1 | 1 | 3 | 0.6667 | NA | NA | 1 | 1 | 5 | 0.8 | NA | NA | NA | NA | NA | NA | 1 | 1 | 1 | 1 |
| 3_1173690 | 1 | 1 | 3 | 0 | NA | NA | 5 | 0.4 | 9 | 0.3333 | NA | NA | 2 | 0.5 | 4 | 0.5 | 1 | 1 | 7 | 0.5714 |
| 3_1173691 | 1 | 1 | 3 | 0.3333 | 1 | 0 | 4 | 1 | 9 | 0.6667 | 1 | 0 | 2 | 0.5 | 3 | 0.3333 | 2 | 1 | 8 | 0.5 |
| 3_1173695 | 3 | 0.3333 | 5 | 0.2 | 1 | 0 | 5 | 0 | 14 | 0.1429 | 4 | 0 | 1 | 0 | 4 | 0.25 | 3 | 0.3333 | 12 | 0.1667 |
| 3_1174193 | 4 | 1 | 6 | 0.6667 | 1 | 1 | 6 | 0 | 17 | 0.5294 | 3 | 0 | 3 | 0 | 4 | 0.25 | 4 | 0.5 | 14 | 0.2143 |
| 3_1174804 | 6 | 0.5 | 5 | 0.4 | 3 | 0.6667 | 7 | 0.5714 | 21 | 0.5238 | 4 | 1 | 4 | 0.5 | 6 | 0.8333 | 4 | 0.5 | 18 | 0.7222 |
| 3_1174806 | 5 | 0 | 6 | 0.1667 | 3 | 0 | 6 | 0 | 20 | 0.05 | 3 | 0.3333 | 4 | 0 | 5 | 0 | 4 | 0.25 | 16 | 0.125 |
| 3_1174807 | 6 | 0.1667 | 7 | 0.2857 | 4 | 0 | 7 | 0 | 24 | 0.125 | 3 | 0.3333 | 4 | 0.25 | 4 | 0 | 4 | 0 | 15 | 0.1333 |
| 3_1174810 | 4 | 0 | 3 | 0 | 3 | 0 | 4 | 0.25 | 14 | 0.0714 | 3 | 0.3333 | 4 | 0 | 4 | 0 | 2 | 0 | 13 | 0.0769 |
| 3_1175759 | 5 | 0.4 | 4 | 0.25 | 3 | 0 | 1 | 0 | 13 | 0.2308 | 3 | 0.3333 | 3 | 0 | 5 | 0.2 | 3 | 0 | 14 | 0.1429 |
| 3_1195139 | 7 | 0.5714 | 16 | 0.75 | 11 | 0.4545 | 6 | 0.3333 | 40 | 0.575 | 7 | 0.5714 | 12 | 0.5 | 4 | 0.75 | 8 | 0.375 | 31 | 0.5161 |
| 3_1204442 | NA | NA | 2 | 1 | 1 | 1 | NA | NA | 3 | 1 | NA | NA | 3 | 0.6667 | NA | NA | NA | NA | 3 | 0.6667 |
| 3_1204445 | NA | NA | 2 | 0 | 1 | 0 | 1 | 1 | 4 | 0.25 | NA | NA | 3 | 0.3333 | NA | NA | NA | NA | 3 | 0.3333 |
| 3_1204798 | NA | NA | NA | NA | 2 | 0.5 | NA | NA | 2 | 0.5 | NA | NA | NA | NA | NA | NA | 1 | 1 | 1 | 1 |
| 3_1206310 | NA | NA | 1 | 1 | NA | NA | NA | NA | 1 | 1 | NA | NA | 1 | 1 | NA | NA | NA | NA | 1 | 1 |
| 3_1209496 | NA | NA | 1 | 1 | NA | NA | NA | NA | 1 | 1 | 1 | 1 | NA | NA | NA | NA | NA | NA | 1 | 1 |
| 3_1209498 | NA | NA | 1 | 1 | NA | NA | NA | NA | 1 | 1 | 1 | 1 | NA | NA | NA | NA | NA | NA | 1 | 1 |
| 3_1209525 | NA | NA | 1 | 1 | NA | NA | NA | NA | 1 | 1 | 1 | 1 | NA | NA | NA | NA | NA | NA | 1 | 1 |
| 3_1209966 | 8 | 0.125 | 1 | 0 | 5 | 0.2 | 3 | 0 | 17 | 0.1176 | 3 | 0 | 3 | 0.3333 | 7 | 0.2857 | 7 | 0.1429 | 20 | 0.2 |
| 3_1209972 | 7 | 0 | 1 | 0 | 4 | 0.25 | 3 | 0.3333 | 15 | 0.1333 | 2 | 0 | 3 | 0 | 7 | 0.1429 | 5 | 0.2 | 17 | 0.1176 |
| 3_1209973 | 8 | 0 | 1 | 1 | 4 | 0 | 3 | 0 | 16 | 0.0625 | 2 | 0 | 2 | 0 | 7 | 0.1429 | 5 | 0.2 | 16 | 0.125 |
| 3_1209995 | 3 | 0 | 1 | 1 | 2 | 0 | NA | NA | 6 | 0.1667 | NA | NA | 2 | 0 | 5 | 0 | 4 | 0.25 | 11 | 0.0909 |
| 3_1209997 | 3 | 0 | 1 | 0 | 2 | 0.5 | 1 | 1 | 7 | 0.2857 | NA | NA | 2 | 0.5 | 5 | 0 | 4 | 0 | 11 | 0.0909 |
| 3_1209999 | 3 | 0 | 1 | 0 | 2 | 1 | 1 | 1 | 7 | 0.4286 | NA | NA | 2 | 0 | 5 | 0 | 4 | 0.25 | 11 | 0.0909 |
| 3_1209999 | 2 | 0.5 | NA | NA | 2 | 1 | 1 | 0 | 5 | 0.6 | NA | NA | 2 | 0.5 | 5 | 0 | 4 | 0 | 11 | 0.0909 |
| 3_1210000 | 2 | 1 | 1 | 0 | NA | NA | 1 | 1 | 4 | 0.75 | NA | NA | 1 | 1 | 3 | 0 | 4 | 1 | 8 | 0.625 |
| 3_1210001 | NA | NA | 1 | 1 | 2 | 0.5 | 1 | 0 | 4 | 0.5 | NA | NA | 2 | 0.5 | 4 | 0.25 | 4 | 0 | 10 | 0.2 |
| 3_1210003 | 3 | 1 | NA | NA | 2 | 1 | 1 | 1 | 6 | 1 | NA | NA | 2 | 1 | 3 | 0.6667 | 3 | 0.6667 | 8 | 0.75 |
| 3_1210004 | 2 | 1 | NA | NA | 2 | 1 | NA | NA | 4 | 1 | NA | NA | 1 | 1 | 5 | 0.8 | 4 | 1 | 10 | 0.9 |
| 3_1210005 | 3 | 0.3333 | NA | NA | 1 | 1 | 1 | 0 | 5 | 0.4 | NA | NA | 2 | 1 | 4 | 0.25 | 4 | 0.25 | 10 | 0.4 |
| 3_1210006 | 2 | 0 | NA | NA | 2 | 0.5 | 2 | 0 | 6 | 0.1667 | NA | NA | 2 | 0.5 | 5 | 0 | 3 | 0 | 10 | 0.1 |
| 3_1210007 | 3 | 0.6667 | NA | NA | 2 | 1 | 2 | 1 | 7 | 0.8571 | NA | NA | 2 | 1 | 5 | 1 | 3 | 0.6667 | 10 | 0.9 |
| 3_1210009 | 3 | 1 | NA | NA | 1 | 0 | 2 | 0 | 6 | 0.5 | NA | NA | 2 | 0.5 | 5 | 0.4 | 2 | 0.5 | 9 | 0.4444 |
| 3_1210009 | 2 | 1 | NA | NA | 1 | 1 | 1 | 1 | 4 | 1 | NA | NA | 2 | 0 | 4 | 1 | 3 | 1 | 9 | 0.7778 |
| 3_1211554 | 4 | 0.5 | NA | NA | 2 | 0 | 2 | 1 | 8 | 0.5 | 2 | 1 | 1 | 1 | NA | NA | 1 | 1 | 4 | 1 |
| 3_1212513 | 8 | 0.125 | 16 | 0.25 | 12 | 0.3333 | 6 | 0.1667 | 42 | 0.2381 | 12 | 0.0833 | 6 | 0.3333 | 11 | 0.1818 | 4 | 0.25 | 33 | 0.1818 |
| 3_1214303 | NA | NA | NA | NA | 1 | 1 | NA | NA | 1 | 1 | NA | NA | 1 | 1 | 1 | 0 | NA | NA | 2 | 0.5 |
| 3_1215278 | 14 | 0 | 6 | 0 | 8 | 0.25 | 12 | 0 | 40 | 0.05 | 10 | 0.3 | 5 | 0 | 7 | 0 | 6 | 0 | 28 | 0.1071 |
| 3_1215279 | 10 | 0.4 | 5 | 0.4 | 10 | 0.5 | 12 | 0.4167 | 37 | 0.4324 | 10 | 0.5 | 6 | 0.5 | 10 | 0.4 | 6 | 0.6667 | 32 | 0.5 |
| 3_1215280 | 13 | 0.6923 | 6 | 0.5 | 11 | 0.2727 | 12 | 0.8333 | 42 | 0.5952 | 10 | 0.6 | 5 | 0.6 | 9 | 0.6667 | 7 | 0.2857 | 31 | 0.5484 |
| 3_1215280 | 13 | 0.0769 | 6 | 0.1667 | 11 | 0.0909 | 12 | 0.3333 | 42 | 0.1667 | 9 | 0.1111 | 6 | 0.1667 | 10 | 0.3 | 6 | 0.1667 | 31 | 0.1935 |
| 3_1215281 | 14 | 0.2143 | 6 | 0.3333 | 10 | 0.4 | 12 | 0.1667 | 42 | 0.2619 | 8 | 0.25 | 5 | 0.4 | 10 | 0.3 | 4 | 0.25 | 27 | 0.2963 |
| 3_1215282 | 15 | 0.1333 | 5 | 0.2 | 7 | 0.1429 | 12 | 0.1667 | 39 | 0.1538 | 9 | 0.2222 | 5 | 0.2 | 10 | 0.2 | 7 | 0.1429 | 31 | 0.1935 |
| 3_1215528 | 7 | 0 | 10 | 0.1 | 14 | 0 | 10 | 0.2 | 41 | 0.0732 | 7 | 0 | 7 | 0.1429 | 7 | 0.1429 | 10 | 0 | 31 | 0.0645 |
| 3_1215528 | 8 | 0.25 | 10 | 0.1 | 13 | 0.3077 | 10 | 0.1 | 41 | 0.1951 | 7 | 0 | 6 | 0.1667 | 8 | 0.125 | 12 | 0.5833 | 33 | 0.2727 |
| 3_1215530 | 6 | 0.1667 | 8 | 0 | 12 | 0.3333 | 8 | 0.375 | 34 | 0.2353 | 8 | 0.125 | 6 | 0.1667 | 6 | 0 | 8 | 0.125 | 28 | 0.1071 |
| 3_1218078 | 1 | 0 | 2 | 1 | 2 | 0.5 | 1 | 0 | 6 | 0.5 | NA | NA | 1 | 0 | 5 | 0 | 4 | 0.5 | 10 | 0.2 |
| 3_1218078 | 1 | 1 | 1 | 0 | 2 | 0.5 | 1 | 0 | 5 | 0.4 | NA | NA | 1 | 0 | 4 | 0 | 3 | 0.3333 | 8 | 0.125 |
| 3_1218079 | 1 | 0 | 2 | 0.5 | 3 | 0.6667 | 1 | 0 | 7 | 0.4286 | NA | NA | NA | NA | 5 | 0.8 | 4 | 0.75 | 9 | 0.7778 |
| 3_1218080 | 1 | 1 | 3 | 1 | 2 | 1 | 1 | 1 | 7 | 1 | NA | NA | 1 | 1 | 5 | 0.8 | 3 | 1 | 9 | 0.8889 |
| 3_1218080 | 1 | 0 | 3 | 1 | 3 | 0.3333 | 1 | 1 | 8 | 0.625 | NA | NA | 1 | 1 | 5 | 0.6 | 4 | 0.5 | 10 | 0.6 |

| | | | | | | | | | | | | | | | | | | | | |
|---|---|---|---|---|---|---|---|---|---|---|---|---|---|---|---|---|---|---|---|---|
| 3_12180809 | 1 | 0 | 3 | 1 | 3 | 0.6667 | 1 | 1 | 8 | 0.75 | NA | NA | 1 | 1 | 5 | 1 | 4 | 0.5 | 10 | 0.8 |
| 3_12180811 | 1 | 1 | 2 | 1 | 1 | 0 | 1 | 1 | 5 | 0.8 | NA | NA | NA | NA | 5 | 0.8 | 3 | 1 | 8 | 0.875 |
| 3_12180813 | 1 | 1 | 2 | 0 | 2 | 0.5 | 2 | 0 | 7 | 0.2857 | NA | NA | 1 | 0 | 4 | 0.75 | 3 | 0.3333 | 8 | 0.5 |
| 3_12180820 | 1 | 1 | 3 | 0.3333 | 2 | 0.5 | 2 | 1 | 8 | 0.625 | NA | NA | 1 | 0 | 5 | 0.6 | 3 | 1 | 9 | 0.6667 |
| 3_12180830 | 1 | 0 | 3 | 0.6667 | 2 | 0 | 2 | 0.5 | 8 | 0.375 | NA | NA | 1 | 1 | 4 | 0.5 | 3 | 1 | 8 | 0.75 |
| 3_12180832 | 1 | 1 | 2 | 0.5 | 3 | 1 | 2 | 1 | 8 | 0.875 | NA | NA | 1 | 0 | 4 | 1 | 4 | 0.5 | 9 | 0.6667 |
| 3_12180837 | 1 | 1 | 3 | 1 | 3 | 1 | 2 | 1 | 9 | 1 | NA | NA | 1 | 1 | 5 | 1 | 4 | 1 | 10 | 1 |
| 3_12180844 | 1 | 0 | 3 | 0.3333 | 4 | 0.25 | 3 | 0.6667 | 11 | 0.3636 | NA | NA | 1 | 0 | 5 | 0.2 | 3 | 0.6667 | 9 | 0.3333 |
| 3_12180845 | 1 | 0 | 3 | 0 | 3 | 0 | 1 | 1 | 8 | 0.125 | NA | NA | 1 | 0 | 3 | 0 | 3 | 0.3333 | 7 | 0.1429 |
| 3_12180846 | NA | NA | 2 | 0.5 | 2 | 1 | 2 | 0.5 | 6 | 0.6667 | NA | NA | 1 | 1 | 4 | 0.75 | 3 | 1 | 8 | 0.875 |
| 3_12180849 | 1 | 0 | 3 | 0.3333 | 4 | 0 | 3 | 0.3333 | 11 | 0.1818 | NA | NA | 1 | 0 | 4 | 0 | 4 | 0.25 | 9 | 0.1111 |
| 3_12180854 | 1 | 1 | 3 | 0.6667 | 3 | 0.6667 | 3 | 0.6667 | 10 | 0.7 | NA | NA | 1 | 0 | 5 | 0.8 | 4 | 0.75 | 10 | 0.7 |
| 3_12206009 | 8 | 0.25 | 12 | 0.4167 | 10 | 0.4 | 8 | 0.625 | 38 | 0.4211 | 7 | 0.2857 | 5 | 0.2 | 6 | 0.1667 | 5 | 0.2 | 23 | 0.2174 |
| 3_12206011 | 6 | 0 | 13 | 0.0769 | 12 | 0.0833 | 6 | 0.1667 | 37 | 0.0811 | 7 | 0 | 6 | 0.1667 | 4 | 0 | 3 | 0 | 20 | 0.05 |
| 3_12206077 | 2 | 0 | 11 | 0.2727 | 10 | 0 | 6 | 0.1667 | 29 | 0.1379 | 4 | 0 | 5 | 0.2 | 2 | 0 | 4 | 0.25 | 15 | 0.1333 |
| 3_12206081 | 1 | 0 | 8 | 0.5 | 8 | 0.875 | 4 | 0.5 | 21 | 0.619 | 4 | 0.25 | 2 | 0.5 | 1 | 1 | 3 | 0.6667 | 10 | 0.5 |
| 3_12206085 | 2 | 0 | 9 | 0.3333 | 10 | 0 | 6 | 0.1667 | 27 | 0.1481 | 4 | 0 | 5 | 0.2 | 3 | 0 | 3 | 0 | 15 | 0.0667 |
| 3_12206122 | 2 | 0 | 12 | 0 | 9 | 0.1111 | 6 | 0.1667 | 29 | 0.069 | 4 | 0.25 | 5 | 0 | 3 | 0 | 4 | 0.25 | 16 | 0.125 |
| 3_12206273 | 2 | 0 | 13 | 0.1538 | 7 | 0 | 7 | 0 | 29 | 0.069 | 3 | 0 | 5 | 0 | 2 | 0 | 2 | 0.5 | 12 | 0.0833 |
| 3_12206905 | 1 | 0 | 1 | 0 | 2 | 0.5 | 2 | 0 | 6 | 0.1667 | NA | NA | 5 | 0.2 | 2 | 0.5 | 1 | 0 | 8 | 0.25 |
| 3_12206929 | 1 | 1 | 1 | 1 | 3 | 0.6667 | 2 | 1 | 7 | 0.8571 | NA | NA | 3 | 1 | 2 | 1 | 1 | 0 | 6 | 0.8333 |
| 3_12206946 | 1 | 1 | 2 | 1 | 2 | 1 | 2 | 0.5 | 7 | 0.8571 | 1 | 1 | 5 | 1 | 2 | 1 | 2 | 1 | 10 | 1 |
| 3_12206955 | NA | NA | 2 | 1 | 2 | 1 | 2 | 1 | 6 | 1 | 1 | 1 | 2 | 0.5 | 2 | 1 | 2 | 1 | 7 | 0.8571 |
| 3_12206959 | 1 | 0 | 2 | 0 | 3 | 0.6667 | 2 | 0.5 | 8 | 0.375 | 1 | 1 | 4 | 0 | 2 | 0.5 | 2 | 0 | 9 | 0.2222 |
| 3_12206959 | 1 | 0 | 2 | 0 | 3 | 0.3333 | 2 | 0 | 8 | 0.125 | 1 | 0 | 5 | 0.2 | 2 | 0 | 1 | 0 | 9 | 0.1111 |
| 3_12206994 | 1 | 0 | 2 | 0.5 | 2 | 0 | 1 | 0 | 6 | 0.1667 | 1 | 0 | 3 | 0.3333 | 1 | 0 | 2 | 0 | 7 | 0.1429 |
| 3_12207007 | 1 | 0 | 2 | 0.5 | 2 | 0 | 2 | 0 | 7 | 0.1429 | 1 | 0 | 5 | 0.8 | 2 | 0 | 2 | 1 | 10 | 0.6 |
| 3_12207010 | 1 | 0 | 1 | 0 | 1 | 1 | 1 | 0 | 4 | 0.25 | 1 | 1 | 4 | 1 | 1 | 0 | 2 | 1 | 8 | 0.875 |
| 3_12207017 | 1 | 1 | 2 | 0 | 3 | 1 | 2 | 1 | 8 | 0.75 | NA | NA | 5 | 0.8 | 2 | 0.5 | 2 | 1 | 9 | 0.7778 |
| 3_12207022 | 1 | 1 | 2 | 0 | 3 | 0 | 1 | 0 | 7 | 0.1429 | 1 | 0 | 5 | 0.4 | 2 | 0 | 2 | 0.5 | 10 | 0.3 |
| 3_12207029 | NA | NA | 2 | 0.5 | 3 | 0.3333 | 2 | 0.5 | 7 | 0.4286 | 1 | 0 | 4 | 0.5 | 1 | 0 | 2 | 0.5 | 8 | 0.375 |
| 3_12230625 | 2 | 0.5 | NA | NA | NA | NA | 1 | 1 | 3 | 0.6667 | 2 | 0.5 | NA | NA | NA | NA | NA | NA | 2 | 0.5 |
| 3_12230637 | 2 | 1 | NA | NA | NA | NA | 1 | 1 | 3 | 1 | 1 | 1 | NA | NA | NA | NA | NA | NA | 1 | 1 |
| 3_12230652 | 2 | 1 | NA | NA | NA | NA | 1 | 0 | 3 | 0.6667 | 2 | 0.5 | NA | NA | NA | NA | NA | NA | 2 | 0.5 |
| 3_12230659 | 2 | 0.5 | NA | NA | NA | NA | 1 | 0 | 3 | 0.3333 | 3 | 0.3333 | NA | NA | NA | NA | NA | NA | 3 | 0.3333 |
| 3_12233256 | 10 | 0.5 | 7 | 0.4286 | 4 | 0.25 | 7 | 0.4286 | 28 | 0.4286 | 5 | 0.6 | 7 | 0.8571 | 8 | 0.125 | 7 | 0.5714 | 27 | 0.5185 |
| 3_12233268 | 7 | 0.5714 | 8 | 0.125 | 4 | 0 | 3 | 0 | 22 | 0.2273 | 5 | 0.4 | 4 | 0.75 | 6 | 0 | 6 | 0.1667 | 21 | 0.2857 |
| 3_12233275 | 8 | 0.375 | 8 | 0.75 | 5 | 0.4 | 7 | 0.4286 | 28 | 0.5 | 5 | 0.4 | 6 | 0.3333 | 7 | 0.2857 | 8 | 0.375 | 26 | 0.3462 |
| 3_12233338 | 6 | 0 | 3 | 0 | 1 | 1 | 1 | 0 | 11 | 0.0909 | 4 | 0.25 | 2 | 0 | 1 | 0 | 5 | 0 | 12 | 0.0833 |
| 3_12233349 | 4 | 0.75 | 3 | 1 | 1 | 1 | 1 | 1 | 9 | 0.8889 | 3 | 1 | 2 | 1 | 2 | 1 | 4 | 1 | 11 | 1 |
| 3_12233354 | 5 | 0.6 | 1 | 0 | NA | NA | 2 | 0.5 | 8 | 0.5 | 3 | 0 | 2 | 0.5 | 2 | 1 | 3 | 0.6667 | 10 | 0.5 |
| 3_12233364 | 5 | 0.6 | 2 | 0 | 1 | 1 | 1 | 0 | 9 | 0.4444 | 2 | 0.5 | NA | NA | 2 | 0.5 | 2 | 0.5 | 6 | 0.5 |
| 3_12233377 | 4 | 0.75 | 2 | 1 | 1 | 1 | 1 | 1 | 8 | 0.875 | 1 | 1 | 1 | 1 | 1 | 1 | 2 | 0.5 | 5 | 0.8 |
| 3_12233561 | 2 | 0.5 | 1 | 0 | 1 | 0 | 1 | 0 | 5 | 0.2 | 1 | 0 | NA | NA | NA | NA | 1 | 1 | 2 | 0.5 |
| 3_12271918 | 23 | 0.3043 | 32 | 0.2812 | 17 | 0.2353 | 14 | 0.2857 | 86 | 0.2791 | 24 | 0.2083 | 21 | 0.2857 | 20 | 0.35 | 16 | 0.25 | 81 | 0.2716 |
| 3_12271920 | 23 | 0.0435 | 34 | 0.0882 | 19 | 0 | 14 | 0.1429 | 90 | 0.0667 | 22 | 0.0455 | 21 | 0.0952 | 23 | 0.0435 | 14 | 0.0714 | 80 | 0.0625 |
| 3_12271961 | 3 | 0 | 7 | 0.1429 | 1 | 0 | 3 | 0 | 14 | 0.0714 | 4 | 0 | 8 | 0.125 | 4 | 0.25 | 3 | 0.3333 | 19 | 0.1579 |
| 3_12271966 | 3 | 0.3333 | 8 | 0.125 | 1 | 0 | 3 | 0 | 15 | 0.1333 | 4 | 0 | 5 | 0 | 5 | 0.2 | 4 | 0 | 18 | 0.0556 |
| 3_12272542 | 1 | 0 | 2 | 0.5 | 3 | 0 | 1 | 1 | 7 | 0.2857 | 1 | 1 | 1 | 1 | 1 | 1 | 1 | 1 | 4 | 1 |
| 3_12272569 | 1 | 0 | 3 | 0.3333 | 7 | 0.4286 | 5 | 0.6 | 16 | 0.4375 | 3 | 0.3333 | 1 | 0 | 1 | 1 | 1 | 0 | 6 | 0.3333 |
| 3_12272723 | 3 | 0.3333 | 1 | 1 | 3 | 0.6667 | 3 | 0.6667 | 10 | 0.6 | 2 | 0.5 | 1 | 1 | 2 | 0.5 | NA | NA | 5 | 0.6 |
| 3_12272727 | 3 | 0 | 1 | 0 | 4 | 0.25 | 3 | 0.3333 | 11 | 0.1818 | 2 | 1 | 1 | 0 | 2 | 0 | NA | NA | 5 | 0.4 |
| 3_12272731 | 3 | 0 | 1 | 1 | 3 | 0 | 3 | 0.3333 | 10 | 0.2 | 2 | 0 | 1 | 1 | 2 | 0.5 | NA | NA | 5 | 0.4 |
| 3_12272733 | 3 | 0.3333 | 1 | 0 | 3 | 0.3333 | 3 | 0 | 10 | 0.2 | 2 | 1 | 1 | 0 | 2 | 0 | NA | NA | 5 | 0.4 |
| 3_12272740 | 2 | 0.5 | NA | NA | 2 | 1 | 3 | 0.3333 | 7 | 0.5714 | 1 | 1 | 1 | 0 | 1 | 1 | NA | NA | 3 | 0.6667 |
| 3_12272745 | 2 | 0.5 | NA | NA | 4 | 0.75 | 3 | 0.6667 | 9 | 0.6667 | 2 | 0.5 | 1 | 1 | 2 | 1 | NA | NA | 5 | 0.8 |
| 3_12272749 | 2 | 1 | NA | NA | 4 | 0.75 | 2 | 0.5 | 8 | 0.75 | 2 | 1 | 1 | 0 | 2 | 1 | NA | NA | 5 | 0.8 |
| 3_12272752 | 1 | 0 | 1 | 0 | 3 | 0.3333 | 3 | 0 | 8 | 0.125 | 1 | 1 | NA | NA | 1 | 0 | NA | NA | 2 | 0.5 |
| 3_12272753 | 2 | 0 | 1 | 1 | 2 | 0 | 3 | 0.6667 | 8 | 0.375 | 2 | 0 | 1 | 1 | 2 | 0 | NA | NA | 5 | 0.2 |
| 3_12272755 | 1 | 0 | 1 | 0 | 4 | 0.5 | 3 | 1 | 9 | 0.5556 | 2 | 1 | 1 | 0 | 2 | 1 | NA | NA | 5 | 0.8 |
| 3_12272768 | 2 | 0 | 1 | 0 | 4 | 0.25 | 3 | 0.3333 | 10 | 0.2 | 2 | 0.5 | 1 | 0 | 2 | 0.5 | NA | NA | 5 | 0.4 |
| 3_12272773 | 2 | 0 | 1 | 1 | 2 | 0 | 3 | 0.6667 | 8 | 0.375 | 2 | 0.5 | 1 | 0 | 1 | 0 | NA | NA | 4 | 0.25 |
| 3_12272776 | 2 | 1 | 1 | 1 | 4 | 1 | 3 | 1 | 10 | 1 | 2 | 1 | 1 | 1 | 1 | 1 | NA | NA | 4 | 1 |
| 3_12272784 | 2 | 0 | 1 | 0 | 4 | 0.75 | 3 | 0 | 10 | 0.3 | 1 | 1 | 1 | 0 | 1 | 0 | NA | NA | 3 | 0.3333 |
| 3_12272789 | 1 | 1 | 1 | 1 | 4 | 0.75 | 2 | 1 | 8 | 0.875 | 2 | 0 | 1 | 1 | 2 | 1 | NA | NA | 5 | 0.6 |

| | | | | | | | | | | | | | | | | | | | | |
|---|---|---|---|---|---|---|---|---|---|---|---|---|---|---|---|---|---|---|---|---|
| 3_12272792 | 2 | 0.5 | 1 | 1 | 3 | 0.3333 | 2 | 0 | 8 | 0.375 | 2 | 0.5 | 1 | 1 | 2 | 0.5 | NA | NA | 5 | 0.6 |
| 3_12272796 | 2 | 1 | 1 | 1 | 4 | 1 | 3 | 0.6667 | 10 | 0.9 | 2 | 1 | 1 | 1 | 2 | 0.5 | NA | NA | 5 | 0.8 |
| 3_12272797 | 1 | 1 | 1 | 0 | 3 | 0.3333 | 2 | 0 | 7 | 0.2857 | 1 | 0 | 1 | 1 | 2 | 0 | NA | NA | 4 | 0.25 |
| 3_12272799 | 2 | 1 | 1 | 1 | 3 | 0.6667 | 2 | 1 | 8 | 0.875 | 1 | 0 | NA | NA | 1 | 1 | NA | NA | 2 | 0.5 |
| 3_12272809 | 2 | 0 | NA | NA | 2 | 0.5 | 2 | 0 | 6 | 0.1667 | NA | NA | 1 | 0 | 2 | 0.5 | NA | NA | 3 | 0.3333 |
| 3_12272810 | 1 | 1 | NA | NA | 4 | 0.25 | 2 | 0.5 | 7 | 0.4286 | 2 | 1 | 1 | 1 | 2 | 0.5 | NA | NA | 5 | 0.8 |
| 3_12272815 | 1 | 1 | NA | NA | 3 | 0.6667 | 2 | 0.5 | 6 | 0.6667 | 2 | 0.5 | 1 | 1 | 2 | 1 | NA | NA | 5 | 0.8 |
| 3_12274283 | NA | NA | NA | NA | 2 | 0 | 2 | 0.5 | 4 | 0.25 | 2 | 1 | 1 | 0 | 1 | 1 | NA | NA | 4 | 0.75 |
| 3_12274288 | NA | NA | NA | NA | 3 | 0.6667 | 3 | 0.6667 | 6 | 0.6667 | 2 | 0.5 | NA | NA | 1 | 0 | NA | NA | 3 | 0.3333 |
| 3_12274305 | NA | NA | NA | NA | 2 | 1 | 2 | 0.5 | 4 | 0.75 | 2 | 0.5 | 1 | 1 | NA | NA | NA | NA | 3 | 0.6667 |
| 3_12274323 | NA | NA | NA | NA | 2 | 0.5 | 1 | 0 | 3 | 0.3333 | 2 | 0.5 | 1 | 0 | NA | NA | NA | NA | 3 | 0.3333 |
| 3_12274339 | NA | NA | NA | NA | 2 | 0 | 2 | 1 | 4 | 0.5 | 2 | 0 | 1 | 1 | NA | NA | NA | NA | 3 | 0.3333 |
| 3_12276248 | NA | NA | NA | NA | 1 | 1 | 2 | 0.5 | 3 | 0.6667 | 1 | 1 | 1 | 0 | NA | NA | NA | NA | 2 | 0.5 |
| 3_12276251 | NA | NA | NA | NA | 1 | 0 | 2 | 0.5 | 3 | 0.3333 | 1 | 1 | 1 | 0 | NA | NA | NA | NA | 2 | 0.5 |
| 3_12290473 | NA | NA | NA | NA | 1 | 1 | NA | NA | 1 | 1 | NA | NA | NA | NA | 1 | 1 | NA | NA | 1 | 1 |
| 3_12290474 | NA | NA | NA | NA | 1 | 1 | NA | NA | 1 | 1 | NA | NA | NA | NA | 1 | 1 | NA | NA | 1 | 1 |
| 3_12290479 | NA | NA | NA | NA | 1 | 1 | NA | NA | 1 | 1 | NA | NA | NA | NA | 1 | 1 | NA | NA | 1 | 1 |
| 3_12290528 | NA | NA | NA | NA | 1 | 1 | NA | NA | 1 | 1 | NA | NA | NA | NA | 1 | 1 | NA | NA | 1 | 1 |
| 3_12290792 | NA | NA | NA | NA | 1 | 1 | NA | NA | 1 | 1 | NA | NA | NA | NA | 1 | 1 | NA | NA | 1 | 1 |
| 3_12316214 | 2 | 0 | 2 | 0 | 4 | 0.25 | 2 | 0 | 10 | 0.1 | 1 | 0 | 3 | 0.3333 | 2 | 0 | NA | NA | 6 | 0.1667 |
| 3_12316289 | 1 | 0 | 1 | 0 | 2 | 0.5 | NA | NA | 4 | 0.25 | 1 | 0 | 2 | 1 | 1 | 0 | 1 | 1 | 5 | 0.6 |
| 3_12316291 | 1 | 1 | 1 | 0 | 2 | 1 | NA | NA | 4 | 0.75 | 1 | 0 | 2 | 0.5 | 1 | 1 | 1 | 1 | 5 | 0.6 |
| 3_12316318 | 1 | 0 | 1 | 1 | 1 | 0 | NA | NA | 3 | 0.3333 | 1 | 0 | 1 | 0 | 1 | 1 | 1 | 0 | 4 | 0.25 |
| 3_12329994 | 1 | 1 | 2 | 0.5 | 1 | 1 | 2 | 1 | 6 | 0.8333 | 4 | 1 | 1 | 1 | 1 | 0 | NA | NA | 6 | 0.8333 |
| 3_12330000 | 1 | 0 | 2 | 0.5 | 1 | 1 | 1 | 0 | 5 | 0.4 | 5 | 0 | 1 | 0 | 1 | 1 | NA | NA | 7 | 0.1429 |
| 3_12332126 | 1 | 1 | 3 | 0.3333 | 2 | 0 | NA | NA | 6 | 0.3333 | 1 | 0 | NA | NA | 1 | 1 | 1 | 0 | 3 | 0.3333 |
| 3_12332197 | 1 | 1 | 2 | 0 | 2 | 1 | NA | NA | 5 | 0.6 | 1 | 1 | NA | NA | 2 | 1 | 1 | 0 | 4 | 0.75 |
| 3_12346504 | NA | NA | 2 | 1 | 1 | 0 | NA | NA | 3 | 0.6667 | 1 | 1 | NA | NA | NA | NA | NA | NA | 1 | 1 |
| 3_12346506 | NA | NA | 1 | 1 | 1 | 0 | NA | NA | 2 | 0.5 | 1 | 0 | NA | NA | 1 | 1 | NA | NA | 2 | 0.5 |
| 3_12346514 | NA | NA | 2 | 1 | 1 | 0 | NA | NA | 3 | 0.6667 | 1 | 0 | NA | NA | 1 | 1 | NA | NA | 2 | 0.5 |
| 3_12346525 | NA | NA | 2 | 0.5 | 1 | 1 | NA | NA | 3 | 0.6667 | 1 | 1 | NA | NA | 1 | 0 | NA | NA | 2 | 0.5 |
| 3_12346531 | NA | NA | 2 | 0.5 | 1 | 1 | NA | NA | 3 | 0.6667 | 1 | 1 | NA | NA | 1 | 1 | NA | NA | 2 | 1 |
| 3_12346531 | NA | NA | 2 | 0.5 | 1 | 1 | NA | NA | 3 | 0.6667 | 1 | 1 | NA | NA | NA | NA | NA | NA | 1 | 1 |
| 3_12346533 | NA | NA | 2 | 0 | 1 | 1 | NA | NA | 3 | 0.3333 | 1 | 1 | NA | NA | 1 | 0 | NA | NA | 2 | 0.5 |
| 3_12346581 | NA | NA | 2 | 1 | 1 | 1 | NA | NA | 3 | 1 | NA | NA | NA | NA | 1 | 1 | NA | NA | 1 | 1 |
| 3_12346594 | NA | NA | 1 | 1 | 1 | 1 | NA | NA | 2 | 1 | NA | NA | NA | NA | 1 | 1 | NA | NA | 1 | 1 |
| 3_12411504 | 7 | 0.4286 | 3 | 0 | 1 | 0 | 5 | 0.6 | 16 | 0.375 | 4 | 0 | 5 | 0 | 14 | 0.2857 | 7 | 0.1429 | 30 | 0.1667 |
| 3_12411516 | 7 | 0.7143 | 3 | 0.3333 | 1 | 1 | 6 | 0.6667 | 17 | 0.6471 | 4 | 0.75 | 4 | 0.25 | 13 | 0.5385 | 7 | 0.5714 | 28 | 0.5357 |
| 3_12411519 | 5 | 1 | 3 | 1 | 1 | 1 | 4 | 1 | 13 | 1 | 2 | 1 | 5 | 1 | 8 | 1 | 7 | 0.8571 | 22 | 0.9545 |
| 3_12411526 | 5 | 1 | 3 | 1 | 1 | 1 | 5 | 1 | 14 | 1 | 4 | 1 | 3 | 1 | 11 | 0.9091 | 7 | 0.8571 | 25 | 0.92 |
| 3_12411528 | 6 | 0.1667 | 3 | 0 | 1 | 1 | 6 | 0.5 | 16 | 0.3125 | 4 | 0 | 4 | 0 | 14 | 0.2857 | 8 | 0.125 | 30 | 0.1667 |
| 3_12411532 | 7 | 1 | 3 | 1 | NA | NA | 6 | 0.6667 | 16 | 0.875 | 4 | 0.5 | 5 | 0.4 | 12 | 0.75 | 7 | 0.7143 | 28 | 0.6429 |
| 3_12411548 | 7 | 0.2857 | 4 | 0 | 1 | 0 | 3 | 0.3333 | 15 | 0.2 | 2 | 1 | 5 | 0.2 | 14 | 0.2143 | 5 | 0 | 26 | 0.2308 |
| 3_12411562 | 5 | 0.6 | 4 | 0 | 1 | 0 | 5 | 0.8 | 15 | 0.4667 | 4 | 0.75 | 4 | 0.25 | 15 | 0.7333 | 8 | 0.75 | 31 | 0.6774 |
| 3_12411562 | 7 | 0.4286 | 4 | 0.75 | 1 | 1 | 4 | 0.5 | 16 | 0.5625 | 5 | 0.2 | 4 | 0.75 | 13 | 0.5385 | 5 | 0.8 | 27 | 0.5556 |
| 3_12411567 | 8 | 0.875 | 4 | 0.75 | 1 | 1 | 7 | 0.5714 | 20 | 0.75 | 4 | 0.5 | 5 | 1 | 14 | 0.6429 | 9 | 0.8889 | 32 | 0.75 |
| 3_12411577 | 8 | 0.875 | 3 | 1 | 1 | 1 | 7 | 1 | 19 | 0.9474 | 4 | 1 | 5 | 1 | 15 | 1 | 9 | 1 | 33 | 1 |
| 3_12411578 | 6 | 0 | 4 | 0 | 1 | 0 | 6 | 0.1667 | 17 | 0.0588 | 5 | 0.2 | 3 | 0 | 16 | 0.1875 | 7 | 0.1429 | 31 | 0.1613 |
| 3_12411581 | 10 | 0.1 | 4 | 0 | 1 | 0 | 6 | 0.1667 | 21 | 0.0952 | 5 | 0 | 5 | 0 | 15 | 0.1333 | 9 | 0 | 34 | 0.0588 |
| 3_12411589 | 7 | 1 | 4 | 1 | NA | NA | 6 | 0.8333 | 17 | 0.9412 | 4 | 1 | 5 | 1 | 13 | 0.8462 | 7 | 1 | 29 | 0.931 |
| 3_12411634 | 10 | 0 | 6 | 0.3333 | 2 | 0.5 | 8 | 0.125 | 26 | 0.1538 | 7 | 0.4286 | 5 | 0.2 | 17 | 0.4118 | 9 | 0.3333 | 38 | 0.3684 |
| 3_12519794 | 1 | 1 | 3 | 0.3333 | 2 | 1 | NA | NA | 6 | 0.6667 | 2 | 1 | 1 | 1 | NA | NA | 2 | 0.5 | 5 | 0.8 |
| 3_12519803 | 1 | 1 | 1 | 0 | 2 | 1 | NA | NA | 4 | 0.75 | 2 | 1 | 1 | 1 | NA | NA | 1 | 1 | 4 | 1 |
| 3_12523128 | 2 | 1 | 1 | 0 | 1 | 0 | NA | NA | 4 | 0.5 | 2 | 0.5 | NA | NA | NA | NA | 2 | 0 | 4 | 0.25 |
| 3_12523135 | 2 | 1 | 1 | 1 | 2 | 0.5 | NA | NA | 5 | 0.8 | 2 | 1 | 1 | 0 | NA | NA | 2 | 0.5 | 5 | 0.6 |
| 3_12523146 | 2 | 0 | NA | NA | 2 | 1 | NA | NA | 4 | 0.5 | 1 | 1 | 1 | 1 | NA | NA | 1 | 0 | 3 | 0.6667 |
| 3_12523155 | 2 | 0.5 | 1 | 0 | 2 | 0.5 | NA | NA | 5 | 0.4 | 2 | 0.5 | 1 | 0 | NA | NA | 2 | 0 | 5 | 0.2 |
| 3_12523177 | 2 | 0 | 1 | 1 | 3 | 0 | NA | NA | 6 | 0.1667 | 2 | 0.5 | 1 | 0 | NA | NA | 2 | 0 | 5 | 0.2 |
| 3_12523186 | 3 | 1 | 1 | 1 | 2 | 1 | NA | NA | 6 | 1 | 2 | 1 | 1 | 1 | NA | NA | 2 | 1 | 5 | 1 |
| 3_12523194 | 4 | 0.75 | 1 | 1 | 3 | 1 | NA | NA | 8 | 0.875 | 1 | 0 | 1 | 0 | NA | NA | 1 | 1 | 3 | 0.3333 |
| 3_12523198 | 4 | 1 | 1 | 1 | 2 | 1 | NA | NA | 7 | 1 | 2 | 1 | 1 | 0 | NA | NA | 2 | 1 | 5 | 0.8 |
| 3_12523200 | 4 | 1 | 1 | 0 | 3 | 0.6667 | NA | NA | 8 | 0.75 | 2 | 0.5 | 1 | 1 | NA | NA | 2 | 1 | 5 | 0.8 |
| 3_12523207 | 4 | 0.5 | 2 | 1 | 3 | 0.6667 | NA | NA | 9 | 0.6667 | 2 | 0.5 | 1 | 0 | NA | NA | 2 | 1 | 5 | 0.6 |
| 3_12523208 | 3 | 0.6667 | 2 | 0 | 3 | 0 | NA | NA | 8 | 0.25 | 2 | 0 | 1 | 0 | NA | NA | 2 | 1 | 5 | 0.4 |
| 3_12523209 | 4 | 1 | 3 | 0.6667 | 4 | 1 | NA | NA | 11 | 0.9091 | 2 | 1 | 1 | 1 | NA | NA | 2 | 1 | 5 | 1 |

| | | | | | | | | | | | | | | | | | | | | |
|---|---|---|---|---|---|---|---|---|---|---|---|---|---|---|---|---|---|---|---|---|
| 3_12523213 | 3 | 0.3333 | 2 | 0 | 4 | 0.25 | NA | NA | 9 | 0.2222 | 2 | 0 | 1 | 1 | NA | NA | 1 | 0 | 4 | 0.25 |
| 3_12523235 | 2 | 1 | 2 | 0.5 | 3 | 1 | NA | NA | 7 | 0.8571 | 1 | 1 | 1 | 1 | NA | NA | 1 | 1 | 3 | 1 |
| 3_12551851 | 4 | 1 | 7 | 1 | 6 | 0.8333 | NA | NA | 17 | 0.9412 | 5 | 1 | 3 | 1 | NA | NA | 4 | 1 | 12 | 1 |
| 3_12551872 | 2 | 0 | 9 | 0.1111 | 5 | 0.2 | NA | NA | 16 | 0.125 | 3 | 0.3333 | 3 | 0 | 1 | 0 | 3 | 0 | 10 | 0.1 |
| 3_12551893 | 2 | 0 | 7 | 0 | 4 | 0.5 | NA | NA | 13 | 0.1538 | 4 | 0 | 3 | 0.3333 | 1 | 0 | 2 | 0 | 10 | 0.1 |
| 3_12551929 | 3 | 0 | 6 | 0.1667 | 3 | 0 | NA | NA | 12 | 0.0833 | 2 | 0 | 3 | 0.3333 | 1 | 0 | 3 | 0.3333 | 9 | 0.2222 |
| 3_12552108 | 3 | 0 | 15 | 0.1333 | 6 | 0 | NA | NA | 24 | 0.0833 | 6 | 0 | 4 | 0.25 | NA | NA | 6 | 0.1667 | 16 | 0.125 |
| 3_12552147 | 3 | 0 | 18 | 0.2222 | 8 | 0.375 | NA | NA | 29 | 0.2414 | 5 | 0.2 | 7 | 0.5714 | 1 | 0 | 4 | 0.5 | 17 | 0.4118 |
| 3_12552150 | 3 | 0.3333 | 18 | 0 | 8 | 0.25 | 1 | 0 | 30 | 0.1 | 7 | 0.1429 | 7 | 0 | 1 | 0 | 5 | 0.2 | 20 | 0.1 |
| 3_12552174 | 3 | 0 | 15 | 0.1333 | 8 | 0.125 | 1 | 0 | 27 | 0.1111 | 7 | 0.1429 | 6 | 0 | 1 | 0 | 7 | 0.1429 | 21 | 0.0952 |
| 3_12565455 | 1 | 0 | 1 | 1 | NA | NA | NA | NA | 2 | 0.5 | 2 | 0 | NA | NA | 3 | 0.3333 | NA | NA | 5 | 0.2 |
| 3_12566094 | 2 | 1 | 5 | 1 | 8 | 1 | 4 | 1 | 19 | 1 | 5 | 1 | 3 | 1 | 4 | 0.75 | 3 | 1 | 15 | 0.9333 |
| 3_12566116 | 2 | 0 | 5 | 0 | 6 | 0.1667 | 2 | 0.5 | 15 | 0.1333 | 4 | 0.5 | 3 | 0.3333 | 2 | 0 | 3 | 0 | 12 | 0.25 |
| 3_12566541 | 1 | 0 | 5 | 0.2 | 3 | 0 | 3 | 0 | 12 | 0.0833 | 3 | 0.3333 | 3 | 0 | 1 | 0 | 5 | 0 | 12 | 0.0833 |
| 3_12570854 | 2 | 1 | 4 | 1 | 5 | 0.8 | 1 | 1 | 12 | 0.9167 | 2 | 1 | 3 | 0.6667 | 2 | 0.5 | 3 | 1 | 10 | 0.8 |
| 3_12570855 | 2 | 0.5 | 5 | 0.2 | 3 | 0 | 3 | 0 | 13 | 0.1538 | 3 | 0 | 2 | 0 | 3 | 0.6667 | 4 | 0.75 | 12 | 0.4167 |
| 3_12570876 | 2 | 0.5 | 4 | 0.5 | 5 | 0.4 | 3 | 0.6667 | 14 | 0.5 | 3 | 0 | 3 | 0 | 3 | 0.6667 | 3 | 0.3333 | 12 | 0.25 |
| 3_12570880 | 2 | 1 | 4 | 1 | 2 | 0.5 | NA | NA | 8 | 0.875 | 2 | 0.5 | 3 | 0.6667 | 2 | 1 | 3 | 1 | 10 | 0.8 |
| 3_12570892 | 2 | 1 | 5 | 0.8 | 5 | 0.8 | 2 | 1 | 14 | 0.8571 | 3 | 1 | 3 | 1 | 3 | 0.3333 | 4 | 1 | 13 | 0.8462 |
| 3_12570899 | NA | NA | 3 | 0.3333 | 5 | 0 | 2 | 0 | 10 | 0.1 | 3 | 0 | 3 | 0.3333 | 3 | 0 | 3 | 0.3333 | 12 | 0.1667 |
| 3_12639098 | 2 | 0 | 3 | 0 | 2 | 0.5 | 4 | 0 | 11 | 0.0909 | NA | NA | 1 | 0 | 3 | 0 | 2 | 0.5 | 6 | 0.1667 |
| 3_12663654 | 7 | 0.2857 | 14 | 0.0714 | 6 | 0.1667 | 8 | 0.125 | 35 | 0.1429 | 2 | 0 | 2 | 0 | 6 | 0.1667 | 6 | 0.3333 | 16 | 0.1875 |
| 3_12663656 | 7 | 0.2857 | 13 | 0.1538 | 6 | 0 | 7 | 0 | 33 | 0.1212 | 1 | 0 | 3 | 0 | 6 | 0.3333 | 7 | 0 | 17 | 0.1176 |
| 3_12663665 | 7 | 0 | 12 | 0.4167 | 6 | 0.3333 | 6 | 0.3333 | 31 | 0.2903 | 1 | 0 | 4 | 0.25 | 6 | 0 | 6 | 0.1667 | 17 | 0.1176 |
| 3_12663674 | 7 | 1 | 13 | 0.8462 | 7 | 1 | 8 | 1 | 35 | 0.9429 | 3 | 0.6667 | 4 | 0.75 | 6 | 0.8333 | 6 | 0.6667 | 19 | 0.7368 |
| 3_12663677 | 5 | 0 | 14 | 0.1429 | 7 | 0.1429 | 7 | 0 | 33 | 0.0909 | 2 | 0 | 5 | 0.4 | 7 | 0.1429 | 4 | 0.25 | 18 | 0.2222 |
| 3_12663763 | 9 | 0.3333 | 22 | 0.3182 | 8 | 0.25 | 8 | 0.375 | 47 | 0.3191 | 6 | 0.1667 | 7 | 0.4286 | 8 | 0.25 | 8 | 0.375 | 29 | 0.3103 |
| 3_12663770 | 9 | 0 | 17 | 0.0588 | 7 | 0.1429 | 7 | 0 | 40 | 0.05 | 4 | 0 | 7 | 0.1429 | 6 | 0.1667 | 7 | 0.1429 | 24 | 0.125 |
| 3_12663788 | 11 | 0.0909 | 22 | 0.0909 | 7 | 0 | 8 | 0.125 | 48 | 0.0833 | 6 | 0 | 7 | 0.1429 | 8 | 0 | 7 | 0.1429 | 28 | 0.0714 |
| 3_12663848 | 11 | 0.0909 | 22 | 0.1364 | 13 | 0.0769 | 7 | 0.1429 | 53 | 0.1132 | 10 | 0.1 | 10 | 0.1 | 5 | 0.2 | 8 | 0 | 33 | 0.0909 |
| 3_12671567 | 45 | 0.0889 | 45 | 0.1778 | 41 | 0.1463 | 45 | 0.0667 | 176 | 0.1193 | 35 | 0.1429 | 42 | 0.1667 | 34 | 0.1471 | 32 | 0.1562 | 143 | 0.1538 |
| 3_12671582 | 38 | 0.1053 | 38 | 0.1053 | 39 | 0.2308 | 45 | 0.0667 | 160 | 0.125 | 31 | 0.0323 | 39 | 0.0513 | 34 | 0.0882 | 31 | 0.0968 | 135 | 0.0667 |
| 3_12671587 | 38 | 0.3158 | 42 | 0.3333 | 44 | 0.2955 | 47 | 0.1702 | 171 | 0.2749 | 33 | 0.1818 | 37 | 0.2703 | 36 | 0.3056 | 33 | 0.2121 | 139 | 0.2446 |
| 3_12671664 | 27 | 0.5185 | 34 | 0.5882 | 32 | 0.4062 | 32 | 0.7188 | 125 | 0.56 | 18 | 0.6111 | 23 | 0.4348 | 27 | 0.5926 | 21 | 0.7143 | 89 | 0.5843 |
| 3_12671665 | 29 | 0.1724 | 35 | 0.0571 | 33 | 0.0606 | 33 | 0.2424 | 130 | 0.1308 | 23 | 0.2174 | 23 | 0.1739 | 28 | 0.2143 | 24 | 0.0417 | 98 | 0.1633 |
| 3_12671673 | 27 | 0.6667 | 32 | 0.8125 | 35 | 0.7714 | 32 | 0.7812 | 126 | 0.7619 | 19 | 0.7895 | 27 | 0.8519 | 23 | 0.6957 | 23 | 0.7826 | 92 | 0.7826 |
| 3_12671676 | 27 | 0.8889 | 33 | 0.9091 | 30 | 0.9333 | 26 | 0.9231 | 116 | 0.9138 | 18 | 0.8333 | 23 | 0.8261 | 25 | 0.88 | 19 | 0.9474 | 85 | 0.8706 |
| 3_12671679 | 24 | 0.75 | 38 | 0.7368 | 33 | 0.697 | 33 | 0.8182 | 128 | 0.75 | 23 | 0.7391 | 27 | 0.8148 | 27 | 0.8889 | 22 | 0.8636 | 99 | 0.8283 |
| 3_12671687 | 27 | 0.4074 | 38 | 0.2632 | 28 | 0.25 | 31 | 0.3226 | 124 | 0.3065 | 20 | 0.15 | 26 | 0.3462 | 29 | 0.3448 | 20 | 0.55 | 95 | 0.3474 |
| 3_12671689 | 28 | 0.2857 | 39 | 0.1282 | 33 | 0.2727 | 34 | 0.2647 | 134 | 0.2313 | 21 | 0.0476 | 27 | 0.2222 | 29 | 0.2414 | 23 | 0.3043 | 100 | 0.21 |
| 3_12671696 | 24 | 0.625 | 33 | 0.6364 | 27 | 0.5185 | 29 | 0.6207 | 113 | 0.6018 | 16 | 0.3125 | 22 | 0.6364 | 28 | 0.5357 | 16 | 0.5 | 82 | 0.5122 |
| 3_12671704 | 25 | 0.28 | 35 | 0.3714 | 32 | 0.2812 | 30 | 0.2 | 122 | 0.2869 | 18 | 0.3333 | 23 | 0.1739 | 24 | 0.3333 | 18 | 0.1111 | 83 | 0.241 |
| 3_12671710 | 26 | 0.1538 | 37 | 0.027 | 35 | 0.1143 | 32 | 0.0625 | 130 | 0.0846 | 23 | 0.1304 | 26 | 0.0769 | 26 | 0.1538 | 19 | 0.0526 | 94 | 0.1064 |
| 3_12671711 | 26 | 0.1923 | 31 | 0.2581 | 33 | 0.2727 | 31 | 0.2903 | 121 | 0.2562 | 21 | 0.2381 | 21 | 0.2857 | 22 | 0.3182 | 17 | 0.3529 | 81 | 0.2963 |
| 3_12671712 | 26 | 0.3462 | 37 | 0.2432 | 33 | 0.1818 | 32 | 0.25 | 128 | 0.25 | 24 | 0.0417 | 25 | 0.28 | 27 | 0.2222 | 18 | 0.3889 | 94 | 0.2234 |
| 3_12671718 | 21 | 0.9524 | 33 | 0.9394 | 24 | 1 | 26 | 1 | 104 | 0.9712 | 19 | 0.9474 | 20 | 0.95 | 24 | 1 | 16 | 0.9375 | 79 | 0.962 |
| 3_12671725 | 20 | 0.5 | 28 | 0.2857 | 29 | 0.2414 | 27 | 0.3333 | 104 | 0.3269 | 18 | 0.3333 | 24 | 0.375 | 20 | 0.35 | 17 | 0.2353 | 79 | 0.3291 |
| 3_12671727 | 22 | 0.4091 | 28 | 0.25 | 27 | 0.3333 | 24 | 0.5 | 101 | 0.3663 | 19 | 0.2632 | 22 | 0.5 | 19 | 0.4211 | 15 | 0.3333 | 75 | 0.3867 |
| 3_12671733 | 23 | 0.7826 | 31 | 0.8387 | 33 | 0.697 | 26 | 0.6923 | 113 | 0.7522 | 24 | 0.8333 | 25 | 0.76 | 24 | 0.9167 | 18 | 0.7222 | 91 | 0.8132 |
| 3_12671734 | 24 | 0.75 | 30 | 0.9333 | 30 | 0.8 | 31 | 0.7097 | 115 | 0.8 | 24 | 0.6667 | 25 | 0.88 | 25 | 0.64 | 19 | 0.7895 | 93 | 0.7419 |
| 3_12671736 | 23 | 0.913 | 33 | 0.7879 | 31 | 0.8065 | 30 | 0.8333 | 117 | 0.8291 | 19 | 0.7368 | 21 | 0.9048 | 24 | 0.9583 | 17 | 0.8235 | 81 | 0.8642 |
| 3_12671740 | 23 | 0.087 | 31 | 0.129 | 32 | 0 | 29 | 0.1034 | 115 | 0.0783 | 23 | 0 | 22 | 0.0909 | 26 | 0.1923 | 19 | 0 | 90 | 0.0778 |
| 3_12673530 | 6 | 0 | 13 | 0.0769 | 4 | 0 | 11 | 0.1818 | 34 | 0.0882 | 5 | 0.2 | 5 | 0.2 | 6 | 0.1667 | 3 | 0 | 19 | 0.1579 |
| 3_12673534 | 6 | 0 | 12 | 0.0833 | 4 | 0.25 | 13 | 0.0769 | 35 | 0.0857 | 4 | 0 | 5 | 0 | 6 | 0.1667 | 3 | 0 | 18 | 0.0556 |
| 3_12673587 | 4 | 0.25 | 6 | 0.8333 | 3 | 1 | 10 | 0.4 | 23 | 0.5652 | 2 | 0.5 | 5 | 0.6 | 4 | 0.75 | 1 | 1 | 12 | 0.6667 |
| 3_12673601 | 4 | 0.75 | 6 | 0.8333 | 4 | 0.75 | 12 | 0.9167 | 26 | 0.8462 | 4 | 0.75 | 5 | 0.8 | 6 | 0.8333 | NA | NA | 15 | 0.8 |
| 3_12673613 | 2 | 0 | 6 | 0 | 4 | 0.25 | 10 | 0.1 | 22 | 0.0909 | 4 | 0.25 | 5 | 0.2 | 6 | 0.1667 | NA | NA | 15 | 0.2 |
| 3_12673617 | 3 | 1 | 5 | 1 | 4 | 0.5 | 10 | 1 | 22 | 0.9091 | 4 | 0.75 | 4 | 1 | 6 | 1 | 1 | 1 | 15 | 0.9333 |
| 3_12673642 | 3 | 0.3333 | 5 | 0 | 3 | 0.3333 | 11 | 0.1818 | 22 | 0.1818 | 4 | 0 | 5 | 0 | 4 | 0.25 | 1 | 0 | 14 | 0.0714 |
| 3_12673660 | 2 | 0.5 | 5 | 0.2 | 2 | 0 | 9 | 0 | 18 | 0.1111 | 4 | 0 | 3 | 0.3333 | 3 | 0.3333 | 1 | 0 | 11 | 0.1818 |
| 3_12673694 | 2 | 0 | 4 | 0 | 2 | 0 | 10 | 0.1 | 18 | 0.0556 | 4 | 0.25 | 3 | 0 | 3 | 0.3333 | NA | NA | 10 | 0.2 |
| 3_12673699 | 1 | 1 | 3 | 0.6667 | 1 | 0 | 6 | 0.5 | 11 | 0.5455 | 2 | 0 | 2 | 0.5 | 2 | 1 | NA | NA | 6 | 0.5 |
| 3_12673718 | 1 | 1 | 4 | 0 | 2 | 0 | 8 | 0.125 | 15 | 0.1333 | 4 | 0 | 3 | 0.3333 | 3 | 0.3333 | NA | NA | 10 | 0.2 |
| 3_12673726 | NA | NA | 3 | 0.3333 | 2 | 0 | 8 | 0.625 | 13 | 0.4615 | 4 | 0.5 | 3 | 0.3333 | 3 | 0.6667 | NA | NA | 10 | 0.5 |
| 3_12673749 | 1 | 0 | 4 | 0.5 | 2 | 0 | 4 | 0.25 | 11 | 0.2727 | 3 | 0.3333 | 3 | 0.3333 | 2 | 0.5 | NA | NA | 8 | 0.375 |

| | | | | | | | | | | | | | | | | | | | | |
|---|---|---|---|---|---|---|---|---|---|---|---|---|---|---|---|---|---|---|---|---|
| 3_1267375 | 1 | 0 | 4 | 0.5 | 2 | 0 | 5 | 0.2 | 12 | 0.25 | 4 | 0.5 | 3 | 0.3333 | 3 | 0 | NA | NA | 10 | 0.3 |
| 3_1267376 | 1 | 0 | 4 | 0.25 | 2 | 0 | 5 | 0 | 12 | 0.0833 | 4 | 0 | 3 | 0.3333 | 3 | 0 | NA | NA | 10 | 0.1 |
| 3_1267376 | 1 | 0 | 3 | 0.3333 | 2 | 0 | 5 | 0.2 | 11 | 0.1818 | 4 | 0.25 | 3 | 0 | 3 | 0.3333 | NA | NA | 10 | 0.2 |
| 3_1267378 | 1 | 0 | 3 | 0.3333 | 2 | 0.5 | 5 | 0.6 | 11 | 0.4545 | 3 | 0.6667 | 3 | 0 | 3 | 1 | NA | NA | 9 | 0.5556 |
| 3_1267380 | 1 | 0 | 3 | 0 | 2 | 0 | 5 | 0.2 | 11 | 0.0909 | 3 | 0.3333 | 2 | 0 | 3 | 0 | NA | NA | 8 | 0.125 |
| 3_1267381 | 1 | 0 | 2 | 0 | 2 | 0 | 4 | 0.25 | 9 | 0.1111 | 3 | 0.6667 | 2 | 0 | 3 | 0.3333 | NA | NA | 8 | 0.375 |
| 3_1267385 | 1 | 1 | 3 | 0 | 2 | 0 | 4 | 0 | 10 | 0.1 | 2 | 0 | 1 | 0 | 1 | 1 | NA | NA | 4 | 0.25 |
| 3_1267386 | 1 | 0 | 3 | 0.6667 | 1 | 0 | 5 | 0 | 10 | 0.2 | 2 | 0.5 | 1 | 0 | 3 | 0.3333 | NA | NA | 6 | 0.3333 |
| 3_1267398 | NA | NA | 1 | 1 | 2 | 1 | 3 | 0.6667 | 6 | 0.8333 | 1 | 1 | NA | NA | 2 | 1 | NA | NA | 3 | 1 |
| 3_1267402 | NA | NA | 2 | 0 | 2 | 0.5 | 3 | 0 | 7 | 0.1429 | 1 | 0 | NA | NA | 1 | 1 | NA | NA | 2 | 0.5 |
| 3_1267409 | NA | NA | 1 | 1 | 2 | 1 | 3 | 0.3333 | 6 | 0.6667 | 1 | 0 | NA | NA | 2 | 0.5 | NA | NA | 3 | 0.3333 |
| 3_1267587 | 3 | 0.3333 | 7 | 0.4286 | 8 | 0.25 | 6 | 0.1667 | 24 | 0.2917 | 7 | 0.1429 | 8 | 0 | 10 | 0.2 | 2 | 0 | 27 | 0.1111 |
| 3_1273304 | 5 | 0.2 | 6 | 0.3333 | 3 | 0.6667 | 4 | 0.5 | 18 | 0.3889 | 2 | 0.5 | 4 | 0 | 2 | 0 | 5 | 0.8 | 13 | 0.3846 |
| 3_1273305 | 5 | 0.2 | 6 | 0.3333 | 3 | 0 | 4 | 0 | 18 | 0.1667 | 1 | 0 | 4 | 0 | 3 | 0 | 3 | 0.3333 | 11 | 0.0909 |
| 3_1274100 | NA | NA | 1 | 1 | NA | NA | NA | NA | 1 | 1 | NA | NA | 1 | 1 | NA | NA | 1 | 0 | 2 | 0.5 |
| 3_1274103 | NA | NA | 1 | 1 | NA | NA | 1 | 1 | 2 | 1 | NA | NA | 1 | 1 | NA | NA | 1 | 1 | 2 | 1 |
| 3_1275828 | 2 | 1 | NA | NA | NA | NA | NA | NA | 2 | 1 | 1 | 1 | NA | NA | NA | NA | NA | NA | 1 | 1 |
| 3_1275828 | 1 | 1 | NA | NA | NA | NA | NA | NA | 1 | 1 | 1 | 1 | NA | NA | NA | NA | NA | NA | 1 | 1 |
| 3_1275833 | 1 | 1 | NA | NA | NA | NA | NA | NA | 1 | 1 | 1 | 1 | NA | NA | NA | NA | NA | NA | 1 | 1 |
| 3_1275904 | 3 | 0.3333 | 1 | 1 | NA | NA | 1 | 1 | 5 | 0.6 | 3 | 1 | NA | NA | 1 | 1 | 1 | 1 | 5 | 1 |
| 3_1275904 | 2 | 0.5 | NA | NA | NA | NA | NA | NA | 2 | 0.5 | 1 | 0 | NA | NA | 1 | 0 | 2 | 0.5 | 4 | 0.25 |
| 3_1275904 | 3 | 0.3333 | 1 | 0 | NA | NA | 1 | 0 | 5 | 0.2 | 3 | 0.6667 | NA | NA | 1 | 0 | 2 | 0 | 6 | 0.3333 |
| 3_1275905 | 3 | 0.6667 | 1 | 1 | NA | NA | 1 | 1 | 5 | 0.8 | 2 | 1 | NA | NA | 1 | 1 | 2 | 1 | 5 | 1 |
| 3_1275906 | 3 | 1 | 1 | 1 | NA | NA | 1 | 1 | 5 | 1 | 3 | 1 | NA | NA | 1 | 1 | 3 | 1 | 7 | 1 |
| 3_1275907 | 2 | 1 | NA | NA | NA | NA | 1 | 1 | 3 | 1 | 2 | 1 | NA | NA | 1 | 1 | 2 | 1 | 5 | 1 |
| 3_1275908 | 3 | 1 | 1 | 1 | NA | NA | 1 | 1 | 5 | 1 | 3 | 1 | NA | NA | 1 | 1 | 3 | 0.6667 | 7 | 0.8571 |
| 3_1275908 | 2 | 0.5 | 1 | 1 | NA | NA | 1 | 0 | 4 | 0.5 | 3 | 1 | NA | NA | 1 | 1 | 2 | 0.5 | 6 | 0.8333 |
| 3_1275909 | 2 | 1 | 1 | 1 | NA | NA | NA | NA | 3 | 1 | 2 | 1 | NA | NA | 1 | 1 | 3 | 1 | 6 | 1 |
| 3_1275911 | 3 | 0.3333 | 1 | 0 | NA | NA | 1 | 0 | 5 | 0.2 | 2 | 0 | NA | NA | 1 | 0 | 3 | 0.3333 | 6 | 0.1667 |
| 3_1275912 | 3 | 0.3333 | 1 | 0 | NA | NA | 1 | 1 | 5 | 0.4 | 3 | 0.3333 | NA | NA | 1 | 1 | 3 | 0.3333 | 7 | 0.4286 |
| 3_1275913 | 3 | 0 | 1 | 1 | NA | NA | 1 | 0 | 5 | 0.2 | 3 | 0 | NA | NA | 1 | 0 | 3 | 0.3333 | 7 | 0.1429 |
| 3_1275919 | 3 | 0.3333 | NA | NA | NA | NA | NA | NA | 3 | 0.3333 | 2 | 0.5 | NA | NA | 1 | 0 | NA | NA | 3 | 0.3333 |
| 3_1276019 | 12 | 0.5833 | 17 | 0.7059 | 5 | 0.4 | 15 | 0.4667 | 49 | 0.5714 | 7 | 0.7143 | 14 | 0.9286 | 13 | 0.6923 | 11 | 0.4545 | 45 | 0.7111 |
| 3_1276020 | 10 | 0.1 | 12 | 0.0833 | 4 | 0.25 | 12 | 0 | 38 | 0.0789 | 7 | 0.2857 | 10 | 0 | 13 | 0.0769 | 9 | 0 | 39 | 0.0769 |
| 3_1276021 | 10 | 0.2 | 15 | 0.0667 | 5 | 0.2 | 15 | 0.1333 | 45 | 0.1333 | 6 | 0.1667 | 12 | 0.1667 | 17 | 0.0588 | 8 | 0 | 43 | 0.093 |
| 3_1276021 | 8 | 0.75 | 11 | 0.8182 | 5 | 1 | 11 | 0.5455 | 35 | 0.7429 | 6 | 0.8333 | 8 | 0.75 | 15 | 0.8667 | 9 | 0.7778 | 38 | 0.8158 |
| 3_1276644 | 1 | 1 | 1 | 1 | 2 | 1 | NA | NA | 4 | 1 | NA | NA | 1 | 1 | 1 | 0 | 2 | 0 | 4 | 0.25 |
| 3_1276645 | 1 | 0 | 1 | 1 | 2 | 1 | NA | NA | 4 | 0.75 | NA | NA | 1 | 1 | 1 | 1 | 3 | 0.3333 | 5 | 0.6 |
| 3_1294268 | NA | NA | NA | NA | 1 | 1 | NA | NA | 1 | 1 | NA | NA | NA | NA | 1 | 1 | 1 | 0 | 2 | 0.5 |
| 3_1294268 | NA | NA | NA | NA | 1 | 1 | NA | NA | 1 | 1 | NA | NA | NA | NA | 1 | 1 | 1 | 0 | 2 | 0.5 |
| 3_1294272 | NA | NA | 1 | 1 | 1 | 0 | NA | NA | 2 | 0.5 | 1 | 0 | NA | NA | 2 | 0.5 | 1 | 0 | 4 | 0.25 |
| 3_1294273 | NA | NA | 1 | 1 | 1 | 1 | NA | NA | 2 | 1 | 1 | 1 | NA | NA | 2 | 0.5 | 1 | 1 | 4 | 0.75 |
| 3_1294278 | NA | NA | 1 | 1 | 1 | 1 | 1 | 1 | 3 | 1 | 1 | 1 | NA | NA | 2 | 0.5 | 1 | 0 | 4 | 0.5 |
| 3_1294280 | NA | NA | 1 | 1 | NA | NA | 1 | 0 | 2 | 0.5 | 1 | 1 | NA | NA | 2 | 0.5 | 1 | 0 | 4 | 0.5 |
| 3_1294281 | NA | NA | 1 | 1 | 1 | 0 | 1 | 0 | 3 | 0.3333 | 1 | 0 | NA | NA | 2 | 1 | 1 | 0 | 4 | 0.5 |
| 3_1294282 | NA | NA | NA | NA | 1 | 1 | 1 | 1 | 2 | 1 | 1 | 0 | NA | NA | 1 | 0 | 1 | 1 | 3 | 0.3333 |
| 3_1294309 | NA | NA | 3 | 0 | 3 | 0.3333 | NA | NA | 6 | 0.1667 | 3 | 0.3333 | NA | NA | 2 | 0 | 3 | 0 | 8 | 0.125 |
| 3_1294313 | NA | NA | 3 | 0.6667 | 2 | 0.5 | NA | NA | 5 | 0.6 | 3 | 0 | NA | NA | 2 | 0.5 | 2 | 0 | 7 | 0.1429 |
| 3_1294319 | 1 | 0 | 2 | 0 | 3 | 0.3333 | 1 | 0 | 7 | 0.1429 | 4 | 0 | NA | NA | 1 | 1 | 3 | 0 | 8 | 0.125 |
| 3_1296186 | 3 | 1 | 4 | 0.75 | 4 | 0.5 | 2 | 0.5 | 13 | 0.6923 | 3 | 0.3333 | 6 | 0.8333 | 5 | 0.8 | NA | NA | 14 | 0.7143 |
| 3_1296885 | 5 | 0.2 | 1 | 0 | 5 | 0 | 2 | 0 | 13 | 0.0769 | 1 | 1 | 2 | 0 | 5 | 0 | 2 | 0 | 10 | 0.1 |
| 3_1296886 | 6 | 0 | 4 | 0.25 | 5 | 0 | 2 | 0 | 17 | 0.0588 | 2 | 0.5 | 3 | 0.6667 | 10 | 0.1 | 4 | 0.5 | 19 | 0.3158 |
| 3_1296886 | 5 | 0 | 3 | 0.3333 | 4 | 0 | 2 | 0 | 14 | 0.0714 | 2 | 0 | 2 | 0 | 9 | 0.1111 | 4 | 0.25 | 17 | 0.1176 |
| 3_1296886 | 5 | 0 | 4 | 0.25 | 5 | 0 | 2 | 0.5 | 16 | 0.125 | 2 | 0 | 3 | 0 | 10 | 0.1 | 4 | 0 | 19 | 0.0526 |
| 3_1296889 | 3 | 0.3333 | 2 | 0.5 | 2 | 1 | 1 | 1 | 8 | 0.625 | 2 | 1 | 2 | 0.5 | 5 | 0.6 | 3 | 0.3333 | 12 | 0.5833 |
| 3_1297200 | NA | NA | 1 | 0 | NA | NA | 1 | 1 | 2 | 0.5 | NA | NA | NA | NA | 1 | 1 | 1 | 0 | 2 | 0.5 |
| 3_1297770 | 3 | 1 | 9 | 0.8889 | 6 | 1 | 8 | 0.875 | 26 | 0.9231 | 4 | 0.75 | 5 | 0.8 | 10 | 0.7 | 6 | 0.8333 | 25 | 0.76 |
| 3_1297771 | 2 | 1 | 9 | 0.7778 | 7 | 0.8571 | 7 | 1 | 25 | 0.88 | 4 | 0.5 | 5 | 0.8 | 10 | 0.9 | 5 | 0.8 | 24 | 0.7917 |
| 3_1297772 | 3 | 0.6667 | 4 | 0.5 | 6 | 1 | 6 | 1 | 19 | 0.8421 | 4 | 0.75 | 5 | 0.8 | 6 | 1 | 5 | 1 | 20 | 0.9 |
| 3_1297772 | 3 | 1 | 8 | 1 | 6 | 0.8333 | 6 | 1 | 23 | 0.9565 | 4 | 0.25 | 5 | 0.8 | 6 | 0.8333 | 6 | 1 | 21 | 0.7619 |
| 3_1297772 | 3 | 0.3333 | 7 | 0.4286 | 3 | 0.3333 | 8 | 0.125 | 21 | 0.2857 | 4 | 0.25 | 5 | 0.2 | 7 | 0 | 6 | 0.1667 | 22 | 0.1364 |
| 3_1297772 | 4 | 0 | 8 | 0.125 | 4 | 0 | 8 | 0.125 | 24 | 0.0833 | 4 | 0.25 | 6 | 0.1667 | 9 | 0.1111 | 5 | 0 | 24 | 0.125 |
| 3_1297774 | 4 | 0 | 6 | 0.1667 | 3 | 0 | 4 | 0 | 17 | 0.0588 | 3 | 0 | 6 | 0.3333 | 7 | 0 | 3 | 0 | 19 | 0.1053 |
| 3_1297774 | 3 | 0 | 6 | 0.3333 | 1 | 0 | 6 | 0.1667 | 16 | 0.1875 | 2 | 0 | 5 | 0.4 | 4 | 0 | 4 | 0.25 | 15 | 0.2 |
| 3_1297774 | 3 | 0.6667 | 6 | 0.8333 | 4 | 0.75 | 6 | 0.8333 | 19 | 0.7895 | 3 | 0.3333 | 4 | 0 | 5 | 0.8 | 4 | 0.5 | 16 | 0.4375 |

| | | | | | | | | | | | | | | | | | | | | |
|---|---|---|---|---|---|---|---|---|---|---|---|---|---|---|---|---|---|---|---|---|
| 3_1297776 | 4 | 0.5 | 4 | 0 | 2 | 0 | 7 | 0.5714 | 17 | 0.3529 | 3 | 0.3333 | 6 | 0.3333 | 5 | 0.4 | 5 | 0.4 | 19 | 0.3684 |
| 3_1297897 | 3 | 0.6667 | 2 | 0 | 2 | 0 | 6 | 0.1667 | 13 | 0.2308 | 3 | 0 | 3 | 0.3333 | 1 | 0 | 3 | 0 | 10 | 0.1 |
| 3_1298185 | 1 | 1 | 1 | 0 | NA | NA | NA | NA | 2 | 0.5 | NA | NA | 1 | 1 | NA | NA | 1 | 0 | 2 | 0.5 |
| 3_1298185 | 1 | 0 | 1 | 1 | NA | NA | NA | NA | 2 | 0.5 | NA | NA | 1 | 1 | NA | NA | 1 | 1 | 2 | 1 |
| 3_1298349 | NA | NA | 2 | 0 | NA | NA | 1 | 1 | 3 | 0.3333 | NA | NA | 1 | 0 | NA | NA | 2 | 1 | 3 | 0.6667 |
| 3_1299750 | NA | NA | NA | NA | NA | NA | 1 | 1 | 1 | 1 | NA | NA | 2 | 1 | NA | NA | NA | NA | 2 | 1 |
| 3_1299752 | NA | NA | NA | NA | NA | NA | 1 | 1 | 1 | 1 | NA | NA | 2 | 0.5 | NA | NA | NA | NA | 2 | 0.5 |
| 3_1303890 | NA | NA | NA | NA | 1 | 0 | 1 | 1 | 2 | 0.5 | 1 | 1 | 3 | 0.6667 | NA | NA | 4 | 0.75 | 8 | 0.75 |
| 3_1303891 | NA | NA | NA | NA | 1 | 1 | NA | NA | 1 | 1 | NA | NA | 2 | 1 | NA | NA | 4 | 1 | 6 | 1 |
| 3_1303891 | 1 | 1 | NA | NA | 1 | 0 | 1 | 1 | 3 | 0.6667 | 1 | 1 | 3 | 0.6667 | NA | NA | 3 | 0.3333 | 7 | 0.5714 |
| 3_1303896 | 1 | 0 | NA | NA | 1 | 1 | 1 | 0 | 3 | 0.3333 | NA | NA | 3 | 0 | NA | NA | 3 | 0.3333 | 6 | 0.1667 |
| 3_1303897 | 1 | 1 | NA | NA | 1 | 0 | 1 | 0 | 3 | 0.3333 | 1 | 1 | 3 | 1 | NA | NA | 3 | 0.6667 | 7 | 0.8571 |
| 3_1303897 | 1 | 1 | NA | NA | 1 | 1 | 1 | 1 | 3 | 1 | NA | NA | 2 | 1 | NA | NA | 3 | 1 | 5 | 1 |
| 3_1303897 | 1 | 0 | NA | NA | 1 | 1 | 1 | 1 | 3 | 0.6667 | 1 | 1 | 2 | 1 | NA | NA | 2 | 0.5 | 5 | 0.8 |
| 3_1303898 | 1 | 1 | NA | NA | 1 | 1 | 1 | 0 | 3 | 0.6667 | 1 | 1 | 3 | 0.6667 | NA | NA | 4 | 0.75 | 8 | 0.75 |
| 3_1303904 | NA | NA | NA | NA | NA | NA | 2 | 0.5 | 2 | 0.5 | 1 | 0 | 3 | 0 | NA | NA | 4 | 1 | 8 | 0.5 |
| 3_1303909 | 1 | 1 | NA | NA | 1 | 0 | 2 | 0.5 | 4 | 0.5 | 2 | 0 | 4 | 0.25 | NA | NA | 4 | 0 | 10 | 0.1 |
| 3_1303911 | 1 | 1 | NA | NA | 1 | 0 | 1 | 0 | 3 | 0.3333 | 3 | 0.3333 | 4 | 0.75 | NA | NA | 4 | 0 | 11 | 0.3636 |
| 3_1303926 | 1 | 0 | 1 | 1 | 1 | 0 | 2 | 0 | 5 | 0.2 | 2 | 0.5 | 4 | 0 | NA | NA | 1 | 0 | 7 | 0.1429 |
| 3_1303929 | 2 | 0.5 | NA | NA | 2 | 0 | 2 | 0 | 6 | 0.1667 | 3 | 1 | 4 | 0 | NA | NA | 5 | 0 | 12 | 0.25 |
| 3_1304140 | 4 | 0 | 1 | 0 | 4 | 0 | 2 | 1 | 11 | 0.1818 | 2 | 0 | 3 | 0.3333 | 4 | 0 | NA | NA | 9 | 0.1111 |
| 3_1304142 | 3 | 0 | 1 | 0 | 3 | 0 | 2 | 0.5 | 9 | 0.1111 | 2 | 0 | 2 | 0.5 | 4 | 0 | NA | NA | 8 | 0.125 |
| 3_1304195 | 1 | 1 | NA | NA | NA | NA | NA | NA | 1 | 1 | 1 | 1 | NA | NA | NA | NA | NA | NA | 1 | 1 |
| 3_1304283 | 8 | 0.125 | 1 | 1 | 4 | 0 | 3 | 0 | 16 | 0.125 | 1 | 0 | 2 | 0 | 5 | 0.2 | NA | NA | 8 | 0.125 |
| 3_1304289 | 5 | 0.2 | 2 | 0 | 2 | 0 | 5 | 0 | 14 | 0.0714 | NA | NA | 2 | 0.5 | 5 | 0 | NA | NA | 7 | 0.1429 |
| 3_1304897 | 8 | 1 | 7 | 0.7143 | 13 | 0.7692 | 5 | 0.8 | 33 | 0.8182 | 5 | 1 | 6 | 0.6667 | 8 | 0.625 | 4 | 0.75 | 23 | 0.7391 |
| 3_1304898 | 4 | 0.5 | 7 | 0.1429 | 12 | 0.3333 | 5 | 0.4 | 28 | 0.3214 | 8 | 0.125 | 7 | 0 | 10 | 0.3 | 2 | 0 | 27 | 0.1481 |
| 3_1304898 | 2 | 0.5 | 2 | 1 | NA | NA | NA | NA | 4 | 0.75 | NA | NA | 1 | 1 | 1 | 1 | NA | NA | 2 | 1 |
| 3_1304899 | 8 | 0 | 8 | 0.25 | 13 | 0.1538 | 6 | 0.5 | 35 | 0.2 | 9 | 0 | 8 | 0.375 | 8 | 0 | 4 | 0.25 | 29 | 0.1379 |
| 3_1305238 | 2 | 0 | 4 | 0 | 6 | 0.1667 | 1 | 0 | 13 | 0.0769 | 6 | 0.1667 | 5 | 0 | 2 | 0 | 3 | 0 | 16 | 0.0625 |
| 3_1309051 | NA | NA | 1 | 1 | NA | NA | NA | NA | 1 | 1 | NA | NA | NA | NA | NA | NA | 1 | 1 | 1 | 1 |
| 3_1310168 | NA | NA | 1 | 1 | NA | NA | NA | NA | 1 | 1 | NA | NA | 1 | 1 | NA | NA | 1 | 0 | 2 | 0.5 |
| 3_1310171 | NA | NA | 1 | 1 | NA | NA | NA | NA | 1 | 1 | NA | NA | 1 | 0 | NA | NA | 2 | 0.5 | 3 | 0.3333 |
| 3_1310273 | 7 | 0.1429 | 6 | 0 | 6 | 0.1667 | 4 | 0 | 23 | 0.087 | 2 | 0 | 3 | 0 | 4 | 0.25 | 4 | 0 | 13 | 0.0769 |
| 3_1310276 | 7 | 0 | 5 | 0 | 6 | 0.3333 | 5 | 0.2 | 23 | 0.1304 | 2 | 0 | 3 | 0 | 4 | 0.25 | 2 | 0 | 11 | 0.0909 |
| 3_1310420 | 5 | 0 | 2 | 0 | 3 | 0 | 5 | 0.2 | 15 | 0.0667 | 2 | 0 | 3 | 0 | 1 | 0 | 2 | 0.5 | 8 | 0.125 |
| 3_1310823 | 1 | 0 | 4 | 1 | NA | NA | NA | NA | 5 | 0.8 | 1 | 1 | 1 | 1 | NA | NA | 1 | 1 | 3 | 1 |
| 3_1310823 | 1 | 1 | 4 | 0.5 | 1 | 1 | NA | NA | 6 | 0.6667 | 3 | 0.6667 | NA | NA | 2 | 1 | 2 | 0.5 | 7 | 0.7143 |
| 3_1310824 | 1 | 0 | 5 | 0.4 | 1 | 0 | NA | NA | 7 | 0.2857 | 3 | 0.3333 | NA | NA | 1 | 0 | 1 | 0 | 5 | 0.2 |
| 3_1310825 | 1 | 0 | 5 | 0.4 | 1 | 1 | NA | NA | 7 | 0.4286 | 2 | 1 | NA | NA | 2 | 0.5 | 1 | 1 | 5 | 0.8 |
| 3_1310826 | 1 | 1 | 5 | 0 | 1 | 0 | NA | NA | 7 | 0.1429 | 2 | 0 | NA | NA | 2 | 0.5 | 1 | 0 | 5 | 0.2 |
| 3_1310839 | 1 | 0 | NA | NA | 1 | 1 | NA | NA | 2 | 0.5 | 2 | 0.5 | NA | NA | NA | NA | NA | NA | 2 | 0.5 |
| 3_1310844 | 1 | 1 | 1 | 0 | 1 | 1 | NA | NA | 3 | 0.6667 | 1 | 1 | NA | NA | NA | NA | NA | NA | 1 | 1 |
| 3_1310845 | 1 | 1 | 1 | 1 | 1 | 1 | NA | NA | 3 | 1 | 1 | 1 | NA | NA | NA | NA | NA | NA | 1 | 1 |
| 3_1310846 | 1 | 1 | 1 | 1 | NA | NA | NA | NA | 2 | 1 | 1 | 1 | NA | NA | NA | NA | NA | NA | 1 | 1 |
| 3_1310847 | 1 | 1 | 1 | 1 | 1 | 0 | NA | NA | 3 | 0.6667 | 1 | 1 | NA | NA | NA | NA | NA | NA | 1 | 1 |
| 3_1310848 | 1 | 1 | 1 | 0 | 1 | 1 | NA | NA | 3 | 0.6667 | 1 | 1 | NA | NA | NA | NA | NA | NA | 1 | 1 |
| 3_1310871 | 1 | 0 | 3 | 0.3333 | 1 | 0 | 1 | 1 | 6 | 0.3333 | 3 | 0.3333 | NA | NA | NA | NA | NA | NA | 3 | 0.3333 |
| 3_1310887 | NA | NA | 1 | 1 | NA | NA | 1 | 0 | 2 | 0.5 | NA | NA | 1 | 1 | 1 | 1 | NA | NA | 2 | 1 |
| 3_1310889 | NA | NA | 1 | 1 | NA | NA | 1 | 0 | 2 | 0.5 | NA | NA | NA | NA | 1 | 1 | NA | NA | 1 | 1 |
| 3_1310891 | 1 | 1 | 4 | 0.25 | 1 | 1 | 3 | 0.3333 | 9 | 0.4444 | 3 | 0.3333 | NA | NA | 2 | 0.5 | NA | NA | 5 | 0.4 |
| 3_1310894 | NA | NA | 2 | 1 | NA | NA | 2 | 1 | 4 | 1 | NA | NA | NA | NA | 1 | 1 | NA | NA | 1 | 1 |
| 3_1310909 | NA | NA | 1 | 1 | NA | NA | 2 | 0 | 3 | 0.3333 | NA | NA | NA | NA | 1 | 1 | NA | NA | 1 | 1 |
| 3_1310912 | NA | NA | 1 | 0 | NA | NA | 2 | 0.5 | 3 | 0.3333 | NA | NA | NA | NA | 1 | 1 | NA | NA | 1 | 1 |
| 3_1310914 | 1 | 0 | 2 | 0.5 | 1 | 0 | 2 | 0.5 | 6 | 0.3333 | 3 | 0.3333 | NA | NA | 1 | 0 | NA | NA | 4 | 0.25 |
| 3_1310915 | NA | NA | 2 | 0.5 | 1 | 1 | 3 | 1 | 6 | 0.8333 | 2 | 0 | NA | NA | 1 | 1 | NA | NA | 3 | 0.3333 |
| 3_1310915 | NA | NA | 2 | 0.5 | 1 | 0 | 3 | 0.6667 | 6 | 0.5 | 3 | 0.3333 | NA | NA | 1 | 0 | NA | NA | 4 | 0.25 |
| 3_1310916 | NA | NA | 1 | 1 | 1 | 1 | 3 | 0.3333 | 5 | 0.6 | 2 | 0.5 | NA | NA | 1 | 1 | NA | NA | 3 | 0.6667 |
| 3_1310916 | NA | NA | 1 | 0 | 1 | 0 | 3 | 0.6667 | 5 | 0.4 | 2 | 0.5 | NA | NA | 1 | 1 | NA | NA | 3 | 0.6667 |
| 3_1310916 | NA | NA | 2 | 1 | 1 | 1 | 3 | 0.6667 | 6 | 0.8333 | 2 | 1 | NA | NA | 1 | 1 | NA | NA | 3 | 1 |
| 3_1310965 | NA | NA | NA | NA | 1 | 0 | 3 | 0.3333 | 4 | 0.25 | 2 | 0 | NA | NA | 1 | 1 | NA | NA | 3 | 0.3333 |
| 3_1311166 | 1 | 1 | NA | NA | NA | NA | NA | NA | 1 | 1 | NA | NA | NA | NA | 1 | 1 | 1 | 1 | 2 | 1 |
| 3_1311357 | 1 | 1 | NA | NA | NA | NA | NA | NA | 1 | 1 | NA | NA | NA | NA | 1 | 1 | NA | NA | 1 | 1 |
| 3_1311358 | 1 | 1 | NA | NA | NA | NA | NA | NA | 1 | 1 | NA | NA | NA | NA | 1 | 1 | NA | NA | 1 | 1 |
| 3_1313488 | 13 | 0.6923 | 13 | 0.8462 | 16 | 0.75 | 9 | 0.8889 | 51 | 0.7843 | 9 | 1 | 9 | 0.8889 | 18 | 0.8333 | 5 | 0.8 | 41 | 0.878 |

| | | | | | | | | | | | | | | | | | | | | |
|---|---|---|---|---|---|---|---|---|---|---|---|---|---|---|---|---|---|---|---|---|
| 3_13134895 | 12 | 0.6667 | 10 | 0.7 | 15 | 0.4667 | 9 | 0.5556 | 46 | 0.587 | 9 | 0.4444 | 7 | 0.5714 | 17 | 0.3529 | 6 | 0.8333 | 39 | 0.4872 |
| 3_13236631 | 12 | 0 | 13 | 0.1538 | 15 | 0 | 15 | 0.0667 | 55 | 0.0545 | 14 | 0.2143 | 7 | 0.1429 | 15 | 0 | 8 | 0.125 | 44 | 0.1136 |
| 3_13254396 | 1 | 1 | NA | NA | NA | NA | NA | NA | 1 | 1 | 1 | 0 | 1 | 1 | 1 | 1 | NA | NA | 3 | 0.6667 |
| 3_13254402 | 1 | 1 | NA | NA | NA | NA | NA | NA | 1 | 1 | 1 | 0 | 1 | 1 | 1 | 1 | NA | NA | 3 | 0.6667 |
| 3_13254413 | 1 | 1 | NA | NA | NA | NA | 1 | 0 | 2 | 0.5 | 1 | 1 | 1 | 1 | NA | NA | NA | NA | 2 | 1 |
| 3_13254416 | 2 | 1 | NA | NA | NA | NA | NA | NA | 2 | 1 | 1 | 1 | NA | NA | 1 | 1 | NA | NA | 2 | 1 |
| 3_13254434 | 2 | 1 | NA | NA | NA | NA | 1 | 0 | 3 | 0.6667 | 1 | 0 | 1 | 0 | 1 | 1 | NA | NA | 3 | 0.3333 |
| 3_13254435 | 2 | 1 | NA | NA | NA | NA | 1 | 1 | 3 | 1 | 1 | 1 | 1 | 1 | 1 | 1 | NA | NA | 3 | 1 |
| 3_13254441 | NA | NA | NA | NA | NA | NA | 1 | 1 | 1 | 1 | 1 | 0 | 1 | 1 | 1 | 1 | NA | NA | 3 | 0.6667 |
| 3_13254455 | 2 | 1 | NA | NA | NA | NA | 1 | 1 | 3 | 1 | 1 | 1 | 1 | 0 | 1 | 1 | NA | NA | 3 | 0.6667 |
| 3_13254497 | 2 | 0.5 | NA | NA | NA | NA | NA | NA | 2 | 0.5 | 1 | 1 | 1 | 0 | NA | NA | NA | NA | 2 | 0.5 |
| 3_13254513 | 1 | 1 | 1 | 1 | NA | NA | 2 | 0.5 | 4 | 0.75 | NA | NA | 1 | 1 | 1 | 1 | NA | NA | 2 | 1 |
| 3_13254542 | 2 | 1 | 1 | 1 | 1 | 1 | 1 | 1 | 5 | 1 | 2 | 1 | 2 | 1 | 1 | 1 | NA | NA | 5 | 1 |
| 3_13254551 | 2 | 1 | 1 | 1 | 1 | 1 | 2 | 1 | 6 | 1 | 2 | 0 | 3 | 0 | 2 | 0.5 | NA | NA | 7 | 0.1429 |
| 3_13254565 | 2 | 0.5 | 2 | 0 | 1 | 1 | 2 | 1 | 7 | 0.5714 | 2 | 0.5 | 4 | 0.5 | 3 | 0.6667 | NA | NA | 9 | 0.5556 |
| 3_13254568 | 2 | 1 | 2 | 1 | 1 | 1 | 2 | 1 | 7 | 1 | 2 | 0.5 | 3 | 1 | 3 | 1 | NA | NA | 8 | 0.875 |
| 3_13254570 | 2 | 1 | 2 | 1 | 1 | 1 | 2 | 1 | 7 | 1 | 2 | 1 | 3 | 0.6667 | 3 | 1 | NA | NA | 8 | 0.875 |
| 3_13254573 | 2 | 0.5 | 1 | 1 | NA | NA | 2 | 1 | 5 | 0.8 | 2 | 1 | 2 | 0.5 | 2 | 0.5 | NA | NA | 6 | 0.6667 |
| 3_13254583 | 2 | 1 | NA | NA | NA | NA | 1 | 1 | 3 | 1 | 2 | 1 | 3 | 1 | 3 | 1 | NA | NA | 8 | 1 |
| 3_13254611 | 1 | 0 | 2 | 1 | NA | NA | 3 | 0.3333 | 6 | 0.5 | 1 | 1 | 3 | 0.3333 | 2 | 0 | 1 | 1 | 7 | 0.4286 |
| 3_13254624 | 2 | 0 | 2 | 0 | NA | NA | 3 | 0.6667 | 7 | 0.2857 | 2 | 0.5 | 2 | 0 | 3 | 0.6667 | 2 | 0.5 | 9 | 0.4444 |
| 3_13254639 | 2 | 0.5 | 2 | 0 | NA | NA | 3 | 0 | 7 | 0.1429 | 2 | 0 | 3 | 0.3333 | 3 | 0.3333 | 2 | 0.5 | 10 | 0.3 |
| 3_13254665 | 2 | 0 | 1 | 0 | 1 | 0 | 2 | 0.5 | 6 | 0.1667 | 1 | 0 | 2 | 0 | 2 | 0.5 | 2 | 0 | 7 | 0.1429 |
| 3_13254723 | 2 | 0.5 | 3 | 0 | 2 | 0 | 1 | 1 | 8 | 0.25 | 1 | 0 | 2 | 0 | 4 | 0.25 | 3 | 0.3333 | 10 | 0.2 |
| 3_13258784 | 1 | 1 | 5 | 0.6 | 2 | 0 | NA | NA | 8 | 0.5 | 3 | 0.3333 | NA | NA | NA | NA | 2 | 1 | 5 | 0.6 |
| 3_13258831 | NA | NA | 3 | 0.3333 | 1 | 1 | NA | NA | 4 | 0.5 | 2 | 0.5 | NA | NA | NA | NA | NA | NA | 2 | 0.5 |
| 3_13259002 | NA | NA | 3 | 0.3333 | NA | NA | NA | NA | 3 | 0.3333 | 2 | 0.5 | NA | NA | NA | NA | NA | NA | 2 | 0.5 |
| 3_13281490 | 1 | 0 | 1 | 1 | 1 | 1 | NA | NA | 3 | 0.6667 | NA | NA | 1 | 0 | 1 | 1 | 1 | 1 | 3 | 0.6667 |
| 3_13281506 | 2 | 0 | 1 | 0 | 1 | 1 | NA | NA | 4 | 0.25 | 1 | 1 | 1 | 0 | 2 | 0 | 2 | 1 | 6 | 0.5 |
| 3_13281526 | 2 | 0 | 2 | 0.5 | 1 | 0 | NA | NA | 5 | 0.2 | 1 | 1 | 1 | 0 | 2 | 0.5 | 1 | 0 | 5 | 0.4 |
| 3_13301187 | NA | NA | NA | NA | 1 | 1 | NA | NA | 1 | 1 | NA | NA | NA | NA | NA | NA | 1 | 1 | 1 | 1 |
| 3_13301419 | NA | NA | NA | NA | NA | NA | 1 | 1 | 1 | 1 | NA | NA | NA | NA | NA | NA | 2 | 1 | 2 | 1 |
| 3_13301563 | NA | NA | NA | NA | NA | NA | 1 | 1 | 1 | 1 | NA | NA | NA | NA | NA | NA | 3 | 1 | 3 | 1 |
| 3_13301598 | NA | NA | NA | NA | NA | NA | 1 | 1 | 1 | 1 | NA | NA | NA | NA | NA | NA | 3 | 1 | 3 | 1 |
| 3_13301612 | NA | NA | NA | NA | NA | NA | 1 | 1 | 1 | 1 | NA | NA | NA | NA | NA | NA | 3 | 0.3333 | 3 | 0.3333 |
| 3_13308371 | 9 | 0 | 10 | 0 | 5 | 0.2 | 5 | 0.2 | 29 | 0.069 | NA | NA | 7 | 0.1429 | 11 | 0.0909 | 6 | 0 | 24 | 0.0833 |
| 3_13317027 | 5 | 0.4 | 2 | 0 | 3 | 0.3333 | 3 | 0 | 13 | 0.2308 | 2 | 0 | 3 | 0 | 5 | 0.2 | 4 | 0 | 14 | 0.0714 |
| 3_13317032 | 5 | 0 | 1 | 0 | 2 | 0.5 | 3 | 0.6667 | 11 | 0.2727 | 2 | 0 | 1 | 1 | 3 | 0 | 1 | 0 | 7 | 0.1429 |
| 3_13317113 | 4 | 0.75 | 1 | 0 | 3 | 0 | 3 | 0.3333 | 11 | 0.3636 | 2 | 0 | 4 | 0.25 | 3 | 0.3333 | 2 | 0.5 | 11 | 0.2727 |
| 3_13317117 | 3 | 0.3333 | 1 | 0 | 2 | 0 | 2 | 0 | 8 | 0.125 | 2 | 0 | 4 | 0 | 3 | 0.3333 | 2 | 1 | 11 | 0.2727 |
| 3_13317179 | 3 | 0 | 2 | 0 | 3 | 0 | 2 | 0.5 | 10 | 0.1 | 2 | 0.5 | 4 | 0 | 1 | 0 | 2 | 0.5 | 9 | 0.2222 |
| 3_13317205 | 4 | 0.5 | NA | NA | 1 | 1 | 2 | 1 | 7 | 0.7143 | NA | NA | 1 | 0 | 1 | 0 | 2 | 1 | 4 | 0.5 |
| 3_13317213 | 2 | 0 | 1 | 1 | 1 | 1 | 3 | 0 | 7 | 0.2857 | 1 | 0 | 3 | 0 | 1 | 1 | 2 | 0.5 | 7 | 0.2857 |
| 3_13317224 | 4 | 0.5 | 1 | 1 | 1 | 1 | 3 | 1 | 9 | 0.7778 | 1 | 0 | 3 | 1 | 1 | 1 | 2 | 1 | 7 | 0.8571 |
| 3_13317242 | 3 | 0.6667 | 1 | 1 | 1 | 1 | 1 | 0 | 6 | 0.6667 | 1 | 1 | 2 | 0.5 | 1 | 1 | 2 | 0.5 | 6 | 0.6667 |
| 3_13317255 | 3 | 0.6667 | 1 | 1 | 1 | 1 | 3 | 0.3333 | 8 | 0.625 | 1 | 1 | 3 | 0.6667 | NA | NA | 1 | 1 | 5 | 0.8 |
| 3_13317257 | 2 | 0 | 1 | 0 | 1 | 1 | 3 | 0.6667 | 7 | 0.4286 | 1 | 0 | 3 | 0.3333 | 1 | 0 | 2 | 0 | 7 | 0.1429 |
| 3_13317258 | 3 | 0.3333 | 1 | 1 | 1 | 0 | 3 | 0.3333 | 8 | 0.375 | 1 | 1 | 3 | 0.6667 | 1 | 0 | 2 | 0 | 7 | 0.4286 |
| 3_13317262 | 3 | 0.6667 | 1 | 1 | 1 | 1 | 3 | 1 | 8 | 0.875 | 1 | 0 | 2 | 0 | 1 | 0 | 2 | 1 | 6 | 0.3333 |
| 3_13317276 | 1 | 0 | 1 | 1 | NA | NA | NA | NA | 2 | 0.5 | NA | NA | NA | NA | 1 | 1 | 1 | 1 | 2 | 1 |
| 3_13317304 | 2 | 1 | NA | NA | 1 | 1 | 3 | 1 | 6 | 1 | 1 | 1 | 1 | 1 | 1 | 1 | 1 | 1 | 4 | 1 |
| 3_13317306 | 3 | 1 | NA | NA | 1 | 1 | 3 | 1 | 7 | 1 | 1 | 1 | 1 | 1 | 1 | 1 | 2 | 1 | 5 | 1 |
| 3_13317311 | 3 | 0 | 1 | 0 | 1 | 0 | 3 | 0.3333 | 8 | 0.125 | 1 | 0 | 1 | 0 | 1 | 0 | 2 | 0.5 | 5 | 0.2 |
| 3_13317321 | 3 | 0.3333 | 1 | 1 | 1 | 0 | 3 | 0.3333 | 8 | 0.375 | NA | NA | 2 | 0 | 1 | 1 | 2 | 0 | 5 | 0.2 |
| 3_13317340 | 1 | 1 | 1 | 1 | NA | NA | 2 | 0 | 4 | 0.5 | 1 | 0 | 1 | 0 | 1 | 1 | 2 | 1 | 5 | 0.6 |
| 3_13317344 | 1 | 1 | 1 | 1 | NA | NA | 3 | 0.6667 | 5 | 0.8 | NA | NA | 1 | 0 | 1 | 1 | 1 | 0 | 3 | 0.3333 |
| 3_13317346 | 2 | 0.5 | NA | NA | NA | NA | 3 | 0.6667 | 5 | 0.6 | 1 | 0 | 1 | 0 | 1 | 0 | 2 | 0.5 | 5 | 0.2 |
| 3_13317353 | 2 | 1 | 1 | 1 | NA | NA | 2 | 1 | 5 | 1 | 1 | 0 | 2 | 1 | 1 | 1 | 2 | 1 | 6 | 0.8333 |
| 3_13317360 | 1 | 1 | 1 | 1 | NA | NA | 2 | 0 | 4 | 0.5 | 1 | 1 | 2 | 0 | 1 | 1 | 2 | 0 | 6 | 0.3333 |
| 3_13317366 | 1 | 1 | 1 | 1 | NA | NA | 1 | 1 | 3 | 1 | 1 | 1 | 1 | 1 | 1 | 1 | 1 | 1 | 4 | 1 |
| 3_13317510 | 1 | 0 | NA | NA | NA | NA | 2 | 0.5 | 3 | 0.3333 | 1 | 0 | 2 | 0 | 1 | 0 | 2 | 0.5 | 6 | 0.1667 |
| 3_13394550 | 1 | 1 | NA | NA | 1 | 0 | 1 | 0 | 3 | 0.3333 | 2 | 0 | NA | NA | NA | NA | 1 | 1 | 3 | 0.3333 |
| 3_13394568 | 1 | 1 | NA | NA | 1 | 1 | 1 | 1 | 3 | 1 | 1 | 1 | 1 | 1 | NA | NA | NA | NA | 2 | 1 |
| 3_13394602 | 1 | 0 | NA | NA | 2 | 0.5 | 1 | 1 | 4 | 0.5 | 1 | 1 | 1 | 0 | NA | NA | 1 | 1 | 3 | 0.6667 |
| 3_13394603 | 3 | 0.3333 | NA | NA | 3 | 0.6667 | 2 | 0.5 | 8 | 0.5 | 2 | 0.5 | 2 | 0.5 | NA | NA | 1 | 1 | 5 | 0.6 |

| | | | | | | | | | | | | | | | | | | | | |
|---|---|---|---|---|---|---|---|---|---|---|---|---|---|---|---|---|---|---|---|---|
| 3_1339460( | 2 | 0.5 | NA | NA | 3 | 1 | 1 | 1 | 6 | 0.8333 | 1 | 1 | 1 | 0 | NA | NA | 1 | 1 | 3 | 0.6667 |
| 3_1339461: | 3 | 0.3333 | NA | NA | 3 | 1 | 2 | 1 | 8 | 0.75 | 2 | 1 | 2 | 1 | 1 | 0 | 1 | 1 | 6 | 0.8333 |
| 3_1339462: | 3 | 1 | 1 | 1 | 2 | 1 | 2 | 1 | 8 | 1 | 2 | 1 | 2 | 1 | 1 | 0 | 1 | 1 | 6 | 0.8333 |
| 3_1339462! | 2 | 0.5 | 1 | 1 | 3 | 0.6667 | 2 | 1 | 8 | 0.75 | 2 | 1 | 1 | 0 | NA | NA | 1 | 1 | 4 | 0.75 |
| 3_1339464 | 3 | 0.6667 | 1 | 1 | 3 | 0.6667 | 2 | 1 | 9 | 0.7778 | 2 | 0.5 | 2 | 0.5 | 1 | 1 | 1 | 1 | 6 | 0.6667 |
| 3_1339640! | 2 | 1 | 3 | 0.6667 | 2 | 1 | 3 | 0.6667 | 10 | 0.8 | 1 | 1 | 1 | 0 | NA | NA | NA | NA | 2 | 0.5 |
| 3_1339660! | 3 | 0.3333 | 1 | 0 | 2 | 0 | 2 | 0 | 8 | 0.125 | 1 | 1 | 2 | 0.5 | NA | NA | 2 | 0 | 5 | 0.4 |
| 3_1339661! | 2 | 1 | 2 | 0 | 4 | 0 | 3 | 0 | 11 | 0.1818 | 1 | 0 | 2 | 0 | 2 | 0 | 2 | 0.5 | 7 | 0.1429 |
| 3_1339669! | 3 | 0.3333 | 2 | 0 | 4 | 0.25 | 5 | 0 | 14 | 0.1429 | 1 | 1 | 3 | 0 | 4 | 0 | 3 | 0.3333 | 11 | 0.1818 |
| 3_1339670( | 2 | 0 | 1 | 0 | 2 | 0 | 3 | 0.3333 | 8 | 0.125 | 1 | 0 | 3 | 0 | 2 | 0.5 | 4 | 0.25 | 10 | 0.2 |
| 3_1339676: | 3 | 0.3333 | NA | NA | 4 | 0.5 | 3 | 0.3333 | 10 | 0.4 | 2 | 0.5 | 3 | 0 | 5 | 0.6 | 4 | 0.5 | 14 | 0.4286 |
| 3_1339678: | 3 | 0 | NA | NA | 3 | 0.3333 | 3 | 0.3333 | 9 | 0.2222 | 2 | 0 | 2 | 0 | 3 | 0.6667 | 2 | 0 | 9 | 0.2222 |
| 3_1340904( | NA | NA | NA | NA | 1 | 1 | NA | NA | 1 | 1 | NA | NA | 1 | 1 | NA | NA | NA | NA | 1 | 1 |
| 3_1340907: | NA | NA | NA | NA | 1 | 1 | NA | NA | 1 | 1 | NA | NA | 1 | 1 | NA | NA | NA | NA | 1 | 1 |
| 3_1340908( | NA | NA | NA | NA | 1 | 1 | NA | NA | 1 | 1 | NA | NA | 1 | 1 | NA | NA | NA | NA | 1 | 1 |
| 3_1340909( | NA | NA | NA | NA | 1 | 1 | NA | NA | 1 | 1 | NA | NA | 1 | 1 | NA | NA | NA | NA | 1 | 1 |
| 3_1350480( | 1 | 0 | 1 | 1 | NA | NA | NA | NA | 2 | 0.5 | NA | NA | NA | NA | 1 | 0 | 1 | 1 | 2 | 0.5 |
| 3_1351125: | 4 | 0.5 | 2 | 0.5 | 7 | 0.1429 | 7 | 0.2857 | 20 | 0.3 | 2 | 1 | 6 | 0 | 5 | 0 | 10 | 0.1 | 23 | 0.1304 |
| 3_1351126( | 6 | 0.5 | 2 | 0 | 8 | 0.125 | 6 | 0.3333 | 22 | 0.2727 | 2 | 0 | 7 | 0.2857 | 5 | 0.2 | 11 | 0.3636 | 25 | 0.28 |
| 3_1351134 | 3 | 0 | 2 | 0 | 5 | 0.4 | 7 | 0 | 17 | 0.1176 | 1 | 0 | 5 | 0 | 4 | 0 | 8 | 0.25 | 18 | 0.1111 |
| 3_1351135: | 3 | 0 | 1 | 1 | 5 | 0 | 7 | 0 | 16 | 0.0625 | 2 | 0 | 6 | 0.1667 | 5 | 0.4 | 8 | 0.25 | 21 | 0.2381 |
| 3_1351140! | 3 | 0.6667 | 5 | 0.2 | 6 | 0.6667 | 8 | 0.25 | 22 | 0.4091 | 1 | 0 | 8 | 0.25 | 6 | 0.3333 | 8 | 0.25 | 23 | 0.2609 |
| 3_1351141: | 2 | 0 | 4 | 0 | 5 | 0.4 | 6 | 0.5 | 17 | 0.2941 | NA | NA | 7 | 0.1429 | 6 | 0.1667 | 5 | 0.4 | 18 | 0.2222 |
| 3_1351142! | 3 | 0 | 6 | 0.1667 | 6 | 0.3333 | 10 | 0.1 | 25 | 0.16 | 2 | 0.5 | 9 | 0.1111 | 6 | 0 | 6 | 0 | 23 | 0.087 |
| 3_1351710 | 33 | 0.1515 | 31 | 0.0968 | 30 | 0.2333 | 34 | 0.1765 | 128 | 0.1641 | 26 | 0.1923 | 34 | 0.0882 | 33 | 0.2424 | 39 | 0.2051 | 132 | 0.1818 |
| 3_1351711! | 29 | 0.4828 | 27 | 0.4444 | 26 | 0.4615 | 35 | 0.4857 | 117 | 0.4701 | 25 | 0.48 | 34 | 0.4706 | 29 | 0.6552 | 32 | 0.5625 | 120 | 0.5417 |
| 3_1351712( | 40 | 0.125 | 29 | 0.3448 | 32 | 0.3438 | 31 | 0.2581 | 132 | 0.2576 | 26 | 0.2692 | 33 | 0.303 | 32 | 0.1562 | 38 | 0.2895 | 129 | 0.2558 |
| 3_1351712: | 29 | 0.0345 | 20 | 0.05 | 26 | 0.2308 | 30 | 0.1 | 105 | 0.1048 | 22 | 0.2727 | 37 | 0.0541 | 22 | 0.0455 | 34 | 0.0294 | 115 | 0.087 |
| 3_1351713 | 35 | 0.1143 | 25 | 0 | 31 | 0.129 | 35 | 0.1429 | 126 | 0.1032 | 27 | 0.1852 | 35 | 0.2 | 32 | 0.125 | 33 | 0.1212 | 127 | 0.1575 |
| 3_1351720! | 25 | 0.24 | 23 | 0.2609 | 30 | 0.1333 | 35 | 0.1714 | 113 | 0.1947 | 20 | 0.25 | 27 | 0.037 | 22 | 0.1818 | 32 | 0.1562 | 101 | 0.1485 |
| 3_1351721( | 25 | 0.04 | 22 | 0.0909 | 28 | 0.0357 | 36 | 0.1944 | 111 | 0.0991 | 20 | 0.15 | 28 | 0.0714 | 22 | 0.1818 | 32 | 0.1875 | 102 | 0.1471 |
| 3_1352761 | 1 | 1 | NA | NA | 1 | 1 | NA | NA | 2 | 1 | NA | NA | 1 | 1 | NA | NA | NA | NA | 1 | 1 |
| 3_1352762( | 1 | 1 | NA | NA | 1 | 0 | NA | NA | 2 | 0.5 | NA | NA | 1 | 1 | NA | NA | NA | NA | 1 | 1 |
| 3_1352762: | 1 | 1 | NA | NA | 1 | 1 | NA | NA | 2 | 1 | NA | NA | 1 | 1 | NA | NA | NA | NA | 1 | 1 |
| 3_1352764 | 1 | 1 | NA | NA | 1 | 1 | NA | NA | 2 | 1 | NA | NA | 1 | 1 | NA | NA | NA | NA | 1 | 1 |
| 3_1352911: | 2 | 1 | 2 | 1 | 1 | 1 | NA | NA | 5 | 1 | 1 | 1 | 1 | 1 | 1 | 1 | 3 | 1 | 6 | 1 |
| 3_1355937 | NA | NA | NA | NA | NA | NA | 1 | 1 | 1 | 1 | NA | NA | NA | NA | NA | NA | 1 | 1 | 1 | 1 |
| 3_1363759: | 5 | 0 | 5 | 0 | 9 | 0.3333 | 11 | 0.0909 | 30 | 0.1333 | 4 | 0 | 6 | 0.1667 | 10 | 0.3 | 13 | 0.0769 | 33 | 0.1515 |
| 3_1366409: | 76 | 0.3158 | 63 | 0.381 | 91 | 0.3407 | 92 | 0.3587 | 322 | 0.3478 | 40 | 0.4 | 87 | 0.3908 | 86 | 0.3488 | 99 | 0.3434 | 312 | 0.3654 |
| 3_1366409! | 68 | 0.2206 | 60 | 0.2667 | 82 | 0.1585 | 91 | 0.2198 | 301 | 0.2126 | 37 | 0.2703 | 83 | 0.253 | 76 | 0.2632 | 88 | 0.2386 | 284 | 0.2535 |
| 3_1366413 | 86 | 0.0465 | 65 | 0.2154 | 84 | 0.1548 | 91 | 0.0879 | 326 | 0.1196 | 31 | 0.2258 | 87 | 0.1034 | 92 | 0.1413 | 92 | 0.087 | 302 | 0.1225 |
| 3_1366413! | 92 | 0.087 | 73 | 0.0959 | 86 | 0.093 | 106 | 0.066 | 357 | 0.084 | 39 | 0.1026 | 92 | 0.0543 | 89 | 0.0112 | 96 | 0.0729 | 316 | 0.0538 |
| 3_1366414 | 99 | 0.2828 | 74 | 0.3919 | 93 | 0.3548 | 103 | 0.2524 | 369 | 0.3144 | 41 | 0.3171 | 98 | 0.2551 | 102 | 0.3529 | 103 | 0.3981 | 344 | 0.3343 |
| 3_1366420: | 108 | 0.3519 | 88 | 0.3068 | 116 | 0.3621 | 122 | 0.2787 | 434 | 0.3249 | 54 | 0.3333 | 114 | 0.3246 | 129 | 0.3256 | 108 | 0.3611 | 405 | 0.3358 |
| 3_1366428! | 48 | 0.1667 | 51 | 0.3333 | 65 | 0.2615 | 60 | 0.1833 | 224 | 0.2366 | 35 | 0.2571 | 58 | 0.2586 | 59 | 0.2373 | 53 | 0.3774 | 205 | 0.2829 |
| 3_1373318 | NA | NA | NA | NA | 1 | 1 | NA | NA | 1 | 1 | 2 | 0 | 1 | 1 | 1 | 0 | 2 | 0.5 | 6 | 0.3333 |
| 3_1373320! | NA | NA | NA | NA | 1 | 1 | NA | NA | 1 | 1 | 2 | 1 | 1 | 1 | 1 | 0 | 2 | 1 | 6 | 0.8333 |
| 3_1373321: | NA | NA | NA | NA | 1 | 1 | NA | NA | 1 | 1 | 1 | 1 | 1 | 1 | 1 | 1 | 2 | 1 | 5 | 1 |
| 3_1375722: | 46 | 0.3478 | 39 | 0.4872 | 31 | 0.4516 | 34 | 0.4412 | 150 | 0.4267 | 25 | 0.28 | 32 | 0.375 | 38 | 0.5789 | 39 | 0.4872 | 134 | 0.4478 |
| 3_1375722! | 46 | 0.3913 | 43 | 0.4884 | 32 | 0.375 | 28 | 0.3571 | 149 | 0.4094 | 25 | 0.48 | 30 | 0.5 | 32 | 0.5 | 41 | 0.439 | 128 | 0.4766 |
| 3_1375726 | 18 | 0.1667 | 16 | 0.0625 | 14 | 0.0714 | 16 | 0.125 | 64 | 0.1094 | 11 | 0.0909 | 15 | 0.0667 | 20 | 0.05 | 22 | 0.0455 | 68 | 0.0588 |
| 3_1376117 | 1 | 1 | 2 | 0.5 | 3 | 0.6667 | 1 | 1 | 7 | 0.7143 | NA | NA | 1 | 1 | 1 | 0 | NA | NA | 2 | 0.5 |
| 3_1376143 | 2 | 0.5 | 1 | 0 | 3 | 1 | 1 | 1 | 7 | 0.7143 | 1 | 1 | 1 | 1 | 1 | 0 | NA | NA | 3 | 0.6667 |
| 3_1376143! | 1 | 1 | 1 | 1 | 2 | 1 | 1 | 1 | 5 | 1 | NA | NA | 1 | 1 | NA | NA | NA | NA | 1 | 1 |
| 3_1376143( | 1 | 1 | 1 | 1 | 4 | 1 | 1 | 1 | 7 | 1 | NA | NA | 1 | 1 | NA | NA | NA | NA | 1 | 1 |
| 3_1376384! | 2 | 1 | 1 | 1 | 1 | 1 | NA | NA | 4 | 1 | NA | NA | 1 | 1 | NA | NA | NA | NA | 1 | 1 |
| 3_1376388! | NA | NA | 1 | 0 | 1 | 1 | NA | NA | 2 | 0.5 | NA | NA | 1 | 1 | NA | NA | NA | NA | 1 | 1 |
| 3_1377968! | NA | NA | 3 | 0.6667 | 1 | 0 | 2 | 0.5 | 6 | 0.5 | 2 | 0 | NA | NA | 1 | 0 | 3 | 0.6667 | 6 | 0.3333 |
| 3_1378491! | NA | NA | NA | NA | NA | NA | 1 | 1 | 1 | 1 | NA | NA | NA | NA | 1 | 1 | NA | NA | 1 | 1 |
| 3_1378625( | 1 | 1 | NA | NA | NA | NA | NA | NA | 1 | 1 | 1 | 0 | 1 | 1 | 1 | 0 | NA | NA | 3 | 0.3333 |
| 3_1378626 | 1 | 1 | NA | NA | NA | NA | NA | NA | 1 | 1 | NA | NA | 1 | 0 | 1 | 1 | NA | NA | 2 | 0.5 |
| 3_1378627! | 1 | 1 | NA | NA | NA | NA | NA | NA | 1 | 1 | 1 | 1 | 1 | 0 | 1 | 1 | NA | NA | 3 | 0.6667 |
| 3_1378629( | 1 | 1 | NA | NA | NA | NA | NA | NA | 1 | 1 | 1 | 1 | 1 | 0 | 1 | 0 | NA | NA | 3 | 0.3333 |
| 3_1378631: | 1 | 1 | NA | NA | NA | NA | NA | NA | 1 | 1 | 1 | 1 | 1 | 1 | 1 | 1 | NA | NA | 3 | 1 |
| 3_1381567 | 4 | 0.5 | 5 | 0.2 | 5 | 0 | 3 | 0.3333 | 17 | 0.2353 | 8 | 0.125 | 2 | 0 | 5 | 0.2 | 8 | 0.25 | 23 | 0.1739 |

| | | | | | | | | | | | | | | | | | | | | |
|---|---|---|---|---|---|---|---|---|---|---|---|---|---|---|---|---|---|---|---|---|
| 3_13815705 | 4 | 0 | 6 | 0.3333 | 9 | 0.2222 | 4 | 0.5 | 23 | 0.2609 | 8 | 0.375 | 1 | 0 | 4 | 0 | 6 | 0.1667 | 19 | 0.2105 |
| 3_13816099 | 8 | 0.125 | 10 | 0.1 | 12 | 0.25 | 4 | 0 | 34 | 0.1471 | 11 | 0 | 6 | 0 | 8 | 0 | 9 | 0.2222 | 34 | 0.0588 |
| 3_13816547 | 7 | 0.1429 | 6 | 0 | 6 | 0.1667 | 7 | 0.2857 | 26 | 0.1538 | 3 | 0.3333 | 9 | 0.1111 | 4 | 0 | 7 | 0.2857 | 23 | 0.1739 |
| 3_13816564 | 5 | 0.4 | 4 | 0 | 5 | 0.2 | 8 | 0 | 22 | 0.1364 | 4 | 0.5 | 5 | 0 | 3 | 0 | 4 | 0.25 | 16 | 0.1875 |
| 3_13816573 | 7 | 0.1429 | 6 | 0 | 6 | 0.1667 | 9 | 0.3333 | 28 | 0.1786 | 4 | 0.5 | 7 | 0.5714 | 3 | 0 | 6 | 0 | 20 | 0.3 |
| 3_13816591 | 6 | 0 | 7 | 0.1429 | 7 | 0 | 8 | 0.125 | 28 | 0.0714 | 4 | 0 | 5 | 0 | 4 | 0 | 4 | 0.25 | 17 | 0.0588 |
| 3_13816597 | 6 | 0.8333 | 7 | 0.7143 | 7 | 0.7143 | 7 | 0.8571 | 27 | 0.7778 | 2 | 1 | 6 | 1 | 5 | 0.8 | 4 | 0.75 | 17 | 0.8824 |
| 3_13816599 | 7 | 0.5714 | 7 | 0.1429 | 5 | 0.4 | 8 | 0.25 | 27 | 0.3333 | 4 | 0.5 | 7 | 0.2857 | 5 | 0.2 | 4 | 0.75 | 20 | 0.4 |
| 3_13816607 | 5 | 0.4 | 4 | 0.25 | 6 | 0.1667 | 6 | 0.1667 | 21 | 0.2381 | 3 | 0 | 3 | 0.6667 | 3 | 0.3333 | 3 | 0.3333 | 12 | 0.3333 |
| 3_13816616 | 7 | 0.4286 | 6 | 0.3333 | 7 | 0.7143 | 8 | 0.375 | 28 | 0.4643 | 4 | 0.5 | 8 | 0.25 | 5 | 0.4 | 4 | 0.25 | 21 | 0.3333 |
| 3_13825944 | 10 | 0.4 | 21 | 0.1429 | 16 | 0.3125 | 20 | 0.15 | 67 | 0.2239 | 7 | 0.1429 | 8 | 0.25 | 13 | 0.2308 | 12 | 0.1667 | 40 | 0.2 |
| 3_13826117 | 11 | 0.2727 | 23 | 0.2609 | 21 | 0.3333 | 20 | 0.1 | 75 | 0.24 | 7 | 0 | 8 | 0.25 | 12 | 0.3333 | 13 | 0.0769 | 40 | 0.175 |
| 3_13826129 | 13 | 0.0769 | 24 | 0.0833 | 21 | 0.1905 | 22 | 0 | 80 | 0.0875 | 7 | 0.1429 | 12 | 0 | 13 | 0.1538 | 12 | 0.1667 | 44 | 0.1136 |
| 3_13826135 | 11 | 0.5455 | 24 | 0.7917 | 21 | 0.619 | 23 | 0.8261 | 79 | 0.7215 | 10 | 0.6 | 13 | 0.7692 | 14 | 0.5714 | 13 | 0.6154 | 50 | 0.64 |
| 3_13826160 | 13 | 0.5385 | 25 | 0.6 | 20 | 0.7 | 22 | 0.5 | 80 | 0.5875 | 10 | 0.6 | 13 | 0.5385 | 13 | 0.4615 | 14 | 0.5714 | 50 | 0.54 |
| 3_13826166 | 13 | 0.6154 | 22 | 0.5455 | 21 | 0.381 | 21 | 0.4286 | 77 | 0.4805 | 11 | 0.7273 | 12 | 0.5 | 14 | 0.4286 | 13 | 0.3846 | 50 | 0.5 |
| 3_13826201 | 12 | 0.0833 | 18 | 0.2222 | 17 | 0.2353 | 21 | 0.3333 | 68 | 0.2353 | 7 | 0.4286 | 11 | 0.2727 | 15 | 0.0667 | 13 | 0.3077 | 46 | 0.2391 |
| 3_13826246 | 4 | 0.25 | 7 | 0.1429 | 8 | 0.5 | 9 | 0.4444 | 28 | 0.3571 | 3 | 0.3333 | 5 | 0.4 | 6 | 0.1667 | 4 | 0.75 | 18 | 0.3889 |
| 3_13826565 | 1 | 1 | NA | NA | NA | NA | NA | NA | 1 | 1 | 1 | 1 | NA | NA | NA | NA | 1 | 0 | 2 | 0.5 |
| 3_13860823 | 13 | 1 | 18 | 0.7222 | 13 | 0.9231 | 11 | 0.8182 | 55 | 0.8545 | 13 | 0.8462 | 10 | 0.8 | 8 | 1 | 15 | 0.9333 | 46 | 0.8913 |
| 3_13860825 | 9 | 0.2222 | 13 | 0.6923 | 9 | 0.4444 | 8 | 0.25 | 39 | 0.4359 | 8 | 0.375 | 7 | 0.1429 | 5 | 0 | 13 | 0.4615 | 33 | 0.303 |
| 3_13860834 | 13 | 0.4615 | 12 | 0.5 | 13 | 0.8462 | 10 | 0.6 | 48 | 0.6042 | 12 | 0.75 | 10 | 0.7 | 7 | 0.5714 | 17 | 0.7647 | 46 | 0.7174 |
| 3_13860834 | 10 | 0.2 | 7 | 0.2857 | 11 | 0 | 10 | 0 | 38 | 0.1053 | 8 | 0.125 | 6 | 0.1667 | 5 | 0 | 10 | 0 | 29 | 0.069 |
| 3_13860836 | 16 | 0.5 | 16 | 0.625 | 11 | 0.7273 | 12 | 0.5 | 55 | 0.5818 | 10 | 0.7 | 9 | 0.5556 | 6 | 0.3333 | 19 | 0.5789 | 44 | 0.5682 |
| 3_13860839 | 19 | 0.2632 | 19 | 0.3158 | 14 | 0.4286 | 12 | 0.6667 | 64 | 0.3906 | 13 | 0.3077 | 10 | 0.4 | 11 | 0.4545 | 20 | 0.45 | 54 | 0.4074 |
| 3_13860873 | 17 | 0.0588 | 21 | 0.1905 | 12 | 0.25 | 16 | 0.125 | 66 | 0.1515 | 15 | 0.0667 | 9 | 0.2222 | 8 | 0.25 | 19 | 0.1579 | 51 | 0.1569 |
| 3_13860879 | 19 | 0.7368 | 21 | 0.8095 | 14 | 0.9286 | 15 | 0.7333 | 69 | 0.7971 | 17 | 0.7059 | 11 | 0.5455 | 10 | 0.6 | 24 | 0.875 | 62 | 0.7258 |
| 3_13860886 | 21 | 0.4286 | 24 | 0.3333 | 14 | 0.5 | 17 | 0.7059 | 76 | 0.4737 | 17 | 0.4706 | 13 | 0.6923 | 11 | 0.3636 | 23 | 0.4783 | 64 | 0.5 |
| 3_13860996 | 20 | 0.15 | 33 | 0.3333 | 20 | 0.35 | 14 | 0.0714 | 87 | 0.2529 | 20 | 0.3 | 14 | 0.3571 | 16 | 0.25 | 22 | 0.2727 | 72 | 0.2917 |
| 3_13861001 | 28 | 0.0714 | 34 | 0.1765 | 23 | 0.3043 | 18 | 0.3333 | 103 | 0.2039 | 22 | 0.3636 | 15 | 0.3333 | 20 | 0.35 | 27 | 0.3704 | 84 | 0.3571 |
| 3_13863189 | 32 | 0.4062 | 40 | 0.4 | 41 | 0.5854 | 32 | 0.375 | 145 | 0.4483 | 24 | 0.4583 | 18 | 0.2778 | 19 | 0.4211 | 28 | 0.4286 | 89 | 0.4045 |
| 3_13863216 | 26 | 0.0769 | 37 | 0.2162 | 37 | 0.0811 | 30 | 0.1667 | 130 | 0.1385 | 24 | 0.1667 | 19 | 0.1053 | 19 | 0.1579 | 28 | 0.1786 | 90 | 0.1556 |
| 3_13863243 | 22 | 0.1364 | 37 | 0.0541 | 27 | 0.037 | 22 | 0 | 108 | 0.0556 | 17 | 0.1176 | 17 | 0.1176 | 18 | 0.0556 | 23 | 0.087 | 75 | 0.0933 |
| 3_13863315 | 15 | 0.0667 | 26 | 0.1923 | 26 | 0.1538 | 18 | 0.0556 | 85 | 0.1294 | 16 | 0.125 | 16 | 0.1875 | 14 | 0.0714 | 19 | 0.1579 | 65 | 0.1385 |
| 3_13863321 | 15 | 0.4667 | 23 | 0.6087 | 24 | 0.375 | 17 | 0.2353 | 79 | 0.4304 | 14 | 0.5714 | 14 | 0.6429 | 13 | 0.3077 | 17 | 0.4118 | 58 | 0.4828 |
| 3_13863334 | 10 | 0.2 | 16 | 0.1875 | 21 | 0.1905 | 14 | 0.1429 | 61 | 0.1803 | 13 | 0.4615 | 9 | 0.1111 | 12 | 0.1667 | 13 | 0.4615 | 47 | 0.3191 |
| 3_13863336 | 11 | 0.1818 | 23 | 0.5652 | 26 | 0.4615 | 17 | 0.5294 | 77 | 0.4675 | 15 | 0.8 | 11 | 0.9091 | 13 | 0.5385 | 19 | 0.6316 | 58 | 0.7069 |
| 3_13878169 | NA | NA | NA | NA | 2 | 0.5 | NA | NA | 2 | 0.5 | NA | NA | NA | NA | NA | NA | 1 | 1 | 1 | 1 |
| 3_13878191 | NA | NA | NA | NA | 2 | 0.5 | NA | NA | 2 | 0.5 | NA | NA | NA | NA | NA | NA | 1 | 1 | 1 | 1 |
| 3_13878195 | NA | NA | NA | NA | 2 | 0.5 | NA | NA | 2 | 0.5 | NA | NA | NA | NA | NA | NA | 1 | 1 | 1 | 1 |
| 3_14153647 | NA | NA | 2 | 0 | 2 | 0.5 | 1 | 0 | 5 | 0.2 | 1 | 1 | 1 | 0 | NA | NA | 1 | 0 | 3 | 0.3333 |
| 3_14153651 | NA | NA | 3 | 0.3333 | 2 | 0 | 2 | 0 | 7 | 0.1429 | 1 | 1 | 1 | 1 | NA | NA | 1 | 0 | 3 | 0.6667 |
| 3_14153660 | NA | NA | 3 | 0.3333 | 2 | 0 | 2 | 0.5 | 7 | 0.2857 | 1 | 1 | 1 | 0 | NA | NA | 1 | 0 | 3 | 0.3333 |
| 3_14153669 | NA | NA | NA | NA | 1 | 0 | 2 | 0.5 | 3 | 0.3333 | 1 | 1 | 1 | 0 | NA | NA | 1 | 0 | 3 | 0.3333 |
| 3_14153682 | NA | NA | 2 | 0 | 2 | 0.5 | 1 | 0 | 5 | 0.2 | 1 | 1 | 1 | 0 | NA | NA | 1 | 0 | 3 | 0.3333 |
| 3_14154315 | 2 | 0 | 4 | 0.25 | NA | NA | 3 | 0 | 9 | 0.1111 | 3 | 0.6667 | 9 | 0.1111 | 2 | 0 | 2 | 0 | 16 | 0.1875 |
| 3_14154378 | 2 | 0.5 | 3 | 0.3333 | 1 | 0 | 3 | 0 | 9 | 0.2222 | 2 | 0 | 7 | 0.2857 | 2 | 0.5 | 2 | 0 | 13 | 0.2308 |
| 3_14154518 | 2 | 0.5 | 3 | 0 | NA | NA | NA | NA | 5 | 0.2 | 2 | 0 | 4 | 0 | 2 | 0.5 | 2 | 0 | 10 | 0.1 |
| 3_14154582 | 2 | 1 | 3 | 0.3333 | 1 | 0 | 1 | 1 | 7 | 0.5714 | 1 | 1 | 5 | 0.2 | 1 | 0 | 2 | 0.5 | 9 | 0.3333 |
| 3_14154590 | 2 | 1 | 3 | 0.6667 | 1 | 0 | 1 | 1 | 7 | 0.7143 | 1 | 1 | 4 | 1 | 2 | 1 | 2 | 0 | 9 | 0.7778 |
| 3_14154620 | 2 | 0.5 | 3 | 0.6667 | NA | NA | NA | NA | 5 | 0.6 | 1 | 1 | 3 | 1 | 2 | 0 | 2 | 1 | 8 | 0.75 |
| 3_14154628 | 2 | 1 | 3 | 1 | 1 | 0 | 1 | 1 | 7 | 0.8571 | 1 | 1 | 4 | 0.5 | 2 | 0.5 | 1 | 1 | 8 | 0.625 |
| 3_14154635 | NA | NA | 3 | 1 | 1 | 1 | 1 | 1 | 5 | 1 | 1 | 0 | 4 | 0.75 | 2 | 1 | 1 | 1 | 8 | 0.75 |
| 3_14154638 | 2 | 0.5 | 3 | 0.3333 | NA | NA | 1 | 0 | 6 | 0.3333 | 1 | 1 | 4 | 0.25 | 1 | 0 | 2 | 0.5 | 8 | 0.375 |
| 3_14156234 | 1 | 0 | 2 | 0.5 | 1 | 0 | 1 | 0 | 5 | 0.2 | 1 | 0 | 4 | 0.25 | 2 | 0 | 2 | 0 | 9 | 0.1111 |
| 3_14194832 | 4 | 0 | 2 | 0.5 | 3 | 0.3333 | NA | NA | 9 | 0.2222 | 2 | 0.5 | 1 | 1 | 1 | 1 | 1 | 0 | 5 | 0.6 |
| 3_14225253 | NA | NA | 1 | 1 | NA | NA | NA | NA | 1 | 1 | NA | NA | 1 | 0 | 1 | 1 | NA | NA | 2 | 0.5 |
| 3_14225340 | NA | NA | 3 | 0.6667 | NA | NA | NA | NA | 3 | 0.6667 | NA | NA | 1 | 0 | 1 | 1 | NA | NA | 2 | 0.5 |
| 3_14225367 | NA | NA | 3 | 1 | NA | NA | NA | NA | 3 | 1 | NA | NA | 1 | 0 | 1 | 1 | NA | NA | 2 | 0.5 |
| 3_14225367 | NA | NA | 3 | 0.6667 | NA | NA | NA | NA | 3 | 0.6667 | NA | NA | 1 | 0 | 1 | 1 | NA | NA | 2 | 0.5 |
| 3_14225381 | NA | NA | 2 | 1 | NA | NA | NA | NA | 2 | 1 | NA | NA | 1 | 1 | 1 | 1 | NA | NA | 2 | 1 |
| 3_14225420 | NA | NA | 2 | 0.5 | NA | NA | NA | NA | 2 | 0.5 | 1 | 0 | 1 | 0 | 1 | 1 | NA | NA | 3 | 0.3333 |
| 3_14225547 | NA | NA | 3 | 0.3333 | NA | NA | NA | NA | 3 | 0.3333 | 1 | 1 | 1 | 0 | 1 | 0 | NA | NA | 3 | 0.3333 |
| 3_14233921 | NA | NA | 1 | 0 | NA | NA | 1 | 1 | 2 | 0.5 | 1 | 1 | 1 | 1 | NA | NA | 1 | 0 | 3 | 0.6667 |
| 3_14234362 | NA | NA | 1 | 0 | NA | NA | 1 | 1 | 2 | 0.5 | 1 | 0 | 1 | 0 | NA | NA | 1 | 1 | 3 | 0.3333 |

| | | | | | | | | | | | | | | | | | | | | |
|---|---|---|---|---|---|---|---|---|---|---|---|---|---|---|---|---|---|---|---|---|
| 3_1424499 | 3 | 0.6667 | 1 | 0 | NA | NA | 1 | 1 | 5 | 0.6 | 1 | 0 | 1 | 0 | 2 | 0.5 | 1 | 1 | 5 | 0.4 |
| 3_1424499 | 2 | 0.5 | 1 | 0 | NA | NA | 3 | 0.3333 | 6 | 0.3333 | 1 | 0 | 1 | 0 | 3 | 0.3333 | NA | NA | 5 | 0.2 |
| 3_1424872 | 9 | 0.1111 | 4 | 0 | 3 | 0.3333 | 6 | 0 | 22 | 0.0909 | 2 | 0.5 | 4 | 0 | 3 | 0 | 1 | 0 | 10 | 0.1 |
| 3_1424890 | 10 | 0.5 | 2 | 0.5 | 3 | 0.3333 | 9 | 0.4444 | 24 | 0.4583 | 3 | 0 | 4 | 0.5 | 3 | 0.6667 | 2 | 0 | 12 | 0.3333 |
| 3_1424891 | 11 | 0.4545 | 4 | 0.25 | 4 | 1 | 10 | 0.6 | 29 | 0.5517 | 6 | 0.6667 | 6 | 0.5 | 5 | 0.6 | 2 | 0.5 | 19 | 0.5789 |
| 3_1424986 | 3 | 0 | 3 | 0 | 3 | 0.3333 | 1 | 0 | 10 | 0.1 | 1 | 0 | 1 | 1 | 5 | 0 | 1 | 0 | 8 | 0.125 |
| 3_1424988 | 2 | 0.5 | 2 | 0 | 3 | 0 | NA | NA | 7 | 0.1429 | NA | NA | 1 | 1 | 4 | 0 | 1 | 0 | 6 | 0.1667 |
| 3_1424988 | 3 | 1 | 2 | 1 | 2 | 1 | 1 | 1 | 8 | 1 | NA | NA | 1 | 1 | 4 | 1 | 1 | 1 | 6 | 1 |
| 3_1424989 | 3 | 0.3333 | 2 | 0 | 3 | 0 | 1 | 0 | 9 | 0.1111 | NA | NA | 1 | 0 | 5 | 0.2 | 1 | 0 | 7 | 0.1429 |
| 3_1424992 | 2 | 0.5 | NA | NA | 2 | 0 | 2 | 0 | 6 | 0.1667 | NA | NA | 1 | 0 | 4 | 0.25 | 1 | 0 | 6 | 0.1667 |
| 3_1424993 | 2 | 1 | 1 | 0 | 3 | 0.6667 | 2 | 0.5 | 8 | 0.625 | NA | NA | NA | NA | 4 | 0.5 | 1 | 1 | 5 | 0.6 |
| 3_1424994 | 2 | 1 | NA | NA | 2 | 0.5 | 2 | 0 | 6 | 0.5 | NA | NA | NA | NA | 2 | 0.5 | 1 | 1 | 3 | 0.6667 |
| 3_1424995 | 2 | 1 | NA | NA | 2 | 0.5 | 1 | 0 | 5 | 0.6 | NA | NA | NA | NA | 3 | 0.6667 | 1 | 0 | 4 | 0.5 |
| 3_1424995 | 3 | 0.6667 | 1 | 0 | 2 | 0.5 | 1 | 1 | 7 | 0.5714 | NA | NA | NA | NA | 2 | 0.5 | 1 | 0 | 3 | 0.3333 |
| 3_1424996 | 2 | 0.5 | 1 | 1 | 2 | 0 | 2 | 0 | 7 | 0.2857 | NA | NA | NA | NA | 4 | 0.5 | 1 | 1 | 5 | 0.6 |
| 3_1438948 | 25 | 0.4 | 19 | 0.3684 | 24 | 0.5 | 28 | 0.1786 | 96 | 0.3542 | 12 | 0.0833 | 23 | 0.1739 | 24 | 0.2917 | 21 | 0.3333 | 80 | 0.2375 |
| 3_1438948 | 30 | 0.1 | 20 | 0.05 | 29 | 0.0345 | 32 | 0.0312 | 111 | 0.0541 | 16 | 0 | 33 | 0.1212 | 29 | 0.069 | 30 | 0 | 108 | 0.0556 |
| 3_1438948 | 27 | 0.6296 | 20 | 0.45 | 29 | 0.5517 | 34 | 0.7059 | 110 | 0.6 | 13 | 0.5385 | 32 | 0.5312 | 29 | 0.5172 | 31 | 0.7097 | 105 | 0.581 |
| 3_1438948 | 22 | 0.2727 | 18 | 0.2222 | 24 | 0.0833 | 28 | 0.2143 | 92 | 0.1957 | 13 | 0.4615 | 28 | 0.4286 | 30 | 0.1 | 29 | 0.0345 | 100 | 0.22 |
| 3_1438948 | 25 | 0.28 | 20 | 0.2 | 25 | 0.2 | 32 | 0.2812 | 102 | 0.2451 | 16 | 0.25 | 32 | 0.4062 | 32 | 0.25 | 30 | 0.2 | 110 | 0.2818 |
| 3_1441364 | NA | NA | NA | NA | 1 | 1 | NA | NA | 1 | 1 | NA | NA | 1 | 1 | 1 | 0 | 1 | 0 | 3 | 0.3333 |
| 3_1441370 | 1 | 1 | 1 | 0 | 1 | 1 | NA | NA | 3 | 0.6667 | 2 | 0.5 | 2 | 0 | 3 | 0.3333 | 2 | 0 | 9 | 0.2222 |
| 3_1441377 | 2 | 0 | 1 | 0 | 1 | 1 | NA | NA | 4 | 0.25 | 5 | 0.4 | 3 | 0.6667 | 3 | 0 | NA | NA | 11 | 0.3636 |
| 3_1441378 | 2 | 0.5 | NA | NA | 1 | 1 | NA | NA | 3 | 0.6667 | 4 | 0 | 4 | 0.5 | 2 | 0 | NA | NA | 10 | 0.2 |
| 3_1441389 | 1 | 0 | 1 | 0 | 2 | 0.5 | NA | NA | 4 | 0.25 | 3 | 0 | 2 | 0 | 2 | 0.5 | NA | NA | 7 | 0.1429 |
| 3_1441395 | 6 | 0.6667 | 7 | 0.4286 | 7 | 0.7143 | 6 | 0.3333 | 26 | 0.5385 | 8 | 0.375 | 4 | 0.5 | 6 | 0.6667 | 4 | 0.75 | 22 | 0.5455 |
| 3_1441399 | 6 | 0.8333 | 7 | 0.8571 | 7 | 1 | 4 | 1 | 24 | 0.9167 | 7 | 0.8571 | 4 | 1 | 4 | 0.75 | 5 | 1 | 20 | 0.9 |
| 3_1441400 | 5 | 0.4 | 6 | 0.8333 | 7 | 1 | 5 | 0.4 | 23 | 0.6957 | 8 | 0.625 | 4 | 0.5 | 5 | 0.6 | 6 | 0.3333 | 23 | 0.5217 |
| 3_1441401 | 6 | 0.1667 | 6 | 0.1667 | 6 | 0 | 6 | 0.1667 | 24 | 0.125 | 8 | 0.5 | 4 | 0 | 5 | 0.2 | 5 | 0 | 22 | 0.2273 |
| 3_1441401 | 6 | 0.1667 | 6 | 0.6667 | 6 | 0.8333 | 4 | 0.25 | 22 | 0.5 | 8 | 0.125 | 2 | 0 | 5 | 0.6 | 5 | 0.4 | 20 | 0.3 |
| 3_1441402 | 4 | 0.75 | 5 | 0.6 | 4 | 1 | 4 | 1 | 17 | 0.8235 | 3 | 0.6667 | 3 | 0.6667 | 4 | 1 | 4 | 0.75 | 14 | 0.7857 |
| 3_1441402 | 7 | 0.8571 | 7 | 0.7143 | 5 | 0.6 | 4 | 0.75 | 23 | 0.7391 | 5 | 0.8 | 3 | 0.3333 | 4 | 1 | 4 | 1 | 16 | 0.8125 |
| 3_1441402 | 4 | 1 | 5 | 1 | 2 | 1 | 1 | 1 | 12 | 1 | 3 | 1 | NA | NA | 3 | 1 | 4 | 1 | 10 | 1 |
| 3_1441402 | 5 | 0.8 | 6 | 0.5 | 7 | 0.5714 | 4 | 0.75 | 22 | 0.6364 | 5 | 0.4 | 4 | 1 | 4 | 0.5 | 6 | 0.6667 | 19 | 0.6316 |
| 3_1441404 | 7 | 0 | 7 | 0.2857 | 7 | 0.1429 | 6 | 0 | 27 | 0.1111 | 7 | 0 | 4 | 0 | 5 | 0.2 | 6 | 0.1667 | 22 | 0.0909 |
| 3_1442763 | NA | NA | 3 | 1 | 1 | 0 | NA | NA | 4 | 0.75 | 1 | 0 | 1 | 0 | 1 | 1 | NA | NA | 3 | 0.3333 |
| 3_1443033 | 34 | 0.1471 | 51 | 0.1373 | 38 | 0.1316 | 35 | 0.0857 | 158 | 0.1266 | 49 | 0.1633 | 41 | 0.2195 | 49 | 0.1837 | 23 | 0.2174 | 162 | 0.1914 |
| 3_1443941 | 4 | 0 | 4 | 0 | 4 | 0 | 6 | 0.1667 | 18 | 0.0556 | 3 | 0 | 2 | 0 | 10 | 0.1 | 3 | 0 | 18 | 0.0556 |
| 3_1443943 | 2 | 0 | 4 | 0 | 5 | 0.2 | 6 | 0 | 17 | 0.0588 | 3 | 0 | 3 | 0 | 10 | 0.1 | 3 | 0 | 19 | 0.0526 |
| 3_1443943 | 3 | 0 | 4 | 0.25 | 6 | 0.1667 | 6 | 0 | 19 | 0.1053 | 2 | 0 | 3 | 0.3333 | 10 | 0 | 3 | 0 | 18 | 0.0556 |
| 3_1443963 | 1 | 0 | 3 | 0 | 3 | 0 | 3 | 0.3333 | 10 | 0.1 | 3 | 0 | 2 | 0 | 5 | 0 | 1 | 1 | 11 | 0.0909 |
| 3_1443963 | 1 | 0 | 2 | 0.5 | 3 | 0 | 2 | 0 | 8 | 0.125 | 2 | 0 | 3 | 0 | 4 | 0 | 1 | 1 | 10 | 0.1 |
| 3_1443965 | 2 | 0 | 2 | 0 | 3 | 0.3333 | 4 | 0.25 | 11 | 0.1818 | 3 | 0 | 3 | 0 | 5 | 0 | 1 | 1 | 12 | 0.0833 |
| 3_1443968 | 1 | 0 | 3 | 0.3333 | 3 | 0 | 3 | 0 | 10 | 0.1 | 3 | 0.3333 | 2 | 0 | 5 | 0 | 1 | 0 | 11 | 0.0909 |
| 3_1443970 | 7 | 0.4286 | 6 | 0.8333 | 8 | 0.625 | 7 | 0.5714 | 28 | 0.6071 | 8 | 0.375 | 2 | 1 | 8 | 0.75 | 2 | 0.5 | 20 | 0.6 |
| 3_1443972 | 8 | 0.25 | 6 | 0 | 8 | 0.25 | 6 | 0.1667 | 28 | 0.1786 | 8 | 0 | 3 | 0.3333 | 7 | 0.2857 | 4 | 0.25 | 22 | 0.1818 |
| 3_1452822 | NA | NA | 1 | 1 | 1 | 0 | 3 | 0.3333 | 5 | 0.4 | 1 | 1 | 2 | 0.5 | NA | NA | 2 | 0.5 | 5 | 0.6 |
| 3_1452823 | 1 | 0 | 1 | 1 | 1 | 1 | 3 | 0.6667 | 6 | 0.6667 | 1 | 1 | 2 | 1 | 1 | 0 | 2 | 0 | 6 | 0.5 |
| 3_1452823 | 1 | 1 | 1 | 0 | 1 | 0 | 3 | 0 | 6 | 0.1667 | 1 | 1 | 2 | 0 | 1 | 0 | 2 | 0 | 6 | 0.1667 |
| 3_1452825 | 1 | 1 | 1 | 0 | NA | NA | 3 | 0 | 5 | 0.2 | 1 | 1 | 2 | 0.5 | 1 | 0 | 2 | 1 | 6 | 0.6667 |
| 3_1452825 | 1 | 0 | 1 | 0 | 1 | 0 | 2 | 0.5 | 5 | 0.2 | 1 | 0 | 3 | 0.6667 | 1 | 1 | 2 | 0 | 7 | 0.4286 |
| 3_1452832 | NA | NA | 1 | 1 | NA | NA | NA | NA | 1 | 1 | NA | NA | 1 | 1 | NA | NA | NA | NA | 1 | 1 |
| 3_1453097 | 1 | 1 | 4 | 1 | 1 | 1 | 3 | 0.3333 | 9 | 0.7778 | 1 | 1 | 5 | 0.8 | 2 | 1 | 2 | 0.5 | 10 | 0.8 |
| 3_1453099 | 1 | 0 | 3 | 0.3333 | NA | NA | NA | NA | 4 | 0.25 | NA | NA | 1 | 1 | 1 | 0 | NA | NA | 2 | 0.5 |
| 3_1453103 | 1 | 0 | 1 | 1 | NA | NA | NA | NA | 2 | 0.5 | NA | NA | 1 | 1 | 1 | 0 | NA | NA | 2 | 0.5 |
| 3_1453103 | 1 | 1 | 1 | 1 | NA | NA | NA | NA | 2 | 1 | NA | NA | 1 | 1 | 1 | 0 | NA | NA | 2 | 0.5 |
| 3_1453529 | 1 | 1 | 1 | 0 | 1 | 0 | NA | NA | 3 | 0.3333 | 1 | 0 | 2 | 0.5 | NA | NA | NA | NA | 3 | 0.3333 |
| 3_1453532 | 1 | 1 | 1 | 0 | 1 | 0 | NA | NA | 3 | 0.3333 | 1 | 0 | 2 | 1 | NA | NA | NA | NA | 3 | 0.6667 |
| 3_1453532 | 1 | 0 | 1 | 0 | 1 | 1 | NA | NA | 3 | 0.3333 | 1 | 1 | 1 | 1 | NA | NA | NA | NA | 2 | 1 |
| 3_1453533 | 2 | 0.5 | 1 | 1 | 2 | 1 | NA | NA | 5 | 0.8 | 1 | 1 | 1 | 1 | NA | NA | NA | NA | 2 | 1 |
| 3_1453534 | 2 | 1 | 1 | 1 | 2 | 1 | NA | NA | 5 | 1 | NA | NA | 2 | 1 | NA | NA | NA | NA | 2 | 1 |
| 3_1453544 | 2 | 0.5 | 1 | 1 | 1 | 0 | NA | NA | 4 | 0.5 | 1 | 1 | 2 | 0.5 | NA | NA | NA | NA | 3 | 0.6667 |
| 3_1455976 | 4 | 0.25 | 12 | 0.0833 | 8 | 0.125 | 4 | 0 | 28 | 0.1071 | 7 | 0.1429 | 7 | 0 | 9 | 0.2222 | 9 | 0 | 32 | 0.0938 |
| 3_1455976 | 5 | 0 | 11 | 0 | 10 | 0.1 | 4 | 0.25 | 30 | 0.0667 | 8 | 0 | 9 | 0.1111 | 8 | 0.25 | 9 | 0 | 34 | 0.0882 |
| 3_1456362 | 1 | 0 | 3 | 0.3333 | 2 | 0.5 | 2 | 0 | 8 | 0.25 | 5 | 0.6 | NA | NA | 1 | 0 | 2 | 0.5 | 8 | 0.5 |

| | | | | | | | | | | | | | | | | | | | | |
|---|---|---|---|---|---|---|---|---|---|---|---|---|---|---|---|---|---|---|---|---|
| 3_1456388 | NA | NA | 2 | 1 | 1 | 1 | 2 | 0 | 5 | 0.6 | 3 | 0 | 1 | 0 | 1 | 0 | 2 | 0.5 | 7 | 0.1429 |
| 3_1456389 | 1 | 0 | 3 | 1 | 2 | 1 | 2 | 0.5 | 8 | 0.75 | 4 | 1 | 1 | 0 | 2 | 0.5 | 2 | 1 | 9 | 0.7778 |
| 3_1456389 | 1 | 1 | 3 | 0.3333 | 1 | 0 | 2 | 0.5 | 7 | 0.4286 | 4 | 0.75 | 1 | 0 | 2 | 0.5 | 2 | 0.5 | 9 | 0.5556 |
| 3_1456390 | 1 | 1 | 3 | 0 | 2 | 0 | 2 | 0 | 8 | 0.125 | 5 | 0.2 | 1 | 0 | 1 | 0 | 2 | 0 | 9 | 0.1111 |
| 3_1456392 | 1 | 0 | 3 | 0 | 3 | 0.3333 | 2 | 0 | 9 | 0.1111 | 5 | 0 | 1 | 0 | 2 | 0.5 | 2 | 0 | 10 | 0.1 |
| 3_1456393 | NA | NA | 1 | 1 | 3 | 0 | 2 | 0.5 | 6 | 0.3333 | 4 | 0.25 | NA | NA | 2 | 0 | 2 | 0 | 8 | 0.125 |
| 3_1456497 | 3 | 0.6667 | 13 | 0.3077 | 4 | 0.75 | 6 | 0.5 | 26 | 0.4615 | 5 | 0.4 | 3 | 0.3333 | 6 | 0.5 | 6 | 0.6667 | 20 | 0.5 |
| 3_1456497 | 6 | 0.1667 | 13 | 0.3077 | 5 | 0.6 | 7 | 0.2857 | 31 | 0.3226 | 7 | 0.2857 | 3 | 0 | 8 | 0.125 | 6 | 0.3333 | 24 | 0.2083 |
| 3_1456498 | 7 | 0.4286 | 13 | 0.2308 | 6 | 0.5 | 6 | 0.5 | 32 | 0.375 | 7 | 0.1429 | 2 | 0.5 | 5 | 0.4 | 6 | 0.5 | 20 | 0.35 |
| 3_1456498 | 6 | 0.5 | 15 | 0.2667 | 6 | 1 | 9 | 0.5556 | 36 | 0.5 | 7 | 0.7143 | 2 | 0.5 | 6 | 0.1667 | 6 | 0.1667 | 21 | 0.381 |
| 3_1456503 | 2 | 0 | 3 | 0.3333 | 2 | 0 | 2 | 0 | 9 | 0.1111 | 3 | 0 | 1 | 1 | 4 | 0 | 1 | 0 | 9 | 0.1111 |
| 3_1456503 | 2 | 0 | 2 | 0.5 | 2 | 1 | 2 | 0.5 | 8 | 0.5 | 3 | 0 | 1 | 0 | 3 | 0.3333 | 1 | 0 | 8 | 0.125 |
| 3_1457073 | 2 | 1 | NA | NA | 1 | 1 | NA | NA | 3 | 1 | NA | NA | NA | NA | 1 | 1 | NA | NA | 1 | 1 |
| 3_1457075 | 2 | 0 | NA | NA | 1 | 1 | NA | NA | 3 | 0.3333 | NA | NA | NA | NA | 1 | 1 | NA | NA | 1 | 1 |
| 3_1457075 | 2 | 0.5 | NA | NA | NA | NA | NA | NA | 2 | 0.5 | NA | NA | NA | NA | 1 | 1 | NA | NA | 1 | 1 |
| 3_1457078 | 1 | 1 | NA | NA | NA | NA | NA | NA | 1 | 1 | NA | NA | NA | NA | 1 | 1 | NA | NA | 1 | 1 |
| 3_1457255 | 2 | 0.5 | NA | NA | 1 | 0 | NA | NA | 3 | 0.3333 | 1 | 0 | NA | NA | 1 | 1 | 1 | 0 | 3 | 0.3333 |
| 3_1457256 | 2 | 0.5 | NA | NA | 1 | 0 | NA | NA | 3 | 0.3333 | 1 | 0 | NA | NA | NA | NA | 1 | 1 | 2 | 0.5 |
| 3_1457257 | 2 | 1 | NA | NA | 1 | 1 | NA | NA | 3 | 1 | 2 | 0.5 | NA | NA | NA | NA | 1 | 1 | 3 | 0.6667 |
| 3_1457258 | 2 | 1 | NA | NA | NA | NA | NA | NA | 2 | 1 | 2 | 1 | NA | NA | 1 | 1 | 1 | 1 | 4 | 1 |
| 3_1457259 | 2 | 0.5 | NA | NA | 1 | 0 | NA | NA | 3 | 0.3333 | 2 | 0.5 | NA | NA | 1 | 0 | 1 | 1 | 4 | 0.5 |
| 3_1457261 | 1 | 0 | NA | NA | 1 | 1 | NA | NA | 2 | 0.5 | 2 | 0 | NA | NA | 1 | 1 | 1 | 0 | 4 | 0.25 |
| 3_1457262 | 1 | 1 | NA | NA | NA | NA | NA | NA | 1 | 1 | 1 | 0 | NA | NA | NA | NA | 1 | 1 | 2 | 0.5 |
| 3_1457262 | 2 | 1 | NA | NA | 1 | 1 | NA | NA | 3 | 1 | 2 | 1 | NA | NA | 1 | 0 | 1 | 0 | 4 | 0.5 |
| 3_1457264 | 2 | 0.5 | NA | NA | 1 | 0 | NA | NA | 3 | 0.3333 | 2 | 0 | NA | NA | 1 | 0 | 1 | 1 | 4 | 0.25 |
| 3_1457264 | 1 | 1 | NA | NA | 1 | 1 | NA | NA | 2 | 1 | 2 | 1 | NA | NA | NA | NA | 1 | 1 | 3 | 1 |
| 3_1461249 | 3 | 0 | 6 | 0 | 1 | 0 | 8 | 0.25 | 18 | 0.1111 | 2 | 0 | 1 | 0 | 5 | 0.2 | 4 | 0.5 | 12 | 0.25 |
| 3_1461445 | 4 | 0.25 | 9 | 0.2222 | 5 | 0.2 | 12 | 0.25 | 30 | 0.2333 | 3 | 0 | 6 | 0.1667 | 8 | 0.375 | 3 | 0.3333 | 20 | 0.25 |
| 3_1461445 | 4 | 0.25 | 8 | 0.125 | 6 | 0 | 11 | 0.0909 | 29 | 0.1034 | 3 | 0.3333 | 7 | 0.1429 | 8 | 0.375 | 3 | 0.3333 | 21 | 0.2857 |
| 3_1461446 | 1 | 1 | 9 | 0.7778 | 6 | 1 | 11 | 0.8182 | 27 | 0.8519 | 2 | 0.5 | 7 | 0.8571 | 7 | 1 | 4 | 0.75 | 20 | 0.85 |
| 3_1461447 | 4 | 0.25 | 9 | 0.1111 | 6 | 0.1667 | 12 | 0.0833 | 31 | 0.129 | 3 | 0.3333 | 6 | 0.5 | 7 | 0 | 4 | 0 | 20 | 0.2 |
| 3_1461448 | 4 | 1 | 11 | 0.3636 | 7 | 0.8571 | 11 | 0.2727 | 33 | 0.5152 | 3 | 0.6667 | 7 | 0.5714 | 8 | 0.5 | 4 | 0.25 | 22 | 0.5 |
| 3_1461455 | 4 | 0 | 10 | 0.1 | 12 | 0.0833 | 14 | 0 | 40 | 0.05 | 4 | 0 | 5 | 0 | 12 | 0.1667 | 6 | 0.1667 | 27 | 0.1111 |
| 3_1461698 | 1 | 0 | 1 | 0 | 1 | 0 | 1 | 1 | 4 | 0.25 | 2 | 0.5 | 2 | 0 | 2 | 0 | 1 | 0 | 7 | 0.1429 |
| 3_1461698 | 1 | 0 | 1 | 1 | 1 | 0 | 1 | 0 | 4 | 0.25 | 2 | 0 | 2 | 0.5 | 2 | 0 | 1 | 0 | 7 | 0.1429 |
| 3_1461698 | 1 | 1 | 1 | 0 | 1 | 0 | 1 | 1 | 4 | 0.5 | 2 | 0.5 | 2 | 1 | 2 | 1 | 1 | 1 | 7 | 0.8571 |
| 3_1464357 | 59 | 0.0508 | 41 | 0.0976 | 50 | 0.08 | 69 | 0.0725 | 219 | 0.0731 | 42 | 0.1667 | 69 | 0.058 | 66 | 0.0758 | 49 | 0.1429 | 226 | 0.1018 |
| 3_1464359 | 42 | 0.0714 | 33 | 0.1212 | 34 | 0.0882 | 43 | 0.093 | 152 | 0.0921 | 31 | 0.129 | 49 | 0.102 | 47 | 0.0851 | 34 | 0.2059 | 161 | 0.1242 |
| 3_1464361 | 55 | 0.1091 | 35 | 0.0571 | 46 | 0.1957 | 55 | 0.0727 | 191 | 0.1099 | 40 | 0.15 | 62 | 0.1613 | 61 | 0.1148 | 48 | 0.2083 | 211 | 0.1564 |
| 3_1464366 | 4 | 0 | NA | NA | 6 | 0.1667 | 2 | 0 | 12 | 0.0833 | NA | NA | 5 | 0 | 5 | 0.2 | 1 | 0 | 11 | 0.0909 |
| 3_1464370 | 40 | 0.125 | 28 | 0.0357 | 36 | 0.1389 | 51 | 0.1765 | 155 | 0.129 | 30 | 0.2 | 35 | 0.1429 | 46 | 0.1304 | 30 | 0.2667 | 141 | 0.1773 |
| 3_1464375 | 39 | 0.0769 | 27 | 0.037 | 33 | 0.1212 | 44 | 0.0909 | 143 | 0.0839 | 22 | 0.0909 | 34 | 0.2059 | 41 | 0.0976 | 28 | 0.1071 | 125 | 0.128 |
| 3_1464375 | 36 | 0.2778 | 23 | 0.1739 | 34 | 0.3529 | 41 | 0.2439 | 134 | 0.2687 | 21 | 0.4762 | 38 | 0.3158 | 38 | 0.3947 | 24 | 0.375 | 121 | 0.3802 |
| 3_1464377 | 33 | 0.2121 | 24 | 0.25 | 30 | 0.3 | 42 | 0.1667 | 129 | 0.2248 | 22 | 0.2727 | 40 | 0.25 | 42 | 0.3333 | 31 | 0.2258 | 135 | 0.2741 |
| 3_1464378 | 34 | 0.0882 | 23 | 0.0435 | 27 | 0.037 | 41 | 0.1463 | 125 | 0.088 | 15 | 0.0667 | 35 | 0.0857 | 37 | 0.1892 | 26 | 0.1538 | 113 | 0.1327 |
| 3_1464379 | 34 | 0.1765 | 23 | 0.2174 | 27 | 0.2593 | 39 | 0.3333 | 123 | 0.252 | 20 | 0.2 | 39 | 0.3077 | 42 | 0.2857 | 28 | 0.3571 | 129 | 0.2946 |
| 3_1466854 | 7 | 0.2857 | 13 | 0.2308 | 10 | 0.1 | 15 | 0.2 | 45 | 0.2 | 4 | 0 | 5 | 0 | 8 | 0.125 | 5 | 0.2 | 22 | 0.0909 |
| 3_1466855 | 6 | 0 | 10 | 0 | 9 | 0.1111 | 13 | 0.1538 | 38 | 0.0789 | 2 | 0 | 6 | 0.1667 | 10 | 0 | 5 | 0.2 | 23 | 0.087 |
| 3_1466887 | 9 | 0 | 11 | 0.1818 | 8 | 0 | 12 | 0 | 40 | 0.05 | 3 | 0.3333 | 10 | 0.2 | 8 | 0.125 | 10 | 0.1 | 31 | 0.1613 |
| 3_1466889 | 15 | 0.1333 | 17 | 0.0588 | 14 | 0.1429 | 21 | 0.1905 | 67 | 0.1343 | 7 | 0 | 14 | 0.1429 | 17 | 0.0588 | 13 | 0.0769 | 51 | 0.0784 |
| 3_1466890 | 15 | 0.4667 | 16 | 0.625 | 13 | 0.9231 | 19 | 0.7368 | 63 | 0.6825 | 4 | 1 | 16 | 0.6875 | 18 | 0.6111 | 12 | 0.8333 | 50 | 0.72 |
| 3_1466893 | 11 | 0.0909 | 13 | 0.5385 | 13 | 0.3846 | 22 | 0.3182 | 59 | 0.339 | 4 | 0.25 | 15 | 0.4 | 16 | 0.125 | 13 | 0.3846 | 48 | 0.2917 |
| 3_1485217 | 2 | 0.5 | 2 | 0.5 | 3 | 0.3333 | 7 | 0.2857 | 14 | 0.3571 | 2 | 0 | 10 | 0.7 | 3 | 0 | 2 | 0 | 17 | 0.4118 |
| 3_1485221 | 4 | 0.75 | 5 | 0.8 | 4 | 1 | 6 | 0.8333 | 19 | 0.8421 | 2 | 1 | 11 | 1 | 5 | 1 | 1 | 1 | 19 | 1 |
| 3_1485223 | 4 | 0.5 | 5 | 0.2 | 4 | 0 | 5 | 0.2 | 18 | 0.2222 | 2 | 0 | 10 | 0.3 | 4 | 0.75 | 1 | 0 | 17 | 0.3529 |
| 3_1485223 | 5 | 0.8 | 5 | 1 | 4 | 0.5 | 7 | 0.7143 | 21 | 0.7619 | 3 | 1 | 10 | 0.9 | 4 | 1 | 2 | 0.5 | 19 | 0.8947 |
| 3_1485223 | 5 | 0.2 | 4 | 0.5 | 4 | 0.25 | 7 | 0.4286 | 20 | 0.35 | 3 | 0.6667 | 11 | 0.1818 | 5 | 0.4 | 2 | 0.5 | 21 | 0.3333 |
| 3_1485224 | 5 | 0.8 | 5 | 0.8 | 4 | 1 | 7 | 0.8571 | 21 | 0.8571 | 4 | 1 | 11 | 0.7273 | 5 | 0.8 | 2 | 0 | 22 | 0.7273 |
| 3_1485225 | 4 | 0.5 | 3 | 0 | 4 | 0.25 | 6 | 0.1667 | 17 | 0.2353 | 4 | 0.5 | 10 | 0.6 | 6 | 0.3333 | 2 | 0 | 22 | 0.4545 |
| 3_1485226 | 5 | 1 | 3 | 1 | 4 | 1 | 7 | 1 | 19 | 1 | 2 | 1 | 7 | 1 | 4 | 1 | 1 | 1 | 14 | 1 |
| 3_1485226 | 5 | 0.8 | 3 | 1 | 1 | 1 | 5 | 0.6 | 14 | 0.7857 | 3 | 1 | 8 | 0.75 | 6 | 1 | NA | NA | 17 | 0.8824 |
| 3_1485227 | 5 | 0.4 | 3 | 0.3333 | 2 | 0 | 5 | 0.6 | 15 | 0.4 | 4 | 0 | 12 | 0.3333 | 6 | 0 | 2 | 0.5 | 24 | 0.2083 |
| 3_1485228 | 4 | 1 | 2 | 1 | 3 | 0.6667 | 4 | 0.75 | 13 | 0.8462 | 2 | 0.5 | 9 | 0.7778 | 5 | 0.8 | 2 | 1 | 18 | 0.7778 |
| 3_1485229 | 5 | 0.8 | 2 | 1 | 3 | 0.6667 | 6 | 1 | 16 | 0.875 | 3 | 1 | 11 | 1 | 6 | 1 | 2 | 1 | 22 | 1 |
| 3_1485230 | 4 | 0.25 | 1 | 1 | 3 | 0.3333 | 5 | 0.4 | 13 | 0.3846 | 2 | 0.5 | 10 | 0.5 | 6 | 0.6667 | 2 | 0.5 | 20 | 0.55 |

| | | | | | | | | | | | | | | | | | | | | |
|---|---|---|---|---|---|---|---|---|---|---|---|---|---|---|---|---|---|---|---|---|
| 3_14852305 | 2 | 1 | 2 | 0.5 | 3 | 1 | 6 | 0.6667 | 13 | 0.7692 | 2 | 0.5 | 11 | 0.7273 | 6 | 1 | 2 | 1 | 21 | 0.8095 |
| 3_14852308 | 4 | 1 | 3 | 1 | 3 | 1 | 6 | 1 | 16 | 1 | 3 | 1 | 11 | 0.9091 | 7 | 1 | 2 | 1 | 23 | 0.9565 |
| 3_14852314 | 4 | 0 | 3 | 0 | 3 | 0.3333 | 5 | 0.2 | 15 | 0.1333 | 3 | 0.3333 | 12 | 0.1667 | 6 | 0 | 2 | 0 | 23 | 0.1304 |
| 3_14852319 | 4 | 0.5 | 3 | 0 | 2 | 1 | 4 | 0.25 | 13 | 0.3846 | 3 | 1 | 8 | 0.75 | 6 | 0.8333 | 2 | 0.5 | 19 | 0.7895 |
| 3_14852343 | 4 | 0.5 | 2 | 1 | 3 | 1 | 4 | 0.75 | 13 | 0.7692 | 3 | 0.6667 | 9 | 1 | 5 | 0.6 | 2 | 1 | 19 | 0.8421 |
| 3_14852344 | 3 | 1 | 3 | 0.6667 | 3 | 0.6667 | 4 | 0.5 | 13 | 0.6923 | 3 | 0.6667 | 9 | 0.5556 | 6 | 1 | 2 | 1 | 20 | 0.75 |
| 3_14852347 | 3 | 0 | 3 | 0.6667 | 3 | 0 | 4 | 0 | 13 | 0.1538 | 3 | 0.3333 | 9 | 0.3333 | 5 | 0 | NA | NA | 17 | 0.2353 |
| 3_14856486 | NA | NA | NA | NA | 1 | 1 | NA | NA | 1 | 1 | 1 | 1 | 1 | 1 | 1 | 1 | 1 | 1 | 4 | 1 |
| 3_14856504 | NA | NA | 1 | 1 | 1 | 1 | 1 | 1 | 3 | 1 | 1 | 1 | 1 | 1 | 1 | 1 | 1 | 0 | 4 | 0.75 |
| 3_14856508 | NA | NA | 1 | 0 | 1 | 0 | 2 | 0.5 | 4 | 0.25 | 1 | 0 | 1 | 0 | 1 | 0 | 1 | 1 | 4 | 0.25 |
| 3_15122316 | 3 | 0.3333 | 1 | 0 | 1 | 1 | 2 | 0 | 7 | 0.2857 | NA | NA | 1 | 0 | 1 | 1 | 5 | 0.2 | 7 | 0.2857 |
| 3_15125016 | NA | NA | 3 | 0.6667 | 1 | 1 | NA | NA | 4 | 0.75 | NA | NA | NA | NA | 1 | 0 | 1 | 1 | 2 | 0.5 |
| 3_15125041 | 1 | 1 | 3 | 0.6667 | 1 | 1 | NA | NA | 5 | 0.8 | NA | NA | NA | NA | 1 | 1 | 3 | 0.3333 | 4 | 0.5 |
| 3_15179970 | NA | NA | 1 | 1 | NA | NA | NA | NA | 1 | 1 | NA | NA | 1 | 1 | NA | NA | 1 | 1 | 2 | 1 |
| 3_15188989 | 13 | 0 | 20 | 0.15 | 8 | 0.25 | 17 | 0.0588 | 58 | 0.1034 | 9 | 0.1111 | 8 | 0 | 15 | 0.1333 | 9 | 0.1111 | 41 | 0.0976 |
| 3_15189065 | 12 | 0.1667 | 17 | 0.0588 | 9 | 0 | 12 | 0.0833 | 50 | 0.08 | 11 | 0.1818 | 9 | 0.1111 | 13 | 0.0769 | 9 | 0.3333 | 42 | 0.1667 |
| 3_15189066 | 11 | 0.0909 | 19 | 0.1053 | 8 | 0.25 | 11 | 0.0909 | 49 | 0.1224 | 9 | 0 | 8 | 0 | 13 | 0.3077 | 7 | 0 | 37 | 0.1081 |
| 3_15189068 | 9 | 0.1111 | 17 | 0.1765 | 10 | 0.1 | 11 | 0.1818 | 47 | 0.1489 | 9 | 0 | 7 | 0.2857 | 14 | 0.1429 | 10 | 0.2 | 40 | 0.15 |
| 3_15189071 | 10 | 0.1 | 17 | 0.0588 | 9 | 0 | 9 | 0.1111 | 45 | 0.0667 | 8 | 0 | 9 | 0.1111 | 12 | 0.1667 | 9 | 0.2222 | 38 | 0.1316 |
| 3_15189074 | 8 | 1 | 16 | 0.8125 | 7 | 0.8571 | 8 | 0.5 | 39 | 0.7949 | 11 | 1 | 9 | 0.5556 | 13 | 1 | 8 | 1 | 41 | 0.9024 |
| 3_15189089 | 2 | 0 | 4 | 0 | 2 | 0.5 | 2 | 0 | 10 | 0.1 | 1 | 1 | 3 | 0.3333 | 2 | 0 | 2 | 0 | 8 | 0.25 |
| 3_15189211 | 6 | 0 | 15 | 0.0667 | 7 | 0 | 7 | 0.4286 | 35 | 0.1143 | 9 | 0.1111 | 6 | 0 | 14 | 0.2857 | 7 | 0.1429 | 36 | 0.1667 |
| 3_15189212 | 7 | 0.1429 | 15 | 0.2667 | 8 | 0 | 8 | 0.25 | 38 | 0.1842 | 9 | 0.2222 | 9 | 0 | 13 | 0.1538 | 8 | 0.125 | 39 | 0.1282 |
| 3_15189214 | 8 | 0.5 | 14 | 0.6429 | 8 | 0.625 | 9 | 0.4444 | 39 | 0.5641 | 8 | 0.625 | 9 | 0.3333 | 14 | 0.2143 | 7 | 0.5714 | 38 | 0.3947 |
| 3_15189224 | 8 | 1 | 17 | 0.9412 | 7 | 0.8571 | 9 | 0.6667 | 41 | 0.878 | 10 | 0.8 | 9 | 0.8889 | 13 | 0.9231 | 9 | 0.8889 | 41 | 0.878 |
| 3_15189234 | 8 | 1 | 12 | 1 | 7 | 1 | 7 | 1 | 34 | 1 | 8 | 1 | 6 | 1 | 11 | 0.9091 | 7 | 1 | 32 | 0.9688 |
| 3_15189236 | 8 | 0.125 | 16 | 0.1875 | 7 | 0.4286 | 9 | 0 | 40 | 0.175 | 10 | 0 | 6 | 0.1667 | 11 | 0.1818 | 8 | 0.375 | 35 | 0.1714 |
| 3_15189238 | 8 | 0.25 | 17 | 0.2941 | 8 | 0.375 | 9 | 0.6667 | 42 | 0.381 | 9 | 0.4444 | 7 | 0 | 11 | 0.8182 | 8 | 0.375 | 35 | 0.4571 |
| 3_15189239 | 8 | 0.25 | 16 | 0.5 | 7 | 0.2857 | 7 | 0.2857 | 38 | 0.3684 | 9 | 0.3333 | 7 | 0.4286 | 11 | 0.2727 | 8 | 0.5 | 35 | 0.3714 |
| 3_15189921 | 7 | 0.5714 | 12 | 0.6667 | 4 | 0.5 | 7 | 0.1429 | 30 | 0.5 | 5 | 0.6 | 6 | 0.3333 | 10 | 0.7 | 7 | 0.8571 | 28 | 0.6429 |
| 3_15191444 | 1 | 1 | NA | NA | 1 | 0 | 2 | 0.5 | 4 | 0.5 | NA | NA | NA | NA | NA | NA | 2 | 0.5 | 2 | 0.5 |
| 3_15191584 | 1 | 1 | NA | NA | 2 | 0.5 | NA | NA | 3 | 0.6667 | NA | NA | NA | NA | NA | NA | 1 | 1 | 1 | 1 |
| 3_15193805 | 9 | 0.4444 | 8 | 0 | 7 | 0.1429 | 5 | 0.2 | 29 | 0.2069 | 9 | 0.3333 | 8 | 0.5 | 8 | 0.375 | 12 | 0.25 | 37 | 0.3514 |
| 3_15193936 | 2 | 0 | 3 | 0 | 1 | 0 | 4 | 0.25 | 10 | 0.1 | NA | NA | 1 | 0 | 3 | 0.6667 | 4 | 0 | 8 | 0.25 |
| 3_15193970 | 2 | 0.5 | 3 | 0 | 3 | 0 | 5 | 0.2 | 13 | 0.1538 | 1 | 0 | 1 | 0 | 3 | 0.3333 | 4 | 0 | 9 | 0.1111 |
| 3_15194121 | 2 | 0 | 5 | 0 | 3 | 0 | 4 | 0.25 | 14 | 0.0714 | 1 | 1 | 1 | 0 | 3 | 0.3333 | 4 | 0 | 9 | 0.2222 |
| 3_15194202 | 2 | 0 | 7 | 0 | 1 | 0 | 8 | 0.125 | 18 | 0.0556 | 1 | 0 | 2 | 0 | 3 | 0.3333 | 4 | 0 | 10 | 0.1 |
| 3_15194232 | 3 | 0.6667 | 7 | 0.1429 | 2 | 1 | 8 | 0.25 | 20 | 0.35 | 2 | 0.5 | 2 | 0 | 4 | 0.5 | 3 | 0.3333 | 11 | 0.3636 |
| 3_15194296 | 2 | 0 | 1 | 0 | 4 | 0.25 | 3 | 0 | 10 | 0.1 | NA | NA | 3 | 0.3333 | 4 | 0 | 2 | 0 | 9 | 0.1111 |
| 3_15194308 | 2 | 0 | 1 | 0 | 2 | 0.5 | 2 | 0 | 7 | 0.1429 | NA | NA | 3 | 0.3333 | 4 | 0 | 2 | 1 | 9 | 0.3333 |
| 3_15194311 | 2 | 0 | 1 | 0 | 4 | 0.25 | 3 | 0.3333 | 10 | 0.2 | NA | NA | 2 | 0 | 4 | 0.5 | 2 | 0.5 | 8 | 0.375 |
| 3_15194312 | 1 | 0 | 1 | 0 | 4 | 0.25 | 2 | 0 | 8 | 0.125 | NA | NA | 3 | 0.3333 | 4 | 0 | 2 | 0 | 9 | 0.1111 |
| 3_15194318 | 2 | 1 | NA | NA | 3 | 1 | 3 | 1 | 8 | 1 | NA | NA | 3 | 1 | 4 | 0.5 | 1 | 1 | 8 | 0.75 |
| 3_15194329 | NA | NA | NA | NA | 2 | 0.5 | 1 | 0 | 3 | 0.3333 | NA | NA | 1 | 1 | 1 | 1 | NA | NA | 2 | 1 |
| 3_15194343 | 2 | 0 | 1 | 0 | 4 | 0.25 | 3 | 0 | 10 | 0.1 | NA | NA | 3 | 0.3333 | 2 | 0 | 2 | 0 | 7 | 0.1429 |
| 3_15194353 | 2 | 0 | 1 | 1 | 2 | 0 | 3 | 0 | 8 | 0.125 | NA | NA | 3 | 0.6667 | 1 | 0 | 2 | 0.5 | 6 | 0.5 |
| 3_15197926 | 3 | 0.6667 | 1 | 0 | 1 | 0 | 1 | 0 | 6 | 0.3333 | 1 | 0 | 2 | 0.5 | NA | NA | 1 | 0 | 4 | 0.25 |
| 3_15197960 | 12 | 0.1667 | 14 | 0.3571 | 11 | 0.1818 | 17 | 0.2353 | 54 | 0.2407 | 8 | 0.25 | 15 | 0.1333 | 12 | 0.25 | 11 | 0.0909 | 46 | 0.1739 |
| 3_15198014 | 13 | 0.1538 | 11 | 0 | 12 | 0.0833 | 13 | 0.2308 | 49 | 0.1224 | 9 | 0.3333 | 12 | 0.1667 | 10 | 0.2 | 12 | 0.1667 | 43 | 0.2093 |
| 3_15198191 | 4 | 0 | 12 | 0.1667 | 4 | 0.5 | 6 | 0.3333 | 26 | 0.2308 | 5 | 0.2 | 11 | 0.2727 | 8 | 0.625 | 8 | 0.5 | 32 | 0.4062 |
| 3_15198203 | 3 | 0.6667 | 12 | 0.5833 | 3 | 1 | 5 | 0.6 | 23 | 0.6522 | 5 | 0.6 | 12 | 0.5 | 8 | 0.625 | 8 | 0.625 | 33 | 0.5758 |
| 3_15198208 | 4 | 0.5 | 11 | 1 | 5 | 1 | 5 | 0.8 | 25 | 0.88 | 6 | 1 | 12 | 1 | 8 | 1 | 9 | 1 | 35 | 1 |
| 3_15198216 | 4 | 0.75 | 4 | 0.75 | 2 | 1 | 4 | 0.75 | 14 | 0.7857 | 3 | 0.6667 | 10 | 0.8 | 4 | 0.75 | 6 | 0.6667 | 23 | 0.7391 |
| 3_15198222 | 4 | 0.5 | 10 | 0.4 | 5 | 0.8 | 5 | 0.6 | 24 | 0.5417 | 5 | 0.6 | 11 | 0.7273 | 7 | 0.8571 | 9 | 0.5556 | 32 | 0.6875 |
| 3_15198226 | 2 | 0.5 | 10 | 0.7 | 4 | 0.75 | 4 | 1 | 20 | 0.75 | 5 | 0.8 | 12 | 0.5833 | 7 | 0.2857 | 8 | 0.5 | 32 | 0.5312 |
| 3_15198232 | 4 | 0.5 | 10 | 0.6 | 4 | 0.25 | 6 | 0.8333 | 24 | 0.5833 | 6 | 0.5 | 11 | 0.4545 | 6 | 0.6667 | 8 | 0.75 | 31 | 0.5806 |
| 3_15204347 | 2 | 0.5 | 9 | 0.4444 | 3 | 0 | 2 | 0 | 16 | 0.3125 | 3 | 0.3333 | 7 | 0.1429 | 2 | 0 | 2 | 0 | 14 | 0.1429 |
| 3_15207390 | 3 | 0.3333 | 5 | 0 | 11 | 0 | 8 | 0.25 | 27 | 0.1111 | 3 | 0 | 6 | 0 | 7 | 0.1429 | 4 | 0 | 20 | 0.05 |
| 3_15207408 | 4 | 0 | 4 | 0.25 | 10 | 0.1 | 9 | 0.1111 | 27 | 0.1111 | 1 | 0 | 7 | 0.2857 | 8 | 0.125 | 5 | 0.2 | 21 | 0.1905 |
| 3_15207414 | 5 | 0.8 | 5 | 0.4 | 10 | 0.4 | 9 | 0.4444 | 29 | 0.4828 | 3 | 0 | 7 | 0.4286 | 7 | 0.1429 | 5 | 0.4 | 22 | 0.2727 |
| 3_15207419 | 5 | 0.4 | 5 | 0.4 | 10 | 0.6 | 8 | 0.25 | 28 | 0.4286 | 2 | 0.5 | 8 | 0.125 | 8 | 0.375 | 7 | 0.2857 | 25 | 0.28 |
| 3_15207420 | 5 | 0.2 | 4 | 0.25 | 9 | 0.1111 | 6 | 0 | 24 | 0.125 | 3 | 0.3333 | 6 | 0.1667 | 3 | 0 | 5 | 0 | 17 | 0.1176 |
| 3_15208510 | 9 | 0.1111 | 7 | 0 | 14 | 0.1429 | 11 | 0 | 41 | 0.0732 | 4 | 0 | 8 | 0.125 | 10 | 0 | 6 | 0.1667 | 28 | 0.0714 |
| 3_15209801 | 12 | 0.3333 | 12 | 0.4167 | 17 | 0.1176 | 14 | 0.2143 | 55 | 0.2545 | 4 | 0.25 | 10 | 0.1 | 15 | 0.1333 | 10 | 0.3 | 39 | 0.1795 |
| 3_15213295 | NA | NA | 3 | 0.6667 | NA | NA | 3 | 0.3333 | 6 | 0.5 | NA | NA | 1 | 1 | NA | NA | 1 | 1 | 2 | 1 |

| | | | | | | | | | | | | | | | | | | | | |
|---|---|---|---|---|---|---|---|---|---|---|---|---|---|---|---|---|---|---|---|---|
| 3_1521330 | NA | NA | 4 | 0.5 | NA | NA | 1 | 0 | 5 | 0.4 | NA | NA | 1 | 0 | NA | NA | 1 | 1 | 2 | 0.5 |
| 3_1521333 | NA | NA | 3 | 0.3333 | NA | NA | 3 | 0.3333 | 6 | 0.3333 | NA | NA | 1 | 0 | 1 | 0 | 1 | 1 | 3 | 0.3333 |
| 3_1521336 | NA | NA | 4 | 0.5 | NA | NA | 2 | 0.5 | 6 | 0.5 | NA | NA | NA | NA | NA | NA | 1 | 1 | 1 | 1 |
| 3_1521336 | NA | NA | 4 | 0.25 | NA | NA | 3 | 0 | 7 | 0.1429 | NA | NA | 1 | 1 | 1 | 0 | 1 | 1 | 3 | 0.6667 |
| 3_1521337 | NA | NA | 3 | 0.6667 | 1 | 0 | 2 | 1 | 6 | 0.6667 | NA | NA | 1 | 1 | 1 | 1 | 1 | 1 | 3 | 1 |
| 3_1521340 | NA | NA | 3 | 0.3333 | 1 | 1 | 2 | 0.5 | 6 | 0.5 | NA | NA | 1 | 1 | 1 | 1 | 1 | 1 | 3 | 1 |
| 3_1521343 | NA | NA | 3 | 1 | 1 | 1 | 3 | 1 | 7 | 1 | NA | NA | 1 | 0 | 1 | 1 | 1 | 1 | 3 | 0.6667 |
| 3_1521343 | NA | NA | 4 | 1 | 1 | 1 | 3 | 0.6667 | 8 | 0.875 | NA | NA | 1 | 1 | 1 | 1 | 1 | 1 | 3 | 1 |
| 3_1521640 | 3 | 0 | 5 | 0.8 | 1 | 1 | 4 | 0.25 | 13 | 0.4615 | 2 | 0.5 | 1 | 0 | 3 | 0.6667 | 2 | 0 | 8 | 0.375 |
| 3_1521641 | 2 | 0.5 | 4 | 1 | 3 | 1 | 3 | 1 | 12 | 0.9167 | 2 | 0.5 | 1 | 1 | 2 | 0.5 | 2 | 1 | 7 | 0.7143 |
| 3_1521641 | 2 | 0 | 3 | 0.3333 | 1 | 1 | 4 | 0 | 10 | 0.2 | NA | NA | 1 | 0 | 3 | 1 | 2 | 0 | 6 | 0.5 |
| 3_1521645 | 3 | 0 | 5 | 0 | 3 | 0.3333 | 5 | 0.2 | 16 | 0.125 | 2 | 0 | 2 | 0 | 3 | 0 | 2 | 0.5 | 9 | 0.1111 |
| 3_1521657 | NA | NA | NA | NA | NA | NA | 1 | 1 | 1 | 1 | NA | NA | NA | NA | NA | NA | 1 | 1 | 1 | 1 |
| 3_1521658 | NA | NA | NA | NA | NA | NA | 1 | 1 | 1 | 1 | NA | NA | NA | NA | NA | NA | 1 | 1 | 1 | 1 |
| 3_1521659 | NA | NA | NA | NA | NA | NA | 1 | 1 | 1 | 1 | NA | NA | NA | NA | NA | NA | 1 | 1 | 1 | 1 |
| 3_1521660 | NA | NA | NA | NA | NA | NA | 1 | 1 | 1 | 1 | NA | NA | NA | NA | NA | NA | 1 | 1 | 1 | 1 |
| 3_1521687 | 10 | 0.1 | 12 | 0.0833 | 6 | 0.1667 | 6 | 0.1667 | 34 | 0.1176 | 7 | 0 | 4 | 0.5 | 5 | 0.2 | 7 | 0.5714 | 23 | 0.3043 |
| 3_1521688 | 13 | 0.2308 | 13 | 0.3077 | 8 | 0.25 | 5 | 0.2 | 39 | 0.2564 | 10 | 0.1 | 7 | 0.1429 | 6 | 0 | 10 | 0.7 | 33 | 0.2727 |
| 3_1521688 | 13 | 0.1538 | 13 | 0.4615 | 7 | 0.2857 | 6 | 0.1667 | 39 | 0.2821 | 10 | 0.3 | 6 | 0.5 | 6 | 0.5 | 8 | 0.625 | 30 | 0.4667 |
| 3_1521689 | 14 | 0.0714 | 12 | 0.1667 | 9 | 0.2222 | 6 | 0.1667 | 41 | 0.1463 | 10 | 0.3 | 6 | 0 | 5 | 0.2 | 12 | 0.0833 | 33 | 0.1515 |
| 3_1521689 | 9 | 0.4444 | 9 | 0.5556 | 8 | 0.625 | 6 | 0.5 | 32 | 0.5312 | 6 | 0.5 | 4 | 0.75 | 4 | 0.75 | 9 | 0.5556 | 23 | 0.6087 |
| 3_1521690 | 12 | 0.1667 | 12 | 0.1667 | 7 | 0.4286 | 6 | 0.3333 | 37 | 0.2432 | 7 | 0.4286 | 5 | 0 | 4 | 0.25 | 9 | 0.4444 | 25 | 0.32 |
| 3_1521690 | 12 | 0 | 12 | 0.0833 | 7 | 0.1429 | 6 | 0 | 37 | 0.0541 | 8 | 0.125 | 8 | 0.125 | 6 | 0.3333 | 9 | 0.2222 | 31 | 0.1935 |
| 3_1521690 | 12 | 0.1667 | 12 | 0.1667 | 7 | 0 | 6 | 0.1667 | 37 | 0.1351 | 7 | 0 | 7 | 0 | 5 | 0 | 9 | 0.2222 | 28 | 0.0714 |
| 3_1521692 | 9 | 0.3333 | 10 | 0.5 | 7 | 0.4286 | 6 | 0 | 32 | 0.3438 | 6 | 0.6667 | 5 | 0.4 | 5 | 0.2 | 10 | 0.6 | 26 | 0.5 |
| 3_1521692 | 10 | 0.1 | 12 | 0.1667 | 7 | 0.4286 | 6 | 0.1667 | 35 | 0.2 | 8 | 0 | 7 | 0 | 6 | 0.1667 | 11 | 0.3636 | 32 | 0.1562 |
| 3_1521692 | 11 | 0.8182 | 10 | 0.6 | 6 | 0.8333 | 5 | 0.2 | 32 | 0.6562 | 7 | 0.7143 | 8 | 0.375 | 6 | 0.1667 | 11 | 0.8182 | 32 | 0.5625 |
| 3_1521694 | 10 | 0.9 | 7 | 1 | 4 | 1 | 5 | 0.8 | 26 | 0.9231 | 8 | 1 | 7 | 0.8571 | 6 | 1 | 10 | 1 | 31 | 0.9677 |
| 3_1521695 | 9 | 0.6667 | 7 | 0.7143 | 4 | 0.25 | 6 | 0.3333 | 26 | 0.5385 | 6 | 0.3333 | 6 | 0.8333 | 4 | 0.25 | 8 | 0.375 | 24 | 0.4583 |
| 3_1521785 | 10 | 0.2 | 7 | 0.2857 | 2 | 0 | 6 | 0.3333 | 25 | 0.24 | 4 | 0.25 | 4 | 0.5 | 6 | 0.3333 | 10 | 0.1 | 24 | 0.25 |
| 3_1524910 | 3 | 0 | 4 | 0 | 3 | 0 | 5 | 0.2 | 15 | 0.0667 | 5 | 0 | 5 | 0.4 | 4 | 0 | 7 | 0.1429 | 21 | 0.1429 |
| 3_1524911 | 4 | 0 | 4 | 0.5 | 3 | 0 | 3 | 0.3333 | 14 | 0.2143 | 4 | 0.5 | 7 | 0.1429 | 5 | 0.4 | 5 | 0.2 | 21 | 0.2857 |
| 3_1524911 | 4 | 0 | 4 | 0.5 | 4 | 0 | 4 | 0 | 16 | 0.125 | 5 | 0.2 | 6 | 0.5 | 5 | 0.2 | 7 | 0.1429 | 23 | 0.2609 |
| 3_1524912 | 4 | 0.75 | 4 | 1 | 4 | 0.75 | 4 | 1 | 16 | 0.875 | 5 | 1 | 7 | 1 | 5 | 0.8 | 4 | 1 | 21 | 0.9524 |
| 3_1524912 | 3 | 0 | 4 | 0.25 | 4 | 0 | 3 | 0.3333 | 14 | 0.1429 | 5 | 0.4 | 6 | 0.1667 | 5 | 0 | 6 | 0 | 22 | 0.1364 |
| 3_1524913 | 4 | 0 | 4 | 0.25 | 4 | 0 | 4 | 0 | 16 | 0.0625 | 4 | 0.5 | 7 | 0 | 5 | 0 | 5 | 0 | 21 | 0.0952 |
| 3_1524913 | 3 | 0.6667 | 3 | 0.3333 | 4 | 0.25 | 4 | 0.75 | 14 | 0.5 | 3 | 0.6667 | 7 | 0.2857 | 4 | 0.25 | 6 | 0.8333 | 20 | 0.5 |
| 3_1524915 | 1 | 0 | 2 | 1 | 1 | 1 | 3 | 1 | 7 | 0.8571 | 4 | 1 | 1 | 1 | 2 | 1 | 1 | 1 | 8 | 1 |
| 3_1524916 | 1 | 0 | 2 | 0.5 | 1 | 1 | 2 | 0.5 | 6 | 0.5 | 3 | 0.3333 | NA | NA | 2 | 0 | 1 | 0 | 6 | 0.1667 |
| 3_1524918 | 1 | 1 | 2 | 0.5 | 1 | 1 | 3 | 1 | 7 | 0.8571 | 3 | 1 | NA | NA | 3 | 1 | 1 | 1 | 7 | 1 |
| 3_1524919 | 1 | 0 | 2 | 0 | 1 | 1 | 4 | 0.25 | 8 | 0.25 | 4 | 0.25 | 1 | 0 | 3 | 0 | 1 | 0 | 9 | 0.1111 |
| 3_1524920 | 1 | 1 | 2 | 0 | 1 | 0 | 3 | 0 | 7 | 0.1429 | 4 | 0 | 1 | 1 | 3 | 0 | 1 | 0 | 9 | 0.1111 |
| 3_1524926 | 1 | 0 | 2 | 0 | 1 | 0 | 4 | 0.25 | 8 | 0.125 | 4 | 0 | 3 | 0.6667 | 2 | 0 | 1 | 0 | 10 | 0.2 |
| 3_1524927 | NA | NA | NA | NA | 1 | 1 | 3 | 1 | 4 | 1 | 3 | 1 | 1 | 1 | 3 | 1 | 1 | 1 | 8 | 1 |
| 3_1524927 | 1 | 1 | 3 | 1 | NA | NA | 5 | 1 | 9 | 1 | 3 | 1 | 2 | 0.5 | 1 | 1 | 1 | 1 | 7 | 0.8571 |
| 3_1524928 | 1 | 1 | 2 | 1 | 1 | 1 | 4 | 1 | 8 | 1 | 3 | 0.6667 | 2 | 1 | 2 | 1 | 1 | 1 | 8 | 0.875 |
| 3_1524928 | NA | NA | 2 | 1 | 1 | 1 | 4 | 0.75 | 7 | 0.8571 | 3 | 1 | 2 | 1 | 3 | 1 | NA | NA | 8 | 1 |
| 3_1524928 | 1 | 1 | 3 | 0.3333 | 2 | 1 | 5 | 0.4 | 11 | 0.5455 | 4 | 1 | 3 | 0.6667 | 3 | 1 | NA | NA | 10 | 0.9 |
| 3_1524928 | 1 | 0 | 3 | 1 | 2 | 1 | 5 | 1 | 11 | 0.9091 | 3 | 1 | 2 | 1 | 3 | 1 | NA | NA | 8 | 1 |
| 3_1524929 | 1 | 0 | 3 | 0.3333 | 1 | 0 | 5 | 0.2 | 10 | 0.2 | 4 | 0 | 2 | 0 | 2 | 0.5 | 1 | 1 | 9 | 0.2222 |
| 3_1524929 | NA | NA | 2 | 1 | 1 | 1 | 4 | 0.75 | 7 | 0.8571 | 4 | 1 | 2 | 1 | 3 | 0.6667 | 1 | 1 | 10 | 0.9 |
| 3_1524929 | NA | NA | 3 | 1 | 2 | 1 | 5 | 1 | 10 | 1 | 3 | 1 | 3 | 1 | 2 | 1 | 1 | 1 | 9 | 1 |
| 3_1524932 | 2 | 0 | 3 | 0.3333 | 1 | 0 | 7 | 0.1429 | 13 | 0.1538 | 4 | 0.25 | 4 | 0.25 | 4 | 0 | NA | NA | 12 | 0.1667 |
| 3_1524933 | 2 | 0 | 3 | 0 | 1 | 0 | 6 | 0.1667 | 12 | 0.0833 | 2 | 0.5 | 4 | 0 | 4 | 0.5 | 1 | 0 | 11 | 0.2727 |
| 3_1524933 | 3 | 0 | 2 | 0 | 1 | 0 | 7 | 0.1429 | 13 | 0.0769 | 4 | 0.25 | 4 | 0 | 2 | 0 | 1 | 0 | 11 | 0.0909 |
| 3_1524943 | 1 | 0 | 2 | 0 | NA | NA | 6 | 0.1667 | 9 | 0.1111 | 3 | 0.3333 | 2 | 0 | 4 | 0 | 1 | 0 | 10 | 0.1 |
| 3_1524944 | 1 | 0 | 3 | 0 | 1 | 0 | 6 | 0.1667 | 11 | 0.0909 | 5 | 0.2 | 4 | 0.5 | 5 | 0 | 1 | 0 | 15 | 0.2 |
| 3_1524944 | 2 | 0 | 3 | 0.3333 | 1 | 0 | 7 | 0.4286 | 13 | 0.3077 | 6 | 0.3333 | 4 | 0.75 | 3 | 0.3333 | 2 | 0 | 15 | 0.4 |
| 3_1524944 | 2 | 1 | 3 | 1 | 1 | 1 | 7 | 1 | 13 | 1 | 6 | 0.8333 | 2 | 1 | 3 | 0.3333 | 2 | 1 | 13 | 0.7692 |
| 3_1524944 | 2 | 0.5 | 2 | 0.5 | 1 | 0 | 9 | 0.7778 | 14 | 0.6429 | 6 | 0.8333 | 2 | 0.5 | 5 | 0.2 | 2 | 0.5 | 15 | 0.5333 |
| 3_1524944 | 2 | 0 | 2 | 0.5 | 1 | 0 | 7 | 0.5714 | 12 | 0.4167 | 5 | 0.2 | 3 | 0.3333 | 6 | 0.1667 | 1 | 0 | 15 | 0.2 |
| 3_1525238 | NA | NA | 2 | 0 | 1 | 0 | 1 | 1 | 4 | 0.25 | 2 | 0.5 | 2 | 0.5 | 1 | 1 | 1 | 0 | 6 | 0.5 |
| 3_1525267 | 5 | 0.6 | 1 | 1 | 5 | 0.4 | 6 | 1 | 17 | 0.7059 | 7 | 0.8571 | 4 | 0.75 | 3 | 0.6667 | 5 | 0.6 | 19 | 0.7368 |
| 3_1525267 | 6 | 0.6667 | 1 | 0 | 6 | 0.1667 | 4 | 0.5 | 17 | 0.4118 | 8 | 0.625 | 4 | 0.5 | 3 | 0 | 6 | 0.3333 | 21 | 0.4286 |
| 3_1525268 | 5 | 1 | NA | NA | 6 | 1 | 4 | 1 | 15 | 1 | 6 | 0.8333 | 4 | 1 | 3 | 1 | 5 | 1 | 18 | 0.9444 |

| | | | | | | | | | | | | | | | | | | | | |
|---|---|---|---|---|---|---|---|---|---|---|---|---|---|---|---|---|---|---|---|---|
| 3_1525268 | 4 | 1 | NA | NA | 5 | 0.4 | 5 | 0.8 | 14 | 0.7143 | 4 | 0.5 | 4 | 0.75 | 2 | 0 | 3 | 0.6667 | 13 | 0.5385 |
| 3_1525269 | 4 | 0.75 | 1 | 1 | 6 | 0.6667 | 5 | 0.6 | 16 | 0.6875 | 7 | 0.5714 | 4 | 0.75 | 2 | 1 | 5 | 0.6 | 18 | 0.6667 |
| 3_1525270 | 5 | 0.4 | 1 | 1 | 5 | 0.4 | 4 | 0.25 | 15 | 0.4 | 7 | 0.1429 | 4 | 0.25 | 2 | 0 | 5 | 0.6 | 18 | 0.2778 |
| 3_1525270 | 4 | 1 | NA | NA | 5 | 0.4 | 4 | 0.75 | 13 | 0.6923 | 6 | 0.5 | 3 | 0.6667 | 3 | 0 | 4 | 0.5 | 16 | 0.4375 |
| 3_1525270 | 5 | 0.4 | 1 | 1 | 6 | 0.1667 | 4 | 0.75 | 16 | 0.4375 | 7 | 0.4286 | 2 | 0.5 | 3 | 0.3333 | 4 | 0.5 | 16 | 0.4375 |
| 3_1525270 | 3 | 0 | 1 | 0 | 3 | 0.3333 | 1 | 0 | 8 | 0.125 | 6 | 0 | 2 | 0.5 | 2 | 0 | 6 | 0 | 16 | 0.0625 |
| 3_1525270 | 5 | 1 | NA | NA | 3 | 1 | 4 | 1 | 12 | 1 | 7 | 0.8571 | 3 | 1 | 2 | 1 | 5 | 1 | 17 | 0.9412 |
| 3_1525271 | 2 | 0.5 | 1 | 1 | 3 | 0 | 4 | 0 | 10 | 0.2 | 4 | 0.25 | 1 | 0 | 2 | 0 | 2 | 0 | 9 | 0.1111 |
| 3_1525272 | 4 | 0.25 | 1 | 0 | 5 | 0.4 | 4 | 0.75 | 14 | 0.4286 | 7 | 0.4286 | 1 | 0 | 1 | 0 | 5 | 0.6 | 14 | 0.4286 |
| 3_1525272 | 5 | 1 | 1 | 1 | 6 | 1 | 4 | 1 | 16 | 1 | 7 | 0.8571 | 3 | 0.6667 | 2 | 1 | 6 | 1 | 18 | 0.8889 |
| 3_1525272 | 5 | 1 | NA | NA | 5 | 1 | 4 | 1 | 14 | 1 | 7 | 1 | 2 | 0.5 | 2 | 1 | 3 | 1 | 14 | 0.9286 |
| 3_1525273 | 5 | 0 | NA | NA | 6 | 0.1667 | 4 | 0 | 15 | 0.0667 | 7 | 0 | 2 | 0 | 2 | 0 | 6 | 0.1667 | 17 | 0.0588 |
| 3_1529104 | 1 | 1 | 2 | 0.5 | NA | NA | NA | NA | 3 | 0.6667 | 2 | 1 | NA | NA | NA | NA | NA | NA | 2 | 1 |
| 3_1529104 | 1 | 1 | 2 | 0.5 | NA | NA | NA | NA | 3 | 0.6667 | 2 | 1 | NA | NA | NA | NA | NA | NA | 2 | 1 |
| 3_1529106 | 1 | 1 | 2 | 1 | NA | NA | NA | NA | 3 | 1 | 1 | 1 | NA | NA | NA | NA | NA | NA | 1 | 1 |
| 3_1529109 | 1 | 1 | NA | NA | NA | NA | NA | NA | 1 | 1 | 2 | 0.5 | NA | NA | NA | NA | NA | NA | 2 | 0.5 |
| 3_1529109 | 1 | 1 | 2 | 1 | NA | NA | NA | NA | 3 | 1 | 2 | 1 | NA | NA | NA | NA | NA | NA | 2 | 1 |
| 3_1529110 | 1 | 1 | 2 | 0.5 | NA | NA | NA | NA | 3 | 0.6667 | 1 | 1 | NA | NA | NA | NA | NA | NA | 1 | 1 |
| 3_1536281 | 5 | 0.4 | 9 | 0.4444 | 6 | 0.1667 | 9 | 0.1111 | 29 | 0.2759 | 6 | 0 | 5 | 0.4 | 7 | 0.1429 | 4 | 0.25 | 22 | 0.1818 |
| 3_1536282 | 5 | 0 | 9 | 0.2222 | 6 | 0.1667 | 9 | 0.1111 | 29 | 0.1379 | 6 | 0 | 3 | 0.3333 | 9 | 0.2222 | 4 | 0 | 22 | 0.1364 |
| 3_1536282 | 5 | 0 | 8 | 0 | 6 | 0.3333 | 7 | 0.1429 | 26 | 0.1154 | 5 | 0 | 6 | 0.1667 | 8 | 0.125 | 4 | 0.25 | 23 | 0.1304 |
| 3_1536282 | 4 | 0.5 | 8 | 0.25 | 7 | 0.1429 | 7 | 0.1429 | 26 | 0.2308 | 6 | 0.3333 | 5 | 0.4 | 9 | 0 | 4 | 0.5 | 24 | 0.25 |
| 3_1536282 | 3 | 0.3333 | 7 | 0.7143 | 6 | 0.6667 | 8 | 0.75 | 24 | 0.6667 | 6 | 0.5 | 4 | 0.5 | 8 | 0.75 | 3 | 1 | 21 | 0.6667 |
| 3_1536283 | 4 | 0.5 | 9 | 0.2222 | 5 | 0.4 | 5 | 0 | 23 | 0.2609 | 6 | 0 | 3 | 0 | 9 | 0.2222 | 4 | 0.25 | 22 | 0.1364 |
| 3_1539735 | NA | NA | NA | NA | NA | NA | 2 | 1 | 2 | 1 | NA | NA | 1 | 0 | NA | NA | 1 | 1 | 2 | 0.5 |
| 3_1539756 | NA | NA | NA | NA | NA | NA | 3 | 1 | 3 | 1 | NA | NA | 1 | 1 | NA | NA | 1 | 1 | 2 | 1 |
| 3_1539764 | NA | NA | 1 | 0 | NA | NA | 3 | 0.3333 | 4 | 0.25 | NA | NA | 2 | 0.5 | NA | NA | 2 | 0 | 4 | 0.25 |
| 3_1540122 | 1 | 0 | 1 | 1 | NA | NA | 2 | 1 | 4 | 0.75 | NA | NA | 2 | 0.5 | 4 | 0.25 | 1 | 1 | 7 | 0.4286 |
| 3_1540395 | 1 | 1 | NA | NA | NA | NA | NA | NA | 1 | 1 | NA | NA | 1 | 1 | 1 | 1 | NA | NA | 2 | 1 |
| 3_1540397 | 1 | 1 | NA | NA | NA | NA | NA | NA | 1 | 1 | NA | NA | 1 | 1 | 1 | 1 | NA | NA | 2 | 1 |
| 3_1540398 | 1 | 1 | NA | NA | NA | NA | NA | NA | 1 | 1 | NA | NA | 1 | 1 | 1 | 1 | NA | NA | 2 | 1 |
| 3_1540402 | 1 | 1 | NA | NA | NA | NA | NA | NA | 1 | 1 | NA | NA | 1 | 0 | 1 | 1 | NA | NA | 2 | 0.5 |
| 3_1542823 | NA | NA | NA | NA | 2 | 0.5 | NA | NA | 2 | 0.5 | 1 | 1 | NA | NA | 2 | 0 | 1 | 0 | 4 | 0.25 |
| 3_1542837 | NA | NA | NA | NA | 1 | 1 | NA | NA | 1 | 1 | 1 | 1 | NA | NA | 2 | 1 | 1 | 1 | 4 | 1 |
| 3_1542838 | NA | NA | NA | NA | 1 | 1 | NA | NA | 1 | 1 | 1 | 1 | NA | NA | 2 | 0 | 1 | 0 | 4 | 0.25 |
| 3_1542839 | 1 | 1 | NA | NA | 1 | 0 | NA | NA | 2 | 0.5 | 1 | 0 | NA | NA | 2 | 0 | 1 | 1 | 4 | 0.25 |
| 3_1542844 | 1 | 1 | NA | NA | 1 | 0 | NA | NA | 2 | 0.5 | NA | NA | NA | NA | 1 | 0 | 1 | 1 | 2 | 0.5 |
| 3_1542845 | 1 | 1 | NA | NA | 1 | 0 | NA | NA | 2 | 0.5 | NA | NA | NA | NA | 1 | 1 | 1 | 0 | 2 | 0.5 |
| 3_1542846 | 1 | 1 | NA | NA | NA | NA | NA | NA | 1 | 1 | NA | NA | NA | NA | 1 | 0 | 1 | 1 | 2 | 0.5 |
| 3_1542846 | 1 | 1 | NA | NA | 1 | 1 | NA | NA | 2 | 1 | NA | NA | NA | NA | 1 | 0 | 1 | 1 | 2 | 0.5 |
| 3_1542853 | 1 | 1 | NA | NA | 1 | 0 | NA | NA | 2 | 0.5 | NA | NA | NA | NA | 1 | 0 | 1 | 1 | 2 | 0.5 |
| 3_1542855 | 2 | 1 | 1 | 0 | 1 | 1 | 1 | 0 | 5 | 0.6 | NA | NA | 3 | 0.3333 | 1 | 0 | 1 | 1 | 5 | 0.4 |
| 3_1542855 | 2 | 0 | 2 | 0.5 | 1 | 0 | 1 | 0 | 6 | 0.1667 | NA | NA | 4 | 0 | 1 | 0 | 1 | 1 | 6 | 0.1667 |
| 3_1543279 | 2 | 0 | 2 | 0.5 | 4 | 0.25 | 2 | 1 | 10 | 0.4 | 2 | 0.5 | 3 | 0.6667 | 5 | 0.6 | 5 | 0.4 | 15 | 0.5333 |
| 3_1544703 | 1 | 1 | NA | NA | NA | NA | NA | NA | 1 | 1 | NA | NA | NA | NA | 1 | 1 | NA | NA | 1 | 1 |
| 3_1544705 | 1 | 1 | NA | NA | NA | NA | NA | NA | 1 | 1 | NA | NA | NA | NA | 1 | 0 | 1 | 1 | 2 | 0.5 |
| 3_1544705 | 1 | 1 | NA | NA | NA | NA | NA | NA | 1 | 1 | NA | NA | NA | NA | 1 | 1 | NA | NA | 1 | 1 |
| 3_1544727 | 2 | 0 | 1 | 1 | 5 | 0.2 | NA | NA | 8 | 0.25 | NA | NA | 1 | 1 | 3 | 0.6667 | 2 | 0.5 | 6 | 0.6667 |
| 3_1544730 | 2 | 1 | 1 | 1 | 5 | 0.8 | 1 | 1 | 9 | 0.8889 | NA | NA | 1 | 1 | 2 | 1 | 1 | 1 | 4 | 1 |
| 3_1544732 | 2 | 0.5 | 2 | 0 | 5 | 0.2 | 2 | 0.5 | 11 | 0.2727 | NA | NA | 1 | 1 | 3 | 0 | 3 | 0.6667 | 7 | 0.4286 |
| 3_1547911 | 9 | 0.3333 | 2 | 0 | 2 | 0 | 2 | 0.5 | 15 | 0.2667 | 2 | 0 | 4 | 0.5 | 4 | 0.25 | 1 | 0 | 11 | 0.2727 |
| 3_1547911 | 10 | 0.2 | 1 | 0 | 4 | 0.25 | 2 | 0 | 17 | 0.1765 | 5 | 0.4 | 4 | 0.25 | 4 | 0.25 | 2 | 0.5 | 15 | 0.3333 |
| 3_1547917 | 6 | 0.3333 | 1 | 1 | 1 | 0 | 2 | 0 | 10 | 0.3 | 3 | 0 | 2 | 0 | 4 | 0.25 | 2 | 0 | 11 | 0.0909 |
| 3_1547920 | 6 | 0.3333 | NA | NA | 1 | 0 | 2 | 0 | 9 | 0.2222 | 3 | 0.3333 | 2 | 0.5 | 4 | 0.5 | 2 | 0 | 11 | 0.3636 |
| 3_1547921 | 6 | 0 | 1 | 0 | 1 | 0 | 2 | 0.5 | 10 | 0.1 | 3 | 0 | 1 | 0 | 4 | 0 | 1 | 1 | 9 | 0.1111 |
| 3_1547923 | 4 | 0.75 | 1 | 1 | 1 | 1 | 2 | 1 | 8 | 0.875 | 4 | 0.75 | 1 | 1 | 5 | 0.8 | 1 | 1 | 11 | 0.8182 |
| 3_1547924 | 5 | 0 | 1 | 1 | 1 | 1 | 2 | 0 | 9 | 0.2222 | 3 | 0 | 2 | 0 | 5 | 0 | 1 | 1 | 11 | 0.0909 |
| 3_1547924 | 5 | 1 | 1 | 1 | 1 | 1 | 2 | 1 | 9 | 1 | 4 | 1 | 2 | 1 | 4 | 1 | NA | NA | 10 | 1 |
| 3_1547925 | 4 | 0.25 | 1 | 0 | 1 | 0 | 2 | 0 | 8 | 0.125 | 4 | 0.25 | 2 | 0 | 3 | 0.3333 | 1 | 0 | 10 | 0.2 |
| 3_1547925 | 2 | 0.5 | 1 | 1 | 1 | 1 | 2 | 0.5 | 6 | 0.6667 | 4 | 0.5 | 1 | 1 | 4 | 1 | 1 | 1 | 10 | 0.8 |
| 3_1547926 | 5 | 0.2 | 1 | 0 | NA | NA | 2 | 0 | 8 | 0.125 | 3 | 0.3333 | 1 | 0 | 3 | 0.3333 | 1 | 1 | 8 | 0.375 |
| 3_1547926 | 5 | 0.4 | 1 | 0 | 1 | 0 | 2 | 0 | 9 | 0.2222 | 4 | 0 | 2 | 0 | 4 | 0 | 1 | 1 | 11 | 0.0909 |
| 3_1547926 | 5 | 1 | 1 | 1 | 1 | 1 | 2 | 1 | 9 | 1 | 4 | 1 | 2 | 1 | 4 | 1 | 1 | 1 | 11 | 1 |
| 3_1547927 | 4 | 1 | 1 | 1 | 1 | 1 | 1 | 1 | 7 | 1 | 3 | 1 | NA | NA | 4 | 1 | 1 | 1 | 8 | 1 |
| 3_1547928 | 2 | 0.5 | 1 | 0 | 1 | 0 | 2 | 0 | 6 | 0.1667 | 1 | 0 | 1 | 1 | 1 | 0 | NA | NA | 3 | 0.3333 |

| | | | | | | | | | | | | | | | | | | | | |
|---|---|---|---|---|---|---|---|---|---|---|---|---|---|---|---|---|---|---|---|---|
| 3_1547929 | 5 | 1 | 1 | 1 | NA | NA | 1 | 1 | 7 | 1 | 4 | 0.75 | 1 | 0 | 3 | 0 | 1 | 0 | 9 | 0.3333 |
| 3_15479299 | 5 | 0.2 | 1 | 0 | 1 | 0 | 2 | 0 | 9 | 0.1111 | 3 | 0 | 1 | 1 | 3 | 0 | 1 | 1 | 8 | 0.25 |
| 3_1547930 | 5 | 0 | NA | NA | 1 | 0 | 2 | 0.5 | 8 | 0.125 | 2 | 0 | 2 | 0 | 3 | 0.3333 | NA | NA | 7 | 0.1429 |
| 3_1547930 | 5 | 0.4 | NA | NA | 1 | 1 | 2 | 0 | 8 | 0.375 | 3 | 0.6667 | 2 | 0 | 4 | 0.5 | 1 | 1 | 10 | 0.5 |
| 3_1547930 | 5 | 0.2 | NA | NA | 1 | 0 | 2 | 0 | 8 | 0.125 | 2 | 0.5 | 2 | 0 | 4 | 0 | 1 | 0 | 9 | 0.1111 |
| 3_1547931 | 4 | 1 | NA | NA | 1 | 1 | 1 | 1 | 6 | 1 | 3 | 1 | 2 | 1 | 4 | 0.75 | 1 | 1 | 10 | 0.9 |
| 3_1547931 | 3 | 1 | NA | NA | NA | NA | 1 | 1 | 4 | 1 | 3 | 1 | 2 | 1 | 1 | 1 | 1 | 1 | 7 | 1 |
| 3_1547932 | 5 | 1 | NA | NA | 1 | 1 | 2 | 1 | 8 | 1 | 2 | 1 | 1 | 1 | 4 | 1 | NA | NA | 7 | 1 |
| 3_1547932 | 5 | 0 | NA | NA | 1 | 0 | 1 | 1 | 7 | 0.1429 | 3 | 0 | 2 | 0.5 | 4 | 0 | 1 | 1 | 10 | 0.2 |
| 3_1547932 | 2 | 0.5 | NA | NA | NA | NA | 2 | 1 | 4 | 0.75 | 3 | 0.3333 | 2 | 0.5 | 3 | 0.6667 | 1 | 1 | 9 | 0.5556 |
| 3_1547933 | 5 | 0.2 | NA | NA | 1 | 0 | 2 | 0 | 8 | 0.125 | 3 | 0 | 1 | 0 | 3 | 0 | 1 | 1 | 8 | 0.125 |
| 3_1547933 | 5 | 0.2 | NA | NA | 1 | 0 | 2 | 0 | 8 | 0.125 | 3 | 0 | 2 | 0.5 | 4 | 0.5 | 1 | 0 | 10 | 0.3 |
| 3_1547934 | 4 | 0.5 | NA | NA | 1 | 0 | 2 | 0.5 | 7 | 0.4286 | 1 | 0 | 1 | 0 | 4 | 0.25 | 1 | 1 | 7 | 0.2857 |
| 3_1547935 | 5 | 1 | NA | NA | 1 | 1 | 2 | 1 | 8 | 1 | 3 | 1 | 2 | 1 | 4 | 1 | 1 | 1 | 10 | 1 |
| 3_1547935 | 3 | 1 | NA | NA | 1 | 1 | 2 | 1 | 6 | 1 | 2 | 1 | 1 | 1 | 4 | 0.75 | 1 | 1 | 8 | 0.875 |
| 3_1547969 | 1 | 1 | NA | NA | 1 | 1 | NA | NA | 2 | 1 | 1 | 1 | NA | NA | NA | NA | NA | NA | 1 | 1 |
| 3_1570184 | 8 | 0.125 | 12 | 0.0833 | 7 | 0 | 9 | 0.2222 | 36 | 0.1111 | 3 | 0.3333 | 11 | 0 | 6 | 0.1667 | 7 | 0.1429 | 27 | 0.1111 |
| 3_1570185 | 11 | 0.4545 | 10 | 0.4 | 5 | 0.4 | 8 | 0.625 | 34 | 0.4706 | 3 | 0.3333 | 11 | 0.1818 | 4 | 0.5 | 5 | 0.2 | 23 | 0.2609 |
| 3_1570187 | 8 | 0 | 13 | 0.2308 | 7 | 0.2857 | 10 | 0.3 | 38 | 0.2105 | 4 | 0.75 | 14 | 0.2857 | 7 | 0.5714 | 6 | 0 | 31 | 0.3548 |
| 3_1570198 | 9 | 0.4444 | 12 | 0.0833 | 8 | 0.25 | 15 | 0.1333 | 44 | 0.2045 | 3 | 0 | 14 | 0.1429 | 11 | 0.1818 | 8 | 0.375 | 36 | 0.1944 |
| 3_1570201 | 13 | 0.3077 | 15 | 0 | 12 | 0.3333 | 18 | 0.1111 | 58 | 0.1724 | 5 | 0 | 18 | 0.1667 | 12 | 0.3333 | 10 | 0.2 | 45 | 0.2 |
| 3_1570207 | 10 | 0 | 16 | 0.25 | 11 | 0 | 20 | 0.15 | 57 | 0.1228 | 6 | 0 | 14 | 0.2857 | 10 | 0 | 7 | 0.1429 | 37 | 0.1351 |
| 3_1570208 | 13 | 0 | 18 | 0.1667 | 13 | 0 | 22 | 0.1818 | 66 | 0.1061 | 6 | 0.1667 | 25 | 0.04 | 14 | 0.0714 | 11 | 0.1818 | 56 | 0.0893 |
| 3_1570213 | 15 | 0.3333 | 21 | 0.381 | 15 | 0.5333 | 32 | 0.4688 | 83 | 0.4337 | 10 | 0.3 | 23 | 0.6087 | 18 | 0.6111 | 17 | 0.4118 | 68 | 0.5147 |
| 3_1572524 | 8 | 0 | 9 | 0.2222 | 8 | 0.125 | 14 | 0.2143 | 39 | 0.1538 | 7 | 0.2857 | 10 | 0 | 8 | 0.125 | 9 | 0.1111 | 34 | 0.1176 |
| 3_1572527 | 7 | 0.1429 | 13 | 0.2308 | 11 | 0.1818 | 20 | 0.25 | 51 | 0.2157 | 6 | 0 | 9 | 0.1111 | 10 | 0.2 | 12 | 0.1667 | 37 | 0.1351 |
| 3_1572536 | 11 | 0.7273 | 17 | 0.5882 | 9 | 0.7778 | 21 | 0.5238 | 58 | 0.6207 | 7 | 0.8571 | 12 | 0.8333 | 13 | 0.6154 | 14 | 0.7857 | 46 | 0.7609 |
| 3_1572538 | 10 | 0.1 | 13 | 0.1538 | 10 | 0.1 | 20 | 0.3 | 53 | 0.1887 | 7 | 0.2857 | 14 | 0.2143 | 15 | 0.2 | 13 | 0.6154 | 49 | 0.3265 |
| 3_1576002 | 1 | 0 | 2 | 0.5 | NA | NA | 2 | 0.5 | 5 | 0.4 | 1 | 1 | 2 | 1 | NA | NA | NA | NA | 3 | 1 |
| 3_1576534 | 5 | 0.4 | 7 | 0.1429 | 5 | 0.6 | 7 | 0.2857 | 24 | 0.3333 | 3 | 0.6667 | 5 | 0.4 | NA | NA | 6 | 0.1667 | 14 | 0.3571 |
| 3_1576535 | 6 | 0.1667 | 9 | 0.2222 | 4 | 0 | 9 | 0.2222 | 28 | 0.1786 | 6 | 0.1667 | 5 | 0.2 | 2 | 0 | 5 | 0 | 18 | 0.1111 |
| 3_1577002 | 5 | 0.2 | 12 | 0.1667 | 4 | 0 | 12 | 0 | 33 | 0.0909 | 8 | 0.125 | 7 | 0.1429 | 2 | 0 | 9 | 0 | 26 | 0.0769 |
| 3_1577114 | 7 | 0 | 12 | 0.1667 | 7 | 0 | 10 | 0.3 | 36 | 0.1389 | 8 | 0.125 | 8 | 0.375 | 2 | 0 | 8 | 0.125 | 26 | 0.1923 |
| 3_1577115 | 5 | 0.6 | 10 | 0.7 | 7 | 0.2857 | 9 | 0.7778 | 31 | 0.6129 | 7 | 0.4286 | 7 | 0.7143 | 2 | 0 | 9 | 0.7778 | 25 | 0.6 |
| 3_1577176 | 11 | 0.8182 | 8 | 0.5 | 9 | 0.5556 | 14 | 0.6429 | 42 | 0.6429 | 7 | 0.5714 | 4 | 0.25 | 15 | 0.2667 | 11 | 0.2727 | 37 | 0.3243 |
| 3_1577318 | 3 | 0 | 3 | 0 | 2 | 0 | 3 | 0.3333 | 11 | 0.0909 | 1 | 0 | NA | NA | 4 | 0.25 | 7 | 0 | 12 | 0.0833 |
| 3_1577694 | 5 | 0 | 1 | 0 | 2 | 0 | 6 | 0.1667 | 14 | 0.0714 | 2 | 0 | 2 | 0 | 2 | 0 | 2 | 0.5 | 8 | 0.125 |
| 3_1577717 | 4 | 0.25 | 1 | 0 | 1 | 1 | 6 | 0 | 12 | 0.1667 | 2 | 0 | 3 | 0 | 1 | 0 | 1 | 1 | 7 | 0.1429 |
| 3_1577718 | 4 | 0.25 | 1 | 0 | 1 | 0 | 6 | 0 | 12 | 0.0833 | 2 | 0 | 3 | 0 | 1 | 0 | 1 | 1 | 7 | 0.1429 |
| 3_1577719 | 5 | 0 | 1 | 0 | 1 | 0 | 6 | 0.1667 | 13 | 0.0769 | 1 | 1 | 3 | 0 | 1 | 0 | 1 | 1 | 6 | 0.3333 |
| 3_1577720 | 2 | 0.5 | 1 | 1 | 1 | 0 | 5 | 0.2 | 9 | 0.3333 | 2 | 0.5 | 2 | 0 | 1 | 0 | 1 | 0 | 6 | 0.1667 |
| 3_1577721 | 3 | 0.6667 | 1 | 0 | NA | NA | 7 | 0 | 11 | 0.1818 | 2 | 0.5 | 2 | 1 | 2 | 0.5 | NA | NA | 6 | 0.6667 |
| 3_1577722 | 2 | 0.5 | 1 | 1 | 1 | 0 | 7 | 0.2857 | 11 | 0.3636 | 1 | 1 | 3 | 0.3333 | 2 | 0.5 | 1 | 1 | 7 | 0.5714 |
| 3_1577723 | NA | NA | 1 | 1 | 2 | 0 | NA | NA | 3 | 0.3333 | 1 | 0 | NA | NA | 1 | 0 | 1 | 1 | 3 | 0.3333 |
| 3_1577724 | NA | NA | 1 | 1 | 3 | 0.6667 | 1 | 0 | 5 | 0.6 | 1 | 1 | 1 | 0 | 1 | 0 | 2 | 1 | 5 | 0.6 |
| 3_1577727 | NA | NA | 1 | 0 | 3 | 0.3333 | 1 | 0 | 5 | 0.2 | 1 | 0 | 2 | 0 | 1 | 0 | 4 | 0.25 | 8 | 0.125 |
| 3_1577736 | 11 | 0 | 12 | 0.25 | 10 | 0.1 | 11 | 0.1818 | 44 | 0.1364 | 6 | 0 | 13 | 0.0769 | 9 | 0 | 13 | 0.1538 | 41 | 0.0732 |
| 3_1577737 | 6 | 0.1667 | 8 | 0.125 | 7 | 0.1429 | 11 | 0.1818 | 32 | 0.1562 | 5 | 0 | 10 | 0 | 8 | 0.125 | 12 | 0.25 | 35 | 0.1143 |
| 3_1577893 | 6 | 0.1667 | 5 | 0.2 | 6 | 0 | 6 | 0.1667 | 23 | 0.1304 | 3 | 0 | 3 | 0 | 6 | 0.1667 | 7 | 0 | 19 | 0.0526 |
| 3_1577894 | 7 | 0.1429 | 6 | 0.3333 | 6 | 0 | 7 | 0 | 26 | 0.1154 | 3 | 0 | 3 | 0 | 6 | 0.1667 | 7 | 0.1429 | 19 | 0.1053 |
| 3_1577895 | 5 | 0.4 | 4 | 0.25 | 3 | 0 | 6 | 0 | 18 | 0.1667 | 4 | 0 | 4 | 0 | 6 | 0 | 6 | 0.1667 | 20 | 0.05 |
| 3_1578659 | 5 | 0.4 | 4 | 0.75 | 2 | 0 | 6 | 0.6667 | 17 | 0.5294 | 3 | 0.3333 | 4 | 0.25 | 3 | 1 | 5 | 0.6 | 15 | 0.5333 |
| 3_1578659 | 4 | 0.5 | 4 | 0 | 2 | 0 | 5 | 0.2 | 15 | 0.2 | 2 | 0 | 4 | 0.25 | 4 | 0 | 5 | 0 | 15 | 0.0667 |
| 3_1578660 | 5 | 0.2 | 3 | 0 | 2 | 0 | 5 | 0.2 | 15 | 0.1333 | 3 | 0 | 3 | 0.3333 | 4 | 0 | 5 | 0 | 15 | 0.0667 |
| 3_1578660 | 4 | 0.25 | 2 | 0 | 2 | 0.5 | 3 | 0.3333 | 11 | 0.2727 | 2 | 0.5 | 4 | 0 | 4 | 0 | 5 | 0 | 15 | 0.0667 |
| 3_1578660 | 5 | 0.4 | 3 | 1 | 2 | 0.5 | 6 | 0.5 | 16 | 0.5625 | 2 | 0.5 | 4 | 0.75 | 4 | 0.75 | 6 | 0.8333 | 16 | 0.75 |
| 3_1578662 | 4 | 1 | 3 | 1 | NA | NA | 5 | 1 | 12 | 1 | 2 | 1 | 3 | 0.6667 | 3 | 1 | 5 | 0.8 | 13 | 0.8462 |
| 3_1578663 | 4 | 0.75 | 3 | 0.3333 | 1 | 0 | 5 | 0.2 | 13 | 0.3846 | 3 | 0.6667 | 4 | 0.25 | 4 | 0.25 | 5 | 0.4 | 16 | 0.375 |
| 3_1578664 | 2 | 0.5 | 1 | 0 | 1 | 0 | 3 | 0.3333 | 7 | 0.2857 | 3 | 0.3333 | 1 | 1 | 2 | 0 | 3 | 0.3333 | 9 | 0.3333 |
| 3_1578667 | 3 | 0.3333 | 3 | 0.3333 | 1 | 0 | 6 | 0.6667 | 13 | 0.4615 | 2 | 0.5 | 4 | 0 | 4 | 0.5 | 3 | 0.3333 | 13 | 0.3077 |
| 3_1578668 | 4 | 0.25 | 3 | 0 | 1 | 0 | 4 | 0.25 | 12 | 0.1667 | 1 | 0 | 4 | 0.25 | 4 | 0 | 2 | 0.5 | 11 | 0.1818 |
| 3_1578670 | 3 | 0 | 3 | 0.3333 | 1 | 0 | 7 | 0.2857 | 14 | 0.2143 | 2 | 0 | 3 | 0 | 3 | 0 | 3 | 0.3333 | 11 | 0.0909 |
| 3_1578671 | 4 | 1 | 3 | 1 | 1 | 1 | 6 | 1 | 14 | 1 | 2 | 1 | 4 | 1 | 3 | 1 | 2 | 1 | 11 | 1 |
| 3_1578671 | 5 | 0.4 | 3 | 0.6667 | 1 | 0 | 5 | 0.2 | 14 | 0.3571 | 2 | 1 | 4 | 0 | 2 | 0 | 3 | 0 | 11 | 0.1818 |
| 3_1578671 | 5 | 0.4 | 2 | 0.5 | 1 | 0 | 5 | 0.4 | 13 | 0.3846 | 2 | 0.5 | 3 | 0.6667 | 3 | 0.6667 | 3 | 0.3333 | 11 | 0.5455 |

| | | | | | | | | | | | | | | | | | | | | |
|---|---|---|---|---|---|---|---|---|---|---|---|---|---|---|---|---|---|---|---|---|
| 3_15786723 | 4 | 0 | 3 | 0 | 1 | 0 | 5 | 0.2 | 13 | 0.0769 | 2 | 0 | 4 | 0.25 | 2 | 0 | 2 | 0.5 | 10 | 0.2 |
| 3_15786725 | 4 | 0 | 2 | 0.5 | NA | NA | 4 | 0.5 | 10 | 0.3 | 1 | 0 | 3 | 0.3333 | 1 | 0 | 1 | 0 | 6 | 0.1667 |
| 3_15786742 | 4 | 0.75 | 3 | 1 | 1 | 1 | 6 | 1 | 14 | 0.9286 | 2 | 1 | 4 | 0.75 | 1 | 1 | 2 | 1 | 9 | 0.8889 |
| 3_15826685 | 2 | 0.5 | 1 | 0 | 1 | 1 | 1 | 0 | 5 | 0.4 | 1 | 0 | 2 | 0 | 1 | 0 | 2 | 0.5 | 6 | 0.1667 |
| 3_15826698 | 1 | 1 | 1 | 0 | 1 | 1 | 2 | 1 | 5 | 0.8 | NA | NA | 1 | 1 | NA | NA | 2 | 0.5 | 3 | 0.6667 |
| 3_15932877 | NA | NA | NA | NA | NA | NA | 1 | 1 | 1 | 1 | NA | NA | 1 | 1 | NA | NA | NA | NA | 1 | 1 |
| 3_15957864 | 7 | 0 | 6 | 0 | 3 | 0 | 2 | 0.5 | 18 | 0.0556 | 6 | 0 | 6 | 0.1667 | 3 | 0.3333 | 1 | 0 | 16 | 0.125 |
| 3_15957870 | 7 | 0 | 6 | 0 | 3 | 0.3333 | 2 | 0 | 18 | 0.0556 | 6 | 0.3333 | 6 | 0 | 3 | 0 | 2 | 0 | 17 | 0.1176 |
| 3_15964013 | 4 | 0.75 | 7 | 0.7143 | 4 | 1 | 3 | 0.6667 | 18 | 0.7778 | 5 | 0.4 | 5 | 1 | 4 | 0.75 | 3 | 0.3333 | 17 | 0.6471 |
| 4_3549357 | 11 | 0.0909 | 7 | 0 | 9 | 0 | 11 | 0.0909 | 38 | 0.0526 | 6 | 0.1667 | 10 | 0.1 | 10 | 0.2 | 3 | 0 | 29 | 0.1379 |
| 4_3564179 | 8 | 0 | 5 | 0 | 5 | 0.2 | 9 | 0.1111 | 27 | 0.0741 | 3 | 0.3333 | 8 | 0 | 7 | 0 | 2 | 1 | 20 | 0.15 |
| 4_3569582 | 6 | 0.8333 | 2 | 1 | 5 | 0.8 | 9 | 0.6667 | 22 | 0.7727 | 4 | 0.75 | 6 | 0.6667 | 5 | 0.6 | 2 | 1 | 17 | 0.7059 |
| 4_3569651 | 8 | 0.625 | 4 | 0.5 | 5 | 0.6 | 8 | 0.75 | 25 | 0.64 | 4 | 0.75 | 7 | 0.7143 | 4 | 0 | 1 | 0 | 16 | 0.5 |
| 4_3569656 | 8 | 0.125 | 4 | 0.25 | 5 | 0 | 8 | 0 | 25 | 0.08 | 3 | 0 | 6 | 0.1667 | 4 | 0 | 1 | 1 | 14 | 0.1429 |
| 4_3585868 | 1 | 0 | 1 | 0 | 1 | 1 | 5 | 0.6 | 8 | 0.5 | NA | NA | 2 | 0.5 | 1 | 0 | 1 | 0 | 4 | 0.25 |
| 4_3585899 | 1 | 1 | 3 | 1 | 3 | 1 | 5 | 0.8 | 12 | 0.9167 | NA | NA | 2 | 1 | 1 | 1 | 2 | 1 | 5 | 1 |
| 4_3585918 | 1 | 1 | 3 | 0.6667 | 3 | 0.6667 | 5 | 0.4 | 12 | 0.5833 | NA | NA | 2 | 0.5 | 1 | 0 | 2 | 0.5 | 5 | 0.4 |
| 4_3585995 | 1 | 1 | 3 | 1 | 2 | 1 | 5 | 1 | 11 | 1 | NA | NA | 2 | 1 | 1 | 1 | 2 | 1 | 5 | 1 |
| 4_3614905 | 1 | 0 | 4 | 0 | 3 | 0.6667 | 8 | 0 | 16 | 0.125 | 1 | 1 | 2 | 0.5 | 1 | 0 | 3 | 0.3333 | 7 | 0.4286 |
| 4_3615612 | 1 | 0 | 4 | 0 | 3 | 0.3333 | 8 | 0 | 16 | 0.0625 | 1 | 1 | 2 | 0 | 1 | 0 | 3 | 0 | 7 | 0.1429 |
| 4_3789805 | 1 | 0 | 1 | 0 | 2 | 0.5 | 1 | 0 | 5 | 0.2 | 1 | 0 | NA | NA | 2 | 0.5 | 1 | 0 | 4 | 0.25 |
| 4_3790001 | NA | NA | 1 | 0 | 3 | 0.3333 | 1 | 0 | 5 | 0.2 | NA | NA | NA | NA | 1 | 1 | 1 | 0 | 2 | 0.5 |
| 4_3790114 | 1 | 0 | 1 | 0 | 3 | 1 | 2 | 0.5 | 7 | 0.5714 | 1 | 0 | NA | NA | 2 | 0.5 | 1 | 1 | 4 | 0.5 |
| 4_3790179 | NA | NA | 1 | 0 | 3 | 0.6667 | 1 | 0 | 5 | 0.4 | 1 | 0 | NA | NA | 2 | 0 | 1 | 1 | 4 | 0.25 |
| 4_3790193 | NA | NA | 1 | 1 | 3 | 1 | 2 | 1 | 6 | 1 | 1 | 1 | NA | NA | 2 | 1 | 1 | 1 | 4 | 1 |
| 4_3790225 | NA | NA | 1 | 0 | 3 | 0.6667 | 2 | 1 | 6 | 0.6667 | 1 | 1 | NA | NA | 2 | 0.5 | 1 | 1 | 4 | 0.75 |
| 4_3790445 | 2 | 0.5 | 1 | 0 | 3 | 1 | 2 | 1 | 8 | 0.75 | 1 | 1 | 1 | 0 | 2 | 1 | 1 | 1 | 5 | 0.8 |
| 4_3790453 | 3 | 0.6667 | 1 | 1 | 3 | 0.3333 | 2 | 0.5 | 9 | 0.5556 | 1 | 0 | 1 | 1 | 2 | 0.5 | NA | NA | 4 | 0.5 |
| 4_3790481 | 3 | 0.3333 | 1 | 0 | 2 | 1 | 2 | 0.5 | 8 | 0.5 | 1 | 1 | NA | NA | 2 | 0.5 | 1 | 1 | 4 | 0.75 |
| 4_3790548 | 1 | 1 | 1 | 0 | 2 | 1 | 1 | 1 | 5 | 0.8 | NA | NA | 1 | 1 | 1 | 1 | 1 | 1 | 3 | 1 |
| 4_3790720 | 3 | 0.6667 | 1 | 1 | 4 | 0.75 | 2 | 1 | 10 | 0.8 | 1 | 0 | 1 | 1 | 2 | 1 | 1 | 1 | 5 | 0.8 |
| 4_3790730 | 3 | 0 | 1 | 0 | 4 | 0.25 | 2 | 0.5 | 10 | 0.2 | 1 | 1 | 1 | 0 | 2 | 0 | 1 | 0 | 5 | 0.2 |
| 4_3790772 | 3 | 0 | 1 | 0 | 4 | 0.5 | 2 | 1 | 10 | 0.4 | 1 | 1 | 1 | 0 | 1 | 0 | 1 | 1 | 4 | 0.5 |
| 4_3790778 | 3 | 0.3333 | 1 | 0 | 1 | 0 | 2 | 0.5 | 7 | 0.2857 | 1 | 1 | 1 | 1 | 2 | 0.5 | 1 | 0 | 5 | 0.6 |
| 4_3790890 | 3 | 0.3333 | 2 | 0.5 | 4 | 0.75 | 2 | 1 | 11 | 0.6364 | 1 | 1 | 1 | 1 | 2 | 0.5 | 1 | 0 | 5 | 0.6 |
| 4_3792236 | NA | NA | NA | NA | 1 | 1 | NA | NA | 1 | 1 | NA | NA | 1 | 1 | NA | NA | NA | NA | 1 | 1 |
| 4_3950774 | NA | NA | 1 | 0 | 2 | 0.5 | NA | NA | 3 | 0.3333 | 1 | 1 | NA | NA | NA | NA | NA | NA | 1 | 1 |
| 4_4133388 | 1 | 1 | 2 | 0.5 | NA | NA | 3 | 0 | 6 | 0.3333 | NA | NA | 6 | 0.1667 | 3 | 0.3333 | 1 | 0 | 10 | 0.2 |
| 4_4133478 | 2 | 0 | 2 | 0 | 3 | 0 | 4 | 0.25 | 11 | 0.0909 | NA | NA | 8 | 0.125 | 4 | 0 | 1 | 0 | 13 | 0.0769 |
| 4_4133519 | 3 | 0.3333 | 2 | 0 | 3 | 0.6667 | 4 | 0.25 | 12 | 0.3333 | NA | NA | 8 | 0.25 | 5 | 0 | 1 | 0 | 14 | 0.1429 |
| 4_4133820 | 307 | 0.0586 | 122 | 0.0574 | 186 | 0.0806 | 365 | 0.0603 | 980 | 0.0633 | 103 | 0.0388 | 431 | 0.0603 | 424 | 0.0613 | 224 | 0.058 | 1182 | 0.0584 |
| 4_4133969 | 293 | 0.9795 | 102 | 0.9902 | 174 | 0.977 | 332 | 0.9669 | 901 | 0.9756 | 83 | 0.9518 | 400 | 0.9525 | 394 | 0.9772 | 198 | 0.9646 | 1075 | 0.9637 |
| 4_4134137 | 249 | 0.1084 | 91 | 0.0989 | 154 | 0.0779 | 289 | 0.1384 | 783 | 0.1124 | 81 | 0.1235 | 352 | 0.0625 | 339 | 0.1091 | 190 | 0.1053 | 962 | 0.0925 |
| 4_4134389 | 268 | 0.1119 | 97 | 0.0928 | 157 | 0.0955 | 310 | 0.1032 | 832 | 0.1034 | 80 | 0.0625 | 370 | 0.1027 | 358 | 0.1173 | 195 | 0.1128 | 1003 | 0.1067 |
| 4_4134465 | 1 | 1 | NA | NA | NA | NA | NA | NA | 1 | 1 | NA | NA | 2 | 1 | NA | NA | NA | NA | 2 | 1 |
| 4_4766886 | 5 | 0 | 4 | 0 | 3 | 0 | 7 | 0.1429 | 19 | 0.0526 | 5 | 0.2 | 8 | 0 | 9 | 0.3333 | 10 | 0 | 32 | 0.125 |
| 4_4767481 | 3 | 0 | 5 | 0 | 3 | 0 | 5 | 0.4 | 16 | 0.125 | 4 | 0.75 | 5 | 0.2 | 6 | 0.1667 | 9 | 0.2222 | 24 | 0.2917 |
| 4_4767560 | 3 | 1 | 5 | 1 | 4 | 1 | 6 | 1 | 18 | 1 | 3 | 1 | 7 | 1 | 7 | 0.8571 | 10 | 1 | 27 | 0.963 |
| 4_4767639 | 2 | 1 | 5 | 1 | 3 | 1 | 7 | 1 | 17 | 1 | 2 | 1 | 6 | 1 | 7 | 1 | 6 | 1 | 21 | 1 |
| 4_4767710 | 3 | 1 | 5 | 1 | 3 | 1 | 6 | 0.8333 | 17 | 0.9412 | 2 | 1 | 7 | 0.8571 | 8 | 1 | 10 | 1 | 27 | 0.963 |
| 4_4767734 | 3 | 0 | 3 | 0 | 2 | 0 | 6 | 0.1667 | 14 | 0.0714 | 2 | 0.5 | 6 | 0.1667 | 7 | 0.1429 | 9 | 0 | 24 | 0.125 |
| 4_4767779 | 3 | 1 | 3 | 1 | 3 | 1 | 5 | 1 | 14 | 1 | 2 | 0.5 | 5 | 1 | 6 | 1 | 9 | 1 | 22 | 0.9545 |
| 4_4767901 | 3 | 0 | 4 | 0.5 | 2 | 1 | 6 | 0.3333 | 15 | 0.4 | 2 | 0 | 5 | 0.4 | 5 | 0.2 | 8 | 0.625 | 20 | 0.4 |
| 4_4793149 | 1 | 0 | 2 | 0.5 | 2 | 1 | 5 | 0.6 | 10 | 0.6 | 1 | 0 | 3 | 0.3333 | 4 | 0.5 | 5 | 0.4 | 13 | 0.3846 |
| 4_5798357 | 7 | 0.5714 | 3 | 1 | 6 | 0.3333 | 5 | 0.2 | 21 | 0.4762 | 5 | 0.8 | 10 | 0.5 | 7 | 0.2857 | 10 | 0.6 | 32 | 0.5312 |
| 4_5798596 | 12 | 0.3333 | 3 | 0.6667 | 8 | 0.75 | 8 | 0.75 | 31 | 0.5806 | 5 | 0.4 | 11 | 0.5455 | 6 | 0.1667 | 12 | 0.5 | 34 | 0.4412 |
| 4_5798682 | 11 | 0.1818 | 3 | 0.3333 | 9 | 0 | 8 | 0 | 31 | 0.0968 | 7 | 0.7143 | 11 | 0.3636 | 6 | 0.5 | 11 | 0.4545 | 35 | 0.4857 |
| 4_5798724 | 12 | 0.8333 | 3 | 1 | 9 | 0.7778 | 8 | 0.75 | 32 | 0.8125 | 6 | 0.8333 | 11 | 0.9091 | 6 | 1 | 12 | 0.9167 | 35 | 0.9143 |
| 4_5798951 | 14 | 0.0714 | 4 | 0.25 | 9 | 0.1111 | 6 | 0.3333 | 33 | 0.1515 | 6 | 0 | 9 | 0 | 4 | 0.25 | 11 | 0.0909 | 30 | 0.0667 |
| 4_5799011 | 11 | 0.1818 | 4 | 0.25 | 7 | 0 | 7 | 0 | 29 | 0.1034 | 4 | 0 | 12 | 0 | 6 | 0.1667 | 10 | 0.1 | 32 | 0.0625 |
| 4_5799029 | 12 | 0.5833 | 4 | 1 | 8 | 0.125 | 6 | 0.6667 | 30 | 0.5333 | 8 | 0.625 | 11 | 0.5455 | 6 | 0.5 | 13 | 0.3846 | 38 | 0.5 |
| 4_5799185 | 13 | 0 | 4 | 0 | 9 | 0 | 10 | 0.2 | 36 | 0.0556 | 8 | 0 | 11 | 0.1818 | 6 | 0 | 13 | 0 | 38 | 0.0526 |
| 4_5799356 | 14 | 0.3571 | 4 | 0 | 9 | 0.2222 | 10 | 0.2 | 37 | 0.2432 | 9 | 0.7778 | 12 | 0.5 | 6 | 0.3333 | 11 | 0.4545 | 38 | 0.5263 |
| 4_5799365 | 14 | 0.4286 | 4 | 0.25 | 9 | 0.2222 | 10 | 0.3 | 37 | 0.3243 | 9 | 0.1111 | 11 | 0.3636 | 7 | 0 | 11 | 0.2727 | 38 | 0.2105 |
| 4_5799417 | 11 | 0.8182 | 4 | 1 | 6 | 0.5 | 9 | 1 | 30 | 0.8333 | 8 | 0.875 | 10 | 0.9 | 6 | 1 | 9 | 0.6667 | 33 | 0.8485 |

| | | | | | | | | | | | | | | | | | | | | |
|---|---|---|---|---|---|---|---|---|---|---|---|---|---|---|---|---|---|---|---|---|
| 4_5799455 | 13 | 1 | 4 | 1 | 9 | 1 | 11 | 0.9091 | 37 | 0.973 | 9 | 0.8889 | 12 | 0.75 | 7 | 1 | 10 | 1 | 38 | 0.8947 |
| 4_5799476 | 12 | 0 | 4 | 0.25 | 9 | 0 | 9 | 0.1111 | 34 | 0.0588 | 9 | 0 | 12 | 0.25 | 6 | 0 | 8 | 0 | 35 | 0.0857 |
| 4_5799521 | 14 | 0.6429 | 4 | 0.5 | 10 | 0.7 | 11 | 0.9091 | 39 | 0.7179 | 9 | 0.5556 | 12 | 0.6667 | 8 | 0.625 | 12 | 0.6667 | 41 | 0.6341 |
| 4_5799528 | 6 | 0.6667 | 3 | 0.3333 | 8 | 0.25 | 7 | 0.1429 | 24 | 0.3333 | 7 | 0 | 4 | 0.5 | 3 | 0.6667 | 7 | 0 | 21 | 0.1905 |
| 4_5799539 | 12 | 0.9167 | 4 | 1 | 9 | 0.8889 | 11 | 1 | 36 | 0.9444 | 7 | 1 | 9 | 1 | 6 | 1 | 10 | 1 | 32 | 1 |
| 4_5799545 | 6 | 0.1667 | 1 | 0 | 4 | 0.25 | 3 | 0 | 14 | 0.1429 | 3 | 0 | 7 | 0 | 3 | 0.3333 | 5 | 0.4 | 18 | 0.1667 |
| 4_5799551 | 14 | 0.5 | 3 | 0.3333 | 9 | 0.5556 | 11 | 0.5455 | 37 | 0.5135 | 7 | 0.7143 | 12 | 0.4167 | 7 | 0.4286 | 11 | 0.3636 | 37 | 0.4595 |
| 4_5799566 | 13 | 0.3846 | 4 | 0.5 | 11 | 0.2727 | 9 | 0.3333 | 37 | 0.3514 | 7 | 0.2857 | 12 | 0.5833 | 4 | 0.5 | 10 | 0.4 | 33 | 0.4545 |
| 4_5799578 | 13 | 0.1538 | 3 | 0 | 10 | 0 | 10 | 0.3 | 36 | 0.1389 | 8 | 0.25 | 11 | 0.1818 | 6 | 0.1667 | 10 | 0.2 | 35 | 0.2 |
| 4_5799881 | 12 | 0 | 2 | 0 | 10 | 0.2 | 8 | 0.375 | 32 | 0.1562 | 8 | 0.125 | 13 | 0.2308 | 3 | 0 | 10 | 0.3 | 34 | 0.2059 |
| 4_5799936 | 16 | 0.5625 | 3 | 0.6667 | 11 | 0.1818 | 11 | 0.5455 | 41 | 0.4634 | 9 | 0.6667 | 14 | 0.4286 | 8 | 0.375 | 11 | 0.3636 | 42 | 0.4524 |
| 4_5800131 | 5 | 0.4 | 1 | 1 | 3 | 0.6667 | 2 | 0.5 | 11 | 0.5455 | 2 | 0.5 | 4 | 0.75 | 4 | 0.5 | 1 | 1 | 11 | 0.6364 |
| 4_5800153 | 6 | 0.1667 | NA | NA | 2 | 0 | 2 | 0 | 10 | 0.1 | 3 | 0.3333 | 5 | 0 | 4 | 0.25 | 1 | 0 | 13 | 0.1538 |
| 4_5800399 | 7 | 0.1429 | NA | NA | 3 | 0.3333 | 3 | 0.3333 | 13 | 0.2308 | 4 | 0.25 | 6 | 0 | 4 | 0.25 | 1 | 0 | 15 | 0.1333 |
| 4_5801027 | 6 | 0.1667 | 1 | 0 | 3 | 0 | 3 | 0 | 13 | 0.0769 | 4 | 0 | 7 | 0.1429 | 5 | 0 | 1 | 0 | 17 | 0.0588 |
| 4_6215968 | 1 | 0 | 5 | 0 | 4 | 0 | 7 | 0.1429 | 17 | 0.0588 | 1 | 0 | 6 | 0 | 9 | 0 | 3 | 0.3333 | 19 | 0.0526 |
| 4_6216002 | 1 | 0 | 3 | 0 | 4 | 0.25 | 6 | 0 | 14 | 0.0714 | 1 | 0 | 6 | 0 | 8 | 0.125 | 2 | 0 | 17 | 0.0588 |
| 4_6216037 | 1 | 0 | 5 | 0.6 | 3 | 0 | 7 | 0.5714 | 16 | 0.4375 | 1 | 0 | 5 | 0.4 | 9 | 0.3333 | 3 | 0.3333 | 18 | 0.3333 |
| 4_6220656 | 7 | 0.2857 | 7 | 0 | 4 | 0 | 3 | 0 | 21 | 0.0952 | 2 | 0 | 5 | 0.4 | 5 | 0 | 4 | 0 | 16 | 0.125 |
| 4_6221193 | 7 | 0 | 6 | 0 | 6 | 0.3333 | 3 | 0 | 22 | 0.0909 | 2 | 0 | 6 | 0 | 5 | 0.2 | 4 | 0 | 17 | 0.0588 |
| 4_6394773 | 17 | 0 | 28 | 0.1071 | 18 | 0.0556 | 35 | 0.0286 | 98 | 0.051 | 22 | 0.0909 | 42 | 0.0476 | 16 | 0.125 | 32 | 0.0625 | 112 | 0.0714 |
| 4_6394792 | 15 | 0.2 | 23 | 0.3478 | 17 | 0.1176 | 30 | 0.3 | 85 | 0.2588 | 20 | 0.25 | 35 | 0.4 | 15 | 0 | 32 | 0.3125 | 102 | 0.2843 |
| 4_6394954 | 17 | 0.4706 | 24 | 0.3333 | 17 | 0.2941 | 31 | 0.4839 | 89 | 0.4045 | 21 | 0.381 | 35 | 0.4 | 17 | 0.3529 | 34 | 0.4706 | 107 | 0.4112 |
| 4_6395104 | 20 | 0.1 | 32 | 0.1875 | 19 | 0.1579 | 39 | 0.3077 | 110 | 0.2091 | 23 | 0.2174 | 42 | 0.1905 | 21 | 0.2381 | 38 | 0.2105 | 124 | 0.2097 |
| 4_6396000 | NA | NA | 1 | 0 | 4 | 0.25 | NA | NA | 5 | 0.2 | 2 | 0.5 | NA | NA | 2 | 0 | 1 | 0 | 5 | 0.2 |
| 4_6396345 | 3 | 0.6667 | 3 | 1 | 4 | 0.75 | 4 | 0.75 | 14 | 0.7857 | 1 | 1 | 2 | 1 | 2 | 1 | 2 | 1 | 7 | 1 |
| 4_6396408 | 4 | 0.75 | 4 | 0.25 | 2 | 0.5 | 5 | 0.6 | 15 | 0.5333 | 1 | 0 | 2 | 0 | 2 | 0.5 | 2 | 0.5 | 7 | 0.2857 |
| 4_6396453 | 5 | 0.6 | 4 | 0 | 4 | 0.75 | 5 | 0.6 | 18 | 0.5 | 1 | 0 | 2 | 0 | 2 | 0.5 | 1 | 0 | 6 | 0.1667 |
| 4_6396518 | 5 | 0.8 | 3 | 0 | 5 | 0.4 | 4 | 1 | 17 | 0.5882 | 1 | 0 | 2 | 0.5 | 2 | 0 | 2 | 0 | 7 | 0.1429 |
| 4_6396670 | 5 | 0.4 | 2 | 0 | 3 | 0 | 4 | 0.5 | 14 | 0.2857 | NA | NA | 2 | 0.5 | 2 | 0 | 2 | 0 | 6 | 0.1667 |
| 4_6688059 | 2 | 0 | 1 | 1 | NA | NA | 1 | 0 | 4 | 0.25 | 5 | 0.4 | 2 | 0.5 | 1 | 1 | 4 | 0.25 | 12 | 0.4167 |
| 4_6688197 | 2 | 0 | 2 | 0.5 | NA | NA | 1 | 1 | 5 | 0.4 | 5 | 0.4 | 2 | 0 | 1 | 0 | 4 | 0 | 12 | 0.1667 |
| 4_6688228 | 2 | 1 | 1 | 1 | NA | NA | 1 | 1 | 4 | 1 | 5 | 0.8 | 3 | 0.6667 | 1 | 1 | 3 | 1 | 12 | 0.8333 |
| 4_6688303 | 2 | 1 | 2 | 1 | NA | NA | 1 | 0 | 5 | 0.8 | 4 | 0.5 | 3 | 0.3333 | 1 | 1 | 3 | 0.3333 | 11 | 0.4545 |
| 4_6697583 | 1 | 1 | NA | NA | NA | NA | 1 | 1 | 2 | 1 | 3 | 0.6667 | 2 | 0.5 | 1 | 1 | 2 | 1 | 8 | 0.75 |
| 4_8606612 | 36 | 0.2222 | 45 | 0.1778 | 44 | 0.1136 | 36 | 0.1389 | 161 | 0.1615 | 30 | 0.1667 | 39 | 0.1282 | 47 | 0.1915 | 32 | 0.2188 | 148 | 0.1757 |
| 4_8606876 | 30 | 0.0667 | 43 | 0.1395 | 48 | 0.1667 | 29 | 0.069 | 150 | 0.12 | 29 | 0.1724 | 36 | 0.0556 | 41 | 0.0976 | 27 | 0.037 | 133 | 0.0902 |
| 4_8606925 | 31 | 0.129 | 43 | 0.093 | 43 | 0.0465 | 31 | 0.0968 | 148 | 0.0878 | 26 | 0.0769 | 34 | 0.0882 | 42 | 0.0476 | 28 | 0.0714 | 130 | 0.0692 |
| 4_8607006 | 31 | 0.0968 | 45 | 0.1556 | 42 | 0.1667 | 23 | 0.2609 | 141 | 0.1631 | 26 | 0.0769 | 32 | 0.125 | 40 | 0.25 | 30 | 0.1667 | 128 | 0.1641 |
| 4_8607081 | 30 | 0.2667 | 37 | 0.3514 | 46 | 0.2391 | 30 | 0.3333 | 143 | 0.2937 | 30 | 0.4333 | 34 | 0.3529 | 39 | 0.3077 | 32 | 0.375 | 135 | 0.363 |
| 4_8607087 | 33 | 0.0909 | 45 | 0.1111 | 48 | 0.0625 | 32 | 0.125 | 158 | 0.0949 | 27 | 0.0741 | 34 | 0.0588 | 43 | 0.1395 | 35 | 0.0571 | 139 | 0.0863 |
| 4_8865922 | NA | NA | NA | NA | 1 | 1 | NA | NA | 1 | 1 | 1 | 0 | NA | NA | NA | NA | 1 | 1 | 2 | 0.5 |
| 4_8866397 | 1 | 1 | NA | NA | 2 | 1 | 1 | 1 | 4 | 1 | 1 | 1 | NA | NA | NA | NA | 1 | 1 | 2 | 1 |
| 4_8866410 | 1 | 1 | NA | NA | NA | NA | 1 | 1 | 2 | 1 | NA | NA | NA | NA | NA | NA | 1 | 1 | 1 | 1 |
| 4_8866439 | 1 | 1 | NA | NA | 2 | 1 | 1 | 1 | 4 | 1 | 1 | 0 | NA | NA | NA | NA | 1 | 1 | 2 | 0.5 |
| 4_8866515 | 1 | 0 | NA | NA | 1 | 1 | 1 | 0 | 3 | 0.3333 | 1 | 1 | NA | NA | NA | NA | 1 | 1 | 2 | 1 |
| 4_8866549 | 1 | 1 | NA | NA | NA | NA | 1 | 1 | 2 | 1 | 1 | 1 | NA | NA | NA | NA | 1 | 0 | 2 | 0.5 |
| 4_8866554 | 1 | 1 | NA | NA | 1 | 1 | 1 | 1 | 3 | 1 | 1 | 1 | NA | NA | NA | NA | NA | NA | 1 | 1 |
| 4_8866606 | NA | NA | NA | NA | NA | NA | 1 | 1 | 1 | 1 | 1 | 1 | NA | NA | NA | NA | NA | NA | 1 | 1 |
| 4_8866688 | 1 | 1 | NA | NA | 1 | 1 | 1 | 1 | 3 | 1 | 1 | 1 | NA | NA | NA | NA | 1 | 1 | 2 | 1 |
| 4_8867516 | 1 | 0 | NA | NA | 2 | 0.5 | 1 | 1 | 4 | 0.5 | 1 | 0 | 1 | 0 | NA | NA | 2 | 0.5 | 4 | 0.25 |
| 4_9281467 | 2 | 0 | 2 | 0.5 | 3 | 0.3333 | 6 | 0.1667 | 13 | 0.2308 | 4 | 0.5 | 7 | 0.2857 | 4 | 0 | 5 | 0.2 | 20 | 0.25 |
| 4_9281563 | 2 | 0 | 2 | 0.5 | 3 | 0.6667 | 5 | 0.6 | 12 | 0.5 | 5 | 0.4 | 7 | 0.5714 | 5 | 0.6 | 3 | 1 | 20 | 0.6 |
| 4_9281680 | 2 | 0.5 | 2 | 0.5 | 3 | 0.6667 | 5 | 0.8 | 12 | 0.6667 | 4 | 0.25 | 8 | 0.375 | 4 | 0.75 | 4 | 0.5 | 20 | 0.45 |
| 4_9281712 | 2 | 0.5 | 3 | 1 | 3 | 0.6667 | 6 | 0.6667 | 14 | 0.7143 | 4 | 0.75 | 6 | 0.5 | 5 | 0.8 | 5 | 0.6 | 20 | 0.65 |
| 4_9281730 | 2 | 0.5 | 4 | 1 | 3 | 0 | 6 | 0.3333 | 15 | 0.4667 | 5 | 0.6 | 9 | 0.4444 | 5 | 0.6 | 4 | 0.75 | 23 | 0.5652 |
| 4_9281751 | 2 | 0.5 | 4 | 0.25 | 3 | 0.6667 | 6 | 0.3333 | 15 | 0.4 | 5 | 0.6 | 8 | 0.25 | 5 | 0.8 | 4 | 0.25 | 22 | 0.4545 |
| 4_9281833 | 2 | 0.5 | 4 | 0.5 | 4 | 0.5 | 5 | 0.8 | 15 | 0.6 | 5 | 0.2 | 9 | 0.5556 | 5 | 0.4 | 5 | 0.2 | 24 | 0.375 |
| 4_9281868 | 2 | 0 | 3 | 0.3333 | 4 | 0 | 5 | 0 | 14 | 0.0714 | 5 | 0.2 | 8 | 0.25 | 5 | 0.2 | 5 | 0 | 23 | 0.1739 |
| 4_9281874 | 2 | 1 | 4 | 0.5 | 3 | 1 | 5 | 0.6 | 14 | 0.7143 | 3 | 0.3333 | 10 | 0.8 | 5 | 1 | 4 | 1 | 22 | 0.8182 |
| 4_9281886 | 2 | 1 | 4 | 0.5 | 3 | 1 | 5 | 1 | 14 | 0.8571 | 5 | 0.8 | 6 | 0.8333 | 3 | 0.6667 | 2 | 1 | 16 | 0.8125 |
| 4_9429842 | 2 | 0 | 5 | 0 | 4 | 0.25 | 4 | 0.75 | 15 | 0.2667 | 5 | 0.4 | 7 | 0.1429 | 6 | 0.3333 | 5 | 0 | 23 | 0.2174 |
| 4_9429854 | 2 | 0 | 5 | 0.6 | 4 | 0.25 | 6 | 0.3333 | 17 | 0.3529 | 5 | 0.2 | 9 | 0.4444 | 4 | 0 | 5 | 0 | 23 | 0.2174 |
| 4_9449473 | 3 | 0.3333 | 3 | 0.6667 | 3 | 0 | 5 | 0.4 | 14 | 0.3571 | 4 | 0 | 7 | 0.7143 | 5 | 0 | 7 | 0.4286 | 23 | 0.3478 |
| 4_9449546 | 4 | 0.25 | 5 | 0.4 | 5 | 0 | 6 | 0.1667 | 20 | 0.2 | 4 | 0.25 | 10 | 0.2 | 5 | 0.2 | 6 | 0.1667 | 25 | 0.2 |

| | | | | | | | | | | | | | | | | | | | | |
|---|---|---|---|---|---|---|---|---|---|---|---|---|---|---|---|---|---|---|---|---|
| 4_9449599 | 2 | 0.5 | 3 | 0.6667 | 4 | 0.5 | 6 | 0.5 | 15 | 0.5333 | 4 | 0.5 | 10 | 0.3 | 3 | 0 | 6 | 0.3333 | 23 | 0.3043 |
| 4_9449636 | 4 | 0 | 4 | 0.25 | 4 | 0.25 | 6 | 0.1667 | 18 | 0.1667 | 5 | 0 | 10 | 0.1 | 6 | 0.3333 | 5 | 0.4 | 26 | 0.1923 |
| 4_9449818 | NA | NA | 4 | 0.25 | 5 | 0.6 | 4 | 0.25 | 13 | 0.3846 | 6 | 0.3333 | 11 | 0.0909 | 6 | 0.3333 | 7 | 0.2857 | 30 | 0.2333 |
| 4_9450087 | 2 | 0 | 6 | 0 | 5 | 0.2 | 6 | 0.1667 | 19 | 0.1053 | 5 | 0.2 | 11 | 0.2727 | 6 | 0.3333 | 5 | 0.4 | 27 | 0.2963 |
| 4_9452950 | NA | NA | 4 | 0.25 | 1 | 0 | NA | NA | 5 | 0.2 | 3 | 0.3333 | 1 | 1 | NA | NA | 4 | 0.25 | 8 | 0.375 |
| 4_9453073 | NA | NA | 3 | 0.3333 | 1 | 0 | NA | NA | 4 | 0.25 | 2 | 0 | 1 | 0 | NA | NA | 3 | 0.3333 | 6 | 0.1667 |
| 4_9453151 | NA | NA | 1 | 1 | 1 | 0 | NA | NA | 2 | 0.5 | NA | NA | 1 | 0 | NA | NA | 2 | 1 | 3 | 0.6667 |
| 4_9576850 | NA | NA | 1 | 0 | 1 | 1 | 2 | 0 | 4 | 0.25 | 2 | 0.5 | 1 | 1 | NA | NA | 2 | 0 | 5 | 0.4 |
| 4_9621167 | 1 | 0 | NA | NA | NA | NA | 1 | 1 | 2 | 0.5 | 1 | 1 | NA | NA | NA | NA | NA | NA | 1 | 1 |
| 4_9630408 | 7 | 0.1429 | 16 | 0.0625 | 5 | 0 | 7 | 0 | 35 | 0.0571 | 2 | 0 | 3 | 0.3333 | 4 | 0 | 5 | 0.2 | 14 | 0.1429 |
| 4_9630611 | 5 | 0.6 | 17 | 0.6471 | 5 | 0.6 | 7 | 0.4286 | 34 | 0.5882 | 2 | 0 | 3 | 0.6667 | 3 | 1 | 5 | 0.6 | 13 | 0.6154 |
| 4_9630694 | 3 | 1 | 13 | 1 | 4 | 1 | 6 | 1 | 26 | 1 | 2 | 1 | 2 | 0.5 | 5 | 1 | 4 | 1 | 13 | 0.9231 |
| 4_9630708 | 4 | 0.5 | 13 | 0.2308 | 4 | 0.25 | 5 | 0.6 | 26 | 0.3462 | 1 | 0 | 2 | 0.5 | 5 | 0.2 | 4 | 0 | 12 | 0.1667 |
| 4_9630740 | 5 | 0.4 | 15 | 0.3333 | 5 | 0.4 | 6 | 0.5 | 31 | 0.3871 | 2 | 1 | 2 | 1 | 5 | 0.6 | 6 | 1 | 15 | 0.8667 |
| 4_9630791 | 5 | 0 | 17 | 0.1176 | 4 | 0.25 | 6 | 0 | 32 | 0.0938 | 3 | 0 | 1 | 0 | 4 | 0 | 5 | 0.2 | 13 | 0.0769 |
| 4_9630833 | 3 | 0.6667 | 16 | 0.9375 | 3 | 1 | 6 | 1 | 28 | 0.9286 | 3 | 0.3333 | 1 | 1 | 5 | 1 | 3 | 1 | 12 | 0.8333 |
| 4_10874765 | 3 | 0 | 1 | 0 | 4 | 0.25 | 5 | 0 | 13 | 0.0769 | 1 | 1 | 6 | 0.1667 | 3 | 0.3333 | 6 | 0.3333 | 16 | 0.3125 |
| 4_10897987 | 6 | 0.5 | 2 | 0.5 | 4 | 0.5 | 7 | 0.2857 | 19 | 0.4211 | 1 | 0 | 8 | 0.75 | 4 | 0.25 | 5 | 0.6 | 18 | 0.5556 |
| 4_10897998 | 6 | 0 | 2 | 0 | 4 | 0.25 | 8 | 0.125 | 20 | 0.1 | 2 | 0 | 8 | 0.125 | 4 | 0 | 5 | 0 | 19 | 0.0526 |
| 4_10898183 | 5 | 1 | 2 | 1 | 3 | 0.6667 | 8 | 1 | 18 | 0.9444 | 2 | 1 | 9 | 0.8889 | 4 | 0.75 | 6 | 1 | 21 | 0.9048 |
| 4_10898236 | 6 | 0.8333 | 2 | 1 | 4 | 0.75 | 8 | 0.875 | 20 | 0.85 | 2 | 0 | 9 | 0.7778 | 5 | 0.8 | 6 | 0.6667 | 22 | 0.6818 |
| 4_10898289 | 6 | 1 | 1 | 1 | 4 | 1 | 8 | 0.5 | 19 | 0.7895 | 3 | 0.6667 | 8 | 0.5 | 5 | 0.6 | 5 | 0.6 | 21 | 0.5714 |
| 4_10898301 | 6 | 0 | 2 | 0 | 4 | 0 | 7 | 0.1429 | 19 | 0.0526 | 3 | 0 | 9 | 0.2222 | 5 | 0 | 6 | 0.1667 | 23 | 0.1304 |
| 4_10898347 | 6 | 0.6667 | 2 | 1 | 3 | 0.6667 | 6 | 0.6667 | 17 | 0.7059 | 2 | 0.5 | 8 | 0.75 | 5 | 0.6 | 5 | 0.4 | 20 | 0.6 |
| 4_10898474 | 5 | 0.2 | 2 | 0.5 | 4 | 0.25 | 8 | 0 | 19 | 0.1579 | 3 | 0.3333 | 6 | 0 | 4 | 0 | 5 | 0.2 | 18 | 0.1111 |
| 4_10988694 | 1 | 0 | 2 | 0.5 | 1 | 0 | 1 | 0 | 5 | 0.2 | 2 | 0 | 1 | 0 | 2 | 0.5 | 1 | 0 | 6 | 0.1667 |
| 4_10988789 | 1 | 0 | 2 | 0 | 1 | 1 | 1 | 1 | 5 | 0.4 | 2 | 0 | NA | NA | 2 | 0.5 | 1 | 0 | 5 | 0.2 |
| 4_10990425 | 1 | 1 | 2 | 1 | 2 | 1 | 2 | 1 | 7 | 1 | 2 | 1 | NA | NA | 1 | 1 | 1 | 1 | 4 | 1 |
| 4_10990516 | 1 | 1 | 1 | 1 | NA | NA | 2 | 0.5 | 4 | 0.75 | 2 | 1 | NA | NA | NA | NA | 1 | 1 | 3 | 1 |
| 4_10990565 | 1 | 1 | 2 | 1 | NA | NA | 2 | 1 | 5 | 1 | 2 | 1 | NA | NA | 1 | 1 | 1 | 1 | 4 | 1 |
| 4_10990595 | 1 | 1 | 2 | 0 | 1 | 1 | 2 | 1 | 6 | 0.6667 | 2 | 1 | NA | NA | 1 | 0 | 1 | 1 | 4 | 0.75 |
| 4_10990667 | 1 | 1 | 2 | 1 | 1 | 1 | 2 | 1 | 6 | 1 | 2 | 1 | NA | NA | 1 | 1 | 1 | 1 | 4 | 1 |
| 4_10990727 | NA | NA | 2 | 0 | NA | NA | 2 | 0.5 | 4 | 0.25 | 1 | 1 | NA | NA | 1 | 1 | 1 | 0 | 3 | 0.6667 |
| 4_10990886 | NA | NA | 2 | 0.5 | NA | NA | 2 | 1 | 4 | 0.75 | 2 | 1 | NA | NA | 1 | 1 | 1 | 1 | 4 | 1 |
| 4_10991031 | 1 | 1 | 2 | 1 | 2 | 0.5 | 2 | 0.5 | 7 | 0.7143 | 2 | 1 | NA | NA | 1 | 1 | 1 | 0 | 4 | 0.75 |
| 4_10991113 | 1 | 1 | 2 | 0.5 | NA | NA | 2 | 0 | 5 | 0.4 | 2 | 0.5 | NA | NA | 1 | 0 | 1 | 1 | 4 | 0.5 |
| 4_10991214 | 1 | 1 | 2 | 1 | 2 | 0 | 2 | 0.5 | 7 | 0.5714 | 2 | 1 | NA | NA | NA | NA | 1 | 1 | 3 | 1 |
| 4_11164469 | 1 | 1 | 1 | 1 | 1 | 0 | 2 | 1 | 5 | 0.8 | NA | NA | 1 | 1 | 1 | 1 | NA | NA | 2 | 1 |
| 4_11165206 | 1 | 1 | 1 | 0 | 1 | 0 | NA | NA | 3 | 0.3333 | NA | NA | 1 | 1 | NA | NA | NA | NA | 1 | 1 |
| 4_11169827 | 1 | 1 | 1 | 1 | 1 | 0 | 2 | 0.5 | 5 | 0.6 | NA | NA | 1 | 1 | 1 | 0 | NA | NA | 2 | 0.5 |
| 4_11169839 | NA | NA | 1 | 0 | 1 | 0 | 2 | 1 | 4 | 0.5 | NA | NA | 1 | 1 | NA | NA | NA | NA | 1 | 1 |
| 4_11172933 | NA | NA | 3 | 0.3333 | 1 | 0 | 1 | 0 | 5 | 0.2 | 3 | 0.3333 | 3 | 0 | 4 | 0 | NA | NA | 10 | 0.1 |
| 4_11173332 | 1 | 0 | 2 | 0.5 | 1 | 0 | 2 | 0.5 | 6 | 0.3333 | 3 | 0 | 2 | 0.5 | 3 | 0 | 1 | 0 | 9 | 0.1111 |
| 4_11174078 | 1 | 1 | 3 | 0 | 1 | 1 | 3 | 0.6667 | 8 | 0.5 | 3 | 0.3333 | 3 | 0.6667 | 4 | 0.5 | 1 | 0 | 11 | 0.4545 |
| 4_11203500 | NA | NA | NA | NA | 2 | 0.5 | 1 | 0 | 3 | 0.3333 | 1 | 0 | 2 | 0.5 | NA | NA | NA | NA | 3 | 0.3333 |
| 4_11261544 | 5 | 0.2 | 3 | 0 | 3 | 0 | 9 | 0 | 20 | 0.05 | 3 | 0 | 4 | 0.25 | 6 | 0 | 2 | 0 | 15 | 0.0667 |
| 4_11262869 | 4 | 0 | 1 | 0 | 1 | 0 | 7 | 0.1429 | 13 | 0.0769 | 2 | 0.5 | 4 | 0 | 3 | 0.3333 | 1 | 0 | 10 | 0.2 |
| 4_11262906 | 3 | 0.3333 | 1 | 0 | 1 | 0 | 7 | 0.1429 | 12 | 0.1667 | 2 | 0.5 | 4 | 0.25 | 3 | 0 | 1 | 0 | 10 | 0.2 |
| 4_11262924 | 4 | 0 | 1 | 0 | 1 | 1 | 7 | 0.2857 | 13 | 0.2308 | 2 | 0 | 4 | 0.25 | 3 | 0.3333 | 1 | 0 | 10 | 0.2 |
| 4_11265751 | 5 | 0.4 | NA | NA | NA | NA | 7 | 0.1429 | 12 | 0.25 | 1 | 1 | 2 | 0 | 1 | 1 | NA | NA | 4 | 0.5 |
| 4_11265778 | 5 | 0 | NA | NA | NA | NA | 7 | 0.1429 | 12 | 0.0833 | 1 | 1 | 2 | 0 | 1 | 0 | NA | NA | 4 | 0.25 |
| 4_11265830 | 4 | 0.25 | NA | NA | NA | NA | 3 | 0.3333 | 7 | 0.2857 | 1 | 0 | 2 | 0.5 | 1 | 0 | NA | NA | 4 | 0.25 |
| 4_11265872 | 4 | 0 | NA | NA | NA | NA | 4 | 0.25 | 8 | 0.125 | 1 | 1 | 1 | 0 | 1 | 0 | NA | NA | 3 | 0.3333 |
| 4_11513308 | 2 | 1 | 3 | 0.6667 | NA | NA | 1 | 1 | 6 | 0.8333 | NA | NA | 2 | 1 | 6 | 0.8333 | NA | NA | 8 | 0.875 |
| 4_11513424 | 2 | 0.5 | 2 | 0.5 | 1 | 0 | 1 | 0 | 6 | 0.3333 | NA | NA | 2 | 0.5 | 5 | 0.4 | NA | NA | 7 | 0.4286 |
| 4_11513450 | 2 | 0 | 3 | 1 | 1 | 1 | 1 | 1 | 7 | 0.7143 | 1 | 0 | 2 | 0.5 | 5 | 0.6 | NA | NA | 8 | 0.5 |
| 4_11513537 | 1 | 0 | 3 | 0 | 1 | 1 | 1 | 0 | 6 | 0.1667 | 1 | 0 | 3 | 0.3333 | 6 | 0.1667 | NA | NA | 10 | 0.2 |
| 4_11513670 | 2 | 1 | 2 | 1 | 1 | 0 | 1 | 1 | 6 | 0.8333 | 1 | 1 | 2 | 1 | 3 | 1 | NA | NA | 6 | 1 |
| 4_11513755 | 2 | 0.5 | 3 | 0.3333 | NA | NA | 1 | 0 | 6 | 0.3333 | 1 | 0 | NA | NA | 5 | 0.2 | NA | NA | 6 | 0.1667 |
| 4_11513774 | 2 | 0.5 | 2 | 0.5 | 1 | 1 | 1 | 0 | 6 | 0.5 | 1 | 1 | 2 | 0.5 | 5 | 1 | NA | NA | 8 | 0.875 |
| 4_11513851 | 2 | 1 | 3 | 1 | 1 | 1 | 1 | 1 | 7 | 1 | 1 | 1 | 3 | 1 | 6 | 1 | NA | NA | 10 | 1 |
| 4_11513976 | 2 | 1 | 3 | 1 | 1 | 1 | 1 | 1 | 7 | 1 | 1 | 0 | 3 | 0.6667 | 6 | 1 | NA | NA | 10 | 0.8 |
| 4_11514065 | 2 | 0.5 | 2 | 0 | 1 | 0 | 1 | 0 | 6 | 0.1667 | 1 | 0 | 2 | 0.5 | 6 | 0.1667 | NA | NA | 9 | 0.2222 |
| 4_11519090 | 2 | 0.5 | 3 | 0.6667 | 1 | 0 | 1 | 0 | 7 | 0.4286 | 1 | 0 | 2 | 0.5 | 4 | 0.5 | NA | NA | 7 | 0.4286 |
| 4_11519189 | 2 | 0.5 | 3 | 1 | 1 | 1 | 1 | 1 | 7 | 0.8571 | 1 | 1 | 3 | 0.6667 | 6 | 1 | NA | NA | 10 | 0.9 |

| | | | | | | | | | | | | | | | | | | | | |
|---|---|---|---|---|---|---|---|---|---|---|---|---|---|---|---|---|---|---|---|---|
| 4_1151920 | 2 | 1 | 3 | 0.6667 | 1 | 0 | 1 | 0 | 7 | 0.5714 | 1 | 1 | 3 | 0 | 5 | 0.4 | NA | NA | 9 | 0.3333 |
| 4_1151926 | 2 | 1 | 1 | 0 | 1 | 0 | 1 | 0 | 5 | 0.4 | 1 | 0 | 3 | 0.3333 | 5 | 0.6 | NA | NA | 9 | 0.4444 |
| 4_1152665 | 2 | 0.5 | 3 | 0.3333 | 2 | 0.5 | 1 | 1 | 8 | 0.5 | 3 | 1 | 3 | 0.6667 | 6 | 0.5 | 1 | 1 | 13 | 0.6923 |
| 4_1152668 | 2 | 0 | 2 | 0.5 | 2 | 0.5 | 1 | 0 | 7 | 0.2857 | 2 | 1 | 2 | 0.5 | 6 | 0.5 | 1 | 1 | 11 | 0.6364 |
| 4_1152668 | 1 | 0 | 1 | 1 | 2 | 0.5 | NA | NA | 4 | 0.5 | 3 | 0.3333 | 1 | 0 | 3 | 0 | NA | NA | 7 | 0.1429 |
| 4_1152673 | 1 | 1 | 3 | 0.6667 | 1 | 1 | NA | NA | 5 | 0.8 | 3 | 1 | 3 | 0.6667 | 6 | 0.6667 | 1 | 1 | 13 | 0.7692 |
| 4_1152679 | 1 | 1 | 2 | 0 | 2 | 0.5 | 1 | 1 | 6 | 0.5 | 3 | 0 | 2 | 1 | 4 | 0.25 | 1 | 1 | 10 | 0.4 |
| 4_1152683 | 2 | 1 | 3 | 1 | 2 | 1 | 1 | 1 | 8 | 1 | 3 | 0.3333 | 3 | 1 | 5 | 0.8 | 1 | 1 | 12 | 0.75 |
| 4_1152695 | 2 | 1 | 2 | 1 | 1 | 1 | 1 | 0 | 6 | 0.8333 | 2 | 1 | 2 | 1 | 5 | 1 | 1 | 0 | 10 | 0.9 |
| 4_1152700 | 3 | 1 | 3 | 0.3333 | 2 | 1 | 1 | 1 | 9 | 0.7778 | 4 | 0.75 | 3 | 0.6667 | 6 | 0.8333 | 1 | 1 | 14 | 0.7857 |
| 4_1152708 | 2 | 0.5 | 3 | 0.6667 | 1 | 1 | 1 | 1 | 7 | 0.7143 | 4 | 0.75 | 2 | 1 | 6 | 0.5 | 1 | 0 | 13 | 0.6154 |
| 4_1152724 | 1 | 1 | NA | NA | NA | NA | NA | NA | 1 | 1 | NA | NA | 1 | 1 | 1 | 1 | NA | NA | 2 | 1 |
| 4_1152725 | 1 | 1 | NA | NA | NA | NA | NA | NA | 1 | 1 | 1 | 1 | 1 | 0 | NA | NA | NA | NA | 2 | 0.5 |
| 4_1152744 | 1 | 1 | NA | NA | NA | NA | NA | NA | 1 | 1 | 1 | 1 | 1 | 1 | 1 | 1 | NA | NA | 3 | 1 |
| 4_1152745 | 1 | 1 | NA | NA | NA | NA | NA | NA | 1 | 1 | 1 | 1 | 1 | 0 | 1 | 0 | NA | NA | 3 | 0.3333 |
| 4_1152746 | 1 | 1 | NA | NA | NA | NA | NA | NA | 1 | 1 | 1 | 0 | 1 | 1 | 1 | 0 | NA | NA | 3 | 0.3333 |
| 4_1152784 | 2 | 0 | 3 | 0.3333 | 2 | 0.5 | 1 | 0 | 8 | 0.25 | 4 | 0 | 2 | 0 | 8 | 0.125 | NA | NA | 14 | 0.0714 |
| 4_1152865 | 3 | 0 | 3 | 0 | 2 | 0.5 | 1 | 0 | 9 | 0.1111 | 2 | 0 | 2 | 0 | 7 | 0.4286 | 2 | 0 | 13 | 0.2308 |
| 4_1154612 | 3 | 0 | 4 | 0 | 7 | 0.2857 | 2 | 0 | 16 | 0.125 | 6 | 0 | 2 | 0.5 | 9 | 0 | 3 | 0 | 20 | 0.05 |
| 4_1154627 | 4 | 0 | 3 | 0.3333 | 5 | 0 | 3 | 0 | 15 | 0.0667 | 4 | 0 | 2 | 0.5 | 9 | 0 | 2 | 0 | 17 | 0.0588 |
| 4_1154974 | 3 | 1 | 4 | 0.25 | 6 | 0 | 3 | 0.3333 | 16 | 0.3125 | 7 | 0.5714 | 2 | 0 | 10 | 0.2 | 4 | 0.25 | 23 | 0.3043 |
| 4_1154976 | 5 | 0.2 | 4 | 0.25 | 8 | 0.125 | 3 | 0.6667 | 20 | 0.25 | 8 | 0 | 2 | 0 | 11 | 0.0909 | 4 | 0.5 | 25 | 0.12 |
| 4_1155044 | 1 | 0 | 1 | 0 | 2 | 0.5 | 2 | 0.5 | 6 | 0.3333 | 1 | 0 | NA | NA | 2 | 0.5 | 2 | 0 | 5 | 0.2 |
| 4_1155046 | 1 | 1 | 1 | 1 | 3 | 0.3333 | 2 | 1 | 7 | 0.7143 | 3 | 0.3333 | NA | NA | 5 | 0.6 | 2 | 1 | 10 | 0.6 |
| 4_1155048 | 1 | 0 | 2 | 1 | 2 | 0 | 1 | 0 | 6 | 0.3333 | 2 | 1 | NA | NA | 5 | 0.2 | 2 | 0.5 | 9 | 0.4444 |
| 4_1161566 | 2 | 0.5 | NA | NA | NA | NA | NA | NA | 2 | 0.5 | NA | NA | 1 | 1 | NA | NA | NA | NA | 1 | 1 |
| 4_1171355 | NA | NA | 1 | 1 | NA | NA | 1 | 1 | 2 | 1 | 1 | 1 | NA | NA | 2 | 0.5 | 1 | 1 | 4 | 0.75 |
| 4_1171380 | 1 | 1 | 1 | 1 | NA | NA | 1 | 1 | 3 | 1 | 1 | 1 | NA | NA | 2 | 0.5 | 1 | 1 | 4 | 0.75 |
| 4_1171398 | 1 | 1 | 1 | 0 | NA | NA | 1 | 0 | 3 | 0.3333 | 1 | 0 | NA | NA | 2 | 0.5 | 1 | 0 | 4 | 0.25 |
| 4_1195909 | 6 | 0.1667 | 10 | 0.2 | 8 | 0.125 | 6 | 0.1667 | 30 | 0.1667 | 9 | 0.2222 | 10 | 0.2 | 8 | 0.25 | 6 | 0 | 33 | 0.1818 |
| 4_1195980 | 4 | 0 | 8 | 0.25 | 9 | 0.1111 | 6 | 0.3333 | 27 | 0.1852 | 8 | 0.125 | 9 | 0.2222 | 5 | 0.2 | 6 | 0.3333 | 28 | 0.2143 |
| 4_1195999 | 16 | 0.1875 | 16 | 0 | 18 | 0.0556 | 14 | 0 | 64 | 0.0625 | 15 | 0.1333 | 15 | 0 | 12 | 0.0833 | 8 | 0 | 50 | 0.06 |
| 4_1196082 | 14 | 0.2857 | 13 | 0.4615 | 9 | 0 | 10 | 0.3 | 46 | 0.2826 | 11 | 0.1818 | 10 | 0.4 | 9 | 0.1111 | 5 | 0.2 | 35 | 0.2286 |
| 4_1196083 | 16 | 0.1875 | 14 | 0.4286 | 13 | 0.3846 | 10 | 0.3 | 53 | 0.3208 | 12 | 0.5 | 10 | 0.6 | 8 | 0.625 | 7 | 0.5714 | 37 | 0.5676 |
| 4_1196105 | 15 | 0.1333 | 16 | 0.25 | 14 | 0.4286 | 10 | 0.4 | 55 | 0.2909 | 11 | 0.2727 | 10 | 0.3 | 12 | 0.4167 | 8 | 0.125 | 41 | 0.2927 |
| 4_1196110 | 14 | 0.2857 | 15 | 0.2 | 11 | 0 | 9 | 0 | 49 | 0.1429 | 12 | 0.1667 | 8 | 0 | 11 | 0.0909 | 9 | 0 | 40 | 0.075 |
| 4_1196116 | 14 | 0.1429 | 15 | 0.1333 | 13 | 0.1538 | 9 | 0.3333 | 51 | 0.1765 | 11 | 0.1818 | 7 | 0 | 12 | 0.1667 | 7 | 0.1429 | 37 | 0.1351 |
| 4_1196120 | 12 | 0 | 12 | 0.25 | 11 | 0.0909 | 8 | 0.375 | 43 | 0.1628 | 9 | 0.1111 | 4 | 0 | 10 | 0.2 | 5 | 0 | 28 | 0.1071 |
| 4_1196121 | 15 | 0.2667 | 16 | 0.4375 | 13 | 0.4615 | 8 | 0.25 | 52 | 0.3654 | 11 | 0.6364 | 8 | 0.125 | 11 | 0.1818 | 8 | 0.25 | 38 | 0.3158 |
| 4_1196122 | 14 | 0.5 | 14 | 0.6429 | 11 | 0.5455 | 10 | 0.6 | 49 | 0.5714 | 11 | 0.6364 | 9 | 0.4444 | 11 | 0.8182 | 8 | 0.25 | 39 | 0.5641 |
| 4_1196145 | 12 | 1 | 14 | 0.8571 | 11 | 0.8182 | 8 | 0.75 | 45 | 0.8667 | 9 | 0.8889 | 8 | 0.875 | 9 | 0.7778 | 6 | 0.6667 | 32 | 0.8125 |
| 4_1196146 | 12 | 1 | 14 | 1 | 14 | 1 | 9 | 1 | 49 | 1 | 12 | 1 | 9 | 1 | 10 | 1 | 8 | 1 | 39 | 1 |
| 4_1196152 | 11 | 0.3636 | 12 | 0.5 | 11 | 0.5455 | 7 | 0.4286 | 41 | 0.4634 | 10 | 0.2 | 8 | 0.375 | 11 | 0.2727 | 7 | 0.7143 | 36 | 0.3611 |
| 4_1196153 | 10 | 0.8 | 10 | 1 | 10 | 1 | 6 | 0.6667 | 36 | 0.8889 | 10 | 1 | 7 | 1 | 10 | 0.9 | 5 | 0.8 | 32 | 0.9375 |
| 4_1196155 | 11 | 1 | 12 | 1 | 11 | 1 | 8 | 1 | 42 | 1 | 10 | 1 | 8 | 1 | 11 | 1 | 8 | 1 | 37 | 1 |
| 4_1196167 | 11 | 1 | 12 | 1 | 11 | 1 | 7 | 1 | 41 | 1 | 12 | 1 | 8 | 1 | 11 | 1 | 6 | 1 | 37 | 1 |
| 4_1196171 | 11 | 1 | 11 | 1 | 11 | 1 | 7 | 1 | 40 | 1 | 9 | 1 | 9 | 1 | 8 | 1 | 7 | 1 | 33 | 1 |
| 4_1196178 | 9 | 0.5556 | 11 | 0.1818 | 10 | 0.1 | 7 | 0.4286 | 37 | 0.2973 | 11 | 0.3636 | 8 | 0.125 | 10 | 0.5 | 5 | 0.2 | 34 | 0.3235 |
| 4_1196181 | 7 | 0.7143 | 12 | 0.8333 | 9 | 0.8889 | 9 | 0.7778 | 37 | 0.8108 | 10 | 0.8 | 8 | 1 | 10 | 0.9 | 7 | 0.7143 | 35 | 0.8571 |
| 4_1196200 | 11 | 0.8182 | 10 | 0.6 | 8 | 0.875 | 8 | 0.5 | 37 | 0.7027 | 10 | 0.6 | 8 | 0.75 | 7 | 0.7143 | 7 | 0.7143 | 32 | 0.6875 |
| 4_1196214 | 6 | 1 | 9 | 1 | 6 | 1 | 4 | 1 | 25 | 1 | 5 | 1 | 3 | 1 | 8 | 0.875 | 5 | 0.8 | 21 | 0.9048 |
| 4_1199412 | NA | NA | 1 | 1 | 1 | 1 | NA | NA | 2 | 1 | NA | NA | 1 | 1 | 1 | 1 | 1 | 1 | 3 | 1 |
| 4_1199422 | NA | NA | 1 | 1 | 1 | 1 | NA | NA | 2 | 1 | NA | NA | 1 | 1 | 1 | 1 | 1 | 1 | 3 | 1 |
| 4_1203978 | 1 | 1 | 1 | 1 | 1 | 1 | NA | NA | 3 | 1 | 1 | 1 | NA | NA | 3 | 0.3333 | NA | NA | 4 | 0.5 |
| 4_1209340 | NA | NA | 1 | 1 | 1 | 1 | NA | NA | 2 | 1 | 1 | 0 | NA | NA | 1 | 1 | NA | NA | 2 | 0.5 |
| 4_1209355 | NA | NA | 1 | 1 | 2 | 1 | 1 | 1 | 4 | 1 | 1 | 1 | NA | NA | 1 | 1 | 1 | 1 | 3 | 1 |
| 4_1209406 | NA | NA | 1 | 0 | 2 | 0 | 2 | 0.5 | 5 | 0.2 | 1 | 0 | 1 | 1 | 1 | 0 | 1 | 1 | 4 | 0.5 |
| 4_1209414 | NA | NA | 1 | 0 | 2 | 0.5 | 2 | 1 | 5 | 0.6 | 1 | 1 | 1 | 0 | 1 | 1 | 1 | 1 | 4 | 0.75 |
| 4_1209417 | NA | NA | 1 | 1 | 2 | 1 | 2 | 1 | 5 | 1 | 1 | 1 | 1 | 1 | 1 | 1 | 1 | 1 | 4 | 1 |
| 4_1209441 | NA | NA | 1 | 1 | 2 | 0.5 | 2 | 0.5 | 5 | 0.6 | 1 | 1 | 1 | 0 | 1 | 1 | 1 | 0 | 4 | 0.5 |
| 4_1209456 | NA | NA | 1 | 1 | 2 | 1 | 1 | 0 | 4 | 0.75 | 1 | 1 | 1 | 1 | 1 | 1 | 1 | 0 | 4 | 0.75 |
| 4_1209462 | NA | NA | 1 | 0 | 2 | 0.5 | 2 | 1 | 5 | 0.6 | 1 | 1 | 1 | 0 | 1 | 0 | 1 | 0 | 4 | 0.25 |
| 4_1209466 | NA | NA | 1 | 0 | 2 | 0.5 | 2 | 0.5 | 5 | 0.4 | 1 | 0 | 1 | 0 | 1 | 1 | 1 | 0 | 4 | 0.5 |
| 4_1209474 | NA | NA | NA | NA | 2 | 0.5 | 2 | 1 | 4 | 0.75 | 1 | 1 | 1 | 1 | 1 | 1 | 1 | 1 | 4 | 1 |
| 4_1209481 | NA | NA | 1 | 0 | 2 | 0.5 | 2 | 1 | 5 | 0.6 | 1 | 1 | 2 | 0 | 1 | 1 | 1 | 0 | 5 | 0.4 |

| | | | | | | | | | | | | | | | | | | | | |
|---|---|---|---|---|---|---|---|---|---|---|---|---|---|---|---|---|---|---|---|---|
| 4_1209488 | NA | NA | 1 | 0 | 2 | 0.5 | 2 | 0 | 5 | 0.2 | 1 | 0 | 2 | 0.5 | 2 | 0 | NA | NA | 5 | 0.2 |
| 4_1209505 | NA | NA | 1 | 1 | NA | NA | 1 | 1 | 2 | 1 | 1 | 1 | 1 | 0 | 2 | 0.5 | NA | NA | 4 | 0.5 |
| 4_1209510 | NA | NA | 2 | 0 | 1 | 1 | 2 | 0.5 | 5 | 0.4 | 1 | 1 | 2 | 0.5 | 2 | 0.5 | 1 | 1 | 6 | 0.6667 |
| 4_1209512 | NA | NA | 2 | 0.5 | 2 | 1 | 2 | 0.5 | 6 | 0.6667 | NA | NA | 2 | 1 | 2 | 1 | 1 | 1 | 5 | 1 |
| 4_1209516 | NA | NA | 2 | 1 | 2 | 1 | 2 | 1 | 6 | 1 | 1 | 1 | 2 | 1 | 2 | 1 | 1 | 1 | 6 | 1 |
| 4_1209521 | NA | NA | 2 | 0 | 2 | 0.5 | 2 | 0.5 | 6 | 0.3333 | 1 | 1 | 2 | 0.5 | 2 | 1 | 1 | 1 | 6 | 0.8333 |
| 4_1209526 | NA | NA | 2 | 0 | 2 | 0.5 | 2 | 1 | 6 | 0.5 | 1 | 1 | 2 | 0.5 | 2 | 0.5 | NA | NA | 5 | 0.6 |
| 4_1209530 | NA | NA | 2 | 1 | 1 | 1 | 2 | 1 | 5 | 1 | 1 | 1 | 1 | 1 | 1 | 1 | 1 | 1 | 4 | 1 |
| 4_1209546 | NA | NA | 2 | 1 | 1 | 0 | 3 | 0.6667 | 6 | 0.6667 | 1 | 1 | 2 | 0.5 | 1 | 1 | 1 | 1 | 5 | 0.8 |
| 4_1209568 | 1 | 0 | 2 | 0 | 3 | 0.3333 | 4 | 0 | 10 | 0.1 | 2 | 0 | 2 | 0 | 3 | 0.3333 | 2 | 0.5 | 9 | 0.2222 |
| 4_1209587 | 1 | 0 | 1 | 1 | 3 | 0.6667 | 6 | 0.5 | 11 | 0.5455 | 2 | 0.5 | 3 | 0.6667 | 2 | 0.5 | 2 | 1 | 9 | 0.6667 |
| 4_1209610 | 1 | 1 | 1 | 1 | 1 | 0 | 6 | 0.5 | 9 | 0.5556 | 2 | 1 | 3 | 1 | 1 | 0 | 1 | 1 | 7 | 0.8571 |
| 4_1209616 | 2 | 0 | 2 | 0 | 4 | 0.25 | 5 | 0.4 | 13 | 0.2308 | 1 | 1 | 3 | 0 | 2 | 0.5 | 2 | 0 | 8 | 0.25 |
| 4_1214429 | NA | NA | 2 | 1 | NA | NA | 1 | 1 | 3 | 1 | NA | NA | NA | NA | 1 | 1 | NA | NA | 1 | 1 |
| 4_1214443 | NA | NA | 2 | 1 | NA | NA | 1 | 1 | 3 | 1 | NA | NA | NA | NA | 1 | 1 | NA | NA | 1 | 1 |
| 4_1214481 | 2 | 1 | 3 | 0 | 1 | 0 | 2 | 0 | 8 | 0.25 | NA | NA | NA | NA | 2 | 1 | NA | NA | 2 | 1 |
| 4_1214503 | NA | NA | 2 | 1 | 1 | 1 | 2 | 0.5 | 5 | 0.8 | NA | NA | NA | NA | 2 | 1 | NA | NA | 2 | 1 |
| 4_1214506 | 1 | 1 | 3 | 1 | 1 | 0 | 1 | 1 | 6 | 0.8333 | NA | NA | NA | NA | 2 | 1 | NA | NA | 2 | 1 |
| 4_1214529 | 1 | 1 | 2 | 1 | 1 | 0 | 2 | 0.5 | 6 | 0.6667 | NA | NA | NA | NA | 2 | 1 | NA | NA | 2 | 1 |
| 4_1214545 | 1 | 1 | 3 | 0.6667 | NA | NA | 2 | 0.5 | 6 | 0.6667 | NA | NA | NA | NA | 2 | 1 | NA | NA | 2 | 1 |
| 4_1214551 | 1 | 0 | 3 | 0.3333 | 1 | 0 | 2 | 0 | 7 | 0.1429 | NA | NA | NA | NA | 2 | 0.5 | NA | NA | 2 | 0.5 |
| 4_1214563 | 1 | 1 | 1 | 0 | NA | NA | 2 | 0 | 4 | 0.25 | NA | NA | 1 | 1 | 2 | 1 | NA | NA | 3 | 1 |
| 4_1214577 | NA | NA | 3 | 0.3333 | 1 | 0 | 2 | 0 | 6 | 0.1667 | NA | NA | 1 | 0 | 2 | 0.5 | NA | NA | 3 | 0.3333 |
| 4_1214577 | 1 | 0 | 3 | 0.6667 | 1 | 0 | 1 | 0 | 6 | 0.3333 | NA | NA | 1 | 1 | 2 | 0 | NA | NA | 3 | 0.3333 |
| 4_1214594 | 1 | 1 | 3 | 0.3333 | 1 | 1 | 1 | 0 | 6 | 0.5 | NA | NA | 1 | 1 | 2 | 0 | NA | NA | 3 | 0.3333 |
| 4_1214599 | 2 | 0 | 3 | 0 | 1 | 0 | 1 | 1 | 7 | 0.1429 | NA | NA | 1 | 1 | 2 | 0.5 | NA | NA | 3 | 0.6667 |
| 4_1214601 | 2 | 0 | 3 | 1 | 1 | 0 | 1 | 1 | 7 | 0.5714 | NA | NA | 1 | 0 | 2 | 1 | NA | NA | 3 | 0.6667 |
| 4_1214606 | 2 | 1 | 2 | 1 | 1 | 1 | 2 | 1 | 7 | 1 | NA | NA | NA | NA | 1 | 1 | NA | NA | 1 | 1 |
| 4_1214611 | 2 | 0.5 | 1 | 1 | 1 | 1 | 1 | 0 | 5 | 0.6 | NA | NA | 1 | 0 | 1 | 1 | NA | NA | 2 | 0.5 |
| 4_1214615 | 2 | 0.5 | 3 | 1 | 1 | 1 | 2 | 1 | 8 | 0.875 | NA | NA | 1 | 1 | 2 | 0.5 | NA | NA | 3 | 0.6667 |
| 4_1214619 | 2 | 1 | 1 | 1 | 1 | 0 | 2 | 1 | 6 | 0.8333 | NA | NA | 1 | 1 | 2 | 1 | NA | NA | 3 | 1 |
| 4_1214636 | 2 | 0.5 | 3 | 0.6667 | 2 | 0.5 | 1 | 1 | 8 | 0.625 | NA | NA | 1 | 1 | 2 | 0.5 | NA | NA | 3 | 0.6667 |
| 4_1214668 | 2 | 0 | 3 | 0.6667 | 2 | 0.5 | 2 | 0 | 9 | 0.3333 | NA | NA | 1 | 0 | 2 | 0.5 | NA | NA | 3 | 0.3333 |
| 4_1214770 | NA | NA | 1 | 1 | NA | NA | NA | NA | 1 | 1 | NA | NA | 1 | 0 | 1 | 1 | NA | NA | 2 | 0.5 |
| 4_1215122 | NA | NA | NA | NA | NA | NA | 1 | 1 | 1 | 1 | NA | NA | NA | NA | NA | NA | 1 | 1 | 1 | 1 |
| 4_1215133 | NA | NA | NA | NA | NA | NA | 1 | 1 | 1 | 1 | NA | NA | NA | NA | NA | NA | 1 | 1 | 1 | 1 |
| 4_1215279 | NA | NA | NA | NA | NA | NA | 1 | 1 | 1 | 1 | NA | NA | NA | NA | NA | NA | 1 | 1 | 1 | 1 |
| 4_1215378 | 18 | 0.0556 | 8 | 0.125 | 9 | 0 | 13 | 0.1538 | 48 | 0.0833 | 9 | 0.3333 | 9 | 0 | 9 | 0 | 7 | 0 | 34 | 0.0882 |
| 4_1223405 | NA | NA | 1 | 1 | 1 | 1 | NA | NA | 2 | 1 | 6 | 0.1667 | 1 | 1 | 2 | 0 | 2 | 0 | 11 | 0.1818 |
| 4_1223414 | NA | NA | 1 | 1 | 2 | 0.5 | 1 | 1 | 4 | 0.75 | 6 | 0.6667 | 1 | 1 | 3 | 1 | 3 | 0.3333 | 13 | 0.6923 |
| 4_1251491 | 1 | 1 | NA | NA | NA | NA | NA | NA | 1 | 1 | 1 | 1 | NA | NA | NA | NA | NA | NA | 1 | 1 |
| 4_1251509 | 1 | 1 | NA | NA | NA | NA | NA | NA | 1 | 1 | 1 | 1 | NA | NA | NA | NA | NA | NA | 1 | 1 |
| 4_1298057 | 1 | 1 | 1 | 1 | 2 | 0 | NA | NA | 4 | 0.5 | 1 | 1 | 1 | 0 | 3 | 0.6667 | 2 | 0.5 | 7 | 0.5714 |
| 4_1388962 | 1 | 1 | 2 | 1 | 4 | 0 | 3 | 0 | 10 | 0.3 | NA | NA | 5 | 0.2 | 1 | 1 | 3 | 0.3333 | 9 | 0.3333 |
| 4_1388970 | 2 | 1 | 2 | 1 | 3 | 1 | 3 | 1 | 10 | 1 | NA | NA | 5 | 1 | 1 | 1 | 2 | 1 | 8 | 1 |
| 4_1388972 | 1 | 1 | 1 | 1 | 3 | 0.3333 | 3 | 0.6667 | 8 | 0.625 | NA | NA | 2 | 1 | 1 | 1 | 2 | 1 | 5 | 1 |
| 4_1388981 | 2 | 1 | 2 | 1 | 4 | 1 | 3 | 1 | 11 | 1 | NA | NA | 3 | 1 | 1 | 1 | 3 | 1 | 7 | 1 |
| 4_1388990 | 3 | 0.6667 | NA | NA | 4 | 0.75 | 3 | 1 | 10 | 0.8 | NA | NA | 3 | 1 | 1 | 1 | 3 | 1 | 7 | 1 |
| 4_1388991 | 3 | 0.3333 | 2 | 0 | 4 | 0.5 | 3 | 0.3333 | 12 | 0.3333 | NA | NA | 5 | 0.2 | 1 | 1 | 3 | 0.3333 | 9 | 0.3333 |
| 4_1388999 | 2 | 1 | 1 | 1 | 2 | 1 | 2 | 1 | 7 | 1 | NA | NA | 3 | 1 | NA | NA | 2 | 1 | 5 | 1 |
| 4_1389003 | 1 | 0 | 1 | 0 | 3 | 0.3333 | 3 | 0.6667 | 8 | 0.375 | NA | NA | 5 | 0.2 | 1 | 0 | 3 | 1 | 9 | 0.4444 |
| 4_1389005 | 2 | 0 | 2 | 0 | 3 | 0 | 2 | 0.5 | 9 | 0.1111 | NA | NA | 3 | 0 | 1 | 0 | 3 | 0.3333 | 7 | 0.1429 |
| 4_1389008 | 3 | 0 | 2 | 0.5 | 4 | 0.75 | 3 | 0.6667 | 12 | 0.5 | NA | NA | 5 | 0.4 | 1 | 0 | 2 | 0.5 | 8 | 0.375 |
| 4_1389012 | 3 | 0.3333 | 2 | 1 | 3 | 1 | 3 | 1 | 11 | 0.8182 | NA | NA | 4 | 1 | 1 | 1 | 2 | 1 | 7 | 1 |
| 4_1389037 | 3 | 0 | 2 | 0 | 3 | 0.3333 | 2 | 0.5 | 10 | 0.2 | NA | NA | 5 | 0.6 | NA | NA | 2 | 0 | 7 | 0.4286 |
| 4_1389041 | 2 | 0 | 2 | 0 | 4 | 0.25 | 3 | 0.3333 | 11 | 0.1818 | NA | NA | 2 | 0 | 1 | 0 | 2 | 0.5 | 5 | 0.2 |
| 4_1389042 | 2 | 1 | 1 | 0 | 1 | 1 | 1 | 1 | 5 | 0.8 | NA | NA | 3 | 1 | NA | NA | NA | NA | 3 | 1 |
| 4_1389068 | 2 | 0.5 | 3 | 0.3333 | 4 | 1 | 4 | 1 | 13 | 0.7692 | NA | NA | 5 | 0.2 | 1 | 0 | 2 | 0.5 | 8 | 0.25 |
| 4_1389077 | 3 | 0 | 2 | 0 | 3 | 0 | 4 | 0.5 | 12 | 0.1667 | NA | NA | 5 | 0.2 | 1 | 0 | 3 | 0.3333 | 9 | 0.2222 |
| 4_1389078 | 4 | 0 | 3 | 0.3333 | 2 | 0 | 4 | 0 | 13 | 0.0769 | NA | NA | 5 | 0.4 | 1 | 0 | 3 | 0.6667 | 9 | 0.4444 |
| 4_1389168 | 3 | 0 | 3 | 0.3333 | 4 | 0.5 | 4 | 0 | 14 | 0.2143 | NA | NA | 7 | 0.1429 | 1 | 1 | 3 | 0 | 11 | 0.1818 |
| 4_1389169 | 3 | 0 | 3 | 0 | 4 | 0 | 4 | 0.25 | 14 | 0.0714 | NA | NA | 7 | 0.1429 | 1 | 0 | 3 | 0.3333 | 11 | 0.1818 |
| 4_1389331 | 9 | 0.1111 | 3 | 0 | 6 | 0.3333 | 5 | 0 | 23 | 0.1304 | NA | NA | 8 | 0.125 | 4 | 0 | 5 | 0.2 | 17 | 0.1176 |
| 4_1423127 | NA | NA | 1 | 0 | 1 | 1 | NA | NA | 2 | 0.5 | 3 | 0.3333 | 2 | 0 | 1 | 0 | 2 | 0 | 8 | 0.125 |
| 4_1480159 | NA | NA | NA | NA | 1 | 1 | 1 | 0 | 2 | 0.5 | NA | NA | NA | NA | 1 | 1 | NA | NA | 1 | 1 |

| | | | | | | | | | | | | | | | | | | | | |
|---|---|---|---|---|---|---|---|---|---|---|---|---|---|---|---|---|---|---|---|---|
| 4_14810193 | 11 | 0.6364 | 12 | 0.75 | 12 | 0.9167 | 13 | 0.9231 | 48 | 0.8125 | 6 | 0.8333 | 11 | 0.6364 | 6 | 0.8333 | 6 | 1 | 29 | 0.7931 |
| 4_14810212 | 11 | 0.1818 | 11 | 0.0909 | 12 | 0.1667 | 12 | 0.0833 | 46 | 0.1304 | 6 | 0.3333 | 11 | 0.0909 | 6 | 0.1667 | 4 | 0.5 | 27 | 0.2222 |
| 4_14811487 | 7 | 0 | 7 | 0.1429 | 7 | 0.2857 | 8 | 0.25 | 29 | 0.1724 | 6 | 0.3333 | 10 | 0.2 | 6 | 0.5 | 2 | 0.5 | 24 | 0.3333 |
| 4_14811588 | 5 | 0 | 6 | 0.1667 | 7 | 0.1429 | 5 | 0 | 23 | 0.087 | 5 | 0 | 8 | 0.125 | 3 | 0 | 1 | 0 | 17 | 0.0588 |
| 4_14811689 | 7 | 0.2857 | 5 | 0.2 | 8 | 0.125 | 9 | 0.1111 | 29 | 0.1724 | 6 | 0 | 9 | 0 | 5 | 0.4 | 3 | 0.6667 | 23 | 0.1739 |
| 4_14818292 | 4 | 0.5 | 8 | 0.125 | 8 | 0.125 | 7 | 0.2857 | 27 | 0.2222 | 5 | 0.6 | 7 | 0.2857 | 6 | 0.8333 | 3 | 0 | 21 | 0.4762 |
| 4_14818427 | 4 | 0 | 7 | 0.1429 | 8 | 0.125 | 7 | 0.2857 | 26 | 0.1538 | 5 | 0.2 | 7 | 0.1429 | 5 | 0.2 | 3 | 0 | 20 | 0.15 |
| 4_14914008 | 4 | 0 | 5 | 0.2 | 7 | 0.1429 | 6 | 0 | 22 | 0.0909 | 10 | 0.1 | 8 | 0 | 8 | 0.125 | 5 | 0 | 31 | 0.0645 |
| 4_14914064 | 5 | 0.2 | 8 | 0.25 | 9 | 0.2222 | 9 | 0.3333 | 31 | 0.2581 | 10 | 0.2 | 9 | 0.1111 | 9 | 0 | 4 | 0.25 | 32 | 0.125 |
| 4_14914570 | 5 | 0.2 | 9 | 0.1111 | 10 | 0.3 | 7 | 0.2857 | 31 | 0.2258 | 10 | 0.1 | 9 | 0 | 10 | 0 | 6 | 0.1667 | 35 | 0.0571 |
| 4_14914735 | 4 | 0 | 8 | 0 | 9 | 0 | 9 | 0.4444 | 30 | 0.1333 | 9 | 0 | 9 | 0.1111 | 10 | 0.1 | 6 | 0.3333 | 34 | 0.1176 |
| 4_14914825 | 4 | 1 | 8 | 0.875 | 10 | 0.9 | 8 | 0.875 | 30 | 0.9 | 10 | 0.8 | 9 | 1 | 11 | 0.5455 | 6 | 0.8333 | 36 | 0.7778 |
| 4_14914968 | 3 | 1 | 9 | 0.7778 | 6 | 1 | 10 | 0.9 | 28 | 0.8929 | 10 | 1 | 7 | 1 | 11 | 0.7273 | 5 | 1 | 33 | 0.9091 |
| 4_14915097 | 6 | 0 | 6 | 0 | 10 | 0.1 | 10 | 0.2 | 32 | 0.0938 | 9 | 0.3333 | 9 | 0.2222 | 11 | 0.3636 | 7 | 0.1429 | 36 | 0.2778 |
| 4_14952602 | 14 | 0.1429 | 10 | 0 | 16 | 0.0625 | 11 | 0 | 51 | 0.0588 | 9 | 0 | 10 | 0.1 | 20 | 0.1 | 12 | 0 | 51 | 0.0588 |
| 4_14952739 | 1 | 0 | NA | NA | NA | NA | 1 | 1 | 2 | 0.5 | NA | NA | 1 | 1 | NA | NA | NA | NA | 1 | 1 |
| 4_14953068 | 9 | 0 | 9 | 0.3333 | 15 | 0.2 | 7 | 0 | 40 | 0.15 | 7 | 0 | 6 | 0.1667 | 17 | 0.2353 | 11 | 0.1818 | 41 | 0.1707 |
| 4_14953348 | 7 | 0.1429 | 7 | 0 | 12 | 0.1667 | 7 | 0.1429 | 33 | 0.1212 | 5 | 0.2 | 4 | 0 | 16 | 0.125 | 8 | 0.375 | 33 | 0.1818 |
| 4_14953608 | 9 | 0.5556 | 7 | 0.5714 | 10 | 0.3 | 7 | 0.5714 | 33 | 0.4848 | 2 | 0.5 | 4 | 0.75 | 14 | 0.7143 | 10 | 0.6 | 30 | 0.6667 |
| 4_14953642 | 8 | 0.125 | 8 | 0 | 13 | 0 | 7 | 0.1429 | 36 | 0.0556 | 5 | 0 | 5 | 0 | 18 | 0.0556 | 11 | 0.0909 | 39 | 0.0513 |
| 4_14953704 | 9 | 0.6667 | 8 | 0.5 | 14 | 0.1429 | 7 | 0.5714 | 38 | 0.4211 | 5 | 0.6 | 6 | 0.5 | 19 | 0.7368 | 10 | 0.8 | 40 | 0.7 |
| 4_14954031 | 8 | 0 | 8 | 0.25 | 12 | 0.0833 | 8 | 0.125 | 36 | 0.1111 | 4 | 0 | 3 | 0 | 18 | 0.0556 | 8 | 0.125 | 33 | 0.0606 |
| 4_14954131 | 8 | 0.125 | 7 | 0.1429 | 10 | 0.1 | 6 | 0.1667 | 31 | 0.129 | 4 | 0.25 | 3 | 0.3333 | 15 | 0 | 5 | 0 | 27 | 0.0741 |
| 4_14954260 | 7 | 0.4286 | 7 | 0.2857 | 8 | 0.375 | 7 | 0 | 29 | 0.2759 | 3 | 0 | 3 | 0.3333 | 15 | 0.2667 | 5 | 0.6 | 26 | 0.3077 |
| 4_15266196 | 2 | 0 | 3 | 0.3333 | 1 | 1 | 1 | 0 | 7 | 0.2857 | NA | NA | 4 | 0.75 | 2 | 0.5 | 2 | 1 | 8 | 0.75 |
| 4_15266277 | 2 | 0 | 3 | 0 | 1 | 1 | 2 | 0 | 8 | 0.125 | 1 | 0 | 4 | 0.5 | 5 | 0.2 | NA | NA | 10 | 0.3 |
| 4_15281103 | 5 | 0 | 6 | 0 | 2 | 0.5 | 4 | 0.25 | 17 | 0.1176 | 1 | 0 | 5 | 0 | 6 | 0.3333 | 3 | 0 | 15 | 0.1333 |
| 4_15283245 | 5 | 0.8 | 5 | 0.6 | 2 | 1 | 4 | 0.75 | 16 | 0.75 | 2 | 1 | 6 | 0.8333 | 7 | 0.8571 | 3 | 0.6667 | 18 | 0.8333 |
| 4_15283360 | 5 | 0.4 | 4 | 0.75 | 3 | 1 | 3 | 1 | 15 | 0.7333 | 2 | 1 | 6 | 0.8333 | 6 | 0.1667 | 4 | 1 | 18 | 0.6667 |
| 4_15283423 | 6 | 0.3333 | 5 | 0.2 | 3 | 0 | 4 | 0 | 18 | 0.1667 | 2 | 0.5 | 5 | 0.8 | 5 | 0 | 3 | 0.6667 | 15 | 0.4667 |
| 4_15283454 | 5 | 0.2 | 5 | 0.4 | 3 | 0.6667 | 2 | 0 | 15 | 0.3333 | 2 | 0 | 4 | 0.5 | 5 | 0 | 4 | 0.5 | 15 | 0.2667 |
| 4_15283458 | 5 | 0.4 | 6 | 0.1667 | 3 | 0.3333 | 4 | 0.5 | 18 | 0.3333 | 2 | 0.5 | 5 | 0.2 | 6 | 0.3333 | 4 | 0 | 17 | 0.2353 |
| 4_15283613 | 7 | 0.4286 | 5 | 0.4 | 3 | 0.3333 | 6 | 0.3333 | 21 | 0.381 | 2 | 0.5 | 5 | 0.8 | 4 | 0.75 | 5 | 0.4 | 16 | 0.625 |
| 4_15283943 | 6 | 0 | 5 | 0.2 | 3 | 0 | 6 | 0 | 20 | 0.05 | 1 | 0 | 5 | 0 | 6 | 0.1667 | 4 | 0 | 16 | 0.0625 |
| 4_15905177 | 7 | 0.1429 | 18 | 0.1111 | 13 | 0.0769 | 18 | 0.1111 | 56 | 0.1071 | 8 | 0.25 | 13 | 0.1538 | 11 | 0.1818 | 17 | 0.1765 | 49 | 0.1837 |
| 4_15906132 | NA | NA | 5 | 0.4 | 2 | 0.5 | 3 | 0 | 10 | 0.3 | 1 | 0 | 4 | 0.25 | 1 | 0 | 1 | 0 | 7 | 0.1429 |
| 4_15917376 | 1 | 0 | 2 | 0.5 | 1 | 0 | 1 | 0 | 5 | 0.2 | 1 | 0 | 1 | 0 | 1 | 1 | 2 | 0.5 | 5 | 0.4 |
| 4_15917738 | 1 | 0 | 2 | 0 | 2 | 0.5 | 1 | 0 | 6 | 0.1667 | 1 | 0 | 1 | 0 | 2 | 0.5 | 3 | 0 | 7 | 0.1429 |
| 4_15918976 | 6 | 0.5 | 3 | 0 | 3 | 0.3333 | 4 | 0.25 | 16 | 0.3125 | 4 | 0.25 | 5 | 0.2 | 1 | 1 | 5 | 0.2 | 15 | 0.2667 |
| 4_15918995 | 6 | 0 | 3 | 0 | 4 | 0.25 | 4 | 0 | 17 | 0.0588 | 2 | 0 | 5 | 0.2 | 4 | 0 | 4 | 0 | 15 | 0.0667 |
| 4_15979183 | 1 | 0 | 1 | 1 | 1 | 0 | 1 | 0 | 4 | 0.25 | 1 | 0 | NA | NA | 3 | 0 | 3 | 0.3333 | 7 | 0.1429 |
| 4_15981486 | 1 | 1 | 2 | 0.5 | NA | NA | 1 | 1 | 4 | 0.75 | 2 | 0.5 | NA | NA | 3 | 0.3333 | 2 | 1 | 7 | 0.5714 |
| 4_15981493 | 1 | 0 | 1 | 0 | 1 | 0 | 1 | 1 | 4 | 0.25 | 1 | 1 | NA | NA | NA | NA | 3 | 0 | 4 | 0.25 |
| 4_15981536 | 1 | 0 | 1 | 1 | 1 | 1 | NA | NA | 3 | 0.6667 | 1 | 0 | NA | NA | 3 | 0.3333 | 3 | 0.6667 | 7 | 0.4286 |
| 4_15997945 | NA | NA | 5 | 0.8 | 5 | 0.6 | 3 | 0.6667 | 13 | 0.6923 | 3 | 1 | 2 | 1 | NA | NA | 4 | 1 | 9 | 1 |
| 4_15998025 | 1 | 1 | 3 | 0 | 4 | 0.5 | 2 | 0 | 10 | 0.3 | 3 | 0 | 2 | 0 | 1 | 0 | 4 | 0.25 | 10 | 0.1 |
| 4_15998194 | NA | NA | 2 | 0 | 3 | 0.3333 | 2 | 0 | 7 | 0.1429 | 2 | 0 | 2 | 0.5 | NA | NA | 4 | 0 | 8 | 0.125 |
| 4_15998217 | 1 | 0 | NA | NA | 1 | 0 | 2 | 0.5 | 4 | 0.25 | 2 | 0.5 | 1 | 0 | NA | NA | 2 | 0.5 | 5 | 0.4 |
| 4_15998386 | 1 | 0 | 3 | 0.3333 | 1 | 0 | 2 | 1 | 7 | 0.4286 | 2 | 0 | 2 | 0.5 | NA | NA | 3 | 1 | 7 | 0.5714 |
| 4_15998483 | NA | NA | NA | NA | 1 | 0 | 1 | 1 | 2 | 0.5 | 1 | 1 | NA | NA | NA | NA | 3 | 1 | 4 | 1 |
| 4_15998584 | 1 | 1 | 2 | 1 | 1 | 0 | 2 | 1 | 6 | 0.8333 | 2 | 1 | 2 | 1 | NA | NA | 2 | 1 | 6 | 1 |
| 4_15998699 | 1 | 1 | 2 | 1 | 1 | 1 | 2 | 1 | 6 | 1 | 2 | 1 | 2 | 1 | NA | NA | 1 | 1 | 5 | 1 |
| 4_15998788 | 1 | 1 | 1 | 0 | 1 | 1 | 1 | 1 | 4 | 0.75 | 1 | 0 | 1 | 1 | NA | NA | NA | NA | 2 | 0.5 |
| 4_15998845 | 1 | 0 | 3 | 0 | 1 | 0 | 2 | 0.5 | 7 | 0.1429 | 2 | 0 | 2 | 0.5 | NA | NA | 2 | 0.5 | 6 | 0.3333 |
| 4_15998869 | 1 | 1 | 3 | 0.6667 | 1 | 0 | 2 | 0.5 | 7 | 0.5714 | 2 | 0.5 | 2 | 0 | NA | NA | 3 | 0.6667 | 7 | 0.4286 |
| 4_15998880 | 1 | 1 | 3 | 0.3333 | 1 | 0 | 2 | 1 | 7 | 0.5714 | 1 | 1 | 2 | 1 | NA | NA | 3 | 1 | 6 | 1 |
| 4_16123759 | NA | NA | 1 | 1 | NA | NA | NA | NA | 1 | 1 | NA | NA | 1 | 1 | 1 | 0 | NA | NA | 2 | 0.5 |
| 4_18115980 | 1 | 1 | 1 | 1 | NA | NA | 1 | 1 | 3 | 1 | NA | NA | 2 | 1 | NA | NA | 1 | 1 | 3 | 1 |
| 4_18115999 | 1 | 0 | 1 | 1 | NA | NA | 1 | 1 | 3 | 0.6667 | NA | NA | 2 | 1 | NA | NA | 1 | 0 | 3 | 0.6667 |
| 4_18116023 | NA | NA | 1 | 0 | NA | NA | 1 | 1 | 2 | 0.5 | NA | NA | 2 | 1 | NA | NA | 1 | 1 | 3 | 1 |
| 4_18116046 | 1 | 1 | 1 | 0 | NA | NA | 1 | 1 | 3 | 0.6667 | NA | NA | 2 | 0.5 | NA | NA | NA | NA | 2 | 0.5 |
| 4_18116064 | 1 | 0 | 1 | 1 | NA | NA | 1 | 1 | 3 | 0.6667 | NA | NA | 2 | 1 | NA | NA | 1 | 0 | 3 | 0.6667 |
| 4_18116729 | NA | NA | NA | NA | NA | NA | 1 | 1 | 1 | 1 | NA | NA | 1 | 1 | NA | NA | 1 | 1 | 2 | 1 |
| 4_18116839 | 1 | 0 | 1 | 1 | NA | NA | 1 | 0 | 3 | 0.3333 | NA | NA | 2 | 0.5 | NA | NA | 2 | 0 | 4 | 0.25 |
| 4_18116857 | 1 | 0 | 1 | 1 | NA | NA | 1 | 1 | 3 | 0.6667 | NA | NA | 2 | 1 | NA | NA | 2 | 0.5 | 4 | 0.75 |

| | | | | | | | | | | | | | | | | | | | | |
|---|---|---|---|---|---|---|---|---|---|---|---|---|---|---|---|---|---|---|---|---|
| 4_1811686 | 1 | 1 | 1 | 1 | NA | NA | 1 | 1 | 3 | 1 | NA | NA | 2 | 1 | NA | NA | NA | NA | 2 | 1 |
| 4_1811701 | NA | NA | 1 | 1 | NA | NA | 1 | 0 | 2 | 0.5 | NA | NA | 2 | 1 | NA | NA | 2 | 0.5 | 4 | 0.75 |
| 4_1811724 | 1 | 0 | 1 | 1 | NA | NA | 1 | 0 | 3 | 0.3333 | 1 | 0 | 3 | 0.6667 | NA | NA | 2 | 0 | 6 | 0.3333 |
| 4_1960728 | NA | NA | NA | NA | 3 | 0.3333 | NA | NA | 3 | 0.3333 | NA | NA | 1 | 0 | 2 | 0 | 1 | 1 | 4 | 0.25 |
| 4_1960851 | NA | NA | 7 | 0.1429 | 5 | 0 | 1 | 0 | 13 | 0.0769 | 2 | 0.5 | 4 | 0 | 3 | 0 | 3 | 0 | 12 | 0.0833 |
| 4_1960999 | NA | NA | 5 | 0 | 6 | 0 | 1 | 1 | 12 | 0.0833 | 1 | 0 | 2 | 0 | 4 | 0.25 | 4 | 0 | 11 | 0.0909 |
| 4_1961152 | 2 | 0.5 | 5 | 0.6 | 3 | 0.6667 | 3 | 0.3333 | 13 | 0.5385 | 1 | 1 | NA | NA | 4 | 0 | 4 | 0.75 | 9 | 0.4444 |
| 4_1961161 | 2 | 0 | 8 | 0 | 3 | 0 | 4 | 0.25 | 17 | 0.0588 | NA | NA | NA | NA | 4 | 0 | 3 | 0.3333 | 7 | 0.1429 |
| 4_1961171 | 2 | 0.5 | 7 | 0.7143 | 2 | 1 | 5 | 0.2 | 16 | 0.5625 | 1 | 0 | NA | NA | 4 | 0.5 | 3 | 0 | 8 | 0.25 |
| 4_1961172 | 2 | 0 | 6 | 0.1667 | 2 | 0 | 3 | 0 | 13 | 0.0769 | NA | NA | NA | NA | 4 | 0 | 3 | 0.3333 | 7 | 0.1429 |
| 4_1961177 | 1 | 0 | 7 | 0 | 2 | 0.5 | 4 | 0 | 14 | 0.0714 | 1 | 0 | NA | NA | 4 | 0.25 | 4 | 0.25 | 9 | 0.2222 |
| 4_2002878 | NA | NA | 1 | 1 | 1 | 0 | 2 | 0 | 4 | 0.25 | 1 | 0 | NA | NA | 1 | 1 | 1 | 1 | 3 | 0.6667 |
| 4_2011126 | 2 | 0 | 3 | 0 | 1 | 0 | 4 | 0.25 | 10 | 0.1 | 1 | 0 | 1 | 0 | NA | NA | 3 | 0.3333 | 5 | 0.2 |
| 4_2174789 | 3 | 0 | 2 | 0.5 | 1 | 0 | 3 | 0 | 9 | 0.1111 | 2 | 0.5 | 1 | 0 | NA | NA | 3 | 0 | 6 | 0.1667 |
| 4_2174816 | 1 | 0 | 3 | 0.3333 | 2 | 0 | NA | NA | 6 | 0.1667 | NA | NA | 1 | 1 | NA | NA | 1 | 0 | 2 | 0.5 |
| 4_2174824 | NA | NA | 3 | 0.3333 | 2 | 1 | NA | NA | 5 | 0.6 | NA | NA | NA | NA | NA | NA | 1 | 1 | 1 | 1 |
| 4_2174831 | 1 | 0 | 2 | 0 | 2 | 0.5 | NA | NA | 5 | 0.2 | NA | NA | 1 | 0 | NA | NA | 1 | 1 | 2 | 0.5 |
| 4_2174953 | 1 | 0 | 1 | 1 | 1 | 1 | NA | NA | 3 | 0.6667 | 1 | 0 | NA | NA | 1 | 0 | 1 | 1 | 3 | 0.3333 |
| 4_2175757 | 9 | 0.2222 | 2 | 0 | 1 | 0 | 3 | 0.6667 | 15 | 0.2667 | 1 | 0 | 2 | 0.5 | 4 | 0.75 | 1 | 1 | 8 | 0.625 |
| 4_2175766 | 9 | 0.1111 | 2 | 0.5 | 1 | 1 | 2 | 0.5 | 14 | 0.2857 | 1 | 1 | 1 | 0 | 4 | 0.5 | 1 | 0 | 7 | 0.4286 |
| 4_2175772 | 8 | 0.875 | 2 | 1 | 1 | 1 | 2 | 1 | 13 | 0.9231 | 1 | 1 | 2 | 1 | 3 | 1 | 1 | 1 | 7 | 1 |
| 4_2175774 | 9 | 0.4444 | 2 | 0 | 1 | 0 | 2 | 0 | 14 | 0.2857 | 1 | 0 | 2 | 1 | 4 | 0.5 | 1 | 1 | 8 | 0.625 |
| 4_2175775 | 9 | 0.2222 | 2 | 0 | 1 | 0 | 2 | 1 | 14 | 0.2857 | NA | NA | 2 | 0 | 3 | 0.3333 | 1 | 0 | 6 | 0.1667 |
| 4_2175827 | 4 | 0.5 | 2 | 0 | 1 | 1 | 1 | 0 | 8 | 0.375 | 2 | 1 | 2 | 0.5 | 1 | 0 | NA | NA | 5 | 0.6 |
| 4_2175830 | 4 | 0.75 | 3 | 0 | 2 | 0 | 1 | 0 | 10 | 0.3 | 1 | 0 | 3 | 0 | 3 | 0.3333 | NA | NA | 7 | 0.1429 |
| 4_2175838 | 6 | 0.3333 | 3 | 0.6667 | 2 | 0 | 1 | 0 | 12 | 0.3333 | 1 | 0 | 2 | 0.5 | 3 | 0.6667 | NA | NA | 6 | 0.5 |
| 4_2175840 | 7 | 0 | 3 | 0.3333 | 2 | 0 | 1 | 1 | 13 | 0.1538 | 1 | 1 | 3 | 0.3333 | 3 | 0.3333 | NA | NA | 7 | 0.4286 |
| 4_2182480 | 3 | 0.6667 | 3 | 0.3333 | 1 | 1 | 1 | 0 | 8 | 0.5 | 2 | 0.5 | NA | NA | 2 | 0.5 | 2 | 0 | 6 | 0.3333 |
| 4_2182491 | 3 | 0 | 2 | 0.5 | 2 | 0 | 1 | 0 | 8 | 0.125 | 4 | 0 | NA | NA | 2 | 0 | 2 | 0.5 | 8 | 0.125 |
| 4_2182503 | 2 | 0 | 2 | 0 | 2 | 1 | 1 | 0 | 7 | 0.2857 | 2 | 0.5 | NA | NA | 2 | 1 | 1 | 1 | 5 | 0.8 |
| 4_2183586 | 4 | 0 | 4 | 0.25 | 3 | 0 | 4 | 0.5 | 15 | 0.2 | 3 | 0 | NA | NA | 2 | 0 | 8 | 0.125 | 13 | 0.0769 |
| 4_2183747 | 6 | 0.5 | 5 | 0.6 | 4 | 0.25 | 5 | 0.4 | 20 | 0.45 | 4 | 0.75 | 4 | 0 | 3 | 0.3333 | 7 | 0.1429 | 18 | 0.2778 |
| 4_2185915 | 2 | 1 | 4 | 0.25 | 4 | 0.25 | 4 | 0.25 | 14 | 0.3571 | NA | NA | NA | NA | 2 | 0 | 3 | 0.3333 | 5 | 0.2 |
| 4_2187371 | 3 | 0.3333 | 5 | 0.6 | 5 | 0.2 | 4 | 0 | 17 | 0.2941 | 2 | 0.5 | 1 | 1 | 2 | 0 | 7 | 0.1429 | 12 | 0.25 |
| 4_2187431 | 5 | 0 | 12 | 0.0833 | 7 | 0.1429 | 7 | 0 | 31 | 0.0645 | 2 | 0.5 | 5 | 0.2 | 5 | 0.2 | 7 | 0.1429 | 19 | 0.2105 |
| 4_2187441 | 5 | 0 | 12 | 0 | 6 | 0.1667 | 8 | 0.125 | 31 | 0.0645 | 2 | 0 | 5 | 0 | 5 | 0.2 | 6 | 0.1667 | 18 | 0.1111 |
| 4_2187515 | 7 | 0.1429 | 13 | 0 | 8 | 0.375 | 10 | 0.1 | 38 | 0.1316 | 2 | 0 | 8 | 0.25 | 13 | 0.0769 | 11 | 0 | 34 | 0.0882 |
| 4_2187539 | 7 | 0.8571 | 7 | 0.7143 | 6 | 0.8333 | 7 | 0.8571 | 27 | 0.8148 | 2 | 1 | 7 | 0.8571 | 8 | 0.5 | 8 | 0.625 | 25 | 0.68 |
| 4_2187563 | 8 | 0.25 | 9 | 0 | 7 | 0.1429 | 7 | 0.1429 | 31 | 0.129 | 2 | 0.5 | 9 | 0.1111 | 8 | 0 | 8 | 0 | 27 | 0.0741 |
| 4_2191043 | 3 | 0 | NA | NA | 4 | 0.25 | 2 | 0 | 9 | 0.1111 | 2 | 0.5 | 2 | 0.5 | 1 | 0 | NA | NA | 5 | 0.4 |
| 4_2191063 | 3 | 0 | 1 | 0 | 4 | 0 | 2 | 0.5 | 10 | 0.1 | 2 | 0.5 | 2 | 0 | 1 | 0 | NA | NA | 5 | 0.2 |
| 4_2191065 | 4 | 0.5 | 1 | 1 | 4 | 0.5 | 2 | 0.5 | 11 | 0.5455 | 2 | 1 | 2 | 1 | 1 | 0 | NA | NA | 5 | 0.8 |
| 4_2191078 | 1 | 1 | 1 | 1 | 2 | 0 | 1 | 0 | 5 | 0.4 | 3 | 0.6667 | 2 | 0 | NA | NA | NA | NA | 5 | 0.4 |
| 4_2191211 | 1 | 1 | 2 | 0 | 2 | 0 | NA | NA | 5 | 0.2 | 4 | 0 | NA | NA | NA | NA | 1 | 1 | 5 | 0.2 |
| 4_2193161 | NA | NA | 2 | 0.5 | NA | NA | NA | NA | 2 | 0.5 | 1 | 1 | NA | NA | 3 | 0 | 2 | 1 | 6 | 0.5 |
| 4_2199341 | NA | NA | 3 | 0.3333 | NA | NA | 1 | 0 | 4 | 0.25 | 1 | 0 | NA | NA | 2 | 0 | 3 | 0.3333 | 6 | 0.1667 |
| 4_2199345 | NA | NA | 3 | 0.3333 | NA | NA | 1 | 0 | 4 | 0.25 | 1 | 1 | NA | NA | 2 | 1 | 2 | 1 | 5 | 1 |
| 4_2199346 | NA | NA | 2 | 1 | NA | NA | 1 | 1 | 3 | 1 | 1 | 1 | NA | NA | 2 | 1 | 3 | 0.6667 | 6 | 0.8333 |
| 4_2199348 | NA | NA | 2 | 0.5 | NA | NA | 1 | 1 | 3 | 0.6667 | NA | NA | NA | NA | 1 | 0 | 2 | 1 | 3 | 0.6667 |
| 4_2199352 | NA | NA | 3 | 1 | NA | NA | NA | NA | 3 | 1 | 1 | 1 | NA | NA | 2 | 1 | 2 | 1 | 5 | 1 |
| 4_2199357 | NA | NA | 2 | 1 | NA | NA | 1 | 0 | 3 | 0.6667 | NA | NA | NA | NA | NA | NA | 1 | 1 | 1 | 1 |
| 4_2199369 | NA | NA | 3 | 0.6667 | NA | NA | 1 | 1 | 4 | 0.75 | 1 | 1 | NA | NA | 1 | 1 | 2 | 0.5 | 4 | 0.75 |
| 4_2199371 | NA | NA | 3 | 1 | NA | NA | 1 | 1 | 4 | 1 | 1 | 1 | NA | NA | 3 | 0.6667 | 2 | 1 | 6 | 0.8333 |
| 4_2199376 | NA | NA | 3 | 0.3333 | NA | NA | 1 | 0 | 4 | 0.25 | 1 | 1 | NA | NA | 2 | 0 | 1 | 0 | 4 | 0.25 |
| 4_2199377 | NA | NA | 1 | 0 | NA | NA | 1 | 1 | 2 | 0.5 | 1 | 1 | NA | NA | 3 | 0 | 2 | 0.5 | 6 | 0.3333 |
| 4_2199380 | NA | NA | 3 | 0 | NA | NA | 1 | 1 | 4 | 0.25 | 1 | 0 | NA | NA | 3 | 0.3333 | 2 | 0 | 6 | 0.1667 |
| 4_2199439 | NA | NA | 3 | 1 | 1 | 1 | 3 | 1 | 7 | 1 | 1 | 1 | 1 | 1 | 3 | 0.6667 | 4 | 0.75 | 9 | 0.7778 |
| 4_2199447 | NA | NA | 3 | 0.3333 | 1 | 0 | 2 | 0.5 | 6 | 0.3333 | 1 | 1 | 2 | 1 | 4 | 0.25 | 4 | 0.25 | 11 | 0.4545 |
| 4_2199457 | 1 | 0 | 3 | 0.3333 | NA | NA | 3 | 0 | 7 | 0.1429 | NA | NA | 2 | 0 | 2 | 0 | 4 | 0.25 | 8 | 0.125 |
| 4_2199796 | 2 | 0 | 7 | 0.1429 | 4 | 0 | 3 | 0 | 16 | 0.0625 | 3 | 0.3333 | 3 | 0 | 11 | 0.2727 | 10 | 0.1 | 27 | 0.1852 |
| 4_2199820 | 3 | 0.3333 | 7 | 0.4286 | 3 | 0 | 3 | 0.3333 | 16 | 0.3125 | 2 | 0.5 | 5 | 0.2 | 9 | 0.3333 | 9 | 0.4444 | 25 | 0.36 |
| 4_2199895 | 3 | 0 | 3 | 0.3333 | 2 | 0 | 1 | 0 | 9 | 0.1111 | 2 | 0 | 5 | 0.2 | 6 | 0 | 7 | 0 | 20 | 0.05 |
| 4_2200099 | 4 | 0.25 | 4 | 0 | 3 | 0 | 1 | 1 | 12 | 0.1667 | 6 | 0.1667 | 4 | 0.5 | 5 | 0.2 | 12 | 0.25 | 27 | 0.2593 |
| 4_2200116 | 6 | 0.1667 | 6 | 0.3333 | 4 | 0.25 | 3 | 0 | 19 | 0.2105 | 6 | 0 | 7 | 0.1429 | 8 | 0.375 | 11 | 0.0909 | 32 | 0.1562 |
| 4_2200117 | 5 | 0.4 | 6 | 0.6667 | 4 | 0.25 | 3 | 1 | 18 | 0.5556 | 6 | 0.1667 | 7 | 0.8571 | 8 | 0.625 | 12 | 0.5 | 33 | 0.5455 |

| | | | | | | | | | | | | | | | | | | | | |
|---|---|---|---|---|---|---|---|---|---|---|---|---|---|---|---|---|---|---|---|---|
| 4_2200120 | 5 | 0 | 5 | 0.2 | 4 | 0 | 2 | 0 | 16 | 0.0625 | 5 | 0.2 | 7 | 0.1429 | 6 | 0 | 11 | 0.1818 | 29 | 0.1379 |
| 4_2237663 | 1 | 0 | 1 | 0 | 1 | 0 | 2 | 0.5 | 5 | 0.2 | 4 | 0.5 | 2 | 0 | 1 | 1 | 1 | 0 | 8 | 0.375 |
| 4_2237667 | 1 | 1 | 1 | 1 | 2 | 1 | 2 | 0.5 | 6 | 0.8333 | 4 | 0.5 | 2 | 1 | 2 | 1 | 1 | 1 | 9 | 0.7778 |
| 4_2237669 | 1 | 1 | 1 | 1 | NA | NA | 1 | 1 | 3 | 1 | 2 | 0.5 | 1 | 1 | 1 | 1 | 1 | 1 | 5 | 0.8 |
| 4_2237677 | 1 | 0 | NA | NA | 1 | 1 | 2 | 0.5 | 4 | 0.5 | 3 | 0.6667 | 1 | 1 | 2 | 0.5 | 1 | 1 | 7 | 0.7143 |
| 4_2237686 | 1 | 0 | NA | NA | NA | NA | 2 | 1 | 3 | 0.6667 | 4 | 1 | 1 | 0 | 2 | 1 | 1 | 1 | 8 | 0.875 |
| 4_2237693 | 1 | 1 | NA | NA | 1 | 1 | 2 | 0.5 | 4 | 0.75 | 4 | 0.25 | 2 | 1 | 2 | 0.5 | 1 | 1 | 9 | 0.5556 |
| 4_2237694 | 1 | 1 | 1 | 1 | 1 | 0 | 1 | 0 | 4 | 0.5 | 4 | 0.75 | 1 | 1 | 1 | 1 | NA | NA | 6 | 0.8333 |
| 4_2237695 | NA | NA | 1 | 0 | 1 | 1 | 1 | 1 | 3 | 0.6667 | 4 | 0.5 | 1 | 0 | 1 | 0 | NA | NA | 6 | 0.3333 |
| 4_2238605 | 1 | 0 | 1 | 0 | 1 | 0 | 1 | 1 | 4 | 0.25 | 3 | 0.3333 | 1 | 0 | NA | NA | 2 | 0.5 | 6 | 0.3333 |
| 4_2238608 | 1 | 0 | NA | NA | 2 | 0.5 | 2 | 1 | 5 | 0.6 | 3 | 0 | 1 | 0 | 2 | 0.5 | 2 | 0 | 8 | 0.125 |
| 4_2238610 | 1 | 0 | 1 | 1 | 1 | 0 | 2 | 0.5 | 5 | 0.4 | 4 | 0.5 | 2 | 0.5 | 2 | 0.5 | 2 | 0 | 10 | 0.4 |
| 4_2242720 | 1 | 1 | 1 | 1 | 3 | 0.3333 | 3 | 0.3333 | 8 | 0.5 | 4 | 0.5 | 2 | 0 | 1 | 1 | 3 | 0.6667 | 10 | 0.5 |
| 4_2242728 | 1 | 0 | 2 | 0 | 2 | 0 | 3 | 0.3333 | 8 | 0.125 | 4 | 0.5 | 1 | 0 | 2 | 0.5 | 3 | 0.6667 | 10 | 0.5 |
| 4_2242734 | 1 | 0 | 2 | 0 | 2 | 0 | 3 | 0.6667 | 8 | 0.25 | 3 | 0 | 2 | 0 | 2 | 0 | 3 | 0.6667 | 10 | 0.2 |
| 4_2242737 | 1 | 0 | 1 | 1 | 3 | 1 | 3 | 0.3333 | 8 | 0.625 | 4 | 0.5 | 2 | 0 | 1 | 0 | 3 | 0.3333 | 10 | 0.3 |
| 4_2242741 | 1 | 0 | 2 | 0 | 3 | 0 | 4 | 0.25 | 10 | 0.1 | 4 | 0.25 | 2 | 0 | 2 | 0 | 3 | 0 | 11 | 0.0909 |
| 4_2243382 | 1 | 0 | 3 | 0.3333 | 3 | 0 | 4 | 0.25 | 11 | 0.1818 | 5 | 0.4 | 2 | 0 | 2 | 0 | 4 | 0 | 13 | 0.1538 |
| 4_2248458 | 1 | 0 | 1 | 1 | NA | NA | 1 | 1 | 3 | 0.6667 | 1 | 0 | 1 | 1 | 1 | 1 | 3 | 1 | 6 | 0.8333 |
| 4_2248471 | 1 | 0 | 1 | 0 | NA | NA | 1 | 1 | 3 | 0.3333 | 1 | 0 | 1 | 0 | NA | NA | 2 | 0.5 | 4 | 0.25 |
| 4_2248660 | 1 | 1 | NA | NA | NA | NA | NA | NA | 1 | 1 | NA | NA | 1 | 1 | 1 | 1 | 1 | 1 | 3 | 1 |
| 4_2248679 | NA | NA | 1 | 1 | NA | NA | NA | NA | 1 | 1 | NA | NA | NA | NA | NA | NA | 1 | 1 | 1 | 1 |
| 4_2248687 | 1 | 1 | NA | NA | NA | NA | NA | NA | 1 | 1 | NA | NA | 1 | 1 | NA | NA | 1 | 1 | 2 | 1 |
| 4_2248690 | 1 | 1 | NA | NA | NA | NA | NA | NA | 1 | 1 | NA | NA | NA | NA | NA | NA | 2 | 0.5 | 2 | 0.5 |
| 4_2248696 | 1 | 1 | NA | NA | NA | NA | NA | NA | 1 | 1 | NA | NA | 1 | 1 | NA | NA | 2 | 1 | 3 | 1 |
| 4_2248705 | 1 | 1 | NA | NA | NA | NA | NA | NA | 1 | 1 | NA | NA | 1 | 1 | NA | NA | 2 | 1 | 3 | 1 |
| 4_2249296 | NA | NA | 1 | 1 | NA | NA | NA | NA | 1 | 1 | 1 | 0 | NA | NA | 1 | 1 | NA | NA | 2 | 0.5 |
| 4_2490177 | 2 | 1 | 6 | 0.6667 | 5 | 1 | 10 | 0.5 | 23 | 0.6957 | 2 | 1 | 4 | 1 | 5 | 1 | 6 | 0.8333 | 17 | 0.9412 |
| 4_2490180 | 2 | 0 | 6 | 0 | 5 | 0.2 | 6 | 0 | 19 | 0.0526 | 2 | 0.5 | 5 | 0.2 | 5 | 0 | 6 | 0 | 18 | 0.1111 |
| 4_2490182 | 2 | 1 | 6 | 0.6667 | 5 | 0.6 | 7 | 0.7143 | 20 | 0.7 | 1 | 0 | 1 | 1 | 2 | 0.5 | 5 | 0.8 | 9 | 0.6667 |
| 4_2490184 | 2 | 1 | 6 | 0.8333 | 5 | 1 | 6 | 1 | 19 | 0.9474 | 3 | 1 | 5 | 1 | 4 | 1 | 6 | 1 | 18 | 1 |
| 4_2490188 | 2 | 1 | 7 | 1 | 5 | 0.8 | 6 | 0.6667 | 20 | 0.85 | 3 | 1 | 4 | 0.75 | 5 | 0.6 | 5 | 1 | 17 | 0.8235 |
| 4_2490308 | 1 | 0 | 6 | 0.5 | 3 | 0 | 5 | 0.2 | 15 | 0.2667 | 2 | 0.5 | 4 | 0 | 4 | 0.25 | 8 | 0.25 | 18 | 0.2222 |
| 4_2490327 | 2 | 0 | 5 | 0 | 7 | 0.2857 | 8 | 0 | 22 | 0.0909 | 1 | 0 | 6 | 0.3333 | 4 | 0.25 | 9 | 0.1111 | 20 | 0.2 |
| 4_2490341 | 2 | 0 | 6 | 0 | 8 | 0.125 | 9 | 0.1111 | 25 | 0.08 | 3 | 0 | 5 | 0.6 | 4 | 0 | 7 | 0.2857 | 19 | 0.2632 |
| 4_2496258 | NA | NA | NA | NA | 1 | 1 | NA | NA | 1 | 1 | 1 | 1 | NA | NA | NA | NA | NA | NA | 1 | 1 |
| 4_2496551 | NA | NA | NA | NA | 1 | 1 | NA | NA | 1 | 1 | 1 | 1 | NA | NA | NA | NA | NA | NA | 1 | 1 |
| 4_2496575 | NA | NA | NA | NA | 1 | 1 | NA | NA | 1 | 1 | 1 | 1 | NA | NA | NA | NA | NA | NA | 1 | 1 |
| 4_2500818 | NA | NA | NA | NA | 1 | 1 | 3 | 0.3333 | 4 | 0.5 | NA | NA | NA | NA | NA | NA | 1 | 1 | 1 | 1 |
| 4_2500856 | NA | NA | NA | NA | NA | NA | 2 | 1 | 2 | 1 | NA | NA | NA | NA | NA | NA | 1 | 1 | 1 | 1 |
| 4_2500860 | NA | NA | NA | NA | 1 | 1 | 4 | 1 | 5 | 1 | NA | NA | NA | NA | NA | NA | 1 | 1 | 1 | 1 |
| 4_2501233 | NA | NA | 1 | 0 | 3 | 0.3333 | 6 | 0 | 10 | 0.1 | NA | NA | NA | NA | 2 | 0 | 3 | 0.3333 | 5 | 0.2 |
| 4_2501243 | NA | NA | 2 | 1 | 4 | 0.75 | 6 | 0.5 | 12 | 0.6667 | NA | NA | 1 | 1 | NA | NA | 4 | 0.5 | 5 | 0.6 |
| 4_2525032 | 2 | 0 | 4 | 0.25 | 2 | 0 | 2 | 0 | 10 | 0.1 | 1 | 0 | 3 | 0.3333 | 2 | 0.5 | 3 | 0 | 9 | 0.2222 |
| 4_2525045 | 2 | 0.5 | 5 | 0.4 | 2 | 0.5 | 2 | 1 | 11 | 0.5455 | 2 | 0.5 | 5 | 0.4 | 1 | 0 | 9 | 0.2222 | 17 | 0.2941 |
| 4_2525051 | 1 | 1 | 5 | 0 | 2 | 1 | 2 | 0 | 10 | 0.3 | 3 | 0.6667 | 6 | 0.6667 | 3 | 0.6667 | 7 | 0.5714 | 19 | 0.6316 |
| 4_2525053 | 1 | 0 | 5 | 0.6 | 2 | 0.5 | 3 | 0.3333 | 11 | 0.4545 | 3 | 0 | 6 | 0.6667 | 3 | 0.3333 | 9 | 0.4444 | 21 | 0.4286 |
| 4_2525061 | 2 | 0.5 | 3 | 0 | 2 | 0.5 | 3 | 0 | 10 | 0.2 | 3 | 0 | 6 | 0.1667 | 3 | 0 | 8 | 0 | 20 | 0.05 |
| 4_2525064 | 2 | 0.5 | 4 | 0 | 2 | 0.5 | 2 | 0 | 10 | 0.2 | 3 | 0 | 5 | 0.4 | 2 | 0 | 7 | 0 | 17 | 0.1176 |
| 4_2560974 | 11 | 0.0909 | 10 | 0.1 | 6 | 0.3333 | 5 | 0 | 32 | 0.125 | 4 | 0 | 10 | 0.1 | 11 | 0.0909 | 11 | 0.0909 | 36 | 0.0833 |
| 4_2560990 | 10 | 0.1 | 8 | 0 | 5 | 0.2 | 5 | 0 | 28 | 0.0714 | 4 | 0 | 9 | 0.3333 | 11 | 0 | 11 | 0.2727 | 35 | 0.1714 |
| 4_2561059 | 7 | 0.2857 | 6 | 0.6667 | 4 | 0.5 | 5 | 0.4 | 22 | 0.4545 | 2 | 0.5 | 4 | 0 | 8 | 0.5 | 9 | 0.3333 | 23 | 0.3478 |
| 4_2561060 | 9 | 0.4444 | 6 | 0.1667 | 5 | 0.6 | 4 | 0.25 | 24 | 0.375 | 3 | 0.6667 | 5 | 0 | 7 | 0.1429 | 7 | 0 | 22 | 0.1364 |
| 4_2561061 | 9 | 0.4444 | 6 | 0.1667 | 6 | 0.6667 | 4 | 0 | 25 | 0.36 | 3 | 0.3333 | 6 | 0.3333 | 9 | 0.4444 | 8 | 0.25 | 26 | 0.3462 |
| 4_2561062 | 7 | 0.2857 | 6 | 0 | 5 | 0.2 | 5 | 0.2 | 23 | 0.1739 | 4 | 0.25 | 6 | 0 | 10 | 0 | 9 | 0.2222 | 29 | 0.1034 |
| 4_2561064 | 9 | 0.1111 | 6 | 0.1667 | 6 | 0 | 5 | 0 | 26 | 0.0769 | 3 | 0 | 6 | 0 | 10 | 0.1 | 9 | 0.2222 | 28 | 0.1071 |
| 4_2561073 | 4 | 0.25 | 4 | 0.25 | 4 | 0.5 | 4 | 0.5 | 16 | 0.375 | 1 | 0 | 3 | 0 | 6 | 0 | 6 | 0.3333 | 16 | 0.125 |
| 4_2561185 | 8 | 0.125 | 2 | 0 | 4 | 0.25 | 4 | 0 | 18 | 0.1111 | 2 | 0.5 | 4 | 0 | 3 | 0 | 6 | 0 | 15 | 0.0667 |
| 4_2561187 | 6 | 0.3333 | 3 | 0 | 5 | 0.4 | 4 | 0.5 | 18 | 0.3333 | 2 | 1 | 1 | 1 | 3 | 0.3333 | 4 | 0 | 10 | 0.4 |
| 4_2561998 | 1 | 1 | NA | NA | NA | NA | 1 | 0 | 2 | 0.5 | NA | NA | 1 | 1 | NA | NA | NA | NA | 1 | 1 |
| 4_2562035 | 1 | 1 | NA | NA | NA | NA | 1 | 1 | 2 | 1 | NA | NA | 1 | 1 | NA | NA | NA | NA | 1 | 1 |
| 4_2562036 | 2 | 1 | NA | NA | NA | NA | 1 | 1 | 3 | 1 | NA | NA | 1 | 1 | NA | NA | NA | NA | 1 | 1 |
| 4_2562047 | 2 | 1 | NA | NA | NA | NA | 1 | 0 | 3 | 0.6667 | NA | NA | 1 | 1 | NA | NA | NA | NA | 1 | 1 |
| 4_2562058 | 2 | 1 | NA | NA | NA | NA | 1 | 0 | 3 | 0.6667 | NA | NA | 1 | 1 | NA | NA | NA | NA | 1 | 1 |
| 4_2562064 | NA | NA | NA | NA | NA | NA | 1 | 1 | 1 | 1 | NA | NA | 1 | 1 | NA | NA | NA | NA | 1 | 1 |

| | | | | | | | | | | | | | | | | | | | | |
|---|---|---|---|---|---|---|---|---|---|---|---|---|---|---|---|---|---|---|---|---|
| 4_26324632 | 5 | 0.2 | 6 | 0.1667 | 4 | 0 | 10 | 0 | 25 | 0.08 | 7 | 0.1429 | 3 | 0.3333 | 10 | 0.1 | 4 | 0 | 24 | 0.125 |
| 4_26325549 | 3 | 0 | 4 | 0.25 | NA | NA | 9 | 0.1111 | 16 | 0.125 | 4 | 0.25 | 1 | 0 | 6 | 0 | 5 | 0 | 16 | 0.0625 |
| 4_26327581 | 2 | 0.5 | 4 | 0.25 | NA | NA | 6 | 0.5 | 12 | 0.4167 | 3 | 0 | 2 | 0.5 | 5 | 0.6 | 4 | 0 | 14 | 0.2857 |
| 4_26327725 | 3 | 0.3333 | 1 | 0 | NA | NA | 4 | 0.5 | 8 | 0.375 | 2 | 0.5 | 2 | 1 | 5 | 0.4 | 2 | 0.5 | 11 | 0.5455 |
| 4_26327729 | 1 | 0 | 2 | 0.5 | NA | NA | 2 | 1 | 5 | 0.6 | 2 | 0.5 | 1 | 1 | 4 | 1 | 2 | 1 | 9 | 0.8889 |
| 4_26327785 | 3 | 1 | 4 | 1 | NA | NA | 6 | 1 | 13 | 1 | 2 | 1 | 2 | 0.5 | 5 | 1 | 4 | 1 | 13 | 0.9231 |
| 4_26327815 | 2 | 1 | 3 | 1 | NA | NA | 5 | 1 | 10 | 1 | 3 | 1 | 2 | 1 | 5 | 1 | 3 | 1 | 13 | 1 |
| 4_26327916 | 3 | 0 | 2 | 0 | NA | NA | 4 | 0.25 | 9 | 0.1111 | 1 | 1 | 2 | 0 | 5 | 0.4 | 3 | 0.3333 | 11 | 0.3636 |
| 4_26327925 | 3 | 0 | 4 | 0.25 | NA | NA | 3 | 0 | 10 | 0.1 | 3 | 0 | 1 | 1 | 5 | 0 | 3 | 0 | 12 | 0.0833 |
| 4_26327934 | 3 | 0.3333 | 2 | 0.5 | NA | NA | 1 | 1 | 6 | 0.5 | 2 | 0.5 | 2 | 0.5 | 3 | 1 | 3 | 0.6667 | 10 | 0.7 |
| 4_26327997 | 3 | 1 | 3 | 0.6667 | NA | NA | 4 | 0.75 | 10 | 0.8 | 3 | 0.3333 | 1 | 1 | 4 | 0.75 | 2 | 0.5 | 10 | 0.6 |
| 4_26328268 | 2 | 0.5 | 1 | 1 | NA | NA | 4 | 0.5 | 7 | 0.5714 | 1 | 1 | 1 | 1 | 3 | 0 | 2 | 1 | 7 | 0.5714 |
| 4_26328274 | 2 | 0 | 2 | 0 | NA | NA | 4 | 0.25 | 8 | 0.125 | 3 | 0 | 2 | 0 | 5 | 0.2 | 2 | 0 | 12 | 0.0833 |
| 4_26340535 | 3 | 0.6667 | 2 | 0 | NA | NA | 4 | 0.5 | 9 | 0.4444 | 3 | 0.3333 | 2 | 0.5 | 3 | 0.3333 | 1 | 1 | 9 | 0.4444 |
| 4_26340566 | 3 | 0.6667 | 3 | 0.6667 | NA | NA | 5 | 0.2 | 11 | 0.4545 | 3 | 0.3333 | 2 | 0 | 4 | 0 | 1 | 0 | 10 | 0.1 |
| 4_26340676 | 3 | 0 | 2 | 1 | NA | NA | 5 | 0.6 | 10 | 0.5 | 3 | 0.6667 | 1 | 0 | 3 | 0.6667 | 1 | 1 | 8 | 0.625 |
| 4_26340717 | 3 | 1 | 2 | 0 | NA | NA | 4 | 0.5 | 9 | 0.5556 | 2 | 1 | NA | NA | 3 | 1 | 1 | 0 | 6 | 0.8333 |
| 4_26340820 | 2 | 1 | 2 | 0 | NA | NA | 4 | 0 | 8 | 0.25 | 2 | 0.5 | 1 | 0 | 3 | 0.3333 | 1 | 0 | 7 | 0.2857 |
| 4_26340838 | 3 | 1 | 2 | 1 | NA | NA | 5 | 1 | 10 | 1 | 3 | 0.6667 | 1 | 1 | 3 | 1 | 1 | 1 | 8 | 0.875 |
| 4_26340850 | 1 | 1 | 2 | 1 | NA | NA | 4 | 0.25 | 7 | 0.5714 | 3 | 0.6667 | 1 | 0 | 3 | 0.6667 | 1 | 1 | 8 | 0.625 |
| 4_26346735 | 2 | 0.5 | 1 | 0 | NA | NA | 2 | 0 | 5 | 0.2 | 2 | 0 | 1 | 1 | 3 | 0 | 1 | 0 | 7 | 0.1429 |
| 4_28813625 | NA | NA | 1 | 1 | NA | NA | 1 | 1 | 2 | 1 | 1 | 1 | 1 | 1 | NA | NA | NA | NA | 2 | 1 |
| 4_28814152 | NA | NA | 1 | 1 | NA | NA | 1 | 0 | 2 | 0.5 | 1 | 1 | 2 | 0 | NA | NA | NA | NA | 3 | 0.3333 |
| 4_28814181 | NA | NA | 1 | 1 | NA | NA | 1 | 0 | 2 | 0.5 | 1 | 1 | 2 | 0.5 | NA | NA | NA | NA | 3 | 0.6667 |
| 4_28814356 | NA | NA | 1 | 1 | NA | NA | NA | NA | 1 | 1 | 1 | 1 | 1 | 0 | NA | NA | NA | NA | 2 | 0.5 |
| 4_28814671 | NA | NA | 1 | 1 | NA | NA | NA | NA | 1 | 1 | 1 | 0 | 1 | 1 | NA | NA | NA | NA | 2 | 0.5 |
| 4_28821095 | 1 | 1 | 5 | 0.6 | 2 | 0.5 | 3 | 0 | 11 | 0.4545 | 3 | 0.6667 | 2 | 1 | 4 | 0.25 | 10 | 0.7 | 19 | 0.6316 |
| 4_28821188 | 1 | 0 | 5 | 0.2 | 2 | 0.5 | 4 | 0.75 | 12 | 0.4167 | 4 | 0.5 | 2 | 1 | 3 | 0 | 10 | 0.4 | 19 | 0.4211 |
| 4_28821224 | NA | NA | 4 | 0 | 1 | 1 | 2 | 0 | 7 | 0.1429 | 4 | 0 | 3 | 0 | 3 | 0 | 7 | 0.1429 | 17 | 0.0588 |
| 4_28821228 | 1 | 1 | 5 | 0.4 | 2 | 0.5 | 4 | 1 | 12 | 0.6667 | 4 | 0 | 3 | 0.3333 | 3 | 0.3333 | 9 | 0.5556 | 19 | 0.3684 |
| 4_28821276 | 1 | 0 | 5 | 0.6 | 2 | 1 | 4 | 1 | 12 | 0.75 | 3 | 0.3333 | 3 | 0.3333 | 4 | 0.5 | 9 | 0.3333 | 19 | 0.3684 |
| 4_28821339 | 1 | 1 | 4 | 0.5 | 2 | 0.5 | 2 | 1 | 9 | 0.6667 | 2 | 0 | 1 | 0 | 3 | 0.3333 | 8 | 0.375 | 14 | 0.2857 |
| 4_28821361 | 1 | 1 | 5 | 0.2 | 2 | 0 | 4 | 0.5 | 12 | 0.3333 | 4 | 0.5 | 3 | 0.6667 | 4 | 0.75 | 9 | 0.4444 | 20 | 0.55 |
| 4_28821468 | 1 | 0 | 4 | 0.5 | 1 | 1 | 4 | 0.25 | 10 | 0.4 | 4 | 0.5 | 2 | 1 | 2 | 1 | 9 | 0.7778 | 17 | 0.7647 |
| 4_28821494 | 1 | 1 | 5 | 0.4 | 2 | 0.5 | 4 | 0.75 | 12 | 0.5833 | 3 | 1 | 3 | 1 | 2 | 1 | 8 | 0.75 | 16 | 0.875 |
| 4_28821533 | 1 | 1 | 5 | 0 | 2 | 1 | 4 | 0.75 | 12 | 0.5 | 3 | 0.6667 | 3 | 1 | 4 | 0.5 | 9 | 0.3333 | 19 | 0.5263 |
| 4_28871883 | NA | NA | 3 | 0.6667 | 2 | 1 | 5 | 1 | 10 | 0.9 | 4 | 1 | 2 | 1 | 5 | 1 | 10 | 0.8 | 21 | 0.9048 |
| 4_28871940 | 1 | 0 | 5 | 0.2 | 3 | 0.6667 | 4 | 0.5 | 13 | 0.3846 | 4 | 0.75 | 3 | 0.3333 | 4 | 0.5 | 8 | 0.375 | 19 | 0.4737 |
| 4_28946266 | 2 | 0.5 | 5 | 0 | 2 | 0 | 5 | 0 | 14 | 0.0714 | 4 | 0.5 | 2 | 0 | NA | NA | 7 | 0.4286 | 13 | 0.3846 |
| 4_28963953 | 1 | 0 | 3 | 0.6667 | 2 | 0.5 | 2 | 0.5 | 8 | 0.5 | 1 | 0 | 1 | 0 | 2 | 0.5 | NA | NA | 4 | 0.25 |
| 4_28963986 | 1 | 1 | 3 | 0.6667 | 2 | 1 | 3 | 0 | 9 | 0.5556 | NA | NA | 1 | 0 | NA | NA | 1 | 1 | 2 | 0.5 |
| 4_28964003 | NA | NA | 2 | 0.5 | 3 | 0 | 2 | 0 | 7 | 0.1429 | 1 | 0 | 1 | 1 | 2 | 0 | 1 | 0 | 5 | 0.2 |
| 4_28964034 | 1 | 0 | 3 | 1 | 2 | 1 | 2 | 1 | 8 | 0.875 | 1 | 1 | 1 | 1 | 2 | 0.5 | 1 | 1 | 5 | 0.8 |
| 4_28964045 | NA | NA | 1 | 1 | 3 | 0.3333 | 3 | 0.6667 | 7 | 0.5714 | NA | NA | 1 | 0 | 3 | 0.6667 | 1 | 1 | 5 | 0.6 |
| 4_28964145 | 1 | 0 | 3 | 0 | 2 | 0.5 | 2 | 0.5 | 8 | 0.25 | 1 | 1 | 1 | 0 | 3 | 0.6667 | 1 | 1 | 6 | 0.6667 |
| 4_28964151 | 1 | 0 | 3 | 1 | 2 | 1 | 3 | 1 | 9 | 0.8889 | 1 | 1 | NA | NA | 1 | 1 | NA | NA | 2 | 1 |
| 4_28964213 | 1 | 0 | 3 | 0 | 2 | 0.5 | 3 | 0.3333 | 9 | 0.2222 | 1 | 1 | 1 | 0 | 3 | 0 | 1 | 0 | 6 | 0.1667 |
| 4_28964224 | 1 | 1 | 3 | 0.6667 | 3 | 1 | 4 | 1 | 11 | 0.9091 | 1 | 1 | 1 | 1 | 3 | 1 | 1 | 1 | 6 | 1 |
| 4_28964295 | 1 | 1 | 3 | 1 | 3 | 1 | 3 | 1 | 10 | 1 | 1 | 1 | 1 | 1 | 3 | 1 | 1 | 1 | 6 | 1 |
| 4_28964449 | 1 | 0 | 1 | 1 | 2 | 0.5 | 5 | 0.8 | 9 | 0.6667 | 1 | 1 | NA | NA | 4 | 0.5 | 2 | 1 | 7 | 0.7143 |
| 4_28964481 | 1 | 0 | 2 | 0.5 | 2 | 0.5 | 4 | 0.25 | 9 | 0.3333 | 1 | 0 | NA | NA | 4 | 0.5 | 2 | 0 | 7 | 0.2857 |
| 4_28964533 | 1 | 0 | 2 | 0.5 | 3 | 0 | 5 | 0.2 | 11 | 0.1818 | 1 | 0 | NA | NA | 4 | 0.5 | 3 | 0.3333 | 8 | 0.375 |
| 4_28964582 | 1 | 0 | 2 | 1 | 3 | 0.6667 | 5 | 0.8 | 11 | 0.7273 | 1 | 1 | NA | NA | 3 | 0.6667 | 3 | 0.6667 | 7 | 0.7143 |
| 4_28965487 | 2 | 0 | 2 | 0 | 4 | 0.5 | 1 | 0 | 9 | 0.2222 | 2 | 0 | NA | NA | 4 | 0.25 | 2 | 0 | 8 | 0.125 |
| 4_28965584 | 2 | 0 | 2 | 0 | 4 | 0.25 | 4 | 0 | 12 | 0.0833 | 2 | 0 | 1 | 0 | 6 | 0.1667 | 4 | 0 | 13 | 0.0769 |
| 4_28966067 | 4 | 0.25 | 4 | 0.25 | 3 | 0 | 6 | 0 | 17 | 0.1176 | 1 | 0 | 2 | 0 | 6 | 0 | 4 | 0.5 | 13 | 0.1538 |
| 4_28966249 | 1 | 0 | 2 | 0 | 2 | 0.5 | 5 | 0 | 10 | 0.1 | 1 | 0 | 2 | 0 | 3 | 0.3333 | 3 | 0 | 9 | 0.1111 |
| 4_28966693 | 1 | 1 | 3 | 0.3333 | 2 | 0 | 4 | 0.25 | 10 | 0.3 | 1 | 1 | 1 | 0 | 3 | 0 | 4 | 0 | 9 | 0.1111 |
| 4_28967097 | NA | NA | 4 | 0.5 | 1 | 1 | 4 | 0.25 | 9 | 0.4444 | 1 | 0 | 2 | 0 | 4 | 0.25 | 7 | 0 | 14 | 0.0714 |
| 4_28967215 | 1 | 0 | 4 | 0 | 2 | 0 | 6 | 0.1667 | 13 | 0.0769 | 1 | 0 | 2 | 0 | 4 | 0.25 | 7 | 0 | 14 | 0.0714 |
| 4_32019691 | 4 | 0.75 | 2 | 1 | 2 | 1 | 5 | 0.4 | 13 | 0.6923 | 6 | 1 | 11 | 0.9091 | 6 | 0.6667 | 3 | 0.6667 | 26 | 0.8462 |
| 4_32019717 | 4 | 1 | 1 | 1 | 3 | 1 | 5 | 0.6 | 13 | 0.8462 | 6 | 0.6667 | 11 | 0.7273 | 6 | 0.5 | 3 | 0.3333 | 26 | 0.6154 |
| 4_32019931 | 5 | 0.2 | 2 | 0 | 3 | 0.3333 | 4 | 0.25 | 14 | 0.2143 | 6 | 0.1667 | 10 | 0.3 | 5 | 0.2 | 4 | 0.25 | 25 | 0.24 |
| 4_32020036 | 5 | 0.2 | 2 | 0 | 3 | 0.3333 | 5 | 0.2 | 15 | 0.2 | 7 | 0.4286 | 12 | 0.3333 | 7 | 0.1429 | 3 | 0 | 29 | 0.2759 |
| 4_32020114 | 5 | 0.2 | 2 | 0 | 3 | 0 | 5 | 0 | 15 | 0.0667 | 6 | 0.1667 | 11 | 0.0909 | 7 | 0 | 4 | 0 | 28 | 0.0714 |

| | | | | | | | | | | | | | | | | | | | | |
|---|---|---|---|---|---|---|---|---|---|---|---|---|---|---|---|---|---|---|---|---|
| 4_3202021 | 3 | 0 | 2 | 0.5 | 3 | 0 | 4 | 0 | 12 | 0.0833 | 3 | 0.6667 | 12 | 0.1667 | 5 | 0 | 2 | 0.5 | 22 | 0.2273 |
| 4_3202028 | 4 | 0 | 2 | 0 | 3 | 0 | 4 | 0.25 | 13 | 0.0769 | 6 | 0.3333 | 11 | 0.4545 | 6 | 0.5 | 3 | 0.3333 | 26 | 0.4231 |
| 4_3202038 | 5 | 0 | 3 | 0 | 5 | 0.2 | 5 | 0 | 18 | 0.0556 | 6 | 0 | 12 | 0 | 6 | 0.1667 | 4 | 0.25 | 28 | 0.0714 |
| 4_3202162 | 6 | 0 | 3 | 0 | 4 | 0 | 2 | 0.5 | 15 | 0.0667 | 4 | 0 | 10 | 0.1 | 4 | 0 | 2 | 0.5 | 20 | 0.1 |
| 4_3202254 | 5 | 0.2 | 3 | 0.3333 | 4 | 0.25 | 1 | 0 | 13 | 0.2308 | 3 | 0.6667 | 10 | 0.3 | 5 | 0.4 | 3 | 0 | 21 | 0.3333 |
| 4_3202256 | 6 | 0.1667 | 3 | 0.6667 | 4 | 0.25 | 1 | 0 | 14 | 0.2857 | 5 | 0.6 | 9 | 0.5556 | 6 | 0.3333 | 3 | 0.3333 | 23 | 0.4783 |
| 4_3202283 | 6 | 0.1667 | 3 | 0 | 4 | 0 | 1 | 0 | 14 | 0.0714 | 5 | 0 | 9 | 0.3333 | 6 | 0.1667 | 3 | 0.6667 | 23 | 0.2609 |
| 4_3202313 | 6 | 0.1667 | 4 | 0.5 | 3 | 1 | NA | NA | 13 | 0.4615 | 3 | 0.6667 | 8 | 0.375 | 6 | 0.5 | 3 | 0.6667 | 20 | 0.5 |
| 4_3202319 | 5 | 0.6 | 4 | 1 | 5 | 0.6 | 1 | 1 | 15 | 0.7333 | 4 | 0.75 | 8 | 0.875 | 6 | 0.3333 | 4 | 0.75 | 22 | 0.6818 |
| 4_3202328 | 3 | 1 | 3 | 0.6667 | 5 | 0.8 | 1 | 0 | 12 | 0.75 | 6 | 0.8333 | 11 | 0.8182 | 7 | 0.7143 | 4 | 1 | 28 | 0.8214 |
| 4_3202336 | 5 | 0.8 | 3 | 1 | 5 | 0.8 | 1 | 1 | 14 | 0.8571 | 5 | 0.8 | 7 | 0.5714 | 4 | 0.5 | 3 | 0.6667 | 19 | 0.6316 |
| 4_3258017 | 1 | 1 | NA | NA | NA | NA | NA | NA | 1 | 1 | 1 | 1 | NA | NA | NA | NA | NA | NA | 1 | 1 |
| 4_3258023 | 1 | 1 | NA | NA | NA | NA | NA | NA | 1 | 1 | 1 | 1 | NA | NA | NA | NA | NA | NA | 1 | 1 |
| 4_3258024 | 1 | 1 | NA | NA | NA | NA | NA | NA | 1 | 1 | 1 | 1 | NA | NA | NA | NA | NA | NA | 1 | 1 |
| 4_3258059 | 1 | 0 | 1 | 1 | NA | NA | NA | NA | 2 | 0.5 | 1 | 1 | NA | NA | NA | NA | NA | NA | 1 | 1 |
| 4_3258088 | 1 | 1 | 1 | 0 | NA | NA | NA | NA | 2 | 0.5 | 1 | 1 | NA | NA | NA | NA | NA | NA | 1 | 1 |
| 4_3258482 | 1 | 0 | 2 | 0.5 | NA | NA | 1 | 0 | 4 | 0.25 | 1 | 1 | NA | NA | NA | NA | NA | NA | 1 | 1 |
| 4_3258565 | 1 | 1 | 2 | 0.5 | NA | NA | 1 | 1 | 4 | 0.75 | 1 | 1 | NA | NA | NA | NA | NA | NA | 1 | 1 |
| 4_3264046 | NA | NA | 2 | 0 | 1 | 1 | NA | NA | 3 | 0.3333 | NA | NA | 1 | 1 | NA | NA | NA | NA | 1 | 1 |
| 4_3264052 | NA | NA | 2 | 1 | 1 | 0 | NA | NA | 3 | 0.6667 | NA | NA | 1 | 1 | NA | NA | NA | NA | 1 | 1 |
| 4_3264090 | NA | NA | NA | NA | 1 | 1 | NA | NA | 1 | 1 | NA | NA | 1 | 1 | NA | NA | NA | NA | 1 | 1 |
| 4_3264101 | NA | NA | 2 | 1 | 1 | 1 | NA | NA | 3 | 1 | NA | NA | 1 | 1 | NA | NA | NA | NA | 1 | 1 |
| 4_3264122 | NA | NA | 2 | 1 | 1 | 1 | NA | NA | 3 | 1 | NA | NA | 1 | 1 | NA | NA | NA | NA | 1 | 1 |
| 4_3264128 | NA | NA | 2 | 0.5 | 1 | 1 | NA | NA | 3 | 0.6667 | NA | NA | 1 | 1 | NA | NA | NA | NA | 1 | 1 |
| 4_3264132 | NA | NA | 2 | 1 | 1 | 1 | NA | NA | 3 | 1 | NA | NA | 1 | 1 | NA | NA | NA | NA | 1 | 1 |
| 4_3264383 | NA | NA | 3 | 0.3333 | 1 | 0 | NA | NA | 4 | 0.25 | NA | NA | 2 | 1 | NA | NA | NA | NA | 2 | 1 |
| 4_3264458 | NA | NA | 2 | 1 | 1 | 1 | NA | NA | 3 | 1 | NA | NA | 1 | 1 | NA | NA | NA | NA | 1 | 1 |
| 4_3264468 | NA | NA | 3 | 0.6667 | 1 | 0 | NA | NA | 4 | 0.5 | NA | NA | 1 | 1 | NA | NA | NA | NA | 1 | 1 |
| 4_3264565 | NA | NA | 4 | 1 | NA | NA | NA | NA | 4 | 1 | NA | NA | 2 | 1 | NA | NA | NA | NA | 2 | 1 |
| 4_3264629 | NA | NA | 4 | 0.75 | 1 | 1 | NA | NA | 5 | 0.8 | NA | NA | 2 | 0.5 | NA | NA | 1 | 0 | 3 | 0.3333 |
| 4_3264642 | NA | NA | 3 | 0.6667 | 1 | 1 | NA | NA | 4 | 0.75 | NA | NA | 1 | 1 | NA | NA | 1 | 1 | 2 | 1 |
| 4_3264645 | NA | NA | 3 | 0 | 1 | 1 | NA | NA | 4 | 0.25 | NA | NA | 2 | 0.5 | NA | NA | 1 | 1 | 3 | 0.6667 |
| 4_3277420 | NA | NA | 1 | 0 | 1 | 0 | 3 | 0.3333 | 5 | 0.2 | NA | NA | 2 | 0.5 | 3 | 0.3333 | NA | NA | 5 | 0.4 |
| 4_3277423 | NA | NA | 2 | 0.5 | 1 | 1 | 3 | 0 | 6 | 0.3333 | NA | NA | 3 | 0.6667 | 3 | 0.3333 | NA | NA | 6 | 0.5 |
| 4_3277427 | NA | NA | 2 | 0.5 | 1 | 0 | 2 | 0 | 5 | 0.2 | NA | NA | 2 | 0 | 2 | 0.5 | NA | NA | 4 | 0.25 |
| 4_3277433 | NA | NA | 1 | 1 | 1 | 0 | 3 | 0.6667 | 5 | 0.6 | NA | NA | 3 | 0.3333 | 3 | 0.3333 | NA | NA | 6 | 0.3333 |
| 4_3277439 | NA | NA | 2 | 0 | 1 | 0 | 3 | 0.3333 | 6 | 0.1667 | NA | NA | 3 | 0 | 3 | 0.3333 | NA | NA | 6 | 0.1667 |
| 4_3277467 | NA | NA | 2 | 0.5 | 1 | 0 | 3 | 0.3333 | 6 | 0.3333 | NA | NA | 2 | 0 | 3 | 0.6667 | NA | NA | 5 | 0.4 |
| 4_3279121 | NA | NA | 2 | 0.5 | NA | NA | NA | NA | 2 | 0.5 | NA | NA | 2 | 1 | NA | NA | NA | NA | 2 | 1 |
| 4_3279211 | NA | NA | 1 | 1 | NA | NA | NA | NA | 1 | 1 | NA | NA | 2 | 1 | NA | NA | NA | NA | 2 | 1 |
| 4_3280119 | 8 | 0.375 | 5 | 0.2 | 3 | 0.3333 | 2 | 0 | 18 | 0.2778 | 1 | 0 | 5 | 0 | 7 | 0.1429 | 4 | 0 | 17 | 0.0588 |
| 4_3280136 | 7 | 0.2857 | 4 | 0.5 | 2 | 1 | 2 | 0.5 | 15 | 0.4667 | 1 | 0 | 5 | 0.4 | 7 | 0.5714 | 4 | 0.5 | 17 | 0.4706 |
| 4_3280151 | 4 | 0 | 4 | 0.5 | 1 | 1 | 1 | 0 | 10 | 0.3 | NA | NA | 3 | 0.3333 | 4 | 0 | 1 | 0 | 8 | 0.125 |
| 4_3280152 | 5 | 0.2 | 3 | 0.3333 | 2 | 0.5 | 1 | 0 | 11 | 0.2727 | NA | NA | 2 | 0.5 | 4 | 0.75 | 1 | 1 | 7 | 0.7143 |
| 4_3280168 | 4 | 0.25 | 4 | 0 | 2 | 1 | NA | NA | 10 | 0.3 | NA | NA | 4 | 0.25 | 3 | 0 | 2 | 0 | 9 | 0.1111 |
| 4_3280199 | 6 | 0.3333 | 4 | 0 | 2 | 0 | 1 | 0 | 13 | 0.1538 | 1 | 0 | 5 | 0.2 | 4 | 0.5 | 4 | 0 | 14 | 0.2143 |
| 4_3280453 | 3 | 0 | 11 | 0.1818 | 4 | 0.25 | 9 | 0.2222 | 27 | 0.1852 | 2 | 0 | 4 | 0.25 | 1 | 0 | 4 | 0.5 | 11 | 0.2727 |
| 4_3280489 | 2 | 0 | 9 | 0.2222 | 3 | 0.3333 | 7 | 0.2857 | 21 | 0.2381 | 2 | 0 | 3 | 0 | 2 | 1 | 4 | 0.25 | 11 | 0.2727 |
| 4_3280501 | 2 | 0.5 | 4 | 0.25 | 1 | 1 | 6 | 0.3333 | 13 | 0.3846 | 1 | 1 | 4 | 0.25 | NA | NA | 3 | 0.6667 | 8 | 0.5 |
| 4_3280516 | 2 | 0 | 8 | 0.125 | 2 | 0.5 | 7 | 0 | 19 | 0.1053 | NA | NA | 4 | 0 | 2 | 0 | 3 | 0.6667 | 9 | 0.2222 |
| 4_3280519 | 2 | 0 | 8 | 0.125 | 2 | 0 | 6 | 0 | 18 | 0.0556 | NA | NA | 4 | 0 | 2 | 1 | 2 | 0 | 8 | 0.25 |
| 4_3280522 | 1 | 0 | 8 | 1 | 2 | 0.5 | 7 | 0.7143 | 18 | 0.7778 | NA | NA | 4 | 0.75 | 2 | 0.5 | 3 | 1 | 9 | 0.7778 |
| 4_3280524 | 2 | 0.5 | 5 | 0.4 | 2 | 1 | 6 | 0.3333 | 15 | 0.4667 | NA | NA | 3 | 0.6667 | 2 | 0 | 3 | 0.6667 | 8 | 0.5 |
| 4_3280529 | 2 | 0 | 7 | 0.1429 | 1 | 0 | 7 | 0.1429 | 17 | 0.1176 | NA | NA | 3 | 0.3333 | 2 | 0 | 2 | 0 | 7 | 0.1429 |
| 4_3280529 | 1 | 0 | 6 | 0.1667 | 1 | 0 | 7 | 0 | 15 | 0.0667 | NA | NA | 3 | 0.3333 | 1 | 0 | 2 | 0 | 6 | 0.1667 |
| 4_3280532 | 2 | 1 | 8 | 0.5 | 2 | 0.5 | 6 | 0.1667 | 18 | 0.4444 | NA | NA | 4 | 0.25 | 2 | 0 | 3 | 0.3333 | 9 | 0.2222 |
| 4_3280535 | 2 | 1 | 8 | 0.75 | 2 | 1 | 6 | 1 | 18 | 0.8889 | NA | NA | 2 | 1 | 1 | 1 | 2 | 1 | 5 | 1 |
| 4_3280540 | 1 | 1 | 4 | 0 | 1 | 0 | 5 | 0.2 | 11 | 0.1818 | NA | NA | 2 | 0.5 | 1 | 0 | NA | NA | 3 | 0.3333 |
| 4_3280542 | 2 | 0.5 | 7 | 0.2857 | 2 | 0.5 | 7 | 0.2857 | 18 | 0.3333 | NA | NA | 4 | 0.5 | 2 | 0 | 3 | 0 | 9 | 0.2222 |
| 4_3280547 | 1 | 1 | 8 | 0.875 | 2 | 1 | 6 | 1 | 17 | 0.9412 | NA | NA | 2 | 1 | 2 | 1 | 3 | 1 | 7 | 1 |
| 4_3280554 | 1 | 0 | 7 | 0.7143 | 2 | 0 | 6 | 0.6667 | 16 | 0.5625 | NA | NA | 3 | 0.6667 | 2 | 0.5 | 2 | 0.5 | 7 | 0.5714 |
| 4_3280555 | 1 | 1 | 2 | 0.5 | 1 | 0 | 5 | 0.4 | 9 | 0.4444 | NA | NA | 3 | 0.3333 | NA | NA | 1 | 1 | 4 | 0.5 |
| 4_3280560 | 2 | 0.5 | 6 | 0 | 1 | 0 | 5 | 0 | 14 | 0.0714 | NA | NA | 4 | 0.5 | 2 | 0 | 3 | 0 | 9 | 0.2222 |
| 4_3280560 | 2 | 1 | 6 | 1 | 1 | 1 | 5 | 0.8 | 14 | 0.9286 | NA | NA | 4 | 1 | 2 | 1 | 3 | 1 | 9 | 1 |
| 4_3280573 | 2 | 0.5 | 5 | 0.6 | 1 | 1 | 2 | 1 | 10 | 0.7 | 1 | 1 | 4 | 0.5 | 1 | 1 | 3 | 0.3333 | 9 | 0.5556 |

| | | | | | | | | | | | | | | | | | | | | |
|---|---|---|---|---|---|---|---|---|---|---|---|---|---|---|---|---|---|---|---|---|
| 4_3280579 | 2 | 0 | 6 | 0.1667 | 1 | 0 | 4 | 0 | 13 | 0.0769 | 1 | 1 | 4 | 0 | 2 | 1 | 3 | 0 | 10 | 0.3 |
| 4_3280585 | 1 | 1 | 7 | 1 | 1 | 1 | 4 | 1 | 13 | 1 | 1 | 1 | 4 | 1 | 2 | 0.5 | 2 | 1 | 9 | 0.8889 |
| 4_3280586 | 1 | 0 | 7 | 0.2857 | 1 | 1 | 4 | 0.5 | 13 | 0.3846 | 1 | 0 | 4 | 0.25 | 1 | 1 | 2 | 0.5 | 8 | 0.375 |
| 4_3280598 | 2 | 1 | 7 | 0.8571 | NA | NA | 5 | 0.8 | 14 | 0.8571 | NA | NA | 4 | 1 | 2 | 0.5 | 3 | 1 | 9 | 0.8889 |
| 4_3280602 | 2 | 1 | 7 | 1 | NA | NA | 5 | 1 | 14 | 1 | NA | NA | 3 | 1 | 2 | 1 | 2 | 1 | 7 | 1 |
| 4_3280609 | 2 | 0.5 | 7 | 0.4286 | NA | NA | 5 | 0.4 | 14 | 0.4286 | NA | NA | 4 | 0 | 2 | 0 | 3 | 0.3333 | 9 | 0.1111 |
| 4_3280613 | 2 | 0 | 6 | 0.6667 | NA | NA | 4 | 1 | 12 | 0.6667 | NA | NA | 3 | 1 | 2 | 0 | 3 | 0.3333 | 8 | 0.5 |
| 4_3280628 | 2 | 0.5 | 7 | 0.4286 | NA | NA | 5 | 0.2 | 14 | 0.3571 | NA | NA | 3 | 0 | 2 | 0.5 | 3 | 0 | 8 | 0.125 |
| 4_3280644 | 2 | 0 | 6 | 0.1667 | NA | NA | 5 | 0.2 | 13 | 0.1538 | NA | NA | 3 | 0.3333 | 2 | 0 | 3 | 0 | 8 | 0.125 |
| 4_3280651 | 1 | 1 | 6 | 0.5 | NA | NA | 5 | 0.4 | 12 | 0.5 | NA | NA | 3 | 0.6667 | NA | NA | 2 | 1 | 5 | 0.8 |
| 4_3299600 | 2 | 0 | 6 | 0.3333 | 4 | 1 | 17 | 0.3529 | 29 | 0.4138 | 5 | 0.6 | 7 | 0.7143 | 7 | 0.1429 | 5 | 0.8 | 24 | 0.5417 |
| 4_3299602 | 3 | 0.6667 | 4 | 0.75 | 4 | 0 | 14 | 0.3571 | 25 | 0.4 | 4 | 0.25 | 8 | 0.5 | 8 | 0.25 | 5 | 0.4 | 25 | 0.36 |
| 4_3301294 | 4 | 0.25 | 8 | 0 | 6 | 0 | 18 | 0.0556 | 36 | 0.0556 | 8 | 0.25 | 12 | 0.0833 | 10 | 0.1 | 6 | 0 | 36 | 0.1111 |
| 4_3301877 | 2 | 0 | 7 | 0 | 5 | 0 | 18 | 0.1111 | 32 | 0.0625 | 8 | 0 | 10 | 0.2 | 10 | 0.2 | 5 | 0 | 33 | 0.1212 |
| 4_3301880 | 3 | 0.3333 | 7 | 0.1429 | 7 | 0 | 19 | 0.3158 | 36 | 0.2222 | 8 | 0.25 | 12 | 0.1667 | 9 | 0.4444 | 6 | 0 | 35 | 0.2286 |
| 4_3301884 | 2 | 0 | 7 | 0 | 5 | 0 | 15 | 0.1333 | 29 | 0.069 | 8 | 0 | 10 | 0.1 | 9 | 0.1111 | 6 | 0.1667 | 33 | 0.0909 |
| 4_3301888 | 3 | 0 | 7 | 0.1429 | 7 | 0 | 19 | 0.2105 | 36 | 0.1389 | 9 | 0 | 13 | 0 | 11 | 0.1818 | 6 | 0 | 39 | 0.0513 |
| 4_3301903 | 5 | 0 | 7 | 0.2857 | 6 | 0.1667 | 17 | 0.2941 | 35 | 0.2286 | 9 | 0.4444 | 12 | 0.0833 | 11 | 0 | 8 | 0.125 | 40 | 0.15 |
| 4_3301904 | 5 | 0.2 | 6 | 0.1667 | 7 | 0.4286 | 16 | 0.1875 | 34 | 0.2353 | 8 | 0 | 11 | 0.2727 | 10 | 0.4 | 7 | 0.2857 | 36 | 0.25 |
| 4_3301908 | 6 | 0 | 7 | 0.2857 | 6 | 0 | 18 | 0.1667 | 37 | 0.1351 | 9 | 0 | 12 | 0.0833 | 10 | 0 | 8 | 0.125 | 39 | 0.0513 |
| 4_3301913 | 5 | 0 | 9 | 0.3333 | 6 | 0.1667 | 18 | 0.0556 | 38 | 0.1316 | 8 | 0 | 13 | 0.0769 | 10 | 0.1 | 9 | 0 | 40 | 0.05 |
| 4_3301919 | 7 | 0.1429 | 8 | 0.375 | 7 | 0.1429 | 19 | 0.0526 | 41 | 0.1463 | 9 | 0.1111 | 15 | 0.2 | 11 | 0 | 9 | 0.1111 | 44 | 0.1136 |
| 4_3301921 | 7 | 0.4286 | 7 | 0.1429 | 5 | 0.2 | 18 | 0.1667 | 37 | 0.2162 | 9 | 0.1111 | 15 | 0.3333 | 8 | 0 | 6 | 0 | 38 | 0.1579 |
| 4_3302151 | 17 | 0.1176 | 16 | 0.3125 | 10 | 0.2 | 28 | 0.1786 | 71 | 0.1972 | 9 | 0.2222 | 30 | 0.1667 | 20 | 0.15 | 13 | 0.1538 | 72 | 0.1667 |
| 4_3302187 | 17 | 0.1176 | 11 | 0.0909 | 10 | 0.3 | 24 | 0.1667 | 62 | 0.1613 | 13 | 0.0769 | 30 | 0.0667 | 22 | 0.0909 | 14 | 0.0714 | 79 | 0.0759 |
| 4_3302335 | 1 | 1 | 1 | 1 | 1 | 1 | 2 | 0.5 | 5 | 0.8 | 1 | 1 | NA | NA | NA | NA | NA | NA | 1 | 1 |
| 4_3302342 | 2 | 1 | NA | NA | 1 | 1 | 1 | 1 | 4 | 1 | 1 | 1 | NA | NA | NA | NA | NA | NA | 1 | 1 |
| 4_3302446 | 1 | 1 | NA | NA | 1 | 0 | 1 | 1 | 3 | 0.6667 | NA | NA | NA | NA | NA | NA | 1 | 1 | 1 | 1 |
| 4_3304988 | 8 | 0.375 | 6 | 0.5 | 7 | 0 | 8 | 0.375 | 29 | 0.3103 | 1 | 0 | 10 | 0.2 | 4 | 0 | 7 | 0.5714 | 22 | 0.2727 |
| 4_3304994 | 7 | 0.2857 | 5 | 0 | 6 | 0.5 | 11 | 0.8182 | 29 | 0.4828 | 3 | 0.3333 | 8 | 0.375 | 3 | 0.3333 | 8 | 0.375 | 22 | 0.3636 |
| 4_3304997 | 8 | 0.125 | 7 | 0 | 6 | 0.3333 | 11 | 0.1818 | 32 | 0.1562 | 3 | 0 | 10 | 0.1 | 4 | 0 | 9 | 0.2222 | 26 | 0.1154 |
| 4_3304998 | 8 | 0 | 7 | 0 | 6 | 0.1667 | 12 | 0.0833 | 33 | 0.0606 | 2 | 0 | 10 | 0.1 | 4 | 0 | 9 | 0.2222 | 25 | 0.12 |
| 4_3305015 | 9 | 0.3333 | 6 | 0.3333 | 7 | 0.1429 | 10 | 0.6 | 32 | 0.375 | 2 | 0.5 | 9 | 0.2222 | 2 | 0 | 8 | 0.125 | 21 | 0.1905 |
| 4_3309564 | NA | NA | 2 | 1 | NA | NA | NA | NA | 2 | 1 | 1 | 0 | NA | NA | NA | NA | 1 | 1 | 2 | 0.5 |
| 4_3309570 | NA | NA | 3 | 0.6667 | NA | NA | NA | NA | 3 | 0.6667 | 1 | 0 | NA | NA | NA | NA | 1 | 1 | 2 | 0.5 |
| 4_3317402 | NA | NA | 1 | 1 | 1 | 0 | NA | NA | 2 | 0.5 | NA | NA | NA | NA | 1 | 1 | NA | NA | 1 | 1 |
| 4_3323101 | 1 | 0 | 4 | 0.25 | 1 | 0 | 4 | 0 | 10 | 0.1 | NA | NA | 5 | 0.4 | 2 | 0 | 2 | 0 | 9 | 0.2222 |
| 4_3323111 | 1 | 1 | 4 | 0.5 | 2 | 1 | 3 | 0.3333 | 10 | 0.6 | NA | NA | 2 | 0 | 2 | 0 | 1 | 1 | 5 | 0.2 |
| 4_3323116 | 2 | 0 | 3 | 0.6667 | 2 | 0.5 | 4 | 0.5 | 11 | 0.4545 | NA | NA | 4 | 0.25 | 2 | 0 | 2 | 1 | 8 | 0.375 |
| 4_3323128 | NA | NA | 1 | 1 | NA | NA | 1 | 0 | 2 | 0.5 | NA | NA | 3 | 0.6667 | NA | NA | NA | NA | 3 | 0.6667 |
| 4_3323138 | 1 | 0 | 4 | 0.5 | 2 | 0 | 4 | 0.25 | 11 | 0.2727 | NA | NA | 5 | 0.4 | 2 | 0 | 2 | 0 | 9 | 0.2222 |
| 4_3323170 | 1 | 0 | 4 | 1 | 2 | 0.5 | 4 | 0.25 | 11 | 0.5455 | NA | NA | 5 | 0.2 | 3 | 0.3333 | 2 | 0 | 10 | 0.2 |
| 4_3323203 | 4 | 0.5 | 4 | 0.25 | 3 | 1 | 5 | 0 | 16 | 0.375 | NA | NA | 4 | 0.5 | 3 | 0 | 2 | 0.5 | 9 | 0.3333 |
| 4_3323205 | 4 | 0 | 4 | 0 | 2 | 0 | 5 | 0.2 | 15 | 0.0667 | NA | NA | 5 | 0.2 | 2 | 0 | 2 | 0 | 9 | 0.1111 |
| 4_3323213 | 4 | 0.25 | 4 | 0 | 2 | 0 | 5 | 0 | 15 | 0.0667 | NA | NA | 5 | 0.2 | 3 | 0 | 2 | 0 | 10 | 0.1 |
| 4_3324559 | 19 | 0.1053 | 21 | 0.1429 | 12 | 0 | 16 | 0.1875 | 68 | 0.1176 | 10 | 0.2 | 13 | 0.1538 | 23 | 0 | 10 | 0.4 | 56 | 0.1429 |
| 4_3324568 | 17 | 0.0588 | 13 | 0.1538 | 9 | 0.3333 | 12 | 0.3333 | 51 | 0.1961 | 8 | 0 | 9 | 0 | 15 | 0.1333 | 9 | 0.1111 | 41 | 0.0732 |
| 4_3324572 | 20 | 0.05 | 18 | 0.1111 | 12 | 0 | 13 | 0.2308 | 63 | 0.0952 | 9 | 0 | 12 | 0.0833 | 20 | 0.15 | 10 | 0.1 | 51 | 0.098 |
| 4_3324575 | 20 | 0.75 | 18 | 0.7222 | 11 | 0.5455 | 16 | 0.4375 | 65 | 0.6308 | 9 | 0.6667 | 11 | 0.5455 | 20 | 0.65 | 10 | 0.7 | 50 | 0.64 |
| 4_3324586 | 18 | 0.7778 | 20 | 0.9 | 11 | 0.6364 | 15 | 0.6667 | 64 | 0.7656 | 7 | 0.8571 | 9 | 0.5556 | 19 | 0.7368 | 9 | 0.6667 | 44 | 0.7045 |
| 4_3324598 | 17 | 0.1765 | 17 | 0.2941 | 10 | 0.2 | 16 | 0.3125 | 60 | 0.25 | 6 | 0 | 9 | 0.3333 | 18 | 0.1667 | 7 | 0.2857 | 40 | 0.2 |
| 4_3324601 | 16 | 1 | 10 | 1 | 7 | 1 | 13 | 1 | 46 | 1 | 6 | 1 | 7 | 1 | 12 | 1 | 6 | 1 | 31 | 1 |
| 4_3324602 | 18 | 0.0556 | 17 | 0.0588 | 9 | 0.2222 | 15 | 0.2 | 59 | 0.1186 | 6 | 0.1667 | 8 | 0 | 21 | 0.1905 | 9 | 0.2222 | 44 | 0.1591 |
| 4_3324619 | 15 | 0 | 18 | 0.1111 | 9 | 0.2222 | 17 | 0.2353 | 59 | 0.1356 | 7 | 0 | 9 | 0.1111 | 18 | 0.2778 | 7 | 0.1429 | 41 | 0.1707 |
| 4_3324626 | 14 | 0.1429 | 17 | 0.1765 | 9 | 0 | 16 | 0.0625 | 56 | 0.1071 | 6 | 0 | 8 | 0.125 | 18 | 0.0556 | 7 | 0 | 39 | 0.0513 |
| 4_3324627 | 15 | 0.9333 | 12 | 1 | 8 | 1 | 12 | 1 | 47 | 0.9787 | 4 | 1 | 8 | 1 | 14 | 1 | 8 | 1 | 34 | 1 |
| 4_3324634 | 11 | 1 | 13 | 1 | 8 | 1 | 12 | 1 | 44 | 1 | 1 | 1 | 5 | 1 | 14 | 1 | 4 | 1 | 24 | 1 |
| 4_3324640 | 14 | 0.5 | 13 | 0.3077 | 9 | 0.5556 | 15 | 0.5333 | 51 | 0.4706 | 4 | 0.25 | 9 | 0.4444 | 17 | 0.4118 | 7 | 0.4286 | 37 | 0.4054 |
| 4_3324642 | 12 | 0 | 16 | 0.0625 | 8 | 0 | 14 | 0.2143 | 50 | 0.08 | 5 | 0.2 | 6 | 0.1667 | 16 | 0.0625 | 7 | 0 | 34 | 0.0882 |
| 4_3350029 | 2 | 0 | 3 | 0 | 4 | 0.25 | 5 | 0 | 14 | 0.0714 | 2 | 0 | 2 | 0 | 3 | 0.3333 | NA | NA | 7 | 0.1429 |
| 4_3350038 | 1 | 1 | 3 | 1 | 3 | 0.6667 | 4 | 1 | 11 | 0.9091 | 3 | 1 | 2 | 1 | 2 | 1 | NA | NA | 7 | 1 |
| 4_3350054 | 2 | 0 | 3 | 0.3333 | 4 | 0.5 | 4 | 0.75 | 13 | 0.4615 | 2 | 0.5 | 2 | 0 | 3 | 0.3333 | NA | NA | 7 | 0.2857 |
| 4_3350056 | 2 | 1 | 3 | 1 | 5 | 1 | 4 | 1 | 14 | 1 | 3 | 1 | 2 | 1 | 3 | 1 | NA | NA | 8 | 1 |
| 4_3350131 | 1 | 1 | 1 | 1 | 4 | 1 | 4 | 1 | 10 | 1 | 3 | 1 | 3 | 0.6667 | NA | NA | NA | NA | 6 | 0.8333 |
| 4_3350134 | 1 | 0 | 1 | 1 | 2 | 0.5 | 4 | 0.25 | 8 | 0.375 | 3 | 0.6667 | 3 | 1 | 2 | 0 | NA | NA | 8 | 0.625 |

| | | | | | | | | | | | | | | | | | | | | |
|---|---|---|---|---|---|---|---|---|---|---|---|---|---|---|---|---|---|---|---|---|
| 4_3350143 | NA | NA | 2 | 0.5 | 4 | 0.75 | 4 | 0.75 | 10 | 0.7 | 3 | 0.3333 | 3 | 0.3333 | 2 | 0 | NA | NA | 8 | 0.25 |
| 4_3350144 | 1 | 1 | NA | NA | 3 | 0.3333 | 3 | 0.6667 | 7 | 0.5714 | 1 | 0 | 2 | 0.5 | 2 | 0 | NA | NA | 5 | 0.2 |
| 4_3350202 | 1 | 1 | 2 | 0 | 4 | 0 | 3 | 0 | 10 | 0.1 | 1 | 1 | NA | NA | 2 | 0 | NA | NA | 3 | 0.3333 |
| 4_3350221 | 1 | 0 | 2 | 1 | 4 | 0.75 | 4 | 0 | 11 | 0.4545 | 3 | 1 | 3 | 0.6667 | 1 | 0 | NA | NA | 7 | 0.7143 |
| 4_3350236 | 1 | 0 | NA | NA | 4 | 0.75 | 4 | 0.25 | 9 | 0.4444 | 2 | 0.5 | 3 | 0.3333 | 1 | 0 | 1 | 0 | 7 | 0.2857 |
| 4_3350245 | NA | NA | 2 | 0 | 4 | 0.5 | 4 | 0 | 10 | 0.2 | 4 | 0.25 | 2 | 0.5 | 2 | 0 | 1 | 0 | 9 | 0.2222 |
| 4_3394360 | 2 | 1 | 3 | 0.6667 | 3 | 1 | NA | NA | 8 | 0.875 | 5 | 0.4 | 1 | 1 | 3 | 1 | 3 | 0.6667 | 12 | 0.6667 |
| 4_3394369 | 2 | 0 | 5 | 0.2 | 3 | 0 | 1 | 1 | 11 | 0.1818 | 5 | 0.4 | 2 | 0.5 | 5 | 0 | 6 | 0.3333 | 18 | 0.2778 |
| 4_3394372 | 2 | 0 | 5 | 0.2 | 3 | 0 | 1 | 0 | 11 | 0.0909 | 6 | 0 | 2 | 0 | 5 | 0.2 | 5 | 0 | 18 | 0.0556 |
| 4_3394389 | 2 | 0 | 5 | 0.6 | 3 | 0.3333 | 2 | 0 | 12 | 0.3333 | 5 | 0.4 | 2 | 1 | 5 | 0.2 | 6 | 0.5 | 18 | 0.4444 |
| 4_3394392 | 3 | 1 | 7 | 1 | 3 | 1 | 2 | 1 | 15 | 1 | 6 | 1 | 1 | 1 | 5 | 1 | 6 | 0.8333 | 18 | 0.9444 |
| 4_3394414 | 4 | 0.5 | 7 | 0.2857 | 3 | 0.3333 | 2 | 0.5 | 16 | 0.375 | 7 | 0.8571 | 2 | 0.5 | 5 | 0 | 6 | 0 | 20 | 0.35 |
| 4_3394419 | 4 | 0.25 | 3 | 0.3333 | 3 | 1 | 1 | 0 | 11 | 0.4545 | 3 | 1 | 1 | 1 | 3 | 0.3333 | 2 | 1 | 9 | 0.7778 |
| 4_3394434 | 4 | 1 | 7 | 1 | 3 | 1 | 2 | 1 | 16 | 1 | 5 | 1 | 2 | 1 | 6 | 1 | 3 | 1 | 16 | 1 |
| 4_3394438 | 5 | 0.8 | 7 | 0.8571 | 2 | 1 | 2 | 1 | 16 | 0.875 | 5 | 1 | 2 | 1 | 6 | 0.6667 | 4 | 0.5 | 17 | 0.7647 |
| 4_3394441 | 4 | 0.25 | 6 | 0.1667 | 3 | 0 | 2 | 0.5 | 15 | 0.2 | 7 | 0.1429 | 2 | 1 | 6 | 0 | 6 | 0.1667 | 21 | 0.1905 |
| 4_3394462 | 5 | 1 | 10 | 1 | 3 | 1 | 2 | 1 | 20 | 1 | 8 | 1 | 2 | 1 | 7 | 1 | 7 | 1 | 24 | 1 |
| 4_3394473 | 5 | 0 | 7 | 0.4286 | 3 | 0.3333 | 2 | 0.5 | 17 | 0.2941 | 8 | 0.125 | 2 | 0 | 8 | 0.125 | 6 | 0 | 24 | 0.0833 |
| 4_3394478 | 5 | 0.4 | 8 | 0.5 | 3 | 0.6667 | 2 | 1 | 18 | 0.5556 | 4 | 0.5 | 1 | 0 | 6 | 0.6667 | 8 | 0.5 | 19 | 0.5263 |
| 4_3394479 | 5 | 0 | 7 | 0.1429 | 3 | 0.3333 | 1 | 0 | 16 | 0.125 | 8 | 0.125 | 1 | 0 | 7 | 0.1429 | 7 | 0.2857 | 23 | 0.1739 |
| 4_3394482 | 5 | 1 | 10 | 1 | 3 | 1 | 2 | 1 | 20 | 1 | 8 | 1 | 2 | 1 | 8 | 1 | 8 | 1 | 26 | 1 |
| 4_3394483 | 4 | 0.25 | 8 | 0.125 | 2 | 0 | 2 | 0.5 | 16 | 0.1875 | 7 | 0.2857 | 2 | 0.5 | 7 | 0.2857 | 7 | 0.1429 | 23 | 0.2609 |
| 4_3394489 | 5 | 1 | 9 | 1 | 2 | 1 | 1 | 1 | 17 | 1 | 8 | 1 | 2 | 1 | 7 | 1 | 7 | 1 | 24 | 1 |
| 4_3394490 | 5 | 0.8 | 10 | 0.7 | 3 | 0.6667 | 1 | 1 | 19 | 0.7368 | 8 | 0.5 | 2 | 1 | 8 | 0.875 | 7 | 0.8571 | 25 | 0.76 |
| 4_3394515 | 5 | 1 | 14 | 1 | 5 | 1 | 4 | 1 | 28 | 1 | 9 | 1 | 2 | 1 | 7 | 0.8571 | 6 | 0.8333 | 24 | 0.9167 |
| 4_3394573 | 6 | 0.1667 | 17 | 0.1176 | 8 | 0.125 | 5 | 0.2 | 36 | 0.1389 | 9 | 0 | 2 | 0 | 10 | 0.4 | 11 | 0 | 32 | 0.125 |
| 4_3394576 | 6 | 0.6667 | 16 | 0.5625 | 8 | 0.5 | 5 | 0.8 | 35 | 0.6 | 9 | 0.3333 | 2 | 0.5 | 11 | 0.7273 | 11 | 0.3636 | 33 | 0.4848 |
| 4_3394576 | 6 | 0.3333 | 16 | 0.1875 | 6 | 0.1667 | 4 | 0.75 | 32 | 0.2812 | 8 | 0.125 | 1 | 0 | 11 | 0.4545 | 11 | 0 | 31 | 0.1935 |
| 4_3394579 | 6 | 0.3333 | 14 | 0.2857 | 6 | 0.5 | 3 | 1 | 29 | 0.4138 | 7 | 0.2857 | 1 | 0 | 5 | 0.2 | 6 | 0.3333 | 19 | 0.2632 |
| 4_3394587 | 4 | 0.25 | 16 | 0.1875 | 7 | 0.1429 | 5 | 0.8 | 32 | 0.2812 | 9 | 0.3333 | 2 | 0.5 | 10 | 0.4 | 10 | 0.2 | 31 | 0.3226 |
| 4_3394624 | 6 | 0.1667 | 19 | 0.0526 | 8 | 0.125 | 5 | 0 | 38 | 0.0789 | 9 | 0 | 2 | 0 | 13 | 0.0769 | 9 | 0.1111 | 33 | 0.0606 |
| 4_3394735 | 9 | 0.1111 | 24 | 0.125 | 11 | 0 | 8 | 0 | 52 | 0.0769 | 10 | 0.3 | 3 | 0 | 10 | 0.1 | 14 | 0.0714 | 37 | 0.1351 |
| 4_3394745 | 11 | 0.0909 | 28 | 0.1429 | 11 | 0 | 8 | 0.125 | 58 | 0.1034 | 10 | 0 | 4 | 0 | 16 | 0.1875 | 18 | 0.1667 | 48 | 0.125 |
| 4_3394821 | 8 | 0.375 | 26 | 0.1923 | 12 | 0.0833 | 8 | 0.125 | 54 | 0.1852 | 10 | 0.4 | 5 | 0.6 | 13 | 0 | 17 | 0.2353 | 45 | 0.2444 |
| 4_3455107 | 10 | 0.8 | 12 | 0.5833 | 10 | 0.3 | 8 | 0.625 | 40 | 0.575 | 6 | 1 | 14 | 0.7857 | 14 | 0.9286 | 8 | 0.875 | 42 | 0.881 |
| 4_3456642 | 19 | 0.4211 | 15 | 0.4 | 10 | 0.2 | 11 | 0.6364 | 55 | 0.4182 | 9 | 0.7778 | 15 | 0.4667 | 17 | 0.4118 | 10 | 0.2 | 51 | 0.451 |
| 4_3456672 | 19 | 0.1053 | 15 | 0.2 | 11 | 0.1818 | 10 | 0 | 55 | 0.1273 | 10 | 0.2 | 13 | 0.2308 | 15 | 0.1333 | 10 | 0 | 48 | 0.1458 |
| 4_3456937 | 2 | 0.5 | 1 | 0 | NA | NA | NA | NA | 3 | 0.3333 | NA | NA | NA | NA | NA | NA | 2 | 0.5 | 2 | 0.5 |
| 4_3457005 | 5 | 0 | NA | NA | 2 | 0.5 | NA | NA | 7 | 0.1429 | NA | NA | 1 | 0 | 1 | 0 | 3 | 0.3333 | 5 | 0.2 |
| 4_3457098 | 4 | 0.75 | 5 | 0 | 2 | 0 | 3 | 0 | 14 | 0.2143 | 5 | 0 | 3 | 0 | 8 | 0 | 3 | 0.6667 | 19 | 0.1053 |
| 4_3457966 | 1 | 1 | 2 | 0.5 | 4 | 0.25 | NA | NA | 7 | 0.4286 | 2 | 0 | 1 | 0 | 2 | 0.5 | 2 | 0.5 | 7 | 0.2857 |
| 4_3457998 | 1 | 0 | 3 | 0 | 4 | 0.25 | NA | NA | 8 | 0.125 | 2 | 0.5 | 1 | 0 | 2 | 0 | 2 | 0.5 | 7 | 0.2857 |
| 4_3458050 | 2 | 0.5 | 4 | 0 | 5 | 0.6 | NA | NA | 11 | 0.3636 | 2 | 0.5 | 1 | 0 | 2 | 0.5 | 2 | 0.5 | 7 | 0.4286 |
| 4_3458052 | 2 | 0 | 4 | 0.25 | 4 | 0 | NA | NA | 10 | 0.1 | 2 | 0 | 1 | 0 | 2 | 0.5 | 2 | 0 | 7 | 0.1429 |
| 4_3466390 | 2 | 0 | NA | NA | NA | NA | 1 | 1 | 3 | 0.3333 | 1 | 0 | 4 | 0.25 | 6 | 0.3333 | 3 | 0.6667 | 14 | 0.3571 |
| 4_3466407 | 2 | 0.5 | NA | NA | 1 | 1 | 1 | 1 | 4 | 0.75 | 2 | 1 | 4 | 1 | 6 | 1 | 4 | 0.75 | 16 | 0.9375 |
| 4_3476743 | NA | NA | NA | NA | 1 | 1 | NA | NA | 1 | 1 | NA | NA | NA | NA | 1 | 1 | NA | NA | 1 | 1 |
| 4_3476773 | NA | NA | NA | NA | 1 | 1 | NA | NA | 1 | 1 | NA | NA | NA | NA | 1 | 1 | NA | NA | 1 | 1 |
| 4_3476841 | NA | NA | NA | NA | 1 | 1 | NA | NA | 1 | 1 | NA | NA | NA | NA | 1 | 1 | NA | NA | 1 | 1 |
| 4_3476893 | 4 | 1 | 4 | 0.75 | 4 | 1 | 3 | 1 | 15 | 0.9333 | 6 | 0.8333 | 4 | 0.75 | 6 | 0.8333 | 2 | 1 | 18 | 0.8333 |
| 4_3476899 | 2 | 0.5 | 4 | 0.25 | 5 | 0.8 | 4 | 0.75 | 15 | 0.6 | 6 | 0.6667 | 1 | 0 | 6 | 0.8333 | 2 | 0.5 | 15 | 0.6667 |
| 4_3476905 | 2 | 0 | 5 | 0 | 5 | 0 | 4 | 0.25 | 16 | 0.0625 | 6 | 0 | 4 | 0.25 | 6 | 0.1667 | 3 | 0 | 19 | 0.1053 |
| 4_3480331 | 3 | 0.6667 | 2 | 0.5 | 1 | 0 | 3 | 0.6667 | 9 | 0.5556 | NA | NA | 3 | 0.3333 | 1 | 0 | 1 | 1 | 5 | 0.4 |
| 4_3480359 | 2 | 1 | 2 | 1 | 3 | 0.3333 | 2 | 1 | 9 | 0.7778 | NA | NA | 3 | 0.6667 | 1 | 0 | 2 | 1 | 6 | 0.6667 |
| 4_3480420 | 8 | 0.25 | 3 | 0.6667 | 5 | 0 | 5 | 0.2 | 21 | 0.2381 | NA | NA | 8 | 0.5 | 4 | 0.5 | 6 | 0.3333 | 18 | 0.4444 |
| 4_3480446 | 5 | 0 | 3 | 0.3333 | 5 | 0.2 | 5 | 0 | 18 | 0.1111 | NA | NA | 7 | 0 | 1 | 0 | 6 | 0.5 | 14 | 0.2143 |
| 4_3480448 | 4 | 0.5 | 3 | 0.3333 | 6 | 0.1667 | 5 | 0.4 | 18 | 0.3333 | NA | NA | 8 | 0.125 | 4 | 0.25 | 6 | 0.3333 | 18 | 0.2222 |
| 4_3480478 | 6 | 0.8333 | 2 | 1 | 5 | 1 | 4 | 0.75 | 17 | 0.8824 | NA | NA | 7 | 1 | 4 | 1 | 6 | 1 | 17 | 1 |
| 4_3480480 | 4 | 0.75 | 2 | 1 | 4 | 0.75 | 3 | 1 | 13 | 0.8462 | NA | NA | 6 | 1 | 3 | 0.6667 | 6 | 1 | 15 | 0.9333 |
| 4_3480495 | 5 | 0 | 1 | 0 | 4 | 0.25 | 3 | 0 | 13 | 0.0769 | NA | NA | 6 | 0 | 3 | 0.3333 | 3 | 0 | 12 | 0.0833 |
| 4_3480501 | 5 | 1 | 2 | 1 | 5 | 1 | 4 | 1 | 16 | 1 | NA | NA | 7 | 0.8571 | 4 | 1 | 5 | 1 | 16 | 0.9375 |
| 4_3480504 | 4 | 0.5 | 2 | 0 | 3 | 0.3333 | 1 | 0 | 10 | 0.3 | NA | NA | 5 | 0.2 | 1 | 1 | 4 | 0.75 | 10 | 0.5 |
| 4_3480504 | 1 | 1 | NA | NA | 1 | 0 | 2 | 0.5 | 4 | 0.5 | NA | NA | 1 | 1 | 1 | 1 | NA | NA | 2 | 1 |
| 4_3480506 | 6 | 0.1667 | 2 | 0 | 5 | 0 | 4 | 0 | 17 | 0.0588 | NA | NA | 7 | 0.1429 | 4 | 0 | 5 | 0.4 | 16 | 0.1875 |
| 4_3480511 | 7 | 0.7143 | 2 | 1 | 5 | 1 | 3 | 0.6667 | 17 | 0.8235 | NA | NA | 6 | 0.8333 | 3 | 0.6667 | 5 | 0.8 | 14 | 0.7857 |

| | | | | | | | | | | | | | | | | | | | | |
|---|---|---|---|---|---|---|---|---|---|---|---|---|---|---|---|---|---|---|---|---|
| 4_3480519 | 4 | 1 | 2 | 0.5 | 5 | 1 | 3 | 1 | 14 | 0.9286 | NA | NA | 5 | 1 | 2 | 1 | 5 | 1 | 12 | 1 |
| 4_3480524 | 7 | 0.1429 | 2 | 0 | 4 | 0.25 | 2 | 0 | 15 | 0.1333 | NA | NA | 6 | 0 | 2 | 0.5 | 5 | 0 | 13 | 0.0769 |
| 4_3480529 | 7 | 0.5714 | 2 | 1 | 3 | 1 | 4 | 0.5 | 16 | 0.6875 | NA | NA | 6 | 0.8333 | 3 | 0.3333 | 5 | 1 | 14 | 0.7857 |
| 4_3480530 | 6 | 0.1667 | 2 | 0.5 | 5 | 0.6 | 4 | 0 | 17 | 0.2941 | NA | NA | 5 | 0.4 | 3 | 0.3333 | 5 | 0.4 | 13 | 0.3846 |
| 4_3480538 | 3 | 0.6667 | 1 | 0 | 5 | 0.2 | 4 | 0.25 | 13 | 0.3077 | NA | NA | 6 | 0 | 2 | 0 | 5 | 0.2 | 13 | 0.0769 |
| 4_3480548 | 3 | 0.3333 | 1 | 0 | 3 | 0.3333 | 4 | 0.25 | 11 | 0.2727 | NA | NA | 4 | 0.5 | 2 | 0.5 | 4 | 0.5 | 10 | 0.5 |
| 4_3480552 | 4 | 0.25 | NA | NA | 5 | 0.2 | 3 | 0 | 12 | 0.1667 | NA | NA | 4 | 0 | 3 | 0.3333 | 2 | 0.5 | 9 | 0.2222 |
| 4_3480556 | 5 | 0.4 | 2 | 0.5 | 5 | 0.2 | 4 | 0.25 | 16 | 0.3125 | NA | NA | 4 | 0.5 | 3 | 0 | 5 | 0.4 | 12 | 0.3333 |
| 4_3480591 | 5 | 0.4 | 1 | 1 | 3 | 0.6667 | 2 | 1 | 11 | 0.6364 | NA | NA | 5 | 0.8 | 2 | 0.5 | 4 | 0.25 | 11 | 0.5455 |
| 4_3480613 | 5 | 0 | 1 | 0 | 4 | 0.25 | 3 | 0 | 13 | 0.0769 | NA | NA | 4 | 0.25 | 2 | 0 | 4 | 0 | 10 | 0.1 |
| 4_3480635 | 3 | 1 | NA | NA | 4 | 0.75 | 3 | 1 | 10 | 0.9 | NA | NA | 5 | 1 | 2 | 1 | 3 | 1 | 10 | 1 |
| 4_3480638 | 3 | 0 | 1 | 0 | 5 | 0.4 | 3 | 0.6667 | 12 | 0.3333 | NA | NA | 5 | 0.2 | 3 | 0.3333 | 3 | 0.3333 | 11 | 0.2727 |
| 4_3480659 | 4 | 1 | 1 | 1 | 4 | 1 | 3 | 0.6667 | 12 | 0.9167 | NA | NA | 4 | 1 | 2 | 1 | 3 | 1 | 9 | 1 |
| 4_3480664 | 4 | 0.5 | 1 | 0 | 4 | 0 | 3 | 0 | 12 | 0.1667 | NA | NA | 4 | 0.25 | 2 | 0 | 3 | 0 | 9 | 0.1111 |
| 4_3480679 | 3 | 0 | 1 | 0 | 3 | 0.3333 | 3 | 0.3333 | 10 | 0.2 | NA | NA | 4 | 0.5 | 3 | 0 | 1 | 0 | 8 | 0.25 |
| 4_3480681 | 2 | 1 | 1 | 0 | 4 | 0.25 | 2 | 1 | 9 | 0.5556 | NA | NA | 1 | 1 | 2 | 1 | 3 | 1 | 6 | 1 |
| 4_3480707 | 3 | 0.3333 | 1 | 0 | 3 | 0 | 1 | 0 | 8 | 0.125 | NA | NA | 3 | 0.6667 | 2 | 0.5 | 3 | 0.6667 | 8 | 0.625 |
| 4_3480725 | 1 | 1 | 1 | 0 | 2 | 0 | 3 | 0 | 7 | 0.1429 | NA | NA | 3 | 0.3333 | 2 | 0 | 2 | 0 | 7 | 0.1429 |
| 4_3480758 | 2 | 0 | 1 | 1 | 2 | 0 | 2 | 0.5 | 7 | 0.2857 | NA | NA | 1 | 1 | 3 | 0 | 1 | 0 | 5 | 0.2 |
| 4_3480760 | NA | NA | 1 | 1 | 2 | 1 | 2 | 0.5 | 5 | 0.8 | NA | NA | NA | NA | 3 | 1 | NA | NA | 3 | 1 |
| 4_3480783 | 1 | 1 | 1 | 1 | 2 | 0 | 2 | 0.5 | 6 | 0.5 | NA | NA | 1 | 0 | 1 | 0 | 1 | 1 | 3 | 0.3333 |
| 4_3480784 | NA | NA | 1 | 0 | 2 | 1 | 2 | 0.5 | 5 | 0.6 | NA | NA | 1 | 0 | 1 | 0 | 1 | 1 | 3 | 0.3333 |
| 4_3480799 | NA | NA | 1 | 1 | 2 | 1 | 2 | 0 | 5 | 0.6 | NA | NA | 1 | 1 | NA | NA | 1 | 1 | 2 | 1 |
| 4_3480836 | NA | NA | 1 | 1 | 1 | 1 | 1 | 1 | 3 | 1 | NA | NA | 1 | 1 | NA | NA | 1 | 1 | 2 | 1 |
| 4_3480848 | NA | NA | 1 | 1 | 1 | 1 | 1 | 1 | 3 | 1 | NA | NA | 1 | 1 | NA | NA | 1 | 1 | 2 | 1 |
| 4_3480861 | NA | NA | 1 | 0 | 1 | 0 | 1 | 1 | 3 | 0.3333 | NA | NA | 1 | 1 | NA | NA | 1 | 0 | 2 | 0.5 |
| 4_3480886 | NA | NA | 1 | 0 | 1 | 0 | 1 | 1 | 3 | 0.3333 | NA | NA | 1 | 0 | NA | NA | 1 | 1 | 2 | 0.5 |
| 4_3480926 | NA | NA | 1 | 1 | 1 | 0 | NA | NA | 2 | 0.5 | NA | NA | 1 | 1 | NA | NA | NA | NA | 1 | 1 |
| 4_3480998 | NA | NA | 1 | 1 | NA | NA | NA | NA | 1 | 1 | NA | NA | 1 | 1 | NA | NA | NA | NA | 1 | 1 |
| 4_3481126 | NA | NA | 1 | 1 | NA | NA | NA | NA | 1 | 1 | NA | NA | 1 | 1 | NA | NA | NA | NA | 1 | 1 |
| 4_3495377 | NA | NA | 1 | 1 | NA | NA | NA | NA | 1 | 1 | NA | NA | NA | NA | 1 | 1 | NA | NA | 1 | 1 |
| 4_3508399 | NA | NA | 1 | 1 | NA | NA | NA | NA | 1 | 1 | NA | NA | NA | NA | 1 | 1 | NA | NA | 1 | 1 |
| 4_3519271 | 2 | 0 | 4 | 0.25 | 1 | 1 | 5 | 0 | 12 | 0.1667 | 3 | 0.3333 | 2 | 0.5 | 2 | 0.5 | NA | NA | 7 | 0.4286 |
| 4_3519289 | 3 | 0 | 4 | 0.25 | 1 | 1 | 3 | 0 | 11 | 0.1818 | 4 | 0.25 | 2 | 0.5 | 2 | 0.5 | NA | NA | 8 | 0.375 |
| 4_3521852 | 3 | 0.6667 | 4 | 1 | 1 | 1 | 3 | 1 | 11 | 0.9091 | 2 | 1 | 2 | 1 | 2 | 1 | NA | NA | 6 | 1 |
| 4_3521868 | 2 | 0.5 | 2 | 0.5 | 1 | 0 | 1 | 0 | 6 | 0.3333 | NA | NA | NA | NA | 1 | 1 | NA | NA | 1 | 1 |
| 4_3521871 | 3 | 0.3333 | 4 | 0.5 | NA | NA | 3 | 0.3333 | 10 | 0.4 | 1 | 0 | 2 | 0 | 2 | 0.5 | NA | NA | 5 | 0.2 |
| 4_3570787 | NA | NA | 1 | 0 | 2 | 1 | NA | NA | 3 | 0.6667 | NA | NA | 1 | 1 | 1 | 1 | NA | NA | 2 | 1 |
| 4_3570800 | NA | NA | 1 | 1 | 3 | 1 | NA | NA | 4 | 1 | NA | NA | 1 | 1 | NA | NA | NA | NA | 1 | 1 |
| 4_3570809 | NA | NA | 1 | 1 | 1 | 1 | NA | NA | 2 | 1 | NA | NA | 1 | 1 | 1 | 0 | NA | NA | 2 | 0.5 |
| 4_3570824 | NA | NA | 1 | 0 | 2 | 0.5 | NA | NA | 3 | 0.3333 | NA | NA | 1 | 1 | NA | NA | NA | NA | 1 | 1 |
| 4_3570836 | NA | NA | 1 | 1 | 1 | 1 | NA | NA | 2 | 1 | NA | NA | 1 | 1 | 1 | 1 | NA | NA | 2 | 1 |
| 4_3570854 | NA | NA | NA | NA | 2 | 0.5 | NA | NA | 2 | 0.5 | NA | NA | 1 | 1 | NA | NA | NA | NA | 1 | 1 |
| 4_3570959 | NA | NA | 1 | 1 | 2 | 1 | NA | NA | 3 | 1 | NA | NA | 1 | 1 | NA | NA | NA | NA | 1 | 1 |
| 4_3570971 | NA | NA | 1 | 1 | 1 | 1 | NA | NA | 2 | 1 | NA | NA | 1 | 1 | NA | NA | NA | NA | 1 | 1 |
| 4_3570979 | NA | NA | 1 | 1 | 1 | 1 | NA | NA | 2 | 1 | NA | NA | 1 | 0 | 1 | 1 | NA | NA | 2 | 0.5 |
| 4_3698716 | 1 | 1 | 1 | 1 | NA | NA | NA | NA | 2 | 1 | 2 | 0.5 | NA | NA | 1 | 1 | NA | NA | 3 | 0.6667 |
| 4_3698743 | 1 | 1 | 1 | 1 | NA | NA | NA | NA | 2 | 1 | 2 | 1 | NA | NA | 1 | 1 | NA | NA | 3 | 1 |
| 4_4019793 | 4 | 0.5 | 5 | 0.6 | 5 | 0.8 | 3 | 1 | 17 | 0.7059 | 4 | 0.75 | NA | NA | 2 | 0 | NA | NA | 6 | 0.5 |
| 4_4019809 | 4 | 0.25 | 5 | 0 | 4 | 0 | 3 | 0 | 16 | 0.0625 | 3 | 0 | 2 | 0.5 | 3 | 0 | NA | NA | 8 | 0.125 |
| 4_4019810 | 4 | 0 | 5 | 0.2 | 4 | 0 | 4 | 0 | 17 | 0.0588 | 4 | 0 | 2 | 0 | 3 | 0.3333 | NA | NA | 9 | 0.1111 |
| 4_4019828 | 4 | 0 | 5 | 0 | 4 | 0.5 | 3 | 0.3333 | 16 | 0.1875 | 4 | 0 | 2 | 0.5 | 3 | 0.3333 | NA | NA | 9 | 0.2222 |
| 4_4019866 | NA | NA | NA | NA | 1 | 1 | 1 | 1 | 2 | 1 | NA | NA | 2 | 1 | NA | NA | NA | NA | 2 | 1 |
| 4_4019867 | NA | NA | NA | NA | 1 | 1 | NA | NA | 1 | 1 | NA | NA | 2 | 1 | NA | NA | NA | NA | 2 | 1 |
| 4_4020212 | NA | NA | NA | NA | NA | NA | 1 | 1 | 1 | 1 | NA | NA | 1 | 1 | NA | NA | NA | NA | 1 | 1 |
| 4_4020254 | NA | NA | NA | NA | NA | NA | 1 | 1 | 1 | 1 | NA | NA | 1 | 1 | NA | NA | NA | NA | 1 | 1 |
| 4_4025963 | 4 | 0.25 | 2 | 0 | 1 | 0 | 4 | 0 | 11 | 0.0909 | 5 | 0.2 | 2 | 0.5 | 5 | 0 | 2 | 0 | 14 | 0.1429 |
| 4_4025966 | 4 | 0.25 | 2 | 0 | 1 | 1 | 2 | 0 | 9 | 0.2222 | 4 | 0.25 | 3 | 0.3333 | 5 | 0.2 | 3 | 0.3333 | 15 | 0.2667 |
| 4_4025980 | 2 | 1 | 3 | 0 | NA | NA | 4 | 0 | 9 | 0.2222 | 6 | 0.1667 | 3 | 0.3333 | 5 | 0.4 | 3 | 0.3333 | 17 | 0.2941 |
| 4_4026009 | 3 | 0.6667 | 3 | 1 | NA | NA | 3 | 0 | 9 | 0.5556 | 3 | 0.6667 | 1 | 1 | 2 | 1 | 2 | 1 | 8 | 0.875 |
| 4_4026009 | 1 | 1 | NA | NA | NA | NA | 1 | 1 | 2 | 1 | 1 | 1 | NA | NA | 1 | 1 | NA | NA | 2 | 1 |
| 4_4026017 | 3 | 1 | 3 | 1 | NA | NA | 4 | 1 | 10 | 1 | 5 | 1 | 2 | 1 | 3 | 1 | 3 | 1 | 13 | 1 |
| 4_4026019 | 3 | 1 | 2 | 1 | NA | NA | 4 | 1 | 9 | 1 | 4 | 1 | 2 | 1 | 4 | 1 | 3 | 1 | 13 | 1 |
| 4_4026022 | 3 | 0.6667 | 1 | 1 | NA | NA | 2 | 0 | 6 | 0.5 | 4 | 1 | 2 | 0.5 | 3 | 0.6667 | 2 | 0.5 | 11 | 0.7273 |
| 4_4026033 | 2 | 1 | 2 | 1 | NA | NA | 4 | 1 | 8 | 1 | 4 | 1 | 2 | 1 | 3 | 0.6667 | 2 | 0.5 | 11 | 0.8182 |

| | | | | | | | | | | | | | | | | | | | | |
|---|---|---|---|---|---|---|---|---|---|---|---|---|---|---|---|---|---|---|---|---|
| 4_4026035 | 3 | 0 | 2 | 0.5 | NA | NA | 4 | 0 | 9 | 0.1111 | 5 | 0.4 | 2 | 0 | 4 | 0.5 | 2 | 0 | 13 | 0.3077 |
| 4_4026037 | 3 | 0.6667 | 2 | 0.5 | NA | NA | 3 | 0.3333 | 8 | 0.5 | 5 | 0.2 | 2 | 0.5 | 1 | 0 | 2 | 0 | 10 | 0.2 |
| 4_4026039 | 3 | 0.3333 | 2 | 0.5 | NA | NA | 4 | 0.25 | 9 | 0.3333 | 5 | 0.4 | 2 | 0 | 4 | 0.25 | 3 | 0.6667 | 14 | 0.3571 |
| 4_4026041 | 3 | 1 | 2 | 1 | NA | NA | 4 | 1 | 9 | 1 | 4 | 1 | 2 | 1 | 2 | 1 | 3 | 1 | 11 | 1 |
| 4_4026042 | 2 | 0.5 | 2 | 0 | NA | NA | 4 | 0.5 | 8 | 0.375 | 4 | 0.5 | 1 | 1 | 4 | 0.5 | 3 | 0.3333 | 12 | 0.5 |
| 4_4026066 | 3 | 0.6667 | 2 | 0.5 | NA | NA | 4 | 0 | 9 | 0.3333 | 4 | 0.25 | 2 | 0 | 3 | 0 | 3 | 0 | 12 | 0.0833 |
| 4_4026073 | 2 | 0 | 2 | 0.5 | NA | NA | 4 | 0.25 | 8 | 0.25 | 3 | 0.3333 | 2 | 0.5 | 4 | 0 | 3 | 0 | 12 | 0.1667 |
| 4_4026082 | 2 | 1 | 1 | 1 | NA | NA | 3 | 1 | 6 | 1 | 3 | 0.6667 | NA | NA | 3 | 1 | 3 | 1 | 9 | 0.8889 |
| 4_4026095 | 2 | 1 | 2 | 0.5 | NA | NA | 3 | 0.3333 | 7 | 0.5714 | 3 | 0 | 1 | 0 | 2 | 1 | 3 | 0.3333 | 9 | 0.3333 |
| 4_4026108 | 2 | 0.5 | 2 | 0 | NA | NA | 4 | 0.25 | 8 | 0.25 | 3 | 0 | 1 | 1 | 3 | 0 | 3 | 0.3333 | 10 | 0.2 |
| 4_4026111 | 2 | 0.5 | 2 | 1 | NA | NA | 4 | 0.75 | 8 | 0.75 | 3 | 0.3333 | 1 | 0 | 4 | 0.75 | 3 | 1 | 11 | 0.6364 |
| 4_4026114 | 2 | 1 | 2 | 0.5 | NA | NA | 4 | 0.5 | 8 | 0.625 | 2 | 0.5 | 1 | 0 | 4 | 0.75 | 2 | 0.5 | 9 | 0.5556 |
| 4_4026130 | 1 | 1 | 2 | 0 | NA | NA | 4 | 0 | 7 | 0.1429 | 3 | 0 | 1 | 1 | 4 | 0 | 2 | 0 | 10 | 0.1 |
| 4_4026135 | 2 | 0.5 | 2 | 0.5 | NA | NA | 3 | 0 | 7 | 0.2857 | 3 | 0 | 1 | 0 | 3 | 0.3333 | 2 | 0 | 9 | 0.1111 |
| 4_4026141 | 2 | 0 | 2 | 0 | NA | NA | 3 | 0.3333 | 7 | 0.1429 | 3 | 0.6667 | 1 | 0 | 3 | 0 | 2 | 0 | 9 | 0.2222 |
| 4_4026165 | 2 | 0.5 | 2 | 0 | NA | NA | 4 | 0.25 | 8 | 0.25 | 3 | 0 | 1 | 0 | 3 | 0.3333 | 2 | 0 | 9 | 0.1111 |
| 4_4026179 | 2 | 1 | 1 | 1 | NA | NA | 3 | 1 | 6 | 1 | 3 | 1 | 1 | 1 | 2 | 1 | 1 | 1 | 7 | 1 |
| 4_4026180 | 2 | 1 | 1 | 0 | NA | NA | 4 | 0.25 | 7 | 0.4286 | 3 | 0.3333 | NA | NA | 3 | 0.3333 | 2 | 0.5 | 8 | 0.375 |
| 4_4026180 | 2 | 0.5 | 2 | 0.5 | NA | NA | 3 | 0 | 7 | 0.2857 | 2 | 0 | 1 | 0 | 3 | 0.3333 | 2 | 1 | 8 | 0.375 |
| 4_4026187 | 2 | 0.5 | 2 | 0.5 | NA | NA | 3 | 0.6667 | 7 | 0.5714 | 3 | 0.3333 | NA | NA | 3 | 0.6667 | 2 | 0.5 | 8 | 0.5 |
| 4_4026201 | 2 | 0.5 | 1 | 0 | NA | NA | 3 | 0 | 6 | 0.1667 | 3 | 0 | 1 | 0 | 2 | 0.5 | 2 | 0 | 8 | 0.125 |
| 4_4026223 | NA | NA | 2 | 1 | NA | NA | 2 | 1 | 4 | 1 | 2 | 1 | NA | NA | 1 | 1 | 1 | 1 | 4 | 1 |
| 4_4026225 | 2 | 0 | 2 | 1 | NA | NA | 3 | 0 | 7 | 0.2857 | 1 | 0 | NA | NA | 2 | 0.5 | 1 | 0 | 4 | 0.25 |
| 4_4026236 | 1 | 1 | 1 | 1 | NA | NA | 2 | 0.5 | 4 | 0.75 | 2 | 0.5 | NA | NA | 3 | 1 | 1 | 0 | 6 | 0.6667 |
| 4_4026249 | 2 | 1 | 1 | 1 | NA | NA | 3 | 0.6667 | 6 | 0.8333 | 2 | 1 | NA | NA | 2 | 1 | 1 | 1 | 5 | 1 |
| 4_4026260 | 2 | 1 | 2 | 0 | NA | NA | 3 | 0 | 7 | 0.2857 | 1 | 1 | NA | NA | 2 | 0.5 | 1 | 0 | 4 | 0.5 |
| 4_4026273 | 1 | 1 | 1 | 0 | NA | NA | 3 | 0 | 5 | 0.2 | 1 | 0 | NA | NA | 2 | 0.5 | 1 | 1 | 4 | 0.5 |
| 4_4026304 | 2 | 1 | 2 | 0.5 | NA | NA | 2 | 0.5 | 6 | 0.6667 | 1 | 1 | NA | NA | 2 | 0.5 | NA | NA | 3 | 0.6667 |
| 4_4027014 | 5 | 0.4 | 1 | 0 | 2 | 0.5 | 3 | 0 | 11 | 0.2727 | 2 | 0 | 1 | 0 | 5 | 0.4 | 2 | 0.5 | 10 | 0.3 |
| 4_4047425 | NA | NA | NA | NA | NA | NA | 2 | 1 | 2 | 1 | NA | NA | NA | NA | NA | NA | 1 | 1 | 1 | 1 |
| 4_4047447 | NA | NA | NA | NA | NA | NA | 2 | 1 | 2 | 1 | NA | NA | NA | NA | NA | NA | 1 | 1 | 1 | 1 |
| 4_4047466 | NA | NA | NA | NA | NA | NA | 2 | 0.5 | 2 | 0.5 | NA | NA | NA | NA | NA | NA | 1 | 1 | 1 | 1 |
| 4_4047484 | NA | NA | NA | NA | NA | NA | 1 | 1 | 1 | 1 | NA | NA | NA | NA | NA | NA | 1 | 1 | 1 | 1 |
| 4_4068629 | 2 | 0.5 | 1 | 0 | 3 | 0.3333 | 2 | 0.5 | 8 | 0.375 | 3 | 0.3333 | 2 | 0 | 2 | 0.5 | 5 | 0 | 12 | 0.1667 |
| 4_4068632 | 2 | 0 | 1 | 0 | 3 | 0.3333 | 3 | 0 | 9 | 0.1111 | 3 | 0 | 2 | 0 | 2 | 0 | 5 | 0.4 | 12 | 0.1667 |
| 4_4068634 | 2 | 0 | 1 | 0 | 3 | 0.3333 | 3 | 0 | 9 | 0.1111 | 2 | 0 | 2 | 0.5 | 2 | 0 | 5 | 0.2 | 11 | 0.1818 |
| 4_4068641 | 2 | 0.5 | 1 | 0 | 3 | 1 | 3 | 0.6667 | 9 | 0.6667 | 1 | 1 | 1 | 0 | 2 | 0.5 | 3 | 1 | 7 | 0.7143 |
| 4_4068649 | 2 | 0 | 1 | 0 | 2 | 0.5 | 2 | 0.5 | 7 | 0.2857 | 1 | 0 | 4 | 0 | 2 | 0 | 5 | 0.6 | 12 | 0.25 |
| 4_4068667 | 2 | 0.5 | 2 | 0 | 2 | 0.5 | 3 | 0.3333 | 9 | 0.3333 | 2 | 0 | 4 | 0 | 1 | 1 | 4 | 0.25 | 11 | 0.1818 |
| 4_4069504 | 2 | 0.5 | 1 | 0 | 3 | 0.3333 | 3 | 0.6667 | 9 | 0.4444 | 2 | 1 | 4 | 0 | 1 | 0 | 4 | 0.75 | 11 | 0.4545 |
| 4_4072110 | NA | NA | 1 | 1 | NA | NA | NA | NA | 1 | 1 | NA | NA | 1 | 1 | NA | NA | NA | NA | 1 | 1 |
| 4_4073281 | 49 | 0.2653 | 59 | 0.2881 | 40 | 0.325 | 51 | 0.3137 | 199 | 0.2965 | 39 | 0.4359 | 55 | 0.2909 | 36 | 0.25 | 50 | 0.28 | 180 | 0.3111 |
| 4_4073283 | 49 | 0.2245 | 51 | 0.2549 | 38 | 0.2632 | 47 | 0.2128 | 185 | 0.2378 | 37 | 0.3514 | 48 | 0.3542 | 34 | 0.1765 | 45 | 0.2889 | 164 | 0.2988 |
| 4_4073284 | 54 | 0.4815 | 59 | 0.4407 | 40 | 0.375 | 51 | 0.3725 | 204 | 0.4216 | 42 | 0.381 | 52 | 0.4808 | 41 | 0.561 | 48 | 0.5 | 183 | 0.4809 |
| 4_4073299 | 58 | 0.6552 | 59 | 0.5932 | 38 | 0.5789 | 49 | 0.5714 | 204 | 0.6029 | 43 | 0.5814 | 54 | 0.6852 | 34 | 0.6176 | 50 | 0.56 | 181 | 0.6133 |
| 4_4075573 | 6 | 0.1667 | 5 | 0.2 | 4 | 0 | 4 | 0 | 19 | 0.1053 | 5 | 0 | 6 | 0.1667 | 8 | 0.125 | 4 | 0 | 23 | 0.087 |
| 4_4077529 | 4 | 0.25 | 4 | 0 | 3 | 0 | 1 | 0 | 12 | 0.0833 | 1 | 1 | 1 | 0 | 3 | 0 | 2 | 0 | 7 | 0.1429 |
| 4_4093617 | 2 | 0 | 2 | 0 | 1 | 1 | 1 | 0 | 6 | 0.1667 | NA | NA | NA | NA | 3 | 0.6667 | 1 | 1 | 4 | 0.75 |
| 4_4093627 | 2 | 0.5 | 3 | 0.6667 | 4 | 0.5 | 2 | 0.5 | 11 | 0.5455 | NA | NA | 1 | 0 | 4 | 0.5 | NA | NA | 5 | 0.4 |
| 4_4093641 | 2 | 0.5 | 3 | 1 | 5 | 0.8 | 2 | 1 | 12 | 0.8333 | NA | NA | 2 | 0 | 5 | 0.4 | 1 | 0 | 8 | 0.25 |
| 4_4096462 | 15 | 0.1333 | 17 | 0.0588 | 11 | 0.1818 | 14 | 0 | 57 | 0.0877 | 16 | 0.125 | 17 | 0.1176 | 16 | 0.0625 | 9 | 0 | 58 | 0.0862 |
| 4_4096469 | 17 | 0.1176 | 17 | 0 | 13 | 0.2308 | 14 | 0 | 61 | 0.082 | 17 | 0.1765 | 20 | 0.15 | 19 | 0.2105 | 10 | 0.1 | 66 | 0.1667 |
| 4_4096503 | 22 | 0.1364 | 19 | 0.2632 | 18 | 0.1111 | 17 | 0.0588 | 76 | 0.1447 | 15 | 0.2667 | 22 | 0.0909 | 22 | 0.2273 | 11 | 0.0909 | 70 | 0.1714 |
| 4_4096516 | 12 | 0 | 13 | 0 | 14 | 0.1429 | 13 | 0.0769 | 52 | 0.0577 | 13 | 0.1538 | 25 | 0 | 18 | 0.1111 | 7 | 0 | 63 | 0.0635 |
| 4_4097686 | 2 | 0 | 5 | 0 | NA | NA | 4 | 0.5 | 11 | 0.1818 | 2 | 0.5 | 3 | 0 | 5 | 0 | 4 | 0 | 14 | 0.0714 |
| 4_4097687 | 1 | 0 | 4 | 0 | NA | NA | 5 | 0.2 | 10 | 0.1 | 1 | 0 | 2 | 0 | 5 | 0.2 | 4 | 0 | 12 | 0.0833 |
| 4_4101161 | NA | NA | 1 | 1 | 1 | 0 | 1 | 1 | 3 | 0.6667 | NA | NA | 1 | 0 | 1 | 0 | 1 | 1 | 3 | 0.3333 |
| 4_4101194 | NA | NA | 1 | 1 | 1 | 1 | 2 | 1 | 4 | 1 | NA | NA | 1 | 1 | NA | NA | 1 | 1 | 2 | 1 |
| 4_4101208 | 4 | 0.5 | 6 | 0.5 | 4 | 0 | 6 | 0.8333 | 20 | 0.5 | 3 | 0 | 6 | 0.1667 | 11 | 0.6364 | 5 | 0.2 | 25 | 0.36 |
| 4_4101209 | 5 | 0.6 | 6 | 0.1667 | 5 | 0.4 | 8 | 0.5 | 24 | 0.4167 | 3 | 0.3333 | 7 | 0.4286 | 13 | 0.2308 | 5 | 0.4 | 28 | 0.3214 |
| 4_4101210 | 4 | 0.25 | 6 | 0.5 | 5 | 0.8 | 8 | 0.625 | 23 | 0.5652 | 3 | 0.6667 | 7 | 0.5714 | 11 | 0.1818 | 5 | 0.8 | 26 | 0.4615 |
| 4_4101212 | 5 | 0.8 | 5 | 0.8 | 5 | 0.8 | 6 | 0.8333 | 21 | 0.8095 | 3 | 0.6667 | 6 | 1 | 13 | 0.7692 | 5 | 0.8 | 27 | 0.8148 |
| 4_4101215 | 4 | 0.5 | 8 | 0.625 | 4 | 0.75 | 4 | 0.5 | 20 | 0.6 | 3 | 0.3333 | 5 | 1 | 9 | 0.6667 | 4 | 0.25 | 21 | 0.619 |
| 4_4101256 | 5 | 0.2 | 7 | 0.1429 | 7 | 0.1429 | 7 | 0.1429 | 26 | 0.1538 | 3 | 0.3333 | 7 | 0.1429 | 13 | 0.3846 | 5 | 0 | 28 | 0.25 |
| 4_4101529 | 1 | 0 | NA | NA | NA | NA | 4 | 0.25 | 5 | 0.2 | 2 | 0 | 2 | 0 | 2 | 0.5 | 2 | 0 | 8 | 0.125 |

| | | | | | | | | | | | | | | | | | | | | |
|---|---|---|---|---|---|---|---|---|---|---|---|---|---|---|---|---|---|---|---|---|
| 4_41024922 | 1 | 1 | NA | NA | NA | NA | 4 | 0 | 5 | 0.2 | 2 | 0 | 2 | 0.5 | 2 | 1 | 4 | 0.5 | 10 | 0.5 |
| 4_4102574 | NA | NA | NA | NA | 1 | 0 | 2 | 0.5 | 3 | 0.3333 | 1 | 0 | 3 | 0.3333 | 3 | 0 | 3 | 0.3333 | 10 | 0.2 |
| 4_4111454 | 12 | 0.0833 | 6 | 0.1667 | 12 | 0.4167 | 13 | 0.3077 | 43 | 0.2558 | 9 | 0.2222 | 15 | 0.0667 | 14 | 0.2857 | 11 | 0.0909 | 49 | 0.1633 |
| 4_41114708 | 13 | 0.3846 | 9 | 0.1111 | 10 | 0.3 | 10 | 0.1 | 42 | 0.2381 | 8 | 0.25 | 12 | 0.1667 | 16 | 0.125 | 11 | 0.3636 | 47 | 0.2128 |
| 4_4111508 | 13 | 0.6923 | 6 | 0.8333 | 11 | 0.9091 | 10 | 0.7 | 40 | 0.775 | 8 | 0.875 | 8 | 0.875 | 14 | 0.6429 | 8 | 0.625 | 38 | 0.7368 |
| 4_4111513 | 13 | 0.6154 | 7 | 0.8571 | 10 | 0.8 | 12 | 0.6667 | 42 | 0.7143 | 6 | 0.6667 | 12 | 0.6667 | 19 | 0.7368 | 8 | 0.75 | 45 | 0.7111 |
| 4_4111534 | 10 | 0.2 | 6 | 0 | 7 | 0.1429 | 8 | 0.125 | 31 | 0.129 | 7 | 0.1429 | 7 | 0.4286 | 11 | 0 | 6 | 0 | 31 | 0.129 |
| 4_4111536 | 11 | 0.8182 | 5 | 0.8 | 9 | 0.6667 | 10 | 0.7 | 35 | 0.7429 | 8 | 0.75 | 12 | 0.6667 | 15 | 0.4667 | 7 | 0.7143 | 42 | 0.619 |
| 4_4111542 | 11 | 0.9091 | 6 | 0.6667 | 6 | 1 | 8 | 0.625 | 31 | 0.8065 | 8 | 0.875 | 11 | 0.5455 | 11 | 0.5455 | 4 | 0.75 | 34 | 0.6471 |
| 4_4112037 | 3 | 0.3333 | NA | NA | 4 | 0.25 | 5 | 0 | 12 | 0.1667 | 3 | 0.3333 | 5 | 0 | 5 | 0 | 3 | 0 | 16 | 0.0625 |
| 4_4113595 | 8 | 0.375 | 5 | 0.4 | 4 | 0.25 | 8 | 0.125 | 25 | 0.28 | 5 | 0.2 | 7 | 0.4286 | 9 | 0.1111 | 7 | 0 | 28 | 0.1786 |
| 4_4113603 | 7 | 0.2857 | 2 | 0.5 | 6 | 0.1667 | 5 | 0.4 | 20 | 0.3 | 2 | 0 | 6 | 0.1667 | 6 | 0.5 | 5 | 0.4 | 19 | 0.3158 |
| 4_4119091 | 8 | 0.625 | 11 | 0.5455 | 14 | 0.5 | 7 | 0.4286 | 40 | 0.525 | 6 | 0.6667 | 7 | 0.4286 | 14 | 0.2143 | 10 | 0.3 | 37 | 0.3514 |
| 4_4119095 | 14 | 0.0714 | 13 | 0.0769 | 18 | 0.0556 | 14 | 0 | 59 | 0.0508 | 8 | 0.25 | 12 | 0.0833 | 22 | 0 | 14 | 0 | 56 | 0.0536 |
| 4_4119098 | 13 | 1 | 13 | 0.9231 | 15 | 1 | 14 | 1 | 55 | 0.9818 | 7 | 1 | 8 | 1 | 21 | 0.9048 | 9 | 1 | 45 | 0.9556 |
| 4_4119107 | 3 | 0 | 7 | 0 | 6 | 0 | 4 | 0.5 | 20 | 0.1 | 4 | 0 | 5 | 0 | 6 | 0 | 9 | 0.2222 | 24 | 0.0833 |
| 4_4119117 | 16 | 0.3125 | 15 | 0.2667 | 15 | 0.2667 | 15 | 0.0667 | 61 | 0.2295 | 9 | 0.5556 | 13 | 0.2308 | 20 | 0 | 15 | 0.1333 | 57 | 0.1754 |
| 4_4119132 | 15 | 0.2 | 13 | 0.1538 | 18 | 0.1667 | 15 | 0.2 | 61 | 0.1803 | 7 | 0 | 9 | 0.3333 | 14 | 0 | 16 | 0.125 | 46 | 0.1087 |
| 4_4119146 | 19 | 0.3158 | 15 | 0.3333 | 17 | 0.3529 | 19 | 0.3684 | 70 | 0.3429 | 9 | 0.6667 | 13 | 0.3846 | 21 | 0.5238 | 17 | 0.3529 | 60 | 0.4667 |
| 4_4119453 | 7 | 0.1429 | 3 | 0 | 4 | 0.75 | 7 | 0.4286 | 21 | 0.3333 | 5 | 0.4 | 6 | 0.1667 | 3 | 0.6667 | 2 | 0 | 16 | 0.3125 |
| 4_4119481 | 7 | 0.5714 | 2 | 0.5 | 3 | 1 | 6 | 0.6667 | 18 | 0.6667 | 2 | 0.5 | 4 | 0.25 | 2 | 1 | 2 | 1 | 10 | 0.6 |
| 4_4119515 | 5 | 0 | 4 | 0 | 3 | 0 | 6 | 0.1667 | 18 | 0.0556 | 3 | 0.3333 | 5 | 0 | 2 | 0 | 2 | 0 | 12 | 0.0833 |
| 4_4119515 | 6 | 0.1667 | 3 | 0.3333 | 1 | 0 | 5 | 0 | 15 | 0.1333 | 2 | 0.5 | 6 | 0.3333 | 2 | 0 | 2 | 0 | 12 | 0.25 |
| 4_4120691 | 2 | 1 | NA | NA | 2 | 0 | 5 | 0.2 | 9 | 0.3333 | 3 | 0.6667 | 1 | 0 | 1 | 0 | NA | NA | 5 | 0.4 |
| 4_4129222 | NA | NA | 8 | 0 | 4 | 0.25 | 7 | 0 | 19 | 0.0526 | 3 | 0 | 2 | 0.5 | 5 | 0 | 2 | 0 | 12 | 0.0833 |
| 4_4129306 | 2 | 0.5 | 12 | 0.5 | 5 | 0.6 | 8 | 0.625 | 27 | 0.5556 | 6 | 0.8333 | 4 | 0.75 | 7 | 0.8571 | 3 | 0.6667 | 20 | 0.8 |
| 4_4129318 | 2 | 0 | 9 | 0.1111 | 5 | 0 | 5 | 0.8 | 21 | 0.2381 | 7 | 0.2857 | 4 | 0.25 | 5 | 0.2 | 3 | 0.3333 | 19 | 0.2632 |
| 4_4131328 | 2 | 0 | 8 | 0.125 | 3 | 0 | 4 | 0.25 | 17 | 0.1176 | 3 | 0 | 3 | 0.3333 | 3 | 0 | 4 | 0 | 13 | 0.0769 |
| 4_4137910 | 3 | 0 | 3 | 0 | 3 | 0 | 2 | 1 | 11 | 0.1818 | 2 | 0.5 | 3 | 0 | 4 | 0 | 3 | 0.6667 | 12 | 0.25 |
| 4_4137924 | 3 | 0.3333 | 3 | 0.3333 | 3 | 0 | 3 | 0 | 12 | 0.1667 | 2 | 0 | 3 | 0.3333 | 4 | 0.25 | 3 | 0 | 12 | 0.1667 |
| 4_4137927 | 3 | 0.6667 | 3 | 0 | 3 | 0 | 3 | 0 | 12 | 0.1667 | 2 | 0.5 | 3 | 0.3333 | 5 | 0.4 | 3 | 0 | 13 | 0.3077 |
| 4_4137933 | 3 | 0.3333 | 2 | 0 | 4 | 0.25 | 3 | 0 | 12 | 0.1667 | 1 | 0 | 3 | 0.3333 | 4 | 0.5 | 3 | 0.3333 | 11 | 0.3636 |
| 4_4137935 | 3 | 1 | 3 | 0.6667 | 4 | 0.75 | 3 | 0.6667 | 13 | 0.7692 | 2 | 0.5 | 3 | 0.6667 | 5 | 0.4 | 3 | 1 | 13 | 0.6154 |
| 4_4137937 | 1 | 0 | 3 | 1 | NA | NA | 3 | 0.6667 | 7 | 0.7143 | 2 | 0.5 | 1 | 1 | 4 | 0.75 | 3 | 0.6667 | 10 | 0.7 |
| 4_4137945 | 4 | 0.25 | 3 | 0.6667 | 3 | 1 | 3 | 0.3333 | 13 | 0.5385 | 1 | 0 | 3 | 0.6667 | 5 | 0.2 | 3 | 0 | 12 | 0.25 |
| 4_4137960 | 3 | 0.6667 | 3 | 0.6667 | 4 | 1 | 2 | 1 | 12 | 0.8333 | 2 | 0.5 | 4 | 0.25 | 5 | 0.8 | 3 | 0 | 14 | 0.4286 |
| 4_4137973 | 4 | 0.75 | 4 | 0.25 | 4 | 0.5 | 3 | 0 | 15 | 0.4 | 2 | 0 | 4 | 0 | 4 | 0 | 2 | 0.5 | 12 | 0.0833 |
| 4_4137976 | 2 | 1 | 3 | 0.6667 | 2 | 1 | 2 | 0.5 | 9 | 0.7778 | 1 | 1 | 4 | 0.75 | 5 | 0.8 | 3 | 0.3333 | 13 | 0.6923 |
| 4_4138873 | 5 | 0.8 | 4 | 0.25 | 3 | 0.3333 | 3 | 0.6667 | 15 | 0.5333 | 2 | 0.5 | 3 | 0.6667 | 6 | 0.3333 | 4 | 0 | 15 | 0.3333 |
| 4_4138873 | 7 | 0.4286 | 5 | 0.2 | 4 | 0.75 | 3 | 0.6667 | 19 | 0.4737 | 3 | 0.3333 | 4 | 0 | 5 | 0.6 | 4 | 0.25 | 16 | 0.3125 |
| 4_4138877 | 7 | 0.7143 | 3 | 1 | 4 | 0.5 | 3 | 0.6667 | 17 | 0.7059 | 3 | 0.3333 | 3 | 0.6667 | 3 | 0.3333 | 4 | 0.75 | 13 | 0.5385 |
| 4_4138890 | 9 | 0.5556 | 4 | 0.5 | 6 | 0.3333 | 1 | 1 | 20 | 0.5 | 3 | 0.3333 | 5 | 0.8 | 4 | 0.5 | 4 | 0 | 16 | 0.4375 |
| 4_4138895 | 7 | 0.2857 | 4 | 0 | 6 | 0 | 3 | 0 | 20 | 0.1 | 4 | 0.25 | 5 | 0 | 5 | 0.2 | 3 | 0 | 17 | 0.1176 |
| 4_4138918 | 11 | 0.4545 | 3 | 0 | 5 | 0 | 4 | 0 | 23 | 0.2174 | 4 | 0.25 | 4 | 0 | 3 | 0 | 6 | 0 | 17 | 0.0588 |
| 4_4138923 | 10 | 0.3 | 3 | 0.3333 | 5 | 0 | 4 | 0.25 | 22 | 0.2273 | 4 | 0.25 | 4 | 0.75 | 3 | 0 | 6 | 0.3333 | 17 | 0.3529 |
| 4_4138941 | 12 | 0.4167 | 2 | 0 | 5 | 0.2 | 4 | 0.25 | 23 | 0.3043 | 3 | 1 | 4 | 0 | 3 | 1 | 7 | 0.2857 | 17 | 0.4706 |
| 4_4148029 | 3 | 0 | 3 | 0 | 3 | 0.3333 | 1 | 0 | 10 | 0.1 | 2 | 0 | 5 | 0.2 | 3 | 0 | 6 | 0.1667 | 16 | 0.125 |
| 4_4148041 | 3 | 0.3333 | 2 | 1 | 2 | 1 | 2 | 0 | 9 | 0.5556 | 2 | 0.5 | 5 | 0.8 | 4 | 0.5 | 6 | 0.6667 | 17 | 0.6471 |
| 4_4148053 | 4 | 0.5 | 4 | 0.5 | 2 | 0.5 | 1 | 0 | 11 | 0.4545 | 2 | 0 | 5 | 0.6 | 3 | 0.3333 | 7 | 0.2857 | 17 | 0.3529 |
| 4_4148076 | 3 | 0.3333 | 3 | 0.6667 | 2 | 0.5 | 2 | 1 | 10 | 0.6 | 2 | 0.5 | 4 | 0.5 | 4 | 0.5 | 6 | 0.1667 | 16 | 0.375 |
| 4_4148089 | 4 | 0 | 4 | 0 | 2 | 0.5 | 2 | 0 | 12 | 0.0833 | 3 | 0.3333 | 5 | 0 | 4 | 0 | 7 | 0 | 19 | 0.0526 |
| 4_4148094 | NA | NA | NA | NA | 1 | 1 | NA | NA | 1 | 1 | 1 | 1 | NA | NA | NA | NA | NA | NA | 1 | 1 |
| 4_4148210 | NA | NA | 1 | 1 | NA | NA | NA | NA | 1 | 1 | 1 | 1 | NA | NA | NA | NA | NA | NA | 1 | 1 |
| 4_4149628 | 7 | 0.1429 | 3 | 0 | 1 | 0 | 1 | 0 | 12 | 0.0833 | 2 | 0 | 2 | 0.5 | NA | NA | 2 | 0 | 6 | 0.1667 |
| 4_4149629 | 11 | 0.1818 | 7 | 0.1429 | 5 | 0 | 4 | 0 | 27 | 0.1111 | 3 | 0.3333 | 5 | 0.2 | 4 | 0.25 | 4 | 0.25 | 16 | 0.25 |
| 4_4149641 | 4 | 0.25 | 5 | 0 | 5 | 0.2 | 3 | 0.3333 | 17 | 0.1765 | 4 | 0 | 3 | 0.3333 | 3 | 0.3333 | 3 | 0 | 13 | 0.1538 |
| 4_4149652 | 8 | 0.125 | 6 | 0 | 2 | 0 | 2 | 0 | 18 | 0.0556 | 3 | 0 | 4 | 0.25 | 3 | 0 | 1 | 0 | 11 | 0.0909 |
| 4_4149659 | 9 | 0.8889 | 5 | 1 | 6 | 0.6667 | 2 | 1 | 22 | 0.8636 | 4 | 0.75 | 5 | 0.8 | 4 | 1 | 1 | 1 | 14 | 0.8571 |
| 4_4149661 | 8 | 0.5 | 5 | 0.8 | 3 | 0 | 2 | 0.5 | 18 | 0.5 | 3 | 1 | 6 | 0.1667 | 4 | 0.75 | 3 | 1 | 16 | 0.625 |
| 4_4149664 | 9 | 0.1111 | 5 | 0 | 5 | 0 | 2 | 0.5 | 21 | 0.0952 | 4 | 0.25 | 4 | 0 | 4 | 0 | 4 | 0 | 16 | 0.0625 |
| 4_4149668 | 8 | 0 | 4 | 0.25 | 3 | 0 | 1 | 0 | 16 | 0.0625 | 4 | 0 | 6 | 0 | 3 | 0 | 3 | 0.3333 | 16 | 0.0625 |
| 4_4149670 | 8 | 0.875 | 5 | 0.8 | 5 | 0.8 | 1 | 1 | 19 | 0.8421 | 3 | 1 | 6 | 0.8333 | 4 | 1 | 3 | 1 | 16 | 0.9375 |
| 4_4149672 | 6 | 0.8333 | 3 | 0.6667 | 6 | 0.3333 | 1 | 1 | 16 | 0.625 | 4 | 0.5 | 4 | 0 | 4 | 0.5 | 2 | 0 | 14 | 0.2857 |
| 4_4149681 | 6 | 0 | 4 | 0.25 | 4 | 0 | 2 | 0 | 16 | 0.0625 | 4 | 0.25 | 5 | 0.4 | 4 | 0.5 | 3 | 0 | 16 | 0.3125 |
| 4_4149693 | 7 | 0.8571 | 5 | 0.2 | 3 | 0 | 2 | 0.5 | 17 | 0.4706 | 3 | 0.3333 | 5 | 0.4 | 4 | 0.5 | 3 | 0.3333 | 15 | 0.4 |

| | | | | | | | | | | | | | | | | | | | | |
|---|---|---|---|---|---|---|---|---|---|---|---|---|---|---|---|---|---|---|---|---|
| 4_4149729 | 8 | 0.875 | 3 | 0.6667 | 6 | 0.8333 | 1 | 1 | 18 | 0.8333 | 3 | 1 | 5 | 0.8 | 3 | 0.3333 | 2 | 0.5 | 13 | 0.6923 |
| 4_4149734 | 8 | 0.875 | 4 | 1 | 4 | 0.75 | 2 | 1 | 18 | 0.8889 | 4 | 1 | 5 | 0.8 | 3 | 1 | 2 | 1 | 14 | 0.9286 |
| 4_4149739 | 6 | 0 | 4 | 0.25 | 4 | 0.25 | 2 | 0 | 16 | 0.125 | 3 | 0 | 5 | 0.2 | 2 | 0 | 1 | 0 | 11 | 0.0909 |
| 4_4149742 | 8 | 1 | 4 | 1 | 5 | 1 | 2 | 1 | 19 | 1 | 3 | 1 | 5 | 0.8 | 3 | 1 | 2 | 1 | 13 | 0.9231 |
| 4_4149746 | 7 | 0.1429 | 4 | 0 | 6 | 0 | 1 | 0 | 18 | 0.0556 | 4 | 0 | 5 | 0.4 | 3 | 0.3333 | 2 | 0 | 14 | 0.2143 |
| 4_4149755 | 7 | 1 | 4 | 0.75 | 5 | 0.8 | 2 | 0.5 | 18 | 0.8333 | 3 | 1 | 4 | 1 | 3 | 1 | 3 | 0.3333 | 13 | 0.8462 |
| 4_4149765 | 8 | 0.25 | 3 | 0.3333 | 6 | 0.5 | 2 | 0.5 | 19 | 0.3684 | 3 | 0.3333 | 5 | 0.2 | 2 | 0 | 2 | 0.5 | 12 | 0.25 |
| 4_4149767 | 6 | 0.8333 | 3 | 0.6667 | 4 | 0.75 | 1 | 1 | 14 | 0.7857 | 1 | 0 | 5 | 1 | 2 | 0.5 | 2 | 0 | 10 | 0.6 |
| 4_4149807 | 4 | 0.75 | 3 | 0.3333 | 4 | 0.25 | 2 | 0 | 13 | 0.3846 | NA | NA | 4 | 0.25 | 2 | 0 | 1 | 0 | 7 | 0.1429 |
| 4_4149810 | 6 | 0.3333 | 3 | 0 | 6 | 0 | 2 | 0 | 17 | 0.1176 | 2 | 0 | 4 | 0 | 2 | 0.5 | 1 | 0 | 9 | 0.1111 |
| 4_4149821 | 5 | 0.8 | 3 | 1 | 5 | 1 | 2 | 1 | 15 | 0.9333 | 2 | 1 | 5 | 1 | 2 | 0.5 | 2 | 1 | 11 | 0.9091 |
| 4_4149829 | 6 | 0.6667 | 3 | 0.6667 | 5 | 0.2 | 1 | 1 | 15 | 0.5333 | 1 | 0 | 4 | 0.5 | 2 | 1 | 2 | 1 | 9 | 0.6667 |
| 4_4149833 | 4 | 1 | 2 | 1 | 4 | 1 | 1 | 1 | 11 | 1 | 1 | 1 | 5 | 1 | 2 | 1 | 1 | 1 | 9 | 1 |
| 4_4149850 | 5 | 0 | 1 | 1 | 3 | 0 | 2 | 0.5 | 11 | 0.1818 | 2 | 0 | 3 | 0.3333 | 2 | 0 | 2 | 0 | 9 | 0.1111 |
| 4_4149864 | 4 | 0.75 | 3 | 0.3333 | 4 | 0.5 | 1 | 1 | 12 | 0.5833 | 1 | 1 | 2 | 0 | 2 | 0.5 | 1 | 0 | 6 | 0.3333 |
| 4_4149886 | 3 | 0.3333 | 2 | 0 | 4 | 1 | NA | NA | 9 | 0.5556 | 1 | 0 | 4 | 0.75 | 2 | 0.5 | 2 | 0.5 | 9 | 0.5556 |
| 4_4149917 | 2 | 1 | 2 | 1 | 2 | 1 | NA | NA | 6 | 1 | 1 | 1 | 2 | 1 | 2 | 1 | 2 | 1 | 7 | 1 |
| 4_4149918 | 3 | 0.6667 | 3 | 0.3333 | 3 | 0.6667 | NA | NA | 9 | 0.5556 | NA | NA | 3 | 0.6667 | 2 | 0.5 | 1 | 1 | 6 | 0.6667 |
| 4_4149935 | 3 | 0.3333 | 2 | 0 | 3 | 0.3333 | NA | NA | 8 | 0.25 | NA | NA | 3 | 0.3333 | 2 | 0 | 2 | 0 | 7 | 0.1429 |
| 4_4149944 | 3 | 1 | 2 | 1 | 3 | 1 | NA | NA | 8 | 1 | NA | NA | 3 | 1 | 2 | 1 | 1 | 1 | 6 | 1 |
| 4_4151883 | 16 | 0.25 | 21 | 0.2381 | 26 | 0.1538 | 10 | 0.1 | 73 | 0.1918 | 10 | 0.1 | 17 | 0.2941 | 10 | 0.2 | 15 | 0.1333 | 52 | 0.1923 |
| 4_4151888 | 17 | 0.2941 | 24 | 0.3333 | 25 | 0.28 | 11 | 0.5455 | 77 | 0.3377 | 10 | 0.5 | 16 | 0.375 | 9 | 0.1111 | 15 | 0.4 | 50 | 0.36 |
| 4_4151908 | 15 | 0.6 | 27 | 0.7037 | 23 | 0.5652 | 13 | 0.5385 | 78 | 0.6154 | 7 | 0.5714 | 16 | 0.6875 | 8 | 0.5 | 17 | 0.7059 | 48 | 0.6458 |
| 4_4152032 | 13 | 0.2308 | 23 | 0.0435 | 19 | 0.1053 | 12 | 0.0833 | 67 | 0.1045 | 6 | 0.1667 | 14 | 0.0714 | 8 | 0.125 | 11 | 0.0909 | 39 | 0.1026 |
| 4_4163822 | 13 | 0.1538 | 10 | 0.6 | 9 | 0.4444 | 20 | 0.2 | 52 | 0.3077 | 13 | 0.1538 | 11 | 0.1818 | 11 | 0.4545 | 8 | 0.5 | 43 | 0.3023 |
| 4_4163839 | 13 | 0.8462 | 14 | 0.8571 | 14 | 0.9286 | 21 | 0.9048 | 62 | 0.8871 | 16 | 0.75 | 10 | 0.5 | 14 | 0.7857 | 12 | 0.9167 | 52 | 0.75 |
| 4_4163858 | 11 | 0.3636 | 12 | 0.5833 | 14 | 0.5714 | 21 | 0.381 | 58 | 0.4655 | 14 | 0.6429 | 10 | 0.6 | 14 | 0.5714 | 11 | 0.7273 | 49 | 0.6327 |
| 4_4163895 | 8 | 0.25 | 11 | 0.0909 | 12 | 0.1667 | 15 | 0.2 | 46 | 0.1739 | 9 | 0.1111 | 7 | 0 | 9 | 0.1111 | 12 | 0.0833 | 37 | 0.0811 |
| 4_4163899 | 11 | 0.2727 | 10 | 0 | 11 | 0 | 15 | 0.1333 | 47 | 0.1064 | 12 | 0.0833 | 11 | 0.2727 | 12 | 0.0833 | 13 | 0 | 48 | 0.1042 |
| 4_4165787 | 6 | 0.1667 | 14 | 0.0714 | 12 | 0.0833 | 9 | 0 | 41 | 0.0732 | 12 | 0 | 15 | 0 | 17 | 0.2941 | 13 | 0 | 57 | 0.0877 |
| 4_4165805 | 4 | 0.5 | 14 | 0.3571 | 11 | 0.4545 | 9 | 0.5556 | 38 | 0.4474 | 12 | 0.0833 | 14 | 0.2857 | 14 | 0.2143 | 13 | 0.3846 | 53 | 0.2453 |
| 4_4171212 | 6 | 1 | 4 | 1 | 5 | 1 | 1 | 1 | 16 | 1 | 7 | 1 | 6 | 1 | 4 | 1 | 3 | 1 | 20 | 1 |
| 4_4171214 | 3 | 1 | 5 | 1 | 5 | 1 | 3 | 1 | 16 | 1 | 6 | 1 | 4 | 1 | 4 | 1 | 4 | 1 | 18 | 1 |
| 4_4171216 | 6 | 1 | 4 | 0.75 | 3 | 1 | 2 | 0.5 | 15 | 0.8667 | 7 | 1 | 4 | 0.75 | 4 | 0.75 | 5 | 0.8 | 20 | 0.85 |
| 4_4171219 | 6 | 1 | 4 | 0.25 | 3 | 0.3333 | 1 | 1 | 14 | 0.6429 | 3 | 0.6667 | 4 | 0.75 | 3 | 0.3333 | 4 | 0.75 | 14 | 0.6429 |
| 4_4171226 | 6 | 0.8333 | 7 | 0.4286 | 4 | 0.75 | 3 | 0.6667 | 20 | 0.65 | 5 | 0.6 | 5 | 0.4 | 5 | 0.4 | 5 | 0.8 | 20 | 0.55 |
| 4_4171239 | 7 | 0.4286 | 5 | 0.2 | 5 | 0.6 | 2 | 1 | 19 | 0.4737 | 4 | 0.25 | 5 | 0 | 5 | 0 | 2 | 0 | 16 | 0.0625 |
| 4_4171240 | 7 | 0 | 5 | 0.2 | 5 | 0 | 3 | 1 | 20 | 0.2 | 6 | 0.1667 | 6 | 0 | 6 | 0 | 3 | 0.3333 | 21 | 0.0952 |
| 4_4171246 | 5 | 1 | 4 | 1 | 5 | 0.8 | 2 | 0.5 | 16 | 0.875 | 4 | 1 | 4 | 1 | 5 | 0.8 | 4 | 1 | 17 | 0.9412 |
| 4_4171246 | 6 | 0.1667 | 4 | 0 | 5 | 0 | 3 | 0 | 18 | 0.0556 | 5 | 0 | 5 | 0.2 | 6 | 0.1667 | 4 | 0.25 | 20 | 0.15 |
| 4_4171249 | 5 | 0.6 | 6 | 0.1667 | 5 | 0.6 | 3 | 0.3333 | 19 | 0.4211 | 6 | 0.3333 | 5 | 0.6 | 4 | 0.25 | 4 | 0.5 | 19 | 0.4211 |
| 4_4171266 | 4 | 0 | 4 | 0.25 | 5 | 0 | 3 | 0 | 16 | 0.0625 | 5 | 0.2 | 5 | 0 | 4 | 0 | 3 | 0 | 17 | 0.0588 |
| 4_4173904 | 1 | 0 | 10 | 0.1 | 8 | 0 | NA | NA | 19 | 0.0526 | 6 | 0 | 6 | 0.1667 | 5 | 0.2 | 5 | 0.2 | 22 | 0.1364 |
| 4_4173915 | 2 | 1 | 9 | 0.5556 | 8 | 0.5 | NA | NA | 19 | 0.5789 | 6 | 0.5 | 9 | 0.6667 | 7 | 0.1429 | 7 | 0.2857 | 29 | 0.4138 |
| 4_4173921 | 2 | 0.5 | 9 | 0.8889 | 8 | 0.625 | NA | NA | 19 | 0.7368 | 6 | 0.6667 | 8 | 0.875 | 6 | 1 | 7 | 0.7143 | 27 | 0.8148 |
| 4_4173927 | 2 | 0.5 | 6 | 0.6667 | 5 | 0.4 | NA | NA | 13 | 0.5385 | 5 | 0.6 | 8 | 0.625 | 6 | 0.8333 | 5 | 1 | 24 | 0.75 |
| 4_4173932 | 2 | 0 | 8 | 0.125 | 7 | 0 | NA | NA | 17 | 0.0588 | 6 | 0 | 8 | 0 | 7 | 0.2857 | 7 | 0.2857 | 28 | 0.1429 |
| 4_4173944 | 2 | 0.5 | 9 | 0.4444 | 7 | 0.5714 | NA | NA | 18 | 0.5 | 5 | 0.6 | 9 | 0.6667 | 6 | 0.6667 | 6 | 0.5 | 26 | 0.6154 |
| 4_4175834 | NA | NA | NA | NA | 1 | 1 | 1 | 0 | 2 | 0.5 | 2 | 0.5 | 1 | 1 | NA | NA | NA | NA | 3 | 0.6667 |
| 4_4175837 | NA | NA | 1 | 1 | 1 | 0 | 1 | 0 | 3 | 0.3333 | 2 | 0 | 1 | 1 | NA | NA | NA | NA | 3 | 0.3333 |
| 4_4175839 | NA | NA | 1 | 1 | 1 | 0 | 1 | 0 | 3 | 0.3333 | 2 | 0.5 | 1 | 1 | NA | NA | NA | NA | 3 | 0.6667 |
| 4_4175858 | 10 | 0.4 | 10 | 0.6 | 7 | 0.2857 | 7 | 0.1429 | 34 | 0.3824 | 7 | 0.2857 | 11 | 0.3636 | 5 | 0.2 | 7 | 0.1429 | 30 | 0.2667 |
| 4_4175863 | 10 | 0.3 | 10 | 0.3 | 6 | 0.1667 | 7 | 0.1429 | 33 | 0.2424 | 7 | 0.1429 | 11 | 0.2727 | 6 | 0.1667 | 7 | 0.1429 | 31 | 0.1935 |
| 4_4175866 | 7 | 0.1429 | 8 | 0.25 | 4 | 0 | 7 | 0 | 26 | 0.1154 | 5 | 0 | 11 | 0.1818 | 6 | 0 | 5 | 0 | 27 | 0.0741 |
| 4_4175871 | 2 | 1 | 3 | 1 | 1 | 1 | NA | NA | 6 | 1 | 1 | 1 | 4 | 1 | NA | NA | 2 | 1 | 7 | 1 |
| 4_4175875 | 3 | 1 | 4 | 0 | 1 | 0 | 2 | 0.5 | 10 | 0.4 | 1 | 1 | 4 | 0 | NA | NA | 2 | 0.5 | 7 | 0.2857 |
| 4_4175877 | 2 | 1 | 4 | 0.75 | 1 | 1 | 2 | 1 | 9 | 0.8889 | NA | NA | 3 | 0.3333 | NA | NA | 2 | 1 | 5 | 0.6 |
| 4_4175902 | 1 | 0 | 3 | 0.3333 | 1 | 0 | 2 | 0 | 7 | 0.1429 | NA | NA | 3 | 0.3333 | NA | NA | 2 | 0 | 5 | 0.2 |
| 4_4175904 | 1 | 0 | 3 | 0.3333 | 1 | 1 | 2 | 0.5 | 7 | 0.4286 | NA | NA | 3 | 0 | NA | NA | 1 | 1 | 4 | 0.25 |
| 4_4176916 | 8 | 0.375 | 1 | 0 | 1 | 0 | 1 | 0 | 11 | 0.2727 | 4 | 0 | 4 | 0.25 | 6 | 0.3333 | 2 | 0.5 | 16 | 0.25 |
| 4_4176927 | 5 | 0.6 | NA | NA | 1 | 0 | NA | NA | 6 | 0.5 | 2 | 0 | 2 | 0.5 | 4 | 0.25 | 2 | 0.5 | 10 | 0.3 |
| 4_4176956 | 9 | 0 | 19 | 0.0526 | 15 | 0.2 | 6 | 0 | 49 | 0.0816 | 13 | 0.1538 | 14 | 0.0714 | 17 | 0 | 11 | 0.1818 | 55 | 0.0909 |
| 4_4186989 | NA | NA | NA | NA | NA | NA | 2 | 0.5 | 2 | 0.5 | NA | NA | NA | NA | NA | NA | 1 | 1 | 1 | 1 |
| 4_4187044 | NA | NA | NA | NA | NA | NA | 1 | 1 | 1 | 1 | NA | NA | 1 | 1 | NA | NA | 1 | 1 | 2 | 1 |
| 4_4187058 | NA | NA | NA | NA | NA | NA | 2 | 1 | 2 | 1 | NA | NA | 1 | 1 | 1 | 1 | 2 | 1 | 4 | 1 |

| | | | | | | | | | | | | | | | | | | | | |
|---|---|---|---|---|---|---|---|---|---|---|---|---|---|---|---|---|---|---|---|---|
| 4_4187063 | NA | NA | NA | NA | NA | NA | 2 | 0.5 | 2 | 0.5 | NA | NA | 1 | 1 | NA | NA | 2 | 1 | 3 | 1 |
| 4_4187069 | NA | NA | NA | NA | NA | NA | 2 | 0.5 | 2 | 0.5 | NA | NA | 1 | 1 | 1 | 1 | 2 | 1 | 4 | 1 |
| 4_4187069 | NA | NA | NA | NA | NA | NA | 2 | 0.5 | 2 | 0.5 | NA | NA | 1 | 0 | 1 | 1 | 1 | 0 | 3 | 0.3333 |
| 4_4187071 | NA | NA | NA | NA | NA | NA | 2 | 0.5 | 2 | 0.5 | NA | NA | 1 | 0 | 1 | 1 | 2 | 0 | 4 | 0.25 |
| 4_4187082 | NA | NA | NA | NA | NA | NA | 2 | 0.5 | 2 | 0.5 | NA | NA | 1 | 1 | 1 | 1 | 2 | 1 | 4 | 1 |
| 4_4187086 | NA | NA | NA | NA | NA | NA | 2 | 1 | 2 | 1 | NA | NA | 1 | 1 | 1 | 1 | 2 | 1 | 4 | 1 |
| 4_4187138 | NA | NA | NA | NA | NA | NA | 2 | 0.5 | 2 | 0.5 | NA | NA | 1 | 0 | 1 | 0 | 2 | 0.5 | 4 | 0.25 |
| 4_4188435 | NA | NA | NA | NA | 1 | 1 | NA | NA | 1 | 1 | NA | NA | NA | NA | 1 | 1 | NA | NA | 1 | 1 |
| 4_4188476 | NA | NA | NA | NA | 1 | 1 | NA | NA | 1 | 1 | NA | NA | NA | NA | 1 | 1 | NA | NA | 1 | 1 |
| 4_4188932 | NA | NA | NA | NA | 2 | 0.5 | 2 | 0.5 | 4 | 0.5 | NA | NA | 1 | 1 | NA | NA | NA | NA | 1 | 1 |
| 4_4188954 | NA | NA | NA | NA | 2 | 1 | 1 | 1 | 3 | 1 | NA | NA | 1 | 1 | NA | NA | 1 | 1 | 2 | 1 |
| 4_4188976 | NA | NA | NA | NA | 1 | 1 | 2 | 1 | 3 | 1 | NA | NA | 1 | 1 | NA | NA | 1 | 1 | 2 | 1 |
| 4_4189002 | NA | NA | 1 | 0 | 2 | 0.5 | 2 | 1 | 5 | 0.6 | NA | NA | 1 | 0 | NA | NA | 1 | 1 | 2 | 0.5 |
| 4_4189050 | NA | NA | NA | NA | 2 | 0.5 | 2 | 1 | 4 | 0.75 | NA | NA | 1 | 1 | NA | NA | 1 | 0 | 2 | 0.5 |
| 4_4189087 | NA | NA | NA | NA | 2 | 0.5 | 1 | 0 | 3 | 0.3333 | NA | NA | 1 | 1 | NA | NA | 1 | 0 | 2 | 0.5 |
| 4_4189113 | NA | NA | NA | NA | 2 | 0 | 2 | 0.5 | 4 | 0.25 | NA | NA | NA | NA | NA | NA | 1 | 1 | 1 | 1 |
| 4_4189116 | NA | NA | NA | NA | 2 | 0.5 | 2 | 0 | 4 | 0.25 | NA | NA | 1 | 1 | NA | NA | 1 | 0 | 2 | 0.5 |
| 4_4189141 | NA | NA | NA | NA | 2 | 0.5 | 2 | 0 | 4 | 0.25 | NA | NA | 1 | 0 | NA | NA | 1 | 1 | 2 | 0.5 |
| 4_4189145 | NA | NA | NA | NA | 2 | 0.5 | 1 | 0 | 3 | 0.3333 | NA | NA | 1 | 0 | NA | NA | 1 | 1 | 2 | 0.5 |
| 4_4189161 | NA | NA | NA | NA | 2 | 0.5 | 2 | 0 | 4 | 0.25 | NA | NA | 1 | 1 | NA | NA | 1 | 1 | 2 | 1 |
| 4_4189167 | NA | NA | NA | NA | 1 | 1 | 2 | 0.5 | 3 | 0.6667 | NA | NA | 1 | 1 | NA | NA | 1 | 1 | 2 | 1 |
| 4_4189220 | NA | NA | NA | NA | 2 | 1 | 1 | 1 | 3 | 1 | NA | NA | 1 | 1 | NA | NA | 1 | 1 | 2 | 1 |
| 4_4189247 | NA | NA | NA | NA | 2 | 0.5 | 2 | 0.5 | 4 | 0.5 | NA | NA | 1 | 1 | NA | NA | 1 | 0 | 2 | 0.5 |
| 4_4189263 | NA | NA | NA | NA | 2 | 0.5 | 2 | 0.5 | 4 | 0.5 | 1 | 1 | 1 | 1 | NA | NA | 1 | 1 | 3 | 1 |
| 4_4189269 | NA | NA | NA | NA | NA | NA | 2 | 1 | 2 | 1 | NA | NA | 1 | 1 | NA | NA | NA | NA | 1 | 1 |
| 4_4189289 | NA | NA | NA | NA | 2 | 0.5 | 2 | 0.5 | 4 | 0.5 | NA | NA | 1 | 0 | NA | NA | 1 | 1 | 2 | 0.5 |
| 4_4196836 | NA | NA | NA | NA | 1 | 1 | 1 | 1 | 2 | 1 | 1 | 1 | NA | NA | NA | NA | NA | NA | 1 | 1 |
| 4_4196866 | NA | NA | NA | NA | 1 | 1 | 1 | 0 | 2 | 0.5 | 1 | 1 | NA | NA | NA | NA | NA | NA | 1 | 1 |
| 4_4196880 | NA | NA | NA | NA | 1 | 1 | 1 | 1 | 2 | 1 | 1 | 1 | NA | NA | NA | NA | NA | NA | 1 | 1 |
| 4_4196886 | NA | NA | NA | NA | 1 | 1 | 1 | 1 | 2 | 1 | 1 | 1 | NA | NA | NA | NA | NA | NA | 1 | 1 |
| 4_4196891 | NA | NA | NA | NA | 1 | 0 | 1 | 1 | 2 | 0.5 | 1 | 1 | NA | NA | NA | NA | NA | NA | 1 | 1 |
| 4_4196891 | NA | NA | NA | NA | 1 | 1 | 1 | 0 | 2 | 0.5 | 1 | 1 | NA | NA | NA | NA | NA | NA | 1 | 1 |
| 4_4196897 | NA | NA | NA | NA | 1 | 1 | 1 | 1 | 2 | 1 | 1 | 1 | NA | NA | NA | NA | NA | NA | 1 | 1 |
| 4_4196901 | NA | NA | NA | NA | 1 | 1 | 1 | 0 | 2 | 0.5 | 1 | 1 | NA | NA | NA | NA | NA | NA | 1 | 1 |
| 4_4196930 | NA | NA | NA | NA | 1 | 1 | 1 | 1 | 2 | 1 | 1 | 1 | NA | NA | NA | NA | NA | NA | 1 | 1 |
| 4_4196988 | NA | NA | NA | NA | 2 | 1 | NA | NA | 2 | 1 | 1 | 1 | NA | NA | NA | NA | 1 | 1 | 2 | 1 |
| 4_4196997 | NA | NA | NA | NA | 2 | 1 | 1 | 1 | 3 | 1 | 1 | 1 | NA | NA | NA | NA | 1 | 1 | 2 | 1 |
| 4_4197034 | NA | NA | NA | NA | 3 | 0.3333 | 2 | 0 | 5 | 0.2 | 1 | 1 | 1 | 0 | NA | NA | 1 | 1 | 3 | 0.6667 |
| 4_4197036 | NA | NA | NA | NA | 3 | 0.6667 | 2 | 0 | 5 | 0.4 | 1 | 1 | 1 | 0 | NA | NA | 1 | 0 | 3 | 0.3333 |
| 4_4197117 | NA | NA | NA | NA | 3 | 1 | 2 | 0.5 | 5 | 0.8 | 1 | 0 | 1 | 1 | NA | NA | 1 | 1 | 3 | 0.6667 |
| 4_4197152 | NA | NA | NA | NA | 3 | 1 | 2 | 1 | 5 | 1 | 1 | 1 | 1 | 1 | NA | NA | 1 | 1 | 3 | 1 |
| 4_4202773 | 2 | 0.5 | 1 | 0 | 2 | 0.5 | NA | NA | 5 | 0.4 | 1 | 0 | 1 | 0 | 1 | 1 | 1 | 0 | 4 | 0.25 |
| 4_4202799 | 1 | 1 | NA | NA | 2 | 0 | NA | NA | 3 | 0.3333 | 1 | 0 | NA | NA | 1 | 1 | 2 | 0 | 4 | 0.25 |
| 4_4202808 | 2 | 0.5 | 1 | 0 | 2 | 0 | NA | NA | 5 | 0.2 | 1 | 0 | 2 | 0 | 1 | 1 | 2 | 0 | 6 | 0.1667 |
| 4_4202868 | 2 | 1 | 1 | 1 | 1 | 1 | NA | NA | 4 | 1 | NA | NA | 2 | 1 | 1 | 1 | 1 | 1 | 4 | 1 |
| 4_4202920 | 2 | 0.5 | 1 | 1 | NA | NA | NA | NA | 3 | 0.6667 | 1 | 0 | 2 | 0.5 | 1 | 0 | 1 | 0 | 5 | 0.2 |
| 4_4202921 | 2 | 1 | 1 | 1 | NA | NA | NA | NA | 3 | 1 | NA | NA | 2 | 0.5 | 1 | 1 | 1 | 1 | 4 | 0.75 |
| 4_4202940 | 2 | 1 | NA | NA | NA | NA | NA | NA | 2 | 1 | 1 | 0 | 2 | 0 | 1 | 1 | 1 | 0 | 5 | 0.2 |
| 4_4203096 | 2 | 0.5 | NA | NA | NA | NA | NA | NA | 2 | 0.5 | NA | NA | 2 | 0 | 1 | 1 | NA | NA | 3 | 0.3333 |
| 4_4203112 | 2 | 0.5 | NA | NA | NA | NA | NA | NA | 2 | 0.5 | NA | NA | 2 | 1 | 1 | 0 | 1 | 1 | 4 | 0.75 |
| 4_4203116 | 2 | 0.5 | NA | NA | NA | NA | NA | NA | 2 | 0.5 | NA | NA | 2 | 0.5 | 1 | 0 | 1 | 0 | 4 | 0.25 |
| 4_4203124 | 1 | 1 | NA | NA | NA | NA | NA | NA | 1 | 1 | 1 | 0 | 2 | 0 | 1 | 1 | 1 | 1 | 5 | 0.4 |
| 4_4203136 | 3 | 1 | 1 | 1 | NA | NA | 2 | 1 | 6 | 1 | 2 | 0.5 | 2 | 1 | 3 | 1 | 1 | 0 | 8 | 0.75 |
| 4_4203156 | 2 | 0.5 | NA | NA | NA | NA | 1 | 0 | 3 | 0.3333 | 2 | 0 | 2 | 0.5 | 3 | 0 | 1 | 1 | 8 | 0.25 |
| 4_4203186 | 1 | 1 | 1 | 0 | NA | NA | 2 | 0 | 4 | 0.25 | 2 | 0 | 3 | 0.3333 | 3 | 0 | 1 | 0 | 9 | 0.1111 |
| 4_4203187 | 1 | 0 | 1 | 1 | NA | NA | 2 | 1 | 4 | 0.75 | 2 | 0 | 3 | 1 | 4 | 1 | 1 | 1 | 10 | 0.8 |
| 4_4203243 | 1 | 1 | 1 | 1 | NA | NA | 2 | 1 | 4 | 1 | 2 | 1 | 1 | 1 | 3 | 0.6667 | 1 | 1 | 7 | 0.8571 |
| 4_4203244 | 1 | 1 | 1 | 1 | NA | NA | 2 | 1 | 4 | 1 | 2 | 1 | 1 | 1 | 4 | 1 | 1 | 1 | 8 | 1 |
| 4_4203256 | 1 | 1 | 1 | 0 | NA | NA | 2 | 0 | 4 | 0.25 | 2 | 0 | 2 | 0 | 4 | 0 | 1 | 1 | 9 | 0.1111 |
| 4_4203266 | 1 | 1 | 1 | 1 | NA | NA | 2 | 0.5 | 4 | 0.75 | 2 | 1 | 2 | 0 | 4 | 1 | 1 | 0 | 9 | 0.6667 |
| 4_4203374 | 1 | 1 | 1 | 1 | NA | NA | 1 | 1 | 3 | 1 | 2 | 0.5 | 2 | 0.5 | 3 | 1 | 1 | 1 | 8 | 0.75 |
| 4_4203398 | 1 | 1 | 1 | 1 | NA | NA | 1 | 1 | 3 | 1 | 2 | 1 | 2 | 0.5 | 3 | 0.6667 | 1 | 1 | 8 | 0.75 |
| 4_4203444 | 1 | 1 | 1 | 1 | NA | NA | NA | NA | 2 | 1 | 2 | 0.5 | 2 | 1 | 3 | 0.6667 | 1 | 1 | 8 | 0.75 |
| 4_4203486 | 1 | 1 | 1 | 1 | NA | NA | NA | NA | 2 | 1 | NA | NA | 2 | 1 | 3 | 1 | NA | NA | 5 | 1 |
| 4_4203489 | 1 | 1 | 1 | 0 | NA | NA | NA | NA | 2 | 0.5 | 2 | 0 | 2 | 0 | 3 | 0.6667 | 1 | 0 | 8 | 0.25 |

| | | | | | | | | | | | | | | | | | | | | |
|---|---|---|---|---|---|---|---|---|---|---|---|---|---|---|---|---|---|---|---|---|
| 4_4203503 | 1 | 1 | 1 | 1 | NA | NA | NA | NA | 2 | 1 | 2 | 1 | 2 | 1 | 3 | 1 | 1 | 0 | 8 | 0.875 |
| 4_4203509 | 1 | 0 | 1 | 1 | NA | NA | NA | NA | 2 | 0.5 | 2 | 0.5 | 2 | 0.5 | 3 | 0.3333 | 1 | 1 | 8 | 0.5 |
| 4_4203513 | 1 | 1 | 1 | 1 | NA | NA | NA | NA | 2 | 1 | 2 | 1 | 2 | 1 | 3 | 1 | NA | NA | 7 | 1 |
| 4_4203527 | 1 | 1 | 1 | 1 | NA | NA | NA | NA | 2 | 1 | 2 | 1 | 2 | 0 | 3 | 0.3333 | 1 | 1 | 8 | 0.5 |
| 4_4221969 | NA | NA | NA | NA | NA | NA | 2 | 0.5 | 2 | 0.5 | 1 | 1 | NA | NA | NA | NA | NA | NA | 1 | 1 |
| 4_4221982 | NA | NA | 1 | 1 | NA | NA | 2 | 1 | 3 | 1 | 1 | 1 | NA | NA | NA | NA | NA | NA | 1 | 1 |
| 4_4221987 | NA | NA | 1 | 1 | NA | NA | 2 | 1 | 3 | 1 | 1 | 1 | NA | NA | NA | NA | NA | NA | 1 | 1 |
| 4_4221993 | NA | NA | 1 | 1 | NA | NA | 2 | 1 | 3 | 1 | 1 | 1 | NA | NA | NA | NA | NA | NA | 1 | 1 |
| 4_4222007 | NA | NA | NA | NA | NA | NA | 2 | 1 | 2 | 1 | 1 | 1 | NA | NA | NA | NA | NA | NA | 1 | 1 |
| 4_4222008 | NA | NA | 1 | 0 | NA | NA | 1 | 1 | 2 | 0.5 | 1 | 1 | NA | NA | NA | NA | NA | NA | 1 | 1 |
| 4_4222010 | NA | NA | 1 | 1 | NA | NA | 2 | 1 | 3 | 1 | 1 | 1 | NA | NA | NA | NA | NA | NA | 1 | 1 |
| 4_4222029 | NA | NA | 1 | 0 | NA | NA | 3 | 0.3333 | 4 | 0.25 | 1 | 1 | NA | NA | NA | NA | NA | NA | 1 | 1 |
| 4_4222063 | NA | NA | 1 | 1 | NA | NA | 3 | 0 | 4 | 0.25 | 2 | 0.5 | NA | NA | NA | NA | NA | NA | 2 | 0.5 |
| 4_4222175 | NA | NA | 1 | 1 | 1 | 0 | 3 | 0 | 5 | 0.2 | 3 | 0.3333 | NA | NA | NA | NA | 2 | 0 | 5 | 0.2 |
| 4_4223233 | 1 | 1 | NA | NA | 1 | 0 | NA | NA | 2 | 0.5 | 1 | 1 | NA | NA | NA | NA | 1 | 0 | 2 | 0.5 |
| 4_4223234 | 1 | 1 | NA | NA | 1 | 1 | NA | NA | 2 | 1 | NA | NA | NA | NA | NA | NA | 1 | 1 | 1 | 1 |
| 4_4223282 | 1 | 1 | NA | NA | 1 | 1 | NA | NA | 2 | 1 | 1 | 1 | NA | NA | NA | NA | NA | NA | 1 | 1 |
| 4_4223326 | 2 | 1 | NA | NA | NA | NA | NA | NA | 2 | 1 | 1 | 1 | NA | NA | NA | NA | NA | NA | 1 | 1 |
| 4_4223466 | 2 | 0.5 | NA | NA | NA | NA | 1 | 0 | 3 | 0.3333 | 1 | 1 | NA | NA | NA | NA | NA | NA | 1 | 1 |
| 4_4223490 | 2 | 0.5 | NA | NA | 1 | 1 | 1 | 1 | 4 | 0.75 | 1 | 1 | NA | NA | NA | NA | 1 | 0 | 2 | 0.5 |
| 4_4223511 | 2 | 0 | NA | NA | 1 | 1 | 1 | 0 | 4 | 0.25 | 1 | 0 | NA | NA | NA | NA | 1 | 1 | 2 | 0.5 |
| 4_4223516 | 2 | 1 | NA | NA | 1 | 1 | 1 | 1 | 4 | 1 | 1 | 1 | NA | NA | NA | NA | 1 | 1 | 2 | 1 |
| 4_4223521 | 2 | 0.5 | NA | NA | 1 | 0 | 1 | 0 | 4 | 0.25 | 1 | 1 | NA | NA | NA | NA | NA | NA | 1 | 1 |
| 4_4223528 | 2 | 0 | NA | NA | 1 | 1 | 1 | 0 | 4 | 0.25 | 1 | 1 | NA | NA | NA | NA | 1 | 0 | 2 | 0.5 |
| 4_4223570 | 2 | 1 | NA | NA | NA | NA | 1 | 1 | 3 | 1 | 1 | 1 | NA | NA | NA | NA | 1 | 1 | 2 | 1 |
| 4_4223613 | 2 | 1 | NA | NA | 1 | 1 | 1 | 0 | 4 | 0.75 | NA | NA | NA | NA | NA | NA | 1 | 1 | 1 | 1 |
| 4_4223646 | 2 | 0.5 | 1 | 0 | 2 | 0.5 | 1 | 0 | 6 | 0.3333 | 1 | 1 | NA | NA | NA | NA | 1 | 1 | 2 | 1 |
| 4_4223688 | NA | NA | 1 | 1 | 2 | 0.5 | 1 | 0 | 4 | 0.5 | NA | NA | NA | NA | NA | NA | 1 | 1 | 1 | 1 |
| 4_4223753 | 1 | 1 | 2 | 1 | 1 | 0 | 1 | 1 | 5 | 0.8 | NA | NA | NA | NA | NA | NA | 1 | 1 | 1 | 1 |
| 4_4223808 | NA | NA | 2 | 0.5 | 1 | 0 | 1 | 0 | 4 | 0.25 | NA | NA | NA | NA | NA | NA | 1 | 1 | 1 | 1 |
| 4_4223857 | NA | NA | 3 | 0.6667 | 2 | 0.5 | 1 | 0 | 6 | 0.5 | NA | NA | NA | NA | NA | NA | 1 | 1 | 1 | 1 |
| 4_4223878 | 1 | 0 | 1 | 1 | 2 | 0.5 | NA | NA | 4 | 0.5 | NA | NA | NA | NA | NA | NA | 2 | 1 | 2 | 1 |
| 4_4223879 | 1 | 0 | 3 | 0.6667 | 2 | 0 | 1 | 0 | 7 | 0.2857 | NA | NA | NA | NA | NA | NA | 2 | 0.5 | 2 | 0.5 |
| 4_4223900 | 1 | 0 | 2 | 1 | 1 | 1 | 1 | 1 | 5 | 0.8 | NA | NA | NA | NA | NA | NA | 2 | 1 | 2 | 1 |
| 4_4223933 | 1 | 1 | 3 | 0.6667 | 2 | 0 | 1 | 0 | 7 | 0.4286 | NA | NA | NA | NA | NA | NA | 2 | 1 | 2 | 1 |
| 4_4223946 | NA | NA | 3 | 1 | 1 | 1 | NA | NA | 4 | 1 | NA | NA | NA | NA | NA | NA | 2 | 1 | 2 | 1 |
| 4_4223951 | 1 | 0 | 2 | 0 | 1 | 1 | NA | NA | 4 | 0.25 | NA | NA | NA | NA | NA | NA | 2 | 0.5 | 2 | 0.5 |
| 4_4223975 | 1 | 0 | 3 | 0.6667 | 1 | 0 | NA | NA | 5 | 0.4 | NA | NA | NA | NA | NA | NA | 2 | 0.5 | 2 | 0.5 |
| 4_4224011 | 1 | 1 | 2 | 0 | 1 | 1 | NA | NA | 4 | 0.5 | NA | NA | NA | NA | NA | NA | 2 | 0.5 | 2 | 0.5 |
| 4_4224012 | 1 | 0 | 3 | 0.3333 | 1 | 0 | NA | NA | 5 | 0.2 | NA | NA | NA | NA | NA | NA | 2 | 0.5 | 2 | 0.5 |
| 4_4224016 | 1 | 1 | 3 | 1 | 2 | 0 | NA | NA | 6 | 0.6667 | NA | NA | NA | NA | NA | NA | 2 | 1 | 2 | 1 |
| 4_4224037 | 1 | 0 | 3 | 0 | 1 | 1 | NA | NA | 5 | 0.2 | NA | NA | NA | NA | NA | NA | 2 | 0.5 | 2 | 0.5 |
| 4_4224040 | 1 | 1 | 3 | 1 | 1 | 1 | NA | NA | 5 | 1 | NA | NA | NA | NA | NA | NA | 2 | 1 | 2 | 1 |
| 4_4224065 | 1 | 1 | 3 | 0.6667 | 2 | 1 | NA | NA | 6 | 0.8333 | 1 | 0 | NA | NA | NA | NA | 2 | 0.5 | 3 | 0.3333 |
| 4_4224069 | 1 | 1 | 3 | 1 | 1 | 0 | NA | NA | 5 | 0.8 | 1 | 0 | NA | NA | NA | NA | 1 | 1 | 2 | 0.5 |
| 4_4224091 | 1 | 1 | 3 | 1 | 1 | 1 | NA | NA | 5 | 1 | 2 | 0.5 | NA | NA | NA | NA | 1 | 1 | 3 | 0.6667 |
| 4_4224095 | 1 | 1 | 1 | 0 | 3 | 0 | NA | NA | 5 | 0.2 | 2 | 0.5 | NA | NA | NA | NA | 1 | 1 | 3 | 0.6667 |
| 4_4224144 | 2 | 0.5 | 3 | 0.6667 | 2 | 1 | NA | NA | 7 | 0.7143 | 2 | 1 | NA | NA | NA | NA | 2 | 1 | 4 | 1 |
| 4_4224165 | 2 | 0.5 | 3 | 0 | 3 | 1 | NA | NA | 8 | 0.5 | 2 | 1 | NA | NA | NA | NA | 1 | 0 | 3 | 0.6667 |
| 4_4224167 | 2 | 0.5 | 4 | 0.25 | 3 | 0 | NA | NA | 9 | 0.2222 | 2 | 0 | NA | NA | NA | NA | 2 | 1 | 4 | 0.5 |
| 4_4224187 | 2 | 1 | 4 | 0.75 | 2 | 1 | NA | NA | 8 | 0.875 | 1 | 1 | NA | NA | NA | NA | 2 | 1 | 3 | 1 |
| 4_4224221 | 2 | 0 | 4 | 0.5 | 2 | 1 | NA | NA | 8 | 0.5 | 1 | 0 | NA | NA | NA | NA | 2 | 0.5 | 3 | 0.3333 |
| 4_4224233 | NA | NA | 4 | 0.75 | 2 | 0.5 | NA | NA | 6 | 0.6667 | 2 | 0.5 | NA | NA | NA | NA | NA | NA | 2 | 0.5 |
| 4_4224262 | NA | NA | 2 | 0.5 | 1 | 1 | NA | NA | 3 | 0.6667 | 2 | 1 | NA | NA | NA | NA | NA | NA | 2 | 1 |
| 4_4224303 | 1 | 0 | 2 | 0 | 1 | 1 | NA | NA | 4 | 0.25 | 2 | 0.5 | NA | NA | NA | NA | NA | NA | 2 | 0.5 |
| 4_4224327 | 1 | 1 | 3 | 0.6667 | 1 | 1 | NA | NA | 5 | 0.8 | 3 | 0.6667 | NA | NA | NA | NA | NA | NA | 3 | 0.6667 |
| 4_4224352 | 1 | 1 | 3 | 0.6667 | NA | NA | NA | NA | 4 | 0.75 | 2 | 0.5 | NA | NA | NA | NA | NA | NA | 2 | 0.5 |
| 4_4224357 | NA | NA | 3 | 0.6667 | 1 | 1 | NA | NA | 4 | 0.75 | 3 | 0.3333 | NA | NA | NA | NA | NA | NA | 3 | 0.3333 |
| 4_4224373 | 1 | 0 | 3 | 0.3333 | 1 | 1 | NA | NA | 5 | 0.4 | 2 | 0.5 | NA | NA | NA | NA | NA | NA | 2 | 0.5 |
| 4_4224378 | 1 | 1 | 3 | 1 | 1 | 1 | NA | NA | 5 | 1 | 2 | 1 | NA | NA | NA | NA | NA | NA | 2 | 1 |
| 4_4224431 | 1 | 1 | 3 | 1 | 1 | 1 | NA | NA | 5 | 1 | 3 | 1 | NA | NA | NA | NA | NA | NA | 3 | 1 |
| 4_4224453 | NA | NA | 2 | 0.5 | 1 | 0 | NA | NA | 3 | 0.3333 | 2 | 0.5 | NA | NA | NA | NA | NA | NA | 2 | 0.5 |
| 4_4224474 | NA | NA | 3 | 0.6667 | 1 | 0 | NA | NA | 4 | 0.5 | 2 | 0.5 | NA | NA | NA | NA | NA | NA | 2 | 0.5 |
| 4_4224500 | 1 | 0 | 3 | 0.3333 | 1 | 0 | 1 | 1 | 6 | 0.3333 | 2 | 0.5 | NA | NA | NA | NA | NA | NA | 2 | 0.5 |
| 4_4225647 | 2 | 0 | 2 | 0 | 2 | 0 | 4 | 0.5 | 10 | 0.2 | 3 | 0 | 2 | 0.5 | 1 | 0 | 2 | 0 | 8 | 0.125 |

| | | | | | | | | | | | | | | | | | | | | |
|---|---|---|---|---|---|---|---|---|---|---|---|---|---|---|---|---|---|---|---|---|
| 4_4225652 | 2 | 0 | 2 | 0.5 | 4 | 0.25 | 3 | 0 | 11 | 0.1818 | 3 | 0 | 3 | 0.3333 | 2 | 0 | 2 | 0 | 10 | 0.1 |
| 4_4225670 | 2 | 0 | 2 | 0.5 | 4 | 0.25 | 5 | 0.4 | 13 | 0.3077 | 2 | 1 | 3 | 0.6667 | 2 | 0.5 | 2 | 0 | 9 | 0.5556 |
| 4_4232209 | 1 | 1 | NA | NA | NA | NA | NA | NA | 1 | 1 | NA | NA | NA | NA | 1 | 1 | NA | NA | 1 | 1 |
| 4_4232219 | 2 | 0.5 | NA | NA | NA | NA | NA | NA | 2 | 0.5 | NA | NA | NA | NA | 1 | 1 | NA | NA | 1 | 1 |
| 4_4232255 | 2 | 0.5 | NA | NA | NA | NA | NA | NA | 2 | 0.5 | NA | NA | NA | NA | 1 | 1 | NA | NA | 1 | 1 |
| 4_4232257 | 2 | 0.5 | NA | NA | NA | NA | NA | NA | 2 | 0.5 | NA | NA | NA | NA | 1 | 1 | NA | NA | 1 | 1 |
| 4_4232333 | 2 | 0.5 | NA | NA | NA | NA | NA | NA | 2 | 0.5 | NA | NA | NA | NA | 1 | 1 | NA | NA | 1 | 1 |
| 4_4232339 | 1 | 1 | NA | NA | NA | NA | NA | NA | 1 | 1 | NA | NA | 1 | 1 | 1 | 1 | NA | NA | 2 | 1 |
| 4_4232374 | 1 | 1 | NA | NA | NA | NA | NA | NA | 1 | 1 | NA | NA | NA | NA | 1 | 1 | NA | NA | 1 | 1 |
| 4_4238090 | NA | NA | 1 | 1 | 2 | 0.5 | NA | NA | 3 | 0.6667 | 1 | 1 | 1 | 1 | 1 | 1 | 1 | 1 | 4 | 1 |
| 4_4238318 | NA | NA | NA | NA | 1 | 1 | NA | NA | 1 | 1 | 1 | 0 | NA | NA | 1 | 1 | 1 | 0 | 3 | 0.3333 |
| 4_4238357 | NA | NA | NA | NA | 1 | 1 | NA | NA | 1 | 1 | NA | NA | 1 | 1 | 1 | 0 | 1 | 1 | 3 | 0.6667 |
| 4_4238408 | NA | NA | NA | NA | 1 | 1 | NA | NA | 1 | 1 | NA | NA | 1 | 1 | NA | NA | NA | NA | 1 | 1 |
| 4_4245901 | 1 | 1 | 1 | 1 | 2 | 1 | NA | NA | 4 | 1 | 1 | 1 | NA | NA | NA | NA | NA | NA | 1 | 1 |
| 4_4245915 | 1 | 1 | 1 | 1 | 1 | 1 | NA | NA | 3 | 1 | 1 | 1 | NA | NA | NA | NA | NA | NA | 1 | 1 |
| 4_4245919 | 1 | 0 | 1 | 1 | 2 | 0.5 | NA | NA | 4 | 0.5 | 1 | 1 | NA | NA | NA | NA | NA | NA | 1 | 1 |
| 4_4245925 | 1 | 1 | 1 | 0 | 2 | 1 | NA | NA | 4 | 0.75 | 1 | 1 | NA | NA | NA | NA | NA | NA | 1 | 1 |
| 4_4245964 | 1 | 1 | 1 | 1 | 2 | 1 | NA | NA | 4 | 1 | 1 | 1 | NA | NA | NA | NA | NA | NA | 1 | 1 |
| 4_4245985 | 1 | 1 | 1 | 1 | 3 | 1 | NA | NA | 5 | 1 | 1 | 1 | NA | NA | NA | NA | NA | NA | 1 | 1 |
| 4_4246025 | NA | NA | 1 | 1 | 2 | 0.5 | NA | NA | 3 | 0.6667 | 1 | 1 | NA | NA | NA | NA | NA | NA | 1 | 1 |
| 4_4246030 | 1 | 1 | 1 | 0 | 3 | 0.6667 | NA | NA | 5 | 0.6 | 1 | 1 | NA | NA | NA | NA | NA | NA | 1 | 1 |
| 4_4246054 | 1 | 0 | 1 | 1 | 3 | 0.6667 | NA | NA | 5 | 0.6 | 1 | 1 | NA | NA | NA | NA | NA | NA | 1 | 1 |
| 4_4246162 | 1 | 0 | 1 | 1 | 4 | 0 | NA | NA | 6 | 0.1667 | 1 | 1 | NA | NA | 1 | 1 | NA | NA | 2 | 1 |
| 4_4246184 | 1 | 1 | 1 | 1 | 4 | 0.75 | NA | NA | 6 | 0.8333 | 1 | 0 | NA | NA | 1 | 1 | NA | NA | 2 | 0.5 |
| 4_4246241 | 1 | 0 | 1 | 1 | 5 | 0.2 | 1 | 1 | 8 | 0.375 | 1 | 1 | 2 | 1 | 1 | 1 | 1 | 1 | 5 | 1 |
| 4_4246242 | 1 | 1 | 1 | 0 | 5 | 0 | 1 | 0 | 8 | 0.125 | 1 | 0 | 2 | 0 | 1 | 0 | 1 | 1 | 5 | 0.2 |
| 4_4246292 | 1 | 1 | 1 | 1 | 5 | 0.8 | NA | NA | 7 | 0.8571 | 1 | 1 | 2 | 1 | 1 | 1 | 1 | 1 | 5 | 1 |
| 4_4246298 | NA | NA | 1 | 0 | 2 | 0.5 | NA | NA | 3 | 0.3333 | 2 | 0 | 1 | 1 | 1 | 1 | 2 | 0 | 6 | 0.3333 |
| 4_4246492 | NA | NA | NA | NA | 2 | 0.5 | NA | NA | 2 | 0.5 | 2 | 0 | 1 | 0 | 2 | 0 | 2 | 0.5 | 7 | 0.1429 |
| 4_4246560 | NA | NA | NA | NA | 3 | 1 | 1 | 0 | 4 | 0.75 | 2 | 1 | 1 | 1 | 3 | 1 | 2 | 1 | 8 | 1 |
| 4_4246620 | NA | NA | NA | NA | 3 | 0.3333 | 2 | 0.5 | 5 | 0.4 | 2 | 0 | 1 | 0 | 3 | 0.3333 | 2 | 0 | 8 | 0.125 |
| 4_4246629 | NA | NA | NA | NA | 3 | 0.3333 | NA | NA | 3 | 0.3333 | 1 | 0 | 1 | 0 | 2 | 0 | 2 | 0.5 | 6 | 0.1667 |
| 4_4246655 | NA | NA | NA | NA | 1 | 0 | 2 | 0.5 | 3 | 0.3333 | 4 | 0.5 | 1 | 0 | 4 | 0 | 2 | 0.5 | 11 | 0.2727 |
| 4_4246870 | NA | NA | NA | NA | NA | NA | 2 | 1 | 2 | 1 | 1 | 1 | NA | NA | 1 | 1 | 1 | 1 | 3 | 1 |
| 4_4246889 | NA | NA | NA | NA | NA | NA | 1 | 1 | 1 | 1 | NA | NA | NA | NA | 1 | 1 | NA | NA | 1 | 1 |
| 4_4246892 | NA | NA | NA | NA | NA | NA | 2 | 0.5 | 2 | 0.5 | 1 | 1 | NA | NA | 1 | 0 | 1 | 0 | 3 | 0.3333 |
| 4_4247051 | NA | NA | NA | NA | NA | NA | 2 | 1 | 2 | 1 | 1 | 1 | NA | NA | 1 | 1 | 1 | 1 | 3 | 1 |
| 4_4247073 | NA | NA | NA | NA | NA | NA | 2 | 0.5 | 2 | 0.5 | 1 | 0 | NA | NA | NA | NA | 1 | 1 | 2 | 0.5 |
| 4_4247090 | NA | NA | NA | NA | NA | NA | 2 | 1 | 2 | 1 | 1 | 1 | NA | NA | NA | NA | 1 | 0 | 2 | 0.5 |
| 4_4247099 | NA | NA | NA | NA | NA | NA | 2 | 1 | 2 | 1 | 1 | 0 | NA | NA | NA | NA | 1 | 1 | 2 | 0.5 |
| 4_4252276 | 1 | 1 | 2 | 1 | NA | NA | 1 | 1 | 4 | 1 | NA | NA | 1 | 1 | NA | NA | NA | NA | 1 | 1 |
| 4_4252311 | 1 | 0 | 2 | 1 | NA | NA | NA | NA | 3 | 0.6667 | NA | NA | 1 | 1 | NA | NA | NA | NA | 1 | 1 |
| 4_4252352 | 1 | 1 | 2 | 0 | NA | NA | 1 | 0 | 4 | 0.25 | NA | NA | 1 | 1 | NA | NA | NA | NA | 1 | 1 |
| 4_4252379 | NA | NA | 1 | 0 | NA | NA | 1 | 1 | 2 | 0.5 | NA | NA | 1 | 1 | NA | NA | NA | NA | 1 | 1 |
| 4_4252395 | NA | NA | NA | NA | NA | NA | 1 | 1 | 1 | 1 | NA | NA | 1 | 1 | NA | NA | NA | NA | 1 | 1 |
| 4_4265591 | 1 | 1 | NA | NA | NA | NA | NA | NA | 1 | 1 | 2 | 0.5 | NA | NA | NA | NA | NA | NA | 2 | 0.5 |
| 4_4271499 | 1 | 0 | NA | NA | 1 | 1 | 1 | 0 | 3 | 0.3333 | NA | NA | NA | NA | 1 | 1 | NA | NA | 1 | 1 |
| 4_4271500 | 1 | 1 | NA | NA | 1 | 1 | 1 | 1 | 3 | 1 | NA | NA | NA | NA | 1 | 1 | NA | NA | 1 | 1 |
| 4_4271508 | 2 | 1 | NA | NA | 1 | 1 | 1 | 1 | 4 | 1 | NA | NA | NA | NA | 1 | 1 | NA | NA | 1 | 1 |
| 4_4271516 | 2 | 1 | NA | NA | 1 | 1 | 1 | 1 | 4 | 1 | NA | NA | NA | NA | 1 | 1 | NA | NA | 1 | 1 |
| 4_4271517 | 2 | 0.5 | NA | NA | 1 | 1 | 1 | 1 | 4 | 0.75 | NA | NA | NA | NA | 1 | 1 | NA | NA | 1 | 1 |
| 4_4271521 | 2 | 1 | NA | NA | 1 | 1 | 1 | 1 | 4 | 1 | NA | NA | NA | NA | 1 | 1 | NA | NA | 1 | 1 |
| 4_4271536 | NA | NA | NA | NA | 1 | 1 | NA | NA | 1 | 1 | NA | NA | NA | NA | 1 | 1 | NA | NA | 1 | 1 |
| 4_4271580 | NA | NA | NA | NA | 1 | 1 | NA | NA | 1 | 1 | NA | NA | NA | NA | 1 | 1 | NA | NA | 1 | 1 |
| 4_4272656 | 1 | 1 | 1 | 0 | NA | NA | NA | NA | 2 | 0.5 | NA | NA | NA | NA | 1 | 1 | NA | NA | 1 | 1 |
| 4_4282985 | NA | NA | 1 | 1 | NA | NA | NA | NA | 1 | 1 | 1 | 1 | NA | NA | 2 | 0.5 | 1 | 1 | 4 | 0.75 |
| 4_4287316 | 1 | 1 | NA | NA | NA | NA | NA | NA | 1 | 1 | 1 | 0 | 1 | 1 | 1 | 1 | NA | NA | 3 | 0.6667 |
| 4_4293819 | 9 | 0.3333 | 10 | 0.2 | 2 | 0 | 12 | 0.25 | 33 | 0.2424 | 5 | 0.2 | 6 | 0.3333 | 7 | 0.1429 | 7 | 0 | 25 | 0.16 |
| 4_4293960 | 9 | 0.1111 | 11 | 0 | 2 | 0.5 | 7 | 0.2857 | 29 | 0.1379 | 3 | 0.3333 | 7 | 0 | 6 | 0.1667 | 8 | 0.25 | 24 | 0.1667 |
| 4_4293969 | 9 | 0.7778 | 11 | 1 | 2 | 1 | 11 | 0.7273 | 33 | 0.8485 | 7 | 0.8571 | 11 | 0.7273 | 8 | 1 | 9 | 0.7778 | 35 | 0.8286 |
| 4_4293970 | 10 | 0.6 | 14 | 0.3571 | 3 | 1 | 11 | 0.6364 | 38 | 0.5526 | 7 | 0.7143 | 12 | 0.6667 | 9 | 0.3333 | 11 | 0.4545 | 39 | 0.5385 |
| 4_4294006 | 10 | 0.1 | 16 | 0.0625 | 1 | 0 | 10 | 0 | 37 | 0.0541 | 8 | 0.375 | 13 | 0.3077 | 10 | 0.2 | 12 | 0.1667 | 43 | 0.2558 |
| 4_4294018 | 9 | 0 | 14 | 0 | 3 | 0 | 11 | 0.2727 | 37 | 0.0811 | 7 | 0.1429 | 10 | 0 | 9 | 0 | 13 | 0.1538 | 39 | 0.0769 |
| 4_4294031 | 11 | 0.1818 | 16 | 0.3125 | 4 | 0.25 | 12 | 0.0833 | 43 | 0.2093 | 7 | 0.1429 | 11 | 0.4545 | 11 | 0.0909 | 11 | 0.1818 | 40 | 0.225 |
| 4_4294056 | 8 | 0.25 | 16 | 0.5 | 4 | 0.75 | 13 | 0.4615 | 41 | 0.4634 | 8 | 0.125 | 11 | 0.2727 | 10 | 0.7 | 9 | 0.4444 | 38 | 0.3947 |

| | | | | | | | | | | | | | | | | | | | | |
|---|---|---|---|---|---|---|---|---|---|---|---|---|---|---|---|---|---|---|---|---|
| 4_42940719 | 13 | 0.2308 | 16 | 0 | 4 | 0 | 13 | 0 | 46 | 0.0652 | 7 | 0.2857 | 16 | 0 | 12 | 0 | 11 | 0.0909 | 46 | 0.0652 |
| 4_42940797 | 13 | 0.1538 | 16 | 0.4375 | 4 | 0.75 | 13 | 0.3077 | 46 | 0.3478 | 8 | 0.375 | 15 | 0.2 | 12 | 0.3333 | 11 | 0.3636 | 46 | 0.3043 |
| 4_42940806 | 13 | 0.4615 | 17 | 0.4706 | 4 | 0.75 | 13 | 0.4615 | 47 | 0.4894 | 7 | 0.5714 | 14 | 0.4286 | 11 | 0.3636 | 12 | 0 | 44 | 0.3182 |
| 4_42940919 | 12 | 0.1667 | 14 | 0.1429 | 4 | 0 | 14 | 0.0714 | 44 | 0.1136 | 9 | 0.1111 | 15 | 0.3333 | 12 | 0 | 14 | 0.3571 | 50 | 0.22 |
| 4_42940961 | 12 | 0.5 | 12 | 0.25 | 5 | 0.6 | 12 | 0.3333 | 41 | 0.3902 | 7 | 0.4286 | 12 | 0.6667 | 12 | 0.5 | 13 | 0.3846 | 44 | 0.5 |
| 4_42941004 | 14 | 0.2857 | 16 | 0.3125 | 5 | 0.2 | 11 | 0.3636 | 46 | 0.3043 | 8 | 0.375 | 15 | 0.4 | 12 | 0.4167 | 13 | 0.2308 | 48 | 0.3542 |
| 4_42941150 | 15 | 0 | 16 | 0 | 6 | 0.1667 | 14 | 0.2143 | 51 | 0.0784 | 8 | 0 | 16 | 0.1875 | 15 | 0 | 13 | 0.1538 | 52 | 0.0962 |
| 4_42941175 | 15 | 0 | 15 | 0.2 | 6 | 0.1667 | 11 | 0.0909 | 47 | 0.1064 | 9 | 0.1111 | 16 | 0.1875 | 13 | 0.0769 | 16 | 0.0625 | 54 | 0.1111 |
| 4_42941228 | 13 | 0.1538 | 12 | 0.1667 | 5 | 0.2 | 13 | 0.2308 | 43 | 0.186 | 9 | 0.2222 | 15 | 0.4667 | 15 | 0.2 | 12 | 0.5833 | 51 | 0.3725 |
| 4_42941289 | 14 | 0.0714 | 16 | 0 | 5 | 0 | 13 | 0.1538 | 48 | 0.0625 | 8 | 0.125 | 16 | 0.3125 | 13 | 0.0769 | 16 | 0.1875 | 53 | 0.1887 |
| 4_42941345 | 13 | 0.2308 | 12 | 0 | 7 | 0.4286 | 13 | 0.3846 | 45 | 0.2444 | 12 | 0 | 14 | 0.2857 | 15 | 0.0667 | 15 | 0.2 | 56 | 0.1429 |
| 4_42941458 | 15 | 0.0667 | 17 | 0.1176 | 7 | 0.4286 | 15 | 0.0667 | 54 | 0.1296 | 13 | 0.0769 | 16 | 0.125 | 17 | 0.1176 | 13 | 0.3846 | 59 | 0.1695 |
| 4_42941477 | 14 | 0.9286 | 15 | 1 | 8 | 1 | 13 | 1 | 50 | 0.98 | 10 | 1 | 14 | 1 | 15 | 0.9333 | 13 | 0.8462 | 52 | 0.9423 |
| 4_42941545 | 17 | 0.3529 | 15 | 0.6 | 7 | 0.4286 | 14 | 0.5 | 53 | 0.4717 | 13 | 0.7692 | 16 | 0.6875 | 17 | 0.5294 | 13 | 0.5385 | 59 | 0.6271 |
| 4_42941711 | 7 | 1 | 12 | 0.9167 | 5 | 1 | 10 | 1 | 34 | 0.9706 | 11 | 1 | 15 | 0.9333 | 8 | 0.875 | 11 | 1 | 45 | 0.9556 |
| 4_42941742 | 9 | 0.6667 | 11 | 0.5455 | 4 | 1 | 10 | 0.5 | 34 | 0.6176 | 11 | 0.8182 | 15 | 0.7333 | 12 | 0.6667 | 12 | 0.5833 | 50 | 0.7 |
| 4_42941865 | 10 | 1 | 13 | 1 | 5 | 1 | 11 | 1 | 39 | 1 | 12 | 0.9167 | 16 | 0.9375 | 12 | 0.9167 | 12 | 1 | 52 | 0.9423 |
| 4_42941947 | 10 | 0.7 | 12 | 0.5833 | 6 | 0.3333 | 11 | 0.5455 | 39 | 0.5641 | 11 | 0.8182 | 17 | 0.5882 | 13 | 0.7692 | 12 | 0.3333 | 53 | 0.6226 |
| 4_42942108 | 10 | 0.6 | 14 | 0.5 | 8 | 0.25 | 11 | 0.6364 | 43 | 0.5116 | 8 | 0 | 17 | 0.3529 | 13 | 0.6154 | 10 | 0.4 | 48 | 0.375 |
| 4_42942142 | 8 | 0.625 | 10 | 0.8 | 7 | 1 | 9 | 0.6667 | 34 | 0.7647 | 9 | 0.7778 | 13 | 0.9231 | 14 | 0.7143 | 8 | 0.75 | 44 | 0.7955 |
| 4_42942487 | 10 | 0.1 | 10 | 0.4 | 8 | 0 | 11 | 0.1818 | 39 | 0.1795 | 11 | 0.2727 | 16 | 0.1875 | 14 | 0.1429 | 13 | 0.3846 | 54 | 0.2407 |
| 4_42942666 | 9 | 0.2222 | 12 | 0.25 | 8 | 0.375 | 13 | 0.3077 | 42 | 0.2857 | 10 | 0.5 | 16 | 0.5625 | 13 | 0.4615 | 14 | 0.7143 | 53 | 0.566 |
| 4_42942696 | 9 | 0.6667 | 12 | 0.4167 | 9 | 0.5556 | 15 | 0.6 | 45 | 0.5556 | 11 | 0.3636 | 15 | 0.3333 | 15 | 0.2 | 15 | 0.4667 | 56 | 0.3393 |
| 4_42942907 | 11 | 0.3636 | 10 | 0.3 | 8 | 0.75 | 15 | 0.2667 | 44 | 0.3864 | 10 | 0.6 | 18 | 0.5556 | 14 | 0.2857 | 15 | 0.7333 | 57 | 0.5439 |
| 4_42942982 | 7 | 0.4286 | 12 | 0.5 | 5 | 0 | 6 | 0.5 | 30 | 0.4 | 8 | 0.375 | 8 | 0.375 | 8 | 0.625 | 6 | 0.3333 | 30 | 0.4333 |
| 4_42943073 | 9 | 0.3333 | 14 | 0.3571 | 9 | 0.3333 | 16 | 0.0625 | 48 | 0.25 | 11 | 0.2727 | 19 | 0.4211 | 15 | 0.1333 | 19 | 0.2632 | 64 | 0.2812 |
| 4_42943135 | 10 | 0.9 | 13 | 1 | 10 | 1 | 14 | 1 | 47 | 0.9787 | 11 | 0.9091 | 20 | 0.95 | 14 | 0.9286 | 19 | 0.9474 | 64 | 0.9375 |
| 4_42943150 | 11 | 0.2727 | 14 | 0.2143 | 10 | 0.1 | 16 | 0.1875 | 51 | 0.1961 | 12 | 0.3333 | 20 | 0.4 | 16 | 0.25 | 18 | 0.5 | 66 | 0.3788 |
| 4_42943271 | 10 | 0.1 | 11 | 0.0909 | 10 | 0 | 12 | 0.0833 | 43 | 0.0698 | 8 | 0.25 | 17 | 0.1176 | 14 | 0 | 15 | 0.1333 | 54 | 0.1111 |
| 4_42943280 | 8 | 0.375 | 11 | 0.3636 | 7 | 0.7143 | 12 | 0.25 | 38 | 0.3947 | 10 | 0.3 | 17 | 0.3529 | 10 | 0.1 | 13 | 0.3077 | 50 | 0.28 |
| 4_42943335 | 9 | 0.5556 | 10 | 0.6 | 10 | 0.4 | 14 | 0.2857 | 43 | 0.4419 | 10 | 0.5 | 18 | 0.3889 | 15 | 0.5333 | 17 | 0.7647 | 60 | 0.55 |
| 4_42944633 | 1 | 1 | 3 | 1 | 1 | 1 | NA | NA | 5 | 1 | 1 | 1 | 1 | 0 | 3 | 1 | 4 | 1 | 9 | 0.8889 |
| 4_42953068 | 1 | 0 | NA | NA | 1 | 0 | 1 | 1 | 3 | 0.3333 | 1 | 0 | NA | NA | NA | NA | 2 | 0.5 | 3 | 0.3333 |
| 4_42956592 | 4 | 0.5 | 11 | 0.2727 | 5 | 0.2 | 6 | 0.3333 | 26 | 0.3077 | 3 | 0.6667 | 3 | 0.3333 | 8 | 0.25 | 9 | 0.2222 | 23 | 0.3043 |
| 4_42956721 | 5 | 0.6 | 11 | 0.6364 | 7 | 0.8571 | 7 | 0.7143 | 30 | 0.7 | 5 | 1 | 2 | 0.5 | 8 | 0.375 | 9 | 0.7778 | 24 | 0.6667 |
| 4_42956817 | 5 | 0.2 | 12 | 0.1667 | 6 | 0.1667 | 7 | 0 | 30 | 0.1333 | 6 | 0.3333 | 3 | 0 | 9 | 0 | 9 | 0.2222 | 27 | 0.1481 |
| 4_42956921 | 6 | 0.5 | 9 | 0.4444 | 6 | 0.5 | 7 | 0.4286 | 28 | 0.4643 | 3 | 1 | 3 | 0.3333 | 9 | 0.5556 | 9 | 0.4444 | 24 | 0.5417 |
| 4_42956937 | 5 | 0.4 | 11 | 0.5455 | 7 | 0.1429 | 7 | 0 | 30 | 0.3 | 7 | 0.2857 | 3 | 0.3333 | 10 | 0.1 | 9 | 0.2222 | 29 | 0.2069 |
| 4_42957340 | 2 | 0 | 8 | 0.125 | 6 | 0.1667 | 4 | 0 | 20 | 0.1 | 3 | 0 | 2 | 0 | 6 | 0.1667 | 7 | 0 | 18 | 0.0556 |
| 4_42957413 | 3 | 1 | 5 | 1 | 2 | 0.5 | 4 | 1 | 14 | 0.9286 | 5 | 0.6 | NA | NA | 3 | 0.6667 | 5 | 1 | 13 | 0.7692 |
| 4_42957428 | 7 | 0.2857 | 8 | 0.25 | 7 | 0.5714 | 7 | 0.1429 | 29 | 0.3103 | 7 | 0.2857 | 5 | 0 | 8 | 0.375 | 11 | 0.1818 | 31 | 0.2258 |
| 4_42957513 | 8 | 0.125 | 12 | 0.0833 | 9 | 0.1111 | 8 | 0 | 37 | 0.0811 | 8 | 0.125 | 5 | 0 | 12 | 0 | 10 | 0.2 | 35 | 0.0857 |
| 4_42957617 | 6 | 0.3333 | 10 | 0.4 | 10 | 0.5 | 8 | 0.125 | 34 | 0.3529 | 9 | 0.3333 | 4 | 0 | 10 | 0.1 | 9 | 0.5556 | 32 | 0.2812 |
| 4_42957638 | 9 | 0.5556 | 10 | 0.6 | 12 | 0.5 | 8 | 0.75 | 39 | 0.5897 | 9 | 0.6667 | 5 | 0.4 | 10 | 0.3 | 10 | 0.6 | 34 | 0.5 |
| 4_42957682 | 9 | 0.4444 | 14 | 0.5714 | 12 | 0.5833 | 8 | 0.125 | 43 | 0.4651 | 8 | 0.5 | 5 | 0.6 | 16 | 0.375 | 10 | 0.6 | 39 | 0.4872 |
| 4_42957823 | 9 | 0.4444 | 15 | 0.6667 | 13 | 0.5385 | 12 | 0.5833 | 49 | 0.5714 | 8 | 0.5 | 6 | 0.5 | 16 | 0.5625 | 11 | 0.5455 | 41 | 0.5366 |
| 4_42958078 | 9 | 0.1111 | 18 | 0.2222 | 9 | 0 | 12 | 0 | 48 | 0.1042 | 9 | 0 | 6 | 0.3333 | 16 | 0.25 | 12 | 0.0833 | 43 | 0.1628 |
| 4_42958598 | 10 | 0.1 | 17 | 0.0588 | 14 | 0 | 18 | 0.0556 | 59 | 0.0508 | 9 | 0.1111 | 8 | 0.125 | 16 | 0.0625 | 13 | 0.0769 | 46 | 0.087 |
| 4_42980492 | 49 | 0.2245 | 46 | 0.1957 | 35 | 0.2 | 33 | 0.2727 | 163 | 0.2209 | 38 | 0.1579 | 38 | 0.1316 | 32 | 0.1875 | 50 | 0.22 | 158 | 0.1772 |
| 4_43002867 | NA | NA | 2 | 0.5 | 1 | 1 | 2 | 0.5 | 5 | 0.6 | 2 | 0 | 1 | 0 | 1 | 0 | 3 | 0.6667 | 7 | 0.2857 |
| 4_43002944 | NA | NA | 2 | 0.5 | 1 | 1 | 2 | 0 | 5 | 0.4 | 2 | 1 | 1 | 0 | NA | NA | 3 | 1 | 6 | 0.8333 |
| 4_43002960 | NA | NA | NA | NA | 1 | 1 | 2 | 1 | 3 | 1 | 2 | 0.5 | 1 | 0 | NA | NA | 2 | 1 | 5 | 0.6 |
| 4_43017651 | 2 | 0 | 3 | 0.3333 | 1 | 0 | 4 | 0 | 10 | 0.1 | 3 | 0.3333 | 6 | 0 | 3 | 1 | 1 | 0 | 13 | 0.3077 |
| 4_43017685 | 4 | 0.5 | 4 | 0.5 | 2 | 0.5 | 4 | 1 | 14 | 0.6429 | 3 | 1 | 6 | 0.6667 | 3 | 0.6667 | 3 | 0.6667 | 15 | 0.7333 |
| 4_43017768 | 4 | 1 | 4 | 1 | 2 | 1 | 3 | 1 | 13 | 1 | 3 | 1 | 5 | 1 | 3 | 0.6667 | 2 | 1 | 13 | 0.9231 |
| 4_43017784 | 3 | 1 | 4 | 0.75 | 3 | 1 | 4 | 0.5 | 14 | 0.7857 | 3 | 1 | 5 | 0.8 | 3 | 0.6667 | 2 | 1 | 13 | 0.8462 |
| 4_43017821 | 4 | 0.75 | 4 | 0.5 | 1 | 1 | 4 | 0 | 13 | 0.4615 | 3 | 0.3333 | 5 | 0.6 | 3 | 0 | 3 | 0.6667 | 14 | 0.4286 |
| 4_43017864 | 4 | 0.5 | 4 | 0.75 | 2 | 0.5 | 3 | 0.3333 | 13 | 0.5385 | 2 | 0.5 | 3 | 0 | 3 | 1 | 3 | 0.3333 | 11 | 0.4545 |
| 4_43017996 | 3 | 0.3333 | 2 | 0.5 | NA | NA | 4 | 0 | 9 | 0.2222 | 2 | 0 | 3 | 0.3333 | 2 | 0 | 2 | 0 | 9 | 0.1111 |
| 4_43020471 | 2 | 0 | 2 | 0.5 | 2 | 0.5 | 4 | 0 | 10 | 0.2 | 2 | 0.5 | 3 | 0.3333 | 2 | 0.5 | 3 | 0.3333 | 10 | 0.4 |
| 4_43020582 | NA | NA | 3 | 0.6667 | 1 | 1 | 4 | 0.5 | 8 | 0.625 | 2 | 1 | 2 | 0 | 2 | 1 | 3 | 0.6667 | 9 | 0.6667 |
| 4_43020588 | 2 | 0 | 2 | 0.5 | 2 | 0.5 | 4 | 0 | 10 | 0.2 | 2 | 0 | 3 | 0 | 2 | 0.5 | 3 | 0 | 10 | 0.1 |
| 4_43021388 | 2 | 1 | 3 | 0.6667 | 2 | 0 | 1 | 0 | 8 | 0.5 | 2 | 0.5 | 2 | 0.5 | NA | NA | 1 | 0 | 5 | 0.4 |
| 4_43021434 | 2 | 0.5 | 3 | 0.3333 | 1 | 1 | 1 | 1 | 7 | 0.5714 | 2 | 0.5 | 1 | 0 | 1 | 1 | 1 | 0 | 5 | 0.4 |
| 4_43024832 | 2 | 0.5 | 2 | 0 | 1 | 0 | 1 | 0 | 6 | 0.1667 | 2 | 0 | 1 | 1 | NA | NA | 1 | 0 | 4 | 0.25 |

| | | | | | | | | | | | | | | | | | | | | |
|---|---|---|---|---|---|---|---|---|---|---|---|---|---|---|---|---|---|---|---|---|
| 4_4302540 | 1 | 1 | 1 | 1 | NA | NA | 1 | 1 | 3 | 1 | 2 | 1 | 1 | 1 | NA | NA | NA | NA | 3 | 1 |
| 4_4302777 | 1 | 1 | NA | NA | NA | NA | NA | NA | 1 | 1 | NA | NA | NA | NA | NA | NA | 1 | 1 | 1 | 1 |
| 4_4302778 | 1 | 1 | NA | NA | NA | NA | NA | NA | 1 | 1 | NA | NA | NA | NA | NA | NA | 1 | 1 | 1 | 1 |
| 4_4302806 | 12 | 0.0833 | 16 | 0.0625 | 7 | 0.1429 | 9 | 0.1111 | 44 | 0.0909 | 9 | 0.2222 | 8 | 0 | 7 | 0 | 5 | 0 | 29 | 0.069 |
| 4_4303269 | 4 | 0 | 5 | 0.2 | 2 | 0 | 4 | 0 | 15 | 0.0667 | 4 | 0.5 | 2 | 0 | 7 | 0.2857 | 1 | 1 | 14 | 0.3571 |
| 4_4303357 | 2 | 0 | 3 | 0 | 1 | 1 | 2 | 0 | 8 | 0.125 | 4 | 0.25 | 1 | 0 | 5 | 0.2 | 3 | 0 | 13 | 0.1538 |
| 4_4303358 | 1 | 1 | 3 | 0 | 1 | 0 | 2 | 0 | 7 | 0.1429 | 3 | 0.3333 | 1 | 1 | 5 | 0.4 | 2 | 0 | 11 | 0.3636 |
| 4_4303625 | 1 | 0 | 1 | 1 | 1 | 1 | 2 | 1 | 5 | 0.8 | 3 | 0.6667 | 1 | 1 | 3 | 0.3333 | 2 | 0.5 | 9 | 0.5556 |
| 4_4303632 | NA | NA | 1 | 1 | NA | NA | 2 | 0.5 | 3 | 0.6667 | 2 | 0 | NA | NA | 2 | 1 | 2 | 0.5 | 6 | 0.5 |
| 4_4303640 | NA | NA | 1 | 1 | NA | NA | 2 | 0.5 | 3 | 0.6667 | 3 | 1 | 1 | 0 | 3 | 0.6667 | 1 | 1 | 8 | 0.75 |
| 4_4303641 | NA | NA | NA | NA | 1 | 1 | 2 | 0 | 3 | 0.3333 | 2 | 1 | 1 | 0 | 2 | 0.5 | 1 | 1 | 6 | 0.6667 |
| 4_4303643 | 1 | 0 | 1 | 0 | 1 | 1 | 1 | 1 | 4 | 0.5 | 2 | 0.5 | 1 | 1 | 2 | 1 | 2 | 1 | 7 | 0.8571 |
| 4_4326339 | 2 | 1 | 1 | 1 | 2 | 0 | 1 | 0 | 6 | 0.5 | NA | NA | NA | NA | 1 | 1 | 1 | 1 | 2 | 1 |
| 4_4326374 | 2 | 0.5 | 1 | 1 | 3 | 0.3333 | NA | NA | 6 | 0.5 | NA | NA | NA | NA | 1 | 1 | 1 | 1 | 2 | 1 |
| 4_4326397 | 2 | 0 | 1 | 0 | 3 | 0.6667 | 2 | 0 | 8 | 0.25 | NA | NA | NA | NA | NA | NA | 1 | 1 | 1 | 1 |
| 4_4326408 | 2 | 1 | NA | NA | 3 | 0.3333 | 2 | 0 | 7 | 0.4286 | NA | NA | NA | NA | 1 | 1 | 1 | 0 | 2 | 0.5 |
| 4_4326436 | 3 | 0.6667 | 1 | 0 | 3 | 0 | 2 | 0 | 9 | 0.2222 | NA | NA | NA | NA | 1 | 1 | 1 | 1 | 2 | 1 |
| 4_4326458 | 1 | 1 | 1 | 0 | 3 | 0.6667 | NA | NA | 5 | 0.6 | NA | NA | NA | NA | 1 | 1 | NA | NA | 1 | 1 |
| 4_4326461 | 2 | 0 | NA | NA | 3 | 1 | 1 | 0 | 6 | 0.5 | NA | NA | NA | NA | 1 | 1 | 1 | 1 | 2 | 1 |
| 4_4326468 | 3 | 0.3333 | 1 | 1 | 4 | 0.25 | 2 | 0 | 10 | 0.3 | NA | NA | 1 | 0 | 1 | 1 | 1 | 1 | 3 | 0.6667 |
| 4_4326469 | 2 | 0 | 1 | 1 | 2 | 0 | 2 | 0 | 7 | 0.1429 | NA | NA | NA | NA | 1 | 1 | 1 | 0 | 2 | 0.5 |
| 4_4326474 | 3 | 1 | 1 | 1 | 4 | 0.5 | 2 | 0 | 10 | 0.6 | NA | NA | NA | NA | NA | NA | 1 | 1 | 1 | 1 |
| 4_4341471 | 4 | 0.25 | 1 | 0 | 1 | 1 | 3 | 0.3333 | 9 | 0.3333 | 1 | 1 | NA | NA | 1 | 1 | 1 | 1 | 3 | 1 |
| 4_4341474 | 5 | 0.6 | 1 | 0 | 1 | 0 | 4 | 0.5 | 11 | 0.4545 | 1 | 1 | NA | NA | 1 | 1 | NA | NA | 2 | 1 |
| 4_4341475 | 5 | 0.4 | 2 | 0 | 1 | 1 | 4 | 0 | 12 | 0.25 | 1 | 0 | NA | NA | 2 | 0 | 1 | 1 | 4 | 0.25 |
| 4_4341489 | 6 | 0.3333 | 2 | 0.5 | 1 | 0 | 3 | 0 | 12 | 0.25 | 1 | 1 | NA | NA | 2 | 1 | 1 | 0 | 4 | 0.75 |
| 4_4341505 | 6 | 0.6667 | 2 | 1 | 1 | 1 | 3 | 0.6667 | 12 | 0.75 | 1 | 1 | NA | NA | 2 | 1 | 1 | 0 | 4 | 0.75 |
| 4_4341524 | 6 | 0.1667 | 2 | 0 | 1 | 0 | 4 | 0 | 13 | 0.0769 | 1 | 0 | NA | NA | 2 | 0.5 | 1 | 0 | 4 | 0.25 |
| 4_4341526 | 6 | 0.1667 | 2 | 0 | 1 | 0 | 3 | 0.3333 | 12 | 0.1667 | 1 | 0 | NA | NA | 3 | 0 | 1 | 1 | 5 | 0.2 |
| 4_4341533 | 5 | 0.6 | 1 | 1 | 1 | 1 | 2 | 1 | 9 | 0.7778 | 1 | 1 | NA | NA | 3 | 1 | 1 | 1 | 5 | 1 |
| 4_4341568 | 6 | 0.1667 | 4 | 0 | 1 | 0 | 6 | 0.3333 | 17 | 0.1765 | 1 | 1 | NA | NA | 4 | 0.25 | 1 | 0 | 6 | 0.3333 |
| 4_4341590 | 6 | 0.8333 | 1 | 1 | 1 | 1 | 5 | 0.6 | 13 | 0.7692 | 2 | 1 | NA | NA | 2 | 1 | NA | NA | 4 | 1 |
| 4_4341653 | 7 | 0.7143 | 4 | 0.25 | 3 | 0.6667 | 8 | 0.75 | 22 | 0.6364 | 2 | 0 | NA | NA | 2 | 1 | 1 | 1 | 5 | 0.6 |
| 4_4341662 | 6 | 0.5 | 5 | 0 | 4 | 0 | 10 | 0.2 | 25 | 0.2 | 2 | 1 | NA | NA | 4 | 0.25 | 1 | 0 | 7 | 0.4286 |
| 4_4342190 | 8 | 0.375 | 5 | 0.4 | 5 | 0.4 | 11 | 0.3636 | 29 | 0.3793 | 3 | 0.3333 | NA | NA | 4 | 0.25 | 3 | 0.3333 | 10 | 0.3 |
| 4_4342542 | 10 | 0.3 | 4 | 0.5 | 8 | 0.75 | 10 | 0.6 | 32 | 0.5312 | 4 | 0.75 | 2 | 1 | 7 | 0.4286 | 3 | 0.3333 | 16 | 0.5625 |
| 4_4342547 | 9 | 0.3333 | 5 | 0.4 | 8 | 0.25 | 9 | 0.2222 | 31 | 0.2903 | 5 | 0.2 | 1 | 0 | 5 | 0.2 | 5 | 0.2 | 16 | 0.1875 |
| 4_4342556 | 13 | 0.0769 | 4 | 0 | 8 | 0 | 10 | 0.1 | 35 | 0.0571 | 6 | 0.1667 | 2 | 0 | 6 | 0 | 5 | 0.2 | 19 | 0.1053 |
| 4_4342557 | 10 | 0.4 | 4 | 1 | 8 | 1 | 9 | 0.7778 | 31 | 0.7419 | 5 | 0.8 | 2 | 1 | 5 | 0.6 | 5 | 0.6 | 17 | 0.7059 |
| 4_4342596 | 14 | 0.2857 | 6 | 0 | 10 | 0.3 | 12 | 0 | 42 | 0.1667 | 8 | 0.125 | 4 | 0.5 | 12 | 0.0833 | 5 | 0.2 | 29 | 0.1724 |
| 4_4342639 | 5 | 0.2 | 1 | 0 | 6 | 0 | 5 | 0 | 17 | 0.0588 | 4 | 0 | 2 | 0 | 3 | 0.3333 | 2 | 0 | 11 | 0.0909 |
| 4_4342640 | 5 | 0.4 | 1 | 1 | 6 | 0.8333 | 5 | 0.2 | 17 | 0.5294 | 3 | 0.3333 | 1 | 1 | 3 | 0.6667 | 1 | 0 | 8 | 0.5 |
| 4_4342649 | 10 | 0.2 | 7 | 0.1429 | 13 | 0.0769 | 13 | 0.3077 | 43 | 0.186 | 8 | 0 | 3 | 0 | 10 | 0.3 | 5 | 0.2 | 26 | 0.1538 |
| 4_4342660 | 12 | 0.25 | 6 | 0.1667 | 12 | 0.5 | 13 | 0.2308 | 43 | 0.3023 | 8 | 0.375 | 3 | 0.6667 | 8 | 0.375 | 6 | 0.3333 | 25 | 0.4 |
| 4_4342667 | 15 | 0.1333 | 5 | 0 | 14 | 0.5 | 13 | 0.2308 | 47 | 0.2553 | 8 | 0.25 | 5 | 0.4 | 11 | 0.3636 | 5 | 0.2 | 29 | 0.3103 |
| 4_4342669 | 14 | 0.1429 | 8 | 0.125 | 12 | 0.0833 | 13 | 0.1538 | 47 | 0.1277 | 8 | 0 | 4 | 0 | 11 | 0.1818 | 6 | 0 | 29 | 0.069 |
| 4_4342685 | 10 | 0.1 | 4 | 0.25 | 9 | 0.1111 | 11 | 0 | 34 | 0.0882 | 8 | 0.125 | 6 | 0 | 8 | 0 | 7 | 0.1429 | 29 | 0.069 |
| 4_4344290 | NA | NA | NA | NA | 1 | 1 | 2 | 1 | 3 | 1 | 2 | 1 | 1 | 1 | 2 | 1 | NA | NA | 5 | 1 |
| 4_4344673 | 6 | 0 | 10 | 0.2 | 10 | 0 | 8 | 0.125 | 34 | 0.0882 | 10 | 0.1 | 16 | 0.0625 | 9 | 0.1111 | 9 | 0 | 44 | 0.0682 |
| 4_4344680 | 6 | 0.3333 | 11 | 0.1818 | 10 | 0.1 | 6 | 0 | 33 | 0.1515 | 8 | 0.125 | 12 | 0.0833 | 12 | 0.25 | 6 | 0.3333 | 38 | 0.1842 |
| 4_4344696 | 9 | 0.6667 | 12 | 0.4167 | 12 | 0.5 | 9 | 0.4444 | 42 | 0.5 | 14 | 0.5714 | 16 | 0.5625 | 13 | 0.6923 | 7 | 0.4286 | 50 | 0.58 |
| 4_4344702 | 8 | 1 | 8 | 1 | 10 | 1 | 9 | 1 | 35 | 1 | 7 | 1 | 14 | 1 | 10 | 1 | 7 | 1 | 38 | 1 |
| 4_4344709 | 10 | 0 | 14 | 0 | 10 | 0.1 | 9 | 0.2222 | 43 | 0.0698 | 11 | 0.0909 | 15 | 0.0667 | 12 | 0.1667 | 9 | 0.2222 | 47 | 0.1277 |
| 4_4344714 | 7 | 0.4286 | 14 | 0.4286 | 12 | 0.1667 | 7 | 0.1429 | 40 | 0.3 | 13 | 0.3077 | 18 | 0.2222 | 13 | 0.3846 | 10 | 0.2 | 54 | 0.2778 |
| 4_4344726 | 8 | 0.875 | 12 | 0.8333 | 10 | 1 | 9 | 0.7778 | 39 | 0.8718 | 13 | 1 | 15 | 0.8 | 14 | 0.6429 | 7 | 0.8571 | 49 | 0.8163 |
| 4_4344740 | 4 | 1 | 10 | 0.9 | 9 | 0.8889 | 6 | 1 | 29 | 0.931 | 11 | 1 | 5 | 0.8 | 9 | 1 | 8 | 1 | 33 | 0.9697 |
| 4_4344755 | 8 | 1 | 14 | 0.9286 | 12 | 0.9167 | 8 | 1 | 42 | 0.9524 | 11 | 1 | 16 | 1 | 15 | 0.8667 | 10 | 1 | 52 | 0.9615 |
| 4_4344757 | 9 | 0.7778 | 12 | 1 | 13 | 0.9231 | 8 | 1 | 42 | 0.9286 | 8 | 0.875 | 13 | 1 | 7 | 1 | 11 | 0.9091 | 39 | 0.9487 |
| 4_4344764 | 9 | 1 | 15 | 0.8667 | 11 | 0.9091 | 8 | 0.875 | 43 | 0.907 | 11 | 1 | 19 | 0.9474 | 14 | 1 | 9 | 0.6667 | 53 | 0.9245 |
| 4_4344769 | 7 | 0.2857 | 14 | 0.2857 | 11 | 0.2727 | 8 | 0.125 | 40 | 0.25 | 10 | 0.2 | 16 | 0.375 | 11 | 0.0909 | 10 | 0.1 | 47 | 0.2128 |
| 4_4344772 | 10 | 0.8 | 13 | 0.7692 | 12 | 0.9167 | 11 | 1 | 46 | 0.8696 | 11 | 0.9091 | 18 | 0.7222 | 16 | 0.75 | 11 | 0.8182 | 56 | 0.7857 |
| 4_4344776 | 7 | 0 | 11 | 0.0909 | 10 | 0 | 8 | 0.375 | 36 | 0.1111 | 8 | 0 | 11 | 0 | 14 | 0.1429 | 8 | 0.125 | 41 | 0.0732 |
| 4_4353192 | 1 | 1 | 4 | 0.5 | 4 | 1 | 4 | 0.5 | 13 | 0.6923 | 6 | 0.8333 | 2 | 1 | 2 | 1 | 1 | 0 | 11 | 0.8182 |
| 4_4355951 | 3 | 0.6667 | 5 | 1 | 1 | 1 | 2 | 1 | 11 | 0.9091 | 2 | 0 | 3 | 1 | 4 | 0.75 | 5 | 0.4 | 14 | 0.5714 |
| 4_4355960 | 3 | 0 | 4 | 0.25 | 1 | 1 | 2 | 0 | 10 | 0.2 | 1 | 0 | 3 | 0 | 5 | 0 | 5 | 0.2 | 14 | 0.0714 |

| | | | | | | | | | | | | | | | | | | | | |
|---|---|---|---|---|---|---|---|---|---|---|---|---|---|---|---|---|---|---|---|---|
| 4_4356274 | 5 | 0.2 | 5 | 0.2 | 5 | 0.2 | 5 | 0 | 20 | 0.15 | 3 | 0 | 9 | 0 | 5 | 0.2 | 5 | 0.2 | 22 | 0.0909 |
| 4_4356293 | 4 | 0 | 5 | 0.2 | 6 | 0.1667 | 5 | 0 | 20 | 0.1 | 2 | 0 | 9 | 0.2222 | 6 | 0.3333 | 4 | 0.75 | 21 | 0.3333 |
| 4_4356306 | 6 | 0 | 5 | 0 | 4 | 0.25 | 5 | 0 | 20 | 0.05 | 2 | 0 | 7 | 0 | 6 | 0 | 5 | 0.2 | 20 | 0.05 |
| 4_4356506 | 1 | 1 | NA | NA | NA | NA | 1 | 1 | 2 | 1 | NA | NA | NA | NA | NA | NA | 1 | 1 | 1 | 1 |
| 4_4356655 | 7 | 0.1429 | 7 | 0.1429 | 6 | 0.6667 | 5 | 0.4 | 25 | 0.32 | 2 | 0 | 9 | 0.3333 | 6 | 0.3333 | 7 | 0.2857 | 24 | 0.2917 |
| 4_4356657 | 8 | 0.5 | 5 | 0.4 | 6 | 0.1667 | 5 | 0 | 24 | 0.2917 | 4 | 0 | 10 | 0.3 | 7 | 0.2857 | 6 | 0.1667 | 27 | 0.2222 |
| 4_4356683 | 9 | 0.4444 | 5 | 0.6 | 7 | 0.4286 | 6 | 0.8333 | 27 | 0.5556 | 2 | 0.5 | 12 | 0.5833 | 6 | 0.3333 | 6 | 0.5 | 26 | 0.5 |
| 4_4356690 | 8 | 0.125 | 5 | 0.4 | 4 | 0 | 4 | 1 | 21 | 0.3333 | 3 | 0 | 6 | 0.1667 | 4 | 0.25 | 3 | 0.3333 | 16 | 0.1875 |
| 4_4356699 | 8 | 0 | 4 | 0.25 | 4 | 0.25 | 4 | 0 | 20 | 0.1 | 2 | 0 | 7 | 0.1429 | 3 | 0 | 4 | 0 | 16 | 0.0625 |
| 4_4356702 | 6 | 0.3333 | 2 | 1 | 1 | 1 | 1 | 1 | 10 | 0.6 | 2 | 0.5 | 4 | 0.5 | 2 | 0.5 | 4 | 0.5 | 12 | 0.5 |
| 4_4356710 | 16 | 0.8125 | 14 | 0.6429 | 8 | 0.5 | 13 | 0.6154 | 51 | 0.6667 | 2 | 1 | 4 | 0.75 | 6 | 0.8333 | 13 | 0.7692 | 25 | 0.8 |
| 4_4356715 | 15 | 0.4 | 8 | 0.5 | 3 | 0.3333 | 9 | 0.4444 | 35 | 0.4286 | 2 | 0 | 3 | 1 | 3 | 1 | 10 | 0.5 | 18 | 0.6111 |
| 4_4356722 | 16 | 0.25 | 13 | 0.0769 | 7 | 0.1429 | 13 | 0.0769 | 49 | 0.1429 | 2 | 0 | 2 | 0 | 5 | 0.2 | 12 | 0.4167 | 21 | 0.2857 |
| 4_4356724 | 15 | 0.7333 | 11 | 0.7273 | 5 | 0.8 | 11 | 0.4545 | 42 | 0.6667 | 2 | 1 | 2 | 0.5 | 6 | 0.5 | 10 | 0.6 | 20 | 0.6 |
| 4_4356850 | 14 | 0 | 8 | 0.125 | 7 | 0 | 11 | 0.0909 | 40 | 0.05 | 1 | 0 | 1 | 0 | 4 | 0.25 | 10 | 0 | 16 | 0.0625 |
| 4_4356962 | 10 | 0 | 5 | 0.4 | 6 | 0.1667 | 6 | 0.3333 | 27 | 0.1852 | 1 | 0 | 2 | 0 | 2 | 0.5 | 6 | 0 | 11 | 0.0909 |
| 4_4357035 | 7 | 0 | 3 | 0 | 4 | 0.25 | 5 | 0 | 19 | 0.0526 | NA | NA | NA | NA | 2 | 0 | 5 | 0.2 | 7 | 0.1429 |
| 4_4357036 | 7 | 0 | 4 | 0.25 | 6 | 0.3333 | 6 | 0.1667 | 23 | 0.1739 | 1 | 1 | 1 | 0 | 3 | 0.6667 | 5 | 0.2 | 10 | 0.4 |
| 4_4357038 | 8 | 0 | 4 | 0 | 6 | 0.3333 | 6 | 0.1667 | 24 | 0.125 | 1 | 1 | 1 | 0 | 3 | 0 | 7 | 0 | 12 | 0.0833 |
| 4_4357039 | 7 | 0.2857 | 1 | 0 | 5 | 0 | 4 | 0 | 17 | 0.1176 | 1 | 0 | NA | NA | 2 | 0.5 | 7 | 0.1429 | 10 | 0.2 |
| 4_4357048 | 1 | 0 | NA | NA | 1 | 1 | NA | NA | 2 | 0.5 | 1 | 0 | NA | NA | NA | NA | 1 | 1 | 2 | 0.5 |
| 4_4357081 | 7 | 0.1429 | 3 | 0 | 5 | 0.2 | 6 | 0 | 21 | 0.0952 | 1 | 0 | NA | NA | 1 | 1 | 8 | 0.125 | 10 | 0.2 |
| 4_4357091 | 6 | 0.1667 | 1 | 0 | 4 | 0 | 5 | 0 | 16 | 0.0625 | NA | NA | NA | NA | 1 | 0 | 5 | 0.2 | 6 | 0.1667 |
| 4_4357818 | 5 | 0.8 | 2 | 1 | 2 | 1 | 3 | 1 | 12 | 0.9167 | NA | NA | NA | NA | 1 | 1 | 7 | 1 | 8 | 1 |
| 4_4357824 | 6 | 0.1667 | 2 | 0 | 3 | 0.6667 | 4 | 0.25 | 15 | 0.2667 | NA | NA | 1 | 0 | 1 | 0 | 6 | 0.1667 | 8 | 0.125 |
| 4_4357850 | 5 | 0 | 3 | 0 | 3 | 0.3333 | 3 | 0 | 14 | 0.0714 | NA | NA | 1 | 0 | 1 | 0 | 4 | 0.5 | 6 | 0.3333 |
| 4_4363216 | 1 | 1 | 1 | 0 | 2 | 0 | 1 | 0 | 5 | 0.2 | NA | NA | 1 | 1 | NA | NA | NA | NA | 1 | 1 |
| 4_4363237 | 1 | 1 | 1 | 1 | 2 | 0.5 | 1 | 1 | 5 | 0.8 | NA | NA | 1 | 1 | NA | NA | NA | NA | 1 | 1 |
| 4_4363264 | 1 | 1 | NA | NA | 2 | 0.5 | 1 | 1 | 4 | 0.75 | NA | NA | 1 | 1 | NA | NA | NA | NA | 1 | 1 |
| 4_4363360 | 1 | 0 | 3 | 0.3333 | 2 | 0.5 | NA | NA | 6 | 0.3333 | NA | NA | 2 | 0.5 | NA | NA | NA | NA | 2 | 0.5 |
| 4_4365096 | 1 | 0 | 8 | 0 | 5 | 0.2 | 2 | 0 | 16 | 0.0625 | 2 | 0 | 6 | 0.1667 | 2 | 0 | 4 | 0 | 14 | 0.0714 |
| 4_4365096 | 1 | 0 | 6 | 0 | 4 | 0 | 1 | 1 | 12 | 0.0833 | 2 | 0 | 5 | 0.2 | 2 | 0 | 4 | 0.25 | 13 | 0.1538 |
| 4_4366419 | NA | NA | NA | NA | 1 | 1 | NA | NA | 1 | 1 | 1 | 1 | NA | NA | NA | NA | NA | NA | 1 | 1 |
| 4_4366466 | NA | NA | 1 | 1 | 1 | 1 | NA | NA | 2 | 1 | 1 | 1 | NA | NA | NA | NA | NA | NA | 1 | 1 |
| 4_4366508 | NA | NA | NA | NA | 1 | 1 | NA | NA | 1 | 1 | 1 | 1 | NA | NA | NA | NA | NA | NA | 1 | 1 |
| 4_4366515 | NA | NA | NA | NA | 1 | 1 | NA | NA | 1 | 1 | 1 | 1 | NA | NA | NA | NA | NA | NA | 1 | 1 |
| 4_4367453 | 1 | 1 | 3 | 0 | 1 | 0 | 1 | 0 | 6 | 0.1667 | 4 | 0.25 | 1 | 0 | 1 | 0 | 1 | 0 | 7 | 0.1429 |
| 4_4368206 | 1 | 0 | 3 | 0.3333 | 2 | 0 | 1 | 0 | 7 | 0.1429 | 3 | 0.3333 | NA | NA | 1 | 1 | 1 | 0 | 5 | 0.4 |
| 4_4368975 | 2 | 0.5 | 2 | 0 | 3 | 0 | 2 | 0 | 9 | 0.1111 | 4 | 0.25 | 3 | 0 | NA | NA | 3 | 0 | 10 | 0.1 |
| 4_4369002 | 2 | 0.5 | 3 | 0.3333 | 1 | 0 | 2 | 0.5 | 8 | 0.375 | 4 | 0.25 | 1 | 0 | NA | NA | 2 | 0 | 7 | 0.1429 |
| 4_4369014 | 2 | 0.5 | 2 | 0 | 2 | 0.5 | 2 | 1 | 8 | 0.5 | 4 | 0.25 | 4 | 0 | NA | NA | 3 | 0.3333 | 11 | 0.1818 |
| 4_4369015 | 3 | 0.3333 | 3 | 1 | 2 | 0.5 | 2 | 1 | 10 | 0.7 | 4 | 0.5 | 3 | 0 | 1 | 0 | 3 | 0 | 11 | 0.1818 |
| 4_4369023 | 4 | 0.75 | 4 | 0.75 | 2 | 0 | NA | NA | 10 | 0.6 | 3 | 0.6667 | 2 | 0.5 | 1 | 0 | 2 | 1 | 8 | 0.625 |
| 4_4369029 | 2 | 0.5 | 4 | 0.5 | 3 | 0.6667 | 2 | 0.5 | 11 | 0.5455 | 3 | 0.6667 | 2 | 0 | 1 | 0 | 3 | 1 | 9 | 0.5556 |
| 4_4369031 | 1 | 1 | 5 | 0.6 | 2 | 1 | 2 | 1 | 10 | 0.8 | 4 | 0.75 | 4 | 0.5 | NA | NA | 1 | 1 | 9 | 0.6667 |
| 4_4369033 | 4 | 0.5 | 3 | 0.3333 | 1 | 0 | 2 | 0.5 | 10 | 0.4 | 4 | 0 | 4 | 0 | NA | NA | 2 | 0.5 | 10 | 0.1 |
| 4_4369039 | 4 | 0.5 | 5 | 0.2 | 3 | 0 | 2 | 0 | 14 | 0.2143 | 5 | 0.2 | 3 | 1 | 1 | 1 | 2 | 0.5 | 11 | 0.5455 |
| 4_4369044 | 5 | 0.2 | 5 | 0.2 | 4 | 0 | 2 | 0 | 16 | 0.125 | 4 | 0.25 | 5 | 0 | 1 | 1 | 3 | 0.3333 | 13 | 0.2308 |
| 4_4369048 | 4 | 0.25 | 5 | 0.2 | 4 | 0 | 1 | 0 | 14 | 0.1429 | 4 | 0.25 | 5 | 0 | NA | NA | 3 | 0 | 12 | 0.0833 |
| 4_4369052 | 5 | 0 | 5 | 0 | 3 | 0.3333 | 2 | 0 | 15 | 0.0667 | 5 | 0.2 | 6 | 0 | NA | NA | 3 | 0 | 14 | 0.0714 |
| 4_4369059 | 4 | 0.25 | 3 | 0.3333 | 2 | 0 | 1 | 0 | 10 | 0.2 | 4 | 0.25 | 3 | 0 | NA | NA | 3 | 0.3333 | 10 | 0.2 |
| 4_4369061 | 5 | 0.2 | 5 | 0 | 4 | 0 | 2 | 0 | 16 | 0.0625 | 5 | 0 | 5 | 0.2 | 1 | 0 | 3 | 0 | 14 | 0.0714 |
| 4_4369083 | 5 | 0.2 | 7 | 0 | 4 | 0 | 2 | 0 | 18 | 0.0556 | 5 | 0 | 6 | 0.1667 | NA | NA | 3 | 0 | 14 | 0.0714 |
| 4_4369096 | 5 | 0.2 | 8 | 0 | 3 | 0 | 2 | 0 | 18 | 0.0556 | 6 | 0.1667 | 7 | 0 | 1 | 0 | 3 | 0 | 17 | 0.0588 |
| 4_4369099 | 6 | 0 | 7 | 0 | 4 | 0.25 | 2 | 0 | 19 | 0.0526 | 7 | 0.1429 | 6 | 0 | NA | NA | 3 | 0 | 16 | 0.0625 |
| 4_4369163 | 4 | 0.25 | 8 | 0 | 4 | 0.25 | 4 | 0.25 | 20 | 0.15 | 8 | 0.125 | 7 | 0.1429 | 1 | 0 | 2 | 0.5 | 18 | 0.1667 |
| 4_4369201 | 7 | 0.1429 | 8 | 0 | 4 | 0.25 | 4 | 0 | 23 | 0.087 | 8 | 0.125 | 5 | 0 | 2 | 0 | 3 | 0.3333 | 18 | 0.1111 |
| 4_4372780 | NA | NA | 1 | 1 | 1 | 0 | NA | NA | 2 | 0.5 | NA | NA | NA | NA | 2 | 1 | NA | NA | 2 | 1 |
| 4_4372793 | NA | NA | 1 | 1 | 1 | 1 | NA | NA | 2 | 1 | NA | NA | NA | NA | 2 | 1 | NA | NA | 2 | 1 |
| 4_4372805 | 1 | 1 | 1 | 1 | 1 | 1 | NA | NA | 3 | 1 | NA | NA | NA | NA | 2 | 1 | NA | NA | 2 | 1 |
| 4_4394335 | 1 | 1 | NA | NA | 3 | 0.6667 | 2 | 0.5 | 6 | 0.6667 | NA | NA | NA | NA | NA | NA | 1 | 1 | 1 | 1 |
| 4_4394342 | 1 | 1 | NA | NA | 3 | 0.3333 | 1 | 1 | 5 | 0.6 | NA | NA | NA | NA | NA | NA | 2 | 0.5 | 2 | 0.5 |
| 4_4394343 | 1 | 0 | 1 | 1 | 3 | 0.6667 | 1 | 1 | 6 | 0.6667 | NA | NA | NA | NA | NA | NA | 2 | 0.5 | 2 | 0.5 |
| 4_4394388 | 1 | 1 | 1 | 0 | 3 | 0 | 2 | 0.5 | 7 | 0.2857 | NA | NA | NA | NA | NA | NA | 3 | 0.3333 | 3 | 0.3333 |
| 4_4394392 | 1 | 0 | 1 | 1 | 2 | 0 | 2 | 0 | 6 | 0.1667 | NA | NA | NA | NA | NA | NA | 2 | 0.5 | 2 | 0.5 |

| | | | | | | | | | | | | | | | | | | | | |
|---|---|---|---|---|---|---|---|---|---|---|---|---|---|---|---|---|---|---|---|---|
| 4_4397835 | 1 | 0 | 1 | 0 | NA | NA | 1 | 1 | 3 | 0.3333 | 2 | 0.5 | NA | NA | NA | NA | NA | NA | 2 | 0.5 |
| 4_4403248 | 51 | 0.1176 | 51 | 0.1765 | 49 | 0.102 | 44 | 0.1591 | 195 | 0.1385 | 42 | 0.119 | 45 | 0.1111 | 55 | 0.2727 | 46 | 0.1957 | 188 | 0.1809 |
| 4_4403266 | 52 | 0.5 | 65 | 0.3692 | 50 | 0.42 | 43 | 0.4419 | 210 | 0.4286 | 42 | 0.4286 | 49 | 0.4082 | 50 | 0.32 | 43 | 0.3256 | 184 | 0.3696 |
| 4_4403439 | 1 | 1 | 1 | 0 | 1 | 0 | NA | NA | 3 | 0.3333 | 1 | 0 | 3 | 0.3333 | 1 | 0 | 2 | 0 | 7 | 0.1429 |
| 4_4403456 | 1 | 1 | 1 | 0 | NA | NA | NA | NA | 2 | 0.5 | NA | NA | 4 | 0.5 | 3 | 0.6667 | 2 | 0.5 | 9 | 0.5556 |
| 4_4403493 | 1 | 1 | 1 | 0 | 1 | 0 | NA | NA | 3 | 0.3333 | NA | NA | 4 | 0.25 | 3 | 0 | 2 | 0 | 9 | 0.1111 |
| 4_4403496 | 1 | 1 | 1 | 0 | NA | NA | NA | NA | 2 | 0.5 | NA | NA | 3 | 0.6667 | 2 | 0.5 | 1 | 1 | 6 | 0.6667 |
| 4_4403508 | 1 | 1 | 1 | 0 | NA | NA | NA | NA | 2 | 0.5 | NA | NA | 2 | 0.5 | 3 | 0 | 1 | 1 | 6 | 0.3333 |
| 4_4403519 | 1 | 1 | NA | NA | NA | NA | NA | NA | 1 | 1 | NA | NA | 3 | 0.3333 | 1 | 1 | NA | NA | 4 | 0.5 |
| 4_4403523 | 1 | 1 | 1 | 0 | NA | NA | NA | NA | 2 | 0.5 | NA | NA | 3 | 0.3333 | 2 | 0.5 | 1 | 1 | 6 | 0.5 |
| 4_4403530 | NA | NA | 1 | 0 | 1 | 1 | NA | NA | 2 | 0.5 | NA | NA | 3 | 0.3333 | 2 | 1 | 1 | 1 | 6 | 0.6667 |
| 4_4403574 | NA | NA | 1 | 1 | 1 | 1 | NA | NA | 2 | 1 | NA | NA | 2 | 1 | 1 | 1 | 1 | 1 | 4 | 1 |
| 4_4403593 | NA | NA | 1 | 0 | 1 | 1 | NA | NA | 2 | 0.5 | NA | NA | 2 | 1 | 1 | 0 | 1 | 1 | 4 | 0.75 |
| 4_4403600 | NA | NA | 1 | 1 | 1 | 0 | NA | NA | 2 | 0.5 | NA | NA | 3 | 0.6667 | NA | NA | 1 | 0 | 4 | 0.5 |
| 4_4403600 | NA | NA | 1 | 1 | 1 | 0 | NA | NA | 2 | 0.5 | NA | NA | 2 | 0.5 | 1 | 0 | 1 | 0 | 4 | 0.25 |
| 4_4403624 | NA | NA | 1 | 0 | 1 | 1 | NA | NA | 2 | 0.5 | NA | NA | 3 | 0.3333 | 1 | 0 | 1 | 1 | 5 | 0.4 |
| 4_4403629 | NA | NA | 1 | 0 | 1 | 1 | NA | NA | 2 | 0.5 | NA | NA | 3 | 0.6667 | 1 | 1 | NA | NA | 4 | 0.75 |
| 4_4403691 | 3 | 0.3333 | 6 | 0.1667 | 4 | 0.5 | NA | NA | 13 | 0.3077 | 3 | 0 | 5 | 0.2 | 3 | 0.3333 | 2 | 0 | 13 | 0.1538 |
| 4_4403709 | 3 | 0.6667 | 4 | 0.5 | 3 | 0.6667 | NA | NA | 10 | 0.6 | 3 | 0.6667 | 4 | 0.25 | 2 | 0.5 | 2 | 0.5 | 11 | 0.4545 |
| 4_4403709 | 2 | 0.5 | 5 | 0.6 | 3 | 0 | NA | NA | 10 | 0.4 | 3 | 0.3333 | 4 | 0.75 | 3 | 1 | 1 | 1 | 11 | 0.7273 |
| 4_4403710 | 3 | 0.6667 | 5 | 0.6 | 3 | 0.3333 | NA | NA | 11 | 0.5455 | 2 | 0.5 | 4 | 0 | 4 | 0.5 | 1 | 0 | 11 | 0.2727 |
| 4_4403714 | 3 | 0.3333 | 4 | 0.75 | 3 | 1 | NA | NA | 10 | 0.7 | 2 | 0.5 | 3 | 0.6667 | 4 | 1 | 2 | 1 | 11 | 0.8182 |
| 4_4403717 | 3 | 0 | 6 | 0.1667 | 3 | 0 | NA | NA | 12 | 0.0833 | 3 | 0.3333 | 5 | 0 | 4 | 0.25 | 2 | 0 | 14 | 0.1429 |
| 4_4403718 | 2 | 0.5 | 6 | 0 | 4 | 0 | NA | NA | 12 | 0.0833 | 3 | 0 | 3 | 0.3333 | 4 | 0.25 | NA | NA | 10 | 0.2 |
| 4_4403733 | 3 | 0 | 5 | 0.2 | 4 | 0.25 | NA | NA | 12 | 0.1667 | 2 | 0.5 | 4 | 0 | 4 | 0 | 2 | 0 | 12 | 0.0833 |
| 4_4405987 | 2 | 0 | 2 | 0.5 | 3 | 0.3333 | NA | NA | 7 | 0.2857 | 3 | 0 | 4 | 0 | 4 | 0.25 | 2 | 0 | 13 | 0.0769 |
| 4_4405993 | 1 | 0 | 2 | 0.5 | 3 | 0.3333 | NA | NA | 6 | 0.3333 | 3 | 0.6667 | 2 | 0.5 | 2 | 0.5 | 1 | 1 | 8 | 0.625 |
| 4_4405995 | 2 | 0.5 | 3 | 0.3333 | 2 | 0.5 | NA | NA | 7 | 0.4286 | 3 | 0 | 4 | 0 | 4 | 0 | 2 | 0.5 | 13 | 0.0769 |
| 4_4405996 | 2 | 1 | 3 | 1 | 3 | 0.6667 | NA | NA | 8 | 0.875 | 2 | 1 | 4 | 0.75 | 3 | 1 | 2 | 0.5 | 11 | 0.8182 |
| 4_4405997 | 2 | 0 | 3 | 0.3333 | 2 | 0.5 | NA | NA | 7 | 0.2857 | 3 | 0.3333 | 4 | 0.5 | 4 | 0.5 | 2 | 0 | 13 | 0.3846 |
| 4_4406001 | 1 | 1 | 2 | 0 | 2 | 0 | NA | NA | 5 | 0.2 | 3 | 0.3333 | 3 | 0 | 3 | 0 | 2 | 0 | 11 | 0.0909 |
| 4_4412694 | 2 | 0.5 | 6 | 0.5 | 7 | 0.2857 | 2 | 0 | 17 | 0.3529 | 3 | 0.6667 | 3 | 0.3333 | 3 | 0.3333 | 1 | 0 | 10 | 0.4 |
| 4_4412947 | 1 | 1 | 4 | 0.25 | 3 | 0.3333 | 1 | 1 | 9 | 0.4444 | 2 | 0 | 1 | 1 | 1 | 1 | 1 | 1 | 5 | 0.6 |
| 4_4412952 | NA | NA | 3 | 0 | 4 | 0.25 | 1 | 0 | 8 | 0.125 | 1 | 0 | 1 | 1 | NA | NA | 1 | 0 | 3 | 0.3333 |
| 4_4412953 | 1 | 1 | 4 | 1 | 4 | 1 | 1 | 0 | 10 | 0.9 | 2 | 1 | 1 | 1 | 1 | 1 | NA | NA | 4 | 1 |
| 4_4493158 | 4 | 0.5 | 3 | 0.3333 | 6 | 0.5 | 4 | 0.75 | 17 | 0.5294 | 2 | 0.5 | 4 | 0.25 | 3 | 0.3333 | 3 | 0.3333 | 12 | 0.3333 |
| 4_4493166 | 4 | 0 | 3 | 0 | 6 | 0.3333 | 4 | 0 | 17 | 0.1176 | 2 | 0.5 | 3 | 0 | 2 | 0 | 2 | 0 | 9 | 0.1111 |
| 4_4493172 | 5 | 0.6 | 3 | 1 | 6 | 0.8333 | 3 | 0.3333 | 17 | 0.7059 | 1 | 1 | 4 | 0.25 | 3 | 0.6667 | 4 | 0.75 | 12 | 0.5833 |
| 4_4493179 | 5 | 0.6 | 3 | 0.6667 | 5 | 0.6 | 5 | 0.4 | 18 | 0.5556 | 2 | 0.5 | 4 | 0.25 | 2 | 0 | 4 | 0.5 | 12 | 0.3333 |
| 4_4493187 | 3 | 0 | 2 | 0 | 7 | 0.2857 | 4 | 0.5 | 16 | 0.25 | 2 | 0 | 4 | 0 | 3 | 0 | 3 | 0.3333 | 12 | 0.0833 |
| 4_4493192 | 4 | 0.75 | 3 | 1 | 7 | 0.7143 | 6 | 1 | 20 | 0.85 | 2 | 0 | 4 | 0.75 | 3 | 1 | 3 | 0.6667 | 12 | 0.6667 |
| 4_4493194 | 5 | 0.2 | 2 | 0 | 7 | 0 | 5 | 0 | 19 | 0.0526 | 2 | 0 | 3 | 0 | 1 | 1 | 4 | 0 | 10 | 0.1 |
| 4_4493200 | 4 | 0 | NA | NA | 7 | 0.1429 | 5 | 0 | 16 | 0.0625 | 2 | 0 | 4 | 0.75 | 2 | 0 | 3 | 0 | 11 | 0.2727 |
| 4_4493202 | 5 | 0.6 | 3 | 1 | 7 | 0.7143 | 7 | 0.7143 | 22 | 0.7273 | 2 | 0.5 | 4 | 0 | 2 | 1 | 3 | 0.6667 | 11 | 0.4545 |
| 4_4493205 | 4 | 0.5 | 3 | 0.3333 | 7 | 0.1429 | 7 | 0.2857 | 21 | 0.2857 | 2 | 0 | 4 | 0.25 | 3 | 0.3333 | 4 | 0.25 | 13 | 0.2308 |
| 4_4499055 | 5 | 0.2 | 2 | 0.5 | 4 | 0 | 6 | 0.5 | 17 | 0.2941 | 1 | 1 | 1 | 0 | 3 | 0 | 5 | 0 | 10 | 0.1 |
| 4_4499055 | 5 | 1 | 2 | 1 | 4 | 1 | 7 | 0.5714 | 18 | 0.8333 | 1 | 0 | NA | NA | 6 | 0.5 | 6 | 0.8333 | 13 | 0.6154 |
| 4_4499140 | NA | NA | 2 | 0.5 | 1 | 0 | 1 | 1 | 4 | 0.5 | NA | NA | 1 | 0 | 2 | 0.5 | NA | NA | 3 | 0.3333 |
| 4_4499169 | NA | NA | NA | NA | 1 | 1 | 1 | 0 | 2 | 0.5 | NA | NA | NA | NA | 2 | 0.5 | NA | NA | 2 | 0.5 |
| 4_4499189 | NA | NA | 1 | 0 | 1 | 1 | NA | NA | 2 | 0.5 | NA | NA | 1 | 0 | 2 | 1 | NA | NA | 3 | 0.6667 |
| 4_4499323 | NA | NA | NA | NA | 1 | 1 | NA | NA | 1 | 1 | NA | NA | 1 | 1 | 1 | 1 | NA | NA | 2 | 1 |
| 4_4499333 | NA | NA | NA | NA | 1 | 1 | NA | NA | 1 | 1 | NA | NA | 1 | 0 | 1 | 1 | NA | NA | 2 | 0.5 |
| 4_4499390 | NA | NA | NA | NA | 1 | 1 | NA | NA | 1 | 1 | NA | NA | NA | NA | 1 | 1 | NA | NA | 1 | 1 |
| 4_4499408 | NA | NA | 1 | 1 | 1 | 1 | NA | NA | 2 | 1 | NA | NA | 1 | 1 | 1 | 1 | NA | NA | 2 | 1 |
| 4_4499418 | NA | NA | NA | NA | 1 | 1 | NA | NA | 1 | 1 | NA | NA | NA | NA | 1 | 1 | NA | NA | 1 | 1 |
| 4_4499493 | NA | NA | NA | NA | 1 | 1 | NA | NA | 1 | 1 | NA | NA | 1 | 1 | 1 | 1 | NA | NA | 2 | 1 |
| 4_4499530 | NA | NA | 1 | 1 | 1 | 1 | NA | NA | 2 | 1 | NA | NA | NA | NA | 1 | 1 | NA | NA | 1 | 1 |
| 4_4503216 | NA | NA | 1 | 1 | NA | NA | NA | NA | 1 | 1 | NA | NA | 1 | 1 | 2 | 1 | 1 | 0 | 4 | 0.75 |
| 4_4503479 | 6 | 0 | 2 | 0 | 6 | 0.3333 | 5 | 0.4 | 19 | 0.2105 | 1 | 0 | 7 | 0.5714 | 4 | 0.5 | 2 | 0.5 | 14 | 0.5 |
| 4_4503482 | 6 | 0.1667 | 2 | 0 | 6 | 0.1667 | 6 | 0 | 20 | 0.1 | 1 | 0 | 7 | 0.1429 | 4 | 0 | 2 | 0 | 14 | 0.0714 |
| 4_4503529 | 5 | 0 | NA | NA | 5 | 0.2 | 4 | 0.25 | 14 | 0.1429 | 1 | 0 | 6 | 0.3333 | 4 | 0 | 2 | 0 | 13 | 0.1538 |
| 4_4503533 | 4 | 0.25 | NA | NA | 6 | 0.6667 | 4 | 0.5 | 14 | 0.5 | 1 | 0 | 6 | 0.5 | 4 | 0.5 | 2 | 0.5 | 13 | 0.4615 |
| 4_4503538 | 4 | 0.5 | NA | NA | 6 | 0.8333 | 2 | 1 | 12 | 0.75 | NA | NA | 4 | 0.75 | 4 | 0.25 | 2 | 0.5 | 10 | 0.5 |
| 4_4503539 | 4 | 0.5 | NA | NA | 4 | 0.25 | 4 | 0.25 | 12 | 0.3333 | NA | NA | 4 | 0.5 | 4 | 0.25 | 2 | 0.5 | 10 | 0.4 |
| 4_4503547 | 4 | 0 | NA | NA | 5 | 0 | 3 | 0.6667 | 12 | 0.1667 | 1 | 0 | 5 | 0.2 | 3 | 0.6667 | 1 | 0 | 10 | 0.3 |

| | | | | | | | | | | | | | | | | | | | | |
|---|---|---|---|---|---|---|---|---|---|---|---|---|---|---|---|---|---|---|---|---|
| 4_4503550( | 1 | 0 | NA | NA | 2 | 0 | 2 | 1 | 5 | 0.4 | 1 | 0 | 4 | 0.5 | 3 | 0.3333 | 1 | 1 | 9 | 0.4444 |
| 4_4503554 | 5 | 0.4 | NA | NA | 4 | 0.25 | 3 | 0.6667 | 12 | 0.4167 | 1 | 0 | 5 | 0.2 | 3 | 0 | 1 | 0 | 10 | 0.1 |
| 4_4503570! | 4 | 0 | NA | NA | 2 | 0.5 | 3 | 0.6667 | 9 | 0.3333 | NA | NA | 4 | 0.25 | 4 | 0 | NA | NA | 8 | 0.125 |
| 4_4503571 | 4 | 0.5 | NA | NA | 3 | 0.3333 | 5 | 1 | 12 | 0.6667 | 1 | 0 | 5 | 0.8 | 3 | 1 | 2 | 1 | 11 | 0.8182 |
| 4_4503576 | 4 | 0 | NA | NA | 3 | 0 | 6 | 0.1667 | 13 | 0.0769 | NA | NA | 5 | 0.4 | 1 | 0 | 2 | 1 | 8 | 0.5 |
| 4_4503584 | 4 | 0 | NA | NA | 4 | 0.75 | 3 | 0 | 11 | 0.2727 | 1 | 0 | 5 | 0.2 | 2 | 0 | 2 | 0 | 10 | 0.1 |
| 4_4505881 | NA | NA | NA | NA | 1 | 1 | 1 | 1 | 2 | 1 | NA | NA | 1 | 1 | NA | NA | NA | NA | 1 | 1 |
| 4_4505886 | NA | NA | NA | NA | 1 | 1 | 1 | 0 | 2 | 0.5 | NA | NA | 1 | 1 | NA | NA | NA | NA | 1 | 1 |
| 4_4505893( | NA | NA | NA | NA | NA | NA | 1 | 1 | 1 | 1 | NA | NA | 1 | 1 | NA | NA | NA | NA | 1 | 1 |
| 4_4510546 | 10 | 0.8 | 11 | 0.8182 | 9 | 0.4444 | 9 | 0.8889 | 39 | 0.7436 | 8 | 0.625 | 14 | 0.8571 | 9 | 0.8889 | 8 | 0.75 | 39 | 0.7949 |
| 4_4510547( | 14 | 0.7857 | 12 | 0.4167 | 12 | 0.3333 | 11 | 0.2727 | 49 | 0.4694 | 10 | 0.5 | 17 | 0.5882 | 10 | 0.7 | 8 | 0.5 | 45 | 0.5778 |
| 4_4510550 | 10 | 1 | 8 | 0.875 | 11 | 0.7273 | 6 | 0.8333 | 35 | 0.8571 | 5 | 1 | 14 | 0.8571 | 6 | 1 | 7 | 0.5714 | 32 | 0.8438 |
| 4_4527904 | NA | NA | 2 | 0.5 | NA | NA | NA | NA | 2 | 0.5 | 1 | 0 | 2 | 1 | NA | NA | 1 | 1 | 4 | 0.75 |
| 4_4527925! | NA | NA | 2 | 0.5 | 1 | 0 | NA | NA | 3 | 0.3333 | 1 | 1 | 2 | 1 | NA | NA | 1 | 1 | 4 | 1 |
| 4_4527933 | NA | NA | 1 | 1 | 1 | 0 | NA | NA | 2 | 0.5 | NA | NA | NA | NA | NA | NA | 1 | 1 | 1 | 1 |
| 4_4527934( | NA | NA | 1 | 1 | 1 | 1 | NA | NA | 2 | 1 | NA | NA | 1 | 1 | NA | NA | 1 | 0 | 2 | 0.5 |
| 4_4527938( | NA | NA | 2 | 1 | 1 | 1 | NA | NA | 3 | 1 | 1 | 1 | 2 | 1 | NA | NA | 1 | 1 | 4 | 1 |
| 4_4527943 | NA | NA | 2 | 0.5 | NA | NA | NA | NA | 2 | 0.5 | 1 | 1 | NA | NA | NA | NA | 1 | 1 | 2 | 1 |
| 4_4528384( | NA | NA | 2 | 0.5 | 1 | 1 | NA | NA | 3 | 0.6667 | 1 | 1 | 2 | 0 | 1 | 1 | 1 | 1 | 5 | 0.6 |
| 4_4528419 | NA | NA | 2 | 1 | NA | NA | NA | NA | 2 | 1 | 1 | 1 | 2 | 0.5 | 1 | 1 | 2 | 1 | 6 | 0.8333 |
| 4_4528446 | NA | NA | 2 | 0 | 1 | 1 | NA | NA | 3 | 0.3333 | 1 | 0 | 2 | 0.5 | NA | NA | 1 | 0 | 4 | 0.25 |
| 4_4528447( | NA | NA | 2 | 0.5 | 1 | 1 | NA | NA | 3 | 0.6667 | NA | NA | 2 | 1 | 1 | 1 | 2 | 1 | 5 | 1 |
| 4_4528453 | NA | NA | 2 | 0.5 | 1 | 1 | NA | NA | 3 | 0.6667 | 1 | 1 | 2 | 0.5 | 1 | 0 | 1 | 1 | 5 | 0.6 |
| 4_4528465 | NA | NA | 2 | 0.5 | 1 | 0 | NA | NA | 3 | 0.3333 | NA | NA | 1 | 0 | 1 | 0 | 2 | 0.5 | 4 | 0.25 |
| 4_4528472! | NA | NA | 2 | 1 | 1 | 1 | NA | NA | 3 | 1 | 1 | 1 | 2 | 0.5 | 1 | 0 | 2 | 1 | 6 | 0.6667 |
| 4_4528476 | NA | NA | 2 | 0.5 | 1 | 1 | NA | NA | 3 | 0.6667 | 1 | 1 | 2 | 0.5 | 1 | 1 | 2 | 0.5 | 6 | 0.6667 |
| 4_4528509 | NA | NA | 2 | 0.5 | 1 | 0 | NA | NA | 3 | 0.3333 | 2 | 0 | 2 | 0 | 1 | 1 | 1 | 0 | 6 | 0.1667 |
| 4_4528517 | NA | NA | 2 | 0.5 | 1 | 0 | NA | NA | 3 | 0.3333 | 2 | 0 | 3 | 0 | 1 | 1 | 1 | 0 | 7 | 0.1429 |
| 4_4528532( | NA | NA | 2 | 0.5 | 1 | 0 | NA | NA | 3 | 0.3333 | 2 | 0 | 2 | 0.5 | 2 | 0 | 2 | 0.5 | 8 | 0.25 |
| 4_4528533( | NA | NA | 2 | 0.5 | 1 | 0 | NA | NA | 3 | 0.3333 | 2 | 0.5 | 1 | 0 | 2 | 0.5 | NA | NA | 5 | 0.4 |
| 4_4528539 | NA | NA | 2 | 0.5 | 1 | 0 | NA | NA | 3 | 0.3333 | 2 | 0.5 | 4 | 1 | 2 | 1 | 1 | 1 | 9 | 0.8889 |
| 4_4528553 | NA | NA | 2 | 1 | NA | NA | NA | NA | 2 | 1 | 2 | 1 | 3 | 1 | 2 | 1 | 2 | 1 | 9 | 1 |
| 4_4528561 | NA | NA | 2 | 1 | 1 | 0 | NA | NA | 3 | 0.6667 | 2 | 1 | 3 | 1 | 1 | 1 | 2 | 1 | 8 | 1 |
| 4_4528573( | NA | NA | 2 | 0 | 2 | 0.5 | NA | NA | 4 | 0.25 | 2 | 0 | 4 | 0 | 2 | 0 | 2 | 0.5 | 10 | 0.1 |
| 4_4528574 | NA | NA | 1 | 1 | 2 | 0.5 | NA | NA | 3 | 0.6667 | 2 | 1 | 4 | 0.5 | 2 | 0.5 | NA | NA | 8 | 0.625 |
| 4_4528579 | NA | NA | 2 | 0.5 | 2 | 0 | NA | NA | 4 | 0.25 | 2 | 0 | 4 | 0.25 | 1 | 0 | 1 | 0 | 8 | 0.125 |
| 4_4528592( | NA | NA | 2 | 0 | 2 | 0.5 | NA | NA | 4 | 0.25 | 2 | 0.5 | 3 | 0.3333 | 1 | 1 | 2 | 0.5 | 8 | 0.5 |
| 4_4529742( | NA | NA | NA | NA | NA | NA | 1 | 1 | 1 | 1 | 1 | 1 | NA | NA | NA | NA | NA | NA | 1 | 1 |
| 4_4531517 | 2 | 1 | 6 | 0.1667 | 5 | 0.6 | 4 | 0 | 17 | 0.3529 | 3 | 0.3333 | 1 | 0 | 1 | 1 | 1 | 0 | 6 | 0.3333 |
| 4_4531525 | 2 | 0.5 | 7 | 0.2857 | 5 | 0.2 | 3 | 0.3333 | 17 | 0.2941 | 3 | 0.3333 | 1 | 0 | 1 | 0 | 1 | 0 | 6 | 0.1667 |
| 4_4531679 | 3 | 0.6667 | NA | NA | 1 | 0 | 4 | 0 | 8 | 0.25 | NA | NA | 2 | 0.5 | 1 | 0 | 3 | 0 | 6 | 0.1667 |
| 4_4534249( | NA | NA | 1 | 1 | NA | NA | NA | NA | 1 | 1 | 2 | 0.5 | NA | NA | 1 | 0 | 1 | 1 | 4 | 0.5 |
| 4_4534261 | 1 | 0 | 2 | 0.5 | NA | NA | 1 | 0 | 4 | 0.25 | 2 | 0.5 | 2 | 1 | 2 | 0 | NA | NA | 6 | 0.5 |
| 4_4537402 | 2 | 1 | 2 | 1 | NA | NA | 2 | 1 | 6 | 1 | 3 | 1 | 2 | 1 | 2 | 1 | 3 | 1 | 10 | 1 |
| 4_4537412 | 2 | 1 | 2 | 1 | NA | NA | 1 | 1 | 5 | 1 | 1 | 1 | 1 | 1 | NA | NA | 3 | 1 | 5 | 1 |
| 4_4537430 | 1 | 0 | 3 | 0.6667 | NA | NA | NA | NA | 4 | 0.5 | 2 | 1 | 2 | 0.5 | 1 | 1 | 2 | 0.5 | 7 | 0.7143 |
| 4_4537440 | 2 | 0.5 | 4 | 0.5 | NA | NA | 1 | 1 | 7 | 0.5714 | 2 | 1 | 2 | 0.5 | 2 | 1 | 3 | 1 | 9 | 0.8889 |
| 4_4537463 | 2 | 0.5 | 2 | 1 | NA | NA | 1 | 0 | 5 | 0.6 | 2 | 1 | 2 | 1 | 1 | 0 | 2 | 1 | 7 | 0.8571 |
| 4_4537469( | 2 | 0 | 4 | 0.25 | NA | NA | 1 | 1 | 7 | 0.2857 | 1 | 0 | 1 | 0 | 2 | 0.5 | 4 | 0.25 | 8 | 0.25 |
| 4_4537477( | 1 | 0 | 2 | 0.5 | NA | NA | 1 | 0 | 4 | 0.25 | NA | NA | 1 | 0 | NA | NA | 3 | 0.3333 | 4 | 0.25 |
| 4_4537898 | 1 | 1 | 5 | 0 | 1 | 0 | 2 | 0.5 | 9 | 0.2222 | NA | NA | NA | NA | NA | NA | 2 | 0.5 | 2 | 0.5 |
| 4_4539626 | 8 | 0.25 | 16 | 0.125 | 10 | 0.1 | 13 | 0.1538 | 47 | 0.1489 | 5 | 0.2 | 10 | 0 | 17 | 0.1176 | 9 | 0.1111 | 41 | 0.0976 |
| 4_4539662 | 6 | 0.1667 | 13 | 0.3846 | 9 | 0.4444 | 8 | 0.375 | 36 | 0.3611 | 3 | 0.3333 | 8 | 0 | 13 | 0.5385 | 8 | 0.75 | 32 | 0.4375 |
| 4_4539663( | 7 | 0 | 16 | 0 | 9 | 0.2222 | 12 | 0.1667 | 44 | 0.0909 | 3 | 0 | 8 | 0 | 17 | 0.1765 | 8 | 0 | 36 | 0.0833 |
| 4_4539732 | 4 | 0.25 | 10 | 0 | 3 | 0.6667 | 7 | 0.2857 | 24 | 0.2083 | 3 | 0 | 4 | 0 | 9 | 0.4444 | 7 | 0.2857 | 23 | 0.2609 |
| 4_4539734 | 4 | 0.25 | 11 | 0.1818 | 6 | 0.3333 | 10 | 0.4 | 31 | 0.2903 | 3 | 0.6667 | 6 | 0.3333 | 9 | 0.2222 | 7 | 0.2857 | 25 | 0.32 |
| 4_4539845! | 11 | 0.2727 | 10 | 0.3 | 9 | 0.2222 | 9 | 0.1111 | 39 | 0.2308 | 3 | 0.3333 | 7 | 0.1429 | 11 | 0.1818 | 6 | 0.1667 | 27 | 0.1852 |
| 4_4539870 | 9 | 0 | 7 | 0 | 7 | 0.4286 | 7 | 0.1429 | 30 | 0.1333 | 3 | 0.3333 | 3 | 0 | 9 | 0 | 5 | 0 | 20 | 0.05 |
| 4_4539887 | 11 | 0.6364 | 10 | 0.5 | 7 | 0.1429 | 7 | 0.7143 | 35 | 0.5143 | 3 | 0.3333 | 6 | 0.3333 | 8 | 0.375 | 6 | 0.3333 | 23 | 0.3478 |
| 4_4539912 | 9 | 0 | 10 | 0.1 | 7 | 0 | 6 | 0.1667 | 32 | 0.0625 | 4 | 0.5 | 6 | 0 | 6 | 0 | 7 | 0.2857 | 23 | 0.1739 |
| 4_4539935( | 8 | 0.375 | 6 | 0.5 | 7 | 0.5714 | 5 | 0.6 | 26 | 0.5 | 4 | 0.25 | 6 | 0.6667 | 5 | 0.4 | 5 | 0.6 | 20 | 0.5 |
| 4_4539936 | 7 | 0.7143 | 7 | 0.7143 | 7 | 0.4286 | 6 | 0.8333 | 27 | 0.6667 | 3 | 1 | 6 | 0.3333 | 5 | 0.6 | 7 | 0.7143 | 21 | 0.619 |
| 4_4539966 | 6 | 0.8333 | 3 | 1 | 6 | 0.6667 | 3 | 1 | 18 | 0.8333 | 2 | 0.5 | 4 | 1 | 1 | 1 | 3 | 1 | 10 | 0.9 |
| 4_4539968 | 5 | 0.2 | 6 | 0 | 6 | 0.1667 | 5 | 0.4 | 22 | 0.1818 | 3 | 0 | 5 | 0.2 | 3 | 0 | 4 | 0 | 15 | 0.0667 |
| 4_4539969! | 7 | 0.2857 | 6 | 0.3333 | 6 | 0.5 | 5 | 0.6 | 24 | 0.4167 | 2 | 0.5 | 6 | 0.5 | 4 | 0.5 | 6 | 0.8333 | 18 | 0.6111 |

| | | | | | | | | | | | | | | | | | | | | |
|---|---|---|---|---|---|---|---|---|---|---|---|---|---|---|---|---|---|---|---|---|
| 4_45399708 | 8 | 1 | 5 | 1 | 7 | 0.7143 | 6 | 0.8333 | 26 | 0.8846 | 2 | 1 | 6 | 1 | 4 | 1 | 5 | 1 | 17 | 1 |
| 4_45399827 | 4 | 1 | 6 | 0.8333 | 8 | 0.875 | 5 | 1 | 23 | 0.913 | 2 | 0.5 | 5 | 1 | 4 | 1 | 4 | 1 | 15 | 0.9333 |
| 4_45400042 | 6 | 0.1667 | 4 | 0.5 | 7 | 0 | 2 | 0 | 19 | 0.1579 | 2 | 0 | 6 | 0.1667 | 3 | 0.3333 | 1 | 0 | 12 | 0.1667 |
| 4_45400079 | 7 | 1 | 5 | 0.8 | 7 | 0.8571 | 5 | 1 | 24 | 0.9167 | 2 | 1 | 6 | 1 | 3 | 1 | 3 | 1 | 14 | 1 |
| 4_45400130 | 7 | 0.2857 | 4 | 0.5 | 7 | 0.4286 | 4 | 0.25 | 22 | 0.3636 | 2 | 0 | 5 | 0.2 | 4 | 0.75 | 3 | 0.3333 | 14 | 0.3571 |
| 4_45423425 | 7 | 0.1429 | 5 | 0 | 2 | 0 | 3 | 0 | 17 | 0.0588 | 1 | 0 | 3 | 0 | 4 | 0.25 | 2 | 0 | 10 | 0.1 |
| 4_45423547 | 6 | 0 | 5 | 0.2 | 1 | 1 | 4 | 0.25 | 16 | 0.1875 | 1 | 0 | 3 | 0.6667 | 4 | 0 | 2 | 0 | 10 | 0.2 |
| 4_45423948 | 6 | 0.6667 | 4 | 0.75 | 1 | 1 | 3 | 1 | 14 | 0.7857 | 2 | 1 | 3 | 0.6667 | 5 | 0.6 | 2 | 1 | 12 | 0.75 |
| 4_45529926 | 3 | 0.6667 | 3 | 1 | NA | NA | NA | NA | 6 | 0.8333 | NA | NA | 3 | 0.6667 | 2 | 1 | 2 | 1 | 7 | 0.8571 |
| 4_45530184 | 3 | 0 | 4 | 0.25 | NA | NA | NA | NA | 7 | 0.1429 | 2 | 0.5 | 2 | 0.5 | 3 | 0.3333 | 2 | 0 | 9 | 0.3333 |
| 4_45530215 | 3 | 0.3333 | 3 | 0 | NA | NA | 1 | 0 | 7 | 0.1429 | 2 | 0 | 2 | 0 | 3 | 0.3333 | 2 | 0 | 9 | 0.1111 |
| 4_45530340 | 2 | 1 | 4 | 0 | NA | NA | 1 | 1 | 7 | 0.4286 | 2 | 1 | 1 | 1 | 2 | 0 | 2 | 0.5 | 7 | 0.5714 |
| 4_45530392 | 2 | 0 | 3 | 0.3333 | NA | NA | 1 | 1 | 6 | 0.3333 | 2 | 0 | 1 | 0 | 2 | 0.5 | 1 | 1 | 6 | 0.3333 |
| 4_45530398 | 2 | 0.5 | 3 | 0.3333 | NA | NA | 1 | 0 | 6 | 0.3333 | 2 | 0.5 | 2 | 0 | 2 | 0.5 | 2 | 0 | 8 | 0.25 |
| 4_45802626 | NA | NA | NA | NA | 2 | 1 | 1 | 0 | 3 | 0.6667 | 2 | 0.5 | NA | NA | 1 | 0 | NA | NA | 3 | 0.3333 |
| 4_45803026 | NA | NA | NA | NA | 1 | 0 | 1 | 1 | 2 | 0.5 | 2 | 1 | NA | NA | 1 | 1 | NA | NA | 3 | 1 |
| 4_45803442 | NA | NA | NA | NA | 3 | 0.6667 | 1 | 1 | 4 | 0.75 | 2 | 0.5 | NA | NA | 1 | 1 | NA | NA | 3 | 0.6667 |
| 4_45803484 | NA | NA | NA | NA | 3 | 0.6667 | 1 | 0 | 4 | 0.5 | 1 | 1 | NA | NA | 1 | 1 | NA | NA | 2 | 1 |
| 4_45803533 | NA | NA | NA | NA | 2 | 0.5 | 1 | 1 | 3 | 0.6667 | 2 | 0 | NA | NA | 1 | 1 | NA | NA | 3 | 0.3333 |
| 4_45803600 | NA | NA | NA | NA | 3 | 0.3333 | 1 | 0 | 4 | 0.25 | 2 | 0.5 | NA | NA | 1 | 0 | NA | NA | 3 | 0.3333 |
| 4_45803988 | NA | NA | NA | NA | 3 | 0.3333 | 1 | 1 | 4 | 0.5 | 2 | 0.5 | NA | NA | NA | NA | NA | NA | 2 | 0.5 |
| 4_45804399 | NA | NA | NA | NA | 2 | 1 | 2 | 1 | 4 | 1 | 2 | 1 | NA | NA | 1 | 1 | 1 | 1 | 4 | 1 |
| 4_45804424 | NA | NA | NA | NA | 3 | 0.3333 | 2 | 1 | 5 | 0.6 | 2 | 0.5 | NA | NA | 1 | 1 | 1 | 0 | 4 | 0.5 |
| 4_45804489 | NA | NA | NA | NA | 3 | 0.6667 | 2 | 0.5 | 5 | 0.6 | 2 | 1 | NA | NA | 1 | 1 | 1 | 1 | 4 | 1 |
| 4_45804499 | NA | NA | NA | NA | 3 | 0.6667 | 1 | 0 | 4 | 0.5 | 2 | 0.5 | NA | NA | 1 | 1 | 1 | 0 | 4 | 0.5 |
| 4_46005358 | 1 | 0 | 1 | 1 | NA | NA | 2 | 0.5 | 4 | 0.5 | 1 | 1 | 4 | 0.75 | 5 | 0.8 | 1 | 1 | 11 | 0.8182 |
| 4_46005430 | 1 | 0 | NA | NA | NA | NA | 2 | 0.5 | 3 | 0.3333 | 1 | 0 | 2 | 0.5 | 3 | 0.6667 | 1 | 0 | 7 | 0.4286 |
| 4_46005610 | 1 | 1 | 1 | 0 | NA | NA | 3 | 0 | 5 | 0.2 | NA | NA | 5 | 0.2 | 5 | 0.6 | 1 | 0 | 11 | 0.3636 |
| 4_46025916 | 1 | 0 | 3 | 0 | NA | NA | 5 | 0.2 | 9 | 0.1111 | 2 | 1 | 4 | 0.75 | 6 | 0.1667 | 1 | 1 | 13 | 0.5385 |
| 4_46025944 | 1 | 0 | 2 | 1 | NA | NA | 3 | 1 | 6 | 0.8333 | 1 | 1 | 3 | 0.3333 | 6 | 0.6667 | NA | NA | 10 | 0.6 |
| 4_46025980 | 2 | 0 | 3 | 0 | NA | NA | 5 | 0.2 | 10 | 0.1 | 2 | 0 | 5 | 0.2 | 6 | 0 | 1 | 0 | 14 | 0.0714 |
| 4_46114406 | 8 | 0.125 | 10 | 0.2 | 10 | 0 | 11 | 0.1818 | 39 | 0.1282 | 6 | 0 | 8 | 0.25 | 13 | 0.2308 | 8 | 0.125 | 35 | 0.1714 |
| 4_46114516 | 8 | 0.75 | 10 | 0.4 | 11 | 0.5455 | 12 | 0.5 | 41 | 0.5366 | 6 | 0.6667 | 9 | 0.5556 | 14 | 0.5 | 8 | 0.375 | 37 | 0.5135 |
| 4_46114527 | 8 | 0.5 | 12 | 0.8333 | 10 | 0.9 | 12 | 1 | 42 | 0.8333 | 6 | 0.8333 | 9 | 0.6667 | 14 | 0.7143 | 8 | 0.875 | 37 | 0.7568 |
| 4_46114808 | 7 | 0 | 11 | 0.2727 | 11 | 0.0909 | 11 | 0.1818 | 40 | 0.15 | 7 | 0 | 8 | 0.25 | 14 | 0 | 11 | 0.1818 | 40 | 0.1 |
| 4_46114842 | 7 | 0.1429 | 15 | 0.0667 | 11 | 0.1818 | 10 | 0 | 43 | 0.093 | 8 | 0.125 | 8 | 0.125 | 14 | 0 | 8 | 0.25 | 38 | 0.1053 |
| 4_46114934 | 2 | 0 | 1 | 0 | 4 | 0.25 | NA | NA | 7 | 0.1429 | 4 | 0.25 | NA | NA | 2 | 0.5 | 7 | 0 | 13 | 0.1538 |
| 4_46115005 | 3 | 0 | 1 | 0 | 3 | 0.3333 | NA | NA | 7 | 0.1429 | 4 | 0.5 | NA | NA | 2 | 0 | 7 | 0 | 13 | 0.1538 |
| 4_46115243 | 6 | 0 | 14 | 0.1429 | 12 | 0.1667 | 12 | 0 | 44 | 0.0909 | 8 | 0 | 8 | 0.125 | 15 | 0.0667 | 9 | 0.2222 | 40 | 0.1 |
| 4_46115397 | 3 | 0.3333 | 1 | 0 | 3 | 0.3333 | NA | NA | 7 | 0.2857 | 3 | 0.3333 | 2 | 0 | 1 | 0 | 5 | 0.4 | 11 | 0.2727 |
| 4_46115829 | 10 | 0.5 | 15 | 0.2 | 14 | 0.2857 | 12 | 0.5 | 51 | 0.3529 | 10 | 0.6 | 8 | 0.625 | 15 | 0.4667 | 13 | 0.3077 | 46 | 0.4783 |
| 4_46115844 | 8 | 0.625 | 15 | 0.5333 | 12 | 0.3333 | 14 | 0.2143 | 49 | 0.4082 | 11 | 0.2727 | 6 | 0.5 | 14 | 0.3571 | 11 | 0.3636 | 42 | 0.3571 |
| 4_46115938 | 9 | 0.1111 | 16 | 0.1875 | 17 | 0.0588 | 13 | 0.0769 | 55 | 0.1091 | 11 | 0.2727 | 8 | 0.125 | 19 | 0.2105 | 12 | 0.25 | 50 | 0.22 |
| 4_46116687 | 2 | 0 | 1 | 1 | 2 | 0 | NA | NA | 5 | 0.2 | 2 | 0.5 | 2 | 0.5 | 1 | 1 | 3 | 0.3333 | 8 | 0.5 |
| 4_46116694 | 2 | 0.5 | 1 | 1 | 2 | 0.5 | NA | NA | 5 | 0.6 | 1 | 1 | 2 | 0.5 | 1 | 0 | 5 | 0.2 | 9 | 0.3333 |
| 4_46116748 | 2 | 0.5 | 1 | 1 | 2 | 1 | NA | NA | 5 | 0.8 | 2 | 0.5 | 2 | 0.5 | 1 | 0 | 5 | 0.2 | 10 | 0.3 |
| 4_46116918 | 2 | 1 | 1 | 1 | 2 | 0.5 | NA | NA | 5 | 0.8 | 2 | 1 | 2 | 1 | 1 | 1 | 5 | 1 | 10 | 1 |
| 4_46175267 | 1 | 1 | 3 | 0 | 3 | 0 | 2 | 0 | 9 | 0.1111 | NA | NA | 3 | 0.3333 | 5 | 0.2 | 1 | 0 | 9 | 0.2222 |
| 4_46175288 | 1 | 1 | 4 | 0.5 | 2 | 1 | 2 | 0 | 9 | 0.5556 | 1 | 0 | 4 | 0.75 | 5 | 0 | NA | NA | 10 | 0.3 |
| 4_46379943 | NA | NA | 1 | 1 | 1 | 1 | NA | NA | 2 | 1 | NA | NA | 1 | 1 | NA | NA | 1 | 1 | 2 | 1 |
| 4_46380172 | NA | NA | 1 | 1 | 1 | 0 | NA | NA | 2 | 0.5 | NA | NA | 1 | 0 | NA | NA | 1 | 1 | 2 | 0.5 |
| 4_46382179 | NA | NA | NA | NA | 1 | 1 | NA | NA | 1 | 1 | NA | NA | 1 | 0 | NA | NA | 1 | 1 | 2 | 0.5 |
| 4_46472182 | 7 | 0.1429 | 17 | 0.3529 | 11 | 0.2727 | 9 | 0.1111 | 44 | 0.25 | 8 | 0.375 | 17 | 0.4706 | 21 | 0.1429 | 8 | 0.25 | 54 | 0.2963 |
| 4_46472201 | 6 | 0.8333 | 16 | 0.8125 | 11 | 0.8182 | 12 | 0.8333 | 45 | 0.8222 | 8 | 0.75 | 20 | 0.75 | 21 | 0.7619 | 9 | 0.6667 | 58 | 0.7414 |
| 4_46472392 | 10 | 0.2 | 20 | 0.15 | 12 | 0.1667 | 12 | 0.25 | 54 | 0.1852 | 9 | 0 | 21 | 0.1429 | 23 | 0.087 | 9 | 0.1111 | 62 | 0.0968 |
| 4_46502918 | NA | NA | NA | NA | 1 | 1 | NA | NA | 1 | 1 | NA | NA | NA | NA | NA | NA | 1 | 1 | 1 | 1 |
| 4_46537245 | 2 | 0 | 4 | 0 | 1 | 0 | 2 | 0.5 | 9 | 0.1111 | 3 | 0.6667 | 3 | 0.6667 | 3 | 0 | 4 | 0.5 | 13 | 0.4615 |
| 4_46537976 | 1 | 1 | 2 | 0 | 1 | 1 | 1 | 1 | 5 | 0.6 | 1 | 1 | 1 | 0 | NA | NA | 4 | 0 | 6 | 0.1667 |
| 4_46538655 | 1 | 1 | 3 | 0.3333 | NA | NA | 1 | 0 | 5 | 0.4 | 1 | 1 | 1 | 0 | 1 | 1 | 4 | 0.25 | 7 | 0.4286 |
| 4_46538962 | 1 | 0 | 3 | 0.3333 | NA | NA | 1 | 0 | 5 | 0.2 | NA | NA | 2 | 0 | 1 | 1 | 3 | 0.3333 | 6 | 0.3333 |
| 4_46539145 | NA | NA | 2 | 0 | 1 | 1 | NA | NA | 3 | 0.3333 | NA | NA | 2 | 0 | 1 | 1 | 4 | 0.5 | 7 | 0.4286 |
| 4_46662702 | 15 | 0.0667 | 30 | 0.1667 | 17 | 0.1765 | 14 | 0 | 76 | 0.1184 | 16 | 0.25 | 7 | 0.1429 | 9 | 0.1111 | 14 | 0.2857 | 46 | 0.2174 |
| 4_46663505 | 13 | 0.8462 | 23 | 0.5217 | 11 | 0.7273 | 10 | 0.8 | 57 | 0.6842 | 13 | 0.7692 | 7 | 0.8571 | 6 | 0.6667 | 14 | 0.7143 | 40 | 0.75 |
| 4_46663561 | 13 | 0.1538 | 21 | 0.2857 | 10 | 0.5 | 9 | 0.3333 | 53 | 0.3019 | 15 | 0.4 | 7 | 0.2857 | 3 | 0.6667 | 11 | 0.4545 | 36 | 0.4167 |
| 4_46663627 | 13 | 0.3077 | 21 | 0.1429 | 12 | 0.5 | 7 | 0.2857 | 53 | 0.283 | 14 | 0.2143 | 7 | 0.4286 | 6 | 0.6667 | 16 | 0.25 | 43 | 0.3256 |

| | | | | | | | | | | | | | | | | | | | | |
|---|---|---|---|---|---|---|---|---|---|---|---|---|---|---|---|---|---|---|---|---|
| 4_4666390 | 14 | 0.1429 | 19 | 0.1053 | 11 | 0.6364 | 6 | 0.5 | 50 | 0.28 | 13 | 0.3846 | 5 | 0.2 | 6 | 0.5 | 13 | 0.3077 | 37 | 0.3514 |
| 4_4666414 | 9 | 0.8889 | 17 | 1 | 9 | 0.8889 | 5 | 1 | 40 | 0.95 | 9 | 0.8889 | 6 | 1 | 5 | 1 | 9 | 0.8889 | 29 | 0.931 |
| 4_4666425 | 11 | 0.3636 | 21 | 0.0476 | 10 | 0 | 7 | 0 | 49 | 0.102 | 9 | 0.2222 | 6 | 0 | 5 | 0 | 9 | 0 | 29 | 0.069 |
| 4_4666448 | 11 | 0.8182 | 21 | 0.4286 | 9 | 0.1111 | 7 | 0.5714 | 48 | 0.4792 | 11 | 0.3636 | 6 | 0.3333 | 4 | 0.75 | 11 | 0.5455 | 32 | 0.4688 |
| 4_4666451 | 10 | 0.8 | 15 | 0.6667 | 6 | 0.5 | 5 | 0.4 | 36 | 0.6389 | 9 | 0.7778 | 5 | 0.6 | 3 | 1 | 9 | 1 | 26 | 0.8462 |
| 4_4666451 | 7 | 0.1429 | 12 | 0.0833 | 7 | 0 | 4 | 0.5 | 30 | 0.1333 | 8 | 0.125 | 5 | 0.2 | 3 | 0 | 9 | 0.2222 | 25 | 0.16 |
| 4_4666452 | 13 | 0.7692 | 18 | 0.7778 | 10 | 0.9 | 6 | 0.6667 | 47 | 0.7872 | 10 | 0.6 | 6 | 0.8333 | 5 | 0.8 | 11 | 0.8182 | 32 | 0.75 |
| 4_4666457 | 9 | 0.5556 | 15 | 0.2667 | 6 | 0.3333 | 6 | 0.6667 | 36 | 0.4167 | 8 | 0.625 | 4 | 0.25 | 4 | 0.75 | 11 | 0.4545 | 27 | 0.5185 |
| 4_4666464 | 13 | 1 | 20 | 0.95 | 10 | 1 | 5 | 1 | 48 | 0.9792 | 10 | 1 | 6 | 0.8333 | 3 | 1 | 11 | 0.9091 | 30 | 0.9333 |
| 4_4666467 | 11 | 0.5455 | 17 | 0.2353 | 8 | 0.5 | 4 | 0.25 | 40 | 0.375 | 7 | 0.5714 | 5 | 0.6 | 5 | 0.6 | 11 | 0.6364 | 28 | 0.6071 |
| 4_4666475 | 10 | 1 | 13 | 0.8462 | 7 | 0.8571 | 4 | 0.75 | 34 | 0.8824 | 8 | 0.875 | 4 | 0.75 | 5 | 0.8 | 7 | 1 | 24 | 0.875 |
| 4_4666482 | 7 | 0.5714 | 14 | 0.2143 | 6 | 0.3333 | 6 | 0.5 | 33 | 0.3636 | 9 | 0.6667 | 3 | 0.3333 | 3 | 0 | 8 | 0.375 | 23 | 0.4348 |
| 4_4701571 | NA | NA | 1 | 0 | 4 | 0.25 | 3 | 0.3333 | 8 | 0.25 | 2 | 0.5 | 1 | 0 | NA | NA | 2 | 0 | 5 | 0.2 |
| 4_4701588 | NA | NA | 1 | 1 | 3 | 0.6667 | 3 | 0.6667 | 7 | 0.7143 | 1 | 1 | 1 | 1 | NA | NA | 2 | 0.5 | 4 | 0.75 |
| 4_4701594 | NA | NA | 1 | 0 | 2 | 0.5 | 1 | 0 | 4 | 0.25 | 1 | 0 | NA | NA | NA | NA | 2 | 0.5 | 3 | 0.3333 |
| 4_4701614 | NA | NA | NA | NA | 3 | 0.6667 | 2 | 0.5 | 5 | 0.6 | 1 | 1 | 1 | 1 | NA | NA | 2 | 0.5 | 4 | 0.75 |
| 4_4701618 | NA | NA | 1 | 1 | 4 | 1 | 2 | 0.5 | 7 | 0.8571 | NA | NA | 1 | 1 | NA | NA | 2 | 0.5 | 3 | 0.6667 |
| 4_4701619 | NA | NA | 1 | 0 | 4 | 0.25 | 2 | 0 | 7 | 0.1429 | 1 | 1 | 1 | 0 | NA | NA | 2 | 0 | 4 | 0.25 |
| 4_4701629 | NA | NA | 1 | 0 | 4 | 0.5 | 2 | 0.5 | 7 | 0.4286 | 1 | 1 | 1 | 0 | NA | NA | 1 | 1 | 3 | 0.6667 |
| 4_4701635 | NA | NA | 1 | 1 | 3 | 1 | 2 | 0.5 | 6 | 0.8333 | 2 | 0.5 | 1 | 0 | NA | NA | 2 | 0.5 | 5 | 0.4 |
| 4_4701646 | NA | NA | NA | NA | 3 | 1 | 2 | 1 | 5 | 1 | 2 | 0.5 | 1 | 1 | NA | NA | 2 | 0.5 | 5 | 0.6 |
| 4_4701654 | NA | NA | NA | NA | 4 | 0.25 | 2 | 0.5 | 6 | 0.3333 | 2 | 0.5 | 1 | 0 | NA | NA | 2 | 0 | 5 | 0.2 |
| 4_4701665 | NA | NA | 1 | 1 | 1 | 1 | 2 | 1 | 4 | 1 | 2 | 1 | 1 | 1 | NA | NA | 1 | 1 | 4 | 1 |
| 4_4712252 | 1 | 1 | 2 | 1 | NA | NA | NA | NA | 3 | 1 | NA | NA | NA | NA | NA | NA | 1 | 1 | 1 | 1 |
| 4_4712261 | 1 | 1 | 2 | 1 | NA | NA | NA | NA | 3 | 1 | NA | NA | NA | NA | NA | NA | 1 | 1 | 1 | 1 |
| 4_4741116 | 2 | 0.5 | 1 | 0 | 2 | 0.5 | 2 | 0.5 | 7 | 0.4286 | 2 | 0.5 | 2 | 0.5 | NA | NA | 1 | 1 | 5 | 0.6 |
| 4_4741419 | 3 | 0.3333 | 6 | 0.3333 | 5 | 0.2 | 2 | 0 | 16 | 0.25 | 4 | 0.5 | 3 | 0.6667 | 1 | 0 | 1 | 1 | 9 | 0.5556 |
| 4_4741434 | 3 | 0.3333 | 6 | 0.5 | 5 | 0.4 | 2 | 0 | 16 | 0.375 | 3 | 0.3333 | 2 | 0 | 1 | 0 | 2 | 0 | 8 | 0.125 |
| 4_4741477 | 4 | 0 | 6 | 0.1667 | 6 | 0.3333 | 2 | 0 | 18 | 0.1667 | 4 | 0.25 | 2 | 0 | 1 | 0 | 3 | 0 | 10 | 0.1 |
| 4_4746479 | NA | NA | NA | NA | 2 | 0.5 | NA | NA | 2 | 0.5 | NA | NA | 1 | 0 | 1 | 0 | 2 | 0.5 | 4 | 0.25 |
| 4_4746489 | NA | NA | NA | NA | 1 | 1 | NA | NA | 1 | 1 | NA | NA | NA | NA | NA | NA | 2 | 0.5 | 2 | 0.5 |
| 4_4746535 | NA | NA | NA | NA | 1 | 1 | NA | NA | 1 | 1 | NA | NA | NA | NA | NA | NA | 1 | 1 | 1 | 1 |
| 4_4746536 | NA | NA | NA | NA | 1 | 1 | NA | NA | 1 | 1 | NA | NA | NA | NA | NA | NA | 2 | 0.5 | 2 | 0.5 |
| 4_4746541 | NA | NA | NA | NA | 1 | 1 | NA | NA | 1 | 1 | NA | NA | NA | NA | NA | NA | 1 | 1 | 1 | 1 |
| 4_4746573 | NA | NA | NA | NA | 1 | 1 | NA | NA | 1 | 1 | NA | NA | NA | NA | NA | NA | 2 | 0.5 | 2 | 0.5 |
| 4_4746591 | NA | NA | NA | NA | 1 | 1 | NA | NA | 1 | 1 | NA | NA | NA | NA | NA | NA | 2 | 0.5 | 2 | 0.5 |
| 4_4746620 | NA | NA | NA | NA | 1 | 1 | NA | NA | 1 | 1 | NA | NA | NA | NA | NA | NA | 2 | 0.5 | 2 | 0.5 |
| 4_4746627 | NA | NA | NA | NA | 1 | 1 | NA | NA | 1 | 1 | NA | NA | NA | NA | NA | NA | 2 | 0.5 | 2 | 0.5 |
| 4_4746652 | NA | NA | NA | NA | 1 | 1 | NA | NA | 1 | 1 | NA | NA | NA | NA | NA | NA | 2 | 1 | 2 | 1 |
| 4_4746669 | NA | NA | NA | NA | 1 | 1 | NA | NA | 1 | 1 | NA | NA | NA | NA | NA | NA | 2 | 0.5 | 2 | 0.5 |
| 4_4746680 | NA | NA | NA | NA | 1 | 1 | NA | NA | 1 | 1 | 1 | 0 | NA | NA | 2 | 0 | 2 | 0.5 | 5 | 0.2 |
| 4_4747013 | 19 | 0.7368 | 13 | 0.8462 | 17 | 0.9412 | 22 | 0.8182 | 71 | 0.831 | 16 | 0.9375 | 23 | 0.6957 | 14 | 0.7143 | 19 | 0.6842 | 72 | 0.75 |
| 4_4747015 | 15 | 0.6667 | 16 | 0.6875 | 19 | 0.7895 | 23 | 0.6087 | 73 | 0.6849 | 17 | 0.4118 | 23 | 0.913 | 16 | 0.5 | 17 | 0.8235 | 73 | 0.6849 |
| 4_4747031 | 15 | 0.4 | 12 | 0.3333 | 18 | 0.1667 | 22 | 0.4091 | 67 | 0.3284 | 15 | 0.3333 | 24 | 0.4583 | 13 | 0.5385 | 17 | 0.4118 | 69 | 0.4348 |
| 4_4747038 | 18 | 0.5556 | 14 | 0.4286 | 20 | 0.45 | 21 | 0.5714 | 73 | 0.5068 | 16 | 0.5625 | 24 | 0.625 | 13 | 0.6154 | 18 | 0.6667 | 71 | 0.6197 |
| 4_4747040 | 17 | 1 | 15 | 1 | 19 | 1 | 22 | 1 | 73 | 1 | 14 | 1 | 24 | 1 | 16 | 1 | 16 | 1 | 70 | 1 |
| 4_4747051 | 12 | 0.1667 | 11 | 0.4545 | 9 | 0.4444 | 14 | 0.5714 | 46 | 0.413 | 8 | 0.25 | 13 | 0.4615 | 8 | 0.125 | 10 | 0.4 | 39 | 0.3333 |
| 4_4747062 | 16 | 0.6875 | 15 | 0.9333 | 17 | 0.7647 | 23 | 0.7391 | 71 | 0.7746 | 14 | 0.8571 | 24 | 0.7917 | 14 | 0.7857 | 17 | 1 | 69 | 0.8551 |
| 4_4747065 | 10 | 0.4 | 12 | 0.25 | 17 | 0.2941 | 18 | 0.4444 | 57 | 0.3509 | 12 | 0.5833 | 20 | 0.25 | 7 | 0.2857 | 11 | 0.4545 | 50 | 0.38 |
| 4_4747189 | 15 | 0.4 | 8 | 0.25 | 11 | 0.1818 | 15 | 0.3333 | 49 | 0.3061 | 7 | 0.2857 | 15 | 0.4667 | 10 | 0.5 | 12 | 0.5 | 44 | 0.4545 |
| 4_4747193 | 16 | 0 | 10 | 0.1 | 12 | 0.0833 | 17 | 0.0588 | 55 | 0.0545 | 8 | 0.125 | 16 | 0.0625 | 12 | 0.0833 | 12 | 0.25 | 48 | 0.125 |
| 4_4747218 | 14 | 0.2857 | 11 | 0.0909 | 10 | 0.2 | 16 | 0.125 | 51 | 0.1765 | 7 | 0 | 17 | 0.2353 | 9 | 0.2222 | 13 | 0.1538 | 46 | 0.1739 |
| 4_4747406 | 9 | 0.6667 | 7 | 0.2857 | 10 | 0.3 | 13 | 0.2308 | 39 | 0.359 | 6 | 0.1667 | 10 | 0.3 | 8 | 0 | 11 | 0.4545 | 35 | 0.2571 |
| 4_4807109 | NA | NA | 1 | 1 | 1 | 1 | 1 | 1 | 3 | 1 | 1 | 1 | NA | NA | 1 | 1 | NA | NA | 2 | 1 |
| 4_4808323 | NA | NA | 2 | 0.5 | 3 | 0.6667 | NA | NA | 5 | 0.6 | NA | NA | NA | NA | 1 | 1 | NA | NA | 1 | 1 |
| 4_4808329 | NA | NA | 2 | 0.5 | 2 | 1 | 1 | 1 | 5 | 0.8 | 1 | 1 | NA | NA | 1 | 1 | NA | NA | 2 | 1 |
| 4_4808348 | NA | NA | 2 | 1 | 3 | 0.6667 | 1 | 1 | 6 | 0.8333 | 1 | 1 | NA | NA | 1 | 1 | NA | NA | 2 | 1 |
| 4_4808416 | 2 | 0 | 2 | 0 | 5 | 0.2 | NA | NA | 9 | 0.1111 | 2 | 0 | NA | NA | 1 | 0 | 1 | 1 | 4 | 0.25 |
| 4_4808428 | 1 | 1 | 1 | 0 | 5 | 0.2 | 1 | 0 | 8 | 0.25 | 4 | 0 | 1 | 0 | 2 | 0.5 | 1 | 0 | 8 | 0.125 |
| 4_4808544 | 1 | 0 | 2 | 0 | 6 | 0.1667 | 2 | 0 | 11 | 0.0909 | 3 | 0 | 3 | 0 | 3 | 0 | 1 | 1 | 10 | 0.1 |
| 4_4808597 | 2 | 0 | 3 | 0 | 4 | 0.25 | 4 | 0 | 13 | 0.0769 | 5 | 0.4 | 4 | 0.25 | 2 | 0 | 1 | 1 | 12 | 0.3333 |
| 4_4808606 | 2 | 0 | 3 | 0 | 5 | 0 | 4 | 0.5 | 14 | 0.1429 | 5 | 0.4 | 4 | 0.25 | 3 | 0 | 1 | 1 | 13 | 0.3077 |
| 4_4808618 | 2 | 0 | 4 | 0 | 5 | 0 | 3 | 0.6667 | 14 | 0.1429 | 6 | 0 | 4 | 0.25 | 3 | 0.3333 | NA | NA | 13 | 0.1538 |
| 4_4818214 | 1 | 1 | 1 | 0 | 2 | 0.5 | 2 | 0 | 6 | 0.3333 | 2 | 0.5 | NA | NA | 2 | 0.5 | 2 | 0 | 6 | 0.3333 |
| 4_4818351 | 1 | 0 | NA | NA | 1 | 1 | 2 | 0 | 4 | 0.25 | 1 | 1 | NA | NA | 1 | 0 | 1 | 1 | 3 | 0.6667 |

| | | | | | | | | | | | | | | | | | | | | |
|---|---|---|---|---|---|---|---|---|---|---|---|---|---|---|---|---|---|---|---|---|
| 4_48184187 | 1 | 0 | 1 | 0 | 2 | 0.5 | 3 | 0 | 7 | 0.1429 | 1 | 0 | NA | NA | 1 | 0 | 2 | 0.5 | 4 | 0.25 |
| 4_48185138 | 1 | 1 | 1 | 0 | 2 | 1 | 3 | 0.6667 | 7 | 0.7143 | 2 | 0.5 | NA | NA | 2 | 0.5 | 2 | 1 | 6 | 0.6667 |
| 4_48185221 | 1 | 1 | 1 | 1 | 2 | 0 | 3 | 1 | 7 | 0.7143 | 2 | 1 | NA | NA | 2 | 1 | 2 | 1 | 6 | 1 |
| 4_48185503 | 2 | 0.5 | 1 | 0 | 2 | 0 | 3 | 0.3333 | 8 | 0.25 | 2 | 0.5 | NA | NA | 3 | 0 | 2 | 0.5 | 7 | 0.2857 |
| 4_48185630 | 1 | 1 | 1 | 1 | 2 | 1 | 3 | 1 | 7 | 1 | 2 | 1 | NA | NA | 3 | 0.6667 | 2 | 1 | 7 | 0.8571 |
| 4_48185678 | 1 | 0 | 1 | 0 | 2 | 0 | 2 | 1 | 6 | 0.3333 | 2 | 0.5 | NA | NA | 4 | 0.25 | 2 | 0 | 8 | 0.25 |
| 4_48185749 | 2 | 0 | 1 | 0 | 2 | 0.5 | 3 | 0.3333 | 8 | 0.25 | 2 | 0.5 | NA | NA | 4 | 0.5 | 2 | 0 | 8 | 0.375 |
| 4_48185775 | 2 | 0 | 1 | 0 | 1 | 0 | 3 | 0.6667 | 7 | 0.2857 | 2 | 0.5 | NA | NA | 4 | 0.25 | 2 | 1 | 8 | 0.5 |
| 4_48190837 | 1 | 0 | NA | NA | 1 | 0 | 2 | 0.5 | 4 | 0.25 | NA | NA | 2 | 1 | 1 | 0 | NA | NA | 3 | 0.6667 |
| 4_48192400 | 1 | 1 | NA | NA | 1 | 1 | 2 | 0.5 | 4 | 0.75 | NA | NA | 1 | 1 | 1 | 0 | NA | NA | 2 | 0.5 |
| 4_48192561 | 1 | 1 | NA | NA | NA | NA | 2 | 0.5 | 3 | 0.6667 | NA | NA | 1 | 1 | 1 | 0 | NA | NA | 2 | 0.5 |
| 4_48192620 | 1 | 1 | NA | NA | 1 | 0 | 2 | 0.5 | 4 | 0.5 | NA | NA | 1 | 1 | 1 | 0 | NA | NA | 2 | 0.5 |
| 4_48194353 | 2 | 0 | 2 | 0 | 2 | 0 | 3 | 0.3333 | 9 | 0.1111 | NA | NA | 2 | 0.5 | 4 | 0 | NA | NA | 6 | 0.1667 |
| 4_48194503 | 4 | 0.25 | 4 | 0 | 6 | 0 | 5 | 0 | 19 | 0.0526 | 3 | 0.3333 | 8 | 0.25 | 9 | 0 | 6 | 0.1667 | 26 | 0.1538 |
| 4_48421723 | 1 | 1 | NA | NA | NA | NA | NA | NA | 1 | 1 | NA | NA | NA | NA | 1 | 1 | NA | NA | 1 | 1 |
| 4_48421891 | 1 | 1 | NA | NA | NA | NA | NA | NA | 1 | 1 | NA | NA | NA | NA | 2 | 0.5 | NA | NA | 2 | 0.5 |
| 4_48421992 | 1 | 1 | NA | NA | NA | NA | NA | NA | 1 | 1 | NA | NA | NA | NA | 2 | 1 | NA | NA | 2 | 1 |
| 4_48422021 | 1 | 1 | NA | NA | NA | NA | NA | NA | 1 | 1 | NA | NA | NA | NA | 2 | 0.5 | NA | NA | 2 | 0.5 |
| 4_48429794 | 1 | 1 | NA | NA | NA | NA | 2 | 1 | 3 | 1 | NA | NA | 1 | 0 | 1 | 1 | NA | NA | 2 | 0.5 |
| 4_48429893 | 1 | 0 | NA | NA | NA | NA | 2 | 0.5 | 3 | 0.3333 | NA | NA | 1 | 0 | 1 | 1 | NA | NA | 2 | 0.5 |
| 4_48429916 | NA | NA | NA | NA | NA | NA | 2 | 0.5 | 2 | 0.5 | NA | NA | 1 | 1 | 1 | 1 | NA | NA | 2 | 1 |
| 4_48429965 | 1 | 0 | NA | NA | NA | NA | 1 | 1 | 2 | 0.5 | NA | NA | 1 | 0 | 1 | 1 | NA | NA | 2 | 0.5 |
| 4_48430154 | 1 | 1 | NA | NA | NA | NA | 2 | 0.5 | 3 | 0.6667 | NA | NA | 1 | 1 | 1 | 1 | NA | NA | 2 | 1 |
| 4_48430902 | 1 | 0 | 2 | 0.5 | 1 | 1 | NA | NA | 4 | 0.5 | 2 | 0.5 | 3 | 0.3333 | 5 | 0.6 | 2 | 0.5 | 12 | 0.5 |
| 4_48468580 | NA | NA | 1 | 1 | NA | NA | NA | NA | 1 | 1 | 1 | 1 | 1 | 1 | 3 | 1 | 1 | 0 | 6 | 0.8333 |
| 4_48468620 | NA | NA | 1 | 1 | NA | NA | NA | NA | 1 | 1 | NA | NA | 1 | 1 | 3 | 1 | 1 | 1 | 5 | 1 |
| 4_48468698 | NA | NA | 1 | 1 | NA | NA | NA | NA | 1 | 1 | 1 | 1 | 1 | 1 | 3 | 1 | 1 | 1 | 6 | 1 |
| 4_48468783 | NA | NA | 1 | 1 | NA | NA | NA | NA | 1 | 1 | 1 | 0 | 2 | 0.5 | 2 | 1 | 1 | 1 | 6 | 0.6667 |
| 4_48552533 | 1 | 1 | 2 | 1 | NA | NA | NA | NA | 3 | 1 | 2 | 0.5 | 2 | 1 | 2 | 0.5 | 1 | 1 | 7 | 0.7143 |
| 4_48552554 | 1 | 1 | 2 | 1 | NA | NA | NA | NA | 3 | 1 | 2 | 0 | 2 | 1 | 2 | 0 | NA | NA | 6 | 0.3333 |
| 4_48552609 | NA | NA | 1 | 1 | NA | NA | NA | NA | 1 | 1 | 2 | 0.5 | 2 | 1 | 2 | 0.5 | 1 | 0 | 7 | 0.5714 |
| 4_48560869 | 2 | 0.5 | 1 | 1 | NA | NA | NA | NA | 3 | 0.6667 | 1 | 1 | NA | NA | 1 | 0 | 1 | 0 | 3 | 0.3333 |
| 4_48561194 | 2 | 1 | 1 | 1 | NA | NA | NA | NA | 3 | 1 | 2 | 1 | 2 | 1 | 2 | 1 | 1 | 1 | 7 | 1 |
| 4_48561255 | 1 | 1 | NA | NA | NA | NA | 1 | 1 | 2 | 1 | 2 | 1 | 2 | 1 | 2 | 1 | 1 | 1 | 7 | 1 |
| 4_48561339 | 2 | 0 | 2 | 0.5 | NA | NA | 1 | 0 | 5 | 0.2 | 2 | 0.5 | 2 | 0 | 2 | 0.5 | 1 | 0 | 7 | 0.2857 |
| 4_48561775 | 3 | 0.6667 | 1 | 1 | NA | NA | NA | NA | 4 | 0.75 | 2 | 0.5 | 3 | 0 | 1 | 0 | 1 | 0 | 7 | 0.1429 |
| 4_48585606 | 6 | 0.1667 | 6 | 0.1667 | 4 | 0 | 3 | 0.3333 | 19 | 0.1579 | 3 | 0.3333 | 7 | 0 | 3 | 0.3333 | 3 | 0 | 16 | 0.125 |
| 4_48658815 | 11 | 0.1818 | 15 | 0.1333 | 11 | 0.0909 | 16 | 0.0625 | 53 | 0.1132 | 7 | 0 | 8 | 0 | 7 | 0.1429 | 11 | 0.0909 | 33 | 0.0606 |
| 4_48662093 | 12 | 0.4167 | 17 | 0.4118 | 13 | 0.4615 | 16 | 0.5 | 58 | 0.4483 | 6 | 0.6667 | 8 | 0.75 | 7 | 0.2857 | 13 | 0.6154 | 34 | 0.5882 |
| 4_48662183 | 11 | 0.3636 | 19 | 0.6316 | 8 | 0.625 | 16 | 0.4375 | 54 | 0.5185 | 5 | 0.6 | 10 | 0.7 | 4 | 0.5 | 10 | 0.7 | 29 | 0.6552 |
| 4_48662323 | 13 | 0.2308 | 20 | 0 | 12 | 0.1667 | 15 | 0.0667 | 60 | 0.1 | 6 | 0 | 9 | 0.1111 | 8 | 0.125 | 13 | 0.0769 | 36 | 0.0833 |
| 4_48662370 | 10 | 0 | 17 | 0.0588 | 11 | 0.0909 | 18 | 0.1111 | 56 | 0.0714 | 6 | 0 | 11 | 0 | 6 | 0 | 11 | 0.1818 | 34 | 0.0588 |
| 4_48662437 | 12 | 0.25 | 17 | 0.1765 | 12 | 0.0833 | 17 | 0.1765 | 58 | 0.1724 | 3 | 0.3333 | 9 | 0 | 8 | 0.25 | 12 | 0.3333 | 32 | 0.2188 |
| 4_48662489 | 14 | 0.1429 | 20 | 0.25 | 13 | 0.0769 | 18 | 0.1111 | 65 | 0.1538 | 8 | 0.25 | 10 | 0 | 8 | 0.25 | 12 | 0.1667 | 38 | 0.1579 |
| 4_48662983 | 19 | 0.0526 | 27 | 0.1111 | 18 | 0.1667 | 17 | 0.0588 | 81 | 0.0988 | 8 | 0.125 | 8 | 0.125 | 8 | 0 | 12 | 0 | 36 | 0.0556 |
| 4_49055229 | NA | NA | NA | NA | 1 | 1 | NA | NA | 1 | 1 | NA | NA | 1 | 1 | 1 | 0 | NA | NA | 2 | 0.5 |
| 4_49055337 | NA | NA | NA | NA | 1 | 1 | NA | NA | 1 | 1 | NA | NA | NA | NA | 1 | 1 | NA | NA | 1 | 1 |
| 4_49057038 | 1 | 1 | NA | NA | NA | NA | NA | NA | 1 | 1 | NA | NA | 1 | 1 | NA | NA | NA | NA | 1 | 1 |
| 4_49057400 | 1 | 1 | NA | NA | NA | NA | NA | NA | 1 | 1 | NA | NA | 1 | 1 | NA | NA | NA | NA | 1 | 1 |
| 4_49338480 | 13 | 0.2308 | 7 | 0 | 3 | 0 | 18 | 0.0556 | 41 | 0.0976 | 5 | 0.4 | 20 | 0.2 | 15 | 0.1333 | 8 | 0 | 48 | 0.1667 |
| 4_49340118 | 1 | 1 | 3 | 0 | 1 | 0 | 5 | 0.2 | 10 | 0.2 | 1 | 0 | 4 | 0 | 3 | 0 | 3 | 0.3333 | 11 | 0.0909 |
| 4_49512688 | 2 | 0 | NA | NA | 5 | 0 | 9 | 0.1111 | 16 | 0.0625 | 2 | 0.5 | 5 | 0 | 6 | 0 | 6 | 0 | 19 | 0.0526 |
| 4_49512716 | 1 | 1 | 2 | 0.5 | 5 | 0.2 | 7 | 0.5714 | 15 | 0.4667 | 1 | 1 | 5 | 0.8 | 5 | 0.4 | 5 | 0.2 | 16 | 0.5 |
| 4_49512809 | 1 | 1 | 2 | 0 | 4 | 0.25 | 9 | 0 | 16 | 0.125 | 2 | 0 | 5 | 0.2 | 5 | 0 | 8 | 0 | 20 | 0.05 |
| 4_49513515 | 1 | 0 | NA | NA | 5 | 0 | 7 | 0.1429 | 13 | 0.0769 | NA | NA | 5 | 0 | 4 | 0.25 | 7 | 0 | 16 | 0.0625 |
| 4_49513571 | 1 | 1 | NA | NA | 5 | 1 | 7 | 0.8571 | 13 | 0.9231 | NA | NA | 5 | 0.8 | 5 | 0.8 | 8 | 0.75 | 18 | 0.7778 |
| 4_49513682 | 1 | 0 | NA | NA | 4 | 0 | 7 | 0.1429 | 12 | 0.0833 | NA | NA | 5 | 0.2 | 4 | 0 | 7 | 0 | 16 | 0.0625 |
| 4_49513753 | 1 | 0 | NA | NA | 5 | 0 | 5 | 0.2 | 11 | 0.0909 | NA | NA | 5 | 0 | 5 | 0.4 | 8 | 0 | 18 | 0.1111 |
| 4_49513814 | 1 | 1 | NA | NA | 3 | 0.3333 | 7 | 0.4286 | 11 | 0.4545 | NA | NA | 5 | 0 | 5 | 0.2 | 7 | 0.1429 | 17 | 0.1176 |
| 4_49513909 | 1 | 0 | NA | NA | 4 | 0.25 | 6 | 0 | 11 | 0.0909 | NA | NA | 5 | 0 | 4 | 0.25 | 8 | 0 | 17 | 0.0588 |
| 4_49513960 | 1 | 0 | NA | NA | 4 | 0 | 6 | 0.1667 | 11 | 0.0909 | NA | NA | 3 | 0.3333 | 4 | 0.25 | 5 | 0.2 | 12 | 0.25 |
| 4_49514106 | 4 | 0 | NA | NA | 4 | 0.25 | 9 | 0 | 17 | 0.0588 | NA | NA | 6 | 0.1667 | 4 | 0 | 7 | 0 | 17 | 0.0588 |
| 4_49514118 | 4 | 0.5 | 1 | 1 | 5 | 0.2 | 9 | 0.7778 | 19 | 0.5789 | NA | NA | 6 | 0.5 | 5 | 0.4 | 7 | 0.4286 | 18 | 0.4444 |
| 4_49514134 | 4 | 0.5 | 1 | 0 | 4 | 0.5 | 8 | 0.5 | 17 | 0.4706 | NA | NA | 4 | 0.25 | 5 | 0 | 6 | 0.6667 | 15 | 0.3333 |
| 4_49514174 | 2 | 0 | 1 | 0 | 5 | 0 | 9 | 0.1111 | 17 | 0.0588 | 1 | 0 | 6 | 0.1667 | 5 | 0 | 7 | 0.1429 | 19 | 0.1053 |

| | | | | | | | | | | | | | | | | | | | | |
|---|---|---|---|---|---|---|---|---|---|---|---|---|---|---|---|---|---|---|---|---|
| 4_4951424 | 3 | 1 | 1 | 1 | 5 | 1 | 7 | 0.8571 | 16 | 0.9375 | NA | NA | 4 | 1 | 3 | 0.6667 | 6 | 0.8333 | 13 | 0.8462 |
| 4_4951437 | 3 | 1 | 2 | 0.5 | 5 | 0.8 | 8 | 0.875 | 18 | 0.8333 | NA | NA | 5 | 1 | 5 | 0.8 | 6 | 1 | 16 | 0.9375 |
| 4_4951438 | 3 | 0.6667 | 2 | 1 | 6 | 1 | 9 | 1 | 20 | 0.95 | NA | NA | 5 | 1 | 4 | 1 | 5 | 1 | 14 | 1 |
| 4_4951441 | 2 | 0 | 2 | 0 | 5 | 0 | 9 | 0.2222 | 18 | 0.1111 | NA | NA | 3 | 0.3333 | 5 | 0.2 | 5 | 0.2 | 13 | 0.2308 |
| 4_4952915 | 1 | 1 | 1 | 1 | NA | NA | NA | NA | 2 | 1 | NA | NA | 3 | 1 | NA | NA | NA | NA | 3 | 1 |
| 4_4952920 | 1 | 0 | 1 | 1 | NA | NA | 1 | 1 | 3 | 0.6667 | NA | NA | 3 | 1 | NA | NA | NA | NA | 3 | 1 |
| 4_4952923 | 1 | 1 | 1 | 0 | NA | NA | 1 | 0 | 3 | 0.3333 | NA | NA | 3 | 0.3333 | NA | NA | NA | NA | 3 | 0.3333 |
| 4_4952926 | 1 | 0 | 1 | 0 | NA | NA | 1 | 1 | 3 | 0.3333 | NA | NA | 3 | 0.3333 | NA | NA | NA | NA | 3 | 0.3333 |
| 4_4952942 | 2 | 0.5 | NA | NA | NA | NA | 1 | 0 | 3 | 0.3333 | NA | NA | 2 | 0.5 | NA | NA | NA | NA | 2 | 0.5 |
| 4_4952963 | 2 | 1 | 1 | 1 | NA | NA | 2 | 1 | 5 | 1 | NA | NA | 3 | 1 | NA | NA | NA | NA | 3 | 1 |
| 4_4952992 | 2 | 0.5 | 1 | 1 | NA | NA | 2 | 0 | 5 | 0.4 | NA | NA | 4 | 0.25 | NA | NA | NA | NA | 4 | 0.25 |
| 4_4953012 | 2 | 0.5 | 1 | 1 | NA | NA | 2 | 1 | 5 | 0.8 | NA | NA | 3 | 0.6667 | NA | NA | NA | NA | 3 | 0.6667 |
| 4_4953016 | 2 | 0 | 1 | 0 | NA | NA | 2 | 0.5 | 5 | 0.2 | NA | NA | 4 | 0.25 | NA | NA | NA | NA | 4 | 0.25 |
| 4_4953022 | 1 | 1 | 1 | 1 | NA | NA | 2 | 1 | 4 | 1 | NA | NA | 4 | 1 | NA | NA | NA | NA | 4 | 1 |
| 4_4953037 | 2 | 1 | 1 | 1 | NA | NA | 1 | 1 | 4 | 1 | NA | NA | 4 | 0.75 | 1 | 1 | NA | NA | 5 | 0.8 |
| 4_4953049 | 3 | 1 | 1 | 1 | NA | NA | 3 | 0.6667 | 7 | 0.8571 | NA | NA | 4 | 1 | NA | NA | NA | NA | 4 | 1 |
| 4_4953056 | 3 | 0.3333 | 1 | 0 | NA | NA | 3 | 0.3333 | 7 | 0.2857 | NA | NA | 4 | 0.25 | 1 | 1 | NA | NA | 5 | 0.4 |
| 4_4953088 | 4 | 0.5 | 1 | 0 | 2 | 0.5 | 3 | 0.3333 | 10 | 0.4 | NA | NA | 4 | 0.75 | 1 | 0 | NA | NA | 5 | 0.6 |
| 4_4953113 | 4 | 0.5 | 1 | 0 | 2 | 1 | 3 | 0 | 10 | 0.4 | NA | NA | 4 | 0.25 | 1 | 0 | NA | NA | 5 | 0.2 |
| 4_4953335 | 2 | 0.5 | NA | NA | NA | NA | 1 | 1 | 3 | 0.6667 | NA | NA | 1 | 1 | NA | NA | NA | NA | 1 | 1 |
| 4_4958547 | 8 | 0 | 15 | 0.1333 | 11 | 0.1818 | 11 | 0.4545 | 45 | 0.2 | 9 | 0.2222 | 17 | 0.3529 | 10 | 0.6 | 6 | 0.6667 | 42 | 0.4286 |
| 4_4958554 | 8 | 0.25 | 13 | 0 | 10 | 0 | 6 | 0 | 37 | 0.0541 | 8 | 0.125 | 16 | 0.125 | 8 | 0.125 | 4 | 0.25 | 36 | 0.1389 |
| 4_4958556 | 4 | 0.5 | 5 | 0.6 | 4 | 0.25 | 5 | 0.4 | 18 | 0.4444 | 4 | 0.75 | 4 | 0 | 2 | 0.5 | 1 | 0 | 11 | 0.3636 |
| 4_4958559 | 7 | 0.5714 | 12 | 0.75 | 8 | 0.75 | 9 | 0.8889 | 36 | 0.75 | 8 | 0.75 | 14 | 0.6429 | 9 | 0.6667 | 6 | 0.8333 | 37 | 0.7027 |
| 4_4958562 | 9 | 0.8889 | 14 | 0.8571 | 8 | 0.875 | 11 | 0.9091 | 42 | 0.881 | 8 | 0.75 | 16 | 0.9375 | 8 | 1 | 6 | 1 | 38 | 0.9211 |
| 4_4958579 | 6 | 0.3333 | 11 | 0.3636 | 7 | 0.1429 | 10 | 0.1 | 34 | 0.2353 | 8 | 0.125 | 13 | 0.1538 | 8 | 0.125 | 6 | 0.1667 | 35 | 0.1429 |
| 4_4958582 | 8 | 0.5 | 11 | 0.6364 | 8 | 0.5 | 11 | 0.7273 | 38 | 0.6053 | 7 | 0.7143 | 14 | 0.5714 | 9 | 0.5556 | 5 | 0.6 | 35 | 0.6 |
| 4_4958607 | 9 | 1 | 9 | 0.8889 | 7 | 1 | 8 | 0.875 | 33 | 0.9394 | 7 | 1 | 11 | 0.9091 | 5 | 1 | 5 | 1 | 28 | 0.9643 |
| 4_4958612 | 9 | 0.5556 | 10 | 0.7 | 7 | 0.8571 | 9 | 0.6667 | 35 | 0.6857 | 7 | 0.5714 | 10 | 0.7 | 7 | 0.7143 | 5 | 0.6 | 29 | 0.6552 |
| 4_4958622 | 6 | 1 | 8 | 1 | 7 | 1 | 5 | 1 | 26 | 1 | 4 | 1 | 10 | 1 | 4 | 1 | 6 | 1 | 24 | 1 |
| 4_4958624 | 6 | 1 | 10 | 0.9 | 8 | 1 | 7 | 1 | 31 | 0.9677 | 6 | 1 | 11 | 1 | 7 | 1 | 5 | 1 | 29 | 1 |
| 4_4958651 | 6 | 0 | 8 | 0.125 | 7 | 0.1429 | 7 | 0.2857 | 28 | 0.1429 | 2 | 0 | 11 | 0.0909 | 2 | 0.5 | 4 | 0.5 | 19 | 0.2105 |
| 4_4958655 | 6 | 0.3333 | 8 | 0.5 | 7 | 0.5714 | 7 | 0.4286 | 28 | 0.4643 | 3 | 0.6667 | 9 | 0.3333 | 5 | 0.6 | 4 | 0.75 | 21 | 0.5238 |
| 4_4958659 | 6 | 0.5 | 8 | 0.375 | 5 | 1 | 5 | 0.6 | 24 | 0.5833 | 3 | 0 | 12 | 0.25 | 5 | 0.4 | 3 | 0.3333 | 23 | 0.2609 |
| 4_4958662 | 4 | 0.5 | 6 | 0.5 | 6 | 0 | 4 | 0.5 | 20 | 0.35 | 2 | 0 | 11 | 0.5455 | 5 | 0 | 3 | 0.6667 | 21 | 0.381 |
| 4_4958697 | 2 | 1 | 8 | 1 | 6 | 0.8333 | 5 | 1 | 21 | 0.9524 | 2 | 1 | 10 | 0.9 | 4 | 1 | 4 | 1 | 20 | 0.95 |
| 4_4958706 | 1 | 1 | 5 | 0.4 | 5 | 0.6 | 6 | 0.8333 | 17 | 0.6471 | 1 | 0 | 7 | 0.7143 | 4 | 1 | 2 | 1 | 14 | 0.7857 |
| 4_4958719 | 2 | 0.5 | 6 | 0 | 5 | 0.2 | 6 | 0.1667 | 19 | 0.1579 | 2 | 0 | 8 | 0.125 | 4 | 0 | 2 | 0 | 16 | 0.0625 |
| 4_4963874 | 3 | 0.3333 | 2 | 0 | 1 | 0 | 2 | 0 | 8 | 0.125 | 1 | 1 | 4 | 0.25 | 2 | 0 | 4 | 0 | 11 | 0.1818 |
| 4_4964224 | 1 | 1 | NA | NA | 1 | 1 | NA | NA | 2 | 1 | NA | NA | 1 | 1 | NA | NA | NA | NA | 1 | 1 |
| 4_4964225 | 1 | 1 | NA | NA | 1 | 1 | NA | NA | 2 | 1 | NA | NA | 1 | 1 | NA | NA | NA | NA | 1 | 1 |
| 4_4964389 | 2 | 0.5 | NA | NA | 1 | 1 | NA | NA | 3 | 0.6667 | NA | NA | 1 | 1 | NA | NA | 1 | 0 | 2 | 0.5 |
| 4_4965021 | 4 | 0.25 | 5 | 0 | 2 | 0 | 2 | 0 | 13 | 0.0769 | 1 | 1 | 2 | 0 | 2 | 0.5 | 7 | 0 | 12 | 0.1667 |
| 4_4965456 | 7 | 0.2857 | 7 | 0.1429 | 6 | 0.3333 | 2 | 0.5 | 22 | 0.2727 | 3 | 0 | 4 | 0 | 4 | 0 | 7 | 0.2857 | 18 | 0.1111 |
| 4_4965599 | 9 | 1 | 6 | 0.6667 | 6 | 0.6667 | 2 | 1 | 23 | 0.8261 | 2 | 1 | 4 | 1 | 5 | 1 | 6 | 0.8333 | 17 | 0.9412 |
| 4_4965608 | 5 | 0.4 | 7 | 0 | 7 | 0 | 3 | 0 | 22 | 0.0909 | 3 | 0.6667 | 2 | 0 | 5 | 0.4 | 4 | 0.25 | 14 | 0.3571 |
| 4_4965613 | 9 | 0.1111 | 8 | 0 | 9 | 0.1111 | 4 | 0.25 | 30 | 0.1 | 3 | 0 | 4 | 0.25 | 4 | 0.25 | 6 | 0.3333 | 17 | 0.2353 |
| 4_4965614 | 9 | 0.3333 | 7 | 0.5714 | 9 | 0.4444 | 4 | 1 | 29 | 0.5172 | 3 | 0.3333 | 3 | 0.3333 | 5 | 0.8 | 7 | 0.5714 | 18 | 0.5556 |
| 4_4965615 | 6 | 0 | 7 | 0.1429 | 7 | 0.1429 | 4 | 0.75 | 24 | 0.2083 | 3 | 0.3333 | 3 | 0 | 4 | 0.25 | 7 | 0.2857 | 17 | 0.2353 |
| 4_4965624 | 8 | 0.125 | 6 | 0 | 6 | 0.1667 | 5 | 0 | 25 | 0.08 | 4 | 0.25 | 4 | 0 | 5 | 0 | 7 | 0.1429 | 20 | 0.1 |
| 4_4965653 | 8 | 0.25 | 7 | 0.1429 | 5 | 0 | 5 | 0.2 | 25 | 0.16 | 4 | 0 | 3 | 0 | 4 | 0.25 | 5 | 0.2 | 16 | 0.125 |
| 4_4966176 | 7 | 0.2857 | 4 | 0 | 3 | 0.3333 | 4 | 0.5 | 18 | 0.2778 | 3 | 0 | 4 | 0.5 | 5 | 0.2 | NA | NA | 12 | 0.25 |
| 4_4966227 | 7 | 0.1429 | 3 | 0.3333 | 2 | 0 | 3 | 0.3333 | 15 | 0.2 | 1 | 0 | 4 | 0.25 | 4 | 0.25 | NA | NA | 9 | 0.2222 |
| 4_4966255 | 6 | 0.1667 | 4 | 0 | 2 | 0 | 3 | 0 | 15 | 0.0667 | 3 | 0.3333 | 3 | 0 | 5 | 0 | NA | NA | 11 | 0.0909 |
| 4_4966316 | 7 | 0.5714 | 3 | 0.3333 | 3 | 0.6667 | 3 | 0.3333 | 16 | 0.5 | 3 | 0.6667 | NA | NA | 5 | 0.4 | 1 | 1 | 9 | 0.5556 |
| 4_4966344 | 7 | 0.2857 | 2 | 0 | 2 | 0 | 3 | 0 | 14 | 0.1429 | 2 | 0 | 3 | 0.3333 | 5 | 0 | 2 | 0 | 12 | 0.0833 |
| 4_4966524 | 3 | 1 | 2 | 0.5 | NA | NA | 2 | 1 | 7 | 0.8571 | NA | NA | 1 | 1 | 3 | 0.6667 | 2 | 0.5 | 6 | 0.6667 |
| 4_4966534 | 3 | 0.3333 | 2 | 0.5 | NA | NA | 2 | 0 | 7 | 0.2857 | 1 | 1 | 2 | 1 | 3 | 0.6667 | 1 | 1 | 7 | 0.8571 |
| 4_4966535 | 2 | 0 | 2 | 0 | NA | NA | 2 | 0.5 | 6 | 0.1667 | 1 | 0 | 2 | 1 | 3 | 0.3333 | NA | NA | 6 | 0.5 |
| 4_4966537 | 3 | 0.6667 | 2 | 1 | NA | NA | 2 | 0.5 | 7 | 0.7143 | 1 | 0 | 1 | 1 | 3 | 0.3333 | 1 | 0 | 6 | 0.3333 |
| 4_4966544 | 3 | 0.6667 | 2 | 0.5 | NA | NA | 2 | 1 | 7 | 0.7143 | 1 | 1 | NA | NA | 3 | 0.6667 | 1 | 0 | 5 | 0.6 |
| 4_4966547 | 1 | 1 | 1 | 0 | NA | NA | 2 | 1 | 4 | 0.75 | 1 | 1 | 1 | 0 | 3 | 0.3333 | 1 | 0 | 6 | 0.3333 |
| 4_4966559 | 1 | 1 | NA | NA | NA | NA | 1 | 1 | 2 | 1 | 1 | 0 | NA | NA | 1 | 1 | 1 | 0 | 3 | 0.3333 |
| 4_4977055 | 3 | 0.6667 | 1 | 1 | NA | NA | 1 | 1 | 5 | 0.8 | 1 | 0 | 1 | 1 | 3 | 1 | 1 | 1 | 6 | 0.8333 |
| 4_4977074 | 3 | 0.6667 | NA | NA | NA | NA | 2 | 0.5 | 5 | 0.6 | 1 | 0 | 2 | 1 | 2 | 0.5 | 1 | 0 | 6 | 0.5 |

| | | | | | | | | | | | | | | | | | | | | |
|---|---|---|---|---|---|---|---|---|---|---|---|---|---|---|---|---|---|---|---|---|
| 4_4977079 | 1 | 0 | 1 | 1 | NA | NA | 2 | 0.5 | 4 | 0.5 | 1 | 1 | 2 | 1 | 3 | 0.3333 | NA | NA | 6 | 0.6667 |
| 4_4977093 | 3 | 0.6667 | NA | NA | NA | NA | 2 | 1 | 5 | 0.8 | NA | NA | 1 | 1 | 2 | 1 | NA | NA | 3 | 1 |
| 4_4977099 | 2 | 0 | 1 | 0 | NA | NA | 2 | 1 | 5 | 0.4 | 1 | 1 | 1 | 0 | NA | NA | 1 | 0 | 3 | 0.3333 |
| 4_4977099 | 3 | 1 | 1 | 1 | NA | NA | 2 | 1 | 6 | 1 | 1 | 0 | 2 | 1 | 1 | 1 | NA | NA | 4 | 0.75 |
| 4_4977112 | 3 | 0.6667 | 1 | 1 | NA | NA | 2 | 1 | 6 | 0.8333 | 1 | 1 | NA | NA | 1 | 0 | 1 | 1 | 3 | 0.6667 |
| 4_4977129 | 3 | 1 | 1 | 1 | NA | NA | 2 | 1 | 6 | 1 | 1 | 1 | 1 | 0 | 1 | 1 | NA | NA | 3 | 0.6667 |
| 4_4977133 | 3 | 0.6667 | 1 | 1 | NA | NA | 1 | 0 | 5 | 0.6 | 1 | 1 | 2 | 1 | NA | NA | 1 | 1 | 4 | 1 |
| 4_4979247 | 3 | 0.3333 | 1 | 0 | NA | NA | 2 | 0 | 6 | 0.1667 | 1 | 0 | 1 | 0 | 1 | 1 | NA | NA | 3 | 0.3333 |
| 4_4979250 | 3 | 0.3333 | 1 | 1 | NA | NA | 2 | 0 | 6 | 0.3333 | 1 | 1 | 1 | 1 | 1 | 1 | NA | NA | 3 | 1 |
| 4_4979263 | 2 | 0.5 | NA | NA | NA | NA | 2 | 0.5 | 4 | 0.5 | 1 | 1 | 1 | 0 | 1 | 1 | NA | NA | 3 | 0.6667 |
| 4_4979265 | 2 | 1 | 1 | 1 | NA | NA | 2 | 0 | 5 | 0.6 | 1 | 1 | 1 | 1 | 1 | 1 | 1 | 0 | 4 | 0.75 |
| 4_4979274 | 2 | 0 | 1 | 0 | NA | NA | 2 | 0.5 | 5 | 0.2 | 1 | 1 | 1 | 0 | 1 | 0 | NA | NA | 3 | 0.3333 |
| 4_4979277 | 2 | 0.5 | 1 | 0 | NA | NA | 2 | 0 | 5 | 0.2 | 1 | 0 | 1 | 1 | 1 | 1 | 1 | 1 | 4 | 0.75 |
| 4_4979281 | 2 | 0.5 | 1 | 1 | NA | NA | 2 | 1 | 5 | 0.8 | 1 | 1 | 1 | 1 | 1 | 0 | 1 | 1 | 4 | 0.75 |
| 4_4984445 | 2 | 0.5 | 1 | 0 | NA | NA | 1 | 1 | 4 | 0.5 | 1 | 1 | NA | NA | 1 | 0 | NA | NA | 2 | 0.5 |
| 4_5248770 | 1 | 0 | NA | NA | NA | NA | 3 | 0.6667 | 4 | 0.5 | NA | NA | 1 | 1 | NA | NA | 1 | 0 | 2 | 0.5 |
| 4_5301517 | 5 | 0 | 2 | 0 | 2 | 0.5 | NA | NA | 9 | 0.1111 | 3 | 0.6667 | 1 | 0 | 1 | 0 | 1 | 0 | 6 | 0.3333 |
| 4_5303225 | 4 | 1 | 1 | 1 | NA | NA | 6 | 1 | 11 | 1 | 2 | 1 | 2 | 1 | 1 | 1 | 4 | 0.5 | 9 | 0.7778 |
| 4_5303228 | 2 | 0.5 | 1 | 0 | NA | NA | 4 | 0.5 | 7 | 0.4286 | 2 | 0.5 | 2 | 1 | 1 | 0 | 3 | 0.3333 | 8 | 0.5 |
| 4_5303236 | 4 | 0.5 | 1 | 0 | NA | NA | 7 | 0.2857 | 12 | 0.3333 | 2 | 0 | 2 | 1 | 1 | 0 | 4 | 0.25 | 9 | 0.3333 |
| 4_5303238 | 4 | 0.25 | 1 | 0 | NA | NA | 4 | 0 | 9 | 0.1111 | 2 | 0 | NA | NA | 1 | 0 | 4 | 0.25 | 7 | 0.1429 |
| 4_5303281 | 4 | 0.5 | 1 | 0 | NA | NA | 6 | 0.5 | 11 | 0.4545 | 1 | 0 | 1 | 0 | NA | NA | 4 | 0.25 | 6 | 0.1667 |
| 4_5303318 | 4 | 0.5 | 1 | 0 | NA | NA | 6 | 0.3333 | 11 | 0.3636 | 1 | 0 | 1 | 0 | 1 | 0 | 4 | 0.5 | 7 | 0.2857 |
| 4_5303333 | 3 | 0 | 1 | 0 | NA | NA | 5 | 0.2 | 9 | 0.1111 | NA | NA | 1 | 1 | 1 | 0 | 4 | 0.25 | 6 | 0.3333 |
| 4_5303340 | 3 | 0 | 1 | 0 | NA | NA | 5 | 0.2 | 9 | 0.1111 | 1 | 0 | 1 | 0 | NA | NA | 3 | 0.3333 | 5 | 0.2 |
| 4_5303365 | 4 | 1 | 1 | 1 | NA | NA | 5 | 0.8 | 10 | 0.9 | NA | NA | 1 | 0 | 1 | 0 | 3 | 0.6667 | 5 | 0.4 |
| 4_5303367 | 3 | 0.6667 | 1 | 0 | NA | NA | 3 | 0.6667 | 7 | 0.5714 | NA | NA | NA | NA | 1 | 1 | 1 | 0 | 2 | 0.5 |
| 4_5303371 | 4 | 1 | 1 | 1 | NA | NA | 5 | 0.6 | 10 | 0.8 | NA | NA | 1 | 1 | 1 | 1 | 2 | 1 | 4 | 1 |
| 4_5303378 | 4 | 0.5 | 1 | 0 | NA | NA | 5 | 0.6 | 10 | 0.5 | NA | NA | 1 | 0 | 1 | 0 | 2 | 0.5 | 4 | 0.25 |
| 4_5303381 | 3 | 1 | 1 | 0 | NA | NA | 6 | 0.5 | 10 | 0.6 | NA | NA | 1 | 1 | 1 | 1 | 2 | 0.5 | 4 | 0.75 |
| 4_5303383 | 4 | 0.5 | NA | NA | NA | NA | 6 | 0.5 | 10 | 0.5 | NA | NA | 1 | 0 | 1 | 0 | 3 | 0.3333 | 5 | 0.2 |
| 4_5303401 | 2 | 1 | 1 | 1 | NA | NA | 4 | 0.75 | 7 | 0.8571 | NA | NA | 1 | 1 | NA | NA | NA | NA | 1 | 1 |
| 4_5303403 | 3 | 0.3333 | 1 | 1 | NA | NA | 6 | 0.8333 | 10 | 0.7 | NA | NA | NA | NA | 1 | 0 | 3 | 0.3333 | 4 | 0.25 |
| 4_5303550 | 3 | 0.3333 | 1 | 0 | NA | NA | 4 | 0.25 | 8 | 0.25 | NA | NA | 1 | 0 | 1 | 0 | 3 | 0.3333 | 5 | 0.2 |
| 4_5356756 | 2 | 0.5 | 2 | 1 | 5 | 1 | 2 | 1 | 11 | 0.9091 | 1 | 0 | 6 | 0.8333 | NA | NA | 2 | 1 | 9 | 0.7778 |
| 4_5356761 | 3 | 1 | 4 | 1 | 3 | 1 | 2 | 1 | 12 | 1 | NA | NA | 5 | 0.8 | NA | NA | 2 | 1 | 7 | 0.8571 |
| 4_5356770 | 3 | 0 | 2 | 1 | 3 | 0.3333 | 1 | 0 | 9 | 0.3333 | 2 | 0.5 | 4 | 0.25 | NA | NA | 2 | 0 | 8 | 0.25 |
| 4_5362163 | 1 | 0 | 7 | 0.4286 | 6 | 0.5 | 4 | 0.25 | 18 | 0.3889 | 10 | 0.2 | 1 | 0 | 2 | 0 | 3 | 0 | 16 | 0.125 |
| 4_5362192 | 4 | 0 | 6 | 0.1667 | 7 | 0.2857 | 4 | 0.25 | 21 | 0.1905 | 8 | 0.25 | 2 | 0.5 | 2 | 0 | 4 | 0 | 16 | 0.1875 |
| 4_5370117 | NA | NA | 3 | 1 | NA | NA | 1 | 1 | 4 | 1 | NA | NA | NA | NA | NA | NA | 2 | 0.5 | 2 | 0.5 |
| 4_5370266 | NA | NA | 3 | 0.3333 | NA | NA | 2 | 0.5 | 5 | 0.4 | 2 | 0 | NA | NA | NA | NA | 3 | 0.3333 | 5 | 0.2 |
| 4_5370578 | 1 | 1 | 3 | 0.3333 | 1 | 0 | 1 | 0 | 6 | 0.3333 | 3 | 0.3333 | NA | NA | NA | NA | 4 | 0.25 | 7 | 0.2857 |
| 4_5370583 | 2 | 0.5 | 3 | 0.3333 | 1 | 1 | 2 | 0.5 | 8 | 0.5 | 3 | 0 | NA | NA | NA | NA | 6 | 0.3333 | 9 | 0.2222 |
| 4_5370612 | 2 | 0.5 | 3 | 0 | 1 | 1 | 2 | 0.5 | 8 | 0.375 | 2 | 0.5 | 1 | 0 | NA | NA | 6 | 0.3333 | 9 | 0.3333 |
| 4_5370641 | 2 | 0.5 | 3 | 0.3333 | 1 | 1 | 2 | 1 | 8 | 0.625 | 4 | 0.25 | 2 | 1 | NA | NA | 6 | 0.1667 | 12 | 0.3333 |
| 4_5370668 | 1 | 1 | 3 | 0.3333 | 1 | 0 | 2 | 0.5 | 7 | 0.4286 | 4 | 0.25 | 1 | 0 | NA | NA | 6 | 0 | 11 | 0.0909 |
| 4_5376120 | 1 | 1 | NA | NA | 1 | 1 | 1 | 1 | 3 | 1 | NA | NA | 1 | 0 | 1 | 0 | 2 | 1 | 4 | 0.5 |
| 4_5376125 | 1 | 0 | NA | NA | 2 | 0.5 | 1 | 0 | 4 | 0.25 | NA | NA | 1 | 0 | 1 | 1 | 2 | 0 | 4 | 0.25 |
| 4_5376129 | 1 | 0 | NA | NA | 2 | 0 | 1 | 1 | 4 | 0.25 | NA | NA | 1 | 1 | 1 | 0 | 1 | 0 | 3 | 0.3333 |
| 4_5376130 | 1 | 1 | NA | NA | 2 | 1 | 1 | 1 | 4 | 1 | NA | NA | 1 | 1 | 1 | 1 | 1 | 1 | 3 | 1 |
| 4_5376139 | 1 | 1 | NA | NA | 1 | 0 | NA | NA | 2 | 0.5 | NA | NA | NA | NA | 1 | 1 | 2 | 0.5 | 3 | 0.6667 |
| 4_5376143 | 1 | 1 | NA | NA | 2 | 1 | 1 | 1 | 4 | 1 | 1 | 1 | 1 | 1 | 1 | 1 | 2 | 1 | 5 | 1 |
| 4_5376150 | 1 | 0 | NA | NA | 2 | 0 | 1 | 1 | 4 | 0.25 | 1 | 1 | 1 | 1 | 1 | 0 | 2 | 1 | 5 | 0.8 |
| 4_5376166 | 1 | 0 | NA | NA | 2 | 1 | 1 | 0 | 4 | 0.5 | 1 | 1 | 1 | 0 | 1 | 0 | 2 | 1 | 5 | 0.6 |
| 4_5376187 | 1 | 0 | NA | NA | 2 | 0 | 2 | 0.5 | 5 | 0.2 | 1 | 1 | 1 | 0 | 1 | 0 | 2 | 0 | 5 | 0.2 |
| 4_5376202 | 2 | 1 | NA | NA | 2 | 0.5 | 1 | 0 | 5 | 0.6 | 1 | 1 | NA | NA | 1 | 0 | 1 | 1 | 3 | 0.6667 |
| 4_5376398 | 7 | 0.1429 | NA | NA | 3 | 0 | 1 | 0 | 11 | 0.0909 | 1 | 0 | 3 | 0 | 4 | 0.25 | 2 | 0 | 10 | 0.1 |
| 4_5376405 | 6 | 0.1667 | NA | NA | 3 | 0 | 1 | 0 | 10 | 0.1 | 1 | 0 | 2 | 0.5 | 4 | 0 | 1 | 0 | 8 | 0.125 |
| 4_5376484 | 5 | 0.4 | 1 | 1 | 3 | 0 | 4 | 0 | 13 | 0.2308 | 1 | 0 | 4 | 0.25 | 7 | 0 | 3 | 0.3333 | 15 | 0.1333 |
| 4_5376558 | 7 | 0 | 1 | 0 | 6 | 0.1667 | 3 | 0 | 17 | 0.0588 | 1 | 0 | 5 | 0.2 | 9 | 0 | 4 | 0 | 19 | 0.0526 |
| 4_5386009 | 2 | 0.5 | 2 | 0.5 | 3 | 0 | 1 | 0 | 8 | 0.25 | NA | NA | 2 | 0 | 1 | 0 | 2 | 0.5 | 5 | 0.2 |
| 4_5386023 | 2 | 0 | 2 | 0 | 2 | 0.5 | 1 | 0 | 7 | 0.1429 | 1 | 0 | 2 | 0 | 2 | 0.5 | 2 | 0.5 | 7 | 0.2857 |
| 4_5386116 | NA | NA | 2 | 0.5 | NA | NA | 3 | 0 | 5 | 0.2 | 1 | 1 | 1 | 0 | 1 | 0 | 3 | 0.3333 | 6 | 0.3333 |
| 4_5386124 | 1 | 1 | 2 | 0 | NA | NA | 2 | 0 | 5 | 0.2 | 1 | 0 | 1 | 0 | 1 | 1 | 2 | 0 | 5 | 0.2 |
| 4_5386136 | 1 | 1 | 2 | 0.5 | NA | NA | 2 | 0 | 5 | 0.4 | 1 | 0 | 1 | 0 | 1 | 1 | 3 | 0.3333 | 6 | 0.3333 |

| | | | | | | | | | | | | | | | | | | | | |
|---|---|---|---|---|---|---|---|---|---|---|---|---|---|---|---|---|---|---|---|---|
| 4_55050293 | 2 | 0 | 1 | 0 | NA | NA | 2 | 0.5 | 5 | 0.2 | NA | NA | NA | NA | 1 | 1 | NA | NA | 1 | 1 |
| 4_55060153 | 2 | 0.5 | 2 | 0 | NA | NA | 3 | 0.3333 | 7 | 0.2857 | 1 | 0 | NA | NA | 1 | 1 | NA | NA | 2 | 0.5 |
| 4_55080785 | 4 | 0.5 | 2 | 1 | 1 | 1 | 3 | 0.6667 | 10 | 0.7 | 2 | 0.5 | NA | NA | 1 | 1 | NA | NA | 3 | 0.6667 |
| 4_55080908 | 4 | 1 | 2 | 0.5 | 1 | 1 | 4 | 1 | 11 | 0.9091 | 2 | 0.5 | NA | NA | 1 | 1 | NA | NA | 3 | 0.6667 |
| 4_55080956 | 3 | 0.3333 | 1 | 0 | NA | NA | 4 | 0 | 8 | 0.125 | 2 | 0.5 | NA | NA | NA | NA | NA | NA | 2 | 0.5 |
| 4_55080999 | 4 | 0.5 | 2 | 1 | NA | NA | 5 | 1 | 11 | 0.8182 | 2 | 0.5 | 1 | 1 | 1 | 1 | NA | NA | 4 | 0.75 |
| 4_55081004 | 2 | 1 | 2 | 1 | NA | NA | 3 | 0.6667 | 7 | 0.8571 | 2 | 0.5 | NA | NA | NA | NA | NA | NA | 2 | 0.5 |
| 4_55081194 | 2 | 1 | 2 | 1 | NA | NA | 4 | 1 | 8 | 1 | 2 | 0.5 | 1 | 1 | NA | NA | NA | NA | 3 | 0.6667 |
| 4_55081221 | 3 | 0.3333 | 2 | 0.5 | NA | NA | 4 | 0 | 9 | 0.2222 | 1 | 0 | 1 | 0 | 1 | 1 | NA | NA | 3 | 0.3333 |
| 4_55081370 | 3 | 0.6667 | 1 | 1 | NA | NA | 3 | 1 | 7 | 0.8571 | 1 | 1 | 1 | 1 | 1 | 1 | NA | NA | 3 | 1 |
| 4_55081480 | 2 | 1 | 2 | 1 | NA | NA | 4 | 0.75 | 8 | 0.875 | 1 | 0 | 1 | 0 | 1 | 1 | NA | NA | 3 | 0.3333 |
| 4_55081488 | 3 | 0.6667 | 2 | 1 | NA | NA | 3 | 0.3333 | 8 | 0.625 | 1 | 1 | 1 | 0 | NA | NA | NA | NA | 2 | 0.5 |
| 4_55081634 | 3 | 0.6667 | 2 | 0.5 | NA | NA | 4 | 0.25 | 9 | 0.4444 | 1 | 0 | 1 | 0 | 1 | 1 | NA | NA | 3 | 0.3333 |
| 4_55082981 | 8 | 0.625 | 5 | 0.8 | NA | NA | 6 | 0.8333 | 19 | 0.7368 | 2 | 0.5 | 4 | 0.5 | 3 | 0.3333 | 5 | 0.8 | 14 | 0.5714 |
| 4_55083307 | 9 | 0.1111 | 6 | 0.3333 | NA | NA | 8 | 0.25 | 23 | 0.2174 | 5 | 0.2 | 4 | 0 | 5 | 0 | 6 | 0.3333 | 20 | 0.15 |
| 4_55389910 | 12 | 0.5 | 19 | 0.5263 | 16 | 0.4375 | 21 | 0.2381 | 68 | 0.4118 | 10 | 0.5 | 13 | 0.6154 | 11 | 0.4545 | 14 | 0.2857 | 48 | 0.4583 |
| 4_55389981 | 15 | 0.1333 | 19 | 0 | 14 | 0.0714 | 23 | 0.0435 | 71 | 0.0563 | 10 | 0.2 | 12 | 0.3333 | 9 | 0.1111 | 15 | 0 | 46 | 0.1522 |
| 4_55389993 | 17 | 0.1765 | 20 | 0.15 | 14 | 0.0714 | 21 | 0.0476 | 72 | 0.1111 | 10 | 0.1 | 16 | 0.0625 | 12 | 0.0833 | 16 | 0.0625 | 54 | 0.0741 |
| 4_55390511 | 22 | 0.2273 | 27 | 0.2963 | 19 | 0.2632 | 27 | 0.1852 | 95 | 0.2421 | 13 | 0.3077 | 19 | 0.3158 | 20 | 0.3 | 20 | 0.25 | 72 | 0.2917 |
| 4_55391339 | 4 | 0.5 | 7 | 0.8571 | 6 | 0.3333 | 5 | 0.4 | 22 | 0.5455 | 2 | 0.5 | 5 | 0.6 | 7 | 0.4286 | 3 | 0.3333 | 17 | 0.4706 |
| 4_55391350 | 4 | 0.25 | 8 | 0 | 9 | 0 | 6 | 0.3333 | 27 | 0.1111 | 4 | 0 | 5 | 0.6 | 7 | 0.4286 | 6 | 0.3333 | 22 | 0.3636 |
| 4_55391456 | 2 | 1 | 9 | 0.1111 | 9 | 0.1111 | 5 | 0 | 25 | 0.16 | 4 | 0 | 5 | 0.2 | 10 | 0.4 | 6 | 0 | 25 | 0.2 |
| 4_55391464 | 2 | 0.5 | 9 | 0 | 9 | 0.1111 | 4 | 0 | 24 | 0.0833 | 5 | 0 | 4 | 0.25 | 7 | 0.4286 | 6 | 0.1667 | 22 | 0.2273 |
| 4_55391993 | 1 | 1 | 2 | 0 | 4 | 0 | NA | NA | 7 | 0.1429 | NA | NA | 2 | 0 | 3 | 0.6667 | 3 | 0.3333 | 8 | 0.375 |
| 4_55527782 | 2 | 0.5 | 3 | 0 | 1 | 0 | 5 | 0.4 | 11 | 0.2727 | NA | NA | 1 | 1 | 1 | 0 | 1 | 0 | 3 | 0.3333 |
| 4_55527804 | 2 | 1 | 3 | 1 | NA | NA | 4 | 1 | 9 | 1 | NA | NA | 1 | 1 | 1 | 1 | 1 | 1 | 3 | 1 |
| 4_55527873 | 1 | 1 | 2 | 1 | NA | NA | 4 | 0.25 | 7 | 0.5714 | NA | NA | NA | NA | 1 | 1 | 1 | 1 | 2 | 1 |
| 4_55527923 | 2 | 0.5 | 2 | 0 | 1 | 0 | 4 | 0 | 9 | 0.1111 | NA | NA | NA | NA | 1 | 1 | 1 | 0 | 2 | 0.5 |
| 4_55527943 | 2 | 0 | 2 | 0 | NA | NA | 5 | 0.2 | 9 | 0.1111 | NA | NA | 1 | 1 | 1 | 0 | 1 | 0 | 3 | 0.3333 |
| 4_55528239 | 2 | 1 | 2 | 1 | 1 | 0 | 5 | 0.8 | 10 | 0.8 | NA | NA | NA | NA | 1 | 1 | NA | NA | 1 | 1 |
| 4_55530112 | 2 | 1 | 2 | 0.5 | 1 | 1 | 5 | 1 | 10 | 0.9 | NA | NA | NA | NA | 1 | 1 | 1 | 1 | 2 | 1 |
| 4_55530129 | 2 | 1 | 1 | 0 | 1 | 0 | 4 | 0.75 | 8 | 0.625 | NA | NA | NA | NA | NA | NA | 1 | 1 | 1 | 1 |
| 4_55530155 | 2 | 1 | NA | NA | 1 | 1 | 5 | 1 | 8 | 1 | NA | NA | 1 | 1 | NA | NA | 1 | 1 | 2 | 1 |
| 4_55530225 | 2 | 0 | 2 | 0 | 1 | 1 | 4 | 0 | 9 | 0.1111 | NA | NA | NA | NA | NA | NA | 1 | 1 | 1 | 1 |
| 4_55530493 | 1 | 1 | 2 | 1 | 1 | 0 | 4 | 0.75 | 8 | 0.75 | NA | NA | 1 | 1 | NA | NA | 1 | 0 | 2 | 0.5 |
| 4_55530553 | 2 | 1 | 2 | 1 | 1 | 1 | 2 | 0.5 | 7 | 0.8571 | NA | NA | 1 | 0 | NA | NA | 1 | 1 | 2 | 0.5 |
| 4_55530799 | 2 | 1 | 2 | 0 | 1 | 1 | 3 | 0.6667 | 8 | 0.625 | NA | NA | NA | NA | NA | NA | 1 | 1 | 1 | 1 |
| 4_55530805 | 2 | 0.5 | 2 | 1 | 1 | 1 | 3 | 1 | 8 | 0.875 | NA | NA | NA | NA | NA | NA | 1 | 1 | 1 | 1 |
| 4_56751015 | 3 | 0 | 6 | 0 | 5 | 0 | 13 | 0.2308 | 27 | 0.1111 | 4 | 0 | 4 | 0.75 | 7 | 0.1429 | 4 | 0 | 19 | 0.2105 |
| 4_56751248 | 4 | 0.5 | 5 | 0.6 | 7 | 0.1429 | 14 | 0.2143 | 30 | 0.3 | 5 | 0.4 | 4 | 0.25 | 8 | 0.75 | 5 | 0.6 | 22 | 0.5455 |
| 4_56751278 | 4 | 0.5 | 10 | 0.8 | 7 | 0.8571 | 14 | 0.5714 | 35 | 0.6857 | 3 | 1 | 3 | 0.6667 | 6 | 0.6667 | 5 | 0.8 | 17 | 0.7647 |
| 4_56751379 | 4 | 0.5 | 9 | 0.7778 | 7 | 1 | 16 | 0.75 | 36 | 0.7778 | 4 | 1 | 4 | 0.75 | 9 | 0.8889 | 6 | 0.6667 | 23 | 0.8261 |
| 4_56751422 | 4 | 0 | 9 | 0.1111 | 7 | 0.2857 | 16 | 0.4375 | 36 | 0.2778 | 4 | 0 | 5 | 0.6 | 8 | 0.5 | 7 | 0.2857 | 24 | 0.375 |
| 4_56751485 | 3 | 0 | 8 | 0.25 | 6 | 0.3333 | 14 | 0.3571 | 31 | 0.2903 | 4 | 0 | 5 | 1 | 5 | 0.6 | 5 | 0.4 | 19 | 0.5263 |
| 4_56751580 | 4 | 0 | 9 | 0.1111 | 7 | 0.1429 | 15 | 0.1333 | 35 | 0.1143 | 3 | 0 | 5 | 0.4 | 8 | 0.375 | 6 | 0 | 22 | 0.2273 |
| 4_56806488 | 2 | 1 | 1 | 0 | NA | NA | NA | NA | 3 | 0.6667 | 2 | 0.5 | NA | NA | 1 | 0 | NA | NA | 3 | 0.3333 |
| 4_56806695 | 2 | 1 | 1 | 0 | NA | NA | NA | NA | 3 | 0.6667 | 2 | 1 | NA | NA | 2 | 1 | NA | NA | 4 | 1 |
| 4_56806722 | 2 | 0.5 | 1 | 0 | NA | NA | NA | NA | 3 | 0.3333 | 2 | 0 | NA | NA | 1 | 1 | NA | NA | 3 | 0.3333 |
| 4_56806923 | 2 | 0 | 2 | 0.5 | NA | NA | NA | NA | 4 | 0.25 | 2 | 0 | NA | NA | 3 | 0.3333 | NA | NA | 5 | 0.2 |
| 4_56807473 | 2 | 0.5 | 3 | 0 | NA | NA | NA | NA | 5 | 0.2 | 2 | 0.5 | NA | NA | 3 | 0.6667 | NA | NA | 5 | 0.6 |
| 4_56807503 | 2 | 0 | 3 | 0.3333 | NA | NA | NA | NA | 5 | 0.2 | 2 | 0 | NA | NA | 3 | 0.3333 | NA | NA | 5 | 0.2 |
| 4_56807594 | 2 | 0.5 | 4 | 0.75 | NA | NA | NA | NA | 6 | 0.6667 | 2 | 0 | NA | NA | 1 | 1 | NA | NA | 3 | 0.3333 |
| 4_56807767 | 2 | 0 | 4 | 0.25 | NA | NA | NA | NA | 6 | 0.1667 | 2 | 0 | NA | NA | 2 | 0.5 | NA | NA | 4 | 0.25 |
| 4_56808897 | 2 | 0 | 6 | 0.1667 | NA | NA | 2 | 0 | 10 | 0.1 | 2 | 0.5 | 1 | 0 | 4 | 0.25 | NA | NA | 7 | 0.2857 |
| 4_56809569 | 3 | 0 | 6 | 0.1667 | NA | NA | 2 | 0 | 11 | 0.0909 | 2 | 0 | 1 | 1 | 4 | 0 | NA | NA | 7 | 0.1429 |
| 4_56811650 | 2 | 0.5 | 2 | 1 | 3 | 0.3333 | 2 | 0 | 9 | 0.4444 | 2 | 0.5 | 3 | 1 | 2 | 0.5 | 1 | 1 | 8 | 0.75 |
| 4_56811933 | 1 | 1 | 2 | 1 | 3 | 1 | 2 | 1 | 8 | 1 | 2 | 1 | 2 | 1 | 1 | 1 | 1 | 1 | 6 | 1 |
| 4_56812134 | 1 | 0 | 2 | 0 | 3 | 0 | 2 | 0.5 | 8 | 0.125 | 1 | 1 | 3 | 0.3333 | 2 | 0 | 1 | 0 | 7 | 0.2857 |
| 4_56812192 | 1 | 0 | 2 | 0.5 | 4 | 0.75 | 2 | 0 | 9 | 0.4444 | NA | NA | 2 | 1 | 2 | 0.5 | 1 | 0 | 5 | 0.6 |
| 4_56867469 | 3 | 0 | 3 | 0.3333 | 3 | 0 | 1 | 1 | 10 | 0.2 | 3 | 0.3333 | 3 | 0 | 1 | 0 | 2 | 0 | 9 | 0.1111 |
| 4_56867502 | 3 | 0 | 5 | 0.2 | 4 | 0 | 2 | 0 | 14 | 0.0714 | 3 | 0.3333 | 3 | 0 | 3 | 0 | 2 | 0 | 11 | 0.0909 |
| 4_56867721 | 4 | 0 | 4 | 0.25 | 4 | 0.25 | 2 | 0.5 | 14 | 0.2143 | 4 | 0 | 2 | 0 | 3 | 0 | 2 | 0.5 | 11 | 0.0909 |
| 4_56957535 | 22 | 0 | 25 | 0.12 | 22 | 0.1364 | 19 | 0 | 88 | 0.0682 | 21 | 0 | 19 | 0.1053 | 17 | 0.0588 | 35 | 0.0571 | 92 | 0.0543 |
| 4_56957580 | 22 | 0.6364 | 21 | 0.5238 | 20 | 0.4 | 20 | 0.45 | 83 | 0.506 | 19 | 0.5789 | 22 | 0.6364 | 20 | 0.6 | 32 | 0.5 | 93 | 0.5699 |
| 4_56957670 | 23 | 0.2609 | 22 | 0.2273 | 23 | 0.4348 | 22 | 0.1364 | 90 | 0.2667 | 19 | 0.2105 | 19 | 0.4211 | 19 | 0.2632 | 30 | 0.2 | 87 | 0.2644 |

| | | | | | | | | | | | | | | | | | | | | |
|---|---|---|---|---|---|---|---|---|---|---|---|---|---|---|---|---|---|---|---|---|
| 4_5695770 | 17 | 0.5882 | 21 | 0.4286 | 19 | 0.5789 | 18 | 0.5556 | 75 | 0.5333 | 15 | 0.4 | 21 | 0.5238 | 16 | 0.5625 | 30 | 0.5667 | 82 | 0.5244 |
| 4_5695900 | 14 | 0.3571 | 19 | 0.3158 | 12 | 0.3333 | 17 | 0.4706 | 62 | 0.371 | 11 | 0.1818 | 18 | 0.2222 | 12 | 0.1667 | 21 | 0.381 | 62 | 0.2581 |
| 4_5695901 | 13 | 0.0769 | 17 | 0.0588 | 11 | 0.1818 | 18 | 0 | 59 | 0.0678 | 11 | 0.1818 | 16 | 0.125 | 13 | 0.1538 | 21 | 0.0476 | 61 | 0.1148 |
| 4_5695921 | 12 | 0 | 17 | 0 | 9 | 0.1111 | 17 | 0.2353 | 55 | 0.0909 | 12 | 0.0833 | 13 | 0.1538 | 11 | 0.0909 | 20 | 0 | 56 | 0.0714 |
| 4_5696047 | 7 | 0.1429 | 8 | 0 | 4 | 0 | 9 | 0.1111 | 28 | 0.0714 | 7 | 0 | 6 | 0.1667 | 7 | 0.1429 | 10 | 0 | 30 | 0.0667 |
| 4_5696109 | 8 | 0 | 9 | 0.1111 | 7 | 0.1429 | 10 | 0.2 | 34 | 0.1176 | 6 | 0.3333 | 8 | 0.125 | 6 | 0.1667 | 10 | 0.1 | 30 | 0.1667 |
| 4_5696110 | 6 | 0 | 10 | 0.2 | 7 | 0.1429 | 7 | 0.2857 | 30 | 0.1667 | 2 | 0 | 7 | 0 | 5 | 0.2 | 5 | 0 | 19 | 0.0526 |
| 4_5696246 | 6 | 0.1667 | 7 | 0.2857 | 5 | 0.2 | 8 | 0.375 | 26 | 0.2692 | 3 | 0 | 3 | 0.3333 | 3 | 0.6667 | 6 | 0.3333 | 15 | 0.3333 |
| 4_5696257 | 5 | 0.2 | 4 | 0 | 3 | 0.3333 | 7 | 0.1429 | 19 | 0.1579 | 3 | 0 | 3 | 0 | 3 | 0 | 7 | 0.1429 | 16 | 0.0625 |
| 4_5696271 | 6 | 0.5 | 6 | 0.1667 | 4 | 0.75 | 7 | 0.2857 | 23 | 0.3913 | 2 | 0 | 2 | 0 | 2 | 1 | 5 | 0.2 | 11 | 0.2727 |
| 4_5696277 | 3 | 0.3333 | 4 | 0.25 | 3 | 0 | 7 | 0 | 17 | 0.1176 | 2 | 0.5 | 3 | 0 | 1 | 1 | 6 | 0 | 12 | 0.1667 |
| 4_5696278 | 3 | 0.3333 | 5 | 0 | 5 | 0.2 | 5 | 0.4 | 18 | 0.2222 | 1 | 0 | 3 | 0 | 3 | 0.6667 | 3 | 0.3333 | 10 | 0.3 |
| 4_5696289 | 3 | 0 | 5 | 0.4 | 4 | 0.25 | 6 | 0.3333 | 18 | 0.2778 | 2 | 0 | 2 | 0.5 | 2 | 0.5 | 5 | 0.4 | 11 | 0.3636 |
| 4_5696292 | 3 | 0 | 6 | 0 | 5 | 0 | 9 | 0.2222 | 23 | 0.087 | 2 | 0 | 3 | 0 | 3 | 0.3333 | 5 | 0 | 13 | 0.0769 |
| 4_5696296 | 4 | 1 | 5 | 0.4 | 2 | 1 | 7 | 1 | 18 | 0.8333 | 1 | 1 | 3 | 1 | 3 | 1 | 5 | 0.8 | 12 | 0.9167 |
| 4_5696308 | 3 | 0.6667 | 4 | 0.5 | 4 | 0.25 | 7 | 0.2857 | 18 | 0.3889 | 2 | 0 | 2 | 0.5 | 2 | 0 | 4 | 0 | 10 | 0.1 |
| 4_5696313 | 4 | 0.75 | 2 | 0 | 2 | 0 | 8 | 0.5 | 16 | 0.4375 | 2 | 0.5 | 3 | 0.3333 | 1 | 1 | 4 | 0.5 | 10 | 0.5 |
| 4_5699295 | 1 | 0 | 2 | 0 | 2 | 0 | 4 | 0.25 | 9 | 0.1111 | 3 | 0 | 5 | 0.4 | 2 | 0 | NA | NA | 10 | 0.2 |
| 4_5699394 | NA | NA | 3 | 0 | NA | NA | 3 | 0.3333 | 6 | 0.1667 | 1 | 0 | 3 | 0.6667 | 2 | 0 | NA | NA | 6 | 0.3333 |
| 4_5699574 | NA | NA | 1 | 1 | NA | NA | 2 | 0 | 3 | 0.3333 | 1 | 0 | 2 | 0.5 | 2 | 0 | NA | NA | 5 | 0.2 |
| 4_5719340 | 2 | 0.5 | 2 | 0 | 2 | 0.5 | 1 | 0 | 7 | 0.2857 | 1 | 1 | NA | NA | 1 | 1 | 1 | 1 | 3 | 1 |
| 4_5719345 | 2 | 0 | 2 | 0 | 1 | 1 | 1 | 0 | 6 | 0.1667 | NA | NA | NA | NA | 1 | 0 | 1 | 1 | 2 | 0.5 |
| 4_5719360 | 2 | 0 | 2 | 0 | 2 | 0.5 | 1 | 0 | 7 | 0.1429 | 1 | 0 | NA | NA | 2 | 0 | 1 | 1 | 4 | 0.25 |
| 4_5719361 | 2 | 0 | 2 | 0.5 | 2 | 0 | 1 | 1 | 7 | 0.2857 | 1 | 0 | NA | NA | 2 | 0.5 | 1 | 0 | 4 | 0.25 |
| 4_5719380 | 1 | 0 | 2 | 0 | 1 | 1 | 1 | 1 | 5 | 0.4 | 1 | 0 | NA | NA | 1 | 0 | 1 | 1 | 3 | 0.3333 |
| 4_5719404 | 2 | 1 | 2 | 0.5 | 2 | 1 | 1 | 1 | 7 | 0.8571 | NA | NA | NA | NA | 2 | 1 | 1 | 1 | 3 | 1 |
| 4_5719411 | 2 | 1 | 2 | 1 | 2 | 1 | 1 | 1 | 7 | 1 | NA | NA | NA | NA | 1 | 1 | 1 | 1 | 2 | 1 |
| 4_5719755 | 2 | 0 | 1 | 0 | 2 | 0.5 | 1 | 0 | 6 | 0.1667 | 1 | 1 | NA | NA | 2 | 0 | 1 | 0 | 4 | 0.25 |
| 4_5770967 | 1 | 1 | 1 | 1 | 1 | 1 | NA | NA | 3 | 1 | NA | NA | NA | NA | 1 | 1 | 1 | 1 | 2 | 1 |
| 4_5770971 | 2 | 1 | 1 | 1 | 1 | 1 | NA | NA | 4 | 1 | NA | NA | NA | NA | 1 | 1 | 1 | 1 | 2 | 1 |
| 4_5770992 | 2 | 0.5 | 1 | 0 | 1 | 0 | NA | NA | 4 | 0.25 | NA | NA | NA | NA | 1 | 0 | 1 | 1 | 2 | 0.5 |
| 4_5771018 | 2 | 1 | 1 | 1 | 1 | 0 | NA | NA | 4 | 0.75 | NA | NA | NA | NA | 1 | 0 | 1 | 1 | 2 | 0.5 |
| 4_5771024 | 2 | 1 | 1 | 1 | 1 | 1 | NA | NA | 4 | 1 | NA | NA | NA | NA | 1 | 1 | 1 | 1 | 2 | 1 |
| 4_5771028 | 2 | 0.5 | 2 | 1 | 1 | 0 | NA | NA | 5 | 0.6 | NA | NA | NA | NA | 1 | 0 | 1 | 1 | 2 | 0.5 |
| 4_5771040 | 1 | 1 | 1 | 1 | 1 | 1 | NA | NA | 3 | 1 | NA | NA | NA | NA | 1 | 1 | 1 | 0 | 2 | 0.5 |
| 4_5771047 | 2 | 1 | 2 | 1 | 1 | 1 | NA | NA | 5 | 1 | NA | NA | NA | NA | 1 | 1 | 1 | 1 | 2 | 1 |
| 4_5771051 | 1 | 1 | 2 | 0.5 | 1 | 0 | NA | NA | 4 | 0.5 | NA | NA | NA | NA | 1 | 1 | 1 | 1 | 2 | 1 |
| 4_5771052 | 2 | 1 | 2 | 1 | 1 | 1 | NA | NA | 5 | 1 | NA | NA | NA | NA | 1 | 1 | 1 | 1 | 2 | 1 |
| 4_5771075 | 2 | 1 | 2 | 0.5 | NA | NA | NA | NA | 4 | 0.75 | 1 | 1 | NA | NA | NA | NA | 1 | 1 | 2 | 1 |
| 4_5771090 | NA | NA | 2 | 0.5 | 1 | 0 | NA | NA | 3 | 0.3333 | 1 | 0 | NA | NA | 1 | 0 | 1 | 1 | 3 | 0.3333 |
| 4_5771096 | 2 | 0.5 | 2 | 0.5 | 1 | 0 | NA | NA | 5 | 0.4 | 1 | 1 | NA | NA | 1 | 0 | 1 | 0 | 3 | 0.3333 |
| 4_5771098 | 2 | 1 | 2 | 0.5 | 1 | 0 | NA | NA | 5 | 0.6 | 1 | 1 | NA | NA | 1 | 0 | 1 | 0 | 3 | 0.3333 |
| 4_5771469 | 2 | 0 | 3 | 0 | 3 | 0.3333 | NA | NA | 8 | 0.125 | 4 | 0.25 | 3 | 0.3333 | 1 | 0 | 3 | 0 | 11 | 0.1818 |
| 4_5771491 | 2 | 0.5 | 3 | 0.6667 | 4 | 0.5 | NA | NA | 9 | 0.5556 | 3 | 0.3333 | 3 | 0.6667 | 1 | 0 | 3 | 1 | 10 | 0.6 |
| 4_5771504 | 3 | 0.6667 | 3 | 0 | 4 | 0 | NA | NA | 10 | 0.2 | 4 | 0 | 2 | 0.5 | NA | NA | 4 | 0.25 | 10 | 0.2 |
| 4_5771683 | 4 | 0.25 | 4 | 0 | 4 | 0 | 1 | 0 | 13 | 0.0769 | 5 | 0 | 7 | 0.1429 | 3 | 0 | 5 | 0 | 20 | 0.05 |
| 4_5771704 | 5 | 0.2 | 4 | 0 | 5 | 0 | 1 | 0 | 15 | 0.0667 | 5 | 0 | 7 | 0 | 3 | 0.3333 | 5 | 0 | 20 | 0.05 |
| 4_5771712 | 4 | 0 | 4 | 0 | 6 | 0.3333 | 2 | 0 | 16 | 0.125 | 6 | 0 | 7 | 0.1429 | 3 | 0 | 4 | 0 | 20 | 0.05 |
| 4_5789305 | NA | NA | NA | NA | 1 | 1 | NA | NA | 1 | 1 | NA | NA | NA | NA | 1 | 1 | NA | NA | 1 | 1 |
| 4_5789307 | NA | NA | NA | NA | 1 | 1 | NA | NA | 1 | 1 | NA | NA | NA | NA | 1 | 1 | NA | NA | 1 | 1 |
| 4_5789325 | NA | NA | NA | NA | 1 | 1 | NA | NA | 1 | 1 | NA | NA | NA | NA | 1 | 1 | NA | NA | 1 | 1 |
| 4_5789328 | NA | NA | NA | NA | 1 | 1 | NA | NA | 1 | 1 | NA | NA | NA | NA | 1 | 1 | NA | NA | 1 | 1 |
| 4_5789661 | 2 | 0.5 | 6 | 0.5 | 3 | 1 | 5 | 0.2 | 16 | 0.5 | 3 | 0 | 3 | 0 | 7 | 0.2857 | 5 | 0.2 | 18 | 0.1667 |
| 4_5837327 | NA | NA | 1 | 1 | 1 | 1 | NA | NA | 2 | 1 | NA | NA | 1 | 1 | NA | NA | NA | NA | 1 | 1 |
| 4_5837333 | NA | NA | 1 | 1 | 1 | 1 | NA | NA | 2 | 1 | NA | NA | 1 | 1 | NA | NA | NA | NA | 1 | 1 |
| 4_5837358 | NA | NA | 1 | 1 | 1 | 1 | NA | NA | 2 | 1 | NA | NA | 1 | 1 | NA | NA | NA | NA | 1 | 1 |
| 4_5837359 | NA | NA | 1 | 1 | 1 | 1 | NA | NA | 2 | 1 | NA | NA | 1 | 1 | NA | NA | NA | NA | 1 | 1 |
| 4_5837375 | NA | NA | 1 | 1 | 1 | 1 | NA | NA | 2 | 1 | NA | NA | 1 | 1 | NA | NA | NA | NA | 1 | 1 |
| 4_5837413 | NA | NA | 1 | 1 | 1 | 1 | NA | NA | 2 | 1 | NA | NA | 1 | 1 | NA | NA | NA | NA | 1 | 1 |
| 4_5837427 | NA | NA | 1 | 1 | 1 | 0 | NA | NA | 2 | 0.5 | NA | NA | 1 | 1 | NA | NA | NA | NA | 1 | 1 |
| 4_5837484 | NA | NA | 1 | 0 | 1 | 1 | NA | NA | 2 | 0.5 | NA | NA | 1 | 1 | NA | NA | NA | NA | 1 | 1 |
| 4_5837485 | NA | NA | 1 | 1 | 1 | 0 | NA | NA | 2 | 0.5 | NA | NA | 1 | 1 | NA | NA | NA | NA | 1 | 1 |
| 4_5843531 | 2 | 0.5 | 6 | 0 | 9 | 0.2222 | 5 | 0.4 | 22 | 0.2273 | 4 | 0 | 3 | 0.3333 | 2 | 0 | 4 | 0 | 13 | 0.0769 |
| 4_5843535 | 2 | 1 | 5 | 0.6 | 6 | 0.6667 | 5 | 0.8 | 18 | 0.7222 | 3 | 0.6667 | 3 | 1 | 2 | 0.5 | 3 | 0.6667 | 11 | 0.7273 |
| 4_5843535 | 2 | 0.5 | 6 | 0 | 8 | 0.625 | 5 | 0.4 | 21 | 0.381 | 4 | 0 | 2 | 0 | 1 | 0 | 4 | 0.25 | 11 | 0.0909 |

| | | | | | | | | | | | | | | | | | | | | |
|---|---|---|---|---|---|---|---|---|---|---|---|---|---|---|---|---|---|---|---|---|
| 4_5843547 | 1 | 0 | 6 | 0 | 9 | 0.2222 | 5 | 0.4 | 21 | 0.1905 | 4 | 0 | 3 | 0.3333 | 1 | 0 | 4 | 0 | 12 | 0.0833 |
| 4_5843568 | 1 | 1 | 6 | 0.3333 | 8 | 0.625 | 4 | 0.75 | 19 | 0.5789 | 3 | 0 | 3 | 0 | 1 | 0 | 4 | 0.25 | 11 | 0.0909 |
| 4_5843578 | NA | NA | 6 | 0 | 9 | 0.1111 | 5 | 0 | 20 | 0.05 | 3 | 0 | 2 | 0 | 1 | 0 | 4 | 0.25 | 10 | 0.1 |
| 4_5843582 | 1 | 0 | 6 | 0.1667 | 8 | 0.125 | 5 | 0.2 | 20 | 0.15 | 2 | 0.5 | 3 | 0.3333 | 1 | 0 | 4 | 0 | 10 | 0.2 |
| 4_5843642 | 1 | 1 | 5 | 0.2 | 7 | 0.5714 | 2 | 1 | 15 | 0.5333 | 2 | 0.5 | 3 | 0.6667 | 1 | 1 | 3 | 0.3333 | 9 | 0.5556 |
| 4_5843650 | 1 | 1 | 5 | 0 | 6 | 0.3333 | 3 | 0.3333 | 15 | 0.2667 | 2 | 0.5 | 3 | 0.3333 | 1 | 1 | 3 | 0.3333 | 9 | 0.4444 |
| 4_5843691 | 1 | 1 | 4 | 0.75 | 3 | 1 | 2 | 1 | 10 | 0.9 | 2 | 1 | 3 | 0.6667 | 1 | 1 | 4 | 0.75 | 10 | 0.8 |
| 4_5843697 | 1 | 0 | 4 | 0.25 | 4 | 0.25 | 2 | 0 | 11 | 0.1818 | 2 | 0 | 3 | 0.3333 | 1 | 0 | 3 | 0 | 9 | 0.1111 |
| 4_5843707 | 1 | 0 | 3 | 0.6667 | 4 | 0.5 | 2 | 0 | 10 | 0.4 | 2 | 0.5 | 3 | 0.6667 | 1 | 0 | 3 | 0.3333 | 9 | 0.4444 |
| 4_5843710 | 1 | 1 | 4 | 1 | 4 | 1 | 2 | 1 | 11 | 1 | 2 | 1 | 3 | 1 | 1 | 1 | 4 | 1 | 10 | 1 |
| 4_5843713 | 1 | 0 | 3 | 0.3333 | 3 | 0 | 2 | 1 | 9 | 0.3333 | 2 | 0 | 2 | 0 | 1 | 0 | 3 | 0.6667 | 8 | 0.25 |
| 4_5843719 | 1 | 1 | 4 | 1 | 4 | 1 | 2 | 1 | 11 | 1 | 2 | 1 | 3 | 1 | 1 | 1 | 4 | 1 | 10 | 1 |
| 4_5843724 | 1 | 1 | 4 | 1 | 4 | 1 | 2 | 1 | 11 | 1 | 2 | 1 | 2 | 1 | 1 | 1 | 4 | 0.75 | 9 | 0.8889 |
| 4_5848656 | 1 | 0 | 3 | 1 | 3 | 0.6667 | 2 | 0.5 | 9 | 0.6667 | 2 | 1 | 3 | 0.6667 | 1 | 1 | 4 | 0.5 | 10 | 0.7 |
| 4_5848665 | 1 | 1 | 2 | 1 | 3 | 0.6667 | 1 | 1 | 7 | 0.8571 | 1 | 1 | 3 | 1 | 1 | 1 | 3 | 1 | 8 | 1 |
| 4_5848671 | 1 | 1 | 3 | 1 | 3 | 1 | 1 | 1 | 8 | 1 | 1 | 1 | 2 | 1 | NA | NA | 4 | 0.75 | 7 | 0.8571 |
| 4_5848675 | 1 | 1 | 1 | 1 | 3 | 1 | 2 | 0.5 | 7 | 0.8571 | 2 | 1 | 2 | 0 | 1 | 1 | 4 | 1 | 9 | 0.7778 |
| 4_5848692 | 1 | 1 | 3 | 0.6667 | 3 | 0.3333 | NA | NA | 7 | 0.5714 | 2 | 1 | 3 | 1 | 1 | 1 | 4 | 0.5 | 10 | 0.8 |
| 4_5848692 | 1 | 1 | 3 | 0.3333 | 3 | 0 | 2 | 1 | 9 | 0.4444 | 2 | 0.5 | 2 | 0 | 1 | 0 | 4 | 0.5 | 9 | 0.3333 |
| 4_5848694 | NA | NA | 2 | 0.5 | 3 | 0 | 2 | 0.5 | 7 | 0.2857 | 1 | 0 | 3 | 0.3333 | 1 | 0 | 4 | 0.5 | 9 | 0.3333 |
| 4_5848698 | 1 | 0 | 3 | 0.3333 | 3 | 0.3333 | 2 | 0 | 9 | 0.2222 | 2 | 0 | 3 | 0 | 1 | 0 | 3 | 0.3333 | 9 | 0.1111 |
| 4_5848710 | 1 | 1 | 2 | 1 | 3 | 0.6667 | 2 | 0.5 | 8 | 0.75 | 1 | 1 | 3 | 1 | NA | NA | 4 | 0.5 | 8 | 0.75 |
| 4_5848713 | 1 | 1 | 3 | 0 | 2 | 0 | 2 | 0 | 8 | 0.125 | 1 | 0 | 3 | 0.3333 | NA | NA | 2 | 0 | 6 | 0.1667 |
| 4_5879731 | 1 | 0 | 3 | 0 | 4 | 0.25 | 3 | 0 | 11 | 0.0909 | 2 | 0.5 | 2 | 0 | 4 | 0 | 2 | 0.5 | 10 | 0.2 |
| 4_5879907 | 2 | 0 | 6 | 0.3333 | 2 | 0 | 7 | 0.2857 | 17 | 0.2353 | 3 | 0.3333 | 3 | 0 | 5 | 0.2 | 5 | 0 | 16 | 0.125 |
| 4_5879928 | 2 | 0 | 6 | 0.1667 | 2 | 0 | 7 | 0 | 17 | 0.0588 | 2 | 0 | 3 | 0 | 5 | 0.2 | 5 | 0 | 15 | 0.0667 |
| 4_5879992 | 2 | 0 | 3 | 0 | 2 | 0.5 | 6 | 0.3333 | 13 | 0.2308 | 2 | 1 | 2 | 0.5 | 5 | 0 | 4 | 0.25 | 13 | 0.3077 |
| 4_5880028 | 8 | 0.125 | 7 | 0.1429 | 7 | 0 | 8 | 0 | 30 | 0.0667 | 5 | 0.2 | 9 | 0 | 7 | 0.1429 | 6 | 0 | 27 | 0.0741 |
| 4_5891272 | NA | NA | NA | NA | 1 | 1 | NA | NA | 1 | 1 | NA | NA | 1 | 1 | 1 | 1 | NA | NA | 2 | 1 |
| 4_5891286 | NA | NA | NA | NA | 1 | 1 | NA | NA | 1 | 1 | NA | NA | 1 | 1 | 1 | 1 | NA | NA | 2 | 1 |
| 4_5891316 | NA | NA | NA | NA | 1 | 1 | NA | NA | 1 | 1 | NA | NA | 1 | 1 | 1 | 1 | NA | NA | 2 | 1 |
| 4_5891320 | NA | NA | NA | NA | 1 | 1 | NA | NA | 1 | 1 | NA | NA | 1 | 1 | NA | NA | NA | NA | 1 | 1 |
| 4_5891379 | NA | NA | NA | NA | 1 | 1 | NA | NA | 1 | 1 | NA | NA | 1 | 0 | 1 | 1 | NA | NA | 2 | 0.5 |
| 4_5891428 | NA | NA | NA | NA | 1 | 1 | NA | NA | 1 | 1 | NA | NA | 1 | 0 | 1 | 1 | NA | NA | 2 | 0.5 |
| 4_5891444 | NA | NA | NA | NA | 1 | 1 | NA | NA | 1 | 1 | NA | NA | 1 | 0 | 1 | 1 | NA | NA | 2 | 0.5 |
| 4_5891466 | NA | NA | NA | NA | 1 | 1 | NA | NA | 1 | 1 | NA | NA | 1 | 1 | 1 | 0 | NA | NA | 2 | 0.5 |
| 4_5891509 | NA | NA | NA | NA | 1 | 1 | NA | NA | 1 | 1 | NA | NA | 2 | 1 | 1 | 1 | NA | NA | 3 | 1 |
| 4_5891514 | NA | NA | NA | NA | 1 | 1 | NA | NA | 1 | 1 | NA | NA | 2 | 1 | 1 | 1 | NA | NA | 3 | 1 |
| 4_5891541 | NA | NA | NA | NA | 1 | 1 | NA | NA | 1 | 1 | NA | NA | 2 | 1 | 1 | 1 | NA | NA | 3 | 1 |
| 4_5891548 | NA | NA | NA | NA | 1 | 1 | NA | NA | 1 | 1 | NA | NA | 2 | 1 | 1 | 0 | NA | NA | 3 | 0.6667 |
| 4_5891570 | NA | NA | NA | NA | 1 | 1 | NA | NA | 1 | 1 | NA | NA | 2 | 0.5 | 1 | 0 | NA | NA | 3 | 0.3333 |
| 4_5891590 | NA | NA | NA | NA | 1 | 1 | NA | NA | 1 | 1 | NA | NA | 2 | 0.5 | 1 | 1 | NA | NA | 3 | 0.6667 |
| 4_5891603 | NA | NA | NA | NA | 1 | 1 | NA | NA | 1 | 1 | NA | NA | 2 | 0.5 | 1 | 1 | NA | NA | 3 | 0.6667 |
| 4_5891612 | NA | NA | NA | NA | 1 | 1 | NA | NA | 1 | 1 | NA | NA | NA | NA | 1 | 1 | NA | NA | 1 | 1 |
| 4_5894610 | 2 | 0.5 | 1 | 1 | NA | NA | 1 | 1 | 4 | 0.75 | 2 | 0 | 1 | 1 | 2 | 0 | 2 | 1 | 7 | 0.4286 |
| 4_5894630 | 2 | 0 | 1 | 1 | NA | NA | 2 | 0 | 5 | 0.2 | 2 | 0 | 1 | 0 | 3 | 0.3333 | 2 | 0 | 8 | 0.125 |
| 4_5894636 | 2 | 0.5 | 1 | 0 | NA | NA | 2 | 0 | 5 | 0.2 | 2 | 0 | 1 | 0 | 3 | 0 | 2 | 0.5 | 8 | 0.125 |
| 4_5895664 | 2 | 0.5 | 2 | 0 | 2 | 0.5 | 3 | 0.6667 | 9 | 0.4444 | 3 | 0.3333 | NA | NA | 4 | 0.5 | 2 | 0 | 9 | 0.3333 |
| 4_5895667 | 2 | 0 | 2 | 0 | 2 | 0 | 3 | 0.3333 | 9 | 0.1111 | 1 | 1 | NA | NA | 4 | 0 | 1 | 0 | 6 | 0.1667 |
| 4_5895686 | 2 | 0 | 2 | 0.5 | 2 | 0.5 | 2 | 0.5 | 8 | 0.375 | 3 | 0.6667 | NA | NA | 5 | 0.2 | 2 | 0 | 10 | 0.3 |
| 4_5895689 | 2 | 0 | 2 | 0 | 2 | 0.5 | 3 | 0.3333 | 9 | 0.2222 | 3 | 0 | NA | NA | 5 | 0.2 | 2 | 0.5 | 10 | 0.2 |
| 4_5895745 | 2 | 0 | 1 | 0 | 2 | 0.5 | 3 | 0 | 8 | 0.125 | 3 | 0 | NA | NA | 5 | 0.4 | 1 | 1 | 9 | 0.3333 |
| 4_5895751 | 2 | 1 | 2 | 1 | 2 | 0.5 | 3 | 1 | 9 | 0.8889 | 3 | 1 | NA | NA | 5 | 0.8 | 2 | 1 | 10 | 0.9 |
| 4_5895752 | 2 | 1 | 2 | 1 | 2 | 1 | 3 | 1 | 9 | 1 | 2 | 1 | NA | NA | 4 | 1 | 2 | 1 | 8 | 1 |
| 4_5895756 | 1 | 0 | 3 | 1 | 2 | 1 | 3 | 1 | 9 | 0.8889 | 3 | 1 | NA | NA | 5 | 0.8 | 1 | 0 | 9 | 0.7778 |
| 4_5895769 | 2 | 0.5 | 2 | 0 | 2 | 0 | 3 | 0.3333 | 9 | 0.2222 | 2 | 0.5 | 1 | 0 | 5 | 0.4 | 2 | 0 | 10 | 0.3 |
| 4_5895773 | 2 | 1 | 2 | 1 | 2 | 1 | 2 | 1 | 8 | 1 | 3 | 1 | NA | NA | 3 | 1 | 2 | 1 | 8 | 1 |
| 4_5895775 | 2 | 0 | 3 | 0 | 2 | 1 | 3 | 0 | 10 | 0.2 | 3 | 0 | 1 | 0 | 4 | 0 | 2 | 0.5 | 10 | 0.1 |
| 4_5895780 | 2 | 0.5 | 2 | 0.5 | 2 | 1 | 3 | 0.6667 | 9 | 0.6667 | 2 | 0.5 | 1 | 1 | 4 | 0.25 | 2 | 0.5 | 9 | 0.4444 |
| 4_5895781 | 2 | 0.5 | 3 | 0.3333 | 2 | 0.5 | 2 | 0 | 9 | 0.3333 | 3 | 0.3333 | 1 | 0 | 4 | 0.5 | 2 | 0.5 | 10 | 0.4 |
| 4_5895785 | 2 | 1 | 3 | 1 | 2 | 1 | 2 | 0.5 | 9 | 0.8889 | 3 | 1 | NA | NA | 5 | 1 | 2 | 0.5 | 10 | 0.9 |
| 4_5895797 | 1 | 1 | 3 | 0.3333 | 3 | 0.3333 | 3 | 0.3333 | 10 | 0.4 | 3 | 0.3333 | 2 | 0 | 4 | 0.25 | 2 | 0.5 | 11 | 0.2727 |
| 4_5895803 | NA | NA | 3 | 1 | 2 | 1 | 3 | 1 | 8 | 1 | 3 | 0.6667 | 2 | 1 | 5 | 1 | 1 | 1 | 11 | 0.9091 |
| 4_5895814 | 2 | 0 | 3 | 0 | 3 | 0 | 3 | 0.3333 | 11 | 0.0909 | 4 | 0.25 | 2 | 0 | 5 | 0 | 2 | 0 | 13 | 0.0769 |
| 4_5895820 | 2 | 0 | 3 | 0 | 3 | 0.6667 | 3 | 0.3333 | 11 | 0.2727 | 4 | 0.75 | 1 | 0 | 5 | 0.4 | 2 | 1 | 12 | 0.5833 |

| | | | | | | | | | | | | | | | | | | | | |
|---|---|---|---|---|---|---|---|---|---|---|---|---|---|---|---|---|---|---|---|---|
| 4_58958216 | 1 | 1 | 3 | 1 | 2 | 1 | 2 | 1 | 8 | 1 | 3 | 1 | 1 | 1 | 3 | 1 | 2 | 1 | 9 | 1 |
| 4_58958285 | 2 | 1 | 3 | 0.6667 | 3 | 1 | 3 | 1 | 11 | 0.9091 | 4 | 1 | 2 | 1 | 3 | 1 | 2 | 1 | 11 | 1 |
| 4_58958332 | 2 | 0.5 | 3 | 0 | 2 | 0 | 3 | 0.3333 | 10 | 0.2 | 4 | 0 | 2 | 0.5 | 4 | 0.5 | 2 | 0.5 | 12 | 0.3333 |
| 4_58958446 | 2 | 0.5 | 3 | 0 | 3 | 0 | 3 | 0 | 11 | 0.0909 | 4 | 0.25 | 2 | 0 | 5 | 0.2 | 2 | 0 | 13 | 0.1538 |
| 4_58958653 | 3 | 0.6667 | 3 | 0.3333 | 3 | 0.3333 | 3 | 0.3333 | 12 | 0.4167 | 4 | 0.25 | 2 | 1 | 5 | 0.4 | 2 | 0 | 13 | 0.3846 |
| 4_58958788 | 3 | 0 | 3 | 0 | 4 | 0.25 | 3 | 0.3333 | 13 | 0.1538 | 4 | 0 | 1 | 1 | 6 | 0.3333 | 2 | 0 | 13 | 0.2308 |
| 4_58959169 | 3 | 0.6667 | 3 | 0.3333 | 4 | 0.5 | 3 | 0.6667 | 13 | 0.5385 | 3 | 1 | 2 | 0 | 7 | 0.8571 | 4 | 0.75 | 16 | 0.75 |
| 4_58959247 | 2 | 0.5 | 4 | 0.25 | 4 | 0.25 | 3 | 1 | 13 | 0.4615 | 3 | 0.6667 | 2 | 0 | 7 | 0.5714 | 3 | 0.3333 | 15 | 0.4667 |
| 4_58959365 | 2 | 0 | 4 | 0.25 | 4 | 0.75 | 4 | 0 | 14 | 0.2857 | 4 | 0.25 | 1 | 0 | 8 | 0.375 | 3 | 0 | 16 | 0.25 |
| 4_58960155 | 3 | 0.3333 | 6 | 0.1667 | 6 | 0.1667 | 9 | 0.1111 | 24 | 0.1667 | 4 | 0.25 | 2 | 0 | 9 | 0 | 4 | 0 | 19 | 0.0526 |
| 4_58962126 | 4 | 0.75 | 6 | 0.6667 | 3 | 1 | 9 | 0.7778 | 22 | 0.7727 | 3 | 0.6667 | 2 | 0.5 | 4 | 0.75 | 4 | 1 | 13 | 0.7692 |
| 4_58962212 | 6 | 0.8333 | 9 | 0.8889 | 8 | 0.75 | 12 | 0.8333 | 35 | 0.8286 | 6 | 1 | 5 | 0.8 | 12 | 0.9167 | 5 | 1 | 28 | 0.9286 |
| 4_58962308 | 7 | 0.4286 | 9 | 0.4444 | 7 | 0.5714 | 9 | 0.3333 | 32 | 0.4375 | 6 | 0.3333 | 4 | 0.75 | 8 | 0.875 | 5 | 0.6 | 23 | 0.6522 |
| 4_58962387 | 6 | 1 | 7 | 1 | 7 | 0.8571 | 11 | 0.9091 | 31 | 0.9355 | 6 | 1 | 6 | 0.8333 | 10 | 0.7 | 4 | 1 | 26 | 0.8462 |
| 4_59020297 | 2 | 0.5 | 1 | 0 | 1 | 1 | NA | NA | 4 | 0.5 | 1 | 1 | 1 | 1 | 1 | 1 | NA | NA | 3 | 1 |
| 4_59020415 | 1 | 1 | 1 | 0 | NA | NA | 1 | 1 | 3 | 0.6667 | 1 | 0 | 1 | 1 | 1 | 0 | NA | NA | 3 | 0.3333 |
| 4_59022914 | 2 | 0.5 | 2 | 1 | 1 | 1 | 1 | 1 | 6 | 0.8333 | NA | NA | 1 | 1 | NA | NA | NA | NA | 1 | 1 |
| 4_59041336 | 17 | 0 | 17 | 0.1176 | 15 | 0.0667 | 19 | 0.1579 | 68 | 0.0882 | 15 | 0 | 13 | 0.1538 | 29 | 0.069 | 8 | 0.125 | 65 | 0.0769 |
| 4_59220337 | 2 | 0 | 7 | 0 | 5 | 0.2 | 6 | 0.1667 | 20 | 0.1 | 1 | 0 | 5 | 0.6 | 1 | 0 | 8 | 0.25 | 15 | 0.3333 |
| 4_59220407 | 2 | 0 | 9 | 0.1111 | 5 | 0.2 | 7 | 0.1429 | 23 | 0.1304 | NA | NA | 5 | 0 | 1 | 0 | 7 | 0.1429 | 13 | 0.0769 |
| 4_59220458 | 2 | 0.5 | 9 | 0.2222 | 3 | 0.3333 | 6 | 0.3333 | 20 | 0.3 | 1 | 0 | 5 | 0.4 | 1 | 1 | 7 | 0.7143 | 14 | 0.5714 |
| 4_59221776 | 4 | 0 | 9 | 0 | 5 | 0 | 10 | 0.2 | 28 | 0.0714 | 2 | 0 | 7 | 0 | 2 | 0.5 | 13 | 0.0769 | 24 | 0.0833 |
| 4_59315070 | 1 | 1 | 1 | 1 | NA | NA | NA | NA | 2 | 1 | NA | NA | NA | NA | NA | NA | 2 | 0.5 | 2 | 0.5 |
| 4_59315085 | 1 | 1 | 1 | 1 | NA | NA | NA | NA | 2 | 1 | NA | NA | NA | NA | 1 | 0 | 2 | 0.5 | 3 | 0.3333 |
| 4_59315093 | 1 | 0 | 2 | 0.5 | NA | NA | NA | NA | 3 | 0.3333 | NA | NA | NA | NA | 1 | 0 | 2 | 0.5 | 3 | 0.3333 |
| 4_59315172 | 1 | 0 | 1 | 1 | NA | NA | NA | NA | 2 | 0.5 | NA | NA | NA | NA | 1 | 1 | 2 | 0 | 3 | 0.3333 |
| 4_59315235 | 1 | 0 | 2 | 1 | NA | NA | NA | NA | 3 | 0.6667 | NA | NA | NA | NA | 1 | 1 | 2 | 1 | 3 | 1 |
| 4_59472408 | 3 | 0.3333 | 5 | 0 | NA | NA | 3 | 0 | 11 | 0.0909 | 2 | 0.5 | NA | NA | 1 | 0 | 2 | 0.5 | 5 | 0.4 |
| 4_59476653 | 1 | 1 | 2 | 1 | NA | NA | 1 | 0 | 4 | 0.75 | NA | NA | NA | NA | 1 | 1 | NA | NA | 1 | 1 |
| 4_59476754 | 2 | 0.5 | 1 | 0 | NA | NA | NA | NA | 3 | 0.3333 | NA | NA | NA | NA | 1 | 1 | NA | NA | 1 | 1 |
| 4_59476821 | 2 | 1 | 2 | 0.5 | 2 | 0.5 | NA | NA | 6 | 0.6667 | 1 | 1 | NA | NA | 1 | 1 | NA | NA | 2 | 1 |
| 4_59476852 | 2 | 0 | 2 | 0.5 | 1 | 1 | NA | NA | 5 | 0.4 | 1 | 1 | NA | NA | 1 | 0 | NA | NA | 2 | 0.5 |
| 4_59610816 | 8 | 0.75 | 3 | 0.3333 | 7 | 0.5714 | 5 | 0.8 | 23 | 0.6522 | 5 | 0.4 | 5 | 0 | 4 | 0.25 | 4 | 0.75 | 18 | 0.3333 |
| 4_59610868 | 6 | 0 | 2 | 0 | 6 | 0 | 6 | 0.1667 | 20 | 0.05 | 5 | 0.2 | 5 | 0 | 4 | 0 | 3 | 0 | 17 | 0.0588 |
| 4_59617770 | 7 | 0 | 3 | 0.3333 | 9 | 0.1111 | 5 | 0 | 24 | 0.0833 | 5 | 0 | 5 | 0 | 4 | 0 | 5 | 0.2 | 19 | 0.0526 |
| 4_59617835 | 6 | 0.3333 | 3 | 0 | 9 | 0.1111 | 6 | 0 | 24 | 0.125 | 5 | 0 | 5 | 0.2 | 5 | 0 | 3 | 0 | 18 | 0.0556 |
| 4_59618106 | 5 | 0 | 4 | 0.5 | 7 | 0 | 7 | 0 | 23 | 0.087 | 3 | 0 | 4 | 0.25 | 6 | 0 | 5 | 0.2 | 18 | 0.1111 |
| 4_59618132 | 5 | 0 | 4 | 0 | 10 | 0.1 | 7 | 0.2857 | 26 | 0.1154 | 3 | 0 | 5 | 0.2 | 6 | 0 | 4 | 0 | 18 | 0.0556 |
| 4_59618173 | 3 | 0 | 4 | 0.5 | 7 | 0 | 5 | 0 | 19 | 0.1053 | 3 | 0.3333 | 4 | 0 | 6 | 0.1667 | 3 | 0 | 16 | 0.125 |
| 4_59618291 | 3 | 0 | 2 | 0 | 8 | 0 | 4 | 0.25 | 17 | 0.0588 | 2 | 0 | 3 | 0.3333 | 3 | 0.3333 | 3 | 0 | 11 | 0.1818 |
| 4_59690502 | 1 | 1 | 1 | 1 | NA | NA | NA | NA | 2 | 1 | 1 | 1 | 1 | 1 | NA | NA | NA | NA | 2 | 1 |
| 4_59690520 | 1 | 0 | 1 | 1 | NA | NA | NA | NA | 2 | 0.5 | 1 | 0 | 1 | 1 | NA | NA | NA | NA | 2 | 0.5 |
| 4_59690637 | 1 | 0 | 1 | 1 | NA | NA | NA | NA | 2 | 0.5 | 1 | 1 | 1 | 1 | NA | NA | NA | NA | 2 | 1 |
| 4_59690643 | 1 | 1 | 1 | 0 | NA | NA | NA | NA | 2 | 0.5 | NA | NA | 1 | 1 | NA | NA | NA | NA | 1 | 1 |
| 4_59690858 | 1 | 0 | 1 | 1 | NA | NA | NA | NA | 2 | 0.5 | 1 | 1 | 1 | 0 | NA | NA | NA | NA | 2 | 0.5 |
| 4_59690885 | 1 | 0 | 1 | 1 | NA | NA | NA | NA | 2 | 0.5 | 1 | 0 | 1 | 1 | NA | NA | NA | NA | 2 | 0.5 |
| 4_59729056 | 1 | 0 | 1 | 1 | 1 | 0 | NA | NA | 3 | 0.3333 | NA | NA | NA | NA | 2 | 0.5 | 1 | 0 | 3 | 0.3333 |
| 4_59752267 | 1 | 0 | 1 | 1 | NA | NA | NA | NA | 2 | 0.5 | 1 | 1 | 1 | 0 | NA | NA | NA | NA | 2 | 0.5 |
| 4_59752415 | NA | NA | 1 | 1 | NA | NA | NA | NA | 1 | 1 | 1 | 1 | 1 | 0 | NA | NA | NA | NA | 2 | 0.5 |
| 4_59752599 | 1 | 1 | 1 | 1 | NA | NA | NA | NA | 2 | 1 | 1 | 1 | 1 | 1 | NA | NA | NA | NA | 2 | 1 |
| 4_59752680 | 1 | 0 | 1 | 1 | NA | NA | NA | NA | 2 | 0.5 | 1 | 1 | 1 | 1 | NA | NA | NA | NA | 2 | 1 |
| 4_59752760 | NA | NA | 1 | 1 | NA | NA | NA | NA | 1 | 1 | 1 | 1 | NA | NA | NA | NA | NA | NA | 1 | 1 |
| 4_59752873 | 1 | 1 | 1 | 1 | NA | NA | NA | NA | 2 | 1 | 1 | 1 | 1 | 1 | NA | NA | NA | NA | 2 | 1 |
| 4_59752975 | NA | NA | 1 | 1 | NA | NA | NA | NA | 1 | 1 | 1 | 1 | 1 | 1 | NA | NA | NA | NA | 2 | 1 |
| 4_59753074 | 1 | 1 | 1 | 1 | NA | NA | NA | NA | 2 | 1 | 1 | 1 | 1 | 1 | NA | NA | NA | NA | 2 | 1 |
| 4_59753134 | NA | NA | 1 | 1 | NA | NA | NA | NA | 1 | 1 | 1 | 0 | 1 | 1 | NA | NA | NA | NA | 2 | 0.5 |
| 4_59753166 | 1 | 1 | 1 | 1 | NA | NA | NA | NA | 2 | 1 | 1 | 1 | NA | NA | NA | NA | NA | NA | 1 | 1 |
| 4_59753637 | 2 | 1 | 1 | 1 | NA | NA | NA | NA | 3 | 1 | 1 | 1 | NA | NA | NA | NA | NA | NA | 1 | 1 |
| 4_59771714 | 3 | 0.3333 | 1 | 0 | 1 | 0 | 1 | 0 | 6 | 0.1667 | 2 | 0 | 3 | 0 | 3 | 0.3333 | 6 | 0 | 14 | 0.0714 |
| 4_59771904 | 3 | 0.6667 | 1 | 1 | 1 | 0 | NA | NA | 5 | 0.6 | 2 | 0 | 3 | 0.3333 | 2 | 0.5 | 6 | 0.5 | 13 | 0.3846 |
| 4_59772075 | 3 | 0.6667 | NA | NA | 1 | 1 | NA | NA | 4 | 0.75 | 2 | 1 | 3 | 1 | NA | NA | 6 | 1 | 11 | 1 |
| 4_59772098 | 3 | 1 | NA | NA | 1 | 1 | NA | NA | 4 | 1 | 2 | 1 | 2 | 1 | 2 | 1 | 5 | 1 | 11 | 1 |
| 4_59772122 | 2 | 1 | 1 | 0 | 1 | 0 | NA | NA | 4 | 0.5 | 2 | 1 | 2 | 1 | 2 | 0.5 | 6 | 1 | 12 | 0.9167 |
| 4_59772134 | 1 | 1 | 1 | 1 | 1 | 1 | NA | NA | 3 | 1 | 2 | 1 | 3 | 1 | 2 | 1 | 6 | 1 | 13 | 1 |
| 4_59772155 | 1 | 1 | 1 | 1 | 1 | 1 | NA | NA | 3 | 1 | 2 | 0.5 | 3 | 0.6667 | 2 | 0.5 | 6 | 1 | 13 | 0.7692 |

| | | | | | | | | | | | | | | | | | | | | |
|---|---|---|---|---|---|---|---|---|---|---|---|---|---|---|---|---|---|---|---|---|
| 4_5977227 | 2 | 1 | 1 | 1 | 1 | 0 | NA | NA | 4 | 0.75 | 2 | 1 | 2 | 1 | 2 | 0.5 | 5 | 0.6 | 11 | 0.7273 |
| 4_5977228 | 2 | 1 | NA | NA | 1 | 1 | NA | NA | 3 | 1 | 1 | 1 | NA | NA | 2 | 0.5 | 5 | 1 | 8 | 0.875 |
| 4_5977229 | 1 | 1 | 1 | 1 | NA | NA | NA | NA | 2 | 1 | 2 | 1 | 1 | 1 | 2 | 1 | 2 | 1 | 7 | 1 |
| 4_5977233 | 1 | 1 | NA | NA | 1 | 1 | NA | NA | 2 | 1 | 2 | 0.5 | 3 | 1 | 2 | 1 | 6 | 0.6667 | 13 | 0.7692 |
| 4_5977446 | 2 | 0 | 2 | 0.5 | 1 | 0 | 1 | 0 | 6 | 0.1667 | 4 | 0.5 | NA | NA | 1 | 0 | 2 | 0.5 | 7 | 0.4286 |
| 4_5977451 | 1 | 0 | 1 | 1 | 1 | 0 | 1 | 0 | 4 | 0.25 | 4 | 0.5 | NA | NA | 1 | 0 | 2 | 0.5 | 7 | 0.4286 |
| 4_5977519 | 2 | 0.5 | 2 | 0 | 1 | 0 | 1 | 1 | 6 | 0.3333 | 3 | 0.6667 | NA | NA | NA | NA | 2 | 0 | 5 | 0.4 |
| 4_5989898 | NA | NA | 1 | 1 | NA | NA | NA | NA | 1 | 1 | NA | NA | 2 | 1 | NA | NA | NA | NA | 2 | 1 |
| 4_5989914 | NA | NA | 1 | 1 | NA | NA | NA | NA | 1 | 1 | NA | NA | 2 | 1 | NA | NA | NA | NA | 2 | 1 |
| 4_5989926 | NA | NA | 1 | 0 | 1 | 1 | 1 | 1 | 3 | 0.6667 | NA | NA | 2 | 1 | NA | NA | NA | NA | 2 | 1 |
| 4_5989928 | NA | NA | 1 | 1 | NA | NA | 1 | 1 | 2 | 1 | NA | NA | 1 | 1 | NA | NA | NA | NA | 1 | 1 |
| 4_5989952 | NA | NA | 1 | 0 | 2 | 1 | 1 | 1 | 4 | 0.75 | NA | NA | 1 | 1 | 1 | 1 | NA | NA | 2 | 1 |
| 4_5989967 | NA | NA | 1 | 1 | 2 | 0.5 | NA | NA | 3 | 0.6667 | NA | NA | 2 | 0.5 | 1 | 1 | NA | NA | 3 | 0.6667 |
| 4_5989973 | NA | NA | 1 | 0 | 2 | 0.5 | NA | NA | 3 | 0.3333 | NA | NA | 1 | 0 | 1 | 1 | NA | NA | 2 | 0.5 |
| 4_5989988 | NA | NA | 1 | 0 | 2 | 0.5 | NA | NA | 3 | 0.3333 | NA | NA | 2 | 0.5 | 1 | 0 | NA | NA | 3 | 0.3333 |
| 4_5990187 | 1 | 0 | 2 | 0.5 | 5 | 0.2 | NA | NA | 8 | 0.25 | NA | NA | 2 | 0.5 | 1 | 0 | 1 | 0 | 4 | 0.25 |
| 4_5990355 | 3 | 0.3333 | 2 | 0 | 5 | 0.4 | NA | NA | 10 | 0.3 | NA | NA | 2 | 0.5 | 4 | 0 | 4 | 0 | 10 | 0.1 |
| 4_5990367 | 3 | 0.6667 | 2 | 0.5 | 5 | 1 | NA | NA | 10 | 0.8 | NA | NA | 2 | 1 | 4 | 0.75 | 4 | 0.75 | 10 | 0.8 |
| 4_5990392 | 3 | 0 | 2 | 0.5 | 4 | 0.5 | 1 | 0 | 10 | 0.3 | NA | NA | 2 | 0 | 4 | 0 | 4 | 0.5 | 10 | 0.2 |
| 4_5990398 | 4 | 0 | 1 | 0 | 4 | 0.25 | 1 | 0 | 10 | 0.1 | NA | NA | 2 | 0.5 | 5 | 0.2 | 4 | 0 | 11 | 0.1818 |
| 4_5990403 | 1 | 0 | 2 | 0 | 3 | 0.6667 | NA | NA | 6 | 0.3333 | NA | NA | 1 | 0 | 2 | 0 | 3 | 1 | 6 | 0.5 |
| 4_5990412 | 3 | 0 | 3 | 0.3333 | 5 | 0.2 | NA | NA | 11 | 0.1818 | NA | NA | 2 | 0 | 4 | 0.25 | 3 | 0 | 9 | 0.1111 |
| 4_5990414 | 4 | 0.25 | 2 | 1 | 5 | 0.6 | 1 | 1 | 12 | 0.5833 | NA | NA | 3 | 0 | 5 | 0 | 4 | 0.25 | 12 | 0.0833 |
| 4_5990416 | 3 | 0 | 3 | 0.3333 | 4 | 0.25 | 1 | 0 | 11 | 0.1818 | NA | NA | 1 | 0 | 5 | 0 | 4 | 0.5 | 10 | 0.2 |
| 4_5990422 | 3 | 0.6667 | 3 | 0.6667 | 5 | 0.8 | NA | NA | 11 | 0.7273 | NA | NA | 3 | 0.6667 | 4 | 0.25 | 2 | 1 | 9 | 0.5556 |
| 4_5990438 | 5 | 0.4 | 3 | 0.6667 | 5 | 0.6 | 2 | 1 | 15 | 0.6 | NA | NA | 2 | 0 | 5 | 0.2 | 3 | 0.3333 | 10 | 0.2 |
| 4_6210894 | 1 | 0 | 2 | 0.5 | 2 | 0.5 | 2 | 0.5 | 7 | 0.4286 | NA | NA | 2 | 0 | 1 | 1 | 1 | 0 | 4 | 0.25 |
| 4_6210938 | 2 | 1 | 2 | 1 | 1 | 1 | 1 | 1 | 6 | 1 | NA | NA | 3 | 0.6667 | NA | NA | 1 | 1 | 4 | 0.75 |
| 4_6210960 | 2 | 1 | 1 | 1 | NA | NA | 1 | 0 | 4 | 0.75 | NA | NA | 4 | 0.75 | 1 | 1 | NA | NA | 5 | 0.8 |
| 4_6210975 | 2 | 1 | NA | NA | 1 | 1 | 2 | 0.5 | 5 | 0.8 | NA | NA | 4 | 0.75 | 2 | 1 | 1 | 1 | 7 | 0.8571 |
| 4_6210977 | 2 | 1 | 1 | 1 | 1 | 1 | 2 | 1 | 6 | 1 | NA | NA | 4 | 1 | NA | NA | 1 | 1 | 5 | 1 |
| 4_6210980 | 2 | 1 | 1 | 1 | 1 | 1 | 2 | 1 | 6 | 1 | NA | NA | 5 | 1 | 2 | 1 | 1 | 1 | 8 | 1 |
| 4_6210985 | 1 | 1 | 1 | 1 | 1 | 1 | 2 | 1 | 5 | 1 | NA | NA | 3 | 1 | 2 | 1 | 1 | 1 | 6 | 1 |
| 4_6210989 | 2 | 0.5 | 1 | 0 | NA | NA | 1 | 1 | 4 | 0.5 | NA | NA | 4 | 0.25 | 1 | 0 | 1 | 0 | 6 | 0.1667 |
| 4_6211002 | NA | NA | 1 | 1 | 1 | 0 | 2 | 0 | 4 | 0.25 | NA | NA | 3 | 0 | 1 | 0 | 1 | 1 | 5 | 0.2 |
| 4_6211003 | NA | NA | 1 | 0 | 1 | 1 | 2 | 0.5 | 4 | 0.5 | NA | NA | 4 | 0.5 | 2 | 0 | 1 | 1 | 7 | 0.4286 |
| 4_6211020 | NA | NA | 1 | 1 | 1 | 0 | NA | NA | 2 | 0.5 | NA | NA | 5 | 0.2 | 2 | 0 | 1 | 0 | 8 | 0.125 |
| 4_6211022 | NA | NA | 1 | 1 | 1 | 1 | 2 | 1 | 4 | 1 | NA | NA | 5 | 1 | 2 | 1 | 1 | 1 | 8 | 1 |
| 4_6211024 | NA | NA | 1 | 0 | 1 | 1 | 2 | 0.5 | 4 | 0.5 | NA | NA | 5 | 0.2 | 2 | 0 | 1 | 0 | 8 | 0.125 |
| 4_6211027 | NA | NA | 1 | 1 | NA | NA | 2 | 0.5 | 3 | 0.6667 | NA | NA | 4 | 0.75 | 1 | 0 | NA | NA | 5 | 0.6 |
| 4_6211036 | NA | NA | 1 | 1 | 1 | 1 | 2 | 1 | 4 | 1 | NA | NA | 4 | 1 | 1 | 1 | 1 | 1 | 6 | 1 |
| 4_6211037 | NA | NA | 1 | 1 | NA | NA | 2 | 1 | 3 | 1 | NA | NA | 3 | 1 | 2 | 1 | NA | NA | 5 | 1 |
| 4_6211071 | NA | NA | 1 | 1 | 1 | 1 | 2 | 1 | 4 | 1 | NA | NA | 3 | 0.3333 | 2 | 1 | 1 | 1 | 6 | 0.6667 |
| 4_6211073 | NA | NA | 1 | 1 | 1 | 1 | 2 | 0 | 4 | 0.5 | NA | NA | 4 | 0.5 | 1 | 0 | 1 | 0 | 6 | 0.3333 |
| 4_6211076 | NA | NA | 1 | 1 | 1 | 1 | 1 | 0 | 3 | 0.6667 | NA | NA | 2 | 0.5 | 1 | 1 | NA | NA | 3 | 0.6667 |
| 4_6221537 | 1 | 0 | 2 | 1 | 1 | 1 | NA | NA | 4 | 0.75 | 1 | 1 | NA | NA | 1 | 1 | 1 | 0 | 3 | 0.6667 |
| 4_6221543 | NA | NA | 1 | 1 | 1 | 0 | NA | NA | 2 | 0.5 | 3 | 0.3333 | NA | NA | NA | NA | 1 | 1 | 4 | 0.5 |
| 4_6221550 | 1 | 1 | 2 | 0.5 | 1 | 0 | NA | NA | 4 | 0.5 | 3 | 0.6667 | NA | NA | 1 | 1 | 1 | 1 | 5 | 0.8 |
| 4_6221568 | 1 | 1 | 2 | 1 | 1 | 1 | NA | NA | 4 | 1 | 2 | 0.5 | NA | NA | 1 | 1 | 1 | 1 | 4 | 0.75 |
| 4_6221578 | 1 | 1 | 2 | 0.5 | 1 | 1 | NA | NA | 4 | 0.75 | 3 | 0 | NA | NA | 1 | 1 | 1 | 1 | 5 | 0.4 |
| 4_6221592 | NA | NA | 2 | 1 | 1 | 1 | NA | NA | 3 | 1 | 3 | 0.6667 | NA | NA | NA | NA | NA | NA | 3 | 0.6667 |
| 4_6221596 | 1 | 0 | 2 | 0.5 | 1 | 0 | NA | NA | 4 | 0.25 | 2 | 0.5 | NA | NA | 1 | 0 | 2 | 0 | 5 | 0.2 |
| 4_6221601 | 1 | 1 | 2 | 1 | NA | NA | NA | NA | 3 | 1 | 4 | 1 | NA | NA | 1 | 1 | 2 | 1 | 7 | 1 |
| 4_6221613 | 1 | 0 | 2 | 0.5 | 1 | 0 | NA | NA | 4 | 0.25 | 4 | 0.25 | NA | NA | 1 | 0 | 2 | 0 | 7 | 0.1429 |
| 4_6221631 | 1 | 0 | 2 | 0.5 | 1 | 1 | NA | NA | 4 | 0.5 | 4 | 0.25 | NA | NA | 1 | 0 | 2 | 0 | 7 | 0.1429 |
| 4_6221870 | 5 | 0.2 | 4 | 0.25 | 2 | 0 | 2 | 0 | 13 | 0.1538 | 3 | 0.3333 | 1 | 1 | 2 | 0 | 3 | 0.6667 | 9 | 0.4444 |
| 4_6221873 | 5 | 0.6 | 4 | 0 | 2 | 0 | 2 | 0 | 13 | 0.2308 | 2 | 0 | 1 | 0 | 3 | 0.3333 | NA | NA | 6 | 0.1667 |
| 4_6221874 | 4 | 0 | 4 | 0.25 | 2 | 0 | 2 | 0 | 12 | 0.0833 | 4 | 0.5 | NA | NA | 3 | 0.6667 | 2 | 0.5 | 9 | 0.5556 |
| 4_6221879 | 5 | 0.2 | 4 | 0.5 | 1 | 1 | 2 | 1 | 12 | 0.5 | 4 | 0.5 | 1 | 1 | 2 | 0.5 | 3 | 0.3333 | 10 | 0.5 |
| 4_6221892 | 2 | 0 | 4 | 0.25 | 1 | 0 | 3 | 0 | 10 | 0.1 | 3 | 0.3333 | NA | NA | 3 | 0 | 1 | 0 | 7 | 0.1429 |
| 4_6221895 | 3 | 0.3333 | 3 | 0.3333 | 2 | 0 | 2 | 0.5 | 10 | 0.3 | 4 | 0.25 | NA | NA | 1 | 0 | 3 | 0.3333 | 8 | 0.25 |
| 4_6221904 | 2 | 0 | 2 | 0.5 | 1 | 1 | 3 | 0 | 8 | 0.25 | 3 | 0.3333 | 1 | 0 | 2 | 1 | 3 | 0.3333 | 9 | 0.4444 |
| 4_6221971 | 1 | 1 | 2 | 0.5 | 2 | 0.5 | 1 | 1 | 6 | 0.6667 | 2 | 0.5 | 2 | 0.5 | 1 | 1 | NA | NA | 5 | 0.6 |
| 4_6222004 | NA | NA | 2 | 0.5 | 4 | 0.25 | 2 | 0 | 8 | 0.25 | 2 | 0.5 | 2 | 0 | 2 | 0 | NA | NA | 6 | 0.1667 |
| 4_6222028 | NA | NA | 2 | 0 | 2 | 0.5 | 2 | 0.5 | 6 | 0.3333 | 2 | 0.5 | 2 | 0 | 1 | 1 | NA | NA | 5 | 0.4 |

| | | | | | | | | | | | | | | | | | | | | |
|---|---|---|---|---|---|---|---|---|---|---|---|---|---|---|---|---|---|---|---|---|
| 4_62221278 | NA | NA | 2 | 0.5 | 1 | 0 | 2 | 0.5 | 5 | 0.4 | 1 | 1 | 1 | 0 | 2 | 0 | NA | NA | 4 | 0.25 |
| 4_62221292 | NA | NA | 1 | 0 | 1 | 1 | 1 | 1 | 3 | 0.6667 | 1 | 1 | NA | NA | 2 | 0.5 | NA | NA | 3 | 0.6667 |
| 4_62222443 | 2 | 0 | 4 | 0.25 | 2 | 0 | 4 | 0 | 12 | 0.0833 | 4 | 0.25 | NA | NA | 5 | 0 | 3 | 0 | 12 | 0.0833 |
| 4_62222584 | 2 | 0 | 6 | 0.1667 | 1 | 0 | 4 | 0 | 13 | 0.0769 | 3 | 0 | 1 | 0 | 5 | 0.2 | 3 | 0 | 12 | 0.0833 |
| 4_62222621 | 2 | 0 | 5 | 0 | 2 | 0 | 3 | 0.3333 | 12 | 0.0833 | 3 | 0.3333 | 1 | 1 | 4 | 0.5 | 2 | 0 | 10 | 0.4 |
| 4_62222682 | 1 | 0 | 5 | 0.2 | 2 | 0 | 2 | 0.5 | 10 | 0.2 | 2 | 0 | 1 | 0 | 5 | 0.2 | 3 | 0 | 11 | 0.0909 |
| 4_62222752 | 2 | 1 | 6 | 0.6667 | 2 | 1 | 2 | 0.5 | 12 | 0.75 | 3 | 0.6667 | 1 | 1 | 4 | 1 | 3 | 0.6667 | 11 | 0.8182 |
| 4_62222766 | 2 | 0 | 5 | 0.4 | 2 | 1 | 2 | 1 | 11 | 0.5455 | 3 | 0 | 1 | 0 | 5 | 0.4 | 1 | 0 | 10 | 0.2 |
| 4_62222851 | 2 | 1 | 4 | 0.25 | 2 | 0.5 | 2 | 0.5 | 10 | 0.5 | 3 | 1 | 1 | 1 | 5 | 0.8 | 3 | 1 | 12 | 0.9167 |
| 4_62307043 | 1 | 1 | 2 | 0.5 | 1 | 0 | 2 | 0.5 | 6 | 0.5 | 1 | 0 | NA | NA | 2 | 0.5 | NA | NA | 3 | 0.3333 |
| 4_62307087 | 1 | 1 | 3 | 1 | 1 | 1 | 2 | 0.5 | 7 | 0.8571 | 1 | 1 | NA | NA | NA | NA | NA | NA | 1 | 1 |
| 4_62307293 | 1 | 0 | 3 | 0 | 1 | 0 | 2 | 0.5 | 7 | 0.1429 | 2 | 0.5 | NA | NA | 2 | 1 | NA | NA | 4 | 0.75 |
| 4_62307702 | 1 | 0 | 2 | 0.5 | NA | NA | 2 | 0 | 5 | 0.2 | 2 | 0.5 | NA | NA | 3 | 0 | NA | NA | 5 | 0.2 |
| 4_62312295 | 2 | 0.5 | 4 | 0.5 | 2 | 1 | NA | NA | 8 | 0.625 | 1 | 1 | NA | NA | 1 | 1 | 2 | 1 | 4 | 1 |
| 4_62312375 | 2 | 1 | 2 | 1 | 2 | 1 | NA | NA | 6 | 1 | 1 | 1 | 1 | 1 | 1 | 1 | 1 | 1 | 4 | 1 |
| 4_62312396 | 2 | 1 | 4 | 0.5 | 3 | 1 | 1 | 0 | 10 | 0.7 | 1 | 1 | 1 | 1 | 2 | 0.5 | 2 | 0.5 | 6 | 0.6667 |
| 4_62312497 | 2 | 1 | 5 | 0.6 | 2 | 0.5 | NA | NA | 9 | 0.6667 | 1 | 1 | 1 | 1 | 2 | 0.5 | 2 | 0.5 | 6 | 0.6667 |
| 4_62312504 | 1 | 0 | 3 | 0.6667 | 1 | 1 | 1 | 0 | 6 | 0.5 | 1 | 1 | 1 | 0 | 1 | 0 | 2 | 0.5 | 5 | 0.4 |
| 4_62312861 | 4 | 0.25 | 7 | 0.2857 | 4 | 0 | 6 | 0 | 21 | 0.1429 | 3 | 0 | 3 | 0.3333 | 4 | 0 | 3 | 0 | 13 | 0.0769 |
| 4_62312876 | 5 | 0 | 6 | 0 | 4 | 0.25 | 7 | 0.1429 | 22 | 0.0909 | 2 | 0 | 3 | 0 | 4 | 0.25 | 3 | 0.3333 | 12 | 0.1667 |
| 4_62312954 | 5 | 0 | 8 | 0.125 | 8 | 0 | 7 | 0.1429 | 28 | 0.0714 | 4 | 0.25 | 4 | 0.25 | 3 | 0 | 4 | 0.5 | 15 | 0.2667 |
| 4_62313049 | 5 | 0.8 | 9 | 0.8889 | 8 | 0.5 | 7 | 0.8571 | 29 | 0.7586 | 3 | 1 | 4 | 1 | 5 | 0.6 | 4 | 1 | 16 | 0.875 |
| 4_62313058 | 5 | 0.2 | 9 | 0 | 8 | 0 | 7 | 0.1429 | 29 | 0.069 | 4 | 0 | 3 | 0.3333 | 5 | 0 | 4 | 0 | 16 | 0.0625 |
| 4_62313095 | 5 | 0.2 | 8 | 0 | 6 | 0.3333 | 7 | 0.1429 | 26 | 0.1538 | 4 | 0.25 | 3 | 0.6667 | 5 | 0.4 | 4 | 0.25 | 16 | 0.375 |
| 4_62341575 | 2 | 1 | 3 | 0.3333 | NA | NA | 1 | 1 | 6 | 0.6667 | NA | NA | 1 | 1 | NA | NA | 2 | 1 | 3 | 1 |
| 4_62341665 | 3 | 1 | 1 | 1 | NA | NA | 2 | 1 | 6 | 1 | 1 | 1 | 1 | 1 | 1 | 1 | 2 | 1 | 5 | 1 |
| 4_62341672 | 3 | 0.6667 | 3 | 0.3333 | NA | NA | 2 | 0.5 | 8 | 0.5 | 1 | 0 | 1 | 1 | 1 | 0 | 2 | 0.5 | 5 | 0.4 |
| 4_62341861 | 2 | 0.5 | 3 | 1 | NA | NA | 2 | 1 | 7 | 0.8571 | 1 | 1 | 1 | 1 | 1 | 0 | 1 | 1 | 4 | 0.75 |
| 4_62341879 | 3 | 0.6667 | 2 | 0.5 | NA | NA | 2 | 1 | 7 | 0.7143 | 1 | 1 | 1 | 1 | 1 | 1 | 2 | 1 | 5 | 1 |
| 4_62341993 | 3 | 0.3333 | 3 | 0 | NA | NA | 2 | 0.5 | 8 | 0.25 | 1 | 1 | 1 | 0 | NA | NA | 2 | 0 | 4 | 0.25 |
| 4_62342101 | 3 | 0.3333 | 3 | 0 | NA | NA | 1 | 1 | 7 | 0.2857 | 1 | 0 | 1 | 1 | 1 | 0 | 2 | 0 | 5 | 0.2 |
| 4_62342134 | 3 | 0.6667 | 3 | 0.3333 | NA | NA | 2 | 1 | 8 | 0.625 | 1 | 1 | 1 | 0 | 1 | 0 | 2 | 0 | 5 | 0.2 |
| 4_62342166 | 2 | 1 | 3 | 0.3333 | NA | NA | 1 | 1 | 6 | 0.6667 | 1 | 0 | 1 | 1 | 1 | 0 | 2 | 0 | 5 | 0.2 |
| 4_62342272 | 3 | 0 | 2 | 0.5 | NA | NA | 2 | 0.5 | 7 | 0.2857 | 1 | 0 | 1 | 1 | 1 | 0 | 2 | 0 | 5 | 0.2 |
| 4_62342324 | 3 | 1 | 3 | 1 | NA | NA | 2 | 1 | 8 | 1 | 1 | 1 | 1 | 1 | 1 | 1 | 2 | 0.5 | 5 | 0.8 |
| 4_62342354 | 3 | 0.6667 | 3 | 0.3333 | NA | NA | 2 | 0.5 | 8 | 0.5 | 1 | 1 | NA | NA | NA | NA | 2 | 0.5 | 3 | 0.6667 |
| 4_62342417 | 1 | 1 | 3 | 0.6667 | NA | NA | 2 | 1 | 6 | 0.8333 | 1 | 1 | NA | NA | NA | NA | 1 | 0 | 2 | 0.5 |
| 4_62342498 | 3 | 1 | 3 | 0.6667 | NA | NA | 3 | 1 | 9 | 0.8889 | 2 | 1 | 1 | 1 | 1 | 1 | 2 | 0.5 | 6 | 0.8333 |
| 4_62342505 | 2 | 1 | 3 | 0.6667 | NA | NA | 1 | 1 | 6 | 0.8333 | NA | NA | NA | NA | NA | NA | 1 | 1 | 1 | 1 |
| 4_62367936 | NA | NA | NA | NA | 2 | 0.5 | NA | NA | 2 | 0.5 | NA | NA | NA | NA | NA | NA | 1 | 1 | 1 | 1 |
| 4_62368166 | NA | NA | NA | NA | 1 | 1 | NA | NA | 1 | 1 | NA | NA | NA | NA | NA | NA | 1 | 1 | 1 | 1 |
| 4_62368648 | NA | NA | NA | NA | 1 | 1 | NA | NA | 1 | 1 | NA | NA | NA | NA | NA | NA | 1 | 1 | 1 | 1 |
| 4_62368734 | NA | NA | NA | NA | 1 | 1 | NA | NA | 1 | 1 | NA | NA | NA | NA | NA | NA | 1 | 1 | 1 | 1 |
| 4_62372044 | 1 | 1 | NA | NA | NA | NA | NA | NA | 1 | 1 | NA | NA | 1 | 1 | NA | NA | 2 | 0 | 3 | 0.3333 |
| 4_62417491 | 1 | 0 | 1 | 1 | 1 | 0 | 1 | 1 | 4 | 0.5 | 2 | 0.5 | 1 | 1 | 1 | 0 | NA | NA | 4 | 0.5 |
| 4_62417522 | 1 | 1 | 1 | 1 | 1 | 0 | NA | NA | 3 | 0.6667 | 2 | 1 | 1 | 1 | 1 | 1 | NA | NA | 4 | 1 |
| 4_62417700 | NA | NA | 1 | 1 | NA | NA | 1 | 0 | 2 | 0.5 | 1 | 1 | 1 | 1 | 1 | 0 | NA | NA | 3 | 0.6667 |
| 4_62417719 | 1 | 1 | 1 | 1 | NA | NA | 1 | 1 | 3 | 1 | 2 | 1 | 1 | 1 | 1 | 1 | NA | NA | 4 | 1 |
| 4_62417864 | 1 | 1 | NA | NA | NA | NA | NA | NA | 1 | 1 | 2 | 0.5 | 1 | 0 | 1 | 1 | NA | NA | 4 | 0.5 |
| 4_62427410 | 5 | 1 | 3 | 0.6667 | 2 | 0.5 | 3 | 0.3333 | 13 | 0.6923 | 1 | 1 | 3 | 1 | 5 | 0.8 | 1 | 1 | 10 | 0.9 |
| 4_62427471 | 5 | 0.6 | 3 | 0.3333 | 3 | 0.3333 | 3 | 0.3333 | 14 | 0.4286 | 1 | 1 | 4 | 0.5 | 4 | 0.5 | 5 | 0.8 | 14 | 0.6429 |
| 4_62440942 | 4 | 0 | 4 | 0 | 5 | 0 | 5 | 0.2 | 18 | 0.0556 | 5 | 0.2 | 1 | 0 | 2 | 0 | 1 | 0 | 9 | 0.1111 |
| 4_62440991 | 4 | 0 | 4 | 0 | 5 | 0.4 | 4 | 0 | 17 | 0.1176 | 4 | 0 | 1 | 1 | 2 | 0 | 1 | 0 | 8 | 0.125 |
| 4_62441347 | 5 | 0 | 2 | 0 | 5 | 0.2 | 5 | 0.2 | 17 | 0.1176 | 3 | 0.6667 | 1 | 0 | 2 | 0 | 1 | 0 | 7 | 0.2857 |
| 4_62441553 | 5 | 0.2 | 2 | 0 | 5 | 0 | 5 | 0 | 17 | 0.0588 | 3 | 0 | 1 | 0 | 1 | 1 | 1 | 0 | 6 | 0.1667 |
| 4_62441760 | 5 | 0 | 2 | 0 | 4 | 0.25 | 4 | 0 | 15 | 0.0667 | 2 | 0.5 | 1 | 0 | 1 | 0 | 1 | 0 | 5 | 0.2 |
| 4_62442032 | 3 | 1 | NA | NA | 2 | 1 | 2 | 1 | 7 | 1 | 2 | 1 | 1 | 1 | NA | NA | 1 | 1 | 4 | 1 |
| 4_62442127 | 5 | 0 | 2 | 0 | 5 | 0.2 | 4 | 0 | 16 | 0.0625 | 3 | 0 | 3 | 0 | 3 | 0 | 2 | 0.5 | 11 | 0.0909 |
| 4_62442143 | 5 | 0.4 | 2 | 0.5 | 5 | 0 | 4 | 0.5 | 16 | 0.3125 | 2 | 0.5 | 3 | 0 | 1 | 0 | 2 | 0 | 8 | 0.125 |
| 4_62442163 | 5 | 0.8 | 2 | 0.5 | 5 | 0.6 | 4 | 0.5 | 16 | 0.625 | 3 | 0.3333 | 3 | 0.6667 | 2 | 1 | 2 | 0.5 | 10 | 0.6 |
| 4_62442226 | 5 | 0.2 | 2 | 0 | 4 | 0 | 4 | 0.25 | 15 | 0.1333 | 2 | 0.5 | 3 | 0.3333 | 2 | 0.5 | 2 | 0 | 9 | 0.3333 |
| 4_62608037 | 1 | 1 | 1 | 1 | 1 | 0 | NA | NA | 3 | 0.6667 | NA | NA | 1 | 1 | NA | NA | NA | NA | 1 | 1 |
| 4_62608179 | 2 | 0 | 1 | 1 | 1 | 0 | NA | NA | 4 | 0.25 | NA | NA | 1 | 1 | NA | NA | NA | NA | 1 | 1 |
| 4_62608340 | 2 | 0 | 1 | 0 | 1 | 1 | NA | NA | 4 | 0.25 | NA | NA | 1 | 1 | NA | NA | NA | NA | 1 | 1 |
| 4_62608609 | 2 | 1 | 2 | 1 | 1 | 0 | NA | NA | 5 | 0.8 | NA | NA | 1 | 1 | NA | NA | NA | NA | 1 | 1 |

| | | | | | | | | | | | | | | | | | | | | |
|---|---|---|---|---|---|---|---|---|---|---|---|---|---|---|---|---|---|---|---|---|
| 4_6260880 | 2 | 0.5 | 2 | 0 | NA | NA | NA | NA | 4 | 0.25 | NA | NA | 1 | 1 | NA | NA | NA | NA | 1 | 1 |
| 4_6262040 | 4 | 1 | 2 | 1 | 2 | 0.5 | NA | NA | 8 | 0.875 | NA | NA | 3 | 1 | 1 | 1 | 1 | 1 | 5 | 1 |
| 4_6262044 | 5 | 0.4 | 3 | 0 | 2 | 0.5 | 1 | 0 | 11 | 0.2727 | 1 | 0 | 3 | 0 | 1 | 0 | 1 | 1 | 6 | 0.1667 |
| 4_6262048 | 4 | 0 | 3 | 0 | 2 | 0.5 | 1 | 0 | 10 | 0.1 | 1 | 0 | 3 | 0.3333 | 1 | 0 | 1 | 0 | 6 | 0.1667 |
| 4_6262060 | 4 | 1 | 3 | 0.6667 | 2 | 1 | NA | NA | 9 | 0.8889 | 1 | 1 | 3 | 1 | 1 | 1 | 1 | 1 | 6 | 1 |
| 4_6262063 | 5 | 1 | 3 | 1 | 2 | 1 | 1 | 1 | 11 | 1 | 1 | 1 | 3 | 1 | 1 | 1 | 2 | 1 | 7 | 1 |
| 4_6262066 | 5 | 0.2 | 3 | 0 | 1 | 1 | 1 | 0 | 10 | 0.2 | 1 | 1 | 3 | 0 | 1 | 0 | 2 | 0.5 | 7 | 0.2857 |
| 4_6262078 | 5 | 0.4 | 3 | 0.3333 | 1 | 0 | 1 | 0 | 10 | 0.3 | 1 | 0 | 2 | 0.5 | NA | NA | 2 | 0.5 | 5 | 0.4 |
| 4_6262081 | 5 | 0.4 | 3 | 0.3333 | 2 | 0.5 | 1 | 0 | 11 | 0.3636 | 1 | 1 | 1 | 0 | 1 | 0 | NA | NA | 3 | 0.3333 |
| 4_6262082 | 1 | 0 | 1 | 0 | 2 | 0 | 1 | 1 | 5 | 0.2 | 1 | 0 | 3 | 0.3333 | 1 | 0 | 2 | 0 | 7 | 0.1429 |
| 4_6262095 | 5 | 0.6 | 2 | 1 | 2 | 0.5 | NA | NA | 9 | 0.6667 | 1 | 1 | 2 | 0.5 | 1 | 0 | 1 | 0 | 5 | 0.4 |
| 4_6262111 | 4 | 0.25 | 3 | 0 | 1 | 1 | 1 | 0 | 9 | 0.2222 | 1 | 1 | 2 | 0 | 1 | 0 | NA | NA | 4 | 0.25 |
| 4_6305208 | NA | NA | 1 | 1 | NA | NA | NA | NA | 1 | 1 | NA | NA | 1 | 1 | NA | NA | NA | NA | 1 | 1 |
| 4_6305256 | NA | NA | 1 | 1 | NA | NA | NA | NA | 1 | 1 | 1 | 1 | 1 | 1 | NA | NA | NA | NA | 2 | 1 |
| 4_6305275 | NA | NA | 1 | 1 | NA | NA | NA | NA | 1 | 1 | NA | NA | 2 | 0.5 | NA | NA | NA | NA | 2 | 0.5 |
| 4_6305295 | NA | NA | 1 | 1 | NA | NA | NA | NA | 1 | 1 | 1 | 1 | 2 | 0 | NA | NA | NA | NA | 3 | 0.3333 |
| 4_6305319 | NA | NA | 1 | 1 | NA | NA | NA | NA | 1 | 1 | 1 | 1 | 1 | 0 | NA | NA | NA | NA | 2 | 0.5 |
| 4_6305322 | NA | NA | 1 | 1 | NA | NA | NA | NA | 1 | 1 | 1 | 0 | 2 | 0.5 | NA | NA | NA | NA | 3 | 0.3333 |
| 4_6305345 | NA | NA | 1 | 1 | NA | NA | NA | NA | 1 | 1 | 1 | 0 | 2 | 0.5 | NA | NA | NA | NA | 3 | 0.3333 |
| 4_6305358 | NA | NA | 1 | 1 | NA | NA | NA | NA | 1 | 1 | 1 | 1 | 1 | 0 | NA | NA | NA | NA | 2 | 0.5 |
| 4_6305361 | NA | NA | 1 | 1 | NA | NA | NA | NA | 1 | 1 | 1 | 1 | 2 | 1 | NA | NA | NA | NA | 3 | 1 |
| 4_6305369 | NA | NA | 1 | 1 | NA | NA | NA | NA | 1 | 1 | 1 | 0 | 1 | 1 | NA | NA | NA | NA | 2 | 0.5 |
| 4_6305435 | NA | NA | 1 | 1 | NA | NA | NA | NA | 1 | 1 | NA | NA | 1 | 1 | NA | NA | NA | NA | 1 | 1 |
| 4_6305503 | NA | NA | 1 | 1 | NA | NA | NA | NA | 1 | 1 | 1 | 1 | 2 | 1 | NA | NA | NA | NA | 3 | 1 |
| 4_6305514 | NA | NA | 1 | 1 | NA | NA | NA | NA | 1 | 1 | 1 | 1 | 2 | 1 | NA | NA | NA | NA | 3 | 1 |
| 4_6305517 | NA | NA | 1 | 1 | NA | NA | NA | NA | 1 | 1 | NA | NA | 2 | 0.5 | NA | NA | NA | NA | 2 | 0.5 |
| 4_6305581 | NA | NA | 1 | 1 | NA | NA | NA | NA | 1 | 1 | 1 | 1 | 1 | 1 | NA | NA | NA | NA | 2 | 1 |
| 4_6305582 | NA | NA | 1 | 1 | NA | NA | NA | NA | 1 | 1 | 1 | 0 | 2 | 0.5 | NA | NA | NA | NA | 3 | 0.3333 |
| 4_6305587 | NA | NA | 1 | 1 | NA | NA | NA | NA | 1 | 1 | 1 | 0 | 2 | 1 | NA | NA | NA | NA | 3 | 0.6667 |
| 4_6305596 | NA | NA | 1 | 1 | NA | NA | NA | NA | 1 | 1 | 1 | 1 | 2 | 1 | NA | NA | NA | NA | 3 | 1 |
| 4_6324957 | NA | NA | 1 | 1 | NA | NA | NA | NA | 1 | 1 | NA | NA | NA | NA | NA | NA | 1 | 1 | 1 | 1 |
| 4_6325152 | NA | NA | 1 | 1 | NA | NA | NA | NA | 1 | 1 | NA | NA | NA | NA | NA | NA | 2 | 0.5 | 2 | 0.5 |
| 4_6325159 | NA | NA | 1 | 1 | NA | NA | NA | NA | 1 | 1 | NA | NA | NA | NA | NA | NA | 2 | 0.5 | 2 | 0.5 |
| 4_6333342 | 3 | 0.3333 | 3 | 0 | 2 | 0.5 | 4 | 0.5 | 12 | 0.3333 | 1 | 1 | 2 | 1 | 2 | 0.5 | 1 | 1 | 6 | 0.8333 |
| 4_6333387 | 3 | 0 | 3 | 0 | 1 | 0 | 2 | 1 | 9 | 0.2222 | 2 | 0 | 4 | 0 | 1 | 1 | NA | NA | 7 | 0.1429 |
| 4_6333663 | 2 | 0.5 | 3 | 0.6667 | 3 | 1 | 5 | 0.8 | 13 | 0.7692 | 2 | 0.5 | 3 | 0.3333 | 2 | 1 | 1 | 1 | 8 | 0.625 |
| 4_6333664 | 2 | 0 | 1 | 1 | 3 | 0 | 5 | 0 | 11 | 0.0909 | 1 | 1 | 3 | 0 | 2 | 0 | NA | NA | 6 | 0.1667 |
| 4_6341402 | NA | NA | 1 | 1 | NA | NA | 3 | 0.3333 | 4 | 0.5 | 2 | 0.5 | 1 | 0 | NA | NA | NA | NA | 3 | 0.3333 |
| 4_6341406 | 1 | 0 | 1 | 1 | NA | NA | 3 | 0 | 5 | 0.2 | 2 | 1 | 1 | 0 | NA | NA | NA | NA | 3 | 0.6667 |
| 4_6346833 | 27 | 0.037 | 20 | 0 | 21 | 0.1905 | 32 | 0.0312 | 100 | 0.06 | 19 | 0.1053 | 13 | 0.0769 | 25 | 0.04 | 21 | 0.1429 | 78 | 0.0897 |
| 4_6346847 | 16 | 0.375 | 18 | 0.2222 | 21 | 0.2857 | 24 | 0.2917 | 79 | 0.2911 | 18 | 0.2222 | 12 | 0.0833 | 19 | 0.1579 | 20 | 0.15 | 69 | 0.1594 |
| 4_6346852 | 17 | 0.1765 | 18 | 0.1667 | 21 | 0.0952 | 24 | 0.2083 | 80 | 0.1625 | 17 | 0.2941 | 11 | 0.1818 | 18 | 0.1667 | 21 | 0.1429 | 67 | 0.194 |
| 4_6350212 | 2 | 0.5 | 1 | 1 | 1 | 1 | 1 | 0 | 5 | 0.6 | 1 | 1 | 1 | 1 | 1 | 0 | NA | NA | 3 | 0.6667 |
| 4_6350223 | 2 | 0 | 1 | 0 | 1 | 0 | 1 | 1 | 5 | 0.2 | 1 | 0 | 1 | 1 | 1 | 0 | NA | NA | 3 | 0.3333 |
| 4_6350230 | 2 | 1 | 1 | 0 | 3 | 0.6667 | 1 | 0 | 7 | 0.5714 | 1 | 0 | 1 | 1 | 2 | 0.5 | NA | NA | 4 | 0.5 |
| 4_6350278 | 2 | 1 | 1 | 1 | 2 | 0.5 | 2 | 0 | 7 | 0.5714 | 2 | 0.5 | 2 | 1 | 3 | 1 | NA | NA | 7 | 0.8571 |
| 4_6350285 | 2 | 1 | 3 | 0.3333 | 3 | 0.6667 | 2 | 0.5 | 10 | 0.6 | 2 | 0.5 | 2 | 0.5 | 3 | 0.3333 | NA | NA | 7 | 0.4286 |
| 4_6350287 | 2 | 0 | 3 | 0.3333 | 3 | 0 | 2 | 0 | 10 | 0.1 | 2 | 0 | 2 | 0 | 4 | 0.25 | NA | NA | 8 | 0.125 |
| 4_6350288 | 2 | 0 | 3 | 0 | 3 | 0.6667 | 2 | 0 | 10 | 0.2 | 1 | 1 | 2 | 0 | 4 | 0.25 | NA | NA | 7 | 0.2857 |
| 4_6350291 | 2 | 0 | 3 | 0.3333 | 3 | 0 | 2 | 0 | 10 | 0.1 | 2 | 0 | 2 | 0.5 | 4 | 0 | NA | NA | 8 | 0.125 |
| 4_6350291 | 2 | 1 | 3 | 1 | 2 | 1 | 2 | 1 | 9 | 1 | 2 | 0 | 2 | 0.5 | 4 | 0.75 | NA | NA | 8 | 0.5 |
| 4_6350293 | 2 | 1 | 3 | 1 | 3 | 1 | 2 | 0.5 | 10 | 0.9 | 2 | 0.5 | 2 | 1 | 4 | 0.75 | NA | NA | 8 | 0.75 |
| 4_6350304 | 2 | 0 | 3 | 1 | 2 | 0.5 | 2 | 0.5 | 9 | 0.5556 | 2 | 0.5 | 2 | 0.5 | 4 | 0.25 | 1 | 0 | 9 | 0.3333 |
| 4_6350305 | 2 | 1 | 3 | 1 | 2 | 1 | 2 | 1 | 9 | 1 | 1 | 1 | 2 | 1 | 4 | 1 | 1 | 1 | 8 | 1 |
| 4_6350308 | 2 | 1 | 3 | 1 | 3 | 1 | 3 | 1 | 11 | 1 | 1 | 1 | 2 | 1 | 3 | 1 | 1 | 1 | 7 | 1 |
| 4_6350320 | 2 | 0.5 | 3 | 0.3333 | 3 | 0.3333 | 3 | 0.6667 | 11 | 0.4545 | 2 | 1 | 2 | 0 | 4 | 0.5 | 1 | 1 | 9 | 0.5556 |
| 4_6350326 | 2 | 0.5 | 3 | 0.3333 | 3 | 0.6667 | 3 | 0.3333 | 11 | 0.4545 | 2 | 1 | 1 | 1 | 4 | 0.25 | 1 | 1 | 8 | 0.625 |
| 4_6350338 | 1 | 0 | 1 | 0 | 1 | 0 | 2 | 0.5 | 5 | 0.2 | 2 | 0.5 | 1 | 0 | 2 | 0 | NA | NA | 5 | 0.2 |
| 4_6350353 | NA | NA | 1 | 0 | 3 | 0.3333 | 3 | 0 | 7 | 0.1429 | 2 | 0.5 | 1 | 0 | 2 | 0 | 1 | 0 | 6 | 0.1667 |
| 4_6350390 | 1 | 1 | 1 | 1 | 2 | 0 | 3 | 0 | 7 | 0.2857 | 3 | 0.6667 | 1 | 0 | 2 | 0 | 1 | 0 | 7 | 0.2857 |
| 4_6387812 | 1 | 1 | NA | NA | NA | NA | NA | NA | 1 | 1 | NA | NA | NA | NA | NA | NA | 1 | 1 | 1 | 1 |
| 4_6529931 | 2 | 0 | 3 | 0.6667 | 3 | 0.3333 | 3 | 0 | 11 | 0.2727 | 6 | 0.3333 | 3 | 0.3333 | 2 | 0 | 2 | 0 | 13 | 0.2308 |
| 4_6529931 | 2 | 1 | 4 | 0.75 | 3 | 0 | 4 | 0 | 13 | 0.3846 | 6 | 0 | 3 | 0.6667 | 2 | 0 | 2 | 0 | 13 | 0.1538 |
| 4_6529934 | 2 | 0 | 2 | 0.5 | 3 | 0.3333 | 2 | 0 | 9 | 0.2222 | 3 | 0.3333 | 2 | 0 | 1 | 0 | 2 | 0.5 | 8 | 0.25 |
| 4_6529952 | 1 | 1 | 2 | 1 | 3 | 1 | 4 | 1 | 10 | 1 | 5 | 0.4 | 3 | 1 | 2 | 1 | 1 | 1 | 11 | 0.7273 |

| | | | | | | | | | | | | | | | | | | | | |
|---|---|---|---|---|---|---|---|---|---|---|---|---|---|---|---|---|---|---|---|---|
| 4_6529953 | 1 | 1 | 2 | 0.5 | 3 | 0.3333 | 3 | 0.6667 | 9 | 0.5556 | 2 | 1 | 3 | 0.6667 | 2 | 0.5 | 1 | 0 | 8 | 0.625 |
| 4_6529955 | 1 | 0 | 2 | 1 | 3 | 0.3333 | 4 | 0.25 | 10 | 0.4 | 5 | 0.4 | 3 | 0.6667 | 2 | 0.5 | 2 | 1 | 12 | 0.5833 |
| 4_6529976 | NA | NA | 2 | 0 | 2 | 0 | 3 | 0.3333 | 7 | 0.1429 | 4 | 0.25 | 2 | 0 | 1 | 0 | 1 | 0 | 8 | 0.125 |
| 4_6553189 | 13 | 0.4615 | 14 | 0.1429 | 9 | 0.6667 | 5 | 0.4 | 41 | 0.3902 | 12 | 0.5 | 11 | 0.2727 | 8 | 0.25 | 8 | 0.125 | 39 | 0.3077 |
| 4_6553192 | 15 | 0.2667 | 14 | 0.4286 | 7 | 0.7143 | 6 | 0.3333 | 42 | 0.4048 | 11 | 0.0909 | 9 | 0.3333 | 8 | 0.25 | 7 | 0.4286 | 35 | 0.2571 |
| 4_6553216 | 17 | 0.1765 | 18 | 0.2222 | 10 | 0.1 | 7 | 0 | 52 | 0.1538 | 14 | 0.0714 | 8 | 0 | 10 | 0.2 | 10 | 0.2 | 42 | 0.119 |
| 4_6553229 | 16 | 0.3125 | 18 | 0 | 10 | 0 | 7 | 0 | 51 | 0.098 | 15 | 0.0667 | 11 | 0.2727 | 11 | 0.0909 | 12 | 0.0833 | 49 | 0.1224 |
| 4_6553234 | 16 | 0.75 | 15 | 0.8 | 9 | 0.7778 | 7 | 1 | 47 | 0.8085 | 13 | 0.7692 | 10 | 1 | 11 | 0.9091 | 10 | 1 | 44 | 0.9091 |
| 4_6553242 | 15 | 1 | 13 | 0.9231 | 10 | 1 | 8 | 1 | 46 | 0.9783 | 13 | 1 | 11 | 1 | 12 | 1 | 13 | 1 | 49 | 1 |
| 4_6553258 | 20 | 0.15 | 16 | 0.25 | 14 | 0.0714 | 10 | 0 | 60 | 0.1333 | 14 | 0 | 13 | 0.2308 | 12 | 0.0833 | 16 | 0.0625 | 55 | 0.0909 |
| 4_6553276 | 12 | 1 | 14 | 1 | 12 | 0.9167 | 7 | 1 | 45 | 0.9778 | 11 | 1 | 8 | 1 | 11 | 0.9091 | 10 | 0.8 | 40 | 0.925 |
| 4_6553290 | 19 | 0.8947 | 15 | 1 | 9 | 0.8889 | 10 | 0.8 | 53 | 0.9057 | 11 | 1 | 13 | 0.8462 | 14 | 1 | 14 | 0.7857 | 52 | 0.9038 |
| 4_6553294 | 20 | 0.1 | 19 | 0.0526 | 12 | 0 | 9 | 0 | 60 | 0.05 | 16 | 0.125 | 16 | 0.0625 | 14 | 0 | 16 | 0.125 | 62 | 0.0806 |
| 4_6553307 | 12 | 0.25 | 5 | 0.6 | 4 | 0 | 7 | 0.2857 | 28 | 0.2857 | 9 | 0.1111 | 5 | 0 | 7 | 0 | 9 | 0.3333 | 30 | 0.1333 |
| 4_6553335 | 21 | 1 | 23 | 1 | 14 | 1 | 9 | 1 | 67 | 1 | 18 | 1 | 17 | 1 | 17 | 1 | 18 | 1 | 70 | 1 |
| 4_6553379 | 25 | 0.4 | 25 | 0.52 | 15 | 0.4667 | 10 | 0.4 | 75 | 0.4533 | 17 | 0.2353 | 18 | 0.3889 | 18 | 0.3889 | 18 | 0.5 | 71 | 0.3803 |
| 4_6553401 | 25 | 0 | 25 | 0.32 | 20 | 0.2 | 14 | 0.1429 | 84 | 0.1667 | 27 | 0.1481 | 25 | 0.16 | 22 | 0.1364 | 18 | 0.1667 | 92 | 0.1522 |
| 4_6553408 | 22 | 0.1364 | 27 | 0.1111 | 21 | 0.0476 | 14 | 0.2857 | 84 | 0.131 | 24 | 0.0833 | 25 | 0.16 | 23 | 0.087 | 19 | 0.1579 | 91 | 0.1209 |
| 4_6553411 | 21 | 0.1905 | 21 | 0 | 15 | 0.0667 | 12 | 0.25 | 69 | 0.1159 | 25 | 0.12 | 18 | 0.1667 | 21 | 0.0952 | 18 | 0.0556 | 82 | 0.1098 |
| 4_6553418 | 26 | 0.1154 | 23 | 0.087 | 17 | 0.0588 | 16 | 0.0625 | 82 | 0.0854 | 25 | 0.12 | 21 | 0.0476 | 23 | 0.087 | 18 | 0.1111 | 87 | 0.092 |
| 4_6553436 | 27 | 0.1111 | 26 | 0.0385 | 26 | 0.0769 | 12 | 0 | 91 | 0.0659 | 29 | 0.0345 | 28 | 0.0357 | 25 | 0.16 | 22 | 0 | 104 | 0.0577 |
| 4_6553652 | 1 | 1 | 1 | 1 | NA | NA | NA | NA | 2 | 1 | 2 | 0 | 2 | 0.5 | 1 | 0 | 1 | 0 | 6 | 0.1667 |
| 4_6553685 | 1 | 1 | 1 | 0 | 1 | 1 | 1 | 0 | 4 | 0.5 | 1 | 0 | 3 | 0 | 1 | 0 | 1 | 1 | 6 | 0.1667 |
| 4_6553755 | 1 | 0 | 1 | 1 | NA | NA | 1 | 1 | 3 | 0.6667 | 1 | 0 | 3 | 0.3333 | 1 | 1 | NA | NA | 5 | 0.4 |
| 4_6553758 | NA | NA | 1 | 1 | 2 | 1 | 1 | 0 | 4 | 0.75 | 2 | 0 | 2 | 0.5 | 1 | 1 | 1 | 0 | 6 | 0.3333 |
| 4_6675757 | NA | NA | 2 | 1 | 1 | 1 | NA | NA | 3 | 1 | NA | NA | NA | NA | NA | NA | 1 | 1 | 1 | 1 |
| 4_6675764 | NA | NA | 2 | 1 | 1 | 1 | NA | NA | 3 | 1 | NA | NA | NA | NA | NA | NA | 1 | 1 | 1 | 1 |
| 4_6675793 | NA | NA | 2 | 0.5 | 1 | 0 | NA | NA | 3 | 0.3333 | NA | NA | NA | NA | NA | NA | 1 | 1 | 1 | 1 |
| 4_6675804 | NA | NA | 1 | 1 | 1 | 1 | NA | NA | 2 | 1 | NA | NA | NA | NA | NA | NA | 1 | 1 | 1 | 1 |
| 4_6675858 | NA | NA | 1 | 1 | 1 | 1 | NA | NA | 2 | 1 | NA | NA | NA | NA | NA | NA | 1 | 1 | 1 | 1 |
| 4_6675876 | NA | NA | 1 | 1 | 1 | 1 | NA | NA | 2 | 1 | NA | NA | NA | NA | NA | NA | 1 | 1 | 1 | 1 |
| 4_6675887 | NA | NA | 2 | 1 | 1 | 1 | NA | NA | 3 | 1 | NA | NA | NA | NA | NA | NA | 1 | 1 | 1 | 1 |
| 4_6675933 | NA | NA | 2 | 1 | 1 | 1 | NA | NA | 3 | 1 | NA | NA | NA | NA | NA | NA | 1 | 1 | 1 | 1 |
| 4_6675941 | NA | NA | 2 | 0.5 | 1 | 1 | NA | NA | 3 | 0.6667 | NA | NA | NA | NA | NA | NA | 1 | 1 | 1 | 1 |
| 4_6675959 | NA | NA | 2 | 0.5 | 1 | 1 | NA | NA | 3 | 0.6667 | NA | NA | NA | NA | NA | NA | 1 | 1 | 1 | 1 |
| 4_6675971 | NA | NA | 2 | 1 | 1 | 1 | NA | NA | 3 | 1 | NA | NA | NA | NA | NA | NA | 1 | 1 | 1 | 1 |
| 4_6675979 | NA | NA | 2 | 0.5 | 1 | 1 | NA | NA | 3 | 0.6667 | NA | NA | NA | NA | NA | NA | 1 | 1 | 1 | 1 |
| 4_6675983 | NA | NA | 2 | 1 | 1 | 1 | NA | NA | 3 | 1 | NA | NA | NA | NA | NA | NA | 1 | 1 | 1 | 1 |
| 4_6675991 | NA | NA | 1 | 1 | 1 | 0 | NA | NA | 2 | 0.5 | NA | NA | NA | NA | NA | NA | 1 | 1 | 1 | 1 |
| 4_6676007 | NA | NA | 1 | 1 | 1 | 1 | NA | NA | 2 | 1 | NA | NA | NA | NA | NA | NA | 2 | 1 | 2 | 1 |
| 4_6676067 | NA | NA | 2 | 1 | NA | NA | NA | NA | 2 | 1 | 1 | 1 | NA | NA | NA | NA | 2 | 1 | 3 | 1 |
| 4_6676073 | NA | NA | 2 | 0.5 | 1 | 1 | NA | NA | 3 | 0.6667 | 1 | 1 | 1 | 0 | NA | NA | 2 | 0.5 | 4 | 0.5 |
| 4_6676309 | NA | NA | 1 | 1 | 1 | 1 | NA | NA | 2 | 1 | 1 | 1 | NA | NA | NA | NA | NA | NA | 1 | 1 |
| 4_6868002 | 18 | 0.1111 | 28 | 0.2143 | 10 | 0.3 | 13 | 0.2308 | 69 | 0.2029 | 16 | 0.125 | 16 | 0.0625 | 20 | 0.05 | 7 | 0.1429 | 59 | 0.0847 |
| 4_6868007 | 11 | 0 | 15 | 0.1333 | 9 | 0.1111 | 6 | 0 | 41 | 0.0732 | 11 | 0.0909 | 10 | 0.1 | 12 | 0 | 9 | 0.2222 | 42 | 0.0952 |
| 4_6868014 | 26 | 0.0385 | 27 | 0 | 14 | 0.0714 | 13 | 0.2308 | 80 | 0.0625 | 19 | 0.1053 | 20 | 0.2 | 21 | 0.0952 | 12 | 0.0833 | 72 | 0.125 |
| 4_6868014 | 24 | 0 | 30 | 0.0667 | 14 | 0.0714 | 13 | 0.1538 | 81 | 0.0617 | 20 | 0.05 | 21 | 0.1429 | 21 | 0.1429 | 11 | 0 | 73 | 0.0959 |
| 4_6868016 | 17 | 0.2941 | 22 | 0.1364 | 14 | 0.2857 | 12 | 0.4167 | 65 | 0.2615 | 15 | 0.2667 | 21 | 0.3333 | 16 | 0.25 | 10 | 0.2 | 62 | 0.2742 |
| 4_6868022 | 20 | 1 | 26 | 1 | 11 | 1 | 10 | 1 | 67 | 1 | 14 | 1 | 17 | 1 | 18 | 1 | 10 | 1 | 59 | 1 |
| 4_6868023 | 22 | 0.7273 | 26 | 0.8077 | 13 | 0.8462 | 14 | 0.7143 | 75 | 0.7733 | 19 | 0.7895 | 21 | 0.7619 | 20 | 0.8 | 11 | 0.9091 | 71 | 0.8028 |
| 4_6868030 | 21 | 0.5714 | 28 | 0.3214 | 9 | 0.4444 | 14 | 0.2143 | 72 | 0.3889 | 18 | 0.2222 | 21 | 0.4286 | 18 | 0.2778 | 12 | 0.5833 | 69 | 0.3623 |
| 4_6868037 | 21 | 0.8571 | 28 | 0.9643 | 12 | 0.9167 | 13 | 1 | 74 | 0.9324 | 19 | 0.9474 | 17 | 0.9412 | 21 | 1 | 10 | 0.9 | 67 | 0.9552 |
| 4_6868043 | 20 | 0.4 | 27 | 0.3333 | 12 | 0.4167 | 13 | 0.4615 | 72 | 0.3889 | 17 | 0.1765 | 18 | 0.2778 | 22 | 0.4545 | 12 | 0.25 | 69 | 0.3043 |
| 4_6868043 | 14 | 1 | 26 | 0.9615 | 9 | 0.8889 | 12 | 0.9167 | 61 | 0.9508 | 15 | 0.9333 | 19 | 0.9474 | 19 | 0.9474 | 9 | 0.8889 | 62 | 0.9355 |
| 4_6868061 | 20 | 0.25 | 26 | 0.3462 | 11 | 0.1818 | 11 | 0.0909 | 68 | 0.25 | 15 | 0.1333 | 18 | 0.2778 | 22 | 0.4091 | 12 | 0.1667 | 67 | 0.2687 |
| 4_6868063 | 18 | 0.6667 | 22 | 0.7727 | 11 | 0.9091 | 10 | 0.3 | 61 | 0.6885 | 14 | 0.8571 | 17 | 0.8235 | 17 | 0.7059 | 10 | 0.5 | 58 | 0.7414 |
| 4_6868066 | 19 | 0.3158 | 24 | 0.375 | 10 | 0.5 | 11 | 0.1818 | 64 | 0.3438 | 15 | 0.4 | 17 | 0.4706 | 21 | 0.381 | 12 | 0.3333 | 65 | 0.4 |
| 4_6868068 | 16 | 0.75 | 24 | 0.8333 | 9 | 0.8889 | 8 | 0.875 | 57 | 0.8246 | 12 | 0.9167 | 15 | 0.8667 | 15 | 0.9333 | 10 | 0.8 | 52 | 0.8846 |
| 4_6868070 | 17 | 0.1176 | 20 | 0.3 | 8 | 0.25 | 9 | 0.1111 | 54 | 0.2037 | 12 | 0.1667 | 15 | 0.0667 | 21 | 0.1429 | 8 | 0.125 | 56 | 0.125 |
| 4_6868070 | 2 | 0 | 4 | 0.25 | 2 | 0.5 | 1 | 0 | 9 | 0.2222 | NA | NA | 3 | 0.3333 | NA | NA | 1 | 0 | 4 | 0.25 |
| 4_6868072 | 18 | 1 | 16 | 0.9375 | 9 | 1 | 7 | 1 | 50 | 0.98 | 8 | 1 | 13 | 1 | 17 | 1 | 11 | 1 | 49 | 1 |
| 4_6868080 | 14 | 0.0714 | 13 | 0.2308 | 11 | 0 | 6 | 0.1667 | 44 | 0.1136 | 10 | 0.1 | 13 | 0.1538 | 13 | 0.0769 | 5 | 0.2 | 41 | 0.122 |
| 4_6868085 | 18 | 0.7778 | 24 | 0.875 | 8 | 0.75 | 10 | 0.9 | 60 | 0.8333 | 14 | 0.8571 | 18 | 0.9444 | 21 | 0.7143 | 12 | 0.9167 | 65 | 0.8462 |
| 4_6868092 | 18 | 0.8333 | 23 | 1 | 9 | 1 | 8 | 1 | 58 | 0.9483 | 15 | 1 | 17 | 0.9412 | 21 | 1 | 10 | 1 | 63 | 0.9841 |
| 4_6868106 | 16 | 0.5625 | 19 | 0.5789 | 5 | 0.2 | 6 | 0.6667 | 46 | 0.5435 | 12 | 0.5 | 14 | 0.5 | 14 | 0.5714 | 11 | 0.6364 | 51 | 0.549 |

| | | | | | | | | | | | | | | | | | | | | |
|---|---|---|---|---|---|---|---|---|---|---|---|---|---|---|---|---|---|---|---|---|
| 4_68681075 | 20 | 0.4 | 23 | 0.4348 | 7 | 0.4286 | 10 | 0.3 | 60 | 0.4 | 12 | 0.5 | 18 | 0.5556 | 18 | 0.6111 | 10 | 0.6 | 58 | 0.569 |
| 4_68681084 | 18 | 0.1111 | 23 | 0.1739 | 8 | 0.125 | 9 | 0 | 58 | 0.1207 | 11 | 0.1818 | 16 | 0.25 | 17 | 0.2353 | 11 | 0.1818 | 55 | 0.2182 |
| 4_68681129 | 12 | 1 | 19 | 0.9474 | 6 | 1 | 8 | 0.75 | 45 | 0.9333 | 10 | 1 | 13 | 0.9231 | 14 | 1 | 9 | 0.8889 | 46 | 0.9565 |
| 4_68716955 | 13 | 0 | 21 | 0.0476 | 7 | 0.2857 | 8 | 0.375 | 49 | 0.1224 | 11 | 0.1818 | 12 | 0.0833 | 15 | 0.0667 | 7 | 0.2857 | 45 | 0.1333 |
| 4_68823187 | 10 | 0.1 | 13 | 0.0769 | 4 | 0.25 | 7 | 0 | 34 | 0.0882 | 8 | 0 | 9 | 0.1111 | 8 | 0.125 | 6 | 0.1667 | 31 | 0.0968 |
| 4_68872421 | 7 | 0 | 9 | 0.1111 | 6 | 0.1667 | 8 | 0.125 | 30 | 0.1 | 7 | 0.1429 | 8 | 0.375 | 8 | 0.25 | 4 | 0.25 | 27 | 0.2593 |
| 4_69843816 | 5 | 0.6 | 8 | 0.375 | 4 | 0.75 | 2 | 0.5 | 19 | 0.5263 | 7 | 0.7143 | 5 | 0.4 | NA | NA | 4 | 0.25 | 16 | 0.5 |
| 4_69843850 | 3 | 0 | 5 | 0 | 3 | 0.3333 | 2 | 0 | 13 | 0.0769 | 5 | 0.2 | 4 | 0 | 1 | 0 | 5 | 0 | 15 | 0.0667 |
| 4_70135253 | NA | NA | NA | NA | 1 | 1 | 1 | 1 | 2 | 1 | 1 | 1 | 1 | 1 | NA | NA | 1 | 1 | 3 | 1 |
| 4_70135369 | NA | NA | NA | NA | 1 | 1 | 1 | 1 | 2 | 1 | 1 | 1 | 1 | 1 | NA | NA | 1 | 1 | 3 | 1 |
| 4_70135471 | NA | NA | NA | NA | 1 | 1 | 1 | 0 | 2 | 0.5 | NA | NA | 1 | 0 | NA | NA | 1 | 1 | 2 | 0.5 |
| 4_70135561 | NA | NA | NA | NA | 1 | 1 | 1 | 0 | 2 | 0.5 | 1 | 0 | 1 | 0 | NA | NA | 1 | 1 | 3 | 0.3333 |
| 4_70135730 | NA | NA | NA | NA | 1 | 1 | 1 | 0 | 2 | 0.5 | 1 | 1 | 1 | 1 | NA | NA | NA | NA | 2 | 1 |
| 4_70182762 | NA | NA | NA | NA | 1 | 1 | NA | NA | 1 | 1 | NA | NA | NA | NA | 1 | 1 | 1 | 1 | 2 | 1 |
| 4_70185145 | NA | NA | 1 | 0 | 1 | 1 | NA | NA | 2 | 0.5 | NA | NA | NA | NA | 1 | 0 | 1 | 1 | 2 | 0.5 |
| 4_70199624 | NA | NA | 1 | 0 | 1 | 1 | NA | NA | 2 | 0.5 | NA | NA | NA | NA | 1 | 1 | 1 | 0 | 2 | 0.5 |
| 4_70199668 | NA | NA | 1 | 1 | 1 | 0 | NA | NA | 2 | 0.5 | NA | NA | NA | NA | 1 | 0 | 1 | 1 | 2 | 0.5 |
| 4_70347444 | 11 | 0.2727 | 15 | 0.2 | 10 | 0.2 | 6 | 0.1667 | 42 | 0.2143 | 6 | 0.1667 | 10 | 0 | 12 | 0.25 | 13 | 0.0769 | 41 | 0.122 |
| 4_70348305 | 10 | 0.1 | 10 | 0.2 | 8 | 0 | 7 | 0.1429 | 35 | 0.1143 | 7 | 0.1429 | 7 | 0.1429 | 9 | 0 | 12 | 0 | 35 | 0.0571 |
| 4_70349167 | 7 | 0.2857 | 9 | 0.3333 | 7 | 0.1429 | 6 | 0 | 29 | 0.2069 | 6 | 0.3333 | 5 | 0 | 9 | 0 | 3 | 0 | 23 | 0.087 |
| 4_70350048 | 5 | 0.4 | 7 | 0.4286 | 6 | 0.1667 | 2 | 0 | 20 | 0.3 | 3 | 0.6667 | 6 | 0.3333 | 8 | 0.625 | 4 | 0.25 | 21 | 0.4762 |
| 4_70350195 | 6 | 0.3333 | 7 | 0.2857 | 6 | 0.3333 | 2 | 0 | 21 | 0.2857 | 3 | 0 | 6 | 0 | 8 | 0.25 | 5 | 0.2 | 22 | 0.1364 |
| 4_70350210 | 4 | 0.25 | 7 | 0.1429 | 3 | 0 | 2 | 0.5 | 16 | 0.1875 | 3 | 0.3333 | 4 | 0 | 7 | 0.4286 | 3 | 0.3333 | 17 | 0.2941 |
| 4_70350435 | 3 | 0 | 6 | 0.1667 | 5 | 0 | 2 | 0.5 | 16 | 0.125 | 3 | 0.3333 | 5 | 0 | 6 | 0.5 | 3 | 0 | 17 | 0.2353 |
| 4_70350472 | 4 | 0.25 | 6 | 0.1667 | 5 | 0 | 2 | 0.5 | 17 | 0.1765 | 3 | 0.3333 | 6 | 0 | 7 | 0.4286 | 5 | 0.6 | 21 | 0.3333 |
| 4_70350968 | 4 | 0.75 | 6 | 1 | 5 | 1 | 2 | 1 | 17 | 0.9412 | 2 | 1 | 6 | 1 | 6 | 1 | 2 | 1 | 16 | 1 |
| 4_70351027 | 3 | 1 | 6 | 1 | 4 | 1 | 2 | 1 | 15 | 1 | 2 | 1 | 6 | 1 | 6 | 1 | 2 | 1 | 16 | 1 |
| 4_70351122 | 4 | 1 | 7 | 1 | 5 | 1 | 2 | 1 | 18 | 1 | 1 | 1 | 5 | 1 | 5 | 1 | 2 | 1 | 13 | 1 |
| 4_70351192 | 1 | 1 | 3 | 1 | 5 | 1 | 2 | 1 | 11 | 1 | 1 | 1 | 3 | 1 | 4 | 1 | 1 | 1 | 9 | 1 |
| 4_70351453 | 4 | 0 | 7 | 0 | 5 | 0.2 | 2 | 0.5 | 18 | 0.1111 | 1 | 0 | 4 | 0.5 | 4 | 0.25 | 2 | 0.5 | 11 | 0.3636 |
| 4_70351472 | 4 | 0.5 | 3 | 0.6667 | 5 | 0.8 | 2 | 0.5 | 14 | 0.6429 | NA | NA | 4 | 0.5 | 3 | 0.6667 | 1 | 1 | 8 | 0.625 |
| 4_70351491 | 3 | 0.3333 | 5 | 0.2 | 4 | 0.25 | 2 | 0.5 | 14 | 0.2857 | 1 | 1 | 4 | 0.5 | 4 | 0.75 | 2 | 0 | 11 | 0.5455 |
| 4_70351571 | 2 | 0 | 3 | 0 | 4 | 0.5 | 1 | 0 | 10 | 0.2 | 1 | 1 | 2 | 0 | 4 | 0.25 | 1 | 0 | 8 | 0.25 |
| 4_70351595 | 4 | 1 | 7 | 1 | 5 | 1 | 1 | 1 | 17 | 1 | 1 | 1 | 4 | 1 | 4 | 1 | 2 | 1 | 11 | 1 |
| 4_70351655 | 3 | 1 | 4 | 0.75 | 4 | 0.75 | 1 | 1 | 12 | 0.8333 | 1 | 1 | 4 | 0.75 | 4 | 0.75 | 1 | 1 | 10 | 0.8 |
| 4_70353591 | 2 | 0 | 5 | 0.2 | 4 | 0 | 1 | 0 | 12 | 0.0833 | 1 | 1 | 3 | 0 | 3 | 0.6667 | 2 | 0 | 9 | 0.3333 |
| 4_70353666 | 3 | 0 | 4 | 0.5 | 3 | 0 | 1 | 0 | 11 | 0.1818 | 1 | 0 | 4 | 0.5 | 3 | 0 | 2 | 0 | 10 | 0.2 |
| 4_72031180 | 1 | 0 | 1 | 1 | NA | NA | 1 | 0 | 3 | 0.3333 | NA | NA | NA | NA | 2 | 0.5 | NA | NA | 2 | 0.5 |
| 4_72036165 | 3 | 0.6667 | 4 | 0.75 | 1 | 1 | 5 | 0.6 | 13 | 0.6923 | NA | NA | 5 | 0.4 | 8 | 0.5 | 8 | 0.5 | 21 | 0.4762 |
| 4_72036320 | 4 | 0.75 | 3 | 0.3333 | NA | NA | 4 | 1 | 11 | 0.7273 | NA | NA | 4 | 1 | 5 | 0.4 | 4 | 0.5 | 13 | 0.6154 |
| 4_72036335 | 3 | 0.3333 | 3 | 0.6667 | NA | NA | 4 | 0.25 | 10 | 0.4 | NA | NA | 4 | 0 | 8 | 0.25 | 6 | 0.1667 | 18 | 0.1667 |
| 4_72036380 | 3 | 1 | 1 | 0 | 1 | 1 | 5 | 0.8 | 10 | 0.8 | 1 | 1 | 4 | 0.75 | 4 | 1 | 6 | 0.6667 | 15 | 0.8 |
| 4_72036434 | 2 | 0 | 1 | 1 | NA | NA | 2 | 0.5 | 5 | 0.4 | 1 | 0 | 1 | 0 | 2 | 0 | 2 | 1 | 6 | 0.3333 |
| 4_72036555 | 4 | 0.25 | 4 | 0.5 | 1 | 0 | 4 | 0.75 | 13 | 0.4615 | 1 | 1 | 4 | 0.5 | 6 | 0.3333 | 7 | 0.2857 | 18 | 0.3889 |
| 4_72076162 | 4 | 0.75 | NA | NA | 1 | 1 | 3 | 0.3333 | 8 | 0.625 | 2 | 1 | 1 | 0 | 2 | 0.5 | 5 | 0.6 | 10 | 0.6 |
| 4_72076204 | 7 | 0 | NA | NA | 1 | 0 | 2 | 0.5 | 10 | 0.1 | 3 | 0.3333 | 1 | 0 | 3 | 0 | 5 | 0 | 12 | 0.0833 |
| 4_72076215 | 7 | 0 | NA | NA | NA | NA | 2 | 0.5 | 9 | 0.1111 | 3 | 0.3333 | 1 | 0 | 3 | 0 | 4 | 0 | 11 | 0.0909 |
| 4_72076404 | 4 | 0 | NA | NA | 1 | 0 | 1 | 1 | 6 | 0.1667 | 2 | 0.5 | 1 | 1 | 2 | 0 | 4 | 0.25 | 9 | 0.3333 |
| 4_72076580 | 8 | 0.5 | 2 | 0 | 1 | 1 | 1 | 0 | 12 | 0.4167 | 3 | 0 | 2 | 0 | 6 | 0 | 6 | 0.3333 | 17 | 0.1176 |
| 4_72118444 | 5 | 0.4 | 1 | 0 | NA | NA | 2 | 0.5 | 8 | 0.375 | 3 | 0 | 1 | 1 | 3 | 0.3333 | 3 | 0.3333 | 10 | 0.3 |
| 4_72118928 | 5 | 0.8 | 2 | 0 | NA | NA | 1 | 0 | 8 | 0.5 | 2 | 0 | 1 | 1 | 2 | 0.5 | 3 | 0.3333 | 8 | 0.375 |
| 4_72119636 | NA | NA | NA | NA | 1 | 1 | 1 | 1 | 2 | 1 | NA | NA | NA | NA | 3 | 1 | NA | NA | 3 | 1 |
| 4_72119691 | NA | NA | 1 | 1 | 1 | 1 | 1 | 1 | 3 | 1 | 1 | 0 | NA | NA | 3 | 0.6667 | NA | NA | 4 | 0.5 |
| 4_72120098 | 1 | 1 | 1 | 1 | 3 | 1 | 2 | 1 | 7 | 1 | 1 | 0 | NA | NA | 3 | 1 | NA | NA | 4 | 0.75 |
| 4_72120230 | 1 | 1 | NA | NA | 3 | 0.6667 | 1 | 1 | 5 | 0.8 | 1 | 1 | NA | NA | 2 | 0.5 | NA | NA | 3 | 0.6667 |
| 4_72120319 | 1 | 1 | 1 | 0 | 3 | 0.3333 | NA | NA | 5 | 0.4 | 1 | 1 | NA | NA | 1 | 1 | NA | NA | 2 | 1 |
| 4_72120824 | 1 | 0 | NA | NA | 2 | 0.5 | NA | NA | 3 | 0.3333 | NA | NA | NA | NA | 1 | 1 | NA | NA | 1 | 1 |
| 4_72120994 | NA | NA | NA | NA | 2 | 0.5 | NA | NA | 2 | 0.5 | NA | NA | NA | NA | 1 | 1 | NA | NA | 1 | 1 |
| 4_72121369 | 1 | 1 | NA | NA | 2 | 0 | NA | NA | 3 | 0.3333 | NA | NA | NA | NA | 2 | 0.5 | NA | NA | 2 | 0.5 |
| 4_74105752 | NA | NA | 1 | 1 | NA | NA | NA | NA | 1 | 1 | 1 | 1 | 1 | 1 | NA | NA | NA | NA | 2 | 1 |
| 4_74105804 | 1 | 1 | 1 | 1 | NA | NA | NA | NA | 2 | 1 | 1 | 1 | NA | NA | NA | NA | NA | NA | 1 | 1 |
| 4_74105960 | 1 | 1 | 1 | 1 | NA | NA | NA | NA | 2 | 1 | 1 | 1 | 1 | 1 | NA | NA | NA | NA | 2 | 1 |
| 4_74323291 | 1 | 1 | 4 | 0 | 1 | 1 | 1 | 1 | 7 | 0.4286 | 4 | 0.25 | 6 | 0.1667 | 3 | 0.6667 | 3 | 0.3333 | 16 | 0.3125 |
| 4_74323314 | 3 | 0.3333 | 3 | 0.3333 | NA | NA | 1 | 0 | 7 | 0.2857 | 3 | 0.3333 | 6 | 0.1667 | 3 | 0.6667 | 3 | 0.6667 | 15 | 0.4 |
| 4_75192925 | NA | NA | NA | NA | 1 | 1 | 1 | 1 | 2 | 1 | NA | NA | 1 | 1 | NA | NA | 1 | 1 | 2 | 1 |

| | | | | | | | | | | | | | | | | | | | | |
|---|---|---|---|---|---|---|---|---|---|---|---|---|---|---|---|---|---|---|---|---|
| 4_7519295 | NA | NA | NA | NA | NA | NA | 1 | 1 | 1 | 1 | NA | NA | 1 | 1 | NA | NA | 1 | 1 | 2 | 1 |
| 4_7519300 | NA | NA | NA | NA | 1 | 1 | 1 | 1 | 2 | 1 | NA | NA | 1 | 1 | NA | NA | 1 | 1 | 2 | 1 |
| 4_7519320 | NA | NA | NA | NA | 1 | 1 | 1 | 1 | 2 | 1 | NA | NA | 1 | 1 | NA | NA | 1 | 1 | 2 | 1 |
| 4_7519322 | NA | NA | NA | NA | 1 | 1 | 1 | 1 | 2 | 1 | NA | NA | 2 | 1 | NA | NA | 1 | 1 | 3 | 1 |
| 4_7519336 | NA | NA | NA | NA | 1 | 1 | 1 | 0 | 2 | 0.5 | NA | NA | 1 | 0 | NA | NA | 1 | 1 | 2 | 0.5 |
| 4_7519341 | NA | NA | NA | NA | 1 | 1 | 1 | 1 | 2 | 1 | NA | NA | 1 | 1 | NA | NA | 1 | 1 | 2 | 1 |
| 4_7519486 | NA | NA | NA | NA | 1 | 1 | NA | NA | 1 | 1 | NA | NA | 1 | 1 | NA | NA | 1 | 1 | 2 | 1 |
| 4_7519574 | 6 | 0.5 | 9 | 0.4444 | 7 | 0 | 11 | 0.5455 | 33 | 0.3939 | 3 | 0 | 4 | 0.25 | 5 | 0.2 | 9 | 0.5556 | 21 | 0.3333 |
| 4_7519582 | 6 | 0.5 | 10 | 0.6 | 7 | 0.4286 | 10 | 0.8 | 33 | 0.6061 | 3 | 0.6667 | 5 | 0.8 | 5 | 0.8 | 8 | 0.875 | 21 | 0.8095 |
| 4_7519583 | 5 | 0.4 | 8 | 0.125 | 8 | 0 | 11 | 0.3636 | 32 | 0.2188 | 2 | 0 | 4 | 0 | 4 | 0 | 7 | 0.1429 | 17 | 0.0588 |
| 4_7585970 | 3 | 0 | 9 | 0.6667 | 11 | 0.4545 | 7 | 0.5714 | 30 | 0.5 | 5 | 0.2 | 6 | 0.1667 | 1 | 1 | 5 | 0.6 | 17 | 0.3529 |
| 4_7586003 | 3 | 0 | 9 | 0.5556 | 9 | 0 | 7 | 0.1429 | 28 | 0.2143 | 5 | 0 | 6 | 0.1667 | NA | NA | 5 | 0.2 | 16 | 0.125 |
| 4_7586276 | 5 | 0.8 | 7 | 0.8571 | 7 | 1 | 4 | 0.75 | 23 | 0.8696 | 1 | 1 | 6 | 1 | NA | NA | 1 | 1 | 8 | 1 |
| 4_7586283 | 6 | 0.5 | 7 | 0.7143 | 5 | 0.6 | 5 | 0.6 | 23 | 0.6087 | 1 | 1 | 5 | 0.8 | NA | NA | 2 | 0.5 | 8 | 0.75 |
| 4_7586286 | 4 | 0 | 5 | 0 | 5 | 0.2 | 4 | 0 | 18 | 0.0556 | 1 | 0 | 4 | 0 | NA | NA | 3 | 0.3333 | 8 | 0.125 |
| 4_7586287 | 6 | 0 | 9 | 0.2222 | 8 | 0 | 6 | 0 | 29 | 0.069 | 1 | 0 | 6 | 0.1667 | NA | NA | 3 | 0 | 10 | 0.1 |
| 4_7586294 | 6 | 0 | 8 | 0.125 | 10 | 0.2 | 5 | 0 | 29 | 0.1034 | NA | NA | 2 | 0.5 | NA | NA | 3 | 0.3333 | 5 | 0.4 |
| 4_7586300 | 6 | 0.6667 | 7 | 0.8571 | 8 | 0.875 | 6 | 0.3333 | 27 | 0.7037 | 1 | 0 | 6 | 0.6667 | NA | NA | 4 | 1 | 11 | 0.7273 |
| 4_7586301 | 3 | 0.6667 | 8 | 0.375 | 6 | 0.3333 | 2 | 0.5 | 19 | 0.4211 | NA | NA | 5 | 0.6 | NA | NA | 2 | 1 | 7 | 0.7143 |
| 4_7586302 | 4 | 0 | 6 | 0.3333 | 6 | 0 | 5 | 0.2 | 21 | 0.1429 | NA | NA | 6 | 0.1667 | NA | NA | NA | NA | 6 | 0.1667 |
| 4_7586302 | 3 | 0 | 1 | 0 | 3 | 0 | 3 | 0.3333 | 10 | 0.1 | NA | NA | 1 | 1 | NA | NA | NA | NA | 1 | 1 |
| 4_7586304 | NA | NA | 2 | 0.5 | 3 | 0 | 3 | 0 | 8 | 0.125 | NA | NA | 2 | 0.5 | NA | NA | 2 | 0.5 | 4 | 0.5 |
| 4_7586306 | 3 | 0 | 6 | 0 | 5 | 0.4 | 4 | 0.25 | 18 | 0.1667 | 1 | 1 | 3 | 0.3333 | NA | NA | 1 | 0 | 5 | 0.4 |
| 4_7586307 | 5 | 0.8 | 8 | 0.875 | 6 | 1 | 5 | 0.8 | 24 | 0.875 | 1 | 1 | 5 | 0.8 | NA | NA | 4 | 1 | 10 | 0.9 |
| 4_7587554 | 3 | 0 | 3 | 0 | 5 | 0.2 | 3 | 0.3333 | 14 | 0.1429 | NA | NA | 3 | 0.3333 | NA | NA | NA | NA | 3 | 0.3333 |
| 4_7604680 | 1 | 0 | 1 | 1 | 1 | 0 | 2 | 0 | 5 | 0.2 | 1 | 0 | 1 | 1 | NA | NA | 1 | 0 | 3 | 0.3333 |
| 4_7604689 | 2 | 0.5 | 2 | 1 | 1 | 0 | 3 | 1 | 8 | 0.75 | NA | NA | 2 | 0.5 | NA | NA | 2 | 1 | 4 | 0.75 |
| 4_7604723 | 1 | 0 | 1 | 1 | NA | NA | 2 | 1 | 4 | 0.75 | 1 | 1 | 1 | 0 | NA | NA | 1 | 1 | 3 | 0.6667 |
| 4_7813092 | NA | NA | NA | NA | 2 | 0.5 | 1 | 0 | 3 | 0.3333 | NA | NA | NA | NA | 1 | 1 | NA | NA | 1 | 1 |
| 4_7813237 | NA | NA | NA | NA | 2 | 1 | 1 | 1 | 3 | 1 | NA | NA | NA | NA | 1 | 1 | NA | NA | 1 | 1 |
| 4_7813244 | NA | NA | NA | NA | 2 | 1 | 1 | 1 | 3 | 1 | NA | NA | NA | NA | 1 | 1 | NA | NA | 1 | 1 |
| 4_7813258 | NA | NA | NA | NA | 2 | 1 | 1 | 1 | 3 | 1 | NA | NA | NA | NA | 1 | 1 | NA | NA | 1 | 1 |
| 4_8082894 | 1 | 0 | 1 | 0 | 1 | 1 | NA | NA | 3 | 0.3333 | NA | NA | NA | NA | 1 | 0 | 3 | 0.3333 | 4 | 0.25 |
| 4_8082922 | 1 | 1 | NA | NA | NA | NA | NA | NA | 1 | 1 | NA | NA | NA | NA | 1 | 1 | 3 | 1 | 4 | 1 |
| 4_8082947 | 1 | 0 | 2 | 0.5 | 1 | 0 | NA | NA | 4 | 0.25 | 1 | 0 | NA | NA | 2 | 0.5 | 4 | 0 | 7 | 0.1429 |
| 4_8087203 | NA | NA | 5 | 1 | 1 | 1 | NA | NA | 6 | 1 | 1 | 1 | NA | NA | 2 | 1 | 4 | 1 | 7 | 1 |
| 4_8087208 | 1 | 1 | 5 | 0.4 | 1 | 0 | NA | NA | 7 | 0.4286 | 2 | 0 | 1 | 0 | 1 | 0 | 6 | 0.3333 | 10 | 0.2 |
| 4_8087211 | 1 | 1 | 5 | 1 | 1 | 0 | NA | NA | 7 | 0.8571 | 2 | 1 | 1 | 1 | 2 | 1 | 4 | 0.75 | 9 | 0.8889 |
| 4_8087219 | 1 | 1 | 5 | 1 | 1 | 1 | NA | NA | 7 | 1 | 2 | 1 | 2 | 1 | 2 | 1 | 5 | 1 | 11 | 1 |
| 4_8119749 | NA | NA | 4 | 0.5 | 2 | 0 | 2 | 0 | 8 | 0.25 | 4 | 0.75 | 7 | 0.5714 | NA | NA | 3 | 0.3333 | 14 | 0.5714 |
| 4_8119760 | NA | NA | 4 | 0.25 | 1 | 0 | 2 | 0 | 7 | 0.1429 | 4 | 0.25 | 6 | 0.1667 | NA | NA | 2 | 0 | 12 | 0.1667 |
| 4_8119779 | NA | NA | 4 | 0.5 | 1 | 0 | 2 | 0 | 7 | 0.2857 | 3 | 0 | 6 | 0.1667 | NA | NA | 3 | 0.3333 | 12 | 0.1667 |
| 4_8119785 | NA | NA | 3 | 0.3333 | 1 | 0 | 3 | 0 | 7 | 0.1429 | 3 | 0.3333 | 6 | 0.3333 | NA | NA | 2 | 0 | 11 | 0.2727 |
| 4_8220895 | 4 | 0.5 | 6 | 0.3333 | 4 | 0.5 | 4 | 0.5 | 18 | 0.4444 | 5 | 0.6 | 5 | 0.4 | 5 | 0.2 | 10 | 0.2 | 25 | 0.32 |
| 4_8220902 | 4 | 0.5 | 6 | 0.3333 | 4 | 0.5 | 3 | 0.3333 | 17 | 0.4118 | 4 | 0.25 | 7 | 0.4286 | 4 | 0 | 9 | 0.5556 | 24 | 0.375 |
| 4_8220915 | 4 | 0.5 | 6 | 0.1667 | 3 | 0.3333 | 4 | 0.5 | 17 | 0.3529 | 5 | 0.2 | 7 | 0.2857 | 4 | 0 | 9 | 0.1111 | 25 | 0.16 |
| 4_8220923 | NA | NA | 2 | 0 | 1 | 1 | 1 | 0 | 4 | 0.25 | 1 | 1 | 2 | 0 | 1 | 0 | 1 | 0 | 5 | 0.2 |
| 4_8220929 | 2 | 0 | 6 | 0.1667 | 4 | 0.25 | 4 | 0 | 16 | 0.125 | 5 | 0.2 | 7 | 0.1429 | 4 | 0.5 | 9 | 0.2222 | 25 | 0.24 |
| 4_8220933 | 2 | 1 | 6 | 0.5 | 4 | 0.5 | 4 | 1 | 16 | 0.6875 | 5 | 0.8 | 7 | 0.8571 | 4 | 1 | 9 | 0.5556 | 25 | 0.76 |
| 4_8220944 | 2 | 0 | 6 | 0.3333 | 3 | 0.3333 | 3 | 0.3333 | 14 | 0.2857 | 5 | 0 | 7 | 0.2857 | 3 | 0 | 9 | 0.1111 | 24 | 0.125 |
| 4_8221013 | 2 | 0 | 3 | 0.3333 | 3 | 0.3333 | 1 | 1 | 9 | 0.3333 | 3 | 1 | 5 | 0.4 | 3 | 0.6667 | 7 | 0.5714 | 18 | 0.6111 |
| 4_8221026 | 2 | 0 | 2 | 0 | 2 | 0.5 | NA | NA | 6 | 0.1667 | 3 | 0 | 4 | 0.5 | 1 | 0 | 4 | 0 | 12 | 0.1667 |
| 4_8221037 | 2 | 0.5 | 3 | 0 | 4 | 0.25 | 1 | 0 | 10 | 0.2 | 3 | 0 | 5 | 0 | 3 | 0 | 5 | 0.2 | 16 | 0.0625 |
| 4_8221038 | 2 | 0.5 | 3 | 0 | 2 | 0 | 1 | 0 | 8 | 0.125 | 3 | 0 | 6 | 0 | 2 | 0 | 6 | 0.5 | 17 | 0.1765 |
| 4_8221475 | NA | NA | NA | NA | NA | NA | 1 | 1 | 1 | 1 | NA | NA | 1 | 0 | 1 | 1 | 2 | 0.5 | 4 | 0.5 |
| 4_8221500 | NA | NA | NA | NA | NA | NA | 1 | 1 | 1 | 1 | NA | NA | 1 | 0 | 2 | 1 | 2 | 1 | 5 | 0.8 |
| 4_8241676 | NA | NA | NA | NA | NA | NA | 1 | 1 | 1 | 1 | NA | NA | NA | NA | 2 | 1 | 1 | 1 | 3 | 1 |
| 4_8242318 | NA | NA | NA | NA | NA | NA | 1 | 1 | 1 | 1 | NA | NA | NA | NA | 2 | 0 | 1 | 1 | 3 | 0.3333 |
| 4_8246534 | NA | NA | 1 | 0 | 1 | 1 | 1 | 0 | 3 | 0.3333 | NA | NA | 1 | 1 | NA | NA | NA | NA | 1 | 1 |
| 4_8246536 | NA | NA | 1 | 1 | 1 | 1 | 1 | 0 | 3 | 0.6667 | NA | NA | 1 | 1 | NA | NA | NA | NA | 1 | 1 |
| 4_8271716 | NA | NA | 3 | 0 | 5 | 0.2 | 1 | 0 | 9 | 0.1111 | 2 | 0 | 2 | 0 | 4 | 0.25 | 3 | 0 | 11 | 0.0909 |
| 4_8272439 | NA | NA | 1 | 0 | 1 | 1 | 1 | 1 | 3 | 0.6667 | NA | NA | 1 | 0 | NA | NA | 2 | 0.5 | 3 | 0.3333 |
| 4_8313901 | 4 | 0 | 4 | 0.25 | 5 | 0.2 | 3 | 0.3333 | 16 | 0.1875 | 2 | 0.5 | 2 | 0 | 7 | 0.1429 | 3 | 0 | 14 | 0.1429 |
| 4_8313924 | 3 | 0 | 4 | 0.25 | 4 | 0 | 6 | 0.3333 | 17 | 0.1765 | 1 | 0 | 2 | 0 | 7 | 0.1429 | 4 | 0 | 14 | 0.0714 |
| 4_8313952 | 4 | 0.25 | 3 | 0 | 5 | 0.2 | 4 | 0 | 16 | 0.125 | 1 | 0 | 1 | 0 | 7 | 0.1429 | 2 | 0 | 11 | 0.0909 |

| | | | | | | | | | | | | | | | | | | | | |
|---|---|---|---|---|---|---|---|---|---|---|---|---|---|---|---|---|---|---|---|---|
| 4_83139608 | 3 | 0 | 4 | 0 | 4 | 0.25 | 4 | 0 | 15 | 0.0667 | NA | NA | 1 | 0 | 5 | 0 | 1 | 1 | 7 | 0.1429 |
| 4_83139654 | 4 | 0.25 | 4 | 0.5 | 5 | 0 | 5 | 0.2 | 18 | 0.2222 | 1 | 0 | 1 | 0 | 7 | 0.1429 | 2 | 0.5 | 11 | 0.1818 |
| 4_83140583 | 3 | 0 | 3 | 0 | 7 | 0.5714 | 5 | 0.6 | 18 | 0.3889 | 1 | 0 | 2 | 1 | 4 | 0.5 | 1 | 0 | 8 | 0.5 |
| 4_83142413 | 1 | 0 | 2 | 0 | 3 | 0.3333 | 3 | 0 | 9 | 0.1111 | 1 | 0 | 1 | 0 | 3 | 0.3333 | NA | NA | 5 | 0.2 |
| 4_83142455 | 1 | 0 | 2 | 0.5 | 3 | 0 | 4 | 0 | 10 | 0.1 | 1 | 0 | 1 | 0 | 4 | 0.25 | 1 | 0 | 7 | 0.1429 |
| 4_83145190 | 1 | 0 | NA | NA | 1 | 1 | 1 | 0 | 3 | 0.3333 | 1 | 0 | 1 | 1 | NA | NA | 1 | 1 | 3 | 0.6667 |
| 4_83145328 | 1 | 1 | NA | NA | 1 | 1 | 1 | 1 | 3 | 1 | 1 | 1 | 1 | 1 | NA | NA | NA | NA | 2 | 1 |
| 4_83364145 | NA | NA | 1 | 1 | NA | NA | 2 | 0 | 3 | 0.3333 | NA | NA | 1 | 1 | NA | NA | NA | NA | 1 | 1 |
| 4_83364283 | NA | NA | NA | NA | NA | NA | 1 | 1 | 1 | 1 | NA | NA | 1 | 1 | NA | NA | NA | NA | 1 | 1 |
| 4_83364543 | NA | NA | 1 | 0 | NA | NA | 1 | 1 | 2 | 0.5 | NA | NA | 1 | 1 | NA | NA | NA | NA | 1 | 1 |
| 4_83364737 | NA | NA | 1 | 1 | NA | NA | 2 | 0 | 3 | 0.3333 | NA | NA | 1 | 1 | NA | NA | NA | NA | 1 | 1 |
| 4_83365234 | NA | NA | 1 | 1 | NA | NA | 2 | 0 | 3 | 0.3333 | NA | NA | 1 | 1 | NA | NA | NA | NA | 1 | 1 |
| 4_83365268 | NA | NA | 1 | 1 | NA | NA | 2 | 1 | 3 | 1 | NA | NA | 1 | 1 | NA | NA | NA | NA | 1 | 1 |
| 4_83365339 | NA | NA | 1 | 1 | NA | NA | 1 | 0 | 2 | 0.5 | NA | NA | 1 | 1 | NA | NA | NA | NA | 1 | 1 |
| 4_83365380 | NA | NA | 1 | 1 | NA | NA | 2 | 1 | 3 | 1 | NA | NA | 1 | 1 | NA | NA | NA | NA | 1 | 1 |
| 4_83365564 | NA | NA | NA | NA | NA | NA | 1 | 1 | 1 | 1 | NA | NA | 1 | 1 | NA | NA | NA | NA | 1 | 1 |
| 4_83365835 | NA | NA | 1 | 0 | NA | NA | 2 | 0.5 | 3 | 0.3333 | NA | NA | 1 | 1 | NA | NA | NA | NA | 1 | 1 |
| 4_83365928 | NA | NA | 1 | 1 | NA | NA | 1 | 1 | 2 | 1 | NA | NA | 1 | 1 | NA | NA | NA | NA | 1 | 1 |
| 4_83366401 | NA | NA | 1 | 1 | NA | NA | 2 | 0 | 3 | 0.3333 | NA | NA | 1 | 1 | NA | NA | NA | NA | 1 | 1 |
| 4_83366413 | NA | NA | NA | NA | NA | NA | 1 | 1 | 1 | 1 | NA | NA | 1 | 1 | NA | NA | NA | NA | 1 | 1 |
| 4_83367265 | NA | NA | 1 | 1 | NA | NA | 1 | 1 | 2 | 1 | NA | NA | 1 | 1 | NA | NA | NA | NA | 1 | 1 |
| 4_83367348 | NA | NA | 1 | 0 | NA | NA | 2 | 0.5 | 3 | 0.3333 | NA | NA | 1 | 1 | NA | NA | NA | NA | 1 | 1 |
| 4_83371313 | 2 | 0 | 4 | 0 | 2 | 0 | 7 | 0.1429 | 15 | 0.0667 | 2 | 0 | 4 | 0 | 6 | 0 | 8 | 0.125 | 20 | 0.05 |
| 4_83374075 | 10 | 0.1 | 10 | 0 | 11 | 0.0909 | 11 | 0.1818 | 42 | 0.0952 | 2 | 0.5 | 5 | 0.2 | 9 | 0.1111 | 9 | 0.1111 | 25 | 0.16 |
| 4_83375210 | 7 | 0.1429 | 9 | 0 | 12 | 0 | 9 | 0.1111 | 37 | 0.0541 | 3 | 0.3333 | 5 | 0 | 6 | 0 | 6 | 0.1667 | 20 | 0.1 |
| 4_83378348 | 3 | 0.3333 | NA | NA | NA | NA | NA | NA | 3 | 0.3333 | NA | NA | NA | NA | 1 | 1 | NA | NA | 1 | 1 |
| 4_83380501 | 17 | 0.2941 | 14 | 0.1429 | 15 | 0.1333 | 27 | 0.1852 | 73 | 0.1918 | 8 | 0.25 | 22 | 0.2273 | 16 | 0.125 | 17 | 0.4706 | 63 | 0.2698 |
| 4_83380525 | 16 | 0.75 | 17 | 0.8824 | 10 | 0.8 | 25 | 0.84 | 68 | 0.8235 | 8 | 1 | 24 | 0.8333 | 15 | 0.8667 | 17 | 0.7059 | 64 | 0.8281 |
| 4_83380582 | 18 | 0.9444 | 16 | 0.9375 | 15 | 0.8 | 26 | 0.9615 | 75 | 0.92 | 9 | 1 | 20 | 0.95 | 18 | 0.9444 | 19 | 0.9474 | 66 | 0.9545 |
| 4_83380591 | 20 | 0.35 | 19 | 0.1579 | 17 | 0.0588 | 30 | 0.2667 | 86 | 0.2209 | 6 | 0.1667 | 24 | 0.375 | 16 | 0.4375 | 18 | 0.2778 | 64 | 0.3438 |
| 4_83380643 | 21 | 0.1905 | 18 | 0.1111 | 14 | 0.1429 | 28 | 0.2143 | 81 | 0.1728 | 7 | 0 | 21 | 0.2381 | 19 | 0.1053 | 14 | 0.2143 | 61 | 0.1639 |
| 4_83380794 | 6 | 0 | 3 | 0 | 3 | 0.3333 | 5 | 0 | 17 | 0.0588 | 1 | 1 | 9 | 0.3333 | 1 | 1 | 5 | 0.6 | 16 | 0.5 |
| 4_83380996 | 7 | 0.1429 | 3 | 0 | 5 | 0.2 | 8 | 0.125 | 23 | 0.1304 | 1 | 1 | 6 | 0.1667 | 2 | 0.5 | 4 | 0.5 | 13 | 0.3846 |
| 4_83381427 | 6 | 0.1667 | 3 | 0 | 3 | 0 | 7 | 0.1429 | 19 | 0.1053 | 1 | 1 | 5 | 0 | 1 | 0 | 3 | 0 | 10 | 0.1 |
| 4_83381641 | 6 | 0 | 2 | 0 | 3 | 0 | 6 | 0.1667 | 17 | 0.0588 | 1 | 0 | 6 | 0.3333 | 2 | 0 | 3 | 0 | 12 | 0.1667 |
| 4_83381732 | 7 | 0.2857 | 2 | 0.5 | 4 | 0 | 6 | 0 | 19 | 0.1579 | 1 | 0 | 6 | 0 | 2 | 0.5 | 3 | 0 | 12 | 0.0833 |
| 4_83783440 | 1 | 1 | NA | NA | NA | NA | NA | NA | 1 | 1 | NA | NA | NA | NA | NA | NA | 2 | 0.5 | 2 | 0.5 |
| 4_83784324 | 1 | 1 | NA | NA | NA | NA | NA | NA | 1 | 1 | NA | NA | NA | NA | NA | NA | 1 | 1 | 1 | 1 |
| 4_85049126 | NA | NA | NA | NA | 2 | 0.5 | NA | NA | 2 | 0.5 | 1 | 0 | NA | NA | NA | NA | 2 | 0.5 | 3 | 0.3333 |
| 4_85049300 | NA | NA | NA | NA | 2 | 1 | NA | NA | 2 | 1 | 1 | 1 | NA | NA | NA | NA | 1 | 0 | 2 | 0.5 |
| 4_85305908 | 7 | 0.4286 | 13 | 0.3846 | 9 | 0.3333 | 4 | 0.5 | 33 | 0.3939 | 10 | 0.1 | 10 | 0.2 | 9 | 0.6667 | 11 | 0.5455 | 40 | 0.375 |
| 4_85306301 | 15 | 0.0667 | 24 | 0.0833 | 21 | 0.0952 | 9 | 0 | 69 | 0.0725 | 16 | 0 | 15 | 0.2 | 19 | 0.0526 | 17 | 0.1176 | 67 | 0.0896 |
| 4_86260855 | NA | NA | NA | NA | 1 | 1 | NA | NA | 1 | 1 | NA | NA | NA | NA | 1 | 1 | NA | NA | 1 | 1 |
| 4_86260999 | NA | NA | NA | NA | 1 | 1 | NA | NA | 1 | 1 | NA | NA | NA | NA | 1 | 1 | NA | NA | 1 | 1 |
| 4_86261277 | NA | NA | NA | NA | 1 | 1 | NA | NA | 1 | 1 | NA | NA | NA | NA | 1 | 1 | NA | NA | 1 | 1 |
| 4_86261298 | NA | NA | NA | NA | 1 | 1 | NA | NA | 1 | 1 | NA | NA | NA | NA | 1 | 1 | NA | NA | 1 | 1 |
| 4_86344228 | NA | NA | 1 | 1 | NA | NA | NA | NA | 1 | 1 | NA | NA | NA | NA | 1 | 1 | NA | NA | 1 | 1 |
| 4_86345125 | NA | NA | 1 | 0 | 1 | 1 | NA | NA | 2 | 0.5 | NA | NA | 1 | 1 | 1 | 0 | NA | NA | 2 | 0.5 |
| 4_86345483 | NA | NA | 1 | 0 | 1 | 1 | NA | NA | 2 | 0.5 | NA | NA | 1 | 0 | 1 | 1 | NA | NA | 2 | 0.5 |
| 4_86345885 | NA | NA | 1 | 0 | 1 | 1 | NA | NA | 2 | 0.5 | NA | NA | 1 | 1 | 1 | 0 | NA | NA | 2 | 0.5 |
| 4_86494331 | 15 | 0.0667 | 23 | 0.087 | 9 | 0 | 17 | 0.1176 | 64 | 0.0781 | 9 | 0 | 16 | 0.125 | 13 | 0.0769 | 11 | 0.0909 | 49 | 0.0816 |
| 4_86494340 | 17 | 0.9412 | 20 | 1 | 11 | 0.8182 | 17 | 1 | 65 | 0.9538 | 11 | 0.9091 | 17 | 0.8824 | 14 | 1 | 16 | 0.875 | 58 | 0.9138 |
| 4_86494355 | 18 | 0.2222 | 22 | 0.0909 | 11 | 0.0909 | 18 | 0.1667 | 69 | 0.1449 | 14 | 0.1429 | 19 | 0.2632 | 14 | 0.1429 | 15 | 0.1333 | 62 | 0.1774 |
| 4_86494391 | 16 | 0.25 | 23 | 0.2174 | 12 | 0.0833 | 16 | 0.125 | 67 | 0.1791 | 13 | 0.1538 | 17 | 0.0588 | 13 | 0.1538 | 15 | 0 | 58 | 0.0862 |
| 4_86494449 | 16 | 0.875 | 20 | 0.95 | 11 | 0.9091 | 17 | 0.8235 | 64 | 0.8906 | 13 | 0.8462 | 16 | 0.75 | 13 | 0.7692 | 14 | 0.7857 | 56 | 0.7857 |
| 4_86494724 | 16 | 0.4375 | 28 | 0.5357 | 13 | 0.4615 | 16 | 0.4375 | 73 | 0.4795 | 13 | 0.6923 | 16 | 0.5625 | 13 | 0.6154 | 16 | 0.5 | 58 | 0.5862 |
| 4_86494961 | 12 | 0 | 19 | 0 | 8 | 0.125 | 14 | 0.2857 | 53 | 0.0943 | 8 | 0.125 | 10 | 0 | 13 | 0.0769 | 10 | 0.1 | 41 | 0.0732 |
| 4_86495040 | 18 | 0.1667 | 24 | 0.2917 | 18 | 0.0556 | 18 | 0.1667 | 78 | 0.1795 | 15 | 0.3333 | 18 | 0.2222 | 15 | 0.2 | 16 | 0.3125 | 64 | 0.2656 |
| 4_86500219 | NA | NA | 2 | 0.5 | NA | NA | NA | NA | 2 | 0.5 | NA | NA | 1 | 0 | NA | NA | 2 | 1 | 3 | 0.6667 |
| 4_86501208 | NA | NA | 2 | 0.5 | 1 | 1 | NA | NA | 3 | 0.6667 | NA | NA | 1 | 1 | NA | NA | 1 | 1 | 2 | 1 |
| 4_86501276 | NA | NA | 2 | 0.5 | 1 | 1 | NA | NA | 3 | 0.6667 | NA | NA | 1 | 0 | NA | NA | 2 | 1 | 3 | 0.6667 |
| 4_86501329 | NA | NA | 2 | 1 | 1 | 1 | NA | NA | 3 | 1 | NA | NA | 1 | 1 | NA | NA | 1 | 0 | 2 | 0.5 |
| 4_86501699 | NA | NA | 1 | 1 | NA | NA | NA | NA | 1 | 1 | NA | NA | 1 | 1 | NA | NA | 2 | 1 | 3 | 1 |
| 4_86743439 | NA | NA | 1 | 1 | 2 | 0.5 | 1 | 1 | 4 | 0.75 | NA | NA | 1 | 1 | NA | NA | 1 | 0 | 2 | 0.5 |
| 4_86743484 | NA | NA | 1 | 1 | 1 | 1 | 1 | 1 | 3 | 1 | NA | NA | 2 | 1 | NA | NA | 1 | 0 | 3 | 0.6667 |

| | | | | | | | | | | | | | | | | | | | | |
|---|---|---|---|---|---|---|---|---|---|---|---|---|---|---|---|---|---|---|---|---|
| 4_8674359 | NA | NA | 1 | 1 | 2 | 0.5 | 1 | 1 | 4 | 0.75 | NA | NA | 2 | 0 | NA | NA | 1 | 1 | 3 | 0.3333 |
| 4_8674375 | NA | NA | 1 | 0 | 3 | 1 | 1 | 1 | 5 | 0.8 | NA | NA | 2 | 0.5 | NA | NA | 1 | 1 | 3 | 0.6667 |
| 4_8674386 | NA | NA | NA | NA | 2 | 0.5 | NA | NA | 2 | 0.5 | NA | NA | 1 | 1 | NA | NA | 1 | 1 | 2 | 1 |
| 4_8674402 | NA | NA | 1 | 0 | 3 | 0.3333 | 1 | 0 | 5 | 0.2 | NA | NA | 1 | 0 | NA | NA | 1 | 1 | 2 | 0.5 |
| 4_8674406 | NA | NA | NA | NA | 2 | 0.5 | 1 | 0 | 3 | 0.3333 | NA | NA | 1 | 0 | NA | NA | 1 | 1 | 2 | 0.5 |
| 4_8674419 | NA | NA | 1 | 1 | 3 | 0.6667 | 1 | 0 | 5 | 0.6 | NA | NA | 2 | 0.5 | NA | NA | 1 | 1 | 3 | 0.6667 |
| 4_8676702 | NA | NA | 2 | 0.5 | 4 | 0.25 | 2 | 0.5 | 8 | 0.375 | NA | NA | 4 | 0.75 | 2 | 0 | 2 | 0.5 | 8 | 0.5 |
| 4_8676719 | NA | NA | 2 | 1 | 4 | 0.75 | 1 | 1 | 7 | 0.8571 | NA | NA | 3 | 0.3333 | 2 | 0.5 | 2 | 1 | 7 | 0.5714 |
| 4_8676724 | NA | NA | 2 | 0 | 4 | 0.75 | 2 | 0.5 | 8 | 0.5 | NA | NA | 3 | 1 | 2 | 1 | 2 | 1 | 7 | 1 |
| 4_8676727 | NA | NA | 2 | 1 | 4 | 1 | 2 | 1 | 8 | 1 | NA | NA | 3 | 1 | NA | NA | 2 | 1 | 5 | 1 |
| 4_8676734 | NA | NA | 2 | 0 | 4 | 0.25 | 2 | 0 | 8 | 0.125 | NA | NA | 3 | 0 | 2 | 0.5 | 2 | 0 | 7 | 0.1429 |
| 4_8676750 | NA | NA | 2 | 0 | 4 | 0.25 | 2 | 0.5 | 8 | 0.25 | NA | NA | 4 | 0.25 | 1 | 1 | 1 | 0 | 6 | 0.3333 |
| 4_8677282 | 4 | 0.25 | 4 | 0 | 2 | 0 | 6 | 0.3333 | 16 | 0.1875 | 6 | 0.3333 | 4 | 0.75 | 3 | 0.3333 | 3 | 0.6667 | 16 | 0.5 |
| 4_8685034 | 1 | 1 | 2 | 0 | 1 | 0 | 1 | 1 | 5 | 0.4 | 2 | 0.5 | 2 | 0 | 2 | 0.5 | 4 | 0 | 10 | 0.2 |
| 4_8685052 | 1 | 1 | 4 | 1 | 1 | 1 | 1 | 1 | 7 | 1 | 2 | 1 | 2 | 1 | 1 | 1 | 5 | 1 | 10 | 1 |
| 4_8685055 | 1 | 0 | 3 | 0.3333 | 1 | 0 | NA | NA | 5 | 0.2 | 1 | 0 | 2 | 0 | 2 | 0 | 3 | 0.6667 | 8 | 0.25 |
| 4_8685057 | 1 | 0 | 3 | 0.3333 | 1 | 0 | 1 | 0 | 6 | 0.1667 | 2 | 0 | 2 | 0.5 | 2 | 0.5 | 5 | 0.8 | 11 | 0.5455 |
| 4_8685071 | 1 | 0 | 3 | 0.3333 | 1 | 0 | 1 | 0 | 6 | 0.1667 | 2 | 0 | 2 | 0 | 2 | 0 | 5 | 0.4 | 11 | 0.1818 |
| 4_8685071 | NA | NA | 3 | 0.6667 | 1 | 1 | 1 | 1 | 5 | 0.8 | 2 | 1 | 2 | 1 | 2 | 1 | 5 | 0.8 | 11 | 0.9091 |
| 4_8685085 | 1 | 0 | 3 | 0.6667 | 1 | 0 | 1 | 0 | 6 | 0.3333 | 2 | 0 | 2 | 1 | 1 | 1 | 4 | 1 | 9 | 0.7778 |
| 4_8690278 | 18 | 0.1111 | 29 | 0.3448 | 23 | 0.087 | 12 | 0.3333 | 82 | 0.2195 | 13 | 0.3077 | 14 | 0.2143 | 18 | 0.1111 | 22 | 0.2273 | 67 | 0.209 |
| 4_8690334 | 18 | 0.1111 | 26 | 0.0769 | 17 | 0.1765 | 11 | 0.3636 | 72 | 0.1528 | 9 | 0.1111 | 12 | 0.1667 | 13 | 0.1538 | 19 | 0.2105 | 53 | 0.1698 |
| 4_8690376 | 18 | 0.0556 | 24 | 0.25 | 16 | 0.1875 | 9 | 0 | 67 | 0.1493 | 9 | 0.1111 | 11 | 0.0909 | 13 | 0.1538 | 17 | 0.0588 | 50 | 0.1 |
| 4_8690401 | 17 | 0.1176 | 21 | 0.0476 | 16 | 0.0625 | 7 | 0.1429 | 61 | 0.082 | 8 | 0 | 12 | 0.1667 | 13 | 0.2308 | 17 | 0 | 50 | 0.1 |
| 4_8690457 | 15 | 0.0667 | 16 | 0.125 | 13 | 0.2308 | 6 | 0.1667 | 50 | 0.14 | 6 | 0.5 | 12 | 0.3333 | 12 | 0.25 | 15 | 0.1333 | 45 | 0.2667 |
| 4_8690460 | 11 | 0.0909 | 13 | 0 | 14 | 0.3571 | 6 | 0.1667 | 44 | 0.1591 | 6 | 0.6667 | 13 | 0.1538 | 12 | 0.4167 | 15 | 0.2 | 46 | 0.3043 |
| 4_8690465 | 15 | 0.1333 | 14 | 0.2857 | 12 | 0.25 | 6 | 0.1667 | 47 | 0.2128 | 6 | 0.8333 | 11 | 0.5455 | 11 | 0.4545 | 16 | 0.25 | 44 | 0.4545 |
| 4_8690470 | 10 | 0.1 | 10 | 0.1 | 9 | 0.1111 | 6 | 0 | 35 | 0.0857 | 5 | 0 | 9 | 0.1111 | 6 | 0.1667 | 14 | 0 | 34 | 0.0588 |
| 4_8690731 | 3 | 0.3333 | 5 | 0 | 5 | 0 | 2 | 0 | 15 | 0.0667 | NA | NA | 5 | 0.2 | 5 | 0 | 3 | 0 | 13 | 0.0769 |
| 4_8690750 | 3 | 0.6667 | 4 | 0 | 4 | 0.5 | 2 | 0 | 13 | 0.3077 | NA | NA | 2 | 0 | 4 | 0.25 | 1 | 0 | 7 | 0.1429 |
| 4_8690783 | 3 | 0 | 5 | 0.2 | 4 | 0 | 1 | 0 | 13 | 0.0769 | NA | NA | 3 | 0 | 4 | 0.25 | 2 | 0 | 9 | 0.1111 |
| 4_8690802 | 2 | 0.5 | 5 | 0 | 4 | 0.25 | 2 | 0.5 | 13 | 0.2308 | NA | NA | 4 | 0.25 | 4 | 0.25 | 2 | 0 | 10 | 0.2 |
| 4_8690929 | 2 | 0.5 | 5 | 0.8 | 3 | 0.6667 | 1 | 1 | 11 | 0.7273 | NA | NA | 4 | 0.25 | 3 | 0.3333 | 4 | 0.5 | 11 | 0.3636 |
| 4_8690936 | 6 | 0.3333 | 6 | 0.5 | 2 | 0 | 1 | 1 | 15 | 0.4 | NA | NA | 3 | 0.3333 | 4 | 0.5 | 4 | 0.5 | 11 | 0.4545 |
| 4_8690938 | 5 | 0.8 | 5 | 0.6 | 2 | 0.5 | 1 | 1 | 13 | 0.6923 | NA | NA | 1 | 1 | 2 | 0.5 | 2 | 0.5 | 5 | 0.6 |
| 4_8690942 | 2 | 1 | 5 | 1 | 3 | 1 | 1 | 1 | 11 | 1 | NA | NA | 4 | 1 | 4 | 1 | 3 | 1 | 11 | 1 |
| 4_8690961 | 6 | 0 | 7 | 0.1429 | 3 | 0.3333 | 1 | 0 | 17 | 0.1176 | NA | NA | 3 | 0.3333 | 4 | 0 | 4 | 0 | 11 | 0.0909 |
| 4_8690977 | 6 | 0.6667 | 7 | 0.7143 | 1 | 1 | 1 | 0 | 15 | 0.6667 | NA | NA | 3 | 1 | 4 | 0.75 | 3 | 1 | 10 | 0.9 |
| 4_8714548 | 5 | 1 | 3 | 1 | NA | NA | 1 | 0 | 9 | 0.8889 | NA | NA | 1 | 1 | 1 | 1 | 1 | 1 | 3 | 1 |
| 4_8714554 | 2 | 0 | 3 | 1 | NA | NA | NA | NA | 5 | 0.6 | NA | NA | 1 | 1 | 2 | 1 | NA | NA | 3 | 1 |
| 4_8714556 | 4 | 0.25 | 3 | 0 | 1 | 0 | 1 | 0 | 9 | 0.1111 | NA | NA | 1 | 0 | 2 | 0.5 | 2 | 1 | 5 | 0.6 |
| 4_8714568 | 6 | 0.5 | 2 | 0.5 | 1 | 0 | 1 | 0 | 10 | 0.4 | NA | NA | 1 | 0 | 1 | 0 | 3 | 0.6667 | 5 | 0.4 |
| 4_8714571 | 5 | 0.2 | 3 | 0 | NA | NA | NA | NA | 8 | 0.125 | NA | NA | NA | NA | 1 | 1 | 3 | 0 | 4 | 0.25 |
| 4_8714578 | 4 | 0 | 2 | 0.5 | NA | NA | 1 | 0 | 7 | 0.1429 | NA | NA | 1 | 0 | 1 | 1 | 3 | 0.3333 | 5 | 0.4 |
| 4_8714579 | 3 | 0.6667 | 2 | 1 | NA | NA | 1 | 0 | 6 | 0.6667 | NA | NA | 1 | 1 | 1 | 1 | 2 | 1 | 4 | 1 |
| 4_8714586 | 3 | 1 | 2 | 0 | 1 | 0 | 1 | 1 | 7 | 0.5714 | NA | NA | 1 | 0 | 1 | 1 | 1 | 1 | 3 | 0.6667 |
| 4_8714586 | 4 | 0 | 3 | 0 | 1 | 1 | 1 | 0 | 9 | 0.1111 | NA | NA | 1 | 0 | 1 | 0 | 3 | 0.3333 | 5 | 0.2 |
| 4_8714595 | 2 | 1 | 2 | 1 | 1 | 1 | 1 | 0 | 6 | 0.8333 | NA | NA | NA | NA | NA | NA | 2 | 0.5 | 2 | 0.5 |
| 4_8768873 | 6 | 0.6667 | NA | NA | 1 | 0 | 3 | 1 | 10 | 0.7 | NA | NA | 3 | 1 | 1 | 0 | 1 | 1 | 5 | 0.8 |
| 4_8769082 | 4 | 0.25 | 1 | 0 | NA | NA | 3 | 0 | 8 | 0.125 | NA | NA | 1 | 1 | NA | NA | 1 | 1 | 2 | 1 |
| 4_8769087 | 3 | 1 | 2 | 1 | NA | NA | 3 | 1 | 8 | 1 | NA | NA | 1 | 1 | NA | NA | 1 | 1 | 2 | 1 |
| 4_8769109 | 5 | 0.4 | 2 | 0 | 1 | 1 | 5 | 0 | 13 | 0.2308 | 2 | 0.5 | 1 | 1 | NA | NA | 4 | 0.25 | 7 | 0.4286 |
| 4_8769121 | 5 | 0.6 | 2 | 0 | 1 | 1 | 5 | 0.8 | 13 | 0.6154 | 3 | 0.6667 | 1 | 0 | NA | NA | 4 | 0.75 | 8 | 0.625 |
| 4_8769136 | 5 | 0.8 | 2 | 0 | NA | NA | 5 | 0.4 | 12 | 0.5 | 2 | 0.5 | 1 | 0 | NA | NA | 2 | 0.5 | 5 | 0.4 |
| 4_8769148 | 5 | 0.2 | 3 | 1 | NA | NA | 5 | 0.6 | 13 | 0.5385 | 3 | 0.3333 | NA | NA | 1 | 0 | 4 | 0.5 | 8 | 0.375 |
| 4_8769155 | 5 | 0.8 | 3 | 0 | NA | NA | 5 | 0.4 | 13 | 0.4615 | 3 | 1 | 1 | 1 | NA | NA | 3 | 0.3333 | 7 | 0.7143 |
| 4_8769166 | 6 | 0.6667 | 2 | 0.5 | NA | NA | 6 | 0.6667 | 14 | 0.6429 | 4 | 0.5 | 1 | 0 | 1 | 0 | 3 | 1 | 9 | 0.5556 |
| 4_8769175 | 6 | 0.6667 | 3 | 0.3333 | NA | NA | 5 | 0.2 | 14 | 0.4286 | 4 | 0.25 | 1 | 0 | 1 | 1 | 4 | 0.5 | 10 | 0.4 |
| 4_8769188 | 2 | 1 | NA | NA | NA | NA | 3 | 0 | 5 | 0.4 | 3 | 0.3333 | NA | NA | NA | NA | 1 | 1 | 4 | 0.5 |
| 4_8769204 | 2 | 1 | 3 | 1 | NA | NA | 6 | 0.8333 | 11 | 0.9091 | 5 | 0.8 | NA | NA | 1 | 1 | 3 | 1 | 9 | 0.8889 |
| 4_8769217 | 1 | 0 | 2 | 0 | NA | NA | 4 | 0.25 | 7 | 0.1429 | 5 | 0.2 | 3 | 0 | 2 | 0 | 3 | 0 | 13 | 0.0769 |
| 4_8769222 | 3 | 0.3333 | NA | NA | NA | NA | 1 | 1 | 4 | 0.5 | 1 | 0 | 3 | 0.3333 | 1 | 1 | 1 | 1 | 6 | 0.5 |
| 4_8769230 | 3 | 0 | 1 | 0 | NA | NA | 6 | 0.1667 | 10 | 0.1 | 2 | 1 | 3 | 0.3333 | 2 | 0.5 | 2 | 0.5 | 9 | 0.5556 |
| 4_8769235 | 3 | 0.6667 | 2 | 0 | NA | NA | 3 | 0.6667 | 8 | 0.5 | 2 | 0.5 | 1 | 0 | 2 | 1 | 1 | 1 | 6 | 0.6667 |
| 4_8775916 | 1 | 0 | 1 | 0 | NA | NA | 2 | 0.5 | 4 | 0.25 | 1 | 1 | 1 | 1 | 2 | 1 | 2 | 0 | 6 | 0.6667 |

| | | | | | | | | | | | | | | | | | | | | |
|---|---|---|---|---|---|---|---|---|---|---|---|---|---|---|---|---|---|---|---|---|
| 4_8775941 | 1 | 1 | 1 | 1 | NA | NA | 1 | 0 | 3 | 0.6667 | 1 | 0 | 1 | 1 | 2 | 0.5 | 1 | 1 | 5 | 0.6 |
| 4_8832890 | 7 | 0.1429 | 5 | 0 | 7 | 0.1429 | 4 | 0 | 23 | 0.087 | 3 | 0 | 2 | 0.5 | 6 | 0 | 2 | 0 | 13 | 0.0769 |
| 4_8832899 | 5 | 0.8 | 5 | 0.8 | 7 | 1 | 4 | 0.5 | 21 | 0.8095 | 3 | 0.6667 | 2 | 1 | 6 | 0.8333 | 1 | 0 | 12 | 0.75 |
| 4_8832912 | 7 | 0.1429 | 6 | 0.3333 | 7 | 0.1429 | 4 | 0.5 | 24 | 0.25 | 2 | 0 | 1 | 0 | 8 | 0.375 | 2 | 0.5 | 13 | 0.3077 |
| 4_8832913 | 6 | 0.1667 | 3 | 0 | 6 | 0 | 5 | 0.2 | 20 | 0.1 | 2 | 0 | 2 | 0.5 | 7 | 0 | 2 | 0 | 13 | 0.0769 |
| 4_8832913 | 7 | 0.1429 | 6 | 0.1667 | 7 | 0 | 6 | 0 | 26 | 0.0769 | 3 | 0 | 1 | 0 | 8 | 0.125 | 2 | 0 | 14 | 0.0714 |
| 4_8832915 | 5 | 0.6 | 6 | 1 | 6 | 0.6667 | 5 | 0.6 | 22 | 0.7273 | 3 | 0 | 2 | 0.5 | 5 | 0.4 | 2 | 1 | 12 | 0.4167 |
| 4_8863689 | 21 | 0.0952 | 23 | 0.087 | 20 | 0.15 | 13 | 0.0769 | 77 | 0.1039 | 11 | 0 | 17 | 0.1765 | 19 | 0.0526 | 22 | 0.0909 | 69 | 0.087 |
| 4_8863844 | 8 | 0 | 14 | 0.2143 | 9 | 0.2222 | 8 | 0.25 | 39 | 0.1795 | 9 | 0.2222 | 11 | 0.2727 | 10 | 0.1 | 16 | 0.0625 | 46 | 0.1522 |
| 4_8863853 | 9 | 0.1111 | 10 | 0.1 | 13 | 0.1538 | 10 | 0.1 | 42 | 0.119 | 9 | 0 | 13 | 0.0769 | 13 | 0.0769 | 14 | 0.1429 | 49 | 0.0816 |
| 4_8863860 | 9 | 0 | 8 | 0.25 | 13 | 0.0769 | 7 | 0.1429 | 37 | 0.1081 | 10 | 0 | 12 | 0.0833 | 10 | 0.2 | 13 | 0 | 45 | 0.0667 |
| 4_8863866 | 10 | 1 | 11 | 1 | 12 | 1 | 9 | 1 | 42 | 1 | 9 | 1 | 12 | 1 | 14 | 1 | 14 | 1 | 49 | 1 |
| 4_8863872 | 9 | 0.1111 | 7 | 0.4286 | 11 | 0.0909 | 7 | 0.5714 | 34 | 0.2647 | 9 | 0.1111 | 9 | 0.3333 | 11 | 0 | 11 | 0.1818 | 40 | 0.15 |
| 4_8863880 | 11 | 0.1818 | 11 | 0.0909 | 10 | 0 | 9 | 0.2222 | 41 | 0.122 | 10 | 0.1 | 12 | 0 | 12 | 0.0833 | 13 | 0.1538 | 47 | 0.0851 |
| 4_8863882 | 9 | 0.8889 | 11 | 1 | 11 | 0.9091 | 9 | 0.8889 | 40 | 0.925 | 8 | 0.75 | 11 | 1 | 11 | 1 | 12 | 0.9167 | 42 | 0.9286 |
| 4_8863882 | 11 | 0.0909 | 10 | 0.3 | 12 | 0.25 | 10 | 0.5 | 43 | 0.2791 | 10 | 0.1 | 11 | 0.0909 | 12 | 0.1667 | 14 | 0.1429 | 47 | 0.1277 |
| 4_8863887 | 11 | 0.2727 | 10 | 0.2 | 12 | 0.0833 | 9 | 0.2222 | 42 | 0.1905 | 9 | 0.5556 | 10 | 0 | 13 | 0.1538 | 12 | 0.25 | 44 | 0.2273 |
| 4_8863891 | 7 | 1 | 9 | 1 | 10 | 1 | 8 | 0.5 | 34 | 0.8824 | 8 | 0.75 | 9 | 0.8889 | 9 | 1 | 10 | 0.9 | 36 | 0.8889 |
| 4_8863900 | 9 | 0.7778 | 9 | 0.4444 | 10 | 0.9 | 10 | 0.9 | 38 | 0.7632 | 8 | 0.875 | 8 | 0.375 | 11 | 0.8182 | 11 | 0.5455 | 38 | 0.6579 |
| 4_8863904 | 9 | 1 | 9 | 0.8889 | 11 | 1 | 8 | 0.875 | 37 | 0.9459 | 9 | 1 | 11 | 0.9091 | 12 | 1 | 14 | 1 | 46 | 0.9783 |
| 4_8863936 | 9 | 0.6667 | 8 | 0.375 | 9 | 0.2222 | 8 | 0.5 | 34 | 0.4412 | 5 | 0.6 | 7 | 0.5714 | 9 | 0.4444 | 12 | 0.3333 | 33 | 0.4545 |
| 4_8863937 | 9 | 1 | 8 | 0.375 | 9 | 0.3333 | 8 | 0.25 | 34 | 0.5 | 5 | 0.8 | 11 | 0.6364 | 9 | 1 | 10 | 0.1 | 35 | 0.6 |
| 4_8863945 | 10 | 0.3 | 8 | 0.5 | 8 | 0.25 | 8 | 0.125 | 34 | 0.2941 | 8 | 0.5 | 11 | 0.2727 | 9 | 0.4444 | 11 | 0.4545 | 39 | 0.4103 |
| 4_8863945 | 8 | 0.375 | 9 | 0 | 10 | 0.3 | 6 | 0.5 | 33 | 0.2727 | 7 | 0.1429 | 9 | 0.1111 | 7 | 0.1429 | 12 | 0 | 35 | 0.0857 |
| 4_8863947 | 9 | 0.2222 | 8 | 0 | 9 | 0.5556 | 8 | 0.375 | 34 | 0.2941 | 7 | 0.1429 | 10 | 0.2 | 10 | 0.3 | 9 | 0.1111 | 36 | 0.1944 |
| 4_8863954 | 9 | 0.8889 | 9 | 1 | 10 | 0.9 | 8 | 0.75 | 36 | 0.8889 | 8 | 0.875 | 10 | 1 | 11 | 1 | 12 | 0.9167 | 41 | 0.9512 |
| 4_8863960 | 8 | 0.5 | 8 | 0.375 | 9 | 0.4444 | 9 | 0.6667 | 34 | 0.5 | 7 | 0.4286 | 9 | 0.2222 | 8 | 0.25 | 12 | 0.5 | 36 | 0.3611 |
| 4_8863970 | 8 | 1 | 8 | 0.875 | 10 | 0.8 | 7 | 1 | 33 | 0.9091 | 8 | 0.875 | 9 | 0.8889 | 9 | 0.7778 | 10 | 0.9 | 36 | 0.8611 |
| 4_8863978 | 6 | 0.5 | 6 | 0.1667 | 6 | 0.1667 | 7 | 0.2857 | 25 | 0.28 | 3 | 0.6667 | 6 | 0.1667 | 7 | 0.7143 | 4 | 0.25 | 20 | 0.45 |
| 4_8863986 | 7 | 0.8571 | 8 | 0.75 | 7 | 0.4286 | 7 | 0.5714 | 29 | 0.6552 | 8 | 0.875 | 9 | 0.7778 | 10 | 0.7 | 11 | 0.6364 | 38 | 0.7368 |
| 4_8863988 | 8 | 1 | 5 | 1 | 9 | 0.8889 | 7 | 1 | 29 | 0.9655 | 8 | 1 | 8 | 1 | 9 | 1 | 11 | 1 | 36 | 1 |
| 4_8863989 | 5 | 0.6 | 5 | 0.8 | 6 | 0.5 | 6 | 0.3333 | 22 | 0.5455 | 6 | 0.6667 | 8 | 0.625 | 10 | 0.6 | 7 | 0.5714 | 31 | 0.6129 |
| 4_8863989 | NA | NA | NA | NA | 1 | 1 | NA | NA | 1 | 1 | NA | NA | NA | NA | NA | NA | 1 | 1 | 1 | 1 |
| 4_8864017 | 6 | 0.3333 | 7 | 0 | 7 | 0.2857 | 9 | 0.3333 | 29 | 0.2414 | 7 | 0.2857 | 9 | 0.1111 | 8 | 0.25 | 10 | 0.2 | 34 | 0.2059 |
| 4_8909785 | 2 | 1 | 2 | 1 | 1 | 0 | NA | NA | 5 | 0.8 | NA | NA | 4 | 0.75 | 3 | 0.6667 | 2 | 1 | 9 | 0.7778 |
| 4_8909789 | 3 | 0.3333 | 2 | 0 | 1 | 0 | NA | NA | 6 | 0.1667 | NA | NA | 5 | 0.2 | 3 | 0.6667 | 2 | 0 | 10 | 0.3 |
| 4_8909796 | 3 | 0.3333 | 2 | 0 | NA | NA | NA | NA | 5 | 0.2 | NA | NA | 5 | 0.6 | 3 | 0.6667 | 1 | 1 | 9 | 0.6667 |
| 4_8909803 | 3 | 1 | 2 | 1 | 1 | 1 | NA | NA | 6 | 1 | NA | NA | 5 | 1 | 3 | 1 | 2 | 1 | 10 | 1 |
| 4_8909805 | 3 | 0.6667 | 3 | 0.3333 | NA | NA | NA | NA | 6 | 0.5 | NA | NA | 5 | 0.2 | 3 | 0.3333 | 2 | 0.5 | 10 | 0.3 |
| 4_8909817 | 3 | 0.3333 | 2 | 0 | 1 | 0 | NA | NA | 6 | 0.1667 | NA | NA | 5 | 0.6 | 2 | 1 | 2 | 0 | 9 | 0.5556 |
| 4_8909833 | 2 | 1 | 3 | 1 | 1 | 1 | NA | NA | 6 | 1 | NA | NA | 4 | 1 | 2 | 1 | 1 | 1 | 7 | 1 |
| 4_8909834 | 2 | 1 | 3 | 1 | 1 | 0 | NA | NA | 6 | 0.8333 | NA | NA | 3 | 0.3333 | 1 | 1 | 1 | 1 | 5 | 0.6 |
| 4_8909838 | 2 | 0 | 3 | 0.3333 | 1 | 0 | NA | NA | 6 | 0.1667 | NA | NA | 4 | 0 | 2 | 1 | 2 | 0 | 8 | 0.25 |
| 4_8909902 | 1 | 0 | 3 | 0.6667 | NA | NA | NA | NA | 4 | 0.5 | NA | NA | 2 | 0 | 2 | 0.5 | 2 | 0 | 6 | 0.1667 |
| 4_8922468 | NA | NA | 1 | 1 | 1 | 1 | NA | NA | 2 | 1 | NA | NA | NA | NA | 1 | 1 | NA | NA | 1 | 1 |
| 4_8922490 | NA | NA | 1 | 1 | NA | NA | NA | NA | 1 | 1 | NA | NA | NA | NA | 1 | 1 | NA | NA | 1 | 1 |
| 4_8922496 | NA | NA | 1 | 1 | NA | NA | NA | NA | 1 | 1 | NA | NA | NA | NA | 1 | 1 | NA | NA | 1 | 1 |
| 4_8922526 | NA | NA | 1 | 1 | NA | NA | NA | NA | 1 | 1 | NA | NA | NA | NA | 1 | 1 | NA | NA | 1 | 1 |
| 4_9113965 | 7 | 0 | 8 | 0.125 | 13 | 0 | 11 | 0.0909 | 39 | 0.0513 | 4 | 0 | 4 | 0 | 6 | 0 | 6 | 0.1667 | 20 | 0.05 |
| 4_9114003 | 7 | 0 | 7 | 0 | 14 | 0.1429 | 12 | 0.0833 | 40 | 0.075 | 3 | 0.3333 | 2 | 0 | 5 | 0.2 | 6 | 0.1667 | 16 | 0.1875 |
| 4_9114120 | 5 | 0.2 | 6 | 0 | 10 | 0 | 9 | 0.1111 | 30 | 0.0667 | 3 | 0.3333 | 2 | 0 | 3 | 0 | 4 | 0.25 | 12 | 0.1667 |
| 4_9114127 | 5 | 0 | 6 | 0.1667 | 10 | 0.2 | 10 | 0 | 31 | 0.0968 | 3 | 0 | 2 | 0 | 3 | 0 | 3 | 0.3333 | 11 | 0.0909 |
| 4_9114141 | 5 | 0 | 7 | 0.1429 | 11 | 0.0909 | 10 | 0.1 | 33 | 0.0909 | 3 | 0 | 2 | 0.5 | 3 | 0 | 4 | 0 | 12 | 0.0833 |
| 4_9114150 | 5 | 0.2 | 7 | 0 | 11 | 0.2727 | 7 | 0.2857 | 30 | 0.2 | 3 | 0.3333 | 1 | 0 | 2 | 0 | 4 | 0.5 | 10 | 0.3 |
| 4_9114158 | 4 | 0.5 | 5 | 0.8 | 9 | 0.5556 | 5 | 0.4 | 23 | 0.5652 | 3 | 1 | 1 | 1 | 3 | 0.3333 | 3 | 0.6667 | 10 | 0.7 |
| 4_9114159 | 5 | 0.4 | 5 | 0.6 | 9 | 0.6667 | 9 | 0.4444 | 28 | 0.5357 | 3 | 0.3333 | 1 | 1 | 3 | 0.3333 | 4 | 1 | 11 | 0.6364 |
| 4_9114162 | 5 | 0.2 | 6 | 0.3333 | 11 | 0.3636 | 7 | 0.2857 | 29 | 0.3103 | 3 | 0.6667 | 1 | 0 | 3 | 0.6667 | 4 | 0.5 | 11 | 0.5455 |
| 4_9126434 | 2 | 0.5 | 2 | 0.5 | 2 | 0.5 | 1 | 0 | 7 | 0.4286 | NA | NA | 1 | 0 | 2 | 1 | 1 | 1 | 4 | 0.75 |
| 4_9322261 | NA | NA | 4 | 0.75 | 3 | 0.6667 | 2 | 0.5 | 9 | 0.6667 | 1 | 0 | 1 | 0 | 5 | 0.8 | 5 | 0.4 | 12 | 0.5 |
| 4_9322271 | 1 | 0 | 4 | 0.5 | 3 | 0 | 2 | 0.5 | 10 | 0.3 | 1 | 0 | 2 | 0.5 | 4 | 0.25 | 5 | 0.8 | 12 | 0.5 |
| 4_9322276 | NA | NA | 2 | 0 | NA | NA | 2 | 0.5 | 4 | 0.25 | NA | NA | 1 | 1 | 1 | 1 | 1 | 1 | 3 | 1 |
| 4_9322278 | 1 | 0 | 4 | 0.5 | 2 | 0 | 2 | 0 | 9 | 0.2222 | 1 | 0 | 3 | 0.3333 | 5 | 0 | 4 | 0 | 13 | 0.0769 |
| 4_9322293 | 1 | 0 | 2 | 0.5 | 2 | 0 | 1 | 0 | 6 | 0.1667 | 1 | 0 | 3 | 0.3333 | 3 | 0.6667 | 2 | 0 | 9 | 0.3333 |
| 4_9322309 | 1 | 1 | 3 | 0.6667 | 2 | 1 | NA | NA | 6 | 0.8333 | NA | NA | 1 | 1 | 3 | 0.3333 | 1 | 1 | 5 | 0.6 |
| 4_9438932 | NA | NA | NA | NA | 2 | 1 | 2 | 1 | 4 | 1 | NA | NA | 1 | 1 | NA | NA | 2 | 0.5 | 3 | 0.6667 |

| | | | | | | | | | | | | | | | | | | | | |
|---|---|---|---|---|---|---|---|---|---|---|---|---|---|---|---|---|---|---|---|---|
| 4_9438942 | NA | NA | NA | NA | 2 | 1 | 2 | 1 | 4 | 1 | NA | NA | 1 | 1 | NA | NA | 2 | 1 | 3 | 1 |
| 4_9443962 | NA | NA | NA | NA | 2 | 0 | 1 | 1 | 3 | 0.3333 | NA | NA | 2 | 0.5 | 1 | 0 | 2 | 0 | 5 | 0.2 |
| 4_9445593 | 3 | 0 | 3 | 0 | NA | NA | 2 | 0.5 | 8 | 0.125 | 2 | 0.5 | 2 | 0 | 3 | 0 | NA | NA | 7 | 0.1429 |
| 4_9445608 | 3 | 0.3333 | 3 | 0.6667 | NA | NA | 1 | 0 | 7 | 0.4286 | 1 | 1 | 3 | 1 | 3 | 1 | NA | NA | 7 | 1 |
| 4_9445772 | 5 | 0.8 | 4 | 0.5 | 3 | 0.3333 | 6 | 0.8333 | 18 | 0.6667 | 7 | 0.4286 | 5 | 0.4 | 5 | 0.8 | 4 | 0.25 | 21 | 0.4762 |
| 4_9445782 | 5 | 0 | 3 | 0 | 4 | 0.25 | 5 | 0 | 17 | 0.0588 | 7 | 0.1429 | 4 | 0 | 5 | 0.4 | 4 | 0 | 20 | 0.15 |
| 4_9445784 | 5 | 0.2 | 2 | 0.5 | 3 | 0.6667 | 5 | 0.2 | 15 | 0.3333 | 7 | 0.4286 | 4 | 0.25 | 4 | 0.25 | 3 | 0.3333 | 18 | 0.3333 |
| 4_9445785 | 5 | 0 | 3 | 0 | 4 | 0 | 6 | 0.1667 | 18 | 0.0556 | 7 | 0 | 4 | 0 | 5 | 0.2 | 4 | 0 | 20 | 0.05 |
| 4_9445788 | 3 | 0.6667 | 2 | 0 | 4 | 0.25 | 6 | 0.1667 | 15 | 0.2667 | 6 | 0.1667 | 3 | 0 | 4 | 0.75 | 4 | 0.5 | 17 | 0.3529 |
| 4_9445790 | 3 | 0 | 3 | 0.3333 | 2 | 0.5 | 4 | 0.75 | 12 | 0.4167 | 7 | 1 | 3 | 0 | 4 | 0.75 | 4 | 0.25 | 18 | 0.6111 |
| 4_9445795 | 5 | 0.2 | 3 | 0 | 3 | 0 | 5 | 0 | 16 | 0.0625 | 7 | 0 | 4 | 0 | 4 | 0 | 4 | 0.25 | 19 | 0.0526 |
| 4_9445798 | 5 | 1 | 3 | 1 | 3 | 1 | 6 | 1 | 17 | 1 | 6 | 1 | 4 | 1 | 4 | 1 | 4 | 1 | 18 | 1 |
| 4_9476241 | 2 | 0 | 1 | 1 | 1 | 0 | 2 | 0 | 6 | 0.1667 | NA | NA | NA | NA | 1 | 1 | 1 | 1 | 2 | 1 |
| 4_9482434 | 2 | 0 | 2 | 0 | 3 | 1 | NA | NA | 7 | 0.4286 | 1 | 1 | NA | NA | NA | NA | NA | NA | 1 | 1 |
| 4_9483491 | 1 | 1 | NA | NA | 3 | 0.3333 | 1 | 0 | 5 | 0.4 | NA | NA | NA | NA | NA | NA | 1 | 1 | 1 | 1 |
| 4_9483524 | 1 | 0 | NA | NA | 2 | 0.5 | 1 | 0 | 4 | 0.25 | NA | NA | 1 | 1 | NA | NA | NA | NA | 1 | 1 |
| 4_9493804 | 6 | 0.5 | 8 | 0.25 | 3 | 0 | 6 | 0 | 23 | 0.2174 | 5 | 0 | 4 | 0 | NA | NA | 2 | 0.5 | 11 | 0.0909 |
| 4_9493805 | 6 | 0 | 6 | 0 | 3 | 0.6667 | 7 | 0 | 22 | 0.0909 | 4 | 0.25 | 4 | 0 | NA | NA | 2 | 0 | 10 | 0.1 |
| 4_9493813 | 7 | 0.7143 | 7 | 0.8571 | 3 | 1 | 7 | 1 | 24 | 0.875 | 5 | 0.6 | 4 | 0.75 | NA | NA | 2 | 1 | 11 | 0.7273 |
| 4_9493819 | 7 | 0.4286 | 6 | 0.5 | 2 | 1 | 6 | 1 | 21 | 0.6667 | 4 | 1 | 4 | 1 | NA | NA | 2 | 1 | 10 | 1 |
| 4_9493820 | 7 | 0.8571 | 5 | 0.4 | 2 | 1 | 6 | 0.6667 | 20 | 0.7 | 4 | 0.5 | 3 | 0 | NA | NA | 2 | 1 | 9 | 0.4444 |
| 4_9493823 | 7 | 1 | 6 | 1 | 2 | 1 | 6 | 1 | 21 | 1 | 3 | 1 | 4 | 1 | NA | NA | 2 | 1 | 9 | 1 |
| 4_9493827 | 5 | 0.8 | 5 | 1 | 2 | 1 | 5 | 0.8 | 17 | 0.8824 | 4 | 1 | 4 | 1 | NA | NA | 2 | 1 | 10 | 1 |
| 4_9493837 | 6 | 1 | 4 | 1 | 2 | 1 | 4 | 1 | 16 | 1 | 2 | 1 | 4 | 0.75 | NA | NA | 1 | 1 | 7 | 0.8571 |
| 4_9493843 | 5 | 0.6 | 5 | 0.2 | 1 | 0 | 5 | 0.2 | 16 | 0.3125 | 2 | 0.5 | 4 | 0.75 | NA | NA | 1 | 1 | 7 | 0.7143 |
| 4_9493844 | 7 | 0.8571 | 5 | 0.6 | 1 | 1 | 4 | 1 | 17 | 0.8235 | 2 | 1 | 4 | 0.75 | NA | NA | 1 | 0 | 7 | 0.7143 |
| 4_9493849 | 4 | 1 | 4 | 1 | NA | NA | 5 | 1 | 13 | 1 | 2 | 0.5 | 3 | 1 | NA | NA | 1 | 1 | 6 | 0.8333 |
| 4_9493855 | 3 | 0 | 4 | 0.5 | 1 | 0 | 6 | 0.3333 | 14 | 0.2857 | 2 | 1 | 3 | 0 | NA | NA | 1 | 1 | 6 | 0.5 |
| 4_9493859 | 4 | 0 | 3 | 0.3333 | 1 | 0 | 4 | 0.25 | 12 | 0.1667 | 1 | 0 | 2 | 0.5 | NA | NA | 1 | 0 | 4 | 0.25 |
| 4_9493914 | 3 | 1 | 2 | 0.5 | NA | NA | 4 | 1 | 9 | 0.8889 | 1 | 0 | 2 | 0.5 | NA | NA | 1 | 0 | 4 | 0.25 |
| 4_9493951 | 2 | 0 | 3 | 0.3333 | NA | NA | 5 | 0.2 | 10 | 0.2 | 2 | 0 | 4 | 0.25 | NA | NA | 1 | 0 | 7 | 0.1429 |
| 4_9493962 | 3 | 0.6667 | 2 | 1 | NA | NA | 2 | 1 | 7 | 0.8571 | 1 | 0 | 3 | 0.6667 | NA | NA | NA | NA | 4 | 0.5 |
| 4_9493968 | 2 | 0 | 3 | 0 | NA | NA | 3 | 0.3333 | 8 | 0.125 | 1 | 0 | 3 | 0.6667 | NA | NA | 1 | 1 | 5 | 0.6 |
| 4_9493982 | 2 | 0.5 | 3 | 0.3333 | NA | NA | 4 | 0.75 | 9 | 0.5556 | 2 | 0.5 | 3 | 0 | NA | NA | NA | NA | 5 | 0.2 |
| 4_9494004 | 2 | 1 | 3 | 0 | NA | NA | 2 | 0.5 | 7 | 0.4286 | 2 | 1 | 3 | 0.6667 | NA | NA | 1 | 1 | 6 | 0.8333 |
| 4_9494008 | 2 | 1 | 3 | 0.3333 | NA | NA | 2 | 0 | 7 | 0.4286 | 2 | 0.5 | 4 | 0.25 | NA | NA | 1 | 1 | 7 | 0.4286 |
| 4_9494032 | 2 | 0 | 3 | 0 | NA | NA | 3 | 1 | 8 | 0.375 | 2 | 0 | 4 | 0.5 | NA | NA | NA | NA | 6 | 0.3333 |
| 4_9494033 | 2 | 0 | 3 | 0.3333 | NA | NA | 3 | 0 | 8 | 0.125 | 2 | 0.5 | 4 | 0 | NA | NA | 1 | 1 | 7 | 0.2857 |
| 4_9584871 | NA | NA | 2 | 1 | NA | NA | NA | NA | 2 | 1 | 1 | 1 | NA | NA | NA | NA | NA | NA | 1 | 1 |
| 4_9591046 | 2 | 0 | 1 | 0 | 4 | 0 | 3 | 0.3333 | 10 | 0.1 | 3 | 0 | 1 | 0 | 3 | 0.3333 | 2 | 0 | 9 | 0.1111 |
| 4_9591062 | 1 | 1 | NA | NA | 4 | 0 | 2 | 0 | 7 | 0.1429 | 2 | 0 | 1 | 0 | 3 | 0.3333 | 2 | 0 | 8 | 0.125 |
| 4_9591071 | 1 | 1 | 1 | 1 | 1 | 1 | 1 | 1 | 4 | 1 | 3 | 0 | NA | NA | 2 | 1 | 2 | 1 | 7 | 0.5714 |
| 4_9591139 | 2 | 0 | 1 | 0 | 4 | 0.25 | 3 | 0 | 10 | 0.1 | 3 | 0 | 1 | 1 | 3 | 0.3333 | 3 | 0.6667 | 10 | 0.4 |
| 4_9591247 | 3 | 0.3333 | 2 | 0 | 5 | 0 | 2 | 0 | 12 | 0.0833 | 4 | 0 | 1 | 0 | 3 | 0.3333 | 4 | 0 | 12 | 0.0833 |
| 4_9591331 | 6 | 0.1667 | 2 | 0 | 7 | 0.2857 | 2 | 0 | 17 | 0.1765 | 4 | 0.25 | 2 | 0 | 3 | 0.3333 | 4 | 0.25 | 13 | 0.2308 |
| 4_9602928 | NA | NA | 1 | 1 | NA | NA | NA | NA | 1 | 1 | 1 | 1 | 1 | 1 | NA | NA | 1 | 1 | 3 | 1 |
| 4_9640455 | 1 | 0 | 2 | 0 | 1 | 1 | 3 | 0 | 7 | 0.1429 | NA | NA | NA | NA | 2 | 0.5 | NA | NA | 2 | 0.5 |
| 4_9640538 | NA | NA | 1 | 0 | 1 | 1 | NA | NA | 2 | 0.5 | NA | NA | NA | NA | 1 | 1 | NA | NA | 1 | 1 |
| 4_9640576 | 1 | 1 | 2 | 0 | 2 | 0.5 | 1 | 1 | 6 | 0.5 | NA | NA | NA | NA | 4 | 0.5 | NA | NA | 4 | 0.5 |
| 4_9640596 | 2 | 0.5 | 4 | 0.25 | 5 | 0 | 3 | 0.6667 | 14 | 0.2857 | 2 | 0 | 2 | 0 | 7 | 0.2857 | 2 | 0 | 13 | 0.1538 |
| 4_9644163 | 2 | 0.5 | 2 | 0.5 | 3 | 0.6667 | 3 | 0.6667 | 10 | 0.6 | NA | NA | 1 | 0 | 3 | 0.3333 | 1 | 0 | 5 | 0.2 |
| 4_9645677 | 7 | 0.4286 | 4 | 0.5 | 5 | 0.2 | 3 | 0.3333 | 19 | 0.3684 | 6 | 0 | 3 | 0.6667 | 4 | 0.25 | NA | NA | 13 | 0.2308 |
| 4_9645684 | 5 | 0.4 | 4 | 0.25 | 2 | 0 | 2 | 1 | 13 | 0.3846 | 4 | 0.5 | 3 | 0.3333 | 3 | 0 | NA | NA | 10 | 0.3 |
| 4_9645685 | 7 | 0.7143 | 4 | 0.75 | 5 | 0.6 | 3 | 1 | 19 | 0.7368 | 5 | 0.2 | 3 | 0.6667 | 3 | 0.3333 | NA | NA | 11 | 0.3636 |
| 4_9645687 | 7 | 0 | 4 | 0.5 | 5 | 0.2 | 3 | 0 | 19 | 0.1579 | 5 | 0.2 | 3 | 0 | 3 | 0 | NA | NA | 11 | 0.0909 |
| 4_9646582 | NA | NA | NA | NA | NA | NA | 1 | 1 | 1 | 1 | NA | NA | NA | NA | NA | NA | 1 | 1 | 1 | 1 |
| 4_9745680 | 1 | 1 | 2 | 1 | 1 | 1 | 1 | 1 | 5 | 1 | 1 | 1 | 4 | 0.75 | 1 | 1 | 2 | 1 | 8 | 0.875 |
| 4_9745682 | 1 | 1 | 2 | 1 | 1 | 1 | 2 | 0.5 | 6 | 0.8333 | 2 | 0.5 | 5 | 0.8 | 1 | 0 | 2 | 1 | 10 | 0.7 |
| 4_9745690 | 1 | 1 | 1 | 1 | 1 | 0 | NA | NA | 3 | 0.6667 | 1 | 1 | 4 | 0.5 | 1 | 1 | NA | NA | 6 | 0.6667 |
| 4_9745697 | 1 | 1 | 2 | 1 | 1 | 1 | 2 | 1 | 6 | 1 | 2 | 1 | 5 | 0.8 | NA | NA | 2 | 0.5 | 9 | 0.7778 |
| 4_9745699 | NA | NA | 2 | 0.5 | 1 | 0 | 2 | 0.5 | 5 | 0.4 | 2 | 0.5 | 5 | 0.2 | 1 | 0 | 1 | 0 | 9 | 0.2222 |
| 4_9745706 | 1 | 0 | 2 | 0 | 1 | 0 | 2 | 0.5 | 6 | 0.1667 | 2 | 0 | 5 | 0.4 | 1 | 1 | 2 | 0 | 10 | 0.3 |
| 4_9767146 | NA | NA | NA | NA | 2 | 0.5 | NA | NA | 2 | 0.5 | 1 | 1 | NA | NA | NA | NA | NA | NA | 1 | 1 |
| 4_9767168 | NA | NA | NA | NA | 2 | 0.5 | NA | NA | 2 | 0.5 | 2 | 0 | NA | NA | 1 | 1 | 1 | 0 | 4 | 0.25 |
| 4_9799972 | NA | NA | 1 | 0 | 1 | 1 | NA | NA | 2 | 0.5 | 3 | 1 | NA | NA | 1 | 1 | 3 | 0.6667 | 7 | 0.8571 |

| | | | | | | | | | | | | | | | | | | | | |
|---|---|---|---|---|---|---|---|---|---|---|---|---|---|---|---|---|---|---|---|---|
| 4_9799984 | NA | NA | 1 | 1 | 3 | 0.6667 | NA | NA | 4 | 0.75 | 3 | 0.6667 | NA | NA | 1 | 0 | 3 | 0.6667 | 7 | 0.5714 |
| 4_9799998 | NA | NA | 1 | 0 | 3 | 0.3333 | NA | NA | 4 | 0.25 | 2 | 0.5 | NA | NA | 1 | 0 | 3 | 0.3333 | 6 | 0.3333 |
| 4_9800014 | NA | NA | 1 | 0 | 2 | 0.5 | NA | NA | 3 | 0.3333 | 3 | 0.6667 | 2 | 0.5 | 1 | 0 | 3 | 0.3333 | 9 | 0.4444 |
| 4_9800553 | 2 | 0 | 3 | 0.3333 | 1 | 0 | 3 | 0 | 9 | 0.1111 | 2 | 0.5 | 1 | 0 | NA | NA | 3 | 0.3333 | 6 | 0.3333 |
| 4_9800570 | 3 | 0 | 5 | 0.2 | 1 | 0 | 3 | 0 | 12 | 0.0833 | 3 | 0 | 3 | 0.6667 | 4 | 0.25 | 3 | 0.6667 | 13 | 0.3846 |
| 4_9800609 | 3 | 0.3333 | 4 | 0.5 | 2 | 0.5 | 3 | 0 | 12 | 0.3333 | 3 | 0 | 2 | 0.5 | 5 | 0.6 | 4 | 0.5 | 14 | 0.4286 |
| 4_9800611 | 4 | 0 | 5 | 0.2 | 2 | 0 | 2 | 0 | 13 | 0.0769 | 4 | 0.25 | 3 | 0 | 5 | 0.2 | 4 | 0.25 | 16 | 0.1875 |
| 4_9860626 | 2 | 0.5 | 2 | 1 | NA | NA | 1 | 1 | 5 | 0.8 | 2 | 1 | NA | NA | 3 | 1 | 1 | 1 | 6 | 1 |
| 4_9860627 | 2 | 0 | 2 | 0.5 | NA | NA | 2 | 0.5 | 6 | 0.3333 | 2 | 0 | NA | NA | 4 | 0.25 | 2 | 1 | 8 | 0.375 |
| 4_9860632 | 2 | 0 | 1 | 1 | NA | NA | 2 | 0 | 5 | 0.2 | 2 | 0.5 | NA | NA | 3 | 0 | 1 | 0 | 6 | 0.1667 |
| 4_9860636 | 2 | 0.5 | 1 | 1 | NA | NA | 1 | 1 | 4 | 0.75 | 2 | 0.5 | NA | NA | 2 | 1 | 2 | 0.5 | 6 | 0.6667 |
| 4_9860640 | 1 | 0 | 1 | 0 | NA | NA | 2 | 0.5 | 4 | 0.25 | 1 | 1 | NA | NA | 4 | 0.5 | 1 | 1 | 6 | 0.6667 |
| 4_9860659 | 1 | 1 | 2 | 0 | NA | NA | 1 | 0 | 4 | 0.25 | 2 | 0 | NA | NA | 2 | 0 | 1 | 1 | 5 | 0.2 |
| 4_9860661 | 1 | 1 | 2 | 1 | NA | NA | 1 | 0 | 4 | 0.75 | 2 | 1 | NA | NA | 2 | 1 | 1 | 1 | 5 | 1 |
| 4_9860674 | 1 | 1 | 2 | 1 | NA | NA | 1 | 0 | 4 | 0.75 | 2 | 1 | NA | NA | 2 | 0.5 | 1 | 0 | 5 | 0.6 |
| 4_9860756 | 1 | 0 | 1 | 1 | NA | NA | 1 | 0 | 3 | 0.3333 | 2 | 0.5 | NA | NA | 3 | 0 | 1 | 0 | 6 | 0.1667 |
| 4_9881943 | 3 | 0.3333 | NA | NA | 1 | 0 | 2 | 0.5 | 6 | 0.3333 | 1 | 0 | 2 | 0.5 | 3 | 0.3333 | 4 | 0.5 | 10 | 0.4 |
| 4_9881964 | 3 | 0 | NA | NA | 1 | 0 | 3 | 0.3333 | 7 | 0.1429 | 1 | 0 | 1 | 0 | 3 | 0 | 4 | 0.25 | 9 | 0.1111 |
| 4_9882232 | 3 | 0.3333 | NA | NA | 1 | 0 | 4 | 0 | 8 | 0.125 | 1 | 0 | 2 | 0 | 3 | 0.3333 | 4 | 0 | 10 | 0.1 |
| 4_9882236 | 3 | 0 | NA | NA | 1 | 0 | 5 | 0.2 | 9 | 0.1111 | 1 | 1 | 2 | 0 | 3 | 0 | 4 | 0 | 10 | 0.1 |
| 4_9882248 | 3 | 0.3333 | NA | NA | NA | NA | 5 | 0.2 | 8 | 0.25 | 1 | 0 | 2 | 0.5 | 3 | 0 | 5 | 0.2 | 11 | 0.1818 |
| 4_9882254 | 3 | 0.3333 | NA | NA | NA | NA | 2 | 0 | 5 | 0.2 | 2 | 0.5 | 2 | 1 | 2 | 1 | 5 | 0.4 | 11 | 0.6364 |
| 4_9882268 | 4 | 1 | 1 | 1 | NA | NA | 5 | 0.8 | 10 | 0.9 | 2 | 1 | 2 | 1 | 3 | 1 | 4 | 1 | 11 | 1 |
| 4_9882269 | 4 | 0.75 | 1 | 0 | 1 | 1 | 4 | 0.75 | 10 | 0.7 | 2 | 0.5 | 2 | 0.5 | 3 | 0.3333 | 4 | 0.5 | 11 | 0.4545 |
| 4_9882275 | 4 | 0.5 | 1 | 0 | 1 | 0 | 5 | 0 | 11 | 0.1818 | 2 | 0.5 | 2 | 0.5 | 3 | 0.3333 | 5 | 0 | 12 | 0.25 |
| 4_9882278 | 4 | 0.25 | 1 | 0 | 1 | 0 | 5 | 0.2 | 11 | 0.1818 | 2 | 0 | 2 | 0 | 3 | 0 | 5 | 0.4 | 12 | 0.1667 |
| 4_9882280 | 3 | 0.6667 | NA | NA | 1 | 0 | 4 | 0.5 | 8 | 0.5 | 2 | 1 | 2 | 1 | 2 | 0.5 | 4 | 0.5 | 10 | 0.7 |
| 4_9882286 | 4 | 1 | 1 | 1 | 1 | 1 | 5 | 1 | 11 | 1 | 2 | 1 | 1 | 0 | 3 | 1 | 5 | 1 | 11 | 0.9091 |
| 4_9882296 | 4 | 0.5 | 1 | 1 | 1 | 1 | 5 | 0.8 | 11 | 0.7273 | 2 | 1 | 1 | 1 | 3 | 1 | 5 | 0.8 | 11 | 0.9091 |
| 4_9882308 | 3 | 0.3333 | 2 | 0.5 | 2 | 0.5 | 5 | 0.2 | 12 | 0.3333 | 2 | 0.5 | NA | NA | 2 | 0.5 | 4 | 0.25 | 8 | 0.375 |
| 4_9882542 | 6 | 0.3333 | 1 | 1 | 1 | 0 | 3 | 0.6667 | 11 | 0.4545 | 2 | 0 | 2 | 0.5 | 1 | 0 | 3 | 0.6667 | 8 | 0.375 |
| 4_9900887 | 2 | 0.5 | NA | NA | 1 | 0 | NA | NA | 3 | 0.3333 | 1 | 0 | NA | NA | NA | NA | 2 | 0.5 | 3 | 0.3333 |
| 4_9900946 | 2 | 0.5 | NA | NA | 2 | 0 | NA | NA | 4 | 0.25 | 2 | 0 | NA | NA | 1 | 0 | 3 | 0.6667 | 6 | 0.3333 |
| 4_9900955 | 1 | 1 | NA | NA | 1 | 1 | NA | NA | 2 | 1 | 2 | 1 | NA | NA | 1 | 1 | 2 | 1 | 5 | 1 |
| 4_9900987 | 2 | 1 | NA | NA | 2 | 0.5 | NA | NA | 4 | 0.75 | 1 | 1 | NA | NA | 1 | 1 | 1 | 1 | 3 | 1 |
| 4_9901661 | NA | NA | NA | NA | 1 | 1 | NA | NA | 1 | 1 | NA | NA | 1 | 1 | NA | NA | NA | NA | 1 | 1 |
| 4_9901780 | NA | NA | NA | NA | 1 | 1 | NA | NA | 1 | 1 | NA | NA | NA | NA | 1 | 1 | NA | NA | 1 | 1 |
| 4_9901788 | NA | NA | NA | NA | 1 | 1 | NA | NA | 1 | 1 | NA | NA | 1 | 0 | 1 | 1 | NA | NA | 2 | 0.5 |
| 4_9901789 | NA | NA | NA | NA | 1 | 1 | NA | NA | 1 | 1 | NA | NA | 1 | 1 | 1 | 1 | NA | NA | 2 | 1 |
| 4_9901794 | NA | NA | NA | NA | 1 | 1 | NA | NA | 1 | 1 | NA | NA | NA | NA | 1 | 1 | NA | NA | 1 | 1 |
| 4_9901801 | NA | NA | NA | NA | 1 | 1 | NA | NA | 1 | 1 | NA | NA | 1 | 1 | 1 | 1 | NA | NA | 2 | 1 |
| 4_9908244 | NA | NA | 4 | 0.25 | 2 | 0 | 2 | 0 | 8 | 0.125 | 1 | 0 | 2 | 0.5 | 2 | 0 | NA | NA | 5 | 0.2 |
| 4_9908268 | 2 | 0.5 | 3 | 0.3333 | 3 | 0.3333 | 3 | 0 | 11 | 0.2727 | 2 | 0 | 1 | 1 | 1 | 1 | NA | NA | 4 | 0.5 |
| 4_9910948 | 2 | 0.5 | 5 | 0.6 | 3 | 0.3333 | 3 | 0.6667 | 13 | 0.5385 | 2 | 1 | 2 | 0.5 | 2 | 1 | NA | NA | 6 | 0.8333 |
| 4_9910953 | 2 | 0 | 4 | 0.25 | 5 | 0.2 | 3 | 0.3333 | 14 | 0.2143 | 2 | 0.5 | 2 | 0 | 2 | 0 | NA | NA | 6 | 0.1667 |
| 4_9910955 | 2 | 0 | 5 | 0.2 | 6 | 0 | 3 | 0 | 16 | 0.0625 | 2 | 0 | 2 | 0 | 2 | 0.5 | 1 | 0 | 7 | 0.1429 |
| 4_9910973 | 2 | 0.5 | 5 | 0.2 | 4 | 0.25 | 3 | 0 | 14 | 0.2143 | 2 | 0.5 | 2 | 0 | 2 | 0 | 1 | 0 | 7 | 0.1429 |
| 4_9914669 | 2 | 1 | 1 | 1 | 7 | 0.7143 | 3 | 0.6667 | 13 | 0.7692 | 4 | 0.75 | 3 | 1 | NA | NA | 3 | 1 | 10 | 0.9 |
| 4_9914681 | 3 | 0.3333 | 5 | 0.2 | 7 | 0 | 3 | 0 | 18 | 0.1111 | 4 | 0 | 3 | 0 | 1 | 1 | 2 | 0.5 | 10 | 0.2 |
| 4_9914681 | 1 | 0 | 5 | 0.8 | 7 | 0.7143 | 3 | 0.3333 | 16 | 0.625 | 4 | 0.75 | 1 | 1 | 2 | 1 | 3 | 0.6667 | 10 | 0.8 |
| 4_9914684 | 3 | 0.3333 | 2 | 0.5 | 3 | 0.3333 | 1 | 1 | 9 | 0.4444 | 2 | 0 | 3 | 0 | 1 | 0 | 2 | 0.5 | 8 | 0.125 |
| 4_9914688 | 2 | 1 | 5 | 0.8 | 7 | 0.7143 | 3 | 0.6667 | 17 | 0.7647 | 4 | 0.75 | 2 | 0.5 | 2 | 0.5 | 3 | 0.6667 | 11 | 0.6364 |
| 4_9914693 | 2 | 1 | 2 | 0.5 | 3 | 0.6667 | NA | NA | 7 | 0.7143 | 1 | 1 | 1 | 1 | 1 | 1 | 1 | 1 | 4 | 1 |
| 4_9914696 | 3 | 1 | 2 | 0.5 | 6 | 1 | 3 | 1 | 14 | 0.9286 | 4 | 1 | 3 | 0.6667 | 2 | 1 | 3 | 1 | 12 | 0.9167 |
| 4_9914704 | 3 | 0.6667 | 5 | 0.4 | 7 | 0.5714 | 3 | 0 | 18 | 0.4444 | 3 | 1 | 3 | 0.3333 | 2 | 0.5 | 4 | 0.5 | 12 | 0.5833 |
| 4_9914719 | 3 | 0.3333 | 6 | 0.3333 | 6 | 0.1667 | 3 | 0 | 18 | 0.2222 | 4 | 0 | 3 | 0 | 2 | 0.5 | 4 | 0 | 13 | 0.0769 |
| 4_9914724 | 2 | 1 | 5 | 1 | 7 | 1 | 2 | 1 | 16 | 1 | 4 | 1 | 3 | 1 | 1 | 1 | 1 | 1 | 9 | 1 |
| 4_9914726 | 3 | 0 | 5 | 0.2 | 6 | 0 | 2 | 0 | 16 | 0.0625 | 3 | 0 | 3 | 0 | NA | NA | 4 | 0.25 | 10 | 0.1 |
| 4_9914727 | 3 | 0.6667 | 6 | 0.5 | 6 | 0.6667 | 2 | 0.5 | 17 | 0.5882 | 4 | 0.5 | 3 | 1 | 2 | 1 | 4 | 1 | 13 | 0.8462 |
| 4_9914758 | 3 | 0 | 5 | 0.2 | 5 | 0 | 4 | 0 | 17 | 0.0588 | 3 | 0 | 3 | 0 | 2 | 0 | 4 | 0.25 | 12 | 0.0833 |
| 4_9914773 | 5 | 0.2 | 8 | 0.125 | 7 | 0 | 7 | 0 | 27 | 0.0741 | 4 | 0 | 2 | 0 | 3 | 0.3333 | 4 | 0 | 13 | 0.0769 |
| 4_9965572 | NA | NA | 1 | 0 | NA | NA | 1 | 1 | 2 | 0.5 | 1 | 1 | NA | NA | NA | NA | NA | NA | 1 | 1 |
| 4_9978800 | NA | NA | NA | NA | NA | NA | 1 | 1 | 1 | 1 | 1 | 1 | NA | NA | NA | NA | 2 | 0.5 | 3 | 0.6667 |
| 4_9978805 | NA | NA | NA | NA | NA | NA | 1 | 1 | 1 | 1 | 1 | 1 | NA | NA | NA | NA | 2 | 1 | 3 | 1 |
| 4_9982193 | 5 | 0.2 | 4 | 0.25 | 11 | 0 | 7 | 0 | 27 | 0.0741 | 7 | 0.1429 | 9 | 0.1111 | 8 | 0 | 9 | 0 | 33 | 0.0606 |

| | | | | | | | | | | | | | | | | | | | | |
|---|---|---|---|---|---|---|---|---|---|---|---|---|---|---|---|---|---|---|---|---|
| 4_9984399 | 7 | 0.2857 | 6 | 0.1667 | 16 | 0 | 10 | 0 | 39 | 0.0769 | 10 | 0.1 | 12 | 0.0833 | 12 | 0.0833 | 11 | 0.0909 | 45 | 0.0889 |
| 4_9984402 | 8 | 0 | 6 | 0.1667 | 16 | 0.0625 | 12 | 0.1667 | 42 | 0.0952 | 11 | 0.2727 | 12 | 0.0833 | 13 | 0.1538 | 13 | 0.1538 | 49 | 0.1633 |
| 4_9984406 | 9 | 0.3333 | 6 | 1 | 15 | 0.6 | 12 | 0.75 | 42 | 0.6429 | 12 | 0.6667 | 14 | 0.5714 | 15 | 0.6 | 14 | 0.7857 | 55 | 0.6545 |
| 4_1008604 | NA | NA | 2 | 1 | NA | NA | 3 | 1 | 5 | 1 | NA | NA | NA | NA | NA | NA | 1 | 1 | 1 | 1 |
| 4_1008605 | NA | NA | 2 | 1 | NA | NA | 2 | 0.5 | 4 | 0.75 | NA | NA | NA | NA | NA | NA | 1 | 1 | 1 | 1 |
| 4_1008606 | NA | NA | 3 | 1 | NA | NA | 3 | 1 | 6 | 1 | NA | NA | 1 | 1 | NA | NA | 2 | 1 | 3 | 1 |
| 4_1008606 | NA | NA | 2 | 1 | NA | NA | 3 | 0.3333 | 5 | 0.6 | NA | NA | 1 | 1 | NA | NA | 2 | 1 | 3 | 1 |
| 4_1008607 | NA | NA | 1 | 1 | NA | NA | 2 | 1 | 3 | 1 | NA | NA | NA | NA | NA | NA | 1 | 1 | 1 | 1 |
| 4_1008607 | NA | NA | NA | NA | NA | NA | 1 | 1 | 1 | 1 | NA | NA | NA | NA | NA | NA | 2 | 0.5 | 2 | 0.5 |
| 4_1008608 | NA | NA | 1 | 1 | NA | NA | 3 | 0.6667 | 4 | 0.75 | NA | NA | 1 | 1 | NA | NA | 2 | 0.5 | 3 | 0.6667 |
| 4_1010077 | 2 | 1 | 1 | 0 | NA | NA | 3 | 0.3333 | 6 | 0.5 | 2 | 0 | 3 | 0 | 1 | 0 | 1 | 1 | 7 | 0.1429 |
| 4_1010093 | 4 | 0 | 3 | 0.3333 | NA | NA | 3 | 0.6667 | 10 | 0.3 | 3 | 0.3333 | 4 | 0 | 3 | 0 | NA | NA | 10 | 0.1 |
| 4_1010101 | 10 | 0.1 | 17 | 0.1176 | 14 | 0.0714 | 9 | 0 | 50 | 0.08 | 9 | 0 | 12 | 0.0833 | 12 | 0.1667 | 10 | 0 | 43 | 0.0698 |
| 4_1010102 | 7 | 0.7143 | 15 | 0.3333 | 12 | 0.5 | 8 | 0.5 | 42 | 0.4762 | 7 | 0.4286 | 11 | 0.1818 | 11 | 0.5455 | 11 | 0.2727 | 40 | 0.35 |
| 4_1010103 | 9 | 0.8889 | 17 | 0.5882 | 13 | 0.5385 | 10 | 0.6 | 49 | 0.6327 | 8 | 0.5 | 12 | 0.6667 | 12 | 0.8333 | 11 | 0.4545 | 43 | 0.6279 |
| 4_1010103 | 6 | 0.1667 | 15 | 0.7333 | 12 | 0.3333 | 7 | 0.7143 | 40 | 0.525 | 7 | 0.5714 | 10 | 0.5 | 8 | 0.5 | 7 | 0.1429 | 32 | 0.4375 |
| 4_1010104 | 9 | 0.2222 | 18 | 0.3889 | 12 | 0.0833 | 7 | 0.5714 | 46 | 0.3043 | 7 | 0.1429 | 13 | 0.0769 | 12 | 0.3333 | 11 | 0.0909 | 43 | 0.1628 |
| 4_1010104 | 9 | 0.3333 | 13 | 0.6154 | 10 | 0.3 | 9 | 0.3333 | 41 | 0.4146 | 5 | 0.4 | 8 | 0.25 | 8 | 0.75 | 8 | 0.5 | 29 | 0.4828 |
| 4_1010106 | 6 | 0 | 12 | 0.0833 | 12 | 0 | 7 | 0.1429 | 37 | 0.0541 | 7 | 0 | 10 | 0.1 | 9 | 0 | 10 | 0.1 | 36 | 0.0556 |
| 4_1010285 | 2 | 0 | 3 | 0.3333 | 10 | 0 | 4 | 0 | 19 | 0.0526 | 3 | 0 | 4 | 0.25 | 6 | 0 | 4 | 0 | 17 | 0.0588 |
| 4_1010367 | 2 | 0 | 3 | 0.3333 | 10 | 0 | 3 | 0 | 18 | 0.0556 | 2 | 0 | 4 | 0 | 4 | 0.25 | 4 | 0.25 | 14 | 0.1429 |
| 4_1012038 | NA | NA | NA | NA | 3 | 1 | 2 | 0.5 | 5 | 0.8 | 1 | 1 | 1 | 0 | 3 | 0.6667 | NA | NA | 5 | 0.6 |
| 4_1012039 | NA | NA | NA | NA | 2 | 0.5 | 2 | 0 | 4 | 0.25 | NA | NA | 1 | 1 | 2 | 0 | NA | NA | 3 | 0.3333 |
| 4_1012039 | NA | NA | NA | NA | 3 | 0.3333 | 1 | 1 | 4 | 0.5 | 1 | 0 | 1 | 1 | 2 | 0 | NA | NA | 4 | 0.25 |
| 4_1012777 | NA | NA | 1 | 1 | NA | NA | NA | NA | 1 | 1 | 1 | 1 | NA | NA | NA | NA | NA | NA | 1 | 1 |
| 4_1012777 | NA | NA | 1 | 1 | NA | NA | NA | NA | 1 | 1 | 1 | 1 | NA | NA | NA | NA | NA | NA | 1 | 1 |
| 4_1013211 | 1 | 1 | 2 | 1 | 1 | 1 | 1 | 1 | 5 | 1 | 2 | 0.5 | 1 | 1 | 1 | 1 | NA | NA | 4 | 0.75 |
| 4_1013213 | 1 | 1 | NA | NA | 1 | 1 | NA | NA | 2 | 1 | 2 | 0.5 | 2 | 0 | 1 | 1 | 1 | 0 | 6 | 0.3333 |
| 4_1013229 | 2 | 0 | 2 | 0.5 | 1 | 0 | 1 | 0 | 6 | 0.1667 | 2 | 0 | 2 | 0.5 | NA | NA | 1 | 0 | 5 | 0.2 |
| 4_1013238 | 2 | 0 | 4 | 0.5 | 1 | 1 | 3 | 0.3333 | 10 | 0.4 | 2 | 0 | 3 | 0.3333 | 1 | 0 | 1 | 0 | 7 | 0.1429 |
| 4_1014757 | 8 | 0.125 | 6 | 0.3333 | 6 | 0 | 12 | 0.25 | 32 | 0.1875 | 5 | 0.2 | 9 | 0.4444 | 3 | 0.3333 | 7 | 0.2857 | 24 | 0.3333 |
| 4_1014758 | 8 | 0.5 | 9 | 0.6667 | 8 | 0.75 | 13 | 0.4615 | 38 | 0.5789 | 5 | 0.6 | 11 | 0.5455 | 5 | 0.6 | 11 | 0.4545 | 32 | 0.5312 |
| 4_1014758 | 8 | 0.125 | 10 | 0 | 9 | 0.1111 | 13 | 0 | 40 | 0.05 | 6 | 0 | 9 | 0.3333 | 3 | 0 | 10 | 0.1 | 28 | 0.1429 |
| 4_1014976 | 12 | 0.3333 | 9 | 0.2222 | 10 | 0.5 | 11 | 0.1818 | 42 | 0.3095 | 10 | 0.5 | 10 | 0.3 | 6 | 0 | 12 | 0.25 | 38 | 0.2895 |
| 4_1014978 | 13 | 0.2308 | 17 | 0.2353 | 11 | 0.3636 | 20 | 0.2 | 61 | 0.2459 | 9 | 0.3333 | 15 | 0.2667 | 13 | 0.1538 | 14 | 0.3571 | 51 | 0.2745 |
| 4_1014978 | 13 | 0.6923 | 14 | 0.5714 | 12 | 0.5833 | 22 | 0.5455 | 61 | 0.5902 | 7 | 0.5714 | 16 | 0.5 | 12 | 0.6667 | 11 | 0.9091 | 46 | 0.6522 |
| 4_1014979 | 17 | 0.5882 | 14 | 0.6429 | 11 | 0.3636 | 17 | 0.2941 | 59 | 0.4746 | 10 | 0.6 | 16 | 0.4375 | 10 | 0.8 | 13 | 0.7692 | 49 | 0.6327 |
| 4_1014979 | 19 | 0.0526 | 17 | 0.0588 | 15 | 0.0667 | 21 | 0.0952 | 72 | 0.0694 | 10 | 0.1 | 17 | 0.1176 | 13 | 0.0769 | 16 | 0 | 56 | 0.0714 |
| 4_1014980 | 17 | 0.2941 | 19 | 0.1579 | 16 | 0.125 | 25 | 0.16 | 77 | 0.1818 | 10 | 0.1 | 16 | 0.1875 | 12 | 0.0833 | 12 | 0.25 | 50 | 0.16 |
| 4_1014981 | 19 | 0.8421 | 19 | 0.7895 | 13 | 0.8462 | 22 | 0.6818 | 73 | 0.7808 | 8 | 0.75 | 16 | 0.6875 | 12 | 1 | 17 | 0.7647 | 53 | 0.7925 |
| 4_1014984 | 20 | 0.05 | 15 | 0.1333 | 15 | 0.1333 | 23 | 0 | 73 | 0.0685 | 8 | 0.125 | 15 | 0.0667 | 14 | 0.1429 | 18 | 0.1111 | 55 | 0.1091 |
| 4_1014990 | 24 | 0.4167 | 18 | 0.3889 | 16 | 0.375 | 27 | 0.2963 | 85 | 0.3647 | 13 | 0.2308 | 16 | 0.3125 | 17 | 0.2941 | 20 | 0.45 | 66 | 0.3333 |
| 4_1015151 | 3 | 0 | 10 | 0.1 | 2 | 0 | 3 | 0 | 18 | 0.0556 | 2 | 0 | 1 | 0 | 1 | 0 | 5 | 0.2 | 9 | 0.1111 |
| 4_1015153 | 3 | 0 | 12 | 0.1667 | 3 | 0 | 4 | 0 | 22 | 0.0909 | 2 | 0 | 2 | 0.5 | 1 | 0 | 5 | 0 | 10 | 0.1 |
| 4_1015160 | 4 | 0.5 | 12 | 0.1667 | 3 | 0 | 5 | 0 | 24 | 0.1667 | 2 | 0 | 3 | 0.3333 | 3 | 0 | 4 | 0 | 12 | 0.0833 |
| 4_1015161 | 4 | 0 | 8 | 0.125 | 2 | 0 | 5 | 0.2 | 19 | 0.1053 | 3 | 0.3333 | 1 | 0 | 2 | 0 | 3 | 0 | 9 | 0.1111 |
| 4_1015163 | 2 | 1 | 7 | 0.1429 | NA | NA | 5 | 0.8 | 14 | 0.5 | 3 | 0.3333 | 2 | 1 | 1 | 1 | 4 | 0.5 | 10 | 0.6 |
| 4_1015163 | 4 | 0.25 | 14 | 0.1429 | 4 | 0.5 | 7 | 0 | 29 | 0.1724 | 3 | 0 | 1 | 1 | 3 | 0.3333 | 5 | 0 | 12 | 0.1667 |
| 4_1024469 | NA | NA | 4 | 1 | 1 | 1 | 5 | 0.4 | 10 | 0.7 | 1 | 1 | 2 | 1 | 3 | 0.3333 | 2 | 0.5 | 8 | 0.625 |
| 4_1024470 | 2 | 0.5 | 4 | 0.5 | 1 | 0 | 6 | 0.8333 | 13 | 0.6154 | 1 | 1 | 2 | 0.5 | 4 | 0.75 | 4 | 0.75 | 11 | 0.7273 |
| 4_1024472 | 2 | 0.5 | 4 | 0 | 2 | 1 | 5 | 0.2 | 13 | 0.3077 | 1 | 0 | 1 | 0 | 4 | 0.25 | 4 | 0.5 | 10 | 0.3 |
| 4_1024473 | 3 | 0.3333 | 4 | 0.5 | 3 | 1 | 7 | 0.5714 | 17 | 0.5882 | 1 | 1 | 4 | 0.5 | 5 | 0.4 | 5 | 0.6 | 15 | 0.5333 |
| 4_1024622 | 19 | 0.7368 | 14 | 0.7857 | 10 | 0.6 | 22 | 0.8636 | 65 | 0.7692 | 6 | 0.8333 | 17 | 0.5294 | 19 | 0.6842 | 10 | 0.5 | 52 | 0.6154 |
| 4_1024623 | 19 | 0.7895 | 16 | 1 | 12 | 1 | 21 | 0.7619 | 68 | 0.8676 | 8 | 1 | 16 | 0.875 | 22 | 0.8182 | 8 | 0.875 | 54 | 0.8704 |
| 4_1024623 | 18 | 0.9444 | 15 | 0.8667 | 11 | 0.9091 | 20 | 0.8 | 64 | 0.875 | 7 | 0.8571 | 13 | 0.9231 | 16 | 0.75 | 7 | 0.7143 | 43 | 0.814 |
| 4_1024623 | 16 | 0.3125 | 12 | 0.5 | 11 | 0.1818 | 19 | 0.4737 | 58 | 0.3793 | 8 | 0.25 | 15 | 0.4 | 19 | 0.3684 | 10 | 0.1 | 52 | 0.3077 |
| 4_1024624 | 18 | 1 | 15 | 0.9333 | 12 | 1 | 23 | 1 | 68 | 0.9853 | 6 | 1 | 19 | 1 | 20 | 1 | 11 | 1 | 56 | 1 |
| 4_1024624 | 20 | 0.95 | 13 | 1 | 11 | 0.9091 | 21 | 0.9524 | 65 | 0.9538 | 8 | 0.75 | 16 | 0.9375 | 23 | 0.913 | 10 | 1 | 57 | 0.9123 |
| 4_1024625 | 16 | 1 | 13 | 0.9231 | 12 | 0.9167 | 18 | 0.8889 | 59 | 0.9322 | 5 | 1 | 17 | 0.9412 | 12 | 1 | 8 | 0.875 | 42 | 0.9524 |
| 4_1024626 | 17 | 0.5294 | 13 | 0.6923 | 10 | 0.7 | 17 | 0.5882 | 57 | 0.614 | 8 | 0.625 | 18 | 0.7222 | 20 | 0.5 | 6 | 0.6667 | 52 | 0.6154 |
| 4_1024628 | 22 | 0.8182 | 14 | 0.7857 | 11 | 0.7273 | 22 | 0.9091 | 69 | 0.8261 | 8 | 1 | 19 | 0.8421 | 22 | 0.8636 | 12 | 0.9167 | 61 | 0.8852 |
| 4_1024628 | 21 | 0.5714 | 16 | 0.5 | 10 | 0.5 | 20 | 0.6 | 67 | 0.5522 | 8 | 0.75 | 13 | 0.4615 | 21 | 0.6667 | 11 | 0.5455 | 53 | 0.6038 |
| 4_1024628 | 17 | 0.5882 | 13 | 0.5385 | 10 | 0.4 | 23 | 0.6957 | 63 | 0.5873 | 7 | 0.5714 | 15 | 0.8667 | 17 | 0.7059 | 11 | 0.5455 | 50 | 0.7 |
| 4_1024630 | 21 | 0.0952 | 16 | 0.0625 | 15 | 0.1333 | 26 | 0.2308 | 78 | 0.141 | 8 | 0.25 | 18 | 0.2222 | 22 | 0.0909 | 11 | 0 | 59 | 0.1356 |
| 4_1024631 | 19 | 0.1053 | 18 | 0.1111 | 15 | 0 | 25 | 0.08 | 77 | 0.0779 | 8 | 0 | 17 | 0.1765 | 22 | 0.1364 | 11 | 0 | 58 | 0.1034 |

| | | | | | | | | | | | | | | | | | | | | |
|---|---|---|---|---|---|---|---|---|---|---|---|---|---|---|---|---|---|---|---|---|
| 4_1024640 | 13 | 0.3077 | 17 | 0.1176 | 16 | 0.25 | 23 | 0.1304 | 69 | 0.1884 | 12 | 0.0833 | 14 | 0.0714 | 22 | 0.1364 | 9 | 0 | 57 | 0.0877 |
| 4_1024641 | 17 | 0.7059 | 19 | 0.7368 | 18 | 0.7778 | 27 | 0.6296 | 81 | 0.7037 | 17 | 0.3529 | 22 | 0.6364 | 22 | 0.4545 | 13 | 0.6154 | 74 | 0.5135 |
| 4_1024644 | 20 | 0.15 | 21 | 0 | 25 | 0.04 | 34 | 0.1176 | 100 | 0.08 | 18 | 0.1667 | 22 | 0.1364 | 30 | 0.0333 | 14 | 0.1429 | 84 | 0.1071 |
| 4_1026176 | 2 | 0 | 5 | 0 | 4 | 0.25 | 1 | 0 | 12 | 0.0833 | 1 | 0 | 6 | 0 | 7 | 0.1429 | 4 | 0.25 | 18 | 0.1111 |
| 4_1026176 | 1 | 0 | 5 | 0 | 3 | 0.3333 | 1 | 0 | 10 | 0.1 | 1 | 0 | 7 | 0 | 7 | 0.1429 | 4 | 0 | 19 | 0.0526 |
| 4_1027782 | 1 | 1 | 3 | 0 | 5 | 0.2 | 2 | 0 | 11 | 0.1818 | 2 | 0 | 3 | 0.6667 | 7 | 0.2857 | 4 | 0 | 16 | 0.25 |
| 4_1027783 | 1 | 0 | 4 | 0 | 4 | 0.25 | 3 | 0.3333 | 12 | 0.1667 | 1 | 0 | 5 | 0 | 7 | 0.1429 | 5 | 0 | 18 | 0.0556 |
| 4_1027786 | 1 | 1 | 5 | 0 | 4 | 0 | 4 | 0 | 14 | 0.0714 | 2 | 0.5 | 7 | 0.1429 | 7 | 0.1429 | 5 | 0 | 21 | 0.1429 |
| 4_1028016 | NA | NA | NA | NA | 1 | 1 | NA | NA | 1 | 1 | NA | NA | NA | NA | NA | NA | 1 | 1 | 1 | 1 |
| 4_1028018 | NA | NA | NA | NA | 1 | 1 | NA | NA | 1 | 1 | NA | NA | NA | NA | NA | NA | 1 | 1 | 1 | 1 |
| 4_1028273 | 9 | 0.6667 | 14 | 0.5714 | 8 | 0.625 | 13 | 0.4615 | 44 | 0.5682 | 4 | 0.75 | 13 | 0.6154 | 13 | 0.5385 | 5 | 0.6 | 35 | 0.6 |
| 4_1028273 | 9 | 0.3333 | 15 | 0.5333 | 10 | 0.4 | 11 | 0.3636 | 45 | 0.4222 | 4 | 0.5 | 17 | 0.4118 | 13 | 0.5385 | 7 | 0.8571 | 41 | 0.5366 |
| 4_1028273 | 6 | 0.3333 | 13 | 0.1538 | 9 | 0.3333 | 12 | 0.3333 | 40 | 0.275 | 3 | 0 | 16 | 0.1875 | 13 | 0.1538 | 6 | 0.1667 | 38 | 0.1579 |
| 4_1028274 | 9 | 0.1111 | 14 | 0.2143 | 10 | 0.1 | 12 | 0 | 45 | 0.1111 | 4 | 0.25 | 19 | 0.1053 | 14 | 0.0714 | 6 | 0.3333 | 43 | 0.1395 |
| 4_1028329 | 1 | 0 | 5 | 0 | 4 | 0.25 | 3 | 0.3333 | 13 | 0.1538 | NA | NA | 2 | 0 | 2 | 0.5 | 2 | 0 | 6 | 0.1667 |
| 4_1028340 | 3 | 0.3333 | 6 | 0.3333 | 6 | 0 | 6 | 0.1667 | 21 | 0.1905 | 3 | 0 | 5 | 0 | 4 | 0.25 | 5 | 0.4 | 17 | 0.1765 |
| 4_1028342 | 2 | 0 | 8 | 0.25 | 5 | 0 | 5 | 0.4 | 20 | 0.2 | 1 | 0 | 5 | 0 | 4 | 0.25 | 5 | 0 | 15 | 0.0667 |
| 4_1028343 | 3 | 0.6667 | 7 | 0.1429 | 6 | 0.1667 | 6 | 0.5 | 22 | 0.3182 | 3 | 0 | 5 | 0.2 | 4 | 0.25 | 5 | 0.2 | 17 | 0.1765 |
| 4_1028343 | 4 | 0.25 | 7 | 0 | 6 | 0.1667 | 6 | 0.6667 | 23 | 0.2609 | 3 | 0 | 6 | 0 | 5 | 0.4 | 6 | 0 | 20 | 0.1 |
| 4_1028921 | NA | NA | 1 | 0 | 1 | 1 | 1 | 0 | 3 | 0.3333 | NA | NA | NA | NA | NA | NA | 2 | 0.5 | 2 | 0.5 |
| 4_1029662 | NA | NA | NA | NA | 1 | 1 | NA | NA | 1 | 1 | 1 | 1 | NA | NA | NA | NA | 1 | 1 | 2 | 1 |
| 4_1029883 | NA | NA | 1 | 1 | NA | NA | NA | NA | 1 | 1 | NA | NA | NA | NA | NA | NA | 1 | 1 | 1 | 1 |
| 4_1030196 | 2 | 1 | 5 | 0.6 | 3 | 1 | 3 | 0.6667 | 13 | 0.7692 | 2 | 1 | 5 | 1 | 2 | 1 | 1 | 0 | 10 | 0.9 |
| 4_1030197 | 1 | 1 | 4 | 1 | 3 | 1 | 3 | 1 | 11 | 1 | 2 | 1 | 3 | 1 | 2 | 1 | NA | NA | 7 | 1 |
| 4_1030197 | 4 | 0 | 4 | 0 | 3 | 0.3333 | 3 | 0 | 14 | 0.0714 | 1 | 1 | 5 | 0 | 2 | 0 | 1 | 0 | 9 | 0.1111 |
| 4_1030197 | 4 | 0 | 5 | 0.2 | 1 | 0 | 3 | 0 | 13 | 0.0769 | 2 | 0 | 5 | 0.4 | 2 | 0 | 1 | 1 | 10 | 0.3 |
| 4_1030197 | 3 | 0.6667 | 5 | 0.6 | 3 | 0.6667 | 3 | 0.6667 | 14 | 0.6429 | 1 | 0 | 5 | 1 | 3 | 1 | 1 | 1 | 10 | 0.9 |
| 4_1030198 | 3 | 0 | 5 | 0.4 | 3 | 0.6667 | 3 | 0.6667 | 14 | 0.4286 | 2 | 0.5 | 5 | 0.6 | 3 | 0.3333 | NA | NA | 10 | 0.5 |
| 4_1030198 | 3 | 0 | 4 | 0.25 | 3 | 0.3333 | 3 | 0 | 13 | 0.1538 | NA | NA | 5 | 0.2 | 2 | 0 | 1 | 0 | 8 | 0.125 |
| 4_1030201 | 4 | 0 | 5 | 0.2 | 3 | 0.3333 | 3 | 0 | 15 | 0.1333 | 2 | 0 | 6 | 0 | 3 | 0.6667 | 1 | 0 | 12 | 0.1667 |
| 4_1030207 | 6 | 0 | 5 | 0 | 2 | 0.5 | 4 | 0 | 17 | 0.0588 | 2 | 0 | 6 | 0.1667 | 3 | 0 | 2 | 0 | 13 | 0.0769 |
| 4_1030279 | 2 | 0 | 1 | 1 | 1 | 0 | 2 | 0 | 6 | 0.1667 | 1 | 0 | 1 | 0 | 1 | 0 | 2 | 0.5 | 5 | 0.2 |
| 4_1030281 | 4 | 0.5 | 1 | 0 | 1 | 0 | 3 | 0.3333 | 9 | 0.3333 | 1 | 0 | 2 | 0 | 1 | 0 | 2 | 1 | 6 | 0.3333 |
| 4_1030294 | 2 | 0.5 | NA | NA | NA | NA | 3 | 0 | 5 | 0.2 | NA | NA | 2 | 0 | 1 | 1 | 1 | 0 | 4 | 0.25 |
| 4_1030297 | 2 | 0 | NA | NA | NA | NA | 3 | 0.6667 | 5 | 0.4 | NA | NA | 2 | 0 | 1 | 0 | 1 | 1 | 4 | 0.25 |
| 4_1030298 | 2 | 0 | NA | NA | NA | NA | 3 | 1 | 5 | 0.6 | NA | NA | 2 | 0.5 | 1 | 1 | 1 | 1 | 4 | 0.75 |
| 4_1030298 | 2 | 0 | NA | NA | NA | NA | 3 | 0.3333 | 5 | 0.2 | NA | NA | 2 | 0 | 1 | 1 | 1 | 0 | 4 | 0.25 |
| 4_1030301 | 1 | 0 | NA | NA | NA | NA | 3 | 0.3333 | 4 | 0.25 | NA | NA | 1 | 0 | 1 | 0 | 1 | 1 | 3 | 0.3333 |
| 4_1030307 | 1 | 1 | 1 | 0 | NA | NA | 2 | 0 | 4 | 0.25 | NA | NA | 2 | 0.5 | 1 | 0 | 2 | 0 | 5 | 0.2 |
| 4_1030307 | 1 | 0 | 1 | 0 | NA | NA | 2 | 1 | 4 | 0.5 | NA | NA | 2 | 1 | 1 | 0 | 2 | 0 | 5 | 0.4 |
| 4_1031763 | NA | NA | NA | NA | 1 | 1 | 2 | 1 | 3 | 1 | 1 | 0 | 2 | 1 | NA | NA | NA | NA | 3 | 0.6667 |
| 4_1031764 | NA | NA | NA | NA | 1 | 0 | 2 | 0.5 | 3 | 0.3333 | 1 | 0 | 2 | 0.5 | NA | NA | 1 | 0 | 4 | 0.25 |
| 4_1031764 | NA | NA | NA | NA | 1 | 1 | 2 | 1 | 3 | 1 | 1 | 1 | 2 | 1 | NA | NA | 1 | 0 | 4 | 0.75 |
| 4_1031764 | NA | NA | NA | NA | NA | NA | 2 | 1 | 2 | 1 | 1 | 1 | 2 | 1 | NA | NA | 1 | 1 | 4 | 1 |
| 4_1031789 | NA | NA | 1 | 1 | NA | NA | NA | NA | 1 | 1 | 1 | 1 | 2 | 0.5 | NA | NA | 1 | 0 | 4 | 0.5 |
| 4_1032233 | 2 | 1 | 2 | 0 | 1 | 1 | 3 | 0 | 8 | 0.375 | 1 | 0 | 2 | 0 | NA | NA | 1 | 1 | 4 | 0.25 |
| 4_1032234 | 2 | 1 | 2 | 0.5 | 1 | 1 | 3 | 0.6667 | 8 | 0.75 | 1 | 1 | 2 | 1 | NA | NA | 1 | 1 | 4 | 1 |
| 4_1034210 | 1 | 1 | 3 | 1 | 2 | 1 | 4 | 0.75 | 10 | 0.9 | 5 | 1 | 1 | 0 | 1 | 0 | 2 | 1 | 9 | 0.7778 |
| 4_1034217 | 3 | 0.6667 | 2 | 0.5 | 2 | 1 | 4 | 1 | 11 | 0.8182 | 4 | 0.5 | NA | NA | NA | NA | 3 | 0.3333 | 7 | 0.4286 |
| 4_1034218 | 4 | 0.5 | 4 | 0.25 | 3 | 0.3333 | 6 | 0 | 17 | 0.2353 | 7 | 0 | 1 | 0 | NA | NA | 4 | 0.25 | 12 | 0.0833 |
| 4_1045849 | 1 | 1 | 6 | 0.8333 | 2 | 0.5 | 7 | 0.8571 | 16 | 0.8125 | 6 | 1 | 2 | 1 | 3 | 1 | 1 | 1 | 12 | 1 |
| 4_1045850 | 4 | 0.5 | 8 | 0.25 | 3 | 0.3333 | 9 | 0.3333 | 24 | 0.3333 | 6 | 0.1667 | 2 | 0.5 | 4 | 0.25 | 1 | 1 | 13 | 0.3077 |
| 4_1045988 | 6 | 0.1667 | 10 | 0.1 | 5 | 0.2 | 11 | 0 | 32 | 0.0938 | 6 | 0 | 3 | 0 | 2 | 0 | 3 | 0.3333 | 14 | 0.0714 |
| 4_1045989 | 4 | 0 | 14 | 0.0714 | 2 | 0 | 10 | 0.1 | 30 | 0.0667 | 6 | 0 | 3 | 0 | 4 | 0.25 | 1 | 0 | 14 | 0.0714 |
| 4_1045989 | 4 | 0.25 | 9 | 0.2222 | 3 | 0 | 9 | 0.1111 | 25 | 0.16 | 7 | 0.1429 | 5 | 0 | 2 | 0 | 3 | 0.3333 | 17 | 0.1176 |
| 4_1045989 | 6 | 0 | 10 | 0.1 | 4 | 0 | 8 | 0.125 | 28 | 0.0714 | 4 | 0.25 | 3 | 0 | 3 | 0 | 2 | 0 | 12 | 0.0833 |
| 4_1045989 | 6 | 0 | 12 | 0.0833 | 6 | 0.1667 | 9 | 0 | 33 | 0.0606 | 6 | 0.1667 | 4 | 0 | 3 | 0 | 1 | 0 | 14 | 0.0714 |
| 4_1045989 | 5 | 0.2 | 10 | 0.2 | 5 | 0.2 | 11 | 0.1818 | 31 | 0.1935 | 7 | 0.1429 | 5 | 0 | 3 | 0 | 3 | 0.3333 | 18 | 0.1111 |
| 4_1045989 | 7 | 0.2857 | 11 | 0.0909 | 4 | 0.25 | 10 | 0.1 | 32 | 0.1562 | 7 | 0.1429 | 5 | 0.4 | 4 | 0.5 | 3 | 0.3333 | 19 | 0.3158 |
| 4_1045990 | 6 | 0.1667 | 9 | 0 | 3 | 0.3333 | 9 | 0.4444 | 27 | 0.2222 | 7 | 0.1429 | 3 | 0 | 4 | 0.25 | 3 | 0 | 17 | 0.1176 |
| 4_1045990 | 7 | 0 | 11 | 0.1818 | 5 | 0 | 8 | 0.125 | 31 | 0.0968 | 7 | 0.1429 | 5 | 0 | 4 | 0 | 3 | 0 | 19 | 0.0526 |
| 4_1045990 | 8 | 0.125 | 13 | 0 | 6 | 0.1667 | 10 | 0.1 | 37 | 0.0811 | 7 | 0 | 4 | 0.25 | 4 | 0 | 2 | 0 | 17 | 0.0588 |
| 4_1045991 | 8 | 0.625 | 11 | 0.6364 | 6 | 0.5 | 8 | 0.25 | 33 | 0.5152 | 9 | 0.4444 | 7 | 0.5714 | 6 | 0.8333 | 2 | 0.5 | 24 | 0.5833 |
| 4_1045991 | 9 | 0 | 13 | 0.0769 | 6 | 0.1667 | 11 | 0.0909 | 39 | 0.0769 | 8 | 0.125 | 8 | 0 | 6 | 0.1667 | 3 | 0 | 25 | 0.08 |
| 4_1045991 | 9 | 0.5556 | 13 | 0.7692 | 6 | 1 | 11 | 0.7273 | 39 | 0.7436 | 9 | 0.7778 | 6 | 0.6667 | 6 | 0.6667 | 3 | 0.6667 | 24 | 0.7083 |

| | | | | | | | | | | | | | | | | | | | | |
|---|---|---|---|---|---|---|---|---|---|---|---|---|---|---|---|---|---|---|---|---|
| 4_1045993 | 8 | 0.5 | 13 | 0.3077 | 5 | 0.8 | 9 | 0.4444 | 35 | 0.4571 | 9 | 0.2222 | 7 | 0.2857 | 4 | 0.25 | 4 | 0.25 | 24 | 0.25 |
| 4_1045994 | 7 | 0.1429 | 11 | 0.1818 | 7 | 0 | 11 | 0.1818 | 36 | 0.1389 | 8 | 0 | 8 | 0 | 5 | 0.2 | 4 | 0.25 | 25 | 0.08 |
| 4_1045995 | 9 | 0.7778 | 14 | 0.7143 | 7 | 0.5714 | 11 | 0.8182 | 41 | 0.7317 | 9 | 0.7778 | 7 | 0.8571 | 6 | 1 | 4 | 0.75 | 26 | 0.8462 |
| 4_1045996 | 8 | 0.875 | 12 | 0.8333 | 5 | 1 | 11 | 1 | 36 | 0.9167 | 8 | 0.875 | 6 | 1 | 6 | 1 | 3 | 1 | 23 | 0.9565 |
| 4_1045998 | 8 | 0 | 11 | 0 | 4 | 0 | 12 | 0.25 | 35 | 0.0857 | 8 | 0.125 | 7 | 0 | 5 | 0 | 5 | 0.2 | 25 | 0.08 |
| 4_1046003 | 6 | 0.3333 | 8 | 0.375 | 6 | 0.1667 | 7 | 0 | 27 | 0.2222 | 8 | 0.125 | 7 | 0.1429 | 8 | 0.375 | 6 | 0 | 29 | 0.1724 |
| 4_1046004 | 8 | 0.125 | 9 | 0.1111 | 5 | 0 | 9 | 0 | 31 | 0.0645 | 8 | 0.375 | 8 | 0 | 6 | 0.3333 | 6 | 0.1667 | 28 | 0.2143 |
| 4_1046012 | 7 | 0 | 12 | 0.0833 | 10 | 0 | 9 | 0.1111 | 38 | 0.0526 | 8 | 0 | 8 | 0 | 11 | 0.0909 | 8 | 0.125 | 35 | 0.0571 |
| 4_1046016 | 10 | 0.2 | 11 | 0.0909 | 10 | 0 | 9 | 0 | 40 | 0.075 | 11 | 0.0909 | 11 | 0 | 11 | 0.0909 | 4 | 0.25 | 37 | 0.0811 |
| 4_1046018 | 11 | 0.0909 | 8 | 0.125 | 10 | 0 | 11 | 0.0909 | 40 | 0.075 | 10 | 0.3 | 9 | 0.1111 | 9 | 0 | 4 | 0.25 | 32 | 0.1562 |
| 4_1046019 | 11 | 0.0909 | 8 | 0.25 | 9 | 0.1111 | 12 | 0.0833 | 40 | 0.125 | 10 | 0.2 | 7 | 0 | 9 | 0.3333 | 6 | 0.1667 | 32 | 0.1875 |
| 4_1048874 | 1 | 0 | 3 | 0.3333 | NA | NA | 3 | 0.3333 | 7 | 0.2857 | 4 | 0.25 | 6 | 0.6667 | 3 | 0.6667 | 4 | 0.25 | 17 | 0.4706 |
| 4_1048927 | NA | NA | 2 | 0.5 | NA | NA | NA | NA | 2 | 0.5 | 2 | 0 | 3 | 0.3333 | 2 | 0 | 2 | 0 | 9 | 0.1111 |
| 4_1050148 | 15 | 0.8 | 9 | 0.6667 | 15 | 0.6667 | 7 | 0.5714 | 46 | 0.6957 | 8 | 0.125 | 11 | 0.5455 | 6 | 0.6667 | 11 | 0.6364 | 36 | 0.5 |
| 4_1050881 | 24 | 0.0417 | 26 | 0.1154 | 24 | 0.25 | 16 | 0.25 | 90 | 0.1556 | 13 | 0.1538 | 20 | 0.1 | 9 | 0.1111 | 22 | 0.0909 | 64 | 0.1094 |
| 4_1050882 | 27 | 0.1481 | 28 | 0.0714 | 29 | 0.1724 | 19 | 0.1579 | 103 | 0.1359 | 17 | 0.1176 | 24 | 0.125 | 18 | 0.1667 | 27 | 0.1111 | 86 | 0.1279 |
| 4_1050883 | 26 | 0.2308 | 26 | 0.1538 | 26 | 0.2692 | 19 | 0.4211 | 97 | 0.2577 | 13 | 0.0769 | 24 | 0.2083 | 15 | 0.2 | 22 | 0.2727 | 74 | 0.2027 |
| 4_1050886 | 37 | 0.0811 | 32 | 0.2812 | 31 | 0.2258 | 22 | 0.2727 | 122 | 0.2049 | 16 | 0.25 | 26 | 0.1538 | 18 | 0.1111 | 29 | 0.1724 | 89 | 0.1685 |
| 4_1050888 | 36 | 0.1389 | 27 | 0.0741 | 27 | 0.1111 | 19 | 0.0526 | 109 | 0.1009 | 16 | 0.0625 | 22 | 0.0455 | 15 | 0 | 23 | 0.087 | 76 | 0.0526 |
| 4_1050889 | 38 | 0.0526 | 36 | 0.0556 | 35 | 0.2286 | 28 | 0.1429 | 137 | 0.1168 | 18 | 0.0556 | 25 | 0.08 | 16 | 0.0625 | 32 | 0.0938 | 91 | 0.0769 |
| 4_1050893 | 40 | 0.025 | 38 | 0.0526 | 39 | 0.1026 | 31 | 0.0968 | 148 | 0.0676 | 23 | 0 | 30 | 0.2 | 22 | 0.0455 | 34 | 0.1471 | 109 | 0.1101 |
| 4_1062961 | 4 | 0.5 | 6 | 0.6667 | 5 | 0.2 | 3 | 1 | 18 | 0.5556 | 6 | 0.5 | NA | NA | 2 | 0.5 | 4 | 0.25 | 12 | 0.4167 |
| 4_1062962 | 4 | 0 | 6 | 0 | 6 | 0.1667 | 3 | 0 | 19 | 0.0526 | 6 | 0 | 1 | 0 | 3 | 0 | 5 | 0.2 | 15 | 0.0667 |
| 4_1062962 | 4 | 0.25 | 5 | 0 | 4 | 0 | 3 | 0 | 16 | 0.0625 | 6 | 0 | 1 | 0 | 3 | 0.6667 | 5 | 0 | 15 | 0.1333 |
| 4_1062962 | 4 | 1 | 3 | 0.6667 | 4 | 0.5 | 3 | 1 | 14 | 0.7857 | 3 | 0.6667 | 1 | 1 | 3 | 0.6667 | 4 | 1 | 11 | 0.8182 |
| 4_1062963 | 4 | 0.25 | 6 | 0 | 5 | 0.2 | 3 | 0 | 18 | 0.1111 | 6 | 0.1667 | 1 | 0 | 3 | 0 | 5 | 0 | 15 | 0.0667 |
| 4_1062965 | 5 | 0 | 6 | 0.1667 | 4 | 0.25 | 4 | 0.25 | 19 | 0.1579 | 6 | 0.1667 | 1 | 0 | 4 | 0 | 6 | 0 | 17 | 0.0588 |
| 4_1062973 | 2 | 1 | 6 | 0.3333 | 4 | 0 | 4 | 0.5 | 16 | 0.375 | 6 | 0.5 | 1 | 1 | 3 | 0.6667 | 4 | 0.5 | 14 | 0.5714 |
| 4_1062973 | 3 | 0.3333 | 6 | 0.6667 | 4 | 0.25 | 5 | 0.2 | 18 | 0.3889 | 6 | 0.3333 | 1 | 0 | 3 | 0.3333 | 4 | 0 | 14 | 0.2143 |
| 4_1062975 | 3 | 0.6667 | 7 | 0.7143 | 5 | 0.4 | 5 | 0.2 | 20 | 0.5 | 6 | 0.6667 | 1 | 1 | 3 | 1 | 2 | 0 | 12 | 0.6667 |
| 4_1062976 | 3 | 0 | 7 | 0.1429 | 4 | 0 | 3 | 0 | 17 | 0.0588 | 4 | 0 | 1 | 0 | 3 | 0.3333 | 2 | 0 | 10 | 0.1 |
| 4_1062976 | 2 | 0 | 7 | 0.2857 | 4 | 0 | 4 | 0 | 17 | 0.1176 | 6 | 0.5 | 1 | 1 | 3 | 0.3333 | 4 | 0.25 | 14 | 0.4286 |
| 4_1062976 | 3 | 0.3333 | 6 | 0.1667 | 7 | 0 | 5 | 0.2 | 21 | 0.1429 | 6 | 0.1667 | 2 | 0.5 | 2 | 0.5 | 4 | 0.5 | 14 | 0.3571 |
| 4_1062979 | 3 | 1 | 7 | 1 | 7 | 1 | 4 | 0.75 | 21 | 0.9524 | 7 | 1 | 3 | 1 | 3 | 1 | 3 | 1 | 16 | 1 |
| 4_1062979 | 2 | 0.5 | 5 | 0.8 | 7 | 0.1429 | 2 | 0 | 16 | 0.375 | 5 | 0.4 | 1 | 1 | 1 | 1 | 4 | 0.75 | 11 | 0.6364 |
| 4_1062981 | 2 | 1 | 7 | 0.8571 | 5 | 1 | 4 | 0.5 | 18 | 0.8333 | 6 | 0.3333 | 3 | 1 | 3 | 1 | 4 | 0.25 | 16 | 0.5625 |
| 4_1062981 | 4 | 1 | 5 | 0.6 | 6 | 0.8333 | 4 | 1 | 19 | 0.8421 | 5 | 1 | 2 | 1 | 3 | 1 | 4 | 1 | 14 | 1 |
| 4_1062981 | 3 | 0.6667 | 6 | 0.6667 | 8 | 0.625 | 5 | 0.6 | 22 | 0.6364 | 6 | 0.8333 | 2 | 1 | 3 | 1 | 4 | 0.25 | 15 | 0.7333 |
| 4_1062982 | 4 | 0.25 | 7 | 0.2857 | 8 | 0.125 | 3 | 0.3333 | 22 | 0.2273 | 7 | 0.8571 | 4 | 0.75 | 2 | 0 | 4 | 0.25 | 17 | 0.5882 |
| 4_1062983 | 6 | 0.1667 | 7 | 0.2857 | 6 | 0 | 5 | 0.4 | 24 | 0.2083 | 7 | 0.1429 | 4 | 0.25 | 3 | 0.3333 | 4 | 0.25 | 18 | 0.2222 |
| 4_1062997 | 2 | 0.5 | NA | NA | 1 | 0 | 1 | 0 | 4 | 0.25 | NA | NA | NA | NA | 1 | 1 | 1 | 1 | 2 | 1 |
| 4_1062998 | 2 | 0.5 | 1 | 1 | 1 | 0 | 1 | 1 | 5 | 0.6 | NA | NA | NA | NA | 1 | 1 | 1 | 1 | 2 | 1 |
| 4_1062999 | 2 | 0.5 | NA | NA | 1 | 0 | 1 | 1 | 4 | 0.5 | NA | NA | NA | NA | 1 | 0 | 1 | 1 | 2 | 0.5 |
| 4_1062999 | 2 | 1 | 1 | 0 | 1 | 0 | 1 | 1 | 5 | 0.6 | NA | NA | NA | NA | 1 | 1 | 1 | 1 | 2 | 1 |
| 4_1062999 | 2 | 0.5 | 1 | 0 | 1 | 0 | 1 | 0 | 5 | 0.2 | NA | NA | NA | NA | 1 | 1 | 1 | 0 | 2 | 0.5 |
| 4_1063001 | 2 | 0.5 | 1 | 1 | 1 | 1 | NA | NA | 4 | 0.75 | NA | NA | NA | NA | NA | NA | 1 | 1 | 1 | 1 |
| 4_1063002 | 2 | 0 | 1 | 0 | 1 | 1 | NA | NA | 4 | 0.25 | NA | NA | NA | NA | 1 | 0 | 1 | 1 | 2 | 0.5 |
| 4_1063002 | 2 | 1 | 1 | 1 | 1 | 1 | NA | NA | 4 | 1 | NA | NA | NA | NA | NA | NA | 1 | 1 | 1 | 1 |
| 4_1063003 | 1 | 1 | 1 | 1 | 1 | 1 | NA | NA | 3 | 1 | NA | NA | NA | NA | 1 | 1 | 1 | 1 | 2 | 1 |
| 4_1063004 | 2 | 0.5 | NA | NA | 1 | 1 | NA | NA | 3 | 0.6667 | NA | NA | NA | NA | 1 | 0 | 1 | 1 | 2 | 0.5 |
| 4_1064439 | 5 | 0.6 | 7 | 0.4286 | 5 | 0.8 | 3 | 1 | 20 | 0.65 | 4 | 0.75 | 3 | 1 | 5 | 0.6 | 5 | 0.6 | 17 | 0.7059 |
| 4_1064441 | 9 | 0.3333 | 10 | 0.2 | 4 | 0.5 | 4 | 0.25 | 27 | 0.2963 | 5 | 0.2 | 3 | 0.6667 | 4 | 0 | 6 | 0 | 18 | 0.1667 |
| 4_1064441 | 9 | 0.3333 | 9 | 0.3333 | 4 | 0.25 | 3 | 0 | 25 | 0.28 | 5 | 0.2 | 1 | 1 | 5 | 0.4 | 6 | 0.5 | 17 | 0.4118 |
| 4_1064441 | 5 | 0.6 | 6 | 0.3333 | 2 | 0.5 | 3 | 1 | 16 | 0.5625 | 3 | 0.6667 | 3 | 0.6667 | 2 | 0.5 | 2 | 0 | 10 | 0.5 |
| 4_1064441 | 4 | 0.5 | 5 | 0.8 | 5 | 0.4 | 2 | 1 | 16 | 0.625 | 3 | 1 | 2 | 0.5 | 3 | 1 | 6 | 0.5 | 14 | 0.7143 |
| 4_1064445 | 6 | 0.1667 | 10 | 0.3 | 5 | 0.2 | 4 | 0.25 | 25 | 0.24 | 5 | 0.4 | 3 | 0.3333 | 6 | 0.6667 | 8 | 0.625 | 22 | 0.5455 |
| 4_1064446 | 9 | 1 | 6 | 0.6667 | 5 | 1 | 4 | 0.75 | 24 | 0.875 | 4 | 1 | 3 | 1 | 1 | 1 | 6 | 0.8333 | 14 | 0.9286 |
| 4_1064447 | 9 | 0.3333 | 10 | 0.2 | 4 | 0.75 | 4 | 0.25 | 27 | 0.3333 | 7 | 0.2857 | 2 | 0 | 7 | 0.4286 | 10 | 0.3 | 26 | 0.3077 |
| 4_1064450 | 11 | 0 | 12 | 0.0833 | 7 | 0 | 5 | 0.2 | 35 | 0.0571 | 8 | 0.125 | 3 | 0 | 5 | 0 | 11 | 0.0909 | 27 | 0.0741 |
| 4_1064451 | 10 | 0.1 | 11 | 0 | 7 | 0 | 4 | 0.25 | 32 | 0.0625 | 9 | 0.1111 | 3 | 0 | 5 | 0 | 9 | 0.1111 | 26 | 0.0769 |
| 4_1065091 | NA | NA | NA | NA | NA | NA | 2 | 0.5 | 2 | 0.5 | NA | NA | 1 | 1 | NA | NA | 1 | 0 | 2 | 0.5 |
| 4_1065101 | NA | NA | NA | NA | NA | NA | 2 | 0.5 | 2 | 0.5 | NA | NA | 1 | 0 | NA | NA | 1 | 1 | 2 | 0.5 |
| 4_1065102 | NA | NA | NA | NA | NA | NA | 2 | 0.5 | 2 | 0.5 | NA | NA | 1 | 1 | NA | NA | 1 | 0 | 2 | 0.5 |
| 4_1065107 | NA | NA | NA | NA | NA | NA | 2 | 0.5 | 2 | 0.5 | 1 | 0 | 1 | 0 | NA | NA | 2 | 0.5 | 4 | 0.25 |
| 4_1065107 | NA | NA | NA | NA | NA | NA | 3 | 0.6667 | 3 | 0.6667 | 1 | 1 | 1 | 1 | NA | NA | 2 | 0.5 | 4 | 0.75 |

| | | | | | | | | | | | | | | | | | | | | |
|---|---|---|---|---|---|---|---|---|---|---|---|---|---|---|---|---|---|---|---|---|
| 4_10651079 | NA | NA | NA | NA | NA | NA | 3 | 0.3333 | 3 | 0.3333 | 1 | 1 | 1 | 0 | NA | NA | 1 | 0 | 3 | 0.3333 |
| 4_10651095 | NA | NA | NA | NA | NA | NA | 3 | 1 | 3 | 1 | 1 | 1 | NA | NA | NA | NA | 2 | 1 | 3 | 1 |
| 4_10651111 | NA | NA | NA | NA | NA | NA | 1 | 1 | 1 | 1 | NA | NA | 1 | 1 | NA | NA | 2 | 1 | 3 | 1 |
| 4_10651118 | NA | NA | NA | NA | NA | NA | 3 | 1 | 3 | 1 | 1 | 1 | 1 | 1 | NA | NA | 2 | 1 | 4 | 1 |
| 4_10651122 | NA | NA | NA | NA | NA | NA | 3 | 0.3333 | 3 | 0.3333 | 1 | 1 | 1 | 1 | NA | NA | 2 | 0.5 | 4 | 0.75 |
| 4_10651159 | NA | NA | NA | NA | NA | NA | 3 | 1 | 3 | 1 | 1 | 1 | 1 | 1 | NA | NA | 2 | 1 | 4 | 1 |
| 4_10651165 | NA | NA | NA | NA | NA | NA | 3 | 0.3333 | 3 | 0.3333 | NA | NA | 1 | 1 | NA | NA | 2 | 0 | 3 | 0.3333 |
| 4_10651168 | NA | NA | NA | NA | NA | NA | 3 | 0.3333 | 3 | 0.3333 | NA | NA | 1 | 1 | NA | NA | 1 | 1 | 2 | 1 |
| 4_10651172 | NA | NA | NA | NA | NA | NA | 2 | 1 | 2 | 1 | 1 | 1 | NA | NA | NA | NA | 2 | 0.5 | 3 | 0.6667 |
| 4_10651197 | NA | NA | 1 | 1 | NA | NA | 3 | 1 | 4 | 1 | 1 | 0 | NA | NA | NA | NA | 2 | 1 | 3 | 0.6667 |
| 4_10651979 | 3 | 1 | 8 | 0.875 | 3 | 1 | 10 | 0.8 | 24 | 0.875 | 1 | 1 | 4 | 0.75 | 4 | 0.75 | 7 | 0.8571 | 16 | 0.8125 |
| 4_10651980 | 3 | 0.6667 | 7 | 0.7143 | 1 | 0 | 7 | 0.7143 | 18 | 0.6667 | 2 | 1 | 4 | 0.5 | 1 | 0 | 5 | 0.6 | 12 | 0.5833 |
| 4_10651981 | 2 | 1 | 7 | 0.8571 | 2 | 1 | 8 | 1 | 19 | 0.9474 | NA | NA | 3 | 1 | 2 | 1 | 6 | 1 | 11 | 1 |
| 4_10651984 | 3 | 1 | 8 | 0.375 | 3 | 0.3333 | 9 | 0.3333 | 23 | 0.4348 | 2 | 0.5 | 3 | 0.6667 | 4 | 0.5 | 5 | 0 | 14 | 0.3571 |
| 4_10651990 | 2 | 1 | 7 | 1 | 3 | 1 | 9 | 1 | 21 | 1 | 2 | 0.5 | 4 | 1 | 4 | 1 | 7 | 1 | 17 | 0.9412 |
| 4_10652000 | 3 | 0.6667 | 7 | 0.5714 | 2 | 0.5 | 8 | 0.625 | 20 | 0.6 | 2 | 0.5 | 4 | 0.25 | 3 | 0.6667 | 6 | 0.8333 | 15 | 0.6 |
| 4_10652002 | 3 | 1 | 8 | 1 | 2 | 1 | 9 | 1 | 22 | 1 | 2 | 1 | 3 | 1 | 3 | 1 | 7 | 1 | 15 | 1 |
| 4_10652018 | 1 | 0 | 9 | 0.3333 | 2 | 0 | 9 | 0.4444 | 21 | 0.3333 | NA | NA | 3 | 0 | 4 | 0.25 | 6 | 0.1667 | 13 | 0.1538 |
| 4_10652019 | 3 | 0.3333 | 9 | 0.4444 | 2 | 0.5 | 10 | 0.3 | 24 | 0.375 | 1 | 0 | 4 | 0 | 3 | 1 | 7 | 0.2857 | 15 | 0.3333 |
| 4_10652023 | 2 | 1 | 9 | 0.8889 | 1 | 1 | 10 | 1 | 22 | 0.9545 | 1 | 1 | 4 | 0.75 | 4 | 0.75 | 7 | 1 | 16 | 0.875 |
| 4_10652028 | 2 | 1 | 7 | 0.8571 | 2 | 1 | 8 | 0.625 | 19 | 0.7895 | 1 | 1 | 4 | 0.75 | 3 | 1 | 5 | 0.8 | 13 | 0.8462 |
| 4_10660193 | 6 | 0.1667 | 8 | 0 | 3 | 0 | 3 | 0 | 20 | 0.05 | 4 | 0.25 | 5 | 0 | 7 | 0 | 4 | 0 | 20 | 0.05 |
| 4_10660243 | 3 | 0 | 9 | 0.1111 | 2 | 0 | 2 | 0 | 16 | 0.0625 | 3 | 0 | 2 | 1 | 5 | 0 | 5 | 0.2 | 15 | 0.2 |
| 4_10660258 | 5 | 0 | 8 | 0.125 | 4 | 0 | 3 | 0 | 20 | 0.05 | 4 | 0 | 4 | 0.25 | 5 | 0.4 | 4 | 0 | 17 | 0.1765 |
| 4_10660291 | 5 | 0 | 4 | 0 | 2 | 0.5 | 3 | 0 | 14 | 0.0714 | 2 | 0.5 | 3 | 0.3333 | 4 | 0.75 | 3 | 0 | 12 | 0.4167 |
| 4_10660298 | 5 | 0.2 | 7 | 0 | 2 | 0 | 4 | 0.25 | 18 | 0.1111 | 4 | 0.25 | 3 | 0.3333 | 5 | 0.2 | 3 | 0 | 15 | 0.2 |
| 4_10660319 | 4 | 0.25 | 6 | 0.1667 | 1 | 0 | 3 | 0 | 14 | 0.1429 | 2 | 0.5 | 3 | 0.6667 | 4 | 0.25 | 1 | 0 | 10 | 0.4 |
| 4_10660321 | 4 | 0 | 6 | 0.3333 | 1 | 0 | 4 | 0.25 | 15 | 0.2 | 2 | 0 | 3 | 0 | 4 | 0.25 | 2 | 0 | 11 | 0.0909 |
| 4_10660358 | 3 | 0 | 6 | 0 | 2 | 0 | 4 | 0.25 | 15 | 0.0667 | 2 | 0 | 3 | 0 | 4 | 0.25 | 2 | 0.5 | 11 | 0.1818 |
| 4_10660361 | 3 | 0 | 6 | 0.1667 | 1 | 0 | 4 | 0.25 | 14 | 0.1429 | 2 | 0 | 2 | 0 | 4 | 0.25 | 2 | 0 | 10 | 0.1 |
| 4_10660413 | 2 | 0 | 4 | 0.25 | 1 | 0 | 2 | 0.5 | 9 | 0.2222 | 2 | 0 | 3 | 0.3333 | 3 | 0.3333 | 1 | 0 | 9 | 0.2222 |
| 4_10660426 | 2 | 0 | 2 | 0.5 | 1 | 0 | 4 | 0.25 | 9 | 0.2222 | NA | NA | 3 | 0 | 3 | 0 | 1 | 1 | 7 | 0.1429 |
| 4_10660430 | 2 | 0.5 | 3 | 0.6667 | 1 | 0 | 4 | 0.5 | 10 | 0.5 | 1 | 1 | 2 | 0 | 2 | 0 | 1 | 0 | 6 | 0.1667 |
| 4_10660436 | 2 | 0.5 | 2 | 0 | NA | NA | 4 | 0.25 | 8 | 0.25 | NA | NA | NA | NA | 1 | 1 | 1 | 0 | 2 | 0.5 |
| 4_10660445 | 2 | 0.5 | 2 | 0 | 1 | 0 | 4 | 0 | 9 | 0.1111 | 1 | 0 | 2 | 0 | 3 | 0.3333 | 1 | 0 | 7 | 0.1429 |
| 4_10660448 | 2 | 0.5 | 2 | 0 | 2 | 0 | 3 | 0 | 9 | 0.1111 | 1 | 0 | 3 | 0.6667 | 3 | 0 | NA | NA | 7 | 0.2857 |
| 4_10660457 | 2 | 1 | 2 | 1 | 1 | 0 | 2 | 1 | 7 | 0.8571 | 1 | 0 | 2 | 1 | 3 | 1 | 1 | 0 | 7 | 0.7143 |
| 4_10660458 | 1 | 0 | 2 | 1 | 1 | 1 | 3 | 0 | 7 | 0.4286 | 1 | 1 | 3 | 0.6667 | 1 | 1 | 1 | 1 | 6 | 0.8333 |
| 4_10660467 | 1 | 1 | 1 | 1 | 2 | 1 | 2 | 1 | 6 | 1 | 1 | 1 | 2 | 0.5 | 1 | 1 | 1 | 1 | 5 | 0.8 |
| 4_10660498 | 2 | 0.5 | 1 | 0 | 1 | 1 | 2 | 0.5 | 6 | 0.5 | 2 | 0.5 | 1 | 1 | 2 | 1 | 1 | 1 | 6 | 0.8333 |
| 4_10660499 | 3 | 1 | 1 | 0 | 1 | 1 | 3 | 1 | 8 | 0.875 | 2 | 1 | NA | NA | 2 | 1 | 1 | 1 | 5 | 1 |
| 4_10660501 | 3 | 1 | 1 | 1 | 1 | 1 | 4 | 0.75 | 9 | 0.8889 | 2 | 0.5 | 1 | 1 | 3 | 0.6667 | 1 | 1 | 7 | 0.7143 |
| 4_10660506 | 3 | 0.3333 | 1 | 0 | 1 | 0 | 4 | 0 | 9 | 0.1111 | 1 | 0 | 1 | 1 | 3 | 0.3333 | 1 | 1 | 6 | 0.5 |
| 4_10660517 | 3 | 0.6667 | 1 | 1 | NA | NA | 3 | 0.6667 | 7 | 0.7143 | 1 | 1 | 1 | 1 | 3 | 0.6667 | 1 | 1 | 6 | 0.8333 |
| 4_10660520 | 3 | 0.6667 | 1 | 0 | NA | NA | 2 | 0 | 6 | 0.3333 | NA | NA | 1 | 0 | 3 | 0.6667 | NA | NA | 4 | 0.5 |
| 4_10660521 | 2 | 1 | 1 | 1 | NA | NA | 3 | 1 | 6 | 1 | 1 | 1 | 1 | 1 | 3 | 1 | 1 | 1 | 6 | 1 |
| 4_10660526 | 3 | 0.3333 | NA | NA | NA | NA | 2 | 1 | 5 | 0.6 | NA | NA | 1 | 1 | 3 | 0.3333 | 1 | 1 | 5 | 0.6 |
| 4_10660534 | 3 | 1 | NA | NA | NA | NA | 3 | 1 | 6 | 1 | 1 | 1 | 1 | 1 | 2 | 1 | 1 | 1 | 5 | 1 |
| 4_10660547 | 3 | 0.6667 | NA | NA | NA | NA | 1 | 1 | 4 | 0.75 | 1 | 1 | 1 | 0 | 2 | 0.5 | 1 | 1 | 5 | 0.6 |
| 4_10660556 | 3 | 0.6667 | NA | NA | NA | NA | 3 | 0.3333 | 6 | 0.5 | 1 | 0 | 1 | 1 | 1 | 0 | 1 | 0 | 4 | 0.25 |
| 4_10690544 | 9 | 0.7778 | 8 | 0.25 | 19 | 0.6316 | 16 | 0.4375 | 52 | 0.5385 | 12 | 0.6667 | 18 | 0.7778 | 10 | 0.5 | 11 | 0.7273 | 51 | 0.6863 |
| 4_10690551 | 8 | 0.75 | 9 | 0.4444 | 18 | 0.5 | 15 | 0.5333 | 50 | 0.54 | 12 | 0.4167 | 17 | 0.4118 | 11 | 0.5455 | 10 | 0.3 | 50 | 0.42 |
| 4_10690587 | 10 | 0.3 | 6 | 0.1667 | 18 | 0.3889 | 17 | 0.4706 | 51 | 0.3725 | 9 | 0.3333 | 14 | 0.5 | 11 | 0.2727 | 12 | 0.6667 | 46 | 0.4565 |
| 4_10690588 | 9 | 0.8889 | 7 | 0.8571 | 14 | 0.8571 | 18 | 0.8333 | 48 | 0.8542 | 12 | 0.9167 | 14 | 0.7857 | 12 | 0.8333 | 8 | 0.75 | 46 | 0.8261 |
| 4_10690593 | 9 | 0 | 7 | 0.1429 | 18 | 0.0556 | 14 | 0.1429 | 48 | 0.0833 | 9 | 0.4444 | 16 | 0.1875 | 13 | 0.0769 | 8 | 0.375 | 46 | 0.2391 |
| 4_10690607 | 8 | 0.25 | 4 | 0 | 13 | 0.0769 | 10 | 0.2 | 35 | 0.1429 | 8 | 0.25 | 12 | 0.25 | 8 | 0.125 | 5 | 0.2 | 33 | 0.2121 |
| 4_10690640 | 12 | 0.5 | 6 | 0.8333 | 22 | 0.7273 | 16 | 0.5625 | 56 | 0.6429 | 14 | 0.5 | 13 | 0.5385 | 13 | 0.6154 | 6 | 0.6667 | 46 | 0.5652 |
| 4_10690641 | 10 | 0.5 | 7 | 0.7143 | 20 | 0.55 | 12 | 0.5 | 49 | 0.551 | 10 | 0.3 | 12 | 0.5833 | 13 | 0.4615 | 8 | 0.375 | 43 | 0.4419 |
| 4_10690642 | 13 | 0.0769 | 7 | 0.1429 | 18 | 0.2222 | 14 | 0.1429 | 52 | 0.1538 | 13 | 0.3077 | 13 | 0.0769 | 12 | 0.25 | 8 | 0 | 46 | 0.1739 |
| 4_10690646 | 13 | 0.3077 | 7 | 0.4286 | 24 | 0.125 | 15 | 0.4 | 59 | 0.2712 | 11 | 0.1818 | 14 | 0.0714 | 12 | 0.3333 | 10 | 0.3 | 47 | 0.2128 |
| 4_10690650 | 12 | 0.0833 | 7 | 0.1429 | 21 | 0.2381 | 16 | 0.125 | 56 | 0.1607 | 13 | 0.1538 | 16 | 0.25 | 13 | 0.1538 | 10 | 0.1 | 52 | 0.1731 |
| 4_10691321 | 4 | 0.75 | 9 | 0.1111 | 5 | 0.4 | 3 | 0.6667 | 21 | 0.381 | 7 | 0.5714 | 2 | 0 | 7 | 0.1429 | 6 | 0.5 | 22 | 0.3636 |
| 4_10692370 | 3 | 0.3333 | 5 | 0.2 | 1 | 0 | 3 | 0 | 12 | 0.1667 | 4 | 0.25 | NA | NA | 6 | 0.3333 | 2 | 0 | 12 | 0.25 |
| 4_10692385 | 5 | 0 | 4 | 0.25 | 2 | 0.5 | 4 | 0.5 | 15 | 0.2667 | 4 | 0.25 | NA | NA | 5 | 0.2 | 1 | 0 | 10 | 0.2 |
| 4_10692392 | 5 | 0.4 | 5 | 0 | 2 | 0.5 | 4 | 0.5 | 16 | 0.3125 | 3 | 0 | NA | NA | 6 | 0.5 | NA | NA | 9 | 0.3333 |

| | | | | | | | | | | | | | | | | | | | | |
|---|---|---|---|---|---|---|---|---|---|---|---|---|---|---|---|---|---|---|---|---|
| 4_1069414( | 2 | 1 | NA | NA | 2 | 1 | NA | NA | 4 | 1 | 1 | 1 | 1 | 1 | NA | NA | NA | NA | 2 | 1 |
| 4_1069415( | 2 | 1 | NA | NA | 2 | 0.5 | NA | NA | 4 | 0.75 | NA | NA | 1 | 1 | NA | NA | NA | NA | 1 | 1 |
| 4_1069416: | 1 | 1 | NA | NA | 2 | 0.5 | NA | NA | 3 | 0.6667 | 1 | 1 | 1 | 1 | NA | NA | NA | NA | 2 | 1 |
| 4_1069416! | 1 | 1 | NA | NA | 2 | 0 | NA | NA | 3 | 0.3333 | 1 | 0 | 1 | 1 | NA | NA | NA | NA | 2 | 0.5 |
| 4_1069418 | NA | NA | NA | NA | 2 | 1 | NA | NA | 2 | 1 | 1 | 1 | 1 | 0 | NA | NA | NA | NA | 2 | 0.5 |
| 4_1069964! | NA | NA | NA | NA | 1 | 1 | NA | NA | 1 | 1 | 1 | 1 | 1 | 1 | NA | NA | 1 | 1 | 3 | 1 |
| 4_1069965: | NA | NA | NA | NA | 1 | 1 | NA | NA | 1 | 1 | 1 | 1 | 2 | 1 | NA | NA | 1 | 1 | 4 | 1 |
| 4_1070577 | 50 | 0.18 | 31 | 0.2258 | 35 | 0.1429 | 56 | 0.125 | 172 | 0.1628 | 34 | 0.0882 | 45 | 0.1333 | 39 | 0.0513 | 46 | 0.1957 | 164 | 0.122 |
| 4_1070577 | 44 | 0.0682 | 25 | 0.08 | 35 | 0.0286 | 46 | 0.087 | 150 | 0.0667 | 29 | 0.0345 | 38 | 0.0526 | 35 | 0.0286 | 37 | 0.0811 | 139 | 0.0504 |
| 4_1070578 | 49 | 0.2245 | 29 | 0.3103 | 37 | 0.3243 | 56 | 0.1964 | 171 | 0.2515 | 31 | 0.2581 | 47 | 0.2766 | 38 | 0.3947 | 41 | 0.2439 | 157 | 0.293 |
| 4_1070909 | 4 | 0.75 | 3 | 0.6667 | 4 | 0.5 | 3 | 0.3333 | 14 | 0.5714 | 3 | 0.6667 | 1 | 1 | 3 | 1 | 1 | 0 | 8 | 0.75 |
| 4_1070910 | 7 | 0.8571 | 4 | 0.75 | 6 | 0.6667 | 4 | 0.75 | 21 | 0.7619 | 6 | 0.6667 | 4 | 0.75 | 5 | 0.8 | 3 | 1 | 18 | 0.7778 |
| 4_1070911 | 6 | 0.3333 | 2 | 0.5 | 2 | 0 | 4 | 0.25 | 14 | 0.2857 | 5 | 0.4 | 3 | 0 | 6 | 0 | 3 | 0.3333 | 17 | 0.1765 |
| 4_1071047 | 4 | 0.25 | 1 | 0 | NA | NA | 2 | 0 | 7 | 0.1429 | 5 | 0 | 4 | 0.25 | 1 | 0 | 1 | 0 | 11 | 0.0909 |
| 4_1071048 | 5 | 1 | NA | NA | 1 | 1 | 1 | 0 | 7 | 0.8571 | 6 | 0.8333 | 2 | 1 | 1 | 1 | 1 | 1 | 10 | 0.9 |
| 4_1071403( | NA | NA | NA | NA | 1 | 1 | NA | NA | 1 | 1 | 1 | 1 | NA | NA | NA | NA | NA | NA | 1 | 1 |
| 4_1072712 | 4 | 0.25 | NA | NA | NA | NA | 5 | 0 | 9 | 0.1111 | 4 | 0.25 | 2 | 0 | NA | NA | 3 | 0.3333 | 9 | 0.2222 |
| 4_1072713( | 3 | 0 | NA | NA | NA | NA | 5 | 0.8 | 8 | 0.5 | 4 | 1 | 2 | 0.5 | 1 | 1 | 3 | 1 | 10 | 0.9 |
| 4_1072713: | 4 | 0.5 | NA | NA | NA | NA | 4 | 0.5 | 8 | 0.5 | 3 | 0.3333 | 1 | 0 | 1 | 1 | 3 | 1 | 8 | 0.625 |
| 4_1072713 | 4 | 0.75 | NA | NA | NA | NA | 5 | 1 | 9 | 0.8889 | 4 | 0.25 | 2 | 0 | 1 | 1 | 3 | 1 | 10 | 0.5 |
| 4_1072714 | 3 | 0.6667 | NA | NA | NA | NA | 3 | 0 | 6 | 0.3333 | 3 | 0.3333 | 2 | 0 | NA | NA | 2 | 0 | 7 | 0.1429 |
| 4_1072715! | 2 | 0 | NA | NA | NA | NA | 4 | 0.5 | 6 | 0.3333 | 2 | 0 | 1 | 0 | 1 | 1 | 3 | 0.6667 | 7 | 0.4286 |
| 4_1072728! | 2 | 0.5 | 1 | 0 | 2 | 0 | 5 | 0.2 | 10 | 0.2 | 4 | 0.25 | 1 | 0 | 3 | 0.3333 | 3 | 0.3333 | 11 | 0.2727 |
| 4_1072729( | NA | NA | 1 | 0 | 1 | 0 | 5 | 0.2 | 7 | 0.1429 | 4 | 0 | 1 | 0 | 3 | 0.3333 | 3 | 0 | 11 | 0.0909 |
| 4_1072731 | 2 | 0 | 2 | 1 | 1 | 1 | 6 | 0.3333 | 11 | 0.4545 | 4 | 0 | 1 | 1 | 2 | 1 | 3 | 0 | 10 | 0.3 |
| 4_1072733 | 2 | 0 | 1 | 0 | 1 | 0 | 6 | 0.3333 | 10 | 0.2 | 2 | 0 | 1 | 0 | 3 | 0.3333 | 2 | 0 | 8 | 0.125 |
| 4_1072733! | 2 | 0 | 2 | 0 | 2 | 0 | 6 | 0.1667 | 12 | 0.0833 | 3 | 0 | 1 | 0 | 3 | 0.6667 | 3 | 0 | 10 | 0.2 |
| 4_1076633: | 2 | 0.5 | NA | NA | 2 | 0.5 | 1 | 1 | 5 | 0.6 | NA | NA | 3 | 1 | NA | NA | 1 | 0 | 4 | 0.75 |
| 4_1076634 | 1 | 0 | NA | NA | 1 | 1 | NA | NA | 2 | 0.5 | NA | NA | 3 | 0.3333 | NA | NA | NA | NA | 3 | 0.3333 |
| 4_1076634! | 2 | 0.5 | NA | NA | 2 | 0 | 1 | 0 | 5 | 0.2 | NA | NA | 3 | 0.6667 | NA | NA | 1 | 0 | 4 | 0.5 |
| 4_1076643: | NA | NA | NA | NA | NA | NA | 1 | 1 | 1 | 1 | NA | NA | 2 | 1 | NA | NA | NA | NA | 2 | 1 |
| 4_1077297 | NA | NA | 2 | 1 | NA | NA | 1 | 0 | 3 | 0.6667 | NA | NA | 2 | 0.5 | NA | NA | NA | NA | 2 | 0.5 |
| 4_1077313 | NA | NA | 3 | 0.3333 | 1 | 0 | 1 | 1 | 5 | 0.4 | 1 | 1 | NA | NA | 2 | 0.5 | 3 | 0.6667 | 6 | 0.6667 |
| 4_1077313 | NA | NA | 4 | 0.25 | 1 | 1 | 1 | 1 | 6 | 0.5 | 1 | 1 | NA | NA | 2 | 0 | 3 | 0.3333 | 6 | 0.3333 |
| 4_1077313 | NA | NA | 3 | 0.6667 | 1 | 0 | 1 | 0 | 5 | 0.4 | 1 | 0 | NA | NA | 2 | 0.5 | 3 | 0.3333 | 6 | 0.3333 |
| 4_1077313( | NA | NA | 4 | 0.5 | 1 | 1 | 1 | 0 | 6 | 0.5 | 1 | 1 | NA | NA | 1 | 1 | 3 | 0.3333 | 5 | 0.6 |
| 4_1077315 | NA | NA | 4 | 0.25 | 1 | 0 | NA | NA | 5 | 0.2 | 1 | 0 | NA | NA | 1 | 0 | 3 | 0.3333 | 5 | 0.2 |
| 4_1077316 | NA | NA | 5 | 0.6 | 1 | 1 | 1 | 1 | 7 | 0.7143 | 1 | 1 | NA | NA | 2 | 0.5 | 1 | 0 | 4 | 0.5 |
| 4_1077316! | NA | NA | 4 | 0.75 | 1 | 0 | NA | NA | 5 | 0.6 | NA | NA | NA | NA | 1 | 1 | 3 | 0.3333 | 4 | 0.5 |
| 4_1077318 | NA | NA | 5 | 0.2 | 1 | 1 | NA | NA | 6 | 0.3333 | 1 | 0 | NA | NA | 2 | 0 | 3 | 0.3333 | 6 | 0.1667 |
| 4_1077320: | NA | NA | 5 | 0.2 | 1 | 0 | 1 | 0 | 7 | 0.1429 | NA | NA | NA | NA | 2 | 0 | 2 | 0.5 | 4 | 0.25 |
| 4_1077320! | NA | NA | 3 | 0.6667 | NA | NA | 1 | 1 | 4 | 0.75 | NA | NA | NA | NA | 2 | 0.5 | 1 | 0 | 3 | 0.3333 |
| 4_1077320! | NA | NA | 3 | 0.3333 | 1 | 1 | 1 | 0 | 5 | 0.4 | 1 | 0 | NA | NA | 2 | 0.5 | 3 | 1 | 6 | 0.6667 |
| 4_1077321 | NA | NA | 5 | 0.2 | 1 | 0 | 1 | 0 | 7 | 0.1429 | 1 | 0 | NA | NA | 2 | 0 | 3 | 0.3333 | 6 | 0.1667 |
| 4_1077321( | NA | NA | 5 | 0.8 | 1 | 0 | 1 | 0 | 7 | 0.5714 | 1 | 0 | NA | NA | 2 | 0.5 | 3 | 0.6667 | 6 | 0.5 |
| 4_1077323( | NA | NA | 4 | 0.5 | 1 | 0 | NA | NA | 5 | 0.4 | 2 | 1 | NA | NA | 2 | 0.5 | 2 | 1 | 6 | 0.8333 |
| 4_1077324! | NA | NA | 5 | 0.4 | 1 | 0 | 1 | 0 | 7 | 0.2857 | 2 | 0 | NA | NA | 2 | 0 | 3 | 0.3333 | 7 | 0.1429 |
| 4_1077325: | NA | NA | 6 | 0.3333 | 1 | 1 | 1 | 0 | 8 | 0.375 | 1 | 0 | NA | NA | 2 | 0 | 3 | 0.3333 | 6 | 0.1667 |
| 4_1077328 | NA | NA | 6 | 1 | 1 | 1 | 1 | 1 | 8 | 1 | 2 | 1 | NA | NA | 2 | 1 | 3 | 1 | 7 | 1 |
| 4_1077329: | NA | NA | 6 | 0.3333 | 1 | 0 | 1 | 0 | 8 | 0.25 | 2 | 0 | NA | NA | 2 | 0.5 | 2 | 0.5 | 6 | 0.3333 |
| 4_1077330: | NA | NA | 4 | 0.5 | 1 | 0 | NA | NA | 5 | 0.4 | 2 | 0.5 | NA | NA | 2 | 0 | 3 | 0.3333 | 7 | 0.2857 |
| 4_1077330 | NA | NA | 6 | 0.8333 | 1 | 1 | 1 | 1 | 8 | 0.875 | 2 | 1 | NA | NA | 2 | 1 | 3 | 0.6667 | 7 | 0.8571 |
| 4_1077333 | NA | NA | 5 | 0.2 | 1 | 0 | 1 | 1 | 7 | 0.2857 | 2 | 0.5 | NA | NA | 2 | 0 | 3 | 0 | 7 | 0.1429 |
| 4_1077335: | NA | NA | 6 | 0.5 | 1 | 1 | 1 | 1 | 8 | 0.625 | 3 | 0.6667 | 3 | 1 | 2 | 0 | 3 | 0.6667 | 11 | 0.6364 |
| 4_1077336 | NA | NA | 6 | 0.1667 | 1 | 0 | NA | NA | 7 | 0.1429 | 2 | 0 | 2 | 0.5 | 2 | 0 | 3 | 0.3333 | 9 | 0.2222 |
| 4_1077336! | NA | NA | 5 | 0.4 | 1 | 0 | 2 | 0 | 8 | 0.25 | 3 | 1 | 4 | 0.75 | 1 | 0 | 3 | 0.3333 | 11 | 0.6364 |
| 4_1077337: | NA | NA | 6 | 0.1667 | 1 | 0 | 2 | 1 | 9 | 0.3333 | 3 | 0.3333 | 4 | 0 | 2 | 0 | 3 | 0.3333 | 12 | 0.1667 |
| 4_1077444! | 7 | 0.5714 | 10 | 0.5 | 7 | 0.2857 | 8 | 0.625 | 32 | 0.5 | 6 | 0.1667 | 6 | 0.5 | 9 | 0.2222 | 6 | 0.6667 | 27 | 0.3704 |
| 4_1077540( | 8 | 0 | 3 | 0 | 5 | 0.2 | 2 | 0 | 18 | 0.0556 | 6 | 0.5 | 6 | 0.1667 | 7 | 0.1429 | 6 | 0.5 | 25 | 0.32 |
| 4_1077540 | 5 | 0 | 1 | 1 | 3 | 0 | 3 | 0 | 12 | 0.0833 | 3 | 0 | 5 | 0.2 | 6 | 0 | 5 | 0 | 19 | 0.0526 |
| 4_1077541 | 5 | 0.6 | 3 | 1 | 6 | 0.6667 | 2 | 0 | 16 | 0.625 | 6 | 0.3333 | 6 | 0.5 | 7 | 0.2857 | 5 | 0.4 | 24 | 0.375 |
| 4_1077541 | 8 | 0.375 | 3 | 0.3333 | 6 | 0.3333 | 3 | 0 | 20 | 0.3 | 5 | 0.2 | 5 | 0 | 8 | 0.125 | 5 | 0.2 | 23 | 0.1304 |
| 4_1077541( | 8 | 0 | 3 | 0 | 6 | 0.1667 | 3 | 0.3333 | 20 | 0.1 | 4 | 0.25 | 6 | 0.1667 | 7 | 0.1429 | 6 | 0.1667 | 23 | 0.1739 |
| 4_1077543( | 1 | 1 | NA | NA | 3 | 1 | 1 | 1 | 5 | 1 | 1 | 1 | 1 | 1 | 2 | 1 | 1 | 1 | 5 | 1 |
| 4_1077547 | 2 | 1 | NA | NA | 3 | 1 | 3 | 0.6667 | 8 | 0.875 | 2 | 1 | NA | NA | 2 | 0.5 | 1 | 1 | 5 | 0.8 |

| | | | | | | | | | | | | | | | | | | | | |
|---|---|---|---|---|---|---|---|---|---|---|---|---|---|---|---|---|---|---|---|---|
| 4_1077548 | 1 | 0 | NA | NA | 3 | 0.6667 | 2 | 0 | 6 | 0.3333 | 1 | 0 | 2 | 0 | 2 | 0.5 | 1 | 0 | 6 | 0.1667 |
| 4_1077548 | 2 | 0.5 | NA | NA | 1 | 1 | 2 | 0 | 5 | 0.4 | 2 | 0.5 | 2 | 0.5 | 1 | 1 | 1 | 1 | 6 | 0.6667 |
| 4_1077612 | 2 | 0.5 | 2 | 1 | 2 | 1 | 2 | 1 | 8 | 0.875 | 1 | 1 | 1 | 1 | NA | NA | NA | NA | 2 | 1 |
| 4_1077642 | 1 | 1 | 2 | 1 | 2 | 1 | 2 | 1 | 7 | 1 | 1 | 1 | 1 | 1 | NA | NA | NA | NA | 2 | 1 |
| 4_1077645 | NA | NA | 2 | 0.5 | 1 | 1 | 2 | 1 | 5 | 0.8 | 1 | 1 | 1 | 0 | NA | NA | NA | NA | 2 | 0.5 |
| 4_1077648 | 1 | 1 | 2 | 1 | NA | NA | 2 | 1 | 5 | 1 | 1 | 1 | 1 | 1 | NA | NA | NA | NA | 2 | 1 |
| 4_1077650 | 1 | 1 | 2 | 1 | 1 | 1 | 2 | 0.5 | 6 | 0.8333 | 1 | 0 | 1 | 1 | NA | NA | NA | NA | 2 | 0.5 |
| 4_1077650 | 1 | 1 | 1 | 1 | 1 | 1 | 2 | 1 | 5 | 1 | 1 | 1 | 1 | 1 | NA | NA | NA | NA | 2 | 1 |
| 4_1077651 | 1 | 1 | 2 | 1 | 1 | 0 | 2 | 0 | 6 | 0.5 | 1 | 0 | 1 | 1 | NA | NA | NA | NA | 2 | 0.5 |
| 4_1079015 | 14 | 0.1429 | 14 | 0.0714 | 14 | 0.2143 | 14 | 0.2857 | 56 | 0.1786 | 8 | 0 | 16 | 0.125 | 10 | 0.1 | 20 | 0.25 | 54 | 0.1481 |
| 4_1079016 | 21 | 0.0476 | 17 | 0.1765 | 17 | 0 | 16 | 0.0625 | 71 | 0.0704 | 15 | 0 | 21 | 0.1429 | 14 | 0 | 22 | 0.1364 | 72 | 0.0833 |
| 4_1079017 | 24 | 0.5 | 25 | 0.36 | 26 | 0.1923 | 23 | 0.2609 | 98 | 0.3265 | 23 | 0.4348 | 29 | 0.1379 | 23 | 0.3478 | 28 | 0.3571 | 103 | 0.3107 |
| 4_1080362 | NA | NA | NA | NA | 2 | 0.5 | NA | NA | 2 | 0.5 | NA | NA | NA | NA | 2 | 0.5 | 1 | 1 | 3 | 0.6667 |
| 4_1080363 | NA | NA | NA | NA | 1 | 1 | NA | NA | 1 | 1 | NA | NA | NA | NA | 1 | 1 | 2 | 1 | 3 | 1 |
| 4_1080871 | 18 | 0.1111 | 14 | 0.5 | 12 | 0.0833 | 18 | 0.1111 | 62 | 0.1935 | 7 | 0.2857 | 17 | 0.2941 | 15 | 0.4 | 10 | 0.2 | 49 | 0.3061 |
| 4_1080872 | 18 | 0.1111 | 14 | 0 | 13 | 0.1538 | 19 | 0 | 64 | 0.0625 | 8 | 0 | 16 | 0 | 16 | 0.25 | 13 | 0 | 53 | 0.0755 |
| 4_1080931 | 2 | 0 | 1 | 1 | 2 | 0.5 | 2 | 0 | 7 | 0.2857 | 5 | 0.2 | 6 | 0.1667 | 1 | 0 | 3 | 0.6667 | 15 | 0.2667 |
| 4_1080932 | 3 | 1 | 1 | 1 | 2 | 0.5 | 3 | 1 | 9 | 0.8889 | 5 | 1 | 5 | 1 | 1 | 1 | 3 | 1 | 14 | 1 |
| 4_1081230 | 3 | 0.6667 | 1 | 0 | 1 | 1 | 3 | 0.3333 | 8 | 0.5 | 4 | 0 | 6 | 0.5 | 1 | 1 | 2 | 0.5 | 13 | 0.3846 |
| 4_1081231 | 2 | 0 | 1 | 0 | 1 | 0 | 3 | 0.3333 | 7 | 0.1429 | 4 | 0 | 4 | 0.25 | 1 | 0 | 1 | 0 | 10 | 0.1 |
| 4_1081231 | 2 | 1 | 1 | 1 | NA | NA | 3 | 0.6667 | 6 | 0.8333 | 4 | 1 | 5 | 1 | NA | NA | 2 | 1 | 11 | 1 |
| 4_1081233 | 3 | 0.6667 | 1 | 1 | 1 | 0 | 3 | 0.6667 | 8 | 0.625 | 4 | 0.5 | 4 | 0.5 | 1 | 0 | 2 | 0.5 | 11 | 0.4545 |
| 4_1081233 | 2 | 0 | 1 | 0 | 1 | 0 | 3 | 0.3333 | 7 | 0.1429 | 4 | 0.5 | 4 | 0.25 | 1 | 0 | 2 | 0 | 11 | 0.2727 |
| 4_1081234 | 2 | 0 | 1 | 1 | NA | NA | 3 | 0 | 6 | 0.1667 | 4 | 0 | 4 | 0.5 | 1 | 1 | 2 | 0 | 11 | 0.2727 |
| 4_1081235 | NA | NA | NA | NA | NA | NA | 1 | 1 | 1 | 1 | 1 | 1 | 1 | 1 | NA | NA | 1 | 1 | 3 | 1 |
| 4_1081235 | 2 | 0.5 | 1 | 1 | NA | NA | 3 | 0 | 6 | 0.3333 | 4 | 0.25 | 3 | 0.3333 | 1 | 0 | 2 | 0 | 10 | 0.2 |
| 4_1081236 | 1 | 1 | 1 | 1 | NA | NA | 3 | 0.6667 | 5 | 0.8 | 3 | 0.6667 | 4 | 0.5 | 1 | 1 | 2 | 1 | 10 | 0.7 |
| 4_1081954 | 1 | 1 | 3 | 1 | NA | NA | 1 | 0 | 5 | 0.8 | 2 | 0 | 1 | 0 | 4 | 0.5 | 2 | 0.5 | 9 | 0.3333 |
| 4_1081954 | 1 | 1 | 2 | 1 | NA | NA | 1 | 1 | 4 | 1 | 3 | 0.3333 | 1 | 0 | 4 | 0.5 | 2 | 1 | 10 | 0.5 |
| 4_1081954 | 2 | 0.5 | 3 | 0.3333 | NA | NA | 2 | 1 | 7 | 0.5714 | 3 | 1 | NA | NA | 5 | 0.8 | 2 | 1 | 10 | 0.9 |
| 4_1081955 | 2 | 1 | 3 | 0.6667 | NA | NA | 2 | 1 | 7 | 0.8571 | 3 | 1 | 1 | 1 | 4 | 1 | 2 | 1 | 10 | 1 |
| 4_1081957 | 2 | 0.5 | 2 | 0 | NA | NA | 1 | 0 | 5 | 0.2 | 2 | 1 | 1 | 0 | 6 | 0.1667 | 1 | 1 | 10 | 0.4 |
| 4_1081959 | NA | NA | 1 | 1 | NA | NA | NA | NA | 1 | 1 | NA | NA | NA | NA | 2 | 1 | NA | NA | 2 | 1 |
| 4_1081960 | NA | NA | 1 | 1 | NA | NA | NA | NA | 1 | 1 | NA | NA | NA | NA | 1 | 1 | NA | NA | 1 | 1 |
| 4_1081964 | NA | NA | 1 | 1 | NA | NA | 1 | 1 | 2 | 1 | NA | NA | NA | NA | 2 | 1 | 1 | 0 | 3 | 0.6667 |
| 4_1081965 | NA | NA | 1 | 1 | NA | NA | 1 | 1 | 2 | 1 | NA | NA | NA | NA | 2 | 0.5 | 1 | 0 | 3 | 0.3333 |
| 4_1081988 | 1 | 1 | 1 | 0 | 1 | 0 | 1 | 0 | 4 | 0.25 | NA | NA | NA | NA | 2 | 0.5 | 1 | 0 | 3 | 0.3333 |
| 4_1083362 | NA | NA | NA | NA | 3 | 0.6667 | NA | NA | 3 | 0.6667 | NA | NA | NA | NA | NA | NA | 1 | 1 | 1 | 1 |
| 4_1083365 | NA | NA | NA | NA | 3 | 0.3333 | NA | NA | 3 | 0.3333 | NA | NA | NA | NA | NA | NA | 2 | 0.5 | 2 | 0.5 |
| 4_1083367 | NA | NA | NA | NA | 3 | 0.6667 | NA | NA | 3 | 0.6667 | NA | NA | NA | NA | NA | NA | 2 | 0.5 | 2 | 0.5 |
| 4_1083436 | NA | NA | NA | NA | 3 | 0.6667 | NA | NA | 3 | 0.6667 | NA | NA | NA | NA | 1 | 0 | 3 | 1 | 4 | 0.75 |
| 4_1083703 | NA | NA | NA | NA | 2 | 0.5 | NA | NA | 2 | 0.5 | NA | NA | NA | NA | 2 | 0.5 | 1 | 1 | 3 | 0.6667 |
| 4_1083704 | NA | NA | NA | NA | 3 | 1 | NA | NA | 3 | 1 | NA | NA | NA | NA | 2 | 1 | 3 | 1 | 5 | 1 |
| 4_1083704 | NA | NA | NA | NA | 3 | 0.3333 | NA | NA | 3 | 0.3333 | NA | NA | NA | NA | 1 | 1 | 3 | 1 | 4 | 1 |
| 4_1083704 | NA | NA | NA | NA | 3 | 0.6667 | NA | NA | 3 | 0.6667 | NA | NA | NA | NA | 2 | 0.5 | 3 | 0 | 5 | 0.2 |
| 4_1083705 | NA | NA | NA | NA | 3 | 0.6667 | NA | NA | 3 | 0.6667 | NA | NA | NA | NA | 2 | 0 | 3 | 0.3333 | 5 | 0.2 |
| 4_1083705 | NA | NA | NA | NA | 2 | 0.5 | NA | NA | 2 | 0.5 | NA | NA | NA | NA | 2 | 0.5 | 3 | 1 | 5 | 0.8 |
| 4_1083705 | NA | NA | NA | NA | 3 | 0.6667 | NA | NA | 3 | 0.6667 | NA | NA | NA | NA | 1 | 1 | 2 | 1 | 3 | 1 |
| 4_1083706 | NA | NA | NA | NA | 2 | 1 | NA | NA | 2 | 1 | NA | NA | NA | NA | 1 | 1 | 3 | 0.6667 | 4 | 0.75 |
| 4_1084127 | 1 | 0 | 1 | 0 | 3 | 0.3333 | 1 | 0 | 6 | 0.1667 | NA | NA | 1 | 1 | 3 | 0 | 4 | 0 | 8 | 0.125 |
| 4_1084160 | 2 | 0 | 2 | 0 | 4 | 0 | 4 | 0.25 | 12 | 0.0833 | NA | NA | 1 | 0 | 3 | 0.3333 | 4 | 0 | 8 | 0.125 |
| 4_1084161 | 2 | 0 | 2 | 0 | 3 | 0.6667 | 4 | 0.25 | 11 | 0.2727 | 1 | 0 | 1 | 1 | 4 | 0.25 | 4 | 0 | 10 | 0.2 |
| 4_1084161 | 2 | 0 | 2 | 0 | 4 | 0.25 | 4 | 0 | 12 | 0.0833 | 1 | 0 | 1 | 0 | 4 | 0 | 3 | 0.3333 | 9 | 0.1111 |
| 4_1084224 | 8 | 0.25 | 4 | 0 | 3 | 0 | 3 | 0 | 18 | 0.1111 | 3 | 0 | 3 | 0 | 6 | 0 | 1 | 1 | 13 | 0.0769 |
| 4_1084910 | NA | NA | NA | NA | 3 | 1 | 1 | 1 | 4 | 1 | 3 | 0.6667 | 1 | 0 | NA | NA | 2 | 1 | 6 | 0.6667 |
| 4_1084910 | NA | NA | 1 | 1 | 4 | 0.5 | 1 | 1 | 6 | 0.6667 | 3 | 1 | 1 | 1 | NA | NA | 4 | 0.5 | 8 | 0.75 |
| 4_1084911 | NA | NA | 1 | 0 | 4 | 0.25 | 1 | 0 | 6 | 0.1667 | 4 | 0.25 | 1 | 0 | 1 | 0 | 4 | 0 | 10 | 0.1 |
| 4_1084911 | NA | NA | 1 | 0 | 5 | 0.6 | 2 | 0.5 | 8 | 0.5 | 4 | 0.5 | 1 | 1 | 1 | 1 | 4 | 0 | 10 | 0.4 |
| 4_1084912 | NA | NA | 1 | 0 | 3 | 0.6667 | 3 | 0.6667 | 7 | 0.5714 | 4 | 0.5 | 1 | 0 | 1 | 0 | 2 | 1 | 8 | 0.5 |
| 4_1084919 | 1 | 0 | 1 | 0 | 1 | 0 | 2 | 0.5 | 5 | 0.2 | 1 | 1 | 1 | 0 | NA | NA | NA | NA | 2 | 0.5 |
| 4_1084960 | 1 | 0 | 4 | 0.25 | 4 | 0 | 3 | 0.3333 | 12 | 0.1667 | 6 | 0 | 5 | 0.2 | 5 | 0 | 4 | 0.5 | 20 | 0.15 |
| 4_1084963 | NA | NA | 1 | 0 | 3 | 0.3333 | 1 | 0 | 5 | 0.2 | 2 | 0 | 1 | 0 | 3 | 0 | 2 | 0.5 | 8 | 0.125 |
| 4_1084964 | NA | NA | 1 | 0 | 3 | 0.6667 | 1 | 1 | 5 | 0.6 | 1 | 1 | 2 | 1 | 2 | 1 | 2 | 0.5 | 7 | 0.8571 |
| 4_1085755 | NA | NA | NA | NA | 1 | 1 | NA | NA | 1 | 1 | NA | NA | NA | NA | NA | NA | 2 | 1 | 2 | 1 |
| 4_1085760 | NA | NA | NA | NA | 1 | 1 | NA | NA | 1 | 1 | NA | NA | NA | NA | NA | NA | 2 | 1 | 2 | 1 |

| | | | | | | | | | | | | | | | | | | | | |
|---|---|---|---|---|---|---|---|---|---|---|---|---|---|---|---|---|---|---|---|---|
| 4_1085765 | NA | NA | NA | NA | 1 | 1 | NA | NA | 1 | 1 | NA | NA | NA | NA | NA | NA | 2 | 0.5 | 2 | 0.5 |
| 4_1085766 | NA | NA | NA | NA | 1 | 1 | NA | NA | 1 | 1 | NA | NA | NA | NA | NA | NA | 2 | 0.5 | 2 | 0.5 |
| 4_1085767 | NA | NA | NA | NA | 1 | 1 | NA | NA | 1 | 1 | NA | NA | NA | NA | NA | NA | 2 | 1 | 2 | 1 |
| 4_1085768 | NA | NA | NA | NA | 1 | 1 | NA | NA | 1 | 1 | NA | NA | NA | NA | NA | NA | 2 | 0.5 | 2 | 0.5 |
| 4_1086721 | 3 | 0 | 4 | 0.25 | 4 | 0 | 2 | 0 | 13 | 0.0769 | 3 | 0 | 3 | 0 | 1 | 0 | 4 | 0.25 | 11 | 0.0909 |
| 4_1086733 | 5 | 0 | 9 | 0 | 10 | 0.1 | 5 | 0.4 | 29 | 0.1034 | 6 | 0.1667 | 12 | 0 | 5 | 0 | 9 | 0.1111 | 32 | 0.0625 |
| 4_1086740 | 12 | 0.4167 | 13 | 0.7692 | 13 | 0.4615 | 6 | 0.6667 | 44 | 0.5682 | 6 | 0.8333 | 26 | 0.6538 | 12 | 0.75 | 11 | 0.4545 | 55 | 0.6545 |
| 4_1087053 | 1 | 1 | 1 | 0 | 1 | 0 | 3 | 0.6667 | 6 | 0.5 | NA | NA | 3 | 0.6667 | NA | NA | NA | NA | 3 | 0.6667 |
| 4_1087054 | 2 | 0 | NA | NA | 2 | 0 | 3 | 0.3333 | 7 | 0.1429 | NA | NA | 3 | 0.3333 | NA | NA | NA | NA | 3 | 0.3333 |
| 4_1087056 | 2 | 0.5 | NA | NA | 1 | 1 | 2 | 1 | 5 | 0.8 | NA | NA | 3 | 0.6667 | NA | NA | 1 | 0 | 4 | 0.5 |
| 4_1087060 | 3 | 0 | 1 | 0 | 2 | 0 | 3 | 0.3333 | 9 | 0.1111 | NA | NA | 2 | 0.5 | NA | NA | 1 | 0 | 3 | 0.3333 |
| 4_1087061 | 3 | 0.3333 | 1 | 0 | 2 | 0 | 3 | 0 | 9 | 0.1111 | NA | NA | 3 | 0.3333 | NA | NA | 1 | 0 | 4 | 0.25 |
| 4_1087061 | 3 | 0.3333 | 1 | 0 | 2 | 0.5 | 4 | 0.25 | 10 | 0.3 | NA | NA | 3 | 0.3333 | NA | NA | 2 | 0.5 | 5 | 0.4 |
| 4_1087062 | 3 | 1 | 1 | 1 | 2 | 1 | 3 | 0.6667 | 9 | 0.8889 | NA | NA | 4 | 0.5 | NA | NA | 1 | 0 | 5 | 0.4 |
| 4_1087063 | 1 | 0 | 1 | 0 | 2 | 0 | 3 | 0.3333 | 7 | 0.1429 | NA | NA | 4 | 0 | NA | NA | 1 | 1 | 5 | 0.2 |
| 4_1087065 | 3 | 0.3333 | 1 | 0 | 3 | 0.3333 | 5 | 0.4 | 12 | 0.3333 | NA | NA | 4 | 0.25 | NA | NA | 2 | 0 | 6 | 0.1667 |
| 4_1087188 | 8 | 0.125 | 4 | 0.25 | 5 | 0.4 | 4 | 0 | 21 | 0.1905 | 3 | 0 | 6 | 0.3333 | 4 | 0.5 | 3 | 0 | 16 | 0.25 |
| 4_1087188 | 8 | 0.125 | 5 | 0.2 | 5 | 0.2 | 3 | 0.3333 | 21 | 0.1905 | 3 | 0.3333 | 4 | 0.5 | 4 | 0.5 | 4 | 0.5 | 15 | 0.4667 |
| 4_1087188 | 8 | 0.375 | 5 | 0.4 | 5 | 0 | 4 | 0 | 22 | 0.2273 | 3 | 0 | 5 | 0.2 | 2 | 0 | 4 | 0 | 14 | 0.0714 |
| 4_1087190 | 8 | 0.375 | 5 | 0.4 | 6 | 0 | 5 | 0.4 | 24 | 0.2917 | 3 | 0.6667 | 6 | 0.3333 | 5 | 0.8 | 4 | 0.25 | 18 | 0.5 |
| 4_1087979 | 19 | 0.1053 | 23 | 0.087 | 20 | 0.05 | 30 | 0.0667 | 92 | 0.0761 | 13 | 0.1538 | 25 | 0.12 | 12 | 0 | 10 | 0.1 | 60 | 0.1 |
| 4_1087979 | 20 | 0.05 | 21 | 0.2381 | 20 | 0 | 26 | 0.0385 | 87 | 0.0805 | 16 | 0 | 21 | 0.1429 | 13 | 0.1538 | 10 | 0 | 60 | 0.0833 |
| 4_1087982 | 19 | 0.3158 | 23 | 0.4783 | 23 | 0.3478 | 30 | 0.5667 | 95 | 0.4421 | 20 | 0.55 | 25 | 0.44 | 16 | 0.5 | 13 | 0.4615 | 74 | 0.4865 |
| 4_1087986 | 1 | 0 | NA | NA | 4 | 0.25 | 1 | 0 | 6 | 0.1667 | 3 | 0.3333 | 3 | 0.3333 | NA | NA | 1 | 0 | 7 | 0.2857 |
| 4_1087994 | 2 | 0 | NA | NA | 4 | 0.25 | 1 | 1 | 7 | 0.2857 | 3 | 0.6667 | 3 | 0 | NA | NA | 2 | 0 | 8 | 0.25 |
| 4_1088000 | 2 | 0 | 1 | 1 | 4 | 0 | 1 | 0 | 8 | 0.125 | 2 | 0 | 3 | 0.3333 | NA | NA | 3 | 0 | 8 | 0.125 |
| 4_1088582 | 4 | 0.5 | 4 | 0.5 | 1 | 0 | 3 | 0.3333 | 12 | 0.4167 | 2 | 0 | 3 | 0 | 3 | 0.3333 | NA | NA | 8 | 0.125 |
| 4_1089183 | 12 | 0.1667 | 15 | 0.2 | 19 | 0.1053 | 10 | 0.1 | 56 | 0.1429 | 10 | 0.3 | 8 | 0.375 | 7 | 0.2857 | 8 | 0.125 | 33 | 0.2727 |
| 4_1089190 | 1 | 0 | 2 | 0.5 | 1 | 0 | NA | NA | 4 | 0.25 | 3 | 0.3333 | 2 | 0.5 | 1 | 1 | 2 | 1 | 8 | 0.625 |
| 4_1089191 | 1 | 1 | 2 | 1 | 1 | 1 | NA | NA | 4 | 1 | 3 | 1 | 3 | 1 | 2 | 1 | 2 | 1 | 10 | 1 |
| 4_1089191 | 1 | 0 | 1 | 1 | 1 | 0 | NA | NA | 3 | 0.3333 | 3 | 0.3333 | 3 | 0.3333 | 1 | 1 | 2 | 0 | 9 | 0.3333 |
| 4_1089191 | NA | NA | 2 | 0.5 | 1 | 1 | NA | NA | 3 | 0.6667 | 3 | 1 | 3 | 1 | NA | NA | 2 | 1 | 8 | 1 |
| 4_1089191 | 4 | 0 | 4 | 1 | 7 | 0.1429 | 5 | 0.4 | 20 | 0.35 | 5 | 0.4 | 4 | 0.75 | 9 | 0.1111 | 2 | 0.5 | 20 | 0.35 |
| 4_1089192 | 5 | 0.2 | 7 | 0.1429 | 9 | 0.1111 | 9 | 0.1111 | 30 | 0.1333 | 5 | 0.2 | 4 | 0.5 | 8 | 0 | 3 | 0 | 20 | 0.15 |
| 4_1089192 | 1 | 1 | 1 | 1 | 1 | 0 | NA | NA | 3 | 0.6667 | 3 | 0.3333 | NA | NA | 2 | 0 | NA | NA | 5 | 0.2 |
| 4_1089193 | 5 | 0.4 | 3 | 0 | 8 | 0 | 7 | 0.5714 | 23 | 0.2609 | 7 | 0.2857 | 3 | 0.3333 | 8 | 0.125 | 2 | 1 | 20 | 0.3 |
| 4_1089193 | 9 | 0.5556 | 8 | 0.5 | 11 | 0.8182 | 8 | 0.875 | 36 | 0.6944 | 6 | 0.6667 | 6 | 0.6667 | 9 | 0.4444 | 2 | 1 | 23 | 0.6087 |
| 4_1089194 | 2 | 1 | 2 | 1 | 1 | 1 | NA | NA | 5 | 1 | 3 | 0.6667 | 3 | 1 | 1 | 1 | 2 | 1 | 9 | 0.8889 |
| 4_1089196 | 2 | 0.5 | 1 | 0 | 2 | 0 | NA | NA | 5 | 0.2 | 3 | 0 | 3 | 0.3333 | 2 | 0 | 2 | 0 | 10 | 0.1 |
| 4_1089197 | 2 | 1 | 1 | 0 | 2 | 0.5 | NA | NA | 5 | 0.6 | 3 | 0 | 2 | 1 | 2 | 1 | 3 | 0.6667 | 10 | 0.6 |
| 4_1089198 | 2 | 0 | 2 | 0.5 | 2 | 0 | NA | NA | 6 | 0.1667 | 3 | 0.6667 | 3 | 0.3333 | 2 | 1 | 1 | 1 | 9 | 0.6667 |
| 4_1089199 | 2 | 0.5 | 2 | 0.5 | 2 | 0.5 | 1 | 1 | 7 | 0.5714 | 3 | 0.3333 | 3 | 0.3333 | 1 | 0 | 3 | 0.3333 | 10 | 0.3 |
| 4_1089199 | 2 | 1 | 1 | 1 | 2 | 0.5 | 1 | 0 | 6 | 0.6667 | 3 | 1 | 2 | 0 | 2 | 1 | 3 | 0.6667 | 10 | 0.7 |
| 4_1089200 | 2 | 1 | 2 | 1 | 2 | 1 | 1 | 0 | 7 | 0.8571 | 1 | 0 | 3 | 0.6667 | 2 | 1 | 3 | 1 | 9 | 0.7778 |
| 4_1089200 | 2 | 0.5 | 2 | 0 | 2 | 0 | NA | NA | 6 | 0.1667 | 3 | 0 | 3 | 0 | 2 | 0.5 | 3 | 1 | 11 | 0.3636 |
| 4_1089205 | 2 | 1 | 1 | 1 | 2 | 1 | 1 | 1 | 6 | 1 | 3 | 0.6667 | 3 | 0.6667 | 2 | 1 | 3 | 0.6667 | 11 | 0.7273 |
| 4_1089206 | 2 | 0.5 | 2 | 0 | 2 | 0 | 1 | 0 | 7 | 0.1429 | 3 | 0.3333 | 3 | 0.3333 | 2 | 0.5 | 3 | 0.3333 | 11 | 0.3636 |
| 4_1092426 | 13 | 0.6923 | 14 | 0.7857 | 8 | 0.625 | 16 | 0.75 | 51 | 0.7255 | 5 | 0.4 | 12 | 0.6667 | 14 | 0.8571 | 17 | 0.9412 | 48 | 0.7917 |
| 4_1092427 | 9 | 0.3333 | 11 | 0.2727 | 8 | 0.375 | 13 | 0.3846 | 41 | 0.3415 | 6 | 0.1667 | 11 | 0.3636 | 11 | 0.1818 | 17 | 0.1176 | 45 | 0.2 |
| 4_1092433 | 11 | 0.0909 | 19 | 0.2632 | 11 | 0.0909 | 16 | 0.375 | 57 | 0.2281 | 7 | 0.1429 | 11 | 0.0909 | 23 | 0.2174 | 18 | 0.2222 | 59 | 0.1864 |
| 4_1093114 | 29 | 0.4483 | 24 | 0.4583 | 27 | 0.5185 | 28 | 0.4643 | 108 | 0.4722 | 23 | 0.6522 | 20 | 0.15 | 25 | 0.24 | 23 | 0.3478 | 91 | 0.3516 |
| 4_1093115 | 28 | 0.2143 | 31 | 0.129 | 33 | 0.0303 | 35 | 0.2286 | 127 | 0.1496 | 29 | 0.2069 | 27 | 0.1111 | 27 | 0.2222 | 23 | 0.1304 | 106 | 0.1698 |
| 4_1093115 | 27 | 0.7778 | 28 | 0.7857 | 29 | 0.8621 | 31 | 0.7419 | 115 | 0.7913 | 25 | 0.8 | 22 | 0.7727 | 24 | 0.8333 | 23 | 0.8261 | 94 | 0.8085 |
| 4_1093115 | 31 | 0.1935 | 29 | 0.1724 | 29 | 0.1034 | 32 | 0.1875 | 121 | 0.1653 | 30 | 0.0667 | 27 | 0.1852 | 26 | 0 | 24 | 0.2917 | 107 | 0.1308 |
| 4_1093115 | 31 | 0.7419 | 30 | 0.7667 | 34 | 0.8529 | 33 | 0.9091 | 128 | 0.8203 | 31 | 0.8387 | 27 | 0.7778 | 28 | 0.8571 | 23 | 0.7826 | 109 | 0.8165 |
| 4_1100627 | 3 | 0 | 5 | 0 | 4 | 0.25 | 5 | 0 | 17 | 0.0588 | 2 | 0 | 2 | 0.5 | 2 | 0 | 6 | 0 | 12 | 0.0833 |
| 4_1100634 | 3 | 1 | 4 | 0.75 | 3 | 0.6667 | 1 | 1 | 11 | 0.8182 | 3 | 1 | 1 | 1 | 1 | 1 | 5 | 1 | 10 | 1 |
| 4_1100634 | 4 | 0.5 | 4 | 0.25 | 4 | 0.25 | 4 | 0 | 16 | 0.25 | 3 | 0 | 1 | 0 | NA | NA | 5 | 0.2 | 9 | 0.1111 |
| 4_1100634 | 4 | 0.25 | 4 | 0 | 5 | 0.2 | 4 | 0.5 | 17 | 0.2353 | 3 | 0 | NA | NA | 2 | 0 | 5 | 0.2 | 10 | 0.1 |
| 4_1100634 | 3 | 0.3333 | 4 | 0.75 | 2 | 0 | 2 | 0 | 11 | 0.3636 | NA | NA | 1 | 0 | 2 | 0.5 | 4 | 0 | 7 | 0.1429 |
| 4_1100638 | 4 | 0 | 3 | 0.3333 | 3 | 0.3333 | 2 | 0 | 12 | 0.1667 | 2 | 1 | NA | NA | NA | NA | 3 | 0.3333 | 5 | 0.6 |
| 4_1100669 | 4 | 0 | 4 | 0.25 | 4 | 0.25 | 4 | 0 | 16 | 0.125 | 2 | 0.5 | 1 | 0 | 1 | 0 | 5 | 0.2 | 9 | 0.2222 |
| 4_1100687 | 4 | 0.25 | 4 | 0.25 | 3 | 0 | 4 | 0 | 15 | 0.1333 | 2 | 0.5 | 1 | 0 | 1 | 0 | 5 | 0 | 9 | 0.1111 |
| 4_1112905 | 2 | 1 | 1 | 0 | 1 | 1 | 4 | 0.75 | 8 | 0.75 | 2 | 1 | 1 | 0 | 2 | 0.5 | 1 | 1 | 6 | 0.6667 |
| 4_1112905 | 2 | 1 | 1 | 1 | 1 | 1 | 3 | 1 | 7 | 1 | 2 | 1 | NA | NA | 2 | 1 | 3 | 1 | 7 | 1 |

| | | | | | | | | | | | | | | | | | | | | |
|---|---|---|---|---|---|---|---|---|---|---|---|---|---|---|---|---|---|---|---|---|
| 4_11129062 | 2 | 0 | 1 | 0 | 1 | 0 | 4 | 0.25 | 8 | 0.125 | 2 | 0 | 1 | 1 | 2 | 0 | 3 | 0.3333 | 8 | 0.25 |
| 4_11131700 | 2 | 0.5 | 1 | 1 | NA | NA | 4 | 0.25 | 7 | 0.4286 | 2 | 0 | 1 | 0 | 3 | 0 | 3 | 0.6667 | 9 | 0.2222 |
| 4_11131729 | 1 | 1 | NA | NA | NA | NA | 1 | 0 | 2 | 0.5 | NA | NA | 1 | 1 | NA | NA | NA | NA | 1 | 1 |
| 4_11131741 | NA | NA | NA | NA | NA | NA | 1 | 1 | 1 | 1 | NA | NA | 1 | 1 | NA | NA | NA | NA | 1 | 1 |
| 4_11168557 | NA | NA | 2 | 0.5 | NA | NA | NA | NA | 2 | 0.5 | NA | NA | 3 | 0.6667 | NA | NA | NA | NA | 3 | 0.6667 |
| 4_11481835 | 23 | 0.087 | 27 | 0.037 | 15 | 0 | 13 | 0.1538 | 78 | 0.0641 | 15 | 0.1333 | 13 | 0 | 12 | 0 | 17 | 0.0588 | 57 | 0.0526 |
| 4_11481842 | 23 | 0.2609 | 25 | 0.32 | 14 | 0.2143 | 13 | 0.1538 | 75 | 0.2533 | 12 | 0.25 | 14 | 0.0714 | 13 | 0.0769 | 16 | 0.3125 | 55 | 0.1818 |
| 4_11481847 | 23 | 0.087 | 28 | 0.0357 | 14 | 0.0714 | 14 | 0.0714 | 79 | 0.0633 | 15 | 0 | 14 | 0.1429 | 12 | 0 | 17 | 0.1176 | 58 | 0.069 |
| 4_11484473 | NA | NA | NA | NA | 1 | 1 | NA | NA | 1 | 1 | NA | NA | NA | NA | 1 | 1 | NA | NA | 1 | 1 |
| 4_11496466 | 1 | 1 | 5 | 0 | 1 | 0 | 1 | 0 | 8 | 0.125 | NA | NA | 1 | 0 | 1 | 1 | 2 | 0 | 4 | 0.25 |
| 4_11496500 | 1 | 1 | 3 | 0.3333 | 1 | 1 | 1 | 1 | 6 | 0.6667 | NA | NA | 1 | 0 | 1 | 1 | 2 | 0 | 4 | 0.25 |
| 4_11496504 | 1 | 1 | 4 | 1 | 1 | 1 | 1 | 1 | 7 | 1 | NA | NA | 1 | 1 | 1 | 1 | 1 | 0 | 3 | 0.6667 |
| 4_11496510 | 1 | 1 | 5 | 0.4 | 1 | 1 | 1 | 1 | 8 | 0.625 | NA | NA | 1 | 1 | 1 | 1 | 1 | 1 | 3 | 1 |
| 4_11496516 | 1 | 0 | 5 | 0.2 | 1 | 1 | 1 | 1 | 8 | 0.375 | NA | NA | 1 | 0 | 1 | 1 | 2 | 1 | 4 | 0.75 |
| 4_11496517 | 1 | 1 | 5 | 1 | 1 | 1 | 1 | 1 | 8 | 1 | NA | NA | 1 | 1 | 1 | 1 | 1 | 1 | 3 | 1 |
| 4_11496523 | 1 | 1 | 5 | 1 | 1 | 1 | 1 | 1 | 8 | 1 | NA | NA | 1 | 1 | NA | NA | 2 | 1 | 3 | 1 |
| 4_11496534 | 1 | 0 | 5 | 0.4 | 1 | 0 | 1 | 0 | 8 | 0.25 | NA | NA | 1 | 1 | 1 | 0 | 2 | 0 | 4 | 0.25 |
| 4_11496537 | 1 | 0 | 4 | 0 | 1 | 0 | 1 | 1 | 7 | 0.1429 | NA | NA | 1 | 0 | 1 | 0 | 2 | 0.5 | 4 | 0.25 |
| 4_11496546 | 1 | 0 | 5 | 0.2 | 1 | 0 | 1 | 0 | 8 | 0.125 | NA | NA | 1 | 1 | 1 | 0 | 2 | 0 | 4 | 0.25 |
| 4_11496561 | 1 | 0 | 4 | 0.75 | NA | NA | NA | NA | 5 | 0.6 | NA | NA | 1 | 0 | 1 | 1 | 1 | 0 | 3 | 0.3333 |
| 4_11496571 | 1 | 0 | 5 | 0.2 | NA | NA | 1 | 1 | 7 | 0.2857 | NA | NA | 1 | 1 | 1 | 1 | 2 | 1 | 4 | 1 |
| 4_11496577 | 1 | 1 | 4 | 1 | NA | NA | NA | NA | 5 | 1 | NA | NA | 1 | 1 | 1 | 1 | 2 | 1 | 4 | 1 |
| 4_11496589 | 1 | 1 | 5 | 0.2 | NA | NA | 1 | 1 | 7 | 0.4286 | NA | NA | 1 | 0 | 1 | 0 | 1 | 1 | 3 | 0.3333 |
| 4_11559522 | 2 | 0.5 | 1 | 1 | NA | NA | NA | NA | 3 | 0.6667 | 1 | 0 | 3 | 0.3333 | 2 | 0.5 | 1 | 1 | 7 | 0.4286 |
| 4_11559552 | 2 | 0.5 | 3 | 0.3333 | NA | NA | 2 | 0.5 | 7 | 0.4286 | 1 | 0 | 6 | 0.6667 | 2 | 0.5 | 3 | 0.6667 | 12 | 0.5833 |
| 4_11559581 | 2 | 0 | 2 | 0.5 | NA | NA | 1 | 0 | 5 | 0.2 | 1 | 0 | 6 | 0 | 3 | 0.3333 | 3 | 0.3333 | 13 | 0.1538 |
| 4_11563109 | 1 | 0 | 4 | 0 | NA | NA | 4 | 0.25 | 9 | 0.1111 | 1 | 0 | 8 | 0 | 3 | 0.3333 | 3 | 0 | 15 | 0.0667 |
| 4_11563124 | 2 | 1 | 2 | 0 | NA | NA | 5 | 0.8 | 9 | 0.6667 | 1 | 1 | 9 | 0.3333 | 3 | 0 | 2 | 0.5 | 15 | 0.3333 |
| 4_11563127 | 2 | 1 | 2 | 0.5 | 1 | 0 | 5 | 0 | 10 | 0.3 | 1 | 0 | 8 | 0.125 | 3 | 0.3333 | 2 | 0 | 14 | 0.1429 |
| 4_11563129 | 2 | 0 | 2 | 0 | 1 | 1 | 5 | 0 | 10 | 0.1 | 1 | 0 | 8 | 0 | 3 | 0.3333 | 2 | 0 | 14 | 0.0714 |
| 4_11563247 | 3 | 0 | 4 | 0.25 | 2 | 0 | 3 | 0 | 12 | 0.0833 | 3 | 0 | 7 | 0.1429 | 3 | 0 | 6 | 0 | 19 | 0.0526 |
| 4_11563334 | 7 | 0.1429 | 7 | 0 | 3 | 0.3333 | 6 | 0 | 23 | 0.087 | 2 | 0.5 | 8 | 0 | 4 | 0 | 5 | 0.2 | 19 | 0.1053 |
| 4_11566828 | 5 | 0.6 | 9 | 0.3333 | 15 | 0.2 | 15 | 0.6 | 44 | 0.4091 | 4 | 0.25 | 12 | 0.3333 | 10 | 0.5 | 7 | 0.2857 | 33 | 0.3636 |
| 4_11566834 | 5 | 1 | 11 | 0.5455 | 14 | 0.7143 | 13 | 0.8462 | 43 | 0.7442 | 3 | 0 | 12 | 0.6667 | 11 | 0.7273 | 8 | 0.75 | 34 | 0.6471 |
| 4_11566836 | 3 | 0.6667 | 10 | 0.3 | 11 | 0.3636 | 13 | 0.2308 | 37 | 0.3243 | 3 | 0.3333 | 10 | 0.4 | 9 | 0.6667 | 7 | 0.5714 | 29 | 0.5172 |
| 4_11566838 | 5 | 0.6 | 11 | 0.5455 | 14 | 0.2857 | 14 | 0.5 | 44 | 0.4545 | 3 | 0.6667 | 11 | 0.3636 | 9 | 0.3333 | 6 | 0.6667 | 29 | 0.4483 |
| 4_11566854 | 5 | 0.2 | 8 | 0.375 | 12 | 0.4167 | 13 | 0.6154 | 38 | 0.4474 | 3 | 0.3333 | 11 | 0.4545 | 12 | 0.5 | 7 | 0.1429 | 33 | 0.3939 |
| 4_11566871 | 6 | 0 | 12 | 0.1667 | 12 | 0.0833 | 15 | 0 | 45 | 0.0667 | 3 | 0 | 11 | 0.0909 | 11 | 0.1818 | 7 | 0 | 32 | 0.0938 |
| 4_11566893 | 6 | 0.3333 | 12 | 0.1667 | 13 | 0.0769 | 14 | 0.2143 | 45 | 0.1778 | 4 | 0.25 | 14 | 0.0714 | 12 | 0.1667 | 10 | 0 | 40 | 0.1 |
| 4_11566903 | 6 | 0.1667 | 13 | 0.0769 | 14 | 0 | 14 | 0.0714 | 47 | 0.0638 | 4 | 0 | 11 | 0.1818 | 12 | 0.0833 | 6 | 0 | 33 | 0.0909 |
| 4_11567097 | 2 | 0 | 12 | 0.0833 | 1 | 0 | 4 | 0 | 19 | 0.0526 | 1 | 1 | 8 | 0.375 | 7 | 0 | 5 | 0 | 21 | 0.1905 |
| 4_11568866 | NA | NA | NA | NA | NA | NA | 1 | 1 | 1 | 1 | NA | NA | NA | NA | NA | NA | 1 | 1 | 1 | 1 |
| 4_11569309 | NA | NA | NA | NA | NA | NA | 1 | 1 | 1 | 1 | NA | NA | 1 | 0 | NA | NA | 1 | 1 | 2 | 0.5 |
| 4_11569322 | NA | NA | NA | NA | NA | NA | 2 | 0.5 | 2 | 0.5 | NA | NA | NA | NA | NA | NA | 1 | 1 | 1 | 1 |
| 4_11570546 | 1 | 0 | NA | NA | 2 | 0 | 1 | 1 | 4 | 0.25 | 1 | 0 | NA | NA | 2 | 0 | 1 | 1 | 4 | 0.25 |
| 4_11573457 | NA | NA | 1 | 0 | 3 | 0.3333 | NA | NA | 4 | 0.25 | NA | NA | NA | NA | NA | NA | 1 | 1 | 1 | 1 |
| 4_11573516 | 2 | 0 | 2 | 1 | 3 | 0.3333 | 1 | 0 | 8 | 0.375 | 2 | 0.5 | 1 | 0 | 1 | 0 | 3 | 1 | 7 | 0.5714 |
| 4_11573523 | NA | NA | NA | NA | 2 | 0.5 | 1 | 1 | 3 | 0.6667 | 2 | 0 | 1 | 1 | 2 | 0.5 | 2 | 1 | 7 | 0.5714 |
| 4_11573536 | 1 | 0 | 1 | 0 | 3 | 0.3333 | 1 | 0 | 6 | 0.1667 | 2 | 0 | NA | NA | 2 | 0.5 | 3 | 0 | 7 | 0.1429 |
| 4_11573546 | 2 | 0 | 2 | 1 | 3 | 0 | 1 | 1 | 8 | 0.375 | NA | NA | 1 | 0 | 2 | 0.5 | 3 | 0.3333 | 6 | 0.3333 |
| 4_11573563 | 2 | 0 | 2 | 0 | 5 | 0 | 2 | 0.5 | 11 | 0.0909 | 2 | 0 | 1 | 0 | 3 | 0 | 3 | 0.3333 | 9 | 0.1111 |
| 4_11573618 | 2 | 0.5 | 3 | 1 | 5 | 0.6 | 2 | 0.5 | 12 | 0.6667 | 3 | 0.6667 | 2 | 0.5 | 3 | 0.6667 | 3 | 0.6667 | 11 | 0.6364 |
| 4_11573624 | 2 | 0 | 3 | 0.3333 | 4 | 0.25 | 2 | 0.5 | 11 | 0.2727 | 3 | 0.3333 | 2 | 0 | 3 | 0.3333 | 4 | 0.25 | 12 | 0.25 |
| 4_11585396 | 1 | 0 | 2 | 0 | 4 | 0.25 | 3 | 0 | 10 | 0.1 | 1 | 1 | 1 | 0 | NA | NA | 1 | 0 | 3 | 0.3333 |
| 4_11585403 | NA | NA | 3 | 0.6667 | 3 | 0.6667 | 1 | 0 | 7 | 0.5714 | 1 | 0 | 1 | 0 | 1 | 1 | 2 | 0 | 5 | 0.2 |
| 4_11585489 | NA | NA | 3 | 0.6667 | 1 | 1 | 1 | 1 | 5 | 0.8 | NA | NA | 1 | 1 | NA | NA | 2 | 1 | 3 | 1 |
| 4_11585513 | 4 | 0.25 | 8 | 0 | 3 | 0 | 2 | 0 | 17 | 0.0588 | 3 | 0 | 1 | 0 | 1 | 0 | 5 | 0.2 | 10 | 0.1 |
| 4_11585518 | 3 | 1 | 8 | 0.75 | 3 | 0.3333 | 2 | 0.5 | 16 | 0.6875 | 3 | 0.3333 | 1 | 0 | NA | NA | 5 | 0.4 | 9 | 0.3333 |
| 4_11585524 | 4 | 0.5 | 9 | 0.7778 | 2 | 0.5 | 2 | 0 | 17 | 0.5882 | 3 | 0.6667 | 1 | 0 | 1 | 0 | 6 | 0.6667 | 11 | 0.5455 |
| 4_11585535 | 4 | 0 | 8 | 0.25 | 2 | 0 | 2 | 0 | 16 | 0.125 | 3 | 0.3333 | 1 | 0 | NA | NA | 6 | 0 | 10 | 0.1 |
| 4_11585536 | 2 | 0 | 5 | 0.4 | 2 | 0 | 1 | 1 | 10 | 0.3 | 2 | 0.5 | 1 | 1 | NA | NA | 3 | 0 | 6 | 0.3333 |
| 4_11585542 | 6 | 0.1667 | 8 | 0.625 | 4 | 0.25 | 2 | 0.5 | 20 | 0.4 | 3 | 0.3333 | 1 | 0 | 1 | 1 | 6 | 0.5 | 11 | 0.4545 |
| 4_11585550 | 5 | 1 | 6 | 1 | 4 | 1 | 2 | 1 | 17 | 1 | 3 | 0.6667 | 1 | 1 | 1 | 1 | 5 | 1 | 10 | 0.9 |
| 4_11585553 | 6 | 0.3333 | 9 | 0.6667 | 4 | 0.5 | 2 | 0.5 | 21 | 0.5238 | 3 | 0.6667 | 1 | 0 | 1 | 1 | 6 | 0.5 | 11 | 0.5455 |
| 4_11585555 | 6 | 0.6667 | 9 | 0.8889 | 3 | 0.6667 | 1 | 1 | 19 | 0.7895 | 3 | 0.3333 | NA | NA | NA | NA | 4 | 0.75 | 7 | 0.5714 |

| | | | | | | | | | | | | | | | | | | | | |
|---|---|---|---|---|---|---|---|---|---|---|---|---|---|---|---|---|---|---|---|---|
| 4_1158905 | 4 | 0.25 | 5 | 0.4 | 3 | 0.3333 | NA | NA | 12 | 0.3333 | NA | NA | 1 | 1 | 2 | 0 | 3 | 0 | 6 | 0.1667 |
| 4_1158912 | 2 | 0 | 3 | 0.3333 | 4 | 0 | NA | NA | 9 | 0.1111 | NA | NA | 2 | 0 | 2 | 0 | 1 | 1 | 5 | 0.2 |
| 4_1158913 | 2 | 1 | 3 | 1 | 1 | 1 | NA | NA | 6 | 1 | NA | NA | 2 | 1 | 1 | 1 | 3 | 1 | 6 | 1 |
| 4_1159055 | 2 | 0.5 | 2 | 0.5 | 1 | 0 | NA | NA | 5 | 0.4 | NA | NA | 2 | 0 | 2 | 0 | 3 | 0.6667 | 7 | 0.2857 |
| 4_1159056 | 2 | 1 | 2 | 0.5 | 1 | 0 | NA | NA | 5 | 0.6 | NA | NA | 2 | 0 | 2 | 0 | 3 | 0.3333 | 7 | 0.1429 |
| 4_1159242 | 94 | 0.0532 | 109 | 0.055 | 78 | 0.0641 | 95 | 0.0632 | 376 | 0.0585 | 70 | 0.1 | 86 | 0.0465 | 84 | 0.0714 | 70 | 0.0571 | 310 | 0.0677 |
| 4_1159458 | 5 | 0.4 | 5 | 0.6 | 7 | 0.4286 | 4 | 0.75 | 21 | 0.5238 | 2 | 0.5 | 2 | 0.5 | 3 | 0.3333 | 4 | 0.5 | 11 | 0.4545 |
| 4_1159462 | 4 | 1 | 3 | 1 | 7 | 0.7143 | 4 | 1 | 18 | 0.8889 | 2 | 1 | 2 | 1 | 3 | 1 | 4 | 0.75 | 11 | 0.9091 |
| 4_1159465 | 6 | 0 | 5 | 0 | 7 | 0.1429 | 6 | 0.1667 | 24 | 0.0833 | 2 | 0 | 2 | 0.5 | 4 | 0.25 | 5 | 0.2 | 13 | 0.2308 |
| 4_1159465 | 6 | 0.8333 | 6 | 0.6667 | 8 | 0.875 | 6 | 1 | 26 | 0.8462 | 2 | 1 | 2 | 0.5 | 2 | 1 | 5 | 1 | 11 | 0.9091 |
| 4_1159536 | 7 | 0.8571 | 9 | 0.7778 | 9 | 0.8889 | 8 | 0.875 | 33 | 0.8485 | 2 | 1 | 4 | 1 | 6 | 0.6667 | 6 | 0.8333 | 18 | 0.8333 |
| 4_1159537 | 7 | 0.8571 | 6 | 0.8333 | 8 | 0.75 | 6 | 0.8333 | 27 | 0.8148 | 3 | 0.6667 | 4 | 1 | 7 | 0.5714 | 6 | 1 | 20 | 0.8 |
| 4_1159940 | 1 | 0 | NA | NA | NA | NA | 2 | 1 | 3 | 0.6667 | 1 | 0 | 2 | 1 | 2 | 0 | 1 | 1 | 6 | 0.5 |
| 4_1159940 | NA | NA | NA | NA | NA | NA | 2 | 1 | 2 | 1 | 1 | 1 | 2 | 1 | 2 | 1 | 1 | 1 | 6 | 1 |
| 4_1159943 | 1 | 0 | NA | NA | NA | NA | 2 | 0.5 | 3 | 0.3333 | 1 | 1 | 2 | 0.5 | 1 | 1 | 1 | 1 | 5 | 0.8 |
| 4_1159945 | 1 | 0 | NA | NA | NA | NA | 2 | 0.5 | 3 | 0.3333 | 1 | 1 | 1 | 0 | 1 | 0 | NA | NA | 3 | 0.3333 |
| 4_1159945 | 1 | 1 | NA | NA | NA | NA | 2 | 1 | 3 | 1 | 1 | 0 | 1 | 0 | NA | NA | 1 | 1 | 3 | 0.3333 |
| 4_1159946 | NA | NA | NA | NA | NA | NA | 1 | 1 | 1 | 1 | NA | NA | 2 | 0.5 | 1 | 0 | 1 | 1 | 4 | 0.5 |
| 4_1159946 | 1 | 1 | NA | NA | NA | NA | 2 | 1 | 3 | 1 | 1 | 1 | 1 | 1 | 1 | 1 | 1 | 1 | 4 | 1 |
| 4_1159946 | 1 | 1 | NA | NA | NA | NA | 2 | 1 | 3 | 1 | 1 | 1 | 1 | 1 | 2 | 0 | 1 | 0 | 5 | 0.4 |
| 4_1159947 | 1 | 1 | NA | NA | NA | NA | 2 | 1 | 3 | 1 | 1 | 1 | 2 | 0.5 | 2 | 0 | 1 | 1 | 6 | 0.5 |
| 4_1159947 | 1 | 1 | NA | NA | NA | NA | 1 | 1 | 2 | 1 | 1 | 1 | 2 | 1 | 2 | 1 | 1 | 1 | 6 | 1 |
| 4_1159947 | 1 | 1 | NA | NA | NA | NA | 2 | 1 | 3 | 1 | 1 | 1 | 2 | 1 | 1 | 1 | 1 | 1 | 5 | 1 |
| 4_1159948 | 1 | 1 | NA | NA | NA | NA | 2 | 1 | 3 | 1 | 1 | 0 | 1 | 0 | 2 | 0.5 | 1 | 1 | 5 | 0.4 |
| 4_1159948 | 1 | 1 | NA | NA | NA | NA | 2 | 1 | 3 | 1 | 1 | 0 | NA | NA | 2 | 0.5 | 1 | 0 | 4 | 0.25 |
| 4_1160082 | 5 | 0 | 5 | 0.2 | 3 | 0 | 2 | 0 | 15 | 0.0667 | 5 | 0.2 | 4 | 0.25 | 2 | 0 | 4 | 0 | 15 | 0.1333 |
| 4_1160082 | 3 | 0 | 4 | 0 | 3 | 0.3333 | 3 | 0 | 13 | 0.0769 | 4 | 0 | 2 | 0 | 3 | 0.3333 | 4 | 0 | 13 | 0.0769 |
| 4_1160164 | 11 | 0.1818 | 9 | 0.2222 | 10 | 0.4 | 6 | 0.1667 | 36 | 0.25 | 6 | 0.1667 | 7 | 0.7143 | 6 | 0.1667 | 7 | 0.4286 | 26 | 0.3846 |
| 4_1160168 | 11 | 0.1818 | 8 | 0 | 10 | 0 | 5 | 0 | 34 | 0.0588 | 6 | 0 | 9 | 0.1111 | 7 | 0 | 9 | 0.1111 | 31 | 0.0645 |
| 4_1161570 | NA | NA | 2 | 0.5 | NA | NA | NA | NA | 2 | 0.5 | 1 | 0 | NA | NA | 1 | 0 | 2 | 0.5 | 4 | 0.25 |
| 4_1161571 | NA | NA | 2 | 0.5 | NA | NA | NA | NA | 2 | 0.5 | NA | NA | NA | NA | NA | NA | 2 | 1 | 2 | 1 |
| 4_1161572 | NA | NA | 2 | 1 | NA | NA | NA | NA | 2 | 1 | 1 | 1 | NA | NA | 1 | 1 | 2 | 1 | 4 | 1 |
| 4_1161635 | NA | NA | NA | NA | NA | NA | 1 | 1 | 1 | 1 | NA | NA | 1 | 0 | NA | NA | 2 | 0.5 | 3 | 0.3333 |
| 4_1161636 | NA | NA | NA | NA | NA | NA | 1 | 1 | 1 | 1 | NA | NA | 1 | 0 | NA | NA | 2 | 0.5 | 3 | 0.3333 |
| 4_1161638 | NA | NA | NA | NA | NA | NA | 1 | 1 | 1 | 1 | NA | NA | 1 | 1 | NA | NA | 2 | 0 | 3 | 0.3333 |
| 4_1161640 | 7 | 0 | 9 | 0 | 7 | 0.1429 | 9 | 0.1111 | 32 | 0.0625 | 10 | 0.1 | 11 | 0.1818 | 8 | 0 | 9 | 0.2222 | 38 | 0.1316 |
| 4_1161640 | 8 | 0.875 | 10 | 0.7 | 7 | 1 | 8 | 0.5 | 33 | 0.7576 | 11 | 0.7273 | 11 | 0.5455 | 7 | 0.7143 | 9 | 1 | 38 | 0.7368 |
| 4_1161640 | 8 | 0.875 | 10 | 0.6 | 5 | 0.4 | 9 | 1 | 32 | 0.75 | 9 | 0.8889 | 9 | 0.8889 | 7 | 0.8571 | 8 | 0.75 | 33 | 0.8485 |
| 4_1161641 | 8 | 0.875 | 8 | 0.875 | 7 | 1 | 8 | 0.875 | 31 | 0.9032 | 11 | 0.9091 | 11 | 1 | 7 | 1 | 9 | 0.8889 | 38 | 0.9474 |
| 4_1161642 | 8 | 0 | 8 | 0.25 | 5 | 0.2 | 8 | 0.125 | 29 | 0.1379 | 8 | 0.25 | 6 | 0 | 6 | 0.1667 | 9 | 0.1111 | 29 | 0.1379 |
| 4_1161642 | 6 | 0.3333 | 6 | 0.5 | 7 | 0 | 7 | 0.2857 | 26 | 0.2692 | 9 | 0.5556 | 6 | 0.3333 | 7 | 0 | 9 | 0.4444 | 31 | 0.3548 |
| 4_1161643 | 5 | 0.2 | 7 | 0.2857 | 7 | 0 | 4 | 0.25 | 23 | 0.1739 | 8 | 0.25 | 9 | 0 | 5 | 0.4 | 7 | 0.4286 | 29 | 0.2414 |
| 4_1161644 | 7 | 0.4286 | 6 | 0 | 7 | 0.4286 | 6 | 0.1667 | 26 | 0.2692 | 10 | 0.3 | 10 | 0.1 | 6 | 0.3333 | 9 | 0.2222 | 35 | 0.2286 |
| 4_1161644 | 6 | 0 | 7 | 0.5714 | 6 | 0.5 | 8 | 0.125 | 27 | 0.2963 | 6 | 0.3333 | 7 | 0.5714 | 4 | 0 | 9 | 0.4444 | 26 | 0.3846 |
| 4_1161645 | 5 | 0.2 | 6 | 0 | 4 | 0.75 | 5 | 0.4 | 20 | 0.3 | 8 | 0 | 6 | 0.1667 | 3 | 0.3333 | 8 | 0.25 | 25 | 0.16 |
| 4_1161645 | 5 | 0.2 | 6 | 0.5 | 5 | 0.6 | 7 | 0.4286 | 23 | 0.4348 | 10 | 0.5 | 10 | 0.5 | 5 | 0.2 | 9 | 0.2222 | 34 | 0.3824 |
| 4_1161647 | 4 | 0.5 | 7 | 0.5714 | 3 | 1 | 8 | 1 | 22 | 0.7727 | 6 | 0.8333 | 9 | 0.8889 | 3 | 0.3333 | 6 | 0.6667 | 24 | 0.75 |
| 4_1161649 | 7 | 0.1429 | 8 | 0.125 | 4 | 0.25 | 8 | 0.125 | 27 | 0.1481 | 8 | 0.125 | 10 | 0.1 | 6 | 0 | 5 | 0.4 | 29 | 0.1379 |
| 4_1161650 | 3 | 0.3333 | 4 | 0.5 | 2 | 1 | 4 | 0.75 | 13 | 0.6154 | 4 | 0.5 | 6 | 0.8333 | 3 | 0.6667 | 6 | 0.8333 | 19 | 0.7368 |
| 4_1161650 | 5 | 0.6 | 5 | 0.2 | 3 | 0 | 7 | 0.4286 | 20 | 0.35 | 7 | 0 | 7 | 0.2857 | 4 | 0.5 | 4 | 0.25 | 22 | 0.2273 |
| 4_1161652 | 2 | 0 | 4 | 0 | 3 | 0.3333 | 7 | 0 | 16 | 0.0625 | 4 | 0.25 | 7 | 0.1429 | 5 | 0.2 | 7 | 0 | 23 | 0.1304 |
| 4_1161654 | 3 | 0 | 7 | 0.2857 | 3 | 0 | 4 | 0.5 | 17 | 0.2353 | 7 | 0.1429 | 2 | 0 | 4 | 0.25 | 5 | 0.4 | 18 | 0.2222 |
| 4_1161677 | 1 | 0 | 5 | 0 | 2 | 0.5 | 4 | 0 | 12 | 0.0833 | 7 | 0 | 4 | 0 | 2 | 0.5 | 5 | 0 | 18 | 0.0556 |
| 4_1161689 | NA | NA | 5 | 0 | 1 | 1 | 3 | 0 | 9 | 0.1111 | 4 | 0 | 3 | 0 | 1 | 0 | 4 | 0.25 | 12 | 0.0833 |
| 4_1161701 | NA | NA | 4 | 0 | 1 | 0 | 2 | 0.5 | 7 | 0.1429 | 3 | 0.6667 | NA | NA | 1 | 0 | 2 | 0 | 6 | 0.3333 |
| 4_1163951 | 2 | 1 | NA | NA | 1 | 1 | NA | NA | 3 | 1 | NA | NA | 1 | 1 | 3 | 1 | 1 | 1 | 5 | 1 |
| 4_1163952 | 1 | 0 | NA | NA | 2 | 0.5 | NA | NA | 3 | 0.3333 | 1 | 1 | NA | NA | 3 | 0.3333 | 1 | 0 | 5 | 0.4 |
| 4_1164000 | NA | NA | NA | NA | NA | NA | 1 | 1 | 1 | 1 | NA | NA | 1 | 1 | NA | NA | NA | NA | 1 | 1 |
| 4_1164001 | NA | NA | NA | NA | NA | NA | 1 | 1 | 1 | 1 | NA | NA | 1 | 1 | NA | NA | NA | NA | 1 | 1 |
| 4_1164088 | 3 | 0.6667 | NA | NA | 1 | 0 | 1 | 0 | 5 | 0.4 | 1 | 1 | NA | NA | 3 | 0 | NA | NA | 4 | 0.25 |
| 4_1164089 | 3 | 0.3333 | NA | NA | 1 | 0 | 1 | 0 | 5 | 0.2 | 1 | 1 | NA | NA | 2 | 0.5 | NA | NA | 3 | 0.6667 |
| 4_1164103 | 2 | 0 | NA | NA | 2 | 0 | 4 | 0.25 | 8 | 0.125 | 1 | 1 | 1 | 1 | 4 | 0.25 | NA | NA | 6 | 0.5 |
| 4_1164104 | 2 | 0 | NA | NA | 2 | 0 | 4 | 0.25 | 8 | 0.125 | 1 | 0 | 1 | 1 | 4 | 0.25 | NA | NA | 6 | 0.3333 |
| 4_1164108 | 2 | 0 | NA | NA | 1 | 0 | 4 | 0.25 | 7 | 0.1429 | 1 | 0 | 1 | 0 | 3 | 0.3333 | NA | NA | 5 | 0.2 |
| 4_1164108 | 2 | 1 | NA | NA | 2 | 0 | 3 | 0 | 7 | 0.2857 | 1 | 0 | 1 | 0 | 3 | 0.6667 | NA | NA | 5 | 0.4 |

| | | | | | | | | | | | | | | | | | | | | |
|---|---|---|---|---|---|---|---|---|---|---|---|---|---|---|---|---|---|---|---|---|
| 4_1164501( | 5 | 0.2 | 3 | 0 | 3 | 0.3333 | 6 | 0 | 17 | 0.1176 | 4 | 0 | 3 | 0 | 5 | 0.2 | 2 | 0.5 | 14 | 0.1429 |
| 4_1164504 | 5 | 0 | 3 | 0 | 5 | 0.4 | 5 | 0 | 18 | 0.1111 | 4 | 0.25 | 4 | 0 | 5 | 0 | 4 | 0 | 17 | 0.0588 |
| 4_1164505 | 6 | 0 | 3 | 0 | 5 | 0.2 | 5 | 0 | 19 | 0.0526 | 4 | 0 | 3 | 0.3333 | 5 | 0.2 | 5 | 0 | 17 | 0.1176 |
| 4_1164526 | NA | NA | 1 | 1 | 1 | 0 | NA | NA | 2 | 0.5 | 3 | 0.3333 | NA | NA | NA | NA | NA | NA | 3 | 0.3333 |
| 4_1164529 | NA | NA | 1 | 0 | 2 | 1 | NA | NA | 3 | 0.6667 | 3 | 0.3333 | NA | NA | NA | NA | NA | NA | 3 | 0.3333 |
| 4_1164531 | NA | NA | NA | NA | 2 | 1 | NA | NA | 2 | 1 | 2 | 1 | NA | NA | NA | NA | NA | NA | 2 | 1 |
| 4_1164532 | NA | NA | 1 | 1 | 3 | 1 | 1 | 1 | 5 | 1 | 3 | 0.6667 | NA | NA | NA | NA | NA | NA | 3 | 0.6667 |
| 4_1164532( | NA | NA | 1 | 1 | 3 | 1 | 1 | 1 | 5 | 1 | 3 | 1 | NA | NA | NA | NA | NA | NA | 3 | 1 |
| 4_1164534 | NA | NA | 1 | 0 | 2 | 0.5 | NA | NA | 3 | 0.3333 | 3 | 1 | NA | NA | NA | NA | NA | NA | 3 | 1 |
| 4_1164536 | NA | NA | 1 | 1 | 1 | 1 | NA | NA | 2 | 1 | 1 | 1 | NA | NA | NA | NA | NA | NA | 1 | 1 |
| 4_1164538 | NA | NA | 1 | 0 | 1 | 1 | NA | NA | 2 | 0.5 | 2 | 0.5 | NA | NA | NA | NA | NA | NA | 2 | 0.5 |
| 4_1164539 | NA | NA | NA | NA | 1 | 1 | NA | NA | 1 | 1 | 2 | 1 | NA | NA | NA | NA | NA | NA | 2 | 1 |
| 4_1164539 | NA | NA | 1 | 1 | NA | NA | NA | NA | 1 | 1 | 2 | 1 | NA | NA | NA | NA | NA | NA | 2 | 1 |
| 4_1164544 | NA | NA | NA | NA | 1 | 1 | NA | NA | 1 | 1 | 1 | 1 | NA | NA | NA | NA | NA | NA | 1 | 1 |
| 4_1164546 | NA | NA | 2 | 0.5 | NA | NA | NA | NA | 2 | 0.5 | 2 | 0.5 | NA | NA | NA | NA | NA | NA | 2 | 0.5 |
| 4_1165567 | 1 | 0 | 2 | 0.5 | 4 | 1 | 4 | 0.5 | 11 | 0.6364 | 2 | 1 | 1 | 0 | 1 | 1 | 5 | 0 | 9 | 0.3333 |
| 4_1165568 | 2 | 0 | 2 | 0.5 | 3 | 0 | 5 | 0 | 12 | 0.0833 | 1 | 0 | NA | NA | 1 | 0 | 4 | 0.25 | 6 | 0.1667 |
| 4_1165571 | 3 | 0 | 2 | 0 | 4 | 0.25 | 5 | 0 | 14 | 0.0714 | 3 | 0 | 1 | 1 | 1 | 0 | 5 | 0 | 10 | 0.1 |
| 4_1165592 | 3 | 1 | NA | NA | 2 | 1 | NA | NA | 5 | 1 | 1 | 0 | 3 | 0.3333 | 1 | 1 | 1 | 1 | 6 | 0.5 |
| 4_1165608 | NA | NA | NA | NA | 2 | 1 | NA | NA | 2 | 1 | 2 | 0.5 | 4 | 0.25 | 2 | 0 | NA | NA | 8 | 0.25 |
| 4_1165608 | NA | NA | NA | NA | 3 | 0.6667 | NA | NA | 3 | 0.6667 | 3 | 1 | 5 | 0.2 | 2 | 0 | NA | NA | 10 | 0.4 |
| 4_1165609 | NA | NA | NA | NA | 3 | 0.3333 | NA | NA | 3 | 0.3333 | 2 | 0 | 5 | 0.2 | 2 | 0 | NA | NA | 9 | 0.1111 |
| 4_1166599 | 1 | 1 | NA | NA | NA | NA | NA | NA | 1 | 1 | 2 | 0.5 | NA | NA | 1 | 1 | NA | NA | 3 | 0.6667 |
| 4_1166601 | 2 | 1 | NA | NA | NA | NA | NA | NA | 2 | 1 | 1 | 1 | NA | NA | 1 | 1 | NA | NA | 2 | 1 |
| 4_1166602 | 2 | 0.5 | 1 | 1 | NA | NA | NA | NA | 3 | 0.6667 | 2 | 0.5 | NA | NA | 1 | 0 | NA | NA | 3 | 0.3333 |
| 4_1166603 | 2 | 1 | 1 | 1 | NA | NA | NA | NA | 3 | 1 | 2 | 0.5 | NA | NA | 1 | 1 | NA | NA | 3 | 0.6667 |
| 4_1166603( | 2 | 1 | NA | NA | NA | NA | NA | NA | 2 | 1 | 2 | 1 | NA | NA | 1 | 1 | NA | NA | 3 | 1 |
| 4_1166605 | 2 | 0.5 | 1 | 1 | 1 | 1 | NA | NA | 4 | 0.75 | 2 | 0 | NA | NA | 1 | 1 | NA | NA | 3 | 0.3333 |
| 4_1166606 | 2 | 1 | 1 | 1 | NA | NA | NA | NA | 3 | 1 | 2 | 1 | NA | NA | 1 | 1 | NA | NA | 3 | 1 |
| 4_1166606 | 2 | 1 | 1 | 1 | 1 | 0 | NA | NA | 4 | 0.75 | 2 | 1 | NA | NA | NA | NA | NA | NA | 2 | 1 |
| 4_1166607 | 2 | 0.5 | NA | NA | 1 | 0 | NA | NA | 3 | 0.3333 | 2 | 0.5 | NA | NA | 1 | 1 | NA | NA | 3 | 0.6667 |
| 4_1166609 | 2 | 1 | NA | NA | 1 | 1 | NA | NA | 3 | 1 | 2 | 0.5 | NA | NA | 1 | 1 | NA | NA | 3 | 0.6667 |
| 4_1166611 | 2 | 0 | NA | NA | 1 | 1 | NA | NA | 3 | 0.3333 | 1 | 1 | NA | NA | 1 | 1 | NA | NA | 2 | 1 |
| 4_1166613 | 2 | 0 | 2 | 0 | 5 | 0.2 | NA | NA | 9 | 0.1111 | 1 | 0 | NA | NA | 4 | 0.75 | NA | NA | 5 | 0.6 |
| 4_1166615 | 2 | 1 | 1 | 1 | 5 | 1 | NA | NA | 8 | 1 | NA | NA | NA | NA | 4 | 0.5 | NA | NA | 4 | 0.5 |
| 4_1166616 | 2 | 0 | 2 | 0 | 4 | 0.25 | NA | NA | 8 | 0.125 | NA | NA | NA | NA | 4 | 0.5 | NA | NA | 4 | 0.5 |
| 4_1166617 | 2 | 1 | 2 | 0.5 | 4 | 1 | NA | NA | 8 | 0.875 | NA | NA | NA | NA | 4 | 1 | NA | NA | 4 | 1 |
| 4_1166618 | 2 | 1 | 2 | 0.5 | 4 | 1 | NA | NA | 8 | 0.875 | NA | NA | NA | NA | 3 | 0.6667 | NA | NA | 3 | 0.6667 |
| 4_1166621( | 2 | 0.5 | 2 | 0 | 2 | 0.5 | NA | NA | 6 | 0.3333 | NA | NA | NA | NA | 3 | 0.3333 | NA | NA | 3 | 0.3333 |
| 4_1167321 | 1 | 1 | 2 | 0.5 | NA | NA | 1 | 1 | 4 | 0.75 | NA | NA | NA | NA | 1 | 1 | NA | NA | 1 | 1 |
| 4_1167335 | 1 | 1 | 2 | 0 | NA | NA | 1 | 0 | 4 | 0.25 | NA | NA | NA | NA | 1 | 1 | NA | NA | 1 | 1 |
| 4_1167339 | NA | NA | 1 | 1 | NA | NA | 1 | 1 | 2 | 1 | NA | NA | NA | NA | 1 | 1 | NA | NA | 1 | 1 |
| 4_1167341 | NA | NA | NA | NA | NA | NA | 1 | 1 | 1 | 1 | NA | NA | NA | NA | 1 | 1 | NA | NA | 1 | 1 |
| 4_1168438( | 1 | 1 | NA | NA | NA | NA | 2 | 1 | 3 | 1 | 1 | 1 | NA | NA | 2 | 1 | 1 | 1 | 4 | 1 |
| 4_1168438 | 1 | 0 | NA | NA | NA | NA | 3 | 0.3333 | 4 | 0.25 | 1 | 0 | NA | NA | 1 | 1 | 1 | 0 | 3 | 0.3333 |
| 4_1168439 | 1 | 0 | NA | NA | NA | NA | 3 | 0.3333 | 4 | 0.25 | 1 | 1 | NA | NA | 1 | 0 | 1 | 1 | 3 | 0.6667 |
| 4_1168441 | 1 | 1 | NA | NA | NA | NA | 2 | 1 | 3 | 1 | 1 | 1 | NA | NA | 3 | 1 | 1 | 1 | 5 | 1 |
| 4_1168442 | NA | NA | NA | NA | NA | NA | 3 | 0.6667 | 3 | 0.6667 | NA | NA | 1 | 0 | 3 | 0.6667 | 2 | 0.5 | 6 | 0.5 |
| 4_1168444( | 2 | 0 | NA | NA | NA | NA | 3 | 0.6667 | 5 | 0.4 | 1 | 0 | 2 | 0 | 2 | 1 | 2 | 0 | 7 | 0.2857 |
| 4_1168445 | 2 | 0.5 | NA | NA | 1 | 0 | 2 | 0 | 5 | 0.2 | 1 | 1 | 1 | 1 | 3 | 0.6667 | 2 | 1 | 7 | 0.8571 |
| 4_1168445 | 2 | 1 | NA | NA | 1 | 1 | 3 | 1 | 6 | 1 | 1 | 1 | 2 | 1 | 3 | 1 | 2 | 1 | 8 | 1 |
| 4_1168446 | 2 | 1 | NA | NA | 1 | 1 | 3 | 1 | 6 | 1 | 1 | 1 | 1 | 1 | 2 | 1 | 1 | 1 | 5 | 1 |
| 4_1168446 | 2 | 0 | NA | NA | 1 | 0 | 3 | 0.3333 | 6 | 0.1667 | 1 | 0 | 1 | 0 | 2 | 0.5 | 2 | 0 | 6 | 0.1667 |
| 4_1168447 | NA | NA | NA | NA | 1 | 0 | 3 | 0.6667 | 4 | 0.5 | 1 | 0 | 2 | 0.5 | 3 | 0 | 2 | 0.5 | 8 | 0.25 |
| 4_1168450 | 1 | 1 | NA | NA | 1 | 1 | 4 | 0.5 | 6 | 0.6667 | 1 | 1 | 4 | 0.75 | 2 | 1 | 2 | 1 | 9 | 0.8889 |
| 4_1168464( | NA | NA | 1 | 0 | 1 | 0 | 3 | 0.3333 | 5 | 0.2 | 2 | 0 | 4 | 0.25 | 3 | 0 | 4 | 0 | 13 | 0.0769 |
| 4_1168475 | 12 | 0.1667 | 5 | 0.4 | 10 | 0.1 | 9 | 0.2222 | 36 | 0.1944 | 3 | 0 | 8 | 0.5 | 6 | 0.1667 | 8 | 0.25 | 25 | 0.28 |
| 4_1168485 | 3 | 0.3333 | 1 | 1 | NA | NA | 1 | 0 | 5 | 0.4 | NA | NA | 1 | 0 | 1 | 1 | NA | NA | 2 | 0.5 |
| 4_1168486 | 12 | 0.25 | 6 | 0.3333 | 7 | 0 | 9 | 0.2222 | 34 | 0.2059 | 5 | 0.6 | 6 | 0.1667 | 7 | 0.4286 | 5 | 0.2 | 23 | 0.3478 |
| 4_1168486 | 13 | 0.3077 | 6 | 0.5 | 8 | 0.125 | 8 | 0.25 | 35 | 0.2857 | 4 | 0.5 | 6 | 0.6667 | 6 | 0.1667 | 4 | 0 | 20 | 0.35 |
| 4_1168486 | 13 | 0.4615 | 7 | 0.2857 | 7 | 0.1429 | 10 | 0.3 | 37 | 0.3243 | 5 | 0.6 | 5 | 0.4 | 8 | 0.625 | 5 | 0.2 | 23 | 0.4783 |
| 4_1168499 | 13 | 0.3077 | 6 | 0.1667 | 7 | 0.1429 | 10 | 0.4 | 36 | 0.2778 | 5 | 0 | 6 | 0.1667 | 6 | 0.3333 | 5 | 0.4 | 22 | 0.2273 |
| 4_1168608( | 4 | 1 | 3 | 0.6667 | 1 | 1 | 2 | 1 | 10 | 0.9 | 1 | 1 | 5 | 1 | 6 | 1 | 2 | 1 | 14 | 1 |
| 4_1168609 | 7 | 0.1429 | 4 | 0 | 1 | 0 | 2 | 0 | 14 | 0.0714 | 1 | 0 | 4 | 0 | 6 | 0.5 | 2 | 0 | 13 | 0.2308 |
| 4_1168610 | 7 | 0.4286 | 4 | 0.25 | 2 | 1 | 2 | 0 | 15 | 0.4 | 1 | 0 | 5 | 0.2 | 5 | 0.6 | 2 | 0.5 | 13 | 0.3846 |

| | | | | | | | | | | | | | | | | | | | | |
|---|---|---|---|---|---|---|---|---|---|---|---|---|---|---|---|---|---|---|---|---|
| 4_11686120 | 7 | 0 | 4 | 0.25 | 2 | 0.5 | 1 | 0 | 14 | 0.1429 | 1 | 0 | 6 | 0.1667 | 4 | 0 | 2 | 0 | 13 | 0.0769 |
| 4_11686152 | NA | NA | 2 | 0.5 | NA | NA | NA | NA | 2 | 0.5 | NA | NA | 3 | 0 | 4 | 0.25 | 2 | 0 | 9 | 0.1111 |
| 4_11692371 | 1 | 0 | 2 | 0.5 | 1 | 0 | NA | NA | 4 | 0.25 | 1 | 0 | 1 | 0 | NA | NA | 1 | 1 | 3 | 0.3333 |
| 4_11697206 | 4 | 0.25 | NA | NA | NA | NA | 1 | 0 | 5 | 0.2 | 1 | 0 | NA | NA | 2 | 0.5 | NA | NA | 3 | 0.3333 |
| 4_11697699 | 2 | 0.5 | 2 | 0.5 | 2 | 0.5 | 2 | 1 | 8 | 0.625 | 2 | 1 | 1 | 1 | 1 | 1 | 1 | 1 | 5 | 1 |
| 4_11697717 | 2 | 0.5 | 3 | 0 | 1 | 0 | 3 | 0 | 9 | 0.1111 | 1 | 0 | 2 | 0.5 | 1 | 1 | 4 | 1 | 8 | 0.75 |
| 4_11697719 | 1 | 0 | 2 | 0 | 1 | 1 | 3 | 0 | 7 | 0.1429 | 1 | 0 | 2 | 0 | 1 | 1 | 4 | 0.75 | 8 | 0.5 |
| 4_11697732 | 2 | 0.5 | 3 | 0.3333 | 2 | 1 | 3 | 0.6667 | 10 | 0.6 | 2 | 0.5 | 2 | 0.5 | 1 | 1 | 4 | 0.75 | 9 | 0.6667 |
| 4_11697740 | 2 | 0.5 | 2 | 0 | 2 | 1 | 2 | 0 | 8 | 0.375 | 2 | 1 | 1 | 1 | 1 | 1 | 4 | 1 | 8 | 1 |
| 4_11697750 | 2 | 0.5 | 2 | 0.5 | 2 | 1 | 3 | 0.6667 | 9 | 0.6667 | 2 | 0.5 | 1 | 0 | 1 | 1 | 4 | 1 | 8 | 0.75 |
| 4_11697771 | 1 | 0 | 2 | 0.5 | 1 | 0 | 3 | 0.3333 | 7 | 0.2857 | 2 | 0.5 | 1 | 0 | 1 | 0 | 4 | 0.25 | 8 | 0.25 |
| 4_11697778 | 2 | 0 | 1 | 0 | 2 | 0.5 | 2 | 0 | 7 | 0.1429 | 2 | 0 | 1 | 0 | 1 | 0 | 4 | 0.5 | 8 | 0.25 |
| 4_11697780 | 2 | 0.5 | 2 | 0.5 | 2 | 1 | 3 | 0.6667 | 9 | 0.6667 | 2 | 0.5 | 1 | 1 | 1 | 1 | 4 | 0.5 | 8 | 0.625 |
| 4_11697791 | 1 | 1 | 1 | 1 | 2 | 0.5 | 2 | 0.5 | 6 | 0.6667 | 2 | 0 | NA | NA | 1 | 0 | 4 | 0.75 | 7 | 0.4286 |
| 4_11697794 | 2 | 0.5 | 1 | 1 | 2 | 0.5 | 3 | 0.3333 | 8 | 0.5 | 1 | 0 | NA | NA | 1 | 1 | 4 | 0.75 | 6 | 0.6667 |
| 4_11697809 | NA | NA | 2 | 1 | 1 | 1 | 1 | 1 | 4 | 1 | NA | NA | NA | NA | 1 | 1 | 1 | 1 | 2 | 1 |
| 4_11697834 | NA | NA | 1 | 1 | NA | NA | 2 | 0 | 3 | 0.3333 | NA | NA | NA | NA | 1 | 0 | 2 | 0.5 | 3 | 0.3333 |
| 4_11697838 | NA | NA | 1 | 0 | NA | NA | 2 | 0.5 | 3 | 0.3333 | NA | NA | NA | NA | 1 | 0 | 2 | 0.5 | 3 | 0.3333 |
| 4_11698594 | NA | NA | 1 | 0 | 2 | 0 | 3 | 0.3333 | 6 | 0.1667 | 1 | 0 | NA | NA | 2 | 0.5 | 1 | 1 | 4 | 0.5 |
| 4_11712564 | NA | NA | 2 | 1 | NA | NA | 2 | 1 | 4 | 1 | NA | NA | 1 | 1 | NA | NA | 1 | 1 | 2 | 1 |
| 4_11712567 | NA | NA | 1 | 1 | NA | NA | 2 | 0.5 | 3 | 0.6667 | NA | NA | 1 | 1 | NA | NA | NA | NA | 1 | 1 |
| 4_11712917 | 19 | 0.2105 | 31 | 0.0645 | 31 | 0.1935 | 20 | 0.1 | 101 | 0.1386 | 15 | 0.2667 | 15 | 0.0667 | 28 | 0.1429 | 22 | 0.1364 | 80 | 0.15 |
| 4_11712931 | 15 | 0.0667 | 25 | 0.12 | 25 | 0.08 | 13 | 0 | 78 | 0.0769 | 12 | 0.1667 | 12 | 0.1667 | 26 | 0.1154 | 14 | 0.2857 | 64 | 0.1719 |
| 4_11714557 | NA | NA | 3 | 0.3333 | 5 | 0.2 | 3 | 0 | 11 | 0.1818 | NA | NA | NA | NA | 2 | 0 | 2 | 0.5 | 4 | 0.25 |
| 4_11714577 | NA | NA | 3 | 0.3333 | 4 | 0.25 | 1 | 0 | 8 | 0.25 | 1 | 1 | NA | NA | 1 | 0 | 1 | 0 | 3 | 0.3333 |
| 4_11714581 | NA | NA | 2 | 0.5 | 4 | 0.75 | 1 | 1 | 7 | 0.7143 | 1 | 1 | NA | NA | 1 | 0 | 1 | 0 | 3 | 0.3333 |
| 4_11714590 | NA | NA | 3 | 0.3333 | 3 | 0.3333 | 1 | 1 | 7 | 0.4286 | 1 | 0 | NA | NA | NA | NA | 1 | 1 | 2 | 0.5 |
| 4_11714591 | NA | NA | 4 | 0.25 | 3 | 0 | 1 | 0 | 8 | 0.125 | NA | NA | NA | NA | 1 | 1 | 1 | 1 | 2 | 1 |
| 4_11714598 | NA | NA | 2 | 0 | 2 | 0.5 | NA | NA | 4 | 0.25 | 1 | 0 | NA | NA | NA | NA | 1 | 1 | 2 | 0.5 |
| 4_11714599 | NA | NA | 2 | 0.5 | 3 | 0.6667 | 1 | 1 | 6 | 0.6667 | NA | NA | NA | NA | 1 | 1 | 1 | 1 | 2 | 1 |
| 4_11714603 | NA | NA | 3 | 1 | 3 | 1 | 1 | 1 | 7 | 1 | 1 | 1 | NA | NA | 1 | 1 | 1 | 1 | 3 | 1 |
| 4_11740757 | 1 | 0 | 3 | 0.3333 | 1 | 0 | 1 | 0 | 6 | 0.1667 | NA | NA | 3 | 0.3333 | 3 | 0.3333 | 5 | 0 | 11 | 0.1818 |
| 4_11742205 | 16 | 0.0625 | 20 | 0 | 6 | 0.1667 | 15 | 0.0667 | 57 | 0.0526 | 10 | 0.1 | 12 | 0.0833 | 15 | 0.0667 | 19 | 0.0526 | 56 | 0.0714 |
| 4_11753188 | NA | NA | 2 | 0.5 | 1 | 1 | 2 | 0.5 | 5 | 0.6 | NA | NA | 3 | 1 | NA | NA | 3 | 0.6667 | 6 | 0.8333 |
| 4_11753228 | 4 | 0 | 3 | 0 | 3 | 0.3333 | 4 | 0.25 | 14 | 0.1429 | 1 | 1 | 8 | 0.25 | 2 | 0 | 8 | 0.125 | 19 | 0.2105 |
| 4_11772530 | NA | NA | 6 | 0.5 | 7 | 0.2857 | 7 | 0.5714 | 20 | 0.45 | 3 | 0 | NA | NA | 2 | 0.5 | 1 | 0 | 6 | 0.1667 |
| 4_11772537 | 3 | 0.3333 | 6 | 0 | 9 | 0 | 7 | 0.1429 | 25 | 0.08 | 4 | 0 | 1 | 0 | 6 | 0.1667 | 2 | 0 | 13 | 0.0769 |
| 4_11772542 | 3 | 0.6667 | 5 | 1 | 10 | 0.8 | 7 | 0.8571 | 25 | 0.84 | 2 | 0.5 | NA | NA | 4 | 0.75 | 2 | 0.5 | 8 | 0.625 |
| 4_11772549 | 2 | 1 | 3 | 0.3333 | 11 | 0.5455 | 7 | 0.5714 | 23 | 0.5652 | 3 | 0 | 1 | 1 | 5 | 0.2 | 2 | 0 | 11 | 0.1818 |
| 4_11772550 | 3 | 0.3333 | 3 | 1 | 7 | 0.5714 | 5 | 0.2 | 18 | 0.5 | 3 | 0.3333 | NA | NA | 5 | 0.4 | 1 | 1 | 9 | 0.4444 |
| 4_11772560 | 1 | 1 | 4 | 0.5 | 8 | 0 | 5 | 0.6 | 18 | 0.3333 | 4 | 0.25 | 1 | 0 | 3 | 0 | 2 | 0 | 10 | 0.1 |
| 4_11772591 | 2 | 0.5 | 5 | 0.2 | 11 | 0.5455 | 7 | 0.8571 | 25 | 0.56 | 6 | 0.3333 | 1 | 1 | 6 | 0.6667 | 3 | 0.3333 | 16 | 0.5 |
| 4_11772614 | 2 | 0 | 4 | 0.25 | 9 | 0.2222 | 6 | 0 | 21 | 0.1429 | 4 | 0 | NA | NA | 7 | 0.1429 | 4 | 0 | 15 | 0.0667 |
| 4_11773048 | 8 | 0.25 | 7 | 0.7143 | 6 | 0.1667 | 6 | 0.6667 | 27 | 0.4444 | 3 | 0 | 10 | 0.5 | 5 | 0.2 | 6 | 0.1667 | 24 | 0.2917 |
| 4_11773091 | 10 | 0 | 8 | 0.125 | 10 | 0.1 | 7 | 0.2857 | 35 | 0.1143 | 6 | 0.3333 | 11 | 0.0909 | 5 | 0.2 | 8 | 0.25 | 30 | 0.2 |
| 4_11773093 | 9 | 0.6667 | 5 | 0.4 | 7 | 0.7143 | 4 | 0.75 | 25 | 0.64 | 2 | 1 | 11 | 0.3636 | 3 | 1 | 7 | 0.7143 | 23 | 0.6087 |
| 4_11773094 | 8 | 0.75 | 7 | 0.8571 | 11 | 0.8182 | 7 | 0.4286 | 33 | 0.7273 | 5 | 0.8 | 10 | 0.8 | 5 | 1 | 7 | 0.5714 | 27 | 0.7778 |
| 4_11773104 | 5 | 0.4 | 6 | 0.5 | 7 | 0.7143 | 5 | 0.6 | 23 | 0.5652 | 6 | 0.3333 | 8 | 0.625 | 6 | 0.3333 | 8 | 0.625 | 28 | 0.5 |
| 4_11773108 | 4 | 0.25 | 5 | 0 | 9 | 0.1111 | 5 | 0.6 | 23 | 0.2174 | 3 | 0 | 9 | 0.1111 | 2 | 0 | 5 | 0.2 | 19 | 0.1053 |
| 4_11773108 | 9 | 0.6667 | 4 | 0.75 | 8 | 0.5 | 5 | 0.4 | 26 | 0.5769 | 5 | 1 | 9 | 0.6667 | 6 | 0.8333 | 5 | 0.8 | 25 | 0.8 |
| 4_11773880 | 5 | 0.6 | 2 | 0 | 4 | 0.25 | 3 | 0 | 14 | 0.2857 | 1 | 1 | 6 | 0.1667 | 3 | 0 | 3 | 0.6667 | 13 | 0.3077 |
| 4_11774161 | 62 | 0.0806 | 59 | 0.0339 | 52 | 0.0769 | 41 | 0.122 | 214 | 0.0748 | 47 | 0.1064 | 48 | 0.0833 | 47 | 0.0638 | 42 | 0.119 | 184 | 0.0924 |
| 4_11774167 | 63 | 0.0476 | 56 | 0.0536 | 49 | 0.1224 | 41 | 0.0732 | 209 | 0.0718 | 47 | 0.0851 | 47 | 0.0426 | 44 | 0.0682 | 41 | 0.0488 | 179 | 0.0615 |
| 4_11774618 | 1 | 1 | 1 | 1 | 2 | 0.5 | 2 | 1 | 6 | 0.8333 | 1 | 1 | 4 | 0.75 | 3 | 1 | NA | NA | 8 | 0.875 |
| 4_11774627 | 1 | 0 | 1 | 0 | 1 | 1 | 1 | 1 | 4 | 0.5 | 1 | 0 | 2 | 0 | 2 | 0.5 | NA | NA | 5 | 0.2 |
| 4_11774660 | NA | NA | 1 | 0 | 1 | 1 | 1 | 0 | 3 | 0.3333 | NA | NA | 3 | 0.3333 | 1 | 1 | NA | NA | 4 | 0.5 |
| 4_11774746 | 1 | 0 | 1 | 1 | 1 | 0 | NA | NA | 3 | 0.3333 | NA | NA | 2 | 0.5 | 1 | 1 | NA | NA | 3 | 0.6667 |
| 4_11774757 | 1 | 1 | 1 | 1 | 1 | 1 | 1 | 1 | 4 | 1 | NA | NA | 3 | 0.6667 | 1 | 1 | NA | NA | 4 | 0.75 |
| 4_11774761 | 1 | 0 | 1 | 0 | 1 | 1 | 1 | 0 | 4 | 0.25 | NA | NA | 3 | 0.3333 | 1 | 1 | NA | NA | 4 | 0.5 |
| 4_11774770 | 1 | 0 | NA | NA | 1 | 1 | 1 | 1 | 3 | 0.6667 | NA | NA | 3 | 0.3333 | 2 | 1 | NA | NA | 5 | 0.6 |
| 4_11775169 | 8 | 0.25 | 13 | 0.5385 | 6 | 0.1667 | 7 | 0.2857 | 34 | 0.3529 | 5 | 0.6 | 7 | 0.4286 | 10 | 0.2 | 8 | 0.375 | 30 | 0.3667 |
| 4_11775208 | 9 | 0.2222 | 11 | 0.1818 | 9 | 0.4444 | 7 | 0.1429 | 36 | 0.25 | 4 | 0.75 | 9 | 0.6667 | 14 | 0.4286 | 6 | 0.3333 | 33 | 0.5152 |
| 4_11776903 | 4 | 0.75 | 5 | 0.2 | 4 | 0.5 | 2 | 0.5 | 15 | 0.4667 | NA | NA | 4 | 0.5 | 6 | 0.5 | 2 | 0.5 | 12 | 0.5 |
| 4_11776907 | 4 | 0.5 | 5 | 0.8 | 4 | 0.75 | 2 | 1 | 15 | 0.7333 | NA | NA | 4 | 0.75 | 7 | 0.4286 | 3 | 0.6667 | 14 | 0.5714 |
| 4_11776919 | 3 | 0.6667 | 4 | 0.75 | 4 | 0.5 | 1 | 1 | 12 | 0.6667 | NA | NA | 4 | 0.75 | 5 | 1 | 4 | 1 | 13 | 0.9231 |

| | | | | | | | | | | | | | | | | | | | | |
|---|---|---|---|---|---|---|---|---|---|---|---|---|---|---|---|---|---|---|---|---|
| 4_11776921 | 3 | 0.3333 | 3 | 0.6667 | 3 | 0 | 1 | 0 | 10 | 0.3 | NA | NA | 4 | 0.5 | 5 | 0.4 | 2 | 0.5 | 11 | 0.4545 |
| 4_11776933 | 3 | 0.6667 | 3 | 0.6667 | 3 | 0.6667 | 1 | 1 | 10 | 0.7 | NA | NA | 2 | 0.5 | 5 | 0.6 | 2 | 1 | 9 | 0.6667 |
| 4_11776943 | 3 | 0 | 3 | 0 | 3 | 0.6667 | 1 | 1 | 10 | 0.3 | NA | NA | 2 | 1 | 6 | 0.5 | 2 | 1 | 10 | 0.7 |
| 4_11777151 | 2 | 0.5 | 2 | 0 | 2 | 0.5 | 1 | 0 | 7 | 0.2857 | NA | NA | 3 | 0.3333 | 4 | 0 | 1 | 0 | 8 | 0.125 |
| 4_11777185 | 2 | 0.5 | 2 | 1 | 2 | 0.5 | 1 | 1 | 7 | 0.7143 | NA | NA | 2 | 0 | 3 | 0 | 2 | 0.5 | 7 | 0.1429 |
| 4_11777187 | 1 | 1 | 2 | 0 | 2 | 0 | 1 | 1 | 6 | 0.3333 | NA | NA | 2 | 0 | 5 | 0.2 | 2 | 0.5 | 9 | 0.2222 |
| 4_11778942 | 2 | 0 | 7 | 0.1429 | 4 | 0 | 5 | 0 | 18 | 0.0556 | 10 | 0 | 8 | 0.125 | 7 | 0.1429 | 8 | 0 | 33 | 0.0606 |
| 4_11778947 | 2 | 0 | 7 | 0.1429 | 4 | 0.5 | 4 | 0.5 | 17 | 0.2941 | 8 | 0.375 | 8 | 0 | 6 | 0.3333 | 8 | 0.5 | 30 | 0.3 |
| 4_11778950 | 2 | 0.5 | 5 | 0.8 | 4 | 1 | 4 | 0.75 | 15 | 0.8 | 8 | 0.75 | 6 | 0.6667 | 6 | 0.5 | 5 | 0.6 | 25 | 0.64 |
| 4_11778959 | 2 | 0.5 | 6 | 0.3333 | 4 | 0.75 | 4 | 0.5 | 16 | 0.5 | 9 | 0.5556 | 6 | 0.5 | 6 | 0.3333 | 7 | 0.7143 | 28 | 0.5357 |
| 4_11778961 | 1 | 0 | 3 | 0 | 4 | 0.25 | 4 | 0 | 12 | 0.0833 | 10 | 0 | 4 | 0 | 6 | 0.1667 | 9 | 0.3333 | 29 | 0.1379 |
| 4_11778982 | 2 | 0 | 8 | 0.125 | 4 | 0 | 4 | 0 | 18 | 0.0556 | 10 | 0 | 7 | 0 | 6 | 0.1667 | 9 | 0.1111 | 32 | 0.0625 |
| 4_11778998 | 1 | 0 | NA | NA | 1 | 1 | NA | NA | 2 | 0.5 | 3 | 0.3333 | 1 | 0 | 1 | 0 | 4 | 0.5 | 9 | 0.3333 |
| 4_11779008 | 2 | 1 | 5 | 1 | 4 | 0.75 | 2 | 0 | 13 | 0.7692 | 6 | 0.6667 | 6 | 0.1667 | 5 | 0.4 | 8 | 0.625 | 25 | 0.48 |
| 4_11779014 | 2 | 0.5 | 5 | 0 | 4 | 0.25 | 2 | 0 | 13 | 0.1538 | 6 | 0.3333 | 6 | 0 | 5 | 0.2 | 8 | 0.5 | 25 | 0.28 |
| 4_11779016 | 2 | 1 | 4 | 0.75 | 2 | 1 | 2 | 1 | 10 | 0.9 | 5 | 1 | 3 | 0.6667 | 3 | 1 | 5 | 0.8 | 16 | 0.875 |
| 4_11796485 | 1 | 1 | 2 | 0 | 1 | 0 | 2 | 0 | 6 | 0.1667 | 4 | 0.25 | 5 | 0 | 4 | 0 | 5 | 0 | 18 | 0.0556 |
| 4_11799422 | 5 | 0.4 | 5 | 0.4 | 4 | 0.25 | 5 | 0.6 | 19 | 0.4211 | 2 | 0.5 | 2 | 0.5 | 2 | 1 | 2 | 0.5 | 8 | 0.625 |
| 4_11799462 | 8 | 0.125 | 3 | 0.3333 | 2 | 1 | 3 | 0.6667 | 16 | 0.375 | 1 | 1 | 3 | 0 | 1 | 0 | 2 | 0 | 7 | 0.1429 |
| 4_11799471 | 6 | 0 | 1 | 0 | 2 | 0.5 | 3 | 0 | 12 | 0.0833 | 1 | 0 | 3 | 0.3333 | 1 | 0 | 2 | 0.5 | 7 | 0.2857 |
| 4_11799523 | 5 | 0.4 | 2 | 0.5 | 2 | 0 | 3 | 0 | 12 | 0.25 | 1 | 1 | 3 | 0 | 1 | 1 | 1 | 1 | 6 | 0.5 |
| 4_11799533 | 4 | 0.5 | 2 | 0.5 | 1 | 0 | 3 | 0 | 10 | 0.3 | NA | NA | 2 | 0.5 | 1 | 0 | 1 | 1 | 4 | 0.5 |
| 4_11799534 | 4 | 0.75 | 1 | 1 | 1 | 1 | 2 | 1 | 8 | 0.875 | 1 | 1 | 2 | 0.5 | 1 | 1 | 2 | 1 | 6 | 0.8333 |
| 4_11799534 | 3 | 1 | 1 | 0 | 1 | 1 | 2 | 1 | 7 | 0.8571 | 1 | 1 | 2 | 1 | 1 | 1 | 1 | 1 | 5 | 1 |
| 4_11806576 | 3 | 0.6667 | 11 | 0.6364 | 6 | 0.6667 | 1 | 1 | 21 | 0.6667 | 3 | 0.3333 | 3 | 0.6667 | 9 | 0.4444 | 7 | 0.2857 | 22 | 0.4091 |
| 4_11806659 | 1 | 0 | 5 | 0.6 | 6 | 0.1667 | NA | NA | 12 | 0.3333 | 3 | 0.6667 | 3 | 0.3333 | 9 | 0.5556 | 7 | 0.2857 | 22 | 0.4545 |
| 4_11806667 | 1 | 0 | 5 | 0.2 | 5 | 0 | 1 | 0 | 12 | 0.0833 | 3 | 0.3333 | 3 | 0 | 8 | 0.375 | 6 | 0.1667 | 20 | 0.25 |
| 4_11806682 | 1 | 0 | 5 | 0.8 | 3 | 0.6667 | 1 | 1 | 10 | 0.7 | 2 | 1 | 3 | 1 | 9 | 1 | 6 | 0.8333 | 20 | 0.95 |
| 4_11806687 | 1 | 0 | 2 | 0.5 | 4 | 0 | 1 | 0 | 8 | 0.125 | 1 | 0 | 1 | 1 | 7 | 0 | 6 | 0 | 15 | 0.0667 |
| 4_11808091 | 1 | 0 | 4 | 0.25 | 3 | 0 | NA | NA | 8 | 0.125 | 2 | 0 | 2 | 0 | 5 | 0.2 | 4 | 0 | 13 | 0.0769 |
| 4_11808325 | 1 | 1 | NA | NA | NA | NA | NA | NA | 1 | 1 | NA | NA | NA | NA | 1 | 1 | NA | NA | 1 | 1 |
| 4_11809323 | 1 | 0 | 1 | 1 | NA | NA | NA | NA | 2 | 0.5 | 1 | 0 | 1 | 1 | 3 | 0.6667 | NA | NA | 5 | 0.6 |
| 4_11809362 | 1 | 1 | 1 | 1 | NA | NA | NA | NA | 2 | 1 | 1 | 0 | NA | NA | 2 | 1 | NA | NA | 3 | 0.6667 |
| 4_11821998 | 1 | 0 | 2 | 0.5 | 2 | 0 | 1 | 0 | 6 | 0.1667 | 1 | 1 | 2 | 0 | NA | NA | 2 | 0.5 | 5 | 0.4 |
| 4_11821999 | 2 | 0.5 | 3 | 0 | 3 | 0.3333 | 1 | 0 | 9 | 0.2222 | 1 | 0 | 3 | 0.3333 | NA | NA | 1 | 0 | 5 | 0.2 |
| 4_11822011 | 2 | 0.5 | 2 | 1 | 2 | 1 | NA | NA | 6 | 0.8333 | 1 | 1 | 2 | 1 | NA | NA | 1 | 1 | 4 | 1 |
| 4_11822018 | 1 | 0 | 2 | 0.5 | 1 | 0 | 1 | 0 | 5 | 0.2 | 1 | 1 | 3 | 1 | NA | NA | 1 | 1 | 5 | 1 |
| 4_11822020 | 1 | 0 | 3 | 1 | 2 | 1 | 1 | 1 | 7 | 0.8571 | 1 | 0 | 2 | 1 | NA | NA | 2 | 0 | 5 | 0.4 |
| 4_11822033 | 1 | 1 | 3 | 0.3333 | 3 | 0.6667 | NA | NA | 7 | 0.5714 | 1 | 1 | 2 | 0 | NA | NA | 1 | 0 | 4 | 0.25 |
| 4_11822040 | 2 | 1 | 2 | 0.5 | 3 | 1 | 1 | 1 | 8 | 0.875 | 1 | 1 | 2 | 1 | NA | NA | 2 | 0.5 | 5 | 0.8 |
| 4_11822044 | 2 | 0 | 3 | 0.3333 | 3 | 0 | 1 | 0 | 9 | 0.1111 | NA | NA | 3 | 0 | NA | NA | 2 | 0.5 | 5 | 0.2 |
| 4_11822045 | 2 | 0.5 | 3 | 0.6667 | 3 | 0 | 1 | 1 | 9 | 0.4444 | 1 | 1 | 3 | 0.6667 | NA | NA | 2 | 1 | 6 | 0.8333 |
| 4_11822046 | 1 | 1 | 2 | 0.5 | 3 | 0.3333 | 1 | 0 | 7 | 0.4286 | 1 | 0 | 3 | 0 | NA | NA | 1 | 1 | 5 | 0.2 |
| 4_11822051 | 2 | 1 | 3 | 0.6667 | 2 | 0.5 | NA | NA | 7 | 0.7143 | 1 | 1 | 2 | 0.5 | NA | NA | 2 | 0.5 | 5 | 0.6 |
| 4_11822053 | 1 | 0 | 3 | 0.3333 | 3 | 0 | 1 | 0 | 8 | 0.125 | 1 | 1 | 2 | 0 | NA | NA | 2 | 0.5 | 5 | 0.4 |
| 4_11827207 | 10 | 0.5 | 6 | 0.3333 | 5 | 1 | 5 | 0.6 | 26 | 0.5769 | 1 | 1 | 5 | 0.6 | 7 | 0.4286 | 4 | 0.75 | 17 | 0.5882 |
| 4_11827213 | 11 | 0.1818 | 7 | 0.4286 | 6 | 0.3333 | 6 | 0.3333 | 30 | 0.3 | 2 | 0 | 5 | 0.4 | 6 | 0.6667 | 4 | 0.5 | 17 | 0.4706 |
| 4_11827250 | 1 | 0 | NA | NA | 1 | 1 | NA | NA | 2 | 0.5 | 1 | 0 | 1 | 0 | NA | NA | 1 | 1 | 3 | 0.3333 |
| 4_11828935 | 2 | 0 | 6 | 0.8333 | 7 | 0.8571 | 4 | 1 | 19 | 0.7895 | 5 | 1 | 5 | 1 | 3 | 1 | 5 | 0.6 | 18 | 0.8889 |
| 4_11828942 | 2 | 0.5 | 5 | 0.8 | 8 | 1 | 4 | 0.75 | 19 | 0.8421 | 5 | 1 | 4 | 1 | 2 | 1 | 5 | 1 | 16 | 1 |
| 4_11828943 | 2 | 0.5 | 6 | 1 | 7 | 0.8571 | 4 | 0.5 | 19 | 0.7895 | 5 | 0.6 | 3 | 0.6667 | 3 | 1 | 5 | 0.4 | 16 | 0.625 |
| 4_11828954 | 2 | 0 | 4 | 0.5 | 6 | 0.3333 | 3 | 0.6667 | 15 | 0.4 | 4 | 0.5 | 4 | 0.25 | 2 | 0.5 | 4 | 0 | 14 | 0.2857 |
| 4_11828956 | 1 | 1 | 5 | 0.8 | 8 | 0.75 | 3 | 0.6667 | 17 | 0.7647 | 4 | 1 | 5 | 1 | 3 | 0.6667 | 4 | 1 | 16 | 0.9375 |
| 4_11828960 | 2 | 0 | 5 | 0 | 6 | 0.5 | 4 | 0 | 17 | 0.1765 | 3 | 0 | 4 | 0 | 2 | 0 | 3 | 0.3333 | 12 | 0.0833 |
| 4_11828968 | 2 | 1 | 4 | 1 | 8 | 1 | 3 | 0.6667 | 17 | 0.9412 | 3 | 0.6667 | 4 | 0.75 | 3 | 1 | 4 | 1 | 14 | 0.8571 |
| 4_11832865 | 2 | 0 | 2 | 1 | NA | NA | NA | NA | 4 | 0.5 | NA | NA | NA | NA | 2 | 0.5 | 2 | 0 | 4 | 0.25 |
| 4_11839758 | NA | NA | NA | NA | 1 | 1 | NA | NA | 1 | 1 | 2 | 0 | 1 | 1 | NA | NA | 1 | 0 | 4 | 0.25 |
| 4_11839761 | NA | NA | NA | NA | 1 | 1 | NA | NA | 1 | 1 | 2 | 0.5 | 1 | 1 | NA | NA | 1 | 0 | 4 | 0.5 |
| 4_11839789 | NA | NA | NA | NA | 1 | 1 | NA | NA | 1 | 1 | 1 | 0 | 1 | 1 | NA | NA | NA | NA | 2 | 0.5 |
| 4_11839826 | 1 | 0 | 1 | 1 | 2 | 0 | 1 | 1 | 5 | 0.4 | 1 | 0 | 1 | 0 | 1 | 1 | 1 | 0 | 4 | 0.25 |
| 4_11839835 | 1 | 1 | 1 | 1 | 3 | 1 | 1 | 1 | 6 | 1 | 2 | 0.5 | 2 | 1 | NA | NA | 1 | 1 | 5 | 0.8 |
| 4_11839844 | NA | NA | 1 | 1 | 2 | 1 | NA | NA | 3 | 1 | 3 | 0.3333 | 2 | 0.5 | 1 | 0 | 1 | 1 | 7 | 0.4286 |
| 4_11839851 | 1 | 0 | 1 | 1 | 3 | 0.3333 | 1 | 0 | 6 | 0.3333 | 3 | 0 | 2 | 0.5 | 1 | 1 | 1 | 0 | 7 | 0.2857 |
| 4_11839857 | 1 | 1 | 1 | 1 | 2 | 1 | 1 | 1 | 5 | 1 | 2 | 1 | 1 | 1 | NA | NA | 1 | 1 | 4 | 1 |
| 4_11839872 | NA | NA | NA | NA | 3 | 0.6667 | 1 | 1 | 4 | 0.75 | 2 | 0.5 | 2 | 0.5 | 1 | 0 | 1 | 0 | 6 | 0.3333 |

| | | | | | | | | | | | | | | | | | | | | |
|---|---|---|---|---|---|---|---|---|---|---|---|---|---|---|---|---|---|---|---|---|
| 4_1183987 | NA | NA | NA | NA | 3 | 0.3333 | NA | NA | 3 | 0.3333 | 2 | 0 | 1 | 1 | NA | NA | NA | NA | 3 | 0.3333 |
| 4_1183988 | NA | NA | NA | NA | 2 | 0.5 | 1 | 1 | 3 | 0.6667 | 2 | 1 | 2 | 1 | 1 | 1 | NA | NA | 5 | 1 |
| 4_1183989 | NA | NA | NA | NA | 3 | 1 | 1 | 1 | 4 | 1 | 2 | 1 | 1 | 1 | 1 | 0 | 1 | 1 | 5 | 0.8 |
| 4_1183991 | NA | NA | NA | NA | 1 | 1 | NA | NA | 1 | 1 | 1 | 1 | 1 | 1 | 1 | 1 | 1 | 1 | 4 | 1 |
| 4_1184842 | NA | NA | 1 | 1 | 2 | 0 | 1 | 0 | 4 | 0.25 | 1 | 0 | 2 | 0 | 5 | 0 | 2 | 0.5 | 10 | 0.1 |
| 4_1184848 | 1 | 1 | NA | NA | 1 | 1 | 1 | 0 | 3 | 0.6667 | 2 | 1 | 3 | 0.6667 | 3 | 0.6667 | 2 | 0.5 | 10 | 0.7 |
| 4_1184849 | 1 | 1 | 1 | 1 | 2 | 1 | 1 | 1 | 5 | 1 | 2 | 1 | 3 | 0.6667 | 6 | 0.5 | 2 | 0.5 | 13 | 0.6154 |
| 4_1189936 | 34 | 0.2647 | 31 | 0.3226 | 14 | 0.0714 | 27 | 0.3333 | 106 | 0.2736 | 22 | 0.2727 | 22 | 0.0455 | 18 | 0.2778 | 20 | 0.05 | 82 | 0.1585 |
| 4_1189938 | 39 | 0.9744 | 33 | 0.9394 | 15 | 1 | 31 | 0.9677 | 118 | 0.9661 | 27 | 0.963 | 30 | 0.9 | 27 | 0.963 | 24 | 1 | 108 | 0.9537 |
| 4_1189938 | 37 | 0.1081 | 34 | 0.0588 | 15 | 0.2667 | 28 | 0.25 | 114 | 0.1491 | 25 | 0.16 | 34 | 0.2059 | 26 | 0.1923 | 25 | 0.2 | 110 | 0.1909 |
| 4_1189939 | 43 | 0.1163 | 35 | 0.1714 | 16 | 0.25 | 32 | 0.125 | 126 | 0.1508 | 32 | 0.0938 | 36 | 0.0833 | 29 | 0.0345 | 23 | 0 | 120 | 0.0583 |
| 4_1189939 | 42 | 0.119 | 38 | 0.1579 | 16 | 0.1875 | 32 | 0.0938 | 128 | 0.1328 | 31 | 0.1613 | 34 | 0.1471 | 29 | 0.1379 | 23 | 0.2174 | 117 | 0.1624 |
| 4_1189940 | 40 | 0.65 | 39 | 0.5641 | 16 | 0.5625 | 31 | 0.5484 | 126 | 0.5873 | 29 | 0.5172 | 36 | 0.5278 | 30 | 0.6333 | 25 | 0.28 | 120 | 0.5 |
| 4_1189940 | 43 | 0.907 | 37 | 0.7838 | 17 | 0.7647 | 34 | 0.8235 | 131 | 0.8321 | 32 | 0.9375 | 37 | 0.8378 | 31 | 0.871 | 21 | 0.8095 | 121 | 0.8678 |
| 4_1189940 | 35 | 0.9143 | 28 | 0.8929 | 11 | 1 | 23 | 1 | 97 | 0.9381 | 21 | 1 | 27 | 0.963 | 21 | 0.9524 | 14 | 0.9286 | 83 | 0.9639 |
| 4_1189941 | 34 | 0.1765 | 33 | 0.2727 | 11 | 0 | 25 | 0.32 | 103 | 0.2233 | 30 | 0.2333 | 32 | 0.2188 | 26 | 0.2692 | 16 | 0.25 | 104 | 0.2404 |
| 4_1189941 | 42 | 0.8095 | 38 | 0.8421 | 17 | 0.8824 | 37 | 0.6216 | 134 | 0.7761 | 32 | 0.9375 | 33 | 0.7576 | 29 | 0.7241 | 24 | 0.8333 | 118 | 0.8136 |
| 4_1189942 | 41 | 0.0244 | 37 | 0.0811 | 17 | 0.1765 | 33 | 0.0909 | 128 | 0.0781 | 27 | 0.037 | 36 | 0.0278 | 26 | 0.1538 | 24 | 0.0417 | 113 | 0.0619 |
| 4_1189943 | 47 | 0.3191 | 40 | 0.4 | 18 | 0.3889 | 35 | 0.3429 | 140 | 0.3571 | 31 | 0.3226 | 39 | 0.4103 | 32 | 0.1562 | 24 | 0.2917 | 126 | 0.3016 |
| 4_1189943 | 48 | 0.25 | 36 | 0.3611 | 18 | 0.2778 | 35 | 0.2286 | 137 | 0.2774 | 30 | 0.2333 | 35 | 0.2857 | 27 | 0.3704 | 24 | 0.1667 | 116 | 0.2672 |
| 4_1189943 | 50 | 0.84 | 38 | 0.7105 | 18 | 0.7222 | 34 | 0.7647 | 140 | 0.7714 | 31 | 0.7097 | 37 | 0.8649 | 28 | 0.6786 | 22 | 0.7727 | 118 | 0.7627 |
| 4_1189944 | 46 | 0.1739 | 36 | 0.3889 | 16 | 0.25 | 32 | 0.1875 | 130 | 0.2462 | 32 | 0.2188 | 39 | 0.2051 | 30 | 0.4 | 25 | 0.44 | 126 | 0.3016 |
| 4_1189944 | 41 | 0.0732 | 36 | 0.1389 | 17 | 0.1176 | 34 | 0.2059 | 128 | 0.1328 | 31 | 0.1613 | 35 | 0.1143 | 29 | 0.2759 | 23 | 0.1304 | 118 | 0.1695 |
| 4_1189947 | 4 | 0.25 | 1 | 1 | 1 | 1 | 1 | 0 | 7 | 0.4286 | 1 | 0 | 2 | 0.5 | NA | NA | NA | NA | 3 | 0.3333 |
| 4_1190582 | 5 | 0 | 10 | 0.4 | 6 | 0 | 9 | 0.1111 | 30 | 0.1667 | 3 | 0 | 4 | 0 | 1 | 0 | 2 | 0.5 | 10 | 0.1 |
| 4_1190582 | 6 | 0.3333 | 9 | 0.2222 | 6 | 0 | 11 | 0.0909 | 32 | 0.1562 | 3 | 0 | 4 | 0.25 | NA | NA | 3 | 0 | 10 | 0.1 |
| 4_1190582 | 5 | 0.8 | 9 | 0.6667 | 7 | 0.8571 | 11 | 0.4545 | 32 | 0.6562 | 3 | 0.6667 | 4 | 0.5 | NA | NA | 3 | 0.6667 | 10 | 0.6 |
| 4_1190583 | 5 | 0.2 | 9 | 0.4444 | 5 | 0.2 | 8 | 0.125 | 27 | 0.2593 | 4 | 0 | 3 | 0.3333 | 1 | 0 | 2 | 0 | 10 | 0.1 |
| 4_1190583 | 5 | 0.2 | 9 | 0.1111 | 7 | 0.2857 | 7 | 0.2857 | 28 | 0.2143 | 4 | 0.25 | 4 | 0 | 1 | 0 | 3 | 0.6667 | 12 | 0.25 |
| 4_1190625 | 2 | 0 | NA | NA | NA | NA | 1 | 1 | 3 | 0.3333 | NA | NA | NA | NA | NA | NA | 1 | 1 | 1 | 1 |
| 4_1190625 | 2 | 0.5 | NA | NA | NA | NA | 1 | 0 | 3 | 0.3333 | NA | NA | NA | NA | NA | NA | 1 | 1 | 1 | 1 |
| 4_1190626 | 2 | 0 | NA | NA | NA | NA | 1 | 1 | 3 | 0.3333 | NA | NA | NA | NA | NA | NA | 1 | 1 | 1 | 1 |
| 4_1190780 | 3 | 0.6667 | 1 | 0 | 1 | 0 | 2 | 0 | 7 | 0.2857 | NA | NA | 5 | 0 | 1 | 1 | 3 | 0.6667 | 9 | 0.3333 |
| 4_1190801 | 2 | 1 | 1 | 1 | 2 | 1 | NA | NA | 5 | 1 | 1 | 0 | 2 | 0.5 | NA | NA | 2 | 0 | 5 | 0.2 |
| 4_1190833 | 2 | 1 | 1 | 1 | NA | NA | 1 | 1 | 4 | 1 | NA | NA | NA | NA | NA | NA | 1 | 1 | 1 | 1 |
| 4_1190834 | 2 | 1 | NA | NA | NA | NA | 4 | 0.5 | 6 | 0.6667 | 1 | 0 | NA | NA | 2 | 0.5 | 1 | 1 | 4 | 0.5 |
| 4_1190834 | 2 | 1 | 1 | 0 | NA | NA | 4 | 0.5 | 7 | 0.5714 | 1 | 0 | NA | NA | 2 | 0.5 | 1 | 1 | 4 | 0.5 |
| 4_1190835 | 2 | 1 | 1 | 1 | NA | NA | 4 | 0.75 | 7 | 0.8571 | 1 | 1 | NA | NA | 2 | 1 | 1 | 1 | 4 | 1 |
| 4_1190836 | 2 | 0.5 | NA | NA | NA | NA | 3 | 0.3333 | 5 | 0.4 | 1 | 0 | NA | NA | 2 | 0 | 1 | 1 | 4 | 0.25 |
| 4_1191052 | 1 | 1 | NA | NA | 1 | 1 | 2 | 0.5 | 4 | 0.75 | 1 | 1 | 2 | 1 | 2 | 1 | NA | NA | 5 | 1 |
| 4_1191054 | 1 | 1 | NA | NA | 1 | 0 | 1 | 0 | 3 | 0.3333 | 2 | 0 | 1 | 0 | 2 | 0.5 | NA | NA | 5 | 0.2 |
| 4_1191352 | 41 | 0.0488 | 38 | 0.0526 | 31 | 0.0323 | 46 | 0.1087 | 156 | 0.0641 | 36 | 0 | 38 | 0.0526 | 54 | 0.0741 | 33 | 0.1212 | 161 | 0.0621 |
| 4_1191354 | 31 | 0.0968 | 34 | 0.0294 | 28 | 0.0714 | 35 | 0.0857 | 128 | 0.0703 | 27 | 0.1481 | 31 | 0 | 43 | 0.0465 | 31 | 0.129 | 132 | 0.0758 |
| 4_1191572 | 1 | 0 | 5 | 0.2 | 1 | 0 | 3 | 0 | 10 | 0.1 | 1 | 0 | 2 | 0 | 1 | 0 | 6 | 0.3333 | 10 | 0.2 |
| 4_1191811 | 1 | 0 | NA | NA | NA | NA | 1 | 1 | 2 | 0.5 | NA | NA | 2 | 0 | 2 | 0.5 | NA | NA | 4 | 0.25 |
| 4_1192757 | 1 | 1 | 3 | 0.3333 | 2 | 1 | NA | NA | 6 | 0.6667 | 1 | 0 | 1 | 0 | 3 | 0.6667 | 1 | 1 | 6 | 0.5 |
| 4_1192757 | 1 | 1 | 3 | 0.3333 | 2 | 1 | NA | NA | 6 | 0.6667 | 1 | 1 | 2 | 0.5 | 2 | 0.5 | 2 | 0.5 | 7 | 0.5714 |
| 4_1192759 | 1 | 1 | 3 | 1 | 2 | 1 | NA | NA | 6 | 1 | NA | NA | NA | NA | 3 | 1 | 1 | 1 | 4 | 1 |
| 4_1192760 | 1 | 1 | 2 | 1 | 2 | 1 | NA | NA | 5 | 1 | 1 | 1 | 1 | 1 | 3 | 1 | 1 | 1 | 6 | 1 |
| 4_1192761 | NA | NA | 2 | 1 | 2 | 0 | NA | NA | 4 | 0.5 | NA | NA | 1 | 1 | 2 | 1 | 1 | 1 | 4 | 1 |
| 4_1192761 | 1 | 1 | 3 | 0.6667 | 1 | 1 | NA | NA | 5 | 0.8 | 1 | 1 | NA | NA | 1 | 1 | 2 | 1 | 4 | 1 |
| 4_1192762 | NA | NA | 3 | 1 | NA | NA | NA | NA | 3 | 1 | NA | NA | 1 | 0 | 2 | 0.5 | NA | NA | 3 | 0.3333 |
| 4_1192764 | 1 | 0 | 2 | 1 | 1 | 0 | NA | NA | 4 | 0.5 | 1 | 1 | 1 | 0 | 3 | 0.6667 | NA | NA | 5 | 0.6 |
| 4_1193234 | 2 | 0 | 1 | 0 | NA | NA | 1 | 1 | 4 | 0.25 | 1 | 0 | 2 | 0.5 | 4 | 0 | 2 | 0 | 9 | 0.1111 |
| 4_1193243 | NA | NA | NA | NA | NA | NA | 1 | 1 | 1 | 1 | NA | NA | 1 | 1 | 3 | 1 | 1 | 1 | 5 | 1 |
| 4_1193244 | 2 | 0.5 | NA | NA | NA | NA | 1 | 1 | 3 | 0.6667 | 1 | 1 | 1 | 0 | 3 | 0.3333 | 1 | 0 | 6 | 0.3333 |
| 4_1193244 | 2 | 1 | NA | NA | NA | NA | 1 | 0 | 3 | 0.6667 | 1 | 0 | 1 | 1 | 3 | 1 | 1 | 1 | 6 | 0.8333 |
| 4_1193244 | 2 | 1 | NA | NA | NA | NA | 1 | 1 | 3 | 1 | 1 | 1 | 1 | 1 | 1 | 1 | NA | NA | 3 | 1 |
| 4_1193245 | 2 | 1 | NA | NA | NA | NA | 1 | 1 | 3 | 1 | 1 | 1 | 1 | 1 | 3 | 1 | 1 | 1 | 6 | 1 |
| 4_1193245 | 2 | 1 | NA | NA | NA | NA | 1 | 1 | 3 | 1 | NA | NA | 1 | 1 | 2 | 1 | NA | NA | 3 | 1 |
| 4_1193247 | 2 | 0.5 | NA | NA | NA | NA | NA | NA | 2 | 0.5 | 1 | 1 | 1 | 0 | 3 | 0.3333 | 1 | 1 | 6 | 0.5 |
| 4_1193249 | 2 | 1 | NA | NA | NA | NA | NA | NA | 2 | 1 | 1 | 1 | 1 | 1 | 2 | 1 | 1 | 1 | 5 | 1 |
| 4_1193250 | 2 | 1 | NA | NA | NA | NA | NA | NA | 2 | 1 | 1 | 1 | 1 | 1 | 2 | 1 | 1 | 1 | 5 | 1 |
| 4_1193251 | 2 | 0.5 | NA | NA | NA | NA | NA | NA | 2 | 0.5 | NA | NA | 1 | 1 | 2 | 1 | 1 | 1 | 4 | 1 |
| 4_1193255 | 2 | 1 | NA | NA | NA | NA | NA | NA | 2 | 1 | 1 | 1 | 1 | 1 | 1 | 1 | 1 | 1 | 4 | 1 |

| | | | | | | | | | | | | | | | | | | | | |
|---|---|---|---|---|---|---|---|---|---|---|---|---|---|---|---|---|---|---|---|---|
| 4_1193951( | NA | NA | 1 | 1 | NA | NA | NA | NA | 1 | 1 | NA | NA | NA | NA | 1 | 1 | 3 | 1 | 4 | 1 |
| 4_1193959! | NA | NA | 2 | 1 | NA | NA | 1 | 1 | 3 | 1 | NA | NA | NA | NA | NA | NA | 1 | 1 | 1 | 1 |
| 4_1193960: | NA | NA | 2 | 0.5 | NA | NA | 1 | 1 | 3 | 0.6667 | NA | NA | NA | NA | NA | NA | 1 | 1 | 1 | 1 |
| 4_1193961' | NA | NA | 2 | 0.5 | NA | NA | 1 | 1 | 3 | 0.6667 | NA | NA | NA | NA | NA | NA | 1 | 1 | 1 | 1 |
| 4_1193961‹ | NA | NA | 2 | 0.5 | NA | NA | NA | NA | 2 | 0.5 | NA | NA | NA | NA | NA | NA | 1 | 1 | 1 | 1 |
| 4_1193962‹ | NA | NA | 1 | 0 | NA | NA | 1 | 1 | 2 | 0.5 | NA | NA | NA | NA | NA | NA | 1 | 1 | 1 | 1 |
| 4_1193962( | NA | NA | 1 | 1 | NA | NA | NA | NA | 1 | 1 | NA | NA | NA | NA | NA | NA | 1 | 1 | 1 | 1 |
| 4_1193962( | NA | NA | 2 | 0.5 | NA | NA | 1 | 0 | 3 | 0.3333 | NA | NA | NA | NA | NA | NA | 1 | 1 | 1 | 1 |
| 4_1193962! | NA | NA | 2 | 1 | 1 | 1 | 1 | 1 | 4 | 1 | NA | NA | NA | NA | NA | NA | 1 | 1 | 1 | 1 |
| 4_1193966: | NA | NA | 1 | 1 | NA | NA | NA | NA | 1 | 1 | NA | NA | NA | NA | NA | NA | 1 | 1 | 1 | 1 |
| 4_1194307' | 3 | 0.3333 | 1 | 1 | NA | NA | 1 | 0 | 5 | 0.4 | NA | NA | 1 | 1 | 1 | 0 | 1 | 0 | 3 | 0.3333 |
| 4_1197300( | 2 | 0 | 2 | 0.5 | 1 | 0 | 2 | 0 | 7 | 0.1429 | 2 | 0 | 3 | 0 | 1 | 0 | 4 | 0.75 | 10 | 0.3 |
| 4_1197300‹ | 1 | 0 | 2 | 1 | 1 | 0 | 2 | 0 | 6 | 0.3333 | 2 | 0.5 | 1 | 0 | NA | NA | 4 | 0 | 7 | 0.1429 |
| 4_1197300: | 2 | 0.5 | 2 | 1 | NA | NA | 2 | 0.5 | 6 | 0.6667 | 1 | 1 | 3 | 0.6667 | NA | NA | 3 | 1 | 7 | 0.8571 |
| 4_1197302! | 1 | 1 | 1 | 1 | 2 | 0.5 | 1 | 1 | 5 | 0.8 | 2 | 0.5 | 2 | 0.5 | 1 | 1 | 2 | 0 | 7 | 0.4286 |
| 4_1199891: | 1 | 0 | NA | NA | 1 | 1 | NA | NA | 2 | 0.5 | 1 | 0 | NA | NA | 2 | 0 | 1 | 1 | 4 | 0.25 |
| 4_1199892! | 1 | 1 | 1 | 1 | 1 | 1 | NA | NA | 3 | 1 | 1 | 0 | NA | NA | 1 | 1 | 1 | 1 | 3 | 0.6667 |
| 4_1199894! | 1 | 1 | 1 | 0 | 1 | 0 | NA | NA | 3 | 0.3333 | NA | NA | NA | NA | 2 | 0.5 | NA | NA | 2 | 0.5 |
| 4_1199895! | 1 | 1 | 1 | 0 | 1 | 1 | NA | NA | 3 | 0.6667 | NA | NA | NA | NA | 1 | 1 | NA | NA | 1 | 1 |
| 4_1199896( | 1 | 0 | 1 | 0 | 1 | 1 | 1 | 0 | 4 | 0.25 | 1 | 0 | NA | NA | 2 | 0 | 1 | 1 | 4 | 0.25 |
| 4_1199897: | 1 | 0 | 1 | 0 | 1 | 0 | 1 | 1 | 4 | 0.25 | 1 | 1 | NA | NA | 2 | 0 | 1 | 0 | 4 | 0.25 |
| 4_1199910' | 1 | 0 | 1 | 1 | 1 | 1 | 1 | 0 | 4 | 0.5 | 1 | 0 | NA | NA | 2 | 1 | 1 | 1 | 4 | 0.75 |
| 4_1199910! | 1 | 1 | 1 | 0 | 1 | 0 | 1 | 1 | 4 | 0.5 | 1 | 0 | NA | NA | 1 | 0 | 1 | 1 | 3 | 0.3333 |
| 4_1199912' | 1 | 1 | 1 | 0 | 1 | 0 | 1 | 0 | 4 | 0.25 | 1 | 0 | NA | NA | 1 | 0 | 1 | 1 | 3 | 0.3333 |
| 4_1199912‹ | NA | NA | 1 | 1 | 1 | 1 | 1 | 1 | 3 | 1 | 1 | 1 | NA | NA | 2 | 1 | NA | NA | 3 | 1 |
| 4_1199913! | 1 | 1 | 1 | 1 | 1 | 1 | 1 | 1 | 4 | 1 | 1 | 1 | 1 | 1 | 1 | 1 | 1 | 1 | 4 | 1 |
| 4_1199914( | 1 | 0 | 1 | 1 | 1 | 1 | 1 | 0 | 4 | 0.5 | 1 | 0 | NA | NA | 2 | 0.5 | 1 | 0 | 4 | 0.25 |
| 4_1199914: | 1 | 1 | 1 | 0 | 1 | 0 | 1 | 0 | 4 | 0.25 | NA | NA | 1 | 0 | 2 | 0.5 | 1 | 0 | 4 | 0.25 |
| 4_1199914! | 1 | 0 | 1 | 1 | 1 | 0 | 1 | 1 | 4 | 0.5 | 1 | 0 | 1 | 0 | 2 | 0.5 | 1 | 1 | 5 | 0.4 |
| 4_1199915( | NA | NA | 1 | 1 | 1 | 1 | 1 | 0 | 3 | 0.6667 | 1 | 0 | 1 | 1 | 2 | 1 | 1 | 0 | 5 | 0.6 |
| 4_1199916( | 1 | 1 | 1 | 0 | 1 | 0 | 1 | 0 | 4 | 0.25 | 1 | 0 | 1 | 1 | 1 | 0 | 1 | 0 | 4 | 0.25 |
| 4_1199919: | 1 | 1 | 1 | 1 | 1 | 0 | 1 | 1 | 4 | 0.75 | 1 | 0 | 2 | 0.5 | 2 | 1 | 1 | 1 | 6 | 0.6667 |
| 4_1199920( | 1 | 0 | 1 | 1 | 1 | 0 | 1 | 0 | 4 | 0.25 | 1 | 0 | 1 | 1 | 2 | 0.5 | 1 | 0 | 5 | 0.4 |
| 4_1199921: | 1 | 1 | 1 | 1 | 1 | 0 | 1 | 1 | 4 | 0.75 | 1 | 0 | 3 | 1 | 2 | 0.5 | 1 | 0 | 7 | 0.5714 |
| 4_1199922! | 1 | 1 | 1 | 0 | 1 | 0 | 1 | 0 | 4 | 0.25 | 1 | 0 | 3 | 0.6667 | 2 | 0 | 1 | 1 | 7 | 0.4286 |
| 4_1199932( | 1 | 1 | 2 | 0.5 | 2 | 0 | 1 | 1 | 6 | 0.5 | 1 | 0 | 3 | 0.6667 | 2 | 0.5 | 1 | 0 | 7 | 0.4286 |
| 4_1199934: | 1 | 0 | 2 | 0 | 2 | 0 | 1 | 1 | 6 | 0.1667 | NA | NA | 1 | 0 | 2 | 0.5 | NA | NA | 3 | 0.3333 |
| 4_1199940! | 1 | 0 | 3 | 0.6667 | 2 | 0.5 | 1 | 0 | 7 | 0.4286 | 1 | 0 | 3 | 0.3333 | 2 | 0.5 | 1 | 0 | 7 | 0.2857 |
| 4_1199949! | 2 | 0.5 | 2 | 0 | 3 | 0 | 1 | 0 | 8 | 0.125 | 3 | 0.6667 | 3 | 0.3333 | 2 | 1 | 1 | 1 | 9 | 0.6667 |
| 4_1201251! | 2 | 1 | 3 | 1 | 3 | 1 | 3 | 1 | 11 | 1 | 3 | 1 | 4 | 1 | 2 | 1 | 1 | 1 | 10 | 1 |
| 4_1201252' | 2 | 1 | 2 | 1 | 3 | 1 | 3 | 1 | 10 | 1 | 2 | 1 | 4 | 1 | 2 | 1 | 1 | 1 | 9 | 1 |
| 4_1201252( | 1 | 1 | 3 | 0.3333 | 3 | 0.3333 | 2 | 0.5 | 9 | 0.4444 | 2 | 0.5 | 5 | 0.6 | NA | NA | 1 | 1 | 8 | 0.625 |
| 4_1201256: | 1 | 1 | 2 | 0.5 | 1 | 0 | 3 | 0.3333 | 7 | 0.4286 | 1 | 0 | 3 | 0.6667 | 1 | 1 | 1 | 1 | 6 | 0.6667 |
| 4_1201257( | 1 | 1 | 3 | 1 | 3 | 0.6667 | 2 | 0.5 | 9 | 0.7778 | 2 | 1 | 4 | 0.75 | 1 | 1 | 1 | 1 | 8 | 0.875 |
| 4_1201262: | 1 | 1 | 3 | 1 | 4 | 0.75 | 1 | 1 | 9 | 0.8889 | 1 | 1 | 3 | 1 | 1 | 1 | 1 | 1 | 6 | 1 |
| 4_1201441: | 1 | 1 | 2 | 0.5 | 2 | 0.5 | 3 | 0.3333 | 8 | 0.5 | 1 | 0 | 3 | 0.3333 | 1 | 0 | 1 | 1 | 6 | 0.3333 |
| 4_1201442: | 1 | 0 | 2 | 0.5 | 3 | 0 | 3 | 0 | 9 | 0.1111 | NA | NA | 3 | 0.3333 | 1 | 0 | 1 | 1 | 5 | 0.4 |
| 4_1201442! | NA | NA | 2 | 1 | 2 | 1 | 3 | 0.6667 | 7 | 0.8571 | NA | NA | 3 | 1 | 1 | 1 | 1 | 1 | 5 | 1 |
| 4_1201443! | 1 | 0 | 1 | 0 | 2 | 0 | 3 | 0.6667 | 7 | 0.2857 | NA | NA | 3 | 0.3333 | 1 | 1 | 1 | 0 | 5 | 0.4 |
| 4_1201444( | 1 | 0 | 1 | 0 | 1 | 1 | 2 | 1 | 5 | 0.6 | NA | NA | 2 | 0.5 | 1 | 1 | 1 | 0 | 4 | 0.5 |
| 4_1203865: | 4 | 1 | 6 | 0.8333 | 3 | 1 | 7 | 1 | 20 | 0.95 | 1 | 1 | 6 | 0.6667 | 7 | 0.8571 | 2 | 0.5 | 16 | 0.75 |
| 4_1203865! | 4 | 0 | 5 | 0.2 | 2 | 0 | 5 | 0.4 | 16 | 0.1875 | NA | NA | 6 | 0.5 | 4 | 0 | 1 | 0 | 11 | 0.2727 |
| 4_1203867! | 5 | 0 | 6 | 0.1667 | 3 | 0 | 7 | 0.2857 | 21 | 0.1429 | 1 | 0 | 5 | 0 | 7 | 0.1429 | 3 | 0.3333 | 16 | 0.125 |
| 4_1203868! | 5 | 0.4 | 8 | 0 | 3 | 0.3333 | 7 | 0.1429 | 23 | 0.1739 | 1 | 0 | 6 | 0.1667 | 7 | 0.1429 | 3 | 0 | 17 | 0.1176 |
| 4_1203869: | 4 | 0.25 | 9 | 0.4444 | 3 | 0.6667 | 8 | 0.625 | 24 | 0.5 | 1 | 1 | 4 | 0.5 | 8 | 0.5 | 3 | 0.6667 | 16 | 0.5625 |
| 4_1203870! | 5 | 0.6 | 7 | 0.4286 | 3 | 0 | 9 | 0.5556 | 24 | 0.4583 | 1 | 0 | 5 | 0.2 | 6 | 0.3333 | 2 | 0 | 14 | 0.2143 |
| 4_1203976: | 2 | 0 | 4 | 0.5 | 3 | 0 | 3 | 0 | 12 | 0.1667 | 7 | 0.5714 | 7 | 0.1429 | 4 | 0 | 2 | 0 | 20 | 0.25 |
| 4_1205239' | 2 | 1 | 1 | 0 | 1 | 0 | NA | NA | 4 | 0.5 | 1 | 0 | NA | NA | 2 | 1 | 1 | 1 | 4 | 0.75 |
| 4_1205245( | 3 | 1 | NA | NA | NA | NA | NA | NA | 3 | 1 | 1 | 1 | 3 | 1 | 2 | 0 | 1 | 1 | 7 | 0.7143 |
| 4_1205246! | 3 | 0 | 1 | 1 | 1 | 0 | NA | NA | 5 | 0.2 | 1 | 0 | 3 | 0.3333 | 2 | 0.5 | 1 | 1 | 7 | 0.4286 |
| 4_1205247‹ | 3 | 0.6667 | 1 | 1 | 1 | 0 | NA | NA | 5 | 0.6 | NA | NA | 3 | 0.6667 | 3 | 0.6667 | 1 | 1 | 7 | 0.7143 |
| 4_1205546: | NA | NA | 2 | 0.5 | NA | NA | NA | NA | 2 | 0.5 | 1 | 1 | 1 | 0 | NA | NA | NA | NA | 2 | 0.5 |
| 4_1205550: | NA | NA | 2 | 1 | 1 | 1 | NA | NA | 3 | 1 | 1 | 1 | 1 | 1 | NA | NA | NA | NA | 2 | 1 |
| 4_1205550! | NA | NA | 1 | 1 | 1 | 1 | NA | NA | 2 | 1 | 1 | 1 | 1 | 1 | NA | NA | NA | NA | 2 | 1 |
| 4_1206146( | 5 | 0.2 | 10 | 0.2 | 8 | 0.375 | 7 | 0.4286 | 30 | 0.3 | 5 | 0.2 | 3 | 0.3333 | 4 | 0 | 3 | 0.6667 | 15 | 0.2667 |

| | | | | | | | | | | | | | | | | | | | | |
|---|---|---|---|---|---|---|---|---|---|---|---|---|---|---|---|---|---|---|---|---|
| 4_12061475 | 6 | 0.6667 | 9 | 0.8889 | 10 | 0.6 | 7 | 0.8571 | 32 | 0.75 | 5 | 0.6 | 3 | 1 | 4 | 1 | 3 | 0.6667 | 15 | 0.8 |
| 4_12061477 | 7 | 0.8571 | 9 | 0.8889 | 10 | 0.5 | 7 | 0.2857 | 33 | 0.6364 | 6 | 0.5 | 4 | 1 | 4 | 1 | 3 | 1 | 17 | 0.8235 |
| 4_12061483 | 3 | 1 | 6 | 1 | 6 | 1 | 6 | 0.6667 | 21 | 0.9048 | 6 | 0.8333 | 3 | 0.6667 | 1 | 1 | 3 | 1 | 13 | 0.8462 |
| 4_12061522 | 6 | 0.5 | 7 | 0.8571 | 5 | 0.8 | 6 | 0.8333 | 24 | 0.75 | 5 | 0.8 | 4 | 0.25 | 4 | 0.5 | 3 | 0.3333 | 16 | 0.5 |
| 4_12061527 | 6 | 0.8333 | 8 | 0.875 | 8 | 0.75 | 6 | 0.6667 | 28 | 0.7857 | 5 | 1 | 4 | 1 | 3 | 1 | 2 | 1 | 14 | 1 |
| 4_12061528 | 8 | 0.125 | 10 | 0.1 | 9 | 0.1111 | 6 | 0.3333 | 33 | 0.1515 | 6 | 0.1667 | 3 | 0 | 4 | 0 | 3 | 0 | 16 | 0.0625 |
| 4_12061530 | 8 | 0.25 | 9 | 0 | 9 | 0.1111 | 5 | 0 | 31 | 0.0968 | 6 | 0 | 4 | 0 | 3 | 0 | 3 | 0.3333 | 16 | 0.0625 |
| 4_12061534 | 6 | 0.5 | 10 | 0.2 | 9 | 0.2222 | 6 | 0.1667 | 31 | 0.2581 | 6 | 0.1667 | 4 | 0.25 | 4 | 0.5 | 3 | 0.3333 | 17 | 0.2941 |
| 4_12061537 | 7 | 0 | 10 | 0.2 | 8 | 0.25 | 6 | 0.1667 | 31 | 0.1613 | 5 | 0 | 3 | 0 | 4 | 0.5 | 3 | 0 | 15 | 0.1333 |
| 4_12061538 | 6 | 0 | 9 | 0.1111 | 8 | 0.125 | 6 | 0 | 29 | 0.069 | 6 | 0.1667 | 3 | 0.3333 | 4 | 0.25 | 2 | 0 | 15 | 0.2 |
| 4_12074860 | NA | NA | 1 | 1 | 1 | 0 | 2 | 0 | 4 | 0.25 | NA | NA | 2 | 0 | 3 | 0.6667 | NA | NA | 5 | 0.4 |
| 4_12074870 | NA | NA | 1 | 0 | 1 | 1 | NA | NA | 2 | 0.5 | 1 | 0 | 2 | 0.5 | 2 | 1 | NA | NA | 5 | 0.6 |
| 4_12074900 | NA | NA | 1 | 0 | 1 | 1 | 1 | 1 | 3 | 0.6667 | 1 | 1 | 2 | 1 | 4 | 1 | NA | NA | 7 | 1 |
| 4_12074995 | NA | NA | 1 | 1 | 1 | 0 | 2 | 1 | 4 | 0.75 | 1 | 1 | 2 | 1 | 3 | 0.6667 | NA | NA | 6 | 0.8333 |
| 4_12075072 | NA | NA | 4 | 0.25 | 1 | 0 | 3 | 0.6667 | 8 | 0.375 | 1 | 1 | 4 | 0.5 | 4 | 0 | NA | NA | 9 | 0.3333 |
| 4_12075106 | NA | NA | 3 | 0.3333 | 1 | 0 | 2 | 0 | 6 | 0.1667 | 1 | 1 | 7 | 0.2857 | 5 | 0.4 | 1 | 0 | 14 | 0.3571 |
| 4_12075136 | 1 | 0 | 5 | 0.6 | 1 | 1 | 4 | 0.75 | 11 | 0.6364 | 1 | 1 | 7 | 0.5714 | 4 | 0.5 | 1 | 1 | 13 | 0.6154 |
| 4_12075141 | 1 | 0 | 2 | 0.5 | 1 | 1 | 3 | 0 | 7 | 0.2857 | 1 | 0 | 5 | 0.8 | 5 | 0.4 | 1 | 1 | 12 | 0.5833 |
| 4_12075148 | 1 | 0 | 3 | 0.3333 | 1 | 0 | 5 | 0.4 | 10 | 0.3 | 1 | 1 | 8 | 0.375 | 5 | 0.6 | NA | NA | 14 | 0.5 |
| 4_12075160 | 1 | 0 | 1 | 0 | NA | NA | 5 | 0.2 | 7 | 0.1429 | 1 | 0 | 7 | 0.2857 | 5 | 0.4 | 1 | 0 | 14 | 0.2857 |
| 4_12075163 | 1 | 0 | 3 | 0 | 1 | 1 | 5 | 0 | 10 | 0.1 | NA | NA | 4 | 0 | 2 | 0.5 | 1 | 0 | 7 | 0.1429 |
| 4_12075174 | 2 | 0 | 4 | 0.5 | 1 | 0 | 6 | 0.1667 | 13 | 0.2308 | 1 | 0 | 7 | 0.2857 | 5 | 0.4 | 1 | 0 | 14 | 0.2857 |
| 4_12075193 | 2 | 0.5 | 4 | 1 | 1 | 1 | 6 | 1 | 13 | 0.9231 | NA | NA | 10 | 0.9 | 5 | 1 | 1 | 1 | 16 | 0.9375 |
| 4_12075205 | 1 | 0 | 5 | 0.6 | 1 | 0 | 5 | 1 | 12 | 0.6667 | 1 | 1 | 6 | 0.3333 | 5 | 0.6 | 1 | 1 | 13 | 0.5385 |
| 4_12075215 | 3 | 0 | 3 | 0.3333 | 1 | 0 | 3 | 0.3333 | 10 | 0.2 | 1 | 1 | 7 | 0.1429 | 2 | 0 | 1 | 0 | 11 | 0.1818 |
| 4_12075221 | 2 | 0 | 5 | 0.8 | 1 | 1 | 6 | 0.3333 | 14 | 0.5 | 1 | 1 | 10 | 0.4 | 5 | 0.8 | 2 | 0.5 | 18 | 0.5556 |
| 4_12075244 | 4 | 0.5 | 5 | 0.4 | 2 | 1 | 5 | 0.6 | 16 | 0.5625 | NA | NA | 7 | 0.8571 | 4 | 1 | 2 | 0.5 | 13 | 0.8462 |
| 4_12075260 | 4 | 0.5 | 7 | 0.1429 | 3 | 0.3333 | 7 | 0 | 21 | 0.1905 | 1 | 0 | 10 | 0.2 | 5 | 0 | 2 | 0 | 18 | 0.1111 |
| 4_12075270 | 1 | 0 | 5 | 0.4 | 3 | 0 | 8 | 0.375 | 17 | 0.2941 | 1 | 0 | 6 | 0 | 3 | 0.3333 | 2 | 0 | 12 | 0.0833 |
| 4_12075286 | 4 | 0.25 | 7 | 0.1429 | 4 | 0 | 7 | 0 | 22 | 0.0909 | 1 | 0 | 10 | 0.2 | 4 | 0 | 2 | 0 | 17 | 0.1176 |
| 4_12075314 | 2 | 0.5 | 1 | 0 | 3 | 0 | 6 | 0.1667 | 12 | 0.1667 | 1 | 0 | 9 | 0.1111 | 3 | 0.3333 | 1 | 0 | 14 | 0.1429 |
| 4_12075326 | 5 | 0 | 7 | 0.2857 | 5 | 0.2 | 8 | 0.125 | 25 | 0.16 | 2 | 0.5 | 10 | 0 | 5 | 0.4 | 2 | 0 | 19 | 0.1579 |
| 4_12077865 | 2 | 1 | 1 | 1 | NA | NA | 2 | 1 | 5 | 1 | NA | NA | NA | NA | NA | NA | 1 | 1 | 1 | 1 |
| 4_12077896 | 2 | 1 | NA | NA | NA | NA | 2 | 1 | 4 | 1 | NA | NA | NA | NA | NA | NA | 1 | 1 | 1 | 1 |
| 4_12077904 | 1 | 1 | NA | NA | NA | NA | 2 | 1 | 3 | 1 | NA | NA | NA | NA | NA | NA | 1 | 1 | 1 | 1 |
| 4_12077911 | 2 | 1 | NA | NA | NA | NA | 1 | 1 | 3 | 1 | NA | NA | NA | NA | NA | NA | 1 | 1 | 1 | 1 |
| 4_12077916 | 1 | 0 | NA | NA | NA | NA | 2 | 1 | 3 | 0.6667 | NA | NA | NA | NA | NA | NA | 1 | 1 | 1 | 1 |
| 4_12077922 | 1 | 1 | NA | NA | NA | NA | 2 | 1 | 3 | 1 | NA | NA | NA | NA | NA | NA | 1 | 1 | 1 | 1 |
| 4_12077952 | NA | NA | NA | NA | NA | NA | 2 | 1 | 2 | 1 | NA | NA | NA | NA | NA | NA | 1 | 1 | 1 | 1 |
| 4_12077959 | 1 | 1 | NA | NA | NA | NA | 2 | 1 | 3 | 1 | NA | NA | NA | NA | NA | NA | 1 | 1 | 1 | 1 |
| 4_12077979 | 1 | 0 | NA | NA | NA | NA | 1 | 1 | 2 | 0.5 | NA | NA | NA | NA | NA | NA | 1 | 1 | 1 | 1 |
| 4_12082566 | 27 | 0.037 | 15 | 0.2 | 16 | 0.375 | 15 | 0.0667 | 73 | 0.1507 | 18 | 0.0556 | 16 | 0.1875 | 9 | 0 | 18 | 0.1667 | 61 | 0.1148 |
| 4_12082581 | 28 | 0.1429 | 16 | 0.0625 | 17 | 0.1765 | 13 | 0.0769 | 74 | 0.1216 | 20 | 0.05 | 18 | 0 | 14 | 0.0714 | 23 | 0.1304 | 75 | 0.0667 |
| 4_12082606 | 22 | 0.1818 | 13 | 0.3846 | 15 | 0.4667 | 11 | 0.4545 | 61 | 0.3443 | 17 | 0.2353 | 16 | 0.375 | 14 | 0.2857 | 23 | 0.3478 | 70 | 0.3143 |
| 4_12082636 | 20 | 0.05 | 12 | 0.3333 | 11 | 0.0909 | 13 | 0.2308 | 56 | 0.1607 | 16 | 0.25 | 12 | 0.0833 | 13 | 0.0769 | 21 | 0.2381 | 62 | 0.1774 |
| 4_12082655 | 22 | 0.0455 | 11 | 0.2727 | 12 | 0.0833 | 11 | 0 | 56 | 0.0893 | 16 | 0 | 14 | 0.0714 | 12 | 0.0833 | 20 | 0.2 | 62 | 0.0968 |
| 4_12082676 | 19 | 0.1053 | 16 | 0.0625 | 11 | 0.2727 | 13 | 0.1538 | 59 | 0.1356 | 16 | 0.1875 | 16 | 0.0625 | 9 | 0.2222 | 18 | 0.1111 | 59 | 0.1356 |
| 4_12082680 | 10 | 0.5 | 6 | 0.6667 | 9 | 0.4444 | 4 | 0.5 | 29 | 0.5172 | 7 | 0.1429 | 5 | 0.2 | 6 | 0.1667 | 5 | 0.8 | 23 | 0.3043 |
| 4_12082680 | 18 | 0.0556 | 11 | 0.1818 | 12 | 0.0833 | 11 | 0 | 52 | 0.0769 | 13 | 0.0769 | 12 | 0.1667 | 7 | 0 | 14 | 0.0714 | 46 | 0.087 |
| 4_12082913 | 15 | 0.2 | 12 | 0 | 11 | 0 | 10 | 0 | 48 | 0.0625 | 14 | 0.0714 | 15 | 0.0667 | 9 | 0.1111 | 16 | 0 | 54 | 0.0556 |
| 4_12091233 | 3 | 0 | 1 | 0 | 1 | 1 | 1 | 0 | 6 | 0.1667 | 1 | 1 | NA | NA | NA | NA | 1 | 1 | 2 | 1 |
| 4_12091646 | 1 | 0 | 4 | 0.5 | 8 | 0.5 | 4 | 0.5 | 17 | 0.4706 | 2 | 0 | 6 | 0.1667 | 4 | 0 | 3 | 0.6667 | 15 | 0.2 |
| 4_12091651 | 2 | 0 | 4 | 0 | 7 | 0.2857 | 6 | 0 | 19 | 0.1053 | 1 | 0 | 6 | 0 | 4 | 0 | 4 | 0.75 | 15 | 0.2 |
| 4_12091662 | 2 | 1 | 4 | 1 | 9 | 1 | 6 | 0.8333 | 21 | 0.9524 | 1 | 1 | 3 | 1 | 3 | 0.6667 | 5 | 1 | 12 | 0.9167 |
| 4_12091680 | 2 | 0 | 3 | 0.3333 | 9 | 0.2222 | 5 | 0 | 19 | 0.1579 | 2 | 0 | 6 | 0 | 4 | 0 | 5 | 0.2 | 17 | 0.0588 |
| 4_12091817 | 2 | 0 | 2 | 0 | 6 | 0 | 6 | 0.1667 | 16 | 0.0625 | 2 | 0.5 | 3 | 0 | 5 | 0 | 5 | 0.2 | 15 | 0.1333 |
| 4_12091819 | 2 | 0.5 | 3 | 0.3333 | 5 | 0.8 | 4 | 0.25 | 14 | 0.5 | 1 | 1 | 4 | 0.25 | 5 | 0.8 | 4 | 0.75 | 14 | 0.6429 |
| 4_12091827 | 2 | 0 | 2 | 0.5 | 7 | 0.2857 | 6 | 0.5 | 17 | 0.3529 | 3 | 0 | 3 | 0.3333 | 5 | 0.6 | 6 | 0.3333 | 17 | 0.3529 |
| 4_12091833 | 1 | 1 | 2 | 0.5 | 8 | 0.75 | 6 | 0.6667 | 17 | 0.7059 | 3 | 0.6667 | 3 | 0.6667 | 5 | 0.6 | 5 | 1 | 16 | 0.75 |
| 4_12100332 | 3 | 0 | 3 | 0 | 2 | 0 | 2 | 0.5 | 10 | 0.1 | 1 | 0 | 2 | 0 | 2 | 0.5 | 2 | 0.5 | 7 | 0.2857 |
| 4_12100350 | 3 | 1 | 3 | 0.6667 | 1 | 1 | 2 | 1 | 9 | 0.8889 | 2 | 0.5 | NA | NA | 2 | 1 | 2 | 1 | 6 | 0.8333 |
| 4_12100360 | 3 | 1 | 3 | 0.6667 | 2 | 1 | 2 | 1 | 10 | 0.9 | 1 | 1 | 2 | 0.5 | 3 | 0.6667 | 2 | 1 | 8 | 0.75 |
| 4_12100367 | 2 | 1 | 3 | 0 | 1 | 1 | 2 | 0.5 | 8 | 0.5 | 1 | 0 | 1 | 1 | 3 | 0.3333 | 1 | 0 | 6 | 0.3333 |
| 4_12100368 | 3 | 1 | 2 | 1 | 2 | 1 | 2 | 1 | 9 | 1 | 2 | 1 | 2 | 1 | 3 | 1 | 2 | 1 | 9 | 1 |
| 4_12100370 | 2 | 0 | 2 | 0.5 | 2 | 0.5 | 2 | 0 | 8 | 0.25 | 1 | 1 | 1 | 1 | 3 | 0.3333 | 2 | 1 | 7 | 0.7143 |

| | | | | | | | | | | | | | | | | | | | | |
|---|---|---|---|---|---|---|---|---|---|---|---|---|---|---|---|---|---|---|---|---|
| 4_12100377 | 2 | 0 | 3 | 0.3333 | 2 | 0 | 2 | 0 | 9 | 0.1111 | 2 | 0 | 2 | 0.5 | 2 | 0.5 | 2 | 0 | 8 | 0.25 |
| 4_12100387 | 2 | 0.5 | 2 | 0 | 2 | 1 | 2 | 1 | 8 | 0.625 | 1 | 0 | 1 | 0 | 1 | 0 | 1 | 1 | 4 | 0.25 |
| 4_12100393 | 2 | 0.5 | 3 | 0 | 2 | 0.5 | 1 | 0 | 8 | 0.25 | 1 | 1 | 2 | 0 | 3 | 0.3333 | 2 | 1 | 8 | 0.5 |
| 4_12100400 | 2 | 0 | 2 | 0.5 | 2 | 0 | 1 | 0 | 7 | 0.1429 | 1 | 0 | 1 | 0 | 3 | 0.3333 | 2 | 0 | 7 | 0.1429 |
| 4_12100415 | 2 | 1 | 1 | 1 | 2 | 1 | 1 | 1 | 6 | 1 | 1 | 1 | 2 | 1 | 3 | 1 | 1 | 1 | 7 | 1 |
| 4_12100417 | 2 | 0.5 | 1 | 0 | 2 | 1 | 1 | 0 | 6 | 0.5 | 1 | 0 | 1 | 0 | 3 | 0.3333 | 1 | 1 | 6 | 0.3333 |
| 4_12100423 | 2 | 0.5 | 1 | 0 | 2 | 0 | 1 | 0 | 6 | 0.1667 | 1 | 1 | 2 | 0.5 | 3 | 1 | 1 | 0 | 7 | 0.7143 |
| 4_12100428 | 2 | 0 | 1 | 0 | 1 | 1 | 1 | 0 | 5 | 0.2 | 1 | 0 | 2 | 0 | 3 | 0.3333 | 1 | 0 | 7 | 0.1429 |
| 4_12100458 | 1 | 0 | 1 | 0 | 2 | 0 | 1 | 1 | 5 | 0.2 | NA | NA | 2 | 1 | 2 | 1 | 1 | 0 | 5 | 0.8 |
| 4_12100482 | 2 | 1 | 1 | 1 | 1 | 1 | 1 | 1 | 5 | 1 | NA | NA | 2 | 1 | 2 | 1 | 1 | 1 | 5 | 1 |
| 4_12100491 | 2 | 1 | 1 | 1 | 2 | 1 | 1 | 1 | 6 | 1 | 1 | 1 | 2 | 1 | 2 | 1 | 1 | 1 | 6 | 1 |
| 4_12100517 | 1 | 1 | 1 | 1 | 2 | 1 | 1 | 1 | 5 | 1 | 1 | 1 | 1 | 1 | 2 | 0 | NA | NA | 4 | 0.5 |
| 4_12100522 | 1 | 1 | 1 | 1 | 2 | 1 | 1 | 1 | 5 | 1 | 1 | 1 | 2 | 1 | 2 | 1 | NA | NA | 5 | 1 |
| 4_12100529 | 1 | 1 | 1 | 0 | 2 | 0.5 | NA | NA | 4 | 0.5 | 1 | 0 | 2 | 0.5 | 1 | 1 | NA | NA | 4 | 0.5 |
| 4_12100533 | 1 | 1 | NA | NA | 2 | 1 | NA | NA | 3 | 1 | 1 | 1 | 2 | 0.5 | 1 | 1 | NA | NA | 4 | 0.75 |
| 4_12100545 | 1 | 1 | 1 | 1 | 2 | 1 | NA | NA | 4 | 1 | 1 | 1 | 2 | 1 | 1 | 1 | NA | NA | 4 | 1 |
| 4_12100556 | 1 | 1 | 1 | 1 | 2 | 0.5 | NA | NA | 4 | 0.75 | 1 | 0 | NA | NA | 1 | 1 | NA | NA | 2 | 0.5 |
| 4_12100560 | 1 | 1 | 1 | 1 | 1 | 0 | NA | NA | 3 | 0.6667 | 1 | 0 | 1 | 0 | 1 | 1 | NA | NA | 3 | 0.3333 |
| 4_12100563 | 1 | 0 | 1 | 0 | 2 | 0.5 | NA | NA | 4 | 0.25 | 1 | 0 | 1 | 0 | 1 | 1 | NA | NA | 3 | 0.3333 |
| 4_12100564 | NA | NA | 1 | 1 | 1 | 0 | NA | NA | 2 | 0.5 | 1 | 0 | 1 | 0 | 1 | 1 | NA | NA | 3 | 0.3333 |
| 4_12100565 | 1 | 1 | 1 | 0 | 2 | 0.5 | NA | NA | 4 | 0.5 | 1 | 1 | 1 | 1 | 1 | 1 | NA | NA | 3 | 1 |
| 4_12100570 | 1 | 1 | 1 | 1 | 1 | 1 | NA | NA | 3 | 1 | 1 | 1 | 1 | 0 | 1 | 1 | NA | NA | 3 | 0.6667 |
| 4_12100610 | NA | NA | 1 | 1 | 2 | 1 | NA | NA | 3 | 1 | NA | NA | 1 | 1 | NA | NA | NA | NA | 1 | 1 |
| 4_12100614 | 1 | 1 | 1 | 1 | 2 | 1 | NA | NA | 4 | 1 | NA | NA | 1 | 1 | 1 | 1 | NA | NA | 2 | 1 |
| 4_12100634 | 1 | 1 | 1 | 0 | 1 | 0 | NA | NA | 3 | 0.3333 | NA | NA | 1 | 1 | 1 | 0 | NA | NA | 2 | 0.5 |
| 4_12100647 | 1 | 0 | 1 | 1 | 2 | 0.5 | NA | NA | 4 | 0.5 | NA | NA | 1 | 0 | 1 | 1 | NA | NA | 2 | 0.5 |
| 4_12100655 | 1 | 1 | 1 | 0 | 2 | 1 | NA | NA | 4 | 0.75 | NA | NA | 1 | 1 | 1 | 1 | NA | NA | 2 | 1 |
| 4_12100676 | 1 | 0 | 1 | 0 | 2 | 0.5 | NA | NA | 4 | 0.25 | NA | NA | 1 | 0 | 1 | 1 | NA | NA | 2 | 0.5 |
| 4_12100704 | 1 | 1 | 1 | 1 | 2 | 1 | NA | NA | 4 | 1 | NA | NA | 1 | 0 | 1 | 1 | NA | NA | 2 | 0.5 |
| 4_12100731 | NA | NA | 1 | 1 | 2 | 1 | NA | NA | 3 | 1 | NA | NA | 1 | 1 | 1 | 1 | NA | NA | 2 | 1 |
| 4_12100777 | 1 | 0 | 1 | 1 | 2 | 0.5 | NA | NA | 4 | 0.5 | NA | NA | 1 | 0 | 1 | 1 | NA | NA | 2 | 0.5 |
| 4_12100789 | 1 | 1 | 1 | 1 | 2 | 1 | NA | NA | 4 | 1 | NA | NA | 1 | 1 | 1 | 1 | NA | NA | 2 | 1 |
| 4_12100801 | 1 | 1 | 1 | 1 | 2 | 1 | NA | NA | 4 | 1 | NA | NA | 1 | 1 | 1 | 1 | NA | NA | 2 | 1 |
| 4_12184272 | 7 | 0.1429 | 12 | 0.4167 | 12 | 0.4167 | 10 | 0.3 | 41 | 0.3415 | 19 | 0.2105 | 25 | 0.2 | 11 | 0.2727 | 9 | 0 | 64 | 0.1875 |
| 4_12223566 | 11 | 0.2727 | 7 | 0.1429 | 7 | 0.2857 | 9 | 0.2222 | 34 | 0.2353 | 13 | 0.0769 | 13 | 0.0769 | 13 | 0 | 4 | 0.75 | 43 | 0.1163 |
| 4_12274346 | NA | NA | NA | NA | 2 | 0.5 | NA | NA | 2 | 0.5 | 1 | 0 | 1 | 1 | 1 | 1 | NA | NA | 3 | 0.6667 |
| 4_12274353 | NA | NA | NA | NA | 2 | 1 | NA | NA | 2 | 1 | 1 | 1 | 1 | 1 | 1 | 1 | NA | NA | 3 | 1 |
| 4_12274356 | NA | NA | NA | NA | 2 | 1 | NA | NA | 2 | 1 | 1 | 0 | 1 | 1 | 1 | 0 | NA | NA | 3 | 0.3333 |
| 4_12274359 | NA | NA | NA | NA | 2 | 0.5 | NA | NA | 2 | 0.5 | 1 | 1 | 1 | 1 | 1 | 1 | NA | NA | 3 | 1 |
| 4_12274375 | NA | NA | 1 | 1 | 2 | 1 | NA | NA | 3 | 1 | 1 | 1 | 1 | 1 | 1 | 1 | NA | NA | 3 | 1 |
| 4_12275150 | 10 | 0.4 | 9 | 0.2222 | 4 | 0.25 | 8 | 0.375 | 31 | 0.3226 | 7 | 0.4286 | 9 | 0.3333 | 6 | 0.3333 | 3 | 0.3333 | 25 | 0.36 |
| 4_12275188 | 8 | 0.25 | 9 | 0.4444 | 6 | 0.1667 | 2 | 1 | 25 | 0.36 | 7 | 0.1429 | 9 | 0.3333 | 3 | 0.3333 | 4 | 0 | 23 | 0.2174 |
| 4_12275204 | 8 | 0 | 7 | 0 | 8 | 0.25 | 3 | 0 | 26 | 0.0769 | 7 | 0.1429 | 9 | 0.2222 | 4 | 0.5 | 4 | 0.25 | 24 | 0.25 |
| 4_12275254 | 11 | 0.0909 | 8 | 0.25 | 7 | 0.2857 | 2 | 0 | 28 | 0.1786 | 6 | 0.1667 | 4 | 0 | 1 | 0 | 3 | 0 | 14 | 0.0714 |
| 4_12275303 | 42 | 0.2381 | 38 | 0.2895 | 40 | 0.225 | 29 | 0.1379 | 149 | 0.2282 | 36 | 0.2778 | 40 | 0.1 | 44 | 0.2045 | 34 | 0.1765 | 154 | 0.1883 |
| 4_12275311 | 42 | 0.3095 | 33 | 0.3333 | 45 | 0.4222 | 31 | 0.3871 | 151 | 0.3642 | 35 | 0.5714 | 43 | 0.4651 | 42 | 0.5238 | 34 | 0.6176 | 154 | 0.539 |
| 4_12275319 | 41 | 0.2439 | 31 | 0 | 38 | 0.1579 | 32 | 0.1562 | 142 | 0.1479 | 31 | 0.0645 | 40 | 0.075 | 37 | 0.2162 | 30 | 0.1333 | 138 | 0.1232 |
| 4_12275342 | 39 | 0.1795 | 33 | 0.1818 | 39 | 0.1795 | 28 | 0.0714 | 139 | 0.1583 | 35 | 0.1714 | 40 | 0.125 | 40 | 0.25 | 31 | 0.3226 | 146 | 0.2123 |
| 4_12275375 | 28 | 0.1429 | 25 | 0.2 | 31 | 0.0968 | 16 | 0.125 | 100 | 0.14 | 23 | 0.2174 | 25 | 0.04 | 23 | 0.087 | 20 | 0.25 | 91 | 0.1429 |
| 4_12275382 | 34 | 0.1471 | 28 | 0.1071 | 29 | 0.0345 | 20 | 0.1 | 111 | 0.0991 | 26 | 0.1923 | 38 | 0.1053 | 32 | 0.0625 | 23 | 0 | 119 | 0.0924 |
| 4_12275396 | 28 | 0.1071 | 26 | 0.1154 | 33 | 0.1212 | 22 | 0.1818 | 109 | 0.1284 | 30 | 0.1333 | 34 | 0.0588 | 28 | 0.0714 | 25 | 0.24 | 117 | 0.1197 |
| 4_12275425 | 1 | 1 | NA | NA | NA | NA | NA | NA | 1 | 1 | NA | NA | NA | NA | NA | NA | 1 | 1 | 1 | 1 |
| 4_12275437 | 1 | 1 | NA | NA | NA | NA | NA | NA | 1 | 1 | 1 | 1 | NA | NA | NA | NA | 1 | 1 | 2 | 1 |
| 4_12275446 | 1 | 1 | NA | NA | NA | NA | NA | NA | 1 | 1 | NA | NA | NA | NA | NA | NA | 2 | 0.5 | 2 | 0.5 |
| 4_12275447 | 1 | 1 | NA | NA | NA | NA | NA | NA | 1 | 1 | NA | NA | NA | NA | NA | NA | 2 | 1 | 2 | 1 |
| 4_12275455 | 1 | 1 | NA | NA | NA | NA | NA | NA | 1 | 1 | NA | NA | NA | NA | NA | NA | 1 | 1 | 1 | 1 |
| 4_12275472 | 1 | 1 | NA | NA | NA | NA | NA | NA | 1 | 1 | NA | NA | NA | NA | NA | NA | 2 | 1 | 2 | 1 |
| 4_12275474 | 2 | 1 | NA | NA | NA | NA | NA | NA | 2 | 1 | NA | NA | NA | NA | NA | NA | 2 | 1 | 2 | 1 |
| 4_12275481 | 2 | 1 | NA | NA | NA | NA | NA | NA | 2 | 1 | NA | NA | NA | NA | NA | NA | 3 | 1 | 3 | 1 |
| 4_12275482 | 2 | 1 | NA | NA | NA | NA | NA | NA | 2 | 1 | NA | NA | NA | NA | NA | NA | 1 | 1 | 1 | 1 |
| 4_12284074 | 7 | 0.1429 | 7 | 0.4286 | 8 | 0.5 | 10 | 0.2 | 32 | 0.3125 | 11 | 0.1818 | 7 | 0.2857 | 8 | 0.375 | 5 | 0.6 | 31 | 0.3226 |
| 4_12284077 | 7 | 0.1429 | 5 | 0 | 6 | 0 | 10 | 0.1 | 28 | 0.0714 | 10 | 0 | 6 | 0.1667 | 8 | 0.125 | 5 | 0 | 29 | 0.069 |
| 4_12284079 | 7 | 0.4286 | 7 | 0.2857 | 7 | 0.8571 | 10 | 0.9 | 31 | 0.6452 | 9 | 0.7778 | 7 | 1 | 8 | 0.625 | 5 | 0.6 | 29 | 0.7586 |
| 4_12284088 | 6 | 0.5 | 6 | 0 | 8 | 0 | 10 | 0.2 | 30 | 0.1667 | 11 | 0.2727 | 7 | 0 | 7 | 0.1429 | 5 | 0.4 | 30 | 0.2 |
| 4_12284091 | 6 | 0.1667 | 4 | 0.5 | 6 | 0 | 7 | 0.1429 | 23 | 0.1739 | 6 | 0 | 4 | 0.5 | 6 | 0.3333 | 4 | 0 | 20 | 0.2 |

| | | | | | | | | | | | | | | | | | | | | |
|---|---|---|---|---|---|---|---|---|---|---|---|---|---|---|---|---|---|---|---|---|
| 4_1228409 | 6 | 0 | 6 | 0.1667 | 7 | 0 | 9 | 0.1111 | 28 | 0.0714 | 10 | 0.1 | 6 | 0.1667 | 6 | 0.3333 | 5 | 0.2 | 27 | 0.1852 |
| 4_1228410 | 5 | 0.8 | 4 | 0.5 | 7 | 0.4286 | 6 | 0.6667 | 22 | 0.5909 | 7 | 0.4286 | 6 | 0.5 | 8 | 0.25 | 4 | 0.5 | 25 | 0.4 |
| 4_1228944 | 1 | 1 | 2 | 1 | NA | NA | NA | NA | 3 | 1 | 1 | 1 | 3 | 1 | 2 | 1 | 1 | 1 | 7 | 1 |
| 4_1228947 | 1 | 1 | 2 | 0.5 | NA | NA | NA | NA | 3 | 0.6667 | NA | NA | 2 | 1 | 2 | 1 | 1 | 1 | 5 | 1 |
| 4_1228948 | 1 | 1 | 2 | 0.5 | NA | NA | NA | NA | 3 | 0.6667 | 1 | 0 | 2 | 0 | 2 | 0.5 | 1 | 0 | 6 | 0.1667 |
| 4_1228949 | NA | NA | 2 | 1 | NA | NA | NA | NA | 2 | 1 | 1 | 0 | 2 | 1 | 2 | 1 | 1 | 1 | 6 | 0.8333 |
| 4_1228950 | 1 | 1 | 2 | 1 | NA | NA | NA | NA | 3 | 1 | 1 | 0 | 3 | 0.6667 | 2 | 1 | 1 | 1 | 7 | 0.7143 |
| 4_1228950 | 1 | 1 | 2 | 1 | NA | NA | NA | NA | 3 | 1 | 1 | 0 | 3 | 1 | 2 | 1 | 1 | 1 | 7 | 0.8571 |
| 4_1228951 | 1 | 1 | 2 | 1 | NA | NA | NA | NA | 3 | 1 | 1 | 0 | 2 | 0.5 | 2 | 1 | NA | NA | 5 | 0.6 |
| 4_1228951 | 1 | 1 | 2 | 1 | NA | NA | NA | NA | 3 | 1 | 1 | 0 | 1 | 1 | 2 | 1 | 1 | 0 | 5 | 0.6 |
| 4_1228952 | 1 | 0 | 3 | 1 | NA | NA | NA | NA | 4 | 0.75 | NA | NA | 2 | 1 | 4 | 0.5 | 1 | 1 | 7 | 0.7143 |
| 4_1228953 | 1 | 0 | 3 | 0.6667 | NA | NA | NA | NA | 4 | 0.5 | 1 | 0 | 3 | 0.3333 | 3 | 0.3333 | 1 | 1 | 8 | 0.375 |
| 4_1228953 | 1 | 0 | 3 | 0.3333 | NA | NA | NA | NA | 4 | 0.25 | 1 | 0 | 3 | 0.3333 | 3 | 0.3333 | NA | NA | 7 | 0.2857 |
| 4_1228957 | 1 | 1 | 2 | 1 | NA | NA | NA | NA | 3 | 1 | 1 | 1 | 1 | 1 | 3 | 1 | 1 | 1 | 6 | 1 |
| 4_1228957 | 2 | 1 | 3 | 0 | NA | NA | NA | NA | 5 | 0.4 | 1 | 0 | 3 | 0.6667 | 3 | 0.6667 | 1 | 0 | 8 | 0.5 |
| 4_1228957 | 1 | 0 | 2 | 1 | NA | NA | NA | NA | 3 | 0.6667 | 1 | 1 | 2 | 1 | 3 | 0.6667 | 1 | 1 | 7 | 0.8571 |
| 4_1228959 | 2 | 1 | 2 | 1 | NA | NA | NA | NA | 4 | 1 | 2 | 1 | 3 | 1 | 4 | 1 | 1 | 1 | 10 | 1 |
| 4_1228959 | 2 | 1 | 3 | 1 | NA | NA | NA | NA | 5 | 1 | 2 | 1 | 3 | 1 | 4 | 1 | 1 | 1 | 10 | 1 |
| 4_1228959 | 2 | 1 | 2 | 0.5 | NA | NA | NA | NA | 4 | 0.75 | 1 | 1 | 1 | 1 | 2 | 1 | NA | NA | 4 | 1 |
| 4_1228959 | 2 | 0.5 | 3 | 0.6667 | NA | NA | NA | NA | 5 | 0.6 | 1 | 1 | 2 | 1 | 3 | 0.3333 | 1 | 0 | 7 | 0.5714 |
| 4_1228962 | 1 | 1 | 3 | 0 | NA | NA | NA | NA | 4 | 0.25 | 2 | 0.5 | 2 | 0 | 3 | 0 | 2 | 0 | 9 | 0.1111 |
| 4_1229479 | 1 | 0 | 4 | 0.75 | NA | NA | 4 | 0.5 | 9 | 0.5556 | NA | NA | 3 | 1 | 2 | 0 | 3 | 0.3333 | 8 | 0.5 |
| 4_1229480 | 1 | 1 | 4 | 0.25 | NA | NA | 5 | 0.4 | 10 | 0.4 | NA | NA | 5 | 0.4 | 2 | 1 | 3 | 0.6667 | 10 | 0.6 |
| 4_1229481 | NA | NA | 5 | 1 | NA | NA | 4 | 1 | 9 | 1 | NA | NA | 5 | 0.8 | 1 | 1 | 3 | 0.6667 | 9 | 0.7778 |
| 4_1229485 | 1 | 0 | 3 | 0.3333 | NA | NA | 3 | 0.3333 | 7 | 0.2857 | NA | NA | 5 | 0.6 | 2 | 0.5 | 1 | 0 | 8 | 0.5 |
| 4_1230209 | 3 | 0 | 2 | 0 | 1 | 0 | 2 | 0.5 | 8 | 0.125 | NA | NA | 3 | 0.6667 | 1 | 0 | 1 | 1 | 5 | 0.6 |
| 4_1230848 | 29 | 0.069 | 37 | 0.0541 | 34 | 0.0294 | 20 | 0.1 | 120 | 0.0583 | 30 | 0.0667 | 34 | 0.1176 | 32 | 0.1562 | 39 | 0.1026 | 135 | 0.1111 |
| 4_1230849 | 25 | 0.48 | 42 | 0.5238 | 33 | 0.5455 | 20 | 0.6 | 120 | 0.5333 | 30 | 0.4333 | 33 | 0.6364 | 32 | 0.4375 | 45 | 0.5111 | 140 | 0.5071 |
| 4_1230851 | 31 | 0.1935 | 41 | 0.1463 | 31 | 0.2903 | 22 | 0.2727 | 125 | 0.216 | 27 | 0.2222 | 33 | 0.2121 | 30 | 0.3333 | 42 | 0.1667 | 132 | 0.2273 |
| 4_1230853 | 34 | 0.0294 | 38 | 0.0789 | 37 | 0.0811 | 29 | 0.069 | 138 | 0.0652 | 31 | 0.0645 | 36 | 0.1111 | 35 | 0.1143 | 45 | 0.1333 | 147 | 0.1088 |
| 4_1230854 | 30 | 0.1667 | 41 | 0.1951 | 43 | 0.186 | 28 | 0.1429 | 142 | 0.1761 | 30 | 0.2333 | 36 | 0.1389 | 30 | 0.3333 | 49 | 0.102 | 145 | 0.1862 |
| 4_1230857 | 33 | 0.1515 | 43 | 0.186 | 42 | 0.1667 | 35 | 0.0571 | 153 | 0.1438 | 35 | 0.0286 | 39 | 0.1795 | 44 | 0.1591 | 51 | 0.1765 | 169 | 0.142 |
| 4_1230860 | 35 | 0 | 52 | 0.1154 | 49 | 0.0612 | 44 | 0.0455 | 180 | 0.0611 | 44 | 0.0909 | 45 | 0.0667 | 46 | 0.0435 | 59 | 0.1186 | 194 | 0.0825 |
| 4_1231134 | NA | NA | 1 | 1 | NA | NA | NA | NA | 1 | 1 | NA | NA | NA | NA | 1 | 1 | NA | NA | 1 | 1 |
| 4_1231138 | NA | NA | 1 | 1 | NA | NA | NA | NA | 1 | 1 | 1 | 1 | NA | NA | 1 | 1 | 1 | 1 | 3 | 1 |
| 4_1231139 | NA | NA | 1 | 1 | NA | NA | NA | NA | 1 | 1 | 1 | 1 | NA | NA | 1 | 1 | 1 | 1 | 3 | 1 |
| 4_1231141 | NA | NA | 1 | 1 | NA | NA | NA | NA | 1 | 1 | NA | NA | NA | NA | 1 | 0 | 1 | 1 | 2 | 0.5 |
| 4_1231150 | 1 | 0 | 2 | 0.5 | 1 | 1 | 1 | 1 | 5 | 0.6 | NA | NA | NA | NA | 1 | 0 | 1 | 1 | 2 | 0.5 |
| 4_1231407 | 1 | 1 | NA | NA | 1 | 1 | NA | NA | 2 | 1 | NA | NA | NA | NA | 2 | 1 | NA | NA | 2 | 1 |
| 4_1231409 | 1 | 1 | NA | NA | NA | NA | NA | NA | 1 | 1 | NA | NA | NA | NA | 2 | 0.5 | NA | NA | 2 | 0.5 |
| 4_1231414 | 1 | 0 | NA | NA | 1 | 1 | NA | NA | 2 | 0.5 | NA | NA | 1 | 1 | 2 | 1 | NA | NA | 3 | 1 |
| 4_1231414 | 1 | 0 | NA | NA | 1 | 1 | NA | NA | 2 | 0.5 | NA | NA | 1 | 1 | 2 | 0.5 | NA | NA | 3 | 0.6667 |
| 4_1231415 | 1 | 0 | NA | NA | 1 | 1 | NA | NA | 2 | 0.5 | NA | NA | 1 | 0 | 2 | 0.5 | NA | NA | 3 | 0.3333 |
| 4_1231767 | 3 | 0.3333 | NA | NA | 1 | 0 | 1 | 0 | 5 | 0.2 | 3 | 0 | 1 | 0 | 2 | 0 | 2 | 0.5 | 8 | 0.125 |
| 4_1231769 | 3 | 0.3333 | 1 | 0 | NA | NA | 4 | 0.25 | 8 | 0.25 | 2 | 0 | 1 | 1 | 3 | 0.3333 | 2 | 0 | 8 | 0.25 |
| 4_1232435 | 14 | 0.2857 | 11 | 0.2727 | 14 | 0.2143 | 11 | 0.2727 | 50 | 0.26 | 6 | 0 | 15 | 0.2667 | 9 | 0 | 8 | 0.125 | 38 | 0.1316 |
| 4_1232437 | 14 | 0.2143 | 15 | 0.2667 | 17 | 0.0588 | 12 | 0.0833 | 58 | 0.1552 | 6 | 0.3333 | 14 | 0 | 11 | 0.2727 | 8 | 0.25 | 39 | 0.1795 |
| 4_1232447 | 16 | 0.4375 | 12 | 0.5833 | 14 | 0.4286 | 14 | 0.3571 | 56 | 0.4464 | 9 | 0.1111 | 10 | 0.5 | 14 | 0.3571 | 9 | 0.1111 | 42 | 0.2857 |
| 4_1233259 | 2 | 0 | 3 | 0.3333 | 1 | 0 | 1 | 1 | 7 | 0.2857 | 1 | 0 | NA | NA | 2 | 0.5 | 1 | 1 | 4 | 0.5 |
| 4_1233259 | 2 | 0.5 | 2 | 0.5 | 1 | 0 | 1 | 0 | 6 | 0.3333 | 1 | 0 | 1 | 0 | 1 | 1 | 1 | 0 | 4 | 0.25 |
| 4_1233260 | 2 | 1 | 3 | 0.3333 | 1 | 0 | 1 | 1 | 7 | 0.5714 | 1 | 1 | 1 | 0 | 1 | 1 | 1 | 1 | 4 | 0.75 |
| 4_1233261 | 2 | 0.5 | 3 | 0.3333 | 1 | 0 | 1 | 0 | 7 | 0.2857 | 1 | 0 | 1 | 0 | 1 | 0 | 1 | 1 | 4 | 0.25 |
| 4_1233269 | 1 | 1 | 2 | 1 | 1 | 0 | NA | NA | 4 | 0.75 | 1 | 1 | 1 | 1 | 1 | 1 | 1 | 0 | 4 | 0.75 |
| 4_1233269 | 1 | 0 | 2 | 0.5 | 1 | 0 | NA | NA | 4 | 0.25 | 1 | 1 | 1 | 1 | 1 | 0 | 1 | 1 | 4 | 0.75 |
| 4_1233286 | 1 | 1 | 2 | 0 | 1 | 0 | 1 | 1 | 5 | 0.4 | NA | NA | 1 | 1 | 1 | 0 | 1 | 1 | 3 | 0.6667 |
| 4_1236293 | 6 | 0.1667 | 11 | 0.1818 | 12 | 0.4167 | 9 | 0.1111 | 38 | 0.2368 | 7 | 0.2857 | 10 | 0.3 | 14 | 0.2857 | 6 | 0.3333 | 37 | 0.2973 |
| 4_1236293 | 6 | 0.1667 | 16 | 0 | 16 | 0.125 | 12 | 0.0833 | 50 | 0.08 | 8 | 0 | 10 | 0.1 | 11 | 0.2727 | 7 | 0.1429 | 36 | 0.1389 |
| 4_1236298 | 5 | 0 | 11 | 0.0909 | 14 | 0.0714 | 10 | 0 | 40 | 0.05 | 7 | 0 | 6 | 0 | 11 | 0.1818 | 5 | 0.2 | 29 | 0.1034 |
| 4_1236299 | 6 | 1 | 11 | 0.9091 | 13 | 0.8462 | 10 | 0.9 | 40 | 0.9 | 6 | 1 | 9 | 0.8889 | 15 | 0.9333 | 7 | 1 | 37 | 0.9459 |
| 4_1236300 | 6 | 0.3333 | 12 | 0.25 | 14 | 0.2857 | 12 | 0.25 | 44 | 0.2727 | 7 | 0.2857 | 9 | 0.4444 | 14 | 0.4286 | 7 | 0.1429 | 37 | 0.3514 |
| 4_1236301 | 6 | 0.5 | 12 | 0.1667 | 14 | 0.2143 | 11 | 0.1818 | 43 | 0.2326 | 6 | 0.5 | 7 | 0 | 13 | 0.3077 | 8 | 0 | 34 | 0.2059 |
| 4_1236304 | 5 | 0.4 | 8 | 0.25 | 12 | 0.3333 | 10 | 0.2 | 35 | 0.2857 | 5 | 0.8 | 6 | 0.3333 | 11 | 0.2727 | 5 | 0.2 | 27 | 0.3704 |
| 4_1236304 | NA | NA | 2 | 0.5 | NA | NA | NA | NA | 2 | 0.5 | 3 | 0 | 1 | 1 | 2 | 0 | NA | NA | 6 | 0.1667 |
| 4_1236304 | 4 | 0.5 | 12 | 0.25 | 12 | 0.3333 | 10 | 0.8 | 38 | 0.4474 | 8 | 0.625 | 7 | 0.5714 | 14 | 0.2857 | 5 | 0.6 | 34 | 0.4706 |
| 4_1236305 | 5 | 0.8 | 10 | 0.2 | 12 | 0.1667 | 9 | 0.2222 | 36 | 0.2778 | 8 | 0.25 | 5 | 0.6 | 13 | 0.0769 | 7 | 0.1429 | 33 | 0.2121 |

| | | | | | | | | | | | | | | | | | | | | |
|---|---|---|---|---|---|---|---|---|---|---|---|---|---|---|---|---|---|---|---|---|
| 4_12379719 | NA | NA | NA | NA | NA | NA | 1 | 1 | 1 | 1 | NA | NA | NA | NA | NA | NA | 1 | 1 | 1 | 1 |
| 4_12379749 | NA | NA | NA | NA | NA | NA | 1 | 1 | 1 | 1 | NA | NA | NA | NA | NA | NA | 1 | 1 | 1 | 1 |
| 4_12379762 | NA | NA | NA | NA | NA | NA | 1 | 1 | 1 | 1 | NA | NA | NA | NA | NA | NA | 1 | 1 | 1 | 1 |
| 4_12381367 | 4 | 0 | 8 | 0.25 | 5 | 0 | 5 | 0 | 22 | 0.0909 | 2 | 0 | 7 | 0.1429 | 10 | 0.1 | 11 | 0.0909 | 30 | 0.1 |
| 4_12381376 | 5 | 0.4 | 8 | 0.5 | 6 | 0.1667 | 5 | 0.2 | 24 | 0.3333 | 2 | 0 | 8 | 0.25 | 10 | 0.4 | 13 | 0.2308 | 33 | 0.2727 |
| 4_12381484 | 4 | 0 | 7 | 0.4286 | 6 | 0.1667 | 5 | 0.4 | 22 | 0.2727 | 2 | 0 | 7 | 0.5714 | 11 | 0.3636 | 13 | 0.3846 | 33 | 0.3939 |
| 4_12382955 | 6 | 0.6667 | 9 | 0.5556 | 10 | 0.7 | 5 | 0.4 | 30 | 0.6 | 6 | 1 | 11 | 0.6364 | 13 | 0.6154 | 13 | 0.4615 | 43 | 0.6279 |
| 4_12382968 | 6 | 0.8333 | 9 | 0.6667 | 11 | 0.4545 | 6 | 0.8333 | 32 | 0.6562 | 7 | 0.7143 | 10 | 0.6 | 14 | 0.6429 | 14 | 0.7143 | 45 | 0.6667 |
| 4_12382969 | 7 | 1 | 9 | 0.8889 | 8 | 0.875 | 6 | 0.8333 | 30 | 0.9 | 9 | 0.8889 | 11 | 1 | 14 | 0.9286 | 15 | 0.7333 | 49 | 0.8776 |
| 4_12382975 | 7 | 1 | 9 | 0.8889 | 11 | 0.9091 | 6 | 1 | 33 | 0.9394 | 8 | 1 | 10 | 1 | 14 | 1 | 15 | 1 | 47 | 1 |
| 4_12382981 | 6 | 0.3333 | 5 | 0 | 5 | 0.2 | 4 | 0 | 20 | 0.15 | 5 | 0 | 7 | 0.1429 | 15 | 0.2 | 12 | 0 | 39 | 0.1026 |
| 4_12382982 | 6 | 0.6667 | 6 | 0.5 | 11 | 0.3636 | 4 | 0.25 | 27 | 0.4444 | 7 | 0.4286 | 11 | 0.3636 | 14 | 0.7143 | 12 | 0.75 | 44 | 0.5909 |
| 4_12382987 | 8 | 1 | 9 | 0.8889 | 8 | 1 | 5 | 1 | 30 | 0.9667 | 9 | 1 | 11 | 1 | 15 | 1 | 15 | 1 | 50 | 1 |
| 4_12382993 | 7 | 0.4286 | 7 | 0.1429 | 9 | 0 | 6 | 0.1667 | 29 | 0.1724 | 9 | 0.6667 | 11 | 0.2727 | 14 | 0.3571 | 14 | 0.4286 | 48 | 0.4167 |
| 4_12383007 | 8 | 0.625 | 8 | 0.875 | 9 | 0.7778 | 6 | 0.8333 | 31 | 0.7742 | 8 | 0.625 | 10 | 1 | 16 | 0.6875 | 16 | 0.75 | 50 | 0.76 |
| 4_12383017 | 9 | 0.2222 | 8 | 0 | 11 | 0.1818 | 6 | 0 | 34 | 0.1176 | 8 | 0 | 8 | 0.125 | 15 | 0.4 | 14 | 0.2857 | 45 | 0.2444 |
| 4_12383018 | 7 | 0.8571 | 9 | 0.8889 | 12 | 0.8333 | 5 | 0.8 | 33 | 0.8485 | 10 | 1 | 9 | 0.8889 | 15 | 0.9333 | 14 | 0.9286 | 48 | 0.9375 |
| 4_12383021 | 8 | 0.125 | 8 | 0.25 | 12 | 0.25 | 5 | 0.4 | 33 | 0.2424 | 10 | 0.3 | 10 | 0.5 | 14 | 0.6429 | 15 | 0.3333 | 49 | 0.449 |
| 4_12383021 | 9 | 0.5556 | 9 | 0 | 11 | 0.3636 | 5 | 0 | 34 | 0.2647 | 8 | 0.125 | 10 | 0.2 | 16 | 0.25 | 16 | 0.125 | 50 | 0.18 |
| 4_12383066 | 13 | 0.2308 | 11 | 0 | 13 | 0 | 8 | 0.25 | 45 | 0.1111 | 10 | 0.1 | 12 | 0.0833 | 16 | 0 | 16 | 0.125 | 54 | 0.0741 |
| 4_12383067 | 14 | 0.5714 | 12 | 0.4167 | 12 | 0.8333 | 7 | 0.5714 | 45 | 0.6 | 12 | 0.6667 | 12 | 0.5 | 17 | 0.7059 | 17 | 0.6471 | 58 | 0.6379 |
| 4_12383069 | 14 | 0.5 | 11 | 0.4545 | 12 | 0.5 | 7 | 0.4286 | 44 | 0.4773 | 11 | 0.3636 | 12 | 0.5 | 15 | 0.6667 | 18 | 0.6667 | 56 | 0.5714 |
| 4_12441713 | NA | NA | NA | NA | NA | NA | 2 | 0.5 | 2 | 0.5 | NA | NA | 1 | 1 | 1 | 0 | NA | NA | 2 | 0.5 |
| 4_12441954 | NA | NA | NA | NA | NA | NA | 1 | 1 | 1 | 1 | NA | NA | NA | NA | 1 | 1 | NA | NA | 1 | 1 |
| 4_12455212 | 1 | 0 | NA | NA | NA | NA | 1 | 1 | 2 | 0.5 | 1 | 1 | 2 | 1 | 2 | 0.5 | 1 | 0 | 6 | 0.6667 |
| 4_12455214 | 1 | 0 | NA | NA | NA | NA | 2 | 1 | 3 | 0.6667 | 1 | 1 | 2 | 1 | 2 | 1 | 1 | 1 | 6 | 1 |
| 4_12455232 | 1 | 1 | NA | NA | NA | NA | 1 | 0 | 2 | 0.5 | 1 | 0 | 2 | 0.5 | 3 | 0.3333 | 1 | 0 | 7 | 0.2857 |
| 4_12455248 | 1 | 0 | NA | NA | NA | NA | 1 | 1 | 2 | 0.5 | 1 | 1 | NA | NA | 3 | 0.6667 | 1 | 1 | 5 | 0.8 |
| 4_12455285 | 1 | 1 | NA | NA | NA | NA | 3 | 1 | 4 | 1 | 1 | 1 | 2 | 1 | 4 | 1 | 1 | 1 | 8 | 1 |
| 4_12455291 | 1 | 1 | NA | NA | NA | NA | 2 | 1 | 3 | 1 | 1 | 1 | 2 | 1 | 5 | 1 | 1 | 1 | 9 | 1 |
| 4_12455299 | 1 | 1 | NA | NA | NA | NA | 3 | 1 | 4 | 1 | NA | NA | 2 | 1 | 3 | 1 | 1 | 1 | 6 | 1 |
| 4_12455331 | NA | NA | NA | NA | NA | NA | 3 | 0.3333 | 3 | 0.3333 | 1 | 0 | 2 | 0 | 4 | 0.25 | 1 | 0 | 8 | 0.125 |
| 4_12455388 | 2 | 0.5 | NA | NA | NA | NA | 3 | 0.6667 | 5 | 0.6 | 2 | 0 | 3 | 0.3333 | 4 | 0.25 | 2 | 0.5 | 11 | 0.2727 |
| 4_12462611 | 15 | 0.1333 | 8 | 0.375 | 4 | 0 | 7 | 0.2857 | 34 | 0.2059 | 4 | 0.5 | 10 | 0 | 13 | 0.2308 | 5 | 0.2 | 32 | 0.1875 |
| 4_12468708 | 4 | 0.25 | 2 | 0 | 2 | 0 | 3 | 0 | 11 | 0.0909 | 1 | 0 | 2 | 1 | 3 | 0.3333 | NA | NA | 6 | 0.5 |
| 4_12469468 | 4 | 0 | 4 | 0.25 | 1 | 0 | 4 | 0.5 | 13 | 0.2308 | 1 | 0 | 1 | 0 | 5 | 0.4 | 1 | 0 | 8 | 0.25 |
| 4_12469479 | 7 | 0 | 7 | 0.1429 | 4 | 0 | 4 | 0.25 | 22 | 0.0909 | 1 | 0 | 3 | 0 | 6 | 0.1667 | 1 | 0 | 11 | 0.0909 |
| 4_12469483 | 6 | 0.3333 | 7 | 0.4286 | 3 | 0.6667 | 4 | 0.75 | 20 | 0.5 | NA | NA | 3 | 0.3333 | 4 | 0.25 | 1 | 0 | 8 | 0.25 |
| 4_12469490 | 6 | 0.1667 | 6 | 0 | 3 | 0.3333 | 5 | 0 | 20 | 0.1 | 1 | 0 | 3 | 0 | 6 | 0.1667 | 1 | 0 | 11 | 0.0909 |
| 4_12469499 | 7 | 0.1429 | 4 | 0 | 3 | 0.6667 | 4 | 0.25 | 18 | 0.2222 | 1 | 1 | 2 | 0.5 | 4 | 0.25 | NA | NA | 7 | 0.4286 |
| 4_12469504 | 7 | 0.5714 | 7 | 0 | 4 | 0.5 | 7 | 0.2857 | 25 | 0.32 | 1 | 0 | 3 | 0.3333 | 5 | 0.4 | NA | NA | 9 | 0.3333 |
| 4_12469906 | NA | NA | 1 | 0 | 1 | 1 | NA | NA | 2 | 0.5 | NA | NA | 1 | 0 | 1 | 1 | NA | NA | 2 | 0.5 |
| 4_12469907 | NA | NA | 1 | 1 | 1 | 1 | NA | NA | 2 | 1 | NA | NA | 1 | 0 | 1 | 1 | NA | NA | 2 | 0.5 |
| 4_12473802 | NA | NA | NA | NA | 1 | 1 | NA | NA | 1 | 1 | NA | NA | NA | NA | 2 | 0.5 | NA | NA | 2 | 0.5 |
| 4_12473817 | NA | NA | NA | NA | 1 | 1 | NA | NA | 1 | 1 | 1 | 1 | NA | NA | 2 | 0 | NA | NA | 3 | 0.3333 |
| 4_12473854 | NA | NA | NA | NA | 1 | 1 | 1 | 0 | 2 | 0.5 | 1 | 0 | NA | NA | 2 | 0.5 | NA | NA | 3 | 0.3333 |
| 4_12473859 | NA | NA | NA | NA | 1 | 1 | NA | NA | 1 | 1 | 1 | 1 | NA | NA | 1 | 1 | NA | NA | 2 | 1 |
| 4_12474292 | NA | NA | 1 | 1 | 6 | 0.8333 | 1 | 0 | 8 | 0.75 | 2 | 1 | 3 | 0.3333 | 4 | 0.5 | 1 | 1 | 10 | 0.6 |
| 4_12474360 | 20 | 0.85 | 16 | 0.9375 | 14 | 0.8571 | 13 | 0.7692 | 63 | 0.8571 | 12 | 1 | 14 | 1 | 9 | 1 | 10 | 1 | 45 | 1 |
| 4_12474420 | 8 | 0.25 | 10 | 0.1 | 10 | 0.1 | 8 | 0.25 | 36 | 0.1667 | 9 | 0 | 10 | 0.1 | 4 | 0.25 | 7 | 0 | 30 | 0.0667 |
| 4_12474438 | 3 | 0 | 6 | 0 | 3 | 0 | 3 | 0.3333 | 15 | 0.0667 | NA | NA | 3 | 0.3333 | 2 | 0.5 | NA | NA | 5 | 0.4 |
| 4_12474652 | 1 | 1 | 1 | 1 | NA | NA | 1 | 0 | 3 | 0.6667 | NA | NA | NA | NA | NA | NA | 2 | 0.5 | 2 | 0.5 |
| 4_12474668 | 1 | 1 | 1 | 1 | NA | NA | 1 | 0 | 3 | 0.6667 | NA | NA | NA | NA | NA | NA | 2 | 1 | 2 | 1 |
| 4_12474670 | NA | NA | 1 | 1 | NA | NA | 1 | 0 | 2 | 0.5 | NA | NA | NA | NA | NA | NA | 2 | 1 | 2 | 1 |
| 4_12474906 | 16 | 0.1875 | 16 | 0.0625 | 6 | 0.1667 | 10 | 0.1 | 48 | 0.125 | 6 | 0 | 6 | 0.3333 | 11 | 0.0909 | 10 | 0 | 33 | 0.0909 |
| 4_12474909 | 15 | 0.3333 | 11 | 0.2727 | 4 | 0.25 | 9 | 0.7778 | 39 | 0.4103 | 5 | 0.2 | 6 | 0.3333 | 9 | 0.6667 | 6 | 0.3333 | 26 | 0.4231 |
| 4_12474913 | 16 | 0.1875 | 15 | 0 | 6 | 0 | 9 | 0.1111 | 46 | 0.087 | 6 | 0 | 7 | 0.1429 | 12 | 0.25 | 10 | 0 | 35 | 0.1143 |
| 4_12474925 | 10 | 0.2 | 5 | 0.4 | 4 | 0.25 | 5 | 0.4 | 24 | 0.2917 | 2 | 0 | 5 | 0 | 6 | 0.3333 | 6 | 0 | 19 | 0.1053 |
| 4_12474929 | 15 | 0.6 | 16 | 0.5 | 5 | 0.4 | 9 | 0.6667 | 45 | 0.5556 | 6 | 0.3333 | 7 | 0.8571 | 11 | 0.6364 | 10 | 0.3 | 34 | 0.5294 |
| 4_12474934 | 4 | 0.5 | 8 | 0.375 | 3 | 0.3333 | 5 | 0.4 | 20 | 0.4 | 2 | 0 | 4 | 0.5 | 8 | 0.5 | 7 | 0.2857 | 21 | 0.381 |
| 4_12474937 | 10 | 0.9 | 12 | 1 | 5 | 1 | 6 | 0.8333 | 33 | 0.9394 | 5 | 1 | 7 | 0.8571 | 8 | 1 | 7 | 1 | 27 | 0.963 |
| 4_12474938 | 14 | 0.1429 | 15 | 0.0667 | 4 | 0 | 9 | 0 | 42 | 0.0714 | 6 | 0.1667 | 9 | 0.1111 | 11 | 0 | 10 | 0 | 36 | 0.0556 |
| 4_12474946 | 11 | 0.0909 | 14 | 0 | 4 | 0.25 | 9 | 0 | 38 | 0.0526 | 7 | 0 | 9 | 0.1111 | 11 | 0.0909 | 9 | 0.1111 | 36 | 0.0833 |
| 4_12474958 | 13 | 0.0769 | 13 | 0.1538 | 4 | 0 | 8 | 0.125 | 38 | 0.1053 | 6 | 0 | 7 | 0 | 10 | 0.1 | 8 | 0.25 | 31 | 0.0968 |
| 4_12474997 | 10 | 0.2 | 9 | 0.1111 | 3 | 0.3333 | 8 | 0.25 | 30 | 0.2 | 3 | 0 | 9 | 0.2222 | 11 | 0.1818 | 10 | 0.1 | 33 | 0.1515 |

| | | | | | | | | | | | | | | | | | | | | |
|---|---|---|---|---|---|---|---|---|---|---|---|---|---|---|---|---|---|---|---|---|
| 4_1247499 | 11 | 0.1818 | 13 | 0.2308 | 4 | 0.25 | 8 | 0.25 | 36 | 0.2222 | 4 | 0 | 9 | 0 | 12 | 0.1667 | 10 | 0.3 | 35 | 0.1429 |
| 4_1247500 | 10 | 0.2 | 14 | 0.2143 | 4 | 0.25 | 7 | 0.2857 | 35 | 0.2286 | 4 | 0.5 | 9 | 0.6667 | 11 | 0.3636 | 9 | 0.4444 | 33 | 0.4848 |
| 4_1247500 | 5 | 0.4 | 7 | 0.2857 | 2 | 0.5 | 4 | 0.25 | 18 | 0.3333 | 2 | 0.5 | 5 | 0.4 | 6 | 0.5 | 5 | 0.2 | 18 | 0.3889 |
| 4_1247502 | 8 | 0.125 | 10 | 0 | 4 | 0.25 | 8 | 0.125 | 30 | 0.1 | 3 | 0 | 7 | 0.1429 | 10 | 0.2 | 9 | 0.3333 | 29 | 0.2069 |
| 4_1247502 | 7 | 1 | 10 | 1 | 4 | 1 | 7 | 1 | 28 | 1 | 3 | 0.6667 | 7 | 1 | 9 | 1 | 10 | 1 | 29 | 0.9655 |
| 4_1247502 | 8 | 0.25 | 10 | 0.3 | 4 | 0 | 6 | 0.1667 | 28 | 0.2143 | 3 | 0.3333 | 7 | 0.2857 | 10 | 0 | 9 | 0.2222 | 29 | 0.1724 |
| 4_1247503 | 6 | 0.1667 | 9 | 0.4444 | 3 | 0.6667 | 8 | 0.5 | 26 | 0.4231 | 1 | 1 | 5 | 0.6 | 8 | 0.75 | 7 | 0.7143 | 21 | 0.7143 |
| 4_1247503 | 6 | 0.1667 | 11 | 0.1818 | 2 | 1 | 5 | 0.2 | 24 | 0.25 | 3 | 0 | 5 | 0.2 | 11 | 0.2727 | 9 | 0.2222 | 28 | 0.2143 |
| 4_1247503 | 6 | 1 | 11 | 0.9091 | 3 | 1 | 8 | 1 | 28 | 0.9643 | 3 | 1 | 8 | 1 | 11 | 1 | 9 | 1 | 31 | 1 |
| 4_1247504 | 6 | 0.8333 | 8 | 1 | 2 | 1 | 8 | 1 | 24 | 0.9583 | 4 | 1 | 7 | 1 | 10 | 0.9 | 8 | 0.875 | 29 | 0.931 |
| 4_1247505 | 5 | 0.8 | 8 | 0.25 | 2 | 1 | 7 | 0.5714 | 22 | 0.5455 | 3 | 0.6667 | 7 | 0.7143 | 8 | 0.5 | 10 | 0.5 | 28 | 0.5714 |
| 4_1247506 | 2 | 0 | 6 | 0.1667 | NA | NA | 3 | 0.3333 | 11 | 0.1818 | 2 | 0 | 3 | 0.3333 | 7 | 0.2857 | 6 | 0.8333 | 18 | 0.4444 |
| 4_1247506 | 3 | 0 | 9 | 0.1111 | 1 | 0 | 7 | 0.1429 | 20 | 0.1 | 3 | 0 | 5 | 0 | 7 | 0.1429 | 8 | 0.125 | 23 | 0.087 |
| 4_1247507 | 5 | 0 | 10 | 0.4 | 1 | 0 | 8 | 0.125 | 24 | 0.2083 | 4 | 0.25 | 7 | 0.2857 | 10 | 0.4 | 10 | 0.5 | 31 | 0.3871 |
| 4_1247507 | 5 | 0.6 | 7 | 0.7143 | 1 | 0 | 6 | 0.5 | 19 | 0.5789 | 2 | 1 | 3 | 0.6667 | 5 | 0.6 | 6 | 0.6667 | 16 | 0.6875 |
| 4_1247555 | 1 | 0 | 6 | 0 | NA | NA | 5 | 0.2 | 12 | 0.0833 | 1 | 1 | 6 | 0 | 5 | 0.2 | 7 | 0.1429 | 19 | 0.1579 |
| 4_1247555 | 1 | 0 | 5 | 0.2 | NA | NA | 6 | 0.1667 | 12 | 0.1667 | 1 | 0 | 2 | 0 | 5 | 0.2 | 7 | 0 | 15 | 0.0667 |
| 4_1248382 | 2 | 0.5 | NA | NA | 2 | 0 | NA | NA | 4 | 0.25 | 1 | 1 | NA | NA | 2 | 0 | 1 | 0 | 4 | 0.25 |
| 4_1249462 | 8 | 0.125 | 3 | 0.3333 | 4 | 0 | 4 | 0 | 19 | 0.1053 | 3 | 0.3333 | 8 | 0.25 | 2 | 0 | 7 | 0 | 20 | 0.15 |
| 4_1249463 | 8 | 0.25 | 3 | 0.3333 | 4 | 0 | 6 | 0.3333 | 21 | 0.2381 | 3 | 0 | 6 | 0.5 | 2 | 0.5 | 7 | 0.5714 | 18 | 0.4444 |
| 4_1249463 | 8 | 0.125 | 3 | 0 | 4 | 0 | 5 | 0.2 | 20 | 0.1 | 3 | 0.3333 | 8 | 0.25 | 2 | 0 | 7 | 0.4286 | 20 | 0.3 |
| 4_1249651 | 2 | 1 | NA | NA | 1 | 1 | 2 | 1 | 5 | 1 | 1 | 1 | NA | NA | 2 | 1 | NA | NA | 3 | 1 |
| 4_1249652 | 2 | 1 | NA | NA | 1 | 0 | 1 | 1 | 4 | 0.75 | 1 | 0 | NA | NA | 2 | 1 | 1 | 1 | 4 | 0.75 |
| 4_1249652 | 2 | 1 | NA | NA | 1 | 1 | 2 | 1 | 5 | 1 | 1 | 1 | NA | NA | 2 | 1 | 1 | 1 | 4 | 1 |
| 4_1249665 | 1 | 1 | 1 | 1 | 2 | 1 | 1 | 1 | 5 | 1 | 2 | 1 | 1 | 1 | 2 | 1 | 1 | 1 | 6 | 1 |
| 4_1249665 | 2 | 1 | 1 | 1 | 2 | 1 | 1 | 1 | 6 | 1 | 1 | 1 | 1 | 1 | 2 | 1 | NA | NA | 4 | 1 |
| 4_1249666 | 2 | 0 | 1 | 1 | 1 | 1 | 2 | 0 | 6 | 0.3333 | 2 | 0.5 | 1 | 1 | 2 | 0.5 | 2 | 0 | 7 | 0.4286 |
| 4_1249668 | 1 | 1 | 1 | 1 | 2 | 1 | 1 | 1 | 5 | 1 | 1 | 1 | 1 | 1 | 2 | 1 | 1 | 1 | 5 | 1 |
| 4_1249669 | 2 | 1 | 1 | 0 | 2 | 0.5 | 2 | 0 | 7 | 0.4286 | 1 | 0 | NA | NA | 1 | 1 | 1 | 1 | 3 | 0.6667 |
| 4_1249669 | 2 | 1 | NA | NA | 2 | 1 | 1 | 1 | 5 | 1 | 2 | 1 | NA | NA | 1 | 1 | NA | NA | 3 | 1 |
| 4_1249670 | 2 | 0.5 | 2 | 0 | 2 | 1 | 1 | 0 | 7 | 0.4286 | 2 | 0.5 | NA | NA | 2 | 1 | 1 | 1 | 5 | 0.8 |
| 4_1249672 | 2 | 1 | 1 | 0 | 1 | 0 | 3 | 0.3333 | 7 | 0.4286 | 2 | 0.5 | 1 | 0 | 2 | 1 | 1 | 0 | 6 | 0.5 |
| 4_1249672 | 2 | 0 | 2 | 0 | 2 | 0 | 3 | 0.3333 | 9 | 0.1111 | NA | NA | 1 | 0 | 2 | 0.5 | 1 | 0 | 4 | 0.25 |
| 4_1249672 | 2 | 1 | 2 | 0.5 | 2 | 1 | 4 | 0.75 | 10 | 0.8 | 2 | 1 | NA | NA | 3 | 0.6667 | NA | NA | 5 | 0.8 |
| 4_1249674 | 2 | 0.5 | 1 | 0 | 1 | 1 | 3 | 0.3333 | 7 | 0.4286 | 2 | 0.5 | 1 | 0 | 1 | 0 | 1 | 1 | 5 | 0.4 |
| 4_1249676 | 2 | 0.5 | 2 | 0 | 2 | 0 | 4 | 0.25 | 10 | 0.2 | 2 | 0.5 | 3 | 0.3333 | 3 | 0.3333 | 1 | 0 | 9 | 0.3333 |
| 4_1250031 | 19 | 0.1579 | 11 | 0 | 4 | 0.25 | 11 | 0 | 45 | 0.0889 | 11 | 0.0909 | 14 | 0.0714 | 10 | 0 | 15 | 0.0667 | 50 | 0.06 |
| 4_1250032 | 14 | 0.0714 | 11 | 0.0909 | 6 | 0.1667 | 10 | 0.1 | 41 | 0.0976 | 11 | 0.1818 | 12 | 0.1667 | 10 | 0 | 13 | 0.3077 | 46 | 0.1739 |
| 4_1250032 | 18 | 0.6111 | 9 | 0.6667 | 5 | 0.4 | 7 | 0.5714 | 39 | 0.5897 | 10 | 0.5 | 10 | 0.5 | 7 | 0 | 12 | 0.5 | 39 | 0.4103 |
| 4_1250043 | 9 | 0.3333 | 3 | 0.3333 | 4 | 0 | 1 | 1 | 17 | 0.2941 | 2 | 0.5 | 5 | 0.2 | 4 | 0 | 5 | 0.6 | 16 | 0.3125 |
| 4_1250058 | 8 | 0.125 | 3 | 0 | 5 | 0.2 | 1 | 0 | 17 | 0.1176 | 2 | 0 | 8 | 0 | 3 | 0 | 5 | 0.2 | 18 | 0.0556 |
| 4_1256052 | NA | NA | NA | NA | 1 | 0 | 3 | 0.3333 | 4 | 0.25 | 2 | 0.5 | NA | NA | 4 | 0 | 1 | 0 | 7 | 0.1429 |
| 4_1256054 | NA | NA | NA | NA | 2 | 0.5 | 3 | 0.3333 | 5 | 0.4 | 2 | 0.5 | NA | NA | 3 | 0 | 2 | 0 | 7 | 0.1429 |
| 4_1256054 | NA | NA | 1 | 1 | 1 | 0 | 1 | 1 | 3 | 0.6667 | 4 | 0.75 | NA | NA | 3 | 1 | 2 | 0.5 | 9 | 0.7778 |
| 4_1256055 | NA | NA | 2 | 0.5 | 2 | 0 | 4 | 0.75 | 8 | 0.5 | 2 | 0 | NA | NA | 4 | 0.5 | 1 | 1 | 7 | 0.4286 |
| 4_1256055 | NA | NA | 2 | 0 | 1 | 1 | 4 | 0.5 | 7 | 0.4286 | 4 | 0 | NA | NA | 2 | 0.5 | 2 | 0 | 8 | 0.125 |
| 4_1256056 | NA | NA | 3 | 0.3333 | 2 | 0.5 | 5 | 0.4 | 10 | 0.4 | 5 | 0.2 | NA | NA | 4 | 0.25 | 2 | 0.5 | 11 | 0.2727 |
| 4_1256059 | 1 | 1 | 3 | 0.3333 | 2 | 1 | 6 | 0.6667 | 12 | 0.6667 | 5 | 0.8 | NA | NA | 5 | 0.6 | 6 | 0.5 | 16 | 0.625 |
| 4_1256060 | 1 | 1 | 3 | 1 | 2 | 0.5 | 2 | 0.5 | 8 | 0.75 | 5 | 0.6 | 1 | 0 | 5 | 0.8 | 4 | 1 | 15 | 0.7333 |
| 4_1256076 | 3 | 0.6667 | 4 | 0 | 4 | 0 | 7 | 0.1429 | 18 | 0.1667 | 6 | 0.1667 | 3 | 0 | 9 | 0.3333 | 9 | 0.4444 | 27 | 0.2963 |
| 4_1256077 | 3 | 0.6667 | 4 | 0 | 5 | 0.2 | 10 | 0.1 | 22 | 0.1818 | 4 | 0.5 | 3 | 0.3333 | 12 | 0.25 | 11 | 0.1818 | 30 | 0.2667 |
| 4_1257821 | 1 | 1 | NA | NA | NA | NA | NA | NA | 1 | 1 | NA | NA | NA | NA | 1 | 1 | NA | NA | 1 | 1 |
| 4_1257831 | 1 | 1 | NA | NA | NA | NA | NA | NA | 1 | 1 | NA | NA | NA | NA | 1 | 1 | NA | NA | 1 | 1 |
| 4_1259375 | NA | NA | NA | NA | 1 | 1 | NA | NA | 1 | 1 | 2 | 0.5 | NA | NA | NA | NA | NA | NA | 2 | 0.5 |
| 4_1259376 | NA | NA | NA | NA | 1 | 1 | 1 | 1 | 2 | 1 | 2 | 1 | NA | NA | NA | NA | NA | NA | 2 | 1 |
| 4_1259378 | NA | NA | NA | NA | 1 | 0 | 1 | 1 | 2 | 0.5 | 2 | 0.5 | NA | NA | NA | NA | NA | NA | 2 | 0.5 |
| 4_1259459 | 2 | 0 | 1 | 1 | NA | NA | 2 | 0 | 5 | 0.2 | 1 | 0 | NA | NA | NA | NA | 1 | 1 | 2 | 0.5 |
| 4_1259822 | 1 | 1 | 4 | 0.75 | 1 | 1 | 4 | 0.25 | 10 | 0.6 | NA | NA | NA | NA | 4 | 0.75 | 1 | 1 | 5 | 0.8 |
| 4_1259822 | 2 | 0.5 | 4 | 1 | 2 | 1 | 6 | 0.8333 | 14 | 0.8571 | NA | NA | NA | NA | 4 | 0.75 | 2 | 1 | 6 | 0.8333 |
| 4_1259822 | 2 | 0 | 1 | 0 | 2 | 0.5 | 3 | 0 | 8 | 0.125 | NA | NA | NA | NA | 1 | 1 | 1 | 1 | 2 | 1 |
| 4_1259823 | 2 | 0.5 | 4 | 0.5 | 1 | 0 | 6 | 1 | 13 | 0.6923 | NA | NA | NA | NA | 4 | 0.75 | 2 | 1 | 6 | 0.8333 |
| 4_1259824 | 1 | 0 | 2 | 0.5 | NA | NA | 1 | 1 | 4 | 0.5 | NA | NA | NA | NA | 1 | 1 | 2 | 0.5 | 3 | 0.6667 |
| 4_1259824 | NA | NA | 1 | 1 | NA | NA | NA | NA | 1 | 1 | NA | NA | NA | NA | NA | NA | 1 | 1 | 1 | 1 |
| 4_1259825 | NA | NA | 2 | 0.5 | NA | NA | 1 | 0 | 3 | 0.3333 | NA | NA | NA | NA | 1 | 1 | 2 | 0.5 | 3 | 0.6667 |
| 4_1259826 | 1 | 1 | 1 | 1 | NA | NA | 1 | 1 | 3 | 1 | NA | NA | NA | NA | 1 | 1 | 2 | 1 | 3 | 1 |

| | | | | | | | | | | | | | | | | | | | | |
|---|---|---|---|---|---|---|---|---|---|---|---|---|---|---|---|---|---|---|---|---|
| 4_1259827 | 1 | 1 | 1 | 1 | NA | NA | 1 | 1 | 3 | 1 | NA | NA | NA | NA | 1 | 1 | 1 | 0 | 2 | 0.5 |
| 4_1259828 | 1 | 1 | 1 | 1 | NA | NA | 1 | 1 | 3 | 1 | NA | NA | NA | NA | 1 | 1 | 2 | 0.5 | 3 | 0.6667 |
| 4_1259844 | NA | NA | 1 | 1 | NA | NA | 1 | 0 | 2 | 0.5 | NA | NA | NA | NA | 1 | 1 | 2 | 0 | 3 | 0.3333 |
| 4_1259856 | NA | NA | 1 | 1 | NA | NA | 1 | 1 | 2 | 1 | NA | NA | NA | NA | 2 | 1 | NA | NA | 2 | 1 |
| 4_1259856 | NA | NA | 1 | 0 | 1 | 1 | NA | NA | 2 | 0.5 | NA | NA | NA | NA | 2 | 1 | 2 | 1 | 4 | 1 |
| 4_1259857 | NA | NA | 1 | 1 | 2 | 0.5 | 2 | 1 | 5 | 0.8 | NA | NA | NA | NA | 2 | 1 | 2 | 1 | 4 | 1 |
| 4_1259858 | NA | NA | 3 | 0.3333 | 1 | 0 | 1 | 1 | 5 | 0.4 | NA | NA | NA | NA | 2 | 0 | 2 | 1 | 4 | 0.5 |
| 4_1259916 | 2 | 0 | NA | NA | NA | NA | 5 | 0.2 | 7 | 0.1429 | 1 | 0 | 2 | 0.5 | 1 | 0 | 2 | 0 | 6 | 0.1667 |
| 4_1259917 | 3 | 0.6667 | NA | NA | NA | NA | 5 | 0.6 | 8 | 0.625 | 1 | 1 | 2 | 0 | 1 | 1 | 2 | 1 | 6 | 0.6667 |
| 4_1259918 | 2 | 1 | NA | NA | NA | NA | 4 | 0.5 | 6 | 0.6667 | NA | NA | 3 | 1 | 1 | 1 | 3 | 1 | 7 | 1 |
| 4_1259919 | 1 | 0 | NA | NA | NA | NA | 4 | 0.25 | 5 | 0.2 | 1 | 0 | 2 | 0 | 1 | 0 | 4 | 0.75 | 8 | 0.375 |
| 4_1259920 | 3 | 1 | NA | NA | NA | NA | 4 | 1 | 7 | 1 | 1 | 1 | 3 | 1 | 1 | 1 | 4 | 0.75 | 9 | 0.8889 |
| 4_1259920 | 3 | 0.3333 | NA | NA | NA | NA | 5 | 1 | 8 | 0.75 | 1 | 0 | 2 | 1 | NA | NA | 4 | 1 | 7 | 0.8571 |
| 4_1259921 | 3 | 0.3333 | NA | NA | NA | NA | 4 | 0.75 | 7 | 0.5714 | 1 | 1 | 3 | 0.6667 | 1 | 1 | 3 | 0.6667 | 8 | 0.75 |
| 4_1259921 | 2 | 0 | NA | NA | NA | NA | 5 | 0.4 | 7 | 0.2857 | 1 | 0 | 2 | 1 | 1 | 0 | 4 | 0 | 8 | 0.25 |
| 4_1259921 | 4 | 0.5 | NA | NA | NA | NA | 5 | 0.6 | 9 | 0.5556 | 1 | 0 | 2 | 0.5 | 1 | 1 | 4 | 0.75 | 8 | 0.625 |
| 4_1259921 | 4 | 0.25 | NA | NA | NA | NA | 5 | 0.6 | 9 | 0.4444 | 1 | 0 | 3 | 0 | NA | NA | 3 | 0.3333 | 7 | 0.1429 |
| 4_1259922 | 3 | 0 | NA | NA | NA | NA | 3 | 0.3333 | 6 | 0.1667 | 1 | 0 | 2 | 0.5 | 1 | 1 | 2 | 1 | 6 | 0.6667 |
| 4_1259923 | 4 | 0.5 | NA | NA | NA | NA | 5 | 0.2 | 9 | 0.3333 | NA | NA | 2 | 0.5 | 1 | 0 | 3 | 0.3333 | 6 | 0.3333 |
| 4_1260327 | 2 | 0.5 | 2 | 0 | 1 | 0 | 3 | 0.6667 | 8 | 0.375 | 2 | 0.5 | 2 | 0 | 2 | 0.5 | 3 | 0 | 9 | 0.2222 |
| 4_1260329 | 2 | 1 | 2 | 1 | 2 | 0.5 | 3 | 1 | 9 | 0.8889 | 2 | 1 | 2 | 1 | 2 | 1 | 3 | 1 | 9 | 1 |
| 4_1260336 | 1 | 0 | 2 | 0 | 3 | 0.3333 | 5 | 0 | 11 | 0.0909 | 2 | 0.5 | 3 | 0 | 3 | 0 | 4 | 0.25 | 12 | 0.1667 |
| 4_1260338 | 2 | 0 | 3 | 1 | 2 | 0.5 | 6 | 0.5 | 13 | 0.5385 | 2 | 0.5 | 2 | 0.5 | 4 | 0.25 | 4 | 0.25 | 12 | 0.3333 |
| 4_1260347 | 1 | 0 | 3 | 0 | 1 | 0 | 7 | 0.1429 | 12 | 0.0833 | 3 | 0.3333 | 2 | 0 | 4 | 0 | 4 | 0 | 13 | 0.0769 |
| 4_1260435 | NA | NA | 1 | 1 | 1 | 1 | NA | NA | 2 | 1 | NA | NA | 2 | 0.5 | 1 | 1 | 1 | 1 | 4 | 0.75 |
| 4_1260588 | 12 | 0.1667 | 11 | 0.0909 | 4 | 0.25 | 6 | 0.1667 | 33 | 0.1515 | 6 | 0 | 9 | 0.1111 | 12 | 0.25 | 9 | 0.2222 | 36 | 0.1667 |
| 4_1260589 | 12 | 0.8333 | 12 | 0.8333 | 4 | 1 | 10 | 1 | 38 | 0.8947 | 6 | 0.8333 | 10 | 0.7 | 11 | 0.7273 | 9 | 0.8889 | 36 | 0.7778 |
| 4_1260590 | 11 | 0.1818 | 10 | 0.2 | 6 | 0.1667 | 7 | 0.1429 | 34 | 0.1765 | 5 | 0 | 10 | 0 | 12 | 0.0833 | 8 | 0.125 | 35 | 0.0571 |
| 4_1260590 | 9 | 0.1111 | 9 | 0.3333 | 4 | 0 | 5 | 0.8 | 27 | 0.2963 | 5 | 0.2 | 8 | 0.125 | 12 | 0.4167 | 6 | 0.5 | 31 | 0.3226 |
| 4_1260592 | 9 | 0.7778 | 6 | 1 | 4 | 1 | 8 | 0.75 | 27 | 0.8519 | 4 | 0.75 | 7 | 1 | 10 | 1 | 10 | 1 | 31 | 0.9677 |
| 4_1260592 | 7 | 0.1429 | 7 | 0.1429 | 4 | 0.25 | 8 | 0.75 | 26 | 0.3462 | 4 | 0.5 | 6 | 0.5 | 7 | 0.4286 | 7 | 0.1429 | 24 | 0.375 |
| 4_1260655 | 8 | 0 | 4 | 0 | 5 | 0 | 8 | 0.25 | 25 | 0.08 | 2 | 0 | 7 | 0.1429 | 7 | 0 | 4 | 0.25 | 20 | 0.1 |
| 4_1260716 | 6 | 0.1667 | 1 | 0 | 3 | 0.3333 | 5 | 0 | 15 | 0.1333 | NA | NA | 4 | 0.25 | 5 | 0.2 | 3 | 0 | 12 | 0.1667 |
| 4_1260720 | 5 | 0.4 | 1 | 0 | 3 | 0 | 5 | 0 | 14 | 0.1429 | NA | NA | 4 | 0 | 4 | 0.25 | 3 | 0.6667 | 11 | 0.2727 |
| 4_1260722 | 6 | 0.6667 | 2 | 0.5 | 3 | 0.6667 | 4 | 0.75 | 15 | 0.6667 | NA | NA | 4 | 1 | 4 | 0.75 | 2 | 1 | 10 | 0.9 |
| 4_1260741 | 4 | 1 | 1 | 0 | 3 | 0.3333 | 3 | 1 | 11 | 0.7273 | NA | NA | 2 | 1 | 4 | 1 | 1 | 1 | 7 | 1 |
| 4_1260743 | 3 | 1 | 1 | 1 | 2 | 0.5 | 4 | 0.75 | 10 | 0.8 | NA | NA | 2 | 0.5 | 4 | 0.75 | 2 | 0.5 | 8 | 0.625 |
| 4_1260744 | 3 | 0.3333 | NA | NA | 2 | 0 | 4 | 0.25 | 9 | 0.2222 | NA | NA | 2 | 0 | 4 | 0.5 | 2 | 0.5 | 8 | 0.375 |
| 4_1261263 | 14 | 0 | 38 | 0.0526 | 21 | 0.0952 | 23 | 0.1304 | 96 | 0.0729 | 16 | 0.1875 | 32 | 0.2188 | 25 | 0.16 | 17 | 0.1176 | 90 | 0.1778 |
| 4_1261263 | 15 | 0.4 | 40 | 0.4 | 25 | 0.44 | 22 | 0.4091 | 102 | 0.4118 | 17 | 0.5882 | 34 | 0.3824 | 26 | 0.4615 | 18 | 0.4444 | 95 | 0.4526 |
| 4_1261263 | 14 | 0.2857 | 44 | 0.1818 | 25 | 0.44 | 22 | 0.1818 | 105 | 0.2571 | 17 | 0.3529 | 31 | 0.2903 | 23 | 0.4348 | 18 | 0.2222 | 89 | 0.3258 |
| 4_1261501 | 2 | 0.5 | 1 | 0 | 1 | 0 | 1 | 0 | 5 | 0.2 | NA | NA | 2 | 0 | 1 | 1 | NA | NA | 3 | 0.3333 |
| 4_1261690 | 6 | 0.1667 | 7 | 0.5714 | 5 | 0.6 | 4 | 0 | 22 | 0.3636 | 6 | 0.5 | 6 | 0.3333 | 3 | 0.6667 | 3 | 0.3333 | 18 | 0.4444 |
| 4_1261691 | 6 | 0.3333 | 7 | 0.2857 | 7 | 0.7143 | 3 | 0 | 23 | 0.3913 | 6 | 0.1667 | 6 | 0.3333 | 3 | 0.3333 | 3 | 0.3333 | 18 | 0.2778 |
| 4_1261691 | 5 | 0.8 | 7 | 1 | 7 | 1 | 4 | 1 | 23 | 0.9565 | 4 | 0.5 | 6 | 0.8333 | 3 | 0.6667 | 3 | 0.6667 | 16 | 0.6875 |
| 4_1261691 | 6 | 0.1667 | 7 | 0.1429 | 7 | 0.1429 | 5 | 0 | 25 | 0.12 | 6 | 0 | 7 | 0 | 3 | 0.3333 | 3 | 0.3333 | 19 | 0.1053 |
| 4_1261693 | 6 | 0.8333 | 6 | 0.8333 | 8 | 0.75 | 4 | 0.75 | 24 | 0.7917 | 7 | 0.7143 | 8 | 0.75 | 4 | 1 | 5 | 0.4 | 24 | 0.7083 |
| 4_1262100 | 1 | 0 | 1 | 1 | NA | NA | 3 | 0.6667 | 5 | 0.6 | 10 | 0.5 | 3 | 0.3333 | 4 | 1 | 5 | 0.4 | 22 | 0.5455 |
| 4_1262100 | 4 | 0 | 3 | 0.3333 | NA | NA | 3 | 0.3333 | 10 | 0.2 | 9 | 0.2222 | 3 | 1 | 3 | 0 | 5 | 0.4 | 20 | 0.35 |
| 4_1262102 | 2 | 0 | 3 | 0 | NA | NA | 3 | 0.3333 | 8 | 0.125 | 7 | 0.4286 | 1 | 1 | 2 | 0.5 | 3 | 0 | 13 | 0.3846 |
| 4_1262102 | 4 | 0 | 2 | 1 | NA | NA | 3 | 0.3333 | 9 | 0.3333 | 10 | 0.9 | 2 | 0.5 | 3 | 0.6667 | 5 | 0.2 | 20 | 0.65 |
| 4_1262103 | 3 | 1 | 3 | 1 | NA | NA | 4 | 0.75 | 10 | 0.9 | 10 | 0.8 | 2 | 1 | 3 | 1 | 4 | 0.75 | 19 | 0.8421 |
| 4_1262103 | 4 | 0.25 | 3 | 0.3333 | NA | NA | 3 | 0.3333 | 10 | 0.3 | 9 | 0.7778 | 2 | 1 | 4 | 0.75 | 5 | 0.8 | 20 | 0.8 |
| 4_1262103 | 4 | 0 | 3 | 0 | NA | NA | 4 | 0.5 | 11 | 0.1818 | 9 | 0.3333 | 2 | 0 | 4 | 0 | 4 | 0.25 | 19 | 0.2105 |
| 4_1262104 | 3 | 0.3333 | 2 | 0 | NA | NA | 4 | 0 | 9 | 0.1111 | 10 | 0.1 | 2 | 0.5 | 3 | 0 | 5 | 0 | 20 | 0.1 |
| 4_1262105 | 4 | 0.5 | 1 | 1 | NA | NA | 3 | 0.3333 | 8 | 0.5 | 8 | 0.375 | 2 | 0.5 | 4 | 0.5 | 5 | 0.4 | 19 | 0.4211 |
| 4_1262105 | 3 | 0.6667 | 3 | 1 | NA | NA | 3 | 0.3333 | 9 | 0.6667 | 7 | 0.2857 | 2 | 0.5 | 3 | 0.6667 | 5 | 0.2 | 17 | 0.3529 |
| 4_1262266 | 1 | 0 | 1 | 0 | 1 | 1 | 3 | 0.6667 | 6 | 0.5 | 2 | 0 | 3 | 0.3333 | 2 | 0.5 | 1 | 0 | 8 | 0.25 |
| 4_1262270 | 1 | 0 | 1 | 1 | 1 | 0 | 4 | 0 | 7 | 0.1429 | 1 | 1 | 4 | 0 | 1 | 0 | 1 | 0 | 7 | 0.1429 |
| 4_1262272 | NA | NA | 1 | 0 | 1 | 0 | 4 | 0.25 | 6 | 0.1667 | 1 | 1 | 5 | 0.2 | 1 | 0 | 1 | 1 | 8 | 0.375 |
| 4_1262272 | 1 | 0 | 2 | 0 | 1 | 0 | 4 | 0.25 | 8 | 0.125 | 1 | 1 | 4 | 0 | 2 | 0 | 1 | 0 | 8 | 0.125 |
| 4_1262275 | 3 | 0 | 1 | 0 | 3 | 0.3333 | 3 | 0 | 10 | 0.1 | 1 | 0 | 6 | 0.1667 | 3 | 0.3333 | 2 | 0 | 12 | 0.1667 |
| 4_1262277 | 3 | 0 | 2 | 0 | 2 | 0.5 | 4 | 0 | 11 | 0.0909 | 1 | 1 | 6 | 0 | 3 | 0.3333 | 2 | 0 | 12 | 0.1667 |
| 4_1263291 | 3 | 0.3333 | 3 | 0 | 7 | 0.2857 | 2 | 0.5 | 15 | 0.2667 | 2 | 0 | 4 | 0 | 5 | 0.2 | 5 | 0 | 16 | 0.0625 |
| 4_1263294 | 5 | 0 | 2 | 0 | 10 | 0.3 | 2 | 0 | 19 | 0.1579 | 2 | 0 | 3 | 0.3333 | 4 | 0.25 | 2 | 0.5 | 11 | 0.2727 |

| | | | | | | | | | | | | | | | | | | | | |
|---|---|---|---|---|---|---|---|---|---|---|---|---|---|---|---|---|---|---|---|---|
| 4_12632987 | 4 | 0.5 | 1 | 1 | 5 | 0.8 | 1 | 0 | 11 | 0.6364 | 2 | 0 | 4 | 0.75 | 3 | 0.6667 | 2 | 0 | 11 | 0.4545 |
| 4_12633103 | 7 | 0.1429 | 2 | 0 | 5 | 0 | 2 | 0 | 16 | 0.0625 | 1 | 0 | 3 | 0 | NA | NA | 2 | 0.5 | 6 | 0.1667 |
| 4_12633122 | 7 | 0 | 1 | 0 | 8 | 0.25 | 2 | 0 | 18 | 0.1111 | 1 | 0 | 4 | 0.75 | 3 | 0 | 3 | 0.3333 | 11 | 0.3636 |
| 4_12633211 | 6 | 0 | 2 | 0 | 6 | 0.1667 | 2 | 0.5 | 16 | 0.125 | NA | NA | 4 | 0.5 | 1 | 0 | 1 | 0 | 6 | 0.3333 |
| 4_12633297 | 3 | 0.6667 | 2 | 1 | 3 | 0 | 2 | 0.5 | 10 | 0.5 | NA | NA | 2 | 0.5 | NA | NA | NA | NA | 2 | 0.5 |
| 4_12633312 | 3 | 0.3333 | 1 | 0 | 3 | 0 | 2 | 0 | 9 | 0.1111 | NA | NA | 2 | 0.5 | NA | NA | NA | NA | 2 | 0.5 |
| 4_12638469 | NA | NA | NA | NA | NA | NA | 2 | 0.5 | 2 | 0.5 | 1 | 1 | 2 | 1 | NA | NA | NA | NA | 3 | 1 |
| 4_12638472 | 1 | 1 | NA | NA | NA | NA | 2 | 1 | 3 | 1 | 1 | 1 | 2 | 1 | NA | NA | NA | NA | 3 | 1 |
| 4_12638480 | NA | NA | NA | NA | NA | NA | 2 | 0.5 | 2 | 0.5 | 1 | 1 | 1 | 1 | NA | NA | NA | NA | 2 | 1 |
| 4_12638487 | 1 | 1 | NA | NA | NA | NA | 2 | 1 | 3 | 1 | 1 | 1 | 1 | 1 | NA | NA | NA | NA | 2 | 1 |
| 4_12638488 | 1 | 0 | NA | NA | NA | NA | 1 | 1 | 2 | 0.5 | 1 | 1 | 2 | 1 | NA | NA | NA | NA | 3 | 1 |
| 4_12638494 | 1 | 0 | NA | NA | NA | NA | 2 | 0.5 | 3 | 0.3333 | 1 | 0 | 2 | 1 | NA | NA | NA | NA | 3 | 0.6667 |
| 4_12638506 | 1 | 0 | NA | NA | NA | NA | 2 | 0.5 | 3 | 0.3333 | 1 | 0 | 2 | 0.5 | NA | NA | NA | NA | 3 | 0.3333 |
| 4_12638633 | 1 | 1 | NA | NA | NA | NA | 2 | 1 | 3 | 1 | NA | NA | 1 | 1 | NA | NA | NA | NA | 1 | 1 |
| 4_12638638 | 1 | 1 | NA | NA | NA | NA | 2 | 0.5 | 3 | 0.6667 | NA | NA | 2 | 0.5 | NA | NA | NA | NA | 2 | 0.5 |
| 4_12638650 | 1 | 1 | NA | NA | NA | NA | 2 | 1 | 3 | 1 | NA | NA | 1 | 1 | NA | NA | NA | NA | 1 | 1 |
| 4_12638674 | 1 | 1 | NA | NA | NA | NA | 1 | 1 | 2 | 1 | NA | NA | 1 | 1 | NA | NA | NA | NA | 1 | 1 |
| 4_12638675 | NA | NA | NA | NA | NA | NA | 1 | 1 | 1 | 1 | NA | NA | 1 | 1 | NA | NA | NA | NA | 1 | 1 |
| 4_12650745 | 1 | 0 | 2 | 0 | 2 | 0.5 | 3 | 0 | 8 | 0.125 | 3 | 0.6667 | NA | NA | 4 | 0.5 | 4 | 0 | 11 | 0.3636 |
| 4_12650754 | 2 | 0 | 3 | 0.6667 | 1 | 0 | 4 | 1 | 10 | 0.6 | 3 | 1 | 1 | 1 | 4 | 0.5 | 4 | 0.25 | 12 | 0.5833 |
| 4_12650769 | 2 | 0.5 | 3 | 0.3333 | 2 | 0.5 | 3 | 0.6667 | 10 | 0.5 | 3 | 1 | 1 | 1 | 4 | 0.75 | 6 | 0.6667 | 14 | 0.7857 |
| 4_12650808 | 4 | 0.25 | 4 | 0.25 | 3 | 0 | 6 | 0.3333 | 17 | 0.2353 | 3 | 0 | 2 | 0 | 4 | 0.25 | 7 | 0 | 16 | 0.0625 |
| 4_12663757 | 77 | 0.0779 | 108 | 0.0648 | 107 | 0.0467 | 101 | 0.0198 | 393 | 0.0509 | 89 | 0.0449 | 124 | 0.0726 | 161 | 0.0373 | 92 | 0.0652 | 466 | 0.0536 |
| 4_12663788 | 87 | 0.0345 | 142 | 0.0915 | 124 | 0.0484 | 100 | 0.08 | 453 | 0.0662 | 105 | 0.0762 | 152 | 0.0724 | 179 | 0.0615 | 107 | 0.0561 | 543 | 0.0663 |
| 4_12663840 | 56 | 0.1607 | 104 | 0.2019 | 80 | 0.175 | 78 | 0.1538 | 318 | 0.1761 | 72 | 0.1528 | 103 | 0.165 | 125 | 0.184 | 79 | 0.2025 | 379 | 0.1768 |
| 4_12663846 | NA | NA | 1 | 1 | NA | NA | 2 | 0 | 3 | 0.3333 | 1 | 0 | 1 | 1 | NA | NA | 1 | 0 | 3 | 0.3333 |
| 4_12663854 | 60 | 0.0667 | 95 | 0.0842 | 72 | 0.0972 | 67 | 0.0597 | 294 | 0.0782 | 66 | 0.0303 | 110 | 0.0545 | 127 | 0.0709 | 80 | 0.0625 | 383 | 0.0574 |
| 4_12664419 | 37 | 0.7027 | 55 | 0.5273 | 40 | 0.45 | 45 | 0.4889 | 177 | 0.5367 | 41 | 0.5122 | 66 | 0.5758 | 81 | 0.4815 | 40 | 0.575 | 228 | 0.5307 |
| 4_12664439 | 34 | 0.2647 | 55 | 0.1455 | 37 | 0.2432 | 42 | 0.2143 | 168 | 0.2083 | 38 | 0.2368 | 65 | 0.1692 | 72 | 0.3056 | 42 | 0.2857 | 217 | 0.2488 |
| 4_12664443 | 26 | 0.0385 | 38 | 0.0789 | 25 | 0.16 | 29 | 0.069 | 118 | 0.0847 | 26 | 0.1154 | 47 | 0.0851 | 51 | 0.0784 | 23 | 0.1304 | 147 | 0.0952 |
| 4_12664446 | 34 | 0.1765 | 52 | 0.2115 | 37 | 0.1622 | 42 | 0.2619 | 165 | 0.2061 | 38 | 0.2895 | 58 | 0.2241 | 74 | 0.2297 | 36 | 0.1389 | 206 | 0.2233 |
| 4_12664471 | 1 | 1 | NA | NA | NA | NA | NA | NA | 1 | 1 | NA | NA | NA | NA | 2 | 0.5 | NA | NA | 2 | 0.5 |
| 4_12664676 | 4 | 0.25 | 11 | 0.1818 | 5 | 0.2 | 7 | 0 | 27 | 0.1481 | 4 | 0 | 3 | 0.3333 | 10 | 0.1 | 4 | 0.25 | 21 | 0.1429 |
| 4_12664703 | 4 | 0.5 | 13 | 0 | 8 | 0.25 | 8 | 0.25 | 33 | 0.1818 | 9 | 0 | 9 | 0.1111 | 9 | 0 | 6 | 0.1667 | 33 | 0.0606 |
| 4_12668562 | 1 | 0 | NA | NA | 2 | 0 | 2 | 0.5 | 5 | 0.2 | 3 | 0.3333 | 4 | 0.25 | 1 | 0 | 2 | 0 | 10 | 0.2 |
| 4_12668570 | 1 | 1 | NA | NA | 2 | 0 | 2 | 0.5 | 5 | 0.4 | 4 | 0.75 | 5 | 0.6 | 1 | 1 | 2 | 1 | 12 | 0.75 |
| 4_12668579 | 1 | 1 | NA | NA | NA | NA | 1 | 1 | 2 | 1 | 4 | 1 | 4 | 0.25 | NA | NA | 1 | 1 | 9 | 0.6667 |
| 4_12668590 | 1 | 0 | NA | NA | 1 | 0 | 2 | 0.5 | 4 | 0.25 | 3 | 0.3333 | 5 | 0.6 | 1 | 1 | 2 | 0.5 | 11 | 0.5455 |
| 4_12674728 | 7 | 0.8571 | 5 | 0.8 | 4 | 0.5 | 5 | 0.8 | 21 | 0.7619 | 6 | 0.6667 | 8 | 0.625 | 5 | 0.4 | 10 | 0.6 | 29 | 0.5862 |
| 4_12674762 | 7 | 0.5714 | 5 | 0.6 | 3 | 0 | 6 | 0.6667 | 21 | 0.5238 | 4 | 0.75 | 9 | 0.4444 | 4 | 0.5 | 9 | 0.5556 | 26 | 0.5385 |
| 4_12674778 | 8 | 0.125 | 6 | 0.1667 | 5 | 0.2 | 7 | 0.4286 | 26 | 0.2308 | 6 | 0 | 8 | 0.375 | 5 | 0.2 | 11 | 0.2727 | 30 | 0.2333 |
| 4_12674779 | 6 | 0.1667 | 5 | 0 | 4 | 0 | 5 | 0.2 | 20 | 0.1 | 5 | 0.4 | 7 | 0.2857 | 3 | 0 | 7 | 0.1429 | 22 | 0.2273 |
| 4_12674797 | 7 | 0.5714 | 7 | 0.5714 | 5 | 0.4 | 8 | 0.375 | 27 | 0.4815 | 6 | 0.1667 | 9 | 0.2222 | 6 | 0.1667 | 10 | 0.2 | 31 | 0.1935 |
| 4_12674807 | 8 | 0.375 | 6 | 0.1667 | 5 | 0.2 | 9 | 0.1111 | 28 | 0.2143 | 6 | 0.3333 | 9 | 0.3333 | 6 | 0.1667 | 10 | 0.3 | 31 | 0.2903 |
| 4_12675768 | 3 | 0 | 6 | 0 | 4 | 0.25 | 5 | 0.4 | 18 | 0.1667 | NA | NA | 6 | 0.3333 | 2 | 0 | 6 | 0 | 14 | 0.1429 |
| 4_12675776 | 3 | 1 | 8 | 1 | 4 | 1 | 3 | 1 | 18 | 1 | NA | NA | 6 | 1 | 1 | 1 | 4 | 1 | 11 | 1 |
| 4_12675792 | NA | NA | 6 | 0.5 | 3 | 0.3333 | 2 | 1 | 11 | 0.5455 | NA | NA | 3 | 1 | 1 | 1 | 4 | 0.75 | 8 | 0.875 |
| 4_12675795 | 3 | 0 | 8 | 0.375 | 2 | 0 | 3 | 0.3333 | 16 | 0.25 | NA | NA | 3 | 0.6667 | 1 | 1 | 4 | 0.75 | 8 | 0.75 |
| 4_12675799 | 1 | 0 | 8 | 0.125 | 3 | 0 | 2 | 0 | 14 | 0.0714 | NA | NA | 3 | 0.3333 | 1 | 0 | 4 | 0.25 | 8 | 0.25 |
| 4_12675801 | 3 | 0 | 7 | 0.7143 | 3 | 0 | 3 | 0.6667 | 16 | 0.4375 | NA | NA | 3 | 0.6667 | 1 | 0 | 5 | 0.4 | 9 | 0.4444 |
| 4_12680468 | NA | NA | 1 | 1 | 1 | 1 | 1 | 0 | 3 | 0.6667 | NA | NA | 2 | 1 | NA | NA | 1 | 0 | 3 | 0.6667 |
| 4_12680470 | NA | NA | 1 | 1 | NA | NA | NA | NA | 1 | 1 | NA | NA | 2 | 1 | NA | NA | 1 | 0 | 3 | 0.6667 |
| 4_12681949 | NA | NA | 1 | 1 | 1 | 1 | NA | NA | 2 | 1 | NA | NA | 1 | 1 | NA | NA | 1 | 1 | 2 | 1 |
| 4_12681960 | 1 | 1 | 1 | 0 | 1 | 0 | NA | NA | 3 | 0.3333 | NA | NA | 1 | 1 | NA | NA | 1 | 0 | 2 | 0.5 |
| 4_12681968 | NA | NA | 1 | 1 | NA | NA | NA | NA | 1 | 1 | NA | NA | 1 | 1 | NA | NA | NA | NA | 1 | 1 |
| 4_12691568 | 6 | 0.1667 | 5 | 0 | 3 | 0 | 3 | 0.3333 | 17 | 0.1176 | 1 | 1 | 11 | 0.0909 | 10 | 0.1 | 5 | 0 | 27 | 0.1111 |
| 4_12691672 | 11 | 0.2727 | 16 | 0.125 | 6 | 0.1667 | 7 | 0.2857 | 40 | 0.2 | 13 | 0 | 15 | 0.0667 | 24 | 0.125 | 10 | 0 | 62 | 0.0645 |
| 4_12691719 | 10 | 0.1 | 13 | 0.3846 | 7 | 0.1429 | 7 | 0.1429 | 37 | 0.2162 | 8 | 0.375 | 13 | 0.2308 | 16 | 0.1875 | 8 | 0.125 | 45 | 0.2222 |
| 4_12692323 | NA | NA | 1 | 1 | NA | NA | NA | NA | 1 | 1 | NA | NA | NA | NA | 2 | 0 | 2 | 0.5 | 4 | 0.25 |
| 4_12692359 | NA | NA | NA | NA | 2 | 0.5 | NA | NA | 2 | 0.5 | NA | NA | NA | NA | 2 | 1 | 3 | 0.3333 | 5 | 0.6 |
| 4_12692455 | 12 | 0 | 14 | 0 | 11 | 0.0909 | 13 | 0.1538 | 50 | 0.06 | 11 | 0 | 17 | 0.1176 | 13 | 0.0769 | 10 | 0.1 | 51 | 0.0784 |
| 4_12692742 | 2 | 0 | 2 | 0 | 4 | 0.25 | 3 | 0 | 11 | 0.0909 | 1 | 0 | 2 | 0.5 | 3 | 0 | 3 | 0 | 9 | 0.1111 |
| 4_12694148 | NA | NA | 4 | 1 | 1 | 1 | NA | NA | 5 | 1 | NA | NA | 1 | 1 | 2 | 1 | NA | NA | 3 | 1 |
| 4_12694149 | NA | NA | 2 | 0.5 | 1 | 0 | NA | NA | 3 | 0.3333 | NA | NA | 1 | 0 | 2 | 1 | NA | NA | 3 | 0.6667 |
| 4_12694150 | NA | NA | 2 | 0.5 | 1 | 1 | NA | NA | 3 | 0.6667 | NA | NA | 1 | 1 | 1 | 0 | NA | NA | 2 | 0.5 |

| | | | | | | | | | | | | | | | | | | | | |
|---|---|---|---|---|---|---|---|---|---|---|---|---|---|---|---|---|---|---|---|---|
| 4_1269415 | NA | NA | 3 | 1 | 1 | 0 | NA | NA | 4 | 0.75 | NA | NA | NA | NA | 2 | 0.5 | NA | NA | 2 | 0.5 |
| 4_1269415 | NA | NA | 4 | 0.75 | NA | NA | NA | NA | 4 | 0.75 | NA | NA | 1 | 1 | NA | NA | NA | NA | 1 | 1 |
| 4_1269416 | 3 | 0.3333 | 2 | 0.5 | 3 | 0.6667 | 4 | 0 | 12 | 0.3333 | 1 | 1 | NA | NA | 1 | 0 | 4 | 0.25 | 6 | 0.3333 |
| 4_1269417 | 3 | 0.3333 | 2 | 0.5 | 3 | 0.3333 | 4 | 0.5 | 12 | 0.4167 | 2 | 0.5 | 1 | 0 | 3 | 0.3333 | 4 | 0.25 | 10 | 0.3 |
| 4_1269419 | NA | NA | 3 | 0.6667 | 1 | 0 | NA | NA | 4 | 0.5 | NA | NA | 1 | 1 | 2 | 0 | NA | NA | 3 | 0.3333 |
| 4_1269419 | 3 | 0.3333 | 1 | 0 | 2 | 1 | 3 | 0.3333 | 9 | 0.4444 | 2 | 0 | 1 | 1 | 1 | 1 | 4 | 0.5 | 8 | 0.5 |
| 4_1269419 | NA | NA | 3 | 1 | 1 | 1 | NA | NA | 4 | 1 | NA | NA | 1 | 1 | 2 | 0.5 | NA | NA | 3 | 0.6667 |
| 4_1269419 | 3 | 0.3333 | 2 | 0.5 | 3 | 0 | 3 | 0 | 11 | 0.1818 | 2 | 0 | 1 | 1 | 2 | 1 | 4 | 0.25 | 9 | 0.4444 |
| 4_1269419 | NA | NA | 3 | 0.6667 | 1 | 0 | NA | NA | 4 | 0.5 | NA | NA | 1 | 1 | 1 | 1 | NA | NA | 2 | 1 |
| 4_1269420 | NA | NA | 2 | 1 | 1 | 0 | NA | NA | 3 | 0.6667 | NA | NA | 1 | 0 | 2 | 0.5 | NA | NA | 3 | 0.3333 |
| 4_1269420 | 2 | 1 | NA | NA | 2 | 1 | 4 | 1 | 8 | 1 | NA | NA | 1 | 1 | 1 | 1 | 2 | 0.5 | 4 | 0.75 |
| 4_1269422 | NA | NA | 4 | 0.5 | NA | NA | NA | NA | 4 | 0.5 | NA | NA | 1 | 1 | 2 | 0.5 | NA | NA | 3 | 0.6667 |
| 4_1269422 | 2 | 0.5 | 2 | 0.5 | 3 | 0 | 3 | 0.3333 | 10 | 0.3 | 2 | 0.5 | NA | NA | 2 | 0.5 | 2 | 0.5 | 6 | 0.5 |
| 4_1269422 | 3 | 0.3333 | 2 | 0 | 3 | 0 | 4 | 0 | 12 | 0.0833 | 2 | 0 | 1 | 0 | 3 | 0 | 4 | 0.25 | 10 | 0.1 |
| 4_1269423 | NA | NA | 4 | 0.75 | NA | NA | NA | NA | 4 | 0.75 | NA | NA | 1 | 1 | 2 | 0.5 | NA | NA | 3 | 0.6667 |
| 4_1269424 | NA | NA | 4 | 1 | NA | NA | NA | NA | 4 | 1 | NA | NA | 1 | 1 | 1 | 1 | NA | NA | 2 | 1 |
| 4_1269424 | NA | NA | 3 | 0.6667 | NA | NA | NA | NA | 3 | 0.6667 | NA | NA | 1 | 1 | 2 | 0 | NA | NA | 3 | 0.3333 |
| 4_1269424 | 3 | 0.6667 | 2 | 1 | 3 | 1 | 4 | 0.5 | 12 | 0.75 | 2 | 1 | 1 | 1 | 4 | 0.5 | 4 | 0.5 | 11 | 0.6364 |
| 4_1269425 | 3 | 0 | 2 | 0 | 3 | 0.3333 | 3 | 0 | 11 | 0.0909 | 2 | 0 | 1 | 0 | 3 | 0.3333 | 4 | 0.25 | 10 | 0.2 |
| 4_1269425 | 3 | 1 | 2 | 1 | 3 | 1 | 4 | 1 | 12 | 1 | 2 | 1 | 1 | 1 | 2 | 1 | 4 | 1 | 9 | 1 |
| 4_1269426 | 3 | 0 | NA | NA | 3 | 0 | 4 | 0.25 | 10 | 0.1 | 2 | 0.5 | 1 | 0 | 3 | 0 | 3 | 0.3333 | 9 | 0.2222 |
| 4_1269427 | 2 | 0.5 | 2 | 0.5 | 3 | 0 | 4 | 0.25 | 11 | 0.2727 | 1 | 0 | 1 | 0 | 4 | 0.5 | 4 | 0 | 10 | 0.2 |
| 4_1269427 | NA | NA | 3 | 0.6667 | NA | NA | NA | NA | 3 | 0.6667 | NA | NA | NA | NA | 2 | 1 | NA | NA | 2 | 1 |
| 4_1269428 | NA | NA | 3 | 0.6667 | NA | NA | NA | NA | 3 | 0.6667 | NA | NA | 1 | 1 | 2 | 1 | NA | NA | 3 | 1 |
| 4_1269428 | 3 | 0 | 2 | 0 | 1 | 1 | 3 | 0 | 9 | 0.1111 | 2 | 0 | 1 | 1 | 3 | 0.3333 | 3 | 0 | 9 | 0.2222 |
| 4_1269428 | 3 | 0.3333 | 2 | 0 | 3 | 0 | 3 | 0 | 11 | 0.0909 | 2 | 0 | 1 | 0 | 4 | 0.25 | 2 | 0 | 9 | 0.1111 |
| 4_1269428 | NA | NA | 4 | 1 | NA | NA | NA | NA | 4 | 1 | NA | NA | 1 | 1 | 2 | 0.5 | NA | NA | 3 | 0.6667 |
| 4_1269430 | 3 | 0.3333 | 2 | 0.5 | 3 | 0.6667 | 4 | 0.5 | 12 | 0.5 | 2 | 1 | 1 | 1 | 5 | 0.4 | 5 | 0.2 | 13 | 0.4615 |
| 4_1269430 | NA | NA | 3 | 1 | NA | NA | NA | NA | 3 | 1 | NA | NA | NA | NA | 1 | 1 | NA | NA | 1 | 1 |
| 4_1269430 | 3 | 0 | 2 | 0.5 | 3 | 0 | 3 | 0.6667 | 11 | 0.2727 | 2 | 0.5 | 1 | 0 | 5 | 0.2 | 6 | 0 | 14 | 0.1429 |
| 4_1269430 | NA | NA | 4 | 0.25 | NA | NA | NA | NA | 4 | 0.25 | NA | NA | 1 | 1 | 2 | 0 | NA | NA | 3 | 0.3333 |
| 4_1269431 | 2 | 1 | 1 | 1 | 2 | 1 | 3 | 1 | 8 | 1 | 1 | 1 | 1 | 1 | 2 | 1 | 3 | 1 | 7 | 1 |
| 4_1269431 | 3 | 1 | 2 | 1 | 3 | 1 | 3 | 1 | 11 | 1 | 2 | 1 | 1 | 1 | 4 | 1 | 6 | 1 | 13 | 1 |
| 4_1269431 | NA | NA | 2 | 1 | NA | NA | NA | NA | 2 | 1 | NA | NA | NA | NA | 2 | 1 | NA | NA | 2 | 1 |
| 4_1269432 | NA | NA | 3 | 0.6667 | NA | NA | NA | NA | 3 | 0.6667 | NA | NA | 1 | 0 | 2 | 1 | NA | NA | 3 | 0.6667 |
| 4_1269432 | 1 | 1 | 1 | 1 | 2 | 1 | 3 | 1 | 7 | 1 | 1 | 1 | 1 | 1 | 2 | 1 | 2 | 1 | 6 | 1 |
| 4_1269432 | 1 | 1 | 1 | 1 | 2 | 1 | 3 | 1 | 7 | 1 | 1 | 1 | 1 | 1 | 4 | 1 | 4 | 1 | 10 | 1 |
| 4_1269433 | NA | NA | 2 | 0.5 | NA | NA | NA | NA | 2 | 0.5 | NA | NA | 1 | 0 | 2 | 1 | NA | NA | 3 | 0.6667 |
| 4_1269434 | 1 | 0 | 1 | 1 | 1 | 0 | 2 | 0 | 5 | 0.2 | 1 | 0 | 1 | 0 | 4 | 0 | 4 | 0.75 | 10 | 0.3 |
| 4_1269435 | 1 | 1 | 1 | 1 | 2 | 1 | 3 | 1 | 7 | 1 | 1 | 1 | NA | NA | 5 | 1 | 4 | 1 | 10 | 1 |
| 4_1269435 | 1 | 1 | 1 | 1 | 2 | 1 | 3 | 0.3333 | 7 | 0.7143 | 1 | 1 | 1 | 1 | 6 | 0.8333 | 4 | 0.5 | 12 | 0.75 |
| 4_1269436 | NA | NA | 3 | 0.3333 | NA | NA | NA | NA | 3 | 0.3333 | NA | NA | 1 | 0 | 1 | 1 | NA | NA | 2 | 0.5 |
| 4_1269436 | 1 | 1 | 1 | 1 | 2 | 0 | 2 | 0.5 | 6 | 0.5 | 1 | 1 | 1 | 1 | 5 | 0.8 | 3 | 0.6667 | 10 | 0.8 |
| 4_1269437 | 1 | 1 | 1 | 0 | 2 | 1 | 2 | 0.5 | 6 | 0.6667 | 1 | 1 | 1 | 1 | 5 | 0.6 | 5 | 1 | 12 | 0.8333 |
| 4_1269446 | NA | NA | NA | NA | 2 | 0.5 | 1 | 0 | 3 | 0.3333 | NA | NA | 1 | 0 | 3 | 0 | 2 | 0.5 | 6 | 0.1667 |
| 4_1269446 | NA | NA | 1 | 0 | 3 | 0.6667 | 3 | 0.6667 | 7 | 0.5714 | NA | NA | 1 | 0 | 4 | 0.5 | 5 | 0.4 | 10 | 0.4 |
| 4_1269446 | NA | NA | 1 | 1 | 2 | 0 | 3 | 0.3333 | 6 | 0.3333 | NA | NA | 1 | 0 | 4 | 0 | 5 | 0.2 | 10 | 0.1 |
| 4_1269990 | 2 | 0 | 2 | 0 | NA | NA | 1 | 1 | 5 | 0.2 | NA | NA | 1 | 1 | NA | NA | 1 | 1 | 2 | 1 |
| 4_1270162 | 2 | 0 | 3 | 0.3333 | 1 | 1 | 2 | 0 | 8 | 0.25 | NA | NA | 2 | 0 | 2 | 0.5 | 3 | 0.3333 | 7 | 0.2857 |
| 4_1270162 | 2 | 0.5 | 3 | 0 | 1 | 0 | 2 | 0 | 8 | 0.125 | NA | NA | 2 | 0.5 | NA | NA | 2 | 0 | 4 | 0.25 |
| 4_1270163 | 2 | 0 | 3 | 0.6667 | 1 | 1 | 3 | 1 | 9 | 0.6667 | NA | NA | 2 | 0 | 2 | 0.5 | 3 | 0 | 7 | 0.1429 |
| 4_1270163 | 2 | 1 | 1 | 1 | NA | NA | 3 | 0.6667 | 6 | 0.8333 | NA | NA | 2 | 0.5 | 2 | 0.5 | 3 | 0.3333 | 7 | 0.4286 |
| 4_1270166 | 1 | 1 | 3 | 0.6667 | 1 | 1 | 3 | 0.6667 | 8 | 0.75 | NA | NA | 2 | 0.5 | 2 | 0.5 | 3 | 0.6667 | 7 | 0.5714 |
| 4_1270169 | 3 | 0.3333 | 3 | 0 | 1 | 0 | 2 | 0.5 | 9 | 0.2222 | NA | NA | 1 | 0 | 2 | 0.5 | 2 | 0 | 5 | 0.2 |
| 4_1270170 | 2 | 0.5 | 1 | 0 | 1 | 1 | 3 | 0.6667 | 7 | 0.5714 | NA | NA | 2 | 0 | 2 | 1 | 2 | 0.5 | 6 | 0.5 |
| 4_1270170 | 2 | 0 | 3 | 0.3333 | 1 | 0 | 2 | 0 | 8 | 0.125 | NA | NA | 2 | 0.5 | 2 | 0 | 1 | 0 | 5 | 0.2 |
| 4_1270171 | 3 | 0.6667 | 3 | 0.3333 | 1 | 1 | 3 | 0.3333 | 10 | 0.5 | NA | NA | 1 | 0 | 2 | 0 | 2 | 0.5 | 5 | 0.2 |
| 4_1270172 | 3 | 0.3333 | 4 | 1 | 1 | 1 | 2 | 1 | 10 | 0.8 | NA | NA | 2 | 1 | 2 | 1 | 3 | 0.6667 | 7 | 0.8571 |
| 4_1270173 | 3 | 0 | 3 | 0 | 1 | 1 | 3 | 0.3333 | 10 | 0.2 | NA | NA | 2 | 0.5 | 2 | 0.5 | 4 | 0 | 8 | 0.25 |
| 4_1270175 | 3 | 0.6667 | 4 | 1 | 1 | 1 | 2 | 0.5 | 10 | 0.8 | NA | NA | 1 | 1 | 2 | 1 | 3 | 0.6667 | 6 | 0.8333 |
| 4_1270175 | 3 | 0.3333 | 3 | 0 | 1 | 0 | 3 | 0 | 10 | 0.1 | NA | NA | 2 | 0.5 | 2 | 0 | 3 | 0.3333 | 7 | 0.2857 |
| 4_1270175 | 2 | 0.5 | 4 | 1 | 1 | 1 | 3 | 1 | 10 | 0.9 | NA | NA | 2 | 1 | 2 | 1 | 3 | 1 | 7 | 1 |
| 4_1270176 | 3 | 0.6667 | 4 | 0.75 | 1 | 1 | 3 | 0.6667 | 11 | 0.7273 | NA | NA | 2 | 0.5 | 2 | 0.5 | 4 | 0.5 | 8 | 0.5 |
| 4_1270176 | 3 | 0.3333 | 3 | 0.3333 | 1 | 0 | 3 | 0.3333 | 10 | 0.3 | NA | NA | 2 | 0 | 2 | 0.5 | 2 | 0 | 6 | 0.1667 |
| 4_1270177 | 2 | 0 | 4 | 0.25 | 1 | 1 | 3 | 0 | 10 | 0.2 | NA | NA | 2 | 0 | 2 | 0.5 | 3 | 0.3333 | 7 | 0.2857 |

| | | | | | | | | | | | | | | | | | | | | |
|---|---|---|---|---|---|---|---|---|---|---|---|---|---|---|---|---|---|---|---|---|
| 4_1270178( | 4 | 0 | 4 | 0.5 | 1 | 1 | 3 | 0 | 12 | 0.25 | NA | NA | 2 | 0.5 | 2 | 0 | 3 | 0.3333 | 7 | 0.2857 |
| 4_1270178 | 4 | 0.5 | 3 | 0.3333 | 1 | 1 | 2 | 0.5 | 10 | 0.5 | NA | NA | 2 | 0 | 2 | 0.5 | 4 | 0.5 | 8 | 0.375 |
| 4_1270178 | 4 | 0.75 | 4 | 1 | 1 | 1 | 3 | 1 | 12 | 0.9167 | NA | NA | 2 | 0.5 | 2 | 1 | 4 | 0.5 | 8 | 0.625 |
| 4_1270179 | 3 | 1 | 4 | 0.75 | 1 | 1 | 2 | 1 | 10 | 0.9 | NA | NA | 2 | 1 | 1 | 1 | 2 | 1 | 5 | 1 |
| 4_1270623 | 3 | 0.6667 | 2 | 0.5 | 1 | 1 | 1 | 1 | 7 | 0.7143 | 1 | 1 | 2 | 0.5 | 2 | 0.5 | NA | NA | 5 | 0.6 |
| 4_1270625 | 8 | 0.125 | 8 | 0.125 | 6 | 0.1667 | 2 | 0 | 24 | 0.125 | 2 | 0.5 | 4 | 0 | 6 | 0.1667 | NA | NA | 12 | 0.1667 |
| 4_1270885 | 20 | 0.5 | 20 | 0.45 | 20 | 0.6 | 7 | 0.5714 | 67 | 0.5224 | 14 | 0.6429 | 22 | 0.5 | 25 | 0.6 | 11 | 0.3636 | 72 | 0.5417 |
| 4_1270885 | 22 | 0.3636 | 19 | 0.1579 | 19 | 0.1579 | 5 | 0.2 | 65 | 0.2308 | 16 | 0.3125 | 20 | 0.15 | 27 | 0.3333 | 12 | 0.4167 | 75 | 0.2933 |
| 4_1270887 | 21 | 0.8571 | 17 | 0.7647 | 18 | 0.7778 | 7 | 0.7143 | 63 | 0.7937 | 15 | 0.7333 | 22 | 0.7727 | 25 | 0.56 | 10 | 0.8 | 72 | 0.6944 |
| 4_1270887 | 22 | 0.3182 | 19 | 0.3684 | 23 | 0.3478 | 7 | 0.1429 | 71 | 0.3239 | 18 | 0.4444 | 25 | 0.28 | 27 | 0.4815 | 13 | 0.3846 | 83 | 0.3976 |
| 4_1270888 | 23 | 0.6522 | 22 | 0.6364 | 23 | 0.6087 | 7 | 0.4286 | 75 | 0.6133 | 19 | 0.7895 | 25 | 0.92 | 27 | 0.7778 | 12 | 0.75 | 83 | 0.8193 |
| 4_1270889 | 14 | 0.5714 | 18 | 0.5 | 16 | 0.375 | 6 | 0.5 | 54 | 0.4815 | 17 | 0.4706 | 19 | 0.3158 | 26 | 0.5385 | 10 | 0.3 | 72 | 0.4306 |
| 4_1270890 | 21 | 0.7619 | 18 | 0.6667 | 19 | 0.6842 | 5 | 1 | 63 | 0.7302 | 17 | 0.7059 | 23 | 0.8261 | 24 | 0.625 | 10 | 0.6 | 74 | 0.7027 |
| 4_1270891 | 21 | 0.5714 | 22 | 0.8636 | 22 | 0.7727 | 8 | 0.75 | 73 | 0.7397 | 16 | 0.4375 | 25 | 0.64 | 29 | 0.6897 | 11 | 0.9091 | 81 | 0.6543 |
| 4_1270891 | 21 | 0.2381 | 22 | 0.3182 | 21 | 0.381 | 8 | 0.5 | 72 | 0.3333 | 20 | 0.3 | 18 | 0.2222 | 22 | 0.3636 | 11 | 0.3636 | 71 | 0.3099 |
| 4_1270892 | 17 | 0.9412 | 19 | 1 | 13 | 1 | 7 | 0.8571 | 56 | 0.9643 | 15 | 1 | 17 | 0.8235 | 23 | 0.913 | 11 | 0.9091 | 66 | 0.9091 |
| 4_1270892 | 15 | 0.7333 | 16 | 0.5 | 13 | 0.6923 | 5 | 0.8 | 49 | 0.6531 | 15 | 0.6667 | 18 | 0.6111 | 17 | 0.5294 | 8 | 0.625 | 58 | 0.6034 |
| 4_1270892 | 21 | 0.9524 | 21 | 0.8095 | 19 | 0.8421 | 11 | 1 | 72 | 0.8889 | 21 | 0.9524 | 21 | 0.9524 | 26 | 0.8462 | 10 | 0.9 | 78 | 0.9103 |
| 4_1270893 | 20 | 0.8 | 22 | 0.5 | 19 | 0.6842 | 7 | 0.5714 | 68 | 0.6471 | 18 | 0.5556 | 22 | 0.5909 | 23 | 0.5652 | 8 | 0.625 | 71 | 0.5775 |
| 4_1270894 | 23 | 0.3478 | 25 | 0.52 | 23 | 0.3478 | 14 | 0.5714 | 85 | 0.4353 | 20 | 0.6 | 23 | 0.4348 | 33 | 0.3939 | 12 | 0.1667 | 88 | 0.4205 |
| 4_1271277 | 30 | 0.0667 | 39 | 0.1538 | 37 | 0.0811 | 18 | 0 | 124 | 0.0887 | 30 | 0.1667 | 37 | 0.0811 | 51 | 0.0588 | 22 | 0.0909 | 140 | 0.0929 |
| 4_1271300 | 27 | 0.1481 | 43 | 0.1163 | 31 | 0.1613 | 18 | 0.1111 | 119 | 0.1345 | 26 | 0.1538 | 30 | 0.2 | 41 | 0.1951 | 19 | 0.1579 | 116 | 0.181 |
| 4_1271301 | 32 | 0.0938 | 49 | 0.1429 | 38 | 0.0263 | 19 | 0 | 138 | 0.0797 | 31 | 0.0645 | 39 | 0.0513 | 47 | 0.0213 | 24 | 0.1667 | 141 | 0.0638 |
| 4_1271303 | 28 | 0.5 | 37 | 0.5405 | 30 | 0.4667 | 18 | 0.5 | 113 | 0.5044 | 23 | 0.3913 | 35 | 0.5429 | 39 | 0.5641 | 20 | 0.4 | 117 | 0.4957 |
| 4_1271306 | 37 | 0.4054 | 58 | 0.4138 | 49 | 0.4082 | 24 | 0.2917 | 168 | 0.3929 | 39 | 0.3333 | 49 | 0.4286 | 63 | 0.3968 | 29 | 0.1724 | 180 | 0.3556 |
| 4_1271306 | 36 | 0.3333 | 50 | 0.34 | 43 | 0.1628 | 23 | 0.087 | 152 | 0.25 | 36 | 0.3056 | 46 | 0.2174 | 59 | 0.2034 | 27 | 0.1852 | 168 | 0.2262 |
| 4_1271306 | 30 | 0.1 | 44 | 0.0909 | 35 | 0.1429 | 20 | 0.15 | 129 | 0.1163 | 32 | 0 | 39 | 0.1026 | 54 | 0.2037 | 30 | 0.0333 | 155 | 0.1032 |
| 4_1271308 | 19 | 0.5263 | 26 | 0.5 | 21 | 0.381 | 13 | 0.3846 | 79 | 0.4557 | 24 | 0.4167 | 37 | 0.5135 | 37 | 0.5676 | 18 | 0.5 | 116 | 0.5086 |
| 4_1271309 | NA | NA | 1 | 1 | NA | NA | NA | NA | 1 | 1 | 1 | 1 | NA | NA | NA | NA | NA | NA | 1 | 1 |
| 4_1271313 | NA | NA | 1 | 1 | NA | NA | NA | NA | 1 | 1 | 1 | 1 | NA | NA | NA | NA | NA | NA | 1 | 1 |
| 4_1271408 | 9 | 0.1111 | 4 | 0.75 | 10 | 0.6 | 5 | 0.4 | 28 | 0.4286 | 6 | 0.3333 | 6 | 0.1667 | 8 | 0.375 | 6 | 0.1667 | 26 | 0.2692 |
| 4_1271409 | 9 | 1 | 6 | 1 | 13 | 1 | 5 | 1 | 33 | 1 | 7 | 0.8571 | 7 | 0.8571 | 9 | 1 | 7 | 1 | 30 | 0.9333 |
| 4_1271409 | 6 | 0.1667 | 3 | 0.3333 | 6 | 0.1667 | 4 | 0 | 19 | 0.1579 | 4 | 0 | 5 | 0.6 | 5 | 0.2 | 5 | 0.2 | 19 | 0.2632 |
| 4_1271409 | 7 | 0.8571 | 3 | 0.3333 | 11 | 0.6364 | 5 | 0.6 | 26 | 0.6538 | 6 | 0.6667 | 5 | 0.6 | 8 | 0.625 | 6 | 0.5 | 25 | 0.6 |
| 4_1271410 | 6 | 0.1667 | 5 | 0 | 12 | 0.25 | 5 | 0 | 28 | 0.1429 | 7 | 0 | 7 | 0.1429 | 8 | 0.25 | 7 | 0 | 29 | 0.1034 |
| 4_1271411 | 8 | 0.625 | 6 | 0.3333 | 13 | 0.6154 | 5 | 0.6 | 32 | 0.5625 | 4 | 0.75 | 5 | 0.8 | 10 | 0.6 | 5 | 0.6 | 24 | 0.6667 |
| 4_1271412 | 9 | 0.4444 | 6 | 0.3333 | 12 | 0.4167 | 7 | 0.4286 | 34 | 0.4118 | 7 | 0.5714 | 6 | 0.6667 | 8 | 0.625 | 8 | 0.375 | 29 | 0.5517 |
| 4_1271412 | 9 | 0.8889 | 7 | 0.8571 | 13 | 1 | 5 | 0.8 | 34 | 0.9118 | 5 | 1 | 7 | 1 | 11 | 0.9091 | 4 | 1 | 27 | 0.963 |
| 4_1271415 | 8 | 0 | 7 | 0 | 12 | 0.1667 | 7 | 0 | 34 | 0.0588 | 8 | 0.5 | 5 | 0.2 | 10 | 0 | 9 | 0.1111 | 32 | 0.1875 |
| 4_1284594 | 1 | 0 | 2 | 0.5 | NA | NA | 2 | 0 | 5 | 0.2 | NA | NA | 1 | 0 | 2 | 0.5 | NA | NA | 3 | 0.3333 |
| 4_1284596 | 2 | 0 | 3 | 0.3333 | NA | NA | 2 | 0 | 7 | 0.1429 | NA | NA | 1 | 1 | 3 | 0 | NA | NA | 4 | 0.25 |
| 4_1284596 | 2 | 0 | 3 | 0.3333 | NA | NA | 2 | 0.5 | 7 | 0.2857 | NA | NA | 1 | 0 | 3 | 0.3333 | NA | NA | 4 | 0.25 |
| 4_1284597 | 1 | 1 | 3 | 0 | NA | NA | 2 | 0.5 | 6 | 0.3333 | NA | NA | 1 | 0 | 3 | 0.6667 | NA | NA | 4 | 0.5 |
| 4_1284597 | 2 | 1 | 3 | 1 | NA | NA | 2 | 1 | 7 | 1 | NA | NA | 1 | 1 | 2 | 1 | NA | NA | 3 | 1 |
| 4_1284603 | 1 | 0 | 4 | 0.75 | 1 | 0 | 2 | 0 | 8 | 0.375 | NA | NA | 1 | 1 | 3 | 0 | NA | NA | 4 | 0.25 |
| 4_1284613 | 1 | 0 | 4 | 0.25 | 1 | 0 | 3 | 0.3333 | 9 | 0.2222 | 1 | 0 | 1 | 1 | 3 | 0 | 1 | 0 | 6 | 0.1667 |
| 4_1284779 | NA | NA | NA | NA | 3 | 0.3333 | NA | NA | 3 | 0.3333 | NA | NA | 1 | 1 | NA | NA | NA | NA | 1 | 1 |
| 4_1284791 | NA | NA | NA | NA | 3 | 0.3333 | NA | NA | 3 | 0.3333 | NA | NA | 1 | 1 | NA | NA | NA | NA | 1 | 1 |
| 4_1284792 | NA | NA | NA | NA | 1 | 1 | 1 | 1 | 2 | 1 | NA | NA | 1 | 1 | NA | NA | NA | NA | 1 | 1 |
| 4_1284793 | NA | NA | NA | NA | 3 | 0.3333 | 1 | 0 | 4 | 0.25 | NA | NA | 1 | 1 | NA | NA | NA | NA | 1 | 1 |
| 4_1284794 | NA | NA | NA | NA | 3 | 1 | 1 | 1 | 4 | 1 | NA | NA | 1 | 1 | NA | NA | NA | NA | 1 | 1 |
| 4_1284797 | NA | NA | NA | NA | 2 | 0.5 | NA | NA | 2 | 0.5 | NA | NA | 1 | 1 | NA | NA | NA | NA | 1 | 1 |
| 4_1284797 | NA | NA | NA | NA | 3 | 1 | NA | NA | 3 | 1 | NA | NA | 1 | 1 | NA | NA | NA | NA | 1 | 1 |
| 4_1286418 | 3 | 0.3333 | 7 | 0.2857 | 4 | 0 | 5 | 0 | 19 | 0.1579 | 3 | 0 | 15 | 0.0667 | 7 | 0.1429 | 5 | 0 | 30 | 0.0667 |
| 4_1286419 | 3 | 1 | 6 | 0.1667 | 4 | 0.5 | 3 | 0 | 16 | 0.375 | 3 | 0 | 13 | 0.1538 | 9 | 0 | 6 | 0.1667 | 31 | 0.0968 |
| 4_1286454 | 4 | 1 | 12 | 0.9167 | 4 | 1 | 7 | 1 | 27 | 0.963 | 4 | 0.5 | 15 | 0.8667 | 13 | 0.8462 | 4 | 1 | 36 | 0.8333 |
| 4_1286455 | 4 | 1 | 14 | 1 | 5 | 1 | 7 | 1 | 30 | 1 | 3 | 1 | 12 | 1 | 9 | 0.8889 | 6 | 1 | 30 | 0.9667 |
| 4_1286460 | 3 | 0.3333 | 15 | 0.2 | 6 | 0.5 | 7 | 0.2857 | 31 | 0.2903 | 6 | 0.1667 | 13 | 0.0769 | 15 | 0 | 6 | 0 | 40 | 0.05 |
| 4_1286460 | 5 | 0 | 17 | 0.0588 | 5 | 0.2 | 9 | 0.2222 | 36 | 0.1111 | 6 | 0 | 16 | 0.1875 | 16 | 0.125 | 7 | 0 | 45 | 0.1111 |
| 4_1286461 | 5 | 0 | 14 | 0.2857 | 4 | 0.25 | 5 | 0.6 | 28 | 0.2857 | 5 | 0.2 | 13 | 0.2308 | 8 | 0 | 3 | 0.3333 | 29 | 0.1724 |
| 4_1286463 | 5 | 0.2 | 17 | 0.3529 | 8 | 0.375 | 8 | 0.125 | 38 | 0.2895 | 7 | 0.4286 | 14 | 0.4286 | 13 | 0.1538 | 8 | 0.5 | 42 | 0.3571 |
| 4_1286463 | 5 | 0 | 16 | 0.25 | 8 | 0.25 | 11 | 0.1818 | 40 | 0.2 | 7 | 0.1429 | 17 | 0.2941 | 15 | 0.1333 | 8 | 0.25 | 47 | 0.2128 |
| 4_1286466 | 4 | 0.5 | 15 | 0.4 | 11 | 0.5455 | 11 | 0.5455 | 41 | 0.4878 | 9 | 0.4444 | 16 | 0.5 | 18 | 0.4444 | 9 | 0.3333 | 52 | 0.4423 |
| 4_1286671 | 2 | 0 | 2 | 0 | 3 | 0 | 8 | 0.125 | 15 | 0.0667 | 3 | 0.3333 | 4 | 0 | 6 | 0 | 3 | 0 | 16 | 0.0625 |
| 4_1286672 | 2 | 0 | 3 | 0 | 3 | 0 | 8 | 0.375 | 16 | 0.1875 | 2 | 0.5 | 4 | 0 | 6 | 0 | 3 | 0 | 15 | 0.0667 |

| | | | | | | | | | | | | | | | | | | | | |
|---|---|---|---|---|---|---|---|---|---|---|---|---|---|---|---|---|---|---|---|---|
| 4_1286674 | 2 | 0 | 2 | 0 | 3 | 0.3333 | 7 | 0.1429 | 14 | 0.1429 | NA | NA | 4 | 0.25 | 5 | 0.2 | 3 | 0 | 12 | 0.1667 |
| 4_1286675 | 1 | 1 | 2 | 1 | 2 | 0.5 | 7 | 0.1429 | 12 | 0.4167 | 1 | 1 | 4 | 0.75 | 2 | 0.5 | 2 | 0 | 9 | 0.5556 |
| 4_1286676 | 2 | 1 | 2 | 1 | 3 | 0.6667 | 7 | 0.8571 | 14 | 0.8571 | 1 | 0 | 4 | 1 | 3 | 0.6667 | 3 | 0.3333 | 11 | 0.6364 |
| 4_1286678 | 1 | 0 | NA | NA | 2 | 0 | 5 | 0.4 | 8 | 0.25 | NA | NA | 1 | 0 | 4 | 0.5 | 2 | 0 | 7 | 0.2857 |
| 4_1286679 | 1 | 1 | 2 | 1 | 3 | 0.3333 | 7 | 0.5714 | 13 | 0.6154 | 2 | 1 | 2 | 1 | 2 | 0.5 | 3 | 0.6667 | 9 | 0.7778 |
| 4_1286679 | 1 | 1 | 2 | 0.5 | 2 | 1 | 4 | 1 | 9 | 0.8889 | 2 | 1 | 3 | 1 | 3 | 1 | 2 | 1 | 10 | 1 |
| 4_1286680 | 1 | 0 | 2 | 0.5 | 4 | 0 | 7 | 0.1429 | 14 | 0.1429 | 1 | 0 | 3 | 0.6667 | 2 | 0 | 3 | 0.3333 | 9 | 0.3333 |
| 4_1286681 | 1 | 1 | 2 | 1 | 1 | 1 | 6 | 1 | 10 | 1 | 1 | 1 | 3 | 1 | 2 | 1 | 3 | 0.6667 | 9 | 0.8889 |
| 4_1287740 | 1 | 0 | 2 | 1 | NA | NA | 1 | 1 | 4 | 0.75 | NA | NA | 1 | 1 | NA | NA | 2 | 1 | 3 | 1 |
| 4_1287777 | 1 | 0 | NA | NA | 1 | 1 | NA | NA | 2 | 0.5 | 1 | 1 | 1 | 1 | NA | NA | NA | NA | 2 | 1 |
| 4_1287780 | 1 | 1 | NA | NA | 2 | 0.5 | 2 | 0 | 5 | 0.4 | 1 | 1 | 2 | 0 | NA | NA | NA | NA | 3 | 0.3333 |
| 4_1287782 | 2 | 0 | NA | NA | 2 | 0 | 2 | 0.5 | 6 | 0.1667 | NA | NA | 2 | 0 | 2 | 0.5 | NA | NA | 4 | 0.25 |
| 4_1288029 | 2 | 0.5 | NA | NA | 1 | 0 | 4 | 0.25 | 7 | 0.2857 | 1 | 0 | 1 | 0 | 2 | 1 | 2 | 0.5 | 6 | 0.5 |
| 4_1288030 | 2 | 0 | 1 | 0 | 2 | 0.5 | 4 | 0 | 9 | 0.1111 | 1 | 0 | 4 | 0.25 | 3 | 0 | 1 | 0 | 9 | 0.1111 |
| 4_1288031 | 1 | 0 | 1 | 0 | 2 | 0.5 | 3 | 0.3333 | 7 | 0.2857 | 1 | 1 | 3 | 1 | 3 | 0 | 1 | 0 | 8 | 0.5 |
| 4_1288032 | 2 | 1 | 1 | 0 | 2 | 1 | 3 | 0.6667 | 8 | 0.75 | NA | NA | 2 | 0 | 3 | 1 | 2 | 1 | 7 | 0.7143 |
| 4_1288034 | 3 | 1 | 1 | 1 | 1 | 0 | 4 | 0.75 | 9 | 0.7778 | 1 | 1 | 6 | 0.8333 | 4 | 1 | 2 | 1 | 13 | 0.9231 |
| 4_1288034 | 3 | 1 | NA | NA | 3 | 1 | 4 | 1 | 10 | 1 | 1 | 1 | 5 | 1 | 4 | 1 | 2 | 1 | 12 | 1 |
| 4_1288034 | 2 | 0 | 1 | 0 | 2 | 0.5 | 2 | 0 | 7 | 0.1429 | 1 | 0 | 5 | 0.2 | 4 | 0.25 | 1 | 0 | 11 | 0.1818 |
| 4_1288035 | 4 | 0.25 | 1 | 1 | 3 | 0 | 4 | 0 | 12 | 0.1667 | 1 | 0 | 6 | 0.3333 | 6 | 0.3333 | 2 | 0.5 | 15 | 0.3333 |
| 4_1288037 | 2 | 1 | NA | NA | 2 | 0.5 | 4 | 0.5 | 8 | 0.625 | NA | NA | 3 | 0.3333 | 5 | 0.6 | 2 | 1 | 10 | 0.6 |
| 4_1288038 | 2 | 1 | 1 | 1 | 2 | 1 | 2 | 0.5 | 7 | 0.8571 | 1 | 1 | 5 | 1 | 3 | 0.6667 | 1 | 1 | 10 | 0.9 |
| 4_1288039 | 5 | 0.8 | 1 | 0 | 3 | 0.3333 | 4 | 0.75 | 13 | 0.6154 | 1 | 1 | 6 | 0.5 | 6 | 1 | 2 | 1 | 15 | 0.8 |
| 4_1288040 | 4 | 0 | NA | NA | 3 | 0.3333 | 4 | 0 | 11 | 0.0909 | 1 | 0 | 5 | 0.2 | 6 | 0 | 2 | 0.5 | 14 | 0.1429 |
| 4_1288040 | 2 | 0.5 | 1 | 0 | 2 | 0 | 2 | 0 | 7 | 0.1429 | NA | NA | 5 | 0.2 | 5 | 0 | 1 | 0 | 11 | 0.0909 |
| 4_1288041 | 5 | 0.8 | 1 | 1 | 3 | 1 | 4 | 0.5 | 13 | 0.7692 | NA | NA | 5 | 0.8 | 5 | 0.8 | 2 | 1 | 12 | 0.8333 |
| 4_1288042 | 4 | 0.5 | 1 | 0 | 4 | 0 | 4 | 0 | 13 | 0.1538 | 1 | 0 | 5 | 0 | 6 | 0 | 3 | 0.3333 | 15 | 0.0667 |
| 4_1288043 | 5 | 0.4 | 1 | 0 | 4 | 0.25 | 4 | 0.25 | 14 | 0.2857 | 1 | 0 | 6 | 0.3333 | 6 | 0.3333 | 2 | 0 | 15 | 0.2667 |
| 4_1288043 | 5 | 0 | 1 | 0 | 3 | 0.3333 | 4 | 0.25 | 13 | 0.1538 | 1 | 0 | 6 | 0 | 6 | 0.3333 | 3 | 0 | 16 | 0.125 |
| 4_1288045 | 5 | 0.2 | 1 | 0 | 4 | 0 | 5 | 0 | 15 | 0.0667 | 2 | 0 | 7 | 0.1429 | 7 | 0 | 3 | 0 | 19 | 0.0526 |
| 4_1288045 | 3 | 0.3333 | 1 | 1 | 4 | 1 | 3 | 0.6667 | 11 | 0.7273 | 2 | 0.5 | 7 | 0.4286 | 6 | 0.3333 | 2 | 0.5 | 17 | 0.4118 |
| 4_1288045 | 5 | 0.4 | NA | NA | 4 | 0.25 | 4 | 0.5 | 13 | 0.3846 | 1 | 0 | 5 | 0.6 | 6 | 0.6667 | 2 | 1 | 14 | 0.6429 |
| 4_1288046 | 4 | 0.25 | 1 | 0 | 4 | 0.5 | 4 | 0.25 | 13 | 0.3077 | 2 | 0.5 | 7 | 0.2857 | 7 | 0.5714 | 2 | 0 | 18 | 0.3889 |
| 4_1288047 | 3 | 1 | NA | NA | 4 | 1 | 4 | 1 | 11 | 1 | 1 | 1 | 7 | 1 | 7 | 1 | 4 | 1 | 19 | 1 |
| 4_1288047 | 5 | 1 | 1 | 1 | 4 | 0.5 | 3 | 0.3333 | 13 | 0.6923 | 2 | 0.5 | 7 | 0.8571 | 7 | 0.7143 | 4 | 0.75 | 20 | 0.75 |
| 4_1288048 | 5 | 0.6 | 1 | 0 | 3 | 0.3333 | 4 | 0.75 | 13 | 0.5385 | 1 | 0 | 7 | 0.4286 | 7 | 0.5714 | 3 | 0.3333 | 18 | 0.4444 |
| 4_1288048 | 4 | 0.75 | NA | NA | 4 | 0.75 | 4 | 0.75 | 12 | 0.75 | 1 | 0 | 5 | 1 | 6 | 0.3333 | 3 | 1 | 15 | 0.6667 |
| 4_1288049 | 6 | 1 | NA | NA | 4 | 1 | 5 | 1 | 15 | 1 | 1 | 1 | 6 | 1 | 8 | 1 | 3 | 1 | 18 | 1 |
| 4_1288241 | NA | NA | NA | NA | 1 | 1 | 1 | 1 | 2 | 1 | NA | NA | NA | NA | 1 | 1 | NA | NA | 1 | 1 |
| 4_1288242 | NA | NA | NA | NA | 1 | 1 | 1 | 1 | 2 | 1 | NA | NA | NA | NA | 1 | 1 | NA | NA | 1 | 1 |
| 4_1288259 | 2 | 0.5 | NA | NA | 7 | 0 | 2 | 0 | 11 | 0.0909 | 5 | 0.2 | 3 | 0 | 3 | 0 | 3 | 0 | 14 | 0.0714 |
| 4_1288260 | 2 | 1 | 1 | 0 | 5 | 0.8 | 3 | 0.6667 | 11 | 0.7273 | 5 | 0.6 | 3 | 0.6667 | 3 | 0.6667 | 2 | 0.5 | 13 | 0.6154 |
| 4_1289020 | 2 | 0 | 3 | 0 | 4 | 0.25 | NA | NA | 9 | 0.1111 | NA | NA | 3 | 0 | 3 | 0.3333 | NA | NA | 6 | 0.1667 |
| 4_1289047 | 2 | 0.5 | 5 | 0 | 1 | 0 | 1 | 0 | 9 | 0.1111 | 2 | 0.5 | 2 | 1 | 6 | 0.1667 | 2 | 0 | 12 | 0.3333 |
| 4_1289049 | 2 | 0.5 | 5 | 0.4 | 1 | 0 | 1 | 0 | 9 | 0.3333 | 2 | 0.5 | 1 | 1 | 5 | 1 | 1 | 0 | 9 | 0.7778 |
| 4_1289050 | 2 | 0.5 | 5 | 0.2 | 1 | 0 | 1 | 0 | 9 | 0.2222 | 2 | 0.5 | 1 | 1 | 6 | 0.3333 | 1 | 0 | 10 | 0.4 |
| 4_1289052 | 2 | 0 | 4 | 0.5 | 1 | 0 | 2 | 0.5 | 9 | 0.3333 | 2 | 0 | 1 | 1 | 6 | 0.3333 | 1 | 0 | 10 | 0.3 |
| 4_1289520 | NA | NA | 2 | 0.5 | 3 | 0.6667 | NA | NA | 5 | 0.6 | NA | NA | 2 | 0.5 | 1 | 0 | 1 | 1 | 4 | 0.5 |
| 4_1289776 | 2 | 1 | 6 | 0.3333 | 8 | 0.25 | NA | NA | 16 | 0.375 | 2 | 0 | 9 | 0.3333 | 3 | 0.3333 | 7 | 0.1429 | 21 | 0.2381 |
| 4_1289777 | 1 | 1 | 3 | 0 | 4 | 0.25 | NA | NA | 8 | 0.25 | 1 | 0 | 4 | 0.25 | 2 | 0 | 4 | 0 | 11 | 0.0909 |
| 4_1289777 | 2 | 0.5 | 7 | 1 | 7 | 0.8571 | 1 | 1 | 17 | 0.8824 | 2 | 1 | 8 | 1 | 3 | 1 | 6 | 0.6667 | 19 | 0.8947 |
| 4_1289777 | 2 | 0.5 | 6 | 0.1667 | 7 | 0.1429 | 1 | 0 | 16 | 0.1875 | 3 | 0.3333 | 10 | 0.3 | 3 | 0 | 5 | 0.8 | 21 | 0.381 |
| 4_1289777 | 2 | 0 | 6 | 0.1667 | 6 | 0 | 1 | 0 | 15 | 0.0667 | 3 | 0.6667 | 9 | 0.1111 | 3 | 0 | 9 | 0.1111 | 24 | 0.1667 |
| 4_1289779 | 2 | 0.5 | 4 | 0.5 | 5 | 0.8 | 1 | 0 | 12 | 0.5833 | 2 | 0.5 | 9 | 0.6667 | 4 | 0 | 5 | 0.4 | 20 | 0.45 |
| 4_1289780 | 1 | 0 | 7 | 0 | 7 | 0.1429 | 1 | 0 | 16 | 0.0625 | 3 | 0 | 8 | 0.25 | 1 | 0 | 7 | 0 | 19 | 0.1053 |
| 4_1290057 | 118 | 0.1102 | 87 | 0.1034 | 128 | 0.0312 | 119 | 0.1008 | 452 | 0.0841 | 79 | 0.0506 | 165 | 0.1212 | 198 | 0.0909 | 109 | 0.0734 | 551 | 0.0907 |
| 4_1290061 | 132 | 0.1364 | 91 | 0.1429 | 121 | 0.124 | 130 | 0.1231 | 474 | 0.1308 | 90 | 0.1556 | 173 | 0.1503 | 197 | 0.1269 | 106 | 0.1415 | 566 | 0.1413 |
| 4_1290124 | 88 | 0.5568 | 55 | 0.5091 | 78 | 0.6282 | 93 | 0.5806 | 314 | 0.5732 | 61 | 0.5246 | 133 | 0.5564 | 147 | 0.5986 | 79 | 0.6835 | 420 | 0.5905 |
| 4_1290125 | 93 | 0.172 | 58 | 0.1379 | 76 | 0.2632 | 97 | 0.1443 | 324 | 0.179 | 57 | 0.2281 | 127 | 0.189 | 130 | 0.2462 | 69 | 0.2609 | 383 | 0.2272 |
| 4_1290365 | 8 | 0.25 | 6 | 0.3333 | 7 | 0.5714 | 10 | 0.2 | 31 | 0.3226 | 8 | 0.5 | 5 | 0.4 | 8 | 0.375 | 11 | 0.6364 | 32 | 0.5 |
| 4_1290366 | 5 | 0.2 | 6 | 0.6667 | 7 | 0.4286 | 10 | 0.6 | 28 | 0.5 | 7 | 0.5714 | 3 | 0 | 8 | 0.25 | 6 | 0.3333 | 24 | 0.3333 |
| 4_1290366 | 9 | 0.8889 | 5 | 1 | 8 | 1 | 12 | 1 | 34 | 0.9706 | 7 | 1 | 4 | 1 | 8 | 1 | 10 | 1 | 29 | 1 |
| 4_1290367 | 8 | 0.625 | 4 | 1 | 7 | 0.4286 | 12 | 0.5 | 31 | 0.5806 | 8 | 0.125 | 4 | 0 | 6 | 0.3333 | 9 | 0.4444 | 27 | 0.2593 |
| 4_1290367 | 9 | 0.2222 | 5 | 0.2 | 7 | 0.1429 | 11 | 0.4545 | 32 | 0.2812 | 8 | 0.75 | 5 | 0.6 | 7 | 0.1429 | 8 | 0.25 | 28 | 0.4286 |
| 4_1290368 | 8 | 0.875 | 4 | 1 | 7 | 1 | 11 | 0.9091 | 30 | 0.9333 | 8 | 1 | 4 | 1 | 5 | 0.8 | 10 | 1 | 27 | 0.963 |

| | | | | | | | | | | | | | | | | | | | | |
|---|---|---|---|---|---|---|---|---|---|---|---|---|---|---|---|---|---|---|---|---|
| 4_1290369 | 6 | 0.1667 | 6 | 0.1667 | 7 | 0 | 8 | 0.375 | 27 | 0.1852 | 7 | 0.5714 | 5 | 0 | 8 | 0 | 6 | 0.1667 | 26 | 0.1923 |
| 4_1290370 | 8 | 0.25 | 5 | 0 | 6 | 0.1667 | 9 | 0.2222 | 28 | 0.1786 | 9 | 0.3333 | 5 | 0.8 | 5 | 0 | 10 | 0.2 | 29 | 0.3103 |
| 4_1290371 | 10 | 0.7 | 7 | 0.5714 | 10 | 0.7 | 12 | 0.8333 | 39 | 0.7179 | 10 | 0.9 | 7 | 1 | 8 | 0.75 | 10 | 0.8 | 35 | 0.8571 |
| 4_1290372 | 8 | 1 | 6 | 0.8333 | 10 | 0.9 | 10 | 0.9 | 34 | 0.9118 | 8 | 1 | 7 | 0.8571 | 7 | 1 | 11 | 1 | 33 | 0.9697 |
| 4_1290372 | 8 | 0.125 | 5 | 0.2 | 9 | 0.3333 | 11 | 0.3636 | 33 | 0.2727 | 9 | 0.4444 | 8 | 0.125 | 8 | 0.25 | 10 | 0.3 | 35 | 0.2857 |
| 4_1290372 | 9 | 1 | 6 | 0.8333 | 7 | 1 | 11 | 0.8182 | 33 | 0.9091 | 9 | 0.4444 | 7 | 1 | 8 | 0.75 | 12 | 1 | 36 | 0.8056 |
| 4_1290373 | 10 | 0.9 | 6 | 0.8333 | 10 | 0.9 | 12 | 0.9167 | 38 | 0.8947 | 9 | 0.7778 | 7 | 1 | 10 | 0.8 | 13 | 0.7692 | 39 | 0.8205 |
| 4_1290374 | 8 | 1 | 7 | 1 | 10 | 0.9 | 12 | 1 | 37 | 0.973 | 9 | 1 | 9 | 1 | 10 | 1 | 12 | 1 | 40 | 1 |
| 4_1290374 | 9 | 0 | 7 | 0 | 10 | 0 | 12 | 0.1667 | 38 | 0.0526 | 8 | 0.125 | 7 | 0 | 10 | 0 | 12 | 0.0833 | 37 | 0.0541 |
| 4_1290376 | 9 | 0 | 6 | 0 | 10 | 0.3 | 11 | 0 | 36 | 0.0833 | 10 | 0.3 | 8 | 0 | 7 | 0 | 10 | 0.2 | 35 | 0.1429 |
| 4_1290376 | 9 | 0.8889 | 6 | 0.8333 | 9 | 0.7778 | 11 | 1 | 35 | 0.8857 | 9 | 0.8889 | 8 | 0.75 | 10 | 1 | 9 | 1 | 36 | 0.9167 |
| 4_1290378 | 10 | 0.2 | 8 | 0.125 | 14 | 0.1429 | 13 | 0.1538 | 45 | 0.1556 | 13 | 0.2308 | 9 | 0.3333 | 12 | 0.4167 | 14 | 0.2143 | 48 | 0.2917 |
| 4_1290379 | 8 | 0.25 | 6 | 0 | 11 | 0.3636 | 13 | 0.0769 | 38 | 0.1842 | 14 | 0.1429 | 11 | 0.1818 | 12 | 0.3333 | 15 | 0.3333 | 52 | 0.25 |
| 4_1290379 | 8 | 0.75 | 8 | 0.375 | 8 | 1 | 9 | 0.6667 | 33 | 0.697 | 8 | 0.75 | 8 | 0.5 | 11 | 0.3636 | 12 | 0.5833 | 39 | 0.5385 |
| 4_1290379 | 8 | 0.875 | 8 | 0.75 | 12 | 1 | 13 | 1 | 41 | 0.9268 | 11 | 0.9091 | 11 | 0.9091 | 9 | 1 | 14 | 1 | 45 | 0.9556 |
| 4_1290381 | 9 | 0.4444 | 8 | 0.625 | 12 | 0.6667 | 9 | 0.5556 | 38 | 0.5789 | 11 | 0.5455 | 10 | 0.6 | 10 | 0.6 | 11 | 0.4545 | 42 | 0.5476 |
| 4_1290383 | 9 | 0 | 10 | 0 | 16 | 0.1875 | 15 | 0.1333 | 50 | 0.1 | 13 | 0 | 14 | 0.0714 | 14 | 0 | 15 | 0.2 | 56 | 0.0714 |
| 4_1290431 | 6 | 0.1667 | 7 | 0 | 3 | 0 | 9 | 0.1111 | 25 | 0.08 | 3 | 0 | 6 | 0.3333 | 5 | 0 | 7 | 0.4286 | 21 | 0.2381 |
| 4_1290756 | 2 | 0.5 | 3 | 0 | 1 | 0 | 2 | 0.5 | 8 | 0.25 | 2 | 0 | NA | NA | 2 | 0.5 | 3 | 0 | 7 | 0.1429 |
| 4_1290758 | 4 | 0.25 | 4 | 0.25 | 2 | 0.5 | 4 | 0 | 14 | 0.2143 | 2 | 0 | NA | NA | 2 | 0 | 3 | 0.3333 | 7 | 0.1429 |
| 4_1290759 | 4 | 0.75 | 4 | 0.75 | 2 | 0.5 | 2 | 0.5 | 12 | 0.6667 | 1 | 1 | NA | NA | 1 | 1 | 2 | 0.5 | 4 | 0.75 |
| 4_1290759 | 4 | 0.5 | 4 | 0.5 | 2 | 0.5 | 4 | 0.75 | 14 | 0.5714 | 2 | 0.5 | NA | NA | 2 | 0.5 | 2 | 1 | 6 | 0.6667 |
| 4_1290759 | 4 | 1 | 4 | 1 | 2 | 1 | 4 | 1 | 14 | 1 | 2 | 1 | NA | NA | 2 | 0.5 | 3 | 1 | 7 | 0.8571 |
| 4_1290760 | 4 | 0.25 | 4 | 0 | 2 | 0 | 4 | 0 | 14 | 0.0714 | 2 | 0 | NA | NA | 2 | 0.5 | 2 | 0.5 | 6 | 0.3333 |
| 4_1290761 | 4 | 0 | 3 | 0 | 2 | 0.5 | 3 | 0.3333 | 12 | 0.1667 | 2 | 0 | NA | NA | 1 | 0 | 3 | 0.3333 | 6 | 0.1667 |
| 4_1291124 | 13 | 0.5385 | 11 | 0.5455 | 16 | 0.25 | 9 | 0.2222 | 49 | 0.3878 | 11 | 0.4545 | 15 | 0.5333 | 8 | 0.375 | 11 | 0.4545 | 45 | 0.4667 |
| 4_1291124 | 15 | 0.6667 | 14 | 0.5714 | 20 | 0.85 | 11 | 0.7273 | 60 | 0.7167 | 11 | 0.7273 | 18 | 0.6111 | 9 | 0.5556 | 12 | 0.4167 | 50 | 0.58 |
| 4_1291124 | 14 | 0.4286 | 10 | 0.5 | 15 | 0.6667 | 10 | 0.5 | 49 | 0.5306 | 10 | 0.7 | 16 | 0.625 | 6 | 0.3333 | 12 | 0.5 | 44 | 0.5682 |
| 4_1291125 | 15 | 0.2 | 16 | 0.1875 | 19 | 0.1053 | 10 | 0.1 | 60 | 0.15 | 11 | 0.2727 | 20 | 0.15 | 8 | 0.125 | 13 | 0.4615 | 52 | 0.25 |
| 4_1291125 | 15 | 0 | 16 | 0.25 | 21 | 0.0952 | 10 | 0.3 | 62 | 0.1452 | 10 | 0.3 | 16 | 0.0625 | 8 | 0.25 | 14 | 0.2143 | 48 | 0.1875 |
| 4_1291125 | 15 | 0.2 | 16 | 0.125 | 20 | 0.05 | 9 | 0 | 60 | 0.1 | 12 | 0 | 18 | 0.0556 | 9 | 0.2222 | 9 | 0.1111 | 48 | 0.0833 |
| 4_1291126 | 13 | 0.2308 | 14 | 0.0714 | 17 | 0.0588 | 10 | 0.4 | 54 | 0.1667 | 10 | 0.3 | 15 | 0.1333 | 8 | 0.125 | 12 | 0 | 45 | 0.1333 |
| 4_1291128 | 17 | 0.3529 | 17 | 0.4706 | 20 | 0.4 | 12 | 0.5 | 66 | 0.4242 | 12 | 0.25 | 17 | 0.3529 | 10 | 0.4 | 16 | 0.5 | 55 | 0.3818 |
| 4_1291128 | 15 | 0.8667 | 16 | 0.625 | 16 | 0.6875 | 8 | 0.875 | 55 | 0.7455 | 12 | 0.8333 | 19 | 0.8947 | 8 | 0.75 | 17 | 0.8824 | 56 | 0.8571 |
| 4_1291129 | 12 | 1 | 13 | 1 | 15 | 0.9333 | 11 | 1 | 51 | 0.9804 | 13 | 0.8462 | 16 | 1 | 8 | 0.625 | 13 | 0.8462 | 50 | 0.86 |
| 4_1291129 | 16 | 0.375 | 16 | 0.1875 | 18 | 0.1667 | 13 | 0.0769 | 63 | 0.2063 | 14 | 0.0714 | 21 | 0.1429 | 9 | 0 | 16 | 0.0625 | 60 | 0.0833 |
| 4_1291145 | NA | NA | 1 | 1 | 1 | 0 | 2 | 0.5 | 4 | 0.5 | NA | NA | 2 | 0.5 | 1 | 1 | NA | NA | 3 | 0.6667 |
| 4_1291151 | NA | NA | 1 | 1 | NA | NA | 2 | 0 | 3 | 0.3333 | NA | NA | 2 | 0.5 | NA | NA | NA | NA | 2 | 0.5 |
| 4_1291156 | NA | NA | 1 | 0 | 1 | 1 | 1 | 1 | 3 | 0.6667 | NA | NA | 2 | 1 | NA | NA | 1 | 1 | 3 | 1 |
| 4_1291160 | NA | NA | 1 | 1 | NA | NA | 1 | 0 | 2 | 0.5 | NA | NA | 2 | 0.5 | NA | NA | 1 | 0 | 3 | 0.3333 |
| 4_1291162 | NA | NA | 1 | 1 | NA | NA | 1 | 1 | 2 | 1 | NA | NA | 2 | 1 | NA | NA | NA | NA | 2 | 1 |
| 4_1291170 | NA | NA | 1 | 1 | 1 | 1 | 1 | 1 | 3 | 1 | NA | NA | NA | NA | NA | NA | 1 | 1 | 1 | 1 |
| 4_1291173 | NA | NA | 1 | 1 | NA | NA | 1 | 0 | 2 | 0.5 | NA | NA | 1 | 1 | NA | NA | 1 | 0 | 2 | 0.5 |
| 4_1292038 | 22 | 0 | 20 | 0.25 | 21 | 0.2381 | 25 | 0.04 | 88 | 0.125 | 25 | 0 | 25 | 0.12 | 22 | 0.0909 | 27 | 0.0741 | 99 | 0.0707 |
| 4_1292407 | 4 | 0.75 | NA | NA | 5 | 0.6 | 2 | 0.5 | 11 | 0.6364 | NA | NA | 2 | 0.5 | 4 | 1 | 1 | 1 | 7 | 0.8571 |
| 4_1292478 | NA | NA | NA | NA | 2 | 1 | 2 | 0.5 | 4 | 0.75 | 1 | 1 | 1 | 0 | NA | NA | NA | NA | 2 | 0.5 |
| 4_1292481 | NA | NA | NA | NA | 1 | 1 | 2 | 1 | 3 | 1 | 1 | 0 | 1 | 1 | NA | NA | NA | NA | 2 | 0.5 |
| 4_1292541 | NA | NA | NA | NA | 1 | 1 | NA | NA | 1 | 1 | NA | NA | 1 | 1 | NA | NA | NA | NA | 1 | 1 |
| 4_1292542 | NA | NA | NA | NA | 1 | 1 | 1 | 1 | 2 | 1 | NA | NA | 1 | 1 | NA | NA | NA | NA | 1 | 1 |
| 4_1292542 | NA | NA | NA | NA | 1 | 1 | 1 | 1 | 2 | 1 | NA | NA | 1 | 1 | NA | NA | NA | NA | 1 | 1 |
| 4_1293208 | 1 | 1 | 1 | 0 | NA | NA | NA | NA | 2 | 0.5 | NA | NA | NA | NA | 1 | 1 | NA | NA | 1 | 1 |
| 4_1293210 | 1 | 1 | 2 | 1 | NA | NA | NA | NA | 3 | 1 | NA | NA | 1 | 1 | 1 | 0 | NA | NA | 2 | 0.5 |
| 4_1293210 | 1 | 0 | 2 | 0.5 | NA | NA | NA | NA | 3 | 0.3333 | NA | NA | 1 | 1 | 1 | 1 | NA | NA | 2 | 1 |
| 4_1293210 | 1 | 0 | 1 | 1 | NA | NA | NA | NA | 2 | 0.5 | NA | NA | 1 | 1 | 1 | 1 | NA | NA | 2 | 1 |
| 4_1293211 | 1 | 1 | 2 | 1 | NA | NA | NA | NA | 3 | 1 | NA | NA | 1 | 1 | 1 | 1 | NA | NA | 2 | 1 |
| 4_1293211 | 1 | 1 | 2 | 0.5 | NA | NA | NA | NA | 3 | 0.6667 | NA | NA | 1 | 1 | 1 | 1 | NA | NA | 2 | 1 |
| 4_1293212 | 1 | 1 | 1 | 1 | NA | NA | NA | NA | 2 | 1 | NA | NA | 1 | 1 | 1 | 1 | NA | NA | 2 | 1 |
| 4_1293213 | 1 | 0 | 1 | 1 | NA | NA | NA | NA | 2 | 0.5 | NA | NA | 1 | 1 | 1 | 1 | NA | NA | 2 | 1 |
| 4_1293213 | 1 | 1 | 1 | 1 | NA | NA | NA | NA | 2 | 1 | NA | NA | 1 | 1 | NA | NA | NA | NA | 1 | 1 |
| 4_1293263 | 1 | 1 | NA | NA | NA | NA | NA | NA | 1 | 1 | NA | NA | 1 | 1 | 1 | 1 | NA | NA | 2 | 1 |
| 4_1293264 | 1 | 1 | 1 | 0 | NA | NA | NA | NA | 2 | 0.5 | NA | NA | 1 | 1 | 1 | 1 | NA | NA | 2 | 1 |
| 4_1293265 | NA | NA | 1 | 1 | NA | NA | NA | NA | 1 | 1 | NA | NA | 1 | 1 | NA | NA | NA | NA | 1 | 1 |
| 4_1293266 | NA | NA | 1 | 1 | NA | NA | NA | NA | 1 | 1 | NA | NA | 1 | 1 | NA | NA | NA | NA | 1 | 1 |
| 4_1293677 | 8 | 0 | 12 | 0.25 | 4 | 0.25 | 6 | 0.1667 | 30 | 0.1667 | 4 | 0.5 | 2 | 0 | 7 | 0.1429 | 11 | 0.0909 | 24 | 0.1667 |
| 4_1293678 | 6 | 0.5 | 6 | 0 | 4 | 0 | 5 | 0.4 | 21 | 0.2381 | 3 | 0.3333 | 2 | 0 | 5 | 0 | 10 | 0.2 | 20 | 0.15 |

| | | | | | | | | | | | | | | | | | | | | |
|---|---|---|---|---|---|---|---|---|---|---|---|---|---|---|---|---|---|---|---|---|
| 4_12936794 | 7 | 0.1429 | 12 | 0.1667 | 4 | 0 | 6 | 0.1667 | 29 | 0.1379 | 4 | 0 | 3 | 0 | 6 | 0 | 11 | 0.2727 | 24 | 0.125 |
| 4_12936797 | 8 | 0.125 | 10 | 0.1 | 4 | 0.5 | 6 | 0.3333 | 28 | 0.2143 | 4 | 0.25 | 3 | 0 | 6 | 0.1667 | 10 | 0.2 | 23 | 0.1739 |
| 4_12938093 | 8 | 0.375 | 12 | 0.1667 | 4 | 0.25 | 7 | 0.5714 | 31 | 0.3226 | 4 | 0.5 | 4 | 0 | 6 | 0.1667 | 12 | 0.0833 | 26 | 0.1538 |
| 4_12938098 | 9 | 0.5556 | 11 | 0.6364 | 3 | 0.6667 | 7 | 0.5714 | 30 | 0.6 | 4 | 0.5 | 4 | 1 | 7 | 0.5714 | 9 | 0.5556 | 24 | 0.625 |
| 4_12938110 | 10 | 0.3 | 12 | 0.4167 | 4 | 0.5 | 7 | 0.1429 | 33 | 0.3333 | 4 | 0.5 | 2 | 0.5 | 7 | 0.4286 | 11 | 0.4545 | 24 | 0.4583 |
| 4_12938119 | 9 | 0.3333 | 12 | 0.3333 | 4 | 0 | 8 | 0 | 33 | 0.2121 | 4 | 0 | 4 | 0 | 7 | 0.2857 | 11 | 0.1818 | 26 | 0.1538 |
| 4_12938128 | 9 | 0.1111 | 9 | 0.3333 | 4 | 0 | 6 | 0 | 28 | 0.1429 | 4 | 0 | 4 | 0.25 | 5 | 0.4 | 11 | 0.1818 | 24 | 0.2083 |
| 4_12938129 | 8 | 0.5 | 9 | 0.4444 | 3 | 0.3333 | 4 | 0.5 | 24 | 0.4583 | 4 | 0 | 4 | 0.25 | 4 | 0 | 9 | 0.2222 | 21 | 0.1429 |
| 4_12938130 | 7 | 0.7143 | 9 | 0.7778 | 4 | 0.75 | 4 | 0.75 | 24 | 0.75 | 4 | 1 | 2 | 1 | 5 | 1 | 8 | 0.375 | 19 | 0.7368 |
| 4_12938136 | 10 | 0.9 | 11 | 0.7273 | 2 | 1 | 8 | 1 | 31 | 0.871 | 4 | 1 | 3 | 1 | 7 | 0.8571 | 10 | 1 | 24 | 0.9583 |
| 4_12938145 | 7 | 0.1429 | 12 | 0 | 4 | 0 | 9 | 0.2222 | 32 | 0.0938 | 2 | 0 | 4 | 0.25 | 7 | 0.1429 | 10 | 0.1 | 23 | 0.1304 |
| 4_12938147 | 9 | 0.2222 | 8 | 0.25 | 1 | 0 | 7 | 0.2857 | 25 | 0.24 | 3 | 0.3333 | 3 | 0.3333 | 5 | 0.2 | 9 | 0.4444 | 20 | 0.35 |
| 4_12938168 | 9 | 0.4444 | 13 | 0.3077 | 3 | 0 | 9 | 0.3333 | 34 | 0.3235 | 4 | 0.25 | 4 | 0.75 | 7 | 0.5714 | 10 | 0.4 | 25 | 0.48 |
| 4_12938181 | 10 | 0 | 13 | 0.0769 | 4 | 0 | 9 | 0.3333 | 36 | 0.1111 | 5 | 0 | 4 | 0.5 | 6 | 0.5 | 11 | 0.3636 | 26 | 0.3462 |
| 4_12938184 | 9 | 0.3333 | 10 | 0.3 | 3 | 0.6667 | 6 | 0.5 | 28 | 0.3929 | 4 | 0 | 3 | 0.6667 | 6 | 0.1667 | 10 | 0.3 | 23 | 0.2609 |
| 4_12938187 | 8 | 0 | 11 | 0.1818 | 4 | 0.25 | 9 | 0 | 32 | 0.0938 | 6 | 0.3333 | 3 | 0 | 6 | 0.3333 | 7 | 0.1429 | 22 | 0.2273 |
| 4_12938195 | 10 | 0.3 | 10 | 0.7 | 3 | 0.6667 | 8 | 0.375 | 31 | 0.4839 | 4 | 1 | 4 | 0.75 | 5 | 1 | 10 | 0.5 | 23 | 0.7391 |
| 4_12940903 | 10 | 0.1 | 29 | 0.1724 | 18 | 0.0556 | 7 | 0 | 64 | 0.1094 | 23 | 0 | 10 | 0.1 | 5 | 0.4 | 8 | 0.25 | 46 | 0.1087 |
| 4_12940906 | 8 | 0.5 | 29 | 0.7586 | 16 | 0.5625 | 8 | 0.875 | 61 | 0.6885 | 19 | 0.5263 | 13 | 0.8462 | 6 | 0.6667 | 10 | 0.7 | 48 | 0.6667 |
| 4_12940909 | 7 | 0.2857 | 28 | 0.2143 | 15 | 0.2 | 7 | 0.2857 | 57 | 0.2281 | 18 | 0.1667 | 9 | 0 | 4 | 0.25 | 10 | 0.2 | 41 | 0.1463 |
| 4_12940948 | 9 | 0.1111 | 32 | 0.1875 | 16 | 0.1875 | 6 | 0.3333 | 63 | 0.1905 | 23 | 0.1739 | 13 | 0.1538 | 4 | 0 | 10 | 0.1 | 50 | 0.14 |
| 4_12941676 | 1 | 0 | 3 | 0 | 2 | 0 | 2 | 0.5 | 8 | 0.125 | 3 | 0.3333 | 1 | 0 | 3 | 0 | 2 | 0 | 9 | 0.1111 |
| 4_12944249 | NA | NA | NA | NA | NA | NA | 3 | 0.6667 | 3 | 0.6667 | 1 | 1 | NA | NA | 1 | 0 | NA | NA | 2 | 0.5 |
| 4_12944905 | NA | NA | NA | NA | NA | NA | 1 | 1 | 1 | 1 | 1 | 1 | NA | NA | NA | NA | NA | NA | 1 | 1 |
| 4_12944923 | NA | NA | NA | NA | NA | NA | 1 | 1 | 1 | 1 | 1 | 1 | NA | NA | NA | NA | NA | NA | 1 | 1 |
| 4_12944936 | NA | NA | NA | NA | NA | NA | 2 | 0.5 | 2 | 0.5 | 2 | 0.5 | NA | NA | 1 | 0 | NA | NA | 3 | 0.3333 |
| 4_12948579 | 17 | 0.0588 | 21 | 0.0476 | 15 | 0 | 20 | 0.15 | 73 | 0.0685 | 18 | 0.0556 | 18 | 0.0556 | 22 | 0.2273 | 24 | 0.125 | 82 | 0.122 |
| 4_12948915 | 19 | 0.0526 | 23 | 0.0435 | 12 | 0.0833 | 17 | 0.0588 | 71 | 0.0563 | 15 | 0 | 17 | 0 | 14 | 0.0714 | 18 | 0.1667 | 64 | 0.0625 |
| 4_12950194 | 10 | 0.7 | 15 | 0.8 | 12 | 0.6667 | 12 | 0.8333 | 49 | 0.7551 | 11 | 0.7273 | 11 | 0.5455 | 11 | 0.5455 | 13 | 0.6923 | 46 | 0.6304 |
| 4_12950203 | 9 | 0.3333 | 18 | 0.7222 | 13 | 0.3846 | 12 | 0.5833 | 52 | 0.5385 | 13 | 0.5385 | 9 | 0.4444 | 10 | 0.7 | 13 | 0.6154 | 45 | 0.5778 |
| 4_12950216 | 7 | 0.4286 | 16 | 0.4375 | 11 | 0.2727 | 12 | 0.4167 | 46 | 0.3913 | 14 | 0.3571 | 10 | 0.2 | 15 | 0.4 | 12 | 0.1667 | 51 | 0.2941 |
| 4_12950224 | 9 | 0 | 18 | 0.1111 | 14 | 0.1429 | 15 | 0 | 56 | 0.0714 | 12 | 0.1667 | 11 | 0 | 14 | 0 | 12 | 0.0833 | 49 | 0.0612 |
| 4_12950225 | 6 | 0.1667 | 9 | 0 | 10 | 0.1 | 6 | 0.1667 | 31 | 0.0968 | 9 | 0.3333 | 7 | 0.2857 | 10 | 0.4 | 11 | 0.1818 | 37 | 0.2973 |
| 4_12950956 | NA | NA | 3 | 0.6667 | 1 | 0 | NA | NA | 4 | 0.5 | NA | NA | NA | NA | 1 | 1 | 3 | 0 | 4 | 0.25 |
| 4_12950960 | NA | NA | 3 | 0.6667 | 1 | 0 | NA | NA | 4 | 0.5 | NA | NA | NA | NA | 1 | 0 | 3 | 1 | 4 | 0.75 |
| 4_12950969 | NA | NA | 3 | 1 | NA | NA | NA | NA | 3 | 1 | NA | NA | 1 | 1 | 1 | 1 | 2 | 0.5 | 4 | 0.75 |
| 4_12950978 | NA | NA | 3 | 0.6667 | 1 | 0 | NA | NA | 4 | 0.5 | NA | NA | 1 | 0 | 1 | 0 | 3 | 1 | 5 | 0.6 |
| 4_12950997 | NA | NA | 3 | 1 | 1 | 0 | NA | NA | 4 | 0.75 | NA | NA | 1 | 1 | NA | NA | 3 | 1 | 4 | 1 |
| 4_12951048 | NA | NA | 3 | 0.3333 | NA | NA | NA | NA | 3 | 0.3333 | NA | NA | 1 | 1 | NA | NA | 3 | 0 | 4 | 0.25 |
| 4_12951070 | NA | NA | 2 | 0.5 | NA | NA | NA | NA | 2 | 0.5 | NA | NA | 1 | 0 | NA | NA | 3 | 0.3333 | 4 | 0.25 |
| 4_12951444 | 1 | 1 | NA | NA | NA | NA | NA | NA | 1 | 1 | NA | NA | 2 | 0 | 1 | 1 | NA | NA | 3 | 0.3333 |
| 4_12951482 | 4 | 0.25 | 2 | 0 | 4 | 0 | 3 | 0 | 13 | 0.0769 | 3 | 0.3333 | 8 | 0 | 4 | 0 | 4 | 0 | 19 | 0.0526 |
| 4_12951643 | 2 | 0 | 2 | 0 | 4 | 0.25 | 2 | 0 | 10 | 0.1 | 4 | 0 | 8 | 0 | 4 | 0.25 | 5 | 0.4 | 21 | 0.1429 |
| 4_12951666 | NA | NA | 1 | 1 | NA | NA | 1 | 1 | 2 | 1 | NA | NA | NA | NA | 1 | 0 | 2 | 1 | 3 | 0.6667 |
| 4_12951668 | NA | NA | NA | NA | 1 | 1 | NA | NA | 1 | 1 | NA | NA | NA | NA | 1 | 0 | 2 | 0.5 | 3 | 0.3333 |
| 4_12951715 | NA | NA | NA | NA | 1 | 1 | NA | NA | 1 | 1 | NA | NA | NA | NA | 1 | 0 | 2 | 0.5 | 3 | 0.3333 |
| 4_12951991 | 4 | 0.25 | 4 | 0 | 7 | 0 | 3 | 0.3333 | 18 | 0.1111 | 5 | 0 | 2 | 0 | 4 | 0 | 2 | 0.5 | 13 | 0.0769 |
| 4_12952132 | 3 | 0 | 5 | 0.2 | 3 | 0 | 2 | 0 | 13 | 0.0769 | 3 | 0.3333 | 2 | 0.5 | 5 | 0.2 | 5 | 0.2 | 15 | 0.2667 |
| 4_12952151 | 3 | 0 | 6 | 0.1667 | 5 | 0 | 2 | 0 | 16 | 0.0625 | 3 | 0 | 3 | 0.3333 | 4 | 0 | 5 | 0 | 15 | 0.0667 |
| 4_12952161 | 4 | 0.5 | 6 | 0.5 | 3 | 0.6667 | 3 | 0.3333 | 16 | 0.5 | 2 | 0 | 3 | 0.6667 | 6 | 0.1667 | 4 | 0.5 | 15 | 0.3333 |
| 4_12952225 | 4 | 0.25 | 6 | 0.6667 | 4 | 0.25 | 4 | 1 | 18 | 0.5556 | 1 | 0 | 2 | 0.5 | 6 | 0.6667 | 4 | 0.75 | 13 | 0.6154 |
| 4_12952227 | 3 | 1 | 7 | 0.8571 | 4 | 0.75 | 4 | 0.5 | 18 | 0.7778 | 4 | 0.75 | 2 | 1 | 8 | 0.75 | 5 | 0.8 | 19 | 0.7895 |
| 4_12952236 | 4 | 0.25 | 6 | 0 | 5 | 0.2 | 4 | 0 | 19 | 0.1053 | 4 | 0.25 | 2 | 0 | 8 | 0 | 5 | 0 | 19 | 0.0526 |
| 4_12952243 | 4 | 0.5 | 7 | 0.1429 | 5 | 0.2 | 4 | 0.5 | 20 | 0.3 | 4 | 0 | 3 | 0.3333 | 8 | 0.125 | 4 | 0.25 | 19 | 0.1579 |
| 4_12952245 | 3 | 1 | 6 | 0.8333 | 4 | 1 | 2 | 1 | 15 | 0.9333 | 1 | 1 | 1 | 1 | 4 | 1 | 4 | 1 | 10 | 1 |
| 4_12952257 | 2 | 1 | 7 | 1 | 4 | 1 | 4 | 1 | 17 | 1 | 3 | 0.6667 | 2 | 1 | 7 | 0.5714 | 4 | 1 | 16 | 0.75 |
| 4_12952264 | 4 | 0.25 | 7 | 0.4286 | 4 | 0.5 | 4 | 0 | 19 | 0.3158 | 3 | 0.3333 | 1 | 1 | 8 | 0.375 | 4 | 0.25 | 16 | 0.375 |
| 4_12952272 | 3 | 0 | 6 | 0.1667 | 5 | 0 | 3 | 0 | 17 | 0.0588 | 2 | 0 | 2 | 0 | 6 | 0 | 5 | 0.4 | 15 | 0.1333 |
| 4_12952280 | 2 | 0.5 | 6 | 0.3333 | 3 | 0.3333 | 3 | 0 | 14 | 0.2857 | 2 | 0.5 | 2 | 0 | 8 | 0.5 | 4 | 0.5 | 16 | 0.4375 |
| 4_12952285 | 2 | 0.5 | 6 | 0.3333 | 3 | 0.6667 | 4 | 0.5 | 15 | 0.4667 | 2 | 0.5 | 1 | 0 | 5 | 0.2 | 5 | 0.4 | 13 | 0.3077 |
| 4_12952287 | 2 | 0.5 | 4 | 0.25 | 5 | 0.2 | 2 | 0 | 13 | 0.2308 | 2 | 0.5 | NA | NA | 4 | 0.25 | 5 | 0.2 | 11 | 0.2727 |
| 4_12952287 | 4 | 0.5 | 7 | 0.4286 | 4 | 0.5 | 3 | 0.6667 | 18 | 0.5 | 3 | 0 | 2 | 0.5 | 6 | 0.3333 | 5 | 0.2 | 16 | 0.25 |
| 4_12952599 | 2 | 0 | 2 | 0 | 5 | 0.2 | 3 | 0 | 12 | 0.0833 | 2 | 0 | NA | NA | 6 | 0 | 4 | 0.25 | 12 | 0.0833 |
| 4_12953799 | 9 | 0.1111 | 6 | 0 | 6 | 0.1667 | 10 | 0.2 | 31 | 0.129 | 9 | 0.5556 | 9 | 0.2222 | 9 | 0.3333 | 11 | 0.2727 | 38 | 0.3421 |
| 4_12953821 | 6 | 0.5 | 3 | 0 | 4 | 0.75 | 7 | 0.1429 | 20 | 0.35 | 9 | 0.8889 | 4 | 0.5 | 4 | 0.75 | 6 | 0.5 | 23 | 0.6957 |

| | | | | | | | | | | | | | | | | | | | | |
|---|---|---|---|---|---|---|---|---|---|---|---|---|---|---|---|---|---|---|---|---|
| 4_12953829 | 7 | 0.7143 | 4 | 0.5 | 5 | 0.8 | 10 | 0.5 | 26 | 0.6154 | 9 | 0.7778 | 8 | 0.625 | 8 | 0.75 | 7 | 0.4286 | 32 | 0.6562 |
| 4_12953850 | 6 | 0 | 7 | 0.1429 | 3 | 0 | 8 | 0.125 | 24 | 0.0833 | 6 | 0 | 5 | 0.2 | 6 | 0.1667 | 6 | 0 | 23 | 0.087 |
| 4_12953852 | 8 | 0.125 | 5 | 0 | 6 | 0.1667 | 10 | 0.1 | 29 | 0.1034 | 6 | 0.1667 | 7 | 0 | 6 | 0.1667 | 6 | 0 | 25 | 0.08 |
| 4_12954049 | 4 | 0 | 1 | 0 | 2 | 0.5 | 4 | 0 | 11 | 0.0909 | 4 | 0.25 | 8 | 0.125 | 3 | 0 | 5 | 0.4 | 20 | 0.2 |
| 4_12954061 | 4 | 0 | 4 | 0 | 2 | 0 | 6 | 0.1667 | 16 | 0.0625 | 3 | 0 | 5 | 0 | 4 | 0 | 5 | 0.2 | 17 | 0.0588 |
| 4_12954148 | 6 | 1 | 6 | 0.8333 | 1 | 1 | 6 | 1 | 19 | 0.9474 | 3 | 1 | 4 | 1 | 3 | 1 | 3 | 1 | 13 | 1 |
| 4_12954157 | 5 | 0 | 6 | 0.1667 | 1 | 0 | 6 | 0.3333 | 18 | 0.1667 | 3 | 0 | 4 | 0 | 4 | 0.25 | 3 | 0.3333 | 14 | 0.1429 |
| 4_12954159 | 5 | 0 | 3 | 0 | 1 | 0 | 5 | 0.2 | 14 | 0.0714 | 3 | 0 | 3 | 0 | 4 | 0 | 4 | 0.25 | 14 | 0.0714 |
| 4_12954160 | 3 | 0 | 4 | 0 | 1 | 0 | 5 | 0.2 | 13 | 0.0769 | 2 | 0 | 3 | 0 | 3 | 0 | 4 | 0.5 | 12 | 0.1667 |
| 4_12954160 | 5 | 1 | 6 | 0.8333 | 1 | 1 | 4 | 1 | 16 | 0.9375 | 2 | 1 | 2 | 1 | 3 | 1 | 5 | 0.4 | 12 | 0.75 |
| 4_12954166 | 6 | 0.8333 | 5 | 0.8 | 1 | 0 | 6 | 0.6667 | 18 | 0.7222 | 3 | 1 | 3 | 1 | 4 | 1 | 3 | 1 | 13 | 1 |
| 4_12954308 | 4 | 0.5 | 3 | 0.3333 | 1 | 1 | 6 | 0.1667 | 14 | 0.3571 | 3 | 0 | 2 | 0.5 | 2 | 0.5 | 5 | 0 | 12 | 0.1667 |
| 4_12954312 | 4 | 0.25 | 4 | 0 | NA | NA | 6 | 0 | 14 | 0.0714 | 3 | 0 | 2 | 0 | 2 | 0 | 5 | 0.2 | 12 | 0.0833 |
| 4_12958064 | 1 | 0 | NA | NA | 1 | 1 | 3 | 0.6667 | 5 | 0.6 | NA | NA | NA | NA | 1 | 1 | 1 | 1 | 2 | 1 |
| 4_12971420 | 4 | 0.25 | 11 | 0.0909 | 6 | 0.5 | 5 | 0.2 | 26 | 0.2308 | 4 | 0 | 14 | 0.1429 | 4 | 0.25 | 2 | 0.5 | 24 | 0.1667 |
| 4_12971421 | 3 | 0 | 8 | 0 | 6 | 0 | 3 | 0.3333 | 20 | 0.05 | 1 | 0 | 11 | 0.1818 | 3 | 0 | 3 | 0 | 18 | 0.1111 |
| 4_12973291 | 5 | 0.4 | 16 | 0 | 10 | 0.1 | 4 | 0 | 35 | 0.0857 | 6 | 0 | 14 | 0.1429 | 5 | 0.4 | 8 | 0.25 | 33 | 0.1818 |
| 4_12973298 | 8 | 0.125 | 23 | 0.6957 | 14 | 0.2143 | 9 | 0.3333 | 54 | 0.4259 | 8 | 0.125 | 19 | 0.4211 | 7 | 0 | 10 | 0.3 | 44 | 0.2727 |
| 4_12973302 | 7 | 0.1429 | 23 | 0.0435 | 15 | 0.0667 | 10 | 0 | 55 | 0.0545 | 8 | 0.125 | 18 | 0.1111 | 8 | 0 | 11 | 0 | 45 | 0.0667 |
| 4_12973328 | 9 | 0 | 22 | 0.1364 | 13 | 0 | 10 | 0 | 54 | 0.0556 | 7 | 0.2857 | 16 | 0.3125 | 7 | 0.1429 | 14 | 0.1429 | 44 | 0.2273 |
| 4_12974165 | 9 | 0.3333 | 25 | 0.56 | 12 | 0.5833 | 10 | 0.4 | 56 | 0.5 | 8 | 0.625 | 20 | 0.45 | 10 | 0.6 | 19 | 0.5789 | 57 | 0.5439 |
| 4_12974330 | 13 | 0.0769 | 23 | 0.087 | 20 | 0.05 | 11 | 0.0909 | 67 | 0.0746 | 8 | 0.125 | 16 | 0.0625 | 13 | 0.1538 | 17 | 0.3529 | 54 | 0.1852 |
| 4_12985524 | 1 | 0 | 1 | 1 | NA | NA | NA | NA | 2 | 0.5 | 1 | 1 | NA | NA | 2 | 1 | NA | NA | 3 | 1 |
| 4_12985527 | 2 | 1 | 1 | 1 | NA | NA | NA | NA | 3 | 1 | 1 | 1 | NA | NA | 2 | 1 | NA | NA | 3 | 1 |
| 4_12985529 | 1 | 0 | 1 | 1 | NA | NA | NA | NA | 2 | 0.5 | 1 | 0 | NA | NA | 2 | 0.5 | NA | NA | 3 | 0.3333 |
| 4_12985531 | 1 | 1 | 1 | 1 | NA | NA | NA | NA | 2 | 1 | 1 | 0 | NA | NA | 2 | 1 | NA | NA | 3 | 0.6667 |
| 4_12985533 | 2 | 1 | 1 | 1 | NA | NA | NA | NA | 3 | 1 | NA | NA | NA | NA | 2 | 1 | NA | NA | 2 | 1 |
| 4_12985543 | NA | NA | 1 | 1 | NA | NA | NA | NA | 1 | 1 | NA | NA | NA | NA | 2 | 0.5 | NA | NA | 2 | 0.5 |
| 4_12985546 | 2 | 0.5 | 1 | 0 | NA | NA | NA | NA | 3 | 0.3333 | 1 | 0 | NA | NA | 2 | 1 | NA | NA | 3 | 0.6667 |
| 4_12985724 | 2 | 0 | 7 | 0.1429 | 1 | 0 | NA | NA | 10 | 0.1 | NA | NA | 4 | 0 | 1 | 1 | 1 | 1 | 6 | 0.3333 |
| 4_12985743 | 2 | 0.5 | 4 | 0.75 | NA | NA | NA | NA | 6 | 0.6667 | NA | NA | 2 | 1 | 1 | 1 | NA | NA | 3 | 1 |
| 4_12985744 | NA | NA | 3 | 0.3333 | NA | NA | NA | NA | 3 | 0.3333 | NA | NA | 1 | 1 | NA | NA | 1 | 1 | 2 | 1 |
| 4_12985744 | NA | NA | 1 | 1 | NA | NA | NA | NA | 1 | 1 | NA | NA | 1 | 1 | NA | NA | NA | NA | 1 | 1 |
| 4_12985836 | 2 | 0 | 4 | 0.25 | 1 | 1 | NA | NA | 7 | 0.2857 | 2 | 0 | 4 | 0.25 | NA | NA | NA | NA | 6 | 0.1667 |
| 4_12985872 | 4 | 0 | 7 | 0.1429 | 1 | 0 | NA | NA | 12 | 0.0833 | 2 | 0 | 4 | 0.25 | 1 | 0 | 1 | 1 | 8 | 0.25 |
| 4_12985875 | 4 | 0.75 | 6 | 1 | 1 | 1 | NA | NA | 11 | 0.9091 | 2 | 0.5 | 4 | 1 | 1 | 1 | 1 | 1 | 8 | 0.875 |
| 4_12988585 | 6 | 0.1667 | 10 | 0.2 | 3 | 0 | 3 | 0 | 22 | 0.1364 | 4 | 0.25 | 10 | 0 | 6 | 0 | 11 | 0.0909 | 31 | 0.0645 |
| 4_12988599 | 4 | 0 | 5 | 0.4 | 5 | 0.6 | 3 | 0.6667 | 17 | 0.4118 | 7 | 0.4286 | 8 | 0.125 | 6 | 0.3333 | 10 | 0.3 | 31 | 0.2903 |
| 4_12988602 | 7 | 0.2857 | 8 | 0.375 | 5 | 0.2 | 3 | 0 | 23 | 0.2609 | 5 | 0.4 | 11 | 0.6364 | 6 | 0.5 | 11 | 0.2727 | 33 | 0.4545 |
| 4_12988612 | 4 | 0.5 | 10 | 0.5 | 4 | 0 | 3 | 0.3333 | 21 | 0.381 | 6 | 0.6667 | 9 | 0.2222 | 5 | 0.2 | 9 | 0.7778 | 29 | 0.4828 |
| 4_12988630 | 6 | 1 | 11 | 0.5455 | 6 | 0.5 | 2 | 0.5 | 25 | 0.64 | 5 | 0.4 | 12 | 0.5 | 8 | 0.625 | 11 | 0.6364 | 36 | 0.5556 |
| 4_12991270 | 27 | 0.4815 | 28 | 0.6429 | 15 | 0.5333 | 32 | 0.5938 | 102 | 0.5686 | 25 | 0.52 | 26 | 0.6923 | 26 | 0.7308 | 27 | 0.5556 | 104 | 0.625 |
| 4_12991274 | 24 | 0.625 | 26 | 0.6154 | 18 | 0.4444 | 32 | 0.625 | 100 | 0.59 | 23 | 0.5652 | 27 | 0.4815 | 24 | 0.5833 | 28 | 0.5714 | 102 | 0.549 |
| 4_12991276 | 25 | 0.92 | 26 | 0.8077 | 19 | 0.8947 | 32 | 0.875 | 102 | 0.8725 | 26 | 0.9231 | 31 | 0.8065 | 25 | 0.84 | 30 | 0.7667 | 112 | 0.8304 |
| 4_12991607 | 14 | 0.2143 | 25 | 0.12 | 14 | 0 | 31 | 0.0645 | 84 | 0.0952 | 18 | 0.0556 | 20 | 0.1 | 17 | 0.0588 | 25 | 0.12 | 80 | 0.0875 |
| 4_12991614 | 26 | 0.8462 | 33 | 0.9697 | 22 | 0.7727 | 41 | 0.9268 | 122 | 0.8934 | 32 | 0.875 | 33 | 0.9394 | 30 | 0.8333 | 37 | 0.8919 | 132 | 0.8864 |
| 4_12991619 | 19 | 0.4211 | 29 | 0.5862 | 15 | 0.5333 | 28 | 0.4286 | 91 | 0.4945 | 22 | 0.5 | 20 | 0.5 | 22 | 0.3636 | 27 | 0.2593 | 91 | 0.3956 |
| 4_12991624 | 29 | 0.8966 | 34 | 0.9412 | 23 | 0.913 | 46 | 0.9348 | 132 | 0.9242 | 32 | 0.9688 | 35 | 0.8286 | 29 | 0.8621 | 36 | 0.8889 | 132 | 0.8864 |
| 4_12991627 | 31 | 0.1935 | 34 | 0.2353 | 23 | 0.087 | 47 | 0.2128 | 135 | 0.1926 | 36 | 0.1944 | 37 | 0.2703 | 32 | 0.1562 | 36 | 0.25 | 141 | 0.2199 |
| 4_13002124 | 12 | 0.75 | 13 | 0.8462 | 19 | 0.8947 | 10 | 0.5 | 54 | 0.7778 | 7 | 0.2857 | 10 | 0.8 | 17 | 0.7647 | 10 | 0.6 | 44 | 0.6591 |
| 4_13002138 | 14 | 0.2143 | 13 | 0.1538 | 19 | 0.1053 | 9 | 0 | 55 | 0.1273 | 5 | 0 | 9 | 0.2222 | 15 | 0.0667 | 10 | 0.1 | 39 | 0.1026 |
| 4_13002158 | 13 | 0.1538 | 11 | 0.2727 | 15 | 0.1333 | 9 | 0.4444 | 48 | 0.2292 | 7 | 0.1429 | 7 | 0.2857 | 16 | 0.3125 | 10 | 0.3 | 40 | 0.275 |
| 4_13002161 | 14 | 0.0714 | 12 | 0 | 17 | 0.0588 | 11 | 0.1818 | 54 | 0.0741 | 6 | 0.1667 | 8 | 0.25 | 17 | 0.1176 | 11 | 0 | 42 | 0.119 |
| 4_13002169 | 10 | 0.9 | 12 | 1 | 19 | 0.9474 | 11 | 0.9091 | 52 | 0.9423 | 7 | 0.8571 | 10 | 0.9 | 17 | 0.8824 | 12 | 0.8333 | 46 | 0.8696 |
| 4_13002175 | 10 | 0.1 | 6 | 0 | 14 | 0.0714 | 6 | 0.1667 | 36 | 0.0833 | 7 | 0.2857 | 10 | 0.4 | 11 | 0.1818 | 9 | 0.1111 | 37 | 0.2432 |
| 4_13002190 | 13 | 0.1538 | 10 | 0.2 | 19 | 0.0526 | 10 | 0.1 | 52 | 0.1154 | 7 | 0 | 10 | 0.1 | 18 | 0.1111 | 12 | 0.0833 | 47 | 0.0851 |
| 4_13005965 | 2 | 0 | 2 | 1 | 1 | 1 | 3 | 0.3333 | 8 | 0.5 | NA | NA | 2 | 0.5 | 2 | 0 | NA | NA | 4 | 0.25 |
| 4_13005975 | 3 | 0 | NA | NA | NA | NA | 4 | 0.25 | 7 | 0.1429 | 2 | 0.5 | 3 | 0.6667 | 2 | 0.5 | NA | NA | 7 | 0.5714 |
| 4_13005987 | 3 | 0 | 2 | 0.5 | 1 | 0 | 4 | 0 | 10 | 0.1 | 2 | 1 | 2 | 0 | 2 | 0 | NA | NA | 6 | 0.3333 |
| 4_13014758 | NA | NA | 2 | 0.5 | NA | NA | NA | NA | 2 | 0.5 | 1 | 1 | NA | NA | 1 | 1 | NA | NA | 2 | 1 |
| 4_13020983 | 5 | 0.6 | 5 | 0.6 | 2 | 0.5 | 3 | 0.6667 | 15 | 0.6 | 2 | 1 | 4 | 0.25 | 3 | 0.6667 | 3 | 0.6667 | 12 | 0.5833 |
| 4_13020994 | 14 | 0.5 | 13 | 0.7692 | 6 | 0.5 | 12 | 0.5833 | 45 | 0.6 | 11 | 0.4545 | 13 | 0.6154 | 11 | 0.4545 | 15 | 0.5333 | 50 | 0.52 |
| 4_13021001 | 21 | 0.7143 | 13 | 0.6923 | 11 | 0.5455 | 13 | 0.4615 | 58 | 0.6207 | 9 | 0.6667 | 15 | 0.7333 | 13 | 0.6923 | 15 | 0.5333 | 52 | 0.6538 |
| 4_13021005 | 20 | 0.5 | 13 | 0.4615 | 10 | 0.5 | 12 | 0.5833 | 55 | 0.5091 | 12 | 0.5 | 14 | 0.9286 | 11 | 0.7273 | 14 | 0.5 | 51 | 0.6667 |
| 4_13021036 | 20 | 0.55 | 13 | 0.4615 | 8 | 0.5 | 13 | 0.3846 | 54 | 0.4815 | 9 | 0.4444 | 10 | 0.6 | 9 | 0.6667 | 15 | 0.6667 | 43 | 0.6047 |

| | | | | | | | | | | | | | | | | | | | | |
|---|---|---|---|---|---|---|---|---|---|---|---|---|---|---|---|---|---|---|---|---|
| 4_13021040 | 18 | 0.8333 | 13 | 0.7692 | 7 | 0.8571 | 12 | 1 | 50 | 0.86 | 8 | 0.75 | 11 | 1 | 9 | 1 | 14 | 0.8571 | 42 | 0.9048 |
| 4_13028363 | 4 | 0.25 | 2 | 0.5 | 6 | 0.1667 | 2 | 0 | 14 | 0.2143 | 4 | 0.25 | 3 | 0.3333 | 3 | 0 | 1 | 0 | 11 | 0.1818 |
| 4_13033950 | 1 | 0 | 5 | 0.8 | 2 | 0.5 | 1 | 1 | 9 | 0.6667 | 2 | 1 | NA | NA | NA | NA | 3 | 0.6667 | 5 | 0.8 |
| 4_13033952 | NA | NA | 5 | 0 | 3 | 0.6667 | NA | NA | 8 | 0.25 | 2 | 0.5 | NA | NA | 1 | 1 | 3 | 0 | 6 | 0.3333 |
| 4_13033954 | 1 | 0 | 4 | 0.75 | 1 | 1 | 1 | 1 | 7 | 0.7143 | 1 | 1 | NA | NA | 1 | 1 | 3 | 0.6667 | 5 | 0.8 |
| 4_13034006 | 2 | 0 | 7 | 0.4286 | 1 | 1 | 1 | 1 | 11 | 0.4545 | 8 | 0.25 | 1 | 0 | NA | NA | 3 | 0 | 12 | 0.1667 |
| 4_13034051 | 2 | 0.5 | 4 | 0 | 3 | 0 | 1 | 0 | 10 | 0.1 | 8 | 0 | 1 | 0 | 1 | 0 | 5 | 0.2 | 15 | 0.0667 |
| 4_13039629 | 2 | 0.5 | 3 | 1 | 2 | 0.5 | NA | NA | 7 | 0.7143 | 2 | 0 | 3 | 0 | 5 | 0.6 | 3 | 0.3333 | 13 | 0.3077 |
| 4_13039649 | 2 | 0 | 2 | 0 | 1 | 0 | 1 | 1 | 6 | 0.1667 | 2 | 0 | 3 | 0 | 4 | 0.5 | 3 | 0 | 12 | 0.1667 |
| 4_13045781 | 2 | 0 | 2 | 0 | 3 | 0 | 5 | 0.2 | 12 | 0.0833 | 1 | 0 | 3 | 0 | 5 | 0.4 | 4 | 0 | 13 | 0.1538 |
| 4_13047995 | 2 | 0.5 | 6 | 0 | 4 | 0 | 5 | 0.2 | 17 | 0.1176 | 1 | 0 | 5 | 0.2 | 9 | 0 | 3 | 0.3333 | 18 | 0.1111 |
| 4_13048135 | 2 | 0 | 5 | 0.2 | 4 | 0 | 4 | 0.25 | 15 | 0.1333 | 2 | 0.5 | 5 | 0 | 6 | 0 | 3 | 0 | 16 | 0.0625 |
| 4_13048139 | 2 | 0 | 7 | 0.1429 | 5 | 0.4 | 4 | 0 | 18 | 0.1667 | 2 | 0 | 5 | 0 | 7 | 0.2857 | 3 | 0 | 17 | 0.1176 |
| 4_13048142 | 2 | 0 | 7 | 0.1429 | 5 | 0 | 4 | 0.25 | 18 | 0.1111 | 1 | 0 | 5 | 0 | 9 | 0.2222 | 3 | 0.3333 | 18 | 0.1667 |
| 4_13050717 | 1 | 1 | 3 | 0 | 2 | 1 | 1 | 1 | 7 | 0.5714 | NA | NA | 3 | 1 | 2 | 1 | 1 | 1 | 6 | 1 |
| 4_13050721 | 3 | 1 | 13 | 0.1538 | 5 | 0.2 | 5 | 0 | 26 | 0.2308 | NA | NA | 5 | 0 | 7 | 0.1429 | 5 | 0.4 | 17 | 0.1765 |
| 4_13050753 | 3 | 0.6667 | 11 | 0.0909 | 3 | 0 | 6 | 0.1667 | 23 | 0.1739 | 2 | 0 | 5 | 0.2 | 14 | 0.1429 | 5 | 0 | 26 | 0.1154 |
| 4_13050794 | 8 | 0 | 17 | 0.0588 | 5 | 0.2 | 6 | 0.1667 | 36 | 0.0833 | 5 | 0.2 | 6 | 0 | 15 | 0.0667 | 6 | 0 | 32 | 0.0625 |
| 4_13050796 | 7 | 0 | 21 | 0.1905 | 10 | 0.1 | 11 | 0 | 49 | 0.102 | 5 | 0 | 6 | 0 | 18 | 0.1111 | 7 | 0 | 36 | 0.0556 |
| 4_13050866 | 8 | 0.25 | 23 | 0.087 | 12 | 0 | 14 | 0.1429 | 57 | 0.1053 | 7 | 0.1429 | 13 | 0.1538 | 19 | 0 | 10 | 0.1 | 49 | 0.0816 |
| 4_13050872 | 8 | 0.375 | 21 | 0 | 11 | 0 | 14 | 0.2143 | 54 | 0.1111 | 8 | 0 | 10 | 0 | 20 | 0.05 | 11 | 0.1818 | 49 | 0.0612 |
| 4_13054600 | 2 | 0.5 | 5 | 0.2 | 6 | 0.1667 | 1 | 0 | 14 | 0.2143 | NA | NA | NA | NA | 3 | 0 | 2 | 0.5 | 5 | 0.2 |
| 4_13054602 | 2 | 0 | 5 | 0.4 | 7 | 0.7143 | 1 | 0 | 15 | 0.4667 | 1 | 0 | NA | NA | 3 | 0.3333 | 3 | 0.3333 | 7 | 0.2857 |
| 4_13054625 | 1 | 1 | 3 | 0.6667 | 5 | 0.8 | NA | NA | 9 | 0.7778 | 1 | 1 | 1 | 1 | 4 | 0.75 | 1 | 1 | 7 | 0.8571 |
| 4_13054626 | NA | NA | NA | NA | NA | NA | 1 | 1 | 1 | 1 | 1 | 1 | NA | NA | NA | NA | NA | NA | 1 | 1 |
| 4_13054636 | 2 | 1 | 4 | 0.5 | 7 | 0.7143 | 1 | 1 | 14 | 0.7143 | 2 | 1 | 1 | 1 | 4 | 0.75 | 2 | 0 | 9 | 0.6667 |
| 4_13054875 | 3 | 0.3333 | 6 | 0.3333 | 6 | 0.3333 | 1 | 1 | 16 | 0.375 | 4 | 0.5 | 1 | 1 | 4 | 0.5 | 3 | 0.3333 | 12 | 0.5 |
| 4_13054885 | 3 | 0.6667 | 5 | 0.6 | 6 | 0.6667 | 1 | 1 | 15 | 0.6667 | 3 | 1 | 2 | 1 | 3 | 0.6667 | 3 | 0.3333 | 11 | 0.7273 |
| 4_13055026 | 5 | 1 | 3 | 0.6667 | 6 | 1 | 2 | 1 | 16 | 0.9375 | 3 | 1 | 3 | 0.6667 | 3 | 1 | 5 | 0.8 | 14 | 0.8571 |
| 4_13055030 | 5 | 0.8 | 6 | 0.3333 | 7 | 0.8571 | 2 | 1 | 20 | 0.7 | 4 | 0.75 | 4 | 0.75 | 4 | 0.75 | 6 | 0.5 | 18 | 0.6667 |
| 4_13055037 | 6 | 0.8333 | 6 | 0.5 | 8 | 0.25 | 2 | 1 | 22 | 0.5455 | 3 | 0.3333 | 2 | 0.5 | 3 | 0.6667 | 7 | 0.2857 | 15 | 0.4 |
| 4_13055135 | 8 | 0.25 | 8 | 0 | 9 | 0 | 5 | 0.2 | 30 | 0.1 | 5 | 0.2 | 8 | 0.25 | 6 | 0.1667 | 8 | 0 | 27 | 0.1481 |
| 4_13055147 | 9 | 0.2222 | 7 | 0 | 10 | 0.1 | 5 | 0 | 31 | 0.0968 | 4 | 1 | 7 | 0.2857 | 7 | 0.4286 | 8 | 0.375 | 26 | 0.4615 |
| 4_13055172 | 10 | 0.1 | 11 | 0.1818 | 9 | 0 | 7 | 0 | 37 | 0.0811 | 7 | 0 | 7 | 0.1429 | 5 | 0 | 9 | 0.1111 | 28 | 0.0714 |
| 4_13055288 | 17 | 0.0588 | 15 | 0.1333 | 15 | 0 | 9 | 0 | 56 | 0.0536 | 10 | 0 | 9 | 0.1111 | 17 | 0.2353 | 15 | 0 | 51 | 0.098 |
| 4_13066195 | 4 | 0.5 | 6 | 0.3333 | 4 | 0.25 | 3 | 0 | 17 | 0.2941 | 7 | 0.1429 | 4 | 0.5 | 1 | 0 | 1 | 0 | 13 | 0.2308 |
| 4_13149625 | 4 | 0.75 | 2 | 0 | 1 | 1 | NA | NA | 7 | 0.5714 | 1 | 1 | 1 | 1 | 1 | 0 | 2 | 0.5 | 5 | 0.6 |
| 4_13149635 | 3 | 0 | 2 | 0.5 | NA | NA | 1 | 0 | 6 | 0.1667 | 1 | 0 | 1 | 0 | 2 | 0.5 | 1 | 0 | 5 | 0.2 |
| 4_13149638 | 3 | 0.6667 | 1 | 0 | NA | NA | 1 | 1 | 5 | 0.6 | 1 | 1 | NA | NA | 1 | 1 | 1 | 1 | 3 | 1 |
| 4_13149642 | 2 | 1 | 2 | 1 | NA | NA | 1 | 1 | 5 | 1 | 1 | 1 | 1 | 1 | 2 | 1 | NA | NA | 4 | 1 |
| 4_13149656 | 2 | 1 | 2 | 0.5 | NA | NA | 1 | 1 | 5 | 0.8 | 1 | 1 | 1 | 0 | 2 | 0 | 1 | 0 | 5 | 0.2 |
| 4_13149662 | 3 | 0.3333 | 2 | 1 | NA | NA | 1 | 1 | 6 | 0.6667 | 1 | 1 | 1 | 1 | 2 | 0.5 | 1 | 1 | 5 | 0.8 |
| 4_13149668 | 3 | 0.6667 | 2 | 0.5 | NA | NA | 1 | 1 | 6 | 0.6667 | 1 | 1 | 1 | 0 | 1 | 0 | NA | NA | 3 | 0.3333 |
| 4_13149670 | 3 | 0.3333 | 2 | 0 | NA | NA | 1 | 0 | 6 | 0.1667 | 1 | 0 | 1 | 0 | 2 | 0.5 | 1 | 0 | 5 | 0.2 |
| 4_13159499 | NA | NA | 7 | 0.4286 | 4 | 0.5 | 2 | 0 | 13 | 0.3846 | 1 | 0 | 2 | 0 | 3 | 0.3333 | 1 | 1 | 7 | 0.2857 |
| 4_13162805 | 7 | 0 | 9 | 0.5556 | 6 | 0 | 6 | 0.1667 | 28 | 0.2143 | 4 | 0 | 12 | 0.1667 | 8 | 0.125 | 8 | 0 | 32 | 0.0938 |
| 4_13162809 | 6 | 0.1667 | 9 | 0.2222 | 6 | 0.1667 | 5 | 0 | 26 | 0.1538 | 5 | 0.2 | 14 | 0.1429 | 8 | 0.25 | 8 | 0.125 | 35 | 0.1714 |
| 4_13162814 | 5 | 0 | 7 | 0 | 4 | 0.25 | 5 | 0.2 | 21 | 0.0952 | 4 | 0 | 10 | 0.1 | 8 | 0.125 | 7 | 0 | 29 | 0.069 |
| 4_13162840 | 3 | 0.3333 | 7 | 0.8571 | 3 | 0.6667 | 4 | 0.75 | 17 | 0.7059 | 6 | 1 | 8 | 0.875 | 4 | 1 | 5 | 0.6 | 23 | 0.8696 |
| 4_13162848 | 6 | 0.1667 | 3 | 0 | 5 | 0 | 5 | 0.4 | 19 | 0.1579 | 5 | 0.4 | 12 | 0 | 8 | 0.5 | 6 | 0.5 | 31 | 0.2903 |
| 4_13162860 | 7 | 0.8571 | 8 | 0.875 | 7 | 0.7143 | 5 | 0.6 | 27 | 0.7778 | 7 | 0.7143 | 12 | 0.5833 | 9 | 1 | 6 | 0.6667 | 34 | 0.7353 |
| 4_13162880 | 6 | 1 | 9 | 0.8889 | 5 | 0.8 | 8 | 0.875 | 28 | 0.8929 | 8 | 0.75 | 13 | 0.7692 | 9 | 1 | 10 | 1 | 40 | 0.875 |
| 4_13162899 | 6 | 0.6667 | 9 | 0.6667 | 7 | 0.8571 | 6 | 0.5 | 28 | 0.6786 | 3 | 0.3333 | 13 | 0.9231 | 6 | 0.8333 | 8 | 0.625 | 30 | 0.7667 |
| 4_13164095 | 1 | 1 | NA | NA | NA | NA | NA | NA | 1 | 1 | NA | NA | NA | NA | 1 | 1 | NA | NA | 1 | 1 |
| 4_13164103 | 1 | 1 | NA | NA | NA | NA | NA | NA | 1 | 1 | NA | NA | NA | NA | 1 | 1 | NA | NA | 1 | 1 |
| 4_13164569 | NA | NA | 1 | 0 | 1 | 1 | NA | NA | 2 | 0.5 | NA | NA | NA | NA | 1 | 1 | 1 | 1 | 2 | 1 |
| 4_13164625 | NA | NA | 1 | 1 | 1 | 0 | NA | NA | 2 | 0.5 | NA | NA | NA | NA | 1 | 0 | 1 | 1 | 2 | 0.5 |
| 4_13164706 | NA | NA | NA | NA | 1 | 1 | NA | NA | 1 | 1 | NA | NA | NA | NA | 1 | 1 | NA | NA | 1 | 1 |
| 4_13164742 | NA | NA | 1 | 1 | 1 | 0 | NA | NA | 2 | 0.5 | NA | NA | NA | NA | 1 | 1 | NA | NA | 1 | 1 |
| 4_13164754 | NA | NA | NA | NA | 1 | 1 | NA | NA | 1 | 1 | NA | NA | NA | NA | 1 | 1 | NA | NA | 1 | 1 |
| 4_13164776 | NA | NA | 1 | 1 | NA | NA | NA | NA | 1 | 1 | NA | NA | NA | NA | 1 | 1 | NA | NA | 1 | 1 |
| 4_13164830 | NA | NA | 1 | 1 | 1 | 1 | NA | NA | 2 | 1 | NA | NA | NA | NA | 1 | 1 | NA | NA | 1 | 1 |
| 4_13164973 | 2 | 1 | NA | NA | NA | NA | NA | NA | 2 | 1 | NA | NA | 1 | 1 | 3 | 1 | NA | NA | 4 | 1 |
| 4_13164984 | 1 | 1 | NA | NA | NA | NA | NA | NA | 1 | 1 | NA | NA | 1 | 1 | 4 | 1 | NA | NA | 5 | 1 |
| 4_13164987 | 2 | 1 | NA | NA | NA | NA | 1 | 1 | 3 | 1 | NA | NA | 1 | 0 | 3 | 0.3333 | NA | NA | 4 | 0.25 |

| | | | | | | | | | | | | | | | | | | | | |
|---|---|---|---|---|---|---|---|---|---|---|---|---|---|---|---|---|---|---|---|---|
| 4_1316499 | 1 | 1 | 1 | 0 | NA | NA | 1 | 1 | 3 | 0.6667 | NA | NA | NA | NA | 2 | 0.5 | NA | NA | 2 | 0.5 |
| 4_1316499 | 2 | 0 | 1 | 0 | NA | NA | 1 | 1 | 4 | 0.25 | NA | NA | 2 | 0 | 4 | 0.25 | NA | NA | 6 | 0.1667 |
| 4_1316501 | 3 | 0.6667 | 1 | 1 | NA | NA | 1 | 1 | 5 | 0.8 | NA | NA | 3 | 1 | 4 | 1 | NA | NA | 7 | 1 |
| 4_1316504 | 2 | 1 | 1 | 1 | 2 | 1 | NA | NA | 5 | 1 | NA | NA | 4 | 0.25 | 4 | 0.75 | NA | NA | 8 | 0.5 |
| 4_1316508 | 2 | 0 | 1 | 0 | 1 | 1 | 1 | 1 | 5 | 0.4 | NA | NA | 5 | 0.6 | 5 | 0.2 | NA | NA | 10 | 0.4 |
| 4_1316512 | 2 | 0.5 | NA | NA | 1 | 0 | 1 | 0 | 4 | 0.25 | NA | NA | 5 | 0 | 5 | 0.6 | NA | NA | 10 | 0.3 |
| 4_1316513 | 1 | 0 | 1 | 0 | 3 | 0 | 5 | 0.2 | 10 | 0.1 | 3 | 0 | 3 | 0.3333 | 4 | 0 | 2 | 0 | 12 | 0.0833 |
| 4_1316515 | 1 | 1 | 1 | 0 | 3 | 1 | 4 | 0.25 | 9 | 0.5556 | 3 | 0 | 3 | 0.3333 | 4 | 0.5 | 1 | 0 | 11 | 0.2727 |
| 4_1316516 | 1 | 1 | 1 | 0 | 2 | 0.5 | 2 | 0 | 6 | 0.3333 | 2 | 0 | 3 | 0.6667 | 3 | 0 | 1 | 0 | 9 | 0.2222 |
| 4_1316524 | 1 | 0 | 1 | 1 | 3 | 1 | 3 | 1 | 8 | 0.875 | 3 | 0.6667 | 1 | 0 | 2 | 1 | 1 | 1 | 7 | 0.7143 |
| 4_1316525 | 1 | 1 | 1 | 0 | 3 | 1 | 3 | 0.6667 | 8 | 0.75 | 3 | 0.6667 | 3 | 0.6667 | 3 | 1 | NA | NA | 9 | 0.7778 |
| 4_1316526 | 1 | 1 | 1 | 0 | 1 | 1 | 1 | 1 | 4 | 0.75 | 1 | 0 | 3 | 1 | 2 | 1 | NA | NA | 6 | 0.8333 |
| 4_1316527 | 1 | 0 | 1 | 0 | 3 | 0.3333 | 1 | 0 | 6 | 0.1667 | 1 | 0 | 3 | 0.3333 | 3 | 0.3333 | 1 | 0 | 8 | 0.25 |
| 4_1316528 | 1 | 1 | 1 | 1 | 1 | 0 | 1 | 1 | 4 | 0.75 | 1 | 1 | NA | NA | 3 | 1 | 1 | 1 | 5 | 1 |
| 4_1316528 | NA | NA | NA | NA | 2 | 0.5 | 1 | 1 | 3 | 0.6667 | 1 | 1 | 2 | 1 | 2 | 0 | 1 | 0 | 6 | 0.5 |
| 4_1316528 | 1 | 0 | 1 | 0 | 3 | 0.3333 | 1 | 0 | 6 | 0.1667 | 1 | 0 | 3 | 0.3333 | 3 | 0.3333 | 1 | 0 | 8 | 0.25 |
| 4_1316528 | 1 | 1 | 1 | 0 | 3 | 0.3333 | 1 | 0 | 6 | 0.3333 | 1 | 0 | 3 | 0 | 3 | 0.3333 | 1 | 0 | 8 | 0.125 |
| 4_1316529 | 1 | 0 | 1 | 0 | 3 | 0 | 1 | 1 | 6 | 0.1667 | 1 | 0 | 3 | 0 | 3 | 0.3333 | 1 | 0 | 8 | 0.125 |
| 4_1316529 | 1 | 1 | 1 | 1 | 3 | 0 | 1 | 0 | 6 | 0.3333 | 1 | 0 | 3 | 0 | 2 | 0 | 1 | 1 | 7 | 0.1429 |
| 4_1316529 | 1 | 0 | 1 | 0 | 3 | 0.3333 | NA | NA | 5 | 0.2 | 1 | 1 | 3 | 0.3333 | 3 | 0 | 1 | 0 | 8 | 0.25 |
| 4_1316530 | 1 | 1 | 1 | 1 | 3 | 1 | NA | NA | 5 | 1 | 1 | 0 | 3 | 0.6667 | 2 | 0.5 | 1 | 1 | 7 | 0.5714 |
| 4_1316530 | 1 | 1 | 1 | 1 | 3 | 1 | 1 | 1 | 6 | 1 | 1 | 1 | 2 | 1 | 1 | 1 | NA | NA | 4 | 1 |
| 4_1316530 | 1 | 1 | 1 | 1 | 3 | 0.6667 | 1 | 1 | 6 | 0.8333 | 1 | 1 | 3 | 0.6667 | 2 | 1 | 1 | 0 | 7 | 0.7143 |
| 4_1316530 | NA | NA | 1 | 1 | 3 | 1 | 1 | 1 | 5 | 1 | 1 | 1 | 3 | 0.6667 | 2 | 1 | 1 | 1 | 7 | 0.8571 |
| 4_1316531 | 1 | 1 | 1 | 0 | 3 | 0 | 1 | 0 | 6 | 0.1667 | NA | NA | 3 | 0 | 2 | 0.5 | 1 | 0 | 6 | 0.1667 |
| 4_1316766 | 1 | 0 | 4 | 0 | 2 | 0 | 1 | 1 | 8 | 0.125 | NA | NA | 2 | 0 | NA | NA | 1 | 1 | 3 | 0.3333 |
| 4_1316767 | 2 | 0.5 | 2 | 0.5 | 1 | 0 | 1 | 1 | 6 | 0.5 | NA | NA | 2 | 1 | NA | NA | 1 | 1 | 3 | 1 |
| 4_1316768 | 2 | 0.5 | 2 | 0 | 1 | 1 | 1 | 0 | 6 | 0.3333 | NA | NA | 1 | 0 | NA | NA | 1 | 1 | 2 | 0.5 |
| 4_1316798 | 1 | 0 | 3 | 0.3333 | 2 | 0 | 1 | 1 | 7 | 0.2857 | 1 | 1 | 2 | 1 | NA | NA | 2 | 0 | 5 | 0.6 |
| 4_1316800 | 1 | 0 | 2 | 0 | 1 | 1 | 2 | 0 | 6 | 0.1667 | 1 | 1 | 1 | 0 | 1 | 0 | 2 | 0 | 5 | 0.2 |
| 4_1316801 | 1 | 1 | 2 | 0 | 1 | 0 | 1 | 0 | 5 | 0.2 | 1 | 1 | 1 | 0 | 1 | 1 | 2 | 0 | 5 | 0.4 |
| 4_1316809 | 1 | 1 | 3 | 0.6667 | 1 | 0 | 2 | 1 | 7 | 0.7143 | 1 | 1 | 1 | 1 | 1 | 1 | 2 | 0.5 | 5 | 0.8 |
| 4_1316811 | 1 | 1 | 3 | 0.3333 | 1 | 1 | 2 | 0.5 | 7 | 0.5714 | 1 | 1 | 2 | 1 | 1 | 0 | 2 | 0 | 6 | 0.5 |
| 4_1316812 | 1 | 1 | 3 | 1 | 1 | 1 | 2 | 1 | 7 | 1 | 1 | 1 | NA | NA | NA | NA | 1 | 1 | 2 | 1 |
| 4_1316812 | 1 | 1 | 3 | 0.3333 | 1 | 1 | 2 | 1 | 7 | 0.7143 | 1 | 1 | 1 | 1 | 1 | 1 | 2 | 0.5 | 5 | 0.8 |
| 4_1316812 | 1 | 1 | 2 | 0 | NA | NA | 2 | 0 | 5 | 0.2 | 1 | 1 | 1 | 0 | 1 | 0 | 2 | 0 | 5 | 0.2 |
| 4_1316814 | 1 | 1 | 2 | 0 | 1 | 0 | 1 | 1 | 5 | 0.4 | 1 | 1 | 1 | 0 | 1 | 0 | 1 | 1 | 4 | 0.5 |
| 4_1316823 | 1 | 0 | 2 | 0.5 | 1 | 0 | 2 | 0.5 | 6 | 0.3333 | 1 | 1 | 1 | 1 | NA | NA | 1 | 0 | 3 | 0.6667 |
| 4_1316849 | 1 | 0 | 3 | 0.3333 | NA | NA | NA | NA | 4 | 0.25 | 1 | 1 | 1 | 1 | 2 | 0 | 3 | 0 | 7 | 0.2857 |
| 4_1316850 | 1 | 0 | 3 | 0 | NA | NA | 1 | 1 | 5 | 0.2 | 1 | 0 | 1 | 1 | 2 | 0.5 | 3 | 0 | 7 | 0.2857 |
| 4_1317345 | 1 | 0 | 2 | 0.5 | NA | NA | NA | NA | 3 | 0.3333 | 1 | 1 | NA | NA | 2 | 0 | 2 | 0.5 | 5 | 0.4 |
| 4_1317345 | 1 | 1 | 2 | 0.5 | NA | NA | NA | NA | 3 | 0.6667 | 1 | 0 | NA | NA | 2 | 0.5 | 2 | 0 | 5 | 0.2 |
| 4_1318328 | NA | NA | 1 | 1 | NA | NA | 1 | 1 | 2 | 1 | 1 | 1 | 2 | 1 | NA | NA | 1 | 1 | 4 | 1 |
| 4_1318405 | NA | NA | NA | NA | 1 | 0 | 1 | 1 | 2 | 0.5 | 1 | 0 | NA | NA | NA | NA | 1 | 1 | 2 | 0.5 |
| 4_1318406 | NA | NA | NA | NA | 1 | 1 | 1 | 1 | 2 | 1 | 1 | 1 | NA | NA | NA | NA | 2 | 0.5 | 3 | 0.6667 |
| 4_1318406 | NA | NA | NA | NA | NA | NA | 1 | 1 | 1 | 1 | 1 | 1 | NA | NA | NA | NA | 2 | 1 | 3 | 1 |
| 4_1318407 | NA | NA | NA | NA | NA | NA | 1 | 1 | 1 | 1 | 1 | 1 | NA | NA | NA | NA | 2 | 1 | 3 | 1 |
| 4_1318409 | NA | NA | NA | NA | 1 | 0 | 1 | 1 | 2 | 0.5 | 1 | 1 | NA | NA | NA | NA | 2 | 0 | 3 | 0.3333 |
| 4_1318409 | NA | NA | NA | NA | 1 | 0 | 1 | 1 | 2 | 0.5 | NA | NA | NA | NA | NA | NA | 1 | 1 | 1 | 1 |
| 4_1318410 | NA | NA | NA | NA | 1 | 1 | 1 | 1 | 2 | 1 | 1 | 1 | NA | NA | NA | NA | 1 | 1 | 2 | 1 |
| 4_1319140 | 4 | 0.25 | 5 | 0.8 | 4 | 0.75 | 7 | 0.2857 | 20 | 0.5 | 6 | 0.3333 | 7 | 0.5714 | 4 | 1 | 6 | 0.6667 | 23 | 0.6087 |
| 4_1319140 | 4 | 0 | 5 | 0 | 4 | 0.25 | 6 | 0 | 19 | 0.0526 | 7 | 0.1429 | 7 | 0.2857 | 5 | 0.2 | 6 | 0 | 25 | 0.16 |
| 4_1319141 | 4 | 0.25 | 5 | 0.8 | 4 | 0.5 | 7 | 0.5714 | 20 | 0.55 | 6 | 0.5 | 8 | 0.875 | 5 | 0.6 | 6 | 0.5 | 25 | 0.64 |
| 4_1319141 | 2 | 0 | 5 | 0.4 | 1 | 0 | 6 | 0 | 14 | 0.1429 | 5 | 0.2 | 7 | 0 | 5 | 0.4 | 5 | 0.2 | 22 | 0.1818 |
| 4_1319141 | 4 | 1 | 5 | 1 | 3 | 1 | 7 | 1 | 19 | 1 | 7 | 0.8571 | 8 | 1 | 5 | 1 | 6 | 1 | 26 | 0.9615 |
| 4_1319314 | 3 | 0 | 4 | 0 | 2 | 0 | 5 | 0.2 | 14 | 0.0714 | 7 | 0 | 8 | 0.125 | 4 | 0 | 4 | 0.25 | 23 | 0.087 |
| 4_1319315 | 3 | 0.6667 | 3 | 1 | 1 | 1 | 5 | 0.8 | 12 | 0.8333 | 5 | 1 | 5 | 0.8 | 5 | 0.6 | 4 | 1 | 19 | 0.8421 |
| 4_1319315 | 3 | 0.3333 | 4 | 0 | 2 | 0 | 4 | 0.25 | 13 | 0.1538 | 5 | 0 | 6 | 0.1667 | 2 | 0 | 1 | 0 | 14 | 0.0714 |
| 4_1319316 | 3 | 0.6667 | 4 | 1 | 2 | 1 | 5 | 0.8 | 14 | 0.8571 | 7 | 0.5714 | 6 | 0.3333 | 5 | 0.6 | 5 | 1 | 23 | 0.6087 |
| 4_1319317 | 2 | 0.5 | 4 | 0.75 | 2 | 1 | 5 | 0.8 | 13 | 0.7692 | 6 | 0.8333 | 6 | 1 | 4 | 0.75 | 5 | 1 | 21 | 0.9048 |
| 4_1319317 | 1 | 1 | 3 | 1 | 2 | 0.5 | 4 | 1 | 10 | 0.9 | 4 | 1 | 6 | 0.8333 | 4 | 0.5 | 5 | 0.8 | 19 | 0.7895 |
| 4_1319318 | 2 | 1 | 3 | 1 | 2 | 1 | 5 | 1 | 12 | 1 | 5 | 1 | 6 | 1 | 5 | 1 | 5 | 1 | 21 | 1 |
| 4_1319319 | 2 | 0 | 3 | 0 | 1 | 0 | 5 | 0.4 | 11 | 0.1818 | 4 | 0 | 6 | 0 | 3 | 0.3333 | 5 | 0.2 | 18 | 0.1111 |
| 4_1319319 | 3 | 0.3333 | 2 | 0 | 2 | 0 | 5 | 0.2 | 12 | 0.1667 | 5 | 0.4 | 6 | 0.1667 | 4 | 0 | 5 | 0.2 | 20 | 0.2 |
| 4_1319320 | 2 | 1 | 2 | 0.5 | 1 | 1 | NA | NA | 5 | 0.8 | 3 | 0.6667 | 3 | 0.6667 | 4 | 0.75 | 3 | 1 | 13 | 0.7692 |

| | | | | | | | | | | | | | | | | | | | | |
|---|---|---|---|---|---|---|---|---|---|---|---|---|---|---|---|---|---|---|---|---|
| 4_1319320 | 2 | 1 | 2 | 1 | 2 | 0.5 | 4 | 0.75 | 10 | 0.8 | 5 | 0.2 | 6 | 0.1667 | 3 | 1 | 4 | 0.75 | 18 | 0.4444 |
| 4_1319322 | NA | NA | 1 | 0 | 2 | 0.5 | 4 | 0.75 | 7 | 0.5714 | 4 | 0.5 | 5 | 0.2 | 3 | 0 | 4 | 0.75 | 16 | 0.375 |
| 4_1319323 | 2 | 1 | 1 | 1 | 1 | 0 | 5 | 0.6 | 9 | 0.6667 | 4 | 1 | 6 | 0.8333 | 4 | 1 | 4 | 0.75 | 18 | 0.8889 |
| 4_1319326 | 3 | 0 | 1 | 0 | 2 | 0 | 5 | 0.2 | 11 | 0.0909 | 5 | 0.2 | 6 | 0.1667 | 3 | 0.3333 | 5 | 0 | 19 | 0.1579 |
| 4_1319326 | 3 | 0.6667 | 1 | 0 | 2 | 1 | 4 | 0 | 10 | 0.4 | 5 | 0.2 | 5 | 0.2 | 4 | 0.25 | 6 | 0.1667 | 20 | 0.2 |
| 4_1319326 | 2 | 1 | 1 | 1 | 2 | 1 | 2 | 1 | 7 | 1 | 4 | 1 | 4 | 1 | 4 | 1 | 5 | 1 | 17 | 1 |
| 4_1319327 | 2 | 0.5 | 1 | 1 | 1 | 1 | 3 | 0.6667 | 7 | 0.7143 | 4 | 1 | 4 | 0.75 | 4 | 1 | 4 | 1 | 16 | 0.9375 |
| 4_1319327 | 3 | 0.3333 | NA | NA | 1 | 1 | 3 | 0 | 7 | 0.2857 | 3 | 0 | 5 | 0.2 | 2 | 0 | 3 | 0.3333 | 13 | 0.1538 |
| 4_1319328 | 3 | 1 | 1 | 0 | 2 | 0.5 | 5 | 1 | 11 | 0.8182 | 4 | 0.75 | 4 | 0.5 | 3 | 1 | 5 | 0.8 | 16 | 0.75 |
| 4_1319512 | 1 | 0 | 1 | 1 | 1 | 0 | NA | NA | 3 | 0.3333 | 1 | 1 | 3 | 0.6667 | NA | NA | NA | NA | 4 | 0.75 |
| 4_1319522 | NA | NA | 1 | 1 | NA | NA | NA | NA | 1 | 1 | 1 | 1 | 2 | 1 | NA | NA | NA | NA | 3 | 1 |
| 4_1319522 | 1 | 0 | 1 | 1 | NA | NA | NA | NA | 2 | 0.5 | 1 | 1 | 2 | 1 | NA | NA | NA | NA | 3 | 1 |
| 4_1319523 | 1 | 0 | 1 | 1 | 1 | 1 | NA | NA | 3 | 0.6667 | 1 | 1 | 2 | 0 | NA | NA | NA | NA | 3 | 0.3333 |
| 4_1319528 | NA | NA | 1 | 1 | 1 | 0 | NA | NA | 2 | 0.5 | NA | NA | 1 | 1 | NA | NA | NA | NA | 1 | 1 |
| 4_1319528 | 1 | 1 | 1 | 1 | NA | NA | NA | NA | 2 | 1 | 1 | 1 | NA | NA | NA | NA | NA | NA | 1 | 1 |
| 4_1319529 | 1 | 0 | 1 | 1 | 1 | 1 | 1 | 1 | 4 | 0.75 | 1 | 1 | 1 | 1 | NA | NA | NA | NA | 2 | 1 |
| 4_1319530 | 1 | 1 | 1 | 1 | 1 | 1 | 1 | 1 | 4 | 1 | NA | NA | 1 | 1 | NA | NA | NA | NA | 1 | 1 |
| 4_1319531 | 1 | 1 | 1 | 1 | 2 | 1 | 1 | 1 | 5 | 1 | NA | NA | 1 | 1 | NA | NA | NA | NA | 1 | 1 |
| 4_1319534 | 1 | 1 | 1 | 0 | 1 | 1 | 1 | 0 | 4 | 0.5 | NA | NA | 1 | 1 | NA | NA | NA | NA | 1 | 1 |
| 4_1319534 | 1 | 0 | 1 | 1 | 1 | 1 | NA | NA | 3 | 0.6667 | NA | NA | 2 | 1 | NA | NA | NA | NA | 2 | 1 |
| 4_1319537 | 2 | 0.5 | 1 | 0 | 1 | 0 | 1 | 0 | 5 | 0.2 | NA | NA | 3 | 0.3333 | NA | NA | NA | NA | 3 | 0.3333 |
| 4_1320199 | 3 | 0 | 1 | 0 | 3 | 0.3333 | 3 | 0 | 10 | 0.1 | 4 | 0.25 | 6 | 0.3333 | 3 | 0.3333 | 4 | 0.5 | 17 | 0.3529 |
| 4_1320200 | 4 | 0 | 1 | 0 | 2 | 0 | 4 | 0.25 | 11 | 0.0909 | 3 | 0.3333 | 3 | 0 | 2 | 0 | 4 | 0 | 12 | 0.0833 |
| 4_1320201 | 3 | 0 | 1 | 0 | 2 | 0.5 | 5 | 0.6 | 11 | 0.3636 | 3 | 0.3333 | 6 | 0.5 | 3 | 0.6667 | 5 | 0.4 | 17 | 0.4706 |
| 4_1320203 | 3 | 0 | 2 | 0 | 5 | 0.4 | 5 | 0.2 | 15 | 0.2 | 6 | 0.3333 | 7 | 0.4286 | 3 | 0.6667 | 3 | 0 | 19 | 0.3684 |
| 4_1320215 | 1 | 1 | NA | NA | NA | NA | NA | NA | 1 | 1 | 1 | 0 | 1 | 0 | 1 | 1 | NA | NA | 3 | 0.3333 |
| 4_1320216 | 1 | 1 | NA | NA | 1 | 1 | NA | NA | 2 | 1 | 1 | 1 | 1 | 1 | NA | NA | NA | NA | 2 | 1 |
| 4_1320216 | 1 | 1 | NA | NA | 1 | 1 | NA | NA | 2 | 1 | 1 | 1 | 1 | 0 | 1 | 1 | NA | NA | 3 | 0.6667 |
| 4_1320216 | 1 | 1 | NA | NA | NA | NA | NA | NA | 1 | 1 | 1 | 1 | 1 | 1 | 1 | 0 | NA | NA | 3 | 0.6667 |
| 4_1320392 | 6 | 0 | 3 | 0 | 4 | 0.25 | 3 | 0 | 16 | 0.0625 | 2 | 0 | 1 | 1 | 3 | 0.3333 | 11 | 0.2727 | 17 | 0.2941 |
| 4_1320594 | 1 | 0 | 1 | 1 | 1 | 1 | 1 | 1 | 4 | 0.75 | 2 | 1 | 1 | 1 | NA | NA | 1 | 1 | 4 | 1 |
| 4_1320595 | 1 | 0 | 1 | 1 | 1 | 0 | 2 | 1 | 5 | 0.6 | 2 | 0 | 1 | 1 | NA | NA | 1 | 0 | 4 | 0.25 |
| 4_1320597 | 1 | 0 | 1 | 0 | 1 | 1 | 2 | 1 | 5 | 0.6 | 2 | 0.5 | 1 | 0 | NA | NA | 1 | 0 | 4 | 0.25 |
| 4_1320597 | 1 | 0 | 1 | 1 | 1 | 1 | 2 | 0 | 5 | 0.4 | 1 | 1 | 1 | 1 | NA | NA | 1 | 1 | 3 | 1 |
| 4_1320598 | 1 | 0 | 1 | 0 | 1 | 1 | 1 | 0 | 4 | 0.25 | 2 | 0.5 | 1 | 0 | NA | NA | 1 | 0 | 4 | 0.25 |
| 4_1320602 | 1 | 0 | NA | NA | 1 | 0 | 1 | 1 | 3 | 0.3333 | NA | NA | 1 | 0 | NA | NA | 1 | 1 | 2 | 0.5 |
| 4_1320833 | 2 | 0.5 | 4 | 0 | 2 | 0.5 | 2 | 0 | 10 | 0.2 | 1 | 0 | 2 | 0 | 2 | 0.5 | 2 | 0 | 7 | 0.1429 |
| 4_1320847 | 2 | 0.5 | 1 | 1 | 1 | 0 | 2 | 0.5 | 6 | 0.5 | 1 | 0 | 1 | 1 | NA | NA | 1 | 0 | 3 | 0.3333 |
| 4_1320851 | 2 | 0.5 | 3 | 0.6667 | 1 | 1 | 2 | 0.5 | 8 | 0.625 | 2 | 1 | 1 | 1 | NA | NA | 1 | 1 | 4 | 1 |
| 4_1320853 | 1 | 0 | 3 | 0.3333 | 1 | 0 | 2 | 0.5 | 7 | 0.2857 | 1 | 0 | NA | NA | NA | NA | 1 | 1 | 2 | 0.5 |
| 4_1320981 | 2 | 0.5 | 2 | 0 | 1 | 0 | 2 | 1 | 7 | 0.4286 | 1 | 1 | NA | NA | NA | NA | 1 | 0 | 2 | 0.5 |
| 4_1321866 | 1 | 1 | 1 | 1 | NA | NA | 2 | 0.5 | 4 | 0.75 | NA | NA | 2 | 1 | 1 | 1 | NA | NA | 3 | 1 |
| 4_1321866 | 1 | 0 | 1 | 0 | 1 | 0 | 3 | 0.6667 | 6 | 0.3333 | NA | NA | 2 | 0.5 | 1 | 1 | NA | NA | 3 | 0.6667 |
| 4_1321866 | 1 | 1 | 1 | 1 | 1 | 1 | 1 | 0 | 4 | 0.75 | NA | NA | 2 | 0.5 | 1 | 1 | NA | NA | 3 | 0.6667 |
| 4_1321868 | 1 | 1 | 1 | 0 | 1 | 1 | 2 | 1 | 5 | 0.8 | NA | NA | 2 | 1 | 1 | 1 | NA | NA | 3 | 1 |
| 4_1321869 | 1 | 1 | 1 | 1 | 1 | 1 | 2 | 1 | 5 | 1 | NA | NA | 2 | 1 | 1 | 1 | NA | NA | 3 | 1 |
| 4_1321869 | 1 | 1 | 1 | 1 | 1 | 1 | 2 | 0.5 | 5 | 0.8 | NA | NA | 2 | 1 | 1 | 1 | NA | NA | 3 | 1 |
| 4_1321870 | 1 | 1 | 1 | 1 | 1 | 1 | 2 | 0 | 5 | 0.6 | NA | NA | 1 | 1 | 1 | 1 | NA | NA | 2 | 1 |
| 4_1321870 | 1 | 1 | 1 | 0 | 1 | 1 | 1 | 1 | 4 | 0.75 | NA | NA | 2 | 0.5 | 1 | 1 | NA | NA | 3 | 0.6667 |
| 4_1321871 | 1 | 0 | 1 | 0 | 1 | 0 | 2 | 0.5 | 5 | 0.2 | NA | NA | 2 | 1 | 1 | 0 | NA | NA | 3 | 0.6667 |
| 4_1321871 | 1 | 1 | 1 | 1 | 1 | 1 | 2 | 1 | 5 | 1 | NA | NA | 2 | 1 | NA | NA | NA | NA | 2 | 1 |
| 4_1321872 | 1 | 0 | NA | NA | 1 | 1 | 1 | 0 | 3 | 0.3333 | NA | NA | 1 | 1 | 1 | 0 | NA | NA | 2 | 0.5 |
| 4_1322198 | 2 | 0 | 1 | 0 | NA | NA | 1 | 1 | 4 | 0.25 | 1 | 1 | 2 | 0 | NA | NA | 2 | 0 | 5 | 0.2 |
| 4_1322198 | 3 | 0.3333 | NA | NA | NA | NA | 1 | 0 | 4 | 0.25 | 1 | 0 | 2 | 0 | NA | NA | 1 | 1 | 4 | 0.25 |
| 4_1322199 | 3 | 0.3333 | 1 | 0 | NA | NA | 1 | 0 | 5 | 0.2 | 1 | 0 | 1 | 0 | NA | NA | 2 | 0.5 | 4 | 0.25 |
| 4_1322200 | 3 | 0.3333 | 1 | 0 | NA | NA | 1 | 0 | 5 | 0.2 | 1 | 1 | 2 | 0 | NA | NA | 1 | 0 | 4 | 0.25 |
| 4_1322205 | 3 | 0.6667 | 1 | 0 | NA | NA | 1 | 0 | 5 | 0.4 | 1 | 1 | 1 | 0 | NA | NA | 1 | 0 | 3 | 0.3333 |
| 4_1322206 | 3 | 1 | NA | NA | NA | NA | 1 | 1 | 4 | 1 | 1 | 1 | 1 | 1 | NA | NA | 1 | 0 | 3 | 0.6667 |
| 4_1322206 | 2 | 0.5 | NA | NA | NA | NA | 1 | 1 | 3 | 0.6667 | 1 | 0 | 1 | 1 | NA | NA | 1 | 0 | 3 | 0.3333 |
| 4_1322206 | 2 | 1 | NA | NA | NA | NA | 1 | 1 | 3 | 1 | NA | NA | 1 | 1 | NA | NA | 1 | 0 | 2 | 0.5 |
| 4_1322208 | 3 | 1 | NA | NA | NA | NA | 1 | 1 | 4 | 1 | 1 | 1 | 1 | 1 | NA | NA | NA | NA | 2 | 1 |
| 4_1322208 | 3 | 0.6667 | NA | NA | NA | NA | 1 | 0 | 4 | 0.5 | 1 | 0 | 1 | 1 | NA | NA | 1 | 1 | 3 | 0.6667 |
| 4_1322208 | 2 | 1 | NA | NA | NA | NA | 1 | 1 | 3 | 1 | 1 | 1 | 1 | 1 | NA | NA | 1 | 1 | 3 | 1 |
| 4_1322209 | 3 | 0 | NA | NA | NA | NA | 1 | 1 | 4 | 0.25 | 1 | 1 | 1 | 1 | NA | NA | 1 | 0 | 3 | 0.6667 |
| 4_1322209 | 3 | 0.6667 | NA | NA | NA | NA | 1 | 1 | 4 | 0.75 | 1 | 1 | 1 | 1 | NA | NA | 1 | 1 | 3 | 1 |

| | | | | | | | | | | | | | | | | | | | | |
|---|---|---|---|---|---|---|---|---|---|---|---|---|---|---|---|---|---|---|---|---|
| 4_13228049 | 19 | 1 | 22 | 0.6818 | 11 | 0.5455 | 12 | 0.8333 | 64 | 0.7812 | 9 | 0.8889 | 10 | 0.7 | 21 | 0.5238 | 20 | 0.75 | 60 | 0.6833 |
| 4_13228067 | 17 | 0 | 26 | 0.1154 | 14 | 0.2143 | 14 | 0.0714 | 71 | 0.0986 | 11 | 0 | 10 | 0.2 | 22 | 0.0455 | 20 | 0.1 | 63 | 0.0794 |
| 4_13244867 | NA | NA | 1 | 1 | 1 | 1 | 1 | 1 | 3 | 1 | NA | NA | 3 | 0.3333 | 1 | 1 | 2 | 0.5 | 6 | 0.5 |
| 4_13244868 | NA | NA | 1 | 0 | 1 | 1 | 3 | 0.3333 | 5 | 0.4 | NA | NA | 3 | 0.3333 | 1 | 0 | 2 | 0.5 | 6 | 0.3333 |
| 4_13244873 | NA | NA | 1 | 0 | 1 | 1 | 3 | 0 | 5 | 0.2 | NA | NA | 3 | 0.3333 | 2 | 0 | 3 | 0 | 8 | 0.125 |
| 4_13244883 | NA | NA | 1 | 0 | 1 | 1 | 3 | 0 | 5 | 0.2 | NA | NA | 2 | 0 | 2 | 0 | 3 | 0.3333 | 7 | 0.1429 |
| 4_13244902 | NA | NA | 1 | 1 | NA | NA | NA | NA | 1 | 1 | NA | NA | 2 | 0.5 | 3 | 1 | 3 | 1 | 8 | 0.875 |
| 4_13244906 | NA | NA | 1 | 1 | 1 | 1 | 2 | 0 | 4 | 0.5 | NA | NA | 2 | 0 | 3 | 0.6667 | 3 | 0.3333 | 8 | 0.375 |
| 4_13244924 | NA | NA | 1 | 0 | NA | NA | 4 | 0.5 | 5 | 0.4 | NA | NA | 3 | 0 | 4 | 0.75 | 3 | 0 | 10 | 0.3 |
| 4_13244932 | NA | NA | 2 | 0.5 | 1 | 1 | 4 | 0.5 | 7 | 0.5714 | NA | NA | 3 | 0 | 4 | 0.75 | 3 | 0.3333 | 10 | 0.4 |
| 4_13244942 | NA | NA | 2 | 0 | 1 | 1 | 3 | 0.3333 | 6 | 0.3333 | NA | NA | 3 | 0.3333 | 4 | 0.25 | 4 | 0 | 11 | 0.1818 |
| 4_13244951 | NA | NA | 1 | 1 | 1 | 1 | 3 | 0.3333 | 5 | 0.6 | NA | NA | 2 | 0.5 | 1 | 0 | 3 | 0.6667 | 6 | 0.5 |
| 4_13245023 | NA | NA | NA | NA | 1 | 1 | 3 | 0 | 4 | 0.25 | NA | NA | 3 | 0.3333 | 1 | 0 | 3 | 0 | 7 | 0.1429 |
| 4_13245125 | 2 | 0 | 2 | 0 | 1 | 1 | 4 | 0.25 | 9 | 0.2222 | 1 | 1 | 6 | 0.3333 | 1 | 0 | 4 | 0 | 12 | 0.25 |
| 4_13245140 | 2 | 0 | 2 | 0 | 1 | 0 | 4 | 0.25 | 9 | 0.1111 | 1 | 1 | 6 | 0.1667 | 1 | 0 | 4 | 0 | 12 | 0.1667 |
| 4_13245150 | 2 | 0.5 | 2 | 0 | 1 | 0 | 4 | 0 | 9 | 0.1111 | 1 | 1 | 4 | 0.5 | 1 | 0 | 3 | 0.3333 | 9 | 0.4444 |
| 4_13245191 | 1 | 1 | NA | NA | 1 | 1 | 3 | 0.3333 | 5 | 0.6 | 1 | 1 | 4 | 0.5 | 2 | 0.5 | 4 | 1 | 11 | 0.7273 |
| 4_13245246 | NA | NA | NA | NA | 1 | 1 | 1 | 0 | 2 | 0.5 | 1 | 0 | NA | NA | 1 | 1 | NA | NA | 2 | 0.5 |
| 4_13245260 | 1 | 1 | NA | NA | 1 | 0 | 1 | 0 | 3 | 0.3333 | NA | NA | NA | NA | 1 | 1 | NA | NA | 1 | 1 |
| 4_13245397 | 1 | 1 | NA | NA | 1 | 1 | NA | NA | 2 | 1 | 1 | 1 | NA | NA | NA | NA | NA | NA | 1 | 1 |
| 4_13246030 | NA | NA | 2 | 0.5 | NA | NA | 5 | 1 | 7 | 0.8571 | NA | NA | 1 | 1 | NA | NA | 1 | 1 | 2 | 1 |
| 4_13246035 | NA | NA | 2 | 0.5 | NA | NA | 4 | 0.25 | 6 | 0.3333 | NA | NA | 1 | 1 | NA | NA | 1 | 1 | 2 | 1 |
| 4_13246044 | NA | NA | 2 | 1 | NA | NA | 5 | 1 | 7 | 1 | NA | NA | 1 | 1 | NA | NA | 1 | 1 | 2 | 1 |
| 4_13246070 | NA | NA | 2 | 1 | NA | NA | 5 | 0.8 | 7 | 0.8571 | NA | NA | 1 | 0 | NA | NA | 1 | 1 | 2 | 0.5 |
| 4_13246085 | NA | NA | 2 | 0.5 | NA | NA | 4 | 0.5 | 6 | 0.5 | NA | NA | 1 | 1 | NA | NA | 1 | 0 | 2 | 0.5 |
| 4_13246102 | NA | NA | NA | NA | NA | NA | 3 | 0.3333 | 3 | 0.3333 | NA | NA | 1 | 0 | NA | NA | 1 | 1 | 2 | 0.5 |
| 4_13246106 | NA | NA | 1 | 0 | NA | NA | 4 | 1 | 5 | 0.8 | NA | NA | 1 | 1 | NA | NA | 1 | 1 | 2 | 1 |
| 4_13250527 | 5 | 0.2 | 1 | 0 | NA | NA | 3 | 0.3333 | 9 | 0.2222 | 4 | 0 | 2 | 0 | 4 | 0.25 | 4 | 0.5 | 14 | 0.2143 |
| 4_13250528 | 5 | 0.4 | 1 | 0 | NA | NA | 3 | 0.3333 | 9 | 0.3333 | 2 | 0.5 | 1 | 1 | 5 | 0.6 | 5 | 0.6 | 13 | 0.6154 |
| 4_13250532 | 6 | 0.1667 | 1 | 0 | NA | NA | 3 | 0.3333 | 10 | 0.2 | 2 | 0 | 2 | 0 | 4 | 0.25 | 5 | 0 | 13 | 0.0769 |
| 4_13250540 | 5 | 0.8 | 1 | 0 | NA | NA | 3 | 0.3333 | 9 | 0.5556 | 4 | 0.5 | 1 | 1 | 4 | 0.5 | 5 | 0.6 | 14 | 0.5714 |
| 4_13255490 | 11 | 0.0909 | 3 | 0 | 6 | 0.3333 | 5 | 0.2 | 25 | 0.16 | 6 | 0.1667 | 7 | 0.2857 | 9 | 0.1111 | 6 | 0.3333 | 28 | 0.2143 |
| 4_13255531 | 8 | 0.875 | 2 | 1 | 3 | 1 | 6 | 1 | 19 | 0.9474 | 4 | 0.75 | 6 | 0.8333 | 7 | 0.8571 | 4 | 1 | 21 | 0.8571 |
| 4_13255537 | 9 | 0.3333 | 2 | 0.5 | 3 | 0 | 5 | 0.4 | 19 | 0.3158 | 3 | 0.6667 | 5 | 0 | 7 | 0.4286 | 5 | 0.2 | 20 | 0.3 |
| 4_13255543 | 6 | 1 | 2 | 1 | 3 | 1 | 3 | 1 | 14 | 1 | 3 | 0.3333 | 7 | 0.7143 | 4 | 0.5 | 4 | 0.75 | 18 | 0.6111 |
| 4_13255545 | 9 | 0.3333 | 2 | 1 | 1 | 0 | 3 | 0 | 15 | 0.3333 | 4 | 0.25 | 5 | 0 | 6 | 0 | 5 | 0 | 20 | 0.05 |
| 4_13258093 | 2 | 0 | 8 | 0.125 | 3 | 0 | 5 | 0.2 | 18 | 0.1111 | 2 | 0 | 2 | 0 | 5 | 0 | 7 | 0.1429 | 16 | 0.0625 |
| 4_13258100 | NA | NA | 8 | 0.875 | 4 | 0.5 | 5 | 0.8 | 17 | 0.7647 | 2 | 1 | 2 | 1 | 5 | 1 | 6 | 0.6667 | 15 | 0.8667 |
| 4_13258111 | 1 | 0 | 8 | 0.625 | 4 | 0.25 | 5 | 0.4 | 18 | 0.4444 | 2 | 0 | 2 | 0.5 | 5 | 0.6 | 7 | 0.2857 | 16 | 0.375 |
| 4_13258122 | NA | NA | 5 | 0.2 | 4 | 0 | 5 | 0 | 14 | 0.0714 | 2 | 0 | 2 | 0 | 4 | 0 | 6 | 0.1667 | 14 | 0.0714 |
| 4_13258217 | 15 | 0.2 | 17 | 0.1765 | 12 | 0.25 | 14 | 0.3571 | 58 | 0.2414 | 11 | 0.0909 | 18 | 0.3889 | 16 | 0.25 | 15 | 0.5333 | 60 | 0.3333 |
| 4_13258235 | 18 | 0.0556 | 20 | 0.1 | 13 | 0.0769 | 14 | 0.0714 | 65 | 0.0769 | 9 | 0 | 24 | 0.125 | 20 | 0.15 | 19 | 0 | 72 | 0.0833 |
| 4_13258270 | 13 | 0.0769 | 14 | 0.2857 | 8 | 0.25 | 12 | 0.3333 | 47 | 0.234 | 6 | 0.1667 | 20 | 0.3 | 19 | 0.3684 | 16 | 0.375 | 61 | 0.3279 |
| 4_13262685 | 9 | 0 | 7 | 0 | 12 | 0.0833 | 7 | 0.1429 | 35 | 0.0571 | 3 | 0 | 12 | 0.0833 | 9 | 0.3333 | 14 | 0 | 38 | 0.1053 |
| 4_13262705 | 8 | 0.5 | 7 | 0.2857 | 11 | 0.4545 | 6 | 0.6667 | 32 | 0.4688 | 3 | 0 | 11 | 0.4545 | 9 | 0.7778 | 14 | 0.2143 | 37 | 0.4054 |
| 4_13262727 | 8 | 0.125 | 5 | 0 | 10 | 0.1 | 7 | 0.1429 | 30 | 0.1 | 2 | 0 | 11 | 0.0909 | 8 | 0.25 | 14 | 0 | 35 | 0.0857 |
| 4_13262838 | 5 | 0 | 4 | 0 | 6 | 0 | 7 | 0.4286 | 22 | 0.1364 | 2 | 0 | 7 | 0.1429 | 8 | 0.125 | 9 | 0.5556 | 26 | 0.2692 |
| 4_13278879 | 1 | 1 | NA | NA | NA | NA | 1 | 0 | 2 | 0.5 | 2 | 0 | 4 | 0.25 | NA | NA | 1 | 0 | 7 | 0.1429 |
| 4_13278914 | 1 | 1 | 1 | 1 | NA | NA | 1 | 1 | 3 | 1 | 1 | 1 | 4 | 1 | NA | NA | 1 | 1 | 6 | 1 |
| 4_13278919 | NA | NA | 1 | 1 | NA | NA | NA | NA | 1 | 1 | 1 | 0 | 4 | 1 | NA | NA | NA | NA | 5 | 0.8 |
| 4_13278935 | 1 | 1 | 1 | 1 | 2 | 0.5 | NA | NA | 4 | 0.75 | 1 | 1 | 4 | 0.5 | 1 | 1 | 1 | 0 | 7 | 0.5714 |
| 4_13278949 | 1 | 1 | NA | NA | 2 | 0.5 | 1 | 1 | 4 | 0.75 | 2 | 1 | 4 | 1 | 1 | 1 | NA | NA | 7 | 1 |
| 4_13278976 | NA | NA | 1 | 1 | 2 | 0 | 1 | 0 | 4 | 0.25 | 1 | 0 | 4 | 0.5 | 1 | 0 | 1 | 0 | 7 | 0.2857 |
| 4_13278985 | 1 | 1 | 1 | 1 | 1 | 1 | NA | NA | 3 | 1 | 2 | 1 | 4 | 1 | 1 | 1 | NA | NA | 7 | 1 |
| 4_13278997 | NA | NA | 1 | 1 | 1 | 0 | 1 | 0 | 3 | 0.3333 | NA | NA | 4 | 0.5 | 1 | 1 | 1 | 0 | 6 | 0.5 |
| 4_13279057 | 1 | 0 | 4 | 0 | 1 | 1 | NA | NA | 6 | 0.1667 | 4 | 0.25 | 4 | 0 | 2 | 0 | 1 | 1 | 11 | 0.1818 |
| 4_13279066 | 2 | 1 | 3 | 0.6667 | NA | NA | 1 | 1 | 6 | 0.8333 | 2 | 0.5 | 4 | 0.75 | 1 | 1 | 1 | 1 | 8 | 0.75 |
| 4_13279070 | 2 | 1 | 3 | 1 | 1 | 1 | 1 | 1 | 7 | 1 | 4 | 1 | 4 | 1 | 1 | 1 | 1 | 1 | 10 | 1 |
| 4_13279082 | 1 | 1 | 3 | 1 | 2 | 1 | 1 | 1 | 7 | 1 | 2 | 1 | 3 | 1 | 1 | 1 | 1 | 1 | 7 | 1 |
| 4_13279099 | 2 | 0.5 | 4 | 0.75 | 2 | 0 | 2 | 1 | 10 | 0.6 | 2 | 1 | 4 | 1 | 1 | 1 | 3 | 0.6667 | 10 | 0.9 |
| 4_13279112 | 2 | 1 | 5 | 0.8 | 2 | 0.5 | 3 | 1 | 12 | 0.8333 | 4 | 0.5 | 3 | 1 | 1 | 1 | 3 | 1 | 11 | 0.8182 |
| 4_13279113 | 1 | 0 | 4 | 0.75 | 1 | 1 | 2 | 0 | 8 | 0.5 | 4 | 0.25 | 4 | 0 | 2 | 0.5 | 2 | 0 | 12 | 0.1667 |
| 4_13279130 | 2 | 1 | 5 | 1 | 1 | 1 | 2 | 1 | 10 | 1 | 4 | 1 | 3 | 1 | 1 | 1 | 2 | 1 | 10 | 1 |
| 4_13279135 | 2 | 0.5 | 4 | 0.25 | 1 | 0 | 3 | 0 | 10 | 0.2 | 4 | 0.25 | 3 | 0.6667 | 1 | 0 | 3 | 0 | 11 | 0.2727 |
| 4_13279155 | 2 | 0.5 | 5 | 0.6 | 2 | 0 | 3 | 0.6667 | 12 | 0.5 | 3 | 0.6667 | 4 | 0.5 | 1 | 1 | 1 | 1 | 9 | 0.6667 |

| | | | | | | | | | | | | | | | | | | | | |
|---|---|---|---|---|---|---|---|---|---|---|---|---|---|---|---|---|---|---|---|---|
| 4_1327915 | 2 | 0 | 5 | 0.2 | 2 | 0 | 3 | 0.3333 | 12 | 0.1667 | 3 | 0.3333 | 4 | 0.25 | 1 | 0 | 3 | 0 | 11 | 0.1818 |
| 4_1327917 | 2 | 1 | 5 | 1 | 2 | 1 | 2 | 1 | 11 | 1 | 4 | 1 | 4 | 1 | 1 | 1 | 3 | 1 | 12 | 1 |
| 4_1327917 | 2 | 1 | 5 | 0.8 | 1 | 1 | 3 | 1 | 11 | 0.9091 | 4 | 0.75 | 4 | 1 | 1 | 1 | 2 | 0.5 | 11 | 0.8182 |
| 4_1327919 | 1 | 0 | 4 | 0.5 | 2 | 0 | 3 | 0 | 10 | 0.2 | 3 | 0 | 2 | 0 | 1 | 0 | 3 | 0.3333 | 9 | 0.1111 |
| 4_1327920 | 1 | 0 | 5 | 1 | 2 | 1 | 3 | 0.6667 | 11 | 0.8182 | 2 | 1 | 4 | 1 | NA | NA | 3 | 1 | 9 | 1 |
| 4_1327920 | 1 | 0 | 1 | 1 | 1 | 0 | 1 | 1 | 4 | 0.5 | 2 | 0.5 | 4 | 0.75 | 1 | 1 | 2 | 1 | 9 | 0.7778 |
| 4_1327921 | 1 | 0 | 4 | 0.25 | 1 | 0 | 2 | 0 | 8 | 0.125 | 4 | 0.25 | 3 | 0.6667 | 1 | 0 | 3 | 0.3333 | 11 | 0.3636 |
| 4_1327923 | 2 | 0.5 | 5 | 1 | 2 | 1 | 4 | 1 | 13 | 0.9231 | 4 | 1 | 5 | 1 | 1 | 0 | 3 | 1 | 13 | 0.9231 |
| 4_1327924 | 2 | 0 | 5 | 0 | 3 | 0 | 4 | 0.25 | 14 | 0.0714 | 4 | 0.5 | 6 | 0 | 2 | 0.5 | 3 | 0 | 15 | 0.2 |
| 4_1327926 | NA | NA | 3 | 1 | 3 | 1 | 2 | 1 | 8 | 1 | 5 | 1 | 3 | 1 | 2 | 1 | 1 | 1 | 11 | 1 |
| 4_1327927 | 2 | 0.5 | 5 | 0.4 | 5 | 0.2 | 4 | 0.25 | 16 | 0.3125 | 5 | 0.6 | 6 | 0.5 | 3 | 0.6667 | 4 | 0.25 | 18 | 0.5 |
| 4_1327928 | 1 | 1 | 4 | 0.25 | 4 | 0.25 | 4 | 0.25 | 13 | 0.3077 | 5 | 0.6 | 3 | 0.3333 | 4 | 0.5 | 3 | 0 | 15 | 0.4 |
| 4_1327929 | 3 | 0.3333 | 4 | 0.25 | 5 | 0.6 | 4 | 0.5 | 16 | 0.4375 | 3 | 0.6667 | 5 | 0.2 | 2 | 0 | 4 | 0.5 | 14 | 0.3571 |
| 4_1327930 | 4 | 1 | 4 | 0.5 | 5 | 0.6 | 4 | 0.75 | 17 | 0.7059 | 5 | 0.8 | 5 | 1 | 3 | 1 | 4 | 0.75 | 17 | 0.8824 |
| 4_1327932 | 4 | 0.25 | 4 | 0.25 | 5 | 0.2 | 4 | 0.5 | 17 | 0.2941 | 5 | 0.6 | 5 | 0.2 | 2 | 1 | 3 | 0 | 15 | 0.4 |
| 4_1327933 | 5 | 0.8 | 4 | 1 | 5 | 1 | 4 | 1 | 18 | 0.9444 | 5 | 1 | 5 | 0.8 | 3 | 1 | 3 | 1 | 16 | 0.9375 |
| 4_1327933 | 3 | 0 | 2 | 0 | 4 | 0 | 3 | 0.3333 | 12 | 0.0833 | 5 | 0.2 | 5 | 0 | 3 | 0.3333 | 3 | 0 | 16 | 0.125 |
| 4_1327933 | 5 | 0.8 | 3 | 0.3333 | 5 | 0.8 | 3 | 0.6667 | 16 | 0.6875 | 5 | 0.4 | 7 | 1 | 3 | 0.6667 | 4 | 0.75 | 19 | 0.7368 |
| 4_1327934 | 4 | 1 | 5 | 0.8 | 4 | 0.5 | 4 | 1 | 17 | 0.8235 | 5 | 0.8 | 6 | 0.5 | 3 | 1 | 5 | 0.6 | 19 | 0.6842 |
| 4_1327935 | 5 | 0.2 | 4 | 0.5 | 3 | 0 | 3 | 0.3333 | 15 | 0.2667 | 3 | 0.3333 | 6 | 0.5 | 3 | 0 | 5 | 0.4 | 17 | 0.3529 |
| 4_1328046 | 4 | 0 | 4 | 0 | 6 | 0.1667 | 5 | 0 | 19 | 0.0526 | 6 | 0.1667 | 8 | 0.125 | 4 | 0 | 5 | 0 | 23 | 0.087 |
| 4_1328256 | 11 | 0 | 16 | 0.0625 | 7 | 0.1429 | 6 | 0 | 40 | 0.05 | 7 | 0.1429 | 3 | 0.3333 | 7 | 0 | 12 | 0.0833 | 29 | 0.1034 |
| 4_1328257 | 4 | 0.25 | 11 | 0.0909 | 2 | 0 | 6 | 0 | 23 | 0.087 | 6 | 0.5 | 4 | 0.25 | 6 | 0.1667 | 10 | 0.4 | 26 | 0.3462 |
| 4_1328257 | 10 | 0.1 | 13 | 0.1538 | 8 | 0 | 3 | 0 | 34 | 0.0882 | 4 | 0 | 4 | 0.5 | 6 | 0.1667 | 11 | 0.1818 | 25 | 0.2 |
| 4_1328257 | 9 | 0 | 15 | 0.0667 | 8 | 0.125 | 6 | 0 | 38 | 0.0526 | 5 | 0.2 | 4 | 0 | 7 | 0 | 14 | 0.0714 | 30 | 0.0667 |
| 4_1328260 | 12 | 0.5 | 19 | 0.5263 | 9 | 0.6667 | 5 | 0.2 | 45 | 0.5111 | 9 | 0.6667 | 4 | 0.75 | 6 | 0.5 | 14 | 0.7857 | 33 | 0.697 |
| 4_1328261 | 12 | 0.0833 | 15 | 0.1333 | 9 | 0.1111 | 4 | 0.25 | 40 | 0.125 | 8 | 0.125 | 3 | 0 | 7 | 0 | 16 | 0.125 | 34 | 0.0882 |
| 4_1328262 | 12 | 0.0833 | 15 | 0.0667 | 10 | 0 | 5 | 0.6 | 42 | 0.119 | 10 | 0.2 | 4 | 0 | 5 | 0 | 14 | 0.1429 | 33 | 0.1212 |
| 4_1328262 | 11 | 0.7273 | 16 | 0.625 | 9 | 0.7778 | 6 | 0.8333 | 42 | 0.7143 | 10 | 0.9 | 3 | 1 | 4 | 0.75 | 17 | 1 | 34 | 0.9412 |
| 4_1328263 | 10 | 0.1 | 17 | 0.0588 | 10 | 0 | 6 | 0.1667 | 43 | 0.0698 | 9 | 0.1111 | 3 | 0.3333 | 6 | 0.1667 | 15 | 0.0667 | 33 | 0.1212 |
| 4_1328263 | 10 | 0.5 | 15 | 0.3333 | 8 | 0.25 | 6 | 0.6667 | 39 | 0.4103 | 9 | 0.2222 | 3 | 0.6667 | 7 | 0.4286 | 15 | 0.6 | 34 | 0.4706 |
| 4_1328264 | 11 | 0.1818 | 20 | 0.1 | 11 | 0.0909 | 7 | 0.2857 | 49 | 0.1429 | 10 | 0.2 | 4 | 0.25 | 6 | 0.3333 | 18 | 0.2222 | 38 | 0.2368 |
| 4_1328267 | 10 | 0.2 | 18 | 0.1667 | 7 | 0 | 9 | 0.1111 | 44 | 0.1364 | 15 | 0.6 | 5 | 0.4 | 7 | 0.5714 | 14 | 0.2143 | 41 | 0.439 |
| 4_1328269 | 14 | 0.1429 | 19 | 0.2105 | 11 | 0.1818 | 9 | 0.2222 | 53 | 0.1887 | 14 | 0.2143 | 6 | 0.5 | 4 | 0.75 | 17 | 0.2353 | 41 | 0.3171 |
| 4_1328271 | 10 | 0.9 | 17 | 0.9412 | 10 | 0.8 | 9 | 0.7778 | 46 | 0.8696 | 12 | 1 | 6 | 0.8333 | 6 | 1 | 15 | 0.9333 | 39 | 0.9487 |
| 4_1328271 | 15 | 0.0667 | 20 | 0.15 | 11 | 0.0909 | 8 | 0.375 | 54 | 0.1481 | 15 | 0.1333 | 7 | 0.2857 | 8 | 0.125 | 16 | 0.25 | 46 | 0.1957 |
| 4_1328271 | 14 | 0.2143 | 21 | 0.1905 | 10 | 0.1 | 9 | 0.3333 | 54 | 0.2037 | 15 | 0.2667 | 5 | 0.4 | 8 | 0 | 16 | 0.125 | 44 | 0.1818 |
| 4_1328271 | 14 | 0.1429 | 18 | 0.2222 | 9 | 0.2222 | 9 | 0.2222 | 50 | 0.2 | 12 | 0.4167 | 4 | 0.25 | 7 | 0.1429 | 15 | 0.2667 | 38 | 0.2895 |
| 4_1328273 | 17 | 0.2941 | 21 | 0.1905 | 11 | 0.1818 | 9 | 0.2222 | 58 | 0.2241 | 16 | 0.25 | 5 | 0.4 | 7 | 0.5714 | 17 | 0.0588 | 45 | 0.2444 |
| 4_1328275 | 14 | 0.7143 | 18 | 0.7778 | 10 | 0.8 | 10 | 0.6 | 52 | 0.7308 | 14 | 0.8571 | 7 | 0.8571 | 8 | 0.875 | 16 | 0.8125 | 45 | 0.8444 |
| 4_1328275 | 15 | 0.1333 | 18 | 0.0556 | 11 | 0 | 10 | 0.1 | 54 | 0.0741 | 15 | 0 | 6 | 0 | 8 | 0.125 | 17 | 0.1176 | 46 | 0.0652 |
| 4_1328277 | 16 | 0 | 22 | 0.0909 | 13 | 0 | 11 | 0.1818 | 62 | 0.0645 | 19 | 0.1053 | 7 | 0 | 10 | 0 | 21 | 0.0476 | 57 | 0.0526 |
| 4_1328277 | 16 | 0.0625 | 19 | 0.3158 | 13 | 0.1538 | 12 | 0.25 | 60 | 0.2 | 20 | 0.3 | 8 | 0.25 | 10 | 0.3 | 21 | 0.3333 | 59 | 0.3051 |
| 4_1328278 | 16 | 0.5 | 21 | 0.5714 | 12 | 0.4167 | 13 | 0.7692 | 62 | 0.5645 | 17 | 0.8235 | 7 | 1 | 10 | 0.5 | 15 | 0.6667 | 49 | 0.7347 |
| 4_1328281 | 17 | 0.1765 | 28 | 0.1429 | 14 | 0.0714 | 15 | 0.3333 | 74 | 0.1757 | 20 | 0.05 | 9 | 0.1111 | 10 | 0.4 | 22 | 0.1818 | 61 | 0.1639 |
| 4_1328281 | 16 | 0.3125 | 23 | 0.2609 | 14 | 0.0714 | 15 | 0.3333 | 68 | 0.25 | 17 | 0.2941 | 8 | 0 | 9 | 0.1111 | 19 | 0.5263 | 53 | 0.3019 |
| 4_1328281 | 18 | 0.3333 | 23 | 0.3913 | 11 | 0.5455 | 13 | 0.6923 | 65 | 0.4615 | 18 | 0.4444 | 8 | 0.75 | 8 | 0.5 | 20 | 0.6 | 54 | 0.5556 |
| 4_1328282 | 17 | 0.2941 | 24 | 0.125 | 11 | 0.1818 | 11 | 0.2727 | 63 | 0.2063 | 18 | 0.2778 | 9 | 0.2222 | 10 | 0.2 | 19 | 0.3158 | 56 | 0.2679 |
| 4_1328283 | 17 | 0.2353 | 27 | 0.1481 | 15 | 0.0667 | 10 | 0.1 | 69 | 0.1449 | 20 | 0.1 | 10 | 0.3 | 11 | 0.3636 | 18 | 0.1111 | 59 | 0.1864 |
| 4_1328285 | 18 | 0.1111 | 23 | 0.0435 | 15 | 0.0667 | 15 | 0.0667 | 71 | 0.0704 | 25 | 0.08 | 11 | 0.0909 | 12 | 0.0833 | 20 | 0.1 | 68 | 0.0882 |
| 4_1329232 | NA | NA | 1 | 1 | NA | NA | 1 | 0 | 2 | 0.5 | 1 | 0 | 2 | 0 | 3 | 0.3333 | 1 | 0 | 7 | 0.1429 |
| 4_1329232 | 1 | 1 | 1 | 1 | NA | NA | 1 | 1 | 3 | 1 | 1 | 1 | 1 | 1 | 3 | 0.6667 | 1 | 1 | 6 | 0.8333 |
| 4_1329237 | NA | NA | 1 | 1 | NA | NA | 1 | 1 | 2 | 1 | 1 | 1 | 3 | 1 | 3 | 0.6667 | NA | NA | 7 | 0.8571 |
| 4_1329261 | 4 | 0.25 | 1 | 0 | 1 | 0 | 2 | 0 | 8 | 0.125 | 1 | 0 | 5 | 0.2 | 2 | 0 | 1 | 0 | 9 | 0.1111 |
| 4_1329263 | 5 | 0 | 2 | 0 | 1 | 0 | 3 | 0.3333 | 11 | 0.0909 | 1 | 0 | 4 | 0 | 4 | 0 | 1 | 1 | 10 | 0.1 |
| 4_1329801 | NA | NA | 3 | 0 | 2 | 0 | 3 | 0.3333 | 8 | 0.125 | 1 | 1 | NA | NA | NA | NA | NA | NA | 1 | 1 |
| 4_1329936 | 13 | 0 | 17 | 0.2353 | 14 | 0.2143 | 7 | 0 | 51 | 0.1373 | 12 | 0.25 | 9 | 0.1111 | 19 | 0.1053 | 11 | 0.1818 | 51 | 0.1569 |
| 4_1329937 | 11 | 0.3636 | 18 | 0.2222 | 13 | 0.2308 | 8 | 0.125 | 50 | 0.24 | 15 | 0.2667 | 9 | 0.2222 | 18 | 0.2778 | 11 | 0.5455 | 53 | 0.3208 |
| 4_1329940 | 10 | 0.1 | 11 | 0.1818 | 12 | 0.1667 | 7 | 0 | 40 | 0.125 | 12 | 0.3333 | 7 | 0.1429 | 16 | 0.25 | 11 | 0 | 46 | 0.1957 |
| 4_1329940 | 12 | 0.3333 | 14 | 0.3571 | 11 | 0.4545 | 8 | 0.375 | 45 | 0.3778 | 13 | 0.3077 | 8 | 0.25 | 18 | 0.5 | 12 | 0.25 | 51 | 0.3529 |
| 4_1330200 | 11 | 0 | 9 | 0 | 12 | 0.1667 | 6 | 0 | 38 | 0.0526 | 9 | 0.2222 | 7 | 0 | 11 | 0.0909 | 10 | 0 | 37 | 0.0811 |
| 4_1330207 | 6 | 0.3333 | 9 | 0 | 9 | 0.2222 | 5 | 0.2 | 29 | 0.1724 | 11 | 0.1818 | 5 | 0 | 10 | 0.2 | 6 | 0.1667 | 32 | 0.1562 |
| 4_1330211 | 4 | 0.75 | 6 | 0.6667 | 6 | 1 | 3 | 0.6667 | 19 | 0.7895 | 6 | 1 | 3 | 1 | 8 | 0.625 | 6 | 1 | 23 | 0.8696 |
| 4_1330213 | 6 | 0 | 7 | 0.1429 | 6 | 0.1667 | 3 | 0.3333 | 22 | 0.1364 | 9 | 0 | 5 | 0 | 8 | 0.25 | 5 | 0 | 27 | 0.0741 |
| 4_1330213 | 6 | 0.1667 | 7 | 0.2857 | 6 | 0.5 | 2 | 0 | 21 | 0.2857 | 8 | 0.5 | 4 | 0.25 | 8 | 0.125 | 3 | 0.3333 | 23 | 0.3043 |

| | | | | | | | | | | | | | | | | | | | | |
|---|---|---|---|---|---|---|---|---|---|---|---|---|---|---|---|---|---|---|---|---|
| 4_1330213 | 4 | 0.25 | 6 | 0.6667 | 5 | 0.2 | 3 | 0.6667 | 18 | 0.4444 | 7 | 0.4286 | 3 | 0 | 7 | 0.2857 | 6 | 0.3333 | 23 | 0.3043 |
| 4_1330215 | 5 | 0.2 | 5 | 0 | 4 | 0 | NA | NA | 14 | 0.0714 | 6 | 0 | 3 | 1 | 7 | 0.2857 | 5 | 0.2 | 21 | 0.2857 |
| 4_1330215 | 3 | 0 | 6 | 0 | 4 | 0 | 2 | 1 | 15 | 0.1333 | 5 | 0.4 | 4 | 0.25 | 9 | 0.4444 | 4 | 0.5 | 22 | 0.4091 |
| 4_1330973 | 1 | 1 | NA | NA | 1 | 1 | 3 | 1 | 5 | 1 | NA | NA | 1 | 0 | 1 | 1 | NA | NA | 2 | 0.5 |
| 4_1330976 | NA | NA | NA | NA | 1 | 1 | 1 | 1 | 2 | 1 | NA | NA | 2 | 1 | 1 | 1 | NA | NA | 3 | 1 |
| 4_1330977 | 1 | 0 | NA | NA | 1 | 0 | 2 | 1 | 4 | 0.5 | NA | NA | 2 | 1 | 2 | 0.5 | NA | NA | 4 | 0.75 |
| 4_1330979 | NA | NA | NA | NA | NA | NA | 2 | 1 | 2 | 1 | 2 | 1 | 2 | 1 | 1 | 1 | NA | NA | 5 | 1 |
| 4_1330981 | 1 | 1 | NA | NA | 2 | 0.5 | 3 | 0.6667 | 6 | 0.6667 | 1 | 1 | 2 | 0.5 | 2 | 0.5 | NA | NA | 5 | 0.6 |
| 4_1331394 | 1 | 0 | NA | NA | 2 | 0 | 3 | 0.3333 | 6 | 0.1667 | 3 | 0 | 4 | 0.5 | 1 | 0 | NA | NA | 8 | 0.25 |
| 4_1331395 | 2 | 0 | NA | NA | 1 | 0 | 4 | 0.25 | 7 | 0.1429 | 3 | 0 | 4 | 0.5 | 2 | 0.5 | NA | NA | 9 | 0.3333 |
| 4_1331395 | 2 | 0 | NA | NA | NA | NA | 4 | 0.5 | 6 | 0.3333 | 3 | 0.6667 | 5 | 0.4 | 2 | 0.5 | NA | NA | 10 | 0.5 |
| 4_1331495 | 9 | 0.1111 | 3 | 0 | 3 | 0 | 3 | 0 | 18 | 0.0556 | 5 | 0 | 7 | 0 | 2 | 0.5 | 1 | 0 | 15 | 0.0667 |
| 4_1331495 | 11 | 0.1818 | 4 | 0.25 | 3 | 0 | 4 | 0 | 22 | 0.1364 | 5 | 0 | 6 | 0.1667 | 3 | 0.3333 | 3 | 0 | 17 | 0.1176 |
| 4_1331496 | 11 | 0.7273 | 4 | 1 | 3 | 1 | 3 | 1 | 21 | 0.8571 | 4 | 0.75 | 6 | 1 | 4 | 1 | NA | NA | 14 | 0.9286 |
| 4_1331496 | 8 | 0.5 | 4 | 0.25 | 4 | 0.25 | 3 | 0.3333 | 19 | 0.3684 | 3 | 0.3333 | 4 | 0 | 3 | 0 | 2 | 0.5 | 12 | 0.1667 |
| 4_1331498 | 8 | 0.5 | 2 | 1 | 3 | 0.6667 | 3 | 0.3333 | 16 | 0.5625 | 4 | 0.75 | 2 | 0 | 3 | 0.3333 | 1 | 0 | 10 | 0.4 |
| 4_1331500 | 9 | 0.3333 | 4 | 0.25 | 4 | 0.5 | 3 | 0 | 20 | 0.3 | 3 | 0 | 5 | 0.6 | 3 | 0.3333 | 2 | 0.5 | 13 | 0.3846 |
| 4_1331502 | 10 | 0.5 | 7 | 0.4286 | 6 | 0.3333 | 4 | 0 | 27 | 0.3704 | 4 | 0.25 | 8 | 0.25 | 6 | 0.5 | 2 | 0 | 20 | 0.3 |
| 4_1331503 | 10 | 0.4 | 3 | 0.6667 | 4 | 0.5 | 2 | 0.5 | 19 | 0.4737 | 4 | 0 | 7 | 0.4286 | 4 | 0.5 | 2 | 0.5 | 17 | 0.3529 |
| 4_1332186 | 30 | 0.1 | 43 | 0.1628 | 61 | 0.2295 | 28 | 0.3214 | 162 | 0.2037 | 45 | 0.2222 | 52 | 0.1923 | 31 | 0.3226 | 28 | 0.1786 | 156 | 0.2244 |
| 4_1332188 | 30 | 0.3333 | 41 | 0.2439 | 55 | 0.4 | 27 | 0.4444 | 153 | 0.3529 | 38 | 0.2632 | 44 | 0.2045 | 29 | 0.3793 | 23 | 0.2609 | 134 | 0.2687 |
| 4_1332188 | 30 | 0.2 | 42 | 0.2143 | 54 | 0.4074 | 29 | 0.3103 | 155 | 0.2968 | 36 | 0.2222 | 50 | 0.3 | 32 | 0.3438 | 26 | 0.2308 | 144 | 0.2778 |
| 4_1332189 | 28 | 0.1786 | 43 | 0.186 | 47 | 0.1915 | 31 | 0.2258 | 149 | 0.1946 | 37 | 0.0541 | 51 | 0.1569 | 31 | 0.0323 | 26 | 0.1923 | 145 | 0.1103 |
| 4_1332190 | 30 | 0.4 | 46 | 0.3261 | 49 | 0.4286 | 32 | 0.4062 | 157 | 0.3885 | 39 | 0.4872 | 52 | 0.4423 | 26 | 0.4231 | 28 | 0.3214 | 145 | 0.4276 |
| 4_1332191 | 23 | 0.1304 | 34 | 0.1765 | 44 | 0.2727 | 24 | 0.2917 | 125 | 0.224 | 31 | 0.2258 | 44 | 0.3636 | 24 | 0.1667 | 23 | 0.2174 | 122 | 0.2623 |
| 4_1332600 | 16 | 0.0625 | 22 | 0.0909 | 17 | 0.0588 | 9 | 0 | 64 | 0.0625 | 18 | 0.1111 | 18 | 0 | 16 | 0.0625 | 19 | 0.0526 | 71 | 0.0563 |
| 4_1333054 | NA | NA | 1 | 1 | NA | NA | 1 | 0 | 2 | 0.5 | NA | NA | 2 | 1 | 1 | 0 | 1 | 0 | 4 | 0.5 |
| 4_1333054 | NA | NA | 1 | 1 | NA | NA | 2 | 0 | 3 | 0.3333 | NA | NA | 2 | 1 | 1 | 0 | 1 | 0 | 4 | 0.5 |
| 4_1333110 | NA | NA | 1 | 1 | NA | NA | 2 | 0.5 | 3 | 0.6667 | NA | NA | 2 | 0.5 | 1 | 0 | 1 | 1 | 4 | 0.5 |
| 4_1333110 | NA | NA | 1 | 1 | NA | NA | 2 | 0.5 | 3 | 0.6667 | NA | NA | 2 | 0.5 | NA | NA | 1 | 0 | 3 | 0.3333 |
| 4_1333111 | NA | NA | 1 | 1 | NA | NA | 2 | 1 | 3 | 1 | NA | NA | 3 | 0.6667 | 1 | 0 | 1 | 1 | 5 | 0.6 |
| 4_1333111 | NA | NA | 1 | 1 | NA | NA | 2 | 0 | 3 | 0.3333 | NA | NA | 3 | 0.3333 | 1 | 0 | 1 | 0 | 5 | 0.2 |
| 4_1333111 | NA | NA | 1 | 0 | NA | NA | 2 | 1 | 3 | 0.6667 | NA | NA | 3 | 0.6667 | NA | NA | 1 | 1 | 4 | 0.75 |
| 4_1333112 | NA | NA | 1 | 1 | NA | NA | 2 | 1 | 3 | 1 | NA | NA | 2 | 1 | 1 | 1 | 1 | 1 | 4 | 1 |
| 4_1333113 | NA | NA | NA | NA | NA | NA | 1 | 1 | 1 | 1 | NA | NA | 2 | 0.5 | NA | NA | 2 | 0.5 | 4 | 0.5 |
| 4_1333113 | NA | NA | NA | NA | NA | NA | 2 | 1 | 2 | 1 | NA | NA | 3 | 1 | 1 | 1 | 1 | 1 | 5 | 1 |
| 4_1333119 | 1 | 1 | 2 | 0.5 | NA | NA | NA | NA | 3 | 0.6667 | 2 | 0.5 | 2 | 1 | 1 | 0 | 1 | 0 | 6 | 0.5 |
| 4_1333120 | 1 | 1 | 2 | 0.5 | NA | NA | 1 | 0 | 4 | 0.5 | 2 | 1 | 1 | 1 | 1 | 0 | NA | NA | 4 | 0.75 |
| 4_1333136 | 1 | 1 | NA | NA | NA | NA | NA | NA | 1 | 1 | 1 | 1 | NA | NA | NA | NA | NA | NA | 1 | 1 |
| 4_1333141 | 1 | 1 | NA | NA | NA | NA | NA | NA | 1 | 1 | 1 | 1 | NA | NA | NA | NA | NA | NA | 1 | 1 |
| 4_1333188 | 1 | 1 | 3 | 0.3333 | NA | NA | 1 | 0 | 5 | 0.4 | 2 | 0.5 | 2 | 1 | 3 | 0.6667 | NA | NA | 7 | 0.7143 |
| 4_1333188 | 1 | 1 | 3 | 1 | NA | NA | 1 | 0 | 5 | 0.8 | 2 | 1 | 3 | 0.6667 | 2 | 0 | 1 | 1 | 8 | 0.625 |
| 4_1333189 | 2 | 0.5 | 3 | 0.3333 | NA | NA | 2 | 0 | 7 | 0.2857 | 1 | 0 | 3 | 0.3333 | 3 | 0.3333 | NA | NA | 7 | 0.2857 |
| 4_1333280 | 1 | 1 | 1 | 1 | NA | NA | 1 | 1 | 3 | 1 | 1 | 1 | NA | NA | 2 | 1 | NA | NA | 3 | 1 |
| 4_1333282 | 1 | 1 | 1 | 1 | NA | NA | 1 | 1 | 3 | 1 | 1 | 1 | NA | NA | 2 | 1 | NA | NA | 3 | 1 |
| 4_1333282 | 1 | 1 | 1 | 1 | NA | NA | 1 | 0 | 3 | 0.6667 | 1 | 1 | NA | NA | 2 | 1 | NA | NA | 3 | 1 |
| 4_1333285 | 1 | 0 | 1 | 1 | NA | NA | 1 | 1 | 3 | 0.6667 | 1 | 1 | NA | NA | 2 | 0.5 | NA | NA | 3 | 0.6667 |
| 4_1333293 | NA | NA | 1 | 1 | NA | NA | 1 | 1 | 2 | 1 | NA | NA | NA | NA | 2 | 0.5 | NA | NA | 2 | 0.5 |
| 4_1333294 | NA | NA | 1 | 0 | NA | NA | 1 | 1 | 2 | 0.5 | NA | NA | NA | NA | 2 | 1 | NA | NA | 2 | 1 |
| 4_1333295 | NA | NA | NA | NA | NA | NA | 1 | 1 | 1 | 1 | NA | NA | NA | NA | 1 | 1 | NA | NA | 1 | 1 |
| 4_1333312 | NA | NA | 1 | 1 | NA | NA | 1 | 1 | 2 | 1 | NA | NA | NA | NA | 1 | 1 | NA | NA | 1 | 1 |
| 4_1333316 | NA | NA | 1 | 1 | NA | NA | 1 | 1 | 2 | 1 | NA | NA | NA | NA | 1 | 1 | NA | NA | 1 | 1 |
| 4_1333327 | 7 | 0 | 5 | 0.2 | 7 | 0 | 6 | 0.3333 | 25 | 0.12 | 2 | 0 | 6 | 0 | 3 | 0.6667 | 3 | 0 | 14 | 0.1429 |
| 4_1333355 | 6 | 0 | 1 | 1 | 1 | 0 | 5 | 0.2 | 13 | 0.1538 | NA | NA | 6 | 0 | NA | NA | 2 | 0.5 | 8 | 0.125 |
| 4_1333356 | 5 | 0.4 | 1 | 1 | NA | NA | 4 | 1 | 10 | 0.7 | NA | NA | 6 | 0.6667 | NA | NA | 1 | 0 | 7 | 0.5714 |
| 4_1333356 | 5 | 0.2 | 1 | 1 | 1 | 1 | 3 | 0.3333 | 10 | 0.4 | NA | NA | 3 | 0.3333 | NA | NA | 1 | 0 | 4 | 0.25 |
| 4_1333358 | 4 | 0.25 | 1 | 0 | 1 | 0 | 4 | 0.5 | 10 | 0.3 | NA | NA | 5 | 0.8 | NA | NA | 1 | 1 | 6 | 0.8333 |
| 4_1333897 | NA | NA | NA | NA | 2 | 0.5 | 2 | 0.5 | 4 | 0.5 | 1 | 1 | 1 | 1 | NA | NA | NA | NA | 2 | 1 |
| 4_1333898 | NA | NA | NA | NA | NA | NA | 1 | 1 | 1 | 1 | 2 | 1 | 2 | 1 | NA | NA | NA | NA | 4 | 1 |
| 4_1333898 | NA | NA | NA | NA | 3 | 0.6667 | 2 | 1 | 5 | 0.8 | 2 | 1 | 2 | 1 | NA | NA | NA | NA | 4 | 1 |
| 4_1333899 | NA | NA | NA | NA | 3 | 1 | 2 | 1 | 5 | 1 | 2 | 1 | 2 | 1 | NA | NA | NA | NA | 4 | 1 |
| 4_1333899 | NA | NA | NA | NA | 3 | 0 | 2 | 0.5 | 5 | 0.2 | 1 | 0 | 2 | 0.5 | NA | NA | NA | NA | 3 | 0.3333 |
| 4_1333922 | NA | NA | NA | NA | 2 | 1 | 1 | 1 | 3 | 1 | 1 | 1 | NA | NA | NA | NA | NA | NA | 1 | 1 |
| 4_1334072 | 8 | 0.25 | 11 | 0 | 12 | 0.1667 | 11 | 0.1818 | 42 | 0.1429 | 14 | 0.1429 | 15 | 0 | 14 | 0 | 12 | 0.25 | 55 | 0.0909 |
| 4_1334072 | 7 | 0 | 8 | 0.125 | 11 | 0.2727 | 10 | 0.2 | 36 | 0.1667 | 11 | 0.2727 | 14 | 0.1429 | 13 | 0.1538 | 10 | 0.2 | 48 | 0.1875 |

| | | | | | | | | | | | | | | | | | | | | |
|---|---|---|---|---|---|---|---|---|---|---|---|---|---|---|---|---|---|---|---|---|
| 4_13340728 | 7 | 0.1429 | 7 | 0.1429 | 8 | 0.25 | 9 | 0 | 31 | 0.129 | 10 | 0.1 | 12 | 0.1667 | 11 | 0.0909 | 7 | 0.1429 | 40 | 0.125 |
| 4_13340738 | 9 | 0.3333 | 9 | 0.3333 | 12 | 0.1667 | 11 | 0.0909 | 41 | 0.2195 | 10 | 0.4 | 13 | 0.2308 | 12 | 0.0833 | 10 | 0.2 | 45 | 0.2222 |
| 4_13340758 | 6 | 0.3333 | 8 | 0.25 | 10 | 0.4 | 12 | 0.5 | 36 | 0.3889 | 10 | 0.4 | 12 | 0.4167 | 13 | 0.1538 | 10 | 0.3 | 45 | 0.3111 |
| 4_13340758 | 7 | 0 | 6 | 0.1667 | 12 | 0.0833 | 12 | 0.25 | 37 | 0.1351 | 10 | 0.3 | 13 | 0.1538 | 10 | 0 | 9 | 0.2222 | 42 | 0.1667 |
| 4_13340759 | 7 | 0.1429 | 8 | 0 | 11 | 0.3636 | 12 | 0 | 38 | 0.1316 | 10 | 0.3 | 14 | 0.0714 | 13 | 0 | 11 | 0.0909 | 48 | 0.1042 |
| 4_13340763 | 7 | 0.1429 | 8 | 0.25 | 10 | 0.2 | 5 | 0 | 30 | 0.1667 | 10 | 0.1 | 14 | 0.3571 | 12 | 0.3333 | 7 | 0.7143 | 43 | 0.3488 |
| 4_13340769 | 4 | 1 | 2 | 0.5 | 6 | 1 | 3 | 1 | 15 | 0.9333 | 5 | 1 | 7 | 1 | 3 | 1 | 7 | 1 | 22 | 1 |
| 4_13340772 | 7 | 0.5714 | 8 | 0.625 | 8 | 0.625 | 8 | 0.5 | 31 | 0.5806 | 9 | 0.8889 | 9 | 0.7778 | 10 | 0.9 | 6 | 0.8333 | 34 | 0.8529 |
| 4_13340776 | 6 | 0.1667 | 7 | 0 | 11 | 0.0909 | 11 | 0.1818 | 35 | 0.1143 | 9 | 0 | 12 | 0.25 | 11 | 0.1818 | 9 | 0.4444 | 41 | 0.2195 |
| 4_13340781 | 5 | 1 | 6 | 1 | 7 | 1 | 8 | 1 | 26 | 1 | 9 | 1 | 8 | 1 | 7 | 1 | 4 | 1 | 28 | 1 |
| 4_13340782 | 4 | 0 | 6 | 0.3333 | 9 | 0.1111 | 9 | 0.3333 | 28 | 0.2143 | 8 | 0 | 12 | 0.1667 | 10 | 0 | 6 | 0 | 36 | 0.0556 |
| 4_13340789 | 6 | 0 | 8 | 0.375 | 10 | 0.1 | 7 | 0.2857 | 31 | 0.1935 | 10 | 0.2 | 11 | 0.1818 | 9 | 0.2222 | 10 | 0.4 | 40 | 0.25 |
| 4_13340796 | 4 | 1 | 4 | 1 | 7 | 1 | 6 | 1 | 21 | 1 | 6 | 0.8333 | 12 | 0.9167 | 5 | 0.6 | 6 | 0.6667 | 29 | 0.7931 |
| 4_13340802 | 5 | 0.4 | 9 | 0.2222 | 10 | 0.5 | 9 | 0.4444 | 33 | 0.3939 | 7 | 0.4286 | 12 | 0.4167 | 5 | 0.2 | 9 | 0.2222 | 33 | 0.3333 |
| 4_13340803 | 5 | 0 | 9 | 0.3333 | 11 | 0.5455 | 7 | 0.5714 | 32 | 0.4062 | 7 | 0.7143 | 13 | 0.3846 | 7 | 0.1429 | 9 | 0.2222 | 36 | 0.3611 |
| 4_13340810 | 5 | 0.8 | 6 | 1 | 10 | 0.9 | 8 | 0.875 | 29 | 0.8966 | 9 | 1 | 12 | 0.9167 | 8 | 1 | 10 | 0.8 | 39 | 0.9231 |
| 4_13340811 | 4 | 0.25 | 8 | 0.75 | 11 | 0.2727 | 10 | 0.7 | 33 | 0.5152 | 9 | 0.4444 | 11 | 0.3636 | 7 | 0.2857 | 8 | 0.5 | 35 | 0.4 |
| 4_13340812 | 4 | 0 | 3 | 0 | 11 | 0.0909 | 9 | 0.2222 | 27 | 0.1111 | 8 | 0.5 | 12 | 0.1667 | 9 | 0.2222 | 10 | 0.5 | 39 | 0.3333 |
| 4_13340819 | 5 | 0.2 | 9 | 0.2222 | 11 | 0.3636 | 9 | 0.4444 | 34 | 0.3235 | 9 | 0.3333 | 12 | 0.0833 | 9 | 0.4444 | 9 | 0.4444 | 39 | 0.3077 |
| 4_13340823 | 4 | 1 | 5 | 1 | 9 | 1 | 8 | 1 | 26 | 1 | 7 | 1 | 13 | 1 | 7 | 1 | 9 | 0.8889 | 36 | 0.9722 |
| 4_13340828 | 5 | 1 | 8 | 0.875 | 10 | 0.9 | 8 | 0.75 | 31 | 0.871 | 8 | 0.75 | 13 | 0.7692 | 8 | 0.875 | 8 | 1 | 37 | 0.8378 |
| 4_13340834 | 5 | 1 | 9 | 1 | 10 | 0.9 | 8 | 0.875 | 32 | 0.9375 | 9 | 0.8889 | 13 | 1 | 9 | 1 | 9 | 1 | 40 | 0.975 |
| 4_13340836 | 4 | 0 | 8 | 0 | 10 | 0.2 | 7 | 0 | 29 | 0.069 | 9 | 0.1111 | 12 | 0.1667 | 9 | 0.1111 | 6 | 0 | 36 | 0.1111 |
| 4_13340838 | 5 | 0.8 | 8 | 0.875 | 10 | 0.6 | 7 | 0.5714 | 30 | 0.7 | 9 | 1 | 12 | 0.5 | 9 | 0.5556 | 7 | 0.5714 | 37 | 0.6486 |
| 4_13340839 | 4 | 0 | 3 | 0 | 8 | 0.25 | 5 | 0 | 20 | 0.1 | 6 | 0.1667 | 8 | 0.125 | 5 | 0 | 5 | 0 | 24 | 0.0833 |
| 4_13340841 | 4 | 0 | 7 | 0 | 11 | 0.0909 | 7 | 0.1429 | 29 | 0.069 | 6 | 0 | 12 | 0.1667 | 9 | 0 | 5 | 0.2 | 32 | 0.0938 |
| 4_13340858 | 4 | 0.75 | 7 | 1 | 11 | 1 | 7 | 0.8571 | 29 | 0.931 | 9 | 1 | 11 | 1 | 8 | 0.75 | 9 | 1 | 37 | 0.9459 |
| 4_13340862 | 4 | 0 | 7 | 0.1429 | 11 | 0.1818 | 6 | 0 | 28 | 0.1071 | 7 | 0.2857 | 9 | 0.2222 | 9 | 0.1111 | 8 | 0.125 | 33 | 0.1818 |
| 4_13340873 | 4 | 0.75 | 5 | 1 | 11 | 1 | 6 | 1 | 26 | 0.9615 | 3 | 1 | 9 | 0.8889 | 5 | 1 | 8 | 0.875 | 25 | 0.92 |
| 4_13340882 | 5 | 0.2 | 7 | 0.4286 | 8 | 0.5 | 9 | 0.5556 | 29 | 0.4483 | 6 | 0.3333 | 8 | 0.5 | 7 | 0.5714 | 7 | 0.1429 | 28 | 0.3929 |
| 4_13340893 | 3 | 0 | 7 | 0.2857 | 10 | 0 | 8 | 0 | 28 | 0.0714 | 8 | 0 | 10 | 0.2 | 9 | 0 | 7 | 0 | 34 | 0.0588 |
| 4_13340902 | 3 | 1 | 6 | 1 | 7 | 1 | 8 | 1 | 24 | 1 | 7 | 1 | 11 | 1 | 9 | 1 | 8 | 1 | 35 | 1 |
| 4_13340911 | 3 | 0.3333 | 6 | 0.3333 | 9 | 0.4444 | 6 | 0.1667 | 24 | 0.3333 | 9 | 0.2222 | 11 | 0.2727 | 9 | 0.3333 | 8 | 0.25 | 37 | 0.2703 |
| 4_13340926 | 3 | 0 | 5 | 0.2 | 8 | 0.375 | 4 | 0 | 20 | 0.2 | 6 | 0.6667 | 5 | 0.2 | 6 | 0.3333 | 7 | 0.4286 | 24 | 0.4167 |
| 4_13340927 | 3 | 0.3333 | 6 | 0.3333 | 8 | 0.375 | 7 | 0.1429 | 24 | 0.2917 | 8 | 0.625 | 10 | 0 | 7 | 0.4286 | 8 | 0.25 | 33 | 0.303 |
| 4_13341259 | 1 | 0 | 1 | 0 | 5 | 0.6 | 3 | 0.3333 | 10 | 0.4 | 4 | 0.5 | 7 | 0.1429 | 4 | 0.25 | 3 | 0 | 18 | 0.2222 |
| 4_13341259 | 2 | 0.5 | 3 | 0 | 6 | 0 | 3 | 0.3333 | 14 | 0.1429 | 1 | 0 | 8 | 0.125 | 6 | 0 | 6 | 0.1667 | 21 | 0.0952 |
| 4_13341266 | 1 | 0 | 3 | 0.3333 | 5 | 0 | 5 | 0 | 14 | 0.0714 | 3 | 0 | 8 | 0.25 | 7 | 0 | 5 | 0.4 | 23 | 0.1739 |
| 4_13341267 | 1 | 0 | 3 | 0.3333 | 5 | 0.6 | 4 | 0.5 | 13 | 0.4615 | 3 | 0 | 9 | 0.2222 | 7 | 0.2857 | 3 | 0 | 22 | 0.1818 |
| 4_13341275 | 2 | 0 | 2 | 0 | 4 | 0.25 | 2 | 0 | 10 | 0.1 | 2 | 0 | 6 | 0.5 | 5 | 0.2 | 4 | 0.25 | 17 | 0.2941 |
| 4_13348171 | NA | NA | 1 | 1 | NA | NA | NA | NA | 1 | 1 | NA | NA | 1 | 1 | NA | NA | NA | NA | 1 | 1 |
| 4_13348193 | NA | NA | 1 | 1 | NA | NA | NA | NA | 1 | 1 | NA | NA | 1 | 1 | NA | NA | NA | NA | 1 | 1 |
| 4_13348199 | NA | NA | 1 | 1 | NA | NA | NA | NA | 1 | 1 | NA | NA | 1 | 1 | NA | NA | NA | NA | 1 | 1 |
| 4_13348225 | NA | NA | 1 | 1 | NA | NA | NA | NA | 1 | 1 | NA | NA | 1 | 1 | NA | NA | NA | NA | 1 | 1 |
| 4_13348227 | NA | NA | 1 | 1 | NA | NA | NA | NA | 1 | 1 | NA | NA | 1 | 1 | NA | NA | NA | NA | 1 | 1 |
| 4_13348264 | NA | NA | 1 | 1 | NA | NA | NA | NA | 1 | 1 | NA | NA | 1 | 1 | NA | NA | NA | NA | 1 | 1 |
| 4_13348305 | NA | NA | 2 | 1 | NA | NA | NA | NA | 2 | 1 | NA | NA | 1 | 1 | NA | NA | NA | NA | 1 | 1 |
| 4_13348305 | NA | NA | 2 | 0.5 | NA | NA | NA | NA | 2 | 0.5 | NA | NA | 1 | 1 | NA | NA | NA | NA | 1 | 1 |
| 4_13348306 | NA | NA | 2 | 0.5 | NA | NA | NA | NA | 2 | 0.5 | NA | NA | 1 | 1 | NA | NA | NA | NA | 1 | 1 |
| 4_13348321 | NA | NA | 2 | 0.5 | NA | NA | NA | NA | 2 | 0.5 | NA | NA | 1 | 1 | NA | NA | NA | NA | 1 | 1 |
| 4_13348329 | NA | NA | 2 | 0.5 | NA | NA | NA | NA | 2 | 0.5 | NA | NA | 1 | 1 | NA | NA | NA | NA | 1 | 1 |
| 4_13348330 | NA | NA | 2 | 0.5 | NA | NA | NA | NA | 2 | 0.5 | NA | NA | 1 | 1 | NA | NA | NA | NA | 1 | 1 |
| 4_13348350 | NA | NA | 1 | 1 | NA | NA | NA | NA | 1 | 1 | NA | NA | 1 | 1 | NA | NA | NA | NA | 1 | 1 |
| 4_13348492 | NA | NA | 1 | 1 | NA | NA | NA | NA | 1 | 1 | NA | NA | NA | NA | NA | NA | 1 | 1 | 1 | 1 |
| 4_13357486 | 1 | 0 | 1 | 0 | 4 | 0.25 | 2 | 0 | 8 | 0.125 | 2 | 0 | 1 | 0 | 2 | 0.5 | NA | NA | 5 | 0.2 |
| 4_13357493 | 1 | 0 | 1 | 0 | 3 | 0.6667 | 1 | 0 | 6 | 0.3333 | 1 | 0 | 1 | 1 | 2 | 0 | NA | NA | 4 | 0.25 |
| 4_13357517 | NA | NA | NA | NA | 3 | 0.6667 | 2 | 1 | 5 | 0.8 | 2 | 1 | NA | NA | NA | NA | NA | NA | 2 | 1 |
| 4_13357530 | NA | NA | 1 | 0 | 4 | 0 | 2 | 0.5 | 7 | 0.1429 | 2 | 0 | 1 | 0 | 1 | 1 | NA | NA | 4 | 0.25 |
| 4_13357532 | NA | NA | 1 | 0 | 3 | 0.3333 | 1 | 0 | 5 | 0.2 | 1 | 0 | 1 | 0 | 1 | 1 | NA | NA | 3 | 0.3333 |
| 4_13369239 | 39 | 0.0769 | 19 | 0.1053 | 20 | 0.05 | 25 | 0.12 | 103 | 0.0874 | 15 | 0 | 39 | 0.1538 | 32 | 0.125 | 19 | 0.1053 | 105 | 0.1143 |
| 4_13369240 | 40 | 0.075 | 20 | 0.2 | 20 | 0.2 | 25 | 0.08 | 105 | 0.1238 | 18 | 0.1667 | 40 | 0.125 | 32 | 0.0938 | 22 | 0.0909 | 112 | 0.1161 |
| 4_13369246 | 40 | 0.1 | 20 | 0.15 | 19 | 0.1053 | 29 | 0.1379 | 108 | 0.1204 | 19 | 0.1053 | 40 | 0.075 | 31 | 0.0645 | 23 | 0.1304 | 113 | 0.0885 |
| 4_13369248 | 38 | 0.0789 | 19 | 0.0526 | 20 | 0.1 | 27 | 0.037 | 104 | 0.0673 | 17 | 0.1176 | 39 | 0.1282 | 29 | 0.0345 | 24 | 0.0417 | 109 | 0.0826 |
| 4_13369646 | 4 | 0.25 | 3 | 0.3333 | 4 | 0.25 | 5 | 0 | 16 | 0.1875 | 3 | 0 | 4 | 0 | 7 | 0.1429 | 6 | 0 | 20 | 0.05 |
| 4_13369704 | 3 | 0.3333 | 4 | 0.25 | 5 | 0 | 3 | 0 | 15 | 0.1333 | 3 | 0 | 6 | 0.1667 | 4 | 0 | 5 | 0.2 | 18 | 0.1111 |

| | | | | | | | | | | | | | | | | | | | | |
|---|---|---|---|---|---|---|---|---|---|---|---|---|---|---|---|---|---|---|---|---|
| 4_13369749 | 4 | 0.5 | 3 | 0.6667 | 3 | 0.3333 | 3 | 0.3333 | 13 | 0.4615 | 4 | 0.25 | 1 | 1 | 3 | 1 | 5 | 0.8 | 13 | 0.6923 |
| 4_13369774 | 4 | 0.25 | 3 | 0 | 4 | 0.25 | 4 | 0 | 15 | 0.1333 | 2 | 0 | 5 | 0.2 | 4 | 0.25 | 3 | 0.3333 | 14 | 0.2143 |
| 4_13369815 | 3 | 1 | 7 | 0.5714 | 2 | 1 | 6 | 1 | 18 | 0.8333 | 4 | 0.75 | 7 | 1 | 5 | 0.8 | 2 | 1 | 18 | 0.8889 |
| 4_13369837 | 5 | 0.4 | 5 | 0.2 | 3 | 0.6667 | 5 | 0 | 18 | 0.2778 | 4 | 0.5 | 6 | 0.3333 | 5 | 0 | 2 | 0 | 17 | 0.2353 |
| 4_13369842 | 5 | 0.2 | 3 | 0 | 3 | 0.3333 | 7 | 0 | 18 | 0.1111 | 4 | 0 | 7 | 0.1429 | 5 | 0 | 2 | 0 | 18 | 0.0556 |
| 4_13369844 | 5 | 0.2 | 4 | 0 | 3 | 0 | 8 | 0.125 | 20 | 0.1 | 4 | 0.5 | 6 | 0.1667 | 5 | 0 | 3 | 0.3333 | 18 | 0.2222 |
| 4_13385396 | 1 | 1 | NA | NA | NA | NA | 2 | 0.5 | 3 | 0.6667 | NA | NA | NA | NA | 3 | 1 | NA | NA | 3 | 1 |
| 4_13385474 | 39 | 0.1026 | 38 | 0.1053 | 36 | 0.1389 | 36 | 0.1111 | 149 | 0.1141 | 34 | 0.0588 | 46 | 0.087 | 39 | 0.0513 | 43 | 0.0698 | 162 | 0.0679 |
| 4_13385510 | 30 | 0.1333 | 26 | 0.2692 | 21 | 0.1429 | 26 | 0 | 103 | 0.1359 | 22 | 0.1364 | 30 | 0.1 | 25 | 0.08 | 27 | 0.1111 | 104 | 0.1058 |
| 4_13385511 | 36 | 0.2778 | 32 | 0.1562 | 31 | 0.2903 | 31 | 0.2581 | 130 | 0.2462 | 29 | 0.1379 | 33 | 0.1818 | 26 | 0.0769 | 33 | 0.0909 | 121 | 0.124 |
| 4_13385839 | 1 | 1 | 3 | 0.6667 | 1 | 1 | 1 | 0 | 6 | 0.6667 | 1 | 1 | 1 | 0 | 3 | 0.6667 | 1 | 0 | 6 | 0.5 |
| 4_13385857 | 1 | 1 | 2 | 1 | 1 | 1 | NA | NA | 4 | 1 | 1 | 1 | 1 | 1 | 3 | 0.6667 | 2 | 1 | 7 | 0.8571 |
| 4_13385860 | 1 | 0 | 1 | 1 | 1 | 0 | 1 | 0 | 4 | 0.25 | 1 | 0 | NA | NA | 1 | 1 | 2 | 0 | 4 | 0.25 |
| 4_13385861 | 1 | 1 | 2 | 0 | 1 | 0 | 1 | 0 | 5 | 0.2 | 1 | 0 | 1 | 0 | 2 | 0.5 | 2 | 0 | 6 | 0.1667 |
| 4_13385865 | 1 | 1 | 2 | 0.5 | 1 | 1 | 1 | 1 | 5 | 0.8 | 1 | 0 | 1 | 0 | 2 | 0.5 | 2 | 0.5 | 6 | 0.3333 |
| 4_13388843 | NA | NA | 2 | 1 | 2 | 0.5 | 3 | 0.6667 | 7 | 0.7143 | NA | NA | 2 | 0.5 | 3 | 0.6667 | 2 | 0 | 7 | 0.4286 |
| 4_13388927 | 1 | 0 | 1 | 0 | 2 | 1 | 1 | 0 | 5 | 0.4 | 2 | 0 | 2 | 0 | 3 | 0.6667 | 2 | 0 | 9 | 0.2222 |
| 4_13388958 | 1 | 0 | 1 | 0 | 2 | 0 | 1 | 1 | 5 | 0.2 | 2 | 0 | 2 | 0.5 | 3 | 0 | 2 | 0.5 | 9 | 0.2222 |
| 4_13388978 | NA | NA | 1 | 1 | 2 | 1 | 1 | 1 | 4 | 1 | 3 | 1 | 2 | 1 | 4 | 1 | 2 | 1 | 11 | 1 |
| 4_13388988 | 1 | 1 | 2 | 0 | 2 | 0.5 | 1 | 0 | 6 | 0.3333 | 3 | 0.3333 | 2 | 0.5 | 4 | 0.5 | 2 | 0.5 | 11 | 0.4545 |
| 4_13388991 | NA | NA | 1 | 0 | 2 | 0.5 | 1 | 1 | 4 | 0.5 | 3 | 0 | 2 | 0 | 4 | 0.5 | 2 | 0 | 11 | 0.1818 |
| 4_13388998 | 1 | 1 | 1 | 0 | 2 | 0 | 1 | 0 | 5 | 0.2 | 3 | 0.3333 | 2 | 0 | 4 | 1 | 2 | 0 | 11 | 0.4545 |
| 4_13389006 | 1 | 1 | 2 | 0 | 2 | 0 | 1 | 0 | 6 | 0.1667 | 3 | 0 | 1 | 0 | 4 | 0 | 2 | 0.5 | 10 | 0.1 |
| 4_13389006 | 1 | 0 | 2 | 0 | 2 | 0.5 | 1 | 0 | 6 | 0.1667 | 3 | 0 | 1 | 1 | 3 | 0.3333 | 2 | 0 | 9 | 0.2222 |
| 4_13389015 | 1 | 1 | 2 | 0 | 2 | 0.5 | 1 | 0 | 6 | 0.3333 | 2 | 0 | 2 | 0 | 3 | 0.6667 | 2 | 1 | 9 | 0.4444 |
| 4_13389016 | 1 | 1 | 2 | 0 | 1 | 0 | 1 | 0 | 5 | 0.2 | 2 | 0 | 2 | 0 | 4 | 0 | 2 | 1 | 10 | 0.2 |
| 4_13389025 | NA | NA | 1 | 0 | 2 | 0 | 1 | 1 | 4 | 0.25 | 2 | 0 | 2 | 0 | 4 | 0.25 | 2 | 0 | 10 | 0.1 |
| 4_13389028 | 1 | 1 | 2 | 0 | 2 | 0 | NA | NA | 5 | 0.2 | 3 | 0 | 2 | 1 | 2 | 0.5 | 2 | 0 | 9 | 0.3333 |
| 4_13389030 | 1 | 1 | 2 | 0 | 2 | 1 | 1 | 1 | 6 | 0.6667 | 3 | 0.6667 | 2 | 0.5 | 4 | 0.75 | 2 | 1 | 11 | 0.7273 |
| 4_13389033 | 1 | 0 | 2 | 0 | 2 | 0 | 1 | 1 | 6 | 0.1667 | 3 | 0 | 2 | 0.5 | 3 | 0 | 2 | 0 | 10 | 0.1 |
| 4_13389058 | 1 | 0 | 2 | 0.5 | 2 | 0 | 1 | 1 | 6 | 0.3333 | 2 | 0 | 2 | 0 | 3 | 0.3333 | 2 | 0 | 9 | 0.1111 |
| 4_13389063 | NA | NA | 2 | 0.5 | 2 | 0.5 | 1 | 1 | 5 | 0.6 | 3 | 0 | 2 | 0 | 4 | 0 | 3 | 0.3333 | 12 | 0.0833 |
| 4_13389069 | 1 | 1 | 2 | 0 | 2 | 0 | 1 | 0 | 6 | 0.1667 | 3 | 0.6667 | 2 | 1 | 4 | 0.5 | 3 | 0.3333 | 12 | 0.5833 |
| 4_13389093 | 2 | 0 | 4 | 0.25 | 4 | 0.75 | 3 | 0 | 13 | 0.3077 | 3 | 0.3333 | 2 | 0.5 | 4 | 0.75 | 2 | 0.5 | 11 | 0.5455 |
| 4_13389121 | 3 | 1 | 4 | 1 | 5 | 0.8 | 3 | 1 | 15 | 0.9333 | 3 | 1 | 2 | 1 | 5 | 1 | 3 | 1 | 13 | 1 |
| 4_13389130 | 4 | 0.25 | 3 | 0 | 3 | 0 | 2 | 0 | 12 | 0.0833 | 2 | 0.5 | 1 | 0 | 5 | 0 | 2 | 0 | 10 | 0.1 |
| 4_13389134 | 4 | 0 | 3 | 0 | 5 | 0.4 | 3 | 0 | 15 | 0.1333 | 3 | 0.3333 | 2 | 0 | 5 | 0 | 3 | 0 | 13 | 0.0769 |
| 4_13389167 | 4 | 0 | 4 | 0.25 | 4 | 0 | 3 | 0 | 15 | 0.0667 | 2 | 0.5 | 3 | 0 | 4 | 0.25 | 2 | 0 | 11 | 0.1818 |
| 4_13397196 | 1 | 1 | NA | NA | 1 | 0 | NA | NA | 2 | 0.5 | 3 | 0.6667 | NA | NA | 1 | 0 | 2 | 0.5 | 6 | 0.5 |
| 4_13397197 | 3 | 0.6667 | NA | NA | 1 | 0 | NA | NA | 4 | 0.5 | 4 | 0.5 | NA | NA | 2 | 0.5 | 1 | 0 | 7 | 0.4286 |
| 4_13397211 | 4 | 0.25 | NA | NA | 2 | 0 | NA | NA | 6 | 0.1667 | 4 | 0 | NA | NA | 1 | 0 | 2 | 0.5 | 7 | 0.1429 |
| 4_13397847 | 1 | 1 | 2 | 0.5 | 3 | 1 | 1 | 1 | 7 | 0.8571 | 1 | 1 | NA | NA | NA | NA | NA | NA | 1 | 1 |
| 4_13397853 | NA | NA | 2 | 0.5 | 4 | 1 | 1 | 1 | 7 | 0.8571 | 1 | 1 | NA | NA | NA | NA | NA | NA | 1 | 1 |
| 4_13397881 | 1 | 0 | 1 | 1 | 4 | 0.5 | NA | NA | 6 | 0.5 | 1 | 1 | NA | NA | NA | NA | NA | NA | 1 | 1 |
| 4_13397884 | NA | NA | 1 | 1 | 4 | 0.5 | NA | NA | 5 | 0.6 | 1 | 1 | NA | NA | NA | NA | NA | NA | 1 | 1 |
| 4_13397909 | 1 | 1 | 1 | 1 | 3 | 1 | NA | NA | 5 | 1 | 1 | 1 | NA | NA | NA | NA | NA | NA | 1 | 1 |
| 4_13397920 | NA | NA | NA | NA | 3 | 1 | NA | NA | 3 | 1 | 1 | 1 | NA | NA | NA | NA | NA | NA | 1 | 1 |
| 4_13400458 | 1 | 1 | 1 | 1 | NA | NA | 4 | 1 | 6 | 1 | 1 | 1 | 1 | 1 | NA | NA | 1 | 1 | 3 | 1 |
| 4_13400473 | NA | NA | 1 | 1 | NA | NA | 4 | 0.5 | 5 | 0.6 | 1 | 1 | 1 | 0 | NA | NA | 1 | 1 | 3 | 0.6667 |
| 4_13400570 | 3 | 0.6667 | 1 | 1 | NA | NA | 2 | 0 | 6 | 0.5 | 1 | 1 | 1 | 0 | NA | NA | 1 | 1 | 3 | 0.6667 |
| 4_13400573 | 2 | 1 | 1 | 1 | NA | NA | 4 | 1 | 7 | 1 | 1 | 1 | 2 | 1 | NA | NA | 1 | 1 | 4 | 1 |
| 4_13400574 | 2 | 1 | NA | NA | NA | NA | 3 | 1 | 5 | 1 | 1 | 1 | 1 | 1 | NA | NA | 1 | 1 | 3 | 1 |
| 4_13400584 | 2 | 1 | 1 | 1 | NA | NA | 4 | 1 | 7 | 1 | 1 | 1 | 2 | 1 | NA | NA | 1 | 1 | 4 | 1 |
| 4_13400589 | 2 | 1 | NA | NA | NA | NA | 4 | 1 | 6 | 1 | NA | NA | 1 | 1 | NA | NA | 1 | 1 | 2 | 1 |
| 4_13400594 | 3 | 0.3333 | 1 | 1 | NA | NA | 3 | 0.3333 | 7 | 0.4286 | 1 | 0 | 1 | 1 | NA | NA | 1 | 0 | 3 | 0.3333 |
| 4_13400599 | 3 | 0.6667 | 1 | 0 | NA | NA | 4 | 0 | 8 | 0.25 | 1 | 0 | NA | NA | NA | NA | 1 | 1 | 2 | 0.5 |
| 4_13400601 | 4 | 1 | 1 | 1 | NA | NA | 4 | 0.75 | 9 | 0.8889 | 1 | 1 | 1 | 1 | NA | NA | 1 | 1 | 3 | 1 |
| 4_13400604 | 3 | 1 | 1 | 1 | NA | NA | 4 | 1 | 8 | 1 | 1 | 1 | 1 | 1 | NA | NA | 1 | 1 | 3 | 1 |
| 4_13400609 | 4 | 0.5 | 1 | 1 | NA | NA | 4 | 0.25 | 9 | 0.4444 | NA | NA | 1 | 1 | NA | NA | 1 | 0 | 2 | 0.5 |
| 4_13400617 | 2 | 1 | 1 | 1 | NA | NA | 4 | 1 | 7 | 1 | 1 | 1 | 1 | 1 | NA | NA | 1 | 1 | 3 | 1 |
| 4_13400618 | 4 | 1 | 1 | 0 | NA | NA | 4 | 0.75 | 9 | 0.7778 | 1 | 1 | 1 | 1 | NA | NA | 1 | 1 | 3 | 1 |
| 4_13400629 | 3 | 1 | 1 | 1 | NA | NA | 3 | 1 | 7 | 1 | 1 | 1 | 1 | 1 | NA | NA | 1 | 1 | 3 | 1 |
| 4_13401500 | 1 | 1 | NA | NA | NA | NA | NA | NA | 1 | 1 | NA | NA | NA | NA | 1 | 1 | NA | NA | 1 | 1 |
| 4_13408143 | NA | NA | 2 | 0.5 | NA | NA | NA | NA | 2 | 0.5 | NA | NA | NA | NA | 1 | 1 | NA | NA | 1 | 1 |
| 4_13408373 | 3 | 0 | 6 | 0.1667 | 2 | 0 | 4 | 0.5 | 15 | 0.2 | 5 | 0.6 | 4 | 0.25 | 3 | 0.3333 | 2 | 0.5 | 14 | 0.4286 |

| | | | | | | | | | | | | | | | | | | | | |
|---|---|---|---|---|---|---|---|---|---|---|---|---|---|---|---|---|---|---|---|---|
| 4_1340838 | 5 | 0 | 10 | 0.1 | 4 | 0.25 | 4 | 0 | 23 | 0.087 | 5 | 0 | 4 | 0 | 2 | 0.5 | 1 | 0 | 12 | 0.0833 |
| 4_1340838 | 6 | 0 | 8 | 0.125 | 2 | 0 | 2 | 0 | 18 | 0.0556 | 3 | 0 | 3 | 0 | 1 | 1 | 1 | 1 | 8 | 0.25 |
| 4_1340839 | 5 | 0.4 | 10 | 0.5 | 3 | 1 | 3 | 1 | 21 | 0.619 | 5 | 1 | 5 | 0.4 | 2 | 1 | 2 | 1 | 14 | 0.7857 |
| 4_1340839 | 4 | 0.25 | 6 | 0.3333 | 4 | 0 | 3 | 0.3333 | 17 | 0.2353 | 5 | 0.4 | 4 | 0.5 | 3 | 0.6667 | 2 | 0.5 | 14 | 0.5 |
| 4_1340840 | 4 | 1 | 9 | 1 | 4 | 1 | 4 | 1 | 21 | 1 | 5 | 1 | 5 | 1 | 3 | 1 | 1 | 1 | 14 | 1 |
| 4_1340840 | 3 | 1 | 6 | 1 | 4 | 1 | 4 | 1 | 17 | 1 | 2 | 1 | 2 | 1 | 1 | 1 | 1 | 1 | 6 | 1 |
| 4_1340841 | 6 | 0.8333 | 9 | 0.5556 | 5 | 0.8 | 3 | 0.6667 | 23 | 0.6957 | 5 | 1 | 5 | 0.6 | 2 | 0.5 | 2 | 0.5 | 14 | 0.7143 |
| 4_1340842 | 5 | 0.6 | 9 | 0.7778 | 5 | 1 | 5 | 1 | 24 | 0.8333 | 4 | 1 | 4 | 1 | 2 | 1 | 1 | 1 | 11 | 1 |
| 4_1340842 | 7 | 0.4286 | 10 | 0.6 | 2 | 0.5 | 6 | 0.3333 | 25 | 0.48 | 5 | 0.4 | 5 | 0.2 | 3 | 0.3333 | 2 | 0.5 | 15 | 0.3333 |
| 4_1340843 | 5 | 0.2 | 8 | 0.625 | 2 | 0.5 | 5 | 0.2 | 20 | 0.4 | 5 | 0.2 | 5 | 0.4 | 3 | 0.3333 | 2 | 0 | 15 | 0.2667 |
| 4_1340844 | 7 | 0.7143 | 9 | 0.3333 | 2 | 0 | 6 | 0.5 | 24 | 0.4583 | 4 | 0 | 5 | 0.8 | 3 | 0.3333 | 2 | 0.5 | 14 | 0.4286 |
| 4_1340845 | 6 | 0.1667 | 8 | 0.375 | 2 | 0.5 | 6 | 0.3333 | 22 | 0.3182 | 4 | 0.25 | 4 | 0 | 3 | 0 | 2 | 0 | 13 | 0.0769 |
| 4_1340845 | 6 | 0.6667 | 7 | 0.4286 | 1 | 1 | 5 | 0.4 | 19 | 0.5263 | 3 | 0.3333 | 3 | 0.3333 | 3 | 1 | 2 | 1 | 11 | 0.6364 |
| 4_1340845 | 3 | 1 | 5 | 1 | 1 | 1 | 5 | 1 | 14 | 1 | 2 | 0.5 | 2 | 0.5 | 2 | 0.5 | NA | NA | 6 | 0.5 |
| 4_1340849 | 7 | 0.2857 | 7 | 0.1429 | NA | NA | 6 | 0 | 20 | 0.15 | 3 | 0.3333 | 4 | 0 | 2 | 0 | 2 | 0 | 11 | 0.0909 |
| 4_1340850 | 6 | 0 | 5 | 0 | 1 | 0 | 5 | 0.6 | 17 | 0.1765 | 4 | 0 | 4 | 0.75 | 3 | 0.3333 | 1 | 1 | 12 | 0.4167 |
| 4_1340850 | 3 | 0 | 7 | 0.1429 | 1 | 0 | 6 | 0.1667 | 17 | 0.1176 | 4 | 0.25 | 4 | 0.25 | 2 | 0 | 1 | 1 | 11 | 0.2727 |
| 4_1340984 | 2 | 0.5 | 5 | 0.6 | 1 | 0 | 3 | 0.6667 | 11 | 0.5455 | NA | NA | 1 | 1 | 2 | 0.5 | 2 | 1 | 5 | 0.8 |
| 4_1340985 | 3 | 0.6667 | 5 | 0.4 | 1 | 0 | 4 | 0.5 | 13 | 0.4615 | NA | NA | 1 | 1 | 2 | 0 | 1 | 1 | 4 | 0.5 |
| 4_1340985 | 1 | 0 | 4 | 0.25 | 1 | 0 | 3 | 0 | 9 | 0.1111 | 1 | 0 | 1 | 1 | 2 | 0.5 | 2 | 0.5 | 6 | 0.5 |
| 4_1340985 | 2 | 0.5 | 2 | 1 | NA | NA | 2 | 1 | 6 | 0.8333 | 1 | 1 | 1 | 1 | 2 | 0.5 | 1 | 0 | 5 | 0.6 |
| 4_1340988 | 3 | 0 | 2 | 0 | NA | NA | 4 | 0.25 | 9 | 0.1111 | 1 | 0 | 1 | 0 | 1 | 0 | 2 | 0.5 | 5 | 0.2 |
| 4_1340990 | 1 | 1 | NA | NA | NA | NA | NA | NA | 1 | 1 | NA | NA | 2 | 1 | NA | NA | 1 | 1 | 3 | 1 |
| 4_1340990 | 1 | 1 | NA | NA | NA | NA | NA | NA | 1 | 1 | NA | NA | 2 | 1 | NA | NA | 1 | 0 | 3 | 0.6667 |
| 4_1340991 | 3 | 0.3333 | 1 | 0 | 1 | 0 | 2 | 0 | 7 | 0.1429 | NA | NA | 1 | 0 | 1 | 1 | 2 | 0.5 | 4 | 0.5 |
| 4_1340992 | 2 | 1 | NA | NA | 1 | 0 | 1 | 1 | 4 | 0.75 | NA | NA | NA | NA | 1 | 0 | 2 | 1 | 3 | 0.6667 |
| 4_1340993 | 1 | 1 | NA | NA | NA | NA | NA | NA | 1 | 1 | NA | NA | 2 | 0.5 | NA | NA | 1 | 1 | 3 | 0.6667 |
| 4_1340993 | 2 | 0 | 1 | 1 | NA | NA | 2 | 0 | 5 | 0.2 | NA | NA | 1 | 0 | NA | NA | 1 | 1 | 2 | 0.5 |
| 4_1340993 | 3 | 0.3333 | 1 | 1 | NA | NA | 2 | 0 | 6 | 0.3333 | NA | NA | NA | NA | 1 | 1 | 1 | 1 | 2 | 1 |
| 4_1340994 | 1 | 1 | NA | NA | NA | NA | NA | NA | 1 | 1 | NA | NA | 2 | 1 | NA | NA | 1 | 1 | 3 | 1 |
| 4_1340995 | 1 | 1 | NA | NA | NA | NA | NA | NA | 1 | 1 | NA | NA | 2 | 1 | NA | NA | 1 | 1 | 3 | 1 |
| 4_1340995 | 1 | 0 | 1 | 1 | NA | NA | 2 | 0.5 | 4 | 0.5 | NA | NA | 1 | 1 | 1 | 0 | 1 | 0 | 3 | 0.3333 |
| 4_1340995 | 1 | 1 | NA | NA | NA | NA | NA | NA | 1 | 1 | NA | NA | 2 | 0.5 | NA | NA | 1 | 0 | 3 | 0.3333 |
| 4_1340996 | 1 | 0 | 1 | 1 | NA | NA | 2 | 0.5 | 4 | 0.5 | NA | NA | 1 | 1 | NA | NA | 1 | 1 | 2 | 1 |
| 4_1341000 | 2 | 1 | 2 | 1 | NA | NA | NA | NA | 4 | 1 | NA | NA | 2 | 1 | NA | NA | NA | NA | 2 | 1 |
| 4_1341010 | 1 | 0 | 1 | 1 | NA | NA | NA | NA | 2 | 0.5 | NA | NA | 2 | 0 | NA | NA | 1 | 1 | 3 | 0.3333 |
| 4_1341530 | NA | NA | 1 | 1 | NA | NA | NA | NA | 1 | 1 | NA | NA | 1 | 0 | 1 | 1 | 1 | 1 | 3 | 0.6667 |
| 4_1341531 | NA | NA | 1 | 1 | NA | NA | NA | NA | 1 | 1 | NA | NA | 2 | 0.5 | 1 | 0 | 2 | 0 | 5 | 0.2 |
| 4_1341531 | NA | NA | 1 | 1 | NA | NA | NA | NA | 1 | 1 | NA | NA | 2 | 0.5 | 1 | 0 | 2 | 1 | 5 | 0.6 |
| 4_1341534 | NA | NA | 1 | 1 | NA | NA | NA | NA | 1 | 1 | NA | NA | 2 | 1 | 1 | 1 | 2 | 0 | 5 | 0.6 |
| 4_1341536 | NA | NA | 1 | 1 | NA | NA | NA | NA | 1 | 1 | NA | NA | 2 | 0.5 | 1 | 1 | 1 | 1 | 4 | 0.75 |
| 4_1341541 | NA | NA | 2 | 0.5 | NA | NA | NA | NA | 2 | 0.5 | NA | NA | 2 | 0.5 | 1 | 1 | 2 | 1 | 5 | 0.8 |
| 4_1341542 | NA | NA | 2 | 1 | NA | NA | NA | NA | 2 | 1 | NA | NA | 2 | 0.5 | 1 | 0 | 2 | 1 | 5 | 0.6 |
| 4_1342007 | 84 | 0.0952 | 119 | 0.1092 | 82 | 0.0854 | 70 | 0.2 | 355 | 0.1183 | 99 | 0.0808 | 74 | 0.1081 | 52 | 0.1346 | 109 | 0.1009 | 334 | 0.1018 |
| 4_1342007 | 99 | 0.4141 | 152 | 0.4342 | 96 | 0.4271 | 82 | 0.4756 | 429 | 0.4359 | 117 | 0.4615 | 86 | 0.5 | 66 | 0.3788 | 125 | 0.48 | 394 | 0.4619 |
| 4_1342343 | NA | NA | 2 | 1 | 2 | 1 | 2 | 1 | 6 | 1 | 1 | 1 | 3 | 1 | 2 | 1 | 1 | 1 | 7 | 1 |
| 4_1342343 | 1 | 1 | 2 | 0 | 3 | 0.3333 | 3 | 0 | 9 | 0.2222 | 1 | 0 | 4 | 0 | 2 | 0.5 | 1 | 0 | 8 | 0.125 |
| 4_1342344 | 1 | 1 | 2 | 0.5 | 2 | 1 | 3 | 1 | 8 | 0.875 | 1 | 1 | 4 | 0.75 | 3 | 0.6667 | 1 | 0 | 9 | 0.6667 |
| 4_1342345 | NA | NA | 2 | 1 | 3 | 1 | 2 | 1 | 7 | 1 | NA | NA | 4 | 1 | 3 | 1 | NA | NA | 7 | 1 |
| 4_1342347 | 1 | 1 | 2 | 1 | 4 | 0.75 | 3 | 1 | 10 | 0.9 | 1 | 1 | 4 | 0.75 | 1 | 1 | 1 | 1 | 7 | 0.8571 |
| 4_1342349 | 1 | 1 | 4 | 1 | 4 | 1 | 2 | 1 | 11 | 1 | 2 | 1 | 5 | 1 | 4 | 1 | 2 | 1 | 13 | 1 |
| 4_1342350 | 1 | 0 | 4 | 0.5 | 4 | 0.5 | 4 | 0.25 | 13 | 0.3846 | 2 | 0.5 | 4 | 0.5 | 2 | 0 | 1 | 1 | 9 | 0.4444 |
| 4_1342352 | 2 | 0.5 | 6 | 0.1667 | 4 | 0.5 | 3 | 0.3333 | 15 | 0.3333 | 2 | 0.5 | 4 | 0 | 4 | 0 | 2 | 1 | 12 | 0.25 |
| 4_1342352 | 3 | 1 | 5 | 0.8 | 4 | 1 | 3 | 1 | 15 | 0.9333 | 2 | 1 | 6 | 0.8333 | 5 | 1 | 2 | 1 | 15 | 0.9333 |
| 4_1342354 | 3 | 0.6667 | 3 | 1 | 2 | 1 | 6 | 1 | 14 | 0.9286 | 2 | 1 | 7 | 1 | 4 | 1 | 3 | 0.6667 | 16 | 0.9375 |
| 4_1342369 | 1 | 0 | NA | NA | 2 | 0.5 | 2 | 0 | 5 | 0.2 | 4 | 0 | 3 | 0.3333 | 1 | 0 | 3 | 0 | 11 | 0.0909 |
| 4_1342529 | 10 | 0.2 | 11 | 0.1818 | 15 | 0.1333 | 11 | 0.3636 | 47 | 0.2128 | 5 | 0 | 11 | 0 | 13 | 0.2308 | 11 | 0.0909 | 40 | 0.1 |
| 4_1342530 | 7 | 0.1429 | 8 | 0.125 | 9 | 0.1111 | 6 | 0.1667 | 30 | 0.1333 | 3 | 0.6667 | 3 | 0.3333 | 12 | 0.4167 | 7 | 0.1429 | 25 | 0.36 |
| 4_1342533 | 7 | 0.2857 | 7 | 0.4286 | 9 | 0.1111 | 10 | 0.1 | 33 | 0.2121 | 3 | 0.3333 | 8 | 0.25 | 9 | 0.1111 | 9 | 0.2222 | 29 | 0.2069 |
| 4_1342536 | 10 | 0 | 12 | 0.1667 | 8 | 0.125 | 11 | 0.0909 | 41 | 0.0976 | 6 | 0.1667 | 6 | 0 | 7 | 0.1429 | 11 | 0.0909 | 30 | 0.1 |
| 4_1342536 | 10 | 0 | 10 | 0.3 | 7 | 0.2857 | 9 | 0.1111 | 36 | 0.1667 | 5 | 0 | 8 | 0.125 | 8 | 0.125 | 9 | 0.1111 | 30 | 0.1 |
| 4_1342537 | 9 | 0.3333 | 9 | 0.1111 | 8 | 0.125 | 8 | 0.375 | 34 | 0.2353 | 5 | 0.2 | 8 | 0.125 | 9 | 0 | 9 | 0 | 31 | 0.0645 |
| 4_1342537 | 8 | 0.375 | 10 | 0.1 | 9 | 0.2222 | 10 | 0.2 | 37 | 0.2162 | 6 | 0.5 | 9 | 0.4444 | 4 | 0.5 | 9 | 0.2222 | 28 | 0.3929 |
| 4_1342537 | 9 | 0.2222 | 10 | 0.4 | 10 | 0.5 | 9 | 0 | 38 | 0.2895 | 6 | 0.5 | 8 | 0.375 | 7 | 0.8571 | 10 | 0.5 | 31 | 0.5484 |
| 4_1342565 | 4 | 0.25 | 8 | 0.25 | 8 | 0 | 6 | 0 | 26 | 0.1154 | 4 | 0 | 6 | 0.1667 | 7 | 0.1429 | 8 | 0.375 | 25 | 0.2 |

| | | | | | | | | | | | | | | | | | | | | |
|---|---|---|---|---|---|---|---|---|---|---|---|---|---|---|---|---|---|---|---|---|
| 4_13426539 | 4 | 0 | 2 | 0.5 | NA | NA | 1 | 0 | 7 | 0.1429 | 7 | 0.1429 | 5 | 0 | 2 | 0 | 3 | 0 | 17 | 0.0588 |
| 4_13426548 | 5 | 0 | 3 | 0.3333 | NA | NA | 1 | 0 | 9 | 0.1111 | 6 | 0.1667 | 4 | 0 | 2 | 0 | 3 | 0.3333 | 15 | 0.1333 |
| 4_13426584 | 3 | 0 | 2 | 0.5 | NA | NA | 1 | 0 | 6 | 0.1667 | 5 | 0.4 | 4 | 0 | 1 | 0 | 1 | 0 | 11 | 0.1818 |
| 4_13426606 | 3 | 0 | 2 | 0.5 | NA | NA | NA | NA | 5 | 0.2 | 5 | 0.2 | 2 | 0.5 | NA | NA | 1 | 0 | 8 | 0.25 |
| 4_13426609 | 5 | 0.6 | 3 | 0.3333 | NA | NA | NA | NA | 8 | 0.5 | 7 | 0.8571 | 5 | 0.6 | 1 | 1 | 2 | 1 | 15 | 0.8 |
| 4_13426616 | 4 | 0 | 2 | 0.5 | NA | NA | NA | NA | 6 | 0.1667 | 6 | 0.3333 | 3 | 0.3333 | 1 | 0 | 1 | 0 | 11 | 0.2727 |
| 4_13426658 | 5 | 0.2 | NA | NA | NA | NA | NA | NA | 5 | 0.2 | 5 | 0.8 | 4 | 0.5 | 1 | 1 | 1 | 1 | 11 | 0.7273 |
| 4_13426660 | 5 | 0.8 | 1 | 1 | NA | NA | NA | NA | 6 | 0.8333 | 5 | 0.4 | 4 | 0.5 | 1 | 0 | 2 | 0 | 12 | 0.3333 |
| 4_13426662 | 5 | 0.2 | NA | NA | NA | NA | NA | NA | 5 | 0.2 | 4 | 0 | 2 | 0.5 | NA | NA | 2 | 0 | 8 | 0.125 |
| 4_13426671 | 5 | 0.6 | NA | NA | NA | NA | NA | NA | 5 | 0.6 | 6 | 0.8333 | 3 | 1 | 1 | 1 | 2 | 0.5 | 12 | 0.8333 |
| 4_13426684 | 5 | 0.2 | NA | NA | NA | NA | NA | NA | 5 | 0.2 | 5 | 0.2 | 1 | 0 | 1 | 0 | 2 | 0 | 9 | 0.1111 |
| 4_13426691 | 4 | 0.25 | 1 | 0 | NA | NA | NA | NA | 5 | 0.2 | 4 | 0.25 | 2 | 0 | 1 | 1 | 2 | 0 | 9 | 0.2222 |
| 4_13426691 | 5 | 0.2 | 1 | 0 | NA | NA | NA | NA | 6 | 0.1667 | 5 | 0.2 | 2 | 0 | 1 | 0 | 2 | 0 | 10 | 0.1 |
| 4_13426696 | 3 | 0.3333 | NA | NA | NA | NA | NA | NA | 3 | 0.3333 | 4 | 0.5 | 1 | 1 | 1 | 1 | 2 | 1 | 8 | 0.75 |
| 4_13429030 | 1 | 0 | 2 | 0.5 | NA | NA | 1 | 1 | 4 | 0.5 | 2 | 0.5 | 2 | 0.5 | 2 | 0.5 | 1 | 0 | 7 | 0.4286 |
| 4_13429075 | 4 | 0.5 | 4 | 0.75 | 3 | 0.6667 | 5 | 0.6 | 16 | 0.625 | 3 | 1 | 3 | 1 | 7 | 0.7143 | 2 | 0.5 | 15 | 0.8 |
| 4_13429089 | 2 | 0.5 | 3 | 0.3333 | 3 | 0.6667 | 5 | 0 | 13 | 0.3077 | 3 | 0.6667 | 4 | 0.75 | 4 | 0.5 | 2 | 0.5 | 13 | 0.6154 |
| 4_13429104 | 2 | 0 | 4 | 0.25 | 2 | 0 | 4 | 0 | 12 | 0.0833 | 3 | 0.3333 | 3 | 0 | 4 | 0.25 | 2 | 0 | 12 | 0.1667 |
| 4_13429109 | 3 | 0 | 4 | 0 | 3 | 0 | 4 | 0.25 | 14 | 0.0714 | 3 | 0 | 3 | 0 | 6 | 0 | 2 | 0.5 | 14 | 0.0714 |
| 4_13443031 | 3 | 0.3333 | 1 | 0 | 2 | 0.5 | 2 | 0.5 | 8 | 0.375 | 1 | 0 | 2 | 0.5 | 4 | 0.75 | NA | NA | 7 | 0.5714 |
| 4_13443060 | 3 | 0.3333 | 1 | 1 | 1 | 0 | 2 | 0 | 7 | 0.2857 | 1 | 1 | 2 | 0.5 | 3 | 0.3333 | NA | NA | 6 | 0.5 |
| 4_13443083 | 3 | 0.3333 | 1 | 1 | 1 | 1 | 2 | 0 | 7 | 0.4286 | 1 | 1 | 2 | 1 | 3 | 1 | NA | NA | 6 | 1 |
| 4_13443084 | 3 | 0 | NA | NA | 1 | 1 | 1 | 0 | 5 | 0.2 | 1 | 1 | 2 | 0.5 | 3 | 0.6667 | NA | NA | 6 | 0.6667 |
| 4_13443095 | 3 | 0.3333 | 1 | 0 | 1 | 0 | 2 | 0.5 | 7 | 0.2857 | 1 | 1 | 2 | 0.5 | 3 | 0.6667 | NA | NA | 6 | 0.6667 |
| 4_13443117 | 3 | 0.3333 | 1 | 0 | 1 | 1 | 1 | 1 | 6 | 0.5 | 1 | 0 | 1 | 1 | 2 | 0 | NA | NA | 4 | 0.25 |
| 4_13453113 | 2 | 0 | 2 | 0 | 1 | 0 | 5 | 0.4 | 10 | 0.2 | 2 | 0 | 6 | 0.5 | 2 | 0.5 | 5 | 0.4 | 15 | 0.4 |
| 4_13453147 | 8 | 0.875 | 5 | 0.8 | 5 | 1 | 9 | 1 | 27 | 0.9259 | 4 | 1 | 8 | 1 | 7 | 0.8571 | 4 | 1 | 23 | 0.9565 |
| 4_13453157 | 8 | 1 | 5 | 1 | 5 | 1 | 10 | 0.9 | 28 | 0.9643 | 2 | 1 | 8 | 0.875 | 6 | 1 | 4 | 1 | 20 | 0.95 |
| 4_13453171 | 5 | 0.8 | 5 | 0.8 | 5 | 0.6 | 11 | 0.7273 | 26 | 0.7308 | 2 | 1 | 7 | 0.8571 | 6 | 0.6667 | 1 | 1 | 16 | 0.8125 |
| 4_13453184 | 7 | 0.1429 | 5 | 0.2 | 4 | 0 | 10 | 0.3 | 26 | 0.1923 | 2 | 0.5 | 8 | 0.25 | 6 | 0.3333 | 4 | 0.25 | 20 | 0.3 |
| 4_13455536 | 3 | 0.6667 | 4 | 0.25 | 3 | 0.6667 | 5 | 0.6 | 15 | 0.5333 | 1 | 0 | 4 | 0.5 | 3 | 0.6667 | 3 | 0.6667 | 11 | 0.5455 |
| 4_13455537 | 4 | 0.75 | 4 | 0 | 4 | 0.75 | 7 | 0.5714 | 19 | 0.5263 | 1 | 1 | 3 | 1 | 3 | 0.6667 | 2 | 0.5 | 9 | 0.7778 |
| 4_13455553 | 4 | 1 | 4 | 1 | 3 | 0.6667 | 6 | 0.8333 | 17 | 0.8824 | 1 | 1 | 3 | 1 | 2 | 1 | 2 | 1 | 8 | 1 |
| 4_13455563 | 4 | 0.75 | 4 | 0.5 | 2 | 1 | 4 | 1 | 14 | 0.7857 | 1 | 0 | 3 | 0.6667 | 2 | 0.5 | 1 | 1 | 7 | 0.5714 |
| 4_13455569 | 4 | 0.25 | 3 | 0 | 3 | 0 | 4 | 0 | 14 | 0.0714 | 1 | 1 | 3 | 0.3333 | 2 | 1 | NA | NA | 6 | 0.6667 |
| 4_13455574 | 3 | 0.3333 | 4 | 0.5 | 4 | 0.5 | 7 | 0.4286 | 18 | 0.4444 | 1 | 0 | 4 | 0 | 2 | 0.5 | NA | NA | 7 | 0.1429 |
| 4_13462536 | 32 | 0.125 | 21 | 0.1905 | 23 | 0 | 20 | 0.15 | 96 | 0.1146 | 24 | 0.125 | 20 | 0.05 | 17 | 0.1765 | 17 | 0.1765 | 78 | 0.1282 |
| 4_13465289 | 5 | 0.2 | 12 | 0.3333 | 7 | 0.2857 | 5 | 0.2 | 29 | 0.2759 | 3 | 0.3333 | 9 | 0.2222 | 6 | 0.1667 | 6 | 0.3333 | 24 | 0.25 |
| 4_13465291 | 2 | 0 | 11 | 0.1818 | 8 | 0.125 | 5 | 0.2 | 26 | 0.1538 | 3 | 0 | 10 | 0.2 | 5 | 0 | 7 | 0.2857 | 25 | 0.16 |
| 4_13465434 | 24 | 0.5833 | 28 | 0.5357 | 37 | 0.6216 | 30 | 0.5 | 119 | 0.563 | 19 | 0.6316 | 29 | 0.5862 | 31 | 0.4194 | 26 | 0.4615 | 105 | 0.5143 |
| 4_13465436 | 12 | 0.4167 | 21 | 0.381 | 20 | 0.45 | 24 | 0.4167 | 77 | 0.4156 | 13 | 0.5385 | 24 | 0.5 | 19 | 0.4737 | 13 | 0.4615 | 69 | 0.4928 |
| 4_13465438 | 22 | 0.1818 | 32 | 0.2188 | 35 | 0.2571 | 34 | 0.1176 | 123 | 0.1951 | 20 | 0.1 | 28 | 0.0714 | 29 | 0.0345 | 20 | 0.15 | 97 | 0.0825 |
| 4_13465448 | 25 | 0.88 | 33 | 0.8485 | 34 | 0.8235 | 34 | 0.8824 | 126 | 0.8571 | 21 | 0.9048 | 24 | 1 | 30 | 0.7333 | 22 | 0.7273 | 97 | 0.8351 |
| 4_13465452 | 28 | 0.1786 | 33 | 0.2121 | 44 | 0.1364 | 38 | 0.1579 | 143 | 0.1678 | 23 | 0.2174 | 31 | 0.1613 | 33 | 0.0909 | 28 | 0.2143 | 115 | 0.1652 |
| 4_13465479 | 2 | 0 | 5 | 0.2 | 6 | 0 | 4 | 0.25 | 17 | 0.1176 | 3 | 0.3333 | 4 | 0 | 4 | 0.5 | 5 | 0.4 | 16 | 0.3125 |
| 4_13466470 | 8 | 0.125 | 5 | 0 | 6 | 0.3333 | 12 | 0.5 | 31 | 0.2903 | 4 | 0.25 | 10 | 0.1 | 12 | 0.4167 | 4 | 0.25 | 30 | 0.2667 |
| 4_13466472 | 11 | 0.3636 | 8 | 0.125 | 6 | 0.5 | 12 | 0.5 | 37 | 0.3784 | 5 | 0.2 | 13 | 0.1538 | 9 | 0.1111 | 4 | 0 | 31 | 0.129 |
| 4_13466477 | 13 | 0.0769 | 9 | 0.2222 | 6 | 0 | 13 | 0.1538 | 41 | 0.122 | 5 | 0.4 | 10 | 0 | 13 | 0.2308 | 4 | 0 | 32 | 0.1562 |
| 4_13494152 | 16 | 0.125 | 10 | 0.1 | 10 | 0.2 | 13 | 0.0769 | 49 | 0.1224 | 5 | 0.2 | 6 | 0.3333 | 11 | 0.3636 | 6 | 0.1667 | 28 | 0.2857 |
| 4_13494189 | 14 | 0.2143 | 9 | 0 | 9 | 0 | 11 | 0.0909 | 43 | 0.093 | 4 | 0 | 6 | 0.1667 | 11 | 0.0909 | 4 | 0.25 | 25 | 0.12 |
| 4_13494211 | 13 | 0.2308 | 9 | 0.4444 | 10 | 0.2 | 9 | 0.5556 | 41 | 0.3415 | 5 | 0.4 | 4 | 0.25 | 12 | 0.5 | 4 | 0.25 | 25 | 0.4 |
| 4_13494378 | 9 | 0 | 8 | 0.375 | 10 | 0.1 | 7 | 0 | 34 | 0.1176 | 2 | 0 | 6 | 0.3333 | 9 | 0 | 3 | 0.3333 | 20 | 0.15 |
| 4_13494431 | 7 | 0 | 6 | 0 | 7 | 0.2857 | 6 | 0 | 26 | 0.0769 | 2 | 0 | 5 | 0 | 8 | 0.125 | 3 | 0.3333 | 18 | 0.1111 |
| 4_13494435 | 8 | 0.75 | 7 | 0.7143 | 7 | 0.7143 | 10 | 0.5 | 32 | 0.6562 | 2 | 1 | 5 | 0.6 | 7 | 0.4286 | 3 | 0.6667 | 17 | 0.5882 |
| 4_13494442 | 9 | 0.7778 | 7 | 0.4286 | 9 | 0.6667 | 9 | 0.6667 | 34 | 0.6471 | 2 | 0 | 5 | 0.8 | 6 | 0.6667 | 3 | 0.6667 | 16 | 0.625 |
| 4_13507260 | 1 | 0 | NA | NA | NA | NA | 2 | 1 | 3 | 0.6667 | 1 | 0 | 2 | 0.5 | 2 | 0 | 4 | 0 | 9 | 0.1111 |
| 4_13514006 | 2 | 0.5 | 2 | 0 | 3 | 0 | 3 | 0 | 10 | 0.1 | 2 | 0 | 3 | 0 | 2 | 0.5 | 3 | 0 | 10 | 0.1 |
| 4_13514037 | 5 | 0.2 | 4 | 0 | 4 | 0 | 3 | 0 | 16 | 0.0625 | 3 | 0 | 5 | 0.2 | 3 | 0 | 5 | 0 | 16 | 0.0625 |
| 4_13514308 | 1 | 0 | 3 | 0 | 1 | 1 | 2 | 0 | 7 | 0.1429 | 2 | 1 | 2 | 0 | 1 | 1 | NA | NA | 5 | 0.6 |
| 4_13514356 | 1 | 1 | NA | NA | 1 | 1 | 1 | 1 | 3 | 1 | 2 | 0.5 | NA | NA | 1 | 1 | 1 | 1 | 4 | 0.75 |
| 4_13514374 | 2 | 1 | 2 | 0.5 | 1 | 1 | 2 | 1 | 7 | 0.8571 | 2 | 1 | 1 | 1 | 1 | 1 | 2 | 1 | 6 | 1 |
| 4_13517643 | 9 | 0.2222 | 11 | 0.2727 | 7 | 0.5714 | 7 | 0.4286 | 34 | 0.3529 | 13 | 0.3077 | 18 | 0.2778 | 13 | 0.3846 | 10 | 0.1 | 54 | 0.2778 |
| 4_13517772 | 6 | 1 | 6 | 1 | 2 | 1 | 6 | 1 | 20 | 1 | 7 | 1 | 10 | 0.9 | 7 | 1 | 6 | 1 | 30 | 0.9667 |
| 4_13517777 | 8 | 0.125 | 4 | 0.25 | 3 | 0 | 7 | 0.2857 | 22 | 0.1818 | 10 | 0.2 | 10 | 0.6 | 6 | 0.1667 | 7 | 0.1429 | 33 | 0.303 |
| 4_13517800 | 4 | 0.25 | 5 | 0.2 | 3 | 0 | 5 | 0 | 17 | 0.1176 | 10 | 0 | 8 | 0 | 9 | 0.3333 | 5 | 0.2 | 32 | 0.125 |

| | | | | | | | | | | | | | | | | | | | | |
|---|---|---|---|---|---|---|---|---|---|---|---|---|---|---|---|---|---|---|---|---|
| 4_13517807 | 6 | 0.1667 | 2 | 0 | 3 | 0.3333 | 5 | 0.4 | 16 | 0.25 | 10 | 0.3 | 9 | 0.2222 | 7 | 0.2857 | 6 | 0.1667 | 32 | 0.25 |
| 4_13517808 | 6 | 0.1667 | 4 | 0.25 | 2 | 0.5 | 7 | 0.4286 | 19 | 0.3158 | 9 | 0.1111 | 8 | 0 | 8 | 0.125 | 8 | 0 | 33 | 0.0606 |
| 4_13517810 | 6 | 0.3333 | 4 | 0.5 | 3 | 0.6667 | 4 | 0.5 | 17 | 0.4706 | 10 | 0.5 | 9 | 0.2222 | 8 | 0.25 | 8 | 0.625 | 35 | 0.4 |
| 4_13517833 | 5 | 0.4 | 3 | 0.3333 | 3 | 0.6667 | 5 | 0.6 | 16 | 0.5 | 8 | 0.375 | 6 | 0 | 4 | 0.75 | 8 | 0.625 | 26 | 0.4231 |
| 4_13517844 | 5 | 0.4 | 4 | 0.25 | 3 | 0 | 6 | 0.1667 | 18 | 0.2222 | 9 | 0.2222 | 7 | 0.2857 | 6 | 0.5 | 8 | 0.375 | 30 | 0.3333 |
| 4_13517852 | 5 | 0 | 4 | 0.25 | 2 | 0 | 5 | 0.2 | 16 | 0.125 | 9 | 0 | 9 | 0 | 4 | 0.25 | 8 | 0.125 | 30 | 0.0667 |
| 4_13517854 | 5 | 0.4 | 3 | 0 | 2 | 0 | 3 | 0.6667 | 13 | 0.3077 | 7 | 0.2857 | 4 | 0.75 | 5 | 0.4 | 8 | 0.125 | 24 | 0.3333 |
| 4_13517902 | 1 | 0 | 3 | 0.3333 | 1 | 0 | 2 | 0 | 7 | 0.1429 | 6 | 0.3333 | 5 | 0.4 | 2 | 0.5 | 7 | 0.5714 | 20 | 0.45 |
| 4_13517904 | 1 | 0 | 1 | 1 | NA | NA | 4 | 0 | 6 | 0.1667 | 5 | 0.2 | 4 | 0 | 1 | 0 | 4 | 0.25 | 14 | 0.1429 |
| 4_13517910 | 3 | 0.3333 | 4 | 0.5 | 2 | 0 | 6 | 0 | 15 | 0.2 | 8 | 0.25 | 8 | 0.125 | 3 | 0.3333 | 6 | 0.1667 | 25 | 0.2 |
| 4_13517917 | 1 | 1 | 2 | 1 | 2 | 1 | 6 | 1 | 11 | 1 | 6 | 0.8333 | 6 | 0.6667 | 3 | 1 | 6 | 1 | 21 | 0.8571 |
| 4_13517917 | 1 | 1 | 3 | 1 | 1 | 1 | 2 | 1 | 7 | 1 | 6 | 0.6667 | 6 | 0.1667 | 3 | 0.3333 | 6 | 1 | 21 | 0.5714 |
| 4_13517924 | 2 | 1 | 4 | 1 | 2 | 1 | 6 | 1 | 14 | 1 | 7 | 1 | 7 | 0.8571 | 4 | 0.75 | 7 | 1 | 25 | 0.92 |
| 4_13517924 | 1 | 1 | 2 | 1 | 1 | 1 | 3 | 0.6667 | 7 | 0.8571 | 3 | 0.6667 | 5 | 0.6 | 3 | 0.3333 | 6 | 0.6667 | 17 | 0.5882 |
| 4_13517936 | 3 | 0 | 4 | 0.5 | 1 | 0 | 5 | 0.4 | 13 | 0.3077 | 8 | 0.25 | 7 | 0.2857 | 4 | 0 | 7 | 0.2857 | 26 | 0.2308 |
| 4_13517942 | 2 | 1 | 3 | 1 | 2 | 1 | 2 | 1 | 9 | 1 | 4 | 1 | 4 | 1 | 2 | 1 | 6 | 1 | 16 | 1 |
| 4_13517952 | 2 | 0.5 | 4 | 0.25 | 2 | 0.5 | 4 | 0.25 | 12 | 0.3333 | 6 | 0.3333 | 6 | 0.1667 | 4 | 0.5 | 5 | 0.8 | 21 | 0.4286 |
| 4_13517958 | NA | NA | 3 | 1 | 2 | 1 | 2 | 1 | 7 | 1 | 4 | 1 | 4 | 1 | 2 | 1 | 3 | 1 | 13 | 1 |
| 4_13517962 | 2 | 0.5 | 4 | 0.5 | 2 | 0.5 | 3 | 0.6667 | 11 | 0.5455 | 4 | 0.5 | 6 | 0.3333 | 3 | 0.6667 | 3 | 0.3333 | 16 | 0.4375 |
| 4_13517985 | 2 | 0.5 | 3 | 0 | 2 | 0.5 | 5 | 0.2 | 12 | 0.25 | 5 | 0.2 | 4 | 0.25 | 3 | 0 | 5 | 0.4 | 17 | 0.2353 |
| 4_13517987 | 2 | 0 | 4 | 0 | 2 | 0 | 5 | 0.2 | 13 | 0.0769 | 4 | 0 | 7 | 0 | 3 | 0 | 6 | 0.1667 | 20 | 0.05 |
| 4_13521848 | 1 | 0 | 2 | 0 | NA | NA | 3 | 0.3333 | 6 | 0.1667 | 3 | 0 | 3 | 0.3333 | 3 | 0 | 2 | 0.5 | 11 | 0.1818 |
| 4_13554191 | 1 | 1 | NA | NA | 2 | 1 | NA | NA | 3 | 1 | NA | NA | 2 | 0.5 | NA | NA | NA | NA | 2 | 0.5 |
| 4_13554224 | 1 | 1 | NA | NA | 2 | 1 | NA | NA | 3 | 1 | NA | NA | 2 | 0.5 | NA | NA | NA | NA | 2 | 0.5 |
| 4_13554235 | 1 | 1 | NA | NA | 2 | 0 | NA | NA | 3 | 0.3333 | NA | NA | 2 | 0.5 | 1 | 0 | NA | NA | 3 | 0.3333 |
| 4_13554250 | 1 | 1 | NA | NA | 1 | 0 | NA | NA | 2 | 0.5 | NA | NA | 2 | 0.5 | NA | NA | NA | NA | 2 | 0.5 |
| 4_13554272 | 1 | 1 | NA | NA | 2 | 0.5 | NA | NA | 3 | 0.6667 | NA | NA | 2 | 1 | NA | NA | NA | NA | 2 | 1 |
| 4_13554285 | NA | NA | NA | NA | 1 | 1 | NA | NA | 1 | 1 | NA | NA | 2 | 0.5 | NA | NA | NA | NA | 2 | 0.5 |
| 4_13559177 | 5 | 0.2 | 4 | 0 | 3 | 0 | 6 | 0 | 18 | 0.0556 | 3 | 0 | 3 | 0 | 5 | 0.4 | 5 | 0.2 | 16 | 0.1875 |
| 4_13559208 | 5 | 0 | 3 | 0.3333 | 4 | 0 | 6 | 0.1667 | 18 | 0.1111 | 4 | 0.5 | 3 | 0.3333 | 6 | 0 | 7 | 0.1429 | 20 | 0.2 |
| 4_13559271 | 5 | 0.2 | 6 | 0.1667 | 6 | 0 | 7 | 0.1429 | 24 | 0.125 | 4 | 0 | 5 | 0 | 5 | 0 | 10 | 0.2 | 24 | 0.0833 |
| 4_13559451 | 1 | 0 | 2 | 0.5 | 2 | 0 | 1 | 0 | 6 | 0.1667 | NA | NA | 3 | 0.3333 | 3 | 0 | 4 | 0 | 10 | 0.1 |
| 4_13559458 | 1 | 1 | 2 | 0.5 | 4 | 0.5 | 1 | 1 | 8 | 0.625 | NA | NA | 3 | 0.3333 | 3 | 0.3333 | 5 | 0.6 | 11 | 0.4545 |
| 4_13559467 | 1 | 0 | 2 | 0 | 6 | 0.3333 | 1 | 1 | 10 | 0.3 | 1 | 0 | 2 | 0 | 4 | 0 | 4 | 0.25 | 11 | 0.0909 |
| 4_13559472 | 1 | 0 | 2 | 0.5 | 5 | 0 | 1 | 0 | 9 | 0.1111 | 1 | 0 | 3 | 0 | 4 | 0 | 2 | 0.5 | 10 | 0.1 |
| 4_13559474 | 1 | 0 | 2 | 0.5 | 5 | 0.8 | 1 | 1 | 9 | 0.6667 | 1 | 0 | 3 | 0.3333 | 3 | 0 | 4 | 0.25 | 11 | 0.1818 |
| 4_13559557 | 1 | 1 | 3 | 0 | 7 | 0.1429 | 1 | 0 | 12 | 0.1667 | 1 | 1 | 5 | 0 | 5 | 0.2 | 3 | 0 | 14 | 0.1429 |
| 4_13559849 | 14 | 0.1429 | 5 | 0.2 | 10 | 0.2 | 16 | 0.4375 | 45 | 0.2667 | 6 | 0.1667 | 9 | 0 | 11 | 0.2727 | 5 | 0.2 | 31 | 0.1613 |
| 4_13559871 | 12 | 0.3333 | 5 | 0.2 | 10 | 0.5 | 16 | 0.3125 | 43 | 0.3488 | 5 | 0.4 | 7 | 0.4286 | 7 | 0.1429 | 3 | 0.3333 | 22 | 0.3182 |
| 4_13559936 | 10 | 0.1 | 5 | 0 | 9 | 0 | 11 | 0.0909 | 35 | 0.0571 | 5 | 0.2 | 7 | 0.1429 | 9 | 0.1111 | 4 | 0.25 | 25 | 0.16 |
| 4_13559947 | 10 | 0.8 | 6 | 0.8333 | 8 | 0.875 | 14 | 0.9286 | 38 | 0.8684 | 6 | 0.8333 | 7 | 0.7143 | 8 | 0.75 | 4 | 1 | 25 | 0.8 |
| 4_13559960 | 8 | 0.375 | 5 | 0.4 | 8 | 0.625 | 12 | 0.6667 | 33 | 0.5455 | 4 | 0.75 | 8 | 0.5 | 7 | 0.4286 | 2 | 1 | 21 | 0.5714 |
| 4_13564574 | 1 | 1 | NA | NA | NA | NA | NA | NA | 1 | 1 | 1 | 1 | NA | NA | NA | NA | NA | NA | 1 | 1 |
| 4_13564676 | 1 | 1 | NA | NA | NA | NA | NA | NA | 1 | 1 | 1 | 1 | NA | NA | NA | NA | NA | NA | 1 | 1 |
| 4_13564808 | 5 | 0 | 3 | 0 | 1 | 1 | 3 | 0 | 12 | 0.0833 | 4 | 0 | NA | NA | 4 | 0.25 | NA | NA | 8 | 0.125 |
| 4_13564810 | 6 | 0 | 4 | 0 | 1 | 1 | 3 | 0.3333 | 14 | 0.1429 | 5 | 0.6 | NA | NA | 6 | 0.6667 | NA | NA | 11 | 0.6364 |
| 4_13564819 | 6 | 0.1667 | 4 | 0.5 | 2 | 0.5 | 3 | 0.6667 | 15 | 0.4 | 5 | 0.6 | NA | NA | 8 | 0.25 | 1 | 0 | 14 | 0.3571 |
| 4_13564820 | 7 | 0.5714 | 3 | 0.6667 | 3 | 0.6667 | 1 | 0 | 14 | 0.5714 | 5 | 0.8 | NA | NA | 7 | 0.4286 | 1 | 0 | 13 | 0.5385 |
| 4_13564843 | 7 | 0 | 5 | 0 | 4 | 0.5 | 1 | 0 | 17 | 0.1176 | 5 | 0.2 | NA | NA | 5 | 0.4 | 1 | 0 | 11 | 0.2727 |
| 4_13566649 | 7 | 0.4286 | 7 | 0.5714 | 4 | 0.5 | 5 | 0.2 | 23 | 0.4348 | 5 | 0.2 | NA | NA | 7 | 0.1429 | 4 | 0.25 | 16 | 0.1875 |
| 4_13566652 | 6 | 0.5 | 6 | 0.1667 | 4 | 0.25 | 6 | 0.1667 | 22 | 0.2727 | 5 | 0.2 | NA | NA | 8 | 0.375 | 4 | 0.25 | 17 | 0.2941 |
| 4_13566654 | 7 | 0.2857 | 7 | 0.1429 | 4 | 0.25 | 6 | 0.3333 | 24 | 0.25 | 7 | 0.2857 | NA | NA | 8 | 0.5 | 2 | 0.5 | 17 | 0.4118 |
| 4_13566654 | 4 | 0.25 | 2 | 0.5 | 3 | 0 | 3 | 0 | 12 | 0.1667 | 7 | 0.1429 | NA | NA | 2 | 0 | 2 | 0.5 | 11 | 0.1818 |
| 4_13568948 | 6 | 0.1667 | 3 | 0 | 7 | 0.1429 | 5 | 0 | 21 | 0.0952 | 6 | 0 | 9 | 0 | 3 | 0 | 5 | 0.4 | 23 | 0.087 |
| 4_13568977 | 6 | 0.5 | 6 | 0.3333 | 5 | 0.4 | 4 | 0.75 | 21 | 0.4762 | 6 | 0.3333 | 8 | 0.75 | 3 | 0.6667 | 4 | 0.75 | 21 | 0.619 |
| 4_13569601 | 12 | 0.1667 | 13 | 0.2308 | 10 | 0 | 10 | 0.1 | 45 | 0.1333 | 7 | 0.1429 | 8 | 0 | 12 | 0.0833 | 12 | 0.0833 | 39 | 0.0769 |
| 4_13569652 | 12 | 0.0833 | 11 | 0 | 10 | 0.1 | 6 | 0 | 39 | 0.0513 | 6 | 0.1667 | 11 | 0 | 7 | 0.1429 | 9 | 0.2222 | 33 | 0.1212 |
| 4_13570315 | 16 | 0.1875 | 12 | 0 | 17 | 0.1765 | 16 | 0.0625 | 61 | 0.1148 | 11 | 0 | 12 | 0.0833 | 18 | 0.0556 | 10 | 0.3 | 51 | 0.098 |
| 4_13570349 | 14 | 0.3571 | 12 | 0.3333 | 17 | 0.2941 | 13 | 0.3846 | 56 | 0.3393 | 11 | 0.2727 | 9 | 0.3333 | 14 | 0.5714 | 9 | 0.2222 | 43 | 0.3721 |
| 4_13570364 | 14 | 0.2857 | 12 | 0.25 | 14 | 0.2143 | 15 | 0.2667 | 55 | 0.2545 | 9 | 0.1111 | 12 | 0 | 16 | 0.125 | 13 | 0.0769 | 50 | 0.08 |
| 4_13573094 | 7 | 0.2857 | 3 | 0 | 3 | 0.3333 | 3 | 0.3333 | 16 | 0.25 | 1 | 0 | 1 | 1 | 1 | 0 | NA | NA | 3 | 0.3333 |
| 4_13573196 | 5 | 0 | 3 | 0.3333 | 3 | 0 | 2 | 0 | 13 | 0.0769 | 1 | 0 | NA | NA | 2 | 0.5 | NA | NA | 3 | 0.3333 |
| 4_13573273 | 5 | 0 | 5 | 0 | 4 | 0.5 | 4 | 0 | 18 | 0.1111 | NA | NA | 1 | 0 | 3 | 0.3333 | NA | NA | 4 | 0.25 |
| 4_13573280 | 5 | 0 | 5 | 0.2 | 3 | 0 | 3 | 0 | 16 | 0.0625 | NA | NA | 2 | 0.5 | 3 | 0 | NA | NA | 5 | 0.2 |
| 4_13573282 | 4 | 0.5 | 4 | 0.75 | 4 | 1 | 2 | 0.5 | 14 | 0.7143 | NA | NA | 1 | 1 | 2 | 1 | NA | NA | 3 | 1 |

| | | | | | | | | | | | | | | | | | | | | |
|---|---|---|---|---|---|---|---|---|---|---|---|---|---|---|---|---|---|---|---|---|
| 4_1357376 | 4 | 0.75 | 4 | 1 | 3 | 0.6667 | 4 | 1 | 15 | 0.8667 | NA | NA | 2 | 0.5 | 3 | 1 | NA | NA | 5 | 0.8 |
| 4_1357376 | 4 | 1 | 4 | 0.75 | 3 | 0.3333 | 4 | 1 | 15 | 0.8 | NA | NA | 2 | 1 | 2 | 1 | NA | NA | 4 | 1 |
| 4_1357378 | 4 | 0.25 | 2 | 0 | 3 | 0 | 4 | 0 | 13 | 0.0769 | NA | NA | 2 | 0.5 | 2 | 0 | NA | NA | 4 | 0.25 |
| 4_1357379 | 2 | 0 | 4 | 0.75 | 2 | 1 | 4 | 0.5 | 12 | 0.5833 | NA | NA | 1 | 1 | 2 | 0.5 | NA | NA | 3 | 0.6667 |
| 4_1357383 | 2 | 0.5 | 3 | 0 | 3 | 0 | 3 | 0.3333 | 11 | 0.1818 | NA | NA | 1 | 1 | 3 | 0 | NA | NA | 4 | 0.25 |
| 4_1357771 | 1 | 1 | 1 | 0 | 1 | 1 | NA | NA | 3 | 0.6667 | 1 | 0 | 1 | 1 | 1 | 0 | 1 | 1 | 4 | 0.5 |
| 4_1358723 | 19 | 0.2105 | 18 | 0.2778 | 8 | 0.25 | 5 | 0.2 | 50 | 0.24 | 15 | 0.4 | 12 | 0.3333 | 11 | 0.1818 | 15 | 0.2667 | 53 | 0.3019 |
| 4_1358724 | 23 | 0.6957 | 19 | 0.6842 | 12 | 0.25 | 9 | 0.4444 | 63 | 0.5714 | 17 | 0.5882 | 14 | 0.4286 | 13 | 0.5385 | 18 | 0.6667 | 62 | 0.5645 |
| 4_1358725 | 18 | 0.2222 | 20 | 0.2 | 10 | 0.1 | 7 | 0.1429 | 55 | 0.1818 | 16 | 0.375 | 10 | 0.6 | 11 | 0.0909 | 13 | 0.1538 | 50 | 0.3 |
| 4_1360179 | 2 | 1 | NA | NA | 1 | 1 | NA | NA | 3 | 1 | 2 | 1 | NA | NA | NA | NA | 3 | 1 | 5 | 1 |
| 4_1360181 | 2 | 1 | NA | NA | NA | NA | 1 | 1 | 3 | 1 | 2 | 0 | 1 | 0 | NA | NA | 4 | 0.25 | 7 | 0.1429 |
| 4_1360185 | 2 | 1 | NA | NA | 1 | 1 | NA | NA | 3 | 1 | 2 | 1 | 2 | 0.5 | NA | NA | 5 | 1 | 9 | 0.8889 |
| 4_1360186 | 1 | 1 | NA | NA | 1 | 1 | NA | NA | 2 | 1 | 2 | 0.5 | 1 | 0 | NA | NA | 3 | 1 | 6 | 0.6667 |
| 4_1360188 | 3 | 0.6667 | NA | NA | 1 | 0 | 1 | 1 | 5 | 0.6 | 3 | 0.3333 | 2 | 1 | NA | NA | 6 | 0.6667 | 11 | 0.6364 |
| 4_1360191 | 4 | 0.5 | 1 | 0 | 2 | 0.5 | 2 | 0.5 | 9 | 0.4444 | 3 | 0 | 3 | 0 | 1 | 0 | 6 | 0.3333 | 13 | 0.1538 |
| 4_1360192 | 4 | 1 | 1 | 1 | 2 | 1 | 2 | 1 | 9 | 1 | 3 | 1 | 3 | 0.6667 | 1 | 1 | 6 | 1 | 13 | 0.9231 |
| 4_1360192 | 3 | 0.6667 | 1 | 1 | 1 | 0 | 2 | 1 | 7 | 0.7143 | 4 | 0.75 | 3 | 0.6667 | 1 | 1 | 5 | 0.6 | 13 | 0.6923 |
| 4_1360193 | 4 | 0.25 | 1 | 1 | 2 | 0 | 2 | 0.5 | 9 | 0.3333 | 3 | 0 | 3 | 0.6667 | 1 | 0 | 6 | 0.3333 | 13 | 0.3077 |
| 4_1360193 | 4 | 0.75 | 1 | 1 | NA | NA | 1 | 1 | 6 | 0.8333 | 3 | 0.6667 | 3 | 1 | NA | NA | 4 | 0.75 | 10 | 0.8 |
| 4_1360193 | 4 | 0.75 | 1 | 1 | 2 | 1 | 2 | 0.5 | 9 | 0.7778 | 4 | 0.25 | 3 | 1 | 1 | 1 | 4 | 0.75 | 12 | 0.6667 |
| 4_1360196 | 4 | 0.25 | 1 | 1 | 1 | 1 | 1 | 0 | 7 | 0.4286 | 3 | 0.3333 | 3 | 0.3333 | 1 | 1 | 7 | 0.5714 | 14 | 0.5 |
| 4_1360197 | 4 | 0.5 | 1 | 0 | 2 | 0 | 2 | 0.5 | 9 | 0.3333 | 3 | 0 | 3 | 0.3333 | 1 | 1 | 6 | 0.5 | 13 | 0.3846 |
| 4_1360197 | 3 | 0.6667 | 2 | 0.5 | 2 | 0.5 | 2 | 1 | 9 | 0.6667 | 4 | 0.25 | 3 | 0.6667 | 1 | 1 | 7 | 0.7143 | 15 | 0.6 |
| 4_1360197 | 4 | 1 | 1 | 0 | 1 | 1 | NA | NA | 6 | 0.8333 | 3 | 1 | 3 | 0.6667 | 1 | 0 | 5 | 0.2 | 12 | 0.5 |
| 4_1360198 | 3 | 1 | 2 | 0.5 | 2 | 0.5 | 2 | 1 | 9 | 0.7778 | 4 | 0.75 | 2 | 1 | 1 | 1 | 7 | 1 | 14 | 0.9286 |
| 4_1360198 | 3 | 0.3333 | 2 | 0 | 3 | 0.3333 | 1 | 1 | 9 | 0.3333 | 4 | 0.25 | 3 | 0.3333 | 1 | 0 | 7 | 0.2857 | 15 | 0.2667 |
| 4_1360200 | 3 | 0 | 1 | 0 | 3 | 0.3333 | 2 | 1 | 9 | 0.3333 | 4 | 0.25 | 2 | 0 | 1 | 0 | 6 | 0.3333 | 13 | 0.2308 |
| 4_1360202 | 4 | 0 | 2 | 0 | 4 | 0 | 2 | 1 | 12 | 0.1667 | 5 | 0.2 | 3 | 0 | 1 | 0 | 8 | 0 | 17 | 0.0588 |
| 4_1360203 | 4 | 0.25 | 3 | 0 | 3 | 0.6667 | 1 | 1 | 11 | 0.3636 | 5 | 0.6 | 3 | 1 | 1 | 0 | 8 | 0.625 | 17 | 0.6471 |
| 4_1360668 | 11 | 0.1818 | 4 | 0.5 | 7 | 0.1429 | 6 | 0.1667 | 28 | 0.2143 | 4 | 0.5 | 4 | 0.25 | 6 | 0.1667 | 4 | 0.5 | 18 | 0.3333 |
| 4_1360670 | 11 | 0.3636 | 3 | 0.6667 | 3 | 0.3333 | 5 | 0 | 22 | 0.3182 | 4 | 0 | 4 | 0.5 | 6 | 0 | 1 | 0 | 15 | 0.1333 |
| 4_1360672 | 15 | 0.5333 | 2 | 0 | 7 | 0.1429 | 4 | 0.5 | 28 | 0.3929 | 3 | 0.6667 | 7 | 0.7143 | 4 | 0.25 | 6 | 0.5 | 20 | 0.55 |
| 4_1360672 | 15 | 0.4667 | 3 | 1 | 9 | 0.3333 | 9 | 0.5556 | 36 | 0.5 | 5 | 0.6 | 4 | 0.25 | 8 | 0.5 | 6 | 0.6667 | 23 | 0.5217 |
| 4_1360672 | 17 | 0.1176 | 3 | 0 | 8 | 0 | 8 | 0 | 36 | 0.0556 | 6 | 0.3333 | 5 | 0 | 5 | 0.2 | 4 | 0.25 | 20 | 0.2 |
| 4_1360672 | 18 | 0.1111 | 5 | 0 | 10 | 0.1 | 11 | 0.1818 | 44 | 0.1136 | 4 | 0 | 7 | 0 | 8 | 0.125 | 7 | 0.1429 | 26 | 0.0769 |
| 4_1360673 | 16 | 0.5 | 4 | 0.5 | 10 | 0.5 | 9 | 0.4444 | 39 | 0.4872 | 6 | 0.1667 | 7 | 0.1429 | 8 | 0.375 | 6 | 0.5 | 27 | 0.2963 |
| 4_1360674 | 18 | 0.1667 | 5 | 0.4 | 10 | 0.1 | 12 | 0.1667 | 45 | 0.1778 | 8 | 0.5 | 8 | 0.125 | 7 | 0 | 7 | 0.4286 | 30 | 0.2667 |
| 4_1360675 | 12 | 0.9167 | 2 | 1 | 8 | 0.75 | 10 | 0.9 | 32 | 0.875 | 6 | 0.8333 | 5 | 1 | 8 | 0.875 | 2 | 1 | 21 | 0.9048 |
| 4_1360676 | 21 | 0.4762 | 5 | 0.4 | 8 | 0.25 | 12 | 0.5 | 46 | 0.4348 | 9 | 0.5556 | 8 | 0.625 | 9 | 0.3333 | 7 | 0.5714 | 33 | 0.5152 |
| 4_1360676 | 23 | 0 | 5 | 0.4 | 9 | 0 | 13 | 0.2308 | 50 | 0.1 | 8 | 0.25 | 9 | 0.1111 | 9 | 0 | 6 | 0 | 32 | 0.0938 |
| 4_1360676 | 17 | 0.5294 | 5 | 0.6 | 9 | 0.2222 | 10 | 0.5 | 41 | 0.4634 | 8 | 0.5 | 9 | 0.4444 | 5 | 0.4 | 6 | 0.6667 | 28 | 0.5 |
| 4_1361693 | NA | NA | 1 | 1 | 2 | 1 | 1 | 0 | 4 | 0.75 | 1 | 1 | 1 | 1 | 2 | 1 | 1 | 1 | 5 | 1 |
| 4_1361693 | NA | NA | 1 | 1 | 2 | 0 | 1 | 0 | 4 | 0.25 | NA | NA | NA | NA | 2 | 0 | 1 | 1 | 3 | 0.3333 |
| 4_1361694 | NA | NA | 1 | 0 | 2 | 1 | 1 | 1 | 4 | 0.75 | 1 | 1 | 1 | 0 | 1 | 0 | 1 | 1 | 4 | 0.5 |
| 4_1361694 | NA | NA | 1 | 1 | 2 | 0.5 | 1 | 1 | 4 | 0.75 | 1 | 1 | 1 | 1 | NA | NA | 1 | 0 | 3 | 0.6667 |
| 4_1361697 | NA | NA | 1 | 1 | 2 | 1 | NA | NA | 3 | 1 | 1 | 1 | 2 | 1 | 1 | 1 | 1 | 1 | 5 | 1 |
| 4_1361700 | 1 | 1 | 1 | 1 | 2 | 1 | 1 | 1 | 5 | 1 | 1 | 1 | 2 | 1 | 2 | 1 | NA | NA | 5 | 1 |
| 4_1361700 | 1 | 1 | 1 | 1 | 2 | 0.5 | 1 | 1 | 5 | 0.8 | 1 | 1 | 2 | 1 | NA | NA | 1 | 1 | 4 | 1 |
| 4_1361700 | NA | NA | 1 | 0 | 2 | 0.5 | 1 | 0 | 4 | 0.25 | 1 | 0 | 2 | 0.5 | 2 | 0.5 | 1 | 0 | 6 | 0.3333 |
| 4_1361701 | NA | NA | 1 | 1 | 2 | 1 | 1 | 1 | 4 | 1 | 1 | 1 | 2 | 1 | 1 | 1 | NA | NA | 4 | 1 |
| 4_1361706 | 1 | 1 | 1 | 1 | 3 | 1 | 1 | 1 | 6 | 1 | 1 | 1 | 2 | 1 | 1 | 1 | 1 | 1 | 5 | 1 |
| 4_1361707 | 2 | 1 | 1 | 1 | 3 | 1 | 1 | 1 | 7 | 1 | 1 | 1 | 2 | 0.5 | 2 | 1 | 1 | 1 | 6 | 0.8333 |
| 4_1361707 | 2 | 1 | 2 | 1 | 2 | 1 | NA | NA | 6 | 1 | 1 | 0 | 2 | 1 | 2 | 1 | 1 | 1 | 6 | 0.8333 |
| 4_1361708 | 2 | 1 | 2 | 0.5 | 3 | 1 | 1 | 1 | 8 | 0.875 | 1 | 1 | 2 | 1 | 2 | 1 | 2 | 1 | 7 | 1 |
| 4_1361708 | 1 | 1 | 2 | 0 | 3 | 0.6667 | 1 | 0 | 7 | 0.4286 | 1 | 1 | 2 | 0 | 2 | 0.5 | 1 | 0 | 6 | 0.3333 |
| 4_1361709 | 2 | 0.5 | 2 | 1 | 3 | 1 | 1 | 1 | 8 | 0.875 | 1 | 1 | 1 | 1 | 3 | 0.3333 | 1 | 1 | 6 | 0.6667 |
| 4_1361709 | 2 | 0 | 2 | 0 | 3 | 0.3333 | 1 | 0 | 8 | 0.125 | 1 | 0 | 1 | 0 | 3 | 0.6667 | 2 | 0 | 7 | 0.2857 |
| 4_1361711 | 2 | 1 | 2 | 1 | 2 | 0.5 | 1 | 1 | 7 | 0.8571 | 1 | 1 | 1 | 1 | 3 | 1 | 2 | 1 | 7 | 1 |
| 4_1361714 | 2 | 0 | 2 | 0 | 1 | 1 | 2 | 0 | 7 | 0.1429 | 1 | 0 | 1 | 0 | 4 | 0.25 | 2 | 0 | 8 | 0.125 |
| 4_1361715 | NA | NA | 2 | 0.5 | 1 | 1 | 2 | 0.5 | 5 | 0.6 | NA | NA | 1 | 1 | 4 | 0.75 | 2 | 0.5 | 7 | 0.7143 |
| 4_1361716 | 2 | 0 | 2 | 0 | 1 | 1 | 3 | 0 | 8 | 0.125 | 1 | 0 | 1 | 1 | 4 | 0.25 | 2 | 0 | 8 | 0.25 |
| 4_1361754 | NA | NA | NA | NA | 1 | 1 | NA | NA | 1 | 1 | 1 | 1 | 1 | 0 | 1 | 1 | NA | NA | 3 | 0.6667 |
| 4_1361755 | NA | NA | NA | NA | 1 | 1 | NA | NA | 1 | 1 | 1 | 1 | 1 | 0 | 1 | 1 | NA | NA | 3 | 0.6667 |
| 4_1361758 | NA | NA | NA | NA | 1 | 1 | NA | NA | 1 | 1 | 2 | 0.5 | 1 | 0 | 1 | 0 | NA | NA | 4 | 0.25 |
| 4_1361759 | NA | NA | NA | NA | 1 | 1 | NA | NA | 1 | 1 | 2 | 1 | NA | NA | 1 | 1 | NA | NA | 3 | 1 |

| | | | | | | | | | | | | | | | | | | | | |
|---|---|---|---|---|---|---|---|---|---|---|---|---|---|---|---|---|---|---|---|---|
| 4_1361760 | NA | NA | NA | NA | 1 | 1 | NA | NA | 1 | 1 | 2 | 1 | 1 | 0 | NA | NA | NA | NA | 3 | 0.6667 |
| 4_1362689 | NA | NA | 1 | 1 | NA | NA | NA | NA | 1 | 1 | 1 | 1 | NA | NA | 1 | 0 | 1 | 1 | 3 | 0.6667 |
| 4_1362690 | 1 | 1 | NA | NA | NA | NA | NA | NA | 1 | 1 | 1 | 0 | NA | NA | 2 | 0 | 2 | 1 | 5 | 0.4 |
| 4_1362692 | 1 | 0 | 1 | 1 | NA | NA | NA | NA | 2 | 0.5 | 1 | 0 | NA | NA | 2 | 0 | 2 | 0.5 | 5 | 0.2 |
| 4_1362695 | 1 | 0 | 1 | 1 | NA | NA | NA | NA | 2 | 0.5 | 1 | 1 | NA | NA | 1 | 1 | 2 | 1 | 4 | 1 |
| 4_1362695 | NA | NA | 1 | 1 | NA | NA | NA | NA | 1 | 1 | 1 | 1 | NA | NA | 2 | 1 | 1 | 1 | 4 | 1 |
| 4_1362699 | 1 | 1 | 1 | 0 | NA | NA | NA | NA | 2 | 0.5 | NA | NA | 1 | 0 | 2 | 0.5 | 2 | 0 | 5 | 0.2 |
| 4_1362701 | 1 | 1 | 2 | 0.5 | NA | NA | NA | NA | 3 | 0.6667 | 1 | 1 | 2 | 1 | 3 | 1 | 1 | 1 | 7 | 1 |
| 4_1362705 | 1 | 1 | 3 | 0.6667 | NA | NA | 1 | 1 | 5 | 0.8 | 1 | 0 | 3 | 0.6667 | 4 | 0.5 | 2 | 1 | 10 | 0.6 |
| 4_1362706 | 2 | 0.5 | 4 | 0.75 | NA | NA | 1 | 1 | 7 | 0.7143 | NA | NA | 3 | 1 | 4 | 1 | 2 | 0.5 | 9 | 0.8889 |
| 4_1362726 | 2 | 1 | 5 | 0.4 | 2 | 0.5 | 1 | 1 | 10 | 0.6 | 1 | 0 | 3 | 0.3333 | 4 | 0.25 | 3 | 0.3333 | 11 | 0.2727 |
| 4_1362726 | 2 | 1 | 4 | 0.5 | 2 | 1 | 1 | 1 | 9 | 0.7778 | 2 | 1 | 3 | 0.6667 | 4 | 0.5 | 3 | 0.3333 | 12 | 0.5833 |
| 4_1362726 | 2 | 1 | 4 | 0.75 | 2 | 0.5 | 1 | 1 | 9 | 0.7778 | 2 | 0.5 | 3 | 0.6667 | 4 | 0.5 | 2 | 0.5 | 11 | 0.5455 |
| 4_1362727 | 2 | 0 | 5 | 0.6 | 2 | 0 | 2 | 0 | 11 | 0.2727 | 2 | 0 | 3 | 0.3333 | 4 | 0.25 | 3 | 0.3333 | 12 | 0.25 |
| 4_1362727 | 2 | 1 | 4 | 0 | 2 | 0 | 2 | 1 | 10 | 0.4 | 2 | 1 | 3 | 1 | 4 | 0.75 | 3 | 0.6667 | 12 | 0.8333 |
| 4_1362728 | 2 | 1 | 6 | 1 | 2 | 0.5 | 2 | 0.5 | 12 | 0.8333 | 2 | 1 | 2 | 1 | 1 | 1 | 3 | 0.6667 | 8 | 0.875 |
| 4_1362728 | 2 | 1 | 6 | 1 | 2 | 1 | 2 | 1 | 12 | 1 | 2 | 1 | 3 | 1 | 4 | 0.75 | 3 | 1 | 12 | 0.9167 |
| 4_1362729 | 2 | 1 | 6 | 0.8333 | 2 | 0.5 | 2 | 1 | 12 | 0.8333 | 2 | 1 | 3 | 0.6667 | 4 | 0.5 | 3 | 1 | 12 | 0.75 |
| 4_1362731 | 2 | 0.5 | 6 | 0.3333 | 2 | 0 | 2 | 0 | 12 | 0.25 | 2 | 0.5 | 3 | 0.6667 | 4 | 0.25 | 3 | 0.3333 | 12 | 0.4167 |
| 4_1362731 | 2 | 0.5 | 5 | 0 | 1 | 0 | 3 | 0.3333 | 11 | 0.1818 | 2 | 0.5 | 3 | 0.3333 | 1 | 0 | 2 | 0.5 | 8 | 0.375 |
| 4_1362732 | 2 | 0.5 | 3 | 0.3333 | 2 | 0.5 | 3 | 0 | 10 | 0.3 | 2 | 0.5 | 3 | 0.3333 | 1 | 0 | 3 | 0.3333 | 9 | 0.3333 |
| 4_1362733 | 2 | 1 | 4 | 0.5 | 1 | 1 | 3 | 0.3333 | 10 | 0.6 | 2 | 1 | 3 | 0.6667 | 4 | 0.25 | 2 | 1 | 11 | 0.6364 |
| 4_1362734 | 2 | 0.5 | 6 | 0.5 | 2 | 0.5 | 4 | 0 | 14 | 0.3571 | 2 | 0.5 | 4 | 0 | 3 | 0 | 2 | 0.5 | 11 | 0.1818 |
| 4_1362735 | 2 | 0 | 6 | 0.1667 | 1 | 0 | 4 | 0 | 13 | 0.0769 | 2 | 0.5 | 4 | 0.75 | 3 | 0 | 3 | 0.3333 | 12 | 0.4167 |
| 4_1362735 | 2 | 0 | 5 | 0.2 | 2 | 0 | 3 | 0 | 12 | 0.0833 | 1 | 1 | 4 | 0.5 | 3 | 0.3333 | 3 | 0.3333 | 11 | 0.4545 |
| 4_1362735 | 2 | 0 | 6 | 0 | 2 | 1 | 4 | 0 | 14 | 0.1429 | 2 | 0 | 4 | 0.25 | 4 | 0.25 | 3 | 0 | 13 | 0.1538 |
| 4_1362736 | 2 | 1 | 4 | 1 | 2 | 0 | 4 | 0.75 | 12 | 0.75 | 1 | 1 | 4 | 0.25 | 3 | 1 | 2 | 1 | 10 | 0.7 |
| 4_1362736 | 2 | 0.5 | 3 | 0.6667 | 2 | 0.5 | 3 | 0.3333 | 10 | 0.5 | 1 | 1 | 4 | 0.25 | 2 | 1 | 2 | 1 | 9 | 0.6667 |
| 4_1362737 | 2 | 0.5 | 6 | 0.5 | 2 | 0.5 | 2 | 0.5 | 12 | 0.5 | 1 | 1 | 5 | 0.4 | 3 | 0.3333 | 3 | 1 | 12 | 0.5833 |
| 4_1362738 | 2 | 0.5 | 6 | 0.6667 | 1 | 0 | 4 | 1 | 13 | 0.6923 | 1 | 1 | 5 | 0.6 | 2 | 0 | 3 | 0.6667 | 11 | 0.5455 |
| 4_1362738 | 1 | 1 | 5 | 0.2 | 2 | 0 | 4 | 0.25 | 12 | 0.25 | 2 | 1 | 5 | 0.2 | 3 | 0 | 2 | 0 | 12 | 0.25 |
| 4_1362739 | 2 | 0.5 | 7 | 1 | 2 | 1 | 3 | 1 | 14 | 0.9286 | 1 | 1 | 5 | 0.6 | 4 | 1 | 4 | 0.75 | 14 | 0.7857 |
| 4_1362746 | 3 | 0 | 7 | 0.1429 | 2 | 0 | 6 | 0 | 18 | 0.0556 | 3 | 0.6667 | 8 | 0 | 2 | 0 | 4 | 0 | 17 | 0.1176 |
| 4_1362752 | 2 | 0.5 | 9 | 0.3333 | 2 | 1 | 9 | 0 | 22 | 0.2727 | 5 | 0.6 | 10 | 0.5 | 4 | 0.25 | 7 | 0.5714 | 26 | 0.5 |
| 4_1362754 | 2 | 0.5 | 12 | 0.0833 | 2 | 0 | 9 | 0 | 25 | 0.08 | 6 | 0 | 10 | 0.1 | 5 | 0 | 7 | 0.1429 | 28 | 0.0714 |
| 4_1362764 | 2 | 0 | 15 | 0.0667 | 6 | 0 | 11 | 0.1818 | 34 | 0.0882 | 9 | 0 | 12 | 0.1667 | 8 | 0 | 13 | 0.1538 | 42 | 0.0952 |
| 4_1362779 | 8 | 1 | 6 | 1 | 6 | 1 | 4 | 0.75 | 24 | 0.9583 | 6 | 0.6667 | 7 | 0.8571 | 11 | 0.7273 | 2 | 1 | 26 | 0.7692 |
| 4_1362780 | 8 | 1 | 7 | 1 | 10 | 1 | 6 | 0.8333 | 31 | 0.9677 | 6 | 1 | 7 | 0.8571 | 10 | 0.8 | 4 | 1 | 27 | 0.8889 |
| 4_1362781 | 7 | 0 | 6 | 0 | 9 | 0.2222 | 7 | 0 | 29 | 0.069 | 4 | 0 | 5 | 0.2 | 9 | 0 | 2 | 0 | 20 | 0.05 |
| 4_1363814 | NA | NA | NA | NA | NA | NA | 1 | 1 | 1 | 1 | NA | NA | NA | NA | NA | NA | 1 | 1 | 1 | 1 |
| 4_1363814 | NA | NA | NA | NA | NA | NA | 1 | 1 | 1 | 1 | NA | NA | NA | NA | NA | NA | 1 | 1 | 1 | 1 |
| 4_1363816 | 1 | 1 | NA | NA | 1 | 0 | NA | NA | 2 | 0.5 | NA | NA | NA | NA | 1 | 1 | 1 | 1 | 2 | 1 |
| 4_1363817 | 2 | 1 | NA | NA | NA | NA | NA | NA | 2 | 1 | NA | NA | NA | NA | 1 | 1 | NA | NA | 1 | 1 |
| 4_1363818 | 1 | 1 | NA | NA | 1 | 0 | NA | NA | 2 | 0.5 | NA | NA | NA | NA | 2 | 0.5 | NA | NA | 2 | 0.5 |
| 4_1363820 | 2 | 1 | NA | NA | 1 | 1 | NA | NA | 3 | 1 | NA | NA | NA | NA | 2 | 1 | NA | NA | 2 | 1 |
| 4_1363821 | 1 | 1 | NA | NA | 1 | 0 | NA | NA | 2 | 0.5 | NA | NA | NA | NA | 2 | 0.5 | NA | NA | 2 | 0.5 |
| 4_1366074 | 8 | 0.25 | 18 | 0.0556 | 15 | 0.0667 | 14 | 0.1429 | 55 | 0.1091 | 17 | 0.0588 | 7 | 0.1429 | 13 | 0.0769 | 10 | 0.2 | 47 | 0.1064 |
| 4_1366075 | 11 | 0.0909 | 18 | 0.0556 | 15 | 0 | 11 | 0.2727 | 55 | 0.0909 | 16 | 0.25 | 8 | 0.25 | 14 | 0.1429 | 9 | 0.1111 | 47 | 0.1915 |
| 4_1366075 | 12 | 0.1667 | 19 | 0.0526 | 14 | 0.0714 | 14 | 0.0714 | 59 | 0.0847 | 17 | 0.2353 | 8 | 0 | 14 | 0.0714 | 10 | 0 | 49 | 0.102 |
| 4_1366076 | 5 | 0.4 | 7 | 0.4286 | 7 | 0.2857 | 5 | 0.2 | 24 | 0.3333 | 10 | 0.5 | 5 | 0.8 | 12 | 0.1667 | 7 | 0 | 34 | 0.3235 |
| 4_1366077 | 9 | 0.2222 | 14 | 0.1429 | 14 | 0.0714 | 14 | 0.1429 | 51 | 0.1373 | 16 | 0.1875 | 7 | 0.1429 | 14 | 0.2143 | 9 | 0 | 46 | 0.1522 |
| 4_1366077 | 9 | 0.4444 | 16 | 0.5 | 15 | 0.2667 | 13 | 0.6923 | 53 | 0.4717 | 14 | 0.5714 | 7 | 0.5714 | 13 | 0.2308 | 9 | 0.4444 | 43 | 0.4419 |
| 4_1366079 | 8 | 0.125 | 14 | 0.5 | 14 | 0.2857 | 14 | 0.2857 | 50 | 0.32 | 14 | 0.4286 | 8 | 0.375 | 13 | 0.3077 | 8 | 0.5 | 43 | 0.3953 |
| 4_1366079 | 9 | 0.1111 | 17 | 0.0588 | 15 | 0.0667 | 13 | 0 | 54 | 0.0556 | 14 | 0.0714 | 8 | 0.125 | 15 | 0.0667 | 8 | 0 | 45 | 0.0667 |
| 4_1366080 | 7 | 0.8571 | 15 | 0.4667 | 12 | 0.6667 | 13 | 0.4615 | 47 | 0.5745 | 15 | 0.4 | 8 | 0.375 | 12 | 0.6667 | 7 | 0.4286 | 42 | 0.4762 |
| 4_1366081 | 11 | 0.1818 | 17 | 0.2941 | 13 | 0.1538 | 13 | 0 | 54 | 0.1667 | 16 | 0.125 | 7 | 0.1429 | 12 | 0.1667 | 8 | 0.25 | 43 | 0.1628 |
| 4_1366081 | 11 | 0.2727 | 14 | 0.3571 | 12 | 0.0833 | 15 | 0.0667 | 52 | 0.1923 | 12 | 0.5 | 5 | 0 | 8 | 0.25 | 8 | 0.25 | 33 | 0.303 |
| 4_1366171 | 16 | 0.3125 | 10 | 0.3 | 7 | 0.4286 | 12 | 0.4167 | 45 | 0.3556 | 22 | 0.4545 | 5 | 0.6 | 9 | 0.3333 | 9 | 0.2222 | 45 | 0.4 |
| 4_1366172 | 10 | 0.1 | 5 | 0 | 7 | 0 | 10 | 0.1 | 32 | 0.0625 | 15 | 0.0667 | 4 | 0 | 9 | 0 | 10 | 0.1 | 38 | 0.0526 |
| 4_1366174 | 14 | 0.5714 | 6 | 0.3333 | 5 | 0.8 | 12 | 0.5833 | 37 | 0.5676 | 15 | 0.4667 | 4 | 0.5 | 8 | 0.25 | 12 | 0.5833 | 39 | 0.4615 |
| 4_1366177 | 11 | 0.1818 | 10 | 0 | 5 | 0.2 | 11 | 0.1818 | 37 | 0.1351 | 13 | 0.0769 | 5 | 0 | 7 | 0 | 9 | 0.1111 | 34 | 0.0588 |
| 4_1366178 | 10 | 0.3 | 9 | 0.3333 | 8 | 0 | 11 | 0 | 38 | 0.1579 | 16 | 0.0625 | 6 | 0.1667 | 6 | 0.1667 | 9 | 0.1111 | 37 | 0.1081 |
| 4_1366197 | 12 | 0.5 | 7 | 0.2857 | 8 | 0.125 | 12 | 0.5 | 39 | 0.3846 | 14 | 0.2143 | 5 | 0.6 | 6 | 0.3333 | 8 | 0 | 33 | 0.2424 |
| 4_1366198 | 11 | 0.2727 | 5 | 0.2 | 7 | 0.1429 | 11 | 0.2727 | 34 | 0.2353 | 17 | 0.0588 | 6 | 0.3333 | 6 | 0.3333 | 9 | 0.2222 | 38 | 0.1842 |
| 4_1366233 | 14 | 0.6429 | 13 | 0.9231 | 19 | 0.6316 | 11 | 0.6364 | 57 | 0.7018 | 23 | 0.7391 | 8 | 1 | 9 | 0.7778 | 13 | 0.6154 | 53 | 0.7547 |

| | | | | | | | | | | | | | | | | | | | | |
|---|---|---|---|---|---|---|---|---|---|---|---|---|---|---|---|---|---|---|---|---|
| 4_1366234 | 13 | 0.4615 | 9 | 0.4444 | 15 | 0.4667 | 8 | 0.25 | 45 | 0.4222 | 17 | 0.5294 | 4 | 0.25 | 3 | 0.3333 | 9 | 0.2222 | 33 | 0.3939 |
| 4_1366235 | 13 | 0.0769 | 15 | 0.2 | 21 | 0 | 11 | 0 | 60 | 0.0667 | 20 | 0.05 | 7 | 0.1429 | 8 | 0.125 | 12 | 0 | 47 | 0.0638 |
| 4_1366236 | 11 | 0.9091 | 13 | 1 | 18 | 0.8333 | 9 | 0.8889 | 51 | 0.902 | 19 | 0.9474 | 6 | 0.8333 | 8 | 0.875 | 12 | 0.8333 | 45 | 0.8889 |
| 4_1366238 | 9 | 0.2222 | 12 | 0.1667 | 21 | 0.1429 | 9 | 0.2222 | 51 | 0.1765 | 19 | 0 | 7 | 0.4286 | 7 | 0.2857 | 12 | 0.5 | 45 | 0.2444 |
| 4_1366239 | 7 | 0.8571 | 13 | 0.8462 | 17 | 0.8235 | 9 | 0.8889 | 46 | 0.8478 | 19 | 0.8947 | 6 | 0.8333 | 8 | 0.875 | 11 | 0.8182 | 44 | 0.8636 |
| 4_1366240 | 9 | 0.1111 | 13 | 0.1538 | 20 | 0.1 | 9 | 0.2222 | 51 | 0.1373 | 19 | 0.1579 | 6 | 0 | 7 | 0.1429 | 12 | 0.0833 | 44 | 0.1136 |
| 4_1366577 | NA | NA | NA | NA | 2 | 1 | 2 | 0.5 | 4 | 0.75 | NA | NA | 2 | 0.5 | 3 | 0.6667 | NA | NA | 5 | 0.6 |
| 4_1366581 | NA | NA | NA | NA | 1 | 1 | 2 | 0 | 3 | 0.3333 | NA | NA | 1 | 0 | 3 | 0.3333 | NA | NA | 4 | 0.25 |
| 4_1366582 | NA | NA | NA | NA | 1 | 1 | 2 | 0.5 | 3 | 0.6667 | NA | NA | 1 | 1 | 3 | 0.3333 | NA | NA | 4 | 0.5 |
| 4_1367072 | 4 | 1 | 1 | 1 | 1 | 1 | 1 | 1 | 7 | 1 | NA | NA | NA | NA | 1 | 1 | 4 | 1 | 5 | 1 |
| 4_1367073 | 4 | 1 | 1 | 0 | 2 | 0.5 | 3 | 1 | 10 | 0.8 | NA | NA | NA | NA | 1 | 0 | 3 | 0.6667 | 4 | 0.5 |
| 4_1367073 | 4 | 0.5 | 1 | 0 | 2 | 0 | 3 | 0.3333 | 10 | 0.3 | NA | NA | NA | NA | 1 | 1 | 3 | 0.6667 | 4 | 0.75 |
| 4_1367076 | 3 | 0.3333 | 1 | 1 | 2 | 0 | 3 | 0.3333 | 9 | 0.3333 | NA | NA | NA | NA | NA | NA | 4 | 0.5 | 4 | 0.5 |
| 4_1367076 | 3 | 0.3333 | 1 | 0 | 2 | 0 | 3 | 0 | 9 | 0.1111 | NA | NA | NA | NA | NA | NA | 3 | 1 | 3 | 1 |
| 4_1367079 | 3 | 1 | 1 | 0 | 1 | 1 | 3 | 1 | 8 | 0.875 | NA | NA | NA | NA | 1 | 1 | 3 | 1 | 4 | 1 |
| 4_1367081 | 3 | 1 | 1 | 1 | 2 | 1 | 3 | 1 | 9 | 1 | NA | NA | NA | NA | 1 | 1 | 2 | 1 | 3 | 1 |
| 4_1367082 | 2 | 1 | NA | NA | 2 | 1 | 2 | 1 | 6 | 1 | NA | NA | NA | NA | 1 | 1 | 4 | 0.75 | 5 | 0.8 |
| 4_1367082 | 3 | 0.6667 | 1 | 1 | 1 | 1 | 2 | 0.5 | 7 | 0.7143 | NA | NA | NA | NA | 1 | 1 | 3 | 0.6667 | 4 | 0.75 |
| 4_1367082 | 3 | 0.3333 | 1 | 0 | 2 | 0.5 | 3 | 0.3333 | 9 | 0.3333 | NA | NA | NA | NA | 1 | 0 | 4 | 0.5 | 5 | 0.4 |
| 4_1367086 | 3 | 1 | NA | NA | 2 | 1 | 2 | 1 | 7 | 1 | NA | NA | NA | NA | NA | NA | 4 | 0.75 | 4 | 0.75 |
| 4_1367086 | 2 | 1 | NA | NA | 2 | 0.5 | 3 | 0.6667 | 7 | 0.7143 | NA | NA | NA | NA | NA | NA | 4 | 1 | 4 | 1 |
| 4_1367088 | 2 | 1 | NA | NA | 1 | 0 | 2 | 0.5 | 5 | 0.6 | NA | NA | NA | NA | NA | NA | 2 | 1 | 2 | 1 |
| 4_1367091 | 3 | 0 | NA | NA | 1 | 1 | 1 | 0 | 5 | 0.2 | NA | NA | NA | NA | NA | NA | 2 | 0.5 | 2 | 0.5 |
| 4_1367099 | 3 | 1 | NA | NA | 1 | 1 | NA | NA | 4 | 1 | NA | NA | NA | NA | NA | NA | 2 | 1 | 2 | 1 |
| 4_1367101 | 2 | 1 | NA | NA | 2 | 1 | NA | NA | 4 | 1 | NA | NA | NA | NA | NA | NA | 2 | 1 | 2 | 1 |
| 4_1370471 | 37 | 0.0811 | 21 | 0.0952 | 26 | 0.1538 | 22 | 0.1364 | 106 | 0.1132 | 29 | 0.1379 | 22 | 0.0455 | 32 | 0.0938 | 15 | 0.2 | 98 | 0.1122 |
| 4_1370480 | 30 | 0.1 | 13 | 0.1538 | 23 | 0.0435 | 21 | 0.0476 | 87 | 0.0805 | 22 | 0.0909 | 19 | 0.1053 | 28 | 0.0357 | 13 | 0.0769 | 82 | 0.0732 |
| 4_1370488 | NA | NA | 2 | 0 | NA | NA | 1 | 1 | 3 | 0.3333 | 1 | 0 | 2 | 0.5 | 5 | 0 | 1 | 1 | 9 | 0.2222 |
| 4_1373442 | NA | NA | 1 | 1 | 1 | 0 | NA | NA | 2 | 0.5 | NA | NA | NA | NA | NA | NA | 1 | 1 | 1 | 1 |
| 4_1373834 | 4 | 0.25 | 8 | 0 | 3 | 0 | 4 | 0 | 19 | 0.0526 | 2 | 0 | 5 | 0 | 4 | 0 | 5 | 0.2 | 16 | 0.0625 |
| 4_1373835 | 4 | 0 | 9 | 0.1111 | 3 | 0 | 6 | 0.3333 | 22 | 0.1364 | 4 | 0.25 | 5 | 0.4 | 6 | 0.1667 | 7 | 0.4286 | 22 | 0.3182 |
| 4_1373835 | 4 | 0 | 8 | 0.25 | 2 | 0 | 6 | 0.6667 | 20 | 0.3 | 4 | 0.25 | 5 | 0.4 | 6 | 0 | 7 | 0.2857 | 22 | 0.2273 |
| 4_1373835 | 4 | 1 | 6 | 1 | 3 | 1 | 6 | 0.6667 | 19 | 0.8947 | 4 | 1 | 5 | 0.8 | 6 | 1 | 7 | 1 | 22 | 0.9545 |
| 4_1373836 | 4 | 0.25 | 8 | 0.5 | 3 | 0 | 4 | 0.5 | 19 | 0.3684 | 4 | 0.5 | 4 | 0 | 7 | 0.2857 | 8 | 0.25 | 23 | 0.2609 |
| 4_1373837 | 4 | 0.5 | 6 | 0.6667 | 2 | 1 | 4 | 0.5 | 16 | 0.625 | 3 | 0 | 4 | 1 | 6 | 0.8333 | 8 | 0.875 | 21 | 0.7619 |
| 4_1373838 | 4 | 0 | 6 | 0 | 2 | 0 | 4 | 0.5 | 16 | 0.125 | 3 | 0 | 4 | 0 | 4 | 0.25 | 7 | 0.2857 | 18 | 0.1667 |
| 4_1373839 | 4 | 0.75 | 7 | 0.8571 | 1 | 0 | 4 | 0.25 | 16 | 0.625 | 3 | 0.3333 | 4 | 0.75 | 7 | 0.7143 | 8 | 0.75 | 22 | 0.6818 |
| 4_1373840 | 2 | 1 | 8 | 0.875 | 2 | 1 | 3 | 1 | 15 | 0.9333 | 2 | 1 | 3 | 0.6667 | 5 | 1 | 8 | 1 | 18 | 0.9444 |
| 4_1373842 | 2 | 0.5 | 5 | 0.2 | 1 | 1 | 3 | 0 | 11 | 0.2727 | 3 | 0.3333 | 4 | 0.5 | 7 | 0.7143 | 6 | 0.6667 | 20 | 0.6 |
| 4_1373842 | 3 | 0.6667 | 5 | 0.8 | 1 | 0 | 1 | 0 | 10 | 0.6 | 3 | 0.6667 | 3 | 0.6667 | 7 | 0.5714 | 6 | 0.8333 | 19 | 0.6842 |
| 4_1373844 | 3 | 1 | 8 | 0.625 | 2 | 0.5 | 3 | 0 | 16 | 0.5625 | 3 | 0.6667 | 3 | 0.6667 | 8 | 0.875 | 9 | 0.6667 | 23 | 0.7391 |
| 4_1373845 | 4 | 0.5 | 8 | 0.125 | 2 | 0.5 | 3 | 0 | 17 | 0.2353 | 1 | 0 | 5 | 0.4 | 10 | 0 | 9 | 0.2222 | 25 | 0.16 |
| 4_1373846 | 5 | 0.4 | 7 | 0.2857 | 3 | 0 | 4 | 0 | 19 | 0.2105 | 3 | 0 | 4 | 0.25 | 9 | 0.4444 | 10 | 0.2 | 26 | 0.2692 |
| 4_1373846 | 5 | 0.8 | 7 | 1 | 2 | 0.5 | 3 | 0.6667 | 17 | 0.8235 | 3 | 1 | 4 | 0.75 | 9 | 0.7778 | 11 | 0.6364 | 27 | 0.7407 |
| 4_1373846 | 5 | 0.2 | 7 | 0 | 3 | 0.3333 | 4 | 0.25 | 19 | 0.1579 | 3 | 0.6667 | 5 | 0.6 | 9 | 0.2222 | 11 | 0.0909 | 28 | 0.2857 |
| 4_1373847 | 5 | 0.4 | 7 | 0.2857 | 2 | 0.5 | 4 | 0.25 | 18 | 0.3333 | 3 | 0 | 4 | 0.25 | 7 | 0.1429 | 8 | 0.375 | 22 | 0.2273 |
| 4_1373852 | 5 | 0.8 | 8 | 0.875 | 3 | 1 | 4 | 1 | 20 | 0.9 | 2 | 1 | 6 | 0.6667 | 12 | 0.9167 | 8 | 1 | 28 | 0.8929 |
| 4_1373854 | 8 | 0.75 | 8 | 0.5 | 4 | 0.5 | 3 | 1 | 23 | 0.6522 | 3 | 0 | 4 | 0.25 | 8 | 0.625 | 11 | 0.7273 | 26 | 0.5385 |
| 4_1374513 | 4 | 0 | 2 | 0 | 2 | 0 | 9 | 0.1111 | 17 | 0.0588 | 1 | 1 | 2 | 0 | 8 | 0 | 4 | 0 | 15 | 0.0667 |
| 4_1374513 | 4 | 0.25 | 2 | 0 | 2 | 0 | 7 | 0 | 15 | 0.0667 | 1 | 1 | 2 | 0.5 | 7 | 0 | 4 | 0.25 | 14 | 0.2143 |
| 4_1374517 | 5 | 0 | 1 | 0 | 3 | 0 | 7 | 0.1429 | 16 | 0.0625 | NA | NA | 4 | 0.25 | 7 | 0 | 1 | 0 | 12 | 0.0833 |
| 4_1374564 | 40 | 0.85 | 35 | 0.7714 | 29 | 0.6207 | 48 | 0.7292 | 152 | 0.75 | 15 | 1 | 38 | 0.7368 | 61 | 0.8197 | 37 | 0.7027 | 151 | 0.7881 |
| 4_1374564 | 30 | 0.1333 | 28 | 0 | 26 | 0.0769 | 35 | 0 | 119 | 0.0504 | 13 | 0 | 28 | 0 | 49 | 0.0816 | 30 | 0.1333 | 120 | 0.0667 |
| 4_1374566 | 41 | 0.439 | 30 | 0.4667 | 31 | 0.3226 | 46 | 0.3043 | 148 | 0.3784 | 14 | 0.4286 | 41 | 0.3659 | 59 | 0.5254 | 33 | 0.4848 | 147 | 0.4626 |
| 4_1374567 | 44 | 0.3636 | 34 | 0.2941 | 32 | 0.3438 | 48 | 0.5 | 158 | 0.3861 | 12 | 0.3333 | 42 | 0.3571 | 55 | 0.4364 | 36 | 0.5 | 145 | 0.4207 |
| 4_1374695 | 2 | 0.5 | NA | NA | NA | NA | NA | NA | 2 | 0.5 | NA | NA | NA | NA | 1 | 1 | NA | NA | 1 | 1 |
| 4_1376901 | 4 | 0.75 | 3 | 0.6667 | NA | NA | NA | NA | 7 | 0.7143 | 3 | 0.6667 | 2 | 0 | NA | NA | 1 | 0 | 6 | 0.3333 |
| 4_1376901 | 5 | 0.2 | 2 | 0 | NA | NA | NA | NA | 7 | 0.1429 | 3 | 0.3333 | 3 | 0.3333 | NA | NA | 1 | 0 | 7 | 0.2857 |
| 4_1376902 | 2 | 1 | 1 | 1 | NA | NA | NA | NA | 3 | 1 | 3 | 1 | 1 | 1 | NA | NA | NA | NA | 4 | 1 |
| 4_1376904 | 6 | 0.1667 | 3 | 0.3333 | NA | NA | NA | NA | 9 | 0.2222 | 4 | 0 | 3 | 0.6667 | NA | NA | 1 | 0 | 8 | 0.25 |
| 4_1376904 | 4 | 0.25 | 1 | 1 | NA | NA | NA | NA | 5 | 0.4 | 3 | 0.6667 | 2 | 0 | NA | NA | 1 | 0 | 6 | 0.3333 |
| 4_1376922 | 6 | 0.8333 | 4 | 0.5 | 2 | 1 | 1 | 0 | 13 | 0.6923 | 4 | 0.5 | 3 | 1 | 2 | 0.5 | 2 | 0.5 | 11 | 0.6364 |
| 4_1376924 | 6 | 0.3333 | 5 | 0.4 | 3 | 0.3333 | 2 | 0.5 | 16 | 0.375 | 4 | 0.5 | 4 | 0.5 | 2 | 0 | 2 | 0.5 | 12 | 0.4167 |
| 4_1378532 | 3 | 0.3333 | 6 | 0.3333 | 4 | 0.25 | 2 | 0 | 15 | 0.2667 | 3 | 0.3333 | 1 | 0 | 2 | 0 | 2 | 0 | 8 | 0.125 |
| 4_1378533 | 3 | 0.6667 | 4 | 0.75 | 5 | 0.4 | 1 | 0 | 13 | 0.5385 | 2 | 0.5 | 2 | 0 | 1 | 0 | 2 | 0.5 | 7 | 0.2857 |

| | | | | | | | | | | | | | | | | | | | | |
|---|---|---|---|---|---|---|---|---|---|---|---|---|---|---|---|---|---|---|---|---|
| 4_1378533 | 1 | 0 | 5 | 0.4 | 6 | 0.1667 | 1 | 1 | 13 | 0.3077 | 3 | 0.3333 | 1 | 0 | 3 | 0.3333 | 2 | 0.5 | 9 | 0.3333 |
| 4_1378535 | 3 | 0.3333 | 5 | 0.4 | 3 | 0.3333 | 2 | 0.5 | 13 | 0.3846 | 2 | 0.5 | 1 | 0 | 3 | 0.6667 | 1 | 0 | 7 | 0.4286 |
| 4_1378536 | 2 | 0.5 | 5 | 0.6 | 5 | 0.8 | 2 | 1 | 14 | 0.7143 | 1 | 1 | 1 | 1 | 4 | 0.5 | 2 | 1 | 8 | 0.75 |
| 4_1378536 | 2 | 0 | 5 | 0.2 | 5 | 0.2 | 2 | 0 | 14 | 0.1429 | 2 | 0 | 2 | 0 | 5 | 0 | 2 | 0.5 | 11 | 0.0909 |
| 4_1378538 | 3 | 0.6667 | 3 | 0.3333 | 7 | 0.7143 | 3 | 0.3333 | 16 | 0.5625 | 2 | 0.5 | 1 | 0 | 4 | 0.5 | 2 | 0.5 | 9 | 0.4444 |
| 4_1378538 | 3 | 0.3333 | 5 | 0 | 7 | 0 | 3 | 0 | 18 | 0.0556 | 3 | 0.6667 | 2 | 0 | 4 | 0 | 2 | 0 | 11 | 0.1818 |
| 4_1379333 | 12 | 0.0833 | 16 | 0.125 | 12 | 0.0833 | 9 | 0.1111 | 49 | 0.102 | 8 | 0 | 7 | 0.1429 | 10 | 0.1 | 6 | 0 | 31 | 0.0645 |
| 4_1379334 | 13 | 0.5385 | 15 | 0.2 | 10 | 0.3 | 9 | 0.2222 | 47 | 0.3191 | 8 | 0.75 | 9 | 0.2222 | 8 | 0.25 | 5 | 0.4 | 30 | 0.4 |
| 4_1379335 | 12 | 0.5833 | 14 | 0.3571 | 8 | 0.25 | 9 | 0.5556 | 43 | 0.4419 | 8 | 0.375 | 7 | 0.2857 | 7 | 0.2857 | 4 | 0.25 | 26 | 0.3077 |
| 4_1379337 | 10 | 0.7 | 11 | 0.9091 | 11 | 0.8182 | 10 | 0.5 | 42 | 0.7381 | 6 | 0.8333 | 6 | 0.6667 | 9 | 0.6667 | 1 | 0 | 22 | 0.6818 |
| 4_1379338 | 13 | 0.3846 | 12 | 0.0833 | 12 | 0.25 | 9 | 0.3333 | 46 | 0.2609 | 10 | 0.3 | 8 | 0.375 | 7 | 0.1429 | 3 | 0 | 28 | 0.25 |
| 4_1379338 | 13 | 0.3077 | 12 | 0.0833 | 9 | 0.4444 | 9 | 0.2222 | 43 | 0.2558 | 8 | 0.25 | 9 | 0.4444 | 7 | 0.4286 | 3 | 0 | 27 | 0.3333 |
| 4_1379341 | 14 | 0.1429 | 16 | 0.125 | 15 | 0.0667 | 10 | 0.2 | 55 | 0.1273 | 8 | 0 | 6 | 0.3333 | 9 | 0.2222 | 5 | 0 | 28 | 0.1429 |
| 4_1379343 | 15 | 0.0667 | 16 | 0 | 14 | 0.0714 | 8 | 0.125 | 53 | 0.0566 | 12 | 0.0833 | 9 | 0.1111 | 9 | 0 | 6 | 0.1667 | 36 | 0.0833 |
| 4_1379683 | NA | NA | NA | NA | 1 | 1 | NA | NA | 1 | 1 | NA | NA | 1 | 1 | 2 | 0 | 1 | 0 | 4 | 0.25 |
| 4_1379691 | 14 | 0.2143 | 19 | 0.3158 | 16 | 0.375 | 24 | 0.5417 | 73 | 0.3836 | 18 | 0.3333 | 28 | 0.3571 | 25 | 0.64 | 26 | 0.5769 | 97 | 0.4845 |
| 4_1379691 | 17 | 0.0588 | 19 | 0.0526 | 14 | 0 | 23 | 0.2609 | 73 | 0.1096 | 19 | 0.0526 | 31 | 0.129 | 23 | 0.0435 | 26 | 0.0385 | 99 | 0.0707 |
| 4_1379692 | 16 | 0.4375 | 17 | 0.7059 | 13 | 0.1538 | 19 | 0.5263 | 65 | 0.4769 | 16 | 0.375 | 24 | 0.4583 | 21 | 0.4286 | 19 | 0.2632 | 80 | 0.3875 |
| 4_1379692 | 17 | 0 | 22 | 0.0909 | 14 | 0 | 23 | 0.087 | 76 | 0.0526 | 19 | 0.1053 | 29 | 0.1034 | 22 | 0.0909 | 25 | 0.12 | 95 | 0.1053 |
| 4_1379694 | 18 | 0 | 21 | 0.1429 | 17 | 0.0588 | 25 | 0.2 | 81 | 0.1111 | 19 | 0 | 29 | 0.2414 | 21 | 0.0952 | 24 | 0.2083 | 93 | 0.1505 |
| 4_1379695 | 18 | 0.1111 | 19 | 0.1053 | 15 | 0 | 24 | 0.0417 | 76 | 0.0658 | 14 | 0.0714 | 28 | 0.0357 | 20 | 0.15 | 23 | 0.087 | 85 | 0.0824 |
| 4_1379697 | 17 | 0 | 21 | 0.0476 | 17 | 0.0588 | 26 | 0.1154 | 81 | 0.0617 | 23 | 0.087 | 27 | 0.2593 | 23 | 0.1739 | 25 | 0.16 | 98 | 0.1735 |
| 4_1379697 | 21 | 0.1429 | 22 | 0.2727 | 17 | 0.2353 | 25 | 0.28 | 85 | 0.2353 | 24 | 0.2083 | 33 | 0.3333 | 25 | 0.32 | 27 | 0.3333 | 109 | 0.3028 |
| 4_1379703 | 25 | 0.72 | 23 | 0.7391 | 24 | 0.75 | 30 | 0.8 | 102 | 0.7549 | 24 | 0.7083 | 31 | 0.8065 | 30 | 0.7 | 25 | 0.84 | 110 | 0.7636 |
| 4_1379704 | 21 | 0.1429 | 28 | 0.1071 | 24 | 0.0417 | 28 | 0.0714 | 101 | 0.0891 | 23 | 0.0435 | 27 | 0.1111 | 29 | 0.069 | 25 | 0.04 | 104 | 0.0673 |
| 4_1379864 | 9 | 0.5556 | 10 | 0.6 | 11 | 0.3636 | 12 | 0.75 | 42 | 0.5714 | 2 | 0.5 | 8 | 0.125 | 13 | 0.2308 | 11 | 0.4545 | 34 | 0.2941 |
| 4_1379864 | 8 | 0.5 | 9 | 0.6667 | 6 | 0.5 | 10 | 0.7 | 33 | 0.6061 | 2 | 1 | 6 | 0.6667 | 12 | 0.4167 | 9 | 0.5556 | 29 | 0.5517 |
| 4_1379865 | 7 | 0.8571 | 8 | 0.875 | 6 | 0.1667 | 13 | 0.6923 | 34 | 0.6765 | 3 | 1 | 9 | 0.5556 | 12 | 0.6667 | 11 | 0.8182 | 35 | 0.7143 |
| 4_1379865 | 9 | 0.3333 | 11 | 0.6364 | 9 | 0.6667 | 13 | 0.3077 | 42 | 0.4762 | 3 | 0.6667 | 8 | 0.625 | 13 | 0.6154 | 11 | 0.6364 | 35 | 0.6286 |
| 4_1379867 | 10 | 0.1 | 11 | 0.0909 | 12 | 0.0833 | 12 | 0 | 45 | 0.0667 | 2 | 0 | 11 | 0.0909 | 14 | 0.0714 | 10 | 0.1 | 37 | 0.0811 |
| 4_1379868 | 7 | 0.4286 | 9 | 0.7778 | 10 | 0.4 | 12 | 0.5833 | 38 | 0.5526 | 3 | 1 | 9 | 0.8889 | 13 | 0.5385 | 8 | 0.625 | 33 | 0.697 |
| 4_1379868 | 9 | 0.6667 | 11 | 0.7273 | 15 | 0.7333 | 14 | 0.7857 | 49 | 0.7347 | 2 | 1 | 11 | 0.4545 | 15 | 0.6667 | 11 | 0.5455 | 39 | 0.5897 |
| 4_1379870 | 7 | 0.1429 | 12 | 0.25 | 11 | 0.4545 | 14 | 0.0714 | 44 | 0.2273 | 3 | 0.6667 | 10 | 0.6 | 13 | 0.3846 | 10 | 0.4 | 36 | 0.4722 |
| 4_1379873 | 8 | 0.125 | 12 | 0.25 | 15 | 0.2667 | 17 | 0.1765 | 52 | 0.2115 | 3 | 0.6667 | 11 | 0.1818 | 17 | 0.2353 | 9 | 0.2222 | 40 | 0.25 |
| 4_1379874 | 8 | 0.125 | 11 | 0.4545 | 15 | 0.1333 | 14 | 0.2143 | 48 | 0.2292 | 3 | 0.3333 | 10 | 0.1 | 11 | 0.1818 | 10 | 0.1 | 34 | 0.1471 |
| 4_1379874 | 9 | 0.2222 | 12 | 0.0833 | 15 | 0.0667 | 15 | 0.0667 | 51 | 0.098 | 2 | 0.5 | 11 | 0.0909 | 16 | 0.0625 | 11 | 0.1818 | 40 | 0.125 |
| 4_1379876 | 8 | 0.125 | 7 | 0 | 12 | 0 | 8 | 0.125 | 35 | 0.0571 | 4 | 0.25 | 9 | 0 | 11 | 0.1818 | 13 | 0 | 37 | 0.0811 |
| 4_1379880 | 7 | 0.4286 | 11 | 0 | 16 | 0 | 7 | 0 | 41 | 0.0732 | 2 | 0.5 | 7 | 0.2857 | 15 | 0 | 9 | 0 | 33 | 0.0909 |
| 4_1380053 | 2 | 0.5 | NA | NA | 1 | 0 | NA | NA | 3 | 0.3333 | NA | NA | 1 | 1 | NA | NA | 1 | 0 | 2 | 0.5 |
| 4_1380053 | 2 | 0.5 | NA | NA | 1 | 1 | NA | NA | 3 | 0.6667 | NA | NA | NA | NA | NA | NA | 1 | 1 | 1 | 1 |
| 4_1380053 | 2 | 0.5 | NA | NA | 1 | 1 | NA | NA | 3 | 0.6667 | NA | NA | 1 | 0 | NA | NA | 1 | 1 | 2 | 0.5 |
| 4_1380154 | NA | NA | NA | NA | 2 | 0.5 | NA | NA | 2 | 0.5 | NA | NA | NA | NA | 1 | 1 | NA | NA | 1 | 1 |
| 4_1380197 | NA | NA | NA | NA | 1 | 1 | NA | NA | 1 | 1 | NA | NA | 1 | 1 | NA | NA | NA | NA | 1 | 1 |
| 4_1380211 | 3 | 0.6667 | NA | NA | 4 | 0.5 | NA | NA | 7 | 0.5714 | NA | NA | 1 | 0 | 3 | 0.3333 | 4 | 0.75 | 8 | 0.5 |
| 4_1380288 | 2 | 0.5 | 1 | 1 | 4 | 0 | NA | NA | 7 | 0.2857 | NA | NA | 2 | 0.5 | 4 | 0.5 | 3 | 0 | 9 | 0.3333 |
| 4_1380289 | 3 | 0.3333 | 1 | 1 | 5 | 0.8 | NA | NA | 9 | 0.6667 | NA | NA | 2 | 1 | 4 | 1 | 3 | 1 | 9 | 1 |
| 4_1380289 | 4 | 0.25 | 1 | 1 | 6 | 0.6667 | NA | NA | 11 | 0.5455 | NA | NA | 2 | 1 | 3 | 0.3333 | 4 | 0.5 | 9 | 0.5556 |
| 4_1380291 | 4 | 1 | 2 | 0.5 | 7 | 0.4286 | NA | NA | 13 | 0.6154 | 1 | 1 | 3 | 1 | 4 | 0.25 | 4 | 1 | 12 | 0.75 |
| 4_1380291 | 4 | 0.25 | 2 | 0 | 6 | 0.1667 | NA | NA | 12 | 0.1667 | 1 | 0 | 3 | 0.6667 | 3 | 0.3333 | 4 | 0.25 | 11 | 0.3636 |
| 4_1380291 | 4 | 0 | 2 | 0.5 | 6 | 0.3333 | NA | NA | 12 | 0.25 | 1 | 0 | 3 | 0.6667 | 4 | 0.5 | 4 | 0.25 | 12 | 0.4167 |
| 4_1380292 | 2 | 0 | 2 | 0.5 | 5 | 0.2 | NA | NA | 9 | 0.2222 | 1 | 0 | 3 | 0.3333 | 2 | 0 | 2 | 0 | 8 | 0.125 |
| 4_1380362 | NA | NA | NA | NA | 1 | 1 | NA | NA | 1 | 1 | NA | NA | NA | NA | 1 | 1 | 1 | 1 | 2 | 1 |
| 4_1380376 | NA | NA | NA | NA | 1 | 1 | NA | NA | 1 | 1 | NA | NA | NA | NA | 1 | 1 | 1 | 0 | 2 | 0.5 |
| 4_1380394 | 17 | 0.5882 | 13 | 0.6923 | 13 | 0.6923 | 21 | 0.7619 | 64 | 0.6875 | 9 | 0.7778 | 13 | 0.7692 | 15 | 0.8667 | 21 | 0.9524 | 58 | 0.8621 |
| 4_1380395 | 6 | 0.5 | 7 | 0.4286 | 5 | 0.4 | 8 | 0.25 | 26 | 0.3846 | 4 | 0.25 | 4 | 0.25 | 10 | 0.4 | 9 | 0.2222 | 27 | 0.2963 |
| 4_1380396 | 6 | 0.1667 | 9 | 0.2222 | 6 | 0.3333 | 8 | 0 | 29 | 0.1724 | 5 | 0.4 | 5 | 0.4 | 10 | 0 | 9 | 0.1111 | 29 | 0.1724 |
| 4_1380397 | 6 | 0.1667 | 9 | 0.5556 | 8 | 0.25 | 7 | 0.1429 | 30 | 0.3 | 4 | 0.25 | 4 | 0.5 | 11 | 0.5455 | 9 | 0.4444 | 28 | 0.4643 |
| 4_1380407 | 26 | 0 | 37 | 0.0811 | 27 | 0.0741 | 33 | 0.0606 | 123 | 0.0569 | 31 | 0.1613 | 29 | 0.1724 | 34 | 0.0588 | 21 | 0.0476 | 115 | 0.113 |
| 4_1380409 | 47 | 0.1489 | 47 | 0.2766 | 51 | 0.1765 | 43 | 0.2326 | 188 | 0.2074 | 41 | 0.2195 | 34 | 0.1471 | 46 | 0.2826 | 33 | 0.1818 | 154 | 0.2143 |
| 4_1380412 | 47 | 0.0638 | 49 | 0.0612 | 40 | 0.05 | 37 | 0.0541 | 173 | 0.0578 | 41 | 0.1463 | 38 | 0.0789 | 41 | 0.0732 | 29 | 0.0345 | 149 | 0.0872 |
| 4_1380412 | 42 | 0.8571 | 47 | 0.8298 | 40 | 0.85 | 39 | 0.7179 | 168 | 0.8155 | 41 | 0.8537 | 33 | 0.8788 | 47 | 0.9149 | 26 | 0.7692 | 147 | 0.8639 |
| 4_1380414 | 45 | 0.0444 | 47 | 0.0638 | 37 | 0.0541 | 42 | 0.0476 | 171 | 0.0526 | 40 | 0.05 | 35 | 0.0571 | 44 | 0.0455 | 30 | 0.0667 | 149 | 0.0537 |
| 4_1381109 | NA | NA | 1 | 1 | NA | NA | NA | NA | 1 | 1 | NA | NA | 1 | 1 | NA | NA | NA | NA | 1 | 1 |
| 4_1381110 | NA | NA | 1 | 1 | NA | NA | NA | NA | 1 | 1 | NA | NA | 1 | 1 | NA | NA | NA | NA | 1 | 1 |
| 4_1381215 | 2 | 1 | 8 | 1 | 3 | 1 | 1 | 1 | 14 | 1 | 4 | 1 | 3 | 1 | 2 | 0.5 | 4 | 1 | 13 | 0.9231 |

| | | | | | | | | | | | | | | | | | | | | |
|---|---|---|---|---|---|---|---|---|---|---|---|---|---|---|---|---|---|---|---|---|
| 4_1381216( | 3 | 1 | 9 | 0.3333 | 2 | 0.5 | NA | NA | 14 | 0.5 | 4 | 0.5 | 4 | 0.75 | 2 | 0.5 | 4 | 0.25 | 14 | 0.5 |
| 4_1381216: | 1 | 0 | 7 | 0.1429 | 2 | 0.5 | 1 | 0 | 11 | 0.1818 | 3 | 0.3333 | 3 | 0 | 1 | 0 | 2 | 0 | 9 | 0.1111 |
| 4_1381216 | 2 | 0 | 8 | 0.5 | 2 | 0 | 1 | 1 | 13 | 0.3846 | 4 | 0.75 | 4 | 0.5 | 1 | 0 | 4 | 0.75 | 13 | 0.6154 |
| 4_1381217( | 2 | 1 | 9 | 0.7778 | 3 | 1 | 1 | 1 | 15 | 0.8667 | 3 | 1 | 4 | 1 | 1 | 1 | 4 | 0.75 | 12 | 0.9167 |
| 4_1381217( | 2 | 0 | 8 | 0.375 | 1 | 0 | 1 | 0 | 12 | 0.25 | 3 | 0.6667 | 3 | 0.6667 | NA | NA | 3 | 0.3333 | 9 | 0.5556 |
| 4_1381217! | 2 | 0.5 | 8 | 0.75 | 2 | 1 | NA | NA | 12 | 0.75 | 4 | 1 | 4 | 0.5 | 1 | 1 | 2 | 0.5 | 11 | 0.7273 |
| 4_1381218: | 2 | 1 | 8 | 0.875 | 3 | 0.6667 | NA | NA | 13 | 0.8462 | 4 | 1 | 4 | 1 | 1 | 1 | 2 | 1 | 11 | 1 |
| 4_1381218( | 2 | 0 | 5 | 0.4 | 2 | 0.5 | NA | NA | 9 | 0.3333 | 3 | 0.6667 | 2 | 0.5 | 1 | 0 | 1 | 0 | 7 | 0.4286 |
| 4_1381222! | 2 | 1 | 6 | 0.5 | 2 | 1 | NA | NA | 10 | 0.7 | 3 | 1 | 3 | 1 | 1 | 1 | 3 | 0.6667 | 10 | 0.9 |
| 4_1381223 | 2 | 0.5 | 4 | 1 | 3 | 1 | 1 | 1 | 10 | 0.9 | 3 | 1 | 3 | 1 | 1 | 1 | 2 | 1 | 9 | 1 |
| 4_1381224! | 3 | 0 | 8 | 0.125 | 3 | 0.6667 | NA | NA | 14 | 0.2143 | 1 | 0 | 3 | 0.3333 | 1 | 0 | 2 | 0.5 | 7 | 0.2857 |
| 4_1381227: | 2 | 0.5 | 6 | 0.1667 | 1 | 0 | NA | NA | 9 | 0.2222 | 1 | 1 | 2 | 0 | NA | NA | 2 | 0.5 | 5 | 0.4 |
| 4_1381229! | 1 | 1 | 6 | 0.1667 | 1 | 0 | NA | NA | 8 | 0.25 | 1 | 0 | 2 | 0 | 1 | 0 | 2 | 0.5 | 6 | 0.1667 |
| 4_1381231 | 3 | 0 | 5 | 0 | 1 | 1 | NA | NA | 9 | 0.1111 | 1 | 0 | 3 | 0 | NA | NA | 2 | 0.5 | 6 | 0.1667 |
| 4_1381231! | 2 | 1 | 5 | 0.4 | NA | NA | NA | NA | 7 | 0.5714 | 1 | 1 | 3 | 0 | 1 | 0 | 2 | 0.5 | 7 | 0.2857 |
| 4_1381233( | 2 | 0.5 | 5 | 0 | 1 | 0 | NA | NA | 8 | 0.125 | 1 | 0 | 3 | 0.3333 | 1 | 0 | 1 | 0 | 6 | 0.1667 |
| 4_1381234: | 2 | 1 | 5 | 1 | 1 | 0 | NA | NA | 8 | 0.875 | 1 | 1 | 3 | 1 | 1 | 1 | 2 | 0.5 | 7 | 0.8571 |
| 4_1381236 | 2 | 0.5 | 4 | 0.5 | 1 | 1 | NA | NA | 7 | 0.5714 | 1 | 0 | 3 | 0 | 1 | 1 | 2 | 1 | 7 | 0.4286 |
| 4_1381238 | NA | NA | 1 | 1 | NA | NA | 1 | 1 | 2 | 1 | NA | NA | NA | NA | 1 | 1 | NA | NA | 1 | 1 |
| 4_1381242 | NA | NA | 1 | 1 | NA | NA | 2 | 0.5 | 3 | 0.6667 | NA | NA | NA | NA | 1 | 1 | NA | NA | 1 | 1 |
| 4_1381662! | 3 | 0 | 10 | 0.2 | 4 | 0.25 | 7 | 0 | 24 | 0.125 | 3 | 0 | 3 | 0 | 5 | 0.6 | 3 | 0.6667 | 14 | 0.3571 |
| 4_1381663! | 3 | 0.6667 | 10 | 0.6 | 5 | 0.8 | 8 | 0.75 | 26 | 0.6923 | 3 | 0.6667 | 4 | 1 | 5 | 1 | 2 | 1 | 14 | 0.9286 |
| 4_1381665: | 4 | 1 | 9 | 1 | 7 | 0.8571 | 6 | 1 | 26 | 0.9615 | 3 | 1 | 3 | 1 | 4 | 1 | 2 | 1 | 12 | 1 |
| 4_1381665 | 4 | 0.75 | 7 | 0.1429 | 5 | 0.8 | 7 | 0.2857 | 23 | 0.4348 | 4 | 0.5 | 2 | 0.5 | 5 | 0.6 | 2 | 0.5 | 13 | 0.5385 |
| 4_1381666( | 5 | 1 | 10 | 1 | 6 | 0.8333 | 8 | 1 | 29 | 0.9655 | 4 | 1 | 4 | 1 | 5 | 1 | 2 | 1 | 15 | 1 |
| 4_1381668 | 5 | 0.6 | 8 | 0.5 | 7 | 0.2857 | 6 | 0.6667 | 26 | 0.5 | 4 | 0.5 | 4 | 0.25 | 5 | 0.8 | 4 | 0.25 | 17 | 0.4706 |
| 4_1381670 | 6 | 0.1667 | 11 | 0 | 6 | 0.1667 | 10 | 0.2 | 33 | 0.1212 | 4 | 0 | 4 | 0.25 | 4 | 0 | 5 | 0 | 17 | 0.0588 |
| 4_1381671: | 5 | 0.4 | 10 | 0.1 | 8 | 0.375 | 12 | 0.3333 | 35 | 0.2857 | 5 | 0.4 | 3 | 0 | 6 | 0.3333 | 6 | 0.5 | 20 | 0.35 |
| 4_1381671: | 3 | 1 | 11 | 1 | 7 | 1 | 8 | 1 | 29 | 1 | 5 | 1 | 3 | 1 | 6 | 1 | 6 | 1 | 20 | 1 |
| 4_1381671! | 4 | 1 | 10 | 0.7 | 5 | 1 | 11 | 0.8182 | 30 | 0.8333 | 4 | 1 | 3 | 0.6667 | 6 | 1 | 5 | 1 | 18 | 0.9444 |
| 4_1381671! | 5 | 0.2 | 11 | 0 | 7 | 0.2857 | 12 | 0.0833 | 35 | 0.1143 | 4 | 0.75 | 4 | 0 | 7 | 0.4286 | 4 | 0.25 | 19 | 0.3684 |
| 4_1381673 | NA | NA | 3 | 0 | 5 | 0.2 | 4 | 0 | 12 | 0.0833 | 2 | 0.5 | 3 | 0 | 4 | 0 | 3 | 0 | 12 | 0.0833 |
| 4_1381679 | NA | NA | 4 | 0 | 5 | 0 | 6 | 0.1667 | 15 | 0.0667 | 4 | 0 | 5 | 0 | 3 | 0 | 3 | 0.3333 | 15 | 0.0667 |
| 4_1381681 | NA | NA | 4 | 0.25 | 5 | 0 | 7 | 0.1429 | 16 | 0.125 | 4 | 0.25 | 5 | 0.4 | 4 | 0.25 | 3 | 0 | 16 | 0.25 |
| 4_1381682 | NA | NA | 2 | 0.5 | 5 | 0 | 5 | 0.4 | 12 | 0.25 | 3 | 0.3333 | 3 | 0.3333 | 4 | 0.25 | 3 | 0 | 13 | 0.2308 |
| 4_1381684 | NA | NA | 4 | 0 | 5 | 0 | 5 | 0.2 | 14 | 0.0714 | 4 | 0.5 | 5 | 0.2 | 6 | 0.3333 | 4 | 0 | 19 | 0.2632 |
| 4_1381691 | NA | NA | 2 | 0 | 6 | 0.1667 | 8 | 0.125 | 16 | 0.125 | 7 | 0.2857 | 5 | 0.2 | 4 | 0.25 | 5 | 0.2 | 21 | 0.2381 |
| 4_1381693 | 1 | 0 | 4 | 0.5 | 8 | 0.125 | 8 | 0.375 | 21 | 0.2857 | 6 | 0.5 | 4 | 0.25 | 6 | 0.1667 | 5 | 0.2 | 21 | 0.2857 |
| 4_1381694( | NA | NA | 2 | 0 | 5 | 0.2 | 6 | 0.1667 | 13 | 0.1538 | 3 | 0 | 5 | 0.4 | 6 | 0.3333 | 2 | 0 | 16 | 0.25 |
| 4_1381715 | 1 | 1 | NA | NA | NA | NA | NA | NA | 1 | 1 | NA | NA | NA | NA | 1 | 1 | NA | NA | 1 | 1 |
| 4_1381718( | 1 | 1 | NA | NA | NA | NA | NA | NA | 1 | 1 | NA | NA | NA | NA | 1 | 1 | NA | NA | 1 | 1 |
| 4_1381718 | 2 | 1 | NA | NA | NA | NA | NA | NA | 2 | 1 | NA | NA | NA | NA | 1 | 1 | NA | NA | 1 | 1 |
| 4_1381719 | 1 | 1 | NA | NA | NA | NA | NA | NA | 1 | 1 | NA | NA | NA | NA | 1 | 1 | NA | NA | 1 | 1 |
| 4_1381723 | 1 | 1 | NA | NA | NA | NA | NA | NA | 1 | 1 | NA | NA | NA | NA | 1 | 1 | NA | NA | 1 | 1 |
| 4_1381725 | 1 | 1 | NA | NA | NA | NA | NA | NA | 1 | 1 | NA | NA | NA | NA | 1 | 1 | NA | NA | 1 | 1 |
| 4_1381820! | 46 | 0.0435 | 41 | 0.0732 | 34 | 0.0294 | 26 | 0.1154 | 147 | 0.0612 | 43 | 0.093 | 19 | 0 | 30 | 0.0667 | 31 | 0.0968 | 123 | 0.0732 |
| 4_1381821 | 40 | 0.15 | 40 | 0.075 | 27 | 0.1481 | 31 | 0 | 138 | 0.0942 | 39 | 0.0513 | 16 | 0.0625 | 29 | 0.0345 | 27 | 0.0741 | 111 | 0.0541 |
| 4_1381841 | 73 | 0.5616 | 66 | 0.5606 | 45 | 0.5333 | 52 | 0.5769 | 236 | 0.5593 | 64 | 0.5469 | 33 | 0.6364 | 54 | 0.537 | 42 | 0.7381 | 193 | 0.601 |
| 4_1381841! | 70 | 0.1714 | 60 | 0.1 | 45 | 0.0889 | 52 | 0.0962 | 227 | 0.1189 | 60 | 0.1833 | 38 | 0.0526 | 56 | 0.0536 | 42 | 0.0714 | 196 | 0.0969 |
| 4_1381847! | 164 | 0.0671 | 160 | 0.0375 | 127 | 0.0551 | 141 | 0.0567 | 592 | 0.0541 | 126 | 0.1111 | 117 | 0.0513 | 141 | 0.0355 | 110 | 0.0818 | 494 | 0.0688 |
| 4_1381848 | 127 | 0.8583 | 136 | 0.7574 | 107 | 0.8224 | 123 | 0.6992 | 493 | 0.783 | 109 | 0.8349 | 98 | 0.7347 | 120 | 0.75 | 99 | 0.8081 | 426 | 0.7817 |
| 4_1381848! | 187 | 0.0802 | 182 | 0.0659 | 156 | 0.0962 | 158 | 0.1203 | 683 | 0.0893 | 144 | 0.125 | 137 | 0.0657 | 165 | 0.0303 | 125 | 0.152 | 571 | 0.0893 |
| 4_1381849! | 194 | 0.1082 | 190 | 0.1053 | 151 | 0.0993 | 166 | 0.0843 | 701 | 0.0999 | 143 | 0.1469 | 136 | 0.0735 | 165 | 0.1394 | 120 | 0.1333 | 564 | 0.1241 |
| 4_1381849( | 183 | 0.1257 | 183 | 0.0546 | 153 | 0.1046 | 164 | 0.0915 | 683 | 0.0937 | 127 | 0.0787 | 131 | 0.1374 | 157 | 0.121 | 114 | 0.1228 | 529 | 0.1153 |
| 4_1381850! | 185 | 0.8757 | 181 | 0.8287 | 148 | 0.8378 | 155 | 0.8452 | 669 | 0.8475 | 137 | 0.8832 | 132 | 0.8333 | 155 | 0.8452 | 113 | 0.8673 | 537 | 0.8566 |
| 4_1381854! | 4 | 0.75 | 2 | 0 | 3 | 0 | NA | NA | 9 | 0.3333 | 1 | 0 | NA | NA | 6 | 0.1667 | 2 | 0.5 | 9 | 0.2222 |
| 4_1381856( | 5 | 1 | 3 | 0.6667 | 3 | 1 | NA | NA | 11 | 0.9091 | 1 | 1 | NA | NA | 5 | 1 | 2 | 1 | 8 | 1 |
| 4_1381856! | 6 | 0.6667 | 4 | 0 | 3 | 0.3333 | NA | NA | 13 | 0.3846 | 1 | 0 | NA | NA | 5 | 0.2 | 2 | 0 | 8 | 0.125 |
| 4_1381857( | 6 | 0.1667 | 4 | 0.25 | 4 | 0.5 | NA | NA | 14 | 0.2857 | 1 | 0 | NA | NA | 4 | 0.25 | 2 | 0.5 | 7 | 0.2857 |
| 4_1381857 | 5 | 1 | 3 | 1 | 4 | 1 | NA | NA | 12 | 1 | 1 | 0 | NA | NA | 4 | 1 | 1 | 1 | 6 | 0.8333 |
| 4_1381857: | 5 | 0 | 4 | 0 | 4 | 0.25 | NA | NA | 13 | 0.0769 | 1 | 0 | NA | NA | 5 | 0.2 | 1 | 0 | 7 | 0.1429 |
| 4_1381857( | 5 | 0 | 3 | 0.3333 | 4 | 0.25 | NA | NA | 12 | 0.1667 | 1 | 0 | NA | NA | 5 | 0 | 2 | 0.5 | 8 | 0.125 |
| 4_1381858( | 4 | 0.25 | 4 | 0.75 | 3 | 0 | NA | NA | 11 | 0.3636 | 1 | 0 | NA | NA | 5 | 0.4 | 2 | 0 | 8 | 0.25 |
| 4_1381859: | 4 | 1 | 3 | 0.3333 | 4 | 1 | NA | NA | 11 | 0.8182 | 1 | 1 | NA | NA | 5 | 0.6 | 2 | 1 | 8 | 0.75 |
| 4_1381865( | 6 | 0.5 | 6 | 0.3333 | 2 | 0.5 | 5 | 0.4 | 19 | 0.4211 | 2 | 0 | 2 | 0 | 5 | 0.4 | 2 | 0 | 11 | 0.1818 |

| | | | | | | | | | | | | | | | | | | | | |
|---|---|---|---|---|---|---|---|---|---|---|---|---|---|---|---|---|---|---|---|---|
| 4_13818652 | 9 | 0 | 8 | 0.125 | 5 | 0 | 7 | 0.1429 | 29 | 0.069 | 3 | 0 | 2 | 0.5 | 6 | 0 | 4 | 0 | 15 | 0.0667 |
| 4_13818663 | 9 | 0.7778 | 9 | 0.8889 | 5 | 0.8 | 7 | 0.7143 | 30 | 0.8 | 5 | 0.6 | 2 | 0 | 7 | 0.4286 | 5 | 0.4 | 19 | 0.4211 |
| 4_13818687 | 11 | 0.1818 | 9 | 0.2222 | 9 | 0.1111 | 7 | 0 | 36 | 0.1389 | 6 | 0.1667 | 3 | 0 | 7 | 0.1429 | 8 | 0.125 | 24 | 0.125 |
| 4_13818694 | 11 | 0.1818 | 6 | 0 | 8 | 0.125 | 5 | 0 | 30 | 0.1 | 5 | 0.2 | 3 | 0.3333 | 7 | 0.2857 | 6 | 0.3333 | 21 | 0.2857 |
| 4_13818710 | 13 | 0.3077 | 7 | 0.5714 | 9 | 0.1111 | 6 | 0.3333 | 35 | 0.3143 | 5 | 0.2 | 3 | 0.6667 | 7 | 0.1429 | 8 | 0.125 | 23 | 0.2174 |
| 4_13818720 | 12 | 0.5833 | 8 | 0.875 | 7 | 0.5714 | 7 | 0.8571 | 34 | 0.7059 | 7 | 0.4286 | 1 | 1 | 6 | 0.3333 | 5 | 1 | 19 | 0.5789 |
| 4_13818739 | 12 | 0.0833 | 9 | 0.1111 | 5 | 0.4 | 7 | 0 | 33 | 0.1212 | 7 | 0.1429 | 4 | 0 | 8 | 0.125 | 8 | 0 | 27 | 0.0741 |
| 4_13829646 | 1 | 1 | NA | NA | NA | NA | NA | NA | 1 | 1 | NA | NA | 1 | 1 | NA | NA | NA | NA | 1 | 1 |
| 4_13862287 | NA | NA | NA | NA | 2 | 1 | NA | NA | 2 | 1 | NA | NA | NA | NA | 1 | 1 | NA | NA | 1 | 1 |
| 4_13862304 | NA | NA | 1 | 1 | 2 | 1 | NA | NA | 3 | 1 | NA | NA | 1 | 1 | NA | NA | NA | NA | 1 | 1 |
| 4_13862312 | NA | NA | 1 | 0 | 2 | 1 | NA | NA | 3 | 0.6667 | NA | NA | 1 | 1 | NA | NA | NA | NA | 1 | 1 |
| 4_13878551 | 1 | 1 | 1 | 0 | NA | NA | NA | NA | 2 | 0.5 | NA | NA | 1 | 1 | 1 | 0 | NA | NA | 2 | 0.5 |
| 4_13878581 | NA | NA | 1 | 1 | NA | NA | NA | NA | 1 | 1 | NA | NA | 1 | 1 | 1 | 1 | NA | NA | 2 | 1 |
| 4_13878582 | 1 | 1 | 1 | 0 | NA | NA | NA | NA | 2 | 0.5 | NA | NA | NA | NA | 1 | 1 | NA | NA | 1 | 1 |
| 4_13878584 | 1 | 1 | NA | NA | NA | NA | NA | NA | 1 | 1 | NA | NA | 1 | 1 | 1 | 1 | NA | NA | 2 | 1 |
| 4_13878622 | 1 | 1 | 1 | 0 | NA | NA | NA | NA | 2 | 0.5 | NA | NA | 1 | 1 | 1 | 0 | NA | NA | 2 | 0.5 |
| 4_13878704 | NA | NA | NA | NA | 1 | 0 | 2 | 0.5 | 3 | 0.3333 | 1 | 1 | 3 | 0.6667 | 4 | 0.5 | 2 | 0.5 | 10 | 0.6 |
| 4_13878715 | 1 | 1 | NA | NA | 1 | 1 | 3 | 0.6667 | 5 | 0.8 | 1 | 1 | 3 | 0.3333 | 3 | 0.6667 | 2 | 1 | 9 | 0.6667 |
| 4_13878873 | 1 | 1 | NA | NA | 2 | 1 | 2 | 1 | 5 | 1 | 1 | 1 | NA | NA | 4 | 0.75 | NA | NA | 5 | 0.8 |
| 4_13878878 | 1 | 1 | NA | NA | 3 | 1 | 2 | 1 | 6 | 1 | 2 | 1 | 2 | 1 | 4 | 1 | 2 | 1 | 10 | 1 |
| 4_13878890 | 1 | 1 | NA | NA | 2 | 1 | 3 | 0.6667 | 6 | 0.8333 | 2 | 0.5 | 2 | 1 | 3 | 0.6667 | 1 | 1 | 8 | 0.75 |
| 4_13878898 | 2 | 1 | NA | NA | 1 | 1 | 3 | 1 | 6 | 1 | NA | NA | 2 | 1 | 2 | 0.5 | 1 | 1 | 5 | 0.8 |
| 4_13878907 | 1 | 1 | NA | NA | 2 | 1 | 2 | 1 | 5 | 1 | 1 | 0 | 2 | 1 | 3 | 1 | 2 | 0 | 8 | 0.625 |
| 4_13880155 | 1 | 1 | NA | NA | 3 | 0 | 3 | 0.6667 | 7 | 0.4286 | 1 | 0 | 3 | 0.6667 | 2 | 0.5 | 1 | 1 | 7 | 0.5714 |
| 4_13880156 | 1 | 1 | NA | NA | 3 | 0.6667 | 3 | 0.6667 | 7 | 0.7143 | 2 | 0.5 | 3 | 1 | 1 | 1 | 1 | 1 | 7 | 0.8571 |
| 4_13880170 | 1 | 0 | NA | NA | 2 | 0.5 | 2 | 0 | 5 | 0.2 | 1 | 1 | 3 | 1 | 2 | 0 | 1 | 0 | 7 | 0.5714 |
| 4_13880176 | 1 | 0 | NA | NA | 2 | 0 | 1 | 1 | 4 | 0.25 | 1 | 0 | 3 | 0.6667 | 2 | 0 | 1 | 1 | 7 | 0.4286 |
| 4_13880185 | 1 | 0 | NA | NA | 1 | 0 | 2 | 0.5 | 4 | 0.25 | 1 | 0 | 3 | 0.3333 | 2 | 0.5 | 1 | 1 | 7 | 0.4286 |
| 4_13880231 | 1 | 1 | NA | NA | 1 | 1 | 1 | 1 | 3 | 1 | 1 | 1 | 2 | 1 | 2 | 1 | 1 | 1 | 6 | 1 |
| 4_13880246 | 1 | 1 | NA | NA | 1 | 1 | 1 | 1 | 3 | 1 | 1 | 1 | 2 | 1 | 1 | 1 | NA | NA | 4 | 1 |
| 4_13880350 | NA | NA | 1 | 1 | 1 | 1 | NA | NA | 2 | 1 | NA | NA | 2 | 1 | NA | NA | NA | NA | 2 | 1 |
| 4_13880404 | NA | NA | 1 | 1 | 1 | 1 | NA | NA | 2 | 1 | NA | NA | 2 | 1 | NA | NA | 1 | 1 | 3 | 1 |
| 4_13880411 | NA | NA | 1 | 1 | 1 | 0 | NA | NA | 2 | 0.5 | NA | NA | 2 | 0.5 | NA | NA | 1 | 0 | 3 | 0.3333 |
| 4_13880557 | NA | NA | 1 | 1 | 1 | 0 | 2 | 0.5 | 4 | 0.5 | NA | NA | 2 | 0.5 | NA | NA | 2 | 0.5 | 4 | 0.5 |
| 4_13880561 | NA | NA | 1 | 0 | 1 | 1 | 2 | 0.5 | 4 | 0.5 | NA | NA | 2 | 0.5 | NA | NA | 2 | 0.5 | 4 | 0.5 |
| 4_13880567 | NA | NA | NA | NA | 1 | 1 | 2 | 1 | 3 | 1 | NA | NA | 2 | 0.5 | NA | NA | 2 | 0.5 | 4 | 0.5 |
| 4_13880617 | NA | NA | 1 | 0 | 1 | 1 | 2 | 0 | 4 | 0.25 | 1 | 1 | 1 | 0 | NA | NA | 2 | 0 | 4 | 0.25 |
| 4_13880634 | NA | NA | 1 | 0 | 1 | 1 | 1 | 1 | 3 | 0.6667 | 1 | 1 | 1 | 1 | NA | NA | 3 | 1 | 5 | 1 |
| 4_13880650 | NA | NA | 1 | 1 | 1 | 1 | 2 | 0.5 | 4 | 0.75 | 1 | 0 | NA | NA | NA | NA | 3 | 0.3333 | 4 | 0.25 |
| 4_13881043 | 6 | 0.3333 | 10 | 0.1 | 6 | 0.3333 | 3 | 0 | 25 | 0.2 | 4 | 0.5 | 4 | 0 | 3 | 0.6667 | 1 | 0 | 12 | 0.3333 |
| 4_13881054 | 6 | 0.8333 | 11 | 0.6364 | 6 | 0.6667 | 3 | 0.6667 | 26 | 0.6923 | 3 | 0.6667 | 3 | 0.3333 | 2 | 0.5 | 2 | 1 | 10 | 0.6 |
| 4_13881066 | 3 | 0.6667 | 9 | 0.2222 | 6 | 0.3333 | 2 | 0.5 | 20 | 0.35 | 3 | 0.3333 | 3 | 0.6667 | 3 | 0.6667 | 2 | 0.5 | 11 | 0.5455 |
| 4_13881075 | 4 | 0 | 9 | 0 | 4 | 0.25 | 3 | 0 | 20 | 0.05 | 2 | 0 | 4 | 0 | 3 | 0 | 2 | 0.5 | 11 | 0.0909 |
| 4_13882932 | 3 | 1 | NA | NA | 6 | 0.3333 | 4 | 0.5 | 13 | 0.5385 | 3 | 0.3333 | 3 | 0.6667 | 2 | 0.5 | 3 | 0.6667 | 11 | 0.5455 |
| 4_13882936 | 4 | 0 | NA | NA | 6 | 0.1667 | 4 | 0 | 14 | 0.0714 | 3 | 0.3333 | 3 | 0 | 2 | 0 | 3 | 0.3333 | 11 | 0.1818 |
| 4_13882951 | 4 | 0.25 | NA | NA | 6 | 0.1667 | 3 | 0 | 13 | 0.1538 | 4 | 0 | 3 | 1 | 2 | 0.5 | 2 | 0.5 | 11 | 0.4545 |
| 4_13882952 | 4 | 0.75 | NA | NA | 5 | 0.6 | 2 | 1 | 11 | 0.7273 | 3 | 1 | 3 | 0.6667 | 2 | 0.5 | 3 | 1 | 11 | 0.8182 |
| 4_13882957 | 2 | 0.5 | NA | NA | 4 | 0.5 | 2 | 0.5 | 8 | 0.5 | 2 | 1 | 1 | 0 | 2 | 0.5 | 2 | 0 | 7 | 0.4286 |
| 4_13883008 | 4 | 0 | NA | NA | 6 | 0.1667 | 2 | 0 | 12 | 0.0833 | 4 | 0.25 | 3 | 0 | 2 | 0 | 2 | 0 | 11 | 0.0909 |
| 4_13883037 | 4 | 0 | NA | NA | 4 | 0 | 1 | 1 | 9 | 0.1111 | 2 | 0 | 3 | 0 | 2 | 0 | 2 | 0.5 | 9 | 0.1111 |
| 4_13883079 | 3 | 0.3333 | NA | NA | 5 | 0.4 | 2 | 0 | 10 | 0.3 | 3 | 0.6667 | NA | NA | 1 | 0 | 2 | 0 | 6 | 0.3333 |
| 4_13883083 | 3 | 0 | NA | NA | 6 | 0.1667 | NA | NA | 9 | 0.1111 | 2 | 0 | 2 | 0.5 | 2 | 0 | 3 | 0 | 9 | 0.1111 |
| 4_13901875 | 27 | 0.037 | 31 | 0.0645 | 21 | 0.0952 | 30 | 0.0333 | 109 | 0.055 | 25 | 0.12 | 32 | 0.0312 | 22 | 0.0455 | 30 | 0.0667 | 109 | 0.0642 |
| 4_13902592 | 1 | 0 | NA | NA | NA | NA | 1 | 1 | 2 | 0.5 | 2 | 0.5 | NA | NA | NA | NA | 1 | 1 | 3 | 0.6667 |
| 4_13904556 | 9 | 0.8889 | 4 | 0.25 | 4 | 0.5 | 3 | 0.6667 | 20 | 0.65 | 6 | 0.8333 | 8 | 0.75 | 5 | 1 | 4 | 0.5 | 23 | 0.7826 |
| 4_13904572 | 7 | 1 | 3 | 0.6667 | 3 | 0.6667 | 4 | 0.75 | 17 | 0.8235 | 7 | 0.8571 | 7 | 0.8571 | 5 | 0.6 | 3 | 0.6667 | 22 | 0.7727 |
| 4_13904573 | 4 | 0 | 2 | 0.5 | 2 | 0 | 2 | 0 | 10 | 0.1 | 4 | 0.5 | 3 | 0 | 4 | 0 | 3 | 0 | 14 | 0.1429 |
| 4_13907484 | 5 | 0.8 | 5 | 0.8 | 4 | 0.25 | 4 | 1 | 18 | 0.7222 | 5 | 1 | 8 | 0.5 | 6 | 1 | 3 | 1 | 22 | 0.8182 |
| 4_13907488 | 6 | 0.5 | 5 | 0.8 | 5 | 0.6 | 5 | 0.2 | 21 | 0.5238 | 5 | 0.8 | 8 | 0.625 | 8 | 0.625 | 2 | 0 | 23 | 0.6087 |
| 4_13907495 | 5 | 0 | 3 | 0.3333 | 3 | 0 | 6 | 0.6667 | 17 | 0.2941 | 5 | 0 | 6 | 0.1667 | 8 | 0.25 | 2 | 0 | 21 | 0.1429 |
| 4_13908155 | NA | NA | 5 | 0 | 3 | 0.6667 | 3 | 0.3333 | 11 | 0.2727 | 2 | 0 | 2 | 1 | 3 | 0 | 1 | 0 | 8 | 0.25 |
| 4_13910287 | 4 | 0.25 | 13 | 0.3077 | 10 | 0.2 | 8 | 0.375 | 35 | 0.2857 | 7 | 0.5714 | 6 | 0.3333 | 6 | 0.1667 | 8 | 0.125 | 27 | 0.2963 |
| 4_13910289 | 4 | 0.5 | 16 | 0.75 | 7 | 1 | 11 | 0.9091 | 38 | 0.8158 | 7 | 0.8571 | 10 | 1 | 8 | 1 | 7 | 1 | 32 | 0.9688 |
| 4_13910299 | 2 | 0.5 | 10 | 0.3 | 5 | 0.2 | 3 | 0.3333 | 20 | 0.3 | 2 | 0.5 | 8 | 0.375 | 4 | 0.5 | 5 | 1 | 19 | 0.5789 |
| 4_13910317 | 6 | 0.1667 | 17 | 0.1765 | 11 | 0.6364 | 11 | 0.1818 | 45 | 0.2889 | 7 | 0.2857 | 11 | 0.4545 | 11 | 0.1818 | 9 | 0.3333 | 38 | 0.3158 |

| | | | | | | | | | | | | | | | | | | | | |
|---|---|---|---|---|---|---|---|---|---|---|---|---|---|---|---|---|---|---|---|---|
| 4_1391046 | 1 | 1 | 9 | 0.2222 | 1 | 0 | 7 | 0.2857 | 18 | 0.2778 | 3 | 0.6667 | 3 | 0 | 3 | 0.3333 | 5 | 0.2 | 14 | 0.2857 |
| 4_1391046 | 2 | 0 | 9 | 0.1111 | 1 | 0 | 7 | 0.2857 | 19 | 0.1579 | 4 | 0.25 | 5 | 0.6 | 4 | 0 | 7 | 0 | 20 | 0.2 |
| 4_1391052 | 2 | 0.5 | NA | NA | NA | NA | NA | NA | 2 | 0.5 | 2 | 0.5 | 1 | 0 | 1 | 0 | 2 | 0 | 6 | 0.1667 |
| 4_1391056 | 3 | 0.3333 | 1 | 0 | 1 | 0 | NA | NA | 5 | 0.2 | 2 | 0 | 2 | 0.5 | 1 | 0 | 2 | 0 | 7 | 0.1429 |
| 4_1392167 | 14 | 0.1429 | 12 | 0.25 | 11 | 0.2727 | 9 | 0.1111 | 46 | 0.1957 | 7 | 0.1429 | 8 | 0.25 | 10 | 0.3 | 14 | 0.2857 | 39 | 0.2564 |
| 4_1392168 | 11 | 0.0909 | 15 | 0.1333 | 13 | 0.2308 | 12 | 0.25 | 51 | 0.1765 | 6 | 0 | 8 | 0 | 8 | 0.25 | 20 | 0.1 | 42 | 0.0952 |
| 4_1393473 | 1 | 0 | 2 | 0 | 4 | 0.25 | 1 | 0 | 8 | 0.125 | NA | NA | 2 | 0.5 | 1 | 0 | 3 | 0.6667 | 6 | 0.5 |
| 4_1393644 | NA | NA | 1 | 0 | 3 | 0 | 3 | 0.3333 | 7 | 0.1429 | NA | NA | 2 | 0.5 | 1 | 0 | NA | NA | 3 | 0.3333 |
| 4_1393755 | 2 | 0 | 1 | 0 | 6 | 0 | 5 | 0.2 | 14 | 0.0714 | NA | NA | 3 | 0 | 3 | 0 | 4 | 0.25 | 10 | 0.1 |
| 4_1393756 | 2 | 1 | 1 | 1 | 6 | 0.5 | 4 | 1 | 13 | 0.7692 | 1 | 0 | 2 | 1 | 2 | 1 | 3 | 0.6667 | 8 | 0.75 |
| 4_1393756 | 2 | 1 | 1 | 0 | 6 | 0.1667 | 5 | 0.2 | 14 | 0.2857 | 1 | 0 | 4 | 0.25 | 3 | 0.3333 | 3 | 0 | 11 | 0.1818 |
| 4_1393758 | 3 | 0.6667 | 1 | 0 | 6 | 0.3333 | 5 | 0.8 | 15 | 0.5333 | 1 | 1 | 7 | 0.5714 | 3 | 0.3333 | 4 | 0 | 15 | 0.4 |
| 4_1393759 | 4 | 0 | 1 | 1 | 6 | 0.3333 | 5 | 0.6 | 16 | 0.375 | 1 | 0 | 7 | 0.5714 | 3 | 0.3333 | 4 | 0 | 15 | 0.3333 |
| 4_1393759 | 3 | 0.3333 | 1 | 0 | 6 | 0.6667 | 5 | 0.8 | 15 | 0.6 | 1 | 1 | 7 | 0.4286 | 2 | 1 | 2 | 0.5 | 12 | 0.5833 |
| 4_1393760 | 3 | 0.3333 | 1 | 0 | 6 | 0 | 5 | 0.6 | 15 | 0.2667 | 1 | 1 | 7 | 0.2857 | 3 | 0 | 4 | 0.75 | 15 | 0.4 |
| 4_1393763 | 3 | 0.6667 | 2 | 1 | 5 | 0.6 | 5 | 0.8 | 15 | 0.7333 | 1 | 1 | 6 | 0.5 | 2 | 1 | 3 | 0 | 12 | 0.5 |
| 4_1393763 | 2 | 0.5 | 2 | 0 | 4 | 0.5 | 6 | 0.5 | 14 | 0.4286 | 1 | 0 | 8 | 0 | 3 | 0.3333 | 4 | 0.25 | 16 | 0.125 |
| 4_1393764 | 4 | 0 | 2 | 0 | 5 | 0.2 | 6 | 0 | 17 | 0.0588 | 1 | 0 | 8 | 0.25 | 3 | 0.3333 | 3 | 0 | 15 | 0.2 |
| 4_1393766 | 4 | 0.5 | 1 | 0 | 7 | 0 | 6 | 0.1667 | 18 | 0.1667 | NA | NA | 5 | 0 | 3 | 0.6667 | 1 | 1 | 9 | 0.3333 |
| 4_1396905 | NA | NA | 1 | 1 | 3 | 0.6667 | 1 | 0 | 5 | 0.6 | 1 | 0 | NA | NA | 1 | 0 | 1 | 1 | 3 | 0.3333 |
| 4_1396906 | NA | NA | 1 | 1 | 3 | 0.6667 | 1 | 0 | 5 | 0.6 | 1 | 1 | NA | NA | 1 | 0 | 1 | 1 | 3 | 0.6667 |
| 4_1396910 | NA | NA | 1 | 1 | 2 | 0.5 | NA | NA | 3 | 0.6667 | 1 | 1 | 1 | 0 | NA | NA | 1 | 1 | 3 | 0.6667 |
| 4_1396914 | NA | NA | 1 | 1 | 2 | 0.5 | NA | NA | 3 | 0.6667 | 1 | 1 | NA | NA | NA | NA | NA | NA | 1 | 1 |
| 4_1396921 | NA | NA | 1 | 1 | NA | NA | NA | NA | 1 | 1 | 1 | 1 | 1 | 0 | NA | NA | NA | NA | 2 | 0.5 |
| 4_1396922 | NA | NA | 1 | 1 | NA | NA | NA | NA | 1 | 1 | 1 | 1 | NA | NA | NA | NA | NA | NA | 1 | 1 |
| 4_1396923 | NA | NA | 1 | 1 | NA | NA | NA | NA | 1 | 1 | 1 | 1 | NA | NA | NA | NA | NA | NA | 1 | 1 |
| 4_1396929 | NA | NA | 1 | 1 | 1 | 1 | NA | NA | 2 | 1 | 1 | 1 | NA | NA | NA | NA | NA | NA | 1 | 1 |
| 4_1396931 | NA | NA | 1 | 1 | 1 | 1 | NA | NA | 2 | 1 | 1 | 1 | NA | NA | NA | NA | NA | NA | 1 | 1 |
| 4_1396931 | NA | NA | 1 | 1 | 1 | 0 | NA | NA | 2 | 0.5 | 1 | 1 | NA | NA | NA | NA | NA | NA | 1 | 1 |
| 4_1396936 | NA | NA | 1 | 1 | 1 | 0 | NA | NA | 2 | 0.5 | 1 | 1 | NA | NA | NA | NA | NA | NA | 1 | 1 |
| 4_1397618 | 6 | 0.5 | 11 | 0.3636 | 8 | 0.125 | 4 | 0.25 | 29 | 0.3103 | 16 | 0.5 | 9 | 0.6667 | 8 | 0.5 | 6 | 0.6667 | 39 | 0.5641 |
| 4_1397619 | 4 | 0 | 5 | 0.4 | 3 | 0 | 3 | 0.3333 | 15 | 0.2 | 7 | 1 | 6 | 0.6667 | 5 | 0.6 | 5 | 1 | 23 | 0.8261 |
| 4_1397620 | 7 | 0.1429 | 11 | 0.1818 | 5 | 0.2 | 3 | 0 | 26 | 0.1538 | 15 | 0 | 6 | 0 | 6 | 0.5 | 5 | 0.2 | 32 | 0.125 |
| 4_1399744 | NA | NA | 1 | 1 | NA | NA | NA | NA | 1 | 1 | 1 | 1 | NA | NA | NA | NA | NA | NA | 1 | 1 |
| 4_1401321 | NA | NA | NA | NA | 1 | 1 | NA | NA | 1 | 1 | NA | NA | NA | NA | NA | NA | 1 | 1 | 1 | 1 |
| 4_1402419 | 7 | 0.2857 | 3 | 0.6667 | NA | NA | 4 | 0.25 | 14 | 0.3571 | 3 | 0.3333 | 3 | 0.3333 | 5 | 0 | 2 | 0.5 | 13 | 0.2308 |
| 4_1402422 | 8 | 0.75 | 6 | 0.8333 | 1 | 0 | 5 | 0.8 | 20 | 0.75 | 1 | 0 | 4 | 1 | 6 | 0.8333 | 1 | 1 | 12 | 0.8333 |
| 4_1402422 | 9 | 0.7778 | 6 | 0.6667 | 2 | 1 | 3 | 1 | 20 | 0.8 | 4 | 0.5 | 5 | 0.8 | 7 | 0.7143 | 1 | 0 | 17 | 0.6471 |
| 4_1402422 | 9 | 0.8889 | 5 | 1 | 2 | 1 | 5 | 1 | 21 | 0.9524 | 2 | 1 | 2 | 1 | 6 | 0.5 | 1 | 0 | 11 | 0.6364 |
| 4_1402423 | 7 | 0.2857 | 4 | 0 | NA | NA | 4 | 0.75 | 15 | 0.3333 | 4 | 0.5 | 5 | 0.4 | 6 | 0 | 1 | 0 | 16 | 0.25 |
| 4_1402423 | 7 | 0.1429 | 3 | 0 | 2 | 0.5 | 5 | 0.2 | 17 | 0.1765 | 4 | 0 | 5 | 0.2 | 7 | 0.1429 | 1 | 0 | 17 | 0.1176 |
| 4_1402424 | 6 | 1 | 4 | 1 | 2 | 1 | 3 | 1 | 15 | 1 | 1 | 1 | 4 | 1 | 2 | 1 | NA | NA | 7 | 1 |
| 4_1402425 | 6 | 0.6667 | 3 | 0.3333 | 2 | 0 | 5 | 0.8 | 16 | 0.5625 | 4 | 0.25 | 2 | 1 | 5 | 0.2 | NA | NA | 11 | 0.3636 |
| 4_1402427 | 7 | 0.1429 | 3 | 0.3333 | 2 | 0 | 5 | 0.2 | 17 | 0.1765 | 4 | 0.5 | NA | NA | 5 | 0 | 1 | 0 | 10 | 0.2 |
| 4_1402428 | 5 | 0 | 4 | 0 | 2 | 0 | 4 | 0.25 | 15 | 0.0667 | 4 | 0 | 3 | 0.3333 | 6 | 0 | 1 | 0 | 14 | 0.0714 |
| 4_1404485 | 6 | 0 | 5 | 0.2 | 3 | 0 | 7 | 0.2857 | 21 | 0.1429 | 4 | 0 | NA | NA | 8 | 0.125 | 2 | 0 | 14 | 0.0714 |
| 4_1404488 | 6 | 0 | 6 | 0.1667 | 4 | 0 | 8 | 0.125 | 24 | 0.0833 | 4 | 0.75 | 1 | 0 | 7 | 0.4286 | 4 | 0 | 16 | 0.375 |
| 4_1404491 | 6 | 0.1667 | 5 | 0 | 4 | 0 | 8 | 0.125 | 23 | 0.087 | 4 | 0 | 1 | 0 | 4 | 0.25 | 4 | 0 | 13 | 0.0769 |
| 4_1406772 | 2 | 0 | 2 | 0.5 | 4 | 0.25 | 1 | 1 | 9 | 0.3333 | 2 | 0.5 | 1 | 0 | 2 | 0.5 | 1 | 0 | 6 | 0.3333 |
| 4_1406779 | 1 | 0 | 1 | 0 | 3 | 0 | 1 | 1 | 6 | 0.1667 | 1 | 1 | NA | NA | 1 | 0 | 1 | 0 | 3 | 0.3333 |
| 4_1406779 | 3 | 0 | 2 | 0.5 | 1 | 0 | NA | NA | 6 | 0.1667 | 1 | 1 | NA | NA | NA | NA | NA | NA | 1 | 1 |
| 4_1406794 | 6 | 0 | 2 | 0 | 4 | 0.25 | 3 | 0 | 15 | 0.0667 | 4 | 0 | 3 | 0.3333 | 5 | 0 | 2 | 0.5 | 14 | 0.1429 |
| 4_1406794 | 6 | 0 | 2 | 0 | 4 | 0.25 | 3 | 0.6667 | 15 | 0.2 | 4 | 0 | 3 | 0 | 5 | 0 | 2 | 1 | 14 | 0.1429 |
| 4_1406795 | 6 | 0.1667 | 2 | 0.5 | 4 | 0.5 | 3 | 0 | 15 | 0.2667 | 4 | 0.25 | 3 | 0.3333 | 5 | 0.2 | 2 | 0 | 14 | 0.2143 |
| 4_1407344 | 17 | 0.8235 | 20 | 0.85 | 11 | 0.2727 | 17 | 0.6471 | 65 | 0.6923 | 7 | 0.8571 | 22 | 0.7727 | 15 | 0.8 | 12 | 0.5 | 56 | 0.7321 |
| 4_1407344 | 10 | 0.4 | 16 | 0.375 | 12 | 0.25 | 11 | 0.1818 | 49 | 0.3061 | 6 | 0.8333 | 13 | 0.2308 | 11 | 0.4545 | 9 | 0.1111 | 39 | 0.359 |
| 4_1407344 | 16 | 0.4375 | 18 | 0.3333 | 14 | 0.2143 | 19 | 0.4211 | 67 | 0.3582 | 9 | 0.4444 | 17 | 0.5294 | 13 | 0.3846 | 9 | 0.6667 | 48 | 0.5 |
| 4_1407345 | 15 | 0.3333 | 19 | 0.4211 | 15 | 0.4 | 18 | 0.2778 | 67 | 0.3582 | 8 | 0.5 | 17 | 0.3529 | 13 | 0.3846 | 9 | 0.6667 | 47 | 0.4468 |
| 4_1407345 | 1 | 0 | 1 | 1 | NA | NA | NA | NA | 2 | 0.5 | NA | NA | 1 | 0 | 1 | 1 | NA | NA | 2 | 0.5 |
| 4_1407345 | 18 | 0.1667 | 21 | 0.0476 | 17 | 0.0588 | 21 | 0 | 77 | 0.0649 | 9 | 0 | 25 | 0.16 | 16 | 0 | 13 | 0.3077 | 63 | 0.127 |
| 4_1407347 | 4 | 0.5 | NA | NA | 1 | 1 | NA | NA | 5 | 0.6 | NA | NA | 2 | 1 | 1 | 1 | NA | NA | 3 | 1 |
| 4_1407347 | 4 | 0.5 | NA | NA | 1 | 1 | NA | NA | 5 | 0.6 | NA | NA | 1 | 1 | NA | NA | NA | NA | 1 | 1 |
| 4_1407443 | 5 | 0.2 | 6 | 0 | 5 | 0 | NA | NA | 16 | 0.0625 | 2 | 0.5 | 3 | 0 | 3 | 0 | 2 | 0 | 10 | 0.1 |
| 4_1407939 | 5 | 0.2 | 4 | 0.25 | 10 | 0.1 | 5 | 0.2 | 24 | 0.1667 | 8 | 0.125 | 4 | 0.25 | 7 | 0.1429 | 5 | 0 | 24 | 0.125 |
| 4_1407943 | 4 | 0.25 | 3 | 0 | 11 | 0.2727 | 4 | 0.75 | 22 | 0.3182 | 6 | 0.3333 | 3 | 0 | 5 | 0.2 | 7 | 0.4286 | 21 | 0.2857 |

| | | | | | | | | | | | | | | | | | | | | |
|---|---|---|---|---|---|---|---|---|---|---|---|---|---|---|---|---|---|---|---|---|
| 4_14079448 | 5 | 0.4 | 3 | 0.6667 | 11 | 0.2727 | 4 | 0.75 | 23 | 0.4348 | 8 | 0.125 | 5 | 1 | 7 | 0.5714 | 6 | 0.3333 | 26 | 0.4615 |
| 4_14079449 | 5 | 0.6 | 2 | 0.5 | 9 | 0.1111 | 3 | 0 | 19 | 0.2632 | 4 | 0.5 | 4 | 0.75 | 5 | 0 | 6 | 0.5 | 19 | 0.4211 |
| 4_14084113 | 6 | 0.1667 | 6 | 0 | 3 | 0 | 3 | 0 | 18 | 0.0556 | 4 | 0 | 4 | 0 | 3 | 0.3333 | 5 | 0 | 16 | 0.0625 |
| 4_14084137 | 5 | 0 | 4 | 0 | 3 | 0 | 4 | 0.25 | 16 | 0.0625 | 4 | 0 | 4 | 0 | 4 | 0 | 6 | 0.1667 | 18 | 0.0556 |
| 4_14084198 | 3 | 0.6667 | 3 | 0.3333 | 3 | 0.6667 | 4 | 1 | 13 | 0.6923 | 3 | 0.6667 | 3 | 0 | 3 | 0.6667 | 2 | 1 | 11 | 0.5455 |
| 4_14084279 | 5 | 0.2 | 3 | 0.3333 | 2 | 0.5 | 4 | 0 | 14 | 0.2143 | 2 | 0.5 | 3 | 0 | 4 | 0.25 | 3 | 0 | 12 | 0.1667 |
| 4_14084296 | 4 | 0.25 | 4 | 0.25 | 2 | 0.5 | 3 | 0.3333 | 13 | 0.3077 | 2 | 0 | 3 | 0.3333 | 5 | 0.2 | 3 | 0 | 13 | 0.1538 |
| 4_14084298 | 3 | 1 | 4 | 0.75 | 2 | 1 | 3 | 1 | 12 | 0.9167 | 2 | 1 | 2 | 0.5 | 6 | 1 | 2 | 1 | 12 | 0.9167 |
| 4_14092697 | 1 | 1 | 1 | 0 | 1 | 1 | NA | NA | 3 | 0.6667 | 1 | 0 | NA | NA | 2 | 0 | 1 | 1 | 4 | 0.25 |
| 4_14092706 | 2 | 1 | 1 | 1 | 2 | 1 | 1 | 0 | 6 | 0.8333 | 2 | 0.5 | NA | NA | 5 | 1 | 1 | 0 | 8 | 0.75 |
| 4_14092715 | 2 | 1 | 1 | 1 | 2 | 1 | 1 | 1 | 6 | 1 | 2 | 0.5 | NA | NA | 4 | 1 | 1 | 1 | 7 | 0.8571 |
| 4_14092720 | 2 | 0.5 | 1 | 1 | 2 | 0.5 | 1 | 1 | 6 | 0.6667 | 1 | 1 | NA | NA | 5 | 0.6 | 1 | 1 | 7 | 0.7143 |
| 4_14092723 | 3 | 1 | 1 | 1 | 2 | 0 | 1 | 1 | 7 | 0.7143 | 3 | 0.3333 | NA | NA | 6 | 0 | NA | NA | 9 | 0.1111 |
| 4_14092728 | 2 | 0.5 | 1 | 1 | 1 | 1 | NA | NA | 4 | 0.75 | 1 | 1 | NA | NA | 4 | 1 | 1 | 1 | 6 | 1 |
| 4_14092753 | 3 | 0 | 1 | 1 | 2 | 0 | 1 | 0 | 7 | 0.1429 | 3 | 0 | 1 | 0 | 7 | 0 | 1 | 1 | 12 | 0.0833 |
| 4_14092764 | 2 | 1 | 1 | 1 | 2 | 1 | 1 | 1 | 6 | 1 | 4 | 1 | 1 | 1 | 5 | 1 | 1 | 1 | 11 | 1 |
| 4_14092767 | 2 | 0.5 | 1 | 0 | 2 | 0 | 1 | 0 | 6 | 0.1667 | 4 | 0 | 1 | 0 | 4 | 0.25 | 1 | 0 | 10 | 0.1 |
| 4_14092767 | 3 | 0.3333 | 1 | 0 | 1 | 1 | 1 | 0 | 6 | 0.3333 | 3 | 0.3333 | 1 | 0 | 5 | 0.6 | 1 | 1 | 10 | 0.5 |
| 4_14092780 | 3 | 0.3333 | 1 | 1 | 1 | 1 | 1 | 1 | 6 | 0.6667 | 3 | 0.6667 | 1 | 1 | 4 | 0.75 | 1 | 0 | 9 | 0.6667 |
| 4_14092784 | 3 | 0.3333 | 2 | 0 | 2 | 0 | 1 | 0 | 8 | 0.125 | 4 | 0.25 | 1 | 0 | 6 | 0.1667 | 1 | 0 | 12 | 0.1667 |
| 4_14092793 | 3 | 0 | 2 | 0.5 | 1 | 0 | NA | NA | 6 | 0.1667 | 4 | 0 | NA | NA | 5 | 0 | 1 | 1 | 10 | 0.1 |
| 4_14092794 | 1 | 1 | 1 | 1 | 1 | 1 | 1 | 1 | 4 | 1 | 4 | 1 | 1 | 1 | 2 | 1 | 1 | 1 | 8 | 1 |
| 4_14092794 | 3 | 0.6667 | 2 | 1 | 2 | 0.5 | 1 | 1 | 8 | 0.75 | 4 | 1 | 1 | 0 | 7 | 0.4286 | 1 | 0 | 13 | 0.5385 |
| 4_14092798 | 3 | 0 | 1 | 0 | 2 | 0.5 | 1 | 1 | 7 | 0.2857 | 3 | 0 | 1 | 0 | 5 | 0.4 | 1 | 1 | 10 | 0.3 |
| 4_14092835 | 2 | 0 | 1 | 1 | 1 | 0 | 1 | 0 | 5 | 0.2 | 1 | 0 | NA | NA | 6 | 0.3333 | 1 | 0 | 8 | 0.25 |
| 4_14092938 | 3 | 0.3333 | 2 | 0 | 2 | 0 | 2 | 0 | 9 | 0.1111 | 1 | 0 | 2 | 0 | 6 | 0.1667 | 1 | 0 | 10 | 0.1 |
| 4_14092957 | 3 | 0 | 2 | 0 | 3 | 0.3333 | 1 | 0 | 9 | 0.1111 | 2 | 0 | 2 | 0 | 6 | 0.1667 | 1 | 0 | 11 | 0.0909 |
| 4_14092971 | 2 | 0 | 1 | 0 | 1 | 0 | 1 | 1 | 5 | 0.2 | 3 | 0.3333 | 4 | 0.25 | 4 | 0 | 1 | 0 | 12 | 0.1667 |
| 4_14092985 | 3 | 0.3333 | 2 | 0 | 3 | 0 | 1 | 0 | 9 | 0.1111 | 3 | 0 | 4 | 0.5 | 7 | 0.1429 | 1 | 0 | 15 | 0.2 |
| 4_14093006 | 3 | 0.3333 | 2 | 0 | 3 | 0 | 2 | 0.5 | 10 | 0.2 | 3 | 0 | 2 | 0.5 | 7 | 0.2857 | 1 | 0 | 13 | 0.2308 |
| 4_14093019 | 2 | 0 | 1 | 0 | 3 | 0.3333 | 2 | 0 | 8 | 0.125 | 3 | 0 | 3 | 0 | 5 | 0.2 | 1 | 0 | 12 | 0.0833 |
| 4_14093027 | 3 | 0 | 2 | 0 | 3 | 0 | 2 | 0.5 | 10 | 0.1 | 2 | 0 | 4 | 0.25 | 5 | 0.2 | NA | NA | 11 | 0.1818 |
| 4_14093070 | 3 | 0 | 2 | 0.5 | 2 | 0 | 3 | 0 | 10 | 0.1 | 3 | 0.3333 | 5 | 0.4 | 7 | 0.1429 | 1 | 0 | 16 | 0.25 |
| 4_14093071 | 3 | 0.3333 | 2 | 0.5 | 3 | 0.3333 | 1 | 1 | 9 | 0.4444 | 1 | 0 | 4 | 0.5 | 7 | 0.5714 | 1 | 0 | 13 | 0.4615 |
| 4_14093080 | 4 | 0 | 1 | 0 | 4 | 0 | 2 | 0.5 | 11 | 0.0909 | 3 | 0 | 4 | 0 | 7 | 0.1429 | 1 | 0 | 15 | 0.0667 |
| 4_14093091 | 4 | 0.25 | 1 | 0 | 4 | 0 | 3 | 0.3333 | 12 | 0.1667 | 3 | 0 | 5 | 0 | 7 | 0.1429 | 1 | 0 | 16 | 0.0625 |
| 4_14093102 | 3 | 0 | 2 | 0 | 2 | 0 | 1 | 1 | 8 | 0.125 | 4 | 0 | 5 | 0.2 | 9 | 0.2222 | 1 | 0 | 19 | 0.1579 |
| 4_14093136 | 3 | 0 | 2 | 0 | 4 | 0 | 2 | 0.5 | 11 | 0.0909 | 3 | 0 | 4 | 0 | 6 | 0.1667 | 1 | 0 | 14 | 0.0714 |
| 4_14098448 | 3 | 0.6667 | 4 | 0.5 | 1 | 0 | NA | NA | 8 | 0.5 | 2 | 0.5 | 3 | 0.3333 | 2 | 0 | 3 | 0.6667 | 10 | 0.4 |
| 4_14098453 | 2 | 0.5 | 2 | 0 | 1 | 0 | NA | NA | 5 | 0.2 | 2 | 0 | 3 | 0 | 3 | 0 | 2 | 0.5 | 10 | 0.1 |
| 4_14098469 | 2 | 1 | 2 | 0.5 | 1 | 1 | NA | NA | 5 | 0.8 | 1 | 0 | 3 | 0.6667 | 3 | 0.6667 | 2 | 0.5 | 9 | 0.5556 |
| 4_14111111 | NA | NA | NA | NA | 1 | 1 | 1 | 1 | 2 | 1 | 2 | 1 | NA | NA | 2 | 1 | NA | NA | 4 | 1 |
| 4_14111113 | NA | NA | NA | NA | 1 | 1 | 1 | 1 | 2 | 1 | 2 | 1 | NA | NA | 2 | 1 | 1 | 1 | 5 | 1 |
| 4_14111119 | NA | NA | NA | NA | 1 | 1 | NA | NA | 1 | 1 | 2 | 0 | NA | NA | 3 | 0.3333 | NA | NA | 5 | 0.2 |
| 4_14111129 | NA | NA | NA | NA | 1 | 1 | 1 | 1 | 2 | 1 | 1 | 1 | NA | NA | 2 | 1 | NA | NA | 3 | 1 |
| 4_14111131 | NA | NA | NA | NA | 1 | 1 | 1 | 1 | 2 | 1 | 1 | 1 | NA | NA | 3 | 1 | NA | NA | 4 | 1 |
| 4_14111132 | NA | NA | NA | NA | 1 | 1 | NA | NA | 1 | 1 | 1 | 1 | NA | NA | 3 | 1 | NA | NA | 4 | 1 |
| 4_14116038 | 2 | 0 | 1 | 1 | 2 | 0 | 1 | 0 | 6 | 0.1667 | 1 | 0 | 2 | 0.5 | 3 | 0 | NA | NA | 6 | 0.1667 |
| 4_14116050 | 2 | 1 | 2 | 1 | 3 | 1 | 2 | 1 | 9 | 1 | 1 | 1 | 2 | 1 | 2 | 1 | 1 | 1 | 6 | 1 |
| 4_14116056 | 2 | 1 | 2 | 0.5 | 2 | 0 | 2 | 0.5 | 8 | 0.5 | NA | NA | 1 | 0 | 1 | 0 | 1 | 1 | 3 | 0.3333 |
| 4_14116057 | 2 | 1 | 1 | 1 | 3 | 1 | 3 | 1 | 9 | 1 | 1 | 1 | 2 | 1 | 2 | 1 | 1 | 1 | 6 | 1 |
| 4_14116070 | 2 | 0 | 2 | 0.5 | 3 | 0.3333 | 3 | 0.3333 | 10 | 0.3 | 1 | 0 | 2 | 0.5 | 2 | 0 | 1 | 0 | 6 | 0.1667 |
| 4_14119187 | NA | NA | NA | NA | 2 | 0.5 | 1 | 0 | 3 | 0.3333 | 4 | 0 | 3 | 0.3333 | 3 | 0 | NA | NA | 10 | 0.1 |
| 4_14119221 | 3 | 0.3333 | 3 | 0.3333 | 2 | 0.5 | 2 | 0.5 | 10 | 0.4 | 5 | 0 | 1 | 0 | 4 | 0.5 | NA | NA | 10 | 0.2 |
| 4_14119428 | 4 | 0.5 | 4 | 0.5 | 3 | 0.3333 | 5 | 0.2 | 16 | 0.375 | 5 | 0.4 | 2 | 0 | 3 | 0.3333 | 2 | 1 | 12 | 0.4167 |
| 4_14119436 | 3 | 0 | 4 | 0.5 | 2 | 0.5 | 6 | 0.3333 | 15 | 0.3333 | 6 | 0 | 3 | 0.3333 | 4 | 0.5 | 2 | 0.5 | 15 | 0.2667 |
| 4_14119445 | 3 | 0.3333 | 4 | 0.25 | 3 | 0.3333 | 4 | 0.25 | 14 | 0.2857 | 5 | 0.6 | 4 | 0.75 | 3 | 0.3333 | NA | NA | 12 | 0.5833 |
| 4_14119448 | 4 | 0.25 | 4 | 0.5 | 2 | 0 | 6 | 0.5 | 16 | 0.375 | 5 | 0.4 | 4 | 0.25 | 5 | 0.4 | 2 | 0 | 16 | 0.3125 |
| 4_14119522 | 4 | 0 | 2 | 0 | 5 | 0.2 | 8 | 0.125 | 19 | 0.1053 | 6 | 0 | 9 | 0 | 10 | 0.3 | 3 | 0.3333 | 28 | 0.1429 |
| 4_14119527 | 6 | 0.1667 | 3 | 0 | 4 | 0.5 | 10 | 0.2 | 23 | 0.2174 | 6 | 0.1667 | 9 | 0.1111 | 10 | 0.2 | 2 | 0 | 27 | 0.1481 |
| 4_14119530 | 6 | 0.1667 | 3 | 0.6667 | 6 | 0 | 10 | 0.3 | 25 | 0.24 | 4 | 0.25 | 8 | 0.25 | 9 | 0.4444 | 3 | 0 | 24 | 0.2917 |
| 4_14119536 | 10 | 0.3 | 3 | 0.6667 | 6 | 0.6667 | 8 | 0.75 | 27 | 0.5556 | 4 | 0.5 | 8 | 0.375 | 4 | 0.5 | 2 | 0.5 | 18 | 0.4444 |
| 4_14119543 | 6 | 0.3333 | 3 | 0.3333 | 4 | 0.25 | 9 | 0.5556 | 22 | 0.4091 | 4 | 0.25 | 8 | 0 | 11 | 0.0909 | NA | NA | 23 | 0.087 |
| 4_14119552 | 8 | 0.5 | 5 | 0.6 | 6 | 0.6667 | 8 | 0.625 | 27 | 0.5926 | 6 | 0.1667 | 10 | 0.7 | 8 | 0.75 | 3 | 0.6667 | 27 | 0.5926 |
| 4_14119557 | 7 | 0.1429 | 4 | 0.5 | 5 | 0.6 | 7 | 0.5714 | 23 | 0.4348 | 5 | 0.2 | 5 | 0.2 | 9 | 0.3333 | 2 | 0 | 21 | 0.2381 |

| | | | | | | | | | | | | | | | | | | | | |
|---|---|---|---|---|---|---|---|---|---|---|---|---|---|---|---|---|---|---|---|---|
| 4_1411957 | 8 | 0.25 | 3 | 0.3333 | 5 | 0.4 | 6 | 0.1667 | 22 | 0.2727 | 5 | 0.4 | 10 | 0.1 | 8 | 0.375 | 2 | 0.5 | 25 | 0.28 |
| 4_1411957 | 6 | 1 | 3 | 1 | 3 | 0.3333 | 5 | 0.8 | 17 | 0.8235 | 5 | 0.8 | 8 | 0.875 | 10 | 0.6 | 2 | 1 | 25 | 0.76 |
| 4_1411957 | 8 | 0.875 | 3 | 1 | 4 | 0.25 | 5 | 1 | 20 | 0.8 | 6 | 0.8333 | 10 | 1 | 10 | 0.8 | 3 | 0.3333 | 29 | 0.8276 |
| 4_1411958 | 7 | 0.5714 | 3 | 1 | 4 | 1 | 5 | 0.8 | 19 | 0.7895 | 6 | 0.6667 | 9 | 0.4444 | 10 | 0.6 | 1 | 1 | 26 | 0.5769 |
| 4_1411958 | 7 | 1 | 3 | 1 | 5 | 1 | 5 | 1 | 20 | 1 | 5 | 1 | 8 | 1 | 7 | 0.7143 | 3 | 1 | 23 | 0.913 |
| 4_1411958 | 6 | 0.3333 | 3 | 0.6667 | 2 | 0.5 | 6 | 0.6667 | 17 | 0.5294 | 4 | 0.75 | 6 | 0.5 | 7 | 1 | 2 | 1 | 19 | 0.7895 |
| 4_1411958 | 5 | 0.8 | 2 | 1 | 5 | 0.6 | 5 | 0.4 | 17 | 0.6471 | 2 | 1 | 8 | 0.875 | 8 | 0.625 | 2 | 0.5 | 20 | 0.75 |
| 4_1411959 | 6 | 0.8333 | 2 | 1 | 4 | 0.75 | 6 | 1 | 18 | 0.8889 | 5 | 0.6 | 7 | 0.8571 | 9 | 0.8889 | 3 | 1 | 24 | 0.8333 |
| 4_1411959 | 4 | 1 | 2 | 1 | 4 | 1 | 5 | 1 | 15 | 1 | 4 | 1 | 9 | 1 | 8 | 1 | 1 | 1 | 22 | 1 |
| 4_1411960 | 5 | 0.2 | 4 | 0.75 | 3 | 0 | 4 | 0.5 | 16 | 0.375 | 3 | 0 | 7 | 0.5714 | 9 | 0.3333 | 3 | 0.3333 | 22 | 0.3636 |
| 4_1411960 | 6 | 0 | 2 | 0 | 5 | 0.2 | 5 | 0 | 18 | 0.0556 | 3 | 0.3333 | 5 | 0 | 7 | 0 | 1 | 0 | 16 | 0.0625 |
| 4_1411961 | 2 | 1 | 1 | 0 | 4 | 0.25 | 3 | 0 | 10 | 0.3 | 2 | 0 | 2 | 0.5 | 2 | 0.5 | NA | NA | 6 | 0.3333 |
| 4_1411975 | 4 | 0 | 2 | 0.5 | 2 | 0 | 4 | 0 | 12 | 0.0833 | 5 | 0 | 4 | 0 | 7 | 0.1429 | 3 | 0.3333 | 19 | 0.1053 |
| 4_1411977 | 5 | 0.6 | 2 | 0 | 3 | 0 | 5 | 0 | 15 | 0.2 | 4 | 0.25 | 3 | 0.3333 | 8 | 0 | 2 | 0 | 17 | 0.1176 |
| 4_1411989 | 3 | 0.3333 | 1 | 0 | 2 | 0.5 | 4 | 0 | 10 | 0.2 | 4 | 0.25 | 4 | 0 | 5 | 0 | 1 | 1 | 14 | 0.1429 |
| 4_1411990 | 3 | 0.3333 | 1 | 0 | 1 | 0 | 4 | 0.25 | 9 | 0.2222 | 4 | 0.5 | 3 | 0.3333 | 4 | 0.75 | NA | NA | 11 | 0.5455 |
| 4_1411994 | 3 | 0 | 1 | 0 | 1 | 0 | 3 | 0.3333 | 8 | 0.125 | 4 | 0.25 | 4 | 0.25 | 1 | 0 | NA | NA | 9 | 0.2222 |
| 4_1411996 | 1 | 1 | NA | NA | NA | NA | NA | NA | 1 | 1 | 3 | 0.3333 | 2 | 0.5 | 2 | 0.5 | NA | NA | 7 | 0.4286 |
| 4_1411996 | 3 | 0.6667 | 1 | 0 | 1 | 0 | 2 | 0.5 | 7 | 0.4286 | 3 | 0.3333 | 3 | 0.3333 | 2 | 0 | NA | NA | 8 | 0.25 |
| 4_1411999 | 3 | 0 | 1 | 0 | 1 | 1 | 2 | 0 | 7 | 0.1429 | 3 | 0.3333 | 4 | 0 | 2 | 0 | NA | NA | 9 | 0.1111 |
| 4_1412000 | 3 | 0.6667 | 1 | 1 | 1 | 0 | 2 | 1 | 7 | 0.7143 | 3 | 1 | 3 | 0 | 2 | 0.5 | NA | NA | 8 | 0.5 |
| 4_1412001 | 3 | 1 | 1 | 1 | 1 | 1 | NA | NA | 5 | 1 | 3 | 1 | 3 | 1 | 2 | 1 | NA | NA | 8 | 1 |
| 4_1412003 | 2 | 1 | NA | NA | 1 | 1 | NA | NA | 3 | 1 | 3 | 0 | 2 | 0 | 2 | 0 | 1 | 1 | 8 | 0.125 |
| 4_1412004 | 3 | 0.3333 | 1 | 0 | NA | NA | NA | NA | 4 | 0.25 | 3 | 0.6667 | 3 | 0.3333 | 2 | 0.5 | NA | NA | 8 | 0.5 |
| 4_1412005 | 2 | 0 | 1 | 0 | 1 | 1 | NA | NA | 4 | 0.25 | 2 | 0 | 3 | 0.3333 | 2 | 0 | 1 | 0 | 8 | 0.125 |
| 4_1412007 | 2 | 1 | NA | NA | 1 | 1 | NA | NA | 3 | 1 | 1 | 0 | 4 | 0.25 | 3 | 0.6667 | 1 | 1 | 9 | 0.4444 |
| 4_1412009 | 2 | 0.5 | 1 | 0 | 1 | 0 | NA | NA | 4 | 0.25 | NA | NA | 4 | 0 | 3 | 0.3333 | NA | NA | 7 | 0.1429 |
| 4_1412010 | 2 | 1 | 1 | 0 | 1 | 1 | NA | NA | 4 | 0.75 | 1 | 1 | 4 | 0.75 | 3 | 1 | NA | NA | 8 | 0.875 |
| 4_1412010 | 2 | 0.5 | 1 | 1 | 1 | 1 | NA | NA | 4 | 0.75 | 1 | 1 | 4 | 1 | 2 | 1 | NA | NA | 7 | 1 |
| 4_1412015 | 2 | 1 | NA | NA | 1 | 1 | NA | NA | 3 | 1 | 1 | 1 | 3 | 0.6667 | 2 | 0.5 | NA | NA | 6 | 0.6667 |
| 4_1412017 | 1 | 0 | 1 | 1 | NA | NA | NA | NA | 2 | 0.5 | 1 | 1 | 2 | 0 | NA | NA | NA | NA | 3 | 0.3333 |
| 4_1412018 | 1 | 1 | 1 | 0 | 1 | 0 | NA | NA | 3 | 0.3333 | 1 | 0 | 1 | 1 | 1 | 0 | NA | NA | 3 | 0.3333 |
| 4_1412019 | 1 | 0 | NA | NA | 1 | 1 | NA | NA | 2 | 0.5 | 1 | 1 | NA | NA | 1 | 0 | NA | NA | 2 | 0.5 |
| 4_1412739 | 2 | 0.5 | NA | NA | 1 | 1 | NA | NA | 3 | 0.6667 | NA | NA | 1 | 1 | NA | NA | NA | NA | 1 | 1 |
| 4_1413462 | 3 | 1 | 3 | 0.6667 | 2 | 0 | 1 | 0 | 9 | 0.5556 | 2 | 1 | NA | NA | 1 | 0 | 1 | 1 | 4 | 0.75 |
| 4_1413472 | NA | NA | 2 | 0.5 | NA | NA | 1 | 0 | 3 | 0.3333 | 3 | 0 | NA | NA | 2 | 0.5 | NA | NA | 5 | 0.2 |
| 4_1413472 | 1 | 0 | 2 | 0.5 | NA | NA | 1 | 1 | 4 | 0.5 | 3 | 0.6667 | NA | NA | 2 | 1 | NA | NA | 5 | 0.8 |
| 4_1413473 | 1 | 0 | 2 | 0.5 | NA | NA | 1 | 1 | 4 | 0.5 | 3 | 0.6667 | NA | NA | 2 | 0.5 | NA | NA | 5 | 0.6 |
| 4_1413476 | 1 | 0 | 1 | 0 | NA | NA | 1 | 1 | 3 | 0.3333 | 2 | 0 | NA | NA | 2 | 0.5 | NA | NA | 4 | 0.25 |
| 4_1413537 | 9 | 0.2222 | 8 | 0.125 | 8 | 0 | 8 | 0.25 | 33 | 0.1515 | 5 | 0 | 8 | 0.125 | 7 | 0.1429 | 6 | 0 | 26 | 0.0769 |
| 4_1413583 | 5 | 0.4 | 6 | 0 | 7 | 0 | 5 | 0 | 23 | 0.087 | 1 | 0 | 5 | 0 | 3 | 0.3333 | 3 | 0 | 12 | 0.0833 |
| 4_1413586 | 5 | 0.8 | 6 | 0.5 | 8 | 0.5 | 6 | 0.5 | 25 | 0.56 | 1 | 0 | 5 | 0.8 | 5 | 1 | 4 | 0.5 | 15 | 0.7333 |
| 4_1413590 | 5 | 1 | 3 | 1 | 7 | 0.7143 | 7 | 0.5714 | 22 | 0.7727 | 1 | 1 | 5 | 1 | 4 | 1 | 3 | 0.6667 | 13 | 0.9231 |
| 4_1413591 | 5 | 0 | 4 | 0.25 | 7 | 0.1429 | 6 | 0.1667 | 22 | 0.1364 | 1 | 0 | 5 | 0 | 6 | 0.1667 | 3 | 0.3333 | 15 | 0.1333 |
| 4_1413594 | 5 | 0.8 | 4 | 0.75 | 5 | 0.8 | 6 | 1 | 20 | 0.85 | 1 | 1 | 2 | 1 | 3 | 0.3333 | 3 | 1 | 9 | 0.7778 |
| 4_1413595 | 5 | 0.4 | 4 | 0.25 | 5 | 0.4 | 6 | 0.3333 | 20 | 0.35 | NA | NA | 4 | 0 | 3 | 0.3333 | 4 | 0.25 | 11 | 0.1818 |
| 4_1413595 | 4 | 0.25 | 3 | 0.3333 | 5 | 0.2 | 7 | 0.2857 | 19 | 0.2632 | NA | NA | 4 | 0 | 4 | 0.25 | 4 | 0 | 12 | 0.0833 |
| 4_1413596 | 4 | 0.25 | 4 | 0.25 | 3 | 0 | 5 | 0.4 | 16 | 0.25 | NA | NA | 4 | 0.75 | 3 | 0.6667 | 3 | 1 | 10 | 0.8 |
| 4_1413596 | 3 | 0 | 4 | 0.5 | 4 | 0.5 | 7 | 0.2857 | 18 | 0.3333 | NA | NA | 3 | 0.6667 | 2 | 0 | 2 | 0.5 | 7 | 0.4286 |
| 4_1413597 | 4 | 0.75 | 4 | 0.25 | 5 | 0.6 | 5 | 0.4 | 18 | 0.5 | NA | NA | 3 | 0.3333 | 3 | 0 | 3 | 0.6667 | 9 | 0.3333 |
| 4_1413597 | 3 | 1 | 2 | 0.5 | 5 | 0.8 | 4 | 1 | 14 | 0.8571 | NA | NA | 4 | 1 | 3 | 1 | 3 | 0.6667 | 10 | 0.9 |
| 4_1414050 | 9 | 0.4444 | 5 | 0.4 | NA | NA | 7 | 0.2857 | 21 | 0.381 | 2 | 0 | 2 | 1 | 4 | 0 | 2 | 0.5 | 10 | 0.3 |
| 4_1414051 | 9 | 0.2222 | 6 | 0.1667 | 2 | 0 | 7 | 0 | 24 | 0.125 | 2 | 0 | 2 | 0.5 | 8 | 0.25 | 2 | 0 | 14 | 0.2143 |
| 4_1414052 | 9 | 0.2222 | 5 | 0.4 | 2 | 0.5 | 8 | 0.25 | 24 | 0.2917 | NA | NA | 2 | 0.5 | 8 | 0.375 | 2 | 0.5 | 12 | 0.4167 |
| 4_1414066 | 3 | 0 | 1 | 1 | 1 | 0 | 3 | 0 | 8 | 0.125 | 1 | 0 | 1 | 0 | 3 | 0.3333 | 2 | 0 | 7 | 0.1429 |
| 4_1414089 | 3 | 0.3333 | 1 | 0 | NA | NA | 3 | 0 | 7 | 0.1429 | 1 | 0 | NA | NA | 1 | 0 | 1 | 1 | 3 | 0.3333 |
| 4_1414105 | 3 | 1 | NA | NA | NA | NA | NA | NA | 3 | 1 | 1 | 1 | 1 | 1 | 2 | 0.5 | 1 | 1 | 5 | 0.8 |
| 4_1414108 | 3 | 0.3333 | NA | NA | NA | NA | NA | NA | 3 | 0.3333 | NA | NA | 1 | 0 | 1 | 0 | 1 | 1 | 3 | 0.3333 |
| 4_1414111 | 4 | 1 | NA | NA | 1 | 1 | NA | NA | 5 | 1 | NA | NA | 6 | 1 | 3 | 1 | 1 | 1 | 10 | 1 |
| 4_1414112 | 4 | 0 | 1 | 1 | 1 | 1 | NA | NA | 6 | 0.3333 | 1 | 0 | 4 | 0.5 | 3 | 0 | 2 | 0 | 10 | 0.2 |
| 4_1414113 | 4 | 0.5 | 1 | 1 | 2 | 1 | NA | NA | 7 | 0.7143 | 1 | 1 | 6 | 0.8333 | 3 | 1 | 2 | 1 | 12 | 0.9167 |
| 4_1414114 | 1 | 1 | NA | NA | 2 | 0.5 | NA | NA | 3 | 0.6667 | 1 | 0 | 5 | 0.6 | 1 | 1 | 1 | 1 | 8 | 0.625 |
| 4_1414115 | 3 | 0.6667 | 1 | 1 | 2 | 0.5 | NA | NA | 6 | 0.6667 | NA | NA | 5 | 0.8 | 3 | 1 | 2 | 1 | 10 | 0.9 |
| 4_1414116 | 1 | 1 | NA | NA | NA | NA | NA | NA | 1 | 1 | 1 | 0 | 3 | 1 | 2 | 1 | 2 | 1 | 8 | 0.875 |
| 4_1414116 | 3 | 0.6667 | NA | NA | 2 | 0 | NA | NA | 5 | 0.4 | NA | NA | 4 | 0 | 2 | 0 | 2 | 0.5 | 8 | 0.125 |

| | | | | | | | | | | | | | | | | | | | | |
|---|---|---|---|---|---|---|---|---|---|---|---|---|---|---|---|---|---|---|---|---|
| 4_1414117 | 4 | 0.25 | NA | NA | 2 | 1 | NA | NA | 6 | 0.5 | 1 | 1 | 4 | 0.5 | 3 | 0.6667 | 2 | 0.5 | 10 | 0.6 |
| 4_1414118 | 4 | 1 | NA | NA | 1 | 1 | NA | NA | 5 | 1 | NA | NA | 3 | 1 | 2 | 1 | 2 | 1 | 7 | 1 |
| 4_1414120 | 4 | 0.25 | NA | NA | 1 | 0 | NA | NA | 5 | 0.2 | 1 | 0 | 3 | 0.6667 | 3 | 0.6667 | 2 | 0.5 | 9 | 0.5556 |
| 4_1414121 | 3 | 0.3333 | NA | NA | 1 | 0 | NA | NA | 4 | 0.25 | 1 | 0 | 4 | 0 | 1 | 1 | 1 | 0 | 7 | 0.1429 |
| 4_1414122 | 4 | 1 | NA | NA | 1 | 1 | NA | NA | 5 | 1 | 1 | 1 | 3 | 1 | 3 | 1 | 2 | 1 | 9 | 1 |
| 4_1414123 | 4 | 0.5 | NA | NA | 1 | 1 | NA | NA | 5 | 0.6 | 1 | 1 | 3 | 0 | 3 | 0.6667 | 2 | 0.5 | 9 | 0.4444 |
| 4_1414127 | 3 | 0.3333 | NA | NA | 1 | 0 | NA | NA | 4 | 0.25 | 1 | 0 | 3 | 0.3333 | 3 | 0.3333 | 1 | 0 | 8 | 0.25 |
| 4_1414127 | 4 | 0.75 | NA | NA | 1 | 1 | NA | NA | 5 | 0.8 | 1 | 0 | 2 | 1 | 3 | 0 | 2 | 0.5 | 8 | 0.375 |
| 4_1414875 | 4 | 0.25 | 6 | 0.1667 | 2 | 0 | 4 | 0 | 16 | 0.125 | 3 | 0 | 7 | 0.1429 | 2 | 0 | 3 | 0 | 15 | 0.0667 |
| 4_1414877 | 5 | 0.2 | 5 | 0.2 | 2 | 0.5 | 4 | 0.5 | 16 | 0.3125 | 2 | 0 | 7 | 0.1429 | NA | NA | 1 | 0 | 10 | 0.1 |
| 4_1414879 | 6 | 0.1667 | 5 | 0 | 1 | 0 | 3 | 0 | 15 | 0.0667 | 5 | 0 | 8 | 0.125 | 1 | 0 | 2 | 0 | 16 | 0.0625 |
| 4_1414887 | 3 | 0 | 1 | 0 | NA | NA | 3 | 0.3333 | 7 | 0.1429 | 5 | 0.4 | 4 | 0 | 1 | 0 | 1 | 0 | 11 | 0.1818 |
| 4_1414888 | 3 | 0.3333 | 3 | 0.3333 | NA | NA | 3 | 1 | 9 | 0.5556 | 4 | 0.75 | 3 | 0.3333 | 1 | 1 | NA | NA | 8 | 0.625 |
| 4_1414893 | 1 | 1 | 5 | 0.4 | 1 | 1 | 2 | 1 | 9 | 0.6667 | 2 | 1 | 2 | 0.5 | 3 | 0.6667 | 2 | 1 | 9 | 0.7778 |
| 4_1414894 | 2 | 0 | 6 | 0.1667 | 1 | 0 | 2 | 0 | 11 | 0.0909 | 3 | 0.3333 | 2 | 0 | 3 | 0 | 3 | 0 | 11 | 0.0909 |
| 4_1414894 | 2 | 0 | 6 | 0.1667 | 1 | 1 | 3 | 0 | 12 | 0.1667 | 2 | 0 | 2 | 0 | 3 | 0 | 2 | 0.5 | 9 | 0.1111 |
| 4_1414894 | 2 | 1 | 6 | 1 | 1 | 1 | NA | NA | 9 | 1 | 3 | 1 | 2 | 1 | 3 | 1 | 2 | 1 | 10 | 1 |
| 4_1414894 | 2 | 0.5 | 6 | 0.6667 | 1 | 0 | 1 | 1 | 10 | 0.6 | 3 | 1 | 2 | 1 | 3 | 1 | 3 | 0.6667 | 11 | 0.9091 |
| 4_1414895 | 2 | 1 | 4 | 1 | 1 | 1 | 2 | 1 | 9 | 1 | 3 | 0.6667 | 2 | 0.5 | 2 | 1 | 3 | 1 | 10 | 0.8 |
| 4_1414897 | 2 | 1 | 6 | 0.6667 | 1 | 1 | 2 | 1 | 11 | 0.8182 | 2 | 1 | 2 | 1 | 3 | 1 | 1 | 1 | 8 | 1 |
| 4_1414897 | 2 | 0 | 6 | 0.1667 | 1 | 0 | 2 | 0 | 11 | 0.0909 | 2 | 0.5 | 2 | 0 | 2 | 1 | 2 | 0.5 | 8 | 0.5 |
| 4_1414897 | 1 | 1 | 4 | 1 | 1 | 1 | 1 | 1 | 7 | 1 | 2 | 0.5 | 1 | 1 | 1 | 1 | NA | NA | 4 | 0.75 |
| 4_1414898 | 1 | 1 | 5 | 0.8 | 1 | 1 | 1 | 1 | 8 | 0.875 | 1 | 1 | 2 | 1 | 1 | 1 | 1 | 1 | 5 | 1 |
| 4_1414898 | 1 | 0 | 6 | 0.6667 | 1 | 0 | 1 | 0 | 9 | 0.4444 | 2 | 0 | 2 | 0 | 2 | 0.5 | 2 | 0.5 | 8 | 0.25 |
| 4_1414898 | 1 | 0 | 6 | 0.3333 | 1 | 0 | 2 | 0 | 10 | 0.2 | NA | NA | 2 | 0.5 | 2 | 0 | 2 | 0 | 6 | 0.1667 |
| 4_1414899 | 1 | 0 | 5 | 0 | 1 | 0 | 2 | 0.5 | 9 | 0.1111 | 1 | 0 | 2 | 0.5 | 1 | 0 | 1 | 0 | 5 | 0.2 |
| 4_1414906 | 1 | 0 | 4 | 0 | 1 | 0 | 1 | 1 | 7 | 0.1429 | 2 | 1 | 1 | 1 | 2 | 0.5 | 1 | 0 | 6 | 0.6667 |
| 4_1414908 | 1 | 1 | 5 | 1 | 1 | 1 | 1 | 1 | 8 | 1 | 1 | 1 | 1 | 1 | 2 | 1 | 2 | 1 | 6 | 1 |
| 4_1414909 | 1 | 1 | 3 | 0.3333 | 1 | 0 | NA | NA | 5 | 0.4 | 2 | 0 | NA | NA | 2 | 0 | 1 | 1 | 5 | 0.2 |
| 4_1414909 | 1 | 1 | 4 | 0.75 | 1 | 0 | 1 | 1 | 7 | 0.7143 | 2 | 1 | 1 | 1 | 2 | 1 | 2 | 0 | 7 | 0.7143 |
| 4_1414910 | 1 | 1 | 3 | 1 | NA | NA | 1 | 1 | 5 | 1 | 1 | 0 | NA | NA | 2 | 1 | 2 | 1 | 5 | 0.8 |
| 4_1414910 | 1 | 1 | 4 | 0 | 1 | 0 | 1 | 1 | 7 | 0.2857 | 1 | 1 | 1 | 0 | 2 | 0.5 | 2 | 0 | 6 | 0.3333 |
| 4_1415410 | 2 | 0.5 | NA | NA | NA | NA | 1 | 1 | 3 | 0.6667 | NA | NA | 1 | 0 | 1 | 1 | 1 | 0 | 3 | 0.3333 |
| 4_1415410 | 2 | 1 | NA | NA | NA | NA | 1 | 1 | 3 | 1 | 1 | 1 | 1 | 1 | NA | NA | 2 | 0.5 | 4 | 0.75 |
| 4_1415412 | 3 | 0.6667 | NA | NA | NA | NA | 2 | 0.5 | 5 | 0.6 | 2 | 0 | 2 | 1 | 1 | 0 | 2 | 0 | 7 | 0.2857 |
| 4_1415413 | 2 | 1 | NA | NA | NA | NA | 2 | 1 | 4 | 1 | 2 | 0.5 | NA | NA | 1 | 1 | 2 | 1 | 5 | 0.8 |
| 4_1415414 | 3 | 0.6667 | NA | NA | NA | NA | 2 | 1 | 5 | 0.8 | 2 | 0.5 | NA | NA | 1 | 1 | 2 | 0 | 5 | 0.4 |
| 4_1415415 | 3 | 0.3333 | NA | NA | NA | NA | 2 | 0.5 | 5 | 0.4 | 1 | 1 | 2 | 0 | 1 | 0 | 2 | 0 | 6 | 0.1667 |
| 4_1415416 | 2 | 0.5 | NA | NA | NA | NA | 1 | 0 | 3 | 0.3333 | NA | NA | 1 | 1 | 1 | 0 | 1 | 0 | 3 | 0.3333 |
| 4_1415416 | 3 | 0.3333 | NA | NA | NA | NA | 2 | 0.5 | 5 | 0.4 | 2 | 0 | 1 | 0 | 1 | 1 | 2 | 0 | 6 | 0.1667 |
| 4_1415859 | 37 | 0.1081 | 40 | 0.05 | 28 | 0.0357 | 19 | 0.0526 | 124 | 0.0645 | 38 | 0.0263 | 18 | 0.1111 | 15 | 0 | 24 | 0.0833 | 95 | 0.0526 |
| 4_1415862 | 31 | 0.1613 | 31 | 0.129 | 26 | 0.1923 | 15 | 0.1333 | 103 | 0.1553 | 28 | 0.1429 | 15 | 0.0667 | 13 | 0.2308 | 20 | 0.05 | 76 | 0.1184 |
| 4_1415866 | 25 | 0.32 | 25 | 0.16 | 20 | 0.15 | 11 | 0.1818 | 81 | 0.2099 | 26 | 0.1923 | 9 | 0.2222 | 10 | 0.3 | 20 | 0.15 | 65 | 0.2 |
| 4_1416022 | NA | NA | NA | NA | 1 | 1 | 2 | 0.5 | 3 | 0.6667 | NA | NA | NA | NA | NA | NA | 1 | 1 | 1 | 1 |
| 4_1416025 | NA | NA | NA | NA | 2 | 1 | 2 | 0.5 | 4 | 0.75 | NA | NA | NA | NA | 1 | 1 | 1 | 1 | 2 | 1 |
| 4_1416027 | NA | NA | NA | NA | 2 | 0.5 | 2 | 0 | 4 | 0.25 | NA | NA | NA | NA | NA | NA | 1 | 1 | 1 | 1 |
| 4_1416027 | NA | NA | NA | NA | 2 | 1 | 1 | 1 | 3 | 1 | NA | NA | NA | NA | 1 | 0 | 1 | 1 | 2 | 0.5 |
| 4_1416028 | NA | NA | NA | NA | 2 | 0.5 | 2 | 0.5 | 4 | 0.5 | NA | NA | NA | NA | 1 | 1 | NA | NA | 1 | 1 |
| 4_1416036 | NA | NA | NA | NA | 2 | 0 | 3 | 0.3333 | 5 | 0.2 | NA | NA | NA | NA | NA | NA | 1 | 1 | 1 | 1 |
| 4_1416039 | NA | NA | NA | NA | 2 | 0 | 3 | 0.3333 | 5 | 0.2 | NA | NA | NA | NA | NA | NA | 1 | 1 | 1 | 1 |
| 4_1416040 | NA | NA | NA | NA | 2 | 0 | 3 | 0.6667 | 5 | 0.4 | NA | NA | NA | NA | 1 | 0 | 1 | 1 | 2 | 0.5 |
| 4_1417585 | 2 | 1 | 1 | 1 | 1 | 1 | 1 | 1 | 5 | 1 | NA | NA | 1 | 1 | 1 | 0 | 2 | 0.5 | 4 | 0.5 |
| 4_1417586 | 2 | 1 | 1 | 1 | 1 | 1 | 1 | 1 | 5 | 1 | NA | NA | 1 | 1 | 1 | 1 | 2 | 1 | 4 | 1 |
| 4_1417587 | 2 | 1 | 1 | 0 | 1 | 0 | 1 | 1 | 5 | 0.6 | NA | NA | 1 | 1 | 1 | 1 | 2 | 1 | 4 | 1 |
| 4_1417589 | 2 | 1 | 1 | 1 | NA | NA | 1 | 1 | 4 | 1 | NA | NA | 1 | 1 | NA | NA | 1 | 1 | 2 | 1 |
| 4_1417589 | 2 | 1 | 1 | 1 | NA | NA | 1 | 1 | 4 | 1 | NA | NA | 1 | 1 | 1 | 1 | 2 | 0.5 | 4 | 0.75 |
| 4_1417590 | 1 | 1 | 1 | 0 | NA | NA | 1 | 0 | 3 | 0.3333 | NA | NA | 1 | 0 | 1 | 1 | 2 | 0 | 4 | 0.25 |
| 4_1417591 | 2 | 1 | 1 | 1 | NA | NA | 1 | 0 | 4 | 0.75 | NA | NA | 1 | 1 | 1 | 1 | 2 | 0.5 | 4 | 0.75 |
| 4_1417591 | 2 | 0.5 | 1 | 1 | NA | NA | 1 | 0 | 4 | 0.5 | NA | NA | 1 | 0 | 1 | 0 | 2 | 0.5 | 4 | 0.25 |
| 4_1417593 | 2 | 1 | 1 | 0 | NA | NA | 1 | 1 | 4 | 0.75 | NA | NA | 1 | 1 | 1 | 1 | 2 | 1 | 4 | 1 |
| 4_1417651 | 2 | 0.5 | 1 | 1 | NA | NA | 1 | 0 | 4 | 0.5 | NA | NA | 1 | 1 | 1 | 1 | NA | NA | 2 | 1 |
| 4_1418297 | 3 | 0.3333 | 3 | 0 | 1 | 0 | 3 | 0 | 10 | 0.1 | 2 | 0.5 | NA | NA | 2 | 0 | 2 | 0 | 6 | 0.1667 |
| 4_1418297 | 3 | 0 | 3 | 0 | 1 | 0 | 3 | 0.3333 | 10 | 0.1 | 2 | 0 | NA | NA | 2 | 0.5 | 1 | 0 | 5 | 0.2 |
| 4_1418299 | 2 | 0.5 | 4 | 0.5 | 1 | 0 | 3 | 0 | 10 | 0.3 | 2 | 0.5 | NA | NA | 2 | 0 | 4 | 0.25 | 8 | 0.25 |
| 4_1418302 | 1 | 1 | 3 | 0.3333 | 1 | 0 | 3 | 0.3333 | 8 | 0.375 | 2 | 1 | NA | NA | 2 | 0.5 | 4 | 0 | 8 | 0.375 |

| | | | | | | | | | | | | | | | | | | | | |
|---|---|---|---|---|---|---|---|---|---|---|---|---|---|---|---|---|---|---|---|---|
| 4_14183164 | NA | NA | 3 | 0.6667 | NA | NA | 1 | 0 | 4 | 0.5 | 1 | 0 | NA | NA | 1 | 1 | NA | NA | 2 | 0.5 |
| 4_14183167 | NA | NA | 2 | 1 | NA | NA | 1 | 0 | 3 | 0.6667 | 1 | 1 | NA | NA | NA | NA | NA | NA | 1 | 1 |
| 4_14183193 | NA | NA | 2 | 1 | NA | NA | 1 | 0 | 3 | 0.6667 | 1 | 1 | NA | NA | NA | NA | NA | NA | 1 | 1 |
| 4_14186667 | 30 | 0.1 | 9 | 0 | 19 | 0.0526 | 21 | 0.1429 | 79 | 0.0886 | 14 | 0.1429 | 24 | 0.25 | 23 | 0.087 | 12 | 0.0833 | 73 | 0.1507 |
| 4_14186825 | 15 | 0.4 | 8 | 0.625 | 13 | 0.6154 | 15 | 0.6 | 51 | 0.549 | 12 | 0.6667 | 16 | 0.75 | 15 | 0.6667 | 8 | 0.625 | 51 | 0.6863 |
| 4_14186850 | 13 | 0.6923 | 10 | 0.5 | 13 | 0.3846 | 14 | 0.4286 | 50 | 0.5 | 11 | 0.3636 | 12 | 0.25 | 16 | 0.4375 | 7 | 0.7143 | 46 | 0.413 |
| 4_14186894 | 12 | 0.9167 | 8 | 0.75 | 12 | 0.8333 | 13 | 0.6923 | 45 | 0.8 | 11 | 0.8182 | 9 | 0.7778 | 15 | 0.3333 | 4 | 0.75 | 39 | 0.6154 |
| 4_14283441 | 4 | 0.75 | 1 | 1 | 4 | 0.5 | 3 | 1 | 12 | 0.75 | NA | NA | 2 | 0.5 | 5 | 0.4 | 7 | 0.2857 | 14 | 0.3571 |
| 4_14283468 | 3 | 0 | NA | NA | 5 | 0.4 | 8 | 0.25 | 16 | 0.25 | NA | NA | 3 | 0 | 4 | 0.25 | 7 | 0 | 14 | 0.0714 |
| 4_14283479 | 3 | 0 | NA | NA | 5 | 0 | 8 | 0.125 | 16 | 0.0625 | NA | NA | 2 | 0 | 3 | 0 | 7 | 0.1429 | 12 | 0.0833 |
| 4_14283491 | 3 | 0.6667 | NA | NA | 5 | 0.2 | 7 | 0.1429 | 15 | 0.2667 | NA | NA | 3 | 0 | 5 | 0.2 | 7 | 0 | 15 | 0.0667 |
| 4_14283493 | 2 | 0 | NA | NA | 4 | 0.5 | 6 | 0.1667 | 12 | 0.25 | NA | NA | 2 | 0.5 | 4 | 0.25 | 5 | 0 | 11 | 0.1818 |
| 4_14283557 | 2 | 1 | NA | NA | 3 | 1 | 7 | 0.8571 | 12 | 0.9167 | NA | NA | 2 | 0 | 4 | 1 | 3 | 1 | 9 | 0.7778 |
| 4_14283570 | 3 | 0.3333 | NA | NA | 4 | 1 | 8 | 0.5 | 15 | 0.6 | NA | NA | 2 | 0 | 4 | 0.75 | 5 | 0.6 | 11 | 0.5455 |
| 4_14285416 | NA | NA | 1 | 1 | NA | NA | NA | NA | 1 | 1 | 1 | 1 | NA | NA | NA | NA | 1 | 0 | 2 | 0.5 |
| 4_14285828 | 3 | 0 | NA | NA | 6 | 0.3333 | 6 | 0.1667 | 15 | 0.2 | NA | NA | 2 | 1 | 2 | 0 | 5 | 0 | 9 | 0.2222 |
| 4_14285833 | 3 | 0 | NA | NA | 5 | 0 | 6 | 0.1667 | 14 | 0.0714 | NA | NA | 2 | 0.5 | 1 | 0 | 4 | 0.25 | 7 | 0.2857 |
| 4_14285841 | 3 | 0 | NA | NA | 6 | 0 | 6 | 0.3333 | 15 | 0.1333 | NA | NA | 3 | 0 | 1 | 0 | 4 | 0.25 | 8 | 0.125 |
| 4_14285841 | 3 | 0 | NA | NA | 6 | 0.1667 | 5 | 0 | 14 | 0.0714 | NA | NA | 2 | 0.5 | 1 | 0 | 4 | 0 | 7 | 0.1429 |
| 4_14285844 | 3 | 0.3333 | NA | NA | 6 | 0.1667 | 6 | 0 | 15 | 0.1333 | NA | NA | 2 | 1 | NA | NA | 4 | 0 | 6 | 0.3333 |
| 4_14285897 | 3 | 0.3333 | NA | NA | 5 | 0.8 | 3 | 0 | 11 | 0.4545 | NA | NA | 2 | 1 | 1 | 0 | 4 | 1 | 7 | 0.8571 |
| 4_14285911 | 2 | 1 | NA | NA | 5 | 1 | 3 | 0.6667 | 10 | 0.9 | NA | NA | 2 | 0.5 | 1 | 0 | 4 | 0.75 | 7 | 0.5714 |
| 4_14285933 | 3 | 0.6667 | NA | NA | 3 | 1 | 3 | 1 | 9 | 0.8889 | NA | NA | 2 | 1 | 1 | 1 | 3 | 0.3333 | 6 | 0.6667 |
| 4_14285987 | 3 | 0.6667 | NA | NA | 3 | 1 | 4 | 0.25 | 10 | 0.6 | NA | NA | 3 | 0.3333 | 1 | 0 | 4 | 0.75 | 8 | 0.5 |
| 4_14285992 | 1 | 1 | NA | NA | 4 | 1 | 4 | 0.5 | 9 | 0.7778 | NA | NA | 2 | 1 | NA | NA | 3 | 1 | 5 | 1 |
| 4_14286052 | 2 | 0.5 | NA | NA | 2 | 0 | 1 | 1 | 5 | 0.4 | NA | NA | 2 | 1 | 1 | 0 | 2 | 0 | 5 | 0.4 |
| 4_14286053 | 2 | 0.5 | NA | NA | 2 | 1 | 1 | 0 | 5 | 0.6 | NA | NA | 2 | 0 | 1 | 0 | 2 | 1 | 5 | 0.4 |
| 4_14286101 | 2 | 0 | NA | NA | 2 | 0.5 | 1 | 0 | 5 | 0.2 | NA | NA | NA | NA | 1 | 1 | 2 | 0.5 | 3 | 0.6667 |
| 4_14286104 | 2 | 0.5 | NA | NA | 2 | 1 | 1 | 1 | 5 | 0.8 | NA | NA | 1 | 0 | 1 | 0 | 2 | 0.5 | 4 | 0.25 |
| 4_14286120 | 2 | 0.5 | NA | NA | 3 | 0.3333 | NA | NA | 5 | 0.4 | NA | NA | 1 | 0 | 1 | 0 | 2 | 0.5 | 4 | 0.25 |
| 4_14286134 | 1 | 1 | NA | NA | 2 | 1 | 1 | 0 | 4 | 0.75 | NA | NA | 1 | 0 | NA | NA | 2 | 0.5 | 3 | 0.3333 |
| 4_14286138 | 2 | 0.5 | NA | NA | 2 | 0 | 1 | 1 | 5 | 0.4 | NA | NA | 1 | 1 | NA | NA | 1 | 0 | 2 | 0.5 |
| 4_14286148 | 2 | 1 | NA | NA | 1 | 0 | 1 | 1 | 4 | 0.75 | NA | NA | 1 | 1 | NA | NA | 2 | 1 | 3 | 1 |
| 4_14286150 | 1 | 1 | NA | NA | 1 | 1 | 1 | 1 | 3 | 1 | NA | NA | 1 | 1 | NA | NA | 2 | 1 | 3 | 1 |
| 4_14286163 | 2 | 1 | NA | NA | 1 | 1 | 1 | 1 | 4 | 1 | NA | NA | 1 | 1 | NA | NA | 2 | 1 | 3 | 1 |
| 4_14286182 | 2 | 0.5 | NA | NA | NA | NA | 1 | 0 | 3 | 0.3333 | NA | NA | 1 | 0 | NA | NA | 2 | 0.5 | 3 | 0.3333 |
| 4_14299408 | 7 | 0.2857 | 4 | 0.5 | 3 | 0 | 2 | 0 | 16 | 0.25 | 2 | 0.5 | NA | NA | 2 | 0.5 | NA | NA | 4 | 0.5 |
| 4_14299410 | 7 | 0.1429 | 5 | 0.2 | 3 | 0.3333 | NA | NA | 15 | 0.2 | 2 | 0 | 1 | 0 | 2 | 0 | 1 | 1 | 6 | 0.1667 |
| 4_14299427 | 5 | 0.6 | 2 | 1 | 4 | 0.5 | 1 | 1 | 12 | 0.6667 | 3 | 0.3333 | 1 | 1 | 1 | 1 | 1 | 1 | 6 | 0.6667 |
| 4_14299429 | 7 | 0.8571 | 3 | 1 | 4 | 0.75 | 1 | 1 | 15 | 0.8667 | 2 | 1 | 1 | 1 | 1 | 1 | 1 | 1 | 5 | 1 |
| 4_14299441 | 6 | 0.6667 | 3 | 0.3333 | 4 | 0.75 | 1 | 1 | 14 | 0.6429 | 3 | 0.6667 | 1 | 1 | 1 | 1 | 1 | 1 | 6 | 0.8333 |
| 4_14299467 | 6 | 0.1667 | 3 | 0 | 3 | 0 | 1 | 0 | 13 | 0.0769 | 3 | 0.3333 | 1 | 0 | 1 | 0 | NA | NA | 5 | 0.2 |
| 4_14299473 | 6 | 0.5 | 4 | 0.25 | 3 | 0 | 1 | 0 | 14 | 0.2857 | 3 | 0 | NA | NA | 1 | 1 | 1 | 0 | 5 | 0.2 |
| 4_14299477 | 5 | 0.2 | 3 | 0 | 3 | 0 | 1 | 0 | 12 | 0.0833 | 3 | 0 | NA | NA | 1 | 1 | 1 | 0 | 5 | 0.2 |
| 4_14309741 | 1 | 1 | NA | NA | 1 | 1 | NA | NA | 2 | 1 | NA | NA | NA | NA | 1 | 1 | NA | NA | 1 | 1 |
| 4_14314328 | 1 | 1 | NA | NA | NA | NA | NA | NA | 1 | 1 | 3 | 0.6667 | 1 | 0 | NA | NA | NA | NA | 4 | 0.5 |
| 4_14314403 | 1 | 1 | NA | NA | NA | NA | NA | NA | 1 | 1 | 3 | 1 | 1 | 1 | NA | NA | NA | NA | 4 | 1 |
| 4_14314433 | 1 | 1 | NA | NA | NA | NA | NA | NA | 1 | 1 | 3 | 0.3333 | 1 | 0 | NA | NA | 1 | 0 | 5 | 0.2 |
| 4_14314436 | 1 | 1 | NA | NA | NA | NA | NA | NA | 1 | 1 | 3 | 0.3333 | 2 | 0 | NA | NA | NA | NA | 5 | 0.2 |
| 4_14314441 | 1 | 1 | NA | NA | NA | NA | NA | NA | 1 | 1 | 3 | 0 | 2 | 0 | NA | NA | 1 | 1 | 6 | 0.1667 |
| 4_14314546 | 2 | 0.5 | NA | NA | NA | NA | NA | NA | 2 | 0.5 | 3 | 1 | 2 | 0.5 | NA | NA | NA | NA | 5 | 0.8 |
| 4_14314552 | 2 | 1 | NA | NA | NA | NA | NA | NA | 2 | 1 | 3 | 1 | 2 | 1 | NA | NA | NA | NA | 5 | 1 |
| 4_14314554 | 2 | 1 | NA | NA | NA | NA | NA | NA | 2 | 1 | 4 | 0.5 | 2 | 1 | NA | NA | 1 | 0 | 7 | 0.5714 |
| 4_14314562 | 2 | 1 | NA | NA | NA | NA | NA | NA | 2 | 1 | 4 | 1 | 2 | 1 | NA | NA | 1 | 0 | 7 | 0.8571 |
| 4_14314569 | 2 | 0.5 | NA | NA | NA | NA | NA | NA | 2 | 0.5 | 4 | 0.5 | 2 | 0 | NA | NA | 1 | 0 | 7 | 0.2857 |
| 4_14314570 | 2 | 0.5 | NA | NA | NA | NA | NA | NA | 2 | 0.5 | 3 | 0.3333 | 2 | 0 | NA | NA | 1 | 0 | 6 | 0.1667 |
| 4_14314572 | 2 | 1 | NA | NA | NA | NA | NA | NA | 2 | 1 | 4 | 1 | 2 | 1 | NA | NA | 1 | 1 | 7 | 1 |
| 4_14314580 | 2 | 0.5 | NA | NA | NA | NA | NA | NA | 2 | 0.5 | 4 | 0 | 2 | 0.5 | NA | NA | 1 | 0 | 7 | 0.1429 |
| 4_14314585 | 2 | 0.5 | NA | NA | 1 | 0 | NA | NA | 3 | 0.3333 | 4 | 0.75 | 2 | 0 | NA | NA | 1 | 0 | 7 | 0.4286 |
| 4_14314595 | 1 | 1 | NA | NA | 1 | 1 | NA | NA | 2 | 1 | 2 | 1 | 1 | 1 | NA | NA | NA | NA | 3 | 1 |
| 4_14314602 | 2 | 0.5 | NA | NA | 1 | 0 | NA | NA | 3 | 0.3333 | 4 | 0 | 1 | 0 | NA | NA | 1 | 1 | 6 | 0.1667 |
| 4_14314611 | 1 | 1 | NA | NA | 1 | 1 | NA | NA | 2 | 1 | 4 | 1 | 2 | 1 | NA | NA | NA | NA | 6 | 1 |
| 4_14314614 | 2 | 0.5 | NA | NA | 1 | 1 | NA | NA | 3 | 0.6667 | 3 | 0.3333 | 2 | 0.5 | NA | NA | 1 | 1 | 6 | 0.5 |
| 4_14314616 | 2 | 1 | NA | NA | 1 | 0 | NA | NA | 3 | 0.6667 | 4 | 0.75 | 1 | 0 | NA | NA | NA | NA | 5 | 0.6 |
| 4_14314645 | 2 | 0.5 | NA | NA | 1 | 1 | 1 | 0 | 4 | 0.5 | 4 | 0.75 | 3 | 0.6667 | NA | NA | 1 | 0 | 8 | 0.625 |

| | | | | | | | | | | | | | | | | | | | | |
|---|---|---|---|---|---|---|---|---|---|---|---|---|---|---|---|---|---|---|---|---|
| 4_14314685 | 2 | 0.5 | NA | NA | 1 | 0 | 2 | 0.5 | 5 | 0.4 | 4 | 0.75 | 2 | 1 | NA | NA | 1 | 1 | 7 | 0.8571 |
| 4_14314699 | 2 | 0 | NA | NA | 1 | 0 | 1 | 1 | 4 | 0.25 | 3 | 0.3333 | 3 | 0.3333 | NA | NA | NA | NA | 6 | 0.3333 |
| 4_14314755 | 1 | 0 | NA | NA | 2 | 0.5 | 3 | 0.3333 | 6 | 0.3333 | 5 | 0 | 3 | 0.6667 | NA | NA | NA | NA | 8 | 0.25 |
| 4_14465375 | 7 | 0 | 2 | 1 | 3 | 0 | 6 | 0 | 18 | 0.1111 | 3 | 0.3333 | 2 | 0 | 2 | 0 | 1 | 0 | 8 | 0.125 |
| 4_14465378 | 7 | 0 | 2 | 0 | 1 | 1 | 6 | 0.1667 | 16 | 0.125 | 3 | 0 | 2 | 0 | 1 | 0 | 1 | 1 | 7 | 0.1429 |
| 4_14465389 | 6 | 0.8333 | 1 | 1 | 4 | 1 | 6 | 1 | 17 | 0.9412 | 3 | 1 | 2 | 1 | 2 | 0.5 | 3 | 1 | 10 | 0.9 |
| 4_14465394 | 6 | 0.3333 | 2 | 0.5 | 4 | 0.5 | 6 | 0.3333 | 18 | 0.3889 | 2 | 0 | 2 | 0 | 2 | 0 | 3 | 0.3333 | 9 | 0.1111 |
| 4_14465396 | 5 | 0.6 | 2 | 1 | 4 | 0.75 | 4 | 1 | 15 | 0.8 | 3 | 1 | 2 | 1 | 1 | 1 | 2 | 1 | 8 | 1 |
| 4_14469925 | 4 | 0.25 | 3 | 0 | 1 | 0 | 4 | 0.25 | 12 | 0.1667 | 4 | 0.5 | 4 | 0.5 | 9 | 0.1111 | 3 | 0 | 20 | 0.25 |
| 4_14469953 | 3 | 1 | 3 | 0.3333 | 1 | 1 | 3 | 0.6667 | 10 | 0.7 | 4 | 0.5 | 4 | 0.5 | 9 | 0.1111 | 2 | 1 | 19 | 0.3684 |
| 4_14469961 | 3 | 1 | 4 | 0.75 | 1 | 1 | 3 | 1 | 11 | 0.9091 | 3 | 1 | 4 | 0.75 | 9 | 0.7778 | 3 | 1 | 19 | 0.8421 |
| 4_14470023 | 3 | 0 | 3 | 0 | 1 | 0 | 3 | 0.3333 | 10 | 0.1 | 4 | 0 | 4 | 0 | 7 | 0.1429 | 3 | 0 | 18 | 0.0556 |
| 4_14470033 | 2 | 0 | NA | NA | NA | NA | 2 | 0.5 | 4 | 0.25 | 2 | 0 | 4 | 0.25 | 3 | 0 | 2 | 1 | 11 | 0.2727 |
| 4_14470131 | 2 | 0.5 | NA | NA | NA | NA | NA | NA | 2 | 0.5 | NA | NA | 1 | 1 | 3 | 0.3333 | 2 | 0.5 | 6 | 0.5 |
| 4_14470164 | 5 | 0.6 | 3 | 0 | 3 | 0 | 1 | 0 | 12 | 0.25 | 5 | 0.2 | 3 | 0 | 4 | 0 | 2 | 0 | 14 | 0.0714 |
| 4_14470171 | 4 | 0.25 | 2 | 0.5 | 1 | 0 | 1 | 0 | 8 | 0.25 | 4 | 0.25 | 3 | 1 | 3 | 0.3333 | 1 | 1 | 11 | 0.5455 |
| 4_14470174 | 5 | 0.4 | 3 | 0.3333 | 3 | 0 | NA | NA | 11 | 0.2727 | 5 | 0.4 | 3 | 0.6667 | 4 | 0 | 2 | 1 | 14 | 0.4286 |
| 4_14470176 | 5 | 0.2 | 2 | 0 | 3 | 0 | 1 | 0 | 11 | 0.0909 | 5 | 0 | 3 | 0 | 2 | 0 | 2 | 0.5 | 12 | 0.0833 |
| 4_14470178 | 4 | 0.25 | 3 | 0 | 2 | 0.5 | 1 | 0 | 10 | 0.2 | 3 | 0.3333 | 3 | 0 | 3 | 0 | 2 | 0 | 11 | 0.0909 |
| 4_14487502 | NA | NA | 2 | 0.5 | NA | NA | NA | NA | 2 | 0.5 | NA | NA | 1 | 1 | NA | NA | NA | NA | 1 | 1 |
| 4_14487555 | NA | NA | 2 | 0.5 | NA | NA | NA | NA | 2 | 0.5 | NA | NA | 1 | 1 | NA | NA | NA | NA | 1 | 1 |
| 4_14487561 | NA | NA | 2 | 1 | NA | NA | NA | NA | 2 | 1 | NA | NA | 1 | 1 | NA | NA | NA | NA | 1 | 1 |
| 4_14488130 | NA | NA | 1 | 1 | NA | NA | NA | NA | 1 | 1 | NA | NA | 1 | 1 | NA | NA | NA | NA | 1 | 1 |
| 4_14488138 | NA | NA | 1 | 1 | NA | NA | NA | NA | 1 | 1 | NA | NA | 1 | 1 | NA | NA | NA | NA | 1 | 1 |
| 4_14488205 | NA | NA | 1 | 1 | NA | NA | NA | NA | 1 | 1 | NA | NA | 1 | 1 | NA | NA | NA | NA | 1 | 1 |
| 4_14488222 | NA | NA | 1 | 1 | NA | NA | NA | NA | 1 | 1 | NA | NA | 1 | 1 | NA | NA | NA | NA | 1 | 1 |
| 4_14488279 | NA | NA | 1 | 1 | NA | NA | NA | NA | 1 | 1 | NA | NA | 1 | 1 | NA | NA | NA | NA | 1 | 1 |
| 4_14555591 | 2 | 0.5 | NA | NA | NA | NA | NA | NA | 2 | 0.5 | NA | NA | 1 | 1 | NA | NA | NA | NA | 1 | 1 |
| 4_14555612 | 1 | 1 | NA | NA | NA | NA | NA | NA | 1 | 1 | NA | NA | 1 | 1 | NA | NA | NA | NA | 1 | 1 |
| 4_14726288 | NA | NA | 1 | 1 | NA | NA | NA | NA | 1 | 1 | NA | NA | NA | NA | 1 | 1 | NA | NA | 1 | 1 |
| 4_14726297 | NA | NA | 1 | 1 | NA | NA | NA | NA | 1 | 1 | NA | NA | NA | NA | 1 | 1 | NA | NA | 1 | 1 |
| 4_14726443 | NA | NA | 1 | 1 | NA | NA | 1 | 1 | 2 | 1 | NA | NA | NA | NA | 2 | 0.5 | NA | NA | 2 | 0.5 |
| 4_14726449 | NA | NA | 1 | 0 | NA | NA | 1 | 1 | 2 | 0.5 | NA | NA | NA | NA | 1 | 1 | NA | NA | 1 | 1 |
| 4_14726482 | NA | NA | 1 | 1 | NA | NA | 1 | 1 | 2 | 1 | NA | NA | NA | NA | 2 | 0.5 | NA | NA | 2 | 0.5 |
| 4_14790632 | 1 | 0 | 1 | 1 | NA | NA | NA | NA | 2 | 0.5 | NA | NA | 2 | 1 | 1 | 1 | 2 | 0.5 | 5 | 0.8 |
| 4_14790691 | 1 | 1 | 1 | 0 | NA | NA | NA | NA | 2 | 0.5 | NA | NA | 2 | 0 | NA | NA | 2 | 0.5 | 4 | 0.25 |
| 4_14790907 | NA | NA | 2 | 1 | 1 | 0 | 1 | 0 | 4 | 0.5 | NA | NA | NA | NA | 3 | 0.3333 | 4 | 0.25 | 7 | 0.2857 |
| 4_14790918 | NA | NA | 2 | 0.5 | 1 | 0 | 1 | 0 | 4 | 0.25 | NA | NA | NA | NA | 3 | 0.3333 | 4 | 0 | 7 | 0.1429 |
| 4_14790989 | NA | NA | 2 | 0 | 1 | 1 | 2 | 0 | 5 | 0.2 | NA | NA | 1 | 1 | 3 | 1 | 2 | 0.5 | 6 | 0.8333 |
| 4_14790998 | NA | NA | 2 | 1 | 1 | 0 | 2 | 0 | 5 | 0.4 | NA | NA | 1 | 1 | 3 | 0.6667 | 2 | 1 | 6 | 0.8333 |
| 4_14791002 | NA | NA | 2 | 0 | 1 | 0 | 2 | 0.5 | 5 | 0.2 | NA | NA | 1 | 1 | 2 | 1 | 2 | 0 | 5 | 0.6 |
| 4_14791020 | NA | NA | 2 | 0.5 | 1 | 0 | 2 | 0 | 5 | 0.2 | NA | NA | 1 | 0 | 3 | 0.6667 | 2 | 0.5 | 6 | 0.5 |
| 4_14791036 | NA | NA | 2 | 0 | 1 | 0 | 2 | 0.5 | 5 | 0.2 | NA | NA | 1 | 0 | 2 | 0.5 | 1 | 0 | 4 | 0.25 |
| 4_14791038 | NA | NA | 2 | 1 | NA | NA | 2 | 1 | 4 | 1 | NA | NA | 1 | 1 | 3 | 1 | 1 | 1 | 5 | 1 |
| 4_14791063 | NA | NA | 2 | 1 | 1 | 1 | 2 | 1 | 5 | 1 | NA | NA | 1 | 1 | 3 | 1 | 2 | 1 | 6 | 1 |
| 4_14791080 | NA | NA | 1 | 0 | 1 | 0 | 1 | 1 | 3 | 0.3333 | NA | NA | 1 | 1 | 3 | 0 | 2 | 1 | 6 | 0.5 |
| 4_14791105 | NA | NA | 1 | 0 | 1 | 0 | 2 | 0.5 | 4 | 0.25 | NA | NA | 1 | 1 | 2 | 1 | 2 | 1 | 5 | 1 |
| 4_14791120 | NA | NA | 2 | 0.5 | 1 | 0 | 2 | 0 | 5 | 0.2 | NA | NA | 1 | 0 | 2 | 0.5 | 2 | 0 | 5 | 0.2 |
| 4_14791126 | NA | NA | 2 | 1 | 1 | 1 | 2 | 1 | 5 | 1 | NA | NA | 1 | 1 | 2 | 1 | 2 | 1 | 5 | 1 |
| 4_14794716 | NA | NA | 2 | 1 | 1 | 0 | 2 | 0.5 | 5 | 0.6 | 1 | 0 | 1 | 0 | 2 | 0.5 | 2 | 0.5 | 6 | 0.3333 |
| 4_14795029 | NA | NA | NA | NA | 2 | 0.5 | 2 | 0.5 | 4 | 0.5 | 1 | 0 | 1 | 0 | 3 | 0.6667 | 1 | 1 | 6 | 0.5 |
| 4_14795816 | 26 | 0.0769 | 31 | 0.0968 | 20 | 0.1 | 17 | 0.0588 | 94 | 0.0851 | 14 | 0.0714 | 22 | 0.1364 | 21 | 0.0952 | 20 | 0.1 | 77 | 0.1039 |
| 4_14795823 | 27 | 0.1481 | 29 | 0.4483 | 20 | 0.2 | 17 | 0.1765 | 93 | 0.2581 | 13 | 0.2308 | 24 | 0.2917 | 18 | 0.3333 | 18 | 0.3889 | 73 | 0.3151 |
| 4_14795849 | 24 | 0.1667 | 27 | 0.1852 | 15 | 0 | 8 | 0 | 74 | 0.1216 | 12 | 0 | 18 | 0.1111 | 17 | 0.0588 | 17 | 0.1176 | 64 | 0.0781 |
| 4_14795870 | 21 | 0.1905 | 27 | 0.3704 | 16 | 0.3125 | 10 | 0.2 | 74 | 0.2838 | 12 | 0.0833 | 15 | 0.1333 | 17 | 0.0588 | 15 | 0.1333 | 59 | 0.1017 |
| 4_14795890 | 19 | 0.1053 | 26 | 0.0385 | 14 | 0.0714 | 12 | 0.0833 | 71 | 0.0704 | 11 | 0.2727 | 20 | 0 | 14 | 0.0714 | 15 | 0.1333 | 60 | 0.1 |
| 4_14795892 | 22 | 0.8182 | 26 | 0.8846 | 15 | 0.9333 | 10 | 0.8 | 73 | 0.863 | 10 | 0.9 | 17 | 0.8824 | 17 | 0.8824 | 13 | 0.8462 | 57 | 0.8772 |
| 4_14795918 | 19 | 0.1579 | 25 | 0.08 | 15 | 0.2 | 7 | 0.1429 | 66 | 0.1364 | 9 | 0 | 20 | 0.2 | 14 | 0 | 14 | 0.2143 | 57 | 0.1228 |
| 4_14795924 | 19 | 0.1053 | 24 | 0.0833 | 15 | 0.1333 | 7 | 0 | 65 | 0.0923 | 10 | 0.1 | 18 | 0.0556 | 16 | 0 | 12 | 0.0833 | 56 | 0.0536 |
| 4_14795931 | 17 | 0.2353 | 21 | 0.0952 | 16 | 0.25 | 9 | 0.1111 | 63 | 0.1746 | 9 | 0.1111 | 17 | 0.2353 | 14 | 0.2143 | 11 | 0.4545 | 51 | 0.2549 |
| 4_14795941 | 17 | 0.3529 | 23 | 0.6087 | 14 | 0.7857 | 7 | 0.7143 | 61 | 0.5902 | 8 | 0.625 | 16 | 0.6875 | 13 | 0.5385 | 14 | 0.5714 | 51 | 0.6078 |
| 4_14795948 | 14 | 1 | 16 | 0.9375 | 11 | 0.9091 | 8 | 0.625 | 49 | 0.898 | 9 | 0.7778 | 14 | 0.8571 | 8 | 0.75 | 13 | 0.9231 | 44 | 0.8409 |
| 4_14795949 | 1 | 1 | 1 | 1 | NA | NA | NA | NA | 2 | 1 | NA | NA | 1 | 1 | 1 | 1 | NA | NA | 2 | 1 |
| 4_14795956 | 15 | 0.9333 | 18 | 0.8889 | 10 | 0.8 | 7 | 0.5714 | 50 | 0.84 | 9 | 0.6667 | 12 | 0.75 | 12 | 0.9167 | 11 | 0.6364 | 44 | 0.75 |
| 4_14795967 | 7 | 0.7143 | 21 | 0.8571 | 9 | 0.5556 | 5 | 0.6 | 42 | 0.7381 | 3 | 0.3333 | 14 | 0.6429 | 12 | 0.3333 | 10 | 0.7 | 39 | 0.5385 |

| | | | | | | | | | | | | | | | | | | | | |
|---|---|---|---|---|---|---|---|---|---|---|---|---|---|---|---|---|---|---|---|---|
| 4_1479596 | 16 | 1 | 20 | 0.8 | 11 | 0.8182 | 7 | 1 | 54 | 0.8889 | 7 | 1 | 15 | 0.9333 | 12 | 0.8333 | 12 | 0.9167 | 46 | 0.913 |
| 4_1479665 | 11 | 0.1818 | 17 | 0.1765 | 13 | 0.1538 | 6 | 0.1667 | 47 | 0.1702 | 4 | 0.25 | 15 | 0.2 | 9 | 0.2222 | 12 | 0.25 | 40 | 0.225 |
| 4_1479946 | 2 | 0 | 3 | 0.6667 | NA | NA | 4 | 0.25 | 9 | 0.3333 | 3 | 0.3333 | 2 | 1 | NA | NA | NA | NA | 5 | 0.6 |
| 4_1479947 | 2 | 1 | 3 | 1 | 1 | 1 | 5 | 1 | 11 | 1 | 3 | 0.6667 | 2 | 1 | NA | NA | NA | NA | 5 | 0.8 |
| 4_1479949 | 3 | 0 | 3 | 0 | 1 | 0 | 4 | 0.25 | 11 | 0.0909 | 3 | 0.3333 | 1 | 0 | NA | NA | NA | NA | 4 | 0.25 |
| 4_1479949 | 3 | 0.3333 | 3 | 0 | 1 | 0 | 5 | 0 | 12 | 0.0833 | 3 | 0 | 2 | 0.5 | NA | NA | NA | NA | 5 | 0.2 |
| 4_1479950 | 3 | 0.3333 | 2 | 1 | NA | NA | 5 | 0.6 | 10 | 0.6 | 3 | 0.3333 | 2 | 1 | NA | NA | NA | NA | 5 | 0.6 |
| 4_1479950 | 2 | 0.5 | 2 | 0.5 | NA | NA | 3 | 0 | 7 | 0.2857 | 2 | 0 | 2 | 1 | NA | NA | NA | NA | 4 | 0.5 |
| 4_1480258 | 1 | 0 | 2 | 1 | NA | NA | 2 | 0 | 5 | 0.4 | 1 | 1 | 1 | 0 | 1 | 0 | 2 | 0.5 | 5 | 0.4 |
| 4_1480259 | 1 | 0 | 2 | 0.5 | NA | NA | 2 | 0 | 5 | 0.2 | NA | NA | 3 | 0.3333 | 1 | 1 | 2 | 0.5 | 6 | 0.5 |
| 4_1480260 | NA | NA | 3 | 1 | NA | NA | 3 | 0.3333 | 6 | 0.6667 | 1 | 1 | 3 | 0.6667 | 1 | 1 | 1 | 1 | 6 | 0.8333 |
| 4_1480260 | 2 | 0 | 2 | 1 | NA | NA | 3 | 0.6667 | 7 | 0.5714 | NA | NA | 3 | 0.6667 | 1 | 1 | 2 | 0.5 | 6 | 0.6667 |
| 4_1480261 | NA | NA | 2 | 1 | NA | NA | 2 | 1 | 4 | 1 | NA | NA | 1 | 1 | 1 | 1 | 2 | 1 | 4 | 1 |
| 4_1480262 | 4 | 0 | 2 | 0.5 | NA | NA | 3 | 0.3333 | 9 | 0.2222 | NA | NA | 3 | 0.6667 | 1 | 1 | 1 | 0 | 5 | 0.6 |
| 4_1480263 | 2 | 0.5 | 2 | 0 | NA | NA | 3 | 0 | 7 | 0.1429 | 1 | 0 | 3 | 0.3333 | 1 | 0 | 2 | 0.5 | 7 | 0.2857 |
| 4_1480264 | 4 | 0.75 | 3 | 0.6667 | NA | NA | 3 | 0.3333 | 10 | 0.6 | 1 | 1 | 3 | 1 | 1 | 0 | 2 | 0 | 7 | 0.5714 |
| 4_1480297 | 3 | 1 | 3 | 0.6667 | 2 | 1 | 2 | 1 | 10 | 0.9 | 1 | 1 | 3 | 1 | 1 | 1 | 1 | 1 | 6 | 1 |
| 4_1480311 | 6 | 0.3333 | 5 | 0.2 | 2 | 0 | 3 | 0 | 16 | 0.1875 | 3 | 0.6667 | 2 | 0.5 | 1 | 0 | 2 | 0 | 8 | 0.375 |
| 4_1480311 | 1 | 1 | 4 | 0.5 | 2 | 1 | 2 | 1 | 9 | 0.7778 | 1 | 1 | 3 | 0.3333 | NA | NA | 2 | 0.5 | 6 | 0.5 |
| 4_1480312 | 5 | 0.8 | 5 | 1 | 2 | 1 | 3 | 0.6667 | 15 | 0.8667 | 3 | 1 | 2 | 0.5 | 1 | 1 | 3 | 1 | 9 | 0.8889 |
| 4_1480314 | 4 | 1 | 4 | 1 | NA | NA | 2 | 1 | 10 | 1 | 3 | 1 | 3 | 0.3333 | 1 | 1 | 3 | 0.6667 | 10 | 0.7 |
| 4_1480889 | 12 | 0.4167 | 12 | 0.5833 | 8 | 0.75 | 13 | 0.8462 | 45 | 0.6444 | 9 | 0.6667 | 7 | 0.5714 | 6 | 0.8333 | 6 | 0.5 | 28 | 0.6429 |
| 4_1480890 | 8 | 0.5 | 12 | 0.3333 | 6 | 0.3333 | 12 | 0.6667 | 38 | 0.4737 | 7 | 0.1429 | 8 | 0.25 | 7 | 0 | 6 | 0.8333 | 28 | 0.2857 |
| 4_1480892 | 9 | 0 | 11 | 0.0909 | 6 | 0.1667 | 11 | 0.0909 | 37 | 0.0811 | 7 | 0.1429 | 8 | 0.125 | 6 | 0 | 4 | 0 | 25 | 0.08 |
| 4_1480895 | 7 | 0.1429 | 8 | 0.25 | 7 | 0.1429 | 11 | 0.2727 | 33 | 0.2121 | 6 | 0.3333 | 6 | 0.3333 | 4 | 0.75 | 4 | 0.25 | 20 | 0.4 |
| 4_1480899 | 8 | 0.25 | 6 | 0.5 | 7 | 0.4286 | 11 | 0.1818 | 32 | 0.3125 | 6 | 0.5 | 8 | 0.375 | 5 | 0 | 2 | 0 | 21 | 0.2857 |
| 4_1480900 | 8 | 0.25 | 10 | 0.6 | 6 | 0.3333 | 12 | 0.25 | 36 | 0.3611 | 4 | 0.5 | 9 | 0.3333 | 6 | 0.5 | 3 | 0.3333 | 22 | 0.4091 |
| 4_1480902 | 9 | 0.1111 | 10 | 0.6 | 5 | 0.2 | 10 | 0.2 | 34 | 0.2941 | 6 | 0.3333 | 9 | 0.1111 | 6 | 0.3333 | 4 | 0.25 | 25 | 0.24 |
| 4_1480904 | 8 | 0 | 7 | 0.5714 | 5 | 0 | 10 | 0.2 | 30 | 0.2 | 7 | 0.2857 | 7 | 0.1429 | 6 | 0 | 3 | 0 | 23 | 0.1304 |
| 4_1480905 | 6 | 0 | 7 | 0 | 5 | 0 | 11 | 0.1818 | 29 | 0.069 | 7 | 0.1429 | 6 | 0 | 5 | 0 | 3 | 0.3333 | 21 | 0.0952 |
| 4_1480905 | 5 | 0 | 8 | 0.375 | 6 | 0.1667 | 5 | 0 | 24 | 0.1667 | 5 | 0.2 | 5 | 0.2 | 5 | 0 | 3 | 0.6667 | 18 | 0.2222 |
| 4_1480906 | 5 | 0.4 | 7 | 0.4286 | 8 | 0.375 | 11 | 0.2727 | 31 | 0.3548 | 6 | 0.5 | 7 | 0.2857 | 7 | 0.1429 | 3 | 0 | 23 | 0.2609 |
| 4_1480907 | 5 | 0.2 | 8 | 0.125 | 7 | 0.2857 | 11 | 0.0909 | 31 | 0.1613 | 7 | 0.1429 | 6 | 0 | 7 | 0.1429 | 3 | 0.3333 | 23 | 0.1304 |
| 4_1480908 | 3 | 0.6667 | 6 | 0.6667 | 5 | 0.8 | 9 | 0.8889 | 23 | 0.7826 | 6 | 0.5 | 4 | 0.5 | 5 | 1 | 2 | 0.5 | 17 | 0.6471 |
| 4_1480908 | 6 | 1 | 9 | 0.8889 | 5 | 1 | 11 | 0.9091 | 31 | 0.9355 | 7 | 0.5714 | 4 | 0.75 | 7 | 1 | 3 | 1 | 21 | 0.8095 |
| 4_1480909 | 6 | 0.8333 | 9 | 0.8889 | 4 | 1 | 11 | 0.8182 | 30 | 0.8667 | 5 | 1 | 5 | 0.8 | 6 | 0.8333 | 2 | 1 | 18 | 0.8889 |
| 4_1480910 | 6 | 0.1667 | 9 | 0 | 4 | 0.25 | 11 | 0.0909 | 30 | 0.1 | 7 | 0 | 5 | 0.2 | 5 | 0 | 3 | 0 | 20 | 0.05 |
| 4_1481399 | 1 | 0 | 1 | 1 | 1 | 0 | 2 | 1 | 5 | 0.6 | 1 | 1 | NA | NA | 1 | 0 | NA | NA | 2 | 0.5 |
| 4_1481402 | 1 | 1 | 1 | 1 | 1 | 0 | 2 | 1 | 5 | 0.8 | 1 | 1 | NA | NA | 1 | 1 | NA | NA | 2 | 1 |
| 4_1481403 | NA | NA | NA | NA | NA | NA | 2 | 1 | 2 | 1 | 1 | 1 | NA | NA | NA | NA | NA | NA | 1 | 1 |
| 4_1481404 | 1 | 0 | 1 | 0 | 1 | 0 | 2 | 0.5 | 5 | 0.2 | 1 | 0 | NA | NA | 1 | 1 | NA | NA | 2 | 0.5 |
| 4_1481405 | 1 | 1 | 1 | 1 | 1 | 1 | 2 | 1 | 5 | 1 | 1 | 1 | NA | NA | 1 | 1 | NA | NA | 2 | 1 |
| 4_1481412 | 1 | 0 | 1 | 1 | 1 | 0 | 1 | 1 | 4 | 0.5 | 1 | 0 | NA | NA | 1 | 0 | 1 | 1 | 3 | 0.3333 |
| 4_1481413 | 1 | 0 | 1 | 0 | 1 | 1 | 1 | 0 | 4 | 0.25 | 1 | 0 | NA | NA | 2 | 0 | 1 | 1 | 4 | 0.25 |
| 4_1481414 | 1 | 0 | NA | NA | 1 | 1 | 1 | 0 | 3 | 0.3333 | 1 | 1 | NA | NA | 3 | 0.6667 | 1 | 1 | 5 | 0.8 |
| 4_1481414 | 1 | 0 | 1 | 1 | NA | NA | 1 | 0 | 3 | 0.3333 | 1 | 1 | NA | NA | 2 | 0.5 | 1 | 1 | 4 | 0.75 |
| 4_1481415 | 1 | 1 | 1 | 1 | 1 | 0 | 1 | 0 | 4 | 0.5 | 1 | 1 | NA | NA | 3 | 0.3333 | NA | NA | 4 | 0.5 |
| 4_1481416 | 1 | 0 | 1 | 1 | 1 | 0 | 1 | 0 | 4 | 0.25 | NA | NA | NA | NA | 2 | 1 | 1 | 0 | 3 | 0.6667 |
| 4_1481423 | 1 | 0 | 2 | 1 | NA | NA | 1 | 1 | 4 | 0.75 | 1 | 1 | NA | NA | 3 | 0 | 1 | 0 | 5 | 0.2 |
| 4_1481425 | 1 | 0 | 2 | 1 | 2 | 0 | 2 | 0.5 | 7 | 0.4286 | 1 | 1 | NA | NA | 1 | 0 | 1 | 0 | 3 | 0.3333 |
| 4_1481427 | 1 | 0 | 2 | 0.5 | 2 | 0.5 | 2 | 0 | 7 | 0.2857 | 1 | 0 | NA | NA | 4 | 0.75 | 1 | 0 | 6 | 0.5 |
| 4_1481427 | 1 | 0 | 2 | 1 | 2 | 1 | 2 | 0.5 | 7 | 0.7143 | 1 | 0 | NA | NA | 5 | 0.4 | 1 | 0 | 7 | 0.2857 |
| 4_1481427 | 1 | 0 | 2 | 1 | 2 | 0.5 | 2 | 0 | 7 | 0.4286 | 1 | 1 | NA | NA | 3 | 1 | 1 | 1 | 5 | 1 |
| 4_1481427 | NA | NA | 2 | 0.5 | 2 | 0 | 2 | 0 | 6 | 0.1667 | 1 | 0 | NA | NA | 5 | 0.4 | 1 | 0 | 7 | 0.2857 |
| 4_1481428 | 1 | 0 | 2 | 0.5 | 2 | 0 | 2 | 0 | 7 | 0.1429 | 1 | 0 | NA | NA | 5 | 0.2 | 1 | 0 | 7 | 0.1429 |
| 4_1481428 | 1 | 0 | 2 | 1 | 2 | 0 | 2 | 0.5 | 7 | 0.4286 | 1 | 1 | NA | NA | 4 | 0.25 | 1 | 0 | 6 | 0.3333 |
| 4_1481431 | 1 | 1 | 2 | 1 | 2 | 0.5 | 1 | 1 | 6 | 0.8333 | 2 | 0.5 | NA | NA | 5 | 0.4 | NA | NA | 7 | 0.4286 |
| 4_1481433 | 1 | 1 | 2 | 0.5 | 3 | 0.3333 | 2 | 0 | 8 | 0.375 | 2 | 0 | NA | NA | 3 | 0.3333 | 1 | 0 | 6 | 0.1667 |
| 4_1481433 | 1 | 1 | 2 | 0.5 | 3 | 0.6667 | 2 | 0.5 | 8 | 0.625 | 2 | 0.5 | NA | NA | 4 | 0.25 | 1 | 0 | 7 | 0.2857 |
| 4_1481435 | 1 | 1 | 1 | 1 | 3 | 1 | 1 | 1 | 6 | 1 | 1 | 1 | NA | NA | 5 | 1 | 1 | 1 | 7 | 1 |
| 4_1481437 | 2 | 0.5 | NA | NA | 2 | 0.5 | 2 | 0 | 6 | 0.3333 | 2 | 0.5 | NA | NA | 4 | 0.5 | 1 | 1 | 7 | 0.5714 |
| 4_1481437 | 2 | 1 | 1 | 1 | 3 | 0.6667 | 1 | 0 | 7 | 0.7143 | 2 | 0.5 | 1 | 1 | 5 | 0.4 | 2 | 1 | 10 | 0.6 |
| 4_1481438 | 2 | 1 | 2 | 0.5 | 3 | 0.3333 | 1 | 1 | 8 | 0.625 | 2 | 0 | 1 | 1 | 4 | 0.25 | 2 | 1 | 9 | 0.4444 |
| 4_1481439 | 2 | 0.5 | 2 | 0.5 | 3 | 0.3333 | 2 | 0.5 | 9 | 0.4444 | 2 | 0 | 1 | 1 | 5 | 0 | 2 | 0.5 | 10 | 0.2 |
| 4_1481648 | NA | NA | 1 | 1 | NA | NA | 2 | 0 | 3 | 0.3333 | NA | NA | NA | NA | 1 | 1 | NA | NA | 1 | 1 |

| | | | | | | | | | | | | | | | | | | | | |
|---|---|---|---|---|---|---|---|---|---|---|---|---|---|---|---|---|---|---|---|---|
| 4_1482304 | 5 | 0 | 3 | 0 | 4 | 0.25 | 2 | 0 | 14 | 0.0714 | 2 | 0 | 7 | 0.2857 | 2 | 1 | 8 | 0.125 | 19 | 0.2632 |
| 4_1482305 | 4 | 0.25 | 3 | 0.3333 | 3 | 0 | 2 | 0 | 12 | 0.1667 | 2 | 0.5 | 5 | 0.2 | 3 | 0 | 8 | 0.5 | 18 | 0.3333 |
| 4_1482376 | 12 | 0.0833 | 16 | 0.0625 | 12 | 0 | 15 | 0.0667 | 55 | 0.0545 | 17 | 0.1176 | 12 | 0.0833 | 16 | 0.0625 | 13 | 0 | 58 | 0.069 |
| 4_1482392 | NA | NA | NA | NA | 1 | 1 | NA | NA | 1 | 1 | 4 | 0.25 | 2 | 0 | 5 | 0.2 | 1 | 0 | 12 | 0.1667 |
| 4_1482492 | 8 | 0.125 | 7 | 0.2857 | 11 | 0.1818 | 9 | 0.2222 | 35 | 0.2 | 14 | 0.1429 | 4 | 0 | 6 | 0.1667 | 9 | 0.1111 | 33 | 0.1212 |
| 4_1482507 | 15 | 0.3333 | 12 | 0.1667 | 16 | 0.3125 | 15 | 0.1333 | 58 | 0.2414 | 20 | 0.25 | 14 | 0.2857 | 13 | 0.0769 | 15 | 0.2 | 62 | 0.2097 |
| 4_1483224 | 2 | 0.5 | 2 | 0.5 | 2 | 0.5 | NA | NA | 6 | 0.5 | 1 | 1 | NA | NA | NA | NA | 1 | 0 | 2 | 0.5 |
| 4_1483229 | 2 | 0.5 | 2 | 0 | 2 | 0 | NA | NA | 6 | 0.1667 | 1 | 1 | NA | NA | 1 | 0 | 1 | 0 | 3 | 0.3333 |
| 4_1483230 | 2 | 1 | 2 | 0.5 | 2 | 0.5 | NA | NA | 6 | 0.6667 | 1 | 1 | NA | NA | 1 | 0 | NA | NA | 2 | 0.5 |
| 4_1483237 | 1 | 1 | 2 | 0.5 | 1 | 0 | NA | NA | 4 | 0.5 | 1 | 1 | NA | NA | 1 | 0 | NA | NA | 2 | 0.5 |
| 4_1483238 | 2 | 1 | 2 | 0 | 1 | 0 | NA | NA | 5 | 0.4 | 1 | 1 | NA | NA | 1 | 0 | NA | NA | 2 | 0.5 |
| 4_1483241 | 3 | 1 | 2 | 1 | 1 | 0 | 4 | 1 | 10 | 0.9 | 1 | 1 | NA | NA | 1 | 1 | 1 | 1 | 3 | 1 |
| 4_1483241 | 3 | 1 | 2 | 0.5 | 1 | 0 | 4 | 0.75 | 10 | 0.7 | 1 | 0 | 1 | 1 | 1 | 1 | 1 | 1 | 4 | 0.75 |
| 4_1483242 | 1 | 1 | 1 | 0 | 1 | 1 | 4 | 1 | 7 | 0.8571 | 1 | 1 | 1 | 0 | 1 | 1 | 1 | 1 | 4 | 0.75 |
| 4_1483245 | 2 | 1 | 2 | 0.5 | 1 | 0 | 4 | 0.25 | 9 | 0.4444 | 1 | 0 | 1 | 1 | 1 | 0 | NA | NA | 3 | 0.3333 |
| 4_1483246 | 1 | 1 | 2 | 1 | 1 | 0 | 2 | 1 | 6 | 0.8333 | 1 | 1 | 1 | 1 | 1 | 1 | NA | NA | 3 | 1 |
| 4_1483246 | 2 | 0 | 2 | 1 | 1 | 1 | 3 | 0.3333 | 8 | 0.5 | 1 | 1 | 1 | 0 | 1 | 1 | NA | NA | 3 | 0.6667 |
| 4_1483246 | 2 | 1 | 2 | 0.5 | 1 | 1 | 3 | 0.3333 | 8 | 0.625 | NA | NA | 1 | 0 | 1 | 1 | NA | NA | 2 | 0.5 |
| 4_1483247 | 1 | 0 | 2 | 0.5 | 1 | 0 | 3 | 0.3333 | 7 | 0.2857 | 1 | 0 | 1 | 0 | 1 | 1 | NA | NA | 3 | 0.3333 |
| 4_1483249 | 2 | 0 | 2 | 0.5 | 1 | 0 | 2 | 0 | 7 | 0.1429 | 1 | 1 | 1 | 0 | 1 | 0 | NA | NA | 3 | 0.3333 |
| 4_1483249 | 2 | 0.5 | 2 | 0 | 1 | 0 | NA | NA | 5 | 0.2 | 2 | 0.5 | 1 | 0 | 1 | 1 | NA | NA | 4 | 0.5 |
| 4_1483250 | 3 | 0.6667 | 4 | 0.25 | 2 | 0.5 | 3 | 0.3333 | 12 | 0.4167 | 4 | 0 | 4 | 0.25 | 1 | 0 | 1 | 0 | 10 | 0.1 |
| 4_1483250 | 1 | 0 | 3 | 0.3333 | 1 | 1 | 1 | 1 | 6 | 0.5 | 4 | 0.5 | 1 | 1 | NA | NA | 1 | 0 | 6 | 0.5 |
| 4_1483251 | 6 | 0.5 | 7 | 0.7143 | 1 | 1 | 3 | 1 | 17 | 0.7059 | 6 | 0.6667 | 3 | 0.6667 | 3 | 1 | 3 | 0.3333 | 15 | 0.6667 |
| 4_1483252 | 6 | 0.8333 | 7 | 0.5714 | 3 | 1 | 4 | 1 | 20 | 0.8 | 6 | 0.8333 | 4 | 0.75 | 3 | 0.6667 | 4 | 0.75 | 17 | 0.7647 |
| 4_1483254 | 6 | 0.5 | 6 | 1 | 3 | 0.6667 | 5 | 1 | 20 | 0.8 | 4 | 0.5 | 4 | 0.75 | 3 | 1 | 4 | 1 | 15 | 0.8 |
| 4_1483254 | 5 | 0.6 | 6 | 0.3333 | 3 | 1 | 4 | 1 | 18 | 0.6667 | 4 | 0.25 | 4 | 0.25 | 3 | 0 | 3 | 1 | 14 | 0.3571 |
| 4_1483254 | 3 | 1 | 3 | 0.3333 | 2 | 1 | 4 | 0.5 | 12 | 0.6667 | 2 | 0 | 3 | 0.6667 | 3 | 1 | 2 | 1 | 10 | 0.7 |
| 4_1483256 | 5 | 1 | 6 | 0.5 | 3 | 1 | 3 | 1 | 17 | 0.8235 | 5 | 0.8 | 4 | 0.75 | 3 | 1 | 3 | 1 | 15 | 0.8667 |
| 4_1483256 | 5 | 0 | 6 | 0 | 3 | 0 | 4 | 0.25 | 18 | 0.0556 | 4 | 0 | 3 | 0 | 3 | 0.3333 | 4 | 0 | 14 | 0.0714 |
| 4_1483256 | 4 | 0 | 5 | 0.2 | 3 | 0 | 5 | 0.4 | 17 | 0.1765 | 4 | 0.25 | 3 | 0 | 3 | 0.3333 | 3 | 0 | 13 | 0.1538 |
| 4_1483256 | 4 | 0.5 | 6 | 0.1667 | 3 | 1 | 4 | 0.5 | 17 | 0.4706 | 4 | 0.5 | 3 | 0.6667 | 3 | 0.3333 | 3 | 0.3333 | 13 | 0.4615 |
| 4_1483257 | 4 | 0.75 | 6 | 1 | 3 | 1 | 5 | 1 | 18 | 0.9444 | 4 | 1 | 2 | 0.5 | 3 | 0.6667 | 1 | 1 | 10 | 0.8 |
| 4_1483258 | 1 | 1 | 5 | 0.6 | 1 | 1 | 5 | 1 | 12 | 0.8333 | 3 | 0.6667 | 2 | 0.5 | 1 | 1 | 2 | 1 | 8 | 0.75 |
| 4_1483258 | 4 | 0.25 | 6 | 0.1667 | 3 | 0 | 2 | 0 | 15 | 0.1333 | 4 | 0.25 | 2 | 0.5 | 2 | 0 | 2 | 0 | 10 | 0.2 |
| 4_1483560 | 2 | 0.5 | 2 | 1 | 1 | 1 | NA | NA | 5 | 0.8 | 2 | 0 | 1 | 0 | 1 | 1 | NA | NA | 4 | 0.25 |
| 4_1483560 | NA | NA | 1 | 1 | 1 | 1 | NA | NA | 2 | 1 | 1 | 1 | NA | NA | 1 | 0 | NA | NA | 2 | 0.5 |
| 4_1483563 | 2 | 1 | 2 | 1 | 1 | 0 | 1 | 1 | 6 | 0.8333 | 2 | 0 | NA | NA | 1 | 1 | NA | NA | 3 | 0.3333 |
| 4_1483564 | 2 | 1 | 2 | 0.5 | NA | NA | 1 | 1 | 5 | 0.8 | 2 | 1 | NA | NA | 1 | 1 | NA | NA | 3 | 1 |
| 4_1483567 | 2 | 1 | NA | NA | 1 | 1 | NA | NA | 3 | 1 | 1 | 1 | NA | NA | 1 | 1 | NA | NA | 2 | 1 |
| 4_1485191 | 2 | 0.5 | 2 | 0.5 | 2 | 0.5 | 1 | 1 | 7 | 0.5714 | NA | NA | 2 | 0.5 | 1 | 0 | NA | NA | 3 | 0.3333 |
| 4_1485191 | NA | NA | 1 | 1 | 3 | 0.3333 | 1 | 1 | 5 | 0.6 | NA | NA | 1 | 0 | 1 | 0 | 1 | 1 | 3 | 0.3333 |
| 4_1485196 | NA | NA | 1 | 1 | 2 | 0 | 1 | 1 | 4 | 0.5 | NA | NA | 1 | 0 | 1 | 0 | 1 | 1 | 3 | 0.3333 |
| 4_1485196 | NA | NA | 2 | 0 | 2 | 0.5 | 1 | 1 | 5 | 0.4 | NA | NA | 1 | 0 | 1 | 1 | NA | NA | 2 | 0.5 |
| 4_1485197 | NA | NA | 1 | 1 | 3 | 0 | 1 | 1 | 5 | 0.4 | NA | NA | 1 | 1 | 1 | 0 | 1 | 0 | 3 | 0.3333 |
| 4_1485197 | NA | NA | 1 | 0 | 3 | 0.3333 | 1 | 1 | 5 | 0.4 | NA | NA | 1 | 1 | 1 | 0 | NA | NA | 2 | 0.5 |
| 4_1485205 | NA | NA | 1 | 1 | 3 | 1 | 1 | 1 | 5 | 1 | NA | NA | 1 | 1 | 1 | 1 | NA | NA | 2 | 1 |
| 4_1485206 | NA | NA | 1 | 1 | 3 | 1 | 1 | 1 | 5 | 1 | NA | NA | 1 | 1 | NA | NA | 1 | 0 | 2 | 0.5 |
| 4_1486413 | 5 | 0.4 | 7 | 0.4286 | 6 | 0.3333 | 2 | 0 | 20 | 0.35 | 7 | 0.7143 | 4 | 0.25 | 9 | 0.6667 | 6 | 0.5 | 26 | 0.5769 |
| 4_1486413 | 4 | 0.75 | 8 | 0.25 | 7 | 0.7143 | 6 | 0.6667 | 25 | 0.56 | 5 | 0.8 | 7 | 0.5714 | 10 | 0.5 | 4 | 0.5 | 26 | 0.5769 |
| 4_1486413 | 9 | 0 | 8 | 0.25 | 8 | 0 | 6 | 0 | 31 | 0.0645 | 8 | 0 | 6 | 0 | 11 | 0.1818 | 6 | 0 | 31 | 0.0645 |
| 4_1486414 | 9 | 1 | 8 | 0.875 | 7 | 0.7143 | 5 | 0.6 | 29 | 0.8276 | 9 | 0.7778 | 6 | 0.3333 | 10 | 0.8 | 6 | 1 | 31 | 0.7419 |
| 4_1486414 | 8 | 0.25 | 7 | 0.4286 | 8 | 0.625 | 6 | 0.1667 | 29 | 0.3793 | 9 | 0.3333 | 5 | 0.4 | 11 | 0.5455 | 4 | 0.25 | 29 | 0.4138 |
| 4_1486414 | 7 | 0.5714 | 9 | 0.3333 | 9 | 0.5556 | 5 | 0.4 | 30 | 0.4667 | 8 | 0.25 | 7 | 0 | 11 | 0.1818 | 7 | 0.1429 | 33 | 0.1515 |
| 4_1486415 | 8 | 0.5 | 9 | 0.5556 | 9 | 0.4444 | 6 | 0.8333 | 32 | 0.5625 | 7 | 0.5714 | 7 | 0.4286 | 10 | 0.6 | 7 | 0.7143 | 31 | 0.5806 |
| 4_1486416 | 7 | 0 | 11 | 0.0909 | 8 | 0.25 | 6 | 0 | 32 | 0.0938 | 7 | 0 | 7 | 0 | 9 | 0.2222 | 7 | 0.1429 | 30 | 0.1 |
| 4_1486416 | 5 | 0.4 | 8 | 0 | 7 | 0.2857 | 6 | 0 | 26 | 0.1538 | 8 | 0.125 | 7 | 0.1429 | 10 | 0.2 | 7 | 0.1429 | 32 | 0.1562 |
| 4_1486417 | 8 | 0.5 | 11 | 0.6364 | 11 | 0.2727 | 7 | 0.5714 | 37 | 0.4865 | 7 | 0.4286 | 8 | 0.5 | 12 | 0.4167 | 8 | 0.125 | 35 | 0.3714 |
| 4_1486418 | 9 | 0.6667 | 11 | 0.6364 | 13 | 0.8462 | 7 | 0.7143 | 40 | 0.725 | 8 | 0.75 | 5 | 0.4 | 10 | 0.9 | 7 | 0.7143 | 30 | 0.7333 |
| 4_1486419 | 10 | 0.4 | 10 | 0.5 | 13 | 0.5385 | 7 | 0.5714 | 40 | 0.5 | 9 | 0.5556 | 7 | 0.4286 | 11 | 0.4545 | 8 | 0.625 | 35 | 0.5143 |
| 4_1486420 | 10 | 0 | 12 | 0.0833 | 11 | 0.0909 | 6 | 0.3333 | 39 | 0.1026 | 9 | 0.1111 | 8 | 0.125 | 12 | 0.1667 | 8 | 0.25 | 37 | 0.1622 |
| 4_1486472 | 5 | 0 | 4 | 0 | 2 | 0 | 1 | 1 | 12 | 0.0833 | 2 | 0.5 | 2 | 0 | 3 | 0 | 5 | 0 | 12 | 0.0833 |
| 4_1486604 | 1 | 1 | NA | NA | 1 | 1 | 2 | 0.5 | 4 | 0.75 | NA | NA | 2 | 1 | 2 | 1 | 3 | 0.6667 | 7 | 0.8571 |
| 4_1486605 | 2 | 1 | NA | NA | NA | NA | 2 | 1 | 4 | 1 | 1 | 0 | 2 | 1 | 1 | 1 | 2 | 1 | 6 | 0.8333 |
| 4_1486605 | 2 | 1 | NA | NA | 1 | 1 | 1 | 0 | 4 | 0.75 | 1 | 0 | 1 | 1 | 1 | 0 | 2 | 0 | 5 | 0.2 |

| | | | | | | | | | | | | | | | | | | | | |
|---|---|---|---|---|---|---|---|---|---|---|---|---|---|---|---|---|---|---|---|---|
| 4_14866058 | 2 | 0.5 | NA | NA | 1 | 0 | 2 | 0 | 5 | 0.2 | 1 | 0 | 2 | 0 | 2 | 0 | 3 | 0.3333 | 8 | 0.125 |
| 4_14866077 | 2 | 1 | NA | NA | 2 | 0.5 | 1 | 0 | 5 | 0.6 | 2 | 0 | 3 | 0 | 2 | 0 | 3 | 0.3333 | 10 | 0.1 |
| 4_14866080 | 3 | 0 | 1 | 1 | NA | NA | 2 | 0.5 | 6 | 0.3333 | 1 | 0 | 3 | 0 | 1 | 0 | 3 | 0.3333 | 8 | 0.125 |
| 4_14866098 | 3 | 0.3333 | NA | NA | 2 | 0 | 2 | 0 | 7 | 0.1429 | 2 | 0.5 | 3 | 0 | 2 | 0 | 4 | 0.5 | 11 | 0.2727 |
| 4_14867886 | 17 | 0.1176 | 11 | 0.3636 | 8 | 0 | 16 | 0 | 52 | 0.1154 | 14 | 0.0714 | 12 | 0.1667 | 15 | 0 | 12 | 0.0833 | 53 | 0.0755 |
| 4_14867894 | 15 | 0.7333 | 11 | 0.8182 | 10 | 0.4 | 19 | 0.6316 | 55 | 0.6545 | 15 | 0.4667 | 12 | 0.5 | 16 | 0.6875 | 9 | 0.7778 | 52 | 0.5962 |
| 4_14868388 | NA | NA | 1 | 1 | 1 | 0 | 2 | 0 | 4 | 0.25 | NA | NA | 1 | 0 | NA | NA | 1 | 1 | 2 | 0.5 |
| 4_14869827 | NA | NA | 4 | 0.25 | NA | NA | 6 | 0 | 10 | 0.1 | 1 | 0 | 2 | 0 | 6 | 0.1667 | 3 | 0 | 12 | 0.0833 |
| 4_14869829 | NA | NA | 5 | 0.2 | NA | NA | 5 | 0 | 10 | 0.1 | 1 | 0 | 2 | 0.5 | 7 | 0 | 4 | 0 | 14 | 0.0714 |
| 4_14870014 | 7 | 0 | 10 | 0 | 4 | 0.5 | 8 | 0.125 | 29 | 0.1034 | 4 | 0 | 5 | 0 | 7 | 0.1429 | 4 | 0 | 20 | 0.05 |
| 4_14870041 | 7 | 0.2857 | 10 | 0.1 | 4 | 0 | 7 | 0.1429 | 28 | 0.1429 | 4 | 0 | 5 | 0.2 | 7 | 0.1429 | 4 | 0 | 20 | 0.1 |
| 4_14870052 | 7 | 0.2857 | 9 | 0.4444 | 3 | 0.3333 | 7 | 0.4286 | 26 | 0.3846 | 4 | 0.25 | 2 | 0 | 8 | 0.125 | 2 | 0.5 | 16 | 0.1875 |
| 4_14870063 | 6 | 0.1667 | 8 | 0 | 4 | 0.5 | 7 | 0.1429 | 25 | 0.16 | 4 | 0 | 4 | 0 | 7 | 0.1429 | 4 | 0 | 19 | 0.0526 |
| 4_14870112 | 6 | 0.5 | 8 | 0.25 | 4 | 0.25 | 5 | 0.2 | 23 | 0.3043 | 3 | 0 | 4 | 0.5 | 4 | 0.5 | 3 | 0.3333 | 14 | 0.3571 |
| 4_14870114 | 5 | 0 | 7 | 0.1429 | 4 | 0 | 3 | 0 | 19 | 0.0526 | 3 | 0 | 4 | 0 | 4 | 0.25 | 3 | 0 | 14 | 0.0714 |
| 4_14870119 | 6 | 0.3333 | 7 | 0.4286 | 4 | 0 | 3 | 0.3333 | 20 | 0.3 | 5 | 0 | 3 | 0.6667 | 3 | 0.3333 | 7 | 0.1429 | 18 | 0.2222 |
| 4_14870124 | 8 | 0.625 | 7 | 0.8571 | 4 | 0 | 5 | 0.4 | 24 | 0.5417 | 5 | 0 | 5 | 0.8 | 5 | 0.6 | 7 | 0.4286 | 22 | 0.4545 |
| 4_14870149 | 6 | 0 | 4 | 0 | 8 | 0.25 | 7 | 0 | 25 | 0.08 | 8 | 0 | 5 | 0.4 | 8 | 0 | 10 | 0.1 | 31 | 0.0968 |
| 4_14870152 | 3 | 0.3333 | 5 | 1 | 4 | 1 | 8 | 0.5 | 20 | 0.7 | 8 | 0.5 | 5 | 0.6 | 6 | 0.6667 | 7 | 0.7143 | 26 | 0.6154 |
| 4_14870155 | 6 | 0.5 | 6 | 0.3333 | 10 | 0.3 | 9 | 0.2222 | 31 | 0.3226 | 5 | 0.2 | 7 | 0.2857 | 11 | 0.1818 | 11 | 0.1818 | 34 | 0.2059 |
| 4_14870169 | 6 | 0.3333 | 5 | 0 | 10 | 0.4 | 11 | 0.1818 | 32 | 0.25 | 6 | 0 | 6 | 0.3333 | 9 | 0.3333 | 11 | 0.2727 | 32 | 0.25 |
| 4_14870171 | 7 | 0.2857 | 4 | 0.75 | 9 | 0.6667 | 7 | 0.7143 | 27 | 0.5926 | 7 | 0.5714 | 6 | 0.5 | 9 | 0.5556 | 10 | 0.1 | 32 | 0.4062 |
| 4_14870181 | 5 | 0 | 4 | 0 | 5 | 0.2 | 9 | 0.1111 | 23 | 0.087 | 5 | 0 | 4 | 0 | 6 | 0.3333 | 6 | 0.1667 | 21 | 0.1429 |
| 4_14870184 | 7 | 0.5714 | 5 | 0.8 | 8 | 0.75 | 10 | 0.9 | 30 | 0.7667 | 8 | 0.5 | 4 | 0.75 | 8 | 0.625 | 10 | 0.7 | 30 | 0.6333 |
| 4_14870192 | 6 | 0.6667 | 4 | 1 | 7 | 0.8571 | 10 | 1 | 27 | 0.8889 | 7 | 0.5714 | 6 | 0.6667 | 8 | 0.875 | 12 | 0.9167 | 33 | 0.7879 |
| 4_14870246 | 3 | 0 | 5 | 0.2 | 4 | 0.25 | 11 | 0.1818 | 23 | 0.1739 | 7 | 0.2857 | 8 | 0 | 8 | 0.125 | 7 | 0.1429 | 30 | 0.1333 |
| 4_14870302 | 3 | 0 | 2 | 0.5 | NA | NA | 3 | 0.6667 | 8 | 0.375 | 1 | 1 | 2 | 0.5 | 3 | 0.3333 | 1 | 0 | 7 | 0.4286 |
| 4_14870315 | 7 | 0 | 6 | 0.1667 | NA | NA | 4 | 0 | 17 | 0.0588 | 3 | 0 | 3 | 0 | 5 | 0.2 | 5 | 0 | 16 | 0.0625 |
| 4_14903325 | NA | NA | 2 | 0 | 1 | 0 | 2 | 0.5 | 5 | 0.2 | 4 | 0.25 | 2 | 0 | 2 | 0 | 2 | 0 | 10 | 0.1 |
| 4_14903606 | NA | NA | 1 | 1 | NA | NA | NA | NA | 1 | 1 | NA | NA | NA | NA | NA | NA | 1 | 1 | 1 | 1 |
| 4_14903608 | NA | NA | NA | NA | 1 | 0 | 1 | 1 | 2 | 0.5 | 3 | 1 | NA | NA | 1 | 0 | 2 | 0.5 | 6 | 0.6667 |
| 4_14903614 | NA | NA | NA | NA | 1 | 0 | 2 | 0.5 | 3 | 0.3333 | 3 | 1 | 2 | 0 | 1 | 1 | 1 | 0 | 7 | 0.5714 |
| 4_14903639 | NA | NA | NA | NA | NA | NA | 2 | 1 | 2 | 1 | 2 | 0.5 | NA | NA | NA | NA | NA | NA | 2 | 0.5 |
| 4_14903650 | NA | NA | 1 | 1 | NA | NA | NA | NA | 1 | 1 | NA | NA | NA | NA | NA | NA | 1 | 1 | 1 | 1 |
| 4_14903666 | NA | NA | 1 | 1 | NA | NA | NA | NA | 1 | 1 | NA | NA | NA | NA | NA | NA | 1 | 1 | 1 | 1 |
| 4_14903697 | NA | NA | 1 | 1 | NA | NA | NA | NA | 1 | 1 | NA | NA | 1 | 0 | NA | NA | 1 | 1 | 2 | 0.5 |
| 4_14903705 | NA | NA | 1 | 1 | NA | NA | NA | NA | 1 | 1 | NA | NA | 1 | 1 | NA | NA | 1 | 1 | 2 | 1 |
| 4_14903746 | NA | NA | 1 | 1 | NA | NA | NA | NA | 1 | 1 | NA | NA | 1 | 1 | NA | NA | 1 | 1 | 2 | 1 |
| 4_14903802 | 1 | 1 | 1 | 0 | NA | NA | NA | NA | 2 | 0.5 | NA | NA | 1 | 1 | NA | NA | 1 | 0 | 2 | 0.5 |
| 4_14903921 | 1 | 1 | 2 | 1 | NA | NA | NA | NA | 3 | 1 | NA | NA | 1 | 1 | NA | NA | 1 | 1 | 2 | 1 |
| 4_14904529 | 4 | 0 | 5 | 0 | 9 | 0.2222 | 8 | 0 | 26 | 0.0769 | 4 | 0 | 3 | 0 | 6 | 0.1667 | 4 | 0.25 | 17 | 0.1176 |
| 4_14904575 | 3 | 0.3333 | 5 | 0 | 7 | 0.1429 | 7 | 0.2857 | 22 | 0.1818 | 4 | 0 | 2 | 0.5 | 3 | 0.3333 | 1 | 1 | 10 | 0.3 |
| 4_14904581 | 3 | 0.6667 | 5 | 0.4 | 6 | 0.5 | 6 | 0.1667 | 20 | 0.4 | 4 | 0.25 | 2 | 0.5 | 3 | 0.6667 | 2 | 0 | 11 | 0.3636 |
| 4_14920240 | 2 | 0.5 | 5 | 0 | NA | NA | NA | NA | 7 | 0.1429 | NA | NA | 2 | 0.5 | 2 | 1 | NA | NA | 4 | 0.75 |
| 4_14920266 | 2 | 1 | 5 | 0.4 | NA | NA | NA | NA | 7 | 0.5714 | NA | NA | 1 | 0 | 1 | 1 | NA | NA | 2 | 0.5 |
| 4_14920270 | 2 | 0 | 6 | 0.3333 | NA | NA | NA | NA | 8 | 0.25 | NA | NA | 2 | 0 | 2 | 0.5 | NA | NA | 4 | 0.25 |
| 4_14920358 | 4 | 0.5 | 6 | 1 | 1 | 1 | NA | NA | 11 | 0.8182 | 1 | 1 | 2 | 0 | 4 | 1 | 4 | 1 | 11 | 0.8182 |
| 4_14920360 | 2 | 1 | 5 | 0.6 | 1 | 1 | NA | NA | 8 | 0.75 | 2 | 0.5 | NA | NA | 4 | 1 | 2 | 0.5 | 8 | 0.75 |
| 4_14920373 | 3 | 0.3333 | 7 | 0.7143 | 1 | 0 | NA | NA | 11 | 0.5455 | 2 | 0.5 | 2 | 0.5 | 5 | 0.4 | 3 | 0.3333 | 12 | 0.4167 |
| 4_14920378 | 1 | 0 | 4 | 0.25 | 1 | 0 | NA | NA | 6 | 0.1667 | 1 | 0 | 2 | 0 | 1 | 1 | 2 | 0 | 6 | 0.1667 |
| 4_14920966 | 2 | 1 | 3 | 1 | 1 | 1 | NA | NA | 6 | 1 | 4 | 0.5 | 4 | 1 | NA | NA | 3 | 1 | 11 | 0.8182 |
| 4_14921300 | 1 | 1 | NA | NA | NA | NA | NA | NA | 1 | 1 | 1 | 1 | NA | NA | NA | NA | NA | NA | 1 | 1 |
| 4_14923461 | 10 | 0.4 | 10 | 0.2 | 4 | 0.25 | 2 | 0 | 26 | 0.2692 | 1 | 0 | 6 | 0.3333 | 5 | 0.2 | 11 | 0.3636 | 23 | 0.3043 |
| 4_14923477 | 9 | 0.5556 | 6 | 0.6667 | 3 | 0.3333 | 2 | 0.5 | 20 | 0.55 | 1 | 0 | 5 | 0.2 | 2 | 0.5 | 6 | 0.5 | 14 | 0.3571 |
| 4_14923484 | 8 | 0.125 | 9 | 0 | 4 | 0.25 | 2 | 0.5 | 23 | 0.1304 | 1 | 0 | 5 | 0.2 | 2 | 0.5 | 9 | 0.2222 | 17 | 0.2353 |
| 4_14923486 | 9 | 0.1111 | 10 | 0 | 4 | 0.25 | 1 | 1 | 24 | 0.125 | 1 | 0 | 5 | 0.2 | 2 | 0 | 10 | 0.1 | 18 | 0.1111 |
| 4_14923492 | 9 | 0 | 9 | 0.2222 | 4 | 0 | 3 | 0 | 25 | 0.08 | 1 | 1 | 6 | 0.1667 | 2 | 0 | 11 | 0.0909 | 20 | 0.15 |
| 4_14923506 | 6 | 0.3333 | 6 | 0.3333 | 1 | 0 | NA | NA | 13 | 0.3077 | NA | NA | 3 | 0.3333 | 2 | 0 | 4 | 0.25 | 9 | 0.2222 |
| 4_14923518 | 7 | 0.8571 | 8 | 0.375 | 4 | 0.75 | 1 | 1 | 20 | 0.65 | 1 | 1 | 5 | 0.6 | 1 | 1 | 10 | 0.7 | 17 | 0.7059 |
| 4_14926184 | 36 | 0.0833 | 45 | 0.0889 | 39 | 0.1026 | 32 | 0.25 | 152 | 0.125 | 26 | 0.2308 | 27 | 0.037 | 35 | 0.0571 | 49 | 0.102 | 137 | 0.1022 |
| 4_14926188 | 25 | 0.28 | 30 | 0.2333 | 32 | 0.125 | 28 | 0.2857 | 115 | 0.2261 | 22 | 0.4545 | 21 | 0.1905 | 29 | 0.2759 | 37 | 0.2162 | 109 | 0.2752 |
| 4_14926307 | 26 | 0 | 26 | 0.1154 | 26 | 0.0385 | 22 | 0.0455 | 100 | 0.05 | 12 | 0.0833 | 14 | 0.2857 | 22 | 0.0455 | 27 | 0.1111 | 75 | 0.12 |
| 4_14926322 | 25 | 0.04 | 24 | 0.125 | 25 | 0.12 | 23 | 0 | 97 | 0.0722 | 13 | 0.0769 | 16 | 0.0625 | 20 | 0.05 | 27 | 0.0741 | 76 | 0.0658 |
| 4_14926325 | 24 | 0.2083 | 17 | 0.0588 | 17 | 0.0588 | 19 | 0.0526 | 77 | 0.1039 | 9 | 0.3333 | 14 | 0.0714 | 17 | 0.2353 | 25 | 0.12 | 65 | 0.1692 |
| 4_14926329 | 24 | 0.5833 | 23 | 0.6957 | 25 | 0.64 | 19 | 0.4737 | 91 | 0.6044 | 11 | 0.2727 | 15 | 0.6 | 22 | 0.6364 | 25 | 0.68 | 73 | 0.589 |

| | | | | | | | | | | | | | | | | | | | | |
|---|---|---|---|---|---|---|---|---|---|---|---|---|---|---|---|---|---|---|---|---|
| 4_1492633 | 24 | 0.3333 | 22 | 0.1364 | 17 | 0.2941 | 22 | 0.3182 | 85 | 0.2706 | 10 | 0.1 | 15 | 0.2667 | 20 | 0.3 | 24 | 0.3333 | 69 | 0.2754 |
| 4_1492634 | 27 | 0.5556 | 23 | 0.4348 | 26 | 0.4615 | 22 | 0.3636 | 98 | 0.4592 | 12 | 0.4167 | 13 | 0.5385 | 18 | 0.6111 | 26 | 0.3462 | 69 | 0.4638 |
| 4_1492636 | 23 | 0.5652 | 19 | 0.3158 | 25 | 0.44 | 23 | 0.4348 | 90 | 0.4444 | 11 | 0.1818 | 13 | 0.4615 | 18 | 0.3333 | 25 | 0.4 | 67 | 0.3582 |
| 4_1492636 | 12 | 0.5 | 18 | 0.3889 | 16 | 0.6875 | 15 | 0.6 | 61 | 0.541 | 10 | 0.3 | 10 | 0.5 | 15 | 0.6 | 16 | 0.5 | 51 | 0.4902 |
| 4_1492639 | 19 | 0.2105 | 20 | 0.2 | 20 | 0.25 | 23 | 0.1739 | 82 | 0.2073 | 12 | 0.0833 | 13 | 0.0769 | 17 | 0.1765 | 26 | 0.1538 | 68 | 0.1324 |
| 4_1492640 | 21 | 0.1429 | 20 | 0.1 | 20 | 0.1 | 21 | 0.0476 | 82 | 0.0976 | 12 | 0 | 12 | 0.0833 | 18 | 0.2222 | 25 | 0.08 | 67 | 0.1045 |
| 4_1492641 | 20 | 0.15 | 21 | 0.0952 | 18 | 0.2222 | 19 | 0 | 78 | 0.1154 | 11 | 0 | 11 | 0.0909 | 16 | 0.0625 | 25 | 0.08 | 63 | 0.0635 |
| 4_1493183 | 2 | 0.5 | 2 | 0.5 | NA | NA | 1 | 0 | 5 | 0.4 | 1 | 1 | 1 | 0 | 1 | 0 | 1 | 1 | 4 | 0.5 |
| 4_1493200 | 2 | 0.5 | 2 | 0.5 | NA | NA | NA | NA | 4 | 0.5 | 1 | 1 | NA | NA | 1 | 1 | 2 | 1 | 4 | 1 |
| 4_1493202 | 2 | 0.5 | 2 | 0 | NA | NA | 1 | 0 | 5 | 0.2 | 1 | 1 | NA | NA | 1 | 1 | 2 | 0.5 | 4 | 0.75 |
| 4_1493202 | 2 | 1 | 2 | 0.5 | NA | NA | NA | NA | 4 | 0.75 | 1 | 0 | NA | NA | 1 | 1 | 1 | 1 | 3 | 0.6667 |
| 4_1493202 | 2 | 0.5 | 2 | 1 | NA | NA | NA | NA | 4 | 0.75 | 1 | 1 | NA | NA | 1 | 1 | 1 | 1 | 3 | 1 |
| 4_1493204 | 2 | 1 | 2 | 1 | NA | NA | 1 | 1 | 5 | 1 | 1 | 0 | NA | NA | 1 | 0 | 2 | 0.5 | 4 | 0.25 |
| 4_1493205 | 1 | 1 | 2 | 1 | NA | NA | 1 | 1 | 4 | 1 | 1 | 1 | NA | NA | 1 | 1 | 1 | 1 | 3 | 1 |
| 4_1493205 | 2 | 1 | NA | NA | NA | NA | 1 | 1 | 3 | 1 | 1 | 1 | NA | NA | 1 | 1 | NA | NA | 2 | 1 |
| 4_1493205 | 2 | 0.5 | 2 | 0 | NA | NA | 1 | 0 | 5 | 0.2 | 1 | 1 | NA | NA | 1 | 0 | 2 | 0 | 4 | 0.25 |
| 4_1493207 | 2 | 1 | 2 | 1 | NA | NA | 1 | 1 | 5 | 1 | 1 | 1 | NA | NA | 1 | 1 | 1 | 1 | 3 | 1 |
| 4_1493207 | 2 | 0 | 2 | 1 | NA | NA | 1 | 0 | 5 | 0.4 | 1 | 1 | NA | NA | 1 | 0 | 1 | 1 | 3 | 0.6667 |
| 4_1493207 | 2 | 0 | 2 | 0.5 | NA | NA | 1 | 0 | 5 | 0.2 | 1 | 1 | NA | NA | 1 | 1 | 1 | 1 | 3 | 1 |
| 4_1493214 | 1 | 0 | 2 | 0.5 | NA | NA | 1 | 1 | 4 | 0.5 | 1 | 1 | NA | NA | NA | NA | 1 | 1 | 2 | 1 |
| 4_1493215 | 1 | 1 | 2 | 0 | NA | NA | 1 | 1 | 4 | 0.5 | 1 | 1 | NA | NA | NA | NA | 1 | 0 | 2 | 0.5 |
| 4_1493220 | 1 | 1 | 1 | 0 | NA | NA | 1 | 1 | 3 | 0.6667 | 1 | 1 | NA | NA | NA | NA | 1 | 0 | 2 | 0.5 |
| 4_1493225 | 1 | 1 | 1 | 1 | NA | NA | 1 | 1 | 3 | 1 | 1 | 1 | NA | NA | NA | NA | 1 | 1 | 2 | 1 |
| 4_1493225 | 1 | 1 | 2 | 1 | NA | NA | 1 | 1 | 4 | 1 | NA | NA | NA | NA | NA | NA | 1 | 1 | 1 | 1 |
| 4_1493226 | 1 | 1 | 1 | 1 | NA | NA | 1 | 1 | 3 | 1 | 1 | 1 | NA | NA | NA | NA | 1 | 1 | 2 | 1 |
| 4_1494129 | 11 | 0 | 11 | 0.0909 | 11 | 0.0909 | 12 | 0.1667 | 45 | 0.0889 | 11 | 0.2727 | 11 | 0.1818 | 13 | 0.0769 | 9 | 0.1111 | 44 | 0.1591 |
| 4_1494129 | 6 | 0.3333 | 8 | 0.5 | 12 | 0.25 | 11 | 0.5455 | 37 | 0.4054 | 8 | 0.5 | 10 | 0.3 | 15 | 0.4 | 7 | 0.7143 | 40 | 0.45 |
| 4_1494133 | 9 | 0 | 9 | 0.2222 | 12 | 0.0833 | 12 | 0.0833 | 42 | 0.0952 | 10 | 0.1 | 11 | 0.2727 | 10 | 0.1 | 8 | 0.25 | 39 | 0.1795 |
| 4_1494137 | 10 | 0.7 | 11 | 0.5455 | 10 | 0.5 | 14 | 0.6429 | 45 | 0.6 | 7 | 0.8571 | 17 | 0.7647 | 10 | 0.7 | 9 | 0.3333 | 43 | 0.6744 |
| 4_1494137 | 11 | 0 | 11 | 0 | 14 | 0.0714 | 16 | 0.25 | 52 | 0.0962 | 8 | 0.125 | 17 | 0.2353 | 10 | 0.2 | 9 | 0.1111 | 44 | 0.1818 |
| 4_1494138 | 8 | 0.25 | 11 | 0.2727 | 13 | 0.3077 | 16 | 0.375 | 48 | 0.3125 | 7 | 0.1429 | 17 | 0.2353 | 8 | 0.125 | 9 | 0.2222 | 41 | 0.1951 |
| 4_1494196 | 6 | 0.1667 | 8 | 0.125 | 6 | 0 | 11 | 0.0909 | 31 | 0.0968 | 4 | 0 | 5 | 0.2 | 3 | 0 | 6 | 0 | 18 | 0.0556 |
| 4_1495104 | 3 | 0 | 2 | 0 | 2 | 0 | 3 | 0.3333 | 10 | 0.1 | 1 | 0 | 3 | 0 | 2 | 0.5 | 2 | 0 | 8 | 0.125 |
| 4_1495104 | 3 | 0 | 2 | 0 | 2 | 0.5 | 2 | 0 | 9 | 0.1111 | NA | NA | 3 | 0 | 1 | 1 | 1 | 0 | 5 | 0.2 |
| 4_1495104 | 3 | 0 | 1 | 0 | 1 | 0 | 3 | 0.6667 | 8 | 0.25 | NA | NA | 2 | 0.5 | 1 | 0 | 2 | 0 | 5 | 0.2 |
| 4_1495197 | 6 | 0.3333 | 5 | 0 | 1 | 0 | 1 | 0 | 13 | 0.1538 | 1 | 1 | NA | NA | NA | NA | NA | NA | 1 | 1 |
| 4_1495198 | 6 | 0.3333 | 4 | 0.75 | 1 | 0 | 2 | 0.5 | 13 | 0.4615 | 1 | 1 | NA | NA | NA | NA | NA | NA | 1 | 1 |
| 4_1495199 | 5 | 0.2 | 4 | 0.75 | 1 | 0 | 1 | 0 | 11 | 0.3636 | 1 | 1 | NA | NA | NA | NA | NA | NA | 1 | 1 |
| 4_1495202 | 5 | 0.4 | 4 | 0.25 | 1 | 0 | 1 | 0 | 11 | 0.2727 | 1 | 1 | NA | NA | NA | NA | NA | NA | 1 | 1 |
| 4_1495208 | 5 | 0.4 | 3 | 0.6667 | 1 | 1 | 2 | 1 | 11 | 0.6364 | 1 | 1 | NA | NA | NA | NA | NA | NA | 1 | 1 |
| 4_1495421 | 11 | 0 | 6 | 0.1667 | 4 | 0 | 8 | 0.25 | 29 | 0.1034 | 6 | 0 | 7 | 0 | 3 | 0.3333 | 7 | 0.1429 | 23 | 0.087 |
| 4_1495520 | 1 | 1 | NA | NA | NA | NA | 1 | 0 | 2 | 0.5 | NA | NA | NA | NA | 1 | 1 | 1 | 0 | 2 | 0.5 |
| 4_1495522 | 1 | 1 | NA | NA | NA | NA | 1 | 0 | 2 | 0.5 | NA | NA | NA | NA | 1 | 0 | 1 | 1 | 2 | 0.5 |
| 4_1495529 | 1 | 1 | NA | NA | NA | NA | 1 | 1 | 2 | 1 | NA | NA | NA | NA | 1 | 1 | 1 | 1 | 2 | 1 |
| 4_1495530 | 1 | 1 | NA | NA | 1 | 1 | 1 | 0 | 3 | 0.6667 | 1 | 0 | NA | NA | 1 | 0 | 2 | 1 | 4 | 0.5 |
| 4_1495531 | 1 | 0 | 1 | 0 | 2 | 1 | 1 | 1 | 5 | 0.6 | 1 | 1 | NA | NA | 1 | 1 | 1 | 1 | 3 | 1 |
| 4_1495531 | 1 | 1 | 1 | 1 | 2 | 1 | NA | NA | 4 | 1 | NA | NA | NA | NA | 1 | 1 | 1 | 1 | 2 | 1 |
| 4_1495533 | NA | NA | 1 | 1 | 2 | 0.5 | NA | NA | 3 | 0.6667 | 1 | 1 | NA | NA | 1 | 1 | 2 | 0.5 | 4 | 0.75 |
| 4_1495700 | 1 | 1 | NA | NA | NA | NA | NA | NA | 1 | 1 | 1 | 1 | 1 | 0 | NA | NA | NA | NA | 2 | 0.5 |
| 4_1495806 | 1 | 1 | 2 | 0.5 | NA | NA | NA | NA | 3 | 0.6667 | 1 | 0 | 1 | 0 | 2 | 0.5 | NA | NA | 4 | 0.25 |
| 4_1495806 | NA | NA | 3 | 0.6667 | NA | NA | NA | NA | 3 | 0.6667 | 1 | 1 | NA | NA | 3 | 1 | NA | NA | 4 | 1 |
| 4_1495807 | 1 | 0 | 3 | 0.3333 | NA | NA | NA | NA | 4 | 0.25 | 1 | 0 | NA | NA | 3 | 0.3333 | NA | NA | 4 | 0.25 |
| 4_1495808 | 1 | 1 | 2 | 0.5 | NA | NA | NA | NA | 3 | 0.6667 | 1 | 0 | 1 | 0 | 3 | 0.3333 | NA | NA | 5 | 0.2 |
| 4_1495808 | 1 | 0 | 3 | 0.6667 | NA | NA | NA | NA | 4 | 0.5 | 1 | 1 | 1 | 1 | 2 | 1 | NA | NA | 4 | 1 |
| 4_1495808 | 1 | 1 | 2 | 0.5 | NA | NA | NA | NA | 3 | 0.6667 | NA | NA | 1 | 1 | 1 | 0 | NA | NA | 2 | 0.5 |
| 4_1495809 | 1 | 1 | 2 | 0 | NA | NA | NA | NA | 3 | 0.3333 | 1 | 0 | NA | NA | 3 | 0.3333 | NA | NA | 4 | 0.25 |
| 4_1495819 | 1 | 1 | NA | NA | NA | NA | NA | NA | 1 | 1 | NA | NA | NA | NA | 1 | 1 | NA | NA | 1 | 1 |
| 4_1496489 | 2 | 0.5 | 3 | 0.6667 | 3 | 1 | 3 | 1 | 11 | 0.8182 | 5 | 0.8 | 2 | 1 | 8 | 0.875 | 5 | 0.6 | 20 | 0.8 |
| 4_1496503 | 1 | 0 | 5 | 0 | 2 | 0.5 | 5 | 0 | 13 | 0.0769 | 4 | 0.25 | 4 | 0 | 7 | 0 | 4 | 0 | 19 | 0.0526 |
| 4_1496504 | NA | NA | 3 | 0.6667 | 1 | 0 | 3 | 1 | 7 | 0.7143 | 2 | 1 | 3 | 0.6667 | 6 | 0.8333 | 2 | 0.5 | 13 | 0.7692 |
| 4_1496505 | NA | NA | 4 | 0.5 | 2 | 0 | 5 | 0.2 | 11 | 0.2727 | 4 | 0.25 | 3 | 0.3333 | 6 | 0 | 4 | 0.25 | 17 | 0.1765 |
| 4_1497320 | 68 | 0.5588 | 93 | 0.5484 | 64 | 0.4688 | 63 | 0.5556 | 288 | 0.5347 | 41 | 0.5122 | 56 | 0.5714 | 73 | 0.4795 | 66 | 0.5303 | 236 | 0.5212 |
| 4_1497321 | 71 | 0.1127 | 90 | 0.0778 | 63 | 0.1429 | 70 | 0.0714 | 294 | 0.0986 | 41 | 0.0976 | 57 | 0.0702 | 74 | 0.0676 | 65 | 0.2 | 237 | 0.1097 |
| 4_1497322 | 73 | 0.2466 | 102 | 0.1667 | 64 | 0.1875 | 71 | 0.1972 | 310 | 0.1968 | 47 | 0.3191 | 72 | 0.1944 | 93 | 0.1935 | 73 | 0.3151 | 285 | 0.2456 |
| 4_1497323 | 79 | 0.3291 | 98 | 0.3469 | 69 | 0.3043 | 72 | 0.2222 | 318 | 0.305 | 49 | 0.3265 | 70 | 0.2857 | 86 | 0.3372 | 73 | 0.3562 | 278 | 0.3273 |

| | | | | | | | | | | | | | | | | | | | | |
|---|---|---|---|---|---|---|---|---|---|---|---|---|---|---|---|---|---|---|---|---|
| 4_1497323 | 79 | 0.5949 | 102 | 0.4314 | 73 | 0.4658 | 77 | 0.4286 | 331 | 0.4773 | 50 | 0.5 | 69 | 0.5652 | 88 | 0.4886 | 76 | 0.4868 | 283 | 0.5088 |
| 4_1497325 | 105 | 0.0667 | 118 | 0.0678 | 80 | 0.05 | 90 | 0.0667 | 393 | 0.0636 | 63 | 0.0635 | 86 | 0.0581 | 111 | 0.0541 | 92 | 0.0435 | 352 | 0.054 |
| 4_1497327 | 95 | 0.2526 | 108 | 0.2315 | 78 | 0.1923 | 85 | 0.3294 | 366 | 0.2514 | 67 | 0.209 | 89 | 0.3258 | 107 | 0.3084 | 86 | 0.2674 | 349 | 0.2837 |
| 4_1497328 | 77 | 0.6104 | 77 | 0.4805 | 45 | 0.4 | 66 | 0.4242 | 265 | 0.4906 | 53 | 0.5849 | 67 | 0.4627 | 83 | 0.5181 | 65 | 0.5385 | 268 | 0.5224 |
| 4_1497329 | 91 | 0.6923 | 110 | 0.5909 | 80 | 0.6625 | 82 | 0.622 | 363 | 0.6391 | 64 | 0.5312 | 83 | 0.5904 | 97 | 0.5979 | 85 | 0.6 | 329 | 0.5836 |
| 4_1497329 | 91 | 0.2198 | 115 | 0.1652 | 74 | 0.1892 | 87 | 0.2529 | 367 | 0.2044 | 66 | 0.3182 | 86 | 0.2558 | 102 | 0.2353 | 80 | 0.2125 | 334 | 0.2515 |
| 4_1497331 | 93 | 0.4301 | 100 | 0.29 | 66 | 0.3636 | 78 | 0.3974 | 337 | 0.368 | 60 | 0.4667 | 78 | 0.3718 | 91 | 0.4615 | 69 | 0.4058 | 298 | 0.4262 |
| 4_1497332 | 121 | 0.157 | 136 | 0.1029 | 93 | 0.0968 | 99 | 0.101 | 449 | 0.1158 | 77 | 0.1299 | 101 | 0.1683 | 126 | 0.1032 | 95 | 0.1368 | 399 | 0.1328 |
| 4_1497340 | 4 | 1 | 4 | 1 | 2 | 1 | 1 | 1 | 11 | 1 | 1 | 1 | 1 | 1 | NA | NA | 1 | 1 | 3 | 1 |
| 4_1497340 | 4 | 0.75 | 4 | 0.5 | 2 | 0 | 1 | 1 | 11 | 0.5455 | 2 | 0 | 1 | 1 | NA | NA | 1 | 0 | 4 | 0.25 |
| 4_1497341 | 3 | 0.3333 | 3 | 0 | 1 | 1 | 1 | 0 | 8 | 0.25 | 1 | 1 | 1 | 0 | NA | NA | 1 | 1 | 3 | 0.6667 |
| 4_1497343 | 3 | 0.3333 | 3 | 0.6667 | 1 | 1 | 1 | 1 | 8 | 0.625 | 1 | 1 | 1 | 0 | 1 | 0 | 1 | 1 | 4 | 0.5 |
| 4_1497343 | 3 | 0.6667 | 4 | 0.5 | 1 | 0 | 1 | 0 | 9 | 0.4444 | 1 | 0 | 1 | 0 | NA | NA | 1 | 1 | 3 | 0.3333 |
| 4_1497344 | 2 | 1 | 1 | 1 | NA | NA | NA | NA | 3 | 1 | NA | NA | 1 | 1 | NA | NA | 1 | 1 | 2 | 1 |
| 4_1497349 | 2 | 0.5 | 2 | 0.5 | NA | NA | NA | NA | 4 | 0.5 | 2 | 0.5 | 1 | 0 | NA | NA | 1 | 0 | 4 | 0.25 |
| 4_1497349 | 2 | 0 | 2 | 0.5 | NA | NA | NA | NA | 4 | 0.25 | NA | NA | 1 | 0 | NA | NA | 1 | 1 | 2 | 0.5 |
| 4_1497350 | 1 | 1 | NA | NA | NA | NA | NA | NA | 1 | 1 | 2 | 0.5 | 1 | 1 | NA | NA | 1 | 0 | 4 | 0.5 |
| 4_1498001 | 2 | 0 | 1 | 0 | 5 | 0.4 | 5 | 0.2 | 13 | 0.2308 | 1 | 1 | 4 | 0 | 4 | 0 | 5 | 0 | 14 | 0.0714 |
| 4_1498007 | 2 | 0 | 1 | 0 | 3 | 0.3333 | 5 | 0 | 11 | 0.0909 | 1 | 0 | 2 | 0 | 1 | 0 | 5 | 0.2 | 9 | 0.1111 |
| 4_1498007 | 2 | 1 | 1 | 1 | 4 | 0.25 | 5 | 0.2 | 12 | 0.4167 | 1 | 0 | 2 | 1 | 1 | 1 | 5 | 0.6 | 9 | 0.6667 |
| 4_1498007 | NA | NA | 1 | 1 | 4 | 0.5 | 4 | 0.5 | 9 | 0.5556 | 1 | 0 | 1 | 1 | NA | NA | 5 | 0.8 | 7 | 0.7143 |
| 4_1498009 | 2 | 1 | 1 | 1 | 2 | 0.5 | 2 | 1 | 7 | 0.8571 | 1 | 1 | 2 | 0.5 | NA | NA | 5 | 0.8 | 8 | 0.75 |
| 4_1498009 | 2 | 0.5 | NA | NA | 2 | 0.5 | 3 | 0.3333 | 7 | 0.4286 | NA | NA | 1 | 1 | 1 | 1 | 1 | 0 | 3 | 0.6667 |
| 4_1498009 | 2 | 1 | 1 | 1 | 4 | 1 | 5 | 1 | 12 | 1 | 1 | 1 | 2 | 1 | NA | NA | 4 | 1 | 7 | 1 |
| 4_1498011 | NA | NA | 2 | 0.5 | 4 | 0 | 5 | 0.4 | 11 | 0.2727 | 1 | 0 | 2 | 0 | 1 | 1 | 4 | 0.5 | 8 | 0.375 |
| 4_1498012 | NA | NA | 1 | 1 | 2 | 1 | 5 | 0.6 | 8 | 0.75 | 1 | 1 | 1 | 1 | 1 | 1 | 4 | 0.5 | 7 | 0.7143 |
| 4_1498012 | NA | NA | 2 | 0.5 | 3 | 0 | 4 | 0 | 9 | 0.1111 | 1 | 1 | 1 | 0 | 1 | 0 | 1 | 0 | 4 | 0.25 |
| 4_1498012 | NA | NA | 2 | 1 | 4 | 1 | 5 | 0.8 | 11 | 0.9091 | 1 | 0 | 2 | 1 | 1 | 1 | 4 | 1 | 8 | 0.875 |
| 4_1498100 | 3 | 0 | 6 | 0.5 | 5 | 0.2 | 6 | 0.1667 | 20 | 0.25 | 2 | 0 | 6 | 0.5 | 5 | 0.2 | 5 | 0.2 | 18 | 0.2778 |
| 4_1498110 | 3 | 0.6667 | 5 | 0.4 | 5 | 0.2 | 5 | 0 | 18 | 0.2778 | 2 | 0 | 4 | 0 | 7 | 0.1429 | 6 | 0.1667 | 19 | 0.1053 |
| 4_1498112 | 3 | 0.3333 | 4 | 0.25 | 4 | 0.25 | 5 | 0 | 16 | 0.1875 | 1 | 0 | 3 | 0 | 6 | 0.1667 | 5 | 0 | 15 | 0.0667 |
| 4_1498258 | NA | NA | 1 | 1 | NA | NA | NA | NA | 1 | 1 | NA | NA | NA | NA | NA | NA | 1 | 1 | 1 | 1 |
| 4_1498260 | NA | NA | 1 | 1 | NA | NA | NA | NA | 1 | 1 | NA | NA | NA | NA | NA | NA | 1 | 1 | 1 | 1 |
| 4_1498261 | NA | NA | 1 | 1 | NA | NA | NA | NA | 1 | 1 | NA | NA | NA | NA | NA | NA | 1 | 1 | 1 | 1 |
| 4_1498262 | NA | NA | 1 | 1 | NA | NA | NA | NA | 1 | 1 | NA | NA | NA | NA | NA | NA | 1 | 1 | 1 | 1 |
| 4_1498267 | NA | NA | 1 | 1 | NA | NA | NA | NA | 1 | 1 | NA | NA | NA | NA | NA | NA | 1 | 1 | 1 | 1 |
| 4_1498273 | NA | NA | 1 | 1 | NA | NA | NA | NA | 1 | 1 | NA | NA | NA | NA | NA | NA | 1 | 1 | 1 | 1 |
| 4_1498277 | NA | NA | 1 | 1 | NA | NA | NA | NA | 1 | 1 | NA | NA | NA | NA | NA | NA | 1 | 1 | 1 | 1 |
| 4_1498285 | 3 | 0.3333 | 2 | 0 | 1 | 0 | 2 | 0.5 | 8 | 0.25 | NA | NA | 1 | 0 | 2 | 0.5 | 1 | 0 | 4 | 0.25 |
| 4_1498289 | 13 | 0.4615 | 5 | 0.6 | 8 | 0.625 | 11 | 0.2727 | 37 | 0.4595 | 4 | 0.25 | 9 | 0.5556 | 11 | 0.7273 | 4 | 0.5 | 28 | 0.5714 |
| 4_1498290 | 12 | 1 | 7 | 1 | 8 | 1 | 15 | 1 | 42 | 1 | 4 | 1 | 9 | 1 | 11 | 1 | 3 | 1 | 27 | 1 |
| 4_1498291 | 11 | 0.3636 | 6 | 0.3333 | 4 | 0.5 | 10 | 0.4 | 31 | 0.3871 | 5 | 0.2 | 10 | 0.2 | 11 | 0.5455 | 4 | 0.25 | 30 | 0.3333 |
| 4_1498291 | 16 | 0.625 | 7 | 0.2857 | 7 | 0.7143 | 12 | 0.1667 | 42 | 0.4524 | 4 | 0 | 9 | 0.1111 | 5 | 0.2 | 5 | 0.8 | 23 | 0.2609 |
| 4_1498292 | 13 | 0.6923 | 5 | 0.8 | 5 | 1 | 13 | 0.8462 | 36 | 0.8056 | 2 | 0.5 | 10 | 0.6 | 11 | 0.9091 | 5 | 0.8 | 28 | 0.75 |
| 4_1498335 | 8 | 0 | 5 | 0.4 | 5 | 0.4 | 10 | 0.2 | 28 | 0.2143 | 4 | 0.5 | 7 | 0.4286 | 9 | 0.2222 | 4 | 0.5 | 24 | 0.375 |
| 4_1499808 | 3 | 0.3333 | 1 | 0 | 2 | 1 | 3 | 0.3333 | 9 | 0.4444 | 7 | 1 | 2 | 0.5 | 1 | 0 | 3 | 0.3333 | 13 | 0.6923 |
| 4_1499814 | 3 | 0 | NA | NA | 2 | 0 | 2 | 0.5 | 7 | 0.1429 | 7 | 0.1429 | 2 | 0 | NA | NA | 3 | 0.3333 | 12 | 0.1667 |
| 4_1499817 | 4 | 0 | 1 | 0 | 2 | 0.5 | 2 | 0 | 9 | 0.1111 | 7 | 0.2857 | 1 | 0 | NA | NA | 3 | 0.3333 | 11 | 0.2727 |
| 4_1499820 | 3 | 0 | NA | NA | 2 | 0 | 1 | 1 | 6 | 0.1667 | 8 | 0.125 | 2 | 0 | NA | NA | 2 | 0 | 12 | 0.0833 |
| 4_1499822 | 4 | 0.5 | NA | NA | 2 | 1 | 2 | 0.5 | 8 | 0.625 | 6 | 0.3333 | 2 | 0 | NA | NA | 2 | 0 | 10 | 0.2 |
| 4_1499822 | 3 | 0.6667 | NA | NA | 2 | 0 | 2 | 1 | 7 | 0.5714 | 6 | 0.8333 | 1 | 0 | NA | NA | 3 | 0.6667 | 10 | 0.7 |
| 4_1499822 | 4 | 0.25 | NA | NA | 2 | 0 | 2 | 0.5 | 8 | 0.25 | 8 | 0.25 | 2 | 0 | NA | NA | 3 | 0.3333 | 13 | 0.2308 |
| 4_1499824 | 3 | 0 | NA | NA | 2 | 0 | 1 | 1 | 6 | 0.1667 | 7 | 0.4286 | 2 | 0 | NA | NA | 3 | 0.3333 | 12 | 0.3333 |
| 4_1499825 | 3 | 1 | NA | NA | 2 | 1 | 2 | 1 | 7 | 1 | 7 | 1 | 2 | 1 | NA | NA | 2 | 1 | 11 | 1 |
| 4_1499826 | 2 | 0 | NA | NA | 2 | 0 | 2 | 0.5 | 6 | 0.1667 | 7 | 0.1429 | 2 | 0 | NA | NA | 2 | 0 | 11 | 0.0909 |
| 4_1499827 | 3 | 0.3333 | NA | NA | 1 | 0 | 2 | 0 | 6 | 0.1667 | 6 | 0.3333 | 2 | 1 | NA | NA | 2 | 1 | 10 | 0.6 |
| 4_1499908 | 3 | 0.6667 | NA | NA | 2 | 0.5 | 2 | 0.5 | 7 | 0.5714 | 6 | 0.1667 | 2 | 0.5 | NA | NA | 2 | 0.5 | 10 | 0.3 |
| 4_1499909 | 2 | 1 | NA | NA | 2 | 1 | 2 | 0.5 | 6 | 0.8333 | 5 | 1 | 2 | 1 | NA | NA | 2 | 1 | 9 | 1 |
| 4_1499909 | 1 | 1 | NA | NA | 2 | 1 | NA | NA | 3 | 1 | 4 | 0.75 | 1 | 0 | NA | NA | 2 | 0.5 | 7 | 0.5714 |
| 4_1499910 | 3 | 0.3333 | NA | NA | 1 | 1 | 2 | 0.5 | 6 | 0.5 | 6 | 0.8333 | 2 | 0.5 | NA | NA | 1 | 1 | 9 | 0.7778 |
| 4_1499910 | 2 | 0.5 | NA | NA | 2 | 0 | 2 | 0 | 6 | 0.1667 | 4 | 0 | 1 | 0 | NA | NA | 2 | 0.5 | 7 | 0.1429 |
| 4_1499910 | 2 | 0.5 | NA | NA | 2 | 0 | 2 | 0 | 6 | 0.1667 | 3 | 0.3333 | 1 | 0 | NA | NA | 2 | 0 | 6 | 0.1667 |
| 4_1499911 | 3 | 0.6667 | NA | NA | 2 | 1 | 1 | 1 | 6 | 0.8333 | 2 | 0.5 | NA | NA | NA | NA | 2 | 1 | 4 | 0.75 |
| 4_1499912 | 1 | 0 | NA | NA | 2 | 1 | 2 | 1 | 5 | 0.8 | 6 | 1 | 1 | 1 | NA | NA | 2 | 1 | 9 | 1 |
| 4_1499915 | 3 | 0.6667 | NA | NA | 2 | 0.5 | 2 | 1 | 7 | 0.7143 | 5 | 0.4 | NA | NA | NA | NA | 2 | 1 | 7 | 0.5714 |

| | | | | | | | | | | | | | | | | | | | | |
|---|---|---|---|---|---|---|---|---|---|---|---|---|---|---|---|---|---|---|---|---|
| 4_1499915 | 2 | 0.5 | NA | NA | 1 | 0 | 2 | 0 | 5 | 0.2 | 6 | 0.3333 | 1 | 1 | NA | NA | 1 | 0 | 8 | 0.375 |
| 4_1500644 | 1 | 0 | 2 | 0.5 | NA | NA | 1 | 0 | 4 | 0.25 | NA | NA | NA | NA | 1 | 1 | 1 | 1 | 2 | 1 |
| 4_1500653 | NA | NA | 1 | 1 | NA | NA | 1 | 1 | 2 | 1 | NA | NA | NA | NA | NA | NA | 1 | 1 | 1 | 1 |
| 4_1500655 | NA | NA | 1 | 1 | NA | NA | 1 | 0 | 2 | 0.5 | NA | NA | NA | NA | 1 | 0 | 1 | 1 | 2 | 0.5 |
| 4_1500656 | NA | NA | 1 | 1 | NA | NA | 1 | 1 | 2 | 1 | NA | NA | NA | NA | 1 | 1 | 1 | 1 | 2 | 1 |
| 4_1500657 | NA | NA | 1 | 1 | NA | NA | 1 | 0 | 2 | 0.5 | NA | NA | NA | NA | 1 | 1 | NA | NA | 1 | 1 |
| 4_1500657 | NA | NA | 1 | 1 | NA | NA | 1 | 0 | 2 | 0.5 | NA | NA | NA | NA | 1 | 1 | 1 | 1 | 2 | 1 |
| 4_1500658 | NA | NA | 1 | 1 | NA | NA | 1 | 1 | 2 | 1 | NA | NA | NA | NA | 1 | 1 | 1 | 1 | 2 | 1 |
| 4_1500659 | NA | NA | 1 | 1 | NA | NA | 1 | 1 | 2 | 1 | NA | NA | NA | NA | 1 | 1 | 1 | 1 | 2 | 1 |
| 4_1500661 | NA | NA | 1 | 1 | NA | NA | 1 | 0 | 2 | 0.5 | NA | NA | NA | NA | 1 | 1 | 1 | 1 | 2 | 1 |
| 4_1500663 | NA | NA | 1 | 1 | NA | NA | 1 | 0 | 2 | 0.5 | NA | NA | NA | NA | 1 | 0 | 1 | 1 | 2 | 0.5 |
| 4_1500669 | NA | NA | 1 | 1 | NA | NA | NA | NA | 1 | 1 | NA | NA | NA | NA | NA | NA | 1 | 1 | 1 | 1 |
| 4_1500671 | NA | NA | 1 | 1 | NA | NA | NA | NA | 1 | 1 | NA | NA | NA | NA | 1 | 1 | 1 | 1 | 2 | 1 |
| 4_1500805 | NA | NA | NA | NA | 1 | 1 | NA | NA | 1 | 1 | NA | NA | NA | NA | NA | NA | 1 | 1 | 1 | 1 |
| 4_1500805 | NA | NA | NA | NA | 1 | 1 | NA | NA | 1 | 1 | NA | NA | NA | NA | NA | NA | 1 | 1 | 1 | 1 |
| 4_1500806 | NA | NA | NA | NA | 1 | 1 | NA | NA | 1 | 1 | NA | NA | NA | NA | NA | NA | 1 | 1 | 1 | 1 |
| 4_1502275 | NA | NA | NA | NA | NA | NA | 2 | 0.5 | 2 | 0.5 | NA | NA | 1 | 1 | 1 | 1 | 1 | 0 | 3 | 0.6667 |
| 4_1502277 | 1 | 1 | NA | NA | NA | NA | 2 | 1 | 3 | 1 | NA | NA | 1 | 0 | NA | NA | 1 | 1 | 2 | 0.5 |
| 4_1502280 | 1 | 1 | NA | NA | NA | NA | 2 | 0.5 | 3 | 0.6667 | NA | NA | 1 | 1 | NA | NA | NA | NA | 1 | 1 |
| 4_1502280 | 1 | 0 | NA | NA | NA | NA | 2 | 0.5 | 3 | 0.3333 | NA | NA | 1 | 1 | NA | NA | 1 | 1 | 2 | 1 |
| 4_1502281 | 1 | 0 | NA | NA | NA | NA | 2 | 0.5 | 3 | 0.3333 | NA | NA | 1 | 0 | NA | NA | 1 | 1 | 2 | 0.5 |
| 4_1506493 | 1 | 1 | NA | NA | NA | NA | NA | NA | 1 | 1 | NA | NA | 1 | 1 | NA | NA | NA | NA | 1 | 1 |
| 4_1506504 | 1 | 1 | 1 | 1 | NA | NA | 1 | 1 | 3 | 1 | NA | NA | 1 | 1 | 1 | 0 | NA | NA | 2 | 0.5 |
| 4_1506509 | 1 | 1 | 1 | 1 | NA | NA | NA | NA | 2 | 1 | NA | NA | 2 | 1 | 1 | 1 | NA | NA | 3 | 1 |
| 4_1506511 | 1 | 1 | 1 | 1 | NA | NA | 1 | 1 | 3 | 1 | NA | NA | 2 | 0.5 | 1 | 0 | NA | NA | 3 | 0.3333 |
| 4_1506511 | 1 | 1 | 1 | 1 | NA | NA | 2 | 0.5 | 4 | 0.75 | NA | NA | 2 | 1 | NA | NA | NA | NA | 2 | 1 |
| 4_1506513 | 1 | 1 | 1 | 0 | NA | NA | 2 | 0.5 | 4 | 0.5 | NA | NA | 2 | 0.5 | 1 | 1 | NA | NA | 3 | 0.6667 |
| 4_1506514 | 1 | 1 | 1 | 1 | NA | NA | 2 | 0.5 | 4 | 0.75 | NA | NA | 2 | 0.5 | 1 | 0 | NA | NA | 3 | 0.3333 |
| 4_1506516 | 1 | 0 | 1 | 1 | NA | NA | 2 | 1 | 4 | 0.75 | NA | NA | 1 | 1 | 1 | 1 | NA | NA | 2 | 1 |
| 4_1506522 | 1 | 1 | NA | NA | NA | NA | 2 | 0.5 | 3 | 0.6667 | NA | NA | 1 | 0 | 1 | 1 | NA | NA | 2 | 0.5 |
| 4_1506989 | 2 | 0 | 2 | 0.5 | 1 | 0 | 1 | 0 | 6 | 0.1667 | 2 | 0.5 | 1 | 0 | 1 | 0 | 1 | 0 | 5 | 0.2 |
| 4_1506990 | 3 | 0 | 4 | 0.5 | 1 | 0 | 1 | 1 | 9 | 0.3333 | 1 | 1 | 3 | 0.3333 | 1 | 1 | 2 | 0.5 | 7 | 0.5714 |
| 4_1506992 | 1 | 0 | 2 | 0 | 1 | 1 | 2 | 0 | 6 | 0.1667 | 3 | 0.3333 | 3 | 0 | 1 | 0 | 3 | 0 | 10 | 0.1 |
| 4_1507004 | 2 | 0 | 5 | 0 | 2 | 0 | 5 | 0.2 | 14 | 0.0714 | 5 | 0 | 6 | 0.1667 | 3 | 0.3333 | 6 | 0 | 20 | 0.1 |
| 4_1507013 | 1 | 0 | 3 | 1 | 2 | 1 | 1 | 0 | 7 | 0.7143 | 2 | 1 | 3 | 1 | 2 | 0.5 | 4 | 1 | 11 | 0.9091 |
| 4_1507016 | 2 | 0 | 3 | 0.3333 | 4 | 0 | 7 | 0.1429 | 16 | 0.125 | 4 | 0.5 | 6 | 0 | 5 | 0.4 | 8 | 0.25 | 23 | 0.2609 |
| 4_1507017 | 4 | 0.75 | 5 | 0.8 | 3 | 0.6667 | 7 | 0.8571 | 19 | 0.7895 | 3 | 0.6667 | 6 | 1 | 5 | 1 | 7 | 0.7143 | 21 | 0.8571 |
| 4_1507017 | 4 | 0.25 | 5 | 0.8 | 4 | 0.5 | 6 | 0.5 | 19 | 0.5263 | 4 | 0.75 | 4 | 0.5 | 3 | 1 | 9 | 0.6667 | 20 | 0.7 |
| 4_1507040 | 3 | 1 | 6 | 0.6667 | 5 | 1 | 7 | 0.5714 | 21 | 0.7619 | 5 | 0.6 | 7 | 0.7143 | 6 | 1 | 10 | 0.7 | 28 | 0.75 |
| 4_1507041 | 5 | 0.2 | 6 | 0.1667 | 4 | 0.25 | 8 | 0.5 | 23 | 0.3043 | 6 | 0.1667 | 7 | 0.1429 | 4 | 0.25 | 12 | 0.25 | 29 | 0.2069 |
| 4_1507050 | 6 | 0.1667 | 7 | 0.1429 | 8 | 0.125 | 12 | 0.1667 | 33 | 0.1515 | 6 | 0.3333 | 13 | 0.0769 | 10 | 0.1 | 15 | 0.3333 | 44 | 0.2045 |
| 4_1507053 | 5 | 0.6 | 6 | 0.6667 | 8 | 0.875 | 9 | 0.5556 | 28 | 0.6786 | 5 | 0.6 | 8 | 0.5 | 7 | 0.7143 | 14 | 0.8571 | 34 | 0.7059 |
| 4_1507060 | 11 | 0.0909 | 10 | 0 | 7 | 0 | 14 | 0.1429 | 42 | 0.0714 | 10 | 0.1 | 19 | 0.1053 | 13 | 0.0769 | 14 | 0 | 56 | 0.0714 |
| 4_1507061 | 11 | 0.3636 | 9 | 1 | 11 | 0.8182 | 11 | 0.4545 | 42 | 0.6429 | 10 | 0.5 | 18 | 0.6111 | 11 | 0.5455 | 15 | 0.4667 | 54 | 0.537 |
| 4_1507062 | 8 | 0.125 | 9 | 0.5556 | 9 | 0.4444 | 13 | 0.0769 | 39 | 0.2821 | 12 | 0.5 | 15 | 0.2 | 14 | 0.2857 | 13 | 0.1538 | 54 | 0.2778 |
| 4_1507139 | 2 | 1 | 5 | 1 | 7 | 1 | 4 | 1 | 18 | 1 | NA | NA | 3 | 1 | 3 | 0.6667 | 3 | 1 | 9 | 0.8889 |
| 4_1507146 | 1 | 0 | 1 | 1 | NA | NA | NA | NA | 2 | 0.5 | 2 | 0 | 1 | 1 | 1 | 0 | 1 | 0 | 5 | 0.2 |
| 4_1507225 | 4 | 0.25 | 5 | 0.4 | 4 | 0.5 | 4 | 0.25 | 17 | 0.3529 | 1 | 0 | 5 | 0.4 | 4 | 0.5 | 5 | 0.4 | 15 | 0.4 |
| 4_1507226 | 5 | 0 | 5 | 0.2 | 4 | 0 | 4 | 0.25 | 18 | 0.1111 | 1 | 0 | 6 | 0.3333 | 3 | 0 | 5 | 0 | 15 | 0.1333 |
| 4_1507227 | 2 | 1 | 3 | 1 | 3 | 1 | 4 | 1 | 12 | 1 | 1 | 1 | 3 | 1 | 3 | 0.6667 | 3 | 0.6667 | 10 | 0.8 |
| 4_1507227 | 3 | 0.3333 | 5 | 0.2 | 3 | 0.3333 | 4 | 0.25 | 15 | 0.2667 | 1 | 0 | 2 | 0 | 4 | 0 | 5 | 0.2 | 12 | 0.0833 |
| 4_1507228 | 1 | 1 | 4 | 0.5 | 3 | 0.6667 | 3 | 0.3333 | 11 | 0.5455 | 1 | 1 | 3 | 0.6667 | 4 | 0.25 | 5 | 0.6 | 13 | 0.5385 |
| 4_1507229 | 3 | 0.3333 | 5 | 0 | 3 | 0 | 4 | 0.25 | 15 | 0.1333 | NA | NA | 3 | 0 | 3 | 0.3333 | 4 | 0.5 | 10 | 0.3 |
| 4_1507270 | 3 | 0 | 4 | 0 | 2 | 0 | 3 | 0.6667 | 12 | 0.1667 | 1 | 1 | 2 | 0 | 2 | 0.5 | 3 | 0 | 8 | 0.25 |
| 4_1507285 | 3 | 0 | 2 | 0.5 | 1 | 1 | 3 | 0.3333 | 9 | 0.3333 | 1 | 0 | 1 | 1 | 2 | 0.5 | 2 | 1 | 6 | 0.6667 |
| 4_1507285 | 3 | 0.6667 | 4 | 0.25 | 2 | 0 | 3 | 0 | 12 | 0.25 | 1 | 0 | 2 | 0 | 2 | 0 | 2 | 0.5 | 7 | 0.1429 |
| 4_1507285 | 3 | 0.3333 | 3 | 1 | 1 | 0 | 2 | 0.5 | 9 | 0.5556 | 1 | 1 | 2 | 0 | 1 | 0 | 2 | 1 | 6 | 0.5 |
| 4_1507287 | 2 | 0.5 | 2 | 1 | 2 | 0.5 | 3 | 0.3333 | 9 | 0.5556 | NA | NA | 1 | 1 | 1 | 0 | 1 | 1 | 3 | 0.6667 |
| 4_1509508 | 4 | 0 | 4 | 0.25 | 3 | 0.3333 | 2 | 0 | 13 | 0.1538 | 2 | 0 | 5 | 0.4 | 1 | 1 | 1 | 0 | 9 | 0.3333 |
| 4_1509509 | 5 | 0.6 | 3 | 1 | 2 | 1 | 2 | 0.5 | 12 | 0.75 | 2 | 0.5 | 5 | 0.8 | NA | NA | 1 | 1 | 8 | 0.75 |
| 4_1509510 | 5 | 0 | 5 | 0.2 | 3 | 0 | 2 | 0 | 15 | 0.0667 | 2 | 0 | 5 | 0.2 | 1 | 0 | 1 | 0 | 9 | 0.1111 |
| 4_1509511 | 4 | 1 | 5 | 0.8 | 3 | 1 | 2 | 1 | 14 | 0.9286 | 2 | 1 | 5 | 1 | NA | NA | 1 | 1 | 8 | 1 |
| 4_1509511 | 5 | 0.4 | 5 | 0.4 | 3 | 0 | 2 | 0 | 15 | 0.2667 | 2 | 0 | 5 | 0.4 | 1 | 0 | 1 | 1 | 9 | 0.3333 |
| 4_1509512 | 4 | 0.25 | 3 | 0.3333 | 4 | 0 | 2 | 0 | 13 | 0.1538 | 2 | 0 | 5 | 0.2 | 1 | 0 | 1 | 0 | 9 | 0.1111 |
| 4_1509513 | 5 | 0 | 5 | 0 | 4 | 0.25 | 2 | 0 | 16 | 0.0625 | 2 | 0.5 | 5 | 0.6 | 2 | 0.5 | 1 | 0 | 10 | 0.5 |

| | | | | | | | | | | | | | | | | | | | | |
|---|---|---|---|---|---|---|---|---|---|---|---|---|---|---|---|---|---|---|---|---|
| 4_15095140 | 5 | 0.8 | 3 | 0.6667 | 5 | 0.6 | NA | NA | 13 | 0.6923 | 3 | 0.3333 | 5 | 0.8 | 3 | 0.6667 | 1 | 0 | 12 | 0.5833 |
| 4_15095160 | 3 | 0.3333 | 6 | 0.5 | 4 | 0.25 | 2 | 1 | 15 | 0.4667 | 3 | 0.6667 | 6 | 0.8333 | 2 | 1 | 1 | 0 | 12 | 0.75 |
| 4_15095178 | 6 | 1 | 6 | 0.6667 | 5 | 1 | 3 | 1 | 20 | 0.9 | 3 | 0.6667 | 5 | 1 | 3 | 1 | 2 | 1 | 13 | 0.9231 |
| 4_15095193 | 7 | 0.1429 | 6 | 0 | 6 | 0.1667 | 2 | 0 | 21 | 0.0952 | 3 | 0 | 6 | 0.1667 | 2 | 0 | 2 | 0 | 13 | 0.0769 |
| 4_15095203 | 6 | 0 | 5 | 0.4 | 5 | 0.6 | 1 | 0 | 17 | 0.2941 | 4 | 0.5 | 6 | 0 | 3 | 0.3333 | 2 | 0.5 | 15 | 0.2667 |
| 4_15095206 | 6 | 0.6667 | 6 | 0.6667 | 6 | 1 | 3 | 1 | 21 | 0.8095 | 2 | 0.5 | 6 | 0.8333 | 3 | 0.6667 | 2 | 1 | 13 | 0.7692 |
| 4_15095257 | 8 | 0 | 7 | 0 | 4 | 0.25 | 4 | 0.25 | 23 | 0.087 | 2 | 0 | 5 | 0 | 2 | 0.5 | 2 | 0 | 11 | 0.0909 |
| 4_15095258 | 8 | 0 | 4 | 0.25 | 4 | 0.25 | 3 | 0 | 19 | 0.1053 | 4 | 0.5 | 4 | 0.25 | 3 | 0 | 2 | 0 | 13 | 0.2308 |
| 4_15095319 | 9 | 0.1111 | 7 | 0.4286 | 5 | 0 | 3 | 0 | 24 | 0.1667 | 2 | 0.5 | 8 | 0.125 | 3 | 0.3333 | 2 | 0 | 15 | 0.2 |
| 4_15098177 | 29 | 0.3103 | 28 | 0.2143 | 25 | 0.16 | 26 | 0.1154 | 108 | 0.2037 | 28 | 0.3571 | 22 | 0.2727 | 27 | 0.3704 | 23 | 0.3913 | 100 | 0.35 |
| 4_15108817 | 5 | 0.2 | 3 | 0.3333 | 1 | 1 | 3 | 0 | 12 | 0.25 | 3 | 0 | 3 | 0 | 6 | 0 | 5 | 0.2 | 17 | 0.0588 |
| 4_15108879 | 3 | 0.3333 | 3 | 0 | 1 | 0 | 3 | 0.6667 | 10 | 0.3 | 3 | 0.6667 | 3 | 0 | 5 | 0.2 | 5 | 0.4 | 16 | 0.3125 |
| 4_15108888 | 3 | 0 | 3 | 0.3333 | 1 | 0 | 2 | 0 | 9 | 0.1111 | 3 | 0 | 3 | 0.3333 | 7 | 0 | 5 | 0 | 18 | 0.0556 |
| 4_15108891 | 1 | 1 | 2 | 0 | 1 | 0 | 2 | 0 | 6 | 0.1667 | 2 | 0 | 1 | 0 | 5 | 0.2 | 4 | 0 | 12 | 0.0833 |
| 4_15108894 | 2 | 0.5 | 3 | 0 | 1 | 0 | 3 | 0 | 9 | 0.1111 | 3 | 0 | 3 | 0 | 7 | 0.2857 | 5 | 0 | 18 | 0.1111 |
| 4_15108961 | 2 | 0.5 | 1 | 0 | NA | NA | 2 | 0 | 5 | 0.2 | 3 | 0 | 3 | 0.3333 | 5 | 0 | 5 | 0.4 | 16 | 0.1875 |
| 4_15108963 | 2 | 1 | 2 | 0 | NA | NA | 3 | 1 | 7 | 0.7143 | 3 | 1 | 3 | 1 | 5 | 1 | 5 | 0.6 | 16 | 0.875 |
| 4_15108973 | 2 | 1 | 2 | 0.5 | NA | NA | 3 | 1 | 7 | 0.8571 | 2 | 1 | 4 | 0.75 | 5 | 1 | 4 | 1 | 15 | 0.9333 |
| 4_15108973 | 2 | 0.5 | 2 | 0.5 | NA | NA | 2 | 1 | 6 | 0.6667 | 2 | 0.5 | 2 | 1 | 6 | 0.6667 | 3 | 1 | 13 | 0.7692 |
| 4_15108978 | 2 | 1 | 2 | 0 | NA | NA | 3 | 0 | 7 | 0.2857 | 3 | 0 | 4 | 0 | 6 | 0 | 4 | 0.25 | 17 | 0.0588 |
| 4_15108984 | 2 | 0.5 | 2 | 0 | NA | NA | 2 | 0 | 6 | 0.1667 | 3 | 0 | 2 | 0 | 3 | 0 | 3 | 0.3333 | 11 | 0.0909 |
| 4_15108988 | NA | NA | 1 | 0 | NA | NA | 2 | 1 | 3 | 0.6667 | 3 | 0.6667 | 3 | 1 | 4 | 0.75 | 4 | 0.75 | 14 | 0.7857 |
| 4_15109002 | 2 | 0.5 | 2 | 0 | NA | NA | 3 | 0 | 7 | 0.1429 | 3 | 0 | 3 | 0 | 3 | 0.6667 | 3 | 0.6667 | 12 | 0.3333 |
| 4_15109023 | 1 | 1 | 2 | 0.5 | NA | NA | 1 | 1 | 4 | 0.75 | 2 | 0.5 | 2 | 1 | 3 | 0.6667 | 2 | 0.5 | 9 | 0.6667 |
| 4_15109025 | 1 | 1 | 2 | 0 | NA | NA | 3 | 0.3333 | 6 | 0.3333 | 2 | 0.5 | 3 | 0.3333 | 4 | 0.25 | 3 | 0.3333 | 12 | 0.3333 |
| 4_15109029 | 1 | 1 | 2 | 0.5 | NA | NA | 3 | 0.3333 | 6 | 0.5 | 3 | 0.3333 | 4 | 0.25 | 4 | 0.5 | 3 | 0.6667 | 14 | 0.4286 |
| 4_15109684 | 1 | 1 | 1 | 0 | NA | NA | 1 | 0 | 3 | 0.3333 | 1 | 1 | 3 | 0.6667 | 3 | 0.3333 | 3 | 0.3333 | 10 | 0.5 |
| 4_15109692 | 1 | 1 | NA | NA | NA | NA | 1 | 0 | 2 | 0.5 | 1 | 0 | 2 | 0.5 | 3 | 0.3333 | 3 | 0 | 9 | 0.2222 |
| 4_15113310 | 4 | 0.25 | 2 | 0 | 3 | 0 | 8 | 0 | 17 | 0.0588 | 4 | 0.25 | 5 | 0 | 3 | 0 | 4 | 0.25 | 16 | 0.125 |
| 4_15113324 | 4 | 0.5 | 1 | 0 | 4 | 0 | 8 | 0.375 | 17 | 0.2941 | 4 | 0 | 4 | 0.25 | 5 | 0.4 | 4 | 0.25 | 17 | 0.2353 |
| 4_15113325 | 5 | 0.8 | 2 | 0 | 3 | 0.3333 | 6 | 0.3333 | 16 | 0.4375 | 3 | 0.6667 | 5 | 0.4 | 1 | 0 | 1 | 0 | 10 | 0.4 |
| 4_15114631 | 8 | 0.375 | 15 | 0.2 | 9 | 0.3333 | 6 | 0.1667 | 38 | 0.2632 | 4 | 0.5 | 11 | 0.3636 | 12 | 0.5833 | 8 | 0.5 | 35 | 0.4857 |
| 4_15114681 | 10 | 0 | 19 | 0.1053 | 11 | 0.0909 | 8 | 0.125 | 48 | 0.0833 | 6 | 0 | 11 | 0.0909 | 11 | 0 | 8 | 0.125 | 36 | 0.0556 |
| 4_15114686 | 8 | 0.25 | 9 | 0.3333 | 6 | 0 | 5 | 0.2 | 28 | 0.2143 | 3 | 0.3333 | 8 | 0.25 | 8 | 0.375 | 2 | 0 | 21 | 0.2857 |
| 4_15114690 | 8 | 1 | 17 | 1 | 11 | 1 | 9 | 0.8889 | 45 | 0.9778 | 6 | 0.8333 | 11 | 1 | 7 | 1 | 6 | 0.8333 | 30 | 0.9333 |
| 4_15114697 | 7 | 0.2857 | 17 | 0.6471 | 7 | 0.8571 | 6 | 0.5 | 37 | 0.5946 | 4 | 0.25 | 8 | 0.625 | 8 | 0.625 | 5 | 0.4 | 25 | 0.52 |
| 4_15114701 | 7 | 0.5714 | 14 | 0.5 | 9 | 0.3333 | 8 | 0.5 | 38 | 0.4737 | 5 | 0.2 | 7 | 0.1429 | 8 | 0.625 | 3 | 0.6667 | 23 | 0.3913 |
| 4_15114705 | 9 | 0.8889 | 17 | 1 | 10 | 1 | 9 | 0.8889 | 45 | 0.9556 | 6 | 1 | 10 | 1 | 9 | 1 | 8 | 1 | 33 | 1 |
| 4_15114720 | 8 | 0 | 17 | 0.1176 | 10 | 0.4 | 9 | 0.2222 | 44 | 0.1818 | 6 | 0.5 | 10 | 0.1 | 8 | 0 | 8 | 0.125 | 32 | 0.1562 |
| 4_15114723 | 5 | 1 | 15 | 0.8 | 8 | 0.75 | 8 | 0.625 | 36 | 0.7778 | 4 | 1 | 6 | 0.5 | 7 | 0.2857 | 7 | 0.5714 | 24 | 0.5417 |
| 4_15114724 | 7 | 0.1429 | 17 | 0.0588 | 8 | 0 | 9 | 0.1111 | 41 | 0.0732 | 6 | 0 | 9 | 0.1111 | 9 | 0.2222 | 8 | 0.125 | 32 | 0.125 |
| 4_15116839 | NA | NA | 5 | 0.2 | 1 | 0 | 3 | 0.6667 | 9 | 0.3333 | 1 | 1 | 4 | 0 | 4 | 0 | 2 | 0.5 | 11 | 0.1818 |
| 4_15122820 | NA | NA | 1 | 1 | 2 | 0 | 1 | 0 | 4 | 0.25 | NA | NA | 2 | 1 | NA | NA | NA | NA | 2 | 1 |
| 4_15122853 | NA | NA | 1 | 0 | 2 | 1 | 1 | 1 | 4 | 0.75 | NA | NA | 1 | 1 | NA | NA | NA | NA | 1 | 1 |
| 4_15122859 | NA | NA | 1 | 1 | 1 | 1 | 1 | 1 | 3 | 1 | NA | NA | 1 | 1 | NA | NA | NA | NA | 1 | 1 |
| 4_15122865 | NA | NA | 1 | 1 | 2 | 1 | 1 | 1 | 4 | 1 | NA | NA | 1 | 1 | NA | NA | NA | NA | 1 | 1 |
| 4_15122905 | NA | NA | 1 | 0 | 2 | 1 | 1 | 0 | 4 | 0.5 | NA | NA | 1 | 1 | NA | NA | NA | NA | 1 | 1 |
| 4_15122938 | NA | NA | NA | NA | 2 | 1 | NA | NA | 2 | 1 | NA | NA | 1 | 1 | NA | NA | NA | NA | 1 | 1 |
| 4_15186514 | NA | NA | NA | NA | 1 | 1 | NA | NA | 1 | 1 | 1 | 1 | 1 | 0 | NA | NA | NA | NA | 2 | 0.5 |
| 4_15186519 | NA | NA | NA | NA | 1 | 1 | NA | NA | 1 | 1 | 1 | 0 | 1 | 1 | NA | NA | NA | NA | 2 | 0.5 |
| 4_15206401 | 2 | 0 | 6 | 0 | 6 | 0.1667 | 5 | 0 | 19 | 0.0526 | 4 | 0 | 5 | 0 | 6 | 0.1667 | 4 | 0.25 | 19 | 0.1053 |
| 4_15206526 | 4 | 0 | 7 | 0 | 8 | 0.25 | 7 | 0.1429 | 26 | 0.1154 | 3 | 0 | 7 | 0.1429 | 6 | 0 | 5 | 0.2 | 21 | 0.0952 |
| 4_15206532 | 4 | 1 | 7 | 0.8571 | 8 | 0.5 | 7 | 0.5714 | 26 | 0.6923 | 4 | 0.5 | 5 | 0.8 | 4 | 0.75 | 2 | 1 | 15 | 0.7333 |
| 4_15206536 | 6 | 1 | 7 | 0.7143 | 7 | 1 | 5 | 0.6 | 25 | 0.84 | 4 | 1 | 7 | 0.5714 | 7 | 1 | 5 | 0.8 | 23 | 0.8261 |
| 4_15206549 | 6 | 0.8333 | 6 | 0.6667 | 5 | 0.6 | 3 | 0.3333 | 20 | 0.65 | 4 | 0.75 | 7 | 0.1429 | 6 | 0.8333 | 5 | 1 | 22 | 0.6364 |
| 4_15206551 | 4 | 1 | 8 | 0.875 | 7 | 0.8571 | 8 | 0.875 | 27 | 0.8889 | 4 | 0.75 | 8 | 0.875 | 6 | 1 | 5 | 0.8 | 23 | 0.8696 |
| 4_15206559 | 6 | 0 | 8 | 0.125 | 7 | 0.1429 | 7 | 0.2857 | 28 | 0.1429 | 5 | 0.2 | 8 | 0.25 | 7 | 0.2857 | 4 | 0.5 | 24 | 0.2917 |
| 4_15206563 | 5 | 0.8 | 8 | 0.875 | 8 | 0.875 | 6 | 0.8333 | 27 | 0.8519 | 5 | 1 | 7 | 1 | 6 | 0.6667 | 4 | 1 | 22 | 0.9091 |
| 4_15206564 | 3 | 1 | 7 | 0.7143 | 7 | 1 | 7 | 0.8571 | 24 | 0.875 | 2 | 1 | 8 | 0.875 | 7 | 1 | 5 | 0.8 | 22 | 0.9091 |
| 4_15206566 | 3 | 0 | 7 | 0.2857 | 4 | 0.75 | 8 | 0.375 | 22 | 0.3636 | 4 | 0.25 | 9 | 0.3333 | 7 | 0.2857 | 4 | 0.25 | 24 | 0.2917 |
| 4_15206567 | 3 | 0.6667 | 3 | 1 | 4 | 1 | 1 | 1 | 11 | 0.9091 | 3 | 0.6667 | 3 | 0.6667 | 3 | 1 | 2 | 0.5 | 11 | 0.7273 |
| 4_15206575 | 3 | 0 | 3 | 0.3333 | 3 | 0.3333 | 2 | 0.5 | 11 | 0.2727 | 4 | 0.25 | 1 | 0 | 3 | 0 | 2 | 0.5 | 10 | 0.2 |
| 4_15206584 | 3 | 0.6667 | 4 | 0.25 | 3 | 0 | 3 | 0.3333 | 13 | 0.3077 | 4 | 0 | 3 | 0 | 3 | 0.3333 | 3 | 0.3333 | 13 | 0.1538 |
| 4_15206636 | 3 | 0 | 5 | 0 | 5 | 0.2 | 4 | 0.25 | 17 | 0.1176 | 5 | 0.2 | 3 | 0.3333 | 3 | 0 | 2 | 0.5 | 13 | 0.2308 |
| 4_15206755 | 2 | 0 | 3 | 0 | 5 | 0.6 | 4 | 0.5 | 14 | 0.3571 | NA | NA | 3 | 0.3333 | 2 | 0.5 | 1 | 0 | 6 | 0.3333 |

| | | | | | | | | | | | | | | | | | | | | |
|---|---|---|---|---|---|---|---|---|---|---|---|---|---|---|---|---|---|---|---|---|
| 4_1520675( | 2 | 0.5 | 3 | 0 | 5 | 0.4 | 4 | 0.5 | 14 | 0.3571 | 1 | 0 | 2 | 0.5 | 1 | 1 | 1 | 1 | 5 | 0.6 |
| 4_1520675! | 1 | 0 | 4 | 0.25 | 3 | 0.6667 | 4 | 0.25 | 12 | 0.3333 | 1 | 1 | 2 | 0.5 | 2 | 0 | NA | NA | 5 | 0.4 |
| 4_1520676 | 1 | 1 | 2 | 1 | 1 | 0 | 1 | 1 | 5 | 0.8 | NA | NA | NA | NA | 2 | 0.5 | 1 | 0 | 3 | 0.3333 |
| 4_1520676! | 2 | 0.5 | 4 | 0.25 | 5 | 0.6 | 5 | 0.6 | 16 | 0.5 | 1 | 1 | 2 | 0.5 | 2 | 1 | 1 | 1 | 6 | 0.8333 |
| 4_1520758( | NA | NA | 1 | 1 | NA | NA | 2 | 1 | 3 | 1 | NA | NA | NA | NA | 1 | 1 | NA | NA | 1 | 1 |
| 4_1520758( | NA | NA | 1 | 1 | NA | NA | 2 | 0.5 | 3 | 0.6667 | NA | NA | NA | NA | 1 | 1 | NA | NA | 1 | 1 |
| 4_1520759; | NA | NA | 1 | 1 | NA | NA | 1 | 0 | 2 | 0.5 | NA | NA | NA | NA | 1 | 1 | NA | NA | 1 | 1 |
| 4_1520759( | NA | NA | 2 | 1 | NA | NA | 2 | 1 | 4 | 1 | NA | NA | NA | NA | 1 | 1 | NA | NA | 1 | 1 |
| 4_1520760' | NA | NA | 2 | 1 | NA | NA | 1 | 1 | 3 | 1 | NA | NA | NA | NA | 1 | 1 | NA | NA | 1 | 1 |
| 4_1520760; | NA | NA | 2 | 1 | NA | NA | 2 | 0.5 | 4 | 0.75 | NA | NA | NA | NA | 1 | 1 | NA | NA | 1 | 1 |
| 4_1520970! | 1 | 0 | 1 | 0 | NA | NA | 1 | 1 | 3 | 0.3333 | 1 | 0 | NA | NA | NA | NA | 1 | 1 | 2 | 0.5 |
| 4_1521001( | 1 | 0 | NA | NA | NA | NA | 1 | 1 | 2 | 0.5 | 1 | 1 | NA | NA | 1 | 0 | NA | NA | 2 | 0.5 |
| 4_1521002; | 1 | 1 | NA | NA | NA | NA | 1 | 1 | 2 | 1 | 2 | 0.5 | NA | NA | 1 | 1 | 1 | 0 | 4 | 0.5 |
| 4_1521005- | 1 | 0 | 1 | 0 | 1 | 1 | 2 | 0 | 5 | 0.2 | 1 | 0 | NA | NA | 1 | 1 | 1 | 0 | 3 | 0.3333 |
| 4_1521017! | 2 | 0 | 4 | 0.25 | 1 | 0 | 3 | 0 | 10 | 0.1 | 3 | 0.3333 | NA | NA | 3 | 0.3333 | 2 | 0 | 8 | 0.25 |
| 4_1521019; | 2 | 0 | 4 | 0 | 1 | 0 | 5 | 0.2 | 12 | 0.0833 | 3 | 0.6667 | 2 | 0 | 4 | 0.25 | 2 | 0 | 11 | 0.2727 |
| 4_1521042! | NA | NA | 1 | 0 | 1 | 1 | NA | NA | 2 | 0.5 | NA | NA | 3 | 0 | 2 | 0.5 | 1 | 1 | 6 | 0.3333 |
| 4_1521043; | NA | NA | 1 | 1 | 1 | 1 | NA | NA | 2 | 1 | NA | NA | 2 | 0.5 | 2 | 0.5 | 2 | 0.5 | 6 | 0.5 |
| 4_1521109( | NA | NA | 1 | 1 | NA | NA | NA | NA | 1 | 1 | NA | NA | 1 | 0 | NA | NA | 1 | 1 | 2 | 0.5 |
| 4_1521110! | NA | NA | 1 | 1 | NA | NA | NA | NA | 1 | 1 | NA | NA | 1 | 0 | NA | NA | 1 | 1 | 2 | 0.5 |
| 4_1521112( | NA | NA | 1 | 1 | NA | NA | NA | NA | 1 | 1 | NA | NA | 1 | 1 | NA | NA | 1 | 1 | 2 | 1 |
| 4_1521195' | 1 | 1 | NA | NA | NA | NA | NA | NA | 1 | 1 | NA | NA | 1 | 1 | NA | NA | NA | NA | 1 | 1 |
| 4_1521196( | 1 | 1 | NA | NA | NA | NA | NA | NA | 1 | 1 | NA | NA | 1 | 1 | NA | NA | NA | NA | 1 | 1 |
| 4_1521198' | 1 | 1 | NA | NA | NA | NA | 1 | 0 | 2 | 0.5 | NA | NA | 1 | 1 | NA | NA | NA | NA | 1 | 1 |
| 4_1521429- | 3 | 1 | 3 | 0.6667 | NA | NA | 3 | 0.6667 | 9 | 0.7778 | NA | NA | 2 | 1 | 1 | 1 | 3 | 1 | 6 | 1 |
| 4_1521429! | 2 | 0.5 | 3 | 0.6667 | NA | NA | 4 | 0 | 9 | 0.3333 | NA | NA | 2 | 0.5 | 2 | 0 | 3 | 0.3333 | 7 | 0.2857 |
| 4_1521430! | 3 | 1 | 2 | 0 | NA | NA | 2 | 1 | 7 | 0.7143 | NA | NA | 2 | 1 | 2 | 1 | 2 | 1 | 6 | 1 |
| 4_1521431- | 2 | 0.5 | 3 | 1 | 2 | 0.5 | 3 | 1 | 10 | 0.8 | NA | NA | 2 | 0 | NA | NA | 2 | 1 | 4 | 0.5 |
| 4_1521431( | 2 | 0.5 | 4 | 0.25 | 2 | 0.5 | 4 | 0 | 12 | 0.25 | NA | NA | 2 | 0 | 2 | 0.5 | 4 | 0 | 8 | 0.125 |
| 4_1521431! | 3 | 1 | 2 | 1 | 3 | 1 | 4 | 0.25 | 12 | 0.75 | NA | NA | 2 | 1 | 2 | 1 | 4 | 0.5 | 8 | 0.75 |
| 4_1521433' | 3 | 0.6667 | 4 | 0.5 | 3 | 0.3333 | 4 | 0.75 | 14 | 0.5714 | NA | NA | 2 | 0.5 | 2 | 1 | 3 | 0.3333 | 7 | 0.5714 |
| 4_1521433- | 2 | 0.5 | 4 | 0.5 | 3 | 0.3333 | 4 | 1 | 13 | 0.6154 | NA | NA | 2 | 0 | 2 | 0.5 | 4 | 1 | 8 | 0.625 |
| 4_1521433( | 3 | 0.3333 | 4 | 0.5 | 3 | 0.3333 | 3 | 0 | 13 | 0.3077 | NA | NA | 2 | 1 | 2 | 0.5 | 3 | 0.6667 | 7 | 0.7143 |
| 4_1521434' | 3 | 1 | 3 | 1 | 1 | 1 | 2 | 1 | 9 | 1 | NA | NA | 2 | 0.5 | 2 | 1 | 2 | 1 | 6 | 0.8333 |
| 4_1521434( | 3 | 1 | 3 | 1 | 3 | 1 | 3 | 1 | 12 | 1 | NA | NA | 2 | 1 | 2 | 1 | 4 | 1 | 8 | 1 |
| 4_1521436( | 3 | 1 | 3 | 1 | 3 | 1 | 4 | 0.75 | 13 | 0.9231 | NA | NA | 2 | 1 | 2 | 1 | 4 | 1 | 8 | 1 |
| 4_1521437; | 3 | 1 | 4 | 1 | 3 | 1 | 3 | 1 | 13 | 1 | NA | NA | 2 | 1 | 3 | 1 | 3 | 1 | 8 | 1 |
| 4_1521438; | 3 | 0.6667 | 4 | 0.5 | 3 | 0.3333 | 3 | 1 | 13 | 0.6154 | NA | NA | 2 | 1 | 3 | 1 | 3 | 1 | 8 | 1 |
| 4_1521438! | 3 | 0.3333 | 4 | 0.25 | 3 | 0.6667 | 3 | 0.3333 | 13 | 0.3846 | NA | NA | 2 | 0.5 | 2 | 0 | 4 | 0.25 | 8 | 0.25 |
| 4_1521439; | 3 | 0 | 4 | 0.25 | 3 | 0.3333 | 4 | 0 | 14 | 0.1429 | NA | NA | 2 | 0 | 3 | 0.6667 | 3 | 0.3333 | 8 | 0.375 |
| 4_1521440( | 3 | 0.6667 | 1 | 0 | 2 | 1 | 2 | 0.5 | 8 | 0.625 | NA | NA | 1 | 1 | 2 | 0.5 | 3 | 0.3333 | 6 | 0.5 |
| 4_1521442- | 3 | 0.3333 | 2 | 0 | 2 | 1 | 2 | 0 | 9 | 0.3333 | NA | NA | 2 | 0 | 3 | 0 | 4 | 0.5 | 9 | 0.2222 |
| 4_1521443- | 3 | 0 | 3 | 0.3333 | 3 | 0 | 3 | 0 | 12 | 0.0833 | NA | NA | 2 | 0 | 3 | 0.3333 | 4 | 0.25 | 9 | 0.2222 |
| 4_1521443! | 2 | 0 | 1 | 0 | 3 | 0 | 4 | 0.75 | 10 | 0.3 | 1 | 0 | NA | NA | 3 | 0 | 3 | 0.3333 | 7 | 0.1429 |
| 4_1521984! | 1 | 0 | 10 | 0.7 | 2 | 1 | 5 | 1 | 18 | 0.7778 | 4 | 1 | 5 | 0.6 | 4 | 0.75 | 4 | 0.75 | 17 | 0.7647 |
| 4_1521984! | 2 | 1 | 12 | 0.4167 | 2 | 1 | 5 | 0.4 | 21 | 0.5238 | 4 | 0.75 | 5 | 0.6 | 4 | 0.5 | 5 | 0.6 | 18 | 0.6111 |
| 4_1521987( | 2 | 1 | 14 | 0.8571 | 2 | 0.5 | 7 | 0.8571 | 25 | 0.84 | 4 | 1 | 4 | 1 | 6 | 1 | 4 | 0.75 | 18 | 0.9444 |
| 4_1521987- | 2 | 1 | 11 | 0.4545 | 4 | 0.5 | 7 | 0.5714 | 24 | 0.5417 | 4 | 0.5 | 5 | 0.6 | 6 | 1 | 5 | 0.6 | 20 | 0.7 |
| 4_1521988; | 2 | 0 | 13 | 0.1538 | 3 | 0 | 7 | 0 | 25 | 0.08 | 5 | 0 | 5 | 0 | 5 | 0.2 | 2 | 0 | 17 | 0.0588 |
| 4_1521989( | 3 | 0 | 8 | 0.25 | 3 | 0.6667 | 8 | 0.375 | 22 | 0.3182 | 4 | 0.5 | 6 | 0.8333 | 4 | 0.5 | 4 | 0.75 | 18 | 0.6667 |
| 4_1521995( | 3 | 1 | 12 | 0.6667 | 1 | 1 | 6 | 0.5 | 22 | 0.6818 | 5 | 0.8 | 6 | 0.5 | 5 | 0.8 | 2 | 0.5 | 18 | 0.6667 |
| 4_1521996( | 4 | 0.25 | 13 | 0.6154 | 6 | 0.6667 | 6 | 0.5 | 29 | 0.5517 | 4 | 0.5 | 4 | 0.25 | 4 | 0.75 | 3 | 1 | 15 | 0.6 |
| 4_1521998; | 4 | 0.25 | 12 | 0.8333 | 5 | 0.8 | 8 | 0.375 | 29 | 0.6207 | 5 | 0.8 | 5 | 0.2 | 4 | 0.75 | 4 | 0.75 | 18 | 0.6111 |
| 4_1522037! | NA | NA | 1 | 0 | 1 | 0 | 1 | 1 | 3 | 0.3333 | NA | NA | 1 | 1 | 2 | 0 | NA | NA | 3 | 0.3333 |
| 4_1522043; | NA | NA | 1 | 1 | 1 | 0 | 1 | 1 | 3 | 0.6667 | NA | NA | 1 | 0 | 2 | 0.5 | 1 | 0 | 4 | 0.25 |
| 4_1522045- | NA | NA | 1 | 1 | 1 | 1 | 2 | 1 | 4 | 1 | NA | NA | 1 | 1 | 2 | 1 | 1 | 0 | 4 | 0.75 |
| 4_1523540( | NA | NA | 1 | 1 | NA | NA | 1 | 0 | 2 | 0.5 | NA | NA | 1 | 1 | NA | NA | NA | NA | 1 | 1 |
| 4_1523562- | 46 | 0.0435 | 68 | 0.0588 | 68 | 0.1029 | 45 | 0.1333 | 227 | 0.0837 | 48 | 0.125 | 37 | 0.1081 | 37 | 0.1081 | 55 | 0.0727 | 177 | 0.1017 |
| 4_1523646! | 1 | 0 | 1 | 0 | 1 | 1 | NA | NA | 3 | 0.3333 | NA | NA | 1 | 1 | 1 | 0 | NA | NA | 2 | 0.5 |
| 4_1523674; | 1 | 1 | 1 | 1 | 2 | 1 | 1 | 1 | 5 | 1 | NA | NA | NA | NA | 2 | 1 | 1 | 1 | 3 | 1 |
| 4_1523676- | NA | NA | 1 | 1 | 2 | 1 | 1 | 1 | 4 | 1 | NA | NA | 1 | 1 | 1 | 1 | 1 | 1 | 3 | 1 |
| 4_1523680; | 1 | 1 | 2 | 1 | 1 | 1 | 1 | 1 | 5 | 1 | 1 | 1 | 1 | 1 | NA | NA | 1 | 1 | 3 | 1 |
| 4_1523681( | 1 | 1 | 2 | 1 | 2 | 1 | NA | NA | 5 | 1 | 1 | 1 | 1 | 1 | 3 | 0.6667 | NA | NA | 5 | 0.8 |
| 4_1523682- | 1 | 1 | 1 | 0 | 2 | 1 | NA | NA | 4 | 0.75 | NA | NA | 1 | 1 | 2 | 0.5 | 1 | 1 | 4 | 0.75 |
| 4_1523682- | 1 | 1 | 2 | 1 | NA | NA | 1 | 1 | 4 | 1 | NA | NA | NA | NA | 2 | 0.5 | NA | NA | 2 | 0.5 |

| | | | | | | | | | | | | | | | | | | | | |
|---|---|---|---|---|---|---|---|---|---|---|---|---|---|---|---|---|---|---|---|---|
| 4_15236830 | 1 | 0 | 2 | 0.5 | 2 | 0.5 | NA | NA | 5 | 0.4 | 1 | 1 | 1 | 0 | 3 | 0.3333 | 1 | 1 | 6 | 0.5 |
| 4_15236857 | 1 | 0 | 2 | 0 | 3 | 0.3333 | 1 | 0 | 7 | 0.1429 | 1 | 0 | 1 | 0 | 2 | 0.5 | 1 | 0 | 5 | 0.2 |
| 4_15236860 | 1 | 1 | 2 | 1 | 3 | 1 | 1 | 1 | 7 | 1 | 1 | 0 | 1 | 1 | 2 | 1 | 1 | 0 | 5 | 0.6 |
| 4_15236873 | 2 | 0 | 2 | 0 | 3 | 0.3333 | 1 | 0 | 8 | 0.125 | 1 | 0 | 3 | 0 | 3 | 0.3333 | 1 | 0 | 8 | 0.125 |
| 4_15236879 | 1 | 0 | 3 | 0.6667 | 3 | 0 | 1 | 0 | 8 | 0.25 | 1 | 0 | 3 | 0 | 4 | 0.25 | 2 | 0 | 10 | 0.1 |
| 4_15236892 | 2 | 0 | 3 | 1 | 3 | 0.6667 | 2 | 1 | 10 | 0.7 | 1 | 0 | 3 | 1 | 3 | 1 | 2 | 1 | 9 | 0.8889 |
| 4_15236940 | 3 | 0.6667 | 3 | 0.3333 | 3 | 0.6667 | 3 | 0.6667 | 12 | 0.5833 | 1 | 1 | 2 | 0.5 | 4 | 1 | 2 | 0.5 | 9 | 0.7778 |
| 4_15238768 | 2 | 1 | 1 | 0 | 1 | 1 | NA | NA | 4 | 0.75 | 2 | 0.5 | 3 | 0.6667 | 2 | 0.5 | 2 | 1 | 9 | 0.6667 |
| 4_15238771 | 2 | 1 | 1 | 1 | 1 | 0 | NA | NA | 4 | 0.75 | 2 | 0.5 | 3 | 1 | 3 | 1 | 2 | 1 | 10 | 0.9 |
| 4_15238771 | NA | NA | NA | NA | 1 | 1 | NA | NA | 1 | 1 | 1 | 1 | 1 | 0 | 1 | 1 | NA | NA | 3 | 0.6667 |
| 4_15238780 | 2 | 0.5 | 1 | 0 | 1 | 0 | NA | NA | 4 | 0.25 | 2 | 0.5 | 3 | 0.3333 | 3 | 0.3333 | 3 | 0.3333 | 11 | 0.3636 |
| 4_15238781 | 1 | 1 | 1 | 0 | 1 | 0 | NA | NA | 3 | 0.3333 | 2 | 0 | 3 | 0 | 2 | 0 | 3 | 0.3333 | 10 | 0.1 |
| 4_15238799 | 2 | 1 | 1 | 1 | 1 | 1 | NA | NA | 4 | 1 | 2 | 1 | 3 | 1 | 2 | 1 | 2 | 1 | 9 | 1 |
| 4_15238815 | 3 | 0.6667 | 1 | 0 | 1 | 1 | NA | NA | 5 | 0.6 | 2 | 0.5 | 3 | 0.3333 | 3 | 0 | 3 | 1 | 11 | 0.4545 |
| 4_15238823 | 3 | 1 | 1 | 0 | 1 | 1 | 1 | 0 | 6 | 0.6667 | 2 | 0.5 | 2 | 0 | 2 | 1 | 3 | 1 | 9 | 0.6667 |
| 4_15238902 | 3 | 0.3333 | 2 | 0 | 2 | 0 | 2 | 0 | 9 | 0.1111 | 4 | 0.25 | 4 | 0 | 2 | 0 | 4 | 0.25 | 14 | 0.1429 |
| 4_15238935 | 2 | 0 | 3 | 0.3333 | 2 | 0 | 2 | 0 | 9 | 0.1111 | 4 | 0 | 3 | 0 | 2 | 0 | 4 | 0.25 | 13 | 0.0769 |
| 4_15238947 | 3 | 0 | 3 | 0 | 3 | 0 | 2 | 0.5 | 11 | 0.0909 | 4 | 0 | 5 | 0.4 | 1 | 0 | 5 | 0 | 15 | 0.1333 |
| 4_15238997 | 3 | 0 | 4 | 0 | 3 | 0.3333 | 2 | 0 | 12 | 0.0833 | 4 | 0 | 5 | 0 | 3 | 0 | 7 | 0.1429 | 19 | 0.0526 |
| 4_15239089 | 3 | 0 | 5 | 0.2 | 4 | 0 | 2 | 0 | 14 | 0.0714 | 5 | 0 | 6 | 0.1667 | 4 | 0 | 7 | 0.1429 | 22 | 0.0909 |
| 4_15239120 | 4 | 0.25 | 4 | 0 | 4 | 0 | 4 | 0 | 16 | 0.0625 | 5 | 0.2 | 6 | 0.1667 | 3 | 0 | 8 | 0.5 | 22 | 0.2727 |
| 4_15239144 | 4 | 0.25 | 6 | 0.5 | 5 | 0.4 | 4 | 0.75 | 19 | 0.4737 | 4 | 0.25 | 6 | 0.5 | 3 | 0.3333 | 9 | 0.5556 | 22 | 0.4545 |
| 4_15241697 | 3 | 0.3333 | 5 | 0 | 3 | 0 | 6 | 0.1667 | 17 | 0.1176 | 2 | 0.5 | 2 | 0 | 5 | 0 | 6 | 0 | 15 | 0.0667 |
| 4_15241699 | 5 | 0.2 | 5 | 0.2 | 3 | 0.3333 | 6 | 0.5 | 19 | 0.3158 | 3 | 0 | 2 | 1 | 6 | 0.3333 | 5 | 0.2 | 16 | 0.3125 |
| 4_15241703 | 5 | 0 | 6 | 0 | 3 | 0.6667 | 5 | 0.4 | 19 | 0.2105 | 2 | 0 | 4 | 0.25 | 6 | 0 | 7 | 0.2857 | 19 | 0.1579 |
| 4_15241820 | 10 | 0 | 11 | 0.3636 | 7 | 0.2857 | 9 | 0 | 37 | 0.1622 | 6 | 0 | 8 | 0.375 | 9 | 0.1111 | 10 | 0.1 | 33 | 0.1515 |
| 4_15247148 | 6 | 0 | 5 | 0 | 4 | 0.25 | 1 | 0 | 16 | 0.0625 | 2 | 0 | 3 | 0.3333 | 6 | 0.1667 | 3 | 0.3333 | 14 | 0.2143 |
| 4_15247160 | 6 | 0.3333 | 3 | 1 | 6 | 0.8333 | 1 | 0 | 16 | 0.625 | 2 | 1 | 2 | 1 | 6 | 0.6667 | 2 | 0.5 | 12 | 0.75 |
| 4_15247160 | 6 | 0.5 | 5 | 0.6 | 6 | 0.5 | 1 | 1 | 18 | 0.5556 | 2 | 0.5 | 3 | 0.3333 | 6 | 0.8333 | 2 | 0.5 | 13 | 0.6154 |
| 4_15247187 | 4 | 0.25 | 4 | 1 | 5 | 0.6 | 2 | 0 | 15 | 0.5333 | 2 | 1 | 2 | 0.5 | 4 | 0.75 | 4 | 0.75 | 12 | 0.75 |
| 4_15247190 | 3 | 0.3333 | 4 | 0.25 | 5 | 0.4 | 2 | 0.5 | 14 | 0.3571 | 2 | 1 | 2 | 0.5 | 3 | 1 | 4 | 0.5 | 11 | 0.7273 |
| 4_15247205 | 4 | 0.5 | 4 | 0.75 | 3 | 0.3333 | 2 | 1 | 13 | 0.6154 | 1 | 1 | 2 | 1 | 3 | 0.6667 | 2 | 1 | 8 | 0.875 |
| 4_15247212 | 5 | 0 | 3 | 0.3333 | 4 | 0.25 | 2 | 0 | 14 | 0.1429 | 2 | 0 | 2 | 0 | 4 | 0.25 | 3 | 0.3333 | 11 | 0.1818 |
| 4_15247213 | 3 | 0.6667 | 2 | 0 | 3 | 0 | 2 | 0 | 10 | 0.2 | 2 | 0.5 | NA | NA | 3 | 0 | 3 | 0 | 8 | 0.125 |
| 4_15247216 | 5 | 1 | 5 | 1 | 5 | 1 | 2 | 0.5 | 17 | 0.9412 | 2 | 1 | 2 | 1 | 4 | 0.75 | 4 | 0.75 | 12 | 0.8333 |
| 4_15247224 | 6 | 0 | 5 | 0 | 5 | 0.2 | 1 | 0 | 17 | 0.0588 | 2 | 0.5 | 1 | 0 | 3 | 0.3333 | 3 | 0 | 9 | 0.2222 |
| 4_15247287 | 7 | 0 | 3 | 0 | 5 | 0.4 | 2 | 0 | 17 | 0.1176 | 3 | 0 | 2 | 0 | 5 | 0.2 | 4 | 0 | 14 | 0.0714 |
| 4_15247429 | 11 | 0.2727 | 13 | 0.0769 | 15 | 0.1333 | 7 | 0.1429 | 46 | 0.1522 | 9 | 0.1111 | 6 | 0.3333 | 11 | 0.0909 | 6 | 0.1667 | 32 | 0.1562 |
| 4_15247431 | 11 | 0.0909 | 14 | 0.1429 | 20 | 0.1 | 9 | 0.2222 | 54 | 0.1296 | 10 | 0 | 6 | 0.5 | 11 | 0 | 7 | 0.5714 | 34 | 0.2059 |
| 4_15247440 | 12 | 0.0833 | 15 | 0.1333 | 19 | 0 | 10 | 0 | 56 | 0.0536 | 7 | 0 | 6 | 0.1667 | 10 | 0.1 | 9 | 0 | 32 | 0.0625 |
| 4_15247526 | 48 | 0.1875 | 61 | 0.1475 | 64 | 0.1875 | 34 | 0.2353 | 207 | 0.1836 | 54 | 0.1111 | 32 | 0.125 | 51 | 0.0588 | 37 | 0.1622 | 174 | 0.1092 |
| 4_15247670 | 28 | 0.25 | 37 | 0.3243 | 29 | 0.1379 | 10 | 0.2 | 104 | 0.2404 | 31 | 0.1935 | 22 | 0.2727 | 31 | 0.2258 | 21 | 0.1429 | 105 | 0.2095 |
| 4_15247678 | 29 | 0.2759 | 33 | 0.5152 | 31 | 0.3226 | 12 | 0.1667 | 105 | 0.3524 | 30 | 0.2667 | 22 | 0.4545 | 29 | 0.2414 | 24 | 0.4583 | 105 | 0.3429 |
| 4_15247703 | 25 | 0.6 | 34 | 0.5588 | 30 | 0.5667 | 10 | 0.7 | 99 | 0.5859 | 25 | 0.56 | 17 | 0.5882 | 29 | 0.6207 | 21 | 0.5238 | 92 | 0.5761 |
| 4_15247719 | 16 | 0.1875 | 23 | 0.2174 | 22 | 0.2273 | 10 | 0.2 | 71 | 0.2113 | 20 | 0.1 | 15 | 0.2667 | 26 | 0.1923 | 14 | 0.1429 | 75 | 0.1733 |
| 4_15247725 | 22 | 0.4091 | 32 | 0.3438 | 31 | 0.3548 | 11 | 0.2727 | 96 | 0.3542 | 26 | 0.3077 | 21 | 0.381 | 28 | 0.3214 | 20 | 0.4 | 95 | 0.3474 |
| 4_15247746 | 22 | 0.5909 | 19 | 0.5789 | 27 | 0.6296 | 9 | 0.5556 | 77 | 0.5974 | 20 | 0.6 | 16 | 0.625 | 19 | 0.5263 | 17 | 0.7059 | 72 | 0.6111 |
| 4_15247750 | 23 | 0.9565 | 29 | 0.9655 | 24 | 0.9167 | 11 | 1 | 87 | 0.954 | 24 | 0.9583 | 18 | 0.9444 | 21 | 1 | 23 | 0.8696 | 86 | 0.9419 |
| 4_15256204 | 1 | 0 | 1 | 1 | 1 | 1 | NA | NA | 3 | 0.6667 | NA | NA | NA | NA | 1 | 1 | 2 | 0.5 | 3 | 0.6667 |
| 4_15256207 | 1 | 1 | 1 | 1 | 1 | 1 | NA | NA | 3 | 1 | NA | NA | NA | NA | 1 | 1 | 2 | 0.5 | 3 | 0.6667 |
| 4_15264746 | 1 | 0 | 1 | 0 | 2 | 0 | 6 | 0.5 | 10 | 0.3 | NA | NA | 1 | 0 | 2 | 0.5 | 3 | 0.3333 | 6 | 0.3333 |
| 4_15264748 | 1 | 0 | 2 | 0.5 | 2 | 0 | 6 | 0.3333 | 11 | 0.2727 | NA | NA | 1 | 0 | 2 | 0.5 | 3 | 0.3333 | 6 | 0.3333 |
| 4_15341997 | NA | NA | 1 | 1 | NA | NA | NA | NA | 1 | 1 | NA | NA | NA | NA | NA | NA | 1 | 1 | 1 | 1 |
| 4_15345772 | 4 | 0 | 6 | 0.3333 | 8 | 0.375 | 5 | 0.2 | 23 | 0.2609 | 5 | 0.4 | 8 | 0.5 | 8 | 0.625 | 5 | 0.4 | 26 | 0.5 |
| 4_15345777 | 2 | 0.5 | 5 | 0 | 9 | 0.2222 | 5 | 0 | 21 | 0.1429 | 4 | 0.25 | 7 | 0 | 11 | 0 | 5 | 0.2 | 27 | 0.0741 |
| 4_15345784 | 2 | 0 | 4 | 0.5 | 6 | 0.3333 | 5 | 0.2 | 17 | 0.2941 | 5 | 0 | 8 | 0.25 | 8 | 0.125 | 7 | 0.2857 | 28 | 0.1786 |
| 4_15345795 | 4 | 0 | 7 | 0.1429 | 7 | 0.2857 | 7 | 0.1429 | 25 | 0.16 | 8 | 0.125 | 8 | 0 | 7 | 0.2857 | 8 | 0.125 | 31 | 0.129 |
| 4_15345836 | 4 | 0.5 | 6 | 0.6667 | 5 | 0.8 | 4 | 0.75 | 19 | 0.6842 | 4 | 0.75 | 6 | 0.6667 | 9 | 0.6667 | 8 | 0.625 | 27 | 0.6667 |
| 4_15345845 | 5 | 0.2 | 4 | 0.25 | 7 | 0 | 6 | 0.1667 | 22 | 0.1364 | 7 | 0 | 6 | 0 | 8 | 0.5 | 9 | 0.1111 | 30 | 0.1667 |
| 4_15345847 | 1 | 1 | 4 | 0.75 | 7 | 0.7143 | 5 | 0.8 | 17 | 0.7647 | 4 | 0.5 | 4 | 1 | 8 | 0.5 | 9 | 0.6667 | 25 | 0.64 |
| 4_15345850 | 4 | 0.5 | 5 | 0.4 | 6 | 0.5 | 6 | 0.1667 | 21 | 0.381 | 6 | 0.1667 | 3 | 0.3333 | 8 | 0.25 | 8 | 0.25 | 25 | 0.24 |
| 4_15345853 | 2 | 0.5 | 4 | 0.25 | 5 | 0.4 | 4 | 0.75 | 15 | 0.4667 | 4 | 0.5 | 5 | 0.6 | 8 | 0.75 | 6 | 0.8333 | 23 | 0.6957 |
| 4_15345863 | 4 | 0.25 | 5 | 0.6 | 7 | 0.1429 | 6 | 0.3333 | 22 | 0.3182 | 7 | 0.2857 | 6 | 0.3333 | 9 | 0.4444 | 9 | 0.4444 | 31 | 0.3871 |
| 4_15345870 | 4 | 0.5 | 3 | 0 | 6 | 0 | 3 | 0.3333 | 16 | 0.1875 | 4 | 0.25 | 4 | 0 | 4 | 0 | 8 | 0 | 20 | 0.05 |
| 4_15345876 | 5 | 0.2 | 4 | 0.25 | 6 | 0.1667 | 5 | 0 | 20 | 0.15 | 5 | 0.2 | 5 | 0 | 8 | 0.125 | 8 | 0 | 26 | 0.0769 |

| | | | | | | | | | | | | | | | | | | | | |
|---|---|---|---|---|---|---|---|---|---|---|---|---|---|---|---|---|---|---|---|---|
| 4_15351663 | 3 | 0.3333 | 3 | 0.6667 | 4 | 0.5 | 4 | 0.5 | 14 | 0.5 | 5 | 0.6 | 2 | 1 | 3 | 0.3333 | 3 | 0.3333 | 13 | 0.5385 |
| 4_15351680 | 4 | 1 | 3 | 1 | 4 | 1 | 4 | 0.5 | 15 | 0.8667 | 4 | 0.75 | 2 | 1 | 5 | 0.8 | 8 | 0.875 | 19 | 0.8421 |
| 4_15351691 | 2 | 0.5 | 2 | 1 | 5 | 0.8 | 4 | 0.5 | 13 | 0.6923 | 3 | 0.6667 | 1 | 1 | 2 | 1 | 4 | 0.75 | 10 | 0.8 |
| 4_15351698 | 1 | 1 | 2 | 0.5 | 1 | 1 | 1 | 0 | 5 | 0.6 | 3 | 0 | 1 | 0 | 4 | 0.5 | 3 | 0 | 11 | 0.1818 |
| 4_15351700 | 4 | 0.5 | 2 | 1 | 2 | 0.5 | 3 | 0.6667 | 11 | 0.6364 | 1 | 0 | 1 | 0 | 4 | 0.5 | 5 | 0.4 | 11 | 0.3636 |
| 4_15351702 | 4 | 0.5 | 2 | 0.5 | 4 | 0.5 | 3 | 0 | 13 | 0.3846 | 3 | 1 | 1 | 0 | 5 | 0.6 | 7 | 0.2857 | 16 | 0.5 |
| 4_15351707 | 4 | 0.25 | 2 | 0.5 | 5 | 0.2 | 4 | 0.75 | 15 | 0.4 | 3 | 0.3333 | 2 | 1 | 4 | 0.5 | 7 | 0.5714 | 16 | 0.5625 |
| 4_15351721 | 3 | 0.3333 | 2 | 0.5 | 3 | 0 | 3 | 0 | 11 | 0.1818 | 2 | 0.5 | 1 | 0 | 4 | 0 | 5 | 0.4 | 12 | 0.25 |
| 4_15356664 | 1 | 1 | NA | NA | NA | NA | NA | NA | 1 | 1 | NA | NA | NA | NA | NA | NA | 1 | 1 | 1 | 1 |
| 4_15356723 | 1 | 1 | NA | NA | NA | NA | NA | NA | 1 | 1 | NA | NA | NA | NA | NA | NA | 1 | 1 | 1 | 1 |
| 4_15356776 | 1 | 1 | NA | NA | NA | NA | NA | NA | 1 | 1 | NA | NA | NA | NA | NA | NA | 1 | 1 | 1 | 1 |
| 4_15356783 | 1 | 1 | NA | NA | NA | NA | NA | NA | 1 | 1 | NA | NA | NA | NA | NA | NA | 1 | 1 | 1 | 1 |
| 4_15404501 | 5 | 0 | 4 | 0.25 | 2 | 0 | 4 | 0 | 15 | 0.0667 | 5 | 0.2 | 2 | 0 | 7 | 0 | 4 | 0 | 18 | 0.0556 |
| 4_15404507 | 5 | 0.2 | 4 | 0 | 2 | 0 | 3 | 0.3333 | 14 | 0.1429 | 5 | 0.2 | 1 | 1 | 7 | 0.2857 | 4 | 0 | 17 | 0.2353 |
| 4_15404509 | 4 | 0.75 | 4 | 0.25 | 2 | 0.5 | 4 | 0.25 | 14 | 0.4286 | 5 | 0.8 | 1 | 1 | 6 | 0.6667 | 4 | 0.25 | 16 | 0.625 |
| 4_15404515 | 3 | 0.6667 | 2 | 1 | 1 | 0 | 2 | 0.5 | 8 | 0.625 | 5 | 0.2 | 2 | 0 | 5 | 0.2 | 3 | 0 | 15 | 0.1333 |
| 4_15404533 | 4 | 0 | 4 | 0.5 | 2 | 0.5 | 2 | 0.5 | 12 | 0.3333 | 5 | 0.4 | 2 | 0.5 | 7 | 0 | 3 | 0.3333 | 17 | 0.2353 |
| 4_15404537 | 4 | 0.25 | 4 | 0.5 | 1 | 0 | 3 | 0.3333 | 12 | 0.3333 | 5 | 0.2 | 2 | 0 | 5 | 0 | 4 | 0.25 | 16 | 0.125 |
| 4_15404587 | NA | NA | NA | NA | NA | NA | 1 | 1 | 1 | 1 | NA | NA | NA | NA | NA | NA | 1 | 1 | 1 | 1 |
| 4_15404610 | 5 | 0 | 5 | 0.2 | 4 | 0 | 2 | 0 | 16 | 0.0625 | 6 | 0.1667 | 2 | 0 | 7 | 0 | 3 | 0 | 18 | 0.0556 |
| 4_15404612 | 4 | 0.5 | 4 | 0.5 | 3 | 0.6667 | 4 | 0.5 | 15 | 0.5333 | 6 | 0.3333 | 2 | 1 | 6 | 0.3333 | 4 | 0.75 | 18 | 0.5 |
| 4_15404620 | 5 | 0 | 5 | 0.2 | 4 | 0 | 4 | 0 | 18 | 0.0556 | 6 | 0.3333 | 2 | 0.5 | 7 | 0.1429 | 3 | 0 | 18 | 0.2222 |
| 4_15404625 | 4 | 0.25 | 4 | 0.5 | 3 | 0 | 2 | 1 | 13 | 0.3846 | 5 | 0 | 2 | 1 | 4 | 0.5 | 4 | 0 | 15 | 0.2667 |
| 4_15404628 | 4 | 0.25 | 5 | 0.8 | 3 | 1 | 4 | 0.5 | 16 | 0.625 | 6 | 0.8333 | 2 | 1 | 6 | 1 | 3 | 0.6667 | 17 | 0.8824 |
| 4_15404932 | 3 | 0.3333 | NA | NA | 1 | 0 | 1 | 0 | 5 | 0.2 | NA | NA | NA | NA | 1 | 1 | NA | NA | 1 | 1 |
| 4_15404937 | 3 | 0.6667 | NA | NA | 2 | 0 | 1 | 0 | 6 | 0.3333 | NA | NA | NA | NA | 3 | 0.6667 | NA | NA | 3 | 0.6667 |
| 4_15404948 | 3 | 0.6667 | NA | NA | 1 | 1 | 1 | 1 | 5 | 0.8 | NA | NA | NA | NA | 3 | 0.3333 | NA | NA | 3 | 0.3333 |
| 4_15404958 | 2 | 1 | NA | NA | 2 | 0.5 | 1 | 1 | 5 | 0.8 | NA | NA | NA | NA | 3 | 0.3333 | NA | NA | 3 | 0.3333 |
| 4_15409124 | 3 | 0 | 1 | 0 | NA | NA | 6 | 0.1667 | 10 | 0.1 | 3 | 0 | 1 | 0 | 4 | 0.25 | 2 | 1 | 10 | 0.3 |
| 4_15409139 | 3 | 0 | 2 | 0 | 1 | 0 | 6 | 0.5 | 12 | 0.25 | 1 | 1 | NA | NA | 4 | 0.25 | 2 | 0 | 7 | 0.2857 |
| 4_15409233 | 5 | 0.2 | 1 | 0 | 2 | 0 | 4 | 0.25 | 12 | 0.1667 | 2 | 0 | 1 | 0 | 5 | 0.2 | 2 | 0 | 10 | 0.1 |
| 4_15409252 | 6 | 0.1667 | 2 | 0 | 2 | 0 | 4 | 0 | 14 | 0.0714 | 3 | 0 | 1 | 0 | 6 | 0 | 2 | 0.5 | 12 | 0.0833 |
| 4_15409485 | NA | NA | 1 | 1 | NA | NA | NA | NA | 1 | 1 | NA | NA | NA | NA | 1 | 1 | NA | NA | 1 | 1 |
| 4_15409491 | NA | NA | 1 | 1 | NA | NA | NA | NA | 1 | 1 | NA | NA | NA | NA | 1 | 1 | NA | NA | 1 | 1 |
| 4_15409665 | 1 | 1 | NA | NA | NA | NA | NA | NA | 1 | 1 | 1 | 1 | 1 | 0 | NA | NA | 1 | 1 | 3 | 0.6667 |
| 4_15410522 | 2 | 0.5 | 2 | 0.5 | 1 | 0 | NA | NA | 5 | 0.4 | 3 | 0.6667 | 2 | 0.5 | NA | NA | NA | NA | 5 | 0.6 |
| 4_15410527 | 4 | 0.75 | 1 | 0 | NA | NA | NA | NA | 5 | 0.6 | 3 | 0.6667 | NA | NA | 1 | 0 | 1 | 1 | 5 | 0.6 |
| 4_15410528 | 3 | 0.6667 | 2 | 0.5 | NA | NA | NA | NA | 5 | 0.6 | 3 | 1 | NA | NA | 1 | 1 | 1 | 1 | 5 | 1 |
| 4_15410531 | 1 | 1 | 1 | 1 | NA | NA | 1 | 0 | 3 | 0.6667 | 3 | 0.6667 | 4 | 0.5 | 1 | 0 | 1 | 1 | 9 | 0.5556 |
| 4_15410535 | 3 | 0.6667 | 2 | 0 | NA | NA | NA | NA | 5 | 0.4 | 3 | 0.3333 | NA | NA | 1 | 0 | NA | NA | 4 | 0.25 |
| 4_15410537 | 4 | 1 | 2 | 0 | NA | NA | NA | NA | 6 | 0.6667 | 2 | 1 | NA | NA | 1 | 1 | NA | NA | 3 | 1 |
| 4_15410549 | 4 | 1 | 1 | 1 | 3 | 1 | 1 | 1 | 9 | 1 | 4 | 1 | 3 | 0.6667 | 1 | 1 | 2 | 1 | 10 | 0.9 |
| 4_15410559 | 3 | 1 | 2 | 0 | 4 | 1 | 1 | 1 | 10 | 0.8 | 4 | 0.5 | 4 | 0.75 | 2 | 1 | 3 | 1 | 13 | 0.7692 |
| 4_15410568 | 4 | 0.25 | 2 | 0 | 3 | 0 | 1 | 0 | 10 | 0.1 | 3 | 0.6667 | 3 | 0 | 2 | 0 | 3 | 0.6667 | 11 | 0.3636 |
| 4_15410602 | 4 | 0.5 | 2 | 0 | NA | NA | NA | NA | 6 | 0.3333 | 2 | 1 | NA | NA | 1 | 0 | NA | NA | 3 | 0.6667 |
| 4_15410681 | 4 | 0.25 | 1 | 0 | NA | NA | NA | NA | 5 | 0.2 | 1 | 1 | NA | NA | 1 | 0 | NA | NA | 2 | 0.5 |
| 4_15410713 | 3 | 0.3333 | 2 | 0 | NA | NA | NA | NA | 5 | 0.2 | 2 | 0.5 | NA | NA | 1 | 1 | NA | NA | 3 | 0.6667 |
| 4_15410715 | 2 | 0.5 | 1 | 0 | NA | NA | NA | NA | 3 | 0.3333 | 2 | 0 | NA | NA | 1 | 1 | NA | NA | 3 | 0.3333 |
| 4_15410717 | 3 | 0.6667 | 1 | 1 | NA | NA | NA | NA | 4 | 0.75 | 1 | 0 | NA | NA | 1 | 1 | NA | NA | 2 | 0.5 |
| 4_15410772 | 2 | 0.5 | 1 | 0 | NA | NA | NA | NA | 3 | 0.3333 | 1 | 1 | NA | NA | 1 | 0 | NA | NA | 2 | 0.5 |
| 4_15424100 | 4 | 0.75 | 4 | 0 | 5 | 0 | 3 | 0 | 16 | 0.1875 | 7 | 0 | 3 | 0.6667 | 4 | 0.5 | 8 | 0.25 | 22 | 0.2727 |
| 4_15424105 | 3 | 0.3333 | 4 | 0 | 4 | 0 | 1 | 0 | 12 | 0.0833 | 3 | 0.3333 | 2 | 0 | 2 | 0.5 | 5 | 0 | 12 | 0.1667 |
| 4_15424222 | 35 | 0.0571 | 26 | 0.0769 | 21 | 0.0952 | 33 | 0.1515 | 115 | 0.0957 | 21 | 0.1905 | 26 | 0.1923 | 27 | 0.1852 | 18 | 0.1667 | 92 | 0.1848 |
| 4_15424256 | 32 | 0.2188 | 25 | 0.2 | 21 | 0.0952 | 32 | 0.25 | 110 | 0.2 | 25 | 0.24 | 27 | 0.1481 | 29 | 0.1034 | 24 | 0.125 | 105 | 0.1524 |
| 4_15424269 | 28 | 0.4286 | 23 | 0.3043 | 19 | 0.5263 | 31 | 0.4516 | 101 | 0.4257 | 21 | 0.3333 | 24 | 0.2917 | 27 | 0.2963 | 23 | 0.4783 | 95 | 0.3474 |
| 4_15424279 | 30 | 0.2 | 18 | 0.1111 | 18 | 0.2222 | 26 | 0.1538 | 92 | 0.1739 | 18 | 0.2222 | 20 | 0.15 | 20 | 0.3 | 22 | 0.1818 | 80 | 0.2125 |
| 4_15424443 | 1 | 1 | 1 | 1 | NA | NA | NA | NA | 2 | 1 | NA | NA | NA | NA | 1 | 1 | 2 | 0 | 3 | 0.3333 |
| 4_15425258 | NA | NA | NA | NA | NA | NA | 1 | 1 | 1 | 1 | NA | NA | NA | NA | 1 | 1 | NA | NA | 1 | 1 |
| 4_15435846 | NA | NA | 1 | 1 | NA | NA | NA | NA | 1 | 1 | 1 | 1 | 3 | 0.6667 | NA | NA | NA | NA | 4 | 0.75 |
| 4_15435853 | NA | NA | 1 | 1 | NA | NA | NA | NA | 1 | 1 | NA | NA | 3 | 0.6667 | 1 | 1 | NA | NA | 4 | 0.75 |
| 4_15435858 | NA | NA | 1 | 1 | NA | NA | NA | NA | 1 | 1 | 1 | 1 | 2 | 1 | 1 | 0 | NA | NA | 4 | 0.75 |
| 4_15435863 | NA | NA | 1 | 1 | NA | NA | NA | NA | 1 | 1 | 1 | 0 | 3 | 0.6667 | 1 | 0 | NA | NA | 5 | 0.4 |
| 4_15435869 | NA | NA | 1 | 1 | NA | NA | NA | NA | 1 | 1 | NA | NA | 2 | 1 | 1 | 1 | NA | NA | 3 | 1 |
| 4_15435882 | NA | NA | 1 | 1 | NA | NA | NA | NA | 1 | 1 | 1 | 0 | 3 | 0.6667 | 1 | 1 | NA | NA | 5 | 0.6 |
| 4_15440313 | NA | NA | 2 | 0.5 | 1 | 1 | 1 | 0 | 4 | 0.5 | 1 | 0 | 1 | 1 | 1 | 0 | 1 | 1 | 4 | 0.5 |

| | | | | | | | | | | | | | | | | | | | | |
|---|---|---|---|---|---|---|---|---|---|---|---|---|---|---|---|---|---|---|---|---|
| 4_1544031 | NA | NA | 1 | 0 | 1 | 1 | 1 | 1 | 3 | 0.6667 | 1 | 1 | 1 | 0 | 1 | 1 | 1 | 1 | 4 | 0.75 |
| 4_1544032 | NA | NA | 2 | 0 | NA | NA | 1 | 1 | 3 | 0.3333 | 1 | 0 | 1 | 0 | NA | NA | 1 | 1 | 3 | 0.3333 |
| 4_1544033 | NA | NA | 1 | 1 | 1 | 0 | 1 | 1 | 3 | 0.6667 | 1 | 0 | 1 | 0 | 1 | 1 | 1 | 0 | 4 | 0.25 |
| 4_1544033 | NA | NA | 1 | 1 | 1 | 0 | 1 | 0 | 3 | 0.3333 | 1 | 0 | 1 | 0 | NA | NA | 1 | 1 | 3 | 0.3333 |
| 4_1544037 | NA | NA | 2 | 0.5 | NA | NA | 1 | 1 | 3 | 0.6667 | 1 | 0 | 1 | 1 | 1 | 1 | NA | NA | 3 | 0.6667 |
| 4_1544040 | NA | NA | 2 | 1 | NA | NA | NA | NA | 2 | 1 | 1 | 1 | 1 | 1 | NA | NA | 1 | 1 | 3 | 1 |
| 4_1544041 | NA | NA | 2 | 1 | NA | NA | NA | NA | 2 | 1 | 1 | 1 | 1 | 1 | NA | NA | NA | NA | 2 | 1 |
| 4_1544041 | NA | NA | 2 | 1 | NA | NA | NA | NA | 2 | 1 | 1 | 1 | 1 | 1 | NA | NA | 1 | 1 | 3 | 1 |
| 4_1544043 | NA | NA | 2 | 1 | NA | NA | NA | NA | 2 | 1 | 1 | 1 | 1 | 1 | NA | NA | 1 | 1 | 3 | 1 |
| 4_1544045 | NA | NA | 1 | 1 | NA | NA | NA | NA | 1 | 1 | 1 | 0 | 1 | 1 | NA | NA | 1 | 0 | 3 | 0.3333 |
| 4_1544046 | NA | NA | 1 | 1 | NA | NA | NA | NA | 1 | 1 | NA | NA | 1 | 1 | NA | NA | NA | NA | 1 | 1 |
| 4_1544265 | NA | NA | 1 | 1 | NA | NA | NA | NA | 1 | 1 | NA | NA | 1 | 1 | NA | NA | NA | NA | 1 | 1 |
| 4_1547158 | 5 | 0 | 3 | 0.6667 | 3 | 0 | 1 | 0 | 12 | 0.1667 | 5 | 0.2 | 2 | 0 | 5 | 0 | 2 | 0 | 14 | 0.0714 |
| 4_1547184 | 5 | 0.2 | 2 | 0 | 1 | 1 | NA | NA | 8 | 0.25 | 5 | 0.4 | 1 | 0 | 3 | 0.6667 | 2 | 0 | 11 | 0.3636 |
| 4_1547184 | 4 | 0.75 | 2 | 1 | 1 | 0 | NA | NA | 7 | 0.7143 | 5 | 0.2 | 1 | 0 | 3 | 0.6667 | 1 | 0 | 10 | 0.3 |
| 4_1547185 | 3 | 0.6667 | 2 | 0 | 1 | 0 | NA | NA | 6 | 0.3333 | 4 | 0.25 | 1 | 0 | 2 | 0.5 | 1 | 0 | 8 | 0.25 |
| 4_1547186 | 4 | 0.75 | 2 | 1 | 1 | 1 | NA | NA | 7 | 0.8571 | 3 | 0.3333 | 1 | 1 | 2 | 1 | 1 | 1 | 7 | 0.7143 |
| 4_1547186 | 5 | 1 | 2 | 1 | 1 | 1 | NA | NA | 8 | 1 | 5 | 1 | 1 | 1 | 3 | 1 | 1 | 1 | 10 | 1 |
| 4_1547186 | 5 | 0.8 | 2 | 1 | 1 | 1 | NA | NA | 8 | 0.875 | 4 | 0.25 | 1 | 0 | 2 | 0 | 1 | 0 | 8 | 0.125 |
| 4_1547187 | 5 | 1 | 2 | 1 | 1 | 1 | NA | NA | 8 | 1 | 5 | 1 | 1 | 1 | 3 | 1 | 1 | 1 | 10 | 1 |
| 4_1547188 | 4 | 0.75 | 2 | 0.5 | 1 | 0 | NA | NA | 7 | 0.5714 | 5 | 0.4 | 1 | 0 | 3 | 0.3333 | 1 | 0 | 10 | 0.3 |
| 4_1547188 | 5 | 0.2 | 2 | 0 | 1 | 0 | NA | NA | 8 | 0.125 | 5 | 0.2 | NA | NA | 3 | 0.3333 | 1 | 1 | 9 | 0.3333 |
| 4_1547188 | 5 | 1 | 1 | 1 | 1 | 1 | NA | NA | 7 | 1 | 5 | 1 | 1 | 1 | 2 | 1 | 1 | 1 | 9 | 1 |
| 4_1547188 | 4 | 0.25 | 2 | 0 | NA | NA | NA | NA | 6 | 0.1667 | 4 | 0.5 | 1 | 0 | 1 | 0 | 1 | 0 | 7 | 0.2857 |
| 4_1547189 | 5 | 0.8 | 2 | 1 | 1 | 1 | NA | NA | 8 | 0.875 | 5 | 0.8 | 1 | 0 | 3 | 0.6667 | 1 | 1 | 10 | 0.7 |
| 4_1547189 | 5 | 0.8 | 2 | 1 | 1 | 1 | NA | NA | 8 | 0.875 | 5 | 1 | 1 | 1 | 3 | 1 | 1 | 1 | 10 | 1 |
| 4_1547191 | 5 | 0.8 | 2 | 1 | 1 | 1 | NA | NA | 8 | 0.875 | 4 | 0.75 | 1 | 1 | 3 | 1 | 1 | 1 | 9 | 0.8889 |
| 4_1547193 | 4 | 1 | 2 | 1 | 1 | 1 | NA | NA | 7 | 1 | 4 | 1 | 1 | 1 | 1 | 1 | 1 | 1 | 7 | 1 |
| 4_1547193 | 5 | 1 | 1 | 1 | NA | NA | NA | NA | 6 | 1 | 5 | 1 | 1 | 1 | 3 | 0.6667 | 1 | 0 | 10 | 0.8 |
| 4_1547194 | 5 | 0.4 | 2 | 0 | 1 | 0 | NA | NA | 8 | 0.25 | 4 | 0 | NA | NA | 3 | 0.3333 | 1 | 0 | 8 | 0.125 |
| 4_1547195 | 3 | 0.6667 | NA | NA | 1 | 1 | NA | NA | 4 | 0.75 | 3 | 0.3333 | 1 | 0 | 3 | 0.3333 | NA | NA | 7 | 0.2857 |
| 4_1547197 | 4 | 0.25 | 2 | 0.5 | 1 | 0 | NA | NA | 7 | 0.2857 | 4 | 0.5 | 1 | 0 | 2 | 0.5 | 1 | 0 | 8 | 0.375 |
| 4_1547197 | 4 | 0.5 | 2 | 0.5 | 1 | 0 | NA | NA | 7 | 0.4286 | 5 | 0.2 | 1 | 0 | 2 | 0 | 1 | 1 | 9 | 0.2222 |
| 4_1547198 | 4 | 0.75 | 2 | 1 | NA | NA | NA | NA | 6 | 0.8333 | 3 | 0.6667 | 1 | 1 | 3 | 1 | NA | NA | 7 | 0.8571 |
| 4_1547200 | 3 | 0.3333 | 1 | 0 | 1 | 0 | NA | NA | 5 | 0.2 | 5 | 0.2 | NA | NA | 3 | 0 | NA | NA | 8 | 0.125 |
| 4_1547203 | 4 | 0.25 | 2 | 0.5 | NA | NA | NA | NA | 6 | 0.3333 | 3 | 0.6667 | NA | NA | 2 | 0.5 | NA | NA | 5 | 0.6 |
| 4_1547205 | 4 | 0.25 | 2 | 0 | 1 | 0 | NA | NA | 7 | 0.1429 | 3 | 0 | NA | NA | 3 | 0.3333 | NA | NA | 6 | 0.1667 |
| 4_1547205 | 3 | 0.3333 | 2 | 1 | 1 | 1 | NA | NA | 6 | 0.6667 | 3 | 0.6667 | NA | NA | 3 | 0.6667 | NA | NA | 6 | 0.6667 |
| 4_1547215 | NA | NA | NA | NA | 1 | 1 | NA | NA | 1 | 1 | NA | NA | 1 | 0 | 2 | 0.5 | NA | NA | 3 | 0.3333 |
| 4_1547215 | NA | NA | NA | NA | 1 | 1 | NA | NA | 1 | 1 | NA | NA | NA | NA | 2 | 0.5 | NA | NA | 2 | 0.5 |
| 4_1547261 | 1 | 0 | NA | NA | 1 | 1 | NA | NA | 2 | 0.5 | NA | NA | NA | NA | NA | NA | 2 | 0.5 | 2 | 0.5 |
| 4_1547277 | NA | NA | NA | NA | 1 | 1 | NA | NA | 1 | 1 | NA | NA | NA | NA | NA | NA | 1 | 1 | 1 | 1 |
| 4_1547299 | 1 | 1 | 2 | 0.5 | 2 | 0 | NA | NA | 5 | 0.4 | 1 | 1 | 1 | 0 | 1 | 0 | 2 | 0 | 5 | 0.2 |
| 4_1547299 | 1 | 1 | 2 | 0 | 1 | 0 | NA | NA | 4 | 0.25 | 1 | 0 | 1 | 1 | 1 | 0 | 2 | 0 | 5 | 0.2 |
| 4_1549457 | NA | NA | 1 | 1 | NA | NA | 2 | 0 | 3 | 0.3333 | 2 | 1 | NA | NA | 1 | 1 | 1 | 1 | 4 | 1 |
| 4_1549458 | NA | NA | 1 | 1 | NA | NA | 1 | 1 | 2 | 1 | 1 | 0 | NA | NA | 1 | 1 | 1 | 1 | 3 | 0.6667 |
| 4_1549459 | NA | NA | NA | NA | NA | NA | 1 | 1 | 1 | 1 | NA | NA | NA | NA | 1 | 1 | 1 | 1 | 2 | 1 |
| 4_1549459 | NA | NA | 1 | 1 | NA | NA | 1 | 1 | 2 | 1 | NA | NA | NA | NA | 1 | 1 | 1 | 1 | 2 | 1 |
| 4_1549501 | NA | NA | NA | NA | NA | NA | 1 | 1 | 1 | 1 | 1 | 1 | NA | NA | NA | NA | NA | NA | 1 | 1 |
| 4_1549503 | NA | NA | NA | NA | NA | NA | 1 | 1 | 1 | 1 | 1 | 1 | NA | NA | NA | NA | NA | NA | 1 | 1 |
| 4_1549507 | NA | NA | NA | NA | NA | NA | 1 | 1 | 1 | 1 | 1 | 1 | NA | NA | NA | NA | NA | NA | 1 | 1 |
| 4_1549805 | NA | NA | 1 | 0 | NA | NA | 2 | 1 | 3 | 0.6667 | 3 | 0 | NA | NA | 2 | 0 | 2 | 0.5 | 7 | 0.1429 |
| 4_1549806 | NA | NA | 1 | 0 | NA | NA | 2 | 1 | 3 | 0.6667 | 3 | 0.6667 | NA | NA | 1 | 1 | 1 | 1 | 5 | 0.8 |
| 4_1549807 | NA | NA | 1 | 1 | NA | NA | 2 | 0.5 | 3 | 0.6667 | 3 | 0.3333 | NA | NA | 1 | 1 | 2 | 1 | 6 | 0.6667 |
| 4_1549807 | NA | NA | 1 | 1 | NA | NA | 2 | 0 | 3 | 0.3333 | 3 | 0.3333 | NA | NA | 1 | 0 | 2 | 0 | 6 | 0.1667 |
| 4_1549807 | NA | NA | 1 | 0 | NA | NA | 2 | 0.5 | 3 | 0.3333 | 3 | 0.6667 | NA | NA | 1 | 1 | 2 | 0 | 6 | 0.5 |
| 4_1549809 | 18 | 0.3889 | 15 | 0.0667 | 15 | 0.0667 | 20 | 0.2 | 68 | 0.1912 | 16 | 0.3125 | 25 | 0.28 | 18 | 0.2778 | 14 | 0.2143 | 73 | 0.274 |
| 4_1550463 | 4 | 0.25 | 10 | 0.1 | 5 | 0.2 | 7 | 0 | 26 | 0.1154 | 1 | 0 | 5 | 0.2 | 4 | 0 | 1 | 0 | 11 | 0.0909 |
| 4_1550464 | 4 | 0.75 | 10 | 1 | 5 | 1 | 8 | 0.875 | 27 | 0.9259 | 2 | 1 | 5 | 1 | 4 | 1 | 2 | 1 | 13 | 1 |
| 4_1550465 | 5 | 0.6 | 10 | 0.6 | 6 | 0.6667 | 6 | 0.8333 | 27 | 0.6667 | 1 | 1 | 6 | 0.3333 | 3 | 0.6667 | 2 | 0 | 12 | 0.4167 |
| 4_1550467 | 4 | 0 | 9 | 0.2222 | 8 | 0 | 6 | 0.1667 | 27 | 0.1111 | 2 | 0 | 4 | 0.25 | 6 | 0.1667 | 1 | 0 | 13 | 0.1538 |
| 4_1550495 | 1 | 1 | 1 | 1 | NA | NA | NA | NA | 2 | 1 | NA | NA | NA | NA | NA | NA | 1 | 1 | 1 | 1 |
| 4_1550496 | 1 | 1 | 1 | 1 | NA | NA | NA | NA | 2 | 1 | 1 | 1 | NA | NA | NA | NA | 1 | 1 | 2 | 1 |
| 4_1550652 | 3 | 1 | 3 | 0.6667 | 7 | 0.5714 | 6 | 0.8333 | 19 | 0.7368 | 6 | 0.3333 | 5 | 0.6 | 7 | 0.8571 | 1 | 1 | 19 | 0.6316 |
| 4_1550652 | 6 | 0.1667 | 5 | 0 | 10 | 0.3 | 6 | 0.3333 | 27 | 0.2222 | 7 | 0.2857 | 4 | 0.25 | 8 | 0.125 | 2 | 0 | 21 | 0.1905 |

| | | | | | | | | | | | | | | | | | | | | |
|---|---|---|---|---|---|---|---|---|---|---|---|---|---|---|---|---|---|---|---|---|
| 4_1550652 | 7 | 0.7143 | 6 | 0.8333 | 10 | 0.8 | 4 | 0.75 | 27 | 0.7778 | 7 | 0.5714 | 5 | 0.8 | 10 | 0.9 | 1 | 1 | 23 | 0.7826 |
| 4_1550653 | 7 | 0.2857 | 6 | 0.5 | 10 | 0.5 | 5 | 0.6 | 28 | 0.4643 | 6 | 0 | 5 | 0 | 8 | 0.375 | 3 | 0.6667 | 22 | 0.2273 |
| 4_1550653 | 7 | 0.1429 | 5 | 0.6 | 9 | 0.4444 | 6 | 0.5 | 27 | 0.4074 | 6 | 0.1667 | 5 | 0.8 | 11 | 0.3636 | 2 | 1 | 24 | 0.4583 |
| 4_1550680 | 4 | 0.25 | 5 | 0 | 3 | 0 | 1 | 0 | 13 | 0.0769 | 3 | 0.6667 | 2 | 0.5 | 3 | 0.6667 | 5 | 0.4 | 13 | 0.5385 |
| 4_1550682 | 5 | 1 | 9 | 1 | 5 | 1 | 3 | 1 | 22 | 1 | 6 | 1 | 3 | 1 | 3 | 1 | 6 | 1 | 18 | 1 |
| 4_1550684 | 4 | 0.75 | 8 | 0.375 | 5 | 0.4 | 3 | 1 | 20 | 0.55 | 4 | 0.75 | 3 | 0.6667 | 3 | 0.3333 | 7 | 0.2857 | 17 | 0.4706 |
| 4_1550686 | 5 | 1 | 8 | 1 | 5 | 1 | 4 | 1 | 22 | 1 | 5 | 1 | 3 | 1 | 3 | 1 | 6 | 1 | 17 | 1 |
| 4_1550689 | 3 | 1 | 7 | 0.7143 | 5 | 0.6 | 4 | 0.75 | 19 | 0.7368 | 4 | 0.75 | 3 | 1 | 1 | 0 | 7 | 0.4286 | 15 | 0.6 |
| 4_1550689 | 5 | 1 | 7 | 0.4286 | 5 | 1 | 4 | 0.75 | 21 | 0.7619 | 5 | 1 | 3 | 0.6667 | 1 | 0 | 4 | 1 | 13 | 0.8462 |
| 4_1550691 | 4 | 0 | 8 | 0.125 | 5 | 0.2 | 3 | 0.3333 | 20 | 0.15 | 4 | 0 | 3 | 0 | 1 | 0 | 6 | 0.3333 | 14 | 0.1429 |
| 4_1550691 | 3 | 1 | 8 | 0.75 | 5 | 0.6 | 4 | 0.25 | 20 | 0.65 | 4 | 0.75 | 2 | 0.5 | 1 | 1 | 6 | 0.6667 | 13 | 0.6923 |
| 4_1550692 | 5 | 0.2 | 8 | 0.125 | 5 | 0 | 4 | 0 | 22 | 0.0909 | 4 | 0.25 | 2 | 0 | 1 | 0 | 5 | 0 | 12 | 0.0833 |
| 4_1550692 | 4 | 0.25 | 8 | 0.125 | 4 | 0 | 1 | 0 | 17 | 0.1176 | 3 | 0.3333 | 2 | 0 | 1 | 0 | 6 | 0.3333 | 12 | 0.25 |
| 4_1550711 | 1 | 1 | 5 | 0.8 | 1 | 0 | 3 | 0 | 10 | 0.5 | 3 | 0.3333 | NA | NA | NA | NA | 4 | 0.25 | 7 | 0.2857 |
| 4_1550914 | 1 | 0 | 2 | 0 | 1 | 1 | 1 | 0 | 5 | 0.2 | 2 | 0.5 | 1 | 0 | NA | NA | 1 | 0 | 4 | 0.25 |
| 4_1551562 | 3 | 0.6667 | NA | NA | 2 | 0 | NA | NA | 5 | 0.4 | NA | NA | 2 | 0.5 | NA | NA | 2 | 0 | 4 | 0.25 |
| 4_1551586 | 3 | 0 | 3 | 0.3333 | 7 | 0 | 1 | 0 | 14 | 0.0714 | 5 | 0 | 2 | 0 | 4 | 0.25 | 1 | 0 | 12 | 0.0833 |
| 4_1551590 | 13 | 0.4615 | 14 | 0.2143 | 15 | 0.3333 | 12 | 0.5 | 54 | 0.3704 | 16 | 0.5 | 9 | 0.4444 | 11 | 0.5455 | 7 | 0.2857 | 43 | 0.4651 |
| 4_1551591 | 13 | 0.1538 | 10 | 0.4 | 14 | 0.1429 | 12 | 0.25 | 49 | 0.2245 | 16 | 0.25 | 7 | 0 | 11 | 0 | 10 | 0.2 | 44 | 0.1364 |
| 4_1551592 | 16 | 0 | 14 | 0.1429 | 18 | 0 | 12 | 0.0833 | 60 | 0.05 | 15 | 0.0667 | 17 | 0.1176 | 13 | 0 | 14 | 0 | 59 | 0.0508 |
| 4_1551593 | 19 | 0.2632 | 16 | 0.125 | 17 | 0.1765 | 17 | 0.2353 | 69 | 0.2029 | 17 | 0.1765 | 17 | 0.0588 | 20 | 0.15 | 13 | 0.1538 | 67 | 0.1343 |
| 4_1551593 | 10 | 0.2 | 14 | 0.3571 | 17 | 0.4706 | 14 | 0.2143 | 55 | 0.3273 | 17 | 0.4706 | 15 | 0.2 | 14 | 0.2143 | 8 | 0.125 | 54 | 0.2778 |
| 4_1552385 | 3 | 0.3333 | 4 | 0 | 2 | 0 | 1 | 0 | 10 | 0.1 | 6 | 0.3333 | 3 | 0.6667 | 4 | 0.5 | 5 | 0.2 | 18 | 0.3889 |
| 4_1552397 | 8 | 0.25 | 16 | 0.3125 | 10 | 0.7 | 9 | 0.5556 | 43 | 0.4419 | 11 | 0.4545 | 12 | 0.5 | 16 | 0.625 | 9 | 0.2222 | 48 | 0.4792 |
| 4_1552401 | 15 | 0.5333 | 21 | 0.381 | 16 | 0.375 | 19 | 0.7368 | 71 | 0.507 | 16 | 0.75 | 19 | 0.4211 | 23 | 0.3043 | 18 | 0.6111 | 76 | 0.5 |
| 4_1552403 | 13 | 0.4615 | 20 | 0.6 | 13 | 0.6154 | 17 | 0.6471 | 63 | 0.5873 | 12 | 0.6667 | 16 | 0.6875 | 23 | 0.3478 | 17 | 0.5294 | 68 | 0.5294 |
| 4_1552403 | 16 | 0.875 | 21 | 0.9048 | 15 | 0.8 | 19 | 0.7895 | 71 | 0.8451 | 15 | 0.7333 | 17 | 1 | 23 | 0.913 | 18 | 0.8333 | 73 | 0.8767 |
| 4_1552406 | 12 | 0.5 | 17 | 0.5882 | 6 | 0.8333 | 10 | 0.6 | 45 | 0.6 | 7 | 0.5714 | 10 | 0.8 | 21 | 0.5238 | 13 | 0.7692 | 51 | 0.6471 |
| 4_1552406 | 11 | 1 | 16 | 0.9375 | 11 | 0.8182 | 12 | 0.9167 | 50 | 0.92 | 8 | 1 | 13 | 1 | 18 | 1 | 12 | 1 | 51 | 1 |
| 4_1552406 | 10 | 0 | 18 | 0.0556 | 13 | 0 | 12 | 0.1667 | 53 | 0.0566 | 10 | 0.2 | 14 | 0 | 21 | 0.0476 | 12 | 0.0833 | 57 | 0.0702 |
| 4_1552408 | 12 | 0.0833 | 18 | 0.1111 | 14 | 0.0714 | 15 | 0 | 59 | 0.0678 | 13 | 0.1538 | 15 | 0.1333 | 21 | 0 | 18 | 0.0556 | 67 | 0.0746 |
| 4_1552408 | 10 | 0.5 | 14 | 0.2143 | 11 | 0.3636 | 14 | 0.5 | 49 | 0.3878 | 12 | 0.75 | 13 | 0.6923 | 18 | 0.4444 | 15 | 0.4667 | 58 | 0.569 |
| 4_1552409 | 10 | 0.1 | 20 | 0.25 | 10 | 0 | 14 | 0.1429 | 54 | 0.1481 | 11 | 0.2727 | 13 | 0.1538 | 20 | 0.1 | 15 | 0.1333 | 59 | 0.1525 |
| 4_1552412 | 6 | 0.6667 | 17 | 0.6471 | 9 | 0.5556 | 13 | 0.4615 | 45 | 0.5778 | 11 | 0.5455 | 12 | 0.5833 | 16 | 0.6875 | 15 | 0.6667 | 54 | 0.6296 |
| 4_1552413 | 10 | 0.5 | 16 | 0.5 | 10 | 0.3 | 12 | 0.5833 | 48 | 0.4792 | 11 | 0.5455 | 9 | 0.6667 | 18 | 0.6667 | 15 | 0.4 | 53 | 0.566 |
| 4_1552414 | 8 | 0.5 | 11 | 0.4545 | 8 | 0.375 | 8 | 0.5 | 35 | 0.4571 | 4 | 0.75 | 9 | 0.3333 | 12 | 0.4167 | 15 | 0.5333 | 40 | 0.475 |
| 4_1552414 | 7 | 0.8571 | 13 | 0.8462 | 3 | 0.6667 | 7 | 0.7143 | 30 | 0.8 | 9 | 0.8889 | 6 | 1 | 11 | 0.9091 | 13 | 0.7692 | 39 | 0.8718 |
| 4_1552414 | 9 | 0.3333 | 16 | 0.75 | 8 | 0.625 | 11 | 0.6364 | 44 | 0.6136 | 9 | 0.7778 | 9 | 0.5556 | 14 | 0.7143 | 14 | 0.7143 | 46 | 0.6957 |
| 4_1552414 | 9 | 0.6667 | 20 | 0.9 | 10 | 0.9 | 12 | 0.5 | 51 | 0.7647 | 10 | 1 | 12 | 0.8333 | 19 | 0.7368 | 15 | 0.7333 | 56 | 0.8036 |
| 4_1552415 | 8 | 0.125 | 15 | 0.2 | 9 | 0.2222 | 11 | 0.2727 | 43 | 0.2093 | 9 | 0.2222 | 8 | 0 | 13 | 0 | 8 | 0.375 | 38 | 0.1316 |
| 4_1552415 | 9 | 1 | 17 | 0.9412 | 8 | 0.875 | 12 | 1 | 46 | 0.9565 | 9 | 1 | 12 | 1 | 19 | 1 | 13 | 1 | 53 | 1 |
| 4_1552417 | 7 | 1 | 15 | 0.6667 | 10 | 0.8 | 8 | 0.875 | 40 | 0.8 | 10 | 0.8 | 8 | 0.625 | 16 | 0.8125 | 12 | 0.75 | 46 | 0.7609 |
| 4_1553061 | 4 | 1 | 6 | 0.8333 | 6 | 0.8333 | 6 | 1 | 22 | 0.9091 | 7 | 1 | 3 | 1 | 13 | 0.9231 | 9 | 1 | 32 | 0.9688 |
| 4_1553061 | 3 | 0.3333 | 3 | 0.3333 | 6 | 0.5 | 4 | 0.75 | 16 | 0.5 | 7 | 0.7143 | 3 | 1 | 9 | 0.6667 | 7 | 0.7143 | 26 | 0.7308 |
| 4_1553066 | 3 | 0.6667 | 6 | 0 | 5 | 0 | 5 | 0.2 | 19 | 0.1579 | 5 | 0.2 | 3 | 0 | 11 | 0.3636 | 7 | 0 | 26 | 0.1923 |
| 4_1553066 | 3 | 0.3333 | 4 | 0 | 5 | 0 | 4 | 0 | 16 | 0.0625 | 4 | 0.25 | 2 | 0.5 | 5 | 0 | 7 | 0.4286 | 18 | 0.2778 |
| 4_1553348 | NA | NA | 2 | 1 | NA | NA | 2 | 1 | 4 | 1 | NA | NA | NA | NA | 3 | 0.6667 | 1 | 0 | 4 | 0.5 |
| 4_1553350 | 5 | 0 | 3 | 0 | 3 | 0 | 8 | 0.125 | 19 | 0.0526 | NA | NA | 2 | 0 | 6 | 0 | 6 | 0.1667 | 14 | 0.0714 |
| 4_1553350 | 3 | 0 | 3 | 0.3333 | 3 | 0 | 8 | 0 | 17 | 0.0588 | NA | NA | 3 | 0.3333 | 6 | 0.1667 | 6 | 0 | 15 | 0.1333 |
| 4_1553352 | 5 | 0.2 | 3 | 0.3333 | 3 | 0.6667 | 9 | 0 | 20 | 0.2 | NA | NA | 3 | 0.3333 | 7 | 0.2857 | 6 | 0.6667 | 16 | 0.4375 |
| 4_1553354 | 5 | 0.4 | 2 | 0 | 3 | 0.3333 | 6 | 0.1667 | 16 | 0.25 | NA | NA | 3 | 0 | 5 | 0 | 5 | 0.2 | 13 | 0.0769 |
| 4_1553407 | 6 | 0.6667 | 4 | 0.5 | 2 | 0 | 9 | 0.3333 | 21 | 0.4286 | NA | NA | 4 | 0 | 8 | 0.25 | 7 | 0.5714 | 19 | 0.3158 |
| 4_1553407 | 6 | 0.6667 | 4 | 0.75 | 3 | 1 | 8 | 1 | 21 | 0.8571 | 1 | 1 | 3 | 1 | 8 | 0.875 | 7 | 0.7143 | 19 | 0.8421 |
| 4_1553408 | 6 | 1 | 4 | 1 | 3 | 1 | 10 | 1 | 23 | 1 | 1 | 1 | 4 | 1 | 7 | 1 | 6 | 1 | 18 | 1 |
| 4_1553410 | 7 | 1 | 3 | 0.3333 | 4 | 1 | 9 | 0.5556 | 23 | 0.7391 | 1 | 0 | 4 | 0.75 | 6 | 0.3333 | 5 | 0.8 | 16 | 0.5625 |
| 4_1553445 | 20 | 0 | 17 | 0.0588 | 23 | 0.1739 | 24 | 0.0417 | 84 | 0.0714 | 9 | 0.2222 | 19 | 0 | 18 | 0.1111 | 15 | 0 | 61 | 0.0656 |
| 4_1553449 | 21 | 0.1429 | 11 | 0.1818 | 23 | 0.087 | 20 | 0.15 | 75 | 0.1333 | 9 | 0.1111 | 16 | 0.0625 | 18 | 0.1667 | 16 | 0 | 59 | 0.0847 |
| 4_1553462 | 14 | 1 | 11 | 0.8182 | 17 | 1 | 14 | 0.9286 | 56 | 0.9464 | 7 | 1 | 12 | 0.9167 | 13 | 1 | 12 | 0.9167 | 44 | 0.9545 |
| 4_1553462 | 12 | 0.0833 | 11 | 0 | 18 | 0.1111 | 13 | 0.1538 | 54 | 0.0926 | 5 | 0 | 10 | 0.1 | 11 | 0.4545 | 11 | 0.1818 | 37 | 0.2162 |
| 4_1553464 | 10 | 0 | 8 | 0 | 16 | 0.1875 | 13 | 0 | 47 | 0.0638 | 7 | 0 | 10 | 0.2 | 8 | 0 | 11 | 0 | 36 | 0.0556 |
| 4_1553465 | 25 | 0.08 | 23 | 0.0435 | 31 | 0.0323 | 34 | 0.1471 | 113 | 0.0796 | 27 | 0 | 32 | 0.0312 | 24 | 0.0833 | 25 | 0.16 | 108 | 0.0648 |
| 4_1553466 | 20 | 0.9 | 18 | 0.9444 | 21 | 0.9048 | 30 | 0.9 | 89 | 0.9101 | 22 | 0.7727 | 27 | 0.8889 | 16 | 0.8125 | 19 | 0.8947 | 84 | 0.8452 |
| 4_1553466 | 9 | 0 | 7 | 0.2857 | 12 | 0.4167 | 15 | 0.2 | 43 | 0.2326 | 14 | 0.0714 | 13 | 0.3077 | 8 | 0.125 | 9 | 0.3333 | 44 | 0.2045 |
| 4_1553466 | 1 | 0 | NA | NA | 1 | 1 | 1 | 0 | 3 | 0.3333 | 1 | 0 | 1 | 0 | 1 | 1 | NA | NA | 3 | 0.3333 |
| 4_1553466 | 21 | 0 | 17 | 0.0588 | 21 | 0.0952 | 24 | 0.125 | 83 | 0.0723 | 20 | 0.1 | 26 | 0.1154 | 22 | 0 | 25 | 0.16 | 93 | 0.0968 |

| | | | | | | | | | | | | | | | | | | | | |
|---|---|---|---|---|---|---|---|---|---|---|---|---|---|---|---|---|---|---|---|---|
| 4_15534668 | 24 | 0.25 | 16 | 0.0625 | 27 | 0.1111 | 24 | 0.0833 | 91 | 0.1319 | 20 | 0.15 | 21 | 0.2381 | 18 | 0.0556 | 20 | 0.1 | 79 | 0.1392 |
| 4_15534674 | 24 | 0.375 | 19 | 0.2632 | 30 | 0.4 | 34 | 0.4706 | 107 | 0.3925 | 26 | 0.3077 | 32 | 0.4688 | 22 | 0.4545 | 27 | 0.4444 | 107 | 0.4206 |
| 4_15534677 | 24 | 0.0417 | 22 | 0.0455 | 29 | 0.1379 | 30 | 0.0333 | 105 | 0.0667 | 27 | 0.0741 | 27 | 0.037 | 19 | 0.1579 | 28 | 0.1429 | 101 | 0.099 |
| 4_15534686 | 26 | 0.4615 | 22 | 0.4091 | 30 | 0.1333 | 33 | 0.1818 | 111 | 0.2793 | 27 | 0.1111 | 33 | 0.1515 | 21 | 0.1429 | 27 | 0.2963 | 108 | 0.1759 |
| 4_15534687 | 24 | 0.6667 | 20 | 0.95 | 29 | 0.5862 | 32 | 0.875 | 105 | 0.7619 | 25 | 0.76 | 32 | 0.75 | 22 | 0.8636 | 24 | 0.6667 | 103 | 0.7573 |
| 4_15546968 | 15 | 0.3333 | 9 | 0.3333 | 12 | 0.3333 | 17 | 0.4706 | 53 | 0.3774 | 5 | 0.2 | 22 | 0.3182 | 19 | 0.5789 | 5 | 0.4 | 51 | 0.4118 |
| 4_15546976 | 18 | 0.6667 | 9 | 0.6667 | 11 | 0.9091 | 17 | 0.8235 | 55 | 0.7636 | 6 | 0.6667 | 19 | 0.5789 | 20 | 0.85 | 5 | 1 | 50 | 0.74 |
| 4_15546979 | 14 | 0.8571 | 11 | 0.6364 | 13 | 0.6154 | 20 | 0.55 | 58 | 0.6552 | 5 | 0.6 | 18 | 0.5 | 18 | 0.6111 | 4 | 1 | 45 | 0.6 |
| 4_15546996 | 18 | 0 | 13 | 0.2308 | 10 | 0.1 | 16 | 0 | 57 | 0.0702 | 8 | 0 | 21 | 0.0476 | 17 | 0.1176 | 3 | 0 | 49 | 0.0612 |
| 4_15564176 | 1 | 1 | NA | NA | NA | NA | NA | NA | 1 | 1 | 1 | 0 | NA | NA | 1 | 1 | NA | NA | 2 | 0.5 |
| 4_15564201 | 40 | 0.1 | 52 | 0.1731 | 49 | 0.1429 | 41 | 0.1463 | 182 | 0.1429 | 40 | 0.225 | 41 | 0.1951 | 34 | 0.0882 | 37 | 0.0811 | 152 | 0.1513 |
| 4_15564203 | 49 | 0.0612 | 63 | 0.0476 | 57 | 0.1579 | 45 | 0.0889 | 214 | 0.0888 | 46 | 0.1087 | 53 | 0.0755 | 41 | 0.1951 | 45 | 0.0667 | 185 | 0.1081 |
| 4_15564270 | 68 | 0.1471 | 78 | 0.1923 | 65 | 0.3077 | 60 | 0.2 | 271 | 0.2103 | 64 | 0.1406 | 67 | 0.2388 | 59 | 0.2203 | 57 | 0.2982 | 247 | 0.2227 |
| 4_15564305 | 80 | 0.1 | 102 | 0.0588 | 71 | 0.0282 | 70 | 0.0143 | 323 | 0.0526 | 67 | 0.0597 | 70 | 0.0714 | 61 | 0.0492 | 65 | 0.0462 | 263 | 0.057 |
| 4_15564340 | 77 | 0.2078 | 93 | 0.2151 | 72 | 0.2222 | 56 | 0.25 | 298 | 0.2215 | 69 | 0.1884 | 71 | 0.2817 | 62 | 0.2097 | 66 | 0.2576 | 268 | 0.2351 |
| 4_15564350 | 82 | 0.2073 | 103 | 0.1845 | 82 | 0.1951 | 73 | 0.1507 | 340 | 0.1853 | 74 | 0.1757 | 73 | 0.1781 | 76 | 0.1711 | 75 | 0.2 | 298 | 0.1812 |
| 4_15564351 | 88 | 0.3864 | 115 | 0.313 | 89 | 0.3034 | 75 | 0.3067 | 367 | 0.327 | 76 | 0.4605 | 85 | 0.3765 | 79 | 0.4557 | 78 | 0.4487 | 318 | 0.434 |
| 4_15564353 | 82 | 0.2317 | 106 | 0.1792 | 71 | 0.1831 | 73 | 0.2192 | 332 | 0.2018 | 71 | 0.1831 | 75 | 0.2267 | 64 | 0.2344 | 65 | 0.2154 | 275 | 0.2145 |
| 4_15567191 | 5 | 0 | 3 | 0.3333 | 2 | 0 | 3 | 0 | 13 | 0.0769 | 5 | 0.2 | 10 | 0.2 | 4 | 0 | 6 | 0.1667 | 25 | 0.16 |
| 4_15567221 | 7 | 0 | 4 | 0.25 | 3 | 0 | 5 | 0 | 19 | 0.0526 | 7 | 0 | 10 | 0.2 | 6 | 0.1667 | 7 | 0 | 30 | 0.1 |
| 4_15567384 | 9 | 0.2222 | 3 | 0.3333 | 4 | 0.5 | 4 | 0.25 | 20 | 0.3 | 7 | 0.2857 | 9 | 0.4444 | 6 | 0 | 9 | 0 | 31 | 0.1935 |
| 4_15567454 | 8 | 0.125 | 7 | 0.1429 | 4 | 0.25 | 4 | 0 | 23 | 0.1304 | 8 | 0 | 12 | 0.5 | 4 | 0.25 | 8 | 0.125 | 32 | 0.25 |
| 4_15567486 | 10 | 0 | 5 | 0.2 | 2 | 0 | 3 | 0 | 20 | 0.05 | 8 | 0.125 | 7 | 0 | 2 | 0 | 3 | 0 | 20 | 0.05 |
| 4_15567501 | 12 | 0.0833 | 9 | 0.2222 | 4 | 0.25 | 6 | 0.3333 | 31 | 0.1935 | 10 | 0.4 | 12 | 0.4167 | 6 | 0 | 10 | 0.1 | 38 | 0.2632 |
| 4_15567511 | 12 | 0.0833 | 9 | 0 | 4 | 0.25 | 7 | 0 | 32 | 0.0625 | 10 | 0.1 | 12 | 0.1667 | 7 | 0 | 9 | 0 | 38 | 0.0789 |
| 4_15567520 | 10 | 0.2 | 8 | 0 | 4 | 0.25 | 7 | 0.1429 | 29 | 0.1379 | 10 | 0.1 | 11 | 0.1818 | 4 | 0 | 10 | 0.1 | 35 | 0.1143 |
| 4_15567531 | 13 | 0.2308 | 9 | 0 | 4 | 0 | 8 | 0.25 | 34 | 0.1471 | 10 | 0.2 | 10 | 0.2 | 7 | 0 | 10 | 0.2 | 37 | 0.1622 |
| 4_15569435 | NA | NA | 1 | 0 | 2 | 0.5 | NA | NA | 3 | 0.3333 | 2 | 0 | NA | NA | 2 | 0.5 | 3 | 0.3333 | 7 | 0.2857 |
| 4_15570311 | 2 | 1 | 1 | 0 | 2 | 1 | 2 | 0 | 7 | 0.5714 | 2 | 1 | NA | NA | 3 | 1 | 2 | 0 | 7 | 0.7143 |
| 4_15570318 | 1 | 1 | 1 | 1 | 2 | 1 | 2 | 1 | 6 | 1 | 2 | 1 | NA | NA | 3 | 1 | 2 | 1 | 7 | 1 |
| 4_15570331 | 2 | 1 | 1 | 1 | 2 | 1 | 2 | 1 | 7 | 1 | 2 | 1 | NA | NA | 3 | 1 | 3 | 1 | 8 | 1 |
| 4_15570334 | 2 | 1 | 1 | 0 | 2 | 0.5 | 2 | 0.5 | 7 | 0.5714 | 1 | 0 | NA | NA | 2 | 0.5 | 3 | 0.3333 | 6 | 0.3333 |
| 4_15570347 | 2 | 0.5 | 1 | 1 | 2 | 1 | 2 | 0.5 | 7 | 0.7143 | 2 | 0.5 | 1 | 0 | 3 | 0 | 3 | 0.3333 | 9 | 0.2222 |
| 4_15570353 | 2 | 0 | 1 | 0 | 2 | 1 | 2 | 0.5 | 7 | 0.4286 | 2 | 0.5 | 1 | 0 | 2 | 0 | 3 | 0 | 8 | 0.125 |
| 4_15570354 | 2 | 0.5 | 1 | 0 | 2 | 0 | 2 | 0 | 7 | 0.1429 | 2 | 0 | 1 | 0 | 2 | 0 | 4 | 0.25 | 9 | 0.1111 |
| 4_15570355 | 1 | 1 | 1 | 1 | 2 | 1 | 2 | 1 | 6 | 1 | 2 | 0.5 | NA | NA | 3 | 1 | 3 | 1 | 8 | 0.875 |
| 4_15570357 | 2 | 1 | 1 | 1 | 2 | 1 | 2 | 1 | 7 | 1 | 2 | 1 | 1 | 1 | 3 | 1 | 4 | 0.75 | 10 | 0.9 |
| 4_15570361 | 2 | 0 | NA | NA | 2 | 0 | 2 | 0.5 | 6 | 0.1667 | 2 | 0.5 | 1 | 1 | 3 | 0.3333 | 4 | 0.5 | 10 | 0.5 |
| 4_15570363 | 2 | 1 | 1 | 1 | 2 | 1 | 2 | 1 | 7 | 1 | 2 | 1 | 1 | 1 | 2 | 0.5 | 4 | 0.75 | 9 | 0.7778 |
| 4_15570368 | 2 | 1 | 1 | 1 | 2 | 1 | 1 | 1 | 6 | 1 | 1 | 1 | 1 | 1 | 2 | 1 | 3 | 1 | 7 | 1 |
| 4_15570394 | 3 | 0.6667 | 1 | 1 | 2 | 1 | 2 | 1 | 8 | 0.875 | 2 | 1 | 2 | 1 | 3 | 1 | 4 | 0.75 | 11 | 0.9091 |
| 4_15570409 | 2 | 0 | 1 | 0 | 1 | 1 | 2 | 0.5 | 6 | 0.3333 | 2 | 1 | 2 | 0 | 3 | 1 | 3 | 0 | 10 | 0.5 |
| 4_15570419 | 3 | 0.3333 | 1 | 0 | 2 | 0 | 3 | 0.3333 | 9 | 0.2222 | 2 | 0.5 | 2 | 1 | 3 | 0 | 4 | 0 | 11 | 0.2727 |
| 4_15570448 | 3 | 0.3333 | 1 | 0 | 2 | 0 | 3 | 0.3333 | 9 | 0.2222 | 1 | 0 | 3 | 0 | 3 | 0.3333 | 5 | 0.4 | 12 | 0.25 |
| 4_15570454 | 3 | 0 | 2 | 0.5 | 2 | 0.5 | 1 | 0 | 8 | 0.25 | 2 | 0 | 3 | 0 | 1 | 0 | 5 | 0.2 | 11 | 0.0909 |
| 4_15570477 | 3 | 0 | 3 | 0.3333 | 2 | 0 | 3 | 0 | 11 | 0.0909 | 1 | 0 | 2 | 0 | 3 | 0.3333 | 4 | 0 | 10 | 0.1 |
| 4_15570480 | 3 | 0.3333 | 3 | 0.3333 | 2 | 0 | 3 | 0 | 11 | 0.1818 | 1 | 0 | 1 | 1 | 3 | 0 | 4 | 0 | 9 | 0.1111 |
| 4_15570489 | 3 | 0.6667 | 3 | 0.3333 | 1 | 0 | 3 | 0 | 10 | 0.3 | 1 | 0 | 3 | 0 | 2 | 0 | 5 | 0.2 | 11 | 0.0909 |
| 4_15570503 | 3 | 0 | 2 | 0 | 2 | 0 | 3 | 0.6667 | 10 | 0.2 | 1 | 0 | 3 | 0.3333 | 3 | 0.6667 | 4 | 0.75 | 11 | 0.5455 |
| 4_15570700 | 9 | 0 | 4 | 0.25 | 4 | 0 | 5 | 0.2 | 22 | 0.0909 | 4 | 0.25 | 6 | 0.1667 | 6 | 0 | 7 | 0 | 23 | 0.087 |
| 4_15570748 | 7 | 0.1429 | 4 | 0.25 | 2 | 0 | 6 | 0.1667 | 19 | 0.1579 | 2 | 0 | 4 | 0 | 7 | 0 | 6 | 0.1667 | 19 | 0.0526 |
| 4_15573913 | 2 | 1 | 3 | 0.6667 | 4 | 1 | 1 | 1 | 10 | 0.9 | 2 | 1 | 3 | 0.6667 | 1 | 1 | NA | NA | 6 | 0.8333 |
| 4_15574293 | 2 | 0.5 | 1 | 1 | NA | NA | 1 | 0 | 4 | 0.5 | NA | NA | 1 | 1 | 1 | 0 | 1 | 1 | 3 | 0.6667 |
| 4_15574312 | 2 | 0.5 | NA | NA | NA | NA | 1 | 1 | 3 | 0.6667 | NA | NA | 2 | 1 | 1 | 0 | 1 | 1 | 4 | 0.75 |
| 4_15574342 | 4 | 0.25 | 4 | 0 | 1 | 0 | 5 | 0 | 14 | 0.0714 | NA | NA | 5 | 0.2 | 1 | 0 | 3 | 0 | 9 | 0.1111 |
| 4_15575330 | 1 | 0 | 3 | 0.3333 | NA | NA | 3 | 0.3333 | 7 | 0.2857 | 1 | 0 | 4 | 0 | 2 | 0.5 | 1 | 0 | 8 | 0.125 |
| 4_15577975 | 2 | 0.5 | 5 | 0.2 | 2 | 0.5 | 2 | 0 | 11 | 0.2727 | 1 | 0 | 2 | 0.5 | 1 | 0 | 1 | 0 | 5 | 0.2 |
| 4_15577976 | 2 | 1 | 5 | 0.8 | 3 | 1 | 2 | 0.5 | 12 | 0.8333 | 1 | 0 | 2 | 1 | 2 | 1 | 1 | 1 | 6 | 0.8333 |
| 4_15577982 | 2 | 0.5 | 5 | 0.8 | 3 | 0 | 2 | 0 | 12 | 0.4167 | 1 | 0 | 2 | 0.5 | 2 | 0 | 1 | 1 | 6 | 0.3333 |
| 4_15577983 | 2 | 1 | 4 | 0.5 | 3 | 0.6667 | 2 | 0 | 11 | 0.5455 | 1 | 0 | 2 | 0.5 | 1 | 0 | NA | NA | 4 | 0.25 |
| 4_15577996 | 2 | 0.5 | 5 | 1 | 2 | 0.5 | 1 | 0 | 10 | 0.7 | 1 | 1 | 2 | 0.5 | 1 | 0 | 1 | 1 | 5 | 0.6 |
| 4_15577997 | 2 | 1 | 3 | 1 | 3 | 0.6667 | 3 | 0.3333 | 11 | 0.7273 | 1 | 0 | 1 | 1 | NA | NA | 1 | 1 | 3 | 0.6667 |
| 4_15578016 | 3 | 0.6667 | 4 | 0.75 | 4 | 0.5 | 3 | 0 | 14 | 0.5 | 1 | 0 | 2 | 1 | 2 | 0.5 | 1 | 0 | 6 | 0.5 |
| 4_15578024 | 1 | 0 | 6 | 0.3333 | 4 | 0.25 | 2 | 0 | 13 | 0.2308 | NA | NA | 2 | 0 | 3 | 0.3333 | 1 | 0 | 6 | 0.1667 |
| 4_15578077 | 2 | 1 | 6 | 0 | 6 | 0.5 | 3 | 0.6667 | 17 | 0.4118 | 2 | 0.5 | 4 | 0 | 3 | 0 | 1 | 0 | 10 | 0.1 |

| | | | | | | | | | | | | | | | | | | | | |
|---|---|---|---|---|---|---|---|---|---|---|---|---|---|---|---|---|---|---|---|---|
| 4_15580006 | 18 | 0.6111 | 19 | 0.5263 | 12 | 0.5833 | 17 | 0.5294 | 66 | 0.5606 | 17 | 0.5882 | 11 | 0.5455 | 17 | 0.7059 | 16 | 0.5 | 61 | 0.5902 |
| 4_15580113 | 2 | 1 | NA | NA | NA | NA | NA | NA | 2 | 1 | 2 | 1 | NA | NA | NA | NA | 2 | 1 | 4 | 1 |
| 4_15580547 | 1 | 1 | NA | NA | NA | NA | 2 | 0.5 | 3 | 0.6667 | NA | NA | 1 | 1 | NA | NA | NA | NA | 1 | 1 |
| 4_15582409 | 5 | 0 | 1 | 0 | 1 | 1 | 1 | 0 | 8 | 0.125 | 6 | 0 | NA | NA | 5 | 0.4 | 1 | 1 | 12 | 0.25 |
| 4_15582431 | 4 | 0.25 | 1 | 0 | 3 | 0.3333 | 2 | 0 | 10 | 0.2 | 5 | 0.2 | 1 | 1 | 7 | 0.1429 | 2 | 0 | 15 | 0.2 |
| 4_15582446 | 7 | 0.2857 | 1 | 1 | 3 | 0 | 2 | 1 | 13 | 0.3846 | 3 | 0 | 1 | 0 | 7 | 0.7143 | 2 | 0 | 13 | 0.3846 |
| 4_15582449 | 6 | 0.3333 | 1 | 0 | 1 | 0 | 2 | 1 | 10 | 0.4 | 5 | 0.2 | 2 | 0.5 | 5 | 0.8 | 1 | 1 | 13 | 0.5385 |
| 4_15582452 | 7 | 0.5714 | 1 | 1 | 1 | 0 | 5 | 0.8 | 14 | 0.6429 | 5 | 0.8 | 1 | 1 | 6 | 0.6667 | 1 | 1 | 13 | 0.7692 |
| 4_15582469 | 7 | 0.2857 | 3 | 0.3333 | 7 | 0.2857 | 4 | 0 | 21 | 0.2381 | 7 | 0.1429 | 1 | 0 | 7 | 0.1429 | 3 | 0.6667 | 18 | 0.2222 |
| 4_15582510 | 2 | 0 | 2 | 0 | 3 | 0.6667 | 1 | 0 | 8 | 0.25 | 3 | 0 | 2 | 0.5 | 5 | 0.6 | NA | NA | 10 | 0.4 |
| 4_15582511 | 2 | 0 | 2 | 0 | 3 | 0.3333 | 1 | 0 | 8 | 0.125 | 3 | 0 | 2 | 0 | 5 | 0.2 | NA | NA | 10 | 0.1 |
| 4_15582527 | 2 | 0.5 | 1 | 1 | 4 | 0.75 | 3 | 0.6667 | 10 | 0.7 | 2 | 1 | 2 | 0.5 | 5 | 0.8 | 1 | 1 | 10 | 0.8 |
| 4_15582531 | 3 | 0 | 2 | 0 | 4 | 0.5 | 2 | 0 | 11 | 0.1818 | 3 | 0.6667 | 1 | 0 | 6 | 0.1667 | 1 | 1 | 11 | 0.3636 |
| 4_15582533 | 3 | 1 | 2 | 1 | 4 | 1 | 3 | 0.6667 | 12 | 0.9167 | 2 | 0.5 | 2 | 1 | 5 | 0.8 | 1 | 1 | 10 | 0.8 |
| 4_15582540 | 2 | 0 | 2 | 0 | 4 | 0 | 3 | 0.3333 | 11 | 0.0909 | 3 | 0 | 2 | 0 | 6 | 0.1667 | 1 | 0 | 12 | 0.0833 |
| 4_15582546 | 1 | 1 | 2 | 0 | 4 | 0.25 | 3 | 0 | 10 | 0.2 | 3 | 0.3333 | 2 | 0 | 5 | 0.2 | 1 | 0 | 11 | 0.1818 |
| 4_15582563 | 2 | 0 | NA | NA | 4 | 0.75 | 3 | 0.3333 | 9 | 0.4444 | 3 | 0.3333 | 2 | 0.5 | 6 | 0.5 | NA | NA | 11 | 0.4545 |
| 4_15585380 | 5 | 0 | 2 | 0.5 | 2 | 0 | 2 | 0 | 11 | 0.0909 | 2 | 0 | 4 | 0.25 | 1 | 0 | 3 | 0 | 10 | 0.1 |
| 4_15585383 | 4 | 0.25 | 2 | 0 | 2 | 0 | 5 | 0 | 13 | 0.0769 | 2 | 1 | 3 | 1 | 1 | 0 | 2 | 0 | 8 | 0.625 |
| 4_15585439 | 4 | 0.25 | 3 | 0.3333 | 3 | 0 | 4 | 0.25 | 14 | 0.2143 | 4 | 0.25 | 4 | 0.5 | NA | NA | 4 | 0.5 | 12 | 0.4167 |
| 4_15585450 | 4 | 0.5 | 3 | 0 | 2 | 0.5 | 4 | 0.5 | 13 | 0.3846 | 3 | 0.6667 | 1 | 1 | NA | NA | 3 | 0.6667 | 7 | 0.7143 |
| 4_15585460 | 6 | 0 | 3 | 0 | 2 | 0.5 | 3 | 0 | 14 | 0.0714 | 4 | 0 | 3 | 0.3333 | NA | NA | 4 | 0 | 11 | 0.0909 |
| 4_15585473 | 7 | 0.5714 | 2 | 0 | 2 | 0.5 | 2 | 1 | 13 | 0.5385 | 4 | 0.5 | 3 | 1 | NA | NA | 2 | 1 | 9 | 0.7778 |
| 4_15585478 | 5 | 0.8 | 3 | 0 | 3 | 0.3333 | 2 | 0.5 | 13 | 0.4615 | 4 | 0.5 | 2 | 0.5 | NA | NA | 3 | 0.6667 | 9 | 0.5556 |
| 4_15585480 | 7 | 0.1429 | 3 | 0.6667 | 3 | 0.3333 | 3 | 0 | 16 | 0.25 | 4 | 0.5 | 1 | 0 | NA | NA | 3 | 0 | 8 | 0.25 |
| 4_15585483 | 7 | 0.1429 | 3 | 0 | 3 | 0 | 2 | 0 | 15 | 0.0667 | 3 | 0.3333 | 2 | 0.5 | NA | NA | 3 | 0 | 8 | 0.25 |
| 4_15585490 | 6 | 0.5 | 1 | 0 | 3 | 0.6667 | 3 | 0.3333 | 13 | 0.4615 | 3 | 0.6667 | 3 | 0.6667 | NA | NA | 2 | 0 | 8 | 0.5 |
| 4_15585491 | 6 | 1 | 3 | 1 | 2 | 1 | 3 | 0.3333 | 14 | 0.8571 | 4 | 1 | 2 | 1 | NA | NA | 3 | 1 | 9 | 1 |
| 4_15585500 | 5 | 0 | 2 | 0 | 2 | 1 | NA | NA | 9 | 0.2222 | 2 | 0.5 | 3 | 0 | NA | NA | 2 | 0.5 | 7 | 0.2857 |
| 4_15585503 | 7 | 0.2857 | 3 | 0.6667 | 3 | 0 | 3 | 0 | 16 | 0.25 | 4 | 0 | 3 | 0.3333 | NA | NA | 3 | 0 | 10 | 0.1 |
| 4_15585539 | 6 | 0.6667 | 2 | 0.5 | 3 | 0.3333 | 3 | 0.6667 | 14 | 0.5714 | 4 | 0.5 | 2 | 0.5 | NA | NA | 2 | 1 | 8 | 0.625 |
| 4_15585743 | 5 | 0.6 | 1 | 1 | 1 | 0 | NA | NA | 7 | 0.5714 | 2 | 0.5 | 2 | 0 | NA | NA | NA | NA | 4 | 0.25 |
| 4_15586801 | NA | NA | 5 | 0.2 | 5 | 0.2 | 5 | 0.2 | 15 | 0.2 | 1 | 0 | 5 | 0.4 | 4 | 0.25 | 1 | 0 | 11 | 0.2727 |
| 4_15586808 | 1 | 0 | 7 | 0.1429 | 5 | 0 | 5 | 0 | 18 | 0.0556 | 2 | 0 | 6 | 0.1667 | 4 | 0.25 | 2 | 0 | 14 | 0.1429 |
| 4_15586813 | 1 | 1 | 4 | 0 | 4 | 0.25 | 5 | 0.4 | 14 | 0.2857 | 2 | 0 | 5 | 0 | 4 | 0.75 | 2 | 0.5 | 13 | 0.3077 |
| 4_15586834 | NA | NA | 4 | 0.25 | 3 | 0 | 3 | 0 | 10 | 0.1 | 1 | 0 | 4 | 0 | 4 | 0.25 | 1 | 0 | 10 | 0.1 |
| 4_15586837 | 1 | 0 | 6 | 0.5 | 4 | 0.25 | 4 | 0.25 | 15 | 0.3333 | 1 | 0 | 5 | 0.2 | 3 | 0 | 2 | 0.5 | 11 | 0.1818 |
| 4_15586866 | 1 | 0 | 5 | 0.2 | 3 | 0.3333 | 3 | 0.3333 | 12 | 0.25 | 1 | 0 | 4 | 0 | 4 | 0.25 | 2 | 0 | 11 | 0.0909 |
| 4_15586875 | 1 | 1 | 5 | 0.6 | 4 | 0.25 | 3 | 0.6667 | 13 | 0.5385 | 1 | 0 | 4 | 0.75 | 4 | 0.75 | 2 | 1 | 11 | 0.7273 |
| 4_15589312 | 11 | 0.4545 | 19 | 0.4737 | 18 | 0.4444 | 18 | 0.5556 | 66 | 0.4848 | 10 | 0.6 | 13 | 0.3846 | 19 | 0.3158 | 19 | 0.5789 | 61 | 0.459 |
| 4_15589315 | 16 | 0.5 | 22 | 0.3182 | 17 | 0.4118 | 18 | 0.5 | 73 | 0.4247 | 11 | 0.2727 | 13 | 0.4615 | 21 | 0.5238 | 15 | 0.4 | 60 | 0.4333 |
| 4_15589646 | NA | NA | 1 | 1 | NA | NA | NA | NA | 1 | 1 | NA | NA | NA | NA | 2 | 0.5 | NA | NA | 2 | 0.5 |
| 4_15589648 | 1 | 1 | 1 | 0 | NA | NA | NA | NA | 2 | 0.5 | NA | NA | NA | NA | 1 | 1 | NA | NA | 1 | 1 |
| 4_15589670 | 1 | 1 | NA | NA | NA | NA | NA | NA | 1 | 1 | NA | NA | NA | NA | 2 | 1 | NA | NA | 2 | 1 |
| 4_15590302 | 3 | 0 | 6 | 0.1667 | 6 | 0 | 4 | 0 | 19 | 0.0526 | 3 | 0 | 5 | 0 | 5 | 0.4 | 7 | 0 | 20 | 0.1 |
| 4_15590311 | 3 | 0 | 7 | 0.1429 | 3 | 0 | 4 | 0 | 17 | 0.0588 | 4 | 0 | 6 | 0.1667 | 9 | 0.1111 | 5 | 0 | 24 | 0.0833 |
| 4_15590344 | 1 | 0 | 8 | 0.25 | 8 | 0.125 | 3 | 0.3333 | 20 | 0.2 | 3 | 0 | 6 | 0.1667 | 7 | 0 | 6 | 0.1667 | 22 | 0.0909 |
| 4_15590418 | 2 | 0 | 7 | 0.7143 | 7 | 0.2857 | 6 | 0.3333 | 22 | 0.4091 | 6 | 0.3333 | 5 | 0.4 | 7 | 0.5714 | 7 | 0.4286 | 25 | 0.44 |
| 4_15590426 | 2 | 1 | 6 | 0 | 4 | 0.25 | 6 | 0.1667 | 18 | 0.2222 | 7 | 0.2857 | 7 | 0.4286 | 3 | 0 | 5 | 0.2 | 22 | 0.2727 |
| 4_15590429 | 1 | 1 | 5 | 0.8 | 5 | 0.4 | 6 | 0.8333 | 17 | 0.7059 | 6 | 0.6667 | 6 | 0.8333 | 9 | 0.6667 | 7 | 0.4286 | 28 | 0.6429 |
| 4_15590436 | 2 | 0 | 9 | 0.2222 | 6 | 0 | 4 | 0 | 21 | 0.0952 | 7 | 0.4286 | 7 | 0.4286 | 10 | 0.2 | 7 | 0.2857 | 31 | 0.3226 |
| 4_15590437 | 2 | 0 | 8 | 0.375 | 6 | 0.1667 | 7 | 0.1429 | 23 | 0.2174 | 7 | 0.1429 | 6 | 0.3333 | 10 | 0.3 | 7 | 0 | 30 | 0.2 |
| 4_15590438 | 1 | 0 | 6 | 0 | 6 | 0 | 5 | 0.2 | 18 | 0.0556 | 5 | 0.4 | 2 | 0 | 8 | 0.125 | 6 | 0 | 21 | 0.1429 |
| 4_15590445 | 2 | 0 | 9 | 0.1111 | 6 | 0.1667 | 4 | 0 | 21 | 0.0952 | 7 | 0.4286 | 6 | 0.1667 | 11 | 0.1818 | 6 | 0 | 30 | 0.2 |
| 4_15590794 | 4 | 0 | 12 | 0.6667 | 8 | 0.25 | 8 | 0.25 | 32 | 0.375 | 9 | 0.5556 | 8 | 0.625 | 13 | 0.3077 | 7 | 0.4286 | 37 | 0.4595 |
| 4_15590802 | 5 | 1 | 12 | 0.6667 | 8 | 0.625 | 9 | 0.6667 | 34 | 0.7059 | 8 | 0.5 | 9 | 0.8889 | 14 | 0.4286 | 10 | 0.7 | 41 | 0.6098 |
| 4_15590813 | 5 | 0.4 | 8 | 0.375 | 7 | 0.2857 | 7 | 0 | 27 | 0.2593 | 8 | 0.125 | 9 | 0.3333 | 11 | 0.2727 | 9 | 0.3333 | 37 | 0.2703 |
| 4_15590825 | 6 | 0.6667 | 10 | 0.7 | 8 | 1 | 9 | 0.6667 | 33 | 0.7576 | 10 | 0.8 | 7 | 0.5714 | 14 | 0.9286 | 9 | 0.7778 | 40 | 0.8 |
| 4_15590844 | 8 | 0.25 | 10 | 0.1 | 6 | 0.6667 | 6 | 0 | 30 | 0.2333 | 9 | 0 | 9 | 0.2222 | 14 | 0.1429 | 11 | 0.1818 | 43 | 0.1395 |
| 4_15591624 | 4 | 0.25 | 5 | 0 | 3 | 0.3333 | 1 | 0 | 13 | 0.1538 | 3 | 0 | 6 | 0.1667 | 6 | 0.1667 | 3 | 0 | 18 | 0.1111 |
| 4_15591629 | 4 | 0.25 | 5 | 0 | 3 | 0.3333 | 1 | 0 | 13 | 0.1538 | 3 | 0 | 6 | 0.1667 | 6 | 0.3333 | 3 | 0 | 18 | 0.1667 |
| 4_15591650 | 17 | 0 | 15 | 0.0667 | 13 | 0.1538 | 18 | 0.1667 | 63 | 0.0952 | 11 | 0.0909 | 18 | 0.0556 | 12 | 0 | 13 | 0.0769 | 54 | 0.0556 |
| 4_15591656 | NA | NA | 1 | 1 | NA | NA | NA | NA | 1 | 1 | NA | NA | 1 | 1 | 1 | 0 | NA | NA | 2 | 0.5 |
| 4_15591703 | 17 | 0.1176 | 16 | 0.125 | 16 | 0.25 | 19 | 0.0526 | 68 | 0.1324 | 11 | 0.2727 | 19 | 0.1053 | 15 | 0.1333 | 12 | 0.0833 | 57 | 0.1404 |
| 4_15591727 | NA | NA | 1 | 1 | NA | NA | 1 | 0 | 2 | 0.5 | NA | NA | 1 | 1 | NA | NA | NA | NA | 1 | 1 |

| | | | | | | | | | | | | | | | | | | | | |
|---|---|---|---|---|---|---|---|---|---|---|---|---|---|---|---|---|---|---|---|---|
| 4_1559173 | 18 | 0.1667 | 17 | 0.1765 | 17 | 0.0588 | 22 | 0.0455 | 74 | 0.1081 | 13 | 0.0769 | 19 | 0.1053 | 20 | 0 | 15 | 0.0667 | 67 | 0.0597 |
| 4_1559174 | 16 | 0.0625 | 17 | 0.0588 | 19 | 0.0526 | 22 | 0.0455 | 74 | 0.0541 | 12 | 0.1667 | 18 | 0.1111 | 21 | 0 | 14 | 0 | 65 | 0.0615 |
| 4_1559175 | 11 | 0.8182 | 11 | 0.9091 | 15 | 0.4667 | 17 | 0.4706 | 54 | 0.6296 | 12 | 0.8333 | 15 | 0.8667 | 18 | 0.6667 | 12 | 0.75 | 57 | 0.7719 |
| 4_1559277 | NA | NA | 3 | 0.3333 | 1 | 0 | NA | NA | 4 | 0.25 | 2 | 0 | NA | NA | 1 | 1 | NA | NA | 3 | 0.3333 |
| 4_1559279 | NA | NA | 3 | 0.6667 | 1 | 0 | NA | NA | 4 | 0.5 | 3 | 0.3333 | NA | NA | 1 | 0 | NA | NA | 4 | 0.25 |
| 4_1559323 | 2 | 0 | 5 | 0.4 | 2 | 0 | 6 | 0.1667 | 15 | 0.2 | 3 | 0 | 3 | 0 | 3 | 0.3333 | 2 | 0.5 | 11 | 0.1818 |
| 4_1559388 | 3 | 0 | 5 | 0.4 | 4 | 0 | 4 | 0 | 16 | 0.125 | 5 | 0 | 3 | 0 | 3 | 0.3333 | 2 | 0 | 13 | 0.0769 |
| 4_1559427 | 2 | 0.5 | 9 | 0.2222 | 7 | 0.1429 | 7 | 0.1429 | 25 | 0.2 | 4 | 0 | 9 | 0.4444 | 3 | 0.6667 | 3 | 0 | 19 | 0.3158 |
| 4_1559428 | 4 | 0 | 9 | 0.4444 | 7 | 0.2857 | 10 | 0.3 | 30 | 0.3 | 6 | 0.1667 | 10 | 0.3 | 3 | 0.3333 | 3 | 0.3333 | 22 | 0.2727 |
| 4_1559432 | 4 | 0 | 9 | 0.1111 | 9 | 0.3333 | 10 | 0.2 | 32 | 0.1875 | 4 | 0.25 | 10 | 0.2 | 3 | 0.3333 | 5 | 0.2 | 22 | 0.2273 |
| 4_1559498 | 2 | 0.5 | 4 | 0 | 3 | 0 | 3 | 0.3333 | 12 | 0.1667 | 5 | 0 | 2 | 0.5 | 6 | 0.3333 | 1 | 0 | 14 | 0.2143 |
| 4_1559498 | 2 | 0.5 | 4 | 0 | 2 | 0.5 | 2 | 0 | 10 | 0.2 | 5 | 0 | 2 | 0.5 | 3 | 0 | 1 | 1 | 11 | 0.1818 |
| 4_1559499 | 1 | 0 | 4 | 0 | 3 | 0 | 3 | 0.3333 | 11 | 0.0909 | 7 | 0.2857 | 2 | 0 | 5 | 0 | 1 | 1 | 15 | 0.2 |
| 4_1559499 | 2 | 0.5 | 4 | 0 | 3 | 0.3333 | 3 | 1 | 12 | 0.4167 | 7 | 0.2857 | 2 | 0.5 | 5 | 0.2 | 1 | 0 | 15 | 0.2667 |
| 4_1559499 | 2 | 1 | 3 | 0 | 3 | 0 | 3 | 0.6667 | 11 | 0.3636 | 7 | 0.1429 | 2 | 0 | 6 | 0.1667 | 1 | 0 | 16 | 0.125 |
| 4_1559500 | 2 | 0.5 | 3 | 0.3333 | 2 | 0 | 3 | 1 | 10 | 0.5 | 5 | 0.4 | 2 | 0 | 4 | 0 | 1 | 0 | 12 | 0.1667 |
| 4_1559501 | 2 | 0.5 | 4 | 0 | 3 | 0.3333 | 3 | 0.3333 | 12 | 0.25 | 6 | 0.5 | 1 | 0 | 4 | 0.25 | 1 | 0 | 12 | 0.3333 |
| 4_1559504 | 2 | 0.5 | 4 | 0.75 | 3 | 0 | 3 | 1 | 12 | 0.5833 | 6 | 0.5 | 1 | 0 | 5 | 0.2 | 1 | 1 | 13 | 0.3846 |
| 4_1559505 | 2 | 1 | 4 | 1 | 3 | 1 | 3 | 1 | 12 | 1 | 6 | 1 | 1 | 1 | 4 | 1 | 1 | 1 | 12 | 1 |
| 4_1559506 | 2 | 0.5 | 1 | 0 | 2 | 0.5 | 2 | 1 | 7 | 0.5714 | 4 | 0.75 | 1 | 0 | 5 | 0.6 | 1 | 1 | 11 | 0.6364 |
| 4_1559506 | 2 | 0.5 | 3 | 0.3333 | 3 | 0 | 2 | 0.5 | 10 | 0.3 | 6 | 0.3333 | 1 | 0 | 4 | 0 | 1 | 0 | 12 | 0.1667 |
| 4_1559507 | 2 | 1 | 4 | 1 | 3 | 1 | 2 | 1 | 11 | 1 | 5 | 0.8 | 1 | 1 | 5 | 1 | 1 | 1 | 12 | 0.9167 |
| 4_1559507 | 1 | 1 | 4 | 1 | 2 | 1 | 2 | 1 | 9 | 1 | 5 | 1 | 1 | 1 | 5 | 0.8 | 1 | 1 | 12 | 0.9167 |
| 4_1559508 | 1 | 1 | 4 | 1 | 2 | 1 | 1 | 1 | 8 | 1 | 4 | 1 | 1 | 1 | 5 | 0.8 | 1 | 1 | 11 | 0.9091 |
| 4_1559511 | 1 | 1 | 4 | 0.75 | 2 | 1 | 2 | 1 | 9 | 0.8889 | 5 | 1 | 1 | 1 | 3 | 1 | 1 | 1 | 10 | 1 |
| 4_1559511 | 1 | 1 | 4 | 0 | 2 | 0 | 2 | 0.5 | 9 | 0.2222 | 5 | 0.2 | 1 | 0 | 2 | 0 | 1 | 0 | 9 | 0.1111 |
| 4_1559512 | 1 | 0 | 3 | 0.3333 | 1 | 0 | 2 | 0 | 7 | 0.1429 | 5 | 0 | 1 | 0 | 1 | 0 | 1 | 1 | 8 | 0.125 |
| 4_1559515 | 1 | 1 | 3 | 0 | 1 | 0 | 2 | 0.5 | 7 | 0.2857 | 5 | 0.2 | 1 | 0 | 2 | 0.5 | 1 | 1 | 9 | 0.3333 |
| 4_1559517 | 1 | 0 | 3 | 0 | 2 | 0 | 2 | 0.5 | 8 | 0.125 | 5 | 0.4 | 1 | 0 | 2 | 0 | 1 | 0 | 9 | 0.2222 |
| 4_1559714 | 1 | 0 | 1 | 1 | 3 | 0 | 3 | 0 | 8 | 0.125 | 3 | 0 | 3 | 0.3333 | 5 | 0 | 2 | 0 | 13 | 0.0769 |
| 4_1559734 | NA | NA | 2 | 1 | NA | NA | 1 | 0 | 3 | 0.6667 | 1 | 0 | 1 | 1 | 4 | 0.5 | 1 | 1 | 7 | 0.5714 |
| 4_1559750 | NA | NA | 2 | 0.5 | 1 | 0 | 1 | 0 | 4 | 0.25 | 1 | 1 | 2 | 0.5 | 5 | 0.2 | 1 | 0 | 9 | 0.3333 |
| 4_1559763 | NA | NA | 1 | 1 | NA | NA | NA | NA | 1 | 1 | NA | NA | 1 | 1 | NA | NA | 1 | 1 | 2 | 1 |
| 4_1559877 | 9 | 0.1111 | 8 | 0 | 11 | 0.1818 | 5 | 0 | 33 | 0.0909 | 4 | 0 | 11 | 0.0909 | 4 | 0.25 | 4 | 0.25 | 23 | 0.1304 |
| 4_1559882 | 5 | 0.2 | 8 | 0.125 | 10 | 0.1 | 3 | 0 | 26 | 0.1154 | 3 | 0 | 12 | 0.1667 | 7 | 0.2857 | 6 | 0.1667 | 28 | 0.1786 |
| 4_1559902 | NA | NA | 1 | 0 | NA | NA | 1 | 1 | 2 | 0.5 | 1 | 0 | 1 | 0 | NA | NA | 2 | 0.5 | 4 | 0.25 |
| 4_1559904 | 4 | 0.25 | 10 | 0.2 | 11 | 0.2727 | 5 | 0.4 | 30 | 0.2667 | 7 | 0.5714 | 13 | 0.1538 | 5 | 0.2 | 13 | 0.2308 | 38 | 0.2632 |
| 4_1559905 | 10 | 0.3 | 12 | 0.0833 | 13 | 0.2308 | 8 | 0 | 43 | 0.1628 | 9 | 0 | 14 | 0.2143 | 10 | 0.2 | 14 | 0.2857 | 47 | 0.1915 |
| 4_1559906 | 3 | 0 | 8 | 0.125 | 12 | 0.1667 | 8 | 0.25 | 31 | 0.1613 | 9 | 0.2222 | 17 | 0.1765 | 9 | 0.2222 | 12 | 0.0833 | 47 | 0.1702 |
| 4_1559912 | 9 | 0.4444 | 10 | 0.1 | 13 | 0.0769 | 7 | 0 | 39 | 0.1538 | 10 | 0.1 | 15 | 0.3333 | 9 | 0.3333 | 11 | 0.0909 | 45 | 0.2222 |
| 4_1559912 | 10 | 0.3 | 9 | 0.2222 | 11 | 0.1818 | 5 | 0.2 | 35 | 0.2286 | 9 | 0.2222 | 13 | 0.1538 | 8 | 0.25 | 12 | 0.1667 | 42 | 0.1905 |
| 4_1559915 | 6 | 0.8333 | 11 | 0.7273 | 13 | 0.7692 | 6 | 0.5 | 36 | 0.7222 | 8 | 0.75 | 15 | 0.6667 | 7 | 1 | 11 | 0.7273 | 41 | 0.7561 |
| 4_1560308 | 5 | 0 | 6 | 0.1667 | 1 | 0 | 4 | 0 | 16 | 0.0625 | 2 | 0.5 | 1 | 0 | NA | NA | 4 | 0 | 7 | 0.1429 |
| 4_1560417 | 18 | 0.7778 | 10 | 1 | 3 | 0.6667 | 8 | 0.875 | 39 | 0.8462 | 3 | 1 | 4 | 0.75 | 7 | 0.8571 | 5 | 0.8 | 19 | 0.8421 |
| 4_1560417 | 17 | 0.2353 | 10 | 0.3 | 3 | 0 | 8 | 0 | 38 | 0.1842 | 3 | 0.3333 | 5 | 0.4 | 7 | 0.2857 | 5 | 0.6 | 20 | 0.4 |
| 4_1560418 | 16 | 0.6875 | 10 | 0.7 | 2 | 1 | 5 | 0.6 | 33 | 0.697 | 4 | 0.5 | 6 | 0.6667 | 8 | 0.625 | 5 | 0.4 | 23 | 0.5652 |
| 4_1560419 | 19 | 0.8421 | 10 | 0.9 | 3 | 0.6667 | 9 | 0.8889 | 41 | 0.8537 | 5 | 1 | 8 | 0.875 | 8 | 1 | 5 | 0.8 | 26 | 0.9231 |
| 4_1560419 | 16 | 0.1875 | 10 | 0 | 3 | 0 | 7 | 0.1429 | 36 | 0.1111 | 4 | 0 | 8 | 0.25 | 6 | 0.5 | 4 | 0.25 | 22 | 0.2727 |
| 4_1560419 | 17 | 0.7059 | 10 | 0.4 | 3 | 0.6667 | 7 | 0.4286 | 37 | 0.5676 | 6 | 0.5 | 8 | 0.625 | 8 | 0.5 | 3 | 0.3333 | 25 | 0.52 |
| 4_1560420 | 14 | 0 | 6 | 0.3333 | 4 | 0.25 | 8 | 0.25 | 32 | 0.1562 | 5 | 0 | 8 | 0.25 | 6 | 0 | 4 | 0 | 23 | 0.087 |
| 4_1560508 | 3 | 0.6667 | 1 | 0 | 4 | 0 | 4 | 0.25 | 12 | 0.25 | 1 | 0 | 5 | 0.4 | 4 | 0 | NA | NA | 10 | 0.2 |
| 4_1560510 | 3 | 0 | NA | NA | 4 | 0.5 | 5 | 0.8 | 12 | 0.5 | 1 | 0 | 5 | 0.2 | 4 | 0.5 | NA | NA | 10 | 0.3 |
| 4_1560741 | 1 | 0 | NA | NA | 1 | 0 | 1 | 1 | 3 | 0.3333 | 1 | 0 | 2 | 0.5 | NA | NA | NA | NA | 3 | 0.3333 |
| 4_1560751 | 1 | 0 | 1 | 1 | 2 | 1 | 1 | 1 | 5 | 0.8 | 1 | 0 | 2 | 1 | 1 | 1 | 1 | 1 | 5 | 0.8 |
| 4_1560755 | 3 | 0 | 1 | 1 | 6 | 0 | 1 | 1 | 11 | 0.1818 | 3 | 0 | 4 | 0.25 | 1 | 0 | 1 | 0 | 9 | 0.1111 |
| 4_1560756 | 2 | 0 | 1 | 0 | 7 | 0.1429 | 1 | 0 | 11 | 0.0909 | 3 | 0 | 4 | 0 | 1 | 1 | 1 | 1 | 9 | 0.2222 |
| 4_1560759 | 3 | 1 | 1 | 1 | 5 | 1 | 1 | 1 | 10 | 1 | 2 | 1 | 3 | 1 | 1 | 1 | NA | NA | 6 | 1 |
| 4_1560759 | 3 | 1 | 1 | 1 | 5 | 0.8 | 1 | 1 | 10 | 0.9 | 2 | 1 | 3 | 0.3333 | 1 | 1 | 1 | 1 | 7 | 0.7143 |
| 4_1560759 | 3 | 1 | 1 | 1 | 5 | 1 | 1 | 1 | 10 | 1 | 2 | 1 | 2 | 1 | 1 | 1 | 1 | 1 | 6 | 1 |
| 4_1560759 | 3 | 1 | 1 | 1 | 4 | 0.75 | 1 | 1 | 9 | 0.8889 | 2 | 1 | 3 | 1 | 1 | 1 | 1 | 1 | 7 | 1 |
| 4_1560760 | 3 | 0.6667 | 1 | 1 | 5 | 1 | 1 | 0 | 10 | 0.8 | 2 | 0.5 | 3 | 0.6667 | 1 | 1 | 1 | 1 | 7 | 0.7143 |
| 4_1560761 | 3 | 1 | 1 | 1 | 2 | 1 | 1 | 1 | 7 | 1 | NA | NA | 2 | 1 | 1 | 1 | 1 | 1 | 4 | 1 |
| 4_1560762 | 2 | 1 | 1 | 1 | 3 | 1 | 1 | 1 | 7 | 1 | NA | NA | 2 | 1 | 1 | 1 | 1 | 1 | 4 | 1 |
| 4_1560764 | 2 | 0 | 1 | 0 | 5 | 0.6 | 1 | 0 | 9 | 0.3333 | 1 | 1 | 3 | 1 | 1 | 1 | 1 | 0 | 6 | 0.8333 |
| 4_1560765 | 3 | 0 | 1 | 0 | 4 | 0.25 | 1 | 0 | 9 | 0.1111 | 1 | 1 | 2 | 0.5 | 1 | 0 | 1 | 0 | 5 | 0.4 |

| | | | | | | | | | | | | | | | | | | | | |
|---|---|---|---|---|---|---|---|---|---|---|---|---|---|---|---|---|---|---|---|---|
| 4_15607664 | 3 | 0.6667 | 1 | 1 | 4 | 1 | 1 | 1 | 9 | 0.8889 | 1 | 1 | 2 | 1 | 1 | 1 | 1 | 0 | 5 | 0.8 |
| 4_15607673 | 3 | 0 | 1 | 0 | 4 | 0.5 | 1 | 1 | 9 | 0.3333 | 1 | 0 | 3 | 0.6667 | 1 | 0 | 1 | 1 | 6 | 0.5 |
| 4_15608087 | 20 | 0.1 | 16 | 0 | 23 | 0.1304 | 20 | 0.1 | 79 | 0.0886 | 14 | 0.3571 | 13 | 0.2308 | 14 | 0.2857 | 17 | 0.2353 | 58 | 0.2759 |
| 4_15608090 | 23 | 0.0435 | 16 | 0 | 21 | 0.0952 | 20 | 0.2 | 80 | 0.0875 | 11 | 0.1818 | 12 | 0.0833 | 14 | 0.0714 | 14 | 0.0714 | 51 | 0.098 |
| 4_15609437 | 30 | 0.9 | 20 | 1 | 28 | 0.8929 | 26 | 0.9615 | 104 | 0.9327 | 14 | 0.9286 | 20 | 0.95 | 17 | 1 | 22 | 0.8182 | 73 | 0.9178 |
| 4_15609477 | 3 | 0.6667 | 1 | 0 | 2 | 1 | 6 | 0.6667 | 12 | 0.6667 | 1 | 1 | 2 | 0.5 | 2 | 0 | 3 | 0.3333 | 8 | 0.375 |
| 4_15609658 | 5 | 0.4 | 7 | 0 | 9 | 0 | 13 | 0.1538 | 34 | 0.1176 | 2 | 0 | 2 | 0 | 4 | 0 | 9 | 0.1111 | 17 | 0.0588 |
| 4_15620992 | 3 | 0.3333 | 2 | 0 | 1 | 0 | 1 | 1 | 7 | 0.2857 | 1 | 1 | 1 | 0 | 1 | 0 | 1 | 0 | 4 | 0.25 |
| 4_15621007 | 3 | 1 | 3 | 1 | NA | NA | 1 | 1 | 7 | 1 | 1 | 1 | 1 | 1 | 1 | 1 | 1 | 1 | 4 | 1 |
| 4_15621017 | 3 | 1 | 2 | 0.5 | 1 | 1 | 1 | 1 | 7 | 0.8571 | 1 | 0 | 1 | 1 | 1 | 1 | 2 | 1 | 5 | 0.8 |
| 4_15621018 | 3 | 1 | 3 | 1 | 1 | 1 | NA | NA | 7 | 1 | 1 | 1 | 1 | 1 | NA | NA | 2 | 1 | 4 | 1 |
| 4_15621024 | 2 | 0 | 3 | 0.6667 | 1 | 1 | 1 | 0 | 7 | 0.4286 | 1 | 0 | 1 | 1 | 1 | 0 | 2 | 0.5 | 5 | 0.4 |
| 4_15621030 | 3 | 0.6667 | 3 | 1 | 1 | 1 | 1 | 1 | 8 | 0.875 | 1 | 1 | 1 | 1 | 1 | 1 | 2 | 0 | 5 | 0.6 |
| 4_15621051 | 3 | 0.6667 | 2 | 0.5 | 1 | 1 | 1 | 0 | 7 | 0.5714 | 1 | 1 | 2 | 1 | 1 | 0 | NA | NA | 4 | 0.75 |
| 4_15621059 | 3 | 0.3333 | 2 | 0.5 | NA | NA | 1 | 0 | 6 | 0.3333 | 1 | 1 | 2 | 0.5 | NA | NA | 1 | 0 | 4 | 0.5 |
| 4_15621066 | 3 | 0 | 3 | 0.3333 | 1 | 0 | 2 | 0.5 | 9 | 0.2222 | 2 | 0.5 | 2 | 0 | 1 | 0 | 1 | 0 | 6 | 0.1667 |
| 4_15621083 | 3 | 0 | 3 | 0.3333 | 1 | 0 | 2 | 0 | 9 | 0.1111 | 2 | 0 | 3 | 0.6667 | 1 | 0 | 1 | 0 | 7 | 0.2857 |
| 4_15621086 | 3 | 0 | 3 | 0.3333 | 1 | 0 | 2 | 0 | 9 | 0.1111 | 2 | 0 | 3 | 0 | 1 | 1 | 1 | 0 | 7 | 0.1429 |
| 4_15621100 | 4 | 0 | 3 | 0 | 1 | 0 | 2 | 0.5 | 10 | 0.1 | 2 | 0 | 1 | 0 | NA | NA | 2 | 0.5 | 5 | 0.2 |
| 4_15621166 | 4 | 0.75 | 2 | 0.5 | 1 | 1 | 2 | 0.5 | 9 | 0.6667 | 2 | 1 | 1 | 1 | 1 | 1 | 1 | 0 | 5 | 0.8 |
| 4_15621704 | NA | NA | 1 | 0 | 1 | 1 | 1 | 1 | 3 | 0.6667 | NA | NA | NA | NA | 1 | 1 | NA | NA | 1 | 1 |
| 4_15621757 | NA | NA | 1 | 0 | 1 | 1 | 1 | 0 | 3 | 0.3333 | NA | NA | NA | NA | 1 | 1 | NA | NA | 1 | 1 |
| 4_15624995 | 11 | 0 | 8 | 0.125 | 3 | 0 | 11 | 0.0909 | 33 | 0.0606 | 6 | 0 | 14 | 0.0714 | 13 | 0.1538 | 6 | 0.3333 | 39 | 0.1282 |
| 4_15625041 | 11 | 0.0909 | 6 | 0 | 3 | 0 | 9 | 0.1111 | 29 | 0.069 | 4 | 0 | 11 | 0 | 9 | 0.1111 | 3 | 0.3333 | 27 | 0.0741 |
| 4_15625078 | 11 | 0.3636 | 6 | 0.5 | 3 | 0 | 7 | 0.4286 | 27 | 0.3704 | 4 | 0.5 | 12 | 0.5833 | 10 | 0.6 | 3 | 0 | 29 | 0.5172 |
| 4_15625080 | 12 | 0.0833 | 8 | 0.125 | 2 | 0.5 | 8 | 0 | 30 | 0.1 | 4 | 0.25 | 12 | 0.0833 | 9 | 0.1111 | 3 | 0 | 28 | 0.1071 |
| 4_15625085 | 12 | 0.5 | 7 | 0.1429 | 3 | 0.6667 | 9 | 0.2222 | 31 | 0.3548 | 1 | 0 | 13 | 0.4615 | 10 | 0.6 | 4 | 0.5 | 28 | 0.5 |
| 4_15625543 | 6 | 0 | 3 | 0.3333 | 1 | 0 | 6 | 0 | 16 | 0.0625 | 1 | 0 | 2 | 0 | 5 | 0.2 | 3 | 0 | 11 | 0.0909 |
| 4_15625673 | 6 | 0 | 4 | 0.25 | 2 | 0 | 5 | 0 | 17 | 0.0588 | 2 | 0 | 3 | 0.3333 | 4 | 0 | 3 | 0 | 12 | 0.0833 |
| 4_15627017 | 1 | 0 | NA | NA | NA | NA | 2 | 1 | 3 | 0.6667 | NA | NA | 1 | 1 | 2 | 0.5 | NA | NA | 3 | 0.6667 |
| 4_15628398 | 3 | 0.6667 | 1 | 0 | NA | NA | NA | NA | 4 | 0.5 | NA | NA | 1 | 1 | NA | NA | 1 | 0 | 2 | 0.5 |
| 4_15628422 | 3 | 0 | 1 | 1 | NA | NA | NA | NA | 4 | 0.25 | NA | NA | 1 | 1 | NA | NA | 1 | 0 | 2 | 0.5 |
| 4_15628430 | 3 | 1 | NA | NA | NA | NA | NA | NA | 3 | 1 | NA | NA | 1 | 1 | NA | NA | 1 | 1 | 2 | 1 |
| 4_15629262 | NA | NA | 2 | 0.5 | NA | NA | NA | NA | 2 | 0.5 | 1 | 0 | NA | NA | NA | NA | 1 | 1 | 2 | 0.5 |
| 4_15629286 | 1 | 1 | 1 | 1 | NA | NA | NA | NA | 2 | 1 | NA | NA | NA | NA | NA | NA | 1 | 1 | 1 | 1 |
| 4_15629486 | 1 | 1 | NA | NA | NA | NA | 1 | 1 | 2 | 1 | NA | NA | NA | NA | 1 | 1 | 2 | 0.5 | 3 | 0.6667 |
| 4_15629936 | NA | NA | NA | NA | 1 | 1 | NA | NA | 1 | 1 | NA | NA | NA | NA | NA | NA | 1 | 1 | 1 | 1 |
| 4_15631383 | 4 | 0 | 2 | 0.5 | 1 | 0 | NA | NA | 7 | 0.1429 | NA | NA | 5 | 0.2 | 4 | 0.25 | 3 | 0 | 12 | 0.1667 |
| 4_15631383 | 6 | 0.1667 | 2 | 0 | 4 | 0 | NA | NA | 12 | 0.0833 | NA | NA | 4 | 0.25 | 6 | 0.3333 | 3 | 0.3333 | 13 | 0.3077 |
| 4_15631395 | 6 | 0.6667 | 5 | 0.6 | 4 | 0.5 | NA | NA | 15 | 0.6 | NA | NA | 6 | 0.8333 | 11 | 0.4545 | 3 | 0.3333 | 20 | 0.55 |
| 4_15631415 | 6 | 0.1667 | 5 | 0.4 | 4 | 0 | NA | NA | 15 | 0.2 | NA | NA | 2 | 0.5 | 9 | 0.2222 | 2 | 0 | 13 | 0.2308 |
| 4_15631419 | 4 | 0.25 | 5 | 0 | 4 | 0.25 | NA | NA | 13 | 0.1538 | NA | NA | 5 | 0 | 9 | 0.3333 | 2 | 0 | 16 | 0.1875 |
| 4_15631428 | 4 | 0.5 | 4 | 0.5 | 4 | 0 | NA | NA | 12 | 0.3333 | NA | NA | 5 | 0.4 | 8 | 0.375 | 2 | 0 | 15 | 0.3333 |
| 4_15631448 | 5 | 0.4 | 5 | 0 | 4 | 0.5 | NA | NA | 14 | 0.2857 | NA | NA | 6 | 0.3333 | 10 | 0.2 | 2 | 0.5 | 18 | 0.2778 |
| 4_15632185 | 1 | 0 | 1 | 0 | 2 | 0 | 5 | 0.4 | 9 | 0.2222 | 4 | 0 | 7 | 0.1429 | 3 | 0 | 3 | 0 | 17 | 0.0588 |
| 4_15633151 | 8 | 0 | 11 | 0.0909 | 1 | 0 | 5 | 0.2 | 25 | 0.08 | 6 | 0 | 17 | 0.1765 | 4 | 0 | 4 | 0.25 | 31 | 0.129 |
| 4_15633162 | 9 | 0.2222 | 12 | 0 | 5 | 0.2 | 5 | 0 | 31 | 0.0968 | 5 | 0 | 15 | 0.0667 | 7 | 0 | 5 | 0.2 | 32 | 0.0625 |
| 4_15633171 | 7 | 0.8571 | 11 | 0.6364 | 4 | 1 | 5 | 0.8 | 27 | 0.7778 | 5 | 0.6 | 14 | 0.8571 | 10 | 0.7 | 7 | 0.5714 | 36 | 0.7222 |
| 4_15633196 | 8 | 0.25 | 8 | 0 | 1 | 0 | 4 | 0.5 | 21 | 0.1905 | 6 | 0 | 9 | 0 | 7 | 0.1429 | 8 | 0.25 | 30 | 0.1 |
| 4_15640981 | 1 | 0 | 1 | 1 | 1 | 0 | 2 | 0.5 | 5 | 0.4 | NA | NA | 1 | 0 | 3 | 0.6667 | 2 | 0.5 | 6 | 0.5 |
| 4_15640998 | 1 | 1 | NA | NA | 1 | 0 | 2 | 0.5 | 4 | 0.5 | NA | NA | 1 | 0 | 3 | 0.3333 | 2 | 0 | 6 | 0.1667 |
| 4_15641022 | 1 | 0 | 1 | 1 | 1 | 0 | 2 | 0.5 | 5 | 0.4 | NA | NA | NA | NA | 3 | 0.3333 | 1 | 0 | 4 | 0.25 |
| 4_15641025 | 1 | 1 | 1 | 1 | 1 | 0 | 1 | 1 | 4 | 0.75 | NA | NA | 1 | 0 | 1 | 1 | NA | NA | 2 | 0.5 |
| 5_3619176 | NA | NA | 3 | 0 | 10 | 0.1 | 3 | 0.6667 | 16 | 0.1875 | 4 | 0 | 4 | 0.25 | 6 | 0.1667 | 7 | 0.1429 | 21 | 0.1429 |
| 5_3619187 | NA | NA | 2 | 1 | 11 | 0.8182 | 3 | 0.6667 | 16 | 0.8125 | 3 | 1 | 4 | 0.75 | 8 | 0.625 | 7 | 0.8571 | 22 | 0.7727 |
| 5_3619277 | NA | NA | 3 | 0 | 9 | 0.1111 | 2 | 0.5 | 14 | 0.1429 | 4 | 0 | 3 | 0.6667 | 8 | 0 | 7 | 0.2857 | 22 | 0.1818 |
| 5_3619616 | NA | NA | 4 | 0.5 | 10 | 0.4 | 4 | 0.75 | 18 | 0.5 | 3 | 0.3333 | 5 | 0.2 | 7 | 0.2857 | 7 | 0.4286 | 22 | 0.3182 |
| 5_3619642 | NA | NA | 3 | 0 | 5 | 0 | 1 | 1 | 9 | 0.1111 | 4 | 0 | 2 | 0 | 3 | 0.3333 | 4 | 0 | 13 | 0.0769 |
| 5_3619693 | NA | NA | 3 | 0.3333 | 10 | 0.2 | 4 | 0.25 | 17 | 0.2353 | 3 | 0.3333 | 3 | 0.3333 | 8 | 0.125 | 4 | 0.25 | 18 | 0.2222 |
| 5_3619866 | 1 | 0 | 5 | 0.2 | 10 | 0.1 | 5 | 0 | 21 | 0.0952 | 4 | 0.25 | 4 | 0 | 8 | 0.25 | 6 | 0 | 22 | 0.1364 |
| 5_3634425 | NA | NA | 1 | 1 | 3 | 0.6667 | 1 | 0 | 5 | 0.6 | 3 | 0.3333 | 3 | 0.3333 | 1 | 1 | 2 | 0.5 | 9 | 0.4444 |
| 5_3634604 | NA | NA | 1 | 0 | 3 | 0.3333 | 1 | 1 | 5 | 0.4 | 3 | 0.3333 | 2 | 0.5 | NA | NA | 2 | 0.5 | 7 | 0.4286 |
| 5_3634640 | NA | NA | NA | NA | 3 | 0.6667 | 1 | 0 | 4 | 0.5 | 2 | 0.5 | 3 | 0.6667 | 1 | 1 | 1 | 1 | 7 | 0.7143 |
| 5_3640310 | NA | NA | 1 | 1 | 2 | 1 | NA | NA | 3 | 1 | 1 | 0 | 2 | 0.5 | 1 | 1 | 2 | 0.5 | 6 | 0.5 |
| 5_3640354 | NA | NA | 1 | 1 | 3 | 0 | NA | NA | 4 | 0.25 | 1 | 0 | 2 | 1 | 1 | 0 | 1 | 1 | 5 | 0.6 |

| | | | | | | | | | | | | | | | | | | | | |
|---|---|---|---|---|---|---|---|---|---|---|---|---|---|---|---|---|---|---|---|---|
| 5_3640361 | NA | NA | 1 | 0 | 3 | 0.3333 | NA | NA | 4 | 0.25 | 1 | 1 | 3 | 0.3333 | NA | NA | 2 | 0 | 6 | 0.3333 |
| 5_3640374 | NA | NA | 1 | 0 | 3 | 1 | NA | NA | 4 | 0.75 | 1 | 0 | 3 | 0 | 1 | 0 | 2 | 0.5 | 7 | 0.1429 |
| 5_3640479 | NA | NA | 1 | 1 | 1 | 0 | NA | NA | 2 | 0.5 | 1 | 1 | 2 | 0.5 | 1 | 1 | 1 | 0 | 5 | 0.6 |
| 5_3640491 | NA | NA | 1 | 1 | 2 | 1 | NA | NA | 3 | 1 | 1 | 1 | 1 | 1 | NA | NA | 2 | 1 | 4 | 1 |
| 5_3640541 | NA | NA | 1 | 1 | 1 | 1 | NA | NA | 2 | 1 | NA | NA | 2 | 1 | NA | NA | 2 | 1 | 4 | 1 |
| 5_3640572 | NA | NA | 1 | 1 | 3 | 0.6667 | NA | NA | 4 | 0.75 | 1 | 1 | 2 | 0.5 | 1 | 1 | 1 | 1 | 5 | 0.8 |
| 5_3640584 | NA | NA | 1 | 1 | 3 | 0.3333 | NA | NA | 4 | 0.5 | NA | NA | 1 | 0 | 1 | 1 | 2 | 0.5 | 4 | 0.5 |
| 5_3640671 | NA | NA | 1 | 0 | 3 | 0.3333 | NA | NA | 4 | 0.25 | 1 | 1 | 2 | 0 | 1 | 1 | 2 | 0 | 6 | 0.3333 |
| 5_3655765 | NA | NA | NA | NA | 4 | 0.25 | NA | NA | 4 | 0.25 | NA | NA | 1 | 1 | NA | NA | NA | NA | 1 | 1 |
| 5_3655836 | NA | NA | NA | NA | 4 | 0.5 | NA | NA | 4 | 0.5 | NA | NA | 2 | 1 | NA | NA | NA | NA | 2 | 1 |
| 5_3655978 | NA | NA | 1 | 1 | 4 | 0.75 | NA | NA | 5 | 0.8 | NA | NA | 2 | 1 | NA | NA | NA | NA | 2 | 1 |
| 5_3680729 | 1 | 0 | NA | NA | NA | NA | 1 | 1 | 2 | 0.5 | NA | NA | NA | NA | NA | NA | 1 | 1 | 1 | 1 |
| 5_3680880 | 1 | 0 | NA | NA | NA | NA | 1 | 1 | 2 | 0.5 | NA | NA | 1 | 1 | NA | NA | 1 | 0 | 2 | 0.5 |
| 5_3681474 | 1 | 0 | NA | NA | NA | NA | 1 | 1 | 2 | 0.5 | NA | NA | 1 | 0 | NA | NA | 1 | 1 | 2 | 0.5 |
| 5_3681514 | NA | NA | NA | NA | NA | NA | 1 | 1 | 1 | 1 | NA | NA | 1 | 1 | NA | NA | NA | NA | 1 | 1 |
| 5_3681527 | 1 | 1 | NA | NA | NA | NA | 1 | 1 | 2 | 1 | NA | NA | 1 | 1 | NA | NA | NA | NA | 1 | 1 |
| 5_3681614 | 1 | 0 | NA | NA | NA | NA | 1 | 1 | 2 | 0.5 | NA | NA | 1 | 1 | NA | NA | 1 | 1 | 2 | 1 |
| 5_3681625 | 1 | 0 | NA | NA | NA | NA | 1 | 1 | 2 | 0.5 | NA | NA | 1 | 1 | NA | NA | 1 | 0 | 2 | 0.5 |
| 5_3683504 | NA | NA | NA | NA | 1 | 0 | 1 | 1 | 2 | 0.5 | NA | NA | NA | NA | 1 | 1 | 2 | 1 | 3 | 1 |
| 5_3683590 | 1 | 1 | NA | NA | 1 | 1 | 1 | 1 | 3 | 1 | NA | NA | NA | NA | 1 | 1 | 2 | 1 | 3 | 1 |
| 5_3686829 | NA | NA | 2 | 0.5 | 5 | 0 | 2 | 0 | 9 | 0.1111 | 1 | 1 | 4 | 0 | 2 | 0 | 1 | 0 | 8 | 0.125 |
| 5_3687812 | NA | NA | NA | NA | 1 | 1 | 1 | 0 | 2 | 0.5 | NA | NA | NA | NA | 1 | 1 | NA | NA | 1 | 1 |
| 5_3690880 | 8 | 0.25 | 7 | 0 | 9 | 0 | 6 | 0.3333 | 30 | 0.1333 | 9 | 0.2222 | 14 | 0.0714 | 10 | 0 | 6 | 0 | 39 | 0.0769 |
| 5_3691081 | 11 | 0 | 8 | 0.25 | 12 | 0 | 8 | 0.125 | 39 | 0.0769 | 12 | 0.1667 | 16 | 0 | 14 | 0.2857 | 10 | 0.1 | 52 | 0.1346 |
| 5_3691370 | 10 | 0.6 | 3 | 0.6667 | 14 | 0.6429 | 7 | 0.5714 | 34 | 0.6176 | 11 | 0.9091 | 16 | 0.4375 | 14 | 0.7857 | 10 | 0.8 | 51 | 0.7059 |
| 5_3691438 | 9 | 0.1111 | 7 | 0.1429 | 14 | 0.0714 | 7 | 0.2857 | 37 | 0.1351 | 10 | 0.2 | 17 | 0.0588 | 16 | 0.125 | 9 | 0.2222 | 52 | 0.1346 |
| 5_3691449 | 10 | 0.9 | 7 | 0.7143 | 14 | 0.5714 | 6 | 0.8333 | 37 | 0.7297 | 12 | 0.9167 | 18 | 0.7778 | 17 | 0.7059 | 8 | 1 | 55 | 0.8182 |
| 5_3740072 | 3 | 0 | 5 | 0 | NA | NA | 4 | 0.5 | 12 | 0.1667 | 2 | 0 | 2 | 1 | 1 | 0 | 3 | 0 | 8 | 0.25 |
| 5_3741361 | 5 | 0.2 | 4 | 0.25 | 5 | 0.2 | 9 | 0.2222 | 23 | 0.2174 | 4 | 0 | 3 | 0 | NA | NA | 8 | 0.125 | 15 | 0.0667 |
| 5_3741899 | 5 | 0 | 4 | 0.5 | 5 | 0.4 | 8 | 0 | 22 | 0.1818 | 3 | 0 | 3 | 0.6667 | NA | NA | 7 | 0.1429 | 13 | 0.2308 |
| 5_3742019 | 5 | 0 | 3 | 0.3333 | 5 | 0.4 | 7 | 0.1429 | 20 | 0.2 | 3 | 0 | 3 | 0.3333 | NA | NA | 7 | 0 | 13 | 0.0769 |
| 5_3742056 | 5 | 0.2 | 3 | 0.3333 | 5 | 0.4 | 7 | 0.1429 | 20 | 0.25 | 3 | 0.3333 | 3 | 0.3333 | NA | NA | 6 | 0.3333 | 12 | 0.3333 |
| 5_3742164 | 2 | 0 | 2 | 0 | 5 | 0 | 7 | 0.1429 | 16 | 0.0625 | 3 | 0 | 3 | 0.3333 | NA | NA | 6 | 0.1667 | 12 | 0.1667 |
| 5_3742594 | 3 | 0.3333 | 3 | 0 | 4 | 0 | 4 | 0 | 14 | 0.0714 | 3 | 0.3333 | 3 | 0.6667 | NA | NA | 6 | 0.3333 | 12 | 0.4167 |
| 5_3742665 | 4 | 0.5 | 3 | 0.3333 | 5 | 0.6 | 6 | 0.3333 | 18 | 0.4444 | 2 | 0 | 3 | 0.6667 | NA | NA | 6 | 0 | 11 | 0.1818 |
| 5_3742686 | 5 | 0.4 | 3 | 0.6667 | 5 | 0.2 | 6 | 0.6667 | 19 | 0.4737 | 3 | 0.6667 | 3 | 1 | NA | NA | 7 | 0.7143 | 13 | 0.7692 |
| 5_3742755 | 3 | 1 | 1 | 1 | 4 | 1 | 5 | 1 | 13 | 1 | 1 | 1 | 3 | 1 | NA | NA | 6 | 1 | 10 | 1 |
| 5_3742856 | 5 | 0.2 | 2 | 0 | 4 | 0 | 6 | 0.1667 | 17 | 0.1176 | 3 | 0.3333 | 3 | 0.3333 | NA | NA | 5 | 0.4 | 11 | 0.3636 |
| 5_3742964 | 5 | 0 | 2 | 0 | 5 | 0 | 6 | 0.1667 | 18 | 0.0556 | 3 | 0.6667 | 3 | 0.6667 | NA | NA | 6 | 0.1667 | 12 | 0.4167 |
| 5_3742981 | 5 | 0.8 | 2 | 0.5 | 4 | 0.5 | 3 | 0.3333 | 14 | 0.5714 | 2 | 0.5 | 3 | 0.6667 | NA | NA | 6 | 0.6667 | 11 | 0.6364 |
| 5_3742988 | 4 | 0.75 | 1 | 0 | 4 | 0.75 | 6 | 0.6667 | 15 | 0.6667 | 1 | 0 | 3 | 0.3333 | NA | NA | 5 | 0.8 | 9 | 0.5556 |
| 5_3743087 | 5 | 0.4 | 2 | 0 | 4 | 0 | 5 | 0.2 | 16 | 0.1875 | 3 | 0 | 2 | 0.5 | NA | NA | 5 | 0.2 | 10 | 0.2 |
| 5_3743129 | 5 | 0.2 | 2 | 0 | 2 | 0.5 | 4 | 0 | 13 | 0.1538 | 3 | 0.3333 | 2 | 0.5 | NA | NA | 4 | 0.25 | 9 | 0.3333 |
| 5_3743144 | 5 | 0.6 | 2 | 0 | 4 | 0.25 | 6 | 0.1667 | 17 | 0.2941 | 3 | 0 | 2 | 0.5 | NA | NA | 5 | 0.8 | 10 | 0.5 |
| 5_3743159 | 3 | 0.3333 | 1 | 0 | 4 | 0.25 | 5 | 0.4 | 13 | 0.3077 | 2 | 0 | 2 | 0 | NA | NA | 3 | 0.3333 | 7 | 0.1429 |
| 5_3743165 | 4 | 0.75 | 2 | 0 | 4 | 0 | 5 | 0.4 | 15 | 0.3333 | 2 | 0.5 | 2 | 1 | NA | NA | 3 | 0.3333 | 7 | 0.5714 |
| 5_3743225 | 4 | 0.25 | 2 | 0.5 | 3 | 1 | 5 | 0.2 | 14 | 0.4286 | 3 | 0.3333 | 2 | 0.5 | NA | NA | 5 | 0.8 | 10 | 0.6 |
| 5_3743240 | 4 | 0.25 | 2 | 0 | 4 | 0 | 6 | 0.3333 | 16 | 0.1875 | 3 | 0 | 2 | 0 | NA | NA | 5 | 0.6 | 10 | 0.3 |
| 5_3743246 | 4 | 0.25 | 1 | 0 | 4 | 0.25 | 6 | 0.3333 | 15 | 0.2667 | 3 | 0.3333 | 2 | 0.5 | NA | NA | 5 | 0.4 | 10 | 0.4 |
| 5_3743257 | 5 | 0.4 | 2 | 0 | 4 | 0.5 | 3 | 0.3333 | 14 | 0.3571 | 3 | 0.3333 | 2 | 0.5 | NA | NA | 5 | 0.2 | 10 | 0.3 |
| 5_3743270 | 2 | 0.5 | 2 | 0 | 2 | 0 | 6 | 0.5 | 12 | 0.3333 | 2 | 0.5 | 2 | 0 | NA | NA | 4 | 0 | 8 | 0.125 |
| 5_3743339 | 3 | 1 | 2 | 1 | 4 | 1 | 6 | 1 | 15 | 1 | 3 | 1 | 2 | 1 | NA | NA | 5 | 1 | 10 | 1 |
| 5_3743382 | 5 | 0.2 | 1 | 0 | 4 | 0 | 6 | 0.3333 | 16 | 0.1875 | 3 | 0 | 2 | 0.5 | NA | NA | 4 | 0.25 | 9 | 0.2222 |
| 5_3743399 | 4 | 0.75 | 2 | 1 | 4 | 0.75 | 5 | 0.8 | 15 | 0.8 | 3 | 1 | 2 | 1 | NA | NA | 5 | 0.8 | 10 | 0.9 |
| 5_3819728 | 1 | 1 | NA | NA | NA | NA | 3 | 0 | 4 | 0.25 | 2 | 0 | NA | NA | NA | NA | 2 | 0.5 | 4 | 0.25 |
| 5_3853302 | 1 | 0 | 3 | 0 | 2 | 0.5 | 3 | 0.6667 | 9 | 0.3333 | NA | NA | 2 | 1 | NA | NA | NA | NA | 2 | 1 |
| 5_3853324 | 2 | 0 | 2 | 0 | 1 | 0 | 3 | 0.3333 | 8 | 0.125 | NA | NA | 1 | 1 | 2 | 0 | 2 | 0 | 5 | 0.2 |
| 5_3853441 | 1 | 0 | 1 | 1 | 2 | 0 | 3 | 0 | 7 | 0.1429 | NA | NA | 1 | 0 | 1 | 0 | 1 | 1 | 3 | 0.3333 |
| 5_3853655 | NA | NA | 1 | 0 | NA | NA | 1 | 1 | 2 | 0.5 | NA | NA | 1 | 1 | 1 | 0 | NA | NA | 2 | 0.5 |
| 5_3863138 | 4 | 0 | 4 | 0 | 4 | 0 | 6 | 0.1667 | 18 | 0.0556 | NA | NA | 4 | 0 | 3 | 0.3333 | 5 | 0.2 | 12 | 0.1667 |
| 5_3892144 | NA | NA | NA | NA | NA | NA | 1 | 1 | 1 | 1 | NA | NA | 1 | 1 | 1 | 1 | NA | NA | 2 | 1 |
| 5_3892393 | 1 | 0 | NA | NA | NA | NA | 1 | 1 | 2 | 0.5 | 1 | 0 | 1 | 1 | 1 | 0 | NA | NA | 3 | 0.3333 |
| 5_3894399 | 1 | 0 | NA | NA | 1 | 1 | 1 | 1 | 3 | 0.6667 | NA | NA | 1 | 0 | 1 | 1 | NA | NA | 2 | 0.5 |
| 5_3895203 | 2 | 1 | NA | NA | NA | NA | 1 | 1 | 3 | 1 | 1 | 1 | NA | NA | NA | NA | NA | NA | 1 | 1 |
| 5_3895215 | 2 | 1 | NA | NA | NA | NA | 1 | 0 | 3 | 0.6667 | 1 | 1 | NA | NA | NA | NA | NA | NA | 1 | 1 |

| | | | | | | | | | | | | | | | | | | | | |
|---|---|---|---|---|---|---|---|---|---|---|---|---|---|---|---|---|---|---|---|---|
| 5_3940696 | NA | NA | NA | NA | NA | NA | 1 | 1 | 1 | 1 | NA | NA | 1 | 1 | 1 | 0 | NA | NA | 2 | 0.5 |
| 5_3940923 | NA | NA | NA | NA | NA | NA | 1 | 1 | 1 | 1 | NA | NA | 1 | 1 | 1 | 1 | NA | NA | 2 | 1 |
| 5_3941061 | NA | NA | NA | NA | NA | NA | 1 | 1 | 1 | 1 | NA | NA | 1 | 1 | 1 | 1 | NA | NA | 2 | 1 |
| 5_3941115 | NA | NA | NA | NA | NA | NA | 1 | 1 | 1 | 1 | NA | NA | 1 | 1 | NA | NA | NA | NA | 1 | 1 |
| 5_3941336 | NA | NA | NA | NA | NA | NA | 1 | 1 | 1 | 1 | NA | NA | 1 | 1 | 1 | 0 | NA | NA | 2 | 0.5 |
| 5_3941356 | NA | NA | NA | NA | NA | NA | 1 | 1 | 1 | 1 | NA | NA | 1 | 1 | NA | NA | NA | NA | 1 | 1 |
| 5_4096123 | NA | NA | 1 | 0 | NA | NA | 1 | 1 | 2 | 0.5 | NA | NA | 2 | 0.5 | 1 | 0 | 1 | 0 | 4 | 0.25 |
| 5_4096187 | NA | NA | 1 | 1 | NA | NA | 1 | 1 | 2 | 1 | NA | NA | 2 | 1 | 1 | 0 | 1 | 0 | 4 | 0.5 |
| 5_4096714 | NA | NA | 2 | 0.5 | 1 | 0 | NA | NA | 3 | 0.3333 | NA | NA | 2 | 0.5 | 2 | 0 | NA | NA | 4 | 0.25 |
| 5_4096815 | NA | NA | 2 | 0.5 | 1 | 1 | 1 | 1 | 4 | 0.75 | NA | NA | 2 | 1 | 2 | 1 | 1 | 1 | 5 | 1 |
| 5_4100189 | NA | NA | 2 | 0 | 1 | 1 | 1 | 0 | 4 | 0.25 | NA | NA | 3 | 0.3333 | 2 | 0 | 1 | 0 | 6 | 0.1667 |
| 5_4100263 | NA | NA | 1 | 1 | 1 | 1 | 2 | 1 | 4 | 1 | NA | NA | 3 | 0.3333 | 2 | 1 | 1 | 1 | 6 | 0.6667 |
| 5_4100545 | NA | NA | 2 | 0 | 1 | 1 | 2 | 0 | 5 | 0.2 | NA | NA | 4 | 0 | 3 | 0.6667 | 1 | 0 | 8 | 0.25 |
| 5_4114369 | NA | NA | 7 | 0.4286 | 4 | 0.5 | 4 | 0.25 | 15 | 0.4 | NA | NA | 4 | 0.75 | 5 | 0.4 | 2 | 0 | 11 | 0.4545 |
| 5_4114424 | NA | NA | 8 | 1 | 3 | 0.6667 | 3 | 1 | 14 | 0.9286 | NA | NA | 4 | 1 | 5 | 0.8 | 2 | 1 | 11 | 0.9091 |
| 5_4114437 | NA | NA | 5 | 0.2 | 4 | 0.25 | 4 | 0.75 | 13 | 0.3846 | NA | NA | 3 | 0.3333 | 5 | 0.6 | 2 | 0.5 | 10 | 0.5 |
| 5_4114451 | NA | NA | 8 | 0 | 2 | 0.5 | 4 | 0.25 | 14 | 0.1429 | NA | NA | 2 | 0 | 5 | 0.2 | 2 | 0 | 9 | 0.1111 |
| 5_4114489 | NA | NA | 6 | 0.5 | 4 | 0.25 | 4 | 0.75 | 14 | 0.5 | NA | NA | 4 | 0.5 | 4 | 0.25 | 2 | 0.5 | 10 | 0.4 |
| 5_4115311 | 1 | 0 | 10 | 0.1 | 5 | 0 | 4 | 0.25 | 20 | 0.1 | NA | NA | 4 | 0.25 | 8 | 0 | 4 | 0 | 16 | 0.0625 |
| 5_4130148 | 6 | 0 | 11 | 0 | 6 | 0.3333 | 8 | 0 | 31 | 0.0645 | 2 | 0 | 9 | 0.2222 | 10 | 0.1 | 7 | 0 | 28 | 0.1071 |
| 5_4132369 | 7 | 0 | 4 | 0 | 4 | 0.25 | 3 | 0.3333 | 18 | 0.1111 | 3 | 0 | 3 | 0.3333 | 6 | 0.1667 | 4 | 0.25 | 16 | 0.1875 |
| 5_4132804 | 9 | 0.4444 | 7 | 0.5714 | 11 | 0.4545 | 7 | 0.5714 | 34 | 0.5 | 6 | 0.3333 | 5 | 0.6 | 7 | 0.7143 | 7 | 0.5714 | 25 | 0.56 |
| 5_4132984 | 9 | 0.4444 | 8 | 0 | 8 | 0 | 6 | 0.1667 | 31 | 0.1613 | 5 | 0.4 | 4 | 0.5 | 7 | 0.1429 | 7 | 0.1429 | 23 | 0.2609 |
| 5_4133092 | 9 | 0.1111 | 6 | 0.1667 | 10 | 0 | 6 | 0 | 31 | 0.0645 | 5 | 0 | 4 | 0.25 | 7 | 0.1429 | 8 | 0.125 | 24 | 0.125 |
| 5_4133256 | 12 | 0.3333 | 8 | 0.375 | 6 | 0.3333 | 6 | 0.3333 | 32 | 0.3438 | 5 | 0.2 | 4 | 0.5 | 7 | 0.7143 | 6 | 0 | 22 | 0.3636 |
| 5_4804268 | 2 | 0.5 | 1 | 0 | 1 | 0 | NA | NA | 4 | 0.25 | 2 | 0 | NA | NA | 1 | 1 | NA | NA | 3 | 0.3333 |
| 5_4804289 | NA | NA | 1 | 1 | NA | NA | NA | NA | 1 | 1 | 2 | 1 | NA | NA | 1 | 1 | NA | NA | 3 | 1 |
| 5_4804312 | NA | NA | 1 | 0 | 1 | 1 | NA | NA | 2 | 0.5 | 2 | 0.5 | NA | NA | 1 | 1 | NA | NA | 3 | 0.6667 |
| 5_4805538 | 1 | 1 | 1 | 1 | NA | NA | NA | NA | 2 | 1 | 2 | 1 | NA | NA | NA | NA | NA | NA | 2 | 1 |
| 5_4805644 | NA | NA | 1 | 1 | NA | NA | NA | NA | 1 | 1 | 1 | 1 | NA | NA | NA | NA | NA | NA | 1 | 1 |
| 5_4805800 | NA | NA | 1 | 1 | NA | NA | NA | NA | 1 | 1 | 2 | 1 | NA | NA | NA | NA | NA | NA | 2 | 1 |
| 5_4805908 | 1 | 0 | 1 | 1 | NA | NA | NA | NA | 2 | 0.5 | 2 | 0.5 | NA | NA | NA | NA | NA | NA | 2 | 0.5 |
| 5_4806221 | NA | NA | 1 | 1 | NA | NA | NA | NA | 1 | 1 | 2 | 1 | NA | NA | NA | NA | NA | NA | 2 | 1 |
| 5_4806712 | 1 | 1 | NA | NA | NA | NA | NA | NA | 1 | 1 | 2 | 0.5 | NA | NA | NA | NA | NA | NA | 2 | 0.5 |
| 5_4807219 | NA | NA | 1 | 1 | NA | NA | NA | NA | 1 | 1 | 1 | 1 | NA | NA | NA | NA | NA | NA | 1 | 1 |
| 5_4853553 | 14 | 0.0714 | 25 | 0.16 | 16 | 0.125 | 8 | 0 | 63 | 0.1111 | 17 | 0.0588 | 4 | 0 | 11 | 0.0909 | 10 | 0.1 | 42 | 0.0714 |
| 5_4853683 | 14 | 0.2143 | 22 | 0.0909 | 14 | 0.2857 | 7 | 0.2857 | 57 | 0.193 | 13 | 0.1538 | 4 | 0.25 | 9 | 0.2222 | 7 | 0.4286 | 33 | 0.2424 |
| 5_4855679 | 7 | 0.1429 | 11 | 0.0909 | 4 | 0 | 6 | 0 | 28 | 0.0714 | 5 | 0 | 2 | 0 | 4 | 0.25 | 5 | 0.2 | 16 | 0.125 |
| 5_4856190 | 7 | 0.2857 | 10 | 0.1 | 4 | 0 | 6 | 0.1667 | 27 | 0.1481 | 4 | 0.5 | 2 | 0 | 3 | 0.3333 | 7 | 0.1429 | 16 | 0.25 |
| 5_4856209 | 2 | 0 | 9 | 0.1111 | 4 | 0 | 5 | 0.2 | 20 | 0.1 | 5 | 0 | 2 | 0 | 3 | 0.3333 | 7 | 0 | 17 | 0.0588 |
| 5_4856510 | 6 | 0.5 | 13 | 0.4615 | 5 | 0.2 | 6 | 0.1667 | 30 | 0.3667 | 3 | 0.6667 | 2 | 0.5 | 4 | 0.5 | 6 | 0.1667 | 15 | 0.4 |
| 5_4856520 | 6 | 0.3333 | 11 | 0.1818 | 5 | 0.2 | 5 | 0 | 27 | 0.1852 | 4 | 0.25 | 2 | 0.5 | 5 | 0.2 | 6 | 0.1667 | 17 | 0.2353 |
| 5_4856574 | 6 | 0 | 12 | 0 | 5 | 0.2 | 7 | 0.1429 | 30 | 0.0667 | 5 | 0.4 | 2 | 0 | 5 | 0 | 7 | 0 | 19 | 0.1053 |
| 5_5299062 | 5 | 0 | 6 | 0 | 2 | 0 | 3 | 0.3333 | 16 | 0.0625 | 1 | 1 | 1 | 0 | 1 | 0 | 4 | 0 | 7 | 0.1429 |
| 5_5555630 | 3 | 0 | 4 | 0.25 | 5 | 0.2 | 1 | 0 | 13 | 0.1538 | 4 | 0 | 7 | 0.2857 | 7 | 0.1429 | 8 | 0 | 26 | 0.1154 |
| 5_5556214 | 3 | 0 | 4 | 0.25 | 5 | 0 | 1 | 0 | 13 | 0.0769 | 4 | 0.5 | 7 | 0 | 8 | 0 | 8 | 0 | 27 | 0.0741 |
| 5_5557213 | 2 | 0.5 | 4 | 0.25 | 2 | 0 | 1 | 1 | 9 | 0.3333 | 3 | 0.3333 | 3 | 0 | 1 | 0 | 7 | 0.2857 | 14 | 0.2143 |
| 5_5557421 | 2 | 0 | 4 | 0.5 | 1 | 0 | 1 | 0 | 8 | 0.25 | 3 | 0.3333 | 4 | 0 | 2 | 0 | 7 | 0.2857 | 16 | 0.1875 |
| 5_5557620 | 2 | 0 | 4 | 0.25 | 1 | 0 | 1 | 0 | 8 | 0.125 | 4 | 0.25 | 4 | 0 | 2 | 0 | 6 | 0.1667 | 16 | 0.125 |
| 5_5557676 | 1 | 1 | 2 | 0.5 | 1 | 0 | NA | NA | 4 | 0.5 | 3 | 0 | 3 | 0 | NA | NA | 6 | 0.1667 | 12 | 0.0833 |
| 5_5557864 | 2 | 0.5 | 4 | 0.25 | 1 | 0 | 1 | 0 | 8 | 0.25 | 3 | 0.3333 | 3 | 0 | 1 | 0 | 7 | 0.1429 | 14 | 0.1429 |
| 5_5557911 | 1 | 0 | 3 | 0.3333 | 1 | 0 | 1 | 1 | 6 | 0.3333 | 3 | 1 | 3 | 0.6667 | 2 | 0 | 7 | 0.5714 | 15 | 0.6 |
| 5_5558015 | 2 | 0.5 | 4 | 1 | 1 | 1 | 1 | 1 | 8 | 0.875 | 3 | 1 | 3 | 1 | 2 | 1 | 6 | 1 | 14 | 1 |
| 5_5558063 | 1 | 1 | 4 | 0.5 | 1 | 1 | NA | NA | 6 | 0.6667 | 3 | 0.3333 | 3 | 0.3333 | 1 | 1 | 5 | 0.8 | 12 | 0.5833 |
| 5_5558181 | 2 | 0.5 | 4 | 0.25 | NA | NA | 1 | 0 | 7 | 0.2857 | 3 | 0.6667 | 3 | 0.3333 | 2 | 0.5 | 5 | 0.4 | 13 | 0.4615 |
| 5_5558229 | 2 | 1 | 4 | 1 | 1 | 1 | 1 | 1 | 8 | 1 | 3 | 1 | 3 | 1 | 2 | 0.5 | 6 | 1 | 14 | 0.9286 |
| 5_5558344 | 2 | 1 | 2 | 1 | 1 | 1 | NA | NA | 5 | 1 | 3 | 1 | 2 | 0.5 | 2 | 1 | 4 | 1 | 11 | 0.9091 |
| 5_5558454 | 2 | 0 | 1 | 1 | NA | NA | NA | NA | 3 | 0.3333 | NA | NA | NA | NA | NA | NA | 1 | 1 | 1 | 1 |
| 5_5589026 | 1 | 1 | NA | NA | 2 | 1 | NA | NA | 3 | 1 | 1 | 1 | NA | NA | 2 | 1 | 1 | 1 | 4 | 1 |
| 5_5589036 | 1 | 0 | NA | NA | 2 | 0.5 | NA | NA | 3 | 0.3333 | 1 | 1 | NA | NA | 2 | 0 | 1 | 0 | 4 | 0.25 |
| 5_5589242 | 1 | 1 | NA | NA | 1 | 1 | NA | NA | 2 | 1 | 1 | 1 | NA | NA | NA | NA | NA | NA | 1 | 1 |
| 5_5631277 | NA | NA | NA | NA | NA | NA | 1 | 1 | 1 | 1 | 1 | 0 | 2 | 0 | 3 | 0.3333 | 1 | 0 | 7 | 0.1429 |
| 5_5719067 | 1 | 1 | 1 | 0 | 2 | 0.5 | NA | NA | 4 | 0.5 | 1 | 1 | NA | NA | 2 | 1 | NA | NA | 3 | 1 |
| 5_5719163 | 1 | 1 | 1 | 0 | 2 | 0.5 | NA | NA | 4 | 0.5 | 1 | 1 | NA | NA | 2 | 0 | NA | NA | 3 | 0.3333 |
| 5_5727600 | NA | NA | NA | NA | 1 | 1 | 1 | 1 | 2 | 1 | NA | NA | 1 | 0 | 1 | 1 | 1 | 1 | 3 | 0.6667 |

| | | | | | | | | | | | | | | | | | | | | |
|---|---|---|---|---|---|---|---|---|---|---|---|---|---|---|---|---|---|---|---|---|
| 5_5727729 | NA | NA | NA | NA | 1 | 1 | 1 | 1 | 2 | 1 | NA | NA | 1 | 0 | 1 | 1 | 1 | 0 | 3 | 0.3333 |
| 5_5732610 | NA | NA | NA | NA | 1 | 1 | 1 | 1 | 2 | 1 | NA | NA | 1 | 0 | 1 | 0 | 1 | 1 | 3 | 0.3333 |
| 5_5732718 | 1 | 1 | NA | NA | 1 | 0 | 1 | 1 | 3 | 0.6667 | NA | NA | 1 | 0 | 1 | 1 | 1 | 1 | 3 | 0.6667 |
| 5_5732738 | NA | NA | NA | NA | 1 | 1 | 1 | 1 | 2 | 1 | NA | NA | 1 | 1 | 1 | 0 | 1 | 1 | 3 | 0.6667 |
| 5_5832681 | 1 | 0 | 6 | 0 | 3 | 0.3333 | 5 | 0 | 15 | 0.0667 | 2 | 0.5 | 2 | 0 | 3 | 0 | 5 | 0.4 | 12 | 0.25 |
| 5_6814125 | 1 | 1 | NA | NA | NA | NA | NA | NA | 1 | 1 | 1 | 1 | NA | NA | NA | NA | NA | NA | 1 | 1 |
| 5_8117689 | 9 | 0.1111 | 12 | 0.1667 | 8 | 0.25 | 10 | 0 | 39 | 0.1282 | 5 | 0.2 | 6 | 0.1667 | 8 | 0 | 12 | 0.0833 | 31 | 0.0968 |
| 5_8122905 | 14 | 0.2143 | 26 | 0.1538 | 15 | 0 | 23 | 0.1304 | 78 | 0.1282 | 8 | 0 | 13 | 0.1538 | 13 | 0.0769 | 21 | 0.1429 | 55 | 0.1091 |
| 5_8123677 | 11 | 0.4545 | 16 | 0.125 | 14 | 0.2143 | 16 | 0.125 | 57 | 0.2105 | 6 | 0.1667 | 12 | 0.25 | 10 | 0.3 | 17 | 0.2941 | 45 | 0.2667 |
| 5_8123737 | 9 | 0 | 17 | 0.0588 | 14 | 0.2857 | 14 | 0.0714 | 54 | 0.1111 | 7 | 0.2857 | 12 | 0.3333 | 11 | 0.2727 | 17 | 0.1176 | 47 | 0.234 |
| 5_8124058 | 8 | 0.125 | 15 | 0.2 | 11 | 0.2727 | 14 | 0.2143 | 48 | 0.2083 | 7 | 0.1429 | 11 | 0.2727 | 10 | 0.2 | 15 | 0.2667 | 43 | 0.2326 |
| 5_8124855 | 9 | 0.1111 | 12 | 0.1667 | 10 | 0 | 9 | 0.1111 | 40 | 0.1 | 4 | 0 | 10 | 0.2 | 7 | 0 | 7 | 0 | 28 | 0.0714 |
| 5_8125724 | 5 | 0.2 | 7 | 0 | 8 | 0.125 | 7 | 0.2857 | 27 | 0.1481 | 4 | 0.25 | 6 | 0.1667 | 4 | 0.25 | 6 | 0.1667 | 20 | 0.2 |
| 5_8125733 | 4 | 0.25 | 9 | 0.2222 | 8 | 0.25 | 6 | 0.1667 | 27 | 0.2222 | 4 | 0 | 3 | 0.3333 | 2 | 0.5 | 6 | 0.1667 | 15 | 0.2 |
| 5_8125798 | 5 | 0 | 8 | 0.125 | 6 | 0.3333 | 7 | 0 | 26 | 0.1154 | 4 | 0 | 5 | 0 | 4 | 0.25 | 5 | 0 | 18 | 0.0556 |
| 5_8126009 | 4 | 0.25 | 7 | 0.4286 | 8 | 0.125 | 6 | 0.1667 | 25 | 0.24 | 4 | 0 | 4 | 0.25 | 3 | 0.3333 | 4 | 0 | 15 | 0.1333 |
| 5_8126072 | 4 | 0.25 | 7 | 0.1429 | 8 | 0 | 6 | 0 | 25 | 0.08 | 5 | 0.4 | 3 | 0.3333 | 2 | 0 | 5 | 0 | 15 | 0.2 |
| 5_8126110 | 4 | 0 | 5 | 0.2 | 7 | 0 | 4 | 0 | 20 | 0.05 | 4 | 0 | 3 | 0.3333 | 2 | 0 | 4 | 0 | 13 | 0.0769 |
| 5_8126979 | 4 | 0.75 | 5 | 0.6 | 4 | 1 | 3 | 0.6667 | 16 | 0.75 | 3 | 0.6667 | 3 | 0.6667 | 2 | 1 | 5 | 0.4 | 13 | 0.6154 |
| 5_8127153 | 4 | 0.5 | 5 | 0 | 5 | 0 | 4 | 0 | 18 | 0.1111 | 3 | 0.3333 | 1 | 0 | 2 | 0 | 6 | 0 | 12 | 0.0833 |
| 5_8127260 | 3 | 0 | 4 | 0 | 4 | 0.25 | 5 | 0 | 16 | 0.0625 | 2 | 0 | 1 | 0 | 1 | 0 | 6 | 0.1667 | 10 | 0.1 |
| 5_8127272 | 3 | 0.6667 | 5 | 0.2 | 4 | 0.5 | 3 | 0.3333 | 15 | 0.4 | 2 | 0.5 | 2 | 0.5 | 2 | 0.5 | 5 | 0.4 | 11 | 0.4545 |
| 5_8127398 | 3 | 0.3333 | 2 | 0 | 3 | 0.3333 | 4 | 0 | 12 | 0.1667 | 2 | 0 | 1 | 0 | 2 | 0 | 4 | 0.5 | 9 | 0.2222 |
| 5_8127424 | 3 | 0.6667 | 3 | 1 | 4 | 1 | 1 | 1 | 11 | 0.9091 | 2 | 1 | 2 | 0.5 | 2 | 1 | 6 | 0.6667 | 12 | 0.75 |
| 5_8127466 | 3 | 0.6667 | 2 | 0 | 4 | 1 | 3 | 0.3333 | 12 | 0.5833 | 3 | 0.3333 | 2 | 0.5 | 2 | 0.5 | 5 | 1 | 12 | 0.6667 |
| 5_8127535 | 3 | 1 | 3 | 0.6667 | 3 | 1 | 3 | 0.6667 | 12 | 0.8333 | 3 | 1 | 2 | 0.5 | 2 | 1 | 2 | 1 | 9 | 0.8889 |
| 5_8127541 | 3 | 0.3333 | NA | NA | 4 | 0.75 | 3 | 0.3333 | 10 | 0.5 | 3 | 1 | 2 | 0.5 | 2 | 0.5 | 6 | 0.8333 | 13 | 0.7692 |
| 5_8127665 | 2 | 0.5 | 2 | 0 | 4 | 0.25 | 2 | 0 | 10 | 0.2 | 2 | 0 | NA | NA | 2 | 0 | 5 | 0.6 | 9 | 0.3333 |
| 5_8127801 | 3 | 0 | 3 | 0 | 4 | 0.5 | 2 | 0 | 12 | 0.1667 | 2 | 0 | 1 | 0 | 3 | 0.3333 | 5 | 0 | 11 | 0.0909 |
| 5_8128016 | 2 | 0.5 | 4 | 0.25 | 5 | 0 | 2 | 0 | 13 | 0.1538 | 2 | 0.5 | NA | NA | 2 | 0 | 5 | 0 | 9 | 0.1111 |
| 5_8128057 | 3 | 0 | 4 | 0 | 7 | 0.2857 | 2 | 0 | 16 | 0.125 | 3 | 0.3333 | 1 | 0 | 3 | 0 | 7 | 0.1429 | 14 | 0.1429 |
| 5_8128154 | 2 | 0.5 | 3 | 0.6667 | 5 | 0.6 | 1 | 0 | 11 | 0.5455 | 3 | 0.6667 | 1 | 1 | 1 | 1 | 4 | 0.75 | 9 | 0.7778 |
| 5_8128318 | 2 | 0.5 | 3 | 0.6667 | 6 | 0.1667 | 1 | 0 | 12 | 0.3333 | 3 | 0.3333 | 1 | 1 | 1 | 1 | 5 | 0.4 | 10 | 0.5 |
| 5_8128533 | 2 | 1 | 5 | 1 | 4 | 0.75 | 2 | 0.5 | 13 | 0.8462 | 2 | 0.5 | 2 | 0 | 1 | 1 | 3 | 0.6667 | 8 | 0.5 |
| 5_8128656 | 4 | 0.25 | 7 | 0.1429 | 7 | 0.2857 | 2 | 0.5 | 20 | 0.25 | 3 | 0.3333 | 2 | 1 | 2 | 1 | 3 | 0 | 10 | 0.5 |
| 5_8141903 | NA | NA | 1 | 1 | NA | NA | NA | NA | 1 | 1 | 1 | 1 | NA | NA | 1 | 1 | NA | NA | 2 | 1 |
| 5_8447444 | NA | NA | 1 | 1 | NA | NA | NA | NA | 1 | 1 | 1 | 1 | 1 | 1 | NA | NA | 1 | 1 | 3 | 1 |
| 5_8503311 | 1 | 1 | 1 | 1 | NA | NA | 2 | 0.5 | 4 | 0.75 | NA | NA | 3 | 1 | 2 | 1 | 1 | 0 | 6 | 0.8333 |
| 5_8540622 | 11 | 0.6364 | 5 | 0.4 | 8 | 0.125 | 7 | 0 | 31 | 0.3226 | 4 | 0.25 | 1 | 0 | 8 | 0.25 | 8 | 0.5 | 21 | 0.3333 |
| 5_8542090 | 6 | 0 | 3 | 0 | 5 | 0.4 | 6 | 0 | 20 | 0.1 | 4 | 0 | 1 | 0 | 7 | 0.1429 | 7 | 0 | 19 | 0.0526 |
| 5_8542440 | 3 | 0.3333 | 3 | 0 | 4 | 0.5 | 6 | 0.6667 | 16 | 0.4375 | 2 | 0 | 1 | 0 | 4 | 0.5 | 7 | 0.1429 | 14 | 0.2143 |
| 5_8542455 | 4 | 0.25 | 3 | 0 | 4 | 0 | 6 | 0 | 17 | 0.0588 | 2 | 0 | 1 | 0 | 4 | 0 | 6 | 0.1667 | 13 | 0.0769 |
| 5_8797562 | NA | NA | 3 | 1 | 2 | 0.5 | 2 | 0.5 | 7 | 0.7143 | 1 | 0 | 2 | 0.5 | 4 | 0.5 | 2 | 0 | 9 | 0.3333 |
| 5_9009176 | NA | NA | 2 | 0.5 | 1 | 1 | NA | NA | 3 | 0.6667 | 1 | 1 | NA | NA | NA | NA | NA | NA | 1 | 1 |
| 5_9016849 | 9 | 0.2222 | 4 | 0.5 | 9 | 0 | 7 | 0.1429 | 29 | 0.1724 | 4 | 0 | 2 | 0 | 7 | 0.4286 | 6 | 0.1667 | 19 | 0.2105 |
| 5_9151257 | 2 | 0 | 1 | 1 | 2 | 0 | 3 | 0.3333 | 8 | 0.25 | 5 | 0 | 2 | 0.5 | 4 | 0 | 2 | 0 | 13 | 0.0769 |
| 5_9151276 | 2 | 0 | NA | NA | 2 | 0 | 3 | 0.3333 | 7 | 0.1429 | 4 | 0 | 2 | 0.5 | 4 | 0 | 2 | 0.5 | 12 | 0.1667 |
| 5_9151325 | 2 | 0.5 | 1 | 0 | 2 | 0 | 3 | 0.3333 | 8 | 0.25 | 5 | 0 | 2 | 0.5 | 3 | 0.3333 | 2 | 0 | 12 | 0.1667 |
| 5_9168476 | 2 | 0.5 | 1 | 0 | 2 | 0 | 3 | 0 | 8 | 0.125 | 5 | 0 | 3 | 0.3333 | 3 | 0.3333 | 2 | 0 | 13 | 0.1538 |
| 5_9168604 | 2 | 0.5 | 1 | 1 | 2 | 0.5 | 2 | 0.5 | 7 | 0.5714 | 4 | 0.25 | 4 | 0.5 | 3 | 0.3333 | 2 | 0.5 | 13 | 0.3846 |
| 5_9168880 | 4 | 0 | 3 | 0.3333 | 2 | 0 | 5 | 0 | 14 | 0.0714 | 5 | 0 | 4 | 0.25 | 3 | 0 | 2 | 0 | 14 | 0.0714 |
| 5_9169991 | 2 | 0 | 2 | 0.5 | 3 | 0 | 5 | 0.2 | 12 | 0.1667 | 4 | 0 | 5 | 0 | 5 | 0 | 3 | 0.3333 | 17 | 0.0588 |
| 5_9170039 | 1 | 1 | 3 | 0.3333 | 3 | 0 | 5 | 0.4 | 12 | 0.3333 | 4 | 0.75 | 6 | 0.3333 | 7 | 0.2857 | 2 | 0 | 19 | 0.3684 |
| 5_9170048 | 2 | 0.5 | 3 | 0.6667 | 3 | 1 | 6 | 0.8333 | 14 | 0.7857 | 4 | 1 | 5 | 1 | 8 | 0.75 | 2 | 1 | 19 | 0.8947 |
| 5_9171179 | NA | NA | 3 | 0.6667 | 1 | 0 | 3 | 0.6667 | 7 | 0.5714 | 2 | 0 | 5 | 0.2 | 3 | 0.3333 | 1 | 0 | 11 | 0.1818 |
| 5_9171187 | 1 | 1 | 3 | 0.6667 | 3 | 0.3333 | 4 | 1 | 11 | 0.7273 | 3 | 0.6667 | 5 | 0.8 | 4 | 0.5 | 2 | 1 | 14 | 0.7143 |
| 5_9171249 | 2 | 0.5 | 3 | 0 | 3 | 0 | 5 | 0.4 | 13 | 0.2308 | 3 | 0 | 5 | 0 | 5 | 0 | 2 | 0.5 | 15 | 0.0667 |
| 5_9171284 | 2 | 0.5 | 3 | 0 | 3 | 0 | 4 | 0 | 12 | 0.0833 | 4 | 0 | 5 | 0.2 | 6 | 0 | 2 | 0.5 | 17 | 0.1176 |
| 5_9210811 | NA | NA | NA | NA | NA | NA | 2 | 0.5 | 2 | 0.5 | NA | NA | 1 | 1 | 2 | 0.5 | 1 | 1 | 4 | 0.75 |
| 5_9211060 | NA | NA | NA | NA | NA | NA | 2 | 0.5 | 2 | 0.5 | NA | NA | 1 | 0 | 2 | 0 | 1 | 1 | 4 | 0.25 |
| 5_9211075 | NA | NA | NA | NA | NA | NA | 2 | 0.5 | 2 | 0.5 | NA | NA | 1 | 1 | 2 | 1 | 1 | 1 | 4 | 1 |
| 5_9513045 | 1 | 1 | NA | NA | NA | NA | 1 | 1 | 2 | 1 | NA | NA | 2 | 0 | NA | NA | 1 | 1 | 3 | 0.3333 |
| 5_9513292 | 2 | 1 | NA | NA | NA | NA | 1 | 1 | 3 | 1 | NA | NA | 2 | 0.5 | NA | NA | 1 | 1 | 3 | 0.6667 |
| 5_9513371 | 2 | 1 | NA | NA | NA | NA | 1 | 0 | 3 | 0.6667 | NA | NA | 3 | 0.6667 | NA | NA | 1 | 0 | 4 | 0.5 |
| 5_9529260 | 1 | 0 | 1 | 0 | 2 | 0 | 4 | 0.25 | 8 | 0.125 | NA | NA | 1 | 0 | 3 | 0 | 4 | 0.5 | 8 | 0.25 |

| | | | | | | | | | | | | | | | | | | | | |
|---|---|---|---|---|---|---|---|---|---|---|---|---|---|---|---|---|---|---|---|---|
| 5_9529280 | 1 | 1 | 1 | 0 | 3 | 0.3333 | 4 | 0.25 | 9 | 0.3333 | NA | NA | 1 | 0 | 3 | 0.3333 | 4 | 0 | 8 | 0.125 |
| 5_9529292 | 1 | 1 | 1 | 0 | 3 | 1 | 4 | 0.75 | 9 | 0.7778 | NA | NA | 1 | 1 | 3 | 1 | 4 | 0.75 | 8 | 0.875 |
| 5_9530348 | NA | NA | 3 | 0.3333 | 3 | 0 | 3 | 0 | 9 | 0.1111 | NA | NA | 1 | 0 | 4 | 0.25 | 4 | 0 | 9 | 0.1111 |
| 5_9531035 | NA | NA | 2 | 0 | 2 | 0.5 | 2 | 0.5 | 6 | 0.3333 | NA | NA | NA | NA | 3 | 0.3333 | 1 | 0 | 4 | 0.25 |
| 5_9531184 | NA | NA | 2 | 0 | 2 | 0.5 | 2 | 1 | 6 | 0.5 | NA | NA | NA | NA | 3 | 0.6667 | 2 | 0 | 5 | 0.4 |
| 5_9531227 | NA | NA | 2 | 0.5 | 1 | 0 | 2 | 0.5 | 5 | 0.4 | NA | NA | NA | NA | 3 | 0.3333 | 2 | 0 | 5 | 0.2 |
| 5_9535615 | 4 | 0.5 | 11 | 0.5455 | 13 | 0.2308 | 5 | 0.2 | 33 | 0.3636 | 4 | 0.5 | 9 | 0.4444 | 12 | 0.5 | 5 | 0.8 | 30 | 0.5333 |
| 5_9535790 | 1 | 0 | 6 | 0.1667 | 8 | 0 | 3 | 0.3333 | 18 | 0.1111 | 4 | 0 | 7 | 0.2857 | 7 | 0 | 2 | 0.5 | 20 | 0.15 |
| 5_9561625 | 4 | 0.25 | 9 | 0 | 9 | 0.1111 | 6 | 0 | 28 | 0.0714 | 5 | 0 | 9 | 0.1111 | 11 | 0 | 6 | 0.3333 | 31 | 0.0968 |
| 5_9561718 | 5 | 0.2 | 8 | 0.125 | 10 | 0 | 5 | 0.2 | 28 | 0.1071 | 4 | 0.5 | 8 | 0.25 | 11 | 0.0909 | 6 | 0.3333 | 29 | 0.2414 |
| 5_9561858 | 4 | 0.5 | 9 | 0.1111 | 5 | 0.2 | 5 | 0.8 | 23 | 0.3478 | 5 | 0.6 | 6 | 0.3333 | 10 | 0.2 | 4 | 0.5 | 25 | 0.36 |
| 5_9561901 | 4 | 1 | 10 | 0.5 | 9 | 0.4444 | 4 | 0.5 | 27 | 0.5556 | 5 | 0.6 | 8 | 0.25 | 11 | 0.7273 | 5 | 0.2 | 29 | 0.4828 |
| 5_9561964 | 1 | 0 | 9 | 0.2222 | 7 | 0.1429 | 5 | 0.2 | 22 | 0.1818 | 5 | 0.2 | 8 | 0 | 10 | 0.3 | 5 | 0.4 | 28 | 0.2143 |
| 5_9562010 | 3 | 1 | 9 | 1 | 3 | 1 | 4 | 0.75 | 19 | 0.9474 | 4 | 0.75 | 7 | 0.8571 | 11 | 1 | 5 | 1 | 27 | 0.9259 |
| 5_9562063 | 3 | 0.6667 | 7 | 0.8571 | 7 | 0.7143 | 4 | 1 | 21 | 0.8095 | 4 | 0.75 | 7 | 0.8571 | 9 | 0.6667 | 5 | 0.8 | 25 | 0.76 |
| 5_9562176 | 3 | 0.3333 | 8 | 0.125 | 7 | 0.1429 | 4 | 0 | 22 | 0.1364 | 5 | 0.2 | 8 | 0.125 | 7 | 0.1429 | 6 | 0.1667 | 26 | 0.1538 |
| 5_9562344 | 2 | 1 | 10 | 0.8 | 7 | 0.8571 | 4 | 0.5 | 23 | 0.7826 | 3 | 0.6667 | 5 | 0.6 | 7 | 0.5714 | 5 | 0.2 | 20 | 0.5 |
| 5_9562365 | 3 | 1 | 10 | 1 | 5 | 1 | 5 | 1 | 23 | 1 | 4 | 0.75 | 8 | 1 | 7 | 1 | 4 | 0.75 | 23 | 0.913 |
| 5_9562384 | 2 | 1 | 10 | 0.8 | 6 | 0.8333 | 4 | 0.75 | 22 | 0.8182 | 3 | 0.6667 | 8 | 0.875 | 6 | 0.8333 | 4 | 1 | 21 | 0.8571 |
| 5_9620088 | 3 | 0.3333 | 9 | 0.1111 | 5 | 0 | 4 | 0 | 21 | 0.0952 | 5 | 0 | 4 | 1 | 7 | 0 | 5 | 0 | 21 | 0.1905 |
| 5_9620185 | 3 | 0 | 10 | 0.2 | 5 | 0 | 4 | 0 | 22 | 0.0909 | 4 | 0 | 7 | 0 | 7 | 0.1429 | 6 | 0.1667 | 24 | 0.0833 |
| 5_9620404 | 2 | 0 | 11 | 0.0909 | 6 | 0.5 | 4 | 0 | 23 | 0.1739 | 4 | 0.5 | 9 | 0 | 6 | 0 | 5 | 0 | 24 | 0.0833 |
| 5_9620570 | 2 | 0 | 8 | 0 | 6 | 0.1667 | 4 | 0 | 20 | 0.05 | 4 | 0 | 5 | 0.2 | 6 | 0 | 5 | 0 | 20 | 0.05 |
| 5_9620581 | 1 | 1 | 8 | 1 | 6 | 1 | 4 | 0.75 | 19 | 0.9474 | 3 | 1 | 6 | 1 | 6 | 1 | 3 | 1 | 18 | 1 |
| 5_9620612 | 2 | 0 | 8 | 0 | 5 | 0.2 | 2 | 0.5 | 17 | 0.1176 | 1 | 0 | 7 | 0.2857 | 4 | 0.5 | 4 | 0.25 | 16 | 0.3125 |
| 5_9620675 | 2 | 0.5 | 6 | 0.3333 | 6 | 0.3333 | 2 | 1 | 16 | 0.4375 | 3 | 0 | 5 | 0.2 | 6 | 0.5 | 4 | 0.75 | 18 | 0.3889 |
| 5_9639739 | 2 | 0.5 | 7 | 0.4286 | 7 | 0.5714 | 3 | 0.3333 | 19 | 0.4737 | 2 | 0.5 | 2 | 0.5 | 4 | 0.75 | 5 | 0.4 | 13 | 0.5385 |
| 5_9639799 | 2 | 1 | 8 | 0.5 | 7 | 0.4286 | 4 | 1 | 21 | 0.619 | 3 | 0.6667 | 1 | 0 | 4 | 0.75 | 5 | 0.6 | 13 | 0.6154 |
| 5_9639884 | 2 | 0.5 | 6 | 0.1667 | 5 | 0.4 | 3 | 0.3333 | 16 | 0.3125 | 2 | 0 | 1 | 1 | 4 | 1 | 3 | 1 | 10 | 0.8 |
| 5_9639931 | 1 | 0 | 6 | 0 | 5 | 0.4 | 4 | 0.5 | 16 | 0.25 | 3 | 0.3333 | 3 | 0 | 3 | 0.3333 | 5 | 0.2 | 14 | 0.2143 |
| 5_9640085 | 1 | 0 | 8 | 0.5 | 5 | 0.6 | 4 | 0.5 | 18 | 0.5 | 3 | 0.6667 | 3 | 0 | 3 | 0.6667 | 4 | 0.75 | 13 | 0.5385 |
| 5_13171304 | NA | NA | 4 | 0.25 | 1 | 0 | 2 | 0 | 7 | 0.1429 | 1 | 0 | 2 | 0 | 2 | 0.5 | NA | NA | 5 | 0.2 |
| 5_13649230 | 1 | 0 | 4 | 0.25 | NA | NA | 1 | 0 | 6 | 0.1667 | NA | NA | 1 | 0 | 1 | 1 | 1 | 0 | 3 | 0.3333 |
| 5_13649280 | 1 | 0 | 4 | 0.25 | NA | NA | 1 | 1 | 6 | 0.3333 | NA | NA | 1 | 0 | 1 | 1 | 1 | 1 | 3 | 0.6667 |
| 5_13649536 | 1 | 1 | 4 | 0.5 | NA | NA | NA | NA | 5 | 0.6 | NA | NA | 1 | 1 | 1 | 0 | NA | NA | 2 | 0.5 |
| 5_13649607 | 1 | 1 | 3 | 1 | NA | NA | 1 | 1 | 5 | 1 | NA | NA | 1 | 1 | 1 | 1 | 1 | 1 | 3 | 1 |
| 5_13649623 | 1 | 1 | 3 | 1 | NA | NA | NA | NA | 4 | 1 | NA | NA | 1 | 0 | 1 | 0 | 2 | 1 | 4 | 0.5 |
| 5_13649648 | 1 | 1 | 1 | 1 | NA | NA | NA | NA | 2 | 1 | NA | NA | 1 | 1 | 1 | 1 | NA | NA | 2 | 1 |
| 5_13649724 | 1 | 1 | 3 | 1 | NA | NA | 1 | 1 | 5 | 1 | NA | NA | 1 | 0 | 1 | 1 | 2 | 1 | 4 | 0.75 |
| 5_13649801 | 1 | 1 | 4 | 1 | NA | NA | 1 | 1 | 6 | 1 | NA | NA | 1 | 0 | 1 | 1 | 1 | 1 | 3 | 0.6667 |
| 5_13649884 | NA | NA | 3 | 1 | NA | NA | NA | NA | 3 | 1 | NA | NA | 1 | 1 | NA | NA | 1 | 1 | 2 | 1 |
| 5_13649929 | NA | NA | 4 | 0.75 | NA | NA | NA | NA | 4 | 0.75 | NA | NA | 1 | 1 | NA | NA | 2 | 1 | 3 | 1 |
| 5_13650042 | 1 | 1 | 5 | 0.8 | NA | NA | 1 | 1 | 7 | 0.8571 | NA | NA | 1 | 0 | 1 | 1 | 2 | 1 | 4 | 0.75 |
| 5_13650075 | 2 | 0.5 | 5 | 0 | NA | NA | 1 | 1 | 8 | 0.25 | NA | NA | 1 | 0 | 1 | 1 | 2 | 0 | 4 | 0.25 |
| 5_14075735 | 1 | 1 | 1 | 0 | 1 | 1 | 1 | 1 | 4 | 0.75 | NA | NA | 1 | 1 | 2 | 0.5 | 1 | 1 | 4 | 0.75 |
| 5_14075868 | 1 | 1 | 1 | 1 | NA | NA | 1 | 1 | 3 | 1 | NA | NA | 2 | 0.5 | 2 | 0.5 | 1 | 1 | 5 | 0.6 |
| 5_14302383 | 2 | 1 | 2 | 0.5 | 2 | 1 | 3 | 0.6667 | 9 | 0.7778 | NA | NA | 2 | 0 | 3 | 0.6667 | 1 | 1 | 6 | 0.5 |
| 5_14302427 | 3 | 0.6667 | 2 | 1 | 2 | 1 | 3 | 1 | 10 | 0.9 | NA | NA | 3 | 0.6667 | 4 | 1 | 1 | 1 | 8 | 0.875 |
| 5_14302442 | 3 | 0 | 2 | 0 | 2 | 0.5 | 3 | 0 | 10 | 0.1 | NA | NA | 3 | 1 | 4 | 0 | 1 | 1 | 8 | 0.5 |
| 5_14302450 | 3 | 0 | 2 | 0.5 | 1 | 0 | 3 | 0.6667 | 9 | 0.3333 | NA | NA | 3 | 0.3333 | 4 | 0.25 | 1 | 0 | 8 | 0.25 |
| 5_14302530 | 2 | 1 | 2 | 1 | 2 | 1 | 3 | 1 | 9 | 1 | NA | NA | 3 | 1 | 4 | 1 | 1 | 1 | 8 | 1 |
| 5_14302546 | 3 | 0 | 2 | 0 | 2 | 0.5 | 3 | 0.3333 | 10 | 0.2 | NA | NA | 3 | 0.3333 | 4 | 0.75 | 1 | 0 | 8 | 0.5 |
| 5_14302592 | 3 | 1 | 2 | 0 | 1 | 0 | 2 | 0.5 | 8 | 0.5 | NA | NA | 2 | 1 | 4 | 1 | 1 | 1 | 7 | 1 |
| 5_14302831 | 3 | 0.3333 | 2 | 0.5 | 2 | 0 | 2 | 0.5 | 9 | 0.3333 | NA | NA | 3 | 0.3333 | 3 | 0.3333 | 1 | 1 | 7 | 0.4286 |
| 5_14302875 | NA | NA | NA | NA | 1 | 1 | 3 | 1 | 4 | 1 | NA | NA | 2 | 1 | 3 | 1 | 1 | 1 | 6 | 1 |
| 5_14302912 | 2 | 0.5 | NA | NA | NA | NA | 1 | 1 | 3 | 0.6667 | NA | NA | 1 | 0 | 4 | 0.75 | 1 | 1 | 6 | 0.6667 |
| 5_14302971 | 3 | 0.6667 | 2 | 0.5 | 2 | 0.5 | 2 | 0.5 | 9 | 0.5556 | NA | NA | 3 | 0.6667 | 5 | 0.4 | NA | NA | 8 | 0.5 |
| 5_14303333 | 3 | 0.3333 | 2 | 0 | 2 | 0.5 | 2 | 0.5 | 9 | 0.3333 | NA | NA | 2 | 0 | 3 | 0.3333 | 1 | 0 | 6 | 0.1667 |
| 5_14306583 | 2 | 0.5 | 1 | 0 | 1 | 0 | 2 | 0 | 6 | 0.1667 | NA | NA | 1 | 1 | 3 | 0.3333 | 1 | 0 | 5 | 0.4 |
| 5_14729488 | 1 | 1 | NA | NA | NA | NA | NA | NA | 1 | 1 | NA | NA | 1 | 0 | NA | NA | 1 | 1 | 2 | 0.5 |
| 5_14729649 | 1 | 1 | NA | NA | NA | NA | NA | NA | 1 | 1 | NA | NA | 1 | 1 | NA | NA | 1 | 1 | 2 | 1 |
| 5_14729917 | 1 | 1 | 1 | 0 | NA | NA | NA | NA | 2 | 0.5 | NA | NA | NA | NA | NA | NA | 1 | 1 | 1 | 1 |
| 5_14729939 | 1 | 1 | 1 | 1 | NA | NA | NA | NA | 2 | 1 | NA | NA | 2 | 1 | NA | NA | 1 | 1 | 3 | 1 |
| 5_14730822 | 1 | 1 | 2 | 1 | NA | NA | 2 | 1 | 5 | 1 | NA | NA | 2 | 1 | NA | NA | NA | NA | 2 | 1 |
| 5_14731259 | 1 | 1 | 3 | 0.6667 | NA | NA | 1 | 1 | 5 | 0.8 | NA | NA | 2 | 1 | NA | NA | 1 | 1 | 3 | 1 |

| | | | | | | | | | | | | | | | | | | | | |
|---|---|---|---|---|---|---|---|---|---|---|---|---|---|---|---|---|---|---|---|---|
| 5_1473143 | 1 | 0 | 3 | 0.3333 | NA | NA | 2 | 0 | 6 | 0.1667 | NA | NA | 2 | 0.5 | 1 | 1 | 1 | 0 | 4 | 0.5 |
| 5_1473161 | 1 | 1 | 3 | 1 | NA | NA | 2 | 1 | 6 | 1 | NA | NA | 2 | 1 | 1 | 1 | 1 | 0 | 4 | 0.75 |
| 5_1473165 | 1 | 1 | 3 | 1 | NA | NA | 2 | 0 | 6 | 0.6667 | NA | NA | 3 | 0.6667 | 1 | 1 | 1 | 0 | 5 | 0.6 |
| 5_1473196 | NA | NA | 1 | 1 | NA | NA | 1 | 0 | 2 | 0.5 | NA | NA | 2 | 1 | 1 | 0 | 1 | 1 | 4 | 0.75 |
| 5_1473205 | NA | NA | 3 | 0.3333 | NA | NA | 1 | 1 | 4 | 0.5 | NA | NA | 3 | 0.3333 | 3 | 0 | 1 | 0 | 7 | 0.1429 |
| 5_1476260 | 1 | 0 | 4 | 0.25 | NA | NA | 2 | 0.5 | 7 | 0.2857 | NA | NA | 4 | 0.5 | 5 | 0 | 1 | 1 | 10 | 0.3 |
| 5_1476280 | 2 | 1 | 3 | 1 | NA | NA | 2 | 1 | 7 | 1 | NA | NA | 4 | 0.25 | 5 | 0.8 | 2 | 0.5 | 11 | 0.5455 |
| 5_1476289 | 2 | 0.5 | 4 | 0.75 | NA | NA | 2 | 0.5 | 8 | 0.625 | NA | NA | 3 | 0.6667 | 6 | 0.6667 | 2 | 0.5 | 11 | 0.6364 |
| 5_1476308 | 2 | 0.5 | 4 | 0.75 | NA | NA | 3 | 0.6667 | 9 | 0.6667 | NA | NA | 4 | 0.25 | 5 | 0.4 | 2 | 0.5 | 11 | 0.3636 |
| 5_1476334 | 3 | 0.3333 | 4 | 0.25 | NA | NA | 3 | 0 | 10 | 0.2 | NA | NA | 4 | 0 | 6 | 0 | 3 | 0.3333 | 13 | 0.0769 |
| 5_1476342 | 3 | 0 | 4 | 0.25 | NA | NA | 3 | 0 | 10 | 0.1 | NA | NA | 6 | 0 | 6 | 0.3333 | 3 | 0 | 15 | 0.1333 |
| 5_1476347 | 3 | 0.3333 | 3 | 0 | NA | NA | 2 | 0 | 8 | 0.125 | NA | NA | 6 | 0.1667 | 5 | 0 | 3 | 0 | 14 | 0.0714 |
| 5_1476348 | 2 | 1 | 3 | 0.3333 | NA | NA | 3 | 1 | 8 | 0.75 | NA | NA | 5 | 0.8 | 4 | 1 | 3 | 1 | 12 | 0.9167 |
| 5_1476358 | 3 | 0 | 4 | 0 | NA | NA | 3 | 0.3333 | 10 | 0.1 | NA | NA | 5 | 0 | 6 | 0.1667 | 3 | 0 | 14 | 0.0714 |
| 5_1476359 | 3 | 0.3333 | 4 | 0.25 | NA | NA | 2 | 1 | 9 | 0.4444 | NA | NA | 6 | 0.1667 | 6 | 0.3333 | 3 | 0.3333 | 15 | 0.2667 |
| 5_1476361 | 2 | 0 | 4 | 0.25 | NA | NA | 2 | 0 | 8 | 0.125 | NA | NA | 6 | 0.1667 | 6 | 0 | 3 | 0 | 15 | 0.0667 |
| 5_1476378 | 4 | 0.5 | 4 | 0.5 | NA | NA | 3 | 0.3333 | 11 | 0.4545 | NA | NA | 6 | 0.6667 | 6 | 0.5 | 3 | 0.3333 | 15 | 0.5333 |
| 5_1476381 | 3 | 0.6667 | 5 | 0.4 | NA | NA | 3 | 0.3333 | 11 | 0.4545 | NA | NA | 5 | 0 | 6 | 0.3333 | 2 | 0.5 | 13 | 0.2308 |
| 5_1476390 | 4 | 0 | 5 | 0 | NA | NA | 3 | 0.3333 | 12 | 0.0833 | NA | NA | 6 | 0.1667 | 7 | 0 | 3 | 0 | 16 | 0.0625 |
| 5_1476394 | 3 | 0.6667 | 5 | 0.6 | NA | NA | 3 | 0 | 11 | 0.4545 | NA | NA | 7 | 0.2857 | 7 | 0.4286 | 4 | 0.75 | 18 | 0.4444 |
| 5_1476395 | 3 | 0 | 5 | 0.2 | NA | NA | 3 | 0 | 11 | 0.0909 | NA | NA | 6 | 0.1667 | 7 | 0.1429 | 4 | 0 | 17 | 0.1176 |
| 5_1476410 | 3 | 0.3333 | 5 | 0.8 | NA | NA | 3 | 1 | 11 | 0.7273 | NA | NA | 6 | 0.8333 | 6 | 0.8333 | 2 | 0.5 | 14 | 0.7857 |
| 5_1476432 | 4 | 0.25 | 5 | 0.2 | NA | NA | 3 | 0.3333 | 12 | 0.25 | NA | NA | 6 | 0.1667 | 5 | 0.4 | 2 | 0.5 | 13 | 0.3077 |
| 5_1476444 | 4 | 0.5 | 6 | 0.3333 | NA | NA | 5 | 0.4 | 15 | 0.4 | NA | NA | 5 | 0.4 | 6 | 0.8333 | 4 | 0.75 | 15 | 0.6667 |
| 5_1476454 | 3 | 0 | 9 | 0.4444 | 1 | 1 | 4 | 0 | 17 | 0.2941 | NA | NA | 7 | 0.2857 | 6 | 0.6667 | 3 | 0.6667 | 16 | 0.5 |
| 5_1484199 | 5 | 0 | 14 | 0 | 4 | 0 | 9 | 0.2222 | 32 | 0.0625 | 2 | 0 | 7 | 0.1429 | 7 | 0.1429 | 6 | 0.1667 | 22 | 0.1364 |
| 5_1484393 | 5 | 0.4 | 14 | 0.2143 | 5 | 0.2 | 6 | 0.3333 | 30 | 0.2667 | 2 | 0.5 | 5 | 0.2 | 6 | 0.1667 | 6 | 0.1667 | 19 | 0.2105 |
| 5_1490887 | NA | NA | 1 | 1 | NA | NA | 1 | 0 | 2 | 0.5 | NA | NA | 1 | 1 | 1 | 1 | 2 | 1 | 4 | 1 |
| 5_1490895 | 1 | 1 | 1 | 1 | 1 | 1 | 1 | 0 | 4 | 0.75 | NA | NA | 1 | 1 | NA | NA | 2 | 1 | 3 | 1 |
| 5_1490905 | 2 | 1 | 1 | 1 | 1 | 1 | 1 | 1 | 5 | 1 | NA | NA | 1 | 1 | 1 | 1 | 2 | 1 | 4 | 1 |
| 5_1490909 | 2 | 1 | 1 | 1 | 1 | 1 | 1 | 1 | 5 | 1 | NA | NA | 1 | 1 | 1 | 1 | 2 | 1 | 4 | 1 |
| 5_1490918 | 2 | 0.5 | 1 | 0 | 1 | 1 | NA | NA | 4 | 0.5 | NA | NA | 1 | 1 | 1 | 0 | 2 | 0.5 | 4 | 0.5 |
| 5_1491298 | 7 | 0 | 3 | 0 | 5 | 0.2 | 2 | 0 | 17 | 0.0588 | 4 | 0 | 3 | 0 | 4 | 0.25 | 2 | 0 | 13 | 0.0769 |
| 5_1657556 | 2 | 0.5 | 8 | 0.5 | 2 | 0 | 3 | 0.6667 | 15 | 0.4667 | 5 | 0.2 | 6 | 0 | 3 | 0 | 4 | 0.5 | 18 | 0.1667 |
| 5_1657561 | 2 | 1 | 7 | 0.7143 | 2 | 1 | 3 | 0.6667 | 14 | 0.7857 | 5 | 1 | 5 | 0.8 | 3 | 1 | 4 | 1 | 17 | 0.9412 |
| 5_1657565 | 2 | 0 | 8 | 0 | 2 | 0 | 3 | 0.3333 | 15 | 0.0667 | 5 | 0.2 | 6 | 0.1667 | 4 | 0.25 | 5 | 0 | 20 | 0.15 |
| 5_1657568 | 2 | 0 | 7 | 0.1429 | 2 | 0 | 3 | 0 | 14 | 0.0714 | 5 | 0.4 | 6 | 0 | 4 | 0 | 5 | 0 | 20 | 0.1 |
| 5_1657577 | 2 | 1 | 5 | 0.8 | 1 | 1 | 2 | 1 | 10 | 0.9 | 5 | 1 | 4 | 1 | 4 | 1 | 4 | 1 | 17 | 1 |
| 5_1657585 | 2 | 1 | 9 | 0.8889 | 2 | 1 | 3 | 0.6667 | 16 | 0.875 | 5 | 0.4 | 5 | 0.6 | 4 | 0.75 | 4 | 1 | 18 | 0.6667 |
| 5_1657586 | 2 | 0.5 | 9 | 0.1111 | 2 | 0 | 3 | 0.3333 | 16 | 0.1875 | 5 | 0.4 | 6 | 0.3333 | 4 | 0.75 | 4 | 0 | 19 | 0.3684 |
| 5_1657590 | 1 | 0 | 8 | 0.5 | 2 | 1 | 2 | 0.5 | 13 | 0.5385 | 5 | 0.4 | 6 | 0.5 | 3 | 0.6667 | 4 | 0.5 | 18 | 0.5 |
| 5_1657594 | 1 | 1 | 7 | 0.7143 | NA | NA | 2 | 0.5 | 10 | 0.7 | 4 | 0.5 | 5 | 0.6 | 3 | 0.6667 | 4 | 0.75 | 16 | 0.625 |
| 5_1657601 | 3 | 0.6667 | 8 | 0 | 2 | 0 | 3 | 0.6667 | 16 | 0.25 | 4 | 0 | 6 | 0.1667 | 4 | 0 | 6 | 0.3333 | 20 | 0.15 |
| 5_1657618 | 3 | 0.3333 | 11 | 0.0909 | 3 | 0 | 4 | 0 | 21 | 0.0952 | 6 | 0 | 7 | 0.1429 | 4 | 0 | 6 | 0.3333 | 23 | 0.1304 |
| 5_1657627 | 2 | 0 | 10 | 0.2 | 3 | 0.6667 | 5 | 0.6 | 20 | 0.35 | 6 | 0.5 | 8 | 0.125 | 3 | 0 | 6 | 0.5 | 23 | 0.3043 |
| 5_1657635 | 3 | 0.3333 | 10 | 0.2 | 2 | 0.5 | 3 | 0.6667 | 18 | 0.3333 | 6 | 0.1667 | 6 | 0.5 | 5 | 0.4 | 8 | 0.25 | 25 | 0.32 |
| 5_1657921 | 15 | 0.0667 | 29 | 0.0345 | 13 | 0.3846 | 13 | 0.0769 | 70 | 0.1143 | 15 | 0.0667 | 16 | 0 | 10 | 0 | 16 | 0.1875 | 57 | 0.0702 |
| 5_1657935 | 15 | 0.0667 | 25 | 0.04 | 14 | 0.0714 | 13 | 0.0769 | 67 | 0.0597 | 13 | 0.0769 | 19 | 0.1053 | 7 | 0 | 16 | 0.0625 | 55 | 0.0727 |
| 5_1793237 | 1 | 1 | 3 | 0.3333 | 1 | 1 | NA | NA | 5 | 0.6 | 2 | 0.5 | NA | NA | 1 | 1 | 1 | 0 | 4 | 0.5 |
| 5_1793241 | 4 | 0.75 | 4 | 0.75 | 1 | 1 | NA | NA | 9 | 0.7778 | 2 | 0.5 | NA | NA | 1 | 1 | 2 | 0.5 | 5 | 0.6 |
| 5_1793245 | 4 | 0.75 | 4 | 0.25 | 1 | 1 | NA | NA | 9 | 0.5556 | 2 | 1 | NA | NA | 1 | 0 | 2 | 0.5 | 5 | 0.6 |
| 5_1793250 | 3 | 0.3333 | 3 | 0 | 1 | 0 | NA | NA | 7 | 0.1429 | 1 | 1 | NA | NA | 1 | 0 | 1 | 0 | 3 | 0.3333 |
| 5_1793251 | 4 | 0 | 3 | 0.6667 | 1 | 0 | NA | NA | 8 | 0.25 | 1 | 0 | NA | NA | 1 | 0 | 2 | 0.5 | 4 | 0.25 |
| 5_1793253 | 4 | 1 | 4 | 1 | 1 | 1 | NA | NA | 9 | 1 | 2 | 1 | NA | NA | 1 | 1 | 1 | 1 | 4 | 1 |
| 5_1793257 | 4 | 1 | 4 | 1 | 1 | 1 | NA | NA | 9 | 1 | 2 | 1 | NA | NA | 1 | 1 | 1 | 1 | 4 | 1 |
| 5_1793268 | 3 | 1 | 3 | 0.6667 | 1 | 1 | NA | NA | 7 | 0.8571 | 2 | 1 | NA | NA | 1 | 1 | 1 | 0 | 4 | 0.75 |
| 5_1793278 | 3 | 0 | 4 | 0.25 | 1 | 0 | NA | NA | 8 | 0.125 | 2 | 0.5 | NA | NA | 1 | 0 | 2 | 0 | 5 | 0.2 |
| 5_1793286 | 4 | 0 | 3 | 0.3333 | 1 | 1 | NA | NA | 8 | 0.25 | 1 | 0 | NA | NA | 1 | 1 | 2 | 0 | 4 | 0.25 |
| 5_1793289 | 2 | 1 | 3 | 1 | 1 | 1 | NA | NA | 6 | 1 | 2 | 1 | NA | NA | 1 | 1 | 2 | 1 | 5 | 1 |
| 5_1793292 | 4 | 0.5 | 4 | 1 | 1 | 1 | NA | NA | 9 | 0.7778 | 2 | 0.5 | NA | NA | 1 | 0 | 2 | 1 | 5 | 0.6 |
| 5_1847048 | 19 | 0.1053 | 44 | 0.0909 | 18 | 0.1111 | 24 | 0.0833 | 105 | 0.0952 | 15 | 0.0667 | 16 | 0.1875 | 15 | 0.0667 | 16 | 0.125 | 62 | 0.1129 |
| 5_1847060 | 14 | 0.0714 | 38 | 0.0263 | 16 | 0.0625 | 21 | 0.0952 | 89 | 0.0562 | 13 | 0 | 19 | 0.1053 | 15 | 0.1333 | 13 | 0.0769 | 60 | 0.0833 |
| 5_1847070 | 17 | 0 | 38 | 0.0526 | 18 | 0.1667 | 20 | 0.1 | 93 | 0.0753 | 15 | 0 | 13 | 0.0769 | 15 | 0.0667 | 14 | 0.1429 | 57 | 0.0702 |
| 5_1847183 | 14 | 0.1429 | 33 | 0.0303 | 18 | 0.1111 | 21 | 0 | 86 | 0.0581 | 14 | 0 | 15 | 0.1333 | 13 | 0.1538 | 13 | 0.0769 | 55 | 0.0909 |
| 5_1847190 | 12 | 0.0833 | 34 | 0.1176 | 16 | 0.25 | 21 | 0.1429 | 83 | 0.1446 | 15 | 0.1333 | 14 | 0.0714 | 13 | 0.0769 | 12 | 0.25 | 54 | 0.1296 |

| | | | | | | | | | | | | | | | | | | | | |
|---|---|---|---|---|---|---|---|---|---|---|---|---|---|---|---|---|---|---|---|---|
| 5_1847203 | 14 | 0.9286 | 29 | 0.7931 | 15 | 1 | 17 | 0.8824 | 75 | 0.88 | 16 | 0.9375 | 13 | 1 | 11 | 1 | 12 | 1 | 52 | 0.9808 |
| 5_1847213 | 12 | 0.0833 | 26 | 0.0385 | 17 | 0.1176 | 17 | 0 | 72 | 0.0556 | 15 | 0.1333 | 12 | 0.0833 | 12 | 0.0833 | 13 | 0 | 52 | 0.0769 |
| 5_1856528 | 8 | 0 | 12 | 0.1667 | 8 | 0.5 | 10 | 0.3 | 38 | 0.2368 | 6 | 0.3333 | 10 | 0 | 10 | 0.2 | 8 | 0.25 | 34 | 0.1765 |
| 5_2067082 | NA | NA | 3 | 0.6667 | 3 | 0.6667 | NA | NA | 6 | 0.6667 | NA | NA | NA | NA | 2 | 1 | 1 | 1 | 3 | 1 |
| 5_2067097 | NA | NA | 3 | 1 | 2 | 1 | 1 | 1 | 6 | 1 | 1 | 0 | 1 | 1 | 3 | 0.3333 | 4 | 1 | 9 | 0.6667 |
| 5_2067104 | NA | NA | 3 | 0.3333 | 3 | 0 | 1 | 1 | 7 | 0.2857 | 1 | 0 | 1 | 1 | 4 | 0.25 | 4 | 0.5 | 10 | 0.4 |
| 5_2067116 | NA | NA | 2 | 0 | 3 | 0 | 1 | 1 | 6 | 0.1667 | 1 | 0 | 1 | 1 | 5 | 0 | 3 | 0.3333 | 10 | 0.2 |
| 5_2090780 | 8 | 0.375 | 7 | 0.4286 | 6 | 0.6667 | 4 | 0.5 | 25 | 0.48 | 2 | 0 | 4 | 0.25 | 7 | 0.1429 | 7 | 0 | 20 | 0.1 |
| 5_2090781 | 9 | 0.6667 | 6 | 0.6667 | 7 | 0.4286 | 4 | 1 | 26 | 0.6538 | 1 | 0 | 4 | 0.5 | 6 | 0.6667 | 5 | 0.6 | 16 | 0.5625 |
| 5_2090789 | 8 | 1 | 6 | 1 | 6 | 1 | 3 | 1 | 23 | 1 | 3 | 0.6667 | 4 | 1 | 8 | 1 | 6 | 0.8333 | 21 | 0.9048 |
| 5_2090797 | 10 | 0.4 | 6 | 0 | 7 | 0 | 5 | 0.6 | 28 | 0.25 | 3 | 0 | 6 | 0.1667 | 8 | 0 | 6 | 0.5 | 23 | 0.1739 |
| 5_2090799 | 9 | 0.6667 | 6 | 0.5 | 6 | 0.3333 | 4 | 0.25 | 25 | 0.48 | 3 | 0 | 5 | 0.2 | 5 | 0.8 | 3 | 0.6667 | 16 | 0.4375 |
| 5_2090799 | 7 | 0.4286 | 7 | 0.8571 | 6 | 0.8333 | 5 | 1 | 25 | 0.76 | 3 | 0.3333 | 5 | 0.6 | 5 | 0.6 | 7 | 0.8571 | 20 | 0.65 |
| 5_2090801 | 9 | 0.7778 | 6 | 0.6667 | 5 | 0.6 | 4 | 0.75 | 24 | 0.7083 | 3 | 1 | 6 | 0.8333 | 7 | 0.8571 | 5 | 0.6 | 21 | 0.8095 |
| 5_2090806 | 10 | 0.1 | 6 | 0 | 7 | 0.4286 | 5 | 0 | 28 | 0.1429 | 3 | 0.6667 | 5 | 0 | 9 | 0.4444 | 7 | 0 | 24 | 0.25 |
| 5_2090816 | 4 | 1 | 4 | 1 | 5 | 0.8 | 4 | 1 | 17 | 0.9412 | 1 | 1 | 6 | 0.5 | 9 | 0.7778 | 5 | 0.8 | 21 | 0.7143 |
| 5_2090911 | 9 | 0.1111 | 8 | 0.125 | 6 | 0.1667 | 7 | 0.1429 | 30 | 0.1333 | 4 | 0.25 | 8 | 0.125 | 9 | 0.1111 | 10 | 0.2 | 31 | 0.1613 |
| 5_2090934 | 9 | 0.1111 | 7 | 0.1429 | 6 | 0.1667 | 8 | 0 | 30 | 0.1 | 5 | 0 | 9 | 0 | 9 | 0.1111 | 9 | 0.2222 | 32 | 0.0938 |
| 5_2090945 | 8 | 0.25 | 8 | 0.125 | 6 | 0 | 5 | 0.2 | 27 | 0.1481 | 4 | 0 | 9 | 0.1111 | 9 | 0.2222 | 6 | 0.1667 | 28 | 0.1429 |
| 5_2090965 | 9 | 0.5556 | 12 | 0.3333 | 9 | 0.2222 | 9 | 0 | 39 | 0.2821 | 5 | 0.2 | 9 | 0.2222 | 11 | 0.5455 | 11 | 0.0909 | 36 | 0.2778 |
| 5_2109122 | 5 | 0.4 | 4 | 0 | 2 | 0 | 6 | 0.5 | 17 | 0.2941 | 2 | 0.5 | 4 | 0.25 | 6 | 0.3333 | 6 | 0.3333 | 18 | 0.3333 |
| 5_2109131 | 6 | 0 | 7 | 0.1429 | 2 | 0 | 6 | 0.1667 | 21 | 0.0952 | 2 | 0 | 4 | 0.25 | 7 | 0.1429 | 4 | 0 | 17 | 0.1176 |
| 5_2109137 | 8 | 0.375 | 6 | 0.3333 | 2 | 0 | 7 | 0.8571 | 23 | 0.4783 | 2 | 0 | 4 | 0 | 8 | 0.125 | 5 | 0.6 | 19 | 0.2105 |
| 5_2109143 | 8 | 0.125 | 7 | 0.1429 | 2 | 0 | 7 | 0.1429 | 24 | 0.125 | 2 | 0 | 4 | 0 | 8 | 0.125 | 6 | 0.1667 | 20 | 0.1 |
| 5_2109152 | 7 | 0.2857 | 7 | 0 | 1 | 0 | 8 | 0.25 | 23 | 0.1739 | 2 | 0 | 5 | 0.2 | 7 | 0.1429 | 5 | 0.2 | 19 | 0.1579 |
| 5_2109153 | 7 | 0.7143 | 7 | 0.8571 | 2 | 1 | 3 | 1 | 19 | 0.8421 | 2 | 1 | 5 | 0.8 | 7 | 1 | 5 | 1 | 19 | 0.9474 |
| 5_2109160 | 7 | 0.7143 | 5 | 0.6 | 3 | 0.6667 | 8 | 0.75 | 23 | 0.6957 | 3 | 0.6667 | 4 | 0.5 | 9 | 0.4444 | 6 | 0.8333 | 22 | 0.5909 |
| 5_2109163 | 6 | 0.8333 | 7 | 1 | 2 | 1 | 6 | 0.8333 | 21 | 0.9048 | 4 | 1 | 2 | 1 | 8 | 1 | 5 | 1 | 19 | 1 |
| 5_2109172 | 8 | 0.75 | 6 | 0.6667 | 2 | 0.5 | 7 | 0.5714 | 23 | 0.6522 | 3 | 0.3333 | 6 | 0.6667 | 11 | 0.8182 | 4 | 0.75 | 24 | 0.7083 |
| 5_2109178 | 8 | 0.375 | 5 | 0.4 | 2 | 0 | 7 | 0.1429 | 22 | 0.2727 | 3 | 0 | 6 | 0.3333 | 8 | 0.5 | 4 | 0.5 | 21 | 0.381 |
| 5_2109980 | NA | NA | 1 | 0 | 2 | 0.5 | 1 | 0 | 4 | 0.25 | 2 | 0.5 | 1 | 0 | NA | NA | NA | NA | 3 | 0.3333 |
| 5_2110117 | NA | NA | 1 | 0 | 2 | 0.5 | 1 | 0 | 4 | 0.25 | 1 | 1 | 1 | 0 | 1 | 0 | 2 | 0 | 5 | 0.2 |
| 5_2110118 | NA | NA | 2 | 1 | 2 | 1 | 2 | 1 | 6 | 1 | NA | NA | 2 | 1 | NA | NA | 2 | 1 | 4 | 1 |
| 5_2110126 | NA | NA | 1 | 0 | 1 | 1 | 1 | 0 | 3 | 0.3333 | 1 | 0 | 2 | 0.5 | 1 | 1 | 2 | 0 | 6 | 0.3333 |
| 5_2110135 | NA | NA | 1 | 1 | 2 | 0.5 | 2 | 1 | 5 | 0.8 | 1 | 1 | 2 | 1 | 1 | 0 | 2 | 1 | 6 | 0.8333 |
| 5_2110139 | NA | NA | 1 | 1 | 2 | 0.5 | NA | NA | 3 | 0.6667 | 1 | 1 | 2 | 0 | NA | NA | 2 | 0.5 | 5 | 0.4 |
| 5_2110141 | NA | NA | NA | NA | 2 | 0.5 | 2 | 1 | 4 | 0.75 | 1 | 1 | 2 | 1 | 1 | 1 | 2 | 1 | 6 | 1 |
| 5_2110151 | NA | NA | 1 | 0 | 2 | 0 | 2 | 0.5 | 5 | 0.2 | 1 | 1 | 2 | 0.5 | 1 | 0 | 2 | 1 | 6 | 0.6667 |
| 5_2110157 | NA | NA | NA | NA | 2 | 0.5 | 2 | 0.5 | 4 | 0.5 | 1 | 1 | 1 | 0 | 1 | 0 | 2 | 0 | 5 | 0.2 |
| 5_2110161 | NA | NA | NA | NA | 2 | 0.5 | 2 | 0.5 | 4 | 0.5 | 1 | 1 | 2 | 1 | NA | NA | 2 | 1 | 5 | 1 |
| 5_2112462 | NA | NA | NA | NA | 1 | 0 | 1 | 1 | 2 | 0.5 | 1 | 0 | 2 | 0.5 | 1 | 1 | 1 | 0 | 5 | 0.4 |
| 5_2112462 | NA | NA | NA | NA | 1 | 1 | 2 | 0.5 | 3 | 0.6667 | 1 | 1 | 1 | 0 | 1 | 0 | 1 | 1 | 4 | 0.5 |
| 5_2112476 | NA | NA | NA | NA | NA | NA | 2 | 1 | 2 | 1 | 1 | 1 | 2 | 1 | 1 | 1 | 1 | 1 | 5 | 1 |
| 5_2112484 | NA | NA | NA | NA | 1 | 0 | 2 | 1 | 3 | 0.6667 | 1 | 0 | 2 | 1 | NA | NA | 1 | 1 | 4 | 0.75 |
| 5_2112490 | NA | NA | NA | NA | NA | NA | 1 | 1 | 1 | 1 | 1 | 1 | 2 | 1 | 1 | 1 | 1 | 1 | 5 | 1 |
| 5_2112493 | NA | NA | NA | NA | 1 | 1 | 2 | 0.5 | 3 | 0.6667 | NA | NA | 1 | 1 | 1 | 1 | NA | NA | 2 | 1 |
| 5_2115666 | NA | NA | 2 | 1 | 2 | 0 | NA | NA | 4 | 0.5 | NA | NA | NA | NA | 1 | 0 | 2 | 0.5 | 3 | 0.3333 |
| 5_2115688 | NA | NA | 1 | 0 | 2 | 1 | 1 | 1 | 4 | 0.75 | NA | NA | NA | NA | 1 | 1 | 2 | 1 | 3 | 1 |
| 5_2115689 | NA | NA | 1 | 1 | 2 | 0 | 1 | 0 | 4 | 0.25 | NA | NA | NA | NA | NA | NA | 2 | 0.5 | 2 | 0.5 |
| 5_2115691 | NA | NA | 1 | 1 | 2 | 1 | 1 | 1 | 4 | 1 | NA | NA | NA | NA | 1 | 1 | 2 | 1 | 3 | 1 |
| 5_2115705 | NA | NA | 2 | 1 | 2 | 0.5 | 1 | 1 | 5 | 0.8 | NA | NA | 2 | 0 | 1 | 1 | 1 | 0 | 4 | 0.25 |
| 5_2115708 | NA | NA | 1 | 1 | 2 | 1 | NA | NA | 3 | 1 | NA | NA | 2 | 0.5 | 1 | 1 | 2 | 1 | 5 | 0.8 |
| 5_2115729 | NA | NA | 2 | 0.5 | 2 | 0 | 1 | 1 | 5 | 0.4 | NA | NA | 2 | 0 | 1 | 1 | 2 | 1 | 5 | 0.6 |
| 5_2115776 | NA | NA | 2 | 1 | 2 | 1 | 1 | 1 | 5 | 1 | NA | NA | 2 | 0 | 1 | 0 | 1 | 1 | 4 | 0.25 |
| 5_2116103 | NA | NA | NA | NA | NA | NA | 1 | 1 | 1 | 1 | NA | NA | NA | NA | 1 | 1 | NA | NA | 1 | 1 |
| 5_2119214 | 4 | 0.5 | 8 | 0.25 | 5 | 0 | 3 | 0 | 20 | 0.2 | 3 | 0 | 5 | 0.2 | 5 | 0 | 5 | 0 | 18 | 0.0556 |
| 5_2119234 | 2 | 0.5 | 3 | 0.6667 | 4 | 0.25 | 3 | 0.6667 | 12 | 0.5 | 2 | 0 | 3 | 1 | 4 | 1 | 4 | 1 | 13 | 0.8462 |
| 5_2119241 | 2 | 0.5 | 7 | 0.1429 | 4 | 0 | 4 | 0 | 17 | 0.1176 | 3 | 0 | 4 | 0.25 | 5 | 0 | 5 | 0 | 17 | 0.0588 |
| 5_2120295 | 3 | 0.3333 | 4 | 1 | 2 | 0.5 | 3 | 0 | 12 | 0.5 | 2 | 1 | 2 | 1 | 2 | 1 | 5 | 0.6 | 11 | 0.8182 |
| 5_2120309 | 2 | 1 | 3 | 0.6667 | 3 | 0.3333 | 2 | 0.5 | 10 | 0.6 | 2 | 0.5 | 2 | 1 | 2 | 1 | 6 | 0.3333 | 12 | 0.5833 |
| 5_2120315 | 2 | 0 | 2 | 0.5 | 3 | 1 | 3 | 0.6667 | 10 | 0.6 | 2 | 1 | 2 | 0.5 | 3 | 0.6667 | 5 | 0.8 | 12 | 0.75 |
| 5_2120318 | 2 | 0 | 2 | 0 | 3 | 0.3333 | 3 | 0 | 10 | 0.1 | 2 | 0.5 | 2 | 0.5 | 2 | 0.5 | 6 | 0.3333 | 12 | 0.4167 |
| 5_2120320 | 2 | 0.5 | 1 | 0 | 3 | 0 | 3 | 0.3333 | 9 | 0.2222 | 2 | 0.5 | 2 | 0 | 3 | 0.3333 | 5 | 0.4 | 12 | 0.3333 |
| 5_2120328 | 1 | 1 | 3 | 1 | 3 | 1 | 3 | 1 | 10 | 1 | 1 | 1 | NA | NA | 3 | 0.6667 | 5 | 0.8 | 9 | 0.7778 |
| 5_2120340 | 1 | 1 | 3 | 0.3333 | 3 | 0.3333 | 3 | 1 | 10 | 0.6 | NA | NA | NA | NA | 3 | 1 | 5 | 0.6 | 8 | 0.75 |

| | | | | | | | | | | | | | | | | | | | | |
|---|---|---|---|---|---|---|---|---|---|---|---|---|---|---|---|---|---|---|---|---|
| 5_2120342 | NA | NA | 2 | 0.5 | 3 | 0 | 1 | 0 | 6 | 0.1667 | NA | NA | NA | NA | 3 | 0.3333 | 5 | 0 | 8 | 0.125 |
| 5_2120353 | 1 | 1 | 3 | 0.3333 | 3 | 1 | 3 | 0.3333 | 10 | 0.6 | 1 | 1 | NA | NA | 2 | 0.5 | 6 | 0.3333 | 9 | 0.4444 |
| 5_2120363 | 1 | 0 | 3 | 0.3333 | 3 | 1 | 3 | 0.3333 | 10 | 0.5 | 1 | 1 | NA | NA | 3 | 0.6667 | 5 | 0.6 | 9 | 0.6667 |
| 5_2157800 | 1 | 1 | 1 | 1 | NA | NA | 1 | 1 | 3 | 1 | 1 | 1 | NA | NA | NA | NA | NA | NA | 1 | 1 |
| 5_2157808 | 1 | 1 | 1 | 1 | NA | NA | 1 | 1 | 3 | 1 | 1 | 1 | NA | NA | NA | NA | NA | NA | 1 | 1 |
| 5_2158055 | 1 | 1 | NA | NA | 1 | 1 | 1 | 1 | 3 | 1 | 1 | 1 | NA | NA | NA | NA | NA | NA | 1 | 1 |
| 5_2158074 | 1 | 1 | NA | NA | 1 | 1 | 1 | 1 | 3 | 1 | 1 | 1 | NA | NA | NA | NA | NA | NA | 1 | 1 |
| 5_2158136 | 1 | 1 | NA | NA | 1 | 1 | 1 | 1 | 3 | 1 | 1 | 1 | NA | NA | NA | NA | NA | NA | 1 | 1 |
| 5_2158327 | 2 | 0 | NA | NA | 2 | 0 | 3 | 0.3333 | 7 | 0.1429 | 1 | 1 | NA | NA | NA | NA | NA | NA | 1 | 1 |
| 5_2168543 | 2 | 1 | 1 | 0 | NA | NA | 3 | 0.6667 | 6 | 0.6667 | 1 | 1 | NA | NA | 2 | 0.5 | NA | NA | 3 | 0.6667 |
| 5_2168547 | 2 | 0.5 | 1 | 1 | NA | NA | 3 | 1 | 6 | 0.8333 | 1 | 1 | NA | NA | 1 | 1 | NA | NA | 2 | 1 |
| 5_2168620 | 2 | 0 | 1 | 1 | NA | NA | 3 | 0.6667 | 6 | 0.5 | 1 | 0 | NA | NA | 1 | 1 | NA | NA | 2 | 0.5 |
| 5_2168671 | 2 | 0 | 3 | 0.3333 | NA | NA | 3 | 0 | 8 | 0.125 | 1 | 0 | NA | NA | 2 | 0.5 | NA | NA | 3 | 0.3333 |
| 5_2169294 | NA | NA | NA | NA | NA | NA | 1 | 1 | 1 | 1 | NA | NA | NA | NA | NA | NA | 1 | 1 | 1 | 1 |
| 5_2169310 | NA | NA | NA | NA | NA | NA | 1 | 1 | 1 | 1 | NA | NA | NA | NA | NA | NA | 2 | 1 | 2 | 1 |
| 5_2186664 | 3 | 1 | 2 | 1 | 2 | 1 | NA | NA | 7 | 1 | NA | NA | 1 | 1 | 1 | 1 | 1 | 1 | 3 | 1 |
| 5_2186687 | 4 | 0.25 | NA | NA | 2 | 0 | 1 | 0 | 7 | 0.1429 | NA | NA | 1 | 0 | 1 | 1 | 1 | 0 | 3 | 0.3333 |
| 5_2186688 | 4 | 0 | 2 | 0.5 | 2 | 0.5 | 1 | 0 | 9 | 0.2222 | NA | NA | 1 | 1 | 1 | 0 | 1 | 0 | 3 | 0.3333 |
| 5_2186690 | 4 | 0.75 | 2 | 1 | 1 | 0 | 1 | 1 | 8 | 0.75 | NA | NA | 1 | 0 | 1 | 1 | 1 | 0 | 3 | 0.3333 |
| 5_2186699 | 5 | 1 | 2 | 0.5 | 2 | 1 | 1 | 1 | 10 | 0.9 | NA | NA | 1 | 1 | NA | NA | 1 | 1 | 2 | 1 |
| 5_2186709 | 5 | 0.8 | 2 | 0 | 1 | 0 | 1 | 1 | 9 | 0.5556 | NA | NA | 1 | 1 | 1 | 0 | 1 | 0 | 3 | 0.3333 |
| 5_2186721 | 5 | 0.8 | 2 | 0.5 | 2 | 1 | 1 | 1 | 10 | 0.8 | NA | NA | 1 | 1 | 1 | 1 | 2 | 1 | 4 | 1 |
| 5_2186746 | 6 | 0.6667 | 1 | 1 | 1 | 1 | 2 | 0.5 | 10 | 0.7 | NA | NA | 2 | 0.5 | 1 | 0 | 1 | 1 | 4 | 0.5 |
| 5_2186749 | 6 | 0.3333 | 2 | 0.5 | 2 | 1 | 1 | 0 | 11 | 0.4545 | NA | NA | NA | NA | 2 | 0.5 | 2 | 0 | 4 | 0.25 |
| 5_2186822 | NA | NA | 3 | 1 | 1 | 1 | NA | NA | 4 | 1 | 2 | 0.5 | NA | NA | NA | NA | NA | NA | 2 | 0.5 |
| 5_2187019 | NA | NA | 2 | 1 | 1 | 1 | NA | NA | 3 | 1 | 1 | 1 | NA | NA | NA | NA | NA | NA | 1 | 1 |
| 5_2187031 | NA | NA | 2 | 0.5 | 1 | 1 | NA | NA | 3 | 0.6667 | 1 | 1 | NA | NA | NA | NA | NA | NA | 1 | 1 |
| 5_2188062 | NA | NA | 2 | 1 | NA | NA | NA | NA | 2 | 1 | 1 | 1 | NA | NA | NA | NA | NA | NA | 1 | 1 |
| 5_2188070 | NA | NA | 1 | 0 | 1 | 1 | NA | NA | 2 | 0.5 | 1 | 1 | NA | NA | NA | NA | NA | NA | 1 | 1 |
| 5_2188074 | NA | NA | 2 | 1 | NA | NA | NA | NA | 2 | 1 | 1 | 1 | NA | NA | NA | NA | NA | NA | 1 | 1 |
| 5_2200862 | 27 | 0.1111 | 16 | 0.0625 | 25 | 0 | 23 | 0.1304 | 91 | 0.0769 | 14 | 0.3571 | 13 | 0.0769 | 16 | 0 | 10 | 0.1 | 53 | 0.1321 |
| 5_2200872 | 23 | 0.0435 | 13 | 0.0769 | 30 | 0.1667 | 22 | 0.0455 | 88 | 0.0909 | 13 | 0.1538 | 12 | 0 | 17 | 0.1765 | 17 | 0.1176 | 59 | 0.1186 |
| 5_2200878 | 12 | 0.25 | 3 | 0.3333 | 12 | 0 | 13 | 0 | 40 | 0.1 | 9 | 0 | 6 | 0.3333 | 7 | 0.2857 | 12 | 0.25 | 34 | 0.2059 |
| 5_2208996 | 3 | 0.3333 | 1 | 1 | NA | NA | 3 | 0 | 7 | 0.2857 | NA | NA | 1 | 0 | 5 | 0 | 3 | 0.6667 | 9 | 0.2222 |
| 5_2209037 | 4 | 1 | 1 | 1 | NA | NA | 2 | 1 | 7 | 1 | NA | NA | 2 | 1 | 4 | 1 | 3 | 0.6667 | 9 | 0.8889 |
| 5_2269183 | 6 | 0.1667 | 5 | 0.2 | 3 | 0 | 2 | 0.5 | 16 | 0.1875 | 3 | 0 | 3 | 0 | 3 | 0 | 5 | 0.2 | 14 | 0.0714 |
| 5_2269185 | 8 | 0.625 | 5 | 0.6 | 3 | 0.6667 | 3 | 0.3333 | 19 | 0.5789 | 3 | 0.3333 | 4 | 1 | 3 | 0.3333 | 6 | 0.3333 | 16 | 0.5 |
| 5_2269214 | 6 | 1 | 5 | 0.8 | 3 | 0.6667 | 3 | 0.6667 | 17 | 0.8235 | 3 | 1 | 4 | 1 | 2 | 0.5 | 6 | 0.6667 | 15 | 0.8 |
| 5_2273326 | 8 | 0.5 | 11 | 0.3636 | 8 | 0.375 | 9 | 0.5556 | 36 | 0.4444 | 5 | 0.8 | 7 | 0.4286 | 6 | 0.6667 | 2 | 0.5 | 20 | 0.6 |
| 5_2281333 | NA | NA | NA | NA | 1 | 1 | NA | NA | 1 | 1 | NA | NA | 1 | 1 | NA | NA | NA | NA | 1 | 1 |
| 5_2281357 | NA | NA | NA | NA | 1 | 1 | NA | NA | 1 | 1 | NA | NA | 1 | 1 | NA | NA | NA | NA | 1 | 1 |
| 5_2281394 | NA | NA | NA | NA | 1 | 1 | NA | NA | 1 | 1 | NA | NA | 1 | 1 | NA | NA | NA | NA | 1 | 1 |
| 5_2294808 | 1 | 1 | NA | NA | NA | NA | NA | NA | 1 | 1 | 1 | 1 | NA | NA | NA | NA | NA | NA | 1 | 1 |
| 5_2294888 | 1 | 1 | NA | NA | NA | NA | NA | NA | 1 | 1 | 1 | 1 | NA | NA | NA | NA | 1 | 1 | 2 | 1 |
| 5_2294934 | 1 | 1 | NA | NA | NA | NA | NA | NA | 1 | 1 | 1 | 1 | NA | NA | NA | NA | 1 | 1 | 2 | 1 |
| 5_2294960 | 1 | 1 | NA | NA | NA | NA | NA | NA | 1 | 1 | 1 | 1 | NA | NA | NA | NA | 1 | 0 | 2 | 0.5 |
| 5_2347851 | 3 | 1 | 9 | 0.7778 | 5 | 0.8 | 3 | 0.6667 | 20 | 0.8 | 2 | 1 | 2 | 1 | 7 | 0.8571 | 6 | 0.6667 | 17 | 0.8235 |
| 5_2347857 | 3 | 0.3333 | 8 | 0.25 | 5 | 0 | 3 | 1 | 19 | 0.3158 | 2 | 0.5 | 2 | 0.5 | 7 | 0.1429 | 3 | 0.3333 | 14 | 0.2857 |
| 5_2347892 | 2 | 0 | 8 | 0.125 | 5 | 0.2 | 3 | 0 | 18 | 0.1111 | 2 | 0 | 2 | 0 | 7 | 0.2857 | 5 | 0.2 | 16 | 0.1875 |
| 5_2363753 | 1 | 0 | 2 | 0 | 2 | 0.5 | 2 | 0.5 | 7 | 0.2857 | 1 | 1 | 2 | 0 | 2 | 0 | NA | NA | 5 | 0.2 |
| 5_2363813 | NA | NA | 1 | 1 | 1 | 1 | NA | NA | 2 | 1 | 1 | 1 | 1 | 1 | 1 | 1 | NA | NA | 3 | 1 |
| 5_2363817 | 1 | 0 | NA | NA | 1 | 1 | 1 | 1 | 3 | 0.6667 | 1 | 1 | 1 | 1 | 1 | 1 | NA | NA | 3 | 1 |
| 5_2363868 | NA | NA | NA | NA | 1 | 1 | NA | NA | 1 | 1 | NA | NA | 1 | 1 | 1 | 1 | NA | NA | 2 | 1 |
| 5_2365522 | 4 | 0 | 3 | 0 | 1 | 1 | 5 | 0 | 13 | 0.0769 | 1 | 1 | 1 | 0 | 4 | 0 | 1 | 0 | 7 | 0.1429 |
| 5_2369052 | 3 | 0.3333 | 2 | 0 | 2 | 0 | 5 | 0.2 | 12 | 0.1667 | 1 | 0 | 1 | 0 | 3 | 0.3333 | 4 | 0.25 | 9 | 0.2222 |
| 5_2369828 | 4 | 0.5 | 4 | 0.25 | 2 | 0.5 | 7 | 0.4286 | 17 | 0.4118 | NA | NA | 1 | 0 | 5 | 0.2 | 3 | 0.3333 | 9 | 0.2222 |
| 5_2369831 | 4 | 0.25 | 3 | 0.3333 | 2 | 0 | 6 | 0.1667 | 15 | 0.2 | 1 | 1 | 1 | 1 | 4 | 0.5 | 2 | 0.5 | 8 | 0.625 |
| 5_2369841 | 4 | 0.25 | 4 | 0.25 | 2 | 0 | 7 | 0.4286 | 17 | 0.2941 | 1 | 1 | 1 | 0 | 4 | 1 | 4 | 0.5 | 10 | 0.7 |
| 5_2370278 | 5 | 0.2 | 6 | 0.1667 | 4 | 0 | 6 | 0.1667 | 21 | 0.1429 | 2 | 0 | 3 | 0 | 5 | 0.2 | 2 | 0 | 12 | 0.0833 |
| 5_2370281 | 3 | 0.6667 | 6 | 1 | 5 | 0.8 | 6 | 0.8333 | 20 | 0.85 | 2 | 1 | 2 | 1 | 5 | 0.4 | 2 | 1 | 11 | 0.7273 |
| 5_2370284 | 4 | 0.5 | 6 | 0 | 4 | 0.75 | 6 | 0.6667 | 20 | 0.45 | 1 | 1 | 3 | 0.6667 | 2 | 0 | 3 | 0.6667 | 9 | 0.5556 |
| 5_2370289 | 4 | 0 | 4 | 0.5 | 5 | 0.2 | 5 | 0.2 | 18 | 0.2222 | 2 | 1 | 2 | 0 | 5 | 0.4 | 2 | 0.5 | 11 | 0.4545 |
| 5_2370296 | 5 | 0.6 | 6 | 0.6667 | 5 | 0.8 | 6 | 0.6667 | 22 | 0.6818 | 2 | 0.5 | 3 | 0.3333 | 6 | 0.5 | 3 | 0.6667 | 14 | 0.5 |
| 5_2370307 | 5 | 0.4 | 6 | 0.6667 | 4 | 0.75 | 5 | 0.8 | 20 | 0.65 | 2 | 0 | 3 | 0.3333 | 6 | 0.3333 | 2 | 0 | 13 | 0.2308 |
| 5_2370323 | 4 | 0.75 | 6 | 0.8333 | 4 | 0.25 | 6 | 0.8333 | 20 | 0.7 | 2 | 0.5 | 4 | 1 | 6 | 1 | 4 | 1 | 16 | 0.9375 |

| | | | | | | | | | | | | | | | | | | | | |
|---|---|---|---|---|---|---|---|---|---|---|---|---|---|---|---|---|---|---|---|---|
| 5_23706659 | 5 | 0.2 | 10 | 0 | 4 | 0.25 | 8 | 0 | 27 | 0.0741 | 2 | 0 | 7 | 0 | 6 | 0.1667 | 7 | 0.1429 | 22 | 0.0909 |
| 5_23706694 | 4 | 0.25 | 9 | 0.2222 | 4 | 0.25 | 6 | 0 | 23 | 0.1739 | 2 | 0 | 6 | 0 | 5 | 0 | 5 | 0.2 | 18 | 0.0556 |
| 5_2370692 | 6 | 0.6667 | 12 | 0.1667 | 4 | 0.5 | 7 | 0.5714 | 29 | 0.4138 | 2 | 0 | 6 | 0.3333 | 5 | 0.6 | 8 | 0.375 | 21 | 0.381 |
| 5_2370693 | 5 | 1 | 11 | 1 | 4 | 1 | 9 | 1 | 29 | 1 | 2 | 1 | 7 | 1 | 6 | 1 | 9 | 0.8889 | 24 | 0.9583 |
| 5_2370694 | 5 | 1 | 12 | 0.6667 | 5 | 0.8 | 9 | 0.4444 | 31 | 0.6774 | 2 | 0 | 6 | 0.5 | 6 | 0.6667 | 7 | 0.8571 | 21 | 0.619 |
| 5_2370701 | 5 | 0 | 14 | 0.0714 | 5 | 0.2 | 8 | 0.125 | 32 | 0.0938 | 2 | 0 | 7 | 0.1429 | 6 | 0 | 9 | 0.1111 | 24 | 0.0833 |
| 5_2370702 | 6 | 1 | 13 | 1 | 6 | 1 | 9 | 0.8889 | 34 | 0.9706 | 2 | 1 | 7 | 0.8571 | 6 | 0.5 | 9 | 1 | 24 | 0.8333 |
| 5_2370714 | 6 | 0.6667 | 14 | 0.2143 | 4 | 0.25 | 8 | 0.625 | 32 | 0.4062 | 3 | 0 | 4 | 0.25 | 7 | 0.1429 | 9 | 0.4444 | 23 | 0.2609 |
| 5_2370718 | 6 | 0.8333 | 12 | 0.8333 | 6 | 0.8333 | 8 | 1 | 32 | 0.875 | 3 | 1 | 9 | 0.8889 | 7 | 1 | 10 | 1 | 29 | 0.9655 |
| 5_2370720 | 6 | 0.5 | 11 | 0.0909 | 5 | 0.4 | 9 | 0.6667 | 31 | 0.3871 | 3 | 0.6667 | 8 | 0.5 | 6 | 0.6667 | 8 | 0.375 | 25 | 0.52 |
| 5_2370725 | 6 | 0.6667 | 14 | 0.4286 | 6 | 0.6667 | 10 | 0.4 | 36 | 0.5 | 3 | 0 | 9 | 0.6667 | 7 | 0.2857 | 10 | 0.7 | 29 | 0.5172 |
| 5_2370755 | 6 | 0.5 | 9 | 0.3333 | 5 | 0.4 | 9 | 0.5556 | 29 | 0.4483 | 3 | 0.6667 | 5 | 0 | 6 | 0.5 | 6 | 0.5 | 20 | 0.4 |
| 5_2370777 | 8 | 0 | 14 | 0.0714 | 7 | 0.1429 | 6 | 0 | 35 | 0.0571 | 3 | 0 | 8 | 0 | 7 | 0.1429 | 7 | 0.1429 | 25 | 0.08 |
| 5_2370778 | 7 | 0 | 16 | 0 | 7 | 0.2857 | 10 | 0.1 | 40 | 0.075 | 3 | 0 | 9 | 0 | 8 | 0.125 | 11 | 0.0909 | 31 | 0.0645 |
| 5_2370785 | 9 | 0.3333 | 15 | 0.3333 | 8 | 0.25 | 10 | 0 | 42 | 0.2381 | 3 | 0 | 9 | 0.3333 | 8 | 0.5 | 11 | 0.4545 | 31 | 0.3871 |
| 5_2370794 | 4 | 0.25 | 7 | 0.4286 | 2 | 0 | 5 | 1 | 18 | 0.5 | 1 | 0 | 5 | 0.4 | 2 | 0.5 | 6 | 0.6667 | 14 | 0.5 |
| 5_2370805 | 9 | 1 | 17 | 1 | 8 | 1 | 12 | 1 | 46 | 1 | 4 | 1 | 9 | 1 | 7 | 1 | 11 | 1 | 31 | 1 |
| 5_2370808 | 9 | 0.4444 | 14 | 0.7143 | 7 | 0.4286 | 9 | 0.3333 | 39 | 0.5128 | 4 | 0.25 | 9 | 0.3333 | 7 | 0.4286 | 8 | 0.5 | 28 | 0.3929 |
| 5_2370852 | 10 | 0.2 | 17 | 0.0588 | 11 | 0.0909 | 11 | 0.0909 | 49 | 0.102 | 5 | 0.2 | 9 | 0.2222 | 10 | 0 | 9 | 0.1111 | 33 | 0.1212 |
| 5_2370854 | 4 | 0.25 | 1 | 0 | NA | NA | 3 | 0.3333 | 8 | 0.25 | 4 | 0 | 2 | 0 | 4 | 0.25 | 2 | 0 | 12 | 0.0833 |
| 5_2370860 | 13 | 0.0769 | 18 | 0.1667 | 11 | 0.0909 | 14 | 0.0714 | 56 | 0.1071 | 5 | 0 | 9 | 0.2222 | 12 | 0.25 | 10 | 0.4 | 36 | 0.25 |
| 5_2371066 | 10 | 0.3 | 7 | 0.2857 | 4 | 0 | 7 | 0.2857 | 28 | 0.25 | 7 | 0.1429 | 6 | 0.1667 | 6 | 0 | 4 | 0 | 23 | 0.087 |
| 5_2371238 | 10 | 0.2 | 18 | 0.2222 | 16 | 0.0625 | 17 | 0.0588 | 61 | 0.1311 | 12 | 0.1667 | 19 | 0.0526 | 12 | 0.0833 | 11 | 0.1818 | 54 | 0.1111 |
| 5_2371242 | 9 | 0.2222 | 18 | 0.2778 | 14 | 0.3571 | 16 | 0.1875 | 57 | 0.2632 | 9 | 0.3333 | 19 | 0.1579 | 13 | 0.5385 | 13 | 0.3077 | 54 | 0.3148 |
| 5_2371248 | 9 | 0.7778 | 17 | 0.8235 | 14 | 1 | 18 | 0.7222 | 58 | 0.8276 | 9 | 0.7778 | 18 | 0.8889 | 12 | 0.8333 | 12 | 0.9167 | 51 | 0.8627 |
| 5_2372918 | 5 | 0.4 | 17 | 0.0588 | 15 | 0.1333 | 14 | 0 | 51 | 0.098 | 9 | 0.1111 | 16 | 0 | 9 | 0.1111 | 12 | 0.0833 | 46 | 0.0652 |
| 5_2372930 | 3 | 0.6667 | 17 | 0.5294 | 14 | 0.4286 | 12 | 0.5 | 46 | 0.5 | 7 | 0.4286 | 14 | 0.5714 | 8 | 0.375 | 10 | 0.5 | 39 | 0.4872 |
| 5_2372931 | 5 | 1 | 18 | 0.6111 | 14 | 0.5 | 13 | 0.4615 | 50 | 0.58 | 8 | 0.75 | 15 | 0.8 | 8 | 0.75 | 11 | 0.7273 | 42 | 0.7619 |
| 5_2372934 | 1 | 1 | 10 | 0.7 | 4 | 0.5 | 6 | 0.6667 | 21 | 0.6667 | 5 | 1 | 8 | 0.625 | 6 | 0.5 | 2 | 1 | 21 | 0.7143 |
| 5_2372942 | 3 | 1 | 15 | 0.1333 | 12 | 0.3333 | 9 | 0.2222 | 39 | 0.2821 | 6 | 0.5 | 14 | 0.1429 | 9 | 0 | 8 | 0.25 | 37 | 0.1892 |
| 5_2372953 | 5 | 0.2 | 16 | 0.3125 | 11 | 0.5455 | 10 | 0.3 | 42 | 0.3571 | 7 | 0.5714 | 14 | 0.4286 | 10 | 0.2 | 9 | 0.1111 | 40 | 0.325 |
| 5_2394559 | 1 | 0 | 1 | 1 | NA | NA | NA | NA | 2 | 0.5 | NA | NA | NA | NA | 1 | 1 | NA | NA | 1 | 1 |
| 5_2394680 | 2 | 0.5 | 5 | 0 | NA | NA | 2 | 0 | 9 | 0.1111 | 1 | 0 | 2 | 0.5 | 1 | 0 | 2 | 0 | 6 | 0.1667 |
| 5_2394688 | 2 | 0.5 | 3 | 0 | NA | NA | 2 | 0 | 7 | 0.1429 | 1 | 0 | 2 | 0.5 | 1 | 0 | 2 | 0.5 | 6 | 0.3333 |
| 5_2394694 | 2 | 0 | 4 | 0 | NA | NA | 2 | 0.5 | 8 | 0.125 | 1 | 0 | 2 | 1 | 1 | 0 | 1 | 1 | 5 | 0.6 |
| 5_2404480 | 40 | 0.1 | 55 | 0.0909 | 33 | 0.0909 | 48 | 0 | 176 | 0.0682 | 45 | 0.0889 | 49 | 0.1429 | 36 | 0.1667 | 45 | 0.0889 | 175 | 0.12 |
| 5_2416630 | 3 | 1 | 6 | 0.5 | 4 | 0.5 | 6 | 0 | 19 | 0.4211 | 3 | 0.6667 | 4 | 0 | 4 | 0.25 | 3 | 0.6667 | 14 | 0.3571 |
| 5_2416697 | 1 | 0 | 4 | 0 | 3 | 0.3333 | 6 | 0.1667 | 14 | 0.1429 | 3 | 0.3333 | 4 | 0 | 3 | 0 | 3 | 0 | 13 | 0.0769 |
| 5_2416711 | 2 | 0.5 | 4 | 0 | 3 | 0 | 5 | 0.2 | 14 | 0.1429 | 3 | 0 | 3 | 0 | 3 | 0.3333 | 3 | 0.3333 | 12 | 0.1667 |
| 5_2416889 | 1 | 0 | 4 | 0 | 2 | 0 | 4 | 0.25 | 11 | 0.0909 | 1 | 0 | 3 | 0 | 3 | 0 | 3 | 0.3333 | 10 | 0.1 |
| 5_2416917 | 1 | 1 | 1 | 0 | 2 | 0 | 3 | 0.6667 | 7 | 0.4286 | 1 | 0 | 3 | 0 | 2 | 0 | 2 | 0.5 | 8 | 0.125 |
| 5_2416978 | 1 | 0 | NA | NA | 2 | 0.5 | 4 | 0.25 | 7 | 0.2857 | NA | NA | 2 | 0 | 1 | 0 | 2 | 0.5 | 5 | 0.2 |
| 5_2416981 | 1 | 0 | 1 | 0 | 2 | 0 | 4 | 0.5 | 8 | 0.25 | NA | NA | 2 | 0 | 1 | 1 | 2 | 0.5 | 5 | 0.4 |
| 5_2416985 | NA | NA | 1 | 1 | 1 | 1 | 2 | 1 | 4 | 1 | NA | NA | 3 | 0.6667 | 1 | 1 | 1 | 1 | 5 | 0.8 |
| 5_2417001 | 1 | 0 | NA | NA | 2 | 0 | 4 | 0.75 | 7 | 0.4286 | NA | NA | 3 | 0 | 1 | 0 | 1 | 1 | 5 | 0.2 |
| 5_2417007 | NA | NA | NA | NA | 2 | 1 | 4 | 1 | 6 | 1 | NA | NA | 3 | 1 | NA | NA | NA | NA | 3 | 1 |
| 5_2417032 | 1 | 0 | NA | NA | 2 | 0 | 4 | 0.5 | 7 | 0.2857 | NA | NA | 2 | 0.5 | NA | NA | 1 | 0 | 3 | 0.3333 |
| 5_2432287 | 8 | 0.125 | 6 | 0.3333 | 2 | 0 | 3 | 0 | 19 | 0.1579 | 7 | 0.1429 | 2 | 0.5 | 6 | 0 | 6 | 0.1667 | 21 | 0.1429 |
| 5_2435419 | 12 | 0.1667 | 10 | 0.2 | 9 | 0.1111 | 16 | 0.125 | 47 | 0.1489 | 5 | 0.2 | 8 | 0 | 12 | 0.3333 | 17 | 0.1765 | 42 | 0.1905 |
| 5_2436471 | 17 | 0.1765 | 18 | 0.1111 | 12 | 0.0833 | 19 | 0.1053 | 66 | 0.1212 | 7 | 0.4286 | 12 | 0.1667 | 16 | 0.1875 | 19 | 0.3158 | 54 | 0.2593 |
| 5_2436474 | 14 | 0.7857 | 18 | 0.7222 | 11 | 1 | 17 | 0.8235 | 60 | 0.8167 | 7 | 0.8571 | 12 | 0.6667 | 15 | 0.8 | 20 | 0.95 | 54 | 0.8333 |
| 5_2436491 | 18 | 0.1667 | 16 | 0.3125 | 12 | 0.0833 | 17 | 0 | 63 | 0.1429 | 7 | 0 | 13 | 0 | 13 | 0.0769 | 21 | 0.1429 | 54 | 0.0741 |
| 5_2436497 | 11 | 0.0909 | 13 | 0.1538 | 9 | 0 | 15 | 0 | 48 | 0.0625 | 7 | 0 | 5 | 0 | 10 | 0.2 | 15 | 0 | 37 | 0.0541 |
| 5_2436527 | 19 | 0.0526 | 17 | 0.0588 | 13 | 0.0769 | 22 | 0.0909 | 71 | 0.0704 | 6 | 0 | 16 | 0 | 15 | 0.2667 | 19 | 0.0526 | 56 | 0.0893 |
| 5_2438707 | 1 | 0 | NA | NA | 1 | 1 | NA | NA | 2 | 0.5 | 4 | 0.25 | 1 | 0 | 1 | 0 | 4 | 1 | 10 | 0.5 |
| 5_2438739 | 2 | 0.5 | NA | NA | 1 | 0 | NA | NA | 3 | 0.3333 | 4 | 0.5 | 1 | 1 | 3 | 0.3333 | 4 | 0.25 | 12 | 0.4167 |
| 5_2438746 | 2 | 0.5 | NA | NA | 1 | 0 | NA | NA | 3 | 0.3333 | 3 | 1 | 1 | 1 | 3 | 0.6667 | 2 | 1 | 9 | 0.8889 |
| 5_2438757 | 1 | 0 | NA | NA | 1 | 1 | NA | NA | 2 | 0.5 | 3 | 0.3333 | NA | NA | 2 | 0 | 4 | 1 | 9 | 0.5556 |
| 5_2438784 | NA | NA | NA | NA | 1 | 1 | NA | NA | 1 | 1 | 2 | 0.5 | 1 | 1 | 2 | 0 | 2 | 0 | 7 | 0.2857 |
| 5_2452471 | 1 | 1 | 6 | 1 | 2 | 1 | 5 | 0.6 | 14 | 0.8571 | 4 | 0.75 | 3 | 0.6667 | 2 | 1 | 5 | 1 | 14 | 0.8571 |
| 5_2452482 | NA | NA | 4 | 0.5 | 3 | 0.3333 | 4 | 0 | 11 | 0.2727 | 2 | 0.5 | 2 | 0.5 | NA | NA | 1 | 1 | 5 | 0.6 |
| 5_2452487 | 3 | 0 | 5 | 0.4 | 1 | 0 | 5 | 0.2 | 14 | 0.2143 | 2 | 0.5 | 3 | 0 | 1 | 1 | 5 | 0.2 | 11 | 0.2727 |
| 5_2452497 | 1 | 1 | 3 | 0.6667 | 1 | 1 | 5 | 0.6 | 10 | 0.7 | 3 | 1 | 3 | 0.6667 | 2 | 1 | 5 | 1 | 13 | 0.9231 |
| 5_2453061 | 1 | 0 | 3 | 0.6667 | 1 | 0 | 4 | 0.25 | 9 | 0.3333 | 2 | 0.5 | 2 | 1 | 1 | 0 | 2 | 0.5 | 7 | 0.5714 |
| 5_2453069 | 1 | 1 | NA | NA | NA | NA | 4 | 1 | 5 | 1 | 2 | 1 | 2 | 1 | NA | NA | 3 | 1 | 7 | 1 |

| | | | | | | | | | | | | | | | | | | | | |
|---|---|---|---|---|---|---|---|---|---|---|---|---|---|---|---|---|---|---|---|---|
| 5_24530818 | 1 | 0 | 2 | 0 | NA | NA | 3 | 0.3333 | 6 | 0.1667 | 2 | 0 | 2 | 0.5 | 1 | 0 | 3 | 0 | 8 | 0.125 |
| 5_24531507 | 1 | 1 | 2 | 0.5 | NA | NA | 3 | 1 | 6 | 0.8333 | 3 | 0.6667 | 1 | 1 | 1 | 0 | 2 | 1 | 7 | 0.7143 |
| 5_24531525 | 1 | 1 | 3 | 0 | NA | NA | 3 | 0 | 7 | 0.1429 | 3 | 0.3333 | 1 | 0 | 1 | 0 | 2 | 0.5 | 7 | 0.2857 |
| 5_24531537 | 1 | 1 | 3 | 0.3333 | NA | NA | 3 | 0.3333 | 7 | 0.4286 | 3 | 0 | 1 | 1 | NA | NA | 2 | 0 | 6 | 0.1667 |
| 5_24531573 | 1 | 1 | 3 | 0.3333 | 1 | 0 | 2 | 0 | 7 | 0.2857 | 3 | 0.3333 | 1 | 0 | 1 | 0 | 2 | 0.5 | 7 | 0.2857 |
| 5_24531672 | 1 | 0 | 3 | 0 | NA | NA | 2 | 0.5 | 6 | 0.1667 | 2 | 0 | 1 | 1 | 1 | 1 | 2 | 1 | 6 | 0.6667 |
| 5_24536927 | NA | NA | 3 | 0.3333 | NA | NA | NA | NA | 3 | 0.3333 | NA | NA | 1 | 0 | 1 | 0 | 2 | 0.5 | 4 | 0.25 |
| 5_24588848 | NA | NA | NA | NA | NA | NA | 3 | 0.6667 | 3 | 0.6667 | 2 | 0.5 | 1 | 0 | NA | NA | 2 | 0.5 | 5 | 0.4 |
| 5_24589216 | NA | NA | NA | NA | NA | NA | 4 | 0.5 | 4 | 0.5 | 3 | 0.6667 | 4 | 1 | NA | NA | 2 | 1 | 9 | 0.8889 |
| 5_24589610 | 1 | 0 | 2 | 0.5 | NA | NA | 2 | 0 | 5 | 0.2 | NA | NA | NA | NA | 1 | 1 | 1 | 0 | 2 | 0.5 |
| 5_24614553 | 10 | 0.2 | 3 | 0 | 2 | 0 | 4 | 0.25 | 19 | 0.1579 | NA | NA | 5 | 0.2 | 5 | 0 | 5 | 0 | 15 | 0.0667 |
| 5_24614673 | 11 | 0.0909 | 5 | 0 | 4 | 0.25 | 4 | 0.5 | 24 | 0.1667 | NA | NA | 6 | 0.6667 | 6 | 0 | 6 | 0.1667 | 18 | 0.2778 |
| 5_24614708 | 11 | 0.0909 | 4 | 0.25 | 4 | 0 | 4 | 0.25 | 23 | 0.1304 | NA | NA | 6 | 0.3333 | 6 | 0.1667 | 8 | 0.125 | 20 | 0.2 |
| 5_24614721 | 13 | 0.6154 | 4 | 0.75 | 5 | 0.2 | 4 | 0.75 | 26 | 0.5769 | NA | NA | 4 | 0.75 | 6 | 0.3333 | 8 | 0.75 | 18 | 0.6111 |
| 5_24614795 | 13 | 0.0769 | 3 | 0.3333 | 5 | 0.2 | 4 | 0.5 | 25 | 0.2 | NA | NA | 4 | 0 | 6 | 0 | 5 | 0.4 | 15 | 0.1333 |
| 5_24614841 | 10 | 0.1 | 5 | 0.4 | 2 | 0.5 | 2 | 0 | 19 | 0.2105 | NA | NA | 2 | 0 | 5 | 0 | 5 | 0.2 | 12 | 0.0833 |
| 5_24616785 | NA | NA | 1 | 1 | NA | NA | NA | NA | 1 | 1 | 1 | 1 | NA | NA | NA | NA | NA | NA | 1 | 1 |
| 5_24623409 | 4 | 0 | 10 | 0.1 | 3 | 0 | 9 | 0.1111 | 26 | 0.0769 | 3 | 0 | 2 | 0 | 2 | 0 | 5 | 0.4 | 12 | 0.1667 |
| 5_24623549 | 4 | 0.25 | 9 | 0.8889 | 4 | 0.25 | 6 | 0.3333 | 23 | 0.5217 | 4 | 1 | 3 | 0.6667 | 1 | 1 | 5 | 0.6 | 13 | 0.7692 |
| 5_24624219 | 16 | 0.375 | 26 | 0.3846 | 20 | 0.35 | 19 | 0.4211 | 81 | 0.3827 | 20 | 0.5 | 16 | 0.375 | 19 | 0.5263 | 10 | 0.8 | 65 | 0.5231 |
| 5_24629064 | 1 | 1 | NA | NA | 1 | 1 | NA | NA | 2 | 1 | 1 | 1 | 1 | 1 | NA | NA | NA | NA | 2 | 1 |
| 5_24640183 | 1 | 0 | NA | NA | 2 | 0 | 1 | 1 | 4 | 0.25 | 2 | 0 | 1 | 1 | 1 | 0 | NA | NA | 4 | 0.25 |
| 5_24641904 | 1 | 0 | NA | NA | 2 | 0 | 5 | 0.4 | 8 | 0.25 | NA | NA | NA | NA | NA | NA | 4 | 0.25 | 4 | 0.25 |
| 5_24643193 | NA | NA | NA | NA | 1 | 0 | 2 | 0.5 | 3 | 0.3333 | NA | NA | 1 | 0 | NA | NA | 4 | 0.25 | 5 | 0.2 |
| 5_24643479 | NA | NA | NA | NA | 2 | 0.5 | 2 | 0.5 | 4 | 0.5 | NA | NA | NA | NA | NA | NA | 3 | 0.6667 | 3 | 0.6667 |
| 5_24643529 | NA | NA | NA | NA | 1 | 0 | 3 | 0.6667 | 4 | 0.5 | NA | NA | 1 | 1 | NA | NA | 3 | 0.3333 | 4 | 0.5 |
| 5_24643550 | NA | NA | NA | NA | 1 | 1 | 2 | 0.5 | 3 | 0.6667 | NA | NA | 1 | 1 | NA | NA | 2 | 0.5 | 3 | 0.6667 |
| 5_24643555 | NA | NA | NA | NA | 1 | 1 | 2 | 0.5 | 3 | 0.6667 | NA | NA | 1 | 0 | NA | NA | 3 | 0.6667 | 4 | 0.5 |
| 5_24643589 | NA | NA | NA | NA | 1 | 1 | 1 | 1 | 2 | 1 | NA | NA | 1 | 1 | NA | NA | 3 | 0.6667 | 4 | 0.75 |
| 5_24643615 | NA | NA | NA | NA | 1 | 0 | 3 | 0.6667 | 4 | 0.5 | NA | NA | 1 | 1 | NA | NA | 3 | 0.6667 | 4 | 0.75 |
| 5_24643674 | NA | NA | NA | NA | 2 | 0.5 | 3 | 0.6667 | 5 | 0.6 | NA | NA | 1 | 1 | NA | NA | 3 | 1 | 4 | 1 |
| 5_24643918 | NA | NA | NA | NA | 2 | 0.5 | 2 | 0.5 | 4 | 0.5 | NA | NA | 1 | 1 | NA | NA | 3 | 1 | 4 | 1 |
| 5_24644026 | NA | NA | NA | NA | 2 | 1 | 3 | 0.6667 | 5 | 0.8 | NA | NA | 1 | 1 | NA | NA | 3 | 1 | 4 | 1 |
| 5_24644170 | NA | NA | NA | NA | NA | NA | 2 | 0.5 | 2 | 0.5 | NA | NA | 1 | 1 | NA | NA | NA | NA | 1 | 1 |
| 5_24644608 | NA | NA | NA | NA | NA | NA | 3 | 0.6667 | 3 | 0.6667 | NA | NA | 1 | 1 | NA | NA | 2 | 1 | 3 | 1 |
| 5_24644627 | NA | NA | NA | NA | 1 | 1 | 3 | 0 | 4 | 0.25 | NA | NA | 1 | 1 | NA | NA | 2 | 0.5 | 3 | 0.6667 |
| 5_24644863 | NA | NA | NA | NA | 1 | 1 | 1 | 0 | 2 | 0.5 | NA | NA | NA | NA | NA | NA | 1 | 1 | 1 | 1 |
| 5_24648125 | 8 | 0.25 | 10 | 0 | 11 | 0.0909 | 13 | 0.1538 | 42 | 0.119 | 7 | 0 | 7 | 0.4286 | 7 | 0 | 12 | 0.0833 | 33 | 0.1212 |
| 5_24648438 | 7 | 0.2857 | 7 | 0.1429 | 7 | 0 | 11 | 0.0909 | 32 | 0.125 | 6 | 0 | 2 | 0 | 6 | 0.1667 | 8 | 0.25 | 22 | 0.1364 |
| 5_24648976 | 6 | 0.3333 | 10 | 0.4 | 8 | 0.125 | 9 | 0.3333 | 33 | 0.303 | 8 | 0.375 | 5 | 0.2 | 6 | 0 | 9 | 0.3333 | 28 | 0.25 |
| 5_24649194 | 7 | 0.1429 | 10 | 0.1 | 8 | 0.125 | 9 | 0.2222 | 34 | 0.1471 | 5 | 0.2 | 5 | 0 | 5 | 0 | 9 | 0.1111 | 24 | 0.0833 |
| 5_24650546 | 7 | 0.7143 | 4 | 1 | 6 | 0.8333 | 2 | 1 | 19 | 0.8421 | 1 | 1 | 2 | 1 | 2 | 1 | 2 | 1 | 7 | 1 |
| 5_24650554 | 6 | 0 | 5 | 0 | 6 | 0.5 | 1 | 0 | 18 | 0.1667 | 1 | 1 | 1 | 1 | 1 | 0 | 2 | 1 | 5 | 0.8 |
| 5_24650705 | 6 | 0.5 | 5 | 0.2 | 5 | 0.8 | 2 | 0 | 18 | 0.4444 | 1 | 1 | 2 | 1 | 2 | 1 | 2 | 0.5 | 7 | 0.8571 |
| 5_24650774 | 6 | 0.6667 | 4 | 0.75 | 4 | 1 | 3 | 0.6667 | 17 | 0.7647 | 1 | 1 | 2 | 0.5 | 2 | 1 | 2 | 0 | 7 | 0.5714 |
| 5_24650879 | 6 | 0.8333 | 5 | 0.6 | 5 | 1 | 3 | 0.6667 | 19 | 0.7895 | 1 | 1 | 2 | 1 | 2 | 1 | 2 | 1 | 7 | 1 |
| 5_24651884 | 4 | 0.75 | 3 | 0.6667 | 2 | 0.5 | 1 | 0 | 10 | 0.6 | NA | NA | 1 | 1 | 2 | 1 | 2 | 0 | 5 | 0.6 |
| 5_24657209 | 13 | 0.0769 | 11 | 0 | 13 | 0.1538 | 9 | 0.2222 | 46 | 0.1087 | 9 | 0 | 21 | 0.0952 | 15 | 0.2 | 11 | 0 | 56 | 0.0893 |
| 5_24687358 | 11 | 0.8182 | 16 | 0.8125 | 17 | 0.8824 | 9 | 0.6667 | 53 | 0.8113 | 16 | 0.9375 | 23 | 0.9565 | 25 | 0.8 | 12 | 0.75 | 76 | 0.8684 |
| 5_24687399 | 19 | 0.8947 | 20 | 0.65 | 22 | 0.9091 | 16 | 0.8125 | 77 | 0.8182 | 17 | 0.6471 | 26 | 0.7308 | 28 | 0.7857 | 15 | 0.8667 | 86 | 0.7558 |
| 5_24687444 | 23 | 0.1739 | 23 | 0.087 | 25 | 0.16 | 17 | 0.1176 | 88 | 0.1364 | 23 | 0.0435 | 32 | 0.1875 | 32 | 0.125 | 17 | 0.2353 | 104 | 0.1442 |
| 5_24687454 | 16 | 0.3125 | 21 | 0.3333 | 26 | 0.4231 | 15 | 0.2 | 78 | 0.3333 | 13 | 0.3846 | 29 | 0.5172 | 26 | 0.2692 | 15 | 0.3333 | 83 | 0.3855 |
| 5_24687463 | 12 | 0.0833 | 20 | 0.1 | 18 | 0.1111 | 16 | 0.0625 | 66 | 0.0909 | 14 | 0.0714 | 22 | 0.0455 | 23 | 0.1304 | 12 | 0 | 71 | 0.0704 |
| 5_24687524 | 21 | 0.1905 | 19 | 0.1053 | 26 | 0 | 14 | 0.0714 | 80 | 0.0875 | 22 | 0.0455 | 25 | 0.08 | 25 | 0.08 | 14 | 0.0714 | 86 | 0.0698 |
| 5_24687701 | 19 | 1 | 28 | 0.8571 | 32 | 0.875 | 17 | 0.9412 | 96 | 0.9062 | 25 | 0.96 | 31 | 0.9355 | 31 | 0.871 | 16 | 0.9375 | 103 | 0.9223 |
| 5_24687807 | 9 | 1 | 14 | 0.8571 | 20 | 0.85 | 14 | 0.7857 | 57 | 0.8596 | 19 | 0.7368 | 20 | 1 | 26 | 0.7692 | 11 | 0.9091 | 76 | 0.8421 |
| 5_24687955 | 21 | 1 | 20 | 1 | 21 | 1 | 15 | 0.9333 | 77 | 0.987 | 24 | 0.9583 | 29 | 0.9655 | 27 | 0.8889 | 16 | 1 | 96 | 0.9479 |
| 5_24706430 | 6 | 0 | 13 | 0.2308 | 5 | 0 | 1 | 0 | 25 | 0.12 | 4 | 0.5 | 9 | 0 | 9 | 0.1111 | 5 | 0.2 | 27 | 0.1481 |
| 5_24706574 | 6 | 0 | 13 | 0.4615 | 4 | 0.5 | 2 | 0 | 25 | 0.32 | 5 | 0.2 | 12 | 0.25 | 10 | 0.1 | 6 | 0.1667 | 33 | 0.1818 |
| 5_24706633 | 6 | 0 | 13 | 0.3846 | 5 | 0.2 | 2 | 0.5 | 26 | 0.2692 | 5 | 0.2 | 12 | 0.25 | 11 | 0.1818 | 7 | 0 | 35 | 0.1714 |
| 5_24706783 | 6 | 0.1667 | 14 | 0.0714 | 4 | 0 | 2 | 0 | 26 | 0.0769 | 5 | 0 | 10 | 0.1 | 12 | 0.25 | 4 | 0.25 | 31 | 0.1613 |
| 5_24770649 | 19 | 0.9474 | 17 | 0.8824 | 10 | 0.9 | 9 | 0.7778 | 55 | 0.8909 | 11 | 0.8182 | 17 | 0.8235 | 17 | 0.8235 | 14 | 0.9286 | 59 | 0.8475 |
| 5_24770701 | 12 | 0.0833 | 13 | 0.3077 | 4 | 0 | 8 | 0 | 37 | 0.1351 | 8 | 0.125 | 16 | 0.25 | 12 | 0.1667 | 10 | 0.2 | 46 | 0.1957 |
| 5_24770788 | 18 | 0.2222 | 15 | 0.2 | 10 | 0 | 8 | 0.125 | 51 | 0.1569 | 11 | 0.2727 | 18 | 0.2222 | 16 | 0.25 | 12 | 0.25 | 57 | 0.2456 |
| 5_24770793 | 13 | 0.1538 | 16 | 0.0625 | 9 | 0.1111 | 7 | 0.2857 | 45 | 0.1333 | 9 | 0.1111 | 17 | 0.1176 | 13 | 0.1538 | 12 | 0 | 51 | 0.098 |

| | | | | | | | | | | | | | | | | | | | | |
|---|---|---|---|---|---|---|---|---|---|---|---|---|---|---|---|---|---|---|---|---|
| 5_2477082 | 15 | 0.0667 | 16 | 0.125 | 8 | 0 | 9 | 0.1111 | 48 | 0.0833 | 11 | 0.0909 | 19 | 0.2105 | 17 | 0.1765 | 13 | 0.1538 | 60 | 0.1667 |
| 5_2477740 | NA | NA | NA | NA | NA | NA | 1 | 1 | 1 | 1 | NA | NA | NA | NA | NA | NA | 1 | 1 | 1 | 1 |
| 5_2477864 | 11 | 0.0909 | 7 | 0 | 5 | 0 | 5 | 0.4 | 28 | 0.1071 | 5 | 0.2 | 10 | 0.2 | 8 | 0 | 6 | 0 | 29 | 0.1034 |
| 5_2478648 | 43 | 0.1163 | 58 | 0.0862 | 38 | 0.0789 | 48 | 0.1042 | 187 | 0.0963 | 41 | 0.0488 | 58 | 0.0517 | 37 | 0.027 | 41 | 0.0976 | 177 | 0.0565 |
| 5_2479202 | 1 | 0 | 1 | 1 | NA | NA | 2 | 0 | 4 | 0.25 | 1 | 1 | 2 | 0.5 | 1 | 1 | NA | NA | 4 | 0.75 |
| 5_2479277 | 2 | 0.5 | 1 | 0 | NA | NA | NA | NA | 3 | 0.3333 | 1 | 0 | 2 | 0.5 | 1 | 0 | NA | NA | 4 | 0.25 |
| 5_2479288 | 2 | 0 | 2 | 0.5 | NA | NA | 1 | 0 | 5 | 0.2 | 2 | 0.5 | 2 | 0 | 1 | 0 | NA | NA | 5 | 0.2 |
| 5_2479292 | 2 | 0 | 2 | 1 | NA | NA | 1 | 0 | 5 | 0.4 | 1 | 1 | 2 | 0.5 | 1 | 0 | NA | NA | 4 | 0.5 |
| 5_2479299 | 2 | 0.5 | 1 | 1 | NA | NA | 1 | 1 | 4 | 0.75 | 2 | 1 | 2 | 0.5 | 1 | 1 | NA | NA | 5 | 0.8 |
| 5_2479309 | 2 | 1 | 3 | 1 | NA | NA | 2 | 0.5 | 7 | 0.8571 | 1 | 1 | 2 | 0.5 | 1 | 1 | NA | NA | 4 | 0.75 |
| 5_2479435 | 2 | 0 | 3 | 0.6667 | 1 | 0 | 3 | 0.3333 | 9 | 0.3333 | 2 | 0 | 2 | 0.5 | 1 | 0 | 1 | 0 | 6 | 0.1667 |
| 5_2479457 | 1 | 1 | 2 | 0 | 1 | 0 | 4 | 0 | 8 | 0.125 | 2 | 0.5 | 2 | 0.5 | 1 | 1 | 2 | 0 | 7 | 0.4286 |
| 5_2479618 | 2 | 0.5 | 5 | 0.6 | 3 | 0 | 4 | 1 | 14 | 0.5714 | 4 | 0.5 | 2 | 0.5 | NA | NA | 2 | 1 | 8 | 0.625 |
| 5_2479627 | 2 | 0.5 | 4 | 1 | 3 | 0.3333 | 3 | 0.3333 | 12 | 0.5833 | 4 | 0.5 | 4 | 0.25 | 1 | 1 | 1 | 0 | 10 | 0.4 |
| 5_2479641 | 1 | 0 | 4 | 0.75 | 1 | 0 | 4 | 0.25 | 10 | 0.4 | 2 | 0.5 | 3 | 0.6667 | 2 | 0 | 1 | 1 | 8 | 0.5 |
| 5_2479661 | 2 | 1 | 5 | 0.8 | 2 | 0.5 | 3 | 0.6667 | 12 | 0.75 | 3 | 0.6667 | 5 | 1 | 3 | 0.3333 | 1 | 0 | 12 | 0.6667 |
| 5_2479681 | 2 | 1 | 5 | 1 | 2 | 1 | 3 | 0.6667 | 12 | 0.9167 | 3 | 0.6667 | 6 | 0.8333 | 4 | 0.75 | 2 | 1 | 15 | 0.8 |
| 5_2479684 | 2 | 1 | 3 | 0.3333 | 3 | 1 | 3 | 0.3333 | 11 | 0.6364 | 3 | 0.6667 | 5 | 0.2 | 3 | 0.3333 | 1 | 0 | 12 | 0.3333 |
| 5_2479684 | 2 | 0.5 | 5 | 0 | 3 | 0 | 3 | 0 | 13 | 0.0769 | 3 | 0.6667 | 5 | 0 | 4 | 0.25 | 2 | 0 | 14 | 0.2143 |
| 5_2479700 | 3 | 1 | 4 | 0.75 | 1 | 1 | 4 | 1 | 12 | 0.9167 | 4 | 0.75 | 4 | 0.75 | 4 | 0.75 | 2 | 1 | 14 | 0.7857 |
| 5_2479707 | 3 | 0.3333 | 5 | 0.2 | 2 | 0 | 3 | 0.3333 | 13 | 0.2308 | 3 | 0 | 4 | 0.25 | 3 | 0 | 2 | 0 | 12 | 0.0833 |
| 5_2479711 | 2 | 1 | 4 | 0.75 | 2 | 1 | 4 | 1 | 12 | 0.9167 | 4 | 1 | 4 | 1 | 3 | 0.3333 | 1 | 1 | 12 | 0.8333 |
| 5_2479723 | 2 | 1 | 5 | 0.8 | 3 | 1 | 5 | 1 | 15 | 0.9333 | 3 | 1 | 5 | 0.8 | 5 | 1 | 2 | 1 | 15 | 0.9333 |
| 5_2479726 | 3 | 1 | 5 | 1 | 4 | 1 | 3 | 1 | 15 | 1 | 4 | 1 | 4 | 1 | 5 | 0.8 | 2 | 1 | 15 | 0.9333 |
| 5_2479733 | 2 | 0 | 2 | 0 | 3 | 0.3333 | 4 | 0 | 11 | 0.0909 | 1 | 0 | 4 | 0 | 5 | 0.2 | 2 | 0 | 12 | 0.0833 |
| 5_2479741 | 3 | 1 | 5 | 1 | 2 | 0.5 | 3 | 1 | 13 | 0.9231 | 3 | 1 | 4 | 1 | 2 | 1 | 2 | 1 | 11 | 1 |
| 5_2479768 | 13 | 0.2308 | 11 | 0.0909 | 12 | 0.1667 | 12 | 0.3333 | 48 | 0.2083 | 9 | 0 | 16 | 0.0625 | 15 | 0.2 | 17 | 0.1765 | 57 | 0.1228 |
| 5_2479832 | 1 | 0 | NA | NA | 2 | 0.5 | 1 | 0 | 4 | 0.25 | NA | NA | NA | NA | 1 | 1 | NA | NA | 1 | 1 |
| 5_2480234 | NA | NA | NA | NA | NA | NA | 1 | 1 | 1 | 1 | 1 | 1 | NA | NA | NA | NA | NA | NA | 1 | 1 |
| 5_2491430 | 12 | 0.3333 | 8 | 0.25 | 7 | 0.4286 | 8 | 0.5 | 35 | 0.3714 | 4 | 0.5 | 9 | 0.7778 | 13 | 0.3077 | 7 | 0.4286 | 33 | 0.4848 |
| 5_2491450 | 12 | 0.25 | 10 | 0.3 | 8 | 0.125 | 8 | 0.25 | 38 | 0.2368 | 5 | 0.2 | 9 | 0.2222 | 11 | 0.2727 | 6 | 0.3333 | 31 | 0.2581 |
| 5_2491457 | 10 | 0.2 | 10 | 0.1 | 6 | 0 | 9 | 0.1111 | 35 | 0.1143 | 2 | 0 | 9 | 0.2222 | 10 | 0.1 | 6 | 0.1667 | 27 | 0.1481 |
| 5_2491468 | 10 | 0.3 | 10 | 0.2 | 5 | 0 | 8 | 0.375 | 33 | 0.2424 | 3 | 1 | 8 | 0.375 | 12 | 0.25 | 6 | 0.3333 | 29 | 0.3793 |
| 5_2491474 | 13 | 0.0769 | 10 | 0 | 8 | 0.125 | 9 | 0 | 40 | 0.05 | 5 | 0 | 10 | 0.1 | 11 | 0.0909 | 6 | 0.1667 | 32 | 0.0938 |
| 5_2491490 | 10 | 0 | 9 | 0 | 6 | 0.1667 | 7 | 0.1429 | 32 | 0.0625 | 4 | 0.25 | 9 | 0.1111 | 9 | 0 | 8 | 0 | 30 | 0.0667 |
| 5_2491495 | 11 | 0.0909 | 7 | 0.1429 | 8 | 0 | 7 | 0 | 33 | 0.0606 | 3 | 0 | 10 | 0.1 | 10 | 0 | 7 | 0.1429 | 30 | 0.0667 |
| 5_2491527 | 10 | 0.6 | 12 | 0.8333 | 10 | 0.7 | 8 | 0.625 | 40 | 0.7 | 4 | 0.25 | 12 | 0.5833 | 11 | 0.5455 | 4 | 0.25 | 31 | 0.4839 |
| 5_2491533 | 11 | 0.1818 | 12 | 0.25 | 12 | 0.1667 | 8 | 0.125 | 43 | 0.186 | 4 | 0.25 | 13 | 0.3846 | 12 | 0.1667 | 7 | 0 | 36 | 0.2222 |
| 5_2500863 | 29 | 0.1379 | 37 | 0.1081 | 27 | 0.1111 | 17 | 0.0588 | 110 | 0.1091 | 25 | 0.08 | 22 | 0.0909 | 24 | 0.0833 | 39 | 0.1026 | 110 | 0.0909 |
| 5_2506865 | 13 | 0.4615 | 11 | 0.5455 | 11 | 0.7273 | 11 | 0.4545 | 46 | 0.5435 | 4 | 0.5 | 10 | 0.8 | 14 | 0.2143 | 11 | 0.6364 | 39 | 0.5128 |
| 5_2506878 | 12 | 0.0833 | 11 | 0 | 11 | 0.0909 | 13 | 0.2308 | 47 | 0.1064 | 3 | 0.3333 | 8 | 0.125 | 19 | 0.2105 | 10 | 0.2 | 40 | 0.2 |
| 5_2544167 | NA | NA | NA | NA | 1 | 1 | NA | NA | 1 | 1 | NA | NA | NA | NA | 1 | 0 | 2 | 1 | 3 | 0.6667 |
| 5_2544191 | NA | NA | NA | NA | 1 | 1 | NA | NA | 1 | 1 | NA | NA | NA | NA | 2 | 1 | 1 | 0 | 3 | 0.6667 |
| 5_2544222 | NA | NA | NA | NA | 1 | 1 | NA | NA | 1 | 1 | NA | NA | NA | NA | 3 | 0.3333 | 1 | 1 | 4 | 0.5 |
| 5_2544225 | NA | NA | NA | NA | 1 | 1 | NA | NA | 1 | 1 | NA | NA | NA | NA | 3 | 1 | 2 | 1 | 5 | 1 |
| 5_2544227 | NA | NA | NA | NA | 1 | 1 | NA | NA | 1 | 1 | NA | NA | NA | NA | 3 | 0.3333 | 2 | 0 | 5 | 0.2 |
| 5_2544232 | NA | NA | NA | NA | 1 | 1 | NA | NA | 1 | 1 | NA | NA | NA | NA | 2 | 1 | NA | NA | 2 | 1 |
| 5_2544233 | NA | NA | NA | NA | 1 | 1 | NA | NA | 1 | 1 | NA | NA | NA | NA | 3 | 1 | 2 | 1 | 5 | 1 |
| 5_2544237 | NA | NA | NA | NA | 1 | 1 | NA | NA | 1 | 1 | NA | NA | NA | NA | 3 | 0.6667 | 2 | 0 | 5 | 0.4 |
| 5_2544237 | NA | NA | NA | NA | 1 | 1 | NA | NA | 1 | 1 | NA | NA | NA | NA | 2 | 0 | 2 | 0.5 | 4 | 0.25 |
| 5_2544339 | 1 | 1 | NA | NA | 1 | 1 | NA | NA | 2 | 1 | NA | NA | NA | NA | 3 | 1 | 2 | 1 | 5 | 1 |
| 5_2545255 | 2 | 0.5 | 4 | 0.25 | 2 | 0.5 | 2 | 0.5 | 10 | 0.4 | 3 | 0.3333 | 2 | 0 | 3 | 0.3333 | NA | NA | 8 | 0.25 |
| 5_2545258 | 2 | 0 | 4 | 0.5 | 1 | 0 | 3 | 0 | 10 | 0.2 | 2 | 0.5 | 2 | 0.5 | 4 | 0.75 | NA | NA | 8 | 0.625 |
| 5_2547030 | 2 | 0 | 6 | 0.1667 | 4 | 0 | 2 | 0.5 | 14 | 0.1429 | 5 | 0.2 | 3 | 0 | 9 | 0.2222 | 1 | 0 | 18 | 0.1667 |
| 5_2547039 | 3 | 0 | 6 | 0.6667 | 3 | 0.3333 | 2 | 0.5 | 14 | 0.4286 | 5 | 0.2 | 2 | 0.5 | 8 | 0.375 | 1 | 1 | 16 | 0.375 |
| 5_2547047 | 3 | 0.3333 | 5 | 0.2 | 2 | 0.5 | 1 | 0 | 11 | 0.2727 | 3 | 1 | 3 | 1 | 8 | 0.5 | NA | NA | 14 | 0.7143 |
| 5_2547069 | 4 | 0 | 8 | 0 | 4 | 0 | 3 | 0.3333 | 19 | 0.0526 | 4 | 0 | 3 | 1 | 9 | 0 | 1 | 0 | 17 | 0.1765 |
| 5_2547071 | 4 | 0.25 | 6 | 0.3333 | 2 | 0 | 4 | 0.25 | 16 | 0.25 | 4 | 0.25 | 1 | 0 | 6 | 0.3333 | 1 | 0 | 12 | 0.25 |
| 5_2547077 | 3 | 0 | 8 | 0 | 5 | 0.2 | 3 | 0 | 19 | 0.0526 | 5 | 0 | 3 | 0 | 9 | 0.2222 | NA | NA | 17 | 0.1176 |
| 5_2547713 | 2 | 0.5 | 2 | 0 | 5 | 0.2 | 8 | 0.125 | 17 | 0.1765 | 3 | 0.3333 | 4 | 0 | 4 | 0.25 | 2 | 0 | 13 | 0.1538 |
| 5_2547735 | 3 | 0 | 5 | 0 | 6 | 0.5 | 11 | 0.0909 | 25 | 0.16 | 3 | 0.3333 | 4 | 0 | 8 | 0.25 | 2 | 0.5 | 17 | 0.2353 |
| 5_2547827 | 2 | 0 | 2 | 0 | 7 | 0.4286 | 10 | 0.5 | 21 | 0.381 | NA | NA | 3 | 0 | 6 | 0.1667 | 2 | 0 | 11 | 0.0909 |
| 5_2547834 | 2 | 0.5 | 4 | 0.25 | 5 | 0 | 8 | 0.375 | 19 | 0.2632 | NA | NA | 3 | 0 | 3 | 0.3333 | 1 | 0 | 7 | 0.1429 |
| 5_2547856 | 2 | 0 | 4 | 0.25 | 3 | 0.3333 | 6 | 0.3333 | 15 | 0.2667 | NA | NA | 3 | 0 | 4 | 0.25 | 2 | 0.5 | 9 | 0.2222 |
| 5_2547866 | 1 | 0 | 4 | 0.5 | 4 | 0.5 | 10 | 0.1 | 19 | 0.2632 | NA | NA | 4 | 0.75 | 3 | 0 | 2 | 0 | 9 | 0.3333 |

| | | | | | | | | | | | | | | | | | | | | |
|---|---|---|---|---|---|---|---|---|---|---|---|---|---|---|---|---|---|---|---|---|
| 5_2547868( | 2 | 0 | 4 | 0 | 4 | 0 | 8 | 0.25 | 18 | 0.1111 | NA | NA | 4 | 0 | 6 | 0.1667 | 2 | 0 | 12 | 0.0833 |
| 5_2548932 | 2 | 0 | 1 | 0 | 2 | 0 | 6 | 0.5 | 11 | 0.2727 | NA | NA | 2 | 0.5 | 4 | 0.25 | 1 | 0 | 7 | 0.2857 |
| 5_2548946( | 2 | 0 | 2 | 0 | 1 | 0 | 5 | 0.6 | 10 | 0.3 | NA | NA | 3 | 0.3333 | 5 | 0.2 | NA | NA | 8 | 0.25 |
| 5_2548958 | 2 | 0.5 | 2 | 1 | 2 | 0.5 | 3 | 0.3333 | 9 | 0.5556 | NA | NA | 3 | 0.6667 | 4 | 0.75 | NA | NA | 7 | 0.7143 |
| 5_2548966 | 2 | 1 | 2 | 1 | 1 | 1 | 1 | 1 | 6 | 1 | NA | NA | 1 | 1 | NA | NA | NA | NA | 1 | 1 |
| 5_2548969 | NA | NA | 1 | 0 | NA | NA | 1 | 1 | 2 | 0.5 | NA | NA | 1 | 1 | NA | NA | NA | NA | 1 | 1 |
| 5_2548976( | 2 | 0.5 | 2 | 0.5 | 1 | 1 | 4 | 0.75 | 9 | 0.6667 | NA | NA | 2 | 0.5 | 2 | 0 | NA | NA | 4 | 0.25 |
| 5_2548992 | 2 | 0 | 2 | 0 | 1 | 0 | 3 | 0.3333 | 8 | 0.125 | NA | NA | 2 | 0.5 | 4 | 0.25 | NA | NA | 6 | 0.3333 |
| 5_2548995( | 2 | 1 | 2 | 1 | 2 | 1 | 4 | 0.5 | 10 | 0.8 | NA | NA | 3 | 1 | 4 | 1 | NA | NA | 7 | 1 |
| 5_2550410 | 1 | 1 | 2 | 1 | 2 | 0.5 | 3 | 0 | 8 | 0.5 | NA | NA | 2 | 0.5 | 3 | 0 | NA | NA | 5 | 0.2 |
| 5_2550443( | 1 | 1 | 2 | 0 | 2 | 0.5 | 3 | 0 | 8 | 0.25 | NA | NA | 2 | 0 | 3 | 0.3333 | NA | NA | 5 | 0.2 |
| 5_2550458 | 1 | 0 | 2 | 0 | 2 | 0.5 | 2 | 0.5 | 7 | 0.2857 | NA | NA | 2 | 0.5 | 3 | 0.3333 | NA | NA | 5 | 0.4 |
| 5_2550473( | NA | NA | 2 | 0.5 | 2 | 0.5 | 2 | 0 | 6 | 0.3333 | NA | NA | 2 | 0 | 2 | 0.5 | NA | NA | 4 | 0.25 |
| 5_2550493 | 1 | 1 | 2 | 0.5 | 1 | 1 | 2 | 1 | 6 | 0.8333 | NA | NA | 2 | 0.5 | 2 | 0.5 | NA | NA | 4 | 0.5 |
| 5_2550494 | 1 | 0 | 2 | 0.5 | 2 | 0.5 | 2 | 0 | 7 | 0.2857 | NA | NA | 2 | 0 | 2 | 0.5 | NA | NA | 4 | 0.25 |
| 5_2550498 | 1 | 1 | 2 | 0 | 2 | 1 | 2 | 0 | 7 | 0.4286 | NA | NA | 2 | 0.5 | 2 | 0.5 | NA | NA | 4 | 0.5 |
| 5_2550521 | 1 | 1 | 2 | 0.5 | 2 | 1 | 2 | 1 | 7 | 0.8571 | NA | NA | 1 | 1 | 2 | 0 | NA | NA | 3 | 0.3333 |
| 5_2550523 | 1 | 0 | 1 | 1 | 1 | 0 | 1 | 0 | 4 | 0.25 | NA | NA | 1 | 0 | 2 | 1 | NA | NA | 3 | 0.6667 |
| 5_2551513 | 1 | 0 | 1 | 0 | 2 | 1 | 2 | 0 | 6 | 0.3333 | NA | NA | 1 | 1 | 1 | 1 | NA | NA | 2 | 1 |
| 5_2551525 | 1 | 1 | 1 | 0 | 2 | 1 | 2 | 1 | 6 | 0.8333 | NA | NA | 1 | 1 | 1 | 0 | NA | NA | 2 | 0.5 |
| 5_2551566 | 1 | 1 | 1 | 1 | 2 | 1 | 2 | 0.5 | 6 | 0.8333 | NA | NA | NA | NA | 1 | 1 | NA | NA | 1 | 1 |
| 5_2551576 | 1 | 1 | 1 | 1 | 1 | 1 | 2 | 0.5 | 5 | 0.8 | NA | NA | NA | NA | 1 | 1 | NA | NA | 1 | 1 |
| 5_2551953 | NA | NA | 1 | 0 | NA | NA | 1 | 1 | 2 | 0.5 | NA | NA | NA | NA | 1 | 1 | NA | NA | 1 | 1 |
| 5_2573541 | NA | NA | 1 | 1 | NA | NA | NA | NA | 1 | 1 | NA | NA | NA | NA | 1 | 1 | NA | NA | 1 | 1 |
| 5_2573565 | NA | NA | 1 | 1 | NA | NA | 1 | 1 | 2 | 1 | NA | NA | 1 | 1 | 1 | 1 | NA | NA | 2 | 1 |
| 5_2589593 | NA | NA | NA | NA | NA | NA | 1 | 1 | 1 | 1 | 2 | 0.5 | 1 | 0 | NA | NA | 1 | 0 | 4 | 0.25 |
| 5_2589616( | NA | NA | NA | NA | NA | NA | 1 | 1 | 1 | 1 | NA | NA | 1 | 1 | NA | NA | NA | NA | 1 | 1 |
| 5_2589750 | NA | NA | NA | NA | NA | NA | 1 | 1 | 1 | 1 | 2 | 1 | 1 | 1 | NA | NA | NA | NA | 3 | 1 |
| 5_2589765 | NA | NA | NA | NA | NA | NA | 1 | 1 | 1 | 1 | 2 | 1 | 1 | 1 | NA | NA | NA | NA | 3 | 1 |
| 5_2589772 | NA | NA | NA | NA | NA | NA | 1 | 1 | 1 | 1 | 2 | 1 | 1 | 1 | NA | NA | NA | NA | 3 | 1 |
| 5_2605526 | 19 | 0.6842 | 23 | 0.3913 | 18 | 0.3889 | 18 | 0.2778 | 78 | 0.4359 | 11 | 0.6364 | 24 | 0.4583 | 23 | 0.6087 | 17 | 0.3529 | 75 | 0.5067 |
| 5_2746720( | 2 | 0.5 | 1 | 0 | 1 | 0 | 6 | 0 | 10 | 0.1 | 3 | 0.6667 | 4 | 0 | 6 | 0.3333 | 1 | 1 | 14 | 0.3571 |
| 5_2793074 | 17 | 0.3529 | 11 | 0.5455 | 12 | 0.4167 | 20 | 0.3 | 60 | 0.3833 | 9 | 0.3333 | 10 | 0.3 | 20 | 0.2 | 13 | 0.3846 | 52 | 0.2885 |
| 5_2793077 | 17 | 0.1765 | 11 | 0 | 13 | 0.0769 | 22 | 0 | 63 | 0.0635 | 10 | 0.1 | 11 | 0.0909 | 23 | 0.0435 | 10 | 0.1 | 54 | 0.0741 |
| 5_2793099( | 14 | 0.8571 | 12 | 1 | 14 | 1 | 22 | 0.8636 | 62 | 0.9194 | 8 | 0.875 | 11 | 0.8182 | 24 | 0.9167 | 15 | 0.8 | 58 | 0.8621 |
| 5_2793102 | 17 | 0.0588 | 12 | 0 | 13 | 0.0769 | 23 | 0.087 | 65 | 0.0615 | 12 | 0 | 11 | 0.0909 | 20 | 0.15 | 12 | 0.0833 | 55 | 0.0909 |
| 5_2793103 | 16 | 0.75 | 11 | 0.8182 | 11 | 1 | 23 | 0.913 | 61 | 0.8689 | 11 | 0.9091 | 12 | 0.9167 | 22 | 0.9091 | 15 | 0.9333 | 60 | 0.9167 |
| 5_2793104( | 15 | 0 | 11 | 0 | 13 | 0.0769 | 23 | 0.1304 | 62 | 0.0645 | 11 | 0 | 13 | 0.1538 | 22 | 0.0455 | 16 | 0.0625 | 62 | 0.0645 |
| 5_2793106( | 16 | 0 | 12 | 0.25 | 15 | 0.1333 | 21 | 0 | 64 | 0.0781 | 11 | 0.0909 | 11 | 0.0909 | 20 | 0.2 | 15 | 0 | 57 | 0.1053 |
| 5_2793140 | 17 | 0.2353 | 11 | 0.3636 | 11 | 0.1818 | 16 | 0.1875 | 55 | 0.2364 | 11 | 0.1818 | 12 | 0.4167 | 20 | 0.3 | 13 | 0.3846 | 56 | 0.3214 |
| 5_2793160( | 9 | 0.1111 | 8 | 0.25 | 5 | 0 | 15 | 0.0667 | 37 | 0.1081 | 3 | 0 | 11 | 0 | 16 | 0 | 11 | 0.4545 | 41 | 0.122 |
| 5_2793231( | 15 | 0.0667 | 12 | 0 | 11 | 0.0909 | 18 | 0.0556 | 56 | 0.0536 | 3 | 0 | 13 | 0.1538 | 16 | 0.0625 | 11 | 0.0909 | 43 | 0.093 |
| 5_2794646( | NA | NA | NA | NA | NA | NA | 1 | 1 | 1 | 1 | NA | NA | NA | NA | 1 | 1 | NA | NA | 1 | 1 |
| 5_2794741 | 7 | 0.1429 | 6 | 0.1667 | 5 | 0 | 7 | 0.4286 | 25 | 0.2 | 2 | 0 | 3 | 0.3333 | 3 | 0.3333 | 1 | 1 | 9 | 0.3333 |
| 5_2796354 | 3 | 0.3333 | 4 | 0 | 1 | 0 | 4 | 0 | 12 | 0.0833 | 1 | 1 | NA | NA | 3 | 0 | NA | NA | 4 | 0.25 |
| 5_2797060 | 5 | 0.2 | 6 | 0 | 3 | 0.3333 | 6 | 0.3333 | 20 | 0.2 | 3 | 0.3333 | 2 | 0.5 | 1 | 0 | 1 | 0 | 7 | 0.2857 |
| 5_2797122 | 3 | 0.3333 | 3 | 0.3333 | 1 | 0 | 5 | 0.2 | 12 | 0.25 | 2 | 0.5 | 1 | 0 | NA | NA | 1 | 0 | 4 | 0.25 |
| 5_2797708 | 1 | 0 | 2 | 0.5 | 2 | 0 | 5 | 0.2 | 10 | 0.2 | 2 | 0 | 1 | 1 | NA | NA | 1 | 1 | 4 | 0.5 |
| 5_2797712 | 1 | 1 | 3 | 0.6667 | 2 | 0.5 | 5 | 0.4 | 11 | 0.5455 | 2 | 0.5 | 1 | 1 | NA | NA | 1 | 1 | 4 | 0.75 |
| 5_2797717 | NA | NA | 3 | 0.3333 | 2 | 0 | 5 | 0.8 | 10 | 0.5 | 2 | 0.5 | 1 | 1 | NA | NA | 1 | 0 | 4 | 0.5 |
| 5_2797721 | 1 | 1 | 3 | 1 | 2 | 1 | 5 | 0.8 | 11 | 0.9091 | 2 | 1 | 1 | 1 | NA | NA | NA | NA | 3 | 1 |
| 5_2797728 | 1 | 0 | 3 | 0.6667 | 1 | 1 | 2 | 0.5 | 7 | 0.5714 | 2 | 1 | 1 | 1 | NA | NA | NA | NA | 3 | 1 |
| 5_2797732 | 1 | 0 | 2 | 0.5 | 2 | 1 | 4 | 0.25 | 9 | 0.4444 | 2 | 0 | 1 | 1 | NA | NA | NA | NA | 3 | 0.3333 |
| 5_2797745( | 1 | 0 | 1 | 0 | 2 | 0.5 | 5 | 0.2 | 9 | 0.2222 | 2 | 0.5 | 1 | 0 | NA | NA | NA | NA | 3 | 0.3333 |
| 5_2797755 | 1 | 0 | 1 | 1 | 2 | 0.5 | 4 | 0.5 | 8 | 0.5 | 2 | 0 | 1 | 1 | NA | NA | NA | NA | 3 | 0.3333 |
| 5_2804730 | 1 | 1 | NA | NA | NA | NA | 1 | 1 | 2 | 1 | NA | NA | NA | NA | NA | NA | 1 | 1 | 1 | 1 |
| 5_2804744 | 1 | 1 | NA | NA | NA | NA | NA | NA | 1 | 1 | 1 | 1 | NA | NA | NA | NA | 1 | 1 | 2 | 1 |
| 5_2804752( | 1 | 1 | NA | NA | NA | NA | 1 | 1 | 2 | 1 | 1 | 0 | NA | NA | NA | NA | 1 | 1 | 2 | 0.5 |
| 5_2804799( | 2 | 1 | NA | NA | NA | NA | 1 | 1 | 3 | 1 | 1 | 1 | NA | NA | NA | NA | 2 | 1 | 3 | 1 |
| 5_2805608 | 2 | 1 | NA | NA | 2 | 0.5 | 1 | 1 | 5 | 0.8 | 1 | 1 | NA | NA | NA | NA | 1 | 1 | 2 | 1 |
| 5_2805614 | 3 | 1 | NA | NA | 2 | 1 | 1 | 1 | 6 | 1 | NA | NA | NA | NA | NA | NA | 1 | 1 | 1 | 1 |
| 5_2805617 | 3 | 1 | NA | NA | 1 | 1 | NA | NA | 4 | 1 | 1 | 1 | NA | NA | NA | NA | 2 | 1 | 3 | 1 |
| 5_2805632 | 2 | 1 | NA | NA | 2 | 1 | 1 | 1 | 5 | 1 | 1 | 1 | NA | NA | NA | NA | 1 | 1 | 2 | 1 |
| 5_2805637 | 2 | 1 | NA | NA | 2 | 1 | 1 | 1 | 5 | 1 | 1 | 1 | NA | NA | NA | NA | 2 | 1 | 3 | 1 |
| 5_2805643 | 3 | 0.6667 | NA | NA | 2 | 1 | 2 | 0.5 | 7 | 0.7143 | 1 | 0 | NA | NA | NA | NA | 2 | 1 | 3 | 0.6667 |

| | | | | | | | | | | | | | | | | | | | | |
|---|---|---|---|---|---|---|---|---|---|---|---|---|---|---|---|---|---|---|---|---|
| 5_2805649 | 2 | 0 | NA | NA | 2 | 0 | 2 | 0.5 | 6 | 0.1667 | 1 | 1 | NA | NA | NA | NA | 1 | 0 | 2 | 0.5 |
| 5_2805660 | 3 | 1 | NA | NA | 3 | 0.6667 | 2 | 1 | 8 | 0.875 | 1 | 1 | NA | NA | NA | NA | 2 | 1 | 3 | 1 |
| 5_2805720 | 3 | 0.3333 | 2 | 0 | 2 | 0.5 | 3 | 0.3333 | 10 | 0.3 | 1 | 0 | 1 | 0 | NA | NA | 1 | 1 | 3 | 0.3333 |
| 5_2805727 | 3 | 0 | 2 | 1 | 3 | 0 | 3 | 0 | 11 | 0.1818 | 1 | 1 | 1 | 0 | NA | NA | 1 | 0 | 3 | 0.3333 |
| 5_2805733 | 3 | 0.3333 | 2 | 0 | 2 | 0.5 | 2 | 0.5 | 9 | 0.3333 | 1 | 0 | 1 | 1 | NA | NA | 1 | 1 | 3 | 0.6667 |
| 5_2805742 | 2 | 0.5 | 2 | 1 | 4 | 0.5 | 1 | 0 | 9 | 0.5556 | 1 | 1 | 1 | 1 | NA | NA | NA | NA | 2 | 1 |
| 5_2805743 | 2 | 1 | 1 | 1 | 2 | 0 | 3 | 0.6667 | 8 | 0.625 | 1 | 0 | 1 | 1 | NA | NA | 1 | 0 | 3 | 0.3333 |
| 5_2805752 | 3 | 1 | 2 | 0.5 | 4 | 0.75 | 2 | 1 | 11 | 0.8182 | 1 | 1 | 1 | 1 | NA | NA | 1 | 0 | 3 | 0.6667 |
| 5_2805754 | 3 | 0.6667 | 2 | 1 | 2 | 0.5 | 3 | 0.3333 | 10 | 0.6 | 2 | 0.5 | 1 | 1 | NA | NA | 2 | 0.5 | 5 | 0.6 |
| 5_2805755 | 3 | 1 | 2 | 1 | 3 | 1 | 2 | 0.5 | 10 | 0.9 | 2 | 1 | 1 | 1 | NA | NA | 1 | 1 | 4 | 1 |
| 5_2805761 | 3 | 0 | 2 | 0.5 | 3 | 0 | 3 | 0 | 11 | 0.0909 | 2 | 0.5 | 1 | 0 | NA | NA | 1 | 0 | 4 | 0.25 |
| 5_2805784 | 2 | 0.5 | 2 | 0 | 3 | 0 | 2 | 0 | 9 | 0.1111 | 2 | 0 | 1 | 1 | NA | NA | 2 | 0 | 5 | 0.2 |
| 5_2805786 | 3 | 0.3333 | 2 | 0.5 | 3 | 0.3333 | 3 | 0 | 11 | 0.2727 | 2 | 0 | NA | NA | NA | NA | 2 | 0.5 | 4 | 0.25 |
| 5_2805788 | 3 | 1 | 2 | 0.5 | 3 | 0.6667 | 3 | 0.3333 | 11 | 0.6364 | 2 | 1 | 1 | 1 | NA | NA | 1 | 0 | 4 | 0.75 |
| 5_2805804 | NA | NA | 2 | 0.5 | 2 | 0.5 | 1 | 1 | 5 | 0.6 | 2 | 1 | 1 | 0 | NA | NA | 1 | 0 | 4 | 0.5 |
| 5_2805808 | 2 | 1 | 2 | 1 | 4 | 0.75 | 3 | 1 | 11 | 0.9091 | 2 | 1 | 1 | 1 | NA | NA | 2 | 1 | 5 | 1 |
| 5_2805810 | 2 | 0.5 | 1 | 1 | 3 | 0.3333 | 3 | 0 | 9 | 0.3333 | 2 | 1 | 1 | 1 | NA | NA | 2 | 0 | 5 | 0.6 |
| 5_2805810 | 2 | 0.5 | 2 | 0.5 | 3 | 0.6667 | 3 | 1 | 10 | 0.7 | 2 | 0 | 1 | 1 | NA | NA | NA | NA | 3 | 0.3333 |
| 5_2805845 | 1 | 1 | NA | NA | 3 | 0 | 2 | 0.5 | 6 | 0.3333 | 3 | 0.3333 | 1 | 0 | NA | NA | 2 | 0.5 | 6 | 0.3333 |
| 5_2805876 | 1 | 1 | NA | NA | 2 | 1 | 1 | 1 | 4 | 1 | 1 | 1 | NA | NA | NA | NA | 1 | 0 | 2 | 0.5 |
| 5_2805879 | 1 | 1 | NA | NA | 2 | 0 | 1 | 0 | 4 | 0.25 | 1 | 1 | NA | NA | NA | NA | 2 | 0.5 | 3 | 0.6667 |
| 5_2805880 | 1 | 0 | NA | NA | 2 | 0.5 | 1 | 1 | 4 | 0.5 | 1 | 0 | NA | NA | NA | NA | 1 | 1 | 2 | 0.5 |
| 5_2805894 | 1 | 0 | NA | NA | 3 | 0 | 1 | 1 | 5 | 0.2 | 1 | 1 | NA | NA | NA | NA | 2 | 1 | 3 | 1 |
| 5_2805907 | 1 | 1 | NA | NA | 2 | 1 | 1 | 1 | 4 | 1 | 1 | 1 | NA | NA | NA | NA | 1 | 1 | 2 | 1 |
| 5_2828185 | 3 | 0 | 9 | 0.1111 | 6 | 0.1667 | 7 | 0.1429 | 25 | 0.12 | 5 | 0.4 | 6 | 0.3333 | 7 | 0.4286 | 6 | 0.5 | 24 | 0.4167 |
| 5_2828192 | 3 | 0 | 8 | 0 | 6 | 0 | 7 | 0.2857 | 24 | 0.0833 | 6 | 0.3333 | 4 | 0 | 7 | 0 | 6 | 0.3333 | 23 | 0.1739 |
| 5_2828193 | 3 | 0.3333 | 10 | 0 | 7 | 0.1429 | 7 | 0.2857 | 27 | 0.1481 | 6 | 0.1667 | 5 | 0 | 7 | 0.1429 | 5 | 0.2 | 23 | 0.1304 |
| 5_2828197 | 3 | 0.3333 | 10 | 0.4 | 7 | 0.2857 | 7 | 0.5714 | 27 | 0.4074 | 6 | 0.3333 | 6 | 0.3333 | 6 | 0.3333 | 6 | 0.3333 | 24 | 0.3333 |
| 5_2828248 | 3 | 0.3333 | 9 | 0.3333 | 7 | 0.2857 | 5 | 0.6 | 24 | 0.375 | 5 | 0.6 | 6 | 0.5 | 8 | 0.625 | 5 | 0.6 | 24 | 0.5833 |
| 5_2828260 | 2 | 1 | 9 | 0.7778 | 7 | 0.8571 | 7 | 0.8571 | 25 | 0.84 | 6 | 0.6667 | 7 | 0.8571 | 8 | 1 | 7 | 1 | 28 | 0.8929 |
| 5_2828270 | 4 | 0.75 | 10 | 0.6 | 9 | 0.7778 | 7 | 0.4286 | 30 | 0.6333 | 3 | 1 | 8 | 0.375 | 9 | 0.5556 | 6 | 0.3333 | 26 | 0.5 |
| 5_2828272 | 3 | 0 | 10 | 0.1 | 7 | 0.1429 | 7 | 0.1429 | 27 | 0.1111 | 6 | 0.1667 | 6 | 0 | 9 | 0.1111 | 6 | 0.1667 | 27 | 0.1111 |
| 5_2828324 | 4 | 0.25 | 14 | 0.1429 | 11 | 0.2727 | 9 | 0.2222 | 38 | 0.2105 | 8 | 0.25 | 6 | 0.3333 | 12 | 0.1667 | 8 | 0 | 34 | 0.1765 |
| 5_2828351 | 6 | 0 | 13 | 0.1538 | 14 | 0 | 9 | 0.2222 | 42 | 0.0952 | 9 | 0.2222 | 11 | 0 | 11 | 0 | 10 | 0.2 | 41 | 0.0976 |
| 5_2854414 | 1 | 1 | NA | NA | NA | NA | NA | NA | 1 | 1 | NA | NA | 2 | 0.5 | 1 | 0 | NA | NA | 3 | 0.3333 |
| 5_2855758 | 1 | 0 | NA | NA | 1 | 1 | NA | NA | 2 | 0.5 | NA | NA | 2 | 0.5 | 1 | 1 | 1 | 0 | 4 | 0.5 |
| 5_2855763 | 1 | 1 | NA | NA | 1 | 1 | NA | NA | 2 | 1 | NA | NA | 2 | 1 | 1 | 0 | NA | NA | 3 | 0.6667 |
| 5_2859924 | 1 | 1 | NA | NA | NA | NA | NA | NA | 1 | 1 | 1 | 0 | 1 | 1 | 1 | 0 | 1 | 1 | 4 | 0.5 |
| 5_2859933 | 1 | 1 | 1 | 0 | NA | NA | 1 | 1 | 3 | 0.6667 | 1 | 1 | 1 | 0 | 1 | 0 | 1 | 1 | 4 | 0.5 |
| 5_2859938 | 1 | 1 | NA | NA | NA | NA | NA | NA | 1 | 1 | 1 | 1 | NA | NA | NA | NA | 1 | 0 | 2 | 0.5 |
| 5_2859943 | 1 | 0 | 1 | 0 | NA | NA | 1 | 1 | 3 | 0.3333 | 1 | 0 | NA | NA | 1 | 1 | 1 | 0 | 3 | 0.3333 |
| 5_2859955 | NA | NA | 1 | 1 | NA | NA | 1 | 1 | 2 | 1 | NA | NA | 1 | 0 | 1 | 0 | 1 | 1 | 3 | 0.3333 |
| 5_2859958 | 1 | 1 | 1 | 1 | NA | NA | 1 | 1 | 3 | 1 | 1 | 1 | 1 | 0 | 1 | 0 | 1 | 1 | 4 | 0.5 |
| 5_2861878 | 1 | 1 | 2 | 0.5 | 1 | 1 | 1 | 1 | 5 | 0.8 | 3 | 0.3333 | 1 | 1 | 1 | 1 | 1 | 0 | 6 | 0.5 |
| 5_2861879 | 2 | 1 | 2 | 1 | 1 | 0 | 2 | 1 | 7 | 0.8571 | 2 | 0.5 | 2 | 1 | 1 | 1 | 1 | 1 | 6 | 0.8333 |
| 5_2861884 | 2 | 1 | 2 | 0.5 | 2 | 1 | 2 | 1 | 8 | 0.875 | 2 | 1 | 2 | 0 | 1 | 0 | 1 | 1 | 6 | 0.5 |
| 5_2861972 | 3 | 0.3333 | 1 | 0 | 2 | 0 | 2 | 0 | 8 | 0.125 | 2 | 0 | 2 | 0 | 1 | 1 | 2 | 0 | 7 | 0.1429 |
| 5_2861983 | 2 | 0.5 | 2 | 0 | 1 | 0 | 1 | 1 | 6 | 0.3333 | 3 | 0 | NA | NA | 1 | 0 | 1 | 1 | 5 | 0.2 |
| 5_2862093 | 3 | 0.6667 | 4 | 0.5 | 3 | 0.6667 | 2 | 0 | 12 | 0.5 | 5 | 0.4 | 1 | 1 | 2 | 0 | 3 | 0.3333 | 11 | 0.3636 |
| 5_2862403 | 6 | 0.1667 | 2 | 0.5 | 4 | 0 | 1 | 0 | 13 | 0.1538 | 2 | 0 | 4 | 0.25 | 4 | 0.25 | 2 | 0 | 12 | 0.1667 |
| 5_2862408 | 6 | 0.3333 | 3 | 0 | 4 | 0.25 | 2 | 0.5 | 15 | 0.2667 | 2 | 0.5 | 5 | 0.4 | 2 | 0.5 | 2 | 0.5 | 11 | 0.4545 |
| 5_2866205 | 1 | 1 | NA | NA | 1 | 0 | 1 | 1 | 3 | 0.6667 | 1 | 1 | NA | NA | 1 | 1 | 1 | 0 | 3 | 0.6667 |
| 5_2866253 | 1 | 0 | NA | NA | 1 | 1 | 1 | 1 | 3 | 0.6667 | 1 | 0 | NA | NA | NA | NA | 1 | 1 | 2 | 0.5 |
| 5_2866269 | 1 | 0 | NA | NA | 1 | 1 | 1 | 0 | 3 | 0.3333 | 1 | 0 | NA | NA | NA | NA | 1 | 1 | 2 | 0.5 |
| 5_2866276 | 1 | 1 | NA | NA | 1 | 1 | 1 | 0 | 3 | 0.6667 | 1 | 0 | NA | NA | NA | NA | 1 | 1 | 2 | 0.5 |
| 5_2866279 | 1 | 1 | NA | NA | 1 | 1 | 1 | 1 | 3 | 1 | 1 | 1 | NA | NA | NA | NA | NA | NA | 1 | 1 |
| 5_2866289 | NA | NA | NA | NA | 1 | 1 | 1 | 0 | 2 | 0.5 | 1 | 0 | NA | NA | NA | NA | 1 | 1 | 2 | 0.5 |
| 5_2866343 | NA | NA | NA | NA | 1 | 0 | 1 | 1 | 2 | 0.5 | 1 | 1 | NA | NA | NA | NA | 1 | 0 | 2 | 0.5 |
| 5_2866351 | NA | NA | NA | NA | 1 | 1 | 1 | 1 | 2 | 1 | 1 | 1 | NA | NA | NA | NA | 1 | 1 | 2 | 1 |
| 5_2942907 | NA | NA | NA | NA | NA | NA | 1 | 1 | 1 | 1 | NA | NA | 2 | 0.5 | NA | NA | NA | NA | 2 | 0.5 |
| 5_2942916 | 1 | 1 | 1 | 0 | NA | NA | 4 | 0.25 | 6 | 0.3333 | 2 | 0 | 1 | 1 | NA | NA | 1 | 0 | 4 | 0.25 |
| 5_2943742 | 2 | 0.5 | 4 | 0.25 | 1 | 0 | 3 | 0.3333 | 10 | 0.3 | 2 | 0.5 | 5 | 0.4 | 5 | 0 | 2 | 0.5 | 14 | 0.2857 |
| 5_2943748 | 2 | 0 | 4 | 0.5 | 2 | 1 | 4 | 0.25 | 12 | 0.4167 | 1 | 0 | 5 | 0.2 | 5 | 0.4 | 1 | 0 | 12 | 0.25 |
| 5_2943754 | 3 | 0.6667 | 4 | 0.5 | 1 | 0 | 4 | 0.75 | 12 | 0.5833 | 2 | 0.5 | 5 | 0.4 | 4 | 0.5 | 3 | 0.3333 | 14 | 0.4286 |
| 5_2943760 | 3 | 0.6667 | 3 | 0.3333 | 2 | 0.5 | 4 | 0.75 | 12 | 0.5833 | 1 | 0 | 5 | 0.6 | 5 | 0.2 | 3 | 0.3333 | 14 | 0.3571 |

| | | | | | | | | | | | | | | | | | | | | |
|---|---|---|---|---|---|---|---|---|---|---|---|---|---|---|---|---|---|---|---|---|
| 5_29437668 | 2 | 0 | 2 | 0 | 2 | 0 | 3 | 0.3333 | 9 | 0.1111 | 2 | 0 | 4 | 0 | 5 | 0 | 3 | 0.6667 | 14 | 0.1429 |
| 5_29437687 | 2 | 0.5 | 3 | 1 | 2 | 0.5 | 3 | 0.3333 | 10 | 0.6 | 2 | 0 | 5 | 0.8 | 5 | 0.2 | 3 | 0.6667 | 15 | 0.4667 |
| 5_29437798 | 3 | 0.3333 | 2 | 0.5 | 2 | 0 | 3 | 0.3333 | 10 | 0.3 | 2 | 0 | 5 | 0.2 | 5 | 0.2 | 3 | 0 | 15 | 0.1333 |
| 5_29437820 | 3 | 0 | 3 | 0 | 2 | 0 | 3 | 0.6667 | 11 | 0.1818 | 2 | 0.5 | 2 | 0 | 5 | 0 | 2 | 0.5 | 11 | 0.1818 |
| 5_29437851 | 2 | 1 | 3 | 0.6667 | 2 | 0.5 | 3 | 1 | 10 | 0.8 | 2 | 0.5 | 3 | 0.3333 | 4 | 0.5 | 3 | 0.3333 | 12 | 0.4167 |
| 5_29437962 | 3 | 0.3333 | 2 | 0 | 2 | 0 | 3 | 0 | 10 | 0.1 | 2 | 0 | 5 | 0.4 | 5 | 0 | 3 | 0.3333 | 15 | 0.2 |
| 5_29437971 | 2 | 0 | 3 | 0 | 2 | 0.5 | 2 | 0 | 9 | 0.1111 | 2 | 0 | 4 | 0 | 5 | 0.2 | 2 | 0 | 13 | 0.0769 |
| 5_29437996 | 3 | 0.3333 | 3 | 1 | 2 | 0.5 | 2 | 1 | 10 | 0.7 | 2 | 1 | 4 | 0.5 | 4 | 1 | 3 | 0 | 13 | 0.6154 |
| 5_29438008 | 3 | 0 | 2 | 0 | 2 | 0 | 3 | 0.3333 | 10 | 0.1 | 1 | 1 | 4 | 0.25 | 5 | 0 | 3 | 0 | 13 | 0.1538 |
| 5_29583130 | NA | NA | 1 | 1 | NA | NA | 1 | 1 | 2 | 1 | NA | NA | 1 | 1 | NA | NA | NA | NA | 1 | 1 |
| 5_29583272 | 1 | 1 | 1 | 1 | 1 | 0 | 2 | 0.5 | 5 | 0.6 | 2 | 0 | 3 | 0.6667 | 1 | 1 | 1 | 1 | 7 | 0.5714 |
| 5_29583286 | NA | NA | 1 | 0 | NA | NA | 1 | 1 | 2 | 0.5 | NA | NA | 1 | 1 | NA | NA | NA | NA | 1 | 1 |
| 5_29583503 | NA | NA | 1 | 1 | NA | NA | 1 | 1 | 2 | 1 | NA | NA | 1 | 1 | NA | NA | NA | NA | 1 | 1 |
| 5_29583539 | NA | NA | 1 | 1 | NA | NA | 1 | 1 | 2 | 1 | NA | NA | 1 | 1 | NA | NA | NA | NA | 1 | 1 |
| 5_29583573 | NA | NA | 1 | 1 | NA | NA | 1 | 1 | 2 | 1 | NA | NA | 1 | 1 | NA | NA | NA | NA | 1 | 1 |
| 5_29583962 | NA | NA | 1 | 1 | NA | NA | 1 | 1 | 2 | 1 | NA | NA | 1 | 1 | NA | NA | NA | NA | 1 | 1 |
| 5_29639847 | NA | NA | 3 | 0.3333 | NA | NA | 2 | 0 | 5 | 0.2 | 1 | 1 | 2 | 0 | NA | NA | 1 | 0 | 4 | 0.25 |
| 5_29639998 | NA | NA | 3 | 0.3333 | NA | NA | 3 | 0.3333 | 6 | 0.3333 | 1 | 0 | 1 | 0 | NA | NA | 2 | 0.5 | 4 | 0.25 |
| 5_29640552 | NA | NA | 2 | 0.5 | NA | NA | 3 | 1 | 5 | 0.8 | 2 | 1 | 2 | 0.5 | NA | NA | 1 | 1 | 5 | 0.8 |
| 5_29640570 | NA | NA | 2 | 0 | NA | NA | 2 | 0.5 | 4 | 0.25 | 1 | 1 | 2 | 0.5 | NA | NA | 1 | 0 | 4 | 0.5 |
| 5_29642733 | NA | NA | 2 | 0.5 | NA | NA | 3 | 0 | 5 | 0.2 | 2 | 0.5 | 2 | 0 | NA | NA | 2 | 0 | 6 | 0.1667 |
| 5_29642877 | NA | NA | NA | NA | NA | NA | 1 | 1 | 1 | 1 | 1 | 1 | NA | NA | NA | NA | NA | NA | 1 | 1 |
| 5_29642900 | NA | NA | NA | NA | NA | NA | 1 | 1 | 1 | 1 | 1 | 1 | NA | NA | NA | NA | NA | NA | 1 | 1 |
| 5_29643149 | NA | NA | NA | NA | NA | NA | 1 | 1 | 1 | 1 | 1 | 1 | NA | NA | NA | NA | NA | NA | 1 | 1 |
| 5_29643317 | NA | NA | NA | NA | NA | NA | 1 | 1 | 1 | 1 | 1 | 1 | NA | NA | NA | NA | NA | NA | 1 | 1 |
| 5_29644898 | NA | NA | 3 | 0.6667 | NA | NA | 2 | 0.5 | 5 | 0.6 | 1 | 1 | 1 | 0 | NA | NA | 2 | 0.5 | 4 | 0.5 |
| 5_29644992 | NA | NA | 3 | 0.3333 | NA | NA | 2 | 0.5 | 5 | 0.4 | 1 | 0 | 2 | 0 | NA | NA | 2 | 1 | 5 | 0.4 |
| 5_29645008 | NA | NA | 3 | 0.3333 | NA | NA | 2 | 0 | 5 | 0.2 | 1 | 0 | 2 | 0.5 | NA | NA | 2 | 1 | 5 | 0.6 |
| 5_29646434 | NA | NA | 2 | 0 | NA | NA | 2 | 0.5 | 4 | 0.25 | 1 | 0 | 2 | 0 | NA | NA | 2 | 0.5 | 5 | 0.2 |
| 5_29806227 | 9 | 0.1111 | 7 | 0.1429 | 3 | 0 | 1 | 0 | 20 | 0.1 | 2 | 0 | 6 | 0 | 3 | 0 | 4 | 0.25 | 15 | 0.0667 |
| 5_29851511 | 6 | 0.1667 | 8 | 0.25 | 5 | 0 | 1 | 0 | 20 | 0.15 | 4 | 0 | 8 | 0 | 3 | 0.3333 | 3 | 0.3333 | 18 | 0.1111 |
| 5_29863378 | 9 | 0 | 11 | 0.0909 | 5 | 0.2 | 1 | 0 | 26 | 0.0769 | 4 | 0.25 | 9 | 0 | 4 | 0.25 | 6 | 0.1667 | 23 | 0.1304 |
| 5_29879582 | 9 | 0.6667 | 14 | 0.5714 | 4 | 0.25 | 3 | 0.3333 | 30 | 0.5333 | 5 | 1 | 8 | 0.375 | 7 | 0.5714 | 7 | 1 | 27 | 0.7037 |
| 5_29879737 | 10 | 0.9 | 13 | 0.9231 | 5 | 0.4 | 3 | 1 | 31 | 0.8387 | 6 | 1 | 8 | 1 | 7 | 0.8571 | 6 | 0.8333 | 27 | 0.9259 |
| 5_29879992 | 9 | 0.2222 | 8 | 0.125 | 5 | 0.2 | 4 | 0 | 26 | 0.1538 | 6 | 0.3333 | 7 | 0.1429 | 7 | 0 | 7 | 0.1429 | 27 | 0.1481 |
| 5_29880120 | 11 | 0.1818 | 12 | 0 | 5 | 0 | 6 | 0.1667 | 34 | 0.0882 | 5 | 0 | 7 | 0.1429 | 11 | 0.1818 | 6 | 0.1667 | 29 | 0.1379 |
| 5_29880150 | 8 | 0.25 | 13 | 0.2308 | 5 | 0.2 | 6 | 0.1667 | 32 | 0.2188 | 5 | 0.4 | 7 | 0.4286 | 9 | 0.7778 | 5 | 0.2 | 26 | 0.5 |
| 5_29880173 | 10 | 0.2 | 13 | 0.2308 | 6 | 0 | 5 | 0.2 | 34 | 0.1765 | 6 | 0 | 8 | 0.25 | 9 | 0.2222 | 6 | 0.3333 | 29 | 0.2069 |
| 5_29880417 | 11 | 0.5455 | 8 | 0.375 | 4 | 0.5 | 4 | 0.75 | 27 | 0.5185 | 4 | 0.25 | 6 | 0.6667 | 8 | 0.625 | 7 | 0.7143 | 25 | 0.6 |
| 5_29880633 | 9 | 0.5556 | 8 | 0.25 | 3 | 0.3333 | 5 | 0.6 | 25 | 0.44 | 8 | 0.5 | 5 | 0.6 | 9 | 0.3333 | 6 | 0.3333 | 28 | 0.4286 |
| 5_29880640 | 11 | 0.1818 | 9 | 0.2222 | 4 | 0.25 | 5 | 0.2 | 29 | 0.2069 | 8 | 0 | 4 | 0.25 | 9 | 0.2222 | 8 | 0.125 | 29 | 0.1379 |
| 5_29880961 | 11 | 0.0909 | 9 | 0.1111 | 4 | 0 | 4 | 0 | 28 | 0.0714 | 8 | 0 | 3 | 0 | 7 | 0.1429 | 7 | 0.1429 | 25 | 0.08 |
| 5_29971492 | 20 | 0.65 | 31 | 0.5484 | 30 | 0.7 | 18 | 0.8889 | 99 | 0.6768 | 17 | 0.5882 | 19 | 0.7368 | 17 | 0.7059 | 24 | 0.75 | 77 | 0.7013 |
| 5_29971518 | 19 | 0.5789 | 30 | 0.3333 | 35 | 0.5143 | 21 | 0.5714 | 105 | 0.4857 | 17 | 0.4118 | 24 | 0.375 | 19 | 0.3684 | 22 | 0.5909 | 82 | 0.439 |
| 5_29971535 | 18 | 0.1111 | 29 | 0.3448 | 34 | 0.2941 | 20 | 0.2 | 101 | 0.2574 | 16 | 0.25 | 21 | 0.1429 | 18 | 0.2222 | 22 | 0.2273 | 77 | 0.2078 |
| 5_29971612 | 16 | 0.125 | 29 | 0.1034 | 32 | 0.1562 | 14 | 0.1429 | 91 | 0.1319 | 16 | 0 | 21 | 0.2381 | 19 | 0.1053 | 22 | 0.0455 | 78 | 0.1026 |
| 5_29971625 | 21 | 0.1429 | 32 | 0.0938 | 37 | 0.1081 | 22 | 0.1818 | 112 | 0.125 | 20 | 0.05 | 25 | 0.12 | 21 | 0 | 23 | 0.1739 | 89 | 0.0899 |
| 5_29971741 | 23 | 0.1304 | 34 | 0.2647 | 38 | 0.2368 | 23 | 0.4348 | 118 | 0.2627 | 22 | 0.3182 | 28 | 0.1071 | 22 | 0.3182 | 23 | 0.2174 | 95 | 0.2316 |
| 5_29971815 | 20 | 0.15 | 35 | 0.2857 | 36 | 0.3611 | 22 | 0.1818 | 113 | 0.2655 | 22 | 0.1364 | 26 | 0.2692 | 21 | 0.3333 | 24 | 0.1667 | 93 | 0.2258 |
| 5_29991065 | 3 | 0 | 4 | 0 | 3 | 0.3333 | 3 | 0 | 13 | 0.0769 | 3 | 0 | 5 | 0.4 | 8 | 0.125 | 5 | 0.2 | 21 | 0.1905 |
| 5_30319388 | 2 | 1 | 2 | 1 | 1 | 1 | NA | NA | 5 | 1 | NA | NA | 1 | 1 | NA | NA | NA | NA | 1 | 1 |
| 5_30319489 | 1 | 1 | 2 | 1 | NA | NA | 1 | 1 | 4 | 1 | NA | NA | NA | NA | 2 | 1 | NA | NA | 2 | 1 |
| 5_30319903 | 1 | 0 | 2 | 0 | 2 | 0.5 | NA | NA | 5 | 0.2 | NA | NA | 1 | 1 | 2 | 0 | NA | NA | 3 | 0.3333 |
| 5_30321336 | 2 | 0.5 | 1 | 0 | 1 | 1 | 1 | 1 | 5 | 0.6 | NA | NA | 4 | 0.5 | 2 | 0 | NA | NA | 6 | 0.3333 |
| 5_30321486 | 2 | 0.5 | 2 | 0.5 | 1 | 1 | 1 | 0 | 6 | 0.5 | 1 | 1 | 3 | 1 | 3 | 0.6667 | NA | NA | 7 | 0.8571 |
| 5_30321762 | 2 | 0 | 2 | 0 | 2 | 0.5 | 1 | 1 | 7 | 0.2857 | 2 | 0.5 | 3 | 0.3333 | 3 | 0.6667 | NA | NA | 8 | 0.5 |
| 5_30321850 | 2 | 1 | 3 | 0.3333 | 2 | 1 | NA | NA | 7 | 0.7143 | 2 | 1 | 4 | 0.5 | 3 | 0.3333 | NA | NA | 9 | 0.5556 |
| 5_30321965 | 1 | 0 | 3 | 0 | 2 | 0.5 | 1 | 0 | 7 | 0.1429 | 1 | 0 | 4 | 0.5 | 2 | 0.5 | NA | NA | 7 | 0.4286 |
| 5_30322361 | 1 | 0 | 2 | 1 | 1 | 1 | NA | NA | 4 | 0.75 | 1 | 0 | 2 | 0.5 | 1 | 0 | NA | NA | 4 | 0.25 |
| 5_30322411 | NA | NA | 2 | 1 | 3 | 0.6667 | NA | NA | 5 | 0.8 | 1 | 0 | 4 | 0.75 | 2 | 0.5 | NA | NA | 7 | 0.5714 |
| 5_30324439 | 7 | 0.4286 | 9 | 0.5556 | 11 | 0.0909 | 8 | 0 | 35 | 0.2571 | 11 | 0.5455 | 11 | 0.1818 | 10 | 0.1 | 7 | 0.1429 | 39 | 0.2564 |
| 5_30324629 | 12 | 0.1667 | 12 | 0.0833 | 11 | 0.3636 | 8 | 0.125 | 43 | 0.186 | 13 | 0.2308 | 13 | 0.1538 | 13 | 0.2308 | 10 | 0.4 | 49 | 0.2449 |
| 5_30324640 | 13 | 0.1538 | 10 | 0.1 | 13 | 0.1538 | 8 | 0 | 44 | 0.1136 | 12 | 0.25 | 13 | 0.1538 | 13 | 0.0769 | 9 | 0 | 47 | 0.1277 |
| 5_30389335 | 7 | 0.4286 | 10 | 0.5 | 13 | 0.4615 | 12 | 0.5 | 42 | 0.4762 | 4 | 0 | 11 | 0.3636 | 9 | 0.4444 | 8 | 0.625 | 32 | 0.4062 |
| 5_30428378 | 1 | 0 | 2 | 0.5 | 2 | 0 | 1 | 0 | 6 | 0.1667 | 1 | 0 | 3 | 0.3333 | 2 | 0 | 1 | 0 | 7 | 0.1429 |

| | | | | | | | | | | | | | | | | | | | | |
|---|---|---|---|---|---|---|---|---|---|---|---|---|---|---|---|---|---|---|---|---|
| 5_3047038 | 1 | 1 | NA | NA | 1 | 0 | 2 | 0 | 4 | 0.25 | 1 | 1 | 2 | 0 | 3 | 0.6667 | 4 | 0.25 | 10 | 0.4 |
| 5_3047195 | 2 | 0.5 | NA | NA | 1 | 0 | 3 | 0 | 6 | 0.1667 | 1 | 0 | 4 | 0 | 5 | 0 | 5 | 0.2 | 15 | 0.0667 |
| 5_3047196 | 2 | 0.5 | NA | NA | 1 | 1 | 3 | 0 | 6 | 0.3333 | 1 | 0 | 4 | 0 | 5 | 0.4 | 5 | 0 | 15 | 0.1333 |
| 5_3047205 | 1 | 1 | NA | NA | 1 | 1 | 3 | 0 | 5 | 0.4 | NA | NA | 1 | 1 | 4 | 1 | 4 | 0.75 | 9 | 0.8889 |
| 5_3047210 | 1 | 1 | NA | NA | 1 | 0 | 1 | 0 | 3 | 0.3333 | 1 | 0 | 4 | 0.5 | 4 | 1 | 5 | 0.8 | 14 | 0.7143 |
| 5_3047214 | 2 | 1 | NA | NA | 1 | 1 | 4 | 1 | 7 | 1 | 1 | 1 | 3 | 1 | 5 | 0.6 | 5 | 1 | 14 | 0.8571 |
| 5_3047220 | 2 | 0 | NA | NA | 2 | 0.5 | 4 | 0 | 8 | 0.125 | 1 | 0 | 4 | 0 | 4 | 0 | 4 | 0.5 | 13 | 0.1538 |
| 5_3047227 | 2 | 0 | NA | NA | 2 | 0 | 5 | 0.2 | 9 | 0.1111 | NA | NA | 5 | 0 | 3 | 0 | 5 | 0.2 | 13 | 0.0769 |
| 5_3047246 | 1 | 0 | NA | NA | 2 | 0.5 | 4 | 0 | 7 | 0.1429 | NA | NA | 5 | 0 | 4 | 0.5 | 6 | 0 | 15 | 0.1333 |
| 5_3047315 | 2 | 1 | NA | NA | 2 | 0.5 | 5 | 0.2 | 9 | 0.4444 | NA | NA | 3 | 0.3333 | 4 | 0.5 | 4 | 0 | 11 | 0.2727 |
| 5_3047380 | 2 | 0 | NA | NA | 2 | 0 | 5 | 0.2 | 9 | 0.1111 | 1 | 1 | 3 | 0 | 1 | 0 | 6 | 0.1667 | 11 | 0.1818 |
| 5_3047406 | 1 | 0 | NA | NA | 3 | 0.3333 | 2 | 0 | 6 | 0.1667 | 3 | 1 | 2 | 0 | 2 | 0 | 2 | 0.5 | 9 | 0.4444 |
| 5_3047485 | 1 | 0 | NA | NA | 3 | 0 | 5 | 0.2 | 9 | 0.1111 | 4 | 0.25 | 2 | 0 | 3 | 0 | 3 | 0.3333 | 12 | 0.1667 |
| 5_3047504 | 1 | 0 | 1 | 0 | 3 | 0.3333 | 6 | 0.3333 | 11 | 0.2727 | 4 | 0.25 | 2 | 0 | 2 | 1 | 3 | 0.6667 | 11 | 0.4545 |
| 5_3047508 | 1 | 0 | 1 | 0 | 3 | 0 | 6 | 0.1667 | 11 | 0.0909 | 4 | 0 | 3 | 0 | 3 | 0 | 3 | 0.3333 | 13 | 0.0769 |
| 5_3047520 | 2 | 0 | 1 | 0 | 3 | 0.6667 | 5 | 0.2 | 11 | 0.2727 | 4 | 0.25 | 3 | 0 | 3 | 0.3333 | 3 | 0.3333 | 13 | 0.2308 |
| 5_3047683 | 1 | 0 | 3 | 0 | 8 | 0.25 | 8 | 0.125 | 20 | 0.15 | 6 | 0.1667 | 4 | 0.25 | 6 | 0.1667 | 4 | 0.25 | 20 | 0.2 |
| 5_3047700 | 1 | 0 | 3 | 0 | 8 | 0.25 | 4 | 0.25 | 16 | 0.1875 | 6 | 0 | 4 | 0 | 6 | 0.1667 | 3 | 0 | 19 | 0.0526 |
| 5_3047702 | 2 | 0.5 | 4 | 0 | 8 | 0 | 9 | 0.2222 | 23 | 0.1304 | 6 | 0.1667 | 4 | 0.25 | 5 | 0.2 | 3 | 0.3333 | 18 | 0.2222 |
| 5_3052456 | 5 | 0.2 | 2 | 0.5 | 5 | 0.4 | 4 | 0.25 | 16 | 0.3125 | 2 | 1 | 5 | 0 | 2 | 0.5 | NA | NA | 9 | 0.3333 |
| 5_3052504 | 4 | 0 | 2 | 0.5 | 3 | 0 | 5 | 0.4 | 14 | 0.2143 | 2 | 0.5 | 5 | 0.2 | 2 | 0.5 | NA | NA | 9 | 0.3333 |
| 5_3052506 | 3 | 1 | 2 | 1 | 4 | 0.5 | 5 | 0.4 | 14 | 0.6429 | 2 | 1 | 4 | 0.5 | 1 | 1 | NA | NA | 7 | 0.7143 |
| 5_3052513 | 1 | 1 | 1 | 1 | 2 | 1 | 2 | 1 | 6 | 1 | 1 | 1 | 2 | 0 | 2 | 1 | NA | NA | 5 | 0.6 |
| 5_3073144 | NA | NA | 4 | 0.5 | 1 | 1 | 3 | 0.6667 | 8 | 0.625 | NA | NA | 2 | 0.5 | NA | NA | NA | NA | 2 | 0.5 |
| 5_3077950 | 3 | 1 | 2 | 1 | 1 | 1 | NA | NA | 6 | 1 | 1 | 1 | NA | NA | 2 | 0.5 | 1 | 1 | 4 | 0.75 |
| 5_3077958 | 2 | 1 | 2 | 0.5 | NA | NA | NA | NA | 4 | 0.75 | 2 | 1 | NA | NA | 2 | 1 | 1 | 0 | 5 | 0.8 |
| 5_3077999 | 3 | 0 | 2 | 0.5 | 1 | 0 | NA | NA | 6 | 0.1667 | 2 | 0.5 | NA | NA | 2 | 0 | 1 | 1 | 5 | 0.4 |
| 5_3078022 | 4 | 0.5 | 2 | 0.5 | 1 | 1 | NA | NA | 7 | 0.5714 | 2 | 0.5 | NA | NA | 1 | 0 | 1 | 0 | 4 | 0.25 |
| 5_3078039 | 4 | 0.5 | 2 | 0.5 | 1 | 0 | NA | NA | 7 | 0.4286 | 2 | 0.5 | NA | NA | 2 | 1 | 1 | 1 | 5 | 0.8 |
| 5_3078044 | 3 | 1 | 1 | 1 | NA | NA | NA | NA | 4 | 1 | 2 | 1 | NA | NA | 1 | 1 | 1 | 1 | 4 | 1 |
| 5_3078105 | 4 | 0.25 | 3 | 0 | 1 | 0 | NA | NA | 8 | 0.125 | 2 | 0.5 | NA | NA | 2 | 0 | 2 | 0 | 6 | 0.1667 |
| 5_3078106 | 4 | 1 | 3 | 0.3333 | 1 | 0 | NA | NA | 8 | 0.625 | 2 | 0.5 | NA | NA | 3 | 0.6667 | 1 | 1 | 6 | 0.6667 |
| 5_3078109 | 4 | 1 | 3 | 0.6667 | 1 | 1 | NA | NA | 8 | 0.875 | NA | NA | NA | NA | 3 | 1 | 1 | 0 | 4 | 0.75 |
| 5_3078159 | 5 | 0.2 | 5 | 0.2 | 2 | 0.5 | NA | NA | 12 | 0.25 | 2 | 0 | NA | NA | 4 | 0 | 1 | 1 | 7 | 0.1429 |
| 5_3078160 | 3 | 0 | 3 | 0.6667 | 2 | 0 | NA | NA | 8 | 0.25 | 1 | 0 | NA | NA | 4 | 0 | 2 | 0.5 | 7 | 0.1429 |
| 5_3078233 | 7 | 0.1429 | 4 | 0 | 2 | 0 | 1 | 0 | 14 | 0.0714 | 2 | 0 | NA | NA | 4 | 0.25 | 4 | 0.5 | 10 | 0.3 |
| 5_3081493 | 2 | 0.5 | 2 | 1 | 4 | 0.75 | 3 | 0.6667 | 11 | 0.7273 | 5 | 1 | 3 | 1 | NA | NA | 1 | 1 | 9 | 1 |
| 5_3081515 | 3 | 1 | 3 | 1 | 3 | 1 | 3 | 1 | 12 | 1 | 4 | 1 | 4 | 1 | 2 | 0.5 | 1 | 1 | 11 | 0.9091 |
| 5_3081517 | 3 | 0.6667 | 3 | 1 | 4 | 0.75 | 2 | 1 | 12 | 0.8333 | 5 | 1 | 4 | 0.75 | 2 | 1 | 1 | 1 | 12 | 0.9167 |
| 5_3081524 | 2 | 1 | 2 | 1 | 3 | 0.3333 | 3 | 0.6667 | 10 | 0.7 | 2 | 1 | 4 | 1 | 2 | 1 | NA | NA | 8 | 1 |
| 5_3081537 | 4 | 0.75 | 5 | 0.2 | 5 | 0.8 | 2 | 0 | 16 | 0.5 | 5 | 0.6 | 4 | 0.5 | 2 | 1 | 2 | 0.5 | 13 | 0.6154 |
| 5_3081543 | 4 | 0.25 | 4 | 0.25 | 5 | 0.4 | 2 | 0 | 15 | 0.2667 | 4 | 0 | 4 | 0 | 2 | 0 | 2 | 0.5 | 12 | 0.0833 |
| 5_3081549 | 2 | 0 | 3 | 0.3333 | 6 | 0.3333 | 2 | 0 | 13 | 0.2308 | 5 | 0.6 | 2 | 0 | 2 | 0 | 2 | 0.5 | 11 | 0.3636 |
| 5_3081552 | 3 | 0 | 5 | 0.2 | 5 | 0 | 2 | 1 | 15 | 0.2 | 3 | 0.3333 | 3 | 0.3333 | 2 | 0.5 | 2 | 0 | 10 | 0.3 |
| 5_3081556 | 3 | 0.3333 | 5 | 0.2 | 6 | 0.3333 | 3 | 0.3333 | 17 | 0.2941 | 4 | 0.5 | 2 | 0.5 | 1 | 1 | 1 | 1 | 8 | 0.625 |
| 5_3081557 | 4 | 0.75 | 5 | 1 | 6 | 1 | 3 | 1 | 18 | 0.9444 | 5 | 1 | 3 | 1 | 2 | 0.5 | NA | NA | 10 | 0.9 |
| 5_3081566 | 4 | 0.5 | 4 | 0 | 4 | 0.75 | 2 | 0.5 | 14 | 0.4286 | 5 | 0.4 | 4 | 0 | 2 | 0 | 2 | 0.5 | 13 | 0.2308 |
| 5_3081585 | 4 | 0.25 | 6 | 0.3333 | 5 | 0.2 | 4 | 0.25 | 19 | 0.2632 | 5 | 0.4 | 5 | 0.4 | 2 | 0.5 | 2 | 0 | 14 | 0.3571 |
| 5_3081692 | NA | NA | 1 | 0 | NA | NA | 2 | 0.5 | 3 | 0.3333 | NA | NA | 1 | 1 | NA | NA | NA | NA | 1 | 1 |
| 5_3095656 | 4 | 0.5 | 1 | 0 | 1 | 0 | 3 | 0.3333 | 9 | 0.3333 | 4 | 0.5 | 2 | 0 | 1 | 0 | 1 | 0 | 8 | 0.25 |
| 5_3102134 | NA | NA | 3 | 0 | 3 | 0.3333 | 2 | 0 | 8 | 0.125 | 1 | 0 | 3 | 0.3333 | 3 | 0 | NA | NA | 7 | 0.1429 |
| 5_3102329 | 40 | 0.05 | 43 | 0.0465 | 36 | 0.0833 | 42 | 0.0476 | 161 | 0.0559 | 34 | 0.0294 | 38 | 0.0789 | 29 | 0.1034 | 36 | 0.0833 | 137 | 0.073 |
| 5_3102364 | NA | NA | NA | NA | 1 | 1 | 1 | 1 | 2 | 1 | 1 | 1 | NA | NA | NA | NA | NA | NA | 1 | 1 |
| 5_3102641 | NA | NA | 2 | 0.5 | 1 | 1 | 1 | 1 | 4 | 0.75 | 2 | 0.5 | NA | NA | 1 | 1 | 1 | 0 | 4 | 0.5 |
| 5_3102652 | NA | NA | 1 | 0 | NA | NA | 1 | 1 | 2 | 0.5 | 2 | 1 | NA | NA | 1 | 1 | 1 | 0 | 4 | 0.75 |
| 5_3102666 | NA | NA | 1 | 1 | 1 | 1 | 1 | 1 | 3 | 1 | 2 | 1 | NA | NA | 1 | 1 | 1 | 0 | 4 | 0.75 |
| 5_3102672 | NA | NA | 2 | 0.5 | 1 | 1 | NA | NA | 3 | 0.6667 | 2 | 0.5 | NA | NA | 1 | 1 | 1 | 1 | 4 | 0.75 |
| 5_3102691 | NA | NA | 2 | 0 | 1 | 1 | 1 | 0 | 4 | 0.25 | 2 | 1 | NA | NA | 1 | 1 | 1 | 0 | 4 | 0.75 |
| 5_3102726 | NA | NA | 1 | 1 | 1 | 1 | NA | NA | 2 | 1 | 2 | 1 | NA | NA | NA | NA | 1 | 0 | 3 | 0.6667 |
| 5_3102773 | NA | NA | 1 | 0 | NA | NA | 1 | 1 | 2 | 0.5 | 2 | 0 | NA | NA | 1 | 0 | 1 | 1 | 4 | 0.25 |
| 5_3102884 | 1 | 0 | 8 | 0 | 2 | 0 | 2 | 0.5 | 13 | 0.0769 | 3 | 0 | 1 | 0 | 2 | 0.5 | 1 | 0 | 7 | 0.1429 |
| 5_3103402 | 2 | 0 | 10 | 0.1 | 2 | 0 | 4 | 0 | 18 | 0.0556 | 8 | 0 | 4 | 0.25 | 5 | 0.2 | 2 | 0 | 19 | 0.1053 |
| 5_3103406 | 1 | 1 | 11 | 0.1818 | 4 | 0 | 4 | 0.25 | 20 | 0.2 | 7 | 0.1429 | 4 | 0.25 | 6 | 0.1667 | 2 | 0.5 | 19 | 0.2105 |
| 5_3103410 | 2 | 1 | 10 | 0.9 | 2 | 1 | 3 | 1 | 17 | 0.9412 | 8 | 0.625 | 2 | 1 | 7 | 0.8571 | 2 | 1 | 19 | 0.7895 |
| 5_3103413 | 2 | 1 | 10 | 0.4 | 4 | 0.25 | 2 | 0.5 | 18 | 0.4444 | 7 | 0.4286 | 4 | 0 | 6 | 0.5 | 2 | 0.5 | 19 | 0.3684 |

| | | | | | | | | | | | | | | | | | | | | |
|---|---|---|---|---|---|---|---|---|---|---|---|---|---|---|---|---|---|---|---|---|
| 5_3103423 | 2 | 0.5 | 13 | 0.0769 | 4 | 0.25 | 4 | 0.25 | 23 | 0.1739 | 8 | 0.125 | 4 | 0 | 7 | 0.4286 | 2 | 0 | 21 | 0.1905 |
| 5_3103427 | 2 | 0.5 | 9 | 0.3333 | 4 | 0 | 3 | 0 | 18 | 0.2222 | 6 | 0.1667 | 4 | 0 | 7 | 0.2857 | 1 | 0 | 18 | 0.1667 |
| 5_3103443 | 1 | 0 | 9 | 0.2222 | 1 | 1 | 3 | 0 | 14 | 0.2143 | 5 | 0.2 | 3 | 0 | 6 | 0.1667 | 2 | 0.5 | 16 | 0.1875 |
| 5_3103446 | 2 | 0.5 | 12 | 0.25 | 3 | 0.6667 | 4 | 0 | 21 | 0.2857 | 7 | 0.2857 | 3 | 0 | 7 | 0.2857 | 2 | 0.5 | 19 | 0.2632 |
| 5_3103455 | 2 | 1 | 12 | 0.8333 | 3 | 0.6667 | 4 | 0.75 | 21 | 0.8095 | 7 | 1 | 4 | 0.75 | 8 | 0.625 | 3 | 0.6667 | 22 | 0.7727 |
| 5_3103456 | 1 | 1 | 6 | 0.8333 | 4 | 1 | 3 | 1 | 14 | 0.9286 | 8 | 0.375 | 5 | 0.6 | 7 | 0.8571 | 2 | 0.5 | 22 | 0.5909 |
| 5_3103458 | 2 | 1 | 9 | 0.4444 | 3 | 0.3333 | 3 | 0.3333 | 17 | 0.4706 | 7 | 0.4286 | 5 | 0.4 | 7 | 0.5714 | 3 | 0.3333 | 22 | 0.4545 |
| 5_3103460 | 2 | 0 | 10 | 0.2 | 4 | 0.25 | 5 | 0 | 21 | 0.1429 | 5 | 0.2 | 5 | 0.2 | 6 | 0 | 1 | 0 | 17 | 0.1176 |
| 5_3103479 | 4 | 0.25 | 12 | 0.25 | 3 | 0.3333 | 5 | 0.2 | 24 | 0.25 | 9 | 0.1111 | 5 | 0.2 | 9 | 0.1111 | 2 | 0.5 | 25 | 0.16 |
| 5_3103480 | 3 | 0.6667 | 11 | 0.3636 | 3 | 0.6667 | 3 | 0.6667 | 20 | 0.5 | 7 | 0.2857 | 4 | 0.5 | 4 | 0.75 | 2 | 0.5 | 17 | 0.4706 |
| 5_3103484 | NA | NA | 1 | 1 | 1 | 1 | NA | NA | 2 | 1 | NA | NA | 1 | 1 | NA | NA | 1 | 1 | 2 | 1 |
| 5_3103646 | NA | NA | 2 | 0 | 1 | 1 | NA | NA | 3 | 0.3333 | NA | NA | 1 | 0 | NA | NA | 2 | 0.5 | 3 | 0.3333 |
| 5_3105063 | 1 | 1 | NA | NA | 1 | 1 | 4 | 0.25 | 6 | 0.5 | NA | NA | 2 | 0.5 | 3 | 0 | NA | NA | 5 | 0.2 |
| 5_3105073 | 1 | 0 | NA | NA | 1 | 1 | 5 | 0 | 7 | 0.1429 | NA | NA | 2 | 0.5 | 3 | 0 | NA | NA | 5 | 0.2 |
| 5_3105074 | 1 | 0 | NA | NA | 2 | 0.5 | 4 | 0.25 | 7 | 0.2857 | NA | NA | 2 | 0.5 | 2 | 0.5 | 1 | 1 | 5 | 0.6 |
| 5_3105325 | 2 | 0 | NA | NA | 3 | 0 | 5 | 0.2 | 10 | 0.1 | NA | NA | 2 | 0.5 | 3 | 0 | 1 | 0 | 6 | 0.1667 |
| 5_3106362 | NA | NA | NA | NA | NA | NA | 1 | 1 | 1 | 1 | NA | NA | 1 | 0 | 1 | 1 | NA | NA | 2 | 0.5 |
| 5_3106370 | NA | NA | NA | NA | NA | NA | 1 | 1 | 1 | 1 | NA | NA | 1 | 1 | 1 | 1 | NA | NA | 2 | 1 |
| 5_3108870 | 1 | 1 | NA | NA | NA | NA | 1 | 0 | 2 | 0.5 | 1 | 1 | NA | NA | NA | NA | 1 | 0 | 2 | 0.5 |
| 5_3108933 | 1 | 0 | NA | NA | NA | NA | 1 | 1 | 2 | 0.5 | 1 | 1 | NA | NA | NA | NA | 1 | 1 | 2 | 1 |
| 5_3109057 | 3 | 0.3333 | 7 | 0.4286 | 1 | 0 | 1 | 0 | 12 | 0.3333 | 8 | 0.25 | 1 | 0 | 5 | 0.6 | 2 | 0.5 | 16 | 0.375 |
| 5_3109091 | 3 | 0.6667 | 7 | 0.1429 | 1 | 0 | 1 | 0 | 12 | 0.25 | 6 | 0 | 1 | 0 | 3 | 0 | 2 | 0.5 | 12 | 0.0833 |
| 5_3109094 | 4 | 0 | 7 | 0.5714 | 1 | 0 | 1 | 0 | 13 | 0.3077 | 6 | 0.5 | 1 | 0 | 4 | 0.25 | 2 | 0.5 | 13 | 0.3846 |
| 5_3109106 | 3 | 0 | 6 | 0 | 1 | 1 | 1 | 0 | 11 | 0.0909 | 6 | 0 | 1 | 0 | 4 | 0.5 | 1 | 0 | 12 | 0.1667 |
| 5_3109112 | 4 | 0 | 7 | 0.2857 | 1 | 0 | 1 | 0 | 13 | 0.1538 | 7 | 0.2857 | 1 | 0 | 4 | 0 | 1 | 0 | 13 | 0.1538 |
| 5_3110917 | 1 | 1 | NA | NA | NA | NA | NA | NA | 1 | 1 | 2 | 1 | 1 | 1 | NA | NA | 2 | 1 | 5 | 1 |
| 5_3110927 | 1 | 1 | NA | NA | NA | NA | NA | NA | 1 | 1 | 1 | 0 | 1 | 1 | NA | NA | 3 | 0.3333 | 5 | 0.4 |
| 5_3110928 | 1 | 1 | NA | NA | NA | NA | NA | NA | 1 | 1 | NA | NA | 1 | 1 | NA | NA | 2 | 1 | 3 | 1 |
| 5_3111002 | NA | NA | 1 | 1 | NA | NA | NA | NA | 1 | 1 | 1 | 1 | 1 | 1 | NA | NA | 2 | 1 | 4 | 1 |
| 5_3111028 | NA | NA | 1 | 1 | NA | NA | NA | NA | 1 | 1 | NA | NA | NA | NA | NA | NA | 3 | 0.6667 | 3 | 0.6667 |
| 5_3111028 | NA | NA | 1 | 1 | NA | NA | NA | NA | 1 | 1 | NA | NA | NA | NA | NA | NA | 3 | 0.3333 | 3 | 0.3333 |
| 5_3111035 | NA | NA | 1 | 1 | NA | NA | NA | NA | 1 | 1 | NA | NA | NA | NA | NA | NA | 3 | 0.6667 | 3 | 0.6667 |
| 5_3111042 | NA | NA | 1 | 1 | NA | NA | NA | NA | 1 | 1 | NA | NA | NA | NA | NA | NA | 2 | 1 | 2 | 1 |
| 5_3111043 | NA | NA | 1 | 1 | NA | NA | NA | NA | 1 | 1 | NA | NA | NA | NA | NA | NA | 2 | 0.5 | 2 | 0.5 |
| 5_3111052 | NA | NA | 1 | 1 | NA | NA | NA | NA | 1 | 1 | NA | NA | NA | NA | NA | NA | 1 | 1 | 1 | 1 |
| 5_3111069 | NA | NA | 1 | 1 | NA | NA | NA | NA | 1 | 1 | NA | NA | NA | NA | NA | NA | 2 | 1 | 2 | 1 |
| 5_3111237 | NA | NA | NA | NA | 1 | 1 | 1 | 1 | 2 | 1 | NA | NA | 1 | 1 | 1 | 1 | 1 | 0 | 3 | 0.6667 |
| 5_3111240 | 1 | 0 | 7 | 0 | 6 | 0.1667 | 10 | 0.1 | 24 | 0.0833 | 6 | 0 | 9 | 0.2222 | 3 | 0 | 10 | 0.1 | 28 | 0.1071 |
| 5_3111255 | 3 | 0.3333 | 6 | 0.3333 | 6 | 0.5 | 11 | 0.3636 | 26 | 0.3846 | 5 | 0.6 | 5 | 0.6 | 4 | 0 | 10 | 0.4 | 24 | 0.4167 |
| 5_3111265 | 3 | 0.6667 | 8 | 1 | 6 | 1 | 8 | 0.875 | 25 | 0.92 | 4 | 1 | 8 | 1 | 4 | 1 | 12 | 0.9167 | 28 | 0.9643 |
| 5_3111267 | 3 | 0 | 7 | 0.2857 | 7 | 0.2857 | 10 | 0.2 | 27 | 0.2222 | 6 | 0.1667 | 10 | 0.3 | 5 | 0 | 12 | 0.25 | 33 | 0.2121 |
| 5_3111279 | 3 | 0.6667 | 5 | 0.6 | 3 | 1 | 6 | 0.8333 | 17 | 0.7647 | 3 | 0.3333 | 7 | 0.5714 | 5 | 0.6 | 6 | 0.8333 | 21 | 0.619 |
| 5_3111289 | 3 | 0 | 7 | 0.7143 | 6 | 0.6667 | 9 | 0.7778 | 25 | 0.64 | 4 | 0.25 | 6 | 0.8333 | 4 | 0.5 | 9 | 0.7778 | 23 | 0.6522 |
| 5_3111295 | 2 | 0 | 7 | 0.1429 | 6 | 0.5 | 8 | 1 | 23 | 0.5217 | 4 | 0.75 | 6 | 0.3333 | 2 | 1 | 10 | 0.5 | 22 | 0.5455 |
| 5_3119361 | 3 | 0 | 4 | 0.25 | 3 | 0 | 4 | 0.25 | 14 | 0.1429 | 5 | 0 | 2 | 0 | 1 | 1 | 3 | 0.3333 | 11 | 0.1818 |
| 5_3119378 | 2 | 0 | 4 | 0.25 | 3 | 0 | 3 | 0 | 12 | 0.0833 | 6 | 0.3333 | 2 | 0.5 | NA | NA | 3 | 0.3333 | 11 | 0.3636 |
| 5_3119380 | 3 | 0 | 4 | 0.5 | 3 | 0.3333 | 4 | 0 | 14 | 0.2143 | 5 | 0.2 | 1 | 0 | NA | NA | 3 | 0 | 9 | 0.1111 |
| 5_3119385 | 3 | 0 | 4 | 0 | 3 | 0 | 4 | 0.25 | 14 | 0.0714 | 6 | 0.1667 | 2 | 0 | NA | NA | 3 | 0 | 11 | 0.0909 |
| 5_3119402 | 3 | 0.6667 | 4 | 0.75 | 3 | 0.6667 | 4 | 0.5 | 14 | 0.6429 | 6 | 0.1667 | 2 | 0.5 | NA | NA | 2 | 1 | 10 | 0.4 |
| 5_3119413 | NA | NA | 2 | 0.5 | 3 | 0 | 2 | 0 | 7 | 0.1429 | 3 | 0.3333 | 2 | 0.5 | NA | NA | 1 | 1 | 6 | 0.5 |
| 5_3120051 | NA | NA | 1 | 1 | NA | NA | NA | NA | 1 | 1 | NA | NA | 1 | 1 | NA | NA | 1 | 1 | 2 | 1 |
| 5_3120060 | NA | NA | 2 | 1 | 1 | 1 | 1 | 1 | 4 | 1 | 1 | 1 | 2 | 1 | NA | NA | 1 | 1 | 4 | 1 |
| 5_3121182 | 10 | 0.7 | 9 | 0.8889 | 4 | 0.5 | 3 | 1 | 26 | 0.7692 | 5 | 1 | 6 | 1 | 11 | 0.5455 | 6 | 0.8333 | 28 | 0.7857 |
| 5_3121189 | 16 | 0.5 | 14 | 0.5714 | 9 | 0.5556 | 11 | 0.6364 | 50 | 0.56 | 11 | 0.6364 | 17 | 0.8824 | 21 | 0.7619 | 8 | 0.625 | 57 | 0.7544 |
| 5_3121191 | 13 | 0.2308 | 16 | 0.5 | 11 | 0.3636 | 11 | 0.5455 | 51 | 0.4118 | 12 | 0.8333 | 17 | 0.5882 | 21 | 0.4762 | 14 | 0.6429 | 64 | 0.6094 |
| 5_3124403 | 2 | 0 | 14 | 0.6429 | 6 | 0.6667 | 2 | 0.5 | 24 | 0.5833 | 18 | 0.5 | 1 | 1 | 1 | 1 | 3 | 1 | 23 | 0.6087 |
| 5_3124406 | 2 | 0 | 14 | 0.5 | 5 | 0.4 | 2 | 0.5 | 23 | 0.4348 | 13 | 0.5385 | 1 | 0 | NA | NA | 4 | 0.75 | 18 | 0.5556 |
| 5_3129815 | 5 | 0 | 8 | 0.125 | 7 | 0 | 7 | 0.1429 | 27 | 0.0741 | 7 | 0 | 1 | 1 | 2 | 0.5 | 10 | 0 | 20 | 0.1 |
| 5_3129872 | 2 | 0 | 7 | 0.2857 | 5 | 0 | 4 | 0 | 18 | 0.1111 | 7 | 0.1429 | 3 | 0.3333 | 2 | 0 | 4 | 0.25 | 16 | 0.1875 |
| 5_3129885 | 3 | 0 | 6 | 0.6667 | 6 | 0 | 5 | 0.4 | 20 | 0.3 | 7 | 0 | 3 | 0 | 1 | 0 | 6 | 0.1667 | 17 | 0.0588 |
| 5_3129898 | 2 | 0 | 6 | 0.3333 | 5 | 0.4 | 5 | 0.6 | 18 | 0.3889 | 7 | 0.4286 | 3 | 0.6667 | 2 | 0.5 | 6 | 0.6667 | 18 | 0.5556 |
| 5_3129903 | 3 | 0.6667 | 6 | 0.6667 | 4 | 0.25 | 6 | 0.5 | 19 | 0.5263 | 5 | 0.4 | 3 | 0.3333 | 2 | 1 | 6 | 0.8333 | 16 | 0.625 |
| 5_3129906 | 3 | 0 | 6 | 0.3333 | 5 | 0.2 | 5 | 0.2 | 19 | 0.2105 | 5 | 0.4 | 3 | 0 | 2 | 0 | 6 | 0.1667 | 16 | 0.1875 |
| 5_3131440 | 2 | 0.5 | 4 | 0.5 | 2 | 1 | 4 | 0.25 | 12 | 0.5 | 2 | 0.5 | 2 | 0.5 | 5 | 0.8 | 3 | 0.3333 | 12 | 0.5833 |
| 5_3131453 | 2 | 0.5 | 1 | 1 | 2 | 0 | 4 | 0.25 | 9 | 0.3333 | 2 | 0.5 | 1 | 1 | 3 | 0.3333 | 2 | 0 | 8 | 0.375 |

| | | | | | | | | | | | | | | | | | | | | |
|---|---|---|---|---|---|---|---|---|---|---|---|---|---|---|---|---|---|---|---|---|
| 5_31314734 | 3 | 0 | 7 | 0.4286 | 3 | 0 | 6 | 0.3333 | 19 | 0.2632 | 6 | 0.3333 | 5 | 0.2 | 7 | 0.2857 | 9 | 0.3333 | 27 | 0.2963 |
| 5_31314806 | 3 | 0.6667 | 6 | 0.8333 | 2 | 0.5 | 4 | 0.75 | 15 | 0.7333 | 4 | 1 | 5 | 0.8 | 7 | 0.7143 | 9 | 0.7778 | 25 | 0.8 |
| 5_31314852 | 3 | 0 | 6 | 0.1667 | 1 | 0 | 4 | 0 | 14 | 0.0714 | 4 | 0 | 5 | 0.6 | 6 | 0 | 8 | 0 | 23 | 0.1304 |
| 5_31314877 | 2 | 1 | 5 | 0.8 | 1 | 1 | 4 | 0.75 | 12 | 0.8333 | NA | NA | 4 | 0.5 | 6 | 0.6667 | 6 | 0.6667 | 16 | 0.625 |
| 5_31315025 | 1 | 1 | 4 | 0.75 | 2 | 0.5 | 3 | 0.6667 | 10 | 0.7 | 1 | 1 | 3 | 0.3333 | 2 | 0.5 | 7 | 0.7143 | 13 | 0.6154 |
| 5_31315089 | 2 | 0.5 | 5 | 0.4 | 2 | 0 | 1 | 1 | 10 | 0.4 | 4 | 0.25 | 4 | 0 | 4 | 0 | 4 | 0.25 | 16 | 0.125 |
| 5_31331160 | NA | NA | NA | NA | 2 | 0.5 | NA | NA | 2 | 0.5 | NA | NA | 1 | 1 | 1 | 1 | 4 | 0.25 | 6 | 0.5 |
| 5_31356065 | 13 | 0.3077 | 16 | 0.625 | 13 | 0.2308 | 17 | 0.2941 | 59 | 0.3729 | 8 | 0.375 | 10 | 0.4 | 14 | 0.5714 | 10 | 0.4 | 42 | 0.4524 |
| 5_31356091 | 13 | 0.2308 | 14 | 0.2143 | 13 | 0.0769 | 17 | 0.0588 | 57 | 0.1404 | 7 | 0.2857 | 8 | 0.25 | 8 | 0.375 | 8 | 0 | 31 | 0.2258 |
| 5_31356399 | 1 | 0 | 1 | 0 | 5 | 0.2 | 3 | 0 | 10 | 0.1 | 1 | 0 | 2 | 0.5 | 4 | 0 | 3 | 0 | 10 | 0.1 |
| 5_31356481 | 1 | 1 | 2 | 0.5 | 3 | 0.6667 | 3 | 0.6667 | 9 | 0.6667 | NA | NA | 2 | 1 | 2 | 1 | 3 | 0.6667 | 7 | 0.8571 |
| 5_31356482 | 1 | 1 | NA | NA | NA | NA | NA | NA | 1 | 1 | NA | NA | NA | NA | 1 | 1 | NA | NA | 1 | 1 |
| 5_31356524 | 1 | 0 | 2 | 1 | 4 | 0.25 | 4 | 0.5 | 11 | 0.4545 | 1 | 0 | 1 | 1 | 4 | 0.5 | 4 | 0.5 | 10 | 0.5 |
| 5_31356545 | 2 | 0.5 | 1 | 1 | 5 | 0 | 4 | 0.75 | 12 | 0.4167 | NA | NA | 1 | 0 | 3 | 0.3333 | 4 | 0.25 | 8 | 0.25 |
| 5_31356596 | 1 | 0 | 2 | 1 | 6 | 0.5 | 3 | 0.3333 | 12 | 0.5 | 1 | 0 | 1 | 1 | 2 | 0.5 | 3 | 1 | 7 | 0.7143 |
| 5_31356664 | 2 | 0.5 | 2 | 0.5 | 3 | 0.3333 | 2 | 0 | 9 | 0.3333 | NA | NA | 1 | 1 | 2 | 0 | 3 | 0.6667 | 6 | 0.5 |
| 5_31356675 | 2 | 0.5 | 2 | 0.5 | 4 | 0.75 | 5 | 0.8 | 13 | 0.6923 | NA | NA | NA | NA | 2 | 0 | 4 | 0.5 | 6 | 0.3333 |
| 5_31356837 | 2 | 1 | 2 | 0.5 | 5 | 0.8 | 4 | 0.5 | 13 | 0.6923 | 1 | 1 | 1 | 1 | 1 | 1 | 4 | 0.75 | 7 | 0.8571 |
| 5_31356918 | 1 | 1 | 2 | 0 | 5 | 0.4 | 3 | 1 | 11 | 0.5455 | 1 | 1 | 2 | 1 | 2 | 0 | 4 | 1 | 9 | 0.7778 |
| 5_31357150 | 2 | 0 | 1 | 0 | 4 | 0.25 | 4 | 0 | 11 | 0.0909 | NA | NA | 2 | 0.5 | 2 | 0 | 3 | 0 | 7 | 0.1429 |
| 5_31357325 | 2 | 1 | 1 | 1 | 3 | 0.6667 | 3 | 0.6667 | 9 | 0.7778 | 1 | 1 | 1 | 0 | 2 | 0.5 | 2 | 0.5 | 6 | 0.5 |
| 5_31357348 | 2 | 0.5 | 1 | 0 | 3 | 0.3333 | 3 | 0.3333 | 9 | 0.3333 | NA | NA | 2 | 0.5 | 2 | 0 | 3 | 0.3333 | 7 | 0.2857 |
| 5_31357428 | 2 | 0 | 1 | 0 | 4 | 0.5 | 3 | 0 | 10 | 0.2 | 1 | 0 | 1 | 1 | 2 | 0 | 3 | 0 | 7 | 0.1429 |
| 5_31360118 | 4 | 0 | 15 | 0.5333 | 19 | 0.3158 | 13 | 0.4615 | 51 | 0.3922 | 3 | 0.3333 | 14 | 0.3571 | 11 | 0.6364 | 10 | 0.4 | 38 | 0.4474 |
| 5_31360133 | 3 | 0.3333 | 9 | 0.1111 | 13 | 0 | 8 | 0 | 33 | 0.0606 | 3 | 0 | 10 | 0 | 11 | 0.0909 | 10 | 0.1 | 34 | 0.0588 |
| 5_31360161 | 6 | 0.8333 | 16 | 0.25 | 16 | 0.375 | 13 | 0.3077 | 51 | 0.3725 | 5 | 0.4 | 12 | 0.1667 | 10 | 0.5 | 9 | 0.2222 | 36 | 0.3056 |
| 5_31360371 | 6 | 1 | 15 | 0.2 | 15 | 0.1333 | 12 | 0.25 | 48 | 0.2917 | 4 | 0.5 | 14 | 0.3571 | 11 | 0.2727 | 11 | 0.6364 | 40 | 0.425 |
| 5_31360378 | 6 | 0 | 17 | 0.1176 | 16 | 0.0625 | 13 | 0.0769 | 52 | 0.0769 | 4 | 0 | 14 | 0.1429 | 11 | 0.2727 | 10 | 0 | 39 | 0.1282 |
| 5_31360398 | 6 | 0.8333 | 17 | 0.8824 | 15 | 0.8667 | 12 | 0.8333 | 50 | 0.86 | 4 | 1 | 14 | 1 | 12 | 1 | 10 | 0.9 | 40 | 0.975 |
| 5_31360429 | 6 | 0.1667 | 15 | 0.1333 | 12 | 0.1667 | 10 | 0.1 | 43 | 0.1395 | 3 | 0 | 11 | 0.1818 | 10 | 0.3 | 8 | 0.125 | 32 | 0.1875 |
| 5_31363543 | 1 | 0 | 10 | 0.4 | 10 | 0.8 | 9 | 1 | 30 | 0.7 | 1 | 1 | 6 | 0.8333 | 6 | 0.8333 | 6 | 1 | 19 | 0.8947 |
| 5_31363626 | 5 | 0 | 11 | 0.0909 | 11 | 0.3636 | 9 | 0.1111 | 36 | 0.1667 | 2 | 0 | 5 | 0 | 4 | 0.5 | 9 | 0.1111 | 20 | 0.15 |
| 5_31363634 | 4 | 0.25 | 9 | 0.2222 | 10 | 0.1 | 9 | 0.1111 | 32 | 0.1562 | 2 | 0 | 6 | 0.1667 | 5 | 0.2 | 9 | 0.3333 | 22 | 0.2273 |
| 5_31363678 | 5 | 0.2 | 11 | 0.5455 | 12 | 0.4167 | 9 | 0.3333 | 37 | 0.4054 | 2 | 0 | 5 | 0.8 | 5 | 0.4 | 9 | 0.5556 | 21 | 0.5238 |
| 5_31363719 | 4 | 0.25 | 11 | 0.6364 | 9 | 0.1111 | 7 | 0.2857 | 31 | 0.3548 | 2 | 0 | 4 | 0 | 2 | 0.5 | 9 | 0.5556 | 17 | 0.3529 |
| 5_31407069 | NA | NA | 1 | 1 | 1 | 1 | NA | NA | 2 | 1 | NA | NA | NA | NA | NA | NA | 1 | 1 | 1 | 1 |
| 5_31407444 | NA | NA | NA | NA | 1 | 1 | NA | NA | 1 | 1 | NA | NA | NA | NA | NA | NA | 1 | 1 | 1 | 1 |
| 5_31410679 | 2 | 0 | 5 | 0 | 3 | 0 | 4 | 0.25 | 14 | 0.0714 | 1 | 0 | 3 | 0 | 7 | 0.1429 | 3 | 0 | 14 | 0.0714 |
| 5_31449929 | 1 | 0 | 2 | 0.5 | 6 | 0.1667 | 2 | 0 | 11 | 0.1818 | 5 | 0 | 4 | 0.25 | 4 | 0 | 4 | 0.25 | 17 | 0.1176 |
| 5_31450147 | 1 | 0 | 2 | 0 | 6 | 0.3333 | 3 | 0.3333 | 12 | 0.25 | 5 | 0.4 | 2 | 0.5 | 2 | 0.5 | 4 | 0.5 | 13 | 0.4615 |
| 5_31450227 | 1 | 0 | 2 | 0 | 4 | 0 | 3 | 0.6667 | 10 | 0.2 | 6 | 0 | 3 | 0.3333 | 4 | 0 | 4 | 0 | 17 | 0.0588 |
| 5_31450261 | 7 | 0.1429 | 10 | 0.3 | 11 | 0.4545 | 10 | 0.5 | 38 | 0.3684 | 13 | 0.6923 | 10 | 0.4 | 6 | 0.6667 | 15 | 0.6 | 44 | 0.5909 |
| 5_31450473 | 7 | 0.1429 | 11 | 0.3636 | 12 | 0.0833 | 9 | 0.2222 | 39 | 0.2051 | 11 | 0.3636 | 10 | 0.1 | 5 | 0 | 15 | 0.2667 | 41 | 0.2195 |
| 5_31450589 | 4 | 0 | 8 | 0.125 | 10 | 0.2 | 5 | 0 | 27 | 0.1111 | 6 | 0 | 6 | 0.1667 | 3 | 0 | 9 | 0.1111 | 24 | 0.0833 |
| 5_31637715 | 3 | 0 | 4 | 0 | 3 | 0 | 3 | 0.3333 | 13 | 0.0769 | 4 | 0.25 | 4 | 0 | 3 | 0 | 5 | 0 | 16 | 0.0625 |
| 5_31638049 | 2 | 0.5 | 4 | 0.25 | 2 | 0 | 2 | 0 | 10 | 0.2 | 5 | 0.2 | 4 | 0 | 4 | 0.25 | 5 | 0.8 | 18 | 0.3333 |
| 5_31638118 | 3 | 0.3333 | 4 | 0.75 | 3 | 0.3333 | 3 | 0.6667 | 13 | 0.5385 | 6 | 0.5 | 4 | 0.25 | 3 | 1 | 5 | 0.6 | 18 | 0.5556 |
| 5_31638213 | 3 | 0 | 4 | 0.25 | 2 | 0 | 3 | 0 | 12 | 0.0833 | 5 | 0 | 3 | 0 | 4 | 0.25 | 5 | 0.4 | 17 | 0.1765 |
| 5_31668306 | 4 | 0.25 | 1 | 1 | 4 | 0 | 4 | 0.5 | 13 | 0.3077 | 2 | 0.5 | 2 | 0.5 | 4 | 0.5 | 6 | 0.5 | 14 | 0.5 |
| 5_31668361 | 6 | 1 | 1 | 1 | 5 | 0.4 | 4 | 1 | 16 | 0.8125 | 2 | 0.5 | 1 | 1 | 4 | 0.5 | 5 | 1 | 12 | 0.75 |
| 5_31668376 | 6 | 0.3333 | 1 | 0 | 5 | 0.4 | 4 | 0 | 16 | 0.25 | 4 | 0 | 2 | 0.5 | 6 | 0.1667 | 5 | 0 | 17 | 0.1176 |
| 5_31668531 | 5 | 0.6 | 1 | 0 | 4 | 0.5 | 3 | 0.6667 | 13 | 0.5385 | 4 | 0.25 | 3 | 0.6667 | 4 | 0.5 | 3 | 0.6667 | 14 | 0.5 |
| 5_31668556 | 6 | 1 | 2 | 1 | 4 | 0.75 | 3 | 1 | 15 | 0.9333 | 4 | 1 | 2 | 0.5 | 5 | 1 | 6 | 0.8333 | 17 | 0.8824 |
| 5_31668589 | 6 | 1 | 1 | 1 | 5 | 1 | 3 | 1 | 15 | 1 | 4 | 1 | 3 | 1 | 5 | 0.8 | 6 | 1 | 18 | 0.9444 |
| 5_31668595 | 6 | 0.1667 | 1 | 0 | 4 | 0 | 3 | 0.3333 | 14 | 0.1429 | 4 | 0.5 | 3 | 0 | 3 | 0.3333 | 4 | 0 | 14 | 0.2143 |
| 5_31668737 | 7 | 0.2857 | 2 | 0.5 | 6 | 0.1667 | 4 | 0.25 | 19 | 0.2632 | 4 | 0 | 2 | 0.5 | 6 | 0 | 6 | 0 | 18 | 0.0556 |
| 5_31668854 | 2 | 0.5 | NA | NA | NA | NA | NA | NA | 2 | 0.5 | 1 | 1 | NA | NA | 2 | 0 | NA | NA | 3 | 0.3333 |
| 5_31670075 | 1 | 1 | NA | NA | NA | NA | NA | NA | 1 | 1 | NA | NA | NA | NA | 2 | 0.5 | NA | NA | 2 | 0.5 |
| 5_31672315 | 1 | 0 | 6 | 0 | 5 | 0 | 5 | 0.2 | 17 | 0.0588 | NA | NA | 3 | 0.3333 | 5 | 0 | 3 | 0 | 11 | 0.0909 |
| 5_31672930 | 1 | 0 | 4 | 1 | 5 | 1 | 4 | 0.5 | 14 | 0.7857 | NA | NA | 6 | 1 | 3 | 0.3333 | 1 | 0 | 10 | 0.7 |
| 5_31672968 | 1 | 1 | 2 | 1 | 3 | 1 | 4 | 1 | 10 | 1 | NA | NA | 6 | 0.8333 | 3 | 1 | 1 | 1 | 10 | 0.9 |
| 5_31673099 | 1 | 0 | 3 | 0 | 5 | 0.4 | 3 | 0 | 12 | 0.1667 | NA | NA | 5 | 0.2 | 3 | 0.3333 | 1 | 1 | 9 | 0.3333 |
| 5_31673125 | 1 | 1 | 4 | 0.5 | 5 | 0.8 | 3 | 0.6667 | 13 | 0.6923 | NA | NA | 6 | 0.8333 | 3 | 1 | 1 | 1 | 10 | 0.9 |
| 5_31680061 | 1 | 1 | 3 | 1 | 3 | 1 | 1 | 1 | 8 | 1 | NA | NA | 5 | 0.6 | 3 | 1 | NA | NA | 8 | 0.75 |
| 5_31680256 | NA | NA | 3 | 1 | 2 | 0.5 | 1 | 0 | 6 | 0.6667 | NA | NA | 5 | 1 | 3 | 1 | NA | NA | 8 | 1 |

| | | | | | | | | | | | | | | | | | | | | |
|---|---|---|---|---|---|---|---|---|---|---|---|---|---|---|---|---|---|---|---|---|
| 5_3168037 | 1 | 0 | 3 | 1 | 3 | 0.3333 | NA | NA | 7 | 0.5714 | NA | NA | 5 | 0.2 | 2 | 0 | NA | NA | 7 | 0.1429 |
| 5_3168053 | 1 | 1 | 3 | 0.6667 | 2 | 0 | 1 | 1 | 7 | 0.5714 | NA | NA | 4 | 0 | 3 | 0.6667 | NA | NA | 7 | 0.2857 |
| 5_3168487 | 2 | 0 | 2 | 0.5 | NA | NA | 1 | 0 | 5 | 0.2 | 1 | 1 | 1 | 0 | NA | NA | 2 | 0.5 | 4 | 0.5 |
| 5_3169350 | 4 | 0 | 3 | 0.3333 | 1 | 0 | 3 | 0 | 11 | 0.0909 | NA | NA | 1 | 0 | NA | NA | 3 | 0.3333 | 4 | 0.25 |
| 5_3171123 | 5 | 0.2 | 3 | 0 | 2 | 0.5 | 7 | 0.1429 | 17 | 0.1765 | 2 | 0 | 3 | 0.6667 | NA | NA | 3 | 0.6667 | 8 | 0.5 |
| 5_3172343 | NA | NA | NA | NA | 1 | 0 | 1 | 1 | 2 | 0.5 | 1 | 0 | 2 | 0.5 | NA | NA | NA | NA | 3 | 0.3333 |
| 5_3177983 | 13 | 0.1538 | 15 | 0.0667 | 9 | 0 | 10 | 0 | 47 | 0.0638 | 12 | 0.0833 | 14 | 0.1429 | 12 | 0.1667 | 6 | 0 | 44 | 0.1136 |
| 5_3224225 | 3 | 0.3333 | 7 | 0 | 4 | 0 | 11 | 0.0909 | 25 | 0.08 | 5 | 0.2 | 4 | 0.25 | 3 | 0.3333 | 8 | 0.25 | 20 | 0.25 |
| 5_3229391 | 1 | 1 | 1 | 0 | NA | NA | NA | NA | 2 | 0.5 | 2 | 0.5 | NA | NA | NA | NA | NA | NA | 2 | 0.5 |
| 5_3231026 | 2 | 0 | 2 | 0.5 | NA | NA | 1 | 0 | 5 | 0.2 | 2 | 0.5 | 1 | 0 | 2 | 0.5 | 1 | 0 | 6 | 0.3333 |
| 5_3231048 | 4 | 0.5 | 2 | 0.5 | NA | NA | 1 | 0 | 7 | 0.4286 | 1 | 0 | 1 | 0 | 3 | 0.3333 | 1 | 0 | 6 | 0.1667 |
| 5_3231049 | 4 | 0.75 | 2 | 1 | NA | NA | 1 | 1 | 7 | 0.8571 | 2 | 1 | 1 | 1 | 3 | 1 | 1 | 1 | 7 | 1 |
| 5_3231050 | 4 | 0.75 | 1 | 0 | NA | NA | 1 | 1 | 6 | 0.6667 | 2 | 0.5 | 1 | 0 | 2 | 0.5 | 1 | 0 | 6 | 0.3333 |
| 5_3231059 | 4 | 0.75 | 3 | 1 | NA | NA | 1 | 1 | 8 | 0.875 | 2 | 1 | 1 | 0 | 4 | 0.75 | 1 | 0 | 8 | 0.625 |
| 5_3231067 | 4 | 0.75 | 2 | 1 | NA | NA | 1 | 1 | 7 | 0.8571 | 2 | 1 | 1 | 1 | 4 | 1 | 1 | 1 | 8 | 1 |
| 5_3231068 | 4 | 1 | 3 | 0.3333 | NA | NA | 1 | 1 | 8 | 0.75 | 1 | 1 | NA | NA | 4 | 0.5 | 1 | 0 | 6 | 0.5 |
| 5_3231071 | 4 | 0.5 | 3 | 0.6667 | NA | NA | 1 | 0 | 8 | 0.5 | 2 | 1 | 1 | 0 | 4 | 0.75 | 1 | 1 | 8 | 0.75 |
| 5_3231077 | 4 | 1 | 3 | 0.6667 | NA | NA | NA | NA | 7 | 0.8571 | 2 | 1 | 1 | 1 | 5 | 0.8 | 1 | 1 | 9 | 0.8889 |
| 5_3231078 | 4 | 1 | 2 | 1 | NA | NA | 1 | 1 | 7 | 1 | 2 | 1 | 1 | 1 | 5 | 1 | NA | NA | 8 | 1 |
| 5_3231090 | 2 | 1 | 3 | 1 | NA | NA | 1 | 1 | 6 | 1 | 3 | 1 | 2 | 1 | 4 | 1 | 2 | 1 | 11 | 1 |
| 5_3231107 | 3 | 0.3333 | 2 | 0 | 1 | 0 | NA | NA | 6 | 0.1667 | 3 | 0 | NA | NA | 5 | 0.2 | 2 | 0.5 | 10 | 0.2 |
| 5_3231114 | 3 | 0 | 1 | 0 | 1 | 1 | NA | NA | 5 | 0.2 | 4 | 0 | 1 | 1 | 5 | 0.2 | NA | NA | 10 | 0.2 |
| 5_3231118 | 2 | 0 | 2 | 0.5 | 1 | 1 | NA | NA | 5 | 0.4 | 4 | 0.25 | 1 | 1 | 5 | 0.8 | 2 | 0 | 12 | 0.5 |
| 5_3231123 | 2 | 0 | 2 | 0 | 1 | 1 | NA | NA | 5 | 0.2 | 3 | 0.3333 | 1 | 1 | 3 | 1 | 2 | 0.5 | 9 | 0.6667 |
| 5_3231241 | 3 | 0 | 2 | 0 | 2 | 0.5 | NA | NA | 7 | 0.1429 | 4 | 0.25 | 1 | 0 | 5 | 0 | 2 | 0 | 12 | 0.0833 |
| 5_3231299 | 6 | 0 | 3 | 0.3333 | 2 | 0 | NA | NA | 11 | 0.0909 | 5 | 0 | 2 | 0 | 6 | 0.1667 | 2 | 0 | 15 | 0.0667 |
| 5_3231358 | 5 | 0 | 7 | 0.4286 | 3 | 0.3333 | NA | NA | 15 | 0.2667 | 5 | 0 | 5 | 0 | 5 | 0.2 | 1 | 0 | 16 | 0.0625 |
| 5_3231365 | 5 | 0 | 7 | 0.1429 | 2 | 0 | NA | NA | 14 | 0.0714 | 3 | 0.3333 | 4 | 0 | 3 | 0 | 2 | 0 | 12 | 0.0833 |
| 5_3231369 | 7 | 0 | 7 | 0.1429 | 2 | 0 | NA | NA | 16 | 0.0625 | 4 | 0 | 5 | 0.2 | 6 | 0 | 2 | 0.5 | 17 | 0.1176 |
| 5_3231374 | 6 | 0 | 6 | 0.3333 | 3 | 0 | NA | NA | 15 | 0.1333 | 4 | 0 | 5 | 0 | 5 | 0 | 2 | 0.5 | 16 | 0.0625 |
| 5_3231439 | 7 | 0.2857 | 8 | 0.125 | 3 | 0.6667 | NA | NA | 18 | 0.2778 | 6 | 0.3333 | 6 | 0.1667 | 6 | 0.1667 | 3 | 0.6667 | 21 | 0.2857 |
| 5_3231450 | 8 | 0 | 7 | 0.1429 | 4 | 0.5 | 1 | 0 | 20 | 0.15 | 7 | 0 | 5 | 0 | 3 | 0.3333 | 3 | 0 | 18 | 0.0556 |
| 5_3231452 | 6 | 0 | 6 | 0.1667 | 4 | 0 | 1 | 0 | 17 | 0.0588 | 5 | 0 | 6 | 0.1667 | 2 | 0.5 | 3 | 0 | 16 | 0.125 |
| 5_3231489 | 7 | 0.1429 | 8 | 0 | 5 | 0.2 | 1 | 0 | 21 | 0.0952 | 10 | 0 | 6 | 0.3333 | 4 | 0.25 | 4 | 0 | 24 | 0.125 |
| 5_3252148 | NA | NA | NA | NA | NA | NA | 1 | 1 | 1 | 1 | NA | NA | NA | NA | 1 | 1 | NA | NA | 1 | 1 |
| 5_3264844 | 32 | 0.5938 | 32 | 0.5938 | 24 | 0.75 | 40 | 0.7 | 128 | 0.6562 | 16 | 0.8125 | 37 | 0.7027 | 41 | 0.6098 | 39 | 0.6667 | 133 | 0.6767 |
| 5_3264944 | 40 | 0.25 | 31 | 0.1935 | 29 | 0.1379 | 50 | 0.22 | 150 | 0.2067 | 19 | 0.3158 | 38 | 0.1579 | 42 | 0.119 | 37 | 0.1892 | 136 | 0.1765 |
| 5_3284214 | 2 | 0.5 | 1 | 1 | NA | NA | 1 | 1 | 4 | 0.75 | NA | NA | NA | NA | 1 | 1 | 1 | 1 | 2 | 1 |
| 5_3284237 | 3 | 0.6667 | 1 | 1 | NA | NA | 1 | 1 | 5 | 0.8 | NA | NA | NA | NA | NA | NA | 1 | 1 | 1 | 1 |
| 5_3284247 | 3 | 0.6667 | 1 | 0 | NA | NA | 1 | 1 | 5 | 0.6 | NA | NA | NA | NA | 1 | 1 | 1 | 1 | 2 | 1 |
| 5_3284248 | 3 | 0 | 1 | 1 | NA | NA | 1 | 0 | 5 | 0.2 | NA | NA | NA | NA | 1 | 0 | 1 | 1 | 2 | 0.5 |
| 5_3284274 | 3 | 0.3333 | 1 | 1 | NA | NA | 1 | 0 | 5 | 0.4 | NA | NA | NA | NA | 1 | 0 | 1 | 1 | 2 | 0.5 |
| 5_3289406 | 13 | 0 | 3 | 0 | 5 | 0.2 | 7 | 0.1429 | 28 | 0.0714 | 8 | 0.375 | 8 | 0.25 | 7 | 0.2857 | 7 | 0 | 30 | 0.2333 |
| 5_3289421 | 13 | 0.2308 | 4 | 0.25 | 6 | 0 | 9 | 0.2222 | 32 | 0.1875 | 8 | 0.375 | 9 | 0.6667 | 8 | 0.25 | 8 | 0.25 | 33 | 0.3939 |
| 5_3289463 | 13 | 0.4615 | 4 | 0.5 | 5 | 0.4 | 7 | 0.7143 | 29 | 0.5172 | 8 | 0.75 | 8 | 0.5 | 7 | 0.2857 | 6 | 0.5 | 29 | 0.5172 |
| 5_3294675 | 2 | 0 | NA | NA | 1 | 1 | 5 | 0.2 | 8 | 0.25 | 2 | 0 | 3 | 0 | 2 | 0.5 | 3 | 0 | 10 | 0.1 |
| 5_3294741 | 4 | 0 | 4 | 0 | 3 | 0.3333 | 9 | 0.1111 | 20 | 0.1 | 4 | 0.25 | 7 | 0.1429 | 4 | 0 | 5 | 0 | 20 | 0.1 |
| 5_3294759 | 3 | 0.3333 | 3 | 0.6667 | 3 | 0 | 2 | 0 | 11 | 0.2727 | 2 | 0 | 7 | 0.5714 | 1 | 0 | 6 | 0.1667 | 16 | 0.3125 |
| 5_3294980 | 2 | 0.5 | NA | NA | 1 | 1 | 4 | 0 | 7 | 0.2857 | 1 | 1 | 3 | 0.3333 | 3 | 0 | 1 | 1 | 8 | 0.375 |
| 5_3294995 | 2 | 0 | NA | NA | 1 | 1 | 3 | 0 | 6 | 0.1667 | 2 | 0 | 3 | 0.3333 | 3 | 0 | 1 | 0 | 9 | 0.1111 |
| 5_3295045 | 2 | 0.5 | NA | NA | 1 | 1 | 3 | 0 | 6 | 0.3333 | 2 | 1 | 4 | 0.75 | 1 | 1 | 1 | 0 | 8 | 0.75 |
| 5_3295047 | 2 | 0.5 | NA | NA | 1 | 1 | 4 | 0.5 | 7 | 0.5714 | 2 | 0.5 | 4 | 0.5 | 1 | 0 | 1 | 0 | 8 | 0.375 |
| 5_3295054 | 2 | 0 | NA | NA | NA | NA | 3 | 0.3333 | 5 | 0.2 | NA | NA | 4 | 0.5 | 1 | 0 | NA | NA | 5 | 0.4 |
| 5_3295086 | 2 | 0 | NA | NA | 1 | 0 | 3 | 0.3333 | 6 | 0.1667 | 2 | 0 | 4 | 0.25 | 1 | 0 | 1 | 0 | 8 | 0.125 |
| 5_3295087 | 1 | 1 | NA | NA | 1 | 1 | 4 | 0.5 | 6 | 0.6667 | NA | NA | 3 | 0.6667 | NA | NA | 1 | 1 | 4 | 0.75 |
| 5_3295089 | 2 | 0.5 | NA | NA | 1 | 1 | 4 | 0.75 | 7 | 0.7143 | 1 | 1 | 2 | 0.5 | 1 | 1 | 1 | 0 | 5 | 0.6 |
| 5_3295095 | 1 | 0 | NA | NA | NA | NA | 3 | 0.6667 | 4 | 0.5 | 1 | 1 | 4 | 0.75 | 1 | 1 | 1 | 0 | 7 | 0.7143 |
| 5_3295099 | 2 | 0.5 | NA | NA | NA | NA | 4 | 0.5 | 6 | 0.5 | 2 | 0.5 | 4 | 0.75 | 1 | 0 | 1 | 0 | 8 | 0.5 |
| 5_3298453 | 2 | 0.5 | NA | NA | NA | NA | 3 | 0.6667 | 5 | 0.6 | 1 | 1 | 1 | 1 | 1 | 0 | 1 | 1 | 4 | 0.75 |
| 5_3298454 | 2 | 1 | NA | NA | NA | NA | 3 | 0.3333 | 5 | 0.6 | 1 | 1 | 1 | 1 | 1 | 1 | 1 | 0 | 4 | 0.75 |
| 5_3298475 | 1 | 1 | NA | NA | NA | NA | 3 | 0.6667 | 4 | 0.75 | 1 | 1 | 1 | 1 | NA | NA | 1 | 1 | 3 | 1 |
| 5_3298482 | 2 | 0.5 | NA | NA | NA | NA | 2 | 1 | 4 | 0.75 | 1 | 1 | 1 | 1 | NA | NA | NA | NA | 2 | 1 |
| 5_3298560 | 2 | 0.5 | NA | NA | NA | NA | 1 | 0 | 3 | 0.3333 | NA | NA | 1 | 1 | NA | NA | NA | NA | 1 | 1 |
| 5_3298585 | 2 | 1 | NA | NA | NA | NA | 1 | 1 | 3 | 1 | NA | NA | 1 | 1 | NA | NA | NA | NA | 1 | 1 |
| 5_3298598 | 1 | 1 | NA | NA | NA | NA | 1 | 0 | 2 | 0.5 | NA | NA | 1 | 1 | NA | NA | NA | NA | 1 | 1 |

| | | | | | | | | | | | | | | | | | | | | |
|---|---|---|---|---|---|---|---|---|---|---|---|---|---|---|---|---|---|---|---|---|
| 5_32985989 | NA | NA | NA | NA | NA | NA | 1 | 1 | 1 | 1 | NA | NA | 1 | 1 | NA | NA | NA | NA | 1 | 1 |
| 5_32986564 | 1 | 1 | NA | NA | NA | NA | 1 | 1 | 2 | 1 | NA | NA | 1 | 1 | NA | NA | NA | NA | 1 | 1 |
| 5_32986978 | 1 | 0 | NA | NA | NA | NA | 1 | 1 | 2 | 0.5 | NA | NA | 1 | 1 | NA | NA | NA | NA | 1 | 1 |
| 5_32986989 | 1 | 1 | NA | NA | NA | NA | 1 | 0 | 2 | 0.5 | NA | NA | 1 | 1 | NA | NA | NA | NA | 1 | 1 |
| 5_32987672 | 1 | 1 | NA | NA | NA | NA | 1 | 0 | 2 | 0.5 | NA | NA | 1 | 1 | NA | NA | NA | NA | 1 | 1 |
| 5_33001580 | NA | NA | NA | NA | NA | NA | 1 | 1 | 1 | 1 | NA | NA | 1 | 1 | NA | NA | NA | NA | 1 | 1 |
| 5_33001651 | NA | NA | NA | NA | NA | NA | 1 | 1 | 1 | 1 | NA | NA | 1 | 1 | NA | NA | NA | NA | 1 | 1 |
| 5_33148349 | 5 | 0.6 | 7 | 0.2857 | 4 | 0.25 | 2 | 0.5 | 18 | 0.3889 | 1 | 0 | 1 | 0 | NA | NA | 5 | 0.2 | 7 | 0.1429 |
| 5_33148487 | 6 | 0.8333 | 7 | 1 | 5 | 1 | 3 | 0.6667 | 21 | 0.9048 | 1 | 1 | 3 | 0.6667 | 1 | 1 | 4 | 1 | 9 | 0.8889 |
| 5_33148537 | 5 | 1 | 5 | 1 | 2 | 1 | 2 | 1 | 14 | 1 | 1 | 1 | 3 | 1 | 1 | 1 | 5 | 0.6 | 10 | 0.8 |
| 5_33148541 | 5 | 0 | 5 | 0 | 3 | 0.6667 | 3 | 0 | 16 | 0.125 | 1 | 1 | 2 | 0 | 1 | 0 | 4 | 0.25 | 8 | 0.25 |
| 5_33184081 | 95 | 0.2842 | 117 | 0.3504 | 98 | 0.2347 | 109 | 0.3303 | 419 | 0.3031 | 72 | 0.3333 | 105 | 0.3619 | 89 | 0.3371 | 101 | 0.3663 | 367 | 0.3515 |
| 5_33184130 | 71 | 0.9859 | 92 | 0.9674 | 71 | 0.9437 | 86 | 0.9884 | 320 | 0.9719 | 52 | 0.9808 | 78 | 0.9872 | 61 | 1 | 75 | 0.9733 | 266 | 0.985 |
| 5_33184229 | 80 | 0.05 | 100 | 0.15 | 83 | 0.1446 | 83 | 0.1205 | 346 | 0.1185 | 59 | 0.1695 | 85 | 0.1176 | 73 | 0.0959 | 82 | 0.1341 | 299 | 0.1271 |
| 5_33184424 | 85 | 0.9529 | 115 | 0.9652 | 78 | 0.9615 | 94 | 0.9787 | 372 | 0.9651 | 70 | 0.9571 | 86 | 0.9651 | 74 | 0.973 | 100 | 0.94 | 330 | 0.9576 |
| 5_33184426 | 2 | 1 | NA | NA | NA | NA | 1 | 1 | 3 | 1 | NA | NA | 1 | 1 | 1 | 1 | NA | NA | 2 | 1 |
| 5_33184451 | 110 | 0.9909 | 133 | 0.9925 | 107 | 0.9907 | 123 | 0.9756 | 473 | 0.9873 | 86 | 0.9884 | 107 | 0.9907 | 97 | 0.9897 | 108 | 0.963 | 398 | 0.9824 |
| 5_33184456 | 1 | 1 | NA | NA | NA | NA | NA | NA | 1 | 1 | 1 | 1 | NA | NA | NA | NA | 1 | 1 | 2 | 1 |
| 5_33184697 | 113 | 0.0796 | 147 | 0.0544 | 117 | 0.1026 | 116 | 0.1293 | 493 | 0.0892 | 91 | 0.1429 | 109 | 0.0734 | 101 | 0.0792 | 120 | 0.075 | 421 | 0.0903 |
| 5_33184718 | 117 | 1 | 142 | 0.993 | 117 | 1 | 116 | 1 | 492 | 0.998 | 96 | 1 | 107 | 1 | 107 | 0.9907 | 120 | 0.9833 | 430 | 0.993 |
| 5_33184972 | 133 | 0.1128 | 163 | 0.0675 | 125 | 0.144 | 130 | 0.1077 | 551 | 0.1053 | 101 | 0.0891 | 121 | 0.0826 | 120 | 0.05 | 133 | 0.1053 | 475 | 0.0821 |
| 5_33185015 | 100 | 0.82 | 106 | 0.8774 | 101 | 0.8416 | 100 | 0.83 | 407 | 0.8428 | 74 | 0.8514 | 94 | 0.8511 | 87 | 0.7586 | 94 | 0.8298 | 349 | 0.8223 |
| 5_33185273 | 116 | 0.0948 | 157 | 0.0892 | 126 | 0.1111 | 121 | 0.124 | 520 | 0.1038 | 100 | 0.08 | 121 | 0.0744 | 124 | 0.0726 | 137 | 0.1022 | 482 | 0.083 |
| 5_33405426 | 33 | 0.1818 | 44 | 0.2045 | 35 | 0.2286 | 37 | 0.2973 | 149 | 0.2282 | 33 | 0.2121 | 36 | 0.1667 | 39 | 0.2564 | 31 | 0.3871 | 139 | 0.2518 |
| 5_33405739 | 30 | 0.3667 | 33 | 0.3939 | 29 | 0.5517 | 33 | 0.697 | 125 | 0.504 | 29 | 0.6207 | 26 | 0.5769 | 34 | 0.4412 | 28 | 0.6429 | 117 | 0.5641 |
| 5_33405788 | 32 | 0.0625 | 38 | 0.1053 | 36 | 0.1389 | 33 | 0.1515 | 139 | 0.1151 | 26 | 0.0385 | 28 | 0.1429 | 35 | 0.0571 | 33 | 0.0303 | 122 | 0.0656 |
| 5_33405891 | 33 | 0.0303 | 40 | 0.075 | 36 | 0.0278 | 30 | 0.1 | 139 | 0.0576 | 27 | 0.0741 | 32 | 0.0312 | 39 | 0.2051 | 34 | 0.1176 | 132 | 0.1136 |
| 5_33407692 | 1 | 0 | 1 | 1 | NA | NA | 1 | 1 | 3 | 0.6667 | NA | NA | 1 | 1 | NA | NA | NA | NA | 1 | 1 |
| 5_33515908 | 9 | 0 | 7 | 0 | 6 | 0.1667 | 13 | 0.0769 | 35 | 0.0571 | 5 | 0.4 | 9 | 0 | 8 | 0 | 2 | 0 | 24 | 0.0833 |
| 5_33529897 | 9 | 0.5556 | 10 | 0.2 | 7 | 0 | 13 | 0.3846 | 39 | 0.3077 | 7 | 0 | 14 | 0.1429 | 13 | 0.3077 | 8 | 0.375 | 42 | 0.2143 |
| 5_33529970 | 10 | 0.2 | 10 | 0.1 | 11 | 0.5455 | 15 | 0.2667 | 46 | 0.2826 | 9 | 0.4444 | 10 | 0.2 | 14 | 0.2143 | 10 | 0.2 | 43 | 0.2558 |
| 5_33530091 | 10 | 1 | 11 | 0.9091 | 9 | 1 | 16 | 1 | 46 | 0.9783 | 8 | 1 | 11 | 1 | 13 | 0.9231 | 7 | 1 | 39 | 0.9744 |
| 5_33530185 | 10 | 1 | 11 | 0.8182 | 7 | 1 | 15 | 0.8667 | 43 | 0.907 | 8 | 0.625 | 14 | 0.9286 | 15 | 0.8 | 8 | 0.5 | 45 | 0.7556 |
| 5_33530215 | 11 | 0.2727 | 11 | 0.3636 | 10 | 0 | 17 | 0.3529 | 49 | 0.2653 | 9 | 0 | 15 | 0.2 | 16 | 0.3125 | 11 | 0 | 51 | 0.1569 |
| 5_33530392 | 12 | 1 | 11 | 0.9091 | 9 | 0.8889 | 15 | 1 | 47 | 0.9574 | 11 | 1 | 10 | 1 | 14 | 0.9286 | 10 | 0.9 | 45 | 0.9556 |
| 5_33530462 | 11 | 0.3636 | 12 | 0.0833 | 7 | 0.4286 | 14 | 0.3571 | 44 | 0.2955 | 11 | 0.4545 | 14 | 0.4286 | 16 | 0.1875 | 9 | 0.1111 | 50 | 0.3 |
| 5_33757894 | 4 | 0 | 5 | 0.2 | 4 | 0.25 | 3 | 0 | 16 | 0.125 | 5 | 0.2 | 3 | 0.3333 | 4 | 0.5 | 6 | 0.5 | 18 | 0.3889 |
| 5_33757913 | 4 | 1 | 5 | 0.6 | 4 | 1 | 3 | 0.3333 | 16 | 0.75 | 4 | 0.5 | 3 | 0.6667 | 4 | 1 | 5 | 0.4 | 16 | 0.625 |
| 5_33757920 | 4 | 0 | 3 | 0.3333 | 3 | 0 | 2 | 0 | 12 | 0.0833 | 6 | 0.3333 | 2 | 0 | 5 | 0 | 6 | 0 | 19 | 0.1053 |
| 5_33757938 | 4 | 0.5 | 5 | 0.4 | 4 | 0.5 | 4 | 0.5 | 17 | 0.4706 | 6 | 0.1667 | 3 | 0.6667 | 4 | 0.25 | 6 | 0.5 | 19 | 0.3684 |
| 5_33757986 | 2 | 0.5 | 3 | 0.6667 | 4 | 0.75 | 3 | 0.6667 | 12 | 0.6667 | 3 | 0.6667 | 2 | 0.5 | 3 | 0.3333 | 4 | 0.5 | 12 | 0.5 |
| 5_33757992 | 4 | 0.5 | 5 | 0.2 | 4 | 0 | 4 | 0 | 17 | 0.1765 | 6 | 0.1667 | 3 | 0 | 5 | 0.4 | 6 | 0.1667 | 20 | 0.2 |
| 5_33758155 | 3 | 0 | 5 | 0 | 3 | 0 | 4 | 0.25 | 15 | 0.0667 | 5 | 0.2 | 2 | 0 | 4 | 0 | 6 | 0 | 17 | 0.0588 |
| 5_33764827 | 2 | 0 | 4 | 0.5 | 2 | 0 | 3 | 0 | 11 | 0.1818 | 5 | 0 | 2 | 1 | 4 | 0 | 4 | 0.25 | 15 | 0.2 |
| 5_33765054 | 2 | 1 | 4 | 1 | 2 | 0.5 | 2 | 0.5 | 10 | 0.8 | 5 | 0.8 | 2 | 0.5 | 4 | 0.25 | 3 | 0.6667 | 14 | 0.5714 |
| 5_33765287 | 2 | 1 | 2 | 1 | 1 | 1 | 2 | 1 | 7 | 1 | 4 | 1 | 2 | 1 | 3 | 1 | 3 | 1 | 12 | 1 |
| 5_33765412 | 1 | 0 | 2 | 0.5 | 2 | 0.5 | 2 | 0 | 7 | 0.2857 | 5 | 0.2 | 2 | 0.5 | 3 | 0.3333 | 1 | 0 | 11 | 0.2727 |
| 5_33797847 | 3 | 0.3333 | 3 | 0 | 3 | 0.3333 | 3 | 0.3333 | 12 | 0.25 | 3 | 0 | 3 | 0 | 2 | 0.5 | 3 | 0.3333 | 11 | 0.1818 |
| 5_33798175 | 2 | 0 | 3 | 1 | 2 | 0.5 | 3 | 1 | 10 | 0.7 | 2 | 0 | 3 | 0.3333 | 1 | 1 | 3 | 0 | 9 | 0.2222 |
| 5_33798186 | 4 | 0.25 | 4 | 0 | 3 | 0 | 3 | 0 | 14 | 0.0714 | 4 | 0 | 3 | 0 | 2 | 0.5 | 4 | 0 | 13 | 0.0769 |
| 5_33798232 | 4 | 0.25 | 2 | 0 | 2 | 0 | 2 | 0 | 10 | 0.1 | 3 | 0 | 3 | 0 | 2 | 0.5 | 3 | 0 | 11 | 0.0909 |
| 5_33811527 | NA | NA | 7 | 0.2857 | 3 | 0 | 2 | 0 | 12 | 0.1667 | 1 | 0 | 7 | 0.2857 | 2 | 0 | 6 | 0.3333 | 16 | 0.25 |
| 5_33811649 | 1 | 1 | 5 | 0.4 | 4 | 0.5 | 2 | 0 | 12 | 0.4167 | 3 | 0.3333 | 5 | 0.2 | 2 | 0.5 | 5 | 0.4 | 15 | 0.3333 |
| 5_33812742 | 1 | 0 | 7 | 0.1429 | 3 | 0.3333 | 1 | 0 | 12 | 0.1667 | 2 | 0 | 5 | 0.8 | 2 | 0.5 | 5 | 0 | 14 | 0.3571 |
| 5_33812779 | 1 | 1 | 4 | 0.5 | 2 | 1 | 1 | 0 | 8 | 0.625 | 2 | 1 | 5 | 0.2 | 1 | 0 | 4 | 0.5 | 12 | 0.4167 |
| 5_33812844 | 1 | 1 | 7 | 0.2857 | 2 | 0.5 | 1 | 0 | 11 | 0.3636 | 1 | 0 | 6 | 0.3333 | 1 | 1 | 5 | 0.8 | 13 | 0.5385 |
| 5_33812848 | 1 | 0 | 6 | 0 | 2 | 0 | 1 | 1 | 10 | 0.1 | 3 | 0 | 4 | 0.25 | 1 | 0 | 4 | 0 | 12 | 0.0833 |
| 5_33812989 | 1 | 0 | 6 | 0.1667 | 2 | 0 | 1 | 0 | 10 | 0.1 | 3 | 0 | 6 | 0.5 | 1 | 0 | 4 | 0.25 | 14 | 0.2857 |
| 5_33815167 | NA | NA | 2 | 0.5 | 1 | 0 | 1 | 0 | 4 | 0.25 | 3 | 0 | 4 | 0 | NA | NA | 4 | 0.25 | 11 | 0.0909 |
| 5_33892951 | 8 | 0.25 | 14 | 0.2143 | 11 | 0.4545 | 10 | 0.2 | 43 | 0.2791 | 7 | 0.4286 | 12 | 0.3333 | 14 | 0.2143 | 11 | 0.4545 | 44 | 0.3409 |
| 5_33892987 | 8 | 0.625 | 14 | 0.4286 | 10 | 0.6 | 10 | 0.1 | 42 | 0.4286 | 8 | 0.375 | 12 | 0.4167 | 15 | 0.4 | 12 | 0.4167 | 47 | 0.4043 |
| 5_33893027 | 7 | 0.1429 | 16 | 0.125 | 11 | 0 | 11 | 0.1818 | 45 | 0.1111 | 8 | 0 | 14 | 0.1429 | 15 | 0.2667 | 12 | 0 | 49 | 0.1224 |
| 5_33893076 | 6 | 0.1667 | 13 | 0.1538 | 10 | 0 | 11 | 0.0909 | 40 | 0.1 | 8 | 0 | 13 | 0 | 14 | 0.2143 | 12 | 0.0833 | 47 | 0.0851 |
| 5_33893203 | 9 | 0.1111 | 14 | 0.2857 | 9 | 0.2222 | 11 | 0.3636 | 43 | 0.2558 | 7 | 0.1429 | 12 | 0.0833 | 16 | 0.25 | 9 | 0.2222 | 44 | 0.1818 |
| 5_33893387 | 8 | 0 | 16 | 0.1875 | 10 | 0.2 | 12 | 0.0833 | 46 | 0.1304 | 7 | 0.1429 | 13 | 0.0769 | 19 | 0 | 11 | 0.0909 | 50 | 0.06 |

| | | | | | | | | | | | | | | | | | | | | |
|---|---|---|---|---|---|---|---|---|---|---|---|---|---|---|---|---|---|---|---|---|
| 5_3389712 | 3 | 0.3333 | 2 | 0 | NA | NA | 3 | 0 | 8 | 0.125 | 2 | 1 | 5 | 0.2 | 4 | 0.5 | 1 | 0 | 12 | 0.4167 |
| 5_3389780 | 3 | 0 | 3 | 0.3333 | NA | NA | 2 | 0.5 | 8 | 0.25 | 2 | 0.5 | 7 | 0.1429 | 3 | 0.3333 | 1 | 0 | 13 | 0.2308 |
| 5_3389844 | 1 | 1 | 3 | 0.3333 | NA | NA | 2 | 0.5 | 6 | 0.5 | 2 | 1 | 7 | 0.5714 | 1 | 1 | 1 | 0 | 11 | 0.6364 |
| 5_3389868 | 1 | 1 | 2 | 1 | NA | NA | 2 | 1 | 5 | 1 | 2 | 1 | 6 | 0.5 | 1 | 1 | 1 | 0 | 10 | 0.6 |
| 5_3389895 | 1 | 1 | 1 | 0 | NA | NA | 2 | 0 | 4 | 0.25 | 1 | 1 | 4 | 0.75 | 2 | 1 | 1 | 0 | 8 | 0.75 |
| 5_3389938 | 1 | 1 | 1 | 0 | NA | NA | NA | NA | 2 | 0.5 | 1 | 1 | 2 | 0.5 | 2 | 1 | 1 | 0 | 6 | 0.6667 |
| 5_3389967 | 8 | 0.125 | 12 | 0.25 | 14 | 0.1429 | 6 | 0.1667 | 40 | 0.175 | 8 | 0.125 | 13 | 0.1538 | 12 | 0.25 | 5 | 0.2 | 38 | 0.1842 |
| 5_3392691 | 5 | 0.2 | 6 | 0.1667 | 2 | 0 | 4 | 0 | 17 | 0.1176 | 4 | 0 | 6 | 0.1667 | 4 | 0 | 3 | 0.6667 | 17 | 0.1765 |
| 5_3392694 | 4 | 1 | 5 | 0.4 | 3 | 1 | 4 | 0.75 | 16 | 0.75 | 4 | 0.75 | 5 | 1 | 3 | 1 | 4 | 0.75 | 16 | 0.875 |
| 5_3392700 | 4 | 0.25 | 4 | 0.25 | 3 | 0 | 4 | 0.5 | 15 | 0.2667 | 5 | 0.2 | 5 | 0 | 4 | 0.75 | 4 | 0.25 | 18 | 0.2778 |
| 5_3392703 | 4 | 0.5 | 4 | 0 | 3 | 0 | 2 | 0 | 13 | 0.1538 | 4 | 0 | 4 | 0.25 | 4 | 0 | 4 | 0 | 16 | 0.0625 |
| 5_3392707 | 4 | 0.25 | 4 | 0.75 | 2 | 1 | 3 | 0.3333 | 13 | 0.5385 | 3 | 0.3333 | 4 | 1 | 2 | 0.5 | 3 | 1 | 12 | 0.75 |
| 5_3400059 | NA | NA | 1 | 1 | 1 | 1 | 1 | 0 | 3 | 0.6667 | 1 | 1 | NA | NA | 2 | 1 | 1 | 1 | 4 | 1 |
| 5_3400081 | NA | NA | 1 | 0 | 1 | 1 | NA | NA | 2 | 0.5 | 1 | 0 | NA | NA | 2 | 1 | 1 | 1 | 4 | 0.75 |
| 5_3401272 | NA | NA | 2 | 1 | NA | NA | 1 | 0 | 3 | 0.6667 | 2 | 0.5 | NA | NA | 2 | 0 | 1 | 1 | 5 | 0.4 |
| 5_3401277 | NA | NA | 2 | 0.5 | 1 | 0 | 2 | 0.5 | 5 | 0.4 | NA | NA | NA | NA | 2 | 0.5 | 1 | 0 | 3 | 0.3333 |
| 5_3401279 | NA | NA | 2 | 1 | NA | NA | 2 | 1 | 4 | 1 | NA | NA | NA | NA | 2 | 1 | 1 | 1 | 3 | 1 |
| 5_3402878 | NA | NA | 2 | 0.5 | 2 | 1 | 1 | 0 | 5 | 0.6 | 1 | 0 | 1 | 0 | 1 | 0 | 1 | 1 | 4 | 0.25 |
| 5_3402887 | NA | NA | 2 | 0.5 | 2 | 0.5 | 2 | 0.5 | 6 | 0.5 | 1 | 0 | 1 | 1 | 1 | 1 | 1 | 0 | 4 | 0.5 |
| 5_3402909 | NA | NA | 2 | 0.5 | 1 | 0 | 1 | 0 | 4 | 0.25 | 1 | 0 | 1 | 1 | 1 | 0 | 1 | 1 | 4 | 0.5 |
| 5_3402969 | NA | NA | 2 | 0 | 2 | 0.5 | 2 | 0 | 6 | 0.1667 | NA | NA | 2 | 0.5 | 1 | 0 | 1 | 0 | 4 | 0.25 |
| 5_3403101 | 1 | 1 | 1 | 1 | 1 | 0 | 4 | 0.25 | 7 | 0.4286 | NA | NA | 2 | 0 | 1 | 1 | NA | NA | 3 | 0.3333 |
| 5_3403304 | 3 | 0.3333 | 4 | 0 | NA | NA | 5 | 0.2 | 12 | 0.1667 | 1 | 0 | 3 | 0 | 2 | 0.5 | 1 | 0 | 7 | 0.1429 |
| 5_3403308 | 3 | 0.3333 | 4 | 0.5 | NA | NA | 6 | 0.6667 | 13 | 0.5385 | 2 | 0 | 2 | 0 | 2 | 0.5 | 1 | 1 | 7 | 0.2857 |
| 5_3403309 | 2 | 0 | 4 | 0 | NA | NA | 5 | 0.2 | 11 | 0.0909 | 1 | 0 | 2 | 0 | 2 | 0.5 | 1 | 1 | 6 | 0.3333 |
| 5_3403324 | 3 | 0.6667 | 5 | 0.2 | NA | NA | 6 | 0.1667 | 14 | 0.2857 | 1 | 0 | 2 | 0 | 2 | 1 | 1 | 0 | 6 | 0.3333 |
| 5_3403399 | 3 | 0.3333 | 5 | 0 | NA | NA | 4 | 0 | 12 | 0.0833 | 3 | 0 | 3 | 0.3333 | 2 | 0.5 | 1 | 0 | 9 | 0.2222 |
| 5_3403401 | 2 | 0.5 | 6 | 0.3333 | 2 | 0 | 5 | 0.4 | 15 | 0.3333 | 6 | 0.3333 | 5 | 0 | 1 | 0 | 2 | 0 | 14 | 0.1429 |
| 5_3403405 | 3 | 0.6667 | 5 | 0 | 1 | 0 | 5 | 0.2 | 14 | 0.2143 | 6 | 0 | 4 | 0.25 | 2 | 0 | 2 | 0 | 14 | 0.0714 |
| 5_3403416 | 2 | 0.5 | 5 | 0.4 | 2 | 0.5 | 3 | 0 | 12 | 0.3333 | 5 | 0.4 | 3 | 0.3333 | 2 | 1 | 2 | 1 | 12 | 0.5833 |
| 5_3403418 | 3 | 0.6667 | 6 | 0.8333 | 2 | 0.5 | 6 | 1 | 17 | 0.8235 | 5 | 1 | 5 | 0.8 | 2 | 1 | 2 | 1 | 14 | 0.9286 |
| 5_3403419 | 3 | 0.6667 | 4 | 0 | 2 | 0 | 5 | 0.4 | 14 | 0.2857 | 4 | 0.25 | 4 | 0.5 | 2 | 0 | 2 | 0.5 | 12 | 0.3333 |
| 5_3403429 | 3 | 0.6667 | 5 | 0 | 2 | 0 | 6 | 0 | 16 | 0.125 | 6 | 0 | 6 | 0.3333 | 2 | 0 | 2 | 0.5 | 16 | 0.1875 |
| 5_3403433 | 3 | 0 | 6 | 0.1667 | 4 | 0.25 | 6 | 0.1667 | 19 | 0.1579 | 4 | 0.25 | 5 | 0.4 | 2 | 0 | 2 | 0.5 | 13 | 0.3077 |
| 5_3403434 | 2 | 0 | 4 | 0 | 3 | 0 | 5 | 0.2 | 14 | 0.0714 | 5 | 0 | 6 | 0.1667 | 1 | 0 | 1 | 1 | 13 | 0.1538 |
| 5_3403441 | 1 | 1 | 5 | 0.6 | 3 | 0.3333 | 6 | 0.3333 | 15 | 0.4667 | 3 | 0.6667 | 6 | 0.6667 | 2 | 1 | 2 | 1 | 13 | 0.7692 |
| 5_3405235 | 1 | 1 | 3 | 1 | NA | NA | NA | NA | 4 | 1 | 1 | 1 | 1 | 0 | 1 | 0 | NA | NA | 3 | 0.3333 |
| 5_3405243 | 1 | 1 | 3 | 0.6667 | NA | NA | NA | NA | 4 | 0.75 | 1 | 1 | 1 | 0 | NA | NA | NA | NA | 2 | 0.5 |
| 5_3405256 | 1 | 1 | 3 | 0.3333 | NA | NA | NA | NA | 4 | 0.5 | 1 | 1 | 1 | 0 | 1 | 0 | NA | NA | 3 | 0.3333 |
| 5_3405444 | 1 | 1 | 1 | 0 | 1 | 0 | NA | NA | 3 | 0.3333 | 2 | 0.5 | NA | NA | 1 | 0 | NA | NA | 3 | 0.3333 |
| 5_3405485 | 1 | 1 | 1 | 1 | 1 | 0 | NA | NA | 3 | 0.6667 | 2 | 0.5 | NA | NA | 1 | 1 | NA | NA | 3 | 0.6667 |
| 5_3405603 | 2 | 0 | 4 | 0.25 | 7 | 0 | 2 | 0 | 15 | 0.0667 | 1 | 0 | 4 | 0.25 | NA | NA | NA | NA | 5 | 0.2 |
| 5_3405614 | 3 | 0 | 4 | 0.5 | 7 | 0 | 2 | 0.5 | 16 | 0.1875 | NA | NA | 5 | 0.6 | NA | NA | NA | NA | 5 | 0.6 |
| 5_3414712 | NA | NA | NA | NA | 1 | 1 | NA | NA | 1 | 1 | 1 | 1 | NA | NA | NA | NA | NA | NA | 1 | 1 |
| 5_3414715 | NA | NA | NA | NA | 1 | 1 | NA | NA | 1 | 1 | 1 | 1 | NA | NA | NA | NA | NA | NA | 1 | 1 |
| 5_3414936 | NA | NA | 1 | 1 | 2 | 1 | NA | NA | 3 | 1 | 1 | 1 | 1 | 1 | 1 | 1 | NA | NA | 3 | 1 |
| 5_3414942 | NA | NA | NA | NA | 2 | 1 | NA | NA | 2 | 1 | 1 | 1 | 1 | 1 | 1 | 0 | NA | NA | 3 | 0.6667 |
| 5_3414944 | NA | NA | 1 | 1 | 1 | 1 | NA | NA | 2 | 1 | 1 | 1 | 1 | 1 | 1 | 1 | NA | NA | 3 | 1 |
| 5_3414947 | NA | NA | 1 | 0 | 2 | 0.5 | NA | NA | 3 | 0.3333 | 1 | 0 | NA | NA | 1 | 1 | NA | NA | 2 | 0.5 |
| 5_3414951 | NA | NA | 1 | 1 | 2 | 1 | NA | NA | 3 | 1 | 1 | 1 | 1 | 1 | NA | NA | NA | NA | 2 | 1 |
| 5_3414973 | NA | NA | 1 | 1 | 2 | 1 | NA | NA | 3 | 1 | 1 | 1 | 1 | 1 | 1 | 1 | NA | NA | 3 | 1 |
| 5_3414977 | NA | NA | 1 | 1 | 2 | 0 | NA | NA | 3 | 0.3333 | 1 | 1 | 1 | 0 | 1 | 0 | NA | NA | 3 | 0.3333 |
| 5_3414987 | NA | NA | 1 | 1 | 2 | 1 | NA | NA | 3 | 1 | 1 | 1 | 1 | 1 | 1 | 1 | NA | NA | 3 | 1 |
| 5_3415003 | NA | NA | 1 | 1 | 1 | 1 | NA | NA | 2 | 1 | NA | NA | 1 | 0 | 1 | 1 | NA | NA | 2 | 0.5 |
| 5_3415014 | NA | NA | 1 | 1 | NA | NA | NA | NA | 1 | 1 | NA | NA | 1 | 1 | 1 | 1 | NA | NA | 2 | 1 |
| 5_3415015 | NA | NA | 1 | 1 | NA | NA | NA | NA | 1 | 1 | NA | NA | 1 | 1 | 1 | 1 | NA | NA | 2 | 1 |
| 5_3415028 | NA | NA | 1 | 1 | NA | NA | NA | NA | 1 | 1 | NA | NA | 1 | 1 | 1 | 1 | NA | NA | 2 | 1 |
| 5_3415029 | NA | NA | 1 | 1 | NA | NA | NA | NA | 1 | 1 | NA | NA | NA | NA | 1 | 1 | NA | NA | 1 | 1 |
| 5_3415030 | NA | NA | 1 | 1 | NA | NA | NA | NA | 1 | 1 | NA | NA | 1 | 0 | 1 | 1 | NA | NA | 2 | 0.5 |
| 5_3415036 | NA | NA | 1 | 1 | NA | NA | NA | NA | 1 | 1 | NA | NA | 1 | 1 | 1 | 1 | NA | NA | 2 | 1 |
| 5_3415181 | 1 | 1 | 1 | 1 | NA | NA | 1 | 1 | 3 | 1 | NA | NA | NA | NA | 1 | 1 | NA | NA | 1 | 1 |
| 5_3415193 | 1 | 1 | 1 | 1 | NA | NA | 1 | 1 | 3 | 1 | NA | NA | NA | NA | 1 | 1 | NA | NA | 1 | 1 |
| 5_3415223 | 1 | 1 | 1 | 1 | NA | NA | 1 | 1 | 3 | 1 | NA | NA | NA | NA | 1 | 1 | NA | NA | 1 | 1 |
| 5_3415227 | NA | NA | 1 | 1 | NA | NA | 1 | 1 | 2 | 1 | NA | NA | NA | NA | 1 | 1 | NA | NA | 1 | 1 |
| 5_3415819 | 9 | 0.1111 | 16 | 0.1875 | 5 | 0.4 | 8 | 0.25 | 38 | 0.2105 | 8 | 0.125 | 8 | 0 | 3 | 0.6667 | 8 | 0.125 | 27 | 0.1481 |

| | | | | | | | | | | | | | | | | | | | | |
|---|---|---|---|---|---|---|---|---|---|---|---|---|---|---|---|---|---|---|---|---|
| 5_34158229 | 11 | 0.4545 | 13 | 0.8462 | 7 | 0.7143 | 6 | 0.3333 | 37 | 0.6216 | 6 | 0.5 | 8 | 0.875 | 4 | 0.75 | 6 | 0.6667 | 24 | 0.7083 |
| 5_34158377 | 10 | 0.9 | 15 | 0.5333 | 5 | 0.8 | 10 | 0.5 | 40 | 0.65 | 7 | 0.4286 | 7 | 0.5714 | 3 | 1 | 6 | 0.8333 | 23 | 0.6522 |
| 5_34158773 | 16 | 0.375 | 14 | 0.3571 | 12 | 0.25 | 9 | 0.1111 | 51 | 0.2941 | 9 | 0.3333 | 8 | 0.5 | 4 | 0.5 | 11 | 0.0909 | 32 | 0.3125 |
| 5_34158782 | 14 | 0.4286 | 11 | 0.3636 | 11 | 0.1818 | 7 | 0.5714 | 43 | 0.3721 | 10 | 0.3 | 5 | 0.6 | 3 | 0.6667 | 8 | 0.25 | 26 | 0.3846 |
| 5_34158823 | 14 | 0.5 | 17 | 0.7647 | 11 | 0.7273 | 9 | 0.5556 | 51 | 0.6471 | 12 | 0.5833 | 9 | 0.6667 | 4 | 0.75 | 11 | 0.6364 | 36 | 0.6389 |
| 5_34158848 | 16 | 0.125 | 15 | 0.3333 | 13 | 0.0769 | 9 | 0 | 53 | 0.1509 | 11 | 0.1818 | 9 | 0 | 3 | 0 | 10 | 0.1 | 33 | 0.0909 |
| 5_34159014 | 13 | 0.4615 | 15 | 0.4667 | 11 | 0.6364 | 9 | 0.5556 | 48 | 0.5208 | 8 | 0.625 | 7 | 0.5714 | 4 | 0.5 | 9 | 0.3333 | 28 | 0.5 |
| 5_34160013 | 18 | 0.0556 | 18 | 0.0556 | 14 | 0.0714 | 14 | 0.1429 | 64 | 0.0781 | 19 | 0.0526 | 14 | 0.0714 | 8 | 0.125 | 13 | 0.0769 | 54 | 0.0741 |
| 5_34160307 | 26 | 0.1538 | 27 | 0.037 | 20 | 0.15 | 19 | 0.0526 | 92 | 0.0978 | 20 | 0.2 | 19 | 0.3158 | 12 | 0.3333 | 14 | 0.0714 | 65 | 0.2308 |
| 5_34160518 | 5 | 0 | 8 | 0.25 | 3 | 0.3333 | 4 | 0 | 20 | 0.15 | 7 | 0.2857 | 3 | 0.6667 | 5 | 0.4 | 3 | 0 | 18 | 0.3333 |
| 5_34161043 | 5 | 0 | 15 | 0 | 5 | 0.2 | 5 | 0.2 | 30 | 0.0667 | 7 | 0.2857 | 4 | 0 | 6 | 0 | 2 | 0 | 19 | 0.1053 |
| 5_34161860 | 8 | 0.25 | 22 | 0.0909 | 9 | 0 | 6 | 0.1667 | 45 | 0.1111 | 10 | 0 | 9 | 0.2222 | 7 | 0.1429 | 3 | 0 | 29 | 0.1034 |
| 5_34311406 | NA | NA | 2 | 0 | 3 | 0.3333 | NA | NA | 5 | 0.2 | NA | NA | 4 | 0 | 1 | 0 | 3 | 0.3333 | 8 | 0.125 |
| 5_34321086 | NA | NA | 3 | 0 | 3 | 0.3333 | NA | NA | 6 | 0.1667 | NA | NA | 4 | 0 | 1 | 0 | 2 | 0.5 | 7 | 0.1429 |
| 5_34323538 | NA | NA | 1 | 1 | 2 | 0 | NA | NA | 3 | 0.3333 | NA | NA | 3 | 0.6667 | 1 | 1 | 2 | 1 | 6 | 0.8333 |
| 5_34325163 | NA | NA | 2 | 0.5 | 1 | 1 | NA | NA | 3 | 0.6667 | NA | NA | 1 | 1 | NA | NA | 1 | 1 | 2 | 1 |
| 5_34325317 | NA | NA | 2 | 0 | 2 | 0.5 | NA | NA | 4 | 0.25 | NA | NA | 1 | 0 | 1 | 1 | 2 | 0.5 | 4 | 0.5 |
| 5_34325402 | NA | NA | 2 | 0.5 | 1 | 0 | NA | NA | 3 | 0.3333 | NA | NA | 3 | 0.6667 | 1 | 1 | 2 | 0.5 | 6 | 0.6667 |
| 5_34325511 | NA | NA | 2 | 0 | 2 | 0.5 | NA | NA | 4 | 0.25 | NA | NA | 2 | 0.5 | 1 | 0 | 2 | 0 | 5 | 0.2 |
| 5_34325575 | NA | NA | 2 | 0.5 | 1 | 0 | NA | NA | 3 | 0.3333 | NA | NA | 2 | 1 | 1 | 1 | 1 | 1 | 4 | 1 |
| 5_34325630 | NA | NA | 2 | 0.5 | 2 | 0 | NA | NA | 4 | 0.25 | NA | NA | 2 | 0.5 | 1 | 0 | 2 | 0 | 5 | 0.2 |
| 5_34325714 | NA | NA | 2 | 0.5 | 2 | 0.5 | NA | NA | 4 | 0.5 | NA | NA | 1 | 0 | 1 | 1 | 1 | 1 | 3 | 0.6667 |
| 5_34325745 | NA | NA | 2 | 1 | 2 | 1 | NA | NA | 4 | 1 | NA | NA | 1 | 1 | 1 | 0 | 1 | 1 | 3 | 0.6667 |
| 5_34331543 | 9 | 0.1111 | 12 | 0 | 7 | 0 | 11 | 0.2727 | 39 | 0.1026 | 5 | 0 | 7 | 0 | 8 | 0.375 | 9 | 0.1111 | 29 | 0.1379 |
| 5_34332628 | 7 | 0 | 14 | 0.0714 | 9 | 0 | 10 | 0.1 | 40 | 0.05 | 5 | 0 | 10 | 0.1 | 7 | 0.2857 | 8 | 0 | 30 | 0.1 |
| 5_34333148 | 4 | 0.25 | 12 | 0.4167 | 8 | 0 | 8 | 0.125 | 32 | 0.2188 | 4 | 0.25 | 12 | 0.1667 | 2 | 0 | 6 | 0 | 24 | 0.125 |
| 5_34333191 | 5 | 0.6 | 11 | 0.3636 | 8 | 0 | 6 | 0 | 30 | 0.2333 | 3 | 0.3333 | 9 | 0.1111 | 4 | 0.25 | 4 | 0 | 20 | 0.15 |
| 5_34333381 | 4 | 0.5 | 9 | 0.6667 | 9 | 0.2222 | 4 | 0 | 26 | 0.3846 | 3 | 0.3333 | 8 | 0.25 | 4 | 0.5 | 6 | 0.3333 | 21 | 0.3333 |
| 5_34333547 | 4 | 0.25 | 10 | 0.1 | 10 | 0.2 | 8 | 0.125 | 32 | 0.1562 | 4 | 0.75 | 8 | 0.125 | 2 | 0 | 7 | 0 | 21 | 0.1905 |
| 5_34333723 | 4 | 0.25 | 13 | 0.3846 | 11 | 0.0909 | 6 | 0.3333 | 34 | 0.2647 | 3 | 0 | 8 | 0 | 3 | 0 | 6 | 0.1667 | 20 | 0.05 |
| 5_34334161 | 1 | 1 | 3 | 0.6667 | 4 | 1 | 3 | 1 | 11 | 0.9091 | 3 | 1 | 4 | 0.5 | 1 | 1 | 3 | 0.3333 | 11 | 0.6364 |
| 5_34334451 | 1 | 0 | 2 | 0.5 | 3 | 0 | 2 | 0 | 8 | 0.125 | 1 | 1 | 2 | 0 | 1 | 1 | 2 | 0 | 6 | 0.3333 |
| 5_34352558 | 6 | 0.5 | 2 | 1 | 4 | 0.25 | 6 | 0.5 | 18 | 0.5 | 4 | 0.5 | 3 | 0.6667 | 7 | 0.1429 | 6 | 0.5 | 20 | 0.4 |
| 5_34352787 | 6 | 0.1667 | 3 | 0 | 3 | 0 | 6 | 0 | 18 | 0.0556 | 2 | 0 | 4 | 0.25 | 6 | 0 | 7 | 0 | 19 | 0.0526 |
| 5_34352989 | 7 | 0.7143 | 4 | 0.5 | 3 | 0.6667 | 6 | 0.6667 | 20 | 0.65 | 3 | 1 | 3 | 0.6667 | 6 | 0.6667 | 6 | 0.5 | 18 | 0.6667 |
| 5_34353046 | 4 | 0.25 | 3 | 0.3333 | 5 | 0.4 | 6 | 0.5 | 18 | 0.3889 | 3 | 0.3333 | 3 | 0.6667 | 6 | 0.3333 | 6 | 0.3333 | 18 | 0.3889 |
| 5_34374058 | NA | NA | 1 | 1 | 3 | 1 | 1 | 0 | 5 | 0.8 | 1 | 0 | NA | NA | NA | NA | 2 | 0.5 | 3 | 0.3333 |
| 5_34374354 | NA | NA | 1 | 0 | 1 | 1 | NA | NA | 2 | 0.5 | 1 | 1 | NA | NA | NA | NA | 3 | 0.3333 | 4 | 0.5 |
| 5_34374367 | NA | NA | 1 | 1 | 3 | 0.6667 | 1 | 1 | 5 | 0.8 | 1 | 1 | NA | NA | NA | NA | 2 | 1 | 3 | 1 |
| 5_34374442 | NA | NA | NA | NA | 2 | 1 | 1 | 1 | 3 | 1 | 1 | 1 | NA | NA | NA | NA | 2 | 1 | 3 | 1 |
| 5_34374489 | NA | NA | 1 | 0 | 3 | 0 | 1 | 1 | 5 | 0.2 | 1 | 0 | NA | NA | NA | NA | 1 | 1 | 2 | 0.5 |
| 5_34374527 | NA | NA | 1 | 0 | 3 | 0 | 1 | 1 | 5 | 0.2 | 1 | 1 | NA | NA | NA | NA | 1 | 1 | 2 | 1 |
| 5_34374981 | NA | NA | NA | NA | 1 | 1 | 1 | 0 | 2 | 0.5 | 1 | 1 | NA | NA | NA | NA | 1 | 1 | 2 | 1 |
| 5_34375023 | NA | NA | 1 | 1 | 1 | 1 | 1 | 1 | 3 | 1 | 1 | 1 | NA | NA | NA | NA | 1 | 1 | 2 | 1 |
| 5_34508757 | 3 | 0.6667 | 6 | 0.8333 | 7 | 0.8571 | 5 | 0.8 | 21 | 0.8095 | 6 | 0.1667 | 3 | 1 | 6 | 0.5 | 5 | 0.6 | 20 | 0.5 |
| 5_34508819 | 3 | 0.6667 | 8 | 0.25 | 8 | 0.5 | 6 | 1 | 25 | 0.56 | 7 | 0.5714 | 6 | 0.8333 | 6 | 0.6667 | 4 | 0.75 | 23 | 0.6957 |
| 5_34509218 | 4 | 0.25 | 8 | 0.25 | 8 | 0.375 | 10 | 0.1 | 30 | 0.2333 | 6 | 0 | 7 | 0.2857 | 9 | 0.4444 | 4 | 0.5 | 26 | 0.3077 |
| 5_34509846 | 4 | 0.25 | 7 | 0.1429 | 12 | 0 | 7 | 0 | 30 | 0.0667 | 8 | 0.125 | 5 | 0 | 10 | 0.3 | 6 | 0 | 29 | 0.1379 |
| 5_34510326 | 5 | 0 | 7 | 0.4286 | 10 | 0.1 | 7 | 0 | 29 | 0.1379 | 5 | 0 | 7 | 0 | 4 | 0.25 | 3 | 0.6667 | 19 | 0.1579 |
| 5_34510425 | 5 | 0 | 11 | 0 | 15 | 0.1333 | 8 | 0.375 | 39 | 0.1282 | 8 | 0.375 | 7 | 0.2857 | 10 | 0.1 | 7 | 0.2857 | 32 | 0.25 |
| 5_34568010 | 1 | 0 | 1 | 0 | 1 | 1 | NA | NA | 3 | 0.3333 | NA | NA | NA | NA | 1 | 0 | 3 | 0.3333 | 4 | 0.25 |
| 5_34618606 | 2 | 0 | NA | NA | NA | NA | 1 | 1 | 3 | 0.3333 | NA | NA | 3 | 0.3333 | 1 | 0 | 1 | 0 | 5 | 0.2 |
| 5_34623120 | 5 | 1 | 3 | 1 | NA | NA | 2 | 1 | 10 | 1 | 2 | 1 | 3 | 1 | 4 | 1 | 6 | 1 | 15 | 1 |
| 5_34623135 | 4 | 0.25 | 3 | 1 | 1 | 0 | NA | NA | 8 | 0.5 | 2 | 0.5 | 3 | 0.6667 | 5 | 0.6 | 5 | 0.8 | 15 | 0.6667 |
| 5_34623253 | 3 | 0 | 3 | 0.3333 | 1 | 0 | 2 | 0 | 9 | 0.1111 | 2 | 0 | 3 | 0.3333 | 4 | 0.5 | 4 | 0 | 13 | 0.2308 |
| 5_34623261 | 2 | 0.5 | 2 | 0 | 1 | 0 | 2 | 0 | 7 | 0.1429 | 2 | 0.5 | 3 | 0.6667 | 3 | 0.6667 | 2 | 0 | 10 | 0.5 |
| 5_34623327 | 4 | 0 | 3 | 0 | 1 | 1 | 1 | 0 | 9 | 0.1111 | 2 | 0.5 | 3 | 0 | 5 | 0 | 5 | 0.4 | 15 | 0.2 |
| 5_34623396 | 3 | 0 | 1 | 1 | 1 | 1 | 1 | 1 | 6 | 0.5 | 1 | 0 | 3 | 1 | 3 | 1 | 4 | 1 | 11 | 0.9091 |
| 5_34643009 | 2 | 0 | 3 | 0.3333 | 1 | 0 | 5 | 0 | 11 | 0.0909 | 2 | 0.5 | 3 | 0 | 2 | 0 | 4 | 0 | 11 | 0.0909 |
| 5_34643034 | 4 | 0 | 3 | 0 | 1 | 0 | 4 | 0.25 | 12 | 0.0833 | 2 | 0 | 3 | 0 | 4 | 0.25 | 7 | 0 | 16 | 0.0625 |
| 5_34643098 | 3 | 1 | 3 | 0.3333 | 1 | 0 | 3 | 0.6667 | 10 | 0.6 | 2 | 1 | 2 | 1 | 4 | 1 | 6 | 1 | 14 | 1 |
| 5_34643237 | 4 | 0 | 4 | 0.25 | NA | NA | 6 | 0.1667 | 14 | 0.1429 | 2 | 0 | 3 | 0.3333 | 5 | 0.6 | 5 | 0.6 | 15 | 0.4667 |
| 5_34643290 | 4 | 0.5 | 4 | 0.25 | 1 | 0 | 5 | 0 | 14 | 0.2143 | 2 | 0.5 | 4 | 0.25 | 5 | 0.2 | 7 | 0.4286 | 18 | 0.3333 |
| 5_34646776 | NA | NA | NA | NA | 1 | 1 | NA | NA | 1 | 1 | NA | NA | 1 | 1 | NA | NA | NA | NA | 1 | 1 |
| 5_34653849 | NA | NA | 2 | 0.5 | 2 | 0.5 | 2 | 0 | 6 | 0.3333 | NA | NA | 1 | 1 | NA | NA | 1 | 0 | 2 | 0.5 |

| | | | | | | | | | | | | | | | | | | | | |
|---|---|---|---|---|---|---|---|---|---|---|---|---|---|---|---|---|---|---|---|---|
| 5_34774169 | 2 | 0 | 9 | 0 | 4 | 0 | 4 | 0.25 | 19 | 0.0526 | 4 | 0 | 6 | 0 | 8 | 0.25 | 3 | 0 | 21 | 0.0952 |
| 5_34788057 | 29 | 0.5517 | 43 | 0.5116 | 21 | 0.4762 | 27 | 0.4815 | 120 | 0.5083 | 17 | 0.7059 | 29 | 0.5862 | 25 | 0.52 | 23 | 0.5652 | 94 | 0.5851 |
| 5_34788153 | 38 | 0.1316 | 54 | 0.1111 | 27 | 0.1111 | 29 | 0.0345 | 148 | 0.1014 | 24 | 0.0417 | 33 | 0.0909 | 29 | 0 | 26 | 0.0769 | 112 | 0.0536 |
| 5_34788582 | 35 | 0.6857 | 40 | 0.525 | 27 | 0.5926 | 29 | 0.4828 | 131 | 0.5725 | 22 | 0.6364 | 33 | 0.6061 | 27 | 0.4444 | 24 | 0.6667 | 106 | 0.5849 |
| 5_34788998 | 45 | 0.0667 | 59 | 0.0678 | 34 | 0.0588 | 38 | 0.0526 | 176 | 0.0625 | 26 | 0.0385 | 40 | 0.025 | 36 | 0.0833 | 35 | 0.0571 | 137 | 0.0511 |
| 5_34789328 | 42 | 0.0952 | 50 | 0.06 | 26 | 0.1538 | 36 | 0.1667 | 154 | 0.1104 | 21 | 0.0476 | 31 | 0.1613 | 32 | 0.1562 | 32 | 0.1562 | 116 | 0.1379 |
| 5_34789449 | 54 | 0.4074 | 58 | 0.4655 | 32 | 0.4375 | 37 | 0.2703 | 181 | 0.4033 | 29 | 0.4828 | 39 | 0.2821 | 40 | 0.325 | 35 | 0.2286 | 143 | 0.3217 |
| 5_34791260 | 2 | 0.5 | 5 | 0.2 | 1 | 0 | 2 | 0 | 10 | 0.2 | 2 | 0 | 1 | 0 | 6 | 0.3333 | 2 | 0 | 11 | 0.1818 |
| 5_34987122 | 1 | 0 | 1 | 1 | NA | NA | NA | NA | 2 | 0.5 | NA | NA | 1 | 1 | NA | NA | 1 | 0 | 2 | 0.5 |
| 5_35066050 | 8 | 0 | 4 | 0.25 | 2 | 0 | 8 | 0.375 | 22 | 0.1818 | 2 | 0.5 | 3 | 0.3333 | 2 | 0 | 7 | 0.1429 | 14 | 0.2143 |
| 5_35066204 | 8 | 0.875 | 4 | 0.75 | 2 | 0.5 | 6 | 0.5 | 20 | 0.7 | 2 | 1 | 3 | 1 | 2 | 0 | 6 | 0.8333 | 13 | 0.7692 |
| 5_35066419 | 7 | 0.4286 | 3 | 0.3333 | 2 | 0 | 4 | 0 | 16 | 0.25 | 1 | 0 | 4 | 0.25 | 3 | 0 | 5 | 0.2 | 13 | 0.1538 |
| 5_35066614 | 9 | 0.3333 | 4 | 0.75 | 2 | 0.5 | 6 | 0.1667 | 21 | 0.381 | 2 | 1 | 4 | 0.75 | 3 | 0.3333 | 7 | 0.8571 | 16 | 0.75 |
| 5_35066975 | 7 | 0 | 2 | 0.5 | 1 | 1 | 4 | 0 | 14 | 0.1429 | 2 | 0 | 5 | 0 | 3 | 0.3333 | 5 | 0 | 15 | 0.0667 |
| 5_35067015 | 5 | 0 | 2 | 0 | 1 | 1 | 5 | 0.6 | 13 | 0.3077 | 4 | 0.25 | 5 | 0.6 | 3 | 0.3333 | 3 | 0.3333 | 15 | 0.4 |
| 5_35067023 | 6 | 0.1667 | 3 | 0 | 1 | 1 | 5 | 0 | 15 | 0.1333 | 4 | 0.5 | 5 | 0 | 2 | 0 | 3 | 0 | 14 | 0.1429 |
| 5_35067027 | 6 | 0 | 3 | 0.3333 | 1 | 0 | 6 | 0.1667 | 16 | 0.125 | 3 | 0 | 3 | 0 | 3 | 0.3333 | 4 | 0 | 13 | 0.0769 |
| 5_35067096 | 7 | 0.1429 | 3 | 0 | NA | NA | 6 | 0 | 16 | 0.0625 | 3 | 0 | 5 | 0.2 | 3 | 0 | 5 | 0 | 16 | 0.0625 |
| 5_35067668 | 9 | 0.1111 | 3 | 0 | 1 | 0 | 5 | 0 | 18 | 0.0556 | 4 | 0 | 4 | 0 | 3 | 0 | 5 | 0.2 | 16 | 0.0625 |
| 5_35067961 | 9 | 0.1111 | 5 | 0 | 3 | 0 | 7 | 0.1429 | 24 | 0.0833 | 4 | 0.25 | 4 | 0 | 3 | 0 | 5 | 0 | 16 | 0.0625 |
| 5_35068020 | 7 | 0 | 5 | 0.2 | 2 | 0 | 7 | 0.5714 | 21 | 0.2381 | 4 | 0.25 | 5 | 0.2 | 4 | 0.25 | 4 | 0 | 17 | 0.1765 |
| 5_35068602 | 10 | 0.1 | 5 | 0.2 | 3 | 0 | 8 | 0 | 26 | 0.0769 | 4 | 0 | 6 | 0 | 4 | 0 | 5 | 0.2 | 19 | 0.0526 |
| 5_35069569 | 5 | 1 | 6 | 0.6667 | 7 | 0.5714 | 4 | 0.5 | 22 | 0.6818 | 3 | 1 | 6 | 0.5 | 5 | 0.8 | 4 | 0.25 | 18 | 0.6111 |
| 5_35069639 | 11 | 0.0909 | 9 | 0.2222 | 7 | 0.1429 | 9 | 0.1111 | 36 | 0.1389 | 4 | 0.25 | 8 | 0.125 | 5 | 0 | 5 | 0.2 | 22 | 0.1364 |
| 5_35069719 | 7 | 0 | 4 | 0 | 6 | 0.3333 | 6 | 0 | 23 | 0.087 | 2 | 0 | 6 | 0 | 3 | 0 | 4 | 0.25 | 15 | 0.0667 |
| 5_35069789 | 9 | 0.4444 | 8 | 0.625 | 3 | 0.6667 | 8 | 0.375 | 28 | 0.5 | 4 | 0.75 | 9 | 0.6667 | 6 | 0.5 | 4 | 0.25 | 23 | 0.5652 |
| 5_35080817 | 2 | 0.5 | 1 | 0 | NA | NA | NA | NA | 3 | 0.3333 | NA | NA | 1 | 1 | 1 | 1 | 2 | 0 | 4 | 0.5 |
| 5_35081030 | 2 | 1 | 1 | 1 | NA | NA | 1 | 1 | 4 | 1 | 1 | 1 | NA | NA | 1 | 1 | 2 | 1 | 4 | 1 |
| 5_35081049 | 1 | 1 | 1 | 1 | NA | NA | 1 | 0 | 3 | 0.6667 | 1 | 1 | 1 | 1 | 1 | 1 | 2 | 0 | 5 | 0.6 |
| 5_35081084 | 2 | 1 | 1 | 1 | NA | NA | NA | NA | 3 | 1 | 1 | 1 | 1 | 1 | NA | NA | 2 | 0.5 | 4 | 0.75 |
| 5_35082596 | 1 | 1 | NA | NA | NA | NA | NA | NA | 1 | 1 | NA | NA | NA | NA | 1 | 1 | NA | NA | 1 | 1 |
| 5_35082958 | 1 | 1 | 1 | 1 | NA | NA | NA | NA | 2 | 1 | NA | NA | NA | NA | 1 | 1 | NA | NA | 1 | 1 |
| 5_35082992 | 1 | 1 | 1 | 1 | NA | NA | NA | NA | 2 | 1 | NA | NA | NA | NA | 1 | 1 | NA | NA | 1 | 1 |
| 5_35083040 | 1 | 1 | 1 | 1 | NA | NA | NA | NA | 2 | 1 | NA | NA | NA | NA | 1 | 1 | NA | NA | 1 | 1 |
| 5_35083505 | 1 | 1 | 1 | 1 | NA | NA | NA | NA | 2 | 1 | NA | NA | NA | NA | 1 | 1 | NA | NA | 1 | 1 |
| 5_35083722 | 1 | 1 | 1 | 1 | NA | NA | NA | NA | 2 | 1 | NA | NA | NA | NA | 1 | 1 | NA | NA | 1 | 1 |
| 5_35083770 | 1 | 1 | 1 | 1 | NA | NA | NA | NA | 2 | 1 | NA | NA | NA | NA | 1 | 1 | NA | NA | 1 | 1 |
| 5_35083958 | 1 | 0 | 1 | 1 | NA | NA | NA | NA | 2 | 0.5 | NA | NA | NA | NA | 1 | 1 | NA | NA | 1 | 1 |
| 5_35084253 | 1 | 0 | 1 | 1 | NA | NA | NA | NA | 2 | 0.5 | NA | NA | NA | NA | 1 | 1 | NA | NA | 1 | 1 |
| 5_35147055 | NA | NA | NA | NA | 1 | 1 | 1 | 1 | 2 | 1 | 1 | 0 | 2 | 0.5 | 2 | 0 | 2 | 0.5 | 7 | 0.2857 |
| 5_35147069 | 1 | 0 | NA | NA | 1 | 1 | 2 | 0 | 4 | 0.25 | 1 | 0 | 2 | 1 | 2 | 0.5 | 2 | 0 | 7 | 0.4286 |
| 5_35189333 | 4 | 0.25 | 4 | 0.5 | 4 | 0.5 | 4 | 0.5 | 16 | 0.4375 | 2 | 0.5 | 10 | 0.3 | 7 | 0.4286 | 2 | 0 | 21 | 0.3333 |
| 5_35189541 | 3 | 0.3333 | 2 | 0 | 2 | 0 | 5 | 0.6 | 12 | 0.3333 | 2 | 0.5 | 9 | 0.4444 | 8 | 0.375 | 2 | 1 | 21 | 0.4762 |
| 5_35189655 | 3 | 1 | 4 | 0.5 | 5 | 1 | 6 | 0.6667 | 18 | 0.7778 | 2 | 0.5 | 10 | 0.8 | 9 | 1 | 2 | 1 | 23 | 0.8696 |
| 5_35189685 | 5 | 0 | 2 | 0 | 3 | 0.3333 | 5 | 0 | 15 | 0.0667 | 2 | 0 | 7 | 0.1429 | 6 | 0 | 1 | 0 | 16 | 0.0625 |
| 5_35189704 | 5 | 0.4 | 5 | 0 | 5 | 0.2 | 6 | 0.1667 | 21 | 0.1905 | 3 | 0 | 9 | 0.4444 | 10 | 0.4 | 2 | 0 | 24 | 0.3333 |
| 5_35189765 | 4 | 0.25 | 4 | 0 | 5 | 0.4 | 7 | 0.1429 | 20 | 0.2 | 3 | 0.3333 | 9 | 0.2222 | 7 | 0.2857 | 2 | 0 | 21 | 0.2381 |
| 5_35189795 | 5 | 0.4 | 4 | 0.25 | 5 | 0.2 | 5 | 0.4 | 19 | 0.3158 | 3 | 0.3333 | 8 | 0.25 | 8 | 0.25 | 2 | 0.5 | 21 | 0.2857 |
| 5_35189833 | 2 | 0.5 | NA | NA | 3 | 0.6667 | 4 | 0.25 | 9 | 0.4444 | 1 | 0 | 5 | 0.2 | 6 | 0.1667 | 2 | 0.5 | 14 | 0.2143 |
| 5_35189994 | 5 | 0.2 | 4 | 0 | 5 | 0 | 6 | 0 | 20 | 0.05 | 4 | 0.25 | 10 | 0.1 | 12 | 0.0833 | 2 | 0 | 28 | 0.1071 |
| 5_35190235 | 5 | 0.4 | 4 | 0 | 5 | 0 | 6 | 0 | 20 | 0.1 | 4 | 0 | 11 | 0.0909 | 13 | 0.1538 | 2 | 0 | 30 | 0.1 |
| 5_35190603 | 5 | 0 | 4 | 0 | 3 | 0 | 5 | 0.4 | 17 | 0.1176 | 4 | 0.25 | 8 | 0 | 10 | 0.1 | 2 | 0 | 24 | 0.0833 |
| 5_35190795 | NA | NA | NA | NA | 1 | 1 | NA | NA | 1 | 1 | NA | NA | NA | NA | 1 | 1 | NA | NA | 1 | 1 |
| 5_35196745 | NA | NA | NA | NA | NA | NA | 1 | 1 | 1 | 1 | NA | NA | 1 | 1 | 1 | 1 | NA | NA | 2 | 1 |
| 5_35196810 | NA | NA | 1 | 0 | NA | NA | 1 | 1 | 2 | 0.5 | NA | NA | 1 | 1 | 1 | 1 | NA | NA | 2 | 1 |
| 5_35236757 | 2 | 0.5 | NA | NA | 1 | 1 | 3 | 0.6667 | 6 | 0.6667 | NA | NA | 1 | 0 | 2 | 0.5 | 3 | 1 | 6 | 0.6667 |
| 5_35236855 | 2 | 0 | NA | NA | 1 | 1 | 2 | 0 | 5 | 0.2 | 1 | 0 | 2 | 0 | 3 | 0 | 3 | 0.6667 | 9 | 0.2222 |
| 5_35236903 | 1 | 1 | NA | NA | 1 | 1 | 2 | 0.5 | 4 | 0.75 | 1 | 0 | 1 | 1 | 2 | 0.5 | 1 | 1 | 5 | 0.6 |
| 5_35236964 | 2 | 0.5 | NA | NA | 1 | 0 | 1 | 0 | 4 | 0.25 | 1 | 1 | 2 | 0 | 3 | 0.3333 | 3 | 0 | 9 | 0.2222 |
| 5_35237093 | 2 | 0 | NA | NA | 1 | 1 | 2 | 0.5 | 5 | 0.4 | 1 | 0 | 2 | 0 | 3 | 0.6667 | 3 | 0.3333 | 9 | 0.3333 |
| 5_35237463 | 2 | 0 | 1 | 0 | 1 | 0 | 3 | 0.3333 | 7 | 0.1429 | 2 | 0.5 | 2 | 0 | 3 | 0.3333 | 2 | 0 | 9 | 0.2222 |
| 5_35244503 | NA | NA | 1 | 1 | NA | NA | NA | NA | 1 | 1 | 1 | 0 | NA | NA | NA | NA | 1 | 1 | 2 | 0.5 |
| 5_35244550 | NA | NA | 1 | 1 | NA | NA | NA | NA | 1 | 1 | 1 | 1 | NA | NA | NA | NA | 1 | 0 | 2 | 0.5 |
| 5_35244763 | NA | NA | 1 | 1 | NA | NA | NA | NA | 1 | 1 | 1 | 0 | NA | NA | NA | NA | 1 | 1 | 2 | 0.5 |
| 5_35248981 | 12 | 0.0833 | 9 | 0.1111 | 14 | 0.0714 | 14 | 0.0714 | 49 | 0.0816 | 8 | 0.125 | 16 | 0.125 | 10 | 0.2 | 19 | 0.2105 | 53 | 0.1698 |

| | | | | | | | | | | | | | | | | | | | | |
|---|---|---|---|---|---|---|---|---|---|---|---|---|---|---|---|---|---|---|---|---|
| 5_35249592 | 10 | 0.1 | 9 | 0.1111 | 14 | 0 | 10 | 0.1 | 43 | 0.0698 | 7 | 0.1429 | 16 | 0.125 | 9 | 0.2222 | 17 | 0 | 49 | 0.102 |
| 5_35250066 | 10 | 0 | 10 | 0.3 | 12 | 0 | 10 | 0 | 42 | 0.0714 | 7 | 0.1429 | 15 | 0.0667 | 9 | 0.1111 | 14 | 0.0714 | 45 | 0.0889 |
| 5_35250138 | 9 | 0.3333 | 8 | 0.375 | 13 | 0.1538 | 7 | 0.1429 | 37 | 0.2432 | 5 | 0.4 | 14 | 0.3571 | 9 | 0.3333 | 11 | 0.3636 | 39 | 0.359 |
| 5_35250237 | 11 | 0.2727 | 9 | 0.1111 | 12 | 0 | 8 | 0.25 | 40 | 0.15 | 5 | 0.2 | 15 | 0.2667 | 11 | 0.0909 | 14 | 0.1429 | 45 | 0.1778 |
| 5_35250363 | 36 | 0.3056 | 21 | 0.1905 | 31 | 0.2903 | 21 | 0.2381 | 109 | 0.2661 | 19 | 0.2105 | 29 | 0.2414 | 25 | 0.4 | 31 | 0.1935 | 104 | 0.2596 |
| 5_35250423 | 35 | 0.8 | 19 | 0.6316 | 27 | 0.6296 | 22 | 0.7727 | 103 | 0.7184 | 15 | 0.6 | 30 | 0.9333 | 26 | 0.7308 | 28 | 0.9643 | 99 | 0.8384 |
| 5_35250428 | 35 | 0.5143 | 16 | 0.625 | 25 | 0.84 | 25 | 0.6 | 101 | 0.6337 | 17 | 0.7059 | 30 | 0.8 | 25 | 0.72 | 35 | 0.6571 | 107 | 0.7196 |
| 5_35250504 | 40 | 0.9 | 20 | 1 | 30 | 1 | 26 | 1 | 116 | 0.9655 | 20 | 0.85 | 34 | 0.9412 | 31 | 1 | 34 | 0.9706 | 119 | 0.9496 |
| 5_35250577 | 37 | 0.1622 | 22 | 0.2273 | 29 | 0.1724 | 26 | 0.0385 | 114 | 0.1491 | 20 | 0.15 | 32 | 0.0938 | 27 | 0.2222 | 36 | 0.1111 | 115 | 0.1391 |
| 5_35250587 | 36 | 0.0278 | 20 | 0.15 | 27 | 0.037 | 26 | 0.1538 | 109 | 0.0826 | 20 | 0.1 | 31 | 0.1613 | 29 | 0.2069 | 35 | 0 | 115 | 0.113 |
| 5_35436526 | 4 | 0.75 | 4 | 0.75 | 1 | 1 | 4 | 0.25 | 13 | 0.6154 | 1 | 1 | 5 | 0.8 | 3 | 0.6667 | 1 | 0 | 10 | 0.7 |
| 5_35436642 | 4 | 0 | 3 | 0 | NA | NA | 3 | 0.3333 | 10 | 0.1 | 3 | 0 | 5 | 0.4 | 4 | 0 | 3 | 0 | 15 | 0.1333 |
| 5_35436833 | 4 | 0.5 | 4 | 0.25 | 1 | 0 | 4 | 0.25 | 13 | 0.3077 | 3 | 0 | 9 | 0.3333 | 7 | 0.2857 | 2 | 0 | 21 | 0.2381 |
| 5_35436864 | 2 | 0.5 | 3 | 0.3333 | NA | NA | 3 | 0.3333 | 8 | 0.375 | 3 | 0 | 5 | 0.4 | 7 | 0.4286 | 2 | 1 | 17 | 0.4118 |
| 5_35436936 | 5 | 0.4 | 4 | 0 | 1 | 1 | 3 | 0.3333 | 13 | 0.3077 | 3 | 0 | 8 | 0.125 | 9 | 0.3333 | 4 | 0.75 | 24 | 0.2917 |
| 5_35436943 | 5 | 0.4 | 5 | 0.6 | 1 | 0 | 3 | 0.6667 | 14 | 0.5 | 3 | 0.3333 | 10 | 0.7 | 9 | 0.2222 | 4 | 0.25 | 26 | 0.4231 |
| 5_35437207 | 7 | 0.1429 | 5 | 0.2 | 1 | 1 | 5 | 0 | 18 | 0.1667 | 4 | 0 | 11 | 0.0909 | 8 | 0.25 | 5 | 0.2 | 28 | 0.1429 |
| 5_35437227 | 7 | 0.2857 | 4 | 1 | 1 | 1 | 5 | 0.2 | 17 | 0.4706 | 4 | 0.25 | 10 | 0.5 | 7 | 0.7143 | 3 | 0 | 24 | 0.4583 |
| 5_35437338 | 9 | 0.5556 | 5 | 0.6 | 1 | 1 | 5 | 0 | 20 | 0.45 | 5 | 0 | 12 | 0.1667 | 9 | 0.3333 | 5 | 0.4 | 31 | 0.2258 |
| 5_35437489 | 8 | 0.375 | 3 | 0 | 2 | 0.5 | 5 | 0.2 | 18 | 0.2778 | 4 | 0.25 | 9 | 0.2222 | 8 | 0 | 4 | 0 | 25 | 0.12 |
| 5_35437519 | 9 | 0.1111 | 5 | 0 | 3 | 0.3333 | 6 | 0.1667 | 23 | 0.1304 | 6 | 0.1667 | 11 | 0.2727 | 8 | 0.375 | 5 | 0 | 30 | 0.2333 |
| 5_35437672 | 8 | 0.375 | 5 | 0 | 2 | 0.5 | 5 | 0.4 | 20 | 0.3 | 4 | 0.5 | 10 | 0.3 | 8 | 0.375 | 6 | 0.3333 | 28 | 0.3571 |
| 5_35438161 | 5 | 0.2 | 6 | 0.1667 | 3 | 0 | 5 | 0.4 | 19 | 0.2105 | 5 | 0.2 | 13 | 0.3846 | 10 | 0.5 | 3 | 0 | 31 | 0.3548 |
| 5_35438180 | 8 | 0.125 | 7 | 0.1429 | 3 | 0.3333 | 4 | 0.25 | 22 | 0.1818 | 5 | 0 | 16 | 0.125 | 9 | 0 | 7 | 0.4286 | 37 | 0.1351 |
| 5_35438192 | 7 | 1 | 8 | 0.875 | 3 | 0.6667 | 3 | 1 | 21 | 0.9048 | 6 | 0.8333 | 16 | 0.8125 | 10 | 0.6 | 4 | 0.5 | 36 | 0.7222 |
| 5_35438465 | 9 | 0.2222 | 8 | 0 | 3 | 0 | 6 | 0 | 26 | 0.0769 | 6 | 0.1667 | 12 | 0.0833 | 11 | 0.5455 | 6 | 0.5 | 35 | 0.3143 |
| 5_35438545 | 10 | 0.9 | 6 | 0.8333 | 3 | 1 | 5 | 1 | 24 | 0.9167 | 6 | 1 | 17 | 0.9412 | 13 | 1 | 8 | 0.875 | 44 | 0.9545 |
| 5_35438623 | 10 | 0.7 | 8 | 0.75 | 4 | 1 | 7 | 0.1429 | 29 | 0.6207 | 6 | 0.6667 | 18 | 0.6111 | 12 | 0.5 | 9 | 0.5556 | 45 | 0.5778 |
| 5_35438629 | 8 | 0.25 | 8 | 0.375 | 4 | 0.5 | 7 | 0.2857 | 27 | 0.3333 | 7 | 0 | 19 | 0.3158 | 10 | 0.5 | 10 | 0.2 | 46 | 0.2826 |
| 5_35438646 | 8 | 0.875 | 7 | 0.5714 | 4 | 0.75 | 8 | 0.875 | 27 | 0.7778 | 7 | 1 | 15 | 0.6667 | 12 | 0.75 | 7 | 0.8571 | 41 | 0.7805 |
| 5_35438656 | 7 | 1 | 4 | 0.75 | 4 | 1 | 4 | 1 | 19 | 0.9474 | 5 | 0.6 | 13 | 1 | 8 | 1 | 6 | 1 | 32 | 0.9375 |
| 5_35438709 | 8 | 1 | 9 | 0.8889 | 5 | 1 | 7 | 1 | 29 | 0.9655 | 7 | 1 | 15 | 0.8667 | 10 | 0.9 | 9 | 0.8889 | 41 | 0.9024 |
| 5_35438756 | 8 | 0.5 | 6 | 0.6667 | 3 | 0.6667 | 7 | 0.5714 | 24 | 0.5833 | 6 | 0.5 | 18 | 0.6667 | 10 | 0.6 | 10 | 0.8 | 44 | 0.6591 |
| 5_35438878 | 3 | 0.3333 | 5 | 0.4 | 4 | 0.5 | 6 | 0.3333 | 18 | 0.3889 | 5 | 0.4 | 10 | 0.4 | 11 | 0.4545 | 5 | 0.6 | 31 | 0.4516 |
| 5_35713777 | NA | NA | 2 | 0.5 | 1 | 1 | NA | NA | 3 | 0.6667 | NA | NA | NA | NA | NA | NA | 1 | 1 | 1 | 1 |
| 5_35713859 | NA | NA | 2 | 1 | 1 | 1 | NA | NA | 3 | 1 | NA | NA | NA | NA | NA | NA | 1 | 1 | 1 | 1 |
| 5_35715199 | NA | NA | 2 | 0.5 | 1 | 0 | NA | NA | 3 | 0.3333 | 2 | 0.5 | NA | NA | NA | NA | NA | NA | 2 | 0.5 |
| 5_35769529 | 3 | 0 | 2 | 0 | 2 | 0.5 | NA | NA | 7 | 0.1429 | 1 | 0 | 2 | 0.5 | 1 | 0 | 1 | 0 | 5 | 0.2 |
| 5_35769695 | 3 | 0.6667 | 1 | 1 | 2 | 1 | NA | NA | 6 | 0.8333 | 2 | 1 | 2 | 1 | 1 | 0 | 1 | 1 | 6 | 0.8333 |
| 5_35769771 | 1 | 1 | 1 | 1 | 1 | 1 | NA | NA | 3 | 1 | 2 | 0.5 | 1 | 1 | 1 | 1 | 1 | 1 | 5 | 0.8 |
| 5_35769893 | 3 | 0.3333 | 1 | 0 | 2 | 0 | NA | NA | 6 | 0.1667 | 2 | 0 | 2 | 0.5 | 1 | 0 | 1 | 1 | 6 | 0.3333 |
| 5_35770045 | 4 | 0.5 | 1 | 0 | 2 | 1 | 1 | 1 | 8 | 0.625 | 2 | 0.5 | 2 | 0.5 | 1 | 0 | 2 | 0.5 | 7 | 0.4286 |
| 5_35770137 | 3 | 0.6667 | 2 | 0 | 1 | 1 | NA | NA | 6 | 0.5 | 2 | 0 | 2 | 1 | 1 | 1 | 2 | 0.5 | 7 | 0.5714 |
| 5_35770415 | 1 | 0 | 2 | 1 | 2 | 0.5 | NA | NA | 5 | 0.6 | NA | NA | 1 | 1 | 1 | 0 | 2 | 0 | 4 | 0.25 |
| 5_35770557 | 3 | 0.3333 | 2 | 0 | 2 | 0.5 | NA | NA | 7 | 0.2857 | 3 | 0 | 2 | 0.5 | 2 | 0 | 2 | 0 | 9 | 0.1111 |
| 5_35770807 | 4 | 0.75 | 1 | 1 | 2 | 1 | NA | NA | 7 | 0.8571 | 2 | 1 | 2 | 0.5 | 2 | 0.5 | 2 | 0.5 | 8 | 0.625 |
| 5_35809595 | 3 | 0.6667 | NA | NA | 1 | 0 | NA | NA | 4 | 0.5 | NA | NA | 1 | 1 | 3 | 1 | NA | NA | 4 | 1 |
| 5_35809704 | 3 | 0.3333 | NA | NA | 1 | 1 | NA | NA | 4 | 0.5 | NA | NA | 1 | 0 | 3 | 0.6667 | NA | NA | 4 | 0.5 |
| 5_35810233 | 3 | 0.6667 | NA | NA | NA | NA | NA | NA | 3 | 0.6667 | NA | NA | 1 | 0 | 3 | 0.6667 | NA | NA | 4 | 0.5 |
| 5_35823422 | 3 | 0.3333 | NA | NA | NA | NA | NA | NA | 3 | 0.3333 | NA | NA | NA | NA | 2 | 0.5 | NA | NA | 2 | 0.5 |
| 5_36040813 | 15 | 0.2667 | 13 | 0.1538 | 18 | 0.0556 | 18 | 0.2222 | 64 | 0.1719 | 14 | 0.1429 | 20 | 0.2 | 21 | 0.1905 | 14 | 0.2143 | 69 | 0.1884 |
| 5_36041159 | 17 | 0 | 14 | 0.1429 | 18 | 0.1111 | 26 | 0.0769 | 75 | 0.08 | 14 | 0 | 20 | 0.2 | 23 | 0.1304 | 13 | 0.0769 | 70 | 0.1143 |
| 5_36041943 | 8 | 0.125 | 8 | 0.375 | 6 | 0.3333 | 10 | 0.2 | 32 | 0.25 | 1 | 1 | 7 | 0.4286 | 11 | 0.0909 | 9 | 0.1111 | 28 | 0.2143 |
| 5_36042177 | 10 | 0 | 11 | 0.0909 | 8 | 0 | 11 | 0.0909 | 40 | 0.05 | 4 | 0.25 | 8 | 0.125 | 13 | 0.0769 | 8 | 0 | 33 | 0.0909 |
| 5_36157252 | NA | NA | 1 | 0 | 3 | 0.3333 | 1 | 1 | 5 | 0.4 | 1 | 1 | 1 | 0 | 2 | 0.5 | NA | NA | 4 | 0.5 |
| 5_36157372 | 1 | 0 | 1 | 0 | 3 | 0 | 1 | 1 | 6 | 0.1667 | 1 | 1 | 1 | 0 | 2 | 1 | NA | NA | 4 | 0.75 |
| 5_36161045 | 3 | 0.3333 | 6 | 0 | 3 | 0 | 1 | 0 | 13 | 0.0769 | 4 | 0 | 1 | 0 | 5 | 0.2 | NA | NA | 10 | 0.1 |
| 5_36170847 | NA | NA | 1 | 1 | NA | NA | NA | NA | 1 | 1 | 1 | 0 | 2 | 0.5 | NA | NA | NA | NA | 3 | 0.3333 |
| 5_36175728 | 6 | 0.3333 | 7 | 0.1429 | 2 | 0 | 2 | 0 | 17 | 0.1765 | 6 | 0 | 3 | 0.3333 | 1 | 0 | 2 | 0 | 12 | 0.0833 |
| 5_36176283 | 4 | 0 | 7 | 0.1429 | 2 | 0.5 | 2 | 0 | 15 | 0.1333 | 9 | 0.1111 | 3 | 1 | 1 | 0 | 1 | 0 | 14 | 0.2857 |
| 5_36176430 | 3 | 0.3333 | 4 | 0 | 2 | 0 | 2 | 0.5 | 11 | 0.1818 | 7 | 0 | 3 | 0.6667 | NA | NA | NA | NA | 10 | 0.2 |
| 5_36176468 | NA | NA | 2 | 0.5 | 1 | 1 | NA | NA | 3 | 0.6667 | 3 | 1 | 1 | 1 | NA | NA | 1 | 1 | 5 | 1 |
| 5_36176492 | 4 | 0.5 | 4 | 1 | 1 | 1 | 2 | 0.5 | 11 | 0.7273 | 7 | 0.7143 | 3 | 0.3333 | 1 | 1 | 1 | 0 | 12 | 0.5833 |
| 5_36176535 | 3 | 0.3333 | 2 | 0 | 1 | 0 | 2 | 0 | 8 | 0.125 | 8 | 0.5 | 3 | 0.3333 | 1 | 0 | 1 | 0 | 13 | 0.3846 |
| 5_36176539 | 3 | 0 | 5 | 0.2 | 1 | 1 | 2 | 0.5 | 11 | 0.2727 | 7 | 0.1429 | 3 | 0.6667 | 1 | 0 | 1 | 0 | 12 | 0.25 |

| | | | | | | | | | | | | | | | | | | | | |
|---|---|---|---|---|---|---|---|---|---|---|---|---|---|---|---|---|---|---|---|---|
| 5_36176653 | 3 | 0 | 4 | 0.25 | 2 | 0.5 | 2 | 0.5 | 11 | 0.2727 | 7 | 0.1429 | 3 | 0 | 1 | 0 | 1 | 0 | 12 | 0.0833 |
| 5_36605077 | NA | NA | NA | NA | 1 | 1 | NA | NA | 1 | 1 | NA | NA | NA | NA | NA | NA | 1 | 1 | 1 | 1 |
| 5_36628614 | 8 | 0.125 | 10 | 0.1 | 9 | 0.2222 | 10 | 0.2 | 37 | 0.1622 | 9 | 0.2222 | 10 | 0.1 | 10 | 0.2 | 9 | 0.2222 | 38 | 0.1842 |
| 5_36628767 | 9 | 0.1111 | 11 | 0.6364 | 9 | 0.6667 | 12 | 0.5833 | 41 | 0.5122 | 10 | 0.6 | 12 | 0.6667 | 7 | 0.4286 | 11 | 0.4545 | 40 | 0.55 |
| 5_36628789 | 8 | 0.125 | 9 | 0.1111 | 4 | 0 | 9 | 0 | 30 | 0.0667 | 12 | 0.1667 | 10 | 0 | 8 | 0 | 9 | 0.1111 | 39 | 0.0769 |
| 5_36628903 | 11 | 0.2727 | 11 | 0.1818 | 8 | 0.375 | 13 | 0.1538 | 43 | 0.2326 | 10 | 0.3 | 12 | 0.25 | 11 | 0.0909 | 10 | 0.4 | 43 | 0.2558 |
| 5_36630229 | NA | NA | 1 | 1 | 1 | 1 | NA | NA | 2 | 1 | NA | NA | NA | NA | NA | NA | 1 | 1 | 1 | 1 |
| 5_36630576 | NA | NA | 1 | 1 | 1 | 1 | NA | NA | 2 | 1 | NA | NA | NA | NA | NA | NA | 1 | 1 | 1 | 1 |
| 5_36630768 | NA | NA | 1 | 1 | 1 | 1 | NA | NA | 2 | 1 | NA | NA | NA | NA | NA | NA | 1 | 1 | 1 | 1 |
| 5_36631151 | NA | NA | 1 | 1 | 1 | 1 | NA | NA | 2 | 1 | NA | NA | NA | NA | NA | NA | 1 | 1 | 1 | 1 |
| 5_36631378 | 3 | 0 | 4 | 0 | 3 | 0 | 8 | 0.125 | 18 | 0.0556 | 6 | 0 | 5 | 0 | 6 | 0.3333 | 5 | 0 | 22 | 0.0909 |
| 5_36631449 | 3 | 0 | 4 | 0.25 | 3 | 0.6667 | 5 | 0 | 15 | 0.2 | 3 | 0.3333 | 5 | 0.4 | 5 | 0.2 | 3 | 0 | 16 | 0.25 |
| 5_36631655 | 2 | 0.5 | 4 | 0.25 | 3 | 0.3333 | 4 | 0.25 | 13 | 0.3077 | 3 | 0 | 5 | 0.2 | 3 | 0.6667 | 5 | 0 | 16 | 0.1875 |
| 5_36631798 | 3 | 0 | 5 | 0.2 | 3 | 0 | 5 | 0.2 | 16 | 0.125 | 5 | 0.2 | 4 | 0.25 | 3 | 0.6667 | 4 | 0 | 16 | 0.25 |
| 5_36631813 | 2 | 0 | 5 | 0.2 | 2 | 0 | 7 | 0.5714 | 16 | 0.3125 | 2 | 0.5 | 4 | 0.5 | 3 | 0.6667 | 3 | 0.6667 | 12 | 0.5833 |
| 5_36631849 | 3 | 0.3333 | 4 | 0 | 3 | 0 | 6 | 0.1667 | 16 | 0.125 | 4 | 0 | 6 | 0 | 1 | 0 | 3 | 0.3333 | 14 | 0.0714 |
| 5_36632298 | 4 | 0.25 | 4 | 0.25 | 4 | 0.25 | 5 | 0.2 | 17 | 0.2353 | 3 | 0.3333 | 5 | 0 | 3 | 0 | 5 | 0 | 16 | 0.0625 |
| 5_36632460 | 4 | 0 | 4 | 0.5 | 3 | 0.6667 | 6 | 0.1667 | 17 | 0.2941 | 3 | 0.3333 | 7 | 0.2857 | 2 | 1 | 4 | 0.25 | 16 | 0.375 |
| 5_36633071 | 2 | 0.5 | 3 | 0.3333 | 1 | 1 | 5 | 0.4 | 11 | 0.4545 | NA | NA | 4 | 0.5 | 4 | 1 | 3 | 1 | 11 | 0.8182 |
| 5_36633159 | 4 | 0.5 | 3 | 0 | 1 | 1 | 4 | 0.25 | 12 | 0.3333 | NA | NA | 5 | 0.4 | 2 | 0.5 | 3 | 1 | 10 | 0.6 |
| 5_36633384 | 3 | 0 | 1 | 0 | 1 | 0 | 3 | 1 | 8 | 0.375 | NA | NA | 5 | 1 | 2 | 0 | 2 | 1 | 9 | 0.7778 |
| 5_36633457 | 1 | 1 | 2 | 0 | 1 | 0 | 4 | 1 | 8 | 0.625 | NA | NA | 4 | 0.25 | 1 | 1 | 3 | 1 | 8 | 0.625 |
| 5_36633466 | 1 | 1 | 2 | 1 | 1 | 1 | 2 | 1 | 6 | 1 | NA | NA | 4 | 1 | 1 | 1 | 3 | 1 | 8 | 1 |
| 5_36633489 | 1 | 1 | 2 | 1 | 1 | 1 | 2 | 1 | 6 | 1 | NA | NA | 6 | 0.8333 | 1 | 1 | 3 | 0.6667 | 10 | 0.8 |
| 5_36633570 | 1 | 1 | 1 | 1 | NA | NA | 1 | 1 | 3 | 1 | NA | NA | 4 | 0.5 | 1 | 0 | 2 | 0.5 | 7 | 0.4286 |
| 5_36633615 | 1 | 0 | 1 | 0 | 1 | 0 | 3 | 0.6667 | 6 | 0.3333 | NA | NA | 3 | 0 | 2 | 0.5 | 1 | 0 | 6 | 0.1667 |
| 5_36633903 | 1 | 1 | 2 | 1 | 1 | 1 | 2 | 0.5 | 6 | 0.8333 | NA | NA | 5 | 1 | 2 | 1 | 3 | 0.6667 | 10 | 0.9 |
| 5_36633915 | 1 | 1 | 1 | 1 | 1 | 0 | 1 | 1 | 4 | 0.75 | NA | NA | 5 | 0.2 | 2 | 0 | 3 | 0 | 10 | 0.1 |
| 5_36634005 | NA | NA | 1 | 1 | 1 | 1 | 2 | 1 | 4 | 1 | NA | NA | 6 | 1 | 2 | 0.5 | 3 | 1 | 11 | 0.9091 |
| 5_36634113 | NA | NA | 1 | 0 | NA | NA | 1 | 1 | 2 | 0.5 | NA | NA | 4 | 1 | 2 | 1 | 3 | 1 | 9 | 1 |
| 5_36648224 | 7 | 0.2857 | 8 | 0.125 | 3 | 0 | 8 | 0.5 | 26 | 0.2692 | 4 | 0.25 | 8 | 0.125 | 8 | 0.25 | 7 | 0.1429 | 27 | 0.1852 |
| 5_36648377 | 6 | 0.3333 | 8 | 0.125 | 6 | 0.3333 | 9 | 0.5556 | 29 | 0.3448 | 5 | 0 | 9 | 0.3333 | 9 | 0.1111 | 7 | 0 | 30 | 0.1333 |
| 5_36648387 | 6 | 0 | 10 | 0.1 | 6 | 0.1667 | 8 | 0.25 | 30 | 0.1333 | 4 | 0 | 8 | 0 | 8 | 0.375 | 6 | 0.5 | 26 | 0.2308 |
| 5_36648555 | 6 | 0.1667 | 7 | 0.1429 | 3 | 0 | 6 | 0.3333 | 22 | 0.1818 | 4 | 0 | 6 | 0.1667 | 7 | 0.1429 | 7 | 0 | 24 | 0.0833 |
| 5_36648594 | 5 | 0 | 7 | 0.1429 | 3 | 0 | 6 | 0.1667 | 21 | 0.0952 | 4 | 0 | 6 | 0.3333 | 8 | 0.5 | 4 | 0.25 | 22 | 0.3182 |
| 5_36648745 | 6 | 0.3333 | 8 | 0 | 5 | 0.2 | 6 | 0.1667 | 25 | 0.16 | 4 | 0 | 9 | 0.1111 | 6 | 0.1667 | 6 | 0 | 25 | 0.08 |
| 5_36649661 | 5 | 0 | 10 | 0.1 | 4 | 0 | 5 | 0.2 | 24 | 0.0833 | 3 | 0.3333 | 8 | 0.125 | 6 | 0 | 5 | 0 | 22 | 0.0909 |
| 5_36649919 | 4 | 0 | 9 | 0.3333 | 4 | 0.75 | 2 | 0 | 19 | 0.3158 | 1 | 0 | 5 | 0.4 | 5 | 0.4 | 4 | 0 | 15 | 0.2667 |
| 5_36650326 | 4 | 0.75 | 5 | 0.8 | 4 | 0.5 | 3 | 0.6667 | 16 | 0.6875 | 2 | 0.5 | 6 | 0.8333 | 4 | 0.75 | 4 | 1 | 16 | 0.8125 |
| 5_36652435 | 4 | 0 | 5 | 0.2 | 4 | 0 | 4 | 0 | 17 | 0.0588 | 2 | 0 | 6 | 0.1667 | 6 | 0 | 5 | 0 | 19 | 0.0526 |
| 5_36652477 | 3 | 0 | 4 | 0.25 | 3 | 0 | 3 | 0 | 13 | 0.0769 | 2 | 0.5 | 5 | 0 | 6 | 0.3333 | 2 | 0 | 15 | 0.2 |
| 5_36652585 | 4 | 0 | 3 | 0 | 4 | 0.25 | 4 | 0 | 15 | 0.0667 | 1 | 0 | 4 | 0.25 | 6 | 0 | 4 | 0 | 15 | 0.0667 |
| 5_36728695 | 1 | 1 | 1 | 0 | 1 | 0 | 1 | 1 | 4 | 0.5 | 1 | 0 | 1 | 1 | 1 | 0 | NA | NA | 3 | 0.3333 |
| 5_36728826 | 1 | 0 | 1 | 1 | 1 | 0 | 2 | 0.5 | 5 | 0.4 | NA | NA | 1 | 0 | 1 | 1 | NA | NA | 2 | 0.5 |
| 5_36728835 | 1 | 1 | 1 | 1 | 1 | 1 | 2 | 0.5 | 5 | 0.8 | NA | NA | 1 | 1 | 1 | 1 | NA | NA | 2 | 1 |
| 5_36728965 | NA | NA | 1 | 1 | 1 | 1 | 1 | 0 | 3 | 0.6667 | 1 | 0 | 1 | 1 | 1 | 0 | NA | NA | 3 | 0.3333 |
| 5_36729168 | NA | NA | 1 | 1 | 1 | 0 | 1 | 1 | 3 | 0.6667 | NA | NA | 1 | 1 | NA | NA | NA | NA | 1 | 1 |
| 5_36758262 | 5 | 0.6 | 7 | 0.2857 | 6 | 0.1667 | 3 | 0.3333 | 21 | 0.3333 | 5 | 0.2 | 6 | 0.1667 | 10 | 0.3 | 3 | 0.6667 | 24 | 0.2917 |
| 5_36758984 | 7 | 0.1429 | 7 | 0 | 5 | 0.6 | 6 | 0 | 25 | 0.16 | 3 | 0.6667 | 8 | 0 | 10 | 0.2 | 2 | 0 | 23 | 0.1739 |
| 5_36759454 | 7 | 0.2857 | 9 | 0.1111 | 5 | 0 | 5 | 0 | 26 | 0.1154 | 4 | 0 | 7 | 0 | 12 | 0.3333 | 1 | 0 | 24 | 0.1667 |
| 5_36759489 | 6 | 0.6667 | 7 | 0.5714 | 4 | 0.75 | 4 | 0.5 | 21 | 0.619 | 4 | 1 | 1 | 0 | 11 | 0.4545 | 2 | 0.5 | 18 | 0.5556 |
| 5_36759608 | 7 | 0.7143 | 6 | 0.3333 | 4 | 1 | 5 | 0.4 | 22 | 0.5909 | 4 | 0.25 | 6 | 0.3333 | 12 | 0.5 | 2 | 0.5 | 24 | 0.4167 |
| 5_36759749 | 5 | 0.6 | 7 | 0 | 4 | 0.5 | 3 | 0 | 19 | 0.2632 | 4 | 0.25 | 6 | 0.3333 | 11 | 0.2727 | 2 | 0.5 | 23 | 0.3043 |
| 5_36759865 | 5 | 0.2 | 6 | 0 | 1 | 0 | 1 | 1 | 13 | 0.1538 | 3 | 0 | 6 | 0.1667 | 6 | 0.1667 | 1 | 0 | 16 | 0.125 |
| 5_36770691 | 4 | 0 | 5 | 0 | 2 | 0 | 3 | 0.3333 | 14 | 0.0714 | 2 | 0 | 3 | 0 | 9 | 0.1111 | 2 | 0 | 16 | 0.0625 |
| 5_36770705 | 5 | 0.6 | 4 | 0.75 | 2 | 0.5 | 3 | 0.6667 | 14 | 0.6429 | 2 | 0.5 | 2 | 0 | 8 | 0.75 | 2 | 0.5 | 14 | 0.5714 |
| 5_36770750 | 3 | 0 | 4 | 0.25 | 3 | 0.6667 | 3 | 0.3333 | 13 | 0.3077 | 2 | 0.5 | 2 | 1 | 9 | 0.4444 | 2 | 0.5 | 15 | 0.5333 |
| 5_36771040 | 2 | 0.5 | 4 | 0.5 | 2 | 1 | 1 | 1 | 9 | 0.6667 | NA | NA | 3 | 0.3333 | 6 | 0.3333 | 2 | 0 | 11 | 0.2727 |
| 5_36771069 | 5 | 1 | 4 | 0.75 | 2 | 0.5 | 3 | 1 | 14 | 0.8571 | 1 | 0 | 1 | 1 | 7 | 0.5714 | 2 | 1 | 11 | 0.6364 |
| 5_36771102 | 5 | 0.4 | 4 | 0.25 | 2 | 0.5 | 3 | 0.6667 | 14 | 0.4286 | 1 | 0 | 2 | 1 | 9 | 0.5556 | 2 | 0.5 | 14 | 0.5714 |
| 5_36771143 | 4 | 0.25 | 4 | 0 | 2 | 0 | 3 | 0.3333 | 13 | 0.1538 | 1 | 0 | 2 | 0.5 | 9 | 0.1111 | 2 | 0 | 14 | 0.1429 |
| 5_36771257 | 5 | 0 | 4 | 0 | 2 | 0 | 3 | 0.3333 | 14 | 0.0714 | 1 | 0 | 1 | 0 | 8 | 0.125 | 2 | 0 | 12 | 0.0833 |
| 5_36771326 | 5 | 0 | 4 | 0.25 | 2 | 0 | 3 | 0 | 14 | 0.0714 | 1 | 0 | 2 | 0 | 6 | 0.1667 | 2 | 0.5 | 11 | 0.1818 |
| 5_36771721 | 3 | 0.3333 | 3 | 1 | 1 | 0 | 2 | 0.5 | 9 | 0.5556 | 1 | 1 | 1 | 1 | 6 | 0.5 | 2 | 0 | 10 | 0.5 |
| 5_36771849 | 2 | 0 | 3 | 0.3333 | 1 | 0 | 2 | 0.5 | 8 | 0.25 | 1 | 1 | 2 | 0.5 | 7 | 0.5714 | 2 | 1 | 12 | 0.6667 |

| | | | | | | | | | | | | | | | | | | | | |
|---|---|---|---|---|---|---|---|---|---|---|---|---|---|---|---|---|---|---|---|---|
| 5_36771858 | 5 | 0.2 | 4 | 0 | 1 | 0 | 2 | 0 | 12 | 0.0833 | 1 | 0 | 2 | 0 | 7 | 0.1429 | 1 | 0 | 11 | 0.0909 |
| 5_36771897 | 3 | 0 | 4 | 0.25 | 1 | 0 | 1 | 0 | 9 | 0.1111 | 1 | 0 | 1 | 0 | 6 | 0.1667 | 1 | 0 | 9 | 0.1111 |
| 5_36771905 | 4 | 0.75 | 4 | 0.25 | 1 | 0 | 2 | 0.5 | 11 | 0.4545 | 1 | 0 | 2 | 0.5 | 7 | 0.7143 | 1 | 1 | 11 | 0.6364 |
| 5_36771977 | 4 | 0.75 | 4 | 0.5 | 1 | 1 | 2 | 0.5 | 11 | 0.6364 | 1 | 1 | 2 | 1 | 4 | 0.75 | 1 | 1 | 8 | 0.875 |
| 5_36772071 | 4 | 0.25 | 4 | 0.5 | 1 | 1 | 2 | 0 | 11 | 0.3636 | NA | NA | 2 | 0.5 | 6 | 0.3333 | NA | NA | 8 | 0.375 |
| 5_36772212 | 4 | 0 | 4 | 0 | 1 | 1 | 1 | 0 | 10 | 0.1 | 1 | 1 | 2 | 0 | 6 | 0.1667 | NA | NA | 9 | 0.2222 |
| 5_36772260 | 3 | 0 | 4 | 0.25 | NA | NA | NA | NA | 7 | 0.1429 | 1 | 1 | 1 | 1 | 6 | 0.1667 | NA | NA | 8 | 0.375 |
| 5_36772285 | 3 | 0 | 4 | 0.5 | 1 | 1 | 1 | 1 | 9 | 0.4444 | 1 | 0 | 1 | 0 | 6 | 0.3333 | NA | NA | 8 | 0.25 |
| 5_36772314 | 4 | 1 | 2 | 1 | 1 | 1 | 1 | 1 | 8 | 1 | NA | NA | 1 | 1 | 6 | 1 | NA | NA | 7 | 1 |
| 5_36772454 | 3 | 0.3333 | 3 | 0.3333 | 1 | 0 | NA | NA | 7 | 0.2857 | 1 | 1 | NA | NA | 4 | 0.5 | NA | NA | 5 | 0.6 |
| 5_36772532 | 3 | 1 | 3 | 1 | 1 | 1 | 1 | 0 | 8 | 0.875 | 1 | 1 | 1 | 1 | 4 | 1 | NA | NA | 6 | 1 |
| 5_36772549 | 4 | 0 | 4 | 0.25 | 1 | 0 | 1 | 1 | 10 | 0.2 | 1 | 1 | 2 | 0.5 | 1 | 0 | NA | NA | 4 | 0.5 |
| 5_36772562 | 4 | 0.25 | 4 | 0 | 1 | 0 | 1 | 0 | 10 | 0.1 | 1 | 0 | 2 | 0 | 5 | 0.2 | NA | NA | 8 | 0.125 |
| 5_36772686 | 1 | 1 | 3 | 1 | 1 | 1 | 1 | 1 | 6 | 1 | NA | NA | 2 | 1 | 3 | 0.6667 | NA | NA | 5 | 0.8 |
| 5_36772873 | 4 | 0.25 | 4 | 0.25 | NA | NA | 1 | 1 | 9 | 0.3333 | 1 | 0 | 2 | 0.5 | 3 | 0.3333 | NA | NA | 6 | 0.3333 |
| 5_36772932 | 3 | 1 | 4 | 1 | 1 | 1 | 1 | 0 | 9 | 0.8889 | NA | NA | 2 | 1 | 3 | 1 | NA | NA | 5 | 1 |
| 5_36773016 | 2 | 0.5 | 2 | 0 | 1 | 0 | 1 | 0 | 6 | 0.1667 | NA | NA | 2 | 0 | 3 | 0.6667 | NA | NA | 5 | 0.4 |
| 5_36773069 | 2 | 0.5 | 3 | 0.3333 | 1 | 0 | 1 | 0 | 7 | 0.2857 | NA | NA | 1 | 1 | 4 | 0.5 | NA | NA | 5 | 0.6 |
| 5_36773222 | 1 | 0 | 3 | 0 | 1 | 1 | 1 | 0 | 6 | 0.1667 | NA | NA | 1 | 1 | 3 | 0 | NA | NA | 4 | 0.25 |
| 5_36773726 | 2 | 0 | 3 | 0.3333 | 1 | 1 | 1 | 0 | 7 | 0.2857 | NA | NA | 1 | 0 | 4 | 0.25 | NA | NA | 5 | 0.2 |
| 5_36905062 | 3 | 0.3333 | 2 | 0.5 | 3 | 0 | 4 | 0.25 | 12 | 0.25 | 2 | 0 | NA | NA | 2 | 0.5 | 2 | 0 | 6 | 0.1667 |
| 5_36905139 | 3 | 0.3333 | 2 | 1 | 1 | 0 | 4 | 0.75 | 10 | 0.6 | 3 | 1 | NA | NA | 2 | 1 | 3 | 0.6667 | 8 | 0.875 |
| 5_36905273 | 2 | 1 | 2 | 0.5 | 1 | 1 | 3 | 1 | 8 | 0.875 | 1 | 1 | NA | NA | NA | NA | 3 | 1 | 4 | 1 |
| 5_36905305 | 2 | 0.5 | 2 | 1 | 1 | 1 | 4 | 0.75 | 9 | 0.7778 | 3 | 1 | NA | NA | 1 | 1 | 3 | 1 | 7 | 1 |
| 5_36905316 | 2 | 0 | 2 | 0.5 | 1 | 1 | 4 | 0.75 | 9 | 0.5556 | 3 | 1 | NA | NA | 1 | 0 | 3 | 1 | 7 | 0.8571 |
| 5_36905379 | 2 | 1 | 1 | 1 | 1 | 1 | 2 | 0.5 | 6 | 0.8333 | 1 | 1 | NA | NA | 1 | 1 | 1 | 0 | 3 | 0.6667 |
| 5_36905386 | 2 | 0 | 1 | 1 | 1 | 0 | 3 | 0 | 7 | 0.1429 | 3 | 0.6667 | NA | NA | 1 | 0 | 2 | 1 | 6 | 0.6667 |
| 5_36905500 | 1 | 0 | 1 | 0 | 1 | 1 | 3 | 0 | 6 | 0.1667 | 2 | 0.5 | NA | NA | 1 | 1 | 3 | 0.3333 | 6 | 0.5 |
| 5_36905667 | 2 | 1 | 2 | 0.5 | 1 | 0 | 3 | 0 | 8 | 0.375 | NA | NA | NA | NA | 1 | 1 | 1 | 0 | 2 | 0.5 |
| 5_36905682 | 2 | 0.5 | 2 | 0.5 | 1 | 0 | 2 | 0 | 7 | 0.2857 | 2 | 0.5 | NA | NA | 1 | 1 | 2 | 0 | 5 | 0.4 |
| 5_36952412 | 98 | 0.2653 | 95 | 0.3158 | 89 | 0.3258 | 94 | 0.2021 | 376 | 0.2766 | 68 | 0.2794 | 98 | 0.2857 | 84 | 0.2262 | 86 | 0.2907 | 336 | 0.2708 |
| 5_36952573 | 84 | 0.131 | 81 | 0.0864 | 71 | 0.0986 | 72 | 0.1389 | 308 | 0.1136 | 60 | 0.1667 | 84 | 0.0595 | 69 | 0.0725 | 76 | 0.1316 | 289 | 0.1038 |
| 5_36952679 | 82 | 0.2927 | 84 | 0.4167 | 73 | 0.3973 | 76 | 0.3947 | 315 | 0.3746 | 57 | 0.3333 | 89 | 0.2921 | 65 | 0.4615 | 63 | 0.4762 | 274 | 0.3832 |
| 5_36952729 | 71 | 0.831 | 79 | 0.8987 | 66 | 0.8485 | 73 | 0.9178 | 289 | 0.8754 | 53 | 0.8491 | 77 | 0.9221 | 71 | 0.8451 | 61 | 0.9344 | 262 | 0.8893 |
| 5_36952756 | 68 | 0.2206 | 79 | 0.3038 | 61 | 0.2623 | 78 | 0.2564 | 286 | 0.2622 | 47 | 0.3404 | 84 | 0.2619 | 66 | 0.2121 | 70 | 0.3714 | 267 | 0.2921 |
| 5_36952883 | 76 | 0.1447 | 82 | 0.0854 | 64 | 0.0938 | 70 | 0.1571 | 292 | 0.1199 | 58 | 0.1379 | 86 | 0.1279 | 75 | 0.1733 | 70 | 0.1286 | 289 | 0.1419 |
| 5_36953756 | 58 | 0.569 | 70 | 0.6 | 58 | 0.6207 | 57 | 0.6842 | 243 | 0.6173 | 44 | 0.5455 | 73 | 0.4658 | 52 | 0.4615 | 56 | 0.6786 | 225 | 0.5333 |
| 5_36964430 | 3 | 0.3333 | 3 | 0.6667 | 2 | 0 | 2 | 0 | 10 | 0.3 | 4 | 0.25 | 1 | 1 | 5 | 0.4 | 4 | 0.25 | 14 | 0.3571 |
| 5_36964437 | 3 | 0.3333 | 2 | 0 | 2 | 0 | 2 | 0.5 | 9 | 0.2222 | 3 | 0.6667 | 1 | 1 | 4 | 0.25 | 4 | 0.5 | 12 | 0.5 |
| 5_36964486 | 2 | 0.5 | 2 | 0.5 | 2 | 0.5 | 2 | 0 | 8 | 0.375 | 3 | 0 | 1 | 0 | 4 | 0 | 4 | 0.25 | 12 | 0.0833 |
| 5_37109862 | 19 | 0.3684 | 25 | 0.36 | 21 | 0.3333 | 17 | 0.4118 | 82 | 0.3659 | 14 | 0.3571 | 23 | 0.4348 | 18 | 0.3889 | 27 | 0.2593 | 82 | 0.3537 |
| 5_37109871 | 19 | 0.6316 | 26 | 0.5769 | 17 | 0.6471 | 16 | 0.75 | 78 | 0.641 | 18 | 0.6667 | 19 | 0.5263 | 14 | 0.5714 | 24 | 0.75 | 75 | 0.64 |
| 5_37110029 | 22 | 0.9545 | 24 | 0.9167 | 21 | 0.8571 | 16 | 1 | 83 | 0.9277 | 18 | 0.8889 | 19 | 0.8947 | 14 | 1 | 23 | 0.913 | 74 | 0.9189 |
| 5_37110055 | 18 | 0.0556 | 26 | 0.1923 | 24 | 0.25 | 18 | 0.1111 | 86 | 0.1628 | 20 | 0.2 | 19 | 0.3158 | 19 | 0.1579 | 25 | 0.16 | 83 | 0.2048 |
| 5_37110217 | 18 | 1 | 35 | 0.8286 | 26 | 0.8846 | 22 | 0.9091 | 101 | 0.8911 | 24 | 0.8333 | 21 | 0.9524 | 21 | 0.8571 | 30 | 0.8667 | 96 | 0.875 |
| 5_37110383 | 24 | 0.7917 | 41 | 0.7073 | 30 | 0.8 | 20 | 0.75 | 115 | 0.7565 | 24 | 0.7083 | 27 | 0.7037 | 25 | 0.64 | 36 | 0.8056 | 112 | 0.7232 |
| 5_37110639 | 29 | 0.3793 | 34 | 0.4706 | 27 | 0.4815 | 23 | 0.5217 | 113 | 0.4602 | 23 | 0.3478 | 31 | 0.6129 | 23 | 0.6087 | 36 | 0.4444 | 113 | 0.5044 |
| 5_37110763 | 37 | 0.0811 | 50 | 0.08 | 39 | 0.0769 | 27 | 0.1111 | 153 | 0.085 | 36 | 0.0278 | 42 | 0.0476 | 39 | 0.1026 | 49 | 0.1224 | 166 | 0.0783 |
| 5_37111217 | 27 | 0.0741 | 38 | 0.0263 | 32 | 0.0312 | 29 | 0.1724 | 126 | 0.0714 | 28 | 0 | 30 | 0.0333 | 26 | 0.0769 | 38 | 0.1316 | 122 | 0.0656 |
| 5_37111260 | 38 | 0.0526 | 51 | 0.098 | 44 | 0.0909 | 37 | 0.1351 | 170 | 0.0941 | 36 | 0.1667 | 40 | 0.125 | 44 | 0.1818 | 52 | 0.0385 | 172 | 0.1221 |
| 5_37111294 | 28 | 0.5 | 42 | 0.5238 | 37 | 0.5676 | 29 | 0.4828 | 136 | 0.5221 | 35 | 0.6286 | 35 | 0.5429 | 32 | 0.5312 | 44 | 0.4318 | 146 | 0.5274 |
| 5_37111360 | 35 | 0.9429 | 43 | 0.9767 | 36 | 0.9722 | 32 | 0.9688 | 146 | 0.9658 | 30 | 1 | 35 | 1 | 31 | 0.9355 | 46 | 0.9783 | 142 | 0.9789 |
| 5_37111493 | 48 | 0.3125 | 48 | 0.4583 | 48 | 0.3958 | 37 | 0.3514 | 181 | 0.3812 | 38 | 0.3421 | 46 | 0.3261 | 41 | 0.561 | 53 | 0.566 | 178 | 0.4551 |
| 5_37111580 | 51 | 0.9804 | 49 | 0.9592 | 49 | 0.9592 | 42 | 1 | 191 | 0.9738 | 40 | 1 | 48 | 0.9792 | 46 | 0.9565 | 57 | 0.9474 | 191 | 0.9686 |
| 5_37111602 | 48 | 0.0833 | 57 | 0.1228 | 58 | 0.3103 | 43 | 0.186 | 206 | 0.1796 | 44 | 0.1591 | 53 | 0.0943 | 52 | 0.1538 | 60 | 0.0833 | 209 | 0.1196 |
| 5_37111897 | 54 | 0.537 | 61 | 0.4754 | 61 | 0.5902 | 47 | 0.5532 | 223 | 0.5381 | 41 | 0.7317 | 50 | 0.6 | 58 | 0.5345 | 64 | 0.5781 | 213 | 0.6009 |
| 5_37111944 | 54 | 0.537 | 57 | 0.5263 | 60 | 0.6333 | 43 | 0.4884 | 214 | 0.5514 | 44 | 0.6591 | 51 | 0.6275 | 56 | 0.5179 | 65 | 0.6769 | 216 | 0.6204 |
| 5_37111985 | 57 | 0.0877 | 62 | 0.1452 | 63 | 0.1746 | 49 | 0.1224 | 231 | 0.1342 | 47 | 0.234 | 57 | 0.2105 | 64 | 0.25 | 68 | 0.2353 | 236 | 0.2331 |
| 5_37112015 | 54 | 0.963 | 61 | 0.9508 | 60 | 0.95 | 51 | 0.9608 | 226 | 0.9558 | 48 | 0.9167 | 52 | 0.9808 | 63 | 0.9683 | 69 | 0.942 | 232 | 0.9526 |
| 5_37112092 | 54 | 0.0185 | 63 | 0.0476 | 64 | 0.0938 | 53 | 0.0566 | 234 | 0.0556 | 53 | 0.0755 | 61 | 0.0656 | 67 | 0.0448 | 72 | 0.0417 | 253 | 0.0553 |
| 5_37112114 | 51 | 0.4118 | 61 | 0.377 | 62 | 0.5161 | 48 | 0.5208 | 222 | 0.455 | 49 | 0.449 | 56 | 0.4286 | 62 | 0.371 | 69 | 0.4783 | 236 | 0.4322 |
| 5_37123567 | 60 | 0.5 | 25 | 0.68 | 32 | 0.6875 | 92 | 0.6196 | 209 | 0.6029 | 22 | 0.7727 | 73 | 0.7397 | 76 | 0.6053 | 34 | 0.5882 | 205 | 0.6683 |
| 5_37123646 | 46 | 0.1304 | 12 | 0.1667 | 18 | 0.2778 | 71 | 0.169 | 147 | 0.1701 | 16 | 0.1875 | 53 | 0.1698 | 57 | 0.2281 | 26 | 0.1538 | 152 | 0.1908 |
| 5_37123780 | 52 | 0.0577 | 23 | 0.1739 | 32 | 0.0938 | 86 | 0.0698 | 193 | 0.0829 | 19 | 0.0526 | 71 | 0.0845 | 72 | 0.0556 | 31 | 0.0645 | 193 | 0.0674 |
| 5_37123899 | 50 | 0.06 | 23 | 0.0435 | 31 | 0.1613 | 88 | 0.1136 | 192 | 0.099 | 18 | 0.1667 | 63 | 0.0952 | 72 | 0.125 | 30 | 0.0333 | 183 | 0.1038 |

| | | | | | | | | | | | | | | | | | | | | |
|---|---|---|---|---|---|---|---|---|---|---|---|---|---|---|---|---|---|---|---|---|
| 5_37123920 | 50 | 0.22 | 23 | 0.1739 | 29 | 0.2069 | 84 | 0.1786 | 186 | 0.1935 | 18 | 0.0556 | 62 | 0.1452 | 64 | 0.2188 | 25 | 0.12 | 169 | 0.1598 |
| 5_37124293 | 44 | 0.9773 | 18 | 0.9444 | 24 | 1 | 82 | 1 | 168 | 0.9881 | 18 | 1 | 53 | 1 | 62 | 0.9677 | 26 | 0.9615 | 159 | 0.9811 |
| 5_37124298 | 28 | 0.3929 | 16 | 0.0625 | 20 | 0.3 | 53 | 0.434 | 117 | 0.3504 | 9 | 0.2222 | 35 | 0.3143 | 35 | 0.2857 | 16 | 0.1875 | 95 | 0.2737 |
| 5_37124488 | 41 | 0.9512 | 18 | 1 | 25 | 1 | 77 | 0.974 | 161 | 0.9752 | 18 | 0.9444 | 50 | 0.94 | 57 | 1 | 25 | 0.88 | 150 | 0.9533 |
| 5_37124783 | 32 | 0.75 | 12 | 0.5 | 19 | 0.7368 | 59 | 0.6271 | 122 | 0.6639 | 14 | 0.9286 | 45 | 0.7111 | 46 | 0.587 | 24 | 0.625 | 129 | 0.6744 |
| 5_37124828 | 41 | 0.0976 | 13 | 0.3077 | 21 | 0.0476 | 71 | 0.1831 | 146 | 0.1507 | 15 | 0.2667 | 46 | 0.1304 | 57 | 0.2105 | 26 | 0.1154 | 144 | 0.1736 |
| 5_37124843 | 32 | 0.0625 | 12 | 0.1667 | 13 | 0.0769 | 59 | 0.0678 | 116 | 0.0776 | 11 | 0.1818 | 41 | 0.0488 | 49 | 0.1224 | 26 | 0.0769 | 127 | 0.0945 |
| 5_37124844 | 2 | 0.5 | NA | NA | NA | NA | NA | NA | 2 | 0.5 | NA | NA | 1 | 0 | 2 | 0 | 1 | 1 | 4 | 0.25 |
| 5_37125044 | 36 | 0.0833 | 12 | 0 | 18 | 0.1667 | 69 | 0.1159 | 135 | 0.1037 | 12 | 0.25 | 45 | 0.1778 | 50 | 0.08 | 20 | 0.2 | 127 | 0.1496 |
| 5_37125106 | 36 | 0.4167 | 10 | 0.8 | 18 | 0.1667 | 64 | 0.5 | 128 | 0.4531 | 12 | 0.3333 | 45 | 0.5778 | 50 | 0.46 | 26 | 0.3846 | 133 | 0.4737 |
| 5_37125258 | 31 | 0.9677 | 11 | 1 | 21 | 0.9524 | 59 | 0.9492 | 122 | 0.959 | 11 | 1 | 41 | 0.9756 | 51 | 0.9804 | 22 | 1 | 125 | 0.984 |
| 5_37125316 | 32 | 0.375 | 10 | 0.3 | 19 | 0.6316 | 55 | 0.5273 | 116 | 0.4828 | 9 | 0.5556 | 37 | 0.6486 | 43 | 0.5116 | 23 | 0.4783 | 112 | 0.5536 |
| 5_37125425 | 32 | 0.2812 | 12 | 0.3333 | 17 | 0.3529 | 54 | 0.2222 | 115 | 0.2696 | 10 | 0.3 | 37 | 0.2703 | 38 | 0.3421 | 18 | 0.2778 | 103 | 0.301 |
| 5_37125454 | 25 | 0.96 | 8 | 1 | 17 | 0.9412 | 49 | 0.8571 | 99 | 0.9091 | 12 | 0.8333 | 36 | 1 | 38 | 0.9737 | 17 | 0.9412 | 103 | 0.9612 |
| 5_37125610 | 34 | 0.9412 | 9 | 0.8889 | 18 | 0.8889 | 50 | 0.98 | 111 | 0.9459 | 12 | 1 | 33 | 1 | 44 | 1 | 17 | 1 | 106 | 1 |
| 5_37125697 | 34 | 0.0882 | 9 | 0.1111 | 18 | 0.0556 | 50 | 0.02 | 111 | 0.0541 | 12 | 0.1667 | 35 | 0.1143 | 45 | 0.0667 | 18 | 0 | 110 | 0.0818 |
| 5_37125766 | 31 | 0.2258 | 9 | 0.4444 | 15 | 0.2667 | 50 | 0.3 | 105 | 0.2857 | 10 | 0.1 | 30 | 0.1667 | 42 | 0.2143 | 18 | 0.3333 | 100 | 0.21 |
| 5_37125784 | 28 | 0.1429 | 7 | 0.1429 | 15 | 0.2 | 42 | 0.0952 | 92 | 0.1304 | 10 | 0 | 26 | 0 | 36 | 0.1111 | 16 | 0.0625 | 88 | 0.0568 |
| 5_37242763 | 7 | 0.1429 | 4 | 0.5 | 3 | 0.6667 | 4 | 0.25 | 18 | 0.3333 | 4 | 0.25 | 3 | 0.3333 | 7 | 0.8571 | 7 | 0.8571 | 21 | 0.6667 |
| 5_37248590 | 6 | 0 | 4 | 0 | 4 | 0 | 4 | 0.25 | 18 | 0.0556 | 3 | 0.3333 | 3 | 0 | 8 | 0.125 | 7 | 0.1429 | 21 | 0.1429 |
| 5_37248729 | 8 | 0.375 | 5 | 0.4 | 2 | 0 | 4 | 0.5 | 19 | 0.3684 | 3 | 0.6667 | 2 | 0.5 | 7 | 0.2857 | 7 | 0.8571 | 19 | 0.5789 |
| 5_37248768 | 10 | 0.7 | 5 | 0.8 | 4 | 1 | 6 | 0.8333 | 25 | 0.8 | 6 | 0.6667 | 5 | 0.8 | 9 | 0.7778 | 7 | 1 | 27 | 0.8148 |
| 5_37248853 | 9 | 0.1111 | 5 | 0.2 | 4 | 0.25 | 5 | 0.2 | 23 | 0.1739 | 6 | 0.1667 | 5 | 0.2 | 9 | 0 | 5 | 0.2 | 25 | 0.12 |
| 5_37282327 | 15 | 0.3333 | 8 | 0.5 | 11 | 0.2727 | 9 | 0.6667 | 43 | 0.4186 | 11 | 0.2727 | 15 | 0.4667 | 14 | 0.1429 | 12 | 0.25 | 52 | 0.2885 |
| 5_37306597 | 3 | 0.3333 | NA | NA | 1 | 1 | NA | NA | 4 | 0.5 | NA | NA | 1 | 0 | NA | NA | 1 | 1 | 2 | 0.5 |
| 5_37306630 | 3 | 0.3333 | NA | NA | 1 | 0 | 1 | 1 | 5 | 0.4 | NA | NA | 1 | 1 | NA | NA | 1 | 0 | 2 | 0.5 |
| 5_37306737 | 3 | 0.6667 | NA | NA | 1 | 1 | 1 | 1 | 5 | 0.8 | NA | NA | 1 | 1 | NA | NA | 1 | 1 | 2 | 1 |
| 5_37306824 | 3 | 1 | NA | NA | 1 | 1 | NA | NA | 4 | 1 | NA | NA | NA | NA | NA | NA | 1 | 1 | 1 | 1 |
| 5_37306866 | 3 | 0.6667 | NA | NA | 1 | 1 | 1 | 1 | 5 | 0.8 | NA | NA | 1 | 1 | NA | NA | 1 | 1 | 2 | 1 |
| 5_37306936 | 2 | 1 | NA | NA | 1 | 1 | NA | NA | 3 | 1 | NA | NA | NA | NA | NA | NA | 1 | 1 | 1 | 1 |
| 5_37307445 | 1 | 1 | NA | NA | 1 | 0 | NA | NA | 2 | 0.5 | 1 | 0 | NA | NA | 1 | 1 | 1 | 0 | 3 | 0.3333 |
| 5_37307712 | 2 | 0.5 | NA | NA | 1 | 0 | NA | NA | 3 | 0.3333 | 1 | 1 | NA | NA | 1 | 1 | 1 | 0 | 3 | 0.6667 |
| 5_37307752 | 2 | 0.5 | NA | NA | 1 | 0 | 1 | 0 | 4 | 0.25 | 1 | 0 | NA | NA | 1 | 1 | 1 | 1 | 3 | 0.6667 |
| 5_37386825 | NA | NA | NA | NA | 2 | 0.5 | 1 | 1 | 3 | 0.6667 | 1 | 0 | 2 | 1 | 1 | 0 | 1 | 1 | 5 | 0.6 |
| 5_37388735 | NA | NA | 1 | 0 | NA | NA | 1 | 1 | 2 | 0.5 | NA | NA | 3 | 0.3333 | NA | NA | NA | NA | 3 | 0.3333 |
| 5_37388846 | NA | NA | 1 | 0 | NA | NA | 1 | 1 | 2 | 0.5 | NA | NA | 3 | 0.6667 | NA | NA | NA | NA | 3 | 0.6667 |
| 5_37388894 | NA | NA | NA | NA | NA | NA | 1 | 1 | 1 | 1 | NA | NA | 2 | 1 | NA | NA | NA | NA | 2 | 1 |
| 5_37389064 | NA | NA | 1 | 1 | NA | NA | 1 | 1 | 2 | 1 | NA | NA | 3 | 0.3333 | NA | NA | NA | NA | 3 | 0.3333 |
| 5_37389210 | NA | NA | 1 | 0 | NA | NA | 1 | 1 | 2 | 0.5 | NA | NA | 3 | 0.6667 | 1 | 1 | NA | NA | 4 | 0.75 |
| 5_37389692 | NA | NA | 1 | 0 | NA | NA | 1 | 1 | 2 | 0.5 | NA | NA | 1 | 0 | 1 | 1 | NA | NA | 2 | 0.5 |
| 5_37389762 | NA | NA | NA | NA | NA | NA | 1 | 1 | 1 | 1 | NA | NA | 3 | 0.6667 | 1 | 1 | NA | NA | 4 | 0.75 |
| 5_37390079 | NA | NA | 1 | 0 | NA | NA | 2 | 0.5 | 3 | 0.3333 | NA | NA | 3 | 0.6667 | 2 | 0.5 | NA | NA | 5 | 0.6 |
| 5_37390145 | NA | NA | 1 | 1 | NA | NA | 2 | 1 | 3 | 1 | 1 | 1 | 3 | 1 | 2 | 1 | NA | NA | 6 | 1 |
| 5_37390257 | NA | NA | 1 | 1 | NA | NA | 1 | 1 | 2 | 1 | 1 | 1 | 2 | 0.5 | 2 | 1 | NA | NA | 5 | 0.8 |
| 5_37390478 | NA | NA | 1 | 0 | NA | NA | 1 | 1 | 2 | 0.5 | 1 | 1 | 2 | 0.5 | 2 | 0 | NA | NA | 5 | 0.4 |
| 5_37390495 | NA | NA | 1 | 0 | NA | NA | 2 | 0.5 | 3 | 0.3333 | 1 | 0 | 3 | 0.3333 | 2 | 0.5 | NA | NA | 6 | 0.3333 |
| 5_37390557 | 1 | 1 | 1 | 1 | NA | NA | 1 | 0 | 3 | 0.6667 | 1 | 1 | 3 | 1 | 1 | 1 | NA | NA | 5 | 1 |
| 5_37390603 | 1 | 0 | 1 | 0 | NA | NA | 2 | 0.5 | 4 | 0.25 | NA | NA | 1 | 0 | 1 | 1 | NA | NA | 2 | 0.5 |
| 5_37390698 | 1 | 1 | NA | NA | NA | NA | 1 | 1 | 2 | 1 | 1 | 1 | 2 | 1 | 2 | 0.5 | NA | NA | 5 | 0.8 |
| 5_37390887 | 1 | 0 | NA | NA | NA | NA | 2 | 1 | 3 | 0.6667 | 1 | 1 | 3 | 0 | 2 | 1 | NA | NA | 6 | 0.5 |
| 5_37390925 | 1 | 1 | NA | NA | NA | NA | 1 | 1 | 2 | 1 | 1 | 0 | 3 | 0.6667 | 2 | 1 | NA | NA | 6 | 0.6667 |
| 5_37390965 | 1 | 1 | NA | NA | NA | NA | 2 | 1 | 3 | 1 | 1 | 1 | 3 | 0.6667 | 2 | 0.5 | NA | NA | 6 | 0.6667 |
| 5_37390993 | 1 | 1 | NA | NA | NA | NA | 2 | 1 | 3 | 1 | 1 | 1 | 3 | 1 | 2 | 1 | NA | NA | 6 | 1 |
| 5_37391167 | 1 | 0 | 1 | 0 | NA | NA | 2 | 1 | 4 | 0.5 | 1 | 1 | 3 | 0.6667 | 2 | 1 | 2 | 0.5 | 8 | 0.75 |
| 5_37392308 | 1 | 0 | NA | NA | 1 | 0 | 3 | 0.3333 | 5 | 0.2 | 1 | 0 | 3 | 0.3333 | 2 | 0.5 | 1 | 0 | 7 | 0.2857 |
| 5_37392535 | 1 | 1 | 1 | 0 | 1 | 0 | 2 | 0.5 | 5 | 0.4 | 1 | 0 | 3 | 0 | 1 | 0 | 2 | 0.5 | 7 | 0.1429 |
| 5_37392678 | 2 | 0.5 | 1 | 0 | 1 | 0 | 3 | 1 | 7 | 0.5714 | 1 | 0 | 3 | 0 | 2 | 0.5 | 1 | 0 | 7 | 0.1429 |
| 5_37392701 | 2 | 0 | NA | NA | 1 | 1 | 3 | 0.3333 | 6 | 0.3333 | 1 | 0 | 1 | 0 | 2 | 1 | 2 | 0 | 6 | 0.3333 |
| 5_37392757 | 1 | 0 | 1 | 1 | 1 | 1 | 1 | 1 | 4 | 0.75 | 1 | 0 | NA | NA | 2 | 1 | 1 | 1 | 4 | 0.75 |
| 5_37392978 | 2 | 0 | 1 | 1 | NA | NA | 3 | 0.6667 | 6 | 0.5 | 2 | 1 | 3 | 0.3333 | 2 | 1 | 1 | 1 | 8 | 0.75 |
| 5_37393060 | 1 | 0 | NA | NA | 1 | 1 | 2 | 0 | 4 | 0.25 | 2 | 0 | 1 | 0 | 2 | 0.5 | 2 | 0.5 | 7 | 0.2857 |
| 5_37393082 | 1 | 0 | 1 | 0 | 1 | 1 | 2 | 0 | 5 | 0.2 | 1 | 1 | 1 | 0 | 2 | 0 | 2 | 0 | 6 | 0.1667 |
| 5_37393562 | 2 | 0.5 | 2 | 0 | 1 | 0 | 1 | 0 | 6 | 0.1667 | 3 | 0 | 1 | 0 | 2 | 0 | 2 | 0.5 | 8 | 0.125 |
| 5_37393874 | 2 | 0.5 | 2 | 0 | 2 | 0 | 1 | 0 | 7 | 0.1429 | 2 | 0.5 | 1 | 1 | 2 | 0 | 2 | 0.5 | 7 | 0.4286 |
| 5_37581884 | 2 | 0.5 | 2 | 0.5 | NA | NA | 2 | 1 | 6 | 0.6667 | NA | NA | NA | NA | NA | NA | 1 | 1 | 1 | 1 |

| | | | | | | | | | | | | | | | | | | | | |
|---|---|---|---|---|---|---|---|---|---|---|---|---|---|---|---|---|---|---|---|---|
| 5_3758201 | 1 | 0 | 1 | 0 | NA | NA | 1 | 1 | 3 | 0.3333 | NA | NA | NA | NA | NA | NA | 1 | 1 | 1 | 1 |
| 5_3760593 | NA | NA | NA | NA | 1 | 1 | 1 | 1 | 2 | 1 | NA | NA | 1 | 1 | NA | NA | NA | NA | 1 | 1 |
| 5_3797804 | 1 | 1 | NA | NA | 1 | 1 | NA | NA | 2 | 1 | NA | NA | 1 | 0 | 1 | 1 | NA | NA | 2 | 0.5 |
| 5_3797825 | 1 | 1 | NA | NA | 1 | 1 | NA | NA | 2 | 1 | NA | NA | 1 | 1 | 1 | 1 | NA | NA | 2 | 1 |
| 5_3797864 | 1 | 1 | NA | NA | 1 | 1 | NA | NA | 2 | 1 | NA | NA | 1 | 1 | 1 | 1 | NA | NA | 2 | 1 |
| 5_3797882 | 1 | 1 | NA | NA | NA | NA | NA | NA | 1 | 1 | NA | NA | 1 | 1 | NA | NA | NA | NA | 1 | 1 |
| 5_3829384 | 9 | 0.2222 | 5 | 0.2 | 1 | 0 | 5 | 0.2 | 20 | 0.2 | 3 | 0 | 8 | 0.375 | 7 | 0.1429 | 6 | 0.3333 | 24 | 0.25 |
| 5_3829391 | 10 | 0.1 | 5 | 0 | 1 | 0 | 3 | 0 | 19 | 0.0526 | 3 | 0 | 7 | 0.1429 | 7 | 0.1429 | 6 | 0 | 23 | 0.087 |
| 5_3829393 | 8 | 0.875 | 7 | 0.5714 | 1 | 0 | 4 | 0.75 | 20 | 0.7 | 1 | 1 | 8 | 0.5 | 6 | 0.8333 | 4 | 1 | 19 | 0.7368 |
| 5_3829411 | 8 | 0.375 | 7 | 0.1429 | 1 | 1 | 4 | 0.5 | 20 | 0.35 | 3 | 0 | 6 | 0.3333 | 7 | 0.1429 | 5 | 0 | 21 | 0.1429 |
| 5_3829585 | 46 | 0.0652 | 39 | 0.1538 | 31 | 0.0323 | 58 | 0.1897 | 174 | 0.1207 | 24 | 0.125 | 58 | 0.1897 | 63 | 0.1905 | 59 | 0.1695 | 204 | 0.1765 |
| 5_3829586 | 48 | 0.0833 | 45 | 0.0667 | 38 | 0.0526 | 65 | 0.0615 | 196 | 0.0663 | 26 | 0.0385 | 59 | 0.0678 | 65 | 0.0923 | 58 | 0.0862 | 208 | 0.0769 |
| 5_3829595 | 47 | 0.8511 | 42 | 0.881 | 35 | 0.8857 | 63 | 0.8889 | 187 | 0.877 | 26 | 0.8077 | 58 | 0.8621 | 58 | 0.8621 | 50 | 0.86 | 192 | 0.8542 |
| 5_3829596 | 43 | 0.8605 | 37 | 0.9189 | 33 | 0.9697 | 59 | 0.9153 | 172 | 0.9128 | 26 | 0.7692 | 60 | 0.95 | 61 | 0.918 | 50 | 0.94 | 197 | 0.9137 |
| 5_3829596 | NA | NA | NA | NA | NA | NA | 1 | 1 | 1 | 1 | NA | NA | NA | NA | NA | NA | 1 | 1 | 1 | 1 |
| 5_3831653 | NA | NA | 1 | 1 | 3 | 0 | NA | NA | 4 | 0.25 | 1 | 0 | 3 | 0 | NA | NA | 1 | 1 | 5 | 0.2 |
| 5_3835780 | 1 | 0 | 1 | 1 | 1 | 1 | NA | NA | 3 | 0.6667 | 1 | 1 | 1 | 0 | 2 | 0.5 | 2 | 0.5 | 6 | 0.5 |
| 5_3835799 | NA | NA | 1 | 1 | 1 | 0 | NA | NA | 2 | 0.5 | 1 | 1 | 1 | 0 | 1 | 0 | 1 | 0 | 4 | 0.25 |
| 5_3835803 | NA | NA | 1 | 1 | NA | NA | NA | NA | 1 | 1 | 1 | 0 | 1 | 1 | 2 | 0.5 | 1 | 1 | 5 | 0.6 |
| 5_3835805 | 1 | 1 | NA | NA | NA | NA | NA | NA | 1 | 1 | NA | NA | 1 | 1 | 2 | 1 | 2 | 1 | 5 | 1 |
| 5_3835812 | 1 | 1 | 1 | 0 | NA | NA | NA | NA | 2 | 0.5 | 1 | 1 | 1 | 1 | NA | NA | 2 | 0.5 | 4 | 0.75 |
| 5_3835827 | NA | NA | 1 | 1 | NA | NA | NA | NA | 1 | 1 | 1 | 1 | NA | NA | 2 | 1 | 1 | 0 | 4 | 0.75 |
| 5_3835843 | 1 | 1 | 1 | 1 | NA | NA | NA | NA | 2 | 1 | 1 | 1 | NA | NA | 2 | 1 | 2 | 1 | 5 | 1 |
| 5_3837103 | NA | NA | 1 | 1 | NA | NA | NA | NA | 1 | 1 | NA | NA | 1 | 0 | 1 | 0 | 1 | 1 | 3 | 0.3333 |
| 5_3837105 | NA | NA | 1 | 1 | NA | NA | NA | NA | 1 | 1 | NA | NA | 1 | 1 | 1 | 1 | 1 | 1 | 3 | 1 |
| 5_3837158 | NA | NA | 1 | 1 | NA | NA | NA | NA | 1 | 1 | NA | NA | 1 | 0 | 1 | 0 | 1 | 1 | 3 | 0.3333 |
| 5_3842651 | 3 | 0 | 2 | 0 | 8 | 0 | 5 | 0.2 | 18 | 0.0556 | 8 | 0.25 | 7 | 0 | 6 | 0 | 3 | 0 | 24 | 0.0833 |
| 5_3842673 | 3 | 0.3333 | 2 | 0 | 4 | 0 | 3 | 0 | 12 | 0.0833 | 8 | 0.125 | 7 | 0.4286 | 5 | 0.4 | 4 | 0 | 24 | 0.25 |
| 5_3847749 | 2 | 0.5 | NA | NA | NA | NA | 1 | 1 | 3 | 0.6667 | 1 | 1 | NA | NA | NA | NA | NA | NA | 1 | 1 |
| 5_3847755 | 2 | 0.5 | NA | NA | NA | NA | 1 | 1 | 3 | 0.6667 | 1 | 1 | NA | NA | NA | NA | NA | NA | 1 | 1 |
| 5_3847763 | 2 | 0.5 | NA | NA | NA | NA | 2 | 1 | 4 | 0.75 | 1 | 1 | NA | NA | NA | NA | NA | NA | 1 | 1 |
| 5_3847793 | 1 | 1 | NA | NA | NA | NA | 1 | 1 | 2 | 1 | NA | NA | 1 | 1 | NA | NA | NA | NA | 1 | 1 |
| 5_3847798 | 1 | 1 | NA | NA | NA | NA | 2 | 1 | 3 | 1 | 1 | 1 | 1 | 1 | NA | NA | NA | NA | 2 | 1 |
| 5_3847808 | 2 | 0.5 | NA | NA | NA | NA | 1 | 1 | 3 | 0.6667 | 1 | 1 | 1 | 1 | NA | NA | NA | NA | 2 | 1 |
| 5_3847816 | 2 | 1 | NA | NA | NA | NA | 2 | 1 | 4 | 1 | 1 | 1 | 1 | 1 | NA | NA | NA | NA | 2 | 1 |
| 5_3847837 | 2 | 1 | NA | NA | NA | NA | 1 | 1 | 3 | 1 | 1 | 1 | NA | NA | NA | NA | NA | NA | 1 | 1 |
| 5_3847949 | 2 | 1 | NA | NA | NA | NA | 2 | 1 | 4 | 1 | 1 | 1 | NA | NA | NA | NA | NA | NA | 1 | 1 |
| 5_3847953 | 3 | 0.3333 | NA | NA | NA | NA | 2 | 1 | 5 | 0.6 | 1 | 1 | 1 | 1 | NA | NA | NA | NA | 2 | 1 |
| 5_3847974 | 3 | 0.6667 | NA | NA | NA | NA | 2 | 1 | 5 | 0.8 | 1 | 0 | 1 | 1 | NA | NA | NA | NA | 2 | 0.5 |
| 5_3868462 | 12 | 0.1667 | 21 | 0.0952 | 14 | 0 | 24 | 0.0417 | 71 | 0.0704 | 12 | 0.1667 | 9 | 0 | 14 | 0.1429 | 8 | 0.125 | 43 | 0.1163 |
| 5_3868465 | 10 | 0.2 | 17 | 0 | 15 | 0.1333 | 22 | 0.0909 | 64 | 0.0938 | 13 | 0.0769 | 11 | 0.1818 | 13 | 0 | 8 | 0 | 45 | 0.0667 |
| 5_3868487 | 11 | 0.2727 | 15 | 0.2 | 12 | 0.0833 | 16 | 0.1875 | 54 | 0.1852 | 10 | 0 | 5 | 0.2 | 9 | 0.1111 | 5 | 0.2 | 29 | 0.1034 |
| 5_3868496 | 7 | 0 | 19 | 0.0526 | 12 | 0 | 17 | 0.1176 | 55 | 0.0545 | 11 | 0.0909 | 8 | 0 | 13 | 0.1538 | 6 | 0 | 38 | 0.0789 |
| 5_3868511 | 10 | 0.5 | 21 | 0.5714 | 11 | 0.4545 | 19 | 0.5789 | 61 | 0.541 | 12 | 0.5833 | 10 | 0.3 | 14 | 0.6429 | 8 | 0.5 | 44 | 0.5227 |
| 5_3868512 | 10 | 0 | 19 | 0.0526 | 11 | 0.3636 | 21 | 0.0952 | 61 | 0.1148 | 14 | 0.1429 | 10 | 0.1 | 13 | 0.1538 | 7 | 0.1429 | 44 | 0.1364 |
| 5_3882600 | 1 | 1 | 4 | 1 | 1 | 1 | NA | NA | 6 | 1 | NA | NA | 1 | 1 | 4 | 1 | NA | NA | 5 | 1 |
| 5_3882759 | 1 | 0 | 2 | 0.5 | NA | NA | NA | NA | 3 | 0.3333 | 1 | 1 | NA | NA | 2 | 0.5 | NA | NA | 3 | 0.6667 |
| 5_3882869 | 1 | 1 | 2 | 0.5 | NA | NA | NA | NA | 3 | 0.6667 | NA | NA | NA | NA | 2 | 1 | NA | NA | 2 | 1 |
| 5_3882878 | NA | NA | 2 | 1 | NA | NA | NA | NA | 2 | 1 | NA | NA | NA | NA | 2 | 1 | NA | NA | 2 | 1 |
| 5_3882894 | 1 | 1 | 1 | 0 | NA | NA | NA | NA | 2 | 0.5 | NA | NA | NA | NA | 2 | 1 | NA | NA | 2 | 1 |
| 5_3977145 | 7 | 0.1429 | 6 | 0 | 7 | 0.1429 | 4 | 0.25 | 24 | 0.125 | 5 | 0 | 4 | 0.25 | 3 | 0 | 4 | 0.25 | 16 | 0.125 |
| 5_3977158 | 6 | 0.3333 | 7 | 0.1429 | 8 | 0.25 | 4 | 0.25 | 25 | 0.24 | 4 | 0.5 | 4 | 0.25 | 3 | 0.6667 | 3 | 0.3333 | 14 | 0.4286 |
| 5_3977168 | 7 | 1 | 8 | 1 | 8 | 1 | 3 | 1 | 26 | 1 | 4 | 1 | 4 | 1 | 3 | 1 | 4 | 1 | 15 | 1 |
| 5_3977172 | 4 | 1 | 5 | 0.8 | 5 | 0.8 | 2 | 1 | 16 | 0.875 | 2 | 1 | 3 | 0.6667 | 3 | 1 | 2 | 0.5 | 10 | 0.8 |
| 5_3977177 | 4 | 1 | 5 | 1 | 4 | 1 | 3 | 1 | 16 | 1 | 1 | 1 | 1 | 1 | 3 | 1 | 1 | 1 | 6 | 1 |
| 5_3977184 | 6 | 0.5 | 7 | 0.7143 | 8 | 0.375 | 3 | 0 | 24 | 0.4583 | 3 | 0.6667 | 2 | 0 | 3 | 0.6667 | 4 | 0.5 | 12 | 0.5 |
| 5_3977185 | 5 | 0.8 | 3 | 1 | 5 | 0.8 | 2 | 0.5 | 15 | 0.8 | 1 | 1 | 1 | 1 | 3 | 1 | 3 | 1 | 8 | 1 |
| 5_3977189 | 3 | 0.3333 | 5 | 0.4 | 5 | 0.8 | 3 | 0.6667 | 16 | 0.5625 | 2 | 0.5 | 2 | 1 | 3 | 0.6667 | 4 | 0.5 | 11 | 0.6364 |
| 5_3977196 | 5 | 1 | 6 | 0.8333 | 8 | 0.875 | 3 | 1 | 22 | 0.9091 | 2 | 1 | 2 | 0.5 | 2 | 1 | 4 | 1 | 10 | 0.9 |
| 5_3977202 | 4 | 1 | 5 | 0.8 | 5 | 1 | 2 | 1 | 16 | 0.9375 | 3 | 0.6667 | 1 | 1 | 2 | 0.5 | 2 | 1 | 8 | 0.75 |
| 5_3977207 | 5 | 1 | 3 | 1 | 5 | 1 | 3 | 1 | 16 | 1 | 2 | 1 | 1 | 1 | 3 | 1 | 3 | 1 | 9 | 1 |
| 5_3977207 | 4 | 0.5 | 6 | 0 | 7 | 0.4286 | 1 | 1 | 18 | 0.3333 | 3 | 0 | 2 | 0 | 3 | 0.3333 | 4 | 0.25 | 12 | 0.1667 |
| 5_3977210 | 5 | 0.2 | 5 | 0.2 | 4 | 0.75 | 3 | 0 | 17 | 0.2941 | 1 | 0 | 1 | 1 | 2 | 0 | 3 | 0.3333 | 7 | 0.2857 |
| 5_3977218 | 6 | 0.8333 | 6 | 0.6667 | 6 | 0.5 | 2 | 0.5 | 20 | 0.65 | 2 | 0.5 | 2 | 0.5 | 3 | 0.3333 | 4 | 0.25 | 11 | 0.3636 |
| 5_3977222 | 4 | 0 | 5 | 0 | 7 | 0.1429 | 2 | 0.5 | 18 | 0.1111 | 2 | 0.5 | 1 | 0 | 2 | 0 | 3 | 0.3333 | 8 | 0.25 |

| | | | | | | | | | | | | | | | | | | | | |
|---|---|---|---|---|---|---|---|---|---|---|---|---|---|---|---|---|---|---|---|---|
| 5_39772358 | 5 | 0.6 | 2 | 1 | 5 | 0.8 | 1 | 0 | 13 | 0.6923 | 3 | 0.3333 | 1 | 0 | 2 | 0.5 | 4 | 0.5 | 10 | 0.4 |
| 5_39772461 | 5 | 0 | 6 | 0 | 6 | 0.3333 | 2 | 0 | 19 | 0.1053 | 2 | 1 | 2 | 0 | 3 | 0.3333 | 2 | 0.5 | 9 | 0.4444 |
| 5_39772511 | 5 | 0.4 | 6 | 0.1667 | 6 | 0.3333 | 3 | 0 | 20 | 0.25 | 1 | 0 | 2 | 0 | 3 | 0.3333 | 3 | 0.3333 | 9 | 0.2222 |
| 5_41782431 | 13 | 0.6154 | 17 | 0.2353 | 12 | 0.5 | 17 | 0.4118 | 59 | 0.4237 | 12 | 0.5 | 12 | 0.25 | 14 | 0.7143 | 13 | 0.5385 | 51 | 0.5098 |
| 5_41782759 | 8 | 0.875 | 13 | 0.9231 | 14 | 0.8571 | 12 | 0.75 | 47 | 0.8511 | 10 | 0.6 | 9 | 0.8889 | 10 | 0.9 | 13 | 0.7692 | 42 | 0.7857 |
| 5_41782883 | 8 | 0.625 | 13 | 0.6154 | 13 | 0.9231 | 13 | 0.6923 | 47 | 0.7234 | 10 | 0.7 | 8 | 0.75 | 11 | 0.8182 | 12 | 0.75 | 41 | 0.7561 |
| 5_41782952 | 11 | 0.3636 | 13 | 0.0769 | 13 | 0.1538 | 13 | 0.0769 | 50 | 0.16 | 9 | 0 | 10 | 0.2 | 8 | 0.125 | 11 | 0.1818 | 38 | 0.1316 |
| 5_41783034 | 9 | 1 | 12 | 0.9167 | 12 | 1 | 13 | 1 | 46 | 0.9783 | 10 | 1 | 9 | 1 | 10 | 1 | 12 | 1 | 41 | 1 |
| 5_41783154 | 7 | 0.8571 | 12 | 0.5833 | 9 | 0.8889 | 10 | 1 | 38 | 0.8158 | 8 | 1 | 5 | 0.6 | 10 | 0.7 | 12 | 0.4167 | 35 | 0.6571 |
| 5_41945546 | 4 | 0 | 3 | 0.3333 | 3 | 0 | 2 | 0 | 12 | 0.0833 | NA | NA | 2 | 0 | 6 | 0.8333 | 3 | 0.3333 | 11 | 0.5455 |
| 5_41946128 | 3 | 0.6667 | 3 | 1 | 3 | 0.6667 | 3 | 1 | 12 | 0.8333 | NA | NA | NA | NA | 6 | 0.8333 | 2 | 0.5 | 8 | 0.75 |
| 5_41946155 | 3 | 0.3333 | 4 | 0.25 | 3 | 0 | 2 | 0.5 | 12 | 0.25 | NA | NA | NA | NA | 6 | 0.3333 | 2 | 0 | 8 | 0.25 |
| 5_41973636 | 1 | 0 | NA | NA | 2 | 0.5 | NA | NA | 3 | 0.3333 | NA | NA | NA | NA | 4 | 0.75 | 1 | 0 | 5 | 0.6 |
| 5_41973665 | 1 | 1 | NA | NA | 2 | 1 | NA | NA | 3 | 1 | NA | NA | NA | NA | 3 | 0.3333 | NA | NA | 3 | 0.3333 |
| 5_41973711 | 1 | 1 | NA | NA | 2 | 1 | NA | NA | 3 | 1 | NA | NA | NA | NA | 3 | 0.3333 | NA | NA | 3 | 0.3333 |
| 5_41973970 | NA | NA | NA | NA | 2 | 1 | NA | NA | 2 | 1 | NA | NA | NA | NA | 4 | 0.75 | 1 | 0 | 5 | 0.6 |
| 5_41974006 | 1 | 1 | NA | NA | 2 | 1 | NA | NA | 3 | 1 | NA | NA | NA | NA | 4 | 1 | NA | NA | 4 | 1 |
| 5_41974039 | 1 | 1 | NA | NA | 1 | 0 | NA | NA | 2 | 0.5 | NA | NA | NA | NA | 3 | 1 | NA | NA | 3 | 1 |
| 5_41974135 | 1 | 1 | NA | NA | 2 | 1 | NA | NA | 3 | 1 | NA | NA | NA | NA | 4 | 1 | NA | NA | 4 | 1 |
| 5_41974362 | 2 | 1 | 1 | 0 | 2 | 1 | NA | NA | 5 | 0.8 | NA | NA | NA | NA | 5 | 0.6 | NA | NA | 5 | 0.6 |
| 5_41974858 | 1 | 1 | 1 | 0 | 2 | 1 | NA | NA | 4 | 0.75 | NA | NA | NA | NA | 5 | 0.8 | NA | NA | 5 | 0.8 |
| 5_41975566 | 1 | 0 | NA | NA | 2 | 0.5 | NA | NA | 3 | 0.3333 | NA | NA | NA | NA | 2 | 0.5 | NA | NA | 2 | 0.5 |
| 5_41976050 | 1 | 1 | NA | NA | 1 | 0 | NA | NA | 2 | 0.5 | NA | NA | NA | NA | 1 | 1 | NA | NA | 1 | 1 |
| 5_41977135 | NA | NA | NA | NA | 1 | 1 | NA | NA | 1 | 1 | NA | NA | NA | NA | 2 | 1 | NA | NA | 2 | 1 |
| 5_41977935 | NA | NA | NA | NA | 1 | 1 | NA | NA | 1 | 1 | NA | NA | NA | NA | 1 | 1 | NA | NA | 1 | 1 |
| 5_41978314 | NA | NA | NA | NA | 1 | 1 | NA | NA | 1 | 1 | NA | NA | NA | NA | 1 | 1 | NA | NA | 1 | 1 |
| 5_41978371 | NA | NA | NA | NA | 1 | 1 | NA | NA | 1 | 1 | NA | NA | NA | NA | 1 | 1 | NA | NA | 1 | 1 |
| 5_43392106 | NA | NA | 1 | 1 | NA | NA | 1 | 1 | 2 | 1 | NA | NA | NA | NA | 2 | 0 | 1 | 1 | 3 | 0.3333 |
| 5_43394773 | NA | NA | 2 | 1 | 3 | 0.3333 | 3 | 0.6667 | 8 | 0.625 | NA | NA | NA | NA | 3 | 0.3333 | 2 | 0 | 5 | 0.2 |
| 5_43443285 | NA | NA | 4 | 0.75 | 3 | 0.3333 | 8 | 0.375 | 15 | 0.4667 | 1 | 0 | NA | NA | 4 | 0.5 | 7 | 0.2857 | 12 | 0.3333 |
| 5_43445072 | 2 | 0 | 4 | 0.25 | 5 | 0.2 | 6 | 0.3333 | 17 | 0.2353 | 1 | 1 | 1 | 0 | 7 | 0 | 9 | 0 | 18 | 0.0556 |
| 5_43446460 | 6 | 0.1667 | 6 | 0.1667 | 7 | 0.1429 | 13 | 0.0769 | 32 | 0.125 | 2 | 0 | 6 | 0 | 13 | 0.0769 | 13 | 0.0769 | 34 | 0.0588 |
| 5_43446857 | 7 | 0.1429 | 10 | 0.3 | 8 | 0.5 | 16 | 0.125 | 41 | 0.2439 | 2 | 0.5 | 7 | 0.2857 | 13 | 0.1538 | 15 | 0.2 | 37 | 0.2162 |
| 5_43877714 | 1 | 0 | 2 | 0.5 | 1 | 0 | 2 | 0 | 6 | 0.1667 | NA | NA | 2 | 0.5 | 1 | 0 | NA | NA | 3 | 0.3333 |
| 5_43902225 | 14 | 0.2143 | 12 | 0.25 | 12 | 0.4167 | 14 | 0.1429 | 52 | 0.25 | 8 | 0 | 19 | 0.3684 | 8 | 0.25 | 10 | 0.4 | 45 | 0.2889 |
| 5_43902486 | 11 | 0.0909 | 9 | 0.4444 | 14 | 0.2143 | 15 | 0 | 49 | 0.1633 | 7 | 0.1429 | 17 | 0.2353 | 8 | 0.125 | 9 | 0.3333 | 41 | 0.2195 |
| 5_43915628 | 8 | 0.375 | 7 | 0.1429 | 12 | 0.1667 | 10 | 0.1 | 37 | 0.1892 | 5 | 0 | 14 | 0.1429 | 4 | 0 | 9 | 0.1111 | 32 | 0.0938 |
| 5_43915666 | 8 | 0.75 | 5 | 0.2 | 11 | 0.2727 | 10 | 0.2 | 34 | 0.3529 | 5 | 0.2 | 12 | 0.1667 | 4 | 0.5 | 8 | 0.25 | 29 | 0.2414 |
| 5_43915688 | 6 | 0.1667 | 6 | 0.1667 | 11 | 0.3636 | 10 | 0.6 | 33 | 0.3636 | 5 | 0.6 | 11 | 0.2727 | 3 | 0.3333 | 7 | 0.1429 | 26 | 0.3077 |
| 5_43915739 | 6 | 0.3333 | 6 | 0.5 | 12 | 0.1667 | 11 | 0.7273 | 35 | 0.4286 | 5 | 0.6 | 12 | 0.5 | 3 | 0.6667 | 9 | 0.3333 | 29 | 0.4828 |
| 5_43915751 | 7 | 0.2857 | 5 | 0.2 | 13 | 0.1538 | 11 | 0.1818 | 36 | 0.1944 | 5 | 0.2 | 12 | 0.0833 | 2 | 0 | 9 | 0.1111 | 28 | 0.1071 |
| 5_43915826 | 8 | 0.375 | 5 | 0.4 | 13 | 0.4615 | 10 | 0.5 | 36 | 0.4444 | 3 | 0.3333 | 12 | 0.4167 | 1 | 0 | 9 | 0.3333 | 25 | 0.36 |
| 5_43915843 | 8 | 0.5 | 6 | 0.3333 | 12 | 0.3333 | 10 | 0.5 | 36 | 0.4167 | 4 | 0.5 | 11 | 0.2727 | 2 | 0.5 | 9 | 0.3333 | 26 | 0.3462 |
| 5_43915991 | 6 | 0.1667 | 6 | 0.3333 | 11 | 0.3636 | 10 | 0.2 | 33 | 0.2727 | 5 | 0.2 | 10 | 0.2 | 1 | 0 | 9 | 0 | 25 | 0.12 |
| 5_43916018 | 7 | 0.7143 | 6 | 0.6667 | 12 | 0.6667 | 9 | 0.6667 | 34 | 0.6765 | 4 | 0.5 | 12 | 0.5833 | NA | NA | 7 | 0.4286 | 23 | 0.5217 |
| 5_43916063 | 6 | 0.6667 | 6 | 0.1667 | 12 | 0.25 | 8 | 0.125 | 32 | 0.2812 | 4 | 0.25 | 10 | 0.2 | 1 | 0 | 7 | 0.1429 | 22 | 0.1818 |
| 5_43916110 | 7 | 0.4286 | 6 | 0.1667 | 12 | 0.4167 | 8 | 0.625 | 33 | 0.4242 | 4 | 0.25 | 13 | 0.3846 | 1 | 0 | 9 | 0.4444 | 27 | 0.3704 |
| 5_43916120 | 6 | 1 | 6 | 0.6667 | 13 | 0.7692 | 7 | 0.8571 | 32 | 0.8125 | 5 | 0.4 | 13 | 0.6923 | 1 | 1 | 9 | 0.3333 | 28 | 0.5357 |
| 5_44069279 | 2 | 0.5 | 2 | 0.5 | NA | NA | 1 | 0 | 5 | 0.4 | 1 | 1 | NA | NA | NA | NA | 2 | 1 | 3 | 1 |
| 5_44136312 | 1 | 1 | NA | NA | 1 | 0 | 2 | 1 | 4 | 0.75 | 1 | 1 | NA | NA | 1 | 1 | NA | NA | 2 | 1 |
| 5_44136510 | 1 | 0 | 1 | 0 | 1 | 1 | 1 | 0 | 4 | 0.25 | 1 | 0 | NA | NA | 1 | 1 | NA | NA | 2 | 0.5 |
| 5_44136968 | 1 | 1 | 1 | 0 | 1 | 0 | 1 | 1 | 4 | 0.5 | 1 | 1 | NA | NA | 1 | 0 | NA | NA | 2 | 0.5 |
| 5_44137056 | NA | NA | 1 | 1 | 1 | 1 | 1 | 0 | 3 | 0.6667 | 1 | 1 | NA | NA | 1 | 1 | NA | NA | 2 | 1 |
| 5_44333101 | 8 | 0.375 | 7 | 0.2857 | 7 | 0.4286 | 5 | 0.6 | 27 | 0.4074 | 6 | 0.3333 | 4 | 0.5 | 5 | 0.8 | 5 | 0.4 | 20 | 0.5 |
| 5_44333274 | 9 | 0.5556 | 7 | 0.4286 | 6 | 0.1667 | 4 | 0 | 26 | 0.3462 | 6 | 0.5 | 3 | 0.3333 | 6 | 0.5 | 3 | 0.3333 | 18 | 0.4444 |
| 5_44333602 | 7 | 0.5714 | 6 | 0.8333 | 6 | 0.6667 | 5 | 0.8 | 24 | 0.7083 | 7 | 0.2857 | 4 | 0.5 | 5 | 0.6 | 3 | 1 | 19 | 0.5263 |
| 5_44333718 | 3 | 0.6667 | 6 | 0.1667 | 5 | 0.4 | 3 | 0 | 17 | 0.2941 | 4 | 0.25 | 1 | 0 | 4 | 0.5 | 2 | 0.5 | 11 | 0.3636 |
| 5_44333768 | 5 | 0.6 | 5 | 0.4 | 6 | 0 | 4 | 0 | 20 | 0.25 | 8 | 0.25 | 4 | 0.25 | 4 | 0 | 3 | 0.6667 | 19 | 0.2632 |
| 5_44333866 | 4 | 0.25 | 6 | 0.3333 | 8 | 0.25 | 3 | 0.6667 | 21 | 0.3333 | 7 | 0.4286 | 4 | 0.25 | 3 | 0 | 3 | 0 | 17 | 0.2353 |
| 5_44333871 | 6 | 0.1667 | 6 | 0.3333 | 6 | 0.3333 | 4 | 0.25 | 22 | 0.2727 | 7 | 0.5714 | 4 | 0.25 | 5 | 0.4 | 4 | 0.25 | 20 | 0.4 |
| 5_44333895 | 5 | 1 | 6 | 0.6667 | 7 | 1 | 4 | 1 | 22 | 0.9091 | 7 | 0.8571 | 4 | 0.75 | 5 | 1 | 3 | 1 | 19 | 0.8947 |
| 5_44333935 | 4 | 0.5 | 6 | 0.3333 | 8 | 0.25 | 3 | 0 | 21 | 0.2857 | 7 | 0.2857 | 3 | 0 | 5 | 0 | 3 | 0 | 18 | 0.1111 |
| 5_44333948 | 5 | 0.8 | 5 | 0.8 | 6 | 0.6667 | 2 | 0.5 | 18 | 0.7222 | 5 | 0.6 | 3 | 0.3333 | 5 | 0.8 | 2 | 1 | 15 | 0.6667 |
| 5_44334078 | 4 | 0.5 | 4 | 0.75 | 4 | 0.25 | 2 | 1 | 14 | 0.5714 | 3 | 1 | 2 | 0.5 | 3 | 0.3333 | 3 | 1 | 11 | 0.7273 |
| 5_44334086 | 6 | 0.1667 | 6 | 0.1667 | 8 | 0.25 | 3 | 0 | 23 | 0.1739 | 7 | 0.2857 | 3 | 0 | 5 | 0.2 | 4 | 0 | 19 | 0.1579 |

| | | | | | | | | | | | | | | | | | | | | |
|---|---|---|---|---|---|---|---|---|---|---|---|---|---|---|---|---|---|---|---|---|
| 5_4433409 | 6 | 0.1667 | 6 | 0.1667 | 8 | 0.125 | 3 | 0.3333 | 23 | 0.1739 | 7 | 0 | 3 | 0.3333 | 4 | 0.25 | 4 | 0.25 | 18 | 0.1667 |
| 5_4433419 | 5 | 0.8 | 3 | 0.3333 | 6 | 0.5 | 4 | 0.75 | 18 | 0.6111 | 5 | 1 | 2 | 0 | 2 | 1 | 1 | 1 | 10 | 0.8 |
| 5_4433423 | 5 | 1 | 5 | 1 | 6 | 1 | 4 | 1 | 20 | 1 | 8 | 1 | 3 | 0.6667 | 4 | 1 | 2 | 1 | 17 | 0.9412 |
| 5_4433428 | 5 | 0.2 | 5 | 0.6 | 8 | 0.25 | 4 | 0.5 | 22 | 0.3636 | 7 | 0.1429 | 3 | 0 | 4 | 0.25 | 2 | 1 | 16 | 0.25 |
| 5_4433429 | 4 | 1 | 5 | 0.8 | 6 | 0.6667 | 3 | 0.6667 | 18 | 0.7778 | 5 | 1 | 3 | 1 | 3 | 1 | 2 | 1 | 13 | 1 |
| 5_4433432 | 6 | 0.3333 | 3 | 0.3333 | 8 | 0.375 | 4 | 0 | 21 | 0.2857 | 7 | 0 | 3 | 0 | 4 | 0.25 | 2 | 1 | 16 | 0.1875 |
| 5_4433434 | 6 | 0.8333 | 5 | 1 | 5 | 0.6 | 4 | 1 | 20 | 0.85 | 7 | 0.8571 | 3 | 0.6667 | 3 | 1 | 1 | 1 | 14 | 0.8571 |
| 5_4433439 | 6 | 0.6667 | 4 | 0.25 | 7 | 0.7143 | 3 | 1 | 20 | 0.65 | 7 | 0.2857 | 3 | 1 | 4 | 0.75 | 2 | 0.5 | 16 | 0.5625 |
| 5_4438445 | NA | NA | 1 | 1 | 1 | 1 | NA | NA | 2 | 1 | NA | NA | 2 | 1 | NA | NA | 2 | 1 | 4 | 1 |
| 5_4438450 | NA | NA | 1 | 0 | 1 | 1 | NA | NA | 2 | 0.5 | NA | NA | 2 | 1 | NA | NA | 4 | 1 | 6 | 1 |
| 5_4438454 | NA | NA | NA | NA | 1 | 1 | NA | NA | 1 | 1 | NA | NA | 2 | 1 | NA | NA | 4 | 1 | 6 | 1 |
| 5_4438458 | NA | NA | 1 | 1 | 1 | 1 | NA | NA | 2 | 1 | NA | NA | 2 | 1 | NA | NA | 4 | 1 | 6 | 1 |
| 5_4438464 | NA | NA | 1 | 0 | 1 | 1 | NA | NA | 2 | 0.5 | NA | NA | 1 | 1 | NA | NA | 5 | 0.8 | 6 | 0.8333 |
| 5_4438465 | NA | NA | 1 | 0 | 1 | 1 | NA | NA | 2 | 0.5 | NA | NA | 2 | 0.5 | NA | NA | 4 | 0 | 6 | 0.1667 |
| 5_4438489 | NA | NA | 1 | 0 | 1 | 1 | NA | NA | 2 | 0.5 | NA | NA | 1 | 1 | NA | NA | 4 | 0.5 | 5 | 0.6 |
| 5_4438506 | NA | NA | 1 | 0 | 1 | 1 | NA | NA | 2 | 0.5 | NA | NA | 2 | 0 | NA | NA | 5 | 0.2 | 7 | 0.1429 |
| 5_4438840 | 1 | 1 | NA | NA | 1 | 0 | 1 | 0 | 3 | 0.3333 | NA | NA | 1 | 1 | NA | NA | NA | NA | 1 | 1 |
| 5_4463029 | NA | NA | NA | NA | 2 | 0 | 2 | 0.5 | 4 | 0.25 | NA | NA | 1 | 0 | 3 | 0.3333 | 1 | 0 | 5 | 0.2 |
| 5_4463036 | NA | NA | NA | NA | 2 | 0 | 2 | 0.5 | 4 | 0.25 | 2 | 1 | 1 | 0 | 3 | 0.3333 | 2 | 0.5 | 8 | 0.5 |
| 5_4559149 | 43 | 0.1163 | 41 | 0.2439 | 55 | 0.2545 | 34 | 0.1176 | 173 | 0.1908 | 38 | 0.1579 | 31 | 0.3226 | 24 | 0.0417 | 29 | 0.1724 | 122 | 0.1803 |
| 5_4559169 | 37 | 0.0541 | 38 | 0.0526 | 44 | 0.0682 | 32 | 0.1875 | 151 | 0.0861 | 33 | 0.0909 | 27 | 0.1481 | 17 | 0.1176 | 20 | 0.1 | 97 | 0.1134 |
| 5_4566940 | 4 | 0.25 | 8 | 0.25 | 9 | 0.1111 | 11 | 0.0909 | 32 | 0.1562 | 11 | 0.3636 | 11 | 0.1818 | 8 | 0.25 | 5 | 0.2 | 35 | 0.2571 |
| 5_4568258 | 22 | 0.2273 | 21 | 0.0952 | 12 | 0.0833 | 17 | 0 | 72 | 0.1111 | 9 | 0.2222 | 14 | 0.4286 | 28 | 0.1429 | 15 | 0.4 | 66 | 0.2727 |
| 5_4568276 | 23 | 0.7826 | 22 | 0.8182 | 13 | 0.6923 | 20 | 0.9 | 78 | 0.8077 | 9 | 0.7778 | 14 | 0.8571 | 30 | 0.9 | 18 | 1 | 71 | 0.9014 |
| 5_4568707 | 1 | 0 | 1 | 1 | 1 | 0 | NA | NA | 3 | 0.3333 | NA | NA | 1 | 0 | 1 | 1 | NA | NA | 2 | 0.5 |
| 5_4568717 | 3 | 0 | 3 | 0.6667 | 3 | 0.6667 | 2 | 0 | 11 | 0.3636 | NA | NA | 6 | 0 | 6 | 0.1667 | 1 | 0 | 13 | 0.0769 |
| 5_4568724 | 2 | 0.5 | 2 | 0.5 | 2 | 0.5 | 2 | 0.5 | 8 | 0.5 | NA | NA | 3 | 0.6667 | 3 | 0 | NA | NA | 6 | 0.3333 |
| 5_4568725 | 2 | 1 | 2 | 1 | 3 | 1 | 2 | 0.5 | 9 | 0.8889 | NA | NA | 5 | 1 | 3 | 1 | 1 | 1 | 9 | 1 |
| 5_4568732 | 2 | 0.5 | 1 | 1 | 3 | 0.6667 | 2 | 0 | 8 | 0.5 | NA | NA | 6 | 0.5 | 5 | 0.2 | 1 | 0 | 12 | 0.3333 |
| 5_4568735 | 2 | 1 | 2 | 1 | 3 | 1 | 2 | 1 | 9 | 1 | NA | NA | 6 | 0.8333 | 6 | 0.8333 | 1 | 1 | 13 | 0.8462 |
| 5_4568743 | 2 | 0 | 3 | 0 | 3 | 0.3333 | 2 | 0 | 10 | 0.1 | NA | NA | 5 | 0.2 | 6 | 0 | 1 | 0 | 12 | 0.0833 |
| 5_4568749 | 2 | 0.5 | 3 | 0.3333 | 3 | 0.3333 | 2 | 0 | 10 | 0.3 | NA | NA | 4 | 0.5 | 6 | 0 | 1 | 1 | 11 | 0.2727 |
| 5_4568750 | 2 | 0.5 | 3 | 0.3333 | 3 | 0.6667 | 2 | 0.5 | 10 | 0.5 | NA | NA | 4 | 0.25 | 5 | 0.4 | 1 | 0 | 10 | 0.3 |
| 5_4568756 | 2 | 1 | 2 | 1 | 2 | 0.5 | 2 | 1 | 8 | 0.875 | NA | NA | 4 | 0.75 | 6 | 0.5 | 1 | 1 | 11 | 0.6364 |
| 5_4568776 | 1 | 0 | 3 | 0 | 3 | 0 | 2 | 0.5 | 9 | 0.1111 | NA | NA | 5 | 0.2 | 5 | 0 | 1 | 0 | 11 | 0.0909 |
| 5_4568777 | 2 | 0 | 3 | 0.3333 | 3 | 0.6667 | 2 | 0 | 10 | 0.3 | NA | NA | 5 | 0 | 4 | 0 | 1 | 1 | 10 | 0.1 |
| 5_4568793 | 2 | 0.5 | 3 | 0.6667 | 2 | 1 | 2 | 0 | 9 | 0.5556 | NA | NA | 4 | 0.25 | 5 | 0.6 | NA | NA | 9 | 0.4444 |
| 5_4568797 | 2 | 0 | 3 | 0.3333 | 3 | 0.6667 | 2 | 0.5 | 10 | 0.4 | NA | NA | 4 | 0.75 | 6 | 0.3333 | 1 | 1 | 11 | 0.5455 |
| 5_4568805 | 2 | 1 | 3 | 0.6667 | 3 | 0.6667 | 2 | 1 | 10 | 0.8 | NA | NA | 4 | 1 | 6 | 0.8333 | 1 | 1 | 11 | 0.9091 |
| 5_4589083 | NA | NA | 1 | 1 | NA | NA | NA | NA | 1 | 1 | 1 | 1 | 1 | 1 | 2 | 0.5 | NA | NA | 4 | 0.75 |
| 5_4589108 | NA | NA | 1 | 1 | NA | NA | NA | NA | 1 | 1 | NA | NA | 1 | 1 | 2 | 0.5 | NA | NA | 3 | 0.6667 |
| 5_4589108 | NA | NA | 1 | 1 | NA | NA | NA | NA | 1 | 1 | 1 | 1 | NA | NA | 2 | 1 | NA | NA | 3 | 1 |
| 5_4589109 | NA | NA | 1 | 1 | NA | NA | NA | NA | 1 | 1 | 1 | 1 | 1 | 0 | 1 | 0 | NA | NA | 3 | 0.3333 |
| 5_4589115 | NA | NA | 1 | 1 | NA | NA | NA | NA | 1 | 1 | NA | NA | 1 | 1 | 2 | 1 | NA | NA | 3 | 1 |
| 5_4589121 | NA | NA | 1 | 1 | NA | NA | NA | NA | 1 | 1 | 1 | 1 | NA | NA | 1 | 1 | NA | NA | 2 | 1 |
| 5_4589121 | NA | NA | 1 | 1 | NA | NA | NA | NA | 1 | 1 | 1 | 0 | NA | NA | 1 | 1 | NA | NA | 2 | 0.5 |
| 5_4589124 | NA | NA | 1 | 1 | NA | NA | NA | NA | 1 | 1 | 1 | 1 | 1 | 1 | 1 | 1 | NA | NA | 3 | 1 |
| 5_4589151 | NA | NA | 1 | 1 | NA | NA | NA | NA | 1 | 1 | 1 | 0 | 1 | 1 | 2 | 0.5 | NA | NA | 4 | 0.5 |
| 5_4589162 | NA | NA | 1 | 1 | NA | NA | NA | NA | 1 | 1 | NA | NA | NA | NA | 1 | 1 | NA | NA | 1 | 1 |
| 5_4589172 | NA | NA | 1 | 1 | NA | NA | NA | NA | 1 | 1 | NA | NA | NA | NA | 2 | 1 | NA | NA | 2 | 1 |
| 5_4589174 | NA | NA | 1 | 1 | NA | NA | NA | NA | 1 | 1 | 1 | 1 | NA | NA | 2 | 1 | NA | NA | 3 | 1 |
| 5_4846137 | 2 | 0.5 | NA | NA | NA | NA | 2 | 1 | 4 | 0.75 | NA | NA | 1 | 0 | 3 | 1 | 1 | 1 | 5 | 0.8 |
| 5_4846147 | 2 | 0.5 | NA | NA | NA | NA | 2 | 0 | 4 | 0.25 | 1 | 0 | 1 | 1 | 3 | 1 | 1 | 1 | 6 | 0.8333 |
| 5_4846164 | 2 | 1 | NA | NA | NA | NA | 2 | 1 | 4 | 1 | 1 | 1 | 2 | 1 | 3 | 1 | 1 | 1 | 7 | 1 |
| 5_4846169 | 2 | 0.5 | NA | NA | NA | NA | 2 | 0 | 4 | 0.25 | 1 | 0 | 1 | 1 | NA | NA | 1 | 0 | 3 | 0.3333 |
| 5_4846169 | 1 | 1 | NA | NA | NA | NA | 3 | 0.6667 | 4 | 0.75 | 1 | 1 | 2 | 1 | 2 | 0.5 | 1 | 1 | 6 | 0.8333 |
| 5_4846172 | 2 | 1 | NA | NA | NA | NA | 3 | 0.6667 | 5 | 0.8 | 1 | 1 | 2 | 0.5 | 3 | 0.6667 | 1 | 1 | 7 | 0.7143 |
| 5_4846191 | 1 | 1 | NA | NA | NA | NA | 3 | 1 | 4 | 1 | NA | NA | 2 | 1 | 3 | 1 | 1 | 1 | 6 | 1 |
| 5_4846198 | 1 | 0 | NA | NA | NA | NA | 2 | 0.5 | 3 | 0.3333 | NA | NA | 2 | 1 | 3 | 0 | 1 | 0 | 6 | 0.3333 |
| 5_4846269 | 3 | 0.6667 | 1 | 0 | NA | NA | 1 | 0 | 5 | 0.4 | 1 | 0 | 3 | 0.6667 | 4 | 0 | 2 | 0.5 | 10 | 0.3 |
| 5_4846449 | 2 | 0 | NA | NA | 1 | 1 | NA | NA | 3 | 0.3333 | NA | NA | 1 | 0 | 2 | 0.5 | 1 | 1 | 4 | 0.5 |
| 5_4846452 | 2 | 1 | NA | NA | 1 | 1 | NA | NA | 3 | 1 | NA | NA | 1 | 0 | 2 | 0.5 | 2 | 0.5 | 5 | 0.4 |
| 5_4846465 | 2 | 1 | 1 | 0 | NA | NA | NA | NA | 3 | 0.6667 | NA | NA | 1 | 1 | 2 | 0 | 2 | 0.5 | 5 | 0.4 |
| 5_4846466 | 1 | 1 | 1 | 0 | 1 | 1 | NA | NA | 3 | 0.6667 | NA | NA | NA | NA | 1 | 1 | 2 | 0.5 | 3 | 0.6667 |
| 5_4854555 | 2 | 1 | 1 | 1 | NA | NA | 2 | 1 | 5 | 1 | 1 | 1 | 3 | 1 | 2 | 0.5 | 2 | 1 | 8 | 0.875 |

| | | | | | | | | | | | | | | | | | | | | |
|---|---|---|---|---|---|---|---|---|---|---|---|---|---|---|---|---|---|---|---|---|
| 5_4854577 | 1 | 1 | 2 | 0.5 | NA | NA | 2 | 0.5 | 5 | 0.6 | 1 | 0 | 2 | 0 | 1 | 0 | 2 | 0.5 | 6 | 0.1667 |
| 5_4854584 | 2 | 1 | 2 | 0.5 | NA | NA | 2 | 1 | 6 | 0.8333 | 1 | 1 | 2 | 0.5 | 1 | 0 | 2 | 1 | 6 | 0.6667 |
| 5_4854784 | 6 | 0.1667 | 4 | 0.25 | 7 | 0 | 8 | 0 | 25 | 0.08 | 8 | 0 | 9 | 0 | 6 | 0 | 8 | 0.375 | 31 | 0.0968 |
| 5_4875706 | 1 | 1 | NA | NA | NA | NA | NA | NA | 1 | 1 | NA | NA | 1 | 1 | NA | NA | 1 | 1 | 2 | 1 |
| 5_5011753 | 4 | 1 | NA | NA | 1 | 1 | 2 | 1 | 7 | 1 | 1 | 1 | 1 | 1 | 2 | 1 | NA | NA | 4 | 1 |
| 5_5011755 | 3 | 1 | NA | NA | NA | NA | 2 | 0 | 5 | 0.6 | NA | NA | 1 | 1 | 2 | 1 | NA | NA | 3 | 1 |
| 5_5011764 | 4 | 0.75 | 1 | 1 | 1 | 1 | 2 | 1 | 8 | 0.875 | 1 | 1 | 1 | 1 | 2 | 0.5 | NA | NA | 4 | 0.75 |
| 5_5011771 | 3 | 1 | 1 | 1 | 1 | 1 | 2 | 1 | 7 | 1 | 1 | 1 | 1 | 1 | 2 | 1 | NA | NA | 4 | 1 |
| 5_5011789 | 4 | 0.75 | 1 | 1 | 1 | 1 | 1 | 1 | 7 | 0.8571 | 1 | 0 | 1 | 1 | 1 | 1 | NA | NA | 3 | 0.6667 |
| 5_5011793 | 4 | 1 | 1 | 1 | 1 | 1 | 2 | 0.5 | 8 | 0.875 | 1 | 1 | 1 | 1 | 2 | 0.5 | NA | NA | 4 | 0.75 |
| 5_5011813 | 4 | 1 | NA | NA | 1 | 1 | 2 | 0.5 | 7 | 0.8571 | 1 | 1 | 1 | 1 | 2 | 1 | NA | NA | 4 | 1 |
| 5_5011814 | 3 | 0.6667 | NA | NA | 1 | 1 | 2 | 0.5 | 6 | 0.6667 | NA | NA | 1 | 1 | 1 | 0 | NA | NA | 2 | 0.5 |
| 5_5011825 | NA | NA | 1 | 1 | NA | NA | NA | NA | 1 | 1 | 1 | 1 | 1 | 1 | 1 | 1 | NA | NA | 3 | 1 |
| 5_5011841 | 3 | 0.3333 | 1 | 0 | 1 | 1 | 1 | 0 | 6 | 0.3333 | NA | NA | 1 | 0 | 2 | 0.5 | NA | NA | 3 | 0.3333 |
| 5_5011850 | 3 | 1 | 1 | 1 | 1 | 1 | 2 | 1 | 7 | 1 | 1 | 1 | NA | NA | 2 | 1 | NA | NA | 3 | 1 |
| 5_5011865 | 4 | 1 | 1 | 0 | 1 | 0 | 1 | 1 | 7 | 0.7143 | 1 | 1 | 1 | 1 | 2 | 0.5 | NA | NA | 4 | 0.75 |
| 5_5011878 | 4 | 0.75 | 1 | 1 | 1 | 0 | 2 | 0.5 | 8 | 0.625 | NA | NA | NA | NA | 1 | 1 | NA | NA | 1 | 1 |
| 5_5161177 | 3 | 0.3333 | 4 | 0 | 5 | 0 | 4 | 0.25 | 16 | 0.125 | 6 | 0.3333 | 2 | 0 | NA | NA | 6 | 0 | 14 | 0.1429 |
| 5_5161211 | 2 | 1 | 3 | 0.6667 | 4 | 0.75 | 3 | 0.6667 | 12 | 0.75 | 6 | 0.5 | 1 | 0 | NA | NA | 6 | 0.5 | 13 | 0.4615 |
| 5_5161242 | 2 | 0 | 4 | 0 | 4 | 0 | 3 | 0.3333 | 13 | 0.0769 | 5 | 0.2 | 1 | 0 | NA | NA | 5 | 0 | 11 | 0.0909 |
| 5_5161247 | 1 | 0 | 4 | 0 | 2 | 0 | 3 | 0.6667 | 10 | 0.2 | 4 | 0.25 | 1 | 1 | NA | NA | 5 | 0.6 | 10 | 0.5 |
| 5_5161261 | 1 | 1 | 3 | 0.3333 | 4 | 0 | 3 | 0.3333 | 11 | 0.2727 | 4 | 0.25 | NA | NA | NA | NA | 5 | 0.6 | 9 | 0.4444 |
| 5_5161276 | 7 | 0 | 4 | 0.5 | 4 | 0 | 7 | 0 | 22 | 0.0909 | 4 | 0 | NA | NA | 1 | 0 | 6 | 0.1667 | 11 | 0.0909 |
| 5_5161433 | 3 | 0.3333 | 1 | 0 | 2 | 0 | 4 | 0 | 10 | 0.1 | 1 | 0 | 2 | 0 | 2 | 0.5 | 3 | 0 | 8 | 0.125 |
| 5_5161495 | 2 | 0 | 2 | 0.5 | 2 | 0.5 | 3 | 0 | 9 | 0.2222 | NA | NA | 1 | 1 | 1 | 0 | 3 | 0.3333 | 5 | 0.4 |
| 5_5161497 | 2 | 0.5 | 2 | 0 | 3 | 0.6667 | 4 | 0.25 | 11 | 0.3636 | NA | NA | NA | NA | NA | NA | 4 | 0.25 | 4 | 0.25 |
| 5_5161512 | 2 | 1 | 2 | 0.5 | 3 | 1 | 4 | 1 | 11 | 0.9091 | NA | NA | 2 | 0.5 | 1 | 1 | 4 | 1 | 7 | 0.8571 |
| 5_5161528 | 3 | 1 | 2 | 0 | 3 | 0.6667 | 4 | 0.25 | 12 | 0.5 | NA | NA | 2 | 1 | NA | NA | 4 | 0.5 | 6 | 0.6667 |
| 5_5161539 | 2 | 0 | 1 | 0 | 2 | 0 | 3 | 0.3333 | 8 | 0.125 | NA | NA | 1 | 0 | 1 | 0 | 2 | 0.5 | 4 | 0.25 |
| 5_5161545 | 2 | 1 | 1 | 1 | 2 | 1 | 2 | 1 | 7 | 1 | NA | NA | 2 | 1 | NA | NA | 3 | 1 | 5 | 1 |
| 5_5161547 | 1 | 1 | 2 | 0.5 | 2 | 0.5 | 4 | 1 | 9 | 0.7778 | NA | NA | 2 | 1 | NA | NA | 3 | 0.6667 | 5 | 0.8 |
| 5_5163107 | 1 | 1 | 1 | 0 | NA | NA | 4 | 0.75 | 6 | 0.6667 | 1 | 1 | 2 | 0.5 | NA | NA | 1 | 1 | 4 | 0.75 |
| 5_5163111 | NA | NA | NA | NA | NA | NA | 4 | 1 | 4 | 1 | 1 | 1 | 2 | 1 | NA | NA | 2 | 1 | 5 | 1 |
| 5_5163132 | 1 | 1 | NA | NA | 1 | 1 | 4 | 1 | 6 | 1 | 1 | 1 | 2 | 1 | NA | NA | 1 | 1 | 4 | 1 |
| 5_5163141 | 1 | 1 | 1 | 0 | 1 | 1 | 2 | 0.5 | 5 | 0.6 | 1 | 0 | 1 | 0 | NA | NA | 1 | 1 | 3 | 0.3333 |
| 5_5163153 | 1 | 0 | 1 | 0 | 1 | 0 | 4 | 0.5 | 7 | 0.2857 | 1 | 0 | 2 | 0 | NA | NA | 3 | 0.3333 | 6 | 0.1667 |
| 5_5230797 | 12 | 0.3333 | 14 | 0.1429 | 10 | 0.1 | 22 | 0.0909 | 58 | 0.1552 | 12 | 0.3333 | 12 | 0.25 | 17 | 0.1176 | 8 | 0.125 | 49 | 0.2041 |
| 5_5230798 | 9 | 0.6667 | 13 | 0.4615 | 7 | 0 | 16 | 0.3125 | 45 | 0.3778 | 10 | 0.2 | 7 | 0.1429 | 14 | 0.2857 | 5 | 0.4 | 36 | 0.25 |
| 5_5230804 | 11 | 0.0909 | 12 | 0.1667 | 10 | 0 | 19 | 0.1053 | 52 | 0.0962 | 11 | 0 | 9 | 0 | 16 | 0 | 10 | 0.3 | 46 | 0.0652 |
| 5_5230805 | 11 | 0 | 16 | 0.0625 | 11 | 0.0909 | 22 | 0.0909 | 60 | 0.0667 | 11 | 0 | 12 | 0.1667 | 18 | 0 | 11 | 0.1818 | 52 | 0.0769 |
| 5_5234200 | 4 | 0 | 5 | 0.2 | 4 | 0 | 7 | 0 | 20 | 0.05 | 7 | 0.1429 | 2 | 0.5 | 10 | 0 | 7 | 0 | 26 | 0.0769 |
| 5_5252570 | 7 | 0.7143 | 2 | 1 | 4 | 0.25 | 1 | 1 | 14 | 0.6429 | 5 | 0.4 | 2 | 0.5 | 2 | 0.5 | 1 | 0 | 10 | 0.4 |
| 5_5252585 | 6 | 0.3333 | 3 | 0.3333 | 1 | 0 | 1 | 0 | 11 | 0.2727 | 3 | 0.3333 | 3 | 0 | 2 | 0 | NA | NA | 8 | 0.125 |
| 5_5252624 | 8 | 0.125 | 3 | 0 | 4 | 0 | 1 | 0 | 16 | 0.0625 | 7 | 0.1429 | 3 | 0 | 1 | 0 | 1 | 0 | 12 | 0.0833 |
| 5_5252649 | 6 | 0.1667 | 2 | 0.5 | 2 | 0 | 1 | 0 | 11 | 0.1818 | 6 | 0.3333 | 3 | 0.3333 | 2 | 0.5 | 1 | 1 | 12 | 0.4167 |
| 5_5253229 | 6 | 0 | 2 | 0.5 | 4 | 0 | 1 | 0 | 13 | 0.0769 | 3 | 0 | 6 | 0.1667 | 6 | 0 | 5 | 0 | 20 | 0.05 |
| 5_5253257 | 5 | 0.2 | 2 | 0 | 4 | 0 | 3 | 0 | 14 | 0.0714 | 1 | 0 | 3 | 0 | 5 | 0 | 4 | 0.25 | 13 | 0.0769 |
| 5_5253258 | 5 | 0 | 2 | 0.5 | 4 | 0 | 3 | 0.3333 | 14 | 0.1429 | 2 | 0 | 3 | 0.3333 | 4 | 0 | 3 | 0 | 12 | 0.0833 |
| 5_5253333 | 3 | 0 | 2 | 0.5 | 2 | 0 | 2 | 0 | 9 | 0.1111 | 1 | 0 | 3 | 1 | 5 | 0.2 | NA | NA | 9 | 0.4444 |
| 5_5253351 | 3 | 1 | 1 | 1 | 1 | 0 | 1 | 1 | 6 | 0.8333 | NA | NA | 3 | 0.3333 | 4 | 1 | 2 | 1 | 9 | 0.7778 |
| 5_5253357 | 4 | 1 | 1 | 1 | 3 | 1 | 1 | 1 | 9 | 1 | 1 | 1 | 3 | 1 | 4 | 1 | 2 | 1 | 10 | 1 |
| 5_5253369 | 2 | 0 | 1 | 0 | 3 | 0.3333 | 1 | 0 | 7 | 0.1429 | 1 | 0 | 3 | 0.3333 | NA | NA | 1 | 0 | 5 | 0.2 |
| 5_5253376 | 2 | 0 | 2 | 0.5 | 2 | 0.5 | NA | NA | 6 | 0.3333 | 1 | 0 | 3 | 0.3333 | 3 | 0 | 2 | 0 | 9 | 0.1111 |
| 5_5253382 | 4 | 1 | 1 | 1 | 3 | 1 | 2 | 1 | 10 | 1 | 1 | 1 | 3 | 1 | 4 | 1 | 3 | 1 | 11 | 1 |
| 5_5253384 | 3 | 0.3333 | 2 | 0 | 2 | 0 | 2 | 0 | 9 | 0.1111 | 1 | 0 | 3 | 0 | 3 | 0.3333 | 3 | 0 | 10 | 0.1 |
| 5_5253389 | 4 | 1 | 2 | 0.5 | 3 | 1 | 2 | 1 | 11 | 0.9091 | 1 | 1 | 3 | 1 | 4 | 0.75 | 3 | 1 | 11 | 0.9091 |
| 5_5269092 | 12 | 0 | 22 | 0.0909 | 22 | 0.0909 | 9 | 0 | 65 | 0.0615 | 4 | 0 | 5 | 0 | 7 | 0 | 10 | 0.2 | 26 | 0.0769 |
| 5_5269098 | 15 | 0.2 | 26 | 0.5769 | 21 | 0.381 | 8 | 0.25 | 70 | 0.4 | 7 | 0.1429 | 6 | 0.1667 | 8 | 0 | 9 | 0.3333 | 30 | 0.1667 |
| 5_5269110 | 14 | 0.6429 | 26 | 0.2308 | 19 | 0.4737 | 5 | 0.2 | 64 | 0.3906 | 6 | 0.3333 | 4 | 0.5 | 6 | 0.3333 | 9 | 0.3333 | 25 | 0.36 |
| 5_5269165 | 9 | 0.1111 | 18 | 0.1111 | 15 | 0.1333 | 5 | 0 | 47 | 0.1064 | 6 | 0 | 5 | 0.2 | 5 | 0 | 4 | 0 | 20 | 0.05 |
| 5_5269192 | 12 | 0.0833 | 16 | 0.0625 | 12 | 0.0833 | 6 | 0 | 46 | 0.0652 | 4 | 0 | 4 | 0 | 5 | 0.2 | 4 | 0 | 17 | 0.0588 |
| 5_5269201 | 11 | 0.5455 | 15 | 0.3333 | 11 | 0.6364 | 5 | 0.6 | 42 | 0.5 | 4 | 0.25 | 4 | 1 | 5 | 0.4 | 3 | 0 | 16 | 0.4375 |
| 5_5269215 | 10 | 0.4 | 14 | 0.2143 | 14 | 0.5 | 5 | 0 | 43 | 0.3256 | 5 | 0.4 | 5 | 0.6 | 5 | 0.2 | 4 | 0.5 | 19 | 0.4211 |
| 5_5269246 | 10 | 0.1 | 12 | 0.3333 | 13 | 0.3077 | 5 | 0.2 | 40 | 0.25 | 4 | 0.25 | 2 | 0.5 | 5 | 0 | 4 | 0.25 | 15 | 0.2 |
| 5_5269249 | 9 | 0.2222 | 13 | 0.1538 | 12 | 0.1667 | 6 | 0 | 40 | 0.15 | 4 | 0.25 | 2 | 0 | 5 | 0.2 | 3 | 0 | 14 | 0.1429 |

| | | | | | | | | | | | | | | | | | | | | |
|---|---|---|---|---|---|---|---|---|---|---|---|---|---|---|---|---|---|---|---|---|
| 5_5269274 | 6 | 0 | 9 | 0.2222 | 10 | 0.1 | 3 | 0 | 28 | 0.1071 | 3 | 0 | 1 | 0 | 3 | 0 | 3 | 0.3333 | 10 | 0.1 |
| 5_5269355 | 5 | 0.2 | 8 | 0.125 | 8 | 0 | 4 | 0 | 25 | 0.08 | 1 | 1 | 2 | 0.5 | 3 | 0.3333 | 3 | 0 | 9 | 0.3333 |
| 5_5269356 | 4 | 0 | 8 | 0.125 | 6 | 0.3333 | 3 | 0.3333 | 21 | 0.1905 | 1 | 0 | 2 | 0 | 3 | 0.3333 | 4 | 0.5 | 10 | 0.3 |
| 5_5269481 | 6 | 0.6667 | 5 | 0.6 | 6 | 0.5 | 3 | 1 | 20 | 0.65 | 1 | 1 | 2 | 0.5 | 1 | 0 | 3 | 1 | 7 | 0.7143 |
| 5_5269484 | 6 | 0 | 5 | 0.2 | 5 | 0.2 | 2 | 0 | 18 | 0.1111 | NA | NA | 2 | 0 | 2 | 0.5 | 4 | 0 | 8 | 0.125 |
| 5_5269533 | 6 | 0.8333 | 8 | 0.5 | 3 | 0.3333 | 4 | 0 | 21 | 0.4762 | NA | NA | 2 | 0.5 | 3 | 0.6667 | 3 | 0.6667 | 8 | 0.625 |
| 5_5269545 | 5 | 0.4 | 5 | 0.2 | 4 | 0 | 3 | 0.3333 | 17 | 0.2353 | NA | NA | 1 | 0 | 4 | 0 | 3 | 0.6667 | 8 | 0.25 |
| 5_5269549 | 5 | 0.2 | 6 | 0.3333 | 5 | 0.4 | 3 | 0.6667 | 19 | 0.3684 | NA | NA | 1 | 1 | 3 | 0.3333 | 2 | 1 | 6 | 0.6667 |
| 5_5269554 | 7 | 0 | 7 | 0.1429 | 6 | 0.1667 | 4 | 0 | 24 | 0.0833 | NA | NA | 1 | 0 | 4 | 0 | 2 | 0.5 | 7 | 0.1429 |
| 5_5269559 | 7 | 0.8571 | 6 | 1 | 2 | 1 | 1 | 1 | 16 | 0.9375 | NA | NA | 2 | 1 | 4 | 1 | 1 | 1 | 7 | 1 |
| 5_5269561 | 7 | 1 | 6 | 0.8333 | 4 | 1 | 2 | 1 | 19 | 0.9474 | NA | NA | 2 | 1 | 4 | 1 | 2 | 1 | 8 | 1 |
| 5_5269570 | 6 | 0.1667 | 6 | 0 | 4 | 0.25 | 3 | 0 | 19 | 0.1053 | NA | NA | 1 | 0 | 3 | 0 | 1 | 1 | 5 | 0.2 |
| 5_5269582 | 7 | 0.8571 | 7 | 1 | 5 | 1 | 4 | 1 | 23 | 0.9565 | NA | NA | 2 | 1 | 3 | 1 | 2 | 1 | 7 | 1 |
| 5_5279752 | 1 | 0 | 3 | 0.3333 | 2 | 0 | 1 | 0 | 7 | 0.1429 | 1 | 1 | 1 | 0 | 3 | 0 | 1 | 0 | 6 | 0.1667 |
| 5_5280106 | 2 | 1 | 4 | 0.5 | NA | NA | 1 | 0 | 7 | 0.5714 | NA | NA | 3 | 0.6667 | 1 | 1 | 1 | 1 | 5 | 0.8 |
| 5_5280112 | 2 | 0.5 | 3 | 0 | NA | NA | 1 | 0 | 6 | 0.1667 | NA | NA | 2 | 0 | 1 | 1 | NA | NA | 3 | 0.3333 |
| 5_5280121 | 2 | 0.5 | 3 | 0.6667 | NA | NA | 1 | 1 | 6 | 0.6667 | NA | NA | 2 | 0 | 1 | 0 | 1 | 1 | 4 | 0.25 |
| 5_5280126 | 1 | 0 | 4 | 0.25 | NA | NA | 1 | 0 | 6 | 0.1667 | NA | NA | 2 | 0.5 | 1 | 0 | 1 | 0 | 4 | 0.25 |
| 5_5280130 | 2 | 1 | 4 | 1 | NA | NA | 1 | 1 | 7 | 1 | NA | NA | 3 | 0.6667 | 1 | 1 | 1 | 1 | 5 | 0.8 |
| 5_5280131 | 2 | 0 | 4 | 0.25 | NA | NA | 1 | 1 | 7 | 0.2857 | NA | NA | 3 | 0.3333 | 1 | 1 | 1 | 1 | 5 | 0.6 |
| 5_5298110 | 1 | 1 | NA | NA | 3 | 0.3333 | 1 | 0 | 5 | 0.4 | NA | NA | NA | NA | 1 | 0 | 1 | 1 | 2 | 0.5 |
| 5_5298116 | NA | NA | NA | NA | 3 | 1 | 2 | 1 | 5 | 1 | NA | NA | NA | NA | 1 | 1 | 1 | 1 | 2 | 1 |
| 5_5298187 | 1 | 0 | NA | NA | 3 | 0.6667 | 2 | 0.5 | 6 | 0.5 | NA | NA | NA | NA | 1 | 0 | 1 | 1 | 2 | 0.5 |
| 5_5302395 | 5 | 0.8 | 6 | 0.8333 | 2 | 1 | 5 | 1 | 18 | 0.8889 | 3 | 1 | 8 | 0.625 | 10 | 0.9 | 5 | 1 | 26 | 0.8462 |
| 5_5302401 | 5 | 1 | 5 | 0.6 | 3 | 0.6667 | 6 | 0.8333 | 19 | 0.7895 | 4 | 0.75 | 9 | 0.7778 | 11 | 0.8182 | 6 | 0.5 | 30 | 0.7333 |
| 5_5326468 | 2 | 0 | 9 | 0.1111 | 2 | 0 | NA | NA | 13 | 0.0769 | 7 | 0.1429 | 1 | 0 | 1 | 0 | 1 | 0 | 10 | 0.1 |
| 5_5326513 | 2 | 0.5 | 6 | 0 | 1 | 0 | NA | NA | 9 | 0.1111 | 6 | 0.1667 | NA | NA | 1 | 0 | NA | NA | 7 | 0.1429 |
| 5_5326527 | 3 | 0 | 4 | 0.5 | 2 | 0 | 1 | 0 | 10 | 0.2 | 6 | 0.3333 | NA | NA | 1 | 0 | 1 | 0 | 8 | 0.25 |
| 5_5335773 | 1 | 1 | 2 | 0.5 | 1 | 1 | NA | NA | 4 | 0.75 | 2 | 0.5 | NA | NA | NA | NA | NA | NA | 2 | 0.5 |
| 5_5343541 | 9 | 1 | 5 | 1 | 8 | 0.875 | 4 | 1 | 26 | 0.9615 | 8 | 0.875 | 3 | 1 | 10 | 0.8 | 5 | 1 | 26 | 0.8846 |
| 5_5343542 | 7 | 0.5714 | 5 | 0.6 | 7 | 0.5714 | 5 | 0.4 | 24 | 0.5417 | 6 | 0.5 | 3 | 0.3333 | 10 | 0.6 | 3 | 0.3333 | 22 | 0.5 |
| 5_5343546 | 9 | 0 | 5 | 0.2 | 9 | 0 | 5 | 0.2 | 28 | 0.0714 | 7 | 0 | 3 | 0.3333 | 9 | 0.1111 | 4 | 0 | 23 | 0.087 |
| 5_5343550 | 6 | 1 | 5 | 0.6 | 7 | 1 | 3 | 0.3333 | 21 | 0.8095 | 6 | 0.6667 | 1 | 1 | 8 | 0.5 | 4 | 1 | 19 | 0.6842 |
| 5_5343558 | 8 | 0.25 | 5 | 0 | 8 | 0.375 | 3 | 0 | 24 | 0.2083 | 5 | 0 | 3 | 0 | 10 | 0.2 | 4 | 0 | 22 | 0.0909 |
| 5_5343562 | 10 | 0.2 | 6 | 0 | 9 | 0 | 5 | 0 | 30 | 0.0667 | 8 | 0.25 | 4 | 0.25 | 9 | 0 | 5 | 0 | 26 | 0.1154 |
| 5_5343571 | 8 | 0.875 | 6 | 0.6667 | 7 | 0.5714 | 4 | 1 | 25 | 0.76 | 8 | 0.625 | 5 | 0.8 | 11 | 0.8182 | 6 | 0.8333 | 30 | 0.7667 |
| 5_5343582 | 8 | 0.25 | 7 | 0.2857 | 8 | 0.125 | 5 | 0.4 | 28 | 0.25 | 6 | 0 | 5 | 0 | 10 | 0.2 | 5 | 0.2 | 26 | 0.1154 |
| 5_5343855 | 2 | 0 | 1 | 0 | 1 | 1 | NA | NA | 4 | 0.25 | 1 | 0 | NA | NA | 1 | 1 | 1 | 1 | 3 | 0.6667 |
| 5_5344050 | 7 | 0.2857 | NA | NA | 1 | 0 | NA | NA | 8 | 0.25 | 3 | 0 | NA | NA | 1 | 1 | 2 | 0 | 6 | 0.1667 |
| 5_5344054 | 7 | 0.4286 | 1 | 0 | 1 | 0 | 1 | 1 | 10 | 0.4 | 2 | 0 | NA | NA | 1 | 0 | 3 | 0.3333 | 6 | 0.1667 |
| 5_5381073 | NA | NA | 3 | 0.6667 | 3 | 0.3333 | 1 | 1 | 7 | 0.5714 | 2 | 1 | NA | NA | NA | NA | 2 | 1 | 4 | 1 |
| 5_5381098 | NA | NA | 3 | 1 | 1 | 1 | 1 | 1 | 5 | 1 | 1 | 1 | 1 | 1 | NA | NA | 2 | 1 | 4 | 1 |
| 5_5381099 | NA | NA | 3 | 0.3333 | 2 | 0.5 | 1 | 0 | 6 | 0.3333 | 2 | 0 | NA | NA | NA | NA | 2 | 1 | 4 | 0.5 |
| 5_5381129 | NA | NA | 3 | 0.6667 | 2 | 1 | 1 | 0 | 6 | 0.6667 | 2 | 1 | NA | NA | 1 | 0 | 2 | 0.5 | 5 | 0.6 |
| 5_5381130 | NA | NA | 2 | 1 | 3 | 0.3333 | 2 | 0.5 | 7 | 0.5714 | 2 | 0.5 | NA | NA | 1 | 1 | 1 | 0 | 4 | 0.5 |
| 5_5381131 | NA | NA | 4 | 0.75 | 3 | 1 | 2 | 1 | 9 | 0.8889 | 1 | 1 | NA | NA | 1 | 0 | 2 | 1 | 4 | 0.75 |
| 5_5381250 | NA | NA | 5 | 0.2 | 2 | 0 | 2 | 0 | 9 | 0.1111 | 2 | 0 | 1 | 0 | 2 | 0 | 4 | 0.5 | 9 | 0.2222 |
| 5_5381261 | NA | NA | 3 | 0.3333 | 2 | 0.5 | 2 | 0 | 7 | 0.2857 | 2 | 0 | 2 | 0 | 2 | 0 | 5 | 0.2 | 11 | 0.0909 |
| 5_5381274 | NA | NA | 5 | 0.4 | 2 | 0 | 2 | 0 | 9 | 0.2222 | 2 | 0 | 1 | 0 | 2 | 0.5 | 5 | 0 | 10 | 0.1 |
| 5_5381321 | NA | NA | 7 | 0.2857 | NA | NA | 2 | 0 | 9 | 0.2222 | 2 | 0 | 1 | 0 | 2 | 0 | 6 | 0.1667 | 11 | 0.0909 |
| 5_5406079 | 3 | 0.3333 | 6 | 0.1667 | 8 | 0.375 | 2 | 0.5 | 19 | 0.3158 | 3 | 0.3333 | 4 | 0 | 4 | 0.75 | 6 | 0.3333 | 17 | 0.3529 |
| 5_5406080 | 4 | 0.75 | 6 | 0.5 | 6 | 0.8333 | 1 | 1 | 17 | 0.7059 | 4 | 0.5 | 3 | 0.6667 | 3 | 1 | 6 | 0.8333 | 16 | 0.75 |
| 5_5406083 | 4 | 0.25 | 6 | 0 | 8 | 0.125 | 2 | 0 | 20 | 0.1 | 4 | 0.25 | 3 | 0.3333 | 3 | 1 | 6 | 0.3333 | 16 | 0.4375 |
| 5_5406090 | 2 | 0.5 | 2 | 0.5 | 7 | 0 | 2 | 0 | 13 | 0.1538 | 2 | 0 | 3 | 0.3333 | 2 | 0 | 6 | 0 | 13 | 0.0769 |
| 5_5406100 | 4 | 0.25 | 5 | 0 | 6 | 0 | 2 | 0 | 17 | 0.0588 | 3 | 0 | 4 | 0.25 | 2 | 0 | 6 | 0 | 15 | 0.0667 |
| 5_5406104 | 4 | 0 | 6 | 0 | 7 | 0.2857 | 1 | 0 | 18 | 0.1111 | 2 | 0.5 | 4 | 0.5 | 2 | 0.5 | 5 | 0.6 | 13 | 0.5385 |
| 5_5427564 | 6 | 0 | 8 | 0.125 | 5 | 0 | 8 | 0.125 | 27 | 0.0741 | 3 | 0 | 6 | 0 | 7 | 0.1429 | 4 | 0.25 | 20 | 0.1 |
| 5_5427575 | 5 | 1 | 8 | 0.875 | 8 | 1 | 5 | 0.8 | 26 | 0.9231 | 3 | 1 | 6 | 1 | 9 | 0.8889 | 5 | 1 | 23 | 0.9565 |
| 5_5427609 | 7 | 0.2857 | 8 | 0.5 | 9 | 0.2222 | 7 | 0.7143 | 31 | 0.4194 | 4 | 0.75 | 6 | 0.3333 | 9 | 0.3333 | 6 | 0.6667 | 25 | 0.48 |
| 5_5427613 | 6 | 1 | 9 | 1 | 10 | 1 | 8 | 0.875 | 33 | 0.9697 | 3 | 1 | 9 | 1 | 7 | 1 | 6 | 1 | 25 | 1 |
| 5_5427618 | 4 | 0 | 8 | 0.25 | 9 | 0 | 9 | 0 | 30 | 0.0667 | 3 | 0 | 8 | 0.125 | 8 | 0.125 | 6 | 0 | 25 | 0.08 |
| 5_5427620 | 7 | 0.7143 | 7 | 0.7143 | 9 | 0.6667 | 8 | 1 | 31 | 0.7742 | 3 | 0.3333 | 7 | 0.7143 | 9 | 0.8889 | 5 | 0.8 | 24 | 0.75 |
| 5_5427626 | 6 | 0.1667 | 7 | 0.2857 | 9 | 0 | 8 | 0.25 | 30 | 0.1667 | 3 | 0 | 7 | 0.4286 | 6 | 0.1667 | 5 | 0.4 | 21 | 0.2857 |
| 5_5427626 | 7 | 0.7143 | 8 | 0.625 | 9 | 0.4444 | 8 | 0.625 | 32 | 0.5938 | 3 | 0.6667 | 7 | 0.4286 | 9 | 0.4444 | 7 | 0.4286 | 26 | 0.4615 |
| 5_5427640 | 9 | 1 | 5 | 1 | 7 | 0.8571 | 8 | 1 | 29 | 0.9655 | 4 | 1 | 6 | 1 | 8 | 1 | 6 | 1 | 24 | 1 |

| | | | | | | | | | | | | | | | | | | | | |
|---|---|---|---|---|---|---|---|---|---|---|---|---|---|---|---|---|---|---|---|---|
| 5_5427642 | 9 | 1 | 5 | 1 | 7 | 1 | 6 | 1 | 27 | 1 | 5 | 1 | 6 | 1 | 6 | 1 | 5 | 1 | 22 | 1 |
| 5_5427650 | 10 | 1 | 9 | 1 | 10 | 1 | 8 | 1 | 37 | 1 | 5 | 1 | 7 | 1 | 10 | 1 | 7 | 1 | 29 | 1 |
| 5_5427651 | 9 | 0.3333 | 9 | 0.3333 | 10 | 0.1 | 8 | 0.375 | 36 | 0.2778 | 5 | 0 | 5 | 0 | 7 | 0.2857 | 4 | 0.25 | 21 | 0.1429 |
| 5_5427653 | 9 | 0.6667 | 8 | 0.5 | 7 | 0.1429 | 7 | 0.5714 | 31 | 0.4839 | 4 | 0.5 | 7 | 0.4286 | 13 | 0.4615 | 7 | 0.4286 | 31 | 0.4516 |
| 5_5427655 | 10 | 0.4 | 8 | 0.5 | 10 | 0.4 | 7 | 0.7143 | 35 | 0.4857 | 4 | 0.75 | 8 | 0.625 | 8 | 0.75 | 7 | 0.4286 | 27 | 0.6296 |
| 5_5427658 | 9 | 0.3333 | 9 | 0.5556 | 9 | 0.5556 | 8 | 0.5 | 35 | 0.4857 | 5 | 0.4 | 9 | 0.6667 | 10 | 0.4 | 6 | 0.5 | 30 | 0.5 |
| 5_5427660 | 6 | 0.6667 | 7 | 0.4286 | 6 | 0.6667 | 8 | 0.625 | 27 | 0.5926 | 4 | 0.5 | 5 | 0.4 | 8 | 0.375 | 6 | 0.5 | 23 | 0.4348 |
| 5_5427660 | 7 | 1 | 6 | 1 | 5 | 1 | 7 | 0.8571 | 25 | 0.96 | 3 | 1 | 6 | 0.8333 | 9 | 1 | 5 | 1 | 23 | 0.9565 |
| 5_5427667 | 10 | 0.4 | 10 | 0.6 | 8 | 0.75 | 10 | 0.7 | 38 | 0.6053 | 5 | 0.6 | 7 | 0.5714 | 10 | 0.4 | 6 | 0.6667 | 28 | 0.5357 |
| 5_5427676 | 9 | 0 | 10 | 0.1 | 11 | 0.0909 | 10 | 0 | 40 | 0.05 | 6 | 0.3333 | 7 | 0 | 15 | 0.1333 | 9 | 0.3333 | 37 | 0.1892 |
| 5_5427690 | 10 | 1 | 8 | 1 | 11 | 1 | 10 | 1 | 39 | 1 | 6 | 1 | 8 | 1 | 11 | 1 | 11 | 0.8182 | 36 | 0.9444 |
| 5_5427692 | 6 | 0.3333 | 4 | 0.25 | 7 | 0 | 5 | 0.2 | 22 | 0.1818 | 7 | 0 | 7 | 0.1429 | 6 | 0.3333 | 8 | 0.5 | 28 | 0.25 |
| 5_5427696 | 10 | 0.4 | 7 | 0.4286 | 12 | 0.5 | 10 | 0.6 | 39 | 0.4872 | 6 | 0.5 | 9 | 0.6667 | 10 | 0.5 | 8 | 0.375 | 33 | 0.5152 |
| 5_5787419 | 2 | 0 | 1 | 0 | 1 | 1 | 1 | 0 | 5 | 0.2 | 2 | 1 | NA | NA | 1 | 1 | 1 | 0 | 4 | 0.75 |
| 5_5787477 | 2 | 0.5 | 1 | 1 | 1 | 1 | 1 | 1 | 5 | 0.8 | 2 | 0.5 | NA | NA | 1 | 0 | NA | NA | 3 | 0.3333 |
| 5_5787521 | 1 | 0 | 2 | 0.5 | 1 | 0 | NA | NA | 4 | 0.25 | 2 | 1 | NA | NA | 1 | 0 | NA | NA | 3 | 0.6667 |
| 5_5787601 | NA | NA | 2 | 1 | 1 | 1 | NA | NA | 3 | 1 | 2 | 1 | NA | NA | NA | NA | NA | NA | 2 | 1 |
| 5_5787614 | NA | NA | 1 | 1 | 1 | 1 | NA | NA | 2 | 1 | 1 | 1 | NA | NA | NA | NA | NA | NA | 1 | 1 |
| 5_5787746 | NA | NA | 1 | 1 | 2 | 1 | 2 | 0 | 5 | 0.6 | NA | NA | NA | NA | NA | NA | 1 | 1 | 1 | 1 |
| 5_5787763 | NA | NA | 1 | 0 | 3 | 1 | 1 | 1 | 5 | 0.8 | NA | NA | NA | NA | NA | NA | 1 | 1 | 1 | 1 |
| 5_5787817 | NA | NA | 1 | 1 | 4 | 1 | 1 | 1 | 6 | 1 | 1 | 1 | NA | NA | NA | NA | 1 | 1 | 2 | 1 |
| 5_5787818 | NA | NA | 1 | 1 | 4 | 0.75 | 2 | 0.5 | 7 | 0.7143 | 1 | 1 | NA | NA | NA | NA | 2 | 1 | 3 | 1 |
| 5_5787825 | NA | NA | 1 | 1 | 4 | 0.75 | 2 | 1 | 7 | 0.8571 | 1 | 0 | NA | NA | NA | NA | 1 | 1 | 2 | 0.5 |
| 5_5787848 | NA | NA | 1 | 1 | 4 | 0.25 | 3 | 0.3333 | 8 | 0.375 | 1 | 0 | NA | NA | NA | NA | 2 | 0.5 | 3 | 0.3333 |
| 5_5787852 | NA | NA | NA | NA | 4 | 0.5 | 3 | 0.6667 | 7 | 0.5714 | 1 | 1 | NA | NA | NA | NA | 1 | 0 | 2 | 0.5 |
| 5_5787856 | NA | NA | 1 | 1 | 2 | 0 | 3 | 0.6667 | 6 | 0.5 | 1 | 0 | NA | NA | NA | NA | 2 | 0.5 | 3 | 0.3333 |
| 5_5787875 | NA | NA | 1 | 0 | 5 | 0.2 | 3 | 0.3333 | 9 | 0.2222 | 1 | 1 | NA | NA | NA | NA | 2 | 0 | 3 | 0.3333 |
| 5_5787877 | NA | NA | 1 | 1 | 4 | 1 | 3 | 1 | 8 | 1 | 1 | 1 | NA | NA | NA | NA | 1 | 1 | 2 | 1 |
| 5_5787878 | NA | NA | 1 | 0 | 4 | 0.25 | 3 | 0 | 8 | 0.125 | 1 | 1 | NA | NA | NA | NA | 2 | 0 | 3 | 0.3333 |
| 5_5787899 | NA | NA | 1 | 1 | 4 | 1 | 3 | 0.6667 | 8 | 0.875 | 1 | 1 | NA | NA | NA | NA | 2 | 0.5 | 3 | 0.6667 |
| 5_5787907 | NA | NA | 1 | 1 | 5 | 0.8 | 3 | 0.3333 | 9 | 0.6667 | 1 | 0 | NA | NA | NA | NA | 2 | 0.5 | 3 | 0.3333 |
| 5_5787920 | NA | NA | 1 | 1 | 5 | 0.8 | 2 | 1 | 8 | 0.875 | 1 | 1 | NA | NA | NA | NA | 2 | 1 | 3 | 1 |
| 5_5787925 | NA | NA | 1 | 0 | 5 | 0.6 | 3 | 0.3333 | 9 | 0.4444 | 1 | 1 | NA | NA | NA | NA | 2 | 0.5 | 3 | 0.6667 |
| 5_5787932 | NA | NA | 1 | 1 | 4 | 0.75 | 2 | 0.5 | 7 | 0.7143 | NA | NA | NA | NA | NA | NA | 2 | 1 | 2 | 1 |
| 5_5828638 | NA | NA | 3 | 0.3333 | 6 | 0 | 3 | 0 | 12 | 0.0833 | 3 | 0.3333 | 2 | 0 | 2 | 0 | 2 | 0 | 9 | 0.1111 |
| 5_5828656 | NA | NA | 3 | 0 | 6 | 0.1667 | 3 | 0.6667 | 12 | 0.25 | 3 | 0 | 2 | 0 | 1 | 1 | 2 | 0 | 8 | 0.125 |
| 5_5828680 | 1 | 0 | 4 | 0.25 | 8 | 0 | 4 | 0 | 17 | 0.0588 | 3 | 0 | 2 | 0 | 2 | 0 | 2 | 0.5 | 9 | 0.1111 |
| 5_5828766 | 1 | 0 | 3 | 0.3333 | 11 | 0.0909 | 6 | 0 | 21 | 0.0952 | 3 | 0.3333 | 4 | 0 | 2 | 0 | 2 | 0 | 11 | 0.0909 |
| 5_5828860 | 1 | 0 | 6 | 0.5 | 4 | 0 | 4 | 0.5 | 15 | 0.3333 | NA | NA | 1 | 0 | NA | NA | 2 | 0.5 | 3 | 0.3333 |
| 5_5828973 | 4 | 0 | 8 | 0.25 | 11 | 0.1818 | 10 | 0 | 33 | 0.1212 | 2 | 0 | 8 | 0 | 4 | 0.25 | 6 | 0.1667 | 20 | 0.1 |
| 5_6275998 | 4 | 0.25 | 10 | 0.3 | 8 | 0 | 3 | 0.3333 | 25 | 0.2 | 5 | 0 | 5 | 0.2 | 4 | 0.25 | 7 | 0.2857 | 21 | 0.1905 |
| 5_6276012 | 4 | 0.25 | 9 | 0.3333 | 5 | 0.2 | 2 | 0 | 20 | 0.25 | 2 | 0 | 5 | 0 | 4 | 0.25 | 5 | 0.4 | 16 | 0.1875 |
| 5_6276086 | 3 | 0.3333 | 9 | 0.1111 | 5 | 0 | 3 | 0 | 20 | 0.1 | 4 | 0 | 3 | 0 | 3 | 0 | 5 | 0.2 | 15 | 0.0667 |
| 5_6276156 | 2 | 1 | 5 | 1 | 3 | 1 | 3 | 0.6667 | 13 | 0.9231 | 4 | 0.75 | 3 | 0.3333 | 2 | 1 | 4 | 1 | 13 | 0.7692 |
| 5_6276169 | 3 | 0.3333 | 4 | 0.25 | 4 | 0.25 | 1 | 0 | 12 | 0.25 | 3 | 0.3333 | 2 | 1 | 3 | 0.3333 | 4 | 0.25 | 12 | 0.4167 |
| 5_6276177 | 3 | 0 | 5 | 0.2 | 4 | 0.25 | 3 | 0 | 15 | 0.1333 | 4 | 0 | 3 | 0.3333 | 3 | 0 | 5 | 0 | 15 | 0.0667 |
| 5_6276187 | 3 | 0 | 4 | 0.25 | 4 | 0 | 3 | 0 | 14 | 0.0714 | 4 | 0 | 2 | 0.5 | 2 | 0.5 | 4 | 0.25 | 12 | 0.25 |
| 5_6276193 | 2 | 1 | 4 | 0.5 | 3 | 1 | 2 | 0.5 | 11 | 0.7273 | 4 | 0.75 | 2 | 1 | 2 | 0.5 | 4 | 0.25 | 12 | 0.5833 |
| 5_6276194 | 3 | 0 | 5 | 0 | 3 | 0.6667 | 3 | 0.3333 | 14 | 0.2143 | 4 | 0 | 2 | 0 | 2 | 0 | 5 | 0.2 | 13 | 0.0769 |
| 5_6288804 | 1 | 1 | 2 | 1 | 2 | 0.5 | 1 | 1 | 6 | 0.8333 | 1 | 1 | NA | NA | 1 | 1 | NA | NA | 2 | 1 |
| 5_6290625 | NA | NA | 1 | 1 | 2 | 1 | 1 | 1 | 4 | 1 | 1 | 1 | NA | NA | NA | NA | 1 | 1 | 2 | 1 |
| 5_6380545 | NA | NA | 1 | 1 | NA | NA | NA | NA | 1 | 1 | 1 | 0 | NA | NA | NA | NA | 1 | 1 | 2 | 0.5 |
| 5_6380588 | NA | NA | 1 | 1 | NA | NA | NA | NA | 1 | 1 | 1 | 0 | NA | NA | NA | NA | 1 | 1 | 2 | 0.5 |
| 5_6396387 | NA | NA | 1 | 1 | NA | NA | NA | NA | 1 | 1 | NA | NA | NA | NA | 1 | 1 | NA | NA | 1 | 1 |
| 5_6396396 | NA | NA | 1 | 1 | NA | NA | NA | NA | 1 | 1 | NA | NA | NA | NA | 1 | 1 | NA | NA | 1 | 1 |
| 5_6396522 | NA | NA | 1 | 1 | NA | NA | 1 | 1 | 2 | 1 | NA | NA | NA | NA | 1 | 1 | 1 | 1 | 2 | 1 |
| 5_6396529 | NA | NA | 1 | 1 | NA | NA | 1 | 0 | 2 | 0.5 | NA | NA | NA | NA | 1 | 1 | 1 | 1 | 2 | 1 |
| 5_6396572 | NA | NA | 1 | 1 | NA | NA | NA | NA | 1 | 1 | NA | NA | NA | NA | 1 | 1 | NA | NA | 1 | 1 |
| 5_6396578 | NA | NA | 1 | 1 | NA | NA | NA | NA | 1 | 1 | NA | NA | NA | NA | NA | NA | 1 | 1 | 1 | 1 |
| 5_6396580 | NA | NA | 1 | 1 | NA | NA | 1 | 1 | 2 | 1 | NA | NA | NA | NA | 1 | 1 | 1 | 1 | 2 | 1 |
| 5_6396599 | NA | NA | 1 | 1 | NA | NA | 1 | 1 | 2 | 1 | NA | NA | NA | NA | NA | NA | 1 | 1 | 1 | 1 |
| 5_6396604 | NA | NA | 1 | 1 | NA | NA | 1 | 1 | 2 | 1 | NA | NA | NA | NA | 1 | 1 | 1 | 1 | 2 | 1 |
| 5_6396619 | NA | NA | 1 | 1 | NA | NA | 1 | 0 | 2 | 0.5 | NA | NA | NA | NA | 1 | 0 | 1 | 1 | 2 | 0.5 |
| 5_6396623 | NA | NA | 1 | 1 | NA | NA | 1 | 0 | 2 | 0.5 | NA | NA | NA | NA | 1 | 1 | NA | NA | 1 | 1 |
| 5_6396634 | NA | NA | 1 | 1 | NA | NA | NA | NA | 1 | 1 | NA | NA | NA | NA | NA | NA | 1 | 1 | 1 | 1 |

| | | | | | | | | | | | | | | | | | | | | |
|---|---|---|---|---|---|---|---|---|---|---|---|---|---|---|---|---|---|---|---|---|
| 5_6396704 | NA | NA | 2 | 0.5 | NA | NA | 1 | 0 | 3 | 0.3333 | NA | NA | NA | NA | 1 | 0 | 1 | 1 | 2 | 0.5 |
| 5_6396715 | NA | NA | 2 | 0.5 | NA | NA | 1 | 1 | 3 | 0.6667 | NA | NA | NA | NA | NA | NA | 1 | 1 | 1 | 1 |
| 5_6396755 | NA | NA | 2 | 0.5 | NA | NA | 1 | 1 | 3 | 0.6667 | 1 | 1 | NA | NA | NA | NA | NA | NA | 1 | 1 |
| 5_6405551 | NA | NA | NA | NA | 1 | 1 | 1 | 0 | 2 | 0.5 | NA | NA | 2 | 0.5 | NA | NA | NA | NA | 2 | 0.5 |
| 5_6405565 | NA | NA | NA | NA | NA | NA | 1 | 1 | 1 | 1 | NA | NA | 1 | 1 | NA | NA | NA | NA | 1 | 1 |
| 5_6405575 | NA | NA | NA | NA | 1 | 1 | 1 | 1 | 2 | 1 | NA | NA | 2 | 0.5 | NA | NA | NA | NA | 2 | 0.5 |
| 5_6405598 | NA | NA | NA | NA | 1 | 1 | 1 | 1 | 2 | 1 | 1 | 1 | 2 | 1 | NA | NA | NA | NA | 3 | 1 |
| 5_6406783 | 1 | 1 | NA | NA | NA | NA | 1 | 1 | 2 | 1 | NA | NA | 2 | 1 | NA | NA | NA | NA | 2 | 1 |
| 5_6406785 | 1 | 1 | NA | NA | NA | NA | 1 | 1 | 2 | 1 | NA | NA | 2 | 1 | 1 | 0 | NA | NA | 3 | 0.6667 |
| 5_6406804 | 1 | 1 | NA | NA | NA | NA | 1 | 1 | 2 | 1 | NA | NA | 2 | 0 | 1 | 1 | NA | NA | 3 | 0.3333 |
| 5_6406808 | 1 | 1 | NA | NA | NA | NA | 1 | 1 | 2 | 1 | NA | NA | 2 | 1 | 1 | 0 | NA | NA | 3 | 0.6667 |
| 5_6406816 | 1 | 1 | NA | NA | NA | NA | 1 | 1 | 2 | 1 | NA | NA | 1 | 1 | 1 | 0 | NA | NA | 2 | 0.5 |
| 5_6406833 | NA | NA | NA | NA | NA | NA | 1 | 1 | 1 | 1 | NA | NA | 1 | 1 | 1 | 1 | NA | NA | 2 | 1 |
| 5_6441426 | NA | NA | 1 | 0 | NA | NA | 1 | 1 | 2 | 0.5 | 1 | 1 | NA | NA | NA | NA | NA | NA | 1 | 1 |
| 5_6450728 | NA | NA | 1 | 0 | NA | NA | 1 | 1 | 2 | 0.5 | 1 | 1 | NA | NA | 1 | 1 | NA | NA | 2 | 1 |
| 5_6450750 | NA | NA | 1 | 1 | NA | NA | 1 | 1 | 2 | 1 | 1 | 1 | NA | NA | 1 | 0 | NA | NA | 2 | 0.5 |
| 5_6464113 | 1 | 0 | 2 | 0.5 | NA | NA | 1 | 0 | 4 | 0.25 | 1 | 0 | 1 | 0 | 1 | 1 | 1 | 1 | 4 | 0.5 |
| 5_6464124 | NA | NA | 2 | 0.5 | NA | NA | NA | NA | 2 | 0.5 | NA | NA | NA | NA | 1 | 1 | 1 | 1 | 2 | 1 |
| 5_6464129 | NA | NA | 2 | 0.5 | NA | NA | NA | NA | 2 | 0.5 | NA | NA | NA | NA | NA | NA | 1 | 1 | 1 | 1 |
| 5_6464143 | NA | NA | 2 | 1 | NA | NA | NA | NA | 2 | 1 | NA | NA | NA | NA | 1 | 1 | 1 | 1 | 2 | 1 |
| 5_6464174 | NA | NA | 2 | 1 | NA | NA | NA | NA | 2 | 1 | NA | NA | NA | NA | 1 | 1 | 1 | 1 | 2 | 1 |
| 5_6464176 | NA | NA | 2 | 0.5 | NA | NA | NA | NA | 2 | 0.5 | NA | NA | NA | NA | 1 | 1 | 1 | 1 | 2 | 1 |
| 5_6464177 | NA | NA | 2 | 0.5 | NA | NA | NA | NA | 2 | 0.5 | NA | NA | NA | NA | 1 | 1 | 1 | 0 | 2 | 0.5 |
| 5_6464178 | NA | NA | 2 | 1 | NA | NA | NA | NA | 2 | 1 | NA | NA | NA | NA | 1 | 1 | 1 | 1 | 2 | 1 |
| 5_6464185 | NA | NA | 2 | 0.5 | NA | NA | NA | NA | 2 | 0.5 | NA | NA | NA | NA | 1 | 1 | 1 | 1 | 2 | 1 |
| 5_6464396 | 1 | 1 | 2 | 0.5 | NA | NA | NA | NA | 3 | 0.6667 | NA | NA | NA | NA | 1 | 1 | NA | NA | 1 | 1 |
| 5_6464490 | 1 | 1 | 1 | 1 | NA | NA | 1 | 0 | 3 | 0.6667 | 1 | 1 | NA | NA | 1 | 0 | NA | NA | 2 | 0.5 |
| 5_6464518 | 1 | 1 | 2 | 1 | NA | NA | 1 | 1 | 4 | 1 | 1 | 1 | NA | NA | 1 | 0 | NA | NA | 2 | 0.5 |
| 5_6464523 | 1 | 1 | 2 | 1 | NA | NA | 2 | 1 | 5 | 1 | 1 | 1 | 1 | 0 | NA | NA | NA | NA | 2 | 0.5 |
| 5_6464525 | 1 | 0 | 2 | 1 | NA | NA | 1 | 1 | 4 | 0.75 | 1 | 1 | 1 | 0 | NA | NA | NA | NA | 2 | 0.5 |
| 5_6464530 | 1 | 0 | 2 | 0.5 | NA | NA | 2 | 0 | 5 | 0.2 | 1 | 1 | 2 | 0 | 1 | 0 | NA | NA | 4 | 0.25 |
| 5_6464550 | 1 | 1 | 2 | 1 | NA | NA | 2 | 0.5 | 5 | 0.8 | 1 | 1 | 2 | 1 | NA | NA | NA | NA | 3 | 1 |
| 5_6464552 | 1 | 0 | 2 | 1 | NA | NA | 2 | 0 | 5 | 0.4 | 1 | 0 | 2 | 1 | 1 | 0 | NA | NA | 4 | 0.5 |
| 5_6464566 | 1 | 1 | 1 | 0 | NA | NA | 1 | 1 | 3 | 0.6667 | 1 | 0 | 2 | 0 | 1 | 1 | NA | NA | 4 | 0.25 |
| 5_6464578 | 1 | 0 | 2 | 0.5 | NA | NA | 2 | 0 | 5 | 0.2 | 1 | 0 | 2 | 0.5 | 1 | 0 | NA | NA | 4 | 0.25 |
| 5_6464579 | 2 | 1 | NA | NA | NA | NA | 2 | 1 | 4 | 1 | 1 | 0 | 1 | 1 | 1 | 1 | NA | NA | 3 | 0.6667 |
| 5_6464582 | 2 | 1 | 1 | 1 | NA | NA | 2 | 1 | 5 | 1 | 1 | 0 | 1 | 1 | 1 | 1 | NA | NA | 3 | 0.6667 |
| 5_6498638 | 1 | 1 | NA | NA | 2 | 0 | NA | NA | 3 | 0.3333 | 2 | 1 | NA | NA | 2 | 1 | NA | NA | 4 | 1 |
| 5_6498639 | 1 | 0 | NA | NA | 3 | 0.3333 | NA | NA | 4 | 0.25 | 2 | 1 | NA | NA | 2 | 0 | 1 | 0 | 5 | 0.4 |
| 5_6498643 | 3 | 0 | NA | NA | 3 | 0.3333 | NA | NA | 6 | 0.1667 | 2 | 0.5 | NA | NA | 3 | 0.6667 | 1 | 0 | 6 | 0.5 |
| 5_6498667 | 3 | 0.3333 | NA | NA | 3 | 0 | NA | NA | 6 | 0.1667 | 2 | 0.5 | 1 | 0 | 3 | 0.6667 | 1 | 0 | 7 | 0.4286 |
| 5_6498674 | 2 | 0.5 | NA | NA | 1 | 0 | NA | NA | 3 | 0.3333 | 2 | 1 | 1 | 0 | 3 | 0.6667 | 1 | 1 | 7 | 0.7143 |
| 5_6498681 | 3 | 0.3333 | NA | NA | 3 | 0 | NA | NA | 6 | 0.1667 | 2 | 0.5 | NA | NA | 2 | 0 | NA | NA | 4 | 0.25 |
| 5_6498692 | 2 | 0.5 | NA | NA | 2 | 0.5 | NA | NA | 4 | 0.5 | 2 | 0 | NA | NA | 2 | 1 | NA | NA | 4 | 0.5 |
| 5_6498749 | 2 | 0.5 | NA | NA | 1 | 0 | 1 | 1 | 4 | 0.5 | 2 | 1 | NA | NA | 1 | 1 | NA | NA | 3 | 1 |
| 5_6498939 | 4 | 0.5 | 1 | 0 | 1 | 0 | 4 | 0.25 | 10 | 0.3 | 2 | 0 | 2 | 0 | 3 | 0 | 2 | 0.5 | 9 | 0.1111 |
| 5_6498945 | 4 | 0 | 1 | 0 | NA | NA | 3 | 0.3333 | 8 | 0.125 | 2 | 0 | 1 | 0 | 1 | 0 | 2 | 0.5 | 6 | 0.1667 |
| 5_6498954 | 4 | 0 | 2 | 0 | 2 | 0 | 4 | 0.25 | 12 | 0.0833 | 2 | 0 | 2 | 1 | 3 | 1 | 1 | 0 | 8 | 0.625 |
| 5_6508226 | NA | NA | NA | NA | NA | NA | 1 | 1 | 1 | 1 | NA | NA | NA | NA | NA | NA | 1 | 1 | 1 | 1 |
| 5_6528886 | NA | NA | NA | NA | NA | NA | 2 | 0.5 | 2 | 0.5 | NA | NA | 2 | 0.5 | 1 | 1 | 1 | 0 | 4 | 0.5 |
| 5_6531467 | 1 | 1 | NA | NA | NA | NA | NA | NA | 1 | 1 | NA | NA | 1 | 0 | 2 | 0.5 | NA | NA | 3 | 0.3333 |
| 5_6532021 | 2 | 0.5 | 1 | 0 | 3 | 0 | 8 | 0 | 14 | 0.0714 | 2 | 0 | 5 | 0 | 9 | 0.2222 | 8 | 0.25 | 24 | 0.1667 |
| 5_6532033 | 3 | 0 | 2 | 0 | 4 | 0 | 9 | 0.1111 | 18 | 0.0556 | 4 | 0.25 | 6 | 0 | 9 | 0.1111 | 9 | 0.1111 | 28 | 0.1071 |
| 5_6532435 | 3 | 0 | 2 | 0 | 4 | 0 | 9 | 0.2222 | 18 | 0.1111 | 4 | 0 | 4 | 0 | 10 | 0.2 | 9 | 0 | 27 | 0.0741 |
| 5_6532440 | 3 | 0.3333 | 1 | 1 | 2 | 0 | 7 | 0.4286 | 13 | 0.3846 | 2 | 0.5 | 2 | 0.5 | 8 | 0.5 | 7 | 0.7143 | 19 | 0.5789 |
| 5_6532462 | 4 | 0 | 2 | 0.5 | 3 | 0.6667 | 9 | 0.1111 | 18 | 0.2222 | 4 | 0 | 6 | 0.5 | 8 | 0.5 | 9 | 0.4444 | 27 | 0.4074 |
| 5_6532474 | 4 | 0.25 | 1 | 0 | 3 | 0 | 9 | 0.2222 | 17 | 0.1765 | 4 | 0.5 | 7 | 0.4286 | 9 | 0.2222 | 10 | 0.6 | 30 | 0.4333 |
| 5_6532486 | 4 | 0 | 2 | 0.5 | 2 | 0 | 7 | 0.2857 | 15 | 0.2 | 4 | 0.25 | 5 | 0 | 9 | 0.3333 | 10 | 0.4 | 28 | 0.2857 |
| 5_6541562 | NA | NA | 2 | 0 | 1 | 1 | 2 | 1 | 5 | 0.6 | 1 | 1 | NA | NA | 1 | 1 | 1 | 1 | 3 | 1 |
| 5_6541714 | 1 | 0 | 2 | 1 | 1 | 0 | 1 | 0 | 5 | 0.4 | 2 | 0.5 | NA | NA | 1 | 0 | 1 | 0 | 4 | 0.25 |
| 5_6541947 | 2 | 0.5 | 1 | 0 | 3 | 0 | 2 | 0 | 8 | 0.125 | 1 | 0 | 2 | 0 | 1 | 0 | 1 | 1 | 5 | 0.2 |
| 5_6542014 | 3 | 0.3333 | 3 | 0 | 4 | 0 | 1 | 0 | 11 | 0.0909 | 1 | 0 | 4 | 0.25 | 1 | 0 | 1 | 0 | 7 | 0.1429 |
| 5_6542038 | 3 | 0 | 3 | 0 | 4 | 0 | 1 | 1 | 11 | 0.0909 | 1 | 0 | 3 | 0.3333 | 1 | 0 | 1 | 0 | 6 | 0.1667 |
| 5_6542353 | 2 | 0 | 3 | 0.3333 | 4 | 0.5 | 1 | 0 | 10 | 0.3 | 1 | 0 | 3 | 0 | 1 | 1 | 1 | 0 | 6 | 0.1667 |
| 5_6554583 | 2 | 0 | NA | NA | 2 | 0.5 | 2 | 0.5 | 6 | 0.3333 | 1 | 0 | 2 | 0 | 2 | 0.5 | 1 | 0 | 6 | 0.1667 |

| | | | | | | | | | | | | | | | | | | | | |
|---|---|---|---|---|---|---|---|---|---|---|---|---|---|---|---|---|---|---|---|---|
| 5_6556638 | 7 | 0.8571 | 5 | 1 | 4 | 1 | 3 | 1 | 19 | 0.9474 | 6 | 1 | 3 | 1 | 14 | 1 | 2 | 1 | 25 | 1 |
| 5_6556648 | 7 | 0.7143 | 5 | 0.8 | 2 | 0.5 | 2 | 1 | 16 | 0.75 | 4 | 0.75 | 3 | 0.3333 | 12 | 0.6667 | 2 | 1 | 21 | 0.6667 |
| 5_6556662 | NA | NA | 2 | 0.5 | 1 | 1 | NA | NA | 3 | 0.6667 | 1 | 1 | NA | NA | NA | NA | NA | NA | 1 | 1 |
| 5_6556726 | NA | NA | 2 | 1 | 1 | 1 | NA | NA | 3 | 1 | 1 | 1 | NA | NA | NA | NA | 1 | 0 | 2 | 0.5 |
| 5_6556746 | NA | NA | 2 | 0.5 | NA | NA | NA | NA | 2 | 0.5 | 1 | 1 | NA | NA | NA | NA | 1 | 0 | 2 | 0.5 |
| 5_6557092 | 5 | 0.2 | 6 | 0.5 | 4 | 0 | 1 | 1 | 16 | 0.3125 | 2 | 0 | 1 | 0 | 5 | 0 | 4 | 0.5 | 12 | 0.1667 |
| 5_6557122 | 3 | 0 | 5 | 0.2 | 4 | 0 | 1 | 0 | 13 | 0.0769 | NA | NA | 1 | 0 | 8 | 0.125 | 5 | 0.2 | 14 | 0.1429 |
| 5_6560425 | 7 | 0.4286 | 13 | 0.2308 | 4 | 0.25 | 10 | 0.2 | 34 | 0.2647 | 8 | 0.375 | 4 | 0 | 6 | 0.1667 | 2 | 0.5 | 20 | 0.25 |
| 5_6560443 | 6 | 0.1667 | 11 | 0.0909 | 5 | 0.2 | 9 | 0.1111 | 31 | 0.129 | 7 | 0.1429 | 3 | 0 | 7 | 0 | 3 | 0.3333 | 20 | 0.1 |
| 5_6560458 | 6 | 0.5 | 9 | 0.2222 | 5 | 0.2 | 8 | 0.625 | 28 | 0.3929 | 7 | 0.1429 | 4 | 0 | 6 | 0.3333 | 2 | 0.5 | 19 | 0.2105 |
| 5_6560653 | 2 | 0.5 | 5 | 0 | 5 | 0 | 4 | 0 | 16 | 0.0625 | 3 | 0 | 2 | 0 | 3 | 0.3333 | 2 | 0.5 | 10 | 0.2 |
| 5_6560655 | 3 | 0.6667 | 6 | 0.8333 | 4 | 0.25 | 5 | 0 | 18 | 0.4444 | 3 | 0.3333 | 3 | 0.6667 | 4 | 0.25 | 1 | 1 | 11 | 0.4545 |
| 5_6560667 | 3 | 0.3333 | 6 | 0.5 | 3 | 0.6667 | 4 | 0 | 16 | 0.375 | 2 | 0 | 3 | 0 | 4 | 0.5 | 1 | 0 | 10 | 0.2 |
| 5_6560671 | 3 | 0.6667 | 6 | 0.1667 | 4 | 0.25 | 5 | 0 | 18 | 0.2222 | 2 | 0 | 3 | 0.3333 | 2 | 0.5 | NA | NA | 7 | 0.2857 |
| 5_6560723 | 32 | 0.1562 | 33 | 0.1212 | 29 | 0.0345 | 35 | 0.1429 | 129 | 0.1163 | 24 | 0.0833 | 19 | 0.0526 | 29 | 0.1724 | 21 | 0.1429 | 93 | 0.1183 |
| 5_6560729 | 33 | 0.0606 | 23 | 0.1304 | 20 | 0.2 | 28 | 0.0357 | 104 | 0.0962 | 22 | 0.0455 | 21 | 0.0476 | 21 | 0.0476 | 19 | 0.1053 | 83 | 0.0602 |
| 5_6560767 | 28 | 0.25 | 27 | 0.2593 | 21 | 0.2381 | 25 | 0.28 | 101 | 0.2574 | 16 | 0.1875 | 14 | 0.0714 | 26 | 0.0769 | 16 | 0.0625 | 72 | 0.0972 |
| 5_6575226 | 19 | 0.1053 | 15 | 0.3333 | 17 | 0.1765 | 18 | 0.3333 | 69 | 0.2319 | 24 | 0.375 | 16 | 0.3125 | 15 | 0.2667 | 10 | 0.3 | 65 | 0.3231 |
| 5_6575242 | 18 | 0.1667 | 17 | 0.1176 | 16 | 0.3125 | 19 | 0.3684 | 70 | 0.2429 | 24 | 0.25 | 14 | 0.2857 | 17 | 0.2353 | 11 | 0.2727 | 66 | 0.2576 |
| 5_6575307 | 7 | 0.1429 | 4 | 0.25 | 6 | 0 | 7 | 0.1429 | 24 | 0.125 | 11 | 0.0909 | 4 | 0.5 | 6 | 0 | 2 | 0 | 23 | 0.1304 |
| 5_6576291 | 1 | 1 | 1 | 0 | NA | NA | 3 | 0.3333 | 5 | 0.4 | NA | NA | 2 | 0.5 | 2 | 0 | 1 | 0 | 5 | 0.2 |
| 5_6576660 | 2 | 0 | 1 | 0 | 6 | 0.1667 | 3 | 0 | 12 | 0.0833 | 3 | 0 | 4 | 0 | 4 | 0.25 | 4 | 0.25 | 15 | 0.1333 |
| 5_6576663 | 1 | 1 | 3 | 0 | 4 | 0 | 3 | 0.6667 | 11 | 0.2727 | 3 | 0.3333 | 4 | 0.5 | 5 | 0 | 2 | 0.5 | 14 | 0.2857 |
| 5_6576710 | 2 | 0.5 | 3 | 0.3333 | 4 | 0.5 | 4 | 0.75 | 13 | 0.5385 | 3 | 0.3333 | 2 | 1 | 5 | 0.6 | 1 | 1 | 11 | 0.6364 |
| 5_6576716 | 2 | 1 | 2 | 0.5 | 5 | 1 | 2 | 1 | 11 | 0.9091 | 3 | 0.6667 | 3 | 0.6667 | 5 | 1 | 2 | 1 | 13 | 0.8462 |
| 5_6576724 | 2 | 1 | 2 | 0 | 6 | 0.3333 | 4 | 0.75 | 14 | 0.5 | 1 | 0 | 4 | 0.5 | 5 | 0.6 | 3 | 0.6667 | 13 | 0.5385 |
| 5_6576731 | 2 | 1 | 3 | 0 | 6 | 0.1667 | 4 | 0.5 | 15 | 0.3333 | 2 | 0 | 4 | 0 | 5 | 0.4 | 2 | 0.5 | 13 | 0.2308 |
| 5_6576735 | 2 | 1 | 2 | 0.5 | 5 | 0.6 | 3 | 1 | 12 | 0.75 | 1 | 1 | 3 | 0.6667 | 4 | 1 | 2 | 0.5 | 10 | 0.8 |
| 5_6576741 | 2 | 0.5 | 2 | 0 | 4 | 0.5 | 3 | 0.3333 | 11 | 0.3636 | 2 | 0 | 4 | 0.75 | 3 | 0.6667 | 2 | 0.5 | 11 | 0.5455 |
| 5_6592052 | 1 | 0 | 2 | 1 | NA | NA | NA | NA | 3 | 0.6667 | 1 | 1 | NA | NA | NA | NA | NA | NA | 1 | 1 |
| 5_6596503 | NA | NA | NA | NA | NA | NA | 1 | 1 | 1 | 1 | NA | NA | 1 | 1 | NA | NA | NA | NA | 1 | 1 |
| 5_6596550 | NA | NA | NA | NA | NA | NA | 1 | 1 | 1 | 1 | NA | NA | 1 | 1 | NA | NA | NA | NA | 1 | 1 |
| 5_6598726 | NA | NA | 2 | 0.5 | 1 | 0 | 3 | 0.3333 | 6 | 0.3333 | 1 | 0 | 2 | 0 | 1 | 1 | 3 | 0 | 7 | 0.1429 |
| 5_6641844 | NA | NA | NA | NA | 1 | 1 | NA | NA | 1 | 1 | NA | NA | 1 | 1 | NA | NA | NA | NA | 1 | 1 |
| 5_6642780 | NA | NA | 1 | 1 | NA | NA | NA | NA | 1 | 1 | NA | NA | NA | NA | 1 | 1 | NA | NA | 1 | 1 |
| 5_6642838 | NA | NA | 1 | 1 | NA | NA | NA | NA | 1 | 1 | NA | NA | NA | NA | 1 | 1 | NA | NA | 1 | 1 |
| 5_6642919 | NA | NA | 1 | 1 | NA | NA | NA | NA | 1 | 1 | NA | NA | NA | NA | 2 | 1 | NA | NA | 2 | 1 |
| 5_6642956 | NA | NA | 1 | 1 | NA | NA | NA | NA | 1 | 1 | NA | NA | NA | NA | 2 | 1 | NA | NA | 2 | 1 |
| 5_6642969 | NA | NA | 1 | 1 | NA | NA | NA | NA | 1 | 1 | NA | NA | NA | NA | 2 | 0.5 | NA | NA | 2 | 0.5 |
| 5_6643004 | NA | NA | 1 | 1 | NA | NA | NA | NA | 1 | 1 | NA | NA | NA | NA | 2 | 1 | NA | NA | 2 | 1 |
| 5_6643032 | NA | NA | 1 | 1 | NA | NA | NA | NA | 1 | 1 | NA | NA | NA | NA | 2 | 1 | NA | NA | 2 | 1 |
| 5_6645300 | NA | NA | 3 | 0.6667 | 1 | 1 | NA | NA | 4 | 0.75 | 2 | 1 | NA | NA | 2 | 1 | NA | NA | 4 | 1 |
| 5_6645310 | NA | NA | 3 | 1 | 1 | 1 | NA | NA | 4 | 1 | 2 | 0.5 | NA | NA | 2 | 1 | NA | NA | 4 | 0.75 |
| 5_6645315 | NA | NA | 3 | 0.3333 | 2 | 0.5 | NA | NA | 5 | 0.4 | 2 | 0 | NA | NA | 2 | 0.5 | NA | NA | 4 | 0.25 |
| 5_6645336 | NA | NA | 2 | 0.5 | 1 | 1 | NA | NA | 3 | 0.6667 | 1 | 1 | NA | NA | 2 | 0.5 | NA | NA | 3 | 0.6667 |
| 5_6645398 | NA | NA | 3 | 0.3333 | 2 | 0.5 | NA | NA | 5 | 0.4 | 2 | 1 | NA | NA | 2 | 0 | NA | NA | 4 | 0.5 |
| 5_6645421 | NA | NA | 2 | 0.5 | 1 | 1 | NA | NA | 3 | 0.6667 | 1 | 1 | NA | NA | 2 | 1 | NA | NA | 3 | 1 |
| 5_6645526 | NA | NA | 2 | 0.5 | 1 | 1 | NA | NA | 3 | 0.6667 | NA | NA | NA | NA | 2 | 0.5 | 1 | 0 | 3 | 0.3333 |
| 5_6645623 | 3 | 0.3333 | 13 | 0.0769 | 7 | 0 | 1 | 0 | 24 | 0.0833 | 5 | 0 | 1 | 0 | 2 | 0 | 7 | 0.1429 | 15 | 0.0667 |
| 5_6645643 | 3 | 0.6667 | 13 | 0.4615 | 6 | 0.6667 | 1 | 0 | 23 | 0.5217 | 4 | 0 | 1 | 1 | 2 | 0.5 | 10 | 0.4 | 17 | 0.3529 |
| 5_6645657 | 3 | 0.3333 | 6 | 0.1667 | 7 | 0.2857 | 1 | 0 | 17 | 0.2353 | 4 | 0.25 | 1 | 1 | 4 | 0.25 | 8 | 0.125 | 17 | 0.2353 |
| 5_6645665 | 3 | 0 | 12 | 0.1667 | 6 | 0.3333 | 1 | 0 | 22 | 0.1818 | 5 | 0 | 1 | 0 | 3 | 0 | 10 | 0.1 | 19 | 0.0526 |
| 5_6645675 | 3 | 0.6667 | 10 | 0.3 | 3 | 0.6667 | 1 | 0 | 17 | 0.4118 | 3 | 0 | 1 | 0 | 2 | 0 | 7 | 0.2857 | 13 | 0.1538 |
| 5_6645683 | 4 | 0.5 | 11 | 0.2727 | 4 | 0.25 | NA | NA | 19 | 0.3158 | 3 | 0.3333 | 2 | 0.5 | 2 | 0 | 9 | 0.1111 | 16 | 0.1875 |
| 5_6645783 | 4 | 0.25 | 8 | 0.5 | 5 | 0.4 | NA | NA | 17 | 0.4118 | 2 | 0 | 1 | 1 | 2 | 0.5 | 5 | 0.8 | 10 | 0.6 |
| 5_6645787 | 4 | 0 | 9 | 0.1111 | 7 | 0.2857 | NA | NA | 20 | 0.15 | 3 | 0 | 1 | 1 | 3 | 1 | 7 | 0 | 14 | 0.2857 |
| 5_6645809 | 4 | 0 | 8 | 0.5 | 6 | 0.3333 | NA | NA | 18 | 0.3333 | 2 | 0.5 | 1 | 0 | 3 | 0 | 7 | 0.4286 | 13 | 0.3077 |
| 5_6645815 | 4 | 0 | 9 | 0.3333 | 6 | 0.5 | NA | NA | 19 | 0.3158 | 2 | 1 | 1 | 1 | 3 | 0.6667 | 6 | 0.6667 | 12 | 0.75 |
| 5_6645826 | 4 | 0.25 | 10 | 0.6 | 5 | 0.8 | NA | NA | 19 | 0.5789 | 1 | 0 | 1 | 0 | 3 | 0.6667 | 5 | 0.8 | 10 | 0.6 |
| 5_6645858 | 3 | 0 | 6 | 0 | 4 | 0.25 | NA | NA | 13 | 0.0769 | 2 | 0.5 | 1 | 0 | 2 | 0 | 5 | 0.2 | 10 | 0.2 |
| 5_6645912 | 1 | 0 | 7 | 0.2857 | 4 | 0 | NA | NA | 12 | 0.1667 | NA | NA | 1 | 0 | 3 | 0.3333 | 5 | 0.2 | 9 | 0.2222 |
| 5_6645918 | 2 | 1 | 8 | 0.875 | 4 | 1 | NA | NA | 14 | 0.9286 | 2 | 0.5 | 1 | 1 | 2 | 1 | 3 | 0.6667 | 8 | 0.75 |
| 5_6645927 | 3 | 1 | 11 | 0.9091 | 5 | 1 | NA | NA | 19 | 0.9474 | 2 | 1 | 4 | 0.75 | 4 | 0.75 | 6 | 1 | 16 | 0.875 |
| 5_6645932 | 2 | 1 | 10 | 0.9 | 6 | 1 | NA | NA | 18 | 0.9444 | 3 | 1 | 3 | 1 | 3 | 1 | 6 | 1 | 15 | 1 |
| 5_6645937 | 1 | 1 | 7 | 0.1429 | 4 | 0.5 | NA | NA | 12 | 0.3333 | 1 | 0 | 3 | 0 | 3 | 0.3333 | 5 | 0.2 | 12 | 0.1667 |

| | | | | | | | | | | | | | | | | | | | | |
|---|---|---|---|---|---|---|---|---|---|---|---|---|---|---|---|---|---|---|---|---|
| 5_66459400 | 2 | 0 | 7 | 0.1429 | 5 | 0.2 | NA | NA | 14 | 0.1429 | 2 | 0 | 2 | 0 | 1 | 1 | 7 | 0 | 12 | 0.0833 |
| 5_66459437 | 1 | 1 | 7 | 0.4286 | 4 | 0.75 | NA | NA | 12 | 0.5833 | 2 | 0.5 | 2 | 1 | 3 | 0.6667 | 6 | 0.1667 | 13 | 0.4615 |
| 5_66459525 | 1 | 1 | 8 | 0.5 | 4 | 0.75 | NA | NA | 13 | 0.6154 | 1 | 0 | 3 | 0.3333 | 3 | 0.6667 | 6 | 0.5 | 13 | 0.4615 |
| 5_66459628 | 2 | 1 | 11 | 0.9091 | 4 | 1 | NA | NA | 17 | 0.9412 | 3 | 0.6667 | 3 | 1 | 3 | 0.6667 | 6 | 1 | 15 | 0.8667 |
| 5_66459633 | 1 | 0 | 10 | 0.4 | 3 | 0.6667 | NA | NA | 14 | 0.4286 | 1 | 1 | 3 | 0.3333 | 2 | 0.5 | 1 | 0 | 7 | 0.4286 |
| 5_66459759 | 2 | 0.5 | 9 | 0.7778 | 6 | 0.6667 | NA | NA | 17 | 0.7059 | 2 | 1 | 2 | 0.5 | 3 | 1 | 7 | 0.5714 | 14 | 0.7143 |
| 5_66459893 | 2 | 1 | 10 | 0.8 | 4 | 0.75 | NA | NA | 16 | 0.8125 | 3 | 0.6667 | 3 | 0.6667 | 2 | 0 | 6 | 0.8333 | 14 | 0.6429 |
| 5_66459925 | 2 | 1 | 10 | 0.5 | 6 | 1 | NA | NA | 18 | 0.7222 | 2 | 0.5 | 2 | 0.5 | 3 | 1 | 6 | 0.8333 | 13 | 0.7692 |
| 5_66609032 | 2 | 0 | 4 | 0.5 | 1 | 0 | NA | NA | 7 | 0.2857 | NA | NA | 1 | 1 | NA | NA | 3 | 0.3333 | 4 | 0.5 |
| 5_66609284 | 2 | 0.5 | 2 | 0 | NA | NA | NA | NA | 4 | 0.25 | NA | NA | 1 | 0 | NA | NA | 3 | 0.3333 | 4 | 0.25 |
| 5_66609429 | 1 | 1 | 4 | 0 | NA | NA | NA | NA | 5 | 0.2 | NA | NA | 1 | 1 | NA | NA | 3 | 0.3333 | 4 | 0.5 |
| 5_66609505 | 1 | 1 | 2 | 0.5 | NA | NA | NA | NA | 3 | 0.6667 | NA | NA | 1 | 1 | NA | NA | 2 | 1 | 3 | 1 |
| 5_67199897 | 1 | 1 | NA | NA | NA | NA | NA | NA | 1 | 1 | NA | NA | 1 | 1 | 2 | 0 | NA | NA | 3 | 0.3333 |
| 5_67200321 | 1 | 1 | NA | NA | NA | NA | NA | NA | 1 | 1 | NA | NA | 1 | 1 | 2 | 1 | NA | NA | 3 | 1 |
| 5_67201169 | 1 | 1 | NA | NA | NA | NA | NA | NA | 1 | 1 | NA | NA | 1 | 1 | NA | NA | NA | NA | 1 | 1 |
| 5_67212523 | 2 | 0 | 2 | 0.5 | 13 | 0.3077 | 3 | 0.6667 | 20 | 0.35 | 5 | 0 | 6 | 0.1667 | 7 | 0.4286 | 2 | 0.5 | 20 | 0.25 |
| 5_67212630 | 4 | 0 | 4 | 0.25 | 13 | 0.1538 | 5 | 0 | 26 | 0.1154 | 4 | 0.25 | 7 | 0.1429 | 6 | 0.1667 | 3 | 0 | 20 | 0.15 |
| 5_67212710 | 3 | 1 | 6 | 1 | 14 | 0.9286 | 4 | 1 | 27 | 0.963 | 6 | 1 | 6 | 1 | 6 | 0.6667 | 2 | 1 | 20 | 0.9 |
| 5_67212748 | 4 | 0.25 | 5 | 0.4 | 14 | 0.2143 | 6 | 0.5 | 29 | 0.3103 | 5 | 0.2 | 8 | 0.5 | 7 | 0.5714 | 3 | 0.3333 | 23 | 0.4348 |
| 5_67212803 | 4 | 0.75 | 6 | 0.3333 | 15 | 0.3333 | 7 | 0.4286 | 32 | 0.4062 | 6 | 0.3333 | 4 | 0.5 | 7 | 0.2857 | 3 | 0.3333 | 20 | 0.35 |
| 5_67212941 | 4 | 1 | 6 | 0.8333 | 11 | 0.7273 | 6 | 0.6667 | 27 | 0.7778 | 7 | 0.7143 | 5 | 0.8 | 4 | 1 | 3 | 0 | 19 | 0.6842 |
| 5_67212956 | 4 | 0.75 | 5 | 0.8 | 14 | 0.7857 | 6 | 1 | 29 | 0.8276 | 6 | 0.6667 | 5 | 0.4 | 6 | 0.5 | 2 | 0.5 | 19 | 0.5263 |
| 5_67213016 | 4 | 1 | 4 | 0.5 | 12 | 0.8333 | 6 | 1 | 26 | 0.8462 | 5 | 0.8 | 7 | 1 | 5 | 0.8 | 1 | 1 | 18 | 0.8889 |
| 5_67213739 | 3 | 0 | 5 | 0 | 14 | 0.1429 | 4 | 0 | 26 | 0.0769 | 8 | 0.125 | 10 | 0 | 9 | 0.4444 | 4 | 0 | 31 | 0.1613 |
| 5_67213978 | 5 | 0.2 | 6 | 0.1667 | 14 | 0.2143 | 4 | 0 | 29 | 0.1724 | 8 | 0.375 | 10 | 0.4 | 10 | 0.3 | 3 | 0 | 31 | 0.3226 |
| 5_67214142 | 5 | 0.4 | 7 | 0 | 14 | 0.0714 | 5 | 0 | 31 | 0.0968 | 8 | 0 | 9 | 0.1111 | 8 | 0.125 | 3 | 0 | 28 | 0.0714 |
| 5_67214166 | 6 | 0 | 8 | 0 | 12 | 0.1667 | 6 | 0.3333 | 32 | 0.125 | 8 | 0.125 | 10 | 0.2 | 6 | 0.1667 | 6 | 0.3333 | 30 | 0.2 |
| 5_67511991 | 13 | 0.0769 | 24 | 0.0833 | 24 | 0.0833 | 11 | 0.0909 | 72 | 0.0833 | 11 | 0.0909 | 21 | 0.1429 | 18 | 0 | 12 | 0.1667 | 62 | 0.0968 |
| 5_67512034 | 11 | 0.0909 | 24 | 0.1667 | 23 | 0.1304 | 13 | 0 | 71 | 0.1127 | 10 | 0.2 | 22 | 0.1364 | 14 | 0.1429 | 13 | 0.0769 | 59 | 0.1356 |
| 5_67512051 | 13 | 0.7692 | 27 | 0.8519 | 25 | 0.8 | 15 | 0.9333 | 80 | 0.8375 | 10 | 1 | 22 | 0.8636 | 18 | 0.9444 | 13 | 0.7692 | 63 | 0.8889 |
| 5_67512057 | 9 | 0 | 25 | 0.08 | 25 | 0.04 | 13 | 0.1538 | 72 | 0.0694 | 10 | 0 | 23 | 0.087 | 19 | 0.0526 | 14 | 0.1429 | 66 | 0.0758 |
| 5_67512095 | 13 | 0.3077 | 21 | 0.4286 | 20 | 0.55 | 15 | 0.3333 | 69 | 0.4203 | 10 | 0.4 | 19 | 0.3158 | 14 | 0.5714 | 12 | 0.5833 | 55 | 0.4545 |
| 5_67512198 | 13 | 0.8462 | 26 | 0.7308 | 24 | 0.625 | 14 | 0.5 | 77 | 0.6753 | 12 | 0.75 | 23 | 0.6087 | 20 | 0.65 | 14 | 1 | 69 | 0.7246 |
| 5_67512363 | 10 | 0.3 | 21 | 0.0952 | 19 | 0.1053 | 7 | 0.1429 | 57 | 0.1404 | 4 | 0 | 16 | 0.0625 | 13 | 0.0769 | 7 | 0.1429 | 40 | 0.075 |
| 5_67512650 | 16 | 0.25 | 29 | 0.2069 | 28 | 0.1429 | 17 | 0.1176 | 90 | 0.1778 | 16 | 0 | 23 | 0.1739 | 23 | 0.2174 | 14 | 0.1429 | 76 | 0.1447 |
| 5_67512733 | 10 | 0.1 | 20 | 0.15 | 22 | 0.3182 | 17 | 0 | 69 | 0.1594 | 17 | 0.1765 | 21 | 0.0952 | 20 | 0.2 | 13 | 0.2308 | 71 | 0.169 |
| 5_67512910 | 11 | 0.3636 | 27 | 0.5926 | 26 | 0.3846 | 13 | 0.7692 | 77 | 0.5195 | 14 | 0.5714 | 17 | 0.3529 | 17 | 0.3529 | 14 | 0.2857 | 62 | 0.3871 |
| 5_67512932 | 17 | 0.5294 | 30 | 0.4667 | 30 | 0.7333 | 19 | 0.4737 | 96 | 0.5625 | 14 | 0.5714 | 25 | 0.36 | 25 | 0.4 | 14 | 0.5714 | 78 | 0.4487 |
| 5_67513158 | 2 | 0 | 2 | 0 | 2 | 0 | 2 | 0.5 | 8 | 0.125 | 3 | 0 | NA | NA | 1 | 0 | 5 | 0.4 | 9 | 0.2222 |
| 5_67514968 | 3 | 0 | 2 | 0 | 2 | 0.5 | 2 | 0 | 9 | 0.1111 | 1 | 1 | 1 | 0 | NA | NA | 5 | 0 | 7 | 0.1429 |
| 5_67515624 | 4 | 0 | 4 | 0.25 | 2 | 0 | 3 | 0.6667 | 13 | 0.2308 | NA | NA | 1 | 1 | NA | NA | 5 | 0.2 | 6 | 0.3333 |
| 5_67775775 | 12 | 0.4167 | 11 | 0.2727 | 12 | 0.0833 | 12 | 0.0833 | 47 | 0.2128 | 7 | 0.1429 | 6 | 0.5 | 9 | 0.1111 | 19 | 0.2632 | 41 | 0.2439 |
| 5_67776060 | 11 | 0.0909 | 10 | 0.1 | 9 | 0.1111 | 10 | 0.2 | 40 | 0.125 | 5 | 0.2 | 5 | 0.4 | 9 | 0 | 15 | 0 | 34 | 0.0882 |
| 5_67777087 | 24 | 0.3333 | 18 | 0.3333 | 16 | 0.1875 | 19 | 0.3158 | 77 | 0.2987 | 15 | 0.4 | 10 | 0.4 | 18 | 0.2778 | 22 | 0.3182 | 65 | 0.3385 |
| 5_67777589 | 21 | 0 | 19 | 0.1579 | 15 | 0.1333 | 20 | 0.05 | 75 | 0.08 | 12 | 0.0833 | 9 | 0.1111 | 16 | 0.125 | 18 | 0.1667 | 55 | 0.1273 |
| 5_67777736 | 19 | 0.1053 | 17 | 0.2353 | 8 | 0 | 18 | 0.2222 | 62 | 0.1613 | 11 | 0.0909 | 7 | 0 | 15 | 0.1333 | 17 | 0 | 50 | 0.06 |
| 5_67777785 | 18 | 0.2778 | 15 | 0.2 | 11 | 0.1818 | 16 | 0 | 60 | 0.1667 | 9 | 0.1111 | 7 | 0 | 15 | 0.2 | 15 | 0.0667 | 46 | 0.1087 |
| 5_67777949 | 15 | 0.2 | 12 | 0.3333 | 12 | 0.1667 | 14 | 0.3571 | 53 | 0.2642 | 8 | 0.125 | 7 | 0.1429 | 14 | 0.2857 | 9 | 0.1111 | 38 | 0.1842 |
| 5_67778358 | 16 | 0.125 | 16 | 0.125 | 9 | 0 | 17 | 0.1176 | 58 | 0.1034 | 8 | 0 | 6 | 0 | 13 | 0.2308 | 15 | 0.0667 | 42 | 0.0952 |
| 5_67779534 | 12 | 0.5 | 14 | 0.7857 | 7 | 0.8571 | 10 | 0.6 | 43 | 0.6744 | 6 | 0.6667 | 2 | 1 | 10 | 0.5 | 11 | 0.7273 | 29 | 0.6552 |
| 5_67779549 | 10 | 1 | 13 | 0.8462 | 4 | 0.75 | 11 | 0.9091 | 38 | 0.8947 | 6 | 0.8333 | 3 | 1 | 14 | 0.7857 | 8 | 0.875 | 31 | 0.8387 |
| 5_67779628 | 9 | 0.1111 | 9 | 0 | 3 | 0 | 7 | 0.1429 | 28 | 0.0714 | 2 | 0 | 3 | 0 | 14 | 0 | 10 | 0.3 | 29 | 0.1034 |
| 5_67779660 | 7 | 0.5714 | 12 | 0.75 | 3 | 0.6667 | 7 | 0.4286 | 29 | 0.6207 | 5 | 0.6 | 1 | 1 | 9 | 0.4444 | 9 | 0.6667 | 24 | 0.5833 |
| 5_67779770 | 11 | 0.6364 | 12 | 0.5833 | 6 | 0.1667 | 11 | 0.4545 | 40 | 0.5 | 3 | 1 | 3 | 0.6667 | 12 | 0.5 | 14 | 0.7143 | 32 | 0.6562 |
| 5_67779791 | 11 | 0.7273 | 14 | 1 | 6 | 1 | 10 | 0.9 | 41 | 0.9024 | 4 | 1 | 3 | 1 | 14 | 0.7857 | 13 | 0.9231 | 34 | 0.8824 |
| 5_69267380 | 6 | 0.1667 | 4 | 0 | 1 | 1 | 5 | 0 | 16 | 0.125 | 2 | 0.5 | 1 | 1 | 6 | 0.5 | 3 | 0 | 12 | 0.4167 |
| 5_69267395 | 6 | 0.1667 | 5 | 0.2 | NA | NA | 2 | 0 | 13 | 0.1538 | 1 | 0 | 2 | 0 | 6 | 0.1667 | 2 | 0 | 11 | 0.0909 |
| 5_69267558 | 5 | 1 | 4 | 0.75 | NA | NA | 5 | 0.8 | 14 | 0.8571 | 2 | 1 | 2 | 0 | 6 | 1 | 2 | 0.5 | 12 | 0.75 |
| 5_69267572 | 5 | 0.6 | 2 | 0 | NA | NA | 4 | 0.25 | 11 | 0.3636 | 2 | 0.5 | NA | NA | 6 | 0.1667 | 2 | 0.5 | 10 | 0.3 |
| 5_69267647 | 6 | 0.5 | 3 | 0.3333 | NA | NA | 5 | 0.6 | 14 | 0.5 | 2 | 0.5 | 1 | 1 | 6 | 0.6667 | 3 | 0 | 12 | 0.5 |
| 5_69267702 | 5 | 1 | 2 | 1 | NA | NA | 3 | 1 | 10 | 1 | NA | NA | 1 | 1 | 5 | 0.8 | 2 | 1 | 8 | 0.875 |
| 5_69267867 | 3 | 1 | 1 | 1 | NA | NA | 2 | 1 | 6 | 1 | 1 | 1 | 2 | 1 | 3 | 1 | 3 | 1 | 9 | 1 |
| 5_69267927 | 4 | 0 | 4 | 0 | NA | NA | 5 | 0.2 | 13 | 0.0769 | 2 | 0 | 2 | 0 | 4 | 0 | 3 | 0.3333 | 11 | 0.0909 |
| 5_69268052 | 5 | 1 | 4 | 1 | NA | NA | 5 | 1 | 14 | 1 | 2 | 0.5 | 2 | 1 | 2 | 1 | 3 | 0.6667 | 9 | 0.7778 |
| 5_69497887 | 4 | 0.75 | 3 | 1 | NA | NA | 3 | 0.6667 | 10 | 0.8 | 1 | 1 | 1 | 1 | 2 | 1 | 2 | 0.5 | 6 | 0.8333 |

| | | | | | | | | | | | | | | | | | | | | |
|---|---|---|---|---|---|---|---|---|---|---|---|---|---|---|---|---|---|---|---|---|
| 5_69498028 | 2 | 0.5 | 3 | 0.6667 | NA | NA | 2 | 0 | 7 | 0.4286 | 1 | 1 | 1 | 1 | 3 | 0.3333 | 2 | 0.5 | 7 | 0.5714 |
| 5_69498145 | 3 | 0.6667 | 4 | 0.75 | NA | NA | 3 | 0 | 10 | 0.5 | 1 | 0 | 1 | 0 | 3 | 0.3333 | 3 | 0 | 8 | 0.125 |
| 5_69498631 | 2 | 0 | 3 | 0.3333 | NA | NA | 2 | 1 | 7 | 0.4286 | NA | NA | 1 | 1 | 1 | 0 | 2 | 0 | 4 | 0.25 |
| 5_69729064 | 4 | 0.5 | 2 | 0.5 | 3 | 0.3333 | 1 | 1 | 10 | 0.5 | NA | NA | 3 | 0.3333 | 3 | 0.6667 | 4 | 0.25 | 10 | 0.4 |
| 5_69733090 | 2 | 0 | 2 | 0 | 1 | 0 | 8 | 0.125 | 13 | 0.0769 | 4 | 0 | 4 | 0.25 | 5 | 0 | 2 | 0 | 15 | 0.0667 |
| 5_70909410 | 4 | 0.25 | 8 | 0.125 | 1 | 0 | 8 | 0.125 | 21 | 0.1429 | 4 | 0 | 9 | 0.1111 | 6 | 0.1667 | 6 | 0.1667 | 25 | 0.12 |
| 5_70910114 | 5 | 0 | 5 | 0.2 | 1 | 0 | 7 | 0.1429 | 18 | 0.1111 | 4 | 0.25 | 6 | 0.1667 | 6 | 0 | 6 | 0 | 22 | 0.0909 |
| 5_70910156 | 4 | 0 | 7 | 0 | 1 | 0 | 8 | 0.25 | 20 | 0.1 | 4 | 0.25 | 8 | 0 | 5 | 0 | 5 | 0.2 | 22 | 0.0909 |
| 5_70910181 | 5 | 0.4 | 6 | 0.5 | 1 | 0 | 8 | 0.25 | 20 | 0.35 | 3 | 0 | 8 | 0.125 | 5 | 0.2 | 5 | 0.4 | 21 | 0.1905 |
| 5_70911444 | 4 | 0.25 | 2 | 0.5 | 3 | 0 | 6 | 0.1667 | 15 | 0.2 | 2 | 0 | 6 | 0 | 3 | 0.3333 | 7 | 0.5714 | 18 | 0.2778 |
| 5_70911501 | 4 | 0 | 3 | 0 | 3 | 0.3333 | 6 | 0 | 16 | 0.0625 | 1 | 0 | 5 | 0.2 | 4 | 0 | 6 | 0 | 16 | 0.0625 |
| 5_70911531 | 4 | 0.5 | 2 | 1 | 3 | 0 | 6 | 0.8333 | 15 | 0.6 | 2 | 1 | 5 | 1 | 4 | 1 | 7 | 1 | 18 | 1 |
| 5_70999580 | 4 | 0.25 | 3 | 0 | 2 | 0.5 | 4 | 0 | 13 | 0.1538 | 1 | 0 | 4 | 0.25 | 2 | 0.5 | 4 | 0.5 | 11 | 0.3636 |
| 5_70999600 | 4 | 0.25 | 2 | 0 | NA | NA | 4 | 0.5 | 10 | 0.3 | 1 | 1 | 3 | 0 | 2 | 0 | 2 | 0 | 8 | 0.125 |
| 5_71115062 | 1 | 1 | 2 | 0 | NA | NA | NA | NA | 3 | 0.3333 | 1 | 1 | 1 | 1 | NA | NA | 1 | 1 | 3 | 1 |
| 5_71116556 | 6 | 0.1667 | 5 | 0 | 2 | 0 | 2 | 0 | 15 | 0.0667 | 1 | 0 | 3 | 0 | 3 | 0 | 6 | 0.1667 | 13 | 0.0769 |
| 5_71117754 | 6 | 0 | 8 | 0.25 | 1 | 0 | 3 | 0.3333 | 18 | 0.1667 | 2 | 1 | 4 | 0.25 | 1 | 0 | 8 | 0.125 | 15 | 0.2667 |
| 5_71117770 | 7 | 0.5714 | 8 | 0.125 | 3 | 0 | 4 | 0.5 | 22 | 0.3182 | 2 | 0.5 | 4 | 0.75 | NA | NA | 7 | 0 | 13 | 0.3077 |
| 5_71118092 | 8 | 0.25 | 8 | 0.25 | 1 | 1 | 4 | 0.5 | 21 | 0.3333 | 2 | 0.5 | 4 | 0.25 | 1 | 1 | 7 | 0.5714 | 14 | 0.5 |
| 5_71118119 | 7 | 0.8571 | 7 | 0.8571 | 2 | 1 | 4 | 1 | 20 | 0.9 | 2 | 1 | 5 | 1 | 1 | 1 | 6 | 0.3333 | 14 | 0.7143 |
| 5_71118443 | 7 | 0 | 5 | 0 | 3 | 0 | 3 | 0.3333 | 18 | 0.0556 | NA | NA | 5 | 0.2 | 1 | 1 | 4 | 0 | 10 | 0.2 |
| 5_71118551 | 6 | 0 | 6 | 0.1667 | 3 | 0 | 3 | 0 | 18 | 0.0556 | 1 | 1 | 4 | 0 | NA | NA | 4 | 0 | 9 | 0.1111 |
| 5_71119138 | 4 | 0 | 5 | 0.4 | 3 | 0 | 4 | 0.5 | 16 | 0.25 | 1 | 1 | 4 | 0 | NA | NA | 3 | 0.6667 | 8 | 0.375 |
| 5_71130886 | 4 | 0.5 | 4 | 0.75 | 3 | 0.3333 | 3 | 1 | 14 | 0.6429 | 1 | 1 | 4 | 0.75 | NA | NA | 3 | 0.6667 | 8 | 0.75 |
| 5_71171868 | 1 | 0 | 2 | 0 | 1 | 1 | 2 | 0 | 6 | 0.1667 | 1 | 0 | 2 | 0 | NA | NA | 3 | 0.3333 | 6 | 0.1667 |
| 5_71723519 | NA | NA | 2 | 0 | 1 | 1 | 1 | 0 | 4 | 0.25 | NA | NA | 1 | 0 | 1 | 1 | 1 | 0 | 3 | 0.3333 |
| 5_71725177 | NA | NA | 2 | 0.5 | 1 | 1 | 1 | 0 | 4 | 0.5 | NA | NA | NA | NA | 1 | 1 | NA | NA | 1 | 1 |
| 5_71725645 | 1 | 1 | 2 | 0 | 1 | 0 | 1 | 0 | 5 | 0.2 | NA | NA | NA | NA | 1 | 1 | 1 | 1 | 2 | 1 |
| 5_71725861 | 1 | 0 | 1 | 1 | 1 | 1 | 1 | 0 | 4 | 0.5 | NA | NA | NA | NA | 1 | 1 | 1 | 1 | 2 | 1 |
| 5_71726741 | 1 | 1 | 1 | 0 | 1 | 0 | 1 | 1 | 4 | 0.5 | NA | NA | NA | NA | 1 | 0 | 1 | 1 | 2 | 0.5 |
| 5_71726778 | 1 | 1 | 1 | 0 | 1 | 0 | 1 | 0 | 4 | 0.25 | NA | NA | NA | NA | 1 | 0 | 1 | 1 | 2 | 0.5 |
| 5_71727360 | 21 | 0.4762 | 13 | 0.8462 | 15 | 0.4 | 21 | 0.4762 | 70 | 0.5286 | 6 | 0.8333 | 24 | 0.8333 | 19 | 0.3684 | 16 | 0.6875 | 65 | 0.6615 |
| 5_71728518 | 16 | 0.4375 | 11 | 0.7273 | 11 | 0.3636 | 22 | 0.5909 | 60 | 0.5333 | 2 | 1 | 18 | 0.4444 | 16 | 0.5625 | 18 | 0.3889 | 54 | 0.4815 |
| 5_71728632 | 17 | 0.2353 | 11 | 0.4545 | 10 | 0.1 | 20 | 0.15 | 58 | 0.2241 | 3 | 0.3333 | 17 | 0.3529 | 15 | 0.4 | 12 | 0.3333 | 47 | 0.3617 |
| 5_71728710 | 18 | 0.3333 | 12 | 0.75 | 10 | 0.3 | 23 | 0.5217 | 63 | 0.4762 | 3 | 0.6667 | 18 | 0.2778 | 13 | 0.6923 | 16 | 0.625 | 50 | 0.52 |
| 5_71728999 | 13 | 0.7692 | 12 | 1 | 10 | 0.9 | 19 | 0.8947 | 54 | 0.8889 | 1 | 1 | 16 | 0.9375 | 12 | 0.8333 | 15 | 0.8667 | 44 | 0.8864 |
| 5_71729073 | 19 | 1 | 12 | 1 | 10 | 0.8 | 20 | 1 | 61 | 0.9672 | 2 | 1 | 17 | 0.9412 | 14 | 0.9286 | 15 | 0.9333 | 48 | 0.9375 |
| 5_71729151 | 17 | 1 | 11 | 0.8182 | 8 | 1 | 20 | 0.95 | 56 | 0.9464 | 2 | 1 | 16 | 0.9375 | 11 | 0.8182 | 13 | 0.9231 | 42 | 0.9048 |
| 5_71729245 | 13 | 0.6923 | 10 | 0.5 | 8 | 0.75 | 17 | 0.7647 | 48 | 0.6875 | 2 | 1 | 16 | 0.6875 | 9 | 0.6667 | 12 | 0.8333 | 39 | 0.7436 |
| 5_71729261 | 19 | 0.7368 | 11 | 0.7273 | 8 | 0.625 | 19 | 0.8421 | 57 | 0.7544 | 2 | 1 | 14 | 0.6429 | 13 | 0.7692 | 13 | 1 | 42 | 0.8095 |
| 5_71729374 | 18 | 0.8889 | 11 | 0.7273 | 7 | 0.8571 | 21 | 0.9048 | 57 | 0.8596 | 2 | 1 | 14 | 0.9286 | 10 | 0.8 | 14 | 0.9286 | 40 | 0.9 |
| 5_71729491 | 11 | 1 | 7 | 1 | 7 | 1 | 17 | 1 | 42 | 1 | 1 | 1 | 13 | 0.8462 | 8 | 0.875 | 10 | 0.9 | 32 | 0.875 |
| 5_71814561 | 9 | 0.1111 | 9 | 0.1111 | 5 | 0 | 13 | 0 | 36 | 0.0556 | 2 | 0 | 6 | 0 | 3 | 0 | 6 | 0.1667 | 17 | 0.0588 |
| 5_72293983 | 1 | 1 | 1 | 1 | 2 | 0.5 | NA | NA | 4 | 0.75 | NA | NA | NA | NA | NA | NA | 2 | 1 | 2 | 1 |
| 5_72294057 | 1 | 1 | 1 | 0 | 2 | 0.5 | 1 | 1 | 5 | 0.6 | NA | NA | NA | NA | NA | NA | 3 | 1 | 3 | 1 |
| 5_72294067 | 1 | 1 | 1 | 0 | 1 | 0 | NA | NA | 3 | 0.3333 | NA | NA | NA | NA | NA | NA | 3 | 0.3333 | 3 | 0.3333 |
| 5_72294516 | 1 | 0 | 1 | 0 | 2 | 1 | NA | NA | 4 | 0.5 | NA | NA | NA | NA | NA | NA | 3 | 0.6667 | 3 | 0.6667 |
| 5_72295317 | 3 | 0.3333 | 1 | 1 | 2 | 1 | NA | NA | 6 | 0.6667 | NA | NA | NA | NA | 1 | 1 | 4 | 0.5 | 5 | 0.6 |
| 5_72295481 | 3 | 0 | 1 | 1 | 2 | 0 | NA | NA | 6 | 0.1667 | NA | NA | NA | NA | 1 | 0 | 4 | 0.25 | 5 | 0.2 |
| 5_72296273 | 2 | 0.5 | 1 | 0 | NA | NA | NA | NA | 3 | 0.3333 | 1 | 1 | NA | NA | NA | NA | 1 | 1 | 2 | 1 |
| 5_72296489 | 2 | 0.5 | 2 | 0 | 2 | 0 | NA | NA | 6 | 0.1667 | 1 | 0 | NA | NA | 1 | 0 | 2 | 0.5 | 4 | 0.25 |
| 5_72297152 | 1 | 0 | 2 | 0.5 | 2 | 0 | NA | NA | 5 | 0.2 | NA | NA | 1 | 0 | 2 | 0.5 | 1 | 1 | 4 | 0.5 |
| 5_72297520 | 3 | 0.3333 | 4 | 0.25 | 2 | 0 | 1 | 0 | 10 | 0.2 | NA | NA | 2 | 0.5 | 3 | 0.3333 | 3 | 0 | 8 | 0.25 |
| 5_72297702 | 2 | 0 | 3 | 0.3333 | NA | NA | 3 | 0.6667 | 8 | 0.375 | NA | NA | 2 | 0.5 | 2 | 0 | 3 | 0 | 7 | 0.1429 |
| 5_72298257 | 1 | 1 | 1 | 0 | NA | NA | 1 | 0 | 3 | 0.3333 | 1 | 0 | 1 | 0 | 3 | 0.3333 | 1 | 0 | 6 | 0.1667 |
| 5_72305177 | 5 | 0.2 | 4 | 0.25 | 9 | 0 | 9 | 0 | 27 | 0.0741 | 1 | 0 | 5 | 0 | 6 | 0.1667 | 6 | 0 | 18 | 0.0556 |
| 5_72305214 | 5 | 0.2 | 5 | 0 | 10 | 0 | 9 | 0.1111 | 29 | 0.069 | 1 | 0 | 5 | 0 | 7 | 0 | 6 | 0.1667 | 19 | 0.0526 |
| 5_72305615 | 3 | 0.3333 | 5 | 0.8 | 10 | 0.6 | 9 | 0.6667 | 27 | 0.6296 | 1 | 0 | 5 | 0.4 | 5 | 0.6 | 6 | 1 | 17 | 0.6471 |
| 5_72305845 | 6 | 0 | 3 | 0.3333 | 11 | 0 | 8 | 0.375 | 28 | 0.1429 | 1 | 0 | 5 | 0.2 | 6 | 0.1667 | 7 | 0.2857 | 19 | 0.2105 |
| 5_72305880 | 5 | 0.2 | 4 | 0 | 11 | 0.0909 | 8 | 0.375 | 28 | 0.1786 | 1 | 1 | 6 | 0 | 7 | 0.1429 | 4 | 0 | 18 | 0.1111 |
| 5_72306080 | 6 | 0.5 | 6 | 0.3333 | 11 | 0.3636 | 7 | 0.4286 | 30 | 0.4 | 1 | 1 | 6 | 0.8333 | 8 | 0.5 | 8 | 0.5 | 23 | 0.6087 |
| 5_72306112 | 4 | 0.5 | 4 | 0.5 | 9 | 0.5556 | 7 | 0.7143 | 24 | 0.5833 | NA | NA | 5 | 0.6 | 6 | 0.5 | 5 | 0.8 | 16 | 0.625 |
| 5_72316615 | 9 | 0.2222 | 9 | 0.1111 | 6 | 0 | 9 | 0.1111 | 33 | 0.1212 | 8 | 0.125 | 7 | 0.2857 | 12 | 0.1667 | 5 | 0 | 32 | 0.1562 |
| 5_72316873 | 9 | 0.7778 | 6 | 0.6667 | 4 | 0.75 | 11 | 0.8182 | 30 | 0.7667 | 3 | 1 | 8 | 0.875 | 14 | 1 | 9 | 1 | 34 | 0.9706 |
| 5_72316951 | 10 | 0.5 | 8 | 0.25 | 6 | 0.5 | 10 | 0.5 | 34 | 0.4412 | 6 | 0.5 | 9 | 0.4444 | 14 | 0.2857 | 9 | 0.2222 | 38 | 0.3421 |

| | | | | | | | | | | | | | | | | | | | | |
|---|---|---|---|---|---|---|---|---|---|---|---|---|---|---|---|---|---|---|---|---|
| 5_72317049 | 8 | 0.125 | 6 | 0.3333 | 6 | 0.3333 | 8 | 0.5 | 28 | 0.3214 | 6 | 0 | 11 | 0.0909 | 11 | 0.0909 | 9 | 0.1111 | 37 | 0.0811 |
| 5_72317064 | 10 | 0.4 | 7 | 0.2857 | 6 | 0.1667 | 8 | 0.5 | 31 | 0.3548 | 6 | 0.3333 | 10 | 0.2 | 11 | 0.7273 | 10 | 0.6 | 37 | 0.4865 |
| 5_72454524 | NA | NA | NA | NA | 1 | 1 | NA | NA | 1 | 1 | NA | NA | NA | NA | 2 | 0.5 | NA | NA | 2 | 0.5 |
| 5_72454730 | NA | NA | NA | NA | 1 | 1 | NA | NA | 1 | 1 | NA | NA | NA | NA | 2 | 0.5 | NA | NA | 2 | 0.5 |
| 5_72913367 | NA | NA | NA | NA | NA | NA | 1 | 1 | 1 | 1 | NA | NA | NA | NA | NA | NA | 1 | 1 | 1 | 1 |
| 5_72913558 | NA | NA | NA | NA | NA | NA | 1 | 1 | 1 | 1 | NA | NA | NA | NA | NA | NA | 1 | 1 | 1 | 1 |
| 5_73132033 | 1 | 1 | 5 | 0 | 2 | 0 | 2 | 0 | 10 | 0.1 | 4 | 0.25 | NA | NA | 2 | 0.5 | 2 | 0.5 | 8 | 0.375 |
| 5_73132095 | 1 | 0 | 5 | 0.2 | 4 | 0 | 2 | 0 | 12 | 0.0833 | 4 | 0 | NA | NA | 2 | 1 | 2 | 0 | 8 | 0.25 |
| 5_73133446 | 1 | 0 | 5 | 1 | 3 | 0 | 2 | 0 | 11 | 0.4545 | 4 | 0.25 | 1 | 1 | 2 | 0.5 | 1 | 0 | 8 | 0.375 |
| 5_73133459 | 1 | 0 | 4 | 0.25 | 3 | 0.6667 | 2 | 0 | 10 | 0.3 | 3 | 0.3333 | 1 | 0 | 2 | 0 | 2 | 0.5 | 8 | 0.25 |
| 5_73133483 | 1 | 1 | 5 | 0.4 | 3 | 0.6667 | 2 | 0 | 11 | 0.4545 | 3 | 0.6667 | 1 | 1 | 1 | 1 | 2 | 1 | 7 | 0.8571 |
| 5_73133497 | 1 | 1 | 4 | 1 | 4 | 1 | 2 | 0.5 | 11 | 0.9091 | 3 | 0.6667 | 1 | 1 | 2 | 1 | 3 | 1 | 9 | 0.8889 |
| 5_73133840 | 1 | 1 | 4 | 0 | 4 | 0 | 2 | 0 | 11 | 0.0909 | 4 | 0 | 2 | 0 | 2 | 0.5 | 2 | 0 | 10 | 0.1 |
| 5_73133923 | 1 | 0 | 4 | 0 | 3 | 1 | 2 | 0 | 10 | 0.3 | 4 | 0.25 | 2 | 0.5 | 2 | 0.5 | 3 | 0.6667 | 11 | 0.4545 |
| 5_73134078 | 1 | 0 | 4 | 0 | 4 | 0.25 | 3 | 0 | 12 | 0.0833 | 4 | 0.5 | 2 | 1 | 2 | 0.5 | 3 | 0 | 11 | 0.4545 |
| 5_73134834 | 1 | 0 | 4 | 0 | 5 | 0 | 4 | 0.25 | 14 | 0.0714 | 4 | 0 | 3 | 0 | 2 | 0 | 3 | 0.3333 | 12 | 0.0833 |
| 5_73134919 | NA | NA | 4 | 0 | 4 | 0.25 | 4 | 0 | 12 | 0.0833 | 4 | 0 | 3 | 0.3333 | NA | NA | 2 | 0 | 9 | 0.1111 |
| 5_73135188 | 3 | 0 | 3 | 0.3333 | 4 | 0.25 | 4 | 0.25 | 14 | 0.2143 | 4 | 0 | 3 | 0 | 2 | 0.5 | 3 | 0.3333 | 12 | 0.1667 |
| 5_73164702 | 2 | 0.5 | NA | NA | NA | NA | NA | NA | 2 | 0.5 | NA | NA | 4 | 0.75 | 5 | 0.8 | 3 | 0.3333 | 12 | 0.6667 |
| 5_73169226 | 1 | 0 | NA | NA | NA | NA | 2 | 1 | 3 | 0.6667 | NA | NA | 5 | 0.2 | 6 | 0.3333 | 4 | 0.5 | 15 | 0.3333 |
| 5_73169243 | 2 | 0.5 | NA | NA | NA | NA | 2 | 0.5 | 4 | 0.5 | NA | NA | 6 | 0.5 | 8 | 0.875 | 6 | 0.8333 | 20 | 0.75 |
| 5_73169249 | 1 | 0 | NA | NA | NA | NA | 2 | 0.5 | 3 | 0.3333 | NA | NA | 5 | 0.4 | 4 | 0.5 | 5 | 0.4 | 14 | 0.4286 |
| 5_73169367 | 2 | 1 | NA | NA | NA | NA | 2 | 1 | 4 | 1 | NA | NA | 6 | 1 | 8 | 1 | 5 | 1 | 19 | 1 |
| 5_73169423 | 1 | 1 | NA | NA | NA | NA | 1 | 0 | 2 | 0.5 | NA | NA | 3 | 0.6667 | 6 | 0.6667 | 6 | 0.6667 | 15 | 0.6667 |
| 5_73169600 | 2 | 1 | NA | NA | NA | NA | 2 | 1 | 4 | 1 | NA | NA | 4 | 0.75 | 6 | 0.6667 | 5 | 0.8 | 15 | 0.7333 |
| 5_73169734 | 2 | 1 | NA | NA | NA | NA | 2 | 0.5 | 4 | 0.75 | NA | NA | 4 | 1 | 9 | 0.5556 | 5 | 0.8 | 18 | 0.7222 |
| 5_73170045 | 3 | 0.3333 | NA | NA | NA | NA | 3 | 0.6667 | 6 | 0.5 | NA | NA | 4 | 0.75 | 8 | 0.625 | 6 | 0.3333 | 18 | 0.5556 |
| 5_73170193 | 3 | 0.3333 | NA | NA | 1 | 0 | 3 | 0.3333 | 7 | 0.2857 | NA | NA | 6 | 0.1667 | 9 | 0 | 5 | 0 | 20 | 0.05 |
| 5_73171483 | 1 | 1 | NA | NA | NA | NA | NA | NA | 1 | 1 | NA | NA | 1 | 1 | 1 | 0 | NA | NA | 2 | 0.5 |
| 5_73177641 | 1 | 0 | 4 | 0.25 | 2 | 0 | 2 | 0 | 9 | 0.1111 | 1 | 0 | 2 | 0.5 | 2 | 0 | NA | NA | 5 | 0.2 |
| 5_73177780 | 2 | 0.5 | 12 | 0.75 | 5 | 0.4 | 2 | 0.5 | 21 | 0.619 | 3 | 1 | 6 | 0.3333 | 3 | 0.3333 | 5 | 0.8 | 17 | 0.5882 |
| 5_73177927 | 3 | 0.6667 | 12 | 0.4167 | 6 | 0.5 | 3 | 0.3333 | 24 | 0.4583 | 3 | 0.3333 | 6 | 0.6667 | 7 | 0.4286 | 4 | 1 | 20 | 0.6 |
| 5_73178127 | 2 | 0.5 | 10 | 0.1 | 6 | 0.1667 | 3 | 0.3333 | 21 | 0.1905 | 2 | 0 | 5 | 0 | 6 | 0 | 6 | 0.3333 | 19 | 0.1053 |
| 5_73178145 | 2 | 1 | 9 | 1 | 5 | 1 | 2 | 1 | 18 | 1 | 3 | 1 | 5 | 1 | 7 | 0.8571 | 4 | 1 | 19 | 0.9474 |
| 5_73178171 | 2 | 0.5 | 10 | 0.9 | 6 | 0.8333 | 2 | 1 | 20 | 0.85 | 2 | 1 | 5 | 1 | 6 | 0.8333 | 6 | 1 | 19 | 0.9474 |
| 5_73178190 | 2 | 1 | 9 | 0.7778 | 6 | 0.8333 | 3 | 1 | 20 | 0.85 | 3 | 0.6667 | 5 | 0.8 | 6 | 0.6667 | 6 | 1 | 20 | 0.8 |
| 5_73178215 | 2 | 0.5 | 12 | 0.6667 | 5 | 0.2 | 3 | 0.3333 | 22 | 0.5 | 3 | 0.3333 | 3 | 0.6667 | 7 | 0.4286 | 6 | 0.6667 | 19 | 0.5263 |
| 5_73178288 | 2 | 0 | 8 | 0.125 | 5 | 0.2 | 3 | 0 | 18 | 0.1111 | 3 | 0 | 4 | 0 | 6 | 0 | 6 | 0.1667 | 19 | 0.0526 |
| 5_73178341 | 2 | 0 | 9 | 0 | 5 | 0.2 | 2 | 0 | 18 | 0.0556 | 3 | 0 | 3 | 0 | 5 | 0 | 5 | 0.2 | 16 | 0.0625 |
| 5_73178440 | 2 | 0.5 | 11 | 0.8182 | 5 | 0.8 | 3 | 0.6667 | 21 | 0.7619 | 3 | 0.3333 | 4 | 0.5 | 6 | 0.6667 | 5 | 0.8 | 18 | 0.6111 |
| 5_73178512 | 2 | 0 | 11 | 0.7273 | 5 | 0.8 | 4 | 0.75 | 22 | 0.6818 | 2 | 1 | 4 | 0.75 | 4 | 0.75 | 4 | 0.75 | 14 | 0.7857 |
| 5_73178880 | 2 | 1 | 8 | 0.75 | 1 | 1 | 3 | 0.3333 | 14 | 0.7143 | 2 | 0.5 | 3 | 1 | 2 | 0.5 | 2 | 0.5 | 9 | 0.6667 |
| 5_73178881 | NA | NA | NA | NA | NA | NA | 1 | 1 | 1 | 1 | NA | NA | 1 | 0 | NA | NA | 1 | 1 | 2 | 0.5 |
| 5_73178882 | NA | NA | NA | NA | 1 | 1 | NA | NA | 1 | 1 | NA | NA | NA | NA | 1 | 1 | NA | NA | 1 | 1 |
| 5_73178894 | 1 | 1 | 6 | 0.8333 | 2 | 1 | 3 | 0.6667 | 12 | 0.8333 | 1 | 1 | 4 | 0.75 | 3 | 0.6667 | 2 | 1 | 10 | 0.8 |
| 5_73178939 | 1 | 0 | 8 | 0.125 | 4 | 0 | 4 | 0.25 | 17 | 0.1176 | 2 | 0 | 4 | 0.25 | 5 | 0 | 3 | 0 | 14 | 0.0714 |
| 5_73179012 | 1 | 1 | 8 | 0.125 | 4 | 0.5 | 3 | 0.3333 | 16 | 0.3125 | 2 | 0 | 2 | 1 | 6 | 0.1667 | 4 | 0.5 | 14 | 0.3571 |
| 5_73179107 | 1 | 0 | 6 | 0.6667 | 4 | 0.25 | 2 | 0.5 | 13 | 0.4615 | 2 | 0 | 2 | 0 | 5 | 0.4 | 4 | 0.25 | 13 | 0.2308 |
| 5_73179117 | 1 | 1 | 7 | 0.5714 | 4 | 0.75 | 3 | 0.6667 | 15 | 0.6667 | 2 | 1 | 3 | 1 | 4 | 0.25 | 4 | 1 | 13 | 0.7692 |
| 5_73179217 | 1 | 0 | 7 | 0.2857 | 2 | 0 | 2 | 0 | 12 | 0.1667 | 1 | 1 | 3 | 0 | 1 | 1 | 3 | 0.6667 | 8 | 0.5 |
| 5_73179235 | 1 | 0 | 7 | 0.4286 | 3 | 0.6667 | 3 | 0.3333 | 14 | 0.4286 | 1 | 0 | 2 | 0 | 3 | 1 | 3 | 0.6667 | 9 | 0.5556 |
| 5_73179257 | 1 | 0 | 6 | 0.1667 | 4 | 0.25 | 3 | 0.6667 | 14 | 0.2857 | 2 | 0.5 | 3 | 0.3333 | 4 | 0.5 | 4 | 0 | 13 | 0.3077 |
| 5_73179271 | 1 | 1 | 6 | 0.8333 | 1 | 1 | 3 | 0.3333 | 11 | 0.7273 | 2 | 0.5 | 3 | 0.3333 | 4 | 0.5 | 4 | 0.25 | 13 | 0.3846 |
| 5_73179292 | 1 | 1 | 5 | 1 | 3 | 1 | 3 | 0.6667 | 12 | 0.9167 | NA | NA | 2 | 1 | 1 | 0 | 1 | 1 | 4 | 0.75 |
| 5_73179312 | 1 | 0 | 6 | 0 | 4 | 0.5 | 3 | 0 | 14 | 0.1429 | 2 | 0 | 3 | 0 | 5 | 0.2 | 4 | 0 | 14 | 0.0714 |
| 5_73179337 | 1 | 1 | 6 | 0.6667 | 3 | 0 | 3 | 0 | 13 | 0.3846 | 2 | 0.5 | 3 | 0.3333 | 3 | 0.3333 | 3 | 0.3333 | 11 | 0.3636 |
| 5_73179420 | 1 | 0 | 6 | 0.3333 | 4 | 0 | 3 | 0.6667 | 14 | 0.2857 | 2 | 0 | 2 | 0 | 5 | 0.2 | 4 | 0 | 13 | 0.0769 |
| 5_73179481 | NA | NA | 5 | 0.8 | 2 | 0.5 | 2 | 1 | 9 | 0.7778 | 2 | 1 | 2 | 0.5 | 5 | 0.4 | 3 | 0.6667 | 12 | 0.5833 |
| 5_73179490 | 1 | 0 | 6 | 0 | 4 | 0 | 3 | 0.3333 | 14 | 0.0714 | 1 | 1 | 2 | 0 | 5 | 0 | 3 | 0 | 11 | 0.0909 |
| 5_73179512 | 1 | 1 | 5 | 0.4 | 4 | 0.5 | 3 | 0.3333 | 13 | 0.4615 | 2 | 0 | 2 | 1 | 5 | 0 | 4 | 0 | 13 | 0.1538 |
| 5_73179558 | 1 | 0 | 6 | 0.1667 | 3 | 0 | 3 | 0 | 13 | 0.0769 | 2 | 0 | 2 | 0.5 | 5 | 0.2 | 4 | 0 | 13 | 0.1538 |
| 5_73179588 | 1 | 0 | 6 | 0.1667 | 4 | 0.25 | 3 | 0 | 14 | 0.1429 | 1 | 0 | 2 | 0 | 5 | 0 | 3 | 0.3333 | 11 | 0.0909 |
| 5_73221778 | NA | NA | 1 | 1 | NA | NA | 1 | 0 | 2 | 0.5 | 1 | 1 | NA | NA | NA | NA | NA | NA | 1 | 1 |
| 5_73222651 | NA | NA | 2 | 1 | NA | NA | NA | NA | 2 | 1 | 2 | 1 | 1 | 1 | NA | NA | NA | NA | 3 | 1 |
| 5_73470906 | 70 | 0.4714 | 57 | 0.5965 | 47 | 0.5532 | 51 | 0.5686 | 225 | 0.5422 | 43 | 0.5349 | 58 | 0.4483 | 39 | 0.4103 | 55 | 0.5818 | 195 | 0.4974 |

| | | | | | | | | | | | | | | | | | | | | |
|---|---|---|---|---|---|---|---|---|---|---|---|---|---|---|---|---|---|---|---|---|
| 5_7347098 | 76 | 0.2237 | 68 | 0.2206 | 61 | 0.2131 | 61 | 0.2951 | 266 | 0.2368 | 44 | 0.3182 | 58 | 0.1897 | 53 | 0.2075 | 65 | 0.1846 | 220 | 0.2182 |
| 5_7347109 | 77 | 0.1818 | 69 | 0.1739 | 61 | 0.1639 | 66 | 0.1818 | 273 | 0.1758 | 54 | 0.2407 | 60 | 0.3 | 53 | 0.2075 | 62 | 0.2419 | 229 | 0.2489 |
| 5_7348033 | 15 | 0.2 | 20 | 0.2 | 13 | 0.1538 | 11 | 0.0909 | 59 | 0.1695 | 12 | 0 | 14 | 0.1429 | 8 | 0.125 | 10 | 0.3 | 44 | 0.1364 |
| 5_7348053 | 13 | 0.2308 | 19 | 0 | 14 | 0.0714 | 11 | 0.0909 | 57 | 0.0877 | 11 | 0.1818 | 14 | 0 | 7 | 0.1429 | 11 | 0 | 43 | 0.0698 |
| 5_7348096 | 13 | 0.0769 | 18 | 0.1111 | 13 | 0.3077 | 11 | 0 | 55 | 0.1273 | 11 | 0.0909 | 14 | 0.2143 | 8 | 0 | 9 | 0.1111 | 42 | 0.119 |
| 5_7348107 | 15 | 0.3333 | 30 | 0.1667 | 23 | 0.1304 | 14 | 0.1429 | 82 | 0.1829 | 9 | 0.6667 | 16 | 0.3125 | 9 | 0.1111 | 13 | 0.7692 | 47 | 0.4681 |
| 5_7354826 | 3 | 0.3333 | 3 | 0 | NA | NA | 3 | 0.6667 | 9 | 0.3333 | 2 | 1 | NA | NA | 2 | 0.5 | 2 | 0.5 | 6 | 0.6667 |
| 5_7354847 | 3 | 1 | 5 | 1 | NA | NA | 3 | 1 | 11 | 1 | 4 | 1 | NA | NA | 2 | 1 | 3 | 1 | 9 | 1 |
| 5_7354854 | 3 | 0 | 5 | 0.2 | NA | NA | 3 | 0.3333 | 11 | 0.1818 | 5 | 0.4 | NA | NA | 2 | 0 | 2 | 0 | 9 | 0.2222 |
| 5_7354870 | 4 | 0 | 4 | 0 | NA | NA | 3 | 0.3333 | 11 | 0.0909 | 4 | 0.5 | NA | NA | 1 | 0 | 3 | 0 | 8 | 0.25 |
| 5_7354872 | 4 | 0 | 5 | 0.2 | NA | NA | 3 | 0 | 12 | 0.0833 | 5 | 0.2 | NA | NA | 2 | 0 | 3 | 0 | 10 | 0.1 |
| 5_7354876 | 2 | 0.5 | 5 | 0.4 | NA | NA | 3 | 0.6667 | 10 | 0.5 | 5 | 0 | NA | NA | 2 | 0 | 2 | 0.5 | 9 | 0.1111 |
| 5_7354877 | 4 | 0.25 | 5 | 0.2 | NA | NA | 3 | 0 | 12 | 0.1667 | 5 | 0.2 | NA | NA | 2 | 0 | 2 | 0.5 | 9 | 0.2222 |
| 5_7354885 | 4 | 0.75 | 5 | 1 | NA | NA | 4 | 0.75 | 13 | 0.8462 | 5 | 1 | NA | NA | 2 | 1 | 3 | 1 | 10 | 1 |
| 5_7354919 | 2 | 0 | 2 | 0.5 | NA | NA | 2 | 0.5 | 6 | 0.3333 | 3 | 0.3333 | NA | NA | 2 | 0 | 1 | 0 | 6 | 0.1667 |
| 5_7354928 | 3 | 0.3333 | 2 | 1 | NA | NA | 2 | 1 | 7 | 0.7143 | 4 | 1 | NA | NA | 2 | 1 | 1 | 1 | 7 | 1 |
| 5_7354930 | 2 | 0.5 | 2 | 0 | NA | NA | 2 | 0 | 6 | 0.1667 | 4 | 0.25 | NA | NA | 2 | 0 | 1 | 0 | 7 | 0.1429 |
| 5_7354947 | 2 | 0.5 | 3 | 1 | NA | NA | 1 | 1 | 6 | 0.8333 | 2 | 1 | NA | NA | 2 | 1 | 2 | 1 | 6 | 1 |
| 5_7354948 | 2 | 0.5 | 2 | 0.5 | NA | NA | 2 | 0 | 6 | 0.3333 | 3 | 0.6667 | NA | NA | 1 | 0 | 2 | 0 | 6 | 0.3333 |
| 5_7354964 | 3 | 0 | 3 | 0.3333 | 2 | 0.5 | 1 | 1 | 9 | 0.3333 | 3 | 0.6667 | NA | NA | 1 | 0 | 1 | 0 | 5 | 0.4 |
| 5_7354965 | 3 | 0.6667 | 3 | 1 | 3 | 1 | 1 | 1 | 10 | 0.9 | 4 | 0.75 | NA | NA | 1 | 1 | 2 | 1 | 7 | 0.8571 |
| 5_7354966 | 2 | 1 | 3 | 1 | 3 | 1 | NA | NA | 8 | 1 | 4 | 1 | NA | NA | 1 | 1 | 2 | 1 | 7 | 1 |
| 5_7354987 | 3 | 1 | 3 | 1 | 3 | 1 | 1 | 1 | 10 | 1 | 2 | 1 | 1 | 1 | 1 | 1 | 2 | 1 | 6 | 1 |
| 5_7371609 | 11 | 0.0909 | 11 | 0 | 8 | 0 | 10 | 0.2 | 40 | 0.075 | 8 | 0 | 8 | 0.25 | 4 | 0 | 10 | 0 | 30 | 0.0667 |
| 5_7371652 | 13 | 0 | 17 | 0.1176 | 10 | 0.2 | 17 | 0.0588 | 57 | 0.0877 | 9 | 0.1111 | 9 | 0.1111 | 7 | 0 | 12 | 0 | 37 | 0.0541 |
| 5_7371681 | 15 | 0 | 19 | 0.0526 | 11 | 0.3636 | 18 | 0.2222 | 63 | 0.1429 | 11 | 0.0909 | 8 | 0.25 | 9 | 0.3333 | 12 | 0.0833 | 40 | 0.175 |
| 5_7371685 | 13 | 0.6154 | 16 | 0.375 | 12 | 0.5833 | 17 | 0.4118 | 58 | 0.4828 | 9 | 0.2222 | 9 | 0.6667 | 9 | 0.2222 | 11 | 0.6364 | 38 | 0.4474 |
| 5_7371690 | 13 | 0.6154 | 18 | 0.6111 | 12 | 0.8333 | 17 | 0.7059 | 60 | 0.6833 | 8 | 0.75 | 10 | 0.7 | 10 | 0.6 | 11 | 0.4545 | 39 | 0.6154 |
| 5_7371698 | 13 | 0.3846 | 18 | 0.2778 | 13 | 0.5385 | 19 | 0.3684 | 63 | 0.381 | 10 | 0.4 | 8 | 0.375 | 11 | 0.1818 | 10 | 0.1 | 39 | 0.2564 |
| 5_7371715 | 13 | 1 | 20 | 0.8 | 7 | 0.8571 | 13 | 0.7692 | 53 | 0.8491 | 7 | 0.8571 | 8 | 0.875 | 7 | 0.7143 | 12 | 0.9167 | 34 | 0.8529 |
| 5_7371717 | 16 | 0.9375 | 21 | 1 | 13 | 1 | 17 | 1 | 67 | 0.9851 | 10 | 0.9 | 11 | 1 | 11 | 0.9091 | 13 | 1 | 45 | 0.9556 |
| 5_7371726 | 12 | 1 | 18 | 0.8333 | 10 | 0.9 | 16 | 0.8125 | 56 | 0.875 | 11 | 0.7273 | 6 | 1 | 8 | 1 | 10 | 0.8 | 35 | 0.8571 |
| 5_7371728 | 11 | 0.6364 | 20 | 0.75 | 13 | 0.8462 | 16 | 0.625 | 60 | 0.7167 | 13 | 0.8462 | 9 | 0.8889 | 9 | 0.6667 | 11 | 0.7273 | 42 | 0.7857 |
| 5_7371733 | 16 | 0.6875 | 21 | 0.6667 | 13 | 0.6154 | 17 | 0.7059 | 67 | 0.6716 | 14 | 0.6429 | 8 | 0.875 | 9 | 0.6667 | 14 | 0.9286 | 45 | 0.7778 |
| 5_7371761 | 14 | 0.8571 | 14 | 1 | 12 | 0.9167 | 12 | 1 | 52 | 0.9423 | 13 | 0.9231 | 11 | 0.9091 | 10 | 0.9 | 11 | 1 | 45 | 0.9333 |
| 5_7371763 | 16 | 0.125 | 17 | 0.1176 | 13 | 0.1538 | 15 | 0.2667 | 61 | 0.1639 | 13 | 0.3846 | 10 | 0 | 10 | 0.2 | 12 | 0.0833 | 45 | 0.1778 |
| 5_7379036 | 2 | 0 | 5 | 0.2 | 1 | 0 | 1 | 0 | 9 | 0.1111 | 2 | 0.5 | 6 | 0 | 6 | 0.6667 | 3 | 0 | 17 | 0.2941 |
| 5_7379055 | 2 | 0 | 5 | 0.2 | NA | NA | 1 | 0 | 8 | 0.125 | 2 | 0 | 5 | 0.2 | 2 | 0 | 2 | 0 | 11 | 0.0909 |
| 5_7379120 | 5 | 0.2 | 8 | 0 | 3 | 0 | 2 | 0 | 18 | 0.0556 | 1 | 0 | 8 | 0.25 | 6 | 0.5 | 5 | 0.2 | 20 | 0.3 |
| 5_7379128 | 5 | 0.2 | 6 | 0 | 3 | 0 | 2 | 0 | 16 | 0.0625 | 1 | 0 | 7 | 0.1429 | 6 | 0 | 5 | 0.2 | 19 | 0.1053 |
| 5_7379284 | 3 | 0 | 4 | 0.25 | 2 | 0 | 3 | 0 | 12 | 0.0833 | 3 | 0.6667 | 5 | 0.4 | 5 | 0.2 | 3 | 0.3333 | 16 | 0.375 |
| 5_7379720 | 3 | 0.3333 | 5 | 0.2 | 3 | 0 | 5 | 0 | 16 | 0.125 | 2 | 0.5 | 5 | 0 | 5 | 0.4 | 4 | 0.25 | 16 | 0.25 |
| 5_7415960 | 16 | 0.5 | 16 | 0.5 | 7 | 0.4286 | 8 | 0.75 | 47 | 0.5319 | 8 | 0.875 | 11 | 0.4545 | 15 | 0.3333 | 13 | 0.7692 | 47 | 0.5745 |
| 5_7416034 | 14 | 0.2857 | 19 | 0.0526 | 5 | 0.2 | 6 | 0.3333 | 44 | 0.1818 | 5 | 0.2 | 8 | 0.125 | 9 | 0.1111 | 11 | 0.1818 | 33 | 0.1515 |
| 5_7416036 | 11 | 0.1818 | 15 | 0.2667 | 4 | 0.25 | 5 | 0 | 35 | 0.2 | 4 | 0.25 | 7 | 0.2857 | 5 | 0.2 | 6 | 0.1667 | 22 | 0.2273 |
| 5_7416045 | 13 | 0.3077 | 16 | 0.1875 | 4 | 0.25 | 5 | 0.4 | 38 | 0.2632 | 3 | 0 | 7 | 0.2857 | 7 | 0.1429 | 8 | 0.125 | 25 | 0.16 |
| 5_7416074 | 16 | 0.125 | 24 | 0.125 | 6 | 0 | 8 | 0.125 | 54 | 0.1111 | 6 | 0.3333 | 13 | 0.1538 | 8 | 0.5 | 17 | 0.1176 | 44 | 0.2273 |
| 5_7416097 | 10 | 0.8 | 7 | 0.7143 | 4 | 0.5 | 2 | 0.5 | 23 | 0.6957 | 4 | 0.75 | 7 | 0.5714 | 4 | 0.5 | 5 | 0.8 | 20 | 0.65 |
| 5_7416105 | 17 | 0.6471 | 18 | 0.8333 | 8 | 0.625 | 7 | 0.5714 | 50 | 0.7 | 7 | 0.5714 | 13 | 0.6923 | 9 | 0.6667 | 16 | 0.75 | 45 | 0.6889 |
| 5_7416109 | 17 | 0.2941 | 21 | 0.0952 | 8 | 0.125 | 7 | 0.2857 | 53 | 0.1887 | 6 | 0.6667 | 13 | 0.2308 | 7 | 0.2857 | 16 | 0.1875 | 42 | 0.2857 |
| 5_7416110 | 19 | 0.3158 | 21 | 0.0952 | 8 | 0.25 | 6 | 0 | 54 | 0.1852 | 5 | 0 | 13 | 0.4615 | 8 | 0.5 | 15 | 0.2 | 41 | 0.3171 |
| 5_7416113 | 19 | 1 | 21 | 1 | 7 | 1 | 7 | 1 | 54 | 1 | 5 | 1 | 12 | 0.9167 | 9 | 1 | 16 | 1 | 42 | 0.9762 |
| 5_7416114 | 13 | 0.6923 | 12 | 0.5833 | 5 | 0.6 | 7 | 0.4286 | 37 | 0.5946 | 4 | 0.25 | 8 | 0.5 | 8 | 0.375 | 8 | 0.375 | 28 | 0.3929 |
| 5_7416121 | 17 | 0.6471 | 19 | 0.6842 | 7 | 0.5714 | 8 | 0.5 | 51 | 0.6275 | 7 | 0.4286 | 13 | 0.6154 | 9 | 0.4444 | 14 | 0.6429 | 43 | 0.5581 |
| 5_7416123 | 18 | 0.1667 | 20 | 0.05 | 7 | 0 | 8 | 0 | 53 | 0.0755 | 6 | 0.1667 | 11 | 0 | 8 | 0.25 | 17 | 0.0588 | 42 | 0.0952 |
| 5_7416123 | 13 | 0.1538 | 14 | 0.0714 | 5 | 0.2 | 7 | 0 | 39 | 0.1026 | 3 | 0 | 8 | 0 | 6 | 0.1667 | 11 | 0.2727 | 28 | 0.1429 |
| 5_7416245 | 14 | 0 | 14 | 0.1429 | 6 | 0.3333 | 3 | 0 | 37 | 0.1081 | 5 | 0 | 11 | 0.2727 | 4 | 0 | 10 | 0 | 30 | 0.1 |
| 5_7416251 | 13 | 0.3846 | 14 | 0.1429 | 6 | 0 | 6 | 0.3333 | 39 | 0.2308 | 5 | 0 | 11 | 0.3636 | 6 | 0.1667 | 13 | 0.0769 | 35 | 0.1714 |
| 5_7416268 | 15 | 0.5333 | 14 | 0.3571 | 5 | 0.4 | 4 | 0.25 | 38 | 0.4211 | 6 | 0.5 | 9 | 0.5556 | 6 | 0.5 | 11 | 0.6364 | 32 | 0.5625 |
| 5_7416269 | 10 | 0.2 | 15 | 0.0667 | 5 | 0 | 3 | 0 | 33 | 0.0909 | 5 | 0.2 | 10 | 0.1 | 6 | 0 | 5 | 0 | 26 | 0.0769 |
| 5_7435885 | 7 | 0.1429 | 8 | 0 | 4 | 0.25 | 7 | 0 | 26 | 0.0769 | 3 | 0 | 5 | 0.2 | 1 | 0 | 1 | 0 | 10 | 0.1 |
| 5_7435887 | 8 | 0.75 | 7 | 0.8571 | 4 | 0.25 | 7 | 0.7143 | 26 | 0.6923 | 2 | 0.5 | 4 | 0.5 | NA | NA | 1 | 1 | 7 | 0.5714 |
| 5_7435898 | 8 | 0.375 | 8 | 0.375 | 4 | 1 | 7 | 0.7143 | 27 | 0.5556 | 3 | 0.3333 | 6 | 0.3333 | 1 | 1 | 1 | 0 | 11 | 0.3636 |
| 5_7435913 | 9 | 0 | 8 | 0.375 | 4 | 0 | 8 | 0.25 | 29 | 0.1724 | 3 | 0 | 5 | 0 | 2 | 0.5 | 1 | 0 | 11 | 0.0909 |
| 5_7435922 | 8 | 1 | 6 | 1 | 4 | 1 | 7 | 1 | 25 | 1 | 3 | 1 | 6 | 1 | 2 | 1 | 1 | 1 | 12 | 1 |

| | | | | | | | | | | | | | | | | | | | | |
|---|---|---|---|---|---|---|---|---|---|---|---|---|---|---|---|---|---|---|---|---|
| 5_74359304 | 10 | 1 | 7 | 1 | 4 | 1 | 8 | 0.875 | 29 | 0.9655 | 3 | 1 | 6 | 0.8333 | 2 | 1 | 1 | 1 | 12 | 0.9167 |
| 5_74359398 | 9 | 0.4444 | 7 | 0.7143 | 4 | 0.5 | 8 | 0.75 | 28 | 0.6071 | 2 | 1 | 5 | 1 | 2 | 1 | 1 | 1 | 10 | 1 |
| 5_74359417 | 10 | 1 | 7 | 1 | 4 | 1 | 9 | 1 | 30 | 1 | 1 | 1 | 6 | 1 | 2 | 1 | 1 | 1 | 10 | 1 |
| 5_74359538 | 10 | 0.1 | 6 | 0.1667 | 5 | 0 | 8 | 0.125 | 29 | 0.1034 | 4 | 0.25 | 6 | 0 | 2 | 0 | NA | NA | 12 | 0.0833 |
| 5_74359589 | 9 | 0.1111 | 7 | 0.2857 | 5 | 0.4 | 8 | 0.25 | 29 | 0.2414 | 3 | 0.6667 | 5 | 0.6 | 2 | 0 | 2 | 0 | 12 | 0.4167 |
| 5_74359663 | 11 | 0.5455 | 7 | 0.4286 | 5 | 0.4 | 8 | 0.25 | 31 | 0.4194 | 4 | 0.5 | 6 | 0.8333 | 2 | 1 | 2 | 1 | 14 | 0.7857 |
| 5_74366798 | 2 | 1 | 1 | 0 | 1 | 1 | 1 | 1 | 5 | 0.8 | 2 | 0.5 | 2 | 0 | 3 | 0 | 1 | 0 | 8 | 0.125 |
| 5_74366803 | 1 | 0 | 1 | 1 | 1 | 1 | 1 | 0 | 4 | 0.5 | 2 | 0.5 | 1 | 1 | 3 | 0.6667 | 1 | 1 | 7 | 0.7143 |
| 5_74691838 | NA | NA | 1 | 1 | NA | NA | 1 | 0 | 2 | 0.5 | NA | NA | NA | NA | 1 | 1 | 1 | 1 | 2 | 1 |
| 5_74696386 | 1 | 0 | 1 | 1 | 1 | 0 | 7 | 0 | 10 | 0.1 | 2 | 0 | 1 | 0 | 5 | 0 | 3 | 0.3333 | 11 | 0.0909 |
| 5_74696430 | NA | NA | NA | NA | NA | NA | 2 | 0.5 | 2 | 0.5 | 1 | 0 | NA | NA | 1 | 1 | NA | NA | 2 | 0.5 |
| 5_74757150 | 1 | 0 | 3 | 0 | 4 | 0 | 9 | 0.1111 | 17 | 0.0588 | 4 | 0 | 1 | 0 | 3 | 0 | 5 | 0.2 | 13 | 0.0769 |
| 5_74757649 | 3 | 0 | 2 | 0 | 4 | 0.25 | 9 | 0 | 18 | 0.0556 | 5 | 0.2 | 1 | 0 | 2 | 0 | 2 | 0.5 | 10 | 0.2 |
| 5_74757733 | 3 | 0.3333 | 3 | 0 | 4 | 0 | 10 | 0 | 20 | 0.05 | 5 | 0 | 1 | 0 | 3 | 0 | 5 | 0.2 | 14 | 0.0714 |
| 5_74757800 | 2 | 0.5 | 4 | 0.25 | 1 | 0 | 7 | 0.1429 | 14 | 0.2143 | 4 | 0.25 | 1 | 0 | 2 | 0 | 2 | 0 | 9 | 0.1111 |
| 5_74757990 | 3 | 0 | 5 | 0 | 1 | 1 | 6 | 0 | 15 | 0.0667 | 3 | 0 | 1 | 0 | 1 | 0 | 2 | 0.5 | 7 | 0.1429 |
| 5_74758480 | 4 | 0.25 | 5 | 0 | 5 | 0.8 | 10 | 0.6 | 24 | 0.4583 | 5 | 0.2 | 1 | 0 | 4 | 0.75 | 5 | 0.4 | 15 | 0.4 |
| 5_74758752 | 4 | 0 | 7 | 0.1429 | 4 | 0 | 11 | 0.2727 | 26 | 0.1538 | 6 | 0.1667 | 1 | 0 | 4 | 0 | 7 | 0.2857 | 18 | 0.1667 |
| 5_74758998 | 3 | 0.3333 | 6 | 0.1667 | 5 | 0 | 10 | 0 | 24 | 0.0833 | 6 | 0.1667 | 1 | 0 | 4 | 0 | 5 | 0 | 16 | 0.0625 |
| 5_74780863 | 2 | 0.5 | 3 | 0.3333 | NA | NA | 6 | 0.3333 | 11 | 0.3636 | 3 | 0 | 1 | 1 | NA | NA | 2 | 0.5 | 6 | 0.3333 |
| 5_74780873 | 2 | 0.5 | 3 | 0 | NA | NA | 4 | 0 | 9 | 0.1111 | 3 | 0 | 1 | 0 | NA | NA | 2 | 1 | 6 | 0.3333 |
| 5_75167353 | 8 | 0.625 | 4 | 0.75 | 3 | 0.3333 | 2 | 0 | 17 | 0.5294 | 4 | 0.25 | 4 | 0.5 | 3 | 0.3333 | 2 | 0 | 13 | 0.3077 |
| 5_75237915 | NA | NA | NA | NA | 1 | 1 | NA | NA | 1 | 1 | NA | NA | 1 | 1 | 1 | 1 | NA | NA | 2 | 1 |
| 5_75353159 | 6 | 0.3333 | 3 | 0.3333 | 1 | 0 | 3 | 0.3333 | 13 | 0.3077 | 3 | 0 | 2 | 1 | 4 | 0.25 | NA | NA | 9 | 0.3333 |
| 5_75353216 | 6 | 0 | 3 | 0.3333 | NA | NA | 2 | 0 | 11 | 0.0909 | 4 | 0 | 2 | 0.5 | 3 | 0 | NA | NA | 9 | 0.1111 |
| 5_75353244 | 6 | 0 | 3 | 0.3333 | 1 | 1 | 3 | 0 | 13 | 0.1538 | 4 | 0.5 | 2 | 0 | 4 | 0 | NA | NA | 10 | 0.2 |
| 5_75355697 | 7 | 0.1429 | 3 | 0 | 1 | 0 | 3 | 0 | 14 | 0.0714 | 4 | 0 | 2 | 0 | 3 | 0.3333 | NA | NA | 9 | 0.1111 |
| 5_75355736 | 6 | 0.3333 | 3 | 0 | 1 | 0 | 3 | 0.3333 | 13 | 0.2308 | 4 | 0.25 | 2 | 0.5 | 4 | 0 | NA | NA | 10 | 0.2 |
| 5_75355748 | 6 | 0.3333 | 3 | 0.6667 | 1 | 1 | 2 | 0.5 | 12 | 0.5 | 4 | 0.25 | 2 | 0.5 | 3 | 0.6667 | NA | NA | 9 | 0.4444 |
| 5_75355853 | 5 | 1 | 2 | 0.5 | 1 | 1 | 3 | 1 | 11 | 0.9091 | 5 | 1 | 2 | 1 | 5 | 1 | NA | NA | 12 | 1 |
| 5_75355916 | 6 | 0.8333 | 2 | 1 | 1 | 1 | 3 | 0.6667 | 12 | 0.8333 | 1 | 0 | 1 | 0 | 4 | 0.75 | NA | NA | 6 | 0.5 |
| 5_75355935 | 2 | 1 | NA | NA | 1 | 1 | NA | NA | 3 | 1 | 2 | 1 | 2 | 1 | 3 | 0.6667 | NA | NA | 7 | 0.8571 |
| 5_75356006 | 6 | 0.6667 | 2 | 0.5 | 2 | 1 | 2 | 0.5 | 12 | 0.6667 | 4 | 1 | 2 | 1 | 5 | 0.8 | NA | NA | 11 | 0.9091 |
| 5_75356017 | 6 | 0.8333 | 3 | 1 | 2 | 1 | 3 | 1 | 14 | 0.9286 | 2 | 1 | 2 | 1 | 4 | 1 | NA | NA | 8 | 1 |
| 5_75356120 | 1 | 1 | NA | NA | NA | NA | 3 | 0.3333 | 4 | 0.5 | 2 | 1 | 1 | 1 | 3 | 0.6667 | NA | NA | 6 | 0.8333 |
| 5_75356127 | 6 | 0 | 2 | 0 | 1 | 0 | 3 | 0.3333 | 12 | 0.0833 | 3 | 0 | 1 | 0 | 1 | 1 | NA | NA | 5 | 0.2 |
| 5_75356168 | 7 | 0.5714 | 3 | 0.3333 | 2 | 1 | 3 | 0.6667 | 15 | 0.6 | 5 | 0.6 | 2 | 1 | 5 | 1 | NA | NA | 12 | 0.8333 |
| 5_75356274 | 6 | 1 | 3 | 1 | 2 | 1 | 2 | 1 | 13 | 1 | 4 | 1 | 2 | 1 | 5 | 1 | NA | NA | 11 | 1 |
| 5_75356825 | 8 | 0.125 | 4 | 0 | 2 | 0 | 3 | 0.6667 | 17 | 0.1765 | 3 | 0 | 3 | 0.3333 | 5 | 0 | NA | NA | 11 | 0.0909 |
| 5_75356842 | 6 | 0.8333 | 3 | 0.6667 | 2 | 0.5 | 1 | 0 | 12 | 0.6667 | 4 | 0.5 | 1 | 0 | 3 | 1 | NA | NA | 8 | 0.625 |
| 5_75356873 | 8 | 0.625 | 4 | 0.25 | 2 | 0 | 3 | 0.6667 | 17 | 0.4706 | 5 | 0.4 | 4 | 0.25 | 6 | 0.3333 | NA | NA | 15 | 0.3333 |
| 5_75356942 | 7 | 0 | 3 | 0 | 1 | 0 | 3 | 0.3333 | 14 | 0.0714 | 4 | 0.75 | 3 | 0 | 5 | 0.2 | NA | NA | 12 | 0.3333 |
| 5_75357004 | 7 | 0.5714 | 3 | 0.3333 | 2 | 0 | 3 | 0.3333 | 15 | 0.4 | 5 | 0.4 | 5 | 0.6 | 6 | 0.5 | NA | NA | 16 | 0.5 |
| 5_75357181 | 8 | 0 | 4 | 0.25 | 3 | 0 | 2 | 0 | 17 | 0.0588 | 2 | 0.5 | 4 | 0 | 5 | 0.4 | NA | NA | 11 | 0.2727 |
| 5_75357621 | 8 | 0.25 | 4 | 0.25 | 2 | 0.5 | 3 | 0 | 17 | 0.2353 | 5 | 0.2 | 5 | 0 | 6 | 0.1667 | NA | NA | 16 | 0.125 |
| 5_75357679 | 7 | 0.1429 | 4 | 0 | 3 | 0 | 3 | 0 | 17 | 0.0588 | 7 | 0.1429 | 5 | 0.2 | 6 | 0.1667 | NA | NA | 18 | 0.1667 |
| 5_75357927 | 5 | 0 | 4 | 0 | 2 | 0 | 3 | 0.3333 | 14 | 0.0714 | 6 | 0 | 6 | 0.1667 | 6 | 0 | NA | NA | 18 | 0.0556 |
| 5_75358185 | 6 | 0 | 4 | 0.25 | 3 | 0 | 3 | 0 | 16 | 0.0625 | 5 | 0.2 | 4 | 0.25 | 5 | 0 | NA | NA | 14 | 0.1429 |
| 5_75358409 | 7 | 0.1429 | 4 | 0 | 3 | 0 | 4 | 0.25 | 18 | 0.1111 | 7 | 0 | 6 | 0.1667 | 5 | 0.2 | NA | NA | 18 | 0.1111 |
| 5_75815430 | 2 | 0 | NA | NA | 1 | 1 | 4 | 0 | 7 | 0.1429 | 3 | 0.3333 | 1 | 0 | 2 | 0.5 | 3 | 1 | 9 | 0.5556 |
| 5_75815484 | 4 | 0.5 | NA | NA | 1 | 1 | 4 | 0 | 9 | 0.3333 | 3 | 0.3333 | NA | NA | 3 | 0.3333 | 3 | 0 | 9 | 0.2222 |
| 5_75815662 | 3 | 0.3333 | NA | NA | 2 | 0 | 5 | 0 | 10 | 0.1 | 4 | 0 | 1 | 0 | 3 | 0.3333 | 3 | 0 | 11 | 0.0909 |
| 5_76094622 | NA | NA | 1 | 1 | 1 | 1 | 2 | 1 | 4 | 1 | NA | NA | NA | NA | NA | NA | 1 | 1 | 1 | 1 |
| 5_76094801 | NA | NA | 1 | 0 | 1 | 1 | 2 | 1 | 4 | 0.75 | NA | NA | NA | NA | NA | NA | 1 | 1 | 1 | 1 |
| 5_76094880 | NA | NA | 1 | 1 | 1 | 1 | 2 | 1 | 4 | 1 | NA | NA | NA | NA | NA | NA | 1 | 1 | 1 | 1 |
| 5_76094943 | NA | NA | 1 | 1 | 1 | 1 | 2 | 0.5 | 4 | 0.75 | NA | NA | NA | NA | NA | NA | 1 | 1 | 1 | 1 |
| 5_76094950 | NA | NA | 1 | 1 | 1 | 1 | 1 | 1 | 3 | 1 | NA | NA | NA | NA | NA | NA | 1 | 1 | 1 | 1 |
| 5_76095152 | NA | NA | 1 | 1 | 1 | 1 | 2 | 1 | 4 | 1 | NA | NA | NA | NA | NA | NA | 1 | 1 | 1 | 1 |
| 5_76302660 | 3 | 0 | 3 | 0 | 2 | 0.5 | 3 | 0.3333 | 11 | 0.1818 | 3 | 0.3333 | 7 | 0.4286 | NA | NA | 4 | 0 | 14 | 0.2857 |
| 5_76302687 | 3 | 0 | 4 | 0.25 | 2 | 0 | 2 | 0.5 | 11 | 0.1818 | 4 | 0.25 | 7 | 0.2857 | 1 | 0 | 5 | 0.2 | 17 | 0.2353 |
| 5_76302693 | 3 | 0 | 4 | 0.25 | 2 | 0.5 | 2 | 0 | 11 | 0.1818 | 3 | 0.3333 | 7 | 0.4286 | 1 | 0 | 4 | 0.5 | 15 | 0.4 |
| 5_76302789 | 3 | 0.3333 | 3 | 0 | 2 | 0 | 3 | 0.3333 | 11 | 0.1818 | 4 | 0.25 | 8 | 0 | NA | NA | 4 | 0.25 | 16 | 0.125 |
| 5_76302848 | 2 | 0 | 4 | 0.25 | 2 | 0 | 3 | 0.3333 | 11 | 0.1818 | 4 | 0.5 | 6 | 0.5 | NA | NA | 4 | 0.75 | 14 | 0.5714 |
| 5_76302877 | 1 | 1 | 4 | 0.25 | 2 | 0 | 2 | 0 | 9 | 0.2222 | 3 | 0 | 5 | 0 | 1 | 0 | 5 | 0.2 | 14 | 0.0714 |
| 5_76356995 | 6 | 0.3333 | 6 | 0 | 5 | 0.4 | 6 | 0 | 23 | 0.1739 | 7 | 0.1429 | 4 | 0.25 | 2 | 0.5 | 3 | 0 | 16 | 0.1875 |

| | | | | | | | | | | | | | | | | | | | | |
|---|---|---|---|---|---|---|---|---|---|---|---|---|---|---|---|---|---|---|---|---|
| 5_7635869 | 1 | 0 | 2 | 0.5 | 3 | 0 | 2 | 0 | 8 | 0.125 | 2 | 0 | 2 | 0 | NA | NA | 8 | 0.375 | 12 | 0.25 |
| 5_7635874 | 1 | 0 | 2 | 0.5 | 3 | 0.6667 | 2 | 0 | 8 | 0.375 | 2 | 0.5 | 2 | 0 | NA | NA | 9 | 0.2222 | 13 | 0.2308 |
| 5_7635977 | 1 | 0 | 1 | 0 | 2 | 0.5 | 2 | 0 | 6 | 0.1667 | 2 | 0 | 2 | 0.5 | 1 | 0 | 8 | 0 | 13 | 0.0769 |
| 5_7636020 | 6 | 0.5 | 6 | 0.3333 | 4 | 0.5 | 6 | 0.6667 | 22 | 0.5 | 3 | 0.3333 | 3 | 0.3333 | 4 | 0.75 | 10 | 0.2 | 20 | 0.35 |
| 5_7636021 | 5 | 0.8 | 5 | 0.8 | 4 | 0.75 | 5 | 0.8 | 19 | 0.7895 | 3 | 1 | 3 | 0.3333 | 4 | 1 | 9 | 0.5556 | 19 | 0.6842 |
| 5_7636036 | 6 | 0 | 6 | 0.1667 | 5 | 0 | 6 | 0.1667 | 23 | 0.087 | 3 | 0 | 3 | 0 | 4 | 0.25 | 9 | 0 | 19 | 0.0526 |
| 5_7636435 | 2 | 0 | 1 | 0 | 1 | 0 | 1 | 1 | 5 | 0.2 | 1 | 0 | NA | NA | 1 | 1 | 2 | 0 | 4 | 0.25 |
| 5_7636446 | 1 | 1 | NA | NA | 1 | 0 | NA | NA | 2 | 0.5 | NA | NA | NA | NA | 1 | 1 | 2 | 0 | 3 | 0.3333 |
| 5_7636458 | 2 | 1 | NA | NA | 1 | 1 | 1 | 1 | 4 | 1 | 1 | 1 | NA | NA | 1 | 1 | 2 | 1 | 4 | 1 |
| 5_7637503 | 2 | 0.5 | NA | NA | 1 | 0 | NA | NA | 3 | 0.3333 | 1 | 1 | NA | NA | 1 | 1 | 1 | 0 | 3 | 0.6667 |
| 5_7671486 | 6 | 0.3333 | 2 | 0 | 4 | 0 | 1 | 0 | 13 | 0.1538 | 1 | 0 | 5 | 0.2 | 4 | 0.25 | 2 | 0.5 | 12 | 0.25 |
| 5_7671757 | 7 | 0.7143 | 1 | 1 | 5 | 0.4 | 3 | 1 | 16 | 0.6875 | 2 | 1 | 2 | 0 | 3 | 0.3333 | 2 | 0.5 | 9 | 0.4444 |
| 5_7671773 | 6 | 0.8333 | 2 | 1 | 4 | 0.75 | 3 | 1 | 15 | 0.8667 | 2 | 1 | 4 | 1 | 3 | 0.6667 | 3 | 1 | 12 | 0.9167 |
| 5_7671786 | 6 | 0.6667 | 2 | 0.5 | 5 | 0.2 | 3 | 0.3333 | 16 | 0.4375 | 2 | 0 | 6 | 0.3333 | 4 | 0 | 2 | 1 | 14 | 0.2857 |
| 5_7671787 | 6 | 0.8333 | 1 | 0 | 5 | 0.6 | 2 | 1 | 14 | 0.7143 | 3 | 1 | 6 | 0.8333 | 4 | 1 | 2 | 1 | 15 | 0.9333 |
| 5_7671792 | 6 | 1 | 1 | 0 | 7 | 0.2857 | 3 | 0.6667 | 17 | 0.5882 | 3 | 0 | 4 | 0.75 | 3 | 0.6667 | 3 | 0.6667 | 13 | 0.5385 |
| 5_7671798 | 8 | 0.875 | 2 | 0 | 5 | 1 | 3 | 1 | 18 | 0.8333 | 2 | 0 | 5 | 0.8 | 3 | 1 | 3 | 0.6667 | 13 | 0.6923 |
| 5_7671809 | 8 | 0 | 2 | 0 | 6 | 0.1667 | 4 | 0 | 20 | 0.05 | 3 | 0.3333 | 6 | 0 | 5 | 0 | 3 | 0 | 17 | 0.0588 |
| 5_7671811 | 7 | 0.1429 | 2 | 0 | 7 | 0 | 2 | 0 | 18 | 0.0556 | 3 | 0 | 6 | 0.1667 | 4 | 0 | 3 | 0 | 16 | 0.0625 |
| 5_7700657 | 1 | 1 | NA | NA | 1 | 1 | 1 | 1 | 3 | 1 | 1 | 1 | NA | NA | NA | NA | 1 | 1 | 2 | 1 |
| 5_7700683 | 1 | 1 | NA | NA | 2 | 0.5 | 1 | 0 | 4 | 0.5 | 1 | 1 | NA | NA | NA | NA | 1 | 1 | 2 | 1 |
| 5_7701874 | 2 | 0.5 | 2 | 0 | 3 | 0.6667 | 2 | 0 | 9 | 0.3333 | 1 | 0 | 1 | 1 | 1 | 0 | NA | NA | 3 | 0.3333 |
| 5_7701917 | 2 | 0.5 | 1 | 0 | 3 | 0.3333 | 4 | 0.25 | 10 | 0.3 | 1 | 0 | 2 | 1 | 1 | 0 | NA | NA | 4 | 0.5 |
| 5_7701938 | 3 | 0 | 3 | 0.3333 | 3 | 0.3333 | 4 | 0.25 | 13 | 0.2308 | 1 | 1 | 3 | 0.6667 | 1 | 0 | NA | NA | 5 | 0.6 |
| 5_7701939 | 3 | 0 | 3 | 0 | 3 | 0.3333 | 4 | 0 | 13 | 0.0769 | 1 | 0 | 3 | 0.3333 | 1 | 0 | NA | NA | 5 | 0.2 |
| 5_7701961 | 3 | 0.3333 | 3 | 0 | 3 | 0 | 5 | 0.4 | 14 | 0.2143 | 1 | 1 | 3 | 1 | 1 | 1 | 1 | 0 | 6 | 0.8333 |
| 5_7702094 | 5 | 0 | 4 | 0 | 3 | 0.3333 | 5 | 0 | 17 | 0.0588 | 1 | 0 | 2 | 0.5 | 2 | 0.5 | 1 | 0 | 6 | 0.3333 |
| 5_7702399 | NA | NA | NA | NA | 1 | 0 | 1 | 1 | 2 | 0.5 | NA | NA | 1 | 0 | 1 | 1 | 1 | 0 | 3 | 0.3333 |
| 5_7703045 | NA | NA | NA | NA | 1 | 1 | 1 | 1 | 2 | 1 | NA | NA | 1 | 1 | NA | NA | NA | NA | 1 | 1 |
| 5_7703049 | NA | NA | NA | NA | 1 | 1 | 1 | 1 | 2 | 1 | NA | NA | 1 | 1 | 1 | 1 | NA | NA | 2 | 1 |
| 5_7706261 | NA | NA | 2 | 0.5 | 2 | 0 | 3 | 0 | 7 | 0.1429 | 1 | 0 | 1 | 0 | 5 | 0.2 | 1 | 0 | 8 | 0.125 |
| 5_7706284 | NA | NA | 2 | 0.5 | 2 | 0.5 | 2 | 1 | 6 | 0.6667 | 1 | 1 | 1 | 1 | 2 | 1 | 1 | 1 | 5 | 1 |
| 5_7706304 | NA | NA | 2 | 0.5 | 2 | 0.5 | 5 | 0.4 | 9 | 0.4444 | NA | NA | 1 | 1 | 5 | 0.8 | 1 | 0 | 7 | 0.7143 |
| 5_7706310 | NA | NA | 2 | 1 | 2 | 1 | 4 | 1 | 8 | 1 | NA | NA | 1 | 1 | 3 | 1 | 1 | 1 | 5 | 1 |
| 5_7706321 | NA | NA | 2 | 1 | 2 | 1 | 5 | 0.6 | 9 | 0.7778 | NA | NA | 1 | 1 | 5 | 0.8 | 1 | 1 | 7 | 0.8571 |
| 5_7711513 | 12 | 0.1667 | 20 | 0.15 | 23 | 0.3478 | 17 | 0.1176 | 72 | 0.2083 | 10 | 0.1 | 15 | 0.0667 | 20 | 0.25 | 23 | 0.1739 | 68 | 0.1618 |
| 5_7714621 | 9 | 0 | 6 | 0 | 7 | 0.1429 | 11 | 0.1818 | 33 | 0.0909 | 12 | 0 | 9 | 0.1111 | 4 | 0 | 10 | 0.1 | 35 | 0.0571 |
| 5_7714626 | 8 | 0.125 | 8 | 0.5 | 8 | 0.25 | 9 | 0.1111 | 33 | 0.2424 | 11 | 0.3636 | 11 | 0.3636 | 7 | 0.5714 | 11 | 0.2727 | 40 | 0.375 |
| 5_7714627 | 6 | 0 | 8 | 0.125 | 7 | 0.2857 | 12 | 0 | 33 | 0.0909 | 11 | 0.1818 | 11 | 0.1818 | 7 | 0 | 10 | 0.2 | 39 | 0.1538 |
| 5_7714637 | 9 | 0.2222 | 9 | 0.2222 | 8 | 0.125 | 11 | 0.2727 | 37 | 0.2162 | 10 | 0.3 | 9 | 0.3333 | 6 | 0.5 | 11 | 0.3636 | 36 | 0.3611 |
| 5_7714662 | 10 | 0 | 8 | 0.125 | 9 | 0.2222 | 11 | 0 | 38 | 0.0789 | 11 | 0 | 10 | 0 | 6 | 0 | 13 | 0.1538 | 40 | 0.05 |
| 5_7714669 | 9 | 0.3333 | 9 | 0.6667 | 9 | 0.6667 | 12 | 0.8333 | 39 | 0.641 | 9 | 0.3333 | 10 | 0.5 | 7 | 0.5714 | 12 | 0.5833 | 38 | 0.5 |
| 5_7714691 | 10 | 0.2 | 8 | 0.25 | 9 | 0 | 13 | 0.1538 | 40 | 0.15 | 11 | 0.0909 | 10 | 0.3 | 6 | 0.1667 | 10 | 0.2 | 37 | 0.1892 |
| 5_7714701 | 8 | 0 | 9 | 0.1111 | 9 | 0 | 13 | 0.0769 | 39 | 0.0513 | 12 | 0 | 11 | 0.1818 | 6 | 0 | 9 | 0.1111 | 38 | 0.0789 |
| 5_7714739 | 7 | 0 | 6 | 0.1667 | 10 | 0.1 | 10 | 0.1 | 33 | 0.0909 | 11 | 0 | 10 | 0.1 | 5 | 0 | 11 | 0.2727 | 37 | 0.1081 |
| 5_7714744 | 7 | 0.2857 | 5 | 0.4 | 10 | 0.4 | 9 | 0.2222 | 31 | 0.3226 | 11 | 0.3636 | 10 | 0.4 | 3 | 0.3333 | 10 | 0.6 | 34 | 0.4412 |
| 5_7723526 | 12 | 0 | 3 | 0 | 7 | 0.2857 | 4 | 0.25 | 26 | 0.1154 | 6 | 0 | 6 | 0.5 | 3 | 0.3333 | 5 | 0 | 20 | 0.2 |
| 5_7723555 | 11 | 0 | 3 | 0.3333 | 10 | 0.1 | 4 | 0.25 | 28 | 0.1071 | 6 | 0 | 6 | 0.1667 | 4 | 0 | 8 | 0.125 | 24 | 0.0833 |
| 5_7723559 | 11 | 0.0909 | 4 | 0.75 | 11 | 0.0909 | 5 | 0.2 | 31 | 0.1935 | 6 | 0 | 7 | 0.1429 | 4 | 0.5 | 7 | 0 | 24 | 0.125 |
| 5_7745704 | 1 | 0 | 2 | 0.5 | 1 | 0 | 2 | 1 | 6 | 0.5 | 2 | 0 | 1 | 1 | 2 | 0 | NA | NA | 5 | 0.2 |
| 5_7745714 | 1 | 0 | 2 | 0.5 | 1 | 1 | 1 | 1 | 5 | 0.6 | 3 | 0.3333 | 1 | 0 | 2 | 0 | NA | NA | 6 | 0.1667 |
| 5_7749614 | 3 | 0 | 6 | 0.1667 | 7 | 0 | 9 | 0.1111 | 25 | 0.08 | 5 | 0.2 | 8 | 0 | 12 | 0.0833 | 15 | 0 | 40 | 0.05 |
| 5_8116856 | 1 | 0 | 1 | 1 | NA | NA | 2 | 0.5 | 4 | 0.5 | 1 | 0 | NA | NA | 1 | 1 | NA | NA | 2 | 0.5 |
| 5_8116874 | 1 | 0 | 1 | 0 | 2 | 0.5 | 2 | 0 | 6 | 0.1667 | 2 | 0 | NA | NA | 1 | 1 | 1 | 0 | 4 | 0.25 |
| 5_8116878 | 1 | 1 | 1 | 0 | 2 | 0.5 | 2 | 0.5 | 6 | 0.5 | 2 | 0.5 | NA | NA | 1 | 1 | NA | NA | 3 | 0.6667 |
| 5_8116892 | 1 | 1 | 1 | 1 | 2 | 0.5 | 1 | 0 | 5 | 0.6 | 2 | 0.5 | 1 | 1 | 1 | 1 | 1 | 1 | 5 | 0.8 |
| 5_8116900 | 1 | 0 | 1 | 0 | 2 | 0.5 | 2 | 0 | 6 | 0.1667 | 2 | 0 | 1 | 1 | NA | NA | 1 | 0 | 4 | 0.25 |
| 5_8165984 | 2 | 0 | 2 | 0 | 1 | 0 | 3 | 0.3333 | 8 | 0.125 | NA | NA | 1 | 0 | 2 | 0.5 | 1 | 1 | 4 | 0.5 |
| 5_8165991 | 2 | 0 | 2 | 0 | 2 | 0.5 | 4 | 0.25 | 10 | 0.2 | 2 | 0 | 1 | 0 | 2 | 0.5 | 2 | 1 | 7 | 0.4286 |
| 5_8165995 | 1 | 0 | 1 | 0 | NA | NA | 3 | 0.3333 | 5 | 0.2 | 1 | 0 | NA | NA | 2 | 0.5 | 2 | 0 | 5 | 0.2 |
| 5_8165997 | 2 | 0 | 2 | 0.5 | 2 | 0 | 4 | 1 | 10 | 0.5 | 2 | 0 | NA | NA | 2 | 0 | 2 | 0.5 | 6 | 0.1667 |
| 5_8166010 | 2 | 0 | 2 | 0.5 | 2 | 0 | 4 | 0.25 | 10 | 0.2 | 2 | 0 | 1 | 1 | 1 | 0 | 2 | 0.5 | 6 | 0.3333 |
| 5_8166011 | 1 | 0 | 2 | 0 | 2 | 0 | 2 | 0.5 | 7 | 0.1429 | 2 | 0 | NA | NA | 2 | 1 | 2 | 0.5 | 6 | 0.5 |
| 5_8166013 | 2 | 0.5 | 2 | 0.5 | 2 | 0 | 3 | 1 | 9 | 0.5556 | 2 | 0.5 | 1 | 1 | 2 | 1 | 1 | 1 | 6 | 0.8333 |
| 5_8166022 | 2 | 0 | 2 | 0.5 | 2 | 0.5 | 4 | 0.75 | 10 | 0.5 | 2 | 1 | 1 | 0 | 1 | 1 | 2 | 1 | 6 | 0.8333 |

| | | | | | | | | | | | | | | | | | | | | |
|---|---|---|---|---|---|---|---|---|---|---|---|---|---|---|---|---|---|---|---|---|
| 5_8166027 | 2 | 0 | 2 | 0 | 2 | 0 | 4 | 0.25 | 10 | 0.1 | 2 | 0.5 | NA | NA | NA | NA | 1 | 0 | 3 | 0.3333 |
| 5_8166030 | 2 | 0 | 2 | 0.5 | 3 | 0.3333 | 4 | 0.25 | 11 | 0.2727 | 2 | 0 | 1 | 0 | 3 | 0.3333 | 1 | 0 | 7 | 0.1429 |
| 5_8166033 | 2 | 0.5 | 2 | 0.5 | 3 | 0.3333 | 3 | 1 | 10 | 0.6 | 2 | 0 | 1 | 1 | 2 | 0 | NA | NA | 5 | 0.2 |
| 5_8166034 | 2 | 1 | 2 | 1 | 3 | 1 | 4 | 1 | 11 | 1 | 2 | 1 | 1 | 1 | 3 | 1 | 1 | 1 | 7 | 1 |
| 5_8166045 | 2 | 0 | 2 | 0.5 | 2 | 0.5 | 3 | 0 | 9 | 0.2222 | 2 | 0.5 | NA | NA | 2 | 0.5 | 2 | 0.5 | 6 | 0.5 |
| 5_8166055 | 2 | 0 | 2 | 0 | 3 | 0 | 4 | 0.25 | 11 | 0.0909 | 2 | 0.5 | 1 | 0 | 3 | 0.3333 | 2 | 0 | 8 | 0.25 |
| 5_8166060 | 2 | 0 | 2 | 0 | 3 | 0.3333 | 4 | 0 | 11 | 0.0909 | 2 | 0.5 | 1 | 0 | 2 | 0.5 | 1 | 0 | 6 | 0.3333 |
| 5_8194223 | 5 | 0.4 | 4 | 0.75 | 2 | 0 | 3 | 0.3333 | 14 | 0.4286 | NA | NA | 1 | 0 | 4 | 0.75 | 1 | 1 | 6 | 0.6667 |
| 5_8194228 | 4 | 1 | 4 | 1 | 2 | 1 | 3 | 1 | 13 | 1 | 1 | 1 | 2 | 1 | 4 | 1 | 1 | 1 | 8 | 1 |
| 5_8194234 | 4 | 0 | 3 | 0 | 3 | 0 | 5 | 0.2 | 15 | 0.0667 | 1 | 0 | 2 | 0 | 4 | 0.25 | 1 | 0 | 8 | 0.125 |
| 5_8194236 | 5 | 0.2 | 3 | 0 | 2 | 0 | 5 | 0.2 | 15 | 0.1333 | 1 | 0 | 2 | 0 | 4 | 0.25 | NA | NA | 7 | 0.1429 |
| 5_8194236 | 5 | 0.6 | 4 | 0.25 | 3 | 0.6667 | 4 | 0.5 | 16 | 0.5 | 1 | 0 | 1 | 0 | 4 | 0.25 | NA | NA | 6 | 0.1667 |
| 5_8194245 | 4 | 1 | 3 | 1 | 1 | 1 | 4 | 0.75 | 12 | 0.9167 | 1 | 1 | 2 | 1 | 4 | 0.5 | 1 | 1 | 8 | 0.75 |
| 5_8194262 | 5 | 1 | 5 | 1 | 3 | 1 | 4 | 1 | 17 | 1 | 1 | 1 | 2 | 1 | 5 | 0.8 | 1 | 1 | 9 | 0.8889 |
| 5_8194267 | 5 | 0.4 | 6 | 0.6667 | 4 | 0.25 | 6 | 0.3333 | 21 | 0.4286 | 1 | 0 | 2 | 0.5 | 5 | 0.4 | 1 | 0 | 9 | 0.3333 |
| 5_8194277 | 5 | 0 | 5 | 0.2 | 4 | 0 | 4 | 0 | 18 | 0.0556 | 1 | 0 | 2 | 0 | 4 | 0.25 | 1 | 0 | 8 | 0.125 |
| 5_8194287 | 5 | 0.6 | 4 | 1 | 4 | 1 | 6 | 1 | 19 | 0.8947 | 1 | 1 | 2 | 1 | 5 | 1 | 1 | 0 | 9 | 0.8889 |
| 5_8194288 | 5 | 0.8 | 6 | 0.8333 | 4 | 0.75 | 7 | 0.5714 | 22 | 0.7273 | 1 | 0 | 2 | 0.5 | 5 | 0.8 | 1 | 1 | 9 | 0.6667 |
| 5_8194294 | 3 | 1 | 6 | 1 | 3 | 1 | 6 | 1 | 18 | 1 | 1 | 1 | 2 | 1 | 5 | 1 | NA | NA | 8 | 1 |
| 5_8194295 | 5 | 0.4 | 5 | 0.8 | 4 | 0.75 | 6 | 0.6667 | 20 | 0.65 | 1 | 0 | 2 | 1 | 4 | 1 | 1 | 0 | 8 | 0.75 |
| 5_8194302 | 5 | 1 | 5 | 1 | 4 | 1 | 5 | 1 | 19 | 1 | 1 | 1 | 2 | 1 | 5 | 0.8 | 1 | 1 | 9 | 0.8889 |
| 5_8194303 | 5 | 0.2 | 6 | 0.6667 | 4 | 0.75 | 5 | 0.4 | 20 | 0.5 | 1 | 1 | 2 | 0.5 | 5 | 0.2 | 1 | 1 | 9 | 0.4444 |
| 5_8194305 | 5 | 0.2 | 6 | 0 | 3 | 0 | 4 | 0 | 18 | 0.0556 | 1 | 0 | 2 | 0.5 | 5 | 0 | 1 | 0 | 9 | 0.1111 |
| 5_8194308 | 5 | 0 | 3 | 0 | 4 | 0 | 5 | 0.2 | 17 | 0.0588 | 1 | 0 | 2 | 0 | 5 | 0.2 | 1 | 0 | 9 | 0.1111 |
| 5_8194314 | 5 | 0.8 | 3 | 0.3333 | 4 | 0.75 | 6 | 0.3333 | 18 | 0.5556 | 1 | 1 | 2 | 0.5 | 4 | 0 | 1 | 1 | 8 | 0.375 |
| 5_8194325 | 6 | 1 | 6 | 1 | 4 | 1 | 6 | 1 | 22 | 1 | 1 | 1 | 2 | 1 | 5 | 0.8 | 1 | 1 | 9 | 0.8889 |
| 5_8194331 | 6 | 1 | 5 | 1 | 4 | 1 | 5 | 1 | 20 | 1 | 1 | 0 | 2 | 1 | 5 | 0.8 | 1 | 1 | 9 | 0.7778 |
| 5_8194338 | 4 | 0.5 | 7 | 0.7143 | 4 | 0.75 | 4 | 1 | 19 | 0.7368 | 2 | 0.5 | 1 | 1 | 1 | 0 | 3 | 0.6667 | 7 | 0.5714 |
| 5_8194353 | 5 | 0.4 | 8 | 0.125 | 5 | 0.6 | 6 | 0.5 | 24 | 0.375 | 3 | 0.6667 | 3 | 0.3333 | 4 | 0.5 | 2 | 0 | 12 | 0.4167 |
| 5_8194354 | 5 | 0 | 9 | 0.1111 | 5 | 0 | 5 | 0.2 | 24 | 0.0833 | 3 | 0 | 3 | 0 | 5 | 0.2 | 3 | 0 | 14 | 0.0714 |
| 5_8194878 | 1 | 0 | 3 | 0.3333 | 3 | 0 | 4 | 0 | 11 | 0.0909 | 1 | 0 | 4 | 0.25 | 4 | 0.25 | 2 | 0 | 11 | 0.1818 |
| 5_8194935 | 2 | 0.5 | 4 | 0 | 3 | 0.3333 | 5 | 0 | 14 | 0.1429 | 4 | 0.25 | 4 | 0 | 5 | 0 | 2 | 0 | 15 | 0.0667 |
| 5_8197283 | 1 | 1 | 1 | 0 | 2 | 1 | 2 | 0 | 6 | 0.5 | NA | NA | 1 | 1 | 2 | 0 | 2 | 0 | 5 | 0.2 |
| 5_8197320 | 2 | 1 | 1 | 1 | 2 | 0 | 2 | 0 | 7 | 0.4286 | NA | NA | 1 | 0 | 2 | 0.5 | 1 | 1 | 4 | 0.5 |
| 5_8197338 | 2 | 0.5 | 1 | 1 | 2 | 0 | 3 | 0 | 8 | 0.25 | NA | NA | 1 | 1 | 2 | 0 | 1 | 0 | 4 | 0.25 |
| 5_8420280 | 4 | 0 | 6 | 0.1667 | 4 | 0 | 2 | 0 | 16 | 0.0625 | 7 | 0 | 3 | 0 | 2 | 0.5 | 2 | 0 | 14 | 0.0714 |
| 5_8420282 | 4 | 0 | 6 | 0.1667 | 4 | 0.25 | 2 | 0.5 | 16 | 0.1875 | 6 | 0.1667 | 2 | 0 | 2 | 0 | 2 | 0.5 | 12 | 0.1667 |
| 5_8420424 | 2 | 0.5 | 3 | 1 | 4 | 0.75 | 1 | 0 | 10 | 0.7 | 3 | 0.6667 | 2 | 1 | 1 | 1 | 1 | 0 | 7 | 0.7143 |
| 5_8420430 | 3 | 0.3333 | 3 | 0 | 5 | 0 | NA | NA | 11 | 0.0909 | 3 | 0.3333 | 3 | 0.3333 | 2 | 0 | 2 | 0 | 10 | 0.2 |
| 5_8420448 | 1 | 1 | 3 | 0.6667 | 5 | 0.8 | 1 | 0 | 10 | 0.7 | 2 | 1 | 2 | 0.5 | 2 | 0.5 | 2 | 1 | 8 | 0.75 |
| 5_8420451 | 2 | 1 | 2 | 0.5 | 4 | 1 | 1 | 1 | 9 | 0.8889 | 2 | 1 | 3 | 0.6667 | 1 | 1 | 2 | 1 | 8 | 0.875 |
| 5_8420579 | 3 | 0 | 3 | 0 | 2 | 0 | 3 | 0.6667 | 11 | 0.1818 | NA | NA | 2 | 0 | 2 | 0.5 | 1 | 0 | 5 | 0.2 |
| 5_8420642 | 3 | 0.6667 | 3 | 0.3333 | 3 | 0.3333 | 3 | 0.6667 | 12 | 0.5 | NA | NA | 1 | 1 | 1 | 0 | 1 | 0 | 3 | 0.3333 |
| 5_8420659 | 3 | 1 | 3 | 0.6667 | 3 | 1 | 3 | 1 | 12 | 0.9167 | NA | NA | 1 | 1 | NA | NA | 1 | 1 | 2 | 1 |
| 5_8421393 | 6 | 0.5 | 7 | 0.4286 | 3 | 0 | 3 | 0 | 19 | 0.3158 | 6 | 0.3333 | 1 | 0 | 2 | 1 | 6 | 0 | 15 | 0.2667 |
| 5_8421398 | 6 | 0 | 6 | 0.1667 | 2 | 0.5 | 4 | 0 | 18 | 0.1111 | 6 | 0 | 1 | 0 | NA | NA | 4 | 0.25 | 11 | 0.0909 |
| 5_8447928 | 2 | 0.5 | 1 | 0 | 1 | 0 | 2 | 0 | 6 | 0.1667 | NA | NA | 1 | 1 | NA | NA | 3 | 0 | 4 | 0.25 |
| 5_8447956 | 2 | 1 | 1 | 1 | 1 | 1 | 2 | 1 | 6 | 1 | NA | NA | 1 | 1 | NA | NA | 3 | 1 | 4 | 1 |
| 5_8447956 | 1 | 1 | NA | NA | 1 | 1 | 2 | 1 | 4 | 1 | NA | NA | 1 | 1 | NA | NA | 3 | 0.6667 | 4 | 0.75 |
| 5_8447959 | 1 | 1 | NA | NA | 1 | 1 | 2 | 1 | 4 | 1 | NA | NA | NA | NA | NA | NA | 2 | 1 | 2 | 1 |
| 5_8447961 | 2 | 0.5 | 1 | 0 | 1 | 0 | 2 | 1 | 6 | 0.5 | NA | NA | 1 | 1 | NA | NA | 3 | 0.6667 | 4 | 0.75 |
| 5_8456432 | 2 | 0.5 | 1 | 0 | 1 | 0 | 2 | 0.5 | 6 | 0.3333 | NA | NA | NA | NA | NA | NA | 3 | 0.3333 | 3 | 0.3333 |
| 5_8456461 | 2 | 1 | 1 | 0 | 1 | 1 | 2 | 1 | 6 | 0.8333 | NA | NA | NA | NA | NA | NA | 3 | 0.6667 | 3 | 0.6667 |
| 5_8618530 | 1 | 1 | 2 | 0.5 | 2 | 1 | 3 | 0.3333 | 8 | 0.625 | 1 | 0 | NA | NA | 1 | 1 | 2 | 0.5 | 4 | 0.5 |
| 5_8618536 | 1 | 0 | 2 | 0.5 | 1 | 1 | 2 | 0 | 6 | 0.3333 | 1 | 1 | 2 | 1 | 1 | 1 | 1 | 1 | 5 | 1 |
| 5_8618539 | 1 | 1 | 2 | 1 | 2 | 1 | 2 | 0.5 | 7 | 0.8571 | 1 | 0 | 2 | 0.5 | 1 | 1 | 2 | 1 | 6 | 0.6667 |
| 5_8625214 | 1 | 1 | NA | NA | NA | NA | NA | NA | 1 | 1 | NA | NA | 2 | 0.5 | 1 | 0 | NA | NA | 3 | 0.3333 |
| 5_8625869 | 1 | 1 | NA | NA | 1 | 0 | 2 | 0 | 4 | 0.25 | NA | NA | 1 | 1 | 2 | 0 | 1 | 0 | 4 | 0.25 |
| 5_8625988 | 3 | 0.3333 | 1 | 0 | 3 | 0 | 2 | 0 | 9 | 0.1111 | 1 | 0 | NA | NA | 2 | 0.5 | 1 | 1 | 4 | 0.5 |
| 5_8626030 | 2 | 1 | 1 | 1 | 2 | 1 | 3 | 0.3333 | 8 | 0.75 | 2 | 1 | 1 | 0 | 4 | 1 | 2 | 1 | 9 | 0.8889 |
| 5_8626036 | 3 | 0.3333 | 1 | 0 | 2 | 0.5 | 2 | 0 | 8 | 0.25 | 2 | 0 | 1 | 0 | 4 | 0 | 3 | 0.3333 | 10 | 0.1 |
| 5_8626038 | 4 | 0.75 | 1 | 0 | 3 | 0.3333 | 2 | 0 | 10 | 0.4 | 2 | 0.5 | NA | NA | 4 | 0.75 | 2 | 0.5 | 8 | 0.625 |
| 5_8626040 | 3 | 0.3333 | 1 | 0 | 3 | 0.3333 | 2 | 0 | 9 | 0.2222 | 2 | 0 | 1 | 0 | 4 | 0.25 | 2 | 0.5 | 9 | 0.2222 |
| 5_8626052 | 4 | 0 | 1 | 1 | 3 | 0.6667 | 2 | 0 | 10 | 0.3 | 2 | 0.5 | 1 | 0 | 3 | 0.3333 | 2 | 0 | 8 | 0.25 |
| 5_8697582 | 2 | 0 | 1 | 0 | 6 | 0 | 11 | 0.0909 | 20 | 0.05 | 2 | 1 | 9 | 0 | 6 | 0 | 6 | 0.1667 | 23 | 0.1304 |

| | | | | | | | | | | | | | | | | | | | | |
|---|---|---|---|---|---|---|---|---|---|---|---|---|---|---|---|---|---|---|---|---|
| 5_86975875 | 2 | 1 | 1 | 1 | 6 | 0.1667 | 10 | 0.5 | 19 | 0.4737 | 2 | 0.5 | 9 | 0.3333 | 7 | 0.5714 | 5 | 0.8 | 23 | 0.5217 |
| 5_86975999 | NA | NA | NA | NA | 1 | 1 | 1 | 1 | 2 | 1 | 1 | 1 | NA | NA | NA | NA | NA | NA | 1 | 1 |
| 5_86976108 | NA | NA | NA | NA | 1 | 1 | 1 | 1 | 2 | 1 | 1 | 1 | NA | NA | NA | NA | 1 | 1 | 2 | 1 |
| 5_86976400 | 1 | 1 | NA | NA | 1 | 1 | 2 | 1 | 4 | 1 | 1 | 1 | NA | NA | NA | NA | 1 | 1 | 2 | 1 |
| 5_86976715 | 1 | 1 | NA | NA | 1 | 1 | 2 | 1 | 4 | 1 | 1 | 1 | NA | NA | NA | NA | 1 | 1 | 2 | 1 |
| 5_86976767 | 1 | 1 | NA | NA | NA | NA | 1 | 1 | 2 | 1 | 1 | 1 | NA | NA | NA | NA | 1 | 1 | 2 | 1 |
| 5_86976830 | NA | NA | NA | NA | 1 | 1 | 2 | 1 | 3 | 1 | 1 | 1 | NA | NA | NA | NA | 1 | 1 | 2 | 1 |
| 5_86976913 | 1 | 1 | NA | NA | NA | NA | 2 | 1 | 3 | 1 | NA | NA | NA | NA | NA | NA | 1 | 1 | 1 | 1 |
| 5_86977030 | 1 | 0 | NA | NA | 1 | 1 | 2 | 1 | 4 | 0.75 | NA | NA | NA | NA | NA | NA | 1 | 1 | 1 | 1 |
| 5_86977088 | 1 | 1 | NA | NA | NA | NA | 2 | 1 | 3 | 1 | NA | NA | NA | NA | NA | NA | 1 | 1 | 1 | 1 |
| 5_86977182 | 1 | 1 | NA | NA | 1 | 1 | 1 | 1 | 3 | 1 | 1 | 1 | NA | NA | NA | NA | 1 | 1 | 2 | 1 |
| 5_86977268 | NA | NA | NA | NA | 1 | 0 | 2 | 1 | 3 | 0.6667 | 1 | 1 | NA | NA | NA | NA | 1 | 1 | 2 | 1 |
| 5_86977280 | 1 | 1 | NA | NA | 1 | 0 | 2 | 1 | 4 | 0.75 | NA | NA | NA | NA | NA | NA | 1 | 1 | 1 | 1 |
| 5_86983730 | 4 | 0.75 | NA | NA | 3 | 0.3333 | 12 | 0.3333 | 19 | 0.4211 | 2 | 0 | 9 | 0 | 7 | 0.2857 | 5 | 0 | 23 | 0.087 |
| 5_86983736 | 3 | 0.3333 | 1 | 0 | 5 | 0.2 | 11 | 0.2727 | 20 | 0.25 | 3 | 0 | 8 | 0 | 10 | 0.3 | 6 | 0.3333 | 27 | 0.1852 |
| 5_86983923 | 5 | 0.2 | 1 | 1 | 6 | 0.1667 | 14 | 0.6429 | 26 | 0.4615 | 3 | 0.6667 | 10 | 0.8 | 8 | 0.375 | 6 | 0 | 27 | 0.4815 |
| 5_88668618 | 1 | 0 | 11 | 0.0909 | 2 | 0 | NA | NA | 14 | 0.0714 | 6 | 0.1667 | NA | NA | NA | NA | 5 | 0 | 11 | 0.0909 |
| 5_88668685 | 1 | 0 | 11 | 0.3636 | 2 | 0 | NA | NA | 14 | 0.2857 | 7 | 0.1429 | NA | NA | NA | NA | 6 | 0.1667 | 13 | 0.1538 |
| 5_88668721 | NA | NA | 12 | 0.1667 | 2 | 0 | NA | NA | 14 | 0.1429 | 7 | 0.2857 | NA | NA | NA | NA | 4 | 0 | 11 | 0.1818 |
| 5_88702831 | 6 | 0 | 14 | 0.0714 | 4 | 0.5 | 4 | 0 | 28 | 0.1071 | 2 | 0 | 4 | 0.25 | 5 | 0 | 5 | 0 | 16 | 0.0625 |
| 5_88702879 | 6 | 1 | 15 | 0.9333 | 4 | 1 | 6 | 1 | 31 | 0.9677 | 2 | 1 | 4 | 0.75 | 5 | 0.8 | 4 | 1 | 15 | 0.8667 |
| 5_88703281 | 5 | 0.6 | 13 | 0.6923 | 3 | 0.3333 | 4 | 0.75 | 25 | 0.64 | 2 | 1 | 5 | 0.4 | 4 | 0.25 | 4 | 0.5 | 15 | 0.4667 |
| 5_88703342 | 6 | 1 | 14 | 0.7857 | 5 | 0.8 | 4 | 0.75 | 29 | 0.8276 | 3 | 0.6667 | 4 | 0.75 | 5 | 0.8 | 5 | 1 | 17 | 0.8235 |
| 5_88703411 | 5 | 0.6 | 13 | 0.6923 | 5 | 0.6 | 6 | 0.8333 | 29 | 0.6897 | 3 | 0.6667 | 5 | 0.4 | 6 | 0.6667 | 4 | 0.75 | 18 | 0.6111 |
| 5_88703472 | 5 | 0.2 | 15 | 0.6 | 5 | 0.2 | 6 | 0.3333 | 31 | 0.4194 | 2 | 1 | 5 | 0.2 | 5 | 0.2 | 6 | 0.5 | 18 | 0.3889 |
| 5_88703538 | 6 | 0.3333 | 15 | 0.3333 | 5 | 0.6 | 5 | 0.2 | 31 | 0.3548 | 2 | 0.5 | 5 | 0 | 7 | 0.4286 | 6 | 0.1667 | 20 | 0.25 |
| 5_88703702 | 5 | 0.8 | 15 | 1 | 3 | 1 | 5 | 0.8 | 28 | 0.9286 | 4 | 1 | 4 | 0.75 | 6 | 1 | 6 | 1 | 20 | 0.95 |
| 5_88703834 | 4 | 0.25 | 11 | 0.2727 | 3 | 0 | 5 | 0 | 23 | 0.1739 | 3 | 0.3333 | 4 | 0.25 | 4 | 0 | 4 | 0.5 | 15 | 0.2667 |
| 5_88788499 | 19 | 0.4737 | 27 | 0.1852 | 21 | 0.3333 | 22 | 0.3182 | 89 | 0.3146 | 13 | 0.3077 | 21 | 0.381 | 18 | 0.3889 | 15 | 0.2667 | 67 | 0.3433 |
| 5_88788565 | 20 | 0.55 | 30 | 0.3 | 21 | 0.2381 | 23 | 0.3043 | 94 | 0.3404 | 18 | 0.1667 | 20 | 0.55 | 14 | 0.2857 | 16 | 0 | 68 | 0.2647 |
| 5_88788625 | NA | NA | NA | NA | NA | NA | 1 | 1 | 1 | 1 | NA | NA | 1 | 1 | NA | NA | NA | NA | 1 | 1 |
| 5_88788652 | 21 | 0.8571 | 33 | 0.7879 | 19 | 0.8947 | 27 | 0.8889 | 100 | 0.85 | 17 | 0.8824 | 22 | 0.7727 | 18 | 0.8889 | 17 | 0.7647 | 74 | 0.8243 |
| 5_88788956 | 16 | 0.1875 | 19 | 0.4211 | 9 | 0.2222 | 11 | 0.1818 | 55 | 0.2727 | 9 | 0.3333 | 14 | 0.1429 | 12 | 0.4167 | 11 | 0.3636 | 46 | 0.3043 |
| 5_88797949 | 2 | 0.5 | 3 | 0.3333 | 1 | 0 | 3 | 0 | 9 | 0.2222 | 2 | 0 | 3 | 0 | 3 | 0.3333 | 3 | 0 | 11 | 0.0909 |
| 5_88798089 | 2 | 1 | 3 | 0.3333 | 2 | 0 | 3 | 0.6667 | 10 | 0.5 | 2 | 0.5 | 4 | 0 | 3 | 0.3333 | 3 | 0.3333 | 12 | 0.25 |
| 5_88798169 | 2 | 0 | 2 | 0 | 3 | 0.3333 | 2 | 0 | 9 | 0.1111 | 2 | 0 | 3 | 0 | 3 | 0 | 2 | 0.5 | 10 | 0.1 |
| 5_88798198 | 2 | 0 | 2 | 0.5 | 2 | 0 | 2 | 0 | 8 | 0.125 | 2 | 0 | 4 | 0 | 2 | 0 | 4 | 0.25 | 12 | 0.0833 |
| 5_88798484 | 2 | 0 | 3 | 0 | 3 | 0 | 3 | 0.3333 | 11 | 0.0909 | 2 | 0 | 4 | 0 | 3 | 0.3333 | 4 | 0 | 13 | 0.0769 |
| 5_88798751 | 3 | 0 | 2 | 0.5 | 3 | 0 | 3 | 0 | 11 | 0.0909 | 2 | 0.5 | 5 | 0 | 3 | 0 | 4 | 0.25 | 14 | 0.1429 |
| 5_88798878 | 3 | 0.6667 | 4 | 0.25 | 3 | 0.3333 | 2 | 0.5 | 12 | 0.4167 | 2 | 0 | 4 | 0.25 | 3 | 0.3333 | 3 | 0 | 12 | 0.1667 |
| 5_88799063 | 3 | 0.3333 | 3 | 0 | 2 | 0 | 1 | 0 | 9 | 0.1111 | NA | NA | 7 | 0 | 3 | 0.3333 | 4 | 0 | 14 | 0.0714 |
| 5_88805748 | NA | NA | 2 | 0.5 | NA | NA | 1 | 1 | 3 | 0.6667 | NA | NA | NA | NA | 1 | 1 | NA | NA | 1 | 1 |
| 5_88807617 | 12 | 0.1667 | 17 | 0.2353 | 18 | 0.2778 | 17 | 0.4706 | 64 | 0.2969 | 7 | 0 | 8 | 0.25 | 13 | 0.3077 | 17 | 0.0588 | 45 | 0.1556 |
| 5_88807688 | 12 | 0.1667 | 14 | 0.0714 | 15 | 0 | 10 | 0 | 51 | 0.0588 | 6 | 0 | 7 | 0.2857 | 10 | 0 | 16 | 0 | 39 | 0.0513 |
| 5_88807925 | 10 | 0.1 | 14 | 0.2143 | 16 | 0.125 | 15 | 0.2 | 55 | 0.1636 | 7 | 0 | 8 | 0.125 | 14 | 0.2857 | 17 | 0.1765 | 46 | 0.1739 |
| 5_88807990 | 10 | 0.5 | 15 | 0.1333 | 14 | 0.0714 | 14 | 0 | 53 | 0.1509 | 6 | 0.1667 | 6 | 0.3333 | 11 | 0.0909 | 17 | 0.1176 | 40 | 0.15 |
| 5_88808074 | 10 | 0.4 | 17 | 0.1765 | 16 | 0.1875 | 13 | 0.0769 | 56 | 0.1964 | 7 | 0 | 8 | 0.25 | 12 | 0.1667 | 17 | 0.1765 | 44 | 0.1591 |
| 5_88808106 | 9 | 0.2222 | 14 | 0.1429 | 15 | 0 | 14 | 0.0714 | 52 | 0.0962 | 7 | 0.1429 | 7 | 0.7143 | 11 | 0 | 15 | 0.1333 | 40 | 0.2 |
| 5_88904024 | 1 | 1 | 4 | 0.25 | 1 | 1 | 1 | 0 | 7 | 0.4286 | NA | NA | 3 | 0.6667 | 3 | 0.3333 | 2 | 0 | 8 | 0.375 |
| 5_88904205 | 1 | 1 | 3 | 0 | 1 | 0 | 1 | 0 | 6 | 0.1667 | NA | NA | 3 | 0.3333 | 3 | 0 | 2 | 0 | 8 | 0.125 |
| 5_88904354 | NA | NA | 4 | 0.25 | 1 | 0 | 1 | 0 | 6 | 0.1667 | NA | NA | 3 | 0.3333 | 3 | 0 | 1 | 0 | 7 | 0.1429 |
| 5_88904370 | 1 | 0 | 4 | 0.5 | 1 | 1 | NA | NA | 6 | 0.5 | NA | NA | 3 | 0.6667 | 3 | 0.3333 | 2 | 0.5 | 8 | 0.5 |
| 5_88904422 | 1 | 1 | 2 | 0 | 1 | 1 | 1 | 0 | 5 | 0.4 | NA | NA | 3 | 0 | 3 | 0.3333 | 2 | 0 | 8 | 0.125 |
| 5_88904487 | NA | NA | 4 | 0.75 | 1 | 1 | 1 | 0 | 6 | 0.6667 | NA | NA | 4 | 0.75 | 3 | 0.3333 | 2 | 0.5 | 9 | 0.5556 |
| 5_88908726 | 3 | 0.3333 | 6 | 0 | 2 | 0 | 1 | 0 | 12 | 0.0833 | 1 | 0 | 6 | 0.1667 | 11 | 0.0909 | 3 | 0 | 21 | 0.0952 |
| 5_88929518 | 4 | 0 | 1 | 0 | 4 | 0 | 2 | 0.5 | 11 | 0.0909 | 3 | 0 | 3 | 0.3333 | 2 | 0 | NA | NA | 8 | 0.125 |
| 5_89385894 | 13 | 0.0769 | 12 | 0 | 13 | 0.0769 | 13 | 0.0769 | 51 | 0.0588 | 9 | 0.1111 | 7 | 0 | 12 | 0.0833 | 14 | 0.0714 | 42 | 0.0714 |
| 5_89823909 | 7 | 0 | 4 | 0 | 8 | 0.25 | 7 | 0.1429 | 26 | 0.1154 | 5 | 0 | 7 | 0.1429 | 5 | 0.2 | 8 | 0.125 | 25 | 0.12 |
| 5_89824009 | 6 | 1 | 3 | 0.6667 | 7 | 0.8571 | 7 | 0.8571 | 23 | 0.8696 | 4 | 0.75 | 7 | 0.8571 | 5 | 1 | 8 | 0.875 | 24 | 0.875 |
| 5_89824124 | 5 | 0.2 | 3 | 0 | 8 | 0 | 6 | 0.1667 | 22 | 0.0909 | 2 | 0 | 6 | 0.1667 | 2 | 0.5 | 7 | 0.1429 | 17 | 0.1765 |
| 5_89824128 | 5 | 1 | 4 | 1 | 7 | 1 | 5 | 1 | 21 | 1 | 4 | 1 | 6 | 1 | 3 | 1 | 7 | 1 | 20 | 1 |
| 5_89824293 | 5 | 0.2 | 4 | 0 | 9 | 0 | 7 | 0.1429 | 25 | 0.08 | 5 | 0.2 | 5 | 0 | 3 | 0.3333 | 8 | 0.125 | 21 | 0.1429 |
| 5_89824309 | 5 | 1 | 4 | 1 | 9 | 0.8889 | 7 | 1 | 25 | 0.96 | 4 | 0.75 | 6 | 0.6667 | 3 | 1 | 8 | 0.875 | 21 | 0.8095 |
| 5_89824347 | 4 | 0 | 2 | 0 | 5 | 0 | 4 | 0.25 | 15 | 0.0667 | 4 | 0 | 5 | 0.2 | 2 | 0 | 8 | 0.125 | 19 | 0.1053 |
| 5_89824394 | 5 | 0.6 | 3 | 0.6667 | 8 | 0.875 | 6 | 0.8333 | 22 | 0.7727 | 5 | 1 | 6 | 1 | 4 | 1 | 8 | 0.625 | 23 | 0.8696 |

| | | | | | | | | | | | | | | | | | | | | |
|---|---|---|---|---|---|---|---|---|---|---|---|---|---|---|---|---|---|---|---|---|
| 5_8982468 | 4 | 0 | 4 | 0.25 | 6 | 0.1667 | 3 | 0.6667 | 17 | 0.2353 | 5 | 0.4 | 6 | 0.1667 | 3 | 0.3333 | 8 | 0.5 | 22 | 0.3636 |
| 5_8982488 | 3 | 0.3333 | 2 | 1 | 5 | 0.2 | 2 | 0.5 | 12 | 0.4167 | 2 | 0.5 | 4 | 0.25 | 1 | 0 | 5 | 0.2 | 12 | 0.25 |
| 5_8982489 | 3 | 1 | 3 | 0.6667 | 6 | 0.8333 | 3 | 0.6667 | 15 | 0.8 | 4 | 1 | 4 | 1 | 3 | 1 | 6 | 0.8333 | 17 | 0.9412 |
| 5_8982499 | 3 | 1 | 3 | 1 | 6 | 0.8333 | 2 | 1 | 14 | 0.9286 | 3 | 1 | 5 | 1 | 1 | 1 | 5 | 0.8 | 14 | 0.9286 |
| 5_8982509 | 4 | 0 | 4 | 0.25 | 7 | 0.4286 | 3 | 0.6667 | 18 | 0.3333 | 3 | 0 | 4 | 0.25 | 2 | 1 | 6 | 0.6667 | 15 | 0.4667 |
| 5_8982518 | 3 | 0 | 2 | 0 | 7 | 0.4286 | 3 | 1 | 15 | 0.4 | 3 | 0.3333 | 2 | 0.5 | 2 | 1 | 5 | 0.6 | 12 | 0.5833 |
| 5_8982521 | 5 | 0.2 | 3 | 0.6667 | 6 | 0.3333 | 3 | 0 | 17 | 0.2941 | 4 | 0.75 | 4 | 0.25 | 2 | 0 | 5 | 1 | 15 | 0.6 |
| 5_8982538 | 2 | 1 | 3 | 1 | 6 | 1 | 1 | 1 | 12 | 1 | 3 | 1 | 4 | 0.75 | 2 | 0.5 | 5 | 1 | 14 | 0.8571 |
| 5_8982541 | 5 | 0.8 | 3 | 0.6667 | 7 | 0.4286 | 1 | 1 | 16 | 0.625 | 3 | 1 | 3 | 1 | 2 | 0.5 | 3 | 1 | 11 | 0.9091 |
| 5_8982554 | 4 | 0.5 | 3 | 0.3333 | 6 | 0.1667 | 1 | 1 | 14 | 0.3571 | 3 | 0.6667 | 3 | 0.6667 | 2 | 0 | 4 | 0.25 | 12 | 0.4167 |
| 5_9000920 | 2 | 0.5 | 1 | 0 | 2 | 0 | NA | NA | 5 | 0.2 | 3 | 0.6667 | 1 | 0 | 1 | 1 | 2 | 0 | 7 | 0.4286 |
| 5_9000950 | 2 | 0 | 1 | 1 | 2 | 0.5 | 1 | 1 | 6 | 0.5 | 3 | 0.3333 | 1 | 0 | 1 | 0 | 2 | 0.5 | 7 | 0.2857 |
| 5_9036508 | NA | NA | NA | NA | 2 | 0.5 | 1 | 0 | 3 | 0.3333 | NA | NA | NA | NA | 1 | 0 | 1 | 1 | 2 | 0.5 |
| 5_9037639 | 3 | 0.3333 | 1 | 0 | 2 | 0 | 3 | 0.3333 | 9 | 0.2222 | 2 | 0.5 | 1 | 0 | 2 | 0 | 3 | 0.3333 | 8 | 0.25 |
| 5_9037688 | 4 | 0.25 | 11 | 0.1818 | 6 | 0 | 12 | 0.0833 | 33 | 0.1212 | 6 | 0.3333 | 4 | 0 | 8 | 0.125 | 9 | 0.2222 | 27 | 0.1852 |
| 5_9037698 | 6 | 0.3333 | 14 | 0.0714 | 7 | 0.2857 | 12 | 0.1667 | 39 | 0.1795 | 9 | 0.1111 | 6 | 0.3333 | 8 | 0.125 | 10 | 0.3 | 33 | 0.2121 |
| 5_9037700 | 5 | 0 | 14 | 0.1429 | 8 | 0 | 10 | 0.2 | 37 | 0.1081 | 9 | 0.2222 | 6 | 0.1667 | 8 | 0 | 9 | 0.1111 | 32 | 0.125 |
| 5_9037700 | 8 | 0.375 | 18 | 0.0556 | 8 | 0.125 | 14 | 0.2857 | 48 | 0.1875 | 9 | 0.4444 | 7 | 0.1429 | 10 | 0.2 | 13 | 0.0769 | 39 | 0.2051 |
| 5_9037703 | 8 | 0.125 | 18 | 0.2222 | 8 | 0.25 | 11 | 0.1818 | 45 | 0.2 | 10 | 0.1 | 6 | 0 | 9 | 0.1111 | 14 | 0.1429 | 39 | 0.1026 |
| 5_9039087 | 4 | 0.5 | 12 | 0.5833 | 6 | 0.5 | 10 | 0.7 | 32 | 0.5938 | 8 | 0.625 | 5 | 1 | 8 | 0.75 | 10 | 0.4 | 31 | 0.6452 |
| 5_9039105 | 4 | 0.5 | 10 | 0.7 | 4 | 0.75 | 9 | 0.3333 | 27 | 0.5556 | 7 | 0.2857 | 4 | 0.25 | 7 | 0.5714 | 6 | 0.6667 | 24 | 0.4583 |
| 5_9039111 | 4 | 0.5 | 10 | 0.5 | 5 | 0.2 | 10 | 0.2 | 29 | 0.3448 | 8 | 0.25 | 4 | 0.25 | 7 | 0.4286 | 9 | 0.1111 | 28 | 0.25 |
| 5_9039119 | 2 | 0 | 10 | 0.1 | 4 | 0.25 | 9 | 0 | 25 | 0.08 | 5 | 0 | 3 | 0.3333 | 6 | 0.1667 | 9 | 0.1111 | 23 | 0.1304 |
| 5_9039127 | 3 | 1 | 11 | 0.7273 | 5 | 0.8 | 10 | 0.7 | 29 | 0.7586 | 6 | 0.8333 | 4 | 0.75 | 6 | 0.6667 | 8 | 0.5 | 24 | 0.6667 |
| 5_9039157 | 2 | 1 | 9 | 0.7778 | 3 | 0 | 10 | 0.4 | 24 | 0.5417 | 4 | 0.25 | 2 | 1 | 3 | 0 | 7 | 0.2857 | 16 | 0.3125 |
| 5_9039186 | 3 | 0 | 8 | 0.25 | 4 | 0.25 | 8 | 0.125 | 23 | 0.1739 | 6 | 0 | 2 | 0 | 4 | 0.25 | 4 | 0 | 16 | 0.0625 |
| 5_9039198 | 3 | 0.6667 | 7 | 0.7143 | 4 | 0.75 | 10 | 0.6 | 24 | 0.6667 | 5 | 0.4 | 2 | 0.5 | 4 | 0.75 | 6 | 0.5 | 17 | 0.5294 |
| 5_9039213 | 3 | 0.3333 | 6 | 0.1667 | 4 | 0 | 11 | 0.1818 | 24 | 0.1667 | 5 | 0.4 | 2 | 0 | 4 | 0.25 | 4 | 0.5 | 15 | 0.3333 |
| 5_9039227 | 2 | 0.5 | 4 | 0.25 | 3 | 1 | 11 | 0.6364 | 20 | 0.6 | 6 | 0.1667 | 2 | 1 | 5 | 0.8 | 5 | 0.6 | 18 | 0.5556 |
| 5_9040153 | 2 | 0 | 3 | 0 | 2 | 0 | 9 | 0.1111 | 16 | 0.0625 | 4 | 0 | 1 | 0 | 5 | 0.2 | 2 | 0 | 12 | 0.0833 |
| 5_9040173 | 1 | 1 | 2 | 0.5 | 3 | 0 | 7 | 0.5714 | 13 | 0.4615 | 3 | 0.3333 | NA | NA | 2 | 0 | 2 | 1 | 7 | 0.4286 |
| 5_9043085 | NA | NA | 1 | 1 | NA | NA | 1 | 1 | 2 | 1 | 1 | 0 | NA | NA | 3 | 0.3333 | NA | NA | 4 | 0.25 |
| 5_9043096 | NA | NA | 1 | 1 | NA | NA | 1 | 1 | 2 | 1 | 1 | 0 | NA | NA | 2 | 0.5 | NA | NA | 3 | 0.3333 |
| 5_9043106 | NA | NA | 1 | 1 | NA | NA | 1 | 1 | 2 | 1 | 1 | 1 | NA | NA | 3 | 1 | NA | NA | 4 | 1 |
| 5_9043113 | NA | NA | 1 | 0 | NA | NA | 1 | 1 | 2 | 0.5 | 1 | 0 | NA | NA | 1 | 1 | NA | NA | 2 | 0.5 |
| 5_9107846 | NA | NA | 1 | 1 | 2 | 1 | 1 | 1 | 4 | 1 | NA | NA | NA | NA | 2 | 1 | NA | NA | 2 | 1 |
| 5_9107853 | NA | NA | 1 | 1 | 1 | 1 | 1 | 1 | 3 | 1 | NA | NA | 1 | 1 | 1 | 1 | NA | NA | 2 | 1 |
| 5_9107860 | NA | NA | 1 | 1 | 2 | 0.5 | 1 | 1 | 4 | 0.75 | NA | NA | NA | NA | 1 | 1 | NA | NA | 1 | 1 |
| 5_9107946 | NA | NA | NA | NA | 1 | 1 | 1 | 1 | 2 | 1 | NA | NA | NA | NA | 2 | 0.5 | NA | NA | 2 | 0.5 |
| 5_9107953 | NA | NA | NA | NA | 1 | 1 | 1 | 0 | 2 | 0.5 | NA | NA | NA | NA | 2 | 1 | NA | NA | 2 | 1 |
| 5_9116825 | 1 | 0 | 2 | 1 | 2 | 0 | NA | NA | 5 | 0.4 | 2 | 0.5 | NA | NA | NA | NA | 5 | 0.6 | 7 | 0.5714 |
| 5_9116875 | 1 | 1 | 2 | 0.5 | 2 | 0 | NA | NA | 5 | 0.4 | 2 | 0.5 | NA | NA | NA | NA | 6 | 0.3333 | 8 | 0.375 |
| 5_9116883 | 1 | 0 | 2 | 0.5 | 2 | 0 | NA | NA | 5 | 0.2 | 2 | 0.5 | NA | NA | NA | NA | 5 | 0 | 7 | 0.1429 |
| 5_9174175 | 2 | 0 | 1 | 1 | NA | NA | NA | NA | 3 | 0.3333 | NA | NA | NA | NA | 1 | 1 | NA | NA | 1 | 1 |
| 5_9177089 | 2 | 0.5 | 2 | 0 | NA | NA | NA | NA | 4 | 0.25 | 3 | 0.3333 | NA | NA | 1 | 0 | 1 | 0 | 5 | 0.2 |
| 5_9177120 | 1 | 1 | 2 | 1 | NA | NA | NA | NA | 3 | 1 | 2 | 1 | NA | NA | 2 | 1 | NA | NA | 4 | 1 |
| 5_9177342 | 4 | 0 | 5 | 0.2 | NA | NA | NA | NA | 9 | 0.1111 | 2 | 0.5 | 2 | 0.5 | 3 | 0 | 6 | 0 | 13 | 0.1538 |
| 5_9177362 | 3 | 0 | 4 | 0.5 | NA | NA | NA | NA | 7 | 0.2857 | 2 | 0 | 2 | 0.5 | 3 | 0 | 5 | 0.2 | 12 | 0.1667 |
| 5_9177367 | 4 | 0.25 | 3 | 0 | NA | NA | NA | NA | 7 | 0.1429 | 2 | 0 | 2 | 0 | 2 | 0 | 4 | 0.25 | 10 | 0.1 |
| 5_9177369 | 2 | 0.5 | 5 | 0.8 | NA | NA | NA | NA | 7 | 0.7143 | 2 | 0.5 | 2 | 1 | 3 | 0 | 5 | 0.4 | 12 | 0.4167 |
| 5_9177390 | 4 | 1 | 5 | 0.8 | NA | NA | NA | NA | 9 | 0.8889 | 2 | 0.5 | 1 | 1 | 2 | 0.5 | 7 | 0.7143 | 12 | 0.6667 |
| 5_9177406 | 3 | 0.3333 | 5 | 0.6 | NA | NA | NA | NA | 8 | 0.5 | 4 | 0 | 2 | 0 | 3 | 0 | 6 | 0.1667 | 15 | 0.0667 |
| 5_9177421 | 4 | 0.25 | 3 | 0 | NA | NA | NA | NA | 7 | 0.1429 | 2 | 0 | 3 | 0.3333 | 3 | 0 | 7 | 0.1429 | 15 | 0.1333 |
| 5_9177444 | 5 | 0.2 | 6 | 0.1667 | 1 | 0 | NA | NA | 12 | 0.1667 | 3 | 0.3333 | 2 | 0 | 3 | 0.6667 | 7 | 0.4286 | 15 | 0.4 |
| 5_9209700 | 1 | 0 | 3 | 0 | 1 | 0 | 1 | 1 | 6 | 0.1667 | 3 | 0.3333 | 2 | 0 | 1 | 0 | 2 | 0 | 8 | 0.125 |
| 5_9209728 | 4 | 0.75 | 6 | 0.1667 | 8 | 0.5 | 3 | 0 | 21 | 0.381 | 6 | 0.5 | 7 | 0.2857 | 5 | 0.4 | 4 | 0.25 | 22 | 0.3636 |
| 5_9209765 | 5 | 0 | 3 | 0.6667 | 6 | 0.1667 | 4 | 0.75 | 18 | 0.3333 | 6 | 0.3333 | 9 | 0 | 5 | 0.2 | 5 | 0.4 | 25 | 0.2 |
| 5_9211079 | NA | NA | NA | NA | NA | NA | 1 | 1 | 1 | 1 | 1 | 0 | 2 | 1 | 1 | 0 | 2 | 0 | 6 | 0.3333 |
| 5_9211754 | 2 | 0.5 | NA | NA | NA | NA | NA | NA | 2 | 0.5 | 1 | 0 | 1 | 1 | NA | NA | NA | NA | 2 | 0.5 |
| 5_9211755 | 2 | 0.5 | NA | NA | NA | NA | NA | NA | 2 | 0.5 | 1 | 1 | 1 | 1 | NA | NA | NA | NA | 2 | 1 |
| 5_9211769 | 2 | 0.5 | NA | NA | NA | NA | NA | NA | 2 | 0.5 | 1 | 0 | 1 | 1 | NA | NA | NA | NA | 2 | 0.5 |
| 5_9211781 | 2 | 0.5 | NA | NA | NA | NA | NA | NA | 2 | 0.5 | 1 | 1 | 1 | 1 | NA | NA | NA | NA | 2 | 1 |
| 5_9211971 | 2 | 0.5 | 1 | 0 | NA | NA | NA | NA | 3 | 0.3333 | 1 | 0 | 2 | 0.5 | NA | NA | NA | NA | 3 | 0.3333 |
| 5_9215430 | 6 | 0.1667 | NA | NA | 4 | 0 | 2 | 0 | 12 | 0.0833 | NA | NA | 3 | 0.3333 | 6 | 0 | 9 | 0 | 18 | 0.0556 |
| 5_9215472 | 7 | 0.2857 | NA | NA | 4 | 0 | 2 | 0 | 13 | 0.1538 | NA | NA | 2 | 0 | 5 | 0.6 | 8 | 0.125 | 15 | 0.2667 |

| | | | | | | | | | | | | | | | | | | | | |
|---|---|---|---|---|---|---|---|---|---|---|---|---|---|---|---|---|---|---|---|---|
| 5_9215538 | 6 | 0.1667 | NA | NA | 2 | 0 | 3 | 0 | 11 | 0.0909 | NA | NA | 2 | 0.5 | 3 | 0 | 6 | 0 | 11 | 0.0909 |
| 5_9215585 | 5 | 0.2 | 1 | 0 | 3 | 0.3333 | 3 | 0 | 12 | 0.1667 | NA | NA | 2 | 0 | 4 | 0 | 5 | 0.4 | 11 | 0.1818 |
| 5_9215592 | 6 | 0.3333 | 1 | 1 | 3 | 0 | 3 | 0.6667 | 13 | 0.3846 | NA | NA | 1 | 0 | 4 | 1 | 3 | 0.3333 | 8 | 0.625 |
| 5_9218509 | 2 | 0 | 1 | 1 | 2 | 1 | 1 | 0 | 6 | 0.5 | NA | NA | 1 | 0 | 1 | 0 | 2 | 0.5 | 4 | 0.25 |
| 5_9220032 | 29 | 0.1034 | 44 | 0.1136 | 28 | 0.1429 | 27 | 0.1111 | 128 | 0.1172 | 20 | 0.05 | 32 | 0.0625 | 32 | 0 | 32 | 0.0938 | 116 | 0.0517 |
| 5_9220033 | 32 | 0.6562 | 47 | 0.7872 | 33 | 0.7879 | 34 | 0.7647 | 146 | 0.7534 | 22 | 0.7273 | 40 | 0.825 | 35 | 0.7714 | 35 | 0.8 | 132 | 0.7879 |
| 5_9220035 | 32 | 0.125 | 48 | 0.0625 | 35 | 0.0857 | 32 | 0.125 | 147 | 0.0952 | 22 | 0.1818 | 43 | 0.2093 | 34 | 0.0294 | 33 | 0.0606 | 132 | 0.1212 |
| 5_9220046 | 31 | 0.0323 | 43 | 0.093 | 34 | 0.0588 | 33 | 0.0303 | 141 | 0.0567 | 18 | 0 | 43 | 0.0698 | 35 | 0.0857 | 35 | 0.0571 | 131 | 0.0611 |
| 5_9220092 | 29 | 0.2759 | 32 | 0.0625 | 29 | 0.2414 | 32 | 0.2812 | 122 | 0.2131 | 16 | 0.25 | 33 | 0.2424 | 31 | 0.2581 | 29 | 0.1034 | 109 | 0.211 |
| 5_9220093 | 29 | 0.2069 | 31 | 0.2903 | 25 | 0.2 | 30 | 0.3 | 115 | 0.2522 | 15 | 0.2 | 34 | 0.2353 | 29 | 0.1724 | 24 | 0.2083 | 102 | 0.2059 |
| 5_9220122 | 28 | 0.0714 | 28 | 0.0357 | 24 | 0.0833 | 29 | 0.0345 | 109 | 0.055 | 10 | 0.1 | 36 | 0.1667 | 31 | 0.129 | 31 | 0.0968 | 108 | 0.1296 |
| 5_9220126 | 22 | 0.5455 | 23 | 0.4348 | 17 | 0.2353 | 24 | 0.5417 | 86 | 0.4535 | 8 | 0.125 | 27 | 0.3704 | 26 | 0.5 | 25 | 0.48 | 86 | 0.4186 |
| 5_9220129 | 24 | 0.0833 | 29 | 0.2069 | 27 | 0.1852 | 30 | 0.0667 | 110 | 0.1364 | 9 | 0 | 35 | 0.2286 | 31 | 0.0968 | 24 | 0.2083 | 99 | 0.1616 |
| 5_9220141 | 20 | 0.35 | 24 | 0.2917 | 22 | 0.3182 | 27 | 0.4444 | 93 | 0.3548 | 8 | 0.375 | 31 | 0.5161 | 25 | 0.4 | 23 | 0.3913 | 87 | 0.4368 |
| 5_9220146 | 20 | 0 | 23 | 0 | 21 | 0.0952 | 26 | 0.1154 | 90 | 0.0556 | 8 | 0.125 | 30 | 0.1 | 28 | 0.1071 | 25 | 0.04 | 91 | 0.0879 |
| 5_9220192 | 16 | 0.1875 | 20 | 0.3 | 24 | 0.1667 | 22 | 0.1818 | 82 | 0.2073 | 6 | 0 | 25 | 0.16 | 21 | 0.0952 | 21 | 0.1905 | 73 | 0.137 |
| 5_9220198 | 23 | 0.3913 | 23 | 0.4783 | 24 | 0.4167 | 27 | 0.7407 | 97 | 0.5155 | 9 | 0.1111 | 31 | 0.4516 | 26 | 0.5 | 21 | 0.4762 | 87 | 0.4368 |
| 5_9220227 | 17 | 0.0588 | 20 | 0.1 | 20 | 0.05 | 27 | 0.037 | 84 | 0.0595 | 7 | 0 | 32 | 0.125 | 26 | 0.1923 | 17 | 0 | 82 | 0.1098 |
| 5_9220235 | 20 | 0.55 | 20 | 0.45 | 18 | 0.6111 | 28 | 0.6429 | 86 | 0.5698 | 7 | 0.2857 | 29 | 0.5517 | 26 | 0.5 | 18 | 0.5556 | 80 | 0.5125 |
| 5_9220237 | 16 | 0.75 | 15 | 0.7333 | 19 | 0.7368 | 23 | 0.8261 | 73 | 0.7671 | 7 | 0.7143 | 26 | 0.8846 | 26 | 0.8462 | 19 | 0.7895 | 78 | 0.8333 |
| 5_9220239 | 18 | 0.1667 | 18 | 0.2778 | 16 | 0.125 | 26 | 0.1923 | 78 | 0.1923 | 5 | 0 | 26 | 0.1538 | 25 | 0.2 | 16 | 0.1875 | 72 | 0.1667 |
| 5_9220241 | 20 | 1 | 18 | 0.7778 | 24 | 0.7917 | 26 | 0.8462 | 88 | 0.8523 | 6 | 0.5 | 32 | 0.7812 | 27 | 0.8148 | 18 | 0.9444 | 83 | 0.8072 |
| 5_9220243 | 20 | 0 | 19 | 0.0526 | 22 | 0.0455 | 29 | 0.1034 | 90 | 0.0556 | 7 | 0 | 31 | 0 | 26 | 0.1538 | 18 | 0.1111 | 82 | 0.0732 |
| 5_9220248 | 20 | 0.85 | 17 | 0.7647 | 24 | 0.7917 | 26 | 0.8077 | 87 | 0.8046 | 7 | 1 | 29 | 0.6552 | 25 | 0.8 | 17 | 0.8824 | 78 | 0.7821 |
| 5_9220265 | 17 | 0.1176 | 19 | 0.2105 | 23 | 0.2609 | 27 | 0.3333 | 86 | 0.2442 | 4 | 0 | 28 | 0.1429 | 25 | 0.24 | 18 | 0.2778 | 75 | 0.2 |
| 5_9220270 | 16 | 0.0625 | 17 | 0.1176 | 22 | 0.0455 | 26 | 0.1923 | 81 | 0.1111 | 4 | 0 | 28 | 0.1071 | 25 | 0.12 | 18 | 0.2778 | 75 | 0.1467 |
| 5_9234984 | 6 | 0.1667 | 5 | 0.2 | 1 | 0 | 6 | 0 | 18 | 0.1111 | 2 | 0 | 4 | 0 | 3 | 0 | 3 | 0.3333 | 12 | 0.0833 |
| 5_9234988 | 13 | 0.4615 | 8 | 0.375 | 3 | 0.3333 | 8 | 0.375 | 32 | 0.4062 | 5 | 0 | 8 | 0.25 | 3 | 0.6667 | 6 | 0 | 22 | 0.1818 |
| 5_9234990 | 15 | 0.3333 | 10 | 0.1 | 4 | 0 | 10 | 0.2 | 39 | 0.2051 | 6 | 0.3333 | 7 | 0.2857 | 8 | 0.375 | 6 | 0.1667 | 27 | 0.2963 |
| 5_9234995 | 11 | 0 | 9 | 0.1111 | 3 | 0 | 7 | 0.2857 | 30 | 0.1 | 4 | 0.25 | 7 | 0 | 6 | 0 | 3 | 0 | 20 | 0.05 |
| 5_9234996 | 14 | 0.1429 | 12 | 0.0833 | 4 | 0 | 9 | 0.2222 | 39 | 0.1282 | 6 | 0.1667 | 9 | 0.1111 | 7 | 0 | 7 | 0.1429 | 29 | 0.1034 |
| 5_9234997 | 11 | 0.1818 | 10 | 0.4 | 4 | 0.5 | 8 | 0.625 | 33 | 0.3939 | 6 | 0.5 | 8 | 0.625 | 8 | 0.625 | 5 | 0.8 | 27 | 0.6296 |
| 5_9234999 | 11 | 0.2727 | 7 | 0.1429 | 4 | 0.5 | 10 | 0.3 | 32 | 0.2812 | 4 | 0.5 | 7 | 0 | 5 | 0.4 | 6 | 0 | 22 | 0.1818 |
| 5_9235021 | 13 | 0.3846 | 9 | 0.3333 | 5 | 0.4 | 12 | 0.5 | 39 | 0.4103 | 6 | 0.1667 | 6 | 0.3333 | 7 | 0.4286 | 8 | 0.625 | 27 | 0.4074 |
| 5_9235029 | 11 | 0.2727 | 12 | 0.1667 | 5 | 0.2 | 9 | 0.1111 | 37 | 0.1892 | 4 | 0 | 8 | 0.125 | 9 | 0.1111 | 6 | 0.3333 | 27 | 0.1481 |
| 5_9235038 | 13 | 0 | 13 | 0.0769 | 5 | 0 | 12 | 0.25 | 43 | 0.093 | 7 | 0 | 10 | 0.1 | 10 | 0.1 | 9 | 0.1111 | 36 | 0.0833 |
| 5_9237470 | 1 | 0 | 1 | 1 | NA | NA | 1 | 1 | 3 | 0.6667 | NA | NA | NA | NA | 1 | 1 | NA | NA | 1 | 1 |
| 5_9240628 | 8 | 0.375 | 8 | 0.25 | 7 | 0.1429 | 8 | 0 | 31 | 0.1935 | 5 | 0.2 | 6 | 0.1667 | 8 | 0.25 | 2 | 0 | 21 | 0.1905 |
| 5_9240648 | 7 | 0 | 9 | 0.1111 | 7 | 0.1429 | 9 | 0 | 32 | 0.0625 | 4 | 0 | 7 | 0 | 8 | 0.25 | 3 | 0 | 22 | 0.0909 |
| 5_9240675 | 8 | 0.25 | 8 | 0.125 | 8 | 0.125 | 8 | 0.375 | 32 | 0.2188 | 6 | 0.3333 | 6 | 0 | 8 | 0.25 | 3 | 0 | 23 | 0.1739 |
| 5_9254053 | NA | NA | NA | NA | 1 | 0 | 1 | 1 | 2 | 0.5 | 2 | 1 | NA | NA | NA | NA | NA | NA | 2 | 1 |
| 5_9254058 | 1 | 0 | NA | NA | 1 | 1 | 1 | 0 | 3 | 0.3333 | 2 | 1 | NA | NA | NA | NA | NA | NA | 2 | 1 |
| 5_9254073 | 1 | 1 | NA | NA | 1 | 1 | 1 | 1 | 3 | 1 | 1 | 1 | NA | NA | NA | NA | NA | NA | 1 | 1 |
| 5_9256225 | 2 | 1 | NA | NA | 2 | 0.5 | 3 | 0.6667 | 7 | 0.7143 | 2 | 0.5 | NA | NA | 1 | 0 | NA | NA | 3 | 0.3333 |
| 5_9256233 | 2 | 0.5 | NA | NA | 1 | 0 | 1 | 1 | 4 | 0.5 | 2 | 0 | NA | NA | 1 | 1 | NA | NA | 3 | 0.3333 |
| 5_9256357 | 3 | 0 | 1 | 0 | NA | NA | 3 | 0.6667 | 7 | 0.2857 | NA | NA | 3 | 0 | 1 | 1 | 1 | 0 | 5 | 0.2 |
| 5_9256405 | 3 | 0.6667 | NA | NA | NA | NA | 3 | 1 | 6 | 0.8333 | 1 | 0 | 3 | 0.6667 | 1 | 1 | 1 | 1 | 6 | 0.6667 |
| 5_9258929 | 16 | 0.4375 | 14 | 0.0714 | 13 | 0.2308 | 14 | 0.2143 | 57 | 0.2456 | 19 | 0.2632 | 12 | 0.25 | 18 | 0.3889 | 11 | 0.4545 | 60 | 0.3333 |
| 5_9259210 | 7 | 0.1429 | 4 | 0.25 | 6 | 0.6667 | 9 | 0.5556 | 26 | 0.4231 | 6 | 0.3333 | 4 | 0.25 | 5 | 0.4 | 4 | 0.5 | 19 | 0.3684 |
| 5_9311253 | NA | NA | 1 | 0 | 1 | 0 | 1 | 1 | 3 | 0.3333 | NA | NA | NA | NA | NA | NA | 1 | 1 | 1 | 1 |
| 5_9318941 | 2 | 0 | 3 | 0.6667 | NA | NA | 1 | 0 | 6 | 0.3333 | 3 | 0 | NA | NA | 2 | 0.5 | 1 | 0 | 6 | 0.1667 |
| 5_9318944 | 2 | 1 | 2 | 1 | NA | NA | 1 | 1 | 5 | 1 | 2 | 1 | NA | NA | 2 | 1 | 1 | 0 | 5 | 0.8 |
| 5_9318959 | 2 | 0 | 3 | 0.6667 | NA | NA | 1 | 0 | 6 | 0.3333 | 2 | 0 | NA | NA | 1 | 1 | 1 | 0 | 4 | 0.25 |
| 5_9318959 | 2 | 0.5 | 2 | 0 | NA | NA | 1 | 0 | 5 | 0.2 | 3 | 0.6667 | NA | NA | 2 | 0.5 | 1 | 0 | 6 | 0.5 |
| 5_9318970 | 1 | 0 | 2 | 1 | NA | NA | 1 | 1 | 4 | 0.75 | 3 | 1 | NA | NA | 2 | 0.5 | 1 | 1 | 6 | 0.8333 |
| 5_9318973 | 1 | 0 | 2 | 1 | NA | NA | 1 | 1 | 4 | 0.75 | 2 | 0.5 | NA | NA | 2 | 0 | 1 | 0 | 5 | 0.2 |
| 5_9318975 | 1 | 1 | 3 | 0.6667 | NA | NA | 1 | 1 | 5 | 0.8 | 3 | 0.6667 | NA | NA | 2 | 0.5 | 1 | 1 | 6 | 0.6667 |
| 5_9318980 | 2 | 1 | 1 | 0 | NA | NA | 1 | 1 | 4 | 0.75 | 2 | 1 | NA | NA | 1 | 0 | NA | NA | 3 | 0.6667 |
| 5_9318980 | 1 | 0 | 3 | 0 | NA | NA | 1 | 1 | 5 | 0.2 | 2 | 1 | NA | NA | 1 | 0 | 1 | 0 | 4 | 0.5 |
| 5_9319004 | 2 | 0 | 2 | 0 | NA | NA | 1 | 1 | 5 | 0.2 | 2 | 0 | NA | NA | 2 | 0.5 | 1 | 0 | 5 | 0.2 |
| 5_9319021 | 2 | 1 | 2 | 1 | NA | NA | NA | NA | 4 | 1 | 3 | 0.3333 | NA | NA | 2 | 1 | 1 | 0 | 6 | 0.5 |
| 5_9319022 | 1 | 1 | 2 | 1 | NA | NA | 1 | 1 | 4 | 1 | 3 | 1 | NA | NA | 2 | 1 | 1 | 1 | 6 | 1 |
| 5_9319030 | 2 | 0.5 | 3 | 1 | NA | NA | 1 | 0 | 6 | 0.6667 | 3 | 0.3333 | NA | NA | 2 | 0.5 | 1 | 0 | 6 | 0.3333 |
| 5_9319043 | 2 | 1 | 3 | 1 | NA | NA | 1 | 1 | 6 | 1 | 2 | 1 | NA | NA | 1 | 1 | 1 | 0 | 4 | 0.75 |
| 5_9319068 | 1 | 1 | 3 | 0 | NA | NA | 1 | 0 | 5 | 0.2 | 1 | 1 | NA | NA | 2 | 0.5 | NA | NA | 3 | 0.6667 |

| | | | | | | | | | | | | | | | | | | | | |
|---|---|---|---|---|---|---|---|---|---|---|---|---|---|---|---|---|---|---|---|---|
| 5_9319071 | 1 | 1 | 3 | 1 | NA | NA | 1 | 1 | 5 | 1 | 1 | 1 | NA | NA | 2 | 0.5 | NA | NA | 3 | 0.6667 |
| 5_9319074 | 1 | 1 | 2 | 1 | NA | NA | NA | NA | 3 | 1 | 1 | 1 | NA | NA | 2 | 1 | 1 | 0 | 4 | 0.75 |
| 5_9319093 | 2 | 1 | 3 | 1 | NA | NA | NA | NA | 5 | 1 | 1 | 1 | NA | NA | 2 | 1 | 1 | 1 | 4 | 1 |
| 5_9319097 | 2 | 1 | 2 | 1 | NA | NA | 1 | 1 | 5 | 1 | 1 | 1 | NA | NA | 2 | 1 | NA | NA | 3 | 1 |
| 5_9319125 | 2 | 1 | 2 | 1 | NA | NA | 1 | 1 | 5 | 1 | 1 | 1 | NA | NA | 2 | 0.5 | NA | NA | 3 | 0.6667 |
| 5_9319151 | NA | NA | 2 | 0.5 | NA | NA | NA | NA | 2 | 0.5 | NA | NA | NA | NA | 2 | 1 | NA | NA | 2 | 1 |
| 5_9319156 | 2 | 0.5 | 3 | 0.3333 | NA | NA | 1 | 0 | 6 | 0.3333 | 1 | 1 | NA | NA | 2 | 0 | NA | NA | 3 | 0.3333 |
| 5_9319161 | 2 | 0 | 2 | 0.5 | NA | NA | 1 | 0 | 5 | 0.2 | 1 | 1 | NA | NA | 2 | 0.5 | NA | NA | 3 | 0.6667 |
| 5_9319248 | 2 | 0.5 | 2 | 0 | NA | NA | NA | NA | 4 | 0.25 | 1 | 1 | NA | NA | 1 | 1 | NA | NA | 2 | 1 |
| 5_9331557 | 2 | 0.5 | NA | NA | 1 | 0 | NA | NA | 3 | 0.3333 | 2 | 0 | NA | NA | NA | NA | 3 | 0.3333 | 5 | 0.2 |
| 5_9331562 | 2 | 1 | NA | NA | 1 | 1 | NA | NA | 3 | 1 | 2 | 0.5 | NA | NA | 1 | 1 | 2 | 1 | 5 | 0.8 |
| 5_9331730 | 2 | 0 | 2 | 0.5 | NA | NA | NA | NA | 4 | 0.25 | 2 | 0.5 | NA | NA | 1 | 0 | 4 | 0 | 7 | 0.1429 |
| 5_9331950 | NA | NA | 3 | 0 | 1 | 0 | 2 | 0.5 | 6 | 0.1667 | 5 | 0.2 | 1 | 1 | NA | NA | 2 | 0 | 8 | 0.25 |
| 5_9332018 | 2 | 0.5 | 3 | 0.3333 | 1 | 0 | 4 | 0.25 | 10 | 0.3 | 5 | 0.2 | 4 | 0 | NA | NA | 3 | 0 | 12 | 0.0833 |
| 5_9332185 | 1 | 0 | 5 | 0 | NA | NA | 5 | 0.4 | 11 | 0.1818 | 2 | 0 | 5 | 0 | 2 | 0 | 1 | 1 | 10 | 0.1 |
| 5_9332190 | 4 | 0 | 6 | 0.1667 | 1 | 0 | 5 | 0.4 | 16 | 0.1875 | 5 | 0.2 | 6 | 0 | 2 | 0 | 6 | 0.1667 | 19 | 0.1053 |
| 5_9332246 | 6 | 0 | 7 | 0.1429 | 2 | 0 | 6 | 0.1667 | 21 | 0.0952 | 5 | 0.2 | 9 | 0 | 3 | 0.3333 | 7 | 0 | 24 | 0.0833 |
| 5_9332331 | 3 | 0 | 4 | 0.25 | 5 | 0 | 6 | 0 | 18 | 0.0556 | 6 | 0.1667 | 10 | 0.1 | 7 | 0.2857 | 6 | 0 | 29 | 0.1379 |
| 5_9332347 | 3 | 0 | 5 | 0 | 5 | 0.2 | 6 | 0.3333 | 19 | 0.1579 | 5 | 0.4 | 15 | 0.1333 | 6 | 0.1667 | 5 | 0.2 | 31 | 0.1935 |
| 5_9332361 | 6 | 0.5 | 5 | 0.2 | 8 | 0.5 | 4 | 0.5 | 23 | 0.4348 | 7 | 0.4286 | 13 | 0.3846 | 8 | 0.125 | 6 | 0.1667 | 34 | 0.2941 |
| 5_9332397 | 7 | 0.1429 | 5 | 0.2 | 9 | 0 | 5 | 0.4 | 26 | 0.1538 | 6 | 0 | 12 | 0 | 6 | 0 | 5 | 0.4 | 29 | 0.069 |
| 5_9332845 | 1 | 1 | 1 | 1 | NA | NA | 1 | 0 | 3 | 0.6667 | NA | NA | 1 | 1 | 1 | 0 | NA | NA | 2 | 0.5 |
| 5_9332894 | 1 | 1 | 1 | 1 | NA | NA | 1 | 0 | 3 | 0.6667 | NA | NA | 1 | 1 | NA | NA | NA | NA | 1 | 1 |
| 5_9333060 | NA | NA | NA | NA | 3 | 0 | 1 | 1 | 4 | 0.25 | NA | NA | NA | NA | 3 | 0.3333 | NA | NA | 3 | 0.3333 |
| 5_9333073 | 32 | 0.0312 | 33 | 0.1818 | 43 | 0.2093 | 23 | 0.2609 | 131 | 0.1679 | 35 | 0.1143 | 27 | 0.1111 | 31 | 0.0968 | 29 | 0.1379 | 122 | 0.1148 |
| 5_9333073 | 30 | 0.2333 | 32 | 0.3438 | 42 | 0.1905 | 26 | 0.2692 | 130 | 0.2538 | 32 | 0.1562 | 29 | 0.2069 | 30 | 0.1333 | 35 | 0.1429 | 126 | 0.1587 |
| 5_9333093 | 25 | 0.6 | 32 | 0.5625 | 40 | 0.425 | 22 | 0.4545 | 119 | 0.5042 | 34 | 0.4706 | 22 | 0.4545 | 29 | 0.3793 | 29 | 0.3448 | 114 | 0.4123 |
| 5_9333109 | 28 | 0.0714 | 31 | 0.1613 | 41 | 0.0732 | 24 | 0.125 | 124 | 0.1048 | 34 | 0.2059 | 25 | 0.12 | 32 | 0.2188 | 33 | 0.0909 | 124 | 0.1613 |
| 5_9333313 | NA | NA | 1 | 1 | 1 | 0 | NA | NA | 2 | 0.5 | 1 | 0 | NA | NA | NA | NA | 2 | 0.5 | 3 | 0.3333 |
| 5_9342330 | 5 | 0.2 | 7 | 0 | 5 | 0 | 14 | 0.0714 | 31 | 0.0645 | 7 | 0 | 18 | 0.1111 | 9 | 0.2222 | 8 | 0.125 | 42 | 0.119 |
| 5_9621850 | 6 | 0.5 | 4 | 0.5 | 1 | 1 | 2 | 0 | 13 | 0.4615 | 3 | 0.6667 | NA | NA | 4 | 0 | 5 | 0 | 12 | 0.1667 |
| 5_9621906 | 4 | 0.25 | 2 | 0.5 | 1 | 1 | 2 | 0.5 | 9 | 0.4444 | 4 | 0.25 | 1 | 1 | 5 | 0.4 | 4 | 0.25 | 14 | 0.3571 |
| 5_9622285 | 1 | 0 | 3 | 0 | 3 | 0.3333 | 3 | 0.6667 | 10 | 0.3 | 3 | 0.3333 | 3 | 0.3333 | 3 | 0 | 1 | 0 | 10 | 0.2 |
| 5_9622499 | 3 | 1 | 5 | 0.8 | 3 | 1 | 2 | 1 | 13 | 0.9231 | 2 | 1 | 2 | 1 | NA | NA | 1 | 1 | 5 | 1 |
| 5_9622509 | 2 | 0 | 5 | 0.8 | 2 | 0.5 | 1 | 1 | 10 | 0.6 | 1 | 0 | 1 | 1 | NA | NA | 1 | 0 | 3 | 0.3333 |
| 5_9622522 | 2 | 1 | 4 | 0.5 | 3 | 0.3333 | 1 | 1 | 10 | 0.6 | 2 | 0.5 | 2 | 0.5 | NA | NA | 1 | 0 | 5 | 0.4 |
| 5_9622791 | 2 | 0.5 | 3 | 0 | 3 | 0.3333 | NA | NA | 8 | 0.25 | NA | NA | 2 | 0 | NA | NA | 1 | 1 | 3 | 0.3333 |
| 5_9641226 | 4 | 0 | 2 | 0 | NA | NA | 4 | 0.25 | 10 | 0.1 | 2 | 0 | NA | NA | 3 | 0 | 1 | 1 | 6 | 0.1667 |
| 5_9641365 | 1 | 0 | NA | NA | NA | NA | 1 | 1 | 2 | 0.5 | 2 | 0 | NA | NA | 1 | 1 | 2 | 0 | 5 | 0.2 |
| 5_9692915 | 2 | 0.5 | NA | NA | 2 | 1 | 5 | 0.4 | 9 | 0.5556 | NA | NA | 3 | 1 | 4 | 0.5 | NA | NA | 7 | 0.7143 |
| 5_9692916 | 1 | 0 | NA | NA | 2 | 0 | 7 | 0.2857 | 10 | 0.2 | NA | NA | 3 | 0.3333 | 4 | 0 | NA | NA | 7 | 0.1429 |
| 5_9692918 | 1 | 0 | NA | NA | 2 | 0.5 | 5 | 0.2 | 8 | 0.25 | NA | NA | 3 | 0 | 3 | 0.3333 | NA | NA | 6 | 0.1667 |
| 5_9692925 | 2 | 0.5 | NA | NA | 2 | 0 | 6 | 0.1667 | 10 | 0.2 | NA | NA | 3 | 0.6667 | 4 | 0 | NA | NA | 7 | 0.2857 |
| 5_9692929 | 2 | 0 | NA | NA | 2 | 0.5 | 7 | 0.1429 | 11 | 0.1818 | NA | NA | 3 | 0 | 4 | 0.25 | NA | NA | 7 | 0.1429 |
| 5_9692941 | 2 | 1 | NA | NA | 2 | 1 | 6 | 1 | 10 | 1 | NA | NA | 4 | 1 | 4 | 1 | NA | NA | 8 | 1 |
| 5_9692949 | 1 | 1 | NA | NA | 2 | 1 | 7 | 0.7143 | 10 | 0.8 | NA | NA | 4 | 0.75 | 2 | 1 | NA | NA | 6 | 0.8333 |
| 5_9692953 | 2 | 0.5 | NA | NA | 2 | 0.5 | 7 | 0.5714 | 11 | 0.5455 | NA | NA | 4 | 1 | 4 | 0.25 | NA | NA | 8 | 0.625 |
| 5_9692957 | 2 | 0.5 | NA | NA | 2 | 0.5 | 5 | 1 | 9 | 0.7778 | NA | NA | 3 | 1 | 2 | 1 | NA | NA | 5 | 1 |
| 5_9693090 | 3 | 0 | 1 | 0 | 2 | 0 | 6 | 0.1667 | 12 | 0.0833 | 3 | 0.3333 | 4 | 0 | 5 | 0.4 | NA | NA | 12 | 0.25 |
| 5_9693181 | 4 | 0.25 | 1 | 0 | 4 | 0 | 8 | 0.125 | 17 | 0.1176 | 3 | 0 | 4 | 0 | 6 | 0.1667 | NA | NA | 13 | 0.0769 |
| 5_9693183 | 3 | 0.3333 | 1 | 1 | 5 | 0.2 | 8 | 0.125 | 17 | 0.2353 | 3 | 0.6667 | 3 | 0 | 6 | 0.3333 | NA | NA | 12 | 0.3333 |
| 5_9693202 | 3 | 0.3333 | 1 | 1 | 5 | 0.6 | 8 | 0.375 | 17 | 0.4706 | 3 | 0.6667 | 3 | 0.3333 | 7 | 0.7143 | NA | NA | 13 | 0.6154 |
| 5_9693215 | 4 | 0.25 | 1 | 0 | 4 | 0 | 8 | 0.125 | 17 | 0.1176 | 1 | 0 | 4 | 0.25 | 6 | 0 | NA | NA | 11 | 0.0909 |
| 5_9693241 | 4 | 0.25 | NA | NA | 6 | 0.3333 | 8 | 0.25 | 18 | 0.2778 | 3 | 1 | 5 | 0.2 | 7 | 0.2857 | NA | NA | 15 | 0.4 |
| 5_9699357 | 3 | 0.6667 | 2 | 0.5 | 2 | 0.5 | 4 | 0.75 | 11 | 0.6364 | 4 | 0.5 | 2 | 0.5 | 2 | 0 | 4 | 0.75 | 12 | 0.5 |
| 5_9721638 | 1 | 0 | 2 | 0 | 2 | 0.5 | 1 | 0 | 6 | 0.1667 | 1 | 1 | 1 | 0 | 2 | 0 | 1 | 1 | 5 | 0.4 |
| 5_9721642 | NA | NA | 1 | 0 | 2 | 0.5 | 1 | 1 | 4 | 0.5 | 1 | 1 | NA | NA | 1 | 1 | 2 | 0.5 | 4 | 0.75 |
| 5_9721648 | 1 | 0 | 1 | 1 | 2 | 1 | 1 | 1 | 5 | 0.8 | 1 | 1 | NA | NA | 1 | 1 | 1 | 0 | 3 | 0.6667 |
| 5_9722969 | NA | NA | NA | NA | NA | NA | 1 | 1 | 1 | 1 | 1 | 1 | NA | NA | NA | NA | NA | NA | 1 | 1 |
| 5_9722981 | NA | NA | 1 | 1 | NA | NA | 1 | 0 | 2 | 0.5 | 1 | 0 | 1 | 0 | NA | NA | 2 | 0.5 | 4 | 0.25 |
| 5_9722987 | NA | NA | 1 | 1 | 1 | 0 | NA | NA | 2 | 0.5 | 1 | 1 | 2 | 0 | NA | NA | 1 | 1 | 4 | 0.5 |
| 5_9722997 | NA | NA | 1 | 1 | 1 | 1 | 1 | 1 | 3 | 1 | 1 | 1 | 1 | 1 | NA | NA | 1 | 0 | 3 | 0.6667 |
| 5_9723019 | NA | NA | 1 | 0 | 1 | 1 | 1 | 1 | 3 | 0.6667 | 1 | 1 | NA | NA | NA | NA | 1 | 1 | 2 | 1 |
| 5_9723069 | 1 | 1 | 1 | 1 | NA | NA | NA | NA | 2 | 1 | NA | NA | 1 | 1 | NA | NA | 2 | 1 | 3 | 1 |
| 5_9723073 | NA | NA | 1 | 1 | NA | NA | 1 | 1 | 2 | 1 | NA | NA | NA | NA | NA | NA | 1 | 1 | 1 | 1 |

| | | | | | | | | | | | | | | | | | | | | |
|---|---|---|---|---|---|---|---|---|---|---|---|---|---|---|---|---|---|---|---|---|
| 5_97230739 | NA | NA | 1 | 0 | NA | NA | 1 | 1 | 2 | 0.5 | NA | NA | 1 | 1 | NA | NA | 1 | 0 | 2 | 0.5 |
| 5_97230895 | NA | NA | 1 | 1 | NA | NA | 1 | 0 | 2 | 0.5 | NA | NA | 1 | 0 | NA | NA | 2 | 0.5 | 3 | 0.3333 |
| 5_97230964 | NA | NA | 1 | 0 | NA | NA | 1 | 1 | 2 | 0.5 | NA | NA | 1 | 0 | NA | NA | 2 | 0.5 | 3 | 0.3333 |
| 5_97231044 | NA | NA | 1 | 1 | NA | NA | 1 | 1 | 2 | 1 | NA | NA | 1 | 1 | NA | NA | 1 | 1 | 2 | 1 |
| 5_97231387 | NA | NA | 1 | 1 | NA | NA | 1 | 1 | 2 | 1 | NA | NA | 1 | 0 | NA | NA | 1 | 1 | 2 | 0.5 |
| 5_97231422 | NA | NA | 1 | 0 | NA | NA | 1 | 1 | 2 | 0.5 | NA | NA | 1 | 1 | NA | NA | 1 | 0 | 2 | 0.5 |
| 5_97231689 | NA | NA | 1 | 1 | NA | NA | 1 | 1 | 2 | 1 | NA | NA | NA | NA | NA | NA | 1 | 1 | 1 | 1 |
| 5_97232162 | NA | NA | 1 | 1 | NA | NA | 1 | 1 | 2 | 1 | NA | NA | 1 | 0 | NA | NA | 1 | 1 | 2 | 0.5 |
| 5_97232271 | NA | NA | NA | NA | NA | NA | 1 | 1 | 1 | 1 | NA | NA | 1 | 1 | NA | NA | 1 | 0 | 2 | 0.5 |
| 5_97232319 | NA | NA | NA | NA | NA | NA | 1 | 1 | 1 | 1 | NA | NA | 1 | 0 | NA | NA | 1 | 1 | 2 | 0.5 |
| 5_97232615 | NA | NA | NA | NA | NA | NA | 1 | 1 | 1 | 1 | NA | NA | 1 | 1 | NA | NA | 1 | 0 | 2 | 0.5 |
| 5_97232950 | NA | NA | 1 | 1 | NA | NA | NA | NA | 1 | 1 | NA | NA | 1 | 1 | NA | NA | 1 | 0 | 2 | 0.5 |
| 5_97233127 | NA | NA | 1 | 1 | NA | NA | NA | NA | 1 | 1 | NA | NA | 1 | 1 | NA | NA | 1 | 1 | 2 | 1 |
| 5_97233270 | NA | NA | 1 | 1 | NA | NA | NA | NA | 1 | 1 | NA | NA | 1 | 1 | NA | NA | NA | NA | 1 | 1 |
| 5_97234914 | NA | NA | 1 | 1 | NA | NA | 1 | 1 | 2 | 1 | NA | NA | 1 | 1 | 1 | 1 | NA | NA | 2 | 1 |
| 5_97235090 | 1 | 1 | 1 | 0 | 2 | 0.5 | NA | NA | 4 | 0.5 | NA | NA | 1 | 0 | 1 | 1 | 1 | 1 | 3 | 0.6667 |
| 5_97235171 | 1 | 1 | 2 | 1 | 1 | 1 | NA | NA | 4 | 1 | 1 | 1 | 1 | 1 | 1 | 1 | 2 | 1 | 5 | 1 |
| 5_97235210 | 1 | 0 | 1 | 0 | 2 | 0.5 | NA | NA | 4 | 0.25 | 1 | 0 | 1 | 1 | 1 | 0 | 1 | 0 | 4 | 0.25 |
| 5_97235250 | NA | NA | 1 | 1 | NA | NA | 1 | 1 | 2 | 1 | NA | NA | 1 | 1 | 1 | 1 | NA | NA | 2 | 1 |
| 5_97235376 | 2 | 0.5 | 1 | 0 | 1 | 0 | NA | NA | 4 | 0.25 | 1 | 0 | 1 | 1 | 1 | 0 | 1 | 0 | 4 | 0.25 |
| 5_97235405 | NA | NA | 1 | 0 | NA | NA | 1 | 1 | 2 | 0.5 | NA | NA | 1 | 1 | 1 | 0 | NA | NA | 2 | 0.5 |
| 5_97235430 | 2 | 0.5 | 2 | 1 | 1 | 1 | NA | NA | 5 | 0.8 | 1 | 1 | 1 | 0 | 1 | 0 | NA | NA | 3 | 0.3333 |
| 5_97235489 | 2 | 1 | 1 | 1 | 1 | 1 | NA | NA | 4 | 1 | 1 | 1 | NA | NA | 1 | 1 | NA | NA | 2 | 1 |
| 5_97235604 | NA | NA | 1 | 1 | NA | NA | 1 | 1 | 2 | 1 | NA | NA | 1 | 1 | 1 | 1 | NA | NA | 2 | 1 |
| 5_97235624 | 1 | 1 | 1 | 1 | NA | NA | NA | NA | 2 | 1 | 1 | 1 | 1 | 1 | NA | NA | 1 | 1 | 3 | 1 |
| 5_97235694 | 2 | 0 | 3 | 0.3333 | 1 | 1 | NA | NA | 6 | 0.3333 | 1 | 1 | 1 | 1 | 1 | 1 | 1 | 0 | 4 | 0.75 |
| 5_97235713 | 2 | 0.5 | 2 | 1 | NA | NA | NA | NA | 4 | 0.75 | 1 | 1 | 1 | 1 | 1 | 1 | NA | NA | 3 | 1 |
| 5_97235721 | NA | NA | 1 | 1 | NA | NA | 1 | 1 | 2 | 1 | NA | NA | 1 | 1 | 1 | 0 | NA | NA | 2 | 0.5 |
| 5_97235754 | NA | NA | NA | NA | NA | NA | 1 | 1 | 1 | 1 | NA | NA | 1 | 1 | 1 | 0 | NA | NA | 2 | 0.5 |
| 5_97235867 | NA | NA | 1 | 1 | NA | NA | 1 | 1 | 2 | 1 | NA | NA | 1 | 1 | 1 | 1 | NA | NA | 2 | 1 |
| 5_97235908 | NA | NA | 1 | 1 | NA | NA | 1 | 1 | 2 | 1 | NA | NA | NA | NA | 1 | 1 | NA | NA | 1 | 1 |
| 5_97235939 | 2 | 1 | 2 | 1 | 1 | 1 | NA | NA | 5 | 1 | NA | NA | 1 | 1 | 1 | 1 | 1 | 1 | 3 | 1 |
| 5_97236009 | NA | NA | NA | NA | NA | NA | 1 | 1 | 1 | 1 | NA | NA | 2 | 1 | 1 | 1 | NA | NA | 3 | 1 |
| 5_97236289 | NA | NA | 1 | 1 | NA | NA | 1 | 1 | 2 | 1 | NA | NA | 1 | 0 | 1 | 1 | NA | NA | 2 | 0.5 |
| 5_97236439 | NA | NA | 1 | 1 | NA | NA | 2 | 1 | 3 | 1 | NA | NA | 2 | 1 | 1 | 1 | NA | NA | 3 | 1 |
| 5_97236503 | NA | NA | 1 | 1 | NA | NA | 2 | 1 | 3 | 1 | NA | NA | 2 | 1 | 1 | 1 | NA | NA | 3 | 1 |
| 5_97238839 | NA | NA | 1 | 1 | NA | NA | 4 | 0 | 5 | 0.2 | NA | NA | 2 | 0 | 1 | 0 | 1 | 1 | 4 | 0.25 |
| 5_97241942 | NA | NA | 1 | 0 | 4 | 0.25 | 1 | 0 | 6 | 0.1667 | 3 | 0 | 3 | 0.3333 | 1 | 0 | NA | NA | 7 | 0.1429 |
| 5_97242830 | 1 | 1 | 1 | 0 | 2 | 0.5 | 1 | 0 | 5 | 0.4 | 2 | 0.5 | 2 | 0.5 | NA | NA | 1 | 0 | 5 | 0.4 |
| 5_97243433 | NA | NA | 1 | 1 | 3 | 0.6667 | 1 | 0 | 5 | 0.6 | 1 | 0 | 3 | 1 | NA | NA | NA | NA | 4 | 0.75 |
| 5_97243575 | NA | NA | 1 | 0 | 3 | 0.3333 | 1 | 0 | 5 | 0.2 | 1 | 1 | 3 | 0.3333 | NA | NA | NA | NA | 4 | 0.5 |
| 5_97243612 | NA | NA | 1 | 1 | 3 | 0.6667 | 1 | 1 | 5 | 0.8 | 1 | 1 | 3 | 1 | NA | NA | NA | NA | 4 | 1 |
| 5_97243777 | 2 | 0 | 3 | 0 | 4 | 0.25 | 1 | 0 | 10 | 0.1 | 2 | 0 | 4 | 0.25 | 1 | 0 | 1 | 0 | 8 | 0.125 |
| 5_98332430 | 1 | 0 | 1 | 0 | 2 | 0.5 | NA | NA | 4 | 0.25 | 2 | 0 | NA | NA | 1 | 0 | 4 | 0.25 | 7 | 0.1429 |
| 5_98332974 | NA | NA | 1 | 1 | 1 | 1 | NA | NA | 2 | 1 | 2 | 1 | NA | NA | 1 | 1 | 4 | 0.5 | 7 | 0.7143 |
| 5_98333219 | 1 | 1 | 1 | 1 | 1 | 1 | NA | NA | 3 | 1 | 2 | 1 | NA | NA | NA | NA | 4 | 1 | 6 | 1 |
| 5_98333241 | 1 | 1 | NA | NA | 2 | 0.5 | NA | NA | 3 | 0.6667 | 1 | 1 | NA | NA | NA | NA | 3 | 1 | 4 | 1 |
| 5_98333557 | 1 | 1 | 1 | 0 | 1 | 1 | NA | NA | 3 | 0.6667 | 2 | 0.5 | NA | NA | 1 | 0 | 3 | 0.6667 | 6 | 0.5 |
| 5_98333860 | NA | NA | NA | NA | 2 | 1 | NA | NA | 2 | 1 | 2 | 0.5 | NA | NA | 1 | 1 | 3 | 0.6667 | 6 | 0.6667 |
| 5_98333875 | NA | NA | 1 | 1 | 1 | 1 | NA | NA | 2 | 1 | 2 | 1 | NA | NA | 1 | 1 | 3 | 1 | 6 | 1 |
| 5_98334039 | 1 | 1 | 1 | 1 | 2 | 1 | NA | NA | 4 | 1 | 2 | 1 | NA | NA | NA | NA | 4 | 0.75 | 6 | 0.8333 |
| 5_98334092 | 1 | 1 | 1 | 1 | 1 | 1 | NA | NA | 3 | 1 | 3 | 1 | NA | NA | 1 | 1 | 4 | 1 | 8 | 1 |
| 5_98334305 | NA | NA | NA | NA | 2 | 1 | NA | NA | 2 | 1 | 3 | 1 | NA | NA | 1 | 0 | 4 | 0.75 | 8 | 0.75 |
| 5_98334420 | 1 | 0 | 1 | 0 | 2 | 0.5 | NA | NA | 4 | 0.25 | 3 | 0.3333 | NA | NA | 1 | 0 | 3 | 0 | 7 | 0.1429 |
| 5_98334510 | 1 | 1 | 1 | 1 | 1 | 1 | NA | NA | 3 | 1 | 3 | 1 | NA | NA | 1 | 1 | 5 | 1 | 9 | 1 |
| 5_98334959 | 1 | 0 | 1 | 0 | 2 | 0.5 | NA | NA | 4 | 0.25 | 4 | 0.25 | NA | NA | 1 | 0 | 5 | 0.4 | 10 | 0.3 |
| 5_98335123 | 2 | 0.5 | 1 | 1 | 1 | 1 | NA | NA | 4 | 0.75 | 4 | 1 | 1 | 0 | 1 | 1 | 5 | 1 | 11 | 0.9091 |
| 5_98623460 | 5 | 0.2 | 5 | 0 | 2 | 0.5 | 4 | 0.25 | 16 | 0.1875 | 2 | 0 | 5 | 0.2 | 3 | 0 | 3 | 0 | 13 | 0.0769 |
| 5_98623521 | NA | NA | 1 | 0 | 2 | 0.5 | NA | NA | 3 | 0.3333 | NA | NA | NA | NA | NA | NA | 1 | 1 | 1 | 1 |
| 5_99079821 | 1 | 1 | NA | NA | 1 | 1 | NA | NA | 2 | 1 | 1 | 1 | 1 | 1 | 1 | 1 | NA | NA | 3 | 1 |
| 5_99365913 | 5 | 0.6 | 7 | 0.2857 | 8 | 0.125 | 2 | 0.5 | 22 | 0.3182 | 2 | 0 | 3 | 0.6667 | 8 | 0.25 | 3 | 0.6667 | 16 | 0.375 |
| 5_99366149 | 10 | 0.1 | 7 | 0.2857 | 4 | 0.25 | 4 | 0 | 25 | 0.16 | 4 | 0.75 | 3 | 0.3333 | 7 | 0.2857 | 2 | 0.5 | 16 | 0.4375 |
| 5_99366161 | 12 | 0.25 | 7 | 0.2857 | 6 | 0.3333 | 5 | 0.2 | 30 | 0.2667 | 6 | 0.5 | 5 | 0 | 9 | 0.2222 | 3 | 0.6667 | 23 | 0.3043 |
| 5_99366271 | 7 | 0.8571 | 7 | 1 | 7 | 0.7143 | 4 | 1 | 25 | 0.88 | 4 | 1 | 4 | 1 | 8 | 1 | 2 | 1 | 18 | 1 |
| 5_99366289 | 9 | 0.7778 | 7 | 0.7143 | 5 | 0.8 | 3 | 0.6667 | 24 | 0.75 | 4 | 0.75 | 2 | 1 | 5 | 0.6 | 1 | 1 | 12 | 0.75 |

| | | | | | | | | | | | | | | | | | | | | |
|---|---|---|---|---|---|---|---|---|---|---|---|---|---|---|---|---|---|---|---|---|
| 5_99366325 | 8 | 1 | 8 | 1 | 7 | 1 | 5 | 1 | 28 | 1 | 6 | 0.8333 | 4 | 1 | 10 | 1 | 2 | 1 | 22 | 0.9545 |
| 5_99366436 | 9 | 0.7778 | 6 | 0.3333 | 7 | 0.2857 | 5 | 1 | 27 | 0.5926 | 6 | 0.5 | 4 | 0.5 | 8 | 0.875 | 3 | 0.6667 | 21 | 0.6667 |
| 5_99366454 | 10 | 1 | 7 | 1 | 6 | 1 | 5 | 1 | 28 | 1 | 5 | 1 | 4 | 1 | 10 | 0.9 | 3 | 1 | 22 | 0.9545 |
| 5_99366462 | 10 | 0.1 | 8 | 0 | 7 | 0.1429 | 5 | 0 | 30 | 0.0667 | 6 | 0 | 5 | 0.2 | 10 | 0.2 | 3 | 0 | 24 | 0.125 |
| 5_99366529 | 9 | 0.8889 | 6 | 0.6667 | 7 | 0.5714 | 4 | 1 | 26 | 0.7692 | 4 | 1 | 4 | 1 | 10 | 0.7 | 2 | 1 | 20 | 0.85 |
| 5_99366574 | 9 | 0.1111 | 6 | 0.1667 | 5 | 0 | 4 | 0 | 24 | 0.0833 | 5 | 0 | 4 | 0.25 | 9 | 0.1111 | 2 | 0.5 | 20 | 0.15 |
| 5_99890204 | 1 | 1 | NA | NA | 1 | 1 | NA | NA | 2 | 1 | 1 | 1 | NA | NA | NA | NA | NA | NA | 1 | 1 |
| 5_99892590 | NA | NA | NA | NA | NA | NA | 1 | 1 | 1 | 1 | NA | NA | NA | NA | NA | NA | 1 | 1 | 1 | 1 |
| 5_99892943 | NA | NA | NA | NA | NA | NA | 1 | 1 | 1 | 1 | NA | NA | NA | NA | NA | NA | 1 | 1 | 1 | 1 |
| 5_10008441 | NA | NA | NA | NA | 1 | 1 | NA | NA | 1 | 1 | NA | NA | NA | NA | NA | NA | 1 | 1 | 1 | 1 |
| 5_10008455 | NA | NA | NA | NA | 1 | 1 | NA | NA | 1 | 1 | NA | NA | NA | NA | NA | NA | 1 | 1 | 1 | 1 |
| 5_10008468 | NA | NA | NA | NA | 1 | 1 | NA | NA | 1 | 1 | NA | NA | NA | NA | NA | NA | 1 | 1 | 1 | 1 |
| 5_10008483 | NA | NA | NA | NA | 1 | 1 | NA | NA | 1 | 1 | NA | NA | NA | NA | NA | NA | 1 | 1 | 1 | 1 |
| 5_10008497 | NA | NA | NA | NA | 1 | 1 | NA | NA | 1 | 1 | NA | NA | NA | NA | NA | NA | 1 | 1 | 1 | 1 |
| 5_10008528 | NA | NA | NA | NA | 1 | 1 | NA | NA | 1 | 1 | NA | NA | NA | NA | NA | NA | 1 | 1 | 1 | 1 |
| 5_10010395 | 4 | 0.5 | 7 | 0.4286 | 6 | 0.3333 | 6 | 0.6667 | 23 | 0.4783 | 2 | 0.5 | 5 | 0.4 | 4 | 0.75 | 2 | 1 | 13 | 0.6154 |
| 5_10010489 | 8 | 0.625 | 9 | 0.4444 | 12 | 0.5833 | 11 | 0.4545 | 40 | 0.525 | 3 | 0.6667 | 13 | 0.5385 | 11 | 0.4545 | 10 | 0.5 | 37 | 0.5135 |
| 5_10010812 | 1 | 0 | 3 | 0 | 3 | 0 | 7 | 0.1429 | 14 | 0.0714 | NA | NA | 6 | 0 | 5 | 0.4 | 4 | 0 | 15 | 0.1333 |
| 5_10010846 | 1 | 1 | 5 | 0 | 4 | 0 | 8 | 0.125 | 18 | 0.1111 | NA | NA | 6 | 0.3333 | 5 | 0.2 | 2 | 0 | 13 | 0.2308 |
| 5_10018158 | 7 | 0.2857 | 3 | 0 | 11 | 0.3636 | 7 | 0.2857 | 28 | 0.2857 | 9 | 0 | 15 | 0.4 | 7 | 0.2857 | 7 | 0.4286 | 38 | 0.2895 |
| 5_10018218 | 9 | 0 | 4 | 0 | 8 | 0.125 | 5 | 0.2 | 26 | 0.0769 | 8 | 0.125 | 14 | 0.0714 | 8 | 0 | 7 | 0.2857 | 37 | 0.1081 |
| 5_10018276 | 14 | 0.5 | 17 | 0.4706 | 25 | 0.52 | 14 | 0.5 | 70 | 0.5 | 17 | 0.5882 | 30 | 0.5333 | 23 | 0.6957 | 23 | 0.7826 | 93 | 0.6452 |
| 5_10018711 | 1 | 1 | 1 | 1 | 2 | 0 | NA | NA | 4 | 0.5 | NA | NA | 3 | 0.6667 | 1 | 1 | 2 | 0.5 | 6 | 0.6667 |
| 5_10018718 | 1 | 0 | 1 | 1 | 2 | 0.5 | NA | NA | 4 | 0.5 | NA | NA | 2 | 0.5 | 1 | 0 | 2 | 0.5 | 5 | 0.4 |
| 5_10018732 | 1 | 0 | 1 | 0 | 1 | 1 | NA | NA | 3 | 0.3333 | NA | NA | 2 | 0.5 | 2 | 0.5 | 1 | 0 | 5 | 0.4 |
| 5_10021590 | 13 | 0.2308 | 4 | 0.5 | 6 | 0 | 7 | 0.2857 | 30 | 0.2333 | 4 | 0.25 | 3 | 0 | 1 | 0 | 3 | 0.3333 | 11 | 0.1818 |
| 5_10021598 | 13 | 0.0769 | 4 | 0 | 6 | 0 | 7 | 0.2857 | 30 | 0.1 | 4 | 0.25 | 4 | 0 | 3 | 0 | 6 | 0 | 17 | 0.0588 |
| 5_10021613 | 13 | 0.2308 | 4 | 0.5 | 7 | 0.4286 | 4 | 0.25 | 28 | 0.3214 | 4 | 0.25 | 3 | 0.3333 | 3 | 0.6667 | 6 | 0.3333 | 16 | 0.375 |
| 5_10021614 | 11 | 0.4545 | 3 | 0 | 6 | 0.1667 | 6 | 0.1667 | 26 | 0.2692 | 3 | 0.6667 | 3 | 0 | 1 | 0 | 3 | 0.3333 | 10 | 0.3 |
| 5_10021621 | 13 | 0.4615 | 5 | 0.6 | 7 | 0.7143 | 7 | 0.2857 | 32 | 0.5 | 5 | 0.2 | 4 | 0.25 | 3 | 0 | 4 | 0.75 | 16 | 0.3125 |
| 5_10021630 | 10 | 0 | 3 | 0 | 5 | 0.4 | 6 | 0 | 24 | 0.0833 | 3 | 0 | 3 | 0.3333 | 2 | 0 | 3 | 0.3333 | 11 | 0.1818 |
| 5_10021634 | 5 | 0 | 3 | 0 | 5 | 0.2 | 5 | 0 | 18 | 0.0556 | 4 | 0 | 4 | 0 | 2 | 0 | 5 | 0.2 | 15 | 0.0667 |
| 5_10022576 | 2 | 0.5 | 10 | 0.2 | 7 | 0 | 3 | 0 | 22 | 0.1364 | 8 | 0.125 | 7 | 0.2857 | 2 | 0.5 | 6 | 0 | 23 | 0.1739 |
| 5_10022577 | 2 | 0 | 9 | 0 | 4 | 0.25 | 2 | 1 | 17 | 0.1765 | 4 | 0.25 | 7 | 0 | 2 | 0 | 4 | 0 | 17 | 0.0588 |
| 5_10022596 | 3 | 0.3333 | 7 | 0 | 7 | 0.1429 | 3 | 0.3333 | 20 | 0.15 | 6 | 0 | 7 | 0.1429 | 2 | 0 | 5 | 0.2 | 20 | 0.1 |
| 5_10022609 | 2 | 0 | 10 | 0.1 | 7 | 0.2857 | 3 | 0.3333 | 22 | 0.1818 | 7 | 0.2857 | 7 | 0 | 2 | 1 | 4 | 0.5 | 20 | 0.3 |
| 5_10022619 | 3 | 0 | 7 | 0.1429 | 5 | 0.4 | 1 | 0 | 16 | 0.1875 | 6 | 0 | 8 | 0.125 | 2 | 0.5 | 4 | 0 | 20 | 0.1 |
| 5_10022636 | NA | NA | 8 | 0.125 | 4 | 0 | 2 | 0 | 14 | 0.0714 | 2 | 0.5 | 6 | 0.3333 | 2 | 0.5 | 2 | 0 | 12 | 0.3333 |
| 5_10022636 | NA | NA | 1 | 0 | 1 | 1 | NA | NA | 2 | 0.5 | 2 | 0.5 | NA | NA | NA | NA | 1 | 1 | 3 | 0.6667 |
| 5_10022664 | 1 | 1 | 10 | 1 | 7 | 1 | 2 | 1 | 20 | 1 | 7 | 1 | 5 | 1 | 2 | 1 | 4 | 0.75 | 18 | 0.9444 |
| 5_10022665 | 2 | 1 | 9 | 1 | 7 | 1 | 2 | 1 | 20 | 1 | 6 | 1 | 6 | 1 | 1 | 1 | 4 | 1 | 17 | 1 |
| 5_10022673 | 2 | 0 | 11 | 0.2727 | 6 | 0.1667 | 1 | 0 | 20 | 0.2 | 4 | 0 | 2 | 0.5 | 2 | 0 | 4 | 0 | 12 | 0.0833 |
| 5_10022687 | 2 | 0 | 9 | 0.1111 | 6 | 0.6667 | 1 | 1 | 18 | 0.3333 | 6 | 0.1667 | 2 | 0 | 1 | 0 | 3 | 0.3333 | 12 | 0.1667 |
| 5_10022688 | 2 | 1 | 8 | 0.75 | 4 | 1 | 1 | 0 | 15 | 0.8 | 4 | 1 | 4 | 0.75 | NA | NA | 3 | 0.6667 | 11 | 0.8182 |
| 5_10022703 | 2 | 0.5 | 7 | 0.8571 | 3 | 1 | NA | NA | 12 | 0.8333 | 6 | 0.6667 | 3 | 0.6667 | 1 | 1 | 2 | 0 | 12 | 0.5833 |
| 5_10022708 | 2 | 1 | 10 | 1 | 6 | 0.8333 | NA | NA | 18 | 0.9444 | 7 | 1 | 4 | 0.75 | 1 | 1 | 2 | 1 | 14 | 0.9286 |
| 5_10022727 | 2 | 0.5 | 10 | 0.4 | 5 | 0.2 | NA | NA | 17 | 0.3529 | 5 | 0.8 | 3 | 0.3333 | 1 | 0 | 2 | 1 | 11 | 0.6364 |
| 5_10022753 | 2 | 0 | 7 | 0 | 4 | 0.25 | 1 | 0 | 14 | 0.0714 | 5 | 0.4 | 4 | 0 | 1 | 1 | 3 | 0 | 13 | 0.2308 |
| 5_10022764 | 2 | 0.5 | 8 | 1 | 5 | 1 | 1 | 1 | 16 | 0.9375 | 6 | 0.8333 | 4 | 1 | 1 | 1 | 3 | 1 | 14 | 0.9286 |
| 5_10022777 | 2 | 0.5 | 7 | 0.2857 | 5 | 0.4 | 1 | 0 | 15 | 0.3333 | 7 | 0.5714 | 4 | 0 | 1 | 0 | NA | NA | 12 | 0.3333 |
| 5_10050976 | 10 | 0.5 | 11 | 0.4545 | 15 | 0.4667 | 15 | 0.6667 | 51 | 0.5294 | 9 | 0.3333 | 8 | 0.625 | 13 | 0.3077 | 8 | 0.375 | 38 | 0.3947 |
| 5_10050979 | 13 | 0.1538 | 11 | 0.1818 | 13 | 0.1538 | 11 | 0 | 48 | 0.125 | 8 | 0.25 | 9 | 0 | 13 | 0 | 5 | 0 | 35 | 0.0571 |
| 5_10050984 | 12 | 0.0833 | 10 | 0.1 | 15 | 0.1333 | 12 | 0.0833 | 49 | 0.102 | 8 | 0.25 | 7 | 0.1429 | 12 | 0.25 | 6 | 0 | 33 | 0.1818 |
| 5_10050985 | 14 | 0.0714 | 12 | 0.1667 | 17 | 0.1176 | 15 | 0.4 | 58 | 0.1897 | 10 | 0.4 | 10 | 0 | 13 | 0.1538 | 6 | 0.1667 | 39 | 0.1795 |
| 5_10051168 | NA | NA | 1 | 1 | 1 | 0 | 1 | 0 | 3 | 0.3333 | 1 | 1 | NA | NA | NA | NA | 1 | 0 | 2 | 0.5 |
| 5_10059040 | 2 | 0 | 3 | 0.3333 | 1 | 1 | 1 | 0 | 7 | 0.2857 | 2 | 0 | 4 | 0 | 1 | 1 | 3 | 0 | 10 | 0.1 |
| 5_10059100 | 1 | 0 | NA | NA | 1 | 1 | 1 | 0 | 3 | 0.3333 | NA | NA | 3 | 0.6667 | 1 | 1 | 1 | 0 | 5 | 0.6 |
| 5_10059188 | NA | NA | 2 | 1 | NA | NA | 1 | 0 | 3 | 0.6667 | 1 | 0 | 4 | 0.75 | 1 | 1 | 2 | 0.5 | 8 | 0.625 |
| 5_10059191 | NA | NA | 2 | 1 | NA | NA | 1 | 0 | 3 | 0.6667 | 1 | 0 | 4 | 1 | 1 | 1 | 2 | 1 | 8 | 0.875 |
| 5_10059192 | NA | NA | 2 | 1 | NA | NA | 1 | 1 | 3 | 1 | 1 | 1 | 4 | 0.75 | NA | NA | 2 | 1 | 7 | 0.8571 |
| 5_10059193 | NA | NA | 2 | 0.5 | NA | NA | 1 | 0 | 3 | 0.3333 | 1 | 1 | 3 | 0 | 1 | 0 | 2 | 0 | 7 | 0.1429 |
| 5_10063327 | NA | NA | 1 | 1 | NA | NA | NA | NA | 1 | 1 | NA | NA | 2 | 1 | 1 | 1 | 1 | 1 | 4 | 1 |
| 5_10063335 | NA | NA | 1 | 1 | NA | NA | NA | NA | 1 | 1 | NA | NA | 2 | 0.5 | 1 | 1 | 1 | 1 | 4 | 0.75 |
| 5_10063363 | NA | NA | 1 | 1 | NA | NA | NA | NA | 1 | 1 | NA | NA | 2 | 0.5 | NA | NA | 1 | 1 | 3 | 0.6667 |
| 5_10069526 | 4 | 0.75 | 9 | 0.1111 | 15 | 0 | 7 | 0 | 35 | 0.1143 | 10 | 0.3 | 14 | 0.0714 | 15 | 0.2 | 9 | 0.2222 | 48 | 0.1875 |

| | | | | | | | | | | | | | | | | | | | | |
|---|---|---|---|---|---|---|---|---|---|---|---|---|---|---|---|---|---|---|---|---|
| 5_1006954 | 5 | 0.4 | 11 | 0.6364 | 13 | 0.3077 | 10 | 0.4 | 39 | 0.4359 | 10 | 0.2 | 13 | 0.3846 | 17 | 0.5294 | 14 | 0.4286 | 54 | 0.4074 |
| 5_1006954 | 6 | 0.1667 | 10 | 0 | 14 | 0.0714 | 9 | 0 | 39 | 0.0513 | 7 | 0.5714 | 15 | 0.0667 | 15 | 0.0667 | 10 | 0 | 47 | 0.1277 |
| 5_1006955 | 7 | 0.1429 | 10 | 0.3 | 16 | 0.1875 | 9 | 0 | 42 | 0.1667 | 11 | 0.0909 | 16 | 0 | 16 | 0.125 | 14 | 0.1429 | 57 | 0.0877 |
| 5_1006955 | 7 | 0.1429 | 11 | 0.0909 | 16 | 0.0625 | 10 | 0.1 | 44 | 0.0909 | 13 | 0.0769 | 17 | 0.1176 | 16 | 0.1875 | 15 | 0.0667 | 61 | 0.1148 |
| 5_1008028 | 5 | 0 | 9 | 0.2222 | 4 | 0.25 | 2 | 0 | 20 | 0.15 | 3 | 0.3333 | 4 | 0 | 5 | 0 | 4 | 0 | 16 | 0.0625 |
| 5_1008029 | 15 | 0.6667 | 16 | 0.875 | 7 | 0.7143 | 7 | 0.8571 | 45 | 0.7778 | 8 | 0.625 | 11 | 0.8182 | 9 | 0.7778 | 12 | 0.75 | 40 | 0.75 |
| 5_1008029 | 19 | 0.1053 | 16 | 0.25 | 9 | 0 | 8 | 0.375 | 52 | 0.1731 | 8 | 0.25 | 11 | 0 | 12 | 0.1667 | 13 | 0.2308 | 44 | 0.1591 |
| 5_1008030 | 16 | 0.4375 | 18 | 0.1667 | 7 | 0.2857 | 7 | 0.2857 | 48 | 0.2917 | 7 | 0.2857 | 10 | 0.4 | 11 | 0.5455 | 12 | 0.25 | 40 | 0.375 |
| 5_1008031 | 17 | 0.0588 | 19 | 0.1053 | 9 | 0 | 8 | 0 | 53 | 0.0566 | 10 | 0 | 13 | 0.1538 | 12 | 0.1667 | 13 | 0.0769 | 48 | 0.1042 |
| 5_1009102 | 2 | 0 | 1 | 1 | NA | NA | NA | NA | 3 | 0.3333 | 1 | 1 | 4 | 0.25 | 1 | 0 | 1 | 1 | 7 | 0.4286 |
| 5_1009162 | NA | NA | 1 | 0 | NA | NA | 1 | 1 | 2 | 0.5 | NA | NA | 2 | 0 | NA | NA | 1 | 1 | 3 | 0.3333 |
| 5_1009532 | NA | NA | 1 | 1 | NA | NA | 1 | 0 | 2 | 0.5 | 1 | 1 | 2 | 0 | 1 | 0 | 1 | 0 | 5 | 0.2 |
| 5_1009542 | NA | NA | 1 | 1 | NA | NA | NA | NA | 1 | 1 | NA | NA | 3 | 1 | 1 | 0 | 1 | 1 | 5 | 0.8 |
| 5_1019695 | 5 | 0.2 | 3 | 0 | 4 | 0 | 2 | 0 | 14 | 0.0714 | NA | NA | 5 | 0.4 | 5 | 0.4 | NA | NA | 10 | 0.4 |
| 5_1019701 | 5 | 0 | 5 | 0.4 | 6 | 0.1667 | 1 | 1 | 17 | 0.2353 | NA | NA | 7 | 0.2857 | 4 | 0.25 | 4 | 0.5 | 15 | 0.3333 |
| 5_1019704 | 6 | 0.3333 | 5 | 0 | 5 | 0.2 | 1 | 0 | 17 | 0.1765 | NA | NA | 5 | 0 | 3 | 0.6667 | 4 | 0.5 | 12 | 0.3333 |
| 5_1019710 | 7 | 0.1429 | 8 | 0.25 | 6 | 0.5 | 1 | 0 | 22 | 0.2727 | 2 | 0 | 7 | 0.2857 | 4 | 0.5 | 5 | 0.4 | 18 | 0.3333 |
| 5_1019711 | 8 | 0.25 | 3 | 0 | 6 | 0.1667 | 1 | 1 | 18 | 0.2222 | 2 | 0 | 8 | 0.25 | 1 | 0 | 8 | 0.375 | 19 | 0.2632 |
| 5_1019712 | 8 | 0.875 | 6 | 0.3333 | 7 | 0.8571 | 1 | 0 | 22 | 0.6818 | 1 | 0 | 9 | 0.5556 | 4 | 0.75 | 6 | 0.6667 | 20 | 0.6 |
| 5_1019713 | 8 | 0.875 | 6 | 0.8333 | 5 | 1 | 1 | 1 | 20 | 0.9 | 2 | 1 | 6 | 0.8333 | 3 | 1 | 6 | 0.8333 | 17 | 0.8824 |
| 5_1019714 | 9 | 0.7778 | 8 | 0.5 | 5 | 0.8 | 1 | 1 | 23 | 0.6957 | 1 | 1 | 10 | 0.4 | 4 | 0.5 | 4 | 0.5 | 19 | 0.4737 |
| 5_1019715 | 6 | 0.1667 | 5 | 0.2 | 6 | 0.1667 | NA | NA | 17 | 0.1765 | 1 | 0 | 6 | 0.3333 | 1 | 0 | 6 | 0 | 14 | 0.1429 |
| 5_1019810 | 15 | 0.6 | 9 | 0.8889 | 13 | 0.3077 | 12 | 0.4167 | 49 | 0.5306 | 8 | 0.125 | 14 | 0.4286 | 10 | 0.5 | 9 | 0.3333 | 41 | 0.3659 |
| 5_1019812 | 17 | 0.0588 | 13 | 0.0769 | 17 | 0.0588 | 13 | 0.0769 | 60 | 0.0667 | 13 | 0 | 17 | 0.1176 | 11 | 0 | 17 | 0.0588 | 58 | 0.0517 |
| 5_1019816 | 12 | 0.0833 | 10 | 0.1 | 9 | 0.2222 | 13 | 0.0769 | 44 | 0.1136 | 6 | 0.1667 | 13 | 0.1538 | 9 | 0.1111 | 12 | 0.0833 | 40 | 0.125 |
| 5_1019824 | 9 | 0 | 8 | 0 | 7 | 0.1429 | 10 | 0.1 | 34 | 0.0588 | 7 | 0 | 12 | 0.3333 | 5 | 0.2 | 9 | 0.1111 | 33 | 0.1818 |
| 5_1019834 | 8 | 0.25 | 9 | 0 | 5 | 0 | 4 | 0 | 26 | 0.0769 | 5 | 0 | 9 | 0.1111 | 7 | 0 | 4 | 0.25 | 25 | 0.08 |
| 5_1019835 | 8 | 0 | 7 | 0.2857 | 5 | 0.2 | 4 | 0.25 | 24 | 0.1667 | 5 | 0.4 | 8 | 0.125 | 7 | 0 | 2 | 0 | 22 | 0.1364 |
| 5_1019836 | 7 | 0.2857 | 8 | 0.125 | 4 | 0 | 3 | 0.6667 | 22 | 0.2273 | 5 | 0.4 | 7 | 0.1429 | 5 | 0.2 | 3 | 0.3333 | 20 | 0.25 |
| 5_1019838 | 3 | 0.3333 | 5 | 0.2 | 1 | 0 | 3 | 0 | 12 | 0.1667 | 3 | 0 | 5 | 0.2 | 5 | 0.2 | NA | NA | 13 | 0.1538 |
| 5_1019839 | 6 | 0.1667 | 8 | 0.25 | 4 | 0.25 | 5 | 0.6 | 23 | 0.3043 | 5 | 0.2 | 7 | 0.2857 | 7 | 0.5714 | 2 | 0.5 | 21 | 0.381 |
| 5_1019840 | 8 | 0.625 | 8 | 0.5 | 5 | 0.8 | 5 | 0.2 | 26 | 0.5385 | 6 | 0.6667 | 7 | 0.7143 | 6 | 0.3333 | 2 | 1 | 21 | 0.619 |
| 5_1019840 | 8 | 1 | 8 | 0.625 | 5 | 0.8 | 4 | 0.75 | 25 | 0.8 | 4 | 1 | 5 | 0.8 | 7 | 1 | 2 | 1 | 18 | 0.9444 |
| 5_1019928 | 7 | 0 | 6 | 0.1667 | 3 | 0 | 5 | 0.2 | 21 | 0.0952 | 2 | 0 | 5 | 0.2 | 4 | 0.25 | NA | NA | 11 | 0.1818 |
| 5_1030402 | 2 | 0.5 | NA | NA | 2 | 0 | NA | NA | 4 | 0.25 | 1 | 1 | NA | NA | NA | NA | NA | NA | 1 | 1 |
| 5_1030403 | 2 | 0.5 | NA | NA | 1 | 0 | NA | NA | 3 | 0.3333 | 1 | 1 | NA | NA | NA | NA | NA | NA | 1 | 1 |
| 5_1030452 | 2 | 1 | NA | NA | 3 | 1 | NA | NA | 5 | 1 | 1 | 1 | NA | NA | NA | NA | NA | NA | 1 | 1 |
| 5_1030452 | 2 | 0.5 | NA | NA | 2 | 0.5 | NA | NA | 4 | 0.5 | 1 | 1 | NA | NA | NA | NA | NA | NA | 1 | 1 |
| 5_1030570 | NA | NA | 9 | 0.1111 | 7 | 0.1429 | 1 | 0 | 17 | 0.1176 | 3 | 0 | 7 | 0.1429 | 4 | 0 | NA | NA | 14 | 0.0714 |
| 5_1030580 | 1 | 0 | 6 | 0 | 5 | 0.2 | 1 | 0 | 13 | 0.0769 | 3 | 0 | 5 | 0.4 | 3 | 0.3333 | NA | NA | 11 | 0.2727 |
| 5_1030599 | 2 | 0.5 | 6 | 0.1667 | 3 | 0.3333 | NA | NA | 11 | 0.2727 | 1 | 0 | 4 | 0 | 2 | 1 | NA | NA | 7 | 0.2857 |
| 5_1037027 | 1 | 1 | 1 | 0 | 1 | 0 | NA | NA | 3 | 0.3333 | NA | NA | NA | NA | 1 | 0 | 1 | 1 | 2 | 0.5 |
| 5_1039320 | 1 | 1 | NA | NA | NA | NA | 1 | 1 | 2 | 1 | NA | NA | 1 | 1 | NA | NA | NA | NA | 1 | 1 |
| 5_1039320 | 1 | 1 | NA | NA | NA | NA | 1 | 1 | 2 | 1 | NA | NA | 1 | 1 | NA | NA | NA | NA | 1 | 1 |
| 5_1039819 | 1 | 0 | NA | NA | NA | NA | 1 | 1 | 2 | 0.5 | NA | NA | 1 | 1 | NA | NA | NA | NA | 1 | 1 |
| 5_1039989 | NA | NA | NA | NA | 1 | 1 | 3 | 1 | 4 | 1 | NA | NA | 1 | 1 | NA | NA | NA | NA | 1 | 1 |
| 5_1039989 | 1 | 1 | NA | NA | 1 | 0 | 3 | 0.3333 | 5 | 0.4 | NA | NA | 1 | 1 | NA | NA | NA | NA | 1 | 1 |
| 5_1039990 | 1 | 1 | NA | NA | 1 | 1 | 4 | 1 | 6 | 1 | NA | NA | 1 | 1 | NA | NA | NA | NA | 1 | 1 |
| 5_1039992 | 1 | 1 | NA | NA | 1 | 1 | 3 | 1 | 5 | 1 | NA | NA | 1 | 1 | NA | NA | NA | NA | 1 | 1 |
| 5_1039994 | NA | NA | NA | NA | 1 | 0 | 4 | 0.5 | 5 | 0.4 | NA | NA | 1 | 1 | NA | NA | NA | NA | 1 | 1 |
| 5_1039995 | 1 | 1 | NA | NA | 1 | 1 | 1 | 0 | 3 | 0.6667 | NA | NA | 1 | 1 | NA | NA | NA | NA | 1 | 1 |
| 5_1040019 | 2 | 0 | 1 | 1 | 1 | 0 | 3 | 0 | 7 | 0.1429 | NA | NA | 2 | 0 | 1 | 1 | NA | NA | 3 | 0.3333 |
| 5_1040023 | 2 | 0 | 2 | 0 | 1 | 0 | 5 | 0.2 | 10 | 0.1 | 1 | 1 | 1 | 0 | 1 | 0 | 1 | 0 | 4 | 0.25 |
| 5_1040029 | 1 | 0 | 3 | 1 | 2 | 1 | 4 | 0.25 | 10 | 0.6 | 1 | 0 | 3 | 0.3333 | 2 | 1 | 2 | 0.5 | 8 | 0.5 |
| 5_1040030 | 1 | 1 | 4 | 0.25 | 2 | 0 | 4 | 0 | 11 | 0.1818 | 1 | 1 | 2 | 0.5 | 2 | 0.5 | 2 | 0 | 7 | 0.4286 |
| 5_1040030 | 1 | 1 | 4 | 1 | 2 | 0 | 4 | 0.25 | 11 | 0.5455 | 2 | 0.5 | 3 | 0.3333 | 2 | 1 | 2 | 0 | 9 | 0.4444 |
| 5_1040102 | 3 | 0 | 4 | 0.25 | 2 | 0 | 4 | 0.25 | 13 | 0.1538 | NA | NA | 1 | 0 | NA | NA | 2 | 0.5 | 3 | 0.3333 |
| 5_1040108 | NA | NA | 4 | 1 | 1 | 1 | 4 | 1 | 9 | 1 | NA | NA | 3 | 0.6667 | 1 | 1 | 3 | 0.6667 | 7 | 0.7143 |
| 5_1040108 | 1 | 1 | 2 | 0 | 3 | 0.3333 | 1 | 0 | 7 | 0.2857 | 1 | 0 | 2 | 0 | 3 | 0 | 3 | 0.3333 | 9 | 0.1111 |
| 5_1040109 | 2 | 1 | 4 | 0.25 | 3 | 0 | 2 | 0.5 | 11 | 0.3636 | 1 | 0 | 3 | 0.3333 | 3 | 0.3333 | 3 | 0.3333 | 10 | 0.3 |
| 5_1040109 | 2 | 0.5 | 3 | 0.3333 | 2 | 0 | 3 | 0.6667 | 10 | 0.4 | 1 | 0 | 2 | 0 | 3 | 0.3333 | 3 | 0.3333 | 9 | 0.2222 |
| 5_1040109 | 2 | 1 | 2 | 1 | 3 | 0.6667 | 3 | 1 | 10 | 0.9 | 1 | 1 | 3 | 0.3333 | 2 | 1 | 3 | 0 | 9 | 0.4444 |
| 5_1040238 | 1 | 0 | 2 | 1 | 3 | 0.6667 | 1 | 0 | 7 | 0.5714 | NA | NA | 2 | 0.5 | 2 | 1 | 1 | 1 | 5 | 0.8 |
| 5_1040239 | 1 | 1 | 2 | 1 | 2 | 1 | 1 | 1 | 6 | 1 | NA | NA | 1 | 1 | 1 | 1 | 1 | 1 | 3 | 1 |
| 5_1040240 | 2 | 0 | 2 | 0.5 | 2 | 0 | 2 | 0 | 8 | 0.125 | NA | NA | 1 | 0 | 1 | 1 | 1 | 1 | 3 | 0.6667 |

| | | | | | | | | | | | | | | | | | | | | |
|---|---|---|---|---|---|---|---|---|---|---|---|---|---|---|---|---|---|---|---|---|
| 5_10403383 | 1 | 1 | 2 | 1 | 2 | 0.5 | 2 | 0.5 | 7 | 0.7143 | NA | NA | 2 | 1 | NA | NA | 1 | 0 | 3 | 0.6667 |
| 5_10413771 | 7 | 0 | 9 | 0.2222 | 6 | 0.3333 | 8 | 0.25 | 30 | 0.2 | 8 | 0.125 | 4 | 0 | 6 | 0.1667 | 7 | 0.7143 | 25 | 0.28 |
| 5_10413773 | 6 | 0.3333 | 10 | 0.5 | 7 | 0.8571 | 6 | 0.6667 | 29 | 0.5862 | 9 | 0.3333 | 5 | 0.6 | 5 | 0.4 | 6 | 0.8333 | 25 | 0.52 |
| 5_10413789 | 7 | 0.8571 | 8 | 0.375 | 7 | 0.7143 | 6 | 1 | 28 | 0.7143 | 7 | 0.2857 | 4 | 0.5 | 6 | 0.8333 | 7 | 0.5714 | 24 | 0.5417 |
| 5_10413820 | 1 | 0 | 3 | 0.3333 | 2 | 0 | 1 | 0 | 7 | 0.1429 | 6 | 0 | 1 | 0 | 1 | 1 | 2 | 0 | 10 | 0.1 |
| 5_10413834 | 1 | 0 | 3 | 0 | 1 | 0 | 1 | 1 | 6 | 0.1667 | 5 | 0.2 | 1 | 0 | NA | NA | 3 | 0 | 9 | 0.1111 |
| 5_10419840 | NA | NA | NA | NA | 1 | 1 | NA | NA | 1 | 1 | NA | NA | NA | NA | NA | NA | 1 | 1 | 1 | 1 |
| 5_10419881 | NA | NA | NA | NA | 1 | 1 | NA | NA | 1 | 1 | NA | NA | NA | NA | NA | NA | 1 | 1 | 1 | 1 |
| 5_10419893 | NA | NA | NA | NA | 1 | 1 | NA | NA | 1 | 1 | 1 | 1 | NA | NA | NA | NA | 1 | 0 | 2 | 0.5 |
| 5_10421303 | 7 | 0.5714 | 4 | 0.25 | 7 | 0.4286 | 7 | 0.4286 | 25 | 0.44 | 4 | 1 | 9 | 0.7778 | 5 | 0.8 | 12 | 0.5833 | 30 | 0.7333 |
| 5_10421314 | 7 | 0.7143 | 9 | 1 | 7 | 0.7143 | 9 | 0.6667 | 32 | 0.7812 | 6 | 0.8333 | 10 | 0.8 | 8 | 0.75 | 15 | 0.7333 | 39 | 0.7692 |
| 5_10424040 | 108 | 0.1852 | 124 | 0.0968 | 103 | 0.0777 | 96 | 0.0938 | 431 | 0.1137 | 84 | 0.131 | 109 | 0.0734 | 115 | 0.113 | 115 | 0.0522 | 423 | 0.0898 |
| 5_10424049 | 105 | 0.5238 | 121 | 0.4463 | 104 | 0.5096 | 102 | 0.5784 | 432 | 0.5116 | 89 | 0.427 | 110 | 0.4364 | 119 | 0.5126 | 119 | 0.5294 | 437 | 0.4805 |
| 5_10424054 | 101 | 0.0198 | 116 | 0.0345 | 108 | 0.0833 | 101 | 0.0792 | 426 | 0.054 | 87 | 0.0805 | 97 | 0.0206 | 116 | 0.0603 | 113 | 0.0442 | 413 | 0.0508 |
| 5_10424061 | 92 | 0.2826 | 108 | 0.2315 | 98 | 0.2755 | 91 | 0.2088 | 389 | 0.2494 | 85 | 0.2471 | 99 | 0.2626 | 113 | 0.3097 | 106 | 0.2925 | 403 | 0.2804 |
| 5_10424093 | 93 | 0.2043 | 97 | 0.1237 | 87 | 0.1149 | 84 | 0.1548 | 361 | 0.1496 | 69 | 0.1884 | 91 | 0.1319 | 103 | 0.2136 | 102 | 0.2451 | 365 | 0.1973 |
| 5_10424109 | 83 | 0.0602 | 87 | 0.0345 | 77 | 0.0519 | 72 | 0.0556 | 319 | 0.0502 | 56 | 0.0893 | 78 | 0.1282 | 90 | 0.1111 | 89 | 0.0562 | 313 | 0.0958 |
| 5_10424115 | 89 | 0.0787 | 87 | 0.0575 | 79 | 0.038 | 78 | 0.0385 | 333 | 0.0541 | 59 | 0.0847 | 82 | 0.0854 | 93 | 0.0753 | 97 | 0.0206 | 331 | 0.0634 |
| 5_10426129 | 55 | 0.0727 | 71 | 0.1127 | 52 | 0.1538 | 65 | 0.0923 | 243 | 0.107 | 43 | 0.0233 | 59 | 0.0678 | 58 | 0.1207 | 65 | 0.1077 | 225 | 0.0844 |
| 5_10426941 | NA | NA | NA | NA | NA | NA | 1 | 1 | 1 | 1 | NA | NA | NA | NA | NA | NA | 1 | 1 | 1 | 1 |
| 5_10426970 | NA | NA | NA | NA | NA | NA | 1 | 1 | 1 | 1 | NA | NA | NA | NA | NA | NA | 1 | 1 | 1 | 1 |
| 5_10426974 | NA | NA | NA | NA | NA | NA | 1 | 1 | 1 | 1 | NA | NA | NA | NA | NA | NA | 1 | 1 | 1 | 1 |
| 5_10435971 | 1 | 1 | 1 | 1 | NA | NA | NA | NA | 2 | 1 | NA | NA | 1 | 1 | NA | NA | NA | NA | 1 | 1 |
| 5_10435972 | 1 | 1 | 1 | 1 | NA | NA | NA | NA | 2 | 1 | NA | NA | 1 | 1 | NA | NA | NA | NA | 1 | 1 |
| 5_10435986 | 1 | 1 | 1 | 1 | 1 | 1 | NA | NA | 3 | 1 | NA | NA | 1 | 1 | NA | NA | NA | NA | 1 | 1 |
| 5_10435988 | NA | NA | 1 | 0 | 1 | 1 | NA | NA | 2 | 0.5 | NA | NA | 1 | 1 | NA | NA | NA | NA | 1 | 1 |
| 5_10436065 | 2 | 0.5 | 2 | 0.5 | 1 | 0 | NA | NA | 5 | 0.4 | 1 | 1 | 1 | 1 | NA | NA | NA | NA | 2 | 1 |
| 5_10436066 | 2 | 0.5 | 2 | 0.5 | 1 | 0 | NA | NA | 5 | 0.4 | 1 | 1 | 1 | 0 | NA | NA | NA | NA | 2 | 0.5 |
| 5_10436081 | 1 | 1 | 2 | 1 | 1 | 1 | NA | NA | 4 | 1 | NA | NA | 1 | 1 | NA | NA | NA | NA | 1 | 1 |
| 5_10436094 | 2 | 0 | 2 | 0.5 | 1 | 0 | NA | NA | 5 | 0.2 | 1 | 1 | 1 | 0 | NA | NA | 1 | 0 | 3 | 0.3333 |
| 5_10436100 | 1 | 1 | 2 | 1 | 1 | 1 | NA | NA | 4 | 1 | 1 | 1 | 1 | 1 | NA | NA | 1 | 0 | 3 | 0.6667 |
| 5_10436101 | 2 | 1 | 1 | 1 | 1 | 0 | NA | NA | 4 | 0.75 | 1 | 0 | 1 | 1 | NA | NA | 1 | 1 | 3 | 0.6667 |
| 5_10436116 | 1 | 1 | 2 | 1 | 1 | 1 | NA | NA | 4 | 1 | 1 | 0 | 1 | 1 | NA | NA | 1 | 1 | 3 | 0.6667 |
| 5_10436125 | 2 | 0.5 | 2 | 0.5 | 1 | 0 | NA | NA | 5 | 0.4 | 1 | 1 | 1 | 0 | NA | NA | 1 | 1 | 3 | 0.6667 |
| 5_10458737 | 1 | 0 | 7 | 0.1429 | 2 | 0 | 3 | 0 | 13 | 0.0769 | 5 | 0.4 | 3 | 0.3333 | 2 | 0 | NA | NA | 10 | 0.3 |
| 5_10458827 | 1 | 0 | 7 | 0.1429 | 2 | 0 | 3 | 0 | 13 | 0.0769 | 5 | 0 | 3 | 0.3333 | 2 | 0.5 | NA | NA | 10 | 0.2 |
| 5_10460356 | 2 | 0 | 1 | 1 | NA | NA | 1 | 1 | 4 | 0.5 | 1 | 1 | NA | NA | NA | NA | 2 | 0 | 3 | 0.3333 |
| 5_10460431 | 2 | 1 | NA | NA | NA | NA | NA | NA | 2 | 1 | 1 | 1 | NA | NA | NA | NA | 1 | 1 | 2 | 1 |
| 5_10460433 | 3 | 0.3333 | NA | NA | NA | NA | 1 | 1 | 4 | 0.5 | NA | NA | NA | NA | 1 | 0 | 1 | 1 | 2 | 0.5 |
| 5_10460463 | 2 | 0.5 | NA | NA | NA | NA | 1 | 0 | 3 | 0.3333 | 1 | 1 | NA | NA | 1 | 0 | 1 | 1 | 3 | 0.6667 |
| 5_10460528 | NA | NA | 1 | 1 | NA | NA | NA | NA | 1 | 1 | NA | NA | NA | NA | 1 | 0 | 1 | 1 | 2 | 0.5 |
| 5_10460556 | 2 | 0.5 | 1 | 1 | NA | NA | NA | NA | 3 | 0.6667 | 1 | 1 | NA | NA | NA | NA | 1 | 1 | 2 | 1 |
| 5_10460691 | 2 | 1 | NA | NA | 1 | 1 | NA | NA | 3 | 1 | 1 | 1 | NA | NA | NA | NA | 1 | 1 | 2 | 1 |
| 5_10460709 | 1 | 1 | NA | NA | 1 | 1 | NA | NA | 2 | 1 | NA | NA | NA | NA | NA | NA | 1 | 1 | 1 | 1 |
| 5_10460719 | 2 | 1 | NA | NA | 1 | 1 | NA | NA | 3 | 1 | 1 | 1 | NA | NA | NA | NA | 1 | 0 | 2 | 0.5 |
| 5_10460735 | 2 | 1 | NA | NA | 1 | 1 | NA | NA | 3 | 1 | 1 | 1 | NA | NA | NA | NA | 1 | 1 | 2 | 1 |
| 5_10460735 | NA | NA | NA | NA | 1 | 1 | NA | NA | 1 | 1 | NA | NA | NA | NA | NA | NA | 1 | 1 | 1 | 1 |
| 5_10463106 | 2 | 0.5 | 4 | 0 | NA | NA | 3 | 0 | 9 | 0.1111 | NA | NA | 2 | 0 | 1 | 0 | 1 | 1 | 4 | 0.25 |
| 5_10465144 | 1 | 1 | 3 | 1 | NA | NA | 3 | 1 | 7 | 1 | 1 | 1 | 2 | 1 | NA | NA | NA | NA | 3 | 1 |
| 5_10465158 | 1 | 0 | 6 | 0.1667 | NA | NA | 5 | 0.2 | 12 | 0.1667 | 1 | 0 | 2 | 0.5 | 1 | 0 | NA | NA | 4 | 0.25 |
| 5_10465313 | 3 | 0.3333 | 6 | 0.3333 | NA | NA | 6 | 0.3333 | 15 | 0.3333 | 1 | 1 | 3 | 0.3333 | 1 | 0 | NA | NA | 5 | 0.4 |
| 5_10469925 | NA | NA | NA | NA | 2 | 0.5 | NA | NA | 2 | 0.5 | NA | NA | NA | NA | 1 | 1 | 1 | 1 | 2 | 1 |
| 5_10469926 | NA | NA | NA | NA | 2 | 0.5 | 1 | 1 | 3 | 0.6667 | NA | NA | NA | NA | 1 | 1 | 1 | 1 | 2 | 1 |
| 5_10473020 | NA | NA | NA | NA | NA | NA | 2 | 1 | 2 | 1 | NA | NA | 2 | 1 | 1 | 1 | 1 | 1 | 4 | 1 |
| 5_10473025 | NA | NA | NA | NA | NA | NA | 2 | 0.5 | 2 | 0.5 | NA | NA | 2 | 0.5 | NA | NA | 1 | 0 | 3 | 0.3333 |
| 5_10473059 | NA | NA | NA | NA | NA | NA | 2 | 0.5 | 2 | 0.5 | NA | NA | 2 | 1 | 2 | 0.5 | 2 | 1 | 6 | 0.8333 |
| 5_10473065 | NA | NA | NA | NA | NA | NA | 2 | 1 | 2 | 1 | NA | NA | 2 | 1 | 2 | 1 | 1 | 1 | 5 | 1 |
| 5_10473076 | NA | NA | NA | NA | NA | NA | 2 | 1 | 2 | 1 | NA | NA | 2 | 1 | 2 | 1 | 2 | 1 | 6 | 1 |
| 5_10473103 | NA | NA | NA | NA | NA | NA | 2 | 1 | 2 | 1 | NA | NA | 2 | 0.5 | 2 | 1 | 2 | 0.5 | 6 | 0.6667 |
| 5_10473111 | NA | NA | NA | NA | NA | NA | 2 | 0.5 | 2 | 0.5 | NA | NA | 2 | 1 | 1 | 1 | NA | NA | 3 | 1 |
| 5_10473118 | NA | NA | NA | NA | NA | NA | 2 | 0.5 | 2 | 0.5 | NA | NA | 2 | 0 | 2 | 0.5 | 1 | 1 | 5 | 0.4 |
| 5_10473120 | NA | NA | NA | NA | NA | NA | 1 | 1 | 1 | 1 | NA | NA | 2 | 0 | 2 | 0.5 | 2 | 0 | 6 | 0.1667 |
| 5_10473135 | NA | NA | NA | NA | NA | NA | 2 | 1 | 2 | 1 | NA | NA | 2 | 0.5 | 2 | 1 | 2 | 0.5 | 6 | 0.6667 |
| 5_10473141 | NA | NA | NA | NA | NA | NA | 2 | 1 | 2 | 1 | NA | NA | 2 | 1 | 2 | 0.5 | 2 | 0.5 | 6 | 0.6667 |
| 5_10473142 | NA | NA | NA | NA | NA | NA | 1 | 1 | 1 | 1 | NA | NA | 2 | 1 | 1 | 1 | 2 | 0.5 | 5 | 0.8 |

| | | | | | | | | | | | | | | | | | | | | |
|---|---|---|---|---|---|---|---|---|---|---|---|---|---|---|---|---|---|---|---|---|
| 5_1047314 | NA | NA | NA | NA | NA | NA | 2 | 0.5 | 2 | 0.5 | NA | NA | 1 | 0 | 2 | 1 | 2 | 0.5 | 5 | 0.6 |
| 5_1047315 | NA | NA | NA | NA | NA | NA | 2 | 1 | 2 | 1 | NA | NA | 2 | 1 | 1 | 1 | 2 | 0.5 | 5 | 0.8 |
| 5_1047315 | NA | NA | NA | NA | NA | NA | 2 | 0.5 | 2 | 0.5 | NA | NA | 2 | 1 | 2 | 1 | 1 | 1 | 5 | 1 |
| 5_1047316 | NA | NA | NA | NA | NA | NA | 2 | 1 | 2 | 1 | 1 | 0 | 2 | 0.5 | 1 | 1 | 2 | 0 | 6 | 0.3333 |
| 5_1047316 | 1 | 1 | NA | NA | NA | NA | 2 | 0.5 | 3 | 0.6667 | 1 | 0 | 2 | 0.5 | 2 | 0.5 | 2 | 0 | 7 | 0.2857 |
| 5_1047317 | 1 | 1 | NA | NA | NA | NA | 2 | 0.5 | 3 | 0.6667 | NA | NA | NA | NA | 1 | 1 | 1 | 1 | 2 | 1 |
| 5_1047317 | 1 | 1 | NA | NA | NA | NA | 2 | 1 | 3 | 1 | 1 | 1 | 2 | 1 | 2 | 1 | 2 | 1 | 7 | 1 |
| 5_1047318 | 1 | 1 | NA | NA | NA | NA | NA | NA | 1 | 1 | NA | NA | 2 | 0.5 | 2 | 1 | 2 | 0.5 | 6 | 0.6667 |
| 5_1047319 | 2 | 1 | NA | NA | NA | NA | 3 | 1 | 5 | 1 | 1 | 1 | 2 | 1 | 3 | 1 | 2 | 1 | 8 | 1 |
| 5_1047320 | 1 | 0 | NA | NA | NA | NA | 3 | 0.6667 | 4 | 0.5 | 1 | 0 | 2 | 0 | 1 | 0 | 2 | 0.5 | 6 | 0.1667 |
| 5_1047321 | 2 | 0.5 | NA | NA | NA | NA | 1 | 1 | 3 | 0.6667 | NA | NA | 2 | 0.5 | NA | NA | 1 | 1 | 3 | 0.6667 |
| 5_1047321 | 2 | 0.5 | NA | NA | NA | NA | 3 | 1 | 5 | 0.8 | 1 | 1 | 2 | 1 | 3 | 0.3333 | 2 | 0.5 | 8 | 0.625 |
| 5_1047322 | 2 | 0 | NA | NA | NA | NA | 3 | 0.6667 | 5 | 0.4 | 1 | 0 | 1 | 0 | 2 | 0.5 | 2 | 0 | 6 | 0.1667 |
| 5_1047877 | 1 | 1 | NA | NA | NA | NA | NA | NA | 1 | 1 | 1 | 1 | NA | NA | NA | NA | NA | NA | 1 | 1 |
| 5_1047881 | 1 | 1 | NA | NA | NA | NA | NA | NA | 1 | 1 | 1 | 1 | NA | NA | NA | NA | NA | NA | 1 | 1 |
| 5_1056278 | 1 | 1 | NA | NA | 1 | 0 | 3 | 1 | 5 | 0.8 | NA | NA | NA | NA | 4 | 1 | 3 | 1 | 7 | 1 |
| 5_1056281 | 2 | 1 | 1 | 0 | 1 | 1 | 3 | 0 | 7 | 0.4286 | NA | NA | NA | NA | 5 | 0 | 3 | 0.6667 | 8 | 0.25 |
| 5_1056281 | 2 | 0.5 | 1 | 0 | 1 | 0 | 3 | 0.3333 | 7 | 0.2857 | NA | NA | NA | NA | 6 | 0.1667 | 3 | 0.3333 | 9 | 0.2222 |
| 5_1056288 | 2 | 0 | 2 | 0 | 1 | 0 | 3 | 0.3333 | 8 | 0.125 | 1 | 0 | 1 | 0 | 6 | 0 | 4 | 0.25 | 12 | 0.0833 |
| 5_1056292 | 2 | 1 | 2 | 0.5 | 2 | 0.5 | 2 | 0.5 | 8 | 0.625 | 1 | 1 | NA | NA | 6 | 0.1667 | 4 | 1 | 11 | 0.5455 |
| 5_1056292 | 2 | 1 | 2 | 0 | 1 | 0 | 2 | 1 | 7 | 0.5714 | 1 | 1 | 1 | 1 | 6 | 0.5 | 4 | 0.5 | 12 | 0.5833 |
| 5_1056293 | 2 | 0.5 | 2 | 0 | 2 | 0 | 3 | 0 | 9 | 0.1111 | 1 | 0 | 1 | 0 | 6 | 0 | 4 | 0.25 | 12 | 0.0833 |
| 5_1056337 | NA | NA | 1 | 1 | 2 | 1 | NA | NA | 3 | 1 | NA | NA | NA | NA | 4 | 1 | 1 | 1 | 5 | 1 |
| 5_1056338 | 1 | 1 | 4 | 0.25 | NA | NA | 3 | 0.6667 | 8 | 0.5 | 1 | 1 | NA | NA | 7 | 0.4286 | 4 | 0.75 | 12 | 0.5833 |
| 5_1056338 | 1 | 1 | 4 | 1 | 2 | 1 | 3 | 1 | 10 | 1 | 1 | 1 | NA | NA | 7 | 1 | 5 | 1 | 13 | 1 |
| 5_1056339 | 1 | 0 | 4 | 0.25 | 1 | 1 | 3 | 0 | 9 | 0.2222 | 1 | 0 | NA | NA | 7 | 0.4286 | 5 | 0.8 | 13 | 0.5385 |
| 5_1056339 | NA | NA | 4 | 1 | 2 | 1 | 2 | 0.5 | 8 | 0.875 | 1 | 1 | NA | NA | 5 | 0.4 | 2 | 1 | 8 | 0.625 |
| 5_1056340 | 1 | 1 | 4 | 1 | 2 | 1 | 2 | 1 | 9 | 1 | 1 | 1 | NA | NA | 7 | 1 | 5 | 1 | 13 | 1 |
| 5_1056340 | 1 | 1 | 4 | 0.5 | 2 | 1 | 3 | 0.6667 | 10 | 0.7 | 1 | 1 | NA | NA | 7 | 0.8571 | 5 | 0.8 | 13 | 0.8462 |
| 5_1056341 | 1 | 0 | 2 | 1 | 2 | 0.5 | 2 | 0 | 7 | 0.4286 | NA | NA | NA | NA | 5 | 0.2 | 3 | 0.3333 | 8 | 0.25 |
| 5_1056342 | 1 | 0 | 4 | 0.25 | 2 | 0 | 3 | 0 | 10 | 0.1 | 2 | 0.5 | 1 | 0 | 6 | 0 | 5 | 0 | 14 | 0.0714 |
| 5_1056345 | 1 | 0 | 4 | 0.5 | 2 | 1 | 4 | 0.5 | 11 | 0.5455 | 2 | 1 | NA | NA | 7 | 0.5714 | 5 | 0.6 | 14 | 0.6429 |
| 5_1056353 | NA | NA | 4 | 0.5 | 3 | 0.3333 | 4 | 0 | 11 | 0.2727 | 2 | 0 | NA | NA | 5 | 0 | 4 | 0.5 | 11 | 0.1818 |
| 5_1056358 | 1 | 0 | 6 | 0.1667 | 3 | 0 | 3 | 0.3333 | 13 | 0.1538 | 1 | 0 | 2 | 0 | 8 | 0.375 | 7 | 0 | 18 | 0.1667 |
| 5_1056358 | 1 | 0 | 5 | 0.6 | 3 | 0.3333 | 4 | 0.25 | 13 | 0.3846 | 2 | 0 | 2 | 0 | 8 | 0.25 | 7 | 0 | 19 | 0.1053 |
| 5_1056361 | 2 | 0 | 5 | 0.2 | 3 | 0 | 4 | 0.25 | 14 | 0.1429 | 3 | 0 | 3 | 0 | 8 | 0.25 | 6 | 0 | 20 | 0.1 |
| 5_1056369 | 4 | 0.25 | 7 | 0.2857 | 4 | 0.25 | 4 | 0 | 19 | 0.2105 | 2 | 0.5 | 6 | 0.5 | 6 | 0.5 | 8 | 0.5 | 22 | 0.5 |
| 5_1056373 | 3 | 0.3333 | 7 | 0 | 4 | 0 | 2 | 0 | 16 | 0.0625 | 1 | 0 | 4 | 0 | 6 | 0 | 6 | 0.1667 | 17 | 0.0588 |
| 5_1057550 | 1 | 1 | 2 | 1 | 1 | 1 | 2 | 1 | 6 | 1 | 1 | 1 | 1 | 1 | 1 | 0 | 1 | 0 | 4 | 0.5 |
| 5_1057551 | NA | NA | 2 | 1 | 1 | 1 | 2 | 1 | 5 | 1 | 1 | 1 | 1 | 1 | 1 | 1 | 1 | 1 | 4 | 1 |
| 5_1057551 | 1 | 1 | 2 | 0.5 | 1 | 1 | 2 | 0 | 6 | 0.5 | 1 | 1 | 1 | 1 | 1 | 0 | 1 | 0 | 4 | 0.5 |
| 5_1057551 | 1 | 1 | 2 | 0 | 1 | 0 | 2 | 0.5 | 6 | 0.3333 | 1 | 0 | 1 | 1 | 1 | 0 | 1 | 1 | 4 | 0.5 |
| 5_1057551 | 1 | 1 | 2 | 1 | 1 | 1 | 2 | 1 | 6 | 1 | 1 | 1 | 1 | 1 | 1 | 1 | NA | NA | 3 | 1 |
| 5_1057553 | 1 | 1 | 1 | 1 | 1 | 1 | 2 | 1 | 5 | 1 | 2 | 0.5 | 1 | 1 | NA | NA | 1 | 1 | 4 | 0.75 |
| 5_1057558 | 2 | 0.5 | 3 | 0 | 1 | 0 | 3 | 0.3333 | 9 | 0.2222 | 3 | 1 | 1 | 0 | 1 | 1 | 2 | 0.5 | 7 | 0.7143 |
| 5_1057561 | 2 | 0 | 3 | 0.3333 | 1 | 1 | 2 | 0.5 | 8 | 0.375 | 3 | 0.6667 | 1 | 0 | 1 | 0 | 2 | 0 | 7 | 0.2857 |
| 5_1057562 | 2 | 0 | 2 | 0.5 | 1 | 1 | 3 | 0.6667 | 8 | 0.5 | 3 | 0.3333 | 1 | 0 | NA | NA | 2 | 0 | 6 | 0.1667 |
| 5_1057562 | 2 | 1 | 2 | 1 | NA | NA | 2 | 1 | 6 | 1 | 2 | 1 | 1 | 1 | 1 | 1 | 2 | 1 | 6 | 1 |
| 5_1057562 | 2 | 1 | 3 | 1 | 1 | 1 | 3 | 1 | 9 | 1 | 3 | 1 | 1 | 1 | 1 | 1 | 2 | 1 | 7 | 1 |
| 5_1057563 | 2 | 1 | 2 | 1 | 1 | 1 | 2 | 1 | 7 | 1 | 3 | 1 | 2 | 1 | 2 | 1 | 2 | 1 | 9 | 1 |
| 5_1057564 | 2 | 0.5 | 3 | 0.3333 | 1 | 0 | 3 | 0 | 9 | 0.2222 | 2 | 0 | 2 | 0 | 2 | 0.5 | 2 | 0 | 8 | 0.125 |
| 5_1057564 | 2 | 0.5 | 3 | 0.6667 | 1 | 0 | 1 | 0 | 7 | 0.4286 | 2 | 0.5 | 2 | 0 | 2 | 0.5 | 1 | 1 | 7 | 0.4286 |
| 5_1057564 | 2 | 0 | 2 | 0 | 1 | 1 | 2 | 0 | 7 | 0.1429 | 2 | 0 | NA | NA | 2 | 0.5 | NA | NA | 4 | 0.25 |
| 5_1057564 | 3 | 0.6667 | 2 | 0 | 1 | 0 | 3 | 0.6667 | 9 | 0.4444 | 3 | 0.6667 | 2 | 0 | 1 | 1 | 2 | 0.5 | 8 | 0.5 |
| 5_1057565 | 3 | 1 | 3 | 0.6667 | 1 | 1 | 2 | 1 | 9 | 0.8889 | 2 | 1 | 3 | 1 | 2 | 1 | 2 | 1 | 9 | 1 |
| 5_1057566 | 3 | 0.6667 | 3 | 0.6667 | 1 | 1 | 2 | 0.5 | 9 | 0.6667 | 3 | 0 | 3 | 0.6667 | 1 | 1 | 1 | 0 | 8 | 0.375 |
| 5_1057566 | 2 | 1 | 3 | 0.3333 | 1 | 1 | 2 | 0.5 | 8 | 0.625 | 3 | 0.6667 | 3 | 1 | 1 | 1 | 2 | 0.5 | 9 | 0.7778 |
| 5_1057571 | 1 | 1 | 3 | 0 | 1 | 0 | 3 | 0.3333 | 8 | 0.25 | 2 | 0.5 | 2 | 0.5 | 3 | 0 | 1 | 0 | 8 | 0.25 |
| 5_1057578 | 1 | 0 | 4 | 0.25 | 1 | 0 | 3 | 0.3333 | 9 | 0.2222 | 2 | 0 | 2 | 0 | 2 | 0 | 2 | 0.5 | 8 | 0.125 |
| 5_1057583 | 1 | 0 | 4 | 0 | 2 | 0.5 | 3 | 0 | 10 | 0.1 | 2 | 0.5 | 2 | 0 | 3 | 0 | 2 | 0 | 9 | 0.1111 |
| 5_1057598 | 2 | 0.5 | 7 | 0.2857 | 1 | 0 | 2 | 0 | 12 | 0.25 | 3 | 0 | 2 | 0 | 3 | 0.3333 | NA | NA | 8 | 0.125 |
| 5_1057604 | 1 | 0 | 3 | 0.3333 | 1 | 0 | 4 | 0 | 9 | 0.1111 | 4 | 0.5 | 3 | 0.3333 | 5 | 0.4 | 1 | 1 | 13 | 0.4615 |
| 5_1057605 | 2 | 1 | 4 | 0.5 | 1 | 1 | 4 | 0.25 | 11 | 0.5455 | 3 | 0 | 2 | 0 | 3 | 0.3333 | 2 | 0.5 | 10 | 0.2 |
| 5_1057607 | 3 | 0.3333 | 4 | 0.75 | 1 | 0 | 2 | 1 | 10 | 0.6 | 5 | 1 | 2 | 0 | 2 | 0.5 | 1 | 0 | 10 | 0.6 |
| 5_1057607 | 3 | 1 | 6 | 1 | NA | NA | 4 | 0.25 | 13 | 0.7692 | 6 | 0.8333 | 3 | 0.6667 | 5 | 1 | 2 | 1 | 16 | 0.875 |

| | | | | | | | | | | | | | | | | | | | | |
|---|---|---|---|---|---|---|---|---|---|---|---|---|---|---|---|---|---|---|---|---|
| 5_10595962 | 2 | 0 | 1 | 1 | 2 | 0.5 | 1 | 0 | 6 | 0.3333 | 2 | 0 | 2 | 0 | 2 | 0.5 | 2 | 0 | 8 | 0.125 |
| 5_10595972 | 2 | 1 | 2 | 0 | 2 | 0.5 | 1 | 1 | 7 | 0.5714 | 3 | 0.3333 | 2 | 0.5 | 2 | 1 | 3 | 1 | 10 | 0.7 |
| 5_10595999 | 3 | 0 | 2 | 0 | 3 | 0.3333 | 1 | 0 | 9 | 0.1111 | 2 | 0 | 3 | 0 | 3 | 0.3333 | 3 | 0 | 11 | 0.0909 |
| 5_10596021 | 1 | 1 | 3 | 0 | 3 | 0 | NA | NA | 7 | 0.1429 | 3 | 0.3333 | 2 | 0 | 3 | 0 | 3 | 0 | 11 | 0.0909 |
| 5_10596025 | 2 | 0 | 3 | 0 | 3 | 0.3333 | 1 | 0 | 9 | 0.1111 | 3 | 0.3333 | 2 | 0 | 3 | 0 | 4 | 0.25 | 12 | 0.1667 |
| 5_10596042 | 3 | 0.3333 | 3 | 0.3333 | 2 | 0 | 1 | 0 | 9 | 0.2222 | 4 | 0 | 2 | 0 | 2 | 0.5 | 4 | 0 | 12 | 0.0833 |
| 5_10596049 | 3 | 1 | 3 | 0.6667 | 3 | 0 | 1 | 1 | 10 | 0.6 | 4 | 0.5 | 5 | 0.6 | 3 | 0.3333 | 4 | 0.75 | 16 | 0.5625 |
| 5_10596054 | 3 | 0 | 2 | 0.5 | 2 | 0 | 1 | 0 | 8 | 0.125 | 3 | 0 | 2 | 0 | 3 | 0.3333 | 2 | 0 | 10 | 0.1 |
| 5_10596057 | 3 | 0 | 3 | 0.3333 | 1 | 1 | 1 | 0 | 8 | 0.25 | 4 | 0.75 | 4 | 0 | 3 | 0 | 3 | 1 | 14 | 0.4286 |
| 5_10596092 | 4 | 0.25 | 2 | 0 | 1 | 1 | 1 | 1 | 8 | 0.375 | 4 | 0 | 4 | 0 | 3 | 0.3333 | 5 | 0.2 | 16 | 0.125 |
| 5_10596096 | 2 | 1 | 2 | 1 | 3 | 1 | 1 | 1 | 8 | 1 | 4 | 1 | 3 | 1 | 3 | 1 | 5 | 1 | 15 | 1 |
| 5_10596097 | 4 | 0 | 3 | 0.3333 | 3 | 0 | 1 | 0 | 11 | 0.0909 | 3 | 0 | 5 | 0.2 | 3 | 0 | 5 | 0 | 16 | 0.0625 |
| 5_10596108 | 2 | 0 | 1 | 1 | 4 | 0.75 | 1 | 0 | 8 | 0.5 | 4 | 0.75 | 3 | 0.3333 | 3 | 0.6667 | 6 | 0.8333 | 16 | 0.6875 |
| 5_10596118 | 3 | 0 | 3 | 0.3333 | 2 | 0 | 2 | 1 | 10 | 0.3 | 3 | 0.3333 | 5 | 0.6 | 3 | 0 | 4 | 0.5 | 15 | 0.4 |
| 5_10596130 | 4 | 0.25 | 1 | 0 | 4 | 0 | 1 | 0 | 10 | 0.1 | 5 | 0.2 | 2 | 0.5 | 3 | 0.3333 | 2 | 0 | 12 | 0.25 |
| 5_10596152 | 4 | 0.25 | 2 | 0.5 | 4 | 0 | 3 | 0 | 13 | 0.1538 | 4 | 0.25 | 3 | 0 | 2 | 0.5 | 6 | 0 | 15 | 0.1333 |
| 5_10596165 | 4 | 0.25 | 2 | 0 | 4 | 0.25 | 3 | 0.3333 | 13 | 0.2308 | 5 | 0.2 | 5 | 0.8 | 3 | 0.6667 | 5 | 0.4 | 18 | 0.5 |
| 5_10596178 | 4 | 0.25 | 3 | 0.3333 | 4 | 0.5 | 3 | 0.6667 | 14 | 0.4286 | 4 | 0.75 | 3 | 1 | 3 | 1 | 7 | 0.2857 | 17 | 0.6471 |
| 5_10596185 | 4 | 0.25 | 3 | 0 | 3 | 0.3333 | 3 | 0.3333 | 13 | 0.2308 | 6 | 0 | 5 | 0 | 2 | 0 | 5 | 0.2 | 18 | 0.0556 |
| 5_10596187 | 4 | 0 | 3 | 0 | 2 | 0 | 3 | 0.3333 | 12 | 0.0833 | 4 | 0 | 3 | 0 | 3 | 0.3333 | 4 | 0 | 14 | 0.0714 |
| 5_10596188 | 4 | 1 | 2 | 1 | 3 | 1 | 3 | 1 | 12 | 1 | 5 | 1 | 5 | 1 | 3 | 1 | 7 | 1 | 20 | 1 |
| 5_10596190 | 4 | 0 | 1 | 0 | 4 | 0.5 | 2 | 0 | 11 | 0.1818 | 5 | 0.2 | 4 | 0 | 3 | 0 | 4 | 0 | 16 | 0.0625 |
| 5_10596192 | 4 | 1 | 2 | 1 | 3 | 1 | 2 | 1 | 11 | 1 | 5 | 1 | 4 | 1 | 2 | 1 | 5 | 0.8 | 16 | 0.9375 |
| 5_10596193 | 4 | 0.25 | 3 | 0.3333 | 4 | 0.25 | 3 | 0.3333 | 14 | 0.2857 | 6 | 0.3333 | 5 | 0.2 | 3 | 0.6667 | 6 | 0.3333 | 20 | 0.35 |
| 5_10596194 | 3 | 1 | 3 | 1 | 3 | 0.6667 | 3 | 1 | 12 | 0.9167 | 6 | 1 | 4 | 1 | 2 | 1 | 6 | 1 | 18 | 1 |
| 5_10596210 | 3 | 0 | 2 | 0 | 4 | 0.25 | 2 | 0 | 11 | 0.0909 | 5 | 0 | 3 | 0 | 3 | 0 | 7 | 0.2857 | 18 | 0.1111 |
| 5_10596217 | 2 | 1 | 4 | 1 | 3 | 1 | 1 | 1 | 10 | 1 | 5 | 1 | 3 | 1 | 3 | 1 | 5 | 1 | 16 | 1 |
| 5_10596220 | 4 | 0.25 | 4 | 0.5 | 2 | 0.5 | 3 | 0.3333 | 13 | 0.3846 | 6 | 0.5 | 5 | 0.2 | 3 | 0 | 7 | 0.4286 | 21 | 0.3333 |
| 5_10596229 | 4 | 1 | 5 | 0.8 | 3 | 1 | 1 | 1 | 13 | 0.9231 | 6 | 0.5 | 5 | 0.6 | 4 | 0.5 | 7 | 0.5714 | 22 | 0.5455 |
| 5_10596258 | 4 | 0.25 | 5 | 0 | 1 | 0 | 4 | 0.25 | 14 | 0.1429 | 4 | 0.25 | 4 | 0.25 | 4 | 0.25 | 6 | 0 | 18 | 0.1667 |
| 5_10596275 | 3 | 1 | 1 | 0 | 1 | 0 | 1 | 0 | 6 | 0.5 | 1 | 0 | 2 | 1 | 3 | 0.3333 | 6 | 0.3333 | 12 | 0.4167 |
| 5_10596283 | 3 | 0 | 1 | 0 | 1 | 1 | 2 | 0.5 | 7 | 0.2857 | 1 | 1 | 2 | 0 | 2 | 0.5 | 6 | 0.3333 | 11 | 0.3636 |
| 5_10596288 | 3 | 0.3333 | 1 | 0 | 1 | 0 | 2 | 0 | 7 | 0.1429 | NA | NA | 2 | 0.5 | 3 | 0.3333 | 6 | 0.6667 | 11 | 0.5455 |
| 5_10601101 | 2 | 0.5 | 3 | 0.3333 | 6 | 0 | 1 | 0 | 12 | 0.1667 | 4 | 0 | 5 | 0 | 2 | 0.5 | 4 | 0 | 15 | 0.0667 |
| 5_10670561 | NA | NA | 1 | 1 | NA | NA | NA | NA | 1 | 1 | NA | NA | 2 | 0.5 | 1 | 1 | NA | NA | 3 | 0.6667 |
| 5_10676490 | 4 | 0 | 4 | 0 | 5 | 0 | 4 | 0.25 | 17 | 0.0588 | 5 | 0 | 7 | 0 | 7 | 0.1429 | 7 | 0.1429 | 26 | 0.0769 |
| 5_10676493 | 4 | 0 | 3 | 1 | 4 | 0.25 | 3 | 0.6667 | 14 | 0.4286 | 5 | 0 | 8 | 0.375 | 7 | 0.1429 | 6 | 0 | 26 | 0.1538 |
| 5_10676567 | 2 | 0 | 5 | 0.4 | 4 | 0.25 | 3 | 0 | 14 | 0.2143 | 6 | 0.3333 | 6 | 0 | 6 | 0.3333 | 5 | 0.2 | 23 | 0.2174 |
| 5_10676598 | 2 | 0.5 | 5 | 0.4 | 4 | 0 | 3 | 0.3333 | 14 | 0.2857 | 5 | 0.4 | 5 | 0 | 6 | 0.3333 | 8 | 0.375 | 24 | 0.2917 |
| 5_10676620 | 2 | 0 | 5 | 1 | 4 | 0.75 | 3 | 1 | 14 | 0.7857 | 6 | 0.8333 | 7 | 0.8571 | 6 | 1 | 7 | 0.4286 | 26 | 0.7692 |
| 5_10676625 | 2 | 0 | 5 | 0 | 4 | 0.25 | 3 | 0 | 14 | 0.0714 | 6 | 0 | 6 | 0 | 6 | 0 | 7 | 0.2857 | 25 | 0.08 |
| 5_10676627 | 1 | 0 | 4 | 0.25 | 3 | 0.3333 | 2 | 0.5 | 10 | 0.3 | 5 | 0 | 7 | 0 | 5 | 0.2 | 8 | 0.125 | 25 | 0.08 |
| 5_10676851 | NA | NA | NA | NA | 1 | 1 | NA | NA | 1 | 1 | NA | NA | NA | NA | 1 | 0 | 1 | 1 | 2 | 0.5 |
| 5_10678271 | 2 | 0 | 4 | 0.25 | 4 | 0.25 | 2 | 0.5 | 12 | 0.25 | 5 | 0.4 | 6 | 0.6667 | 4 | 0.5 | 5 | 0 | 20 | 0.4 |
| 5_10678280 | 2 | 0.5 | 4 | 0 | 4 | 0 | 2 | 0 | 12 | 0.0833 | 6 | 0 | 7 | 0 | 5 | 0.2 | 6 | 0.1667 | 24 | 0.0833 |
| 5_10678286 | 1 | 0 | 3 | 0.6667 | 4 | 0.75 | 1 | 1 | 9 | 0.6667 | 4 | 0.75 | 5 | 1 | 4 | 0.5 | 4 | 0.5 | 17 | 0.7059 |
| 5_10678290 | 2 | 1 | 3 | 0.3333 | 2 | 0 | 1 | 1 | 8 | 0.5 | 6 | 0.3333 | 4 | 0.75 | 3 | 0.3333 | 4 | 0.25 | 17 | 0.4118 |
| 5_10678298 | 2 | 0 | 4 | 0.5 | 4 | 0 | 2 | 0 | 12 | 0.1667 | 6 | 0.3333 | 6 | 0.3333 | 5 | 0.4 | 6 | 0.1667 | 23 | 0.3043 |
| 5_10678305 | 2 | 1 | 4 | 1 | 4 | 0.25 | 2 | 0 | 12 | 0.5833 | 6 | 0.3333 | 5 | 0 | 5 | 0.4 | 4 | 0.75 | 20 | 0.35 |
| 5_10678319 | 1 | 1 | 4 | 0.75 | 3 | 1 | 1 | 1 | 9 | 0.8889 | 6 | 0.8333 | 6 | 1 | 5 | 1 | 3 | 0.6667 | 20 | 0.9 |
| 5_10678360 | 1 | 1 | 3 | 0.3333 | 3 | 0 | 1 | 0 | 8 | 0.25 | 5 | 0.2 | 6 | 0.5 | 4 | 0 | 3 | 0.3333 | 18 | 0.2778 |
| 5_10678369 | 1 | 1 | 2 | 1 | 2 | 1 | NA | NA | 5 | 1 | 5 | 1 | 6 | 1 | 4 | 0.75 | 3 | 1 | 18 | 0.9444 |
| 5_10678384 | 1 | 1 | 3 | 0 | 3 | 0.3333 | NA | NA | 7 | 0.2857 | 5 | 0.2 | 6 | 0 | 5 | 0.2 | 4 | 0 | 20 | 0.1 |
| 5_10678389 | NA | NA | 2 | 0.5 | 3 | 0.3333 | NA | NA | 5 | 0.4 | 3 | 0 | 6 | 0.1667 | 4 | 0.25 | 4 | 0.25 | 17 | 0.1765 |
| 5_10678390 | 1 | 1 | 3 | 0 | 2 | 0 | NA | NA | 6 | 0.1667 | NA | NA | 4 | 0.25 | 5 | 0.2 | 3 | 0.3333 | 12 | 0.25 |
| 5_10678404 | 1 | 1 | 2 | 1 | 2 | 0.5 | NA | NA | 5 | 0.8 | 1 | 1 | 5 | 0.8 | 3 | 1 | 4 | 1 | 13 | 0.9231 |
| 5_10678408 | 1 | 1 | 2 | 0 | 3 | 0.3333 | NA | NA | 6 | 0.3333 | 3 | 0.3333 | 6 | 0.6667 | 4 | 0.5 | 4 | 0.75 | 17 | 0.5882 |
| 5_10678432 | 1 | 0 | 2 | 0 | 2 | 0.5 | NA | NA | 5 | 0.2 | 3 | 0 | 6 | 0 | 5 | 0.4 | 4 | 0 | 18 | 0.1111 |
| 5_10678455 | 1 | 0 | 2 | 1 | 2 | 0.5 | NA | NA | 5 | 0.6 | 3 | 0.6667 | 6 | 1 | 5 | 0.6 | 4 | 1 | 18 | 0.8333 |
| 5_10678458 | 1 | 0 | 2 | 0.5 | 2 | 0.5 | NA | NA | 5 | 0.4 | 2 | 0 | 5 | 0.4 | 5 | 0.4 | 4 | 0.75 | 16 | 0.4375 |
| 5_10678467 | 1 | 1 | 2 | 1 | 1 | 1 | NA | NA | 4 | 1 | 3 | 0.6667 | 5 | 0.8 | 5 | 1 | 4 | 1 | 17 | 0.8824 |
| 5_10678482 | 1 | 1 | 1 | 0 | 2 | 0.5 | NA | NA | 4 | 0.5 | 3 | 0.6667 | 4 | 0.75 | 5 | 0.4 | 4 | 0.5 | 16 | 0.5625 |
| 5_10678517 | 1 | 1 | 2 | 0.5 | 2 | 1 | NA | NA | 5 | 0.8 | 2 | 1 | 4 | 1 | 2 | 1 | 3 | 0.6667 | 11 | 0.9091 |
| 5_10678547 | 1 | 1 | 2 | 1 | 1 | 1 | NA | NA | 4 | 1 | 2 | 1 | 4 | 1 | 4 | 1 | 1 | 1 | 11 | 1 |
| 5_10678552 | 1 | 0 | 2 | 0 | 2 | 0.5 | NA | NA | 5 | 0.2 | 1 | 0 | 1 | 0 | 4 | 0.25 | 2 | 0 | 8 | 0.125 |

| | | | | | | | | | | | | | | | | | | | | |
|---|---|---|---|---|---|---|---|---|---|---|---|---|---|---|---|---|---|---|---|---|
| 5_10678553 | 1 | 0 | 2 | 0.5 | 2 | 0 | NA | NA | 5 | 0.2 | 2 | 0.5 | 3 | 0.6667 | 4 | 0 | 2 | 0.5 | 11 | 0.3636 |
| 5_10678561 | 1 | 0 | 2 | 0.5 | 2 | 0 | NA | NA | 5 | 0.2 | 2 | 0 | 3 | 0 | 4 | 0.25 | 1 | 0 | 10 | 0.1 |
| 5_10678616 | NA | NA | 1 | 0 | 2 | 1 | NA | NA | 3 | 0.6667 | NA | NA | 2 | 1 | 1 | 1 | 2 | 0.5 | 5 | 0.8 |
| 5_10678619 | NA | NA | 1 | 1 | 2 | 1 | NA | NA | 3 | 1 | NA | NA | 2 | 1 | 1 | 1 | 2 | 0.5 | 5 | 0.8 |
| 5_10678638 | NA | NA | 1 | 1 | 2 | 0 | NA | NA | 3 | 0.3333 | NA | NA | 2 | 0.5 | 1 | 0 | 1 | 1 | 4 | 0.5 |
| 5_10713090 | 1 | 1 | NA | NA | NA | NA | NA | NA | 1 | 1 | NA | NA | 2 | 0.5 | NA | NA | NA | NA | 2 | 0.5 |
| 5_10713102 | 1 | 1 | NA | NA | 2 | 1 | NA | NA | 3 | 1 | NA | NA | 2 | 1 | NA | NA | NA | NA | 2 | 1 |
| 5_10713104 | 1 | 1 | NA | NA | 2 | 1 | NA | NA | 3 | 1 | NA | NA | 2 | 0.5 | NA | NA | NA | NA | 2 | 0.5 |
| 5_10713107 | NA | NA | 1 | 1 | 2 | 0.5 | NA | NA | 3 | 0.6667 | NA | NA | 2 | 1 | NA | NA | NA | NA | 2 | 1 |
| 5_10713110 | 1 | 1 | 1 | 1 | 2 | 1 | NA | NA | 4 | 1 | NA | NA | 2 | 1 | NA | NA | NA | NA | 2 | 1 |
| 5_10713114 | 1 | 1 | 1 | 1 | 1 | 0 | NA | NA | 3 | 0.6667 | NA | NA | 2 | 0.5 | NA | NA | NA | NA | 2 | 0.5 |
| 5_10725468 | 3 | 0 | 1 | 0 | 1 | 0 | 2 | 0.5 | 7 | 0.1429 | 1 | 0 | NA | NA | 2 | 0.5 | 3 | 0 | 6 | 0.1667 |
| 5_10725506 | 3 | 0.3333 | 1 | 0 | 1 | 0 | 2 | 0.5 | 7 | 0.2857 | 1 | 0 | NA | NA | 2 | 0 | 3 | 0.3333 | 6 | 0.1667 |
| 5_10725551 | 3 | 0.3333 | 1 | 0 | NA | NA | 2 | 0 | 6 | 0.1667 | 1 | 1 | NA | NA | 2 | 0.5 | 3 | 0 | 6 | 0.3333 |
| 5_10725730 | 2 | 0.5 | NA | NA | NA | NA | 2 | 0 | 4 | 0.25 | 1 | 1 | NA | NA | 1 | 1 | 1 | 0 | 3 | 0.6667 |
| 5_10750677 | 1 | 1 | NA | NA | NA | NA | NA | NA | 1 | 1 | 1 | 1 | NA | NA | NA | NA | NA | NA | 1 | 1 |
| 5_10753339 | 2 | 0 | 3 | 0 | 3 | 0.3333 | 2 | 0 | 10 | 0.1 | NA | NA | 5 | 0 | 2 | 0 | 4 | 0.25 | 11 | 0.0909 |
| 5_10753362 | 2 | 1 | 4 | 0.25 | 4 | 0.25 | 2 | 1 | 12 | 0.5 | NA | NA | 5 | 0.2 | 2 | 0.5 | 3 | 0 | 10 | 0.2 |
| 5_10753397 | 2 | 0 | 3 | 0.3333 | 3 | 0 | 2 | 0 | 10 | 0.1 | NA | NA | 4 | 0.5 | 1 | 0 | 5 | 0 | 10 | 0.2 |
| 5_10753806 | 1 | 1 | 1 | 1 | 2 | 1 | 1 | 1 | 5 | 1 | NA | NA | NA | NA | 1 | 1 | NA | NA | 1 | 1 |
| 5_10753815 | 1 | 1 | 1 | 1 | 2 | 0 | NA | NA | 4 | 0.5 | NA | NA | NA | NA | 1 | 1 | NA | NA | 1 | 1 |
| 5_10753823 | 1 | 0 | 1 | 1 | 2 | 0.5 | 1 | 0 | 5 | 0.4 | NA | NA | NA | NA | 1 | 1 | NA | NA | 1 | 1 |
| 5_10757764 | 7 | 0.2857 | 12 | 0.1667 | 8 | 0.125 | 8 | 0.125 | 35 | 0.1714 | 6 | 0.1667 | 12 | 0.1667 | 14 | 0.0714 | 6 | 0.3333 | 38 | 0.1579 |
| 5_10757765 | 8 | 0 | 7 | 0.2857 | 8 | 0.25 | 4 | 0.5 | 27 | 0.2222 | 6 | 0 | 13 | 0.0769 | 8 | 0.125 | 5 | 0 | 32 | 0.0625 |
| 5_10757771 | 9 | 0.5556 | 10 | 0.9 | 10 | 0.6 | 8 | 0.875 | 37 | 0.7297 | 7 | 1 | 12 | 0.75 | 12 | 0.8333 | 7 | 0.8571 | 38 | 0.8421 |
| 5_10757775 | 8 | 0.375 | 13 | 0.3077 | 11 | 0.5455 | 7 | 0.5714 | 39 | 0.4359 | 6 | 0.5 | 15 | 0.6667 | 15 | 0.3333 | 6 | 0.3333 | 42 | 0.4762 |
| 5_10757775 | 9 | 0.1111 | 11 | 0.1818 | 10 | 0.1 | 7 | 0.1429 | 37 | 0.1351 | 5 | 0.2 | 15 | 0.1333 | 10 | 0.1 | 7 | 0 | 37 | 0.1081 |
| 5_10757780 | 9 | 0.1111 | 13 | 0.0769 | 11 | 0.1818 | 8 | 0.375 | 41 | 0.1707 | 3 | 0 | 13 | 0.3077 | 14 | 0.0714 | 5 | 0 | 35 | 0.1429 |
| 5_10765125 | 2 | 0.5 | 3 | 0.3333 | 3 | 0.3333 | 5 | 0.8 | 13 | 0.5385 | NA | NA | 3 | 1 | 5 | 0.6 | 1 | 1 | 9 | 0.7778 |
| 5_10765133 | 2 | 0 | 3 | 0.3333 | 4 | 0 | 6 | 0.1667 | 15 | 0.1333 | NA | NA | 3 | 0.3333 | 5 | 0 | 2 | 0.5 | 10 | 0.2 |
| 5_10765138 | 1 | 0 | 3 | 0 | 4 | 0 | 5 | 0.4 | 13 | 0.1538 | NA | NA | 2 | 0 | 5 | 0.4 | 2 | 0.5 | 9 | 0.3333 |
| 5_10765142 | 2 | 0 | 4 | 0.25 | 3 | 0.3333 | 6 | 0.1667 | 15 | 0.2 | NA | NA | 3 | 0.6667 | 4 | 0 | 1 | 0 | 8 | 0.25 |
| 5_10765145 | 1 | 0 | 4 | 0 | 3 | 0.3333 | 4 | 0.25 | 12 | 0.1667 | NA | NA | 3 | 0.6667 | 5 | 0.8 | 2 | 0 | 10 | 0.6 |
| 5_10765156 | 2 | 0.5 | 5 | 0.2 | 4 | 0.5 | 6 | 0.1667 | 17 | 0.2941 | NA | NA | 4 | 0.75 | 5 | 0.4 | 2 | 0 | 11 | 0.4545 |
| 5_10765482 | 3 | 0 | 6 | 0.1667 | 2 | 0 | 5 | 0.2 | 16 | 0.125 | 1 | 0 | 5 | 0.2 | 6 | 0.3333 | 3 | 0.3333 | 15 | 0.2667 |
| 5_10765486 | 3 | 0.3333 | 6 | 0.5 | 4 | 0.5 | 6 | 0.8333 | 19 | 0.5789 | 1 | 1 | 5 | 0.4 | 4 | 0.5 | 4 | 0.25 | 14 | 0.4286 |
| 5_10765494 | 4 | 0.75 | 6 | 0.8333 | 4 | 0.5 | 6 | 0.8333 | 20 | 0.75 | 1 | 1 | 5 | 0.6 | 6 | 0.6667 | 4 | 0.5 | 16 | 0.625 |
| 5_10765498 | 3 | 0.3333 | 6 | 0.3333 | 4 | 0.25 | 4 | 0.25 | 17 | 0.2941 | 1 | 0 | 5 | 0.8 | 7 | 0.2857 | 4 | 0 | 17 | 0.3529 |
| 5_10765506 | 4 | 0.25 | 5 | 0 | 3 | 0 | 6 | 0 | 18 | 0.0556 | 1 | 0 | 5 | 0 | 6 | 0.1667 | 3 | 0 | 15 | 0.0667 |
| 5_10765520 | 4 | 1 | 7 | 0.8571 | 3 | 0.6667 | 6 | 0.6667 | 20 | 0.8 | 1 | 1 | 4 | 1 | 6 | 1 | 5 | 0.8 | 16 | 0.9375 |
| 5_10765527 | 4 | 0.25 | 7 | 0.2857 | 4 | 1 | 5 | 0.6 | 20 | 0.5 | 1 | 0 | 5 | 0.6 | 7 | 0.4286 | 5 | 0.4 | 18 | 0.4444 |
| 5_10765531 | 4 | 1 | 7 | 0.8571 | 4 | 0.5 | 5 | 0.8 | 20 | 0.8 | 1 | 0 | 6 | 0.6667 | 7 | 0.8571 | 5 | 0.8 | 19 | 0.7368 |
| 5_10765534 | 2 | 0.5 | 6 | 0.3333 | 4 | 0.75 | 5 | 0.4 | 17 | 0.4706 | 1 | 1 | 3 | 0 | 6 | 0.3333 | 4 | 0.25 | 14 | 0.2857 |
| 5_10765539 | 4 | 0 | 7 | 0 | 3 | 0 | 6 | 0.3333 | 20 | 0.1 | 1 | 0 | 4 | 0.25 | 7 | 0.2857 | 5 | 0 | 17 | 0.1765 |
| 5_10765540 | 2 | 0 | 7 | 0.4286 | 4 | 0.75 | 5 | 0.4 | 18 | 0.4444 | 1 | 0 | 4 | 0.5 | 6 | 0.3333 | 4 | 1 | 15 | 0.5333 |
| 5_10765542 | 4 | 0 | 7 | 0.1429 | 4 | 0.5 | 6 | 0.1667 | 21 | 0.1905 | 1 | 1 | 6 | 0.3333 | 7 | 0.1429 | 4 | 0.25 | 18 | 0.2778 |
| 5_10765564 | 5 | 0.4 | 5 | 0.4 | 3 | 0.3333 | 8 | 0.875 | 21 | 0.5714 | 1 | 1 | 6 | 0.5 | 6 | 0.6667 | 5 | 0.8 | 18 | 0.6667 |
| 5_10765569 | 5 | 0.4 | 6 | 0 | 4 | 0.25 | 8 | 0.125 | 23 | 0.1739 | 1 | 0 | 7 | 0.1429 | 7 | 0.2857 | 5 | 0 | 20 | 0.15 |
| 5_10765575 | 6 | 0.1667 | 6 | 0.1667 | 4 | 0 | 6 | 0.1667 | 22 | 0.1364 | 1 | 0 | 6 | 0.1667 | 5 | 0.6 | 6 | 0 | 18 | 0.2222 |
| 5_10765588 | 6 | 0.3333 | 7 | 0.4286 | 3 | 0 | 5 | 0.2 | 21 | 0.2857 | 1 | 0 | 7 | 0.2857 | 8 | 0 | 6 | 0.3333 | 22 | 0.1818 |
| 5_10765603 | 5 | 0.8 | 7 | 0.5714 | 3 | 0.6667 | 9 | 0.5556 | 24 | 0.625 | 1 | 0 | 9 | 0.6667 | 8 | 0.75 | 6 | 0.5 | 24 | 0.625 |
| 5_10765632 | 8 | 0.375 | 7 | 0.2857 | 3 | 0 | 7 | 0.1429 | 25 | 0.24 | 1 | 0 | 9 | 0 | 8 | 0.625 | 6 | 0.1667 | 24 | 0.25 |
| 5_10765652 | 8 | 0.875 | 7 | 1 | 3 | 0.6667 | 9 | 0.7778 | 27 | 0.8519 | 3 | 0.6667 | 10 | 0.9 | 8 | 1 | 7 | 1 | 28 | 0.9286 |
| 5_10765664 | 7 | 0.4286 | 7 | 0.5714 | 2 | 1 | 8 | 0.25 | 24 | 0.4583 | 3 | 0.6667 | 7 | 0.5714 | 6 | 0.6667 | 8 | 0.375 | 24 | 0.5417 |
| 5_10765680 | 6 | 1 | 10 | 0.9 | 5 | 0.8 | 8 | 0.75 | 29 | 0.8621 | 4 | 1 | 11 | 0.7273 | 6 | 1 | 7 | 1 | 28 | 0.8929 |
| 5_10765685 | 9 | 0.6667 | 8 | 0.375 | 5 | 1 | 8 | 0.5 | 30 | 0.6 | 4 | 0.25 | 10 | 0.6 | 8 | 0.625 | 7 | 0.5714 | 29 | 0.5517 |
| 5_10765689 | 8 | 1 | 9 | 1 | 5 | 1 | 9 | 1 | 31 | 1 | 4 | 1 | 11 | 1 | 6 | 1 | 7 | 1 | 28 | 1 |
| 5_10765708 | 7 | 0.1429 | 3 | 0 | 3 | 0 | 8 | 0.375 | 21 | 0.1905 | 2 | 0 | 7 | 0.4286 | 6 | 0.1667 | 5 | 0.2 | 20 | 0.25 |
| 5_10765723 | 7 | 0.1429 | 3 | 0 | 5 | 0.2 | 9 | 0 | 24 | 0.0833 | 3 | 0.3333 | 8 | 0.5 | 6 | 0.1667 | 5 | 0.2 | 22 | 0.3182 |
| 5_10765731 | 5 | 0 | 3 | 0 | 4 | 0.25 | 9 | 0.2222 | 21 | 0.1429 | 2 | 0 | 7 | 0.1429 | 2 | 0.5 | 4 | 0 | 15 | 0.1333 |
| 5_10765732 | 7 | 0.2857 | 3 | 1 | 5 | 0.8 | 9 | 0.5556 | 24 | 0.5833 | 4 | 0.25 | 8 | 0.875 | 6 | 0.6667 | 4 | 0.25 | 22 | 0.5909 |
| 5_10765743 | 8 | 0 | 4 | 0.25 | 5 | 0 | 9 | 0.2222 | 26 | 0.1154 | 4 | 0.25 | 8 | 0.25 | 7 | 0.2857 | 5 | 0.4 | 24 | 0.2917 |
| 5_10765779 | 10 | 0.1 | 4 | 0 | 6 | 0 | 9 | 0.2222 | 29 | 0.1034 | 8 | 0 | 11 | 0.0909 | 7 | 0.1429 | 5 | 0 | 31 | 0.0645 |
| 5_10765816 | 3 | 0.6667 | 3 | 0 | 5 | 0 | 5 | 0 | 16 | 0.125 | 2 | 0 | 4 | 0.25 | 3 | 0 | 3 | 0.6667 | 12 | 0.25 |
| 5_10765839 | 8 | 1 | 11 | 1 | 4 | 0.75 | 8 | 1 | 31 | 0.9677 | 5 | 1 | 8 | 1 | 5 | 0.8 | 9 | 0.8889 | 27 | 0.9259 |

| | | | | | | | | | | | | | | | | | | | | |
|---|---|---|---|---|---|---|---|---|---|---|---|---|---|---|---|---|---|---|---|---|
| 5_1076584 | 6 | 0.8333 | 6 | 0.6667 | 4 | 0.75 | 5 | 1 | 21 | 0.8095 | 3 | 0.3333 | 5 | 0.8 | 3 | 0.6667 | 4 | 0.5 | 15 | 0.6 |
| 5_1076584 | 11 | 0.0909 | 7 | 0.1429 | 6 | 0.1667 | 7 | 0.1429 | 31 | 0.129 | 5 | 0 | 11 | 0.0909 | 3 | 0 | 9 | 0.1111 | 28 | 0.0714 |
| 5_1076584 | 12 | 0 | 11 | 0.0909 | 7 | 0.1429 | 8 | 0.25 | 38 | 0.1053 | 6 | 0.3333 | 11 | 0 | 5 | 0 | 10 | 0 | 32 | 0.0625 |
| 5_1076584 | 11 | 0.6364 | 8 | 0.375 | 5 | 0 | 6 | 0.6667 | 30 | 0.4667 | 3 | 0.3333 | 8 | 0.375 | 3 | 0 | 8 | 0.5 | 22 | 0.3636 |
| 5_1076585 | 11 | 0.1818 | 7 | 0.2857 | 6 | 0 | 5 | 0.4 | 29 | 0.2069 | 5 | 0 | 11 | 0.2727 | 5 | 0 | 6 | 0.5 | 27 | 0.2222 |
| 5_1076585 | 12 | 1 | 9 | 1 | 7 | 0.7143 | 8 | 1 | 36 | 0.9444 | 6 | 1 | 13 | 1 | 5 | 1 | 10 | 1 | 34 | 1 |
| 5_1076586 | 10 | 0.6 | 10 | 0.1 | 7 | 0.7143 | 7 | 0.4286 | 34 | 0.4412 | 2 | 0 | 13 | 0.4615 | 5 | 0.8 | 10 | 0.5 | 30 | 0.5 |
| 5_1076586 | 16 | 0.125 | 10 | 0 | 8 | 0.125 | 8 | 0.125 | 42 | 0.0952 | 7 | 0.1429 | 16 | 0.0625 | 5 | 0.4 | 11 | 0 | 39 | 0.1026 |
| 5_1076587 | 15 | 0.5333 | 9 | 0.6667 | 8 | 0.625 | 9 | 0.7778 | 41 | 0.6341 | 9 | 0.4444 | 15 | 0.6667 | 6 | 0.5 | 12 | 0.75 | 42 | 0.619 |
| 5_1076634 | 3 | 0 | 1 | 1 | NA | NA | NA | NA | 4 | 0.25 | NA | NA | NA | NA | 1 | 1 | NA | NA | 1 | 1 |
| 5_1076965 | NA | NA | 1 | 1 | NA | NA | NA | NA | 1 | 1 | NA | NA | NA | NA | NA | NA | 1 | 1 | 1 | 1 |
| 5_1077679 | 1 | 1 | 2 | 1 | 1 | 1 | NA | NA | 4 | 1 | 1 | 1 | NA | NA | 1 | 1 | 3 | 1 | 5 | 1 |
| 5_1077680 | 1 | 0 | 4 | 0.25 | 2 | 0.5 | NA | NA | 7 | 0.2857 | 1 | 0 | NA | NA | 2 | 0.5 | 3 | 0 | 6 | 0.1667 |
| 5_1077681 | 1 | 0 | 4 | 0.5 | 1 | 0 | NA | NA | 6 | 0.3333 | NA | NA | NA | NA | 1 | 1 | 3 | 0.3333 | 4 | 0.5 |
| 5_1077682 | 1 | 0 | 3 | 1 | 2 | 1 | NA | NA | 6 | 0.8333 | 1 | 1 | NA | NA | 2 | 1 | 2 | 1 | 5 | 1 |
| 5_1077683 | 1 | 0 | 3 | 0.3333 | 2 | 0.5 | NA | NA | 6 | 0.3333 | 1 | 0 | NA | NA | 2 | 1 | 3 | 0.3333 | 6 | 0.5 |
| 5_1077685 | 1 | 0 | 4 | 0.5 | 2 | 0 | NA | NA | 7 | 0.2857 | 1 | 0 | NA | NA | NA | NA | 1 | 1 | 2 | 0.5 |
| 5_1078927 | 4 | 0 | 4 | 0 | 9 | 0.1111 | 4 | 0.25 | 21 | 0.0952 | 6 | 0.1667 | 2 | 0 | 4 | 0.25 | 3 | 0.6667 | 15 | 0.2667 |
| 5_1078937 | 5 | 0 | 3 | 0.3333 | 8 | 0 | 3 | 0 | 19 | 0.0526 | 2 | 0 | 5 | 0.4 | 1 | 0 | 3 | 0 | 11 | 0.1818 |
| 5_1078955 | 3 | 0 | 4 | 0.25 | 6 | 0 | 3 | 0 | 16 | 0.0625 | 2 | 0.5 | 2 | 0 | 1 | 0 | 2 | 0 | 7 | 0.1429 |
| 5_1078957 | 2 | 1 | 5 | 1 | 4 | 1 | 3 | 1 | 14 | 1 | 2 | 1 | 3 | 0.6667 | 1 | 1 | 1 | 1 | 7 | 0.8571 |
| 5_1078958 | 3 | 1 | 3 | 1 | 4 | 0.75 | 1 | 1 | 11 | 0.9091 | 2 | 1 | 2 | 0.5 | 1 | 0 | 1 | 0 | 6 | 0.5 |
| 5_1078959 | 2 | 1 | 3 | 0.6667 | 5 | 0.6 | 3 | 0.6667 | 13 | 0.6923 | 1 | 1 | 2 | 0.5 | NA | NA | 1 | 0 | 4 | 0.5 |
| 5_1080562 | 5 | 0.2 | 4 | 0.5 | 4 | 0 | 1 | 0 | 14 | 0.2143 | 5 | 0 | 4 | 0.25 | 5 | 0 | 2 | 0 | 16 | 0.0625 |
| 5_1080569 | 4 | 0 | 3 | 0 | 4 | 0.25 | 6 | 0.3333 | 17 | 0.1765 | 2 | 0 | 3 | 0 | 3 | 0.3333 | 1 | 0 | 9 | 0.1111 |
| 5_1080570 | 5 | 0.2 | 4 | 0.25 | 3 | 0.6667 | 8 | 0 | 20 | 0.2 | 2 | 0.5 | 3 | 0.3333 | 4 | 0.25 | 1 | 1 | 10 | 0.4 |
| 5_1080571 | 5 | 0 | 4 | 0 | 3 | 0.3333 | 7 | 0 | 19 | 0.0526 | 2 | 0 | 3 | 0 | 2 | 0.5 | 1 | 0 | 8 | 0.125 |
| 5_1080572 | 4 | 0.75 | 3 | 1 | 3 | 1 | 6 | 1 | 16 | 0.9375 | 2 | 1 | 3 | 1 | 2 | 1 | 1 | 1 | 8 | 1 |
| 5_1080572 | 3 | 1 | 3 | 0.6667 | 3 | 0.6667 | 5 | 0.8 | 14 | 0.7857 | 2 | 1 | 2 | 0.5 | 4 | 1 | NA | NA | 8 | 0.875 |
| 5_1080573 | 4 | 1 | 4 | 0.75 | 4 | 0.75 | 9 | 0.8889 | 21 | 0.8571 | 1 | 1 | 2 | 1 | 3 | 1 | NA | NA | 6 | 1 |
| 5_1080573 | 3 | 0 | 3 | 0 | 5 | 0 | 9 | 0.1111 | 20 | 0.05 | 1 | 0 | 2 | 0 | 2 | 0 | 1 | 1 | 6 | 0.1667 |
| 5_1080574 | 3 | 0.6667 | 4 | 1 | 4 | 0.5 | 10 | 0.6 | 21 | 0.6667 | 1 | 1 | 2 | 0.5 | 3 | 1 | 1 | 1 | 7 | 0.8571 |
| 5_1080575 | 4 | 0.25 | 4 | 0.25 | 5 | 0 | 10 | 0.2 | 23 | 0.1739 | 1 | 0 | 1 | 0 | 2 | 0.5 | 1 | 1 | 5 | 0.4 |
| 5_1080575 | 3 | 0.6667 | 4 | 0.5 | 4 | 0.75 | 9 | 0.3333 | 20 | 0.5 | 1 | 0 | 2 | 0.5 | 1 | 0 | NA | NA | 4 | 0.25 |
| 5_1080575 | 4 | 1 | 2 | 1 | 5 | 1 | 9 | 0.7778 | 20 | 0.9 | NA | NA | 2 | 1 | 3 | 1 | 1 | 1 | 6 | 1 |
| 5_1080575 | 3 | 0.6667 | NA | NA | 5 | 0.8 | 7 | 0.5714 | 15 | 0.6667 | 1 | 0 | 1 | 1 | 3 | 0.6667 | 1 | 1 | 6 | 0.6667 |
| 5_1080575 | 4 | 1 | 4 | 1 | 5 | 0.6 | 10 | 0.9 | 23 | 0.8696 | 1 | 1 | 2 | 1 | 3 | 0.6667 | 1 | 1 | 7 | 0.8571 |
| 5_1080575 | 4 | 1 | 4 | 0.5 | 5 | 0.4 | 10 | 0.4 | 23 | 0.5217 | 1 | 1 | 2 | 0 | 3 | 0.6667 | 1 | 0 | 7 | 0.4286 |
| 5_1080576 | 4 | 0 | 4 | 0 | 4 | 0.25 | 9 | 0.1111 | 21 | 0.0952 | NA | NA | 2 | 0 | 3 | 0.3333 | 1 | 0 | 6 | 0.1667 |
| 5_1080576 | 3 | 0.3333 | 3 | 0 | 4 | 0.25 | 8 | 0.375 | 18 | 0.2778 | NA | NA | 2 | 0 | 3 | 0.3333 | 1 | 1 | 6 | 0.3333 |
| 5_1080577 | 4 | 1 | 4 | 1 | 3 | 1 | 9 | 0.8889 | 20 | 0.95 | 1 | 1 | 1 | 1 | 2 | 0.5 | 1 | 1 | 5 | 0.8 |
| 5_1080578 | 2 | 0.5 | 3 | 1 | 4 | 1 | 6 | 0.8333 | 15 | 0.8667 | 1 | 1 | 1 | 1 | 2 | 1 | 1 | 1 | 5 | 1 |
| 5_1080579 | 4 | 0 | 3 | 0 | 4 | 0.25 | 9 | 0 | 20 | 0.05 | NA | NA | 2 | 0.5 | 2 | 0 | 1 | 0 | 5 | 0.2 |
| 5_1080580 | 4 | 0 | 3 | 0 | 4 | 0 | 9 | 0.1111 | 20 | 0.05 | NA | NA | 1 | 0 | 3 | 0 | 1 | 1 | 5 | 0.2 |
| 5_1082137 | NA | NA | 4 | 0.25 | 2 | 0 | 2 | 0 | 8 | 0.125 | NA | NA | 3 | 0 | 1 | 0 | 3 | 0.3333 | 7 | 0.1429 |
| 5_1082544 | NA | NA | 4 | 0.25 | 3 | 0 | 5 | 0 | 12 | 0.0833 | 2 | 0 | 6 | 0 | 4 | 0 | 5 | 0.2 | 17 | 0.0588 |
| 5_1082545 | NA | NA | 4 | 0.25 | 3 | 0 | 5 | 0 | 12 | 0.0833 | 2 | 0 | 6 | 0 | 5 | 0.6 | 5 | 0.2 | 18 | 0.2222 |
| 5_1082545 | NA | NA | 3 | 0.3333 | 2 | 0 | 5 | 0.4 | 10 | 0.3 | 2 | 0.5 | 4 | 0 | 1 | 0 | 4 | 0.5 | 11 | 0.2727 |
| 5_1082546 | NA | NA | 4 | 0.75 | 3 | 0.6667 | 4 | 0.75 | 11 | 0.7273 | 2 | 0.5 | 5 | 1 | 5 | 0.8 | 5 | 1 | 17 | 0.8824 |
| 5_1082546 | NA | NA | 4 | 0.25 | 4 | 0.5 | 5 | 0.2 | 13 | 0.3077 | 2 | 0 | 4 | 0.25 | 5 | 0.2 | 4 | 0.5 | 15 | 0.2667 |
| 5_1082546 | NA | NA | 3 | 0.3333 | 4 | 0.25 | 4 | 0.5 | 11 | 0.3636 | 2 | 0.5 | 6 | 0.6667 | 3 | 0.3333 | 5 | 0.4 | 16 | 0.5 |
| 5_1082548 | NA | NA | 4 | 0.25 | 4 | 0.25 | 5 | 0.2 | 13 | 0.2308 | 2 | 0.5 | 5 | 0.2 | 6 | 0.3333 | 6 | 0.3333 | 19 | 0.3158 |
| 5_1082548 | NA | NA | 4 | 1 | 4 | 1 | 5 | 1 | 13 | 1 | 2 | 1 | 5 | 1 | 6 | 0.8333 | 6 | 1 | 19 | 0.9474 |
| 5_1082548 | NA | NA | 4 | 0 | 3 | 0 | 4 | 0.25 | 11 | 0.0909 | 2 | 0 | 4 | 0 | 6 | 0.3333 | 5 | 0.2 | 17 | 0.1765 |
| 5_1082550 | NA | NA | 4 | 0 | 4 | 0.25 | 5 | 0 | 13 | 0.0769 | 2 | 0 | 6 | 0.5 | 6 | 0.3333 | 6 | 0.3333 | 20 | 0.35 |
| 5_1082553 | 1 | 0 | 4 | 0.25 | 4 | 0.5 | 4 | 0.75 | 13 | 0.4615 | 2 | 0 | 6 | 0.3333 | 7 | 0.4286 | 5 | 0.4 | 20 | 0.35 |
| 5_1082564 | 1 | 0 | 5 | 0 | 4 | 0 | 4 | 0.25 | 14 | 0.0714 | 3 | 0.3333 | 3 | 0 | 6 | 0.1667 | 6 | 0 | 18 | 0.1111 |
| 5_1082567 | NA | NA | 6 | 0.1667 | 4 | 0 | 3 | 0.3333 | 13 | 0.1538 | 4 | 0 | 4 | 0 | 6 | 0.1667 | 6 | 0.3333 | 20 | 0.15 |
| 5_1082703 | 3 | 0.3333 | 4 | 0.25 | 4 | 0 | 3 | 0.3333 | 14 | 0.2143 | 1 | 0 | 3 | 0.3333 | 4 | 0.5 | 3 | 0 | 11 | 0.2727 |
| 5_1082717 | 3 | 0.3333 | 4 | 0.5 | 2 | 0.5 | 3 | 0.3333 | 12 | 0.4167 | 2 | 0.5 | 3 | 0 | 2 | 0 | 4 | 0.5 | 11 | 0.2727 |
| 5_1082801 | 2 | 0 | 2 | 0.5 | 2 | 0 | 2 | 0 | 8 | 0.125 | 1 | 0 | 4 | 0.5 | 3 | 0 | 2 | 0.5 | 10 | 0.3 |
| 5_1084173 | 7 | 0 | 4 | 0.5 | 7 | 0 | 4 | 0.25 | 22 | 0.1364 | 3 | 0 | 4 | 0.5 | 6 | 0.1667 | 6 | 0 | 19 | 0.1579 |
| 5_1084173 | 6 | 0.3333 | 3 | 0 | 7 | 0.2857 | 4 | 0 | 20 | 0.2 | 4 | 0 | 4 | 0 | 5 | 0 | 9 | 0.4444 | 22 | 0.1818 |
| 5_1084174 | 5 | 0.2 | 3 | 0.6667 | 6 | 0 | 2 | 0.5 | 16 | 0.25 | 3 | 0 | 4 | 0 | 5 | 0 | 7 | 0.2857 | 19 | 0.1053 |
| 5_1084175 | 6 | 1 | 4 | 1 | 7 | 0.8571 | 4 | 1 | 21 | 0.9524 | 4 | 1 | 4 | 1 | 6 | 1 | 9 | 1 | 23 | 1 |

| | | | | | | | | | | | | | | | | | | | | |
|---|---|---|---|---|---|---|---|---|---|---|---|---|---|---|---|---|---|---|---|---|
| 5_1084175 | 7 | 0.4286 | 4 | 0.25 | 6 | 0.3333 | 4 | 0.5 | 21 | 0.381 | 4 | 0.75 | 4 | 0.25 | 6 | 0.5 | 8 | 0.5 | 22 | 0.5 |
| 5_1084266 | 7 | 1 | 3 | 1 | 7 | 0.5714 | 4 | 1 | 21 | 0.8571 | 4 | 0.75 | 4 | 1 | 7 | 1 | 8 | 1 | 23 | 0.9565 |
| 5_1084266 | 8 | 0 | 4 | 0.25 | 6 | 0 | 4 | 0.25 | 22 | 0.0909 | 4 | 0 | 4 | 0 | 8 | 0.25 | 10 | 0 | 26 | 0.0769 |
| 5_1084266 | 8 | 0.375 | 1 | 1 | 7 | 0.2857 | 4 | 0 | 20 | 0.3 | 4 | 0.5 | 2 | 0 | 6 | 0.3333 | 9 | 0.1111 | 21 | 0.2381 |
| 5_1084268 | 6 | 0.3333 | 4 | 0 | 6 | 0.3333 | 2 | 0.5 | 18 | 0.2778 | 4 | 0.75 | 4 | 0.25 | 8 | 0 | 10 | 0 | 26 | 0.1538 |
| 5_1084268 | 8 | 0.125 | 4 | 0.25 | 6 | 0 | 4 | 0 | 22 | 0.0909 | 5 | 0 | 4 | 0 | 7 | 0.1429 | 8 | 0.125 | 24 | 0.0833 |
| 5_1084272 | 7 | 0.2857 | 4 | 0 | 5 | 0 | 3 | 0 | 19 | 0.1053 | 5 | 0.2 | 4 | 0 | 8 | 0 | 11 | 0.0909 | 28 | 0.0714 |
| 5_1084958 | 1 | 1 | 2 | 0.5 | 1 | 1 | 1 | 0 | 5 | 0.6 | 1 | 1 | 1 | 0 | 1 | 0 | 1 | 1 | 4 | 0.5 |
| 5_1084959 | 1 | 1 | 2 | 1 | NA | NA | 1 | 1 | 4 | 1 | 1 | 0 | 1 | 1 | 1 | 0 | 1 | 1 | 4 | 0.5 |
| 5_1084962 | NA | NA | 2 | 0.5 | NA | NA | 2 | 0.5 | 4 | 0.5 | 1 | 0 | 2 | 1 | 1 | 0 | 1 | 1 | 5 | 0.6 |
| 5_1085134 | 9 | 0.3333 | 2 | 0.5 | 3 | 0.3333 | 7 | 0.1429 | 21 | 0.2857 | 4 | 0.25 | 8 | 0.375 | 11 | 0.4545 | 1 | 0 | 24 | 0.375 |
| 5_1085136 | 9 | 0.6667 | 3 | 0.3333 | 4 | 0.5 | 11 | 0.6364 | 27 | 0.5926 | 4 | 0.25 | 8 | 0.625 | 12 | 0.5833 | 3 | 0 | 27 | 0.4815 |
| 5_1085137 | 8 | 0.5 | 3 | 1 | 3 | 0.3333 | 11 | 0.9091 | 25 | 0.72 | 2 | 0.5 | 6 | 0.5 | 13 | 0.6923 | 3 | 0.3333 | 24 | 0.5833 |
| 5_1085138 | 9 | 0.6667 | 2 | 1 | 4 | 0.25 | 12 | 0.5833 | 27 | 0.5926 | 3 | 0.6667 | 6 | 0.8333 | 13 | 0.8462 | 3 | 1 | 25 | 0.84 |
| 5_1085139 | 10 | 0.3 | 5 | 0.2 | 11 | 0.3636 | 13 | 0.4615 | 39 | 0.359 | 4 | 0.25 | 9 | 0.5556 | 19 | 0.5263 | 7 | 0.2857 | 39 | 0.4615 |
| 5_1085139 | 15 | 0.2667 | 5 | 0.2 | 7 | 0.1429 | 14 | 0.0714 | 41 | 0.1707 | 7 | 0 | 7 | 0.1429 | 24 | 0.125 | 7 | 0 | 45 | 0.0889 |
| 5_1085140 | 17 | 0.1176 | 7 | 0.2857 | 14 | 0.1429 | 20 | 0.15 | 58 | 0.1552 | 11 | 0.1818 | 11 | 0.0909 | 28 | 0.25 | 11 | 0.0909 | 61 | 0.1803 |
| 5_1085140 | 18 | 0.1667 | 6 | 0.1667 | 14 | 0 | 19 | 0.2105 | 57 | 0.1404 | 12 | 0 | 10 | 0.2 | 24 | 0 | 10 | 0.2 | 56 | 0.0714 |
| 5_1085140 | 18 | 1 | 7 | 0.8571 | 11 | 0.9091 | 17 | 0.8824 | 53 | 0.9245 | 10 | 0.9 | 10 | 1 | 25 | 1 | 11 | 1 | 56 | 0.9821 |
| 5_1085140 | 15 | 0.3333 | 5 | 0.2 | 12 | 0.4167 | 18 | 0.3889 | 50 | 0.36 | 10 | 0.4 | 9 | 0.4444 | 20 | 0.4 | 11 | 0.3636 | 50 | 0.4 |
| 5_1085141 | 14 | 0.2857 | 4 | 0.25 | 10 | 0.2 | 17 | 0.2353 | 45 | 0.2444 | 9 | 0.3333 | 8 | 0.125 | 23 | 0.3478 | 10 | 0.8 | 50 | 0.4 |
| 5_1085141 | 14 | 0.9286 | 4 | 0.75 | 9 | 0.8889 | 16 | 1 | 43 | 0.9302 | 8 | 0.75 | 5 | 0.8 | 25 | 0.84 | 7 | 1 | 45 | 0.8444 |
| 5_1085142 | 17 | 1 | 7 | 1 | 12 | 1 | 20 | 1 | 56 | 1 | 9 | 1 | 10 | 1 | 26 | 1 | 10 | 1 | 55 | 1 |
| 5_1085143 | 16 | 0.9375 | 5 | 1 | 12 | 0.8333 | 19 | 0.8421 | 52 | 0.8846 | 9 | 1 | 9 | 0.8889 | 22 | 0.8636 | 10 | 1 | 50 | 0.92 |
| 5_1085144 | 8 | 0.625 | 6 | 0.6667 | 11 | 0.4545 | 14 | 0.5714 | 39 | 0.5641 | 8 | 0.375 | 9 | 0.3333 | 19 | 0.3684 | 7 | 0.5714 | 43 | 0.3953 |
| 5_1085145 | 12 | 0.75 | 6 | 0.6667 | 10 | 0.8 | 18 | 0.6667 | 46 | 0.7174 | 8 | 0.625 | 11 | 0.4545 | 22 | 0.7727 | 9 | 0.5556 | 50 | 0.64 |
| 5_1085145 | 4 | 1 | 4 | 1 | 4 | 0.75 | 9 | 1 | 21 | 0.9524 | 3 | 1 | 5 | 1 | 9 | 0.7778 | 3 | 1 | 20 | 0.9 |
| 5_1085147 | 11 | 0.2727 | 7 | 0.4286 | 8 | 0.25 | 11 | 0.2727 | 37 | 0.2973 | 8 | 0 | 7 | 0.2857 | 21 | 0.3333 | 7 | 0.1429 | 43 | 0.2326 |
| 5_1085147 | 13 | 0.2308 | 6 | 0 | 9 | 0.3333 | 16 | 0.125 | 44 | 0.1818 | 9 | 0.3333 | 10 | 0.3 | 22 | 0.3182 | 8 | 0.25 | 49 | 0.3061 |
| 5_1085148 | 8 | 0.5 | 6 | 0.6667 | 8 | 0.625 | 15 | 0.4667 | 37 | 0.5405 | 7 | 0.7143 | 6 | 0.5 | 16 | 0.75 | 9 | 0.4444 | 38 | 0.6316 |
| 5_1085149 | 11 | 0.4545 | 7 | 0.8571 | 6 | 0.3333 | 10 | 0.6 | 34 | 0.5588 | 6 | 0.8333 | 8 | 0.875 | 17 | 0.6471 | 7 | 0.5714 | 38 | 0.7105 |
| 5_1085149 | 9 | 0.6667 | 6 | 0.6667 | 7 | 0.4286 | 13 | 0.6923 | 35 | 0.6286 | 8 | 0.75 | 10 | 0.8 | 20 | 0.7 | 7 | 0.5714 | 45 | 0.7111 |
| 5_1085152 | 9 | 0.7778 | 5 | 0.6 | 8 | 0.5 | 14 | 0.7857 | 36 | 0.6944 | 7 | 0.5714 | 7 | 1 | 17 | 0.7647 | 8 | 0.5 | 39 | 0.7179 |
| 5_1085156 | 5 | 0.2 | 3 | 0.3333 | 2 | 0 | 9 | 0 | 19 | 0.1053 | 6 | 0 | 4 | 0 | 8 | 0.125 | 2 | 0 | 20 | 0.05 |
| 5_1085160 | NA | NA | NA | NA | 3 | 0.3333 | 1 | 0 | 4 | 0.25 | NA | NA | 3 | 0.3333 | 1 | 1 | 3 | 0.3333 | 7 | 0.4286 |
| 5_1085161 | 1 | 0 | 4 | 0 | 5 | 0.6 | 1 | 0 | 11 | 0.2727 | 2 | 0 | 3 | 0.3333 | 1 | 0 | 4 | 0 | 10 | 0.1 |
| 5_1085174 | 5 | 0.4 | 7 | 0 | 5 | 0.4 | 1 | 1 | 18 | 0.2778 | 2 | 0.5 | 3 | 0.6667 | 3 | 0.3333 | 8 | 0.625 | 16 | 0.5625 |
| 5_1085179 | 2 | 0 | 2 | 0 | NA | NA | 1 | 1 | 5 | 0.2 | NA | NA | 2 | 0.5 | 2 | 0 | 2 | 0.5 | 6 | 0.3333 |
| 5_1085179 | 2 | 0 | 3 | 0.3333 | NA | NA | 1 | 1 | 6 | 0.3333 | NA | NA | 2 | 0.5 | 3 | 0.6667 | 2 | 0 | 7 | 0.4286 |
| 5_1085185 | 3 | 1 | 1 | 0 | NA | NA | 1 | 0 | 5 | 0.6 | NA | NA | 2 | 0.5 | 3 | 0.6667 | NA | NA | 5 | 0.6 |
| 5_1085187 | 3 | 0.6667 | 1 | 0 | 2 | 1 | 3 | 0 | 9 | 0.4444 | 1 | 0 | 5 | 0.6 | 3 | 0.6667 | 5 | 0.6 | 14 | 0.5714 |
| 5_1085187 | 7 | 0.1429 | 3 | 0 | 4 | 0.25 | 3 | 0 | 17 | 0.1176 | 2 | 0 | 6 | 0.1667 | 5 | 0 | 7 | 0 | 20 | 0.05 |
| 5_1085188 | 7 | 0 | 2 | 0 | 3 | 0.3333 | 3 | 0.6667 | 15 | 0.2 | 1 | 0 | 6 | 0.1667 | 4 | 0 | 5 | 0 | 16 | 0.0625 |
| 5_1085189 | 7 | 0.5714 | 2 | 0.5 | 4 | 1 | 3 | 0.6667 | 16 | 0.6875 | 2 | 1 | 7 | 0.7143 | 5 | 0.4 | 7 | 0.7143 | 21 | 0.6667 |
| 5_1085190 | 4 | 0.25 | 2 | 0 | 2 | 0.5 | 3 | 0.6667 | 11 | 0.3636 | 1 | 0 | 5 | 0.8 | 4 | 0 | 6 | 0.3333 | 16 | 0.375 |
| 5_1085192 | 5 | 0 | 2 | 0 | 3 | 0.6667 | 3 | 0 | 13 | 0.1538 | 3 | 0 | 7 | 0.1429 | 5 | 0 | 5 | 0 | 20 | 0.05 |
| 5_1085195 | 6 | 0.8333 | 1 | 0 | 4 | 0.75 | 5 | 0.6 | 16 | 0.6875 | 2 | 0.5 | 8 | 0.625 | 4 | 1 | 8 | 0.875 | 22 | 0.7727 |
| 5_1085196 | 7 | 0.1429 | 5 | 0 | 3 | 0 | 5 | 0 | 20 | 0.05 | 3 | 0.3333 | 7 | 0.2857 | 6 | 0 | 7 | 0 | 23 | 0.1304 |
| 5_1085199 | 8 | 0.125 | 5 | 0 | 4 | 0 | 7 | 0.1429 | 24 | 0.0833 | 3 | 0 | 9 | 0.1111 | 5 | 0 | 9 | 0.3333 | 26 | 0.1538 |
| 5_1085204 | 9 | 0.5556 | 9 | 0.6667 | 7 | 0.5714 | 7 | 0.5714 | 32 | 0.5938 | 3 | 0.6667 | 6 | 0.1667 | 7 | 0.4286 | 10 | 0.6 | 26 | 0.4615 |
| 5_1085204 | 10 | 0.9 | 10 | 0.7 | 9 | 0.8889 | 7 | 0.8571 | 36 | 0.8333 | 4 | 1 | 10 | 0.8 | 4 | 0.75 | 10 | 0.7 | 28 | 0.7857 |
| 5_1085205 | 8 | 0.25 | 8 | 0.375 | 10 | 0.5 | 6 | 0.5 | 32 | 0.4062 | 2 | 0 | 10 | 0.4 | 6 | 0.3333 | 7 | 0.7143 | 25 | 0.44 |
| 5_1085205 | 10 | 0 | 8 | 0 | 9 | 0.1111 | 8 | 0.125 | 35 | 0.0571 | 5 | 0 | 12 | 0.0833 | 8 | 0 | 11 | 0.1818 | 36 | 0.0833 |
| 5_1085206 | 10 | 0 | 9 | 0 | 9 | 0.1111 | 8 | 0.125 | 36 | 0.0556 | 5 | 0 | 12 | 0.0833 | 9 | 0 | 11 | 0.0909 | 37 | 0.0541 |
| 5_1085206 | 3 | 0.6667 | 1 | 1 | 1 | 1 | 3 | 0.6667 | 8 | 0.75 | 2 | 1 | 7 | 0.8571 | 2 | 0.5 | 7 | 0.8571 | 18 | 0.8333 |
| 5_1085207 | 3 | 1 | 1 | 1 | 1 | 0 | 4 | 0 | 9 | 0.4444 | 3 | 0 | 5 | 0.2 | 3 | 0 | 5 | 0.4 | 16 | 0.1875 |
| 5_1085214 | 3 | 0.6667 | 1 | 0 | 5 | 0.8 | 6 | 0.3333 | 15 | 0.5333 | 3 | 0.3333 | 6 | 0.3333 | 3 | 0.6667 | 7 | 0.5714 | 19 | 0.4737 |
| 5_1085214 | 3 | 0 | 1 | 0 | 5 | 0.8 | 6 | 0.1667 | 15 | 0.3333 | 4 | 0.75 | 8 | 0.25 | 3 | 0.3333 | 8 | 0.5 | 23 | 0.4348 |
| 5_1085218 | 3 | 0.3333 | NA | NA | 2 | 0.5 | 7 | 0.4286 | 12 | 0.4167 | 2 | 0 | 7 | 0.4286 | 4 | 0 | 6 | 0.3333 | 19 | 0.2632 |
| 5_1085219 | 3 | 0.3333 | 1 | 0 | 4 | 0 | 8 | 0 | 16 | 0.0625 | 2 | 0.5 | 7 | 0.4286 | 4 | 0.5 | 6 | 0.3333 | 19 | 0.4211 |
| 5_1085219 | 2 | 1 | 1 | 0 | 3 | 0.6667 | 7 | 0.1429 | 13 | 0.3846 | 2 | 0.5 | 4 | 1 | 4 | 0.5 | 4 | 0.5 | 14 | 0.6429 |
| 5_1085219 | 3 | 0.6667 | 1 | 0 | 4 | 0.5 | 6 | 0.1667 | 14 | 0.3571 | NA | NA | 5 | 0.6 | 4 | 0.25 | 6 | 0.6667 | 15 | 0.5333 |
| 5_1085220 | 3 | 1 | NA | NA | 3 | 0.6667 | 8 | 0.875 | 14 | 0.8571 | 1 | 0 | 5 | 0.8 | 4 | 0.75 | 7 | 1 | 17 | 0.8235 |
| 5_1085229 | NA | NA | NA | NA | NA | NA | 1 | 1 | 1 | 1 | NA | NA | 2 | 0 | NA | NA | 1 | 1 | 3 | 0.3333 |
| 5_1086482 | 1 | 1 | 4 | 1 | 1 | 1 | 5 | 1 | 11 | 1 | 1 | 1 | 6 | 0.6667 | 4 | 1 | 4 | 1 | 15 | 0.8667 |

| | | | | | | | | | | | | | | | | | | | | |
|---|---|---|---|---|---|---|---|---|---|---|---|---|---|---|---|---|---|---|---|---|
| 5_10864828 | 1 | 1 | 4 | 0.5 | 2 | 0 | 6 | 0.3333 | 13 | 0.3846 | 1 | 1 | 5 | 0.2 | 4 | 0 | 4 | 0.25 | 14 | 0.2143 |
| 5_10864835 | 1 | 1 | 4 | 1 | 2 | 0.5 | 6 | 1 | 13 | 0.9231 | 2 | 1 | 6 | 1 | 4 | 1 | 4 | 1 | 16 | 1 |
| 5_10864837 | NA | NA | 4 | 0.5 | 1 | 1 | 6 | 0.6667 | 11 | 0.6364 | NA | NA | 5 | 0.6 | 3 | 0.6667 | 3 | 0.6667 | 11 | 0.6364 |
| 5_10864841 | 1 | 0 | 4 | 0.25 | 2 | 0 | 6 | 0.5 | 13 | 0.3077 | 2 | 0.5 | 6 | 0.1667 | 4 | 0.25 | 4 | 0.25 | 16 | 0.25 |
| 5_10864882 | 1 | 0 | 3 | 0.6667 | 2 | 0 | 6 | 0.1667 | 12 | 0.25 | 2 | 0 | 6 | 0 | 5 | 0.2 | 3 | 0 | 16 | 0.0625 |
| 5_10864897 | 1 | 0 | 5 | 0.2 | 2 | 0 | 6 | 0 | 14 | 0.0714 | 2 | 0.5 | 5 | 0 | 4 | 0 | 3 | 0 | 14 | 0.0714 |
| 5_10865015 | 4 | 0 | 7 | 0 | 2 | 0.5 | 5 | 0.2 | 18 | 0.1111 | 3 | 0 | 6 | 0.3333 | 4 | 0.25 | 4 | 0.5 | 17 | 0.2941 |
| 5_10865045 | 5 | 0.2 | 4 | 0.25 | 3 | 0.3333 | 2 | 0.5 | 14 | 0.2857 | 2 | 0 | 4 | 0 | 5 | 0.4 | 3 | 0.6667 | 14 | 0.2857 |
| 5_10865062 | 7 | 0 | 7 | 0.2857 | 5 | 0 | 7 | 0 | 26 | 0.0769 | 3 | 0 | 7 | 0 | 7 | 0 | 7 | 0.2857 | 24 | 0.0833 |
| 5_10865131 | 8 | 0.125 | 7 | 0.1429 | 5 | 0 | 8 | 0.125 | 28 | 0.1071 | 3 | 0 | 7 | 0 | 8 | 0.375 | 8 | 0 | 26 | 0.1154 |
| 5_10866673 | 101 | 0.0792 | 160 | 0.0688 | 116 | 0.0431 | 107 | 0.0561 | 484 | 0.062 | 100 | 0.09 | 96 | 0.0625 | 92 | 0.0543 | 83 | 0.0723 | 371 | 0.0701 |
| 5_10866750 | 71 | 0.0563 | 114 | 0.0702 | 87 | 0.0805 | 84 | 0.0714 | 356 | 0.0702 | 72 | 0.0556 | 69 | 0.0435 | 78 | 0.0769 | 60 | 0.0667 | 279 | 0.0609 |
| 5_10866751 | 80 | 0.1 | 122 | 0.0574 | 89 | 0.0449 | 85 | 0.0588 | 376 | 0.0638 | 82 | 0.0366 | 80 | 0.075 | 86 | 0.0698 | 65 | 0.0615 | 313 | 0.0607 |
| 5_10866752 | 72 | 0.4861 | 115 | 0.5565 | 74 | 0.4324 | 81 | 0.4198 | 342 | 0.4825 | 72 | 0.4444 | 72 | 0.4167 | 70 | 0.4143 | 60 | 0.3833 | 274 | 0.4161 |
| 5_10866759 | 64 | 0.0469 | 102 | 0.0882 | 75 | 0.0667 | 76 | 0.0789 | 317 | 0.0726 | 58 | 0.0345 | 67 | 0.0597 | 65 | 0.0923 | 56 | 0.0893 | 246 | 0.0691 |
| 5_10866801 | 62 | 0.2097 | 94 | 0.3085 | 69 | 0.1449 | 72 | 0.2917 | 297 | 0.2458 | 55 | 0.3273 | 57 | 0.2632 | 66 | 0.2576 | 53 | 0.2453 | 231 | 0.2727 |
| 5_10866804 | 58 | 0.2931 | 99 | 0.3434 | 69 | 0.3333 | 74 | 0.2162 | 300 | 0.3 | 61 | 0.2787 | 57 | 0.2105 | 59 | 0.2881 | 56 | 0.5357 | 233 | 0.3262 |
| 5_10871756 | 4 | 1 | 4 | 1 | 13 | 0.9231 | 9 | 1 | 30 | 0.9667 | 9 | 1 | 7 | 0.8571 | 5 | 0.8 | 11 | 0.9091 | 32 | 0.9062 |
| 5_10871760 | 4 | 0.5 | 4 | 0.25 | 9 | 0.7778 | 10 | 0.3 | 27 | 0.4815 | 7 | 0.7143 | 8 | 0.25 | 5 | 0.4 | 9 | 0.4444 | 29 | 0.4483 |
| 5_10871764 | 3 | 1 | 4 | 0.5 | 11 | 0.8182 | 10 | 0.5 | 28 | 0.6786 | 8 | 0.625 | 7 | 0.8571 | 6 | 0.3333 | 8 | 0.875 | 29 | 0.6897 |
| 5_10871780 | 3 | 0.3333 | 3 | 1 | 9 | 0.3333 | 4 | 0.5 | 19 | 0.4737 | 8 | 0.125 | 5 | 0.8 | 6 | 0.5 | 5 | 0.4 | 24 | 0.4167 |
| 5_10871795 | 2 | 1 | NA | NA | 3 | 0.6667 | 2 | 0 | 7 | 0.5714 | 2 | 0 | 1 | 0 | 3 | 0.3333 | 1 | 0 | 7 | 0.1429 |
| 5_10873006 | 2 | 0 | 3 | 0 | 8 | 0 | 6 | 0.1667 | 19 | 0.0526 | 4 | 0 | 3 | 0.3333 | 3 | 0 | 7 | 0 | 17 | 0.0588 |
| 5_10873014 | 3 | 0 | 3 | 0.3333 | 6 | 0 | 7 | 0.2857 | 19 | 0.1579 | 3 | 0 | 5 | 0.2 | 3 | 0.3333 | 7 | 0 | 18 | 0.1111 |
| 5_10873066 | 2 | 0 | 3 | 0 | 7 | 0.1429 | 5 | 0.2 | 17 | 0.1176 | 5 | 0 | 2 | 0 | 3 | 0 | 7 | 0.1429 | 17 | 0.0588 |
| 5_10873073 | 2 | 0.5 | 3 | 0.6667 | 7 | 0.5714 | 6 | 0.8333 | 18 | 0.6667 | 5 | 0.6 | 3 | 1 | 3 | 1 | 8 | 0.875 | 19 | 0.8421 |
| 5_10873083 | 2 | 0 | 3 | 0.3333 | 6 | 0 | 6 | 0.3333 | 17 | 0.1765 | 5 | 0.8 | 4 | 0 | 2 | 0 | 7 | 0.4286 | 18 | 0.3889 |
| 5_10873084 | 2 | 0.5 | 3 | 0 | 7 | 0.5714 | 6 | 0.1667 | 18 | 0.3333 | 5 | 0 | 4 | 0.25 | 3 | 1 | 8 | 0.5 | 20 | 0.4 |
| 5_10873099 | 1 | 0 | 3 | 0.6667 | 7 | 0 | 5 | 0.4 | 16 | 0.25 | 5 | 0.6 | NA | NA | 3 | 0.3333 | 7 | 0.4286 | 15 | 0.4667 |
| 5_10879205 | 6 | 0 | 3 | 0 | 1 | 1 | 1 | 0 | 11 | 0.0909 | 1 | 0 | 5 | 0 | 6 | 0 | 4 | 0.25 | 16 | 0.0625 |
| 5_10879264 | 8 | 0.25 | NA | NA | NA | NA | NA | NA | 8 | 0.25 | NA | NA | 2 | 0.5 | 4 | 0.25 | NA | NA | 6 | 0.3333 |
| 5_10879371 | 1 | 1 | 3 | 0.3333 | 1 | 1 | 1 | 1 | 6 | 0.6667 | 1 | 1 | 4 | 0.25 | 4 | 0 | 4 | 0.5 | 13 | 0.3077 |
| 5_10879388 | 1 | 1 | 3 | 0.6667 | 1 | 1 | 1 | 1 | 6 | 0.8333 | 1 | 1 | 4 | 0.5 | 4 | 0.5 | 4 | 1 | 13 | 0.6923 |
| 5_10879397 | 1 | 1 | NA | NA | 2 | 1 | 1 | 1 | 4 | 1 | 1 | 1 | 2 | 1 | 3 | 1 | 4 | 0.75 | 10 | 0.9 |
| 5_10879404 | 1 | 1 | 3 | 1 | 2 | 1 | 1 | 1 | 7 | 1 | 1 | 0 | 3 | 1 | 4 | 0.75 | 3 | 0.6667 | 11 | 0.7273 |
| 5_10879406 | 1 | 0 | 4 | 0.25 | 2 | 0 | 2 | 0 | 9 | 0.1111 | 1 | 0 | 4 | 0.25 | 4 | 0 | 4 | 0.25 | 13 | 0.1538 |
| 5_10879407 | 1 | 0 | 3 | 0.3333 | 2 | 0 | NA | NA | 6 | 0.1667 | 1 | 1 | 4 | 0.25 | 4 | 0.25 | 3 | 0.3333 | 12 | 0.3333 |
| 5_10879432 | 1 | 0 | 4 | 0.25 | 2 | 0.5 | 3 | 0 | 10 | 0.2 | 1 | 0 | 5 | 0.2 | 5 | 0.4 | 4 | 0 | 15 | 0.2 |
| 5_10879445 | 1 | 1 | 4 | 0.75 | 3 | 0.3333 | 2 | 0.5 | 10 | 0.6 | 1 | 0 | 6 | 0.8333 | 4 | 0.25 | 4 | 0.5 | 15 | 0.5333 |
| 5_10879452 | NA | NA | 2 | 0 | 2 | 0.5 | 2 | 0 | 6 | 0.1667 | 1 | 1 | 3 | 0.3333 | 4 | 0.5 | 4 | 0.25 | 12 | 0.4167 |
| 5_10879456 | NA | NA | NA | NA | 1 | 0 | 1 | 1 | 2 | 0.5 | 2 | 0.5 | 1 | 0 | 3 | 0.3333 | 3 | 0.3333 | 9 | 0.3333 |
| 5_10879485 | 1 | 0 | 1 | 1 | 2 | 0 | 1 | 0 | 5 | 0.2 | 2 | 0.5 | 2 | 0.5 | 3 | 0 | 3 | 0 | 10 | 0.2 |
| 5_10879532 | 2 | 0 | 6 | 0 | 3 | 0 | 4 | 0.25 | 15 | 0.0667 | 1 | 0 | 3 | 0.3333 | 10 | 0.1 | 1 | 0 | 15 | 0.1333 |
| 5_10879602 | 2 | 0.5 | 6 | 0.1667 | 2 | 0.5 | 3 | 0 | 13 | 0.2308 | 1 | 0 | 3 | 0.3333 | 5 | 0.2 | 1 | 0 | 10 | 0.2 |
| 5_10879609 | 1 | 0 | 7 | 0 | 3 | 0 | 4 | 0.25 | 15 | 0.0667 | NA | NA | 3 | 0 | 6 | 0.1667 | NA | NA | 9 | 0.1111 |
| 5_10879650 | 1 | 1 | 5 | 1 | 2 | 0.5 | 4 | 0.75 | 12 | 0.8333 | NA | NA | 3 | 0.3333 | 7 | 0.7143 | NA | NA | 10 | 0.6 |
| 5_10883125 | 1 | 1 | 3 | 0.3333 | NA | NA | 2 | 1 | 6 | 0.6667 | NA | NA | 4 | 0.25 | 2 | 0.5 | NA | NA | 6 | 0.3333 |
| 5_10883149 | 1 | 0 | 2 | 0.5 | 1 | 1 | 1 | 0 | 5 | 0.4 | NA | NA | 4 | 0 | 3 | 0.6667 | NA | NA | 7 | 0.2857 |
| 5_10885290 | NA | NA | 3 | 0 | 4 | 0.25 | 1 | 0 | 8 | 0.125 | 2 | 0 | 1 | 0 | 2 | 0 | 3 | 0.3333 | 8 | 0.125 |
| 5_10885390 | NA | NA | 3 | 0.6667 | 4 | 0.5 | 3 | 0.6667 | 10 | 0.6 | 2 | 1 | 4 | 0 | 2 | 1 | 5 | 0.4 | 13 | 0.4615 |
| 5_10885422 | NA | NA | 5 | 0.6 | 3 | 0.6667 | 3 | 0.3333 | 11 | 0.5455 | 2 | 0 | 4 | 0.75 | 2 | 0.5 | 4 | 0.75 | 12 | 0.5833 |
| 5_10885423 | NA | NA | 4 | 0 | 4 | 0 | 4 | 0.25 | 12 | 0.0833 | 2 | 0 | 4 | 0 | 1 | 0 | 4 | 0.25 | 11 | 0.0909 |
| 5_10885436 | NA | NA | 5 | 0.2 | 4 | 0.25 | 4 | 0 | 13 | 0.1538 | 3 | 0 | 4 | 0 | 2 | 0.5 | 5 | 0.2 | 14 | 0.1429 |
| 5_10982246 | 1 | 1 | NA | NA | 3 | 0 | NA | NA | 4 | 0.25 | 1 | 1 | 1 | 1 | 1 | 1 | NA | NA | 3 | 1 |
| 5_10982249 | 1 | 1 | NA | NA | 3 | 0.6667 | NA | NA | 4 | 0.75 | 1 | 1 | 1 | 0 | 1 | 0 | NA | NA | 3 | 0.3333 |
| 5_10982260 | 1 | 1 | NA | NA | 2 | 0 | NA | NA | 3 | 0.3333 | 1 | 0 | 1 | 1 | 1 | 0 | NA | NA | 3 | 0.3333 |
| 5_10982277 | 1 | 1 | NA | NA | 2 | 0 | NA | NA | 3 | 0.3333 | 1 | 0 | 1 | 0 | 1 | 1 | NA | NA | 3 | 0.3333 |
| 5_10982416 | 1 | 0 | NA | NA | 1 | 1 | NA | NA | 2 | 0.5 | NA | NA | NA | NA | 1 | 1 | NA | NA | 1 | 1 |
| 5_10982483 | 1 | 1 | NA | NA | 1 | 0 | NA | NA | 2 | 0.5 | NA | NA | NA | NA | 1 | 1 | NA | NA | 1 | 1 |
| 5_10982498 | 1 | 1 | NA | NA | 1 | 0 | NA | NA | 2 | 0.5 | NA | NA | NA | NA | 1 | 1 | NA | NA | 1 | 1 |
| 5_10984577 | NA | NA | 1 | 1 | NA | NA | NA | NA | 1 | 1 | NA | NA | 1 | 1 | NA | NA | NA | NA | 1 | 1 |
| 5_10993028 | NA | NA | NA | NA | 1 | 1 | NA | NA | 1 | 1 | NA | NA | 1 | 1 | NA | NA | NA | NA | 1 | 1 |
| 5_11002977 | NA | NA | 1 | 1 | NA | NA | 2 | 0 | 3 | 0.3333 | 1 | 0 | 1 | 1 | 1 | 1 | 4 | 0.75 | 7 | 0.7143 |
| 5_11002980 | NA | NA | 1 | 0 | NA | NA | 2 | 0.5 | 3 | 0.3333 | 1 | 0 | NA | NA | 1 | 0 | 3 | 0.3333 | 5 | 0.2 |
| 5_11002990 | NA | NA | 1 | 0 | NA | NA | 2 | 0.5 | 3 | 0.3333 | 1 | 0 | 1 | 0 | 1 | 0 | 4 | 0.25 | 7 | 0.1429 |

| | | | | | | | | | | | | | | | | | | | | |
|---|---|---|---|---|---|---|---|---|---|---|---|---|---|---|---|---|---|---|---|---|
| 5_1100302 | NA | NA | 1 | 1 | NA | NA | 2 | 0 | 3 | 0.3333 | 1 | 0 | 1 | 0 | 1 | 0 | 4 | 0.25 | 7 | 0.1429 |
| 5_1100306 | NA | NA | 1 | 0 | NA | NA | 2 | 0.5 | 3 | 0.3333 | 1 | 0 | 1 | 0 | 1 | 0 | 3 | 0.3333 | 6 | 0.1667 |
| 5_1100306 | NA | NA | 1 | 1 | NA | NA | 2 | 1 | 3 | 1 | 1 | 1 | 1 | 0 | 1 | 1 | 3 | 0.3333 | 6 | 0.5 |
| 5_1100308 | NA | NA | 1 | 1 | NA | NA | 1 | 1 | 2 | 1 | 1 | 1 | 1 | 1 | 1 | 1 | 2 | 0.5 | 5 | 0.8 |
| 5_1100310 | NA | NA | 1 | 0 | NA | NA | 2 | 0.5 | 3 | 0.3333 | 1 | 0 | 1 | 0 | NA | NA | 3 | 0.3333 | 5 | 0.2 |
| 5_1100310 | NA | NA | 1 | 0 | NA | NA | 2 | 0.5 | 3 | 0.3333 | 1 | 0 | 1 | 0 | 1 | 0 | 3 | 0.3333 | 6 | 0.1667 |
| 5_1100311 | NA | NA | 1 | 1 | NA | NA | 2 | 1 | 3 | 1 | 1 | 1 | 1 | 0 | 1 | 0 | 2 | 0.5 | 5 | 0.4 |
| 5_1100313 | NA | NA | 1 | 0 | NA | NA | 2 | 1 | 3 | 0.6667 | 1 | 1 | 1 | 0 | NA | NA | 3 | 0.6667 | 5 | 0.6 |
| 5_1100313 | NA | NA | 1 | 1 | NA | NA | 2 | 1 | 3 | 1 | 1 | 1 | NA | NA | 1 | 1 | 3 | 0.6667 | 5 | 0.8 |
| 5_1100314 | NA | NA | 1 | 0 | NA | NA | 2 | 0.5 | 3 | 0.3333 | 1 | 0 | 1 | 0 | 1 | 0 | 3 | 0.3333 | 6 | 0.1667 |
| 5_1100314 | NA | NA | 1 | 1 | NA | NA | 1 | 1 | 2 | 1 | 1 | 1 | 1 | 1 | 1 | 1 | 2 | 1 | 5 | 1 |
| 5_1100319 | 1 | 0 | 1 | 0 | NA | NA | 1 | 1 | 3 | 0.3333 | 1 | 0 | NA | NA | 1 | 0 | 2 | 0.5 | 4 | 0.25 |
| 5_1100325 | 1 | 0 | 1 | 0 | NA | NA | 2 | 0.5 | 4 | 0.25 | 1 | 0 | 1 | 0 | 1 | 0 | 3 | 0.6667 | 6 | 0.3333 |
| 5_1100331 | 1 | 1 | 1 | 0 | NA | NA | 1 | 1 | 3 | 0.6667 | 1 | 0 | 1 | 0 | 1 | 1 | 1 | 0 | 4 | 0.25 |
| 5_1100332 | 1 | 1 | NA | NA | NA | NA | 1 | 1 | 2 | 1 | NA | NA | 1 | 0 | 1 | 1 | NA | NA | 2 | 0.5 |
| 5_1100398 | 2 | 0.5 | 1 | 0 | NA | NA | NA | NA | 3 | 0.3333 | 2 | 0.5 | NA | NA | 1 | 1 | NA | NA | 3 | 0.6667 |
| 5_1100398 | 2 | 0.5 | 1 | 0 | NA | NA | NA | NA | 3 | 0.3333 | 2 | 0 | NA | NA | 1 | 1 | NA | NA | 3 | 0.3333 |
| 5_1100401 | 2 | 0.5 | 1 | 0 | NA | NA | NA | NA | 3 | 0.3333 | 1 | 0 | 1 | 1 | 2 | 0.5 | NA | NA | 4 | 0.5 |
| 5_1100901 | 3 | 0.3333 | 2 | 0 | 1 | 0 | 2 | 0 | 8 | 0.125 | NA | NA | 2 | 0 | NA | NA | 3 | 0.3333 | 5 | 0.2 |
| 5_1100905 | 3 | 0.3333 | 2 | 0 | 1 | 0 | 2 | 0 | 8 | 0.125 | NA | NA | 3 | 0 | NA | NA | 5 | 0.4 | 8 | 0.25 |
| 5_1100907 | 3 | 0.3333 | 2 | 0 | NA | NA | 2 | 0.5 | 7 | 0.2857 | NA | NA | 3 | 0.6667 | NA | NA | 5 | 0 | 8 | 0.25 |
| 5_1100908 | 3 | 0.3333 | 2 | 0 | 1 | 0 | 1 | 0 | 7 | 0.1429 | NA | NA | 2 | 0.5 | NA | NA | 5 | 0 | 7 | 0.1429 |
| 5_1100909 | 4 | 0.25 | 2 | 0 | 1 | 0 | 2 | 0 | 9 | 0.1111 | NA | NA | 3 | 0.6667 | NA | NA | 4 | 0 | 7 | 0.2857 |
| 5_1100915 | 2 | 1 | NA | NA | NA | NA | 1 | 1 | 3 | 1 | NA | NA | 2 | 0.5 | NA | NA | 3 | 1 | 5 | 0.8 |
| 5_1100916 | 2 | 1 | 1 | 1 | NA | NA | 2 | 1 | 5 | 1 | NA | NA | 2 | 0.5 | NA | NA | 4 | 0.5 | 6 | 0.5 |
| 5_1100916 | 2 | 0.5 | 1 | 0 | NA | NA | 2 | 0.5 | 5 | 0.4 | NA | NA | 2 | 0.5 | NA | NA | 2 | 0.5 | 4 | 0.5 |
| 5_1100917 | 2 | 1 | 1 | 1 | NA | NA | 2 | 0 | 5 | 0.6 | NA | NA | 2 | 0.5 | NA | NA | 4 | 0.5 | 6 | 0.5 |
| 5_1100917 | 2 | 0.5 | 1 | 0 | NA | NA | 2 | 1 | 5 | 0.6 | NA | NA | 2 | 0.5 | NA | NA | 4 | 0.5 | 6 | 0.5 |
| 5_1100919 | NA | NA | NA | NA | 1 | 0 | 1 | 1 | 2 | 0.5 | NA | NA | NA | NA | NA | NA | 4 | 0.5 | 4 | 0.5 |
| 5_1101580 | 3 | 0.6667 | 1 | 1 | 3 | 0.6667 | 2 | 1 | 9 | 0.7778 | 1 | 1 | 4 | 1 | 3 | 1 | 1 | 1 | 9 | 1 |
| 5_1101581 | 1 | 0 | 1 | 0 | 3 | 0 | 2 | 1 | 7 | 0.2857 | 1 | 1 | 5 | 0.4 | 3 | 0.6667 | 1 | 1 | 10 | 0.6 |
| 5_1101581 | 3 | 0.3333 | NA | NA | 4 | 0 | 2 | 0.5 | 9 | 0.2222 | 1 | 0 | 4 | 0 | 3 | 1 | 1 | 0 | 9 | 0.3333 |
| 5_1101581 | 2 | 0 | 1 | 1 | 4 | 0.5 | 3 | 0.3333 | 10 | 0.4 | 1 | 0 | 5 | 0.8 | 3 | 0.3333 | 1 | 0 | 10 | 0.5 |
| 5_1101582 | 2 | 0.5 | 1 | 1 | 3 | 1 | 3 | 1 | 9 | 0.8889 | 1 | 1 | 4 | 0.25 | 3 | 1 | 1 | 0 | 9 | 0.5556 |
| 5_1101853 | NA | NA | NA | NA | NA | NA | 1 | 1 | 1 | 1 | 1 | 1 | NA | NA | 4 | 1 | 1 | 0 | 6 | 0.8333 |
| 5_1101854 | NA | NA | NA | NA | NA | NA | 1 | 1 | 1 | 1 | NA | NA | NA | NA | 4 | 0.5 | NA | NA | 4 | 0.5 |
| 5_1101854 | NA | NA | NA | NA | NA | NA | 1 | 1 | 1 | 1 | 1 | 1 | NA | NA | 3 | 1 | 1 | 1 | 5 | 1 |
| 5_1101856 | NA | NA | NA | NA | NA | NA | 1 | 1 | 1 | 1 | 1 | 1 | NA | NA | 4 | 1 | 1 | 1 | 6 | 1 |
| 5_1102516 | NA | NA | NA | NA | 1 | 1 | 1 | 0 | 2 | 0.5 | NA | NA | 1 | 1 | NA | NA | NA | NA | 1 | 1 |
| 5_1102518 | 1 | 1 | NA | NA | NA | NA | NA | NA | 1 | 1 | NA | NA | 1 | 1 | NA | NA | NA | NA | 1 | 1 |
| 5_1102520 | 1 | 0 | NA | NA | NA | NA | 2 | 1 | 3 | 0.6667 | 1 | 1 | 1 | 1 | NA | NA | 1 | 1 | 3 | 1 |
| 5_1102524 | 1 | 0 | 1 | 0 | NA | NA | 2 | 0.5 | 4 | 0.25 | 1 | 1 | 1 | 0 | NA | NA | NA | NA | 2 | 0.5 |
| 5_1102527 | 1 | 1 | 2 | 0.5 | 1 | 0 | 1 | 0 | 5 | 0.4 | 1 | 1 | 1 | 0 | NA | NA | 1 | 0 | 3 | 0.3333 |
| 5_1102528 | 1 | 0 | 3 | 0 | 1 | 1 | 2 | 0 | 7 | 0.1429 | 1 | 0 | 1 | 0 | NA | NA | 1 | 1 | 3 | 0.3333 |
| 5_1102532 | 1 | 1 | 3 | 0 | 1 | 0 | 2 | 0 | 7 | 0.1429 | 1 | 0 | 2 | 1 | 1 | 0 | 1 | 1 | 5 | 0.6 |
| 5_1102532 | 1 | 1 | 3 | 0.3333 | 1 | 0 | 2 | 0.5 | 7 | 0.4286 | 1 | 0 | 2 | 0.5 | 1 | 0 | 1 | 1 | 5 | 0.4 |
| 5_1102534 | 1 | 1 | 3 | 0 | 2 | 0 | NA | NA | 6 | 0.1667 | 1 | 0 | 2 | 0.5 | 1 | 0 | 1 | 0 | 5 | 0.2 |
| 5_1102537 | 1 | 1 | 3 | 0.3333 | 1 | 0 | NA | NA | 5 | 0.4 | 1 | 0 | 2 | 0 | 1 | 0 | 1 | 1 | 5 | 0.2 |
| 5_1102537 | 2 | 0.5 | 3 | 0.3333 | 2 | 0 | 2 | 0 | 9 | 0.2222 | 1 | 0 | 2 | 0 | 1 | 1 | 1 | 0 | 5 | 0.2 |
| 5_1102752 | 1 | 1 | NA | NA | 1 | 1 | NA | NA | 2 | 1 | NA | NA | NA | NA | 1 | 1 | 1 | 1 | 2 | 1 |
| 5_1102754 | 1 | 1 | NA | NA | 1 | 1 | NA | NA | 2 | 1 | NA | NA | NA | NA | 1 | 1 | 1 | 1 | 2 | 1 |
| 5_1102756 | 1 | 1 | NA | NA | 2 | 0 | NA | NA | 3 | 0.3333 | NA | NA | NA | NA | 1 | 0 | 1 | 1 | 2 | 0.5 |
| 5_1102762 | 1 | 0 | NA | NA | 2 | 1 | NA | NA | 3 | 0.6667 | NA | NA | NA | NA | 1 | 0 | 1 | 1 | 2 | 0.5 |
| 5_1102763 | 1 | 1 | NA | NA | 1 | 0 | NA | NA | 2 | 0.5 | NA | NA | NA | NA | 1 | 1 | 1 | 1 | 2 | 1 |
| 5_1102764 | 1 | 1 | NA | NA | 1 | 1 | NA | NA | 2 | 1 | NA | NA | NA | NA | NA | NA | 1 | 1 | 1 | 1 |
| 5_1102765 | 1 | 0 | NA | NA | 2 | 0.5 | NA | NA | 3 | 0.3333 | NA | NA | NA | NA | NA | NA | 1 | 1 | 1 | 1 |
| 5_1102765 | 1 | 0 | NA | NA | 2 | 0.5 | NA | NA | 3 | 0.3333 | NA | NA | NA | NA | 1 | 0 | 1 | 1 | 2 | 0.5 |
| 5_1102766 | 1 | 1 | NA | NA | 2 | 1 | NA | NA | 3 | 1 | NA | NA | NA | NA | 1 | 1 | 1 | 1 | 2 | 1 |
| 5_1102767 | 1 | 1 | NA | NA | 2 | 1 | NA | NA | 3 | 1 | NA | NA | NA | NA | 1 | 1 | 1 | 1 | 2 | 1 |
| 5_1102768 | 1 | 0 | NA | NA | 2 | 0.5 | NA | NA | 3 | 0.3333 | NA | NA | NA | NA | 1 | 1 | 1 | 0 | 2 | 0.5 |
| 5_1102773 | 1 | 0 | NA | NA | 2 | 0.5 | 1 | 0 | 4 | 0.25 | NA | NA | NA | NA | 1 | 1 | 1 | 0 | 2 | 0.5 |
| 5_1102777 | 1 | 0 | NA | NA | 2 | 1 | 1 | 1 | 4 | 0.75 | NA | NA | NA | NA | NA | NA | 1 | 1 | 1 | 1 |
| 5_1102797 | NA | NA | 1 | 1 | 1 | 1 | 1 | 0 | 3 | 0.6667 | NA | NA | NA | NA | 1 | 1 | NA | NA | 1 | 1 |
| 5_1103161 | 1 | 1 | NA | NA | NA | NA | NA | NA | 1 | 1 | NA | NA | 2 | 0.5 | NA | NA | NA | NA | 2 | 0.5 |
| 5_1103188 | 8 | 0.125 | 4 | 0.25 | 2 | 0 | 1 | 0 | 15 | 0.1333 | 3 | 0 | 1 | 0 | 2 | 0 | 4 | 0.25 | 10 | 0.1 |

| | | | | | | | | | | | | | | | | | | | | |
|---|---|---|---|---|---|---|---|---|---|---|---|---|---|---|---|---|---|---|---|---|
| 5_11031902 | 7 | 0.2857 | 5 | 0.4 | 4 | 0.25 | 1 | 0 | 17 | 0.2941 | 3 | 0.3333 | 1 | 0 | 2 | 1 | 4 | 0 | 10 | 0.3 |
| 5_11031904 | 5 | 0.4 | 6 | 0.1667 | 4 | 0 | 1 | 0 | 16 | 0.1875 | 1 | 0 | NA | NA | 3 | 0.3333 | 3 | 0 | 7 | 0.1429 |
| 5_11031907 | 6 | 0 | 6 | 0 | 4 | 0 | 2 | 0.5 | 18 | 0.0556 | 2 | 0 | 1 | 0 | 3 | 0 | 5 | 0.2 | 11 | 0.0909 |
| 5_11031932 | 8 | 0 | 7 | 0.1429 | 4 | 0.25 | 2 | 0 | 21 | 0.0952 | 3 | 0 | 2 | 0 | 3 | 0.3333 | 6 | 0 | 14 | 0.0714 |
| 5_11033543 | NA | NA | 2 | 0.5 | NA | NA | 1 | 0 | 3 | 0.3333 | 1 | 0 | 1 | 0 | 1 | 1 | NA | NA | 3 | 0.3333 |
| 5_11033555 | NA | NA | 3 | 1 | NA | NA | 1 | 1 | 4 | 1 | 1 | 0 | 1 | 1 | 1 | 0 | NA | NA | 3 | 0.3333 |
| 5_11033558 | NA | NA | 2 | 0.5 | NA | NA | 1 | 0 | 3 | 0.3333 | 1 | 1 | 1 | 0 | 1 | 0 | NA | NA | 3 | 0.3333 |
| 5_11033759 | NA | NA | 4 | 0.25 | NA | NA | NA | NA | 4 | 0.25 | 1 | 0 | 1 | 0 | 2 | 0.5 | NA | NA | 4 | 0.25 |
| 5_11035274 | 1 | 0 | 6 | 0.1667 | 1 | 0 | 1 | 0 | 9 | 0.1111 | 3 | 0 | 3 | 0 | 3 | 0 | 2 | 0.5 | 11 | 0.0909 |
| 5_11037090 | 4 | 0 | 9 | 0.2222 | 4 | 0 | 4 | 0.5 | 21 | 0.1905 | 4 | 0 | 4 | 0 | 4 | 0.5 | 4 | 0 | 16 | 0.125 |
| 5_11037097 | 6 | 0.3333 | 8 | 0.25 | 4 | 0.25 | 5 | 0.4 | 23 | 0.3043 | 2 | 0 | 3 | 0 | 4 | 0.25 | 4 | 0.25 | 13 | 0.1538 |
| 5_11037106 | 2 | 0.5 | 3 | 0.6667 | 2 | 0 | 3 | 0.3333 | 10 | 0.4 | NA | NA | NA | NA | 1 | 0 | 2 | 0.5 | 3 | 0.3333 |
| 5_11038229 | 1 | 1 | 1 | 1 | 1 | 1 | 2 | 0.5 | 5 | 0.8 | 1 | 0 | 1 | 1 | 1 | 1 | 1 | 0 | 4 | 0.5 |
| 5_11038230 | 1 | 0 | 3 | 0.3333 | 1 | 0 | 2 | 0.5 | 7 | 0.2857 | 1 | 1 | NA | NA | 1 | 0 | 2 | 1 | 4 | 0.75 |
| 5_11038244 | 1 | 1 | 3 | 1 | 1 | 1 | 2 | 1 | 7 | 1 | 1 | 1 | 1 | 1 | 1 | 1 | 2 | 1 | 5 | 1 |
| 5_11038247 | 1 | 0 | 3 | 0.3333 | 1 | 0 | 2 | 0 | 7 | 0.1429 | 1 | 0 | 1 | 0 | 1 | 1 | 2 | 0 | 5 | 0.2 |
| 5_11038249 | 1 | 0 | 3 | 0.3333 | 1 | 0 | 2 | 0 | 7 | 0.1429 | 1 | 0 | 1 | 0 | 2 | 0.5 | 1 | 0 | 5 | 0.2 |
| 5_11040074 | 2 | 0.5 | 3 | 0.6667 | 3 | 0 | 2 | 0 | 10 | 0.3 | 2 | 0.5 | 1 | 0 | 3 | 0 | 1 | 0 | 7 | 0.1429 |
| 5_11040077 | 2 | 0.5 | 5 | 0.8 | 3 | 0.3333 | 3 | 0.3333 | 13 | 0.5385 | 2 | 0.5 | 1 | 0 | 3 | 0.6667 | 2 | 0.5 | 8 | 0.5 |
| 5_11040082 | 2 | 1 | 5 | 0.8 | 2 | 1 | 2 | 1 | 11 | 0.9091 | NA | NA | 1 | 1 | 3 | 0.6667 | 2 | 1 | 6 | 0.8333 |
| 5_11040084 | 2 | 0 | 5 | 0.8 | 3 | 1 | 3 | 0.6667 | 13 | 0.6923 | 2 | 0.5 | 1 | 1 | 4 | 0.75 | 2 | 1 | 9 | 0.7778 |
| 5_11040089 | 2 | 0.5 | 5 | 0.2 | 3 | 0 | 3 | 0.3333 | 13 | 0.2308 | 2 | 0 | 1 | 0 | 4 | 0.25 | 2 | 0.5 | 9 | 0.2222 |
| 5_11040091 | 1 | 0 | 5 | 1 | 3 | 1 | 3 | 1 | 12 | 0.9167 | 2 | 1 | 1 | 0 | 4 | 1 | 2 | 1 | 9 | 0.8889 |
| 5_11040101 | 3 | 0 | 4 | 0.25 | 2 | 0.5 | 3 | 0.3333 | 12 | 0.25 | 2 | 0 | 1 | 0 | 4 | 0.25 | 2 | 0 | 9 | 0.1111 |
| 5_11040234 | 3 | 0.3333 | 2 | 0 | 3 | 0 | 2 | 0 | 10 | 0.1 | 2 | 0.5 | 3 | 0.3333 | 5 | 0 | 3 | 0.3333 | 13 | 0.2308 |
| 5_11040392 | NA | NA | 2 | 0 | 5 | 0.4 | 2 | 0.5 | 9 | 0.3333 | 1 | 0 | 2 | 0 | 4 | 0.5 | 3 | 0.3333 | 10 | 0.3 |
| 5_11040432 | 2 | 0 | 1 | 1 | 5 | 0.4 | 1 | 0 | 9 | 0.3333 | 1 | 0 | 2 | 0.5 | 4 | 0.25 | 2 | 0 | 9 | 0.2222 |
| 5_11040736 | 12 | 0.0833 | 5 | 0.2 | 9 | 0.2222 | 8 | 0.25 | 34 | 0.1765 | 8 | 0 | 8 | 0.25 | 3 | 0.3333 | 3 | 0 | 22 | 0.1364 |
| 5_11040766 | 10 | 0.1 | 6 | 0 | 9 | 0.1111 | 8 | 0.125 | 33 | 0.0909 | 7 | 0 | 6 | 0.3333 | 3 | 0 | 4 | 0.5 | 20 | 0.2 |
| 5_11040813 | 13 | 0.3077 | 9 | 0.3333 | 10 | 0.3 | 8 | 0.125 | 40 | 0.275 | 6 | 0.3333 | 5 | 0.2 | 2 | 0 | 5 | 0.2 | 18 | 0.2222 |
| 5_11040819 | 13 | 0.0769 | 9 | 0 | 8 | 0.125 | 8 | 0 | 38 | 0.0526 | 4 | 0 | 5 | 0 | 2 | 0.5 | 5 | 0.2 | 16 | 0.125 |
| 5_11050969 | 2 | 0 | 3 | 0.6667 | 2 | 1 | 1 | 1 | 8 | 0.625 | 4 | 0.5 | 1 | 1 | 4 | 0.5 | NA | NA | 9 | 0.5556 |
| 5_11051006 | 2 | 0 | 4 | 0.5 | 1 | 0 | 1 | 0 | 8 | 0.25 | 5 | 0.4 | 1 | 1 | 4 | 0.5 | NA | NA | 10 | 0.5 |
| 5_11051008 | 2 | 0 | 3 | 0.3333 | 3 | 0 | 1 | 0 | 9 | 0.1111 | 5 | 0 | 1 | 1 | 2 | 0 | NA | NA | 8 | 0.125 |
| 5_11051031 | 2 | 1 | 8 | 0.125 | 3 | 0 | 6 | 0.6667 | 19 | 0.3684 | 6 | 0.6667 | 2 | 0.5 | 8 | 0.25 | 1 | 0 | 17 | 0.4118 |
| 5_11051043 | 3 | 0.6667 | 8 | 0.25 | 4 | 0.75 | 5 | 0 | 20 | 0.35 | 6 | 0.1667 | 4 | 0.25 | 11 | 0.1818 | NA | NA | 21 | 0.1905 |
| 5_11051052 | 4 | 0.25 | 7 | 0.4286 | 4 | 0.5 | 3 | 0 | 18 | 0.3333 | 8 | 0.25 | 4 | 0 | 11 | 0.3636 | 3 | 0 | 26 | 0.2308 |
| 5_11051080 | 3 | 0.6667 | 4 | 1 | 2 | 1 | 4 | 0.75 | 13 | 0.8462 | 6 | 0.8333 | 4 | 0.75 | 10 | 0.9 | 2 | 0.5 | 22 | 0.8182 |
| 5_11051087 | 3 | 0.3333 | 4 | 1 | 2 | 1 | 6 | 0.3333 | 15 | 0.6 | 8 | 0.5 | 3 | 0.6667 | 12 | 0.4167 | 2 | 0.5 | 25 | 0.48 |
| 5_11051101 | 2 | 1 | 2 | 1 | 1 | 1 | 4 | 0.75 | 9 | 0.8889 | 5 | 0.6 | 3 | 0.6667 | 11 | 0.6364 | 2 | 1 | 21 | 0.6667 |
| 5_11051122 | 3 | 0.6667 | 5 | 0.2 | 3 | 0.3333 | 5 | 0.2 | 16 | 0.3125 | 8 | 0.375 | 2 | 0.5 | 13 | 0.0769 | 2 | 1 | 25 | 0.28 |
| 5_11051126 | 3 | 0.6667 | 5 | 0.8 | 2 | 0.5 | 5 | 0.4 | 15 | 0.6 | 8 | 0.5 | 2 | 0 | 11 | 0.4545 | 2 | 0.5 | 23 | 0.4348 |
| 5_11051130 | 3 | 0.6667 | 5 | 0.4 | 3 | 0 | 5 | 0.4 | 16 | 0.375 | 9 | 0.4444 | 3 | 0 | 9 | 0.7778 | 1 | 0 | 22 | 0.5 |
| 5_11051140 | 3 | 0.6667 | 5 | 0 | 3 | 0.3333 | 4 | 0.25 | 15 | 0.2667 | 7 | 0.1429 | 3 | 0 | 12 | 0 | 2 | 0.5 | 24 | 0.0833 |
| 5_11051160 | 2 | 0.5 | 5 | 0.8 | 2 | 0 | 6 | 0.1667 | 15 | 0.4 | 7 | 0.2857 | 3 | 0 | 11 | 0.6364 | 2 | 0 | 23 | 0.3913 |
| 5_11057794 | 1 | 0 | 2 | 0.5 | NA | NA | 1 | 1 | 4 | 0.5 | 2 | 1 | NA | NA | 1 | 0 | NA | NA | 3 | 0.6667 |
| 5_11057815 | 1 | 1 | 2 | 0.5 | NA | NA | NA | NA | 3 | 0.6667 | 1 | 1 | NA | NA | 1 | 0 | NA | NA | 2 | 0.5 |
| 5_11057856 | 1 | 1 | 2 | 1 | NA | NA | NA | NA | 3 | 1 | 1 | 1 | NA | NA | 1 | 0 | NA | NA | 2 | 0.5 |
| 5_11057868 | 1 | 1 | 2 | 1 | NA | NA | 1 | 1 | 4 | 1 | 1 | 1 | NA | NA | 1 | 0 | NA | NA | 2 | 0.5 |
| 5_11057890 | 1 | 1 | 2 | 1 | NA | NA | 1 | 1 | 4 | 1 | 1 | 1 | NA | NA | 1 | 0 | NA | NA | 2 | 0.5 |
| 5_11057943 | 1 | 1 | 2 | 1 | NA | NA | NA | NA | 3 | 1 | 1 | 1 | NA | NA | 1 | 1 | NA | NA | 2 | 1 |
| 5_11057949 | 1 | 1 | 2 | 1 | NA | NA | NA | NA | 3 | 1 | 1 | 1 | NA | NA | 1 | 0 | NA | NA | 2 | 0.5 |
| 5_11057953 | 1 | 1 | 1 | 1 | NA | NA | NA | NA | 2 | 1 | 1 | 1 | NA | NA | NA | NA | NA | NA | 1 | 1 |
| 5_11076856 | 6 | 1 | 13 | 0.9231 | 4 | 0.75 | 6 | 0.8333 | 29 | 0.8966 | 2 | 1 | 1 | 1 | 1 | 1 | 8 | 1 | 12 | 1 |
| 5_11076857 | 8 | 1 | 12 | 1 | 3 | 0.6667 | 4 | 1 | 27 | 0.963 | 2 | 1 | 1 | 1 | 1 | 1 | 8 | 0.875 | 12 | 0.9167 |
| 5_11076861 | 5 | 0.2 | 10 | 0.1 | 2 | 0 | 4 | 0.25 | 21 | 0.1429 | 2 | 1 | 1 | 0 | 1 | 0 | 8 | 0.25 | 12 | 0.3333 |
| 5_11076869 | 9 | 0.2222 | 12 | 0.5 | 4 | 0.25 | 6 | 0.5 | 31 | 0.3871 | 2 | 0 | 1 | 0 | 1 | 1 | 9 | 0.5556 | 13 | 0.4615 |
| 5_11076872 | 9 | 0 | 13 | 0.1538 | 4 | 0 | 7 | 0 | 33 | 0.0606 | 2 | 0 | 1 | 0 | 1 | 0 | 9 | 0.1111 | 13 | 0.0769 |
| 5_11076876 | 7 | 0 | 9 | 0 | 4 | 0 | 5 | 0.8 | 25 | 0.16 | 3 | 0 | NA | NA | 1 | 0 | 10 | 0.3 | 14 | 0.2143 |
| 5_11076889 | 6 | 0.5 | 10 | 0.8 | 3 | 0.6667 | 4 | 0.75 | 23 | 0.6957 | 3 | 0.6667 | 1 | 1 | 1 | 0 | 5 | 0.6 | 10 | 0.6 |
| 5_11076908 | 9 | 0.4444 | 13 | 0.4615 | 3 | 0.3333 | 6 | 0.1667 | 31 | 0.3871 | 4 | 0.5 | NA | NA | 1 | 0 | 10 | 0.8 | 15 | 0.6667 |
| 5_11080563 | 5 | 0.8 | 2 | 1 | 1 | 1 | 1 | 1 | 9 | 0.8889 | 2 | 1 | 1 | 1 | 1 | 1 | 1 | 1 | 5 | 1 |
| 5_11080571 | 5 | 0 | 2 | 0.5 | 1 | 0 | NA | NA | 8 | 0.125 | 2 | 0.5 | 1 | 0 | 1 | 0 | 1 | 0 | 5 | 0.2 |
| 5_11080572 | 5 | 0.8 | 2 | 1 | 1 | 1 | 1 | 0 | 9 | 0.7778 | 2 | 1 | 1 | 1 | 1 | 1 | 2 | 1 | 6 | 1 |
| 5_11080572 | 3 | 1 | NA | NA | NA | NA | 1 | 1 | 4 | 1 | 1 | 1 | 1 | 1 | 1 | 0 | 2 | 1 | 5 | 0.8 |

| | | | | | | | | | | | | | | | | | | | | |
|---|---|---|---|---|---|---|---|---|---|---|---|---|---|---|---|---|---|---|---|---|
| 5_1108058 | 5 | 0 | 1 | 1 | 1 | 0 | 1 | 0 | 8 | 0.125 | 1 | 0 | NA | NA | 1 | 0 | 1 | 1 | 3 | 0.3333 |
| 5_1108123 | 11 | 0 | 9 | 0.1111 | 2 | 0.5 | 5 | 0 | 27 | 0.0741 | 4 | 0 | 8 | 0 | 9 | 0.2222 | 5 | 0 | 26 | 0.0769 |
| 5_1108130 | 9 | 0.3333 | 5 | 0 | NA | NA | 4 | 0.25 | 18 | 0.2222 | 2 | 0 | 6 | 0.1667 | 6 | 0.3333 | 4 | 0 | 18 | 0.1667 |
| 5_1108134 | 8 | 0.125 | 4 | 0 | 1 | 0 | 3 | 0 | 16 | 0.0625 | 1 | 0 | 7 | 0 | 5 | 0.2 | 4 | 0.5 | 17 | 0.1765 |
| 5_1108135 | 9 | 0.1111 | 3 | 0 | 1 | 0 | 2 | 0 | 15 | 0.0667 | NA | NA | 5 | 0 | 5 | 0 | 4 | 0.25 | 14 | 0.0714 |
| 5_1108135 | 6 | 0.1667 | 4 | 0 | 1 | 0 | 3 | 0.3333 | 14 | 0.1429 | 2 | 0.5 | 7 | 0 | 3 | 0.3333 | 4 | 0 | 16 | 0.125 |
| 5_1108136 | 7 | 1 | 4 | 1 | NA | NA | 2 | 1 | 13 | 1 | 1 | 1 | 5 | 1 | 5 | 1 | 1 | 1 | 12 | 1 |
| 5_1108136 | 7 | 0.8571 | 4 | 0.25 | NA | NA | 3 | 0 | 14 | 0.5 | 1 | 1 | 6 | 0.5 | 5 | 0.4 | 4 | 0.75 | 16 | 0.5625 |
| 5_1109225 | 5 | 1 | 1 | 1 | 1 | 1 | 2 | 1 | 9 | 1 | 1 | 1 | 1 | 0 | 2 | 1 | 1 | 0 | 5 | 0.6 |
| 5_1109229 | 3 | 0.3333 | 1 | 0 | 1 | 1 | 1 | 0 | 6 | 0.3333 | 2 | 0.5 | 1 | 0 | 1 | 0 | 1 | 0 | 5 | 0.2 |
| 5_1109333 | 5 | 0 | 6 | 0.1667 | 6 | 0 | 2 | 0 | 19 | 0.0526 | 8 | 0 | 2 | 0.5 | 5 | 0 | 5 | 0 | 20 | 0.05 |
| 5_1109333 | 4 | 0 | 6 | 0.1667 | 6 | 0 | 2 | 0.5 | 18 | 0.1111 | 5 | 0 | 2 | 0 | 5 | 0 | 4 | 0.25 | 16 | 0.0625 |
| 5_1109334 | 5 | 0 | 5 | 0 | 5 | 0.2 | 1 | 0 | 16 | 0.0625 | 9 | 0.2222 | 1 | 0 | 4 | 0 | 3 | 0 | 17 | 0.1176 |
| 5_1109335 | 5 | 0 | 5 | 0.2 | 6 | 0 | 2 | 0 | 18 | 0.0556 | 7 | 0 | 1 | 0 | 6 | 0.1667 | 3 | 0 | 17 | 0.0588 |
| 5_1109335 | 5 | 0.2 | 5 | 0.2 | 5 | 0.2 | 1 | 0 | 16 | 0.1875 | 8 | 0 | 2 | 0.5 | 3 | 0 | 2 | 0 | 15 | 0.0667 |
| 5_1109335 | 4 | 1 | 6 | 1 | 4 | 0.5 | 2 | 1 | 16 | 0.875 | 7 | 1 | 1 | 1 | 5 | 1 | 4 | 1 | 17 | 1 |
| 5_1109335 | 2 | 0 | 5 | 0.2 | 5 | 0.4 | 1 | 0 | 13 | 0.2308 | 8 | 0.125 | 2 | 0.5 | 6 | 0.6667 | 4 | 0.5 | 20 | 0.4 |
| 5_1109335 | 4 | 1 | 5 | 0.8 | 5 | 0.8 | 1 | 0 | 15 | 0.8 | 7 | 0.4286 | 2 | 1 | 6 | 0.5 | 3 | 0.3333 | 18 | 0.5 |
| 5_1109336 | 5 | 0 | 5 | 0.2 | 4 | 0 | 2 | 0.5 | 16 | 0.125 | 9 | 0.4444 | 1 | 0 | 6 | 0.1667 | 1 | 0 | 17 | 0.2941 |
| 5_1109337 | 4 | 0 | 4 | 0 | 5 | 0.4 | 1 | 0 | 14 | 0.1429 | 9 | 0.1111 | 2 | 0 | 6 | 0.1667 | 3 | 0 | 20 | 0.1 |
| 5_1109338 | 4 | 0.25 | 4 | 0.5 | 4 | 0.75 | 1 | 0 | 13 | 0.4615 | 9 | 0.5556 | NA | NA | 5 | 0.4 | 3 | 0.3333 | 17 | 0.4706 |
| 5_1109338 | 4 | 0.25 | 4 | 0.5 | 5 | 0.4 | 2 | 1 | 15 | 0.4667 | 8 | 0.375 | 2 | 0 | 3 | 0.6667 | 3 | 0.3333 | 16 | 0.375 |
| 5_1109339 | 5 | 0.6 | 5 | 0.2 | 5 | 0.4 | 1 | 0 | 16 | 0.375 | 7 | 0.2857 | 2 | 0.5 | 5 | 0.4 | 3 | 0.6667 | 17 | 0.4118 |
| 5_1109339 | 5 | 0.2 | 5 | 0.2 | 3 | 0.6667 | 1 | 0 | 14 | 0.2857 | 8 | 0.625 | 2 | 0 | 5 | 0.4 | 4 | 0 | 19 | 0.3684 |
| 5_1109339 | 5 | 0.2 | 4 | 0.75 | 5 | 0.4 | 1 | 1 | 15 | 0.4667 | 9 | 0.1111 | 2 | 0 | 5 | 0 | 3 | 0 | 19 | 0.0526 |
| 5_1109577 | NA | NA | NA | NA | 2 | 0.5 | NA | NA | 2 | 0.5 | 2 | 0 | 2 | 0.5 | NA | NA | 1 | 0 | 5 | 0.2 |
| 5_1114343 | 1 | 1 | 2 | 1 | NA | NA | NA | NA | 3 | 1 | NA | NA | 1 | 1 | NA | NA | NA | NA | 1 | 1 |
| 5_1114345 | 1 | 0 | 3 | 1 | NA | NA | NA | NA | 4 | 0.75 | NA | NA | 1 | 1 | NA | NA | NA | NA | 1 | 1 |
| 5_1114348 | 1 | 1 | 3 | 0.6667 | NA | NA | NA | NA | 4 | 0.75 | NA | NA | 1 | 1 | NA | NA | NA | NA | 1 | 1 |
| 5_1114351 | 1 | 0 | 2 | 0.5 | NA | NA | NA | NA | 3 | 0.3333 | NA | NA | 1 | 1 | NA | NA | NA | NA | 1 | 1 |
| 5_1114355 | 1 | 1 | 2 | 0 | NA | NA | NA | NA | 3 | 0.3333 | NA | NA | 1 | 1 | NA | NA | NA | NA | 1 | 1 |
| 5_1114362 | 1 | 1 | 3 | 0.6667 | NA | NA | NA | NA | 4 | 0.75 | NA | NA | 1 | 1 | NA | NA | NA | NA | 1 | 1 |
| 5_1114364 | 1 | 1 | 3 | 0.6667 | NA | NA | NA | NA | 4 | 0.75 | NA | NA | 1 | 1 | NA | NA | NA | NA | 1 | 1 |
| 5_1114364 | NA | NA | 3 | 1 | NA | NA | NA | NA | 3 | 1 | NA | NA | 1 | 1 | NA | NA | NA | NA | 1 | 1 |
| 5_1114374 | 1 | 1 | 4 | 0.5 | NA | NA | 1 | 0 | 6 | 0.5 | NA | NA | 1 | 0 | 1 | 1 | NA | NA | 2 | 0.5 |
| 5_1115348 | 11 | 0.1818 | 12 | 0 | 8 | 0.125 | 9 | 0.2222 | 40 | 0.125 | 9 | 0.1111 | 15 | 0.2 | 8 | 0 | 9 | 0.3333 | 41 | 0.1707 |
| 5_1115348 | 11 | 0.4545 | 11 | 0.3636 | 10 | 0.8 | 8 | 0.25 | 40 | 0.475 | 10 | 0.4 | 10 | 0.2 | 6 | 0.3333 | 6 | 0.6667 | 32 | 0.375 |
| 5_1115360 | 14 | 0.2143 | 17 | 0.4118 | 11 | 0.4545 | 13 | 0.2308 | 55 | 0.3273 | 13 | 0.1538 | 23 | 0.3913 | 10 | 0.3 | 10 | 0.5 | 56 | 0.3393 |
| 5_1115655 | 1 | 1 | NA | NA | NA | NA | NA | NA | 1 | 1 | 1 | 1 | NA | NA | NA | NA | NA | NA | 1 | 1 |
| 5_1115660 | 1 | 1 | NA | NA | NA | NA | 1 | 0 | 2 | 0.5 | 1 | 0 | NA | NA | 1 | 0 | 1 | 1 | 3 | 0.3333 |
| 5_1115660 | 1 | 1 | 2 | 0.5 | NA | NA | 1 | 1 | 4 | 0.75 | 1 | 1 | 1 | 1 | 1 | 1 | 1 | 1 | 4 | 1 |
| 5_1115661 | 1 | 1 | 2 | 1 | NA | NA | NA | NA | 3 | 1 | 1 | 1 | NA | NA | 1 | 1 | 1 | 1 | 3 | 1 |
| 5_1115661 | 1 | 1 | 2 | 0 | NA | NA | NA | NA | 3 | 0.3333 | 1 | 1 | 1 | 0 | 1 | 1 | 1 | 1 | 4 | 0.75 |
| 5_1115665 | 1 | 1 | 2 | 1 | NA | NA | 1 | 0 | 4 | 0.75 | 1 | 1 | 1 | 1 | 1 | 1 | 1 | 1 | 4 | 1 |
| 5_1115667 | 1 | 1 | 2 | 1 | NA | NA | 1 | 1 | 4 | 1 | 1 | 1 | 1 | 1 | 1 | 1 | 1 | 1 | 4 | 1 |
| 5_1115667 | 1 | 0 | 2 | 1 | 1 | 1 | 1 | 0 | 5 | 0.6 | NA | NA | 1 | 0 | NA | NA | 1 | 1 | 2 | 0.5 |
| 5_1115670 | 1 | 1 | 2 | 0.5 | 1 | 1 | 2 | 1 | 6 | 0.8333 | 1 | 1 | 1 | 0 | 1 | 0 | 1 | 1 | 4 | 0.5 |
| 5_1115673 | NA | NA | 1 | 1 | 1 | 0 | 2 | 1 | 4 | 0.75 | 1 | 0 | 1 | 1 | 1 | 1 | NA | NA | 3 | 0.6667 |
| 5_1115675 | 1 | 1 | 2 | 0.5 | 1 | 1 | 2 | 0.5 | 6 | 0.6667 | 1 | 0 | 1 | 1 | 1 | 1 | NA | NA | 3 | 0.6667 |
| 5_1115677 | 1 | 1 | 2 | 1 | 1 | 1 | 2 | 1 | 6 | 1 | 1 | 1 | 1 | 1 | 1 | 1 | NA | NA | 3 | 1 |
| 5_1115679 | 1 | 1 | 2 | 0 | 1 | 0 | 1 | 0 | 5 | 0.2 | 2 | 0.5 | 1 | 0 | 1 | 0 | 1 | 0 | 5 | 0.2 |
| 5_1115681 | 1 | 1 | 1 | 1 | 1 | 1 | 2 | 1 | 5 | 1 | 2 | 1 | 1 | 1 | 1 | 1 | 1 | 0 | 5 | 0.8 |
| 5_1115681 | 1 | 1 | 1 | 1 | NA | NA | 3 | 1 | 5 | 1 | 2 | 1 | 1 | 1 | 1 | 1 | 1 | 0 | 5 | 0.8 |
| 5_1115684 | NA | NA | 2 | 0.5 | 1 | 1 | 4 | 0.75 | 7 | 0.7143 | 2 | 1 | 2 | 0.5 | 1 | 0 | 1 | 0 | 6 | 0.5 |
| 5_1115686 | 1 | 1 | 2 | 0.5 | NA | NA | 5 | 1 | 8 | 0.875 | 1 | 0 | NA | NA | 1 | 1 | 1 | 1 | 3 | 0.6667 |
| 5_1115686 | 1 | 0 | 1 | 0 | 1 | 1 | 6 | 0.1667 | 9 | 0.2222 | 1 | 0 | 2 | 0 | 1 | 1 | 1 | 0 | 5 | 0.2 |
| 5_1115688 | 1 | 1 | 2 | 1 | 1 | 1 | 5 | 1 | 9 | 1 | 2 | 1 | 3 | 1 | 1 | 1 | 1 | 1 | 7 | 1 |
| 5_1115688 | 1 | 1 | 2 | 1 | 1 | 0 | 6 | 0.5 | 10 | 0.6 | 2 | 1 | 3 | 0.3333 | 2 | 0 | 1 | 1 | 8 | 0.5 |
| 5_1115690 | 1 | 0 | 2 | 0.5 | 1 | 0 | 5 | 0.2 | 9 | 0.2222 | 2 | 0 | 3 | 0 | 2 | 0.5 | 2 | 0 | 9 | 0.1111 |
| 5_1115690 | 1 | 1 | 1 | 0 | 1 | 0 | 4 | 0.75 | 7 | 0.5714 | 2 | 1 | 3 | 1 | 2 | 0.5 | 3 | 1 | 10 | 0.9 |
| 5_1115694 | 1 | 1 | 3 | 0 | 1 | 0 | 5 | 0.2 | 10 | 0.2 | 2 | 1 | 3 | 0.3333 | 2 | 0.5 | 3 | 0 | 10 | 0.4 |
| 5_1115694 | 1 | 0 | 3 | 0 | 1 | 1 | 5 | 0.8 | 10 | 0.5 | 1 | 0 | 3 | 0 | 2 | 0.5 | 1 | 0 | 7 | 0.1429 |
| 5_1115696 | 1 | 1 | 2 | 1 | 1 | 0 | 6 | 0.8333 | 10 | 0.8 | 2 | 1 | 3 | 0.6667 | 2 | 0.5 | NA | NA | 7 | 0.7143 |
| 5_1115696 | 1 | 1 | 3 | 0 | 1 | 0 | 6 | 0.3333 | 11 | 0.2727 | 2 | 0 | 3 | 0 | 2 | 0.5 | 2 | 0 | 9 | 0.1111 |
| 5_1116025 | 2 | 0 | 4 | 0 | 1 | 0 | 6 | 0.1667 | 13 | 0.0769 | 2 | 0 | 4 | 0.25 | 3 | 0 | 2 | 0 | 11 | 0.0909 |

| | | | | | | | | | | | | | | | | | | | | |
|---|---|---|---|---|---|---|---|---|---|---|---|---|---|---|---|---|---|---|---|---|
| 5_11160263 | 2 | 1 | 3 | 0.6667 | 1 | 1 | 6 | 1 | 12 | 0.9167 | 1 | 1 | 4 | 0.5 | 3 | 1 | 3 | 1 | 11 | 0.8182 |
| 5_11160271 | 2 | 1 | 5 | 1 | 1 | 1 | 7 | 0.8571 | 15 | 0.9333 | 2 | 1 | 4 | 1 | 4 | 1 | 3 | 1 | 13 | 1 |
| 5_11160303 | NA | NA | 3 | 0 | 1 | 0 | 4 | 0.75 | 8 | 0.375 | 1 | 0 | 2 | 0.5 | 1 | 0 | NA | NA | 4 | 0.25 |
| 5_11160345 | 2 | 0 | 7 | 0.1429 | 3 | 0 | 7 | 0 | 19 | 0.0526 | 5 | 0.2 | 5 | 0.2 | 5 | 0 | 2 | 0 | 17 | 0.1176 |
| 5_11160404 | 3 | 0.3333 | 6 | 0.1667 | 5 | 0 | 6 | 0.1667 | 20 | 0.15 | 6 | 0 | 5 | 0.4 | 4 | 0 | 4 | 0 | 19 | 0.1053 |
| 5_11160450 | 7 | 0 | 12 | 0.0833 | 4 | 0 | 9 | 0.1111 | 32 | 0.0625 | 9 | 0.1111 | 7 | 0.1429 | 8 | 0 | 5 | 0.2 | 29 | 0.1034 |
| 5_11160792 | NA | NA | 3 | 1 | 3 | 0.6667 | 2 | 0.5 | 8 | 0.75 | 1 | 1 | 4 | 0.25 | 2 | 0 | 3 | 0 | 10 | 0.2 |
| 5_11236124 | 3 | 0.3333 | 6 | 0 | 1 | 1 | 2 | 0.5 | 12 | 0.25 | 3 | 0.3333 | 4 | 0.5 | 3 | 0.6667 | 4 | 0.5 | 14 | 0.5 |
| 5_11236143 | 3 | 0.3333 | 5 | 0 | 1 | 0 | 1 | 0 | 10 | 0.1 | 3 | 0.3333 | 5 | 0 | 5 | 0.4 | 4 | 0 | 17 | 0.1765 |
| 5_11236171 | 3 | 0 | 5 | 0.2 | 1 | 0 | 2 | 0 | 11 | 0.0909 | 3 | 0.3333 | 4 | 0.5 | 3 | 0 | 4 | 0.25 | 14 | 0.2857 |
| 5_11236240 | 3 | 0.3333 | 4 | 0.25 | NA | NA | 2 | 0.5 | 9 | 0.3333 | 2 | 0.5 | 3 | 0.3333 | NA | NA | 2 | 0.5 | 7 | 0.4286 |
| 5_11236255 | 2 | 0 | 3 | 0.3333 | NA | NA | 1 | 0 | 6 | 0.1667 | 2 | 0.5 | 3 | 0 | NA | NA | 1 | 0 | 6 | 0.1667 |
| 5_11236258 | 2 | 0.5 | 3 | 0.3333 | NA | NA | 1 | 0 | 6 | 0.3333 | 2 | 0 | 3 | 0.3333 | NA | NA | 1 | 1 | 6 | 0.3333 |
| 5_11236260 | 1 | 0 | 3 | 0.3333 | NA | NA | 2 | 0 | 6 | 0.1667 | 1 | 1 | 3 | 0 | NA | NA | 1 | 1 | 5 | 0.4 |
| 5_11245565 | 2 | 0.5 | 3 | 0.6667 | 4 | 0.5 | 1 | 0 | 10 | 0.5 | 1 | 1 | 1 | 1 | NA | NA | 7 | 0.4286 | 9 | 0.5556 |
| 5_11245584 | 2 | 0.5 | 4 | 0.25 | 5 | 0.2 | NA | NA | 11 | 0.2727 | 1 | 0 | 1 | 0 | NA | NA | 6 | 0.6667 | 8 | 0.5 |
| 5_11245597 | 3 | 1 | 6 | 0.6667 | 4 | 0.75 | 1 | 0 | 14 | 0.7143 | 1 | 1 | 1 | 1 | 2 | 1 | 8 | 1 | 12 | 1 |
| 5_11245612 | 3 | 0.6667 | 6 | 0.5 | 4 | 0.25 | 1 | 1 | 14 | 0.5 | 1 | 1 | 1 | 1 | 2 | 0 | 7 | 0.4286 | 11 | 0.4545 |
| 5_11245626 | 1 | 0 | 5 | 0.4 | 4 | 0.5 | NA | NA | 10 | 0.4 | NA | NA | 1 | 1 | 2 | 0 | 6 | 0.8333 | 9 | 0.6667 |
| 5_11245630 | 3 | 0.3333 | 6 | 0 | 5 | 0.2 | 1 | 1 | 15 | 0.2 | 1 | 0 | 2 | 0.5 | 2 | 0 | 9 | 0 | 14 | 0.0714 |
| 5_11246280 | 2 | 0 | 6 | 0.3333 | 6 | 0 | 2 | 0 | 16 | 0.125 | NA | NA | 1 | 0 | 2 | 0 | 9 | 0.1111 | 12 | 0.0833 |
| 5_11246289 | 2 | 0 | 6 | 0 | 8 | 0.125 | 1 | 0 | 17 | 0.0588 | 1 | 0 | 2 | 0 | 3 | 0.3333 | 7 | 0 | 13 | 0.0769 |
| 5_11246293 | 4 | 0.25 | 7 | 0.1429 | 8 | 0 | 2 | 1 | 21 | 0.1905 | 1 | 1 | 3 | 0 | 3 | 0 | 8 | 0.25 | 15 | 0.2 |
| 5_11246298 | 4 | 1 | 6 | 0.6667 | 4 | 1 | 1 | 1 | 15 | 0.8667 | NA | NA | 3 | 0.6667 | 3 | 1 | 8 | 1 | 14 | 0.9286 |
| 5_11246302 | 2 | 1 | 5 | 0.6 | 6 | 0.6667 | 1 | 1 | 14 | 0.7143 | 2 | 1 | 3 | 0.3333 | 2 | 1 | 6 | 0.8333 | 13 | 0.7692 |
| 5_11246305 | 3 | 0 | 7 | 0 | 8 | 0.25 | 2 | 0.5 | 20 | 0.15 | 3 | 0.3333 | 4 | 0.25 | 3 | 0 | 8 | 0.625 | 18 | 0.3889 |
| 5_11246308 | 2 | 0 | 6 | 0.3333 | 6 | 0.5 | 2 | 0 | 16 | 0.3125 | 2 | 1 | 2 | 0 | 4 | 0.25 | 9 | 0.1111 | 17 | 0.2353 |
| 5_11246309 | 4 | 0.25 | 3 | 0 | 8 | 0.75 | 2 | 0.5 | 17 | 0.4706 | 3 | 0 | 3 | 0 | 3 | 0.6667 | 8 | 0.375 | 17 | 0.2941 |
| 5_11248106 | 3 | 0.3333 | 1 | 1 | 4 | 0.25 | 1 | 0 | 9 | 0.3333 | NA | NA | 2 | 0.5 | 4 | 0.5 | NA | NA | 6 | 0.5 |
| 5_11248132 | 3 | 0 | 2 | 0 | 4 | 0.75 | 1 | 0 | 10 | 0.3 | 2 | 1 | 1 | 0 | 5 | 0.6 | NA | NA | 8 | 0.625 |
| 5_11248350 | 4 | 0.25 | 3 | 0 | 5 | 0 | 2 | 0 | 14 | 0.0714 | 2 | 0 | 1 | 1 | 3 | 0.3333 | NA | NA | 6 | 0.3333 |
| 5_11248362 | NA | NA | 1 | 1 | NA | NA | 1 | 0 | 2 | 0.5 | NA | NA | 1 | 0 | NA | NA | 1 | 1 | 2 | 0.5 |
| 5_11248500 | 5 | 0.6 | 5 | 1 | 4 | 1 | 1 | 1 | 15 | 0.8667 | 3 | 0.6667 | 1 | 1 | 2 | 1 | 2 | 1 | 8 | 0.875 |
| 5_11248501 | 6 | 0.5 | 5 | 0.6 | 7 | 0.8571 | 2 | 1 | 20 | 0.7 | 3 | 1 | 2 | 1 | 5 | 0.8 | 1 | 1 | 11 | 0.9091 |
| 5_11248502 | 5 | 0 | 4 | 0 | 7 | 0.1429 | 1 | 0 | 17 | 0.0588 | 2 | 0 | 1 | 0 | 5 | 0 | 2 | 0.5 | 10 | 0.1 |
| 5_11248505 | NA | NA | 1 | 0 | NA | NA | 1 | 1 | 2 | 0.5 | 1 | 1 | 1 | 0 | 1 | 0 | NA | NA | 3 | 0.3333 |
| 5_11248512 | 5 | 0.2 | 3 | 0 | 7 | 0 | 2 | 0 | 17 | 0.0588 | 3 | 0 | 2 | 0 | 5 | 0.2 | 1 | 0 | 11 | 0.0909 |
| 5_11248520 | 7 | 0 | 3 | 0 | 8 | 0.125 | 1 | 0 | 19 | 0.0526 | 3 | 0 | 2 | 0 | 4 | 0 | 1 | 1 | 10 | 0.1 |
| 5_11248523 | 6 | 0.3333 | 4 | 0 | 8 | 0.125 | NA | NA | 18 | 0.1667 | 3 | 0 | 2 | 0 | 4 | 0 | 2 | 1 | 11 | 0.1818 |
| 5_11248530 | 5 | 0.4 | 3 | 1 | 4 | 1 | 2 | 1 | 14 | 0.7857 | 2 | 1 | 4 | 0.5 | 5 | 0.4 | 2 | 0 | 13 | 0.4615 |
| 5_11248531 | 2 | 0.5 | 8 | 0.875 | 2 | 1 | 1 | 1 | 13 | 0.8462 | 1 | 1 | 7 | 1 | NA | NA | 3 | 0.6667 | 11 | 0.9091 |
| 5_11248539 | 5 | 1 | 4 | 0.75 | 2 | 1 | 2 | 1 | 13 | 0.9231 | 3 | 0.6667 | 3 | 1 | 5 | 0.8 | 1 | 1 | 12 | 0.8333 |
| 5_11248539 | 2 | 1 | 8 | 0.75 | 2 | 0.5 | 1 | 0 | 13 | 0.6923 | 2 | 1 | 7 | 0.4286 | 1 | 0 | 3 | 0.3333 | 13 | 0.4615 |
| 5_11248545 | 2 | 0.5 | 9 | 0.2222 | 2 | 0 | 1 | 0 | 14 | 0.2143 | 1 | 0 | 6 | 0.6667 | 1 | 1 | 3 | 0.3333 | 11 | 0.5455 |
| 5_11248550 | NA | NA | 6 | 0.6667 | 2 | 1 | 1 | 0 | 9 | 0.6667 | NA | NA | 6 | 0.8333 | 1 | 0 | 2 | 1 | 9 | 0.7778 |
| 5_11248553 | 5 | 0.6 | 3 | 1 | 2 | 0 | 2 | 1 | 12 | 0.6667 | 3 | 0 | 3 | 0.6667 | 5 | 0.4 | 1 | 0 | 12 | 0.3333 |
| 5_11248555 | 1 | 0 | 7 | 0 | 2 | 0.5 | 1 | 0 | 11 | 0.0909 | 1 | 0 | 6 | 0.1667 | 1 | 0 | 2 | 0 | 10 | 0.1 |
| 5_11248574 | 5 | 0 | 5 | 0.4 | 4 | 0 | 2 | 0 | 16 | 0.125 | 3 | 0 | 4 | 0 | 6 | 0 | 3 | 0.6667 | 16 | 0.125 |
| 5_11248581 | 5 | 0.8 | 7 | 0.8571 | 4 | 1 | 2 | 1 | 18 | 0.8889 | 3 | 0.6667 | 4 | 0.75 | 5 | 1 | 3 | 1 | 15 | 0.8667 |
| 5_11249199 | NA | NA | 1 | 0 | 1 | 1 | NA | NA | 2 | 0.5 | 1 | 1 | NA | NA | NA | NA | NA | NA | 1 | 1 |
| 5_11249228 | 1 | 1 | 1 | 1 | 1 | 1 | NA | NA | 3 | 1 | 1 | 1 | NA | NA | NA | NA | NA | NA | 1 | 1 |
| 5_11249248 | 1 | 0 | 1 | 1 | 2 | 1 | NA | NA | 4 | 0.75 | 1 | 1 | NA | NA | NA | NA | NA | NA | 1 | 1 |
| 5_11249266 | 1 | 0 | 1 | 0 | 2 | 1 | NA | NA | 4 | 0.5 | 1 | 1 | NA | NA | 2 | 0.5 | NA | NA | 3 | 0.6667 |
| 5_11251799 | 2 | 1 | 3 | 0.3333 | 4 | 0.75 | 1 | 1 | 10 | 0.7 | 1 | 1 | 1 | 1 | 4 | 1 | NA | NA | 6 | 1 |
| 5_11251807 | 2 | 0.5 | 2 | 0.5 | 4 | 0.5 | 1 | 0 | 9 | 0.4444 | NA | NA | 2 | 0.5 | 2 | 0 | NA | NA | 4 | 0.25 |
| 5_11251807 | 3 | 1 | 2 | 0.5 | 4 | 1 | 1 | 0 | 10 | 0.8 | NA | NA | 2 | 1 | 4 | 0.75 | NA | NA | 6 | 0.8333 |
| 5_11251814 | 3 | 0 | 3 | 0.6667 | 4 | 0.5 | 1 | 0 | 11 | 0.3636 | 1 | 1 | 2 | 0.5 | 4 | 0.75 | NA | NA | 7 | 0.7143 |
| 5_11251825 | 3 | 0.6667 | 3 | 1 | 4 | 1 | 1 | 1 | 11 | 0.9091 | 1 | 1 | 2 | 1 | 4 | 0.5 | NA | NA | 7 | 0.7143 |
| 5_11251830 | 3 | 0.3333 | 3 | 0.3333 | 4 | 0.25 | 1 | 0 | 11 | 0.2727 | 1 | 0 | 2 | 0 | 3 | 0.3333 | NA | NA | 6 | 0.1667 |
| 5_11251846 | 3 | 0 | 3 | 0 | 4 | 0.25 | 1 | 0 | 11 | 0.0909 | 2 | 0 | 3 | 0 | 4 | 0.25 | NA | NA | 9 | 0.1111 |
| 5_11251852 | 2 | 0.5 | 3 | 0 | 4 | 0 | 1 | 0 | 10 | 0.1 | 2 | 0.5 | 3 | 0 | 4 | 0.25 | NA | NA | 9 | 0.2222 |
| 5_11251854 | 3 | 0.3333 | 3 | 0.6667 | 3 | 0 | 1 | 1 | 10 | 0.4 | 1 | 0 | 3 | 0.3333 | 3 | 0.6667 | NA | NA | 7 | 0.4286 |
| 5_11252608 | 3 | 0.3333 | 3 | 0.6667 | 4 | 0.5 | 3 | 1 | 13 | 0.6154 | 1 | 1 | 3 | 1 | 6 | 0.6667 | NA | NA | 10 | 0.8 |
| 5_11253332 | 14 | 0.2143 | 11 | 0.0909 | 15 | 0.2 | 17 | 0.2353 | 57 | 0.193 | 7 | 0.1429 | 22 | 0.1818 | 16 | 0.3125 | 21 | 0.2381 | 66 | 0.2273 |
| 5_11253344 | 19 | 0.4737 | 12 | 0.4167 | 17 | 0.4706 | 19 | 0.4737 | 67 | 0.4627 | 3 | 0.3333 | 18 | 0.5 | 18 | 0.7778 | 19 | 0.4211 | 58 | 0.5517 |

| | | | | | | | | | | | | | | | | | | | | |
|---|---|---|---|---|---|---|---|---|---|---|---|---|---|---|---|---|---|---|---|---|
| 5_1125334 | 20 | 0.2 | 12 | 0.0833 | 15 | 0.1333 | 17 | 0.1176 | 64 | 0.1406 | 5 | 0 | 20 | 0.1 | 17 | 0.0588 | 21 | 0.0476 | 63 | 0.0635 |
| 5_1125335 | 21 | 0.3333 | 12 | 0.1667 | 16 | 0.4375 | 18 | 0.2778 | 67 | 0.3134 | 7 | 0.2857 | 18 | 0.3333 | 18 | 0.2222 | 18 | 0.2778 | 61 | 0.2787 |
| 5_1125335 | 16 | 0.5625 | 9 | 0.5556 | 12 | 0.5 | 17 | 0.4706 | 54 | 0.5185 | 5 | 0.8 | 12 | 0.8333 | 13 | 0.6154 | 17 | 0.7059 | 47 | 0.7234 |
| 5_1125336 | 14 | 0.0714 | 11 | 0.2727 | 12 | 0.1667 | 13 | 0.2308 | 50 | 0.18 | 4 | 0 | 12 | 0.1667 | 14 | 0.3571 | 16 | 0.4375 | 46 | 0.3043 |
| 5_1125390 | 20 | 0.35 | 12 | 0.3333 | 14 | 0.3571 | 18 | 0.2222 | 64 | 0.3125 | 6 | 0.1667 | 18 | 0.3333 | 16 | 0.1875 | 18 | 0.3333 | 58 | 0.2759 |
| 5_1125390 | 15 | 0.6667 | 11 | 0.8182 | 15 | 0.6667 | 19 | 0.7368 | 60 | 0.7167 | 5 | 0.8 | 16 | 0.8125 | 14 | 0.9286 | 20 | 0.8 | 55 | 0.8364 |
| 5_1125391 | 18 | 0.0556 | 8 | 0.25 | 12 | 0.3333 | 14 | 0.2143 | 52 | 0.1923 | 3 | 0 | 16 | 0.1875 | 11 | 0 | 14 | 0.3571 | 44 | 0.1818 |
| 5_1125391 | 20 | 0.2 | 9 | 0 | 13 | 0 | 20 | 0.05 | 62 | 0.0806 | 6 | 0.1667 | 20 | 0.3 | 14 | 0.2857 | 19 | 0.3684 | 59 | 0.3051 |
| 5_1125391 | 16 | 0.8125 | 7 | 0.2857 | 12 | 0.9167 | 17 | 0.5882 | 52 | 0.6923 | 5 | 1 | 16 | 0.625 | 13 | 0.6154 | 16 | 0.75 | 50 | 0.7 |
| 5_1125391 | 16 | 0.75 | 10 | 0.5 | 10 | 0.6 | 14 | 0.7857 | 50 | 0.68 | 4 | 0.75 | 15 | 0.8667 | 11 | 0.4545 | 16 | 0.875 | 46 | 0.7609 |
| 5_1125392 | 18 | 0.9444 | 5 | 0.8 | 12 | 0.9167 | 18 | 0.8889 | 53 | 0.9057 | 6 | 0.8333 | 17 | 0.8824 | 12 | 0.75 | 18 | 0.8333 | 53 | 0.8302 |
| 5_1125392 | 17 | 0.1765 | 6 | 0 | 9 | 0.1111 | 15 | 0.1333 | 47 | 0.1277 | 3 | 0 | 16 | 0.1875 | 10 | 0.1 | 13 | 0.1538 | 42 | 0.1429 |
| 5_1125393 | 16 | 0.5 | 9 | 0.4444 | 11 | 0.3636 | 16 | 0.1875 | 52 | 0.3654 | 4 | 0.5 | 18 | 0.4444 | 12 | 0.4167 | 17 | 0.2941 | 51 | 0.3922 |
| 5_1125397 | 14 | 0.8571 | 5 | 0.6 | 12 | 0.9167 | 16 | 0.6875 | 47 | 0.7872 | 5 | 0.8 | 10 | 0.7 | 9 | 0.8889 | 16 | 0.9375 | 40 | 0.85 |
| 5_1125397 | 16 | 0.9375 | 4 | 0.75 | 11 | 0.8182 | 17 | 0.9412 | 48 | 0.8958 | 4 | 1 | 11 | 0.9091 | 10 | 0.8 | 14 | 1 | 39 | 0.9231 |
| 5_1125398 | 12 | 0.75 | 4 | 0.5 | 12 | 0.9167 | 17 | 0.6471 | 45 | 0.7333 | 5 | 0.8 | 11 | 0.7273 | 9 | 1 | 16 | 0.8125 | 41 | 0.8293 |
| 5_1125399 | 4 | 0.75 | 1 | 1 | 1 | 0 | 7 | 0.8571 | 13 | 0.7692 | 1 | 1 | 4 | 1 | 1 | 1 | 6 | 1 | 12 | 1 |
| 5_1125399 | 13 | 0 | 2 | 0 | 10 | 0.1 | 14 | 0.0714 | 39 | 0.0513 | 5 | 0.2 | 10 | 0.1 | 7 | 0 | 13 | 0 | 35 | 0.0571 |
| 5_1125400 | 8 | 0.125 | 1 | 1 | 7 | 0.5714 | 15 | 0.2 | 31 | 0.2903 | 3 | 0 | 9 | 0.5556 | 7 | 0.8571 | 12 | 0.3333 | 31 | 0.4839 |
| 5_1125400 | 10 | 0.6 | 3 | 0 | 7 | 0.4286 | 14 | 0.5714 | 34 | 0.5 | 3 | 0 | 11 | 0.2727 | 7 | 0.1429 | 13 | 0.4615 | 34 | 0.2941 |
| 5_1125670 | 12 | 0.0833 | 15 | 0.0667 | 12 | 0 | 16 | 0.0625 | 55 | 0.0545 | 8 | 0.125 | 9 | 0.1111 | 11 | 0 | 10 | 0 | 38 | 0.0526 |
| 5_1125672 | 12 | 0.5 | 13 | 0.3077 | 11 | 0.3636 | 15 | 0.2667 | 51 | 0.3529 | 8 | 0.625 | 9 | 0.2222 | 12 | 0.75 | 10 | 0.4 | 39 | 0.5128 |
| 5_1125672 | 13 | 0.0769 | 15 | 0.2 | 11 | 0.1818 | 15 | 0 | 54 | 0.1111 | 7 | 0.1429 | 9 | 0 | 12 | 0 | 9 | 0.1111 | 37 | 0.0541 |
| 5_1125672 | 9 | 0.4444 | 12 | 0.1667 | 10 | 0.2 | 15 | 0.4 | 46 | 0.3043 | 7 | 0.1429 | 8 | 0.25 | 9 | 0.4444 | 8 | 0.25 | 32 | 0.2812 |
| 5_1125673 | 12 | 1 | 15 | 0.7333 | 12 | 0.9167 | 15 | 0.8667 | 54 | 0.8704 | 9 | 1 | 9 | 0.8889 | 11 | 1 | 10 | 0.9 | 39 | 0.9487 |
| 5_1125673 | 12 | 0.0833 | 15 | 0.2667 | 12 | 0.1667 | 15 | 0.2 | 54 | 0.1852 | 7 | 0.2857 | 8 | 0.5 | 11 | 0.3636 | 8 | 0.25 | 34 | 0.3529 |
| 5_1125673 | 9 | 0.7778 | 13 | 0.5385 | 7 | 0.8571 | 13 | 0.6923 | 42 | 0.6905 | 5 | 0.4 | 8 | 0.75 | 10 | 0.4 | 7 | 0.5714 | 30 | 0.5333 |
| 5_1125674 | 12 | 0.0833 | 14 | 0.1429 | 13 | 0.0769 | 14 | 0.4286 | 53 | 0.1887 | 7 | 0 | 7 | 0.1429 | 11 | 0.2727 | 9 | 0.2222 | 34 | 0.1765 |
| 5_1125674 | 9 | 0.6667 | 15 | 0.6 | 12 | 0.5833 | 15 | 0.4667 | 51 | 0.5686 | 6 | 0.8333 | 8 | 0.5 | 11 | 0.4545 | 9 | 0.8889 | 34 | 0.6471 |
| 5_1125678 | 8 | 0.25 | 13 | 0.1538 | 12 | 0 | 10 | 0 | 43 | 0.093 | 6 | 0.3333 | 7 | 0 | 10 | 0.1 | 10 | 0 | 33 | 0.0909 |
| 5_1125682 | 7 | 0.2857 | 12 | 0.1667 | 9 | 0.2222 | 7 | 0.1429 | 35 | 0.2 | 4 | 0 | 5 | 0 | 6 | 0.3333 | 9 | 0.3333 | 24 | 0.2083 |
| 5_1125684 | 5 | 0 | 13 | 0.2308 | 10 | 0.3 | 8 | 0.375 | 36 | 0.25 | 5 | 0.2 | 6 | 0 | 6 | 0.3333 | 10 | 0.1 | 27 | 0.1481 |
| 5_1125686 | 5 | 0.2 | 13 | 0.4615 | 9 | 0.5556 | 8 | 0.75 | 35 | 0.5143 | 5 | 0.8 | 6 | 0.1667 | 6 | 0.5 | 10 | 0.7 | 27 | 0.5556 |
| 5_1125689 | 5 | 0.2 | 11 | 0.2727 | 6 | 0 | 7 | 0.2857 | 29 | 0.2069 | 4 | 0 | 5 | 0 | 2 | 0.5 | 9 | 0.3333 | 20 | 0.2 |
| 5_1125691 | 5 | 0 | 11 | 0.2727 | 7 | 0 | 6 | 0.1667 | 29 | 0.1379 | 3 | 0.3333 | 5 | 0.2 | 3 | 0.6667 | 8 | 0.375 | 19 | 0.3684 |
| 5_1125692 | 5 | 0 | 12 | 0.0833 | 5 | 0 | 6 | 0.1667 | 28 | 0.0714 | 3 | 0.3333 | 5 | 0.2 | 3 | 0 | 8 | 0.375 | 19 | 0.2632 |
| 5_1125697 | 3 | 0.3333 | 6 | 0.3333 | 5 | 0.2 | 5 | 0.4 | 19 | 0.3158 | 3 | 0 | 2 | 0.5 | 2 | 0 | 7 | 0.4286 | 14 | 0.2857 |
| 5_1125697 | 4 | 0.5 | 10 | 0.4 | 6 | 0 | 5 | 0.2 | 25 | 0.28 | 3 | 0.6667 | 2 | 0 | 2 | 0 | 7 | 0.2857 | 14 | 0.2857 |
| 5_1125698 | 6 | 0 | 8 | 0.25 | 6 | 0 | 5 | 0.2 | 25 | 0.12 | 2 | 0 | 2 | 0 | 2 | 0.5 | 7 | 0 | 13 | 0.0769 |
| 5_1125698 | 5 | 0 | 10 | 0 | 6 | 0.3333 | 5 | 0.2 | 26 | 0.1154 | 4 | 0.5 | 2 | 0 | 2 | 0 | 7 | 0.2857 | 15 | 0.2667 |
| 5_1125698 | 6 | 0.8333 | 8 | 0.5 | 6 | 0.6667 | 5 | 0.8 | 25 | 0.68 | 2 | 0.5 | 2 | 0.5 | 2 | 1 | 6 | 0.5 | 12 | 0.5833 |
| 5_1125698 | 4 | 0.25 | 10 | 0 | 5 | 0.4 | 5 | 0 | 24 | 0.125 | 3 | 0 | 2 | 0.5 | 2 | 0 | 5 | 0 | 12 | 0.0833 |
| 5_1125764 | 5 | 0 | 7 | 0.1429 | 4 | 0 | 3 | 0.3333 | 19 | 0.1053 | 3 | 0.3333 | 2 | 0 | 2 | 0 | 5 | 0.2 | 12 | 0.1667 |
| 5_1126229 | 1 | 0 | 1 | 0 | 2 | 0.5 | 1 | 0 | 5 | 0.2 | NA | NA | NA | NA | 1 | 0 | 1 | 1 | 2 | 0.5 |
| 5_1126231 | 1 | 0 | 1 | 0 | 1 | 0 | 1 | 1 | 4 | 0.25 | NA | NA | NA | NA | 1 | 0 | 1 | 1 | 2 | 0.5 |
| 5_1127305 | NA | NA | 1 | 1 | 1 | 1 | 1 | 1 | 3 | 1 | NA | NA | 1 | 1 | NA | NA | 1 | 1 | 2 | 1 |
| 5_1127307 | NA | NA | 1 | 0 | 1 | 1 | 1 | 0 | 3 | 0.3333 | NA | NA | 1 | 0 | NA | NA | 1 | 1 | 2 | 0.5 |
| 5_1127310 | NA | NA | 1 | 0 | 1 | 1 | 1 | 0 | 3 | 0.3333 | 1 | 0 | NA | NA | 1 | 1 | 1 | 0 | 3 | 0.3333 |
| 5_1130585 | 7 | 0 | 10 | 0.5 | 9 | 0.1111 | 4 | 0.5 | 30 | 0.2667 | 7 | 0.2857 | 6 | 0 | 4 | 0.25 | 10 | 0.3 | 27 | 0.2222 |
| 5_1130588 | 6 | 0.1667 | 7 | 0.4286 | 6 | 0 | 4 | 0 | 23 | 0.1739 | 7 | 0.1429 | 5 | 0 | 4 | 0 | 9 | 0.3333 | 25 | 0.16 |
| 5_1130589 | 7 | 0.5714 | 9 | 0.6667 | 6 | 0.6667 | 3 | 1 | 25 | 0.68 | 7 | 0.5714 | 4 | 0.5 | 3 | 0.3333 | 10 | 0.7 | 24 | 0.5833 |
| 5_1130591 | 5 | 0.6 | 8 | 0.375 | 6 | 0.5 | 4 | 0.25 | 23 | 0.4348 | 7 | 0.4286 | 5 | 0.4 | 4 | 0.25 | 9 | 0.3333 | 25 | 0.36 |
| 5_1130592 | 5 | 0.6 | 5 | 0.6 | 4 | 0.5 | 3 | 1 | 17 | 0.6471 | 6 | 0.3333 | 3 | 0.6667 | 3 | 0.3333 | 7 | 0.2857 | 19 | 0.3684 |
| 5_1130593 | 5 | 0.2 | 7 | 0.1429 | 6 | 0.1667 | 4 | 0 | 22 | 0.1364 | 5 | 0.2 | 4 | 0.25 | 4 | 0 | 8 | 0.25 | 21 | 0.1905 |
| 5_1130594 | 5 | 0.2 | 6 | 0 | 6 | 0.3333 | 4 | 0 | 21 | 0.1429 | 5 | 0 | 2 | 0.5 | 4 | 0 | 6 | 0.3333 | 17 | 0.1765 |
| 5_1130595 | 4 | 0.5 | 8 | 0.875 | 6 | 0.3333 | 3 | 0.3333 | 21 | 0.5714 | 5 | 0.4 | 3 | 0.6667 | 4 | 0.75 | 5 | 0.6 | 17 | 0.5882 |
| 5_1130595 | 5 | 0.2 | 7 | 0.5714 | 5 | 0.6 | 4 | 0 | 21 | 0.381 | 5 | 0.4 | 3 | 0 | 4 | 0.25 | 5 | 0.2 | 17 | 0.2353 |
| 5_1130596 | 4 | 1 | 7 | 1 | 4 | 0.75 | 3 | 0.6667 | 18 | 0.8889 | 5 | 0.4 | 3 | 0.3333 | 3 | 0.3333 | 6 | 0.6667 | 17 | 0.4706 |
| 5_1130596 | 5 | 0.2 | 6 | 0.1667 | 6 | 0.1667 | 3 | 0.3333 | 20 | 0.2 | 6 | 0.1667 | 3 | 0 | 4 | 0 | 7 | 0.1429 | 20 | 0.1 |
| 5_1130597 | 4 | 0 | 5 | 0.2 | 6 | 0 | 4 | 0 | 19 | 0.0526 | 3 | 0.3333 | 3 | 0 | 3 | 0.3333 | 6 | 0.3333 | 15 | 0.2667 |
| 5_1130598 | 4 | 0 | 4 | 0.25 | 4 | 0 | 3 | 0.3333 | 15 | 0.1333 | 3 | 0 | 3 | 0 | 4 | 0 | 4 | 0.5 | 14 | 0.1429 |
| 5_1130602 | 4 | 0.25 | 4 | 0 | 4 | 0 | 2 | 0 | 14 | 0.0714 | 5 | 0 | 3 | 0 | 4 | 0 | 6 | 0.1667 | 18 | 0.0556 |
| 5_1130613 | 2 | 0.5 | 2 | 0 | 2 | 0 | 1 | 0 | 7 | 0.1429 | 3 | 0 | 1 | 0 | 3 | 0 | 4 | 0.25 | 11 | 0.0909 |
| 5_1130621 | 2 | 0.5 | 1 | 0 | 1 | 0 | NA | NA | 4 | 0.25 | 3 | 0 | 1 | 0 | 2 | 0.5 | 2 | 0 | 8 | 0.125 |
| 5_1130622 | 2 | 0.5 | 2 | 0 | 1 | 0 | 1 | 0 | 6 | 0.1667 | 3 | 0.6667 | 1 | 0 | 1 | 0 | 2 | 0 | 7 | 0.2857 |

| | | | | | | | | | | | | | | | | | | | | |
|---|---|---|---|---|---|---|---|---|---|---|---|---|---|---|---|---|---|---|---|---|
| 5_11306243 | 1 | 0 | 2 | 1 | 1 | 1 | 1 | 1 | 5 | 0.8 | 2 | 1 | 1 | 1 | 2 | 1 | 1 | 1 | 6 | 1 |
| 5_11306252 | 1 | 0 | 2 | 1 | 1 | 1 | 1 | 1 | 5 | 0.8 | 2 | 0.5 | NA | NA | 2 | 0.5 | 2 | 1 | 6 | 0.6667 |
| 5_11306263 | 1 | 1 | 2 | 0.5 | 1 | 1 | 1 | 1 | 5 | 0.8 | 1 | 1 | NA | NA | 2 | 1 | 2 | 0 | 5 | 0.6 |
| 5_11306269 | NA | NA | 2 | 0.5 | 1 | 1 | 1 | 0 | 4 | 0.5 | NA | NA | NA | NA | 2 | 0 | 2 | 0.5 | 4 | 0.25 |
| 5_11306287 | NA | NA | 2 | 0.5 | NA | NA | 1 | 1 | 3 | 0.6667 | 1 | 1 | NA | NA | 1 | 0 | 2 | 0.5 | 4 | 0.5 |
| 5_11323425 | 4 | 0.5 | 1 | 0 | 4 | 0.25 | 4 | 0.25 | 13 | 0.3077 | NA | NA | 3 | 0 | 1 | 0 | 3 | 0.3333 | 7 | 0.1429 |
| 5_11323430 | 4 | 0 | 2 | 0 | 4 | 0 | 4 | 0.25 | 14 | 0.0714 | NA | NA | 2 | 0 | 1 | 0 | 3 | 0.3333 | 6 | 0.1667 |
| 5_11323436 | 3 | 0.3333 | 2 | 0 | 4 | 0.25 | 4 | 0.5 | 13 | 0.3077 | NA | NA | 2 | 0 | 1 | 1 | 3 | 0.3333 | 6 | 0.3333 |
| 5_11323440 | 3 | 0.3333 | 1 | 0 | 4 | 0.5 | 3 | 0.3333 | 11 | 0.3636 | NA | NA | 2 | 0 | NA | NA | 2 | 0.5 | 4 | 0.25 |
| 5_11323443 | 2 | 1 | 2 | 0.5 | 4 | 0.5 | 4 | 1 | 12 | 0.75 | NA | NA | 2 | 1 | NA | NA | 2 | 0.5 | 4 | 0.75 |
| 5_11323444 | 2 | 0.5 | 1 | 0 | 5 | 0 | 4 | 0.25 | 12 | 0.1667 | NA | NA | 3 | 0.3333 | 1 | 0 | 1 | 0 | 5 | 0.2 |
| 5_11323444 | 2 | 0 | 2 | 0 | 5 | 0 | 4 | 0.25 | 13 | 0.0769 | NA | NA | 2 | 0 | 1 | 0 | 2 | 0.5 | 5 | 0.2 |
| 5_11323455 | 1 | 1 | 2 | 0.5 | 5 | 0.6 | 4 | 0.5 | 12 | 0.5833 | NA | NA | 3 | 0.6667 | 1 | 0 | 2 | 0 | 6 | 0.3333 |
| 5_11323481 | NA | NA | 2 | 0.5 | 5 | 0.2 | 3 | 0 | 10 | 0.2 | NA | NA | 2 | 1 | 1 | 1 | 2 | 0.5 | 5 | 0.8 |
| 5_11323493 | NA | NA | 2 | 0 | 4 | 0.25 | 3 | 0.3333 | 9 | 0.2222 | NA | NA | 2 | 0 | 1 | 1 | 2 | 0.5 | 5 | 0.4 |
| 5_11323615 | NA | NA | 2 | 0.5 | 2 | 0.5 | 1 | 0 | 5 | 0.4 | NA | NA | NA | NA | NA | NA | 2 | 0.5 | 2 | 0.5 |
| 5_11323717 | 1 | 0 | 1 | 0 | 2 | 0.5 | 1 | 0 | 5 | 0.2 | NA | NA | 1 | 0 | 1 | 0 | 2 | 0.5 | 4 | 0.25 |
| 5_11323767 | 1 | 1 | 2 | 1 | 2 | 1 | NA | NA | 5 | 1 | NA | NA | 2 | 1 | 1 | 1 | 2 | 1 | 5 | 1 |
| 5_11323794 | 2 | 0.5 | 1 | 1 | 3 | 1 | NA | NA | 6 | 0.8333 | NA | NA | 2 | 1 | 1 | 1 | 2 | 1 | 5 | 1 |
| 5_11323804 | 2 | 1 | 1 | 0 | 3 | 1 | NA | NA | 6 | 0.8333 | NA | NA | 2 | 1 | NA | NA | 1 | 0 | 3 | 0.6667 |
| 5_11323808 | 1 | 1 | 1 | 1 | 3 | 1 | NA | NA | 5 | 1 | NA | NA | 2 | 1 | NA | NA | 2 | 1 | 4 | 1 |
| 5_11323826 | 2 | 1 | 1 | 1 | 1 | 1 | NA | NA | 4 | 1 | NA | NA | 1 | 1 | NA | NA | 1 | 1 | 2 | 1 |
| 5_11323840 | 2 | 0.5 | 1 | 0 | 2 | 0.5 | NA | NA | 5 | 0.4 | NA | NA | 1 | 1 | 1 | 1 | 2 | 0.5 | 4 | 0.75 |
| 5_11323841 | 2 | 1 | 1 | 0 | 1 | 1 | NA | NA | 4 | 0.75 | NA | NA | 1 | 0 | NA | NA | 2 | 1 | 3 | 0.6667 |
| 5_11323856 | 2 | 1 | NA | NA | 2 | 1 | NA | NA | 4 | 1 | NA | NA | 2 | 1 | 1 | 1 | 1 | 1 | 4 | 1 |
| 5_11323865 | 2 | 0.5 | NA | NA | 2 | 0 | NA | NA | 4 | 0.25 | NA | NA | 2 | 1 | 1 | 0 | 1 | 0 | 4 | 0.5 |
| 5_11328580 | 1 | 1 | NA | NA | NA | NA | NA | NA | 1 | 1 | NA | NA | NA | NA | 1 | 1 | NA | NA | 1 | 1 |
| 5_11328591 | 1 | 0 | 1 | 1 | NA | NA | NA | NA | 2 | 0.5 | NA | NA | 1 | 1 | 1 | 0 | NA | NA | 2 | 0.5 |
| 5_11339248 | 2 | 0 | 6 | 0 | 1 | 1 | 4 | 0 | 13 | 0.0769 | 6 | 0 | 11 | 0.1818 | 6 | 0 | 6 | 0.1667 | 29 | 0.1034 |
| 5_11339273 | 1 | 0 | 6 | 0.1667 | 2 | 0 | 3 | 0 | 12 | 0.0833 | 6 | 0 | 12 | 0.1667 | 7 | 0 | 5 | 0 | 30 | 0.0667 |
| 5_11339279 | 1 | 1 | 7 | 0.7143 | 2 | 1 | 3 | 1 | 13 | 0.8462 | 2 | 1 | 11 | 0.8182 | 6 | 1 | 5 | 0.8 | 24 | 0.875 |
| 5_11339280 | 1 | 1 | 6 | 0.1667 | 2 | 0 | 2 | 0 | 11 | 0.1818 | 5 | 0.4 | 11 | 0.0909 | 7 | 0 | 4 | 0 | 27 | 0.1111 |
| 5_11341140 | 1 | 0 | 2 | 1 | 1 | 0 | 2 | 0 | 6 | 0.3333 | 4 | 0 | 5 | 0 | 3 | 0.3333 | 2 | 0.5 | 14 | 0.1429 |
| 5_11341158 | 1 | 0 | 3 | 0.6667 | 1 | 0 | 2 | 0 | 7 | 0.2857 | 4 | 0.25 | 5 | 0 | 5 | 0.2 | 1 | 0 | 15 | 0.1333 |
| 5_11368862 | 3 | 0.6667 | 2 | 1 | 2 | 1 | NA | NA | 7 | 0.8571 | 1 | 1 | 3 | 1 | 4 | 1 | 1 | 0 | 9 | 0.8889 |
| 5_11368865 | 3 | 0.6667 | 2 | 1 | 2 | 1 | 2 | 1 | 9 | 0.8889 | 1 | 1 | 3 | 1 | 4 | 1 | 3 | 1 | 11 | 1 |
| 5_11368871 | 4 | 0 | 2 | 0 | 2 | 0.5 | 2 | 0.5 | 10 | 0.2 | 2 | 0 | 3 | 0 | 4 | 0 | 4 | 0.5 | 13 | 0.1538 |
| 5_11368873 | 4 | 0.75 | 2 | 0.5 | 2 | 1 | 2 | 0.5 | 10 | 0.7 | 1 | 0 | 3 | 0.3333 | 5 | 0.4 | 4 | 0.5 | 13 | 0.3846 |
| 5_11368878 | 4 | 0.25 | 2 | 0 | 1 | 0 | 3 | 0 | 10 | 0.1 | 2 | 0 | 3 | 0.3333 | 5 | 0.2 | 4 | 0 | 14 | 0.1429 |
| 5_11368882 | 3 | 1 | 2 | 1 | 2 | 1 | 3 | 1 | 10 | 1 | 2 | 1 | 3 | 1 | 6 | 1 | 2 | 1 | 13 | 1 |
| 5_11368923 | 4 | 0.5 | 3 | 0.6667 | 2 | 1 | 2 | 1 | 11 | 0.7273 | 3 | 0.3333 | 3 | 0.6667 | 6 | 0.5 | 4 | 0.75 | 16 | 0.5625 |
| 5_11368927 | 4 | 0.5 | 3 | 0.3333 | 3 | 0.6667 | 3 | 0 | 13 | 0.3846 | 2 | 0 | 3 | 0.6667 | 5 | 0.6 | 2 | 0 | 12 | 0.4167 |
| 5_11368929 | 3 | 0 | 4 | 0 | 4 | 0 | 3 | 0.3333 | 14 | 0.0714 | 3 | 0 | 3 | 0 | 6 | 0.1667 | 4 | 0 | 16 | 0.0625 |
| 5_11372590 | 5 | 0.6 | 5 | 0.4 | 6 | 0.1667 | 2 | 0.5 | 18 | 0.3889 | 2 | 0.5 | 6 | 0.1667 | 8 | 0.375 | 2 | 0.5 | 18 | 0.3333 |
| 5_11372596 | 8 | 0.625 | 6 | 0.8333 | 7 | 0.8571 | 3 | 0.3333 | 24 | 0.7083 | 5 | 0.6 | 6 | 0.8333 | 9 | 0.8889 | 5 | 0.8 | 25 | 0.8 |
| 5_11372612 | 9 | 0.2222 | 7 | 0.2857 | 8 | 0.5 | 3 | 0 | 27 | 0.2963 | 6 | 0.1667 | 6 | 0.1667 | 9 | 0.4444 | 7 | 0.2857 | 28 | 0.2857 |
| 5_11372614 | 6 | 0.8333 | 6 | 0.5 | 8 | 0.25 | 2 | 1 | 22 | 0.5455 | 4 | 0.5 | 4 | 0.5 | 7 | 0.5714 | 7 | 0.5714 | 22 | 0.5455 |
| 5_11372622 | 8 | 0.25 | 9 | 0.3333 | 8 | 0.375 | 3 | 0.3333 | 28 | 0.3214 | 5 | 0.4 | 6 | 0.1667 | 8 | 0.5 | 6 | 0.3333 | 25 | 0.36 |
| 5_11372627 | 6 | 1 | 7 | 0.5714 | 6 | 0.8333 | 3 | 1 | 22 | 0.8182 | 4 | 1 | 4 | 0.75 | 5 | 1 | 3 | 1 | 16 | 0.9375 |
| 5_11372634 | 8 | 0 | 7 | 0.2857 | 7 | 0 | 4 | 0 | 26 | 0.0769 | 7 | 0 | 6 | 0 | 9 | 0.2222 | 5 | 0 | 27 | 0.0741 |
| 5_11372646 | 9 | 0.7778 | 8 | 1 | 9 | 1 | 4 | 1 | 30 | 0.9333 | 7 | 1 | 7 | 1 | 9 | 1 | 4 | 1 | 27 | 1 |
| 5_11372649 | 7 | 0.7143 | 8 | 0.75 | 8 | 0.75 | 1 | 1 | 24 | 0.75 | 7 | 0.7143 | 6 | 0.6667 | 10 | 0.5 | 5 | 1 | 28 | 0.6786 |
| 5_11372657 | 10 | 0.1 | 8 | 0.125 | 8 | 0 | 3 | 0 | 29 | 0.069 | 7 | 0.1429 | 6 | 0 | 10 | 0.1 | 4 | 0 | 27 | 0.0741 |
| 5_11372658 | 11 | 0 | 9 | 0 | 8 | 0.125 | 4 | 0.25 | 32 | 0.0625 | 6 | 0 | 8 | 0.125 | 10 | 0.2 | 6 | 0 | 30 | 0.1 |
| 5_11372662 | 9 | 0.1111 | 7 | 0.1429 | 7 | 0.4286 | 4 | 0.25 | 27 | 0.2222 | 7 | 0 | 7 | 0 | 9 | 0.1111 | 5 | 0.4 | 28 | 0.1071 |
| 5_11372663 | 10 | 0.2 | 9 | 0.1111 | 9 | 0.1111 | 4 | 0 | 32 | 0.125 | 7 | 0.4286 | 8 | 0 | 11 | 0.1818 | 6 | 0.3333 | 32 | 0.2188 |
| 5_11372730 | 9 | 0 | 8 | 0.25 | 11 | 0.1818 | 3 | 0 | 31 | 0.129 | 5 | 0.6 | 9 | 0 | 11 | 0 | 6 | 0.1667 | 31 | 0.129 |
| 5_11372746 | 8 | 0.625 | 10 | 0.3 | 13 | 0.3077 | 3 | 0 | 34 | 0.3529 | 7 | 0.7143 | 8 | 0.75 | 10 | 0.3 | 7 | 0.4286 | 32 | 0.5312 |
| 5_11372748 | 10 | 0.9 | 9 | 0.6667 | 15 | 0.6667 | 4 | 0.75 | 38 | 0.7368 | 8 | 0.5 | 12 | 0.9167 | 12 | 0.8333 | 7 | 0.8571 | 39 | 0.7949 |
| 5_11372760 | 10 | 0.6 | 8 | 0.875 | 15 | 0.3333 | 4 | 0.75 | 37 | 0.5676 | 6 | 0.8333 | 11 | 0.6364 | 10 | 0.7 | 10 | 0.8 | 37 | 0.7297 |
| 5_11372875 | NA | NA | NA | NA | 1 | 1 | NA | NA | 1 | 1 | NA | NA | 1 | 1 | NA | NA | 2 | 0 | 3 | 0.3333 |
| 5_11375138 | 1 | 1 | NA | NA | NA | NA | NA | NA | 1 | 1 | NA | NA | 1 | 1 | NA | NA | NA | NA | 1 | 1 |
| 5_11375140 | 1 | 1 | NA | NA | NA | NA | NA | NA | 1 | 1 | NA | NA | 1 | 1 | NA | NA | NA | NA | 1 | 1 |
| 5_11375144 | 1 | 1 | NA | NA | NA | NA | NA | NA | 1 | 1 | NA | NA | 1 | 1 | NA | NA | NA | NA | 1 | 1 |
| 5_11375165 | 1 | 1 | NA | NA | NA | NA | NA | NA | 1 | 1 | NA | NA | 1 | 1 | NA | NA | NA | NA | 1 | 1 |

| | | | | | | | | | | | | | | | | | | | | |
|---|---|---|---|---|---|---|---|---|---|---|---|---|---|---|---|---|---|---|---|---|
| 5_11375172 | 1 | 1 | NA | NA | NA | NA | NA | NA | 1 | 1 | NA | NA | 1 | 1 | NA | NA | NA | NA | 1 | 1 |
| 5_11375183 | 1 | 1 | NA | NA | NA | NA | NA | NA | 1 | 1 | NA | NA | 1 | 1 | NA | NA | NA | NA | 1 | 1 |
| 5_11387629 | 4 | 0.25 | 1 | 0 | 2 | 0 | 2 | 0.5 | 9 | 0.2222 | NA | NA | NA | NA | 1 | 0 | 1 | 1 | 2 | 0.5 |
| 5_11387637 | 4 | 0.75 | 1 | 1 | 2 | 1 | 2 | 0.5 | 9 | 0.7778 | NA | NA | NA | NA | 1 | 1 | 1 | 1 | 2 | 1 |
| 5_11387676 | 3 | 0 | 1 | 0 | 1 | 0 | 2 | 1 | 7 | 0.2857 | NA | NA | NA | NA | 1 | 1 | 1 | 0 | 2 | 0.5 |
| 5_11387690 | 4 | 1 | NA | NA | 2 | 1 | 2 | 1 | 8 | 1 | NA | NA | NA | NA | 1 | 1 | 1 | 1 | 2 | 1 |
| 5_11387695 | 3 | 1 | 1 | 1 | 2 | 0.5 | 2 | 0 | 8 | 0.625 | NA | NA | NA | NA | 1 | 1 | 1 | 1 | 2 | 1 |
| 5_11387710 | 3 | 1 | 1 | 1 | 1 | 1 | 2 | 0.5 | 7 | 0.8571 | NA | NA | NA | NA | 1 | 1 | 2 | 1 | 3 | 1 |
| 5_11387778 | 5 | 0.4 | 1 | 0 | 2 | 0 | 3 | 0 | 11 | 0.1818 | 1 | 0 | NA | NA | 1 | 0 | 2 | 0.5 | 4 | 0.25 |
| 5_11387806 | 4 | 0.75 | NA | NA | 1 | 0 | 3 | 1 | 8 | 0.75 | 2 | 1 | NA | NA | 1 | 1 | 2 | 0.5 | 5 | 0.8 |
| 5_11387819 | 5 | 0.6 | 1 | 0 | 2 | 0 | 3 | 0 | 11 | 0.2727 | 2 | 0 | NA | NA | 1 | 0 | 2 | 0.5 | 5 | 0.2 |
| 5_11387826 | 6 | 0.8333 | 1 | 1 | 2 | 0.5 | 4 | 0.5 | 13 | 0.6923 | 1 | 1 | NA | NA | NA | NA | 2 | 0.5 | 3 | 0.6667 |
| 5_11387830 | 5 | 1 | 1 | 1 | 1 | 0 | 4 | 0.5 | 11 | 0.7273 | 2 | 1 | NA | NA | 1 | 1 | 2 | 0 | 5 | 0.6 |
| 5_11387835 | 4 | 0.25 | 1 | 0 | 1 | 0 | 4 | 0 | 10 | 0.1 | 2 | 0 | NA | NA | 1 | 0 | 2 | 0.5 | 5 | 0.2 |
| 5_11387840 | 7 | 0.4286 | 2 | 0 | 1 | 1 | 4 | 0.25 | 14 | 0.3571 | 1 | 1 | NA | NA | 1 | 0 | 2 | 0.5 | 4 | 0.5 |
| 5_11387845 | 4 | 0.5 | 1 | 0 | NA | NA | 4 | 0 | 9 | 0.2222 | 2 | 0.5 | 1 | 0 | 1 | 0 | 2 | 0.5 | 6 | 0.3333 |
| 5_11387846 | 3 | 1 | NA | NA | 1 | 1 | 2 | 1 | 6 | 1 | 1 | 1 | NA | NA | NA | NA | 1 | 1 | 2 | 1 |
| 5_11388070 | 9 | 0.2222 | 4 | 0.25 | 2 | 0 | 4 | 0.25 | 19 | 0.2105 | 1 | 0 | 5 | 0.2 | 1 | 0 | 5 | 0.4 | 12 | 0.25 |
| 5_11388095 | 7 | 0.1429 | 4 | 0 | 1 | 0 | 5 | 0 | 17 | 0.0588 | NA | NA | 8 | 0.125 | 1 | 0 | 4 | 0.5 | 13 | 0.2308 |
| 5_11388104 | 3 | 0.3333 | 4 | 0 | 1 | 0 | 3 | 0.3333 | 11 | 0.1818 | 1 | 0 | 5 | 0.8 | 2 | 0 | 4 | 0.25 | 12 | 0.4167 |
| 5_11388118 | 6 | 0.8333 | 4 | 1 | 2 | 1 | 4 | 0.75 | 16 | 0.875 | 1 | 1 | 6 | 1 | 2 | 1 | 4 | 1 | 13 | 1 |
| 5_11388371 | 3 | 0 | 4 | 0.25 | 1 | 0 | 4 | 0 | 12 | 0.0833 | 1 | 0 | 3 | 0.3333 | 2 | 0 | 1 | 0 | 7 | 0.1429 |
| 5_11388378 | 2 | 0.5 | 3 | 0.6667 | 1 | 0 | 4 | 0.75 | 10 | 0.6 | 1 | 1 | 3 | 1 | 2 | 0 | 2 | 0.5 | 8 | 0.625 |
| 5_11388390 | 3 | 1 | 2 | 0.5 | 1 | 1 | 5 | 1 | 11 | 0.9091 | NA | NA | 3 | 1 | 2 | 1 | 1 | 1 | 6 | 1 |
| 5_11388393 | 2 | 1 | 3 | 0.3333 | 1 | 0 | 3 | 0 | 9 | 0.3333 | NA | NA | 1 | 0 | 2 | 0.5 | 3 | 0 | 6 | 0.1667 |
| 5_11391616 | 25 | 0.64 | 22 | 0.7273 | 16 | 0.5625 | 25 | 0.52 | 88 | 0.6136 | 21 | 0.7619 | 18 | 0.5 | 23 | 0.6957 | 14 | 0.6429 | 76 | 0.6579 |
| 5_11391621 | 26 | 0.6538 | 23 | 0.4783 | 16 | 0.8125 | 27 | 0.7778 | 92 | 0.6739 | 21 | 0.619 | 19 | 0.5789 | 25 | 0.64 | 13 | 0.7692 | 78 | 0.641 |
| 5_11393188 | 7 | 0.5714 | 8 | 0.625 | 3 | 0.6667 | 2 | 0.5 | 20 | 0.6 | 3 | 0.3333 | 2 | 0 | 5 | 0.6 | 2 | 0.5 | 12 | 0.4167 |
| 5_11393206 | 6 | 0.6667 | 7 | 0.4286 | 3 | 1 | 2 | 0.5 | 18 | 0.6111 | 4 | 0.25 | 3 | 0.3333 | 5 | 0.8 | 1 | 1 | 13 | 0.5385 |
| 5_11393237 | 7 | 0.4286 | 4 | 0.25 | 1 | 0 | 2 | 1 | 14 | 0.4286 | 2 | 0 | NA | NA | 3 | 0.3333 | 1 | 0 | 6 | 0.1667 |
| 5_11393243 | 6 | 0 | 6 | 0.3333 | 3 | 0 | 2 | 0 | 17 | 0.1176 | 3 | 0 | 2 | 0 | 4 | 0.25 | 3 | 0.3333 | 12 | 0.1667 |
| 5_11393245 | 7 | 1 | 8 | 1 | 3 | 1 | 2 | 1 | 20 | 1 | 4 | 1 | 1 | 1 | 4 | 1 | 2 | 1 | 11 | 1 |
| 5_11393256 | 7 | 0.5714 | 7 | 0.2857 | 1 | 0 | 1 | 1 | 16 | 0.4375 | 3 | 0.6667 | 1 | 1 | 4 | 0.5 | 2 | 0 | 10 | 0.5 |
| 5_11393261 | 8 | 0.125 | 7 | 0 | 2 | 0 | 1 | 0 | 18 | 0.0556 | 4 | 0.5 | 2 | 0 | 4 | 0 | 2 | 0 | 12 | 0.1667 |
| 5_11393264 | 7 | 0.1429 | 8 | 0.375 | 2 | 0 | 2 | 0 | 19 | 0.2105 | 3 | 0.3333 | 1 | 0 | 3 | 0 | 2 | 0 | 9 | 0.1111 |
| 5_11393275 | 8 | 1 | 7 | 1 | NA | NA | 1 | 1 | 16 | 1 | 2 | 1 | 1 | 1 | 5 | 1 | 1 | 1 | 9 | 1 |
| 5_11393282 | 5 | 1 | 8 | 0.875 | 1 | 1 | 1 | 1 | 15 | 0.9333 | 4 | 0.75 | 2 | 0.5 | 4 | 1 | 2 | 1 | 12 | 0.8333 |
| 5_11393293 | 8 | 0 | 7 | 0 | 1 | 1 | 1 | 0 | 17 | 0.0588 | 2 | 0 | 2 | 0 | 5 | 0 | 2 | 0.5 | 11 | 0.0909 |
| 5_11393294 | 4 | 1 | 4 | 0.75 | NA | NA | 1 | 1 | 9 | 0.8889 | 1 | 1 | 2 | 0 | 5 | 0.8 | 1 | 1 | 9 | 0.6667 |
| 5_11393327 | 5 | 0 | 5 | 0.2 | 1 | 0 | NA | NA | 11 | 0.0909 | 3 | 0 | 2 | 0.5 | 4 | 0.25 | 2 | 0 | 11 | 0.1818 |
| 5_11393331 | 5 | 0.6 | 5 | 0.4 | 1 | 0 | NA | NA | 11 | 0.4545 | 3 | 0 | 1 | 0 | 4 | 0.5 | 2 | 0 | 10 | 0.2 |
| 5_11393339 | 5 | 0.4 | 5 | 0.6 | 1 | 0 | NA | NA | 11 | 0.4545 | 2 | 1 | 1 | 1 | 3 | 0.6667 | 1 | 0 | 7 | 0.7143 |
| 5_11395687 | 5 | 0.8 | 9 | 0.7778 | 5 | 1 | 2 | 1 | 21 | 0.8571 | 6 | 1 | 11 | 0.9091 | 5 | 0.8 | 4 | 1 | 26 | 0.9231 |
| 5_11395705 | 5 | 0 | 5 | 0 | 4 | 0.25 | 2 | 0 | 16 | 0.0625 | 7 | 0.2857 | 12 | 0 | 6 | 0 | 3 | 0 | 28 | 0.0714 |
| 5_11395709 | 3 | 0 | 4 | 0.5 | 4 | 0.25 | 2 | 0.5 | 13 | 0.3077 | 7 | 0.4286 | 9 | 0 | 5 | 0.2 | 3 | 0 | 24 | 0.1667 |
| 5_11395711 | 4 | 0.25 | 6 | 0.1667 | 5 | 0.2 | 2 | 0.5 | 17 | 0.2353 | 5 | 0 | 11 | 0.0909 | 6 | 0.3333 | 4 | 0.25 | 26 | 0.1538 |
| 5_11395711 | 6 | 0.3333 | 9 | 0.2222 | 4 | 0 | 2 | 0.5 | 21 | 0.2381 | 7 | 0.1429 | 13 | 0.2308 | 6 | 0.5 | 4 | 0 | 30 | 0.2333 |
| 5_11395721 | 6 | 0.8333 | 3 | 0.3333 | 4 | 0.75 | 2 | 0 | 15 | 0.6 | 4 | 0 | 9 | 0.5556 | 4 | 0.75 | 2 | 1 | 19 | 0.5263 |
| 5_11395723 | 5 | 0 | 9 | 0.3333 | 5 | 0.2 | 2 | 0.5 | 21 | 0.2381 | 7 | 0.1429 | 12 | 0.1667 | 6 | 0.1667 | 4 | 0.25 | 29 | 0.1724 |
| 5_11395743 | 3 | 0.6667 | 7 | 0.5714 | 3 | 0.6667 | 1 | 1 | 14 | 0.6429 | 6 | 0.3333 | 10 | 0.5 | 7 | 0.5714 | 4 | 0.5 | 27 | 0.4815 |
| 5_11395754 | 4 | 1 | 9 | 0.8889 | 6 | 1 | 1 | 1 | 20 | 0.95 | 3 | 1 | 10 | 1 | 6 | 1 | 4 | 1 | 23 | 1 |
| 5_11395763 | 3 | 0 | 10 | 0 | 6 | 0.3333 | 1 | 0 | 20 | 0.1 | 6 | 0.1667 | 12 | 0.25 | 6 | 0.1667 | 4 | 0 | 28 | 0.1786 |
| 5_11395766 | 3 | 0.3333 | 10 | 0 | 5 | 0.2 | 2 | 0 | 20 | 0.1 | 6 | 0 | 12 | 0.1667 | 7 | 0 | 5 | 0 | 30 | 0.0667 |
| 5_11395778 | 3 | 0 | 9 | 0 | 6 | 0.1667 | 2 | 0 | 20 | 0.05 | 6 | 0 | 11 | 0.0909 | 6 | 0.1667 | 4 | 0 | 27 | 0.0741 |
| 5_11395787 | 2 | 0 | 10 | 0.1 | 6 | 0 | 1 | 0 | 19 | 0.0526 | 6 | 0 | 11 | 0.1818 | 6 | 0 | 4 | 0 | 27 | 0.0741 |
| 5_11395790 | 2 | 0.5 | 6 | 0.1667 | 6 | 0.1667 | 1 | 0 | 15 | 0.2 | 6 | 0.3333 | 9 | 0.1111 | 5 | 0.2 | 4 | 0.25 | 24 | 0.2083 |
| 5_11395794 | 3 | 0.6667 | 8 | 0.75 | 5 | 0.4 | 2 | 0.5 | 18 | 0.6111 | 6 | 0.3333 | 10 | 0.6 | 6 | 0.5 | 3 | 0 | 25 | 0.44 |
| 5_11395796 | 2 | 1 | 7 | 0.7143 | 5 | 0.6 | 2 | 0.5 | 16 | 0.6875 | 5 | 0.8 | 8 | 0.75 | 4 | 1 | 3 | 0.3333 | 20 | 0.75 |
| 5_11395812 | 3 | 0 | 10 | 0 | 6 | 0.1667 | 2 | 0.5 | 21 | 0.0952 | 5 | 0.2 | 9 | 0.1111 | 6 | 0 | 3 | 0 | 23 | 0.087 |
| 5_11395814 | 1 | 0 | 8 | 0.375 | 5 | 0.4 | 2 | 0.5 | 16 | 0.375 | 4 | 0 | 9 | 0.1111 | 6 | 0.1667 | 3 | 0 | 22 | 0.0909 |
| 5_11395815 | 3 | 0.6667 | 9 | 0.6667 | 4 | 1 | 2 | 1 | 18 | 0.7778 | 4 | 1 | 10 | 0.9 | 6 | 0.5 | 3 | 1 | 23 | 0.8261 |
| 5_11398058 | 10 | 0.1 | 12 | 0 | 10 | 0.3 | 13 | 0.0769 | 45 | 0.1111 | 14 | 0 | 4 | 0 | 7 | 0.1429 | 16 | 0.1875 | 41 | 0.0976 |
| 5_11398065 | 11 | 0.0909 | 15 | 0.1333 | 10 | 0.2 | 14 | 0.2857 | 50 | 0.18 | 11 | 0.2727 | 7 | 0.1429 | 9 | 0.1111 | 15 | 0.2667 | 42 | 0.2143 |
| 5_11398076 | 13 | 0.5385 | 15 | 0.4667 | 10 | 0.6 | 15 | 0.8 | 53 | 0.6038 | 14 | 0.7857 | 7 | 0.5714 | 9 | 0.8889 | 18 | 0.6111 | 48 | 0.7083 |
| 5_11398095 | 9 | 0.4444 | 12 | 0.4167 | 10 | 0.3 | 13 | 0.6923 | 44 | 0.4773 | 11 | 0.2727 | 8 | 0.625 | 7 | 0.8571 | 14 | 0.3571 | 40 | 0.475 |

| | | | | | | | | | | | | | | | | | | | | |
|---|---|---|---|---|---|---|---|---|---|---|---|---|---|---|---|---|---|---|---|---|
| 5_11398174 | 10 | 0 | 13 | 0 | 8 | 0.375 | 9 | 0.1111 | 40 | 0.1 | 10 | 0 | 4 | 0.75 | 8 | 0 | 11 | 0 | 33 | 0.0909 |
| 5_11398176 | 12 | 0.1667 | 13 | 0.3077 | 9 | 0.2222 | 13 | 0.1538 | 47 | 0.2128 | 12 | 0.0833 | 5 | 0.6 | 8 | 0.25 | 14 | 0.0714 | 39 | 0.1795 |
| 5_11398184 | 6 | 0.5 | 6 | 0.1667 | 6 | 0.3333 | 5 | 0.4 | 23 | 0.3478 | 10 | 0.3 | 4 | 0.25 | 5 | 0.6 | 9 | 0.6667 | 28 | 0.4643 |
| 5_11398185 | 10 | 0.1 | 10 | 0.3 | 7 | 0.1429 | 12 | 0.0833 | 39 | 0.1538 | 12 | 0.0833 | 3 | 0 | 8 | 0.125 | 12 | 0.1667 | 35 | 0.1143 |
| 5_11398189 | 8 | 0.125 | 11 | 0.0909 | 8 | 0 | 7 | 0 | 34 | 0.0588 | 11 | 0.0909 | 3 | 0 | 5 | 0 | 11 | 0.0909 | 30 | 0.0667 |
| 5_11398191 | 8 | 0.125 | 11 | 0.1818 | 8 | 0.25 | 10 | 0.2 | 37 | 0.1892 | 12 | 0.0833 | 4 | 0.5 | 7 | 0.2857 | 11 | 0.3636 | 34 | 0.2647 |
| 5_11398214 | 7 | 0.5714 | 11 | 0.7273 | 5 | 0.8 | 6 | 1 | 29 | 0.7586 | 12 | 0.5 | 2 | 0.5 | 6 | 0.8333 | 11 | 0.6364 | 31 | 0.6129 |
| 5_11398219 | 7 | 0 | 10 | 0.1 | 6 | 0 | 9 | 0.1111 | 32 | 0.0625 | 12 | 0.1667 | 3 | 0.3333 | 7 | 0 | 12 | 0.1667 | 34 | 0.1471 |
| 5_11398227 | 6 | 0.8333 | 9 | 1 | 6 | 1 | 7 | 0.8571 | 28 | 0.9286 | 9 | 1 | 3 | 1 | 5 | 1 | 11 | 1 | 28 | 1 |
| 5_11398232 | 5 | 0.6 | 9 | 0.3333 | 5 | 0.4 | 8 | 0.5 | 27 | 0.4444 | 12 | 0.3333 | 3 | 0.6667 | 5 | 1 | 10 | 0.4 | 30 | 0.5 |
| 5_11398234 | 3 | 0 | 9 | 0 | 4 | 0.25 | 8 | 0.125 | 24 | 0.0833 | 10 | 0.2 | 3 | 0.3333 | 6 | 0.3333 | 7 | 0 | 26 | 0.1923 |
| 5_11406790 | NA | NA | NA | NA | NA | NA | 1 | 1 | 1 | 1 | 1 | 1 | NA | NA | NA | NA | NA | NA | 1 | 1 |
| 5_11406809 | NA | NA | NA | NA | NA | NA | 1 | 1 | 1 | 1 | 1 | 1 | NA | NA | NA | NA | NA | NA | 1 | 1 |
| 5_11407541 | NA | NA | 1 | 1 | 1 | 0 | 2 | 0.5 | 4 | 0.5 | NA | NA | NA | NA | 1 | 0 | 3 | 0.3333 | 4 | 0.25 |
| 5_11407762 | NA | NA | 2 | 0.5 | NA | NA | 1 | 0 | 3 | 0.3333 | NA | NA | NA | NA | 2 | 0.5 | 2 | 0 | 4 | 0.25 |
| 5_11407832 | NA | NA | 2 | 0.5 | NA | NA | 2 | 0 | 4 | 0.25 | NA | NA | NA | NA | 1 | 1 | 2 | 0 | 3 | 0.3333 |
| 5_11407850 | NA | NA | 1 | 0 | NA | NA | 1 | 1 | 2 | 0.5 | NA | NA | NA | NA | 1 | 1 | NA | NA | 1 | 1 |
| 5_11407863 | NA | NA | 2 | 1 | NA | NA | 2 | 1 | 4 | 1 | NA | NA | NA | NA | NA | NA | 2 | 0.5 | 2 | 0.5 |
| 5_11407879 | NA | NA | 1 | 1 | NA | NA | 2 | 0.5 | 3 | 0.6667 | NA | NA | 1 | 1 | 1 | 1 | 2 | 1 | 4 | 1 |
| 5_11407882 | NA | NA | 1 | 1 | NA | NA | 1 | 1 | 2 | 1 | NA | NA | 1 | 1 | 1 | 1 | 1 | 1 | 3 | 1 |
| 5_11407915 | NA | NA | NA | NA | NA | NA | 2 | 0.5 | 2 | 0.5 | NA | NA | 1 | 0 | 1 | 1 | 2 | 0.5 | 4 | 0.5 |
| 5_11407959 | 1 | 1 | 1 | 1 | NA | NA | 1 | 1 | 3 | 1 | NA | NA | 1 | 1 | 1 | 1 | 1 | 0 | 3 | 0.6667 |
| 5_11407972 | 1 | 1 | 1 | 1 | NA | NA | 1 | 1 | 3 | 1 | NA | NA | 1 | 1 | 1 | 1 | NA | NA | 2 | 1 |
| 5_11407974 | 2 | 0 | 2 | 0.5 | NA | NA | NA | NA | 4 | 0.25 | NA | NA | 1 | 1 | 1 | 0 | NA | NA | 2 | 0.5 |
| 5_11407990 | 1 | 1 | 1 | 1 | NA | NA | 1 | 1 | 3 | 1 | NA | NA | 1 | 1 | 2 | 1 | NA | NA | 3 | 1 |
| 5_11408015 | 2 | 1 | 2 | 1 | NA | NA | NA | NA | 4 | 1 | 1 | 1 | NA | NA | 2 | 0.5 | NA | NA | 3 | 0.6667 |
| 5_11408017 | 2 | 0 | 2 | 0.5 | NA | NA | NA | NA | 4 | 0.25 | 1 | 0 | NA | NA | 2 | 0.5 | NA | NA | 3 | 0.3333 |
| 5_11408023 | 2 | 1 | 1 | 1 | NA | NA | NA | NA | 3 | 1 | NA | NA | NA | NA | 2 | 1 | NA | NA | 2 | 1 |
| 5_11408030 | 2 | 0 | 2 | 0.5 | NA | NA | NA | NA | 4 | 0.25 | NA | NA | NA | NA | 2 | 0.5 | NA | NA | 2 | 0.5 |
| 5_11408033 | 1 | 1 | NA | NA | NA | NA | NA | NA | 1 | 1 | 1 | 0 | NA | NA | 2 | 1 | NA | NA | 3 | 0.6667 |
| 5_11408037 | 2 | 0.5 | 2 | 0.5 | NA | NA | NA | NA | 4 | 0.5 | 1 | 1 | NA | NA | 2 | 0.5 | NA | NA | 3 | 0.6667 |
| 5_11408071 | 2 | 0.5 | 2 | 1 | NA | NA | NA | NA | 4 | 0.75 | 1 | 1 | NA | NA | 2 | 1 | NA | NA | 3 | 1 |
| 5_11408448 | 1 | 0 | 2 | 1 | NA | NA | NA | NA | 3 | 0.6667 | 1 | 1 | NA | NA | 2 | 0.5 | NA | NA | 3 | 0.6667 |
| 5_11408468 | 1 | 0 | 2 | 0.5 | NA | NA | NA | NA | 3 | 0.3333 | 1 | 1 | NA | NA | 1 | 0 | NA | NA | 2 | 0.5 |
| 5_11408472 | 1 | 0 | 2 | 0.5 | NA | NA | NA | NA | 3 | 0.3333 | 1 | 1 | NA | NA | 2 | 0 | NA | NA | 3 | 0.3333 |
| 5_11416502 | 10 | 0.6 | 18 | 0.5556 | 18 | 0.6667 | 23 | 0.7826 | 69 | 0.6667 | 12 | 0.75 | 21 | 0.7143 | 22 | 0.5909 | 20 | 0.65 | 75 | 0.6667 |
| 5_11416509 | 19 | 0.6316 | 23 | 0.5652 | 23 | 0.5217 | 27 | 0.5926 | 92 | 0.5761 | 12 | 0.5 | 25 | 0.44 | 27 | 0.7037 | 23 | 0.6957 | 87 | 0.5977 |
| 5_11416620 | 18 | 0.3889 | 10 | 0.2 | 15 | 0.2667 | 25 | 0.36 | 68 | 0.3235 | 8 | 0.25 | 15 | 0.3333 | 15 | 0.1333 | 18 | 0.2778 | 56 | 0.25 |
| 5_11416623 | 11 | 0.1818 | 9 | 0.4444 | 9 | 0.3333 | 15 | 0.6 | 44 | 0.4091 | 8 | 0.375 | 15 | 0.6 | 14 | 0.2143 | 12 | 0.5 | 49 | 0.4286 |
| 5_11416633 | 17 | 0.9412 | 10 | 0.7 | 15 | 0.8667 | 23 | 0.7826 | 65 | 0.8308 | 8 | 0.875 | 15 | 0.9333 | 14 | 0.9286 | 13 | 1 | 50 | 0.94 |
| 5_11416636 | 17 | 0.2941 | 10 | 0.2 | 14 | 0.2143 | 25 | 0.4 | 66 | 0.303 | 9 | 0.1111 | 14 | 0.1429 | 15 | 0.3333 | 17 | 0.4118 | 55 | 0.2727 |
| 5_11425964 | 5 | 0.6 | 3 | 0.3333 | 4 | 0.75 | 1 | 1 | 13 | 0.6154 | 3 | 1 | 5 | 0.4 | 3 | 0.3333 | 4 | 0.25 | 15 | 0.4667 |
| 5_11426002 | 3 | 0.3333 | 3 | 0.3333 | 4 | 0.25 | 2 | 0 | 12 | 0.25 | 4 | 0 | 7 | 0 | 4 | 0 | 4 | 0.25 | 19 | 0.0526 |
| 5_11426041 | 4 | 0.25 | 5 | 0.4 | 5 | 0.8 | 3 | 0.3333 | 17 | 0.4706 | 4 | 0.25 | 4 | 0.5 | 4 | 0.25 | 5 | 0.2 | 17 | 0.2941 |
| 5_11426055 | 5 | 0 | 6 | 0.1667 | 5 | 0 | 2 | 0 | 18 | 0.0556 | 4 | 0 | 6 | 0.1667 | 4 | 0 | 5 | 0 | 19 | 0.0526 |
| 5_11426066 | 2 | 0 | 2 | 0 | 4 | 0.5 | 2 | 1 | 10 | 0.4 | 4 | 0 | 6 | 0.1667 | 4 | 0.25 | 6 | 0 | 20 | 0.1 |
| 5_11426073 | 5 | 0.6 | 4 | 0.25 | 4 | 0.5 | 4 | 1 | 17 | 0.5882 | 5 | 0.6 | 7 | 0.2857 | 4 | 0 | 5 | 0.4 | 21 | 0.3333 |
| 5_11426084 | 4 | 0.25 | 5 | 0.6 | 5 | 0.4 | 4 | 0.75 | 18 | 0.5 | 5 | 0.4 | 7 | 0.8571 | 4 | 0.75 | 4 | 0.25 | 20 | 0.6 |
| 5_11426090 | 2 | 1 | 3 | 0.3333 | 3 | 0.6667 | 3 | 1 | 11 | 0.7273 | 2 | 1 | 5 | 0.8 | 2 | 1 | 3 | 1 | 12 | 0.9167 |
| 5_11426099 | 2 | 0 | 2 | 0.5 | 3 | 0.6667 | 4 | 0.5 | 11 | 0.4545 | 1 | 0 | 2 | 0 | 2 | 0.5 | 2 | 0.5 | 7 | 0.2857 |
| 5_11426101 | 3 | 0 | 3 | 0.3333 | 3 | 0 | 4 | 0 | 13 | 0.0769 | 3 | 0.3333 | 5 | 0 | 2 | 0 | 4 | 0 | 14 | 0.0714 |
| 5_11426114 | 4 | 1 | 4 | 0.75 | 3 | 1 | 4 | 0.75 | 15 | 0.8667 | 3 | 0.6667 | 4 | 1 | 1 | 1 | 4 | 0.5 | 12 | 0.75 |
| 5_11426115 | 3 | 0.6667 | 2 | 0.5 | 3 | 0.6667 | 3 | 1 | 11 | 0.7273 | 3 | 0.6667 | 3 | 0.6667 | 2 | 1 | 4 | 0.75 | 12 | 0.75 |
| 5_11426115 | 4 | 0 | 4 | 0.25 | 3 | 0.3333 | 4 | 0 | 15 | 0.1333 | 2 | 0 | 5 | 0.2 | 1 | 1 | 4 | 0 | 12 | 0.1667 |
| 5_11426117 | 4 | 0.5 | 4 | 0.75 | 3 | 0.3333 | 4 | 0.5 | 15 | 0.5333 | 3 | 0.6667 | 4 | 1 | NA | NA | 3 | 0.6667 | 10 | 0.8 |
| 5_11426122 | 3 | 0 | 3 | 0 | 3 | 0.3333 | 4 | 0.25 | 13 | 0.1538 | 2 | 0.5 | 5 | 0 | NA | NA | 4 | 0.25 | 11 | 0.1818 |
| 5_11426131 | 4 | 0.25 | 3 | 0.6667 | 3 | 1 | 4 | 1 | 14 | 0.7143 | 3 | 0.6667 | 4 | 0.5 | 2 | 1 | 4 | 0.75 | 13 | 0.6923 |
| 5_11426150 | 3 | 0.3333 | 3 | 0.3333 | 2 | 0.5 | 3 | 0.3333 | 11 | 0.3636 | 3 | 0.6667 | 5 | 0.2 | 3 | 0.3333 | 2 | 0 | 13 | 0.3077 |
| 5_11427658 | NA | NA | 1 | 1 | NA | NA | NA | NA | 1 | 1 | 1 | 1 | 1 | 1 | NA | NA | NA | NA | 2 | 1 |
| 5_11427663 | NA | NA | 1 | 1 | NA | NA | NA | NA | 1 | 1 | 1 | 1 | 1 | 1 | 1 | 1 | NA | NA | 3 | 1 |
| 5_11438787 | 1 | 1 | NA | NA | NA | NA | 1 | 1 | 2 | 1 | NA | NA | NA | NA | 1 | 1 | NA | NA | 1 | 1 |
| 5_11438826 | 1 | 1 | NA | NA | NA | NA | 1 | 0 | 2 | 0.5 | NA | NA | NA | NA | 1 | 1 | NA | NA | 1 | 1 |
| 5_11438835 | 1 | 0 | NA | NA | NA | NA | 1 | 1 | 2 | 0.5 | NA | NA | NA | NA | 1 | 1 | NA | NA | 1 | 1 |
| 5_11438860 | 1 | 1 | NA | NA | NA | NA | 1 | 1 | 2 | 1 | NA | NA | NA | NA | 1 | 1 | NA | NA | 1 | 1 |
| 5_11450134 | NA | NA | NA | NA | NA | NA | 1 | 1 | 1 | 1 | 1 | 1 | NA | NA | NA | NA | NA | NA | 1 | 1 |

| | | | | | | | | | | | | | | | | | | | | |
|---|---|---|---|---|---|---|---|---|---|---|---|---|---|---|---|---|---|---|---|---|
| 5_1145018 | 10 | 0.1 | 3 | 0.6667 | 2 | 0 | 6 | 0.1667 | 21 | 0.1905 | 13 | 0.2308 | 8 | 0.625 | 12 | 0.5 | 6 | 0.3333 | 39 | 0.4103 |
| 5_1145036 | 3 | 0 | 2 | 0.5 | 1 | 0 | 3 | 0.3333 | 9 | 0.2222 | 8 | 0.125 | 6 | 0.1667 | 9 | 0.1111 | 3 | 0 | 26 | 0.1154 |
| 5_1145038 | 4 | 0 | 2 | 0 | 1 | 0 | 3 | 0.3333 | 10 | 0.1 | 8 | 0 | 8 | 0.375 | 9 | 0.2222 | 3 | 0 | 28 | 0.1786 |
| 5_1145577 | 7 | 0 | 2 | 0.5 | 2 | 0 | 9 | 0.2222 | 20 | 0.15 | 5 | 0 | 4 | 0 | 7 | 0.1429 | 5 | 0.2 | 21 | 0.0952 |
| 5_1145578 | 10 | 0.2 | 9 | 0 | 4 | 0 | 11 | 0 | 34 | 0.0588 | 8 | 0 | 5 | 0 | 10 | 0 | 8 | 0.25 | 31 | 0.0645 |
| 5_1145578 | 11 | 0.5455 | 8 | 0.625 | 7 | 0.7143 | 10 | 0.6 | 36 | 0.6111 | 9 | 0.2222 | 9 | 0.4444 | 12 | 0.5 | 8 | 0.375 | 38 | 0.3947 |
| 5_1145579 | 11 | 0.0909 | 8 | 0.125 | 8 | 0.125 | 10 | 0.1 | 37 | 0.1081 | 7 | 0 | 9 | 0 | 10 | 0.1 | 7 | 0.1429 | 33 | 0.0606 |
| 5_1145584 | 12 | 0.0833 | 10 | 0.1 | 11 | 0.1818 | 11 | 0.2727 | 44 | 0.1591 | 10 | 0.2 | 9 | 0 | 13 | 0.0769 | 9 | 0.1111 | 41 | 0.0976 |
| 5_1145585 | 11 | 0.2727 | 9 | 0.1111 | 13 | 0.2308 | 12 | 0.0833 | 45 | 0.1778 | 9 | 0 | 11 | 0.0909 | 12 | 0 | 9 | 0.3333 | 41 | 0.0976 |
| 5_1145585 | 13 | 0.1538 | 11 | 0.2727 | 11 | 0.2727 | 10 | 0.1 | 45 | 0.2 | 7 | 0.5714 | 10 | 0.3 | 8 | 0.25 | 9 | 0.1111 | 34 | 0.2941 |
| 5_1145591 | 12 | 0.4167 | 6 | 0 | 11 | 0.4545 | 13 | 0.3077 | 42 | 0.3333 | 8 | 0.625 | 10 | 0.2 | 10 | 0.2 | 7 | 0.2857 | 35 | 0.3143 |
| 5_1145695 | 2 | 0 | 1 | 1 | 1 | 0 | 1 | 0 | 5 | 0.2 | 1 | 0 | 1 | 0 | 2 | 0 | 2 | 0.5 | 6 | 0.1667 |
| 5_1145697 | 1 | 0 | 2 | 1 | 2 | 0.5 | NA | NA | 5 | 0.6 | NA | NA | 2 | 0.5 | 4 | 0.25 | 2 | 0.5 | 8 | 0.375 |
| 5_1145698 | 2 | 0.5 | 2 | 0 | 2 | 0 | 1 | 1 | 7 | 0.2857 | 1 | 0 | 2 | 0.5 | 4 | 0.25 | 2 | 0 | 9 | 0.2222 |
| 5_1145700 | 2 | 0 | 1 | 1 | 2 | 0 | 1 | 0 | 6 | 0.1667 | NA | NA | 1 | 0 | 4 | 0.25 | 2 | 1 | 7 | 0.4286 |
| 5_1145704 | 1 | 1 | 2 | 0 | 1 | 0 | 1 | 0 | 5 | 0.2 | 1 | 0 | 1 | 0 | 4 | 0 | 2 | 0.5 | 8 | 0.125 |
| 5_1145708 | 2 | 0.5 | 1 | 1 | 1 | 1 | 1 | 1 | 5 | 0.8 | 1 | 0 | 1 | 0 | 3 | 0.6667 | 1 | 0 | 6 | 0.3333 |
| 5_1145709 | 2 | 0.5 | 1 | 1 | NA | NA | 1 | 0 | 4 | 0.5 | 1 | 1 | 1 | 0 | 4 | 0.75 | 1 | 1 | 7 | 0.7143 |
| 5_1145710 | 2 | 0.5 | 1 | 1 | 2 | 0.5 | 1 | 0 | 6 | 0.5 | NA | NA | 1 | 0 | 2 | 0 | 1 | 1 | 4 | 0.25 |
| 5_1145710 | 2 | 0.5 | 1 | 1 | 2 | 0 | 1 | 0 | 6 | 0.3333 | 1 | 1 | 1 | 0 | 4 | 0.25 | 1 | 1 | 7 | 0.4286 |
| 5_1145711 | 2 | 0.5 | 1 | 0 | 2 | 0 | 1 | 0 | 6 | 0.1667 | 1 | 0 | 1 | 0 | 2 | 0 | 1 | 1 | 5 | 0.2 |
| 5_1145712 | 1 | 1 | 1 | 1 | 2 | 1 | 1 | 1 | 5 | 1 | 1 | 1 | 1 | 0 | 2 | 1 | 1 | 1 | 5 | 0.8 |
| 5_1145712 | 2 | 1 | 1 | 1 | 2 | 0.5 | 1 | 1 | 6 | 0.8333 | 1 | 1 | 1 | 0 | 3 | 0.6667 | 1 | 1 | 6 | 0.6667 |
| 5_1145983 | 1 | 1 | 3 | 0 | 2 | 0.5 | NA | NA | 6 | 0.3333 | 3 | 0.3333 | 3 | 0.6667 | NA | NA | NA | NA | 6 | 0.5 |
| 5_1145984 | NA | NA | 1 | 1 | 1 | 1 | NA | NA | 2 | 1 | 4 | 0.75 | 3 | 1 | NA | NA | NA | NA | 7 | 0.8571 |
| 5_1147059 | 1 | 1 | NA | NA | 1 | 0 | 1 | 1 | 3 | 0.6667 | NA | NA | NA | NA | 2 | 0.5 | NA | NA | 2 | 0.5 |
| 5_1147492 | NA | NA | 2 | 1 | 1 | 1 | 1 | 1 | 4 | 1 | 1 | 1 | 1 | 1 | 2 | 1 | 1 | 1 | 5 | 1 |
| 5_1147492 | NA | NA | 2 | 0 | 1 | 1 | NA | NA | 3 | 0.3333 | 1 | 1 | 1 | 1 | 1 | 0 | 1 | 1 | 4 | 0.75 |
| 5_1147495 | 1 | 1 | 1 | 1 | NA | NA | NA | NA | 2 | 1 | NA | NA | 1 | 1 | 1 | 1 | NA | NA | 2 | 1 |
| 5_1147498 | 2 | 0.5 | 3 | 1 | NA | NA | 1 | 0 | 6 | 0.6667 | 1 | 1 | NA | NA | 3 | 1 | NA | NA | 4 | 1 |
| 5_1147498 | 3 | 1 | 3 | 1 | 1 | 1 | 1 | 1 | 8 | 1 | 2 | 1 | NA | NA | 3 | 1 | NA | NA | 5 | 1 |
| 5_1147499 | 4 | 0.5 | 2 | 0 | 1 | 0 | 1 | 0 | 8 | 0.25 | 2 | 0 | NA | NA | 3 | 0.3333 | 1 | 0 | 6 | 0.1667 |
| 5_1147500 | 2 | 0 | 1 | 0 | 1 | 1 | 1 | 0 | 5 | 0.2 | 2 | 0 | 1 | 0 | 2 | 0.5 | 1 | 0 | 6 | 0.1667 |
| 5_1147501 | 4 | 0.25 | 3 | 0.6667 | 1 | 1 | 1 | 1 | 9 | 0.5556 | 4 | 1 | NA | NA | 3 | 1 | 1 | 1 | 8 | 1 |
| 5_1147501 | 3 | 0 | 2 | 1 | NA | NA | 1 | 1 | 6 | 0.5 | 1 | 0 | 1 | 0 | 2 | 1 | 1 | 1 | 5 | 0.6 |
| 5_1147504 | 3 | 0.3333 | 3 | 0.6667 | 1 | 1 | 1 | 0 | 8 | 0.5 | 5 | 0.2 | 1 | 0 | 4 | 0.25 | 1 | 0 | 11 | 0.1818 |
| 5_1147507 | 4 | 0 | 2 | 0.5 | 2 | 0 | 1 | 0 | 9 | 0.1111 | 5 | 0.2 | NA | NA | 4 | 0.25 | 2 | 0 | 11 | 0.1818 |
| 5_1147508 | 4 | 0 | 2 | 0.5 | 1 | 1 | 2 | 0 | 9 | 0.2222 | 3 | 0 | 1 | 0 | 4 | 0.5 | 2 | 0.5 | 10 | 0.3 |
| 5_1148076 | 3 | 0.6667 | 8 | 0.25 | 8 | 0.375 | 6 | 0.5 | 25 | 0.4 | 4 | 0.25 | 5 | 0.2 | 6 | 0.1667 | 2 | 0 | 17 | 0.1765 |
| 5_1148077 | 4 | 0.5 | 13 | 0.5385 | 8 | 0.5 | 9 | 0.4444 | 34 | 0.5 | 8 | 0.375 | 8 | 0.5 | 6 | 0.6667 | 4 | 0.5 | 26 | 0.5 |
| 5_1148080 | 5 | 0.4 | 9 | 0.8889 | 7 | 0.4286 | 8 | 0.875 | 29 | 0.6897 | 6 | 0.6667 | 4 | 0.25 | 7 | 0.5714 | 4 | 0.75 | 21 | 0.5714 |
| 5_1148081 | 3 | 0 | 8 | 0.125 | 8 | 0.125 | 6 | 0 | 25 | 0.08 | 6 | 0 | 8 | 0.125 | 5 | 0.4 | 3 | 0 | 22 | 0.1364 |
| 5_1148084 | 2 | 0.5 | 8 | 0.25 | 7 | 0.4286 | 5 | 0.4 | 22 | 0.3636 | 7 | 0.5714 | 7 | 0.4286 | 6 | 0.3333 | 4 | 0.25 | 24 | 0.4167 |
| 5_1148657 | 2 | 0 | 1 | 1 | 3 | 1 | 4 | 1 | 10 | 0.8 | 1 | 0 | 1 | 1 | 1 | 1 | 3 | 0.6667 | 6 | 0.6667 |
| 5_1148658 | 2 | 1 | 1 | 1 | 3 | 1 | 4 | 1 | 10 | 1 | 1 | 1 | 2 | 1 | 2 | 1 | 4 | 1 | 9 | 1 |
| 5_1148663 | 1 | 1 | 1 | 1 | 3 | 1 | 3 | 1 | 8 | 1 | NA | NA | 1 | 1 | 2 | 1 | 3 | 1 | 6 | 1 |
| 5_1148668 | 2 | 0.5 | 1 | 1 | 2 | 1 | 1 | 1 | 6 | 0.8333 | 1 | 1 | 2 | 1 | 1 | 0 | 2 | 0.5 | 6 | 0.6667 |
| 5_1148668 | 2 | 0 | 1 | 0 | 3 | 0 | 3 | 0.3333 | 9 | 0.1111 | 1 | 0 | 1 | 0 | 2 | 0.5 | 3 | 0.3333 | 7 | 0.2857 |
| 5_1148681 | 2 | 0.5 | 5 | 0.2 | 2 | 1 | 5 | 0.4 | 14 | 0.4286 | 2 | 1 | 6 | 0 | 1 | 1 | 5 | 0.2 | 14 | 0.2857 |
| 5_1148682 | 2 | 0.5 | 3 | 0.3333 | 2 | 1 | 3 | 0.6667 | 10 | 0.6 | 2 | 1 | 4 | 0.75 | NA | NA | 2 | 1 | 8 | 0.875 |
| 5_1148682 | 2 | 1 | 5 | 0.8 | 2 | 0.5 | 4 | 0.75 | 13 | 0.7692 | 2 | 0.5 | 6 | 0.6667 | 1 | 1 | 3 | 0.6667 | 12 | 0.6667 |
| 5_1149424 | 3 | 0 | 7 | 0.5714 | 5 | 0.6 | 5 | 0.4 | 20 | 0.45 | 4 | 0.25 | 7 | 0.5714 | 7 | 0.4286 | 5 | 0.4 | 23 | 0.4348 |
| 5_1149433 | 5 | 0.2 | 8 | 0.125 | 7 | 0.1429 | 8 | 0 | 28 | 0.1071 | 4 | 0 | 8 | 0.25 | 7 | 0.2857 | 6 | 0.1667 | 25 | 0.2 |
| 5_1149464 | 12 | 0.1667 | 14 | 0.2857 | 21 | 0.0952 | 10 | 0 | 57 | 0.1404 | 16 | 0.1875 | 21 | 0.0952 | 17 | 0 | 17 | 0.2353 | 71 | 0.1268 |
| 5_1149465 | 15 | 0.1333 | 15 | 0.1333 | 21 | 0.0952 | 11 | 0.2727 | 62 | 0.1452 | 17 | 0.2941 | 21 | 0.1905 | 20 | 0.15 | 17 | 0.1765 | 75 | 0.2 |
| 5_1149466 | 14 | 0.1429 | 14 | 0.0714 | 20 | 0 | 10 | 0 | 58 | 0.0517 | 15 | 0.1333 | 23 | 0.0435 | 19 | 0.0526 | 16 | 0.0625 | 73 | 0.0685 |
| 5_1149468 | 9 | 0.3333 | 14 | 0.2143 | 16 | 0.1875 | 8 | 0.125 | 47 | 0.2128 | 14 | 0.1429 | 20 | 0.15 | 16 | 0.25 | 16 | 0.3125 | 66 | 0.2121 |
| 5_1149470 | 9 | 1 | 12 | 0.75 | 17 | 0.9412 | 9 | 1 | 47 | 0.9149 | 10 | 0.9 | 13 | 0.9231 | 16 | 0.875 | 13 | 0.8462 | 52 | 0.8846 |
| 5_1149470 | 9 | 0.6667 | 12 | 0.9167 | 17 | 0.7059 | 8 | 0.625 | 46 | 0.7391 | 13 | 0.6923 | 18 | 0.7222 | 15 | 0.8667 | 14 | 0.7143 | 60 | 0.75 |
| 5_1149472 | 7 | 0.5714 | 12 | 0.5833 | 16 | 0.375 | 7 | 0.5714 | 42 | 0.5 | 11 | 0.8182 | 16 | 0.4375 | 9 | 0.5556 | 15 | 0.6667 | 51 | 0.6078 |
| 5_1149472 | 10 | 0.1 | 13 | 0.1538 | 14 | 0.2143 | 7 | 0.2857 | 44 | 0.1818 | 9 | 0.1111 | 19 | 0.1579 | 12 | 0.0833 | 12 | 0 | 52 | 0.0962 |
| 5_1149473 | 10 | 0 | 14 | 0.1429 | 13 | 0.0769 | 7 | 0 | 44 | 0.0682 | 13 | 0.0769 | 15 | 0 | 13 | 0.0769 | 13 | 0.1538 | 54 | 0.0741 |
| 5_1149601 | 1 | 1 | 3 | 0 | NA | NA | 3 | 0 | 7 | 0.1429 | 3 | 0 | 2 | 1 | 3 | 0 | 3 | 0 | 11 | 0.1818 |
| 5_1149602 | 1 | 1 | 3 | 0 | NA | NA | 3 | 0.3333 | 7 | 0.2857 | 3 | 0.6667 | 2 | 0.5 | 2 | 0 | 4 | 0.25 | 11 | 0.3636 |
| 5_1149605 | 1 | 0 | 6 | 0.1667 | 1 | 1 | 4 | 0 | 12 | 0.1667 | 3 | 0 | 4 | 0 | 4 | 0.25 | 3 | 0 | 14 | 0.0714 |

| | | | | | | | | | | | | | | | | | | | | |
|---|---|---|---|---|---|---|---|---|---|---|---|---|---|---|---|---|---|---|---|---|
| 5_1149605 | 1 | 1 | 5 | 1 | 1 | 1 | 3 | 1 | 10 | 1 | 3 | 1 | 3 | 1 | 3 | 1 | 4 | 0.75 | 13 | 0.9231 |
| 5_1149728 | NA | NA | NA | NA | NA | NA | 1 | 1 | 1 | 1 | NA | NA | NA | NA | NA | NA | 1 | 1 | 1 | 1 |
| 5_1149728 | NA | NA | NA | NA | NA | NA | 1 | 1 | 1 | 1 | NA | NA | NA | NA | NA | NA | 1 | 1 | 1 | 1 |
| 5_1150308 | 1 | 1 | 1 | 1 | NA | NA | NA | NA | 2 | 1 | 1 | 1 | NA | NA | NA | NA | NA | NA | 1 | 1 |
| 5_1150311 | NA | NA | 1 | 1 | NA | NA | NA | NA | 1 | 1 | 1 | 1 | NA | NA | NA | NA | NA | NA | 1 | 1 |
| 5_1150317 | NA | NA | 1 | 1 | NA | NA | NA | NA | 1 | 1 | 1 | 1 | NA | NA | NA | NA | NA | NA | 1 | 1 |
| 5_1150490 | NA | NA | NA | NA | 1 | 1 | NA | NA | 1 | 1 | 1 | 1 | 1 | 1 | NA | NA | NA | NA | 2 | 1 |
| 5_1150493 | 1 | 1 | NA | NA | 1 | 1 | NA | NA | 2 | 1 | 2 | 1 | 1 | 1 | NA | NA | NA | NA | 3 | 1 |
| 5_1150500 | 1 | 1 | NA | NA | 1 | 0 | NA | NA | 2 | 0.5 | 2 | 0.5 | 1 | 1 | NA | NA | 1 | 1 | 4 | 0.75 |
| 5_1150858 | 5 | 0.4 | 17 | 0.5882 | 15 | 0.6667 | 4 | 0.75 | 41 | 0.6098 | 9 | 0.6667 | 17 | 0.5294 | 10 | 0.7 | 5 | 0.6 | 41 | 0.6098 |
| 5_1150859 | 5 | 0 | 19 | 0.1579 | 18 | 0.0556 | 5 | 0 | 47 | 0.0851 | 9 | 0 | 18 | 0.1667 | 9 | 0.1111 | 8 | 0 | 44 | 0.0909 |
| 5_1150860 | 4 | 0.75 | 18 | 0.7222 | 12 | 0.8333 | 3 | 0.6667 | 37 | 0.7568 | 7 | 0.7143 | 12 | 0.8333 | 9 | 0.8889 | 4 | 1 | 32 | 0.8438 |
| 5_1150861 | 3 | 0.6667 | 16 | 0.5625 | 15 | 0.4 | 3 | 0.3333 | 37 | 0.4865 | 8 | 0.25 | 17 | 0.6471 | 10 | 0.2 | 6 | 0.3333 | 41 | 0.4146 |
| 5_1150861 | 4 | 0.25 | 18 | 0.2778 | 20 | 0.55 | 5 | 0.6 | 47 | 0.4255 | 8 | 0.375 | 15 | 0.2667 | 10 | 0.4 | 8 | 0.25 | 41 | 0.3171 |
| 5_1150862 | 3 | 0.6667 | 17 | 1 | 16 | 1 | 5 | 1 | 41 | 0.9756 | 8 | 0.875 | 18 | 1 | 8 | 0.75 | 6 | 1 | 40 | 0.925 |
| 5_1150863 | 4 | 0 | 19 | 0.0526 | 17 | 0.1765 | 5 | 0.4 | 45 | 0.1333 | 9 | 0 | 18 | 0.1111 | 12 | 0.0833 | 7 | 0.1429 | 46 | 0.087 |
| 5_1150868 | 3 | 0 | 1 | 1 | 1 | 1 | NA | NA | 5 | 0.4 | NA | NA | 2 | 1 | NA | NA | 1 | 1 | 3 | 1 |
| 5_1150869 | 3 | 0.6667 | 1 | 0 | 1 | 0 | NA | NA | 5 | 0.4 | NA | NA | 2 | 0.5 | NA | NA | 1 | 1 | 3 | 0.6667 |
| 5_1150870 | 3 | 0 | 1 | 1 | 1 | 0 | NA | NA | 5 | 0.2 | NA | NA | 2 | 0 | NA | NA | 1 | 1 | 3 | 0.3333 |
| 5_1150870 | 2 | 0.5 | 1 | 1 | 1 | 0 | NA | NA | 4 | 0.5 | NA | NA | 2 | 0.5 | NA | NA | 1 | 0 | 3 | 0.3333 |
| 5_1152353 | 9 | 0.7778 | 13 | 0.6923 | 7 | 0.8571 | 10 | 0.9 | 39 | 0.7949 | 6 | 0.8333 | 7 | 0.7143 | 14 | 0.7143 | 5 | 0.4 | 32 | 0.6875 |
| 5_1152353 | 9 | 0.4444 | 14 | 0.5 | 8 | 0.5 | 12 | 0.4167 | 43 | 0.4651 | 6 | 0.6667 | 9 | 0.5556 | 17 | 0.7059 | 7 | 0.5714 | 39 | 0.641 |
| 5_1152354 | 11 | 0.2727 | 11 | 0.6364 | 7 | 0.5714 | 12 | 0.8333 | 41 | 0.5854 | 6 | 0.6667 | 7 | 0.5714 | 16 | 0.625 | 6 | 0.3333 | 35 | 0.5714 |
| 5_1152355 | 12 | 0.9167 | 13 | 1 | 7 | 0.7143 | 10 | 1 | 42 | 0.9286 | 5 | 1 | 7 | 0.8571 | 13 | 1 | 6 | 1 | 31 | 0.9677 |
| 5_1152355 | 10 | 0 | 15 | 0.4 | 7 | 0.1429 | 12 | 0.25 | 44 | 0.2273 | 5 | 0.2 | 9 | 0.1111 | 12 | 0.25 | 5 | 0 | 31 | 0.1613 |
| 5_1152355 | 13 | 0.6154 | 15 | 0.4 | 5 | 0.6 | 13 | 0.5385 | 46 | 0.5217 | 5 | 0.4 | 9 | 0.3333 | 17 | 0.4706 | 6 | 0.1667 | 37 | 0.3784 |
| 5_1152355 | 10 | 0.1 | 13 | 0.1538 | 6 | 0.5 | 12 | 0.0833 | 41 | 0.1707 | 4 | 0 | 5 | 0.2 | 15 | 0.2 | 6 | 0.1667 | 30 | 0.1667 |
| 5_1152356 | 12 | 0.9167 | 10 | 1 | 6 | 1 | 9 | 0.8889 | 37 | 0.9459 | 4 | 1 | 5 | 1 | 14 | 1 | 5 | 1 | 28 | 1 |
| 5_1152356 | 15 | 0.5333 | 10 | 0.1 | 8 | 0.25 | 11 | 0.2727 | 44 | 0.3182 | 5 | 0 | 6 | 0.6667 | 12 | 0.25 | 5 | 0 | 28 | 0.25 |
| 5_1152356 | 14 | 0.5 | 14 | 0.4286 | 7 | 0.4286 | 11 | 0.6364 | 46 | 0.5 | 6 | 0.5 | 8 | 0.375 | 14 | 0.4286 | 7 | 0.1429 | 35 | 0.3714 |
| 5_1152356 | 9 | 0.4444 | 14 | 0.3571 | 7 | 0.5714 | 8 | 0.375 | 38 | 0.4211 | 5 | 0.2 | 7 | 0.5714 | 15 | 0.4667 | 8 | 0.25 | 35 | 0.4 |
| 5_1152357 | 15 | 0.6 | 15 | 0.3333 | 9 | 0.3333 | 13 | 0.3077 | 52 | 0.4038 | 6 | 0.3333 | 11 | 0.1818 | 15 | 0.2667 | 6 | 0.3333 | 38 | 0.2632 |
| 5_1152357 | 8 | 0.25 | 12 | 0.1667 | 7 | 0.4286 | 11 | 0.0909 | 38 | 0.2105 | 4 | 0.25 | 6 | 0 | 11 | 0.1818 | 2 | 0 | 23 | 0.1304 |
| 5_1152358 | 14 | 0.4286 | 15 | 0.2667 | 8 | 0.375 | 13 | 0.5385 | 50 | 0.4 | 6 | 0.3333 | 11 | 0.3636 | 14 | 0.3571 | 7 | 0.2857 | 38 | 0.3421 |
| 5_1152358 | 9 | 0.4444 | 12 | 0.5 | 6 | 0.1667 | 9 | 0.6667 | 36 | 0.4722 | 3 | 0.6667 | 7 | 0.5714 | 13 | 0.3846 | 4 | 0 | 27 | 0.4074 |
| 5_1152360 | 11 | 0.1818 | 13 | 0 | 6 | 0.1667 | 10 | 0 | 40 | 0.075 | 5 | 0.6 | 6 | 0 | 8 | 0 | 5 | 0 | 24 | 0.125 |
| 5_1152360 | 14 | 0.1429 | 15 | 0.0667 | 8 | 0.125 | 14 | 0.0714 | 51 | 0.098 | 5 | 0.2 | 9 | 0 | 11 | 0.1818 | 7 | 0.1429 | 32 | 0.125 |
| 5_1152361 | 14 | 0.9286 | 15 | 0.6667 | 9 | 0.6667 | 10 | 0.7 | 48 | 0.75 | 6 | 0.6667 | 11 | 0.6364 | 14 | 0.7857 | 6 | 0.8333 | 37 | 0.7297 |
| 5_1152361 | 17 | 0 | 18 | 0.1667 | 9 | 0.1111 | 15 | 0 | 59 | 0.0678 | 6 | 0 | 11 | 0.0909 | 13 | 0.0769 | 6 | 0.1667 | 36 | 0.0833 |
| 5_1152361 | 18 | 0.2222 | 18 | 0 | 9 | 0 | 15 | 0.0667 | 60 | 0.0833 | 6 | 0.3333 | 10 | 0.2 | 14 | 0.2857 | 5 | 0.2 | 35 | 0.2571 |
| 5_1152364 | 10 | 0.2 | 7 | 0.1429 | 5 | 0 | 8 | 0.125 | 30 | 0.1333 | 7 | 0.2857 | 5 | 0 | 11 | 0.3636 | 6 | 0.1667 | 29 | 0.2414 |
| 5_1152367 | 8 | 0.25 | 6 | 0 | 5 | 0 | 7 | 0.1429 | 26 | 0.1154 | 6 | 0.1667 | 5 | 0 | 10 | 0.2 | 5 | 0 | 26 | 0.1154 |
| 5_1152488 | NA | NA | 2 | 0 | 4 | 0.5 | 1 | 0 | 7 | 0.2857 | 2 | 0.5 | 1 | 1 | NA | NA | NA | NA | 3 | 0.6667 |
| 5_1152608 | 6 | 0 | 3 | 0.3333 | 4 | 0 | 2 | 0 | 15 | 0.0667 | 1 | 0 | 6 | 0.1667 | 5 | 0.2 | 2 | 0 | 14 | 0.1429 |
| 5_1152620 | 4 | 0.25 | 3 | 0 | 3 | 0 | 2 | 0 | 12 | 0.0833 | 1 | 0 | 6 | 0.1667 | 4 | 0.25 | 1 | 0 | 12 | 0.1667 |
| 5_1152625 | 2 | 0 | 2 | 0.5 | 4 | 0 | 2 | 0 | 10 | 0.1 | 1 | 0 | 5 | 0 | 4 | 0 | 2 | 0.5 | 12 | 0.0833 |
| 5_1152629 | 1 | 0 | 2 | 0 | 2 | 0.5 | 1 | 0 | 6 | 0.1667 | NA | NA | 5 | 0.2 | 3 | 0.3333 | 2 | 0 | 10 | 0.2 |
| 5_1152634 | 2 | 1 | 3 | 0.6667 | 3 | 0.3333 | 1 | 1 | 9 | 0.6667 | NA | NA | 2 | 1 | 3 | 1 | 2 | 1 | 7 | 1 |
| 5_1152813 | 3 | 0.3333 | 3 | 0 | 6 | 0 | 8 | 0 | 20 | 0.05 | 6 | 0 | 8 | 0.25 | 9 | 0 | 5 | 0 | 28 | 0.0714 |
| 5_1152819 | 4 | 0 | 4 | 0 | 4 | 0 | 6 | 0.1667 | 18 | 0.0556 | 4 | 0.5 | 6 | 0.1667 | 8 | 0.125 | 3 | 0 | 21 | 0.1905 |
| 5_1152823 | 4 | 0 | 4 | 0.25 | 4 | 0 | 7 | 0 | 19 | 0.0526 | 5 | 0.4 | 7 | 0 | 11 | 0.0909 | 3 | 0.3333 | 26 | 0.1538 |
| 5_1152824 | 3 | 0.3333 | 4 | 0.5 | 3 | 0.3333 | 7 | 0.2857 | 17 | 0.3529 | 5 | 0.6 | 6 | 0 | 11 | 0.4545 | 3 | 0 | 25 | 0.32 |
| 5_1152824 | 4 | 0.25 | 4 | 0 | 3 | 0.3333 | 6 | 0.1667 | 17 | 0.1765 | 6 | 0 | 2 | 0 | 9 | 0.1111 | 3 | 0 | 20 | 0.05 |
| 5_1152831 | 1 | 0 | 3 | 0 | 2 | 0 | 5 | 0.4 | 11 | 0.1818 | 3 | 0.3333 | 5 | 0.4 | 9 | 0.3333 | 4 | 0.25 | 21 | 0.3333 |
| 5_1152832 | 3 | 0 | 3 | 0 | 4 | 0 | 5 | 0.2 | 15 | 0.0667 | 3 | 0 | 4 | 0 | 10 | 0.1 | 2 | 0.5 | 19 | 0.1053 |
| 5_1152852 | 1 | 0 | NA | NA | NA | NA | 3 | 0.3333 | 4 | 0.25 | NA | NA | 2 | 0 | 2 | 0.5 | 2 | 0.5 | 6 | 0.3333 |
| 5_1152856 | 3 | 0.3333 | NA | NA | 1 | 1 | 2 | 0 | 6 | 0.3333 | NA | NA | 2 | 0.5 | 1 | 1 | 4 | 0.25 | 7 | 0.4286 |
| 5_1152859 | 3 | 0.6667 | NA | NA | 1 | 0 | NA | NA | 4 | 0.5 | NA | NA | 1 | 0 | 1 | 0 | 3 | 0.3333 | 5 | 0.2 |
| 5_1152861 | 2 | 0.5 | NA | NA | NA | NA | 1 | 0 | 3 | 0.3333 | NA | NA | 2 | 0.5 | 2 | 0.5 | 3 | 0 | 7 | 0.2857 |
| 5_1153069 | 17 | 0.9412 | 38 | 0.6842 | 21 | 0.8095 | 14 | 0.8571 | 90 | 0.7889 | 24 | 0.75 | 9 | 0.7778 | 7 | 0.4286 | 18 | 0.7222 | 58 | 0.7069 |
| 5_1153070 | 16 | 0.75 | 35 | 0.7143 | 24 | 0.7083 | 13 | 0.6923 | 88 | 0.7159 | 26 | 0.7308 | 9 | 0.6667 | 8 | 0.75 | 20 | 0.85 | 63 | 0.7619 |
| 5_1153145 | 1 | 1 | NA | NA | NA | NA | NA | NA | 1 | 1 | 1 | 1 | NA | NA | NA | NA | 1 | 0 | 2 | 0.5 |
| 5_1153320 | 14 | 0.1429 | 24 | 0.0417 | 11 | 0.0909 | 9 | 0.1111 | 58 | 0.0862 | 16 | 0 | 7 | 0.2857 | 2 | 0 | 9 | 0 | 34 | 0.0588 |
| 5_1153327 | 1 | 1 | NA | NA | NA | NA | NA | NA | 1 | 1 | 1 | 1 | NA | NA | NA | NA | NA | NA | 1 | 1 |
| 5_1153431 | 1 | 1 | 1 | 1 | NA | NA | NA | NA | 2 | 1 | 1 | 1 | NA | NA | NA | NA | NA | NA | 1 | 1 |

| | | | | | | | | | | | | | | | | | | | | |
|---|---|---|---|---|---|---|---|---|---|---|---|---|---|---|---|---|---|---|---|---|
| 5_1153742 | NA | NA | NA | NA | 1 | 0 | 2 | 0.5 | 3 | 0.3333 | 1 | 1 | NA | NA | NA | NA | NA | NA | 1 | 1 |
| 5_1153745 | NA | NA | NA | NA | 1 | 1 | 2 | 0.5 | 3 | 0.6667 | 2 | 1 | NA | NA | NA | NA | NA | NA | 2 | 1 |
| 5_1153746 | NA | NA | 1 | 1 | 1 | 1 | 1 | 1 | 3 | 1 | 1 | 1 | NA | NA | NA | NA | NA | NA | 1 | 1 |
| 5_1153759 | NA | NA | NA | NA | NA | NA | 2 | 1 | 2 | 1 | 1 | 0 | 1 | 1 | NA | NA | NA | NA | 2 | 0.5 |
| 5_1153759 | NA | NA | NA | NA | NA | NA | 1 | 1 | 1 | 1 | 1 | 0 | 1 | 1 | NA | NA | NA | NA | 2 | 0.5 |
| 5_1153786 | 2 | 0 | NA | NA | 1 | 0 | 1 | 1 | 4 | 0.25 | 2 | 0 | NA | NA | 1 | 1 | 4 | 0.25 | 7 | 0.2857 |
| 5_1153787 | 14 | 0.2857 | 16 | 0.25 | 12 | 0.1667 | 18 | 0.2222 | 60 | 0.2333 | 10 | 0.1 | 10 | 0.3 | 9 | 0 | 11 | 0.1818 | 40 | 0.15 |
| 5_1153788 | 15 | 0.7333 | 18 | 0.7778 | 14 | 0.5714 | 20 | 0.75 | 67 | 0.7164 | 11 | 0.3636 | 11 | 0.8182 | 13 | 0.6154 | 19 | 0.5789 | 54 | 0.5926 |
| 5_1153788 | 13 | 0.6923 | 20 | 0.4 | 12 | 0.5 | 15 | 0.4667 | 60 | 0.5 | 11 | 0.2727 | 12 | 0.5 | 12 | 0.5 | 16 | 0.375 | 51 | 0.4118 |
| 5_1153789 | 19 | 0.0526 | 25 | 0.08 | 16 | 0 | 25 | 0.12 | 85 | 0.0706 | 14 | 0.1429 | 13 | 0.1538 | 15 | 0.0667 | 22 | 0.0909 | 64 | 0.1094 |
| 5_1154225 | 3 | 0.3333 | 3 | 0 | 2 | 0 | 5 | 0.2 | 13 | 0.1538 | 1 | 0 | 5 | 0.4 | 6 | 0.3333 | 1 | 0 | 13 | 0.3077 |
| 5_1154340 | 4 | 0 | 3 | 0.6667 | 2 | 0 | 5 | 0.4 | 14 | 0.2857 | 2 | 0 | 4 | 0.25 | 6 | 0.1667 | 2 | 0 | 14 | 0.1429 |
| 5_1154341 | 4 | 1 | 4 | 0.75 | 2 | 1 | 6 | 0.6667 | 16 | 0.8125 | 1 | 1 | 4 | 0.75 | 7 | 0.7143 | 2 | 1 | 14 | 0.7857 |
| 5_1154342 | 4 | 0.25 | 5 | 0.2 | 2 | 1 | 6 | 0.6667 | 17 | 0.4706 | 1 | 1 | 5 | 0.4 | 6 | 0.5 | NA | NA | 12 | 0.5 |
| 5_1154348 | 5 | 0 | 4 | 0 | 3 | 0 | 5 | 0.2 | 17 | 0.0588 | 1 | 0 | 4 | 0 | 6 | 0.1667 | NA | NA | 11 | 0.0909 |
| 5_1154366 | 91 | 0.8132 | 119 | 0.6807 | 96 | 0.7812 | 91 | 0.7582 | 397 | 0.7531 | 95 | 0.7053 | 77 | 0.6623 | 85 | 0.7176 | 99 | 0.8586 | 356 | 0.7416 |
| 5_1154367 | 87 | 0.6782 | 126 | 0.7619 | 98 | 0.7653 | 86 | 0.814 | 397 | 0.7557 | 105 | 0.7524 | 76 | 0.8289 | 93 | 0.6774 | 104 | 0.8462 | 378 | 0.7751 |
| 5_1154367 | 99 | 0.9293 | 127 | 0.9134 | 103 | 0.9029 | 99 | 0.8586 | 428 | 0.9019 | 105 | 0.9143 | 80 | 0.95 | 103 | 0.9612 | 104 | 0.9712 | 392 | 0.949 |
| 5_1154654 | 7 | 0.2857 | 10 | 0.2 | 6 | 0 | 8 | 0 | 31 | 0.129 | 2 | 0 | 3 | 0.6667 | 9 | 0 | 6 | 0 | 20 | 0.1 |
| 5_1154664 | 7 | 0.1429 | 6 | 0.1667 | 2 | 0 | 4 | 0.25 | 19 | 0.1579 | NA | NA | 3 | 0 | 5 | 0.4 | 5 | 0 | 13 | 0.1538 |
| 5_1154665 | 5 | 0.8 | 4 | 0.75 | 2 | 1 | 2 | 1 | 13 | 0.8462 | NA | NA | 1 | 0 | 1 | 1 | 2 | 0.5 | 4 | 0.5 |
| 5_1154666 | 7 | 0 | 7 | 0.1429 | 2 | 0.5 | 3 | 0.3333 | 19 | 0.1579 | NA | NA | 3 | 0.3333 | 5 | 0.2 | 5 | 0 | 13 | 0.1538 |
| 5_1154673 | 5 | 0 | 3 | 0 | NA | NA | 1 | 1 | 9 | 0.1111 | NA | NA | 2 | 0.5 | 2 | 0.5 | 3 | 0 | 7 | 0.2857 |
| 5_1154710 | 13 | 0.1538 | 23 | 0.2174 | 9 | 0.1111 | 9 | 0.1111 | 54 | 0.1667 | 8 | 0.375 | 8 | 0.25 | 14 | 0.0714 | 12 | 0.0833 | 42 | 0.1667 |
| 5_1154716 | 5 | 0 | 2 | 0 | 2 | 0.5 | 4 | 0 | 13 | 0.0769 | 3 | 0 | 7 | 0.1429 | 4 | 0 | 4 | 0 | 18 | 0.0556 |
| 5_1154717 | 6 | 0 | 2 | 0 | 5 | 0.2 | 2 | 0 | 15 | 0.0667 | 3 | 0.3333 | 8 | 0.125 | 4 | 0 | 5 | 0.2 | 20 | 0.15 |
| 5_1154723 | 3 | 0 | 2 | 0 | 2 | 0.5 | 3 | 0.3333 | 10 | 0.2 | 4 | 0.25 | 8 | 0.375 | 5 | 0.2 | 5 | 0 | 22 | 0.2273 |
| 5_1154723 | 2 | 1 | 1 | 1 | 2 | 1 | 2 | 1 | 7 | 1 | 3 | 0.6667 | 4 | 1 | 5 | 0.6 | 3 | 1 | 15 | 0.8 |
| 5_1154723 | 3 | 0.3333 | 2 | 0 | 2 | 0 | 1 | 0 | 8 | 0.125 | 3 | 0 | 6 | 0 | 4 | 0 | 2 | 0.5 | 15 | 0.0667 |
| 5_1154724 | 4 | 0 | 1 | 0 | 2 | 0 | 3 | 0.3333 | 10 | 0.1 | 4 | 0 | 8 | 0.125 | 3 | 0 | 5 | 0 | 20 | 0.05 |
| 5_1154724 | 4 | 0.25 | NA | NA | 2 | 0.5 | 2 | 0 | 8 | 0.25 | 4 | 0.25 | 8 | 0 | 3 | 0 | 5 | 0 | 20 | 0.05 |
| 5_1154725 | 3 | 0 | 1 | 1 | 2 | 0.5 | 2 | 0.5 | 8 | 0.375 | 3 | 0 | 9 | 0.4444 | 4 | 0.75 | 4 | 0.5 | 20 | 0.45 |
| 5_1154725 | 3 | 0.6667 | 1 | 0 | 1 | 1 | 1 | 1 | 6 | 0.6667 | 3 | 1 | 5 | 1 | 4 | 1 | 3 | 1 | 15 | 1 |
| 5_1154725 | 3 | 0.6667 | 1 | 1 | 1 | 1 | 1 | 0 | 6 | 0.6667 | 4 | 1 | 6 | 0.6667 | 3 | 1 | 5 | 1 | 18 | 0.8889 |
| 5_1154810 | 4 | 0 | 6 | 0.3333 | 4 | 0.25 | 3 | 0.3333 | 17 | 0.2353 | 5 | 0.6 | 2 | 0 | 6 | 0.1667 | 3 | 0 | 16 | 0.25 |
| 5_1154810 | 3 | 0.6667 | 4 | 1 | 3 | 0.6667 | 2 | 0.5 | 12 | 0.75 | 4 | 0.25 | 1 | 0 | 4 | 0.25 | 2 | 1 | 11 | 0.3636 |
| 5_1154810 | 3 | 0.3333 | 4 | 0 | 4 | 0 | 2 | 0.5 | 13 | 0.1538 | 4 | 0.5 | 2 | 0 | 4 | 0.25 | 2 | 0.5 | 12 | 0.3333 |
| 5_1154810 | 3 | 1 | 5 | 0.6 | 4 | 0.75 | 3 | 1 | 15 | 0.8 | 3 | 1 | 2 | 1 | 4 | 0.75 | 3 | 0.3333 | 12 | 0.75 |
| 5_1154811 | 4 | 1 | 6 | 1 | 4 | 1 | 3 | 1 | 17 | 1 | 4 | 1 | 2 | 1 | 6 | 1 | 2 | 1 | 14 | 1 |
| 5_1154811 | 4 | 1 | 5 | 1 | 3 | 1 | 2 | 1 | 14 | 1 | 5 | 1 | 2 | 1 | 5 | 1 | 3 | 1 | 15 | 1 |
| 5_1154811 | 4 | 0.5 | 6 | 0.3333 | 4 | 0.5 | 3 | 0 | 17 | 0.3529 | 6 | 0.3333 | 2 | 0 | 6 | 0.3333 | 3 | 1 | 17 | 0.4118 |
| 5_1154812 | 5 | 0.8 | 6 | 0.8333 | 4 | 0.75 | 3 | 0.6667 | 18 | 0.7778 | 7 | 0.5714 | 2 | 1 | 6 | 0.6667 | 3 | 0.6667 | 18 | 0.6667 |
| 5_1154812 | 4 | 0 | 4 | 0 | 4 | 0.25 | 3 | 0 | 15 | 0.0667 | 6 | 0 | 2 | 0 | 5 | 0.2 | 3 | 0 | 16 | 0.0625 |
| 5_1154812 | 4 | 0 | 4 | 0 | 3 | 0.3333 | 2 | 0 | 13 | 0.0769 | 3 | 0 | 1 | 0 | 5 | 0 | 3 | 0.6667 | 12 | 0.1667 |
| 5_1154812 | 5 | 0.8 | 4 | 0.5 | 2 | 1 | 3 | 0.3333 | 14 | 0.6429 | 6 | 0.5 | 2 | 1 | 7 | 0.7143 | 3 | 0.3333 | 18 | 0.6111 |
| 5_1154813 | 6 | 0.3333 | 5 | 0.2 | 4 | 0.25 | 3 | 0.3333 | 18 | 0.2778 | 5 | 0.6 | 1 | 1 | 4 | 0.25 | 3 | 0 | 13 | 0.3846 |
| 5_1154813 | 6 | 0.1667 | 4 | 0.25 | 3 | 0 | 2 | 0 | 15 | 0.1333 | 5 | 0 | 2 | 0.5 | 5 | 0.2 | 3 | 0.3333 | 15 | 0.2 |
| 5_1154814 | 4 | 0 | 6 | 0.1667 | 4 | 0 | 3 | 0.3333 | 17 | 0.1176 | 7 | 0 | 2 | 0 | 6 | 0.3333 | 3 | 0 | 18 | 0.1111 |
| 5_1154838 | 188 | 0.133 | 199 | 0.1357 | 192 | 0.1094 | 162 | 0.1358 | 741 | 0.1282 | 188 | 0.1436 | 164 | 0.1585 | 160 | 0.1125 | 144 | 0.1389 | 656 | 0.1387 |
| 5_1155555 | 1 | 1 | NA | NA | NA | NA | NA | NA | 1 | 1 | NA | NA | NA | NA | NA | NA | 1 | 1 | 1 | 1 |
| 5_1156163 | 1 | 1 | NA | NA | NA | NA | NA | NA | 1 | 1 | 3 | 0.6667 | 1 | 1 | NA | NA | 2 | 0.5 | 6 | 0.6667 |
| 5_1156207 | 1 | 1 | NA | NA | NA | NA | NA | NA | 1 | 1 | 1 | 1 | 1 | 1 | NA | NA | NA | NA | 2 | 1 |
| 5_1156209 | 1 | 1 | NA | NA | NA | NA | NA | NA | 1 | 1 | 2 | 1 | 1 | 1 | NA | NA | NA | NA | 3 | 1 |
| 5_1156922 | NA | NA | 4 | 0.25 | NA | NA | 2 | 0 | 6 | 0.1667 | 1 | 1 | NA | NA | NA | NA | NA | NA | 1 | 1 |
| 5_1156926 | 1 | 1 | 4 | 0 | 1 | 0 | 2 | 0.5 | 8 | 0.25 | 1 | 0 | 1 | 1 | NA | NA | NA | NA | 2 | 0.5 |
| 5_1156929 | 1 | 1 | 3 | 0.6667 | 2 | 1 | 3 | 0.3333 | 9 | 0.6667 | 2 | 1 | 3 | 1 | NA | NA | NA | NA | 5 | 1 |
| 5_1156930 | 2 | 0.5 | 3 | 0.3333 | 2 | 1 | 3 | 1 | 10 | 0.7 | 2 | 0.5 | 3 | 0 | NA | NA | NA | NA | 5 | 0.2 |
| 5_1156931 | 1 | 0 | 3 | 0 | 3 | 0.3333 | 3 | 0.6667 | 10 | 0.3 | 1 | 0 | 1 | 1 | NA | NA | NA | NA | 2 | 0.5 |
| 5_1157605 | 2 | 0.5 | 8 | 0.625 | 9 | 0.4444 | 3 | 0.6667 | 22 | 0.5455 | 3 | 0 | 7 | 0.2857 | 5 | 0.2 | 4 | 0 | 19 | 0.1579 |
| 5_1157605 | 5 | 0 | 10 | 0 | 9 | 0.3333 | 4 | 0.25 | 28 | 0.1429 | 3 | 0 | 7 | 0.2857 | 6 | 0.1667 | 5 | 0.2 | 21 | 0.1905 |
| 5_1157837 | 6 | 0 | 9 | 0.1111 | 3 | 0.3333 | 2 | 0 | 20 | 0.1 | 5 | 0 | 5 | 0 | 6 | 0 | 10 | 0.2 | 26 | 0.0769 |
| 5_1157838 | 6 | 1 | 9 | 0.8889 | 7 | 0.8571 | 4 | 1 | 26 | 0.9231 | 4 | 1 | 6 | 0.6667 | 7 | 1 | 7 | 0.7143 | 24 | 0.8333 |
| 5_1157838 | 5 | 1 | 5 | 1 | 5 | 1 | 3 | 1 | 18 | 1 | 3 | 1 | 5 | 0.8 | 6 | 0.8333 | 8 | 1 | 22 | 0.9091 |
| 5_1157840 | 6 | 0.6667 | 9 | 0.6667 | 9 | 0.6667 | 3 | 1 | 27 | 0.7037 | 5 | 1 | 6 | 0.6667 | 8 | 0.75 | 10 | 0.6 | 29 | 0.7241 |
| 5_1157840 | 6 | 1 | 8 | 1 | 8 | 1 | 5 | 1 | 27 | 1 | 5 | 1 | 6 | 1 | 4 | 1 | 10 | 1 | 25 | 1 |

| | | | | | | | | | | | | | | | | | | | | |
|---|---|---|---|---|---|---|---|---|---|---|---|---|---|---|---|---|---|---|---|---|
| 5_11578438 | 7 | 0.2857 | 9 | 0.1111 | 8 | 0.5 | 5 | 0.2 | 29 | 0.2759 | 4 | 0.5 | 3 | 0.3333 | 8 | 0.375 | 12 | 0.4167 | 27 | 0.4074 |
| 5_11578440 | 7 | 0.5714 | 10 | 0.6 | 7 | 0.4286 | 6 | 0.5 | 30 | 0.5333 | 5 | 0.6 | 7 | 0.2857 | 7 | 0.5714 | 11 | 0.6364 | 30 | 0.5333 |
| 5_11578451 | 6 | 0.6667 | 10 | 0.6 | 9 | 0.1111 | 7 | 0.4286 | 32 | 0.4375 | 5 | 0.2 | 7 | 0.2857 | 9 | 0.1111 | 11 | 0.7273 | 32 | 0.375 |
| 5_11578458 | 3 | 1 | 10 | 0.7 | 8 | 1 | 3 | 0.6667 | 24 | 0.8333 | 4 | 1 | 6 | 0.5 | 6 | 1 | 8 | 1 | 24 | 0.875 |
| 5_11578464 | 5 | 0.2 | 7 | 0.2857 | 8 | 0.125 | 6 | 0.3333 | 26 | 0.2308 | 4 | 0.25 | 3 | 0 | 3 | 0.3333 | 10 | 0.1 | 20 | 0.15 |
| 5_11578486 | 8 | 0 | 9 | 0.1111 | 10 | 0.1 | 6 | 0 | 33 | 0.0606 | 5 | 0.2 | 6 | 0 | 11 | 0.0909 | 11 | 0.1818 | 33 | 0.1212 |
| 5_11578506 | 5 | 0.2 | 8 | 0.25 | 12 | 0.5833 | 7 | 0.4286 | 32 | 0.4062 | 6 | 0.3333 | 9 | 0.2222 | 10 | 0 | 12 | 0.5833 | 37 | 0.2973 |
| 5_11578507 | 7 | 0.4286 | 10 | 0.1 | 11 | 0.1818 | 7 | 0.1429 | 35 | 0.2 | 6 | 0 | 6 | 0.1667 | 10 | 0.3 | 10 | 0.1 | 32 | 0.1562 |
| 5_11578513 | 6 | 0.3333 | 8 | 0.875 | 12 | 0.4167 | 3 | 0 | 29 | 0.4828 | 7 | 0.2857 | 8 | 0.25 | 6 | 0.3333 | 10 | 0.3 | 31 | 0.2903 |
| 5_11578676 | 2 | 0 | 7 | 0.5714 | 9 | 0 | 8 | 0.25 | 26 | 0.2308 | 9 | 0.3333 | 3 | 0 | 9 | 0.1111 | 6 | 0 | 27 | 0.1481 |
| 5_11586264 | 2 | 0 | 1 | 0 | 3 | 0 | 3 | 0.3333 | 9 | 0.1111 | 2 | 0 | NA | NA | 3 | 0.3333 | 2 | 0 | 7 | 0.1429 |
| 5_11586945 | 2 | 1 | NA | NA | 2 | 0.5 | 2 | 1 | 6 | 0.8333 | 1 | 1 | NA | NA | 1 | 1 | 2 | 1 | 4 | 1 |
| 5_11593970 | 2 | 1 | 2 | 0.5 | 2 | 1 | 1 | 1 | 7 | 0.8571 | 3 | 1 | 2 | 1 | NA | NA | 2 | 1 | 7 | 1 |
| 5_11614407 | 3 | 0.3333 | 7 | 0 | 2 | 0 | 5 | 0.2 | 17 | 0.1176 | 3 | 0.3333 | 10 | 0 | 1 | 0 | 5 | 0 | 19 | 0.0526 |
| 5_11614448 | 3 | 0 | 6 | 0.1667 | 1 | 0 | 4 | 0 | 14 | 0.0714 | 3 | 0 | 8 | 0.125 | 1 | 0 | 3 | 0 | 15 | 0.0667 |
| 5_11614457 | 3 | 1 | 6 | 0.8333 | 1 | 0 | 4 | 0.75 | 14 | 0.7857 | 2 | 0.5 | 6 | 0.5 | 1 | 1 | 4 | 0 | 13 | 0.3846 |
| 5_11614458 | 3 | 0 | 7 | 0.1429 | 2 | 0.5 | 5 | 0.2 | 17 | 0.1765 | 3 | 0.3333 | 9 | 0.1111 | NA | NA | 5 | 0.2 | 17 | 0.1765 |
| 5_11614467 | 3 | 0 | 6 | 0.1667 | 2 | 0 | 4 | 0.25 | 15 | 0.1333 | 2 | 0 | 9 | 0.1111 | 1 | 0 | 4 | 0 | 16 | 0.0625 |
| 5_11614472 | 2 | 0.5 | 7 | 1 | 3 | 1 | 5 | 0.8 | 17 | 0.8824 | 3 | 1 | 10 | 0.7 | 1 | 1 | 5 | 1 | 19 | 0.8421 |
| 5_11614477 | 3 | 1 | 7 | 0.8571 | 3 | 0.6667 | 5 | 0.8 | 18 | 0.8333 | 2 | 1 | 10 | 1 | 3 | 0.6667 | 5 | 1 | 20 | 0.95 |
| 5_11614478 | 2 | 0 | 6 | 0.3333 | 2 | 0 | 4 | 0 | 14 | 0.1429 | 2 | 0 | 9 | 0 | 3 | 0 | 4 | 0.25 | 18 | 0.0556 |
| 5_11614491 | 3 | 0 | 7 | 0.1429 | 3 | 0 | 5 | 0.2 | 18 | 0.1111 | 4 | 0 | 12 | 0.3333 | 6 | 0 | 4 | 0.25 | 26 | 0.1923 |
| 5_11614532 | 2 | 0 | 8 | 0.375 | 4 | 0.75 | 6 | 0.1667 | 20 | 0.35 | 4 | 0.25 | 12 | 0.25 | 7 | 0.1429 | 5 | 0 | 28 | 0.1786 |
| 5_11614537 | 3 | 0 | 8 | 0.25 | 4 | 0.75 | 5 | 0.4 | 20 | 0.35 | 4 | 0 | 12 | 0.0833 | 7 | 0.4286 | 6 | 0.5 | 29 | 0.2414 |
| 5_11614579 | 5 | 0.4 | 8 | 0.75 | 4 | 0.5 | 7 | 0.7143 | 24 | 0.625 | 5 | 0.4 | 10 | 0.4 | 9 | 0.7778 | 7 | 0.8571 | 31 | 0.6129 |
| 5_11614580 | 5 | 0.2 | 10 | 0.1 | 4 | 0.5 | 7 | 0.2857 | 26 | 0.2308 | 4 | 0.25 | 12 | 0.3333 | 9 | 0.1111 | 7 | 0.1429 | 32 | 0.2188 |
| 5_11614583 | 4 | 0 | 10 | 0.3 | 4 | 0.25 | 8 | 0.375 | 26 | 0.2692 | 4 | 0.5 | 13 | 0.4615 | 9 | 0.7778 | 7 | 0.5714 | 33 | 0.5758 |
| 5_11614584 | 6 | 0.6667 | 10 | 0.8 | 4 | 0.5 | 7 | 0.8571 | 27 | 0.7407 | 6 | 0.5 | 14 | 0.9286 | 8 | 0.875 | 6 | 0.8333 | 34 | 0.8235 |
| 5_11614622 | 8 | 0.125 | 8 | 0 | 5 | 0 | 8 | 0.25 | 29 | 0.1034 | 7 | 0.1429 | 17 | 0.1176 | 11 | 0 | 7 | 0.1429 | 42 | 0.0952 |
| 5_11614658 | 6 | 0.8333 | 7 | 0.8571 | 4 | 0.75 | 10 | 0.9 | 27 | 0.8519 | 7 | 0.5714 | 16 | 0.625 | 11 | 0.6364 | 5 | 0.6 | 39 | 0.6154 |
| 5_11614660 | 7 | 0.1429 | 8 | 0.625 | 5 | 0.6 | 11 | 0.2727 | 31 | 0.3871 | 7 | 0.4286 | 18 | 0.1667 | 13 | 0.0769 | 6 | 0.1667 | 44 | 0.1818 |
| 5_11614674 | 6 | 0.1667 | 4 | 0 | 4 | 0.25 | 4 | 0 | 18 | 0.1111 | 2 | 0.5 | 7 | 0.4286 | 9 | 0.1111 | 3 | 0.3333 | 21 | 0.2857 |
| 5_11614688 | 3 | 0.3333 | 2 | 0.5 | 3 | 0 | 2 | 0 | 10 | 0.2 | 4 | 0.25 | 8 | 0.5 | 9 | 0.1111 | 3 | 0 | 24 | 0.25 |
| 5_11615179 | NA | NA | 2 | 0 | 1 | 1 | 1 | 0 | 4 | 0.25 | 2 | 1 | 3 | 0 | 1 | 0 | NA | NA | 6 | 0.3333 |
| 5_11615185 | NA | NA | 3 | 0.6667 | 1 | 1 | 1 | 1 | 5 | 0.8 | 2 | 0.5 | 3 | 1 | 1 | 0 | 2 | 0.5 | 8 | 0.625 |
| 5_11615196 | NA | NA | 3 | 0.6667 | 2 | 0 | 1 | 1 | 6 | 0.5 | 2 | 1 | 2 | 0.5 | 1 | 1 | 2 | 1 | 7 | 0.8571 |
| 5_11615213 | NA | NA | 3 | 0.6667 | 1 | 1 | 1 | 0 | 5 | 0.6 | 2 | 1 | 3 | 0.3333 | 1 | 1 | 2 | 1 | 8 | 0.75 |
| 5_11615216 | NA | NA | 4 | 0.25 | 2 | 0 | 1 | 0 | 7 | 0.1429 | 2 | 0.5 | 1 | 0 | 1 | 0 | 2 | 0 | 6 | 0.1667 |
| 5_11615218 | NA | NA | 3 | 0 | 2 | 0.5 | NA | NA | 5 | 0.2 | 1 | 1 | 3 | 0.3333 | NA | NA | 1 | 0 | 5 | 0.4 |
| 5_11615224 | NA | NA | 2 | 0.5 | 1 | 0 | 1 | 0 | 4 | 0.25 | NA | NA | 2 | 0 | 1 | 0 | 1 | 1 | 4 | 0.25 |
| 5_11615245 | NA | NA | 3 | 1 | 1 | 0 | NA | NA | 4 | 0.75 | 2 | 1 | 2 | 0.5 | 1 | 1 | 1 | 0 | 6 | 0.6667 |
| 5_11615246 | NA | NA | 4 | 0.25 | 1 | 0 | NA | NA | 5 | 0.2 | 2 | 0.5 | 3 | 0.3333 | NA | NA | 1 | 0 | 6 | 0.3333 |
| 5_11615262 | NA | NA | 3 | 0.3333 | 1 | 0 | NA | NA | 4 | 0.25 | 1 | 1 | 2 | 1 | 1 | 1 | 2 | 0.5 | 6 | 0.8333 |
| 5_11626588 | 1 | 1 | NA | NA | NA | NA | NA | NA | 1 | 1 | NA | NA | NA | NA | 1 | 1 | NA | NA | 1 | 1 |
| 5_11626615 | 1 | 1 | NA | NA | NA | NA | NA | NA | 1 | 1 | NA | NA | NA | NA | 1 | 1 | NA | NA | 1 | 1 |
| 5_11626620 | 1 | 1 | NA | NA | NA | NA | NA | NA | 1 | 1 | NA | NA | NA | NA | 1 | 1 | NA | NA | 1 | 1 |
| 5_11654682 | 8 | 0.375 | 5 | 0 | 4 | 0.5 | 1 | 1 | 18 | 0.3333 | 5 | 0.4 | 4 | 0.5 | 2 | 0.5 | 2 | 0 | 13 | 0.3846 |
| 5_11654697 | 8 | 0.625 | 5 | 0.4 | 4 | 0.25 | 1 | 1 | 18 | 0.5 | 5 | 0.6 | 4 | 0.75 | 3 | 0.3333 | 2 | 0.5 | 14 | 0.5714 |
| 5_11654700 | 10 | 0.3 | 5 | 0 | 3 | 0 | 1 | 0 | 19 | 0.1579 | 3 | 0 | 4 | 0.5 | 3 | 0 | 2 | 0.5 | 12 | 0.25 |
| 5_11654710 | 9 | 0.2222 | 5 | 0 | 4 | 0.25 | 1 | 0 | 19 | 0.1579 | 4 | 0 | 4 | 0.25 | 2 | 0.5 | 2 | 0 | 12 | 0.1667 |
| 5_11654734 | 8 | 0.75 | 4 | 0.25 | 3 | 1 | 1 | 0 | 16 | 0.625 | 4 | 0.75 | 4 | 0.75 | 3 | 1 | 2 | 0.5 | 13 | 0.7692 |
| 5_11656042 | 7 | 1 | 2 | 1 | 4 | 1 | 1 | 1 | 14 | 1 | 3 | 0.6667 | 3 | 1 | 3 | 0.3333 | 1 | 1 | 10 | 0.7 |
| 5_11656047 | 6 | 0.1667 | 1 | 1 | 3 | 0.3333 | NA | NA | 10 | 0.3 | 4 | 0 | 2 | 0 | 3 | 0.3333 | 1 | 0 | 10 | 0.1 |
| 5_11656064 | 6 | 0.1667 | 2 | 0 | 4 | 0 | 1 | 0 | 13 | 0.0769 | 3 | 0.3333 | 2 | 0 | 3 | 0.3333 | 1 | 1 | 9 | 0.3333 |
| 5_11656072 | 6 | 0 | 2 | 0.5 | 3 | 0 | 1 | 0 | 12 | 0.0833 | 4 | 0 | 3 | 0.3333 | 1 | 0 | 1 | 0 | 9 | 0.1111 |
| 5_11658016 | NA | NA | NA | NA | NA | NA | 1 | 1 | 1 | 1 | NA | NA | 1 | 1 | NA | NA | 1 | 1 | 2 | 1 |
| 5_11658037 | NA | NA | NA | NA | 1 | 1 | 1 | 1 | 2 | 1 | NA | NA | 1 | 0 | NA | NA | 2 | 1 | 3 | 0.6667 |
| 5_11658068 | NA | NA | NA | NA | 1 | 1 | 1 | 1 | 2 | 1 | NA | NA | 2 | 1 | NA | NA | 1 | 1 | 3 | 1 |
| 5_11658071 | NA | NA | NA | NA | NA | NA | 1 | 1 | 1 | 1 | NA | NA | 2 | 1 | NA | NA | 2 | 1 | 4 | 1 |
| 5_11658075 | NA | NA | NA | NA | 1 | 1 | 1 | 1 | 2 | 1 | NA | NA | 2 | 1 | NA | NA | 2 | 1 | 4 | 1 |
| 5_11658096 | NA | NA | NA | NA | NA | NA | 1 | 1 | 1 | 1 | NA | NA | 2 | 0 | NA | NA | 2 | 0.5 | 4 | 0.25 |
| 5_11722982 | 6 | 0.3333 | 9 | 0.4444 | 6 | 0.5 | 5 | 0.6 | 26 | 0.4615 | 13 | 0.6923 | 8 | 0.25 | 9 | 0.6667 | 3 | 0.6667 | 33 | 0.5758 |
| 5_11722999 | 7 | 1 | 5 | 1 | 5 | 0.8 | 5 | 0.6 | 22 | 0.8636 | 9 | 0.7778 | 7 | 1 | 3 | 1 | 3 | 0.6667 | 22 | 0.8636 |
| 5_11723010 | 7 | 0 | 8 | 0.125 | 5 | 0.4 | 4 | 0.25 | 24 | 0.1667 | 12 | 0.0833 | 7 | 0.1429 | 5 | 0 | 3 | 0 | 27 | 0.0741 |
| 5_11723015 | 5 | 0.8 | 7 | 0.4286 | 4 | 0.5 | 5 | 0.4 | 21 | 0.5238 | 8 | 0.5 | 5 | 0.6 | 6 | 0.5 | 3 | 0.3333 | 22 | 0.5 |

| | | | | | | | | | | | | | | | | | | | | |
|---|---|---|---|---|---|---|---|---|---|---|---|---|---|---|---|---|---|---|---|---|
| 5_11723017 | 7 | 0.1429 | 8 | 0.25 | 7 | 0 | 3 | 0 | 25 | 0.12 | 12 | 0 | 8 | 0 | 6 | 0.3333 | 3 | 0 | 29 | 0.069 |
| 5_11723045 | 5 | 0.4 | 6 | 0.1667 | 3 | 0 | 3 | 0.3333 | 17 | 0.2353 | 8 | 0.5 | 6 | 0.1667 | 4 | 0.5 | 1 | 0 | 19 | 0.3684 |
| 5_11723047 | 7 | 0.4286 | 10 | 0.4 | 3 | 0 | 5 | 0.2 | 25 | 0.32 | 11 | 0.2727 | 6 | 0 | 7 | 0.4286 | 3 | 0.3333 | 27 | 0.2593 |
| 5_11723050 | 4 | 0.25 | 11 | 0.0909 | 7 | 0 | 5 | 0.4 | 27 | 0.1481 | 10 | 0.1 | 7 | 0.2857 | 7 | 0.1429 | 2 | 0 | 26 | 0.1538 |
| 5_11723077 | 6 | 1 | 14 | 0.8571 | 10 | 0.6 | 4 | 1 | 34 | 0.8235 | 14 | 0.7857 | 8 | 0.75 | 6 | 0.8333 | 2 | 1 | 30 | 0.8 |
| 5_11723079 | 5 | 0.2 | 10 | 0.5 | 10 | 0.2 | 5 | 0.6 | 30 | 0.3667 | 12 | 0.25 | 9 | 0.3333 | 4 | 0.25 | 2 | 1 | 27 | 0.3333 |
| 5_11723088 | 6 | 0.3333 | 9 | 0.1111 | 8 | 0.25 | 3 | 0.3333 | 26 | 0.2308 | 12 | 0.4167 | 6 | 0.5 | 5 | 0.8 | 2 | 0 | 25 | 0.48 |
| 5_11723090 | 2 | 0.5 | 12 | 0.5833 | 10 | 0.6 | 4 | 0.25 | 28 | 0.5357 | 8 | 0.75 | 7 | 0.4286 | 5 | 1 | 2 | 0.5 | 22 | 0.6818 |
| 5_11737879 | 1 | 0 | 1 | 1 | 1 | 1 | 2 | 1 | 5 | 0.8 | NA | NA | 3 | 1 | 2 | 1 | NA | NA | 5 | 1 |
| 5_11737903 | 1 | 0 | 2 | 0.5 | NA | NA | 1 | 1 | 4 | 0.5 | NA | NA | 4 | 0.5 | 2 | 0 | NA | NA | 6 | 0.3333 |
| 5_11740398 | 5 | 0 | 3 | 0 | 1 | 0 | 4 | 0.25 | 13 | 0.0769 | 3 | 0.3333 | 6 | 0 | 2 | 0 | 2 | 0 | 13 | 0.0769 |
| 5_11740400 | 5 | 0.4 | 4 | 0.5 | 1 | 1 | 4 | 0.75 | 14 | 0.5714 | 3 | 0.3333 | 5 | 0.4 | 2 | 1 | 2 | 1 | 12 | 0.5833 |
| 5_11740409 | 4 | 0.5 | 3 | 0.6667 | 1 | 1 | 4 | 0.75 | 12 | 0.6667 | 3 | 0 | 6 | 0.8333 | 3 | 1 | 2 | 0.5 | 14 | 0.6429 |
| 5_11740414 | 5 | 0.4 | 4 | 0.5 | 1 | 1 | 4 | 0.5 | 14 | 0.5 | 3 | 0 | 6 | 0.3333 | 3 | 1 | 1 | 0 | 13 | 0.3846 |
| 5_11741181 | 5 | 0 | 6 | 0.1667 | 1 | 0 | 3 | 0 | 15 | 0.0667 | 3 | 0.3333 | 6 | 0.3333 | 4 | 0 | 2 | 0 | 15 | 0.2 |
| 5_11741182 | 4 | 0.25 | 5 | 0.4 | 1 | 0 | 2 | 0.5 | 12 | 0.3333 | 2 | 0.5 | 4 | 0.25 | 2 | 0.5 | 2 | 0.5 | 10 | 0.4 |
| 5_11741186 | 5 | 0.2 | 6 | 0.1667 | 1 | 0 | 4 | 0 | 16 | 0.125 | 3 | 0.3333 | 5 | 0.4 | 4 | 0.25 | 1 | 0 | 13 | 0.3077 |
| 5_11741188 | 4 | 1 | 2 | 1 | NA | NA | 2 | 1 | 8 | 1 | 2 | 1 | 6 | 1 | 4 | 1 | 2 | 0.5 | 14 | 0.9286 |
| 5_11741248 | 7 | 0.1429 | 6 | 0.1667 | 3 | 0.3333 | 3 | 0.3333 | 19 | 0.2105 | 3 | 0.6667 | 8 | 0 | 2 | 0.5 | 1 | 0 | 14 | 0.2143 |
| 5_11741251 | 7 | 0.4286 | 6 | 0.1667 | 3 | 0.6667 | 4 | 0 | 20 | 0.3 | 3 | 0.3333 | 8 | 0.125 | 3 | 0 | 2 | 0.5 | 16 | 0.1875 |
| 5_11741255 | 7 | 0.2857 | 6 | 0 | 3 | 0 | 4 | 0.25 | 20 | 0.15 | 3 | 0 | 8 | 0.125 | 2 | 0 | 2 | 0 | 15 | 0.0667 |
| 5_11741282 | 9 | 0 | 8 | 0.375 | 4 | 0.25 | 4 | 0 | 25 | 0.16 | 3 | 0 | 8 | 0 | 3 | 0.6667 | 4 | 0.25 | 18 | 0.1667 |
| 5_11750883 | 23 | 0.087 | 21 | 0.0952 | 20 | 0.1 | 17 | 0.0588 | 81 | 0.0864 | 17 | 0.1176 | 17 | 0.1176 | 16 | 0.0625 | 14 | 0.2143 | 64 | 0.125 |
| 5_11751560 | 31 | 0.9677 | 39 | 0.9487 | 30 | 1 | 21 | 0.8571 | 121 | 0.9504 | 35 | 0.9143 | 25 | 0.96 | 23 | 0.9565 | 18 | 0.8889 | 101 | 0.9307 |
| 5_11751566 | 31 | 0.9032 | 38 | 0.8947 | 29 | 0.9655 | 22 | 1 | 120 | 0.9333 | 29 | 0.8966 | 30 | 0.9 | 28 | 0.9643 | 19 | 1 | 106 | 0.934 |
| 5_11751569 | 37 | 0.1622 | 47 | 0.1702 | 37 | 0.1351 | 23 | 0.1739 | 144 | 0.1597 | 35 | 0.2286 | 34 | 0.0882 | 31 | 0.1935 | 20 | 0.25 | 120 | 0.1833 |
| 5_11751570 | 34 | 0.3529 | 42 | 0.5714 | 38 | 0.3684 | 23 | 0.3478 | 137 | 0.4234 | 34 | 0.5 | 33 | 0.3939 | 32 | 0.4062 | 19 | 0.6316 | 118 | 0.4661 |
| 5_11751600 | 31 | 0.4516 | 40 | 0.35 | 30 | 0.4 | 27 | 0.3704 | 128 | 0.3906 | 33 | 0.2727 | 33 | 0.3636 | 28 | 0.3571 | 15 | 0.4 | 109 | 0.3394 |
| 5_11751603 | 29 | 0.5172 | 41 | 0.5122 | 32 | 0.5938 | 25 | 0.44 | 127 | 0.5197 | 24 | 0.375 | 30 | 0.4 | 29 | 0.6552 | 13 | 0.5385 | 96 | 0.4896 |
| 5_11751732 | 1 | 1 | 1 | 0 | 1 | 0 | 2 | 0 | 5 | 0.2 | 4 | 0 | 3 | 0.3333 | 2 | 0.5 | 1 | 1 | 10 | 0.3 |
| 5_11751773 | 1 | 1 | 1 | 0 | NA | NA | 2 | 0 | 4 | 0.25 | 3 | 0 | 4 | 0.5 | 2 | 0.5 | 1 | 0 | 10 | 0.3 |
| 5_11751783 | 1 | 1 | 1 | 1 | NA | NA | 2 | 0 | 4 | 0.5 | 3 | 0.3333 | 3 | 0.6667 | 1 | 1 | 1 | 0 | 8 | 0.5 |
| 5_11752839 | 2 | 1 | NA | NA | NA | NA | NA | NA | 2 | 1 | 2 | 1 | 1 | 1 | NA | NA | 1 | 1 | 4 | 1 |
| 5_11753075 | 1 | 1 | NA | NA | NA | NA | NA | NA | 1 | 1 | NA | NA | NA | NA | NA | NA | 1 | 1 | 1 | 1 |
| 5_11753085 | 1 | 1 | NA | NA | NA | NA | NA | NA | 1 | 1 | NA | NA | NA | NA | NA | NA | 1 | 1 | 1 | 1 |
| 5_11753094 | 1 | 1 | NA | NA | NA | NA | NA | NA | 1 | 1 | NA | NA | NA | NA | NA | NA | 1 | 1 | 1 | 1 |
| 5_11753107 | 1 | 0 | 1 | 1 | NA | NA | NA | NA | 2 | 0.5 | NA | NA | NA | NA | NA | NA | 1 | 1 | 1 | 1 |
| 5_11753124 | 1 | 1 | 1 | 1 | NA | NA | NA | NA | 2 | 1 | NA | NA | NA | NA | NA | NA | 1 | 1 | 1 | 1 |
| 5_11784623 | NA | NA | NA | NA | 1 | 1 | NA | NA | 1 | 1 | NA | NA | NA | NA | 2 | 1 | NA | NA | 2 | 1 |
| 5_11784624 | NA | NA | NA | NA | 1 | 1 | NA | NA | 1 | 1 | NA | NA | NA | NA | 2 | 0.5 | NA | NA | 2 | 0.5 |
| 5_11790383 | NA | NA | NA | NA | 1 | 1 | NA | NA | 1 | 1 | NA | NA | 1 | 0 | 2 | 1 | NA | NA | 3 | 0.6667 |
| 5_11790388 | NA | NA | NA | NA | 1 | 1 | NA | NA | 1 | 1 | NA | NA | 1 | 1 | 2 | 0.5 | NA | NA | 3 | 0.6667 |
| 5_11790390 | NA | NA | NA | NA | 1 | 1 | NA | NA | 1 | 1 | NA | NA | 1 | 1 | 2 | 1 | NA | NA | 3 | 1 |
| 5_11790393 | NA | NA | NA | NA | 1 | 1 | NA | NA | 1 | 1 | NA | NA | NA | NA | 2 | 0.5 | NA | NA | 2 | 0.5 |
| 5_11790398 | NA | NA | NA | NA | 1 | 1 | NA | NA | 1 | 1 | NA | NA | 1 | 1 | 2 | 1 | NA | NA | 3 | 1 |
| 5_11791137 | 3 | 0.3333 | NA | NA | 2 | 0 | NA | NA | 5 | 0.2 | 4 | 0.25 | 2 | 0.5 | 2 | 0 | 2 | 1 | 10 | 0.4 |
| 5_11791158 | 4 | 0.25 | 1 | 0 | 2 | 0 | NA | NA | 7 | 0.1429 | 3 | 0.3333 | 2 | 0.5 | 2 | 0 | 4 | 0 | 11 | 0.1818 |
| 5_11791160 | 4 | 0 | 1 | 1 | 2 | 0 | NA | NA | 7 | 0.1429 | 3 | 0.3333 | 3 | 0 | 3 | 0 | 4 | 0.25 | 13 | 0.1538 |
| 5_11809208 | 3 | 0.3333 | 1 | 0 | 2 | 0.5 | 1 | 1 | 7 | 0.4286 | 3 | 0.3333 | 1 | 1 | 1 | 0 | 1 | 0 | 6 | 0.3333 |
| 5_11814619 | 9 | 0.1111 | 12 | 0.4167 | 20 | 0.15 | 17 | 0.1176 | 58 | 0.1897 | 7 | 0.1429 | 19 | 0.2632 | 7 | 0.1429 | 18 | 0.0556 | 51 | 0.1569 |
| 5_11814621 | 9 | 0.1111 | 12 | 0.25 | 19 | 0.2632 | 17 | 0.0588 | 57 | 0.1754 | 7 | 0.4286 | 15 | 0.2 | 4 | 0 | 17 | 0.2941 | 43 | 0.2558 |
| 5_11814624 | 11 | 0.5455 | 10 | 0.5 | 17 | 0.4706 | 18 | 0.3889 | 56 | 0.4643 | 7 | 0.4286 | 19 | 0.3684 | 6 | 0.6667 | 16 | 0.4375 | 48 | 0.4375 |
| 5_11814626 | 8 | 0.625 | 12 | 0.75 | 16 | 0.6875 | 18 | 0.3333 | 54 | 0.5741 | 8 | 0.5 | 16 | 0.4375 | 5 | 0 | 18 | 0.2778 | 47 | 0.3404 |
| 5_11814632 | 11 | 0.3636 | 15 | 0.1333 | 18 | 0.0556 | 19 | 0.0526 | 63 | 0.127 | 8 | 0.25 | 17 | 0.1176 | 7 | 0 | 17 | 0.1176 | 49 | 0.1224 |
| 5_11814636 | 10 | 0.9 | 14 | 0.5714 | 20 | 0.65 | 18 | 0.5556 | 62 | 0.6452 | 7 | 0.5714 | 18 | 0.7778 | 6 | 0.8333 | 18 | 0.6111 | 49 | 0.6939 |
| 5_11814651 | 9 | 0.1111 | 14 | 0.1429 | 18 | 0.1111 | 20 | 0.15 | 61 | 0.1311 | 5 | 0.2 | 18 | 0.2222 | 7 | 0 | 16 | 0.0625 | 46 | 0.1304 |
| 5_11814652 | 11 | 0.1818 | 15 | 0.2667 | 18 | 0.1667 | 20 | 0.2 | 64 | 0.2031 | 8 | 0.25 | 17 | 0.4118 | 9 | 0.1111 | 18 | 0 | 52 | 0.1923 |
| 5_11814659 | 4 | 0.5 | 8 | 0.5 | 6 | 0.5 | 9 | 0.3333 | 27 | 0.4444 | 2 | 0 | 5 | 0 | 4 | 1 | 7 | 0.2857 | 18 | 0.3333 |
| 5_11814710 | 10 | 0.4 | 16 | 0.5625 | 20 | 0.5 | 18 | 0.6111 | 64 | 0.5312 | 8 | 0.625 | 19 | 0.3684 | 6 | 0.8333 | 18 | 0.3889 | 51 | 0.4706 |
| 5_11814725 | 13 | 0.0769 | 15 | 0 | 25 | 0.12 | 23 | 0.0435 | 76 | 0.0658 | 9 | 0.2222 | 20 | 0.1 | 9 | 0.1111 | 19 | 0 | 57 | 0.0877 |
| 5_11814776 | 12 | 0.25 | 16 | 0.1875 | 27 | 0.3704 | 21 | 0.2381 | 76 | 0.2763 | 10 | 0.4 | 23 | 0.4348 | 13 | 0.3846 | 23 | 0.3043 | 69 | 0.3768 |
| 5_11814790 | 16 | 0.0625 | 17 | 0.0588 | 30 | 0.1 | 23 | 0.0435 | 86 | 0.0698 | 9 | 0 | 27 | 0.0741 | 12 | 0.0833 | 22 | 0.0455 | 70 | 0.0571 |
| 5_11814792 | 16 | 0.125 | 19 | 0.2632 | 31 | 0.0968 | 20 | 0.25 | 86 | 0.1744 | 9 | 0.2222 | 25 | 0.32 | 11 | 0.0909 | 24 | 0.25 | 69 | 0.2464 |
| 5_11819898 | NA | NA | 1 | 0 | NA | NA | 1 | 1 | 2 | 0.5 | NA | NA | NA | NA | NA | NA | 2 | 0.5 | 2 | 0.5 |
| 5_11819937 | NA | NA | 1 | 1 | NA | NA | 1 | 1 | 2 | 1 | NA | NA | NA | NA | NA | NA | 2 | 0.5 | 2 | 0.5 |

| | | | | | | | | | | | | | | | | | | | | |
|---|---|---|---|---|---|---|---|---|---|---|---|---|---|---|---|---|---|---|---|---|
| 5_1181998 | 34 | 0.2059 | 28 | 0.25 | 34 | 0.2353 | 35 | 0.2286 | 131 | 0.229 | 22 | 0.0909 | 21 | 0.1429 | 26 | 0.0769 | 15 | 0.0667 | 84 | 0.0952 |
| 5_1181998 | 31 | 0.0323 | 26 | 0.1923 | 31 | 0.0968 | 36 | 0.0833 | 124 | 0.0968 | 20 | 0 | 20 | 0.05 | 28 | 0.1071 | 15 | 0.1333 | 83 | 0.0723 |
| 5_1182054 | 2 | 0 | 4 | 0 | 1 | 1 | 4 | 0 | 11 | 0.0909 | 3 | 0 | 6 | 0.1667 | 5 | 0 | 6 | 0.1667 | 20 | 0.1 |
| 5_1182152 | NA | NA | NA | NA | NA | NA | 1 | 1 | 1 | 1 | 1 | 1 | 1 | 1 | 1 | 1 | 2 | 1 | 5 | 1 |
| 5_1182153 | NA | NA | NA | NA | NA | NA | 1 | 1 | 1 | 1 | 1 | 0 | 1 | 1 | 1 | 1 | 2 | 1 | 5 | 0.8 |
| 5_1182156 | 2 | 0.5 | 2 | 0 | 1 | 0 | 2 | 0 | 7 | 0.1429 | 3 | 0 | 6 | 0.1667 | 2 | 0 | 6 | 0.3333 | 17 | 0.1765 |
| 5_1183251 | NA | NA | 1 | 1 | NA | NA | 1 | 0 | 2 | 0.5 | 6 | 0.1667 | 1 | 1 | 1 | 0 | 1 | 1 | 9 | 0.3333 |
| 5_1183261 | 1 | 0 | 1 | 0 | 1 | 1 | 1 | 0 | 4 | 0.25 | 4 | 0.5 | 2 | 0.5 | 2 | 0 | NA | NA | 8 | 0.375 |
| 5_1183262 | 1 | 0 | 2 | 1 | 1 | 1 | 1 | 0 | 5 | 0.6 | 6 | 0.1667 | 2 | 0.5 | 1 | 0 | NA | NA | 9 | 0.2222 |
| 5_1183262 | 1 | 1 | 1 | 0 | 1 | 0 | 1 | 0 | 4 | 0.25 | 6 | 0.3333 | 2 | 0.5 | 2 | 0.5 | NA | NA | 10 | 0.4 |
| 5_1183288 | 7 | 0.7143 | 12 | 0.4167 | 7 | 0.4286 | 14 | 0.7857 | 40 | 0.6 | 4 | 0 | 9 | 0.8889 | 11 | 0.6364 | 4 | 0.75 | 28 | 0.6429 |
| 5_1183289 | 10 | 0.6 | 13 | 0.2308 | 8 | 0.5 | 17 | 0.3529 | 48 | 0.3958 | 4 | 0.5 | 9 | 0.4444 | 14 | 0.5 | 6 | 1 | 33 | 0.5758 |
| 5_1183289 | 9 | 0.8889 | 13 | 0.6923 | 7 | 1 | 15 | 0.8 | 44 | 0.8182 | 5 | 0.6 | 10 | 0.6 | 12 | 0.75 | 6 | 0.3333 | 33 | 0.6061 |
| 5_1183290 | 12 | 0.25 | 14 | 0.1429 | 8 | 0.125 | 14 | 0.1429 | 48 | 0.1667 | 4 | 0 | 11 | 0.2727 | 13 | 0.2308 | 6 | 0.3333 | 34 | 0.2353 |
| 5_1183290 | 10 | 0.7 | 13 | 0.6154 | 6 | 1 | 15 | 0.6667 | 44 | 0.7045 | 5 | 0.4 | 11 | 0.5455 | 14 | 0.7143 | 5 | 1 | 35 | 0.6571 |
| 5_1183290 | 8 | 0.875 | 10 | 0.6 | 6 | 0.8333 | 12 | 0.6667 | 36 | 0.7222 | 4 | 0.5 | 11 | 0.6364 | 12 | 0.8333 | 5 | 0.6 | 32 | 0.6875 |
| 5_1183290 | 9 | 0.1111 | 11 | 0 | 5 | 0.2 | 14 | 0 | 39 | 0.0513 | 5 | 0 | 11 | 0.2727 | 13 | 0.1538 | 4 | 0 | 33 | 0.1515 |
| 5_1183291 | 9 | 0 | 12 | 0.0833 | 5 | 0.4 | 14 | 0.0714 | 40 | 0.1 | 3 | 0 | 10 | 0.1 | 11 | 0.0909 | 5 | 0.2 | 29 | 0.1034 |
| 5_1183292 | 8 | 0.125 | 10 | 0 | 5 | 0.8 | 10 | 0.2 | 33 | 0.2121 | 4 | 0 | 10 | 0.3 | 10 | 0.2 | 2 | 0.5 | 26 | 0.2308 |
| 5_1183294 | 6 | 0 | 10 | 0 | 3 | 0 | 12 | 0.1667 | 31 | 0.0645 | 4 | 0 | 8 | 0.125 | 9 | 0.1111 | 5 | 0.2 | 26 | 0.1154 |
| 5_1183297 | 6 | 0.3333 | 8 | 0.125 | 2 | 0 | 8 | 0.375 | 24 | 0.25 | 1 | 1 | 8 | 0.625 | 9 | 0.4444 | 4 | 0 | 22 | 0.4545 |
| 5_1183297 | 6 | 0 | 10 | 0.2 | 3 | 0 | 10 | 0.1 | 29 | 0.1034 | 1 | 0 | 8 | 0.25 | 9 | 0 | 5 | 0.2 | 23 | 0.1304 |
| 5_1183300 | 5 | 0.4 | 7 | 0.7143 | 2 | 0.5 | 7 | 0.2857 | 21 | 0.4762 | 1 | 1 | 7 | 0.2857 | 8 | 0.25 | 4 | 0.5 | 20 | 0.35 |
| 5_1183301 | 4 | 0 | 5 | 0 | 2 | 0 | 8 | 0.125 | 19 | 0.0526 | 1 | 0 | 5 | 0 | 7 | 0.1429 | 3 | 0 | 16 | 0.0625 |
| 5_1183302 | 5 | 0.8 | 9 | 1 | 2 | 1 | 9 | 0.8889 | 25 | 0.92 | 1 | 0 | 6 | 0.8333 | 6 | 1 | 3 | 1 | 16 | 0.875 |
| 5_1183302 | 4 | 0 | 8 | 0.5 | 2 | 0 | 5 | 0.4 | 19 | 0.3158 | 1 | 0 | 7 | 0.2857 | 7 | 0.2857 | 3 | 0.6667 | 18 | 0.3333 |
| 5_1183303 | 4 | 0 | 8 | 0 | 2 | 0 | 7 | 0.2857 | 21 | 0.0952 | 1 | 0 | 6 | 0 | 8 | 0.125 | 3 | 0.3333 | 18 | 0.1111 |
| 5_1183304 | 3 | 0.3333 | 8 | 0.125 | 2 | 0 | 4 | 0 | 17 | 0.1176 | 1 | 1 | 5 | 0.2 | 6 | 0.1667 | 3 | 0 | 15 | 0.2 |
| 5_1183304 | 3 | 0 | 8 | 0.375 | 2 | 0 | 6 | 0.5 | 19 | 0.3158 | 1 | 0 | 4 | 0.5 | 7 | 0.1429 | 3 | 0.6667 | 15 | 0.3333 |
| 5_1183305 | 3 | 1 | 7 | 0.8571 | 2 | 1 | 5 | 1 | 17 | 0.9412 | NA | NA | 5 | 1 | 7 | 1 | 3 | 1 | 15 | 1 |
| 5_1183307 | 3 | 0.3333 | 9 | 0.7778 | 2 | 1 | 5 | 0.6 | 19 | 0.6842 | 1 | 1 | 5 | 1 | 6 | 0.8333 | 3 | 0.6667 | 15 | 0.8667 |
| 5_1183308 | 2 | 0 | 7 | 0.5714 | 2 | 0 | 4 | 0.25 | 15 | 0.3333 | 1 | 1 | 5 | 0.2 | 7 | 0.2857 | 3 | 0.3333 | 16 | 0.3125 |
| 5_1183309 | 2 | 0.5 | 4 | 0.5 | 1 | 1 | 5 | 0.8 | 12 | 0.6667 | 1 | 1 | 2 | 1 | 6 | 0.8333 | 2 | 1 | 11 | 0.9091 |
| 5_1183309 | 2 | 0 | 8 | 0.875 | 1 | 1 | 6 | 0.6667 | 17 | 0.7059 | NA | NA | 5 | 1 | 5 | 0.6 | 1 | 1 | 11 | 0.8182 |
| 5_1183309 | 2 | 0 | 8 | 0.5 | 2 | 0 | 4 | 0.25 | 16 | 0.3125 | 1 | 0 | 5 | 0.2 | 5 | 0.2 | 3 | 1 | 14 | 0.3571 |
| 5_1183309 | 2 | 0 | 7 | 0.4286 | 1 | 0 | 3 | 0.6667 | 13 | 0.3846 | 1 | 1 | 5 | 0.8 | 6 | 0.3333 | 3 | 0.6667 | 15 | 0.6 |
| 5_1183311 | 2 | 0 | 7 | 0.1429 | 2 | 0 | 3 | 0 | 14 | 0.0714 | 1 | 0 | 1 | 0 | 3 | 0.3333 | NA | NA | 5 | 0.2 |
| 5_1183312 | 1 | 0 | 6 | 0.5 | 2 | 0.5 | 4 | 1 | 13 | 0.6154 | 1 | 1 | 2 | 1 | 5 | 0.2 | 1 | 0 | 9 | 0.4444 |
| 5_1183314 | 2 | 0.5 | 8 | 0.25 | 1 | 0 | 5 | 0.4 | 16 | 0.3125 | 1 | 1 | 4 | 0.5 | 6 | 0 | 3 | 0.6667 | 14 | 0.3571 |
| 5_1183315 | 2 | 1 | 7 | 0.8571 | 1 | 1 | 5 | 1 | 15 | 0.9333 | 1 | 1 | 5 | 1 | 5 | 0.8 | 3 | 1 | 14 | 0.9286 |
| 5_1183752 | 2 | 0.5 | 3 | 0 | NA | NA | 3 | 0 | 8 | 0.125 | NA | NA | 3 | 0 | 3 | 0 | 2 | 0.5 | 8 | 0.125 |
| 5_1183994 | 2 | 1 | 1 | 1 | 1 | 0 | NA | NA | 4 | 0.75 | 3 | 0.6667 | 1 | 1 | 2 | 0.5 | NA | NA | 6 | 0.6667 |
| 5_1183994 | 2 | 0.5 | 1 | 1 | 1 | 0 | NA | NA | 4 | 0.5 | 3 | 0.6667 | 1 | 0 | 2 | 1 | 2 | 1 | 8 | 0.75 |
| 5_1183996 | NA | NA | 1 | 1 | NA | NA | NA | NA | 1 | 1 | 2 | 0.5 | 1 | 1 | NA | NA | 2 | 1 | 5 | 0.8 |
| 5_1183999 | NA | NA | 1 | 1 | NA | NA | NA | NA | 1 | 1 | 2 | 1 | 1 | 0 | 1 | 1 | 1 | 0 | 5 | 0.6 |
| 5_1184001 | NA | NA | 1 | 1 | NA | NA | NA | NA | 1 | 1 | 1 | 1 | NA | NA | 1 | 1 | NA | NA | 2 | 1 |
| 5_1184004 | NA | NA | 1 | 1 | NA | NA | NA | NA | 1 | 1 | 1 | 1 | NA | NA | NA | NA | 1 | 1 | 2 | 1 |
| 5_1184005 | NA | NA | 1 | 1 | NA | NA | NA | NA | 1 | 1 | 2 | 1 | NA | NA | 1 | 1 | 1 | 1 | 4 | 1 |
| 5_1184006 | NA | NA | 1 | 1 | NA | NA | NA | NA | 1 | 1 | 2 | 1 | NA | NA | 1 | 1 | 1 | 1 | 4 | 1 |
| 5_1184009 | NA | NA | 1 | 1 | NA | NA | NA | NA | 1 | 1 | 2 | 0.5 | NA | NA | 1 | 1 | 1 | 0 | 4 | 0.5 |
| 5_1184010 | NA | NA | 1 | 1 | NA | NA | 1 | 1 | 2 | 1 | 2 | 0.5 | NA | NA | NA | NA | 1 | 1 | 3 | 0.6667 |
| 5_1184036 | 3 | 0.6667 | NA | NA | 1 | 0 | 1 | 1 | 5 | 0.6 | NA | NA | NA | NA | 1 | 1 | 4 | 0.75 | 5 | 0.8 |
| 5_1184037 | 4 | 0 | NA | NA | NA | NA | 1 | 1 | 5 | 0.2 | NA | NA | 2 | 0 | 1 | 0 | 2 | 0.5 | 5 | 0.2 |
| 5_1185410 | NA | NA | 2 | 1 | NA | NA | NA | NA | 2 | 1 | NA | NA | 1 | 1 | NA | NA | NA | NA | 1 | 1 |
| 5_1188662 | 1 | 1 | NA | NA | NA | NA | 1 | 1 | 2 | 1 | 2 | 0.5 | 1 | 0 | NA | NA | NA | NA | 3 | 0.3333 |
| 5_1188663 | 1 | 0 | NA | NA | NA | NA | 1 | 1 | 2 | 0.5 | 2 | 1 | 1 | 1 | NA | NA | NA | NA | 3 | 1 |
| 5_1188666 | 1 | 0 | NA | NA | NA | NA | 1 | 1 | 2 | 0.5 | 2 | 0.5 | 1 | 1 | NA | NA | NA | NA | 3 | 0.6667 |
| 5_1188667 | NA | NA | NA | NA | NA | NA | 1 | 1 | 1 | 1 | 1 | 1 | 1 | 0 | NA | NA | NA | NA | 2 | 0.5 |
| 5_1188800 | 3 | 1 | NA | NA | NA | NA | 1 | 1 | 4 | 1 | 1 | 1 | 1 | 1 | 1 | 1 | 1 | 1 | 4 | 1 |
| 5_1188800 | 4 | 0.75 | NA | NA | NA | NA | 1 | 0 | 5 | 0.6 | 1 | 1 | 3 | 0 | 1 | 1 | 1 | 0 | 6 | 0.3333 |
| 5_1188802 | 3 | 1 | NA | NA | NA | NA | 1 | 1 | 4 | 1 | NA | NA | 3 | 0.6667 | 1 | 1 | 1 | 1 | 5 | 0.8 |
| 5_1188802 | 3 | 1 | NA | NA | NA | NA | NA | NA | 3 | 1 | NA | NA | NA | NA | 1 | 1 | 1 | 0 | 2 | 0.5 |
| 5_1188803 | 3 | 0.3333 | NA | NA | NA | NA | NA | NA | 3 | 0.3333 | 1 | 0 | 1 | 0 | NA | NA | 1 | 1 | 3 | 0.3333 |
| 5_1188804 | 4 | 0.25 | NA | NA | NA | NA | 1 | 0 | 5 | 0.2 | 2 | 0.5 | 3 | 0.3333 | 1 | 0 | 1 | 0 | 7 | 0.2857 |
| 5_1188806 | 4 | 1 | NA | NA | NA | NA | 1 | 1 | 5 | 1 | 1 | 1 | 3 | 1 | 1 | 1 | 1 | 1 | 6 | 1 |

| | | | | | | | | | | | | | | | | | | | | |
|---|---|---|---|---|---|---|---|---|---|---|---|---|---|---|---|---|---|---|---|---|
| 5_1188807 | 4 | 1 | NA | NA | NA | NA | 1 | 1 | 5 | 1 | 1 | 1 | 3 | 0.6667 | 1 | 1 | 1 | 1 | 6 | 0.8333 |
| 5_1189008 | 3 | 0.6667 | NA | NA | 1 | 0 | 2 | 0.5 | 6 | 0.5 | 2 | 0.5 | 6 | 0 | 1 | 0 | 2 | 0.5 | 11 | 0.1818 |
| 5_1189010 | 5 | 1 | 2 | 1 | 1 | 1 | 3 | 1 | 11 | 1 | 5 | 1 | 7 | 1 | 2 | 1 | 3 | 1 | 17 | 1 |
| 5_1189010 | 5 | 1 | 4 | 0.75 | 1 | 1 | 2 | 0.5 | 12 | 0.8333 | 5 | 1 | 6 | 0.6667 | 2 | 1 | 3 | 0.6667 | 16 | 0.8125 |
| 5_1189011 | 4 | 1 | 3 | 1 | NA | NA | 2 | 1 | 9 | 1 | 5 | 1 | 6 | 1 | 2 | 1 | 2 | 1 | 15 | 1 |
| 5_1189011 | 5 | 0.4 | 4 | 0.25 | 1 | 1 | 3 | 0.3333 | 13 | 0.3846 | 4 | 0.25 | 7 | 0 | 2 | 0.5 | 3 | 0.3333 | 16 | 0.1875 |
| 5_1189013 | 4 | 0.5 | 1 | 0 | 4 | 0.25 | 3 | 0.3333 | 12 | 0.3333 | 3 | 0.3333 | 8 | 0.125 | 1 | 1 | 5 | 0 | 17 | 0.1765 |
| 5_1189014 | 5 | 0 | 3 | 0.3333 | 4 | 0 | 2 | 0 | 14 | 0.0714 | 5 | 0 | 6 | 0.1667 | 1 | 0 | 4 | 0 | 16 | 0.0625 |
| 5_1189014 | 5 | 0.2 | 3 | 0.6667 | 3 | 0 | 2 | 0 | 13 | 0.2308 | 6 | 0 | 7 | 0.4286 | 2 | 0.5 | 5 | 0.2 | 20 | 0.25 |
| 5_1189016 | 6 | 0 | 4 | 0.25 | 4 | 0 | 2 | 0 | 16 | 0.0625 | 6 | 0.1667 | 6 | 0.1667 | 2 | 0 | 4 | 0 | 18 | 0.1111 |
| 5_1189017 | 6 | 0.3333 | 3 | 0.3333 | 4 | 0.25 | 2 | 0.5 | 15 | 0.3333 | 6 | 0.5 | 7 | 0.1429 | 2 | 0.5 | 5 | 0.4 | 20 | 0.35 |
| 5_1189018 | 5 | 0.6 | 4 | 1 | 2 | 1 | 2 | 0.5 | 13 | 0.7692 | 6 | 0.8333 | 8 | 0.625 | 2 | 1 | 3 | 1 | 19 | 0.7895 |
| 5_1189018 | 7 | 0.1429 | 4 | 0 | 2 | 0 | 2 | 0 | 15 | 0.0667 | 6 | 0.1667 | 7 | 0.1429 | 2 | 0.5 | 4 | 0.25 | 19 | 0.2105 |
| 5_1189019 | 5 | 0.6 | 4 | 1 | 2 | 1 | 1 | 1 | 12 | 0.8333 | 6 | 0.5 | 8 | 0.75 | 2 | 1 | 5 | 1 | 21 | 0.7619 |
| 5_1189021 | 6 | 0.1667 | 4 | 0.25 | 4 | 0 | 3 | 0 | 17 | 0.1176 | 5 | 0 | 6 | 0.1667 | 1 | 0 | 6 | 0 | 18 | 0.0556 |
| 5_1189021 | 7 | 0.7143 | 4 | 1 | 5 | 0.8 | 3 | 0 | 19 | 0.6842 | 5 | 0.8 | 7 | 0.7143 | 2 | 1 | 6 | 0.5 | 20 | 0.7 |
| 5_1189023 | 7 | 0.1429 | 3 | 0 | 6 | 0 | 2 | 0 | 18 | 0.0556 | 5 | 0 | 7 | 0 | 2 | 0 | 5 | 0.2 | 19 | 0.0526 |
| 5_1189033 | 10 | 0.8 | 8 | 0.875 | 12 | 0.8333 | 10 | 0.8 | 40 | 0.825 | 9 | 0.6667 | 10 | 0.6 | 8 | 0.5 | 10 | 0.7 | 37 | 0.6216 |
| 5_1198190 | 1 | 0 | 1 | 1 | NA | NA | NA | NA | 2 | 0.5 | 2 | 0.5 | NA | NA | NA | NA | NA | NA | 2 | 0.5 |
| 5_1198209 | 2 | 1 | 1 | 1 | 1 | 1 | NA | NA | 4 | 1 | 2 | 1 | NA | NA | NA | NA | NA | NA | 2 | 1 |
| 5_1198209 | 2 | 1 | NA | NA | 1 | 1 | NA | NA | 3 | 1 | 2 | 1 | NA | NA | NA | NA | NA | NA | 2 | 1 |
| 5_1198213 | 1 | 1 | 1 | 0 | 1 | 1 | NA | NA | 3 | 0.6667 | 2 | 1 | NA | NA | NA | NA | NA | NA | 2 | 1 |
| 5_1198214 | 1 | 1 | NA | NA | 1 | 1 | NA | NA | 2 | 1 | 2 | 1 | NA | NA | NA | NA | NA | NA | 2 | 1 |
| 5_1198216 | 1 | 1 | 1 | 1 | 1 | 1 | NA | NA | 3 | 1 | 2 | 0.5 | NA | NA | NA | NA | NA | NA | 2 | 0.5 |
| 5_1203361 | 2 | 0 | 3 | 0.6667 | 1 | 0 | NA | NA | 6 | 0.3333 | 1 | 0 | NA | NA | 1 | 1 | NA | NA | 2 | 0.5 |
| 5_1203362 | 3 | 0.6667 | 3 | 1 | 1 | 0 | NA | NA | 7 | 0.7143 | 1 | 1 | NA | NA | 1 | 1 | NA | NA | 2 | 1 |
| 5_1203363 | 2 | 1 | 1 | 1 | 1 | 0 | NA | NA | 4 | 0.75 | 1 | 1 | NA | NA | 1 | 1 | NA | NA | 2 | 1 |
| 5_1203363 | 3 | 0.6667 | 3 | 0.6667 | 1 | 1 | NA | NA | 7 | 0.7143 | 1 | 1 | NA | NA | 1 | 1 | NA | NA | 2 | 1 |
| 5_1203364 | 2 | 1 | 2 | 0.5 | NA | NA | NA | NA | 4 | 0.75 | 1 | 0 | NA | NA | 1 | 1 | NA | NA | 2 | 0.5 |
| 5_1203366 | 2 | 1 | 3 | 0.6667 | NA | NA | NA | NA | 5 | 0.8 | 1 | 1 | NA | NA | 1 | 1 | NA | NA | 2 | 1 |
| 5_1203366 | 2 | 0 | 3 | 0.3333 | 1 | 1 | NA | NA | 6 | 0.3333 | 1 | 0 | NA | NA | 1 | 1 | NA | NA | 2 | 0.5 |
| 5_1203367 | 3 | 0.3333 | 3 | 0.3333 | 1 | 0 | NA | NA | 7 | 0.2857 | 1 | 0 | NA | NA | 1 | 1 | NA | NA | 2 | 0.5 |
| 5_1203368 | 2 | 1 | 2 | 1 | NA | NA | NA | NA | 4 | 1 | NA | NA | NA | NA | 1 | 1 | NA | NA | 1 | 1 |
| 5_1203369 | 2 | 1 | 2 | 1 | NA | NA | NA | NA | 4 | 1 | 2 | 1 | NA | NA | NA | NA | NA | NA | 2 | 1 |
| 5_1203370 | 2 | 1 | 2 | 1 | NA | NA | NA | NA | 4 | 1 | 2 | 0.5 | NA | NA | 1 | 1 | NA | NA | 3 | 0.6667 |
| 5_1206221 | 2 | 0.5 | 2 | 0 | 2 | 0 | 4 | 0.25 | 10 | 0.2 | 4 | 0 | 4 | 0.25 | 6 | 0.1667 | 1 | 0 | 15 | 0.1333 |
| 5_1206223 | 2 | 0.5 | 3 | 0.3333 | NA | NA | 4 | 0.25 | 9 | 0.3333 | 4 | 0.5 | 3 | 0.3333 | 4 | 0 | NA | NA | 11 | 0.2727 |
| 5_1206232 | 2 | 0.5 | 6 | 0.6667 | 2 | 0 | 1 | 1 | 11 | 0.5455 | 5 | 0.8 | 3 | 0 | 2 | 0 | NA | NA | 10 | 0.4 |
| 5_1206234 | 3 | 1 | 8 | 0.75 | 2 | 0.5 | 2 | 1 | 15 | 0.8 | 8 | 0.625 | 6 | 0.8333 | 3 | 0 | 1 | 0 | 18 | 0.5556 |
| 5_1206234 | NA | NA | 5 | 0.8 | 1 | 1 | 5 | 1 | 11 | 0.9091 | 5 | 0.6 | 4 | 0.75 | 3 | 0.3333 | 2 | 0.5 | 14 | 0.5714 |
| 5_1206236 | 2 | 0.5 | 6 | 0.3333 | 2 | 0 | 5 | 0.6 | 15 | 0.4 | 7 | 0.5714 | 5 | 0.4 | 2 | 0 | 1 | 1 | 15 | 0.4667 |
| 5_1206236 | 1 | 1 | 5 | 0.6 | 3 | 1 | 5 | 0.6 | 14 | 0.7143 | 8 | 0.625 | 5 | 0.6 | 2 | 1 | 1 | 0 | 16 | 0.625 |
| 5_1206237 | 1 | 1 | 6 | 0.8333 | 3 | 0.6667 | 5 | 0.8 | 15 | 0.8 | 8 | 0.875 | 6 | 0.5 | 3 | 0.6667 | 1 | 0 | 18 | 0.6667 |
| 5_1206726 | 5 | 0 | 7 | 0 | 4 | 0 | 4 | 0.25 | 20 | 0.05 | 6 | 0.1667 | 2 | 0 | 4 | 0 | 5 | 0 | 17 | 0.0588 |
| 5_1206740 | 2 | 0.5 | 2 | 0 | NA | NA | 3 | 1 | 7 | 0.5714 | 2 | 0.5 | 2 | 0 | 3 | 0.6667 | 2 | 1 | 9 | 0.5556 |
| 5_1207472 | 1 | 0 | 3 | 0.6667 | 2 | 0 | 1 | 0 | 7 | 0.2857 | 2 | 0 | 2 | 0 | 2 | 0.5 | 1 | 1 | 7 | 0.2857 |
| 5_1207474 | 1 | 0 | 1 | 0 | 2 | 0.5 | NA | NA | 4 | 0.25 | 1 | 1 | 1 | 0 | 2 | 0.5 | 2 | 1 | 6 | 0.6667 |
| 5_1207495 | NA | NA | NA | NA | 1 | 1 | 1 | 0 | 2 | 0.5 | 2 | 1 | 1 | 0 | NA | NA | NA | NA | 3 | 0.6667 |
| 5_1207597 | 4 | 0.25 | 3 | 0 | 2 | 0 | 1 | 0 | 10 | 0.1 | 3 | 0 | 3 | 0.3333 | 3 | 0 | 1 | 0 | 10 | 0.1 |
| 5_1207654 | 4 | 1 | 4 | 0.5 | 4 | 0.75 | 3 | 1 | 15 | 0.8 | 6 | 0.8333 | 6 | 0.6667 | 4 | 1 | 5 | 0.8 | 21 | 0.8095 |
| 5_1207655 | 3 | 0.6667 | 6 | 0.5 | 3 | 0.6667 | 3 | 0.3333 | 15 | 0.5333 | 3 | 0.3333 | 4 | 0.75 | 2 | 0 | 3 | 0.6667 | 12 | 0.5 |
| 5_1207655 | 4 | 1 | 5 | 1 | 4 | 1 | 1 | 1 | 14 | 1 | 5 | 1 | 6 | 1 | 3 | 1 | 3 | 1 | 17 | 1 |
| 5_1207656 | 3 | 1 | 6 | 0.5 | 3 | 1 | 2 | 1 | 14 | 0.7857 | 5 | 0.8 | 6 | 1 | 4 | 0.75 | 5 | 0.6 | 20 | 0.8 |
| 5_1208171 | 4 | 0.25 | 3 | 0 | 2 | 0.5 | 3 | 1 | 12 | 0.4167 | 1 | 1 | 5 | 0.4 | 3 | 0 | 6 | 0.5 | 15 | 0.4 |
| 5_1208172 | 4 | 0 | 3 | 0 | 3 | 0 | 4 | 0.25 | 14 | 0.0714 | 1 | 0 | 9 | 0.2222 | 4 | 0 | 5 | 0 | 19 | 0.1053 |
| 5_1208173 | 5 | 0.8 | 3 | 0.3333 | 3 | 0.6667 | 4 | 0.25 | 15 | 0.5333 | 1 | 1 | 8 | 0.75 | 3 | 1 | 6 | 0.6667 | 18 | 0.7778 |
| 5_1208174 | 5 | 0.8 | 4 | 1 | 3 | 1 | 4 | 1 | 16 | 0.9375 | 1 | 1 | 7 | 1 | 4 | 1 | 6 | 0.8333 | 18 | 0.9444 |
| 5_1208174 | 5 | 0.4 | 4 | 0.25 | 2 | 1 | 4 | 0 | 15 | 0.3333 | 1 | 0 | 7 | 0.2857 | 4 | 0.5 | 7 | 0.2857 | 19 | 0.3158 |
| 5_1208175 | 4 | 0.75 | 5 | 1 | 3 | 1 | 4 | 0.75 | 16 | 0.875 | 1 | 1 | 7 | 1 | 3 | 0.3333 | 7 | 0.7143 | 18 | 0.7778 |
| 5_1208185 | 6 | 0.5 | 5 | 0.8 | 1 | 1 | 8 | 0.75 | 20 | 0.7 | 2 | 0.5 | 2 | 0 | 6 | 0.6667 | 4 | 0.25 | 14 | 0.4286 |
| 5_1208188 | 6 | 0.1667 | 2 | 0 | 1 | 0 | 5 | 0 | 14 | 0.0714 | 2 | 1 | 2 | 0 | 4 | 0.25 | 4 | 0.25 | 12 | 0.3333 |
| 5_1208189 | 5 | 0.2 | 3 | 0 | 1 | 0 | 7 | 0 | 16 | 0.0625 | 3 | 0 | 4 | 0.5 | 5 | 0.2 | 4 | 0 | 16 | 0.1875 |
| 5_1208189 | 6 | 0.5 | 5 | 0.2 | 1 | 0 | 7 | 0.5714 | 19 | 0.4211 | 4 | 0.25 | 5 | 0.2 | 5 | 0.2 | 4 | 0.25 | 18 | 0.2222 |
| 5_1208326 | 3 | 0 | 4 | 0.5 | NA | NA | 6 | 0.5 | 13 | 0.3846 | 1 | 0 | NA | NA | 3 | 0.3333 | 1 | 1 | 5 | 0.4 |
| 5_1208329 | 4 | 0 | 4 | 0 | NA | NA | 6 | 0.5 | 14 | 0.2143 | 1 | 0 | NA | NA | 3 | 0.3333 | 3 | 0.3333 | 7 | 0.2857 |

| | | | | | | | | | | | | | | | | | | | | |
|---|---|---|---|---|---|---|---|---|---|---|---|---|---|---|---|---|---|---|---|---|
| 5_1208330( | 3 | 0.6667 | 3 | 0.3333 | NA | NA | 5 | 0.2 | 11 | 0.3636 | 1 | 0 | NA | NA | 2 | 0 | 1 | 1 | 4 | 0.25 |
| 5_1208486: | 3 | 0.3333 | 2 | 0.5 | NA | NA | 2 | 0.5 | 7 | 0.4286 | NA | NA | NA | NA | 1 | 1 | 1 | 0 | 2 | 0.5 |
| 5_1208486: | 3 | 0.3333 | 1 | 1 | NA | NA | 2 | 0 | 6 | 0.3333 | NA | NA | NA | NA | 2 | 0 | 2 | 0.5 | 4 | 0.25 |
| 5_1208486( | 3 | 0.6667 | 2 | 0 | NA | NA | 3 | 0.6667 | 8 | 0.5 | NA | NA | NA | NA | 2 | 0.5 | 2 | 1 | 4 | 0.75 |
| 5_1208487' | 3 | 0.6667 | 2 | 1 | NA | NA | 1 | 1 | 6 | 0.8333 | NA | NA | NA | NA | 2 | 0.5 | 2 | 0.5 | 4 | 0.5 |
| 5_1208487: | 2 | 0.5 | 2 | 1 | NA | NA | 2 | 0 | 6 | 0.5 | NA | NA | NA | NA | 2 | 0.5 | 2 | 0 | 4 | 0.25 |
| 5_1208487· | 3 | 0.6667 | 2 | 1 | NA | NA | 1 | 0 | 6 | 0.6667 | NA | NA | NA | NA | 2 | 0 | 2 | 0.5 | 4 | 0.25 |
| 5_1210787' | 22 | 0.2273 | 36 | 0.3333 | 37 | 0.2703 | 18 | 0.2222 | 113 | 0.2743 | 24 | 0.1667 | 20 | 0.25 | 23 | 0.2609 | 12 | 0.25 | 79 | 0.2278 |
| 5_1210789( | 20 | 0.1 | 37 | 0.3243 | 29 | 0.3103 | 11 | 0.0909 | 97 | 0.2474 | 25 | 0.16 | 18 | 0.1111 | 19 | 0.1579 | 13 | 0.0769 | 75 | 0.1333 |
| 5_1210789: | 21 | 0.0476 | 34 | 0.0588 | 32 | 0.125 | 15 | 0.1333 | 102 | 0.0882 | 23 | 0.0435 | 19 | 0.0526 | 19 | 0.1579 | 14 | 0.2857 | 75 | 0.12 |
| 5_1210801! | 15 | 0.3333 | 24 | 0.125 | 16 | 0.375 | 13 | 0.3077 | 68 | 0.2647 | 14 | 0.1429 | 13 | 0.2308 | 14 | 0.3571 | 11 | 0.1818 | 52 | 0.2308 |
| 5_1210802: | 15 | 0.0667 | 27 | 0.037 | 19 | 0.1053 | 12 | 0.0833 | 73 | 0.0685 | 19 | 0.2105 | 17 | 0.1176 | 15 | 0 | 10 | 0.1 | 61 | 0.1148 |
| 5_1212700 | 6 | 0.5 | 4 | 0.75 | 5 | 1 | 6 | 0.8333 | 21 | 0.7619 | 4 | 0.75 | 5 | 0.8 | 4 | 0.75 | 8 | 0.875 | 21 | 0.8095 |
| 5_1212700 | 8 | 0.25 | 4 | 0 | 5 | 0.4 | 6 | 0 | 23 | 0.1739 | 4 | 0 | 4 | 0.25 | 4 | 0.5 | 9 | 0.3333 | 21 | 0.2857 |
| 5_1212701( | 7 | 1 | 3 | 1 | 2 | 1 | 5 | 0.8 | 17 | 0.9412 | 4 | 1 | 5 | 1 | 4 | 1 | 7 | 0.8571 | 20 | 0.95 |
| 5_1212702( | 7 | 0.2857 | 4 | 0.25 | 5 | 0.6 | 6 | 0.3333 | 22 | 0.3636 | 4 | 0.25 | 5 | 0.4 | 4 | 0.25 | 9 | 0.2222 | 22 | 0.2727 |
| 5_1212703( | 7 | 0.5714 | 3 | 1 | 3 | 1 | 5 | 0.6 | 18 | 0.7222 | 4 | 0.5 | 4 | 0.75 | 4 | 0.75 | 7 | 0.5714 | 19 | 0.6316 |
| 5_1212707( | 6 | 0.3333 | 3 | 0.3333 | 4 | 0.5 | 4 | 0.25 | 17 | 0.3529 | 3 | 0 | 4 | 0 | 3 | 0.3333 | 5 | 0.2 | 15 | 0.1333 |
| 5_1212712! | 1 | 1 | 2 | 0 | 2 | 0 | 3 | 0.6667 | 8 | 0.375 | 3 | 0.6667 | 1 | 0 | 2 | 0 | NA | NA | 6 | 0.3333 |
| 5_1212719· | 1 | 1 | 1 | 1 | 1 | 1 | NA | NA | 3 | 1 | 2 | 1 | NA | NA | 2 | 0.5 | 2 | 1 | 6 | 0.8333 |
| 5_1212720: | 1 | 0 | 1 | 0 | 1 | 1 | 1 | 0 | 4 | 0.25 | 2 | 0 | NA | NA | 2 | 0.5 | 1 | 0 | 5 | 0.2 |
| 5_1212720: | 1 | 1 | NA | NA | 1 | 0 | 1 | 0 | 3 | 0.3333 | 2 | 1 | NA | NA | 2 | 1 | 2 | 0.5 | 6 | 0.8333 |
| 5_1212721: | 1 | 0 | 1 | 1 | 1 | 0 | 1 | 0 | 4 | 0.25 | 1 | 1 | NA | NA | 2 | 0.5 | 2 | 0.5 | 5 | 0.6 |
| 5_1214869( | 1 | 1 | 6 | 0.5 | 1 | 0 | 4 | 0 | 12 | 0.3333 | 5 | 0.4 | 5 | 0.4 | 2 | 0 | 3 | 0.6667 | 15 | 0.4 |
| 5_1214870' | 2 | 0 | 6 | 0.1667 | 1 | 0 | 4 | 0.25 | 13 | 0.1538 | 5 | 0 | 6 | 0.1667 | 3 | 0 | 4 | 0 | 18 | 0.0556 |
| 5_1214870! | 2 | 0 | 5 | 0 | 1 | 1 | 3 | 0.3333 | 11 | 0.1818 | 5 | 0.6 | 5 | 0.2 | 4 | 0 | 4 | 0.25 | 18 | 0.2778 |
| 5_1214871: | 1 | 1 | 5 | 1 | 1 | 0 | NA | NA | 7 | 0.8571 | 3 | 0.3333 | 6 | 1 | 4 | 1 | 4 | 1 | 17 | 0.8824 |
| 5_1214873: | 2 | 0 | 5 | 0.2 | 1 | 0 | 4 | 0 | 12 | 0.0833 | 5 | 0.2 | 8 | 0.375 | 4 | 0.25 | 5 | 0 | 22 | 0.2273 |
| 5_1214874( | 1 | 0 | 2 | 0 | NA | NA | 3 | 0.6667 | 6 | 0.3333 | 3 | 0.3333 | 4 | 0.75 | 2 | 0 | 2 | 1 | 11 | 0.5455 |
| 5_1214875' | 1 | 1 | 4 | 0.75 | NA | NA | 5 | 0.8 | 10 | 0.8 | 4 | 0.5 | 6 | 0.8333 | 3 | 0.3333 | 4 | 0.75 | 17 | 0.6471 |
| 5_1214875: | 2 | 0.5 | 5 | 0.4 | 1 | 1 | 5 | 0.4 | 13 | 0.4615 | 5 | 0.6 | 7 | 0.7143 | 3 | 0.3333 | 6 | 0.5 | 21 | 0.5714 |
| 5_1214876' | 2 | 1 | 5 | 1 | NA | NA | 4 | 0.75 | 11 | 0.9091 | 4 | 0.75 | 8 | 0.75 | 5 | 1 | 6 | 0.8333 | 23 | 0.8261 |
| 5_1214876 | 1 | 1 | 5 | 0 | 1 | 0 | 4 | 0 | 11 | 0.0909 | 5 | 0 | 8 | 0 | 4 | 0.5 | 5 | 0 | 22 | 0.0909 |
| 5_1215030' | 11 | 0 | 7 | 0 | 10 | 0.1 | 10 | 0.1 | 38 | 0.0526 | 11 | 0.0909 | 13 | 0 | 14 | 0 | 14 | 0.1429 | 52 | 0.0577 |
| 5_1215047! | 4 | 1 | 5 | 1 | 7 | 0.7143 | 6 | 0.8333 | 22 | 0.8636 | 3 | 1 | 11 | 0.8182 | 6 | 0.8333 | 10 | 0.9 | 30 | 0.8667 |
| 5_1215049: | 9 | 0.7778 | 10 | 0.9 | 12 | 0.75 | 15 | 0.7333 | 46 | 0.7826 | 11 | 0.8182 | 12 | 0.8333 | 11 | 0.7273 | 14 | 0.6429 | 48 | 0.75 |
| 5_1215051! | 10 | 0.1 | 11 | 0.1818 | 11 | 0.0909 | 14 | 0.1429 | 46 | 0.1304 | 9 | 0.4444 | 14 | 0.2857 | 14 | 0.2857 | 12 | 0 | 49 | 0.2449 |
| 5_1215051! | 11 | 0.0909 | 10 | 0.2 | 13 | 0.0769 | 17 | 0.1765 | 51 | 0.1373 | 9 | 0.2222 | 11 | 0.1818 | 14 | 0.2857 | 13 | 0.2308 | 47 | 0.234 |
| 5_1215053( | 8 | 0 | 15 | 0.2 | 14 | 0 | 14 | 0.1429 | 51 | 0.098 | 9 | 0.4444 | 14 | 0.0714 | 13 | 0.2308 | 17 | 0 | 53 | 0.1509 |
| 5_1215054' | 11 | 0.0909 | 14 | 0 | 15 | 0 | 19 | 0.1053 | 59 | 0.0508 | 11 | 0 | 13 | 0 | 15 | 0.2 | 16 | 0 | 55 | 0.0545 |
| 5_1215054! | 8 | 0 | 14 | 0.0714 | 14 | 0.1429 | 18 | 0.2222 | 54 | 0.1296 | 9 | 0 | 11 | 0.0909 | 8 | 0.25 | 14 | 0.1429 | 42 | 0.119 |
| 5_1215054! | 11 | 0.1818 | 16 | 0.0625 | 14 | 0.0714 | 18 | 0.1111 | 59 | 0.1017 | 11 | 0.0909 | 16 | 0 | 15 | 0.2 | 16 | 0.125 | 58 | 0.1034 |
| 5_1215055' | 12 | 0.0833 | 14 | 0 | 13 | 0.1538 | 18 | 0 | 57 | 0.0526 | 8 | 0 | 15 | 0 | 13 | 0.1538 | 17 | 0.0588 | 53 | 0.0566 |
| 5_1215056( | 8 | 0.5 | 10 | 0.5 | 11 | 0.3636 | 13 | 0.7692 | 42 | 0.5476 | 9 | 0.7778 | 11 | 0.7273 | 11 | 0.6364 | 16 | 0.625 | 47 | 0.6809 |
| 5_1215056( | 11 | 0.0909 | 18 | 0.1111 | 15 | 0.2 | 15 | 0.4 | 59 | 0.2034 | 11 | 0.3636 | 18 | 0.2222 | 15 | 0.4 | 16 | 0.125 | 60 | 0.2667 |
| 5_1215058: | 11 | 0.7273 | 16 | 0.5 | 13 | 0.1538 | 17 | 0.5882 | 57 | 0.4912 | 10 | 0.8 | 15 | 0.4667 | 15 | 0.8 | 17 | 0.4706 | 57 | 0.614 |
| 5_1215060: | 13 | 0.1538 | 18 | 0.1111 | 16 | 0 | 20 | 0.25 | 67 | 0.1343 | 11 | 0.5455 | 18 | 0.3333 | 18 | 0.2778 | 20 | 0.2 | 67 | 0.3134 |
| 5_1215063( | 16 | 0.8125 | 17 | 0.6471 | 17 | 0.6471 | 22 | 0.8182 | 72 | 0.7361 | 12 | 0.75 | 16 | 0.875 | 18 | 0.8889 | 18 | 0.8889 | 64 | 0.8594 |
| 5_1215099! | 5 | 0.2 | 8 | 0.25 | 9 | 0.5556 | 11 | 0.6364 | 33 | 0.4545 | 3 | 0.6667 | 3 | 1 | 4 | 0.5 | 5 | 0.8 | 15 | 0.7333 |
| 5_1215100! | 8 | 0 | 6 | 0.1667 | 8 | 0.125 | 13 | 0.1538 | 35 | 0.1143 | 4 | 0 | 5 | 0 | 6 | 0 | 5 | 0.2 | 20 | 0.05 |
| 5_1215100! | 6 | 0.3333 | 8 | 0.5 | 7 | 0.2857 | 12 | 0.5833 | 33 | 0.4545 | 4 | 0.5 | 4 | 0.5 | 6 | 0.3333 | 4 | 0.25 | 18 | 0.3889 |
| 5_1215114! | 5 | 0.2 | 5 | 0.2 | 5 | 0 | 6 | 0.1667 | 21 | 0.1429 | 3 | 0 | 3 | 0 | 4 | 0.5 | 3 | 0 | 13 | 0.1538 |
| 5_1215126( | 1 | 1 | 1 | 0 | 2 | 0 | 1 | 0 | 5 | 0.2 | 1 | 1 | 1 | 0 | 1 | 0 | NA | NA | 3 | 0.3333 |
| 5_1215159: | 1 | 0 | 4 | 0.25 | 2 | 0 | 5 | 0 | 12 | 0.0833 | 1 | 0 | 2 | 0 | 5 | 0.4 | 3 | 0 | 11 | 0.1818 |
| 5_1215166' | 3 | 0 | 4 | 0 | 1 | 1 | 8 | 0.125 | 16 | 0.125 | 1 | 0 | 2 | 0 | 4 | 0 | 4 | 0.5 | 11 | 0.1818 |
| 5_1215166 | 4 | 0.25 | 3 | 0 | 1 | 1 | 8 | 0.125 | 16 | 0.1875 | 1 | 1 | 2 | 0 | 3 | 0.3333 | 2 | 0 | 8 | 0.25 |
| 5_1215167: | 3 | 0.3333 | 4 | 0.25 | 2 | 0.5 | 7 | 0.4286 | 16 | 0.375 | 1 | 0 | 2 | 0 | 5 | 0.4 | 4 | 0.25 | 12 | 0.25 |
| 5_1215168( | 4 | 0 | 3 | 0 | 2 | 0 | 4 | 0.25 | 13 | 0.0769 | 1 | 0 | 2 | 0 | 5 | 0 | 4 | 0.25 | 12 | 0.0833 |
| 5_1215774· | 6 | 0 | 6 | 0.1667 | 2 | 0 | 4 | 0 | 18 | 0.0556 | 1 | 0 | 3 | 0 | 4 | 0 | 6 | 0.1667 | 14 | 0.0714 |
| 5_1215774 | 6 | 0 | 7 | 0.4286 | 2 | 0 | 3 | 0 | 18 | 0.1667 | 1 | 0 | 3 | 0.3333 | 4 | 0.25 | 7 | 0 | 15 | 0.1333 |
| 5_1215779' | 5 | 0.4 | 6 | 0.5 | 2 | 0.5 | 4 | 0.25 | 17 | 0.4118 | 1 | 1 | 4 | 0.5 | 4 | 0.5 | 6 | 0.1667 | 15 | 0.4 |
| 5_1215779! | 6 | 0.5 | 6 | 0.6667 | 2 | 0.5 | 3 | 0.6667 | 17 | 0.5882 | 1 | 1 | 4 | 0.5 | 3 | 0.6667 | 7 | 0.2857 | 15 | 0.4667 |
| 5_1215779! | 6 | 0.1667 | 7 | 0 | 2 | 0.5 | 3 | 0 | 18 | 0.1111 | 1 | 0 | 4 | 0 | 4 | 0.25 | 5 | 0.2 | 14 | 0.1429 |
| 5_1215780( | 6 | 0.1667 | 7 | 0.2857 | 2 | 0.5 | 3 | 0.3333 | 18 | 0.2778 | 1 | 1 | 2 | 0 | 4 | 0.25 | 6 | 0.1667 | 13 | 0.2308 |
| 5_1215781' | 6 | 0.5 | 5 | 0.2 | 2 | 0.5 | 3 | 1 | 16 | 0.5 | NA | NA | 2 | 0.5 | 5 | 0.6 | 5 | 0.2 | 12 | 0.4167 |

| | | | | | | | | | | | | | | | | | | | | |
|---|---|---|---|---|---|---|---|---|---|---|---|---|---|---|---|---|---|---|---|---|
| 5_1215781 | 6 | 1 | 7 | 1 | 3 | 1 | 4 | 1 | 20 | 1 | 1 | 1 | 4 | 1 | 5 | 1 | 6 | 1 | 16 | 1 |
| 5_1215782 | 5 | 0.6 | 7 | 0.4286 | 3 | 0.6667 | 4 | 0.5 | 19 | 0.5263 | 1 | 1 | 3 | 0 | 4 | 0.25 | 5 | 0.4 | 13 | 0.3077 |
| 5_1215782 | 4 | 0 | 8 | 0.125 | 3 | 0.6667 | 4 | 0 | 19 | 0.1579 | 1 | 0 | 3 | 0 | 4 | 0.25 | 6 | 0.1667 | 14 | 0.1429 |
| 5_1215783 | 5 | 1 | 8 | 1 | 3 | 1 | 4 | 0.5 | 20 | 0.9 | 2 | 0.5 | 3 | 1 | 4 | 0.75 | 6 | 0.8333 | 15 | 0.8 |
| 5_1215786 | 5 | 0 | 9 | 0 | 4 | 0.5 | 4 | 0 | 22 | 0.0909 | 2 | 0.5 | 3 | 0 | 2 | 0 | 9 | 0 | 16 | 0.0625 |
| 5_1215787 | 5 | 0.4 | 7 | 0.4286 | 4 | 0.75 | 4 | 0.75 | 20 | 0.55 | 2 | 0 | 2 | 0 | 4 | 0 | 8 | 0.25 | 16 | 0.125 |
| 5_1215790 | 4 | 0.5 | 8 | 0.375 | 4 | 0.75 | 4 | 0 | 20 | 0.4 | 2 | 0.5 | 3 | 0.6667 | 4 | 0.5 | 6 | 0.3333 | 15 | 0.4667 |
| 5_1215791 | 3 | 0 | 8 | 0.25 | 4 | 0 | 4 | 0 | 19 | 0.1053 | 2 | 0.5 | 2 | 0 | 4 | 0.25 | 6 | 0 | 14 | 0.1429 |
| 5_1215791 | 3 | 0.3333 | 7 | 0.5714 | 3 | 0.3333 | 3 | 0 | 16 | 0.375 | 1 | 0 | 3 | 0.3333 | 2 | 0.5 | 7 | 0.1429 | 13 | 0.2308 |
| 5_1215792 | 4 | 0.5 | 9 | 0.4444 | 5 | 0.2 | 4 | 0.25 | 22 | 0.3636 | 2 | 0.5 | 3 | 0.6667 | 4 | 0.5 | 7 | 0.2857 | 16 | 0.4375 |
| 5_1215792 | 4 | 0.75 | 8 | 0.5 | 5 | 0.6 | 4 | 0.25 | 21 | 0.5238 | 2 | 0.5 | 3 | 0 | 4 | 0.5 | 5 | 0.6 | 14 | 0.4286 |
| 5_1215795 | 3 | 0.3333 | 8 | 0 | 2 | 0 | 4 | 0 | 17 | 0.0588 | 1 | 0 | 2 | 0 | 5 | 0.2 | 5 | 0 | 13 | 0.0769 |
| 5_1215797 | 4 | 0 | 6 | 0.1667 | 3 | 0 | 1 | 0 | 14 | 0.0714 | 2 | 0.5 | 2 | 0 | 3 | 0 | 1 | 0 | 8 | 0.125 |
| 5_1215801 | 5 | 0.6 | 6 | 0.6667 | 5 | 0.8 | 3 | 0.6667 | 19 | 0.6842 | 2 | 0 | 2 | 0 | 4 | 0.5 | 4 | 0.75 | 12 | 0.4167 |
| 5_1215802 | 6 | 0.1667 | 7 | 0 | 5 | 0.4 | 2 | 0 | 20 | 0.15 | 2 | 0 | 2 | 0 | 5 | 0.6 | 5 | 0.2 | 14 | 0.2857 |
| 5_1215828 | 9 | 0 | 10 | 0.1 | 10 | 0.2 | 18 | 0.1111 | 47 | 0.1064 | 7 | 0.1429 | 18 | 0 | 15 | 0.2 | 18 | 0.0556 | 58 | 0.0862 |
| 5_1215829 | 11 | 0.3636 | 14 | 0.1429 | 16 | 0.1875 | 16 | 0.25 | 57 | 0.2281 | 8 | 0.25 | 23 | 0.3478 | 21 | 0.2381 | 21 | 0.3333 | 73 | 0.3014 |
| 5_1215829 | 12 | 1 | 14 | 0.9286 | 19 | 1 | 20 | 0.85 | 65 | 0.9385 | 9 | 1 | 22 | 1 | 19 | 1 | 20 | 1 | 70 | 1 |
| 5_1215830 | 12 | 0.5833 | 16 | 0.5625 | 18 | 0.5556 | 20 | 0.65 | 66 | 0.5909 | 8 | 0.375 | 23 | 0.4783 | 20 | 0.5 | 22 | 0.5455 | 73 | 0.4932 |
| 5_1215832 | 11 | 1 | 14 | 1 | 17 | 0.8824 | 19 | 0.8421 | 61 | 0.918 | 8 | 0.875 | 20 | 0.8 | 19 | 0.9474 | 19 | 0.8421 | 66 | 0.8636 |
| 5_1215833 | 9 | 0.2222 | 12 | 0.4167 | 10 | 0.6 | 16 | 0.3125 | 47 | 0.383 | 9 | 0.4444 | 15 | 0.3333 | 18 | 0.5556 | 15 | 0.1333 | 57 | 0.3684 |
| 5_1215835 | 9 | 0 | 10 | 0.1 | 11 | 0 | 18 | 0.1111 | 48 | 0.0625 | 8 | 0 | 17 | 0.2353 | 18 | 0.0556 | 18 | 0.2222 | 61 | 0.1475 |
| 5_1215835 | 10 | 0.7 | 11 | 0.7273 | 13 | 0.6923 | 16 | 0.8125 | 50 | 0.74 | 8 | 0.75 | 17 | 0.6471 | 16 | 0.875 | 18 | 0.6667 | 59 | 0.7288 |
| 5_1215912 | 1 | 1 | NA | NA | NA | NA | NA | NA | 1 | 1 | NA | NA | NA | NA | 1 | 1 | NA | NA | 1 | 1 |
| 5_1215927 | 1 | 1 | NA | NA | NA | NA | NA | NA | 1 | 1 | NA | NA | NA | NA | 1 | 1 | NA | NA | 1 | 1 |
| 5_1215927 | 1 | 1 | NA | NA | NA | NA | NA | NA | 1 | 1 | NA | NA | NA | NA | 1 | 1 | NA | NA | 1 | 1 |
| 5_1215934 | 1 | 1 | NA | NA | NA | NA | NA | NA | 1 | 1 | NA | NA | NA | NA | 1 | 1 | NA | NA | 1 | 1 |
| 5_1215939 | 1 | 0 | 1 | 1 | NA | NA | NA | NA | 2 | 0.5 | NA | NA | NA | NA | 2 | 1 | NA | NA | 2 | 1 |
| 5_1215939 | 1 | 1 | 1 | 1 | NA | NA | NA | NA | 2 | 1 | NA | NA | NA | NA | 1 | 1 | NA | NA | 1 | 1 |
| 5_1215940 | 1 | 1 | NA | NA | NA | NA | NA | NA | 1 | 1 | NA | NA | NA | NA | 1 | 1 | NA | NA | 1 | 1 |
| 5_1216571 | 1 | 1 | 2 | 1 | 1 | 1 | NA | NA | 4 | 1 | NA | NA | 1 | 1 | NA | NA | NA | NA | 1 | 1 |
| 5_1217082 | NA | NA | 1 | 0 | 1 | 1 | NA | NA | 2 | 0.5 | NA | NA | 1 | 1 | NA | NA | NA | NA | 1 | 1 |
| 5_1217085 | 1 | 1 | 1 | 0 | 1 | 1 | NA | NA | 3 | 0.6667 | NA | NA | 1 | 1 | NA | NA | NA | NA | 1 | 1 |
| 5_1217366 | NA | NA | NA | NA | 1 | 1 | NA | NA | 1 | 1 | 1 | 1 | NA | NA | NA | NA | 1 | 1 | 2 | 1 |
| 5_1217367 | NA | NA | NA | NA | 1 | 1 | NA | NA | 1 | 1 | 1 | 0 | NA | NA | NA | NA | 1 | 1 | 2 | 0.5 |
| 5_1218033 | 3 | 0.6667 | 2 | 0.5 | 2 | 0 | 4 | 0.75 | 11 | 0.5455 | 4 | 0.75 | 1 | 0 | 2 | 0 | 1 | 0 | 8 | 0.375 |
| 5_1218233 | 3 | 0.3333 | 3 | 0 | 4 | 0 | 4 | 0 | 14 | 0.0714 | 2 | 0 | 4 | 0 | 6 | 0 | 4 | 0.25 | 16 | 0.0625 |
| 5_1218235 | 3 | 0 | 3 | 0 | 4 | 0.25 | 4 | 0 | 14 | 0.0714 | 2 | 0 | 5 | 0 | 6 | 0.1667 | 5 | 0 | 18 | 0.0556 |
| 5_1218235 | 4 | 1 | 3 | 1 | 4 | 1 | 4 | 1 | 15 | 1 | 2 | 1 | 6 | 1 | 4 | 1 | 6 | 1 | 18 | 1 |
| 5_1218235 | 3 | 0.6667 | 3 | 1 | 4 | 0.5 | 2 | 0.5 | 12 | 0.6667 | 2 | 0.5 | 5 | 1 | 6 | 0.5 | 4 | 0.75 | 17 | 0.7059 |
| 5_1218236 | 3 | 0.3333 | 3 | 0.3333 | 4 | 0.5 | 4 | 0.5 | 14 | 0.4286 | 2 | 0.5 | 6 | 0.8333 | 7 | 0.7143 | 6 | 0.6667 | 21 | 0.7143 |
| 5_1218236 | 3 | 1 | 1 | 1 | 2 | 1 | 3 | 1 | 9 | 1 | 1 | 1 | 5 | 0.6 | 5 | 0.8 | 2 | 1 | 13 | 0.7692 |
| 5_1218237 | 3 | 0 | 3 | 0.3333 | 3 | 0 | 4 | 0.5 | 13 | 0.2308 | 2 | 0 | 5 | 0 | 8 | 0 | 3 | 0.3333 | 18 | 0.0556 |
| 5_1218243 | 5 | 0 | 4 | 0.25 | 4 | 0.5 | 5 | 0.4 | 18 | 0.2778 | 2 | 0 | 7 | 0.1429 | 6 | 0.1667 | 6 | 0 | 21 | 0.0952 |
| 5_1218245 | 6 | 0.1667 | 4 | 0.25 | 3 | 0.3333 | 6 | 0 | 19 | 0.1579 | 2 | 0.5 | 7 | 0.2857 | 8 | 0.125 | 6 | 0.1667 | 23 | 0.2174 |
| 5_1218246 | 6 | 0 | 4 | 0.25 | 4 | 0 | 6 | 0 | 20 | 0.05 | 2 | 0 | 7 | 0 | 8 | 0 | 7 | 0.2857 | 24 | 0.0833 |
| 5_1218248 | 4 | 0.75 | 5 | 0 | 5 | 0.2 | 4 | 0.25 | 18 | 0.2778 | 3 | 0 | 6 | 0 | 9 | 0.3333 | 6 | 0.3333 | 24 | 0.2083 |
| 5_1218248 | 4 | 0.25 | 5 | 0.4 | 5 | 0 | 4 | 0 | 18 | 0.1667 | 3 | 0.3333 | 6 | 0.1667 | 8 | 0.125 | 5 | 0 | 22 | 0.1364 |
| 5_1219525 | 6 | 0.3333 | 12 | 0.1667 | 4 | 0 | 7 | 0.2857 | 29 | 0.2069 | 14 | 0.2143 | 6 | 0.3333 | 10 | 0 | 10 | 0.1 | 40 | 0.15 |
| 5_1219525 | 5 | 0.6 | 10 | 0.1 | 4 | 0 | 4 | 0.25 | 23 | 0.2174 | 6 | 0.3333 | 3 | 0.6667 | 5 | 0 | 9 | 0.2222 | 23 | 0.2609 |
| 5_1219526 | 6 | 0.1667 | 13 | 0.3846 | 4 | 0.5 | 6 | 0.5 | 29 | 0.3793 | 13 | 0.5385 | 5 | 0.2 | 10 | 0.7 | 10 | 0.5 | 38 | 0.5263 |
| 5_1219528 | 7 | 0.1429 | 12 | 0.1667 | 4 | 0 | 6 | 0.1667 | 29 | 0.1379 | 12 | 0.25 | 6 | 0.1667 | 10 | 0 | 6 | 0.1667 | 34 | 0.1471 |
| 5_1219542 | 2 | 0 | 1 | 1 | NA | NA | 1 | 1 | 4 | 0.5 | NA | NA | 4 | 0.25 | 1 | 0 | NA | NA | 5 | 0.2 |
| 5_1219550 | 1 | 1 | NA | NA | NA | NA | 2 | 1 | 3 | 1 | 1 | 1 | 3 | 0.6667 | 2 | 1 | NA | NA | 6 | 0.8333 |
| 5_1219553 | 4 | 0.25 | 4 | 0.25 | 1 | 0 | 5 | 0.2 | 14 | 0.2143 | 3 | 0 | 7 | 0.2857 | 6 | 0.1667 | 4 | 0 | 20 | 0.15 |
| 5_1219555 | 4 | 0 | 2 | 0 | NA | NA | 5 | 0.2 | 11 | 0.0909 | 3 | 0.6667 | 7 | 0.2857 | 6 | 0.1667 | 3 | 0.3333 | 19 | 0.3158 |
| 5_1219558 | 3 | 0.6667 | 1 | 0 | NA | NA | 2 | 0.5 | 6 | 0.5 | 2 | 0.5 | 4 | 0.5 | 4 | 0 | 1 | 0 | 11 | 0.2727 |
| 5_1219559 | 4 | 1 | 3 | 1 | 1 | 1 | 3 | 1 | 11 | 1 | 2 | 1 | 4 | 0.75 | 5 | 1 | 2 | 1 | 13 | 0.9231 |
| 5_1219559 | 4 | 1 | 2 | 0.5 | 1 | 0 | 3 | 0.6667 | 10 | 0.7 | 3 | 0.6667 | 4 | 0.5 | 4 | 0.75 | 2 | 0 | 13 | 0.5385 |
| 5_1219559 | 3 | 0.3333 | 2 | 0 | NA | NA | 4 | 0.5 | 9 | 0.3333 | 3 | 0.6667 | 3 | 0.3333 | 5 | 0.4 | 2 | 0.5 | 13 | 0.4615 |
| 5_1219665 | 2 | 0 | 1 | 1 | 1 | 1 | 1 | 1 | 5 | 0.6 | 3 | 0.3333 | 5 | 1 | 2 | 0.5 | 1 | 1 | 11 | 0.7273 |
| 5_1219666 | 2 | 0 | 1 | 1 | 1 | 1 | 1 | 1 | 5 | 0.6 | 3 | 0 | 4 | 0.25 | 3 | 0.3333 | 2 | 0 | 12 | 0.1667 |
| 5_1219910 | 1 | 1 | NA | NA | NA | NA | 2 | 0 | 3 | 0.3333 | NA | NA | NA | NA | 2 | 0.5 | NA | NA | 2 | 0.5 |
| 5_1219910 | NA | NA | NA | NA | NA | NA | 2 | 1 | 2 | 1 | NA | NA | NA | NA | 2 | 1 | 1 | 1 | 3 | 1 |
| 5_1219912 | 2 | 1 | NA | NA | NA | NA | 2 | 1 | 4 | 1 | 1 | 1 | NA | NA | 2 | 1 | 1 | 1 | 4 | 1 |

| | | | | | | | | | | | | | | | | | | | | |
|---|---|---|---|---|---|---|---|---|---|---|---|---|---|---|---|---|---|---|---|---|
| 5_1219912 | 2 | 0.5 | NA | NA | NA | NA | 2 | 1 | 4 | 0.75 | 1 | 1 | NA | NA | 2 | 0 | 1 | 1 | 4 | 0.5 |
| 5_1219913 | 1 | 1 | NA | NA | NA | NA | 2 | 0 | 3 | 0.3333 | NA | NA | NA | NA | 2 | 0.5 | 1 | 0 | 3 | 0.3333 |
| 5_1219913 | 2 | 1 | NA | NA | NA | NA | 2 | 1 | 4 | 1 | 1 | 0 | NA | NA | 2 | 1 | 1 | 1 | 4 | 0.75 |
| 5_1219914 | 2 | 0.5 | NA | NA | NA | NA | 3 | 0.6667 | 5 | 0.6 | 1 | 1 | NA | NA | 2 | 1 | 1 | 0 | 4 | 0.75 |
| 5_1219914 | 3 | 1 | NA | NA | NA | NA | 3 | 1 | 6 | 1 | 1 | 1 | NA | NA | 1 | 1 | NA | NA | 2 | 1 |
| 5_1219915 | 3 | 0.3333 | NA | NA | NA | NA | 3 | 0 | 6 | 0.1667 | 1 | 0 | NA | NA | 1 | 1 | 1 | 0 | 3 | 0.3333 |
| 5_1219915 | 2 | 0 | NA | NA | NA | NA | 2 | 1 | 4 | 0.5 | NA | NA | NA | NA | 1 | 1 | 1 | 1 | 2 | 1 |
| 5_1219915 | 4 | 0.5 | NA | NA | NA | NA | 3 | 0.3333 | 7 | 0.4286 | 1 | 1 | NA | NA | 2 | 0 | 1 | 0 | 4 | 0.25 |
| 5_1219917 | 4 | 1 | NA | NA | NA | NA | 3 | 1 | 7 | 1 | 1 | 1 | NA | NA | 2 | 0.5 | 1 | 1 | 4 | 0.75 |
| 5_1219917 | 3 | 1 | NA | NA | NA | NA | 2 | 1 | 5 | 1 | 1 | 1 | NA | NA | 2 | 1 | 1 | 1 | 4 | 1 |
| 5_1219917 | 4 | 1 | NA | NA | NA | NA | 3 | 1 | 7 | 1 | 1 | 1 | NA | NA | 2 | 1 | 1 | 1 | 4 | 1 |
| 5_1219918 | 4 | 0.25 | NA | NA | NA | NA | 3 | 0 | 7 | 0.1429 | 1 | 0 | NA | NA | 2 | 0.5 | 1 | 0 | 4 | 0.25 |
| 5_1219919 | 4 | 0.5 | NA | NA | 1 | 0 | 3 | 0.3333 | 8 | 0.375 | 1 | 0 | NA | NA | 1 | 1 | 1 | 1 | 3 | 0.6667 |
| 5_1219964 | 2 | 0 | 2 | 0 | 2 | 0.5 | 2 | 0 | 8 | 0.125 | 2 | 0 | 1 | 0 | 7 | 0.1429 | 7 | 0.2857 | 17 | 0.1765 |
| 5_1219965 | 3 | 1 | 3 | 0.6667 | 1 | 1 | 2 | 1 | 9 | 0.8889 | 2 | 1 | 1 | 1 | 5 | 1 | 8 | 1 | 16 | 1 |
| 5_1219968 | 3 | 0.3333 | 2 | 0 | 2 | 0.5 | NA | NA | 7 | 0.2857 | 2 | 0 | 1 | 0 | 7 | 0 | 6 | 0.3333 | 16 | 0.125 |
| 5_1219968 | 2 | 0 | 3 | 0.3333 | 2 | 0.5 | 1 | 1 | 8 | 0.375 | 2 | 1 | 1 | 0 | 6 | 0.1667 | 5 | 0.6 | 14 | 0.4286 |
| 5_1219968 | 3 | 0.3333 | 3 | 0 | 2 | 0.5 | NA | NA | 8 | 0.25 | 1 | 0 | NA | NA | 5 | 0.2 | 6 | 0.3333 | 12 | 0.25 |
| 5_1219970 | 1 | 1 | 2 | 1 | 1 | 1 | 1 | 1 | 5 | 1 | 1 | 1 | 1 | 1 | 4 | 1 | 4 | 1 | 10 | 1 |
| 5_1219970 | 1 | 1 | 1 | 0 | 1 | 0 | 1 | 1 | 4 | 0.5 | 1 | 1 | 1 | 1 | 6 | 0.8333 | 6 | 0.6667 | 14 | 0.7857 |
| 5_1219971 | 2 | 0.5 | 1 | 1 | 2 | 1 | 1 | 1 | 6 | 0.8333 | 2 | 1 | 1 | 1 | 7 | 0.7143 | 5 | 0.6 | 15 | 0.7333 |
| 5_1219971 | 1 | 1 | 1 | 0 | 1 | 1 | 1 | 1 | 4 | 0.75 | 2 | 1 | 1 | 1 | 5 | 0.6 | 5 | 0.6 | 13 | 0.6923 |
| 5_1219971 | 1 | 1 | 2 | 1 | 2 | 1 | 1 | 0 | 6 | 0.8333 | 2 | 0.5 | 1 | 1 | 6 | 0.6667 | 5 | 0.8 | 14 | 0.7143 |
| 5_1219974 | NA | NA | 2 | 0.5 | 1 | 1 | 1 | 1 | 4 | 0.75 | 2 | 0 | 1 | 0 | 5 | 0.2 | 4 | 0.25 | 12 | 0.1667 |
| 5_1219976 | NA | NA | 1 | 1 | NA | NA | 1 | 0 | 2 | 0.5 | 2 | 0.5 | 1 | 0 | 3 | 0.3333 | 4 | 0.25 | 10 | 0.3 |
| 5_1219977 | NA | NA | 2 | 1 | 1 | 1 | NA | NA | 3 | 1 | 2 | 1 | 1 | 1 | 4 | 0.75 | 2 | 0.5 | 9 | 0.7778 |
| 5_1219979 | NA | NA | 1 | 1 | NA | NA | 1 | 1 | 2 | 1 | 2 | 1 | 1 | 0 | 1 | 0 | 2 | 0.5 | 6 | 0.5 |
| 5_1219979 | NA | NA | 2 | 0.5 | NA | NA | 1 | 1 | 3 | 0.6667 | 2 | 1 | 1 | 1 | 4 | 1 | 4 | 1 | 11 | 1 |
| 5_1219980 | NA | NA | 2 | 1 | NA | NA | 1 | 1 | 3 | 1 | 2 | 1 | 1 | 1 | 4 | 0.5 | 4 | 1 | 11 | 0.8182 |
| 5_1219983 | NA | NA | 1 | 0 | NA | NA | 1 | 1 | 2 | 0.5 | 2 | 0 | 2 | 0 | 4 | 0.25 | 3 | 0.6667 | 11 | 0.2727 |
| 5_1219983 | NA | NA | 3 | 0.3333 | NA | NA | NA | NA | 3 | 0.3333 | 1 | 0 | 3 | 0 | 3 | 0 | 4 | 0.25 | 11 | 0.0909 |
| 5_1219983 | NA | NA | 2 | 0.5 | NA | NA | 1 | 0 | 3 | 0.3333 | 1 | 0 | 3 | 0.6667 | 3 | 0.3333 | 2 | 0.5 | 9 | 0.4444 |
| 5_1219985 | NA | NA | 3 | 0.6667 | NA | NA | 1 | 0 | 4 | 0.5 | 1 | 0 | 3 | 0.3333 | 4 | 0.25 | 4 | 0.5 | 12 | 0.3333 |
| 5_1219985 | NA | NA | 3 | 0.6667 | NA | NA | 1 | 1 | 4 | 0.75 | 1 | 1 | 3 | 1 | 4 | 1 | 3 | 1 | 11 | 1 |
| 5_1219986 | NA | NA | 3 | 0.3333 | NA | NA | 1 | 1 | 4 | 0.5 | 1 | 0 | 3 | 0.3333 | 4 | 0.5 | 3 | 0.3333 | 11 | 0.3636 |
| 5_1219987 | NA | NA | 2 | 1 | NA | NA | NA | NA | 2 | 1 | NA | NA | 1 | 1 | 3 | 1 | 3 | 0.6667 | 7 | 0.8571 |
| 5_1219987 | NA | NA | 1 | 1 | NA | NA | 1 | 0 | 2 | 0.5 | 1 | 0 | 2 | 0 | 4 | 0.25 | 3 | 0 | 10 | 0.1 |
| 5_1219989 | NA | NA | 1 | 1 | NA | NA | NA | NA | 1 | 1 | 1 | 1 | 2 | 1 | 2 | 1 | 2 | 0.5 | 7 | 0.8571 |
| 5_1219990 | NA | NA | 3 | 0.3333 | NA | NA | NA | NA | 3 | 0.3333 | 1 | 0 | 2 | 0 | 3 | 0 | 1 | 1 | 7 | 0.1429 |
| 5_1219990 | NA | NA | 2 | 1 | NA | NA | 1 | 1 | 3 | 1 | 1 | 0 | 2 | 1 | 3 | 1 | 2 | 0.5 | 8 | 0.75 |
| 5_1219990 | NA | NA | 3 | 0.3333 | NA | NA | 1 | 1 | 4 | 0.5 | 1 | 1 | 2 | 1 | 3 | 0.6667 | 2 | 0.5 | 8 | 0.75 |
| 5_1219991 | NA | NA | 3 | 1 | NA | NA | 1 | 1 | 4 | 1 | 1 | 1 | 2 | 0 | 3 | 0.6667 | 2 | 1 | 8 | 0.625 |
| 5_1219991 | NA | NA | 3 | 0 | NA | NA | 1 | 1 | 4 | 0.25 | 1 | 0 | 2 | 0 | 3 | 0.3333 | 2 | 0 | 8 | 0.125 |
| 5_1219993 | NA | NA | 3 | 0.3333 | NA | NA | 1 | 1 | 4 | 0.5 | 1 | 0 | 2 | 0.5 | 4 | 0.75 | 1 | 1 | 8 | 0.625 |
| 5_1220071 | NA | NA | 3 | 1 | NA | NA | NA | NA | 3 | 1 | 1 | 1 | NA | NA | 1 | 1 | NA | NA | 2 | 1 |
| 5_1220118 | NA | NA | 1 | 1 | NA | NA | NA | NA | 1 | 1 | 1 | 1 | NA | NA | 1 | 1 | NA | NA | 2 | 1 |
| 5_1220122 | NA | NA | 1 | 1 | NA | NA | NA | NA | 1 | 1 | 1 | 1 | NA | NA | 2 | 1 | NA | NA | 3 | 1 |
| 5_1223070 | 19 | 0.3158 | 20 | 0.2 | 14 | 0.1429 | 14 | 0.0714 | 67 | 0.194 | 9 | 0.2222 | 30 | 0.3667 | 25 | 0.36 | 27 | 0.2963 | 91 | 0.3297 |
| 5_1223122 | 36 | 0.6667 | 34 | 0.7941 | 29 | 0.6897 | 36 | 0.8056 | 135 | 0.7407 | 20 | 0.95 | 46 | 0.7826 | 47 | 0.6596 | 38 | 0.7105 | 151 | 0.7483 |
| 5_1223122 | 36 | 0.1111 | 27 | 0.1111 | 28 | 0.0714 | 31 | 0.1613 | 122 | 0.1148 | 19 | 0.0526 | 42 | 0.1429 | 44 | 0.0682 | 33 | 0.1212 | 138 | 0.1014 |
| 5_1223124 | 37 | 0.0811 | 31 | 0.129 | 26 | 0.1538 | 37 | 0.0541 | 131 | 0.0992 | 16 | 0.0625 | 44 | 0.0682 | 45 | 0.0667 | 35 | 0.1714 | 140 | 0.0929 |
| 5_1223126 | 39 | 0.0769 | 34 | 0.0882 | 29 | 0.1379 | 41 | 0.1463 | 143 | 0.1119 | 23 | 0.087 | 48 | 0.125 | 46 | 0.1304 | 37 | 0.1081 | 154 | 0.1169 |
| 5_1223128 | 26 | 0.5 | 31 | 0.4516 | 16 | 0.5625 | 26 | 0.5769 | 99 | 0.5152 | 14 | 0.4286 | 31 | 0.6774 | 33 | 0.4242 | 24 | 0.5417 | 102 | 0.5294 |
| 5_1223132 | 45 | 0.1556 | 41 | 0.0244 | 28 | 0.1786 | 41 | 0.0976 | 155 | 0.1097 | 22 | 0.0455 | 53 | 0.0755 | 45 | 0.0667 | 36 | 0.0556 | 156 | 0.0641 |
| 5_1223764 | 1 | 1 | NA | NA | NA | NA | NA | NA | 1 | 1 | 1 | 1 | 1 | 1 | 2 | 1 | NA | NA | 4 | 1 |
| 5_1223784 | 1 | 1 | NA | NA | NA | NA | NA | NA | 1 | 1 | 1 | 1 | 2 | 1 | 2 | 1 | 1 | 1 | 6 | 1 |
| 5_1223784 | 1 | 1 | NA | NA | NA | NA | NA | NA | 1 | 1 | 1 | 0 | 3 | 0.3333 | 3 | 0 | 1 | 0 | 8 | 0.125 |
| 5_1223789 | NA | NA | 1 | 1 | NA | NA | 1 | 1 | 2 | 1 | NA | NA | NA | NA | NA | NA | 1 | 1 | 1 | 1 |
| 5_1223795 | NA | NA | NA | NA | 2 | 0 | 1 | 1 | 3 | 0.3333 | 1 | 1 | 2 | 1 | 1 | 1 | 1 | 1 | 5 | 1 |
| 5_1223795 | NA | NA | NA | NA | 2 | 0 | 2 | 1 | 4 | 0.5 | NA | NA | 2 | 1 | 1 | 0 | 1 | 1 | 4 | 0.75 |
| 5_1223796 | NA | NA | NA | NA | 2 | 0.5 | 2 | 1 | 4 | 0.75 | 1 | 1 | 2 | 1 | 2 | 0.5 | 1 | 0 | 6 | 0.6667 |
| 5_1223799 | NA | NA | 1 | 1 | NA | NA | 1 | 0 | 2 | 0.5 | NA | NA | 1 | 0 | 2 | 0.5 | 1 | 0 | 4 | 0.25 |
| 5_1223800 | NA | NA | 1 | 0 | 1 | 0 | 1 | 1 | 3 | 0.3333 | NA | NA | 1 | 1 | 2 | 1 | 1 | 1 | 4 | 1 |
| 5_1223801 | NA | NA | 1 | 1 | NA | NA | 1 | 1 | 2 | 1 | NA | NA | 1 | 1 | 2 | 0.5 | 1 | 0 | 4 | 0.5 |
| 5_1223802 | NA | NA | 1 | 0 | 1 | 0 | 1 | 1 | 3 | 0.3333 | NA | NA | 1 | 1 | NA | NA | 1 | 1 | 2 | 1 |

| | | | | | | | | | | | | | | | | | | | | |
|---|---|---|---|---|---|---|---|---|---|---|---|---|---|---|---|---|---|---|---|---|
| 5_1224228 | 4 | 0.75 | 2 | 0.5 | 1 | 1 | 2 | 0.5 | 9 | 0.6667 | 2 | 1 | 7 | 0.5714 | 6 | 0.5 | 4 | 0.5 | 19 | 0.5789 |
| 5_1224231 | 1 | 1 | NA | NA | NA | NA | NA | NA | 1 | 1 | NA | NA | 1 | 1 | NA | NA | 1 | 1 | 2 | 1 |
| 5_1224235 | 2 | 1 | NA | NA | NA | NA | 1 | 1 | 3 | 1 | NA | NA | 1 | 1 | 1 | 1 | 1 | 1 | 3 | 1 |
| 5_1224262 | 2 | 0 | NA | NA | 1 | 0 | 2 | 0.5 | 5 | 0.2 | NA | NA | 1 | 0 | 1 | 1 | 1 | 0 | 3 | 0.3333 |
| 5_1224265 | 2 | 0 | 1 | 1 | 1 | 0 | 2 | 0.5 | 6 | 0.3333 | NA | NA | 1 | 0 | 1 | 1 | 1 | 0 | 3 | 0.3333 |
| 5_1224462 | 5 | 0 | 3 | 0 | 2 | 0.5 | 1 | 0 | 11 | 0.0909 | 3 | 0 | 6 | 0.3333 | 5 | 0 | 2 | 0 | 16 | 0.125 |
| 5_1224589 | 4 | 1 | 5 | 1 | 4 | 1 | 3 | 1 | 16 | 1 | 5 | 1 | 7 | 0.8571 | 9 | 0.8889 | 6 | 1 | 27 | 0.9259 |
| 5_1224589 | 5 | 0.4 | 5 | 0.4 | 2 | 0 | 2 | 0.5 | 14 | 0.3571 | 4 | 0.5 | 7 | 0.5714 | 9 | 0.2222 | 5 | 0.6 | 25 | 0.44 |
| 5_1224590 | 5 | 0.2 | 5 | 0.4 | 4 | 0.25 | 3 | 0 | 17 | 0.2353 | 2 | 0 | 6 | 0.5 | 7 | 0.2857 | 4 | 0.25 | 19 | 0.3158 |
| 5_1224590 | 5 | 0 | 5 | 0.2 | 3 | 0 | 3 | 0.3333 | 16 | 0.125 | 4 | 0.25 | 8 | 0.125 | 9 | 0.3333 | 6 | 0.3333 | 27 | 0.2593 |
| 5_1224590 | 5 | 0.6 | 4 | 0.25 | 3 | 0 | 3 | 0 | 15 | 0.2667 | 3 | 0.3333 | 7 | 0.1429 | 6 | 0.3333 | 6 | 0.1667 | 22 | 0.2273 |
| 5_1224592 | 4 | 1 | 5 | 1 | 4 | 1 | 4 | 1 | 17 | 1 | 5 | 1 | 6 | 1 | 9 | 0.8889 | 5 | 1 | 25 | 0.96 |
| 5_1224592 | 5 | 0.2 | 5 | 0.4 | 4 | 0.5 | 5 | 0 | 19 | 0.2632 | 4 | 0.25 | 8 | 0.25 | 7 | 0.1429 | 4 | 0.25 | 23 | 0.2174 |
| 5_1224593 | 4 | 0.25 | 6 | 0.1667 | 4 | 0.25 | 4 | 0 | 18 | 0.1667 | 5 | 0.2 | 7 | 0.2857 | 9 | 0.1111 | 7 | 0.1429 | 28 | 0.1786 |
| 5_1224593 | 4 | 1 | 7 | 1 | 4 | 1 | 5 | 1 | 20 | 1 | 6 | 0.8333 | 9 | 1 | 8 | 1 | 6 | 1 | 29 | 0.9655 |
| 5_1224595 | 8 | 0.25 | 5 | 0 | 4 | 0.25 | 5 | 0 | 22 | 0.1364 | 6 | 0 | 8 | 0.375 | 9 | 0.1111 | 5 | 0 | 28 | 0.1429 |
| 5_1224596 | 6 | 0.6667 | 7 | 0.5714 | 4 | 0.5 | 4 | 0.5 | 21 | 0.5714 | 5 | 0.6 | 8 | 0.625 | 5 | 0.4 | 5 | 0.4 | 23 | 0.5217 |
| 5_1224596 | 9 | 0.3333 | 9 | 0.4444 | 5 | 0 | 5 | 0.2 | 28 | 0.2857 | 5 | 1 | 9 | 0.5556 | 9 | 0.3333 | 6 | 0.3333 | 29 | 0.5172 |
| 5_1224596 | 6 | 0.3333 | 2 | 1 | 4 | 0.5 | 4 | 0 | 16 | 0.375 | 5 | 0.6 | 6 | 0.6667 | 7 | 0.2857 | 4 | 0.25 | 22 | 0.4545 |
| 5_1224598 | 3 | 0.3333 | 7 | 0.1429 | 2 | 0.5 | 5 | 0 | 17 | 0.1765 | 3 | 0.3333 | 7 | 0.2857 | 6 | 0.1667 | 4 | 0 | 20 | 0.2 |
| 5_1224600 | 10 | 0 | 7 | 0.1429 | 4 | 0 | 6 | 0.3333 | 27 | 0.1111 | 6 | 0.3333 | 8 | 0.25 | 9 | 0.1111 | 5 | 0 | 28 | 0.1786 |
| 5_1224602 | 6 | 0 | 8 | 0.25 | 4 | 0 | 7 | 0.2857 | 25 | 0.16 | 5 | 0.4 | 9 | 0.3333 | 6 | 0.1667 | 4 | 0 | 24 | 0.25 |
| 5_1224607 | 8 | 0.125 | 9 | 0 | 7 | 0.1429 | 11 | 0 | 35 | 0.0571 | 7 | 0.1429 | 10 | 0.1 | 12 | 0.0833 | 7 | 0 | 36 | 0.0833 |
| 5_1224610 | 8 | 0 | 6 | 0.5 | 7 | 0.1429 | 12 | 0.0833 | 33 | 0.1515 | 7 | 0.1429 | 11 | 0.3636 | 14 | 0 | 7 | 0 | 39 | 0.1282 |
| 5_1224613 | 5 | 0.2 | 11 | 0 | 7 | 0.1429 | 12 | 0.0833 | 35 | 0.0857 | 5 | 0.2 | 13 | 0.1538 | 13 | 0 | 7 | 0 | 38 | 0.0789 |
| 5_1224615 | NA | NA | NA | NA | NA | NA | 1 | 1 | 1 | 1 | 1 | 1 | NA | NA | NA | NA | NA | NA | 1 | 1 |
| 5_1224621 | NA | NA | NA | NA | NA | NA | 1 | 1 | 1 | 1 | 1 | 1 | NA | NA | NA | NA | 1 | 0 | 2 | 0.5 |
| 5_1225002 | 22 | 0.5 | 26 | 0.5385 | 15 | 0.3333 | 24 | 0.4167 | 87 | 0.4598 | 19 | 0.5263 | 16 | 0.4375 | 26 | 0.3846 | 23 | 0.6522 | 84 | 0.5 |
| 5_1225002 | 20 | 0 | 26 | 0.1154 | 14 | 0.1429 | 21 | 0 | 81 | 0.0617 | 18 | 0 | 18 | 0.1111 | 26 | 0.0769 | 24 | 0.2083 | 86 | 0.1047 |
| 5_1225012 | 22 | 0.1364 | 28 | 0.2857 | 17 | 0 | 23 | 0.087 | 90 | 0.1444 | 24 | 0.25 | 12 | 0.1667 | 26 | 0.1538 | 29 | 0.1724 | 91 | 0.1868 |
| 5_1225027 | 15 | 0.2667 | 7 | 0 | 8 | 0.125 | 10 | 0.3 | 40 | 0.2 | 7 | 0.4286 | 13 | 0.3846 | 16 | 0.25 | 13 | 0.2308 | 49 | 0.3061 |
| 5_1225028 | 2 | 0 | 9 | 0.2222 | 4 | 0.25 | 4 | 0 | 19 | 0.1579 | 3 | 0 | 3 | 0 | 6 | 0.1667 | 7 | 0.1429 | 19 | 0.1053 |
| 5_1225029 | 2 | 0.5 | 9 | 0.3333 | 5 | 0.2 | 4 | 0.5 | 20 | 0.35 | 2 | 0 | 3 | 0 | 5 | 0.6 | 8 | 0.375 | 18 | 0.3333 |
| 5_1225029 | 3 | 0.6667 | 9 | 0.8889 | 4 | 0 | 5 | 0.6 | 21 | 0.619 | 3 | 0.6667 | 2 | 1 | 5 | 1 | 9 | 0.7778 | 19 | 0.8421 |
| 5_1225046 | 1 | 1 | NA | NA | NA | NA | 1 | 0 | 2 | 0.5 | NA | NA | NA | NA | 2 | 0 | 1 | 1 | 3 | 0.3333 |
| 5_1225054 | 16 | 0.125 | 11 | 0.0909 | 11 | 0.0909 | 10 | 0 | 48 | 0.0833 | 6 | 0.5 | 11 | 0.0909 | 13 | 0.0769 | 17 | 0.1176 | 47 | 0.1489 |
| 5_1225205 | 4 | 0.25 | 4 | 0.25 | 4 | 0 | 5 | 0 | 17 | 0.1176 | 3 | 0.3333 | 3 | 0.3333 | 8 | 0.25 | 2 | 0 | 16 | 0.25 |
| 5_1225205 | 3 | 0 | 4 | 0.75 | 6 | 0.3333 | 5 | 0.2 | 18 | 0.3333 | 3 | 0.3333 | 3 | 0.6667 | 9 | 0.1111 | 2 | 0 | 17 | 0.2353 |
| 5_1225206 | 4 | 1 | 4 | 1 | 6 | 0.8333 | 5 | 0.6 | 19 | 0.8421 | 4 | 0.75 | 3 | 0.6667 | 8 | 0.625 | 2 | 0.5 | 17 | 0.6471 |
| 5_1225439 | 45 | 0.0667 | 36 | 0.0556 | 24 | 0.0833 | 25 | 0.08 | 130 | 0.0692 | 21 | 0 | 30 | 0 | 28 | 0.1071 | 36 | 0.0833 | 115 | 0.0522 |
| 5_1225819 | 3 | 0.3333 | 2 | 0 | 3 | 0 | 5 | 0 | 13 | 0.0769 | 1 | 0 | NA | NA | 5 | 0.4 | 3 | 0.3333 | 9 | 0.3333 |
| 5_1225821 | 3 | 1 | 2 | 1 | 4 | 1 | 2 | 1 | 11 | 1 | 1 | 1 | 2 | 1 | 6 | 0.6667 | 3 | 0.6667 | 12 | 0.75 |
| 5_1225821 | 3 | 1 | 1 | 1 | 4 | 1 | 5 | 0.8 | 13 | 0.9231 | 1 | 1 | 2 | 0.5 | 5 | 0.8 | 3 | 0.3333 | 11 | 0.6364 |
| 5_1225822 | 5 | 0.4 | 3 | 0 | 4 | 0.25 | 5 | 0.6 | 17 | 0.3529 | 1 | 1 | 2 | 0.5 | 6 | 0.6667 | 3 | 0.6667 | 12 | 0.6667 |
| 5_1225824 | 5 | 0.8 | 3 | 0.3333 | 5 | 0.4 | 5 | 0.8 | 18 | 0.6111 | NA | NA | 3 | 0.6667 | 7 | 0.4286 | 4 | 0.25 | 14 | 0.4286 |
| 5_1225824 | 3 | 0 | 2 | 0 | 4 | 0 | 6 | 0.1667 | 15 | 0.0667 | 1 | 0 | NA | NA | 6 | 0.1667 | 3 | 0 | 10 | 0.1 |
| 5_1225824 | 1 | 1 | 2 | 0 | 4 | 0.25 | 2 | 0.5 | 9 | 0.3333 | 1 | 1 | 1 | 1 | 7 | 0.5714 | 1 | 0 | 10 | 0.6 |
| 5_1225933 | 4 | 0 | 2 | 0 | 9 | 0 | 3 | 0.3333 | 18 | 0.0556 | 1 | 0 | 5 | 0 | 8 | 0.125 | 2 | 0 | 16 | 0.0625 |
| 5_1225944 | 59 | 0.1017 | 47 | 0.0851 | 64 | 0.1094 | 52 | 0.0962 | 222 | 0.0991 | 28 | 0.0714 | 45 | 0.0667 | 58 | 0.0862 | 52 | 0.0385 | 183 | 0.0656 |
| 5_1225951 | 44 | 0.1818 | 26 | 0.1154 | 45 | 0.0889 | 34 | 0.1176 | 149 | 0.1275 | 27 | 0.1111 | 38 | 0.0789 | 45 | 0.1333 | 37 | 0 | 147 | 0.0816 |
| 5_1225952 | 59 | 0.1186 | 45 | 0.1333 | 54 | 0.1111 | 46 | 0.1304 | 204 | 0.1225 | 32 | 0.2188 | 48 | 0.0833 | 50 | 0.08 | 44 | 0.0909 | 174 | 0.1092 |
| 5_1225952 | 50 | 0.1 | 38 | 0.1316 | 44 | 0.1136 | 42 | 0.1429 | 174 | 0.1207 | 30 | 0.1667 | 44 | 0.2045 | 41 | 0.1463 | 38 | 0.2105 | 153 | 0.183 |
| 5_1225952 | 51 | 0.2157 | 43 | 0.2326 | 49 | 0.3061 | 46 | 0.2609 | 189 | 0.254 | 32 | 0.2812 | 45 | 0.3111 | 47 | 0.1915 | 44 | 0.2727 | 168 | 0.2619 |
| 5_1225981 | 41 | 0.0488 | 33 | 0.0606 | 38 | 0.0789 | 29 | 0.069 | 141 | 0.0638 | 22 | 0.2273 | 35 | 0 | 41 | 0.0488 | 32 | 0.0625 | 130 | 0.0692 |
| 5_1226075 | 26 | 0.4615 | 23 | 0.3043 | 22 | 0.4091 | 16 | 0.3125 | 87 | 0.3793 | 16 | 0.25 | 21 | 0.1429 | 21 | 0.3333 | 24 | 0.4167 | 82 | 0.2927 |
| 5_1226397 | 24 | 0.0833 | 18 | 0.0556 | 18 | 0.0556 | 8 | 0 | 68 | 0.0588 | 12 | 0.0833 | 16 | 0 | 21 | 0 | 20 | 0.15 | 69 | 0.058 |
| 5_1226398 | 15 | 0.1333 | 10 | 0.1 | 11 | 0.2727 | 6 | 0 | 42 | 0.1429 | 8 | 0.375 | 12 | 0.1667 | 15 | 0.2667 | 10 | 0.1 | 45 | 0.2222 |
| 5_1226884 | 2 | 0.5 | 3 | 0.6667 | 1 | 0 | 1 | 0 | 7 | 0.4286 | 5 | 0.4 | 1 | 1 | NA | NA | 1 | 1 | 7 | 0.5714 |
| 5_1226884 | 2 | 0 | 2 | 0.5 | 1 | 1 | 1 | 0 | 6 | 0.3333 | 4 | 0.5 | 1 | 1 | NA | NA | 1 | 0 | 6 | 0.5 |
| 5_1226887 | 2 | 0 | 6 | 0.8333 | 1 | 1 | 2 | 0.5 | 11 | 0.6364 | 5 | 0.6 | 1 | 1 | 2 | 0.5 | 1 | 0 | 9 | 0.5556 |
| 5_1226887 | 2 | 0 | 6 | 0.3333 | 1 | 0 | 2 | 0.5 | 11 | 0.2727 | 6 | 0.5 | 1 | 1 | 2 | 0.5 | 1 | 0 | 10 | 0.5 |
| 5_1226890 | 1 | 1 | 6 | 0.3333 | 1 | 0 | 2 | 0 | 10 | 0.3 | 5 | 0.2 | 1 | 0 | 2 | 0.5 | 1 | 0 | 9 | 0.2222 |
| 5_1226891 | 2 | 0.5 | 2 | 0.5 | NA | NA | 1 | 0 | 5 | 0.4 | 3 | 0.6667 | 1 | 1 | 1 | 0 | 1 | 0 | 6 | 0.5 |
| 5_1228693 | 15 | 0.2667 | 22 | 0.2727 | 25 | 0.04 | 15 | 0.1333 | 77 | 0.1688 | 18 | 0.1111 | 15 | 0.1333 | 18 | 0.0556 | 14 | 0.2857 | 65 | 0.1385 |
| 5_1228694 | 16 | 0.5 | 21 | 0.7619 | 16 | 0.4375 | 15 | 0.8667 | 68 | 0.6471 | 17 | 0.7647 | 11 | 0.7273 | 15 | 0.4667 | 12 | 0.5833 | 55 | 0.6364 |

| | | | | | | | | | | | | | | | | | | | | |
|---|---|---|---|---|---|---|---|---|---|---|---|---|---|---|---|---|---|---|---|---|
| 5_12286945 | 17 | 0.2353 | 23 | 0.3913 | 20 | 0.2 | 15 | 0.1333 | 75 | 0.2533 | 15 | 0.3333 | 13 | 0.1538 | 17 | 0.4118 | 14 | 0.4286 | 59 | 0.339 |
| 5_12286965 | 16 | 0.0625 | 20 | 0.1 | 16 | 0.125 | 13 | 0.1538 | 65 | 0.1077 | 16 | 0.0625 | 13 | 0.0769 | 15 | 0.0667 | 12 | 0.25 | 56 | 0.1071 |
| 5_12287040 | 13 | 0.1538 | 18 | 0.2222 | 11 | 0.4545 | 12 | 0.1667 | 54 | 0.2407 | 14 | 0.3571 | 12 | 0.1667 | 12 | 0.1667 | 10 | 0.1 | 48 | 0.2083 |
| 5_12287098 | 13 | 0.3846 | 17 | 0.1765 | 10 | 0.3 | 10 | 0.4 | 50 | 0.3 | 12 | 0.3333 | 9 | 0.1111 | 8 | 0.25 | 9 | 0.3333 | 38 | 0.2632 |
| 5_12287113 | 13 | 0.0769 | 14 | 0.1429 | 9 | 0.1111 | 10 | 0 | 46 | 0.087 | 11 | 0 | 11 | 0 | 8 | 0.125 | 9 | 0.1111 | 39 | 0.0513 |
| 5_12287149 | 12 | 0.4167 | 12 | 0.5 | 6 | 0.5 | 8 | 0.375 | 38 | 0.4474 | 10 | 0.5 | 10 | 0.2 | 7 | 0.4286 | 6 | 0.3333 | 33 | 0.3636 |
| 5_12287179 | 14 | 1 | 10 | 1 | 6 | 0.8333 | 7 | 1 | 37 | 0.973 | 8 | 0.75 | 8 | 1 | 5 | 0.4 | 4 | 1 | 25 | 0.8 |
| 5_12287184 | 14 | 0.2143 | 11 | 0.0909 | 5 | 0.2 | 6 | 0 | 36 | 0.1389 | 9 | 0.1111 | 8 | 0 | 2 | 0 | 3 | 0.3333 | 22 | 0.0909 |
| 5_12287188 | 13 | 0.9231 | 11 | 1 | 6 | 0.8333 | 7 | 1 | 37 | 0.9459 | 9 | 1 | 8 | 1 | 3 | 0.3333 | 2 | 1 | 22 | 0.9091 |
| 5_12287202 | 9 | 0.2222 | 10 | 0.4 | 6 | 0.5 | 6 | 0.1667 | 31 | 0.3226 | 10 | 0.6 | 6 | 0.1667 | 4 | 0.25 | 1 | 0 | 21 | 0.381 |
| 5_12290196 | 5 | 0.2 | 9 | 0 | 5 | 0 | 3 | 0.3333 | 22 | 0.0909 | 5 | 0.4 | 4 | 0.5 | 3 | 0.3333 | NA | NA | 12 | 0.4167 |
| 5_12290205 | 3 | 0.3333 | 9 | 0.2222 | 6 | 0 | 3 | 0 | 21 | 0.1429 | 5 | 0 | 3 | 0.3333 | 3 | 0.3333 | NA | NA | 11 | 0.1818 |
| 5_12290211 | 4 | 0 | 8 | 0.125 | 6 | 0.3333 | 3 | 0.3333 | 21 | 0.1905 | 3 | 0 | 3 | 0 | 4 | 0.25 | NA | NA | 10 | 0.1 |
| 5_12290219 | 4 | 1 | 9 | 0.7778 | 6 | 0.8333 | 3 | 0.6667 | 22 | 0.8182 | 4 | 0.5 | 5 | 0.8 | 3 | 1 | NA | NA | 12 | 0.75 |
| 5_12292586 | 12 | 0.5 | 18 | 0.1667 | 24 | 0 | 21 | 0.0476 | 75 | 0.1333 | 11 | 0.1818 | 18 | 0.1111 | 22 | 0.1364 | 12 | 0.0833 | 63 | 0.127 |
| 5_12300224 | 7 | 0 | 14 | 0 | 6 | 0 | 8 | 0.25 | 35 | 0.0571 | 6 | 0 | 4 | 0 | 8 | 0 | 6 | 0.3333 | 24 | 0.0833 |
| 5_12300228 | 3 | 0.6667 | 8 | 0.5 | 3 | 0.6667 | 4 | 0.75 | 18 | 0.6111 | 5 | 0.4 | 3 | 1 | 6 | 0.8333 | 3 | 0.6667 | 17 | 0.7059 |
| 5_12300230 | 5 | 0.6 | 14 | 0.7143 | 6 | 0.8333 | 8 | 1 | 33 | 0.7879 | 6 | 1 | 3 | 0.6667 | 8 | 1 | 6 | 0.8333 | 23 | 0.913 |
| 5_12300233 | 5 | 0.4 | 14 | 0.1429 | 6 | 0 | 8 | 0 | 33 | 0.1212 | 4 | 0.25 | 4 | 0.25 | 8 | 0.125 | 6 | 0.1667 | 22 | 0.1818 |
| 5_12300242 | 4 | 0 | 10 | 0.1 | 3 | 0 | 5 | 0.2 | 22 | 0.0909 | 5 | 0.2 | 4 | 0 | 7 | 0.2857 | 7 | 0 | 23 | 0.1304 |
| 5_12300631 | 6 | 0 | 14 | 0.0714 | 6 | 0 | 9 | 0.1111 | 35 | 0.0571 | 7 | 0.1429 | 5 | 0 | 8 | 0.125 | 8 | 0 | 28 | 0.0714 |
| 5_12300635 | 7 | 0.5714 | 16 | 0.5 | 7 | 0 | 10 | 0.7 | 40 | 0.475 | 6 | 0.1667 | 5 | 0.4 | 8 | 0.75 | 7 | 0.4286 | 26 | 0.4615 |
| 5_12300648 | 5 | 0.2 | 12 | 0.25 | 8 | 0.125 | 6 | 0.1667 | 31 | 0.1935 | 6 | 0.1667 | 4 | 0 | 7 | 0.2857 | 5 | 0 | 22 | 0.1364 |
| 5_12300900 | 2 | 0 | 2 | 0.5 | 4 | 0 | NA | NA | 8 | 0.125 | 4 | 0 | 2 | 0.5 | 3 | 0 | NA | NA | 9 | 0.1111 |
| 5_12300938 | 2 | 0 | 2 | 0 | 5 | 0.4 | NA | NA | 9 | 0.2222 | 3 | 0.3333 | 1 | 1 | 1 | 1 | NA | NA | 5 | 0.6 |
| 5_12300951 | 1 | 0 | 2 | 0 | 5 | 0.2 | NA | NA | 8 | 0.125 | 2 | 0 | 1 | 0 | 1 | 1 | NA | NA | 4 | 0.25 |
| 5_12300954 | 1 | 1 | 2 | 1 | 5 | 0.8 | NA | NA | 8 | 0.875 | 2 | 1 | 2 | 0.5 | NA | NA | NA | NA | 4 | 0.75 |
| 5_12300965 | 2 | 1 | 2 | 0.5 | 5 | 0.2 | NA | NA | 9 | 0.4444 | 1 | 1 | 2 | 1 | 1 | 1 | NA | NA | 4 | 1 |
| 5_12300966 | 2 | 1 | 1 | 1 | 4 | 0.75 | NA | NA | 7 | 0.8571 | 1 | 1 | 2 | 0.5 | 1 | 1 | NA | NA | 4 | 0.75 |
| 5_12300971 | 1 | 1 | 2 | 1 | 4 | 1 | NA | NA | 7 | 1 | 1 | 1 | 2 | 1 | 1 | 1 | NA | NA | 4 | 1 |
| 5_12300977 | 1 | 0 | 2 | 0.5 | 4 | 1 | NA | NA | 7 | 0.7143 | 1 | 1 | 2 | 1 | 1 | 1 | NA | NA | 4 | 1 |
| 5_12315348 | NA | NA | 3 | 0.3333 | 2 | 0.5 | 3 | 0 | 8 | 0.25 | 1 | 0 | NA | NA | 2 | 0.5 | NA | NA | 3 | 0.3333 |
| 5_12316745 | NA | NA | 2 | 1 | 2 | 1 | 4 | 1 | 8 | 1 | 1 | 1 | NA | NA | 2 | 1 | NA | NA | 3 | 1 |
| 5_12316763 | 1 | 0 | 2 | 1 | 2 | 0.5 | 2 | 0.5 | 7 | 0.5714 | 2 | 1 | NA | NA | 1 | 1 | NA | NA | 3 | 1 |
| 5_12316768 | 1 | 0 | 3 | 1 | 3 | 0 | 5 | 0.8 | 12 | 0.5833 | 3 | 0.6667 | NA | NA | 2 | 1 | NA | NA | 5 | 0.8 |
| 5_12316769 | 1 | 1 | 3 | 1 | 3 | 1 | 5 | 1 | 12 | 1 | 3 | 1 | NA | NA | 2 | 0.5 | NA | NA | 5 | 0.8 |
| 5_12316770 | 1 | 1 | 3 | 1 | 3 | 1 | 4 | 1 | 11 | 1 | 4 | 1 | NA | NA | 2 | 0.5 | NA | NA | 6 | 0.8333 |
| 5_12316779 | 1 | 0 | 2 | 0.5 | 1 | 0 | 3 | 0.3333 | 7 | 0.2857 | 2 | 0 | NA | NA | 1 | 1 | NA | NA | 3 | 0.3333 |
| 5_12325486 | 1 | 0 | NA | NA | 1 | 1 | 1 | 1 | 3 | 0.6667 | NA | NA | 1 | 0 | 3 | 0.6667 | NA | NA | 4 | 0.5 |
| 5_12325497 | 1 | 0 | NA | NA | 1 | 1 | 1 | 0 | 3 | 0.3333 | 1 | 0 | 1 | 1 | 3 | 0.3333 | NA | NA | 5 | 0.4 |
| 5_12325528 | 1 | 1 | NA | NA | 1 | 1 | NA | NA | 2 | 1 | 1 | 0 | 1 | 0 | 3 | 0.3333 | NA | NA | 5 | 0.2 |
| 5_12325555 | NA | NA | NA | NA | 1 | 1 | NA | NA | 1 | 1 | 1 | 1 | NA | NA | 1 | 1 | NA | NA | 2 | 1 |
| 5_12325559 | NA | NA | NA | NA | 1 | 1 | NA | NA | 1 | 1 | 1 | 1 | NA | NA | 1 | 0 | NA | NA | 2 | 0.5 |
| 5_12325618 | NA | NA | NA | NA | 1 | 1 | NA | NA | 1 | 1 | 1 | 1 | NA | NA | 1 | 0 | NA | NA | 2 | 0.5 |
| 5_12325637 | 3 | 1 | 8 | 0.5 | 10 | 0.5 | 6 | 0.8333 | 27 | 0.6296 | 13 | 0.3077 | 2 | 0.5 | 6 | 0.6667 | 4 | 0.5 | 25 | 0.44 |
| 5_12325643 | 3 | 0.6667 | 8 | 0.25 | 11 | 0.0909 | 4 | 0.25 | 26 | 0.2308 | 10 | 0.1 | 1 | 0 | 7 | 0.5714 | 3 | 0.3333 | 21 | 0.2857 |
| 5_12325659 | 3 | 0.3333 | 8 | 0.125 | 9 | 0 | 6 | 0 | 26 | 0.0769 | 6 | 0.1667 | 2 | 0 | 6 | 0.3333 | 3 | 0 | 17 | 0.1765 |
| 5_12325660 | 3 | 0 | 8 | 0.375 | 9 | 0.2222 | 5 | 0.4 | 25 | 0.28 | 8 | 0.375 | 2 | 1 | 6 | 0.1667 | 4 | 0.25 | 20 | 0.35 |
| 5_12325662 | 3 | 0.6667 | 9 | 0.4444 | 11 | 0.4545 | 3 | 0.3333 | 26 | 0.4615 | 10 | 0.7 | 1 | 0 | 5 | 0.4 | 3 | 1 | 19 | 0.6316 |
| 5_12325665 | 3 | 0.6667 | 8 | 1 | 7 | 0.8571 | 4 | 1 | 22 | 0.9091 | 5 | 0.8 | 1 | 1 | 4 | 1 | 2 | 0.5 | 12 | 0.8333 |
| 5_12325678 | 3 | 0 | 7 | 0 | 11 | 0.1818 | 6 | 0.3333 | 27 | 0.1481 | 10 | 0 | 1 | 1 | 6 | 0.3333 | 3 | 0 | 20 | 0.15 |
| 5_12325697 | 1 | 1 | NA | NA | 5 | 0.6 | 3 | 1 | 9 | 0.7778 | 7 | 1 | 1 | 1 | 4 | 0.75 | 2 | 1 | 14 | 0.9286 |
| 5_12325721 | 1 | 1 | 5 | 0.6 | 6 | 0.8333 | NA | NA | 12 | 0.75 | 7 | 0.8571 | NA | NA | 2 | 0.5 | 2 | 0.5 | 11 | 0.7273 |
| 5_12325721 | 1 | 1 | 7 | 0.2857 | 11 | 0.0909 | 3 | 0 | 22 | 0.1818 | 6 | 0.1667 | 1 | 0 | 5 | 0.2 | 3 | 0 | 15 | 0.1333 |
| 5_12325731 | 1 | 0 | 5 | 0 | 9 | 0.1111 | 4 | 0.5 | 19 | 0.1579 | 4 | 0.25 | 1 | 1 | 4 | 0.25 | 2 | 0.5 | 11 | 0.3636 |
| 5_12325740 | 2 | 1 | 2 | 0 | 8 | 0.75 | 2 | 1 | 14 | 0.7143 | 4 | 0.5 | NA | NA | 2 | 0.5 | 1 | 0 | 7 | 0.4286 |
| 5_12325741 | 1 | 0 | 6 | 1 | 9 | 1 | 3 | 1 | 19 | 0.9474 | 7 | 1 | NA | NA | 5 | 1 | 2 | 1 | 14 | 1 |
| 5_12325745 | 2 | 0.5 | 4 | 0 | 9 | 0.2222 | 2 | 0 | 17 | 0.1765 | 6 | 0.5 | NA | NA | 4 | 0 | 2 | 0 | 12 | 0.25 |
| 5_12325746 | 1 | 1 | 5 | 0.6 | 10 | 0.7 | 2 | 0.5 | 18 | 0.6667 | 6 | 0.6667 | NA | NA | 5 | 1 | 2 | 0.5 | 13 | 0.7692 |
| 5_12325747 | 2 | 0 | 5 | 0.2 | 9 | 0 | 2 | 0 | 18 | 0.0556 | 6 | 0.3333 | NA | NA | 5 | 0.2 | 2 | 0 | 13 | 0.2308 |
| 5_12325748 | 2 | 1 | 4 | 0.5 | 8 | 0.375 | 2 | 0.5 | 16 | 0.5 | 6 | 0.5 | NA | NA | 4 | 0.5 | 2 | 0 | 12 | 0.4167 |
| 5_12325749 | 2 | 0.5 | 6 | 0.1667 | 11 | 0.0909 | 3 | 1 | 22 | 0.2727 | 8 | 0.25 | NA | NA | 5 | 0.2 | 2 | 0 | 15 | 0.2 |
| 5_12325752 | 1 | 0 | 5 | 0.4 | 10 | 0.1 | 3 | 0 | 19 | 0.1579 | 6 | 0.3333 | NA | NA | 5 | 0 | 2 | 0.5 | 13 | 0.2308 |
| 5_12328570 | NA | NA | NA | NA | NA | NA | 1 | 1 | 1 | 1 | NA | NA | 1 | 1 | NA | NA | 1 | 0 | 2 | 0.5 |
| 5_12328649 | NA | NA | NA | NA | NA | NA | 1 | 1 | 1 | 1 | NA | NA | 2 | 1 | NA | NA | 1 | 1 | 3 | 1 |

| | | | | | | | | | | | | | | | | | | | | |
|---|---|---|---|---|---|---|---|---|---|---|---|---|---|---|---|---|---|---|---|---|
| 5_12328668 | NA | NA | NA | NA | NA | NA | 1 | 1 | 1 | 1 | NA | NA | 1 | 1 | NA | NA | NA | NA | 1 | 1 |
| 5_12329026 | NA | NA | NA | NA | NA | NA | 2 | 1 | 2 | 1 | NA | NA | 2 | 1 | NA | NA | NA | NA | 2 | 1 |
| 5_12329028 | NA | NA | NA | NA | NA | NA | 2 | 0.5 | 2 | 0.5 | NA | NA | 2 | 0.5 | NA | NA | 1 | 1 | 3 | 0.6667 |
| 5_12329043 | NA | NA | NA | NA | NA | NA | 2 | 1 | 2 | 1 | NA | NA | 2 | 1 | NA | NA | 1 | 1 | 3 | 1 |
| 5_12329062 | NA | NA | NA | NA | NA | NA | 2 | 1 | 2 | 1 | NA | NA | 2 | 0 | NA | NA | 1 | 1 | 3 | 0.3333 |
| 5_12329665 | NA | NA | 1 | 1 | NA | NA | 2 | 1 | 3 | 1 | 1 | 1 | 1 | 1 | NA | NA | 1 | 1 | 3 | 1 |
| 5_12329671 | 1 | 0 | 1 | 0 | NA | NA | 2 | 1 | 4 | 0.5 | 1 | 1 | 2 | 0.5 | NA | NA | 1 | 1 | 4 | 0.75 |
| 5_12329815 | 1 | 1 | 1 | 0 | NA | NA | 1 | 0 | 3 | 0.3333 | NA | NA | 2 | 0.5 | NA | NA | 1 | 1 | 3 | 0.6667 |
| 5_12329829 | 1 | 1 | 1 | 0 | NA | NA | 2 | 0.5 | 4 | 0.5 | 2 | 1 | 2 | 1 | NA | NA | 1 | 0 | 5 | 0.8 |
| 5_12329830 | 1 | 1 | 1 | 0 | NA | NA | 2 | 1 | 4 | 0.75 | 2 | 1 | 2 | 0 | NA | NA | NA | NA | 4 | 0.5 |
| 5_12329834 | 1 | 0 | 1 | 0 | NA | NA | 2 | 0.5 | 4 | 0.25 | 1 | 1 | 2 | 0.5 | NA | NA | NA | NA | 3 | 0.6667 |
| 5_12329874 | 2 | 1 | 1 | 1 | NA | NA | 2 | 1 | 5 | 1 | 2 | 1 | 2 | 1 | NA | NA | 1 | 0 | 5 | 0.8 |
| 5_12329915 | 2 | 1 | 1 | 1 | NA | NA | 2 | 0.5 | 5 | 0.8 | 2 | 0 | 3 | 0.6667 | NA | NA | NA | NA | 5 | 0.4 |
| 5_12330404 | 1 | 0 | 1 | 1 | 1 | 1 | 2 | 0.5 | 5 | 0.6 | 1 | 1 | 1 | 1 | 1 | 1 | NA | NA | 3 | 1 |
| 5_12330411 | 3 | 0 | 1 | 1 | 1 | 0 | 2 | 0 | 7 | 0.1429 | 2 | 0.5 | 6 | 0 | 2 | 0 | 1 | 0 | 11 | 0.0909 |
| 5_12330448 | 3 | 0.3333 | 1 | 0 | 1 | 0 | 2 | 0 | 7 | 0.1429 | 2 | 0.5 | 5 | 0 | 2 | 0 | NA | NA | 9 | 0.1111 |
| 5_12338670 | 3 | 0.6667 | 3 | 0 | 3 | 0 | 7 | 0.5714 | 16 | 0.375 | 2 | 0.5 | 1 | 0 | 5 | 0 | 6 | 0.6667 | 14 | 0.3571 |
| 5_12347089 | NA | NA | NA | NA | NA | NA | 1 | 1 | 1 | 1 | NA | NA | NA | NA | NA | NA | 1 | 1 | 1 | 1 |
| 5_12347187 | NA | NA | NA | NA | NA | NA | 1 | 1 | 1 | 1 | NA | NA | NA | NA | NA | NA | 1 | 1 | 1 | 1 |
| 5_12347249 | 1 | 0 | 1 | 1 | NA | NA | NA | NA | 2 | 0.5 | NA | NA | 1 | 0 | NA | NA | 2 | 0.5 | 3 | 0.3333 |
| 5_12350925 | 4 | 0.25 | 2 | 1 | 6 | 0.6667 | 5 | 0.8 | 17 | 0.6471 | 5 | 0.6 | 3 | 1 | 7 | 0.8571 | 4 | 0.75 | 19 | 0.7895 |
| 5_12350931 | 4 | 0 | 4 | 0 | 6 | 0 | 5 | 0.2 | 19 | 0.0526 | 5 | 0 | 4 | 0 | 6 | 0.3333 | 5 | 0.2 | 20 | 0.15 |
| 5_12350933 | 4 | 1 | 4 | 1 | 8 | 0.875 | 6 | 1 | 22 | 0.9545 | 5 | 0.8 | 4 | 1 | 7 | 1 | 5 | 1 | 21 | 0.9524 |
| 5_12350936 | 2 | 0 | 4 | 0 | 7 | 0.4286 | 4 | 0 | 17 | 0.1765 | 5 | 0 | 4 | 0.25 | 6 | 0.1667 | 5 | 0 | 20 | 0.1 |
| 5_12350937 | 2 | 0.5 | 3 | 0.6667 | 5 | 0.4 | 6 | 0.8333 | 16 | 0.625 | 3 | 0.6667 | 2 | 1 | 7 | 1 | 3 | 0.6667 | 15 | 0.8667 |
| 5_12350971 | 2 | 0.5 | 5 | 0 | 6 | 0.3333 | 5 | 0.2 | 18 | 0.2222 | 5 | 0.2 | 2 | 0 | 8 | 0.125 | 7 | 0 | 22 | 0.0909 |
| 5_12351015 | 3 | 0.3333 | 3 | 0.6667 | 6 | 0.5 | 6 | 0 | 18 | 0.3333 | 5 | 0.4 | 3 | 0 | 5 | 0.4 | 7 | 0.2857 | 20 | 0.3 |
| 5_12351015 | 2 | 1 | 4 | 0 | 7 | 0 | 6 | 0.1667 | 19 | 0.1579 | 2 | 0 | 3 | 0.3333 | 4 | 0.25 | 5 | 0 | 14 | 0.1429 |
| 5_12351016 | 3 | 0.6667 | 7 | 1 | 8 | 1 | 6 | 1 | 24 | 0.9583 | 4 | 0.75 | 3 | 1 | 7 | 1 | 7 | 1 | 21 | 0.9524 |
| 5_12351027 | 4 | 0.5 | 6 | 0.6667 | 8 | 0.375 | 8 | 0.25 | 26 | 0.4231 | 4 | 0.25 | 3 | 0.3333 | 9 | 0.4444 | 9 | 0.4444 | 25 | 0.4 |
| 5_12351034 | 4 | 1 | 7 | 0.8571 | 8 | 0.875 | 8 | 0.875 | 27 | 0.8889 | 3 | 0.6667 | 1 | 1 | 8 | 1 | 9 | 0.8889 | 21 | 0.9048 |
| 5_12351047 | 5 | 0 | 7 | 0.2857 | 10 | 0 | 9 | 0 | 31 | 0.0645 | 5 | 0.2 | 3 | 0 | 9 | 0.2222 | 8 | 0.125 | 25 | 0.16 |
| 5_12351051 | 5 | 0.8 | 3 | 1 | 10 | 0.9 | 8 | 1 | 26 | 0.9231 | 4 | 0.75 | 2 | 1 | 8 | 0.75 | 8 | 0.875 | 22 | 0.8182 |
| 5_12351051 | 4 | 1 | 5 | 1 | 8 | 1 | 9 | 1 | 26 | 1 | 4 | 0.75 | 3 | 1 | 6 | 1 | 7 | 1 | 20 | 0.95 |
| 5_12351057 | 6 | 0.6667 | 7 | 0.1429 | 9 | 0.1111 | 8 | 0.5 | 30 | 0.3333 | 5 | 0.2 | 2 | 0.5 | 8 | 0.625 | 9 | 0.6667 | 24 | 0.5417 |
| 5_12351062 | 5 | 0.4 | 7 | 0.2857 | 8 | 0.375 | 7 | 0 | 27 | 0.2593 | 3 | 0.6667 | 3 | 0 | 8 | 0.25 | 8 | 0.5 | 22 | 0.3636 |
| 5_12351081 | 5 | 0 | 8 | 0.25 | 12 | 0 | 10 | 0.1 | 35 | 0.0857 | 6 | 0 | 4 | 0.25 | 7 | 0.2857 | 10 | 0.3 | 27 | 0.2222 |
| 5_12351094 | 4 | 1 | 9 | 0.7778 | 15 | 0.8667 | 8 | 0.875 | 36 | 0.8611 | 5 | 1 | 4 | 0.75 | 8 | 0.875 | 12 | 0.9167 | 29 | 0.8966 |
| 5_12351106 | 4 | 0.75 | 8 | 0.25 | 15 | 0.3333 | 13 | 0.1538 | 40 | 0.3 | 6 | 0.5 | 5 | 0 | 7 | 0.7143 | 11 | 0.6364 | 29 | 0.5172 |
| 5_12351135 | 6 | 0.6667 | 7 | 1 | 14 | 0.8571 | 12 | 0.8333 | 39 | 0.8462 | 8 | 1 | 5 | 0.8 | 9 | 1 | 12 | 0.9167 | 34 | 0.9412 |
| 5_12351157 | 6 | 0.6667 | 10 | 0.5 | 17 | 0.7647 | 14 | 0.5714 | 47 | 0.6383 | 6 | 0.5 | 5 | 0.8 | 10 | 0.9 | 14 | 0.8571 | 35 | 0.8 |
| 5_12351168 | 8 | 0.5 | 11 | 0.5455 | 18 | 0.5556 | 12 | 0.5833 | 49 | 0.551 | 7 | 0.8571 | 4 | 0.75 | 9 | 0.8889 | 13 | 0.3846 | 33 | 0.6667 |
| 5_12351174 | 7 | 0.8571 | 11 | 0.8182 | 14 | 0.8571 | 11 | 0.9091 | 43 | 0.8605 | 8 | 1 | 4 | 0.75 | 7 | 1 | 11 | 0.7273 | 30 | 0.8667 |
| 5_12351178 | 6 | 0 | 12 | 0.25 | 16 | 0 | 11 | 0.1818 | 45 | 0.1111 | 8 | 0.25 | 3 | 0 | 11 | 0.0909 | 12 | 0 | 34 | 0.0882 |
| 5_12351189 | 8 | 0 | 14 | 0.1429 | 20 | 0.05 | 14 | 0 | 56 | 0.0536 | 8 | 0.125 | 6 | 0 | 11 | 0.1818 | 15 | 0.1333 | 40 | 0.125 |
| 5_12351195 | 8 | 0.25 | 14 | 0.2857 | 21 | 0.4762 | 13 | 0.3077 | 56 | 0.3571 | 9 | 0.5556 | 5 | 0.4 | 11 | 0.5455 | 16 | 0.3125 | 41 | 0.439 |
| 5_12351197 | 8 | 0.125 | 14 | 0.2143 | 19 | 0.1053 | 15 | 0.1333 | 56 | 0.1429 | 10 | 0.1 | 6 | 0.1667 | 10 | 0.4 | 17 | 0.1176 | 43 | 0.186 |
| 5_12359159 | NA | NA | NA | NA | NA | NA | 1 | 1 | 1 | 1 | NA | NA | NA | NA | 1 | 1 | NA | NA | 1 | 1 |
| 5_12359165 | NA | NA | NA | NA | NA | NA | 1 | 1 | 1 | 1 | NA | NA | NA | NA | 1 | 1 | NA | NA | 1 | 1 |
| 5_12359195 | NA | NA | NA | NA | NA | NA | 1 | 1 | 1 | 1 | 1 | 1 | NA | NA | 1 | 1 | NA | NA | 2 | 1 |
| 5_12359220 | NA | NA | NA | NA | NA | NA | 1 | 1 | 1 | 1 | 1 | 1 | NA | NA | 1 | 1 | NA | NA | 2 | 1 |
| 5_12359225 | NA | NA | NA | NA | NA | NA | 2 | 1 | 2 | 1 | 1 | 1 | NA | NA | 1 | 0 | NA | NA | 2 | 0.5 |
| 5_12359253 | NA | NA | NA | NA | NA | NA | 2 | 1 | 2 | 1 | 1 | 1 | NA | NA | 1 | 1 | NA | NA | 2 | 1 |
| 5_12359268 | NA | NA | NA | NA | NA | NA | 2 | 0.5 | 2 | 0.5 | NA | NA | NA | NA | 1 | 1 | NA | NA | 1 | 1 |
| 5_12359307 | 1 | 1 | NA | NA | NA | NA | 2 | 1 | 3 | 1 | NA | NA | NA | NA | 1 | 1 | NA | NA | 1 | 1 |
| 5_12359311 | 1 | 1 | NA | NA | NA | NA | 4 | 1 | 5 | 1 | 1 | 1 | NA | NA | 1 | 1 | NA | NA | 2 | 1 |
| 5_12359328 | 1 | 0 | NA | NA | NA | NA | 3 | 0.6667 | 4 | 0.5 | 2 | 0.5 | NA | NA | 1 | 1 | 1 | 1 | 4 | 0.75 |
| 5_12359343 | 1 | 1 | NA | NA | NA | NA | 4 | 1 | 5 | 1 | 1 | 1 | NA | NA | NA | NA | NA | NA | 1 | 1 |
| 5_12359432 | 1 | 0 | 1 | 0 | 1 | 1 | 4 | 0.5 | 7 | 0.4286 | NA | NA | 1 | 1 | 1 | 0 | 1 | 1 | 3 | 0.6667 |
| 5_12359558 | 1 | 0 | 1 | 1 | 1 | 1 | 4 | 0 | 7 | 0.2857 | NA | NA | 2 | 0 | 2 | 0 | 1 | 1 | 5 | 0.2 |
| 5_12365047 | 4 | 0 | 3 | 0.6667 | 2 | 1 | 2 | 0.5 | 11 | 0.4545 | NA | NA | 1 | 1 | 4 | 0.5 | 1 | 0 | 6 | 0.5 |
| 5_12366609 | 1 | 1 | 3 | 0.6667 | 3 | 0.3333 | 3 | 0.6667 | 10 | 0.6 | NA | NA | 5 | 0.4 | 7 | 0.2857 | 3 | 0.6667 | 15 | 0.4 |
| 5_12366711 | NA | NA | 3 | 0.3333 | 2 | 0.5 | 3 | 0.6667 | 8 | 0.5 | NA | NA | 4 | 0.25 | 5 | 0 | 1 | 0 | 10 | 0.1 |
| 5_12366723 | 4 | 0 | 6 | 0.1667 | 7 | 0.4286 | 8 | 0.125 | 25 | 0.2 | 3 | 0.3333 | 9 | 0.2222 | 9 | 0.2222 | 3 | 0.3333 | 24 | 0.25 |
| 5_12366797 | 8 | 0.375 | 6 | 0.1667 | 4 | 0.5 | 6 | 0.6667 | 24 | 0.4167 | 2 | 0 | 9 | 0.5556 | 10 | 0.5 | 4 | 0.5 | 25 | 0.48 |

| | | | | | | | | | | | | | | | | | | | | |
|---|---|---|---|---|---|---|---|---|---|---|---|---|---|---|---|---|---|---|---|---|
| 5_12366800 | 6 | 0 | 5 | 0.4 | 2 | 0 | 4 | 0.25 | 17 | 0.1765 | NA | NA | 7 | 0.1429 | 7 | 0.2857 | 5 | 0 | 19 | 0.1579 |
| 5_12366811 | 8 | 0.5 | 5 | 0.4 | 4 | 0.5 | 8 | 0.5 | 25 | 0.48 | 2 | 0 | 12 | 0.3333 | 9 | 0.3333 | 3 | 0.3333 | 26 | 0.3077 |
| 5_12366829 | 8 | 1 | 4 | 1 | 3 | 1 | 7 | 1 | 22 | 1 | 2 | 1 | 11 | 0.9091 | 9 | 1 | 5 | 1 | 27 | 0.963 |
| 5_12366832 | 6 | 0.6667 | 4 | 1 | 3 | 1 | 5 | 1 | 18 | 0.8889 | 1 | 0 | 8 | 0.875 | 5 | 0.8 | 4 | 1 | 18 | 0.8333 |
| 5_12366841 | 5 | 1 | 4 | 1 | 1 | 1 | 7 | 0.7143 | 17 | 0.8824 | 2 | 0.5 | 10 | 0.8 | 8 | 0.75 | 3 | 0.6667 | 23 | 0.7391 |
| 5_12366844 | 5 | 0.2 | 5 | 0.6 | 3 | 0.3333 | 7 | 0.2857 | 20 | 0.35 | 2 | 0 | 9 | 0.4444 | 9 | 0.1111 | 4 | 0 | 24 | 0.2083 |
| 5_12366846 | 6 | 0.3333 | 4 | 0.25 | 3 | 0.6667 | 4 | 0.5 | 17 | 0.4118 | 2 | 0.5 | 7 | 0.2857 | 9 | 0.3333 | 5 | 0.8 | 23 | 0.4348 |
| 5_12366862 | 5 | 0.8 | 5 | 1 | 1 | 1 | 5 | 0.8 | 16 | 0.875 | 2 | 0.5 | 7 | 0.7143 | 6 | 0.8333 | 4 | 1 | 19 | 0.7895 |
| 5_12366872 | 5 | 0.6 | 4 | 0.75 | 3 | 0.3333 | 6 | 0.3333 | 18 | 0.5 | 2 | 0 | 9 | 0.7778 | 8 | 0.5 | 4 | 0.5 | 23 | 0.5652 |
| 5_12366873 | 5 | 0.4 | 4 | 0.5 | 3 | 0.3333 | 6 | 0.3333 | 18 | 0.3889 | 1 | 1 | 9 | 0.2222 | 8 | 0.375 | 3 | 0 | 21 | 0.2857 |
| 5_12366876 | 4 | 0.5 | 4 | 0.5 | 3 | 0.3333 | 4 | 0.5 | 15 | 0.4667 | 1 | 1 | 9 | 0.7778 | 7 | 0.4286 | 4 | 0.5 | 21 | 0.619 |
| 5_12366887 | 5 | 0.2 | 5 | 0.2 | 3 | 0 | 8 | 0.25 | 21 | 0.1905 | 2 | 0 | 9 | 0 | 8 | 0 | 5 | 0.4 | 24 | 0.0833 |
| 5_12366891 | 4 | 0.25 | 5 | 0 | 3 | 0 | 4 | 0 | 16 | 0.0625 | 2 | 0.5 | 8 | 0.125 | 6 | 0.5 | 5 | 0.4 | 21 | 0.3333 |
| 5_12366892 | 4 | 1 | 5 | 1 | 3 | 0.6667 | 8 | 0.875 | 20 | 0.9 | 2 | 0.5 | 7 | 1 | 8 | 0.875 | 5 | 0.8 | 22 | 0.8636 |
| 5_12366895 | 3 | 0 | 4 | 0 | 3 | 0 | 5 | 0.2 | 15 | 0.0667 | 1 | 1 | 5 | 0 | 6 | 0.1667 | 4 | 0.5 | 16 | 0.25 |
| 5_12366902 | 2 | 0 | 3 | 0 | 2 | 0.5 | 5 | 0.4 | 12 | 0.25 | 1 | 0 | 7 | 0 | 7 | 0.2857 | 4 | 0 | 19 | 0.1053 |
| 5_12371606 | 10 | 0.2 | 8 | 0.625 | 10 | 0.3 | 9 | 0.3333 | 37 | 0.3514 | 8 | 0.375 | 14 | 0.2143 | 18 | 0.3333 | 6 | 0.6667 | 46 | 0.3478 |
| 5_12371617 | 9 | 0.2222 | 7 | 0.4286 | 11 | 0.3636 | 9 | 0.4444 | 36 | 0.3611 | 8 | 0.5 | 11 | 0.1818 | 19 | 0.2632 | 5 | 0.4 | 43 | 0.3023 |
| 5_12371651 | 8 | 0 | 5 | 0.2 | 8 | 0.125 | 9 | 0.1111 | 30 | 0.1 | 5 | 0 | 12 | 0.1667 | 14 | 0.2857 | 6 | 0 | 37 | 0.1622 |
| 5_12371668 | 10 | 0.1 | 4 | 0.75 | 7 | 0 | 10 | 0.5 | 31 | 0.2903 | 6 | 0.5 | 13 | 0.0769 | 15 | 0.4667 | 5 | 0.4 | 39 | 0.3333 |
| 5_12371671 | 9 | 0.2222 | 4 | 0.25 | 6 | 0.5 | 10 | 0.6 | 29 | 0.4138 | 7 | 0.7143 | 13 | 0.3077 | 16 | 0.3125 | 6 | 0 | 42 | 0.3333 |
| 5_12371672 | 8 | 0.25 | 4 | 0 | 7 | 0.4286 | 9 | 0.3333 | 28 | 0.2857 | 6 | 0.5 | 13 | 0.3846 | 16 | 0.4375 | 6 | 0 | 41 | 0.3659 |
| 5_12371699 | 9 | 0.2222 | 4 | 0.25 | 5 | 0.8 | 9 | 0.2222 | 27 | 0.3333 | 5 | 0.2 | 12 | 0.25 | 15 | 0.0667 | 6 | 0 | 38 | 0.1316 |
| 5_12371703 | 10 | 0.2 | 4 | 0.25 | 7 | 0.4286 | 6 | 0.5 | 27 | 0.3333 | 5 | 0.4 | 12 | 0.0833 | 17 | 0.2353 | 7 | 0.4286 | 41 | 0.2439 |
| 5_12371704 | 9 | 0 | 3 | 0 | 6 | 0.3333 | 7 | 0.1429 | 25 | 0.12 | 6 | 0.1667 | 12 | 0.1667 | 16 | 0.1875 | 6 | 0.1667 | 40 | 0.175 |
| 5_12371709 | 7 | 0.4286 | 3 | 0.6667 | 3 | 0.3333 | 4 | 0.5 | 17 | 0.4706 | 4 | 0.25 | 9 | 0.4444 | 13 | 0.3077 | 6 | 0.5 | 32 | 0.375 |
| 5_12371748 | 9 | 0.7778 | 3 | 1 | 6 | 0.5 | 6 | 0.6667 | 24 | 0.7083 | 7 | 0.8571 | 11 | 0.8182 | 17 | 0.8235 | 4 | 1 | 39 | 0.8462 |
| 5_12371750 | 11 | 0 | 4 | 0 | 6 | 0.3333 | 7 | 0 | 28 | 0.0714 | 9 | 0 | 11 | 0.0909 | 17 | 0.1765 | 7 | 0 | 44 | 0.0909 |
| 5_12376831 | 3 | 0.6667 | 1 | 1 | NA | NA | 1 | 1 | 5 | 0.8 | 2 | 0.5 | 1 | 0 | 3 | 0.3333 | NA | NA | 6 | 0.3333 |
| 5_12376853 | 3 | 0.3333 | NA | NA | 2 | 0 | 4 | 0.25 | 9 | 0.2222 | 4 | 0.25 | 6 | 0 | 5 | 0.2 | 4 | 0 | 19 | 0.1053 |
| 5_12376858 | 1 | 1 | 1 | 0 | 1 | 1 | NA | NA | 3 | 0.6667 | 3 | 0.3333 | 4 | 0.25 | 6 | 0.1667 | 2 | 0 | 15 | 0.2 |
| 5_12376869 | 3 | 1 | 1 | 1 | 2 | 1 | 3 | 1 | 9 | 1 | 4 | 0.75 | 5 | 1 | 8 | 0.625 | 3 | 1 | 20 | 0.8 |
| 5_12376874 | 3 | 0.6667 | NA | NA | 2 | 1 | 3 | 0.3333 | 8 | 0.625 | 5 | 0.6 | 5 | 0.8 | 7 | 0.2857 | 3 | 1 | 20 | 0.6 |
| 5_12376875 | 3 | 0 | NA | NA | 2 | 0.5 | 3 | 0.6667 | 8 | 0.375 | 5 | 0.2 | 6 | 0.1667 | 7 | 0.1429 | 3 | 0 | 21 | 0.1429 |
| 5_12376878 | 2 | 0 | NA | NA | 2 | 0.5 | 3 | 0.3333 | 7 | 0.2857 | 5 | 0.4 | 6 | 0.1667 | 4 | 0.75 | 3 | 0.6667 | 18 | 0.4444 |
| 5_12376901 | 2 | 0.5 | NA | NA | 1 | 0 | 3 | 0 | 6 | 0.1667 | 3 | 0 | 4 | 0 | 5 | 0 | 2 | 0.5 | 14 | 0.0714 |
| 5_12376918 | 3 | 0 | NA | NA | NA | NA | 3 | 0.3333 | 6 | 0.1667 | 3 | 0.3333 | 5 | 0 | 6 | 0 | 2 | 0 | 16 | 0.0625 |
| 5_12376919 | 3 | 0 | NA | NA | 1 | 1 | 3 | 0 | 7 | 0.1429 | 3 | 0 | 5 | 0.2 | 6 | 0 | 1 | 0 | 15 | 0.0667 |
| 5_12379173 | 2 | 0.5 | NA | NA | NA | NA | NA | NA | 2 | 0.5 | 2 | 0 | 5 | 0.4 | 1 | 0 | NA | NA | 8 | 0.25 |
| 5_12379193 | 1 | 1 | NA | NA | NA | NA | NA | NA | 1 | 1 | 2 | 0.5 | 3 | 0.3333 | 1 | 0 | NA | NA | 6 | 0.3333 |
| 5_12383827 | 2 | 0.5 | 7 | 1 | 1 | 1 | 4 | 0.75 | 14 | 0.8571 | 3 | 0.6667 | 2 | 1 | 8 | 1 | 5 | 1 | 18 | 0.9444 |
| 5_12383834 | 2 | 0 | 7 | 0.1429 | 1 | 0 | 4 | 0 | 14 | 0.0714 | 2 | 0 | 2 | 0.5 | 8 | 0 | 5 | 0.2 | 17 | 0.1176 |
| 5_12383841 | 2 | 1 | 7 | 1 | 1 | 1 | 4 | 0.5 | 14 | 0.8571 | 2 | 0.5 | 1 | 1 | 7 | 0.8571 | 4 | 1 | 14 | 0.8571 |
| 5_12383842 | 2 | 0 | 7 | 0.1429 | 1 | 0 | 3 | 0 | 13 | 0.0769 | 1 | 0 | 2 | 0 | 8 | 0.125 | 5 | 0.2 | 16 | 0.125 |
| 5_12383846 | 2 | 0 | 7 | 0.2857 | 1 | 0 | 4 | 0 | 14 | 0.1429 | 2 | 0 | 3 | 0 | 8 | 0.125 | 5 | 0 | 18 | 0.0556 |
| 5_12383850 | 1 | 1 | 7 | 0.4286 | 1 | 0 | 4 | 0.25 | 13 | 0.3846 | 2 | 0 | 2 | 0 | 5 | 0.6 | 5 | 0.4 | 14 | 0.3571 |
| 5_12383854 | 1 | 0 | 7 | 0.1429 | 1 | 0 | 4 | 0 | 13 | 0.0769 | 2 | 0 | 3 | 0 | 8 | 0.125 | 5 | 0 | 18 | 0.0556 |
| 5_12383856 | 2 | 0 | 4 | 0.25 | 1 | 0 | 3 | 0 | 10 | 0.1 | 2 | 0 | 2 | 0 | 5 | 0.2 | 5 | 0 | 14 | 0.0714 |
| 5_12383859 | NA | NA | 2 | 1 | NA | NA | 1 | 1 | 3 | 1 | NA | NA | 1 | 1 | 3 | 0.6667 | 2 | 0 | 6 | 0.5 |
| 5_12383863 | 2 | 0.5 | 2 | 0.5 | NA | NA | 2 | 0.5 | 6 | 0.5 | 1 | 1 | 1 | 0 | 2 | 1 | 3 | 1 | 7 | 0.8571 |
| 5_12383863 | 2 | 0 | 6 | 0.6667 | 1 | 0 | 2 | 0.5 | 11 | 0.4545 | 1 | 1 | 1 | 1 | 7 | 0.2857 | 5 | 0.6 | 14 | 0.5 |
| 5_12383875 | 2 | 0.5 | 7 | 0.1429 | 1 | 0 | 3 | 0.6667 | 13 | 0.3077 | 1 | 1 | 3 | 1 | 8 | 0.875 | 4 | 0.75 | 16 | 0.875 |
| 5_12383876 | 2 | 0 | 7 | 0.1429 | 1 | 0 | 3 | 0.3333 | 13 | 0.1538 | 2 | 0.5 | 3 | 0 | 8 | 0.125 | 5 | 0 | 18 | 0.1111 |
| 5_12383887 | 2 | 0.5 | 6 | 0.5 | 1 | 0 | 4 | 1 | 13 | 0.6154 | 2 | 1 | 2 | 1 | 6 | 0.8333 | 4 | 1 | 14 | 0.9286 |
| 5_12383903 | 1 | 1 | 5 | 0.2 | NA | NA | 4 | 0.25 | 10 | 0.3 | 1 | 0 | 3 | 0.6667 | 6 | 0.1667 | 2 | 0 | 12 | 0.25 |
| 5_12383904 | 2 | 0.5 | 5 | 0.2 | 1 | 0 | 4 | 0.25 | 12 | 0.25 | 1 | 1 | 3 | 0.3333 | 8 | 0 | 4 | 0.25 | 16 | 0.1875 |
| 5_12386703 | 15 | 0.2667 | 7 | 0.5714 | 9 | 0.1111 | 15 | 0.0667 | 46 | 0.2174 | 8 | 0.25 | 12 | 0.0833 | 15 | 0.1333 | 13 | 0.0769 | 48 | 0.125 |
| 5_12386704 | 17 | 0.0588 | 7 | 0.2857 | 8 | 0 | 15 | 0.0667 | 47 | 0.0851 | 8 | 0.25 | 11 | 0 | 15 | 0.0667 | 14 | 0.1429 | 48 | 0.1042 |
| 5_12386714 | 16 | 0.1875 | 7 | 0.1429 | 8 | 0.125 | 14 | 0 | 45 | 0.1111 | 6 | 0.1667 | 13 | 0.0769 | 15 | 0.2 | 15 | 0.0667 | 49 | 0.1224 |
| 5_12386721 | 16 | 0.1875 | 5 | 0.2 | 7 | 0.2857 | 13 | 0.1538 | 41 | 0.1951 | 4 | 0 | 14 | 0.1429 | 12 | 0.25 | 14 | 0.1429 | 44 | 0.1591 |
| 5_12386722 | 16 | 0.6875 | 5 | 0.8 | 4 | 0.75 | 9 | 0.6667 | 34 | 0.7059 | 4 | 0.25 | 10 | 0.8 | 12 | 0.5 | 10 | 0.7 | 36 | 0.6111 |
| 5_12406589 | 1 | 0 | NA | NA | NA | NA | 2 | 0.5 | 3 | 0.3333 | NA | NA | 1 | 1 | NA | NA | NA | NA | 1 | 1 |
| 5_12414035 | 2 | 1 | 9 | 0.6667 | 3 | 0.6667 | 10 | 0.4 | 24 | 0.5833 | 6 | 1 | 3 | 0.3333 | 3 | 0.6667 | 6 | 0.8333 | 18 | 0.7778 |
| 5_12414039 | 4 | 0.5 | 11 | 0.3636 | 3 | 0 | 10 | 0.2 | 28 | 0.2857 | 5 | 0.2 | 5 | 0.2 | 5 | 0.6 | 4 | 0.25 | 19 | 0.3158 |
| 5_12414040 | 3 | 0.6667 | 12 | 0.3333 | 5 | 0.4 | 11 | 0.2727 | 31 | 0.3548 | 8 | 0.25 | 5 | 0.2 | 5 | 0.4 | 6 | 0.3333 | 24 | 0.2917 |

| | | | | | | | | | | | | | | | | | | | | |
|---|---|---|---|---|---|---|---|---|---|---|---|---|---|---|---|---|---|---|---|---|
| 5_12414054 | 4 | 0.5 | 9 | 0.2222 | 5 | 0.2 | 10 | 0.6 | 28 | 0.3929 | 8 | 0.125 | 5 | 0.4 | 6 | 0.5 | 6 | 0.1667 | 25 | 0.28 |
| 5_12414057 | 2 | 0 | 13 | 0.0769 | 2 | 0 | 9 | 0.1111 | 26 | 0.0769 | 8 | 0 | 6 | 0 | 6 | 0 | 7 | 0.2857 | 27 | 0.0741 |
| 5_12414116 | 3 | 0.6667 | 11 | 0.3636 | 2 | 1 | 7 | 0.5714 | 23 | 0.5217 | 3 | 0.6667 | 4 | 0.25 | 1 | 0 | 1 | 0 | 9 | 0.3333 |
| 5_12414224 | NA | NA | NA | NA | 1 | 1 | 1 | 1 | 2 | 1 | NA | NA | 3 | 0.3333 | 2 | 0.5 | 1 | 0 | 6 | 0.3333 |
| 5_12414262 | NA | NA | NA | NA | 1 | 0 | 1 | 1 | 2 | 0.5 | NA | NA | 3 | 0 | 2 | 0.5 | 2 | 0 | 7 | 0.1429 |
| 5_12414273 | 6 | 0.1667 | 6 | 0 | 5 | 0.2 | 7 | 0 | 24 | 0.0833 | 7 | 0.1429 | 4 | 0 | 3 | 0.3333 | 3 | 0 | 17 | 0.1176 |
| 5_12414289 | 6 | 0.1667 | 7 | 0 | 4 | 0 | 8 | 0.375 | 25 | 0.16 | 8 | 0.125 | 5 | 0 | 3 | 0.3333 | 3 | 0.3333 | 19 | 0.1579 |
| 5_12414292 | 6 | 1 | 7 | 1 | 4 | 0.75 | 8 | 0.875 | 25 | 0.92 | 7 | 0.8571 | 4 | 0.75 | 3 | 1 | 3 | 1 | 17 | 0.8824 |
| 5_12414295 | 5 | 1 | 5 | 1 | 2 | 1 | 6 | 1 | 18 | 1 | 7 | 0.8571 | 3 | 1 | 3 | 1 | 3 | 1 | 16 | 0.9375 |
| 5_12414307 | 6 | 0.1667 | 7 | 0.1429 | 3 | 0.3333 | 6 | 0.1667 | 22 | 0.1818 | 4 | 0 | 4 | 0.25 | 3 | 0.3333 | 3 | 0 | 14 | 0.1429 |
| 5_12414340 | 5 | 0.4 | 7 | 0.1429 | 3 | 0.3333 | 6 | 0.1667 | 21 | 0.2381 | 5 | 0.4 | 4 | 0.5 | 2 | 0.5 | 2 | 0 | 13 | 0.3846 |
| 5_12414372 | 5 | 0 | 2 | 0.5 | 1 | 0 | 4 | 0 | 12 | 0.0833 | 1 | 0 | 1 | 0 | 1 | 1 | 2 | 0.5 | 5 | 0.4 |
| 5_12414391 | 5 | 0.8 | 6 | 0.3333 | 2 | 0.5 | 5 | 0.4 | 18 | 0.5 | 4 | 0 | 2 | 0.5 | 1 | 1 | 2 | 0.5 | 9 | 0.3333 |
| 5_12414405 | 4 | 1 | 6 | 1 | 2 | 1 | 6 | 0.8333 | 18 | 0.9444 | 3 | 1 | 2 | 1 | 1 | 1 | 1 | 1 | 7 | 1 |
| 5_12414417 | 4 | 1 | 5 | 0.4 | 2 | 0.5 | 6 | 0.5 | 17 | 0.5882 | 2 | 0.5 | 2 | 0 | 1 | 1 | NA | NA | 5 | 0.4 |
| 5_12414435 | 4 | 0 | 6 | 0.1667 | 1 | 0 | 6 | 0 | 17 | 0.0588 | 3 | 0.3333 | 2 | 0.5 | 1 | 0 | 1 | 0 | 7 | 0.2857 |
| 5_12414447 | 2 | 0.5 | 6 | 0.8333 | 2 | 0.5 | 6 | 0.6667 | 16 | 0.6875 | 4 | 0.5 | 2 | 0 | 1 | 1 | 1 | 0 | 8 | 0.375 |
| 5_12414451 | 3 | 0.3333 | 5 | 0.4 | 2 | 0 | 4 | 0.5 | 14 | 0.3571 | 3 | 1 | 2 | 1 | 1 | 1 | 1 | 1 | 7 | 1 |
| 5_12414467 | 3 | 1 | 4 | 0.5 | 2 | 0 | 4 | 0.5 | 13 | 0.5385 | 3 | 1 | 2 | 1 | NA | NA | 1 | 1 | 6 | 1 |
| 5_12419995 | 14 | 0 | 8 | 0.125 | 11 | 0.0909 | 6 | 0 | 39 | 0.0513 | 4 | 0 | 10 | 0.1 | 11 | 0.0909 | 9 | 0.1111 | 34 | 0.0882 |
| 5_12420011 | 11 | 0.1818 | 6 | 0 | 8 | 0.125 | 7 | 0.2857 | 32 | 0.1562 | 4 | 0 | 11 | 0 | 8 | 0.25 | 9 | 0.2222 | 32 | 0.125 |
| 5_12420018 | 12 | 0.75 | 5 | 0.2 | 8 | 0.625 | 7 | 1 | 32 | 0.6875 | 3 | 0.3333 | 8 | 0.625 | 7 | 0.7143 | 12 | 0.6667 | 30 | 0.6333 |
| 5_12420019 | 10 | 0.5 | 6 | 0 | 12 | 0.3333 | 4 | 0.25 | 32 | 0.3125 | 6 | 0.3333 | 10 | 0.1 | 6 | 0 | 11 | 0.5455 | 33 | 0.2727 |
| 5_12420037 | 13 | 0.4615 | 4 | 0.25 | 10 | 0.8 | 5 | 0.2 | 32 | 0.5 | 6 | 0.8333 | 8 | 0.375 | 7 | 0.5714 | 7 | 0.4286 | 28 | 0.5357 |
| 5_12420040 | 13 | 0.5385 | 6 | 0.1667 | 9 | 0.4444 | 5 | 0.4 | 33 | 0.4242 | 5 | 0 | 8 | 0.25 | 9 | 0.4444 | 9 | 0.1111 | 31 | 0.2258 |
| 5_12420041 | 14 | 0.6429 | 7 | 0.5714 | 11 | 0.4545 | 7 | 0.7143 | 39 | 0.5897 | 6 | 0.6667 | 10 | 0.6 | 10 | 0.4 | 8 | 0.25 | 34 | 0.4706 |
| 5_12420043 | 13 | 0.4615 | 5 | 0.4 | 11 | 0.3636 | 5 | 0.2 | 34 | 0.3824 | 2 | 0.5 | 10 | 0.5 | 10 | 0.2 | 7 | 0.5714 | 29 | 0.4138 |
| 5_12420047 | 11 | 1 | 5 | 1 | 10 | 1 | 4 | 1 | 30 | 1 | 2 | 1 | 7 | 1 | 6 | 1 | 6 | 1 | 21 | 1 |
| 5_12420048 | 9 | 0.5556 | 5 | 0 | 7 | 0.8571 | 4 | 0 | 25 | 0.44 | 5 | 0.6 | 7 | 0.4286 | 8 | 0.75 | 6 | 0.3333 | 26 | 0.5385 |
| 5_12420073 | 9 | 0.2222 | 4 | 0.25 | 12 | 0.25 | 4 | 0.25 | 29 | 0.2414 | 2 | 0 | 8 | 0.125 | 7 | 0.1429 | 8 | 0 | 25 | 0.08 |
| 5_12420077 | 12 | 0.8333 | 3 | 0.3333 | 12 | 0.75 | 7 | 0.5714 | 34 | 0.7059 | 5 | 0.8 | 7 | 0.5714 | 9 | 0.6667 | 6 | 0.5 | 27 | 0.6296 |
| 5_12420088 | 13 | 0.1538 | 6 | 0.1667 | 8 | 0.375 | 7 | 0.4286 | 34 | 0.2647 | 3 | 0 | 8 | 0.25 | 8 | 0.125 | 7 | 0 | 26 | 0.1154 |
| 5_12422117 | 8 | 0 | 2 | 0 | 3 | 0 | 5 | 0.4 | 18 | 0.1111 | NA | NA | 2 | 0 | 3 | 0 | 3 | 0.6667 | 8 | 0.25 |
| 5_12422787 | 4 | 1 | 1 | 1 | 4 | 0.25 | 3 | 1 | 12 | 0.75 | NA | NA | 3 | 0.6667 | 2 | 1 | 1 | 1 | 6 | 0.8333 |
| 5_12422790 | 7 | 0.5714 | 2 | 1 | 4 | 1 | 4 | 0.5 | 17 | 0.7059 | NA | NA | 3 | 0.6667 | 2 | 1 | 1 | 1 | 6 | 0.8333 |
| 5_12422794 | 3 | 0.6667 | 1 | 0 | 3 | 0.3333 | 3 | 0.6667 | 10 | 0.5 | NA | NA | 3 | 0.6667 | 4 | 0.25 | 1 | 1 | 8 | 0.5 |
| 5_12422802 | 4 | 0.75 | 1 | 1 | 3 | 1 | NA | NA | 8 | 0.875 | NA | NA | 2 | 1 | NA | NA | 1 | 1 | 3 | 1 |
| 5_12422809 | 6 | 1 | 2 | 1 | 4 | 1 | 1 | 0 | 13 | 0.9231 | NA | NA | 3 | 1 | 4 | 1 | 1 | 0 | 8 | 0.875 |
| 5_12422813 | 6 | 0.3333 | 2 | 0 | 4 | 0.25 | 2 | 0 | 14 | 0.2143 | NA | NA | 3 | 0.3333 | 4 | 0.25 | NA | NA | 7 | 0.2857 |
| 5_12425221 | 2 | 0 | 1 | 1 | 4 | 0 | NA | NA | 7 | 0.1429 | NA | NA | 2 | 0 | 2 | 0 | 3 | 0.3333 | 7 | 0.1429 |
| 5_12425246 | 6 | 0 | 1 | 0 | 3 | 0.3333 | 3 | 0 | 13 | 0.0769 | 2 | 0 | 2 | 0 | 2 | 0.5 | 3 | 0 | 9 | 0.1111 |
| 5_12425254 | 8 | 0.25 | NA | NA | 4 | 0.25 | 2 | 0 | 14 | 0.2143 | 2 | 0 | 3 | 0 | 4 | 0 | 2 | 0.5 | 11 | 0.0909 |
| 5_12425266 | 8 | 0.25 | 1 | 0 | 3 | 0.6667 | 2 | 0 | 14 | 0.2857 | 1 | 0 | 2 | 0.5 | 4 | 0.25 | 2 | 0.5 | 9 | 0.3333 |
| 5_12425303 | 9 | 0.1111 | 2 | 0 | 6 | 0.1667 | 3 | 0 | 20 | 0.1 | 3 | 0 | 3 | 0.3333 | 4 | 0.25 | 4 | 0 | 14 | 0.1429 |
| 5_12425317 | 6 | 0.1667 | 1 | 0 | 6 | 0.1667 | 1 | 0 | 14 | 0.1429 | 2 | 0 | 2 | 0.5 | 2 | 0 | 1 | 0 | 7 | 0.1429 |
| 5_12425336 | 9 | 0.3333 | 1 | 0 | 8 | 0.125 | 2 | 0 | 20 | 0.2 | 3 | 0.3333 | 2 | 0 | 4 | 0.5 | 2 | 0 | 11 | 0.2727 |
| 5_12425343 | 9 | 0 | 2 | 0.5 | 6 | 0 | 3 | 0 | 20 | 0.05 | 3 | 0.6667 | 2 | 0 | 5 | 0 | 2 | 0 | 12 | 0.1667 |
| 5_12425461 | 6 | 0.1667 | 5 | 0.2 | 7 | 0 | 7 | 0.2857 | 25 | 0.16 | 7 | 0.1429 | 6 | 0 | 6 | 0.1667 | 8 | 0.25 | 27 | 0.1481 |
| 5_12425524 | 7 | 0 | 5 | 0 | 8 | 0.125 | 9 | 0.2222 | 29 | 0.1034 | 10 | 0.2 | 9 | 0 | 8 | 0 | 12 | 0.0833 | 39 | 0.0769 |
| 5_12425704 | 28 | 0.1071 | 31 | 0.0645 | 32 | 0.0938 | 37 | 0.1622 | 128 | 0.1094 | 18 | 0.2222 | 35 | 0.1143 | 29 | 0.069 | 26 | 0.0385 | 108 | 0.1019 |
| 5_12425738 | 31 | 0.1613 | 35 | 0.0571 | 30 | 0.1667 | 33 | 0.1515 | 129 | 0.1318 | 16 | 0 | 35 | 0.1714 | 34 | 0.1765 | 27 | 0.1111 | 112 | 0.1339 |
| 5_12425768 | 28 | 0.4643 | 31 | 0.5161 | 25 | 0.32 | 34 | 0.4118 | 118 | 0.4322 | 14 | 0.4286 | 36 | 0.4167 | 27 | 0.3704 | 27 | 0.5926 | 104 | 0.4519 |
| 5_12425770 | 25 | 0.32 | 35 | 0.2286 | 23 | 0.1739 | 35 | 0.1143 | 118 | 0.2034 | 15 | 0.1333 | 37 | 0.1622 | 30 | 0.0667 | 27 | 0.2222 | 109 | 0.1468 |
| 5_12425779 | 6 | 0 | 11 | 0.4545 | 2 | 0.5 | 9 | 0.1111 | 28 | 0.25 | 6 | 0.3333 | 9 | 0.3333 | 10 | 0.3 | 7 | 0.2857 | 32 | 0.3125 |
| 5_12425779 | 10 | 0.4 | 9 | 0.5556 | 6 | 0.3333 | 10 | 0.6 | 35 | 0.4857 | 3 | 0.3333 | 8 | 0.375 | 6 | 0.3333 | 8 | 0.625 | 25 | 0.44 |
| 5_12425779 | 3 | 0 | 5 | 0.6 | 4 | 0.5 | 4 | 0.25 | 16 | 0.375 | 2 | 0 | NA | NA | 2 | 0.5 | 2 | 0 | 6 | 0.1667 |
| 5_12425780 | 1 | 1 | 2 | 0 | 1 | 0 | 1 | 0 | 5 | 0.2 | NA | NA | 2 | 1 | 1 | 0 | 1 | 0 | 4 | 0.5 |
| 5_12425786 | 16 | 0.125 | 22 | 0.0909 | 20 | 0.15 | 28 | 0.0714 | 86 | 0.1047 | 12 | 0.25 | 31 | 0.1935 | 24 | 0.0417 | 20 | 0.1 | 87 | 0.1379 |
| 5_12425789 | 29 | 0.2069 | 35 | 0.2571 | 23 | 0.087 | 35 | 0.1143 | 122 | 0.1721 | 15 | 0.2 | 37 | 0.2432 | 35 | 0.1714 | 25 | 0.32 | 112 | 0.2321 |
| 5_12425822 | 23 | 0.2174 | 26 | 0.2692 | 15 | 0.2 | 25 | 0.04 | 89 | 0.1798 | 12 | 0.25 | 27 | 0.2222 | 29 | 0.069 | 12 | 0.0833 | 80 | 0.15 |
| 5_12425850 | 33 | 0.4545 | 30 | 0.5333 | 31 | 0.5806 | 33 | 0.5152 | 127 | 0.5197 | 18 | 0.4444 | 44 | 0.5909 | 31 | 0.5484 | 27 | 0.3333 | 120 | 0.5 |
| 5_12425859 | 30 | 0.5667 | 30 | 0.4667 | 30 | 0.7333 | 31 | 0.7097 | 121 | 0.6198 | 20 | 0.5 | 41 | 0.7073 | 34 | 0.4412 | 26 | 0.5385 | 121 | 0.562 |
| 5_12425864 | 35 | 0.2 | 30 | 0.1667 | 28 | 0.1071 | 35 | 0.2 | 128 | 0.1719 | 23 | 0.1739 | 43 | 0.2326 | 33 | 0.2121 | 25 | 0.16 | 124 | 0.2016 |
| 5_12425880 | 32 | 0.125 | 33 | 0.2727 | 30 | 0.1333 | 32 | 0.3125 | 127 | 0.2126 | 21 | 0 | 35 | 0.1429 | 31 | 0.129 | 26 | 0.1923 | 113 | 0.1239 |
| 5_12425881 | 31 | 0.0645 | 32 | 0.1875 | 29 | 0.2069 | 33 | 0.303 | 125 | 0.192 | 20 | 0.15 | 40 | 0.25 | 33 | 0.1212 | 27 | 0.2222 | 120 | 0.1917 |

| | | | | | | | | | | | | | | | | | | | | |
|---|---|---|---|---|---|---|---|---|---|---|---|---|---|---|---|---|---|---|---|---|
| 5_12425889 | 24 | 0.7083 | 29 | 0.5862 | 29 | 0.5862 | 32 | 0.5938 | 114 | 0.614 | 19 | 0.5789 | 34 | 0.6176 | 27 | 0.6667 | 21 | 0.4762 | 101 | 0.5941 |
| 5_12425918 | 26 | 0.0769 | 30 | 0.1 | 29 | 0.069 | 30 | 0.0667 | 115 | 0.0783 | 19 | 0.1579 | 37 | 0.1351 | 28 | 0.0357 | 21 | 0.0952 | 105 | 0.1048 |
| 5_12425921 | 24 | 0.5 | 30 | 0.3667 | 28 | 0.5 | 26 | 0.4231 | 108 | 0.4444 | 17 | 0.5294 | 32 | 0.4375 | 26 | 0.4231 | 23 | 0.5217 | 98 | 0.4694 |
| 5_12425922 | 26 | 1 | 25 | 0.88 | 23 | 0.9565 | 28 | 0.8929 | 102 | 0.9314 | 16 | 0.9375 | 33 | 0.9697 | 25 | 0.76 | 23 | 0.8696 | 97 | 0.8866 |
| 5_12425932 | 20 | 0.95 | 23 | 1 | 25 | 0.92 | 24 | 0.9583 | 92 | 0.9565 | 16 | 0.875 | 26 | 1 | 24 | 0.9583 | 20 | 1 | 86 | 0.9651 |
| 5_12425933 | 26 | 0.8846 | 24 | 0.7083 | 30 | 0.8333 | 26 | 0.8077 | 106 | 0.8113 | 19 | 0.9474 | 37 | 0.8919 | 29 | 0.8966 | 23 | 0.8261 | 108 | 0.8889 |
| 5_12425935 | 25 | 0.36 | 28 | 0.4286 | 28 | 0.6429 | 28 | 0.3929 | 109 | 0.4587 | 18 | 0.5 | 39 | 0.5385 | 24 | 0.375 | 21 | 0.4286 | 102 | 0.4706 |
| 5_12425942 | 20 | 0.8 | 22 | 0.7727 | 25 | 0.8 | 23 | 0.6087 | 90 | 0.7444 | 15 | 0.8 | 31 | 0.8065 | 25 | 0.72 | 19 | 0.8947 | 90 | 0.8 |
| 5_12425944 | 27 | 0.3333 | 24 | 0.4583 | 24 | 0.4167 | 23 | 0.3478 | 98 | 0.3878 | 17 | 0.4118 | 28 | 0.4643 | 21 | 0.2857 | 22 | 0.4545 | 88 | 0.4091 |
| 5_12425952 | 24 | 0.4583 | 23 | 0.3478 | 28 | 0.3571 | 27 | 0.3704 | 102 | 0.3824 | 19 | 0.3158 | 37 | 0.4865 | 25 | 0.44 | 19 | 0.2632 | 100 | 0.4 |
| 5_12432317 | 2 | 1 | NA | NA | NA | NA | NA | NA | 2 | 1 | NA | NA | 1 | 1 | NA | NA | NA | NA | 1 | 1 |
| 5_12432376 | 2 | 1 | NA | NA | NA | NA | NA | NA | 2 | 1 | NA | NA | 1 | 1 | NA | NA | NA | NA | 1 | 1 |
| 5_12432378 | 1 | 1 | NA | NA | NA | NA | NA | NA | 1 | 1 | NA | NA | 1 | 1 | NA | NA | NA | NA | 1 | 1 |
| 5_12432395 | 2 | 0.5 | NA | NA | NA | NA | NA | NA | 2 | 0.5 | NA | NA | 1 | 1 | NA | NA | NA | NA | 1 | 1 |
| 5_12432411 | 2 | 0.5 | NA | NA | NA | NA | NA | NA | 2 | 0.5 | NA | NA | 1 | 1 | NA | NA | NA | NA | 1 | 1 |
| 5_12432444 | 1 | 1 | NA | NA | NA | NA | NA | NA | 1 | 1 | NA | NA | 1 | 1 | NA | NA | NA | NA | 1 | 1 |
| 5_12438923 | 6 | 0.6667 | 3 | 0.6667 | 1 | 1 | 3 | 0.3333 | 13 | 0.6154 | 1 | 0 | 4 | 0.5 | 3 | 1 | 1 | 1 | 9 | 0.6667 |
| 5_12438940 | 7 | 0.1429 | 4 | 0.25 | 3 | 0 | 3 | 0 | 17 | 0.1176 | 2 | 0.5 | 4 | 0 | 4 | 0.25 | 1 | 0 | 11 | 0.1818 |
| 5_12438944 | 7 | 0.5714 | 3 | 1 | 1 | 0 | 3 | 1 | 14 | 0.7143 | 1 | 1 | 4 | 0.5 | 4 | 1 | 1 | 1 | 10 | 0.8 |
| 5_12438978 | 6 | 0.3333 | 3 | 0.3333 | 1 | 0 | 2 | 1 | 12 | 0.4167 | 2 | 0.5 | 4 | 0.5 | 3 | 0.6667 | 1 | 1 | 10 | 0.6 |
| 5_12439305 | 2 | 0 | 2 | 0.5 | 1 | 0 | 2 | 0 | 7 | 0.1429 | 1 | 0 | 1 | 0 | 3 | 0 | 1 | 1 | 6 | 0.1667 |
| 5_12439307 | 3 | 0.3333 | 2 | 0.5 | 1 | 1 | 2 | 1 | 8 | 0.625 | 1 | 0 | 1 | 1 | 4 | 0.5 | NA | NA | 6 | 0.5 |
| 5_12445356 | NA | NA | NA | NA | 1 | 1 | NA | NA | 1 | 1 | NA | NA | NA | NA | 1 | 1 | NA | NA | 1 | 1 |
| 5_12447883 | 1 | 1 | 2 | 0 | 1 | 0 | 2 | 1 | 6 | 0.5 | 1 | 0 | 1 | 1 | 1 | 1 | NA | NA | 3 | 0.6667 |
| 5_12447897 | 1 | 1 | 2 | 1 | 1 | 1 | 2 | 1 | 6 | 1 | 1 | 1 | 1 | 1 | 1 | 1 | NA | NA | 3 | 1 |
| 5_12447898 | 1 | 1 | 1 | 0 | 1 | 0 | NA | NA | 3 | 0.3333 | 1 | 1 | 1 | 1 | 1 | 1 | 1 | 0 | 4 | 0.75 |
| 5_12447940 | 1 | 0 | 2 | 0.5 | 1 | 0 | 2 | 0 | 6 | 0.1667 | NA | NA | 2 | 0.5 | NA | NA | 1 | 0 | 3 | 0.3333 |
| 5_12447949 | 1 | 0 | 2 | 0.5 | 1 | 1 | 2 | 0.5 | 6 | 0.5 | 1 | 0 | 2 | 1 | 1 | 1 | 1 | 1 | 5 | 0.8 |
| 5_12447951 | 1 | 1 | 2 | 0.5 | 2 | 1 | 2 | 1 | 7 | 0.8571 | 1 | 1 | 2 | 1 | 1 | 1 | 1 | 1 | 5 | 1 |
| 5_12447972 | 1 | 1 | 2 | 1 | 1 | 1 | 2 | 1 | 6 | 1 | 1 | 1 | 3 | 0.6667 | 2 | 0.5 | 1 | 0 | 7 | 0.5714 |
| 5_12448372 | 8 | 0 | 9 | 0.2222 | 8 | 0.125 | 4 | 0.75 | 29 | 0.2069 | 13 | 0.1538 | 15 | 0.1333 | 7 | 0.2857 | 10 | 0 | 45 | 0.1333 |
| 5_12448401 | 6 | 0.1667 | 9 | 0 | 7 | 0.4286 | 5 | 0.4 | 27 | 0.2222 | 13 | 0.0769 | 16 | 0 | 9 | 0.4444 | 10 | 0 | 48 | 0.1042 |
| 5_12448402 | 3 | 0.6667 | 11 | 0.6364 | 6 | 0.6667 | 3 | 1 | 23 | 0.6957 | 12 | 0.6667 | 15 | 0.8667 | 9 | 0.4444 | 10 | 0.8 | 46 | 0.7174 |
| 5_12450696 | 9 | 0.3333 | 7 | 0.1429 | 6 | 0.3333 | 6 | 0.1667 | 28 | 0.25 | 5 | 0 | 5 | 0 | 9 | 0.1111 | 4 | 0.25 | 23 | 0.087 |
| 5_12450709 | 9 | 0.8889 | 6 | 0.6667 | 6 | 1 | 8 | 0.75 | 29 | 0.8276 | 5 | 0.6 | 6 | 0.8333 | 10 | 0.8 | 6 | 0.8333 | 27 | 0.7778 |
| 5_12450762 | 8 | 0.25 | 6 | 0 | 4 | 0 | 8 | 0.125 | 26 | 0.1154 | 2 | 0 | 5 | 0.2 | 9 | 0.1111 | 4 | 0.25 | 20 | 0.15 |
| 5_12450920 | 3 | 0 | 3 | 0 | 4 | 0.25 | 7 | 0 | 17 | 0.0588 | 2 | 0.5 | NA | NA | 6 | 0.1667 | 3 | 0 | 11 | 0.1818 |
| 5_12451245 | 10 | 0.3 | 6 | 0.3333 | 7 | 0.2857 | 5 | 0.4 | 28 | 0.3214 | 2 | 0 | 6 | 0.3333 | 8 | 0.625 | 3 | 0.3333 | 19 | 0.4211 |
| 5_12451247 | 8 | 0.5 | 6 | 0.3333 | 5 | 0.8 | 4 | 0.25 | 23 | 0.4783 | 2 | 0.5 | 3 | 0.3333 | 7 | 0.7143 | 3 | 0.6667 | 15 | 0.6 |
| 5_12451252 | 9 | 1 | 7 | 0.7143 | 9 | 0.7778 | 4 | 0.75 | 29 | 0.8276 | 3 | 0.6667 | 6 | 0.8333 | 7 | 1 | 3 | 0.6667 | 19 | 0.8421 |
| 5_12459861 | 3 | 0 | 4 | 0.25 | 5 | 0 | 3 | 0 | 15 | 0.0667 | 2 | 1 | 3 | 0 | 3 | 0 | 4 | 0.25 | 12 | 0.25 |
| 5_12459880 | 2 | 0.5 | 4 | 0.5 | 5 | 0.4 | 4 | 0.5 | 15 | 0.4667 | 3 | 0.6667 | 3 | 0.3333 | 4 | 0.75 | 6 | 0.8333 | 16 | 0.6875 |
| 5_12459898 | 2 | 0.5 | 4 | 0.25 | 5 | 0.4 | 5 | 0.2 | 16 | 0.3125 | 4 | 0.5 | 3 | 0 | 5 | 0 | 5 | 0.4 | 17 | 0.2353 |
| 5_12459908 | 4 | 0 | 4 | 0.25 | 5 | 0 | 5 | 0.2 | 18 | 0.1111 | 4 | 0.25 | 3 | 0 | 5 | 0 | 6 | 0 | 18 | 0.0556 |
| 5_12459912 | 4 | 0 | 3 | 0 | 5 | 0 | 5 | 0.2 | 17 | 0.0588 | 3 | 0 | 1 | 0 | 4 | 0 | 5 | 0.2 | 13 | 0.0769 |
| 5_12459943 | 2 | 0 | 4 | 0 | 4 | 0 | 4 | 0.25 | 14 | 0.0714 | 5 | 0.4 | NA | NA | 4 | 0 | 5 | 0.4 | 14 | 0.2857 |
| 5_12459953 | 4 | 0 | 4 | 0 | 2 | 0 | 5 | 0.2 | 15 | 0.0667 | 5 | 0.4 | 3 | 0 | 4 | 0.25 | 5 | 0.2 | 17 | 0.2353 |
| 5_12460138 | 1 | 1 | NA | NA | 1 | 0 | 1 | 1 | 3 | 0.6667 | 2 | 0.5 | 4 | 0.5 | 1 | 0 | 3 | 0.6667 | 10 | 0.5 |
| 5_12460146 | 3 | 0 | 4 | 0 | 2 | 0.5 | 2 | 0 | 11 | 0.0909 | 2 | 0 | 4 | 0.25 | 2 | 0 | 3 | 0.3333 | 11 | 0.1818 |
| 5_12460156 | 3 | 0.3333 | 2 | 0.5 | 2 | 1 | 1 | 1 | 8 | 0.625 | 1 | 1 | NA | NA | NA | NA | 3 | 0.3333 | 4 | 0.5 |
| 5_12460162 | 3 | 1 | 3 | 0.6667 | 1 | 1 | NA | NA | 7 | 0.8571 | 1 | 1 | 2 | 0.5 | 2 | 1 | 1 | 1 | 6 | 0.8333 |
| 5_12460167 | 2 | 1 | 3 | 1 | 1 | 1 | 1 | 1 | 7 | 1 | 1 | 1 | 2 | 0.5 | NA | NA | 2 | 1 | 5 | 0.8 |
| 5_12460167 | 4 | 0 | 4 | 0.25 | 2 | 0 | 2 | 0.5 | 12 | 0.1667 | 3 | 0.3333 | 4 | 0 | 2 | 0 | 3 | 0 | 12 | 0.0833 |
| 5_12460168 | 3 | 0 | 2 | 0 | 2 | 0.5 | 2 | 0 | 9 | 0.1111 | 2 | 0.5 | 3 | 0 | 1 | 1 | 3 | 0.3333 | 9 | 0.3333 |
| 5_12461625 | 3 | 0.3333 | 1 | 0 | 2 | 0 | NA | NA | 6 | 0.1667 | 3 | 0 | 4 | 0.5 | 2 | 0 | 3 | 0 | 12 | 0.1667 |
| 5_12462836 | 1 | 1 | 2 | 0.5 | 2 | 0.5 | NA | NA | 5 | 0.6 | 1 | 1 | NA | NA | NA | NA | 1 | 0 | 2 | 0.5 |
| 5_12462844 | 1 | 0 | 1 | 1 | NA | NA | NA | NA | 2 | 0.5 | 1 | 1 | 1 | 1 | NA | NA | 2 | 0 | 4 | 0.5 |
| 5_12462856 | NA | NA | 2 | 1 | 2 | 1 | NA | NA | 4 | 1 | 1 | 1 | 1 | 1 | NA | NA | 2 | 1 | 4 | 1 |
| 5_12462857 | 1 | 1 | 2 | 1 | 2 | 1 | 1 | 1 | 6 | 1 | 1 | 0 | 1 | 1 | NA | NA | 2 | 1 | 4 | 0.75 |
| 5_12463126 | 7 | 0.8571 | 7 | 0.7143 | 8 | 0.875 | 10 | 0.9 | 32 | 0.8438 | 7 | 0.8571 | 5 | 0.8 | 4 | 1 | 5 | 1 | 21 | 0.9048 |
| 5_12463133 | 3 | 0 | 2 | 0 | 5 | 0.2 | 2 | 0 | 12 | 0.0833 | 1 | 0 | 2 | 0 | NA | NA | 1 | 1 | 4 | 0.25 |
| 5_12463151 | 7 | 0 | 6 | 0.3333 | 4 | 0 | 9 | 0.1111 | 26 | 0.1154 | 4 | 0.75 | 5 | 0 | 5 | 0 | 7 | 0.1429 | 21 | 0.1905 |
| 5_12463158 | 5 | 0 | 8 | 0 | 4 | 0.25 | 9 | 0.4444 | 26 | 0.1923 | 3 | 0.6667 | 5 | 0.4 | 5 | 0.6 | 7 | 0.5714 | 20 | 0.55 |
| 5_12463166 | 5 | 0.2 | 7 | 0.2857 | 4 | 1 | 7 | 0.8571 | 23 | 0.5652 | 3 | 0.6667 | 3 | 0.6667 | 5 | 0.8 | 6 | 0.6667 | 17 | 0.7059 |
| 5_12463166 | NA | NA | 1 | 1 | NA | NA | 1 | 0 | 2 | 0.5 | NA | NA | 1 | 1 | NA | NA | NA | NA | 1 | 1 |

| | | | | | | | | | | | | | | | | | | | | |
|---|---|---|---|---|---|---|---|---|---|---|---|---|---|---|---|---|---|---|---|---|
| 5_12463167 | 5 | 0.2 | 9 | 0.4444 | 4 | 0.5 | 8 | 1 | 26 | 0.5769 | 4 | 0.75 | 5 | 0.4 | 5 | 0.4 | 7 | 0.7143 | 21 | 0.5714 |
| 5_12463169 | 3 | 0.6667 | 7 | 0.5714 | 3 | 1 | 7 | 1 | 20 | 0.8 | NA | NA | 5 | 0.6 | 4 | 0.75 | 4 | 1 | 13 | 0.7692 |
| 5_12463174 | 4 | 0.25 | 6 | 0 | 4 | 0.5 | 8 | 0.625 | 22 | 0.3636 | 3 | 1 | 4 | 0.5 | 4 | 0.75 | 5 | 0.4 | 16 | 0.625 |
| 5_12463176 | 5 | 0.4 | 8 | 0.375 | 4 | 0.5 | 4 | 0.25 | 21 | 0.381 | 2 | 0 | 5 | 0.4 | 4 | 0.5 | 7 | 0.2857 | 18 | 0.3333 |
| 5_12463390 | 1 | 1 | NA | NA | NA | NA | NA | NA | 1 | 1 | NA | NA | NA | NA | NA | NA | 2 | 0.5 | 2 | 0.5 |
| 5_12463397 | 1 | 1 | NA | NA | NA | NA | NA | NA | 1 | 1 | NA | NA | NA | NA | NA | NA | 2 | 0.5 | 2 | 0.5 |
| 5_12463399 | 1 | 1 | NA | NA | NA | NA | NA | NA | 1 | 1 | NA | NA | NA | NA | NA | NA | 2 | 1 | 2 | 1 |
| 5_12463453 | 1 | 1 | NA | NA | NA | NA | NA | NA | 1 | 1 | NA | NA | NA | NA | NA | NA | 2 | 1 | 2 | 1 |
| 5_12463470 | 1 | 1 | NA | NA | NA | NA | NA | NA | 1 | 1 | NA | NA | NA | NA | NA | NA | 1 | 1 | 1 | 1 |
| 5_12463471 | 1 | 1 | NA | NA | NA | NA | NA | NA | 1 | 1 | NA | NA | NA | NA | NA | NA | 1 | 1 | 1 | 1 |
| 5_12463531 | NA | NA | 1 | 1 | NA | NA | NA | NA | 1 | 1 | NA | NA | NA | NA | NA | NA | 1 | 1 | 1 | 1 |
| 5_12466876 | 5 | 0.4 | 7 | 0.1429 | 4 | 0 | 5 | 0 | 21 | 0.1429 | NA | NA | 2 | 0 | 1 | 0 | 4 | 0.5 | 7 | 0.2857 |
| 5_12468722 | 18 | 0.2778 | 18 | 0.1111 | 20 | 0.15 | 15 | 0 | 71 | 0.1408 | 16 | 0.125 | 20 | 0.05 | 16 | 0 | 19 | 0.1053 | 71 | 0.0704 |
| 5_12468753 | 22 | 0.0909 | 18 | 0.1111 | 22 | 0.0455 | 14 | 0 | 76 | 0.0658 | 16 | 0.125 | 17 | 0.0588 | 17 | 0.1176 | 20 | 0.25 | 70 | 0.1429 |
| 5_12468778 | 26 | 0.0769 | 19 | 0.1579 | 21 | 0.1905 | 18 | 0 | 84 | 0.1071 | 18 | 0.1111 | 19 | 0.1579 | 19 | 0.1053 | 24 | 0.125 | 80 | 0.125 |
| 5_12468783 | 24 | 0.0833 | 18 | 0.0556 | 21 | 0.0952 | 15 | 0 | 78 | 0.0641 | 16 | 0 | 21 | 0.0952 | 19 | 0.1579 | 22 | 0 | 78 | 0.0641 |
| 5_12468816 | 28 | 0.1429 | 21 | 0.0476 | 26 | 0.1538 | 18 | 0.2778 | 93 | 0.1505 | 25 | 0.08 | 22 | 0.1818 | 22 | 0.1818 | 25 | 0.2 | 94 | 0.1596 |
| 5_12470674 | NA | NA | 2 | 1 | NA | NA | 7 | 0.5714 | 9 | 0.6667 | 3 | 1 | 1 | 1 | 2 | 0.5 | NA | NA | 6 | 0.8333 |
| 5_12470677 | 3 | 0 | 5 | 0.2 | NA | NA | 9 | 0 | 17 | 0.0588 | 4 | 0 | 1 | 0 | 4 | 0.25 | 2 | 0.5 | 11 | 0.1818 |
| 5_12470678 | 3 | 0.6667 | 4 | 0.75 | NA | NA | 8 | 0.625 | 15 | 0.6667 | 3 | 0 | 1 | 1 | 4 | 0.75 | NA | NA | 8 | 0.5 |
| 5_12470687 | 2 | 0.5 | 5 | 0 | NA | NA | 9 | 0.2222 | 16 | 0.1875 | 5 | 0 | 1 | 1 | 3 | 0 | 2 | 0 | 11 | 0.0909 |
| 5_12470689 | 3 | 0.6667 | 5 | 1 | NA | NA | 8 | 0.75 | 16 | 0.8125 | 5 | 1 | 1 | 1 | 4 | 1 | 1 | 0 | 11 | 0.9091 |
| 5_12470690 | 3 | 0.6667 | 4 | 1 | NA | NA | 8 | 0.875 | 15 | 0.8667 | 4 | 0.75 | 1 | 1 | 4 | 0.5 | 2 | 1 | 11 | 0.7273 |
| 5_12470766 | NA | NA | 4 | 0 | NA | NA | 3 | 0.6667 | 7 | 0.2857 | 2 | 0.5 | 2 | 0 | 2 | 0 | NA | NA | 6 | 0.1667 |
| 5_12470872 | 4 | 0.75 | 7 | 0.4286 | 1 | 0 | 1 | 0 | 13 | 0.4615 | 5 | 0.4 | 2 | 0.5 | NA | NA | 2 | 0 | 9 | 0.3333 |
| 5_12470891 | 4 | 0.25 | 8 | 0.125 | 4 | 0 | 3 | 0.3333 | 19 | 0.1579 | 4 | 0 | 5 | 0 | 5 | 0.2 | 6 | 0 | 20 | 0.05 |
| 5_12470895 | 6 | 0.1667 | 9 | 0.2222 | 4 | 0 | 3 | 0 | 22 | 0.1364 | 5 | 0.2 | 6 | 0.1667 | 6 | 0 | 7 | 0 | 24 | 0.0833 |
| 5_12470899 | 6 | 0.5 | 8 | 0.375 | 3 | 0 | 3 | 1 | 20 | 0.45 | 5 | 0.4 | 6 | 0.5 | 6 | 0.3333 | 7 | 0.5714 | 24 | 0.4583 |
| 5_12470904 | 5 | 0.4 | 8 | 0.125 | 3 | 0.6667 | 3 | 0.6667 | 19 | 0.3684 | 5 | 0 | 6 | 0.6667 | 6 | 0.3333 | 5 | 0.8 | 22 | 0.4545 |
| 5_12470905 | 6 | 0.1667 | 5 | 0.4 | 2 | 0 | 3 | 0.3333 | 16 | 0.25 | 5 | 0 | 3 | 0.6667 | 4 | 0 | 5 | 0 | 17 | 0.1176 |
| 5_12470913 | 5 | 0.8 | 7 | 1 | 2 | 1 | 3 | 1 | 17 | 0.9412 | 4 | 1 | 5 | 0.8 | 6 | 0.8333 | 4 | 1 | 19 | 0.8947 |
| 5_12470931 | 5 | 0.8 | 8 | 0.625 | 3 | 0.3333 | 3 | 0.3333 | 19 | 0.5789 | 4 | 0.75 | 4 | 0.75 | 5 | 0.4 | 5 | 0.4 | 18 | 0.5556 |
| 5_12470935 | 5 | 0.6 | 6 | 0.3333 | 3 | 0 | 2 | 0.5 | 16 | 0.375 | 4 | 0 | 2 | 0 | 5 | 0.4 | 3 | 0 | 14 | 0.1429 |
| 5_12471515 | 2 | 0.5 | 6 | 0 | 1 | 0 | 2 | 0 | 11 | 0.0909 | 2 | 0 | 3 | 0 | 3 | 0 | 5 | 0.4 | 13 | 0.1538 |
| 5_12473435 | 1 | 1 | 2 | 0.5 | 2 | 0.5 | 2 | 1 | 7 | 0.7143 | NA | NA | 1 | 0 | 4 | 0.5 | 5 | 0.8 | 10 | 0.6 |
| 5_12476527 | 1 | 0 | 2 | 0 | 4 | 0.5 | 1 | 1 | 8 | 0.375 | 3 | 0 | 2 | 0 | 6 | 0.3333 | 3 | 0 | 14 | 0.1429 |
| 5_12476547 | 1 | 0 | NA | NA | 4 | 0.75 | 2 | 1 | 7 | 0.7143 | 2 | 1 | 1 | 1 | 2 | 0 | 2 | 1 | 7 | 0.7143 |
| 5_12476563 | NA | NA | NA | NA | 1 | 1 | 1 | 0 | 2 | 0.5 | 2 | 0.5 | NA | NA | 1 | 1 | 1 | 0 | 4 | 0.5 |
| 5_12476572 | NA | NA | NA | NA | 2 | 0.5 | 1 | 1 | 3 | 0.6667 | 2 | 0.5 | NA | NA | 1 | 0 | NA | NA | 3 | 0.3333 |
| 5_12479890 | 3 | 1 | 3 | 0.3333 | 5 | 0.6 | 9 | 1 | 20 | 0.8 | 8 | 0.875 | 8 | 0.875 | 9 | 0.5556 | 5 | 0.4 | 30 | 0.7 |
| 5_12479894 | 4 | 0.25 | 3 | 0.3333 | 4 | 0.5 | 8 | 0.125 | 19 | 0.2632 | 6 | 0.1667 | 8 | 0.125 | 7 | 0.2857 | 3 | 0.3333 | 24 | 0.2083 |
| 5_12479905 | 3 | 0.6667 | 3 | 0.3333 | 4 | 0.5 | 8 | 0.75 | 18 | 0.6111 | 6 | 0.8333 | 7 | 0.8571 | 7 | 0.7143 | 4 | 0.75 | 24 | 0.7917 |
| 5_12479916 | 4 | 0.25 | 3 | 0 | 4 | 0.25 | 6 | 0.1667 | 17 | 0.1765 | 9 | 0.2222 | 7 | 0 | 8 | 0 | 4 | 0 | 28 | 0.0714 |
| 5_12479929 | 4 | 0.25 | 3 | 0 | 4 | 0.5 | 5 | 0.4 | 16 | 0.3125 | 7 | 0.4286 | 9 | 0.3333 | 6 | 0.5 | 4 | 0.75 | 26 | 0.4615 |
| 5_12479942 | 5 | 0.2 | 2 | 0.5 | 3 | 0 | 8 | 0.25 | 18 | 0.2222 | 11 | 0.1818 | 10 | 0 | 10 | 0.2 | 6 | 0.1667 | 37 | 0.1351 |
| 5_12479954 | 5 | 0.6 | 4 | 0.25 | 1 | 0 | 7 | 0.2857 | 17 | 0.3529 | 11 | 0.4545 | 8 | 0.25 | 11 | 0.3636 | 6 | 0.8333 | 36 | 0.4444 |
| 5_12479963 | 4 | 0.75 | 4 | 0.5 | 5 | 0.8 | 7 | 0.8571 | 20 | 0.75 | 11 | 0.5455 | 11 | 0.7273 | 12 | 0.5833 | 4 | 0.5 | 38 | 0.6053 |
| 5_12479964 | 5 | 0.4 | 2 | 0 | 5 | 0.6 | 5 | 0.6 | 17 | 0.4706 | 9 | 0.4444 | 11 | 0.6364 | 9 | 0.3333 | 6 | 0.1667 | 35 | 0.4286 |
| 5_12479999 | 1 | 0 | 3 | 0.3333 | 3 | 0 | 5 | 0 | 12 | 0.0833 | 8 | 0.25 | 10 | 0 | 7 | 0.1429 | 4 | 0 | 29 | 0.1034 |
| 5_12480028 | 1 | 0 | 2 | 0.5 | 5 | 0.2 | 6 | 0.6667 | 14 | 0.4286 | 6 | 0.3333 | 10 | 0.4 | 7 | 0.5714 | 3 | 0.3333 | 26 | 0.4231 |
| 5_12480079 | 2 | 0 | 3 | 0 | 5 | 0.2 | 4 | 0.75 | 14 | 0.2857 | 9 | 0.4444 | 10 | 0.1 | 8 | 0.25 | 4 | 0.75 | 31 | 0.3226 |
| 5_12480089 | 2 | 0 | 3 | 0.6667 | 5 | 0.2 | 6 | 0.3333 | 16 | 0.3125 | 10 | 0.4 | 10 | 0.2 | 7 | 0.4286 | 5 | 0.2 | 32 | 0.3125 |
| 5_12480089 | 2 | 0 | 3 | 0 | 5 | 0.4 | 6 | 0 | 16 | 0.125 | 9 | 0.2222 | 12 | 0.0833 | 7 | 0 | 5 | 0 | 33 | 0.0909 |
| 5_12480115 | NA | NA | 4 | 0 | 5 | 0.2 | 6 | 0.1667 | 15 | 0.1333 | 10 | 0.1 | 9 | 0 | 8 | 0.125 | 4 | 0 | 31 | 0.0645 |
| 5_12480126 | 3 | 0.3333 | 3 | 0 | 6 | 0.6667 | 3 | 0.3333 | 15 | 0.4 | 8 | 0.875 | 11 | 0.4545 | 8 | 0.5 | 5 | 0.4 | 32 | 0.5625 |
| 5_12480138 | 4 | 0.5 | 4 | 1 | 9 | 0.5556 | 6 | 0.8333 | 23 | 0.6957 | 12 | 0.4167 | 12 | 0.5 | 8 | 0.5 | 5 | 0.6 | 37 | 0.4865 |
| 5_12480145 | 1 | 1 | 5 | 0.2 | 9 | 0.6667 | 7 | 0.2857 | 22 | 0.4545 | 12 | 0.3333 | 12 | 0.3333 | 6 | 0.3333 | 4 | 0.5 | 34 | 0.3529 |
| 5_12491179 | 3 | 1 | 3 | 1 | 8 | 1 | 10 | 0.9 | 24 | 0.9583 | 10 | 0.9 | 8 | 0.875 | 8 | 0.625 | 2 | 0.5 | 28 | 0.7857 |
| 5_12491195 | 7 | 0.7143 | 4 | 0.5 | 8 | 0.875 | 12 | 0.8333 | 31 | 0.7742 | 10 | 0.6 | 8 | 0.75 | 8 | 0.625 | 3 | 0.6667 | 29 | 0.6552 |
| 5_12491200 | 6 | 1 | 3 | 1 | 7 | 1 | 10 | 0.9 | 26 | 0.9615 | 14 | 1 | 7 | 1 | 7 | 1 | 2 | 1 | 30 | 1 |
| 5_12491214 | 10 | 0.1 | 17 | 0.0588 | 19 | 0.1053 | 24 | 0 | 70 | 0.0571 | 19 | 0.0526 | 22 | 0.1818 | 18 | 0.1111 | 12 | 0 | 71 | 0.0986 |
| 5_12491225 | 7 | 1 | 12 | 0.9167 | 14 | 0.8571 | 17 | 0.9412 | 50 | 0.92 | 16 | 0.875 | 18 | 0.7778 | 15 | 0.8667 | 8 | 1 | 57 | 0.8596 |
| 5_12491245 | 11 | 0.2727 | 13 | 0.3846 | 16 | 0.4375 | 20 | 0.3 | 60 | 0.35 | 18 | 0.2778 | 19 | 0.2105 | 17 | 0.5294 | 10 | 0.4 | 64 | 0.3438 |
| 5_12491259 | 6 | 1 | 10 | 1 | 10 | 1 | 8 | 1 | 34 | 1 | 4 | 1 | 13 | 1 | 8 | 1 | 8 | 1 | 33 | 1 |
| 5_12491259 | 9 | 0 | 12 | 0 | 13 | 0 | 16 | 0.3125 | 50 | 0.1 | 16 | 0.1875 | 15 | 0.1333 | 12 | 0 | 9 | 0 | 52 | 0.0962 |

| | | | | | | | | | | | | | | | | | | | | |
|---|---|---|---|---|---|---|---|---|---|---|---|---|---|---|---|---|---|---|---|---|
| 5_1249126( | 7 | 0.8571 | 9 | 1 | 10 | 1 | 17 | 0.9412 | 43 | 0.9535 | 14 | 0.9286 | 13 | 0.9231 | 14 | 1 | 9 | 0.8889 | 50 | 0.94 |
| 5_1249126‹ | 11 | 0.1818 | 14 | 0.1429 | 15 | 0.3333 | 20 | 0.3 | 60 | 0.25 | 18 | 0.2222 | 18 | 0.3333 | 17 | 0.1765 | 11 | 0.3636 | 64 | 0.2656 |
| 5_1249129! | 10 | 0.2 | 11 | 0.0909 | 14 | 0.1429 | 17 | 0.4118 | 52 | 0.2308 | 15 | 0.3333 | 17 | 0.4706 | 14 | 0.1429 | 10 | 0.1 | 56 | 0.2857 |
| 5_1249131! | 8 | 0.125 | 8 | 0.125 | 12 | 0 | 19 | 0.1579 | 47 | 0.1064 | 12 | 0.0833 | 16 | 0.125 | 15 | 0 | 10 | 0.1 | 53 | 0.0755 |
| 5_1249390! | 8 | 0 | 11 | 0.1818 | 13 | 0.0769 | 17 | 0 | 49 | 0.0612 | 10 | 0.1 | 12 | 0.0833 | 15 | 0 | 11 | 0.0909 | 48 | 0.0625 |
| 5_1249621: | 4 | 0.5 | 2 | 1 | 3 | 0.3333 | 2 | 1 | 11 | 0.6364 | 1 | 1 | NA | NA | 1 | 0 | 1 | 1 | 3 | 0.6667 |
| 5_1249621! | 6 | 0.3333 | 3 | 0 | 4 | 0 | 3 | 0 | 16 | 0.125 | 2 | 0 | 1 | 0 | 5 | 0.2 | 3 | 0.3333 | 11 | 0.1818 |
| 5_1249623 | 6 | 0.6667 | 3 | 0.3333 | 4 | 0.5 | 3 | 0 | 16 | 0.4375 | 3 | 0.3333 | 2 | 0 | 5 | 0.4 | 3 | 0.3333 | 13 | 0.3077 |
| 5_1249624‹ | 6 | 0.1667 | 4 | 0 | 4 | 0 | 3 | 0 | 17 | 0.0588 | 2 | 0 | 1 | 0 | 5 | 0.2 | 3 | 0 | 11 | 0.0909 |
| 5_1249625( | 5 | 0.6 | 4 | 0.75 | 4 | 0.75 | 3 | 1 | 16 | 0.75 | 3 | 0.6667 | 2 | 0.5 | 2 | 1 | 2 | 1 | 9 | 0.7778 |
| 5_1249625! | 6 | 0.8333 | 3 | 1 | 3 | 0.6667 | 3 | 0.6667 | 15 | 0.8 | 1 | 1 | NA | NA | 5 | 0.4 | 3 | 0.6667 | 9 | 0.5556 |
| 5_1249625( | 6 | 0.5 | 4 | 1 | 4 | 0.25 | 3 | 1 | 17 | 0.6471 | 2 | 0.5 | 2 | 0.5 | 4 | 0.75 | 3 | 1 | 11 | 0.7273 |
| 5_1249628: | 4 | 0.25 | 3 | 0.6667 | 3 | 0.3333 | 3 | 0.3333 | 13 | 0.3846 | 2 | 0.5 | 1 | 0 | 1 | 0 | 3 | 0.3333 | 7 | 0.2857 |
| 5_1249635‹ | 7 | 0.2857 | 5 | 0.2 | 5 | 0 | 6 | 0 | 23 | 0.1304 | 3 | 0 | 2 | 0 | 6 | 0.1667 | 3 | 0 | 14 | 0.0714 |
| 5_1249636 | 7 | 0.2857 | 3 | 0.3333 | 4 | 0 | 5 | 0 | 19 | 0.1579 | 3 | 0 | 2 | 0 | 6 | 0.1667 | 4 | 0 | 15 | 0.0667 |
| 5_1249640 | 7 | 0.1429 | 6 | 0.3333 | 5 | 0.2 | 6 | 0.1667 | 24 | 0.2083 | 3 | 0 | 3 | 0.3333 | 3 | 0 | 3 | 0 | 12 | 0.0833 |
| 5_1249650( | 11 | 0 | 11 | 0.1818 | 12 | 0.0833 | 12 | 0 | 46 | 0.0652 | 7 | 0 | 5 | 0.2 | 12 | 0.0833 | 8 | 0 | 32 | 0.0625 |
| 5_1249654! | 9 | 0.1111 | 9 | 0.2222 | 9 | 0 | 11 | 0.2727 | 38 | 0.1579 | 8 | 0.125 | 4 | 0.25 | 10 | 0.1 | 8 | 0.125 | 30 | 0.1333 |
| 5_1250892: | 1 | 1 | NA | NA | NA | NA | NA | NA | 1 | 1 | NA | NA | 1 | 1 | NA | NA | NA | NA | 1 | 1 |
| 5_1250943 | 13 | 0.9231 | 20 | 0.9 | 5 | 0.8 | 10 | 0.7 | 48 | 0.8542 | 9 | 1 | 8 | 0.75 | 10 | 0.8 | 12 | 0.9167 | 39 | 0.8718 |
| 5_1250943! | 12 | 1 | 18 | 0.9444 | 5 | 0.8 | 6 | 0.8333 | 41 | 0.9268 | 8 | 0.75 | 9 | 0.8889 | 9 | 0.8889 | 7 | 1 | 33 | 0.8788 |
| 5_1250944! | 11 | 1 | 20 | 0.9 | 6 | 0.8333 | 9 | 0.8889 | 46 | 0.913 | 6 | 1 | 8 | 0.875 | 9 | 0.8889 | 9 | 0.7778 | 32 | 0.875 |
| 5_1250946( | 12 | 0.5833 | 13 | 0.7692 | 4 | 0.5 | 6 | 0.8333 | 35 | 0.6857 | 7 | 0.5714 | 4 | 0.75 | 6 | 0.6667 | 6 | 1 | 23 | 0.7391 |
| 5_1250946( | 12 | 0.75 | 12 | 0.75 | 4 | 0.5 | 6 | 0.8333 | 34 | 0.7353 | 2 | 0.5 | 9 | 1 | 7 | 0.5714 | 10 | 0.5 | 28 | 0.6786 |
| 5_1250946 | 14 | 0.1429 | 12 | 0.5833 | 7 | 0.5714 | 9 | 0.5556 | 42 | 0.4286 | 7 | 0.2857 | 8 | 0.75 | 10 | 0.2 | 8 | 0.25 | 33 | 0.3636 |
| 5_1250947! | 15 | 0.4667 | 15 | 1 | 6 | 0.5 | 8 | 0.5 | 44 | 0.6591 | 8 | 0.75 | 6 | 0.5 | 8 | 0.875 | 7 | 0.5714 | 29 | 0.6897 |
| 5_1250948( | 13 | 1 | 16 | 0.8125 | 5 | 1 | 8 | 1 | 42 | 0.9286 | 7 | 1 | 8 | 0.875 | 10 | 0.9 | 8 | 1 | 33 | 0.9394 |
| 5_1250948: | 12 | 0.1667 | 14 | 0.0714 | 5 | 0 | 8 | 0.25 | 39 | 0.1282 | 6 | 0.1667 | 8 | 0 | 9 | 0.1111 | 8 | 0.125 | 31 | 0.0968 |
| 5_1250948! | 8 | 0 | 13 | 0.0769 | 7 | 0 | 7 | 0.4286 | 35 | 0.1143 | 8 | 0.125 | 8 | 0 | 9 | 0.1111 | 7 | 0.1429 | 32 | 0.0938 |
| 5_1250949: | 10 | 0.2 | 8 | 0 | 2 | 0.5 | 3 | 0.6667 | 23 | 0.2174 | 2 | 0 | 9 | 0.1111 | 6 | 0 | 6 | 0.1667 | 23 | 0.087 |
| 5_1250949! | 8 | 0 | 7 | 0.2857 | 5 | 0.6 | 5 | 0.6 | 25 | 0.32 | 3 | 0.3333 | 2 | 0.5 | 6 | 0.5 | 9 | 0.3333 | 20 | 0.4 |
| 5_1250949( | 11 | 1 | 13 | 0.9231 | 6 | 0.6667 | 9 | 1 | 39 | 0.9231 | 2 | 1 | 7 | 0.8571 | 9 | 1 | 8 | 0.75 | 26 | 0.8846 |
| 5_1250950 | 11 | 0.3636 | 15 | 0.5333 | 4 | 0.5 | 9 | 0.6667 | 39 | 0.5128 | 6 | 0.3333 | 7 | 0.4286 | 10 | 0.3 | 11 | 0.3636 | 34 | 0.3529 |
| 5_1250950‹ | 13 | 0.8462 | 13 | 1 | 3 | 0.6667 | 7 | 0.8571 | 36 | 0.8889 | 6 | 0.5 | 5 | 0.6 | 5 | 0.6 | 8 | 0.875 | 24 | 0.6667 |
| 5_1250951‹ | 12 | 0.1667 | 15 | 0.2 | 4 | 0 | 7 | 0.1429 | 38 | 0.1579 | 6 | 0 | 6 | 0.6667 | 9 | 0.1111 | 8 | 0.375 | 29 | 0.2759 |
| 5_1250952: | 12 | 0.0833 | 9 | 0 | 4 | 0.25 | 6 | 0 | 31 | 0.0645 | 6 | 0 | 6 | 0.3333 | 6 | 0 | 8 | 0 | 26 | 0.0769 |
| 5_1250952‹ | 11 | 0.1818 | 12 | 0.3333 | 2 | 0 | 7 | 0.2857 | 32 | 0.25 | 5 | 0 | 6 | 0.3333 | 5 | 0.2 | 8 | 0.25 | 24 | 0.2083 |
| 5_1250952! | 8 | 0.125 | 13 | 0.3846 | 4 | 0 | 6 | 0.1667 | 31 | 0.2258 | 5 | 0.4 | 4 | 0 | 6 | 0 | 9 | 0.1111 | 24 | 0.125 |
| 5_1251039 | 6 | 0.3333 | 4 | 0.5 | 1 | 1 | 2 | 0.5 | 13 | 0.4615 | 1 | 0 | 4 | 0.25 | 5 | 0.2 | 3 | 0.3333 | 13 | 0.2308 |
| 5_1251251! | 1 | 1 | 1 | 1 | 1 | 1 | NA | NA | 3 | 1 | NA | NA | NA | NA | 1 | 0 | 1 | 1 | 2 | 0.5 |
| 5_1253546! | 7 | 0.8571 | 4 | 0.25 | 4 | 0.75 | 4 | 0.75 | 19 | 0.6842 | 6 | 0.3333 | 8 | 0.375 | 3 | 0.6667 | 8 | 0.5 | 25 | 0.44 |
| 5_1253547: | 7 | 0.4286 | 3 | 0 | 5 | 0.6 | 2 | 0.5 | 17 | 0.4118 | 5 | 0.6 | 4 | 0.5 | 4 | 0.5 | 9 | 0.5556 | 22 | 0.5455 |
| 5_1253547: | 8 | 0.125 | 2 | 0 | 5 | 0.6 | 3 | 0 | 18 | 0.2222 | 6 | 0.1667 | 8 | 0.125 | 4 | 0 | 10 | 0.3 | 28 | 0.1786 |
| 5_1253548‹ | 7 | 0.1429 | 4 | 0.25 | 3 | 0.3333 | 2 | 0 | 16 | 0.1875 | 3 | 0 | 4 | 0.5 | 2 | 1 | 8 | 0.5 | 17 | 0.4706 |
| 5_1254631! | 106 | 0.0566 | 112 | 0.0625 | 103 | 0.0388 | 111 | 0.0811 | 432 | 0.0602 | 78 | 0.0256 | 76 | 0.0526 | 92 | 0.0652 | 95 | 0.0737 | 341 | 0.0557 |
| 5_1254633: | 79 | 0.1646 | 87 | 0.1034 | 86 | 0.1512 | 86 | 0.0814 | 338 | 0.1243 | 61 | 0.1311 | 61 | 0.0984 | 78 | 0.2436 | 73 | 0.1507 | 273 | 0.1612 |
| 5_1254634( | 79 | 0.4937 | 87 | 0.4598 | 84 | 0.4524 | 87 | 0.5057 | 337 | 0.4777 | 60 | 0.4667 | 58 | 0.3966 | 78 | 0.4744 | 68 | 0.4706 | 264 | 0.4545 |
| 5_1254634‹ | 82 | 0.1585 | 88 | 0.0568 | 86 | 0.1744 | 94 | 0.0957 | 350 | 0.12 | 62 | 0.129 | 60 | 0.1 | 77 | 0.1558 | 68 | 0.0735 | 267 | 0.1161 |
| 5_1254635! | 75 | 0.12 | 88 | 0.0568 | 84 | 0.0595 | 93 | 0.0538 | 340 | 0.0706 | 58 | 0.1034 | 57 | 0.1053 | 67 | 0.1194 | 67 | 0.0896 | 249 | 0.1044 |
| 5_1254636: | 75 | 0.3733 | 84 | 0.3571 | 80 | 0.45 | 87 | 0.3218 | 326 | 0.3742 | 56 | 0.3036 | 55 | 0.3091 | 70 | 0.4 | 63 | 0.3492 | 244 | 0.3443 |
| 5_1254636( | 77 | 0.0909 | 85 | 0.0353 | 84 | 0.0357 | 85 | 0.0471 | 331 | 0.0514 | 57 | 0.0702 | 57 | 0.0702 | 71 | 0.0423 | 69 | 0.0435 | 254 | 0.0551 |
| 5_1254638( | 69 | 0.2754 | 77 | 0.1429 | 74 | 0.2432 | 83 | 0.241 | 303 | 0.2244 | 54 | 0.2222 | 52 | 0.2115 | 73 | 0.274 | 65 | 0.2615 | 244 | 0.2459 |
| 5_1254640 | 80 | 0.0875 | 81 | 0.0617 | 79 | 0.0127 | 90 | 0.0556 | 330 | 0.0545 | 53 | 0 | 54 | 0.1667 | 71 | 0.0986 | 75 | 0.08 | 253 | 0.087 |
| 5_1254640! | 83 | 0.2892 | 80 | 0.3 | 76 | 0.3289 | 88 | 0.3182 | 327 | 0.3089 | 55 | 0.3091 | 51 | 0.2941 | 75 | 0.3067 | 71 | 0.2958 | 252 | 0.3016 |
| 5_1254643 | 77 | 0.2987 | 79 | 0.2658 | 72 | 0.3056 | 82 | 0.1951 | 310 | 0.2645 | 47 | 0.1702 | 48 | 0.2917 | 68 | 0.1471 | 67 | 0.2537 | 230 | 0.213 |
| 5_1254645‹ | 77 | 0.2208 | 73 | 0.2055 | 76 | 0.1053 | 78 | 0.2308 | 304 | 0.1908 | 44 | 0.25 | 46 | 0.2391 | 71 | 0.1972 | 61 | 0.2623 | 222 | 0.2342 |
| 5_1254647 | 74 | 0.2703 | 71 | 0.1549 | 68 | 0.1176 | 76 | 0.1447 | 289 | 0.173 | 42 | 0.1429 | 38 | 0.1842 | 60 | 0.1333 | 54 | 0.2037 | 194 | 0.1649 |
| 5_1254650( | 61 | 0.3115 | 58 | 0.3103 | 57 | 0.2105 | 61 | 0.2459 | 237 | 0.27 | 38 | 0.2105 | 37 | 0.1892 | 57 | 0.1754 | 50 | 0.22 | 182 | 0.1978 |
| 5_1254652: | 43 | 0.2093 | 49 | 0.1837 | 55 | 0.1818 | 50 | 0.1 | 197 | 0.1675 | 33 | 0.1212 | 30 | 0.1667 | 46 | 0.1304 | 37 | 0.1081 | 146 | 0.1301 |
| 5_1254918: | NA | NA | 1 | 0 | 1 | 1 | NA | NA | 2 | 0.5 | NA | NA | NA | NA | 1 | 1 | 2 | 1 | 3 | 1 |
| 5_1254918! | NA | NA | 1 | 1 | 1 | 0 | NA | NA | 2 | 0.5 | NA | NA | NA | NA | 1 | 0 | 2 | 1 | 3 | 0.6667 |
| 5_1254919! | NA | NA | 1 | 1 | 1 | 1 | NA | NA | 2 | 1 | NA | NA | NA | NA | 1 | 1 | 2 | 1 | 3 | 1 |
| 5_1254920 | NA | NA | 1 | 1 | 1 | 1 | NA | NA | 2 | 1 | NA | NA | NA | NA | 1 | 1 | 2 | 1 | 3 | 1 |
| 5_1255314! | 6 | 0.3333 | 2 | 0 | 4 | 0 | 10 | 0.5 | 22 | 0.3182 | 8 | 0.375 | 2 | 0.5 | 5 | 0.2 | 4 | 0.25 | 19 | 0.3158 |
| 5_1255317! | 6 | 0.1667 | 4 | 0.25 | 3 | 0.3333 | 8 | 0.125 | 21 | 0.1905 | 8 | 0.5 | 3 | 0 | 4 | 0.5 | 4 | 0 | 19 | 0.3158 |

| | | | | | | | | | | | | | | | | | | | | |
|---|---|---|---|---|---|---|---|---|---|---|---|---|---|---|---|---|---|---|---|---|
| 5_1255318 | 7 | 0.4286 | 4 | 0.25 | 4 | 0.5 | 10 | 0.4 | 25 | 0.4 | 9 | 0.6667 | 3 | 0.3333 | 4 | 0.5 | 5 | 0.4 | 21 | 0.5238 |
| 5_1255318 | 6 | 0.1667 | 4 | 0.25 | 4 | 0 | 10 | 0.1 | 24 | 0.125 | 9 | 0.3333 | 3 | 0 | 5 | 0.4 | 5 | 0 | 22 | 0.2273 |
| 5_1255319 | 6 | 0.8333 | NA | NA | 3 | 1 | 9 | 1 | 18 | 0.9444 | 7 | 1 | 1 | 1 | 3 | 1 | 4 | 1 | 15 | 1 |
| 5_1255320 | 9 | 0.1111 | 5 | 0.2 | 5 | 0 | 7 | 0.2857 | 26 | 0.1538 | 8 | 0.375 | 3 | 0 | 6 | 0.3333 | 5 | 0 | 22 | 0.2273 |
| 5_1255320 | 8 | 0.125 | 5 | 0 | 4 | 0.5 | 7 | 0.1429 | 24 | 0.1667 | 8 | 0.125 | 1 | 0 | 5 | 0 | 3 | 1 | 17 | 0.2353 |
| 5_1255322 | 3 | 0 | 4 | 0.25 | 4 | 0.25 | 6 | 0.1667 | 17 | 0.1765 | 3 | 0.3333 | NA | NA | 2 | 0 | 2 | 0 | 7 | 0.1429 |
| 5_1255322 | 4 | 0 | 4 | 0 | 4 | 0 | 5 | 0.2 | 17 | 0.0588 | 3 | 0 | NA | NA | 3 | 0 | 3 | 0.3333 | 9 | 0.1111 |
| 5_1255323 | 3 | 1 | 4 | 1 | 2 | 1 | 6 | 1 | 15 | 1 | 2 | 1 | NA | NA | 2 | 1 | 2 | 1 | 6 | 1 |
| 5_1255325 | 4 | 1 | 4 | 1 | 5 | 0.8 | 6 | 1 | 19 | 0.9474 | 3 | 1 | NA | NA | 1 | 1 | 2 | 1 | 6 | 1 |
| 5_1255325 | 4 | 0 | 4 | 0.5 | 5 | 0.4 | 5 | 0.6 | 18 | 0.3889 | 3 | 0.6667 | NA | NA | 3 | 0.3333 | 1 | 1 | 7 | 0.5714 |
| 5_1255325 | 3 | 0.6667 | 4 | 0.25 | 5 | 0 | 7 | 0.2857 | 19 | 0.2632 | 2 | 0 | NA | NA | 2 | 0 | 2 | 0.5 | 6 | 0.1667 |
| 5_1255326 | 4 | 0.25 | 4 | 0 | 5 | 0 | 7 | 0.1429 | 20 | 0.1 | 3 | 0.3333 | NA | NA | 2 | 0 | 2 | 0.5 | 7 | 0.2857 |
| 5_1255327 | 2 | 1 | 3 | 0.3333 | 3 | 0.6667 | 5 | 1 | 13 | 0.7692 | 3 | 0.3333 | NA | NA | 1 | 0 | 2 | 0.5 | 6 | 0.3333 |
| 5_1255327 | 4 | 0.25 | 4 | 0.5 | 5 | 0.6 | 5 | 0.4 | 18 | 0.4444 | 2 | 0 | NA | NA | 3 | 0.3333 | 2 | 0.5 | 7 | 0.2857 |
| 5_1255327 | 2 | 1 | 3 | 0.6667 | 1 | 0 | 4 | 0.5 | 10 | 0.6 | 1 | 0 | NA | NA | 2 | 0 | 2 | 1 | 5 | 0.4 |
| 5_1255329 | 5 | 0.4 | 3 | 0.6667 | 4 | 0.75 | 7 | 0 | 19 | 0.3684 | 3 | 0.3333 | NA | NA | 3 | 0 | 1 | 0 | 7 | 0.1429 |
| 5_1255329 | 5 | 0.8 | 4 | 1 | 3 | 0.6667 | 6 | 0.6667 | 18 | 0.7778 | 3 | 0 | 1 | 1 | 3 | 0.6667 | 2 | 1 | 9 | 0.5556 |
| 5_1255331 | 5 | 0 | 3 | 0 | 6 | 0.3333 | 8 | 0 | 22 | 0.0909 | 3 | 0 | 1 | 1 | 3 | 0 | 3 | 0 | 10 | 0.1 |
| 5_1255331 | 5 | 1 | 4 | 1 | 6 | 1 | 8 | 1 | 23 | 1 | 3 | 1 | 1 | 1 | 2 | 1 | 2 | 1 | 8 | 1 |
| 5_1255354 | 9 | 0.3333 | 9 | 0.5556 | 8 | 0.5 | 9 | 0.6667 | 35 | 0.5143 | 3 | 0.3333 | 4 | 0.25 | 6 | 0.5 | 2 | 0.5 | 15 | 0.4 |
| 5_1255355 | 10 | 0 | 11 | 0.0909 | 10 | 0.1 | 9 | 0 | 40 | 0.05 | 4 | 0 | 5 | 0.2 | 6 | 0 | 3 | 0 | 18 | 0.0556 |
| 5_1255361 | 11 | 0.0909 | 14 | 0.2143 | 9 | 0.2222 | 10 | 0 | 44 | 0.1364 | 3 | 0 | 5 | 0 | 7 | 0.2857 | 7 | 0 | 22 | 0.0909 |
| 5_1255368 | 11 | 0.0909 | 17 | 0.1176 | 12 | 0.0833 | 12 | 0.0833 | 52 | 0.0962 | 5 | 0.2 | 5 | 0 | 6 | 0.1667 | 8 | 0.125 | 24 | 0.125 |
| 5_1255368 | 9 | 0.3333 | 16 | 0.0625 | 7 | 0.1429 | 10 | 0.3 | 42 | 0.1905 | 3 | 0 | 4 | 0.25 | 5 | 0.2 | 6 | 0.3333 | 18 | 0.2222 |
| 5_1255933 | 3 | 0.3333 | 3 | 0.3333 | 2 | 0.5 | 3 | 0 | 11 | 0.2727 | 5 | 0.6 | 4 | 0 | 7 | 0.2857 | NA | NA | 16 | 0.3125 |
| 5_1255933 | 1 | 0 | 3 | 0 | 2 | 0.5 | 2 | 0 | 8 | 0.125 | 5 | 0.2 | 4 | 0 | 7 | 0 | NA | NA | 16 | 0.0625 |
| 5_1255934 | 4 | 0 | 3 | 0.3333 | 3 | 0 | 3 | 0.3333 | 13 | 0.1538 | 5 | 0 | 3 | 0.3333 | 7 | 0.2857 | NA | NA | 15 | 0.2 |
| 5_1255934 | 3 | 0 | 3 | 0.6667 | 2 | 0 | 3 | 0 | 11 | 0.1818 | 5 | 0.4 | 4 | 0 | 6 | 0.3333 | NA | NA | 15 | 0.2667 |
| 5_1255934 | 4 | 1 | 3 | 0.6667 | 3 | 1 | 3 | 0.6667 | 13 | 0.8462 | 3 | 1 | 4 | 0.75 | 7 | 0.7143 | NA | NA | 14 | 0.7857 |
| 5_1255934 | 3 | 0 | 2 | 0 | 3 | 0 | 3 | 0.3333 | 11 | 0.0909 | 5 | 0.2 | 4 | 0 | 5 | 0 | NA | NA | 14 | 0.0714 |
| 5_1255935 | 3 | 0 | 4 | 0.25 | 3 | 0.3333 | 3 | 0 | 13 | 0.1538 | 6 | 0.3333 | 4 | 0.5 | 7 | 0.2857 | NA | NA | 17 | 0.3529 |
| 5_1255936 | 4 | 0.5 | 2 | 0 | 4 | 0 | 2 | 0 | 12 | 0.1667 | 3 | 0 | 4 | 0.25 | 6 | 0.1667 | NA | NA | 13 | 0.1538 |
| 5_1255936 | 1 | 0 | 4 | 0 | 3 | 0 | 2 | 0.5 | 10 | 0.1 | 4 | 0 | 3 | 0 | 6 | 0.1667 | NA | NA | 13 | 0.0769 |
| 5_1255938 | 2 | 0.5 | 4 | 0 | 4 | 0 | 2 | 0.5 | 12 | 0.1667 | 5 | 0.2 | 4 | 0.25 | 5 | 0.4 | NA | NA | 14 | 0.2857 |
| 5_1255942 | 4 | 0.25 | 5 | 0.4 | 5 | 0.2 | 3 | 0.3333 | 17 | 0.2941 | 8 | 0.25 | 4 | 0.25 | 7 | 0.4286 | NA | NA | 19 | 0.3158 |
| 5_1255942 | 5 | 0 | 5 | 0.2 | 5 | 0 | 5 | 0 | 20 | 0.05 | 9 | 0.3333 | 4 | 0.25 | 7 | 0.2857 | NA | NA | 20 | 0.3 |
| 5_1258640 | 2 | 0.5 | NA | NA | NA | NA | NA | NA | 2 | 0.5 | NA | NA | 1 | 1 | NA | NA | NA | NA | 1 | 1 |
| 5_1258644 | 3 | 0.6667 | NA | NA | 1 | 0 | NA | NA | 4 | 0.5 | NA | NA | 1 | 1 | NA | NA | NA | NA | 1 | 1 |
| 5_1258647 | 3 | 1 | NA | NA | 1 | 1 | NA | NA | 4 | 1 | NA | NA | 3 | 1 | NA | NA | NA | NA | 3 | 1 |
| 5_1258648 | 3 | 1 | NA | NA | 1 | 1 | NA | NA | 4 | 1 | NA | NA | 3 | 1 | NA | NA | NA | NA | 3 | 1 |
| 5_1258648 | 3 | 1 | NA | NA | 1 | 1 | NA | NA | 4 | 1 | NA | NA | 3 | 1 | NA | NA | NA | NA | 3 | 1 |
| 5_1258649 | 3 | 0.6667 | NA | NA | NA | NA | NA | NA | 3 | 0.6667 | NA | NA | 1 | 1 | NA | NA | NA | NA | 1 | 1 |
| 5_1258649 | 3 | 1 | NA | NA | 1 | 0 | NA | NA | 4 | 0.75 | NA | NA | 2 | 1 | NA | NA | NA | NA | 2 | 1 |
| 5_1258650 | 4 | 0.25 | NA | NA | 1 | 0 | NA | NA | 5 | 0.2 | NA | NA | 5 | 0.6 | NA | NA | NA | NA | 5 | 0.6 |
| 5_1258650 | 4 | 1 | NA | NA | 1 | 0 | NA | NA | 5 | 0.8 | NA | NA | 5 | 0.8 | NA | NA | NA | NA | 5 | 0.8 |
| 5_1258651 | 4 | 1 | NA | NA | 1 | 1 | NA | NA | 5 | 1 | NA | NA | 5 | 1 | NA | NA | NA | NA | 5 | 1 |
| 5_1258651 | 4 | 0.75 | NA | NA | 1 | 1 | NA | NA | 5 | 0.8 | NA | NA | 5 | 0.4 | NA | NA | NA | NA | 5 | 0.4 |
| 5_1258651 | 4 | 0.25 | NA | NA | 1 | 0 | NA | NA | 5 | 0.2 | NA | NA | 5 | 0.2 | NA | NA | NA | NA | 5 | 0.2 |
| 5_1258651 | 3 | 0.6667 | NA | NA | 1 | 0 | NA | NA | 4 | 0.5 | NA | NA | 4 | 1 | NA | NA | NA | NA | 4 | 1 |
| 5_1258653 | 2 | 0 | NA | NA | 2 | 1 | NA | NA | 4 | 0.5 | NA | NA | 5 | 0.6 | NA | NA | 1 | 0 | 6 | 0.5 |
| 5_1258653 | 4 | 0.75 | NA | NA | 2 | 0.5 | NA | NA | 6 | 0.6667 | NA | NA | 5 | 0.6 | NA | NA | 1 | 1 | 6 | 0.6667 |
| 5_1258654 | 5 | 0.2 | NA | NA | 2 | 1 | NA | NA | 7 | 0.4286 | NA | NA | 5 | 0.6 | NA | NA | 1 | 0 | 6 | 0.5 |
| 5_1258655 | 6 | 0.3333 | 1 | 0 | 2 | 0 | NA | NA | 9 | 0.2222 | NA | NA | 4 | 0.25 | NA | NA | 1 | 1 | 5 | 0.4 |
| 5_1258660 | 7 | 0.2857 | 3 | 0 | 2 | 1 | NA | NA | 12 | 0.3333 | 1 | 0 | 5 | 0.2 | NA | NA | 1 | 0 | 7 | 0.1429 |
| 5_1258683 | 10 | 0.1 | 6 | 0 | 7 | 0.1429 | 4 | 0 | 27 | 0.0741 | 5 | 0 | 13 | 0.1538 | 3 | 0 | 3 | 0 | 24 | 0.0833 |
| 5_1258685 | 8 | 0.375 | 7 | 0.1429 | 4 | 0.5 | 5 | 0.4 | 24 | 0.3333 | 5 | 0.2 | 14 | 0.1429 | 4 | 0.5 | 4 | 0.25 | 27 | 0.2222 |
| 5_1258693 | 9 | 0.1111 | 10 | 0.4 | 7 | 0.1429 | 9 | 0 | 35 | 0.1714 | 8 | 0.125 | 15 | 0 | 9 | 0.1111 | 4 | 0.25 | 36 | 0.0833 |
| 5_1258694 | 13 | 0.9231 | 14 | 0.7143 | 11 | 0.7273 | 8 | 0.875 | 46 | 0.8043 | 10 | 0.7 | 17 | 0.8824 | 11 | 0.8182 | 7 | 1 | 45 | 0.8444 |
| 5_1258694 | 12 | 0.25 | 15 | 0.3333 | 12 | 0.1667 | 6 | 0.1667 | 45 | 0.2444 | 11 | 0.2727 | 14 | 0.2857 | 11 | 0 | 8 | 0.375 | 44 | 0.2273 |
| 5_1258695 | 13 | 0 | 15 | 0.0667 | 12 | 0 | 8 | 0.375 | 48 | 0.0833 | 12 | 0.1667 | 20 | 0 | 11 | 0 | 8 | 0.125 | 51 | 0.0588 |
| 5_1276400 | 1 | 1 | NA | NA | NA | NA | NA | NA | 1 | 1 | 2 | 1 | NA | NA | NA | NA | NA | NA | 2 | 1 |
| 5_1276403 | 1 | 1 | NA | NA | NA | NA | NA | NA | 1 | 1 | 2 | 0.5 | NA | NA | NA | NA | NA | NA | 2 | 0.5 |
| 5_1276405 | 1 | 1 | NA | NA | NA | NA | NA | NA | 1 | 1 | 1 | 1 | NA | NA | NA | NA | NA | NA | 1 | 1 |
| 5_1276408 | 1 | 1 | NA | NA | NA | NA | 1 | 1 | 2 | 1 | 1 | 1 | NA | NA | NA | NA | NA | NA | 1 | 1 |
| 5_1276408 | 1 | 1 | NA | NA | NA | NA | 2 | 0 | 3 | 0.3333 | 1 | 1 | NA | NA | NA | NA | NA | NA | 1 | 1 |

| | | | | | | | | | | | | | | | | | | | | |
|---|---|---|---|---|---|---|---|---|---|---|---|---|---|---|---|---|---|---|---|---|
| 5_12764111 | 1 | 1 | NA | NA | NA | NA | 2 | 0.5 | 3 | 0.6667 | 2 | 1 | NA | NA | NA | NA | NA | NA | 2 | 1 |
| 5_12764118 | 1 | 1 | NA | NA | NA | NA | 2 | 1 | 3 | 1 | 1 | 1 | NA | NA | NA | NA | NA | NA | 1 | 1 |
| 5_12764124 | 1 | 1 | NA | NA | NA | NA | 2 | 1 | 3 | 1 | 2 | 1 | NA | NA | NA | NA | NA | NA | 2 | 1 |
| 5_12764155 | 1 | 1 | NA | NA | NA | NA | 2 | 1 | 3 | 1 | 1 | 1 | NA | NA | NA | NA | NA | NA | 1 | 1 |
| 5_12764164 | NA | NA | NA | NA | NA | NA | 2 | 0.5 | 2 | 0.5 | 1 | 1 | NA | NA | NA | NA | NA | NA | 1 | 1 |
| 5_12764219 | 1 | 1 | NA | NA | NA | NA | 2 | 1 | 3 | 1 | 2 | 1 | NA | NA | NA | NA | NA | NA | 2 | 1 |
| 5_12764234 | 1 | 1 | NA | NA | NA | NA | 2 | 1 | 3 | 1 | 2 | 1 | NA | NA | NA | NA | NA | NA | 2 | 1 |
| 5_12767187 | 1 | 1 | NA | NA | NA | NA | NA | NA | 1 | 1 | 1 | 0 | NA | NA | 1 | 1 | NA | NA | 2 | 0.5 |
| 5_12767208 | 1 | 1 | NA | NA | NA | NA | NA | NA | 1 | 1 | NA | NA | NA | NA | 1 | 1 | NA | NA | 1 | 1 |
| 5_12767210 | 1 | 1 | NA | NA | NA | NA | NA | NA | 1 | 1 | 1 | 1 | NA | NA | NA | NA | NA | NA | 1 | 1 |
| 5_12767273 | 6 | 0.3333 | 3 | 0 | 2 | 0 | 1 | 0 | 12 | 0.1667 | 4 | 0 | 2 | 0.5 | 4 | 0 | 4 | 0 | 14 | 0.0714 |
| 5_12767278 | 5 | 0.2 | 4 | 0.25 | 4 | 0.25 | 1 | 0 | 14 | 0.2143 | 7 | 0.1429 | 2 | 0 | 5 | 0 | 2 | 0.5 | 16 | 0.125 |
| 5_12767290 | 7 | 0.1429 | 5 | 0 | 5 | 0.2 | 3 | 0.3333 | 20 | 0.15 | 7 | 0.5714 | 2 | 0 | 4 | 0 | 4 | 0.25 | 17 | 0.2941 |
| 5_12767290 | 6 | 0.1667 | 4 | 0 | 5 | 0.2 | 3 | 0.6667 | 18 | 0.2222 | 6 | 0.1667 | 2 | 0 | 4 | 0.25 | 4 | 0.25 | 16 | 0.1875 |
| 5_12767302 | 7 | 0.2857 | 2 | 0 | 5 | 0 | 2 | 0 | 16 | 0.125 | 3 | 0.3333 | 2 | 0.5 | 6 | 0 | 4 | 0 | 15 | 0.1333 |
| 5_12767303 | 7 | 0.7143 | 4 | 0 | 5 | 0.2 | 3 | 0 | 19 | 0.3158 | 7 | 0.2857 | 4 | 0.5 | 5 | 0 | 4 | 0 | 20 | 0.2 |
| 5_12767314 | 7 | 0.1429 | 3 | 0 | 3 | 0.3333 | 3 | 0.3333 | 16 | 0.1875 | 7 | 0.4286 | 2 | 0.5 | 5 | 0.4 | 3 | 0 | 17 | 0.3529 |
| 5_12767323 | 6 | 1 | 2 | 1 | 2 | 0.5 | 2 | 1 | 12 | 0.9167 | 3 | 1 | 1 | 1 | 5 | 1 | 3 | 0.6667 | 12 | 0.9167 |
| 5_12767355 | 6 | 1 | 2 | 1 | 5 | 0.8 | 3 | 1 | 16 | 0.9375 | 5 | 1 | 1 | 1 | 6 | 1 | 3 | 0.6667 | 15 | 0.9333 |
| 5_12767357 | 4 | 0.75 | 1 | 1 | 5 | 0.6 | 1 | 1 | 11 | 0.7273 | 5 | 0.8 | 1 | 1 | 1 | 1 | 2 | 1 | 9 | 0.8889 |
| 5_12767360 | 5 | 0.2 | 1 | 1 | 4 | 0.75 | 2 | 0.5 | 12 | 0.5 | 5 | 0.4 | 1 | 1 | 5 | 0.2 | 3 | 0.6667 | 14 | 0.4286 |
| 5_12767365 | 5 | 1 | 1 | 1 | 4 | 0.75 | 3 | 1 | 13 | 0.9231 | 4 | 0.75 | 1 | 1 | 5 | 0.6 | 3 | 1 | 13 | 0.7692 |
| 5_12767369 | 6 | 0.1667 | 2 | 0 | 5 | 0 | 3 | 0.6667 | 16 | 0.1875 | 3 | 0 | 2 | 0 | 4 | 0.25 | 3 | 0 | 12 | 0.0833 |
| 5_12767372 | 4 | 0.5 | 1 | 1 | 5 | 0.6 | 3 | 0 | 13 | 0.4615 | 4 | 0.5 | 1 | 0 | 5 | 0 | 2 | 1 | 12 | 0.3333 |
| 5_12778417 | 1 | 1 | NA | NA | NA | NA | NA | NA | 1 | 1 | 1 | 1 | 2 | 1 | 2 | 0 | 1 | 1 | 6 | 0.6667 |
| 5_12778428 | 1 | 1 | NA | NA | NA | NA | NA | NA | 1 | 1 | 1 | 1 | 2 | 1 | 1 | 1 | 1 | 1 | 5 | 1 |
| 5_12778516 | 1 | 0 | 2 | 0.5 | 1 | 0 | 1 | 0 | 5 | 0.2 | NA | NA | 1 | 0 | 1 | 0 | 2 | 0.5 | 4 | 0.25 |
| 5_12778620 | NA | NA | 1 | 0 | 1 | 0 | 2 | 0.5 | 4 | 0.25 | NA | NA | 2 | 0.5 | NA | NA | NA | NA | 2 | 0.5 |
| 5_12785974 | 1 | 0 | 3 | 0.3333 | 2 | 0 | 2 | 0 | 8 | 0.125 | 4 | 0 | 2 | 0.5 | NA | NA | NA | NA | 6 | 0.1667 |
| 5_12785976 | 1 | 1 | 3 | 0.3333 | 2 | 0 | 2 | 0 | 8 | 0.25 | 4 | 0.25 | 2 | 0.5 | NA | NA | NA | NA | 6 | 0.3333 |
| 5_12786070 | NA | NA | 4 | 1 | 1 | 1 | 1 | 1 | 6 | 1 | 2 | 1 | 2 | 1 | NA | NA | NA | NA | 4 | 1 |
| 5_12786073 | NA | NA | 4 | 1 | 1 | 1 | NA | NA | 5 | 1 | 2 | 1 | 2 | 0.5 | NA | NA | NA | NA | 4 | 0.75 |
| 5_12786082 | NA | NA | 3 | 1 | 1 | 1 | NA | NA | 4 | 1 | 1 | 1 | 2 | 0.5 | NA | NA | NA | NA | 3 | 0.6667 |
| 5_12786099 | NA | NA | 4 | 1 | 1 | 1 | NA | NA | 5 | 1 | 1 | 1 | 1 | 1 | NA | NA | NA | NA | 2 | 1 |
| 5_12786108 | NA | NA | NA | NA | 1 | 1 | NA | NA | 1 | 1 | 2 | 1 | 1 | 1 | NA | NA | NA | NA | 3 | 1 |
| 5_12786113 | NA | NA | 4 | 1 | 1 | 1 | NA | NA | 5 | 1 | 1 | 1 | 2 | 1 | NA | NA | NA | NA | 3 | 1 |
| 5_12786119 | NA | NA | 3 | 1 | 1 | 1 | NA | NA | 4 | 1 | 2 | 1 | NA | NA | NA | NA | NA | NA | 2 | 1 |
| 5_12786121 | NA | NA | 4 | 0.75 | 1 | 0 | NA | NA | 5 | 0.6 | 2 | 1 | 2 | 1 | NA | NA | NA | NA | 4 | 1 |
| 5_12786130 | NA | NA | 4 | 1 | 1 | 1 | NA | NA | 5 | 1 | 2 | 1 | 2 | 0.5 | NA | NA | NA | NA | 4 | 0.75 |
| 5_12786131 | NA | NA | 2 | 1 | 1 | 1 | NA | NA | 3 | 1 | 1 | 1 | 2 | 0.5 | NA | NA | NA | NA | 3 | 0.6667 |
| 5_12786144 | NA | NA | 2 | 0.5 | 1 | 1 | NA | NA | 3 | 0.6667 | 2 | 0.5 | 2 | 0.5 | NA | NA | NA | NA | 4 | 0.5 |
| 5_12786147 | NA | NA | 2 | 0 | 1 | 1 | NA | NA | 3 | 0.3333 | 2 | 0 | 2 | 0.5 | NA | NA | NA | NA | 4 | 0.25 |
| 5_12786169 | NA | NA | 2 | 0.5 | NA | NA | NA | NA | 2 | 0.5 | 2 | 1 | 2 | 1 | NA | NA | NA | NA | 4 | 1 |
| 5_12786172 | NA | NA | 2 | 1 | 1 | 1 | NA | NA | 3 | 1 | 2 | 1 | 2 | 1 | NA | NA | NA | NA | 4 | 1 |
| 5_12786183 | NA | NA | 2 | 0.5 | 1 | 1 | NA | NA | 3 | 0.6667 | 1 | 0 | 2 | 0.5 | NA | NA | NA | NA | 3 | 0.3333 |
| 5_12786195 | NA | NA | 1 | 1 | 1 | 0 | NA | NA | 2 | 0.5 | 2 | 0.5 | 2 | 0 | NA | NA | NA | NA | 4 | 0.25 |
| 5_12834584 | 1 | 0 | 1 | 1 | 1 | 1 | NA | NA | 3 | 0.6667 | 1 | 1 | 1 | 0 | NA | NA | NA | NA | 2 | 0.5 |
| 5_12834595 | 1 | 1 | 1 | 1 | NA | NA | NA | NA | 2 | 1 | 1 | 1 | 2 | 1 | NA | NA | NA | NA | 3 | 1 |
| 5_12834631 | 1 | 0 | 1 | 0 | 1 | 1 | NA | NA | 3 | 0.3333 | 1 | 1 | 2 | 0 | NA | NA | NA | NA | 3 | 0.3333 |
| 5_12883603 | 2 | 1 | 8 | 0.375 | 8 | 0.625 | 4 | 0.25 | 22 | 0.5 | 3 | 1 | 8 | 0.375 | 6 | 0.5 | 6 | 0.3333 | 23 | 0.4783 |
| 5_12883608 | 3 | 0.3333 | 10 | 0.5 | 10 | 0.5 | 5 | 0.4 | 28 | 0.4643 | 4 | 0.5 | 8 | 0.25 | 7 | 0.4286 | 8 | 0.625 | 27 | 0.4444 |
| 5_12883630 | NA | NA | 4 | 0.5 | 6 | 0 | 1 | 0 | 11 | 0.1818 | 1 | 0 | 7 | 0.1429 | 4 | 0 | 2 | 0 | 14 | 0.0714 |
| 5_12883674 | NA | NA | 9 | 0.1111 | 5 | 0.2 | 1 | 1 | 15 | 0.2 | NA | NA | 6 | 0.1667 | 5 | 0.2 | 3 | 0 | 14 | 0.1429 |
| 5_12883764 | NA | NA | 5 | 0.6 | 2 | 0 | NA | NA | 7 | 0.4286 | NA | NA | 4 | 0 | 2 | 0 | 3 | 0.3333 | 9 | 0.1111 |
| 5_12883771 | NA | NA | 7 | 0.7143 | 3 | 0.6667 | 1 | 1 | 11 | 0.7273 | NA | NA | 3 | 0.6667 | 5 | 0.2 | 2 | 0.5 | 10 | 0.4 |
| 5_12883781 | NA | NA | 5 | 0.6 | 3 | 0 | 1 | 0 | 9 | 0.3333 | NA | NA | 3 | 0.3333 | 4 | 0.25 | 3 | 0 | 10 | 0.2 |
| 5_12885926 | NA | NA | NA | NA | 1 | 1 | NA | NA | 1 | 1 | NA | NA | NA | NA | 1 | 1 | NA | NA | 1 | 1 |
| 5_12885953 | NA | NA | NA | NA | 1 | 1 | NA | NA | 1 | 1 | NA | NA | NA | NA | 1 | 1 | NA | NA | 1 | 1 |
| 5_12886550 | NA | NA | 4 | 1 | 2 | 1 | 2 | 0 | 8 | 0.75 | 1 | 1 | NA | NA | 1 | 1 | NA | NA | 2 | 1 |
| 5_12887426 | NA | NA | 3 | 0 | 1 | 0 | 4 | 0.25 | 8 | 0.125 | 1 | 0 | NA | NA | 2 | 0.5 | 1 | 0 | 4 | 0.25 |
| 5_12887448 | NA | NA | 4 | 0.5 | 2 | 1 | 4 | 0.5 | 10 | 0.6 | 1 | 1 | NA | NA | 2 | 0 | 1 | 1 | 4 | 0.5 |
| 5_12887450 | NA | NA | 4 | 0.25 | 2 | 0 | 4 | 0.25 | 10 | 0.2 | NA | NA | NA | NA | 2 | 0.5 | NA | NA | 2 | 0.5 |
| 5_12887453 | NA | NA | 4 | 0.75 | 2 | 1 | 4 | 0.75 | 10 | 0.8 | 1 | 1 | NA | NA | 1 | 1 | 1 | 1 | 3 | 1 |
| 5_12887453 | NA | NA | 4 | 0.75 | 1 | 0 | 3 | 0.3333 | 8 | 0.5 | 1 | 0 | NA | NA | 2 | 1 | 1 | 1 | 4 | 0.75 |
| 5_12887472 | NA | NA | 4 | 0.75 | 1 | 1 | 2 | 1 | 7 | 0.8571 | 1 | 1 | NA | NA | 2 | 1 | 1 | 1 | 4 | 1 |

| | | | | | | | | | | | | | | | | | | | | |
|---|---|---|---|---|---|---|---|---|---|---|---|---|---|---|---|---|---|---|---|---|
| 5_1290623 | 3 | 1 | NA | NA | 1 | 1 | NA | NA | 4 | 1 | 2 | 1 | 1 | 1 | 1 | 1 | 2 | 1 | 6 | 1 |
| 5_1290624 | 3 | 1 | NA | NA | 2 | 0.5 | NA | NA | 5 | 0.8 | 2 | 1 | 1 | 1 | 1 | 1 | 2 | 0.5 | 6 | 0.8333 |
| 5_1290624 | 2 | 1 | NA | NA | 2 | 1 | NA | NA | 4 | 1 | 3 | 0.3333 | 1 | 1 | 1 | 1 | 2 | 1 | 7 | 0.7143 |
| 5_1290627 | 2 | 0.5 | NA | NA | 2 | 0 | NA | NA | 4 | 0.25 | 3 | 0.6667 | 1 | 1 | 1 | 0 | 2 | 0 | 7 | 0.4286 |
| 5_1290628 | 2 | 0 | NA | NA | 2 | 1 | NA | NA | 4 | 0.5 | 3 | 0.6667 | 1 | 1 | NA | NA | 1 | 1 | 5 | 0.8 |
| 5_1290628 | 3 | 0.3333 | NA | NA | 2 | 0.5 | NA | NA | 5 | 0.4 | 3 | 1 | 1 | 1 | 1 | 1 | 2 | 0.5 | 7 | 0.8571 |
| 5_1290629 | 2 | 1 | NA | NA | 2 | 0.5 | NA | NA | 4 | 0.75 | 3 | 0.6667 | 1 | 1 | 1 | 1 | 2 | 1 | 7 | 0.8571 |
| 5_1290630 | 3 | 0.6667 | NA | NA | 2 | 0.5 | NA | NA | 5 | 0.6 | 3 | 0.3333 | 1 | 1 | 1 | 1 | 2 | 1 | 7 | 0.7143 |
| 5_1290631 | 3 | 0.6667 | NA | NA | 2 | 1 | NA | NA | 5 | 0.8 | 2 | 0 | 1 | 1 | 1 | 1 | 2 | 0.5 | 6 | 0.5 |
| 5_1290631 | 3 | 0.6667 | NA | NA | 2 | 1 | NA | NA | 5 | 0.8 | 2 | 1 | NA | NA | 1 | 1 | 2 | 0.5 | 5 | 0.8 |
| 5_1290635 | 1 | 1 | NA | NA | NA | NA | NA | NA | 1 | 1 | 1 | 0 | NA | NA | NA | NA | 2 | 1 | 3 | 0.6667 |
| 5_1291000 | 44 | 0.7727 | 46 | 0.8043 | 28 | 0.8929 | 41 | 0.8049 | 159 | 0.8113 | 33 | 0.9394 | 33 | 0.8485 | 39 | 0.8718 | 32 | 0.8438 | 137 | 0.8759 |
| 5_1291000 | 36 | 0.0556 | 34 | 0.0294 | 30 | 0.0333 | 40 | 0.125 | 140 | 0.0643 | 36 | 0.1111 | 26 | 0.1154 | 33 | 0.1212 | 26 | 0.0769 | 121 | 0.1074 |
| 5_1291001 | 42 | 0.3571 | 44 | 0.3409 | 35 | 0.3429 | 43 | 0.4186 | 164 | 0.3659 | 35 | 0.2571 | 30 | 0.3 | 42 | 0.4048 | 31 | 0.3871 | 138 | 0.3406 |
| 5_1291005 | 15 | 0 | 11 | 0.1818 | 10 | 0.1 | 9 | 0 | 45 | 0.0667 | 6 | 0 | 15 | 0.4 | 11 | 0.0909 | 10 | 0.2 | 42 | 0.2143 |
| 5_1292797 | NA | NA | NA | NA | NA | NA | 1 | 1 | 1 | 1 | NA | NA | NA | NA | NA | NA | 1 | 1 | 1 | 1 |
| 5_1292801 | NA | NA | NA | NA | NA | NA | 1 | 1 | 1 | 1 | NA | NA | NA | NA | NA | NA | 1 | 1 | 1 | 1 |
| 5_1292801 | NA | NA | NA | NA | NA | NA | 1 | 1 | 1 | 1 | NA | NA | NA | NA | NA | NA | 1 | 1 | 1 | 1 |
| 5_1292802 | NA | NA | 1 | 1 | NA | NA | 1 | 1 | 2 | 1 | NA | NA | NA | NA | NA | NA | 1 | 1 | 1 | 1 |
| 5_1295792 | 1 | 1 | NA | NA | NA | NA | NA | NA | 1 | 1 | NA | NA | NA | NA | 1 | 1 | NA | NA | 1 | 1 |
| 5_1295795 | 2 | 1 | NA | NA | NA | NA | 1 | 1 | 3 | 1 | NA | NA | NA | NA | 1 | 1 | 1 | 0 | 2 | 0.5 |
| 5_1295795 | 3 | 0 | NA | NA | NA | NA | 1 | 1 | 4 | 0.25 | NA | NA | 1 | 0 | 1 | 1 | 1 | 0 | 3 | 0.3333 |
| 5_1295796 | 3 | 1 | NA | NA | NA | NA | 1 | 1 | 4 | 1 | NA | NA | 1 | 1 | 1 | 1 | 1 | 0 | 3 | 0.6667 |
| 5_1295796 | 1 | 1 | NA | NA | NA | NA | 1 | 0 | 2 | 0.5 | NA | NA | 1 | 1 | 1 | 1 | 1 | 0 | 3 | 0.6667 |
| 5_1295796 | 2 | 0.5 | NA | NA | NA | NA | 1 | 1 | 3 | 0.6667 | NA | NA | 1 | 1 | 1 | 0 | 1 | 0 | 3 | 0.3333 |
| 5_1295796 | 2 | 1 | NA | NA | NA | NA | 1 | 0 | 3 | 0.6667 | NA | NA | 1 | 1 | 1 | 0 | 1 | 0 | 3 | 0.3333 |
| 5_1295797 | 3 | 1 | NA | NA | NA | NA | 1 | 1 | 4 | 1 | NA | NA | 1 | 1 | 1 | 1 | 1 | 1 | 3 | 1 |
| 5_1295798 | 3 | 0.3333 | NA | NA | NA | NA | 1 | 0 | 4 | 0.25 | NA | NA | 1 | 1 | 1 | 0 | 1 | 0 | 3 | 0.3333 |
| 5_1295803 | 1 | 1 | NA | NA | NA | NA | NA | NA | 1 | 1 | NA | NA | 1 | 1 | NA | NA | 1 | 0 | 2 | 0.5 |
| 5_1295803 | 1 | 1 | NA | NA | NA | NA | NA | NA | 1 | 1 | NA | NA | 1 | 1 | NA | NA | 1 | 0 | 2 | 0.5 |
| 5_1296724 | 3 | 0 | 9 | 0.2222 | 4 | 0 | 4 | 0.25 | 20 | 0.15 | 8 | 0.125 | 2 | 1 | 7 | 0.5714 | 4 | 0 | 21 | 0.3333 |
| 5_1296724 | 4 | 0.25 | 7 | 0 | 3 | 0 | 3 | 0 | 17 | 0.0588 | 6 | 0 | 2 | 0 | 6 | 0.1667 | 3 | 0 | 17 | 0.0588 |
| 5_1296827 | 7 | 0.4286 | 13 | 0.3077 | 5 | 0.8 | 9 | 0.4444 | 34 | 0.4412 | 8 | 0.5 | 5 | 0.4 | 8 | 0.5 | 9 | 0.3333 | 30 | 0.4333 |
| 5_1296834 | 11 | 0.2727 | 16 | 0.1875 | 7 | 0.4286 | 12 | 0.25 | 46 | 0.2609 | 8 | 0.625 | 5 | 0 | 10 | 0.4 | 8 | 0.25 | 31 | 0.3548 |
| 5_1296836 | 11 | 0.4545 | 12 | 0.25 | 5 | 0.4 | 11 | 0.5455 | 39 | 0.4103 | 10 | 0.7 | 6 | 0.5 | 9 | 0.2222 | 8 | 0.375 | 33 | 0.4545 |
| 5_1296836 | 13 | 0.8462 | 17 | 0.7647 | 5 | 0.8 | 14 | 0.7143 | 49 | 0.7755 | 12 | 0.9167 | 7 | 0.8571 | 9 | 0.8889 | 9 | 0.6667 | 37 | 0.8378 |
| 5_1296836 | 11 | 1 | 20 | 0.95 | 7 | 1 | 14 | 1 | 52 | 0.9808 | 14 | 1 | 6 | 1 | 11 | 1 | 10 | 1 | 41 | 1 |
| 5_1296837 | 8 | 0.375 | 12 | 0.4167 | 6 | 0.3333 | 7 | 0.1429 | 33 | 0.3333 | 11 | 0.3636 | 6 | 0.5 | 8 | 0.375 | 6 | 0.5 | 31 | 0.4194 |
| 5_1296837 | 11 | 0 | 16 | 0.375 | 6 | 0.8333 | 13 | 0.3077 | 46 | 0.3261 | 13 | 0.3846 | 7 | 0.2857 | 11 | 0.4545 | 10 | 0.3 | 41 | 0.3659 |
| 5_1296844 | 14 | 0.1429 | 14 | 0.7857 | 9 | 0.2222 | 12 | 0.25 | 49 | 0.3673 | 14 | 0.3571 | 8 | 0.5 | 10 | 0.6 | 14 | 0.4286 | 46 | 0.4565 |
| 5_1296854 | 2 | 0 | 3 | 0 | NA | NA | 2 | 0.5 | 7 | 0.1429 | 6 | 0.1667 | NA | NA | 5 | 0.2 | 5 | 0.2 | 16 | 0.1875 |
| 5_1296854 | 3 | 0.3333 | 3 | 0 | 1 | 1 | 3 | 0 | 10 | 0.2 | 5 | 0.2 | NA | NA | 4 | 0.25 | 5 | 0.2 | 14 | 0.2143 |
| 5_1296855 | 3 | 0.6667 | 3 | 1 | NA | NA | 2 | 1 | 8 | 0.875 | 6 | 1 | NA | NA | 5 | 0.6 | 5 | 1 | 16 | 0.875 |
| 5_1296855 | 3 | 0.3333 | 4 | 0 | 1 | 0 | 2 | 0.5 | 10 | 0.2 | 5 | 0 | NA | NA | 5 | 0.2 | 5 | 0.2 | 15 | 0.1333 |
| 5_1296856 | 2 | 1 | 4 | 0.75 | 2 | 1 | 3 | 1 | 11 | 0.9091 | 7 | 0.5714 | 1 | 1 | 5 | 0.8 | 4 | 0.75 | 17 | 0.7059 |
| 5_1296856 | 3 | 0.3333 | 3 | 0.3333 | 2 | 0 | 3 | 0.3333 | 11 | 0.2727 | 7 | 0.1429 | 1 | 0 | 3 | 0.6667 | 5 | 0.4 | 16 | 0.3125 |
| 5_1296858 | 3 | 0.6667 | 4 | 0.75 | 2 | 0.5 | 4 | 0.25 | 13 | 0.5385 | 5 | 0.2 | 1 | 1 | 5 | 0.4 | 4 | 0.5 | 15 | 0.4 |
| 5_1296859 | 2 | 0.5 | 3 | 0.6667 | 2 | 0.5 | 4 | 0.25 | 11 | 0.4545 | 5 | 0 | 1 | 1 | 5 | 0.2 | 4 | 0.5 | 15 | 0.2667 |
| 5_1296859 | NA | NA | 3 | 0.3333 | 1 | 1 | 1 | 0 | 5 | 0.4 | 1 | 1 | 1 | 0 | 1 | 1 | 1 | 1 | 4 | 0.75 |
| 5_1296860 | NA | NA | 3 | 1 | 1 | 1 | 2 | 1 | 6 | 1 | 2 | 1 | 1 | 1 | 2 | 1 | 2 | 1 | 7 | 1 |
| 5_1296865 | 1 | 0 | 3 | 0.3333 | 1 | 0 | 1 | 0 | 6 | 0.1667 | 1 | 1 | NA | NA | 2 | 1 | 2 | 0.5 | 5 | 0.8 |
| 5_1296866 | 1 | 0 | 2 | 1 | 2 | 0.5 | 1 | 0 | 6 | 0.5 | 2 | 0.5 | NA | NA | 2 | 0.5 | 2 | 0.5 | 6 | 0.5 |
| 5_1297334 | NA | NA | 1 | 1 | NA | NA | 1 | 1 | 2 | 1 | 1 | 0 | 3 | 0.6667 | 2 | 0.5 | 2 | 0 | 8 | 0.375 |
| 5_1297336 | NA | NA | 1 | 1 | NA | NA | 1 | 1 | 2 | 1 | NA | NA | 3 | 1 | 2 | 1 | 2 | 1 | 7 | 1 |
| 5_1297337 | NA | NA | 1 | 1 | NA | NA | 1 | 0 | 2 | 0.5 | NA | NA | 3 | 0.3333 | 2 | 0 | 2 | 1 | 7 | 0.4286 |
| 5_1297337 | NA | NA | 1 | 1 | NA | NA | 1 | 1 | 2 | 1 | NA | NA | 2 | 1 | 2 | 1 | 2 | 1 | 6 | 1 |
| 5_1297339 | NA | NA | 1 | 1 | NA | NA | 1 | 1 | 2 | 1 | NA | NA | 3 | 1 | 1 | 1 | 2 | 1 | 6 | 1 |
| 5_1297339 | NA | NA | 1 | 0 | NA | NA | 1 | 1 | 2 | 0.5 | NA | NA | 3 | 0 | 2 | 0.5 | 2 | 1 | 7 | 0.4286 |
| 5_1297352 | NA | NA | NA | NA | NA | NA | 1 | 1 | 1 | 1 | NA | NA | NA | NA | 1 | 1 | 1 | 1 | 2 | 1 |
| 5_1297354 | NA | NA | NA | NA | NA | NA | 1 | 1 | 1 | 1 | NA | NA | NA | NA | 1 | 1 | 1 | 1 | 2 | 1 |
| 5_1297950 | 10 | 0.5 | 12 | 0.0833 | 4 | 0 | 12 | 0.1667 | 38 | 0.2105 | 7 | 0.2857 | 7 | 0 | 12 | 0.3333 | 9 | 0.2222 | 35 | 0.2286 |
| 5_1297951 | 8 | 0 | 11 | 0.3636 | 3 | 0.3333 | 11 | 0.2727 | 33 | 0.2424 | 6 | 0.6667 | 10 | 0.5 | 12 | 0.5 | 9 | 0.6667 | 37 | 0.5676 |
| 5_1297952 | 11 | 0.0909 | 11 | 0.0909 | 6 | 0.3333 | 13 | 0.3846 | 41 | 0.2195 | 8 | 0.375 | 11 | 0.1818 | 12 | 0.5 | 11 | 0.4545 | 42 | 0.381 |
| 5_1297953 | 9 | 0.5556 | 8 | 0.875 | 4 | 1 | 8 | 0.75 | 29 | 0.7586 | 5 | 0.8 | 9 | 0.7778 | 8 | 0.625 | 7 | 0.7143 | 29 | 0.7241 |
| 5_1297954 | 7 | 0.7143 | 8 | 0.875 | 6 | 1 | 7 | 0.8571 | 28 | 0.8571 | 6 | 1 | 8 | 0.875 | 10 | 0.7 | 8 | 0.875 | 32 | 0.8438 |

| | | | | | | | | | | | | | | | | | | | | |
|---|---|---|---|---|---|---|---|---|---|---|---|---|---|---|---|---|---|---|---|---|
| 5_12979842 | 3 | 0.3333 | 1 | 0 | 3 | 0 | 5 | 0 | 12 | 0.0833 | NA | NA | NA | NA | 3 | 0 | 5 | 0.2 | 8 | 0.125 |
| 5_12979863 | 4 | 0 | 1 | 0 | 3 | 0.3333 | 5 | 0 | 13 | 0.0769 | NA | NA | NA | NA | 4 | 0.25 | 5 | 0 | 9 | 0.1111 |
| 5_12983430 | 14 | 0.0714 | 17 | 0.0588 | 12 | 0.0833 | 14 | 0.0714 | 57 | 0.0702 | 5 | 0 | 15 | 0 | 16 | 0.0625 | 15 | 0.1333 | 51 | 0.0588 |
| 5_12983435 | 15 | 0.4667 | 12 | 0.5833 | 13 | 0.5385 | 12 | 0.5 | 52 | 0.5192 | 4 | 0.75 | 12 | 0.4167 | 15 | 0.2 | 13 | 0.1538 | 44 | 0.2955 |
| 5_12983437 | 15 | 0.0667 | 15 | 0.1333 | 9 | 0 | 12 | 0 | 51 | 0.0588 | 5 | 0 | 16 | 0 | 13 | 0.0769 | 11 | 0.1818 | 45 | 0.0667 |
| 5_12983455 | 14 | 0.0714 | 11 | 0.1818 | 13 | 0.0769 | 10 | 0.2 | 48 | 0.125 | 4 | 0 | 13 | 0.2308 | 13 | 0.1538 | 12 | 0.0833 | 42 | 0.1429 |
| 5_12986405 | NA | NA | 2 | 0 | NA | NA | 1 | 1 | 3 | 0.3333 | NA | NA | 3 | 0.3333 | NA | NA | NA | NA | 3 | 0.3333 |
| 5_12986444 | 2 | 0.5 | 1 | 1 | NA | NA | NA | NA | 3 | 0.6667 | NA | NA | NA | NA | 1 | 1 | 1 | 0 | 2 | 0.5 |
| 5_12987476 | 30 | 0.8 | 30 | 0.7333 | 19 | 0.5789 | 27 | 0.7407 | 106 | 0.7264 | 25 | 0.84 | 27 | 0.8148 | 30 | 0.6667 | 33 | 0.697 | 115 | 0.7478 |
| 5_12989149 | 3 | 0 | 2 | 0 | 2 | 0.5 | 3 | 0 | 10 | 0.1 | 2 | 0.5 | NA | NA | 1 | 0 | 3 | 0 | 6 | 0.1667 |
| 5_12989169 | 2 | 0 | 2 | 0 | 1 | 1 | 3 | 0.3333 | 8 | 0.25 | 2 | 0 | NA | NA | 2 | 0.5 | 2 | 0 | 6 | 0.1667 |
| 5_12989170 | 2 | 0.5 | 1 | 1 | 1 | 1 | 3 | 0.3333 | 7 | 0.5714 | 2 | 1 | NA | NA | 1 | 0 | 3 | 0.6667 | 6 | 0.6667 |
| 5_12989172 | 3 | 1 | 2 | 1 | 2 | 1 | 2 | 1 | 9 | 1 | 2 | 0.5 | NA | NA | 1 | 1 | 3 | 0.6667 | 6 | 0.6667 |
| 5_12989174 | 3 | 0 | 2 | 0.5 | 2 | 0 | 3 | 0.3333 | 10 | 0.2 | 1 | 1 | NA | NA | 2 | 0 | 3 | 0.6667 | 6 | 0.5 |
| 5_12989179 | 3 | 1 | 2 | 0 | 2 | 0 | 2 | 1 | 9 | 0.5556 | 2 | 0.5 | NA | NA | 2 | 1 | 3 | 0.6667 | 7 | 0.7143 |
| 5_12989180 | 2 | 0 | 2 | 0 | 2 | 0.5 | 3 | 0 | 9 | 0.1111 | 2 | 0 | NA | NA | 2 | 0 | 3 | 0.3333 | 7 | 0.1429 |
| 5_12989183 | 2 | 1 | 2 | 0.5 | 2 | 0 | 3 | 0 | 9 | 0.3333 | 2 | 0 | NA | NA | 2 | 0.5 | 2 | 0 | 6 | 0.1667 |
| 5_12989184 | 3 | 0 | 2 | 0 | 2 | 0 | 3 | 0.6667 | 10 | 0.2 | 1 | 0 | NA | NA | 1 | 0 | 3 | 0.3333 | 5 | 0.2 |
| 5_12989195 | 2 | 0 | 1 | 0 | 3 | 0 | 3 | 0.3333 | 9 | 0.1111 | 1 | 0 | NA | NA | 1 | 0 | 3 | 0.6667 | 5 | 0.4 |
| 5_12989201 | 2 | 1 | 2 | 1 | 3 | 0.6667 | 2 | 1 | 9 | 0.8889 | 1 | 1 | NA | NA | 2 | 0.5 | 3 | 0.6667 | 6 | 0.6667 |
| 5_12989245 | NA | NA | 3 | 0.3333 | 3 | 0 | 2 | 0 | 8 | 0.125 | NA | NA | 1 | 0 | 4 | 0.25 | 1 | 0 | 6 | 0.1667 |
| 5_12989328 | 2 | 1 | 2 | 0.5 | 2 | 0 | 1 | 0 | 7 | 0.4286 | NA | NA | NA | NA | 4 | 0.75 | NA | NA | 4 | 0.75 |
| 5_12991016 | 156 | 0.0641 | 125 | 0.048 | 110 | 0.0545 | 116 | 0.0517 | 507 | 0.0552 | 107 | 0.0654 | 106 | 0.0755 | 122 | 0.0492 | 118 | 0.0424 | 453 | 0.0574 |
| 5_12993067 | 1 | 1 | NA | NA | NA | NA | 1 | 0 | 2 | 0.5 | NA | NA | 1 | 1 | NA | NA | 1 | 0 | 2 | 0.5 |
| 5_12993075 | 1 | 1 | NA | NA | NA | NA | 1 | 1 | 2 | 1 | NA | NA | 1 | 1 | NA | NA | NA | NA | 1 | 1 |
| 5_12993090 | 1 | 1 | NA | NA | NA | NA | 1 | 0 | 2 | 0.5 | NA | NA | 1 | 1 | NA | NA | 1 | 0 | 2 | 0.5 |
| 5_12993094 | 1 | 1 | NA | NA | NA | NA | NA | NA | 1 | 1 | NA | NA | 1 | 1 | NA | NA | 1 | 0 | 2 | 0.5 |
| 5_12993113 | NA | NA | NA | NA | NA | NA | 1 | 1 | 1 | 1 | 1 | 1 | 2 | 1 | NA | NA | 1 | 0 | 4 | 0.75 |
| 5_12993117 | 1 | 1 | NA | NA | NA | NA | NA | NA | 1 | 1 | 1 | 1 | 1 | 1 | NA | NA | 1 | 1 | 3 | 1 |
| 5_12993158 | 1 | 1 | NA | NA | NA | NA | 1 | 1 | 2 | 1 | 1 | 1 | 1 | 1 | NA | NA | 1 | 1 | 3 | 1 |
| 5_12993172 | 1 | 1 | NA | NA | NA | NA | 1 | 0 | 2 | 0.5 | 1 | 1 | 2 | 1 | NA | NA | NA | NA | 3 | 1 |
| 5_12993190 | 1 | 1 | NA | NA | NA | NA | 1 | 1 | 2 | 1 | 1 | 1 | 1 | 1 | 1 | 1 | 1 | 0 | 4 | 0.75 |
| 5_12993222 | 1 | 1 | NA | NA | NA | NA | NA | NA | 1 | 1 | 1 | 0 | 2 | 0.5 | NA | NA | NA | NA | 3 | 0.3333 |
| 5_12993234 | 1 | 1 | NA | NA | NA | NA | NA | NA | 1 | 1 | 1 | 1 | 1 | 1 | 1 | 1 | 1 | 1 | 4 | 1 |
| 5_12993238 | 1 | 1 | NA | NA | NA | NA | NA | NA | 1 | 1 | 1 | 1 | 2 | 0 | 1 | 0 | 1 | 1 | 5 | 0.4 |
| 5_13001112 | 5 | 0.2 | 10 | 0.6 | 6 | 0.5 | 5 | 0.6 | 26 | 0.5 | 4 | 0 | 3 | 0.3333 | 8 | 0.125 | 5 | 0.6 | 20 | 0.25 |
| 5_13001120 | 5 | 0.4 | 10 | 0.1 | 8 | 0.5 | 6 | 0.1667 | 29 | 0.2759 | 3 | 0 | 4 | 0.5 | 9 | 0 | 8 | 0 | 24 | 0.0833 |
| 5_13001131 | 6 | 0.6667 | 10 | 0.8 | 6 | 0.5 | 4 | 1 | 26 | 0.7308 | 3 | 0.3333 | 4 | 1 | 8 | 0.625 | 7 | 0.8571 | 22 | 0.7273 |
| 5_13001131 | NA | NA | NA | NA | 1 | 1 | NA | NA | 1 | 1 | NA | NA | NA | NA | 1 | 1 | NA | NA | 1 | 1 |
| 5_13001133 | 6 | 0.1667 | 9 | 0.1111 | 6 | 0.3333 | 5 | 0.2 | 26 | 0.1923 | 4 | 0.5 | 4 | 0.25 | 8 | 0.125 | 7 | 0.4286 | 23 | 0.3043 |
| 5_13001143 | 6 | 0.1667 | 11 | 0.1818 | 7 | 0.7143 | 5 | 0.4 | 29 | 0.3448 | 3 | 0.3333 | 4 | 0 | 9 | 0.4444 | 8 | 0.125 | 24 | 0.25 |
| 5_13001174 | 4 | 0 | 3 | 0 | 6 | 0.5 | 4 | 0 | 17 | 0.1765 | 2 | 0 | 3 | 0.6667 | 4 | 0 | 5 | 0.4 | 14 | 0.2857 |
| 5_13001179 | 4 | 0.5 | 3 | 0.3333 | 6 | 0.3333 | 2 | 0.5 | 15 | 0.4 | 1 | 0 | 3 | 0.6667 | 3 | 0.6667 | 4 | 0.25 | 11 | 0.4545 |
| 5_13001183 | 4 | 0.5 | 4 | 0.25 | 3 | 0.3333 | 2 | 0 | 13 | 0.3077 | 2 | 0 | 3 | 0 | 3 | 0.3333 | 4 | 0.25 | 12 | 0.1667 |
| 5_13001192 | 3 | 0 | 3 | 0 | 3 | 0.3333 | 3 | 0 | 12 | 0.0833 | 1 | 1 | 3 | 0 | 3 | 0 | 5 | 0 | 12 | 0.0833 |
| 5_13001193 | 3 | 0 | 3 | 0 | 6 | 0 | 4 | 0.25 | 16 | 0.0625 | 2 | 0 | 4 | 0 | 4 | 0 | 4 | 0.25 | 14 | 0.0714 |
| 5_13001195 | 4 | 0 | 4 | 0 | 5 | 0 | 4 | 0.25 | 17 | 0.0588 | 2 | 0 | 4 | 0 | 3 | 0 | 5 | 0.2 | 14 | 0.0714 |
| 5_13001196 | 3 | 1 | 3 | 0.3333 | 6 | 0.3333 | 4 | 0.25 | 16 | 0.4375 | 2 | 0 | 3 | 0.6667 | 4 | 0.25 | 4 | 0.25 | 13 | 0.3077 |
| 5_13001197 | 3 | 1 | 3 | 0.3333 | 6 | 0.6667 | 4 | 0.5 | 16 | 0.625 | 1 | 0 | 4 | 1 | 2 | 0 | 5 | 0.4 | 12 | 0.5 |
| 5_13001202 | 4 | 1 | 3 | 1 | 6 | 1 | 4 | 0.75 | 17 | 0.9412 | 2 | 1 | 4 | 1 | 4 | 0.75 | 5 | 0.8 | 15 | 0.8667 |
| 5_13001216 | 3 | 0.6667 | 3 | 1 | 4 | 0.75 | 1 | 1 | 11 | 0.8182 | 1 | 1 | 1 | 0 | 4 | 1 | 5 | 1 | 11 | 0.9091 |
| 5_13001228 | 4 | 1 | 3 | 1 | 4 | 1 | 4 | 0.75 | 15 | 0.9333 | 2 | 1 | 4 | 1 | 4 | 1 | 5 | 1 | 15 | 1 |
| 5_13001230 | 4 | 0.5 | 3 | 0.6667 | 5 | 0.6 | 3 | 0.3333 | 15 | 0.5333 | 2 | 1 | 4 | 0.75 | 4 | 0.5 | 5 | 0.6 | 15 | 0.6667 |
| 5_13001231 | 4 | 0.25 | 4 | 0.25 | 6 | 0 | 4 | 0 | 18 | 0.1111 | 2 | 0 | 4 | 0 | 4 | 0 | 5 | 0.6 | 15 | 0.2 |
| 5_13001233 | 4 | 0 | 4 | 0 | 6 | 0 | 5 | 0.2 | 19 | 0.0526 | 2 | 0 | 4 | 0 | 4 | 0.25 | 5 | 0.2 | 15 | 0.1333 |
| 5_13001253 | 4 | 0.5 | 3 | 0.3333 | 6 | 0.5 | 3 | 0.6667 | 16 | 0.5 | 1 | 1 | 4 | 0.25 | 3 | 0.3333 | 3 | 0.6667 | 11 | 0.4545 |
| 5_13001257 | 4 | 0.5 | 3 | 0 | 5 | 0.8 | 3 | 0.3333 | 15 | 0.4667 | 1 | 1 | 4 | 0.5 | 5 | 0 | 5 | 0.6 | 15 | 0.4 |
| 5_13001258 | 4 | 0.75 | 2 | 1 | 6 | 0.5 | 2 | 0.5 | 14 | 0.6429 | 1 | 0 | 4 | 0.5 | 5 | 0.2 | 5 | 0.6 | 15 | 0.4 |
| 5_13001299 | 4 | 0 | 5 | 0 | 7 | 0.1429 | 5 | 0.2 | 21 | 0.0952 | 1 | 0 | 5 | 0 | 5 | 0 | 5 | 0.2 | 16 | 0.0625 |
| 5_13001811 | NA | NA | 4 | 0.25 | 2 | 0 | NA | NA | 6 | 0.1667 | 1 | 0 | 3 | 0.3333 | 1 | 0 | 1 | 0 | 6 | 0.1667 |
| 5_13001813 | NA | NA | 4 | 0.25 | 2 | 1 | NA | NA | 6 | 0.5 | 1 | 0 | 3 | 0.3333 | 1 | 0 | 2 | 1 | 7 | 0.4286 |
| 5_13001820 | NA | NA | 4 | 1 | 2 | 0.5 | NA | NA | 6 | 0.8333 | 1 | 1 | 3 | 0.3333 | 1 | 1 | 2 | 1 | 7 | 0.7143 |
| 5_13001827 | NA | NA | 3 | 0.6667 | 2 | 0.5 | NA | NA | 5 | 0.6 | 1 | 1 | 2 | 0.5 | 2 | 1 | 1 | 0 | 6 | 0.6667 |
| 5_13002531 | NA | NA | NA | NA | 1 | 1 | NA | NA | 1 | 1 | NA | NA | NA | NA | 1 | 1 | NA | NA | 1 | 1 |
| 5_13002537 | NA | NA | NA | NA | 1 | 1 | NA | NA | 1 | 1 | NA | NA | NA | NA | 1 | 1 | NA | NA | 1 | 1 |

| | | | | | | | | | | | | | | | | | | | | |
|---|---|---|---|---|---|---|---|---|---|---|---|---|---|---|---|---|---|---|---|---|
| 5_13002856 | 1 | 0 | 1 | 0 | 1 | 1 | 2 | 0.5 | 5 | 0.4 | 1 | 0 | 3 | 0.3333 | 1 | 1 | 1 | 0 | 6 | 0.3333 |
| 5_13002862 | 1 | 0 | 1 | 0 | 2 | 0.5 | 2 | 0 | 6 | 0.1667 | 1 | 0 | 2 | 0.5 | NA | NA | 1 | 0 | 4 | 0.25 |
| 5_13002862 | 1 | 0 | 1 | 0 | 1 | 1 | 1 | 0 | 4 | 0.25 | 1 | 0 | 3 | 0 | 1 | 0 | 2 | 1 | 7 | 0.2857 |
| 5_13002885 | NA | NA | 2 | 0 | 1 | 0 | 2 | 1 | 5 | 0.4 | 1 | 0 | 6 | 0.1667 | 1 | 0 | 2 | 0.5 | 10 | 0.2 |
| 5_13004050 | NA | NA | 4 | 0.25 | 2 | 0 | 4 | 0 | 10 | 0.1 | 3 | 0 | 2 | 0.5 | 3 | 0 | NA | NA | 8 | 0.125 |
| 5_13008876 | 6 | 1 | 1 | 1 | 3 | 1 | 1 | 1 | 11 | 1 | 2 | 1 | 3 | 1 | 1 | 1 | 1 | 1 | 7 | 1 |
| 5_13008890 | 9 | 0.3333 | 3 | 0.3333 | 2 | 0.5 | 1 | 1 | 15 | 0.4 | 2 | 0.5 | 3 | 0.3333 | 4 | 0 | 1 | 0 | 10 | 0.2 |
| 5_13008898 | 9 | 0.1111 | 4 | 0 | 3 | 0.3333 | 1 | 0 | 17 | 0.1176 | 2 | 0 | 5 | 0.2 | 5 | 0.2 | 1 | 0 | 13 | 0.1538 |
| 5_13008903 | 9 | 0.5556 | 3 | 0.3333 | 4 | 0.75 | 1 | 1 | 17 | 0.5882 | 3 | 0.6667 | 5 | 0.6 | 6 | 0.5 | 1 | 1 | 15 | 0.6 |
| 5_13008943 | 9 | 0 | 4 | 0 | 4 | 0.25 | 1 | 0 | 18 | 0.0556 | 3 | 0 | 5 | 0 | 7 | 0 | 2 | 0.5 | 17 | 0.0588 |
| 5_13013048 | 1 | 0 | 3 | 0 | 6 | 0 | 7 | 0.1429 | 17 | 0.0588 | 4 | 0 | 8 | 0 | 2 | 0 | 2 | 0.5 | 16 | 0.0625 |
| 5_13013055 | 1 | 1 | 3 | 0.3333 | 6 | 0.1667 | 6 | 0.3333 | 16 | 0.3125 | 4 | 0 | 7 | 0.4286 | 1 | 0 | 2 | 0.5 | 14 | 0.2857 |
| 5_13013058 | 1 | 1 | 3 | 0 | 5 | 0 | 6 | 0.3333 | 15 | 0.2 | 3 | 0.3333 | 8 | 0.375 | 3 | 0.3333 | 2 | 0 | 16 | 0.3125 |
| 5_13013066 | 2 | 0 | 3 | 0 | 6 | 0 | 6 | 0.1667 | 17 | 0.0588 | 4 | 0 | 10 | 0.1 | 3 | 0 | 2 | 0 | 19 | 0.0526 |
| 5_13013071 | 1 | 0 | 3 | 0.3333 | 5 | 0.6 | 7 | 0.1429 | 16 | 0.3125 | NA | NA | 9 | 0 | 3 | 0.3333 | 1 | 0 | 13 | 0.0769 |
| 5_13013103 | 2 | 1 | 4 | 0.5 | 6 | 1 | 6 | 0.8333 | 18 | 0.8333 | 4 | 0.75 | 7 | 0.8571 | 2 | 1 | 2 | 1 | 15 | 0.8667 |
| 5_13013114 | 1 | 0 | 4 | 0.25 | 4 | 0.5 | 5 | 0.2 | 14 | 0.2857 | 4 | 0.25 | 6 | 0 | 3 | 0.3333 | 2 | 0.5 | 15 | 0.2 |
| 5_13013116 | 2 | 0.5 | 3 | 1 | 6 | 1 | 6 | 1 | 17 | 0.9412 | 5 | 1 | 9 | 1 | 3 | 1 | 2 | 1 | 19 | 1 |
| 5_13013125 | 2 | 0 | 3 | 0.6667 | 5 | 0.6 | 5 | 0.6 | 15 | 0.5333 | 3 | 0.3333 | 7 | 0.7143 | 2 | 0 | 1 | 1 | 13 | 0.5385 |
| 5_13016421 | 5 | 0.2 | 6 | 0 | 6 | 0 | 7 | 0.1429 | 24 | 0.0833 | 6 | 0 | 11 | 0.1818 | 5 | 0.2 | 2 | 0 | 24 | 0.125 |
| 5_13016424 | 5 | 0.2 | 6 | 0.3333 | 8 | 0.375 | 6 | 0.1667 | 25 | 0.28 | 6 | 0.5 | 15 | 0.5333 | 5 | 0.4 | 2 | 0.5 | 28 | 0.5 |
| 5_13018123 | 1 | 1 | NA | NA | 1 | 1 | NA | NA | 2 | 1 | 1 | 1 | 1 | 1 | NA | NA | NA | NA | 2 | 1 |
| 5_13018135 | 1 | 1 | NA | NA | NA | NA | NA | NA | 1 | 1 | 1 | 1 | 1 | 1 | NA | NA | NA | NA | 2 | 1 |
| 5_13018150 | 1 | 1 | NA | NA | NA | NA | NA | NA | 1 | 1 | 1 | 1 | NA | NA | NA | NA | NA | NA | 1 | 1 |
| 5_13018153 | 1 | 0 | NA | NA | 1 | 1 | NA | NA | 2 | 0.5 | 1 | 0 | 1 | 1 | NA | NA | NA | NA | 2 | 0.5 |
| 5_13018156 | 1 | 1 | NA | NA | 1 | 0 | NA | NA | 2 | 0.5 | 1 | 1 | 1 | 0 | NA | NA | NA | NA | 2 | 0.5 |
| 5_13018164 | 1 | 1 | 1 | 1 | 1 | 1 | NA | NA | 3 | 1 | 2 | 1 | 1 | 1 | NA | NA | NA | NA | 3 | 1 |
| 5_13018194 | 1 | 1 | 1 | 1 | 1 | 1 | NA | NA | 3 | 1 | 2 | 1 | 1 | 1 | NA | NA | NA | NA | 3 | 1 |
| 5_13018378 | 1 | 0 | 1 | 0 | 1 | 1 | NA | NA | 3 | 0.3333 | 1 | 1 | NA | NA | NA | NA | NA | NA | 1 | 1 |
| 5_13021640 | 2 | 0 | 2 | 0 | 3 | 0 | 5 | 0.2 | 12 | 0.0833 | 1 | 0 | 3 | 0 | 2 | 0.5 | 5 | 0 | 11 | 0.0909 |
| 5_13022687 | 5 | 0 | 2 | 0 | 3 | 0 | 5 | 0.2 | 15 | 0.0667 | 1 | 0 | 3 | 0 | 2 | 0.5 | 4 | 0 | 10 | 0.1 |
| 5_13024071 | 5 | 0.2 | 2 | 0 | 2 | 0.5 | 3 | 0 | 12 | 0.1667 | NA | NA | 3 | 0 | 2 | 0.5 | 5 | 0 | 10 | 0.1 |
| 5_13024183 | 5 | 0.8 | 1 | 1 | 4 | 1 | 6 | 1 | 16 | 0.9375 | 2 | 1 | 4 | 0.5 | 2 | 1 | 5 | 1 | 13 | 0.8462 |
| 5_13024184 | 6 | 0.5 | 1 | 0 | 3 | 0.6667 | 7 | 0.5714 | 17 | 0.5294 | 2 | 0.5 | 4 | 0.25 | 2 | 0.5 | 5 | 0.6 | 13 | 0.4615 |
| 5_13024190 | 6 | 1 | 1 | 1 | 4 | 1 | 5 | 0.4 | 16 | 0.8125 | 1 | 1 | 4 | 0.75 | 2 | 0.5 | 6 | 0.6667 | 13 | 0.6923 |
| 5_13024191 | 3 | 1 | 2 | 1 | 3 | 1 | 5 | 1 | 13 | 1 | NA | NA | 4 | 0.75 | 2 | 1 | 5 | 1 | 11 | 0.9091 |
| 5_13024192 | 4 | 0.75 | 1 | 0 | 2 | 1 | 4 | 0.75 | 11 | 0.7273 | 2 | 1 | 3 | 0.6667 | NA | NA | 5 | 0.4 | 10 | 0.6 |
| 5_13024194 | 7 | 0.1429 | 2 | 0.5 | 4 | 0.25 | 7 | 0 | 20 | 0.15 | 2 | 0.5 | 3 | 0.3333 | 2 | 0 | 6 | 0 | 13 | 0.1538 |
| 5_13024198 | 7 | 0.1429 | 2 | 0 | 3 | 0 | 5 | 0 | 17 | 0.0588 | 2 | 0 | 4 | 0.25 | 1 | 0 | 5 | 0 | 12 | 0.0833 |
| 5_13024214 | 6 | 0.8333 | 2 | 1 | 3 | 1 | 4 | 1 | 15 | 0.9333 | 1 | 1 | 4 | 1 | 1 | 1 | 4 | 0.75 | 10 | 0.9 |
| 5_13024214 | 6 | 0.3333 | 2 | 0.5 | 4 | 0.5 | 5 | 0 | 17 | 0.2941 | 1 | 0 | 3 | 0.6667 | 1 | 0 | 5 | 0.6 | 10 | 0.5 |
| 5_13024268 | 6 | 0.6667 | 8 | 0.5 | 5 | 0.6 | 6 | 0.5 | 25 | 0.56 | 2 | 1 | 5 | 0.6 | 4 | 0.5 | 5 | 0.8 | 16 | 0.6875 |
| 5_13024277 | 8 | 0.125 | 6 | 0.1667 | 4 | 0.25 | 8 | 0.25 | 26 | 0.1923 | 2 | 0.5 | 6 | 0 | 3 | 0.3333 | 5 | 0.2 | 16 | 0.1875 |
| 5_13024282 | 8 | 0 | 7 | 0 | 5 | 0.4 | 8 | 0.25 | 28 | 0.1429 | 3 | 0 | 5 | 0 | 4 | 0 | 5 | 0.4 | 17 | 0.1176 |
| 5_13024287 | 7 | 0.2857 | 7 | 0 | 5 | 0.2 | 7 | 0.1429 | 26 | 0.1538 | 2 | 0 | 6 | 0.1667 | 4 | 0 | 5 | 0.2 | 17 | 0.1176 |
| 5_13024288 | 7 | 0 | 7 | 0.2857 | 4 | 0.25 | 5 | 0.2 | 23 | 0.1739 | 3 | 0.3333 | 4 | 0 | 4 | 0.5 | 4 | 0.5 | 15 | 0.3333 |
| 5_13026063 | 1 | 1 | 2 | 0.5 | 2 | 0 | 2 | 0.5 | 7 | 0.4286 | NA | NA | NA | NA | 3 | 0.3333 | 1 | 1 | 4 | 0.5 |
| 5_13026079 | 1 | 1 | 2 | 0 | 2 | 0 | 3 | 0.6667 | 8 | 0.375 | 1 | 0 | 1 | 0 | 4 | 0.25 | 3 | 0.3333 | 9 | 0.2222 |
| 5_13027025 | 2 | 1 | 3 | 1 | 3 | 0.6667 | 4 | 0.75 | 12 | 0.8333 | 1 | 1 | 1 | 0 | 8 | 0.625 | 3 | 1 | 13 | 0.6923 |
| 5_13027027 | 2 | 0.5 | 2 | 0.5 | 3 | 0 | 4 | 0.25 | 11 | 0.2727 | 1 | 0 | 1 | 0 | 7 | 0.1429 | 3 | 0.6667 | 12 | 0.25 |
| 5_13027029 | 1 | 0 | 3 | 0.3333 | 1 | 0 | 3 | 0.6667 | 8 | 0.375 | NA | NA | 1 | 0 | 6 | 0.3333 | 3 | 0 | 10 | 0.2 |
| 5_13027039 | 2 | 0.5 | 2 | 1 | 1 | 1 | 3 | 1 | 8 | 0.875 | NA | NA | NA | NA | 4 | 0.75 | 3 | 1 | 7 | 0.8571 |
| 5_13027050 | 2 | 0.5 | 2 | 0.5 | NA | NA | 1 | 0 | 5 | 0.4 | NA | NA | NA | NA | 4 | 0.25 | 2 | 0 | 6 | 0.1667 |
| 5_13027054 | 1 | 0 | 2 | 0.5 | 1 | 0 | 3 | 0.6667 | 7 | 0.4286 | NA | NA | NA | NA | 2 | 1 | 3 | 0.3333 | 5 | 0.6 |
| 5_13027236 | 4 | 0.25 | 9 | 0.2222 | 2 | 0 | 4 | 0.5 | 19 | 0.2632 | 2 | 0.5 | 1 | 0 | 9 | 0.1111 | 5 | 0.2 | 17 | 0.1765 |
| 5_13027492 | 8 | 0.25 | 2 | 0 | 3 | 0 | 3 | 0 | 16 | 0.125 | 3 | 0 | 2 | 0 | 4 | 0.25 | 1 | 0 | 10 | 0.1 |
| 5_13036963 | 2 | 1 | 2 | 0.5 | 1 | 1 | 1 | 1 | 6 | 0.8333 | 2 | 1 | 3 | 1 | 4 | 0.75 | 2 | 1 | 11 | 0.9091 |
| 5_13036966 | 2 | 1 | 2 | 1 | 1 | 0 | NA | NA | 5 | 0.8 | 1 | 1 | 3 | 1 | 3 | 1 | 2 | 1 | 9 | 1 |
| 5_13036970 | 2 | 0.5 | 2 | 0.5 | 1 | 0 | 1 | 1 | 6 | 0.5 | 1 | 1 | 3 | 0.6667 | 3 | 0.3333 | 1 | 1 | 8 | 0.625 |
| 5_13036985 | 1 | 1 | 2 | 1 | 1 | 0 | NA | NA | 4 | 0.75 | 2 | 1 | 3 | 1 | 3 | 1 | 2 | 0.5 | 10 | 0.9 |
| 5_13036988 | 2 | 0.5 | 2 | 0 | 1 | 1 | NA | NA | 5 | 0.4 | 2 | 0 | 2 | 0 | 2 | 0 | 1 | 1 | 7 | 0.1429 |
| 5_13086831 | 2 | 1 | 5 | 1 | 2 | 1 | 1 | 1 | 10 | 1 | 1 | 0 | 3 | 0.6667 | 7 | 0.5714 | 1 | 1 | 12 | 0.5833 |
| 5_13086834 | 2 | 1 | 5 | 0.8 | 2 | 1 | 1 | 0 | 10 | 0.8 | 1 | 0 | 2 | 1 | 4 | 1 | 1 | 1 | 8 | 0.875 |
| 5_13086838 | 1 | 1 | 5 | 1 | 2 | 1 | 1 | 1 | 9 | 1 | 1 | 0 | 1 | 1 | 7 | 0.5714 | 1 | 1 | 10 | 0.6 |
| 5_13086844 | 2 | 1 | 5 | 1 | 2 | 0.5 | 1 | 1 | 10 | 0.9 | NA | NA | 3 | 1 | 6 | 1 | 1 | 1 | 10 | 1 |

| | | | | | | | | | | | | | | | | | | | | |
|---|---|---|---|---|---|---|---|---|---|---|---|---|---|---|---|---|---|---|---|---|
| 5_1308686 | 1 | 1 | 3 | 1 | NA | NA | NA | NA | 4 | 1 | NA | NA | NA | NA | 5 | 0.6 | NA | NA | 5 | 0.6 |
| 5_1308686 | 1 | 1 | 3 | 0.3333 | 1 | 0 | NA | NA | 5 | 0.4 | NA | NA | NA | NA | 4 | 0.25 | NA | NA | 4 | 0.25 |
| 5_1308687 | 1 | 1 | 3 | 1 | NA | NA | NA | NA | 4 | 1 | NA | NA | NA | NA | 5 | 0.4 | NA | NA | 5 | 0.4 |
| 5_1308687 | NA | NA | 2 | 1 | NA | NA | NA | NA | 2 | 1 | NA | NA | NA | NA | 4 | 1 | NA | NA | 4 | 1 |
| 5_1308688 | 1 | 1 | 3 | 1 | NA | NA | NA | NA | 4 | 1 | NA | NA | NA | NA | 5 | 1 | NA | NA | 5 | 1 |
| 5_1308690 | NA | NA | 4 | 0.25 | NA | NA | NA | NA | 4 | 0.25 | NA | NA | NA | NA | 5 | 0.2 | NA | NA | 5 | 0.2 |
| 5_1308690 | 1 | 1 | 4 | 1 | 1 | 1 | NA | NA | 6 | 1 | NA | NA | NA | NA | 6 | 1 | NA | NA | 6 | 1 |
| 5_1308693 | 1 | 1 | 4 | 1 | NA | NA | NA | NA | 5 | 1 | NA | NA | NA | NA | 4 | 0.75 | NA | NA | 4 | 0.75 |
| 5_1308694 | 1 | 1 | 3 | 0 | 1 | 1 | NA | NA | 5 | 0.4 | NA | NA | 2 | 0.5 | 6 | 0.1667 | NA | NA | 8 | 0.25 |
| 5_1308694 | 1 | 1 | 4 | 0.5 | NA | NA | NA | NA | 5 | 0.6 | NA | NA | 2 | 0.5 | 5 | 0.4 | NA | NA | 7 | 0.4286 |
| 5_1308695 | 1 | 1 | 2 | 1 | NA | NA | NA | NA | 3 | 1 | NA | NA | NA | NA | 1 | 1 | NA | NA | 1 | 1 |
| 5_1308704 | 1 | 1 | 4 | 0.25 | 1 | 0 | NA | NA | 6 | 0.3333 | 2 | 0.5 | 1 | 0 | 2 | 0 | NA | NA | 5 | 0.2 |
| 5_1308706 | NA | NA | 4 | 0.75 | 1 | 1 | NA | NA | 5 | 0.8 | 4 | 0.75 | 1 | 1 | 1 | 0 | NA | NA | 6 | 0.6667 |
| 5_1308711 | 1 | 0 | 4 | 0.5 | 2 | 0 | NA | NA | 7 | 0.2857 | 4 | 0.5 | 1 | 1 | 3 | 0 | 1 | 1 | 9 | 0.4444 |
| 5_1308724 | 3 | 0 | 4 | 0 | 4 | 0 | 2 | 0.5 | 13 | 0.0769 | 4 | 0 | 2 | 0 | 3 | 0 | 2 | 0.5 | 11 | 0.0909 |
| 5_1308731 | 2 | 0.5 | 6 | 0 | 4 | 0 | 2 | 0 | 14 | 0.0714 | 4 | 0.25 | 4 | 0 | 4 | 0 | 4 | 0.25 | 16 | 0.125 |
| 5_1308733 | 1 | 0 | 3 | 0.3333 | 1 | 0 | 1 | 0 | 6 | 0.1667 | 5 | 0.4 | 1 | 0 | 2 | 0.5 | 2 | 0 | 10 | 0.3 |
| 5_1308733 | 3 | 0 | 6 | 0.1667 | 3 | 0 | 3 | 0.3333 | 15 | 0.1333 | 4 | 0.5 | 4 | 0.25 | 4 | 0 | 4 | 0 | 16 | 0.1875 |
| 5_1308736 | 3 | 0.3333 | 7 | 0.1429 | 5 | 0 | 1 | 0 | 16 | 0.125 | 4 | 0.25 | 4 | 0 | 6 | 0 | 1 | 0 | 15 | 0.0667 |
| 5_1308737 | 2 | 0 | 7 | 0.1429 | 5 | 0 | 3 | 0 | 17 | 0.0588 | 4 | 0 | 3 | 0.3333 | 5 | 0 | 4 | 0.25 | 16 | 0.125 |
| 5_1308748 | 4 | 0.25 | 11 | 0 | 5 | 0.4 | 4 | 0 | 24 | 0.125 | 8 | 0.125 | 8 | 0.125 | 10 | 0.2 | 6 | 0.1667 | 32 | 0.1562 |
| 5_1308754 | 5 | 0 | 11 | 0.0909 | 7 | 0.1429 | 4 | 0.25 | 27 | 0.1111 | 10 | 0.1 | 10 | 0.2 | 13 | 0.2308 | 9 | 0.1111 | 42 | 0.1667 |
| 5_1308758 | 26 | 0.6154 | 18 | 0.5 | 18 | 0.2778 | 17 | 0.3529 | 79 | 0.4557 | 18 | 0.5 | 30 | 0.3667 | 23 | 0.4783 | 19 | 0.5263 | 90 | 0.4556 |
| 5_1308760 | 28 | 0.0357 | 21 | 0.1905 | 24 | 0 | 22 | 0.1364 | 95 | 0.0842 | 22 | 0.0455 | 38 | 0.0526 | 32 | 0.2188 | 20 | 0.05 | 112 | 0.0982 |
| 5_1309043 | 1 | 1 | 2 | 0.5 | 2 | 0.5 | 4 | 0.5 | 9 | 0.5556 | 3 | 0.3333 | 4 | 0.75 | 2 | 0.5 | 5 | 0.6 | 14 | 0.5714 |
| 5_1309044 | 1 | 0 | 2 | 0.5 | 3 | 0 | 4 | 0 | 10 | 0.1 | 6 | 0.3333 | 7 | 0.4286 | 2 | 1 | 8 | 0.5 | 23 | 0.4783 |
| 5_1309045 | 1 | 1 | 2 | 1 | 3 | 1 | 1 | 1 | 7 | 1 | 5 | 0.6 | 6 | 0.8333 | 1 | 1 | 4 | 0.75 | 16 | 0.75 |
| 5_1309045 | 1 | 1 | 1 | 1 | 3 | 0.6667 | 4 | 0.75 | 9 | 0.7778 | 5 | 0.8 | 7 | 1 | 1 | 1 | 7 | 0.8571 | 20 | 0.9 |
| 5_1309046 | 1 | 0 | 2 | 0 | 3 | 0 | 3 | 0.3333 | 9 | 0.1111 | 5 | 0 | 6 | 0.1667 | 1 | 0 | 8 | 0 | 20 | 0.05 |
| 5_1309046 | 1 | 1 | 2 | 0 | 3 | 0.3333 | 4 | 0.25 | 10 | 0.3 | 5 | 0.2 | 6 | 0.3333 | 2 | 1 | 8 | 0 | 21 | 0.2381 |
| 5_1309046 | 1 | 0 | 2 | 0 | 3 | 0.3333 | 2 | 0 | 8 | 0.125 | 3 | 0.3333 | 6 | 0.6667 | 2 | 1 | 6 | 0.8333 | 17 | 0.7059 |
| 5_1309047 | 1 | 0 | 2 | 0 | 3 | 0.3333 | 3 | 0 | 9 | 0.1111 | 5 | 0 | 7 | 0.1429 | 2 | 1 | 7 | 0 | 21 | 0.1429 |
| 5_1309047 | 1 | 1 | 1 | 0 | 2 | 0 | 3 | 0 | 7 | 0.1429 | 5 | 0 | 7 | 0.2857 | 2 | 1 | 7 | 0.1429 | 21 | 0.2381 |
| 5_1309050 | 1 | 0 | 1 | 0 | 3 | 0.3333 | 3 | 0.3333 | 8 | 0.25 | 4 | 0.5 | 5 | 0.4 | 1 | 0 | 6 | 0.6667 | 16 | 0.5 |
| 5_1309051 | 1 | 0 | 2 | 0.5 | 1 | 1 | 3 | 0.6667 | 7 | 0.5714 | 4 | 0.75 | 4 | 0.25 | 2 | 0.5 | 6 | 0.8333 | 16 | 0.625 |
| 5_1309051 | 1 | 0 | 2 | 0.5 | 3 | 0.3333 | 3 | 1 | 9 | 0.5556 | 4 | 0.5 | 7 | 0.2857 | 2 | 0.5 | 7 | 0.5714 | 20 | 0.45 |
| 5_1309052 | 1 | 1 | 2 | 1 | 2 | 0.5 | 1 | 1 | 6 | 0.8333 | 1 | 1 | 5 | 0.8 | 1 | 0 | 4 | 1 | 11 | 0.8182 |
| 5_1309052 | 1 | 1 | 1 | 1 | 1 | 1 | 2 | 1 | 5 | 1 | 3 | 0.6667 | 3 | 0.6667 | 2 | 1 | 4 | 0.75 | 12 | 0.75 |
| 5_1313353 | 1 | 1 | 1 | 1 | NA | NA | NA | NA | 2 | 1 | 1 | 1 | 4 | 0.75 | 1 | 0 | 1 | 0 | 7 | 0.5714 |
| 5_1313356 | 1 | 1 | 1 | 1 | NA | NA | 1 | 1 | 3 | 1 | 1 | 1 | 4 | 1 | 2 | 1 | 2 | 1 | 9 | 1 |
| 5_1313357 | 1 | 0 | 1 | 0 | NA | NA | 1 | 1 | 3 | 0.3333 | NA | NA | 4 | 0.25 | 2 | 0.5 | 2 | 1 | 8 | 0.5 |
| 5_1313357 | 1 | 0 | 1 | 1 | NA | NA | 1 | 1 | 3 | 0.6667 | 1 | 0 | 2 | 1 | 2 | 1 | 2 | 0.5 | 7 | 0.7143 |
| 5_1313357 | 1 | 0 | 1 | 0 | NA | NA | 1 | 1 | 3 | 0.3333 | 1 | 0 | 2 | 0 | 1 | 1 | 3 | 0.6667 | 7 | 0.4286 |
| 5_1313364 | NA | NA | NA | NA | NA | NA | 1 | 1 | 1 | 1 | NA | NA | 1 | 0 | 1 | 1 | 1 | 0 | 3 | 0.3333 |
| 5_1314664 | 5 | 0.4 | 8 | 0.375 | 5 | 0.6 | 10 | 0.6 | 28 | 0.5 | 2 | 0.5 | 9 | 0.5556 | 10 | 0.3 | 6 | 0.6667 | 27 | 0.4815 |
| 5_1314668 | 3 | 0.3333 | 8 | 0.5 | 7 | 0.1429 | 9 | 0.1111 | 27 | 0.2593 | 3 | 0 | 6 | 0 | 10 | 0.2 | 3 | 0.3333 | 22 | 0.1364 |
| 5_1314671 | 3 | 0.3333 | 9 | 0.6667 | 7 | 0.4286 | 8 | 0.375 | 27 | 0.4815 | 2 | 0 | 6 | 0.3333 | 10 | 0.3 | 4 | 0.25 | 22 | 0.2727 |
| 5_1314680 | 1 | 0 | 5 | 0 | 4 | 0.75 | 7 | 0.4286 | 17 | 0.3529 | 2 | 0 | 2 | 0.5 | 4 | 0.25 | 3 | 0.3333 | 11 | 0.2727 |
| 5_1314681 | 1 | 0 | 4 | 0 | 3 | 0.3333 | 5 | 0.2 | 13 | 0.1538 | 1 | 1 | 2 | 0.5 | 3 | 0 | 3 | 0.6667 | 9 | 0.4444 |
| 5_1314681 | 1 | 1 | 3 | 1 | 3 | 1 | 4 | 1 | 11 | 1 | 2 | 1 | 2 | 1 | 6 | 1 | 1 | 1 | 11 | 1 |
| 5_1314682 | 1 | 0 | 6 | 1 | 5 | 1 | 6 | 0.6667 | 18 | 0.8333 | 1 | 1 | 2 | 1 | 6 | 0.6667 | 3 | 1 | 12 | 0.8333 |
| 5_1314682 | 1 | 0 | 3 | 0.3333 | 4 | 0.75 | 7 | 0.8571 | 15 | 0.6667 | 1 | 1 | 2 | 1 | 6 | 0.3333 | 3 | 1 | 12 | 0.6667 |
| 5_1314682 | 1 | 0 | 5 | 0.6 | 5 | 0 | 5 | 0.6 | 16 | 0.375 | NA | NA | 2 | 0.5 | 4 | 0.25 | 3 | 0.3333 | 9 | 0.3333 |
| 5_1314682 | 1 | 1 | 5 | 1 | 5 | 1 | 6 | 1 | 17 | 1 | 1 | 1 | 2 | 1 | 4 | 1 | 2 | 1 | 9 | 1 |
| 5_1314683 | 1 | 1 | 5 | 1 | 4 | 1 | 5 | 0.8 | 15 | 0.9333 | 1 | 1 | 2 | 1 | 5 | 0.6 | 3 | 1 | 11 | 0.8182 |
| 5_1314684 | 1 | 0 | 5 | 0.4 | 3 | 0.6667 | 4 | 0.25 | 13 | 0.3846 | 1 | 0 | 2 | 0 | 4 | 0.5 | 3 | 0 | 10 | 0.2 |
| 5_1314684 | 1 | 1 | 3 | 1 | 4 | 1 | 2 | 1 | 10 | 1 | 1 | 1 | 2 | 1 | 3 | 0.6667 | 3 | 1 | 9 | 0.8889 |
| 5_1314685 | 1 | 1 | 4 | 1 | 4 | 1 | 5 | 1 | 14 | 1 | 1 | 1 | 1 | 1 | 4 | 1 | 3 | 1 | 9 | 1 |
| 5_1314685 | 1 | 1 | 5 | 1 | 3 | 1 | 4 | 1 | 13 | 1 | 1 | 1 | 2 | 1 | 4 | 1 | 3 | 1 | 10 | 1 |
| 5_1314686 | 1 | 0 | 4 | 0.25 | 3 | 0.3333 | 4 | 0 | 12 | 0.1667 | 1 | 0 | 2 | 0 | 5 | 0.2 | 3 | 0 | 11 | 0.0909 |
| 5_1314687 | 1 | 1 | 5 | 0.6 | 2 | 0.5 | 4 | 0.75 | 12 | 0.6667 | NA | NA | 2 | 1 | 4 | 0.75 | 3 | 0.6667 | 9 | 0.7778 |
| 5_1314688 | NA | NA | 2 | 1 | 3 | 1 | 2 | 0.5 | 7 | 0.8571 | NA | NA | 2 | 1 | 3 | 0.6667 | 3 | 1 | 8 | 0.875 |
| 5_1314688 | 1 | 0 | 3 | 0.3333 | 3 | 0.3333 | 4 | 0.25 | 11 | 0.2727 | 1 | 1 | 2 | 0 | 4 | 0.25 | 3 | 0.6667 | 10 | 0.4 |
| 5_1314688 | 1 | 0 | 3 | 0.3333 | 1 | 1 | 2 | 0.5 | 7 | 0.4286 | NA | NA | 2 | 0.5 | 3 | 0.3333 | 3 | 0.6667 | 8 | 0.5 |
| 5_1314689 | 1 | 1 | 3 | 1 | 2 | 0 | 3 | 0.3333 | 9 | 0.5556 | 1 | 1 | 2 | 0.5 | 2 | 0.5 | 3 | 1 | 8 | 0.75 |

| | | | | | | | | | | | | | | | | | | | | |
|---|---|---|---|---|---|---|---|---|---|---|---|---|---|---|---|---|---|---|---|---|
| 5_13146893 | NA | NA | 3 | 0.6667 | 1 | 0 | 2 | 0 | 6 | 0.3333 | 1 | 1 | 1 | 1 | 4 | 0.75 | 2 | 0.5 | 8 | 0.75 |
| 5_13146895 | 1 | 1 | 1 | 1 | NA | NA | 3 | 0.6667 | 5 | 0.8 | 1 | 1 | 1 | 1 | 2 | 1 | NA | NA | 4 | 1 |
| 5_13146896 | 1 | 1 | 4 | 0.5 | 1 | 1 | 4 | 0.25 | 10 | 0.5 | 1 | 0 | 1 | 0 | 3 | 0.6667 | 2 | 0.5 | 7 | 0.4286 |
| 5_13146905 | 1 | 0 | 5 | 0.4 | 2 | 0 | 4 | 0.25 | 12 | 0.25 | 1 | 1 | 2 | 0 | 4 | 0.75 | 3 | 0.3333 | 10 | 0.5 |
| 5_13146935 | 1 | 0 | 4 | 0.25 | 2 | 0 | 3 | 0 | 10 | 0.1 | 1 | 1 | 2 | 0.5 | 4 | 0 | 3 | 0 | 10 | 0.2 |
| 5_13416721 | 1 | 1 | 2 | 0.5 | 3 | 0.3333 | 1 | 0 | 7 | 0.4286 | NA | NA | 3 | 0.6667 | 4 | 0.5 | 1 | 1 | 8 | 0.625 |
| 5_13416726 | 1 | 1 | 1 | 0 | 3 | 0.6667 | NA | NA | 5 | 0.6 | NA | NA | 3 | 0.3333 | 5 | 0.2 | 4 | 0.75 | 12 | 0.4167 |
| 5_13416743 | 1 | 0 | 2 | 0 | 4 | 0.25 | 1 | 0 | 8 | 0.125 | NA | NA | 2 | 0 | 6 | 0.5 | 4 | 0.25 | 12 | 0.3333 |
| 5_13416751 | 1 | 0 | 2 | 0 | 3 | 0.3333 | 1 | 0 | 7 | 0.1429 | NA | NA | 1 | 0 | 6 | 0 | 4 | 0.25 | 11 | 0.0909 |
| 5_13417709 | 1 | 1 | 1 | 0 | 1 | 1 | NA | NA | 3 | 0.6667 | 1 | 1 | NA | NA | 2 | 0.5 | 1 | 0 | 4 | 0.5 |
| 5_13417713 | NA | NA | 1 | 1 | 1 | 0 | NA | NA | 2 | 0.5 | 1 | 0 | NA | NA | 2 | 0.5 | 1 | 1 | 4 | 0.5 |
| 5_13417720 | 1 | 1 | 1 | 1 | 2 | 0.5 | NA | NA | 4 | 0.75 | 1 | 1 | NA | NA | 2 | 1 | 1 | 1 | 4 | 1 |
| 5_13417723 | 1 | 1 | 1 | 1 | 1 | 0 | NA | NA | 3 | 0.6667 | 1 | 1 | NA | NA | 2 | 1 | 1 | 1 | 4 | 1 |
| 5_13417737 | 2 | 0.5 | 1 | 0 | 2 | 0.5 | NA | NA | 5 | 0.4 | 1 | 1 | NA | NA | 2 | 1 | 1 | 0 | 4 | 0.75 |
| 5_13417750 | 1 | 0 | 1 | 0 | 2 | 0.5 | NA | NA | 4 | 0.25 | 1 | 0 | NA | NA | 2 | 0.5 | 1 | 1 | 4 | 0.5 |
| 5_13417759 | 1 | 1 | 1 | 0 | 2 | 0.5 | NA | NA | 4 | 0.5 | 1 | 0 | NA | NA | 2 | 0.5 | NA | NA | 3 | 0.3333 |
| 5_13425027 | 1 | 0 | 1 | 0 | NA | NA | 3 | 0.3333 | 5 | 0.2 | 2 | 0.5 | 1 | 0 | 1 | 0 | NA | NA | 4 | 0.25 |
| 5_13425050 | 1 | 0 | 1 | 1 | NA | NA | 3 | 0.6667 | 5 | 0.6 | 1 | 1 | NA | NA | 1 | 1 | NA | NA | 2 | 1 |
| 5_13438671 | 1 | 0 | 1 | 1 | 3 | 0 | 1 | 0 | 6 | 0.1667 | 4 | 0 | 1 | 0 | 4 | 0.25 | 2 | 0 | 11 | 0.0909 |
| 5_13438672 | NA | NA | 2 | 0.5 | 5 | 0 | 2 | 0.5 | 9 | 0.2222 | 4 | 0.25 | 1 | 0 | 6 | 0 | 2 | 0 | 13 | 0.0769 |
| 5_13438738 | NA | NA | NA | NA | 1 | 1 | 1 | 0 | 2 | 0.5 | 2 | 0 | NA | NA | 4 | 0.25 | NA | NA | 6 | 0.1667 |
| 5_13438773 | NA | NA | NA | NA | 1 | 1 | 1 | 0 | 2 | 0.5 | 1 | 1 | NA | NA | 4 | 0.5 | NA | NA | 5 | 0.6 |
| 5_13451858 | 15 | 0.2 | 18 | 0.2222 | 8 | 0.25 | 11 | 0.0909 | 52 | 0.1923 | 12 | 0.25 | 10 | 0.3 | 13 | 0.3846 | 14 | 0.2857 | 49 | 0.3061 |
| 5_13451894 | 15 | 0.2 | 15 | 0.3333 | 5 | 0.2 | 8 | 0.25 | 43 | 0.2558 | 9 | 0.1111 | 7 | 0.2857 | 13 | 0.1538 | 10 | 0.1 | 39 | 0.1538 |
| 5_13451920 | 12 | 0.0833 | 19 | 0.1579 | 10 | 0.2 | 10 | 0 | 51 | 0.1176 | 6 | 0.1667 | 8 | 0.5 | 13 | 0.2308 | 11 | 0.0909 | 38 | 0.2368 |
| 5_13451928 | 10 | 0.2 | 9 | 0.3333 | 6 | 0.1667 | 7 | 0.2857 | 32 | 0.25 | 4 | 0 | 8 | 0.375 | 12 | 0.1667 | 5 | 0.2 | 29 | 0.2069 |
| 5_13451929 | 8 | 0.5 | 15 | 0.5333 | 7 | 0.4286 | 9 | 0.5556 | 39 | 0.5128 | 7 | 0.5714 | 6 | 0.3333 | 12 | 0.6667 | 4 | 1 | 29 | 0.6207 |
| 5_13451932 | 11 | 0.2727 | 14 | 0.1429 | 7 | 0 | 8 | 0 | 40 | 0.125 | 5 | 0.2 | 7 | 0 | 10 | 0.2 | 7 | 0.2857 | 29 | 0.1724 |
| 5_13451936 | 9 | 0.5556 | 13 | 0.3846 | 9 | 0.3333 | 7 | 0.5714 | 38 | 0.4474 | 5 | 0.8 | 3 | 0 | 7 | 0.2857 | 8 | 0.25 | 23 | 0.3478 |
| 5_13451948 | 10 | 0.1 | 17 | 0.2353 | 7 | 0.1429 | 8 | 0.5 | 42 | 0.2381 | 6 | 0.5 | 6 | 0.6667 | 10 | 0.6 | 9 | 0.1111 | 31 | 0.4516 |
| 5_13451952 | 13 | 0.7692 | 17 | 0.8824 | 9 | 0.8889 | 9 | 0.8889 | 48 | 0.8542 | 5 | 1 | 7 | 0.7143 | 13 | 0.8462 | 11 | 0.9091 | 36 | 0.8611 |
| 5_13451954 | 7 | 1 | 15 | 0.7333 | 6 | 0.8333 | 9 | 0.7778 | 37 | 0.8108 | 6 | 1 | 6 | 0.5 | 12 | 0.75 | 7 | 0.7143 | 31 | 0.7419 |
| 5_13453142 | 7 | 0.8571 | 11 | 0.8182 | 6 | 0.6667 | 8 | 0.375 | 32 | 0.6875 | 3 | 0.3333 | 7 | 0.7143 | 4 | 0.75 | 4 | 0.75 | 18 | 0.6667 |
| 5_13453159 | 9 | 0.5556 | 12 | 0.8333 | 7 | 0.8571 | 7 | 0.8571 | 35 | 0.7714 | 5 | 1 | 7 | 0.8571 | 5 | 0.6 | 4 | 0.75 | 21 | 0.8095 |
| 5_13453163 | 9 | 0.3333 | 12 | 0.1667 | 5 | 0 | 7 | 0.2857 | 33 | 0.2121 | 4 | 0 | 7 | 0.2857 | 5 | 0 | 3 | 0 | 19 | 0.1053 |
| 5_13453189 | 8 | 0.625 | 9 | 0.6667 | 5 | 0.6 | 5 | 0.4 | 27 | 0.5926 | 3 | 0 | 3 | 0.6667 | 4 | 0.5 | 3 | 0.3333 | 13 | 0.3846 |
| 5_13453191 | 10 | 0 | 9 | 0.1111 | 4 | 0.25 | 6 | 0 | 29 | 0.069 | 5 | 0 | 7 | 0.1429 | 4 | 0 | 3 | 0.3333 | 19 | 0.1053 |
| 5_13453194 | 9 | 0.7778 | 9 | 0.6667 | 5 | 1 | 6 | 1 | 29 | 0.8276 | 3 | 1 | 5 | 0.4 | 2 | 0.5 | 2 | 1 | 12 | 0.6667 |
| 5_13453196 | 9 | 0.8889 | 9 | 0.7778 | 5 | 0.6 | 4 | 0.75 | 27 | 0.7778 | 4 | 0.5 | 7 | 0.8571 | 4 | 0.5 | 3 | 1 | 18 | 0.7222 |
| 5_13453206 | 7 | 0.2857 | 7 | 0.2857 | 5 | 0.2 | 3 | 0 | 22 | 0.2273 | 2 | 0 | 5 | 0 | 4 | 0.25 | 3 | 0.3333 | 14 | 0.1429 |
| 5_13453210 | 9 | 0 | 10 | 0.3 | 5 | 0.4 | 5 | 0.4 | 29 | 0.2414 | 5 | 0.6 | 7 | 0.2857 | 4 | 0.5 | 2 | 1 | 18 | 0.5 |
| 5_13453212 | 7 | 0 | 6 | 0 | 5 | 0.2 | 3 | 0.3333 | 21 | 0.0952 | 3 | 0 | 4 | 0 | 4 | 0.25 | 1 | 0 | 12 | 0.0833 |
| 5_13453217 | 5 | 0.6 | 8 | 0.75 | 3 | 0 | 4 | 0.5 | 20 | 0.55 | 3 | 0 | 6 | 0.3333 | 2 | 0.5 | 3 | 0.6667 | 14 | 0.3571 |
| 5_13455188 | 5 | 0.6 | 3 | 0.3333 | 1 | 1 | 1 | 0 | 10 | 0.5 | 2 | 0.5 | 3 | 0.6667 | 2 | 0.5 | 2 | 1 | 9 | 0.6667 |
| 5_13462683 | 7 | 0.1429 | 4 | 0.25 | 9 | 0 | 7 | 0.4286 | 27 | 0.1852 | 4 | 0 | 6 | 0.3333 | 2 | 0 | 3 | 0 | 15 | 0.1333 |
| 5_13462687 | 6 | 0.3333 | 6 | 0 | 6 | 0.1667 | 6 | 0 | 24 | 0.125 | 2 | 0.5 | 6 | 0 | 3 | 0 | 4 | 0 | 15 | 0.0667 |
| 5_13462693 | 5 | 0.6 | 4 | 0.5 | 7 | 0.5714 | 4 | 0.75 | 20 | 0.6 | 3 | 0 | 5 | 0.2 | 3 | 0.6667 | 3 | 0.3333 | 14 | 0.2857 |
| 5_13462701 | 5 | 0.8 | 4 | 0.5 | 9 | 0.6667 | 8 | 0.75 | 26 | 0.6923 | 4 | 1 | 6 | 0.5 | 3 | 0.6667 | 4 | 0.75 | 17 | 0.7059 |
| 5_13462711 | 5 | 0.4 | 5 | 0.6 | 7 | 0.4286 | 5 | 0.4 | 22 | 0.4545 | 2 | 0 | 5 | 0.6 | 2 | 0 | 5 | 0.2 | 14 | 0.2857 |
| 5_13468494 | 5 | 0.2 | 8 | 0 | 2 | 0 | 13 | 0.0769 | 28 | 0.0714 | 4 | 0 | 4 | 0 | 5 | 0 | 7 | 0.1429 | 20 | 0.05 |
| 5_13468499 | 2 | 0.5 | 3 | 0.3333 | 2 | 0.5 | 3 | 0 | 10 | 0.3 | 1 | 0 | 2 | 0.5 | 1 | 0 | 2 | 0 | 6 | 0.1667 |
| 5_13468615 | 1 | 0 | 5 | 0.4 | 1 | 0 | 2 | 0 | 9 | 0.2222 | 1 | 0 | 2 | 0.5 | NA | NA | 3 | 0.3333 | 6 | 0.3333 |
| 5_13468645 | 3 | 0.3333 | 7 | 0 | 3 | 0.3333 | 5 | 0 | 18 | 0.1111 | 2 | 0.5 | 3 | 0.3333 | 3 | 0 | 5 | 0 | 13 | 0.1538 |
| 5_13496223 | NA | NA | 2 | 0.5 | NA | NA | 1 | 0 | 3 | 0.3333 | NA | NA | NA | NA | 1 | 1 | NA | NA | 1 | 1 |
| 5_13496971 | 1 | 1 | 2 | 0.5 | NA | NA | 1 | 1 | 4 | 0.75 | NA | NA | NA | NA | 1 | 0 | 1 | 1 | 2 | 0.5 |
| 5_13496972 | NA | NA | 2 | 1 | 1 | 1 | 1 | 1 | 4 | 1 | NA | NA | NA | NA | 1 | 1 | 1 | 1 | 2 | 1 |
| 5_13496974 | NA | NA | 1 | 0 | 1 | 0 | 1 | 1 | 3 | 0.3333 | NA | NA | NA | NA | 1 | 0 | 1 | 1 | 2 | 0.5 |
| 5_13504090 | 5 | 0.8 | 5 | 0.6 | 4 | 1 | 4 | 0.5 | 18 | 0.7222 | 5 | 0.6 | 1 | 1 | 4 | 0.75 | 4 | 1 | 14 | 0.7857 |
| 5_13504094 | 5 | 0.2 | 7 | 0.2857 | 3 | 0.3333 | 3 | 0.6667 | 18 | 0.3333 | 2 | 0 | 1 | 1 | 4 | 0.25 | 4 | 0.25 | 11 | 0.2727 |
| 5_13504101 | 4 | 0 | 3 | 0 | 4 | 0.5 | 3 | 0 | 14 | 0.1429 | 2 | 0.5 | 1 | 1 | 2 | 0 | 3 | 0.6667 | 8 | 0.5 |
| 5_13504101 | 3 | 0.6667 | NA | NA | NA | NA | 3 | 0.3333 | 6 | 0.5 | 1 | 1 | 3 | 1 | 2 | 1 | 1 | 1 | 7 | 1 |
| 5_13504185 | NA | NA | 1 | 0 | 1 | 1 | NA | NA | 2 | 0.5 | NA | NA | NA | NA | NA | NA | 1 | 1 | 1 | 1 |
| 5_13507910 | 2 | 0 | 19 | 0.2632 | 17 | 0.5294 | NA | NA | 38 | 0.3684 | 14 | 0.4286 | 7 | 0.7143 | 1 | 1 | 3 | 0.3333 | 25 | 0.52 |
| 5_13507911 | 2 | 0 | 17 | 0.1176 | 17 | 0.0588 | 1 | 0 | 37 | 0.0811 | 13 | 0.0769 | 8 | 0 | 1 | 0 | 4 | 0.25 | 26 | 0.0769 |
| 5_13507970 | 2 | 0 | 15 | 0 | 17 | 0.1765 | 2 | 0 | 36 | 0.0833 | 15 | 0.0667 | 8 | 0.125 | 4 | 0 | 5 | 0 | 32 | 0.0625 |

| | | | | | | | | | | | | | | | | | | | | |
|---|---|---|---|---|---|---|---|---|---|---|---|---|---|---|---|---|---|---|---|---|
| 5_1350798 | 1 | 1 | 20 | 0.25 | 20 | 0.4 | 3 | 0.3333 | 44 | 0.3409 | 18 | 0.2778 | 8 | 0.375 | 4 | 0.75 | 4 | 0.5 | 34 | 0.3824 |
| 5_1350818 | 1 | 0 | 6 | 0.5 | 1 | 0 | 4 | 0.25 | 12 | 0.3333 | 5 | 0.4 | 7 | 0.2857 | 4 | 0.5 | 5 | 0.6 | 21 | 0.4286 |
| 5_1350820 | 1 | 1 | 6 | 0.1667 | 2 | 1 | 5 | 0.6 | 14 | 0.5 | 6 | 0.5 | 7 | 0.1429 | 4 | 0 | 2 | 0.5 | 19 | 0.2632 |
| 5_1350820 | 3 | 0.3333 | 8 | 0.625 | 1 | 1 | 7 | 0.4286 | 19 | 0.5263 | 5 | 0.4 | 8 | 0.75 | 4 | 1 | 4 | 1 | 21 | 0.7619 |
| 5_1350934 | 6 | 0.1667 | 9 | 0.1111 | 6 | 0 | 3 | 0 | 24 | 0.0833 | 8 | 0 | 10 | 0.1 | 6 | 0 | 6 | 0.1667 | 30 | 0.0667 |
| 5_1350936 | 7 | 0.1429 | 9 | 0.1111 | 5 | 0.2 | 2 | 0 | 23 | 0.1304 | 5 | 0.2 | 11 | 0.1818 | 7 | 0.5714 | 8 | 0.125 | 31 | 0.2581 |
| 5_1350938 | 4 | 0.5 | 11 | 0.0909 | 5 | 0.2 | 2 | 0.5 | 22 | 0.2273 | 7 | 0 | 12 | 0.1667 | 8 | 0.5 | 7 | 0.1429 | 34 | 0.2059 |
| 5_1350938 | 6 | 0 | 8 | 0.25 | 6 | 0 | 3 | 0.3333 | 23 | 0.1304 | 7 | 0.2857 | 10 | 0.1 | 6 | 0 | 8 | 0.25 | 31 | 0.1613 |
| 5_1350938 | 5 | 1 | 9 | 0.8889 | 5 | 0.8 | 3 | 1 | 22 | 0.9091 | 5 | 0.8 | 9 | 0.6667 | 5 | 1 | 9 | 0.8889 | 28 | 0.8214 |
| 5_1350938 | 6 | 0.3333 | 11 | 0.2727 | 3 | 0.3333 | 2 | 0 | 22 | 0.2727 | 8 | 0.375 | 7 | 0.4286 | 6 | 0 | 6 | 0 | 27 | 0.2222 |
| 5_1350938 | 7 | 0 | 10 | 0.1 | 8 | 0 | 4 | 0.25 | 29 | 0.069 | 9 | 0.1111 | 11 | 0 | 8 | 0 | 8 | 0.125 | 36 | 0.0556 |
| 5_1350940 | 5 | 0.6 | 9 | 0.5556 | 6 | 0.5 | 4 | 0.75 | 24 | 0.5833 | 5 | 0.6 | 9 | 0.2222 | 1 | 0 | 5 | 0.2 | 20 | 0.3 |
| 5_1350941 | 8 | 0.375 | 9 | 0.3333 | 6 | 0.1667 | 6 | 0.1667 | 29 | 0.2759 | 6 | 0.1667 | 10 | 0.2 | 5 | 0.2 | 3 | 0 | 24 | 0.1667 |
| 5_1350941 | 8 | 1 | 10 | 1 | 4 | 1 | 4 | 1 | 26 | 1 | 7 | 1 | 9 | 1 | 4 | 1 | 4 | 1 | 24 | 1 |
| 5_1350942 | 3 | 1 | 6 | 1 | 1 | 1 | 1 | 1 | 11 | 1 | 2 | 1 | 7 | 1 | 2 | 1 | 1 | 1 | 12 | 1 |
| 5_1350942 | 1 | 1 | 2 | 0.5 | 1 | 1 | NA | NA | 4 | 0.75 | NA | NA | 4 | 0.75 | 1 | 1 | NA | NA | 5 | 0.8 |
| 5_1350959 | NA | NA | 1 | 1 | NA | NA | NA | NA | 1 | 1 | NA | NA | 2 | 1 | NA | NA | NA | NA | 2 | 1 |
| 5_1350963 | NA | NA | 1 | 1 | NA | NA | NA | NA | 1 | 1 | NA | NA | 2 | 0.5 | 1 | 1 | NA | NA | 3 | 0.6667 |
| 5_1350968 | 1 | 0 | 1 | 1 | NA | NA | NA | NA | 2 | 0.5 | NA | NA | 2 | 0.5 | 1 | 0 | NA | NA | 3 | 0.3333 |
| 5_1351019 | 2 | 0 | 3 | 0 | 2 | 0.5 | 4 | 0 | 11 | 0.0909 | 3 | 0.3333 | 3 | 0 | 5 | 0 | 3 | 0.3333 | 14 | 0.1429 |
| 5_1351021 | 2 | 0.5 | 3 | 0.6667 | 2 | 1 | 3 | 1 | 10 | 0.8 | 4 | 0.5 | 3 | 1 | 4 | 0.75 | 1 | 0 | 12 | 0.6667 |
| 5_1351022 | 2 | 0.5 | 4 | 0 | 1 | 0 | 3 | 1 | 10 | 0.4 | 2 | 0.5 | 3 | 0 | 4 | 0.25 | 2 | 0 | 11 | 0.1818 |
| 5_1351025 | 3 | 1 | 3 | 0.6667 | 1 | 1 | 2 | 1 | 9 | 0.8889 | 3 | 0.3333 | 2 | 0.5 | 3 | 1 | 3 | 0.3333 | 11 | 0.5455 |
| 5_1351881 | 3 | 0.6667 | NA | NA | NA | NA | 1 | 0 | 4 | 0.5 | NA | NA | NA | NA | 3 | 0.6667 | 1 | 1 | 4 | 0.75 |
| 5_1351883 | 3 | 0.6667 | NA | NA | NA | NA | NA | NA | 3 | 0.6667 | NA | NA | NA | NA | 2 | 1 | 1 | 1 | 3 | 1 |
| 5_1351883 | 3 | 1 | NA | NA | NA | NA | 1 | 1 | 4 | 1 | 1 | 1 | NA | NA | 3 | 0.6667 | 1 | 1 | 5 | 0.8 |
| 5_1351883 | 3 | 0.6667 | 1 | 1 | NA | NA | 1 | 1 | 5 | 0.8 | NA | NA | NA | NA | 3 | 0.3333 | 1 | 0 | 4 | 0.25 |
| 5_1351885 | 3 | 1 | 1 | 1 | NA | NA | 1 | 1 | 5 | 1 | 1 | 1 | NA | NA | 1 | 1 | NA | NA | 2 | 1 |
| 5_1351886 | 3 | 0 | 1 | 0 | NA | NA | 1 | 1 | 5 | 0.2 | 1 | 0 | NA | NA | 2 | 0.5 | 1 | 0 | 4 | 0.25 |
| 5_1351887 | 2 | 0.5 | 1 | 0 | NA | NA | 1 | 0 | 4 | 0.25 | NA | NA | NA | NA | 3 | 0.3333 | NA | NA | 3 | 0.3333 |
| 5_1351887 | 3 | 0.3333 | 1 | 0 | NA | NA | 1 | 0 | 5 | 0.2 | NA | NA | NA | NA | 3 | 0.3333 | NA | NA | 3 | 0.3333 |
| 5_1351888 | 3 | 0 | 2 | 0 | NA | NA | 1 | 1 | 6 | 0.1667 | 1 | 0 | NA | NA | 3 | 0.3333 | 1 | 1 | 5 | 0.4 |
| 5_1351888 | 3 | 0.3333 | 2 | 0 | NA | NA | 1 | 0 | 6 | 0.1667 | 1 | 0 | NA | NA | 2 | 0.5 | 1 | 0 | 4 | 0.25 |
| 5_1351889 | 3 | 0.6667 | 2 | 0.5 | 1 | 0 | 1 | 0 | 7 | 0.4286 | 1 | 1 | NA | NA | 2 | 0.5 | 1 | 1 | 4 | 0.75 |
| 5_1351894 | 2 | 0 | 4 | 0.25 | NA | NA | 1 | 0 | 7 | 0.1429 | 1 | 0 | NA | NA | 3 | 0.6667 | 1 | 0 | 5 | 0.4 |
| 5_1351906 | 2 | 0 | 4 | 0.25 | 1 | 0 | 1 | 0 | 8 | 0.125 | 2 | 0.5 | 1 | 0 | 3 | 0 | 2 | 0 | 8 | 0.125 |
| 5_1351907 | 3 | 0.3333 | 4 | 0.5 | NA | NA | 1 | 1 | 8 | 0.5 | 1 | 0 | 1 | 1 | 2 | 0 | 2 | 0.5 | 6 | 0.3333 |
| 5_1351909 | 3 | 0.6667 | 3 | 0 | NA | NA | 1 | 0 | 7 | 0.2857 | 2 | 0 | 1 | 1 | 3 | 0 | 2 | 0.5 | 8 | 0.25 |
| 5_1351910 | 3 | 0.3333 | 4 | 0.25 | 1 | 0 | 1 | 0 | 9 | 0.2222 | 3 | 0 | 1 | 1 | 3 | 0 | 1 | 0 | 8 | 0.125 |
| 5_1351910 | 3 | 1 | 4 | 1 | 1 | 1 | 1 | 1 | 9 | 1 | 3 | 1 | 1 | 1 | 3 | 1 | 2 | 1 | 9 | 1 |
| 5_1351913 | 3 | 0.3333 | 4 | 0 | NA | NA | 1 | 0 | 8 | 0.125 | 1 | 1 | 1 | 1 | 2 | 0 | 2 | 0.5 | 6 | 0.5 |
| 5_1351920 | 2 | 0 | 1 | 1 | NA | NA | 3 | 0 | 6 | 0.1667 | 2 | 0.5 | 1 | 0 | 2 | 0 | 1 | 0 | 6 | 0.1667 |
| 5_1352097 | 5 | 0 | 9 | 0 | 6 | 0.1667 | 8 | 0.125 | 28 | 0.0714 | 6 | 0.3333 | 8 | 0 | 11 | 0.0909 | 6 | 0 | 31 | 0.0968 |
| 5_1352097 | 4 | 0 | 9 | 0.1111 | 7 | 0.2857 | 7 | 0 | 27 | 0.1111 | 6 | 0 | 9 | 0.1111 | 9 | 0 | 6 | 0.1667 | 30 | 0.0667 |
| 5_1352099 | 5 | 0 | 8 | 0 | 7 | 0 | 9 | 0.2222 | 29 | 0.069 | 6 | 0.1667 | 8 | 0.125 | 9 | 0.2222 | 8 | 0 | 31 | 0.129 |
| 5_1352101 | 7 | 0.7143 | 8 | 0.75 | 8 | 0.75 | 7 | 0.8571 | 30 | 0.7667 | 6 | 0.6667 | 8 | 0.75 | 10 | 0.6 | 7 | 0.5714 | 31 | 0.6452 |
| 5_1352101 | 5 | 1 | 7 | 0.8571 | 7 | 1 | 7 | 1 | 26 | 0.9615 | 5 | 1 | 9 | 1 | 10 | 1 | 8 | 1 | 32 | 1 |
| 5_1352102 | 4 | 0.75 | 10 | 0.9 | 7 | 0.7143 | 8 | 1 | 29 | 0.8621 | 6 | 1 | 7 | 1 | 12 | 0.8333 | 9 | 1 | 34 | 0.9412 |
| 5_1352105 | 8 | 0.375 | 10 | 0.5 | 11 | 0.6364 | 11 | 0.5455 | 40 | 0.525 | 8 | 0.375 | 9 | 0.5556 | 13 | 0.5385 | 9 | 0.6667 | 39 | 0.5385 |
| 5_1352455 | 2 | 0.5 | 3 | 0.6667 | 1 | 0 | 1 | 0 | 7 | 0.4286 | NA | NA | 2 | 1 | 1 | 1 | 1 | 1 | 4 | 1 |
| 5_1352456 | 2 | 0 | 3 | 0.3333 | 1 | 1 | NA | NA | 6 | 0.3333 | NA | NA | 2 | 0.5 | 1 | 0 | 1 | 0 | 4 | 0.25 |
| 5_1352456 | 2 | 0 | 1 | 0 | 1 | 1 | 1 | 0 | 5 | 0.2 | NA | NA | 1 | 0 | 1 | 1 | 1 | 0 | 3 | 0.3333 |
| 5_1352457 | 1 | 0 | 2 | 0 | 1 | 1 | 1 | 0 | 5 | 0.2 | NA | NA | 2 | 0 | 1 | 0 | 1 | 1 | 4 | 0.25 |
| 5_1352458 | 2 | 0.5 | 3 | 0.6667 | NA | NA | 1 | 0 | 6 | 0.5 | NA | NA | 3 | 1 | 1 | 1 | 1 | 1 | 5 | 1 |
| 5_1352463 | 2 | 0 | 5 | 0.4 | 1 | 0 | 1 | 0 | 9 | 0.2222 | NA | NA | 2 | 0 | 1 | 0 | 1 | 1 | 4 | 0.25 |
| 5_1352465 | 2 | 0.5 | 5 | 0.2 | 1 | 0 | 2 | 0.5 | 10 | 0.3 | 1 | 1 | 4 | 0.25 | 1 | 1 | 1 | 0 | 7 | 0.4286 |
| 5_1352470 | 2 | 0 | 5 | 0.2 | 2 | 0.5 | 2 | 0.5 | 11 | 0.2727 | 1 | 0 | 5 | 0.2 | 1 | 1 | 1 | 1 | 8 | 0.375 |
| 5_1352730 | 4 | 0.5 | 8 | 0.75 | 4 | 1 | 2 | 0.5 | 18 | 0.7222 | 2 | 0 | 7 | 0.4286 | 4 | 0.5 | 1 | 1 | 14 | 0.4286 |
| 5_1352730 | 6 | 0 | 9 | 0.2222 | 6 | 0 | 2 | 0 | 23 | 0.087 | 3 | 0 | 8 | 0.125 | 4 | 0 | 1 | 0 | 16 | 0.0625 |
| 5_1352731 | 5 | 0 | 8 | 0.125 | 4 | 0 | 1 | 0 | 18 | 0.0556 | 1 | 1 | 7 | 0 | 4 | 0 | 1 | 0 | 13 | 0.0769 |
| 5_1352732 | 3 | 0.6667 | 7 | 0.7143 | 4 | 0.25 | 2 | 1 | 16 | 0.625 | 4 | 0.5 | 6 | 0.3333 | 5 | 0.6 | NA | NA | 15 | 0.4667 |
| 5_1352732 | 7 | 0.1429 | 9 | 0.3333 | 6 | 0.1667 | 3 | 0 | 25 | 0.2 | 5 | 0 | 7 | 0.2857 | 5 | 0.6 | 1 | 1 | 18 | 0.3333 |
| 5_1352732 | NA | NA | 1 | 1 | 2 | 0.5 | 2 | 1 | 5 | 0.8 | 1 | 1 | 2 | 0.5 | 2 | 1 | NA | NA | 5 | 0.8 |
| 5_1352738 | 7 | 0 | 10 | 0.2 | 9 | 0.3333 | 4 | 0.25 | 30 | 0.2 | 6 | 0.1667 | 9 | 0 | 7 | 0.2857 | 3 | 0 | 25 | 0.12 |
| 5_1352739 | 8 | 0.5 | 9 | 0.2222 | 7 | 0.5714 | 5 | 0.2 | 29 | 0.3793 | 8 | 0.5 | 6 | 0.3333 | 7 | 0.5714 | 1 | 0 | 22 | 0.4545 |

| | | | | | | | | | | | | | | | | | | | | |
|---|---|---|---|---|---|---|---|---|---|---|---|---|---|---|---|---|---|---|---|---|
| 5_13527393 | 11 | 0.4545 | 10 | 0.2 | 7 | 0.1429 | 6 | 0.3333 | 34 | 0.2941 | 7 | 0.2857 | 9 | 0.2222 | 7 | 0.2857 | 3 | 0 | 26 | 0.2308 |
| 5_13527401 | 10 | 0.3 | 8 | 0.125 | 8 | 0.125 | 6 | 0 | 32 | 0.1562 | 7 | 0 | 10 | 0.1 | 8 | 0.25 | 3 | 0 | 28 | 0.1071 |
| 5_13527828 | NA | NA | NA | NA | 1 | 1 | NA | NA | 1 | 1 | 2 | 1 | NA | NA | NA | NA | NA | NA | 2 | 1 |
| 5_13527839 | NA | NA | NA | NA | 1 | 1 | NA | NA | 1 | 1 | 1 | 1 | NA | NA | NA | NA | 1 | 1 | 2 | 1 |
| 5_13527846 | NA | NA | NA | NA | 1 | 1 | NA | NA | 1 | 1 | 1 | 1 | NA | NA | NA | NA | NA | NA | 1 | 1 |
| 5_13527870 | NA | NA | NA | NA | 1 | 1 | NA | NA | 1 | 1 | 1 | 0 | NA | NA | NA | NA | 1 | 1 | 2 | 0.5 |
| 5_13527872 | NA | NA | NA | NA | 1 | 1 | NA | NA | 1 | 1 | 2 | 1 | NA | NA | NA | NA | 1 | 1 | 3 | 1 |
| 5_13527887 | NA | NA | NA | NA | 1 | 1 | NA | NA | 1 | 1 | 2 | 0.5 | NA | NA | NA | NA | 1 | 1 | 3 | 0.6667 |
| 5_13527919 | NA | NA | NA | NA | 1 | 1 | NA | NA | 1 | 1 | 2 | 1 | NA | NA | NA | NA | NA | NA | 2 | 1 |
| 5_13540508 | 10 | 0.1 | 11 | 0 | 6 | 0.1667 | 8 | 0 | 35 | 0.0571 | 10 | 0.1 | 6 | 0.3333 | 3 | 0 | 3 | 0 | 22 | 0.1364 |
| 5_13540606 | 8 | 0.25 | 5 | 0 | 4 | 0.25 | 5 | 0 | 22 | 0.1364 | 6 | 0.1667 | 4 | 0 | 4 | 0 | 2 | 0 | 16 | 0.0625 |
| 5_13540631 | 5 | 0.4 | 3 | 0.6667 | 1 | 0 | 4 | 0.75 | 13 | 0.5385 | 5 | 0.6 | 3 | 0.6667 | 3 | 0.6667 | 1 | 1 | 12 | 0.6667 |
| 5_13540670 | 5 | 0.2 | 3 | 0 | NA | NA | 3 | 0 | 11 | 0.0909 | 4 | 0.25 | 3 | 0 | 1 | 0 | 1 | 0 | 9 | 0.1111 |
| 5_13540671 | 8 | 1 | 4 | 1 | 2 | 1 | 6 | 1 | 20 | 1 | 5 | 1 | 4 | 1 | 2 | 1 | 2 | 1 | 13 | 1 |
| 5_13540688 | 8 | 0.25 | 2 | 0 | 2 | 0 | 6 | 0 | 18 | 0.1111 | 5 | 0.4 | 3 | 0.3333 | 2 | 0 | 1 | 1 | 11 | 0.3636 |
| 5_13540989 | 8 | 0.25 | 3 | 0.3333 | 1 | 0 | 7 | 0 | 19 | 0.1579 | 5 | 0 | 1 | 0 | 1 | 0 | 2 | 0.5 | 9 | 0.1111 |
| 5_13540990 | 7 | 0.2857 | 3 | 0.6667 | 2 | 0 | 5 | 0.6 | 17 | 0.4118 | 3 | 0.3333 | 1 | 1 | 1 | 1 | 1 | 1 | 6 | 0.6667 |
| 5_13540992 | 3 | 0.3333 | 2 | 0 | 1 | 0 | 3 | 0.6667 | 9 | 0.3333 | 5 | 0.4 | 3 | 0 | NA | NA | 1 | 1 | 9 | 0.3333 |
| 5_13541006 | 7 | 0.1429 | 2 | 0 | 2 | 0.5 | 3 | 0.3333 | 14 | 0.2143 | 3 | 0.3333 | 3 | 0 | 1 | 0 | 2 | 0.5 | 9 | 0.2222 |
| 5_13541053 | 7 | 0.1429 | 2 | 0.5 | NA | NA | 5 | 0.4 | 14 | 0.2857 | 4 | 0 | 3 | 0.3333 | 1 | 0 | 2 | 0 | 10 | 0.1 |
| 5_13541128 | 7 | 0.1429 | 2 | 0 | 1 | 0 | 2 | 0 | 12 | 0.0833 | 3 | 0.6667 | NA | NA | 1 | 0 | 1 | 0 | 5 | 0.4 |
| 5_13543547 | 4 | 1 | 7 | 0.8571 | 6 | 0.8333 | 1 | 1 | 18 | 0.8889 | 2 | 0 | 6 | 0.3333 | 3 | 0 | 1 | 0 | 12 | 0.1667 |
| 5_13543565 | 3 | 0.3333 | 3 | 0.3333 | 5 | 0.4 | 3 | 0 | 14 | 0.2857 | 1 | 0 | 5 | 0.2 | 2 | 0 | 1 | 0 | 9 | 0.1111 |
| 5_13543571 | 5 | 0.4 | 7 | 0.4286 | 7 | 0.1429 | 3 | 0.6667 | 22 | 0.3636 | 4 | 0 | 5 | 0.6 | 2 | 0.5 | 1 | 0 | 12 | 0.3333 |
| 5_13543579 | 2 | 0 | 7 | 0.1429 | 7 | 0.1429 | 3 | 0 | 19 | 0.1053 | 4 | 0.25 | 6 | 0 | 3 | 0 | 1 | 0 | 14 | 0.0714 |
| 5_13543582 | 5 | 0.4 | 5 | 0.6 | 7 | 0.5714 | 3 | 0 | 20 | 0.45 | 3 | 0 | 5 | 0.6 | 3 | 0 | 1 | 0 | 12 | 0.25 |
| 5_13543591 | 5 | 0 | 6 | 0 | 6 | 0.3333 | 3 | 0 | 20 | 0.1 | 4 | 0.25 | 6 | 0 | 3 | 0 | 1 | 0 | 14 | 0.0714 |
| 5_13543596 | 5 | 0 | 6 | 0.1667 | 6 | 0 | 2 | 0 | 19 | 0.0526 | 4 | 0 | 5 | 0.2 | 4 | 0.25 | 1 | 0 | 14 | 0.1429 |
| 5_13543598 | 5 | 0.2 | 5 | 0.4 | 6 | 0.3333 | 3 | 0.3333 | 19 | 0.3158 | 4 | 0.25 | 6 | 0.1667 | 4 | 0.25 | 1 | 0 | 15 | 0.2 |
| 5_13543646 | 5 | 0 | 4 | 0 | 6 | 0.1667 | 2 | 0 | 17 | 0.0588 | 4 | 0.25 | 5 | 0 | 4 | 0 | 1 | 0 | 14 | 0.0714 |
| 5_13543790 | 2 | 1 | 3 | 0 | 4 | 0.5 | 2 | 0 | 11 | 0.3636 | 4 | 0.5 | 2 | 0.5 | 2 | 0 | NA | NA | 8 | 0.375 |
| 5_13543856 | 2 | 0.5 | 4 | 0.25 | 5 | 0.4 | 2 | 0.5 | 13 | 0.3846 | 3 | 0.3333 | 2 | 0.5 | NA | NA | NA | NA | 5 | 0.4 |
| 5_13543859 | 3 | 0 | 3 | 0 | 4 | 0.25 | 2 | 0.5 | 12 | 0.1667 | 3 | 0.3333 | 2 | 0.5 | 1 | 1 | NA | NA | 6 | 0.5 |
| 5_13543884 | 3 | 0.3333 | 4 | 0.75 | 5 | 0.6 | 2 | 0.5 | 14 | 0.5714 | 3 | 0.6667 | 2 | 1 | 1 | 1 | NA | NA | 6 | 0.8333 |
| 5_13543891 | 3 | 1 | 3 | 1 | 4 | 1 | 2 | 1 | 12 | 1 | 3 | 1 | 2 | 0.5 | NA | NA | NA | NA | 5 | 0.8 |
| 5_13543898 | 3 | 0 | 4 | 0.25 | 4 | 0 | 1 | 0 | 12 | 0.0833 | 4 | 0.25 | 2 | 0 | 1 | 0 | NA | NA | 7 | 0.1429 |
| 5_13543904 | 1 | 1 | 4 | 0.75 | 3 | 1 | 1 | 1 | 9 | 0.8889 | 2 | 1 | 1 | 0 | 1 | 1 | NA | NA | 4 | 0.75 |
| 5_13543910 | 1 | 1 | 3 | 1 | 3 | 0.6667 | 1 | 1 | 8 | 0.875 | 4 | 0.5 | 1 | 1 | 1 | 1 | NA | NA | 6 | 0.6667 |
| 5_13543913 | 2 | 0 | 3 | 0 | 3 | 0 | 1 | 1 | 9 | 0.1111 | 4 | 0.25 | 1 | 0 | 1 | 1 | NA | NA | 6 | 0.3333 |
| 5_13543921 | 3 | 0 | 4 | 0.5 | 4 | 1 | 1 | 0 | 12 | 0.5 | 4 | 0.25 | 1 | 0 | 1 | 0 | NA | NA | 6 | 0.1667 |
| 5_13543930 | 3 | 0 | 4 | 0 | 4 | 0 | 1 | 1 | 12 | 0.0833 | 2 | 0 | 1 | 0 | 1 | 1 | NA | NA | 4 | 0.25 |
| 5_13543962 | 2 | 1 | 2 | 1 | 4 | 0.5 | 1 | 1 | 9 | 0.7778 | 4 | 1 | 1 | 1 | NA | NA | NA | NA | 5 | 1 |
| 5_13543963 | 3 | 0 | 3 | 0.3333 | 3 | 0 | 1 | 0 | 10 | 0.1 | 4 | 0.25 | NA | NA | NA | NA | NA | NA | 4 | 0.25 |
| 5_13543966 | 3 | 0.3333 | 2 | 1 | 4 | 0.75 | 1 | 0 | 10 | 0.6 | 4 | 0.5 | 1 | 0 | 1 | 0 | NA | NA | 6 | 0.3333 |
| 5_13543970 | 3 | 0.3333 | 3 | 0.3333 | 4 | 0.25 | NA | NA | 10 | 0.3 | 4 | 0.5 | 1 | 0 | 1 | 1 | NA | NA | 6 | 0.5 |
| 5_13543976 | 3 | 0.3333 | 2 | 0.5 | 2 | 0 | 1 | 1 | 8 | 0.375 | 1 | 1 | 1 | 0 | 1 | 0 | NA | NA | 3 | 0.3333 |
| 5_13544001 | 3 | 0.6667 | 1 | 0 | 1 | 0 | 1 | 0 | 6 | 0.3333 | 2 | 0.5 | NA | NA | 1 | 0 | NA | NA | 3 | 0.3333 |
| 5_13567226 | 5 | 0 | 8 | 0 | 6 | 0.3333 | 7 | 0 | 26 | 0.0769 | 5 | 0.2 | 11 | 0.2727 | 7 | 0.2857 | 7 | 0.2857 | 30 | 0.2667 |
| 5_13567279 | 10 | 0 | 8 | 0.25 | 7 | 0.4286 | 9 | 0.3333 | 34 | 0.2353 | 8 | 0 | 12 | 0.5 | 12 | 0.4167 | 6 | 0.3333 | 38 | 0.3421 |
| 5_13567286 | 11 | 0.1818 | 7 | 0.1429 | 7 | 0.2857 | 9 | 0.1111 | 34 | 0.1765 | 9 | 0.2222 | 13 | 0 | 10 | 0.1 | 7 | 0 | 39 | 0.0769 |
| 5_13567291 | 10 | 0.4 | 7 | 0.1429 | 5 | 0.2 | 7 | 0.4286 | 29 | 0.3103 | 10 | 0.1 | 12 | 0.75 | 10 | 0.1 | 7 | 0 | 39 | 0.2821 |
| 5_13567318 | 2 | 0.5 | NA | NA | 3 | 1 | NA | NA | 5 | 0.8 | 1 | 1 | 5 | 0.4 | 4 | 0.75 | 2 | 1 | 12 | 0.6667 |
| 5_13567326 | 3 | 0 | NA | NA | 2 | 0.5 | NA | NA | 5 | 0.2 | 1 | 0 | 5 | 0.2 | 5 | 0 | 2 | 0 | 13 | 0.0769 |
| 5_13567428 | 1 | 0 | 1 | 0 | 1 | 1 | NA | NA | 3 | 0.3333 | NA | NA | 2 | 1 | 2 | 0.5 | 3 | 0.3333 | 7 | 0.5714 |
| 5_13567437 | NA | NA | 1 | 1 | 1 | 1 | NA | NA | 2 | 1 | NA | NA | 2 | 0.5 | 1 | 1 | 1 | 1 | 4 | 0.75 |
| 5_13567463 | 1 | 0 | 1 | 0 | 2 | 0.5 | NA | NA | 4 | 0.25 | NA | NA | 2 | 0.5 | 2 | 0.5 | 3 | 0.3333 | 7 | 0.4286 |
| 5_13567467 | 1 | 1 | 1 | 1 | 1 | 1 | NA | NA | 3 | 1 | NA | NA | 1 | 1 | 2 | 1 | 3 | 0.6667 | 6 | 0.8333 |
| 5_13567475 | 2 | 0 | 1 | 1 | 2 | 0 | NA | NA | 5 | 0.2 | NA | NA | 2 | 0 | 2 | 0.5 | 2 | 0 | 6 | 0.1667 |
| 5_13567493 | 2 | 1 | 2 | 1 | 1 | 1 | NA | NA | 5 | 1 | NA | NA | 1 | 1 | 2 | 1 | 2 | 1 | 5 | 1 |
| 5_13567505 | 2 | 1 | 2 | 0.5 | 2 | 0.5 | NA | NA | 6 | 0.6667 | NA | NA | 1 | 1 | 2 | 0.5 | 2 | 1 | 5 | 0.8 |
| 5_13567509 | 2 | 0.5 | 1 | 0 | 2 | 0 | NA | NA | 5 | 0.2 | NA | NA | 2 | 0 | 2 | 0.5 | 3 | 0 | 7 | 0.1429 |
| 5_13567563 | 1 | 1 | NA | NA | 2 | 1 | 1 | 1 | 4 | 1 | NA | NA | 1 | 1 | 1 | 1 | NA | NA | 2 | 1 |
| 5_13567569 | 1 | 1 | NA | NA | 2 | 1 | 1 | 1 | 4 | 1 | NA | NA | 1 | 1 | 1 | 1 | 1 | 1 | 3 | 1 |
| 5_13567573 | 1 | 0 | NA | NA | 2 | 0.5 | 1 | 1 | 4 | 0.5 | NA | NA | 1 | 1 | 1 | 1 | 1 | 1 | 3 | 1 |
| 5_13567582 | 1 | 0 | NA | NA | 2 | 0.5 | NA | NA | 3 | 0.3333 | NA | NA | 1 | 0 | 1 | 0 | 1 | 1 | 3 | 0.3333 |

| | | | | | | | | | | | | | | | | | | | | |
|---|---|---|---|---|---|---|---|---|---|---|---|---|---|---|---|---|---|---|---|---|
| 5_13567654 | 1 | 0 | NA | NA | 2 | 0.5 | NA | NA | 3 | 0.3333 | NA | NA | 3 | 0.3333 | 1 | 0 | 1 | 0 | 5 | 0.2 |
| 5_13574481 | 1 | 0 | 3 | 0.3333 | 3 | 0.3333 | 1 | 0 | 8 | 0.25 | 4 | 0 | NA | NA | 2 | 0.5 | 2 | 0.5 | 8 | 0.25 |
| 5_13576375 | 2 | 0.5 | 15 | 0.2 | 11 | 0.1818 | 8 | 0.25 | 36 | 0.2222 | 9 | 0.3333 | 6 | 0 | 11 | 0.1818 | 7 | 0.5714 | 33 | 0.2727 |
| 5_13576392 | 3 | 0.3333 | 17 | 0.4118 | 11 | 0.2727 | 10 | 0.4 | 41 | 0.3659 | 9 | 0 | 8 | 0.625 | 10 | 0.2 | 8 | 0.375 | 35 | 0.2857 |
| 5_13576402 | 2 | 1 | 16 | 0.375 | 12 | 0.3333 | 13 | 0.3846 | 43 | 0.3953 | 9 | 0.5556 | 6 | 0.3333 | 11 | 0.3636 | 8 | 0.25 | 34 | 0.3824 |
| 5_13576455 | 2 | 0 | 4 | 0.25 | 3 | 0.3333 | 5 | 0.2 | 14 | 0.2143 | 5 | 0.4 | 1 | 0 | 4 | 0.25 | 3 | 0.6667 | 13 | 0.3846 |
| 5_13581904 | 71 | 0.338 | 68 | 0.3235 | 70 | 0.3143 | 74 | 0.3919 | 283 | 0.3428 | 60 | 0.2667 | 72 | 0.3472 | 59 | 0.3729 | 70 | 0.4143 | 261 | 0.3525 |
| 5_13581921 | 65 | 0.0769 | 53 | 0.0189 | 61 | 0.0492 | 67 | 0.0746 | 246 | 0.0569 | 55 | 0.0545 | 66 | 0.0606 | 55 | 0.0727 | 61 | 0.0164 | 237 | 0.0506 |
| 5_13590207 | 4 | 0 | 4 | 0 | 2 | 0 | 5 | 0.6 | 15 | 0.2 | 5 | 0.2 | 5 | 0.6 | 5 | 0 | 5 | 0.2 | 20 | 0.25 |
| 5_13590217 | 2 | 1 | 2 | 1 | 1 | 1 | 5 | 0.8 | 10 | 0.9 | 5 | 0.6 | 4 | 0.5 | 6 | 0.5 | 4 | 1 | 19 | 0.6316 |
| 5_13590230 | 4 | 0.25 | 4 | 0 | 3 | 1 | 5 | 0.4 | 16 | 0.375 | 4 | 0.25 | 3 | 0.3333 | 6 | 0.3333 | 4 | 0.25 | 17 | 0.2941 |
| 5_13590256 | 4 | 0 | 5 | 0 | 6 | 0 | 5 | 0.2 | 20 | 0.05 | 5 | 0.2 | 7 | 0 | 6 | 0.1667 | 5 | 0.2 | 23 | 0.1304 |
| 5_13590269 | 4 | 0 | 8 | 0.125 | 6 | 0.1667 | 5 | 0 | 23 | 0.087 | 5 | 0 | 7 | 0.1429 | 7 | 0.1429 | 5 | 0 | 24 | 0.0833 |
| 5_13590333 | 7 | 0 | 10 | 0.1 | 7 | 0.1429 | 9 | 0.1111 | 33 | 0.0909 | 8 | 0 | 7 | 0.1429 | 9 | 0.1111 | 8 | 0.25 | 32 | 0.125 |
| 5_13590359 | 4 | 0 | 9 | 0.3333 | 6 | 0.3333 | 10 | 0.2 | 29 | 0.2414 | 8 | 0.625 | 7 | 0 | 7 | 0.4286 | 6 | 0.3333 | 28 | 0.3571 |
| 5_13593735 | 103 | 0.2718 | 136 | 0.2132 | 106 | 0.2358 | 101 | 0.1584 | 446 | 0.2197 | 108 | 0.3241 | 91 | 0.2198 | 86 | 0.1628 | 111 | 0.2162 | 396 | 0.2348 |
| 5_13593773 | 82 | 0.0854 | 100 | 0.07 | 70 | 0.0714 | 79 | 0.0759 | 331 | 0.0755 | 77 | 0.1558 | 74 | 0.0541 | 68 | 0.0588 | 90 | 0.1222 | 309 | 0.1003 |
| 5_13593795 | 57 | 0.1228 | 83 | 0.1205 | 55 | 0.1273 | 64 | 0.25 | 259 | 0.1544 | 64 | 0.1094 | 59 | 0.1695 | 55 | 0.0727 | 66 | 0.197 | 244 | 0.1393 |
| 5_13593812 | 54 | 0.2037 | 67 | 0.2388 | 37 | 0.1622 | 48 | 0.2292 | 206 | 0.2136 | 50 | 0.2 | 53 | 0.1698 | 43 | 0.1395 | 59 | 0.1864 | 205 | 0.1756 |
| 5_13593830 | 72 | 0.0556 | 96 | 0.0312 | 65 | 0.1385 | 69 | 0.0725 | 302 | 0.0695 | 74 | 0.0946 | 67 | 0.0299 | 53 | 0.1321 | 86 | 0.093 | 280 | 0.0857 |
| 5_13593890 | 63 | 0.1587 | 84 | 0.1667 | 69 | 0.1739 | 62 | 0.129 | 278 | 0.1583 | 59 | 0.1525 | 70 | 0.2143 | 55 | 0.1818 | 78 | 0.1667 | 262 | 0.1794 |
| 5_13593961 | 48 | 0.8542 | 59 | 0.9153 | 43 | 0.9535 | 43 | 0.8837 | 193 | 0.9016 | 48 | 0.8333 | 55 | 0.9455 | 48 | 0.8542 | 50 | 0.94 | 201 | 0.8955 |
| 5_13593969 | 50 | 0.34 | 56 | 0.2857 | 50 | 0.24 | 42 | 0.381 | 198 | 0.3081 | 51 | 0.2157 | 57 | 0.2982 | 45 | 0.3111 | 51 | 0.3137 | 204 | 0.2843 |
| 5_13593983 | 45 | 0.9778 | 47 | 0.9787 | 38 | 1 | 38 | 1 | 168 | 0.9881 | 43 | 0.9767 | 49 | 1 | 41 | 1 | 45 | 1 | 178 | 0.9944 |
| 5_13593994 | 42 | 0.9762 | 49 | 0.9796 | 38 | 0.8947 | 41 | 0.9268 | 170 | 0.9471 | 43 | 0.9767 | 46 | 0.9565 | 43 | 1 | 42 | 1 | 174 | 0.9828 |
| 5_13593995 | 44 | 0.8409 | 47 | 0.7234 | 37 | 0.8108 | 33 | 0.8788 | 161 | 0.8075 | 41 | 0.7073 | 48 | 0.8333 | 41 | 0.7561 | 45 | 0.8444 | 175 | 0.7886 |
| 5_13594020 | 38 | 1 | 45 | 0.9778 | 33 | 0.8788 | 33 | 0.9697 | 149 | 0.9597 | 38 | 1 | 35 | 1 | 35 | 1 | 35 | 1 | 143 | 1 |
| 5_13594026 | 40 | 0.25 | 40 | 0.25 | 33 | 0.1818 | 35 | 0.1429 | 148 | 0.2095 | 36 | 0.3333 | 35 | 0.2 | 36 | 0.1667 | 37 | 0.2162 | 144 | 0.2292 |
| 5_13594027 | 38 | 0.0789 | 42 | 0.119 | 32 | 0.1562 | 35 | 0.0286 | 147 | 0.0952 | 37 | 0.0811 | 34 | 0.0588 | 37 | 0.027 | 38 | 0.1053 | 146 | 0.0685 |
| 5_13604102 | NA | NA | NA | NA | 1 | 1 | NA | NA | 1 | 1 | 1 | 1 | NA | NA | 1 | 1 | 1 | 1 | 3 | 1 |
| 5_13604117 | 1 | 1 | NA | NA | NA | NA | NA | NA | 1 | 1 | 1 | 1 | NA | NA | 1 | 1 | 1 | 0 | 3 | 0.6667 |
| 5_13604129 | 1 | 0 | NA | NA | 1 | 1 | NA | NA | 2 | 0.5 | 1 | 0 | NA | NA | NA | NA | 1 | 1 | 2 | 0.5 |
| 5_13615176 | 4 | 0 | 11 | 0.3636 | 9 | 0.1111 | 10 | 0.2 | 34 | 0.2059 | 8 | 0.125 | 10 | 0.1 | 13 | 0.3846 | 12 | 0.25 | 43 | 0.2326 |
| 5_13616918 | NA | NA | 1 | 1 | 1 | 1 | NA | NA | 2 | 1 | NA | NA | NA | NA | 2 | 0.5 | 1 | 1 | 3 | 0.6667 |
| 5_13616946 | NA | NA | NA | NA | 2 | 0.5 | NA | NA | 2 | 0.5 | 1 | 1 | NA | NA | 2 | 0.5 | 1 | 0 | 4 | 0.5 |
| 5_13616956 | NA | NA | 1 | 1 | 2 | 0.5 | NA | NA | 3 | 0.6667 | 1 | 0 | NA | NA | 3 | 1 | 2 | 1 | 6 | 0.8333 |
| 5_13616957 | NA | NA | 1 | 1 | 2 | 1 | NA | NA | 3 | 1 | NA | NA | NA | NA | 3 | 1 | 1 | 0 | 4 | 0.75 |
| 5_13616980 | 1 | 1 | 2 | 1 | 2 | 1 | NA | NA | 5 | 1 | NA | NA | NA | NA | 3 | 1 | 2 | 1 | 5 | 1 |
| 5_13616992 | 1 | 0 | 2 | 0 | 2 | 0.5 | NA | NA | 5 | 0.2 | NA | NA | NA | NA | 3 | 0.6667 | 2 | 0 | 5 | 0.4 |
| 5_13616992 | 1 | 0 | 1 | 0 | 2 | 0.5 | NA | NA | 4 | 0.25 | 1 | 0 | NA | NA | 2 | 0.5 | 2 | 0.5 | 5 | 0.4 |
| 5_13617009 | 1 | 0 | 2 | 0 | 2 | 0.5 | NA | NA | 5 | 0.2 | NA | NA | NA | NA | 2 | 0 | 2 | 0.5 | 4 | 0.25 |
| 5_13617029 | 1 | 0 | 2 | 0 | 2 | 0.5 | NA | NA | 5 | 0.2 | NA | NA | NA | NA | 2 | 0.5 | 1 | 0 | 3 | 0.3333 |
| 5_13617642 | 18 | 0.5556 | 10 | 0.6 | 9 | 0.7778 | 10 | 0.4 | 47 | 0.5745 | 7 | 0.7143 | 7 | 0.7143 | 7 | 0.2857 | 5 | 0.8 | 26 | 0.6154 |
| 5_13617644 | 14 | 0.5 | 10 | 0.3 | 9 | 0.4444 | 8 | 0.25 | 41 | 0.3902 | 9 | 0.3333 | 11 | 0.4545 | 7 | 0.5714 | 7 | 0.7143 | 34 | 0.5 |
| 5_13617647 | 5 | 0.2 | 1 | 0 | 1 | 0 | 1 | 0 | 8 | 0.125 | 1 | 0 | 4 | 0.25 | 2 | 0 | 2 | 0 | 9 | 0.1111 |
| 5_13617652 | 12 | 0.3333 | 8 | 0.25 | 6 | 0.3333 | 9 | 0.3333 | 35 | 0.3143 | 4 | 0.5 | 10 | 0.5 | 7 | 0.1429 | 4 | 0 | 25 | 0.32 |
| 5_13617662 | 10 | 0.5 | 8 | 0.375 | 7 | 0.2857 | 9 | 0 | 34 | 0.2941 | 7 | 0.1429 | 11 | 0.2727 | 6 | 0 | 6 | 0.1667 | 30 | 0.1667 |
| 5_13617668 | 13 | 0.6154 | 10 | 0.6 | 7 | 0.2857 | 8 | 0.25 | 38 | 0.4737 | 5 | 0.2 | 10 | 0.3 | 5 | 0.2 | 6 | 0.3333 | 26 | 0.2692 |
| 5_13617672 | 7 | 0.1429 | 8 | 0.125 | 3 | 0 | 6 | 0 | 24 | 0.0833 | 6 | 0 | 7 | 0 | 4 | 0.25 | 1 | 0 | 18 | 0.0556 |
| 5_13617696 | 13 | 0.0769 | 5 | 0 | 8 | 0 | 8 | 0.125 | 34 | 0.0588 | 5 | 0.2 | 11 | 0.0909 | 5 | 0 | 1 | 0 | 22 | 0.0909 |
| 5_13617727 | 13 | 0.2308 | 6 | 0.1667 | 7 | 0 | 4 | 0 | 30 | 0.1333 | 6 | 0.1667 | 11 | 0 | 5 | 0.2 | NA | NA | 22 | 0.0909 |
| 5_13617753 | 11 | 0 | 4 | 0 | 9 | 0.2222 | 5 | 0 | 29 | 0.069 | 6 | 0 | 8 | 0 | 4 | 0.25 | 2 | 0 | 20 | 0.05 |
| 5_13617773 | 7 | 0.4286 | 2 | 0 | 6 | 0 | 6 | 0 | 21 | 0.1429 | 5 | 0.2 | 6 | 0.1667 | 5 | 0.2 | 1 | 0 | 17 | 0.1765 |
| 5_13617813 | 8 | 0 | 4 | 0 | 5 | 0.2 | 7 | 0.1429 | 24 | 0.0833 | 7 | 0.4286 | 8 | 0.125 | 2 | 0 | 1 | 0 | 18 | 0.2222 |
| 5_13617838 | 8 | 1 | 4 | 0.75 | 5 | 0.8 | 5 | 0.6 | 22 | 0.8182 | 6 | 0.8333 | 4 | 1 | 2 | 1 | 1 | 1 | 13 | 0.9231 |
| 5_13617894 | 5 | 1 | 2 | 1 | 4 | 1 | 7 | 1 | 18 | 1 | 4 | 0.5 | 3 | 1 | NA | NA | 1 | 1 | 8 | 0.75 |
| 5_13617899 | 6 | 1 | 3 | 1 | 4 | 1 | 4 | 1 | 17 | 1 | 3 | 1 | 4 | 1 | 1 | 1 | 1 | 1 | 9 | 1 |
| 5_13617912 | 6 | 1 | 1 | 1 | 2 | 1 | 6 | 1 | 15 | 1 | 3 | 1 | 2 | 1 | 1 | 1 | 1 | 1 | 7 | 1 |
| 5_13617918 | 6 | 0.3333 | 2 | 0 | 3 | 0.3333 | 5 | 0 | 16 | 0.1875 | 4 | 0.25 | 3 | 0.3333 | 1 | 0 | 1 | 1 | 9 | 0.3333 |
| 5_13620978 | NA | NA | NA | NA | NA | NA | 1 | 1 | 1 | 1 | NA | NA | 1 | 0 | NA | NA | 1 | 1 | 2 | 0.5 |
| 5_13620999 | 4 | 0.25 | 4 | 0.25 | 1 | 0 | 2 | 0.5 | 11 | 0.2727 | 2 | 0.5 | 5 | 0.2 | 1 | 0 | 4 | 0 | 12 | 0.1667 |
| 5_13621009 | 4 | 0.25 | 6 | 0.5 | NA | NA | 3 | 0.6667 | 13 | 0.4615 | 2 | 1 | 4 | 0.5 | 2 | 0 | 4 | 0.75 | 12 | 0.5833 |
| 5_13621017 | 4 | 0.5 | 5 | 0 | NA | NA | 3 | 0 | 12 | 0.1667 | 2 | 0 | 4 | 0.75 | 2 | 0 | 3 | 0.3333 | 11 | 0.3636 |
| 5_13624710 | 1 | 1 | 1 | 0 | 1 | 0 | 2 | 0 | 5 | 0.2 | 3 | 0.3333 | 2 | 0 | 1 | 0 | 1 | 0 | 7 | 0.1429 |
| 5_13626065 | NA | NA | NA | NA | NA | NA | 2 | 1 | 2 | 1 | 1 | 1 | NA | NA | 1 | 1 | 1 | 0 | 3 | 0.6667 |

| | | | | | | | | | | | | | | | | | | | | |
|---|---|---|---|---|---|---|---|---|---|---|---|---|---|---|---|---|---|---|---|---|
| 5_13629649 | 2 | 0.5 | 2 | 0 | 4 | 0.25 | 3 | 0 | 11 | 0.1818 | 1 | 0 | 3 | 0.6667 | 2 | 0 | 3 | 0.3333 | 9 | 0.3333 |
| 5_13629778 | 2 | 0 | 1 | 0 | 4 | 0 | 3 | 0.3333 | 10 | 0.1 | 1 | 0 | 2 | 0 | 1 | 1 | 2 | 0.5 | 6 | 0.3333 |
| 5_13677179 | NA | NA | NA | NA | 1 | 1 | NA | NA | 1 | 1 | 1 | 1 | NA | NA | NA | NA | NA | NA | 1 | 1 |
| 5_13694075 | 1 | 1 | 1 | 1 | 1 | 0 | 1 | 1 | 4 | 0.75 | NA | NA | NA | NA | 1 | 1 | NA | NA | 1 | 1 |
| 5_13694186 | 6 | 0 | 10 | 0.1 | 12 | 0.0833 | 9 | 0 | 37 | 0.0541 | 9 | 0.1111 | 7 | 0.1429 | 4 | 0 | 11 | 0.0909 | 31 | 0.0968 |
| 5_13694226 | 1 | 1 | NA | NA | 2 | 0 | 1 | 0 | 4 | 0.25 | 1 | 1 | NA | NA | NA | NA | 2 | 0.5 | 3 | 0.6667 |
| 5_13694399 | 1 | 1 | NA | NA | 1 | 0 | NA | NA | 2 | 0.5 | NA | NA | NA | NA | 2 | 1 | 1 | 0 | 3 | 0.6667 |
| 5_13696268 | NA | NA | 2 | 1 | NA | NA | NA | NA | 2 | 1 | 1 | 1 | NA | NA | NA | NA | 1 | 1 | 2 | 1 |
| 5_13696293 | NA | NA | 1 | 1 | NA | NA | 1 | 0 | 2 | 0.5 | 1 | 1 | 1 | 1 | NA | NA | 1 | 1 | 3 | 1 |
| 5_13696444 | 2 | 0.5 | 1 | 0 | NA | NA | 2 | 0 | 5 | 0.2 | 1 | 0 | 1 | 0 | 1 | 0 | 2 | 0.5 | 5 | 0.2 |
| 5_13696455 | 2 | 0 | 1 | 1 | NA | NA | 1 | 0 | 4 | 0.25 | 1 | 0 | 1 | 0 | 1 | 0 | 2 | 1 | 5 | 0.4 |
| 5_13696584 | 2 | 0.5 | 1 | 0 | NA | NA | 3 | 0 | 6 | 0.1667 | 2 | 0.5 | 2 | 0 | 1 | 0 | 1 | 0 | 6 | 0.1667 |
| 5_13701110 | 7 | 0.7143 | 16 | 0.75 | 17 | 0.8235 | 3 | 0.6667 | 43 | 0.7674 | 8 | 0.25 | 17 | 0.4706 | 14 | 0.5714 | 7 | 0.7143 | 46 | 0.5 |
| 5_13701118 | 7 | 0.1429 | 16 | 0.125 | 15 | 0.2667 | 3 | 0.3333 | 41 | 0.1951 | 8 | 0.125 | 16 | 0.125 | 15 | 0.1333 | 8 | 0.125 | 47 | 0.1277 |
| 5_13701408 | 1 | 1 | NA | NA | 1 | 0 | NA | NA | 2 | 0.5 | 1 | 1 | NA | NA | 1 | 0 | NA | NA | 2 | 0.5 |
| 5_13701579 | 1 | 1 | 1 | 0 | NA | NA | NA | NA | 2 | 0.5 | 1 | 0 | 1 | 1 | 1 | 1 | NA | NA | 3 | 0.6667 |
| 5_13701594 | 1 | 1 | 1 | 0 | NA | NA | NA | NA | 2 | 0.5 | 1 | 0 | 1 | 1 | 1 | 0 | NA | NA | 3 | 0.3333 |
| 5_13701598 | 1 | 1 | 1 | 1 | NA | NA | NA | NA | 2 | 1 | 1 | 0 | 1 | 1 | 1 | 0 | NA | NA | 3 | 0.3333 |
| 5_13702516 | 2 | 0 | 3 | 0 | 4 | 0.5 | 2 | 0 | 11 | 0.1818 | 5 | 0 | 6 | 0.1667 | 5 | 0.6 | NA | NA | 16 | 0.25 |
| 5_13702530 | 3 | 0.6667 | 5 | 0.4 | 4 | 0.5 | 1 | 0 | 13 | 0.4615 | 3 | 0.6667 | 6 | 0.6667 | 6 | 0.6667 | NA | NA | 15 | 0.6667 |
| 5_13702532 | 3 | 0.6667 | 5 | 0.6 | 1 | 0 | 2 | 1 | 11 | 0.6364 | 4 | 0.5 | 7 | 0.2857 | 5 | 0.6 | NA | NA | 16 | 0.4375 |
| 5_13706011 | 9 | 0.1111 | 13 | 0.0769 | 9 | 0 | 9 | 0.1111 | 40 | 0.075 | 12 | 0 | 12 | 0.1667 | 9 | 0.2222 | 6 | 0.1667 | 39 | 0.1282 |
| 5_13706027 | 13 | 0.1538 | 16 | 0.125 | 13 | 0.2308 | 13 | 0.0769 | 55 | 0.1455 | 12 | 0.1667 | 13 | 0.0769 | 13 | 0.1538 | 9 | 0.1111 | 47 | 0.1277 |
| 5_13706034 | 11 | 0.9091 | 13 | 0.5385 | 12 | 0.8333 | 14 | 0.7143 | 50 | 0.74 | 10 | 0.8 | 9 | 0.7778 | 10 | 0.6 | 7 | 0.5714 | 36 | 0.6944 |
| 5_13706040 | 9 | 1 | 14 | 0.9286 | 13 | 1 | 17 | 1 | 53 | 0.9811 | 15 | 0.8667 | 14 | 0.9286 | 13 | 1 | 7 | 1 | 49 | 0.9388 |
| 5_13706057 | 12 | 0.1667 | 16 | 0.125 | 7 | 0 | 15 | 0.0667 | 50 | 0.1 | 13 | 0.2308 | 12 | 0.0833 | 13 | 0 | 8 | 0.125 | 46 | 0.1087 |
| 5_13706062 | 13 | 0.4615 | 15 | 0.6 | 12 | 0.5 | 19 | 0.2105 | 59 | 0.4237 | 15 | 0.5333 | 15 | 0.4667 | 15 | 0.3333 | 12 | 0.0833 | 57 | 0.3684 |
| 5_13706109 | 21 | 0.1429 | 27 | 0.1481 | 19 | 0.0526 | 30 | 0.0667 | 97 | 0.1031 | 20 | 0.3 | 20 | 0.1 | 21 | 0.0952 | 19 | 0.0526 | 80 | 0.1375 |
| 5_13706122 | 16 | 0.875 | 24 | 0.875 | 19 | 1 | 34 | 0.8529 | 93 | 0.8925 | 21 | 0.8571 | 19 | 0.8947 | 27 | 0.7778 | 22 | 0.9545 | 89 | 0.8652 |
| 5_13706141 | 28 | 0.7143 | 34 | 0.7353 | 27 | 0.8148 | 39 | 0.8718 | 128 | 0.7891 | 20 | 0.8 | 26 | 0.8846 | 30 | 0.7333 | 31 | 0.871 | 107 | 0.8224 |
| 5_13706150 | 30 | 0.1 | 36 | 0.1111 | 30 | 0.1333 | 38 | 0.1316 | 134 | 0.1194 | 23 | 0.087 | 27 | 0.1481 | 33 | 0.0606 | 32 | 0.125 | 115 | 0.1043 |
| 5_13706186 | 38 | 0.1053 | 43 | 0.0233 | 38 | 0.0789 | 48 | 0.125 | 167 | 0.0838 | 30 | 0.1667 | 33 | 0.0909 | 46 | 0.0217 | 38 | 0.0789 | 147 | 0.0816 |
| 5_13706189 | 26 | 0.0385 | 27 | 0.037 | 28 | 0.1071 | 29 | 0.0345 | 110 | 0.0545 | 22 | 0.0455 | 24 | 0.0417 | 39 | 0.0769 | 32 | 0.125 | 117 | 0.0769 |
| 5_13706220 | 4 | 0.25 | 2 | 0 | 2 | 0 | 2 | 0 | 10 | 0.1 | 1 | 0 | 2 | 0.5 | 2 | 0 | NA | NA | 5 | 0.2 |
| 5_13706248 | 4 | 0.75 | 2 | 0 | 1 | 1 | 4 | 0 | 11 | 0.3636 | 1 | 0 | 2 | 0.5 | NA | NA | NA | NA | 3 | 0.3333 |
| 5_13706385 | 6 | 0.8333 | 9 | 0.5556 | 7 | 0.4286 | 11 | 0.5455 | 33 | 0.5758 | 9 | 0.4444 | 15 | 0.4 | 15 | 0.6667 | 11 | 0.2727 | 50 | 0.46 |
| 5_13706449 | 50 | 0.1 | 58 | 0.1207 | 42 | 0.1667 | 74 | 0.0946 | 224 | 0.1161 | 41 | 0.122 | 58 | 0.0172 | 63 | 0.0635 | 52 | 0.1923 | 214 | 0.0935 |
| 5_13709179 | NA | NA | 2 | 1 | NA | NA | 1 | 0 | 3 | 0.6667 | NA | NA | NA | NA | 1 | 1 | NA | NA | 1 | 1 |
| 5_13709194 | NA | NA | 2 | 1 | NA | NA | 1 | 1 | 3 | 1 | NA | NA | NA | NA | 1 | 1 | NA | NA | 1 | 1 |
| 5_13728342 | 2 | 0.5 | NA | NA | NA | NA | NA | NA | 2 | 0.5 | NA | NA | NA | NA | 2 | 0.5 | NA | NA | 2 | 0.5 |
| 5_13728369 | 2 | 0 | NA | NA | NA | NA | 1 | 1 | 3 | 0.3333 | NA | NA | NA | NA | 2 | 0 | 1 | 1 | 3 | 0.3333 |
| 5_13728371 | 2 | 0.5 | NA | NA | NA | NA | 1 | 0 | 3 | 0.3333 | 1 | 0 | NA | NA | 2 | 1 | 1 | 0 | 4 | 0.5 |
| 5_13728389 | 2 | 0.5 | NA | NA | NA | NA | 1 | 1 | 3 | 0.6667 | 1 | 0 | NA | NA | 1 | 1 | 1 | 1 | 3 | 0.6667 |
| 5_13728395 | 2 | 0.5 | NA | NA | NA | NA | NA | NA | 2 | 0.5 | NA | NA | NA | NA | 2 | 0.5 | 1 | 0 | 3 | 0.3333 |
| 5_13728396 | 2 | 0 | NA | NA | NA | NA | 1 | 1 | 3 | 0.3333 | NA | NA | NA | NA | NA | NA | 1 | 1 | 1 | 1 |
| 5_13728410 | 1 | 1 | NA | NA | 1 | 0 | 1 | 1 | 3 | 0.6667 | NA | NA | NA | NA | 2 | 0.5 | 1 | 1 | 3 | 0.6667 |
| 5_13728411 | NA | NA | NA | NA | 1 | 0 | 1 | 1 | 2 | 0.5 | NA | NA | NA | NA | 1 | 1 | 1 | 1 | 2 | 1 |
| 5_13728422 | 1 | 1 | NA | NA | NA | NA | NA | NA | 1 | 1 | NA | NA | NA | NA | 1 | 0 | 1 | 1 | 2 | 0.5 |
| 5_13728434 | 1 | 1 | NA | NA | 1 | 1 | NA | NA | 2 | 1 | NA | NA | NA | NA | 2 | 1 | 1 | 1 | 3 | 1 |
| 5_13728439 | NA | NA | NA | NA | 1 | 1 | 1 | 0 | 2 | 0.5 | 1 | 0 | NA | NA | 3 | 0.3333 | 1 | 0 | 5 | 0.2 |
| 5_13728461 | NA | NA | NA | NA | 1 | 1 | NA | NA | 1 | 1 | NA | NA | NA | NA | 3 | 0.3333 | NA | NA | 3 | 0.3333 |
| 5_13728474 | NA | NA | 1 | 0 | 1 | 0 | 1 | 1 | 3 | 0.3333 | NA | NA | NA | NA | 2 | 1 | NA | NA | 2 | 1 |
| 5_13728509 | NA | NA | 1 | 1 | 1 | 1 | 1 | 1 | 3 | 1 | NA | NA | NA | NA | 1 | 1 | NA | NA | 1 | 1 |
| 5_13728821 | 1 | 1 | NA | NA | NA | NA | NA | NA | 1 | 1 | 1 | 0 | 1 | 0 | 1 | 1 | 1 | 1 | 4 | 0.5 |
| 5_13728860 | 6 | 0.3333 | 5 | 1 | 7 | 0.8571 | 7 | 0.5714 | 25 | 0.68 | 6 | 0.5 | 5 | 0.8 | 3 | 0.3333 | 8 | 0.75 | 22 | 0.6364 |
| 5_13728867 | 8 | 0.625 | 5 | 0.6 | 6 | 0.8333 | 8 | 0.875 | 27 | 0.7407 | 7 | 0.7143 | 7 | 0.5714 | 4 | 0.75 | 9 | 0.7778 | 27 | 0.7037 |
| 5_13728869 | 8 | 1 | 6 | 1 | 8 | 0.875 | 6 | 1 | 28 | 0.9643 | 6 | 0.6667 | 6 | 1 | 6 | 0.8333 | 9 | 1 | 27 | 0.8889 |
| 5_13728870 | 6 | 0 | 6 | 0.1667 | 8 | 0.25 | 9 | 0.1111 | 29 | 0.1379 | 8 | 0.25 | 8 | 0.125 | 5 | 0 | 8 | 0.25 | 29 | 0.1724 |
| 5_13728883 | 5 | 1 | 3 | 1 | 5 | 1 | 6 | 1 | 19 | 1 | 5 | 1 | 5 | 1 | 4 | 1 | 7 | 1 | 21 | 1 |
| 5_13728885 | 7 | 0.4286 | 4 | 0.75 | 7 | 0 | 8 | 0.375 | 26 | 0.3462 | 7 | 0.1429 | 6 | 0.6667 | 5 | 0 | 8 | 0.375 | 26 | 0.3077 |
| 5_13728898 | 7 | 0.8571 | 5 | 1 | 9 | 0.8889 | 7 | 1 | 28 | 0.9286 | 5 | 1 | 5 | 1 | 3 | 0.6667 | 9 | 0.8889 | 22 | 0.9091 |
| 5_13728922 | 7 | 0.2857 | 6 | 0 | 12 | 0.0833 | 9 | 0.1111 | 34 | 0.1176 | 11 | 0 | 9 | 0.1111 | 6 | 0.3333 | 9 | 0.3333 | 35 | 0.1714 |
| 5_13728930 | 8 | 0.5 | 6 | 0 | 12 | 0.1667 | 12 | 0.3333 | 38 | 0.2632 | 11 | 0.0909 | 9 | 0.4444 | 6 | 0.3333 | 8 | 0.5 | 34 | 0.3235 |
| 5_13728981 | 9 | 0.5556 | 7 | 0 | 11 | 0.2727 | 12 | 0.5 | 39 | 0.359 | 10 | 0.6 | 11 | 0.3636 | 6 | 0.6667 | 10 | 0.3 | 37 | 0.4595 |
| 5_13728991 | 8 | 0.5 | 7 | 0.7143 | 12 | 0.5833 | 11 | 0.8182 | 38 | 0.6579 | 11 | 0.8182 | 10 | 0.3 | 5 | 0.8 | 9 | 0.7778 | 35 | 0.6571 |

| | | | | | | | | | | | | | | | | | | | | |
|---|---|---|---|---|---|---|---|---|---|---|---|---|---|---|---|---|---|---|---|---|
| 5_13728993 | 9 | 0.3333 | 6 | 0.3333 | 12 | 0.4167 | 11 | 0.0909 | 38 | 0.2895 | 10 | 0.5 | 11 | 0.5455 | 6 | 0 | 10 | 0.8 | 37 | 0.5135 |
| 5_13729005 | 9 | 0.2222 | 3 | 0 | 6 | 0.1667 | 7 | 0.1429 | 25 | 0.16 | 7 | 0.1429 | 9 | 0 | 5 | 0 | 5 | 0.4 | 26 | 0.1154 |
| 5_13729007 | 7 | 0.1429 | 7 | 0.2857 | 12 | 0.3333 | 9 | 0.4444 | 35 | 0.3143 | 6 | 0.5 | 11 | 0.3636 | 5 | 0.4 | 8 | 0.25 | 30 | 0.3667 |
| 5_13729020 | 8 | 0.125 | 7 | 0.1429 | 12 | 0.0833 | 10 | 0.3 | 37 | 0.1622 | 10 | 0.2 | 11 | 0.4545 | 7 | 0.2857 | 11 | 0.0909 | 39 | 0.2564 |
| 5_13729271 | 8 | 0.25 | 10 | 0.1 | 13 | 0.0769 | 14 | 0.0714 | 45 | 0.1111 | 12 | 0 | 14 | 0.1429 | 8 | 0.125 | 11 | 0.0909 | 45 | 0.0889 |
| 5_13729318 | 4 | 1 | 1 | 1 | 3 | 1 | 4 | 1 | 12 | 1 | 1 | 1 | 2 | 1 | 1 | 1 | 2 | 1 | 6 | 1 |
| 5_13729327 | 4 | 0.75 | 2 | 1 | 3 | 1 | 4 | 0.75 | 13 | 0.8462 | 1 | 0 | 4 | 1 | 1 | 1 | 2 | 0.5 | 8 | 0.75 |
| 5_13729337 | 4 | 0.5 | 2 | 0 | NA | NA | 2 | 0 | 8 | 0.25 | 1 | 0 | 1 | 0 | NA | NA | 1 | 1 | 3 | 0.3333 |
| 5_13729339 | 3 | 0.3333 | 2 | 0 | 2 | 1 | 4 | 0.25 | 11 | 0.3636 | 1 | 0 | 3 | 0 | 1 | 0 | 2 | 0.5 | 7 | 0.1429 |
| 5_13729357 | 3 | 0.6667 | 1 | 0 | 3 | 0 | 5 | 0 | 12 | 0.1667 | 1 | 0 | 2 | 0 | 1 | 1 | 2 | 0.5 | 6 | 0.3333 |
| 5_13729361 | 4 | 0.25 | 1 | 0 | 4 | 0.75 | 4 | 0.25 | 13 | 0.3846 | 1 | 0 | 3 | 0.3333 | 1 | 0 | 2 | 0 | 7 | 0.1429 |
| 5_13729374 | 4 | 1 | 2 | 0.5 | 4 | 1 | 5 | 0.8 | 15 | 0.8667 | 1 | 1 | 3 | 1 | 1 | 1 | 3 | 0.3333 | 8 | 0.75 |
| 5_13729375 | 5 | 0.6 | 2 | 0 | 4 | 0.25 | 4 | 0.25 | 15 | 0.3333 | 1 | 0 | 3 | 0 | 1 | 0 | 3 | 0.6667 | 8 | 0.25 |
| 5_13731303 | 1 | 0 | 6 | 0.1667 | 6 | 0.3333 | 3 | 0 | 16 | 0.1875 | 1 | 0 | 4 | 0.25 | 2 | 0.5 | 2 | 0.5 | 9 | 0.3333 |
| 5_13731310 | NA | NA | 6 | 0.1667 | 7 | 0 | 3 | 0.3333 | 16 | 0.125 | 2 | 0 | 3 | 0 | 2 | 0.5 | NA | NA | 7 | 0.1429 |
| 5_13731350 | 1 | 0 | 5 | 0.6 | 8 | 0.375 | 3 | 0 | 17 | 0.3529 | 2 | 0.5 | 3 | 1 | 2 | 0.5 | 1 | 1 | 8 | 0.75 |
| 5_13731353 | 1 | 0 | 5 | 0.8 | 7 | 0.4286 | 3 | 1 | 16 | 0.625 | 2 | 0.5 | 2 | 0.5 | 2 | 0.5 | 1 | 0 | 7 | 0.4286 |
| 5_13731374 | 1 | 1 | 4 | 0.25 | 6 | 0.8333 | 1 | 1 | 12 | 0.6667 | 2 | 0 | 1 | 0 | 3 | 1 | NA | NA | 6 | 0.5 |
| 5_13737202 | NA | NA | 1 | 1 | NA | NA | NA | NA | 1 | 1 | NA | NA | 1 | 1 | NA | NA | 1 | 0 | 2 | 0.5 |
| 5_13749976 | 7 | 1 | 4 | 0.75 | 8 | 0.875 | 4 | 1 | 23 | 0.913 | 6 | 1 | 2 | 1 | 4 | 1 | 2 | 1 | 14 | 1 |
| 5_13749988 | 10 | 0 | 4 | 0.5 | 8 | 0.5 | 3 | 0 | 25 | 0.24 | 6 | 0.3333 | 3 | 0.6667 | 3 | 0.6667 | 1 | 1 | 13 | 0.5385 |
| 5_13749990 | 9 | 0.1111 | 4 | 0 | 6 | 0.1667 | 2 | 0 | 21 | 0.0952 | 6 | 0.1667 | 3 | 0 | 4 | 0 | 2 | 0.5 | 15 | 0.1333 |
| 5_13749998 | 7 | 0.1429 | 3 | 0 | 5 | 0.2 | 2 | 0 | 17 | 0.1176 | 4 | 0.25 | 3 | 0.6667 | 3 | 0 | 2 | 0 | 12 | 0.25 |
| 5_13749998 | 7 | 0.8571 | 4 | 0.75 | 5 | 1 | 2 | 1 | 18 | 0.8889 | 5 | 1 | 2 | 1 | 4 | 0.75 | 1 | 1 | 12 | 0.9167 |
| 5_13750020 | 8 | 0.5 | 3 | 0.6667 | 5 | 0.2 | 2 | 0 | 18 | 0.3889 | 4 | 0.5 | 1 | 0 | 4 | 0.25 | 3 | 0.6667 | 12 | 0.4167 |
| 5_13750020 | 8 | 0.5 | 3 | 0 | 4 | 0.5 | NA | NA | 15 | 0.4 | 3 | 1 | 2 | 0.5 | 2 | 0 | 3 | 1 | 10 | 0.7 |
| 5_13750034 | 8 | 0.125 | 3 | 0 | 7 | 0.1429 | 2 | 0 | 20 | 0.1 | 5 | 0 | 3 | 0.3333 | 4 | 0 | 3 | 0 | 15 | 0.0667 |
| 5_13750114 | NA | NA | 1 | 1 | NA | NA | NA | NA | 1 | 1 | NA | NA | 1 | 1 | NA | NA | NA | NA | 1 | 1 |
| 5_13750130 | NA | NA | 1 | 1 | NA | NA | NA | NA | 1 | 1 | NA | NA | 1 | 1 | NA | NA | NA | NA | 1 | 1 |
| 5_13750155 | NA | NA | 1 | 1 | NA | NA | NA | NA | 1 | 1 | NA | NA | 1 | 1 | NA | NA | NA | NA | 1 | 1 |
| 5_13752402 | 4 | 0 | 4 | 0.25 | 4 | 0.25 | 3 | 0.6667 | 15 | 0.2667 | 3 | 0.3333 | 6 | 0.3333 | 1 | 0 | 3 | 0.6667 | 13 | 0.3846 |
| 5_13752444 | 4 | 0.25 | 4 | 0 | 2 | 0 | 5 | 0 | 15 | 0.0667 | 3 | 0 | 5 | 0.2 | 3 | 0 | 3 | 0.3333 | 14 | 0.1429 |
| 5_13752458 | 4 | 0 | 3 | 0 | 4 | 0 | 5 | 0.2 | 16 | 0.0625 | 2 | 0.5 | 6 | 0 | 3 | 0 | 3 | 0 | 14 | 0.0714 |
| 5_13752493 | 4 | 0.5 | 4 | 0 | 4 | 0 | 5 | 0 | 17 | 0.1176 | 3 | 0.3333 | 6 | 0 | 3 | 0 | 5 | 0 | 17 | 0.0588 |
| 5_13752501 | 4 | 0 | 4 | 0 | 4 | 0.25 | 4 | 0 | 16 | 0.0625 | 3 | 0.3333 | 5 | 0 | 3 | 0.3333 | 4 | 0 | 15 | 0.1333 |
| 5_13752578 | 4 | 0.25 | 4 | 0.25 | 3 | 0 | 6 | 0 | 17 | 0.1176 | 6 | 0.5 | 7 | 0 | 3 | 0 | 6 | 0.5 | 22 | 0.2727 |
| 5_13752587 | 4 | 0.5 | 3 | 0.3333 | 2 | 0.5 | 12 | 0.4167 | 21 | 0.4286 | 5 | 0.6 | 6 | 0.5 | 2 | 1 | 4 | 0.25 | 17 | 0.5294 |
| 5_13752588 | 4 | 0.25 | 3 | 0 | 3 | 0 | 9 | 0 | 19 | 0.0526 | 4 | 0.5 | 7 | 0.2857 | 2 | 0 | 6 | 0 | 19 | 0.2105 |
| 5_13752679 | 57 | 0.1053 | 70 | 0.1 | 48 | 0.1458 | 54 | 0.0556 | 229 | 0.1004 | 41 | 0.0732 | 71 | 0.1127 | 63 | 0.1905 | 54 | 0.0556 | 229 | 0.1135 |
| 5_13753033 | NA | NA | 2 | 0 | 1 | 1 | NA | NA | 3 | 0.3333 | 1 | 0 | 3 | 0 | 3 | 0.3333 | NA | NA | 7 | 0.1429 |
| 5_13756773 | 18 | 0.1111 | 25 | 0.08 | 20 | 0.2 | 22 | 0.1818 | 85 | 0.1412 | 9 | 0.1111 | 14 | 0.0714 | 14 | 0.1429 | 16 | 0.0625 | 53 | 0.0943 |
| 5_13758534 | NA | NA | 1 | 0 | 1 | 1 | NA | NA | 2 | 0.5 | NA | NA | NA | NA | NA | NA | 1 | 1 | 1 | 1 |
| 5_13758535 | NA | NA | 1 | 1 | NA | NA | NA | NA | 1 | 1 | NA | NA | NA | NA | NA | NA | 1 | 1 | 1 | 1 |
| 5_13758564 | NA | NA | 2 | 1 | NA | NA | NA | NA | 2 | 1 | NA | NA | NA | NA | NA | NA | 1 | 1 | 1 | 1 |
| 5_13763840 | 2 | 0 | 5 | 0 | 1 | 0 | 3 | 0.3333 | 11 | 0.0909 | 2 | 0 | 4 | 0.25 | 2 | 0 | 3 | 0 | 11 | 0.0909 |
| 5_13763848 | 3 | 1 | 4 | 0.5 | 1 | 1 | 3 | 0 | 11 | 0.5455 | 2 | 0 | 4 | 0.5 | 1 | 0 | 4 | 0.5 | 11 | 0.3636 |
| 5_13763861 | 3 | 0.3333 | 5 | 0.6 | 1 | 1 | 3 | 0.6667 | 12 | 0.5833 | 2 | 1 | 4 | 0.75 | 2 | 0.5 | 4 | 0.5 | 12 | 0.6667 |
| 5_13763869 | 2 | 1 | 3 | 0.3333 | 1 | 0 | 3 | 0.3333 | 9 | 0.4444 | 2 | 0 | 2 | 0 | 2 | 0 | 3 | 0.3333 | 9 | 0.1111 |
| 5_13763874 | 3 | 0 | 3 | 0 | 1 | 1 | 3 | 0 | 10 | 0.1 | 2 | 0 | 4 | 0 | 2 | 0 | 4 | 0.25 | 12 | 0.0833 |
| 5_13763917 | 3 | 0.6667 | 4 | 0.25 | NA | NA | 2 | 0 | 9 | 0.3333 | 2 | 0 | 5 | 0.2 | 1 | 0 | 4 | 0.25 | 12 | 0.1667 |
| 5_13763921 | 4 | 0.75 | 2 | 1 | NA | NA | 3 | 0.6667 | 9 | 0.7778 | 3 | 1 | 5 | 1 | NA | NA | 3 | 1 | 11 | 1 |
| 5_13763943 | 1 | 0 | 3 | 0.3333 | NA | NA | 1 | 0 | 5 | 0.2 | 1 | 1 | 2 | 0.5 | 1 | 0 | 1 | 0 | 5 | 0.4 |
| 5_13763959 | 1 | 0 | 2 | 0.5 | NA | NA | 1 | 0 | 4 | 0.25 | 1 | 0 | 3 | 0.3333 | 1 | 0 | 1 | 0 | 6 | 0.1667 |
| 5_13764103 | 1 | 1 | 3 | 0.6667 | NA | NA | 2 | 1 | 6 | 0.8333 | 1 | 1 | 4 | 1 | 1 | 0 | 1 | 0 | 7 | 0.7143 |
| 5_13764104 | 1 | 0 | 2 | 0 | NA | NA | 2 | 0.5 | 5 | 0.2 | 1 | 0 | 4 | 0 | 1 | 1 | 1 | 0 | 7 | 0.1429 |
| 5_13764930 | 4 | 0 | 6 | 0.1667 | 5 | 0.2 | 6 | 0.1667 | 21 | 0.1429 | 5 | 0.2 | 6 | 0.3333 | 8 | 0.375 | 5 | 0.2 | 24 | 0.2917 |
| 5_13764951 | 4 | 0.75 | 6 | 0.1667 | 4 | 0.25 | 6 | 0.8333 | 20 | 0.5 | 5 | 0.8 | 7 | 0.5714 | 8 | 0.625 | 5 | 0.4 | 25 | 0.6 |
| 5_13764978 | 3 | 0.3333 | 5 | 0 | 5 | 0 | 4 | 0.5 | 17 | 0.1765 | 4 | 0.25 | 6 | 0 | 7 | 0.2857 | 6 | 0 | 23 | 0.1304 |
| 5_13764982 | 3 | 0.3333 | 6 | 0 | 5 | 0 | 2 | 0 | 16 | 0.0625 | 5 | 0 | 6 | 0.1667 | 4 | 0.5 | 5 | 0.6 | 20 | 0.3 |
| 5_13771832 | NA | NA | NA | NA | NA | NA | 3 | 1 | 3 | 1 | NA | NA | NA | NA | NA | NA | 1 | 1 | 1 | 1 |
| 5_13771842 | NA | NA | NA | NA | NA | NA | 3 | 0.6667 | 3 | 0.6667 | NA | NA | NA | NA | NA | NA | 1 | 1 | 1 | 1 |
| 5_13771845 | NA | NA | NA | NA | NA | NA | 1 | 1 | 1 | 1 | NA | NA | NA | NA | NA | NA | 1 | 1 | 1 | 1 |
| 5_13771878 | NA | NA | NA | NA | NA | NA | 2 | 1 | 2 | 1 | NA | NA | NA | NA | NA | NA | 1 | 1 | 1 | 1 |
| 5_13771941 | NA | NA | NA | NA | NA | NA | 1 | 1 | 1 | 1 | NA | NA | NA | NA | NA | NA | 1 | 1 | 1 | 1 |
| 5_13772007 | NA | NA | NA | NA | NA | NA | 2 | 0.5 | 2 | 0.5 | NA | NA | NA | NA | NA | NA | 1 | 1 | 1 | 1 |

| | | | | | | | | | | | | | | | | | | | | |
|---|---|---|---|---|---|---|---|---|---|---|---|---|---|---|---|---|---|---|---|---|
| 5_1377201 | NA | NA | NA | NA | NA | NA | 3 | 0.6667 | 3 | 0.6667 | NA | NA | NA | NA | NA | NA | 1 | 1 | 1 | 1 |
| 5_1377202 | NA | NA | NA | NA | NA | NA | 3 | 1 | 3 | 1 | NA | NA | NA | NA | NA | NA | 1 | 1 | 1 | 1 |
| 5_1377202 | NA | NA | NA | NA | NA | NA | 3 | 0.6667 | 3 | 0.6667 | NA | NA | NA | NA | NA | NA | 1 | 1 | 1 | 1 |
| 5_1377202 | NA | NA | NA | NA | NA | NA | 2 | 0.5 | 2 | 0.5 | NA | NA | NA | NA | NA | NA | 1 | 1 | 1 | 1 |
| 5_1377210 | NA | NA | NA | NA | 1 | 1 | 3 | 0 | 4 | 0.25 | NA | NA | NA | NA | NA | NA | 1 | 1 | 1 | 1 |
| 5_1377299 | 2 | 0.5 | NA | NA | 1 | 1 | 1 | 1 | 4 | 0.75 | 2 | 1 | 1 | 1 | 1 | 1 | 2 | 1 | 6 | 1 |
| 5_1377332 | 2 | 1 | NA | NA | 1 | 1 | NA | NA | 3 | 1 | 3 | 1 | 1 | 1 | NA | NA | 3 | 1 | 7 | 1 |
| 5_1377333 | 2 | 0.5 | NA | NA | 1 | 1 | 1 | 0 | 4 | 0.5 | 3 | 0.6667 | 1 | 0 | NA | NA | 1 | 0 | 5 | 0.4 |
| 5_1377333 | 2 | 0.5 | NA | NA | NA | NA | 1 | 0 | 3 | 0.3333 | 3 | 0.6667 | 1 | 1 | NA | NA | 1 | 1 | 5 | 0.8 |
| 5_1377335 | 1 | 0 | NA | NA | 1 | 1 | 1 | 0 | 3 | 0.3333 | 3 | 0.3333 | 1 | 0 | NA | NA | 2 | 0.5 | 6 | 0.3333 |
| 5_1377336 | 1 | 0 | NA | NA | 1 | 1 | 1 | 0 | 3 | 0.3333 | 3 | 0.3333 | 1 | 1 | NA | NA | 2 | 0.5 | 6 | 0.5 |
| 5_1377337 | 1 | 1 | NA | NA | 1 | 1 | 1 | 1 | 3 | 1 | 1 | 1 | 1 | 1 | NA | NA | 1 | 1 | 3 | 1 |
| 5_1377338 | 1 | 1 | NA | NA | 1 | 0 | 1 | 1 | 3 | 0.6667 | 2 | 1 | 1 | 0 | NA | NA | NA | NA | 3 | 0.6667 |
| 5_1377340 | 1 | 1 | NA | NA | 1 | 0 | 1 | 0 | 3 | 0.3333 | 3 | 0.3333 | 1 | 1 | NA | NA | 2 | 0 | 6 | 0.3333 |
| 5_1377342 | 1 | 1 | NA | NA | 1 | 1 | 1 | 1 | 3 | 1 | 2 | 0.5 | 1 | 0 | NA | NA | 1 | 1 | 4 | 0.5 |
| 5_1377362 | NA | NA | NA | NA | 1 | 1 | NA | NA | 1 | 1 | 3 | 0.6667 | NA | NA | NA | NA | 1 | 1 | 4 | 0.75 |
| 5_1377445 | 3 | 0 | 6 | 0.5 | 1 | 0 | 1 | 0 | 11 | 0.2727 | 2 | 0.5 | 4 | 0 | NA | NA | NA | NA | 6 | 0.1667 |
| 5_1377452 | 3 | 0.6667 | 6 | 0.6667 | 2 | 0 | 1 | 0 | 12 | 0.5 | 2 | 0 | 6 | 0.1667 | NA | NA | NA | NA | 8 | 0.125 |
| 5_1377454 | 3 | 1 | 5 | 0.8 | 3 | 1 | 1 | 1 | 12 | 0.9167 | 2 | 1 | 6 | 1 | NA | NA | NA | NA | 8 | 1 |
| 5_1377459 | 4 | 0.25 | 7 | 0.1429 | 3 | 0 | 1 | 1 | 15 | 0.2 | 2 | 0.5 | 7 | 0.5714 | NA | NA | NA | NA | 9 | 0.5556 |
| 5_1377459 | 3 | 0 | 7 | 0.2857 | 3 | 0 | 1 | 0 | 14 | 0.1429 | 2 | 0 | 7 | 0.5714 | 1 | 1 | NA | NA | 10 | 0.5 |
| 5_1377579 | 36 | 0.1667 | 34 | 0.2941 | 25 | 0.16 | 38 | 0.3158 | 133 | 0.2406 | 31 | 0.1935 | 23 | 0.1739 | 25 | 0.2 | 16 | 0 | 95 | 0.1579 |
| 5_1377773 | 2 | 0.5 | 4 | 0.25 | 6 | 0.5 | 6 | 0.5 | 18 | 0.4444 | 3 | 0.6667 | 2 | 1 | 4 | 0.75 | 2 | 0.5 | 11 | 0.7273 |
| 5_1377777 | 1 | 0 | 3 | 0 | 5 | 0 | 4 | 0.75 | 13 | 0.2308 | 3 | 0 | 2 | 0 | 3 | 0 | 3 | 0.3333 | 11 | 0.0909 |
| 5_1377779 | 2 | 0.5 | 5 | 0 | 6 | 0 | 6 | 0 | 19 | 0.0526 | 5 | 0 | 2 | 0 | 3 | 0 | 4 | 0.25 | 14 | 0.0714 |
| 5_1377780 | 1 | 0 | 2 | 0 | 3 | 0.3333 | 4 | 0 | 10 | 0.1 | 2 | 0.5 | NA | NA | NA | NA | 2 | 0 | 4 | 0.25 |
| 5_1377781 | 2 | 0 | 6 | 0.3333 | 7 | 0 | 7 | 0 | 22 | 0.0909 | 5 | 0 | 2 | 0 | 5 | 0 | 4 | 0.25 | 16 | 0.0625 |
| 5_1377803 | 9 | 0.6667 | 10 | 0.9 | 3 | 0.3333 | 4 | 1 | 26 | 0.7692 | 6 | 0.5 | 7 | 0.5714 | 4 | 0.75 | 6 | 0.8333 | 23 | 0.6522 |
| 5_1377803 | 9 | 0.7778 | 12 | 0.5833 | 3 | 1 | 5 | 0.6 | 29 | 0.6897 | 5 | 0.6 | 8 | 0.625 | 5 | 0.4 | 6 | 0.3333 | 24 | 0.5 |
| 5_1377803 | 10 | 0.1 | 12 | 0 | 3 | 0 | 6 | 0.1667 | 31 | 0.0645 | 6 | 0.1667 | 8 | 0.125 | 5 | 0 | 5 | 0 | 24 | 0.0833 |
| 5_1377804 | 12 | 0.3333 | 9 | 0.1111 | 3 | 0 | 6 | 0.1667 | 30 | 0.2 | 5 | 0.2 | 7 | 0 | 4 | 0.5 | 5 | 0 | 21 | 0.1429 |
| 5_1377804 | 11 | 1 | 10 | 1 | 3 | 1 | 5 | 1 | 29 | 1 | 6 | 0.8333 | 8 | 1 | 4 | 1 | 6 | 1 | 24 | 0.9583 |
| 5_1377806 | 10 | 0.3 | 7 | 0.4286 | 4 | 0.5 | 5 | 0 | 26 | 0.3077 | 5 | 0 | 7 | 0.4286 | 4 | 0 | 5 | 0.4 | 21 | 0.2381 |
| 5_1377828 | 8 | 0.75 | 5 | 0.4 | 2 | 1 | 1 | 1 | 16 | 0.6875 | 4 | 0.5 | 4 | 0.5 | NA | NA | 3 | 0.6667 | 11 | 0.5455 |
| 5_1377828 | 8 | 0.75 | 7 | 1 | 3 | 1 | 2 | 0.5 | 20 | 0.85 | 4 | 1 | 4 | 1 | 1 | 1 | 2 | 1 | 11 | 1 |
| 5_1377828 | 7 | 0.4286 | 6 | 0.8333 | 2 | 1 | 2 | 0 | 17 | 0.5882 | 4 | 0.5 | 4 | 0 | 1 | 0 | 2 | 1 | 11 | 0.3636 |
| 5_1377828 | 5 | 0.8 | 6 | 0.6667 | 3 | 0.6667 | 2 | 0.5 | 16 | 0.6875 | 3 | 0.3333 | 2 | 1 | 1 | 1 | 1 | 1 | 7 | 0.7143 |
| 5_1377829 | 8 | 0.75 | 5 | 0.6 | 2 | 0.5 | 2 | 1 | 17 | 0.7059 | 3 | 1 | 4 | 0.75 | 1 | 1 | 2 | 1 | 10 | 0.9 |
| 5_1377830 | 8 | 0.125 | 6 | 0.1667 | 2 | 0.5 | 2 | 0 | 18 | 0.1667 | 4 | 0.25 | 4 | 0.25 | 1 | 0 | 1 | 0 | 10 | 0.2 |
| 5_1377832 | 6 | 0.6667 | 6 | 0.6667 | 2 | 1 | 2 | 0.5 | 16 | 0.6875 | 3 | 1 | 4 | 1 | 1 | 1 | 1 | 1 | 9 | 1 |
| 5_1377834 | 2 | 1 | 4 | 0.75 | 2 | 1 | 1 | 1 | 9 | 0.8889 | 2 | 1 | 4 | 1 | 1 | 1 | 1 | 1 | 8 | 1 |
| 5_1377835 | 5 | 0.2 | 4 | 0 | 1 | 0 | 2 | 0 | 12 | 0.0833 | 2 | 0 | 4 | 0.25 | 1 | 0 | 1 | 0 | 8 | 0.125 |
| 5_1377836 | 5 | 0.4 | 3 | 0 | 1 | 0 | 2 | 0 | 11 | 0.1818 | 2 | 0.5 | 4 | 0 | 1 | 0 | 1 | 0 | 8 | 0.125 |
| 5_1377838 | 4 | 0.25 | 4 | 0 | 1 | 1 | 2 | 0 | 11 | 0.1818 | 2 | 0.5 | 3 | 0.3333 | NA | NA | NA | NA | 5 | 0.4 |
| 5_1377838 | 4 | 0.5 | 3 | 0.3333 | 1 | 1 | 1 | 1 | 9 | 0.5556 | 2 | 1 | 3 | 0 | NA | NA | NA | NA | 5 | 0.4 |
| 5_1379524 | 1 | 0 | 5 | 0.4 | 3 | 0.3333 | 5 | 0.6 | 14 | 0.4286 | 2 | 0.5 | 2 | 0 | 2 | 0 | 4 | 0 | 10 | 0.1 |
| 5_1379534 | 1 | 0 | 5 | 0.2 | 1 | 0 | 4 | 0.5 | 11 | 0.2727 | 1 | 0 | 3 | 0 | 2 | 0.5 | 3 | 0 | 9 | 0.1111 |
| 5_1379535 | 1 | 1 | 5 | 0.4 | 1 | 1 | 5 | 1 | 12 | 0.75 | 1 | 1 | 3 | 0.6667 | 2 | 1 | 3 | 1 | 9 | 0.8889 |
| 5_1379546 | NA | NA | 5 | 0.4 | 1 | 0 | 5 | 0.6 | 11 | 0.4545 | NA | NA | 3 | 0.3333 | NA | NA | 3 | 0.6667 | 6 | 0.5 |
| 5_1379555 | NA | NA | 3 | 0.3333 | NA | NA | 3 | 0.6667 | 6 | 0.5 | NA | NA | 3 | 0.3333 | NA | NA | 3 | 0.3333 | 6 | 0.3333 |
| 5_1379559 | 1 | 1 | 2 | 1 | 1 | 1 | 3 | 1 | 7 | 1 | NA | NA | 3 | 0.6667 | NA | NA | 2 | 1 | 5 | 0.8 |
| 5_1379561 | NA | NA | 2 | 1 | 1 | 0 | 3 | 1 | 6 | 0.8333 | NA | NA | 1 | 1 | NA | NA | 1 | 1 | 2 | 1 |
| 5_1380912 | 1 | 1 | 2 | 0 | 1 | 0 | 2 | 0 | 6 | 0.1667 | 1 | 1 | NA | NA | 1 | 0 | 1 | 1 | 3 | 0.6667 |
| 5_1380913 | 1 | 1 | 1 | 1 | 1 | 0 | 1 | 0 | 4 | 0.5 | 1 | 0 | NA | NA | 1 | 0 | 1 | 1 | 3 | 0.3333 |
| 5_1380914 | NA | NA | 2 | 1 | 1 | 0 | 1 | 0 | 4 | 0.5 | NA | NA | NA | NA | 1 | 1 | 1 | 0 | 2 | 0.5 |
| 5_1380915 | NA | NA | 2 | 1 | 1 | 0 | 2 | 0.5 | 5 | 0.6 | 1 | 0 | NA | NA | 1 | 1 | 1 | 1 | 3 | 0.6667 |
| 5_1380915 | 1 | 0 | 1 | 0 | 1 | 0 | 2 | 0.5 | 5 | 0.2 | 1 | 1 | NA | NA | 1 | 0 | NA | NA | 2 | 0.5 |
| 5_1380991 | 1 | 1 | 2 | 0.5 | 2 | 0.5 | 1 | 1 | 6 | 0.6667 | 2 | 0 | 2 | 0 | 1 | 0 | 2 | 0.5 | 7 | 0.1429 |
| 5_1380991 | 1 | 1 | 2 | 1 | 2 | 1 | 2 | 1 | 7 | 1 | 2 | 1 | 1 | 0 | 2 | 1 | 2 | 1 | 7 | 0.8571 |
| 5_1380995 | NA | NA | 2 | 0.5 | 2 | 0 | 2 | 0 | 6 | 0.1667 | 2 | 0 | 2 | 0 | NA | NA | 2 | 1 | 6 | 0.3333 |
| 5_1380999 | 2 | 0 | 3 | 0 | 2 | 0.5 | 3 | 0 | 10 | 0.1 | 2 | 0.5 | 2 | 0 | 2 | 0 | 2 | 0 | 8 | 0.125 |
| 5_1380999 | 2 | 0.5 | 3 | 0.6667 | 2 | 0.5 | 2 | 0 | 9 | 0.4444 | 2 | 0.5 | 2 | 0.5 | 1 | 0 | 2 | 1 | 7 | 0.5714 |
| 5_1381001 | 2 | 0.5 | 3 | 0 | 2 | 0 | 2 | 0 | 9 | 0.1111 | 2 | 0.5 | 1 | 0 | 2 | 0 | 2 | 0 | 7 | 0.1429 |
| 5_1381001 | 2 | 0 | 3 | 0.3333 | 2 | 0 | 3 | 0 | 10 | 0.1 | 2 | 0 | 2 | 0 | 2 | 0 | 2 | 1 | 8 | 0.25 |
| 5_1381001 | 2 | 1 | 2 | 1 | 2 | 0.5 | 3 | 1 | 9 | 0.8889 | 2 | 1 | 2 | 0.5 | 2 | 1 | 1 | 1 | 7 | 0.8571 |

| | | | | | | | | | | | | | | | | | | | | |
|---|---|---|---|---|---|---|---|---|---|---|---|---|---|---|---|---|---|---|---|---|
| 5_13810025 | 1 | 1 | 2 | 1 | 2 | 1 | 4 | 1 | 9 | 1 | 2 | 1 | 1 | 1 | 3 | 1 | 2 | 1 | 8 | 1 |
| 5_13810033 | 2 | 1 | 3 | 1 | 1 | 1 | 3 | 1 | 9 | 1 | 2 | 1 | 2 | 1 | 3 | 1 | 2 | 1 | 9 | 1 |
| 5_13810058 | 2 | 0.5 | 3 | 0 | 2 | 0 | 3 | 0 | 10 | 0.1 | 2 | 0 | 2 | 0 | 3 | 0.6667 | 2 | 0 | 9 | 0.2222 |
| 5_13810063 | 2 | 1 | 2 | 1 | 1 | 1 | 2 | 1 | 7 | 1 | 2 | 1 | 2 | 1 | 2 | 1 | 3 | 1 | 9 | 1 |
| 5_13810069 | 2 | 1 | 2 | 0.5 | 2 | 0 | 3 | 0.6667 | 9 | 0.5556 | 3 | 0 | 1 | 0 | 3 | 0.6667 | 2 | 0 | 9 | 0.2222 |
| 5_13810071 | 2 | 1 | 3 | 1 | 2 | 1 | 3 | 1 | 10 | 1 | 3 | 1 | 2 | 1 | 3 | 1 | 3 | 1 | 11 | 1 |
| 5_13810073 | 2 | 1 | 3 | 1 | 2 | 1 | 3 | 1 | 10 | 1 | 3 | 1 | 2 | 1 | 3 | 1 | 2 | 1 | 10 | 1 |
| 5_13810079 | 2 | 1 | 3 | 1 | 2 | 1 | 4 | 1 | 11 | 1 | 3 | 1 | 1 | 1 | 4 | 1 | 3 | 1 | 11 | 1 |
| 5_13810087 | 2 | 0.5 | 2 | 1 | 2 | 1 | 2 | 0.5 | 8 | 0.75 | 3 | 0.6667 | NA | NA | 4 | 1 | 3 | 1 | 10 | 0.9 |
| 5_13810090 | 2 | 0 | 3 | 0.3333 | 3 | 0 | 3 | 0.3333 | 11 | 0.1818 | 3 | 0 | 2 | 0 | 3 | 0.3333 | 3 | 0 | 11 | 0.0909 |
| 5_13810128 | 2 | 0 | 4 | 0 | 3 | 0.3333 | 4 | 0.25 | 13 | 0.1538 | 3 | 0.3333 | 3 | 0 | 3 | 0 | 3 | 0 | 12 | 0.0833 |
| 5_13810146 | 2 | 0 | 2 | 0 | 3 | 0 | 3 | 0.6667 | 10 | 0.2 | 2 | 0 | 2 | 0 | 4 | 0.25 | 3 | 0 | 11 | 0.0909 |
| 5_13810151 | 3 | 0.6667 | 3 | 0 | 3 | 0.3333 | 3 | 1 | 12 | 0.5 | 3 | 1 | 3 | 0.6667 | 3 | 0.6667 | 3 | 1 | 12 | 0.8333 |
| 5_13810152 | 3 | 0.3333 | 4 | 0.25 | 2 | 0.5 | 3 | 0.3333 | 12 | 0.3333 | 2 | 0.5 | 2 | 0 | 4 | 0.25 | 5 | 0.4 | 13 | 0.3077 |
| 5_13810167 | 3 | 0.3333 | 4 | 0.25 | 3 | 0 | 4 | 0.25 | 14 | 0.2143 | 4 | 0.25 | 3 | 0 | 4 | 0 | 5 | 0 | 16 | 0.0625 |
| 5_13810168 | 3 | 0 | 3 | 0 | 2 | 0 | 4 | 0.25 | 12 | 0.0833 | 3 | 0.3333 | 2 | 0 | 4 | 0 | 4 | 0.25 | 13 | 0.1538 |
| 5_13810171 | 2 | 0 | 3 | 0 | 3 | 0.3333 | 4 | 0.5 | 12 | 0.25 | 4 | 0.25 | 2 | 0.5 | 4 | 0.25 | 6 | 0.1667 | 16 | 0.25 |
| 5_13810191 | 3 | 1 | 2 | 0.5 | 3 | 1 | 3 | 0.6667 | 11 | 0.8182 | 4 | 0.5 | 3 | 0.6667 | 3 | 0.6667 | 6 | 1 | 16 | 0.75 |
| 5_13810202 | 3 | 0 | 4 | 0.25 | 3 | 0 | 4 | 0.25 | 14 | 0.1429 | 4 | 0.25 | 7 | 0.1429 | 5 | 0 | 6 | 0.1667 | 22 | 0.1364 |
| 5_13810206 | 3 | 1 | 3 | 0.3333 | 3 | 1 | 4 | 0.75 | 13 | 0.7692 | 3 | 1 | 7 | 1 | 5 | 1 | 4 | 1 | 19 | 1 |
| 5_13810210 | 3 | 0.3333 | 4 | 0.75 | 2 | 0.5 | 4 | 0.75 | 13 | 0.6154 | 4 | 0.5 | 7 | 0.4286 | 5 | 0.4 | 5 | 0.2 | 21 | 0.381 |
| 5_13810216 | 2 | 1 | 4 | 0.75 | 3 | 0.6667 | 4 | 0.5 | 13 | 0.6923 | 2 | 1 | 7 | 0.5714 | 3 | 1 | 5 | 0.8 | 17 | 0.7647 |
| 5_13810230 | 3 | 0 | 4 | 0.25 | 3 | 0.3333 | 4 | 0 | 14 | 0.1429 | 3 | 0.3333 | 6 | 0 | 5 | 0 | 6 | 0 | 20 | 0.05 |
| 5_13810245 | 3 | 0.6667 | 4 | 0.5 | 3 | 1 | 4 | 0.75 | 14 | 0.7143 | 4 | 0.25 | 7 | 0.4286 | 5 | 0 | 6 | 0.6667 | 22 | 0.3636 |
| 5_13810246 | 1 | 0 | 4 | 0 | 3 | 0.3333 | 4 | 0.25 | 12 | 0.1667 | 4 | 0.25 | 7 | 0.5714 | 4 | 0.25 | 6 | 0.1667 | 21 | 0.3333 |
| 5_13810251 | 1 | 0 | 2 | 0 | 1 | 1 | 3 | 0.3333 | 7 | 0.2857 | 2 | 1 | 4 | 0.5 | 3 | 0.6667 | 5 | 0.6 | 14 | 0.6429 |
| 5_13810260 | 4 | 0.25 | 3 | 0.3333 | 3 | 0.3333 | 3 | 0 | 13 | 0.2308 | 4 | 0.75 | 7 | 0.5714 | 4 | 0.75 | 6 | 0.8333 | 21 | 0.7143 |
| 5_13810262 | 3 | 0.3333 | 3 | 0.3333 | 3 | 1 | 3 | 0 | 12 | 0.4167 | 4 | 0.75 | 5 | 0.2 | 2 | 0.5 | 4 | 0 | 15 | 0.3333 |
| 5_13810269 | 2 | 0 | 3 | 0 | 1 | 0 | 4 | 0.25 | 10 | 0.1 | 4 | 0.5 | 5 | 0.4 | 4 | 0 | 3 | 0.3333 | 16 | 0.3125 |
| 5_13810273 | 2 | 0 | 3 | 0.3333 | 2 | 0.5 | 3 | 0 | 10 | 0.2 | 4 | 0.5 | 5 | 0.4 | 4 | 0 | 5 | 0.8 | 18 | 0.4444 |
| 5_13810277 | NA | NA | 1 | 1 | 1 | 1 | 3 | 0.3333 | 5 | 0.6 | NA | NA | 4 | 0.25 | NA | NA | 4 | 0.75 | 8 | 0.5 |
| 5_13810285 | NA | NA | 1 | 1 | 2 | 0.5 | 3 | 0 | 6 | 0.3333 | 2 | 0.5 | 5 | 0 | 1 | 0 | 4 | 0.5 | 12 | 0.25 |
| 5_13810289 | NA | NA | NA | NA | 1 | 1 | 2 | 0 | 3 | 0.3333 | 1 | 0 | 4 | 0.25 | 1 | 0 | 2 | 0 | 8 | 0.125 |
| 5_13810290 | NA | NA | 1 | 0 | 2 | 0.5 | 3 | 0 | 6 | 0.1667 | 1 | 1 | 5 | 0 | 1 | 0 | 4 | 0.5 | 11 | 0.2727 |
| 5_13810359 | NA | NA | 1 | 0 | 2 | 0 | 5 | 0.2 | 8 | 0.125 | 2 | 0 | 3 | 0.3333 | 3 | 0 | 5 | 0 | 13 | 0.0769 |
| 5_13810373 | NA | NA | 1 | 0 | 1 | 0 | 5 | 0.2 | 7 | 0.1429 | 2 | 0.5 | 4 | 0.25 | 3 | 0 | 4 | 0 | 13 | 0.1538 |
| 5_13810377 | NA | NA | 1 | 0 | 2 | 0 | 5 | 0.2 | 8 | 0.125 | 2 | 0 | 4 | 0 | 3 | 0.6667 | 4 | 0.5 | 13 | 0.3077 |
| 5_13810398 | 1 | 0 | 1 | 0 | 2 | 0.5 | 5 | 0 | 9 | 0.1111 | 2 | 0 | 4 | 0 | 3 | 0 | 5 | 0.2 | 14 | 0.0714 |
| 5_13810399 | 1 | 0 | NA | NA | 2 | 0.5 | 5 | 0.4 | 8 | 0.375 | 2 | 0.5 | 4 | 0.25 | 3 | 0.3333 | 5 | 0.2 | 14 | 0.2857 |
| 5_13810409 | 1 | 0 | NA | NA | 3 | 1 | 4 | 0 | 8 | 0.375 | 1 | 0 | 4 | 0.25 | 3 | 0.3333 | 5 | 0.2 | 13 | 0.2308 |
| 5_13810510 | 1 | 0 | 1 | 0 | 4 | 0 | 5 | 0.2 | 11 | 0.0909 | 3 | 0.3333 | 4 | 0 | 3 | 0 | 3 | 0 | 13 | 0.0769 |
| 5_13810521 | 1 | 0 | 3 | 0.3333 | 5 | 0.2 | 4 | 0 | 13 | 0.1538 | 2 | 0.5 | 4 | 0.25 | 3 | 0.6667 | 6 | 0.6667 | 15 | 0.5333 |
| 5_13810546 | 1 | 0 | 2 | 0.5 | 4 | 0 | 4 | 0.25 | 11 | 0.1818 | 3 | 0.3333 | 4 | 0.25 | 4 | 0 | 7 | 0.2857 | 18 | 0.2222 |
| 5_13812394 | NA | NA | NA | NA | 1 | 1 | 2 | 1 | 3 | 1 | 1 | 1 | 2 | 0.5 | NA | NA | NA | NA | 3 | 0.6667 |
| 5_13813137 | NA | NA | 1 | 0 | 1 | 1 | 2 | 1 | 4 | 0.75 | 1 | 1 | 3 | 0.3333 | NA | NA | 1 | 1 | 5 | 0.6 |
| 5_13813172 | NA | NA | 1 | 1 | 1 | 0 | 2 | 0 | 4 | 0.25 | 1 | 0 | 3 | 0.3333 | NA | NA | 1 | 0 | 5 | 0.2 |
| 5_13813181 | NA | NA | 1 | 1 | 1 | 0 | 1 | 0 | 3 | 0.3333 | 1 | 1 | 2 | 0.5 | NA | NA | 1 | 0 | 4 | 0.5 |
| 5_13816186 | 19 | 0.1579 | 27 | 0.1481 | 21 | 0.1905 | 15 | 0.0667 | 82 | 0.1463 | 15 | 0.2 | 21 | 0.0952 | 28 | 0.1429 | 18 | 0.2222 | 82 | 0.1585 |
| 5_13816196 | 1 | 0 | NA | NA | 2 | 0.5 | NA | NA | 3 | 0.3333 | 1 | 0 | NA | NA | 2 | 0.5 | NA | NA | 3 | 0.3333 |
| 5_13816197 | 1 | 0 | NA | NA | 2 | 0.5 | NA | NA | 3 | 0.3333 | 1 | 0 | NA | NA | 2 | 0.5 | NA | NA | 3 | 0.3333 |
| 5_13816212 | 19 | 0.6842 | 30 | 0.4333 | 20 | 0.45 | 16 | 0.5625 | 85 | 0.5176 | 16 | 0.75 | 25 | 0.52 | 31 | 0.6452 | 15 | 0.4667 | 87 | 0.5977 |
| 5_13816218 | 18 | 0.3333 | 32 | 0.25 | 22 | 0.2727 | 12 | 0.0833 | 84 | 0.25 | 17 | 0.4118 | 22 | 0.3636 | 31 | 0.4839 | 17 | 0.2353 | 87 | 0.3908 |
| 5_13816229 | 2 | 0.5 | 1 | 0 | NA | NA | NA | NA | 3 | 0.3333 | 2 | 1 | NA | NA | 1 | 1 | NA | NA | 3 | 1 |
| 5_13816564 | 1 | 1 | NA | NA | 1 | 1 | NA | NA | 2 | 1 | 1 | 0 | 3 | 0.3333 | 1 | 1 | 2 | 0 | 7 | 0.2857 |
| 5_13817657 | NA | NA | 1 | 0 | 2 | 1 | NA | NA | 3 | 0.6667 | NA | NA | NA | NA | 1 | 1 | NA | NA | 1 | 1 |
| 5_13817666 | NA | NA | 2 | 0.5 | 2 | 1 | NA | NA | 4 | 0.75 | 1 | 0 | NA | NA | 2 | 1 | 1 | 1 | 4 | 0.75 |
| 5_13817677 | NA | NA | 2 | 1 | 2 | 1 | NA | NA | 4 | 1 | 1 | 0 | NA | NA | 2 | 1 | NA | NA | 3 | 0.6667 |
| 5_13817680 | NA | NA | 2 | 1 | 2 | 1 | NA | NA | 4 | 1 | 1 | 0 | NA | NA | 1 | 1 | 1 | 1 | 3 | 0.6667 |
| 5_13817683 | NA | NA | 2 | 0.5 | 1 | 1 | NA | NA | 3 | 0.6667 | 1 | 1 | NA | NA | 2 | 1 | NA | NA | 3 | 1 |
| 5_13817683 | NA | NA | 1 | 1 | 2 | 0.5 | NA | NA | 3 | 0.6667 | NA | NA | NA | NA | 1 | 1 | NA | NA | 1 | 1 |
| 5_13817685 | NA | NA | 1 | 1 | 1 | 1 | NA | NA | 2 | 1 | 1 | 0 | NA | NA | 2 | 0.5 | NA | NA | 3 | 0.3333 |
| 5_13817690 | 1 | 0 | 1 | 0 | 2 | 1 | NA | NA | 4 | 0.5 | 1 | 0 | NA | NA | 2 | 0.5 | NA | NA | 3 | 0.3333 |
| 5_13817693 | 5 | 0.2 | 7 | 0.1429 | 2 | 0 | 3 | 0.3333 | 17 | 0.1765 | 2 | 0 | 1 | 0 | 2 | 0.5 | 2 | 0.5 | 7 | 0.2857 |
| 5_13817705 | 5 | 0.2 | 8 | 0.25 | 2 | 0.5 | 1 | 0 | 16 | 0.25 | 2 | 0.5 | 1 | 1 | 2 | 0 | 2 | 0 | 7 | 0.2857 |
| 5_13817708 | 5 | 0.6 | 8 | 0.25 | 2 | 1 | 3 | 0.6667 | 18 | 0.5 | 2 | 0.5 | 1 | 1 | 2 | 1 | 2 | 1 | 7 | 0.8571 |

| | | | | | | | | | | | | | | | | | | | | |
|---|---|---|---|---|---|---|---|---|---|---|---|---|---|---|---|---|---|---|---|---|
| 5_1381777( | 1 | 1 | 1 | 1 | 2 | 0 | 1 | 0 | 5 | 0.4 | NA | NA | NA | NA | 2 | 0.5 | 1 | 0 | 3 | 0.3333 |
| 5_1381912( | 1 | 0 | NA | NA | 3 | 0.3333 | 1 | 1 | 5 | 0.4 | NA | NA | 3 | 0 | NA | NA | 1 | 1 | 4 | 0.25 |
| 5_1381915( | 2 | 1 | NA | NA | 3 | 0.3333 | 2 | 1 | 7 | 0.7143 | NA | NA | 3 | 0.3333 | NA | NA | NA | NA | 3 | 0.3333 |
| 5_1381929! | NA | NA | NA | NA | NA | NA | 1 | 1 | 1 | 1 | 1 | 1 | NA | NA | NA | NA | NA | NA | 1 | 1 |
| 5_1381930' | 1 | 1 | NA | NA | NA | NA | 1 | 1 | 2 | 1 | 1 | 1 | NA | NA | NA | NA | NA | NA | 1 | 1 |
| 5_1381931! | 1 | 1 | NA | NA | NA | NA | 1 | 1 | 2 | 1 | 1 | 1 | NA | NA | NA | NA | NA | NA | 1 | 1 |
| 5_1381936; | 1 | 1 | NA | NA | NA | NA | 1 | 1 | 2 | 1 | 1 | 1 | NA | NA | NA | NA | NA | NA | 1 | 1 |
| 5_1381937! | 1 | 1 | NA | NA | NA | NA | NA | NA | 1 | 1 | 1 | 1 | NA | NA | NA | NA | NA | NA | 1 | 1 |
| 5_1382575( | 9 | 0.3333 | 9 | 0.2222 | 12 | 0.1667 | 5 | 0.2 | 35 | 0.2286 | 14 | 0.3571 | 14 | 0.3571 | 9 | 0.2222 | 11 | 0.3636 | 48 | 0.3333 |
| 5_1382575; | 7 | 0.1429 | 9 | 0.1111 | 10 | 0.2 | 7 | 0.2857 | 33 | 0.1818 | 12 | 0.1667 | 12 | 0.25 | 9 | 0.1111 | 10 | 0.2 | 43 | 0.186 |
| 5_1382580! | 6 | 0 | 10 | 0.2 | 4 | 0 | 9 | 0 | 29 | 0.069 | 6 | 0.1667 | 2 | 0 | 2 | 0.5 | 5 | 0.2 | 15 | 0.2 |
| 5_1382634· | 1 | 1 | NA | NA | NA | NA | NA | NA | 1 | 1 | 1 | 1 | 1 | 1 | NA | NA | NA | NA | 2 | 1 |
| 5_1382639! | 1 | 1 | NA | NA | NA | NA | NA | NA | 1 | 1 | 1 | 1 | 1 | 1 | NA | NA | NA | NA | 2 | 1 |
| 5_1382639! | 1 | 1 | NA | NA | NA | NA | NA | NA | 1 | 1 | 1 | 1 | 1 | 0 | NA | NA | NA | NA | 2 | 0.5 |
| 5_1382641; | NA | NA | 2 | 0.5 | 1 | 0 | 1 | 1 | 4 | 0.5 | NA | NA | 1 | 1 | NA | NA | 1 | 1 | 2 | 1 |
| 5_1382641; | 1 | 1 | NA | NA | NA | NA | NA | NA | 1 | 1 | 1 | 1 | 1 | 1 | NA | NA | NA | NA | 2 | 1 |
| 5_1382643; | 1 | 1 | NA | NA | NA | NA | NA | NA | 1 | 1 | 1 | 0 | 1 | 1 | NA | NA | NA | NA | 2 | 0.5 |
| 5_1382645; | NA | NA | 1 | 0 | 1 | 1 | 1 | 0 | 3 | 0.3333 | NA | NA | 1 | 1 | 1 | 1 | 1 | 0 | 3 | 0.6667 |
| 5_1382646( | NA | NA | 2 | 1 | NA | NA | 1 | 0 | 3 | 0.6667 | NA | NA | 1 | 0 | 1 | 1 | 1 | 0 | 3 | 0.3333 |
| 5_1382654! | NA | NA | NA | NA | 1 | 1 | NA | NA | 1 | 1 | NA | NA | NA | NA | 1 | 1 | NA | NA | 1 | 1 |
| 5_1382655( | NA | NA | NA | NA | 1 | 1 | NA | NA | 1 | 1 | NA | NA | NA | NA | NA | NA | 1 | 1 | 1 | 1 |
| 5_1382657( | NA | NA | NA | NA | 1 | 1 | NA | NA | 1 | 1 | NA | NA | NA | NA | 1 | 0 | 1 | 1 | 2 | 0.5 |
| 5_1382666( | NA | NA | NA | NA | 1 | 1 | NA | NA | 1 | 1 | NA | NA | NA | NA | 1 | 1 | 1 | 0 | 2 | 0.5 |
| 5_1382667; | NA | NA | NA | NA | 1 | 1 | NA | NA | 1 | 1 | NA | NA | NA | NA | 1 | 1 | 1 | 0 | 2 | 0.5 |
| 5_1384543! | NA | NA | 4 | 0.75 | NA | NA | 4 | 0.5 | 8 | 0.625 | 1 | 1 | 1 | 0 | 3 | 0 | 3 | 0.6667 | 8 | 0.375 |
| 5_1384546' | NA | NA | 4 | 1 | 1 | 1 | 4 | 1 | 9 | 1 | 1 | 1 | 1 | 1 | 6 | 1 | 4 | 1 | 12 | 1 |
| 5_1384546( | NA | NA | 3 | 0.6667 | 1 | 0 | 3 | 0.3333 | 7 | 0.4286 | 1 | 1 | 1 | 1 | 6 | 1 | 4 | 1 | 12 | 1 |
| 5_1384547· | NA | NA | 2 | 0 | 1 | 0 | 3 | 0.3333 | 6 | 0.1667 | 1 | 1 | 2 | 0.5 | 4 | 0.25 | 3 | 0.3333 | 10 | 0.4 |
| 5_1384549( | NA | NA | 4 | 0 | 1 | 0 | 3 | 0.3333 | 8 | 0.125 | 1 | 0 | 2 | 0.5 | 6 | 0.1667 | 3 | 0.3333 | 12 | 0.25 |
| 5_1384549' | NA | NA | 4 | 0.25 | 1 | 1 | 4 | 0.5 | 9 | 0.4444 | 1 | 0 | 1 | 1 | 6 | 0.8333 | 3 | 0.3333 | 11 | 0.6364 |
| 5_1384549! | NA | NA | 4 | 1 | 1 | 1 | 4 | 0.75 | 9 | 0.8889 | 1 | 0 | 1 | 1 | 6 | 1 | 4 | 1 | 12 | 0.9167 |
| 5_1384550! | NA | NA | 3 | 0.6667 | 1 | 1 | 3 | 0.6667 | 7 | 0.7143 | 1 | 0 | 2 | 0.5 | 5 | 0.6 | 4 | 0.5 | 12 | 0.5 |
| 5_1384551' | NA | NA | 4 | 1 | 1 | 1 | 2 | 1 | 7 | 1 | NA | NA | 1 | 1 | 5 | 1 | 3 | 1 | 9 | 1 |
| 5_1384552; | NA | NA | 4 | 0.75 | 1 | 1 | 3 | 1 | 8 | 0.875 | 1 | 1 | 2 | 1 | 5 | 1 | 4 | 0.75 | 12 | 0.9167 |
| 5_1384552! | NA | NA | 4 | 0.75 | 1 | 0 | 4 | 0 | 9 | 0.3333 | 1 | 1 | 2 | 1 | 6 | 0.6667 | 4 | 0.5 | 13 | 0.6923 |
| 5_1384553' | NA | NA | 4 | 1 | 1 | 1 | 4 | 0.75 | 9 | 0.8889 | 1 | 0 | 1 | 1 | 5 | 0.8 | 4 | 0.75 | 11 | 0.7273 |
| 5_1384554' | NA | NA | 3 | 1 | NA | NA | 3 | 1 | 6 | 1 | 1 | 1 | 2 | 1 | 6 | 1 | 4 | 1 | 13 | 1 |
| 5_1384554! | NA | NA | 2 | 0.5 | 1 | 0 | 4 | 0.75 | 7 | 0.5714 | 1 | 1 | 2 | 1 | 5 | 0.8 | 4 | 0.5 | 12 | 0.75 |
| 5_1384555( | NA | NA | 3 | 0 | 1 | 1 | 4 | 0.25 | 8 | 0.25 | 1 | 1 | 1 | 0 | 7 | 0 | 4 | 0 | 13 | 0.0769 |
| 5_1384555! | NA | NA | 3 | 1 | NA | NA | 3 | 1 | 6 | 1 | 1 | 1 | NA | NA | 6 | 0.8333 | 4 | 1 | 11 | 0.9091 |
| 5_1384556( | NA | NA | 4 | 0.25 | 1 | 0 | 4 | 0.5 | 9 | 0.3333 | 1 | 0 | 2 | 0.5 | 6 | 0.1667 | 4 | 0.75 | 13 | 0.3846 |
| 5_1384557' | NA | NA | 3 | 0.6667 | 1 | 0 | 3 | 0.6667 | 7 | 0.5714 | 1 | 1 | 2 | 0 | 1 | 0 | 4 | 0.25 | 8 | 0.25 |
| 5_1384557· | NA | NA | 4 | 0.25 | NA | NA | 3 | 0.6667 | 7 | 0.4286 | 1 | 0 | 2 | 0 | 6 | 0 | 4 | 0.5 | 13 | 0.1538 |
| 5_1384557; | NA | NA | 4 | 0.75 | 1 | 0 | 4 | 0.75 | 9 | 0.6667 | 1 | 1 | 2 | 1 | 6 | 0.6667 | 2 | 1 | 11 | 0.8182 |
| 5_1384558' | NA | NA | 4 | 0.5 | 1 | 0 | 4 | 0 | 9 | 0.2222 | 1 | 0 | 2 | 0.5 | 6 | 0.3333 | 4 | 0.5 | 13 | 0.3846 |
| 5_1384558; | NA | NA | 3 | 1 | 1 | 1 | 4 | 1 | 8 | 1 | 1 | 0 | 2 | 1 | 7 | 1 | 4 | 0.75 | 14 | 0.8571 |
| 5_1384559( | NA | NA | 4 | 1 | 1 | 1 | 4 | 1 | 9 | 1 | 1 | 1 | 2 | 1 | 7 | 1 | 4 | 1 | 14 | 1 |
| 5_1385878' | NA | NA | 1 | 1 | NA | NA | NA | NA | 1 | 1 | NA | NA | 2 | 0.5 | 1 | 0 | 2 | 0.5 | 5 | 0.4 |
| 5_1386029( | NA | NA | 4 | 0.5 | 4 | 0.25 | 2 | 0 | 10 | 0.3 | 2 | 0 | 1 | 1 | 1 | 1 | 1 | 1 | 5 | 0.6 |
| 5_1386039; | NA | NA | 4 | 0.5 | 3 | 0 | NA | NA | 7 | 0.2857 | 2 | 0.5 | 1 | 0 | 1 | 0 | 2 | 0.5 | 6 | 0.3333 |
| 5_1386040( | NA | NA | 4 | 0.5 | 3 | 0.6667 | NA | NA | 7 | 0.5714 | 2 | 0.5 | 1 | 0 | 1 | 1 | 2 | 0.5 | 6 | 0.5 |
| 5_1386041; | NA | NA | 2 | 1 | 2 | 0 | 1 | 1 | 5 | 0.6 | 2 | 0.5 | 1 | 0 | 1 | 0 | 2 | 0 | 6 | 0.1667 |
| 5_1386043; | NA | NA | 4 | 0.75 | 2 | 0.5 | 1 | 0 | 7 | 0.5714 | 1 | 0 | 1 | 0 | 1 | 1 | 2 | 0.5 | 5 | 0.4 |
| 5_1386044; | 2 | 0 | 4 | 0.5 | 4 | 0.25 | 3 | 0.3333 | 13 | 0.3077 | 1 | 0 | 6 | 0.3333 | 7 | 0.2857 | 4 | 0.25 | 18 | 0.2778 |
| 5_1386045( | 2 | 0 | 2 | 0 | 3 | 0.3333 | 2 | 0 | 9 | 0.1111 | 1 | 0 | 6 | 0 | 4 | 0.25 | 3 | 0 | 14 | 0.0714 |
| 5_1386045' | 2 | 0 | 3 | 1 | 3 | 0.3333 | 1 | 0 | 9 | 0.4444 | 1 | 0 | 2 | 0 | 4 | 0.5 | 1 | 0 | 8 | 0.25 |
| 5_1386047; | 2 | 0.5 | 4 | 0 | 2 | 0.5 | 3 | 0 | 11 | 0.1818 | 1 | 1 | 5 | 0.2 | 4 | 0.25 | 3 | 0 | 13 | 0.2308 |
| 5_1386048( | 2 | 0 | 3 | 1 | 3 | 0.3333 | 2 | 0.5 | 10 | 0.5 | NA | NA | 4 | 0.25 | 5 | 0 | 4 | 0 | 13 | 0.0769 |
| 5_1386048! | 2 | 0 | 4 | 0.75 | 3 | 0 | 3 | 0.3333 | 12 | 0.3333 | 1 | 0 | 4 | 0.25 | 5 | 0 | 2 | 0 | 12 | 0.0833 |
| 5_1386049! | 2 | 0 | 2 | 0.5 | 2 | 0 | 3 | 0 | 9 | 0.1111 | 1 | 0 | 4 | 0 | 5 | 0 | 3 | 0.3333 | 13 | 0.0769 |
| 5_1386055! | 1 | 0 | 3 | 0.6667 | 3 | 0.3333 | 1 | 0 | 8 | 0.375 | 1 | 0 | 4 | 0.25 | 3 | 0 | 3 | 0 | 11 | 0.0909 |
| 5_1386058' | 1 | 0 | 2 | 0 | 3 | 0.3333 | 2 | 0 | 8 | 0.125 | 1 | 1 | 5 | 0.4 | 2 | 0.5 | 3 | 0 | 11 | 0.3636 |
| 5_1386059' | 1 | 0 | 1 | 0 | 2 | 0.5 | 2 | 0.5 | 6 | 0.3333 | 1 | 1 | 3 | 0.6667 | NA | NA | 2 | 1 | 6 | 0.8333 |
| 5_1386059; | NA | NA | 2 | 0 | 3 | 0 | 2 | 0.5 | 7 | 0.1429 | 1 | 1 | 3 | 0.3333 | 2 | 0 | 3 | 0 | 9 | 0.2222 |
| 5_1386445( | 1 | 1 | NA | NA | NA | NA | 1 | 1 | 2 | 1 | 1 | 1 | NA | NA | 1 | 1 | NA | NA | 2 | 1 |

| | | | | | | | | | | | | | | | | | | | | |
|---|---|---|---|---|---|---|---|---|---|---|---|---|---|---|---|---|---|---|---|---|
| 5_1386446 | 1 | 0 | NA | NA | NA | NA | 2 | 0.5 | 3 | 0.3333 | NA | NA | NA | NA | 1 | 1 | NA | NA | 1 | 1 |
| 5_1386448 | 2 | 0.5 | NA | NA | NA | NA | 1 | 0 | 3 | 0.3333 | 2 | 1 | NA | NA | 1 | 1 | NA | NA | 3 | 1 |
| 5_1386448 | 2 | 0.5 | NA | NA | NA | NA | 1 | 1 | 3 | 0.6667 | 2 | 0.5 | NA | NA | 1 | 0 | NA | NA | 3 | 0.3333 |
| 5_1386452 | 3 | 1 | 1 | 1 | NA | NA | 3 | 1 | 7 | 1 | 2 | 1 | NA | NA | 2 | 1 | NA | NA | 4 | 1 |
| 5_1386460 | 3 | 1 | 3 | 0.3333 | NA | NA | 2 | 0.5 | 8 | 0.625 | 2 | 0.5 | NA | NA | 3 | 0 | NA | NA | 5 | 0.2 |
| 5_1386461 | 3 | 0.6667 | 2 | 0.5 | NA | NA | 4 | 0.25 | 9 | 0.4444 | 2 | 0 | NA | NA | 3 | 1 | NA | NA | 5 | 0.6 |
| 5_1386462 | 3 | 1 | 2 | 1 | NA | NA | 3 | 1 | 8 | 1 | 2 | 1 | NA | NA | 2 | 1 | NA | NA | 4 | 1 |
| 5_1386462 | 2 | 0 | 1 | 0 | NA | NA | 3 | 0.6667 | 6 | 0.3333 | 2 | 0.5 | NA | NA | 3 | 0.3333 | NA | NA | 5 | 0.4 |
| 5_1386464 | 3 | 0 | 2 | 0.5 | NA | NA | 4 | 0.25 | 9 | 0.2222 | 2 | 0.5 | NA | NA | 3 | 0 | NA | NA | 5 | 0.2 |
| 5_1386464 | 3 | 0.6667 | 2 | 1 | NA | NA | 4 | 0.75 | 9 | 0.7778 | 2 | 0 | NA | NA | 2 | 0.5 | NA | NA | 4 | 0.25 |
| 5_1386466 | 3 | 1 | 1 | 1 | NA | NA | 3 | 1 | 7 | 1 | 2 | 1 | NA | NA | 2 | 1 | NA | NA | 4 | 1 |
| 5_1386468 | 4 | 1 | 2 | 1 | 1 | 1 | 4 | 1 | 11 | 1 | 2 | 1 | NA | NA | 4 | 0.75 | NA | NA | 6 | 0.8333 |
| 5_1386468 | 2 | 1 | 2 | 0.5 | 1 | 1 | 4 | 0.75 | 9 | 0.7778 | 3 | 0.6667 | 1 | 1 | 4 | 0.25 | NA | NA | 8 | 0.5 |
| 5_1387404 | 1 | 1 | NA | NA | NA | NA | 2 | 1 | 3 | 1 | NA | NA | 1 | 1 | 2 | 1 | NA | NA | 3 | 1 |
| 5_1387412 | 2 | 1 | NA | NA | NA | NA | 4 | 0.75 | 6 | 0.8333 | 1 | 1 | 2 | 1 | 1 | 1 | NA | NA | 4 | 1 |
| 5_1387414 | 1 | 0 | 3 | 0 | 2 | 0.5 | 4 | 0.25 | 10 | 0.2 | 2 | 0 | 5 | 0 | 2 | 0.5 | 1 | 0 | 10 | 0.1 |
| 5_1387416 | 1 | 0 | 3 | 0 | 2 | 0.5 | 6 | 0 | 12 | 0.0833 | 1 | 0 | 5 | 0 | 4 | 0.25 | 1 | 0 | 11 | 0.0909 |
| 5_1387949 | 6 | 0.1667 | 2 | 0 | 3 | 0 | 9 | 0 | 20 | 0.05 | 4 | 0 | 7 | 0.1429 | 5 | 0.2 | 4 | 0 | 20 | 0.1 |
| 5_1387950 | 6 | 0.3333 | 4 | 0.25 | 4 | 0.5 | 10 | 0.4 | 24 | 0.375 | 6 | 0.6667 | 8 | 0.5 | 6 | 0.5 | 4 | 0.5 | 24 | 0.5417 |
| 5_1387953 | 7 | 0.5714 | 4 | 0.5 | 3 | 0.6667 | 8 | 0.25 | 22 | 0.4545 | 8 | 0.875 | 9 | 0.5556 | 6 | 0.5 | 3 | 0.6667 | 26 | 0.6538 |
| 5_1387953 | 7 | 0.4286 | 4 | 0.25 | 3 | 0 | 8 | 0 | 22 | 0.1818 | 7 | 0 | 7 | 0.1429 | 6 | 0.3333 | 3 | 0 | 23 | 0.1304 |
| 5_1387954 | 7 | 0.8571 | 6 | 1 | 5 | 0.8 | 9 | 1 | 27 | 0.9259 | 7 | 0.8571 | 11 | 0.9091 | 7 | 1 | 3 | 1 | 28 | 0.9286 |
| 5_1387955 | 11 | 1 | 5 | 1 | 4 | 1 | 7 | 0.8571 | 27 | 0.963 | 6 | 0.8333 | 11 | 0.9091 | 8 | 0.875 | 6 | 1 | 31 | 0.9032 |
| 5_1387955 | 11 | 0.5455 | 6 | 0.3333 | 5 | 0.8 | 9 | 0.5556 | 31 | 0.5484 | 7 | 0.8571 | 11 | 0.8182 | 8 | 1 | 6 | 0.8333 | 32 | 0.875 |
| 5_1387956 | 11 | 0 | 6 | 0.1667 | 5 | 0.2 | 10 | 0.1 | 32 | 0.0938 | 8 | 0.125 | 12 | 0.0833 | 9 | 0 | 6 | 0 | 35 | 0.0571 |
| 5_1387957 | 12 | 0 | 6 | 0.1667 | 5 | 0 | 8 | 0.125 | 31 | 0.0645 | 8 | 0.25 | 8 | 0 | 9 | 0.1111 | 4 | 0 | 29 | 0.1034 |
| 5_1387957 | 11 | 0.7273 | 5 | 0.6 | 4 | 1 | 10 | 0.7 | 30 | 0.7333 | 6 | 0.6667 | 11 | 0.7273 | 9 | 0.8889 | 6 | 0.6667 | 32 | 0.75 |
| 5_1389620 | 4 | 0.5 | 3 | 0 | 1 | 1 | NA | NA | 8 | 0.375 | 1 | 0 | NA | NA | 2 | 1 | 1 | 0 | 4 | 0.5 |
| 5_1389622 | 7 | 0.2857 | 7 | 0 | 5 | 0 | 3 | 0.3333 | 22 | 0.1364 | 3 | 0.6667 | NA | NA | 4 | 0 | 1 | 0 | 8 | 0.25 |
| 5_1389624 | 9 | 0.4444 | 10 | 0.1 | 3 | 0.6667 | 6 | 0 | 28 | 0.25 | 3 | 0.3333 | NA | NA | 5 | 0 | 4 | 0.5 | 12 | 0.25 |
| 5_1389624 | 5 | 0.2 | 8 | 0.375 | 1 | 0 | 4 | 0.25 | 18 | 0.2778 | 2 | 0.5 | NA | NA | 2 | 0 | 3 | 0.6667 | 7 | 0.4286 |
| 5_1389625 | 9 | 1 | 9 | 0.8889 | 4 | 1 | 6 | 1 | 28 | 0.9643 | 2 | 1 | NA | NA | 5 | 1 | 3 | 1 | 10 | 1 |
| 5_1389626 | 6 | 1 | 4 | 1 | 2 | 1 | 4 | 1 | 16 | 1 | 3 | 1 | NA | NA | 3 | 1 | 3 | 1 | 9 | 1 |
| 5_1389626 | 8 | 0 | 7 | 0.2857 | 4 | 0.25 | 5 | 0.2 | 24 | 0.1667 | 1 | 1 | NA | NA | 4 | 0.25 | 4 | 0 | 9 | 0.2222 |
| 5_1389627 | 5 | 1 | 8 | 0.75 | 3 | 0.3333 | 3 | 1 | 19 | 0.7895 | 2 | 1 | NA | NA | 3 | 0.6667 | 4 | 0 | 9 | 0.4444 |
| 5_1389628 | 7 | 0.8571 | 7 | 0.7143 | 4 | 0.5 | 6 | 0.8333 | 24 | 0.75 | 2 | 0.5 | NA | NA | 5 | 0.6 | 4 | 0.75 | 11 | 0.6364 |
| 5_1389628 | 5 | 0.2 | 6 | 0.1667 | 4 | 0 | 5 | 0 | 20 | 0.1 | 2 | 0 | NA | NA | 5 | 0.2 | 4 | 0.5 | 11 | 0.2727 |
| 5_1389628 | 5 | 0.6 | 5 | 0.8 | 2 | 1 | 3 | 1 | 15 | 0.8 | 2 | 1 | NA | NA | 1 | 1 | 3 | 0.3333 | 6 | 0.6667 |
| 5_1389810 | 3 | 0 | 5 | 0.2 | 1 | 0 | 4 | 0 | 13 | 0.0769 | 1 | 1 | NA | NA | 2 | 1 | NA | NA | 3 | 1 |
| 5_1389824 | 2 | 1 | NA | NA | NA | NA | 1 | 1 | 3 | 1 | NA | NA | NA | NA | NA | NA | 1 | 1 | 1 | 1 |
| 5_1389827 | 1 | 0 | NA | NA | 1 | 1 | 2 | 1 | 4 | 0.75 | NA | NA | NA | NA | NA | NA | 1 | 1 | 1 | 1 |
| 5_1389829 | 2 | 0 | NA | NA | 1 | 0 | 1 | 1 | 4 | 0.25 | 1 | 1 | NA | NA | NA | NA | 1 | 0 | 2 | 0.5 |
| 5_1389830 | 2 | 1 | NA | NA | 1 | 1 | 2 | 1 | 5 | 1 | 1 | 1 | NA | NA | NA | NA | 1 | 1 | 2 | 1 |
| 5_1389833 | 2 | 1 | NA | NA | 1 | 1 | 3 | 0 | 6 | 0.5 | 1 | 1 | NA | NA | NA | NA | 1 | 1 | 2 | 1 |
| 5_1389836 | 1 | 0 | 1 | 1 | 1 | 1 | 3 | 0.6667 | 6 | 0.6667 | 1 | 0 | NA | NA | NA | NA | 2 | 1 | 3 | 0.6667 |
| 5_1389836 | NA | NA | 1 | 0 | NA | NA | 2 | 0.5 | 3 | 0.3333 | NA | NA | NA | NA | NA | NA | 2 | 0.5 | 2 | 0.5 |
| 5_1389836 | 1 | 1 | 1 | 1 | NA | NA | 3 | 1 | 5 | 1 | 1 | 1 | NA | NA | NA | NA | 1 | 1 | 2 | 1 |
| 5_1389836 | 1 | 1 | 1 | 1 | 1 | 1 | 3 | 0.3333 | 6 | 0.6667 | 1 | 1 | NA | NA | NA | NA | 2 | 0.5 | 3 | 0.6667 |
| 5_1389837 | NA | NA | 1 | 0 | 1 | 1 | 3 | 0 | 5 | 0.2 | 1 | 0 | NA | NA | NA | NA | 2 | 0.5 | 3 | 0.3333 |
| 5_1390031 | 79 | 0.3291 | 85 | 0.2824 | 74 | 0.2973 | 78 | 0.2692 | 316 | 0.2943 | 80 | 0.275 | 63 | 0.3016 | 63 | 0.2381 | 60 | 0.1833 | 266 | 0.2519 |
| 5_1390038 | 79 | 0.1013 | 81 | 0.1235 | 66 | 0.1061 | 62 | 0.0806 | 288 | 0.1042 | 64 | 0.0781 | 61 | 0.0328 | 61 | 0.0656 | 53 | 0.2075 | 239 | 0.0921 |
| 5_1390042 | 63 | 0.8571 | 62 | 0.8548 | 46 | 0.7609 | 53 | 0.7736 | 224 | 0.817 | 50 | 0.84 | 50 | 0.82 | 50 | 0.74 | 41 | 0.9024 | 191 | 0.822 |
| 5_1392343 | 7 | 0.4286 | 8 | 0.375 | 2 | 0 | 2 | 0.5 | 19 | 0.3684 | 3 | 0.6667 | 5 | 0.2 | 4 | 0.75 | 3 | 0.3333 | 15 | 0.4667 |
| 5_1392343 | 8 | 0.625 | 9 | 0.5556 | 2 | 0 | 4 | 0.25 | 23 | 0.4783 | 5 | 0.4 | 5 | 0.2 | 5 | 0.4 | 3 | 0.6667 | 18 | 0.3889 |
| 5_1392345 | 8 | 0.25 | 10 | 0.7 | NA | NA | 4 | 0.5 | 22 | 0.5 | 5 | 0.6 | 5 | 0.8 | 2 | 1 | 4 | 0.75 | 16 | 0.75 |
| 5_1392345 | 8 | 0 | 6 | 0.1667 | 1 | 0 | 5 | 0 | 20 | 0.05 | 5 | 0 | 4 | 0 | 2 | 0.5 | 4 | 0 | 15 | 0.0667 |
| 5_1392347 | 6 | 0.1667 | 8 | 0.375 | 1 | 0 | 4 | 0.5 | 19 | 0.3158 | 7 | 0.5714 | 5 | 0.4 | 3 | 0.3333 | 3 | 0 | 18 | 0.3889 |
| 5_1392347 | 7 | 0.1429 | 6 | 0.1667 | NA | NA | 3 | 0.6667 | 16 | 0.25 | 8 | 0.375 | 7 | 0 | 3 | 0.3333 | 3 | 0 | 21 | 0.1905 |
| 5_1392348 | 9 | 0 | 10 | 0.2 | 3 | 0 | 5 | 0.4 | 27 | 0.1481 | 7 | 0.1429 | 7 | 0.1429 | 4 | 0.25 | 6 | 0.5 | 24 | 0.25 |
| 5_1392548 | 12 | 0.1667 | 10 | 0.1 | 7 | 0.2857 | 11 | 0 | 40 | 0.125 | 5 | 0 | 10 | 0.2 | 7 | 0.1429 | 11 | 0.0909 | 33 | 0.1212 |
| 5_1392549 | 12 | 0.1667 | 15 | 0.0667 | 9 | 0 | 14 | 0 | 50 | 0.06 | 9 | 0 | 11 | 0.0909 | 9 | 0.2222 | 8 | 0 | 37 | 0.0811 |
| 5_1392551 | 6 | 0.3333 | 10 | 0.2 | 4 | 0 | 5 | 0.4 | 25 | 0.24 | 7 | 0.2857 | 10 | 0.3 | 5 | 0.4 | 7 | 0.4286 | 29 | 0.3448 |
| 5_1392552 | 6 | 0.1667 | 10 | 0.4 | 4 | 0 | 5 | 0.4 | 25 | 0.28 | 7 | 0 | 8 | 0.125 | 5 | 0 | 6 | 0.1667 | 26 | 0.0769 |
| 5_1392560 | 1 | 0 | 1 | 1 | 3 | 0 | 1 | 0 | 6 | 0.1667 | NA | NA | 2 | 0 | NA | NA | 2 | 0.5 | 4 | 0.25 |
| 5_1392577 | 30 | 0.2333 | 32 | 0.25 | 11 | 0.2727 | 23 | 0.2174 | 96 | 0.2396 | 16 | 0.1875 | 16 | 0.125 | 16 | 0.3125 | 11 | 0.2727 | 59 | 0.2203 |

| | | | | | | | | | | | | | | | | | | | | |
|---|---|---|---|---|---|---|---|---|---|---|---|---|---|---|---|---|---|---|---|---|
| 5_13925815 | 23 | 0.0435 | 27 | 0.2222 | 15 | 0.0667 | 21 | 0.1429 | 86 | 0.1279 | 13 | 0.1538 | 13 | 0.0769 | 14 | 0.2143 | 12 | 0.0833 | 52 | 0.1346 |
| 5_13925820 | 20 | 0.2 | 28 | 0.2857 | 15 | 0.0667 | 21 | 0.2857 | 84 | 0.2262 | 19 | 0.1053 | 16 | 0.125 | 13 | 0.3077 | 10 | 0.2 | 58 | 0.1724 |
| 5_13925837 | 22 | 0.9545 | 28 | 0.8214 | 13 | 0.6923 | 16 | 0.75 | 79 | 0.8228 | 14 | 0.7857 | 12 | 0.5833 | 12 | 0.8333 | 13 | 0.6154 | 51 | 0.7059 |
| 5_13925841 | 20 | 0.4 | 17 | 0.1765 | 11 | 0.4545 | 12 | 0.25 | 60 | 0.3167 | 12 | 0.25 | 9 | 0.3333 | 12 | 0.5 | 10 | 0.1 | 43 | 0.3023 |
| 5_13925842 | 18 | 0.5556 | 20 | 0.55 | 9 | 0.4444 | 12 | 0.3333 | 59 | 0.4915 | 6 | 0.8333 | 4 | 0.25 | 8 | 1 | 9 | 0.4444 | 27 | 0.6667 |
| 5_13925843 | 26 | 0.0769 | 25 | 0.12 | 12 | 0.0833 | 12 | 0.0833 | 75 | 0.0933 | 13 | 0.0769 | 10 | 0.1 | 14 | 0.0714 | 14 | 0.1429 | 51 | 0.098 |
| 5_13925845 | 22 | 0.8182 | 22 | 0.7727 | 12 | 0.5833 | 14 | 0.7143 | 70 | 0.7429 | 9 | 0.8889 | 9 | 0.8889 | 8 | 0.625 | 10 | 0.8 | 36 | 0.8056 |
| 5_13925846 | 24 | 0.2917 | 28 | 0.25 | 17 | 0.3529 | 18 | 0.1667 | 87 | 0.2644 | 13 | 0.4615 | 12 | 0.25 | 15 | 0.2667 | 15 | 0.4 | 55 | 0.3455 |
| 5_13925849 | 13 | 0.1538 | 10 | 0.4 | 9 | 0.2222 | 11 | 0.4545 | 43 | 0.3023 | 6 | 0.1667 | 6 | 0.1667 | 7 | 0.5714 | 9 | 0.3333 | 28 | 0.3214 |
| 5_13925850 | 23 | 0.1304 | 25 | 0.2 | 18 | 0.2222 | 16 | 0.1875 | 82 | 0.1829 | 16 | 0.125 | 12 | 0.1667 | 14 | 0.2857 | 14 | 0.2143 | 56 | 0.1964 |
| 5_13925861 | 25 | 0.32 | 27 | 0.2963 | 16 | 0.3125 | 19 | 0.4211 | 87 | 0.3333 | 15 | 0.4 | 12 | 0.1667 | 11 | 0.2727 | 11 | 0.0909 | 49 | 0.2449 |
| 5_13925864 | 23 | 0.1739 | 27 | 0.1111 | 16 | 0 | 19 | 0.1053 | 85 | 0.1059 | 16 | 0.1875 | 13 | 0.1538 | 16 | 0.0625 | 12 | 0.1667 | 57 | 0.1404 |
| 5_13925865 | 22 | 0.8636 | 22 | 0.7727 | 11 | 0.9091 | 17 | 0.7059 | 72 | 0.8056 | 13 | 0.9231 | 12 | 0.6667 | 15 | 0.8 | 10 | 0.6 | 50 | 0.76 |
| 5_13925874 | 18 | 0.8333 | 19 | 0.7895 | 10 | 0.7 | 14 | 0.8571 | 61 | 0.8033 | 13 | 0.6923 | 12 | 0.75 | 7 | 0.8571 | 10 | 0.7 | 42 | 0.7381 |
| 5_13925875 | 21 | 0.0952 | 23 | 0.2174 | 9 | 0 | 18 | 0.2222 | 71 | 0.1549 | 12 | 0.1667 | 14 | 0.0714 | 12 | 0 | 11 | 0.0909 | 49 | 0.0816 |
| 5_13932348 | 2 | 0.5 | 1 | 0 | 1 | 0 | 2 | 0 | 6 | 0.1667 | 1 | 1 | 3 | 0 | 2 | 0 | 2 | 0.5 | 8 | 0.25 |
| 5_13932351 | 1 | 1 | 1 | 0 | 1 | 1 | 3 | 0 | 6 | 0.3333 | 1 | 1 | 3 | 0 | 1 | 0 | 1 | 0 | 6 | 0.1667 |
| 5_13932354 | 1 | 1 | 1 | 0 | 1 | 0 | 2 | 0 | 5 | 0.2 | 1 | 1 | 3 | 0 | 2 | 0 | 3 | 0.3333 | 9 | 0.2222 |
| 5_13932357 | 1 | 1 | 2 | 0 | 1 | 1 | 4 | 0.25 | 8 | 0.375 | 2 | 0.5 | 3 | 1 | 3 | 0.6667 | 3 | 1 | 11 | 0.8182 |
| 5_13932360 | 5 | 0.2 | 5 | 0 | 5 | 0 | 3 | 0 | 18 | 0.0556 | 7 | 0 | 5 | 0.4 | 7 | 0 | 5 | 0 | 24 | 0.0833 |
| 5_13932364 | 7 | 1 | 6 | 0.8333 | 4 | 1 | 6 | 1 | 23 | 0.9565 | 8 | 1 | 5 | 0.8 | 12 | 0.75 | 4 | 1 | 29 | 0.8621 |
| 5_13932370 | 7 | 0.4286 | 8 | 0.25 | 4 | 0 | 5 | 0.4 | 24 | 0.2917 | 7 | 0.2857 | 7 | 0.1429 | 10 | 0 | 8 | 0 | 32 | 0.0938 |
| 5_13932380 | 6 | 0 | 10 | 0.1 | 5 | 0 | 5 | 0.2 | 26 | 0.0769 | 9 | 0.1111 | 7 | 0.1429 | 10 | 0.1 | 7 | 0.4286 | 33 | 0.1818 |
| 5_13932381 | 7 | 1 | 9 | 0.8889 | 5 | 0.8 | 4 | 0.75 | 25 | 0.88 | 8 | 0.875 | 7 | 1 | 9 | 0.8889 | 8 | 0.875 | 32 | 0.9062 |
| 5_13932382 | 9 | 0.3333 | 10 | 0.1 | 5 | 0.2 | 6 | 0.3333 | 30 | 0.2333 | 9 | 0.1111 | 7 | 0.4286 | 12 | 0.25 | 7 | 0.2857 | 35 | 0.2571 |
| 5_13932392 | 4 | 0 | 9 | 0.1111 | 4 | 0 | 5 | 0.2 | 22 | 0.0909 | 7 | 0.1429 | 4 | 0.25 | 8 | 0.5 | 5 | 0.4 | 24 | 0.3333 |
| 5_13932392 | 4 | 1 | 10 | 0.9 | 4 | 1 | 5 | 1 | 23 | 0.9565 | 9 | 0.8889 | 7 | 1 | 11 | 0.9091 | 5 | 1 | 32 | 0.9375 |
| 5_13932397 | 4 | 0 | 10 | 0 | 5 | 0.4 | 5 | 0 | 24 | 0.0833 | 7 | 0.1429 | 7 | 0.2857 | 10 | 0 | 4 | 0.25 | 28 | 0.1429 |
| 5_13932397 | 6 | 0 | 9 | 0 | 5 | 0.2 | 5 | 0.2 | 25 | 0.08 | 7 | 0 | 7 | 0.2857 | 12 | 0.25 | 6 | 0 | 32 | 0.1562 |
| 5_13932403 | 4 | 0.5 | 10 | 0.9 | 5 | 0.8 | 6 | 0.8333 | 25 | 0.8 | 7 | 0.7143 | 6 | 0.8333 | 12 | 0.6667 | 7 | 0.8571 | 32 | 0.75 |
| 5_13934572 | NA | NA | NA | NA | 1 | 1 | 1 | 0 | 2 | 0.5 | 5 | 0.6 | NA | NA | 3 | 0.6667 | 2 | 1 | 10 | 0.7 |
| 5_13934587 | NA | NA | 2 | 0.5 | 1 | 0 | 1 | 1 | 4 | 0.5 | 4 | 0 | NA | NA | 3 | 0.3333 | 2 | 0 | 9 | 0.1111 |
| 5_13934609 | NA | NA | 1 | 1 | 1 | 1 | NA | NA | 2 | 1 | 4 | 0.75 | NA | NA | 4 | 0.5 | 2 | 0.5 | 10 | 0.6 |
| 5_13937836 | 3 | 0.3333 | NA | NA | 1 | 0 | NA | NA | 4 | 0.25 | 2 | 0.5 | NA | NA | NA | NA | 3 | 0.3333 | 5 | 0.4 |
| 5_13937850 | 3 | 0.6667 | NA | NA | 2 | 0.5 | NA | NA | 5 | 0.6 | 2 | 1 | NA | NA | NA | NA | 3 | 0.6667 | 5 | 0.8 |
| 5_13937885 | 1 | 0 | NA | NA | 2 | 0.5 | NA | NA | 3 | 0.3333 | 3 | 0.3333 | NA | NA | 1 | 0 | 2 | 0.5 | 6 | 0.3333 |
| 5_13937886 | 2 | 0.5 | NA | NA | 2 | 0 | NA | NA | 4 | 0.25 | 1 | 0 | NA | NA | 1 | 1 | 3 | 0.3333 | 5 | 0.4 |
| 5_13937887 | 3 | 1 | NA | NA | 1 | 0 | NA | NA | 4 | 0.75 | 3 | 0.3333 | NA | NA | 1 | 1 | 3 | 0.6667 | 7 | 0.5714 |
| 5_13937893 | 3 | 1 | NA | NA | 2 | 1 | NA | NA | 5 | 1 | 3 | 1 | NA | NA | 1 | 1 | 2 | 0.5 | 6 | 0.8333 |
| 5_13937898 | 3 | 1 | NA | NA | 1 | 1 | NA | NA | 4 | 1 | 3 | 1 | NA | NA | 1 | 1 | 3 | 1 | 7 | 1 |
| 5_13937902 | NA | NA | NA | NA | 1 | 1 | NA | NA | 1 | 1 | 2 | 0.5 | NA | NA | 1 | 1 | 3 | 1 | 6 | 0.8333 |
| 5_13937906 | 3 | 0 | NA | NA | 2 | 0.5 | NA | NA | 5 | 0.2 | 2 | 0 | NA | NA | 1 | 0 | 3 | 0.3333 | 6 | 0.1667 |
| 5_13937923 | 3 | 0.6667 | NA | NA | 3 | 1 | NA | NA | 6 | 0.8333 | 3 | 0.3333 | NA | NA | 1 | 1 | 2 | 0.5 | 6 | 0.5 |
| 5_13937924 | 3 | 1 | NA | NA | 3 | 1 | NA | NA | 6 | 1 | 3 | 1 | NA | NA | 1 | 1 | 3 | 1 | 7 | 1 |
| 5_13937947 | 3 | 1 | NA | NA | 3 | 0.6667 | NA | NA | 6 | 0.8333 | 2 | 1 | NA | NA | 1 | 1 | 3 | 1 | 6 | 1 |
| 5_13938031 | 3 | 0.3333 | 1 | 0 | 2 | 0 | 1 | 0 | 7 | 0.1429 | 3 | 0.3333 | 3 | 0 | 1 | 0 | 3 | 0.3333 | 10 | 0.2 |
| 5_13938054 | 4 | 0.5 | 1 | 0 | 2 | 0.5 | 1 | 1 | 8 | 0.5 | 3 | 0 | 4 | 0.5 | 1 | 1 | 3 | 0.3333 | 11 | 0.3636 |
| 5_13938106 | 6 | 0.1667 | NA | NA | 1 | 0 | 2 | 0 | 9 | 0.1111 | 4 | 0.25 | 4 | 0 | 1 | 0 | 3 | 0 | 12 | 0.0833 |
| 5_13945704 | 9 | 0.1111 | 13 | 0.0769 | 5 | 0 | 4 | 0.5 | 31 | 0.129 | 11 | 0.0909 | 9 | 0.1111 | 7 | 0 | 7 | 0.1429 | 34 | 0.0882 |
| 5_13945792 | 7 | 0.5714 | 9 | 0.4444 | 3 | 1 | 2 | 0.5 | 21 | 0.5714 | 6 | 0.8333 | 5 | 1 | 4 | 0.5 | 5 | 0.8 | 20 | 0.8 |
| 5_13945795 | 6 | 0.3333 | 10 | 0.3 | 2 | 0 | 2 | 0 | 20 | 0.25 | 5 | 0 | 6 | 0.1667 | 4 | 0 | 5 | 0 | 20 | 0.05 |
| 5_13945797 | 6 | 0.3333 | 7 | 0.1429 | 3 | 0.6667 | 1 | 0 | 17 | 0.2941 | 6 | 0 | 6 | 0.1667 | 4 | 0 | 4 | 0.25 | 20 | 0.1 |
| 5_13945809 | 7 | 0 | 10 | 0.2 | 3 | 0 | 2 | 0 | 22 | 0.0909 | 5 | 0 | 6 | 0.1667 | 4 | 0.25 | 6 | 0 | 21 | 0.0952 |
| 5_13945833 | 6 | 0.3333 | 11 | 0.2727 | 3 | 0.3333 | 2 | 1 | 22 | 0.3636 | 4 | 0 | 6 | 0 | 3 | 0.3333 | 2 | 0 | 15 | 0.0667 |
| 5_13945835 | 6 | 0.8333 | 10 | 0.5 | 3 | 1 | 2 | 1 | 21 | 0.7143 | 6 | 0.6667 | 4 | 1 | 3 | 1 | 3 | 1 | 16 | 0.875 |
| 5_13945838 | 7 | 1 | 9 | 0.4444 | 3 | 1 | 2 | 1 | 21 | 0.7619 | 4 | 1 | 4 | 1 | 2 | 0.5 | 2 | 1 | 12 | 0.9167 |
| 5_13945845 | 7 | 1 | 9 | 1 | 2 | 1 | 1 | 1 | 19 | 1 | 4 | 1 | 6 | 0.8333 | 3 | 1 | 4 | 0.75 | 17 | 0.8824 |
| 5_13945848 | 4 | 0.25 | 11 | 0.1818 | 2 | 0 | 2 | 0 | 19 | 0.1579 | 6 | 0.3333 | 5 | 0 | 3 | 0 | 3 | 0.6667 | 17 | 0.2353 |
| 5_13945860 | 7 | 0 | 9 | 0.1111 | 1 | 0 | 1 | 0 | 18 | 0.0556 | 5 | 0.2 | 5 | 0 | 2 | 0 | 3 | 0 | 15 | 0.0667 |
| 5_13945870 | 5 | 1 | 8 | 0.875 | 2 | 1 | 2 | 1 | 17 | 0.9412 | 5 | 0.8 | 5 | 0.8 | 1 | 1 | 5 | 0.8 | 16 | 0.8125 |
| 5_13945875 | 7 | 0.7143 | 9 | 0.7778 | 2 | 0.5 | 2 | 1 | 20 | 0.75 | 6 | 0.8333 | 5 | 0.8 | 2 | 1 | 5 | 0.6 | 18 | 0.7778 |
| 5_13945893 | 5 | 0.4 | 9 | 0.1111 | 2 | 0 | 2 | 0 | 18 | 0.1667 | 5 | 0.4 | 5 | 0 | 3 | 0 | 3 | 0 | 16 | 0.125 |
| 5_13945894 | 4 | 1 | 7 | 0.8571 | 2 | 0.5 | NA | NA | 13 | 0.8462 | 6 | 0.8333 | 3 | 1 | 3 | 0.6667 | 3 | 1 | 15 | 0.8667 |
| 5_13945907 | 5 | 1 | 8 | 0.75 | 1 | 0 | 1 | 1 | 15 | 0.8 | 5 | 0.6 | 5 | 0.8 | 3 | 1 | 4 | 1 | 17 | 0.8235 |
| 5_13945908 | 3 | 0.3333 | 5 | 0.4 | 1 | 1 | 1 | 0 | 10 | 0.4 | 5 | 0.4 | 3 | 0.6667 | 2 | 0.5 | 3 | 0 | 13 | 0.3846 |

| | | | | | | | | | | | | | | | | | | | | |
|---|---|---|---|---|---|---|---|---|---|---|---|---|---|---|---|---|---|---|---|---|
| 5_1394591( | 5 | 0.8 | 8 | 0.875 | 2 | 1 | 1 | 1 | 16 | 0.875 | 6 | 0.8333 | 4 | 0.25 | 3 | 0.6667 | 3 | 1 | 16 | 0.6875 |
| 5_1394591( | 5 | 0.8 | 7 | 0.8571 | 1 | 1 | 1 | 1 | 14 | 0.8571 | 5 | 1 | 4 | 0.75 | 3 | 1 | 3 | 1 | 15 | 0.9333 |
| 5_1397372( | 7 | 0 | 8 | 0 | 1 | 0 | 9 | 0.2222 | 25 | 0.08 | 3 | 0.3333 | 11 | 0.0909 | 3 | 0.6667 | 6 | 0.1667 | 23 | 0.2174 |
| 5_1397372: | 6 | 0.5 | 7 | 0 | 1 | 0 | 8 | 0.25 | 22 | 0.2273 | 4 | 0 | 15 | 0.4667 | 7 | 0.2857 | 8 | 0.5 | 34 | 0.3824 |
| 5_1397406: | 4 | 0 | 6 | 0.1667 | 1 | 0 | 5 | 0 | 16 | 0.0625 | 2 | 0 | 10 | 0.2 | 2 | 0 | 7 | 0.1429 | 21 | 0.1429 |
| 5_1397859: | 1 | 0 | 2 | 0.5 | NA | NA | NA | NA | 3 | 0.3333 | 1 | 1 | NA | NA | 1 | 0 | 1 | 0 | 3 | 0.3333 |
| 5_1397860: | 1 | 0 | 1 | 1 | NA | NA | 1 | 0 | 3 | 0.3333 | 1 | 0 | NA | NA | 1 | 0 | 1 | 1 | 3 | 0.3333 |
| 5_1397861: | 1 | 0 | 2 | 0.5 | NA | NA | 1 | 1 | 4 | 0.5 | 1 | 0 | NA | NA | 1 | 0 | 1 | 1 | 3 | 0.3333 |
| 5_1397863: | 1 | 1 | 2 | 0 | NA | NA | 1 | 0 | 4 | 0.25 | 1 | 1 | NA | NA | 1 | 0 | 1 | 0 | 3 | 0.3333 |
| 5_1397864: | 1 | 1 | 1 | 1 | NA | NA | 1 | 1 | 3 | 1 | NA | NA | NA | NA | NA | NA | 1 | 1 | 1 | 1 |
| 5_1397865: | 2 | 0 | 1 | 0 | NA | NA | 1 | 1 | 4 | 0.25 | NA | NA | NA | NA | 1 | 0 | 1 | 1 | 2 | 0.5 |
| 5_1397865: | 2 | 1 | 2 | 0.5 | NA | NA | 1 | 0 | 5 | 0.6 | 1 | 0 | NA | NA | 1 | 1 | 1 | 0 | 3 | 0.3333 |
| 5_1397868: | 2 | 0 | 2 | 1 | NA | NA | 1 | 1 | 5 | 0.6 | 1 | 1 | 1 | 0 | 1 | 1 | 1 | 1 | 4 | 0.75 |
| 5_1397871: | 3 | 0.3333 | 1 | 1 | 1 | 1 | 1 | 0 | 6 | 0.5 | 1 | 1 | 3 | 0 | NA | NA | 1 | 0 | 5 | 0.2 |
| 5_1397872: | 3 | 0 | 3 | 0.3333 | 1 | 0 | 1 | 0 | 8 | 0.125 | NA | NA | 3 | 0 | 1 | 1 | 1 | 0 | 5 | 0.2 |
| 5_1397880: | 2 | 0.5 | 4 | 1 | 1 | 1 | 1 | 1 | 8 | 0.875 | 1 | 1 | 2 | 0.5 | 2 | 0 | 2 | 1 | 7 | 0.5714 |
| 5_1397881( | 3 | 0 | 4 | 0 | 2 | 0.5 | 1 | 0 | 10 | 0.1 | 1 | 1 | 2 | 0.5 | 2 | 0 | 2 | 0 | 7 | 0.2857 |
| 5_1397882: | 3 | 1 | 4 | 1 | 1 | 1 | 1 | 1 | 9 | 1 | 1 | 1 | 1 | 0 | 2 | 0.5 | 2 | 0.5 | 6 | 0.5 |
| 5_1398095: | 4 | 0.75 | 2 | 1 | 2 | 0.5 | 5 | 0.8 | 13 | 0.7692 | 2 | 1 | 5 | 1 | 4 | 0.75 | 1 | 1 | 12 | 0.9167 |
| 5_1399570: | NA | NA | 1 | 1 | NA | NA | NA | NA | 1 | 1 | 1 | 1 | NA | NA | NA | NA | NA | NA | 1 | 1 |
| 5_1399584: | NA | NA | 1 | 1 | NA | NA | 1 | 1 | 2 | 1 | 1 | 1 | NA | NA | 1 | 1 | NA | NA | 2 | 1 |
| 5_1399590: | NA | NA | 2 | 0.5 | NA | NA | 1 | 0 | 3 | 0.3333 | NA | NA | NA | NA | NA | NA | 1 | 1 | 1 | 1 |
| 5_1399594: | NA | NA | 3 | 0.6667 | 2 | 0 | 2 | 0.5 | 7 | 0.4286 | NA | NA | 1 | 0 | 1 | 1 | 1 | 0 | 3 | 0.3333 |
| 5_1399595: | NA | NA | 3 | 0 | 2 | 0.5 | 2 | 0.5 | 7 | 0.2857 | 1 | 1 | 1 | 0 | NA | NA | 1 | 0 | 3 | 0.3333 |
| 5_1399595: | NA | NA | 3 | 0.6667 | 2 | 1 | 2 | 1 | 7 | 0.8571 | 1 | 1 | 1 | 1 | 1 | 1 | 1 | 0 | 4 | 0.75 |
| 5_1399596( | NA | NA | 3 | 1 | 2 | 1 | 2 | 1 | 7 | 1 | 1 | 1 | NA | NA | 1 | 1 | 1 | 1 | 3 | 1 |
| 5_1399596( | NA | NA | 3 | 1 | 2 | 1 | 2 | 1 | 7 | 1 | 1 | 1 | 1 | 1 | NA | NA | 1 | 1 | 3 | 1 |
| 5_1399597: | NA | NA | 4 | 0 | 1 | 0 | 2 | 1 | 7 | 0.2857 | 1 | 0 | 1 | 1 | NA | NA | 1 | 0 | 3 | 0.3333 |
| 5_1399597: | NA | NA | 3 | 0 | 2 | 1 | 2 | 0 | 7 | 0.2857 | 1 | 0 | 1 | 0 | 2 | 0 | 1 | 1 | 5 | 0.2 |
| 5_1399599( | NA | NA | 3 | 0.3333 | 1 | 0 | 1 | 0 | 5 | 0.2 | 1 | 0 | 1 | 0 | 3 | 0.3333 | 2 | 0 | 7 | 0.1429 |
| 5_1399599: | NA | NA | 4 | 0.5 | 2 | 0 | 2 | 0 | 8 | 0.25 | NA | NA | NA | NA | 3 | 0.3333 | 2 | 0.5 | 5 | 0.4 |
| 5_1399602: | NA | NA | 4 | 0.25 | 2 | 0.5 | 2 | 0.5 | 8 | 0.375 | NA | NA | 2 | 0 | 5 | 0.2 | 2 | 0.5 | 9 | 0.2222 |
| 5_1399602( | NA | NA | 3 | 0 | 1 | 1 | 3 | 0.3333 | 7 | 0.2857 | 2 | 0.5 | 2 | 0 | 4 | 0.25 | 2 | 0.5 | 10 | 0.3 |
| 5_1399604: | NA | NA | 3 | 0.6667 | 3 | 0 | 4 | 0.25 | 10 | 0.3 | 2 | 0 | 3 | 1 | 5 | 0 | 1 | 0 | 11 | 0.2727 |
| 5_1399949: | 1 | 0 | 6 | 0 | 2 | 0.5 | 1 | 0 | 10 | 0.1 | 2 | 0 | 3 | 0 | 3 | 0.3333 | 3 | 0 | 11 | 0.0909 |
| 5_1399949: | 1 | 0 | 6 | 0 | 2 | 0.5 | 1 | 0 | 10 | 0.1 | 1 | 0 | 2 | 0 | 3 | 0.3333 | 2 | 0 | 8 | 0.125 |
| 5_1403135: | NA | NA | 2 | 1 | NA | NA | 2 | 1 | 4 | 1 | 1 | 1 | 4 | 1 | 1 | 1 | 2 | 1 | 8 | 1 |
| 5_1403136: | NA | NA | NA | NA | NA | NA | 4 | 0.75 | 4 | 0.75 | 2 | 1 | 5 | 1 | 1 | 1 | 2 | 1 | 10 | 1 |
| 5_1403137: | NA | NA | 2 | 1 | NA | NA | 2 | 1 | 4 | 1 | 2 | 1 | 3 | 1 | 1 | 1 | 2 | 1 | 8 | 1 |
| 5_1403137: | NA | NA | 2 | 0 | NA | NA | 4 | 0.5 | 6 | 0.3333 | 2 | 0.5 | 4 | 0.5 | 1 | 0 | 2 | 0 | 9 | 0.3333 |
| 5_1403137: | NA | NA | 2 | 1 | NA | NA | 3 | 1 | 5 | 1 | 2 | 1 | 3 | 1 | 1 | 1 | 2 | 1 | 8 | 1 |
| 5_1403138: | NA | NA | NA | NA | NA | NA | 2 | 1 | 2 | 1 | 1 | 1 | 2 | 1 | NA | NA | NA | NA | 3 | 1 |
| 5_1403138: | NA | NA | 1 | 1 | NA | NA | 4 | 1 | 5 | 1 | 2 | 0.5 | 3 | 0.3333 | 1 | 1 | 2 | 0.5 | 8 | 0.5 |
| 5_1403139( | NA | NA | 1 | 0 | NA | NA | 4 | 0.5 | 5 | 0.4 | 2 | 0 | 4 | 0.5 | NA | NA | 2 | 0.5 | 8 | 0.375 |
| 5_1403142: | NA | NA | 2 | 1 | NA | NA | 3 | 1 | 5 | 1 | 2 | 1 | 2 | 1 | 1 | 0 | 1 | 1 | 6 | 0.8333 |
| 5_1403142( | NA | NA | 2 | 0.5 | NA | NA | 3 | 1 | 5 | 0.8 | 2 | 1 | 4 | 0.75 | 1 | 1 | 2 | 1 | 9 | 0.8889 |
| 5_1403142: | NA | NA | 2 | 0.5 | NA | NA | 3 | 0.3333 | 5 | 0.4 | 2 | 0 | 3 | 0.3333 | 1 | 1 | 2 | 1 | 8 | 0.5 |
| 5_1403142: | NA | NA | 2 | 0 | NA | NA | 2 | 1 | 4 | 0.5 | 2 | 0.5 | 1 | 1 | 1 | 1 | 2 | 0.5 | 6 | 0.6667 |
| 5_1403144: | NA | NA | 1 | 1 | NA | NA | 3 | 0.3333 | 4 | 0.5 | 2 | 0.5 | 3 | 0 | NA | NA | 2 | 1 | 7 | 0.4286 |
| 5_1403168( | NA | NA | NA | NA | NA | NA | 1 | 1 | 1 | 1 | NA | NA | 2 | 0 | 2 | 1 | 1 | 1 | 5 | 0.6 |
| 5_1403235: | NA | NA | 1 | 0 | 2 | 1 | NA | NA | 3 | 0.6667 | 1 | 1 | 1 | 1 | NA | NA | NA | NA | 2 | 1 |
| 5_1403235: | NA | NA | 1 | 1 | 2 | 1 | NA | NA | 3 | 1 | NA | NA | 1 | 1 | NA | NA | NA | NA | 1 | 1 |
| 5_1403237: | NA | NA | NA | NA | 1 | 1 | NA | NA | 1 | 1 | 1 | 0 | 1 | 1 | NA | NA | NA | NA | 2 | 0.5 |
| 5_1403238: | NA | NA | 1 | 1 | 1 | 1 | NA | NA | 2 | 1 | 1 | 1 | 1 | 1 | NA | NA | NA | NA | 2 | 1 |
| 5_1403261( | 3 | 1 | 1 | 0 | 3 | 0.6667 | 1 | 0 | 8 | 0.625 | 1 | 0 | NA | NA | 1 | 1 | 1 | 1 | 3 | 0.6667 |
| 5_1403262: | NA | NA | 1 | 0 | 2 | 0.5 | NA | NA | 3 | 0.3333 | 2 | 0.5 | NA | NA | NA | NA | 1 | 1 | 3 | 0.6667 |
| 5_1403264: | 2 | 0.5 | NA | NA | 1 | 1 | 1 | 0 | 4 | 0.5 | 1 | 0 | NA | NA | 1 | 1 | 1 | 1 | 3 | 0.6667 |
| 5_1403270( | 2 | 0.5 | 1 | 1 | NA | NA | 1 | 1 | 4 | 0.75 | 2 | 1 | NA | NA | NA | NA | NA | NA | 2 | 1 |
| 5_1404287: | 12 | 0.25 | 13 | 0.3846 | 7 | 0.1429 | 10 | 0.4 | 42 | 0.3095 | 8 | 0 | 16 | 0.5 | 10 | 0.5 | 4 | 0.5 | 38 | 0.3947 |
| 5_1404287: | 14 | 0.3571 | 18 | 0.3889 | 11 | 0.0909 | 14 | 0.0714 | 57 | 0.2456 | 11 | 0.2727 | 17 | 0.4706 | 14 | 0.3571 | 12 | 0.3333 | 54 | 0.3704 |
| 5_1404289: | 14 | 0.7857 | 16 | 0.9375 | 11 | 1 | 13 | 0.8462 | 54 | 0.8889 | 11 | 1 | 19 | 0.9474 | 15 | 1 | 7 | 0.7143 | 52 | 0.9423 |
| 5_1404290: | 17 | 0.2353 | 13 | 0.3077 | 11 | 0.0909 | 15 | 0.3333 | 56 | 0.25 | 10 | 0.5 | 15 | 0.1333 | 13 | 0.2308 | 12 | 0.3333 | 50 | 0.28 |
| 5_1405094: | 1 | 1 | NA | NA | 2 | 1 | 2 | 1 | 5 | 1 | 1 | 1 | 1 | 1 | NA | NA | 3 | 1 | 5 | 1 |
| 5_1405095: | 2 | 0.5 | 1 | 0 | 3 | 0.3333 | 2 | 0 | 8 | 0.25 | 1 | 0 | 1 | 0 | NA | NA | 3 | 0.3333 | 5 | 0.2 |
| 5_1405097: | 3 | 1 | 3 | 1 | 3 | 0.3333 | 2 | 1 | 11 | 0.8182 | 2 | 1 | 1 | 1 | 1 | 1 | 4 | 0.75 | 8 | 0.875 |

| | | | | | | | | | | | | | | | | | | | | |
|---|---|---|---|---|---|---|---|---|---|---|---|---|---|---|---|---|---|---|---|---|
| 5_1405098 | 4 | 0.25 | 3 | 0.6667 | 3 | 0.3333 | 2 | 0.5 | 12 | 0.4167 | 2 | 0.5 | NA | NA | NA | NA | 4 | 0.5 | 6 | 0.5 |
| 5_1405098 | 2 | 1 | 2 | 0.5 | 2 | 1 | 1 | 0 | 7 | 0.7143 | 2 | 0.5 | 1 | 0 | 1 | 1 | 4 | 0.5 | 8 | 0.5 |
| 5_1405099 | 2 | 0.5 | 1 | 0 | 3 | 0 | 1 | 1 | 7 | 0.2857 | 2 | 0.5 | 1 | 0 | 1 | 1 | 4 | 0 | 8 | 0.25 |
| 5_1405103 | 4 | 0.25 | 3 | 0 | 4 | 0 | 2 | 0 | 13 | 0.0769 | 1 | 0 | 1 | 0 | 1 | 1 | 3 | 0.3333 | 6 | 0.3333 |
| 5_1405103 | 4 | 1 | 4 | 0.75 | 4 | 0.5 | 1 | 1 | 13 | 0.7692 | 2 | 1 | 1 | 1 | 1 | 1 | 5 | 1 | 9 | 1 |
| 5_1405104 | 4 | 0.5 | 4 | 0.75 | 4 | 0.25 | 2 | 0 | 14 | 0.4286 | 2 | 0 | 1 | 0 | NA | NA | 4 | 0.25 | 7 | 0.1429 |
| 5_1405104 | 4 | 0.25 | 1 | 0 | 3 | 0 | 2 | 0 | 10 | 0.1 | 1 | 0 | 1 | 0 | NA | NA | 4 | 0.25 | 6 | 0.1667 |
| 5_1405105 | 4 | 0.5 | 3 | 0.6667 | 4 | 0 | 1 | 1 | 12 | 0.4167 | 2 | 0.5 | 1 | 1 | 1 | 1 | 3 | 0.6667 | 7 | 0.7143 |
| 5_1405105 | 4 | 1 | 4 | 1 | 4 | 1 | 2 | 0.5 | 14 | 0.9286 | 2 | 1 | 1 | 1 | 1 | 1 | 5 | 1 | 9 | 1 |
| 5_1405106 | 4 | 1 | 3 | 1 | 4 | 1 | 1 | 1 | 12 | 1 | 2 | 0.5 | 1 | 1 | 1 | 1 | 5 | 1 | 9 | 0.8889 |
| 5_1405107 | 3 | 1 | 4 | 0.75 | 4 | 0.75 | 2 | 1 | 13 | 0.8462 | 1 | 1 | 1 | 1 | 1 | 1 | 6 | 1 | 9 | 1 |
| 5_1405107 | 3 | 0.3333 | 4 | 0 | 3 | 0 | 1 | 0 | 11 | 0.0909 | 2 | 0 | NA | NA | 1 | 0 | 5 | 0.2 | 8 | 0.125 |
| 5_1405108 | 4 | 0 | 4 | 0.25 | 4 | 0.25 | 2 | 0 | 14 | 0.1429 | 2 | 0 | 1 | 0 | 1 | 1 | 6 | 0 | 10 | 0.1 |
| 5_1405109 | 5 | 1 | 3 | 1 | 5 | 0.8 | 2 | 0.5 | 15 | 0.8667 | 2 | 0.5 | 1 | 1 | 1 | 1 | 4 | 0.5 | 8 | 0.625 |
| 5_1406008 | 4 | 0 | 7 | 0.1429 | 4 | 0 | 5 | 0 | 20 | 0.05 | 4 | 0 | 4 | 0 | 6 | 0.1667 | 5 | 0 | 19 | 0.0526 |
| 5_1406065 | 36 | 0.3611 | 26 | 0.3077 | 30 | 0.4667 | 35 | 0.4857 | 127 | 0.4094 | 27 | 0.4444 | 26 | 0.5385 | 33 | 0.3333 | 24 | 0.5417 | 110 | 0.4545 |
| 5_1406068 | 37 | 0.1892 | 28 | 0.25 | 28 | 0.1429 | 34 | 0.2647 | 127 | 0.2126 | 27 | 0.1852 | 24 | 0.25 | 29 | 0.2759 | 23 | 0.2174 | 103 | 0.233 |
| 5_1406072 | 30 | 0.2333 | 22 | 0.1364 | 22 | 0.3182 | 30 | 0.3333 | 104 | 0.2596 | 23 | 0.2174 | 22 | 0.2727 | 22 | 0.2273 | 16 | 0.125 | 83 | 0.2169 |
| 5_1406076 | 28 | 0.5 | 18 | 0.5 | 17 | 0.4706 | 29 | 0.5517 | 92 | 0.5109 | 19 | 0.5263 | 18 | 0.3333 | 20 | 0.4 | 16 | 0.4375 | 73 | 0.4247 |
| 5_1406082 | 24 | 0.5833 | 17 | 0.8235 | 11 | 0.4545 | 22 | 0.5909 | 74 | 0.6216 | 19 | 0.5263 | 16 | 0.75 | 17 | 0.5882 | 15 | 0.6 | 67 | 0.6119 |
| 5_1406082 | 24 | 0.125 | 14 | 0.1429 | 16 | 0 | 22 | 0.0909 | 76 | 0.0921 | 18 | 0.2778 | 16 | 0.1875 | 15 | 0.0667 | 13 | 0 | 62 | 0.1452 |
| 5_1406083 | 24 | 0.0833 | 16 | 0.125 | 15 | 0.1333 | 23 | 0.087 | 78 | 0.1026 | 21 | 0.1905 | 18 | 0.1111 | 15 | 0.1333 | 14 | 0.2143 | 68 | 0.1618 |
| 5_1406083 | 28 | 0.2143 | 13 | 0.3846 | 15 | 0.0667 | 21 | 0.2857 | 77 | 0.2338 | 20 | 0.2 | 19 | 0.3684 | 15 | 0.4667 | 15 | 0.2667 | 69 | 0.3188 |
| 5_1406083 | 24 | 0.375 | 10 | 0.4 | 11 | 0.1818 | 18 | 0.3333 | 63 | 0.3333 | 19 | 0.3684 | 16 | 0.3125 | 13 | 0.2308 | 13 | 0.2308 | 61 | 0.2951 |
| 5_1406085 | 20 | 0.25 | 11 | 0.3636 | 13 | 0.1538 | 22 | 0.1818 | 66 | 0.2273 | 17 | 0.1176 | 14 | 0.4286 | 13 | 0 | 9 | 0.3333 | 53 | 0.2075 |
| 5_1406087 | 24 | 0.125 | 13 | 0 | 14 | 0 | 17 | 0.1765 | 68 | 0.0882 | 19 | 0.0526 | 15 | 0.0667 | 9 | 0 | 10 | 0.2 | 53 | 0.0755 |
| 5_1406089 | 12 | 1 | 9 | 0.8889 | 8 | 0.75 | 13 | 0.9231 | 42 | 0.9048 | 11 | 0.8182 | 9 | 0.8889 | 7 | 0.7143 | 8 | 1 | 35 | 0.8571 |
| 5_1406091 | 18 | 0.1111 | 8 | 0.375 | 11 | 0.0909 | 17 | 0.2353 | 54 | 0.1852 | 18 | 0 | 12 | 0 | 9 | 0.2222 | 8 | 0.125 | 47 | 0.0638 |
| 5_1406218 | 1 | 1 | 1 | 0 | NA | NA | 1 | 0 | 3 | 0.3333 | NA | NA | 1 | 0 | 1 | 1 | NA | NA | 2 | 0.5 |
| 5_1406477 | 1 | 1 | NA | NA | NA | NA | NA | NA | 1 | 1 | NA | NA | 1 | 1 | NA | NA | NA | NA | 1 | 1 |
| 5_1406501 | NA | NA | 3 | 0.3333 | 2 | 0 | 1 | 0 | 6 | 0.1667 | 3 | 0 | 2 | 0 | 1 | 1 | 3 | 0.6667 | 9 | 0.3333 |
| 5_1406510 | NA | NA | 1 | 1 | 1 | 1 | NA | NA | 2 | 1 | 2 | 1 | 1 | 1 | 1 | 1 | NA | NA | 4 | 1 |
| 5_1407040 | 1 | 0 | 3 | 0 | 1 | 1 | 1 | 1 | 6 | 0.3333 | NA | NA | NA | NA | 1 | 1 | NA | NA | 1 | 1 |
| 5_1407040 | NA | NA | 4 | 0 | 1 | 1 | 1 | 1 | 6 | 0.3333 | NA | NA | NA | NA | 1 | 1 | NA | NA | 1 | 1 |
| 5_1407045 | 1 | 0 | 5 | 0.6 | 1 | 1 | 2 | 1 | 9 | 0.6667 | NA | NA | NA | NA | 2 | 0.5 | NA | NA | 2 | 0.5 |
| 5_1407048 | 2 | 0.5 | 4 | 0.25 | 1 | 0 | 2 | 0 | 9 | 0.2222 | NA | NA | NA | NA | 1 | 1 | NA | NA | 1 | 1 |
| 5_1407271 | 1 | 0 | NA | NA | 1 | 0 | 2 | 1 | 4 | 0.5 | NA | NA | 1 | 1 | 2 | 0.5 | NA | NA | 3 | 0.6667 |
| 5_1407271 | NA | NA | NA | NA | 1 | 0 | 1 | 1 | 2 | 0.5 | NA | NA | 1 | 1 | 2 | 1 | NA | NA | 3 | 1 |
| 5_1407377 | 1 | 1 | NA | NA | 3 | 0.3333 | 3 | 0.3333 | 7 | 0.4286 | NA | NA | 1 | 0 | 2 | 0.5 | NA | NA | 3 | 0.3333 |
| 5_1407378 | 1 | 1 | NA | NA | 3 | 0.6667 | 4 | 0.75 | 8 | 0.75 | NA | NA | 1 | 1 | 2 | 1 | NA | NA | 3 | 1 |
| 5_1407379 | 2 | 1 | 1 | 0 | 3 | 0.6667 | 4 | 0.75 | 10 | 0.7 | NA | NA | 1 | 0 | 2 | 1 | NA | NA | 3 | 0.6667 |
| 5_1407380 | 2 | 0.5 | 1 | 0 | 3 | 0 | 3 | 0 | 9 | 0.1111 | NA | NA | 1 | 0 | 2 | 0.5 | NA | NA | 3 | 0.3333 |
| 5_1407381 | 2 | 1 | 1 | 1 | 3 | 1 | 3 | 0.6667 | 9 | 0.8889 | NA | NA | 1 | 1 | 2 | 1 | NA | NA | 3 | 1 |
| 5_1407382 | 1 | 1 | 1 | 0 | 3 | 1 | 4 | 1 | 9 | 0.8889 | NA | NA | 1 | 1 | 1 | 0 | NA | NA | 2 | 0.5 |
| 5_1407383 | 1 | 1 | 1 | 1 | 2 | 1 | 4 | 0.75 | 8 | 0.875 | 1 | 1 | 1 | 1 | NA | NA | NA | NA | 2 | 1 |
| 5_1407386 | 1 | 0 | NA | NA | 2 | 0 | 2 | 1 | 5 | 0.4 | 1 | 0 | 1 | 1 | 2 | 1 | NA | NA | 4 | 0.75 |
| 5_1407388 | 1 | 1 | NA | NA | 3 | 0.6667 | 4 | 1 | 8 | 0.875 | NA | NA | 1 | 1 | 2 | 0.5 | NA | NA | 3 | 0.6667 |
| 5_1407445 | 8 | 0.75 | 14 | 0.6429 | 12 | 0.8333 | 9 | 0.7778 | 43 | 0.7442 | 14 | 0.8571 | 5 | 0.8 | 8 | 0.75 | 7 | 0.7143 | 34 | 0.7941 |
| 5_1407445 | 9 | 0.5556 | 15 | 0.7333 | 14 | 0.6429 | 11 | 0.3636 | 49 | 0.5918 | 17 | 0.2353 | 8 | 0.625 | 12 | 0.6667 | 10 | 0.4 | 47 | 0.4468 |
| 5_1407446 | 5 | 0.2 | 12 | 0 | 11 | 0.1818 | 7 | 0.4286 | 35 | 0.1714 | 12 | 0.3333 | 5 | 0.2 | 11 | 0.1818 | 7 | 0.2857 | 35 | 0.2571 |
| 5_1407447 | 9 | 0 | 14 | 0.0714 | 14 | 0.1429 | 11 | 0 | 48 | 0.0625 | 17 | 0.0588 | 8 | 0.25 | 11 | 0 | 11 | 0 | 47 | 0.0638 |
| 5_1407448 | 10 | 0.1 | 14 | 0.0714 | 14 | 0.0714 | 11 | 0 | 49 | 0.0612 | 15 | 0 | 6 | 0 | 11 | 0.0909 | 11 | 0.1818 | 43 | 0.0698 |
| 5_1407449 | 9 | 0.2222 | 13 | 0.1538 | 14 | 0.2857 | 10 | 0.1 | 46 | 0.1957 | 15 | 0.1333 | 5 | 0 | 10 | 0.2 | 11 | 0.1818 | 41 | 0.1463 |
| 5_1407451 | 8 | 0.75 | 11 | 0.4545 | 11 | 0.5455 | 6 | 0.1667 | 36 | 0.5 | 12 | 0.5833 | 3 | 0.6667 | 7 | 0.4286 | 9 | 0.7778 | 31 | 0.6129 |
| 5_1407452 | 6 | 0 | 11 | 0.0909 | 12 | 0.4167 | 8 | 0.125 | 37 | 0.1892 | 15 | 0.2667 | 3 | 0 | 10 | 0.2 | 9 | 0.2222 | 37 | 0.2162 |
| 5_1407452 | 1 | 1 | NA | NA | NA | NA | NA | NA | 1 | 1 | NA | NA | 1 | 1 | NA | NA | NA | NA | 1 | 1 |
| 5_1407454 | 8 | 0 | 9 | 0 | 14 | 0.1429 | 5 | 0 | 36 | 0.0556 | 13 | 0.0769 | 4 | 0 | 8 | 0.375 | 7 | 0 | 32 | 0.125 |
| 5_1407455 | 8 | 0.5 | 10 | 0.1 | 14 | 0.2143 | 7 | 0.1429 | 39 | 0.2308 | 13 | 0.1538 | 3 | 0 | 9 | 0 | 6 | 0.1667 | 31 | 0.0968 |
| 5_1407456 | 8 | 0 | 11 | 0.0909 | 14 | 0.1429 | 8 | 0 | 41 | 0.0732 | 15 | 0.0667 | 3 | 0.3333 | 9 | 0.2222 | 6 | 0 | 33 | 0.1212 |
| 5_1407459 | 6 | 0 | 9 | 0.1111 | 11 | 0.3636 | 7 | 0 | 33 | 0.1515 | 13 | 0.2308 | 2 | 0 | 8 | 0.375 | 4 | 0.25 | 27 | 0.2593 |
| 5_1407460 | 8 | 0.75 | 9 | 0.5556 | 10 | 0.5 | 7 | 0.4286 | 34 | 0.5588 | 11 | 0.8182 | 4 | 0.5 | 9 | 0.6667 | 5 | 1 | 29 | 0.7586 |
| 5_1407460 | 6 | 1 | 10 | 0.8 | 10 | 0.6 | 4 | 0.5 | 30 | 0.7333 | 12 | 0.8333 | 4 | 0.75 | 9 | 0.6667 | 4 | 1 | 29 | 0.7931 |
| 5_1407460 | 8 | 0.375 | 10 | 0.2 | 11 | 0.2727 | 5 | 0.4 | 34 | 0.2941 | 14 | 0.4286 | 4 | 0.25 | 8 | 0.375 | 5 | 0.6 | 31 | 0.4194 |
| 5_1407460 | 8 | 1 | 10 | 1 | 12 | 1 | 7 | 1 | 37 | 1 | 13 | 1 | 3 | 1 | 9 | 1 | 5 | 1 | 30 | 1 |
| 5_1407461 | 8 | 0.625 | 9 | 0.6667 | 9 | 0.8889 | 5 | 1 | 31 | 0.7742 | 12 | 0.75 | 3 | 0.3333 | 6 | 0.8333 | 4 | 1 | 25 | 0.76 |

| | | | | | | | | | | | | | | | | | | | | |
|---|---|---|---|---|---|---|---|---|---|---|---|---|---|---|---|---|---|---|---|---|
| 5_1407462 | 6 | 1 | 8 | 1 | 10 | 0.9 | 7 | 1 | 31 | 0.9677 | 12 | 1 | 4 | 1 | 8 | 1 | 4 | 1 | 28 | 1 |
| 5_1407463 | 5 | 0.2 | 9 | 0.3333 | 8 | 0.25 | 6 | 0.5 | 28 | 0.3214 | 9 | 0.1111 | 2 | 0 | 7 | 0.4286 | 2 | 0.5 | 20 | 0.25 |
| 5_1407463 | 7 | 0.5714 | 7 | 0.5714 | 7 | 0.5714 | 7 | 0.8571 | 28 | 0.6429 | 13 | 0.8462 | 3 | 1 | 9 | 0.7778 | 4 | 0.75 | 29 | 0.8276 |
| 5_1407463 | 8 | 0.25 | 9 | 0.1111 | 10 | 0.1 | 7 | 0 | 34 | 0.1176 | 12 | 0.0833 | 4 | 0 | 6 | 0.3333 | 4 | 0 | 26 | 0.1154 |
| 5_1407463 | 7 | 0.7143 | 9 | 0.8889 | 10 | 0.8 | 6 | 0.8333 | 32 | 0.8125 | 12 | 1 | 4 | 1 | 8 | 0.625 | 5 | 1 | 29 | 0.8966 |
| 5_1407463 | 6 | 0.3333 | 9 | 0 | 9 | 0.2222 | 7 | 0.2857 | 31 | 0.1935 | 11 | 0.0909 | 4 | 0 | 8 | 0.125 | 4 | 0.5 | 27 | 0.1481 |
| 5_1407465 | 7 | 0.4286 | 8 | 0.375 | 6 | 0.5 | 7 | 0.4286 | 28 | 0.4286 | 11 | 0.3636 | 3 | 0.3333 | 6 | 0.1667 | 4 | 1 | 24 | 0.4167 |
| 5_1407466 | 7 | 0.1429 | 5 | 0.4 | 8 | 0 | 6 | 0.1667 | 26 | 0.1538 | 9 | 0.2222 | 3 | 0.3333 | 8 | 0.125 | 5 | 0.2 | 25 | 0.2 |
| 5_1407466 | 1 | 1 | NA | NA | NA | NA | 1 | 1 | 2 | 1 | NA | NA | 1 | 1 | NA | NA | NA | NA | 1 | 1 |
| 5_1424425 | 2 | 1 | 1 | 1 | NA | NA | 1 | 1 | 4 | 1 | 1 | 0 | 1 | 1 | NA | NA | 1 | 0 | 3 | 0.3333 |
| 5_1424426 | 1 | 1 | 1 | 1 | NA | NA | 1 | 1 | 3 | 1 | 1 | 1 | 1 | 1 | NA | NA | 1 | 1 | 3 | 1 |
| 5_1424428 | 1 | 1 | 1 | 0 | NA | NA | NA | NA | 2 | 0.5 | 1 | 1 | 1 | 1 | NA | NA | NA | NA | 2 | 1 |
| 5_1424429 | 1 | 1 | 1 | 1 | NA | NA | NA | NA | 2 | 1 | 1 | 1 | NA | NA | NA | NA | 1 | 1 | 2 | 1 |
| 5_1424429 | NA | NA | 1 | 1 | NA | NA | NA | NA | 1 | 1 | 1 | 0 | 1 | 1 | NA | NA | NA | NA | 2 | 0.5 |
| 5_1424430 | 1 | 0 | 1 | 1 | NA | NA | 1 | 1 | 3 | 0.6667 | 1 | 0 | 1 | 0 | NA | NA | 1 | 1 | 3 | 0.3333 |
| 5_1424430 | 1 | 1 | 1 | 1 | NA | NA | 1 | 1 | 3 | 1 | 1 | 1 | NA | NA | NA | NA | 1 | 1 | 2 | 1 |
| 5_1424430 | 1 | 1 | 1 | 1 | NA | NA | 1 | 1 | 3 | 1 | 1 | 1 | 1 | 0 | NA | NA | 1 | 1 | 3 | 0.6667 |
| 5_1424435 | 1 | 1 | 2 | 0.5 | NA | NA | 1 | 0 | 4 | 0.5 | 1 | 0 | 1 | 1 | NA | NA | 1 | 1 | 3 | 0.6667 |
| 5_1424435 | 1 | 0 | 2 | 0.5 | NA | NA | 1 | 1 | 4 | 0.5 | 1 | 1 | NA | NA | NA | NA | 1 | 1 | 2 | 1 |
| 5_1424436 | 1 | 1 | 2 | 1 | NA | NA | 1 | 1 | 4 | 1 | 1 | 1 | 1 | 1 | NA | NA | 1 | 1 | 3 | 1 |
| 5_1424436 | 1 | 1 | 2 | 0 | NA | NA | 1 | 1 | 4 | 0.5 | 1 | 0 | 1 | 0 | NA | NA | 1 | 1 | 3 | 0.3333 |
| 5_1424438 | 1 | 1 | 2 | 1 | NA | NA | 1 | 1 | 4 | 1 | 1 | 1 | 1 | 1 | 1 | 1 | 2 | 1 | 5 | 1 |
| 5_1424438 | 2 | 1 | 1 | 1 | NA | NA | NA | NA | 3 | 1 | 1 | 1 | 1 | 1 | 1 | 1 | 1 | 1 | 4 | 1 |
| 5_1424440 | 2 | 1 | 3 | 1 | NA | NA | 1 | 1 | 6 | 1 | 1 | 0 | 1 | 1 | NA | NA | 2 | 0.5 | 4 | 0.5 |
| 5_1424440 | 2 | 0.5 | 3 | 1 | NA | NA | NA | NA | 5 | 0.8 | 1 | 1 | 1 | 1 | 1 | 1 | 2 | 1 | 5 | 1 |
| 5_1424441 | 2 | 0.5 | 2 | 0.5 | 1 | 0 | 1 | 1 | 6 | 0.5 | 1 | 0 | 1 | 0 | 1 | 1 | 1 | 1 | 4 | 0.5 |
| 5_1424443 | 2 | 1 | 2 | 0 | 1 | 0 | 1 | 0 | 6 | 0.3333 | NA | NA | 1 | 1 | 1 | 0 | 2 | 0 | 4 | 0.25 |
| 5_1424476 | 4 | 0 | 8 | 0.125 | 4 | 0.25 | 4 | 0 | 20 | 0.1 | 3 | 0.3333 | 3 | 0 | 3 | 0.3333 | 3 | 0 | 12 | 0.1667 |
| 5_1424631 | 2 | 0.5 | 3 | 0 | NA | NA | 2 | 0 | 7 | 0.1429 | 3 | 0.3333 | 1 | 0 | NA | NA | NA | NA | 4 | 0.25 |
| 5_1424632 | 4 | 0.75 | 1 | 1 | NA | NA | 2 | 0.5 | 7 | 0.7143 | 3 | 1 | 1 | 1 | 1 | 0 | NA | NA | 5 | 0.8 |
| 5_1424633 | 4 | 0 | 4 | 0.25 | NA | NA | 3 | 0 | 11 | 0.0909 | 3 | 0 | 1 | 1 | 1 | 1 | NA | NA | 5 | 0.4 |
| 5_1424633 | 3 | 0.6667 | 4 | 0.5 | NA | NA | 2 | 0.5 | 9 | 0.5556 | 4 | 0.75 | 1 | 1 | 1 | 1 | NA | NA | 6 | 0.8333 |
| 5_1424633 | 4 | 0 | 4 | 0.5 | NA | NA | 1 | 0 | 9 | 0.2222 | 2 | 0.5 | 1 | 1 | NA | NA | NA | NA | 3 | 0.6667 |
| 5_1424634 | 3 | 0 | 4 | 0.5 | NA | NA | 3 | 0 | 10 | 0.2 | 4 | 0.25 | 1 | 0 | 2 | 0.5 | NA | NA | 7 | 0.2857 |
| 5_1424640 | 1 | 0 | 2 | 0.5 | NA | NA | 1 | 0 | 4 | 0.25 | 2 | 0 | NA | NA | 2 | 0.5 | NA | NA | 4 | 0.25 |
| 5_1424642 | 1 | 0 | 2 | 0.5 | NA | NA | 1 | 0 | 4 | 0.25 | 2 | 0.5 | NA | NA | 1 | 1 | NA | NA | 3 | 0.6667 |
| 5_1424708 | NA | NA | 5 | 0 | 3 | 0.3333 | NA | NA | 8 | 0.125 | 1 | 1 | 2 | 0 | 1 | 0 | 5 | 0.2 | 9 | 0.2222 |
| 5_1424925 | NA | NA | 1 | 1 | NA | NA | NA | NA | 1 | 1 | 1 | 1 | 1 | 1 | 1 | 1 | 1 | 1 | 4 | 1 |
| 5_1425293 | NA | NA | 1 | 1 | NA | NA | NA | NA | 1 | 1 | 1 | 0 | 1 | 1 | NA | NA | 1 | 1 | 3 | 0.6667 |
| 5_1425293 | NA | NA | 1 | 1 | NA | NA | NA | NA | 1 | 1 | 1 | 1 | 1 | 1 | 1 | 1 | 1 | 1 | 4 | 1 |
| 5_1425294 | NA | NA | 1 | 1 | NA | NA | NA | NA | 1 | 1 | NA | NA | 1 | 0 | 1 | 1 | 1 | 1 | 3 | 0.6667 |
| 5_1425296 | NA | NA | 1 | 1 | NA | NA | NA | NA | 1 | 1 | 1 | 1 | 1 | 1 | NA | NA | 1 | 1 | 3 | 1 |
| 5_1425487 | 1 | 1 | NA | NA | NA | NA | NA | NA | 1 | 1 | NA | NA | 1 | 1 | NA | NA | 3 | 1 | 4 | 1 |
| 5_1425494 | 26 | 0.1538 | 31 | 0.1613 | 32 | 0.0625 | 29 | 0.1034 | 118 | 0.1186 | 34 | 0.2353 | 20 | 0.15 | 24 | 0.125 | 31 | 0.2581 | 109 | 0.2018 |
| 5_1425494 | 23 | 0.0435 | 24 | 0.0833 | 28 | 0.0357 | 29 | 0.2069 | 104 | 0.0962 | 29 | 0.069 | 22 | 0.0455 | 23 | 0.0435 | 24 | 0.0417 | 98 | 0.051 |
| 5_1425494 | 16 | 1 | 19 | 0.7895 | 20 | 0.8 | 14 | 0.8571 | 69 | 0.8551 | 19 | 0.6316 | 16 | 0.8125 | 18 | 0.8889 | 18 | 0.9444 | 71 | 0.8169 |
| 5_1425494 | 24 | 0.9583 | 27 | 0.9259 | 23 | 0.9565 | 27 | 0.9259 | 101 | 0.9406 | 34 | 0.8235 | 21 | 0.9048 | 21 | 0.6667 | 31 | 0.9355 | 107 | 0.8411 |
| 5_1425495 | 26 | 0.0769 | 31 | 0.2903 | 31 | 0.2581 | 33 | 0.1212 | 121 | 0.1901 | 33 | 0.0606 | 25 | 0.2 | 25 | 0.12 | 28 | 0.2143 | 111 | 0.1441 |
| 5_1425502 | 17 | 0 | 15 | 0 | 14 | 0.2143 | 19 | 0.0526 | 65 | 0.0615 | 17 | 0.1176 | 9 | 0.1111 | 15 | 0 | 16 | 0 | 57 | 0.0526 |
| 5_1425503 | 21 | 0.2381 | 29 | 0.2069 | 22 | 0.0909 | 28 | 0.0714 | 100 | 0.15 | 24 | 0.1667 | 21 | 0.1429 | 24 | 0.125 | 23 | 0.087 | 92 | 0.1304 |
| 5_1425504 | 18 | 0.9444 | 29 | 0.931 | 19 | 0.8947 | 29 | 1 | 95 | 0.9474 | 23 | 0.8261 | 19 | 0.8947 | 19 | 0.9474 | 23 | 0.9565 | 84 | 0.9048 |
| 5_1425504 | 21 | 0.9524 | 27 | 0.8889 | 21 | 0.9524 | 29 | 0.931 | 98 | 0.9286 | 25 | 0.96 | 22 | 0.9545 | 23 | 0.8261 | 24 | 0.9167 | 94 | 0.9149 |
| 5_1426527 | 4 | 0.25 | 7 | 0.4286 | 9 | 0.2222 | 6 | 0.5 | 26 | 0.3462 | 7 | 0 | 7 | 0.2857 | 5 | 0.6 | 6 | 0.3333 | 25 | 0.28 |
| 5_1426527 | 4 | 0.5 | 4 | 0.5 | 6 | 0.3333 | 3 | 0.3333 | 17 | 0.4118 | 6 | 0 | 6 | 0.1667 | 4 | 0.5 | 5 | 0 | 21 | 0.1429 |
| 5_1426527 | 4 | 0.75 | 7 | 0.4286 | 9 | 0.5556 | 6 | 0.8333 | 26 | 0.6154 | 6 | 0.8333 | 7 | 0.4286 | 5 | 0.6 | 6 | 0.3333 | 24 | 0.5417 |
| 5_1426528 | 4 | 1 | 8 | 0.375 | 8 | 0.625 | 4 | 1 | 24 | 0.6667 | 8 | 0.875 | 5 | 1 | 5 | 1 | 5 | 0.8 | 23 | 0.913 |
| 5_1426528 | 4 | 1 | 7 | 0.5714 | 8 | 0.625 | 6 | 0.6667 | 25 | 0.68 | 8 | 0.25 | 5 | 0.6 | 5 | 0.8 | 4 | 0.5 | 22 | 0.5 |
| 5_1426529 | 3 | 0.3333 | 8 | 0.125 | 8 | 0.25 | 5 | 0.4 | 24 | 0.25 | 7 | 0.1429 | 7 | 0.4286 | 5 | 0 | 5 | 0.2 | 24 | 0.2083 |
| 5_1426532 | 2 | 0.5 | 7 | 0.1429 | 7 | 0.1429 | 7 | 0.2857 | 23 | 0.2174 | 6 | 0 | 6 | 0.3333 | 6 | 0.3333 | 4 | 0.5 | 22 | 0.2727 |
| 5_1426533 | 1 | 1 | 4 | 0 | 4 | 0.25 | 5 | 0 | 14 | 0.1429 | 4 | 0 | 4 | 0.75 | 5 | 0.6 | 2 | 0.5 | 15 | 0.4667 |
| 5_1426533 | 2 | 0 | 6 | 0 | 9 | 0.1111 | 9 | 0.1111 | 26 | 0.0769 | 5 | 0 | 5 | 0 | 6 | 0 | 3 | 0.3333 | 19 | 0.0526 |
| 5_1426543 | 6 | 0.1667 | 10 | 0 | 12 | 0.1667 | 11 | 0.0909 | 39 | 0.1026 | 7 | 0 | 6 | 0.1667 | 10 | 0.4 | 4 | 0 | 27 | 0.1852 |
| 5_1426553 | 8 | 0.125 | 10 | 0.2 | 12 | 0 | 12 | 0.25 | 42 | 0.1429 | 9 | 0.2222 | 9 | 0.1111 | 9 | 0.4444 | 9 | 0.3333 | 36 | 0.2778 |
| 5_1427081 | NA | NA | 2 | 0.5 | NA | NA | NA | NA | 2 | 0.5 | NA | NA | NA | NA | 1 | 1 | NA | NA | 1 | 1 |
| 5_1427082 | NA | NA | 2 | 0.5 | NA | NA | NA | NA | 2 | 0.5 | NA | NA | NA | NA | 2 | 1 | NA | NA | 2 | 1 |

| | | | | | | | | | | | | | | | | | | | | |
|---|---|---|---|---|---|---|---|---|---|---|---|---|---|---|---|---|---|---|---|---|
| 5_1427113 | 3 | 0 | 5 | 0 | 6 | 0.1667 | 3 | 0.3333 | 17 | 0.1176 | 6 | 0.1667 | 5 | 0.2 | 16 | 0.125 | 5 | 0 | 32 | 0.125 |
| 5_1427113 | 3 | 0.3333 | 5 | 0.2 | 7 | 0.1429 | 3 | 0 | 18 | 0.1667 | 5 | 0.2 | 4 | 0 | 15 | 0.0667 | 5 | 0 | 29 | 0.069 |
| 5_1427171 | NA | NA | 2 | 0.5 | 6 | 0.1667 | 1 | 1 | 9 | 0.3333 | 5 | 0.4 | 5 | 0.2 | 10 | 0.2 | 4 | 0 | 24 | 0.2083 |
| 5_1427173 | NA | NA | 2 | 0.5 | 6 | 0 | 1 | 0 | 9 | 0.1111 | 5 | 0.4 | 5 | 0.2 | 6 | 0 | 3 | 0.3333 | 19 | 0.2105 |
| 5_1427179 | NA | NA | 1 | 1 | 4 | 1 | NA | NA | 5 | 1 | 2 | 1 | 1 | 1 | 4 | 0.5 | 2 | 0.5 | 9 | 0.6667 |
| 5_1427179 | NA | NA | 2 | 0.5 | 4 | 0.25 | NA | NA | 6 | 0.3333 | 2 | 0.5 | 3 | 0.3333 | 6 | 0.3333 | 4 | 0.25 | 15 | 0.3333 |
| 5_1427179 | NA | NA | 2 | 1 | 5 | 1 | NA | NA | 7 | 1 | 4 | 1 | 3 | 1 | 7 | 1 | 4 | 1 | 18 | 1 |
| 5_1427362 | NA | NA | 1 | 1 | 5 | 0.4 | NA | NA | 6 | 0.5 | 1 | 1 | NA | NA | 2 | 0.5 | 1 | 1 | 4 | 0.75 |
| 5_1427412 | NA | NA | 3 | 0.6667 | 5 | 0.4 | NA | NA | 8 | 0.5 | 3 | 0.3333 | NA | NA | 4 | 0.25 | 3 | 0.3333 | 10 | 0.3 |
| 5_1427456 | NA | NA | 3 | 1 | 3 | 0.3333 | NA | NA | 6 | 0.6667 | 3 | 0.3333 | NA | NA | 3 | 1 | 3 | 1 | 9 | 0.7778 |
| 5_1427456 | NA | NA | 3 | 0.6667 | 2 | 0.5 | NA | NA | 5 | 0.6 | 3 | 0.3333 | NA | NA | 3 | 0.3333 | 3 | 0 | 9 | 0.2222 |
| 5_1427573 | NA | NA | 3 | 1 | NA | NA | NA | NA | 3 | 1 | 2 | 0.5 | NA | NA | 2 | 1 | 2 | 1 | 6 | 0.8333 |
| 5_1427574 | NA | NA | 3 | 1 | 2 | 0.5 | NA | NA | 5 | 0.8 | 2 | 1 | NA | NA | 1 | 1 | 2 | 1 | 5 | 1 |
| 5_1427575 | NA | NA | 3 | 0.3333 | 1 | 1 | NA | NA | 4 | 0.5 | 2 | 0 | NA | NA | 2 | 0 | 2 | 1 | 6 | 0.3333 |
| 5_1427576 | NA | NA | 3 | 0.6667 | 1 | 1 | NA | NA | 4 | 0.75 | 2 | 0 | NA | NA | 2 | 0 | 2 | 0.5 | 6 | 0.1667 |
| 5_1427577 | NA | NA | 3 | 0.6667 | 1 | 1 | NA | NA | 4 | 0.75 | 1 | 0 | NA | NA | 1 | 1 | 1 | 1 | 3 | 0.6667 |
| 5_1427729 | NA | NA | 2 | 0.5 | NA | NA | NA | NA | 2 | 0.5 | 2 | 0.5 | NA | NA | NA | NA | 1 | 1 | 3 | 0.6667 |
| 5_1427730 | NA | NA | 1 | 1 | 1 | 1 | NA | NA | 2 | 1 | 1 | 0 | NA | NA | NA | NA | 1 | 1 | 2 | 0.5 |
| 5_1427730 | NA | NA | 2 | 1 | 1 | 1 | NA | NA | 3 | 1 | 1 | 1 | NA | NA | NA | NA | 1 | 1 | 2 | 1 |
| 5_1427733 | NA | NA | 2 | 1 | 1 | 1 | NA | NA | 3 | 1 | 2 | 0.5 | NA | NA | NA | NA | 1 | 1 | 3 | 0.6667 |
| 5_1427734 | NA | NA | NA | NA | 1 | 1 | NA | NA | 1 | 1 | 1 | 1 | NA | NA | NA | NA | NA | NA | 1 | 1 |
| 5_1427738 | NA | NA | 2 | 0.5 | 1 | 0 | NA | NA | 3 | 0.3333 | 1 | 1 | NA | NA | NA | NA | 1 | 0 | 2 | 0.5 |
| 5_1427742 | NA | NA | 1 | 1 | NA | NA | NA | NA | 1 | 1 | 2 | 0.5 | NA | NA | NA | NA | 1 | 1 | 3 | 0.6667 |
| 5_1427743 | NA | NA | 1 | 1 | 1 | 1 | NA | NA | 2 | 1 | 2 | 0.5 | NA | NA | NA | NA | 1 | 1 | 3 | 0.6667 |
| 5_1428528 | 1 | 0 | 1 | 1 | NA | NA | 4 | 0 | 6 | 0.1667 | 3 | 0.3333 | 1 | 1 | 3 | 0 | NA | NA | 7 | 0.2857 |
| 5_1428528 | 2 | 1 | 1 | 1 | 2 | 0.5 | 5 | 0.4 | 10 | 0.6 | 6 | 0.1667 | 1 | 1 | 6 | 0.3333 | 1 | 1 | 14 | 0.3571 |
| 5_1428529 | 1 | 1 | 3 | 1 | 2 | 0 | 4 | 0.5 | 10 | 0.6 | 4 | 0.75 | 2 | 1 | 6 | 0.3333 | 1 | 1 | 13 | 0.6154 |
| 5_1428579 | NA | NA | NA | NA | 1 | 1 | 1 | 1 | 2 | 1 | NA | NA | 1 | 0 | 2 | 1 | NA | NA | 3 | 0.6667 |
| 5_1428715 | NA | NA | NA | NA | 1 | 1 | 1 | 1 | 2 | 1 | NA | NA | 1 | 0 | 1 | 1 | NA | NA | 2 | 0.5 |
| 5_1428716 | NA | NA | NA | NA | 1 | 1 | 1 | 1 | 2 | 1 | NA | NA | 1 | 0 | 1 | 1 | NA | NA | 2 | 0.5 |
| 5_1428717 | NA | NA | NA | NA | 1 | 1 | 1 | 0 | 2 | 0.5 | NA | NA | 1 | 1 | 1 | 0 | NA | NA | 2 | 0.5 |
| 5_1428719 | NA | NA | NA | NA | NA | NA | 1 | 1 | 1 | 1 | NA | NA | 1 | 0 | 2 | 0.5 | NA | NA | 3 | 0.3333 |
| 5_1428722 | NA | NA | NA | NA | NA | NA | 1 | 1 | 1 | 1 | NA | NA | 1 | 0 | 1 | 1 | NA | NA | 2 | 0.5 |
| 5_1428892 | 410 | 0.3756 | 372 | 0.4167 | 347 | 0.3718 | 332 | 0.3825 | 1461 | 0.3867 | 326 | 0.3988 | 360 | 0.3917 | 338 | 0.3846 | 329 | 0.4134 | 1353 | 0.3969 |
| 5_1428893 | 384 | 0.0938 | 346 | 0.0896 | 334 | 0.0838 | 307 | 0.0782 | 1371 | 0.0868 | 297 | 0.1414 | 339 | 0.0855 | 307 | 0.0879 | 316 | 0.0981 | 1259 | 0.1025 |
| 5_1428894 | 346 | 0.1069 | 304 | 0.125 | 285 | 0.1123 | 267 | 0.0936 | 1202 | 0.1098 | 279 | 0.1362 | 285 | 0.1123 | 270 | 0.1259 | 275 | 0.1091 | 1109 | 0.1208 |
| 5_1428895 | 344 | 0.6453 | 302 | 0.6854 | 290 | 0.6724 | 273 | 0.6484 | 1209 | 0.6625 | 265 | 0.6943 | 289 | 0.692 | 265 | 0.6453 | 273 | 0.7619 | 1092 | 0.6987 |
| 5_1428904 | 239 | 0.1967 | 206 | 0.1748 | 202 | 0.2228 | 208 | 0.2163 | 855 | 0.2023 | 180 | 0.2 | 200 | 0.17 | 193 | 0.1554 | 197 | 0.2132 | 770 | 0.1844 |
| 5_1429579 | 25 | 0.16 | 25 | 0.08 | 30 | 0.0667 | 22 | 0.0909 | 102 | 0.098 | 18 | 0.1667 | 27 | 0.1481 | 33 | 0.0909 | 24 | 0.0417 | 102 | 0.1078 |
| 5_1429579 | 29 | 0.2414 | 26 | 0.1538 | 34 | 0.2059 | 25 | 0.08 | 114 | 0.1754 | 18 | 0.1111 | 28 | 0.1071 | 28 | 0.0357 | 28 | 0.1429 | 102 | 0.098 |
| 5_1429580 | 24 | 0.25 | 27 | 0.0741 | 28 | 0.1429 | 24 | 0.125 | 103 | 0.1456 | 21 | 0.1429 | 30 | 0.1333 | 33 | 0 | 23 | 0.0435 | 107 | 0.0748 |
| 5_1429580 | 28 | 0.1071 | 30 | 0.1 | 31 | 0.0645 | 22 | 0.0909 | 111 | 0.0901 | 19 | 0.0526 | 32 | 0.0625 | 33 | 0.0606 | 29 | 0.0345 | 113 | 0.0531 |
| 5_1429581 | 27 | 0.037 | 28 | 0.1429 | 34 | 0.0294 | 25 | 0.28 | 114 | 0.114 | 20 | 0.05 | 30 | 0.2 | 36 | 0.1111 | 28 | 0.1071 | 114 | 0.1228 |
| 5_1429582 | 17 | 0.3529 | 14 | 0.2143 | 28 | 0.25 | 17 | 0.2941 | 76 | 0.2763 | 14 | 0.2143 | 23 | 0.2609 | 26 | 0.3462 | 18 | 0.3889 | 81 | 0.3086 |
| 5_1429769 | 3 | 0.6667 | 8 | 0.5 | 2 | 0 | 6 | 0 | 19 | 0.3158 | 4 | 0 | 3 | 0 | 3 | 0.3333 | 5 | 0.2 | 15 | 0.1333 |
| 5_1429774 | 3 | 0 | 4 | 0 | 2 | 0 | 5 | 0.2 | 14 | 0.0714 | 4 | 0 | 3 | 0 | 1 | 0 | 6 | 0.1667 | 14 | 0.0714 |
| 5_1429784 | 2 | 0 | 6 | 0.3333 | 2 | 0 | 4 | 0 | 14 | 0.1429 | 3 | 0 | 2 | 0.5 | 2 | 0 | 5 | 0.2 | 12 | 0.1667 |
| 5_1430015 | 2 | 0.5 | 1 | 0 | 2 | 0 | 3 | 0 | 8 | 0.125 | 3 | 0 | NA | NA | 2 | 0 | 4 | 0.25 | 9 | 0.1111 |
| 5_1430716 | 1 | 0 | 1 | 0 | 1 | 0 | 2 | 0.5 | 5 | 0.2 | 3 | 0 | NA | NA | 1 | 0 | 3 | 0.3333 | 7 | 0.1429 |
| 5_1430763 | 1 | 0 | 1 | 1 | NA | NA | 2 | 0 | 4 | 0.25 | 3 | 0.3333 | NA | NA | 1 | 0 | 2 | 1 | 6 | 0.5 |
| 5_1431580 | 1 | 1 | 1 | 0 | 1 | 1 | NA | NA | 3 | 0.6667 | 1 | 1 | NA | NA | 1 | 1 | 1 | 1 | 3 | 1 |
| 5_1431581 | NA | NA | 1 | 1 | 2 | 1 | NA | NA | 3 | 1 | NA | NA | NA | NA | 1 | 1 | NA | NA | 1 | 1 |
| 5_1431582 | 1 | 1 | 1 | 1 | 1 | 1 | NA | NA | 3 | 1 | 1 | 1 | 1 | 1 | 1 | 0 | 1 | 1 | 4 | 0.75 |
| 5_1431582 | 1 | 0 | 1 | 0 | 3 | 1 | NA | NA | 5 | 0.6 | NA | NA | 1 | 1 | 1 | 1 | 1 | 0 | 3 | 0.6667 |
| 5_1431584 | 1 | 1 | NA | NA | NA | NA | NA | NA | 1 | 1 | NA | NA | 1 | 1 | 1 | 1 | 1 | 0 | 3 | 0.6667 |
| 5_1431584 | 1 | 1 | NA | NA | 2 | 1 | NA | NA | 3 | 1 | NA | NA | 1 | 1 | 1 | 1 | 1 | 0 | 3 | 0.6667 |
| 5_1431585 | 1 | 1 | NA | NA | 2 | 1 | NA | NA | 3 | 1 | NA | NA | NA | NA | 1 | 1 | 1 | 1 | 2 | 1 |
| 5_1431586 | 1 | 1 | NA | NA | 2 | 1 | NA | NA | 3 | 1 | NA | NA | 1 | 1 | 1 | 0 | 1 | 0 | 3 | 0.3333 |
| 5_1431588 | 1 | 1 | NA | NA | 2 | 0.5 | NA | NA | 3 | 0.6667 | NA | NA | 1 | 1 | 1 | 0 | 1 | 0 | 3 | 0.3333 |
| 5_1431588 | NA | NA | NA | NA | 2 | 1 | NA | NA | 2 | 1 | NA | NA | 1 | 1 | 1 | 0 | 1 | 0 | 3 | 0.3333 |
| 5_1431590 | 1 | 1 | NA | NA | 2 | 0 | NA | NA | 3 | 0.3333 | NA | NA | NA | NA | NA | NA | 1 | 1 | 1 | 1 |
| 5_1431590 | 1 | 1 | NA | NA | 2 | 1 | NA | NA | 3 | 1 | NA | NA | NA | NA | NA | NA | 1 | 1 | 1 | 1 |
| 5_1431597 | 1 | 1 | 1 | 1 | 2 | 1 | NA | NA | 4 | 1 | NA | NA | NA | NA | NA | NA | 1 | 1 | 1 | 1 |
| 5_1431600 | 1 | 1 | 1 | 0 | 2 | 1 | NA | NA | 4 | 0.75 | NA | NA | NA | NA | NA | NA | 1 | 1 | 1 | 1 |
| 5_1431603 | 1 | 1 | 1 | 1 | 2 | 1 | NA | NA | 4 | 1 | NA | NA | NA | NA | NA | NA | 1 | 1 | 1 | 1 |

| | | | | | | | | | | | | | | | | | | | | |
|---|---|---|---|---|---|---|---|---|---|---|---|---|---|---|---|---|---|---|---|---|
| 5_1431606( | 1 | 1 | 1 | 1 | 2 | 1 | NA | NA | 4 | 1 | NA | NA | NA | NA | NA | NA | 1 | 1 | 1 | 1 |
| 5_1432301! | 2 | 1 | NA | NA | NA | NA | NA | NA | 2 | 1 | NA | NA | NA | NA | 1 | 1 | NA | NA | 1 | 1 |
| 5_1432304: | 2 | 0.5 | 1 | 1 | NA | NA | NA | NA | 3 | 0.6667 | NA | NA | NA | NA | 1 | 1 | NA | NA | 1 | 1 |
| 5_1432309! | 2 | 1 | 1 | 0 | NA | NA | NA | NA | 3 | 0.6667 | NA | NA | NA | NA | 1 | 1 | NA | NA | 1 | 1 |
| 5_1432311( | 1 | 1 | 1 | 1 | NA | NA | NA | NA | 2 | 1 | NA | NA | 1 | 0 | 1 | 1 | 1 | 0 | 3 | 0.3333 |
| 5_1432315: | 1 | 1 | 1 | 0 | 1 | 0 | 1 | 0 | 4 | 0.25 | NA | NA | 1 | 1 | 2 | 1 | 1 | 1 | 4 | 1 |
| 5_1432316( | 1 | 1 | 1 | 1 | NA | NA | 1 | 1 | 3 | 1 | NA | NA | NA | NA | 2 | 1 | 1 | 1 | 3 | 1 |
| 5_1432316( | 2 | 1 | 1 | 0 | 1 | 1 | 1 | 0 | 5 | 0.6 | NA | NA | 1 | 1 | 3 | 1 | 1 | 1 | 5 | 1 |
| 5_1432316! | 2 | 0.5 | NA | NA | 1 | 0 | 1 | 0 | 4 | 0.25 | NA | NA | 1 | 0 | 3 | 0 | 1 | 1 | 5 | 0.2 |
| 5_1432317: | 2 | 0 | 1 | 0 | 1 | 1 | 1 | 0 | 5 | 0.2 | NA | NA | NA | NA | 3 | 0.6667 | NA | NA | 3 | 0.6667 |
| 5_1432321: | 2 | 0 | 1 | 1 | 1 | 0 | NA | NA | 4 | 0.25 | NA | NA | 1 | 0 | 3 | 0.3333 | 2 | 0 | 6 | 0.1667 |
| 5_1432321! | NA | NA | 2 | 0.5 | 1 | 0 | 1 | 1 | 4 | 0.5 | NA | NA | 1 | 0 | 3 | 1 | 2 | 0.5 | 6 | 0.6667 |
| 5_1432322: | 2 | 0.5 | 2 | 1 | 1 | 0 | 1 | 1 | 6 | 0.6667 | NA | NA | 1 | 1 | 3 | 0.6667 | 1 | 0 | 5 | 0.6 |
| 5_1432322! | 2 | 1 | 2 | 1 | 1 | 1 | 1 | 1 | 6 | 1 | NA | NA | 1 | 1 | 2 | 0.5 | 2 | 1 | 5 | 0.8 |
| 5_1432323 | 1 | 1 | 2 | 1 | 1 | 1 | NA | NA | 4 | 1 | NA | NA | 1 | 1 | 1 | 1 | 1 | 1 | 3 | 1 |
| 5_1432324 | 2 | 0 | 3 | 0.6667 | 1 | 1 | 2 | 0.5 | 8 | 0.5 | NA | NA | 1 | 1 | 3 | 0 | 2 | 0 | 6 | 0.1667 |
| 5_1432327! | 2 | 0 | 4 | 0.5 | 1 | 1 | 2 | 0.5 | 9 | 0.4444 | NA | NA | 1 | 0 | 3 | 0.6667 | 3 | 1 | 7 | 0.7143 |
| 5_1432329: | 2 | 0.5 | 4 | 0 | 1 | 0 | 2 | 0 | 9 | 0.1111 | NA | NA | 2 | 0 | 3 | 0.3333 | 3 | 0.6667 | 8 | 0.375 |
| 5_1432330! | 2 | 1 | 3 | 1 | 1 | 1 | 2 | 0 | 8 | 0.75 | NA | NA | 2 | 0 | 3 | 0.6667 | 2 | 0.5 | 7 | 0.4286 |
| 5_1432331: | 2 | 1 | 4 | 1 | 1 | 1 | 1 | 1 | 8 | 1 | NA | NA | 3 | 0.6667 | 3 | 1 | 2 | 1 | 8 | 0.875 |
| 5_1432332: | 2 | 0.5 | 4 | 0 | 1 | 0 | 2 | 0 | 9 | 0.1111 | NA | NA | 2 | 0 | 3 | 0 | 4 | 0.25 | 9 | 0.1111 |
| 5_1432333! | 2 | 1 | 3 | 0 | 1 | 0 | 1 | 0 | 7 | 0.2857 | NA | NA | 3 | 0.3333 | 1 | 0 | 3 | 0.3333 | 7 | 0.2857 |
| 5_1432334! | 2 | 1 | 3 | 1 | 1 | 1 | 2 | 1 | 8 | 1 | NA | NA | 4 | 1 | 3 | 1 | 4 | 1 | 11 | 1 |
| 5_1432335 | 1 | 1 | 4 | 1 | 1 | 1 | 1 | 1 | 7 | 1 | NA | NA | 4 | 1 | 3 | 1 | 2 | 1 | 9 | 1 |
| 5_1432335: | 2 | 1 | 4 | 0.75 | 1 | 1 | 2 | 0.5 | 9 | 0.7778 | NA | NA | 4 | 1 | 3 | 0.6667 | 4 | 0.75 | 11 | 0.8182 |
| 5_1432335( | 2 | 0 | 3 | 0 | 1 | 0 | 2 | 0.5 | 8 | 0.125 | NA | NA | 4 | 0.25 | 3 | 0.3333 | 4 | 0 | 11 | 0.1818 |
| 5_1432337( | 1 | 0 | 3 | 0.3333 | NA | NA | 1 | 0 | 5 | 0.2 | NA | NA | 2 | 0.5 | 1 | 0 | 2 | 0 | 5 | 0.2 |
| 5_1432383: | 1 | 0 | 1 | 1 | NA | NA | 1 | 1 | 3 | 0.6667 | 1 | 1 | NA | NA | NA | NA | NA | NA | 1 | 1 |
| 5_1432394: | 1 | 1 | NA | NA | NA | NA | NA | NA | 1 | 1 | 1 | 1 | NA | NA | NA | NA | NA | NA | 1 | 1 |
| 5_1432394( | 1 | 1 | NA | NA | NA | NA | NA | NA | 1 | 1 | 1 | 1 | NA | NA | NA | NA | NA | NA | 1 | 1 |
| 5_1432732( | NA | NA | NA | NA | NA | NA | 1 | 1 | 1 | 1 | 2 | 1 | NA | NA | NA | NA | NA | NA | 2 | 1 |
| 5_1432732 | NA | NA | NA | NA | NA | NA | 1 | 1 | 1 | 1 | 1 | 1 | NA | NA | NA | NA | NA | NA | 1 | 1 |
| 5_1432732: | NA | NA | NA | NA | NA | NA | 1 | 1 | 1 | 1 | 1 | 1 | NA | NA | NA | NA | NA | NA | 1 | 1 |
| 5_1432733! | NA | NA | NA | NA | 1 | 0 | 1 | 1 | 2 | 0.5 | 2 | 1 | NA | NA | NA | NA | NA | NA | 2 | 1 |
| 5_1432734! | NA | NA | NA | NA | NA | NA | 1 | 1 | 1 | 1 | 1 | 1 | NA | NA | NA | NA | NA | NA | 1 | 1 |
| 5_1432754( | NA | NA | NA | NA | 1 | 1 | NA | NA | 1 | 1 | 1 | 1 | NA | NA | NA | NA | NA | NA | 1 | 1 |
| 5_1432870: | 2 | 0 | 2 | 0.5 | 1 | 1 | 3 | 0 | 8 | 0.25 | 5 | 0.2 | 3 | 0.3333 | 7 | 0 | 7 | 0 | 22 | 0.0909 |
| 5_1432874( | 2 | 1 | 2 | 0.5 | 1 | 0 | 4 | 0 | 9 | 0.3333 | 5 | 0.2 | 5 | 0.2 | 6 | 0 | 6 | 0.1667 | 22 | 0.1364 |
| 5_1432874( | 2 | 0 | 2 | 0 | NA | NA | 2 | 0.5 | 6 | 0.1667 | 4 | 0 | 5 | 0 | 5 | 0 | 4 | 0.25 | 18 | 0.0556 |
| 5_1432934: | 2 | 1 | 2 | 1 | 2 | 1 | 2 | 1 | 8 | 1 | 2 | 1 | 3 | 1 | 5 | 1 | 1 | 1 | 11 | 1 |
| 5_1432934! | 2 | 0.5 | 2 | 0 | 2 | 0 | 2 | 0.5 | 8 | 0.25 | 4 | 0.25 | 4 | 0.5 | 7 | 0.1429 | 5 | 0.2 | 20 | 0.25 |
| 5_1432936: | 1 | 1 | 2 | 1 | 1 | 1 | 1 | 1 | 5 | 1 | 4 | 1 | 4 | 0.75 | 7 | 1 | 6 | 0.8333 | 21 | 0.9048 |
| 5_1432936: | NA | NA | 2 | 0.5 | 1 | 0 | 1 | 0 | 4 | 0.25 | 3 | 1 | 4 | 0.75 | 6 | 1 | 4 | 1 | 17 | 0.9412 |
| 5_1432936! | 1 | 1 | 2 | 0.5 | 1 | 1 | 1 | 1 | 5 | 0.8 | 4 | 0.75 | 3 | 0.3333 | 6 | 0.6667 | 5 | 0.8 | 18 | 0.6667 |
| 5_1433001! | 1 | 1 | 2 | 0 | 1 | 0 | 1 | 0 | 5 | 0.2 | 1 | 0 | 2 | 0 | 4 | 0.25 | 5 | 0 | 12 | 0.0833 |
| 5_1433008: | 1 | 1 | 1 | 0 | NA | NA | NA | NA | 2 | 0.5 | 1 | 0 | 1 | 0 | 3 | 0 | 3 | 0.6667 | 8 | 0.25 |
| 5_1433023: | 3 | 0 | 3 | 0.3333 | 4 | 0 | 2 | 0 | 12 | 0.0833 | NA | NA | 3 | 0 | 1 | 0 | 4 | 0.25 | 8 | 0.125 |
| 5_1433144! | NA | NA | NA | NA | NA | NA | 1 | 1 | 1 | 1 | NA | NA | 1 | 1 | NA | NA | NA | NA | 1 | 1 |
| 5_1433145( | NA | NA | NA | NA | NA | NA | 1 | 1 | 1 | 1 | NA | NA | 1 | 1 | NA | NA | NA | NA | 1 | 1 |
| 5_1434066: | 5 | 0 | 4 | 0.25 | 4 | 0 | 6 | 0.3333 | 19 | 0.1579 | 4 | 0.25 | 5 | 0 | 4 | 0.25 | 6 | 0.1667 | 19 | 0.1579 |
| 5_1434068: | 7 | 0 | 4 | 0.25 | 4 | 0 | 8 | 0.125 | 23 | 0.087 | 4 | 0 | 6 | 0 | 5 | 0 | 5 | 0.2 | 20 | 0.05 |
| 5_1434068( | 6 | 0.5 | 5 | 0 | 3 | 0.6667 | 8 | 0.375 | 22 | 0.3636 | 1 | 1 | 6 | 0.3333 | 5 | 0.8 | 4 | 0.25 | 16 | 0.5 |
| 5_1434604( | 3 | 0.3333 | 4 | 0.75 | 1 | 0 | 5 | 0.6 | 13 | 0.5385 | 2 | 0 | NA | NA | 2 | 1 | 1 | 0 | 5 | 0.4 |
| 5_1434886: | 3 | 0.3333 | 4 | 0.75 | 2 | 0.5 | 8 | 0.625 | 17 | 0.5882 | 2 | 0 | 1 | 0 | 3 | 0.6667 | 4 | 0.75 | 10 | 0.5 |
| 5_1434886! | 2 | 0.5 | 2 | 0 | 2 | 0 | 6 | 0.1667 | 12 | 0.1667 | 1 | 0 | 1 | 0 | 2 | 0 | 4 | 0.5 | 8 | 0.25 |
| 5_1434891( | 3 | 0 | 2 | 0 | 2 | 0 | 7 | 0.1429 | 14 | 0.0714 | 5 | 0 | 3 | 0.3333 | 5 | 0 | 4 | 0 | 17 | 0.0588 |
| 5_1434891! | 3 | 1 | 5 | 0.8 | 1 | 1 | 8 | 0.625 | 17 | 0.7647 | 5 | 0.6 | 3 | 0.3333 | 4 | 1 | 3 | 1 | 15 | 0.7333 |
| 5_1434892: | 3 | 0 | 5 | 0.4 | 2 | 0 | 8 | 0.375 | 18 | 0.2778 | 3 | 0.6667 | 3 | 0.3333 | 3 | 0.3333 | 3 | 0.3333 | 12 | 0.4167 |
| 5_1434892: | 1 | 1 | 3 | 0 | 2 | 0 | 6 | 0.3333 | 12 | 0.25 | 5 | 0.2 | 2 | 0.5 | 4 | 0.75 | 3 | 0.6667 | 14 | 0.5 |
| 5_1434893: | 2 | 1 | 5 | 0.2 | 3 | 0 | 8 | 0.375 | 18 | 0.3333 | 4 | 0.5 | 2 | 0.5 | 3 | 0.6667 | 4 | 1 | 13 | 0.6923 |
| 5_1434894: | 3 | 0.3333 | 2 | 1 | 1 | 0 | 5 | 0.6 | 11 | 0.5455 | 4 | 0.5 | 2 | 0.5 | 4 | 0.5 | 4 | 0 | 14 | 0.3571 |
| 5_1434895( | 3 | 0.6667 | 2 | 1 | NA | NA | 6 | 0.6667 | 11 | 0.7273 | 2 | 0.5 | 2 | 1 | 5 | 0.4 | 2 | 1 | 11 | 0.6364 |
| 5_1434896( | 4 | 1 | 2 | 0.5 | 1 | 1 | 5 | 0.8 | 12 | 0.8333 | 3 | 0.6667 | 2 | 0.5 | 5 | 0.6 | 3 | 0.6667 | 13 | 0.6154 |
| 5_1434897: | 4 | 0.75 | 1 | 1 | NA | NA | 5 | 0.6 | 10 | 0.7 | 4 | 0.25 | 1 | 1 | 5 | 0.6 | 3 | 0 | 13 | 0.3846 |
| 5_1434897! | 2 | 0 | 2 | 0 | NA | NA | 1 | 1 | 5 | 0.2 | 5 | 0.2 | 1 | 0 | 5 | 0 | 4 | 0 | 15 | 0.0667 |

| | | | | | | | | | | | | | | | | | | | | |
|---|---|---|---|---|---|---|---|---|---|---|---|---|---|---|---|---|---|---|---|---|
| 5_1434898 | 4 | 0.25 | 1 | 0 | 1 | 0 | 5 | 0 | 11 | 0.0909 | 5 | 0.4 | 2 | 0 | 5 | 0 | 5 | 0 | 17 | 0.1176 |
| 5_1434900 | NA | NA | 1 | 1 | NA | NA | NA | NA | 1 | 1 | NA | NA | 1 | 1 | NA | NA | NA | NA | 1 | 1 |
| 5_1434921 | 56 | 0.2321 | 45 | 0.2667 | 46 | 0.1087 | 50 | 0.12 | 197 | 0.1827 | 40 | 0.15 | 46 | 0.2174 | 45 | 0.1333 | 50 | 0.2 | 181 | 0.1768 |
| 5_1434921 | 65 | 0.1231 | 47 | 0.1277 | 50 | 0.1 | 53 | 0.0943 | 215 | 0.1116 | 40 | 0.025 | 52 | 0.1154 | 53 | 0.0566 | 56 | 0.0893 | 201 | 0.0746 |
| 5_1434923 | 55 | 0.5818 | 40 | 0.6 | 41 | 0.5122 | 48 | 0.6875 | 184 | 0.5978 | 37 | 0.5946 | 45 | 0.5556 | 43 | 0.6744 | 50 | 0.56 | 175 | 0.5943 |
| 5_1434924 | 46 | 0.2609 | 41 | 0.1463 | 37 | 0.1622 | 39 | 0.2051 | 163 | 0.1963 | 26 | 0.1923 | 40 | 0.175 | 40 | 0.125 | 40 | 0.1 | 146 | 0.1438 |
| 5_1434926 | 51 | 0.098 | 38 | 0.0526 | 43 | 0 | 49 | 0.0816 | 181 | 0.0608 | 35 | 0.0857 | 45 | 0.0889 | 42 | 0.0952 | 50 | 0.04 | 172 | 0.0756 |
| 5_1434952 | NA | NA | NA | NA | NA | NA | 1 | 1 | 1 | 1 | NA | NA | 1 | 1 | NA | NA | NA | NA | 1 | 1 |
| 5_1434954 | NA | NA | NA | NA | NA | NA | 1 | 1 | 1 | 1 | NA | NA | 1 | 1 | NA | NA | NA | NA | 1 | 1 |
| 5_1435374 | 2 | 0 | 2 | 0.5 | 1 | 0 | 2 | 1 | 7 | 0.4286 | 1 | 1 | 1 | 0 | NA | NA | 1 | 0 | 3 | 0.3333 |
| 5_1435375 | 2 | 0.5 | 3 | 1 | 1 | 0 | 2 | 1 | 8 | 0.75 | 1 | 0 | 1 | 1 | NA | NA | 1 | 0 | 3 | 0.3333 |
| 5_1435486 | 4 | 0.25 | 3 | 1 | 1 | 1 | 3 | 0.3333 | 11 | 0.5455 | 5 | 0.4 | 2 | 0 | 3 | 0 | 2 | 0 | 12 | 0.1667 |
| 5_1435486 | 4 | 0.5 | 3 | 0.3333 | 2 | 0 | 3 | 0.6667 | 12 | 0.4167 | 5 | 0.4 | 3 | 1 | 3 | 0.3333 | 2 | 1 | 13 | 0.6154 |
| 5_1435487 | 5 | 0 | 5 | 0.2 | 2 | 0 | 2 | 1 | 14 | 0.2143 | 5 | 0.4 | 2 | 1 | 3 | 0.6667 | 2 | 0.5 | 12 | 0.5833 |
| 5_1435489 | 5 | 0.6 | 5 | 0.6 | 3 | 0.3333 | 2 | 0.5 | 15 | 0.5333 | 4 | 0.5 | 4 | 0.5 | 2 | 0.5 | 1 | 1 | 11 | 0.5455 |
| 5_1435490 | 6 | 0.8333 | 3 | 1 | 2 | 1 | 2 | 1 | 13 | 0.9231 | 4 | 1 | 3 | 1 | 2 | 0 | 2 | 1 | 11 | 0.8182 |
| 5_1435490 | 4 | 1 | 7 | 0.7143 | 1 | 1 | 2 | 1 | 14 | 0.8571 | 4 | 1 | 3 | 1 | 3 | 1 | 2 | 1 | 12 | 1 |
| 5_1435491 | 5 | 1 | 4 | 1 | 3 | 1 | 2 | 1 | 14 | 1 | 5 | 1 | NA | NA | 3 | 1 | 2 | 1 | 10 | 1 |
| 5_1435492 | 7 | 0 | 7 | 0 | 2 | 0 | 2 | 0.5 | 18 | 0.0556 | 6 | 0.1667 | 3 | 0 | 3 | 0 | 3 | 0 | 15 | 0.0667 |
| 5_1435494 | 4 | 0.25 | 7 | 0.2857 | 2 | 0.5 | 1 | 1 | 14 | 0.3571 | 7 | 0.4286 | 3 | 0.6667 | 4 | 0.75 | 3 | 0.6667 | 17 | 0.5882 |
| 5_1435495 | 6 | 0.8333 | 5 | 1 | 3 | 1 | NA | NA | 14 | 0.9286 | 6 | 0.6667 | 2 | 0.5 | 4 | 0.5 | 4 | 1 | 16 | 0.6875 |
| 5_1435496 | 7 | 0 | 8 | 0.375 | 4 | 0.25 | 4 | 0 | 23 | 0.1739 | 6 | 0 | 3 | 0.3333 | 5 | 0.4 | 4 | 0 | 18 | 0.1667 |
| 5_1435498 | 4 | 0.75 | 6 | 0.5 | 2 | 0.5 | 3 | 0.6667 | 15 | 0.6 | 5 | 0.2 | 3 | 0.3333 | 3 | 0 | 4 | 0.75 | 15 | 0.3333 |
| 5_1435500 | 7 | 1 | 8 | 1 | 5 | 1 | 5 | 1 | 25 | 1 | 8 | 1 | 4 | 1 | 6 | 1 | 4 | 1 | 22 | 1 |
| 5_1435500 | 7 | 0.2857 | 7 | 0.2857 | 4 | 0.25 | 5 | 0 | 23 | 0.2174 | 7 | 0.1429 | 2 | 0 | 6 | 0 | 4 | 0 | 19 | 0.0526 |
| 5_1435500 | 7 | 0.4286 | 9 | 0.3333 | 5 | 0.4 | 6 | 0.1667 | 27 | 0.3333 | 8 | 0.25 | 3 | 0.3333 | 6 | 0.3333 | 4 | 0 | 21 | 0.2381 |
| 5_1435501 | 6 | 1 | 6 | 1 | 5 | 1 | 3 | 1 | 20 | 1 | 8 | 0.875 | 4 | 1 | 4 | 1 | 4 | 1 | 20 | 0.95 |
| 5_1435501 | 8 | 0.125 | 8 | 0.25 | 5 | 0.2 | 6 | 0.1667 | 27 | 0.1852 | 8 | 0.25 | 3 | 0 | 6 | 0 | 4 | 0.25 | 21 | 0.1429 |
| 5_1435501 | 5 | 0.2 | 5 | 0 | 1 | 0 | 3 | 0.3333 | 14 | 0.1429 | 3 | 0.3333 | 3 | 0 | 3 | 0.3333 | 3 | 0 | 12 | 0.1667 |
| 5_1436936 | 4 | 0.25 | 4 | 0 | 3 | 0 | 1 | 0 | 12 | 0.0833 | 3 | 0.6667 | 2 | 0 | 5 | 0 | 1 | 0 | 11 | 0.1818 |
| 5_1436937 | 4 | 1 | 3 | 1 | 3 | 1 | 1 | 1 | 11 | 1 | 3 | 1 | 2 | 1 | 5 | 0.8 | 1 | 1 | 11 | 0.9091 |
| 5_1436938 | 4 | 0.25 | 4 | 0.25 | 3 | 0 | 1 | 0 | 12 | 0.1667 | 3 | 0 | 2 | 0.5 | 5 | 0.2 | 1 | 0 | 11 | 0.1818 |
| 5_1436938 | 4 | 0 | 4 | 0.5 | 2 | 0.5 | 1 | 0 | 11 | 0.2727 | 3 | 0.3333 | 3 | 1 | 5 | 0.4 | 1 | 0 | 12 | 0.5 |
| 5_1436939 | 4 | 0.5 | 4 | 0.75 | 3 | 0.6667 | 1 | 0 | 12 | 0.5833 | 2 | 0 | 3 | 0.3333 | 3 | 0.6667 | 1 | 1 | 9 | 0.4444 |
| 5_1436941 | 5 | 0 | 4 | 0.5 | 4 | 0.25 | 1 | 0 | 14 | 0.2143 | 2 | 0 | 2 | 0 | 4 | 0.25 | 1 | 0 | 9 | 0.1111 |
| 5_1436952 | 6 | 0 | 5 | 0.2 | 5 | 0 | 1 | 0 | 17 | 0.0588 | 4 | 0 | 5 | 0.2 | 6 | 0 | 1 | 0 | 16 | 0.0625 |
| 5_1436955 | 6 | 0 | 6 | 0.1667 | 6 | 0 | 1 | 0 | 19 | 0.0526 | 3 | 0.3333 | 4 | 0 | 5 | 0 | 3 | 0 | 15 | 0.0667 |
| 5_1436956 | 7 | 0 | 4 | 0 | 7 | 0.1429 | 1 | 0 | 19 | 0.0526 | 4 | 0.25 | 5 | 0 | 4 | 0 | 3 | 0.3333 | 16 | 0.125 |
| 5_1436962 | 5 | 0 | 1 | 0 | 2 | 0.5 | 5 | 0 | 13 | 0.0769 | 3 | 0 | 2 | 0 | 4 | 0.25 | 3 | 0 | 12 | 0.0833 |
| 5_1436963 | 7 | 0 | 1 | 0 | 2 | 0 | 6 | 0.3333 | 16 | 0.125 | 3 | 0 | 2 | 0.5 | 4 | 0.25 | 4 | 0 | 13 | 0.1538 |
| 5_1436964 | 7 | 0.1429 | 1 | 0 | 1 | 1 | 5 | 0 | 14 | 0.1429 | 3 | 0 | 2 | 0 | 2 | 0.5 | 4 | 0 | 11 | 0.0909 |
| 5_1436965 | 6 | 0.1667 | 1 | 0 | 1 | 0 | 5 | 0.4 | 13 | 0.2308 | 3 | 0.3333 | 2 | 0 | 2 | 0.5 | 4 | 0 | 11 | 0.1818 |
| 5_1436965 | 5 | 0 | 1 | 0 | 1 | 0 | 5 | 0.2 | 12 | 0.0833 | 3 | 0.3333 | 2 | 0.5 | 1 | 0 | 4 | 0.25 | 10 | 0.3 |
| 5_1436966 | 7 | 0.2857 | NA | NA | NA | NA | 4 | 0 | 11 | 0.1818 | 2 | 0 | 2 | 0 | 2 | 0.5 | 4 | 0 | 10 | 0.1 |
| 5_1436966 | 5 | 1 | NA | NA | NA | NA | 3 | 1 | 8 | 1 | 2 | 1 | 2 | 0.5 | 1 | 1 | 4 | 1 | 9 | 0.8889 |
| 5_1436967 | 5 | 0.2 | NA | NA | NA | NA | 4 | 0.25 | 9 | 0.2222 | 2 | 0 | 2 | 0 | 1 | 0 | 3 | 0.3333 | 8 | 0.125 |
| 5_1436967 | 6 | 0.8333 | NA | NA | NA | NA | 4 | 0.5 | 10 | 0.7 | 2 | 1 | 2 | 0.5 | 2 | 0 | 3 | 0.6667 | 9 | 0.5556 |
| 5_1436968 | 5 | 0.8 | NA | NA | NA | NA | 4 | 0.5 | 9 | 0.6667 | 2 | 0.5 | 2 | 0 | 2 | 0.5 | 4 | 0.75 | 10 | 0.5 |
| 5_1436968 | 6 | 0.6667 | NA | NA | NA | NA | 2 | 0 | 8 | 0.5 | 2 | 0 | 2 | 0 | 1 | 0 | 3 | 0.3333 | 8 | 0.125 |
| 5_1436969 | 4 | 1 | NA | NA | NA | NA | 3 | 0.3333 | 7 | 0.7143 | 2 | 1 | 2 | 1 | 1 | 1 | 3 | 0.3333 | 8 | 0.75 |
| 5_1436970 | 6 | 0.5 | NA | NA | NA | NA | 2 | 0.5 | 8 | 0.5 | 2 | 0 | 2 | 0.5 | NA | NA | 4 | 0.75 | 8 | 0.5 |
| 5_1436990 | 2 | 1 | NA | NA | NA | NA | NA | NA | 2 | 1 | NA | NA | NA | NA | NA | NA | 1 | 1 | 1 | 1 |
| 5_1436991 | 1 | 1 | NA | NA | NA | NA | NA | NA | 1 | 1 | NA | NA | NA | NA | NA | NA | 1 | 1 | 1 | 1 |
| 5_1436992 | 2 | 0.5 | NA | NA | NA | NA | NA | NA | 2 | 0.5 | NA | NA | NA | NA | NA | NA | 1 | 1 | 1 | 1 |
| 5_1436994 | 5 | 0.6 | NA | NA | NA | NA | 2 | 1 | 7 | 0.7143 | 2 | 0 | 1 | 1 | 1 | 1 | 5 | 0.8 | 9 | 0.6667 |
| 5_1437005 | 4 | 0.25 | NA | NA | NA | NA | 2 | 0 | 6 | 0.1667 | 2 | 0 | 1 | 0 | 1 | 1 | 5 | 0.4 | 9 | 0.3333 |
| 5_1437006 | 5 | 0.6 | NA | NA | NA | NA | 3 | 0.3333 | 8 | 0.5 | 2 | 0 | 1 | 1 | NA | NA | 5 | 0.4 | 8 | 0.375 |
| 5_1437007 | 3 | 1 | NA | NA | NA | NA | 3 | 1 | 6 | 1 | 1 | 1 | 1 | 1 | NA | NA | 5 | 1 | 7 | 1 |
| 5_1437009 | 4 | 1 | NA | NA | NA | NA | 3 | 1 | 7 | 1 | 1 | 1 | 1 | 1 | NA | NA | 3 | 1 | 5 | 1 |
| 5_1437009 | 4 | 1 | NA | NA | NA | NA | 3 | 0.6667 | 7 | 0.8571 | 1 | 0 | 1 | 0 | NA | NA | 3 | 0.6667 | 5 | 0.4 |
| 5_1437011 | 1 | 1 | NA | NA | NA | NA | 2 | 0.5 | 3 | 0.6667 | 1 | 0 | 1 | 0 | NA | NA | 3 | 0.3333 | 5 | 0.2 |
| 5_1437011 | 3 | 0.3333 | NA | NA | NA | NA | 3 | 0.3333 | 6 | 0.3333 | 1 | 0 | 1 | 1 | NA | NA | 3 | 0 | 5 | 0.2 |
| 5_1437012 | 3 | 0.6667 | NA | NA | NA | NA | 1 | 1 | 4 | 0.75 | 1 | 1 | 1 | 0 | NA | NA | 3 | 0.3333 | 5 | 0.4 |
| 5_1437012 | 3 | 1 | NA | NA | NA | NA | NA | NA | 3 | 1 | 1 | 1 | 1 | 1 | NA | NA | 2 | 0 | 4 | 0.5 |
| 5_1437013 | 3 | 0.3333 | NA | NA | NA | NA | 3 | 0.6667 | 6 | 0.5 | NA | NA | 1 | 0 | NA | NA | 2 | 0.5 | 3 | 0.3333 |

| | | | | | | | | | | | | | | | | | | | | |
|---|---|---|---|---|---|---|---|---|---|---|---|---|---|---|---|---|---|---|---|---|
| 5_14370140 | 3 | 0.6667 | NA | NA | NA | NA | 2 | 0.5 | 5 | 0.6 | 1 | 0 | 1 | 1 | NA | NA | 2 | 0 | 4 | 0.25 |
| 5_14370148 | 3 | 0.6667 | NA | NA | NA | NA | 2 | 1 | 5 | 0.8 | 1 | 1 | 1 | 1 | NA | NA | 2 | 1 | 4 | 1 |
| 5_14370154 | 3 | 0.3333 | NA | NA | NA | NA | 2 | 0 | 5 | 0.2 | 1 | 0 | 1 | 0 | NA | NA | 2 | 0.5 | 4 | 0.25 |
| 5_14374424 | NA | NA | NA | NA | 2 | 0.5 | 1 | 1 | 3 | 0.6667 | NA | NA | 2 | 1 | NA | NA | 1 | 1 | 3 | 1 |
| 5_14374429 | NA | NA | NA | NA | 2 | 1 | 1 | 1 | 3 | 1 | NA | NA | 1 | 1 | NA | NA | NA | NA | 1 | 1 |
| 5_14375313 | NA | NA | 1 | 1 | NA | NA | 2 | 0.5 | 3 | 0.6667 | NA | NA | 1 | 1 | NA | NA | NA | NA | 1 | 1 |
| 5_14375337 | NA | NA | 1 | 1 | 2 | 0 | 2 | 0 | 5 | 0.2 | NA | NA | 1 | 1 | NA | NA | NA | NA | 1 | 1 |
| 5_14375347 | NA | NA | 1 | 1 | 2 | 0 | 2 | 0 | 5 | 0.2 | NA | NA | 1 | 1 | NA | NA | NA | NA | 1 | 1 |
| 5_14375375 | NA | NA | 1 | 1 | 1 | 1 | 2 | 0.5 | 4 | 0.75 | NA | NA | 1 | 1 | NA | NA | 1 | 1 | 2 | 1 |
| 5_14383592 | 7 | 0.1429 | 6 | 0.1667 | 6 | 0.3333 | 4 | 0 | 23 | 0.1739 | 8 | 0 | 4 | 0 | 4 | 0.25 | 3 | 0 | 19 | 0.0526 |
| 5_14383876 | 8 | 0.875 | 7 | 1 | 6 | 1 | 4 | 1 | 25 | 0.96 | 9 | 1 | 5 | 1 | 5 | 1 | 4 | 0.75 | 23 | 0.9565 |
| 5_14383885 | 8 | 0.5 | 9 | 0.5556 | 6 | 0.5 | 3 | 1 | 26 | 0.5769 | 8 | 0.25 | 5 | 0.4 | 6 | 0.6667 | 5 | 0.4 | 24 | 0.4167 |
| 5_14383891 | 5 | 0.4 | 9 | 0.6667 | 7 | 0.8571 | 4 | 0.75 | 25 | 0.68 | 9 | 0.7778 | 4 | 0.5 | 5 | 1 | 5 | 0.4 | 23 | 0.6957 |
| 5_14383918 | 7 | 0.5714 | 8 | 0.375 | 8 | 0.5 | 4 | 0.75 | 27 | 0.5185 | 7 | 0.8571 | 3 | 0.3333 | 4 | 0.25 | 5 | 0.4 | 19 | 0.5263 |
| 5_14383954 | 1 | 0 | 1 | 1 | 2 | 0.5 | 1 | 0 | 5 | 0.4 | 1 | 1 | NA | NA | NA | NA | NA | NA | 1 | 1 |
| 5_14383955 | 5 | 0.8 | 4 | 0.5 | 3 | 1 | 3 | 1 | 15 | 0.8 | 7 | 0.8571 | 6 | 0.6667 | 4 | 1 | 4 | 0.5 | 21 | 0.7619 |
| 5_14383956 | 2 | 0 | 1 | 1 | 1 | 0 | 1 | 0 | 5 | 0.2 | 2 | 0.5 | NA | NA | NA | NA | NA | NA | 2 | 0.5 |
| 5_14383967 | 2 | 1 | 1 | 1 | 1 | 1 | 1 | 1 | 5 | 1 | 2 | 1 | NA | NA | NA | NA | NA | NA | 2 | 1 |
| 5_14383972 | 5 | 0.4 | 4 | 0 | 4 | 0 | 3 | 0 | 16 | 0.125 | 6 | 0 | 3 | 0.3333 | 3 | 0.3333 | 4 | 0 | 16 | 0.125 |
| 5_14383973 | 2 | 0 | 1 | 0 | 2 | 0 | 2 | 0.5 | 7 | 0.1429 | 2 | 0.5 | NA | NA | NA | NA | NA | NA | 2 | 0.5 |
| 5_14383980 | 4 | 1 | 3 | 0.6667 | 2 | 1 | 2 | 1 | 11 | 0.9091 | 1 | 1 | 4 | 1 | 2 | 1 | 2 | 1 | 9 | 1 |
| 5_14383993 | 2 | 0.5 | 2 | 0 | 2 | 0 | 1 | 0 | 7 | 0.1429 | 2 | 0.5 | NA | NA | NA | NA | NA | NA | 2 | 0.5 |
| 5_14383998 | 2 | 1 | 1 | 1 | 1 | 1 | 1 | 1 | 5 | 1 | 2 | 0.5 | NA | NA | NA | NA | NA | NA | 2 | 0.5 |
| 5_14384038 | 1 | 0 | 3 | 0.3333 | NA | NA | NA | NA | 4 | 0.25 | 3 | 0.3333 | NA | NA | NA | NA | NA | NA | 3 | 0.3333 |
| 5_14384049 | 1 | 1 | 2 | 0.5 | 1 | 0 | 1 | 0 | 5 | 0.4 | 3 | 0.3333 | NA | NA | NA | NA | NA | NA | 3 | 0.3333 |
| 5_14384064 | 2 | 0 | 2 | 0.5 | 1 | 0 | 1 | 0 | 6 | 0.1667 | 4 | 0.25 | NA | NA | NA | NA | NA | NA | 4 | 0.25 |
| 5_14386235 | NA | NA | NA | NA | NA | NA | 1 | 1 | 1 | 1 | 1 | 1 | NA | NA | NA | NA | NA | NA | 1 | 1 |
| 5_14386250 | NA | NA | NA | NA | 1 | 1 | 1 | 0 | 2 | 0.5 | 1 | 1 | NA | NA | NA | NA | NA | NA | 1 | 1 |
| 5_14386281 | NA | NA | NA | NA | NA | NA | 1 | 1 | 1 | 1 | 1 | 1 | NA | NA | NA | NA | NA | NA | 1 | 1 |
| 5_14386790 | NA | NA | 2 | 0.5 | 1 | 0 | 1 | 0 | 4 | 0.25 | 1 | 0 | 1 | 1 | 1 | 0 | NA | NA | 3 | 0.3333 |
| 5_14386797 | NA | NA | 1 | 0 | 1 | 1 | 1 | 0 | 3 | 0.3333 | 2 | 0.5 | 1 | 1 | NA | NA | NA | NA | 3 | 0.6667 |
| 5_14386821 | NA | NA | 2 | 0 | 1 | 0 | 1 | 1 | 4 | 0.25 | 3 | 0.3333 | 1 | 0 | 1 | 0 | NA | NA | 5 | 0.2 |
| 5_14386823 | NA | NA | 2 | 0.5 | 1 | 0 | NA | NA | 3 | 0.3333 | 3 | 0 | 1 | 1 | NA | NA | NA | NA | 4 | 0.25 |
| 5_14392491 | 8 | 0 | 4 | 0 | 4 | 0.25 | 6 | 0.1667 | 22 | 0.0909 | 7 | 0 | 3 | 0.3333 | 5 | 0 | 8 | 0.25 | 23 | 0.1304 |
| 5_14392493 | 7 | 0.2857 | 3 | 0.3333 | 5 | 0.4 | 8 | 0.125 | 23 | 0.2609 | 7 | 0.2857 | 2 | 0 | 4 | 0.25 | 8 | 0.125 | 21 | 0.1905 |
| 5_14411055 | 4 | 0.5 | 2 | 0 | 1 | 0 | 1 | 1 | 8 | 0.375 | 1 | 1 | 2 | 1 | NA | NA | 1 | 1 | 4 | 1 |
| 5_14411092 | 3 | 0.3333 | NA | NA | 1 | 1 | 2 | 0.5 | 6 | 0.5 | 1 | 1 | 1 | 0 | NA | NA | 1 | 1 | 3 | 0.6667 |
| 5_14411097 | 4 | 0.75 | 3 | 0.3333 | 2 | 1 | 2 | 1 | 11 | 0.7273 | 1 | 1 | 2 | 0.5 | NA | NA | 1 | 1 | 4 | 0.75 |
| 5_14411109 | 3 | 1 | 4 | 0.5 | 3 | 0.6667 | 2 | 0.5 | 12 | 0.6667 | 1 | 1 | 1 | 1 | NA | NA | 1 | 0 | 3 | 0.6667 |
| 5_14411114 | 4 | 0.5 | 4 | 0.25 | 3 | 0 | 2 | 0 | 13 | 0.2308 | 1 | 0 | 1 | 0 | NA | NA | 1 | 1 | 3 | 0.3333 |
| 5_14411120 | 4 | 0.25 | 2 | 0.5 | 2 | 0.5 | 2 | 0 | 10 | 0.3 | 2 | 1 | 1 | 1 | NA | NA | 1 | 1 | 4 | 1 |
| 5_14411130 | 2 | 0 | 3 | 0.6667 | 2 | 0 | 1 | 0 | 8 | 0.25 | 3 | 0.3333 | 2 | 0.5 | NA | NA | 1 | 0 | 6 | 0.3333 |
| 5_14411146 | 3 | 0.6667 | 3 | 0 | 2 | 0 | 2 | 1 | 10 | 0.4 | 3 | 0 | 2 | 0.5 | NA | NA | 1 | 0 | 6 | 0.1667 |
| 5_14411153 | 3 | 0.6667 | 4 | 0.25 | 3 | 1 | 2 | 0 | 12 | 0.5 | 3 | 0.3333 | 2 | 0.5 | NA | NA | 1 | 1 | 6 | 0.5 |
| 5_14411156 | 3 | 0.3333 | 1 | 0 | 2 | 0 | 1 | 0 | 7 | 0.1429 | 2 | 0.5 | 2 | 0 | NA | NA | 1 | 1 | 5 | 0.4 |
| 5_14411159 | 3 | 0.3333 | 3 | 0.3333 | 2 | 0 | 1 | 0 | 9 | 0.2222 | 2 | 0.5 | 2 | 0.5 | NA | NA | NA | NA | 4 | 0.5 |
| 5_14411178 | 3 | 1 | 3 | 1 | 2 | 0.5 | 2 | 0.5 | 10 | 0.8 | 3 | 1 | 3 | 0.3333 | NA | NA | 1 | 1 | 7 | 0.7143 |
| 5_14411203 | 3 | 0 | 5 | 0.4 | 3 | 0.3333 | 2 | 0.5 | 13 | 0.3077 | 3 | 1 | 3 | 0.6667 | NA | NA | 1 | 0 | 7 | 0.7143 |
| 5_14411224 | 2 | 0 | 4 | 0 | 3 | 0 | 3 | 0.6667 | 12 | 0.1667 | 3 | 0.3333 | 4 | 0.25 | NA | NA | 1 | 1 | 8 | 0.375 |
| 5_14411231 | 3 | 0 | 4 | 0.25 | 3 | 1 | 1 | 0 | 11 | 0.3636 | 3 | 0 | 3 | 0.3333 | NA | NA | 1 | 0 | 7 | 0.1429 |
| 5_14411234 | 4 | 0 | 5 | 0.2 | 3 | 0 | 2 | 0 | 14 | 0.0714 | 3 | 0 | 4 | 0.25 | NA | NA | 1 | 0 | 8 | 0.125 |
| 5_14411249 | 4 | 0 | 5 | 0 | 1 | 0 | 3 | 0.3333 | 13 | 0.0769 | 2 | 0.5 | 3 | 0 | NA | NA | 1 | 0 | 6 | 0.1667 |
| 5_14411251 | 3 | 0.3333 | 6 | 0 | 3 | 0 | 3 | 0.3333 | 15 | 0.1333 | 3 | 0.3333 | 4 | 0 | NA | NA | 1 | 0 | 8 | 0.125 |
| 5_14411282 | 4 | 0.25 | 5 | 0 | 3 | 0 | 3 | 0 | 15 | 0.0667 | 3 | 0 | 4 | 0.25 | 1 | 0 | 1 | 0 | 9 | 0.1111 |
| 5_14411283 | 4 | 0.75 | 5 | 0.2 | 2 | 0 | 2 | 0 | 13 | 0.3077 | 2 | 0 | 4 | 0.25 | NA | NA | 1 | 0 | 7 | 0.1429 |
| 5_14411289 | 4 | 1 | 4 | 0.75 | 2 | 1 | 2 | 1 | 12 | 0.9167 | 3 | 1 | 2 | 1 | NA | NA | 1 | 0 | 6 | 0.8333 |
| 5_14411296 | 2 | 1 | 6 | 0.6667 | 3 | 0.6667 | 1 | 1 | 12 | 0.75 | 3 | 1 | 4 | 0.25 | NA | NA | 1 | 0 | 8 | 0.5 |
| 5_14411299 | 4 | 0.25 | 5 | 0.2 | 3 | 0.3333 | 3 | 0 | 15 | 0.2 | 3 | 0 | 4 | 0.25 | 1 | 0 | 1 | 1 | 9 | 0.2222 |
| 5_14411301 | 3 | 0 | 5 | 0.8 | 4 | 1 | 3 | 0 | 15 | 0.5333 | 3 | 0.6667 | 3 | 0.3333 | NA | NA | 1 | 1 | 7 | 0.5714 |
| 5_14411307 | 4 | 0 | 6 | 0.3333 | 4 | 0 | 3 | 0.3333 | 17 | 0.1765 | 3 | 0 | 4 | 0.25 | NA | NA | 1 | 1 | 8 | 0.25 |
| 5_14412003 | 5 | 0.2 | 5 | 0 | 2 | 0.5 | 3 | 0 | 15 | 0.1333 | 4 | 0.25 | 4 | 0 | 2 | 0 | 1 | 0 | 11 | 0.0909 |
| 5_14412155 | 5 | 0.2 | 6 | 0 | 4 | 0.75 | 4 | 0.5 | 19 | 0.3158 | 4 | 0.75 | 4 | 0.5 | 3 | 0.3333 | 1 | 1 | 12 | 0.5833 |
| 5_14412163 | 5 | 0.4 | 7 | 0.1429 | 2 | 0.5 | 4 | 0.5 | 18 | 0.3333 | 4 | 0.5 | 3 | 0.6667 | 4 | 0 | 1 | 0 | 12 | 0.3333 |
| 5_14412164 | 4 | 0.5 | 7 | 0.1429 | 4 | 0 | 4 | 0 | 19 | 0.1579 | 3 | 0 | 4 | 0.5 | 4 | 0 | 1 | 0 | 12 | 0.1667 |
| 5_14412181 | 4 | 0.25 | 7 | 0.4286 | 3 | 0.6667 | 2 | 1 | 16 | 0.5 | 4 | 0.5 | 5 | 0.8 | 4 | 0.25 | 1 | 1 | 14 | 0.5714 |

| | | | | | | | | | | | | | | | | | | | | |
|---|---|---|---|---|---|---|---|---|---|---|---|---|---|---|---|---|---|---|---|---|
| 5_14412196 | 5 | 0 | 8 | 0.125 | 4 | 0.5 | 3 | 0 | 20 | 0.15 | 3 | 0.3333 | 6 | 0 | 4 | 0.25 | 1 | 0 | 14 | 0.1429 |
| 5_14412225 | 9 | 0 | 10 | 0.2 | 3 | 0 | 4 | 0 | 26 | 0.0769 | 3 | 0.3333 | 8 | 0 | 3 | 0 | 2 | 0 | 16 | 0.0625 |
| 5_14412265 | 7 | 0 | 11 | 0.1818 | 7 | 0.1429 | 8 | 0 | 33 | 0.0909 | 4 | 0 | 5 | 0 | 3 | 0 | 3 | 0.6667 | 15 | 0.1333 |
| 5_14412295 | 9 | 0 | 10 | 0.2 | 7 | 0.2857 | 7 | 0 | 33 | 0.1212 | 6 | 0.3333 | 8 | 0.125 | 4 | 0.25 | 3 | 0.6667 | 21 | 0.2857 |
| 5_14412298 | 9 | 0 | 9 | 0 | 7 | 0.2857 | 7 | 0 | 32 | 0.0625 | 4 | 0 | 7 | 0 | 2 | 0 | 3 | 0.3333 | 16 | 0.0625 |
| 5_14412300 | 8 | 0.25 | 10 | 0.4 | 6 | 0.5 | 7 | 0 | 31 | 0.2903 | 5 | 0.2 | 8 | 0.375 | 4 | 0.25 | 3 | 0.6667 | 20 | 0.35 |
| 5_14412301 | 7 | 0.4286 | 9 | 0.1111 | 4 | 1 | 6 | 0.3333 | 26 | 0.3846 | 6 | 0.3333 | 7 | 0 | 3 | 1 | 2 | 1 | 18 | 0.3889 |
| 5_14412312 | 10 | 0.2 | 9 | 0.3333 | 6 | 0.3333 | 7 | 0 | 32 | 0.2188 | 6 | 0.5 | 5 | 0 | 4 | 0.5 | 3 | 0.3333 | 18 | 0.3333 |
| 5_14412319 | 7 | 0.4286 | 9 | 0.3333 | 6 | 0.5 | 3 | 0.3333 | 25 | 0.4 | 6 | 0.5 | 8 | 0.375 | 4 | 0.5 | 3 | 0.6667 | 21 | 0.4762 |
| 5_14412329 | 8 | 0.375 | 10 | 0.4 | 8 | 0.5 | 7 | 0.1429 | 33 | 0.3636 | 7 | 0.4286 | 5 | 0.2 | 4 | 0.5 | 3 | 0.3333 | 19 | 0.3684 |
| 5_14412422 | 11 | 0.0909 | 9 | 0.1111 | 10 | 0.2 | 16 | 0.125 | 46 | 0.1304 | 9 | 0.5556 | 8 | 0.125 | 5 | 0.6 | 4 | 0.5 | 26 | 0.4231 |
| 5_14413158 | 8 | 0.375 | 8 | 0 | 4 | 0.5 | 4 | 0 | 24 | 0.2083 | 8 | 0 | 8 | 0 | 4 | 0 | 8 | 0.25 | 28 | 0.0714 |
| 5_14413161 | 7 | 0 | 7 | 0 | 2 | 0.5 | 3 | 0 | 19 | 0.0526 | 7 | 0 | 7 | 0.1429 | 4 | 0 | 11 | 0.0909 | 29 | 0.069 |
| 5_14413176 | 3 | 0.3333 | 6 | 0 | 4 | 0 | 2 | 0 | 15 | 0.0667 | 5 | 0.4 | 8 | 0.125 | 5 | 0.2 | 9 | 0.2222 | 27 | 0.2222 |
| 5_14413184 | 8 | 0.5 | 10 | 0 | 3 | 0 | 4 | 0 | 25 | 0.16 | 8 | 0 | 8 | 0 | 5 | 0.2 | 10 | 0.1 | 31 | 0.0645 |
| 5_14413185 | 8 | 0.25 | 10 | 0 | 1 | 0 | 4 | 0 | 23 | 0.087 | 6 | 0.3333 | 8 | 0.125 | 4 | 0.25 | 10 | 0.2 | 28 | 0.2143 |
| 5_14413190 | 6 | 0 | 9 | 0.2222 | 2 | 0 | 4 | 0 | 21 | 0.0952 | 7 | 0 | 7 | 0 | 5 | 0.2 | 10 | 0.1 | 29 | 0.069 |
| 5_14413202 | 4 | 0.5 | 2 | 0 | 2 | 0 | 1 | 0 | 9 | 0.2222 | 6 | 0 | 4 | 0.25 | 3 | 0.3333 | 6 | 0.1667 | 19 | 0.1579 |
| 5_14413217 | 4 | 0.25 | 9 | 0.3333 | 1 | 0 | 2 | 0 | 16 | 0.25 | 6 | 0.3333 | 6 | 0.5 | 5 | 0.2 | 7 | 0.1429 | 24 | 0.2917 |
| 5_14413218 | 6 | 0.1667 | 9 | 0 | 1 | 0 | 3 | 0.3333 | 19 | 0.1053 | 8 | 0.125 | 6 | 0.3333 | 4 | 0.25 | 8 | 0 | 26 | 0.1538 |
| 5_14413257 | 6 | 0.3333 | 4 | 0.25 | NA | NA | NA | NA | 10 | 0.3 | 6 | 0.5 | 6 | 0.3333 | 3 | 1 | 4 | 0.25 | 19 | 0.4737 |
| 5_14413268 | 6 | 0.3333 | 6 | 0.6667 | 2 | 0.5 | 1 | 1 | 15 | 0.5333 | 8 | 0.5 | 6 | 0.5 | 5 | 0.2 | 6 | 0.5 | 25 | 0.44 |
| 5_14414705 | 2 | 1 | 2 | 1 | NA | NA | NA | NA | 4 | 1 | 1 | 1 | 2 | 0.5 | 1 | 1 | 2 | 0 | 6 | 0.5 |
| 5_14416222 | NA | NA | 1 | 0 | NA | NA | 1 | 1 | 2 | 0.5 | NA | NA | NA | NA | 1 | 1 | NA | NA | 1 | 1 |
| 5_14420104 | 14 | 0.5714 | 7 | 0.5714 | 13 | 0.5385 | 9 | 0.5556 | 43 | 0.5581 | 12 | 0.5833 | 16 | 0.1875 | 12 | 0.6667 | 9 | 0.4444 | 49 | 0.449 |
| 5_14449319 | 6 | 0.8333 | 6 | 1 | 5 | 0.8 | 4 | 1 | 21 | 0.9048 | 2 | 1 | 6 | 0.6667 | 1 | 1 | NA | NA | 9 | 0.7778 |
| 5_14449325 | 6 | 0.5 | 4 | 0 | 5 | 0 | 4 | 0.5 | 19 | 0.2632 | 1 | 1 | 6 | 0.6667 | 1 | 1 | NA | NA | 8 | 0.75 |
| 5_14449362 | 5 | 0.2 | 11 | 0.2727 | 6 | 0.3333 | 4 | 0.5 | 26 | 0.3077 | NA | NA | 8 | 0.25 | 2 | 0.5 | NA | NA | 10 | 0.3 |
| 5_14449362 | 2 | 0.5 | 5 | 0.2 | 3 | 0 | 3 | 0.3333 | 13 | 0.2308 | 1 | 1 | 3 | 0 | 2 | 0.5 | NA | NA | 6 | 0.3333 |
| 5_14449402 | 7 | 0.2857 | 11 | 0.1818 | 5 | 0 | 4 | 0 | 27 | 0.1481 | NA | NA | 7 | 0.4286 | 3 | 0.3333 | NA | NA | 10 | 0.4 |
| 5_14449406 | 8 | 0.125 | 12 | 0.3333 | 7 | 0.1429 | 5 | 0.2 | 32 | 0.2188 | 1 | 1 | 10 | 0.4 | 3 | 0 | NA | NA | 14 | 0.3571 |
| 5_14467387 | 14 | 0.0714 | 27 | 0.1111 | 19 | 0.1053 | 29 | 0 | 89 | 0.0674 | 9 | 0 | 25 | 0.04 | 25 | 0.12 | 20 | 0.05 | 79 | 0.0633 |
| 5_14467415 | 14 | 0.1429 | 19 | 0.3158 | 13 | 0.1538 | 18 | 0.2778 | 64 | 0.2344 | 4 | 0.25 | 19 | 0.3158 | 19 | 0.2105 | 12 | 0.4167 | 54 | 0.2963 |
| 5_14467424 | 15 | 0 | 24 | 0.0833 | 15 | 0.1333 | 24 | 0 | 78 | 0.0513 | 7 | 0 | 19 | 0.1053 | 19 | 0.1579 | 19 | 0.1053 | 64 | 0.1094 |
| 5_14469210 | 13 | 0.1538 | 17 | 0.3529 | 14 | 0.0714 | 22 | 0.1818 | 66 | 0.197 | 6 | 0.1667 | 21 | 0.1905 | 20 | 0.3 | 15 | 0.2 | 62 | 0.2258 |
| 5_14479555 | 6 | 0.5 | 6 | 0.3333 | 8 | 0.375 | 7 | 0.1429 | 27 | 0.3333 | 6 | 0.3333 | 5 | 0.4 | 5 | 0.4 | 5 | 0 | 21 | 0.2857 |
| 5_14479559 | 5 | 0.6 | 3 | 0.3333 | 3 | 0 | 6 | 0.6667 | 17 | 0.4706 | 7 | 0.1429 | 4 | 0.5 | 4 | 0.25 | 3 | 0.3333 | 18 | 0.2778 |
| 5_14479572 | 6 | 0 | 6 | 0 | 5 | 0.4 | 9 | 0.1111 | 26 | 0.1154 | 6 | 0.1667 | 7 | 0.2857 | 4 | 0 | 6 | 0.3333 | 23 | 0.2174 |
| 5_14479574 | 5 | 0.8 | 5 | 0.6 | 9 | 0.8889 | 9 | 0.5556 | 28 | 0.7143 | 9 | 1 | 7 | 0.5714 | 4 | 1 | 5 | 0.6 | 25 | 0.8 |
| 5_14479596 | 8 | 0.625 | 6 | 0.3333 | 9 | 0.3333 | 10 | 0.5 | 33 | 0.4545 | 10 | 0.4 | 9 | 0.6667 | 6 | 0.6667 | 6 | 0.1667 | 31 | 0.4839 |
| 5_14479601 | 9 | 0.1111 | 7 | 0.7143 | 9 | 0.2222 | 11 | 0.4545 | 36 | 0.3611 | 8 | 0.25 | 8 | 0.5 | 4 | 0.5 | 6 | 0.3333 | 26 | 0.3846 |
| 5_14479605 | 8 | 0.25 | 6 | 0.5 | 9 | 0.1111 | 11 | 0.4545 | 34 | 0.3235 | 9 | 0.1111 | 9 | 0.4444 | 4 | 0.75 | 8 | 0.25 | 30 | 0.3333 |
| 5_14479608 | 7 | 0 | 5 | 0.2 | 9 | 0.1111 | 11 | 0.0909 | 32 | 0.0938 | 9 | 0 | 9 | 0.3333 | 6 | 0 | 7 | 0 | 31 | 0.0968 |
| 5_14479621 | 8 | 0.5 | 5 | 0.4 | 6 | 0.6667 | 8 | 0.875 | 27 | 0.6296 | 9 | 0.8889 | 5 | 0.6 | 4 | 0.75 | 6 | 0.8333 | 24 | 0.7917 |
| 5_14479642 | 9 | 0.1111 | 5 | 0.2 | 8 | 0 | 15 | 0.0667 | 37 | 0.0811 | 9 | 0.1111 | 11 | 0.2727 | 6 | 0.5 | 10 | 0.2 | 36 | 0.25 |
| 5_14481358 | 4 | 0.25 | 2 | 0 | 4 | 0.5 | 1 | 1 | 11 | 0.3636 | 1 | 0 | 5 | 0.6 | 6 | 0.1667 | 1 | 1 | 13 | 0.3846 |
| 5_14481367 | 2 | 1 | 2 | 1 | 1 | 1 | 1 | 1 | 6 | 1 | 1 | 1 | 4 | 1 | 6 | 1 | NA | NA | 11 | 1 |
| 5_14481402 | 3 | 0.6667 | 2 | 0.5 | 4 | 0.75 | NA | NA | 9 | 0.6667 | 1 | 1 | 4 | 0.25 | 5 | 1 | 1 | 0 | 11 | 0.6364 |
| 5_14481421 | 2 | 0.5 | 1 | 1 | 4 | 0.25 | NA | NA | 7 | 0.4286 | 1 | 1 | 4 | 0.5 | 5 | 0.8 | 1 | 0 | 11 | 0.6364 |
| 5_14481439 | 3 | 0.6667 | 2 | 0.5 | 4 | 0.5 | NA | NA | 9 | 0.5556 | 1 | 1 | 4 | 0.5 | 6 | 0.5 | 1 | 1 | 12 | 0.5833 |
| 5_14481439 | 3 | 0.6667 | 2 | 0 | 4 | 0.5 | NA | NA | 9 | 0.4444 | 1 | 0 | 5 | 0.2 | 6 | 0.5 | 1 | 0 | 13 | 0.3077 |
| 5_14481449 | 1 | 0 | 2 | 0 | 3 | 0.3333 | NA | NA | 6 | 0.1667 | 1 | 0 | 4 | 0.25 | 5 | 0.2 | 1 | 0 | 11 | 0.1818 |
| 5_14481462 | 3 | 0.3333 | 2 | 0.5 | 4 | 0.5 | NA | NA | 9 | 0.4444 | NA | NA | 3 | 0 | 5 | 0.2 | 1 | 0 | 9 | 0.1111 |
| 5_14481466 | 1 | 1 | 2 | 0 | 3 | 0.3333 | NA | NA | 6 | 0.3333 | NA | NA | 2 | 0.5 | 5 | 0.4 | 1 | 0 | 8 | 0.375 |
| 5_14481534 | 2 | 1 | 1 | 1 | 4 | 1 | NA | NA | 7 | 1 | NA | NA | 2 | 0 | 2 | 0.5 | 1 | 1 | 5 | 0.4 |
| 5_14481563 | 1 | 1 | NA | NA | 3 | 0.3333 | NA | NA | 4 | 0.5 | NA | NA | 3 | 0 | 2 | 0 | 1 | 1 | 6 | 0.1667 |
| 5_14481601 | 1 | 1 | 1 | 1 | 3 | 1 | NA | NA | 5 | 1 | NA | NA | 1 | 1 | 2 | 1 | 1 | 1 | 4 | 1 |
| 5_14481606 | 1 | 1 | 1 | 0 | 3 | 0 | NA | NA | 5 | 0.2 | NA | NA | 2 | 0 | 1 | 0 | 1 | 1 | 4 | 0.25 |
| 5_14504526 | 45 | 0.0667 | 33 | 0.1212 | 51 | 0.098 | 48 | 0.1042 | 177 | 0.096 | 30 | 0.1333 | 36 | 0.0556 | 52 | 0.0577 | 37 | 0.1622 | 155 | 0.0968 |
| 5_14506495 | 3 | 0 | 4 | 0.25 | 2 | 0 | NA | NA | 9 | 0.1111 | 2 | 0 | 4 | 0 | 2 | 0 | 4 | 0.25 | 12 | 0.0833 |
| 5_14508453 | 6 | 0.5 | 5 | 0.8 | 1 | 0 | 6 | 0.6667 | 18 | 0.6111 | NA | NA | NA | NA | 2 | 0.5 | 1 | 0 | 3 | 0.3333 |
| 5_14508457 | 7 | 1 | 4 | 1 | NA | NA | 5 | 0.8 | 16 | 0.9375 | 1 | 0 | 2 | 1 | 2 | 0.5 | 2 | 1 | 7 | 0.7143 |
| 5_14508462 | 6 | 0.3333 | 4 | 0 | 1 | 0 | 5 | 0.4 | 16 | 0.25 | 2 | 0 | 3 | 0.6667 | 2 | 0.5 | 2 | 0.5 | 9 | 0.4444 |
| 5_14508473 | 18 | 0.3889 | 8 | 0.25 | 8 | 0.125 | 8 | 0.375 | 42 | 0.3095 | 5 | 0 | 11 | 0.1818 | 9 | 0.1111 | 15 | 0.2667 | 40 | 0.175 |
| 5_14508474 | 20 | 0.15 | 8 | 0.25 | 10 | 0.4 | 12 | 0.0833 | 50 | 0.2 | 8 | 0.375 | 13 | 0.2308 | 9 | 0.3333 | 16 | 0.1875 | 46 | 0.2609 |

| | | | | | | | | | | | | | | | | | | | | |
|---|---|---|---|---|---|---|---|---|---|---|---|---|---|---|---|---|---|---|---|---|
| 5_1450847 | 19 | 0.1053 | 8 | 0.125 | 10 | 0.1 | 12 | 0.0833 | 49 | 0.102 | 7 | 0.1429 | 13 | 0.2308 | 10 | 0.3 | 15 | 0.1333 | 45 | 0.2 |
| 5_1450848 | 7 | 0.1429 | 4 | 0 | 1 | 0 | 6 | 0.1667 | 18 | 0.1111 | 1 | 0 | 3 | 0.3333 | 3 | 0.3333 | NA | NA | 7 | 0.2857 |
| 5_1450849 | 8 | 0.5 | 5 | 0.6 | 1 | 1 | 6 | 0.8333 | 20 | 0.65 | 1 | 1 | 3 | 0.6667 | 2 | 0 | NA | NA | 6 | 0.5 |
| 5_1450851 | 5 | 0 | 5 | 0 | NA | NA | 7 | 0.1429 | 17 | 0.0588 | 1 | 0 | 2 | 0.5 | 2 | 0 | 1 | 0 | 6 | 0.1667 |
| 5_1450917 | 2 | 0.5 | 2 | 0.5 | NA | NA | 3 | 0.3333 | 7 | 0.4286 | NA | NA | 2 | 0.5 | 1 | 0 | 1 | 1 | 4 | 0.5 |
| 5_1450917 | 1 | 1 | 2 | 1 | NA | NA | 2 | 1 | 5 | 1 | 1 | 0 | 1 | 1 | NA | NA | 1 | 1 | 3 | 0.6667 |
| 5_1450918 | 2 | 0.5 | 2 | 0 | NA | NA | 2 | 0 | 6 | 0.1667 | NA | NA | 1 | 0 | 1 | 0 | 1 | 1 | 3 | 0.3333 |
| 5_1450919 | 2 | 0.5 | 1 | 1 | NA | NA | 1 | 0 | 4 | 0.5 | NA | NA | 2 | 0.5 | 1 | 1 | 1 | 1 | 4 | 0.75 |
| 5_1450921 | 2 | 0 | NA | NA | NA | NA | 2 | 1 | 4 | 0.5 | NA | NA | 1 | 0 | 1 | 0 | 1 | 1 | 3 | 0.3333 |
| 5_1451019 | 2 | 0.5 | 1 | 0 | NA | NA | 1 | 0 | 4 | 0.25 | NA | NA | 2 | 0.5 | NA | NA | 1 | 0 | 3 | 0.3333 |
| 5_1451181 | 2 | 1 | 1 | 0 | 2 | 1 | 3 | 0.3333 | 8 | 0.625 | NA | NA | 3 | 0.6667 | 2 | 1 | 2 | 1 | 7 | 0.8571 |
| 5_1451181 | 2 | 0 | 2 | 0.5 | 1 | 1 | 2 | 0.5 | 7 | 0.4286 | 2 | 1 | 3 | 1 | 2 | 0.5 | 2 | 0.5 | 9 | 0.7778 |
| 5_1451182 | 2 | 0.5 | NA | NA | 1 | 0 | 1 | 0 | 4 | 0.25 | 1 | 0 | 3 | 0.6667 | 2 | 0 | 1 | 1 | 7 | 0.4286 |
| 5_1451184 | 3 | 0.6667 | 2 | 0.5 | 3 | 0.3333 | 3 | 0.3333 | 11 | 0.4545 | 2 | 1 | 3 | 0.3333 | 3 | 0.6667 | 3 | 0.6667 | 11 | 0.6364 |
| 5_1451185 | 2 | 0.5 | NA | NA | 3 | 0 | 2 | 0.5 | 7 | 0.2857 | 2 | 0.5 | 2 | 0 | 3 | 0.3333 | 2 | 0 | 9 | 0.2222 |
| 5_1451206 | NA | NA | 1 | 1 | 1 | 0 | NA | NA | 2 | 0.5 | 1 | 1 | NA | NA | 1 | 0 | 2 | 0 | 4 | 0.25 |
| 5_1451207 | NA | NA | 1 | 1 | 1 | 0 | NA | NA | 2 | 0.5 | 1 | 1 | NA | NA | 1 | 0 | 2 | 0.5 | 4 | 0.5 |
| 5_1451318 | NA | NA | 1 | 0 | 2 | 0.5 | 2 | 0 | 5 | 0.2 | 1 | 0 | 1 | 0 | NA | NA | 4 | 0.25 | 6 | 0.1667 |
| 5_1451319 | NA | NA | 1 | 0 | 2 | 1 | 2 | 0.5 | 5 | 0.6 | 1 | 0 | NA | NA | NA | NA | 3 | 0.3333 | 4 | 0.25 |
| 5_1451320 | NA | NA | 2 | 0 | 2 | 0.5 | 2 | 0 | 6 | 0.1667 | 1 | 1 | 1 | 0 | NA | NA | 4 | 0.25 | 6 | 0.3333 |
| 5_1451320 | NA | NA | 1 | 1 | 2 | 1 | 2 | 0.5 | 5 | 0.8 | NA | NA | 1 | 1 | NA | NA | 3 | 0.6667 | 4 | 0.75 |
| 5_1451322 | NA | NA | 1 | 0 | 1 | 0 | 2 | 1 | 4 | 0.5 | NA | NA | 1 | 0 | NA | NA | 2 | 0.5 | 3 | 0.3333 |
| 5_1451324 | NA | NA | 1 | 1 | 1 | 1 | 2 | 0.5 | 4 | 0.75 | NA | NA | 2 | 1 | NA | NA | 1 | 0 | 3 | 0.6667 |
| 5_1451326 | NA | NA | 1 | 0 | 1 | 1 | 2 | 0.5 | 4 | 0.5 | NA | NA | 1 | 1 | NA | NA | 3 | 0.6667 | 4 | 0.75 |
| 5_1451326 | NA | NA | 1 | 1 | 1 | 0 | 2 | 1 | 4 | 0.75 | NA | NA | 1 | 1 | NA | NA | 3 | 0.6667 | 4 | 0.75 |
| 5_1451327 | NA | NA | 2 | 1 | 1 | 1 | 2 | 1 | 5 | 1 | NA | NA | 2 | 1 | NA | NA | 3 | 0.6667 | 5 | 0.8 |
| 5_1451328 | NA | NA | 1 | 0 | 1 | 0 | 2 | 0.5 | 4 | 0.25 | NA | NA | 1 | 1 | NA | NA | 3 | 1 | 4 | 1 |
| 5_1451414 | 1 | 1 | NA | NA | NA | NA | NA | NA | 1 | 1 | 1 | 0 | NA | NA | 1 | 1 | 1 | 1 | 3 | 0.6667 |
| 5_1451417 | 1 | 1 | NA | NA | NA | NA | NA | NA | 1 | 1 | 1 | 0 | NA | NA | 1 | 1 | 1 | 1 | 3 | 0.6667 |
| 5_1451552 | 1 | 1 | NA | NA | NA | NA | 1 | 0 | 2 | 0.5 | 3 | 0.3333 | NA | NA | 1 | 1 | 1 | 0 | 5 | 0.4 |
| 5_1451570 | 1 | 1 | 1 | 1 | NA | NA | 1 | 1 | 3 | 1 | 2 | 1 | NA | NA | 1 | 1 | 1 | 1 | 4 | 1 |
| 5_1451570 | NA | NA | 1 | 0 | NA | NA | 2 | 0.5 | 3 | 0.3333 | 3 | 0.3333 | NA | NA | 2 | 1 | 1 | 1 | 6 | 0.6667 |
| 5_1451574 | 1 | 0 | 3 | 0.3333 | NA | NA | 2 | 0.5 | 6 | 0.3333 | 3 | 0.6667 | NA | NA | 3 | 0.6667 | 1 | 0 | 7 | 0.5714 |
| 5_1451576 | 1 | 1 | 1 | 1 | NA | NA | 2 | 1 | 4 | 1 | 2 | 1 | NA | NA | 2 | 0.5 | NA | NA | 4 | 0.75 |
| 5_1451577 | 1 | 0 | 1 | 1 | NA | NA | 3 | 0.3333 | 5 | 0.4 | 2 | 0 | NA | NA | 3 | 0.6667 | 1 | 0 | 6 | 0.3333 |
| 5_1451579 | 1 | 1 | 2 | 0 | NA | NA | 3 | 1 | 6 | 0.6667 | 4 | 0.5 | 1 | 0 | 2 | 0 | 1 | 1 | 8 | 0.375 |
| 5_1451581 | 1 | 1 | 2 | 1 | NA | NA | 3 | 1 | 6 | 1 | 4 | 1 | 1 | 1 | 2 | 0.5 | 1 | 1 | 8 | 0.875 |
| 5_1451582 | 1 | 0 | 2 | 0 | NA | NA | 2 | 0.5 | 5 | 0.2 | 3 | 0.3333 | 1 | 0 | 2 | 0 | 1 | 0 | 7 | 0.1429 |
| 5_1451582 | 1 | 0 | 2 | 0.5 | NA | NA | 3 | 0.3333 | 6 | 0.3333 | 3 | 0.3333 | 1 | 0 | 2 | 1 | 1 | 1 | 7 | 0.5714 |
| 5_1451583 | 1 | 0 | 1 | 0 | NA | NA | 4 | 0.25 | 6 | 0.1667 | 3 | 0.3333 | 1 | 0 | 3 | 0 | 1 | 0 | 8 | 0.125 |
| 5_1451584 | 1 | 1 | 1 | 0 | NA | NA | 3 | 0 | 5 | 0.2 | 4 | 1 | NA | NA | 2 | 0.5 | 1 | 1 | 7 | 0.8571 |
| 5_1451584 | 1 | 0 | 2 | 1 | NA | NA | 3 | 0.6667 | 6 | 0.6667 | 4 | 0.25 | 1 | 0 | 3 | 0.3333 | 1 | 1 | 9 | 0.3333 |
| 5_1451741 | NA | NA | NA | NA | 4 | 1 | NA | NA | 4 | 1 | 1 | 1 | NA | NA | 1 | 1 | 1 | 1 | 3 | 1 |
| 5_1451741 | NA | NA | NA | NA | 2 | 1 | NA | NA | 2 | 1 | 1 | 1 | NA | NA | 2 | 1 | 1 | 0 | 4 | 0.75 |
| 5_1451805 | 5 | 0.4 | 2 | 0 | 1 | 1 | 2 | 0.5 | 10 | 0.4 | 1 | 0 | 1 | 0 | 4 | 0.5 | 1 | 1 | 7 | 0.4286 |
| 5_1451805 | 5 | 0.6 | 1 | 1 | 1 | 0 | 3 | 0 | 10 | 0.4 | 1 | 0 | 1 | 0 | 4 | 0.25 | 2 | 0 | 8 | 0.125 |
| 5_1451807 | 3 | 0.6667 | 2 | 0.5 | 2 | 1 | 3 | 1 | 10 | 0.8 | 1 | 1 | 1 | 0 | 3 | 0.6667 | 2 | 0.5 | 7 | 0.5714 |
| 5_1451807 | 3 | 0.6667 | 2 | 0.5 | 2 | 0.5 | 2 | 0 | 9 | 0.4444 | 1 | 1 | 1 | 0 | 3 | 0.3333 | NA | NA | 5 | 0.4 |
| 5_1451812 | 3 | 0.3333 | 2 | 0 | 2 | 0 | 3 | 0 | 10 | 0.1 | NA | NA | 2 | 0 | 5 | 0.2 | 2 | 0 | 9 | 0.1111 |
| 5_1451812 | 4 | 1 | 2 | 1 | 2 | 0.5 | 1 | 1 | 9 | 0.8889 | NA | NA | 1 | 1 | 4 | 1 | 2 | 1 | 7 | 1 |
| 5_1451812 | 4 | 0.75 | 2 | 0 | 2 | 0 | 3 | 0 | 11 | 0.2727 | 1 | 0 | 2 | 0.5 | 5 | 0.8 | 2 | 0 | 10 | 0.5 |
| 5_1451813 | 5 | 0.8 | 2 | 1 | 2 | 1 | 2 | 1 | 11 | 0.9091 | 1 | 1 | 2 | 1 | 4 | 0.75 | 2 | 1 | 9 | 0.8889 |
| 5_1451814 | 5 | 0.6 | 2 | 0.5 | 2 | 0.5 | 2 | 0 | 11 | 0.4545 | NA | NA | 2 | 0 | 4 | 0.75 | 2 | 1 | 8 | 0.625 |
| 5_1451816 | 5 | 0.6 | 2 | 0.5 | 2 | 0 | 4 | 0.5 | 13 | 0.4615 | 1 | 0 | 2 | 0 | 4 | 0.5 | 2 | 0.5 | 9 | 0.3333 |
| 5_1451817 | 5 | 1 | 1 | 1 | 2 | 1 | 4 | 1 | 12 | 1 | 1 | 1 | 2 | 1 | 2 | 1 | 1 | 0 | 6 | 0.8333 |
| 5_1451817 | 3 | 0.3333 | 2 | 0 | 2 | 0 | 4 | 0.25 | 11 | 0.1818 | 1 | 0 | 2 | 0 | 5 | 0.2 | 2 | 0 | 10 | 0.1 |
| 5_1451817 | 4 | 1 | 2 | 0.5 | 2 | 1 | 4 | 1 | 12 | 0.9167 | 1 | 1 | 2 | 1 | 5 | 1 | 2 | 1 | 10 | 1 |
| 5_1451818 | 5 | 1 | 2 | 1 | 1 | 0 | 2 | 1 | 10 | 0.9 | 1 | 1 | 2 | 1 | 4 | 1 | 2 | 1 | 9 | 1 |
| 5_1451823 | 5 | 0.6 | 2 | 0.5 | 2 | 0 | 4 | 0.5 | 13 | 0.4615 | 1 | 0 | 3 | 0.6667 | 5 | 0.4 | 3 | 0 | 12 | 0.3333 |
| 5_1451827 | 6 | 0.1667 | 2 | 0 | 1 | 0 | 4 | 0 | 13 | 0.0769 | NA | NA | 3 | 0.3333 | 6 | 0 | 3 | 0 | 12 | 0.0833 |
| 5_1452272 | 1 | 0 | 1 | 1 | NA | NA | NA | NA | 2 | 0.5 | 1 | 1 | NA | NA | 3 | 0 | 1 | 0 | 5 | 0.2 |
| 5_1452272 | 1 | 1 | 3 | 0.3333 | 1 | 1 | 1 | 1 | 6 | 0.6667 | 1 | 1 | 1 | 0 | 3 | 0.6667 | 2 | 0.5 | 7 | 0.5714 |
| 5_1452274 | 2 | 0.5 | 4 | 0.25 | 1 | 1 | 1 | 0 | 8 | 0.375 | 1 | 1 | 2 | 1 | 3 | 0.3333 | 2 | 1 | 8 | 0.75 |
| 5_1452275 | 2 | 1 | 3 | 0 | 1 | 1 | 1 | 0 | 7 | 0.4286 | 1 | 0 | 1 | 1 | 2 | 1 | 2 | 0.5 | 6 | 0.6667 |
| 5_1452276 | 2 | 1 | 3 | 0.6667 | 1 | 1 | 1 | 1 | 7 | 0.8571 | NA | NA | 2 | 0.5 | 2 | 1 | 3 | 1 | 7 | 0.8571 |

| | | | | | | | | | | | | | | | | | | | | |
|---|---|---|---|---|---|---|---|---|---|---|---|---|---|---|---|---|---|---|---|---|
| 5_1452280 | 1 | 1 | 4 | 1 | 1 | 1 | 1 | 1 | 7 | 1 | 1 | 1 | 2 | 1 | 2 | 1 | 2 | 1 | 7 | 1 |
| 5_1452285 | 2 | 0.5 | 4 | 0.75 | 2 | 1 | 1 | 0 | 9 | 0.6667 | 1 | 1 | 2 | 1 | 3 | 0.6667 | 1 | 0 | 7 | 0.7143 |
| 5_1452287 | 2 | 0.5 | 4 | 0.25 | 2 | 0 | 1 | 0 | 9 | 0.2222 | 1 | 0 | 2 | 0.5 | 3 | 0 | 1 | 0 | 7 | 0.1429 |
| 5_1452287 | 2 | 1 | 3 | 1 | 2 | 1 | 1 | 0 | 8 | 0.875 | 1 | 1 | 2 | 0.5 | 2 | 1 | 3 | 0.3333 | 8 | 0.625 |
| 5_1452288 | 2 | 0.5 | 3 | 1 | NA | NA | 1 | 0 | 6 | 0.6667 | 2 | 1 | 2 | 1 | 3 | 1 | 2 | 1 | 9 | 1 |
| 5_1452289 | 1 | 1 | 4 | 0.75 | 1 | 1 | 1 | 1 | 7 | 0.8571 | 3 | 1 | 3 | 1 | 2 | 1 | 2 | 1 | 10 | 1 |
| 5_1452289 | 1 | 0 | 3 | 0 | 2 | 0.5 | 1 | 0 | 7 | 0.1429 | 2 | 0 | 2 | 0.5 | 2 | 0.5 | 2 | 0 | 8 | 0.25 |
| 5_1461470 | 7 | 0.4286 | 9 | 0.6667 | 2 | 0.5 | 8 | 0.5 | 26 | 0.5385 | 4 | 0.5 | 7 | 0.1429 | 5 | 0.4 | 7 | 0.7143 | 23 | 0.4348 |
| 5_1461475 | 7 | 0.1429 | 9 | 0.3333 | 3 | 0 | 6 | 0.1667 | 25 | 0.2 | 3 | 0 | 7 | 0.1429 | 5 | 0 | 7 | 0.1429 | 22 | 0.0909 |
| 5_1461485 | 4 | 0.75 | 6 | 0.3333 | 2 | 0 | 2 | 0.5 | 14 | 0.4286 | 2 | 0.5 | 4 | 1 | 6 | 0.1667 | 7 | 0.2857 | 19 | 0.4211 |
| 5_1461485 | 3 | 0.3333 | 5 | 0.6 | 2 | 0.5 | 1 | 1 | 11 | 0.5455 | 2 | 0.5 | 3 | 0.3333 | 5 | 0.6 | 7 | 0.8571 | 17 | 0.6471 |
| 5_1461485 | 4 | 0 | 7 | 0.1429 | 2 | 0 | 1 | 0 | 14 | 0.0714 | 2 | 0 | 4 | 0 | 6 | 0 | 7 | 0.2857 | 19 | 0.1053 |
| 5_1461486 | 2 | 0.5 | 6 | 0.5 | 2 | 0.5 | 2 | 1 | 12 | 0.5833 | 1 | 1 | 3 | 0.3333 | 6 | 0 | 7 | 0.2857 | 17 | 0.2353 |
| 5_1461487 | 3 | 0 | 7 | 0.2857 | 2 | 0.5 | 2 | 0 | 14 | 0.2143 | 1 | 0 | 4 | 0 | 6 | 0.3333 | 7 | 0 | 18 | 0.1111 |
| 5_1461685( | NA | NA | 1 | 0 | 1 | 1 | 1 | 0 | 3 | 0.3333 | NA | NA | 3 | 0.6667 | 3 | 0.6667 | 1 | 1 | 7 | 0.7143 |
| 5_1462385 | 2 | 0 | 1 | 0 | 4 | 0.25 | 3 | 0.3333 | 10 | 0.2 | 1 | 0 | 5 | 0.2 | 5 | 0.6 | 3 | 0 | 14 | 0.2857 |
| 5_1464071 | 1 | 1 | 3 | 0 | NA | NA | 4 | 0.75 | 8 | 0.5 | 1 | 0 | NA | NA | 1 | 0 | 3 | 1 | 5 | 0.6 |
| 5_1464072 | 1 | 1 | 3 | 1 | NA | NA | 5 | 0.2 | 9 | 0.5556 | NA | NA | NA | NA | 1 | 1 | 2 | 0 | 3 | 0.3333 |
| 5_1465193 | 2 | 0 | 2 | 0 | 3 | 0 | 2 | 0.5 | 9 | 0.1111 | NA | NA | 5 | 0.4 | 3 | 0 | 6 | 0 | 14 | 0.1429 |
| 5_1465196 | 5 | 0.2 | 6 | 0.1667 | 5 | 0.6 | 3 | 0 | 19 | 0.2632 | NA | NA | 10 | 0.5 | 3 | 0.3333 | 5 | 0.4 | 18 | 0.4444 |
| 5_1465197 | 3 | 0.6667 | 3 | 0.6667 | 2 | 0.5 | 2 | 0.5 | 10 | 0.6 | NA | NA | 8 | 0.375 | NA | NA | 3 | 0.3333 | 11 | 0.3636 |
| 5_1465197 | 5 | 1 | 4 | 0.75 | 5 | 0.8 | 3 | 1 | 17 | 0.8824 | NA | NA | 11 | 0.7273 | 2 | 0.5 | 5 | 1 | 18 | 0.7778 |
| 5_1465197 | 4 | 0.25 | 4 | 0.5 | 5 | 0.2 | 3 | 0.6667 | 16 | 0.375 | NA | NA | 10 | 0.2 | 2 | 1 | 5 | 0.2 | 17 | 0.2941 |
| 5_1465198 | 5 | 0.4 | 4 | 0 | 5 | 0.4 | 3 | 0.3333 | 17 | 0.2941 | NA | NA | 9 | 0.1111 | 2 | 0.5 | 5 | 0.2 | 16 | 0.1875 |
| 5_1465201 | 4 | 1 | 5 | 1 | 5 | 0.8 | 2 | 1 | 16 | 0.9375 | NA | NA | 9 | 1 | 3 | 1 | 5 | 1 | 17 | 1 |
| 5_1465201 | 4 | 0.25 | 2 | 0 | 4 | 0.25 | 2 | 0.5 | 12 | 0.25 | NA | NA | 4 | 0.25 | 1 | 0 | 4 | 0.25 | 9 | 0.2222 |
| 5_1465203 | 4 | 0 | 4 | 0.25 | 5 | 0 | 2 | 0 | 15 | 0.0667 | NA | NA | 8 | 0.25 | 2 | 0 | 4 | 0.25 | 14 | 0.2143 |
| 5_1465203 | 4 | 0.25 | 5 | 0.2 | 4 | 0 | 1 | 0 | 14 | 0.1429 | NA | NA | 9 | 0.3333 | 3 | 0.3333 | 5 | 0.2 | 17 | 0.2941 |
| 5_1465203 | 3 | 1 | 1 | 1 | 5 | 1 | 1 | 1 | 10 | 1 | NA | NA | 8 | 0.875 | 1 | 1 | 5 | 1 | 14 | 0.9286 |
| 5_1465204 | 4 | 0.25 | 5 | 0 | 4 | 0 | 2 | 0 | 15 | 0.0667 | NA | NA | 9 | 0.1111 | 2 | 0 | 5 | 0 | 16 | 0.0625 |
| 5_1465204 | 4 | 0.5 | 4 | 0 | 5 | 0.8 | 2 | 0 | 15 | 0.4 | NA | NA | 8 | 0.125 | 2 | 0.5 | 5 | 0.2 | 15 | 0.2 |
| 5_1465205 | 4 | 0.75 | 2 | 0.5 | 3 | 1 | 2 | 0.5 | 11 | 0.7273 | NA | NA | 8 | 0.625 | 2 | 1 | 3 | 1 | 13 | 0.7692 |
| 5_1465205 | 4 | 1 | 4 | 1 | 5 | 1 | 2 | 1 | 15 | 1 | NA | NA | 6 | 1 | 3 | 1 | 5 | 1 | 14 | 1 |
| 5_1465206 | 3 | 1 | 4 | 1 | 5 | 1 | 2 | 1 | 14 | 1 | NA | NA | 8 | 1 | 1 | 1 | 5 | 1 | 14 | 1 |
| 5_1465207 | 3 | 1 | 4 | 1 | 5 | 1 | 1 | 1 | 13 | 1 | NA | NA | 6 | 1 | 1 | 1 | 5 | 1 | 12 | 1 |
| 5_1465208 | 2 | 1 | 4 | 0.75 | 4 | 1 | 2 | 0.5 | 12 | 0.8333 | NA | NA | 6 | 1 | 1 | 1 | 5 | 1 | 12 | 1 |
| 5_1466723 | 6 | 0 | 6 | 0.3333 | 3 | 0.3333 | 3 | 0 | 18 | 0.1667 | 4 | 0 | 5 | 0.2 | 5 | 0.2 | 6 | 0 | 20 | 0.1 |
| 5_1466728 | 3 | 0 | 3 | 0 | 2 | 0 | 2 | 0.5 | 10 | 0.1 | 4 | 0 | 5 | 0 | 5 | 0.2 | 5 | 0 | 19 | 0.0526 |
| 5_1466730 | 1 | 0 | 1 | 1 | NA | NA | 2 | 0.5 | 4 | 0.5 | 3 | 0 | 4 | 0.25 | 5 | 0 | 3 | 0 | 15 | 0.0667 |
| 5_1466731 | 3 | 0.3333 | 3 | 0 | 1 | 1 | 2 | 0.5 | 9 | 0.3333 | 3 | 0 | 5 | 0 | 5 | 0.2 | 5 | 0.4 | 18 | 0.1667 |
| 5_1466736 | 2 | 0.5 | 2 | 0 | NA | NA | 2 | 0 | 6 | 0.1667 | 3 | 0 | 3 | 0.3333 | 4 | 0.25 | 5 | 0.2 | 15 | 0.2 |
| 5_1466738 | 2 | 0.5 | 2 | 0.5 | NA | NA | 2 | 1 | 6 | 0.6667 | 2 | 0.5 | 2 | 1 | 2 | 1 | 4 | 0 | 10 | 0.5 |
| 5_1466743 | 2 | 1 | 2 | 1 | 1 | 1 | 2 | 1 | 7 | 1 | 1 | 1 | 2 | 1 | 3 | 1 | 3 | 1 | 9 | 1 |
| 5_1466744 | 2 | 0 | 2 | 1 | 1 | 1 | 2 | 1 | 7 | 0.7143 | 2 | 1 | 1 | 1 | 3 | 1 | 3 | 0.6667 | 9 | 0.8889 |
| 5_1467839( | NA | NA | 4 | 0.5 | 2 | 1 | 1 | 1 | 7 | 0.7143 | 3 | 0.6667 | 2 | 1 | NA | NA | NA | NA | 5 | 0.8 |
| 5_1467841 | NA | NA | 4 | 0.5 | 2 | 0 | 1 | 0 | 7 | 0.2857 | 3 | 0.6667 | 1 | 1 | 1 | 0 | NA | NA | 5 | 0.6 |
| 5_1467842 | NA | NA | 4 | 0.25 | 1 | 0 | NA | NA | 5 | 0.2 | 3 | 0.6667 | 2 | 0.5 | 1 | 0 | NA | NA | 6 | 0.5 |
| 5_1467843( | NA | NA | 4 | 0.5 | 2 | 1 | 1 | 1 | 7 | 0.7143 | 3 | 0.6667 | 2 | 1 | 1 | 1 | NA | NA | 6 | 0.8333 |
| 5_1467843 | NA | NA | 4 | 0.5 | 2 | 1 | 1 | 1 | 7 | 0.7143 | 3 | 0.6667 | 2 | 1 | NA | NA | NA | NA | 5 | 0.8 |
| 5_1468918 | 1 | 1 | 2 | 1 | 2 | 1 | 4 | 0.75 | 9 | 0.8889 | 2 | 1 | NA | NA | 3 | 1 | 1 | 1 | 6 | 1 |
| 5_1468918 | 2 | 0 | 2 | 0.5 | 2 | 0.5 | 4 | 0 | 10 | 0.2 | 2 | 0.5 | 2 | 0 | 3 | 0 | 2 | 0 | 9 | 0.1111 |
| 5_1468919 | 2 | 1 | 1 | 1 | 2 | 1 | 4 | 1 | 9 | 1 | 2 | 1 | 2 | 1 | 3 | 0.6667 | 2 | 1 | 9 | 0.8889 |
| 5_1468920 | 2 | 1 | 1 | 1 | NA | NA | 2 | 1 | 5 | 1 | 1 | 1 | 2 | 1 | 3 | 1 | 1 | 0 | 7 | 0.8571 |
| 5_1468922 | 2 | 1 | 1 | 1 | 2 | 1 | 4 | 0.75 | 9 | 0.8889 | 2 | 1 | 2 | 1 | 2 | 1 | 1 | 1 | 7 | 1 |
| 5_1468923 | 1 | 0 | 1 | 0 | 2 | 0 | 4 | 0.25 | 8 | 0.125 | 2 | 0 | 2 | 0 | 2 | 0 | 2 | 0.5 | 8 | 0.125 |
| 5_1468924 | 2 | 1 | 1 | 1 | 2 | 1 | 3 | 1 | 8 | 1 | 2 | 0.5 | 2 | 0.5 | 3 | 1 | 2 | 1 | 9 | 0.7778 |
| 5_1468924 | 2 | 0 | 1 | 0 | 2 | 0.5 | 4 | 0 | 9 | 0.1111 | 2 | 0 | 2 | 0 | 2 | 0.5 | 2 | 0 | 8 | 0.125 |
| 5_1468925 | 2 | 0 | 1 | 0 | 2 | 0.5 | 4 | 0.5 | 9 | 0.3333 | 2 | 0 | 2 | 0 | 3 | 1 | 2 | 0.5 | 9 | 0.4444 |
| 5_1468925 | 2 | 0 | 1 | 0 | 2 | 0.5 | 3 | 0 | 8 | 0.125 | 1 | 0 | 2 | 0 | 3 | 0 | 2 | 0.5 | 8 | 0.125 |
| 5_1468957 | 2 | 0.5 | 1 | 0 | NA | NA | 2 | 0 | 5 | 0.2 | 1 | 1 | 1 | 0 | 3 | 0 | 1 | 0 | 6 | 0.1667 |
| 5_1468957 | 2 | 0.5 | 1 | 0 | 1 | 0 | 4 | 0.5 | 8 | 0.375 | 1 | 0 | 2 | 1 | 3 | 0.6667 | 2 | 1 | 8 | 0.75 |
| 5_1468958 | 2 | 1 | 1 | 1 | 1 | 1 | 4 | 0.75 | 8 | 0.875 | 1 | 1 | 2 | 1 | 3 | 1 | 1 | 1 | 7 | 1 |
| 5_1469541 | NA | NA | NA | NA | 2 | 1 | NA | NA | 2 | 1 | 1 | 1 | 1 | 0 | NA | NA | 1 | 1 | 3 | 0.6667 |
| 5_1469551 | NA | NA | NA | NA | 1 | 1 | NA | NA | 1 | 1 | NA | NA | 1 | 1 | NA | NA | 1 | 0 | 2 | 0.5 |
| 5_1469551 | NA | NA | NA | NA | 2 | 1 | NA | NA | 2 | 1 | NA | NA | 1 | 1 | NA | NA | NA | NA | 1 | 1 |

| | | | | | | | | | | | | | | | | | | | | |
|---|---|---|---|---|---|---|---|---|---|---|---|---|---|---|---|---|---|---|---|---|
| 5_1469552 | NA | NA | NA | NA | 2 | 0.5 | NA | NA | 2 | 0.5 | NA | NA | 1 | 1 | NA | NA | 1 | 1 | 2 | 1 |
| 5_1469553 | NA | NA | NA | NA | 2 | 1 | NA | NA | 2 | 1 | NA | NA | 1 | 1 | NA | NA | 1 | 1 | 2 | 1 |
| 5_1469554 | NA | NA | NA | NA | 2 | 0.5 | NA | NA | 2 | 0.5 | NA | NA | 1 | 0 | NA | NA | 1 | 1 | 2 | 0.5 |
| 5_1469556 | NA | NA | NA | NA | 1 | 1 | NA | NA | 1 | 1 | NA | NA | 1 | 0 | NA | NA | 1 | 1 | 2 | 0.5 |
| 5_1469556 | NA | NA | NA | NA | 2 | 1 | NA | NA | 2 | 1 | NA | NA | 1 | 1 | NA | NA | 1 | 0 | 2 | 0.5 |
| 5_1470154 | 4 | 0.5 | 6 | 0 | 9 | 0.2222 | 10 | 0.2 | 29 | 0.2069 | 13 | 0.6154 | 9 | 0.1111 | 5 | 0.4 | 7 | 0.4286 | 34 | 0.4118 |
| 5_1470154 | 4 | 0 | 5 | 0.6 | 9 | 0.2222 | 8 | 0 | 26 | 0.1923 | 10 | 0.1 | 10 | 0.2 | 7 | 0 | 9 | 0.1111 | 36 | 0.1111 |
| 5_1470154 | 5 | 0 | 7 | 0.1429 | 9 | 0.1111 | 11 | 0.3636 | 32 | 0.1875 | 11 | 0.0909 | 10 | 0.1 | 5 | 0.2 | 9 | 0.1111 | 35 | 0.1143 |
| 5_1470154 | 5 | 0 | 5 | 0.2 | 8 | 0.25 | 10 | 0.1 | 28 | 0.1429 | 11 | 0.2727 | 11 | 0.4545 | 6 | 0.1667 | 7 | 0.2857 | 35 | 0.3143 |
| 5_1470155 | 6 | 0 | 5 | 0 | 9 | 0.2222 | 9 | 0.2222 | 29 | 0.1379 | 12 | 0.25 | 9 | 0 | 7 | 0 | 11 | 0.1818 | 39 | 0.1282 |
| 5_1470156 | 5 | 1 | 7 | 1 | 9 | 1 | 10 | 1 | 31 | 1 | 11 | 1 | 9 | 1 | 5 | 1 | 11 | 1 | 36 | 1 |
| 5_1470157 | 5 | 0.4 | 6 | 0 | 8 | 0 | 9 | 0.1111 | 28 | 0.1071 | 10 | 0.2 | 10 | 0.3 | 5 | 0.2 | 6 | 0 | 31 | 0.1935 |
| 5_1470478 | 10 | 0.8 | 2 | 0.5 | 16 | 0.8125 | 9 | 1 | 37 | 0.8378 | 6 | 0.8333 | 7 | 0.8571 | 8 | 0.75 | 11 | 0.9091 | 32 | 0.8438 |
| 5_1470479 | 10 | 0 | 4 | 0 | 16 | 0.1875 | 9 | 0.1111 | 39 | 0.1026 | 9 | 0.1111 | 8 | 0.125 | 9 | 0 | 14 | 0 | 40 | 0.05 |
| 5_1474825 | 1 | 0 | 2 | 0.5 | 1 | 1 | 1 | 1 | 5 | 0.6 | 1 | 1 | 2 | 1 | 1 | 1 | 2 | 1 | 6 | 1 |
| 5_1474825 | NA | NA | 2 | 0.5 | NA | NA | 1 | 0 | 3 | 0.3333 | 2 | 0 | 1 | 0 | 1 | 0 | 2 | 0.5 | 6 | 0.1667 |
| 5_1474826 | NA | NA | 2 | 0.5 | NA | NA | 1 | 1 | 3 | 0.6667 | 2 | 0.5 | 1 | 0 | 1 | 0 | 2 | 0 | 6 | 0.1667 |
| 5_1474827 | 1 | 0 | 2 | 0.5 | NA | NA | 1 | 1 | 4 | 0.5 | 2 | 1 | 3 | 1 | 1 | 1 | 3 | 0.3333 | 9 | 0.7778 |
| 5_1474827 | 1 | 0 | 2 | 0 | NA | NA | 1 | 1 | 4 | 0.25 | 2 | 1 | 3 | 0 | 1 | 0 | 4 | 0 | 10 | 0.2 |
| 5_1474828 | 1 | 0 | 2 | 1 | NA | NA | 1 | 1 | 4 | 0.75 | 2 | 1 | 3 | 0.6667 | 1 | 1 | 4 | 1 | 10 | 0.9 |
| 5_1474829 | NA | NA | 2 | 1 | NA | NA | 1 | 1 | 3 | 1 | 2 | 0.5 | 3 | 0.6667 | 1 | 1 | 4 | 1 | 10 | 0.8 |
| 5_1474834 | 2 | 0.5 | 1 | 0 | 2 | 0.5 | 2 | 0.5 | 7 | 0.4286 | 3 | 0.6667 | 4 | 0 | 1 | 0 | 4 | 0.25 | 12 | 0.25 |
| 5_1474835 | 2 | 0.5 | 1 | 0 | 2 | 0.5 | 1 | 0 | 6 | 0.3333 | 3 | 0 | 5 | 0.4 | 1 | 0 | 2 | 0.5 | 11 | 0.2727 |
| 5_1474835 | 3 | 0.6667 | 2 | 1 | 2 | 1 | 2 | 0.5 | 9 | 0.7778 | 2 | 1 | 4 | 0.75 | 2 | 0.5 | 3 | 0.3333 | 11 | 0.6364 |
| 5_1474835 | 3 | 0.6667 | 2 | 1 | 2 | 1 | 2 | 1 | 9 | 0.8889 | 1 | 1 | 5 | 1 | 2 | 1 | 4 | 1 | 12 | 1 |
| 5_1474835 | 3 | 0 | 2 | 0.5 | 2 | 0 | 2 | 0 | 9 | 0.1111 | 1 | 0 | 4 | 0 | 1 | 0 | 3 | 0.3333 | 9 | 0.1111 |
| 5_1474836 | 2 | 0.5 | 1 | 1 | 1 | 1 | 2 | 0.5 | 6 | 0.6667 | 2 | 1 | 3 | 1 | 1 | 1 | 2 | 1 | 8 | 1 |
| 5_1474836 | 3 | 1 | 2 | 0.5 | 2 | 1 | 2 | 1 | 9 | 0.8889 | 3 | 0 | 4 | 0.75 | 2 | 0 | 4 | 0.75 | 13 | 0.4615 |
| 5_1474837 | 3 | 0.3333 | 3 | 0.3333 | 2 | 0 | 3 | 0.3333 | 11 | 0.2727 | 2 | 0.5 | 5 | 0.2 | 2 | 1 | 4 | 0 | 13 | 0.3077 |
| 5_1474838 | 3 | 0.3333 | 3 | 0.6667 | 2 | 0 | 3 | 0.3333 | 11 | 0.3636 | 3 | 0 | 5 | 0.4 | 2 | 0.5 | 4 | 0.5 | 14 | 0.3571 |
| 5_1474838 | 3 | 0.3333 | 3 | 0 | 2 | 0 | 3 | 0.6667 | 11 | 0.2727 | 3 | 0 | 5 | 0.2 | 2 | 0.5 | 3 | 0 | 13 | 0.1538 |
| 5_1474840 | 4 | 1 | 3 | 1 | 2 | 1 | 5 | 0.6 | 14 | 0.8571 | 3 | 1 | 5 | 0.6 | NA | NA | 3 | 1 | 11 | 0.8182 |
| 5_1474840 | 4 | 0.5 | 2 | 0.5 | 2 | 0.5 | 5 | 0.6 | 13 | 0.5385 | 3 | 0.3333 | 5 | 0.4 | 1 | 0 | 4 | 0.5 | 13 | 0.3846 |
| 5_1474841 | 4 | 0.25 | 2 | 0.5 | 2 | 0 | 4 | 0 | 12 | 0.1667 | 3 | 0.3333 | 4 | 0 | 2 | 0 | 4 | 0 | 13 | 0.0769 |
| 5_1474841 | 4 | 0.75 | 2 | 1 | 2 | 1 | 4 | 0.75 | 12 | 0.8333 | 4 | 1 | 5 | 1 | 1 | 1 | 3 | 1 | 13 | 1 |
| 5_1474842 | 3 | 0.6667 | 2 | 1 | 2 | 1 | 5 | 1 | 12 | 0.9167 | 4 | 1 | 3 | 1 | 2 | 0.5 | 4 | 1 | 13 | 0.9231 |
| 5_1474846 | 4 | 0 | 5 | 0 | 3 | 0.3333 | 5 | 0.2 | 17 | 0.1176 | 6 | 0.1667 | 4 | 0 | 3 | 0 | 5 | 0.2 | 18 | 0.1111 |
| 5_1474849 | 5 | 0 | 4 | 0 | 4 | 0.25 | 6 | 0 | 19 | 0.0526 | 4 | 0 | 5 | 0 | 4 | 0.25 | 4 | 0 | 17 | 0.0588 |
| 5_1475722 | 2 | 0 | 4 | 0.5 | NA | NA | NA | NA | 6 | 0.3333 | NA | NA | 5 | 0.4 | NA | NA | NA | NA | 5 | 0.4 |
| 5_1475755 | NA | NA | 3 | 1 | NA | NA | NA | NA | 3 | 1 | 1 | 1 | 1 | 1 | NA | NA | NA | NA | 2 | 1 |
| 5_1475755 | NA | NA | 3 | 0.6667 | NA | NA | NA | NA | 3 | 0.6667 | 1 | 1 | 1 | 0 | NA | NA | NA | NA | 2 | 0.5 |
| 5_1475758 | NA | NA | 3 | 1 | NA | NA | NA | NA | 3 | 1 | 1 | 1 | 1 | 1 | NA | NA | NA | NA | 2 | 1 |
| 5_1475759 | 1 | 1 | 3 | 1 | NA | NA | NA | NA | 4 | 1 | 1 | 1 | 1 | 1 | NA | NA | NA | NA | 2 | 1 |
| 5_1475761 | 2 | 0.5 | 1 | 0 | NA | NA | 1 | 0 | 4 | 0.25 | 1 | 1 | 1 | 0 | 1 | 0 | 1 | 0 | 4 | 0.25 |
| 5_1475762 | 2 | 1 | 2 | 1 | NA | NA | 1 | 1 | 5 | 1 | 1 | 1 | 1 | 1 | 1 | 0 | 2 | 0 | 5 | 0.4 |
| 5_1475762 | 2 | 0.5 | 2 | 1 | NA | NA | 1 | 1 | 5 | 0.8 | 1 | 1 | 1 | 0 | 1 | 0 | 2 | 0 | 5 | 0.2 |
| 5_1476627 | 1 | 0 | NA | NA | NA | NA | 1 | 1 | 2 | 0.5 | 1 | 1 | NA | NA | 1 | 0 | NA | NA | 2 | 0.5 |
| 5_1478153 | 2 | 0.5 | 3 | 0.6667 | 3 | 0.6667 | NA | NA | 8 | 0.625 | NA | NA | 1 | 1 | 2 | 0.5 | 1 | 0 | 4 | 0.5 |
| 5_1478154 | 1 | 1 | 2 | 0.5 | 2 | 0.5 | NA | NA | 5 | 0.6 | NA | NA | NA | NA | 3 | 0.3333 | 4 | 0.25 | 7 | 0.2857 |
| 5_1478155 | 3 | 0.6667 | 2 | 0.5 | 3 | 0 | NA | NA | 8 | 0.375 | NA | NA | 1 | 1 | 3 | 0.6667 | 1 | 0 | 5 | 0.6 |
| 5_1478155 | 3 | 0.3333 | 2 | 0.5 | 3 | 1 | NA | NA | 8 | 0.625 | NA | NA | 1 | 0 | 3 | 0 | 4 | 0.25 | 8 | 0.125 |
| 5_1478155 | 3 | 0.6667 | 2 | 1 | 3 | 0 | NA | NA | 8 | 0.5 | NA | NA | 1 | 1 | 3 | 0.6667 | 3 | 0.6667 | 7 | 0.7143 |
| 5_1478156 | 3 | 0.3333 | 2 | 0 | 3 | 0 | NA | NA | 8 | 0.125 | NA | NA | 1 | 0 | 3 | 0 | 3 | 0.6667 | 7 | 0.2857 |
| 5_1478157 | 3 | 0.3333 | 2 | 0 | 3 | 0 | NA | NA | 8 | 0.125 | NA | NA | 1 | 0 | 2 | 0 | 4 | 0.25 | 7 | 0.1429 |
| 5_1478158 | 3 | 0.3333 | 2 | 0.5 | 3 | 0 | NA | NA | 8 | 0.25 | NA | NA | 1 | 1 | 1 | 0 | 3 | 0.3333 | 5 | 0.4 |
| 5_1478158 | 3 | 0 | 2 | 0.5 | 3 | 0 | NA | NA | 8 | 0.125 | NA | NA | 1 | 0 | 1 | 0 | 4 | 0.25 | 6 | 0.1667 |
| 5_1478231 | 2 | 1 | 2 | 1 | 1 | 1 | NA | NA | 5 | 1 | NA | NA | 1 | 0 | 2 | 0 | 3 | 1 | 6 | 0.5 |
| 5_1478231 | 2 | 1 | 2 | 1 | 1 | 1 | NA | NA | 5 | 1 | NA | NA | 1 | 1 | 2 | 1 | 3 | 0.3333 | 6 | 0.6667 |
| 5_1478231 | NA | NA | 2 | 0.5 | 1 | 1 | NA | NA | 3 | 0.6667 | NA | NA | 1 | 1 | 2 | 0.5 | 2 | 0.5 | 5 | 0.6 |
| 5_1478232 | 2 | 1 | 2 | 1 | 1 | 0 | NA | NA | 5 | 0.8 | NA | NA | 1 | 1 | NA | NA | 2 | 1 | 3 | 1 |
| 5_1478929 | NA | NA | NA | NA | 2 | 0.5 | NA | NA | 2 | 0.5 | NA | NA | NA | NA | 1 | 1 | NA | NA | 1 | 1 |
| 5_1478934 | NA | NA | 1 | 0 | 1 | 1 | NA | NA | 2 | 0.5 | NA | NA | NA | NA | 1 | 1 | NA | NA | 1 | 1 |
| 5_1478935 | NA | NA | 1 | 1 | 1 | 1 | NA | NA | 2 | 1 | NA | NA | NA | NA | 1 | 1 | NA | NA | 1 | 1 |
| 5_1478936 | NA | NA | 1 | 1 | 1 | 1 | NA | NA | 2 | 1 | NA | NA | NA | NA | 1 | 1 | NA | NA | 1 | 1 |
| 5_1478937 | NA | NA | 1 | 1 | 1 | 1 | NA | NA | 2 | 1 | NA | NA | NA | NA | 1 | 1 | NA | NA | 1 | 1 |

| | | | | | | | | | | | | | | | | | | | | |
|---|---|---|---|---|---|---|---|---|---|---|---|---|---|---|---|---|---|---|---|---|
| 5_1478938 | NA | NA | NA | NA | 1 | 1 | NA | NA | 1 | 1 | NA | NA | NA | NA | 1 | 1 | NA | NA | 1 | 1 |
| 5_1478939 | NA | NA | 1 | 1 | 1 | 0 | NA | NA | 2 | 0.5 | NA | NA | NA | NA | 1 | 1 | NA | NA | 1 | 1 |
| 5_1480147 | NA | NA | 1 | 1 | NA | NA | NA | NA | 1 | 1 | 1 | 0 | 1 | 1 | NA | NA | NA | NA | 2 | 0.5 |
| 5_1482501 | 4 | 0 | 4 | 0.25 | NA | NA | NA | NA | 8 | 0.125 | 4 | 0.25 | 5 | 0.2 | 2 | 0 | 2 | 0 | 13 | 0.1538 |
| 5_1482504 | 4 | 0 | 5 | 0.4 | NA | NA | NA | NA | 9 | 0.2222 | 4 | 0.25 | 5 | 0 | 2 | 0 | 2 | 0 | 13 | 0.0769 |
| 5_1482506 | 3 | 0.3333 | 5 | 0.4 | NA | NA | NA | NA | 8 | 0.375 | 4 | 0 | 5 | 0.2 | 2 | 0.5 | 2 | 0 | 13 | 0.1538 |
| 5_1482506 | 3 | 0.3333 | 5 | 0 | NA | NA | NA | NA | 8 | 0.125 | 4 | 0.25 | 7 | 0.1429 | 1 | 1 | 3 | 0.3333 | 15 | 0.2667 |
| 5_1482516 | 4 | 0 | 6 | 0.3333 | NA | NA | NA | NA | 10 | 0.2 | 4 | 0 | 8 | 0.25 | 1 | 0 | 3 | 0.6667 | 16 | 0.25 |
| 5_1482519 | 4 | 0.25 | 6 | 0.1667 | 1 | 0 | NA | NA | 11 | 0.1818 | 4 | 0 | 8 | 0 | 2 | 0 | 3 | 0.3333 | 17 | 0.0588 |
| 5_1482525 | 5 | 0.2 | 6 | 0.1667 | 2 | 0 | NA | NA | 13 | 0.1538 | 7 | 0.4286 | 7 | 0.1429 | 4 | 0.25 | 5 | 0.2 | 23 | 0.2609 |
| 5_1482527 | 4 | 0.5 | 6 | 0.5 | 2 | 1 | NA | NA | 12 | 0.5833 | 7 | 0.7143 | 7 | 0.5714 | 4 | 0.5 | 5 | 0.4 | 23 | 0.5652 |
| 5_1482646 | 7 | 0.1429 | 5 | 0 | 3 | 0 | 1 | 0 | 16 | 0.0625 | 1 | 1 | NA | NA | 1 | 0 | 2 | 0 | 4 | 0.25 |
| 5_1482648 | 6 | 0.3333 | 5 | 0 | 1 | 0 | 1 | 0 | 13 | 0.1538 | 1 | 1 | NA | NA | 1 | 0 | 2 | 0 | 4 | 0.25 |
| 5_1482653 | 6 | 0 | 6 | 0.1667 | 2 | 0 | 3 | 0 | 17 | 0.0588 | 3 | 0 | NA | NA | 1 | 0 | 2 | 0.5 | 6 | 0.1667 |
| 5_1482658 | 2 | 0 | 4 | 0 | 1 | 0 | 3 | 0.3333 | 10 | 0.1 | 2 | 0 | NA | NA | 1 | 1 | 1 | 0 | 4 | 0.25 |
| 5_1482684 | 3 | 0.6667 | 1 | 0 | NA | NA | 2 | 0 | 6 | 0.3333 | 1 | 1 | 1 | 1 | 1 | 0 | NA | NA | 3 | 0.6667 |
| 5_1482684 | 2 | 0.5 | 1 | 0 | NA | NA | 1 | 0 | 4 | 0.25 | 1 | 1 | 1 | 1 | 1 | 0 | NA | NA | 3 | 0.6667 |
| 5_1482685 | 3 | 0.6667 | 1 | 1 | NA | NA | 2 | 1 | 6 | 0.8333 | 1 | 1 | 1 | 1 | 1 | 0 | NA | NA | 3 | 0.6667 |
| 5_1482685 | 3 | 1 | 1 | 1 | NA | NA | 2 | 1 | 6 | 1 | 1 | 1 | 1 | 1 | 1 | 0 | NA | NA | 3 | 0.6667 |
| 5_1482688 | 3 | 0.3333 | 1 | 0 | NA | NA | 2 | 1 | 6 | 0.5 | 1 | 1 | NA | NA | 1 | 0 | NA | NA | 2 | 0.5 |
| 5_1482688 | 3 | 1 | 1 | 1 | NA | NA | 2 | 1 | 6 | 1 | 1 | 1 | NA | NA | 1 | 1 | NA | NA | 2 | 1 |
| 5_1482689 | 2 | 1 | 1 | 0 | NA | NA | 1 | 1 | 4 | 0.75 | 1 | 1 | NA | NA | 1 | 1 | NA | NA | 2 | 1 |
| 5_1482690 | 2 | 0.5 | 1 | 0 | NA | NA | 1 | 0 | 4 | 0.25 | 1 | 1 | NA | NA | 1 | 0 | NA | NA | 2 | 0.5 |
| 5_1482690 | 3 | 1 | 1 | 1 | NA | NA | NA | NA | 4 | 1 | NA | NA | NA | NA | 1 | 1 | NA | NA | 1 | 1 |
| 5_1482692 | 3 | 0 | 1 | 0 | NA | NA | 1 | 1 | 5 | 0.2 | 1 | 1 | NA | NA | 1 | 0 | NA | NA | 2 | 0.5 |
| 5_1484422 | 23 | 0.9565 | 35 | 0.8571 | 31 | 0.9355 | 23 | 0.9565 | 112 | 0.9196 | 17 | 0.9412 | 30 | 0.9 | 29 | 0.931 | 30 | 0.9333 | 106 | 0.9245 |
| 5_1484424 | 39 | 0.9231 | 36 | 0.9167 | 32 | 0.9375 | 39 | 0.9487 | 146 | 0.9315 | 20 | 1 | 35 | 0.9714 | 29 | 1 | 28 | 0.9643 | 112 | 0.9821 |
| 5_1484424 | 36 | 0.4167 | 33 | 0.3636 | 34 | 0.2353 | 39 | 0.2821 | 142 | 0.3239 | 20 | 0.3 | 35 | 0.2286 | 28 | 0.2857 | 31 | 0.2903 | 114 | 0.2719 |
| 5_1484424 | 36 | 0.1944 | 35 | 0.2571 | 33 | 0.3333 | 36 | 0.2778 | 140 | 0.2643 | 19 | 0.2105 | 37 | 0.2162 | 33 | 0.1515 | 29 | 0.3793 | 118 | 0.2373 |
| 5_1484424 | 30 | 0.8 | 31 | 0.6452 | 34 | 0.6176 | 38 | 0.6053 | 133 | 0.6617 | 19 | 0.5263 | 31 | 0.7419 | 31 | 0.7419 | 28 | 0.6429 | 109 | 0.6789 |
| 5_1484425 | 36 | 0.2222 | 34 | 0.5 | 33 | 0.5758 | 33 | 0.6061 | 136 | 0.4706 | 17 | 0.4706 | 36 | 0.4167 | 34 | 0.4706 | 32 | 0.5 | 119 | 0.4622 |
| 5_1484426 | 37 | 0.973 | 32 | 0.9062 | 33 | 0.9091 | 36 | 0.9167 | 138 | 0.9275 | 21 | 0.9524 | 36 | 0.9444 | 34 | 0.9706 | 31 | 0.9355 | 122 | 0.9508 |
| 5_1484427 | 31 | 0.7097 | 29 | 0.7586 | 27 | 0.9259 | 30 | 0.7667 | 117 | 0.7863 | 20 | 0.8 | 30 | 0.7667 | 27 | 0.7407 | 24 | 0.7083 | 101 | 0.7525 |
| 5_1484427 | 33 | 0.2424 | 26 | 0.1538 | 30 | 0.1667 | 34 | 0.1176 | 123 | 0.1707 | 16 | 0.25 | 33 | 0.2424 | 34 | 0.1765 | 26 | 0.1538 | 109 | 0.2018 |
| 5_1484428 | 33 | 0.9697 | 27 | 1 | 25 | 1 | 31 | 1 | 116 | 0.9914 | 18 | 1 | 29 | 0.9655 | 28 | 1 | 26 | 1 | 101 | 0.9901 |
| 5_1484428 | 31 | 1 | 31 | 0.871 | 26 | 0.9231 | 35 | 0.9143 | 123 | 0.9268 | 17 | 0.9412 | 31 | 0.9032 | 34 | 0.7647 | 24 | 1 | 106 | 0.8868 |
| 5_1484428 | 35 | 0.2857 | 30 | 0.3 | 26 | 0.1154 | 35 | 0.3429 | 126 | 0.2698 | 18 | 0.4444 | 33 | 0.3333 | 32 | 0.3438 | 26 | 0.4231 | 109 | 0.3761 |
| 5_1484429 | 32 | 0.125 | 27 | 0.1481 | 29 | 0.1379 | 36 | 0.1111 | 124 | 0.129 | 17 | 0.1176 | 25 | 0.08 | 31 | 0.0968 | 24 | 0.0833 | 97 | 0.0928 |
| 5_1484891 | 21 | 0.1429 | 14 | 0 | 16 | 0 | 24 | 0.0833 | 75 | 0.0667 | 8 | 0 | 20 | 0.1 | 28 | 0.0714 | 20 | 0.2 | 76 | 0.1053 |
| 5_1488087 | 2 | 0.5 | 8 | 0.125 | NA | NA | 4 | 0.5 | 14 | 0.2857 | 1 | 0 | 3 | 0.3333 | 7 | 0.2857 | 2 | 0 | 13 | 0.2308 |
| 5_1488089 | 1 | 0 | 6 | 0.1667 | 1 | 0 | 5 | 0.2 | 13 | 0.1538 | 1 | 0 | 3 | 0.3333 | 6 | 0.1667 | 1 | 1 | 11 | 0.2727 |
| 5_1488119 | 4 | 0.25 | 6 | 0 | 2 | 0 | 2 | 0 | 14 | 0.0714 | 4 | 0.25 | 1 | 0 | 6 | 0 | 4 | 0 | 15 | 0.0667 |
| 5_1488126 | 3 | 0.3333 | 4 | 0.5 | 2 | 0 | 2 | 0 | 11 | 0.2727 | 3 | 0.3333 | 1 | 0 | 4 | 0 | 5 | 0.4 | 13 | 0.2308 |
| 5_1488130 | 2 | 0 | 2 | 1 | 3 | 0.3333 | 1 | 0 | 8 | 0.375 | 3 | 0.6667 | 1 | 1 | 3 | 0.6667 | 4 | 1 | 11 | 0.8182 |
| 5_1488414 | 1 | 0 | 2 | 0 | 3 | 0.3333 | NA | NA | 6 | 0.1667 | 3 | 0 | NA | NA | 2 | 0.5 | 2 | 0 | 7 | 0.1429 |
| 5_1488654 | 2 | 0.5 | 1 | 0 | 2 | 0 | NA | NA | 5 | 0.2 | 1 | 0 | NA | NA | 1 | 0 | 2 | 0.5 | 4 | 0.25 |
| 5_1489362 | NA | NA | 1 | 1 | NA | NA | NA | NA | 1 | 1 | NA | NA | NA | NA | NA | NA | 1 | 1 | 1 | 1 |
| 5_1489846 | 10 | 0.1 | 6 | 0 | 9 | 0.1111 | 12 | 0.25 | 37 | 0.1351 | 6 | 0.1667 | 8 | 0.375 | 7 | 0 | 15 | 0.2 | 36 | 0.1944 |
| 5_1489847 | 8 | 0.25 | 10 | 0 | 8 | 0 | 15 | 0.0667 | 41 | 0.0732 | 7 | 0.1429 | 8 | 0.25 | 7 | 0 | 17 | 0.1176 | 39 | 0.1282 |
| 5_1489847 | 9 | 0.3333 | 6 | 0.1667 | 9 | 0.3333 | 13 | 0 | 37 | 0.1892 | 4 | 0.25 | 6 | 0.1667 | 6 | 0 | 14 | 0.2143 | 30 | 0.1667 |
| 5_1491307 | 3 | 1 | 1 | 1 | 1 | 1 | 2 | 1 | 7 | 1 | 2 | 1 | 1 | 1 | 1 | 1 | 1 | 1 | 5 | 1 |
| 5_1491308 | 3 | 0 | 1 | 1 | 1 | 0 | 3 | 0 | 8 | 0.125 | 1 | 1 | 1 | 0 | 1 | 0 | 2 | 0 | 5 | 0.2 |
| 5_1491310 | 3 | 0.6667 | 1 | 1 | 3 | 0.3333 | 3 | 0.3333 | 10 | 0.5 | 2 | 0 | 1 | 1 | 1 | 0 | 3 | 0.3333 | 7 | 0.2857 |
| 5_1491310 | 3 | 1 | 1 | 1 | 2 | 0.5 | 3 | 1 | 9 | 0.8889 | 2 | 1 | 1 | 1 | 1 | 0 | 3 | 0.3333 | 7 | 0.5714 |
| 5_1491503 | 4 | 0.25 | 1 | 1 | 3 | 0 | 1 | 0 | 9 | 0.2222 | 1 | 0 | NA | NA | 1 | 0 | 2 | 0.5 | 4 | 0.25 |
| 5_1491504 | 4 | 0.5 | 1 | 1 | 3 | 0.3333 | 2 | 0 | 10 | 0.4 | 1 | 0 | NA | NA | 1 | 1 | 3 | 0.6667 | 5 | 0.6 |
| 5_1491505 | 5 | 0.2 | 1 | 0 | 2 | 0 | 2 | 0.5 | 10 | 0.2 | NA | NA | NA | NA | 1 | 0 | 3 | 0.3333 | 4 | 0.25 |
| 5_1491505 | 5 | 0.2 | 1 | 0 | 3 | 0 | 3 | 0 | 12 | 0.0833 | 1 | 0 | NA | NA | 1 | 0 | 3 | 0.3333 | 5 | 0.2 |
| 5_1491506 | 3 | 1 | 1 | 1 | 2 | 0.5 | 3 | 0.6667 | 9 | 0.7778 | 1 | 1 | NA | NA | 1 | 1 | 1 | 1 | 3 | 1 |
| 5_1491507 | 5 | 0.4 | 1 | 0 | 3 | 0.3333 | 3 | 0.3333 | 12 | 0.3333 | 1 | 0 | NA | NA | 1 | 1 | 3 | 0.3333 | 5 | 0.4 |
| 5_1491508 | 6 | 0.8333 | 1 | 1 | 1 | 1 | 3 | 1 | 11 | 0.9091 | NA | NA | NA | NA | NA | NA | 2 | 1 | 2 | 1 |
| 5_1491508 | 4 | 1 | 1 | 1 | 2 | 1 | 3 | 1 | 10 | 1 | 2 | 0.5 | 1 | 1 | 1 | 1 | 2 | 1 | 6 | 0.8333 |
| 5_1491508 | 5 | 0.8 | 1 | 1 | 3 | 0.3333 | 3 | 0.6667 | 12 | 0.6667 | 2 | 0.5 | 1 | 1 | 1 | 0 | 1 | 0 | 5 | 0.4 |
| 5_1491512 | 9 | 0.6667 | NA | NA | 2 | 0.5 | 3 | 0.6667 | 14 | 0.6429 | 2 | 0.5 | 1 | 1 | 2 | 1 | 2 | 0.5 | 7 | 0.7143 |
| 5_1491512 | 7 | 0.1429 | 1 | 0 | 2 | 0 | 3 | 0.3333 | 13 | 0.1538 | 1 | 0 | NA | NA | 2 | 0 | 2 | 0.5 | 5 | 0.2 |

| | | | | | | | | | | | | | | | | | | | | |
|---|---|---|---|---|---|---|---|---|---|---|---|---|---|---|---|---|---|---|---|---|
| 5_14915142 | 9 | 0.1111 | NA | NA | 4 | 0 | 2 | 0 | 15 | 0.0667 | 2 | 0 | NA | NA | 2 | 0.5 | 2 | 0 | 6 | 0.1667 |
| 5_14915159 | 8 | 0.875 | 1 | 1 | 4 | 0.5 | 3 | 1 | 16 | 0.8125 | 2 | 1 | NA | NA | 2 | 1 | 3 | 0.6667 | 7 | 0.8571 |
| 5_14915163 | 9 | 0.1111 | 1 | 0 | 5 | 0 | 3 | 0 | 18 | 0.0556 | 2 | 0.5 | 1 | 0 | 2 | 0.5 | 2 | 0 | 7 | 0.2857 |
| 5_14915184 | 7 | 0 | 3 | 0 | 5 | 0 | 3 | 0.3333 | 18 | 0.0556 | 3 | 0 | 1 | 1 | 1 | 0 | 2 | 0 | 7 | 0.1429 |
| 5_14915186 | 9 | 0.4444 | 3 | 0.6667 | 5 | 0.8 | 3 | 0.6667 | 20 | 0.6 | 3 | 0.6667 | 1 | 0 | 2 | 0.5 | 3 | 0.3333 | 9 | 0.4444 |
| 5_14915192 | 8 | 1 | 3 | 1 | 3 | 1 | 3 | 1 | 17 | 1 | 3 | 1 | 1 | 1 | 2 | 1 | 2 | 1 | 8 | 1 |
| 5_14915194 | 5 | 0.2 | 2 | 0.5 | 4 | 0.5 | 3 | 0.3333 | 14 | 0.3571 | 3 | 0.3333 | 1 | 1 | 1 | 0 | 2 | 0.5 | 7 | 0.4286 |
| 5_14915194 | 5 | 1 | 3 | 1 | 4 | 0.75 | 1 | 0 | 13 | 0.8462 | 2 | 1 | 1 | 1 | 2 | 1 | 1 | 1 | 6 | 1 |
| 5_14915196 | 8 | 0.625 | 3 | 0.6667 | 5 | 1 | 3 | 1 | 19 | 0.7895 | 1 | 1 | 1 | 1 | 2 | 0.5 | 3 | 0.6667 | 7 | 0.7143 |
| 5_14915203 | 9 | 0.1111 | 3 | 0 | 5 | 0.6 | 3 | 0.3333 | 20 | 0.25 | 3 | 0.6667 | 1 | 1 | 1 | 1 | 3 | 0 | 8 | 0.5 |
| 5_14915204 | 5 | 0.8 | 3 | 0.6667 | 2 | 0.5 | 2 | 1 | 12 | 0.75 | 1 | 1 | 1 | 1 | 1 | 0 | 2 | 0.5 | 5 | 0.6 |
| 5_14922468 | 3 | 0.6667 | 1 | 1 | NA | NA | 2 | 0 | 6 | 0.5 | 1 | 1 | NA | NA | NA | NA | NA | NA | 1 | 1 |
| 5_14935338 | NA | NA | 2 | 0.5 | 1 | 0 | 1 | 1 | 4 | 0.5 | NA | NA | NA | NA | 1 | 1 | NA | NA | 1 | 1 |
| 5_14935345 | NA | NA | 1 | 1 | 1 | 1 | NA | NA | 2 | 1 | NA | NA | NA | NA | 1 | 1 | NA | NA | 1 | 1 |
| 5_14935361 | NA | NA | 1 | 1 | 1 | 1 | 1 | 0 | 3 | 0.6667 | 1 | 1 | NA | NA | 2 | 0.5 | NA | NA | 3 | 0.6667 |
| 5_14935366 | NA | NA | 2 | 1 | 1 | 1 | 1 | 1 | 4 | 1 | 1 | 1 | NA | NA | 2 | 1 | NA | NA | 3 | 1 |
| 5_14935372 | NA | NA | 2 | 0.5 | 1 | 0 | 1 | 0 | 4 | 0.25 | 1 | 0 | NA | NA | 2 | 0.5 | NA | NA | 3 | 0.3333 |
| 5_14935494 | NA | NA | 3 | 0.6667 | NA | NA | 1 | 0 | 4 | 0.5 | NA | NA | 1 | 1 | NA | NA | NA | NA | 1 | 1 |
| 5_14954068 | 40 | 0.075 | 36 | 0.1389 | 34 | 0.0588 | 25 | 0.12 | 135 | 0.0963 | 32 | 0.0938 | 32 | 0.1875 | 34 | 0.0588 | 36 | 0.1111 | 134 | 0.1119 |
| 5_14954079 | 40 | 0.05 | 42 | 0.119 | 37 | 0.1351 | 25 | 0.16 | 144 | 0.1111 | 32 | 0.0938 | 34 | 0.2059 | 37 | 0.1351 | 33 | 0.2121 | 136 | 0.1618 |
| 5_14954087 | 42 | 0.119 | 34 | 0.1471 | 31 | 0.3226 | 27 | 0.2593 | 134 | 0.2015 | 30 | 0.2333 | 38 | 0.3421 | 34 | 0.2353 | 34 | 0.1765 | 136 | 0.25 |
| 5_14954090 | 43 | 0.3256 | 35 | 0.3143 | 28 | 0.5357 | 28 | 0.4643 | 134 | 0.3955 | 32 | 0.375 | 33 | 0.5455 | 36 | 0.1944 | 30 | 0.2667 | 131 | 0.3435 |
| 5_14954097 | 27 | 0.3704 | 24 | 0.3333 | 19 | 0.5789 | 19 | 0.3684 | 89 | 0.4045 | 25 | 0.36 | 23 | 0.3043 | 26 | 0.1923 | 26 | 0.3077 | 100 | 0.29 |
| 5_14954098 | 39 | 0.359 | 34 | 0.5882 | 32 | 0.4375 | 24 | 0.375 | 129 | 0.4419 | 34 | 0.4412 | 33 | 0.5758 | 38 | 0.3947 | 32 | 0.5312 | 137 | 0.4818 |
| 5_14968292 | NA | NA | 2 | 0 | 1 | 0 | 2 | 0.5 | 5 | 0.2 | NA | NA | 3 | 0 | 3 | 0.3333 | 2 | 0 | 8 | 0.125 |
| 5_14968308 | NA | NA | 2 | 1 | 2 | 0 | 2 | 0.5 | 6 | 0.5 | NA | NA | 2 | 0 | 4 | 0.25 | 2 | 0.5 | 8 | 0.25 |
| 5_14968315 | NA | NA | 2 | 1 | 1 | 1 | 2 | 1 | 5 | 1 | NA | NA | 2 | 1 | 4 | 0.5 | 2 | 0.5 | 8 | 0.625 |
| 5_14968412 | NA | NA | 2 | 0 | 1 | 0 | 2 | 0.5 | 5 | 0.2 | NA | NA | 3 | 0 | 3 | 0.3333 | 2 | 0 | 8 | 0.125 |
| 5_15032879 | NA | NA | 1 | 0 | 2 | 0.5 | NA | NA | 3 | 0.3333 | 2 | 0.5 | 4 | 0.25 | NA | NA | 1 | 1 | 7 | 0.4286 |
| 5_15036039 | 2 | 0.5 | 2 | 0 | 4 | 0.25 | 2 | 0.5 | 10 | 0.3 | 3 | 0 | 5 | 0.6 | NA | NA | 1 | 1 | 9 | 0.4444 |
| 5_15036046 | 3 | 0.3333 | 2 | 1 | 5 | 0.8 | 3 | 0.6667 | 13 | 0.6923 | 5 | 0.8 | 4 | 0.75 | 1 | 0 | 3 | 0 | 13 | 0.5385 |
| 5_15036058 | 2 | 0 | 2 | 0 | 5 | 0 | 2 | 0.5 | 11 | 0.0909 | 5 | 0.4 | 5 | 0.2 | 1 | 0 | 1 | 1 | 12 | 0.3333 |
| 5_15036066 | 4 | 0.75 | 2 | 0.5 | 6 | 0.6667 | 2 | 0.5 | 14 | 0.6429 | 5 | 0.6 | 6 | 0.5 | 1 | 0 | 4 | 1 | 16 | 0.625 |
| 5_15036078 | 4 | 0 | 2 | 0 | 6 | 0.1667 | 1 | 0 | 13 | 0.0769 | 5 | 0 | 4 | 0.25 | 1 | 0 | 5 | 0 | 15 | 0.0667 |
| 5_15036303 | 4 | 0 | 2 | 0 | 6 | 0.1667 | 3 | 0 | 15 | 0.0667 | 3 | 0 | 6 | 0 | 1 | 0 | 4 | 0.25 | 14 | 0.0714 |
| 5_15039808 | 1 | 1 | NA | NA | NA | NA | NA | NA | 1 | 1 | NA | NA | 1 | 1 | NA | NA | NA | NA | 1 | 1 |
| 5_15039822 | 1 | 1 | NA | NA | NA | NA | NA | NA | 1 | 1 | NA | NA | 1 | 1 | NA | NA | NA | NA | 1 | 1 |
| 5_15039851 | 1 | 1 | NA | NA | NA | NA | NA | NA | 1 | 1 | NA | NA | 1 | 1 | NA | NA | NA | NA | 1 | 1 |
| 5_15039853 | 1 | 1 | NA | NA | NA | NA | NA | NA | 1 | 1 | NA | NA | 1 | 1 | NA | NA | NA | NA | 1 | 1 |
| 5_15041971 | 10 | 0.7 | 10 | 0.9 | 9 | 0.8889 | 6 | 1 | 35 | 0.8571 | 10 | 1 | 17 | 0.8824 | 4 | 1 | 11 | 1 | 42 | 0.9524 |
| 5_15041972 | 11 | 0.8182 | 11 | 0.9091 | 10 | 1 | 6 | 1 | 38 | 0.9211 | 12 | 1 | 12 | 1 | 4 | 1 | 9 | 1 | 37 | 1 |
| 5_15041976 | 14 | 0.5714 | 11 | 0.7273 | 10 | 0.6 | 7 | 0.7143 | 42 | 0.6429 | 11 | 0.5455 | 14 | 0.8571 | 5 | 0.8 | 12 | 0.75 | 42 | 0.7381 |
| 5_15041979 | 16 | 0.0625 | 9 | 0.2222 | 12 | 0 | 6 | 0 | 43 | 0.0698 | 11 | 0 | 12 | 0.0833 | 3 | 0 | 10 | 0.1 | 36 | 0.0556 |
| 5_15041981 | 15 | 0.0667 | 11 | 0 | 13 | 0.2308 | 6 | 0.1667 | 45 | 0.1111 | 12 | 0.3333 | 11 | 0.1818 | 5 | 0 | 10 | 0.1 | 38 | 0.1842 |
| 5_15041983 | 13 | 0.4615 | 10 | 0.4 | 10 | 0.5 | 6 | 0.5 | 39 | 0.4615 | 11 | 0.5455 | 13 | 0.5385 | 5 | 0.2 | 11 | 0.8182 | 40 | 0.575 |
| 5_15041986 | 14 | 0.3571 | 9 | 0.3333 | 15 | 0.2 | 7 | 0.2857 | 45 | 0.2889 | 12 | 0.25 | 15 | 0.1333 | 5 | 0 | 13 | 0.1538 | 45 | 0.1556 |
| 5_15041998 | 16 | 0.75 | 11 | 0.8182 | 14 | 0.9286 | 7 | 0.8571 | 48 | 0.8333 | 13 | 0.7692 | 14 | 0.6429 | 4 | 1 | 11 | 0.9091 | 42 | 0.7857 |
| 5_15042001 | 15 | 0.9333 | 10 | 1 | 13 | 1 | 5 | 1 | 43 | 0.9767 | 14 | 0.9286 | 16 | 1 | 4 | 1 | 13 | 1 | 47 | 0.9787 |
| 5_15042003 | 12 | 0.0833 | 10 | 0 | 15 | 0.2 | 5 | 0 | 42 | 0.0952 | 13 | 0 | 15 | 0.1333 | 5 | 0.2 | 10 | 0.1 | 43 | 0.093 |
| 5_15042008 | 15 | 0.2 | 11 | 0.1818 | 16 | 0.4375 | 8 | 0.25 | 50 | 0.28 | 14 | 0.5 | 15 | 0.3333 | 5 | 0.2 | 12 | 0 | 46 | 0.2826 |
| 5_15042009 | 15 | 0.2 | 10 | 0.2 | 16 | 0.0625 | 8 | 0 | 49 | 0.1224 | 14 | 0 | 16 | 0.1875 | 4 | 0 | 10 | 0 | 44 | 0.0682 |
| 5_15042010 | 14 | 0.8571 | 11 | 0.5455 | 14 | 0.9286 | 7 | 1 | 46 | 0.8261 | 14 | 0.9286 | 13 | 0.7692 | 4 | 1 | 13 | 1 | 44 | 0.9091 |
| 5_15042016 | 14 | 0.0714 | 9 | 0 | 14 | 0.0714 | 7 | 0.1429 | 44 | 0.0682 | 15 | 0.2667 | 15 | 0.0667 | 4 | 0 | 11 | 0.1818 | 45 | 0.1556 |
| 5_15042017 | 16 | 0.3125 | 10 | 0.3 | 16 | 0.3125 | 6 | 0.1667 | 48 | 0.2917 | 14 | 0.0714 | 16 | 0.1875 | 5 | 0 | 13 | 0.2308 | 48 | 0.1458 |
| 5_15042022 | 16 | 0.875 | 10 | 0.9 | 11 | 0.7273 | 6 | 1 | 43 | 0.8605 | 14 | 1 | 16 | 0.9375 | 4 | 1 | 9 | 0.7778 | 43 | 0.9302 |
| 5_15049243 | 1 | 0 | 3 | 0.6667 | NA | NA | NA | NA | 4 | 0.5 | 1 | 1 | NA | NA | NA | NA | 2 | 0.5 | 3 | 0.6667 |
| 5_15049260 | 1 | 1 | 2 | 0 | NA | NA | NA | NA | 3 | 0.3333 | NA | NA | NA | NA | NA | NA | 2 | 1 | 2 | 1 |
| 5_15049292 | 1 | 0 | 3 | 0.6667 | NA | NA | NA | NA | 4 | 0.5 | 1 | 0 | NA | NA | NA | NA | 2 | 0.5 | 3 | 0.3333 |
| 5_15049600 | 5 | 0.2 | 4 | 0 | 1 | 0 | 2 | 0 | 12 | 0.0833 | 3 | 0 | 4 | 0.25 | 7 | 0.1429 | 3 | 0 | 17 | 0.1176 |
| 5_15049607 | 5 | 0.4 | 4 | 0 | 1 | 0 | 3 | 0 | 13 | 0.1538 | 3 | 0 | 3 | 0 | 8 | 0.125 | 3 | 0 | 17 | 0.0588 |
| 5_15049632 | 3 | 0 | 3 | 0 | NA | NA | 3 | 0.3333 | 9 | 0.1111 | 1 | 0 | 2 | 0 | 9 | 0.1111 | 1 | 0 | 13 | 0.0769 |
| 5_15049640 | 4 | 0.25 | 3 | 0.3333 | NA | NA | 3 | 0 | 10 | 0.2 | 2 | 1 | 2 | 0.5 | 9 | 0.3333 | 3 | 0 | 16 | 0.375 |
| 5_15051564 | NA | NA | 1 | 1 | NA | NA | NA | NA | 1 | 1 | NA | NA | 3 | 1 | 1 | 1 | 1 | 0 | 5 | 0.8 |
| 5_15051632 | 1 | 1 | 2 | 1 | NA | NA | NA | NA | 3 | 1 | NA | NA | 2 | 0.5 | NA | NA | NA | NA | 2 | 0.5 |
| 5_15051653 | 1 | 1 | 2 | 1 | NA | NA | NA | NA | 3 | 1 | NA | NA | 1 | 1 | NA | NA | 1 | 1 | 2 | 1 |

| | | | | | | | | | | | | | | | | | | | | |
|---|---|---|---|---|---|---|---|---|---|---|---|---|---|---|---|---|---|---|---|---|
| 5_1505166 | 2 | 0.5 | 2 | 0 | 1 | 0 | NA | NA | 5 | 0.2 | NA | NA | 5 | 0.4 | 1 | 1 | 1 | 0 | 7 | 0.4286 |
| 5_1505168 | 2 | 1 | 2 | 0.5 | 1 | 0 | NA | NA | 5 | 0.6 | NA | NA | 2 | 1 | NA | NA | NA | NA | 2 | 1 |
| 5_1505170 | 2 | 1 | 1 | 1 | NA | NA | NA | NA | 3 | 1 | NA | NA | 1 | 1 | NA | NA | 1 | 1 | 2 | 1 |
| 5_1505171 | 1 | 1 | NA | NA | NA | NA | NA | NA | 1 | 1 | NA | NA | 1 | 0 | NA | NA | 1 | 1 | 2 | 0.5 |
| 5_1505174 | 2 | 1 | NA | NA | NA | NA | NA | NA | 2 | 1 | NA | NA | 1 | 1 | 1 | 1 | 1 | 1 | 3 | 1 |
| 5_1505174 | 1 | 1 | 1 | 1 | NA | NA | NA | NA | 2 | 1 | NA | NA | 1 | 1 | 1 | 1 | 1 | 0 | 3 | 0.6667 |
| 5_1505176 | 2 | 0.5 | 1 | 0 | NA | NA | NA | NA | 3 | 0.3333 | NA | NA | 1 | 1 | 1 | 1 | 1 | 0 | 3 | 0.6667 |
| 5_1505235 | 1 | 1 | 1 | 1 | 1 | 1 | 1 | 1 | 4 | 1 | NA | NA | 4 | 1 | 2 | 1 | 1 | 1 | 7 | 1 |
| 5_1505237 | 1 | 1 | 1 | 1 | 1 | 1 | NA | NA | 3 | 1 | 1 | 1 | 3 | 0.3333 | 1 | 1 | 2 | 0.5 | 7 | 0.5714 |
| 5_1505238 | NA | NA | 1 | 0 | NA | NA | 1 | 1 | 2 | 0.5 | 1 | 1 | 3 | 1 | 1 | 1 | 2 | 1 | 7 | 1 |
| 5_1505239 | 1 | 1 | 1 | 1 | NA | NA | 1 | 1 | 3 | 1 | 1 | 1 | 3 | 1 | 1 | 1 | 2 | 1 | 7 | 1 |
| 5_1505240 | NA | NA | 1 | 0 | NA | NA | 1 | 1 | 2 | 0.5 | 1 | 0 | 2 | 1 | NA | NA | 2 | 0 | 5 | 0.4 |
| 5_1505241 | 1 | 1 | 1 | 0 | NA | NA | 1 | 0 | 3 | 0.3333 | 1 | 1 | 2 | 0 | NA | NA | 2 | 0 | 5 | 0.2 |
| 5_1505241 | 1 | 1 | NA | NA | NA | NA | 1 | 0 | 2 | 0.5 | NA | NA | 2 | 0 | 1 | 1 | 2 | 0 | 5 | 0.2 |
| 5_1505246 | NA | NA | NA | NA | NA | NA | 1 | 1 | 1 | 1 | NA | NA | 1 | 0 | 1 | 1 | 2 | 0.5 | 4 | 0.5 |
| 5_1505246 | NA | NA | NA | NA | NA | NA | 1 | 1 | 1 | 1 | NA | NA | 3 | 0.6667 | 1 | 1 | 2 | 1 | 6 | 0.8333 |
| 5_1505305 | NA | NA | 1 | 1 | NA | NA | NA | NA | 1 | 1 | 1 | 0 | 1 | 1 | NA | NA | NA | NA | 2 | 0.5 |
| 5_1505308 | NA | NA | 1 | 1 | NA | NA | NA | NA | 1 | 1 | 1 | 0 | 1 | 1 | NA | NA | NA | NA | 2 | 0.5 |
| 5_1505309 | NA | NA | 1 | 1 | NA | NA | NA | NA | 1 | 1 | 1 | 0 | 1 | 1 | NA | NA | NA | NA | 2 | 0.5 |
| 5_1505665 | 2 | 0 | 2 | 0 | 1 | 0 | 2 | 0.5 | 7 | 0.1429 | NA | NA | 1 | 1 | NA | NA | 4 | 0 | 5 | 0.2 |
| 5_1505666 | 2 | 0 | 2 | 0.5 | NA | NA | 2 | 0 | 6 | 0.1667 | 2 | 0.5 | 1 | 0 | NA | NA | 6 | 0.1667 | 9 | 0.2222 |
| 5_1505848 | 2 | 0 | 1 | 1 | NA | NA | 1 | 0 | 4 | 0.25 | 2 | 0 | 2 | 0 | NA | NA | 4 | 0.25 | 8 | 0.125 |
| 5_1505848 | 2 | 1 | 1 | 1 | NA | NA | NA | NA | 3 | 1 | 1 | 1 | 1 | 1 | NA | NA | 4 | 0.75 | 6 | 0.8333 |
| 5_1505849 | 2 | 1 | 1 | 0 | NA | NA | 1 | 1 | 4 | 0.75 | 1 | 1 | 2 | 1 | NA | NA | 4 | 0.5 | 7 | 0.7143 |
| 5_1505849 | 2 | 0.5 | 1 | 0 | NA | NA | 1 | 0 | 4 | 0.25 | 1 | 0 | 2 | 0 | NA | NA | 4 | 0.25 | 7 | 0.1429 |
| 5_1505850 | 2 | 0 | 1 | 0 | NA | NA | 1 | 1 | 4 | 0.25 | 1 | 0 | 2 | 0.5 | NA | NA | 3 | 0.6667 | 6 | 0.5 |
| 5_1505850 | 2 | 0.5 | 1 | 0 | NA | NA | 1 | 0 | 4 | 0.25 | 2 | 0 | 2 | 0.5 | NA | NA | 4 | 0.75 | 8 | 0.5 |
| 5_1505852 | 1 | 1 | 1 | 1 | NA | NA | 1 | 0 | 3 | 0.6667 | 1 | 0 | 2 | 0.5 | NA | NA | 2 | 0.5 | 5 | 0.4 |
| 5_1505853 | 1 | 0 | 1 | 1 | NA | NA | 1 | 0 | 3 | 0.3333 | 2 | 0 | 2 | 0.5 | NA | NA | 2 | 0.5 | 6 | 0.3333 |
| 5_1505857 | NA | NA | 1 | 0 | NA | NA | 1 | 1 | 2 | 0.5 | NA | NA | 2 | 0.5 | NA | NA | 3 | 0.6667 | 5 | 0.6 |
| 5_1507241 | 2 | 0 | 2 | 0 | 6 | 0.1667 | 4 | 0 | 14 | 0.0714 | 5 | 0.2 | 5 | 0 | 9 | 0 | 6 | 0.3333 | 25 | 0.12 |
| 5_1507243 | 2 | 0 | 3 | 0 | 5 | 0.6 | 4 | 0 | 14 | 0.2143 | 6 | 0.1667 | 5 | 0.8 | 10 | 0.1 | 8 | 0.125 | 29 | 0.2414 |
| 5_1507313 | 4 | 0.25 | 2 | 0.5 | 3 | 0.6667 | 2 | 0.5 | 11 | 0.4545 | 7 | 0.2857 | 5 | 0.8 | 11 | 0.4545 | 6 | 0.5 | 29 | 0.4828 |
| 5_1507314 | 4 | 0.25 | 4 | 0.75 | 5 | 0.8 | 3 | 0.6667 | 16 | 0.625 | 5 | 0.6 | 6 | 0.6667 | 12 | 0.8333 | 5 | 0.8 | 28 | 0.75 |
| 5_1507314 | 3 | 1 | 4 | 0.75 | 5 | 0.4 | 3 | 1 | 15 | 0.7333 | 4 | 1 | 5 | 1 | 11 | 0.8182 | 7 | 0.8571 | 27 | 0.8889 |
| 5_1507314 | 3 | 1 | 3 | 1 | 7 | 1 | 4 | 1 | 17 | 1 | 4 | 1 | 5 | 0.8 | 11 | 0.8182 | 6 | 1 | 26 | 0.8846 |
| 5_1507314 | 5 | 0 | 5 | 0 | 7 | 0.1429 | 3 | 0 | 20 | 0.05 | 6 | 0 | 5 | 0.2 | 11 | 0 | 8 | 0.125 | 30 | 0.0667 |
| 5_1507315 | 5 | 0 | 3 | 0 | 7 | 0.2857 | 5 | 0.2 | 20 | 0.15 | 6 | 0.1667 | 6 | 0.1667 | 12 | 0.1667 | 8 | 0 | 32 | 0.125 |
| 5_1507315 | 4 | 0.5 | 4 | 0.25 | 6 | 0.3333 | 4 | 0.25 | 18 | 0.3333 | 8 | 0.125 | 5 | 0.6 | 9 | 0.3333 | 7 | 0 | 29 | 0.2414 |
| 5_1507316 | 7 | 0 | 5 | 0 | 7 | 0.4286 | 6 | 0.1667 | 25 | 0.16 | 8 | 0 | 6 | 0 | 12 | 0.0833 | 8 | 0.125 | 34 | 0.0588 |
| 5_1507319 | 7 | 0.1429 | 5 | 0.2 | 7 | 0 | 7 | 0.1429 | 26 | 0.1154 | 8 | 0 | 6 | 0 | 11 | 0 | 10 | 0.2 | 35 | 0.0571 |
| 5_1507331 | 7 | 0.1429 | 4 | 0 | 5 | 0.4 | 7 | 0 | 23 | 0.1304 | 5 | 0.2 | 7 | 0 | 10 | 0.2 | 7 | 0 | 29 | 0.1034 |
| 5_1507337 | 4 | 0 | 4 | 0 | 7 | 0.1429 | 7 | 0.2857 | 22 | 0.1364 | 4 | 0 | 5 | 0 | 5 | 0 | 6 | 0.5 | 20 | 0.15 |
| 5_1508764 | NA | NA | NA | NA | NA | NA | 1 | 1 | 1 | 1 | NA | NA | NA | NA | NA | NA | 1 | 1 | 1 | 1 |
| 5_1509041 | 1 | 1 | NA | NA | NA | NA | 1 | 0 | 2 | 0.5 | NA | NA | NA | NA | NA | NA | 1 | 1 | 1 | 1 |
| 5_1509042 | 1 | 1 | NA | NA | NA | NA | 1 | 1 | 2 | 1 | NA | NA | NA | NA | NA | NA | 1 | 1 | 1 | 1 |
| 5_1509043 | 1 | 1 | NA | NA | NA | NA | 1 | 1 | 2 | 1 | NA | NA | NA | NA | NA | NA | 1 | 1 | 1 | 1 |
| 5_1509061 | 1 | 0 | NA | NA | 1 | 1 | 1 | 0 | 3 | 0.3333 | NA | NA | NA | NA | NA | NA | 1 | 1 | 1 | 1 |
| 5_1509120 | 1 | 1 | NA | NA | 2 | 1 | 1 | 1 | 4 | 1 | NA | NA | NA | NA | NA | NA | 1 | 1 | 1 | 1 |
| 5_1509125 | 1 | 1 | NA | NA | 2 | 1 | 1 | 1 | 4 | 1 | NA | NA | NA | NA | NA | NA | 2 | 1 | 2 | 1 |
| 5_1509126 | 1 | 0 | NA | NA | 2 | 0.5 | 1 | 0 | 4 | 0.25 | NA | NA | NA | NA | NA | NA | 2 | 0.5 | 2 | 0.5 |
| 5_1509127 | 1 | 1 | NA | NA | 2 | 0.5 | 1 | 1 | 4 | 0.75 | NA | NA | NA | NA | NA | NA | 2 | 0.5 | 2 | 0.5 |
| 5_1509127 | 1 | 0 | NA | NA | 2 | 1 | 1 | 0 | 4 | 0.5 | NA | NA | NA | NA | NA | NA | 2 | 0.5 | 2 | 0.5 |
| 5_1509151 | NA | NA | NA | NA | 1 | 1 | 1 | 0 | 2 | 0.5 | NA | NA | NA | NA | NA | NA | 3 | 0.6667 | 3 | 0.6667 |
| 5_1509151 | NA | NA | NA | NA | 2 | 0.5 | NA | NA | 2 | 0.5 | NA | NA | NA | NA | NA | NA | 3 | 0.6667 | 3 | 0.6667 |
| 5_1509151 | NA | NA | NA | NA | 2 | 1 | 1 | 0 | 3 | 0.6667 | NA | NA | NA | NA | NA | NA | 3 | 1 | 3 | 1 |
| 5_1509156 | NA | NA | NA | NA | 2 | 1 | 1 | 1 | 3 | 1 | NA | NA | NA | NA | NA | NA | 3 | 0.6667 | 3 | 0.6667 |
| 5_1509157 | NA | NA | NA | NA | 2 | 1 | 1 | 1 | 3 | 1 | NA | NA | NA | NA | NA | NA | 2 | 1 | 2 | 1 |
| 5_1509158 | NA | NA | NA | NA | 2 | 1 | 1 | 1 | 3 | 1 | NA | NA | NA | NA | NA | NA | 1 | 1 | 1 | 1 |
| 5_1509161 | NA | NA | NA | NA | 1 | 1 | 1 | 0 | 2 | 0.5 | NA | NA | NA | NA | NA | NA | 2 | 0.5 | 2 | 0.5 |
| 5_1509162 | NA | NA | NA | NA | 2 | 0.5 | 1 | 0 | 3 | 0.3333 | NA | NA | NA | NA | NA | NA | 3 | 0.3333 | 3 | 0.3333 |
| 5_1509162 | NA | NA | NA | NA | 2 | 1 | 1 | 1 | 3 | 1 | NA | NA | NA | NA | NA | NA | 2 | 0.5 | 2 | 0.5 |
| 5_1509162 | 1 | 1 | NA | NA | 2 | 0 | 1 | 0 | 4 | 0.25 | NA | NA | NA | NA | NA | NA | 3 | 0.6667 | 3 | 0.6667 |
| 5_1509163 | 1 | 0 | NA | NA | 2 | 0.5 | 1 | 1 | 4 | 0.5 | NA | NA | NA | NA | NA | NA | 2 | 0.5 | 2 | 0.5 |
| 5_1509165 | NA | NA | NA | NA | 2 | 1 | NA | NA | 2 | 1 | NA | NA | NA | NA | NA | NA | 3 | 0.6667 | 3 | 0.6667 |

| | | | | | | | | | | | | | | | | | | | | |
|---|---|---|---|---|---|---|---|---|---|---|---|---|---|---|---|---|---|---|---|---|
| 5_1509165 | 1 | 0 | NA | NA | 2 | 1 | 1 | 1 | 4 | 0.75 | NA | NA | NA | NA | NA | NA | 1 | 1 | 1 | 1 |
| 5_1509167 | 1 | 1 | NA | NA | 3 | 1 | 1 | 1 | 5 | 1 | NA | NA | NA | NA | NA | NA | 3 | 1 | 3 | 1 |
| 5_1509629 | NA | NA | 1 | 1 | 2 | 1 | NA | NA | 3 | 1 | NA | NA | NA | NA | NA | NA | 2 | 1 | 2 | 1 |
| 5_1509686 | NA | NA | 1 | 1 | 1 | 0 | NA | NA | 2 | 0.5 | NA | NA | NA | NA | NA | NA | 2 | 0.5 | 2 | 0.5 |
| 5_1509858 | NA | NA | NA | NA | 1 | 1 | NA | NA | 1 | 1 | NA | NA | NA | NA | NA | NA | 2 | 0.5 | 2 | 0.5 |
| 5_1509859 | NA | NA | NA | NA | 1 | 1 | NA | NA | 1 | 1 | NA | NA | NA | NA | NA | NA | 1 | 1 | 1 | 1 |
| 5_1509860 | NA | NA | NA | NA | 1 | 1 | NA | NA | 1 | 1 | NA | NA | NA | NA | NA | NA | 1 | 1 | 1 | 1 |
| 5_1509864 | NA | NA | NA | NA | 1 | 1 | NA | NA | 1 | 1 | NA | NA | NA | NA | NA | NA | 1 | 1 | 1 | 1 |
| 5_1515663 | NA | NA | 1 | 0 | 1 | 1 | 1 | 0 | 3 | 0.3333 | NA | NA | 4 | 0.25 | 1 | 1 | 2 | 1 | 7 | 0.5714 |
| 5_1515663 | 1 | 0 | 1 | 1 | 1 | 0 | 1 | 0 | 4 | 0.25 | NA | NA | 4 | 0.75 | 1 | 0 | 2 | 0 | 7 | 0.4286 |
| 6_3372630 | 1 | 1 | NA | NA | NA | NA | NA | NA | 1 | 1 | NA | NA | NA | NA | 1 | 1 | NA | NA | 1 | 1 |
| 6_3373139 | NA | NA | NA | NA | 1 | 1 | NA | NA | 1 | 1 | NA | NA | NA | NA | 1 | 1 | NA | NA | 1 | 1 |
| 6_3373758 | NA | NA | NA | NA | 1 | 1 | NA | NA | 1 | 1 | NA | NA | NA | NA | 1 | 1 | NA | NA | 1 | 1 |
| 6_3373766 | NA | NA | NA | NA | 1 | 1 | NA | NA | 1 | 1 | NA | NA | NA | NA | 1 | 1 | NA | NA | 1 | 1 |
| 6_3373994 | NA | NA | NA | NA | 1 | 1 | NA | NA | 1 | 1 | NA | NA | NA | NA | 1 | 1 | NA | NA | 1 | 1 |
| 6_3374290 | NA | NA | NA | NA | 1 | 1 | NA | NA | 1 | 1 | NA | NA | NA | NA | 1 | 1 | NA | NA | 1 | 1 |
| 6_3374367 | NA | NA | NA | NA | 1 | 1 | NA | NA | 1 | 1 | NA | NA | NA | NA | 1 | 1 | NA | NA | 1 | 1 |
| 6_3374642 | NA | NA | NA | NA | 1 | 1 | NA | NA | 1 | 1 | NA | NA | NA | NA | 1 | 1 | NA | NA | 1 | 1 |
| 6_3602783 | 6 | 0 | 10 | 0.3 | 8 | 0.125 | 8 | 0 | 32 | 0.125 | 3 | 0 | 9 | 0 | 10 | 0.1 | 4 | 0.25 | 26 | 0.0769 |
| 6_3602803 | 5 | 0 | 10 | 0.2 | 9 | 0.3333 | 9 | 0.2222 | 33 | 0.2121 | 3 | 0.3333 | 8 | 0.5 | 9 | 0.2222 | 6 | 0.5 | 26 | 0.3846 |
| 6_3602878 | 6 | 0 | 11 | 0.0909 | 10 | 0.1 | 9 | 0.2222 | 36 | 0.1111 | 3 | 0 | 10 | 0 | 11 | 0.0909 | 6 | 0.1667 | 30 | 0.0667 |
| 6_3602899 | 6 | 0.5 | 11 | 0.6364 | 11 | 0.2727 | 8 | 0.25 | 36 | 0.4167 | 2 | 1 | 10 | 0.4 | 13 | 0.6923 | 5 | 0.4 | 30 | 0.5667 |
| 6_3602944 | 6 | 0.3333 | 12 | 0.5 | 12 | 0.5 | 10 | 0.4 | 40 | 0.45 | 3 | 1 | 11 | 0.2727 | 15 | 0.5333 | 6 | 0.3333 | 35 | 0.4571 |
| 6_4004289 | 4 | 0.25 | 5 | 0.2 | 3 | 0 | 3 | 0 | 15 | 0.1333 | 3 | 0 | 1 | 0 | 2 | 0 | 3 | 0.3333 | 9 | 0.1111 |
| 6_4008164 | 1 | 1 | 2 | 1 | 1 | 1 | 2 | 1 | 6 | 1 | 2 | 0.5 | NA | NA | 1 | 1 | 1 | 1 | 4 | 0.75 |
| 6_4008235 | 4 | 0.75 | 5 | 0.6 | 4 | 0.25 | 3 | 0.3333 | 16 | 0.5 | 3 | 0 | 1 | 0 | 2 | 1 | 2 | 0.5 | 8 | 0.375 |
| 6_4008263 | 4 | 0.75 | 2 | 1 | 4 | 0.75 | 2 | 1 | 12 | 0.8333 | 2 | 0.5 | 1 | 0 | 2 | 0.5 | 3 | 1 | 8 | 0.625 |
| 6_4077373 | 5 | 0 | 4 | 0 | 6 | 0.1667 | 6 | 0.3333 | 21 | 0.1429 | 1 | 0 | 3 | 0 | 4 | 0.75 | 3 | 0.6667 | 11 | 0.4545 |
| 6_4540667 | 4 | 0.75 | 3 | 0.3333 | 1 | 0 | 5 | 0 | 13 | 0.3077 | 2 | 0.5 | 1 | 0 | NA | NA | 1 | 1 | 4 | 0.5 |
| 6_4540753 | 6 | 0.6667 | 3 | 0.6667 | 3 | 0.6667 | 6 | 0.3333 | 18 | 0.5556 | 2 | 0.5 | 1 | 0 | NA | NA | 2 | 0.5 | 5 | 0.4 |
| 6_4540799 | 2 | 0 | 2 | 0.5 | 3 | 0.3333 | 4 | 0 | 11 | 0.1818 | 1 | 0 | NA | NA | NA | NA | 2 | 0.5 | 3 | 0.3333 |
| 6_4540936 | 3 | 0.3333 | 2 | 0.5 | 3 | 0.3333 | 6 | 0.3333 | 14 | 0.3571 | 1 | 1 | 1 | 0 | NA | NA | 1 | 0 | 3 | 0.3333 |
| 6_4541002 | 1 | 1 | NA | NA | NA | NA | 1 | 0 | 2 | 0.5 | 2 | 0.5 | 1 | 0 | NA | NA | 1 | 0 | 4 | 0.25 |
| 6_4642112 | 10 | 0.9 | 8 | 0.75 | 6 | 0.8333 | 5 | 0.8 | 29 | 0.8276 | 3 | 1 | 9 | 0.7778 | 7 | 0.8571 | 14 | 0.8571 | 33 | 0.8485 |
| 6_4642141 | 8 | 0.375 | 8 | 0.125 | 5 | 0.8 | 7 | 0.2857 | 28 | 0.3571 | 2 | 0.5 | 4 | 0.75 | 6 | 0.5 | 8 | 0.625 | 20 | 0.6 |
| 6_4642203 | 5 | 1 | 9 | 0.6667 | 5 | 0.8 | 8 | 1 | 27 | 0.8519 | 3 | 1 | 8 | 0.625 | 6 | 0.8333 | 8 | 0.875 | 25 | 0.8 |
| 6_4642218 | 7 | 0.2857 | 5 | 0.4 | 6 | 0.5 | 5 | 0.4 | 23 | 0.3913 | 2 | 0.5 | 8 | 0.25 | 5 | 0 | 11 | 0.0909 | 26 | 0.1538 |
| 6_4642233 | 9 | 0.7778 | 9 | 0.6667 | 6 | 0.5 | 6 | 0.6667 | 30 | 0.6667 | 4 | 1 | 8 | 0.875 | 6 | 1 | 15 | 0.8 | 33 | 0.8788 |
| 6_4642255 | 10 | 0.5 | 9 | 0.2222 | 7 | 0.2857 | 7 | 0.4286 | 33 | 0.3636 | 4 | 0.25 | 9 | 0.2222 | 7 | 0.2857 | 14 | 0.1429 | 34 | 0.2059 |
| 6_4642419 | 10 | 0.3 | 9 | 0.2222 | 7 | 0.2857 | 8 | 0.375 | 34 | 0.2941 | 5 | 0.4 | 7 | 0.5714 | 6 | 0.5 | 13 | 0.4615 | 31 | 0.4839 |
| 6_4642564 | 10 | 0.1 | 10 | 0.2 | 8 | 0.25 | 8 | 0 | 36 | 0.1389 | 5 | 0 | 10 | 0.1 | 8 | 0 | 13 | 0.0769 | 36 | 0.0556 |
| 6_4642612 | 10 | 0.5 | 10 | 0.5 | 8 | 0.5 | 8 | 0.25 | 36 | 0.4444 | 5 | 0.6 | 10 | 0.5 | 8 | 0.375 | 14 | 0.5714 | 37 | 0.5135 |
| 6_4674490 | 3 | 0 | 3 | 0.3333 | 3 | 0 | 8 | 0 | 17 | 0.0588 | 4 | 0 | 7 | 0 | 5 | 0.2 | 4 | 0 | 20 | 0.05 |
| 6_4756736 | 1 | 1 | 2 | 1 | NA | NA | 4 | 1 | 7 | 1 | 1 | 1 | 2 | 1 | NA | NA | NA | NA | 3 | 1 |
| 6_4756789 | 1 | 1 | 2 | 0 | NA | NA | 4 | 0.25 | 7 | 0.2857 | NA | NA | 1 | 1 | NA | NA | NA | NA | 1 | 1 |
| 6_4757304 | 1 | 1 | 3 | 0.6667 | 1 | 1 | 4 | 0.5 | 9 | 0.6667 | 1 | 1 | 2 | 1 | 1 | 1 | NA | NA | 4 | 1 |
| 6_4757319 | 1 | 0 | 2 | 1 | 1 | 0 | 4 | 0.75 | 8 | 0.625 | 1 | 1 | 2 | 1 | 1 | 1 | NA | NA | 4 | 1 |
| 6_4757360 | 1 | 0 | 3 | 0.3333 | 1 | 0 | 3 | 0 | 8 | 0.125 | 2 | 0.5 | 1 | 1 | 1 | 0 | NA | NA | 4 | 0.5 |
| 6_4757528 | 1 | 0 | 3 | 0 | 1 | 0 | 5 | 0.8 | 10 | 0.4 | 2 | 0 | 2 | 0.5 | 1 | 0 | NA | NA | 5 | 0.2 |
| 6_4757571 | 1 | 0 | 3 | 0.6667 | 1 | 1 | 5 | 0.8 | 10 | 0.7 | 1 | 1 | 2 | 1 | NA | NA | 1 | 1 | 4 | 1 |
| 6_4757741 | NA | NA | 1 | 1 | NA | NA | 5 | 0.4 | 6 | 0.5 | 2 | 1 | 1 | 1 | NA | NA | 1 | 1 | 4 | 1 |
| 6_4757745 | 1 | 1 | 2 | 0.5 | 1 | 0 | 5 | 0.2 | 9 | 0.3333 | 2 | 0 | 2 | 0.5 | 1 | 1 | 1 | 0 | 6 | 0.3333 |
| 6_4757794 | NA | NA | 3 | 1 | 1 | 1 | 6 | 1 | 10 | 1 | 1 | 1 | 2 | 1 | 1 | 1 | NA | NA | 4 | 1 |
| 6_4757949 | 1 | 1 | 2 | 1 | 1 | 1 | 5 | 1 | 9 | 1 | 2 | 1 | 2 | 1 | 1 | 1 | 1 | 1 | 6 | 1 |
| 6_4757970 | 1 | 0 | 1 | 1 | 1 | 0 | 5 | 1 | 8 | 0.75 | 2 | 0.5 | 1 | 0 | 1 | 0 | 1 | 1 | 5 | 0.4 |
| 6_4758117 | 1 | 1 | 2 | 0.5 | 1 | 0 | 4 | 1 | 8 | 0.75 | 2 | 1 | 2 | 1 | 1 | 1 | 1 | 1 | 6 | 1 |
| 6_4758254 | 1 | 1 | 1 | 1 | 1 | 1 | 6 | 1 | 9 | 1 | 1 | 1 | 1 | 1 | 1 | 1 | 1 | 1 | 4 | 1 |
| 6_4758290 | 1 | 0 | 3 | 1 | 1 | 1 | 7 | 0.7143 | 12 | 0.75 | 2 | 0.5 | 2 | 0.5 | 1 | 1 | NA | NA | 5 | 0.6 |
| 6_4758369 | 1 | 0 | 3 | 1 | 1 | 1 | 7 | 0.7143 | 12 | 0.75 | 2 | 0.5 | 3 | 0.6667 | 1 | 1 | 1 | 1 | 7 | 0.7143 |
| 6_4758378 | 1 | 0 | 3 | 0.3333 | 1 | 1 | 6 | 1 | 11 | 0.7273 | 3 | 0.6667 | 3 | 0.6667 | 1 | 0 | 1 | 1 | 8 | 0.625 |
| 6_4758435 | 1 | 1 | 2 | 1 | 1 | 1 | 7 | 1 | 11 | 1 | 3 | 1 | 3 | 0.3333 | 1 | 1 | 1 | 1 | 8 | 0.75 |
| 6_4758807 | 1 | 1 | 2 | 1 | 1 | 1 | 7 | 1 | 11 | 1 | 3 | 1 | 3 | 1 | 1 | 1 | 1 | 1 | 8 | 1 |
| 6_4758916 | 1 | 1 | 3 | 1 | 1 | 1 | 8 | 1 | 13 | 1 | 4 | 1 | 2 | 1 | 1 | 1 | 1 | 1 | 8 | 1 |
| 6_4759118 | 1 | 0 | 3 | 0.3333 | 1 | 0 | 8 | 0 | 13 | 0.0769 | 4 | 0.25 | 3 | 0 | 1 | 0 | 1 | 0 | 9 | 0.1111 |
| 6_4759307 | 1 | 0 | 3 | 0.3333 | 2 | 0 | 9 | 0.5556 | 15 | 0.4 | 4 | 0.25 | 3 | 0 | 1 | 0 | 2 | 0 | 10 | 0.1 |

| | | | | | | | | | | | | | | | | | | | | |
|---|---|---|---|---|---|---|---|---|---|---|---|---|---|---|---|---|---|---|---|---|
| 6_4759338 | 1 | 1 | 2 | 0 | 3 | 0.6667 | 7 | 0.4286 | 13 | 0.4615 | 2 | 1 | 3 | 0.3333 | 1 | 0 | 2 | 1 | 8 | 0.625 |
| 6_4759429 | 1 | 0 | 3 | 0 | 3 | 0.3333 | 8 | 0.125 | 15 | 0.1333 | 4 | 0.5 | 3 | 0 | 1 | 0 | 2 | 1 | 10 | 0.4 |
| 6_4759448 | 1 | 0 | 3 | 0.6667 | 2 | 0.5 | 7 | 0 | 13 | 0.2308 | 2 | 0.5 | 3 | 0.3333 | NA | NA | 2 | 0 | 7 | 0.2857 |
| 6_4759459 | NA | NA | 3 | 0.6667 | 3 | 0 | 8 | 0.625 | 14 | 0.5 | 3 | 0.6667 | 3 | 0 | 1 | 0 | 2 | 0.5 | 9 | 0.3333 |
| 6_4759472 | NA | NA | 3 | 0 | 3 | 0 | 9 | 0.5556 | 15 | 0.3333 | 2 | 0.5 | 2 | 0 | 1 | 0 | 2 | 0 | 7 | 0.1429 |
| 6_4905274 | NA | NA | 1 | 1 | NA | NA | 1 | 1 | 2 | 1 | NA | NA | 3 | 0.3333 | 1 | 1 | NA | NA | 4 | 0.5 |
| 6_4905317 | NA | NA | 1 | 1 | NA | NA | 1 | 0 | 2 | 0.5 | NA | NA | 3 | 0.3333 | 1 | 1 | NA | NA | 4 | 0.5 |
| 6_4905951 | NA | NA | NA | NA | NA | NA | 1 | 1 | 1 | 1 | NA | NA | 4 | 0.75 | 1 | 1 | NA | NA | 5 | 0.8 |
| 6_4906606 | 1 | 1 | 3 | 0 | NA | NA | 1 | 0 | 5 | 0.2 | 2 | 0 | 4 | 0.25 | 1 | 0 | 1 | 0 | 8 | 0.125 |
| 6_4906648 | 1 | 1 | 2 | 1 | NA | NA | NA | NA | 3 | 1 | 2 | 0.5 | 4 | 0.5 | 1 | 1 | 1 | 1 | 8 | 0.625 |
| 6_4906687 | 1 | 0 | 3 | 0.3333 | NA | NA | 1 | 0 | 5 | 0.2 | 2 | 1 | 4 | 0.5 | 1 | 1 | 1 | 0 | 8 | 0.625 |
| 6_5158195 | 2 | 0 | 5 | 0.2 | 1 | 0 | 3 | 0 | 11 | 0.0909 | 2 | 0 | 5 | 0 | 2 | 0 | 4 | 0.5 | 13 | 0.1538 |
| 6_5159599 | NA | NA | 4 | 0.75 | 1 | 1 | 2 | 0 | 7 | 0.5714 | 3 | 0.6667 | 5 | 0.4 | 2 | 1 | 2 | 0 | 12 | 0.5 |
| 6_5159881 | 2 | 1 | 5 | 1 | 3 | 1 | 6 | 0.8333 | 16 | 0.9375 | 2 | 1 | 8 | 1 | 3 | 1 | 5 | 1 | 18 | 1 |
| 6_5160057 | 1 | 1 | 5 | 0.8 | 3 | 1 | 8 | 1 | 17 | 0.9412 | 3 | 1 | 6 | 1 | 2 | 0.5 | 4 | 0.75 | 15 | 0.8667 |
| 6_5160112 | 2 | 0.5 | 6 | 0.3333 | 3 | 0.6667 | 6 | 0.3333 | 17 | 0.4118 | 4 | 0.5 | 7 | 0.1429 | 2 | 0.5 | 5 | 0.2 | 18 | 0.2778 |
| 6_5160131 | 1 | 0 | 5 | 0.2 | 1 | 1 | 7 | 0.7143 | 14 | 0.5 | 2 | 0.5 | 7 | 0.7143 | 3 | 1 | 6 | 0.3333 | 18 | 0.6111 |
| 6_5160187 | 2 | 0 | 4 | 0.25 | 3 | 0 | 8 | 0.125 | 17 | 0.1176 | 2 | 0 | 8 | 0.125 | 3 | 0 | 6 | 0.1667 | 19 | 0.1053 |
| 6_5160590 | 3 | 0 | 9 | 0.1111 | 2 | 0 | 8 | 0.125 | 22 | 0.0909 | 4 | 0 | 7 | 0 | 1 | 0 | 7 | 0.1429 | 19 | 0.0526 |
| 6_5160691 | 4 | 0 | 9 | 0.1111 | 3 | 0 | 11 | 0.0909 | 27 | 0.0741 | 4 | 0 | 6 | 0.1667 | 4 | 0 | 7 | 0.1429 | 21 | 0.0952 |
| 6_5160819 | 3 | 0 | 10 | 0.2 | 3 | 0 | 12 | 0 | 28 | 0.0714 | 2 | 0 | 7 | 0 | 2 | 0 | 8 | 0.125 | 19 | 0.0526 |
| 6_5161139 | 3 | 0.6667 | 10 | 0.2 | 4 | 0.5 | 12 | 0.5833 | 29 | 0.4483 | 5 | 0.6 | 9 | 0.4444 | 4 | 0.75 | 8 | 0.875 | 26 | 0.6538 |
| 6_5161180 | 4 | 0.25 | 9 | 0.3333 | 4 | 0.75 | 13 | 0.2308 | 30 | 0.3333 | 3 | 0 | 9 | 0.2222 | 5 | 0.4 | 10 | 0.7 | 27 | 0.4074 |
| 6_5161359 | 4 | 0 | 10 | 0.2 | 5 | 0.2 | 15 | 0.0667 | 34 | 0.1176 | 4 | 0 | 10 | 0.1 | 5 | 0 | 9 | 0.2222 | 28 | 0.1071 |
| 6_5161677 | 4 | 0.75 | 10 | 0.4 | 7 | 0.4286 | 14 | 0.4286 | 35 | 0.4571 | 5 | 0.4 | 9 | 0.3333 | 5 | 0.2 | 11 | 0.5455 | 30 | 0.4 |
| 6_5161689 | 3 | 0.3333 | 11 | 0.0909 | 7 | 0.4286 | 15 | 0.0667 | 36 | 0.1667 | 5 | 0.2 | 9 | 0.1111 | 6 | 0 | 7 | 0 | 27 | 0.0741 |
| 6_5161704 | 3 | 0 | 5 | 0 | 4 | 0.5 | 10 | 0.2 | 22 | 0.1818 | 4 | 0 | 7 | 0.2857 | 4 | 0 | 8 | 0.25 | 23 | 0.1739 |
| 6_5161807 | 3 | 0.6667 | 6 | 0.3333 | 9 | 0.7778 | 12 | 0.9167 | 30 | 0.7333 | 4 | 0.5 | 7 | 0.5714 | 4 | 0.5 | 10 | 0.9 | 25 | 0.68 |
| 6_5162211 | 5 | 0.2 | 9 | 0.4444 | 8 | 0.5 | 16 | 0.1875 | 38 | 0.3158 | 4 | 0.25 | 10 | 0.2 | 6 | 0.1667 | 10 | 0.4 | 30 | 0.2667 |
| 6_5162398 | 7 | 0.2857 | 10 | 0.5 | 9 | 0.4444 | 16 | 0.6875 | 42 | 0.5238 | 5 | 0.4 | 11 | 0.3636 | 8 | 0.125 | 10 | 0.5 | 34 | 0.3529 |
| 6_5162414 | 7 | 0 | 10 | 0.3 | 9 | 0.1111 | 17 | 0.2353 | 43 | 0.186 | 5 | 0.2 | 11 | 0.1818 | 7 | 0.2857 | 9 | 0.3333 | 32 | 0.25 |
| 6_5162467 | 7 | 0.1429 | 11 | 0.0909 | 8 | 0 | 16 | 0.125 | 42 | 0.0952 | 5 | 0.2 | 9 | 0.1111 | 6 | 0.3333 | 10 | 0.1 | 30 | 0.1667 |
| 6_5162617 | 8 | 0 | 11 | 0.0909 | 11 | 0.1818 | 18 | 0.1111 | 48 | 0.1042 | 5 | 0.2 | 10 | 0 | 11 | 0.1818 | 12 | 0.0833 | 38 | 0.1053 |
| 6_5164367 | 10 | 0.2 | 14 | 0.2143 | 10 | 0 | 26 | 0.1538 | 60 | 0.15 | 2 | 0.5 | 20 | 0.15 | 17 | 0.1176 | 10 | 0.4 | 49 | 0.2041 |
| 6_5165277 | 37 | 0.1351 | 27 | 0.0741 | 22 | 0 | 37 | 0.027 | 123 | 0.065 | 6 | 0 | 27 | 0.1111 | 32 | 0.0938 | 23 | 0.1739 | 88 | 0.1136 |
| 6_5165367 | 39 | 0.1282 | 29 | 0.1379 | 26 | 0.0769 | 39 | 0.2051 | 133 | 0.1429 | 8 | 0.375 | 32 | 0.0938 | 34 | 0.0588 | 26 | 0.1538 | 100 | 0.12 |
| 6_5165373 | 36 | 0.3611 | 26 | 0.2308 | 21 | 0.3333 | 38 | 0.3421 | 121 | 0.3223 | 7 | 0.2857 | 27 | 0.2593 | 33 | 0.2121 | 24 | 0.4167 | 91 | 0.2857 |
| 6_5265138 | 5 | 0 | 14 | 0 | 10 | 0.2 | 10 | 0.2 | 39 | 0.1026 | 14 | 0.2857 | 10 | 0.2 | 8 | 0.125 | 6 | 0 | 38 | 0.1842 |
| 6_5265327 | 4 | 0 | 13 | 0.2308 | 8 | 0.125 | 10 | 0.2 | 35 | 0.1714 | 9 | 0.4444 | 10 | 0.2 | 7 | 0.1429 | 4 | 0.25 | 30 | 0.2667 |
| 6_5483586 | 1 | 0 | 1 | 1 | 1 | 0 | 1 | 0 | 4 | 0.25 | 3 | 0 | NA | NA | 4 | 0.25 | 2 | 0 | 9 | 0.1111 |
| 6_5484675 | NA | NA | 2 | 0.5 | 1 | 0 | 1 | 0 | 4 | 0.25 | 2 | 0 | NA | NA | 3 | 0 | 2 | 0.5 | 7 | 0.1429 |
| 6_5485199 | 1 | 1 | 2 | 0 | 1 | 0 | 1 | 1 | 5 | 0.4 | 2 | 0.5 | NA | NA | 3 | 0.6667 | 1 | 1 | 6 | 0.6667 |
| 6_5485206 | 1 | 1 | 2 | 0.5 | 1 | 0 | 1 | 0 | 5 | 0.4 | 2 | 0.5 | NA | NA | 4 | 1 | 1 | 0 | 7 | 0.7143 |
| 6_5485295 | 1 | 1 | 2 | 0 | 1 | 0 | 1 | 1 | 5 | 0.4 | 2 | 1 | NA | NA | 3 | 0.6667 | 2 | 1 | 7 | 0.8571 |
| 6_5485377 | 1 | 1 | 2 | 0.5 | 1 | 1 | 1 | 1 | 5 | 0.8 | 2 | 1 | NA | NA | 4 | 1 | 2 | 1 | 8 | 1 |
| 6_5485507 | 1 | 0 | 2 | 0.5 | 1 | 1 | 1 | 0 | 5 | 0.4 | 2 | 0 | NA | NA | 4 | 0.25 | 2 | 0.5 | 8 | 0.25 |
| 6_5966797 | 3 | 0 | 7 | 0 | 3 | 0.3333 | 6 | 0 | 19 | 0.0526 | 7 | 0 | 6 | 0.1667 | 5 | 0.2 | 5 | 0.4 | 23 | 0.1739 |
| 6_5972186 | 5 | 0 | 7 | 0 | 6 | 0.1667 | 6 | 0.3333 | 24 | 0.125 | 6 | 0.1667 | 7 | 0.1429 | 5 | 0.2 | 7 | 0.1429 | 25 | 0.16 |
| 6_6027530 | 9 | 0 | 9 | 0.1111 | 9 | 0.1111 | 7 | 0 | 34 | 0.0588 | 9 | 0.2222 | 13 | 0 | 12 | 0.1667 | 7 | 0 | 41 | 0.0976 |
| 6_6041698 | 1 | 1 | 1 | 1 | NA | NA | 1 | 1 | 3 | 1 | 1 | 1 | 1 | 0 | NA | NA | NA | NA | 2 | 0.5 |
| 6_6042153 | NA | NA | 1 | 1 | NA | NA | 1 | 1 | 2 | 1 | 1 | 1 | 1 | 0 | NA | NA | NA | NA | 2 | 0.5 |
| 6_6558415 | 19 | 0.1053 | 31 | 0.1613 | 18 | 0.1111 | 16 | 0.3125 | 84 | 0.1667 | 12 | 0.1667 | 17 | 0.1765 | 16 | 0.1875 | 23 | 0.3913 | 68 | 0.25 |
| 6_6808606 | 2 | 0 | 8 | 0.125 | 2 | 0 | 2 | 0 | 14 | 0.0714 | 3 | 0 | 2 | 1 | 7 | 0.1429 | 2 | 0 | 14 | 0.2143 |
| 6_6808885 | 2 | 0 | 8 | 0 | 1 | 1 | 3 | 0 | 14 | 0.0714 | 3 | 0 | 3 | 0 | 5 | 0.2 | 2 | 0 | 13 | 0.0769 |
| 6_6809691 | 1 | 0 | 7 | 0.1429 | 2 | 0 | 2 | 0 | 12 | 0.0833 | NA | NA | 1 | 0 | 2 | 0.5 | 1 | 0 | 4 | 0.25 |
| 6_6820665 | 1 | 1 | 1 | 1 | 2 | 1 | 2 | 0.5 | 6 | 0.8333 | 2 | 0.5 | 4 | 0.75 | 6 | 1 | 1 | 1 | 13 | 0.8462 |
| 6_6821054 | 1 | 1 | 1 | 0 | NA | NA | 1 | 0 | 3 | 0.3333 | 4 | 0.25 | 3 | 0 | 3 | 0 | 1 | 0 | 11 | 0.0909 |
| 6_6823856 | NA | NA | 1 | 1 | 3 | 0.3333 | 1 | 1 | 5 | 0.6 | NA | NA | 4 | 0.25 | 3 | 0.3333 | 1 | 0 | 8 | 0.25 |
| 6_6823983 | NA | NA | 2 | 0.5 | 2 | 0 | 1 | 0 | 5 | 0.2 | NA | NA | 4 | 0 | 5 | 0.2 | 1 | 1 | 10 | 0.2 |
| 6_6824122 | NA | NA | 2 | 1 | 1 | 1 | 1 | 1 | 4 | 1 | NA | NA | 1 | 1 | 4 | 1 | 1 | 1 | 6 | 1 |
| 6_6824176 | NA | NA | 2 | 1 | 2 | 0.5 | 1 | 1 | 5 | 0.8 | NA | NA | 3 | 1 | 3 | 0.6667 | 1 | 1 | 7 | 0.8571 |
| 6_6824218 | NA | NA | NA | NA | 2 | 0.5 | NA | NA | 2 | 0.5 | NA | NA | 2 | 0.5 | 2 | 1 | 1 | 0 | 5 | 0.6 |
| 6_6824318 | NA | NA | NA | NA | 1 | 1 | NA | NA | 1 | 1 | NA | NA | 1 | 1 | 3 | 1 | 1 | 1 | 5 | 1 |
| 6_6824369 | NA | NA | 2 | 0.5 | 1 | 0 | 1 | 0 | 4 | 0.25 | NA | NA | 4 | 0.25 | 4 | 0.25 | 1 | 0 | 9 | 0.2222 |
| 6_6824465 | NA | NA | 2 | 0 | 2 | 0.5 | 1 | 0 | 5 | 0.2 | NA | NA | 4 | 0.25 | 4 | 0.25 | 1 | 0 | 9 | 0.2222 |

| | | | | | | | | | | | | | | | | | | | | |
|---|---|---|---|---|---|---|---|---|---|---|---|---|---|---|---|---|---|---|---|---|
| 6_6824482 | NA | NA | 1 | 0 | 2 | 0.5 | 1 | 0 | 4 | 0.25 | NA | NA | 4 | 0 | 3 | 0.6667 | 1 | 0 | 8 | 0.25 |
| 6_6863423 | NA | NA | NA | NA | NA | NA | 2 | 0.5 | 2 | 0.5 | NA | NA | 2 | 0.5 | 1 | 0 | 2 | 0.5 | 5 | 0.4 |
| 6_6865256 | 5 | 0.4 | 1 | 0 | 2 | 1 | 3 | 0 | 11 | 0.3636 | 1 | 0 | 3 | 0.3333 | NA | NA | 3 | 0 | 7 | 0.1429 |
| 6_6867944 | 6 | 0.6667 | 1 | 1 | 3 | 0.6667 | 4 | 0.75 | 14 | 0.7143 | 1 | 1 | 4 | 0.5 | 4 | 0.5 | 3 | 0.3333 | 12 | 0.5 |
| 6_6867974 | 5 | 1 | 1 | 1 | 3 | 1 | 5 | 1 | 14 | 1 | 1 | 1 | 3 | 1 | 3 | 1 | 2 | 1 | 9 | 1 |
| 6_6868003 | 4 | 1 | NA | NA | 3 | 0.6667 | 5 | 0.6 | 12 | 0.75 | 1 | 1 | 3 | 0.6667 | 3 | 0.3333 | 2 | 1 | 9 | 0.6667 |
| 6_6868219 | 8 | 0.5 | 3 | 0.3333 | 3 | 0 | 8 | 0.125 | 22 | 0.2727 | NA | NA | 7 | 0.5714 | 8 | 0.375 | 4 | 0.5 | 19 | 0.4737 |
| 6_6868292 | 9 | 0.1111 | 3 | 0 | 3 | 0 | 5 | 0.2 | 20 | 0.1 | 1 | 0 | 4 | 0.25 | 5 | 0.2 | 3 | 0.3333 | 13 | 0.2308 |
| 6_6868346 | 7 | 0.5714 | 1 | 0 | 4 | 0.75 | 5 | 0.6 | 17 | 0.5882 | 3 | 0.3333 | 3 | 0 | 7 | 0.4286 | 1 | 1 | 14 | 0.3571 |
| 6_6868461 | 12 | 0.1667 | 3 | 0 | 4 | 0.25 | 7 | 0 | 26 | 0.1154 | 1 | 0 | 7 | 0.1429 | 7 | 0.5714 | 4 | 0.5 | 19 | 0.3684 |
| 6_6877895 | 9 | 0.5556 | 3 | 0.3333 | 5 | 0.6 | 12 | 0.4167 | 29 | 0.4828 | 4 | 0.5 | 10 | 0.2 | 2 | 1 | 10 | 0.5 | 26 | 0.4231 |
| 6_6877942 | 8 | 0.375 | 4 | 0.25 | 5 | 0.8 | 9 | 0.3333 | 26 | 0.4231 | 3 | 0.3333 | 7 | 0.4286 | 3 | 0.3333 | 8 | 0.625 | 21 | 0.4762 |
| 6_6878068 | 6 | 0.1667 | 3 | 0.3333 | 5 | 0.4 | 10 | 0.1 | 24 | 0.2083 | 2 | 0 | 5 | 0 | 2 | 0 | 9 | 0.2222 | 18 | 0.1111 |
| 6_6878128 | 8 | 1 | 4 | 1 | 3 | 1 | 10 | 1 | 25 | 1 | 3 | 1 | 5 | 0.8 | 3 | 1 | 7 | 0.8571 | 18 | 0.8889 |
| 6_6878265 | 6 | 0.1667 | 3 | 0 | 3 | 0 | 8 | 0.125 | 20 | 0.1 | 3 | 0.3333 | 7 | 0 | 1 | 1 | 7 | 0.4286 | 18 | 0.2778 |
| 6_6879541 | 4 | 0.25 | 3 | 0 | 3 | 0 | 10 | 0.3 | 20 | 0.2 | 4 | 0.5 | 6 | 0 | 3 | 0 | 7 | 0.4286 | 20 | 0.25 |
| 6_6879642 | 1 | 0 | 2 | 0 | 2 | 0.5 | 8 | 1 | 13 | 0.6923 | 2 | 1 | 6 | 0.1667 | 2 | 0.5 | 6 | 0.6667 | 16 | 0.5 |
| 6_6956156 | NA | NA | 4 | 0.5 | 4 | 0.25 | 4 | 0.25 | 12 | 0.3333 | NA | NA | 1 | 0 | 4 | 0.25 | 4 | 0.5 | 9 | 0.3333 |
| 6_7038954 | NA | NA | 3 | 0.3333 | 3 | 0.3333 | 3 | 0.3333 | 9 | 0.3333 | NA | NA | 1 | 0 | 2 | 0.5 | 4 | 0.5 | 7 | 0.4286 |
| 6_7038976 | NA | NA | 3 | 0.6667 | 4 | 0.5 | 4 | 0.5 | 11 | 0.5455 | NA | NA | 1 | 1 | 2 | 1 | 3 | 0.3333 | 6 | 0.6667 |
| 6_7039113 | NA | NA | 3 | 1 | 4 | 0.5 | 3 | 0.6667 | 10 | 0.7 | NA | NA | NA | NA | 3 | 0.6667 | 4 | 0.75 | 7 | 0.7143 |
| 6_7039162 | 1 | 0 | 4 | 0.5 | 3 | 0.6667 | 3 | 0.6667 | 11 | 0.5455 | NA | NA | 1 | 1 | 4 | 0.75 | 3 | 1 | 8 | 0.875 |
| 6_7039171 | 1 | 0 | 2 | 0 | 4 | 0.25 | 3 | 0.3333 | 10 | 0.2 | NA | NA | 1 | 1 | 4 | 0 | 2 | 1 | 7 | 0.4286 |
| 6_7039270 | NA | NA | 3 | 0.3333 | 4 | 0.25 | 4 | 0.25 | 11 | 0.2727 | NA | NA | 1 | 1 | 2 | 0.5 | 1 | 1 | 4 | 0.75 |
| 6_7554900 | 1 | 1 | NA | NA | 1 | 0 | 3 | 0 | 5 | 0.2 | NA | NA | 2 | 1 | NA | NA | 5 | 0.2 | 7 | 0.4286 |
| 6_7562712 | 109 | 0.2752 | 58 | 0.2586 | 102 | 0.2843 | 119 | 0.2605 | 388 | 0.2706 | 36 | 0.3056 | 129 | 0.2636 | 141 | 0.2837 | 95 | 0.3474 | 401 | 0.2943 |
| 6_7562756 | 123 | 0.065 | 68 | 0.0441 | 122 | 0.0738 | 141 | 0.0922 | 454 | 0.0727 | 41 | 0.0488 | 150 | 0.12 | 158 | 0.0633 | 115 | 0.0522 | 464 | 0.0776 |
| 6_7672947 | NA | NA | NA | NA | 1 | 0 | 1 | 1 | 2 | 0.5 | NA | NA | 1 | 1 | NA | NA | NA | NA | 1 | 1 |
| 6_7673430 | NA | NA | NA | NA | 1 | 0 | 1 | 1 | 2 | 0.5 | NA | NA | 1 | 1 | NA | NA | 1 | 0 | 2 | 0.5 |
| 6_7674146 | NA | NA | NA | NA | 1 | 0 | 1 | 1 | 2 | 0.5 | NA | NA | 1 | 1 | NA | NA | 1 | 1 | 2 | 1 |
| 6_7674675 | NA | NA | 1 | 1 | 1 | 1 | NA | NA | 2 | 1 | NA | NA | 1 | 1 | NA | NA | 1 | 0 | 2 | 0.5 |
| 6_7674682 | NA | NA | 1 | 0 | 1 | 1 | NA | NA | 2 | 0.5 | NA | NA | 1 | 1 | NA | NA | 1 | 0 | 2 | 0.5 |
| 6_7675232 | 17 | 0.4118 | 19 | 0.4211 | 24 | 0.25 | 16 | 0.4375 | 76 | 0.3684 | 21 | 0.2857 | 24 | 0.375 | 18 | 0.2778 | 11 | 0.4545 | 74 | 0.3378 |
| 6_7675364 | 14 | 0 | 19 | 0.0526 | 20 | 0.05 | 17 | 0.1176 | 70 | 0.0571 | 12 | 0.1667 | 16 | 0.0625 | 11 | 0.0909 | 9 | 0 | 48 | 0.0833 |
| 6_7675401 | 14 | 0.2857 | 18 | 0.2778 | 22 | 0.0909 | 16 | 0.125 | 70 | 0.1857 | 13 | 0.3077 | 22 | 0.1364 | 13 | 0.1538 | 11 | 0.5455 | 59 | 0.2542 |
| 6_7866643 | 4 | 0.5 | 2 | 0 | 1 | 1 | 2 | 1 | 9 | 0.5556 | 1 | 0 | NA | NA | 1 | 0 | 2 | 1 | 4 | 0.5 |
| 6_7871105 | 4 | 0.5 | 2 | 1 | 2 | 0 | 2 | 1 | 10 | 0.6 | 1 | 1 | NA | NA | 2 | 1 | 2 | 1 | 5 | 1 |
| 6_7871265 | 3 | 0.6667 | 2 | 0.5 | 2 | 0 | 1 | 0 | 8 | 0.375 | 1 | 0 | NA | NA | 2 | 0 | 2 | 1 | 5 | 0.4 |
| 6_7871293 | 3 | 1 | 2 | 1 | 2 | 1 | 1 | 1 | 8 | 1 | 1 | 1 | NA | NA | 2 | 1 | 2 | 1 | 5 | 1 |
| 6_7871348 | 3 | 1 | 1 | 1 | 2 | 1 | 1 | 0 | 7 | 0.8571 | 1 | 1 | NA | NA | 2 | 1 | 1 | 1 | 4 | 1 |
| 6_7871379 | 3 | 0.3333 | 2 | 0 | 1 | 0 | 2 | 0 | 8 | 0.125 | NA | NA | NA | NA | 2 | 0.5 | 3 | 0.3333 | 5 | 0.4 |
| 6_7871386 | 4 | 1 | 1 | 1 | 2 | 1 | 2 | 1 | 9 | 1 | NA | NA | NA | NA | 2 | 1 | 1 | 1 | 3 | 1 |
| 6_7871507 | 3 | 1 | 2 | 1 | 2 | 1 | 2 | 1 | 9 | 1 | 1 | 1 | NA | NA | 2 | 1 | 3 | 1 | 6 | 1 |
| 6_7871519 | 3 | 0.6667 | 2 | 0.5 | 1 | 1 | 2 | 1 | 8 | 0.75 | 1 | 1 | NA | NA | 2 | 1 | 2 | 0.5 | 5 | 0.8 |
| 6_7871642 | 4 | 1 | 2 | 1 | 3 | 0.6667 | 2 | 1 | 11 | 0.9091 | 1 | 1 | NA | NA | 2 | 1 | 3 | 0.6667 | 6 | 0.8333 |
| 6_7871713 | 4 | 0.25 | 2 | 0 | 3 | 0 | NA | NA | 9 | 0.1111 | 1 | 0 | NA | NA | 2 | 1 | 2 | 0 | 5 | 0.4 |
| 6_7871826 | 4 | 0.25 | 2 | 0 | 3 | 0 | 2 | 0 | 11 | 0.0909 | 1 | 0 | NA | NA | 2 | 0.5 | 3 | 0.6667 | 6 | 0.5 |
| 6_7873793 | 4 | 0.25 | 3 | 0 | 2 | 0 | 4 | 0 | 13 | 0.0769 | 2 | 0 | NA | NA | 1 | 1 | 4 | 0 | 7 | 0.1429 |
| 6_7873958 | 4 | 0.5 | 4 | 0.5 | 3 | 0.6667 | 3 | 0.6667 | 14 | 0.5714 | 2 | 0 | NA | NA | NA | NA | 3 | 0.6667 | 5 | 0.4 |
| 6_8214783 | 4 | 0.25 | 1 | 1 | 1 | 0 | 1 | 1 | 7 | 0.4286 | 1 | 0 | 1 | 0 | 2 | 0 | 1 | 1 | 5 | 0.2 |
| 6_8214870 | 4 | 0 | NA | NA | 1 | 1 | 1 | 1 | 6 | 0.3333 | 1 | 1 | 1 | 0 | 3 | 0.6667 | 1 | 0 | 6 | 0.5 |
| 6_8214948 | 3 | 0.3333 | 1 | 0 | 1 | 1 | 1 | 1 | 6 | 0.5 | NA | NA | 1 | 0 | 3 | 0.3333 | 1 | 0 | 5 | 0.2 |
| 6_8214975 | 4 | 0.25 | 1 | 0 | 1 | 1 | 1 | 0 | 7 | 0.2857 | 1 | 0 | 1 | 0 | 3 | 0.6667 | NA | NA | 5 | 0.4 |
| 6_8215048 | 3 | 0.6667 | 1 | 1 | 1 | 1 | 1 | 0 | 6 | 0.6667 | 1 | 0 | 1 | 1 | 2 | 1 | 1 | 0 | 5 | 0.6 |
| 6_8215095 | 3 | 0.3333 | 1 | 0 | NA | NA | 1 | 0 | 5 | 0.2 | NA | NA | 1 | 0 | 2 | 0.5 | 1 | 0 | 4 | 0.25 |
| 6_8215221 | 4 | 0.25 | 1 | 1 | 1 | 1 | 1 | 1 | 7 | 0.5714 | 1 | 0 | 1 | 0 | 3 | 0.3333 | 1 | 1 | 6 | 0.3333 |
| 6_8215291 | 3 | 0.3333 | 1 | 1 | NA | NA | 1 | 0 | 5 | 0.4 | 1 | 1 | 1 | 1 | 2 | 0.5 | 1 | 1 | 5 | 0.8 |
| 6_8215589 | 4 | 0.5 | 1 | 1 | 1 | 1 | 1 | 0 | 7 | 0.5714 | 1 | 0 | 1 | 0 | 3 | 0.3333 | 1 | 0 | 6 | 0.1667 |
| 6_8215829 | 4 | 0 | 1 | 0 | NA | NA | 1 | 1 | 6 | 0.1667 | 1 | 0 | 1 | 0 | 3 | 0.3333 | NA | NA | 5 | 0.2 |
| 6_8215841 | 2 | 0.5 | NA | NA | NA | NA | 1 | 1 | 3 | 0.6667 | 1 | 0 | 1 | 0 | 2 | 0.5 | 1 | 1 | 5 | 0.4 |
| 6_8216014 | 3 | 0 | 1 | 1 | 1 | 1 | 1 | 0 | 6 | 0.3333 | 1 | 1 | 1 | 1 | 3 | 1 | NA | NA | 5 | 1 |
| 6_8235467 | 3 | 0 | 2 | 0.5 | 2 | 0.5 | 2 | 0 | 9 | 0.2222 | 1 | 1 | 2 | 1 | 3 | 0.3333 | 3 | 0.3333 | 9 | 0.5556 |
| 6_8235623 | 2 | 1 | 1 | 1 | 1 | 1 | 1 | 0 | 5 | 0.8 | 1 | 1 | 1 | 1 | 2 | 1 | 2 | 0.5 | 6 | 0.8333 |
| 6_8235717 | 2 | 1 | 1 | 1 | 1 | 1 | 1 | 1 | 5 | 1 | NA | NA | 1 | 1 | 2 | 0.5 | 2 | 1 | 5 | 0.8 |
| 6_8236461 | NA | NA | NA | NA | 1 | 1 | NA | NA | 1 | 1 | 1 | 0 | 1 | 1 | 1 | 0 | NA | NA | 3 | 0.3333 |

| | | | | | | | | | | | | | | | | | | | | |
|---|---|---|---|---|---|---|---|---|---|---|---|---|---|---|---|---|---|---|---|---|
| 6_8236700 | NA | NA | NA | NA | 1 | 1 | NA | NA | 1 | 1 | 1 | 0 | 1 | 1 | 1 | 0 | NA | NA | 3 | 0.3333 |
| 6_8236828 | NA | NA | NA | NA | 1 | 1 | NA | NA | 1 | 1 | 1 | 0 | 1 | 1 | 1 | 1 | NA | NA | 3 | 0.6667 |
| 6_8256992 | 2 | 0.5 | NA | NA | 1 | 0 | NA | NA | 3 | 0.3333 | NA | NA | 2 | 1 | NA | NA | NA | NA | 2 | 1 |
| 6_8427440 | 6 | 0.3333 | 5 | 0.2 | 3 | 0.3333 | 3 | 0 | 17 | 0.2353 | NA | NA | 9 | 0.3333 | 4 | 0.25 | 4 | 0 | 17 | 0.2353 |
| 6_8459025 | 2 | 0 | 5 | 0 | 4 | 0.25 | 1 | 0 | 12 | 0.0833 | 1 | 0 | NA | NA | 7 | 0 | 2 | 0.5 | 10 | 0.1 |
| 6_8507899 | NA | NA | NA | NA | 1 | 1 | 1 | 1 | 2 | 1 | 1 | 0 | 1 | 0 | NA | NA | 2 | 1 | 4 | 0.5 |
| 6_8508116 | NA | NA | NA | NA | 1 | 1 | NA | NA | 1 | 1 | 1 | 0 | 1 | 0 | NA | NA | 2 | 1 | 4 | 0.5 |
| 6_8508798 | NA | NA | NA | NA | 1 | 1 | 1 | 1 | 2 | 1 | 1 | 0 | NA | NA | NA | NA | 2 | 0.5 | 3 | 0.3333 |
| 6_8508851 | NA | NA | NA | NA | 1 | 0 | 1 | 1 | 2 | 0.5 | NA | NA | NA | NA | NA | NA | 2 | 0.5 | 2 | 0.5 |
| 6_8509091 | NA | NA | NA | NA | 2 | 0.5 | 1 | 0 | 3 | 0.3333 | 1 | 0 | NA | NA | NA | NA | 2 | 1 | 3 | 0.6667 |
| 6_8509107 | NA | NA | NA | NA | 2 | 1 | 1 | 1 | 3 | 1 | 1 | 1 | NA | NA | NA | NA | 2 | 0.5 | 3 | 0.6667 |
| 6_8509176 | NA | NA | NA | NA | 2 | 0.5 | 1 | 1 | 3 | 0.6667 | 1 | 1 | NA | NA | NA | NA | 2 | 1 | 3 | 1 |
| 6_8509262 | NA | NA | NA | NA | 1 | 1 | 1 | 1 | 2 | 1 | 1 | 0 | NA | NA | NA | NA | 2 | 1 | 3 | 0.6667 |
| 6_8509526 | NA | NA | NA | NA | 1 | 1 | 1 | 1 | 2 | 1 | 1 | 0 | NA | NA | NA | NA | 2 | 1 | 3 | 0.6667 |
| 6_8514195 | 1 | 1 | NA | NA | NA | NA | NA | NA | 1 | 1 | NA | NA | NA | NA | 1 | 1 | NA | NA | 1 | 1 |
| 6_8593044 | 1 | 0 | 2 | 0.5 | NA | NA | 1 | 1 | 4 | 0.5 | NA | NA | 2 | 0.5 | NA | NA | 1 | 1 | 3 | 0.6667 |
| 6_8593203 | 1 | 1 | 1 | 1 | 1 | 1 | 1 | 1 | 4 | 1 | NA | NA | 3 | 1 | NA | NA | 1 | 0 | 4 | 0.75 |
| 6_8593236 | 1 | 1 | 2 | 1 | 1 | 1 | NA | NA | 4 | 1 | NA | NA | 2 | 1 | NA | NA | 1 | 1 | 3 | 1 |
| 6_8593309 | 2 | 0 | 1 | 0 | 1 | 1 | 1 | 0 | 5 | 0.2 | NA | NA | 3 | 0 | NA | NA | 2 | 0.5 | 5 | 0.2 |
| 6_8593333 | 1 | 1 | 2 | 1 | 1 | 1 | 1 | 1 | 5 | 1 | NA | NA | 3 | 0.3333 | NA | NA | 2 | 0.5 | 5 | 0.4 |
| 6_8594686 | 4 | 0 | 2 | 0 | 1 | 0 | 1 | 1 | 8 | 0.125 | NA | NA | 3 | 0 | NA | NA | 5 | 0.2 | 8 | 0.125 |
| 6_8595130 | 3 | 0 | 3 | 0.3333 | 1 | 0 | 4 | 0 | 11 | 0.0909 | 1 | 0 | 3 | 0 | 1 | 0 | 6 | 0.1667 | 11 | 0.0909 |
| 6_8596727 | 5 | 0.2 | 2 | 0 | 2 | 0 | 5 | 0 | 14 | 0.0714 | 3 | 0.3333 | 4 | 0 | 3 | 0 | 8 | 0.125 | 18 | 0.1111 |
| 6_8596794 | 4 | 0.5 | 3 | 0 | 3 | 0.6667 | 5 | 0.4 | 15 | 0.4 | 3 | 0.6667 | 4 | 0 | 4 | 0.25 | 9 | 0.7778 | 20 | 0.5 |
| 6_8596944 | 6 | 0.1667 | 3 | 0.3333 | 3 | 0 | 5 | 0 | 17 | 0.1176 | 3 | 0 | 5 | 0.6 | 3 | 0.3333 | 9 | 0.1111 | 20 | 0.25 |
| 6_8597159 | 4 | 0.5 | 2 | 1 | 3 | 0.6667 | 8 | 0.875 | 17 | 0.7647 | 2 | 0.5 | 4 | 1 | 3 | 0.6667 | 9 | 0.8889 | 18 | 0.8333 |
| 6_8630697 | 8 | 0.125 | 7 | 0 | 4 | 0 | 8 | 0.125 | 27 | 0.0741 | 6 | 0.1667 | 9 | 0.1111 | 12 | 0.1667 | 8 | 0.125 | 35 | 0.1429 |
| 6_8630708 | 6 | 0 | 6 | 0 | 3 | 0 | 8 | 0.25 | 23 | 0.087 | 5 | 0.2 | 7 | 0.2857 | 11 | 0.0909 | 6 | 0 | 29 | 0.1379 |
| 6_9247169 | 2 | 1 | 3 | 1 | NA | NA | 2 | 1 | 7 | 1 | 3 | 1 | 1 | 0 | 1 | 1 | 2 | 1 | 7 | 0.8571 |
| 6_9247185 | 3 | 0.3333 | 3 | 0.6667 | NA | NA | 4 | 0.5 | 10 | 0.5 | 3 | 0.3333 | 3 | 0.6667 | 1 | 0 | 2 | 1 | 9 | 0.5556 |
| 6_9247214 | 2 | 1 | 3 | 1 | NA | NA | 4 | 1 | 9 | 1 | 3 | 0.6667 | 3 | 1 | 1 | 1 | 2 | 1 | 9 | 0.8889 |
| 6_9247248 | 3 | 0 | 4 | 0.25 | NA | NA | 4 | 0.25 | 11 | 0.1818 | 3 | 0.3333 | 3 | 0.3333 | 1 | 0 | 2 | 0 | 9 | 0.2222 |
| 6_9247296 | 3 | 1 | 4 | 1 | NA | NA | 4 | 1 | 11 | 1 | 3 | 1 | 2 | 1 | 1 | 1 | 2 | 1 | 8 | 1 |
| 6_9247308 | 3 | 0.3333 | 3 | 0 | NA | NA | 4 | 0 | 10 | 0.1 | 3 | 0.3333 | 3 | 0.3333 | NA | NA | 2 | 0 | 8 | 0.25 |
| 6_9247364 | 2 | 1 | 4 | 1 | NA | NA | 4 | 1 | 10 | 1 | 3 | 1 | 1 | 1 | 1 | 1 | 2 | 1 | 7 | 1 |
| 6_9247442 | 3 | 0 | 5 | 0.6 | NA | NA | 4 | 0.5 | 12 | 0.4167 | 2 | 1 | 2 | 0 | 1 | 0 | 2 | 0 | 7 | 0.2857 |
| 6_9247472 | 3 | 1 | 6 | 0.3333 | NA | NA | 3 | 0.3333 | 12 | 0.5 | 3 | 0.6667 | 3 | 0.3333 | 1 | 0 | 2 | 0.5 | 9 | 0.4444 |
| 6_9247497 | 2 | 0 | 6 | 0.1667 | NA | NA | 2 | 0 | 10 | 0.1 | 3 | 0.3333 | 3 | 0 | 1 | 0 | 1 | 0 | 8 | 0.125 |
| 6_9247773 | 1 | 1 | 7 | 1 | NA | NA | 4 | 0.75 | 12 | 0.9167 | 2 | 1 | 3 | 1 | 1 | 1 | 2 | 1 | 8 | 1 |
| 6_9247780 | 2 | 0.5 | 5 | 0 | NA | NA | 4 | 0 | 11 | 0.0909 | 2 | 0.5 | 3 | 0 | NA | NA | 1 | 1 | 6 | 0.3333 |
| 6_9247821 | 1 | 1 | 6 | 0.5 | NA | NA | 4 | 0 | 11 | 0.3636 | 2 | 0 | 3 | 0.3333 | 1 | 0 | 2 | 0 | 8 | 0.125 |
| 6_9247865 | 3 | 0.6667 | 5 | 0.6 | NA | NA | 4 | 0.75 | 12 | 0.6667 | 2 | 1 | 3 | 1 | 1 | 1 | 1 | 1 | 7 | 1 |
| 6_9247895 | 3 | 0.6667 | 7 | 0.1429 | NA | NA | 4 | 0.75 | 14 | 0.4286 | 3 | 0.3333 | 4 | 0.25 | 1 | 0 | 2 | 0 | 10 | 0.2 |
| 6_9247966 | 2 | 0.5 | 4 | 0.5 | NA | NA | 1 | 1 | 7 | 0.5714 | 3 | 1 | 4 | 0.5 | 1 | 0 | 2 | 1 | 10 | 0.7 |
| 6_9247991 | 3 | 0 | 7 | 0.1429 | NA | NA | 4 | 0.25 | 14 | 0.1429 | 3 | 0.6667 | 3 | 0 | 1 | 0 | 2 | 0 | 9 | 0.2222 |
| 6_9248169 | 3 | 0.6667 | 7 | 0.7143 | NA | NA | 4 | 0.5 | 14 | 0.6429 | 2 | 0.5 | 4 | 0.25 | 3 | 0.3333 | 2 | 0.5 | 11 | 0.3636 |
| 6_9248186 | 3 | 1 | 8 | 0.75 | NA | NA | 4 | 0.75 | 15 | 0.8 | 2 | 0.5 | 3 | 0.6667 | 3 | 0.3333 | 2 | 0.5 | 10 | 0.5 |
| 6_9248190 | 2 | 0 | 7 | 0.2857 | NA | NA | 4 | 0.75 | 13 | 0.3846 | 2 | 0.5 | 3 | 0.3333 | 3 | 0.3333 | 2 | 0.5 | 10 | 0.4 |
| 6_9248201 | 3 | 0 | 8 | 0.125 | NA | NA | 4 | 0 | 15 | 0.0667 | 3 | 0 | 4 | 0 | 3 | 0.3333 | 2 | 0 | 12 | 0.0833 |
| 6_9248219 | 3 | 0.3333 | 7 | 0.4286 | NA | NA | 4 | 0.5 | 14 | 0.4286 | 3 | 0 | 4 | 0.25 | 3 | 0.3333 | 2 | 0 | 12 | 0.1667 |
| 6_9248409 | 2 | 0 | 6 | 0.3333 | NA | NA | 2 | 0 | 10 | 0.2 | 3 | 0 | 3 | 0.3333 | 1 | 0 | 2 | 0.5 | 9 | 0.2222 |
| 6_9248547 | 3 | 0 | 8 | 0.625 | 1 | 0 | 3 | 0.6667 | 15 | 0.4667 | 3 | 0 | 3 | 0.6667 | 3 | 0.6667 | 2 | 0.5 | 11 | 0.4545 |
| 6_9248571 | 3 | 1 | 8 | 0.875 | 1 | 1 | 4 | 1 | 16 | 0.9375 | 3 | 1 | 4 | 1 | 3 | 1 | 2 | 1 | 12 | 1 |
| 6_9248872 | 3 | 0.3333 | 6 | 0.5 | 2 | 0 | 3 | 0.3333 | 14 | 0.3571 | 3 | 0.3333 | 3 | 0.3333 | 4 | 0 | 1 | 0 | 11 | 0.1818 |
| 6_9248886 | 3 | 0 | 7 | 0.1429 | 2 | 0 | 5 | 0.2 | 17 | 0.1176 | 3 | 0 | 5 | 0.4 | 5 | 0.2 | 3 | 0 | 16 | 0.1875 |
| 6_9249551 | NA | NA | 2 | 1 | 1 | 1 | 1 | 1 | 4 | 1 | 1 | 1 | 1 | 1 | 1 | 1 | NA | NA | 3 | 1 |
| 6_1200705 | 3 | 0.3333 | 5 | 0 | 4 | 0 | 3 | 0.3333 | 15 | 0.1333 | 4 | 0.25 | 2 | 0 | 5 | 0 | 1 | 0 | 12 | 0.0833 |
| 6_1203930 | 1 | 1 | NA | NA | NA | NA | 2 | 0.5 | 3 | 0.6667 | 1 | 0 | 2 | 0.5 | 1 | 0 | NA | NA | 4 | 0.25 |
| 6_1203932 | 1 | 0 | NA | NA | NA | NA | 2 | 0.5 | 3 | 0.3333 | 1 | 1 | 1 | 0 | NA | NA | NA | NA | 2 | 0.5 |
| 6_1203970 | 1 | 1 | 1 | 0 | NA | NA | 3 | 0.6667 | 5 | 0.6 | 1 | 1 | 3 | 1 | NA | NA | NA | NA | 4 | 1 |
| 6_1203979 | 1 | 1 | 1 | 0 | NA | NA | 3 | 0.6667 | 5 | 0.6 | 1 | 1 | 3 | 0.6667 | NA | NA | NA | NA | 4 | 0.75 |
| 6_1203986 | 1 | 1 | NA | NA | NA | NA | 3 | 0.3333 | 4 | 0.5 | 1 | 0 | 3 | 0.3333 | NA | NA | NA | NA | 4 | 0.25 |
| 6_1203991 | 1 | 1 | 2 | 0.5 | NA | NA | 2 | 0 | 5 | 0.4 | 1 | 1 | 3 | 0.6667 | NA | NA | NA | NA | 4 | 0.75 |
| 6_1203997 | 1 | 1 | 2 | 1 | NA | NA | 3 | 0.6667 | 6 | 0.8333 | 1 | 1 | 3 | 0.6667 | NA | NA | NA | NA | 4 | 0.75 |
| 6_1204030 | NA | NA | 2 | 0.5 | NA | NA | 2 | 0 | 4 | 0.25 | 1 | 1 | 2 | 0.5 | NA | NA | NA | NA | 3 | 0.6667 |

| | | | | | | | | | | | | | | | | | | | | |
|---|---|---|---|---|---|---|---|---|---|---|---|---|---|---|---|---|---|---|---|---|
| 6_1204557 | NA | NA | 2 | 0 | 1 | 1 | 1 | 0 | 4 | 0.25 | NA | NA | 3 | 0.6667 | NA | NA | 1 | 0 | 4 | 0.5 |
| 6_1204561 | 2 | 0 | 3 | 0 | 1 | 1 | 1 | 0 | 7 | 0.1429 | NA | NA | 3 | 0 | NA | NA | 1 | 1 | 4 | 0.25 |
| 6_1231175 | 6 | 0.1667 | 1 | 0 | 3 | 0 | 9 | 0.2222 | 19 | 0.1579 | 1 | 0 | 4 | 0.25 | 3 | 0 | 3 | 0.3333 | 11 | 0.1818 |
| 6_1231237 | 6 | 0.3333 | 1 | 0 | 2 | 0 | 7 | 0.2857 | 16 | 0.25 | 2 | 0 | 4 | 0.25 | 1 | 0 | 2 | 0 | 9 | 0.1111 |
| 6_1231377 | 5 | 0 | 1 | 0 | 3 | 0 | 3 | 0.6667 | 12 | 0.1667 | 2 | 0 | 4 | 0 | 4 | 0 | 1 | 1 | 11 | 0.0909 |
| 6_1231383 | 3 | 0 | 1 | 1 | 3 | 0 | 2 | 0 | 9 | 0.1111 | 2 | 0 | 2 | 0 | 4 | 0.25 | NA | NA | 8 | 0.125 |
| 6_1231398 | 5 | 0 | 1 | 0 | 3 | 0.3333 | 3 | 0 | 12 | 0.0833 | 1 | 0 | 2 | 0 | 4 | 0.25 | 1 | 0 | 8 | 0.125 |
| 6_1231413 | 5 | 0.2 | 2 | 0 | 2 | 0 | 3 | 0 | 12 | 0.0833 | 2 | 0 | 4 | 0 | 2 | 0.5 | 1 | 1 | 9 | 0.2222 |
| 6_1231444 | 4 | 0 | 2 | 0 | 3 | 0.3333 | 2 | 0 | 11 | 0.0909 | NA | NA | 4 | 0 | 2 | 0 | 1 | 1 | 7 | 0.1429 |
| 6_1231529 | 4 | 0.25 | 2 | 0 | 3 | 0 | 2 | 0 | 11 | 0.0909 | 1 | 0 | 3 | 0.3333 | 2 | 0 | 1 | 0 | 7 | 0.1429 |
| 6_1231614 | 3 | 0.6667 | 1 | 1 | 2 | 1 | 2 | 1 | 8 | 0.875 | 2 | 1 | 3 | 0.6667 | 2 | 0.5 | 1 | 1 | 8 | 0.75 |
| 6_1231644 | 2 | 0.5 | 1 | 0 | 2 | 0.5 | 1 | 0 | 6 | 0.3333 | 1 | 0 | 2 | 0 | 2 | 0 | 1 | 1 | 6 | 0.1667 |
| 6_1231649 | 2 | 1 | 1 | 1 | 2 | 0.5 | 1 | 1 | 6 | 0.8333 | 1 | 1 | 1 | 0 | 2 | 1 | 1 | 1 | 5 | 0.8 |
| 6_1231670 | 2 | 1 | 1 | 1 | NA | NA | 2 | 0.5 | 5 | 0.8 | 1 | 1 | 2 | 0.5 | 2 | 1 | 1 | 1 | 6 | 0.8333 |
| 6_1231674 | 2 | 1 | 1 | 1 | 1 | 1 | 2 | 1 | 6 | 1 | 1 | 1 | 2 | 0.5 | 2 | 1 | 1 | 1 | 6 | 0.8333 |
| 6_1231675 | 2 | 0.5 | 1 | 1 | 1 | 0 | 2 | 1 | 6 | 0.6667 | 2 | 0 | 2 | 0 | 1 | 1 | 1 | 1 | 6 | 0.3333 |
| 6_1231675 | 2 | 1 | 1 | 0 | 1 | 0 | 2 | 0 | 6 | 0.3333 | 1 | 1 | 2 | 0.5 | 2 | 0.5 | 1 | 0 | 6 | 0.5 |
| 6_1231690 | 1 | 1 | 1 | 0 | 1 | 1 | 2 | 1 | 5 | 0.8 | 2 | 1 | 2 | 0.5 | 2 | 1 | 1 | 1 | 7 | 0.8571 |
| 6_1231698 | NA | NA | 1 | 0 | NA | NA | 2 | 1 | 3 | 0.6667 | 1 | 0 | NA | NA | 1 | 1 | 1 | 0 | 3 | 0.3333 |
| 6_1231700 | NA | NA | 1 | 1 | 1 | 0 | 2 | 0.5 | 4 | 0.5 | 1 | 0 | NA | NA | 2 | 0.5 | 1 | 0 | 4 | 0.25 |
| 6_1231719 | 1 | 0 | 1 | 1 | 1 | 1 | 2 | 0 | 5 | 0.4 | 1 | 0 | 1 | 1 | 2 | 0.5 | 1 | 0 | 5 | 0.4 |
| 6_1231726 | NA | NA | 1 | 0 | 1 | 0 | 2 | 0.5 | 4 | 0.25 | 1 | 0 | 1 | 0 | 1 | 1 | 1 | 0 | 4 | 0.25 |
| 6_1231726 | NA | NA | 1 | 1 | 1 | 0 | 1 | 1 | 3 | 0.6667 | 1 | 1 | 1 | 0 | 1 | 1 | 1 | 0 | 4 | 0.5 |
| 6_1231733 | 1 | 0 | 1 | 1 | NA | NA | 2 | 0.5 | 4 | 0.5 | 2 | 0.5 | 1 | 1 | 2 | 0.5 | 1 | 1 | 6 | 0.6667 |
| 6_1231734 | 1 | 1 | 1 | 1 | 1 | 1 | 1 | 1 | 4 | 1 | NA | NA | 1 | 0 | 2 | 1 | NA | NA | 3 | 0.6667 |
| 6_1231736 | 1 | 1 | 1 | 1 | NA | NA | 2 | 0.5 | 4 | 0.75 | 1 | 0 | 1 | 1 | 1 | 1 | 1 | 0 | 4 | 0.5 |
| 6_1237498 | NA | NA | 1 | 1 | NA | NA | NA | NA | 1 | 1 | 1 | 0 | 1 | 1 | NA | NA | NA | NA | 2 | 0.5 |
| 6_1237499 | NA | NA | 1 | 1 | NA | NA | NA | NA | 1 | 1 | 1 | 1 | 1 | 0 | NA | NA | NA | NA | 2 | 0.5 |
| 6_1237502 | NA | NA | 1 | 1 | NA | NA | NA | NA | 1 | 1 | 1 | 1 | 1 | 0 | NA | NA | NA | NA | 2 | 0.5 |
| 6_1237507 | NA | NA | 1 | 1 | NA | NA | NA | NA | 1 | 1 | 1 | 1 | NA | NA | NA | NA | NA | NA | 1 | 1 |
| 6_1237529 | NA | NA | 1 | 1 | NA | NA | NA | NA | 1 | 1 | NA | NA | 1 | 1 | NA | NA | NA | NA | 1 | 1 |
| 6_1237543 | NA | NA | 1 | 1 | NA | NA | NA | NA | 1 | 1 | NA | NA | 1 | 1 | NA | NA | NA | NA | 1 | 1 |
| 6_1237561 | NA | NA | 1 | 1 | NA | NA | NA | NA | 1 | 1 | NA | NA | 1 | 1 | NA | NA | NA | NA | 1 | 1 |
| 6_1237570 | NA | NA | 1 | 1 | NA | NA | NA | NA | 1 | 1 | NA | NA | 1 | 1 | NA | NA | NA | NA | 1 | 1 |
| 6_1237583 | NA | NA | 1 | 1 | NA | NA | NA | NA | 1 | 1 | NA | NA | 1 | 1 | NA | NA | NA | NA | 1 | 1 |
| 6_1237584 | NA | NA | 1 | 1 | NA | NA | NA | NA | 1 | 1 | NA | NA | 1 | 1 | NA | NA | NA | NA | 1 | 1 |
| 6_1308421 | 6 | 0.1667 | 7 | 0.2857 | 4 | 0.25 | 9 | 0.8889 | 26 | 0.4615 | 8 | 0.375 | 5 | 0 | 6 | 0.8333 | 11 | 0.4545 | 30 | 0.4333 |
| 6_1308432 | 7 | 0.7143 | 7 | 0.4286 | 3 | 0.3333 | 7 | 0.4286 | 24 | 0.5 | 3 | 0.6667 | 7 | 0.8571 | 4 | 0.75 | 9 | 0.6667 | 23 | 0.7391 |
| 6_1308437 | 6 | 0.8333 | 12 | 0.25 | 5 | 0.2 | 9 | 0.3333 | 32 | 0.375 | 7 | 0.4286 | 7 | 0 | 6 | 0.6667 | 11 | 0.3636 | 31 | 0.3548 |
| 6_1308440 | 8 | 0 | 11 | 0.3636 | 4 | 0.5 | 9 | 0.3333 | 32 | 0.2812 | 8 | 0.375 | 8 | 0.375 | 6 | 0.3333 | 12 | 0.5833 | 34 | 0.4412 |
| 6_1308446 | 7 | 0.4286 | 10 | 0.3 | 4 | 0.75 | 8 | 0.5 | 29 | 0.4483 | 5 | 0.8 | 7 | 0.2857 | 6 | 0.5 | 12 | 0.5 | 30 | 0.5 |
| 6_1308474 | 8 | 0.125 | 7 | 0.1429 | 6 | 0.3333 | 9 | 0.1111 | 30 | 0.1667 | 8 | 0.125 | 8 | 0 | 6 | 0.5 | 13 | 0.2308 | 35 | 0.2 |
| 6_1308706 | 17 | 0.1176 | 18 | 0.0556 | 15 | 0.1333 | 17 | 0.0588 | 67 | 0.0896 | 8 | 0.125 | 13 | 0.0769 | 12 | 0.1667 | 11 | 0.0909 | 44 | 0.1136 |
| 6_1358149 | NA | NA | 4 | 0.25 | 2 | 0 | 1 | 0 | 7 | 0.1429 | 3 | 0 | 1 | 0 | 1 | 1 | 1 | 0 | 6 | 0.1667 |
| 6_1358150 | NA | NA | 4 | 0 | 2 | 0.5 | 2 | 0.5 | 8 | 0.25 | 4 | 0.25 | 1 | 0 | 1 | 0 | 1 | 0 | 7 | 0.1429 |
| 6_1358227 | NA | NA | 4 | 0.25 | 2 | 0 | 2 | 0.5 | 8 | 0.25 | 5 | 0.2 | 2 | 0 | NA | NA | 1 | 0 | 8 | 0.125 |
| 6_1358232 | NA | NA | 3 | 0.3333 | 1 | 0 | 2 | 0.5 | 6 | 0.3333 | 5 | 0.6 | 2 | 0 | NA | NA | 1 | 1 | 8 | 0.5 |
| 6_1358246 | NA | NA | 3 | 1 | 1 | 1 | 2 | 1 | 6 | 1 | 2 | 1 | 2 | 1 | NA | NA | 1 | 1 | 5 | 1 |
| 6_1358278 | NA | NA | 4 | 1 | 1 | 1 | 2 | 1 | 7 | 1 | 4 | 1 | 1 | 1 | NA | NA | 1 | 1 | 6 | 1 |
| 6_1358281 | NA | NA | 4 | 0.25 | 1 | 0 | 2 | 0 | 7 | 0.1429 | 3 | 0.3333 | 1 | 0 | NA | NA | 1 | 0 | 5 | 0.2 |
| 6_1358282 | NA | NA | 4 | 1 | NA | NA | 2 | 1 | 6 | 1 | 3 | 0.6667 | 2 | 1 | NA | NA | 1 | 1 | 6 | 0.8333 |
| 6_1358289 | NA | NA | 4 | 0.5 | 1 | 0 | 2 | 0.5 | 7 | 0.4286 | 4 | 0.5 | 2 | 0 | NA | NA | 1 | 0 | 7 | 0.2857 |
| 6_1358306 | NA | NA | 4 | 1 | NA | NA | 2 | 1 | 6 | 1 | 4 | 1 | 1 | 1 | NA | NA | 1 | 1 | 6 | 1 |
| 6_1358310 | NA | NA | 3 | 1 | NA | NA | 2 | 1 | 5 | 1 | 4 | 1 | 2 | 1 | NA | NA | 1 | 1 | 7 | 1 |
| 6_1358312 | NA | NA | 3 | 1 | NA | NA | 1 | 1 | 4 | 1 | 4 | 0.75 | 2 | 1 | NA | NA | 1 | 0 | 7 | 0.7143 |
| 6_1358316 | NA | NA | 4 | 0.25 | NA | NA | 1 | 0 | 5 | 0.2 | 3 | 0.6667 | 1 | 0 | NA | NA | 1 | 1 | 5 | 0.6 |
| 6_1359135 | NA | NA | 4 | 0.25 | NA | NA | 1 | 0 | 5 | 0.2 | 3 | 0.6667 | 2 | 0 | NA | NA | 1 | 0 | 6 | 0.3333 |
| 6_1359141 | NA | NA | 4 | 0.25 | NA | NA | 1 | 0 | 5 | 0.2 | 4 | 0.25 | 2 | 0.5 | NA | NA | 1 | 1 | 7 | 0.4286 |
| 6_1360239 | NA | NA | 3 | 1 | NA | NA | 2 | 0.5 | 5 | 0.8 | 3 | 0.6667 | NA | NA | NA | NA | 1 | 1 | 4 | 0.75 |
| 6_1360250 | NA | NA | 3 | 1 | NA | NA | 2 | 1 | 5 | 1 | NA | NA | NA | NA | NA | NA | 1 | 1 | 1 | 1 |
| 6_1360257 | NA | NA | 2 | 0.5 | NA | NA | 2 | 0.5 | 4 | 0.5 | 1 | 0 | NA | NA | NA | NA | 1 | 1 | 2 | 0.5 |
| 6_1360263 | NA | NA | 3 | 1 | NA | NA | 2 | 1 | 5 | 1 | 1 | 1 | NA | NA | NA | NA | 1 | 1 | 2 | 1 |
| 6_1360288 | NA | NA | 3 | 0.6667 | NA | NA | 2 | 1 | 5 | 0.8 | 1 | 1 | NA | NA | NA | NA | NA | NA | 1 | 1 |
| 6_1362706 | 10 | 0.1 | 2 | 0 | 6 | 0 | 1 | 0 | 19 | 0.0526 | 4 | 0.25 | 4 | 0 | 3 | 0 | 5 | 0.2 | 16 | 0.125 |
| 6_1362795 | 10 | 0.3 | NA | NA | 6 | 0 | 1 | 0 | 17 | 0.1765 | 4 | 0 | 3 | 0.3333 | 3 | 0 | 6 | 0 | 16 | 0.0625 |

| | | | | | | | | | | | | | | | | | | | | |
|---|---|---|---|---|---|---|---|---|---|---|---|---|---|---|---|---|---|---|---|---|
| 6_1362827 | 8 | 0.25 | NA | NA | 6 | 0.5 | 1 | 0 | 15 | 0.3333 | 4 | 0.5 | 3 | 1 | 3 | 0.3333 | 5 | 0.4 | 15 | 0.5333 |
| 6_1362839 | 6 | 0.8333 | NA | NA | 4 | 0.75 | 1 | 1 | 11 | 0.8182 | 2 | 1 | 1 | 1 | 1 | 1 | 3 | 1 | 7 | 1 |
| 6_1362843 | 7 | 0 | NA | NA | 6 | 0.1667 | 1 | 0 | 14 | 0.0714 | 4 | 0 | 1 | 1 | 3 | 0 | 5 | 0 | 13 | 0.0769 |
| 6_1362869 | 9 | 1 | NA | NA | 4 | 0.75 | 1 | 1 | 14 | 0.9286 | 4 | 1 | 1 | 1 | 1 | 0 | 2 | 1 | 8 | 0.875 |
| 6_1362870 | 7 | 1 | NA | NA | 4 | 1 | 1 | 1 | 12 | 1 | 3 | 1 | 1 | 1 | 2 | 1 | 5 | 1 | 11 | 1 |
| 6_1362872 | 8 | 0.125 | NA | NA | 3 | 0 | 1 | 0 | 12 | 0.0833 | 3 | 0.3333 | 1 | 0 | 2 | 0 | 4 | 0 | 10 | 0.1 |
| 6_1362898 | 6 | 0.6667 | NA | NA | 6 | 1 | 1 | 1 | 13 | 0.8462 | 3 | 1 | NA | NA | 3 | 1 | 3 | 0.3333 | 9 | 0.7778 |
| 6_1383512 | 5 | 0.2 | 9 | 0.1111 | 9 | 0 | 4 | 0 | 27 | 0.0741 | 9 | 0.3333 | 3 | 0.3333 | 11 | 0.0909 | 3 | 0 | 26 | 0.1923 |
| 6_1383531 | 7 | 0.4286 | 7 | 0.7143 | 9 | 0.3333 | 3 | 1 | 26 | 0.5385 | 7 | 0.7143 | 3 | 0.6667 | 9 | 0.5556 | 6 | 0.5 | 25 | 0.6 |
| 6_1383533 | 7 | 0.1429 | 7 | 0.1429 | 8 | 0.375 | 2 | 0.5 | 24 | 0.25 | 8 | 0.375 | 3 | 0 | 11 | 0.3636 | 5 | 0.4 | 27 | 0.3333 |
| 6_1383536 | 6 | 0 | 9 | 0.2222 | 9 | 0.3333 | 4 | 0.5 | 28 | 0.25 | 7 | 0.7143 | 4 | 0.5 | 10 | 0.3 | 5 | 0.6 | 26 | 0.5 |
| 6_1383543 | 6 | 1 | 9 | 1 | 9 | 1 | 3 | 1 | 27 | 1 | 7 | 1 | 4 | 0.75 | 9 | 1 | 5 | 1 | 25 | 0.96 |
| 6_1383545 | 2 | 0.5 | 4 | 0.25 | 2 | 0 | NA | NA | 8 | 0.25 | 2 | 0.5 | 1 | 0 | 5 | 0 | 1 | 0 | 9 | 0.1111 |
| 6_1383546 | 1 | 1 | 3 | 0.3333 | 1 | 1 | NA | NA | 5 | 0.6 | 3 | 0.3333 | NA | NA | 2 | 0.5 | 2 | 0 | 7 | 0.2857 |
| 6_1383553 | 3 | 0.6667 | 8 | 0.375 | 9 | 0.3333 | 4 | 0.5 | 24 | 0.4167 | 6 | 0.3333 | 4 | 0.5 | 9 | 0.1111 | 2 | 0.5 | 21 | 0.2857 |
| 6_1383556 | 4 | 0.75 | 7 | 1 | 8 | 0.875 | 3 | 1 | 22 | 0.9091 | 5 | 1 | 4 | 1 | 9 | 1 | 6 | 1 | 24 | 1 |
| 6_1383560 | 5 | 0.2 | 9 | 0.1111 | 8 | 0.5 | 3 | 0.3333 | 25 | 0.28 | 3 | 0.3333 | 3 | 0.6667 | 9 | 0.1111 | 3 | 0.6667 | 18 | 0.3333 |
| 6_1383568 | 6 | 0 | 8 | 0.25 | 6 | 0 | 3 | 0.3333 | 23 | 0.1304 | 6 | 0.3333 | 4 | 0 | 10 | 0.1 | 6 | 0.1667 | 26 | 0.1538 |
| 6_1383575 | 4 | 0 | 6 | 0 | 6 | 0.3333 | 3 | 0 | 19 | 0.1053 | 5 | 0.2 | 4 | 0.25 | 8 | 0 | 5 | 0.2 | 22 | 0.1364 |
| 6_1383575 | 6 | 0.3333 | 8 | 0.375 | 5 | 0.2 | 3 | 0.3333 | 22 | 0.3182 | 6 | 0.8333 | 4 | 0.5 | 8 | 0.25 | 5 | 0.6 | 23 | 0.5217 |
| 6_1383580 | 6 | 0.5 | 7 | 0.8571 | 4 | 0.25 | 2 | 0.5 | 19 | 0.5789 | 5 | 0.4 | 2 | 1 | 7 | 0.5714 | 5 | 0.8 | 19 | 0.6316 |
| 6_1383603 | 4 | 1 | 4 | 1 | 1 | 1 | 2 | 1 | 11 | 1 | 4 | 1 | 3 | 1 | 4 | 1 | 3 | 1 | 14 | 1 |
| 6_1383621 | 4 | 1 | 7 | 1 | 5 | 1 | NA | NA | 16 | 1 | 6 | 1 | 4 | 1 | 6 | 1 | 3 | 1 | 19 | 1 |
| 6_1383622 | 4 | 1 | 6 | 1 | 5 | 1 | 2 | 1 | 17 | 1 | 3 | 1 | 3 | 0.6667 | 4 | 1 | 4 | 0.5 | 14 | 0.7857 |
| 6_1383646 | 6 | 0.8333 | 7 | 1 | 5 | 1 | 3 | 1 | 21 | 0.9524 | 7 | 1 | 4 | 1 | 7 | 1 | 4 | 1 | 22 | 1 |
| 6_1383662 | 5 | 0.2 | 6 | 0.1667 | 5 | 0 | 3 | 0 | 19 | 0.1053 | 7 | 0.2857 | 2 | 0 | 6 | 0.3333 | 3 | 0 | 18 | 0.2222 |
| 6_1383663 | 6 | 1 | 6 | 1 | 5 | 1 | 2 | 1 | 19 | 1 | 7 | 1 | 3 | 1 | 8 | 1 | 4 | 1 | 22 | 1 |
| 6_1383664 | 5 | 1 | 5 | 1 | 5 | 1 | 2 | 1 | 17 | 1 | 5 | 1 | 2 | 1 | 7 | 1 | 4 | 1 | 18 | 1 |
| 6_1383675 | 5 | 0.4 | 6 | 0.5 | 4 | 0.5 | 2 | 0 | 17 | 0.4118 | 7 | 0.2857 | 4 | 0.75 | 7 | 0.7143 | 5 | 0.6 | 23 | 0.5652 |
| 6_1383683 | 4 | 0.25 | 4 | 0.5 | 3 | 1 | 2 | 0.5 | 13 | 0.5385 | 4 | 0.25 | NA | NA | 4 | 0.25 | 4 | 0.25 | 12 | 0.25 |
| 6_1383683 | 5 | 0.4 | 6 | 0.6667 | 5 | 0.6 | 3 | 0.6667 | 19 | 0.5789 | 6 | 0.8333 | 2 | 1 | 6 | 0.1667 | 4 | 0.5 | 18 | 0.5556 |
| 6_1383689 | 5 | 0.2 | 4 | 0.75 | 4 | 0.75 | 3 | 0 | 16 | 0.4375 | 5 | 0.8 | 2 | 0.5 | 6 | 0.6667 | 3 | 0.3333 | 16 | 0.625 |
| 6_1383693 | 4 | 0 | 4 | 0.25 | 4 | 0 | 3 | 0.6667 | 15 | 0.2 | 5 | 0.2 | 2 | 0 | 5 | 0 | 2 | 1 | 14 | 0.2143 |
| 6_1383693 | 5 | 1 | 6 | 1 | 5 | 1 | 3 | 1 | 19 | 1 | 5 | 1 | 3 | 1 | 7 | 1 | 3 | 1 | 18 | 1 |
| 6_1383885 | 1 | 1 | 1 | 1 | NA | NA | NA | NA | 2 | 1 | 3 | 1 | 1 | 1 | 2 | 0.5 | 1 | 1 | 7 | 0.8571 |
| 6_1383886 | 1 | 0 | 1 | 1 | NA | NA | NA | NA | 2 | 0.5 | 3 | 0.6667 | 1 | 1 | 1 | 0 | 1 | 1 | 6 | 0.6667 |
| 6_1387050 | 5 | 0.4 | 9 | 0.5556 | 4 | 0.75 | 2 | 1 | 20 | 0.6 | 5 | 0.4 | 10 | 0.2 | 8 | 0.5 | 2 | 0.5 | 25 | 0.36 |
| 6_1387052 | 5 | 0.8 | 10 | 1 | 4 | 1 | 2 | 1 | 21 | 0.9524 | 6 | 0.8333 | 10 | 1 | 7 | 0.8571 | 2 | 0.5 | 25 | 0.88 |
| 6_1387053 | 6 | 0.1667 | 9 | 0.6667 | 4 | 0.25 | 2 | 0 | 21 | 0.381 | 6 | 0.1667 | 10 | 0.2 | 8 | 0.5 | 2 | 0 | 26 | 0.2692 |
| 6_1387054 | 5 | 0.8 | 11 | 0.8182 | 3 | 0.6667 | 2 | 0 | 21 | 0.7143 | 5 | 0.6 | 9 | 0.6667 | 7 | 0.8571 | 2 | 0 | 23 | 0.6522 |
| 6_1543801 | 4 | 0 | 5 | 0.4 | 5 | 0.2 | 4 | 0 | 18 | 0.1667 | NA | NA | 7 | 0.2857 | 3 | 0 | 4 | 0.25 | 14 | 0.2143 |
| 6_1543802 | 4 | 1 | 4 | 0.5 | 4 | 0.75 | 4 | 1 | 16 | 0.8125 | NA | NA | 7 | 0.8571 | 3 | 0.3333 | 3 | 1 | 13 | 0.7692 |
| 6_1543805 | 4 | 1 | 5 | 0.6 | 5 | 0.8 | 4 | 0.75 | 18 | 0.7778 | NA | NA | 7 | 1 | 3 | 0.6667 | 4 | 0.5 | 14 | 0.7857 |
| 6_1543811 | 3 | 1 | 5 | 1 | 3 | 0.6667 | 4 | 0.5 | 15 | 0.8 | NA | NA | 5 | 1 | 3 | 1 | 4 | 1 | 12 | 1 |
| 6_1543815 | 4 | 0.5 | 4 | 1 | 4 | 0.5 | 4 | 0.5 | 16 | 0.625 | NA | NA | 6 | 0.8333 | 3 | 1 | 4 | 0.75 | 13 | 0.8462 |
| 6_1543830 | 1 | 0 | 6 | 0.5 | 5 | 0 | 3 | 0.3333 | 15 | 0.2667 | NA | NA | 6 | 0.1667 | 1 | 0 | 4 | 0.5 | 11 | 0.2727 |
| 6_1543833 | 4 | 0.75 | 6 | 0.3333 | 4 | 0.75 | 6 | 0.5 | 20 | 0.55 | NA | NA | 8 | 0.875 | 3 | 1 | 4 | 0.5 | 15 | 0.8 |
| 6_1543839 | 2 | 0 | 7 | 0 | 5 | 0.2 | 7 | 0.1429 | 21 | 0.0952 | NA | NA | 7 | 0.2857 | 2 | 0.5 | 4 | 0 | 13 | 0.2308 |
| 6_1543892 | 5 | 1 | 6 | 0.5 | 5 | 0.4 | 7 | 0.5714 | 23 | 0.6087 | NA | NA | 9 | 0.5556 | 3 | 0.3333 | 4 | 0 | 16 | 0.375 |
| 6_1543896 | 5 | 0 | 8 | 0 | 5 | 0.2 | 7 | 0.1429 | 25 | 0.08 | NA | NA | 9 | 0 | 4 | 0 | 5 | 0.2 | 18 | 0.0556 |
| 6_1543898 | 3 | 0.3333 | 8 | 0 | 5 | 0 | 7 | 0.4286 | 23 | 0.1739 | NA | NA | 9 | 0.4444 | 4 | 0 | 5 | 0 | 18 | 0.2222 |
| 6_1543899 | 4 | 0.5 | 4 | 0.25 | 3 | 0.3333 | 7 | 0.5714 | 18 | 0.4444 | NA | NA | 5 | 0.4 | 3 | 1 | 2 | 0.5 | 10 | 0.6 |
| 6_1543917 | 4 | 0.5 | 7 | 0.1429 | 5 | 0 | 6 | 0.1667 | 22 | 0.1818 | NA | NA | 9 | 0.3333 | 4 | 0.25 | 6 | 0 | 19 | 0.2105 |
| 6_1543918 | 3 | 0 | 8 | 0.375 | 5 | 0.2 | 6 | 0 | 22 | 0.1818 | NA | NA | 8 | 0.25 | 4 | 0 | 5 | 0.2 | 17 | 0.1765 |
| 6_1543976 | 5 | 0 | 8 | 0.25 | 4 | 0 | 7 | 0.1429 | 24 | 0.125 | NA | NA | 10 | 0 | 4 | 0.25 | 6 | 0 | 20 | 0.05 |
| 6_1544002 | 5 | 0.4 | 6 | 0.1667 | 3 | 0.3333 | 7 | 0.1429 | 21 | 0.2381 | NA | NA | 10 | 0 | 7 | 0.2857 | 6 | 0.6667 | 23 | 0.2609 |
| 6_1544039 | 5 | 0.4 | 6 | 0.3333 | 3 | 0 | 6 | 0.3333 | 20 | 0.3 | NA | NA | 8 | 0.375 | 10 | 0.5 | 4 | 0.75 | 22 | 0.5 |
| 6_1544136 | 2 | 0.5 | 1 | 0 | 2 | 0 | 4 | 0 | 9 | 0.1111 | 1 | 1 | 2 | 0 | 2 | 0 | 4 | 0.5 | 9 | 0.3333 |
| 6_1544160 | 2 | 0 | 3 | 0 | 1 | 1 | 2 | 1 | 8 | 0.375 | 1 | 1 | 1 | 0 | 1 | 0 | 3 | 0.6667 | 6 | 0.5 |
| 6_1544169 | 1 | 0 | 2 | 0 | 2 | 0.5 | 3 | 0 | 8 | 0.125 | 1 | 0 | 2 | 0.5 | 2 | 0 | 4 | 0 | 9 | 0.1111 |
| 6_1579975 | NA | NA | NA | NA | NA | NA | 2 | 0.5 | 2 | 0.5 | NA | NA | NA | NA | NA | NA | 1 | 1 | 1 | 1 |
| 6_1579977 | NA | NA | NA | NA | NA | NA | 2 | 1 | 2 | 1 | NA | NA | NA | NA | NA | NA | 1 | 1 | 1 | 1 |
| 6_1579981 | NA | NA | NA | NA | NA | NA | 2 | 1 | 2 | 1 | NA | NA | NA | NA | NA | NA | 1 | 1 | 1 | 1 |
| 6_1579994 | NA | NA | NA | NA | NA | NA | 2 | 0.5 | 2 | 0.5 | NA | NA | NA | NA | NA | NA | 1 | 1 | 1 | 1 |
| 6_1580033 | NA | NA | NA | NA | 1 | 1 | 2 | 0 | 3 | 0.3333 | NA | NA | NA | NA | NA | NA | 1 | 1 | 1 | 1 |

| | | | | | | | | | | | | | | | | | | | | |
|---|---|---|---|---|---|---|---|---|---|---|---|---|---|---|---|---|---|---|---|---|
| 6_1580049 | NA | NA | NA | NA | 1 | 1 | 2 | 0.5 | 3 | 0.6667 | NA | NA | NA | NA | NA | NA | 1 | 1 | 1 | 1 |
| 6_1580065 | NA | NA | NA | NA | 1 | 0 | 2 | 1 | 3 | 0.6667 | NA | NA | NA | NA | NA | NA | 1 | 1 | 1 | 1 |
| 6_1728748 | 3 | 0 | 5 | 0.2 | 6 | 0 | 2 | 0 | 16 | 0.0625 | 1 | 0 | 1 | 0 | 1 | 0 | 2 | 0.5 | 5 | 0.2 |
| 6_1733945 | 1 | 1 | 3 | 1 | NA | NA | 2 | 1 | 6 | 1 | 2 | 1 | 1 | 1 | 2 | 1 | 2 | 1 | 7 | 1 |
| 6_1733956 | 1 | 0 | 4 | 0 | NA | NA | 2 | 0.5 | 7 | 0.1429 | 2 | 0.5 | NA | NA | 1 | 0 | 1 | 0 | 4 | 0.25 |
| 6_1733974 | 1 | 1 | 4 | 0.25 | NA | NA | 1 | 1 | 6 | 0.5 | 1 | 0 | NA | NA | 2 | 0.5 | 1 | 0 | 4 | 0.25 |
| 6_1734032 | 1 | 1 | 5 | 0.6 | NA | NA | 2 | 0.5 | 8 | 0.625 | 1 | 0 | 1 | 1 | 2 | 0.5 | 2 | 0.5 | 6 | 0.5 |
| 6_1734046 | 1 | 1 | 6 | 0.3333 | 1 | 1 | 2 | 1 | 10 | 0.6 | 3 | 0.3333 | 1 | 1 | 3 | 0.3333 | 2 | 0 | 9 | 0.3333 |
| 6_1734056 | 1 | 1 | 6 | 0.3333 | 1 | 1 | 2 | 0 | 10 | 0.4 | 3 | 0.3333 | 1 | 0 | 3 | 0 | 2 | 0 | 9 | 0.1111 |
| 6_1734093 | 1 | 0 | 5 | 0.2 | 1 | 0 | 3 | 0 | 10 | 0.1 | 2 | 0.5 | 1 | 0 | 3 | 0 | 2 | 0 | 8 | 0.125 |
| 6_1734100 | 1 | 0 | 5 | 0 | 2 | 0.5 | 2 | 0.5 | 10 | 0.2 | 3 | 0 | 1 | 0 | 2 | 0.5 | 2 | 0 | 8 | 0.125 |
| 6_1734106 | 1 | 0 | 7 | 0.4286 | 2 | 0.5 | 2 | 0.5 | 12 | 0.4167 | 3 | 0.3333 | 1 | 0 | 3 | 0.3333 | 2 | 0 | 9 | 0.2222 |
| 6_1757341 | 1 | 1 | NA | NA | NA | NA | NA | NA | 1 | 1 | NA | NA | 1 | 1 | NA | NA | 1 | 1 | 2 | 1 |
| 6_1757347 | 1 | 1 | NA | NA | NA | NA | NA | NA | 1 | 1 | NA | NA | 1 | 0 | NA | NA | 1 | 1 | 2 | 0.5 |
| 6_1757355 | 1 | 1 | NA | NA | NA | NA | NA | NA | 1 | 1 | NA | NA | 1 | 1 | NA | NA | 1 | 1 | 2 | 1 |
| 6_1766589 | 35 | 0.0571 | 45 | 0.1111 | 32 | 0.0938 | 38 | 0.0526 | 150 | 0.08 | 17 | 0.2353 | 39 | 0.0256 | 38 | 0.0789 | 30 | 0.1 | 124 | 0.0887 |
| 6_1794269 | NA | NA | 2 | 0.5 | NA | NA | 1 | 0 | 3 | 0.3333 | 1 | 0 | 2 | 0.5 | 3 | 0 | 3 | 0.3333 | 9 | 0.2222 |
| 6_1794282 | 1 | 0 | 2 | 0 | NA | NA | 2 | 0.5 | 5 | 0.2 | 2 | 0 | 4 | 0.5 | 4 | 0.25 | 4 | 0.25 | 14 | 0.2857 |
| 6_1794288 | 1 | 1 | 2 | 1 | NA | NA | 1 | 1 | 4 | 1 | 2 | 1 | 3 | 1 | 4 | 1 | 3 | 1 | 12 | 1 |
| 6_1798965 | 1 | 0 | NA | NA | 1 | 1 | 4 | 1 | 6 | 0.8333 | 1 | 1 | 1 | 1 | 3 | 1 | 3 | 1 | 8 | 1 |
| 6_1798974 | 1 | 1 | NA | NA | 1 | 1 | 4 | 1 | 6 | 1 | 1 | 1 | NA | NA | 3 | 1 | 2 | 1 | 6 | 1 |
| 6_1798975 | 1 | 1 | NA | NA | 1 | 1 | 4 | 0.75 | 6 | 0.8333 | 1 | 1 | 1 | 1 | 3 | 1 | 3 | 1 | 8 | 1 |
| 6_1798977 | 1 | 1 | NA | NA | 1 | 1 | 4 | 1 | 6 | 1 | 1 | 1 | 1 | 1 | 2 | 0.5 | 3 | 1 | 7 | 0.8571 |
| 6_1798988 | 1 | 0 | NA | NA | 1 | 0 | 4 | 0.75 | 6 | 0.5 | 1 | 1 | 1 | 1 | 3 | 0.3333 | 3 | 1 | 8 | 0.75 |
| 6_1798999 | 1 | 0 | NA | NA | 1 | 1 | 3 | 0.3333 | 5 | 0.4 | 1 | 0 | NA | NA | 3 | 0.3333 | 2 | 0 | 6 | 0.1667 |
| 6_1832128 | NA | NA | NA | NA | NA | NA | 1 | 1 | 1 | 1 | NA | NA | NA | NA | 1 | 1 | NA | NA | 1 | 1 |
| 6_1832145 | NA | NA | NA | NA | NA | NA | 1 | 1 | 1 | 1 | NA | NA | NA | NA | 1 | 1 | NA | NA | 1 | 1 |
| 6_1836716 | 9 | 0.4444 | 13 | 0.2308 | 5 | 0.2 | 16 | 0.125 | 43 | 0.2326 | 5 | 0.4 | 14 | 0.2857 | 17 | 0.1765 | 11 | 0.1818 | 47 | 0.234 |
| 6_1836723 | 10 | 0.4 | 16 | 0.1875 | 6 | 0.1667 | 17 | 0.1176 | 49 | 0.2041 | 4 | 0 | 13 | 0.0769 | 17 | 0 | 14 | 0.2857 | 48 | 0.1042 |
| 6_1836728 | 9 | 0 | 17 | 0.1765 | 6 | 0.3333 | 18 | 0.0556 | 50 | 0.12 | 4 | 0 | 13 | 0.2308 | 17 | 0 | 12 | 0.0833 | 46 | 0.087 |
| 6_1836739 | 6 | 0 | 14 | 0.3571 | 6 | 0.1667 | 14 | 0.2857 | 40 | 0.25 | 3 | 0 | 13 | 0.4615 | 16 | 0.1875 | 12 | 0.25 | 44 | 0.2727 |
| 6_1836741 | 8 | 0.25 | 14 | 0.5 | 6 | 0.5 | 17 | 0.3529 | 45 | 0.4 | 3 | 0.3333 | 13 | 0.3077 | 15 | 0.6667 | 12 | 0.4167 | 43 | 0.4651 |
| 6_1836755 | 9 | 0.6667 | 10 | 0.4 | 5 | 0.8 | 16 | 0.5 | 40 | 0.55 | 1 | 1 | 10 | 0.6 | 13 | 0.5385 | 11 | 0.4545 | 35 | 0.5429 |
| 6_1838287 | 7 | 0 | 7 | 0.1429 | 3 | 0 | 13 | 0.0769 | 30 | 0.0667 | NA | NA | 10 | 0.2 | 13 | 0.0769 | 8 | 0 | 31 | 0.0968 |
| 6_1840854 | 4 | 0 | 10 | 0.5 | 2 | 0 | 11 | 0.0909 | 27 | 0.2222 | NA | NA | 7 | 0.1429 | 12 | 0.1667 | 7 | 0.2857 | 26 | 0.1923 |
| 6_1843078 | 2 | 0.5 | NA | NA | NA | NA | 1 | 0 | 3 | 0.3333 | NA | NA | 1 | 1 | 3 | 0 | NA | NA | 4 | 0.25 |
| 6_1843395 | 1 | 0 | 1 | 0 | NA | NA | 2 | 0.5 | 4 | 0.25 | NA | NA | NA | NA | 4 | 0.5 | 1 | 0 | 5 | 0.4 |
| 6_1843410 | 2 | 0 | 6 | 0.3333 | NA | NA | 2 | 0 | 10 | 0.2 | NA | NA | 5 | 0.6 | 5 | 1 | 3 | 0.6667 | 13 | 0.7692 |
| 6_1843427 | 3 | 0.6667 | 4 | 0.75 | NA | NA | 1 | 1 | 8 | 0.75 | NA | NA | 4 | 1 | 4 | 1 | 3 | 1 | 11 | 1 |
| 6_1843433 | 2 | 0.5 | 6 | 0.1667 | NA | NA | 1 | 0 | 9 | 0.2222 | NA | NA | 4 | 0.5 | 4 | 0.25 | 3 | 0 | 11 | 0.2727 |
| 6_1843470 | NA | NA | 5 | 0.4 | NA | NA | 1 | 1 | 6 | 0.5 | NA | NA | 4 | 0.75 | 4 | 0.75 | 1 | 1 | 9 | 0.7778 |
| 6_1843488 | NA | NA | 3 | 1 | NA | NA | 1 | 0 | 4 | 0.75 | NA | NA | 3 | 1 | 3 | 0.6667 | 1 | 1 | 7 | 0.8571 |
| 6_1843492 | NA | NA | 4 | 0.25 | NA | NA | 1 | 0 | 5 | 0.2 | NA | NA | 4 | 0.25 | 2 | 0 | 1 | 0 | 7 | 0.1429 |
| 6_1884332 | 1 | 1 | NA | NA | 1 | 1 | 1 | 1 | 3 | 1 | 3 | 0.6667 | 1 | 1 | 2 | 0 | NA | NA | 6 | 0.5 |
| 6_1884406 | 1 | 0 | NA | NA | NA | NA | 2 | 0.5 | 3 | 0.3333 | 5 | 0.2 | 1 | 0 | 2 | 0.5 | NA | NA | 8 | 0.25 |
| 6_1884415 | 1 | 0 | NA | NA | 1 | 1 | 2 | 0.5 | 4 | 0.5 | 4 | 0.5 | 1 | 0 | 2 | 0.5 | NA | NA | 7 | 0.4286 |
| 6_1884424 | 1 | 0 | NA | NA | 1 | 0 | 2 | 0.5 | 4 | 0.25 | 5 | 0.6 | 1 | 0 | 2 | 0.5 | NA | NA | 8 | 0.5 |
| 6_1884438 | 1 | 0 | NA | NA | 1 | 1 | 2 | 1 | 4 | 0.75 | 5 | 1 | 1 | 1 | 1 | 0 | NA | NA | 7 | 0.8571 |
| 6_1884443 | 1 | 1 | NA | NA | 1 | 0 | 2 | 0.5 | 4 | 0.5 | 4 | 0.25 | 1 | 1 | 2 | 0.5 | NA | NA | 7 | 0.4286 |
| 6_1884450 | 1 | 0 | NA | NA | 1 | 0 | 2 | 0.5 | 4 | 0.25 | 5 | 0 | 1 | 1 | 2 | 0 | NA | NA | 8 | 0.125 |
| 6_1884454 | 1 | 0 | NA | NA | 1 | 1 | 2 | 0.5 | 4 | 0.5 | 3 | 0.3333 | 1 | 0 | 2 | 0.5 | NA | NA | 6 | 0.3333 |
| 6_1884467 | NA | NA | NA | NA | 1 | 1 | 2 | 0 | 3 | 0.3333 | 5 | 0.6 | 1 | 0 | 2 | 0.5 | NA | NA | 8 | 0.5 |
| 6_1884479 | 1 | 0 | NA | NA | 1 | 1 | 2 | 0.5 | 4 | 0.5 | 5 | 0.4 | 1 | 1 | 2 | 0.5 | NA | NA | 8 | 0.5 |
| 6_1884490 | 1 | 0 | NA | NA | 1 | 0 | 2 | 0.5 | 4 | 0.25 | 4 | 0.25 | 1 | 1 | 1 | 0 | NA | NA | 6 | 0.3333 |
| 6_1884505 | 1 | 0 | NA | NA | 1 | 1 | 1 | 0 | 3 | 0.3333 | 2 | 0 | 1 | 0 | 1 | 1 | NA | NA | 4 | 0.25 |
| 6_1884509 | 1 | 1 | NA | NA | NA | NA | 2 | 0 | 3 | 0.3333 | 4 | 0.25 | 1 | 1 | 2 | 0 | NA | NA | 7 | 0.2857 |
| 6_1884526 | 1 | 1 | NA | NA | NA | NA | 2 | 1 | 3 | 1 | 4 | 1 | 1 | 1 | 2 | 0.5 | NA | NA | 7 | 0.8571 |
| 6_2172611 | 8 | 0.875 | 4 | 0.75 | 10 | 0.9 | 9 | 0.8889 | 31 | 0.871 | NA | NA | 9 | 1 | 8 | 1 | 8 | 0.75 | 25 | 0.92 |
| 6_2172617 | 9 | 0.1111 | 4 | 0 | 12 | 0.0833 | 9 | 0.1111 | 34 | 0.0882 | NA | NA | 10 | 0.2 | 10 | 0.1 | 8 | 0.125 | 28 | 0.1429 |
| 6_2172720 | 9 | 0.4444 | 3 | 0 | 10 | 0.2 | 12 | 0.1667 | 34 | 0.2353 | NA | NA | 9 | 0.4444 | 10 | 0.3 | 7 | 0.5714 | 26 | 0.4231 |
| 6_2172721 | 6 | 0 | 3 | 0 | 10 | 0.1 | 11 | 0.1818 | 30 | 0.1 | NA | NA | 6 | 0.1667 | 5 | 0 | 6 | 0 | 17 | 0.0588 |
| 6_2172734 | 10 | 0.5 | 6 | 0.8333 | 12 | 0.6667 | 13 | 0.9231 | 41 | 0.7317 | NA | NA | 10 | 0.8 | 8 | 1 | 9 | 0.8889 | 27 | 0.8889 |
| 6_2172737 | 9 | 0.1111 | 6 | 0 | 12 | 0.1667 | 13 | 0.1538 | 40 | 0.125 | NA | NA | 9 | 0.2222 | 10 | 0.2 | 9 | 0.1111 | 28 | 0.1786 |
| 6_2172743 | 11 | 0.4545 | 6 | 0.1667 | 11 | 0.0909 | 12 | 0.1667 | 40 | 0.225 | NA | NA | 10 | 0 | 8 | 0.375 | 7 | 0 | 25 | 0.12 |
| 6_2172752 | 10 | 0.1 | 6 | 0 | 10 | 0.2 | 12 | 0.0833 | 38 | 0.1053 | NA | NA | 9 | 0.1111 | 11 | 0.1818 | 9 | 0.3333 | 29 | 0.2069 |

| | | | | | | | | | | | | | | | | | | | | |
|---|---|---|---|---|---|---|---|---|---|---|---|---|---|---|---|---|---|---|---|---|
| 6_2172755 | 11 | 0.1818 | 6 | 0 | 12 | 0 | 13 | 0.0769 | 42 | 0.0714 | NA | NA | 8 | 0.125 | 11 | 0 | 9 | 0.1111 | 28 | 0.0714 |
| 6_2172757 | 7 | 0.8571 | 3 | 1 | 7 | 0.7143 | 10 | 0.9 | 27 | 0.8519 | NA | NA | 5 | 1 | 9 | 0.8889 | 9 | 0.8889 | 23 | 0.913 |
| 6_2172762 | 10 | 1 | 3 | 1 | 11 | 1 | 11 | 1 | 35 | 1 | NA | NA | 8 | 0.875 | 9 | 0.8889 | 10 | 1 | 27 | 0.9259 |
| 6_2172764 | 8 | 0 | 5 | 0.4 | 10 | 0.1 | 11 | 0.0909 | 34 | 0.1176 | NA | NA | 9 | 0 | 8 | 0.25 | 10 | 0 | 27 | 0.0741 |
| 6_2172766 | 10 | 1 | 5 | 1 | 11 | 1 | 13 | 1 | 39 | 1 | NA | NA | 9 | 1 | 10 | 0.9 | 9 | 1 | 28 | 0.9643 |
| 6_2172770 | 11 | 0.1818 | 4 | 0.25 | 11 | 0 | 12 | 0.25 | 38 | 0.1579 | NA | NA | 8 | 0.125 | 11 | 0.3636 | 10 | 0.1 | 29 | 0.2069 |
| 6_2172774 | 9 | 0.7778 | 5 | 1 | 7 | 0.8571 | 9 | 1 | 30 | 0.9 | NA | NA | 7 | 1 | 4 | 1 | 9 | 1 | 20 | 1 |
| 6_2172780 | 9 | 0.7778 | 6 | 1 | 11 | 1 | 8 | 1 | 34 | 0.9412 | NA | NA | 7 | 1 | 11 | 1 | 10 | 1 | 28 | 1 |
| 6_2172788 | 11 | 0 | 7 | 0.4286 | 11 | 0.0909 | 11 | 0.2727 | 40 | 0.175 | NA | NA | 10 | 0 | 11 | 0.1818 | 11 | 0.2727 | 32 | 0.1562 |
| 6_2172813 | 11 | 0.4545 | 7 | 0.4286 | 10 | 0.2 | 13 | 0.3077 | 41 | 0.3415 | 1 | 0 | 10 | 0.6 | 9 | 0.3333 | 8 | 0.25 | 28 | 0.3929 |
| 6_2172842 | 14 | 0.0714 | 9 | 0.1111 | 14 | 0.0714 | 16 | 0.125 | 53 | 0.0943 | 3 | 0 | 10 | 0 | 10 | 0.3 | 10 | 0 | 33 | 0.0909 |
| 6_2177259 | 1 | 1 | 4 | 1 | 4 | 0.5 | NA | NA | 9 | 0.7778 | 1 | 1 | 1 | 1 | 3 | 0.6667 | 3 | 1 | 8 | 0.875 |
| 6_2177260 | 1 | 1 | 6 | 0.5 | 2 | 0.5 | NA | NA | 9 | 0.5556 | NA | NA | 1 | 0 | 1 | 1 | 2 | 0.5 | 4 | 0.5 |
| 6_2177271 | 1 | 1 | 6 | 1 | 4 | 1 | NA | NA | 11 | 1 | 1 | 1 | 1 | 1 | 2 | 1 | 3 | 1 | 7 | 1 |
| 6_2197370 | 4 | 1 | 2 | 1 | 4 | 0.75 | 1 | 0 | 11 | 0.8182 | NA | NA | 2 | 1 | NA | NA | 1 | 1 | 3 | 1 |
| 6_2197382 | 5 | 0.6 | 2 | 0.5 | 4 | 0.5 | 1 | 1 | 12 | 0.5833 | NA | NA | 2 | 1 | NA | NA | NA | NA | 2 | 1 |
| 6_2197384 | 4 | 0.25 | 2 | 0.5 | 3 | 0.6667 | 1 | 0 | 10 | 0.4 | NA | NA | 2 | 0.5 | NA | NA | 1 | 1 | 3 | 0.6667 |
| 6_2197483 | 4 | 0.5 | 4 | 0 | 2 | 0 | 2 | 0 | 12 | 0.1667 | NA | NA | 1 | 1 | NA | NA | NA | NA | 1 | 1 |
| 6_2230824 | 8 | 0.25 | 10 | 0.2 | 9 | 0.2222 | 4 | 0.75 | 31 | 0.2903 | 6 | 0.3333 | 3 | 0 | 5 | 0.2 | 10 | 0.4 | 24 | 0.2917 |
| 6_2230841 | 4 | 0 | 7 | 0 | 6 | 0.1667 | 7 | 0.1429 | 24 | 0.0833 | 6 | 0 | 3 | 0.3333 | 5 | 0 | 9 | 0.2222 | 23 | 0.1304 |
| 6_2299454 | 4 | 0 | 7 | 0 | 5 | 0 | 6 | 0.3333 | 22 | 0.0909 | 5 | 0.2 | 8 | 0 | 4 | 0 | 5 | 0.2 | 22 | 0.0909 |
| 6_2299925 | 5 | 0.6 | 7 | 0.2857 | 6 | 0.8333 | 6 | 0.5 | 24 | 0.5417 | 5 | 0.4 | 7 | 0.5714 | 4 | 0.25 | 5 | 0.2 | 21 | 0.381 |
| 6_2299932 | 4 | 1 | 7 | 0.7143 | 6 | 1 | 4 | 1 | 21 | 0.9048 | 4 | 1 | 8 | 0.875 | 4 | 0.75 | 8 | 0.75 | 24 | 0.8333 |
| 6_2299934 | 5 | 0 | 6 | 0 | 6 | 0.1667 | 6 | 0.1667 | 23 | 0.087 | 5 | 0.2 | 8 | 0 | 4 | 0.25 | 8 | 0.125 | 25 | 0.12 |
| 6_2299947 | 3 | 0.6667 | 7 | 0.4286 | 4 | 0.5 | 6 | 0.3333 | 20 | 0.45 | 4 | 0.25 | 7 | 0.1429 | 4 | 0 | 6 | 0.5 | 21 | 0.2381 |
| 6_2299948 | 3 | 0.6667 | 4 | 0.5 | 2 | 0.5 | 3 | 0.6667 | 12 | 0.5833 | 4 | 1 | 4 | 0.5 | 2 | 0.5 | 5 | 0.8 | 15 | 0.7333 |
| 6_2299952 | 4 | 0 | 6 | 0 | 5 | 0 | 5 | 0.2 | 20 | 0.05 | 5 | 0.4 | 6 | 0.1667 | 4 | 0 | 7 | 0 | 22 | 0.1364 |
| 6_2299954 | 4 | 0.5 | 5 | 1 | 5 | 0.6 | 5 | 1 | 19 | 0.7895 | 4 | 0.75 | 6 | 0.8333 | 4 | 0.5 | 7 | 0.4286 | 21 | 0.619 |
| 6_2299956 | 4 | 0.5 | 7 | 1 | 4 | 1 | 5 | 1 | 20 | 0.9 | 4 | 0.75 | 5 | 1 | 3 | 0.6667 | 5 | 1 | 17 | 0.8824 |
| 6_2299957 | 5 | 0.6 | 7 | 0.8571 | 7 | 1 | 6 | 1 | 25 | 0.88 | 5 | 1 | 9 | 0.8889 | 4 | 1 | 7 | 0.8571 | 25 | 0.92 |
| 6_2299965 | 5 | 0 | 5 | 0.2 | 4 | 0.5 | 6 | 0 | 20 | 0.15 | 4 | 0.25 | 9 | 0.2222 | 3 | 0 | 8 | 0.25 | 24 | 0.2083 |
| 6_2299968 | 5 | 0 | 7 | 0 | 8 | 0.125 | 6 | 0.5 | 26 | 0.1538 | 5 | 0 | 10 | 0.2 | 4 | 0 | 6 | 0.1667 | 25 | 0.12 |
| 6_2299993 | 4 | 0.25 | 6 | 0.8333 | 8 | 0.75 | 4 | 0.75 | 22 | 0.6818 | 5 | 0.8 | 9 | 0.8889 | 4 | 0.25 | 7 | 0.7143 | 25 | 0.72 |
| 6_2299997 | 5 | 0.2 | 4 | 0.75 | 8 | 0.25 | 6 | 0.6667 | 23 | 0.4348 | 5 | 0 | 10 | 0.6 | 4 | 0.5 | 7 | 0.4286 | 26 | 0.4231 |
| 6_2300005 | 5 | 0.4 | 7 | 0.5714 | 8 | 0.125 | 7 | 0.4286 | 27 | 0.3704 | 5 | 0.2 | 13 | 0.3077 | 4 | 0 | 8 | 0.25 | 30 | 0.2333 |
| 6_2300020 | 2 | 0.5 | 5 | 0.4 | 6 | 0 | 6 | 0.1667 | 19 | 0.2105 | 1 | 0 | 9 | 0.2222 | 2 | 0 | 4 | 0.5 | 16 | 0.25 |
| 6_2300026 | 2 | 0.5 | 5 | 0.2 | 8 | 0.375 | 5 | 0.4 | 20 | 0.35 | 2 | 0.5 | 9 | 0.4444 | 1 | 0 | 5 | 0.4 | 17 | 0.4118 |
| 6_2300027 | 2 | 0.5 | 5 | 0 | 8 | 0.125 | 7 | 0.4286 | 22 | 0.2273 | 2 | 0.5 | 10 | 0.6 | 2 | 0 | 5 | 0.4 | 19 | 0.4737 |
| 6_2300045 | NA | NA | 5 | 0.2 | 6 | 0 | 7 | 0 | 18 | 0.0556 | 1 | 0 | 9 | 0.1111 | 2 | 0 | 2 | 0 | 14 | 0.0714 |
| 6_2300054 | 2 | 0 | 5 | 0.6 | 7 | 0.7143 | 7 | 0.1429 | 21 | 0.4286 | 2 | 0 | 10 | 0.5 | 1 | 1 | 4 | 0.25 | 17 | 0.4118 |
| 6_2300060 | 2 | 0.5 | 5 | 0.4 | 7 | 0.7143 | 9 | 0.7778 | 23 | 0.6522 | 2 | 0.5 | 9 | 0.8889 | 2 | 0.5 | 2 | 0.5 | 15 | 0.7333 |
| 6_2300071 | 1 | 0 | 4 | 0 | 7 | 0.2857 | 9 | 0.5556 | 21 | 0.3333 | 2 | 1 | 9 | 0.4444 | 2 | 0.5 | 4 | 0.5 | 17 | 0.5294 |
| 6_2300097 | 3 | 1 | 4 | 0.5 | 7 | 1 | 9 | 0.8889 | 23 | 0.8696 | 3 | 1 | 9 | 1 | 2 | 1 | 4 | 1 | 18 | 1 |
| 6_2300108 | 2 | 1 | 5 | 0.6 | 7 | 0.7143 | 9 | 0.6667 | 23 | 0.6957 | 2 | 1 | 9 | 0.5556 | 2 | 1 | 5 | 0.8 | 18 | 0.7222 |
| 6_2300109 | 3 | 1 | 4 | 1 | 7 | 0.8571 | 7 | 0.7143 | 21 | 0.8571 | 3 | 1 | 6 | 1 | 1 | 1 | 3 | 0.6667 | 13 | 0.9231 |
| 6_2300111 | 4 | 0.25 | 4 | 0 | 7 | 0.2857 | 9 | 0.3333 | 24 | 0.25 | 3 | 0 | 9 | 0.4444 | 2 | 0.5 | 5 | 0.8 | 19 | 0.4737 |
| 6_2300119 | 4 | 0 | 5 | 0 | 6 | 0 | 9 | 0.3333 | 24 | 0.125 | 3 | 0 | 10 | 0.1 | 2 | 0 | 5 | 0 | 20 | 0.05 |
| 6_2300127 | 4 | 0.5 | 4 | 0 | 5 | 0.4 | 8 | 0.25 | 21 | 0.2857 | 4 | 0.25 | 7 | 0.4286 | 2 | 0.5 | 3 | 0 | 16 | 0.3125 |
| 6_2300134 | 3 | 1 | 5 | 0.8 | 9 | 1 | 10 | 1 | 27 | 0.963 | 4 | 1 | 10 | 1 | 2 | 1 | 6 | 1 | 22 | 1 |
| 6_2300137 | 4 | 0 | 4 | 0.25 | 8 | 0.125 | 9 | 0.2222 | 25 | 0.16 | 4 | 0 | 7 | 0 | 2 | 0.5 | 6 | 0.3333 | 19 | 0.1579 |
| 6_2300139 | NA | NA | NA | NA | NA | NA | 1 | 1 | 1 | 1 | NA | NA | 1 | 1 | NA | NA | NA | NA | 1 | 1 |
| 6_2300152 | 3 | 1 | 5 | 0.8 | 5 | 1 | 7 | 0.7143 | 20 | 0.85 | 4 | 0.75 | 8 | 0.75 | 1 | 0 | 4 | 0.5 | 17 | 0.6471 |
| 6_2300174 | 1 | 0 | 2 | 0 | 4 | 0.75 | 5 | 0.2 | 12 | 0.3333 | 3 | 0 | 5 | 0.2 | 2 | 0 | 5 | 0 | 15 | 0.0667 |
| 6_2300189 | 2 | 0 | 4 | 0.5 | 5 | 0.4 | 8 | 0.375 | 19 | 0.3684 | 3 | 0 | 7 | 0.8571 | 4 | 1 | 7 | 0 | 21 | 0.4762 |
| 6_2300191 | 3 | 0 | 4 | 0.5 | 3 | 0 | 8 | 0.25 | 18 | 0.2222 | 4 | 0 | 10 | 0.1 | 3 | 0 | 8 | 0.125 | 25 | 0.08 |
| 6_2300193 | 4 | 0.75 | 5 | 1 | 7 | 0.8571 | 9 | 0.5556 | 25 | 0.76 | 6 | 1 | 12 | 0.8333 | 5 | 1 | 6 | 0.8333 | 29 | 0.8966 |
| 6_2300206 | 4 | 0 | 3 | 0.3333 | 5 | 0 | 8 | 0.5 | 20 | 0.25 | 6 | 0.1667 | 10 | 0.2 | 5 | 0.2 | 6 | 0 | 27 | 0.1481 |
| 6_2300214 | 4 | 0.5 | 5 | 0.4 | 9 | 0.4444 | 9 | 0.5556 | 27 | 0.4815 | 9 | 0.4444 | 11 | 0.6364 | 4 | 0.5 | 8 | 0.125 | 32 | 0.4375 |
| 6_2300219 | 5 | 0.4 | 5 | 0.4 | 8 | 0.375 | 10 | 0.8 | 28 | 0.5357 | 9 | 0.5556 | 10 | 0.5 | 6 | 0.6667 | 6 | 0.6667 | 31 | 0.5806 |
| 6_2300221 | 3 | 0.6667 | 3 | 0.6667 | 10 | 0.6 | 7 | 0.5714 | 23 | 0.6087 | 9 | 0.6667 | 11 | 0.6364 | 6 | 0.5 | 6 | 0.6667 | 32 | 0.625 |
| 6_2300222 | 5 | 0.2 | 5 | 0 | 9 | 0.2222 | 7 | 0.2857 | 26 | 0.1923 | 8 | 0.25 | 10 | 0.4 | 6 | 0.1667 | 7 | 0.2857 | 31 | 0.2903 |
| 6_2300230 | 4 | 1 | 3 | 0.6667 | 8 | 0.75 | 10 | 0.7 | 25 | 0.76 | 9 | 0.4444 | 11 | 0.6364 | 5 | 0.6 | 7 | 0.8571 | 32 | 0.625 |
| 6_2300230 | 5 | 0.6 | 5 | 0.6 | 9 | 0.7778 | 9 | 0.4444 | 28 | 0.6071 | 8 | 1 | 10 | 0.8 | 6 | 0.5 | 8 | 0.75 | 32 | 0.7812 |
| 6_2300234 | 5 | 0 | 5 | 0.2 | 12 | 0.3333 | 10 | 0.2 | 32 | 0.2188 | 9 | 0 | 10 | 0.2 | 6 | 0.3333 | 8 | 0 | 33 | 0.1212 |
| 6_2300246 | 5 | 0.8 | 3 | 1 | 10 | 1 | 9 | 0.8889 | 27 | 0.9259 | 7 | 0.8571 | 11 | 0.9091 | 6 | 1 | 7 | 1 | 31 | 0.9355 |

| | | | | | | | | | | | | | | | | | | | | |
|---|---|---|---|---|---|---|---|---|---|---|---|---|---|---|---|---|---|---|---|---|
| 6_2300247 | 5 | 0.6 | 4 | 0.5 | 11 | 0.3636 | 9 | 0.1111 | 29 | 0.3448 | 6 | 0.3333 | 9 | 0.3333 | 6 | 0.5 | 7 | 0.1429 | 28 | 0.3214 |
| 6_2300253 | 6 | 0 | 3 | 0 | 11 | 0.1818 | 10 | 0.1 | 30 | 0.1 | 7 | 0 | 10 | 0.1 | 7 | 0.1429 | 7 | 0.1429 | 31 | 0.0968 |
| 6_2300265 | 6 | 0.3333 | 3 | 0 | 6 | 0.1667 | 4 | 0 | 19 | 0.1579 | 8 | 0.125 | 8 | 0 | 4 | 0.25 | 6 | 0.3333 | 26 | 0.1538 |
| 6_2300265 | 6 | 0.1667 | 5 | 0.2 | 11 | 0.0909 | 9 | 0.1111 | 31 | 0.129 | 9 | 0.1111 | 11 | 0.1818 | 6 | 0.1667 | 8 | 0.125 | 34 | 0.1471 |
| 6_2300284 | 5 | 0.4 | 6 | 0.3333 | 7 | 0.2857 | 6 | 0.1667 | 24 | 0.2917 | 6 | 0 | 10 | 0 | 6 | 0.3333 | 6 | 0.3333 | 28 | 0.1429 |
| 6_2300308 | 7 | 0 | 6 | 0 | 7 | 0.4286 | 5 | 0 | 25 | 0.12 | 5 | 0.2 | 11 | 0.1818 | 6 | 0.5 | 8 | 0.125 | 30 | 0.2333 |
| 6_2300319 | 6 | 0.1667 | 6 | 0.3333 | 7 | 0.2857 | 6 | 0 | 25 | 0.2 | 6 | 0.1667 | 11 | 0.0909 | 5 | 0.4 | 8 | 0.625 | 30 | 0.3 |
| 6_2300328 | 7 | 0.5714 | 6 | 0.1667 | 6 | 0.1667 | 6 | 0.1667 | 25 | 0.28 | 6 | 0 | 10 | 0.2 | 6 | 0 | 7 | 0.1429 | 29 | 0.1034 |
| 6_2300330 | 7 | 0.4286 | 5 | 0.2 | 5 | 0 | 5 | 0.4 | 22 | 0.2727 | 5 | 0 | 8 | 0.375 | 7 | 0.2857 | 5 | 0 | 25 | 0.2 |
| 6_2300357 | 7 | 0.4286 | 7 | 0.1429 | 7 | 0.4286 | 5 | 0 | 26 | 0.2692 | 6 | 0.1667 | 11 | 0.0909 | 6 | 0 | 6 | 0.1667 | 29 | 0.1034 |
| 6_2300397 | 7 | 0 | 4 | 0 | 5 | 0 | 4 | 0.25 | 20 | 0.05 | 5 | 0.2 | 9 | 0.1111 | 4 | 0 | 7 | 0.1429 | 25 | 0.12 |
| 6_2300400 | 6 | 0 | 7 | 0.2857 | 4 | 0.25 | 5 | 0.2 | 22 | 0.1818 | 5 | 0.2 | 10 | 0.4 | 3 | 0.3333 | 2 | 0.5 | 20 | 0.35 |
| 6_2305205 | 6 | 0.1667 | 4 | 0 | 6 | 0.1667 | 8 | 0.125 | 24 | 0.125 | 11 | 0 | 11 | 0.0909 | 8 | 0.125 | 11 | 0.0909 | 41 | 0.0732 |
| 6_2326299 | 6 | 0.5 | 11 | 0.0909 | 5 | 0.2 | 1 | 0 | 23 | 0.2174 | 7 | 0 | 2 | 0.5 | 3 | 0 | 8 | 0.25 | 20 | 0.15 |
| 6_2326330 | 4 | 0 | 10 | 0.1 | 3 | 0 | NA | NA | 17 | 0.0588 | 7 | 0.2857 | 3 | 0 | 4 | 0 | 9 | 0 | 23 | 0.087 |
| 6_2326356 | 2 | 0.5 | 5 | 0.4 | 3 | 0.6667 | NA | NA | 10 | 0.5 | 5 | 0.2 | 2 | 0 | 1 | 0 | 6 | 0.1667 | 14 | 0.1429 |
| 6_2326359 | 5 | 0 | 10 | 0.1 | 4 | 0.25 | NA | NA | 19 | 0.1053 | 5 | 0 | 3 | 0 | 3 | 0 | 7 | 0.2857 | 18 | 0.1111 |
| 6_2362668 | 2 | 0.5 | 3 | 0.3333 | 1 | 0 | NA | NA | 6 | 0.3333 | 2 | 0.5 | 2 | 0.5 | 1 | 0 | 6 | 0.1667 | 11 | 0.2727 |
| 6_2383975 | 1 | 1 | NA | NA | NA | NA | NA | NA | 1 | 1 | NA | NA | NA | NA | NA | NA | 1 | 1 | 1 | 1 |
| 6_2383976 | 1 | 1 | NA | NA | NA | NA | NA | NA | 1 | 1 | NA | NA | NA | NA | NA | NA | 1 | 1 | 1 | 1 |
| 6_2383980 | 1 | 1 | NA | NA | NA | NA | NA | NA | 1 | 1 | NA | NA | NA | NA | NA | NA | 1 | 1 | 1 | 1 |
| 6_2383987 | 1 | 1 | NA | NA | NA | NA | NA | NA | 1 | 1 | NA | NA | NA | NA | NA | NA | 1 | 1 | 1 | 1 |
| 6_2445421 | 1 | 1 | NA | NA | NA | NA | 1 | 0 | 2 | 0.5 | NA | NA | 1 | 1 | NA | NA | NA | NA | 1 | 1 |
| 6_2461449 | 13 | 0.2308 | 17 | 0.3529 | 16 | 0.25 | 12 | 0.5 | 58 | 0.3276 | 7 | 0.2857 | 23 | 0.4783 | 13 | 0.3846 | 25 | 0.44 | 68 | 0.4265 |
| 6_2461628 | 8 | 0 | 14 | 0 | 13 | 0.1538 | 8 | 0.375 | 43 | 0.1163 | 4 | 0 | 15 | 0 | 14 | 0.0714 | 20 | 0.1 | 53 | 0.0566 |
| 6_2495190 | 3 | 0.3333 | 5 | 0 | 5 | 0.2 | 8 | 0.5 | 21 | 0.2857 | 5 | 0 | 4 | 0.5 | 5 | 0.4 | 3 | 0 | 17 | 0.2353 |
| 6_2495197 | 4 | 0 | 8 | 0.25 | 5 | 0 | 7 | 0.2857 | 24 | 0.1667 | 8 | 0.125 | 5 | 0.4 | 4 | 0 | 6 | 0 | 23 | 0.1304 |
| 6_2495382 | 2 | 0 | 5 | 0.4 | 8 | 0.5 | 5 | 0.2 | 20 | 0.35 | 7 | 0.2857 | 3 | 0.6667 | 4 | 0 | 7 | 0.4286 | 21 | 0.3333 |
| 6_2495390 | 2 | 0 | 4 | 0.25 | 6 | 0.1667 | 4 | 0.25 | 16 | 0.1875 | 6 | 0.1667 | 3 | 0 | 4 | 0 | 7 | 0.1429 | 20 | 0.1 |
| 6_2495391 | 2 | 0 | 5 | 0 | 3 | 0 | 4 | 0.25 | 14 | 0.0714 | 5 | 0.2 | 3 | 0.3333 | 4 | 0 | 7 | 0 | 19 | 0.1053 |
| 6_2495393 | 2 | 0 | 5 | 0 | 5 | 0 | 5 | 0.4 | 17 | 0.1176 | 6 | 0.1667 | 3 | 0 | 3 | 0 | 7 | 0 | 19 | 0.0526 |
| 6_2495402 | 1 | 0 | 3 | 0.3333 | 8 | 0.375 | 5 | 0.2 | 17 | 0.2941 | 7 | 0.1429 | 1 | 1 | 3 | 0 | 5 | 0.4 | 16 | 0.25 |
| 6_2495403 | 2 | 0 | 4 | 0.25 | 6 | 0 | 6 | 0.1667 | 18 | 0.1111 | 5 | 0.4 | NA | NA | 4 | 0 | 5 | 0.2 | 14 | 0.2143 |
| 6_2495407 | 1 | 1 | 3 | 0.6667 | 6 | 1 | 4 | 0.75 | 14 | 0.8571 | 5 | 0.8 | 2 | 0.5 | 3 | 0 | 3 | 1 | 13 | 0.6154 |
| 6_2495415 | 1 | 0 | 3 | 0 | 7 | 0.4286 | 5 | 0.2 | 16 | 0.25 | 6 | 0.6667 | 3 | 0.6667 | 3 | 0 | 6 | 0.3333 | 18 | 0.4444 |
| 6_2495426 | 2 | 0 | 6 | 0.1667 | 7 | 0 | 5 | 0.2 | 20 | 0.1 | 7 | 0.1429 | 3 | 0.3333 | 4 | 0 | 5 | 0 | 19 | 0.1053 |
| 6_2495451 | 1 | 0 | 3 | 0 | 4 | 0.25 | 4 | 0 | 12 | 0.0833 | 6 | 0.3333 | 3 | 0.3333 | 4 | 0.25 | 3 | 0 | 16 | 0.25 |
| 6_2495462 | 2 | 1 | 5 | 0.4 | 6 | 0.1667 | 4 | 0.25 | 17 | 0.3529 | 6 | 0.3333 | 2 | 0 | 4 | 0 | 3 | 0 | 15 | 0.1333 |
| 6_2495464 | 1 | 1 | 4 | 1 | 3 | 1 | 2 | 1 | 10 | 1 | 2 | 1 | 2 | 1 | 3 | 0.3333 | 2 | 1 | 9 | 0.7778 |
| 6_2495468 | 2 | 0 | 4 | 0.75 | 7 | 0.4286 | 4 | 0.25 | 17 | 0.4118 | 4 | 0.75 | 3 | 0.6667 | 3 | 0 | 3 | 0.3333 | 13 | 0.4615 |
| 6_2495477 | 2 | 1 | 4 | 1 | 5 | 1 | 5 | 1 | 16 | 1 | 6 | 1 | 3 | 1 | 4 | 1 | 2 | 1 | 15 | 1 |
| 6_2495479 | 2 | 1 | 2 | 1 | 7 | 1 | 6 | 1 | 17 | 1 | 5 | 1 | 3 | 1 | 3 | 1 | 2 | 1 | 13 | 1 |
| 6_2495490 | 1 | 0 | 4 | 0.25 | 3 | 0.3333 | 5 | 0.6 | 13 | 0.3846 | 4 | 0 | 2 | 0.5 | NA | NA | 2 | 0.5 | 8 | 0.25 |
| 6_2495490 | NA | NA | 1 | 1 | 1 | 0 | NA | NA | 2 | 0.5 | NA | NA | 1 | 1 | 2 | 0 | NA | NA | 3 | 0.3333 |
| 6_2495497 | 1 | 0 | 2 | 1 | 1 | 1 | 2 | 1 | 6 | 0.8333 | 2 | 1 | 2 | 0.5 | 1 | 1 | 1 | 1 | 6 | 0.8333 |
| 6_2495506 | NA | NA | 2 | 1 | 7 | 1 | 4 | 1 | 13 | 1 | 6 | 1 | 3 | 1 | 3 | 1 | 2 | 1 | 14 | 1 |
| 6_2495511 | 1 | 0 | 3 | 0 | 6 | 0.1667 | 6 | 0.6667 | 16 | 0.3125 | 5 | 0.4 | 2 | 0 | 3 | 0.3333 | 2 | 0.5 | 12 | 0.3333 |
| 6_2495517 | 1 | 1 | 3 | 1 | 6 | 0.8333 | 6 | 1 | 16 | 0.9375 | 4 | 0.75 | 2 | 0.5 | 3 | 1 | 1 | 1 | 10 | 0.8 |
| 6_2495535 | NA | NA | 3 | 0 | 5 | 0.2 | 5 | 0 | 13 | 0.0769 | 5 | 0.2 | 2 | 0 | 3 | 0 | 2 | 0.5 | 12 | 0.1667 |
| 6_2495567 | NA | NA | 2 | 1 | 2 | 1 | 4 | 1 | 8 | 1 | 4 | 1 | 2 | 0.5 | 2 | 0.5 | 1 | 1 | 9 | 0.7778 |
| 6_2495589 | NA | NA | 3 | 0.3333 | 5 | 0.4 | 3 | 0.3333 | 11 | 0.3636 | 5 | 0 | 2 | 0 | 2 | 0 | 1 | 1 | 10 | 0.1 |
| 6_2495593 | NA | NA | 2 | 0.5 | 2 | 1 | 3 | 0.6667 | 7 | 0.7143 | 5 | 0.8 | 2 | 0.5 | 2 | 0 | 1 | 0 | 10 | 0.5 |
| 6_2495622 | NA | NA | NA | NA | 2 | 0.5 | 1 | 0 | 3 | 0.3333 | 1 | 1 | NA | NA | NA | NA | NA | NA | 1 | 1 |
| 6_2566855 | 3 | 0 | 4 | 0 | 2 | 1 | 3 | 0 | 12 | 0.1667 | 1 | 1 | 5 | 0 | 1 | 0 | 4 | 0 | 11 | 0.0909 |
| 6_2566857 | 3 | 0.3333 | 4 | 0 | 3 | 0 | 3 | 0.3333 | 13 | 0.1538 | 2 | 0.5 | 4 | 0.25 | 1 | 0 | 4 | 0.5 | 11 | 0.3636 |
| 6_2566890 | 3 | 1 | 5 | 1 | 5 | 1 | 3 | 1 | 16 | 1 | 3 | 1 | 5 | 1 | 3 | 1 | 7 | 1 | 18 | 1 |
| 6_2566892 | 5 | 0 | 8 | 0.125 | 9 | 0 | 7 | 0.2857 | 29 | 0.1034 | 4 | 0 | 8 | 0.125 | 5 | 0 | 10 | 0.2 | 27 | 0.1111 |
| 6_2566895 | 3 | 0 | 8 | 0.375 | 8 | 0.25 | 3 | 1 | 22 | 0.3636 | 3 | 0 | 6 | 0 | 5 | 0 | 6 | 0.3333 | 20 | 0.1 |
| 6_2566905 | 4 | 1 | 3 | 1 | 7 | 1 | 6 | 1 | 20 | 1 | 3 | 1 | 6 | 1 | 3 | 1 | 8 | 1 | 20 | 1 |
| 6_2566922 | 4 | 0 | 6 | 0.3333 | 9 | 0 | 6 | 0.1667 | 25 | 0.12 | 4 | 0 | 5 | 0 | 4 | 0 | 8 | 0.25 | 21 | 0.0952 |
| 6_2566929 | 5 | 1 | 8 | 0.75 | 7 | 0.8571 | 5 | 1 | 25 | 0.88 | 3 | 1 | 7 | 0.8571 | 4 | 0.75 | 9 | 1 | 23 | 0.913 |
| 6_2566941 | 2 | 1 | 4 | 1 | 3 | 1 | 1 | 1 | 10 | 1 | 3 | 1 | 4 | 1 | 2 | 1 | 6 | 1 | 15 | 1 |
| 6_2566948 | 2 | 1 | 7 | 0.8571 | 7 | 0.8571 | 6 | 1 | 22 | 0.9091 | 3 | 1 | 6 | 0.8333 | 4 | 1 | 10 | 1 | 23 | 0.9565 |
| 6_2566959 | 2 | 1 | 8 | 0.875 | 6 | 1 | 6 | 0.8333 | 22 | 0.9091 | 4 | 1 | 6 | 1 | 4 | 1 | 9 | 0.8889 | 23 | 0.9565 |
| 6_2566973 | 2 | 0 | 6 | 0 | 6 | 0.1667 | 6 | 0.3333 | 20 | 0.15 | 3 | 0 | 5 | 0 | 4 | 0 | 7 | 0.1429 | 19 | 0.0526 |

| | | | | | | | | | | | | | | | | | | | | |
|---|---|---|---|---|---|---|---|---|---|---|---|---|---|---|---|---|---|---|---|---|
| 6_2568821 | 1 | 1 | 7 | 0.8571 | 6 | 0.8333 | 6 | 0.8333 | 20 | 0.85 | 4 | 0.75 | 5 | 1 | 2 | 1 | 10 | 0.9 | 21 | 0.9048 |
| 6_2568831 | 2 | 1 | 8 | 0.625 | 7 | 0.5714 | 5 | 0.6 | 22 | 0.6364 | 3 | 0.6667 | 6 | 0.6667 | 4 | 0.75 | 6 | 0.8333 | 19 | 0.7368 |
| 6_2568844 | 1 | 0 | 6 | 0 | 2 | 0 | 4 | 0.25 | 13 | 0.0769 | 2 | 0 | 5 | 0.2 | 2 | 0 | 5 | 0 | 14 | 0.0714 |
| 6_2568845 | 1 | 1 | 6 | 0.6667 | 5 | 0.8 | 4 | 1 | 16 | 0.8125 | 2 | 0.5 | 5 | 1 | 2 | 1 | 8 | 0.625 | 17 | 0.7647 |
| 6_2568850 | 1 | 1 | 7 | 0.7143 | 7 | 0.4286 | 5 | 0.2 | 20 | 0.5 | 3 | 0.6667 | 6 | 0.1667 | 4 | 0 | 7 | 0.2857 | 20 | 0.25 |
| 6_2568855 | 2 | 0 | 8 | 0 | 5 | 0.4 | 5 | 0.2 | 20 | 0.15 | 2 | 0 | 5 | 0.4 | 4 | 0 | 7 | 0.1429 | 18 | 0.1667 |
| 6_2568855 | 1 | 0 | 8 | 0.25 | 8 | 0.375 | 4 | 0.25 | 21 | 0.2857 | 3 | 0.6667 | 6 | 0.3333 | 3 | 0 | 5 | 0.4 | 17 | 0.3529 |
| 6_2568869 | 2 | 1 | 7 | 0 | 6 | 0 | 6 | 0.1667 | 21 | 0.1429 | 2 | 0 | 5 | 0.4 | 4 | 0 | 8 | 0 | 19 | 0.1053 |
| 6_2568877 | 2 | 0 | 4 | 0.5 | 7 | 0.1429 | 6 | 0 | 19 | 0.1579 | 3 | 0.6667 | 5 | 0.4 | 4 | 0 | 8 | 0.25 | 20 | 0.3 |
| 6_2568883 | 2 | 0.5 | 6 | 0.1667 | 6 | 0.1667 | 4 | 0.25 | 18 | 0.2222 | 3 | 0.3333 | 5 | 0.6 | 3 | 0 | 7 | 0 | 18 | 0.2222 |
| 6_2568887 | 1 | 1 | 7 | 0.1429 | 4 | 0 | 5 | 0.2 | 17 | 0.1765 | 3 | 1 | 2 | 0.5 | 4 | 0 | 6 | 0.1667 | 15 | 0.3333 |
| 6_2568892 | 1 | 0 | 7 | 0.2857 | 5 | 0 | 5 | 0 | 18 | 0.1111 | 3 | 0.3333 | 4 | 0 | 4 | 0 | 6 | 0.1667 | 17 | 0.1176 |
| 6_2568898 | 1 | 0 | 3 | 1 | 3 | 1 | 1 | 1 | 8 | 0.875 | 1 | 1 | 4 | 1 | 1 | 0 | 4 | 0.5 | 10 | 0.7 |
| 6_2568918 | NA | NA | 6 | 0 | 6 | 0 | 4 | 0.5 | 16 | 0.125 | 2 | 0 | 4 | 0.25 | 2 | 0.5 | 6 | 0.3333 | 14 | 0.2857 |
| 6_2568929 | NA | NA | 4 | 0.25 | 4 | 0 | 5 | 0.2 | 13 | 0.1538 | 3 | 0 | 3 | 0.3333 | 1 | 0 | 4 | 0 | 11 | 0.0909 |
| 6_2568940 | NA | NA | 5 | 0.2 | 4 | 0 | 5 | 0.2 | 14 | 0.1429 | 2 | 0 | 2 | 0.5 | 1 | 0 | 6 | 0.3333 | 11 | 0.2727 |
| 6_2568954 | NA | NA | 4 | 0.25 | 3 | 0 | 3 | 0 | 10 | 0.1 | 2 | 0 | 3 | 0.6667 | 1 | 0 | 4 | 0.25 | 10 | 0.3 |
| 6_2568957 | NA | NA | 5 | 1 | 3 | 1 | 4 | 1 | 12 | 1 | 3 | 1 | 3 | 1 | 1 | 1 | 4 | 1 | 11 | 1 |
| 6_2568977 | NA | NA | 4 | 0.75 | 2 | 0 | 3 | 0.3333 | 9 | 0.4444 | 2 | 0 | 1 | 1 | 1 | 0 | 2 | 0 | 6 | 0.1667 |
| 6_2573827 | NA | NA | 1 | 0 | 1 | 1 | NA | NA | 2 | 0.5 | 1 | 1 | NA | NA | 1 | 1 | NA | NA | 2 | 1 |
| 6_2574450 | 2 | 0 | 1 | 0 | 2 | 0 | 2 | 0.5 | 7 | 0.1429 | NA | NA | 1 | 0 | 1 | 0 | 4 | 0.5 | 6 | 0.3333 |
| 6_2574452 | 2 | 0.5 | 1 | 0 | 1 | 0 | 1 | 1 | 5 | 0.4 | NA | NA | 1 | 0 | 1 | 0 | 4 | 0.5 | 6 | 0.3333 |
| 6_2727558 | NA | NA | 1 | 1 | 1 | 1 | 1 | 1 | 3 | 1 | NA | NA | NA | NA | 1 | 1 | NA | NA | 1 | 1 |
| 6_2736334 | 1 | 1 | 1 | 1 | NA | NA | NA | NA | 2 | 1 | NA | NA | NA | NA | 1 | 1 | NA | NA | 1 | 1 |
| 6_2736354 | 1 | 1 | 1 | 0 | NA | NA | NA | NA | 2 | 0.5 | NA | NA | NA | NA | 1 | 1 | NA | NA | 1 | 1 |
| 6_2824151 | NA | NA | 1 | 1 | NA | NA | NA | NA | 1 | 1 | 1 | 0 | 1 | 0 | NA | NA | 1 | 1 | 3 | 0.3333 |
| 6_2824154 | NA | NA | 1 | 1 | NA | NA | NA | NA | 1 | 1 | 1 | 0 | 2 | 0 | NA | NA | 1 | 1 | 4 | 0.25 |
| 6_2824160 | NA | NA | 1 | 1 | NA | NA | NA | NA | 1 | 1 | 1 | 1 | 2 | 0.5 | NA | NA | 1 | 1 | 4 | 0.75 |
| 6_2824305 | 1 | 0 | 2 | 0.5 | 2 | 0.5 | NA | NA | 5 | 0.4 | 1 | 1 | 2 | 0.5 | NA | NA | 3 | 0.3333 | 6 | 0.5 |
| 6_2824307 | 1 | 0 | 2 | 1 | 2 | 1 | NA | NA | 5 | 0.8 | 1 | 1 | 1 | 1 | 1 | 1 | 2 | 1 | 5 | 1 |
| 6_2824312 | 1 | 1 | NA | NA | 2 | 1 | NA | NA | 3 | 1 | 1 | 1 | NA | NA | 1 | 0 | NA | NA | 2 | 0.5 |
| 6_2824347 | 1 | 0 | 2 | 0.5 | 1 | 1 | NA | NA | 4 | 0.5 | 1 | 1 | 1 | 1 | 1 | 0 | 3 | 0.3333 | 6 | 0.5 |
| 6_2824352 | 1 | 1 | 1 | 0 | 2 | 0.5 | NA | NA | 4 | 0.5 | 1 | 0 | 2 | 1 | 1 | 1 | 3 | 1 | 7 | 0.8571 |
| 6_2824360 | 1 | 1 | 1 | 1 | 2 | 1 | NA | NA | 4 | 1 | 1 | 1 | 2 | 1 | 1 | 1 | 3 | 1 | 7 | 1 |
| 6_2824368 | NA | NA | 1 | 1 | 1 | 0 | NA | NA | 2 | 0.5 | 1 | 0 | 2 | 0 | 1 | 0 | 3 | 0.3333 | 7 | 0.1429 |
| 6_2824383 | 1 | 0 | 1 | 0 | 2 | 0.5 | NA | NA | 4 | 0.25 | 1 | 0 | 2 | 1 | 1 | 1 | 2 | 1 | 6 | 0.8333 |
| 6_2824393 | 1 | 0 | 1 | 0 | 2 | 0.5 | NA | NA | 4 | 0.25 | 1 | 1 | 1 | 0 | 1 | 0 | 1 | 1 | 4 | 0.5 |
| 6_2824402 | NA | NA | 1 | 1 | 2 | 1 | NA | NA | 3 | 1 | 1 | 1 | 1 | 1 | NA | NA | 2 | 1 | 4 | 1 |
| 6_2824436 | NA | NA | 1 | 1 | 2 | 1 | NA | NA | 3 | 1 | 1 | 1 | 3 | 0.6667 | NA | NA | NA | NA | 4 | 0.75 |
| 6_2824451 | 1 | 1 | 1 | 1 | 1 | 1 | NA | NA | 3 | 1 | 1 | 1 | 1 | 1 | NA | NA | 1 | 1 | 3 | 1 |
| 6_2824724 | NA | NA | 1 | 0 | 1 | 1 | NA | NA | 2 | 0.5 | 1 | 1 | NA | NA | NA | NA | 1 | 0 | 2 | 0.5 |
| 6_2826236 | 1 | 1 | 1 | 1 | NA | NA | NA | NA | 2 | 1 | NA | NA | 1 | 1 | NA | NA | NA | NA | 1 | 1 |
| 6_2826243 | 1 | 0 | 1 | 1 | 1 | 0 | NA | NA | 3 | 0.3333 | NA | NA | 1 | 1 | NA | NA | NA | NA | 1 | 1 |
| 6_2826275 | 1 | 0 | 1 | 1 | 1 | 0 | NA | NA | 3 | 0.3333 | NA | NA | 1 | 1 | NA | NA | NA | NA | 1 | 1 |
| 6_2826358 | 1 | 1 | 1 | 1 | 1 | 1 | NA | NA | 3 | 1 | NA | NA | 1 | 1 | NA | NA | NA | NA | 1 | 1 |
| 6_2841689 | 6 | 0.1667 | 1 | 0 | 3 | 0.6667 | 2 | 0.5 | 12 | 0.3333 | 4 | 0.5 | 4 | 0.25 | 2 | 0.5 | 3 | 0.3333 | 13 | 0.3846 |
| 6_2841689 | 6 | 0 | 1 | 1 | 3 | 0 | 2 | 0 | 12 | 0.0833 | 4 | 0.25 | 4 | 0.25 | 3 | 0.3333 | 3 | 0 | 14 | 0.2143 |
| 6_2841721 | 4 | 0.5 | 1 | 1 | 1 | 1 | 2 | 1 | 8 | 0.75 | 4 | 0.75 | 3 | 0.6667 | 1 | 0 | 2 | 1 | 10 | 0.7 |
| 6_2841744 | 6 | 0 | 1 | 1 | 3 | 0 | 1 | 0 | 11 | 0.0909 | 3 | 0 | 1 | 0 | 3 | 0.3333 | 2 | 0 | 9 | 0.1111 |
| 6_2841747 | 5 | 0.2 | 1 | 1 | 3 | 0 | 2 | 0 | 11 | 0.1818 | 4 | 0 | 3 | 0.3333 | 3 | 0.3333 | 3 | 0 | 13 | 0.1538 |
| 6_2841787 | 1 | 0 | NA | NA | 3 | 0.3333 | NA | NA | 4 | 0.25 | 2 | 0.5 | 1 | 0 | 3 | 0.6667 | 2 | 0.5 | 8 | 0.5 |
| 6_2841814 | 1 | 0 | NA | NA | 3 | 0 | 1 | 1 | 5 | 0.2 | 3 | 0 | 2 | 0.5 | 2 | 0.5 | NA | NA | 7 | 0.2857 |
| 6_2841818 | 2 | 1 | NA | NA | 2 | 1 | 1 | 1 | 5 | 1 | 3 | 1 | 1 | 1 | 1 | 1 | 1 | 1 | 6 | 1 |
| 6_2841843 | 2 | 0.5 | NA | NA | 3 | 0.6667 | 1 | 0 | 6 | 0.5 | 4 | 0.25 | 2 | 0 | 2 | 0.5 | 1 | 1 | 9 | 0.3333 |
| 6_2841853 | 2 | 1 | NA | NA | 3 | 0.6667 | 1 | 1 | 6 | 0.8333 | 3 | 1 | 2 | 0.5 | 1 | 1 | 1 | 1 | 7 | 0.8571 |
| 6_2841860 | 2 | 0.5 | NA | NA | 2 | 0.5 | NA | NA | 4 | 0.5 | 2 | 0.5 | 2 | 0 | 1 | 1 | 1 | 1 | 6 | 0.5 |
| 6_2841861 | 2 | 1 | NA | NA | 3 | 0.3333 | 1 | 1 | 6 | 0.6667 | 3 | 0.6667 | 2 | 0.5 | 1 | 0 | 1 | 0 | 7 | 0.4286 |
| 6_2841863 | 1 | 0 | NA | NA | 3 | 0.3333 | 1 | 0 | 5 | 0.2 | 2 | 0 | 1 | 0 | 1 | 0 | 1 | 1 | 5 | 0.2 |
| 6_2841866 | 2 | 1 | NA | NA | 1 | 1 | 1 | 1 | 4 | 1 | 2 | 0.5 | 2 | 1 | 1 | 0 | 1 | 1 | 6 | 0.6667 |
| 6_2841868 | 1 | 1 | NA | NA | 2 | 1 | 1 | 1 | 4 | 1 | 3 | 1 | 2 | 1 | 1 | 1 | 1 | 1 | 7 | 1 |
| 6_2841874 | 2 | 1 | NA | NA | 2 | 0.5 | NA | NA | 4 | 0.75 | 2 | 1 | 2 | 1 | 1 | 1 | 1 | 1 | 6 | 1 |
| 6_2841903 | 2 | 0.5 | NA | NA | 3 | 0.3333 | 1 | 0 | 6 | 0.3333 | 2 | 0 | 2 | 0.5 | 1 | 1 | 1 | 0 | 6 | 0.3333 |
| 6_2842072 | 1 | 1 | NA | NA | 2 | 0.5 | NA | NA | 3 | 0.6667 | 2 | 0.5 | 2 | 0.5 | NA | NA | 1 | 0 | 5 | 0.4 |
| 6_2842085 | 1 | 0 | NA | NA | 2 | 0.5 | NA | NA | 3 | 0.3333 | 1 | 0 | 1 | 0 | NA | NA | 1 | 1 | 3 | 0.3333 |
| 6_2842094 | 1 | 0 | NA | NA | 2 | 0.5 | NA | NA | 3 | 0.3333 | 2 | 0 | 2 | 1 | NA | NA | 1 | 1 | 5 | 0.6 |

| | | | | | | | | | | | | | | | | | | | | |
|---|---|---|---|---|---|---|---|---|---|---|---|---|---|---|---|---|---|---|---|---|
| 6_2842151 | NA | NA | NA | NA | 2 | 1 | NA | NA | 2 | 1 | 2 | 0.5 | 1 | 0 | NA | NA | NA | NA | 3 | 0.3333 |
| 6_2842622 | 30 | 0.1 | 34 | 0.0294 | 46 | 0.1087 | 21 | 0 | 131 | 0.0687 | 36 | 0.1111 | 40 | 0.025 | 41 | 0.0244 | 36 | 0.0833 | 153 | 0.0588 |
| 6_2842637 | 31 | 0.2581 | 33 | 0.2121 | 47 | 0.2979 | 25 | 0.2 | 136 | 0.25 | 37 | 0.4054 | 41 | 0.2195 | 40 | 0.175 | 33 | 0.3636 | 151 | 0.2848 |
| 6_2882900 | 2 | 0.5 | 1 | 0 | 3 | 0 | 1 | 0 | 7 | 0.1429 | 2 | 0.5 | 2 | 0 | 2 | 0 | 1 | 0 | 7 | 0.1429 |
| 6_2882957 | 3 | 0.6667 | 3 | 0.6667 | 4 | 0.5 | 3 | 0.3333 | 13 | 0.5385 | 1 | 0 | 3 | 1 | 3 | 1 | 2 | 0.5 | 9 | 0.7778 |
| 6_2882963 | 2 | 0.5 | 1 | 0 | 3 | 0.3333 | 3 | 0.6667 | 9 | 0.4444 | 1 | 1 | 3 | 0.3333 | 2 | 0 | 1 | 0 | 7 | 0.2857 |
| 6_2882968 | 2 | 1 | 2 | 0.5 | 4 | 0.5 | 2 | 0.5 | 10 | 0.6 | 1 | 0 | 3 | 1 | 4 | 0.5 | 2 | 0.5 | 10 | 0.6 |
| 6_2882970 | 2 | 0 | 2 | 0.5 | 4 | 0.5 | 3 | 0.3333 | 11 | 0.3636 | 1 | 1 | 3 | 0.3333 | 3 | 0.3333 | 2 | 0.5 | 9 | 0.4444 |
| 6_2882974 | 2 | 0 | 3 | 0.6667 | 4 | 0.25 | 3 | 0 | 12 | 0.25 | 1 | 0 | 3 | 0.3333 | 4 | 0 | 2 | 0.5 | 10 | 0.2 |
| 6_2882976 | 1 | 0 | 3 | 0.6667 | 3 | 0.6667 | 2 | 0.5 | 9 | 0.5556 | 1 | 1 | 3 | 0.3333 | 3 | 1 | 2 | 1 | 9 | 0.7778 |
| 6_2882992 | 1 | 1 | 1 | 1 | 4 | 1 | 3 | 0.6667 | 9 | 0.8889 | 1 | 1 | 4 | 1 | 4 | 1 | 2 | 1 | 11 | 1 |
| 6_2882997 | 1 | 1 | 1 | 0 | 3 | 0.6667 | 3 | 1 | 8 | 0.75 | NA | NA | 3 | 0.3333 | 3 | 0.6667 | 2 | 0.5 | 8 | 0.5 |
| 6_2883000 | 1 | 0 | 1 | 0 | 3 | 0.3333 | 3 | 0.6667 | 8 | 0.375 | 1 | 1 | 4 | 0.5 | 3 | 0.3333 | 1 | 0 | 9 | 0.4444 |
| 6_2883021 | 1 | 0 | NA | NA | 2 | 1 | 3 | 0.3333 | 6 | 0.5 | 1 | 0 | 3 | 0.6667 | 2 | 0.5 | 1 | 1 | 7 | 0.5714 |
| 6_2883036 | 1 | 1 | NA | NA | 2 | 1 | 3 | 1 | 6 | 1 | 1 | 1 | 4 | 0.75 | 2 | 1 | NA | NA | 7 | 0.8571 |
| 6_2883038 | 1 | 1 | NA | NA | 1 | 1 | 2 | 1 | 4 | 1 | 1 | 1 | 1 | 1 | 2 | 0.5 | NA | NA | 4 | 0.75 |
| 6_2883044 | NA | NA | NA | NA | 2 | 0 | 3 | 1 | 5 | 0.6 | 1 | 1 | 4 | 0.75 | 4 | 1 | NA | NA | 9 | 0.8889 |
| 6_2883049 | 1 | 1 | NA | NA | 1 | 1 | 2 | 1 | 4 | 1 | 1 | 1 | 3 | 1 | 3 | 1 | NA | NA | 7 | 1 |
| 6_2883050 | 1 | 1 | NA | NA | 2 | 1 | 3 | 0.6667 | 6 | 0.8333 | 1 | 1 | 2 | 0.5 | 4 | 0.5 | 1 | 1 | 8 | 0.625 |
| 6_2883070 | 1 | 1 | NA | NA | 2 | 1 | 2 | 1 | 5 | 1 | NA | NA | 3 | 1 | 1 | 1 | 1 | 1 | 5 | 1 |
| 6_2883080 | NA | NA | NA | NA | 2 | 0.5 | 2 | 0.5 | 4 | 0.5 | NA | NA | 3 | 1 | 3 | 0 | NA | NA | 6 | 0.5 |
| 6_2883082 | 1 | 1 | NA | NA | 2 | 1 | 2 | 1 | 5 | 1 | 1 | 1 | 3 | 1 | 2 | 1 | NA | NA | 6 | 1 |
| 6_2883086 | 1 | 1 | NA | NA | NA | NA | 1 | 1 | 2 | 1 | 1 | 0 | 1 | 1 | NA | NA | NA | NA | 2 | 0.5 |
| 6_2883087 | 1 | 1 | NA | NA | 1 | 1 | NA | NA | 2 | 1 | 1 | 1 | 3 | 1 | NA | NA | NA | NA | 4 | 1 |
| 6_2883101 | 1 | 1 | NA | NA | 1 | 1 | 2 | 0.5 | 4 | 0.75 | 1 | 0 | 3 | 1 | 2 | 0.5 | NA | NA | 6 | 0.6667 |
| 6_2883108 | NA | NA | NA | NA | 1 | 1 | 2 | 1 | 3 | 1 | 1 | 1 | 2 | 1 | 2 | 1 | NA | NA | 5 | 1 |
| 6_2883117 | NA | NA | NA | NA | 1 | 1 | 2 | 0 | 3 | 0.3333 | 1 | 1 | 2 | 0 | 3 | 0.3333 | NA | NA | 6 | 0.3333 |
| 6_2883121 | NA | NA | NA | NA | 1 | 1 | 2 | 0 | 3 | 0.3333 | 1 | 0 | 2 | 0.5 | 1 | 0 | NA | NA | 4 | 0.25 |
| 6_2883123 | NA | NA | NA | NA | NA | NA | 2 | 1 | 2 | 1 | 1 | 1 | 2 | 1 | 2 | 1 | NA | NA | 5 | 1 |
| 6_2883129 | NA | NA | NA | NA | 1 | 0 | 2 | 0.5 | 3 | 0.3333 | 1 | 0 | 3 | 0 | 2 | 0.5 | NA | NA | 6 | 0.1667 |
| 6_2883159 | NA | NA | NA | NA | 1 | 1 | 1 | 1 | 2 | 1 | 1 | 1 | 1 | 0 | 2 | 0.5 | NA | NA | 4 | 0.5 |
| 6_2883218 | NA | NA | NA | NA | 1 | 1 | 1 | 1 | 2 | 1 | 1 | 1 | 3 | 0.6667 | NA | NA | NA | NA | 4 | 0.75 |
| 6_2888860 | 11 | 0.3636 | 18 | 0.3333 | 8 | 0 | 13 | 0.1538 | 50 | 0.24 | 6 | 0.1667 | 7 | 0.2857 | 13 | 0.5385 | 8 | 0.25 | 34 | 0.3529 |
| 6_2912429 | 3 | 0.3333 | 2 | 1 | NA | NA | NA | NA | 5 | 0.6 | 3 | 0 | 1 | 0 | 5 | 0.8 | 1 | 0 | 10 | 0.4 |
| 6_2912473 | 4 | 0 | 1 | 0 | 1 | 1 | NA | NA | 6 | 0.1667 | 2 | 0.5 | 1 | 0 | 5 | 0.2 | 1 | 0 | 9 | 0.2222 |
| 6_2912497 | 5 | 0.2 | 1 | 0 | NA | NA | NA | NA | 6 | 0.1667 | 2 | 0 | 1 | 0 | 5 | 0.4 | 1 | 0 | 9 | 0.2222 |
| 6_2912531 | 8 | 1 | 4 | 1 | 2 | 1 | 2 | 0.5 | 16 | 0.9375 | 4 | 0.75 | 4 | 0.5 | 8 | 0.625 | 1 | 1 | 17 | 0.6471 |
| 6_2912537 | 7 | 0.7143 | 4 | 0.5 | 2 | 0 | 2 | 0 | 15 | 0.4667 | 3 | 1 | 3 | 0.6667 | 7 | 0.4286 | 1 | 0 | 14 | 0.5714 |
| 6_2915319 | NA | NA | 1 | 1 | NA | NA | NA | NA | 1 | 1 | 1 | 1 | NA | NA | 2 | 1 | NA | NA | 3 | 1 |
| 6_2915469 | 1 | 1 | NA | NA | NA | NA | NA | NA | 1 | 1 | 1 | 0 | NA | NA | 2 | 1 | NA | NA | 3 | 0.6667 |
| 6_2915478 | 2 | 0.5 | 1 | 0 | NA | NA | 1 | 0 | 4 | 0.25 | 2 | 0 | 1 | 0 | 5 | 0.6 | NA | NA | 8 | 0.375 |
| 6_2915479 | 2 | 0.5 | 1 | 0 | NA | NA | 1 | 0 | 4 | 0.25 | 2 | 0.5 | 2 | 1 | 5 | 0.2 | NA | NA | 9 | 0.4444 |
| 6_2916929 | NA | NA | NA | NA | 1 | 0 | 2 | 1 | 3 | 0.6667 | 3 | 0.6667 | NA | NA | 1 | 1 | NA | NA | 4 | 0.75 |
| 6_2916933 | NA | NA | NA | NA | 1 | 0 | 1 | 1 | 2 | 0.5 | 2 | 1 | NA | NA | NA | NA | NA | NA | 2 | 1 |
| 6_2916983 | NA | NA | NA | NA | 1 | 1 | 1 | 1 | 2 | 1 | 2 | 0.5 | NA | NA | 1 | 1 | NA | NA | 3 | 0.6667 |
| 6_2917000 | NA | NA | NA | NA | 1 | 0 | 1 | 1 | 2 | 0.5 | 2 | 0.5 | NA | NA | 1 | 1 | NA | NA | 3 | 0.6667 |
| 6_2917001 | NA | NA | NA | NA | 1 | 1 | 2 | 1 | 3 | 1 | 3 | 1 | NA | NA | 1 | 1 | 1 | 1 | 5 | 1 |
| 6_2917029 | NA | NA | NA | NA | 1 | 1 | 1 | 1 | 2 | 1 | 3 | 0.6667 | NA | NA | 1 | 1 | NA | NA | 4 | 0.75 |
| 6_2917047 | NA | NA | NA | NA | NA | NA | 1 | 1 | 1 | 1 | 3 | 0.3333 | NA | NA | 1 | 1 | NA | NA | 4 | 0.5 |
| 6_2917144 | NA | NA | NA | NA | NA | NA | 1 | 1 | 1 | 1 | 2 | 1 | NA | NA | NA | NA | NA | NA | 2 | 1 |
| 6_2927498 | 2 | 1 | 3 | 1 | 1 | 1 | 1 | 1 | 7 | 1 | NA | NA | 2 | 1 | 1 | 1 | NA | NA | 3 | 1 |
| 6_2927515 | 2 | 1 | 3 | 1 | 1 | 1 | 1 | 1 | 7 | 1 | NA | NA | 2 | 1 | 1 | 1 | 1 | 1 | 4 | 1 |
| 6_2927516 | 2 | 1 | 3 | 1 | 1 | 1 | 1 | 1 | 7 | 1 | NA | NA | 2 | 0.5 | 1 | 1 | 1 | 0 | 4 | 0.5 |
| 6_2927524 | 2 | 0.5 | 3 | 0 | 1 | 0 | 1 | 0 | 7 | 0.1429 | 1 | 1 | 2 | 0 | 1 | 0 | 1 | 0 | 5 | 0.2 |
| 6_2927526 | 2 | 0.5 | 3 | 0.6667 | 1 | 1 | 1 | 1 | 7 | 0.7143 | 1 | 1 | 2 | 0.5 | 1 | 0 | 1 | 1 | 5 | 0.6 |
| 6_2937458 | 5 | 1 | 7 | 0.7143 | 7 | 0.8571 | 8 | 0.75 | 27 | 0.8148 | 5 | 0.8 | 3 | 1 | 6 | 1 | 10 | 1 | 24 | 0.9583 |
| 6_2937459 | 4 | 0.25 | 6 | 0.8333 | 7 | 0.1429 | 7 | 0.2857 | 24 | 0.375 | 5 | 0.4 | 3 | 0 | 6 | 0.5 | 9 | 0.6667 | 23 | 0.4783 |
| 6_2937547 | 4 | 0 | 6 | 0.1667 | 6 | 0.1667 | 5 | 0 | 21 | 0.0952 | 4 | 0.25 | 1 | 0 | 6 | 0 | 10 | 0.2 | 21 | 0.1429 |
| 6_2940609 | 6 | 0 | 5 | 0 | 4 | 0 | 6 | 0.3333 | 21 | 0.0952 | 5 | 0.2 | 1 | 0 | 3 | 0.3333 | 4 | 0 | 13 | 0.1538 |
| 6_2942344 | 2 | 0 | NA | NA | 1 | 1 | 1 | 0 | 4 | 0.25 | NA | NA | 1 | 0 | NA | NA | 1 | 1 | 2 | 0.5 |
| 6_2942685 | 10 | 0.6 | 12 | 0.4167 | 4 | 0.5 | 9 | 0.3333 | 35 | 0.4571 | 10 | 0.5 | 3 | 0 | 5 | 0.8 | 8 | 0.375 | 26 | 0.4615 |
| 6_2947021 | 39 | 0.5385 | 27 | 0.6667 | 29 | 0.3448 | 18 | 0.3333 | 113 | 0.4867 | 19 | 0.4211 | 32 | 0.4062 | 30 | 0.4 | 35 | 0.5714 | 116 | 0.4569 |
| 6_2954162 | 1 | 0 | 5 | 0.2 | 1 | 0 | 2 | 0 | 9 | 0.1111 | 5 | 0 | 3 | 0 | 6 | 0 | 5 | 0.2 | 19 | 0.0526 |
| 6_2954182 | 1 | 1 | 6 | 0 | 1 | 1 | 2 | 0 | 10 | 0.2 | 4 | 0 | 3 | 0 | 7 | 0.1429 | 5 | 0 | 19 | 0.0526 |
| 6_2971768 | 1 | 1 | 3 | 0.3333 | NA | NA | NA | NA | 4 | 0.5 | NA | NA | 1 | 0 | 2 | 0.5 | NA | NA | 3 | 0.3333 |

| | | | | | | | | | | | | | | | | | | | | |
|---|---|---|---|---|---|---|---|---|---|---|---|---|---|---|---|---|---|---|---|---|
| 6_2971787 | 1 | 1 | 2 | 0.5 | NA | NA | NA | NA | 3 | 0.6667 | 1 | 1 | 1 | 0 | 2 | 1 | NA | NA | 4 | 0.75 |
| 6_2971797 | 2 | 0.5 | 1 | 1 | NA | NA | NA | NA | 3 | 0.6667 | 1 | 0 | 2 | 0 | 2 | 0.5 | NA | NA | 5 | 0.2 |
| 6_2971800 | 2 | 0 | 1 | 1 | NA | NA | NA | NA | 3 | 0.3333 | 1 | 0 | 2 | 0.5 | 2 | 0.5 | NA | NA | 5 | 0.4 |
| 6_2971837 | 2 | 0.5 | 2 | 0 | NA | NA | NA | NA | 4 | 0.25 | 2 | 0.5 | 2 | 0.5 | 3 | 0.6667 | NA | NA | 7 | 0.5714 |
| 6_2976011 | NA | NA | 1 | 1 | 1 | 1 | 5 | 1 | 7 | 1 | NA | NA | 1 | 1 | 3 | 1 | 1 | 1 | 5 | 1 |
| 6_2976027 | NA | NA | NA | NA | 1 | 0 | 4 | 0.25 | 5 | 0.2 | NA | NA | 2 | 1 | 2 | 1 | 1 | 1 | 5 | 1 |
| 6_2976863 | 2 | 1 | 1 | 1 | 1 | 1 | NA | NA | 4 | 1 | 1 | 1 | 2 | 0.5 | 5 | 0.6 | NA | NA | 8 | 0.625 |
| 6_2990984 | 6 | 0.1667 | 14 | 0.1429 | 3 | 0 | 1 | 1 | 24 | 0.1667 | 2 | 0.5 | 4 | 0 | 6 | 0.1667 | 5 | 0 | 17 | 0.1176 |
| 6_2994535 | 1 | 1 | NA | NA | NA | NA | NA | NA | 1 | 1 | NA | NA | NA | NA | 2 | 0.5 | NA | NA | 2 | 0.5 |
| 6_2995675 | 12 | 0.75 | 7 | 1 | 9 | 1 | 13 | 0.8462 | 41 | 0.878 | 9 | 1 | 15 | 0.8667 | 13 | 0.9231 | 12 | 0.75 | 49 | 0.8776 |
| 6_2995698 | 9 | 0.8889 | 4 | 1 | 8 | 0.875 | 8 | 1 | 29 | 0.931 | 4 | 1 | 6 | 1 | 7 | 0.8571 | 4 | 1 | 21 | 0.9524 |
| 6_2995703 | 13 | 0.1538 | 8 | 0.125 | 10 | 0.1 | 11 | 0.2727 | 42 | 0.1667 | 6 | 0.3333 | 11 | 0.0909 | 10 | 0.4 | 9 | 0.3333 | 36 | 0.2778 |
| 6_2995704 | 11 | 0.0909 | 6 | 0 | 9 | 0 | 12 | 0.3333 | 38 | 0.1316 | 7 | 0.1429 | 12 | 0.1667 | 11 | 0.2727 | 11 | 0.0909 | 41 | 0.1707 |
| 6_2995708 | 11 | 0.7273 | 6 | 1 | 10 | 0.9 | 10 | 1 | 37 | 0.8919 | 7 | 1 | 10 | 0.9 | 8 | 0.875 | 8 | 1 | 33 | 0.9394 |
| 6_2995710 | 14 | 0.7857 | 8 | 0.75 | 12 | 0.4167 | 15 | 0.4667 | 49 | 0.5918 | 8 | 0.625 | 14 | 0.7143 | 14 | 0.4286 | 10 | 0.5 | 46 | 0.5652 |
| 6_2995719 | 12 | 0.8333 | 4 | 0.5 | 8 | 0.5 | 9 | 0.6667 | 33 | 0.6667 | 4 | 0.5 | 12 | 0.6667 | 10 | 0.6 | 7 | 0.5714 | 33 | 0.6061 |
| 6_2995719 | 12 | 0.8333 | 6 | 1 | 9 | 0.7778 | 16 | 0.875 | 43 | 0.8605 | 8 | 0.875 | 16 | 0.8125 | 14 | 0.7857 | 11 | 0.9091 | 49 | 0.8367 |
| 6_2995723 | 12 | 0.0833 | 8 | 0.25 | 11 | 0.1818 | 16 | 0.25 | 47 | 0.1915 | 8 | 0.125 | 17 | 0.1765 | 14 | 0.1429 | 12 | 0.0833 | 51 | 0.1373 |
| 6_2995731 | 14 | 0.6429 | 8 | 0.625 | 10 | 0.4 | 14 | 0.3571 | 46 | 0.5 | 9 | 0.4444 | 15 | 0.6 | 12 | 0.6667 | 14 | 0.3571 | 50 | 0.52 |
| 6_2995739 | 16 | 0.9375 | 7 | 0.8571 | 9 | 1 | 13 | 1 | 45 | 0.9556 | 7 | 1 | 14 | 0.8571 | 14 | 0.9286 | 12 | 0.9167 | 47 | 0.9149 |
| 6_2995749 | 16 | 0.8125 | 6 | 0.6667 | 10 | 0.5 | 15 | 0.6667 | 47 | 0.6809 | 7 | 0.8571 | 9 | 0.8889 | 13 | 0.9231 | 7 | 1 | 36 | 0.9167 |
| 6_2995767 | 15 | 0.6 | 8 | 0.75 | 17 | 0.5294 | 14 | 0.5 | 54 | 0.5741 | 7 | 0.4286 | 17 | 0.4706 | 16 | 0.5625 | 9 | 0.5556 | 49 | 0.5102 |
| 6_2995778 | 16 | 0.25 | 8 | 0.125 | 12 | 0.1667 | 15 | 0.4 | 51 | 0.2549 | 7 | 0.2857 | 16 | 0.3125 | 12 | 0.1667 | 13 | 0.2308 | 48 | 0.25 |
| 6_2995787 | 19 | 0 | 11 | 0.0909 | 17 | 0.0588 | 17 | 0.1176 | 64 | 0.0625 | 11 | 0.0909 | 16 | 0.0625 | 15 | 0.0667 | 13 | 0.0769 | 55 | 0.0727 |
| 6_2995792 | 15 | 0.8 | 10 | 0.5 | 14 | 0.7857 | 16 | 0.9375 | 55 | 0.7818 | 11 | 0.8182 | 15 | 0.6667 | 17 | 0.8824 | 15 | 0.8 | 58 | 0.7931 |
| 6_2995814 | 11 | 0.8182 | 8 | 0.875 | 13 | 0.7692 | 11 | 0.9091 | 43 | 0.8372 | 10 | 0.9 | 11 | 0.9091 | 17 | 0.9412 | 13 | 1 | 51 | 0.9412 |
| 6_2995818 | 19 | 0.2632 | 11 | 0.1818 | 14 | 0.1429 | 18 | 0.1667 | 62 | 0.1935 | 11 | 0.1818 | 17 | 0.1765 | 18 | 0.1111 | 13 | 0.2308 | 59 | 0.1695 |
| 6_2995843 | 15 | 0.3333 | 11 | 0.2727 | 16 | 0.5 | 18 | 0.4444 | 60 | 0.4 | 9 | 0.3333 | 16 | 0.4375 | 16 | 0.5 | 13 | 0.4615 | 54 | 0.4444 |
| 6_2995883 | 20 | 0.2 | 12 | 0.25 | 17 | 0.1176 | 23 | 0.3478 | 72 | 0.2361 | 11 | 0.1818 | 19 | 0.2632 | 22 | 0.2273 | 13 | 0.3077 | 65 | 0.2462 |
| 6_2995958 | 32 | 0.0938 | 22 | 0.0909 | 24 | 0.0417 | 30 | 0.0667 | 108 | 0.0741 | 19 | 0.1053 | 24 | 0.1667 | 34 | 0.1176 | 18 | 0.0556 | 95 | 0.1158 |
| 6_3014047 | NA | NA | NA | NA | NA | NA | 1 | 1 | 1 | 1 | NA | NA | NA | NA | NA | NA | 1 | 1 | 1 | 1 |
| 6_3014077 | NA | NA | NA | NA | NA | NA | 1 | 1 | 1 | 1 | NA | NA | NA | NA | NA | NA | 1 | 1 | 1 | 1 |
| 6_3015191 | 1 | 0 | 2 | 1 | 2 | 1 | 2 | 0.5 | 7 | 0.7143 | 1 | 1 | NA | NA | 2 | 0.5 | 1 | 0 | 4 | 0.5 |
| 6_3015208 | 1 | 0 | 1 | 1 | 2 | 1 | 2 | 1 | 6 | 0.8333 | 1 | 1 | NA | NA | 2 | 1 | NA | NA | 3 | 1 |
| 6_3015219 | NA | NA | 1 | 1 | 1 | 1 | 2 | 1 | 4 | 1 | 1 | 0 | NA | NA | 2 | 1 | NA | NA | 3 | 0.6667 |
| 6_3015224 | 1 | 0 | 2 | 0.5 | 2 | 0 | 1 | 0 | 6 | 0.1667 | 1 | 0 | NA | NA | 2 | 0.5 | NA | NA | 3 | 0.3333 |
| 6_3015232 | 1 | 1 | 2 | 1 | 2 | 1 | 2 | 1 | 7 | 1 | NA | NA | NA | NA | 2 | 1 | NA | NA | 2 | 1 |
| 6_3015236 | 1 | 0 | 2 | 0.5 | 1 | 0 | 2 | 0.5 | 6 | 0.3333 | 1 | 0 | NA | NA | 2 | 0.5 | 1 | 0 | 4 | 0.25 |
| 6_3015277 | 1 | 0 | 3 | 0.3333 | 2 | 0.5 | 2 | 0 | 8 | 0.25 | 3 | 0.3333 | NA | NA | 2 | 0.5 | NA | NA | 5 | 0.4 |
| 6_3015281 | 1 | 1 | 4 | 0 | 2 | 0.5 | 2 | 0.5 | 9 | 0.3333 | 3 | 0 | NA | NA | 2 | 0.5 | NA | NA | 5 | 0.2 |
| 6_3015282 | 1 | 1 | 3 | 1 | 1 | 1 | 2 | 1 | 7 | 1 | 3 | 1 | NA | NA | 1 | 1 | NA | NA | 4 | 1 |
| 6_3015286 | 1 | 0 | 4 | 1 | 1 | 1 | 2 | 1 | 8 | 0.875 | 2 | 1 | NA | NA | 2 | 1 | 1 | 1 | 5 | 1 |
| 6_3015300 | 1 | 1 | 4 | 0.5 | 2 | 1 | 2 | 1 | 9 | 0.7778 | 2 | 1 | NA | NA | 2 | 0.5 | 1 | 0 | 5 | 0.6 |
| 6_3021174 | 9 | 0 | 13 | 0.1538 | 14 | 0.0714 | 6 | 0.1667 | 42 | 0.0952 | 10 | 0.1 | 13 | 0.1538 | 12 | 0 | 12 | 0.1667 | 47 | 0.1064 |
| 6_3021307 | 9 | 0.1111 | 8 | 0.125 | 9 | 0.1111 | 6 | 0 | 32 | 0.0938 | 5 | 0 | 14 | 0.1429 | 8 | 0 | 11 | 0 | 38 | 0.0526 |
| 6_3021353 | 4 | 0 | 5 | 0 | 6 | 0.1667 | 2 | 0 | 17 | 0.0588 | NA | NA | 6 | 0.5 | 6 | 0.5 | 11 | 0.1818 | 23 | 0.3478 |
| 6_3021365 | 7 | 0 | 7 | 0.2857 | 6 | 0 | 4 | 0 | 24 | 0.0833 | 3 | 0 | 9 | 0.1111 | 8 | 0 | 10 | 0.1 | 30 | 0.0667 |
| 6_3021375 | 8 | 0.25 | 8 | 0 | 5 | 0 | 6 | 0.1667 | 27 | 0.1111 | 3 | 0 | 10 | 0.1 | 9 | 0.1111 | 8 | 0.25 | 30 | 0.1333 |
| 6_3021397 | 6 | 0 | 9 | 0 | 7 | 0.2857 | 6 | 0.1667 | 28 | 0.1071 | 2 | 0 | 11 | 0 | 8 | 0.125 | 9 | 0.1111 | 30 | 0.0667 |
| 6_3021399 | 7 | 0.1429 | 7 | 0.2857 | 6 | 0 | 4 | 0 | 24 | 0.125 | 3 | 0 | 10 | 0.3 | 7 | 0 | 9 | 0.2222 | 29 | 0.1724 |
| 6_3021400 | 7 | 0 | 7 | 0.4286 | 7 | 0.2857 | 4 | 0 | 25 | 0.2 | 3 | 0.3333 | 11 | 0.3636 | 6 | 0 | 9 | 0 | 29 | 0.1724 |
| 6_3021402 | 4 | 0.5 | 7 | 0.5714 | 7 | 0.5714 | 5 | 0.4 | 23 | 0.5217 | 2 | 0.5 | 6 | 0.6667 | 5 | 0.4 | 8 | 0.75 | 21 | 0.619 |
| 6_3021413 | 6 | 1 | 9 | 0.8889 | 4 | 1 | 4 | 1 | 23 | 0.9565 | 1 | 1 | 5 | 1 | 6 | 1 | 6 | 1 | 18 | 1 |
| 6_3021421 | 6 | 0.3333 | 7 | 0.2857 | 5 | 0.2 | 6 | 1 | 24 | 0.4583 | 2 | 1 | 6 | 0.5 | 5 | 0.6 | 5 | 0.6 | 18 | 0.6111 |
| 6_3021429 | 6 | 0.5 | 7 | 0.7143 | 5 | 0.8 | 5 | 0.8 | 23 | 0.6957 | 1 | 0 | 6 | 0.6667 | 6 | 0.8333 | 5 | 0.4 | 18 | 0.6111 |
| 6_3021441 | 6 | 0 | 8 | 0 | 3 | 0 | 4 | 0.5 | 21 | 0.0952 | 2 | 0 | 7 | 0 | 7 | 0.1429 | 6 | 0.1667 | 22 | 0.0909 |
| 6_3021455 | 2 | 0 | 7 | 0 | 4 | 0 | 4 | 0.25 | 17 | 0.0588 | 1 | 0 | 6 | 0.1667 | 4 | 0 | 5 | 0.2 | 16 | 0.125 |
| 6_3021467 | 5 | 0.6 | 7 | 0.2857 | 4 | 0.75 | 4 | 0.5 | 20 | 0.5 | 2 | 1 | 4 | 0.5 | 6 | 1 | 5 | 0.8 | 17 | 0.8235 |
| 6_3021490 | 7 | 0.4286 | 7 | 0.2857 | 4 | 0 | 4 | 0 | 22 | 0.2273 | 2 | 0 | 6 | 0.6667 | 5 | 0.2 | 6 | 0.5 | 19 | 0.4211 |
| 6_3021493 | 6 | 1 | 7 | 0.8571 | 4 | 0.75 | 4 | 0.75 | 21 | 0.8571 | 1 | 1 | 7 | 1 | 5 | 1 | 4 | 1 | 17 | 1 |
| 6_3021497 | 7 | 0.1429 | 6 | 0 | 4 | 0 | 3 | 0 | 20 | 0.05 | 2 | 0.5 | 5 | 0 | 4 | 0 | 6 | 0 | 17 | 0.0588 |
| 6_3036665 | 6 | 0.8333 | 6 | 0.6667 | 3 | 0.6667 | 5 | 0.4 | 20 | 0.65 | 3 | 0.6667 | 5 | 0.8 | 5 | 0.4 | 5 | 0.2 | 18 | 0.5 |
| 6_3036665 | 5 | 0 | 6 | 0.1667 | 4 | 0 | 5 | 0.2 | 20 | 0.1 | 3 | 0.3333 | 4 | 0.25 | 4 | 0.25 | 4 | 0.5 | 15 | 0.3333 |
| 6_3045054 | 3 | 0.6667 | 2 | 0 | 4 | 0.5 | 3 | 0.3333 | 12 | 0.4167 | 5 | 0.2 | 4 | 0.25 | 2 | 0 | 3 | 1 | 14 | 0.3571 |
| 6_3045058 | 2 | 0 | 4 | 0.25 | 3 | 0.6667 | 1 | 1 | 10 | 0.4 | 4 | 0.25 | 4 | 0.75 | 1 | 1 | 3 | 0.3333 | 12 | 0.5 |

| | | | | | | | | | | | | | | | | | | | | |
|---|---|---|---|---|---|---|---|---|---|---|---|---|---|---|---|---|---|---|---|---|
| 6_3045066 | 2 | 0.5 | 3 | 0.3333 | 2 | 1 | 3 | 0.3333 | 10 | 0.5 | 3 | 0.3333 | 3 | 0.6667 | 2 | 0.5 | 3 | 0.6667 | 11 | 0.5455 |
| 6_3045070 | 3 | 1 | 3 | 0.6667 | 4 | 0.5 | 3 | 1 | 13 | 0.7692 | 4 | 1 | 4 | 1 | 2 | 1 | 3 | 1 | 13 | 1 |
| 6_3045074 | 3 | 0.6667 | 4 | 0.75 | 3 | 0.6667 | 3 | 0.6667 | 13 | 0.6923 | 5 | 0.8 | 2 | 1 | 2 | 1 | 3 | 1 | 12 | 0.9167 |
| 6_3045085 | 3 | 0.3333 | 3 | 0 | 4 | 0.75 | 3 | 0.6667 | 13 | 0.4615 | 5 | 0.6 | 4 | 1 | 2 | 1 | 3 | 0.3333 | 14 | 0.7143 |
| 6_3045121 | 3 | 0 | 2 | 0.5 | 2 | 0 | 2 | 0 | 9 | 0.1111 | 4 | 0 | 2 | 0.5 | 2 | 0.5 | 6 | 0.3333 | 14 | 0.2857 |
| 6_3045248 | 2 | 0 | 4 | 0.25 | 3 | 0 | 3 | 0 | 12 | 0.0833 | 5 | 0 | 2 | 0.5 | 3 | 0 | 5 | 0 | 15 | 0.0667 |
| 6_3048261 | 5 | 1 | 1 | 0 | 2 | 0.5 | 5 | 1 | 13 | 0.8462 | NA | NA | 4 | 0.5 | 2 | 1 | 4 | 0.75 | 10 | 0.7 |
| 6_3048264 | 3 | 0 | NA | NA | 2 | 0 | 5 | 0.2 | 10 | 0.1 | NA | NA | 3 | 0.3333 | 3 | 0 | 3 | 0.3333 | 9 | 0.2222 |
| 6_3048281 | 5 | 1 | 1 | 1 | 2 | 0.5 | 5 | 0.6 | 13 | 0.7692 | NA | NA | 4 | 0.5 | 2 | 0.5 | 4 | 0.75 | 10 | 0.6 |
| 6_3048290 | 4 | 0.25 | 1 | 1 | 3 | 0.6667 | 4 | 1 | 12 | 0.6667 | NA | NA | 4 | 1 | 3 | 1 | 1 | 1 | 8 | 1 |
| 6_3074807 | 2 | 1 | 2 | 0.5 | 1 | 0 | NA | NA | 5 | 0.6 | NA | NA | 2 | 0.5 | 3 | 0.3333 | NA | NA | 5 | 0.4 |
| 6_3074816 | 4 | 0.75 | 2 | 0 | 2 | 1 | 1 | 0 | 9 | 0.5556 | 1 | 0 | 2 | 1 | 3 | 0.6667 | NA | NA | 6 | 0.6667 |
| 6_3074826 | 2 | 0 | 2 | 0 | 1 | 0 | 3 | 0.3333 | 8 | 0.125 | 3 | 0.3333 | 5 | 0 | 5 | 0 | 1 | 0 | 14 | 0.0714 |
| 6_3074875 | 11 | 0.1818 | 11 | 0.0909 | 12 | 0 | 6 | 0 | 40 | 0.075 | 4 | 0 | 6 | 0.5 | 13 | 0.0769 | 7 | 0 | 30 | 0.1333 |
| 6_3079300 | 6 | 0.3333 | 6 | 0.1667 | 6 | 0.3333 | 1 | 0 | 19 | 0.2632 | 3 | 0.6667 | 5 | 0.2 | 10 | 0.3 | 6 | 0.1667 | 24 | 0.2917 |
| 6_3131424 | 8 | 0.375 | 5 | 0.4 | 3 | 0.3333 | 3 | 0 | 19 | 0.3158 | 5 | 0.2 | 5 | 0.2 | 2 | 0.5 | 3 | 0.3333 | 15 | 0.2667 |
| 6_3148459 | 2 | 0 | 8 | 0.375 | 4 | 0.75 | 6 | 0 | 20 | 0.3 | 5 | 0 | 4 | 0.5 | 5 | 0.4 | 4 | 0.25 | 18 | 0.2778 |
| 6_3148460 | 2 | 0 | 7 | 0 | 2 | 0 | 4 | 0.25 | 15 | 0.0667 | 5 | 0 | 4 | 0 | 5 | 0.2 | 4 | 0 | 18 | 0.0556 |
| 6_3148468 | 1 | 1 | 5 | 0.6 | 3 | 0.6667 | 6 | 0.3333 | 15 | 0.5333 | 3 | 0.3333 | 4 | 0.25 | 4 | 0.75 | 5 | 0 | 16 | 0.3125 |
| 6_3149214 | 2 | 0.5 | 5 | 0 | 2 | 0 | 2 | 0 | 11 | 0.0909 | 1 | 0 | 1 | 1 | 4 | 0.25 | 4 | 0 | 10 | 0.2 |
| 6_3149295 | 2 | 0.5 | 4 | 0.25 | 1 | 0 | 2 | 0 | 9 | 0.2222 | 3 | 0.3333 | NA | NA | 4 | 0.25 | 3 | 0.3333 | 10 | 0.3 |
| 6_3149301 | 2 | 0 | 5 | 0 | 1 | 0 | 1 | 1 | 9 | 0.1111 | 2 | 0.5 | 2 | 1 | 2 | 0.5 | 2 | 0.5 | 8 | 0.625 |
| 6_3149302 | 2 | 0.5 | 7 | 0.1429 | 2 | 0 | 3 | 0.6667 | 14 | 0.2857 | 3 | 0.3333 | 2 | 1 | 3 | 0.6667 | 4 | 0.25 | 12 | 0.5 |
| 6_3149311 | NA | NA | 6 | 0.5 | 2 | 0 | 2 | 0.5 | 10 | 0.4 | 3 | 0.3333 | 1 | 1 | 4 | 0.75 | 3 | 0.6667 | 11 | 0.6364 |
| 6_3149324 | 2 | 0.5 | 5 | 0 | 3 | 0 | 4 | 0 | 14 | 0.0714 | 3 | 0 | 2 | 0 | 6 | 0.1667 | 4 | 0.25 | 15 | 0.1333 |
| 6_3149325 | 1 | 0 | 4 | 0.5 | 3 | 0.3333 | 3 | 0.6667 | 11 | 0.4545 | 3 | 0 | 2 | 1 | 6 | 0.3333 | 4 | 0 | 15 | 0.2667 |
| 6_3149338 | 2 | 0 | 7 | 0.1429 | 3 | 0 | 4 | 0 | 16 | 0.0625 | 3 | 0.3333 | 1 | 0 | 6 | 0 | 4 | 0.25 | 14 | 0.1429 |
| 6_3149343 | 2 | 0 | 5 | 0.2 | 2 | 0 | 3 | 0.3333 | 12 | 0.1667 | 3 | 0.3333 | 2 | 0.5 | 5 | 0.2 | 3 | 0.3333 | 13 | 0.3077 |
| 6_3149355 | 2 | 1 | 6 | 0.3333 | 3 | 1 | 3 | 1 | 14 | 0.7143 | 3 | 0.3333 | 1 | 1 | 5 | 0.8 | 4 | 0.75 | 13 | 0.6923 |
| 6_3149673 | 1 | 1 | 3 | 0 | 5 | 0.2 | 1 | 0 | 10 | 0.2 | 6 | 0.8333 | 2 | 0.5 | 2 | 0.5 | NA | NA | 10 | 0.7 |
| 6_3149983 | NA | NA | 1 | 1 | 4 | 1 | NA | NA | 5 | 1 | 2 | 0.5 | NA | NA | 1 | 1 | 1 | 1 | 4 | 0.75 |
| 6_3149985 | NA | NA | NA | NA | 3 | 1 | NA | NA | 3 | 1 | 2 | 1 | NA | NA | NA | NA | NA | NA | 2 | 1 |
| 6_3150565 | NA | NA | NA | NA | 2 | 0.5 | NA | NA | 2 | 0.5 | 1 | 0 | NA | NA | NA | NA | 1 | 1 | 2 | 0.5 |
| 6_3150568 | NA | NA | NA | NA | 2 | 1 | NA | NA | 2 | 1 | NA | NA | NA | NA | NA | NA | 1 | 1 | 1 | 1 |
| 6_3178109 | 1 | 1 | NA | NA | NA | NA | NA | NA | 1 | 1 | NA | NA | NA | NA | 1 | 1 | NA | NA | 1 | 1 |
| 6_3178117 | 1 | 1 | NA | NA | NA | NA | NA | NA | 1 | 1 | NA | NA | NA | NA | 1 | 1 | NA | NA | 1 | 1 |
| 6_3212184 | 8 | 0.125 | 9 | 0.2222 | 6 | 0 | 7 | 0.1429 | 30 | 0.1333 | 2 | 0 | 6 | 0.1667 | 6 | 0.1667 | 3 | 0 | 17 | 0.1176 |
| 6_3212329 | 5 | 0.2 | 9 | 0.1111 | 3 | 0.3333 | 5 | 0.2 | 22 | 0.1818 | 1 | 0 | 4 | 0 | 3 | 0 | 4 | 0.25 | 12 | 0.0833 |
| 6_3212545 | 4 | 1 | 2 | 1 | 1 | 1 | 1 | 1 | 8 | 1 | NA | NA | 1 | 1 | 1 | 0 | 3 | 0.6667 | 5 | 0.6 |
| 6_3212550 | 4 | 0.5 | 2 | 0 | 1 | 0 | 1 | 0 | 8 | 0.25 | NA | NA | 2 | 0 | 3 | 0 | 4 | 0.5 | 9 | 0.2222 |
| 6_3212567 | 3 | 1 | 2 | 0 | 1 | 0 | 1 | 0 | 7 | 0.4286 | NA | NA | 2 | 0.5 | 2 | 0.5 | 3 | 0.6667 | 7 | 0.5714 |
| 6_3212607 | 3 | 1 | NA | NA | NA | NA | NA | NA | 3 | 1 | NA | NA | 1 | 0 | 1 | 0 | 2 | 1 | 4 | 0.5 |
| 6_3212609 | 3 | 0.6667 | 2 | 1 | NA | NA | NA | NA | 5 | 0.8 | NA | NA | 1 | 1 | 2 | 1 | 3 | 1 | 6 | 1 |
| 6_3212611 | 3 | 1 | 1 | 1 | NA | NA | NA | NA | 4 | 1 | NA | NA | 1 | 1 | 2 | 1 | 3 | 1 | 6 | 1 |
| 6_3212630 | 1 | 1 | 2 | 0.5 | NA | NA | NA | NA | 3 | 0.6667 | NA | NA | 1 | 1 | 2 | 1 | 2 | 1 | 5 | 1 |
| 6_3212637 | 2 | 1 | 2 | 1 | NA | NA | NA | NA | 4 | 1 | NA | NA | 1 | 1 | 2 | 1 | 1 | 0 | 4 | 0.75 |
| 6_3212674 | 2 | 1 | 1 | 1 | NA | NA | NA | NA | 3 | 1 | NA | NA | NA | NA | 1 | 0 | 2 | 0.5 | 3 | 0.3333 |
| 6_3212685 | 2 | 0.5 | 2 | 0.5 | NA | NA | NA | NA | 4 | 0.5 | NA | NA | 1 | 0 | 2 | 0.5 | 2 | 0 | 5 | 0.2 |
| 6_3212685 | 2 | 1 | 2 | 1 | NA | NA | NA | NA | 4 | 1 | NA | NA | 1 | 1 | 1 | 1 | 2 | 1 | 4 | 1 |
| 6_3212689 | 2 | 1 | 2 | 0.5 | NA | NA | NA | NA | 4 | 0.75 | NA | NA | 1 | 1 | 2 | 1 | 2 | 1 | 5 | 1 |
| 6_3212707 | 2 | 1 | 1 | 1 | NA | NA | NA | NA | 3 | 1 | NA | NA | 1 | 0 | 2 | 1 | 1 | 0 | 4 | 0.5 |
| 6_3212720 | 2 | 0 | 2 | 0.5 | NA | NA | NA | NA | 4 | 0.25 | NA | NA | 1 | 0 | 1 | 1 | 1 | 0 | 3 | 0.3333 |
| 6_3212721 | NA | NA | 2 | 0.5 | NA | NA | NA | NA | 2 | 0.5 | NA | NA | 1 | 0 | 2 | 1 | 1 | 0 | 4 | 0.5 |
| 6_3212742 | NA | NA | 1 | 1 | NA | NA | NA | NA | 1 | 1 | NA | NA | 1 | 1 | 1 | 1 | NA | NA | 2 | 1 |
| 6_3276915 | 1 | 0 | 4 | 0.25 | 5 | 0 | 4 | 0.25 | 14 | 0.1429 | 1 | 0 | 4 | 0 | 7 | 0.1429 | 3 | 0 | 15 | 0.0667 |
| 6_3276939 | 12 | 0.3333 | 11 | 0.4545 | 16 | 0.1875 | 11 | 0.5455 | 50 | 0.36 | 5 | 0.2 | 12 | 0.1667 | 13 | 0.2308 | 10 | 0.4 | 40 | 0.25 |
| 6_3276946 | 12 | 0.25 | 10 | 0.4 | 13 | 0.3846 | 9 | 0.3333 | 44 | 0.3409 | 5 | 0.4 | 12 | 0.4167 | 14 | 0.6429 | 7 | 0.2857 | 38 | 0.4737 |
| 6_3276949 | 14 | 0.3571 | 10 | 0.3 | 16 | 0.1875 | 12 | 0.1667 | 52 | 0.25 | 4 | 0 | 11 | 0 | 13 | 0.1538 | 10 | 0.2 | 38 | 0.1053 |
| 6_3276963 | 18 | 0.1667 | 9 | 0.2222 | 17 | 0.1765 | 11 | 0.0909 | 55 | 0.1636 | 5 | 0 | 12 | 0 | 15 | 0.1333 | 7 | 0 | 39 | 0.0513 |
| 6_3276971 | 17 | 0.7647 | 12 | 0.9167 | 18 | 0.7778 | 15 | 0.7333 | 62 | 0.7903 | 6 | 0.8333 | 11 | 0.8182 | 15 | 1 | 9 | 0.8889 | 41 | 0.9024 |
| 6_3394901 | 1 | 0 | 4 | 0.5 | 3 | 0.6667 | 1 | 0 | 9 | 0.4444 | 3 | 0.3333 | 2 | 0 | 4 | 0.25 | 3 | 0 | 12 | 0.1667 |
| 6_3394906 | 1 | 0 | 4 | 0.25 | 3 | 0.3333 | 1 | 0 | 9 | 0.2222 | 2 | 0 | 3 | 0.3333 | 4 | 0 | 4 | 0.75 | 13 | 0.3077 |
| 6_3394913 | NA | NA | 4 | 1 | 2 | 0 | 1 | 0 | 7 | 0.5714 | 3 | 1 | 3 | 0.3333 | 4 | 0.5 | 4 | 0.5 | 14 | 0.5714 |
| 6_3394934 | 1 | 0 | 3 | 0.3333 | 4 | 0 | 1 | 0 | 9 | 0.1111 | 3 | 0.6667 | 4 | 0.25 | 6 | 0 | 4 | 0.25 | 17 | 0.2353 |
| 6_3395057 | 2 | 0 | 4 | 0 | 7 | 0 | 2 | 0.5 | 15 | 0.0667 | 4 | 0 | 5 | 0.2 | 9 | 0 | 5 | 0.2 | 23 | 0.087 |

| | | | | | | | | | | | | | | | | | | | | |
|---|---|---|---|---|---|---|---|---|---|---|---|---|---|---|---|---|---|---|---|---|
| 6_33950609 | 3 | 0 | NA | NA | 7 | 0 | 2 | 0.5 | 12 | 0.0833 | 3 | 0.3333 | 4 | 0.5 | 10 | 0.1 | 4 | 0.25 | 21 | 0.2381 |
| 6_33950739 | 4 | 0 | 5 | 0.2 | 9 | 0.1111 | 2 | 0 | 20 | 0.1 | 4 | 0.25 | 8 | 0.125 | 10 | 0.2 | 5 | 0 | 27 | 0.1481 |
| 6_33950798 | 3 | 1 | 1 | 0 | 8 | 0.5 | 2 | 0 | 14 | 0.5 | 3 | 0.3333 | 7 | 0.4286 | 8 | 0.125 | 4 | 0.5 | 22 | 0.3182 |
| 6_33950867 | 1 | 0 | 3 | 0 | 4 | 0.25 | 3 | 0.3333 | 11 | 0.1818 | 2 | 0.5 | 5 | 0.6 | 9 | 0.2222 | 4 | 0.5 | 20 | 0.4 |
| 6_34080110 | NA | NA | 1 | 1 | NA | NA | NA | NA | 1 | 1 | NA | NA | NA | NA | 1 | 1 | NA | NA | 1 | 1 |
| 6_34080600 | NA | NA | 1 | 1 | NA | NA | NA | NA | 1 | 1 | NA | NA | NA | NA | 1 | 1 | NA | NA | 1 | 1 |
| 6_34081036 | NA | NA | 1 | 1 | NA | NA | NA | NA | 1 | 1 | NA | NA | NA | NA | 1 | 1 | NA | NA | 1 | 1 |
| 6_34081141 | NA | NA | 1 | 1 | NA | NA | NA | NA | 1 | 1 | NA | NA | NA | NA | 1 | 1 | NA | NA | 1 | 1 |
| 6_34081457 | NA | NA | 1 | 1 | NA | NA | NA | NA | 1 | 1 | NA | NA | NA | NA | 1 | 1 | NA | NA | 1 | 1 |
| 6_34081540 | NA | NA | 1 | 1 | NA | NA | NA | NA | 1 | 1 | NA | NA | NA | NA | 1 | 1 | NA | NA | 1 | 1 |
| 6_34081616 | NA | NA | 1 | 1 | NA | NA | NA | NA | 1 | 1 | NA | NA | NA | NA | 1 | 1 | NA | NA | 1 | 1 |
| 6_34081634 | NA | NA | 1 | 1 | NA | NA | NA | NA | 1 | 1 | NA | NA | NA | NA | 1 | 1 | NA | NA | 1 | 1 |
| 6_34081726 | NA | NA | 1 | 1 | NA | NA | NA | NA | 1 | 1 | NA | NA | NA | NA | 1 | 1 | NA | NA | 1 | 1 |
| 6_34081786 | NA | NA | 1 | 1 | NA | NA | NA | NA | 1 | 1 | NA | NA | NA | NA | 1 | 1 | NA | NA | 1 | 1 |
| 6_34083130 | NA | NA | 1 | 1 | NA | NA | NA | NA | 1 | 1 | NA | NA | NA | NA | 1 | 1 | NA | NA | 1 | 1 |
| 6_34083493 | NA | NA | 1 | 1 | NA | NA | NA | NA | 1 | 1 | NA | NA | NA | NA | 1 | 1 | NA | NA | 1 | 1 |
| 6_34133066 | 4 | 0.25 | 3 | 0 | 5 | 0 | 3 | 0.3333 | 15 | 0.1333 | 2 | 0 | 2 | 0 | 7 | 0 | 1 | 1 | 12 | 0.0833 |
| 6_34133303 | 3 | 0.3333 | 1 | 0 | 5 | 0.4 | 3 | 0 | 12 | 0.25 | 2 | 0 | 3 | 0 | 7 | 0.1429 | 1 | 0 | 13 | 0.0769 |
| 6_34133326 | 5 | 0 | 2 | 0 | 5 | 0.2 | 3 | 0 | 15 | 0.0667 | 2 | 0 | 3 | 0.3333 | 7 | 0 | 2 | 0 | 14 | 0.0714 |
| 6_34133490 | 10 | 0.1 | 2 | 0 | 8 | 0.125 | 2 | 0 | 22 | 0.0909 | 3 | 0 | 4 | 0 | 6 | 0.3333 | 3 | 0 | 16 | 0.125 |
| 6_34133536 | 9 | 0.1111 | 1 | 0 | 8 | 0.125 | 4 | 0 | 22 | 0.0909 | 2 | 0 | 4 | 0 | 7 | 0 | 4 | 0.25 | 17 | 0.0588 |
| 6_34133670 | 8 | 0.5 | 1 | 0 | 6 | 0.1667 | 3 | 0.6667 | 18 | 0.3889 | 3 | 0.6667 | 4 | 0.25 | 7 | 0.2857 | 4 | 0.75 | 18 | 0.4444 |
| 6_34133731 | 8 | 0.625 | 3 | 0.3333 | 8 | 0.25 | 5 | 0.4 | 24 | 0.4167 | 2 | 0 | 4 | 0.25 | 8 | 0.5 | 4 | 0.5 | 18 | 0.3889 |
| 6_34133771 | 7 | 0.7143 | 3 | 0.6667 | 8 | 0.75 | 5 | 0.6 | 23 | 0.6957 | 2 | 0.5 | 3 | 0.6667 | 5 | 0.6 | 2 | 0.5 | 12 | 0.5833 |
| 6_34133811 | 8 | 0.375 | 1 | 0 | 7 | 0.2857 | 5 | 0.8 | 21 | 0.4286 | 1 | 0 | 4 | 0 | 6 | 0.3333 | 4 | 0.75 | 15 | 0.3333 |
| 6_34133829 | 7 | 0.1429 | 2 | 0.5 | 6 | 0 | 3 | 0 | 18 | 0.1111 | 2 | 0 | 3 | 0 | 6 | 0.5 | 3 | 0.3333 | 14 | 0.2857 |
| 6_34133909 | 8 | 0.5 | 3 | 0.3333 | 8 | 0.25 | 5 | 0.6 | 24 | 0.4167 | 2 | 1 | 4 | 0.25 | 6 | 0.1667 | 4 | 0.25 | 16 | 0.3125 |
| 6_34133920 | 8 | 0.75 | 1 | 1 | 6 | 0.5 | 4 | 1 | 19 | 0.7368 | 2 | 1 | 2 | 0.5 | 6 | 0.5 | 3 | 1 | 13 | 0.6923 |
| 6_34134062 | 6 | 0.6667 | 3 | 1 | 6 | 0.6667 | 4 | 1 | 19 | 0.7895 | 2 | 0.5 | 4 | 0.5 | 5 | 1 | 2 | 1 | 13 | 0.7692 |
| 6_34134068 | 6 | 0.3333 | 3 | 0 | 7 | 0 | 4 | 0.75 | 20 | 0.25 | 2 | 0 | 3 | 0 | 6 | 0.1667 | 1 | 0 | 12 | 0.0833 |
| 6_34134202 | 5 | 0 | 2 | 0 | 6 | 0 | 4 | 0.25 | 17 | 0.0588 | 2 | 0.5 | 4 | 0 | 7 | 0.1429 | 4 | 0.25 | 17 | 0.1765 |
| 6_34134257 | 6 | 1 | 2 | 1 | 5 | 1 | 5 | 1 | 18 | 1 | 2 | 1 | 4 | 1 | 6 | 1 | 4 | 1 | 16 | 1 |
| 6_34134391 | 5 | 0.2 | 3 | 0 | 7 | 0 | 5 | 0.4 | 20 | 0.15 | 1 | 0 | 3 | 0 | 7 | 0.2857 | 4 | 0.25 | 15 | 0.2 |
| 6_34134433 | 7 | 0.8571 | 2 | 1 | 8 | 1 | 5 | 1 | 22 | 0.9545 | 2 | 1 | 4 | 1 | 6 | 1 | 3 | 1 | 15 | 1 |
| 6_34134472 | 6 | 1 | 3 | 1 | 8 | 1 | 5 | 1 | 22 | 1 | 2 | 1 | 4 | 1 | 7 | 1 | 3 | 0.6667 | 16 | 0.9375 |
| 6_34134599 | 5 | 0 | 3 | 0.3333 | 7 | 0 | NA | NA | 15 | 0.0667 | 2 | 0 | 3 | 0 | 3 | 0.3333 | 2 | 0 | 10 | 0.1 |
| 6_34134817 | 6 | 0.3333 | 3 | 0.6667 | 7 | 0.4286 | 4 | 0.5 | 20 | 0.45 | 2 | 0 | 3 | 0.3333 | 5 | 0.6 | 4 | 0.25 | 14 | 0.3571 |
| 6_34134855 | 6 | 1 | 3 | 0.3333 | 8 | 0.75 | 4 | 1 | 21 | 0.8095 | 2 | 0 | 4 | 0.75 | 5 | 0.8 | 3 | 0.6667 | 14 | 0.6429 |
| 6_34135030 | 6 | 0 | 3 | 0.3333 | 7 | 0.2857 | 4 | 0.25 | 20 | 0.2 | 1 | 0 | 4 | 0 | 3 | 0.3333 | 4 | 0 | 12 | 0.0833 |
| 6_34135038 | 5 | 0.4 | 2 | 0 | 7 | 0.1429 | 4 | 0 | 18 | 0.1667 | 1 | 0 | 3 | 0 | 1 | 0 | 4 | 0.5 | 9 | 0.2222 |
| 6_34135172 | 4 | 1 | 3 | 1 | 4 | 1 | 3 | 1 | 14 | 1 | 1 | 0 | 3 | 1 | 2 | 1 | 2 | 1 | 8 | 0.875 |
| 6_34135302 | 6 | 0.3333 | 3 | 0.3333 | 4 | 0.75 | 4 | 1 | 17 | 0.5882 | 1 | 0 | 4 | 0.75 | 3 | 1 | 2 | 0.5 | 10 | 0.7 |
| 6_34135319 | 4 | 0.75 | 3 | 1 | 3 | 1 | 3 | 1 | 13 | 0.9231 | 1 | 1 | 3 | 1 | 1 | 1 | 1 | 1 | 6 | 1 |
| 6_34135356 | 6 | 0.6667 | 2 | 0.5 | 6 | 0.1667 | 4 | 0.25 | 18 | 0.3889 | 1 | 1 | 4 | 0.5 | 2 | 0 | 2 | 0.5 | 9 | 0.4444 |
| 6_34135392 | 3 | 0.3333 | 3 | 0.3333 | 5 | 0.4 | 3 | 0 | 14 | 0.2857 | NA | NA | 3 | 0 | 3 | 0.3333 | 2 | 0 | 8 | 0.125 |
| 6_34281013 | 15 | 0.5333 | 8 | 0.5 | 7 | 0.2857 | 10 | 0.4 | 40 | 0.45 | 7 | 0.2857 | 9 | 0.6667 | 8 | 0.25 | 14 | 0.5 | 38 | 0.4474 |
| 6_34373620 | 2 | 0.5 | 1 | 0 | 3 | 0 | 2 | 0 | 8 | 0.125 | 3 | 0.3333 | 4 | 0 | 2 | 0 | 1 | 0 | 10 | 0.1 |
| 6_34373654 | 2 | 0.5 | 1 | 0 | 4 | 0 | 1 | 0 | 8 | 0.125 | 3 | 0.3333 | 4 | 0 | 2 | 0.5 | 1 | 1 | 10 | 0.3 |
| 6_34373747 | 2 | 0.5 | 1 | 1 | 4 | 0.5 | NA | NA | 7 | 0.5714 | 3 | 0.6667 | 4 | 0.5 | 2 | 1 | NA | NA | 9 | 0.6667 |
| 6_34373829 | 1 | 1 | 1 | 0 | 5 | 0.6 | 2 | 0.5 | 9 | 0.5556 | 2 | 0 | 4 | 0 | 2 | 0.5 | 1 | 0 | 9 | 0.1111 |
| 6_34373838 | 1 | 0 | 1 | 0 | 3 | 0.3333 | 2 | 0 | 7 | 0.1429 | 2 | 0.5 | 3 | 0 | NA | NA | 1 | 0 | 6 | 0.1667 |
| 6_34464299 | 6 | 0.5 | 4 | 0.25 | 8 | 0 | 5 | 0.2 | 23 | 0.2174 | 4 | 0 | NA | NA | 6 | 0.1667 | 4 | 0.5 | 14 | 0.2143 |
| 6_34464314 | 7 | 0.2857 | 4 | 0 | 6 | 0.1667 | 5 | 0.2 | 22 | 0.1818 | 4 | 0 | 1 | 0 | 5 | 0.2 | 4 | 0.25 | 14 | 0.1429 |
| 6_34464371 | 8 | 0.125 | 4 | 0 | 8 | 0 | 5 | 0.2 | 25 | 0.08 | 5 | 0 | 1 | 0 | 5 | 0.2 | 4 | 0 | 15 | 0.0667 |
| 6_34464386 | 6 | 0.5 | 3 | 1 | 6 | 0.5 | 5 | 0.6 | 20 | 0.6 | 5 | 0.8 | 1 | 1 | 6 | 0.8333 | 3 | 0.6667 | 15 | 0.8 |
| 6_34464403 | 7 | 1 | 2 | 1 | 6 | 0.8333 | 5 | 0.8 | 20 | 0.9 | 5 | 1 | 2 | 1 | 5 | 1 | 4 | 1 | 16 | 1 |
| 6_34464776 | 9 | 0.5556 | 3 | 0.6667 | 12 | 0.4167 | 4 | 0 | 28 | 0.4286 | 6 | 0.3333 | 4 | 0.75 | 7 | 0.8571 | 5 | 1 | 22 | 0.7273 |
| 6_34481572 | 14 | 0.9286 | 4 | 0.5 | 10 | 0.8 | 4 | 0.75 | 32 | 0.8125 | 7 | 0.5714 | 2 | 0.5 | 8 | 0.75 | 4 | 1 | 21 | 0.7143 |
| 6_34482008 | 13 | 0.0769 | 7 | 0 | 16 | 0.125 | 5 | 0 | 41 | 0.0732 | 6 | 0 | 4 | 0.5 | 6 | 0 | 5 | 0 | 21 | 0.0952 |
| 6_34482024 | 14 | 0.0714 | 7 | 0.1429 | 19 | 0.1579 | 8 | 0.125 | 48 | 0.125 | 10 | 0 | 5 | 0 | 8 | 0 | 5 | 0.4 | 28 | 0.0714 |
| 6_34762214 | 5 | 0 | 3 | 0.3333 | NA | NA | 1 | 0 | 9 | 0.1111 | 1 | 0 | NA | NA | 2 | 1 | 2 | 0.5 | 5 | 0.6 |
| 6_34762377 | 5 | 0.6 | 3 | 0.6667 | NA | NA | 1 | 0 | 9 | 0.5556 | 1 | 1 | NA | NA | 3 | 0 | 2 | 0 | 6 | 0.1667 |
| 6_34762574 | 4 | 0.25 | 2 | 0 | 1 | 0 | 1 | 0 | 8 | 0.125 | 1 | 1 | NA | NA | 2 | 0.5 | 2 | 0.5 | 5 | 0.6 |
| 6_34762822 | 3 | 0 | 2 | 0.5 | 1 | 1 | 1 | 0 | 7 | 0.2857 | 1 | 1 | NA | NA | 2 | 0 | 2 | 0 | 5 | 0.2 |
| 6_34762971 | 4 | 0.25 | 2 | 0.5 | 1 | 1 | 1 | 1 | 8 | 0.5 | 1 | 0 | NA | NA | 3 | 0.3333 | 2 | 0 | 6 | 0.1667 |

| | | | | | | | | | | | | | | | | | | | | |
|---|---|---|---|---|---|---|---|---|---|---|---|---|---|---|---|---|---|---|---|---|
| 6_3476306 | 3 | 0 | 2 | 0.5 | 1 | 0 | 1 | 0 | 7 | 0.1429 | 1 | 1 | NA | NA | 2 | 0.5 | 2 | 0.5 | 5 | 0.6 |
| 6_3485075 | NA | NA | 1 | 1 | 2 | 1 | NA | NA | 3 | 1 | 2 | 1 | NA | NA | NA | NA | 1 | 1 | 3 | 1 |
| 6_3485082 | NA | NA | 1 | 1 | 1 | 0 | NA | NA | 2 | 0.5 | 2 | 0.5 | NA | NA | NA | NA | 1 | 1 | 3 | 0.6667 |
| 6_3485104 | NA | NA | 1 | 1 | 1 | 0 | NA | NA | 2 | 0.5 | 2 | 1 | NA | NA | NA | NA | NA | NA | 2 | 1 |
| 6_3485104 | NA | NA | 1 | 1 | 2 | 0.5 | NA | NA | 3 | 0.6667 | 2 | 1 | NA | NA | NA | NA | 1 | 1 | 3 | 1 |
| 6_3485109 | NA | NA | 1 | 1 | 2 | 0.5 | NA | NA | 3 | 0.6667 | 2 | 0.5 | NA | NA | NA | NA | NA | NA | 2 | 0.5 |
| 6_3485121 | NA | NA | 1 | 0 | 2 | 0.5 | NA | NA | 3 | 0.3333 | 2 | 0.5 | NA | NA | NA | NA | 1 | 0 | 3 | 0.3333 |
| 6_3485125 | NA | NA | 1 | 1 | 1 | 1 | NA | NA | 2 | 1 | 2 | 1 | NA | NA | NA | NA | 1 | 1 | 3 | 1 |
| 6_3485146 | NA | NA | 1 | 1 | 2 | 1 | NA | NA | 3 | 1 | 1 | 1 | NA | NA | NA | NA | 1 | 1 | 2 | 1 |
| 6_3485467 | 1 | 1 | 1 | 1 | 2 | 1 | 1 | 1 | 5 | 1 | 3 | 1 | 3 | 1 | 2 | 0.5 | 1 | 1 | 9 | 0.8889 |
| 6_3485481 | 1 | 0 | 1 | 1 | 2 | 0 | 1 | 0 | 5 | 0.2 | 3 | 0.3333 | 2 | 0.5 | 2 | 0 | 1 | 0 | 8 | 0.25 |
| 6_3485606 | 2 | 1 | 2 | 1 | 1 | 0 | NA | NA | 5 | 0.8 | NA | NA | 4 | 0.5 | NA | NA | NA | NA | 4 | 0.5 |
| 6_3485640 | 2 | 0.5 | 2 | 1 | 1 | 0 | NA | NA | 5 | 0.6 | NA | NA | 4 | 0.25 | NA | NA | NA | NA | 4 | 0.25 |
| 6_3485640 | 2 | 0.5 | 1 | 1 | 1 | 0 | NA | NA | 4 | 0.5 | NA | NA | 4 | 0.25 | NA | NA | NA | NA | 4 | 0.25 |
| 6_3485734 | 1 | 1 | NA | NA | NA | NA | NA | NA | 1 | 1 | NA | NA | 1 | 1 | NA | NA | NA | NA | 1 | 1 |
| 6_3485744 | NA | NA | 2 | 0.5 | 1 | 0 | 1 | 1 | 4 | 0.5 | NA | NA | 2 | 1 | 2 | 0.5 | 1 | 0 | 5 | 0.6 |
| 6_3485750 | NA | NA | 2 | 1 | 1 | 1 | 1 | 1 | 4 | 1 | NA | NA | 2 | 1 | 2 | 0.5 | 1 | 1 | 5 | 0.8 |
| 6_3485766 | NA | NA | 2 | 0.5 | NA | NA | 1 | 1 | 3 | 0.6667 | NA | NA | 2 | 0.5 | 2 | 0 | 1 | 0 | 5 | 0.2 |
| 6_3485767 | 1 | 1 | 2 | 0.5 | NA | NA | 1 | 0 | 4 | 0.5 | NA | NA | 1 | 1 | 2 | 0.5 | 1 | 1 | 4 | 0.75 |
| 6_3485783 | 1 | 1 | 1 | 0 | NA | NA | 1 | 1 | 3 | 0.6667 | NA | NA | 1 | 0 | 1 | 1 | NA | NA | 2 | 0.5 |
| 6_3500007 | 1 | 1 | 2 | 0 | 1 | 1 | 3 | 0.6667 | 7 | 0.5714 | NA | NA | 4 | 0.75 | 2 | 0.5 | NA | NA | 6 | 0.6667 |
| 6_3500019 | 1 | 0 | 3 | 0 | 1 | 1 | 5 | 0 | 10 | 0.1 | 1 | 0 | 5 | 0.2 | 3 | 0 | 1 | 0 | 10 | 0.1 |
| 6_3500118 | NA | NA | 2 | 0 | 1 | 0 | 4 | 0.5 | 7 | 0.2857 | 1 | 0 | 4 | 0.5 | 2 | 1 | 1 | 0 | 8 | 0.5 |
| 6_3502304 | 1 | 1 | 2 | 1 | NA | NA | 2 | 0.5 | 5 | 0.8 | 1 | 1 | 3 | 0.6667 | 2 | 0 | 1 | 0 | 7 | 0.4286 |
| 6_3502311 | 1 | 1 | 2 | 0 | 1 | 0 | 3 | 0.6667 | 7 | 0.4286 | 1 | 1 | 2 | 1 | 2 | 0 | 2 | 0.5 | 7 | 0.5714 |
| 6_3502316 | 1 | 1 | 2 | 1 | 1 | 1 | 2 | 1 | 6 | 1 | 1 | 1 | 3 | 1 | 1 | 0 | 2 | 1 | 7 | 0.8571 |
| 6_3502318 | NA | NA | 2 | 0.5 | 1 | 0 | 2 | 0.5 | 5 | 0.4 | 1 | 0 | 1 | 0 | 2 | 0.5 | 2 | 0.5 | 6 | 0.3333 |
| 6_3502805 | 2 | 0.5 | NA | NA | 1 | 0 | 4 | 1 | 7 | 0.7143 | 3 | 1 | 6 | 0.6667 | 3 | 1 | 2 | 0.5 | 14 | 0.7857 |
| 6_3502807 | 1 | 0 | 2 | 0 | 1 | 0 | 4 | 0.25 | 8 | 0.125 | 3 | 0 | 6 | 0.1667 | 3 | 0 | 1 | 0 | 13 | 0.0769 |
| 6_3502808 | 2 | 0 | 2 | 0 | 1 | 0 | 3 | 0.3333 | 8 | 0.125 | 3 | 0 | 5 | 0 | 4 | 0.25 | 1 | 0 | 13 | 0.0769 |
| 6_3502835 | 2 | 0 | 2 | 1 | 1 | 0 | 5 | 0.6 | 10 | 0.5 | 4 | 0 | 6 | 0 | 4 | 0.5 | 2 | 1 | 16 | 0.25 |
| 6_3502837 | 2 | 0 | 2 | 0 | 1 | 0 | 5 | 0.6 | 10 | 0.3 | 3 | 0 | 7 | 0.1429 | 3 | 0 | 2 | 0 | 15 | 0.0667 |
| 6_3505709 | NA | NA | 2 | 0 | 1 | 0 | 3 | 0.3333 | 6 | 0.1667 | 3 | 0.3333 | 3 | 0 | 4 | 0.25 | 2 | 0.5 | 12 | 0.25 |
| 6_3522437 | 3 | 0 | 5 | 0.4 | 2 | 0 | 1 | 0 | 11 | 0.1818 | 2 | 0.5 | NA | NA | 2 | 0.5 | 3 | 0.3333 | 7 | 0.4286 |
| 6_3523739 | 7 | 0.1429 | 5 | 0 | 5 | 0.2 | 3 | 0 | 20 | 0.1 | 2 | 0 | 3 | 0 | 4 | 0 | 1 | 1 | 10 | 0.1 |
| 6_3523819 | 6 | 0.1667 | 6 | 0 | 4 | 0.5 | 3 | 0 | 19 | 0.1579 | 4 | 0 | 4 | 0.25 | 4 | 0 | 1 | 0 | 13 | 0.0769 |
| 6_3523841 | 6 | 1 | 5 | 1 | 5 | 1 | 4 | 1 | 20 | 1 | 4 | 1 | 4 | 1 | 4 | 0.75 | 1 | 1 | 13 | 0.9231 |
| 6_3523849 | 7 | 1 | 5 | 0.8 | 6 | 1 | 4 | 1 | 22 | 0.9545 | 4 | 0.75 | 4 | 1 | 3 | 1 | 1 | 1 | 12 | 0.9167 |
| 6_3523853 | 7 | 0.2857 | 4 | 0.5 | 7 | 0.4286 | 3 | 0.3333 | 21 | 0.381 | 3 | 0 | 3 | 0.6667 | 4 | 0.25 | 1 | 1 | 11 | 0.3636 |
| 6_3523879 | 2 | 0.5 | 2 | 0 | 1 | 0 | 2 | 1 | 7 | 0.4286 | NA | NA | 2 | 1 | NA | NA | NA | NA | 2 | 1 |
| 6_3524507 | 3 | 1 | 3 | 0.6667 | NA | NA | 1 | 0 | 7 | 0.7143 | 2 | 0.5 | 4 | 0.25 | 1 | 1 | NA | NA | 7 | 0.4286 |
| 6_3524548 | 1 | 1 | 3 | 0.6667 | NA | NA | 1 | 0 | 5 | 0.6 | 1 | 1 | 2 | 0.5 | NA | NA | NA | NA | 3 | 0.6667 |
| 6_3524575 | 3 | 0.3333 | 4 | 0.25 | NA | NA | 2 | 0 | 9 | 0.2222 | 1 | 0 | 3 | 0.6667 | 1 | 0 | NA | NA | 5 | 0.4 |
| 6_3528450 | 1 | 0 | 2 | 0.5 | NA | NA | NA | NA | 3 | 0.3333 | 2 | 0.5 | 2 | 0 | NA | NA | NA | NA | 4 | 0.25 |
| 6_3529024 | 1 | 1 | NA | NA | 1 | 0 | NA | NA | 2 | 0.5 | NA | NA | NA | NA | 1 | 1 | NA | NA | 1 | 1 |
| 6_3529029 | 1 | 1 | NA | NA | 1 | 0 | NA | NA | 2 | 0.5 | NA | NA | NA | NA | 1 | 1 | NA | NA | 1 | 1 |
| 6_3529044 | 1 | 0 | NA | NA | 1 | 1 | NA | NA | 2 | 0.5 | NA | NA | NA | NA | 1 | 1 | NA | NA | 1 | 1 |
| 6_3548901 | 20 | 0.1 | 19 | 0.0526 | 15 | 0.1333 | 21 | 0 | 75 | 0.0667 | 10 | 0.1 | 19 | 0.0526 | 21 | 0.1429 | 11 | 0 | 61 | 0.082 |
| 6_3548915 | 17 | 0.2353 | 17 | 0.2353 | 14 | 0.2857 | 23 | 0.1304 | 71 | 0.2113 | 11 | 0.3636 | 20 | 0.35 | 20 | 0.35 | 14 | 0.2857 | 65 | 0.3385 |
| 6_3650026 | NA | NA | NA | NA | 2 | 1 | NA | NA | 2 | 1 | 1 | 1 | NA | NA | NA | NA | NA | NA | 1 | 1 |
| 6_3650031 | NA | NA | NA | NA | 2 | 0.5 | NA | NA | 2 | 0.5 | 1 | 1 | NA | NA | NA | NA | NA | NA | 1 | 1 |
| 6_3650037 | NA | NA | NA | NA | 2 | 1 | NA | NA | 2 | 1 | 1 | 1 | NA | NA | NA | NA | NA | NA | 1 | 1 |
| 6_3650042 | NA | NA | NA | NA | 2 | 1 | NA | NA | 2 | 1 | 1 | 1 | NA | NA | NA | NA | NA | NA | 1 | 1 |
| 6_3650045 | NA | NA | NA | NA | 2 | 1 | NA | NA | 2 | 1 | 1 | 1 | NA | NA | NA | NA | NA | NA | 1 | 1 |
| 6_3650065 | NA | NA | NA | NA | 2 | 1 | NA | NA | 2 | 1 | 1 | 1 | NA | NA | NA | NA | NA | NA | 1 | 1 |
| 6_3650130 | 1 | 1 | 1 | 1 | 2 | 0 | NA | NA | 4 | 0.5 | 2 | 1 | NA | NA | 1 | 0 | NA | NA | 3 | 0.6667 |
| 6_3650141 | NA | NA | 1 | 1 | 2 | 1 | NA | NA | 3 | 1 | 2 | 1 | NA | NA | 1 | 1 | NA | NA | 3 | 1 |
| 6_3650149 | 1 | 1 | 1 | 1 | 1 | 1 | NA | NA | 3 | 1 | 2 | 1 | NA | NA | 1 | 1 | NA | NA | 3 | 1 |
| 6_3650152 | 1 | 1 | 1 | 0 | 2 | 0.5 | NA | NA | 4 | 0.5 | 2 | 0.5 | NA | NA | 1 | 0 | NA | NA | 3 | 0.3333 |
| 6_3650154 | 1 | 1 | 1 | 0 | 2 | 0 | NA | NA | 4 | 0.25 | 2 | 0.5 | 1 | 0 | 1 | 0 | NA | NA | 4 | 0.25 |
| 6_3650160 | NA | NA | 1 | 1 | 2 | 0.5 | NA | NA | 3 | 0.6667 | 2 | 0.5 | 1 | 1 | 1 | 1 | NA | NA | 4 | 0.75 |
| 6_3650168 | 1 | 1 | 1 | 1 | 2 | 1 | NA | NA | 4 | 1 | 2 | 1 | 1 | 1 | 1 | 1 | NA | NA | 4 | 1 |
| 6_3650178 | 1 | 1 | 1 | 1 | 2 | 0 | NA | NA | 4 | 0.5 | 2 | 0 | 1 | 1 | 1 | 1 | NA | NA | 4 | 0.5 |
| 6_3650188 | 1 | 0 | 1 | 0 | 1 | 1 | NA | NA | 3 | 0.3333 | 2 | 0.5 | 1 | 0 | 1 | 0 | NA | NA | 4 | 0.25 |
| 6_3650224 | 1 | 1 | 1 | 1 | 2 | 1 | NA | NA | 4 | 1 | 2 | 1 | 1 | 1 | 1 | 1 | NA | NA | 4 | 1 |

| | | | | | | | | | | | | | | | | | | | | |
|---|---|---|---|---|---|---|---|---|---|---|---|---|---|---|---|---|---|---|---|---|
| 6_3650228 | 1 | 1 | 1 | 1 | 2 | 1 | NA | NA | 4 | 1 | 2 | 1 | 1 | 1 | 1 | 1 | NA | NA | 4 | 1 |
| 6_3650238 | 1 | 1 | NA | NA | 2 | 1 | NA | NA | 3 | 1 | 2 | 1 | 1 | 1 | NA | NA | NA | NA | 3 | 1 |
| 6_3650242 | 1 | 0 | 1 | 0 | 1 | 1 | 1 | 0 | 4 | 0.25 | 2 | 0.5 | 1 | 0 | 1 | 1 | NA | NA | 4 | 0.5 |
| 6_3650246 | 1 | 0 | 1 | 0 | 2 | 0.5 | 1 | 1 | 5 | 0.4 | 2 | 0.5 | 1 | 0 | 1 | 1 | NA | NA | 4 | 0.5 |
| 6_3650257 | NA | NA | NA | NA | 2 | 0.5 | 1 | 1 | 3 | 0.6667 | 1 | 1 | 1 | 1 | NA | NA | NA | NA | 2 | 1 |
| 6_3650271 | 1 | 1 | 1 | 0 | 1 | 1 | 1 | 0 | 4 | 0.5 | 2 | 0.5 | 1 | 0 | 1 | 0 | NA | NA | 4 | 0.25 |
| 6_3650278 | 1 | 1 | NA | NA | 2 | 1 | 1 | 0 | 4 | 0.75 | 1 | 1 | NA | NA | NA | NA | NA | NA | 1 | 1 |
| 6_3650423 | 2 | 0.5 | 1 | 0 | 2 | 1 | 2 | 0.5 | 7 | 0.5714 | 3 | 0.3333 | 1 | 0 | 2 | 0 | NA | NA | 6 | 0.1667 |
| 6_3650522 | 3 | 0 | 1 | 0 | 1 | 0 | 2 | 0.5 | 7 | 0.1429 | 3 | 0.3333 | 1 | 0 | 1 | 0 | 1 | 0 | 6 | 0.1667 |
| 6_3681442 | 1 | 1 | 4 | 0.5 | 2 | 0.5 | 5 | 0.4 | 12 | 0.5 | NA | NA | 2 | 0.5 | 3 | 1 | 2 | 1 | 7 | 0.8571 |
| 6_3682396 | NA | NA | 1 | 1 | 1 | 0 | NA | NA | 2 | 0.5 | 1 | 0 | NA | NA | 1 | 1 | 1 | 1 | 3 | 0.6667 |
| 6_3682404 | NA | NA | 2 | 1 | 2 | 1 | NA | NA | 4 | 1 | 2 | 1 | NA | NA | NA | NA | 1 | 1 | 3 | 1 |
| 6_3682414 | NA | NA | 2 | 0.5 | 2 | 0.5 | NA | NA | 4 | 0.5 | 2 | 0.5 | NA | NA | NA | NA | 1 | 1 | 3 | 0.6667 |
| 6_3682431 | NA | NA | 1 | 1 | 2 | 0.5 | NA | NA | 3 | 0.6667 | 2 | 1 | NA | NA | NA | NA | NA | NA | 2 | 1 |
| 6_3730437 | 3 | 0.3333 | 2 | 0.5 | NA | NA | NA | NA | 5 | 0.4 | 2 | 0.5 | 4 | 0.5 | 3 | 0.3333 | 3 | 0.3333 | 12 | 0.4167 |
| 6_3730447 | 3 | 0 | 2 | 0.5 | NA | NA | NA | NA | 5 | 0.2 | 2 | 0.5 | 1 | 0 | 2 | 0.5 | 2 | 0 | 7 | 0.2857 |
| 6_3730849 | 1 | 1 | 1 | 0 | 1 | 0 | NA | NA | 3 | 0.3333 | 1 | 0 | 1 | 0 | 1 | 1 | 1 | 0 | 4 | 0.25 |
| 6_3730855 | 1 | 1 | 1 | 0 | NA | NA | NA | NA | 2 | 0.5 | 1 | 0 | 1 | 0 | 1 | 0 | 1 | 1 | 4 | 0.25 |
| 6_3730896 | NA | NA | 1 | 1 | 1 | 0 | NA | NA | 2 | 0.5 | 1 | 1 | 1 | 1 | 1 | 1 | NA | NA | 3 | 1 |
| 6_3730901 | 1 | 1 | 1 | 0 | 1 | 1 | NA | NA | 3 | 0.6667 | NA | NA | NA | NA | 1 | 1 | NA | NA | 1 | 1 |
| 6_3730904 | 1 | 0 | 1 | 1 | 1 | 0 | NA | NA | 3 | 0.3333 | 1 | 0 | NA | NA | 1 | 1 | 1 | 0 | 3 | 0.3333 |
| 6_3778355 | 14 | 0.1429 | 11 | 0 | 16 | 0.1875 | 8 | 0 | 49 | 0.102 | 13 | 0.0769 | 7 | 0 | 10 | 0 | 11 | 0.1818 | 41 | 0.0732 |
| 6_3778366 | 27 | 0.3333 | 22 | 0.0909 | 23 | 0.2174 | 13 | 0.1538 | 85 | 0.2118 | 19 | 0.0526 | 13 | 0.2308 | 13 | 0.0769 | 16 | 0.1875 | 61 | 0.1311 |
| 6_3778377 | 37 | 0.0541 | 30 | 0.1333 | 30 | 0.0667 | 12 | 0 | 109 | 0.0734 | 27 | 0.037 | 19 | 0.0526 | 19 | 0.0526 | 23 | 0.1304 | 88 | 0.0682 |
| 6_3778388 | 45 | 0.2 | 41 | 0.2683 | 48 | 0.2708 | 23 | 0.1304 | 157 | 0.2293 | 47 | 0.2553 | 30 | 0.0667 | 28 | 0.1071 | 23 | 0.3043 | 128 | 0.1875 |
| 6_3794248 | 1 | 0 | 2 | 0 | 4 | 0.25 | 2 | 0 | 9 | 0.1111 | 1 | 0 | 2 | 0.5 | NA | NA | 2 | 0 | 5 | 0.2 |
| 6_3794255 | 2 | 0.5 | 2 | 0.5 | 4 | 0 | 2 | 0.5 | 10 | 0.3 | 1 | 0 | 2 | 0.5 | NA | NA | 3 | 0.3333 | 6 | 0.3333 |
| 6_3794269 | 2 | 0.5 | 3 | 0.3333 | 3 | 0.3333 | 1 | 0 | 9 | 0.3333 | 1 | 1 | 1 | 1 | NA | NA | 3 | 1 | 5 | 1 |
| 6_3794276 | 1 | 1 | 2 | 1 | 3 | 1 | 2 | 0.5 | 8 | 0.875 | 1 | 1 | 2 | 1 | NA | NA | 3 | 0.6667 | 6 | 0.8333 |
| 6_3794290 | 2 | 0.5 | 2 | 0 | 4 | 0.75 | 1 | 1 | 9 | 0.5556 | 1 | 0 | 2 | 1 | NA | NA | 2 | 1 | 5 | 0.8 |
| 6_3810029 | 4 | 0.5 | 4 | 0.75 | 5 | 0.4 | 2 | 0.5 | 15 | 0.5333 | 1 | 1 | 2 | 0.5 | 1 | 0 | NA | NA | 4 | 0.5 |
| 6_3810036 | 3 | 0.3333 | 4 | 0.5 | 5 | 0.4 | 1 | 0 | 13 | 0.3846 | 1 | 0 | 2 | 0.5 | NA | NA | NA | NA | 3 | 0.3333 |
| 6_3810044 | 3 | 0.3333 | 3 | 0 | 5 | 0.6 | 1 | 1 | 12 | 0.4167 | 1 | 1 | 2 | 0 | 1 | 0 | NA | NA | 4 | 0.25 |
| 6_3810088 | 2 | 0 | 3 | 0 | 3 | 0.3333 | 2 | 0 | 10 | 0.1 | 1 | 1 | 2 | 0.5 | NA | NA | NA | NA | 3 | 0.6667 |
| 6_3810109 | 3 | 0 | 2 | 1 | 4 | 0.25 | 1 | 0 | 10 | 0.3 | NA | NA | 2 | 0.5 | NA | NA | NA | NA | 2 | 0.5 |
| 6_3810382 | 1 | 1 | NA | NA | 2 | 0.5 | 1 | 1 | 4 | 0.75 | NA | NA | 1 | 0 | 2 | 0.5 | NA | NA | 3 | 0.3333 |
| 6_3810390 | 1 | 1 | NA | NA | 2 | 0 | 1 | 0 | 4 | 0.25 | NA | NA | NA | NA | 2 | 0.5 | NA | NA | 2 | 0.5 |
| 6_3810409 | 2 | 0.5 | NA | NA | NA | NA | 1 | 0 | 3 | 0.3333 | NA | NA | 1 | 1 | 1 | 0 | NA | NA | 2 | 0.5 |
| 6_3810468 | 1 | 1 | 2 | 1 | NA | NA | 2 | 0.5 | 5 | 0.8 | 1 | 1 | NA | NA | 3 | 1 | 3 | 0 | 7 | 0.5714 |
| 6_3810482 | 1 | 0 | 2 | 0 | NA | NA | 2 | 0.5 | 5 | 0.2 | 1 | 0 | 2 | 0 | 3 | 0 | 3 | 0.3333 | 9 | 0.1111 |
| 6_3810509 | 1 | 0 | 1 | 1 | NA | NA | 2 | 1 | 4 | 0.75 | 1 | 1 | 1 | 1 | 3 | 0.6667 | 3 | 0.3333 | 8 | 0.625 |
| 6_3810526 | 1 | 0 | 2 | 0.5 | NA | NA | 1 | 1 | 4 | 0.5 | 1 | 1 | NA | NA | 1 | 1 | 3 | 1 | 5 | 1 |
| 6_3810545 | 1 | 0 | 1 | 0 | NA | NA | 2 | 1 | 4 | 0.5 | 1 | 0 | 1 | 1 | 2 | 0.5 | 2 | 1 | 6 | 0.6667 |
| 6_3817150 | NA | NA | NA | NA | NA | NA | 1 | 1 | 1 | 1 | NA | NA | NA | NA | 1 | 1 | 2 | 1 | 3 | 1 |
| 6_3817155 | NA | NA | NA | NA | NA | NA | 1 | 1 | 1 | 1 | NA | NA | NA | NA | 1 | 1 | 1 | 1 | 2 | 1 |
| 6_3817186 | NA | NA | NA | NA | NA | NA | 1 | 1 | 1 | 1 | NA | NA | NA | NA | 1 | 0 | 1 | 1 | 2 | 0.5 |
| 6_3817286 | NA | NA | NA | NA | NA | NA | 1 | 1 | 1 | 1 | NA | NA | NA | NA | 1 | 1 | NA | NA | 1 | 1 |
| 6_3817289 | NA | NA | NA | NA | NA | NA | 1 | 1 | 1 | 1 | NA | NA | NA | NA | 1 | 1 | 1 | 1 | 2 | 1 |
| 6_3826979 | NA | NA | 1 | 1 | NA | NA | NA | NA | 1 | 1 | 1 | 1 | NA | NA | NA | NA | NA | NA | 1 | 1 |
| 6_3826989 | NA | NA | 1 | 1 | NA | NA | NA | NA | 1 | 1 | 1 | 1 | NA | NA | NA | NA | NA | NA | 1 | 1 |
| 6_3827207 | NA | NA | 1 | 1 | NA | NA | NA | NA | 1 | 1 | 1 | 1 | NA | NA | NA | NA | NA | NA | 1 | 1 |
| 6_3828364 | NA | NA | 1 | 1 | 1 | 0 | NA | NA | 2 | 0.5 | NA | NA | NA | NA | 1 | 1 | 1 | 0 | 2 | 0.5 |
| 6_3830966 | NA | NA | NA | NA | 1 | 1 | 1 | 1 | 2 | 1 | NA | NA | NA | NA | 1 | 1 | NA | NA | 1 | 1 |
| 6_3840222 | NA | NA | 1 | 1 | NA | NA | 1 | 1 | 2 | 1 | NA | NA | 2 | 1 | 1 | 1 | 1 | 1 | 4 | 1 |
| 6_3840232 | NA | NA | 2 | 1 | NA | NA | 1 | 0 | 3 | 0.6667 | NA | NA | 1 | 1 | NA | NA | 1 | 1 | 2 | 1 |
| 6_3840414 | NA | NA | 3 | 0.6667 | NA | NA | 2 | 0.5 | 5 | 0.6 | 2 | 0 | 2 | 0.5 | 2 | 0 | 1 | 0 | 7 | 0.1429 |
| 6_3849978 | NA | NA | 1 | 1 | NA | NA | NA | NA | 1 | 1 | 1 | 1 | NA | NA | NA | NA | NA | NA | 1 | 1 |
| 6_3850013 | NA | NA | 1 | 1 | NA | NA | NA | NA | 1 | 1 | 1 | 1 | NA | NA | NA | NA | NA | NA | 1 | 1 |
| 6_3850015 | NA | NA | 1 | 1 | NA | NA | NA | NA | 1 | 1 | 1 | 1 | NA | NA | NA | NA | NA | NA | 1 | 1 |
| 6_3850087 | NA | NA | 1 | 1 | NA | NA | NA | NA | 1 | 1 | 1 | 1 | NA | NA | NA | NA | NA | NA | 1 | 1 |
| 6_3852861 | 3 | 0.3333 | 3 | 0.3333 | 2 | 0.5 | 3 | 0 | 11 | 0.2727 | 1 | 0 | 2 | 0 | 6 | 0.1667 | 1 | 1 | 10 | 0.2 |
| 6_3856956 | 2 | 0 | 6 | 0 | 4 | 0.25 | 5 | 0 | 17 | 0.0588 | 1 | 0 | NA | NA | 7 | 0.1429 | 3 | 0 | 11 | 0.0909 |
| 6_3858027 | 3 | 0.3333 | 8 | 0 | 7 | 0.1429 | 5 | 0.4 | 23 | 0.1739 | 1 | 0 | 3 | 0 | 7 | 0.1429 | 3 | 0 | 14 | 0.0714 |
| 6_3858379 | 9 | 0.3333 | 11 | 0.3636 | 10 | 0.3 | 8 | 0.125 | 38 | 0.2895 | 4 | 0.75 | 5 | 0 | 3 | 1 | 6 | 0.1667 | 18 | 0.3889 |
| 6_3858397 | 10 | 0.1 | 11 | 0.2727 | 11 | 0 | 9 | 0 | 41 | 0.0976 | 5 | 0.2 | 6 | 0 | 3 | 0.3333 | 9 | 0 | 23 | 0.087 |

| | | | | | | | | | | | | | | | | | | | | |
|---|---|---|---|---|---|---|---|---|---|---|---|---|---|---|---|---|---|---|---|---|
| 6_3858539 | NA | NA | 2 | 0.5 | 2 | 0.5 | 1 | 0 | 5 | 0.4 | 1 | 1 | NA | NA | 5 | 0.6 | 1 | 0 | 7 | 0.5714 |
| 6_3865755 | 8 | 0.375 | 6 | 0.8333 | 6 | 0.8333 | 4 | 0.75 | 24 | 0.6667 | 6 | 0.8333 | 4 | 0.25 | 6 | 0.8333 | 5 | 0.4 | 21 | 0.619 |
| 6_3865762 | 10 | 0.4 | 7 | 0.2857 | 7 | 0.4286 | 4 | 0.25 | 28 | 0.3571 | 7 | 0.5714 | 6 | 0.5 | 6 | 0.3333 | 4 | 0.75 | 23 | 0.5217 |
| 6_3866544 | 5 | 0.6 | 6 | 0.1667 | 11 | 0.0909 | 10 | 0.2 | 32 | 0.2188 | 7 | 0.4286 | 7 | 0.1429 | 11 | 0.4545 | 6 | 0.3333 | 31 | 0.3548 |
| 6_3866756 | 4 | 0 | 3 | 0 | 11 | 0.0909 | 8 | 0.125 | 26 | 0.0769 | 3 | 0 | 3 | 0.3333 | 7 | 0 | 1 | 0 | 14 | 0.0714 |
| 6_3866781 | 6 | 0 | 3 | 0 | 11 | 0.1818 | 7 | 0.1429 | 27 | 0.1111 | 2 | 0 | 1 | 0 | 6 | 0.1667 | NA | NA | 9 | 0.1111 |
| 6_3867462 | NA | NA | 1 | 0 | 1 | 0 | 2 | 0.5 | 4 | 0.25 | NA | NA | NA | NA | 1 | 1 | 1 | 0 | 2 | 0.5 |
| 6_3867468 | NA | NA | NA | NA | NA | NA | 2 | 1 | 2 | 1 | NA | NA | NA | NA | NA | NA | 1 | 1 | 1 | 1 |
| 6_3867469 | NA | NA | 1 | 1 | NA | NA | 3 | 0.6667 | 4 | 0.75 | NA | NA | NA | NA | 1 | 1 | 1 | 0 | 2 | 0.5 |
| 6_3867491 | NA | NA | 1 | 0 | 1 | 0 | 3 | 0.3333 | 5 | 0.2 | NA | NA | NA | NA | 1 | 1 | 1 | 1 | 2 | 1 |
| 6_3867497 | NA | NA | 1 | 1 | NA | NA | 2 | 0 | 3 | 0.3333 | NA | NA | NA | NA | 1 | 1 | NA | NA | 1 | 1 |
| 6_3906414 | 2 | 0.5 | 1 | 0 | 1 | 0 | NA | NA | 4 | 0.25 | 1 | 0 | 2 | 0 | NA | NA | 3 | 0.6667 | 6 | 0.3333 |
| 6_3911432 | 5 | 0 | 5 | 0.2 | 4 | 0.25 | 2 | 0 | 16 | 0.125 | 4 | 0.25 | 1 | 0 | NA | NA | 1 | 0 | 6 | 0.1667 |
| 6_3911991 | 2 | 0.5 | 1 | 0 | NA | NA | 1 | 0 | 4 | 0.25 | 1 | 1 | 1 | 0 | NA | NA | NA | NA | 2 | 0.5 |
| 6_3911996 | 2 | 1 | 1 | 1 | NA | NA | 1 | 1 | 4 | 1 | 1 | 1 | 1 | 1 | NA | NA | NA | NA | 2 | 1 |
| 6_3912001 | 2 | 1 | 1 | 1 | NA | NA | 1 | 1 | 4 | 1 | 1 | 1 | 1 | 1 | NA | NA | NA | NA | 2 | 1 |
| 6_3912141 | 2 | 0 | 1 | 1 | NA | NA | 1 | 1 | 4 | 0.5 | 1 | 0 | 1 | 1 | NA | NA | NA | NA | 2 | 0.5 |
| 6_3931305 | 2 | 0 | 5 | 0 | 5 | 0.2 | 4 | 0 | 16 | 0.0625 | 4 | 0.5 | 4 | 0.25 | 4 | 0.25 | 3 | 0 | 15 | 0.2667 |
| 6_3931322 | 1 | 0 | 4 | 0.25 | 4 | 0 | 3 | 0 | 12 | 0.0833 | 3 | 0.3333 | 6 | 0.3333 | 3 | 0 | 3 | 0 | 15 | 0.2 |
| 6_3931333 | 2 | 0 | 3 | 0.3333 | 5 | 0.2 | 2 | 0.5 | 12 | 0.25 | 3 | 0 | 7 | 0.1429 | 3 | 0 | 3 | 0 | 16 | 0.0625 |
| 6_3931348 | 2 | 1 | 4 | 1 | 5 | 0.8 | 3 | 1 | 14 | 0.9286 | 3 | 1 | 6 | 0.8333 | 2 | 0.5 | 2 | 1 | 13 | 0.8462 |
| 6_3931349 | 2 | 1 | 4 | 1 | 4 | 1 | 3 | 1 | 13 | 1 | 3 | 0.6667 | 7 | 1 | 2 | 1 | 3 | 1 | 15 | 0.9333 |
| 6_3931353 | 2 | 0.5 | 4 | 0.5 | 5 | 0.2 | 2 | 0.5 | 13 | 0.3846 | 3 | 0.3333 | 7 | 0.4286 | 3 | 0 | 3 | 0 | 16 | 0.25 |
| 6_3931376 | 2 | 0.5 | 4 | 0.5 | 5 | 0.8 | 2 | 1 | 13 | 0.6923 | 2 | 1 | 4 | 0.25 | 2 | 0.5 | 2 | 0.5 | 10 | 0.5 |
| 6_3931383 | 2 | 1 | 4 | 0.75 | 4 | 0.75 | 2 | 0.5 | 12 | 0.75 | 2 | 1 | 5 | 0.8 | 3 | 0.3333 | 2 | 0.5 | 12 | 0.6667 |
| 6_3931389 | 2 | 0.5 | 4 | 0.75 | 4 | 0.5 | 2 | 0 | 12 | 0.5 | 2 | 0.5 | 6 | 0.5 | 2 | 1 | 2 | 1 | 12 | 0.6667 |
| 6_3931392 | 1 | 0 | 4 | 0.25 | 3 | 0.6667 | 2 | 0 | 10 | 0.3 | 2 | 0 | 6 | 0.1667 | 3 | 0 | 2 | 0 | 13 | 0.0769 |
| 6_3931410 | 2 | 0.5 | 3 | 1 | 5 | 1 | 2 | 1 | 12 | 0.9167 | 2 | 1 | 6 | 1 | 3 | 1 | 2 | 1 | 13 | 1 |
| 6_3937484 | 17 | 0.0588 | 10 | 0 | 18 | 0.1111 | 10 | 0 | 55 | 0.0545 | 10 | 0 | 10 | 0 | 23 | 0.087 | 17 | 0.0588 | 60 | 0.05 |
| 6_3937549 | 13 | 0.1538 | 11 | 0 | 19 | 0.2632 | 8 | 0 | 51 | 0.1373 | 10 | 0.2 | 10 | 0.3 | 20 | 0.15 | 14 | 0.1429 | 54 | 0.1852 |
| 6_3937586 | 12 | 0.75 | 12 | 0.5 | 19 | 0.5263 | 8 | 0.5 | 51 | 0.5686 | 8 | 0.375 | 8 | 0.25 | 15 | 0.6667 | 13 | 0.6923 | 44 | 0.5455 |
| 6_3937618 | 16 | 0.9375 | 14 | 0.9286 | 21 | 1 | 11 | 1 | 62 | 0.9677 | 10 | 0.8 | 9 | 1 | 18 | 0.8333 | 15 | 0.8667 | 52 | 0.8654 |
| 6_3937620 | 16 | 0.75 | 15 | 0.7333 | 19 | 0.5789 | 9 | 0.7778 | 59 | 0.6949 | 10 | 1 | 9 | 1 | 19 | 0.8421 | 14 | 0.7143 | 52 | 0.8654 |
| 6_3937622 | 16 | 0.125 | 15 | 0.2667 | 20 | 0.4 | 10 | 0.1 | 61 | 0.2459 | 7 | 0.2857 | 8 | 0.5 | 17 | 0.2353 | 15 | 0.2667 | 47 | 0.2979 |
| 6_3937803 | 3 | 0.3333 | 4 | 0 | 3 | 0 | 2 | 0 | 12 | 0.0833 | 4 | 0 | 1 | 1 | 2 | 0 | 2 | 0 | 9 | 0.1111 |
| 6_3937813 | 4 | 0 | 4 | 0 | 3 | 0.3333 | 2 | 0 | 13 | 0.0769 | 4 | 0 | 2 | 0.5 | 2 | 0 | 2 | 1 | 10 | 0.3 |
| 6_3938260 | 4 | 0.25 | 1 | 0 | 1 | 0 | NA | NA | 6 | 0.1667 | 2 | 0 | 1 | 0 | 3 | 0.3333 | 1 | 0 | 7 | 0.1429 |
| 6_3938271 | 21 | 0.1905 | 14 | 0.1429 | 20 | 0.15 | 9 | 0.1111 | 64 | 0.1562 | 10 | 0.1 | 6 | 0.1667 | 14 | 0 | 16 | 0.1875 | 46 | 0.1087 |
| 6_3939734 | 3 | 0 | 5 | 0 | 2 | 0 | 2 | 0.5 | 12 | 0.0833 | 2 | 0 | 3 | 0 | 3 | 0 | 5 | 0.2 | 13 | 0.0769 |
| 6_3943949 | 5 | 0.2 | 1 | 0 | 3 | 0.3333 | 4 | 0 | 13 | 0.1538 | 3 | 0.3333 | 4 | 0 | 3 | 0.3333 | 3 | 0 | 13 | 0.1538 |
| 6_3943959 | 5 | 0.2 | 1 | 0 | 2 | 0 | 4 | 0 | 12 | 0.0833 | 3 | 0 | 3 | 0.3333 | 3 | 0 | 3 | 0 | 12 | 0.0833 |
| 6_3943960 | 4 | 0.25 | 1 | 0 | 2 | 0.5 | 2 | 0 | 9 | 0.2222 | 2 | 0.5 | 1 | 0 | 3 | 0.3333 | 3 | 0.3333 | 9 | 0.3333 |
| 6_3943960 | NA | NA | NA | NA | 1 | 1 | 2 | 0 | 3 | 0.3333 | NA | NA | 2 | 0.5 | NA | NA | NA | NA | 2 | 0.5 |
| 6_3943969 | 4 | 0.25 | NA | NA | 1 | 0 | 3 | 0 | 8 | 0.125 | 2 | 0.5 | 3 | 0.3333 | 3 | 0.6667 | 2 | 0.5 | 10 | 0.5 |
| 6_3943989 | 6 | 0.3333 | 1 | 0 | 1 | 1 | 3 | 1 | 11 | 0.5455 | 3 | 1 | 4 | 0.5 | 3 | 0.3333 | 2 | 0.5 | 12 | 0.5833 |
| 6_3943998 | 4 | 1 | 1 | 1 | 2 | 1 | 3 | 0.6667 | 10 | 0.9 | 3 | 0.6667 | 2 | 1 | 3 | 1 | 1 | 1 | 9 | 0.8889 |
| 6_3943999 | 5 | 0.4 | 1 | 0 | 2 | 1 | 3 | 0.3333 | 11 | 0.4545 | 3 | 0.6667 | 3 | 0.6667 | 3 | 0 | 2 | 1 | 11 | 0.5455 |
| 6_3944009 | 5 | 0.2 | NA | NA | 2 | 0.5 | 3 | 0.6667 | 10 | 0.4 | 2 | 1 | 2 | 0.5 | 1 | 0 | 2 | 0.5 | 7 | 0.5714 |
| 6_3944020 | 7 | 1 | 1 | 0 | 3 | 1 | 4 | 0.5 | 15 | 0.8 | 2 | 1 | 4 | 1 | 2 | 1 | 1 | 1 | 9 | 1 |
| 6_3944024 | 7 | 0.1429 | 1 | 0 | 3 | 0.3333 | 5 | 0 | 16 | 0.125 | 2 | 0 | 4 | 0 | 3 | 0.3333 | 1 | 1 | 10 | 0.2 |
| 6_3944029 | 7 | 0 | 1 | 0 | 3 | 0.3333 | 5 | 0.2 | 16 | 0.125 | 2 | 0 | 4 | 0.25 | 3 | 0 | 1 | 0 | 10 | 0.1 |
| 6_3944115 | 6 | 0 | 1 | 0 | 3 | 0.3333 | 3 | 0.6667 | 13 | 0.2308 | 2 | 0 | 3 | 0 | 3 | 0.3333 | 1 | 0 | 9 | 0.1111 |
| 6_3949977 | 2 | 0.5 | 1 | 1 | NA | NA | 3 | 0.6667 | 6 | 0.6667 | 1 | 0 | 2 | 0.5 | 2 | 0.5 | 1 | 1 | 6 | 0.5 |
| 6_3950000 | 3 | 1 | 1 | 0 | 1 | 1 | 3 | 0.6667 | 8 | 0.75 | 1 | 0 | 1 | 1 | 2 | 0.5 | 1 | 1 | 5 | 0.6 |
| 6_3950006 | 1 | 1 | 1 | 0 | 1 | 1 | 3 | 0.3333 | 6 | 0.5 | 1 | 0 | 2 | 0 | 2 | 1 | 1 | 0 | 6 | 0.3333 |
| 6_3950007 | 2 | 0.5 | 1 | 0 | 1 | 0 | 3 | 0 | 7 | 0.1429 | 1 | 0 | 2 | 0.5 | 2 | 0 | 1 | 0 | 6 | 0.1667 |
| 6_3950015 | 2 | 0 | NA | NA | 1 | 1 | 2 | 0 | 5 | 0.2 | 1 | 1 | 1 | 1 | 2 | 0 | NA | NA | 4 | 0.5 |
| 6_3950020 | 2 | 1 | 1 | 0 | 1 | 0 | 3 | 0.6667 | 7 | 0.5714 | 1 | 0 | 2 | 0.5 | 2 | 1 | 1 | 1 | 6 | 0.6667 |
| 6_3950026 | 2 | 0 | 1 | 0 | 1 | 0 | 3 | 0.3333 | 7 | 0.1429 | 1 | 0 | 1 | 1 | 2 | 0 | NA | NA | 4 | 0.25 |
| 6_3952043 | NA | NA | 1 | 1 | NA | NA | NA | NA | 1 | 1 | NA | NA | NA | NA | NA | NA | 1 | 1 | 1 | 1 |
| 6_3952043 | NA | NA | 1 | 1 | NA | NA | NA | NA | 1 | 1 | NA | NA | 1 | 1 | NA | NA | 1 | 1 | 2 | 1 |
| 6_3952083 | NA | NA | 1 | 1 | NA | NA | NA | NA | 1 | 1 | NA | NA | NA | NA | NA | NA | 1 | 1 | 1 | 1 |
| 6_3955106 | NA | NA | 1 | 1 | 2 | 0 | NA | NA | 3 | 0.3333 | 2 | 0 | 4 | 0.5 | 2 | 0 | NA | NA | 8 | 0.25 |
| 6_3955130 | NA | NA | 1 | 1 | 3 | 0.3333 | 1 | 1 | 5 | 0.6 | 4 | 0.5 | 5 | 0 | 2 | 1 | NA | NA | 11 | 0.3636 |
| 6_3955134 | NA | NA | 1 | 0 | 3 | 0.3333 | 2 | 0 | 6 | 0.1667 | 4 | 0.25 | 5 | 0.2 | 2 | 0.5 | NA | NA | 11 | 0.2727 |

| | | | | | | | | | | | | | | | | | | | | |
|---|---|---|---|---|---|---|---|---|---|---|---|---|---|---|---|---|---|---|---|---|
| 6_3955144 | NA | NA | NA | NA | 3 | 0.6667 | 2 | 0.5 | 5 | 0.6 | 4 | 1 | 5 | 0.6 | 2 | 1 | NA | NA | 11 | 0.8182 |
| 6_3955147 | NA | NA | 1 | 1 | 4 | 1 | 1 | 1 | 6 | 1 | 4 | 1 | 4 | 1 | 3 | 0.6667 | NA | NA | 11 | 0.9091 |
| 6_3955165 | NA | NA | 1 | 0 | 3 | 0.6667 | 1 | 1 | 5 | 0.6 | 4 | 0.75 | 6 | 1 | 3 | 0.6667 | NA | NA | 13 | 0.8462 |
| 6_3956451 | 1 | 0 | 3 | 0.3333 | 4 | 0.25 | 2 | 0 | 10 | 0.2 | 3 | 1 | 8 | 0 | 4 | 0.25 | 1 | 0 | 16 | 0.25 |
| 6_3958000 | NA | NA | NA | NA | 1 | 1 | 1 | 0 | 2 | 0.5 | NA | NA | NA | NA | 1 | 1 | NA | NA | 1 | 1 |
| 6_3958054 | 9 | 0.1111 | 12 | 0.3333 | 13 | 0.3077 | 7 | 0.4286 | 41 | 0.2927 | 10 | 0.4 | 19 | 0.3684 | 15 | 0.3333 | 13 | 0.5385 | 57 | 0.4035 |
| 6_3958062 | 9 | 0 | 9 | 0 | 12 | 0.1667 | 6 | 0 | 36 | 0.0556 | 11 | 0.0909 | 15 | 0.2 | 13 | 0.0769 | 11 | 0 | 50 | 0.1 |
| 6_3958064 | 8 | 0.125 | 12 | 0.6667 | 13 | 0.4615 | 6 | 0.6667 | 39 | 0.4872 | 9 | 0.4444 | 17 | 0.6471 | 11 | 0.7273 | 13 | 0.6923 | 50 | 0.64 |
| 6_3958085 | 8 | 0 | 12 | 0.0833 | 11 | 0.1818 | 6 | 0 | 37 | 0.0811 | 11 | 0.1818 | 17 | 0.1176 | 12 | 0.1667 | 11 | 0.2727 | 51 | 0.1765 |
| 6_3958095 | 7 | 0.1429 | 9 | 0.2222 | 9 | 0 | 4 | 0 | 29 | 0.1034 | 11 | 0.0909 | 12 | 0.0833 | 9 | 0.1111 | 6 | 0.1667 | 38 | 0.1053 |
| 6_3958099 | 8 | 0.25 | 8 | 0.625 | 10 | 0.5 | 5 | 0.6 | 31 | 0.4839 | 10 | 0.5 | 15 | 0.7333 | 12 | 0.4167 | 11 | 0.7273 | 48 | 0.6042 |
| 6_3958100 | 1 | 0 | 5 | 0 | 8 | 0.25 | 4 | 0 | 18 | 0.1111 | 5 | 0 | 6 | 0.3333 | 6 | 0.1667 | 10 | 0 | 27 | 0.1111 |
| 6_3958148 | 7 | 0 | 7 | 0.4286 | 10 | 0.2 | 6 | 0 | 30 | 0.1667 | 10 | 0 | 12 | 0 | 10 | 0.3 | 10 | 0.3 | 42 | 0.1429 |
| 6_3958151 | 7 | 0 | 7 | 0.1429 | 9 | 0.2222 | 5 | 0 | 28 | 0.1071 | 9 | 0.2222 | 14 | 0.0714 | 9 | 0.1111 | 8 | 0.125 | 40 | 0.125 |
| 6_3958161 | 7 | 0 | 7 | 0.2857 | 11 | 0.0909 | 6 | 0 | 31 | 0.0968 | 10 | 0.1 | 15 | 0.1333 | 10 | 0.1 | 10 | 0.1 | 45 | 0.1111 |
| 6_3958435 | 4 | 0.5 | 3 | 0 | 2 | 0 | 3 | 0 | 12 | 0.1667 | 3 | 0 | 6 | 0.5 | NA | NA | 6 | 0 | 15 | 0.2 |
| 6_3958438 | 4 | 0.25 | 4 | 0.25 | 2 | 0.5 | 4 | 0.25 | 14 | 0.2857 | 4 | 0.25 | 6 | 0.5 | 1 | 0 | 6 | 0.1667 | 17 | 0.2941 |
| 6_3958451 | 4 | 0.25 | 3 | 0 | 1 | 0 | 2 | 0 | 10 | 0.1 | 3 | 0 | 7 | 0.1429 | 1 | 0 | 6 | 0.1667 | 17 | 0.1176 |
| 6_3958704 | 2 | 0 | 2 | 0.5 | 1 | 1 | 2 | 0.5 | 7 | 0.4286 | 2 | 0.5 | 2 | 0 | NA | NA | 7 | 0.2857 | 11 | 0.2727 |
| 6_3958709 | 2 | 0.5 | 1 | 0 | NA | NA | 1 | 0 | 4 | 0.25 | 2 | 0 | 2 | 0 | NA | NA | 3 | 0.3333 | 7 | 0.1429 |
| 6_3958713 | 2 | 0 | 2 | 0.5 | 1 | 0 | 2 | 0.5 | 7 | 0.2857 | 2 | 0 | 2 | 0 | NA | NA | 5 | 0.2 | 9 | 0.1111 |
| 6_3958717 | 1 | 0 | 2 | 0 | 1 | 1 | 2 | 0.5 | 6 | 0.3333 | 1 | 0 | 1 | 0 | NA | NA | 5 | 0.2 | 7 | 0.1429 |
| 6_3977883 | 1 | 0 | 5 | 0.2 | NA | NA | 3 | 0 | 9 | 0.1111 | NA | NA | 4 | 0 | 2 | 0.5 | NA | NA | 6 | 0.1667 |
| 6_3977912 | 10 | 0 | 10 | 0.2 | 15 | 0.2667 | 14 | 0.0714 | 49 | 0.1429 | 7 | 0 | 17 | 0.2353 | 22 | 0.1818 | 12 | 0.0833 | 58 | 0.1552 |
| 6_3977918 | 19 | 0.1053 | 14 | 0.1429 | 22 | 0.1364 | 19 | 0.1579 | 74 | 0.1351 | 14 | 0.1429 | 21 | 0.0952 | 25 | 0.12 | 17 | 0.1176 | 77 | 0.1169 |
| 6_3977923 | 23 | 0.6522 | 17 | 0.6471 | 23 | 0.8261 | 19 | 0.4737 | 82 | 0.6585 | 14 | 0.5714 | 22 | 0.7273 | 26 | 0.6538 | 16 | 0.625 | 78 | 0.6538 |
| 6_3977928 | 20 | 0.1 | 19 | 0.1579 | 23 | 0 | 19 | 0.0526 | 81 | 0.0741 | 14 | 0 | 23 | 0.087 | 26 | 0.1154 | 17 | 0.1176 | 80 | 0.0875 |
| 6_3977929 | 21 | 0.1905 | 18 | 0.1667 | 19 | 0.1053 | 15 | 0.2 | 73 | 0.1644 | 13 | 0.0769 | 22 | 0.0455 | 26 | 0.0769 | 14 | 0.0714 | 75 | 0.0667 |
| 6_3977933 | 22 | 0.0909 | 20 | 0.25 | 18 | 0.3333 | 18 | 0.1667 | 78 | 0.2051 | 13 | 0.4615 | 23 | 0.3913 | 25 | 0.28 | 17 | 0.4706 | 78 | 0.3846 |
| 6_3977946 | 21 | 0.0476 | 17 | 0 | 20 | 0.05 | 17 | 0.1176 | 75 | 0.0533 | 14 | 0.0714 | 22 | 0.0455 | 24 | 0.1667 | 17 | 0 | 77 | 0.0779 |
| 6_3978237 | 21 | 0.1429 | 13 | 0.0769 | 20 | 0.3 | 16 | 0.125 | 70 | 0.1714 | 13 | 0.0769 | 21 | 0.381 | 24 | 0.1667 | 16 | 0.1875 | 74 | 0.2162 |
| 6_3984957 | NA | NA | NA | NA | NA | NA | 2 | 1 | 2 | 1 | NA | NA | 1 | 1 | NA | NA | NA | NA | 1 | 1 |
| 6_4022334 | 1 | 1 | NA | NA | 1 | 1 | 3 | 1 | 5 | 1 | NA | NA | 1 | 1 | NA | NA | NA | NA | 1 | 1 |
| 6_4022367 | 1 | 1 | NA | NA | 1 | 0 | 3 | 0 | 5 | 0.2 | NA | NA | 1 | 0 | 1 | 1 | NA | NA | 2 | 0.5 |
| 6_4022383 | 1 | 0 | NA | NA | 1 | 0 | 3 | 0.3333 | 5 | 0.2 | 1 | 0 | 1 | 0 | 1 | 1 | NA | NA | 3 | 0.3333 |
| 6_4022410 | 1 | 1 | NA | NA | 2 | 0 | 2 | 0.5 | 5 | 0.4 | 2 | 1 | 1 | 1 | 1 | 0 | NA | NA | 4 | 0.75 |
| 6_4022426 | 1 | 1 | 1 | 0 | 1 | 1 | 3 | 0.6667 | 6 | 0.6667 | 1 | 1 | NA | NA | 1 | 1 | NA | NA | 2 | 1 |
| 6_4022427 | 1 | 1 | 1 | 0 | 2 | 0.5 | 3 | 0 | 7 | 0.2857 | 1 | 1 | 1 | 1 | 1 | 1 | NA | NA | 3 | 1 |
| 6_4022437 | 1 | 1 | 1 | 0 | 2 | 1 | 3 | 0 | 7 | 0.4286 | 2 | 0 | 1 | 1 | 1 | 0 | NA | NA | 4 | 0.25 |
| 6_4022758 | 3 | 0.3333 | 2 | 0.5 | 5 | 0 | 6 | 0 | 16 | 0.125 | 3 | 0 | 4 | 0.25 | 2 | 0 | NA | NA | 9 | 0.1111 |
| 6_4022918 | 4 | 0.25 | 2 | 0 | 6 | 0.3333 | 5 | 0 | 17 | 0.1765 | 2 | 1 | 3 | 0 | 2 | 0.5 | 2 | 0.5 | 9 | 0.4444 |
| 6_4022930 | 5 | 0 | 3 | 0.3333 | 5 | 0 | 6 | 0 | 19 | 0.0526 | 2 | 0.5 | 3 | 0 | 2 | 0 | 2 | 0 | 9 | 0.1111 |
| 6_4022943 | 5 | 0 | 3 | 0 | 6 | 0.1667 | 6 | 0 | 20 | 0.05 | 1 | 1 | 2 | 0.5 | 2 | 0.5 | 2 | 0 | 7 | 0.4286 |
| 6_4022979 | 6 | 0.1667 | 3 | 0.3333 | 8 | 0 | 6 | 0.1667 | 23 | 0.1304 | 2 | 0 | 1 | 0 | 1 | 0 | 3 | 0.3333 | 7 | 0.1429 |
| 6_4023175 | 11 | 0.0909 | 10 | 0.1 | 12 | 0.0833 | 12 | 0.0833 | 45 | 0.0889 | 3 | 0 | 11 | 0.1818 | 7 | 0 | 10 | 0 | 31 | 0.0645 |
| 6_4023178 | 7 | 0.1429 | 8 | 0.375 | 10 | 0.2 | 8 | 0.375 | 33 | 0.2727 | 3 | 0.3333 | 10 | 0.1 | 6 | 0.3333 | 7 | 0.7143 | 26 | 0.3462 |
| 6_4023183 | 11 | 0.0909 | 10 | 0 | 11 | 0.1818 | 12 | 0.1667 | 44 | 0.1136 | 3 | 0.6667 | 11 | 0.1818 | 8 | 0.25 | 8 | 0.25 | 30 | 0.2667 |
| 6_4023184 | 10 | 1 | 11 | 1 | 11 | 0.8182 | 13 | 0.9231 | 45 | 0.9333 | 2 | 1 | 10 | 0.9 | 8 | 1 | 10 | 0.9 | 30 | 0.9333 |
| 6_4023187 | 11 | 0.2727 | 10 | 0.7 | 11 | 0.5455 | 13 | 0.3846 | 45 | 0.4667 | 2 | 1 | 10 | 0.3 | 10 | 0.4 | 10 | 0.4 | 32 | 0.4062 |
| 6_4023189 | 9 | 0.2222 | 9 | 0.2222 | 13 | 0.2308 | 11 | 0.4545 | 42 | 0.2857 | 5 | 0.2 | 11 | 0.3636 | 8 | 0.5 | 8 | 0.25 | 32 | 0.3438 |
| 6_4037343 | 5 | 0 | 2 | 0 | 4 | 0.25 | 4 | 0 | 15 | 0.0667 | 2 | 0 | 2 | 0 | 10 | 0.1 | 3 | 0 | 17 | 0.0588 |
| 6_4038031 | 4 | 0.25 | 6 | 0 | 2 | 0 | 5 | 0 | 17 | 0.0588 | 2 | 0 | 8 | 0.125 | 5 | 0 | 2 | 0 | 17 | 0.0588 |
| 6_4038185 | 1 | 1 | 4 | 0.5 | 2 | 0 | 2 | 0 | 9 | 0.3333 | 2 | 0 | 4 | 0 | 5 | 0.2 | 1 | 1 | 12 | 0.1667 |
| 6_4038266 | NA | NA | 4 | 0.25 | 2 | 0 | 2 | 0 | 8 | 0.125 | 1 | 0 | 4 | 0 | 3 | 0.3333 | 2 | 0 | 10 | 0.1 |
| 6_4038336 | NA | NA | 3 | 1 | 1 | 1 | 2 | 0.5 | 6 | 0.8333 | 2 | 1 | 1 | 1 | 2 | 0 | 2 | 0.5 | 7 | 0.5714 |
| 6_4038338 | NA | NA | 3 | 0 | 1 | 1 | 2 | 0 | 6 | 0.1667 | 2 | 0 | 1 | 0 | 3 | 0 | 2 | 0.5 | 8 | 0.125 |
| 6_4038356 | NA | NA | 3 | 1 | 1 | 1 | 2 | 0.5 | 6 | 0.8333 | 2 | 1 | 1 | 1 | 3 | 1 | 1 | 1 | 7 | 1 |
| 6_4038364 | NA | NA | 1 | 1 | NA | NA | 2 | 0 | 3 | 0.3333 | 2 | 0.5 | 1 | 0 | 3 | 1 | 1 | 0 | 7 | 0.5714 |
| 6_4038365 | NA | NA | 3 | 1 | 1 | 1 | 2 | 1 | 6 | 1 | 2 | 1 | 1 | 1 | 3 | 1 | 2 | 1 | 8 | 1 |
| 6_4038373 | NA | NA | 2 | 0.5 | 1 | 1 | 2 | 0 | 5 | 0.4 | 2 | 0 | 1 | 0 | 3 | 0.3333 | 2 | 0 | 8 | 0.125 |
| 6_4038376 | NA | NA | 2 | 0.5 | NA | NA | 1 | 0 | 3 | 0.3333 | 2 | 0.5 | 1 | 1 | 2 | 0.5 | 2 | 0.5 | 7 | 0.5714 |
| 6_4038380 | NA | NA | 3 | 0.3333 | 1 | 0 | 2 | 0 | 6 | 0.1667 | 2 | 0 | 1 | 0 | 3 | 0.6667 | 1 | 0 | 7 | 0.2857 |
| 6_4038383 | NA | NA | 3 | 0.3333 | 1 | 0 | 2 | 0 | 6 | 0.1667 | 2 | 0 | 1 | 1 | 3 | 0 | 1 | 0 | 7 | 0.1429 |
| 6_4159122 | NA | NA | 1 | 1 | NA | NA | 2 | 0 | 3 | 0.3333 | NA | NA | NA | NA | 1 | 1 | NA | NA | 1 | 1 |
| 6_4159130 | NA | NA | 1 | 0 | NA | NA | 2 | 0.5 | 3 | 0.3333 | NA | NA | 1 | 0 | 1 | 0 | 1 | 1 | 3 | 0.3333 |

| | | | | | | | | | | | | | | | | | | | | |
|---|---|---|---|---|---|---|---|---|---|---|---|---|---|---|---|---|---|---|---|---|
| 6_4159131 | NA | NA | 1 | 1 | NA | NA | 2 | 0.5 | 3 | 0.6667 | NA | NA | 1 | 1 | NA | NA | NA | NA | 1 | 1 |
| 6_4159140 | NA | NA | 1 | 0 | NA | NA | 1 | 1 | 2 | 0.5 | NA | NA | 1 | 1 | 1 | 1 | 1 | 1 | 3 | 1 |
| 6_4159334 | NA | NA | NA | NA | 1 | 1 | 2 | 0 | 3 | 0.3333 | NA | NA | NA | NA | 2 | 0.5 | NA | NA | 2 | 0.5 |
| 6_4159673 | 2 | 0.5 | 3 | 0.3333 | NA | NA | 5 | 0.2 | 10 | 0.3 | 1 | 0 | 1 | 1 | 2 | 0 | 1 | 0 | 5 | 0.2 |
| 6_4159689 | 1 | 1 | 2 | 0 | 2 | 0.5 | 5 | 0.8 | 10 | 0.6 | 2 | 1 | 1 | 0 | 2 | 0.5 | 1 | 0 | 6 | 0.5 |
| 6_4159690 | NA | NA | 3 | 1 | 2 | 0 | 4 | 1 | 9 | 0.7778 | 2 | 1 | 2 | 0 | 3 | 0.3333 | NA | NA | 7 | 0.4286 |
| 6_4159693 | 2 | 0.5 | 4 | 0.5 | 2 | 0 | 6 | 0.5 | 14 | 0.4286 | 2 | 1 | 1 | 0 | 2 | 0.5 | 1 | 0 | 6 | 0.5 |
| 6_4159731 | 3 | 0.3333 | 4 | 0.5 | 2 | 0.5 | 8 | 0.375 | 17 | 0.4118 | 3 | 0.3333 | 2 | 0.5 | 3 | 0.3333 | 1 | 0 | 9 | 0.3333 |
| 6_4159737 | 2 | 1 | 2 | 0 | 1 | 1 | 8 | 0.75 | 13 | 0.6923 | 3 | 0.3333 | 1 | 0 | 3 | 0.6667 | 1 | 0 | 8 | 0.375 |
| 6_4222386 | 5 | 0 | 3 | 0 | 3 | 0.3333 | 1 | 0 | 12 | 0.0833 | 2 | 0 | 6 | 0.1667 | 4 | 0 | 2 | 0 | 14 | 0.0714 |
| 6_4229295 | 3 | 0 | 4 | 0.5 | 1 | 1 | NA | NA | 8 | 0.375 | 1 | 0 | 1 | 1 | 1 | 0 | 2 | 0 | 5 | 0.2 |
| 6_4229305 | 4 | 0 | 5 | 0.4 | 1 | 1 | 1 | 1 | 11 | 0.3636 | 1 | 0 | 1 | 1 | 2 | 0 | 1 | 0 | 5 | 0.2 |
| 6_4229363 | 3 | 0.3333 | 2 | 0 | 1 | 0 | NA | NA | 6 | 0.1667 | NA | NA | NA | NA | 2 | 0.5 | NA | NA | 2 | 0.5 |
| 6_4229392 | 4 | 0.25 | 4 | 0 | 1 | 0 | NA | NA | 9 | 0.1111 | 1 | 0 | 1 | 1 | 2 | 0 | NA | NA | 4 | 0.25 |
| 6_4229393 | 4 | 0.25 | 3 | 0 | NA | NA | NA | NA | 7 | 0.1429 | 1 | 0 | 1 | 0 | 1 | 1 | NA | NA | 3 | 0.3333 |
| 6_4229437 | 1 | 1 | 2 | 1 | NA | NA | NA | NA | 3 | 1 | 1 | 1 | NA | NA | 1 | 1 | 1 | 1 | 3 | 1 |
| 6_4229444 | 1 | 1 | 4 | 0.75 | NA | NA | NA | NA | 5 | 0.8 | 3 | 0.6667 | NA | NA | 1 | 1 | 1 | 1 | 5 | 0.8 |
| 6_4229456 | 2 | 0.5 | 4 | 0.5 | NA | NA | NA | NA | 6 | 0.5 | 3 | 0 | NA | NA | 1 | 1 | 1 | 1 | 5 | 0.4 |
| 6_4229461 | 1 | 0 | 3 | 0.3333 | NA | NA | NA | NA | 4 | 0.25 | 1 | 1 | NA | NA | 1 | 0 | 1 | 0 | 3 | 0.3333 |
| 6_4229463 | 2 | 1 | 4 | 1 | NA | NA | NA | NA | 6 | 1 | 2 | 1 | NA | NA | 1 | 1 | 1 | 1 | 4 | 1 |
| 6_4229480 | 1 | 0 | 5 | 0.2 | 1 | 1 | NA | NA | 7 | 0.2857 | 1 | 1 | 1 | 1 | 1 | 0 | 2 | 0.5 | 5 | 0.6 |
| 6_4229487 | 2 | 0.5 | 4 | 0.5 | 1 | 0 | NA | NA | 7 | 0.4286 | 1 | 0 | 1 | 1 | 1 | 0 | 2 | 0.5 | 5 | 0.4 |
| 6_4229488 | 2 | 0 | 5 | 0.6 | 1 | 0 | NA | NA | 8 | 0.375 | 1 | 0 | NA | NA | 1 | 1 | 2 | 0 | 4 | 0.25 |
| 6_4229488 | 2 | 1 | 3 | 1 | 1 | 1 | NA | NA | 6 | 1 | 1 | 1 | 1 | 1 | 1 | 1 | 1 | 1 | 4 | 1 |
| 6_4229494 | 1 | 1 | 6 | 0.8333 | NA | NA | NA | NA | 7 | 0.8571 | 1 | 1 | 1 | 1 | 1 | 1 | 2 | 1 | 5 | 1 |
| 6_4229499 | 2 | 0 | 6 | 0.3333 | NA | NA | NA | NA | 8 | 0.25 | 1 | 0 | 1 | 1 | 1 | 0 | 2 | 0.5 | 5 | 0.4 |
| 6_4229519 | 1 | 1 | 4 | 0 | NA | NA | NA | NA | 5 | 0.2 | 1 | 0 | 1 | 1 | NA | NA | 2 | 0.5 | 4 | 0.5 |
| 6_4229529 | 2 | 0 | 5 | 0.4 | NA | NA | NA | NA | 7 | 0.2857 | 1 | 0 | 1 | 1 | NA | NA | 2 | 0.5 | 4 | 0.5 |
| 6_4229531 | 1 | 1 | 3 | 1 | NA | NA | NA | NA | 4 | 1 | 1 | 1 | 1 | 1 | NA | NA | 2 | 1 | 4 | 1 |
| 6_4229532 | 2 | 0 | 5 | 0.4 | NA | NA | NA | NA | 7 | 0.2857 | 1 | 0 | 1 | 0 | NA | NA | 2 | 0.5 | 4 | 0.25 |
| 6_4233490 | 7 | 0.1429 | 4 | 0 | 4 | 0 | 3 | 0 | 18 | 0.0556 | 5 | 0.2 | 4 | 0.25 | 4 | 0 | 1 | 0 | 14 | 0.1429 |
| 6_4233526 | 9 | 0.1111 | 4 | 0 | 5 | 0.4 | 3 | 0 | 21 | 0.1429 | 6 | 0.1667 | 4 | 0.25 | 6 | 0 | 1 | 0 | 17 | 0.1176 |
| 6_4264548 | 12 | 0.0833 | 13 | 0 | 14 | 0.1429 | 20 | 0.1 | 59 | 0.0847 | 10 | 0.2 | 20 | 0.1 | 12 | 0.1667 | 16 | 0.1875 | 58 | 0.1552 |
| 6_4264566 | 11 | 0 | 12 | 0.1667 | 11 | 0.1818 | 20 | 0.05 | 54 | 0.0926 | 8 | 0.25 | 22 | 0.1364 | 11 | 0.1818 | 14 | 0.1429 | 55 | 0.1636 |
| 6_4264579 | 11 | 0.1818 | 10 | 0.1 | 12 | 0.3333 | 20 | 0.25 | 53 | 0.2264 | 9 | 0.3333 | 16 | 0.1875 | 11 | 0.4545 | 15 | 0.1333 | 51 | 0.2549 |
| 6_4264645 | 10 | 0.2 | 6 | 0.3333 | 10 | 0.4 | 15 | 0.3333 | 41 | 0.3171 | 8 | 0.25 | 19 | 0.0526 | 9 | 0 | 14 | 0.0714 | 50 | 0.08 |
| 6_4264653 | 10 | 0.1 | 6 | 0 | 10 | 0.2 | 17 | 0.0588 | 43 | 0.093 | 9 | 0.1111 | 18 | 0 | 8 | 0 | 13 | 0.3077 | 48 | 0.1042 |
| 6_4264673 | 9 | 0.1111 | 4 | 0.25 | 9 | 0.3333 | 16 | 0.125 | 38 | 0.1842 | 8 | 0.375 | 21 | 0.1905 | 4 | 0.25 | 13 | 0.1538 | 46 | 0.2174 |
| 6_4264706 | 15 | 0.8 | 15 | 0.9333 | 17 | 0.9412 | 17 | 0.9412 | 64 | 0.9062 | 14 | 0.8571 | 30 | 0.7333 | 16 | 0.9375 | 21 | 0.8095 | 81 | 0.8148 |
| 6_4264707 | 15 | 0.0667 | 12 | 0.0833 | 14 | 0.1429 | 16 | 0.0625 | 57 | 0.0877 | 10 | 0 | 28 | 0.1429 | 11 | 0.0909 | 18 | 0.1111 | 67 | 0.1045 |
| 6_4264729 | NA | NA | 2 | 0.5 | 2 | 0 | 1 | 1 | 5 | 0.4 | 1 | 1 | 3 | 0.3333 | 1 | 1 | 3 | 0.3333 | 8 | 0.5 |
| 6_4264786 | NA | NA | 2 | 0 | 2 | 0 | 1 | 1 | 5 | 0.2 | NA | NA | 4 | 0.25 | 1 | 0 | 3 | 0.3333 | 8 | 0.25 |
| 6_4264988 | NA | NA | 2 | 1 | 3 | 0.6667 | 3 | 0.3333 | 8 | 0.625 | 1 | 1 | NA | NA | NA | NA | 2 | 1 | 3 | 1 |
| 6_4265223 | 7 | 0.1429 | 3 | 0.3333 | 8 | 0.25 | 12 | 0 | 30 | 0.1333 | 4 | 0.25 | 12 | 0.25 | 8 | 0 | 8 | 0 | 32 | 0.125 |
| 6_4265551 | 4 | 0.5 | 7 | 0.5714 | 4 | 0.5 | 8 | 0.5 | 23 | 0.5217 | 5 | 1 | 14 | 0.4286 | 9 | 0.5556 | 11 | 0.8182 | 39 | 0.641 |
| 6_4265558 | 9 | 0.1111 | 8 | 0 | 6 | 0.1667 | 11 | 0.1818 | 34 | 0.1176 | 5 | 0 | 10 | 0.2 | 9 | 0.1111 | 12 | 0.0833 | 36 | 0.1111 |
| 6_4265574 | 9 | 0.5556 | 8 | 0.875 | 6 | 0.6667 | 10 | 0.9 | 33 | 0.7576 | 4 | 0.5 | 11 | 0.1818 | 11 | 0.9091 | 10 | 0.8 | 36 | 0.6111 |
| 6_4265577 | 6 | 0.1667 | 8 | 0.25 | 6 | 0.6667 | 8 | 0.25 | 28 | 0.3214 | 4 | 0.25 | 7 | 0.4286 | 11 | 0.0909 | 10 | 0.4 | 32 | 0.2812 |
| 6_4265589 | 7 | 0.2857 | 6 | 0.3333 | 4 | 0 | 5 | 0.2 | 22 | 0.2273 | 3 | 0.3333 | 7 | 0 | 10 | 0.2 | 9 | 0.2222 | 29 | 0.1724 |
| 6_4265593 | 8 | 0.25 | 7 | 0.1429 | 5 | 0.6 | 7 | 0.1429 | 27 | 0.2593 | 4 | 0.25 | 8 | 0.25 | 9 | 0.1111 | 8 | 0.5 | 29 | 0.2759 |
| 6_4265628 | 6 | 0.6667 | 7 | 0.2857 | 3 | 0.6667 | 6 | 0.6667 | 22 | 0.5455 | 4 | 0.5 | 8 | 0.5 | 7 | 0.5714 | 8 | 0.5 | 27 | 0.5185 |
| 6_4265818 | 2 | 0.5 | 1 | 0 | NA | NA | NA | NA | 3 | 0.3333 | NA | NA | 1 | 1 | NA | NA | NA | NA | 1 | 1 |
| 6_4266347 | 3 | 0 | 5 | 0 | 3 | 0 | 5 | 0.2 | 16 | 0.0625 | 3 | 0.3333 | 5 | 0 | 6 | 0 | 8 | 0.125 | 22 | 0.0909 |
| 6_4266352 | 3 | 0.6667 | 5 | 0.6 | 3 | 0.3333 | 5 | 0.2 | 16 | 0.4375 | 3 | 0.3333 | 6 | 0.5 | 7 | 0.5714 | 6 | 0.6667 | 22 | 0.5455 |
| 6_4266368 | 3 | 1 | 3 | 0.6667 | 4 | 1 | 3 | 1 | 13 | 0.9231 | 3 | 1 | 4 | 1 | 8 | 1 | 8 | 1 | 23 | 1 |
| 6_4266379 | 3 | 0.6667 | 4 | 0.25 | 5 | 0.4 | 3 | 0.6667 | 15 | 0.4667 | 2 | 0.5 | 5 | 0.2 | 5 | 0.6 | 8 | 0.5 | 20 | 0.45 |
| 6_4722139 | 3 | 0 | 3 | 0 | 5 | 0 | 9 | 0.2222 | 20 | 0.1 | 3 | 0.3333 | 4 | 0 | 4 | 0 | 4 | 0.25 | 15 | 0.1333 |
| 6_4727551 | 5 | 0.4 | 5 | 0.2 | 12 | 0.3333 | 15 | 0.1333 | 37 | 0.2432 | 4 | 0 | 11 | 0.2727 | 10 | 0.2 | 9 | 0.2222 | 34 | 0.2059 |
| 6_4727569 | 8 | 0.125 | 8 | 0.5 | 14 | 0.6429 | 16 | 0.5 | 46 | 0.4783 | 5 | 1 | 13 | 0.4615 | 11 | 0.6364 | 7 | 0.5714 | 36 | 0.6111 |
| 6_4727580 | 10 | 0.1 | 7 | 0 | 11 | 0.0909 | 15 | 0.1333 | 43 | 0.093 | 5 | 0.2 | 12 | 0.0833 | 11 | 0.1818 | 7 | 0.2857 | 35 | 0.1714 |
| 6_4727593 | 7 | 0.1429 | 7 | 0.1429 | 17 | 0.0588 | 15 | 0.0667 | 46 | 0.087 | 4 | 0.25 | 11 | 0 | 11 | 0 | 5 | 0.2 | 31 | 0.0645 |
| 6_4727626 | 6 | 0 | 5 | 0 | 11 | 0.2727 | 10 | 0 | 32 | 0.0938 | 7 | 0 | 13 | 0 | 9 | 0.2222 | 7 | 0.1429 | 36 | 0.0833 |
| 6_4727735 | 1 | 0 | 1 | 0 | 1 | 0 | 2 | 0.5 | 5 | 0.2 | 1 | 0 | 2 | 0 | 2 | 0 | 1 | 1 | 6 | 0.1667 |
| 6_4728000 | 3 | 0 | 2 | 0 | 4 | 0.25 | 7 | 0 | 16 | 0.0625 | 1 | 0 | 5 | 0 | 3 | 0.3333 | 2 | 0 | 11 | 0.0909 |
| 6_4728009 | 3 | 0 | 3 | 0 | 4 | 0 | 6 | 0.1667 | 16 | 0.0625 | 2 | 0 | 4 | 0.25 | 3 | 0 | 2 | 0 | 11 | 0.0909 |

| | | | | | | | | | | | | | | | | | | | | |
|---|---|---|---|---|---|---|---|---|---|---|---|---|---|---|---|---|---|---|---|---|
| 6_4740892 | NA | NA | 1 | 1 | 1 | 1 | 2 | 1 | 4 | 1 | NA | NA | NA | NA | NA | NA | 2 | 1 | 2 | 1 |
| 6_4742978 | 1 | 1 | NA | NA | NA | NA | 1 | 1 | 2 | 1 | NA | NA | NA | NA | 3 | 0.3333 | NA | NA | 3 | 0.3333 |
| 6_4742981 | NA | NA | NA | NA | NA | NA | 1 | 1 | 1 | 1 | NA | NA | NA | NA | 2 | 0.5 | NA | NA | 2 | 0.5 |
| 6_4743060 | 1 | 1 | NA | NA | NA | NA | 1 | 1 | 2 | 1 | NA | NA | NA | NA | 3 | 1 | NA | NA | 3 | 1 |
| 6_4743083 | 1 | 1 | NA | NA | NA | NA | 1 | 1 | 2 | 1 | NA | NA | NA | NA | 2 | 1 | NA | NA | 2 | 1 |
| 6_4746169 | 2 | 0 | 10 | 0.2 | 3 | 0 | 5 | 0 | 20 | 0.1 | 5 | 0 | 4 | 0 | 8 | 0 | 3 | 0.3333 | 20 | 0.05 |
| 6_4746171 | 3 | 0 | 10 | 0 | 1 | 0 | 5 | 0.4 | 19 | 0.1053 | 7 | 0.1429 | 4 | 0 | 7 | 0.2857 | 4 | 0 | 22 | 0.1364 |
| 6_4746178 | 3 | 0 | 11 | 0.1818 | 3 | 0.3333 | 6 | 0 | 23 | 0.1304 | 9 | 0.4444 | 5 | 0 | 9 | 0 | 4 | 0.25 | 27 | 0.1852 |
| 6_4746187 | 2 | 0 | 9 | 0.6667 | NA | NA | 4 | 0.5 | 15 | 0.5333 | 6 | 0.1667 | 5 | 0.2 | 6 | 0.1667 | 3 | 1 | 20 | 0.3 |
| 6_4746197 | 3 | 0.3333 | 11 | 0.0909 | 4 | 0 | 6 | 0.1667 | 24 | 0.125 | 9 | 0.3333 | 5 | 0 | 9 | 0.2222 | 4 | 0 | 27 | 0.1852 |
| 6_4750254 | 11 | 0.2727 | 18 | 0.1111 | 12 | 0.3333 | 11 | 0 | 52 | 0.1731 | 12 | 0.1667 | 11 | 0.0909 | 16 | 0.0625 | 10 | 0.2 | 49 | 0.1224 |
| 6_4750266 | 10 | 0.5 | 17 | 0.7647 | 11 | 0.4545 | 13 | 0.6154 | 51 | 0.6078 | 12 | 0.5833 | 10 | 0.6 | 15 | 0.6667 | 10 | 0.5 | 47 | 0.5957 |
| 6_4750272 | 13 | 0.7692 | 19 | 0.8421 | 10 | 0.8 | 12 | 1 | 54 | 0.8519 | 12 | 0.8333 | 11 | 0.6364 | 15 | 0.8667 | 10 | 1 | 48 | 0.8333 |
| 6_4750273 | 12 | 0.8333 | 21 | 0.9524 | 10 | 0.9 | 14 | 0.8571 | 57 | 0.8947 | 12 | 0.75 | 11 | 1 | 17 | 0.6471 | 10 | 0.8 | 50 | 0.78 |
| 6_4750273 | 12 | 0.0833 | 17 | 0.2353 | 11 | 0.4545 | 8 | 0 | 48 | 0.2083 | 11 | 0.2727 | 9 | 0.1111 | 16 | 0.125 | 8 | 0.125 | 44 | 0.1591 |
| 6_4750279 | 12 | 0.25 | 17 | 0.2941 | 12 | 0.1667 | 11 | 0.1818 | 52 | 0.2308 | 12 | 0.3333 | 8 | 0.125 | 18 | 0.2222 | 8 | 0.125 | 46 | 0.2174 |
| 6_4750291 | 14 | 0.2143 | 21 | 0.1905 | 14 | 0.3571 | 15 | 0.4667 | 64 | 0.2969 | 15 | 0.0667 | 12 | 0.0833 | 17 | 0.3529 | 10 | 0.4 | 54 | 0.2222 |
| 6_4763294 | 2 | 1 | 2 | 0.5 | 4 | 0.75 | NA | NA | 8 | 0.75 | 1 | 1 | 1 | 1 | 1 | 0 | 1 | 1 | 4 | 0.75 |
| 6_4763298 | 2 | 1 | 2 | 1 | 3 | 0.6667 | 1 | 1 | 8 | 0.875 | 1 | 0 | NA | NA | 1 | 0 | 1 | 1 | 3 | 0.3333 |
| 6_4763299 | 2 | 1 | 1 | 1 | 3 | 1 | 1 | 1 | 7 | 1 | 1 | 1 | 1 | 1 | 1 | 0 | 1 | 1 | 4 | 0.75 |
| 6_4763302 | 1 | 1 | 2 | 1 | 3 | 0.6667 | 1 | 1 | 7 | 0.8571 | 1 | 1 | 1 | 1 | 1 | 1 | 1 | 0 | 4 | 0.75 |
| 6_4763312 | 2 | 1 | 1 | 1 | 4 | 1 | 2 | 0.5 | 9 | 0.8889 | 1 | 1 | 1 | 0 | 1 | 1 | 1 | 1 | 4 | 0.75 |
| 6_4763322 | 2 | 0.5 | 2 | 0.5 | 4 | 0.5 | 2 | 1 | 10 | 0.6 | 1 | 1 | 1 | 0 | 1 | 1 | 1 | 1 | 4 | 0.75 |
| 6_4763363 | 2 | 1 | 1 | 1 | 3 | 0.6667 | 1 | 1 | 7 | 0.8571 | 1 | 1 | NA | NA | NA | NA | 1 | 1 | 2 | 1 |
| 6_4763374 | 2 | 1 | 1 | 1 | 4 | 1 | 1 | 1 | 8 | 1 | 1 | 0 | NA | NA | NA | NA | 1 | 1 | 2 | 0.5 |
| 6_4763382 | 2 | 1 | 1 | 1 | 3 | 1 | 1 | 1 | 7 | 1 | 1 | 1 | NA | NA | NA | NA | 1 | 1 | 2 | 1 |
| 6_4763394 | 2 | 0.5 | NA | NA | 2 | 0.5 | 1 | 0 | 5 | 0.4 | 2 | 0.5 | NA | NA | NA | NA | 1 | 0 | 3 | 0.3333 |
| 6_4763397 | 2 | 0.5 | 1 | 0 | 3 | 0.3333 | NA | NA | 6 | 0.3333 | 2 | 0.5 | NA | NA | NA | NA | 1 | 1 | 3 | 0.6667 |
| 6_4763725 | 1 | 1 | NA | NA | NA | NA | NA | NA | 1 | 1 | 3 | 1 | NA | NA | NA | NA | 1 | 1 | 4 | 1 |
| 6_4763729 | 1 | 1 | 1 | 1 | 1 | 0 | NA | NA | 3 | 0.6667 | 3 | 1 | NA | NA | NA | NA | 1 | 1 | 4 | 1 |
| 6_4763729 | 1 | 1 | 1 | 1 | 1 | 1 | NA | NA | 3 | 1 | 2 | 1 | NA | NA | NA | NA | 1 | 1 | 3 | 1 |
| 6_4763732 | 1 | 1 | 1 | 1 | 1 | 1 | NA | NA | 3 | 1 | 2 | 0.5 | NA | NA | NA | NA | NA | NA | 2 | 0.5 |
| 6_4763743 | 1 | 1 | 2 | 0.5 | 1 | 1 | NA | NA | 4 | 0.75 | 3 | 1 | NA | NA | NA | NA | 1 | 1 | 4 | 1 |
| 6_4763794 | 1 | 1 | 1 | 0 | NA | NA | NA | NA | 2 | 0.5 | 2 | 1 | NA | NA | NA | NA | NA | NA | 2 | 1 |
| 6_4763804 | NA | NA | 2 | 0.5 | NA | NA | NA | NA | 2 | 0.5 | 2 | 0.5 | NA | NA | NA | NA | 1 | 0 | 3 | 0.3333 |
| 6_4763828 | 1 | 1 | 2 | 0.5 | NA | NA | NA | NA | 3 | 0.6667 | 2 | 0.5 | NA | NA | NA | NA | 1 | 0 | 3 | 0.3333 |
| 6_4763911 | 1 | 1 | NA | NA | NA | NA | NA | NA | 1 | 1 | 1 | 1 | NA | NA | NA | NA | NA | NA | 1 | 1 |
| 6_4764161 | 1 | 1 | NA | NA | NA | NA | NA | NA | 1 | 1 | 1 | 1 | 1 | 1 | 2 | 0.5 | 2 | 1 | 6 | 0.8333 |
| 6_4764166 | 1 | 1 | NA | NA | NA | NA | NA | NA | 1 | 1 | 1 | 1 | 1 | 0 | NA | NA | 1 | 1 | 3 | 0.6667 |
| 6_4764168 | 1 | 0 | NA | NA | NA | NA | 1 | 1 | 2 | 0.5 | 1 | 1 | 1 | 1 | 1 | 1 | NA | NA | 3 | 1 |
| 6_4764179 | 1 | 1 | NA | NA | NA | NA | 1 | 1 | 2 | 1 | 1 | 1 | 1 | 1 | 2 | 1 | 2 | 0.5 | 6 | 0.8333 |
| 6_4764230 | 1 | 1 | NA | NA | NA | NA | NA | NA | 1 | 1 | NA | NA | NA | NA | NA | NA | 1 | 1 | 1 | 1 |
| 6_4764249 | 1 | 1 | NA | NA | NA | NA | NA | NA | 1 | 1 | NA | NA | NA | NA | 1 | 1 | 1 | 0 | 2 | 0.5 |
| 6_4765129 | NA | NA | 1 | 1 | NA | NA | NA | NA | 1 | 1 | NA | NA | 1 | 1 | NA | NA | NA | NA | 1 | 1 |
| 6_4765891 | 1 | 1 | 1 | 1 | NA | NA | NA | NA | 2 | 1 | NA | NA | NA | NA | 1 | 1 | NA | NA | 1 | 1 |
| 6_4765895 | 1 | 1 | 1 | 1 | NA | NA | NA | NA | 2 | 1 | NA | NA | NA | NA | 1 | 1 | NA | NA | 1 | 1 |
| 6_4765909 | 1 | 1 | 1 | 1 | NA | NA | NA | NA | 2 | 1 | NA | NA | NA | NA | 1 | 1 | NA | NA | 1 | 1 |
| 6_4772210 | 3 | 1 | 2 | 1 | 1 | 1 | 3 | 0.6667 | 9 | 0.8889 | 1 | 1 | 4 | 1 | NA | NA | 2 | 0.5 | 7 | 0.8571 |
| 6_4772214 | 3 | 1 | 1 | 1 | 1 | 1 | 2 | 1 | 7 | 1 | NA | NA | 4 | 1 | NA | NA | 2 | 1 | 6 | 1 |
| 6_4772214 | 2 | 1 | 2 | 1 | 1 | 1 | 3 | 1 | 8 | 1 | 1 | 1 | 3 | 1 | NA | NA | 1 | 1 | 5 | 1 |
| 6_4772217 | 2 | 1 | 2 | 0 | 1 | 0 | 2 | 0.5 | 7 | 0.4286 | NA | NA | 1 | 0 | NA | NA | 1 | 1 | 2 | 0.5 |
| 6_4772228 | 3 | 1 | 2 | 0.5 | 1 | 0 | 2 | 1 | 8 | 0.75 | 1 | 1 | 4 | 0.75 | NA | NA | 2 | 0.5 | 7 | 0.7143 |
| 6_4772238 | 3 | 0.6667 | 2 | 1 | 1 | 0 | 3 | 0.3333 | 9 | 0.5556 | 1 | 0 | 5 | 0.2 | NA | NA | 2 | 0 | 8 | 0.125 |
| 6_4772279 | 2 | 1 | 3 | 1 | 1 | 1 | 2 | 1 | 8 | 1 | 1 | 1 | 1 | 1 | 1 | 1 | 2 | 1 | 5 | 1 |
| 6_4772289 | 2 | 1 | 3 | 0.6667 | 1 | 0 | 2 | 0.5 | 8 | 0.625 | 1 | 1 | 3 | 1 | 1 | 1 | 2 | 1 | 7 | 1 |
| 6_4772298 | 2 | 0.5 | 3 | 0.6667 | 1 | 0 | 2 | 0.5 | 8 | 0.5 | 1 | 1 | 3 | 1 | 1 | 1 | 2 | 1 | 7 | 1 |
| 6_4772309 | 2 | 0 | 3 | 0.6667 | 1 | 1 | 1 | 1 | 7 | 0.5714 | 2 | 0 | 3 | 0.6667 | NA | NA | 2 | 0 | 7 | 0.2857 |
| 6_4772312 | 2 | 0.5 | 3 | 0.3333 | 1 | 1 | 2 | 0.5 | 8 | 0.5 | 2 | 0.5 | 3 | 1 | 1 | 1 | 1 | 0 | 7 | 0.7143 |
| 6_4772413 | 1 | 1 | NA | NA | NA | NA | NA | NA | 1 | 1 | NA | NA | NA | NA | NA | NA | 1 | 1 | 1 | 1 |
| 6_4772625 | 2 | 0 | 1 | 1 | 1 | 0 | 1 | 0 | 5 | 0.2 | 2 | 0 | 2 | 0.5 | 3 | 0.3333 | 2 | 0.5 | 9 | 0.3333 |
| 6_4772644 | 1 | 1 | 1 | 1 | NA | NA | NA | NA | 2 | 1 | 3 | 0 | 2 | 1 | 3 | 1 | 2 | 1 | 10 | 0.7 |
| 6_4772648 | 3 | 1 | NA | NA | 1 | 1 | 1 | 1 | 5 | 1 | 3 | 0.6667 | 2 | 1 | 4 | 0.75 | 2 | 1 | 11 | 0.8182 |
| 6_4772649 | 3 | 1 | 1 | 1 | 1 | 1 | NA | NA | 5 | 1 | 3 | 1 | 2 | 1 | 4 | 1 | 2 | 1 | 11 | 1 |
| 6_4772652 | 2 | 1 | NA | NA | 1 | 1 | 1 | 1 | 4 | 1 | 2 | 0 | 1 | 1 | 2 | 0.5 | 2 | 1 | 7 | 0.5714 |
| 6_4772662 | 3 | 1 | 1 | 1 | 2 | 1 | 1 | 1 | 7 | 1 | 3 | 1 | 2 | 0.5 | 3 | 1 | 2 | 1 | 10 | 0.9 |

| | | | | | | | | | | | | | | | | | | | | |
|---|---|---|---|---|---|---|---|---|---|---|---|---|---|---|---|---|---|---|---|---|
| 6_4772673( | 3 | 0.3333 | 1 | 0 | 2 | 0.5 | 1 | 1 | 7 | 0.4286 | 3 | 0.6667 | 1 | 0 | 3 | 0 | 2 | 0 | 9 | 0.2222 |
| 6_4772697! | 2 | 0.5 | NA | NA | NA | NA | NA | NA | 2 | 0.5 | 1 | 0 | 1 | 0 | 1 | 1 | 2 | 0.5 | 5 | 0.4 |
| 6_4772713: | 2 | 1 | NA | NA | NA | NA | NA | NA | 2 | 1 | 1 | 1 | 2 | 1 | 1 | 0 | 1 | 1 | 5 | 0.8 |
| 6_4772723( | 2 | 0.5 | NA | NA | 1 | 1 | NA | NA | 3 | 0.6667 | 1 | 0 | 2 | 0.5 | 1 | 0 | 2 | 0 | 6 | 0.1667 |
| 6_4772731( | 2 | 0.5 | NA | NA | 1 | 0 | NA | NA | 3 | 0.3333 | 1 | 0 | 2 | 0 | 1 | 1 | 2 | 0 | 6 | 0.1667 |
| 6_4772732( | 2 | 1 | NA | NA | 1 | 0 | NA | NA | 3 | 0.6667 | 1 | 0 | 2 | 1 | 1 | 0 | 2 | 1 | 6 | 0.6667 |
| 6_4772747: | 2 | 0.5 | NA | NA | 1 | 0 | NA | NA | 3 | 0.3333 | 1 | 0 | 1 | 0 | 1 | 0 | 2 | 1 | 5 | 0.4 |
| 6_4773232 | NA | NA | NA | NA | NA | NA | 1 | 1 | 1 | 1 | NA | NA | NA | NA | 1 | 1 | NA | NA | 1 | 1 |
| 6_4773561 | NA | NA | NA | NA | 1 | 1 | 1 | 1 | 2 | 1 | 1 | 1 | 1 | 1 | NA | NA | NA | NA | 2 | 1 |
| 6_4773565! | NA | NA | NA | NA | 1 | 1 | NA | NA | 1 | 1 | NA | NA | 1 | 1 | NA | NA | NA | NA | 1 | 1 |
| 6_4773579( | NA | NA | NA | NA | 1 | 1 | 1 | 1 | 2 | 1 | 1 | 0 | 1 | 1 | NA | NA | NA | NA | 2 | 0.5 |
| 6_4774117( | NA | NA | NA | NA | NA | NA | 2 | 1 | 2 | 1 | 1 | 1 | NA | NA | NA | NA | 1 | 1 | 2 | 1 |
| 6_4774122( | NA | NA | 2 | 0.5 | NA | NA | 2 | 1 | 4 | 0.75 | NA | NA | 2 | 0.5 | NA | NA | 2 | 1 | 4 | 0.75 |
| 6_4774122( | NA | NA | 2 | 0.5 | NA | NA | 1 | 1 | 3 | 0.6667 | 1 | 1 | 1 | 1 | NA | NA | 1 | 1 | 3 | 1 |
| 6_4774125: | NA | NA | 2 | 0 | NA | NA | 2 | 0.5 | 4 | 0.25 | 1 | 0 | 2 | 1 | NA | NA | 2 | 0.5 | 5 | 0.6 |
| 6_4774135 | NA | NA | 2 | 0.5 | NA | NA | 2 | 1 | 4 | 0.75 | NA | NA | 2 | 0.5 | NA | NA | 2 | 1 | 4 | 0.75 |
| 6_4774146: | NA | NA | 2 | 0.5 | NA | NA | 1 | 0 | 3 | 0.3333 | NA | NA | 2 | 0.5 | NA | NA | 2 | 0.5 | 4 | 0.5 |
| 6_4774197! | NA | NA | 2 | 1 | NA | NA | 1 | 1 | 3 | 1 | 1 | 1 | 1 | 1 | NA | NA | 2 | 0.5 | 4 | 0.75 |
| 6_4774206 | NA | NA | 1 | 1 | NA | NA | 1 | 1 | 2 | 1 | 1 | 1 | 1 | 0 | NA | NA | 2 | 0.5 | 4 | 0.5 |
| 6_4774551( | NA | NA | 1 | 0 | 1 | 1 | 1 | 1 | 3 | 0.6667 | NA | NA | 1 | 1 | NA | NA | NA | NA | 1 | 1 |
| 6_4774555( | NA | NA | 1 | 1 | 1 | 1 | 1 | 1 | 3 | 1 | NA | NA | 2 | 1 | NA | NA | NA | NA | 2 | 1 |
| 6_4774556 | NA | NA | 1 | 1 | 1 | 1 | 1 | 1 | 3 | 1 | NA | NA | 1 | 1 | NA | NA | NA | NA | 1 | 1 |
| 6_4774559 | NA | NA | 1 | 0 | 1 | 1 | 1 | 1 | 3 | 0.6667 | NA | NA | 2 | 1 | NA | NA | NA | NA | 2 | 1 |
| 6_4774570: | NA | NA | 1 | 1 | 1 | 1 | 1 | 0 | 3 | 0.6667 | NA | NA | 2 | 1 | NA | NA | NA | NA | 2 | 1 |
| 6_4774619 | NA | NA | NA | NA | 1 | 1 | 1 | 1 | 2 | 1 | NA | NA | 2 | 1 | NA | NA | NA | NA | 2 | 1 |
| 6_4774651! | NA | NA | 1 | 1 | 1 | 0 | 1 | 0 | 3 | 0.3333 | NA | NA | 1 | 1 | NA | NA | 1 | 0 | 2 | 0.5 |
| 6_4774655( | NA | NA | 1 | 1 | 1 | 0 | 1 | 1 | 3 | 0.6667 | NA | NA | 2 | 0 | NA | NA | 1 | 1 | 3 | 0.3333 |
| 6_4774991( | 1 | 1 | 3 | 0.6667 | 1 | 0 | 2 | 1 | 7 | 0.7143 | 1 | 1 | 3 | 0.6667 | NA | NA | 2 | 1 | 6 | 0.8333 |
| 6_4774995( | 1 | 1 | 4 | 0.75 | NA | NA | 2 | 0 | 7 | 0.5714 | 1 | 1 | 1 | 1 | 1 | 1 | 2 | 0.5 | 5 | 0.8 |
| 6_4774996 | NA | NA | 3 | 1 | 1 | 1 | 2 | 0.5 | 6 | 0.8333 | 1 | 1 | 3 | 1 | NA | NA | 3 | 1 | 7 | 1 |
| 6_4774999 | NA | NA | 4 | 0.75 | NA | NA | 2 | 0.5 | 6 | 0.6667 | 1 | 1 | 2 | 1 | 1 | 1 | 1 | 0 | 5 | 0.8 |
| 6_4775009! | 1 | 1 | 3 | 0.6667 | 1 | 0 | 2 | 0 | 7 | 0.4286 | 1 | 1 | 3 | 0.6667 | 1 | 1 | 3 | 0.6667 | 8 | 0.75 |
| 6_4775025( | 1 | 1 | 3 | 0.6667 | NA | NA | 1 | 1 | 5 | 0.8 | NA | NA | 2 | 1 | NA | NA | 3 | 1 | 5 | 1 |
| 6_4775034( | NA | NA | 3 | 0.6667 | 1 | 0 | 1 | 1 | 5 | 0.6 | 1 | 0 | 3 | 0 | 1 | 1 | 3 | 0.3333 | 8 | 0.25 |
| 6_4775041( | NA | NA | 3 | 0.6667 | 1 | 0 | 1 | 0 | 5 | 0.4 | 1 | 1 | 3 | 0.3333 | NA | NA | 2 | 0.5 | 6 | 0.5 |
| 6_4775062: | NA | NA | 3 | 0 | 1 | 0 | 2 | 0.5 | 6 | 0.1667 | 2 | 0 | 4 | 0.25 | NA | NA | 3 | 0 | 9 | 0.1111 |
| 6_4775065( | NA | NA | 3 | 1 | 1 | 1 | 2 | 1 | 6 | 1 | 2 | 1 | 3 | 1 | NA | NA | 3 | 0.6667 | 8 | 0.875 |
| 6_4775079( | NA | NA | 3 | 1 | 1 | 1 | 1 | 1 | 5 | 1 | 2 | 1 | 4 | 1 | 1 | 1 | 3 | 1 | 10 | 1 |
| 6_4775090! | NA | NA | 1 | 1 | NA | NA | NA | NA | 1 | 1 | 1 | 1 | NA | NA | NA | NA | NA | NA | 1 | 1 |
| 6_4775090( | 1 | 1 | 2 | 1 | 1 | 1 | 1 | 1 | 5 | 1 | 1 | 1 | 1 | 0 | 1 | 0 | NA | NA | 3 | 0.3333 |
| 6_4775092: | 1 | 1 | 4 | 1 | 1 | 1 | 2 | 1 | 8 | 1 | 2 | 1 | 3 | 1 | 1 | 1 | 2 | 1 | 8 | 1 |
| 6_4775107 | 2 | 0.5 | 3 | 0 | 1 | 1 | 2 | 0.5 | 8 | 0.375 | 2 | 0.5 | 4 | 0.75 | 3 | 1 | 3 | 1 | 12 | 0.8333 |
| 6_4775121! | 3 | 1 | 4 | 1 | 2 | 1 | 2 | 0.5 | 11 | 0.9091 | 2 | 1 | 4 | 0.5 | 3 | 0.6667 | 2 | 1 | 11 | 0.7273 |
| 6_4775140 | 2 | 0 | 4 | 0.25 | 4 | 0.25 | 2 | 0 | 12 | 0.1667 | 3 | 0 | 4 | 0.25 | 3 | 0.6667 | 3 | 0 | 13 | 0.2308 |
| 6_4775140( | 2 | 0.5 | 4 | 0.75 | 3 | 1 | 1 | 1 | 10 | 0.8 | 3 | 1 | 3 | 1 | 1 | 1 | 3 | 1 | 10 | 1 |
| 6_4775142: | 2 | 0 | 3 | 0.6667 | 3 | 0.3333 | 1 | 1 | 9 | 0.4444 | 3 | 0.6667 | 1 | 1 | 3 | 0.6667 | 3 | 1 | 10 | 0.8 |
| 6_4775144 | 3 | 0.6667 | 3 | 0.6667 | 4 | 1 | 2 | 1 | 12 | 0.8333 | 1 | 1 | 3 | 0.6667 | 3 | 0.6667 | 3 | 1 | 10 | 0.8 |
| 6_4775147( | 3 | 0.3333 | 4 | 0.5 | 3 | 0.6667 | 2 | 0 | 12 | 0.4167 | 3 | 0.6667 | 3 | 0.3333 | 1 | 1 | 3 | 0.6667 | 10 | 0.6 |
| 6_4775150! | 3 | 0.6667 | 5 | 1 | 4 | 1 | 2 | 1 | 14 | 0.9286 | 3 | 1 | 4 | 1 | 3 | 1 | 3 | 0.3333 | 13 | 0.8462 |
| 6_4775152: | 3 | 1 | 5 | 1 | 4 | 1 | 2 | 1 | 14 | 1 | 3 | 1 | 4 | 0.75 | 3 | 1 | 3 | 0.6667 | 13 | 0.8462 |
| 6_4775159: | 3 | 0.6667 | 5 | 0.4 | 4 | 0.75 | 2 | 0.5 | 14 | 0.5714 | 3 | 0.6667 | 4 | 0.25 | 3 | 0.3333 | 3 | 0.3333 | 13 | 0.3846 |
| 6_4775174 | 4 | 0 | 3 | 0.3333 | 1 | 0 | 2 | 0.5 | 10 | 0.2 | 1 | 1 | 1 | 0 | 2 | 0.5 | 1 | 1 | 5 | 0.6 |
| 6_4777326( | 10 | 0.6 | 10 | 0.7 | 10 | 0.6 | 5 | 0.6 | 35 | 0.6286 | 12 | 0.6667 | 5 | 1 | 12 | 0.5 | 9 | 0.8889 | 38 | 0.7105 |
| 6_4777332( | 9 | 0.3333 | 10 | 0.4 | 8 | 0.5 | 4 | 0.5 | 31 | 0.4194 | 12 | 0.3333 | 5 | 0.8 | 11 | 0.2727 | 8 | 0.625 | 36 | 0.4444 |
| 6_4777337 | 9 | 0.6667 | 10 | 1 | 8 | 1 | 3 | 1 | 30 | 0.9 | 11 | 1 | 4 | 1 | 13 | 0.7692 | 9 | 0.8889 | 37 | 0.8919 |
| 6_4777343! | 7 | 0.2857 | 9 | 0.2222 | 8 | 0.25 | 2 | 0.5 | 26 | 0.2692 | 11 | 0 | 5 | 0.2 | 11 | 0.0909 | 8 | 0 | 35 | 0.0571 |
| 6_4777346 | 3 | 0.3333 | 6 | 0.1667 | 5 | 0.2 | 1 | 0 | 15 | 0.2 | 8 | 0.25 | 2 | 0 | 8 | 0.125 | 5 | 0.2 | 23 | 0.1739 |
| 6_4777352: | 7 | 0.7143 | 6 | 0.3333 | 8 | 0.625 | 4 | 0.25 | 25 | 0.52 | 10 | 0.3 | 1 | 1 | 10 | 0.2 | 4 | 0.5 | 25 | 0.32 |
| 6_4777353 | 7 | 0.7143 | 9 | 0.6667 | 8 | 0.5 | 4 | 0.25 | 28 | 0.5714 | 11 | 0.6364 | 3 | 0 | 9 | 0.5556 | 8 | 0.75 | 31 | 0.5806 |
| 6_4777363( | 9 | 0 | 10 | 0.2 | 10 | 0.1 | 4 | 0.25 | 33 | 0.1212 | 9 | 0.1111 | 4 | 0.5 | 10 | 0.4 | 8 | 0.375 | 31 | 0.3226 |
| 6_4777373( | 7 | 0.4286 | 9 | 0.3333 | 7 | 0.5714 | 4 | 0.5 | 27 | 0.4444 | 8 | 0.25 | 3 | 0.3333 | 7 | 0.4286 | 7 | 0.4286 | 25 | 0.36 |
| 6_4777376( | 9 | 0.1111 | 10 | 0 | 8 | 0.125 | 4 | 0.25 | 31 | 0.0968 | 8 | 0.125 | 2 | 0 | 10 | 0.3 | 8 | 0 | 28 | 0.1429 |
| 6_4779646( | NA | NA | NA | NA | NA | NA | 1 | 1 | 1 | 1 | NA | NA | 1 | 1 | NA | NA | NA | NA | 1 | 1 |
| 6_4779674! | NA | NA | 1 | 1 | NA | NA | NA | NA | 1 | 1 | NA | NA | 1 | 1 | NA | NA | NA | NA | 1 | 1 |
| 6_4779681 | NA | NA | 1 | 1 | NA | NA | 1 | 1 | 2 | 1 | NA | NA | 1 | 1 | NA | NA | NA | NA | 1 | 1 |

| | | | | | | | | | | | | | | | | | | | | |
|---|---|---|---|---|---|---|---|---|---|---|---|---|---|---|---|---|---|---|---|---|
| 6_4779694 | NA | NA | 1 | 1 | NA | NA | 1 | 1 | 2 | 1 | NA | NA | 1 | 1 | NA | NA | NA | NA | 1 | 1 |
| 6_4779791 | NA | NA | NA | NA | NA | NA | 1 | 1 | 1 | 1 | NA | NA | 1 | 1 | NA | NA | NA | NA | 1 | 1 |
| 6_4779792 | NA | NA | NA | NA | NA | NA | 1 | 1 | 1 | 1 | NA | NA | 1 | 1 | NA | NA | NA | NA | 1 | 1 |
| 6_4779835 | NA | NA | NA | NA | NA | NA | 1 | 1 | 1 | 1 | NA | NA | 1 | 1 | NA | NA | NA | NA | 1 | 1 |
| 6_4779855 | NA | NA | NA | NA | NA | NA | 1 | 1 | 1 | 1 | NA | NA | 1 | 1 | NA | NA | NA | NA | 1 | 1 |
| 6_4788214 | 2 | 0.5 | 1 | 1 | NA | NA | 1 | 1 | 4 | 0.75 | 1 | 1 | 1 | 0 | 1 | 0 | NA | NA | 3 | 0.3333 |
| 6_4788224 | 1 | 1 | NA | NA | NA | NA | 2 | 1 | 3 | 1 | 1 | 1 | 1 | 1 | 2 | 0.5 | 1 | 1 | 5 | 0.8 |
| 6_4788231 | 2 | 1 | 1 | 1 | NA | NA | 2 | 1 | 5 | 1 | 1 | 1 | 1 | 1 | 1 | 1 | 1 | 1 | 4 | 1 |
| 6_4788234 | 2 | 0.5 | 1 | 1 | NA | NA | 2 | 1 | 5 | 0.8 | 1 | 1 | 1 | 1 | NA | NA | 1 | 1 | 3 | 1 |
| 6_4788247 | 1 | 0 | NA | NA | NA | NA | 1 | 1 | 2 | 0.5 | 1 | 1 | 1 | 1 | 1 | 1 | NA | NA | 3 | 1 |
| 6_4788250 | 1 | 0 | 1 | 1 | NA | NA | 2 | 0.5 | 4 | 0.5 | NA | NA | NA | NA | 1 | 1 | 1 | 1 | 2 | 1 |
| 6_4788253 | 1 | 1 | 1 | 0 | NA | NA | 2 | 0 | 4 | 0.25 | 1 | 0 | 1 | 0 | 2 | 0.5 | 1 | 0 | 5 | 0.2 |
| 6_4788256 | 2 | 1 | NA | NA | NA | NA | 2 | 1 | 4 | 1 | 1 | 0 | 1 | 1 | 2 | 1 | 1 | 1 | 5 | 0.8 |
| 6_4788258 | 2 | 1 | 1 | 1 | NA | NA | 1 | 0 | 4 | 0.75 | NA | NA | 1 | 0 | 2 | 1 | 1 | 1 | 4 | 0.75 |
| 6_4788259 | 1 | 1 | 1 | 0 | NA | NA | 1 | 1 | 3 | 0.6667 | 1 | 1 | 1 | 1 | 2 | 1 | 1 | 1 | 5 | 1 |
| 6_4788314 | 2 | 0.5 | 1 | 0 | 2 | 0.5 | 2 | 0 | 7 | 0.2857 | 2 | 0.5 | 2 | 0.5 | 2 | 1 | 1 | 0 | 7 | 0.5714 |
| 6_4788338 | 3 | 0.3333 | 1 | 0 | 2 | 1 | 3 | 0.3333 | 9 | 0.4444 | 3 | 0.3333 | 2 | 0 | 2 | 0 | 1 | 0 | 8 | 0.125 |
| 6_4788341 | 4 | 0.75 | NA | NA | 2 | 0.5 | 3 | 0.6667 | 9 | 0.6667 | 3 | 0.6667 | 1 | 1 | 2 | 0.5 | 1 | 1 | 7 | 0.7143 |
| 6_4788349 | 4 | 0.25 | 1 | 1 | 1 | 0 | 1 | 0 | 7 | 0.2857 | 2 | 0 | 2 | 0 | 2 | 0.5 | 1 | 0 | 7 | 0.1429 |
| 6_4788351 | 4 | 1 | 1 | 1 | 2 | 1 | 2 | 1 | 9 | 1 | 3 | 1 | 2 | 0 | 3 | 1 | 1 | 1 | 9 | 0.7778 |
| 6_4788352 | 2 | 1 | NA | NA | 2 | 0.5 | 2 | 1 | 6 | 0.8333 | 3 | 1 | 2 | 1 | 1 | 1 | 2 | 1 | 8 | 1 |
| 6_4788352 | 4 | 0 | NA | NA | 2 | 0 | 3 | 0.3333 | 9 | 0.1111 | 3 | 0 | 2 | 0.5 | 3 | 0 | 1 | 1 | 9 | 0.2222 |
| 6_4788357 | 4 | 1 | 1 | 1 | 2 | 1 | 3 | 1 | 10 | 1 | 2 | 1 | 2 | 0.5 | 2 | 1 | 1 | 1 | 7 | 0.8571 |
| 6_4788368 | 3 | 0.6667 | 1 | 1 | 2 | 1 | 3 | 0.3333 | 9 | 0.6667 | 3 | 0.3333 | 2 | 0 | 3 | 0 | 2 | 0.5 | 10 | 0.2 |
| 6_4790781 | 3 | 0.6667 | 1 | 1 | 2 | 1 | NA | NA | 6 | 0.8333 | 2 | 1 | 1 | 1 | 2 | 1 | 3 | 0.6667 | 8 | 0.875 |
| 6_4790783 | 3 | 0.6667 | 1 | 0 | 2 | 0 | NA | NA | 6 | 0.3333 | 3 | 0 | 1 | 0 | 2 | 0 | 3 | 0.3333 | 9 | 0.1111 |
| 6_4790784 | 3 | 0.3333 | 1 | 1 | 2 | 0.5 | NA | NA | 6 | 0.5 | 3 | 0.3333 | 1 | 1 | 2 | 1 | 3 | 0.3333 | 9 | 0.5556 |
| 6_4790787 | 3 | 1 | 1 | 1 | 1 | 1 | NA | NA | 5 | 1 | 3 | 1 | 1 | 1 | 2 | 1 | 3 | 1 | 9 | 1 |
| 6_4790792 | 2 | 0 | 1 | 0 | 2 | 1 | NA | NA | 5 | 0.4 | 3 | 0.3333 | 1 | 1 | 2 | 0.5 | 3 | 0.3333 | 9 | 0.4444 |
| 6_4790794 | 3 | 0.3333 | 1 | 1 | 2 | 0.5 | NA | NA | 6 | 0.5 | 3 | 0.3333 | 1 | 1 | 2 | 0 | 3 | 1 | 9 | 0.5556 |
| 6_4790823 | 2 | 0 | NA | NA | 2 | 0.5 | NA | NA | 4 | 0.25 | 1 | 0 | 1 | 1 | 1 | 0 | 1 | 0 | 4 | 0.25 |
| 6_4790833 | 3 | 1 | 1 | 1 | 1 | 1 | NA | NA | 5 | 1 | 2 | 1 | 1 | 1 | 2 | 1 | 3 | 1 | 8 | 1 |
| 6_4790877 | 1 | 1 | 2 | 0.5 | NA | NA | 2 | 1 | 5 | 0.8 | 2 | 0.5 | NA | NA | 6 | 0.8333 | 2 | 1 | 10 | 0.8 |
| 6_4790893 | 3 | 0.6667 | 3 | 0 | 2 | 0.5 | 1 | 1 | 9 | 0.4444 | 3 | 0.3333 | 1 | 0 | 5 | 0.6 | 2 | 0.5 | 11 | 0.4545 |
| 6_4790925 | 3 | 0 | 3 | 0 | 1 | 0 | 1 | 1 | 8 | 0.125 | 4 | 0.25 | 1 | 0 | 5 | 0 | 3 | 0 | 13 | 0.0769 |
| 6_4790934 | 3 | 0 | 3 | 0.3333 | 1 | 0 | NA | NA | 7 | 0.1429 | 4 | 0.5 | 2 | 0 | 3 | 0 | 3 | 0 | 12 | 0.1667 |
| 6_4792898 | 1 | 1 | NA | NA | NA | NA | 1 | 1 | 2 | 1 | NA | NA | NA | NA | 1 | 1 | NA | NA | 1 | 1 |
| 6_4792927 | NA | NA | NA | NA | NA | NA | 1 | 1 | 1 | 1 | NA | NA | NA | NA | 1 | 1 | NA | NA | 1 | 1 |
| 6_4792941 | 1 | 1 | NA | NA | NA | NA | NA | NA | 1 | 1 | NA | NA | NA | NA | 1 | 1 | NA | NA | 1 | 1 |
| 6_4792960 | 1 | 1 | NA | NA | NA | NA | 1 | 1 | 2 | 1 | NA | NA | NA | NA | 1 | 1 | NA | NA | 1 | 1 |
| 6_4794047 | 3 | 0.6667 | 1 | 0 | 4 | 0.25 | NA | NA | 8 | 0.375 | 2 | 0.5 | 1 | 1 | 2 | 0 | 1 | 0 | 6 | 0.3333 |
| 6_4794083 | 3 | 0.3333 | NA | NA | 4 | 0.75 | NA | NA | 7 | 0.5714 | 1 | 1 | 1 | 0 | 2 | 0.5 | NA | NA | 4 | 0.5 |
| 6_4794123 | 1 | 1 | NA | NA | 3 | 0.6667 | 2 | 1 | 6 | 0.8333 | 1 | 1 | 1 | 1 | 1 | 1 | NA | NA | 3 | 1 |
| 6_4794146 | 3 | 1 | NA | NA | 4 | 1 | 2 | 1 | 9 | 1 | 1 | 0 | 1 | 1 | 2 | 0.5 | 3 | 1 | 7 | 0.7143 |
| 6_4794184 | 4 | 0.25 | 1 | 0 | 4 | 0.25 | 2 | 0 | 11 | 0.1818 | 2 | 0.5 | 2 | 1 | 3 | 0 | 3 | 0 | 10 | 0.3 |
| 6_4794199 | 4 | 1 | 1 | 1 | 4 | 0.75 | 4 | 0.75 | 13 | 0.8462 | 2 | 1 | 1 | 1 | 3 | 0.6667 | 3 | 1 | 9 | 0.8889 |
| 6_4800129 | 3 | 0 | 6 | 0.1667 | 2 | 0 | 3 | 0 | 14 | 0.0714 | 5 | 0.2 | 2 | 0 | 1 | 0 | 5 | 0 | 13 | 0.0769 |
| 6_4800147 | 4 | 1 | 5 | 1 | 2 | 0.5 | 8 | 0.875 | 19 | 0.8947 | 3 | 1 | 2 | 0.5 | 1 | 1 | 3 | 1 | 9 | 0.8889 |
| 6_4800149 | 5 | 1 | 5 | 0.8 | 1 | 1 | 9 | 0.6667 | 20 | 0.8 | 5 | 1 | 2 | 0.5 | 2 | 1 | 3 | 1 | 12 | 0.9167 |
| 6_4800158 | 3 | 0.6667 | 4 | 0.25 | 2 | 1 | 9 | 0.4444 | 18 | 0.5 | 4 | 0.5 | 1 | 1 | 2 | 0.5 | 4 | 0.5 | 11 | 0.5455 |
| 6_4800160 | 2 | 0 | 2 | 0 | 1 | 0 | 5 | 0.2 | 10 | 0.1 | 1 | 0 | NA | NA | 1 | 1 | 5 | 0.2 | 7 | 0.2857 |
| 6_4800191 | 3 | 1 | 2 | 1 | 2 | 1 | 6 | 1 | 13 | 1 | 5 | 1 | 1 | 1 | 1 | 1 | 5 | 1 | 12 | 1 |
| 6_4800246 | 4 | 1 | 1 | 0 | 1 | 1 | 5 | 0.4 | 11 | 0.6364 | 2 | 1 | 1 | 1 | 1 | 1 | 4 | 0.5 | 8 | 0.75 |
| 6_4800247 | 4 | 0.25 | 1 | 0 | 1 | 0 | 6 | 0.3333 | 12 | 0.25 | 3 | 0.6667 | 2 | 0.5 | 1 | 0 | 5 | 0.6 | 11 | 0.5455 |
| 6_4800619 | 1 | 1 | 1 | 1 | 1 | 0 | 7 | 0.4286 | 10 | 0.5 | 2 | 0.5 | 1 | 0 | NA | NA | 4 | 0.5 | 7 | 0.4286 |
| 6_4802089 | 1 | 0 | 1 | 0 | 1 | 0 | 5 | 0.4 | 8 | 0.25 | 3 | 0.6667 | NA | NA | NA | NA | 4 | 0.25 | 7 | 0.4286 |
| 6_4802095 | 2 | 0.5 | 1 | 0 | 1 | 0 | 6 | 0.1667 | 10 | 0.2 | 2 | 1 | NA | NA | NA | NA | 4 | 0.5 | 6 | 0.6667 |
| 6_4802098 | 2 | 0.5 | 1 | 0 | 1 | 0 | 3 | 0.3333 | 7 | 0.2857 | 2 | 0.5 | NA | NA | NA | NA | 4 | 0.75 | 6 | 0.6667 |
| 6_4802100 | 1 | 1 | 1 | 1 | 1 | 1 | 4 | 0.75 | 7 | 0.8571 | 2 | 1 | NA | NA | NA | NA | 3 | 0.3333 | 5 | 0.6 |
| 6_4802113 | 2 | 1 | 1 | 1 | NA | NA | 5 | 1 | 8 | 1 | 3 | 0.6667 | NA | NA | NA | NA | 4 | 0.5 | 7 | 0.5714 |
| 6_4802116 | 2 | 0.5 | 1 | 1 | 1 | 0 | 4 | 0 | 8 | 0.25 | 3 | 0.3333 | NA | NA | NA | NA | 3 | 0.3333 | 6 | 0.3333 |
| 6_4802120 | 1 | 0 | 1 | 1 | NA | NA | 5 | 0.4 | 7 | 0.4286 | 3 | 0.3333 | NA | NA | NA | NA | 4 | 0 | 7 | 0.1429 |
| 6_4802121 | 1 | 0 | 1 | 0 | 1 | 1 | 5 | 0 | 8 | 0.125 | 3 | 0 | NA | NA | NA | NA | 4 | 0.5 | 7 | 0.2857 |
| 6_4802138 | 2 | 0.5 | 1 | 0 | 1 | 0 | 5 | 0.4 | 9 | 0.3333 | 2 | 0.5 | NA | NA | NA | NA | 2 | 1 | 4 | 0.75 |
| 6_4802144 | 2 | 0.5 | 1 | 1 | NA | NA | 2 | 0.5 | 5 | 0.6 | 2 | 0 | NA | NA | NA | NA | 4 | 0.5 | 6 | 0.3333 |

| | | | | | | | | | | | | | | | | | | | | |
|---|---|---|---|---|---|---|---|---|---|---|---|---|---|---|---|---|---|---|---|---|
| 6_4802153 | 2 | 1 | NA | NA | 1 | 1 | 2 | 0 | 5 | 0.6 | 2 | 0.5 | NA | NA | NA | NA | 4 | 0 | 6 | 0.1667 |
| 6_4804037 | 1 | 0 | 2 | 0.5 | NA | NA | 2 | 0.5 | 5 | 0.4 | NA | NA | 2 | 0 | 2 | 1 | 2 | 0.5 | 6 | 0.5 |
| 6_4804057 | 1 | 0 | 2 | 0.5 | 1 | 0 | 1 | 0 | 5 | 0.2 | NA | NA | 2 | 0 | 4 | 0.25 | 3 | 0 | 9 | 0.1111 |
| 6_4804062 | 1 | 1 | 2 | 1 | 1 | 1 | 2 | 1 | 6 | 1 | NA | NA | 2 | 0.5 | 4 | 0.75 | 3 | 0.6667 | 9 | 0.6667 |
| 6_4804063 | 1 | 1 | 2 | 1 | 1 | 1 | 2 | 1 | 6 | 1 | NA | NA | 2 | 1 | 2 | 1 | 2 | 1 | 6 | 1 |
| 6_4804076 | 1 | 0 | 2 | 1 | 1 | 1 | 2 | 1 | 6 | 0.8333 | NA | NA | 2 | 0 | 4 | 0.5 | 2 | 0.5 | 8 | 0.375 |
| 6_4804079 | 1 | 1 | 2 | 1 | 1 | 0 | 2 | 1 | 6 | 0.8333 | NA | NA | 2 | 0.5 | 5 | 0.6 | 2 | 0.5 | 9 | 0.5556 |
| 6_4804081 | 1 | 1 | 2 | 1 | 1 | 1 | 2 | 1 | 6 | 1 | NA | NA | 2 | 1 | 5 | 1 | 2 | 1 | 9 | 1 |
| 6_4804082 | NA | NA | 2 | 0.5 | NA | NA | 1 | 0 | 3 | 0.3333 | NA | NA | 1 | 1 | 4 | 0 | NA | NA | 5 | 0.2 |
| 6_4804082 | NA | NA | 2 | 0.5 | NA | NA | 2 | 1 | 4 | 0.75 | NA | NA | 1 | 1 | 3 | 1 | 1 | 1 | 5 | 1 |
| 6_4804085 | 1 | 0 | 3 | 0 | 1 | 1 | 1 | 1 | 6 | 0.3333 | NA | NA | 2 | 1 | 4 | 0.25 | 2 | 0 | 8 | 0.375 |
| 6_4804118 | NA | NA | 1 | 0 | 1 | 0 | 1 | 1 | 3 | 0.3333 | NA | NA | 1 | 1 | 2 | 1 | NA | NA | 3 | 1 |
| 6_4804129 | 1 | 1 | 2 | 1 | 1 | 1 | 1 | 1 | 5 | 1 | NA | NA | 2 | 1 | 4 | 1 | 1 | 1 | 7 | 1 |
| 6_4804143 | 1 | 1 | 2 | 0 | 1 | 1 | 2 | 0 | 6 | 0.3333 | NA | NA | 2 | 0.5 | 3 | 0.6667 | 1 | 0 | 6 | 0.5 |
| 6_4804166 | NA | NA | 2 | 1 | NA | NA | 1 | 1 | 3 | 1 | NA | NA | 1 | 1 | 3 | 0.6667 | 1 | 1 | 5 | 0.8 |
| 6_4804170 | 1 | 0 | 1 | 0 | NA | NA | 1 | 1 | 3 | 0.3333 | NA | NA | 1 | 1 | 3 | 0 | 1 | 0 | 5 | 0.2 |
| 6_4804179 | 1 | 1 | 3 | 1 | NA | NA | 1 | 1 | 5 | 1 | NA | NA | 2 | 1 | 3 | 0.6667 | 1 | 1 | 6 | 0.8333 |
| 6_4839267 | 2 | 1 | 4 | 1 | 2 | 1 | 4 | 1 | 12 | 1 | 2 | 1 | 1 | 0 | 2 | 1 | 1 | 1 | 6 | 0.8333 |
| 6_4839271 | 2 | 0 | 4 | 0.25 | 2 | 0 | 4 | 0 | 12 | 0.0833 | 3 | 0.3333 | 1 | 0 | 1 | 1 | 2 | 0 | 7 | 0.2857 |
| 6_4839282 | 2 | 1 | 4 | 0 | 1 | 1 | 4 | 0.75 | 11 | 0.5455 | 3 | 0 | 1 | 0 | 2 | 0 | 2 | 0.5 | 8 | 0.125 |
| 6_4839284 | 2 | 0.5 | 3 | 0 | 2 | 1 | 3 | 0.3333 | 10 | 0.4 | 3 | 1 | 1 | 1 | 2 | 0 | 1 | 1 | 7 | 0.7143 |
| 6_4839291 | 1 | 0 | 4 | 0.25 | 1 | 0 | 3 | 0 | 9 | 0.1111 | 3 | 0.3333 | 1 | 0 | 2 | 0 | 1 | 0 | 7 | 0.1429 |
| 6_4839293 | 1 | 1 | 4 | 0 | 2 | 0.5 | 4 | 0.5 | 11 | 0.3636 | 3 | 0 | 1 | 0 | 2 | 0 | 2 | 0.5 | 8 | 0.125 |
| 6_4839294 | 2 | 1 | 3 | 0.6667 | 2 | 0.5 | 4 | 0.75 | 11 | 0.7273 | 3 | 1 | 1 | 0 | 2 | 0.5 | 3 | 1 | 9 | 0.7778 |
| 6_4839307 | 2 | 0 | 4 | 0.25 | 1 | 0 | 4 | 0 | 11 | 0.0909 | 3 | 0.3333 | 1 | 0 | 1 | 0 | 3 | 0 | 8 | 0.125 |
| 6_4839324 | 1 | 0 | 2 | 0 | 2 | 0 | 3 | 0.3333 | 8 | 0.125 | 3 | 0.3333 | 1 | 0 | 2 | 0.5 | 3 | 0.3333 | 9 | 0.3333 |
| 6_4839327 | 2 | 1 | 4 | 0.5 | 2 | 1 | 4 | 1 | 12 | 0.8333 | 2 | 0 | 1 | 1 | 2 | 0.5 | 3 | 1 | 8 | 0.625 |
| 6_4839330 | 2 | 1 | 4 | 0.75 | 2 | 1 | 4 | 0.5 | 12 | 0.75 | 2 | 0.5 | 1 | 0 | 1 | 1 | 3 | 0.6667 | 7 | 0.5714 |
| 6_4839332 | 1 | 1 | 2 | 1 | 2 | 0.5 | 2 | 0 | 7 | 0.5714 | 2 | 0.5 | 1 | 0 | 1 | 1 | 2 | 1 | 6 | 0.6667 |
| 6_4839338 | 2 | 0.5 | 4 | 0.25 | 2 | 0.5 | 4 | 0.75 | 12 | 0.5 | 4 | 0.25 | 1 | 0 | 2 | 0.5 | 3 | 0.3333 | 10 | 0.3 |
| 6_4839356 | 2 | 0 | 3 | 0.3333 | 1 | 0 | 4 | 0.25 | 10 | 0.2 | 3 | 0 | 1 | 0 | 2 | 0 | 3 | 0.3333 | 9 | 0.1111 |
| 6_4839370 | 2 | 0.5 | 3 | 0.3333 | NA | NA | 5 | 0 | 10 | 0.2 | 4 | 0.25 | NA | NA | 2 | 0.5 | 3 | 0.6667 | 9 | 0.4444 |
| 6_4839378 | 2 | 1 | 3 | 0.6667 | 1 | 0 | 5 | 0.6 | 11 | 0.6364 | 4 | 0.25 | NA | NA | 2 | 0.5 | 4 | 0.5 | 10 | 0.4 |
| 6_4841390 | NA | NA | 2 | 1 | 3 | 0.3333 | 1 | 1 | 6 | 0.6667 | 1 | 1 | 2 | 1 | 1 | 0 | 3 | 0.3333 | 7 | 0.5714 |
| 6_4841404 | NA | NA | 2 | 1 | 2 | 0 | 1 | 1 | 5 | 0.6 | NA | NA | 4 | 0.5 | 1 | 1 | 3 | 0.6667 | 8 | 0.625 |
| 6_4841444 | NA | NA | 2 | 0 | 4 | 0 | 1 | 1 | 7 | 0.1429 | 1 | 0 | 3 | 0.3333 | 1 | 1 | 2 | 0 | 7 | 0.2857 |
| 6_4841447 | NA | NA | 2 | 0 | 4 | 0 | 1 | 1 | 7 | 0.1429 | 1 | 1 | 4 | 0.25 | 1 | 0 | 2 | 0 | 8 | 0.25 |
| 6_4841459 | 4 | 0 | 2 | 0 | 5 | 0 | 8 | 0.75 | 19 | 0.3158 | 2 | 0.5 | 6 | 0.6667 | 8 | 0.125 | 3 | 0.3333 | 19 | 0.3684 |
| 6_4841463 | 3 | 0.3333 | 2 | 0 | 7 | 0.4286 | 7 | 0.1429 | 19 | 0.2632 | 2 | 0.5 | 4 | 0.5 | 4 | 0.5 | 2 | 0 | 12 | 0.4167 |
| 6_4841466 | 4 | 1 | 3 | 1 | 6 | 1 | 8 | 0.75 | 21 | 0.9048 | 2 | 1 | 7 | 1 | 8 | 0.875 | 3 | 1 | 20 | 0.95 |
| 6_4841468 | 4 | 0 | 3 | 0 | 6 | 0.1667 | 6 | 0 | 19 | 0.0526 | 2 | 0 | 6 | 0.3333 | 7 | 0 | 3 | 0 | 18 | 0.1111 |
| 6_4841475 | 2 | 1 | 3 | 1 | 7 | 0.7143 | 8 | 1 | 20 | 0.9 | 2 | 1 | 5 | 1 | 8 | 0.875 | 3 | 1 | 18 | 0.9444 |
| 6_4841480 | 2 | 0.5 | 3 | 0.6667 | 7 | 1 | 7 | 1 | 19 | 0.8947 | 2 | 1 | 5 | 0.8 | 8 | 0.75 | 2 | 1 | 17 | 0.8235 |
| 6_4841482 | 2 | 0.5 | 3 | 0.3333 | 7 | 0.7143 | 7 | 0.7143 | 19 | 0.6316 | 2 | 0 | 5 | 0.6 | 8 | 0.375 | 3 | 0.6667 | 18 | 0.4444 |
| 6_4841492 | 1 | 0 | 2 | 0 | 5 | 0.2 | 6 | 0 | 14 | 0.0714 | 2 | 0 | 4 | 0.5 | 8 | 0.125 | 2 | 0.5 | 16 | 0.25 |
| 6_4841519 | 2 | 0 | 2 | 0.5 | 4 | 0.25 | 7 | 0.2857 | 15 | 0.2667 | 1 | 0 | 5 | 0.2 | 7 | 0.1429 | 3 | 0.3333 | 16 | 0.1875 |
| 6_4841535 | 2 | 1 | 2 | 1 | 7 | 0.7143 | 6 | 0.8333 | 17 | 0.8235 | 1 | 1 | 5 | 1 | 7 | 0.5714 | 3 | 1 | 16 | 0.8125 |
| 6_4841541 | 2 | 0.5 | NA | NA | 5 | 0.2 | 5 | 0.2 | 12 | 0.25 | 2 | 0 | 1 | 0 | 5 | 0.4 | 3 | 0.3333 | 11 | 0.2727 |
| 6_4841542 | 2 | 1 | 2 | 1 | 6 | 0.8333 | 3 | 1 | 13 | 0.9231 | 1 | 1 | 2 | 1 | 4 | 0.75 | 3 | 1 | 10 | 0.9 |
| 6_4841546 | 2 | 0.5 | 2 | 0 | 6 | 0.5 | 7 | 0.2857 | 17 | 0.3529 | 1 | 0 | 4 | 0.25 | 6 | 0.1667 | 3 | 0.6667 | 14 | 0.2857 |
| 6_4841565 | 2 | 1 | 2 | 1 | 6 | 0.5 | 7 | 0.8571 | 17 | 0.7647 | 1 | 0 | 3 | 0.3333 | 8 | 0.75 | 3 | 0.6667 | 15 | 0.6 |
| 6_4841607 | 2 | 1 | 2 | 1 | 6 | 1 | 7 | 0.5714 | 17 | 0.8235 | 1 | 1 | 4 | 0.75 | 6 | 0.8333 | 3 | 1 | 14 | 0.8571 |
| 6_4841623 | 1 | 1 | 2 | 0.5 | 3 | 0.6667 | 5 | 0.8 | 11 | 0.7273 | 1 | 0 | 3 | 0.3333 | 6 | 0.6667 | 3 | 1 | 13 | 0.6154 |
| 6_4841624 | 1 | 0 | 1 | 0 | 1 | 1 | 3 | 0.6667 | 6 | 0.5 | 1 | 1 | 1 | 1 | 5 | 0.4 | 3 | 0.6667 | 10 | 0.6 |
| 6_4850550 | 2 | 1 | NA | NA | NA | NA | NA | NA | 2 | 1 | 1 | 1 | NA | NA | NA | NA | NA | NA | 1 | 1 |
| 6_4850594 | 1 | 1 | NA | NA | NA | NA | NA | NA | 1 | 1 | 2 | 0.5 | NA | NA | NA | NA | NA | NA | 2 | 0.5 |
| 6_4850595 | 2 | 0.5 | NA | NA | NA | NA | NA | NA | 2 | 0.5 | 2 | 1 | NA | NA | NA | NA | NA | NA | 2 | 1 |
| 6_4850614 | 2 | 1 | NA | NA | NA | NA | NA | NA | 2 | 1 | 2 | 1 | NA | NA | NA | NA | NA | NA | 2 | 1 |
| 6_4850631 | 2 | 1 | NA | NA | NA | NA | NA | NA | 2 | 1 | 2 | 1 | NA | NA | NA | NA | NA | NA | 2 | 1 |
| 6_4850640 | 2 | 1 | NA | NA | NA | NA | NA | NA | 2 | 1 | 1 | 1 | NA | NA | NA | NA | NA | NA | 1 | 1 |
| 6_4850690 | 1 | 1 | NA | NA | NA | NA | NA | NA | 1 | 1 | 2 | 1 | NA | NA | NA | NA | NA | NA | 2 | 1 |
| 6_4850912 | 2 | 0.5 | NA | NA | 1 | 0 | NA | NA | 3 | 0.3333 | 2 | 0.5 | NA | NA | NA | NA | NA | NA | 2 | 0.5 |
| 6_4850925 | 2 | 0 | NA | NA | 1 | 1 | NA | NA | 3 | 0.3333 | 1 | 1 | NA | NA | NA | NA | NA | NA | 1 | 1 |
| 6_4850931 | 2 | 0 | NA | NA | 1 | 1 | NA | NA | 3 | 0.3333 | 2 | 0.5 | NA | NA | NA | NA | NA | NA | 2 | 0.5 |
| 6_4850941 | 2 | 1 | NA | NA | NA | NA | NA | NA | 2 | 1 | 2 | 1 | NA | NA | NA | NA | NA | NA | 2 | 1 |

| | | | | | | | | | | | | | | | | | | | | |
|---|---|---|---|---|---|---|---|---|---|---|---|---|---|---|---|---|---|---|---|---|
| 6_4850948 | 1 | 1 | NA | NA | 1 | 1 | NA | NA | 2 | 1 | 2 | 0.5 | NA | NA | NA | NA | NA | NA | 2 | 0.5 |
| 6_4850950 | 2 | 1 | NA | NA | 1 | 1 | NA | NA | 3 | 1 | 2 | 1 | NA | NA | NA | NA | NA | NA | 2 | 1 |
| 6_4851009 | 2 | 0.5 | NA | NA | 1 | 1 | NA | NA | 3 | 0.6667 | 3 | 0.6667 | NA | NA | NA | NA | NA | NA | 3 | 0.6667 |
| 6_4851011 | 2 | 0.5 | NA | NA | 1 | 0 | NA | NA | 3 | 0.3333 | 3 | 0.6667 | NA | NA | NA | NA | NA | NA | 3 | 0.6667 |
| 6_4851014 | 2 | 1 | NA | NA | 1 | 1 | NA | NA | 3 | 1 | 3 | 0.6667 | NA | NA | NA | NA | NA | NA | 3 | 0.6667 |
| 6_4851145 | 1 | 1 | NA | NA | 1 | 1 | NA | NA | 2 | 1 | 3 | 0.6667 | NA | NA | NA | NA | NA | NA | 3 | 0.6667 |
| 6_4851152 | 2 | 0 | NA | NA | 1 | 1 | NA | NA | 3 | 0.3333 | 3 | 0.3333 | NA | NA | NA | NA | NA | NA | 3 | 0.3333 |
| 6_4851773 | 48 | 0.1667 | 41 | 0.2439 | 38 | 0.2895 | 42 | 0.3095 | 169 | 0.2485 | 45 | 0.2667 | 55 | 0.2 | 36 | 0.1667 | 37 | 0.2703 | 173 | 0.2254 |
| 6_4851775 | 48 | 0.0833 | 41 | 0.0732 | 40 | 0.2 | 43 | 0.1163 | 172 | 0.1163 | 47 | 0.0851 | 52 | 0.0385 | 35 | 0.0857 | 37 | 0.1351 | 171 | 0.0819 |
| 6_4852180 | NA | NA | NA | NA | NA | NA | 3 | 0.3333 | 3 | 0.3333 | 2 | 1 | NA | NA | NA | NA | 1 | 0 | 3 | 0.6667 |
| 6_4852195 | NA | NA | NA | NA | NA | NA | 3 | 1 | 3 | 1 | 2 | 0.5 | NA | NA | NA | NA | NA | NA | 2 | 0.5 |
| 6_4852197 | NA | NA | NA | NA | NA | NA | 3 | 0.3333 | 3 | 0.3333 | 1 | 1 | NA | NA | NA | NA | 1 | 0 | 2 | 0.5 |
| 6_4852213 | NA | NA | NA | NA | NA | NA | 3 | 0.6667 | 3 | 0.6667 | 2 | 1 | NA | NA | NA | NA | 1 | 0 | 3 | 0.6667 |
| 6_4852217 | NA | NA | NA | NA | NA | NA | 3 | 1 | 3 | 1 | 3 | 1 | NA | NA | NA | NA | 1 | 1 | 4 | 1 |
| 6_4854513 | 3 | 0 | 3 | 0.3333 | 2 | 0 | NA | NA | 8 | 0.125 | NA | NA | 2 | 0 | 4 | 0 | 1 | 1 | 7 | 0.1429 |
| 6_4854516 | 3 | 1 | 3 | 0.6667 | 2 | 1 | NA | NA | 8 | 0.875 | NA | NA | 1 | 1 | 4 | 1 | 1 | 1 | 6 | 1 |
| 6_4854531 | 3 | 1 | 3 | 1 | 2 | 1 | NA | NA | 8 | 1 | NA | NA | 2 | 1 | 4 | 1 | 1 | 1 | 7 | 1 |
| 6_4854533 | 3 | 0.6667 | 3 | 1 | 2 | 0.5 | NA | NA | 8 | 0.75 | NA | NA | 2 | 0.5 | 3 | 0.3333 | 1 | 1 | 6 | 0.5 |
| 6_4854547 | 3 | 0.3333 | 3 | 0 | 3 | 0 | NA | NA | 9 | 0.1111 | NA | NA | 2 | 0 | 3 | 0.3333 | 1 | 1 | 6 | 0.3333 |
| 6_4854557 | 3 | 0 | 3 | 0.3333 | 3 | 0 | NA | NA | 9 | 0.1111 | NA | NA | 2 | 0.5 | 5 | 0 | 1 | 1 | 8 | 0.25 |
| 6_4854571 | 3 | 0.3333 | 1 | 0 | 2 | 0 | 1 | 1 | 7 | 0.2857 | NA | NA | 2 | 0.5 | 5 | 0 | 1 | 0 | 8 | 0.125 |
| 6_4854571 | 1 | 1 | 2 | 1 | 1 | 1 | NA | NA | 4 | 1 | NA | NA | 1 | 1 | 2 | 1 | 1 | 1 | 4 | 1 |
| 6_4854585 | 3 | 0 | 3 | 0.3333 | 3 | 0 | 1 | 0 | 10 | 0.1 | NA | NA | 2 | 0 | 5 | 0.4 | 1 | 1 | 8 | 0.375 |
| 6_4854622 | 3 | 0 | 2 | 1 | 3 | 0.6667 | 1 | 1 | 9 | 0.5556 | NA | NA | 1 | 0 | 4 | 0.5 | 1 | 1 | 6 | 0.5 |
| 6_4854720 | 4 | 0 | 5 | 0.2 | 3 | 0.3333 | 1 | 0 | 13 | 0.1538 | NA | NA | 2 | 0.5 | 5 | 0 | 1 | 0 | 8 | 0.125 |
| 6_4854760 | 3 | 1 | 5 | 0.6 | 2 | 1 | 1 | 1 | 11 | 0.8182 | NA | NA | 2 | 0.5 | 4 | 0.5 | 2 | 0.5 | 8 | 0.5 |
| 6_4856724 | 1 | 1 | NA | NA | NA | NA | NA | NA | 1 | 1 | NA | NA | NA | NA | 1 | 1 | NA | NA | 1 | 1 |
| 6_4857379 | 3 | 0.6667 | 3 | 1 | 4 | 1 | 4 | 1 | 14 | 0.9286 | 3 | 1 | 3 | 0.6667 | 4 | 0.25 | 2 | 1 | 12 | 0.6667 |
| 6_4857391 | 2 | 1 | 6 | 1 | 4 | 1 | 3 | 0.6667 | 15 | 0.9333 | 2 | 1 | 3 | 0.6667 | 4 | 0.75 | 2 | 0 | 11 | 0.6364 |
| 6_4857427 | 3 | 0 | 7 | 0 | 5 | 0 | 5 | 0.2 | 20 | 0.05 | 3 | 0 | 3 | 0 | 5 | 0.2 | 2 | 0.5 | 13 | 0.1538 |
| 6_4857440 | 3 | 0 | 7 | 0.2857 | 4 | 0 | 6 | 0.5 | 20 | 0.25 | 3 | 0 | 4 | 0.25 | 4 | 1 | 2 | 0 | 13 | 0.3846 |
| 6_4857456 | 3 | 0.6667 | 7 | 1 | 4 | 1 | 6 | 1 | 20 | 0.95 | 4 | 1 | 4 | 1 | 5 | 1 | 2 | 1 | 15 | 1 |
| 6_4857482 | 4 | 0 | 7 | 0.2857 | 5 | 0.4 | 5 | 0.2 | 21 | 0.2381 | 5 | 0.4 | 4 | 0 | 5 | 0.2 | 2 | 0 | 16 | 0.1875 |
| 6_4857484 | 3 | 0.6667 | 6 | 0 | 4 | 0.75 | 7 | 0.7143 | 20 | 0.5 | 3 | 1 | 3 | 1 | 3 | 1 | 1 | 0 | 10 | 0.9 |
| 6_4857503 | 5 | 1 | 6 | 0.3333 | 6 | 0.8333 | 5 | 1 | 22 | 0.7727 | 3 | 0.3333 | 2 | 1 | 5 | 0.6 | 2 | 1 | 12 | 0.6667 |
| 6_4857557 | 4 | 0.25 | 5 | 0 | 9 | 0.1111 | 7 | 0.1429 | 25 | 0.12 | 5 | 0 | 4 | 0 | 5 | 0.2 | 1 | 1 | 15 | 0.1333 |
| 6_4859628 | 2 | 1 | 1 | 1 | NA | NA | 2 | 0 | 5 | 0.6 | NA | NA | NA | NA | 2 | 0.5 | 1 | 1 | 3 | 0.6667 |
| 6_4859632 | NA | NA | 1 | 1 | NA | NA | 1 | 1 | 2 | 1 | 1 | 1 | NA | NA | 1 | 1 | 1 | 1 | 3 | 1 |
| 6_4859638 | 1 | 1 | 1 | 0 | NA | NA | 2 | 1 | 4 | 0.75 | 1 | 1 | NA | NA | 2 | 1 | 1 | 1 | 4 | 1 |
| 6_4859645 | 2 | 1 | NA | NA | NA | NA | NA | NA | 2 | 1 | 1 | 1 | NA | NA | 2 | 1 | 1 | 1 | 4 | 1 |
| 6_4859664 | 2 | 0 | 1 | 0 | NA | NA | 2 | 0.5 | 5 | 0.2 | 1 | 1 | NA | NA | 1 | 0 | 1 | 0 | 3 | 0.3333 |
| 6_4859692 | 2 | 1 | 1 | 0 | 1 | 0 | 2 | 0 | 6 | 0.3333 | 2 | 1 | NA | NA | 2 | 0.5 | 2 | 0.5 | 6 | 0.6667 |
| 6_4859695 | 3 | 0.6667 | 1 | 0 | 1 | 0 | 2 | 1 | 7 | 0.5714 | 2 | 1 | NA | NA | 3 | 0.3333 | 1 | 0 | 6 | 0.5 |
| 6_4859743 | 3 | 0.3333 | 1 | 0 | 1 | 0 | 2 | 0 | 7 | 0.1429 | 2 | 0.5 | NA | NA | 3 | 0 | 2 | 0.5 | 7 | 0.2857 |
| 6_4859756 | 2 | 1 | 1 | 1 | 1 | 1 | 1 | 0 | 5 | 0.8 | 2 | 1 | NA | NA | 3 | 0.6667 | 1 | 1 | 6 | 0.8333 |
| 6_4859762 | 3 | 1 | 2 | 1 | 1 | 1 | 2 | 1 | 8 | 1 | 2 | 0.5 | NA | NA | 4 | 1 | 3 | 1 | 9 | 0.8889 |
| 6_4859770 | 3 | 0.3333 | 2 | 0.5 | 1 | 1 | 1 | 1 | 7 | 0.5714 | 2 | 0.5 | NA | NA | 2 | 0.5 | 2 | 1 | 6 | 0.6667 |
| 6_4859806 | 3 | 0.6667 | 1 | 0 | 1 | 1 | 1 | 0 | 6 | 0.5 | 2 | 1 | NA | NA | 4 | 0.75 | 1 | 1 | 7 | 0.8571 |
| 6_4859814 | 4 | 0 | 3 | 0 | NA | NA | 2 | 0.5 | 9 | 0.1111 | 2 | 0.5 | NA | NA | 4 | 0.25 | 2 | 1 | 8 | 0.5 |
| 6_4859821 | 3 | 0.3333 | 3 | 0 | NA | NA | 1 | 0 | 7 | 0.1429 | 2 | 0 | NA | NA | 4 | 0.25 | 2 | 0.5 | 8 | 0.25 |
| 6_4859842 | 3 | 0 | 4 | 0.25 | 2 | 0 | 1 | 0 | 10 | 0.1 | NA | NA | NA | NA | 1 | 1 | 1 | 0 | 2 | 0.5 |
| 6_4859847 | 3 | 1 | 3 | 1 | 1 | 1 | NA | NA | 7 | 1 | 2 | 1 | NA | NA | 2 | 1 | 1 | 1 | 5 | 1 |
| 6_4860867 | 1 | 0 | 2 | 0 | 1 | 1 | 1 | 1 | 5 | 0.4 | 1 | 0 | 1 | 0 | 2 | 1 | NA | NA | 4 | 0.5 |
| 6_4860881 | 1 | 0 | 2 | 1 | NA | NA | 3 | 1 | 6 | 0.8333 | NA | NA | 1 | 0 | 5 | 0.6 | NA | NA | 6 | 0.5 |
| 6_4860882 | 1 | 1 | 2 | 0.5 | 1 | 1 | 3 | 1 | 7 | 0.8571 | NA | NA | 1 | 1 | 5 | 1 | NA | NA | 6 | 1 |
| 6_4860885 | 1 | 0 | NA | NA | 1 | 0 | 2 | 1 | 4 | 0.5 | 1 | 0 | 1 | 0 | 5 | 0.8 | NA | NA | 7 | 0.5714 |
| 6_4860893 | 1 | 0 | 2 | 0.5 | NA | NA | 3 | 1 | 6 | 0.6667 | 1 | 1 | NA | NA | 5 | 0.8 | NA | NA | 6 | 0.8333 |
| 6_4860897 | 1 | 0 | 2 | 1 | 1 | 0 | 3 | 0.6667 | 7 | 0.5714 | 1 | 1 | 1 | 1 | 5 | 0.4 | NA | NA | 7 | 0.5714 |
| 6_4860900 | 1 | 1 | 2 | 1 | 1 | 0 | 3 | 0.6667 | 7 | 0.7143 | 1 | 1 | 2 | 0.5 | 4 | 0.5 | NA | NA | 7 | 0.5714 |
| 6_4860904 | 1 | 1 | 3 | 0.3333 | 2 | 0.5 | 3 | 0.3333 | 9 | 0.4444 | 1 | 0 | 1 | 1 | 5 | 0.2 | NA | NA | 7 | 0.2857 |
| 6_4860905 | 1 | 0 | 3 | 0.6667 | 2 | 0.5 | 3 | 0 | 9 | 0.3333 | 1 | 0 | 1 | 1 | 5 | 0 | NA | NA | 7 | 0.1429 |
| 6_4860929 | NA | NA | 1 | 1 | 1 | 0 | 2 | 0.5 | 4 | 0.5 | NA | NA | NA | NA | 2 | 0.5 | NA | NA | 2 | 0.5 |
| 6_4867878 | NA | NA | NA | NA | 1 | 0 | 1 | 1 | 2 | 0.5 | 3 | 0.3333 | NA | NA | 1 | 1 | 1 | 0 | 5 | 0.4 |
| 6_4867881 | NA | NA | NA | NA | 1 | 0 | 1 | 1 | 2 | 0.5 | 3 | 1 | NA | NA | 1 | 1 | 1 | 1 | 5 | 1 |
| 6_4867887 | NA | NA | NA | NA | 1 | 1 | 1 | 1 | 2 | 1 | 2 | 1 | NA | NA | 1 | 1 | 1 | 1 | 4 | 1 |

| | | | | | | | | | | | | | | | | | | | | |
|---|---|---|---|---|---|---|---|---|---|---|---|---|---|---|---|---|---|---|---|---|
| 6_4867893! | NA | NA | NA | NA | 1 | 0 | 1 | 1 | 2 | 0.5 | 3 | 0.3333 | NA | NA | 1 | 0 | 1 | 0 | 5 | 0.2 |
| 6_4867957( | NA | NA | NA | NA | 1 | 1 | 1 | 1 | 2 | 1 | 3 | 0.3333 | NA | NA | 1 | 1 | 1 | 0 | 5 | 0.4 |
| 6_4867959! | NA | NA | NA | NA | 1 | 1 | 1 | 1 | 2 | 1 | 2 | 1 | NA | NA | 1 | 1 | 1 | 0 | 4 | 0.75 |
| 6_4867972( | NA | NA | NA | NA | NA | NA | 1 | 1 | 1 | 1 | NA | NA | NA | NA | 1 | 1 | 1 | 1 | 2 | 1 |
| 6_4881246' | 4 | 0 | 11 | 0.1818 | 9 | 0 | 9 | 0.1111 | 33 | 0.0909 | 6 | 0.1667 | 11 | 0 | 8 | 0.25 | 2 | 0 | 27 | 0.1111 |
| 6_4902482( | NA | NA | 2 | 0.5 | NA | NA | NA | NA | 2 | 0.5 | 1 | 1 | NA | NA | NA | NA | NA | NA | 1 | 1 |
| 6_4903332( | 1 | 1 | 2 | 1 | NA | NA | NA | NA | 3 | 1 | 3 | 0.3333 | NA | NA | NA | NA | NA | NA | 3 | 0.3333 |
| 6_4906150: | 1 | 1 | 1 | 1 | 4 | 0.5 | 2 | 0 | 8 | 0.5 | 3 | 0.3333 | 3 | 0.3333 | 4 | 0.25 | NA | NA | 10 | 0.3 |
| 6_4906157: | 1 | 0 | 2 | 1 | 6 | 0.5 | 2 | 0 | 11 | 0.4545 | 2 | 1 | 2 | 0.5 | 2 | 0 | NA | NA | 6 | 0.5 |
| 6_4922163' | 9 | 0.2222 | 19 | 0.1053 | 10 | 0.2 | 19 | 0.0526 | 57 | 0.1228 | 1 | 0 | 9 | 0.1111 | 15 | 0.0667 | 9 | 0.1111 | 34 | 0.0882 |
| 6_4931092: | 1 | 1 | 1 | 1 | 2 | 0.5 | 2 | 1 | 6 | 0.8333 | NA | NA | 1 | 0 | 1 | 1 | NA | NA | 2 | 0.5 |
| 6_4931105! | 3 | 1 | 1 | 1 | 2 | 1 | 4 | 1 | 10 | 1 | 1 | 1 | 1 | 1 | 2 | 1 | NA | NA | 4 | 1 |
| 6_4931679( | 3 | 0 | 2 | 0 | 2 | 0 | 4 | 0.25 | 11 | 0.0909 | 1 | 0 | 1 | 1 | 1 | 0 | NA | NA | 3 | 0.3333 |
| 6_4931700: | 2 | 0.5 | 3 | 0 | 2 | 0 | 1 | 0 | 8 | 0.125 | 1 | 0 | 1 | 0 | 2 | 0.5 | NA | NA | 4 | 0.25 |
| 6_4931702( | 2 | 1 | 2 | 1 | 1 | 1 | 4 | 1 | 9 | 1 | 1 | 1 | 1 | 1 | 2 | 1 | NA | NA | 4 | 1 |
| 6_4931730! | 2 | 0.5 | 3 | 0 | 2 | 1 | 5 | 0.6 | 12 | 0.5 | 1 | 1 | 1 | 0 | 1 | 0 | NA | NA | 3 | 0.3333 |
| 6_4931731! | 3 | 0.6667 | 3 | 0.6667 | 2 | 0.5 | 5 | 0.8 | 13 | 0.6923 | 1 | 1 | 1 | 0 | 2 | 0.5 | NA | NA | 4 | 0.5 |
| 6_5017351: | 9 | 0.2222 | 3 | 0.6667 | 4 | 0.25 | 4 | 0.25 | 20 | 0.3 | NA | NA | 5 | 0.2 | 5 | 0.2 | 9 | 0.2222 | 19 | 0.2105 |
| 6_5017513! | 10 | 0.3 | 5 | 1 | 4 | 0 | 4 | 0.25 | 23 | 0.3913 | NA | NA | 5 | 0 | 8 | 0.25 | 9 | 0.3333 | 22 | 0.2273 |
| 6_5017521! | 10 | 0.1 | 6 | 0.5 | 4 | 0.25 | 4 | 0.25 | 24 | 0.25 | NA | NA | 5 | 0.2 | 8 | 0.125 | 10 | 0.1 | 23 | 0.1304 |
| 6_5017525' | 11 | 0.2727 | 4 | 0.75 | 3 | 0 | 4 | 0.25 | 22 | 0.3182 | NA | NA | 4 | 0 | 6 | 0.3333 | 8 | 0.125 | 18 | 0.1667 |
| 6_5017526! | 11 | 0.2727 | 5 | 0.8 | 4 | 0 | 4 | 0 | 24 | 0.2917 | NA | NA | 6 | 0 | 5 | 0.2 | 8 | 0.5 | 19 | 0.2632 |
| 6_5017534: | 8 | 0 | 3 | 0.3333 | 2 | 0 | 3 | 0 | 16 | 0.0625 | NA | NA | 4 | 0.25 | 6 | 0.3333 | 10 | 0.4 | 20 | 0.35 |
| 6_5017535: | 3 | 0 | 4 | 0.75 | 1 | 1 | 1 | 1 | 9 | 0.5556 | NA | NA | 2 | 0.5 | 2 | 1 | 5 | 0.4 | 9 | 0.5556 |
| 6_5017631: | 7 | 0 | 2 | 0 | 1 | 0 | 5 | 0.2 | 15 | 0.0667 | NA | NA | 2 | 0.5 | 2 | 0 | 3 | 0 | 7 | 0.1429 |
| 6_5017781: | 5 | 0.2 | 4 | 0 | 2 | 0 | 3 | 0 | 14 | 0.0714 | 1 | 0 | 6 | 0 | 6 | 0.1667 | 4 | 0 | 17 | 0.0588 |
| 6_5017964: | 1 | 0 | 2 | 0.5 | 1 | 0 | NA | NA | 4 | 0.25 | NA | NA | NA | NA | 2 | 0.5 | 1 | 0 | 3 | 0.3333 |
| 6_5018453! | 1 | 0 | 2 | 0 | NA | NA | 2 | 0.5 | 5 | 0.2 | NA | NA | 1 | 1 | 2 | 0 | NA | NA | 3 | 0.3333 |
| 6_5018466: | NA | NA | 1 | 1 | NA | NA | NA | NA | 1 | 1 | NA | NA | 1 | 1 | NA | NA | NA | NA | 1 | 1 |
| 6_5018472: | NA | NA | 1 | 1 | 1 | 0 | NA | NA | 2 | 0.5 | NA | NA | 1 | 1 | NA | NA | NA | NA | 1 | 1 |
| 6_5018485! | NA | NA | 1 | 1 | 1 | 1 | NA | NA | 2 | 1 | NA | NA | 1 | 1 | NA | NA | NA | NA | 1 | 1 |
| 6_5018489: | 2 | 0.5 | 1 | 0 | NA | NA | 3 | 0 | 6 | 0.1667 | NA | NA | 1 | 1 | NA | NA | 1 | 1 | 2 | 1 |
| 6_5018493! | 2 | 0.5 | 2 | 1 | NA | NA | 3 | 0.6667 | 7 | 0.7143 | NA | NA | 1 | 0 | 1 | 1 | 1 | 0 | 3 | 0.3333 |
| 6_5018494( | NA | NA | 1 | 1 | 1 | 0 | NA | NA | 2 | 0.5 | NA | NA | 1 | 1 | NA | NA | NA | NA | 1 | 1 |
| 6_5018502( | 2 | 0 | 1 | 1 | NA | NA | 3 | 0.6667 | 6 | 0.5 | NA | NA | 1 | 1 | 1 | 1 | 1 | 1 | 3 | 1 |
| 6_5018531! | NA | NA | 1 | 1 | 1 | 0 | 1 | 1 | 3 | 0.6667 | NA | NA | 1 | 1 | NA | NA | 1 | 1 | 2 | 1 |
| 6_5027174: | 4 | 0.75 | 2 | 1 | 1 | 0 | 3 | 0.3333 | 10 | 0.6 | 2 | 0 | 4 | 1 | 2 | 0.5 | 2 | 1 | 10 | 0.7 |
| 6_5027209: | 2 | 1 | 2 | 0 | 1 | 0 | 2 | 1 | 7 | 0.5714 | 2 | 1 | 3 | 0.6667 | 2 | 0.5 | NA | NA | 7 | 0.7143 |
| 6_5027216: | 4 | 0 | 2 | 0 | 1 | 0 | 3 | 0.3333 | 10 | 0.1 | 2 | 0 | 5 | 0.2 | 2 | 0 | 2 | 0 | 11 | 0.0909 |
| 6_5027221: | 1 | 1 | NA | NA | 1 | 0 | 1 | 1 | 3 | 0.6667 | 1 | 1 | 1 | 1 | 1 | 1 | NA | NA | 3 | 1 |
| 6_5027246' | NA | NA | 2 | 1 | 1 | 0 | 3 | 1 | 6 | 0.8333 | 1 | 1 | 5 | 1 | 2 | 0.5 | 2 | 1 | 10 | 0.9 |
| 6_5027406: | 1 | 1 | 2 | 0 | 2 | 0 | 4 | 0 | 9 | 0.1111 | 1 | 1 | 3 | 0 | 2 | 0 | 1 | 1 | 7 | 0.2857 |
| 6_5027409( | 2 | 0 | 4 | 0.25 | 2 | 0 | 4 | 0.25 | 12 | 0.1667 | 1 | 0 | 5 | 0.6 | 1 | 0 | 2 | 0 | 9 | 0.3333 |
| 6_5054103: | 4 | 0.5 | 3 | 0.3333 | 1 | 0 | 1 | 0 | 9 | 0.3333 | 3 | 0.3333 | 2 | 0 | 2 | 0 | 1 | 0 | 8 | 0.125 |
| 6_5054198! | 33 | 0.4242 | 43 | 0.3953 | 36 | 0.3333 | 25 | 0.24 | 137 | 0.3577 | 29 | 0.4483 | 32 | 0.25 | 28 | 0.3571 | 25 | 0.4 | 114 | 0.3596 |
| 6_5054210! | 40 | 0.125 | 49 | 0.3265 | 43 | 0.3256 | 29 | 0.3103 | 161 | 0.2733 | 33 | 0.2121 | 37 | 0.2432 | 32 | 0.2188 | 29 | 0.1724 | 131 | 0.2137 |
| 6_5054272( | NA | NA | 1 | 1 | 1 | 1 | NA | NA | 2 | 1 | 1 | 0 | NA | NA | 1 | 1 | NA | NA | 2 | 0.5 |
| 6_5054275: | NA | NA | 1 | 1 | 1 | 1 | NA | NA | 2 | 1 | NA | NA | NA | NA | 1 | 1 | NA | NA | 1 | 1 |
| 6_5054281: | NA | NA | 1 | 1 | 1 | 0 | NA | NA | 2 | 0.5 | 1 | 0 | NA | NA | 1 | 1 | NA | NA | 2 | 0.5 |
| 6_5054283( | NA | NA | 1 | 1 | 1 | 1 | NA | NA | 2 | 1 | 1 | 0 | NA | NA | 1 | 1 | NA | NA | 2 | 0.5 |
| 6_5143467: | NA | NA | NA | NA | 1 | 1 | NA | NA | 1 | 1 | NA | NA | 1 | 1 | NA | NA | NA | NA | 1 | 1 |
| 6_5143476( | NA | NA | NA | NA | 1 | 1 | NA | NA | 1 | 1 | NA | NA | 1 | 1 | NA | NA | NA | NA | 1 | 1 |
| 6_5143477: | NA | NA | NA | NA | 1 | 1 | NA | NA | 1 | 1 | NA | NA | 1 | 1 | NA | NA | NA | NA | 1 | 1 |
| 6_5143504! | NA | NA | 1 | 1 | 1 | 0 | NA | NA | 2 | 0.5 | NA | NA | 1 | 1 | NA | NA | NA | NA | 1 | 1 |
| 6_5143522! | NA | NA | 1 | 0 | 1 | 1 | NA | NA | 2 | 0.5 | NA | NA | 1 | 1 | NA | NA | NA | NA | 1 | 1 |
| 6_5143534' | NA | NA | NA | NA | 1 | 1 | NA | NA | 1 | 1 | NA | NA | 1 | 1 | NA | NA | NA | NA | 1 | 1 |
| 6_5143536: | NA | NA | 1 | 1 | 1 | 0 | NA | NA | 2 | 0.5 | NA | NA | 1 | 1 | NA | NA | NA | NA | 1 | 1 |
| 6_5143545( | NA | NA | 2 | 1 | 1 | 1 | NA | NA | 3 | 1 | NA | NA | 1 | 1 | NA | NA | NA | NA | 1 | 1 |
| 6_5143565: | NA | NA | 2 | 0 | 1 | 1 | NA | NA | 3 | 0.3333 | NA | NA | 1 | 1 | NA | NA | NA | NA | 1 | 1 |
| 6_5143899: | 4 | 0.25 | 20 | 0.1 | 11 | 0 | 16 | 0.0625 | 51 | 0.0784 | 5 | 0 | 10 | 0.1 | 16 | 0.125 | 8 | 0.125 | 39 | 0.1026 |
| 6_5144015: | 45 | 0.0667 | 56 | 0.0714 | 45 | 0.1333 | 51 | 0.0196 | 197 | 0.0711 | 32 | 0.125 | 44 | 0.0682 | 44 | 0.0682 | 38 | 0.0526 | 158 | 0.0759 |
| 6_5156527( | 14 | 0 | 21 | 0.0952 | 13 | 0.0769 | 11 | 0.0909 | 59 | 0.0678 | 12 | 0.0833 | 11 | 0.0909 | 5 | 0 | 17 | 0.0588 | 45 | 0.0667 |
| 6_5156641: | 17 | 0.2941 | 23 | 0.1304 | 21 | 0.1429 | 10 | 0.3 | 71 | 0.1972 | 13 | 0.1538 | 12 | 0.1667 | 8 | 0 | 17 | 0.0588 | 50 | 0.1 |
| 6_5183483( | 4 | 0.5 | 2 | 0 | NA | NA | 1 | 1 | 7 | 0.4286 | 1 | 0 | 1 | 0 | 3 | 0.6667 | 2 | 0 | 7 | 0.2857 |
| 6_5252385: | 4 | 0.25 | 11 | 0.2727 | 11 | 0.0909 | 15 | 0.1333 | 41 | 0.1707 | 11 | 0.0909 | 10 | 0.2 | 5 | 0 | 8 | 0.25 | 34 | 0.1471 |

| | | | | | | | | | | | | | | | | | | | | |
|---|---|---|---|---|---|---|---|---|---|---|---|---|---|---|---|---|---|---|---|---|
| 6_5252393 | 6 | 0 | 13 | 0.2308 | 13 | 0 | 17 | 0.1765 | 49 | 0.1224 | 9 | 0.3333 | 12 | 0.25 | 8 | 0 | 13 | 0.4615 | 42 | 0.2857 |
| 6_5274317 | 26 | 0.0385 | 18 | 0.0556 | 17 | 0.1765 | 18 | 0.1667 | 79 | 0.1013 | 13 | 0.0769 | 10 | 0 | 18 | 0.1111 | 9 | 0.2222 | 50 | 0.1 |
| 6_5274644 | 5 | 0 | 5 | 0 | 1 | 0 | 6 | 0.5 | 17 | 0.1765 | 5 | 0 | 6 | 0.5 | 5 | 0.2 | 3 | 0.3333 | 19 | 0.2632 |
| 6_5274650 | 5 | 0 | 4 | 0 | 1 | 0 | 5 | 0.6 | 15 | 0.2 | 4 | 0.25 | 4 | 0 | 5 | 0 | 3 | 0.3333 | 16 | 0.125 |
| 6_5274717 | 4 | 0.5 | 4 | 0.75 | 1 | 1 | 3 | 1 | 12 | 0.75 | 3 | 1 | 4 | 1 | 4 | 1 | 5 | 0.8 | 16 | 0.9375 |
| 6_5379255 | 3 | 1 | 6 | 0.6667 | 4 | 0.5 | 4 | 0.5 | 17 | 0.6471 | 1 | 0 | 4 | 1 | 2 | 0.5 | 1 | 1 | 8 | 0.75 |
| 6_5379258 | 4 | 0.25 | 6 | 0.5 | 4 | 0 | 4 | 0.25 | 18 | 0.2778 | 2 | 0 | 3 | 0.6667 | 3 | 0.3333 | 1 | 1 | 9 | 0.4444 |
| 6_5379283 | 3 | 0.3333 | 5 | 0.6 | 4 | 0.25 | 4 | 0.5 | 16 | 0.4375 | 2 | 0 | 4 | 0 | 3 | 0.6667 | 1 | 0 | 10 | 0.2 |
| 6_5379309 | 3 | 0.3333 | 4 | 0 | 4 | 0 | 4 | 0.75 | 15 | 0.2667 | 2 | 0.5 | 4 | 0.5 | 1 | 1 | 1 | 1 | 8 | 0.625 |
| 6_5379348 | 2 | 0 | 5 | 0.2 | 4 | 0 | 4 | 0 | 15 | 0.0667 | 2 | 0.5 | 4 | 0 | 4 | 0.25 | 1 | 0 | 11 | 0.1818 |
| 6_5379410 | 3 | 0.3333 | 3 | 0 | 4 | 0 | 4 | 0 | 14 | 0.0714 | 1 | 0 | 4 | 0 | 3 | 0.3333 | NA | NA | 8 | 0.125 |
| 6_5379414 | 2 | 1 | 2 | 0.5 | 3 | 0.6667 | 4 | 0.75 | 11 | 0.7273 | NA | NA | 3 | 0.3333 | 2 | 1 | NA | NA | 5 | 0.6 |
| 6_5379415 | 2 | 0.5 | 3 | 1 | 2 | 1 | 5 | 0.8 | 12 | 0.8333 | 1 | 1 | 3 | 0.6667 | 3 | 1 | NA | NA | 7 | 0.8571 |
| 6_5379416 | 3 | 0.3333 | 5 | 0.4 | 5 | 0.8 | 7 | 0.5714 | 20 | 0.55 | 3 | 0.3333 | 4 | 0.75 | 4 | 0.25 | 2 | 1 | 13 | 0.5385 |
| 6_5379417 | 3 | 0.6667 | 6 | 0 | 3 | 0 | 3 | 0.3333 | 15 | 0.2 | 2 | 0 | 2 | 0 | 3 | 0.3333 | 2 | 0 | 9 | 0.1111 |
| 6_5379427 | 4 | 0 | 5 | 0.2 | 5 | 0.2 | 7 | 0 | 21 | 0.0952 | 2 | 0.5 | 5 | 0 | 4 | 0.25 | 1 | 0 | 12 | 0.1667 |
| 6_5379437 | 3 | 0.6667 | 6 | 0.5 | 5 | 0.8 | 7 | 0.8571 | 21 | 0.7143 | 3 | 0.6667 | 5 | 0.6 | 4 | 0.75 | 2 | 1 | 14 | 0.7143 |
| 6_5379440 | 2 | 0 | 3 | 0 | 3 | 0.3333 | 7 | 0.1429 | 15 | 0.1333 | 3 | 0 | 4 | 0 | 4 | 0 | 2 | 0.5 | 13 | 0.0769 |
| 6_5379449 | 3 | 0.6667 | 5 | 0.8 | 5 | 0.8 | 7 | 0.8571 | 20 | 0.8 | 3 | 0.3333 | 5 | 1 | 4 | 0.5 | NA | NA | 12 | 0.6667 |
| 6_5379451 | 3 | 1 | 5 | 0 | 3 | 0.3333 | 5 | 0.2 | 16 | 0.3125 | 3 | 0 | 4 | 0.25 | 4 | 0.25 | 2 | 0 | 13 | 0.1538 |
| 6_5379457 | 3 | 0.3333 | 4 | 1 | 4 | 1 | 6 | 0.8333 | 17 | 0.8235 | 3 | 0.3333 | 3 | 0.6667 | 4 | 0.5 | 1 | 1 | 11 | 0.5455 |
| 6_5379470 | 2 | 1 | 6 | 0.8333 | 5 | 0.6 | 7 | 0.8571 | 20 | 0.8 | 2 | 0.5 | 4 | 1 | 4 | 0.75 | 2 | 1 | 12 | 0.8333 |
| 6_5379472 | 2 | 1 | 5 | 0.8 | 3 | 0.6667 | 7 | 0.5714 | 17 | 0.7059 | 2 | 0 | 2 | 1 | 3 | 0.6667 | 1 | 1 | 8 | 0.625 |
| 6_5379479 | 3 | 0.6667 | 6 | 0.6667 | 5 | 0.4 | 5 | 0.6 | 19 | 0.5789 | 2 | 0 | 5 | 0.8 | 3 | 0.6667 | 1 | 0 | 11 | 0.5455 |
| 6_5379486 | 3 | 1 | 3 | 1 | 2 | 1 | 6 | 1 | 14 | 1 | 2 | 1 | 4 | 1 | 4 | 1 | 1 | 1 | 11 | 1 |
| 6_5379495 | 3 | 1 | 5 | 1 | 5 | 1 | 6 | 1 | 19 | 1 | 2 | 1 | 3 | 1 | 4 | 1 | 2 | 1 | 11 | 1 |
| 6_5379513 | 3 | 0 | 5 | 0 | 4 | 0 | 5 | 0.2 | 17 | 0.0588 | 2 | 0 | 4 | 0 | 4 | 0.25 | 2 | 0 | 12 | 0.0833 |
| 6_5379526 | 2 | 0.5 | 5 | 0.4 | 5 | 0 | 6 | 0 | 18 | 0.1667 | 4 | 0 | 4 | 0.5 | 4 | 0 | 2 | 0 | 14 | 0.1429 |
| 6_5379545 | 2 | 0 | 4 | 0.5 | 2 | 1 | 3 | 0.6667 | 11 | 0.5455 | 1 | 1 | 2 | 1 | 3 | 0.6667 | NA | NA | 6 | 0.8333 |
| 6_5379548 | 2 | 1 | 5 | 0.6 | 3 | 1 | 4 | 1 | 14 | 0.8571 | 2 | 0.5 | 4 | 1 | 4 | 0.75 | 2 | 1 | 12 | 0.8333 |
| 6_5379559 | 2 | 0.5 | 4 | 0 | 3 | 0.3333 | 5 | 0.2 | 14 | 0.2143 | 2 | 0.5 | 3 | 0.3333 | 3 | 0.3333 | 2 | 0.5 | 10 | 0.4 |
| 6_5379560 | 1 | 1 | 4 | 0.75 | 4 | 1 | 4 | 0.25 | 13 | 0.6923 | 2 | 0.5 | 3 | 1 | 3 | 1 | 2 | 0.5 | 10 | 0.8 |
| 6_5379569 | 1 | 0 | 4 | 0.25 | 3 | 0.3333 | 5 | 0.2 | 13 | 0.2308 | 2 | 0.5 | 4 | 0.75 | 2 | 0.5 | 2 | 1 | 10 | 0.7 |
| 6_5379579 | 2 | 1 | 3 | 1 | 4 | 1 | 5 | 1 | 14 | 1 | 2 | 1 | 4 | 1 | 2 | 0.5 | 2 | 1 | 10 | 0.9 |
| 6_5379583 | NA | NA | 5 | 0.6 | 4 | 0.5 | 3 | 0 | 12 | 0.4167 | 2 | 0.5 | 2 | 0.5 | 3 | 0.3333 | 1 | 1 | 8 | 0.5 |
| 6_5379613 | NA | NA | 4 | 1 | 2 | 0.5 | 4 | 0.5 | 10 | 0.7 | 2 | 0.5 | 4 | 1 | 1 | 0 | 2 | 0 | 9 | 0.5556 |
| 6_5379619 | 1 | 1 | 3 | 1 | 2 | 1 | 3 | 1 | 9 | 1 | 2 | 1 | 3 | 1 | 1 | 1 | 1 | 1 | 7 | 1 |
| 6_5379630 | 1 | 1 | 4 | 0 | 2 | 0 | 2 | 0 | 9 | 0.1111 | 2 | 0 | 4 | 0.5 | 2 | 0.5 | 2 | 0.5 | 10 | 0.4 |
| 6_5379639 | 1 | 1 | 4 | 0.75 | 2 | 1 | 2 | 1 | 9 | 0.8889 | 2 | 1 | 4 | 1 | 2 | 1 | 2 | 1 | 10 | 1 |
| 6_5379708 | 1 | 1 | 2 | 0 | 1 | 0 | 2 | 0 | 6 | 0.1667 | 1 | 0 | 3 | 0.3333 | 2 | 0.5 | 2 | 0 | 8 | 0.25 |
| 6_5379740 | NA | NA | 1 | 0 | 1 | 1 | 2 | 0 | 4 | 0.25 | 1 | 0 | 3 | 0 | 2 | 0.5 | 2 | 0 | 8 | 0.125 |
| 6_5401698 | 2 | 0 | 3 | 0.3333 | 6 | 0.1667 | 4 | 0.25 | 15 | 0.2 | 3 | 0.3333 | 6 | 0.3333 | 9 | 0.2222 | 5 | 0.4 | 23 | 0.3043 |
| 6_5424990 | NA | NA | 3 | 0 | 2 | 0.5 | NA | NA | 5 | 0.2 | 3 | 0.3333 | 3 | 0 | NA | NA | 1 | 0 | 7 | 0.1429 |
| 6_5427707 | 7 | 0.1429 | 11 | 0.2727 | 12 | 0.25 | 7 | 0.1429 | 37 | 0.2162 | 9 | 0.1111 | 15 | 0.4 | 15 | 0.1333 | 9 | 0 | 48 | 0.1875 |
| 6_5427732 | 5 | 0.6 | 10 | 0.4 | 15 | 0.3333 | 6 | 0.1667 | 36 | 0.3611 | 10 | 0.1 | 13 | 0.3077 | 17 | 0.3529 | 8 | 0.375 | 48 | 0.2917 |
| 6_5427734 | 7 | 0.4286 | 11 | 0.4545 | 16 | 0.4375 | 6 | 0.5 | 40 | 0.45 | 9 | 0.6667 | 13 | 0.5385 | 13 | 0.3077 | 7 | 0.4286 | 42 | 0.4762 |
| 6_5427749 | 7 | 0.1429 | 12 | 0 | 11 | 0.0909 | 6 | 0 | 36 | 0.0556 | 8 | 0.125 | 13 | 0.0769 | 15 | 0.1333 | 6 | 0 | 42 | 0.0952 |
| 6_5427804 | 8 | 0.375 | 14 | 0.0714 | 15 | 0.3333 | 9 | 0.2222 | 46 | 0.2391 | 10 | 0.3 | 14 | 0.1429 | 19 | 0.2105 | 6 | 0.1667 | 49 | 0.2041 |
| 6_5427811 | 9 | 0.3333 | 10 | 0.3 | 15 | 0.6 | 9 | 0.5556 | 43 | 0.4651 | 5 | 0.4 | 15 | 0.6667 | 14 | 0.5 | 5 | 0.6 | 39 | 0.5641 |
| 6_5427842 | 9 | 0.1111 | 14 | 0.1429 | 17 | 0.0588 | 10 | 0.1 | 50 | 0.1 | 9 | 0 | 17 | 0 | 18 | 0.1111 | 8 | 0.125 | 52 | 0.0577 |
| 6_5427854 | 10 | 0 | 14 | 0.1429 | 18 | 0.1667 | 11 | 0 | 53 | 0.0943 | 9 | 0.1111 | 19 | 0.4211 | 18 | 0.0556 | 8 | 0 | 54 | 0.1852 |
| 6_5447811 | 11 | 0.0909 | 21 | 0.1905 | 17 | 0.0588 | 12 | 0 | 61 | 0.0984 | 11 | 0.0909 | 17 | 0.1176 | 11 | 0 | 9 | 0 | 48 | 0.0625 |
| 6_5447814 | 14 | 0.1429 | 22 | 0.0455 | 19 | 0.0526 | 14 | 0.0714 | 69 | 0.0725 | 12 | 0.0833 | 15 | 0.1333 | 12 | 0 | 9 | 0.2222 | 48 | 0.1042 |
| 6_5447909 | 20 | 0 | 27 | 0.037 | 23 | 0.087 | 24 | 0.0833 | 94 | 0.0532 | 16 | 0.1875 | 21 | 0.0952 | 17 | 0.0588 | 18 | 0.1667 | 72 | 0.125 |
| 6_5448060 | 26 | 0 | 37 | 0.0541 | 35 | 0.1429 | 32 | 0.0312 | 130 | 0.0615 | 20 | 0.05 | 24 | 0.0833 | 22 | 0.0909 | 17 | 0.1176 | 83 | 0.0843 |
| 6_5448300 | 27 | 0.0741 | 27 | 0.1481 | 28 | 0.1429 | 23 | 0.0435 | 105 | 0.1048 | 12 | 0 | 26 | 0.1154 | 19 | 0.0526 | 22 | 0.0455 | 79 | 0.0633 |
| 6_5448306 | 23 | 0.6522 | 25 | 0.64 | 26 | 0.3846 | 21 | 0.7143 | 95 | 0.5895 | 11 | 0.8182 | 20 | 0.6 | 18 | 0.6111 | 19 | 0.6316 | 68 | 0.6471 |
| 6_5448312 | 20 | 0.55 | 24 | 0.5833 | 23 | 0.6957 | 19 | 0.6842 | 86 | 0.6279 | 11 | 0.7273 | 20 | 0.65 | 17 | 0.6471 | 21 | 0.7619 | 69 | 0.6957 |
| 6_5448314 | 26 | 0.1923 | 25 | 0.32 | 29 | 0.2759 | 21 | 0.381 | 101 | 0.2871 | 13 | 0.3846 | 26 | 0.2692 | 17 | 0.5294 | 23 | 0.4783 | 79 | 0.4051 |
| 6_5448333 | 23 | 0.2609 | 27 | 0.2593 | 28 | 0.1786 | 20 | 0.4 | 98 | 0.2653 | 10 | 0.2 | 25 | 0.24 | 13 | 0.3077 | 22 | 0.2727 | 70 | 0.2571 |
| 6_5448337 | 22 | 0.0909 | 21 | 0.2381 | 25 | 0.12 | 19 | 0.1053 | 87 | 0.1379 | 9 | 0.1111 | 23 | 0.1304 | 15 | 0.2667 | 20 | 0.3 | 67 | 0.209 |
| 6_5448340 | 17 | 0.0588 | 20 | 0.1 | 17 | 0.0588 | 13 | 0.0769 | 67 | 0.0746 | 7 | 0.2857 | 18 | 0 | 12 | 0.1667 | 16 | 0.1875 | 53 | 0.1321 |
| 6_5448343 | 22 | 0 | 24 | 0.0833 | 21 | 0.0476 | 18 | 0.1667 | 85 | 0.0706 | 9 | 0 | 15 | 0.0667 | 13 | 0.0769 | 20 | 0.05 | 57 | 0.0526 |
| 6_5448376 | 25 | 0.2 | 23 | 0.0435 | 23 | 0.087 | 19 | 0.0526 | 90 | 0.1 | 10 | 0.2 | 19 | 0.1579 | 14 | 0.0714 | 24 | 0.1667 | 67 | 0.1493 |
| 6_5448521 | 10 | 0.1 | 10 | 0 | 12 | 0.0833 | 6 | 0 | 38 | 0.0526 | 4 | 0 | 8 | 0 | 5 | 0.2 | 5 | 0.2 | 22 | 0.0909 |

| | | | | | | | | | | | | | | | | | | | | |
|---|---|---|---|---|---|---|---|---|---|---|---|---|---|---|---|---|---|---|---|---|
| 6_5448569 | 16 | 0.25 | 12 | 0.0833 | 13 | 0.0769 | 9 | 0.1111 | 50 | 0.14 | 6 | 0 | 13 | 0.1538 | 10 | 0 | 14 | 0.0714 | 43 | 0.0698 |
| 6_5448590 | 14 | 0.1429 | 7 | 0 | 12 | 0 | 6 | 0 | 39 | 0.0513 | 5 | 0 | 11 | 0.0909 | 7 | 0.1429 | 11 | 0 | 34 | 0.0588 |
| 6_5448595 | 16 | 0.5625 | 10 | 0.5 | 11 | 0.4545 | 7 | 0 | 44 | 0.4318 | 4 | 0.5 | 14 | 0.6429 | 8 | 0.625 | 15 | 0.4 | 41 | 0.5366 |
| 6_5448613 | 14 | 0.0714 | 13 | 0.1538 | 13 | 0 | 12 | 0 | 52 | 0.0577 | 5 | 0.2 | 13 | 0.0769 | 12 | 0 | 15 | 0.0667 | 45 | 0.0667 |
| 6_5448621 | 19 | 0.1579 | 16 | 0.1875 | 16 | 0.1875 | 13 | 0.2308 | 64 | 0.1875 | 6 | 0.3333 | 15 | 0 | 14 | 0.0714 | 14 | 0.2143 | 49 | 0.1224 |
| 6_5448622 | 17 | 0.1765 | 16 | 0.3125 | 15 | 0.4667 | 12 | 0.3333 | 60 | 0.3167 | 5 | 0.6 | 13 | 0.3846 | 13 | 0.3077 | 16 | 0.5 | 47 | 0.4255 |
| 6_5455638 | 2 | 0 | 2 | 1 | NA | NA | 2 | 0.5 | 6 | 0.5 | 1 | 1 | 1 | 0 | 2 | 0 | 2 | 0.5 | 6 | 0.3333 |
| 6_5455651 | NA | NA | 2 | 0.5 | NA | NA | 1 | 0 | 3 | 0.3333 | 1 | 1 | NA | NA | 1 | 1 | 2 | 1 | 4 | 1 |
| 6_5467619 | NA | NA | 1 | 0 | 3 | 1 | NA | NA | 4 | 0.75 | NA | NA | 2 | 1 | 1 | 1 | 1 | 1 | 4 | 1 |
| 6_5467624 | NA | NA | 1 | 0 | 2 | 0.5 | NA | NA | 3 | 0.3333 | NA | NA | 3 | 0.6667 | 1 | 0 | NA | NA | 4 | 0.5 |
| 6_5467642 | 1 | 1 | 1 | 0 | 3 | 1 | NA | NA | 5 | 0.8 | NA | NA | 2 | 1 | 1 | 1 | 1 | 0 | 4 | 0.75 |
| 6_5467836 | NA | NA | 3 | 0.3333 | 8 | 0 | 1 | 0 | 12 | 0.0833 | 1 | 1 | 5 | 0.2 | 2 | 0 | 3 | 0.3333 | 11 | 0.2727 |
| 6_5467890 | 1 | 0 | 4 | 0.5 | 7 | 0.1429 | 1 | 0 | 13 | 0.2308 | 2 | 0 | 5 | 0.2 | 3 | 0 | 4 | 0 | 14 | 0.0714 |
| 6_5467921 | 1 | 0 | 4 | 0 | 9 | 0.1111 | 2 | 0 | 16 | 0.0625 | 2 | 0 | 5 | 0 | 2 | 0 | 5 | 0.2 | 14 | 0.0714 |
| 6_5468026 | 3 | 0 | 4 | 0.25 | 10 | 0.1 | 3 | 0 | 20 | 0.1 | 4 | 0 | 5 | 0 | 4 | 0.75 | 7 | 0 | 20 | 0.15 |
| 6_5468054 | 3 | 0 | 5 | 0.4 | 9 | 0.3333 | 6 | 0 | 23 | 0.2174 | 5 | 0.2 | 7 | 0.2857 | 5 | 0.4 | 7 | 0.1429 | 24 | 0.25 |
| 6_5479254 | 1 | 1 | NA | NA | NA | NA | 1 | 1 | 2 | 1 | NA | NA | 1 | 1 | NA | NA | NA | NA | 1 | 1 |
| 6_5479308 | 1 | 1 | NA | NA | NA | NA | 1 | 1 | 2 | 1 | NA | NA | 1 | 1 | NA | NA | NA | NA | 1 | 1 |
| 6_5479316 | 1 | 1 | NA | NA | NA | NA | 1 | 1 | 2 | 1 | NA | NA | 1 | 1 | NA | NA | NA | NA | 1 | 1 |
| 6_5479317 | 1 | 1 | NA | NA | NA | NA | 1 | 1 | 2 | 1 | NA | NA | 1 | 1 | NA | NA | NA | NA | 1 | 1 |
| 6_5479329 | 1 | 1 | NA | NA | NA | NA | 1 | 1 | 2 | 1 | NA | NA | 1 | 1 | NA | NA | NA | NA | 1 | 1 |
| 6_5486554 | 3 | 0.3333 | 2 | 0.5 | NA | NA | 1 | 0 | 6 | 0.3333 | 2 | 0.5 | NA | NA | 2 | 0.5 | 2 | 1 | 6 | 0.6667 |
| 6_5486561 | 2 | 0.5 | 1 | 0 | NA | NA | 1 | 0 | 4 | 0.25 | 3 | 1 | NA | NA | 2 | 1 | 2 | 0.5 | 7 | 0.8571 |
| 6_5486583 | 5 | 0.2 | 2 | 0 | NA | NA | NA | NA | 7 | 0.1429 | 4 | 0.25 | NA | NA | 2 | 0 | 2 | 0 | 8 | 0.125 |
| 6_5486629 | 4 | 0.25 | 2 | 0.5 | NA | NA | 1 | 0 | 7 | 0.2857 | 4 | 0 | NA | NA | 2 | 0.5 | 2 | 0 | 8 | 0.125 |
| 6_5486644 | 5 | 0.6 | 1 | 0 | NA | NA | 1 | 0 | 7 | 0.4286 | 2 | 0 | NA | NA | 2 | 0.5 | 2 | 0 | 6 | 0.1667 |
| 6_5486646 | 4 | 0.25 | 2 | 0.5 | NA | NA | 1 | 0 | 7 | 0.2857 | 3 | 0 | NA | NA | 1 | 1 | 2 | 0 | 6 | 0.1667 |
| 6_5490100 | NA | NA | 1 | 0 | 2 | 0.5 | NA | NA | 3 | 0.3333 | 2 | 0.5 | NA | NA | NA | NA | NA | NA | 2 | 0.5 |
| 6_5490137 | NA | NA | 1 | 0 | 2 | 0.5 | NA | NA | 3 | 0.3333 | 2 | 0.5 | NA | NA | NA | NA | NA | NA | 2 | 0.5 |
| 6_5490149 | NA | NA | 1 | 0 | 2 | 1 | NA | NA | 3 | 0.6667 | 2 | 1 | NA | NA | NA | NA | NA | NA | 2 | 1 |
| 6_5490175 | NA | NA | NA | NA | 2 | 1 | NA | NA | 2 | 1 | 2 | 0.5 | NA | NA | NA | NA | NA | NA | 2 | 0.5 |
| 6_5492106 | NA | NA | NA | NA | 2 | 1 | NA | NA | 2 | 1 | 1 | 1 | NA | NA | NA | NA | NA | NA | 1 | 1 |
| 6_5496127 | NA | NA | 1 | 1 | 1 | 0 | NA | NA | 2 | 0.5 | NA | NA | 1 | 1 | 1 | 0 | 1 | 1 | 3 | 0.6667 |
| 6_5496175 | NA | NA | 1 | 1 | 1 | 0 | NA | NA | 2 | 0.5 | NA | NA | 1 | 1 | 1 | 0 | 1 | 0 | 3 | 0.3333 |
| 6_5496230 | NA | NA | 1 | 1 | 1 | 0 | NA | NA | 2 | 0.5 | NA | NA | NA | NA | 1 | 1 | 1 | 0 | 2 | 0.5 |
| 6_5496260 | 1 | 1 | 1 | 1 | 1 | 0 | 1 | 1 | 4 | 0.75 | 1 | 0 | NA | NA | 2 | 0.5 | 1 | 1 | 4 | 0.5 |
| 6_5496269 | 6 | 0.3333 | 6 | 0.5 | 8 | 0.625 | 6 | 0.6667 | 26 | 0.5385 | 3 | 1 | 3 | 0.6667 | 3 | 1 | 3 | 0.3333 | 12 | 0.75 |
| 6_5496278 | 6 | 0.1667 | 8 | 0 | 8 | 0 | 10 | 0.3 | 32 | 0.125 | 5 | 0 | 4 | 0.25 | 2 | 0.5 | 6 | 0.1667 | 17 | 0.1765 |
| 6_5496279 | 6 | 0.1667 | 6 | 0.1667 | 7 | 0 | 7 | 0 | 26 | 0.0769 | 4 | 0.25 | 3 | 0.3333 | 1 | 0 | 5 | 0 | 13 | 0.1538 |
| 6_5496283 | 4 | 0.25 | 5 | 1 | 5 | 0.2 | 7 | 0.7143 | 21 | 0.5714 | 3 | 0.3333 | 4 | 1 | 2 | 0.5 | 5 | 0.8 | 14 | 0.7143 |
| 6_5496285 | 5 | 0.6 | 5 | 0.2 | 6 | 0.1667 | 7 | 0.2857 | 23 | 0.3043 | 3 | 0 | 4 | 0.5 | 1 | 0 | 5 | 0 | 13 | 0.1538 |
| 6_5505630 | 18 | 0.1667 | 20 | 0.05 | 17 | 0.0588 | 13 | 0 | 68 | 0.0735 | 15 | 0.0667 | 13 | 0.2308 | 15 | 0.1333 | 16 | 0.0625 | 59 | 0.1186 |
| 6_5505640 | 21 | 0.0952 | 18 | 0.0556 | 21 | 0.1905 | 12 | 0.0833 | 72 | 0.1111 | 20 | 0.05 | 10 | 0.1 | 15 | 0.0667 | 18 | 0.1111 | 63 | 0.0794 |
| 6_5506736 | NA | NA | NA | NA | NA | NA | 1 | 1 | 1 | 1 | NA | NA | NA | NA | 1 | 0 | 1 | 1 | 2 | 0.5 |
| 6_5520116 | 2 | 0.5 | NA | NA | NA | NA | NA | NA | 2 | 0.5 | NA | NA | 1 | 1 | NA | NA | NA | NA | 1 | 1 |
| 6_5529553 | 2 | 0.5 | NA | NA | 1 | 1 | NA | NA | 3 | 0.6667 | NA | NA | 1 | 1 | NA | NA | NA | NA | 1 | 1 |
| 6_5542918 | 2 | 0 | 2 | 0 | 1 | 1 | 2 | 0 | 7 | 0.1429 | 1 | 0 | 3 | 0.3333 | 1 | 1 | 1 | 0 | 6 | 0.3333 |
| 6_5542918 | 2 | 0 | 3 | 0 | 1 | 0 | 2 | 0.5 | 8 | 0.125 | NA | NA | 3 | 0.6667 | 3 | 0 | 1 | 0 | 7 | 0.2857 |
| 6_5547402 | 5 | 0.4 | 5 | 0.2 | 1 | 0 | 3 | 0.3333 | 14 | 0.2857 | 5 | 0.4 | 5 | 0 | 6 | 0 | 6 | 0.1667 | 22 | 0.1364 |
| 6_5547450 | 6 | 0.8333 | 6 | 0.8333 | 2 | 1 | 7 | 1 | 21 | 0.9048 | 7 | 0.8571 | 8 | 0.875 | 5 | 0.8 | 9 | 0.8889 | 29 | 0.8621 |
| 6_5547453 | 6 | 0.6667 | 5 | 0.6 | 1 | 0 | 9 | 0.4444 | 21 | 0.5238 | 7 | 0.5714 | 8 | 0.375 | 7 | 0.2857 | 10 | 0.4 | 32 | 0.4062 |
| 6_5547458 | 6 | 0.6667 | 6 | 0.8333 | 2 | 0.5 | 8 | 1 | 22 | 0.8182 | 7 | 0.8571 | 8 | 1 | 5 | 0.6 | 10 | 0.7 | 30 | 0.8 |
| 6_5547458 | 5 | 0.8 | 5 | 1 | 2 | 1 | 7 | 1 | 19 | 0.9474 | 7 | 1 | 7 | 1 | 8 | 0.75 | 9 | 0.7778 | 31 | 0.871 |
| 6_5547474 | 4 | 1 | 6 | 1 | 2 | 0.5 | 8 | 0.875 | 20 | 0.9 | 4 | 1 | 8 | 1 | 8 | 0.875 | 7 | 0.8571 | 27 | 0.9259 |
| 6_5547486 | 7 | 0.4286 | 3 | 0.3333 | 2 | 0.5 | 10 | 0.3 | 22 | 0.3636 | 6 | 0.3333 | 7 | 0.5714 | 7 | 0.5714 | 8 | 0.5 | 28 | 0.5 |
| 6_5547487 | 7 | 0.2857 | 5 | 0.8 | 1 | 1 | 8 | 0.25 | 21 | 0.4286 | 6 | 0.3333 | 6 | 0.5 | 7 | 0.5714 | 9 | 0.5556 | 28 | 0.5 |
| 6_5547488 | 6 | 0.3333 | 5 | 0 | 2 | 0 | 9 | 0 | 22 | 0.0909 | 7 | 0 | 9 | 0.1111 | 9 | 0.1111 | 10 | 0.1 | 35 | 0.0857 |
| 6_5547511 | 6 | 0 | 5 | 0.2 | 2 | 0 | 9 | 0.3333 | 22 | 0.1818 | 6 | 0 | 12 | 0.4167 | 8 | 0.375 | 8 | 0.625 | 34 | 0.3824 |
| 6_5547518 | 5 | 1 | 5 | 0.6 | 2 | 1 | 9 | 0.8889 | 21 | 0.8571 | 7 | 0.8571 | 12 | 1 | 8 | 0.75 | 7 | 1 | 34 | 0.9118 |
| 6_5547521 | 4 | 0.5 | 5 | 0.4 | 2 | 0.5 | 9 | 0.1111 | 20 | 0.3 | 5 | 0.6 | 12 | 0.8333 | 8 | 0.5 | 5 | 0.4 | 30 | 0.6333 |
| 6_5547523 | 4 | 0 | 4 | 0.25 | 2 | 0 | 8 | 0.25 | 18 | 0.1667 | 8 | 0 | 9 | 0.2222 | 7 | 0 | 5 | 0.2 | 29 | 0.1034 |
| 6_5547538 | 3 | 0.6667 | 5 | 1 | 3 | 0.6667 | 9 | 0.5556 | 20 | 0.7 | 8 | 0.875 | 12 | 0.75 | 7 | 0.5714 | 4 | 0.75 | 31 | 0.7419 |
| 6_5547553 | 3 | 0 | 4 | 0.75 | 3 | 0.3333 | 10 | 0 | 20 | 0.2 | 5 | 0 | 8 | 0.25 | 6 | 0.3333 | 5 | 0 | 24 | 0.1667 |
| 6_5547563 | 7 | 0.1429 | 4 | 0 | 4 | 0 | 12 | 0.0833 | 27 | 0.0741 | 9 | 0 | 12 | 0.0833 | 7 | 0.2857 | 8 | 0 | 36 | 0.0833 |
| 6_5547589 | 6 | 0.1667 | 4 | 0 | 5 | 0.4 | 8 | 0.125 | 23 | 0.1739 | 6 | 0.1667 | 11 | 0.1818 | 6 | 0 | 7 | 0.2857 | 30 | 0.1667 |

| | | | | | | | | | | | | | | | | | | | | |
|---|---|---|---|---|---|---|---|---|---|---|---|---|---|---|---|---|---|---|---|---|
| 6_55476253 | 7 | 0 | 5 | 0.2 | 5 | 0 | 9 | 0.1111 | 26 | 0.0769 | 11 | 0.0909 | 13 | 0 | 9 | 0.3333 | 10 | 0.1 | 43 | 0.1163 |
| 6_55476977 | 8 | 0 | 12 | 0.0833 | 5 | 0.4 | 12 | 0 | 37 | 0.0811 | 12 | 0.3333 | 16 | 0.25 | 13 | 0.1538 | 9 | 0.2222 | 50 | 0.24 |
| 6_55477365 | 11 | 0 | 14 | 0.2143 | 8 | 0.125 | 14 | 0.0714 | 47 | 0.1064 | 13 | 0.2308 | 20 | 0.2 | 12 | 0.1667 | 10 | 0.2 | 55 | 0.2 |
| 6_55477706 | 10 | 0.2 | 16 | 0.75 | 8 | 0.5 | 15 | 0.2667 | 49 | 0.449 | 13 | 0.3077 | 20 | 0.7 | 9 | 0.5556 | 8 | 0.25 | 50 | 0.5 |
| 6_55477978 | 10 | 0.2 | 17 | 0.4118 | 7 | 0.1429 | 13 | 0.1538 | 47 | 0.2553 | 11 | 0.4545 | 21 | 0.4762 | 7 | 0.4286 | 9 | 0.3333 | 48 | 0.4375 |
| 6_55478096 | 14 | 0.2143 | 16 | 0.25 | 10 | 0.3 | 16 | 0.3125 | 56 | 0.2679 | 15 | 0.6667 | 21 | 0.4762 | 10 | 0.4 | 11 | 0.4545 | 57 | 0.5088 |
| 6_55478330 | 14 | 0.1429 | 17 | 0.2353 | 9 | 0 | 14 | 0.2143 | 54 | 0.1667 | 12 | 0.0833 | 21 | 0.0476 | 9 | 0 | 12 | 0.25 | 54 | 0.0926 |
| 6_55655797 | 3 | 0.6667 | 5 | 1 | NA | NA | 2 | 0.5 | 10 | 0.8 | NA | NA | NA | NA | NA | NA | 3 | 0.6667 | 3 | 0.6667 |
| 6_55655966 | 3 | 1 | 6 | 1 | NA | NA | 2 | 1 | 11 | 1 | NA | NA | NA | NA | NA | NA | 3 | 1 | 3 | 1 |
| 6_55656112 | 3 | 1 | 3 | 0.6667 | NA | NA | 1 | 1 | 7 | 0.8571 | NA | NA | NA | NA | NA | NA | 1 | 1 | 1 | 1 |
| 6_55656233 | 2 | 1 | 4 | 0.75 | NA | NA | 2 | 1 | 8 | 0.875 | NA | NA | NA | NA | NA | NA | 1 | 1 | 1 | 1 |
| 6_55656249 | 3 | 1 | 3 | 1 | NA | NA | 1 | 1 | 7 | 1 | NA | NA | NA | NA | NA | NA | 2 | 1 | 2 | 1 |
| 6_55656292 | 3 | 1 | 4 | 1 | NA | NA | 2 | 1 | 9 | 1 | NA | NA | NA | NA | NA | NA | 2 | 0.5 | 2 | 0.5 |
| 6_56083485 | 1 | 0 | NA | NA | 3 | 0.6667 | NA | NA | 4 | 0.5 | 2 | 0.5 | NA | NA | NA | NA | 1 | 1 | 3 | 0.6667 |
| 6_56083497 | 1 | 1 | NA | NA | 2 | 0.5 | NA | NA | 3 | 0.6667 | 2 | 0.5 | NA | NA | 1 | 1 | 1 | 0 | 4 | 0.5 |
| 6_56083754 | 1 | 1 | NA | NA | 1 | 1 | NA | NA | 2 | 1 | 1 | 1 | NA | NA | 1 | 1 | 1 | 1 | 3 | 1 |
| 6_56713569 | NA | NA | 2 | 1 | 2 | 0 | 3 | 0 | 7 | 0.2857 | NA | NA | 3 | 0.3333 | 1 | 1 | NA | NA | 4 | 0.5 |
| 6_56713664 | 1 | 1 | 2 | 1 | 3 | 0 | 1 | 0 | 7 | 0.4286 | NA | NA | 2 | 1 | 2 | 0.5 | NA | NA | 4 | 0.75 |
| 6_56713735 | 2 | 0.5 | 2 | 0.5 | 3 | 0.3333 | 4 | 0.5 | 11 | 0.4545 | NA | NA | 3 | 0.6667 | 2 | 1 | NA | NA | 5 | 0.8 |
| 6_56713828 | 2 | 1 | 2 | 1 | 3 | 1 | 3 | 1 | 10 | 1 | NA | NA | 2 | 1 | 1 | 1 | NA | NA | 3 | 1 |
| 6_56713843 | 1 | 1 | 2 | 1 | 1 | 1 | 3 | 0.3333 | 7 | 0.7143 | NA | NA | 2 | 0.5 | 1 | 1 | NA | NA | 3 | 0.6667 |
| 6_56734491 | NA | NA | 3 | 1 | 2 | 1 | 4 | 1 | 9 | 1 | 1 | 1 | 2 | 1 | 2 | 1 | NA | NA | 5 | 1 |
| 6_56736077 | 1 | 0 | 2 | 0 | 3 | 0.3333 | 9 | 0 | 15 | 0.0667 | 3 | 0.3333 | 5 | 0 | 4 | 0 | 1 | 0 | 13 | 0.0769 |
| 6_56755021 | 10 | 0.3 | 8 | 0.25 | 12 | 0.4167 | 6 | 0.5 | 36 | 0.3611 | 4 | 0.5 | 6 | 0.5 | 10 | 0.4 | 10 | 0.2 | 30 | 0.3667 |
| 6_56755097 | 11 | 0.2727 | 9 | 0.4444 | 12 | 0.5 | 8 | 0.375 | 40 | 0.4 | 7 | 0.7143 | 9 | 0.4444 | 13 | 0.3846 | 7 | 0.5714 | 36 | 0.5 |
| 6_56755366 | 11 | 0.1818 | 10 | 0.1 | 12 | 0.0833 | 9 | 0.1111 | 42 | 0.119 | 6 | 0.1667 | 6 | 0 | 11 | 0.1818 | 8 | 0 | 31 | 0.0968 |
| 6_56755835 | 5 | 0.6 | 2 | 0 | 2 | 1 | 3 | 0.6667 | 12 | 0.5833 | 3 | 0.6667 | 3 | 0.6667 | 4 | 0.5 | 3 | 0.3333 | 13 | 0.5385 |
| 6_56755859 | 7 | 0.5714 | 7 | 0.5714 | 6 | 0.8333 | 7 | 1 | 27 | 0.7407 | 3 | 1 | 6 | 1 | 8 | 0.875 | 8 | 0.75 | 25 | 0.88 |
| 6_56755899 | 6 | 0.8333 | 5 | 1 | 6 | 1 | 7 | 0.8571 | 24 | 0.9167 | 5 | 1 | 6 | 1 | 9 | 0.8889 | 6 | 1 | 26 | 0.9615 |
| 6_56755913 | 7 | 0.1429 | 8 | 0.125 | 6 | 0.3333 | 7 | 0.1429 | 28 | 0.1786 | 3 | 0.3333 | 8 | 0.125 | 9 | 0.1111 | 7 | 0.2857 | 27 | 0.1852 |
| 6_56755980 | 8 | 0 | 8 | 0.375 | 6 | 0 | 6 | 0 | 28 | 0.1071 | 5 | 0.2 | 8 | 0 | 8 | 0.125 | 6 | 0.1667 | 27 | 0.1111 |
| 6_56756014 | 7 | 0.8571 | 8 | 0.875 | 5 | 0.6 | 7 | 1 | 27 | 0.8519 | 5 | 1 | 7 | 0.4286 | 8 | 0.875 | 6 | 1 | 26 | 0.8077 |
| 6_56756064 | 6 | 1 | 5 | 0.6 | 4 | 1 | 7 | 0.8571 | 22 | 0.8636 | 5 | 1 | 8 | 0.625 | 7 | 0.8571 | 5 | 1 | 25 | 0.84 |
| 6_56756243 | 5 | 0.8 | 8 | 0.875 | 3 | 0.3333 | 7 | 0.8571 | 23 | 0.7826 | 4 | 1 | 7 | 0.5714 | 9 | 1 | 3 | 0.6667 | 23 | 0.8261 |
| 6_56756277 | 5 | 0.6 | 6 | 0.6667 | 6 | 0.6667 | 6 | 0.5 | 23 | 0.6087 | 4 | 1 | 5 | 0.4 | 8 | 0.5 | 3 | 0.6667 | 20 | 0.6 |
| 6_56756395 | 3 | 1 | 8 | 0.125 | 5 | 0.2 | 6 | 0.5 | 22 | 0.3636 | 3 | 0.3333 | 5 | 0.4 | 8 | 0.625 | 4 | 0.75 | 20 | 0.55 |
| 6_56756432 | 6 | 0.3333 | 8 | 0.25 | 6 | 0.3333 | 7 | 0.4286 | 27 | 0.3333 | 4 | 0.5 | 5 | 0.4 | 9 | 0.2222 | 6 | 0.8333 | 24 | 0.4583 |
| 6_56756507 | 4 | 1 | 8 | 1 | 5 | 0.8 | 7 | 1 | 24 | 0.9583 | 4 | 1 | 6 | 1 | 6 | 1 | 6 | 1 | 22 | 1 |
| 6_56756543 | 3 | 1 | 7 | 1 | 4 | 1 | 7 | 1 | 21 | 1 | 4 | 0.75 | 6 | 0.8333 | 8 | 1 | 4 | 1 | 22 | 0.9091 |
| 6_56756556 | 3 | 0.3333 | 8 | 0.5 | 5 | 0.6 | 7 | 0.5714 | 23 | 0.5217 | 4 | 0.75 | 5 | 0.8 | 8 | 0.625 | 6 | 0.3333 | 23 | 0.6087 |
| 6_56756645 | 5 | 0.8 | 7 | 0.5714 | 5 | 0.4 | 7 | 0.5714 | 24 | 0.5833 | 4 | 0.75 | 6 | 0.5 | 8 | 0.75 | 5 | 0.8 | 23 | 0.6957 |
| 6_56756699 | 4 | 1 | 6 | 1 | 4 | 1 | 6 | 1 | 20 | 1 | 3 | 1 | 5 | 1 | 7 | 1 | 4 | 1 | 19 | 1 |
| 6_56756711 | 4 | 1 | 7 | 0.8571 | 4 | 1 | 6 | 0.8333 | 21 | 0.9048 | 3 | 1 | 6 | 0.8333 | 6 | 0.8333 | 3 | 1 | 18 | 0.8889 |
| 6_56756718 | 5 | 0.4 | 5 | 0.4 | 5 | 0.4 | 6 | 0.3333 | 21 | 0.381 | 2 | 0.5 | 6 | 0.1667 | 5 | 0.4 | 3 | 0 | 16 | 0.25 |
| 6_56756749 | 4 | 0 | 6 | 0 | 5 | 0 | 5 | 0.4 | 20 | 0.1 | 3 | 0 | 4 | 0.25 | 4 | 0.5 | 3 | 0 | 14 | 0.2143 |
| 6_56756798 | 5 | 0.2 | 5 | 0.2 | 4 | 0.25 | 7 | 0.1429 | 21 | 0.1905 | 4 | 0.75 | 5 | 0 | 8 | 0 | 2 | 0 | 19 | 0.1579 |
| 6_56756807 | 4 | 1 | 5 | 1 | 4 | 1 | 7 | 1 | 20 | 1 | 3 | 1 | 5 | 1 | 8 | 1 | 4 | 1 | 20 | 1 |
| 6_56756945 | 4 | 0.25 | 3 | 0 | 5 | 0.4 | 7 | 0.1429 | 19 | 0.2105 | 3 | 0 | 6 | 0 | 7 | 0.2857 | 4 | 0.25 | 20 | 0.15 |
| 6_56756955 | 4 | 0.5 | 3 | 0 | 5 | 0.2 | 6 | 0.3333 | 18 | 0.2778 | 3 | 0 | 6 | 0.3333 | 7 | 0.1429 | 3 | 0 | 19 | 0.1579 |
| 6_56757199 | 4 | 0.25 | 4 | 0.5 | 4 | 0 | 6 | 0.5 | 18 | 0.3333 | 3 | 0.6667 | 6 | 0.5 | 7 | 0.7143 | 4 | 0 | 20 | 0.5 |
| 6_56757340 | 1 | 1 | 3 | 1 | 2 | 1 | 3 | 1 | 9 | 1 | 2 | 0.5 | 5 | 1 | 6 | 0.8333 | 3 | 1 | 16 | 0.875 |
| 6_56766279 | 3 | 0 | 2 | 0.5 | 4 | 0.5 | 6 | 0 | 15 | 0.2 | 2 | 0 | 5 | 0 | 4 | 0 | 2 | 0.5 | 13 | 0.0769 |
| 6_56766296 | 4 | 0.25 | 2 | 0 | 4 | 0.25 | 6 | 0.3333 | 16 | 0.25 | 2 | 0 | 5 | 0.2 | 3 | 0 | 2 | 0 | 12 | 0.0833 |
| 6_56766308 | 3 | 0.3333 | 2 | 0.5 | 4 | 0.25 | 5 | 0.4 | 14 | 0.3571 | 2 | 0.5 | 5 | 0.8 | 4 | 0.25 | 2 | 1 | 13 | 0.6154 |
| 6_56766317 | 4 | 0.25 | 2 | 0 | 3 | 0 | 6 | 0.6667 | 15 | 0.3333 | 2 | 0 | 6 | 0.1667 | 5 | 0.2 | 2 | 0 | 15 | 0.1333 |
| 6_56766415 | 4 | 1 | 2 | 1 | 3 | 1 | 6 | 1 | 15 | 1 | 2 | 1 | 6 | 1 | 4 | 1 | 2 | 1 | 14 | 1 |
| 6_56766432 | 4 | 0.25 | 2 | 0 | 3 | 0.3333 | 6 | 0 | 15 | 0.1333 | 2 | 0 | 5 | 0 | 5 | 0.2 | 2 | 0 | 14 | 0.0714 |
| 6_56766464 | 3 | 0.6667 | 2 | 0.5 | 3 | 0.3333 | 5 | 0.4 | 13 | 0.4615 | 2 | 0 | 5 | 0.6 | 4 | 0.25 | 1 | 0 | 12 | 0.3333 |
| 6_56766566 | 3 | 0.6667 | 1 | 1 | 2 | 0.5 | 6 | 0.6667 | 12 | 0.6667 | 2 | 1 | 4 | 0.75 | 3 | 0.6667 | 2 | 0 | 11 | 0.6364 |
| 6_56855159 | 2 | 0 | 4 | 0 | 5 | 0.2 | 3 | 0 | 14 | 0.0714 | 2 | 0 | 3 | 0 | 3 | 0.3333 | 7 | 0.1429 | 15 | 0.1333 |
| 6_56855189 | 3 | 1 | 3 | 1 | 4 | 1 | 2 | 1 | 12 | 1 | 2 | 1 | 3 | 0.3333 | 2 | 1 | 6 | 1 | 13 | 0.8462 |
| 6_56855369 | 3 | 0 | 3 | 0.3333 | 5 | 0 | 1 | 0 | 12 | 0.0833 | 2 | 0.5 | 3 | 0 | 2 | 0 | 9 | 0.3333 | 16 | 0.25 |
| 6_56856192 | 5 | 0.2 | 4 | 0 | 6 | 0.1667 | 3 | 0.3333 | 18 | 0.1667 | 6 | 0.1667 | 3 | 0 | 7 | 0.4286 | 6 | 0.1667 | 22 | 0.2273 |
| 6_56859837 | 6 | 0.8333 | 6 | 0.3333 | 5 | 0.4 | 10 | 0.6 | 27 | 0.5556 | 5 | 0.2 | 1 | 0 | 3 | 0.3333 | 8 | 0.5 | 17 | 0.3529 |
| 6_56859873 | 5 | 1 | 5 | 1 | 5 | 1 | 12 | 1 | 27 | 1 | 6 | 0.6667 | 3 | 1 | 4 | 1 | 9 | 0.8889 | 22 | 0.8636 |

| | | | | | | | | | | | | | | | | | | | | |
|---|---|---|---|---|---|---|---|---|---|---|---|---|---|---|---|---|---|---|---|---|
| 6_56860015 | 6 | 0.8333 | 6 | 0.5 | 5 | 0.8 | 8 | 0.875 | 25 | 0.76 | 3 | 1 | 2 | 0.5 | 4 | 0.75 | 8 | 0.75 | 17 | 0.7647 |
| 6_57483767 | 8 | 0.375 | 6 | 0.6667 | 10 | 0.3 | 13 | 0.3077 | 37 | 0.3784 | 8 | 0.375 | 7 | 0.2857 | 5 | 0.4 | 13 | 0.3077 | 33 | 0.3333 |
| 6_57487603 | 2 | 1 | 1 | 0 | 3 | 0.6667 | 5 | 1 | 11 | 0.8182 | 3 | 0.6667 | 3 | 1 | 2 | 0.5 | 1 | 1 | 9 | 0.7778 |
| 6_57487647 | 2 | 0.5 | 1 | 0 | 3 | 1 | 2 | 1 | 8 | 0.75 | 3 | 0.3333 | 2 | 0 | 1 | 0 | 1 | 0 | 7 | 0.1429 |
| 6_57487761 | 2 | 0 | 1 | 1 | 2 | 1 | 4 | 0.75 | 9 | 0.6667 | 2 | 1 | 2 | 1 | 2 | 1 | 1 | 1 | 7 | 1 |
| 6_57501778 | 2 | 0.5 | 1 | 1 | 2 | 0 | 3 | 1 | 8 | 0.625 | 2 | 0.5 | 2 | 0 | NA | NA | NA | NA | 4 | 0.25 |
| 6_57501977 | 2 | 1 | 1 | 1 | 1 | 1 | 2 | 1 | 6 | 1 | 2 | 1 | 1 | 1 | NA | NA | 1 | 1 | 4 | 1 |
| 6_57502184 | 1 | 0 | NA | NA | 2 | 0.5 | 2 | 0.5 | 5 | 0.4 | 1 | 1 | 2 | 0.5 | NA | NA | 1 | 1 | 4 | 0.75 |
| 6_57666096 | NA | NA | 2 | 0 | 5 | 0 | 3 | 0.3333 | 10 | 0.1 | 2 | 0 | 2 | 0.5 | 1 | 0 | 7 | 0 | 12 | 0.0833 |
| 6_57666348 | 3 | 0 | 5 | 0 | 5 | 0 | 7 | 0.1429 | 20 | 0.05 | 7 | 0 | 4 | 0 | 1 | 0 | 6 | 0.1667 | 18 | 0.0556 |
| 6_57666403 | 3 | 0 | 7 | 0.1429 | 2 | 0 | 6 | 0.1667 | 18 | 0.1111 | 5 | 0 | 4 | 0.25 | 1 | 0 | 6 | 0.1667 | 16 | 0.125 |
| 6_57666438 | 1 | 0 | 3 | 0 | 4 | 0.25 | 6 | 0.1667 | 14 | 0.1429 | 6 | 0.1667 | 3 | 0 | 2 | 0.5 | 7 | 0.5714 | 18 | 0.3333 |
| 6_57666513 | 2 | 0 | 6 | 0.3333 | 4 | 0.25 | 7 | 0.2857 | 19 | 0.2632 | 6 | 0.1667 | 5 | 0.4 | 1 | 0 | 7 | 0.4286 | 19 | 0.3158 |
| 6_57666523 | 2 | 0 | 7 | 0.1429 | 4 | 0 | 7 | 0.1429 | 20 | 0.1 | 6 | 0 | 5 | 0.2 | 2 | 0.5 | 8 | 0.125 | 21 | 0.1429 |
| 6_57666586 | 3 | 0.6667 | 3 | 0.3333 | 3 | 0.6667 | 6 | 0.5 | 15 | 0.5333 | 5 | 0.8 | 4 | 0.5 | 2 | 0.5 | 8 | 0.5 | 19 | 0.5789 |
| 6_57666615 | 3 | 0.6667 | 4 | 0.25 | 2 | 0 | 5 | 0.2 | 14 | 0.2857 | 6 | 0.6667 | 5 | 0.6 | 1 | 1 | 7 | 0.2857 | 19 | 0.5263 |
| 6_57679818 | NA | NA | 1 | 0 | NA | NA | 1 | 1 | 2 | 0.5 | NA | NA | 1 | 0 | 1 | 0 | 2 | 1 | 4 | 0.5 |
| 6_57680068 | NA | NA | 2 | 1 | NA | NA | 2 | 1 | 4 | 1 | 1 | 0 | 2 | 1 | 1 | 1 | 3 | 0.3333 | 7 | 0.5714 |
| 6_57714711 | 2 | 0 | 8 | 0.25 | 1 | 0 | 9 | 0.2222 | 20 | 0.2 | 4 | 0.25 | 6 | 0 | 6 | 0.6667 | 6 | 0.3333 | 22 | 0.3182 |
| 6_57714750 | 3 | 0 | 9 | 0 | 1 | 0 | 11 | 0.1818 | 24 | 0.0833 | 5 | 0.2 | 5 | 0 | 6 | 0.1667 | 7 | 0.1429 | 23 | 0.1304 |
| 6_57714842 | 2 | 0 | 9 | 0.3333 | 1 | 0 | 11 | 0.0909 | 23 | 0.1739 | 5 | 0.2 | 5 | 0.2 | 7 | 0.1429 | 4 | 0.25 | 21 | 0.1905 |
| 6_57714876 | 3 | 0 | 10 | 0.1 | 1 | 0 | 11 | 0.0909 | 25 | 0.08 | 5 | 0.2 | 5 | 0 | 7 | 0 | 7 | 0.1429 | 24 | 0.0833 |
| 6_57729605 | NA | NA | 6 | 0 | 5 | 0 | 9 | 0.1111 | 20 | 0.05 | 2 | 0 | 9 | 0.1111 | 9 | 0 | 7 | 0.1429 | 27 | 0.0741 |
| 6_57729610 | 7 | 0 | 16 | 0.5 | 7 | 0.4286 | 12 | 0.0833 | 42 | 0.2857 | 3 | 0.6667 | 11 | 0.0909 | 12 | 0.4167 | 8 | 0.25 | 34 | 0.2941 |
| 6_57730132 | 4 | 0.25 | 13 | 0.3077 | 5 | 0.2 | 8 | 0 | 30 | 0.2 | 2 | 0 | 4 | 0 | 12 | 0.1667 | 3 | 0 | 21 | 0.0952 |
| 6_57731088 | 3 | 0 | 7 | 0 | 1 | 0 | 7 | 0.1429 | 18 | 0.0556 | 2 | 0 | 2 | 0 | 7 | 0.1429 | 5 | 0 | 16 | 0.0625 |
| 6_57731323 | 3 | 0 | 7 | 0.2857 | 1 | 0 | 8 | 0.125 | 19 | 0.1579 | 1 | 0 | 4 | 0.25 | 5 | 0 | 3 | 0.3333 | 13 | 0.1538 |
| 6_57731375 | 3 | 0 | 7 | 0.1429 | NA | NA | 7 | 0 | 17 | 0.0588 | 2 | 0 | 3 | 0 | 4 | 0 | 4 | 0.25 | 13 | 0.0769 |
| 6_57731439 | 3 | 0 | 7 | 0.2857 | 1 | 0 | 6 | 0.6667 | 17 | 0.3529 | 2 | 0 | 2 | 0 | 3 | 0.3333 | 3 | 0.6667 | 10 | 0.3 |
| 6_57731478 | 3 | 0 | 8 | 0.5 | NA | NA | 7 | 0.4286 | 18 | 0.3889 | 1 | 0 | 2 | 0.5 | 4 | 0 | 3 | 0 | 10 | 0.1 |
| 6_58882367 | 53 | 0.1132 | 68 | 0.0882 | 60 | 0.0833 | 61 | 0.0656 | 242 | 0.0868 | 37 | 0.0811 | 59 | 0.0847 | 56 | 0.0893 | 51 | 0.098 | 203 | 0.0887 |
| 6_58882394 | 42 | 0.2143 | 59 | 0.2203 | 55 | 0.2182 | 59 | 0.2034 | 215 | 0.214 | 33 | 0.2727 | 51 | 0.1569 | 50 | 0.2 | 45 | 0.2444 | 179 | 0.2123 |
| 6_58882874 | 43 | 0.2093 | 49 | 0.2041 | 48 | 0.125 | 50 | 0.24 | 190 | 0.1947 | 33 | 0.1212 | 47 | 0.1915 | 44 | 0.1364 | 37 | 0.1622 | 161 | 0.1553 |
| 6_58882928 | 39 | 0.0513 | 54 | 0.0741 | 49 | 0.0612 | 53 | 0.0755 | 195 | 0.0667 | 33 | 0.1212 | 44 | 0.0227 | 45 | 0.0667 | 41 | 0.0488 | 163 | 0.0613 |
| 6_58882962 | 36 | 1 | 46 | 0.9783 | 36 | 1 | 46 | 0.9565 | 164 | 0.9817 | 29 | 0.931 | 38 | 0.9737 | 38 | 0.8684 | 34 | 0.9118 | 139 | 0.9209 |
| 6_58883009 | 36 | 0.1111 | 53 | 0.0189 | 44 | 0.1136 | 53 | 0.1509 | 186 | 0.0968 | 34 | 0.1765 | 43 | 0.093 | 37 | 0.027 | 38 | 0.0789 | 152 | 0.0921 |
| 6_58883031 | 40 | 1 | 53 | 0.9057 | 45 | 0.9778 | 52 | 0.9423 | 190 | 0.9526 | 28 | 0.9643 | 44 | 1 | 39 | 0.9231 | 41 | 0.9268 | 152 | 0.9539 |
| 6_58883115 | 20 | 0.9 | 35 | 0.7714 | 24 | 0.7917 | 32 | 0.7812 | 111 | 0.8018 | 16 | 0.9375 | 23 | 0.8261 | 26 | 0.8846 | 22 | 0.7727 | 87 | 0.8506 |
| 6_58883120 | 33 | 0.5455 | 43 | 0.4186 | 31 | 0.4516 | 41 | 0.4146 | 148 | 0.4527 | 24 | 0.4583 | 36 | 0.5 | 27 | 0.4444 | 31 | 0.4839 | 118 | 0.4746 |
| 6_58883211 | 39 | 0.9744 | 48 | 0.875 | 45 | 0.8222 | 49 | 0.8776 | 181 | 0.884 | 29 | 1 | 38 | 0.9211 | 33 | 0.9697 | 37 | 0.973 | 137 | 0.9635 |
| 6_58883218 | 36 | 0.8056 | 48 | 0.9375 | 43 | 0.9535 | 39 | 0.8974 | 166 | 0.9036 | 28 | 0.8929 | 40 | 0.95 | 30 | 0.9667 | 31 | 0.9677 | 129 | 0.9457 |
| 6_58883223 | 31 | 0.4516 | 47 | 0.4681 | 37 | 0.4595 | 33 | 0.4545 | 148 | 0.4595 | 25 | 0.56 | 32 | 0.375 | 24 | 0.4583 | 30 | 0.3333 | 111 | 0.4234 |
| 6_58883255 | 35 | 0.0857 | 45 | 0.1556 | 42 | 0.119 | 50 | 0.18 | 172 | 0.1395 | 31 | 0.0645 | 38 | 0.0789 | 34 | 0.1765 | 34 | 0.1765 | 137 | 0.1241 |
| 6_58883319 | 36 | 0.7222 | 50 | 0.66 | 40 | 0.775 | 49 | 0.7347 | 175 | 0.72 | 31 | 0.6774 | 46 | 0.5652 | 31 | 0.7097 | 39 | 0.7692 | 147 | 0.6735 |
| 6_58883375 | 38 | 0.5263 | 51 | 0.6078 | 40 | 0.525 | 49 | 0.5102 | 178 | 0.5449 | 32 | 0.375 | 46 | 0.4565 | 34 | 0.4412 | 41 | 0.6098 | 153 | 0.4771 |
| 6_58883424 | 36 | 0.6667 | 47 | 0.7234 | 35 | 0.5714 | 48 | 0.8333 | 166 | 0.7108 | 25 | 0.64 | 42 | 0.619 | 32 | 0.8125 | 35 | 0.7714 | 134 | 0.709 |
| 6_58883595 | 30 | 0.1333 | 50 | 0.06 | 41 | 0.0976 | 50 | 0.06 | 171 | 0.0819 | 24 | 0.0417 | 42 | 0.119 | 37 | 0.0811 | 39 | 0.0256 | 142 | 0.0704 |
| 6_58883789 | 30 | 0.9333 | 49 | 0.9592 | 40 | 0.95 | 46 | 0.913 | 165 | 0.9394 | 27 | 0.8889 | 40 | 0.95 | 30 | 0.9 | 36 | 0.8056 | 133 | 0.8872 |
| 6_58895927 | 8 | 0.5 | 8 | 0 | 4 | 0 | 4 | 0.25 | 24 | 0.2083 | 10 | 0.3 | 4 | 0 | 9 | 0 | 4 | 0 | 27 | 0.1111 |
| 6_58896141 | 7 | 0.1429 | 9 | 0.1111 | 5 | 0 | 7 | 0 | 28 | 0.0714 | 12 | 0.25 | 6 | 0.1667 | 11 | 0.0909 | 5 | 0.2 | 34 | 0.1765 |
| 6_58896371 | 8 | 0.125 | 8 | 0.125 | 5 | 0 | 4 | 0 | 25 | 0.08 | 11 | 0 | 5 | 0 | 10 | 0.2 | 2 | 0 | 28 | 0.0714 |
| 6_58911217 | 3 | 0 | 3 | 0.6667 | 1 | 0 | NA | NA | 7 | 0.2857 | NA | NA | NA | NA | 1 | 0 | 2 | 0.5 | 3 | 0.3333 |
| 6_58911305 | 3 | 0.6667 | 2 | 0.5 | NA | NA | NA | NA | 5 | 0.6 | NA | NA | 1 | 0 | 1 | 0 | 2 | 1 | 4 | 0.5 |
| 6_58911310 | 3 | 1 | 2 | 1 | 1 | 1 | NA | NA | 6 | 1 | NA | NA | 1 | 1 | 1 | 0 | 2 | 1 | 4 | 0.75 |
| 6_58911503 | 3 | 0.3333 | 1 | 0 | 1 | 1 | NA | NA | 5 | 0.4 | NA | NA | NA | NA | 1 | 1 | 2 | 1 | 3 | 1 |
| 6_58911869 | 3 | 0.6667 | 2 | 0.5 | 1 | 0 | NA | NA | 6 | 0.5 | NA | NA | 1 | 1 | 1 | 0 | 2 | 1 | 4 | 0.75 |
| 6_58911991 | 2 | 0.5 | 2 | 1 | 1 | 1 | NA | NA | 5 | 0.8 | NA | NA | 1 | 1 | 1 | 1 | NA | NA | 2 | 1 |
| 6_58912012 | 2 | 0 | 2 | 0.5 | 1 | 1 | NA | NA | 5 | 0.4 | NA | NA | 1 | 1 | 1 | 0 | 1 | 1 | 3 | 0.6667 |
| 6_58912214 | 1 | 1 | 2 | 1 | 1 | 1 | NA | NA | 4 | 1 | NA | NA | 1 | 1 | 1 | 1 | 1 | 1 | 3 | 1 |
| 6_58912237 | 2 | 0.5 | 1 | 1 | NA | NA | NA | NA | 3 | 0.6667 | NA | NA | 1 | 1 | 1 | 1 | NA | NA | 2 | 1 |
| 6_58912255 | 2 | 0.5 | 2 | 0 | 1 | 0 | NA | NA | 5 | 0.2 | NA | NA | 1 | 0 | 1 | 0 | 1 | 1 | 3 | 0.3333 |
| 6_58912293 | 2 | 0 | 2 | 0.5 | 1 | 0 | NA | NA | 5 | 0.2 | NA | NA | 1 | 1 | 1 | 0 | 1 | 1 | 3 | 0.6667 |
| 6_58912331 | 2 | 1 | 2 | 0.5 | 1 | 1 | NA | NA | 5 | 0.8 | NA | NA | 1 | 1 | 1 | 0 | NA | NA | 2 | 0.5 |
| 6_58912365 | 2 | 0 | 2 | 0.5 | 1 | 0 | NA | NA | 5 | 0.2 | NA | NA | 1 | 1 | NA | NA | 1 | 0 | 2 | 0.5 |
| 6_58912395 | 2 | 0.5 | 2 | 0 | 1 | 1 | NA | NA | 5 | 0.4 | NA | NA | 1 | 0 | 1 | 0 | 1 | 1 | 3 | 0.3333 |

| | | | | | | | | | | | | | | | | | | | | |
|---|---|---|---|---|---|---|---|---|---|---|---|---|---|---|---|---|---|---|---|---|
| 6_5891242 | 2 | 0.5 | 2 | 1 | 1 | 0 | NA | NA | 5 | 0.6 | NA | NA | NA | NA | 1 | 1 | 1 | 1 | 2 | 1 |
| 6_5891245 | 1 | 1 | 2 | 0.5 | 1 | 0 | NA | NA | 4 | 0.5 | NA | NA | NA | NA | NA | NA | 1 | 1 | 1 | 1 |
| 6_5891254 | 2 | 0.5 | 2 | 0.5 | 1 | 1 | NA | NA | 5 | 0.6 | NA | NA | 1 | 1 | 1 | 1 | NA | NA | 2 | 1 |
| 6_5891256 | NA | NA | 1 | 1 | 1 | 1 | NA | NA | 2 | 1 | NA | NA | 1 | 1 | 1 | 1 | 1 | 1 | 3 | 1 |
| 6_5918662 | NA | NA | NA | NA | 2 | 1 | NA | NA | 2 | 1 | 1 | 1 | 2 | 1 | 1 | 1 | 2 | 0.5 | 6 | 0.8333 |
| 6_5918676 | 1 | 0 | NA | NA | 2 | 0.5 | NA | NA | 3 | 0.3333 | 1 | 1 | 2 | 1 | NA | NA | 2 | 0.5 | 5 | 0.8 |
| 6_5918701 | 1 | 1 | NA | NA | 2 | 1 | NA | NA | 3 | 1 | 1 | 1 | 2 | 1 | 2 | 1 | 2 | 1 | 7 | 1 |
| 6_5918706 | 1 | 1 | NA | NA | 2 | 1 | NA | NA | 3 | 1 | 1 | 1 | 2 | 1 | 2 | 1 | 2 | 0.5 | 7 | 0.8571 |
| 6_5918712 | 1 | 1 | NA | NA | 2 | 0.5 | NA | NA | 3 | 0.6667 | 1 | 0 | 2 | 0 | NA | NA | 2 | 0.5 | 5 | 0.2 |
| 6_5918722 | 1 | 1 | NA | NA | 1 | 1 | NA | NA | 2 | 1 | 1 | 0 | 2 | 0.5 | 1 | 0 | 2 | 0 | 6 | 0.1667 |
| 6_5918729 | 1 | 0 | NA | NA | 2 | 0.5 | NA | NA | 3 | 0.3333 | 1 | 1 | 2 | 0.5 | 1 | 0 | 2 | 0.5 | 6 | 0.5 |
| 6_5918770 | 1 | 1 | NA | NA | 2 | 1 | NA | NA | 3 | 1 | 3 | 1 | 2 | 0.5 | 2 | 1 | 2 | 1 | 9 | 0.8889 |
| 6_5918772 | 1 | 0 | NA | NA | 2 | 0.5 | NA | NA | 3 | 0.3333 | 2 | 0.5 | 1 | 1 | 1 | 0 | 2 | 0 | 6 | 0.3333 |
| 6_5918774 | 1 | 1 | NA | NA | 2 | 0.5 | NA | NA | 3 | 0.6667 | 3 | 0.6667 | 2 | 0 | 2 | 0 | 2 | 0 | 9 | 0.2222 |
| 6_5918815 | 1 | 1 | NA | NA | 1 | 0 | NA | NA | 2 | 0.5 | 3 | 1 | 2 | 1 | 2 | 1 | 3 | 0.6667 | 10 | 0.9 |
| 6_5918821 | 3 | 0.6667 | NA | NA | NA | NA | NA | NA | 3 | 0.6667 | 2 | 0 | 2 | 0 | 2 | 0 | 3 | 0.6667 | 9 | 0.2222 |
| 6_5918829 | 3 | 0.3333 | NA | NA | 2 | 0 | NA | NA | 5 | 0.2 | 3 | 0.6667 | 2 | 0 | 3 | 0 | 3 | 0 | 11 | 0.1818 |
| 6_5918839 | 3 | 1 | NA | NA | 2 | 0 | NA | NA | 5 | 0.6 | 3 | 0.6667 | 2 | 1 | 2 | 1 | 3 | 1 | 10 | 0.9 |
| 6_5918843 | 2 | 1 | 1 | 1 | 2 | 1 | NA | NA | 5 | 1 | 2 | 0.5 | 2 | 0.5 | 3 | 0.6667 | 3 | 0.3333 | 10 | 0.5 |
| 6_5918845 | 3 | 0.6667 | 1 | 0 | 2 | 0.5 | NA | NA | 6 | 0.5 | 3 | 0.3333 | 2 | 0 | 4 | 0.5 | 3 | 0 | 12 | 0.25 |
| 6_5918854 | 3 | 1 | NA | NA | 2 | 0.5 | NA | NA | 5 | 0.8 | 3 | 1 | 3 | 1 | 4 | 0.5 | 3 | 0.6667 | 13 | 0.7692 |
| 6_5918858 | 2 | 1 | NA | NA | 2 | 1 | NA | NA | 4 | 1 | 3 | 1 | 3 | 1 | 4 | 1 | 2 | 1 | 12 | 1 |
| 6_5918866 | 3 | 0.6667 | 1 | 1 | 2 | 0 | NA | NA | 6 | 0.5 | 3 | 0.3333 | 3 | 0 | 4 | 0.25 | 4 | 0.5 | 14 | 0.2857 |
| 6_5918874 | 2 | 0.5 | 1 | 1 | 2 | 0.5 | NA | NA | 5 | 0.6 | 2 | 0.5 | 2 | 0 | 3 | 0.3333 | 4 | 0.25 | 11 | 0.2727 |
| 6_5931826 | 1 | 0 | 1 | 0 | 3 | 0.3333 | 5 | 0 | 10 | 0.1 | 1 | 0 | 7 | 0.4286 | 4 | 0.25 | 5 | 0.2 | 17 | 0.2941 |
| 6_5931831 | 2 | 1 | 1 | 0 | 3 | 0.6667 | 5 | 0.6 | 11 | 0.6364 | 1 | 0 | 6 | 0.5 | 3 | 0.3333 | 4 | 1 | 14 | 0.5714 |
| 6_5931834 | 2 | 1 | 1 | 0 | 3 | 0.3333 | 6 | 0.6667 | 12 | 0.5833 | 1 | 1 | 5 | 0.4 | 3 | 0.6667 | 4 | 0.5 | 13 | 0.5385 |
| 6_5931836 | 2 | 1 | 1 | 1 | 2 | 1 | 6 | 1 | 11 | 1 | NA | NA | 3 | 0.6667 | 1 | 1 | 4 | 1 | 8 | 0.875 |
| 6_5931837 | 1 | 0 | NA | NA | 3 | 0.3333 | 5 | 0.2 | 9 | 0.2222 | NA | NA | 2 | 0.5 | 2 | 1 | 3 | 0.6667 | 7 | 0.7143 |
| 6_5931838 | 2 | 1 | 1 | 1 | 3 | 1 | 5 | 0.8 | 11 | 0.9091 | NA | NA | 5 | 1 | 2 | 1 | 4 | 1 | 11 | 1 |
| 6_5931840 | 2 | 0 | 1 | 1 | 3 | 0.6667 | 5 | 0.4 | 11 | 0.4545 | NA | NA | 5 | 0.4 | 2 | 0.5 | 5 | 0.6 | 12 | 0.5 |
| 6_5931848 | 2 | 0 | 1 | 0 | 2 | 1 | 5 | 0.2 | 10 | 0.3 | NA | NA | 3 | 0.3333 | 2 | 0 | 2 | 0.5 | 7 | 0.2857 |
| 6_5931857 | 1 | 1 | 1 | 1 | 3 | 0.6667 | 4 | 0.75 | 9 | 0.7778 | NA | NA | 2 | 1 | 2 | 0.5 | 4 | 0.75 | 8 | 0.75 |
| 6_5931891 | 1 | 1 | 1 | 1 | 2 | 0 | 3 | 0 | 7 | 0.2857 | NA | NA | 4 | 0.25 | 3 | 0.3333 | 2 | 0 | 9 | 0.2222 |
| 6_5931905 | 2 | 0 | 1 | 1 | 3 | 0.3333 | 4 | 0.25 | 10 | 0.3 | NA | NA | 4 | 0 | 2 | 0.5 | 3 | 0.6667 | 9 | 0.3333 |
| 6_5931921 | 2 | 0 | 1 | 0 | 3 | 0 | 4 | 0.5 | 10 | 0.2 | NA | NA | 4 | 0.25 | 2 | 0 | 4 | 0.25 | 10 | 0.2 |
| 6_5931922 | 2 | 0.5 | 1 | 0 | 3 | 0.3333 | 2 | 0.5 | 8 | 0.375 | NA | NA | 4 | 0.25 | 3 | 0.6667 | 3 | 0.3333 | 10 | 0.4 |
| 6_5931986 | 1 | 0 | 1 | 1 | 3 | 0.3333 | 3 | 0 | 8 | 0.25 | NA | NA | 1 | 1 | 2 | 1 | 3 | 0.3333 | 6 | 0.6667 |
| 6_5931987 | 2 | 0.5 | NA | NA | 2 | 0 | 4 | 0.25 | 8 | 0.25 | NA | NA | 1 | 0 | 1 | 1 | 2 | 0 | 4 | 0.25 |
| 6_5932019 | 2 | 0.5 | NA | NA | 4 | 0.75 | 2 | 0 | 8 | 0.5 | NA | NA | 1 | 1 | 1 | 1 | 1 | 0 | 3 | 0.6667 |
| 6_5932026 | 2 | 0.5 | NA | NA | 4 | 0.25 | 1 | 0 | 7 | 0.2857 | NA | NA | 1 | 0 | 1 | 1 | 2 | 0 | 4 | 0.25 |
| 6_5932051 | 1 | 1 | NA | NA | 3 | 0.3333 | 1 | 1 | 5 | 0.6 | NA | NA | 1 | 1 | 1 | 0 | 1 | 1 | 3 | 0.6667 |
| 6_5932091 | NA | NA | NA | NA | 3 | 0.3333 | 2 | 0.5 | 5 | 0.4 | NA | NA | 1 | 0 | 1 | 1 | 1 | 0 | 3 | 0.3333 |
| 6_5932092 | NA | NA | NA | NA | 3 | 0.3333 | 2 | 0.5 | 5 | 0.4 | NA | NA | 1 | 1 | NA | NA | 1 | 0 | 2 | 0.5 |
| 6_5932169 | 2 | 1 | 2 | 0.5 | 4 | 0.5 | 2 | 0.5 | 10 | 0.6 | 1 | 1 | 6 | 0.8333 | 5 | 1 | 3 | 1 | 15 | 0.9333 |
| 6_5932181 | 2 | 0.5 | 2 | 0 | 5 | 0.2 | 2 | 0 | 11 | 0.1818 | 1 | 0 | 6 | 0 | 5 | 0.4 | 3 | 0 | 15 | 0.1333 |
| 6_5932199 | 1 | 0 | 2 | 0 | 6 | 0 | 4 | 0.25 | 13 | 0.0769 | 1 | 1 | 7 | 0.2857 | 4 | 0 | 3 | 0 | 15 | 0.2 |
| 6_5932392 | 1 | 0 | NA | NA | 5 | 0.4 | 2 | 0 | 8 | 0.25 | NA | NA | 2 | 0 | 5 | 0.2 | 1 | 0 | 8 | 0.125 |
| 6_5932409 | NA | NA | NA | NA | 5 | 0.8 | 2 | 0.5 | 7 | 0.7143 | NA | NA | 2 | 0.5 | 3 | 0.6667 | 1 | 1 | 6 | 0.6667 |
| 6_5932411 | 1 | 0 | NA | NA | 5 | 0.2 | 2 | 0 | 8 | 0.125 | NA | NA | 2 | 0 | 3 | 0.3333 | 2 | 0.5 | 7 | 0.2857 |
| 6_5932439 | 1 | 0 | NA | NA | 3 | 0.3333 | 2 | 0 | 6 | 0.1667 | 1 | 1 | 2 | 0 | 4 | 0 | 3 | 0.3333 | 10 | 0.2 |
| 6_5932441 | 1 | 0 | NA | NA | 5 | 0.8 | 1 | 0 | 7 | 0.5714 | 1 | 1 | 2 | 0.5 | 4 | 0.75 | 2 | 1 | 9 | 0.7778 |
| 6_5932455 | 1 | 0 | NA | NA | 4 | 0.5 | 3 | 0.6667 | 8 | 0.5 | 1 | 1 | 2 | 0.5 | 3 | 0.6667 | 3 | 0.3333 | 9 | 0.5556 |
| 6_5932463 | 1 | 0 | NA | NA | 5 | 0.6 | 3 | 0.3333 | 9 | 0.4444 | NA | NA | 2 | 0.5 | 2 | 0 | 2 | 0.5 | 6 | 0.3333 |
| 6_5932468 | 1 | 0 | NA | NA | 4 | 0.75 | 2 | 1 | 7 | 0.7143 | NA | NA | 2 | 1 | 3 | 0.6667 | 2 | 1 | 7 | 0.8571 |
| 6_5932472 | 1 | 0 | NA | NA | 3 | 1 | 1 | 0 | 5 | 0.6 | NA | NA | 2 | 0.5 | 3 | 0.6667 | 3 | 1 | 8 | 0.75 |
| 6_5932501 | 1 | 0 | NA | NA | 5 | 0.8 | 3 | 1 | 9 | 0.7778 | NA | NA | 2 | 0.5 | 3 | 1 | 2 | 1 | 7 | 0.8571 |
| 6_5932505 | 1 | 0 | NA | NA | 4 | 0.5 | 2 | 0 | 7 | 0.2857 | NA | NA | 2 | 0.5 | 3 | 0.3333 | 2 | 1 | 7 | 0.5714 |
| 6_5932512 | 1 | 1 | NA | NA | 4 | 0.75 | 1 | 0 | 6 | 0.6667 | NA | NA | 1 | 1 | 2 | 0.5 | 2 | 1 | 5 | 0.8 |
| 6_5932514 | 1 | 0 | NA | NA | 5 | 0.8 | 1 | 1 | 7 | 0.7143 | NA | NA | NA | NA | 2 | 1 | 2 | 0.5 | 4 | 0.75 |
| 6_5932519 | 1 | 0 | NA | NA | 4 | 0.5 | 3 | 0.6667 | 8 | 0.5 | NA | NA | 1 | 0 | 3 | 0 | 2 | 0.5 | 6 | 0.1667 |
| 6_5932520 | 1 | 0 | NA | NA | 4 | 0 | 2 | 1 | 7 | 0.2857 | NA | NA | 1 | 1 | 3 | 0.6667 | 2 | 1 | 6 | 0.8333 |
| 6_5932529 | 1 | 0 | NA | NA | NA | NA | 3 | 0.3333 | 4 | 0.25 | NA | NA | 1 | 0 | 2 | 0.5 | 2 | 0 | 5 | 0.2 |
| 6_5932531 | 1 | 0 | NA | NA | 4 | 0.25 | 2 | 0 | 7 | 0.1429 | NA | NA | NA | NA | 3 | 0 | 2 | 0.5 | 5 | 0.2 |
| 6_5932536 | 1 | 0 | NA | NA | 4 | 0.75 | 1 | 0 | 6 | 0.5 | NA | NA | 1 | 0 | 2 | 0.5 | 2 | 0.5 | 5 | 0.4 |

| | | | | | | | | | | | | | | | | | | | | |
|---|---|---|---|---|---|---|---|---|---|---|---|---|---|---|---|---|---|---|---|---|
| 6_5932543 | 1 | 0 | NA | NA | 5 | 0.8 | 2 | 0.5 | 8 | 0.625 | NA | NA | 1 | 1 | 3 | 0.6667 | 1 | 1 | 5 | 0.8 |
| 6_5932544 | 1 | 0 | NA | NA | 5 | 0.6 | 3 | 0.3333 | 9 | 0.4444 | NA | NA | 1 | 1 | 3 | 0.6667 | 2 | 0.5 | 6 | 0.6667 |
| 6_5932548 | 1 | 0 | NA | NA | 4 | 0.5 | 1 | 0 | 6 | 0.3333 | NA | NA | NA | NA | 2 | 1 | 2 | 0.5 | 4 | 0.75 |
| 6_5932640 | 1 | 1 | NA | NA | 2 | 0 | NA | NA | 3 | 0.3333 | NA | NA | 1 | 1 | 1 | 1 | 2 | 0.5 | 4 | 0.75 |
| 6_5932644 | 1 | 1 | NA | NA | 3 | 1 | NA | NA | 4 | 1 | NA | NA | NA | NA | 1 | 1 | 2 | 1 | 3 | 1 |
| 6_5932722 | 1 | 0 | NA | NA | 2 | 0.5 | NA | NA | 3 | 0.3333 | NA | NA | 1 | 1 | 3 | 0.3333 | 3 | 0.3333 | 7 | 0.4286 |
| 6_5932905 | 1 | 1 | NA | NA | 3 | 0.6667 | NA | NA | 4 | 0.75 | NA | NA | 1 | 1 | 1 | 1 | 2 | 1 | 4 | 1 |
| 6_5932941 | NA | NA | NA | NA | 2 | 0.5 | NA | NA | 2 | 0.5 | NA | NA | 1 | 0 | 2 | 1 | 1 | 0 | 4 | 0.5 |
| 6_5932947 | NA | NA | NA | NA | 3 | 0.3333 | NA | NA | 3 | 0.3333 | NA | NA | 1 | 1 | 2 | 0 | 2 | 0 | 5 | 0.2 |
| 6_5932973 | NA | NA | NA | NA | 2 | 1 | NA | NA | 2 | 1 | NA | NA | 1 | 1 | 2 | 1 | 2 | 1 | 5 | 1 |
| 6_5933000 | NA | NA | NA | NA | 2 | 1 | NA | NA | 2 | 1 | NA | NA | NA | NA | 4 | 1 | 2 | 0.5 | 6 | 0.8333 |
| 6_5933001 | NA | NA | NA | NA | 3 | 1 | NA | NA | 3 | 1 | NA | NA | NA | NA | 4 | 1 | 1 | 1 | 5 | 1 |
| 6_5933020 | NA | NA | NA | NA | 4 | 1 | NA | NA | 4 | 1 | NA | NA | NA | NA | 4 | 0.5 | 2 | 0.5 | 6 | 0.5 |
| 6_5933021 | NA | NA | NA | NA | 4 | 1 | NA | NA | 4 | 1 | NA | NA | NA | NA | 4 | 0.25 | 2 | 1 | 6 | 0.5 |
| 6_5933035 | NA | NA | NA | NA | 2 | 0.5 | NA | NA | 2 | 0.5 | NA | NA | NA | NA | 2 | 0.5 | 2 | 0 | 4 | 0.25 |
| 6_5933047 | NA | NA | NA | NA | 3 | 1 | NA | NA | 3 | 1 | NA | NA | NA | NA | 3 | 0.6667 | 1 | 1 | 4 | 0.75 |
| 6_5933049 | NA | NA | NA | NA | 2 | 1 | NA | NA | 2 | 1 | NA | NA | NA | NA | 2 | 1 | 1 | 1 | 3 | 1 |
| 6_5933055 | NA | NA | NA | NA | 2 | 0.5 | NA | NA | 2 | 0.5 | NA | NA | NA | NA | 3 | 0.6667 | 1 | 0 | 4 | 0.5 |
| 6_5933072 | NA | NA | NA | NA | 2 | 0.5 | NA | NA | 2 | 0.5 | NA | NA | NA | NA | 3 | 0.6667 | 1 | 1 | 4 | 0.75 |
| 6_5933077 | NA | NA | NA | NA | 2 | 0.5 | NA | NA | 2 | 0.5 | NA | NA | NA | NA | 1 | 1 | 1 | 0 | 2 | 0.5 |
| 6_5933087 | NA | NA | NA | NA | 2 | 0.5 | NA | NA | 2 | 0.5 | NA | NA | NA | NA | 3 | 0.6667 | 1 | 1 | 4 | 0.75 |
| 6_5933087 | NA | NA | NA | NA | 2 | 1 | NA | NA | 2 | 1 | NA | NA | NA | NA | 3 | 1 | 1 | 1 | 4 | 1 |
| 6_5933121 | NA | NA | NA | NA | 1 | 1 | NA | NA | 1 | 1 | NA | NA | NA | NA | 2 | 1 | NA | NA | 2 | 1 |
| 6_5933167 | NA | NA | NA | NA | 2 | 0.5 | NA | NA | 2 | 0.5 | NA | NA | NA | NA | 2 | 0.5 | 1 | 0 | 3 | 0.3333 |
| 6_5933170 | NA | NA | NA | NA | 2 | 1 | NA | NA | 2 | 1 | NA | NA | NA | NA | 2 | 1 | NA | NA | 2 | 1 |
| 6_5933195 | NA | NA | NA | NA | 2 | 1 | NA | NA | 2 | 1 | NA | NA | NA | NA | 2 | 1 | 1 | 1 | 3 | 1 |
| 6_5933215 | NA | NA | NA | NA | 2 | 1 | NA | NA | 2 | 1 | NA | NA | NA | NA | 2 | 0.5 | 1 | 1 | 3 | 0.6667 |
| 6_5933229 | NA | NA | NA | NA | 2 | 0.5 | NA | NA | 2 | 0.5 | NA | NA | NA | NA | 3 | 0.3333 | 1 | 0 | 4 | 0.25 |
| 6_5933233 | NA | NA | NA | NA | 2 | 0.5 | NA | NA | 2 | 0.5 | NA | NA | NA | NA | 3 | 0 | 1 | 1 | 4 | 0.25 |
| 6_6070883 | 57 | 0.1053 | 62 | 0.2097 | 52 | 0.2308 | 52 | 0.0577 | 223 | 0.1525 | 53 | 0.1698 | 53 | 0.1887 | 45 | 0.0889 | 62 | 0.1129 | 213 | 0.1408 |
| 6_6128809 | NA | NA | NA | NA | 4 | 0.25 | 1 | 0 | 5 | 0.2 | NA | NA | NA | NA | 2 | 0.5 | NA | NA | 2 | 0.5 |
| 6_6128828 | NA | NA | NA | NA | 4 | 1 | NA | NA | 4 | 1 | NA | NA | NA | NA | 1 | 1 | 1 | 0 | 2 | 0.5 |
| 6_6128869 | NA | NA | NA | NA | 4 | 0.5 | 1 | 0 | 5 | 0.4 | NA | NA | NA | NA | 2 | 1 | 1 | 0 | 3 | 0.6667 |
| 6_6128899 | NA | NA | NA | NA | 3 | 0.3333 | 1 | 0 | 4 | 0.25 | NA | NA | NA | NA | 2 | 0.5 | 1 | 0 | 3 | 0.3333 |
| 6_6235681 | NA | NA | 1 | 1 | 1 | 0 | NA | NA | 2 | 0.5 | NA | NA | NA | NA | NA | NA | 1 | 1 | 1 | 1 |
| 6_6235693 | NA | NA | 1 | 1 | 1 | 1 | 2 | 1 | 4 | 1 | 1 | 1 | NA | NA | NA | NA | 1 | 1 | 2 | 1 |
| 6_6235700 | NA | NA | 1 | 1 | 1 | 0 | 2 | 1 | 4 | 0.75 | 1 | 0 | NA | NA | NA | NA | 1 | 1 | 2 | 0.5 |
| 6_6235701 | NA | NA | 1 | 1 | 1 | 0 | 2 | 0.5 | 4 | 0.5 | 1 | 0 | NA | NA | NA | NA | 1 | 1 | 2 | 0.5 |
| 6_6235709 | NA | NA | NA | NA | 1 | 1 | 2 | 1 | 3 | 1 | 1 | 1 | NA | NA | NA | NA | 1 | 1 | 2 | 1 |
| 6_6235715 | NA | NA | 1 | 1 | 1 | 1 | 2 | 0.5 | 4 | 0.75 | 1 | 1 | NA | NA | NA | NA | 1 | 0 | 2 | 0.5 |
| 6_6235722 | NA | NA | 1 | 1 | 1 | 0 | 2 | 1 | 4 | 0.75 | 1 | 0 | NA | NA | NA | NA | 1 | 1 | 2 | 0.5 |
| 6_6235727 | NA | NA | 1 | 1 | NA | NA | NA | NA | 1 | 1 | 1 | 0 | NA | NA | NA | NA | 1 | 1 | 2 | 0.5 |
| 6_6235735 | NA | NA | 1 | 1 | 1 | 0 | 2 | 0 | 4 | 0.25 | 1 | 0 | NA | NA | NA | NA | 1 | 1 | 2 | 0.5 |
| 6_6235964 | NA | NA | 1 | 0 | 4 | 0 | 3 | 0.3333 | 8 | 0.125 | 1 | 0 | NA | NA | 1 | 0 | 3 | 0.3333 | 5 | 0.2 |
| 6_6464295 | NA | NA | NA | NA | NA | NA | 1 | 1 | 1 | 1 | 2 | 1 | 3 | 1 | NA | NA | 1 | 1 | 6 | 1 |
| 6_6464312 | NA | NA | NA | NA | NA | NA | 1 | 1 | 1 | 1 | 1 | 1 | 2 | 1 | NA | NA | 1 | 1 | 4 | 1 |
| 6_6464511 | 1 | 0 | 2 | 0 | 3 | 0.3333 | 2 | 0 | 8 | 0.125 | 3 | 0 | 3 | 0 | 2 | 0 | 3 | 0.3333 | 11 | 0.0909 |
| 6_6487605 | 2 | 0 | 5 | 0.6 | 3 | 0.3333 | 1 | 0 | 11 | 0.3636 | 1 | 1 | 1 | 0 | NA | NA | NA | NA | 2 | 0.5 |
| 6_6505192 | 2 | 0.5 | 2 | 0.5 | NA | NA | 1 | 1 | 5 | 0.6 | NA | NA | 1 | 0 | 4 | 0.75 | 1 | 1 | 6 | 0.6667 |
| 6_6505197 | 2 | 0 | 2 | 0.5 | NA | NA | 1 | 0 | 5 | 0.2 | NA | NA | 1 | 1 | 4 | 0.75 | 1 | 0 | 6 | 0.6667 |
| 6_6509140 | 2 | 1 | 6 | 0.8333 | NA | NA | 4 | 1 | 12 | 0.9167 | NA | NA | NA | NA | 2 | 1 | 2 | 0.5 | 4 | 0.75 |
| 6_6509141 | 2 | 0 | 6 | 0 | NA | NA | 4 | 0.25 | 12 | 0.0833 | NA | NA | 1 | 1 | 4 | 0.25 | 2 | 0.5 | 7 | 0.4286 |
| 6_6509148 | 2 | 0.5 | 7 | 0.7143 | NA | NA | 4 | 0.75 | 13 | 0.6923 | NA | NA | 1 | 1 | 4 | 0.25 | 2 | 1 | 7 | 0.5714 |
| 6_6543319 | 1 | 0 | 1 | 0 | 2 | 0.5 | NA | NA | 4 | 0.25 | NA | NA | 1 | 1 | 1 | 1 | NA | NA | 2 | 1 |
| 6_6543320 | NA | NA | 1 | 1 | 2 | 1 | NA | NA | 3 | 1 | NA | NA | 3 | 1 | 1 | 1 | NA | NA | 4 | 1 |
| 6_6543346 | 1 | 1 | 1 | 1 | 2 | 1 | NA | NA | 4 | 1 | NA | NA | 3 | 0.6667 | 1 | 1 | NA | NA | 4 | 0.75 |
| 6_6543357 | 1 | 1 | 1 | 0 | 2 | 0.5 | NA | NA | 4 | 0.5 | NA | NA | 2 | 0.5 | 1 | 0 | NA | NA | 3 | 0.3333 |
| 6_6543358 | 1 | 0 | 1 | 0 | 2 | 0.5 | NA | NA | 4 | 0.25 | NA | NA | 1 | 1 | NA | NA | NA | NA | 1 | 1 |
| 6_6543371 | 1 | 0 | NA | NA | 2 | 0.5 | NA | NA | 3 | 0.3333 | NA | NA | 2 | 0.5 | 1 | 0 | NA | NA | 3 | 0.3333 |
| 6_6543414 | NA | NA | NA | NA | 2 | 0.5 | NA | NA | 2 | 0.5 | NA | NA | 1 | 1 | 1 | 0 | NA | NA | 2 | 0.5 |
| 6_6564890 | 1 | 1 | 1 | 1 | 1 | 1 | NA | NA | 3 | 1 | 1 | 0 | 1 | 1 | NA | NA | NA | NA | 2 | 0.5 |
| 6_6568045 | 2 | 1 | 3 | 1 | 2 | 1 | NA | NA | 7 | 1 | 2 | 1 | 1 | 1 | 1 | 1 | NA | NA | 4 | 1 |
| 6_6568047 | 2 | 0.5 | 3 | 0.6667 | 2 | 0.5 | NA | NA | 7 | 0.5714 | 1 | 1 | 1 | 1 | 1 | 1 | NA | NA | 3 | 1 |
| 6_6568058 | 2 | 0.5 | 4 | 1 | 2 | 1 | NA | NA | 8 | 0.875 | 1 | 1 | 1 | 0 | 1 | 0 | NA | NA | 3 | 0.3333 |
| 6_6568061 | 2 | 0.5 | 3 | 1 | 2 | 1 | NA | NA | 7 | 0.8571 | 2 | 0.5 | 1 | 1 | NA | NA | NA | NA | 3 | 0.6667 |

| | | | | | | | | | | | | | | | | | | | | |
|---|---|---|---|---|---|---|---|---|---|---|---|---|---|---|---|---|---|---|---|---|
| 6_6568072 | 1 | 0 | 5 | 0.2 | 2 | 0 | NA | NA | 8 | 0.125 | 2 | 0.5 | 1 | 0 | NA | NA | NA | NA | 3 | 0.3333 |
| 6_6568074 | 2 | 0 | 4 | 0.5 | 2 | 1 | NA | NA | 8 | 0.5 | 2 | 1 | 1 | 1 | 1 | 0 | NA | NA | 4 | 0.75 |
| 6_6568075 | 2 | 0.5 | 4 | 1 | 2 | 0.5 | NA | NA | 8 | 0.75 | 1 | 1 | 1 | 1 | 1 | 1 | NA | NA | 3 | 1 |
| 6_6568123 | 2 | 0 | 6 | 0.1667 | 2 | 0.5 | NA | NA | 10 | 0.2 | 2 | 0.5 | 1 | 1 | 1 | 1 | NA | NA | 4 | 0.75 |
| 6_6568130 | 2 | 0 | 4 | 0.25 | 2 | 0 | NA | NA | 8 | 0.125 | 2 | 0.5 | 1 | 0 | 1 | 1 | NA | NA | 4 | 0.5 |
| 6_6568131 | 2 | 0.5 | 4 | 1 | 1 | 1 | NA | NA | 7 | 0.8571 | 2 | 1 | 1 | 1 | 1 | 1 | NA | NA | 4 | 1 |
| 6_6568136 | 2 | 0.5 | 6 | 0.8333 | 2 | 1 | NA | NA | 10 | 0.8 | 2 | 0.5 | 1 | 1 | 1 | 1 | NA | NA | 4 | 0.75 |
| 6_6568137 | 2 | 1 | 6 | 1 | 2 | 1 | NA | NA | 10 | 1 | 2 | 1 | 1 | 1 | 1 | 1 | NA | NA | 4 | 1 |
| 6_6568145 | 2 | 0 | 6 | 0.1667 | 2 | 0.5 | NA | NA | 10 | 0.2 | 2 | 0.5 | 2 | 0 | NA | NA | NA | NA | 4 | 0.25 |
| 6_6568148 | 2 | 0 | 4 | 0.5 | NA | NA | NA | NA | 6 | 0.3333 | 2 | 0.5 | 1 | 1 | NA | NA | NA | NA | 3 | 0.6667 |
| 6_6568149 | 1 | 1 | 5 | 0.6 | 1 | 1 | NA | NA | 7 | 0.7143 | 2 | 0 | 2 | 1 | 1 | 1 | NA | NA | 5 | 0.6 |
| 6_6568157 | 2 | 0.5 | 8 | 0.875 | 2 | 0.5 | NA | NA | 12 | 0.75 | 2 | 0.5 | 2 | 1 | NA | NA | NA | NA | 4 | 0.75 |
| 6_6568169 | 2 | 1 | 8 | 1 | 2 | 1 | NA | NA | 12 | 1 | 2 | 1 | NA | NA | NA | NA | NA | NA | 2 | 1 |
| 6_6699469 | 3 | 0.3333 | 3 | 0.3333 | 4 | 0 | 10 | 0 | 20 | 0.1 | 5 | 0 | 4 | 0 | 8 | 0.25 | 9 | 0.2222 | 26 | 0.1538 |
| 6_6699613 | NA | NA | 4 | 0 | 6 | 0 | 7 | 0.1429 | 17 | 0.0588 | 4 | 0 | 3 | 0 | 8 | 0 | 10 | 0.2 | 25 | 0.08 |
| 6_6699801 | 3 | 1 | 5 | 0.2 | 7 | 0.1429 | 8 | 0.375 | 23 | 0.3478 | 4 | 0.5 | 6 | 0.3333 | 9 | 0.3333 | 7 | 0.2857 | 26 | 0.3462 |
| 6_6701249 | 1 | 1 | 2 | 0.5 | 3 | 0 | 4 | 0.5 | 10 | 0.4 | NA | NA | 2 | 0.5 | NA | NA | NA | NA | 2 | 0.5 |
| 6_6701261 | 3 | 0 | 2 | 0 | 3 | 0.3333 | 4 | 0.25 | 12 | 0.1667 | NA | NA | 2 | 0.5 | NA | NA | NA | NA | 2 | 0.5 |
| 6_6701262 | 2 | 0.5 | 1 | 0 | 2 | 0 | 4 | 0.25 | 9 | 0.2222 | NA | NA | 2 | 0.5 | NA | NA | NA | NA | 2 | 0.5 |
| 6_6726105 | 26 | 0.9615 | 25 | 1 | 19 | 1 | 20 | 0.95 | 90 | 0.9778 | 16 | 1 | 25 | 1 | 27 | 1 | 20 | 0.95 | 88 | 0.9886 |
| 6_6726108 | 21 | 0.7619 | 24 | 1 | 19 | 0.8947 | 17 | 0.9412 | 81 | 0.9012 | 18 | 0.8889 | 27 | 0.9259 | 29 | 0.931 | 20 | 1 | 94 | 0.9362 |
| 6_6726111 | 21 | 0.2857 | 29 | 0.3103 | 16 | 0.375 | 16 | 0.4375 | 82 | 0.3415 | 15 | 0.5333 | 26 | 0.4615 | 29 | 0.5517 | 20 | 0.45 | 90 | 0.5 |
| 6_6726137 | 9 | 0.2222 | 5 | 0.4 | 2 | 0 | 3 | 0 | 19 | 0.2105 | 4 | 0.25 | 6 | 0.6667 | 6 | 0 | 3 | 0.6667 | 19 | 0.3684 |
| 6_6727193 | 1 | 0 | 1 | 1 | NA | NA | 2 | 0 | 4 | 0.25 | NA | NA | 1 | 0 | 3 | 1 | NA | NA | 4 | 0.75 |
| 6_7067586 | NA | NA | 82 | 0.4268 | 8 | 0.375 | NA | NA | 90 | 0.4222 | NA | NA | 6 | 0.5 | NA | NA | NA | NA | 6 | 0.5 |
| 6_7070358 | 18 | 0.3889 | 194 | 0.3247 | 21 | 0.381 | NA | NA | 233 | 0.3348 | 66 | 0.3485 | 4 | 0.5 | 16 | 0.1875 | 104 | 0.3269 | 190 | 0.3263 |
| 6_7070360 | 19 | 0.1053 | 232 | 0.056 | 20 | 0.05 | 1 | 0 | 272 | 0.0588 | 76 | 0.0921 | 8 | 0.25 | 17 | 0.2353 | 107 | 0.0374 | 208 | 0.0817 |
| 6_7070374 | 16 | 0 | 221 | 0.0724 | 20 | 0.1 | 1 | 0 | 258 | 0.0698 | 74 | 0.1351 | 8 | 0 | 17 | 0.1765 | 111 | 0.0631 | 210 | 0.0952 |
| 6_7074321 | 2 | 0.5 | 4 | 0 | 1 | 0 | NA | NA | 7 | 0.1429 | NA | NA | 2 | 0.5 | 2 | 0 | 2 | 0 | 6 | 0.1667 |
| 6_7086953 | 1 | 1 | 2 | 0.5 | 1 | 0 | NA | NA | 4 | 0.5 | 1 | 0 | 3 | 1 | NA | NA | 3 | 0.6667 | 7 | 0.7143 |
| 6_7086959 | 1 | 0 | 1 | 1 | 1 | 1 | NA | NA | 3 | 0.6667 | 1 | 0 | 1 | 0 | NA | NA | 3 | 0.3333 | 5 | 0.2 |
| 6_7086965 | 1 | 0 | 1 | 0 | 1 | 1 | NA | NA | 3 | 0.3333 | 1 | 1 | 3 | 0.6667 | NA | NA | 2 | 1 | 6 | 0.8333 |
| 6_7086966 | 1 | 1 | 2 | 0 | 1 | 0 | NA | NA | 4 | 0.25 | 1 | 1 | 2 | 0 | NA | NA | 3 | 0.6667 | 6 | 0.5 |
| 6_7086973 | 1 | 1 | 2 | 1 | 1 | 1 | NA | NA | 4 | 1 | 1 | 1 | 3 | 1 | NA | NA | 3 | 1 | 7 | 1 |
| 6_7086991 | NA | NA | 1 | 1 | NA | NA | NA | NA | 1 | 1 | 1 | 1 | 1 | 0 | NA | NA | 2 | 0.5 | 4 | 0.5 |
| 6_7087000 | NA | NA | 1 | 1 | 1 | 0 | NA | NA | 2 | 0.5 | 1 | 1 | 3 | 1 | NA | NA | 2 | 1 | 6 | 1 |
| 6_7087010 | 1 | 0 | 1 | 1 | 1 | 0 | NA | NA | 3 | 0.3333 | 1 | 1 | 2 | 0.5 | NA | NA | 3 | 0.6667 | 6 | 0.6667 |
| 6_7088133 | 1 | 1 | 2 | 0.5 | 3 | 0.6667 | NA | NA | 6 | 0.6667 | 1 | 1 | 2 | 0.5 | 2 | 1 | 2 | 0.5 | 7 | 0.7143 |
| 6_7088136 | 1 | 1 | 2 | 1 | 4 | 1 | 1 | 1 | 8 | 1 | 1 | 1 | 5 | 1 | 2 | 1 | 3 | 1 | 11 | 1 |
| 6_7088139 | 1 | 0 | 4 | 0.25 | 4 | 0 | 1 | 1 | 10 | 0.2 | 1 | 1 | 5 | 0.2 | 2 | 0 | 1 | 0 | 9 | 0.2222 |
| 6_7088142 | 1 | 0 | 3 | 0 | 3 | 0.3333 | 1 | 0 | 8 | 0.125 | 1 | 0 | 3 | 0 | 2 | 0.5 | 3 | 0 | 9 | 0.1111 |
| 6_7088143 | 1 | 1 | 4 | 0.25 | 3 | 0 | 1 | 1 | 9 | 0.3333 | 1 | 0 | 5 | 0.2 | 2 | 0.5 | 3 | 0.3333 | 11 | 0.2727 |
| 6_7088145 | 1 | 0 | 3 | 0.6667 | 4 | 0.25 | 1 | 0 | 9 | 0.3333 | 1 | 0 | 5 | 0.2 | 2 | 0 | 3 | 0 | 11 | 0.0909 |
| 6_7088146 | 1 | 1 | 3 | 0.6667 | 4 | 1 | 1 | 1 | 9 | 0.8889 | 1 | 1 | 5 | 0.6 | 3 | 1 | 3 | 0.3333 | 12 | 0.6667 |
| 6_7088155 | NA | NA | 4 | 0.75 | 3 | 0.6667 | 1 | 0 | 8 | 0.625 | 1 | 0 | 5 | 0.2 | 3 | 1 | 4 | 0 | 13 | 0.3077 |
| 6_7088159 | 1 | 0 | 5 | 0.2 | 4 | 0.25 | 1 | 0 | 11 | 0.1818 | 1 | 0 | 6 | 0.3333 | 3 | 0 | 5 | 0 | 15 | 0.1333 |
| 6_7126106 | 2 | 1 | NA | NA | 5 | 1 | 4 | 0.5 | 11 | 0.8182 | 2 | 1 | 3 | 0.6667 | 4 | 0.5 | 2 | 1 | 11 | 0.7273 |
| 6_7126115 | 3 | 0.6667 | NA | NA | 5 | 0.8 | 4 | 0.75 | 12 | 0.75 | 3 | 0.6667 | 3 | 0.3333 | 2 | 1 | 2 | 1 | 10 | 0.7 |
| 6_7126125 | 3 | 0.6667 | NA | NA | 5 | 0.6 | 3 | 0 | 11 | 0.4545 | 3 | 0 | 3 | 0.6667 | 3 | 0.3333 | 2 | 0 | 11 | 0.2727 |
| 6_7126131 | 3 | 0.3333 | NA | NA | 5 | 0.2 | 5 | 0 | 13 | 0.1538 | 3 | 0.3333 | 3 | 0 | 4 | 0.75 | 2 | 0.5 | 12 | 0.4167 |
| 6_7126142 | 4 | 0 | 1 | 0 | 5 | 0.2 | 5 | 0 | 15 | 0.0667 | 3 | 0.3333 | 3 | 0 | 4 | 0 | 2 | 0 | 12 | 0.0833 |
| 6_7126157 | 4 | 0 | 1 | 0 | 5 | 0.6 | 5 | 0.4 | 15 | 0.3333 | 2 | 1 | 3 | 0 | 3 | 0.6667 | 1 | 1 | 9 | 0.5556 |
| 6_7126165 | 4 | 0.25 | 1 | 0 | 5 | 0.4 | 5 | 0 | 15 | 0.2 | 3 | 0 | 3 | 0 | 4 | 0.25 | 2 | 0 | 12 | 0.0833 |
| 6_7126169 | 4 | 0.5 | NA | NA | 3 | 0.3333 | 4 | 0 | 11 | 0.2727 | 1 | 1 | 3 | 0.3333 | 4 | 0.25 | 1 | 0 | 9 | 0.3333 |
| 6_7126171 | 4 | 0.75 | 1 | 0 | 5 | 0 | 5 | 0.4 | 15 | 0.3333 | 3 | 0.3333 | 3 | 0.6667 | 4 | 0.5 | 2 | 0 | 12 | 0.4167 |
| 6_7126183 | 4 | 0.25 | 1 | 1 | 5 | 0.8 | 4 | 0.75 | 14 | 0.6429 | 2 | 1 | 3 | 0.6667 | 4 | 0.5 | 2 | 1 | 11 | 0.7273 |
| 6_7126185 | 4 | 0.5 | 1 | 0 | 5 | 0 | 4 | 0 | 14 | 0.1429 | 2 | 0 | 2 | 0.5 | 3 | 0.3333 | 2 | 0.5 | 9 | 0.3333 |
| 6_7126200 | 4 | 1 | 1 | 0 | 5 | 0.8 | 5 | 1 | 15 | 0.8667 | 2 | 1 | 2 | 0.5 | 4 | 0.75 | 2 | 0.5 | 10 | 0.7 |
| 6_7126204 | 4 | 0.5 | NA | NA | 5 | 0 | 5 | 0.2 | 14 | 0.2143 | 2 | 0.5 | 3 | 0 | 4 | 0 | 1 | 0 | 10 | 0.1 |
| 6_7136574 | 10 | 0.2 | 9 | 0.2222 | 3 | 0.3333 | 5 | 0.4 | 27 | 0.2593 | 1 | 0 | 4 | 0.75 | 6 | 0.6667 | 9 | 0.3333 | 20 | 0.5 |
| 6_7136624 | 15 | 0.1333 | 8 | 0.125 | 6 | 0.1667 | 8 | 0 | 37 | 0.1081 | 5 | 0.2 | 14 | 0 | 13 | 0.1538 | 15 | 0.0667 | 47 | 0.0851 |
| 6_7136693 | 12 | 0.1667 | 6 | 0 | 4 | 0 | 8 | 0.125 | 30 | 0.1 | 4 | 0 | 7 | 0.1429 | 6 | 0.1667 | 6 | 0 | 23 | 0.087 |
| 6_7136750 | 7 | 0.5714 | 6 | 0.1667 | 6 | 0.6667 | 6 | 0.3333 | 25 | 0.44 | 6 | 0.3333 | 7 | 0.2857 | 4 | 0.5 | 5 | 0.4 | 22 | 0.3636 |
| 6_7136751 | 9 | 0.1111 | 6 | 0 | 6 | 0.1667 | 5 | 0 | 26 | 0.0769 | 5 | 0 | 7 | 0.1429 | 4 | 0 | 4 | 0 | 20 | 0.05 |
| 6_7136778 | 7 | 0.7143 | 6 | 0.3333 | 6 | 0.5 | 6 | 0.3333 | 25 | 0.48 | 5 | 0.6 | 6 | 0.3333 | 4 | 0.75 | 5 | 0 | 20 | 0.4 |

| | | | | | | | | | | | | | | | | | | | | |
|---|---|---|---|---|---|---|---|---|---|---|---|---|---|---|---|---|---|---|---|---|
| 6_7136786 | 8 | 0 | 6 | 0.1667 | 6 | 0.1667 | 6 | 0.3333 | 26 | 0.1538 | 4 | 0 | 6 | 0.5 | 3 | 0 | 5 | 0 | 18 | 0.1667 |
| 6_7136834 | 3 | 0.6667 | 4 | 0.25 | 4 | 0.5 | 3 | 0 | 14 | 0.3571 | 4 | 0.5 | 6 | 0.3333 | 1 | 1 | 3 | 0.3333 | 14 | 0.4286 |
| 6_7136868 | 5 | 1 | 2 | 1 | 3 | 1 | 4 | 1 | 14 | 1 | 4 | 1 | 2 | 1 | 4 | 1 | 3 | 1 | 13 | 1 |
| 6_7136871 | 7 | 0.7143 | 2 | 1 | 4 | 1 | 4 | 0.5 | 17 | 0.7647 | 5 | 1 | 3 | 0.6667 | 3 | 1 | 3 | 1 | 14 | 0.9286 |
| 6_7136876 | 5 | 0.4 | 2 | 0 | 5 | 0.6 | 3 | 0.6667 | 15 | 0.4667 | 4 | 0.75 | 3 | 0.6667 | 2 | 0.5 | 3 | 0 | 12 | 0.5 |
| 6_7136878 | 3 | 0 | 2 | 0.5 | 1 | 0 | 4 | 0 | 10 | 0.1 | 4 | 0 | 3 | 0.3333 | 2 | 0.5 | 4 | 0 | 13 | 0.1538 |
| 6_7136887 | 7 | 0 | 2 | 0 | 5 | 0.4 | 4 | 0 | 18 | 0.1111 | 5 | 0 | 2 | 0 | 3 | 0 | 2 | 0.5 | 12 | 0.0833 |
| 6_7136888 | 5 | 1 | 2 | 1 | 4 | 1 | 4 | 1 | 15 | 1 | 4 | 1 | 3 | 0.6667 | 4 | 1 | 2 | 1 | 13 | 0.9231 |
| 6_7136889 | 6 | 0.1667 | 2 | 0.5 | 5 | 0 | 4 | 0 | 17 | 0.1176 | 3 | 0 | 3 | 0.3333 | 2 | 1 | 3 | 0.6667 | 11 | 0.4545 |
| 6_7136901 | 5 | 0.6 | 1 | 0 | 3 | 0.6667 | 3 | 0.6667 | 12 | 0.5833 | 3 | 0.3333 | 1 | 1 | 3 | 0.3333 | 2 | 0.5 | 9 | 0.4444 |
| 6_7136909 | 4 | 0 | 1 | 0 | 4 | 0.25 | 3 | 0.3333 | 12 | 0.1667 | 4 | 0.25 | 1 | 1 | 3 | 0 | 3 | 0.3333 | 11 | 0.2727 |
| 6_7136972 | 3 | 0 | 1 | 0 | 7 | 0.2857 | 5 | 0 | 16 | 0.125 | 4 | 0.75 | 1 | 1 | 3 | 1 | 1 | 0 | 9 | 0.7778 |
| 6_7137002 | 3 | 1 | 1 | 0 | 6 | 0.8333 | 3 | 0.6667 | 13 | 0.7692 | 3 | 1 | 1 | 1 | 3 | 1 | 3 | 1 | 10 | 1 |
| 6_7137003 | 3 | 1 | 1 | 0 | 4 | 0.75 | 2 | 0.5 | 10 | 0.7 | 4 | 0.5 | 1 | 1 | 1 | 1 | 2 | 0.5 | 8 | 0.625 |
| 6_7137007 | 2 | 1 | 1 | 1 | 5 | 0.8 | 3 | 0.6667 | 11 | 0.8182 | 3 | 1 | 1 | 1 | 3 | 0.6667 | 1 | 0 | 8 | 0.75 |
| 6_7147173 | 8 | 0.375 | 8 | 0.625 | 2 | 0 | 5 | 0.2 | 23 | 0.3913 | 2 | 1 | 8 | 0.375 | 3 | 0.6667 | 6 | 0.5 | 19 | 0.5263 |
| 6_7148601 | 8 | 0.375 | 8 | 0.375 | 3 | 0.3333 | 3 | 0.6667 | 22 | 0.4091 | 2 | 0 | 7 | 0.4286 | 5 | 0.4 | 8 | 0 | 22 | 0.2273 |
| 6_7148618 | 7 | 0.5714 | 7 | 0 | 5 | 0 | 6 | 0.1667 | 25 | 0.2 | 2 | 0 | 8 | 0.25 | 6 | 0 | 5 | 0.6 | 21 | 0.2381 |
| 6_7148620 | 5 | 0.4 | 5 | 0 | 3 | 0 | 2 | 0 | 15 | 0.1333 | 2 | 0.5 | 9 | 0.2222 | 4 | 0 | 6 | 0.3333 | 21 | 0.2381 |
| 6_7148624 | 6 | 0.6667 | 7 | 0.7143 | 4 | 0.75 | 6 | 0.8333 | 23 | 0.7391 | 2 | 0.5 | 6 | 1 | 5 | 0.8 | 6 | 0.8333 | 19 | 0.8421 |
| 6_7148639 | 11 | 1 | 8 | 1 | 6 | 1 | 8 | 1 | 33 | 1 | 2 | 1 | 10 | 1 | 7 | 1 | 7 | 1 | 26 | 1 |
| 6_7148645 | 12 | 0.9167 | 8 | 0.5 | 6 | 0.3333 | 8 | 0.75 | 34 | 0.6765 | 3 | 0.6667 | 11 | 1 | 6 | 0.8333 | 9 | 0.8889 | 29 | 0.8966 |
| 6_7148653 | 10 | 0.2 | 8 | 0.125 | 6 | 0.1667 | 7 | 0.4286 | 31 | 0.2258 | 2 | 0 | 10 | 0.3 | 6 | 0.1667 | 8 | 0.25 | 26 | 0.2308 |
| 6_7148659 | 9 | 0.8889 | 7 | 0.5714 | 6 | 0.8333 | 7 | 0.5714 | 29 | 0.7241 | 5 | 0.4 | 9 | 0.6667 | 5 | 0.2 | 9 | 0.4444 | 28 | 0.4643 |
| 6_7148664 | 10 | 1 | 7 | 0.8571 | 6 | 1 | 8 | 0.875 | 31 | 0.9355 | 4 | 1 | 10 | 1 | 6 | 1 | 9 | 1 | 29 | 1 |
| 6_7148674 | 9 | 1 | 7 | 0.8571 | 6 | 1 | 8 | 1 | 30 | 0.9667 | 6 | 1 | 11 | 0.9091 | 7 | 0.8571 | 8 | 1 | 32 | 0.9375 |
| 6_7148679 | 9 | 1 | 6 | 1 | 6 | 0.6667 | 6 | 1 | 27 | 0.9259 | 4 | 1 | 9 | 1 | 5 | 1 | 6 | 1 | 24 | 1 |
| 6_7148680 | 11 | 0.9091 | 8 | 0.875 | 6 | 1 | 9 | 0.8889 | 34 | 0.9118 | 5 | 1 | 9 | 0.7778 | 7 | 0.8571 | 7 | 0.8571 | 28 | 0.8571 |
| 6_7148682 | 11 | 0.1818 | 7 | 0 | 6 | 0.1667 | 8 | 0.125 | 32 | 0.125 | 4 | 0 | 10 | 0.3 | 7 | 0.1429 | 8 | 0 | 29 | 0.1379 |
| 6_7148689 | 6 | 0.6667 | 7 | 0.5714 | 4 | 1 | 7 | 1 | 24 | 0.7917 | 4 | 0.75 | 6 | 1 | 2 | 0.5 | 8 | 0.875 | 20 | 0.85 |
| 6_7148696 | 10 | 0.2 | 7 | 0.1429 | 6 | 0.1667 | 7 | 0.1429 | 30 | 0.1667 | 4 | 0 | 9 | 0.1111 | 6 | 0 | 8 | 0.25 | 27 | 0.1111 |
| 6_7148698 | 8 | 0.875 | 6 | 0.6667 | 4 | 0.75 | 9 | 0.8889 | 27 | 0.8148 | 3 | 1 | 8 | 0.875 | 3 | 1 | 9 | 1 | 23 | 0.9565 |
| 6_7148700 | 12 | 1 | 7 | 1 | 7 | 1 | 9 | 1 | 35 | 1 | 5 | 1 | 10 | 1 | 6 | 0.8333 | 10 | 1 | 31 | 0.9677 |
| 6_7148718 | 9 | 1 | 7 | 1 | 6 | 1 | 8 | 1 | 30 | 1 | 4 | 1 | 9 | 1 | 4 | 1 | 8 | 1 | 25 | 1 |
| 6_7152097 | 10 | 0.1 | 8 | 0 | 6 | 0 | 13 | 0.3846 | 37 | 0.1622 | 10 | 0 | 15 | 0.0667 | 7 | 0.1429 | 4 | 0 | 36 | 0.0556 |
| 6_7155708 | 13 | 0.2308 | 17 | 0.1765 | 12 | 0.5833 | 18 | 0.3333 | 60 | 0.3167 | 14 | 0.1429 | 23 | 0.2609 | 15 | 0.6667 | 10 | 0.6 | 62 | 0.3871 |
| 6_7155718 | 7 | 0.2857 | 4 | 0 | 7 | 0.4286 | 11 | 0.1818 | 29 | 0.2414 | 8 | 0.125 | 16 | 0 | 4 | 0 | 4 | 0.25 | 32 | 0.0625 |
| 6_7155723 | 10 | 0.1 | 6 | 0.3333 | 9 | 0 | 13 | 0.1538 | 38 | 0.1316 | 9 | 0 | 17 | 0.1176 | 4 | 0.25 | 5 | 0.4 | 35 | 0.1429 |
| 6_7155801 | 11 | 0.1818 | 4 | 0.25 | 6 | 0.3333 | 10 | 0.1 | 31 | 0.1935 | 8 | 0.375 | 18 | 0.2222 | 5 | 0.4 | 6 | 0.3333 | 37 | 0.2973 |
| 6_7155831 | 13 | 0.1538 | 7 | 0.1429 | 7 | 0.1429 | 13 | 0 | 40 | 0.1 | 8 | 0 | 25 | 0.16 | 7 | 0 | 5 | 0.2 | 45 | 0.1111 |
| 6_7156615 | 8 | 0.375 | 5 | 0.4 | 4 | 0 | 8 | 0.5 | 25 | 0.36 | 1 | 0 | 10 | 0.3 | 8 | 0.5 | 6 | 0.3333 | 25 | 0.36 |
| 6_7156632 | 5 | 0.2 | 2 | 0 | 3 | 0 | 5 | 0 | 15 | 0.0667 | NA | NA | 5 | 0.4 | 3 | 0 | 6 | 0 | 14 | 0.1429 |
| 6_7156650 | 8 | 0.125 | 5 | 0 | 3 | 0.3333 | 7 | 0.4286 | 23 | 0.2174 | 1 | 0 | 10 | 0.4 | 8 | 0.25 | 6 | 0.1667 | 25 | 0.28 |
| 6_7156655 | 6 | 0.8333 | 5 | 0.8 | 3 | 0.6667 | 6 | 0.8333 | 20 | 0.8 | 1 | 0 | 10 | 0.8 | 9 | 0.8889 | 6 | 1 | 26 | 0.8462 |
| 6_7158101 | 3 | 0 | 4 | 0 | 1 | 1 | 1 | 1 | 9 | 0.2222 | NA | NA | 4 | 0 | 4 | 0 | 3 | 0.3333 | 11 | 0.0909 |
| 6_7158102 | 2 | 1 | 4 | 0.5 | 1 | 1 | 2 | 0 | 9 | 0.5556 | NA | NA | 5 | 0.2 | 5 | 0.4 | 3 | 0.6667 | 13 | 0.3846 |
| 6_7158870 | 2 | 0.5 | 3 | 0 | NA | NA | 1 | 0 | 6 | 0.1667 | NA | NA | 3 | 0.6667 | 5 | 0.2 | NA | NA | 8 | 0.375 |
| 6_7158871 | 2 | 0.5 | 3 | 1 | 1 | 1 | 1 | 1 | 7 | 0.8571 | NA | NA | 4 | 0.75 | 3 | 0.6667 | 2 | 0.5 | 9 | 0.6667 |
| 6_7158874 | 2 | 0 | 3 | 0.3333 | 1 | 1 | 1 | 1 | 7 | 0.4286 | NA | NA | 5 | 0.4 | 5 | 0.4 | 2 | 0 | 12 | 0.3333 |
| 6_7158881 | 2 | 0 | 2 | 0.5 | 1 | 0 | 1 | 0 | 6 | 0.1667 | NA | NA | 4 | 0.25 | 5 | 0.6 | 2 | 0.5 | 11 | 0.4545 |
| 6_7158886 | 2 | 0 | 3 | 0.6667 | 1 | 0 | 1 | 0 | 7 | 0.2857 | NA | NA | 3 | 0 | 5 | 0.4 | 3 | 0 | 11 | 0.1818 |
| 6_7178496 | 4 | 0.75 | 7 | 0.2857 | 3 | 0 | 2 | 0 | 16 | 0.3125 | 3 | 0 | 8 | 0.25 | 5 | 0 | 8 | 0.25 | 24 | 0.1667 |
| 6_7178499 | 4 | 0.75 | 5 | 0.6 | 4 | 0.75 | 3 | 1 | 16 | 0.75 | 1 | 1 | 6 | 0.8333 | 7 | 1 | 8 | 0.5 | 22 | 0.7727 |
| 6_7178527 | 1 | 0 | 8 | 0.25 | 2 | 0.5 | 4 | 0 | 15 | 0.2 | 4 | 0.75 | 8 | 0.5 | 6 | 0.1667 | 7 | 0.2857 | 25 | 0.4 |
| 6_7178604 | 10 | 0 | 12 | 0.1667 | 4 | 0 | 7 | 0.1429 | 33 | 0.0909 | 7 | 0.1429 | 12 | 0.0833 | 9 | 0.1111 | 11 | 0.1818 | 39 | 0.1282 |
| 6_7178632 | 10 | 0.2 | 14 | 0.0714 | 6 | 0.1667 | 5 | 0 | 35 | 0.1143 | 7 | 0 | 11 | 0.0909 | 12 | 0 | 10 | 0.3 | 40 | 0.1 |
| 6_7178676 | 12 | 0.0833 | 15 | 0.2 | 10 | 0.1 | 9 | 0.3333 | 46 | 0.1739 | 10 | 0.3 | 14 | 0.1429 | 14 | 0.1429 | 14 | 0.3571 | 52 | 0.2308 |
| 6_7178699 | 12 | 0.0833 | 15 | 0 | 9 | 0 | 7 | 0.2857 | 43 | 0.0698 | 9 | 0 | 13 | 0 | 14 | 0.2143 | 11 | 0.0909 | 47 | 0.0851 |
| 6_7178703 | 13 | 0.3077 | 14 | 0.2857 | 9 | 0.4444 | 8 | 0.5 | 44 | 0.3636 | 9 | 0.5556 | 14 | 0.2857 | 15 | 0.4667 | 15 | 0.3333 | 53 | 0.3962 |
| 6_7178735 | NA | NA | 1 | 1 | 1 | 1 | NA | NA | 2 | 1 | NA | NA | NA | NA | NA | NA | 4 | 1 | 4 | 1 |
| 6_7178751 | 14 | 0.2143 | 19 | 0.3158 | 10 | 0.2 | 9 | 0.3333 | 52 | 0.2692 | 11 | 0.3636 | 14 | 0.2857 | 16 | 0.125 | 16 | 0.4375 | 57 | 0.2982 |
| 6_7178758 | 12 | 0.5 | 19 | 0.5789 | 9 | 0.3333 | 10 | 0.6 | 50 | 0.52 | 9 | 0.5556 | 12 | 0.4167 | 12 | 0.5 | 11 | 0.4545 | 44 | 0.4773 |
| 6_7179288 | 3 | 0 | 1 | 0 | 3 | 0 | 4 | 0.25 | 11 | 0.0909 | 4 | 0 | 3 | 0 | 2 | 0.5 | 3 | 0.3333 | 12 | 0.1667 |
| 6_7179299 | 4 | 0.25 | 4 | 0 | 5 | 0.2 | 7 | 0 | 20 | 0.1 | 5 | 0 | 4 | 0 | 4 | 0.25 | 8 | 0.125 | 21 | 0.0952 |
| 6_7185152 | 18 | 0.1111 | 20 | 0.1 | 13 | 0 | 18 | 0.1111 | 69 | 0.087 | 12 | 0.25 | 12 | 0 | 17 | 0.0588 | 15 | 0.1333 | 56 | 0.1071 |

| | | | | | | | | | | | | | | | | | | | | |
|---|---|---|---|---|---|---|---|---|---|---|---|---|---|---|---|---|---|---|---|---|
| 6_7185180 | 19 | 0.2105 | 20 | 0.4 | 14 | 0.3571 | 20 | 0.3 | 73 | 0.3151 | 10 | 0.3 | 10 | 0.2 | 12 | 0.0833 | 12 | 0.5833 | 44 | 0.2955 |
| 6_7185209 | 14 | 0.3571 | 13 | 0.2308 | 13 | 0.0769 | 21 | 0.0952 | 61 | 0.1803 | 11 | 0.3636 | 13 | 0.2308 | 19 | 0.1579 | 16 | 0.125 | 59 | 0.2034 |
| 6_7185771 | 6 | 0 | 8 | 0.25 | 9 | 0.2222 | 4 | 0 | 27 | 0.1481 | 4 | 0 | 2 | 0 | 12 | 0.1667 | 3 | 0 | 21 | 0.0952 |
| 6_7185779 | 8 | 0.125 | 8 | 0.125 | 9 | 0.1111 | 4 | 0 | 29 | 0.1034 | 2 | 0 | 3 | 0 | 12 | 0.0833 | 3 | 0 | 20 | 0.05 |
| 6_7185780 | 9 | 0.2222 | 9 | 0.1111 | 11 | 0.0909 | 5 | 0 | 34 | 0.1176 | 3 | 0 | 3 | 0.3333 | 13 | 0.0769 | 3 | 1 | 22 | 0.2273 |
| 6_7185790 | 8 | 0 | 9 | 0.1111 | 9 | 0.1111 | 4 | 0 | 30 | 0.0667 | 2 | 0 | 2 | 0.5 | 11 | 0.0909 | 3 | 0 | 18 | 0.1111 |
| 6_7185793 | 9 | 0.2222 | 8 | 0.125 | 9 | 0 | 2 | 0.5 | 28 | 0.1429 | 2 | 0 | 3 | 0.3333 | 11 | 0.0909 | 3 | 0.3333 | 19 | 0.1579 |
| 6_7185802 | 8 | 0.25 | 8 | 0.625 | 11 | 0.5455 | 3 | 0.6667 | 30 | 0.5 | 3 | 0.6667 | 3 | 0.3333 | 11 | 0.8182 | 2 | 1 | 19 | 0.7368 |
| 6_7195552 | 4 | 0.75 | 4 | 1 | 5 | 1 | 6 | 0.8333 | 19 | 0.8947 | NA | NA | 1 | 0 | 3 | 1 | 3 | 0.6667 | 7 | 0.7143 |
| 6_7195555 | 4 | 0 | 4 | 0.25 | 6 | 0.1667 | 5 | 0.2 | 19 | 0.1579 | NA | NA | 1 | 0 | 4 | 0 | 4 | 0.25 | 9 | 0.1111 |
| 6_7195575 | 4 | 0 | 2 | 0 | 6 | 0.5 | 4 | 0 | 16 | 0.1875 | NA | NA | 1 | 0 | 3 | 0.3333 | 2 | 1 | 6 | 0.5 |
| 6_7195603 | 3 | 0.3333 | 5 | 0.2 | 4 | 0.5 | 5 | 0.2 | 17 | 0.2941 | NA | NA | 1 | 0 | 3 | 0 | 3 | 0.3333 | 7 | 0.1429 |
| 6_7195605 | 4 | 0 | 4 | 0 | 4 | 0 | 5 | 0.2 | 17 | 0.0588 | NA | NA | 2 | 0 | 3 | 0.3333 | 3 | 0 | 8 | 0.125 |
| 6_7195606 | 4 | 0.25 | 4 | 0.25 | 5 | 0.2 | 6 | 0.3333 | 19 | 0.2632 | NA | NA | 2 | 0.5 | 4 | 0.25 | 4 | 0.25 | 10 | 0.3 |
| 6_7195623 | 4 | 0 | 3 | 0.3333 | 5 | 0.4 | 5 | 0.4 | 17 | 0.2941 | 1 | 1 | 2 | 0.5 | 4 | 0.5 | 3 | 0 | 10 | 0.4 |
| 6_7195624 | 4 | 0.25 | 3 | 0.6667 | 4 | 0 | 4 | 0 | 15 | 0.2 | 1 | 0 | 1 | 1 | 4 | 0.25 | 4 | 0.25 | 10 | 0.3 |
| 6_7195632 | 3 | 0 | 5 | 0.2 | 7 | 0.2857 | 5 | 0.2 | 20 | 0.2 | 1 | 0 | 2 | 0.5 | 4 | 0.25 | 4 | 0.25 | 11 | 0.2727 |
| 6_7196161 | NA | NA | NA | NA | 1 | 1 | NA | NA | 1 | 1 | NA | NA | NA | NA | 1 | 1 | NA | NA | 1 | 1 |
| 6_7207459 | 3 | 0 | 7 | 0.1429 | 5 | 0 | 4 | 0.25 | 19 | 0.1053 | 2 | 0 | 3 | 0 | 6 | 0.1667 | NA | NA | 11 | 0.0909 |
| 6_7211920 | 10 | 0.2 | 32 | 0.3125 | 16 | 0.1875 | 7 | 0.1429 | 65 | 0.2462 | 7 | 0 | 14 | 0.3571 | 14 | 0.2857 | 11 | 0.5455 | 46 | 0.3261 |
| 6_7211935 | 10 | 0.1 | 34 | 0.0588 | 16 | 0.125 | 9 | 0.2222 | 69 | 0.1014 | 8 | 0 | 15 | 0.0667 | 12 | 0.0833 | 11 | 0.0909 | 46 | 0.0652 |
| 6_7211942 | 9 | 0.3333 | 33 | 0.5455 | 18 | 0.2778 | 8 | 0.5 | 68 | 0.4412 | 8 | 0.125 | 13 | 0.4615 | 15 | 0.4 | 11 | 0.7273 | 47 | 0.4468 |
| 6_7213051 | 10 | 0 | 39 | 0.1282 | 21 | 0.1429 | 11 | 0.0909 | 81 | 0.1111 | 10 | 0.1 | 15 | 0 | 19 | 0.0526 | 13 | 0.0769 | 57 | 0.0526 |
| 6_7213058 | 14 | 0.6429 | 38 | 0.3947 | 22 | 0.4545 | 14 | 0.4286 | 88 | 0.4545 | 11 | 0.4545 | 17 | 0.5294 | 17 | 0.4118 | 12 | 0.3333 | 57 | 0.4386 |
| 6_7213063 | 13 | 0.0769 | 40 | 0.15 | 21 | 0.1905 | 14 | 0.0714 | 88 | 0.1364 | 11 | 0 | 17 | 0.0588 | 19 | 0 | 12 | 0.1667 | 59 | 0.0508 |
| 6_7213065 | 12 | 0.25 | 42 | 0.2857 | 22 | 0.5455 | 12 | 0.3333 | 88 | 0.3523 | 8 | 0.375 | 17 | 0.3529 | 17 | 0.4118 | 12 | 0.3333 | 54 | 0.3704 |
| 6_7213072 | 13 | 0.9231 | 34 | 0.9412 | 20 | 1 | 11 | 1 | 78 | 0.9615 | 10 | 1 | 16 | 1 | 17 | 1 | 11 | 1 | 54 | 1 |
| 6_7213076 | 11 | 0.3636 | 37 | 0.1892 | 23 | 0.3478 | 14 | 0.2857 | 85 | 0.2706 | 10 | 0.3 | 19 | 0.3684 | 20 | 0.3 | 12 | 0.1667 | 61 | 0.2951 |
| 6_7213079 | 12 | 0.0833 | 36 | 0.2222 | 17 | 0.1176 | 11 | 0 | 76 | 0.1447 | 12 | 0.0833 | 14 | 0.1429 | 19 | 0.2105 | 12 | 0.25 | 57 | 0.1754 |
| 6_7213091 | 17 | 1 | 40 | 0.9 | 23 | 0.9565 | 14 | 1 | 94 | 0.9468 | 14 | 0.8571 | 17 | 0.8235 | 16 | 1 | 13 | 1 | 60 | 0.9167 |
| 6_7213097 | 14 | 0.5 | 43 | 0.3488 | 22 | 0.4545 | 11 | 0.2727 | 90 | 0.3889 | 11 | 0.6364 | 18 | 0.6111 | 18 | 0.4444 | 13 | 0.5385 | 60 | 0.55 |
| 6_7213108 | 14 | 0.3571 | 41 | 0.4878 | 23 | 0.3913 | 13 | 0.7692 | 91 | 0.4835 | 13 | 0.4615 | 19 | 0.3158 | 18 | 0.5 | 10 | 0.7 | 60 | 0.4667 |
| 6_7213119 | 13 | 0.6923 | 34 | 0.4706 | 20 | 0.6 | 10 | 0.9 | 77 | 0.5974 | 12 | 0.5 | 19 | 0.4737 | 15 | 0.6667 | 13 | 0.5385 | 59 | 0.5424 |
| 6_7218329 | 1 | 0 | 1 | 1 | NA | NA | 1 | 0 | 3 | 0.3333 | 1 | 1 | NA | NA | NA | NA | 1 | 0 | 2 | 0.5 |
| 6_7218349 | NA | NA | 1 | 1 | NA | NA | 1 | 1 | 2 | 1 | 1 | 1 | NA | NA | NA | NA | 1 | 1 | 2 | 1 |
| 6_7218357 | NA | NA | NA | NA | NA | NA | 1 | 1 | 1 | 1 | 1 | 0 | NA | NA | NA | NA | 1 | 1 | 2 | 0.5 |
| 6_7221178 | NA | NA | 1 | 1 | NA | NA | 1 | 1 | 2 | 1 | NA | NA | NA | NA | NA | NA | 1 | 1 | 1 | 1 |
| 6_7221201 | NA | NA | 1 | 1 | NA | NA | 1 | 1 | 2 | 1 | NA | NA | NA | NA | NA | NA | 1 | 1 | 1 | 1 |
| 6_7221209 | NA | NA | 1 | 0 | NA | NA | 1 | 1 | 2 | 0.5 | NA | NA | NA | NA | NA | NA | 1 | 1 | 1 | 1 |
| 6_7229589 | 5 | 0.4 | 3 | 0.3333 | 6 | 0.1667 | 4 | 0.75 | 18 | 0.3889 | 1 | 1 | 3 | 0.3333 | 9 | 0.5556 | 1 | 1 | 14 | 0.5714 |
| 6_7229595 | 8 | 0.75 | 3 | 1 | 4 | 0.5 | 5 | 0.8 | 20 | 0.75 | 2 | 0 | 3 | 0.3333 | 10 | 0.5 | 3 | 0.6667 | 18 | 0.4444 |
| 6_7229609 | 6 | 0.3333 | 3 | 0.3333 | 7 | 0.2857 | 5 | 0.4 | 21 | 0.3333 | 2 | 0.5 | 2 | 0 | 12 | 0.0833 | 3 | 0 | 19 | 0.1053 |
| 6_7229609 | 4 | 0.25 | 2 | 1 | 7 | 0.7143 | 6 | 0.6667 | 19 | 0.6316 | 2 | 1 | 3 | 0.6667 | 8 | 0.875 | 4 | 0.75 | 17 | 0.8235 |
| 6_7232562 | NA | NA | 1 | 0 | 2 | 1 | NA | NA | 3 | 0.6667 | NA | NA | NA | NA | 1 | 0 | 1 | 1 | 2 | 0.5 |
| 6_7232607 | NA | NA | 1 | 0 | 2 | 0.5 | 1 | 0 | 4 | 0.25 | NA | NA | NA | NA | 1 | 0 | 1 | 1 | 2 | 0.5 |
| 6_7232635 | NA | NA | 1 | 0 | 2 | 0.5 | 1 | 1 | 4 | 0.5 | NA | NA | NA | NA | 1 | 0 | 2 | 1 | 3 | 0.6667 |
| 6_7232658 | NA | NA | 1 | 1 | 1 | 0 | 1 | 0 | 3 | 0.3333 | NA | NA | NA | NA | 1 | 0 | 3 | 0.6667 | 4 | 0.5 |
| 6_7232684 | NA | NA | 1 | 0 | 2 | 1 | 1 | 0 | 4 | 0.5 | NA | NA | NA | NA | 1 | 0 | 3 | 0.3333 | 4 | 0.25 |
| 6_7232732 | 1 | 1 | 1 | 0 | 2 | 0.5 | 2 | 0.5 | 6 | 0.5 | NA | NA | NA | NA | 2 | 0.5 | 2 | 0.5 | 4 | 0.5 |
| 6_7232757 | 1 | 0 | 1 | 0 | 2 | 0.5 | 1 | 0 | 5 | 0.2 | 1 | 0 | NA | NA | 1 | 1 | 4 | 0.25 | 6 | 0.3333 |
| 6_7232764 | 1 | 0 | NA | NA | 2 | 0.5 | 2 | 0.5 | 5 | 0.4 | 1 | 0 | NA | NA | 2 | 0.5 | 5 | 0.4 | 8 | 0.375 |
| 6_7232767 | 1 | 1 | 1 | 0 | 1 | 1 | 2 | 0.5 | 5 | 0.6 | 1 | 1 | NA | NA | 2 | 0.5 | 3 | 0.6667 | 6 | 0.6667 |
| 6_7232778 | 1 | 1 | 1 | 0 | 1 | 0 | 1 | 0 | 4 | 0.25 | 1 | 1 | NA | NA | 2 | 0.5 | 4 | 0.5 | 7 | 0.5714 |
| 6_7232778 | 1 | 1 | 1 | 0 | 1 | 0 | 2 | 1 | 5 | 0.6 | 1 | 1 | NA | NA | 2 | 0.5 | 5 | 0.4 | 8 | 0.5 |
| 6_7232797 | 1 | 1 | NA | NA | NA | NA | 2 | 1 | 3 | 1 | 1 | 1 | NA | NA | 2 | 0.5 | 4 | 0.75 | 7 | 0.7143 |
| 6_7232808 | 1 | 0 | 1 | 0 | 2 | 0 | 2 | 0.5 | 6 | 0.1667 | 1 | 0 | NA | NA | 2 | 0.5 | 4 | 0 | 7 | 0.1429 |
| 6_7232812 | 1 | 0 | 1 | 0 | 1 | 0 | 2 | 0.5 | 5 | 0.2 | 1 | 0 | NA | NA | 2 | 0.5 | 4 | 0 | 7 | 0.1429 |
| 6_7232813 | NA | NA | 1 | 1 | 1 | 1 | 1 | 1 | 3 | 1 | NA | NA | NA | NA | 2 | 1 | 5 | 0.6 | 7 | 0.7143 |
| 6_7232898 | 2 | 0.5 | 2 | 0 | 3 | 0 | 3 | 0.3333 | 10 | 0.2 | 1 | 0 | NA | NA | 3 | 0.3333 | 5 | 0 | 9 | 0.1111 |
| 6_7232952 | 3 | 0 | 2 | 0.5 | 3 | 0 | 4 | 0.5 | 12 | 0.25 | 1 | 0 | NA | NA | 2 | 0.5 | 5 | 0 | 8 | 0.125 |
| 6_7233556 | NA | NA | 1 | 1 | NA | NA | 5 | 0 | 6 | 0.1667 | 1 | 0 | 3 | 0.3333 | NA | NA | NA | NA | 4 | 0.25 |
| 6_7233610 | NA | NA | 1 | 1 | NA | NA | 5 | 0.2 | 6 | 0.3333 | 1 | 0 | 3 | 0.3333 | NA | NA | 1 | 1 | 5 | 0.4 |
| 6_7233704 | 10 | 0.3 | 8 | 0.375 | 8 | 0.25 | 11 | 0.1818 | 37 | 0.2703 | 7 | 0.4286 | 10 | 0.4 | 14 | 0.3571 | 8 | 0.375 | 39 | 0.3846 |
| 6_7233725 | 9 | 0 | 8 | 0.125 | 9 | 0.1111 | 16 | 0.25 | 42 | 0.1429 | 6 | 0 | 9 | 0 | 15 | 0.1333 | 10 | 0 | 40 | 0.05 |
| 6_7233735 | 7 | 0.5714 | 6 | 0.3333 | 7 | 0.5714 | 9 | 0.5556 | 29 | 0.5172 | 4 | 0.5 | 7 | 0.4286 | 8 | 0.625 | 7 | 0.4286 | 26 | 0.5 |

| | | | | | | | | | | | | | | | | | | | | |
|---|---|---|---|---|---|---|---|---|---|---|---|---|---|---|---|---|---|---|---|---|
| 6_7234607( | NA | NA | 1 | 0 | NA | NA | 2 | 1 | 3 | 0.6667 | NA | NA | NA | NA | 2 | 1 | NA | NA | 2 | 1 |
| 6_7234609 | NA | NA | 2 | 1 | NA | NA | 2 | 1 | 4 | 1 | NA | NA | 1 | 0 | 3 | 0.6667 | NA | NA | 4 | 0.5 |
| 6_7234621( | NA | NA | 1 | 1 | NA | NA | 1 | 1 | 2 | 1 | NA | NA | NA | NA | 3 | 0.6667 | NA | NA | 3 | 0.6667 |
| 6_7234625 | NA | NA | 2 | 0.5 | NA | NA | 2 | 0 | 4 | 0.25 | NA | NA | NA | NA | 2 | 0.5 | NA | NA | 2 | 0.5 |
| 6_7236232 | 6 | 0.6667 | 1 | 1 | 1 | 0 | 3 | 0.3333 | 11 | 0.5455 | 3 | 0.6667 | 2 | 1 | 1 | 0 | 3 | 0.3333 | 9 | 0.5556 |
| 6_7240723 | NA | NA | 3 | 0.3333 | NA | NA | 3 | 0.6667 | 6 | 0.5 | 2 | 0.5 | 1 | 1 | 1 | 0 | 4 | 0 | 8 | 0.25 |
| 6_7240726 | NA | NA | 3 | 0 | NA | NA | 4 | 0.25 | 7 | 0.1429 | 1 | 0 | 1 | 0 | NA | NA | 4 | 0.25 | 6 | 0.1667 |
| 6_7241134 | 14 | 0.1429 | 23 | 0.087 | 25 | 0 | 19 | 0.1053 | 81 | 0.0741 | 15 | 0.2667 | 20 | 0.1 | 23 | 0.087 | 13 | 0.0769 | 71 | 0.1268 |
| 6_7241635 | 2 | 0.5 | 2 | 1 | 3 | 0 | NA | NA | 7 | 0.4286 | 2 | 0.5 | 2 | 1 | 1 | 0 | 1 | 0 | 5 | 0.6 |
| 6_7241652 | 3 | 0 | 3 | 0.3333 | 3 | 0 | NA | NA | 9 | 0.1111 | 1 | 0 | 2 | 0.5 | 2 | 0 | NA | NA | 5 | 0.2 |
| 6_7241668 | 2 | 0.5 | 1 | 0 | 3 | 0.3333 | NA | NA | 6 | 0.3333 | 3 | 0.3333 | 3 | 0.3333 | 1 | 0 | NA | NA | 7 | 0.2857 |
| 6_7241678 | 2 | 1 | 1 | 1 | 2 | 0.5 | NA | NA | 5 | 0.8 | 2 | 1 | 3 | 0.6667 | 2 | 0.5 | NA | NA | 7 | 0.7143 |
| 6_7241706 | 3 | 0.3333 | 2 | 0.5 | 3 | 0 | NA | NA | 8 | 0.25 | 2 | 1 | 2 | 0.5 | 2 | 0 | 1 | 0 | 7 | 0.4286 |
| 6_7241717 | 3 | 0.6667 | 3 | 0.3333 | 3 | 0 | NA | NA | 9 | 0.3333 | 2 | 0.5 | 3 | 0.3333 | 1 | 0 | 1 | 0 | 7 | 0.2857 |
| 6_7241723 | 3 | 1 | 1 | 1 | NA | NA | NA | NA | 4 | 1 | 1 | 1 | 2 | 0 | 2 | 0.5 | NA | NA | 5 | 0.4 |
| 6_7241733 | 3 | 0 | 4 | 0.25 | 3 | 0 | 1 | 1 | 11 | 0.1818 | 3 | 0.6667 | 3 | 0.3333 | 3 | 0 | 1 | 1 | 10 | 0.4 |
| 6_7241751 | 2 | 0.5 | 2 | 0 | 2 | 0 | 1 | 0 | 7 | 0.1429 | 4 | 0.25 | 3 | 0.3333 | 2 | 0.5 | 1 | 0 | 10 | 0.3 |
| 6_7254313 | 4 | 0 | 4 | 0.25 | NA | NA | 2 | 0.5 | 10 | 0.2 | 1 | 0 | 1 | 0 | 5 | 0 | 4 | 0.25 | 11 | 0.0909 |
| 6_7254318 | 4 | 0 | 6 | 0.5 | NA | NA | 2 | 0 | 12 | 0.25 | 1 | 0 | 1 | 0 | 5 | 0.2 | 4 | 0 | 11 | 0.0909 |
| 6_7254323 | 4 | 0 | 4 | 0.5 | NA | NA | 2 | 0.5 | 10 | 0.3 | 1 | 0 | 1 | 0 | 5 | 0.2 | 3 | 0 | 10 | 0.1 |
| 6_7254323 | 4 | 0.25 | 5 | 0 | NA | NA | 2 | 0 | 11 | 0.0909 | 1 | 0 | 1 | 1 | 5 | 0.4 | 4 | 0 | 11 | 0.2727 |
| 6_7254337 | 3 | 0 | 3 | 0 | NA | NA | 2 | 0.5 | 8 | 0.125 | 1 | 0 | 1 | 0 | 4 | 0.25 | 3 | 0 | 9 | 0.1111 |
| 6_7254337 | 3 | 0.6667 | 4 | 0.75 | NA | NA | 2 | 0.5 | 9 | 0.6667 | 1 | 1 | 1 | 1 | 4 | 0.5 | 3 | 0.3333 | 9 | 0.5556 |
| 6_7254339 | 3 | 0.3333 | 4 | 0.75 | NA | NA | 1 | 1 | 8 | 0.625 | 1 | 0 | 1 | 1 | 5 | 0.8 | 3 | 0.3333 | 10 | 0.6 |
| 6_7254341 | 3 | 0.6667 | 2 | 0 | NA | NA | 2 | 0.5 | 7 | 0.4286 | 1 | 1 | 1 | 1 | 5 | 0.2 | 3 | 0.3333 | 10 | 0.4 |
| 6_7254359 | 4 | 0.5 | 4 | 0.75 | NA | NA | 2 | 0.5 | 10 | 0.6 | 1 | 1 | 1 | 0 | 3 | 0 | 4 | 0.75 | 9 | 0.4444 |
| 6_7254362 | 4 | 0 | 5 | 0.2 | NA | NA | 2 | 0.5 | 11 | 0.1818 | 1 | 0 | 1 | 0 | 4 | 0 | 4 | 0.25 | 10 | 0.1 |
| 6_7257192 | 1 | 0 | 3 | 0.3333 | NA | NA | NA | NA | 4 | 0.25 | 1 | 0 | 1 | 0 | 2 | 0 | 3 | 0.3333 | 7 | 0.1429 |
| 6_7257196 | 1 | 0 | 3 | 0.3333 | NA | NA | NA | NA | 4 | 0.25 | 1 | 0 | 1 | 0 | 2 | 0 | 3 | 0.3333 | 7 | 0.1429 |
| 6_7257488 | 1 | 0 | 1 | 1 | NA | NA | NA | NA | 2 | 0.5 | NA | NA | NA | NA | NA | NA | 1 | 1 | 1 | 1 |
| 6_7257498 | 1 | 1 | 2 | 1 | NA | NA | NA | NA | 3 | 1 | NA | NA | NA | NA | NA | NA | 1 | 1 | 1 | 1 |
| 6_7257853 | NA | NA | 1 | 0 | NA | NA | 1 | 1 | 2 | 0.5 | NA | NA | NA | NA | NA | NA | 1 | 1 | 1 | 1 |
| 6_7258430 | 3 | 0.3333 | 4 | 0 | 1 | 0 | 2 | 0 | 10 | 0.1 | 1 | 0 | 2 | 0.5 | 2 | 0 | 2 | 0 | 7 | 0.1429 |
| 6_7258635 | 7 | 0.1429 | 9 | 0.3333 | 4 | 0 | 7 | 0 | 27 | 0.1481 | 10 | 0 | 10 | 0.2 | 12 | 0.0833 | 8 | 0 | 40 | 0.075 |
| 6_7258853 | 9 | 0.7778 | 7 | 0.4286 | 9 | 0.4444 | 13 | 0.3846 | 38 | 0.5 | 9 | 0.2222 | 7 | 0.5714 | 13 | 0.3077 | 10 | 0.2 | 39 | 0.3077 |
| 6_7258878 | 6 | 0.8333 | 5 | 0.8 | 7 | 0.5714 | 10 | 0.6 | 28 | 0.6786 | 4 | 0.5 | 5 | 0.6 | 7 | 0.8571 | 5 | 0.2 | 21 | 0.5714 |
| 6_7258893 | 8 | 0.25 | 7 | 0.1429 | 8 | 0.125 | 10 | 0.1 | 33 | 0.1515 | 8 | 0.125 | 5 | 0 | 12 | 0.0833 | 8 | 0 | 33 | 0.0606 |
| 6_7258902 | 7 | 1 | 8 | 0.875 | 6 | 0.6667 | 9 | 1 | 30 | 0.9 | 7 | 1 | 5 | 0.8 | 10 | 1 | 7 | 0.5714 | 29 | 0.8621 |
| 6_7259261 | 7 | 0.8571 | 7 | 0.8571 | 7 | 0.8571 | 10 | 0.8 | 31 | 0.8387 | 6 | 0.6667 | 4 | 0.25 | 12 | 0.5833 | 7 | 0.7143 | 29 | 0.5862 |
| 6_7259264 | 8 | 0.25 | 8 | 0.625 | 8 | 0.375 | 10 | 0.3 | 34 | 0.3824 | 8 | 0.125 | 4 | 0.25 | 11 | 0.2727 | 7 | 0.1429 | 30 | 0.2 |
| 6_7259265 | 7 | 0.1429 | 5 | 0.2 | 6 | 0.5 | 10 | 0.2 | 28 | 0.25 | 7 | 0.4286 | 4 | 0 | 11 | 0.4545 | 6 | 0.1667 | 28 | 0.3214 |
| 6_7259282 | 7 | 0.7143 | 7 | 0.7143 | 8 | 0.875 | 9 | 0.7778 | 31 | 0.7742 | 8 | 0.875 | 3 | 1 | 10 | 0.7 | 6 | 0.8333 | 27 | 0.8148 |
| 6_7259285 | 4 | 0 | 4 | 0 | 5 | 0 | 9 | 0.2222 | 22 | 0.0909 | 4 | 0.25 | 1 | 0 | 9 | 0.3333 | 3 | 0.6667 | 17 | 0.3529 |
| 6_7259288 | 7 | 0.4286 | 6 | 0.6667 | 7 | 0 | 9 | 0.3333 | 29 | 0.3448 | 7 | 0.2857 | 3 | 0.6667 | 11 | 0.2727 | 6 | 0.3333 | 27 | 0.3333 |
| 6_7259291 | 8 | 0.125 | 6 | 0.1667 | 6 | 0.1667 | 9 | 0.1111 | 29 | 0.1379 | 8 | 0 | 4 | 0 | 9 | 0.1111 | 5 | 0.6 | 26 | 0.1538 |
| 6_7259331 | 5 | 0 | 6 | 0 | 7 | 0 | 8 | 0.375 | 26 | 0.1154 | 6 | 0.5 | 4 | 0 | 9 | 0.1111 | 5 | 0.2 | 24 | 0.2083 |
| 6_7260423( | NA | NA | NA | NA | NA | NA | 1 | 1 | 1 | 1 | NA | NA | 2 | 0.5 | NA | NA | NA | NA | 2 | 0.5 |
| 6_7281810 | 3 | 0.3333 | 1 | 0 | 3 | 0.3333 | 2 | 0 | 9 | 0.2222 | 2 | 0.5 | 1 | 0 | 1 | 0 | 2 | 0 | 6 | 0.1667 |
| 6_7281835 | 2 | 0 | NA | NA | 2 | 0 | 2 | 1 | 6 | 0.3333 | 3 | 1 | 2 | 0.5 | 1 | 0 | 1 | 0 | 7 | 0.5714 |
| 6_7281841 | 3 | 0.3333 | NA | NA | 2 | 1 | 1 | 1 | 6 | 0.6667 | 4 | 0.5 | 1 | 0 | 1 | 0 | 2 | 0.5 | 8 | 0.375 |
| 6_7281979 | 10 | 0.1 | 9 | 0.3333 | 12 | 0 | 7 | 0.1429 | 38 | 0.1316 | 13 | 0.3077 | 13 | 0.1538 | 11 | 0.0909 | 9 | 0.2222 | 46 | 0.1957 |
| 6_7293441 | 65 | 0.3538 | 73 | 0.2329 | 65 | 0.5077 | 63 | 0.381 | 266 | 0.3647 | 37 | 0.4054 | 57 | 0.4035 | 67 | 0.3433 | 61 | 0.4262 | 222 | 0.3919 |
| 6_7685877 | 10 | 0.1 | 15 | 0.0667 | 17 | 0.1765 | 11 | 0.1818 | 53 | 0.1321 | 10 | 0 | 11 | 0.0909 | 10 | 0.2 | 13 | 0 | 44 | 0.0682 |
| 6_7685878 | 7 | 0.4286 | 11 | 0.3636 | 13 | 0.2308 | 7 | 0.1429 | 38 | 0.2895 | 6 | 0.3333 | 8 | 0.5 | 7 | 0.5714 | 8 | 0.375 | 29 | 0.4483 |
| 6_7685884 | 13 | 0.0769 | 16 | 0.0625 | 17 | 0.0588 | 11 | 0.1818 | 57 | 0.0877 | 9 | 0.1111 | 10 | 0 | 10 | 0.1 | 13 | 0.1538 | 42 | 0.0952 |
| 6_7685940 | 6 | 0.1667 | 12 | 0.0833 | 10 | 0.1 | 10 | 0.1 | 38 | 0.1053 | 6 | 0.3333 | 7 | 0 | 7 | 0.1429 | 10 | 0.1 | 30 | 0.1333 |
| 6_7685944 | 7 | 0 | 12 | 0 | 13 | 0.1538 | 9 | 0.1111 | 41 | 0.0732 | 6 | 0 | 8 | 0.25 | 7 | 0.2857 | 11 | 0.1818 | 32 | 0.1875 |
| 6_7685953 | 8 | 0.75 | 11 | 0.6364 | 11 | 0.8182 | 11 | 0.8182 | 41 | 0.7561 | 6 | 0.6667 | 8 | 1 | 8 | 0.875 | 9 | 0.5556 | 31 | 0.7742 |
| 6_7721983 | 4 | 0.25 | 2 | 0 | NA | NA | 3 | 0.3333 | 9 | 0.2222 | 3 | 0 | 2 | 0.5 | 2 | 0 | 2 | 0.5 | 9 | 0.2222 |
| 6_7722051 | 4 | 1 | 5 | 1 | NA | NA | 5 | 0.8 | 14 | 0.9286 | 4 | 1 | 3 | 0.6667 | 3 | 0.6667 | 3 | 1 | 13 | 0.8462 |
| 6_7722087 | 5 | 0.2 | 6 | 0.1667 | 1 | 0 | 4 | 0.25 | 16 | 0.1875 | 4 | 0 | 1 | 1 | 7 | 0.2857 | 4 | 0 | 16 | 0.1875 |
| 6_7722089 | 5 | 0 | 6 | 0 | 2 | 0.5 | 4 | 0 | 17 | 0.0588 | 4 | 0 | 2 | 0 | 7 | 0.1429 | 4 | 0.5 | 17 | 0.1765 |
| 6_7722092 | 5 | 0.6 | 5 | 0.8 | 2 | 1 | 4 | 0.75 | 16 | 0.75 | 3 | 1 | 2 | 0.5 | 7 | 1 | 4 | 0.5 | 16 | 0.8125 |
| 6_7722097 | 4 | 0.75 | 6 | 0.5 | 2 | 0 | 6 | 0.8333 | 18 | 0.6111 | 4 | 1 | 2 | 1 | 5 | 0.4 | 5 | 0.6 | 16 | 0.6875 |
| 6_7722109 | 5 | 1 | 5 | 1 | 2 | 0.5 | 6 | 0.8333 | 18 | 0.8889 | 4 | 1 | 3 | 0.6667 | 6 | 1 | 4 | 1 | 17 | 0.9412 |

| | | | | | | | | | | | | | | | | | | | | |
|---|---|---|---|---|---|---|---|---|---|---|---|---|---|---|---|---|---|---|---|---|
| 6_77221203 | 4 | 0.25 | 7 | 0.2857 | 2 | 0.5 | 6 | 0.8333 | 19 | 0.4737 | 3 | 1 | 2 | 0.5 | 7 | 0.8571 | 5 | 0.6 | 17 | 0.7647 |
| 6_77221329 | 3 | 1 | 5 | 1 | 2 | 1 | 4 | 1 | 14 | 1 | 3 | 1 | 2 | 1 | 5 | 1 | 4 | 1 | 14 | 1 |
| 6_77221407 | 5 | 0.8 | 6 | 0.6667 | 1 | 1 | 4 | 0 | 16 | 0.5625 | 4 | 0.75 | 3 | 0.6667 | 7 | 0.5714 | 6 | 0.8333 | 20 | 0.7 |
| 6_77221485 | 5 | 0.4 | 7 | 0.2857 | 1 | 1 | 6 | 0.1667 | 19 | 0.3158 | 4 | 0 | 3 | 0.6667 | 7 | 0 | 4 | 0 | 18 | 0.1111 |
| 6_77221494 | 4 | 0.75 | 6 | 1 | 2 | 1 | 5 | 0.8 | 17 | 0.8824 | 4 | 1 | 3 | 1 | 7 | 1 | 6 | 0.6667 | 20 | 0.9 |
| 6_77221695 | 7 | 0.4286 | 6 | 0.3333 | 3 | 1 | 6 | 0 | 22 | 0.3636 | 3 | 0.3333 | 3 | 0.3333 | 5 | 0.4 | 7 | 0.4286 | 18 | 0.3889 |
| 6_77221927 | 8 | 0.125 | 6 | 0 | 4 | 0.25 | 5 | 0 | 23 | 0.087 | 3 | 0 | 3 | 0.3333 | 7 | 0.1429 | 9 | 0.3333 | 22 | 0.2273 |
| 6_77221970 | 8 | 0.25 | 7 | 0.2857 | 4 | 0 | 6 | 0.3333 | 25 | 0.24 | 4 | 0 | 2 | 0 | 9 | 0.1111 | 8 | 0.25 | 23 | 0.1304 |
| 6_77222023 | 6 | 0.1667 | 6 | 0 | 4 | 0 | 7 | 0.2857 | 23 | 0.1304 | 4 | 0 | 2 | 0.5 | 8 | 0 | 6 | 0 | 20 | 0.05 |
| 6_77222186 | 8 | 0.875 | 7 | 1 | 3 | 1 | 5 | 1 | 23 | 0.9565 | 4 | 0.75 | 4 | 1 | 8 | 1 | 9 | 1 | 25 | 0.96 |
| 6_77222215 | 6 | 1 | 4 | 1 | 3 | 1 | 5 | 1 | 18 | 1 | 2 | 1 | 2 | 1 | 7 | 0.7143 | 3 | 1 | 14 | 0.8571 |
| 6_77222221 | 7 | 0.5714 | 7 | 0.7143 | 2 | 0.5 | 5 | 0.8 | 21 | 0.6667 | 2 | 0.5 | 2 | 0.5 | 5 | 0.4 | 5 | 0.4 | 14 | 0.4286 |
| 6_77222245 | 8 | 1 | 7 | 1 | 4 | 1 | 7 | 1 | 26 | 1 | 4 | 1 | 5 | 1 | 9 | 1 | 9 | 1 | 27 | 1 |
| 6_77222250 | 6 | 0.3333 | 6 | 0.1667 | 3 | 0 | 7 | 0 | 22 | 0.1364 | 3 | 0.3333 | 3 | 0 | 7 | 0.4286 | 7 | 0.2857 | 20 | 0.3 |
| 6_77222344 | 8 | 0.75 | 7 | 0.8571 | 3 | 0.3333 | 6 | 0.8333 | 24 | 0.75 | 4 | 1 | 6 | 1 | 9 | 0.8889 | 8 | 1 | 27 | 0.963 |
| 6_77222371 | 6 | 0.6667 | 6 | 0.5 | 3 | 0.6667 | 6 | 1 | 21 | 0.7143 | 4 | 0.5 | 5 | 0.2 | 9 | 0.4444 | 7 | 0.7143 | 25 | 0.48 |
| 6_77222500 | NA | NA | 1 | 1 | NA | NA | NA | NA | 1 | 1 | 1 | 1 | NA | NA | NA | NA | NA | NA | 1 | 1 |
| 6_77234370 | NA | NA | 1 | 1 | NA | NA | 2 | 0.5 | 3 | 0.6667 | 1 | 1 | NA | NA | 1 | 1 | NA | NA | 2 | 1 |
| 6_77234507 | NA | NA | 1 | 1 | NA | NA | 2 | 1 | 3 | 1 | NA | NA | NA | NA | 1 | 1 | NA | NA | 1 | 1 |
| 6_77234520 | NA | NA | 1 | 1 | NA | NA | 2 | 1 | 3 | 1 | 2 | 1 | NA | NA | 1 | 1 | NA | NA | 3 | 1 |
| 6_77234600 | NA | NA | 1 | 0 | NA | NA | 1 | 1 | 2 | 0.5 | 2 | 1 | NA | NA | NA | NA | NA | NA | 2 | 1 |
| 6_77234617 | NA | NA | 1 | 1 | NA | NA | 2 | 0.5 | 3 | 0.6667 | 2 | 1 | NA | NA | 1 | 1 | NA | NA | 3 | 1 |
| 6_77234818 | NA | NA | 1 | 1 | NA | NA | 2 | 1 | 3 | 1 | NA | NA | NA | NA | 1 | 1 | NA | NA | 1 | 1 |
| 6_77234890 | NA | NA | 1 | 1 | NA | NA | 2 | 0.5 | 3 | 0.6667 | 1 | 1 | NA | NA | NA | NA | NA | NA | 1 | 1 |
| 6_77234983 | NA | NA | 1 | 1 | NA | NA | 2 | 1 | 3 | 1 | 2 | 1 | NA | NA | 1 | 1 | 1 | 1 | 4 | 1 |
| 6_79998702 | 2 | 0.5 | 3 | 0.3333 | 3 | 0.6667 | 3 | 0.6667 | 11 | 0.5455 | 3 | 1 | 4 | 1 | 2 | 0.5 | 3 | 0.6667 | 12 | 0.8333 |
| 6_79998765 | 2 | 0 | 3 | 0.6667 | 4 | 0.5 | 3 | 1 | 12 | 0.5833 | 2 | 1 | 5 | 1 | 3 | 0.6667 | 3 | 1 | 13 | 0.9231 |
| 6_79998817 | 2 | 0.5 | 3 | 0.3333 | 5 | 0.8 | 3 | 0.6667 | 13 | 0.6154 | 3 | 0.6667 | 7 | 0.8571 | 3 | 0.6667 | 4 | 0.75 | 17 | 0.7647 |
| 6_79998922 | 2 | 1 | 2 | 1 | 4 | 1 | 3 | 0.6667 | 11 | 0.9091 | 3 | 1 | 6 | 0.8333 | 4 | 1 | 2 | 1 | 15 | 0.9333 |
| 6_79998974 | 2 | 0.5 | 2 | 0.5 | 5 | 0.4 | 2 | 0 | 11 | 0.3636 | 3 | 0.3333 | 5 | 0.6 | 3 | 0.3333 | 4 | 0 | 15 | 0.3333 |
| 6_79999024 | 2 | 0 | 3 | 0.3333 | 6 | 0 | 3 | 0.3333 | 14 | 0.1429 | 2 | 0 | 7 | 0.2857 | 3 | 0 | 5 | 0.2 | 17 | 0.1765 |
| 6_79999066 | 2 | 0 | 3 | 0.3333 | 6 | 0 | 3 | 0 | 14 | 0.0714 | 3 | 0.3333 | 7 | 0.1429 | 4 | 0.25 | 5 | 0 | 19 | 0.1579 |
| 6_79999128 | 2 | 1 | 1 | 1 | 6 | 0.6667 | 2 | 1 | 11 | 0.8182 | 2 | 1 | 7 | 0.5714 | 3 | 0.6667 | 4 | 1 | 16 | 0.75 |
| 6_79999137 | 2 | 0 | 3 | 0 | 5 | 0.2 | 3 | 0 | 13 | 0.0769 | 3 | 0.3333 | 8 | 0 | 4 | 0.25 | 5 | 0.2 | 20 | 0.15 |
| 6_79999143 | 1 | 1 | 1 | 0 | 5 | 0.4 | 3 | 0.3333 | 10 | 0.4 | 3 | 1 | 8 | 0.875 | 3 | 0.3333 | 5 | 0.6 | 19 | 0.7368 |
| 6_79999308 | 2 | 0 | 3 | 0 | 5 | 0 | 3 | 0.3333 | 13 | 0.0769 | 3 | 0 | 8 | 0.125 | 4 | 0 | 5 | 0 | 20 | 0.05 |
| 6_79999343 | 2 | 1 | 3 | 1 | 6 | 0.8333 | 3 | 1 | 14 | 0.9286 | 3 | 1 | 7 | 1 | 4 | 1 | 5 | 1 | 19 | 1 |
| 6_79999478 | 2 | 1 | 3 | 1 | 5 | 0.6 | 3 | 0.3333 | 13 | 0.6923 | 3 | 1 | 7 | 1 | 4 | 1 | 5 | 0.8 | 19 | 0.9474 |
| 6_79999658 | 1 | 1 | 3 | 0.3333 | 6 | 0.3333 | 3 | 0.6667 | 13 | 0.4615 | 3 | 1 | 7 | 1 | 4 | 0.75 | 5 | 0.6 | 19 | 0.8421 |
| 6_80000017 | 2 | 1 | 4 | 0.75 | 6 | 0.6667 | 3 | 0.6667 | 15 | 0.7333 | 3 | 1 | 7 | 0.7143 | 4 | 0.75 | 5 | 0.8 | 19 | 0.7895 |
| 6_80000052 | 2 | 1 | 4 | 0.75 | 6 | 0.6667 | 3 | 1 | 15 | 0.8 | 3 | 1 | 5 | 1 | 4 | 1 | 4 | 1 | 16 | 1 |
| 6_80000067 | 1 | 1 | 3 | 1 | 4 | 1 | 3 | 1 | 11 | 1 | 2 | 1 | 5 | 1 | 4 | 1 | 4 | 1 | 15 | 1 |
| 6_80000075 | 2 | 0 | 4 | 0.25 | 6 | 0.1667 | 2 | 0.5 | 14 | 0.2143 | 3 | 0 | 7 | 0 | 3 | 0.3333 | 3 | 0 | 16 | 0.0625 |
| 6_80000150 | 2 | 0 | 2 | 0.5 | 4 | 0.25 | 2 | 0 | 10 | 0.2 | 2 | 1 | 4 | 1 | 4 | 0.5 | 2 | 0 | 12 | 0.6667 |
| 6_80000167 | 2 | 0 | 2 | 0 | 4 | 0 | 3 | 0.3333 | 11 | 0.0909 | 2 | 0 | 5 | 0.4 | 5 | 0 | 3 | 0 | 15 | 0.1333 |
| 6_80000308 | 2 | 0 | 2 | 0 | 4 | 0 | 3 | 0.3333 | 11 | 0.0909 | 2 | 0.5 | 8 | 0.125 | 4 | 0.5 | 2 | 0 | 16 | 0.25 |
| 6_80000312 | 1 | 1 | 2 | 0.5 | 4 | 0.5 | 4 | 0 | 11 | 0.3636 | 2 | 0 | 5 | 0.4 | 5 | 0.2 | 2 | 0.5 | 14 | 0.2857 |
| 6_80000336 | 2 | 0.5 | 2 | 0 | 4 | 0 | 4 | 0.25 | 12 | 0.1667 | 2 | 0 | 7 | 0.1429 | 5 | 0 | 3 | 0 | 17 | 0.0588 |
| 6_80000587 | 2 | 0.5 | 1 | 1 | 4 | 0.25 | 4 | 0.5 | 11 | 0.4545 | 2 | 0.5 | 6 | 0.8333 | 5 | 0.8 | 3 | 0.6667 | 16 | 0.75 |
| 6_80001284 | 3 | 0.6667 | 2 | 0.5 | 3 | 0.3333 | 5 | 0.8 | 13 | 0.6154 | 2 | 1 | 9 | 0.5556 | 5 | 0.8 | 3 | 1 | 19 | 0.7368 |
| 6_80001519 | 3 | 0 | 2 | 0.5 | 4 | 0 | 5 | 0 | 14 | 0.0714 | 4 | 0 | 9 | 0.1111 | 6 | 0.1667 | 3 | 0 | 22 | 0.0909 |
| 6_80001740 | 3 | 0.3333 | 2 | 1 | 5 | 1 | 5 | 0.6 | 15 | 0.7333 | 5 | 1 | 11 | 0.4545 | 8 | 0.625 | 3 | 0.6667 | 27 | 0.6296 |
| 6_80001749 | 2 | 0.5 | 2 | 0 | 5 | 0.4 | 5 | 0.2 | 14 | 0.2857 | 5 | 0.4 | 11 | 0.3636 | 7 | 0 | 2 | 0.5 | 25 | 0.28 |
| 6_80786315 | 2 | 1 | 1 | 0 | 2 | 0 | 1 | 0 | 6 | 0.3333 | NA | NA | 1 | 1 | 1 | 1 | 1 | 0 | 3 | 0.6667 |
| 6_80786364 | 2 | 0.5 | 1 | 0 | 1 | 0 | 1 | 0 | 5 | 0.2 | NA | NA | 1 | 0 | 1 | 0 | 2 | 0.5 | 4 | 0.25 |
| 6_80786371 | 3 | 0.6667 | 1 | 1 | 2 | 1 | 1 | 1 | 7 | 0.8571 | NA | NA | 1 | 1 | NA | NA | 3 | 0.6667 | 4 | 0.75 |
| 6_80786527 | 3 | 0.3333 | 2 | 0 | 1 | 0 | 1 | 0 | 7 | 0.1429 | NA | NA | 1 | 0 | 2 | 0 | 3 | 0.3333 | 6 | 0.1667 |
| 6_80786849 | 3 | 0.3333 | 2 | 0.5 | 1 | 1 | 4 | 0.5 | 10 | 0.5 | NA | NA | 2 | 0 | 3 | 0.6667 | 3 | 1 | 8 | 0.625 |
| 6_80786923 | 2 | 0.5 | 1 | 0 | 2 | 0.5 | 3 | 0.3333 | 8 | 0.375 | NA | NA | 1 | 0 | 2 | 0.5 | 3 | 0.6667 | 6 | 0.5 |
| 6_80786979 | 3 | 0.3333 | 2 | 0 | 2 | 1 | 3 | 0.3333 | 10 | 0.4 | NA | NA | 2 | 0 | 2 | 0.5 | 3 | 0.6667 | 7 | 0.4286 |
| 6_81935359 | 1 | 1 | 3 | 1 | NA | NA | NA | NA | 4 | 1 | NA | NA | 2 | 0.5 | 1 | 0 | 1 | 0 | 4 | 0.25 |
| 6_81935421 | 1 | 1 | 2 | 1 | NA | NA | NA | NA | 3 | 1 | NA | NA | 1 | 1 | 1 | 1 | 1 | 0 | 3 | 0.6667 |
| 6_81935881 | NA | NA | 4 | 0.5 | 1 | 0 | NA | NA | 5 | 0.4 | NA | NA | 2 | 1 | 1 | 0 | 1 | 0 | 4 | 0.5 |
| 6_81936040 | NA | NA | 3 | 0.3333 | 1 | 0 | NA | NA | 4 | 0.25 | NA | NA | 2 | 0.5 | NA | NA | 1 | 0 | 3 | 0.3333 |
| 6_81936422 | NA | NA | 1 | 1 | NA | NA | 1 | 1 | 2 | 1 | NA | NA | NA | NA | 1 | 0 | 1 | 1 | 2 | 0.5 |

| | | | | | | | | | | | | | | | | | | | | |
|---|---|---|---|---|---|---|---|---|---|---|---|---|---|---|---|---|---|---|---|---|
| 6_81937897 | NA | NA | 1 | 0 | 1 | 1 | NA | NA | 2 | 0.5 | NA | NA | 1 | 1 | NA | NA | 1 | 0 | 2 | 0.5 |
| 6_81938634 | 2 | 0.5 | 1 | 1 | 5 | 0.4 | 3 | 0.3333 | 11 | 0.4545 | 1 | 0 | 4 | 0.25 | 3 | 0 | 2 | 0 | 10 | 0.1 |
| 6_81938853 | 1 | 0 | 2 | 0.5 | 4 | 1 | 1 | 1 | 8 | 0.75 | 1 | 1 | 4 | 0.75 | 1 | 0 | 1 | 1 | 7 | 0.7143 |
| 6_81938858 | 1 | 0 | 3 | 1 | 2 | 0.5 | 2 | 1 | 8 | 0.75 | NA | NA | 3 | 0.3333 | 2 | 0 | 2 | 1 | 7 | 0.4286 |
| 6_81938914 | 1 | 1 | 3 | 1 | 4 | 1 | 3 | 1 | 11 | 1 | 1 | 1 | 5 | 1 | 1 | 1 | 2 | 1 | 9 | 1 |
| 6_81968708 | NA | NA | 1 | 1 | NA | NA | NA | NA | 1 | 1 | NA | NA | NA | NA | 1 | 1 | NA | NA | 1 | 1 |
| 6_82068874 | 3 | 0.6667 | 1 | 0 | 2 | 0.5 | 4 | 0.25 | 10 | 0.4 | 2 | 0 | 4 | 0.25 | 3 | 1 | 1 | 1 | 10 | 0.5 |
| 6_82068955 | 2 | 1 | 2 | 0 | 2 | 0.5 | 1 | 0 | 7 | 0.4286 | 2 | 0.5 | 4 | 0.25 | 4 | 0.25 | 1 | 0 | 11 | 0.2727 |
| 6_82069002 | 3 | 1 | 1 | 1 | 2 | 1 | 3 | 1 | 9 | 1 | 1 | 1 | 3 | 1 | 4 | 1 | 1 | 1 | 9 | 1 |
| 6_82069132 | 3 | 0 | 2 | 0.5 | 2 | 0 | 3 | 0.3333 | 10 | 0.2 | 2 | 0.5 | 3 | 0.3333 | 3 | 0 | 1 | 0 | 9 | 0.2222 |
| 6_82069160 | 3 | 0 | 1 | 0 | 2 | 0 | 3 | 0.3333 | 9 | 0.1111 | 2 | 0 | 3 | 0.3333 | 3 | 0.3333 | 2 | 0 | 10 | 0.2 |
| 6_82069329 | 2 | 0.5 | 2 | 0 | 2 | 0.5 | 4 | 0.5 | 10 | 0.4 | 2 | 0 | 3 | 0.6667 | 4 | 0.25 | 1 | 1 | 10 | 0.4 |
| 6_82069452 | 3 | 0 | 2 | 0 | 2 | 0.5 | 4 | 0.5 | 11 | 0.2727 | 2 | 0 | 3 | 0.3333 | 4 | 0.25 | 2 | 0 | 11 | 0.1818 |
| 6_82069494 | 3 | 0.6667 | 2 | 0 | NA | NA | 4 | 0.75 | 9 | 0.5556 | 2 | 1 | 4 | 0.5 | 3 | 1 | 1 | 1 | 10 | 0.8 |
| 6_82069600 | 3 | 1 | 1 | 1 | 2 | 1 | 3 | 1 | 9 | 1 | 2 | 1 | 4 | 1 | 3 | 0.6667 | 2 | 1 | 11 | 0.9091 |
| 6_82069672 | 3 | 0 | 1 | 1 | 1 | 0 | 2 | 0 | 7 | 0.1429 | 2 | 0 | 1 | 0 | 2 | 0 | 3 | 0.3333 | 8 | 0.125 |
| 6_82069706 | 2 | 0.5 | 3 | 0.3333 | NA | NA | 2 | 0 | 7 | 0.2857 | 2 | 0.5 | 4 | 0 | 2 | 0 | 1 | 0 | 9 | 0.1111 |
| 6_82069710 | 3 | 0 | 3 | 0 | 1 | 0 | 4 | 0.25 | 11 | 0.0909 | NA | NA | 4 | 0 | 3 | 0 | 2 | 0.5 | 9 | 0.1111 |
| 6_82069775 | 3 | 0 | 2 | 0.5 | 2 | 0 | 3 | 0.3333 | 10 | 0.2 | 2 | 0.5 | 4 | 0 | 4 | 0.25 | 3 | 0.3333 | 13 | 0.2308 |
| 6_82069803 | 3 | 0 | 3 | 0.3333 | 2 | 0 | 4 | 0 | 12 | 0.0833 | 1 | 0 | 4 | 0 | 3 | 0.3333 | 3 | 0 | 11 | 0.0909 |
| 6_82069862 | 3 | 0 | 3 | 0 | 2 | 0.5 | 4 | 0.25 | 12 | 0.1667 | 2 | 0 | 4 | 0 | 3 | 0.3333 | 3 | 0.3333 | 12 | 0.1667 |
| 6_82069955 | 4 | 1 | 4 | 0.5 | 2 | 1 | 4 | 0.5 | 14 | 0.7143 | 2 | 0.5 | 4 | 0.25 | 4 | 0.75 | 3 | 0.6667 | 13 | 0.5385 |
| 6_82380426 | 1 | 0 | NA | NA | NA | NA | 1 | 1 | 2 | 0.5 | 1 | 1 | 1 | 0 | NA | NA | NA | NA | 2 | 0.5 |
| 6_82380469 | 1 | 0 | NA | NA | NA | NA | 1 | 1 | 2 | 0.5 | 1 | 1 | 1 | 0 | NA | NA | NA | NA | 2 | 0.5 |
| 6_82380613 | 1 | 1 | NA | NA | NA | NA | 2 | 1 | 3 | 1 | 1 | 1 | 1 | 1 | NA | NA | NA | NA | 2 | 1 |
| 6_82534418 | 1 | 0 | NA | NA | 1 | 0 | 1 | 1 | 3 | 0.3333 | 1 | 1 | 1 | 0 | NA | NA | NA | NA | 2 | 0.5 |
| 6_82534627 | 1 | 0 | NA | NA | NA | NA | 3 | 0.3333 | 4 | 0.25 | 2 | 0.5 | 1 | 0 | NA | NA | NA | NA | 3 | 0.3333 |
| 6_82534703 | 1 | 1 | NA | NA | NA | NA | 2 | 1 | 3 | 1 | 1 | 1 | 1 | 1 | NA | NA | NA | NA | 2 | 1 |
| 6_82534882 | 1 | 0 | NA | NA | 1 | 0 | 3 | 1 | 5 | 0.6 | 2 | 1 | 1 | 1 | 1 | 1 | NA | NA | 4 | 1 |
| 6_82535412 | 1 | 0 | NA | NA | 1 | 0 | 3 | 0.3333 | 5 | 0.2 | 2 | 0.5 | 1 | 0 | 2 | 0 | NA | NA | 5 | 0.2 |
| 6_82535424 | 1 | 1 | NA | NA | NA | NA | 3 | 1 | 4 | 1 | 2 | 1 | 1 | 1 | 1 | 1 | NA | NA | 4 | 1 |
| 6_82535544 | 1 | 1 | 1 | 1 | 1 | 1 | 3 | 0.6667 | 6 | 0.8333 | 2 | 1 | 1 | 1 | 1 | 1 | NA | NA | 4 | 1 |
| 6_82535621 | 1 | 1 | 1 | 1 | 1 | 1 | 3 | 0.6667 | 6 | 0.8333 | 2 | 1 | NA | NA | 2 | 1 | NA | NA | 4 | 1 |
| 6_82535646 | 1 | 0 | 1 | 1 | NA | NA | 2 | 0.5 | 4 | 0.5 | 2 | 1 | 1 | 1 | 2 | 0.5 | NA | NA | 5 | 0.8 |
| 6_82535990 | 1 | 1 | NA | NA | 1 | 1 | 2 | 1 | 4 | 1 | 2 | 1 | 1 | 1 | 1 | 1 | NA | NA | 4 | 1 |
| 6_82536227 | NA | NA | 1 | 1 | 1 | 1 | 4 | 1 | 6 | 1 | 2 | 1 | 1 | 1 | 1 | 1 | NA | NA | 4 | 1 |
| 6_82536245 | 1 | 1 | 1 | 1 | 1 | 0 | 3 | 1 | 6 | 0.8333 | 2 | 0 | 1 | 1 | 2 | 0 | NA | NA | 5 | 0.2 |
| 6_82536360 | 1 | 1 | 1 | 0 | 1 | 0 | 3 | 0 | 6 | 0.1667 | 2 | 0.5 | 1 | 1 | 2 | 0 | NA | NA | 5 | 0.4 |
| 6_82536364 | NA | NA | NA | NA | NA | NA | 3 | 0.6667 | 3 | 0.6667 | 1 | 0 | NA | NA | 1 | 1 | NA | NA | 2 | 0.5 |
| 6_82536379 | 1 | 1 | NA | NA | 1 | 1 | 3 | 1 | 5 | 1 | 2 | 1 | NA | NA | 1 | 1 | NA | NA | 3 | 1 |
| 6_82536476 | 2 | 0 | 1 | 1 | NA | NA | 3 | 0.3333 | 6 | 0.3333 | 1 | 1 | 1 | 0 | 3 | 0 | NA | NA | 5 | 0.2 |
| 6_82536567 | 3 | 0.3333 | NA | NA | 1 | 0 | 4 | 0.5 | 8 | 0.375 | 2 | 0 | 1 | 1 | 3 | 0 | NA | NA | 6 | 0.1667 |
| 6_82536643 | 2 | 1 | 1 | 0 | 1 | 0 | 4 | 0.25 | 8 | 0.375 | 2 | 0.5 | 1 | 0 | 2 | 0 | NA | NA | 5 | 0.2 |
| 6_82536885 | 3 | 0.3333 | 1 | 1 | 2 | 0 | 3 | 0.3333 | 9 | 0.3333 | 2 | 0.5 | 1 | 1 | 3 | 0 | 1 | 0 | 7 | 0.2857 |
| 6_82536903 | 3 | 1 | 1 | 1 | 2 | 1 | 4 | 0.75 | 10 | 0.9 | 2 | 1 | 1 | 1 | 3 | 1 | 1 | 1 | 7 | 1 |
| 6_82536983 | 3 | 0.6667 | 1 | 0 | 1 | 1 | 3 | 0.3333 | 8 | 0.5 | 2 | 0.5 | 1 | 1 | 3 | 0.3333 | 1 | 0 | 7 | 0.4286 |
| 6_82624188 | 4 | 0.25 | 3 | 0.3333 | 1 | 0 | 5 | 0.2 | 13 | 0.2308 | 4 | 0.25 | NA | NA | 6 | 0 | 2 | 0 | 12 | 0.0833 |
| 6_82624646 | 12 | 0.75 | 6 | 0.8333 | 6 | 0.6667 | 7 | 0.5714 | 31 | 0.7097 | 6 | 0.6667 | 2 | 1 | 7 | 0.8571 | 5 | 0.4 | 20 | 0.7 |
| 6_82624673 | 11 | 0.1818 | 6 | 0 | 6 | 0.1667 | 8 | 0.25 | 31 | 0.1613 | 6 | 0.3333 | 2 | 0.5 | 6 | 0.5 | 5 | 0 | 19 | 0.3158 |
| 6_82624740 | 10 | 0.2 | 8 | 0.125 | 6 | 0.5 | 7 | 0.8571 | 31 | 0.3871 | 6 | 0.5 | 5 | 0 | 5 | 0.6 | 6 | 0.1667 | 22 | 0.3182 |
| 6_82624772 | 10 | 1 | 8 | 0.75 | 6 | 1 | 8 | 0.875 | 32 | 0.9062 | 6 | 1 | 5 | 0.8 | 6 | 1 | 6 | 1 | 23 | 0.9565 |
| 6_82702323 | NA | NA | 1 | 1 | 3 | 1 | NA | NA | 4 | 1 | NA | NA | 1 | 1 | NA | NA | 1 | 1 | 2 | 1 |
| 6_82702646 | NA | NA | 1 | 0 | 3 | 0.6667 | NA | NA | 4 | 0.5 | NA | NA | 1 | 1 | NA | NA | 1 | 0 | 2 | 0.5 |
| 6_82702941 | NA | NA | 1 | 1 | 2 | 0.5 | NA | NA | 3 | 0.6667 | NA | NA | 1 | 1 | NA | NA | NA | NA | 1 | 1 |
| 6_82704188 | NA | NA | NA | NA | 1 | 1 | NA | NA | 1 | 1 | NA | NA | 1 | 1 | NA | NA | NA | NA | 1 | 1 |
| 6_82704226 | NA | NA | NA | NA | 1 | 1 | NA | NA | 1 | 1 | NA | NA | 1 | 1 | NA | NA | NA | NA | 1 | 1 |
| 6_82888885 | 3 | 0.3333 | 2 | 0 | 3 | 0.6667 | 2 | 0.5 | 10 | 0.4 | 4 | 0.75 | 3 | 0.3333 | 2 | 0.5 | 3 | 0.3333 | 12 | 0.5 |
| 6_82888962 | 5 | 1 | 5 | 0.8 | 4 | 1 | 1 | 1 | 15 | 0.9333 | 3 | 0.6667 | 4 | 1 | 4 | 0.75 | 5 | 1 | 16 | 0.875 |
| 6_82889114 | 3 | 0.3333 | 2 | 0.5 | 3 | 0.3333 | NA | NA | 8 | 0.375 | 2 | 0.5 | 3 | 0.3333 | 4 | 0 | 4 | 0.25 | 13 | 0.2308 |
| 6_82889294 | 4 | 0.5 | 4 | 0.75 | 3 | 1 | 1 | 1 | 12 | 0.75 | 1 | 1 | 2 | 0.5 | 3 | 0.6667 | 3 | 1 | 9 | 0.7778 |
| 6_82889392 | 4 | 1 | 4 | 0.75 | 2 | 0.5 | 1 | 0 | 11 | 0.7273 | 2 | 0.5 | 3 | 1 | 4 | 1 | 3 | 0.3333 | 12 | 0.75 |
| 6_82889535 | 4 | 0 | 3 | 0 | 3 | 0.3333 | 1 | 0 | 11 | 0.0909 | 2 | 0 | 5 | 0.2 | 3 | 0 | 4 | 0 | 14 | 0.0714 |
| 6_82889574 | 4 | 0.25 | 5 | 0.2 | 2 | 0.5 | 1 | 0 | 12 | 0.25 | 2 | 0.5 | 2 | 0 | 3 | 0 | 2 | 1 | 9 | 0.3333 |
| 6_82889629 | 4 | 0 | 4 | 0 | 2 | 0 | 2 | 0.5 | 12 | 0.0833 | 2 | 0 | 5 | 0 | 3 | 0 | 4 | 0.5 | 14 | 0.1429 |
| 6_82889668 | 3 | 1 | 4 | 1 | 3 | 0.6667 | 1 | 1 | 11 | 0.9091 | 1 | 0 | 2 | 1 | 3 | 1 | 3 | 1 | 9 | 0.8889 |

| | | | | | | | | | | | | | | | | | | | | |
|---|---|---|---|---|---|---|---|---|---|---|---|---|---|---|---|---|---|---|---|---|
| 6_8288979 | 4 | 0.25 | 4 | 0.5 | 3 | 0.3333 | 2 | 0 | 13 | 0.3077 | 2 | 0 | 3 | 0.3333 | 3 | 0.6667 | 4 | 0.75 | 12 | 0.5 |
| 6_8289006 | 4 | 0.5 | 5 | 0.4 | 3 | 1 | 1 | 1 | 13 | 0.6154 | 2 | 0.5 | 5 | 0.4 | 4 | 0.25 | 5 | 0.6 | 16 | 0.4375 |
| 6_8289009 | 4 | 0.25 | 4 | 0 | 3 | 0 | 2 | 0 | 13 | 0.0769 | 1 | 0 | 3 | 0 | 4 | 0.5 | 4 | 0 | 12 | 0.1667 |
| 6_8289020 | 4 | 0.75 | 5 | 0.8 | 3 | 1 | 2 | 0.5 | 14 | 0.7857 | 1 | 1 | 2 | 1 | 4 | 1 | 5 | 0.8 | 12 | 0.9167 |
| 6_8289030 | 4 | 0.25 | 4 | 0 | 3 | 0 | 1 | 1 | 12 | 0.1667 | 2 | 0 | 6 | 0.1667 | 4 | 0.25 | 4 | 0 | 16 | 0.125 |
| 6_8289033 | 3 | 1 | 4 | 0.5 | 3 | 0.6667 | 1 | 1 | 11 | 0.7273 | 2 | 1 | 5 | 1 | 3 | 1 | 3 | 1 | 13 | 1 |
| 6_8289040 | 4 | 0.25 | 5 | 0 | 1 | 0 | 1 | 0 | 11 | 0.0909 | 2 | 0 | 6 | 0 | 4 | 0.25 | 4 | 0.5 | 16 | 0.1875 |
| 6_8289043 | 2 | 1 | 5 | 0.8 | 1 | 1 | 1 | 1 | 9 | 0.8889 | 2 | 1 | 4 | 0.5 | 3 | 0.6667 | 2 | 1 | 11 | 0.7273 |
| 6_8289045 | 4 | 1 | 1 | 1 | 3 | 1 | 1 | 1 | 9 | 1 | 2 | 1 | 3 | 1 | 4 | 1 | 3 | 1 | 12 | 1 |
| 6_8289066 | 4 | 0.25 | 2 | 0 | 2 | 0 | 1 | 0 | 9 | 0.1111 | 2 | 0 | 4 | 0.5 | 3 | 0 | 3 | 0.3333 | 12 | 0.25 |
| 6_8289069 | 3 | 0 | 3 | 0.3333 | 3 | 0.3333 | 2 | 0.5 | 11 | 0.2727 | 2 | 0 | 4 | 0 | 3 | 1 | 3 | 0.6667 | 12 | 0.4167 |
| 6_8289097 | 4 | 0.25 | 3 | 0 | 2 | 0 | 2 | 0 | 11 | 0.0909 | 2 | 0 | 4 | 0.25 | 2 | 0 | 1 | 0 | 9 | 0.1111 |
| 6_8289480 | 3 | 0 | 1 | 1 | 1 | 0 | 2 | 0.5 | 7 | 0.2857 | 1 | 0 | 1 | 0 | 1 | 1 | NA | NA | 3 | 0.3333 |
| 6_8302844 | 3 | 0.3333 | 1 | 1 | 1 | 1 | 5 | 0.4 | 10 | 0.5 | NA | NA | NA | NA | 2 | 0.5 | NA | NA | 2 | 0.5 |
| 6_8302848 | 5 | 0.4 | 2 | 0 | 6 | 0.3333 | 7 | 0.2857 | 20 | 0.3 | 1 | 1 | 1 | 1 | 7 | 0.4286 | NA | NA | 9 | 0.5556 |
| 6_8303035 | 4 | 0.25 | 2 | 0 | 4 | 0 | 7 | 0 | 17 | 0.0588 | 1 | 0 | 1 | 0 | 6 | 0.5 | NA | NA | 8 | 0.375 |
| 6_8303114 | 2 | 0.5 | NA | NA | 4 | 0.75 | 6 | 0.1667 | 12 | 0.4167 | NA | NA | 1 | 1 | 6 | 0.6667 | NA | NA | 7 | 0.7143 |
| 6_8303216 | 3 | 0.6667 | 10 | 0.4 | 10 | 0.4 | 4 | 0.25 | 27 | 0.4074 | 8 | 0.125 | 6 | 0.1667 | 5 | 0.2 | 4 | 0.5 | 23 | 0.2174 |
| 6_8303217 | 3 | 0.3333 | 10 | 0.1 | 9 | 0.2222 | 2 | 0 | 24 | 0.1667 | 7 | 0.4286 | 6 | 0.5 | 8 | 0.5 | 4 | 0.25 | 25 | 0.44 |
| 6_8303455 | 7 | 0 | 13 | 0.0769 | 14 | 0.0714 | 5 | 0 | 39 | 0.0513 | 9 | 0 | 7 | 0.1429 | 12 | 0.0833 | 6 | 0 | 34 | 0.0588 |
| 6_8307921 | 2 | 1 | 1 | 1 | NA | NA | NA | NA | 3 | 1 | NA | NA | NA | NA | NA | NA | 1 | 1 | 1 | 1 |
| 6_8307924 | 2 | 1 | 1 | 1 | NA | NA | NA | NA | 3 | 1 | NA | NA | NA | NA | NA | NA | 1 | 1 | 1 | 1 |
| 6_8308510 | 2 | 0.5 | 1 | 0 | 1 | 0 | NA | NA | 4 | 0.25 | NA | NA | NA | NA | NA | NA | 1 | 1 | 1 | 1 |
| 6_8308540 | 2 | 0.5 | 1 | 1 | 1 | 0 | NA | NA | 4 | 0.5 | NA | NA | NA | NA | NA | NA | 1 | 1 | 1 | 1 |
| 6_8308575 | 1 | 0 | 1 | 1 | NA | NA | NA | NA | 2 | 0.5 | NA | NA | 1 | 0 | NA | NA | 1 | 1 | 2 | 0.5 |
| 6_8308670 | 7 | 0.7143 | 9 | 0.7778 | 7 | 0.7143 | 4 | 0.75 | 27 | 0.7407 | 3 | 1 | 6 | 0.5 | 11 | 0.7273 | 5 | 0.8 | 25 | 0.72 |
| 6_8308677 | 12 | 0.25 | 12 | 0.1667 | 8 | 0 | 6 | 0 | 38 | 0.1316 | 9 | 0.2222 | 11 | 0.3636 | 16 | 0.125 | 8 | 0 | 44 | 0.1818 |
| 6_8309439 | NA | NA | NA | NA | 2 | 1 | 2 | 0.5 | 4 | 0.75 | 2 | 1 | 2 | 1 | 2 | 1 | 2 | 1 | 8 | 1 |
| 6_8309444 | NA | NA | 2 | 0 | 2 | 0.5 | 2 | 0 | 6 | 0.1667 | 3 | 0.3333 | 3 | 0.6667 | 2 | 1 | 2 | 0.5 | 10 | 0.6 |
| 6_8309459 | NA | NA | 2 | 0.5 | 2 | 0 | 2 | 0 | 6 | 0.1667 | 1 | 0 | 3 | 0.3333 | 2 | 1 | 1 | 0 | 7 | 0.4286 |
| 6_8309475 | NA | NA | 2 | 0.5 | 2 | 0 | 2 | 0 | 6 | 0.1667 | 3 | 0 | 3 | 0.3333 | 1 | 0 | 3 | 0 | 10 | 0.1 |
| 6_8309487 | NA | NA | 4 | 0.5 | 2 | 0.5 | 2 | 0 | 8 | 0.375 | 4 | 0 | 3 | 0 | 2 | 1 | 3 | 0 | 12 | 0.1667 |
| 6_8309497 | NA | NA | 4 | 0.25 | 3 | 0.3333 | 2 | 0 | 9 | 0.2222 | 3 | 0 | 3 | 0.6667 | 1 | 0 | 2 | 0 | 9 | 0.2222 |
| 6_8309513 | NA | NA | 2 | 1 | 3 | 1 | 2 | 1 | 7 | 1 | 4 | 0.75 | 3 | 1 | 2 | 1 | 2 | 1 | 11 | 0.9091 |
| 6_8309553 | NA | NA | 5 | 0.6 | 2 | 1 | 2 | 0.5 | 9 | 0.6667 | 4 | 0.5 | 3 | 1 | 4 | 0.75 | 3 | 0.3333 | 14 | 0.6429 |
| 6_8309560 | 1 | 0 | 5 | 0.2 | 4 | 0.25 | 2 | 0 | 12 | 0.1667 | 5 | 0 | 3 | 0.3333 | 3 | 0 | 3 | 0 | 14 | 0.0714 |
| 6_8309565 | NA | NA | 4 | 0.25 | 5 | 0.4 | 2 | 0 | 11 | 0.2727 | 5 | 0 | 3 | 0.3333 | 4 | 0.5 | 3 | 0.3333 | 15 | 0.2667 |
| 6_8309569 | 1 | 0 | 5 | 0.4 | 5 | 0.6 | 2 | 0.5 | 13 | 0.4615 | 5 | 0.6 | 4 | 0.5 | 3 | 0.6667 | 2 | 1 | 14 | 0.6429 |
| 6_8309609 | 6 | 0.8333 | 5 | 0.6 | 4 | 0.5 | 7 | 0.5714 | 22 | 0.6364 | 8 | 0.875 | 7 | 0.7143 | 2 | 0.5 | 1 | 1 | 18 | 0.7778 |
| 6_8309616 | 6 | 0.1667 | 3 | 0 | 4 | 0.25 | 5 | 0 | 18 | 0.1111 | 8 | 0.125 | 3 | 0 | 2 | 0 | NA | NA | 13 | 0.0769 |
| 6_8309660 | 6 | 0.5 | 2 | 0 | 3 | 0.3333 | 5 | 0.4 | 16 | 0.375 | 7 | 0.5714 | 3 | 0.3333 | 2 | 0.5 | 1 | 0 | 13 | 0.4615 |
| 6_8309917 | 6 | 0.5 | 6 | 0.6667 | 4 | 0.25 | 3 | 0 | 19 | 0.4211 | 4 | 0.25 | 4 | 0 | 2 | 0.5 | 4 | 0.25 | 14 | 0.2143 |
| 6_8311500 | NA | NA | 5 | 0.2 | NA | NA | NA | NA | 5 | 0.2 | 1 | 1 | 1 | 1 | 1 | 1 | NA | NA | 3 | 1 |
| 6_8311501 | NA | NA | 5 | 0.2 | NA | NA | NA | NA | 5 | 0.2 | NA | NA | 2 | 0 | 1 | 1 | NA | NA | 3 | 0.3333 |
| 6_8311502 | NA | NA | 5 | 0.2 | NA | NA | NA | NA | 5 | 0.2 | NA | NA | 2 | 0.5 | 1 | 0 | NA | NA | 3 | 0.3333 |
| 6_8311508 | NA | NA | 2 | 0.5 | NA | NA | NA | NA | 2 | 0.5 | 1 | 0 | 2 | 0 | 1 | 1 | NA | NA | 4 | 0.25 |
| 6_8312924 | 8 | 0.75 | 11 | 0.3636 | 7 | 0.2857 | 3 | 0.6667 | 29 | 0.4828 | 7 | 0.5714 | 8 | 0.25 | 7 | 0.2857 | 4 | 0.25 | 26 | 0.3462 |
| 6_8312931 | 1 | 1 | 6 | 0.6667 | 6 | 0.6667 | 2 | 0.5 | 15 | 0.6667 | 6 | 0.6667 | 8 | 0.5 | 7 | 0.7143 | 5 | 1 | 26 | 0.6923 |
| 6_8312944 | 5 | 0 | 7 | 0 | 5 | 0.2 | 1 | 0 | 18 | 0.0556 | 4 | 0.5 | 4 | 0 | 8 | 0 | 3 | 0 | 19 | 0.1053 |
| 6_8313807 | 3 | 0 | 3 | 0.6667 | 3 | 0.3333 | 4 | 0.25 | 13 | 0.3077 | 2 | 0.5 | 1 | 0 | 3 | 0 | 4 | 0.5 | 10 | 0.3 |
| 6_8313814 | 3 | 0 | 3 | 0.3333 | 3 | 0.3333 | 4 | 0 | 13 | 0.1538 | 3 | 0 | 1 | 0 | 4 | 0.25 | 4 | 0 | 12 | 0.0833 |
| 6_8313817 | 3 | 0.3333 | 3 | 0.3333 | 3 | 0.3333 | 5 | 0.2 | 14 | 0.2857 | 3 | 0 | 1 | 0 | 4 | 0 | 5 | 0.2 | 13 | 0.0769 |
| 6_8313829 | 3 | 0.3333 | 2 | 0.5 | 3 | 0 | 3 | 0 | 11 | 0.1818 | 3 | 0.3333 | 1 | 0 | 2 | 0 | 3 | 0.3333 | 9 | 0.2222 |
| 6_8313847 | 2 | 0 | 2 | 0.5 | 2 | 0.5 | 4 | 0 | 10 | 0.2 | 2 | 0.5 | NA | NA | 1 | 0 | 3 | 0.3333 | 6 | 0.3333 |
| 6_8313855 | 2 | 1 | 4 | 0.5 | 2 | 1 | 5 | 0.6 | 13 | 0.6923 | 3 | 0.3333 | 1 | 1 | 3 | 0.3333 | 5 | 0.6 | 12 | 0.5 |
| 6_8313861 | 2 | 0 | 4 | 0 | 3 | 0.3333 | 5 | 0 | 14 | 0.0714 | 3 | 0 | 1 | 0 | 4 | 0 | 4 | 0.25 | 12 | 0.0833 |
| 6_8313891 | 4 | 0.25 | 4 | 0 | 3 | 0 | 5 | 0 | 16 | 0.0625 | 1 | 0 | NA | NA | 3 | 0.3333 | 6 | 0.1667 | 10 | 0.2 |
| 6_8313919 | 3 | 1 | 4 | 1 | 2 | 1 | 5 | 1 | 14 | 1 | 4 | 0.25 | NA | NA | 4 | 0.75 | 3 | 1 | 11 | 0.6364 |
| 6_8313932 | 4 | 0.5 | 4 | 0.25 | 3 | 0 | 6 | 0 | 17 | 0.1765 | 4 | 0 | 1 | 0 | 4 | 0 | 6 | 0.1667 | 15 | 0.0667 |
| 6_8313939 | 4 | 0.25 | 4 | 0 | 4 | 0 | 6 | 0 | 18 | 0.0556 | 4 | 0 | 1 | 0 | 3 | 0 | 5 | 0.2 | 13 | 0.0769 |
| 6_8313951 | 3 | 1 | 4 | 1 | 3 | 1 | 6 | 1 | 16 | 1 | 4 | 1 | 2 | 1 | 2 | 1 | 4 | 1 | 12 | 1 |
| 6_8313956 | 4 | 1 | 4 | 0.25 | 3 | 1 | 4 | 0.75 | 15 | 0.7333 | 4 | 0.5 | 2 | 0.5 | 2 | 0 | 6 | 1 | 14 | 0.6429 |
| 6_8313977 | 4 | 0 | 4 | 0 | 4 | 0 | 6 | 0.1667 | 18 | 0.0556 | 4 | 0 | 3 | 0 | 4 | 0.25 | 6 | 0 | 17 | 0.0588 |
| 6_8313989 | 5 | 0.8 | 4 | 0.75 | 5 | 1 | 6 | 0.6667 | 20 | 0.8 | 5 | 1 | 3 | 0.6667 | 2 | 0 | 6 | 1 | 16 | 0.8125 |
| 6_8313992 | 2 | 0.5 | 1 | 1 | 1 | 1 | 3 | 0.6667 | 7 | 0.7143 | NA | NA | 1 | 1 | 1 | 0 | 3 | 0.3333 | 5 | 0.4 |

| | | | | | | | | | | | | | | | | | | | | |
|---|---|---|---|---|---|---|---|---|---|---|---|---|---|---|---|---|---|---|---|---|
| 6_8313993 | NA | NA | 1 | 0 | 2 | 0.5 | 2 | 0 | 5 | 0.2 | NA | NA | 1 | 1 | 1 | 0 | 2 | 0.5 | 4 | 0.5 |
| 6_8313982 | NA | NA | NA | NA | 1 | 1 | NA | NA | 1 | 1 | NA | NA | 1 | 1 | NA | NA | NA | NA | 1 | 1 |
| 6_8315666 | 5 | 0.2 | 2 | 0 | 3 | 0 | 1 | 0 | 11 | 0.0909 | 1 | 0 | 3 | 0.3333 | NA | NA | NA | NA | 4 | 0.25 |
| 6_8317706 | 37 | 0.4054 | 39 | 0.2821 | 38 | 0.3947 | 41 | 0.2439 | 155 | 0.329 | 27 | 0.3333 | 37 | 0.4324 | 38 | 0.4211 | 30 | 0.3667 | 132 | 0.3939 |
| 6_8328285 | 1 | 1 | 1 | 1 | 2 | 0.5 | NA | NA | 4 | 0.75 | NA | NA | 1 | 1 | 1 | 0 | NA | NA | 2 | 0.5 |
| 6_8328340 | 1 | 1 | 1 | 0 | 1 | 0 | NA | NA | 3 | 0.3333 | NA | NA | 3 | 0.3333 | 2 | 0.5 | 1 | 0 | 6 | 0.3333 |
| 6_8330278 | 1 | 1 | NA | NA | NA | NA | 1 | 1 | 2 | 1 | NA | NA | NA | NA | 1 | 1 | NA | NA | 1 | 1 |
| 6_8330285 | 1 | 1 | NA | NA | NA | NA | 1 | 1 | 2 | 1 | NA | NA | NA | NA | 1 | 1 | NA | NA | 1 | 1 |
| 6_8331730 | 1 | 1 | 3 | 1 | 5 | 1 | 2 | 1 | 11 | 1 | 2 | 1 | NA | NA | 2 | 1 | 1 | 1 | 5 | 1 |
| 6_8331755 | 1 | 0 | 3 | 1 | 4 | 0.75 | 5 | 0.6 | 13 | 0.6923 | 2 | 1 | NA | NA | 2 | 1 | 1 | 1 | 5 | 1 |
| 6_8331789 | 1 | 0 | 2 | 1 | 5 | 0 | 4 | 0.25 | 12 | 0.25 | 1 | 0 | NA | NA | 2 | 0.5 | NA | NA | 3 | 0.3333 |
| 6_8333530 | 10 | 0.3 | 6 | 0.5 | 6 | 0.5 | 10 | 0.2 | 32 | 0.3438 | 8 | 0.375 | 10 | 0.3 | 10 | 0.2 | 5 | 0.4 | 33 | 0.303 |
| 6_8333542 | 1 | 0 | 2 | 0.5 | 1 | 0 | 1 | 1 | 5 | 0.4 | NA | NA | 3 | 0.6667 | NA | NA | NA | NA | 3 | 0.6667 |
| 6_8333942 | 4 | 0 | 3 | 0 | 6 | 0.1667 | 4 | 0 | 17 | 0.0588 | 6 | 0 | 3 | 0 | 4 | 0.25 | 2 | 0 | 15 | 0.0667 |
| 6_8333976 | 5 | 0 | 4 | 0.25 | 5 | 0.2 | 5 | 0 | 19 | 0.1053 | 5 | 0.2 | 5 | 0 | 5 | 0 | 3 | 0 | 18 | 0.0556 |
| 6_8333992 | 6 | 0.3333 | 3 | 0.3333 | 3 | 0 | 5 | 0.2 | 17 | 0.2353 | 5 | 0 | 5 | 0.2 | 4 | 0.75 | 3 | 0.3333 | 17 | 0.2941 |
| 6_8334004 | 6 | 0 | 3 | 0 | 3 | 0.3333 | 3 | 0 | 15 | 0.0667 | 4 | 0 | 4 | 0 | 3 | 0.3333 | 2 | 0 | 13 | 0.0769 |
| 6_8334187 | 5 | 0 | 4 | 0.5 | 1 | 1 | 2 | 0 | 12 | 0.25 | 3 | 0.3333 | 3 | 0 | 1 | 1 | 2 | 0.5 | 9 | 0.3333 |
| 6_8334396 | NA | NA | 1 | 1 | NA | NA | 1 | 1 | 2 | 1 | 1 | 0 | 2 | 0.5 | NA | NA | 2 | 1 | 5 | 0.6 |
| 6_8334486 | NA | NA | 1 | 1 | NA | NA | NA | NA | 1 | 1 | 1 | 1 | 1 | 1 | NA | NA | NA | NA | 2 | 1 |
| 6_8334509 | NA | NA | 1 | 1 | NA | NA | NA | NA | 1 | 1 | NA | NA | NA | NA | NA | NA | 1 | 1 | 1 | 1 |
| 6_8334549 | NA | NA | 1 | 1 | NA | NA | NA | NA | 1 | 1 | 1 | 1 | 1 | 1 | NA | NA | 1 | 1 | 3 | 1 |
| 6_8334560 | NA | NA | 1 | 1 | NA | NA | NA | NA | 1 | 1 | NA | NA | 1 | 1 | NA | NA | NA | NA | 1 | 1 |
| 6_8334576 | NA | NA | 1 | 1 | NA | NA | NA | NA | 1 | 1 | 1 | 1 | 1 | 1 | NA | NA | 1 | 1 | 3 | 1 |
| 6_8334593 | NA | NA | 2 | 1 | NA | NA | NA | NA | 2 | 1 | NA | NA | 1 | 1 | NA | NA | 1 | 1 | 2 | 1 |
| 6_8334617 | NA | NA | 1 | 1 | NA | NA | NA | NA | 1 | 1 | NA | NA | NA | NA | NA | NA | 1 | 1 | 1 | 1 |
| 6_8334620 | NA | NA | 1 | 1 | NA | NA | NA | NA | 1 | 1 | NA | NA | NA | NA | NA | NA | 1 | 1 | 1 | 1 |
| 6_8338025 | NA | NA | 2 | 1 | NA | NA | NA | NA | 2 | 1 | NA | NA | NA | NA | NA | NA | 1 | 1 | 1 | 1 |
| 6_8338032 | NA | NA | 1 | 1 | NA | NA | NA | NA | 1 | 1 | NA | NA | NA | NA | NA | NA | 1 | 1 | 1 | 1 |
| 6_8338078 | NA | NA | 1 | 1 | NA | NA | NA | NA | 1 | 1 | NA | NA | NA | NA | NA | NA | 1 | 1 | 1 | 1 |
| 6_8345736 | 2 | 0.5 | 1 | 1 | 5 | 0.2 | 8 | 0 | 16 | 0.1875 | 2 | 0.5 | 8 | 0.25 | NA | NA | 1 | 0 | 11 | 0.2727 |
| 6_8345738 | 1 | 1 | 2 | 0.5 | 5 | 0 | 7 | 0.2857 | 15 | 0.2667 | 2 | 0 | 8 | 0.25 | 1 | 0 | NA | NA | 11 | 0.1818 |
| 6_8345748 | NA | NA | 1 | 1 | NA | NA | 1 | 1 | 2 | 1 | 1 | 0 | 1 | 1 | NA | NA | 1 | 1 | 3 | 0.6667 |
| 6_8345811 | NA | NA | NA | NA | 1 | 0 | 2 | 0.5 | 3 | 0.3333 | 1 | 0 | 3 | 0.3333 | NA | NA | NA | NA | 4 | 0.25 |
| 6_8352040 | 5 | 0.4 | 7 | 0.4286 | 2 | 0 | 5 | 0.4 | 19 | 0.3684 | NA | NA | 4 | 0.25 | 2 | 0.5 | 6 | 0.5 | 12 | 0.4167 |
| 6_8353874 | 5 | 0.2 | 6 | 0.1667 | 4 | 0.25 | 6 | 0.1667 | 21 | 0.1905 | NA | NA | 5 | 0.2 | 3 | 0 | 5 | 0.2 | 13 | 0.1538 |
| 6_8370929 | NA | NA | NA | NA | 1 | 1 | NA | NA | 1 | 1 | NA | NA | NA | NA | 1 | 1 | NA | NA | 1 | 1 |
| 6_8374811 | 4 | 0.5 | 4 | 0.5 | 3 | 0.3333 | 4 | 0.25 | 15 | 0.4 | 2 | 0 | 3 | 0 | 2 | 1 | 2 | 0 | 9 | 0.2222 |
| 6_8375207 | 4 | 0.5 | 3 | 0.3333 | 3 | 0 | 5 | 0.8 | 15 | 0.4667 | NA | NA | 1 | 1 | 4 | 0 | 1 | 0 | 6 | 0.1667 |
| 6_8377395 | 7 | 0 | 2 | 0 | 4 | 0.25 | 9 | 0.2222 | 22 | 0.1364 | 3 | 0 | 5 | 0.4 | 3 | 0 | 5 | 0.2 | 16 | 0.1875 |
| 6_8377939 | 9 | 0.1111 | 3 | 0 | 6 | 0.5 | 10 | 0.4 | 28 | 0.2857 | 6 | 0.3333 | 7 | 0.4286 | 6 | 0.5 | 9 | 0.4444 | 28 | 0.4286 |
| 6_8377948 | 11 | 0 | 4 | 0.25 | 7 | 0 | 12 | 0.0833 | 34 | 0.0588 | 8 | 0.25 | 10 | 0 | 8 | 0 | 10 | 0.1 | 36 | 0.0833 |
| 6_8378335 | 8 | 0.25 | 11 | 0 | 8 | 0 | 5 | 0 | 32 | 0.0625 | 2 | 0 | 8 | 0.125 | 3 | 0 | 4 | 0.25 | 17 | 0.1176 |
| 6_8378579 | NA | NA | 4 | 0.5 | 4 | 0.25 | 2 | 0.5 | 10 | 0.4 | NA | NA | 2 | 0.5 | 1 | 1 | 1 | 1 | 4 | 0.75 |
| 6_8378692 | 1 | 1 | 2 | 0 | 4 | 0.25 | 5 | 0.4 | 12 | 0.3333 | NA | NA | 2 | 0 | 3 | 0.3333 | 1 | 0 | 6 | 0.1667 |
| 6_8390574 | 3 | 1 | NA | NA | 4 | 1 | 3 | 0.6667 | 10 | 0.9 | NA | NA | NA | NA | NA | NA | 5 | 1 | 5 | 1 |
| 6_8390586 | 3 | 0.6667 | NA | NA | 4 | 0.75 | 2 | 1 | 9 | 0.7778 | NA | NA | 1 | 0 | NA | NA | 5 | 0.6 | 6 | 0.5 |
| 6_8390592 | 3 | 0.6667 | NA | NA | 3 | 0.6667 | 2 | 0.5 | 8 | 0.625 | NA | NA | 1 | 1 | NA | NA | 5 | 0.6 | 6 | 0.6667 |
| 6_8390603 | 2 | 0.5 | NA | NA | 3 | 0.3333 | 3 | 0.6667 | 8 | 0.5 | NA | NA | 1 | 0 | NA | NA | 5 | 0.2 | 6 | 0.1667 |
| 6_8390644 | 3 | 0 | NA | NA | 3 | 0.3333 | 2 | 0.5 | 8 | 0.25 | NA | NA | 1 | 0 | 1 | 1 | 6 | 0.1667 | 8 | 0.25 |
| 6_8390665 | 3 | 1 | NA | NA | 4 | 1 | 3 | 1 | 10 | 1 | 2 | 1 | 1 | 1 | 1 | 1 | 7 | 1 | 11 | 1 |
| 6_8390670 | 3 | 0.3333 | NA | NA | 4 | 0.25 | 1 | 0 | 8 | 0.25 | 2 | 0.5 | 1 | 0 | 1 | 0 | 7 | 0.1429 | 11 | 0.1818 |
| 6_8390673 | 2 | 1 | NA | NA | 4 | 1 | 3 | 0.6667 | 9 | 0.8889 | 1 | 1 | 1 | 1 | 2 | 0.5 | 6 | 0.8333 | 10 | 0.8 |
| 6_8390674 | 3 | 0.3333 | NA | NA | 4 | 0.25 | 3 | 0.6667 | 10 | 0.4 | 2 | 0 | 1 | 1 | 2 | 0 | 7 | 0.4286 | 12 | 0.3333 |
| 6_8390682 | 3 | 0.6667 | NA | NA | 4 | 0.25 | 2 | 0.5 | 9 | 0.4444 | 2 | 0.5 | 1 | 0 | 2 | 0.5 | 7 | 0.2857 | 12 | 0.3333 |
| 6_8390692 | 2 | 1 | NA | NA | 4 | 1 | 3 | 0.6667 | 9 | 0.8889 | 2 | 0 | 1 | 1 | 2 | 0.5 | 5 | 1 | 10 | 0.7 |
| 6_8390696 | 2 | 0 | NA | NA | 4 | 0 | 2 | 0.5 | 8 | 0.125 | 2 | 0 | 1 | 0 | 3 | 0 | 7 | 0.1429 | 13 | 0.0769 |
| 6_8390704 | 3 | 1 | NA | NA | 4 | 0.75 | 3 | 0.6667 | 10 | 0.8 | 2 | 1 | 1 | 1 | 3 | 1 | 5 | 0.8 | 11 | 0.9091 |
| 6_8395336 | 4 | 0.25 | 3 | 0.6667 | 6 | 0.3333 | 5 | 0.2 | 18 | 0.3333 | 2 | 0 | 1 | 1 | 5 | 0.2 | 13 | 0.3077 | 21 | 0.2857 |
| 6_8395339 | 1 | 0 | 3 | 0.3333 | 2 | 0.5 | 4 | 0.25 | 10 | 0.3 | 2 | 0 | 1 | 1 | 5 | 0.2 | 10 | 0.4 | 18 | 0.3333 |
| 6_8395351 | 3 | 1 | 3 | 1 | 5 | 0.8 | 5 | 0.6 | 16 | 0.8125 | 2 | 1 | 1 | 1 | 6 | 1 | 13 | 0.9231 | 22 | 0.9545 |
| 6_8395355 | 4 | 0 | 3 | 0 | 6 | 0.1667 | 4 | 0 | 17 | 0.0588 | 2 | 0 | 2 | 0 | 6 | 0.3333 | 15 | 0.0667 | 25 | 0.12 |
| 6_8395370 | 5 | 0.4 | 3 | 0.6667 | 6 | 0.6667 | 6 | 0.5 | 20 | 0.55 | 1 | 0 | 2 | 0.5 | 3 | 0 | 11 | 0.5455 | 17 | 0.4118 |
| 6_8395401 | 5 | 0.4 | 3 | 0 | 7 | 0.5714 | 5 | 0.6 | 20 | 0.45 | 2 | 0.5 | 1 | 0 | 6 | 0.6667 | 14 | 0.6429 | 23 | 0.6087 |
| 6_8395404 | 3 | 1 | 3 | 1 | 7 | 0.7143 | 5 | 0.8 | 18 | 0.8333 | 2 | 1 | 2 | 1 | 6 | 0.8333 | 14 | 1 | 24 | 0.9583 |

| | | | | | | | | | | | | | | | | | | | | |
|---|---|---|---|---|---|---|---|---|---|---|---|---|---|---|---|---|---|---|---|---|
| 6_8395412 | 2 | 1 | 2 | 0.5 | 5 | 0.8 | 5 | 0.6 | 14 | 0.7143 | 1 | 0 | 1 | 1 | 3 | 0.3333 | 9 | 0.5556 | 14 | 0.5 |
| 6_8395434 | 4 | 0.25 | 3 | 0.3333 | 7 | 0.1429 | 6 | 0.3333 | 20 | 0.25 | 2 | 0.5 | 2 | 0.5 | 6 | 0 | 13 | 0.3077 | 23 | 0.2609 |
| 6_8395611 | 4 | 0.5 | 3 | 1 | 9 | 0.4444 | 6 | 0.5 | 22 | 0.5455 | 1 | 0 | 1 | 0 | 6 | 1 | 12 | 0.9167 | 20 | 0.85 |
| 6_8395631 | 6 | 0.5 | 1 | 0 | 9 | 0.5556 | 7 | 0.4286 | 23 | 0.4783 | 2 | 0 | 3 | 0.6667 | 7 | 0.7143 | 14 | 0.3571 | 26 | 0.4615 |
| 6_8395635 | 5 | 0.2 | 3 | 0 | 7 | 0.2857 | 6 | 0.3333 | 21 | 0.2381 | 2 | 0 | 2 | 0 | 6 | 0.6667 | 14 | 0.2857 | 24 | 0.3333 |
| 6_8395641 | 5 | 0 | 2 | 0 | 8 | 0.25 | 6 | 0 | 21 | 0.0952 | 2 | 0.5 | 3 | 0 | 7 | 0 | 13 | 0.0769 | 25 | 0.08 |
| 6_8395645 | 6 | 0 | 3 | 0 | 9 | 0 | 6 | 0.3333 | 24 | 0.0833 | 2 | 0 | 3 | 0 | 6 | 0 | 15 | 0.2 | 26 | 0.1154 |
| 6_8395659 | 6 | 0.8333 | 2 | 0.5 | 8 | 0.5 | 7 | 0.8571 | 23 | 0.6957 | 3 | 0 | 5 | 0.6 | 7 | 0.2857 | 15 | 0.5333 | 30 | 0.4333 |
| 6_8395662 | 8 | 0.125 | 3 | 0 | 7 | 0.1429 | 7 | 0.1429 | 25 | 0.12 | 3 | 0 | 3 | 0 | 7 | 0.1429 | 16 | 0.0625 | 29 | 0.069 |
| 6_8395679 | 8 | 0 | 3 | 0 | 10 | 0.1 | 6 | 0.1667 | 27 | 0.0741 | 3 | 0 | 5 | 0 | 9 | 0.1111 | 17 | 0.1176 | 34 | 0.0882 |
| 6_8395684 | 8 | 0.125 | 4 | 0 | 8 | 0.25 | 7 | 0 | 27 | 0.1111 | 3 | 0 | 5 | 0.4 | 9 | 0.1111 | 16 | 0.1875 | 33 | 0.1818 |
| 6_8395688 | 8 | 0.375 | 4 | 0.25 | 10 | 0.1 | 7 | 0.5714 | 29 | 0.3103 | 3 | 0 | 6 | 0.1667 | 10 | 0.1 | 14 | 0.1429 | 33 | 0.1212 |
| 6_8459560 | 5 | 0 | 9 | 0.2222 | 7 | 0.1429 | 9 | 0 | 30 | 0.1 | 4 | 0 | 4 | 0 | 1 | 0 | 4 | 0.25 | 13 | 0.0769 |
| 6_8459562 | 7 | 0.1429 | 17 | 0.0588 | 8 | 0 | 8 | 0 | 40 | 0.05 | 6 | 0 | 4 | 0.25 | 2 | 0 | 6 | 0 | 18 | 0.0556 |
| 6_8459591 | 7 | 0.5714 | 16 | 0.75 | 8 | 0.5 | 9 | 0.6667 | 40 | 0.65 | 6 | 0.1667 | 3 | 0.6667 | 2 | 0 | 6 | 0.6667 | 17 | 0.4118 |
| 6_8459596 | 7 | 0.1429 | 17 | 0.0588 | 8 | 0.125 | 9 | 0.1111 | 41 | 0.0976 | 6 | 0 | 4 | 0 | 2 | 0.5 | 7 | 0.2857 | 19 | 0.1579 |
| 6_8459609 | 6 | 0 | 17 | 0.1765 | 8 | 0 | 7 | 0 | 38 | 0.0789 | 5 | 0 | 3 | 0.3333 | 1 | 0 | 6 | 0 | 15 | 0.0667 |
| 6_8459618 | 6 | 0.5 | 16 | 0 | 7 | 0.1429 | 8 | 0.125 | 37 | 0.1351 | 6 | 0.1667 | 3 | 0.3333 | 2 | 0 | 5 | 0 | 16 | 0.125 |
| 6_8459620 | 5 | 0 | 13 | 0.0769 | 6 | 0.1667 | 7 | 0 | 31 | 0.0645 | 4 | 0 | 4 | 0 | 1 | 0 | 4 | 0.25 | 13 | 0.0769 |
| 6_8459620 | 6 | 1 | 15 | 0.8667 | 6 | 1 | 7 | 0.7143 | 34 | 0.8824 | 5 | 0.6 | 3 | 0.3333 | 2 | 0 | 6 | 0.8333 | 16 | 0.5625 |
| 6_8459624 | 5 | 0.6 | 14 | 0.5 | 4 | 0.75 | 7 | 0.5714 | 30 | 0.5667 | 6 | 0.5 | 4 | 0 | 2 | 0 | 5 | 0.4 | 17 | 0.2941 |
| 6_8459629 | 4 | 1 | 13 | 1 | 5 | 0.8 | 8 | 0.875 | 30 | 0.9333 | 6 | 0.8333 | 3 | 1 | 2 | 1 | 5 | 1 | 16 | 0.9375 |
| 6_8459636 | 5 | 0 | 15 | 0 | 5 | 0.2 | 7 | 0.1429 | 32 | 0.0625 | 4 | 0 | 4 | 0 | 2 | 0.5 | 4 | 0 | 14 | 0.0714 |
| 6_8459644 | 4 | 1 | 14 | 0.7857 | 5 | 1 | 7 | 0.8571 | 30 | 0.8667 | 5 | 1 | 3 | 0.6667 | 1 | 1 | 3 | 1 | 12 | 0.9167 |
| 6_8459648 | 5 | 0.2 | 13 | 0.3846 | 5 | 0.6 | 7 | 0.4286 | 30 | 0.4 | 4 | 0.75 | 2 | 0 | 1 | 0 | 5 | 0.2 | 12 | 0.3333 |
| 6_8459652 | 5 | 1 | 11 | 0.9091 | 4 | 1 | 7 | 0.8571 | 27 | 0.9259 | 4 | 1 | 2 | 1 | 1 | 1 | 4 | 0.75 | 11 | 0.9091 |
| 6_8459657 | 4 | 0.75 | 13 | 0.7692 | 5 | 1 | 7 | 1 | 29 | 0.8621 | 5 | 1 | 3 | 0.3333 | 1 | 0 | 4 | 0.75 | 13 | 0.6923 |
| 6_8459666 | 5 | 0.8 | 9 | 0.6667 | 4 | 0.75 | 4 | 0.75 | 22 | 0.7273 | 3 | 0.6667 | 2 | 1 | 1 | 1 | 2 | 1 | 8 | 0.875 |
| 6_8459667 | 5 | 0 | 12 | 0.25 | NA | NA | 5 | 0.2 | 22 | 0.1818 | 4 | 0 | 1 | 0 | 1 | 0 | 2 | 0.5 | 8 | 0.125 |
| 6_8459668 | 5 | 0.4 | 12 | 0.5 | 4 | 0 | 7 | 0.4286 | 28 | 0.3929 | 4 | 0.5 | 3 | 0.6667 | 2 | 0.5 | 4 | 0 | 13 | 0.3846 |
| 6_8459700 | 4 | 0 | 13 | 0.1538 | 5 | 0.4 | 7 | 0 | 29 | 0.1379 | 4 | 0 | 2 | 0.5 | 1 | 0 | 4 | 0.5 | 11 | 0.2727 |
| 6_8459705 | 5 | 1 | 12 | 1 | 5 | 1 | 8 | 0.875 | 30 | 0.9667 | 4 | 1 | 2 | 1 | 2 | 1 | 4 | 0.75 | 12 | 0.9167 |
| 6_8459710 | 3 | 0 | 12 | 0.3333 | 3 | 0.3333 | 8 | 0.125 | 26 | 0.2308 | 2 | 0.5 | 2 | 0 | NA | NA | 4 | 0 | 8 | 0.125 |
| 6_8459714 | 4 | 0 | 13 | 0.1538 | 4 | 0.25 | 7 | 0.1429 | 28 | 0.1429 | 5 | 0.4 | 2 | 0 | NA | NA | 4 | 0 | 11 | 0.1818 |
| 6_8459716 | 5 | 0.2 | 11 | 0.3636 | 3 | 0.3333 | 8 | 0.125 | 27 | 0.2593 | 4 | 0.25 | 2 | 0.5 | 1 | 0 | 3 | 0.3333 | 10 | 0.3 |
| 6_8459717 | 5 | 1 | 13 | 0.7692 | 5 | 1 | 8 | 0.875 | 31 | 0.871 | 5 | 1 | 2 | 1 | 2 | 1 | 4 | 1 | 13 | 1 |
| 6_8459722 | 4 | 0.75 | 12 | 0.6667 | 5 | 0.8 | 7 | 0.8571 | 28 | 0.75 | 4 | 1 | NA | NA | 1 | 1 | 3 | 1 | 8 | 1 |
| 6_8459734 | 5 | 0 | 9 | 0.3333 | 5 | 0.4 | 8 | 0.5 | 27 | 0.3333 | 5 | 0 | 2 | 0.5 | 2 | 0 | 3 | 0 | 12 | 0.0833 |
| 6_8459745 | 4 | 0.5 | 12 | 0.75 | 3 | 0.3333 | 6 | 1 | 25 | 0.72 | 4 | 1 | 2 | 0.5 | 2 | 0.5 | 3 | 1 | 11 | 0.8182 |
| 6_8459839 | 3 | 0 | 10 | 0.3 | 4 | 0.25 | 6 | 0 | 23 | 0.1739 | 3 | 0.6667 | 2 | 0 | 1 | 0 | 3 | 0 | 9 | 0.2222 |
| 6_8511075 | 10 | 0.1 | 9 | 0.1111 | 6 | 0 | 10 | 0.4 | 35 | 0.1714 | 6 | 0.1667 | 6 | 0 | 4 | 0 | 3 | 0.3333 | 19 | 0.1053 |
| 6_8511108 | 10 | 0.1 | 6 | 0.3333 | 7 | 0.2857 | 9 | 0.4444 | 32 | 0.2812 | 6 | 0.1667 | 5 | 0.2 | 4 | 0.25 | 5 | 0.4 | 20 | 0.25 |
| 6_8511109 | 10 | 0.5 | 8 | 0.625 | 7 | 0.8571 | 8 | 0.5 | 33 | 0.6061 | 6 | 0.5 | 5 | 0.4 | 4 | 0.5 | 6 | 0.5 | 21 | 0.4762 |
| 6_8511112 | 10 | 0.5 | 8 | 0.375 | 5 | 1 | 9 | 0.4444 | 32 | 0.5312 | 6 | 0.5 | 5 | 0.2 | 3 | 0.3333 | 5 | 0.6 | 19 | 0.4211 |
| 6_8511116 | 10 | 0.4 | 8 | 0.125 | 4 | 0 | 7 | 0.4286 | 29 | 0.2759 | 5 | 0.2 | 3 | 0.3333 | 4 | 0.5 | 4 | 0 | 16 | 0.25 |
| 6_8511116 | 9 | 0.4444 | 7 | 0.2857 | 6 | 0.3333 | 9 | 0.6667 | 31 | 0.4516 | 6 | 0.3333 | 5 | 0.4 | 4 | 0.75 | 5 | 0.2 | 20 | 0.4 |
| 6_8511394 | 7 | 0 | 6 | 0.5 | 4 | 0.25 | 9 | 0.3333 | 26 | 0.2692 | 5 | 0.2 | 3 | 0.3333 | 4 | 0.5 | 6 | 0.1667 | 18 | 0.2778 |
| 6_8518103 | 1 | 0 | 4 | 0.75 | 1 | 1 | 1 | 1 | 7 | 0.7143 | 1 | 1 | NA | NA | NA | NA | 1 | 0 | 2 | 0.5 |
| 6_8518113 | 1 | 1 | 3 | 0.6667 | 1 | 1 | 1 | 0 | 6 | 0.6667 | 1 | 1 | NA | NA | NA | NA | 1 | 1 | 2 | 1 |
| 6_8518121 | 1 | 1 | 3 | 1 | 1 | 1 | 1 | 1 | 6 | 1 | 1 | 1 | NA | NA | 1 | 1 | 1 | 0 | 3 | 0.6667 |
| 6_8518124 | 1 | 1 | 3 | 0.3333 | 1 | 1 | 1 | 0 | 6 | 0.5 | 1 | 1 | NA | NA | 1 | 0 | 1 | 0 | 3 | 0.3333 |
| 6_8519011 | 19 | 0.1053 | 19 | 0.1579 | 17 | 0.1176 | 14 | 0 | 69 | 0.1014 | 18 | 0.1667 | 16 | 0.0625 | 17 | 0.0588 | 10 | 0.1 | 61 | 0.0984 |
| 6_8519012 | 18 | 0.2778 | 16 | 0 | 13 | 0.1538 | 12 | 0.0833 | 59 | 0.1356 | 16 | 0.25 | 14 | 0.1429 | 15 | 0.1333 | 10 | 0 | 55 | 0.1455 |
| 6_8519039 | 12 | 0.3333 | 13 | 0.1538 | 6 | 0.1667 | 8 | 0.125 | 39 | 0.2051 | 13 | 0.3077 | 12 | 0.25 | 7 | 0 | 5 | 0 | 37 | 0.1892 |
| 6_8519084 | 10 | 0.2 | 14 | 0 | 7 | 0 | 4 | 0 | 35 | 0.0571 | 14 | 0.0714 | 9 | 0 | 11 | 0.0909 | 4 | 0 | 38 | 0.0526 |
| 6_8519096 | 10 | 0.4 | 11 | 0.3636 | 8 | 0.75 | 3 | 0 | 32 | 0.4375 | 14 | 0.2857 | 11 | 0.0909 | 12 | 0.3333 | 6 | 0.5 | 43 | 0.2791 |
| 6_8519105 | 7 | 0.1429 | 11 | 0.0909 | 8 | 0.25 | 3 | 0 | 29 | 0.1379 | 11 | 0.1818 | 11 | 0.1818 | 12 | 0.0833 | 6 | 0.3333 | 40 | 0.175 |
| 6_8519122 | 6 | 0 | 14 | 0.1429 | 8 | 0 | 3 | 0 | 31 | 0.0645 | 11 | 0 | 10 | 0.1 | 12 | 0.0833 | 6 | 0 | 39 | 0.0513 |
| 6_8519133 | 9 | 0.1111 | 14 | 0.4286 | 7 | 0.4286 | 2 | 0.5 | 32 | 0.3438 | 12 | 0.5 | 10 | 0.2 | 12 | 0.3333 | 5 | 0.2 | 39 | 0.3333 |
| 6_8519142 | 6 | 0.5 | 10 | 0.6 | 7 | 0.5714 | 2 | 0 | 25 | 0.52 | 9 | 0.2222 | 9 | 0.5556 | 8 | 0.75 | 3 | 0.3333 | 29 | 0.4828 |
| 6_8519143 | 4 | 0.25 | 8 | 0.125 | 5 | 0 | 3 | 0.3333 | 20 | 0.15 | 5 | 0.4 | 8 | 0.125 | 7 | 0.2857 | 2 | 0 | 22 | 0.2273 |
| 6_8519145 | 8 | 1 | 12 | 0.9167 | 7 | 1 | 2 | 1 | 29 | 0.9655 | 11 | 0.8182 | 9 | 1 | 10 | 0.8 | 4 | 0.5 | 34 | 0.8235 |
| 6_8519152 | 8 | 0.75 | 9 | 1 | 5 | 0.8 | 1 | 1 | 23 | 0.8696 | 12 | 0.9167 | 9 | 0.7778 | 9 | 0.7778 | 5 | 1 | 35 | 0.8571 |
| 6_8519161 | 6 | 0.3333 | 7 | 0.4286 | 4 | 0 | 1 | 0 | 18 | 0.2778 | 7 | 0.7143 | 7 | 0.5714 | 6 | 0.3333 | 3 | 0 | 23 | 0.4783 |
| 6_8519161 | 8 | 0.75 | 8 | 0.625 | 5 | 0.8 | 3 | 0.6667 | 24 | 0.7083 | 10 | 0.7 | 9 | 0.8889 | 7 | 1 | 4 | 1 | 30 | 0.8667 |

| | | | | | | | | | | | | | | | | | | | | |
|---|---|---|---|---|---|---|---|---|---|---|---|---|---|---|---|---|---|---|---|---|
| 6_8519166 | 8 | 0.125 | 9 | 0.1111 | 7 | 0 | 3 | 0.3333 | 27 | 0.1111 | 12 | 0.1667 | 11 | 0.1818 | 7 | 0.1429 | 3 | 0.6667 | 33 | 0.2121 |
| 6_8519238 | 6 | 0.1667 | 6 | 0 | 2 | 0 | 2 | 0 | 16 | 0.0625 | 5 | 0.2 | 9 | 0.3333 | 5 | 0.2 | 2 | 0 | 21 | 0.2381 |
| 6_8530915 | NA | NA | NA | NA | 1 | 1 | NA | NA | 1 | 1 | NA | NA | NA | NA | NA | NA | 1 | 1 | 1 | 1 |
| 6_8530935 | NA | NA | NA | NA | 1 | 1 | NA | NA | 1 | 1 | NA | NA | NA | NA | NA | NA | 1 | 1 | 1 | 1 |
| 6_8530946 | NA | NA | NA | NA | 1 | 1 | NA | NA | 1 | 1 | NA | NA | NA | NA | NA | NA | 1 | 1 | 1 | 1 |
| 6_8530987 | NA | NA | NA | NA | 1 | 1 | NA | NA | 1 | 1 | NA | NA | NA | NA | NA | NA | 1 | 1 | 1 | 1 |
| 6_8531391 | 7 | 0.1429 | 2 | 0.5 | 2 | 0.5 | 5 | 0.2 | 16 | 0.25 | 6 | 0.3333 | 5 | 0.2 | 6 | 0.1667 | 2 | 0 | 19 | 0.2105 |
| 6_8531395 | 5 | 1 | 1 | 1 | 2 | 1 | 5 | 0.8 | 13 | 0.9231 | 4 | 0.25 | 5 | 0.6 | 6 | 1 | 2 | 1 | 17 | 0.7059 |
| 6_8531397 | 5 | 0.4 | 2 | 0.5 | 1 | 1 | 6 | 0.5 | 14 | 0.5 | 6 | 0.6667 | 5 | 0.8 | 7 | 0.5714 | 2 | 0.5 | 20 | 0.65 |
| 6_8531409 | 6 | 0 | 2 | 0 | 1 | 0 | 5 | 0.2 | 14 | 0.0714 | 5 | 0.2 | 4 | 0.25 | 6 | 0 | 2 | 0.5 | 17 | 0.1765 |
| 6_8531876 | 1 | 0 | NA | NA | NA | NA | 1 | 1 | 2 | 0.5 | 1 | 0 | 2 | 0 | 1 | 1 | NA | NA | 4 | 0.25 |
| 6_8531877 | 1 | 1 | NA | NA | NA | NA | 1 | 1 | 2 | 1 | 2 | 1 | 2 | 1 | 1 | 1 | 1 | 1 | 6 | 1 |
| 6_8532510 | 3 | 0.6667 | 1 | 0 | 1 | 0 | 3 | 1 | 8 | 0.625 | 2 | 0 | 3 | 0.3333 | 3 | 1 | NA | NA | 8 | 0.5 |
| 6_8532539 | 3 | 0 | 1 | 1 | 1 | 0 | 3 | 0.3333 | 8 | 0.25 | 1 | 0 | 2 | 0 | 2 | 0.5 | NA | NA | 5 | 0.2 |
| 6_8532543 | 4 | 0.5 | 1 | 0 | 1 | 1 | 3 | 0.3333 | 9 | 0.4444 | 2 | 0 | 2 | 0 | 3 | 0.6667 | NA | NA | 7 | 0.2857 |
| 6_8532582 | 4 | 0.25 | NA | NA | 1 | 1 | 1 | 1 | 6 | 0.5 | 2 | 0.5 | 1 | 0 | 4 | 0.25 | NA | NA | 7 | 0.2857 |
| 6_8532585 | 3 | 0.3333 | NA | NA | 1 | 0 | 1 | 0 | 5 | 0.2 | 1 | 0 | 1 | 0 | 3 | 0.3333 | NA | NA | 5 | 0.2 |
| 6_8542235 | 6 | 0.3333 | 2 | 0 | 4 | 0.75 | 6 | 0.1667 | 18 | 0.3333 | 2 | 0 | 4 | 0.5 | 5 | 0.4 | 1 | 1 | 12 | 0.4167 |
| 6_8542237 | 6 | 0.1667 | 2 | 0 | 4 | 0 | 5 | 0 | 17 | 0.0588 | 3 | 0 | 4 | 0.25 | 4 | 0 | 2 | 0 | 13 | 0.0769 |
| 6_8542243 | 5 | 0.6 | 2 | 1 | 4 | 0.75 | 5 | 1 | 16 | 0.8125 | 2 | 1 | 4 | 0.75 | 3 | 0.6667 | 2 | 1 | 11 | 0.8182 |
| 6_8542256 | 6 | 0.8333 | 2 | 1 | 5 | 1 | 4 | 0.75 | 17 | 0.8824 | 3 | 0.6667 | 3 | 1 | 5 | 1 | 2 | 0.5 | 13 | 0.8462 |
| 6_8542273 | 6 | 0.8333 | 1 | 1 | 4 | 0.5 | 6 | 0.8333 | 17 | 0.7647 | 3 | 1 | 2 | 0.5 | 6 | 0.8333 | 2 | 1 | 13 | 0.8462 |
| 6_8542276 | 3 | 1 | 2 | 1 | 5 | 0.6 | 5 | 1 | 15 | 0.8667 | 3 | 1 | 2 | 1 | 5 | 1 | 2 | 1 | 12 | 1 |
| 6_8542296 | 5 | 0.2 | 2 | 0 | 5 | 0.2 | 6 | 0 | 18 | 0.1111 | 3 | 0 | 4 | 0 | 6 | 0 | 3 | 0.3333 | 16 | 0.0625 |
| 6_8542313 | 7 | 1 | 2 | 0.5 | 5 | 1 | 7 | 0.8571 | 21 | 0.9048 | 3 | 0.6667 | 4 | 0.75 | 6 | 1 | 2 | 1 | 15 | 0.8667 |
| 6_8542325 | 5 | 0.4 | 3 | 0.3333 | 5 | 0.6 | 6 | 0.1667 | 19 | 0.3684 | 3 | 0 | 4 | 0 | 6 | 0.3333 | 2 | 0 | 15 | 0.1333 |
| 6_8542330 | 5 | 0 | 1 | 0 | 5 | 0.2 | 8 | 0.5 | 19 | 0.2632 | 3 | 0.3333 | 3 | 0 | 5 | 0 | 1 | 1 | 12 | 0.1667 |
| 6_8542383 | 2 | 1 | 2 | 1 | 6 | 0.5 | 2 | 0.5 | 12 | 0.6667 | NA | NA | 3 | 0.6667 | NA | NA | 2 | 0 | 5 | 0.4 |
| 6_8542397 | 3 | 0 | 3 | 0 | 6 | 0 | 2 | 0.5 | 14 | 0.0714 | NA | NA | 3 | 0 | NA | NA | 1 | 1 | 4 | 0.25 |
| 6_8542400 | 3 | 1 | 3 | 0.6667 | 4 | 0.75 | 2 | 1 | 12 | 0.8333 | NA | NA | 3 | 0.6667 | NA | NA | 2 | 0.5 | 5 | 0.6 |
| 6_8542412 | 3 | 0 | 3 | 0 | 5 | 0.2 | 2 | 0.5 | 13 | 0.1538 | NA | NA | 2 | 0 | NA | NA | 1 | 1 | 3 | 0.3333 |
| 6_8542417 | 2 | 0.5 | 2 | 0 | 5 | 1 | 1 | 1 | 10 | 0.7 | NA | NA | 1 | 1 | NA | NA | 1 | 0 | 2 | 0.5 |
| 6_8544517 | 38 | 0.1053 | 37 | 0.0811 | 33 | 0.0909 | 30 | 0.1667 | 138 | 0.1087 | 37 | 0.1081 | 30 | 0.0667 | 30 | 0.1667 | 35 | 0.1714 | 132 | 0.1288 |
| 6_8544678 | 9 | 0.1111 | 5 | 0.4 | 8 | 0 | 12 | 0.3333 | 34 | 0.2059 | 4 | 0 | 10 | 0.3 | 19 | 0.0526 | 9 | 0.2222 | 42 | 0.1429 |
| 6_8544678 | 7 | 0 | 4 | 0 | 5 | 0.2 | 13 | 0.0769 | 29 | 0.069 | 2 | 0 | 7 | 0 | 17 | 0 | 7 | 0.2857 | 33 | 0.0606 |
| 6_8544689 | 6 | 0 | 2 | 0 | 7 | 0.1429 | 12 | 0.0833 | 27 | 0.0741 | 4 | 0 | 9 | 0.3333 | 18 | 0.1667 | 7 | 0.1429 | 38 | 0.1842 |
| 6_8544826 | 2 | 0 | 2 | 0 | 3 | 0 | 8 | 0.125 | 15 | 0.0667 | 3 | 0 | 3 | 0 | 10 | 0 | 4 | 0.5 | 20 | 0.1 |
| 6_8544879 | 1 | 0 | 1 | 0 | 2 | 0.5 | 7 | 0.1429 | 11 | 0.1818 | 2 | 0 | 2 | 0.5 | 8 | 0.25 | 2 | 0.5 | 14 | 0.2857 |
| 6_8544925 | 4 | 0 | 2 | 0.5 | 4 | 0 | 7 | 0.1429 | 17 | 0.1176 | 2 | 0.5 | 2 | 0 | 10 | 0.2 | 5 | 0.2 | 19 | 0.2105 |
| 6_8544960 | 5 | 0.4 | 2 | 1 | 2 | 1 | 6 | 0.6667 | 15 | 0.6667 | 2 | 1 | 3 | 0.6667 | 7 | 1 | 2 | 0.5 | 14 | 0.8571 |
| 6_8544981 | 6 | 0.6667 | 2 | 0.5 | 2 | 0.5 | 5 | 0.6 | 15 | 0.6 | NA | NA | 3 | 0.6667 | 8 | 0.5 | 4 | 0.75 | 15 | 0.6 |
| 6_8544984 | 6 | 0.1667 | 2 | 0 | 2 | 0 | 6 | 0.3333 | 16 | 0.1875 | NA | NA | 3 | 0 | 7 | 0.2857 | 4 | 0.25 | 14 | 0.2143 |
| 6_8544985 | 6 | 0.1667 | 2 | 0.5 | 1 | 0 | 6 | 0.1667 | 15 | 0.2 | NA | NA | 3 | 0.3333 | 7 | 0.2857 | 4 | 0.25 | 14 | 0.2857 |
| 6_8544986 | 5 | 0 | 2 | 1 | 2 | 0.5 | 5 | 0.6 | 14 | 0.4286 | NA | NA | 2 | 0.5 | 8 | 0.25 | 3 | 0 | 13 | 0.2308 |
| 6_8544989 | 4 | 0.25 | 2 | 0 | 2 | 1 | 6 | 0.5 | 14 | 0.4286 | NA | NA | 2 | 0.5 | 7 | 0.2857 | 3 | 0.3333 | 12 | 0.3333 |
| 6_8544993 | 3 | 0.6667 | 2 | 1 | 2 | 1 | 6 | 0.8333 | 13 | 0.8462 | NA | NA | 3 | 1 | 7 | 1 | 4 | 1 | 14 | 1 |
| 6_8544995 | 4 | 0.5 | 2 | 0.5 | 2 | 0.5 | 4 | 0.25 | 12 | 0.4167 | NA | NA | 3 | 0.6667 | 7 | 0.5714 | 4 | 0.75 | 14 | 0.6429 |
| 6_8545687 | NA | NA | 2 | 0.5 | 1 | 1 | 3 | 0 | 6 | 0.3333 | NA | NA | 2 | 0 | 2 | 0.5 | 2 | 0 | 6 | 0.1667 |
| 6_8546087 | 1 | 1 | 1 | 0 | 1 | 0 | 4 | 0 | 7 | 0.1429 | NA | NA | 2 | 0 | 2 | 0.5 | 1 | 0 | 5 | 0.2 |
| 6_8546142 | NA | NA | 1 | 1 | 1 | 1 | 1 | 0 | 3 | 0.6667 | NA | NA | 1 | 1 | 1 | 1 | NA | NA | 2 | 1 |
| 6_8546182 | 1 | 0 | 1 | 0 | 1 | 1 | 1 | 0 | 4 | 0.25 | NA | NA | 1 | 1 | 2 | 0 | 1 | 0 | 4 | 0.25 |
| 6_8548814 | 4 | 0 | 3 | 0.6667 | 7 | 0.1429 | NA | NA | 14 | 0.2143 | 4 | 0 | 7 | 0.1429 | 7 | 0 | 1 | 0 | 19 | 0.0526 |
| 6_8548824 | 9 | 0.8889 | 5 | 0.8 | 8 | 0.875 | NA | NA | 22 | 0.8636 | 7 | 0.4286 | 6 | 0.3333 | 7 | 0.7143 | 2 | 1 | 22 | 0.5455 |
| 6_8548834 | 7 | 0.8571 | 5 | 0.8 | 8 | 0.875 | NA | NA | 20 | 0.85 | 5 | 0.8 | 6 | 1 | 7 | 1 | 2 | 1 | 20 | 0.95 |
| 6_8548839 | 5 | 0.8 | 3 | 0.6667 | 4 | 0.75 | NA | NA | 12 | 0.75 | 4 | 0.75 | 6 | 0.8333 | 9 | 0.6667 | 2 | 0.5 | 21 | 0.7143 |
| 6_8548910 | 6 | 0.6667 | 4 | 0.75 | 5 | 0.6 | NA | NA | 15 | 0.6667 | 4 | 0.5 | 5 | 0.2 | 6 | 0.5 | 2 | 0.5 | 17 | 0.4118 |
| 6_8548923 | 5 | 0.2 | 2 | 0 | 6 | 0 | NA | NA | 13 | 0.0769 | 4 | 0 | 5 | 0.2 | 5 | 0.2 | 1 | 0 | 15 | 0.1333 |
| 6_8548925 | 5 | 1 | 2 | 1 | 5 | 0.8 | NA | NA | 12 | 0.9167 | 3 | 1 | 4 | 0.75 | 5 | 0.8 | 2 | 1 | 14 | 0.8571 |
| 6_8548929 | 4 | 0.5 | 3 | 0.3333 | 6 | 0.6667 | NA | NA | 13 | 0.5385 | 4 | 0.75 | 5 | 0.4 | 5 | 0.8 | 2 | 0 | 16 | 0.5625 |
| 6_8548934 | 5 | 0 | 4 | 0 | 6 | 0.1667 | NA | NA | 15 | 0.0667 | 4 | 0.5 | 5 | 0.4 | 6 | 0 | 2 | 0 | 17 | 0.2353 |
| 6_8548935 | 5 | 0.4 | 4 | 0.25 | 6 | 0.1667 | NA | NA | 15 | 0.2667 | 4 | 0 | 5 | 0 | 6 | 0.1667 | 2 | 0.5 | 17 | 0.1176 |
| 6_8548941 | 6 | 0.5 | 2 | 0.5 | 5 | 0.8 | NA | NA | 13 | 0.6154 | 4 | 0.75 | 5 | 0.6 | 5 | 0.4 | 2 | 0.5 | 16 | 0.5625 |
| 6_8548964 | 5 | 0.2 | 3 | 0.6667 | 6 | 0 | NA | NA | 14 | 0.2143 | 3 | 0.3333 | 5 | 0.2 | 5 | 0.4 | 1 | 0 | 14 | 0.2857 |
| 6_8573778 | 1 | 1 | NA | NA | NA | NA | NA | NA | 1 | 1 | NA | NA | 1 | 1 | NA | NA | 1 | 1 | 2 | 1 |
| 6_8573817 | 1 | 1 | NA | NA | NA | NA | NA | NA | 1 | 1 | NA | NA | NA | NA | NA | NA | 1 | 1 | 1 | 1 |

| | | | | | | | | | | | | | | | | | | | | |
|---|---|---|---|---|---|---|---|---|---|---|---|---|---|---|---|---|---|---|---|---|
| 6_8573828 | 1 | 1 | NA | NA | NA | NA | NA | NA | 1 | 1 | NA | NA | 1 | 1 | NA | NA | 1 | 1 | 2 | 1 |
| 6_8573849 | 1 | 1 | NA | NA | NA | NA | NA | NA | 1 | 1 | NA | NA | 1 | 1 | NA | NA | 1 | 1 | 2 | 1 |
| 6_8579470 | 1 | 1 | 1 | 1 | 2 | 1 | NA | NA | 4 | 1 | NA | NA | 1 | 1 | NA | NA | 2 | 0.5 | 3 | 0.6667 |
| 6_8579491 | NA | NA | 1 | 1 | 2 | 1 | NA | NA | 3 | 1 | NA | NA | 1 | 0 | NA | NA | 2 | 0.5 | 3 | 0.3333 |
| 6_8579492 | 1 | 1 | NA | NA | 1 | 1 | NA | NA | 2 | 1 | NA | NA | 1 | 1 | NA | NA | 2 | 0 | 3 | 0.3333 |
| 6_8580763 | NA | NA | 1 | 0 | NA | NA | 1 | 1 | 2 | 0.5 | 1 | 1 | NA | NA | NA | NA | NA | NA | 1 | 1 |
| 6_8580765 | NA | NA | NA | NA | NA | NA | 1 | 1 | 1 | 1 | 1 | 1 | NA | NA | NA | NA | NA | NA | 1 | 1 |
| 6_8580785 | 1 | 1 | 1 | 0 | NA | NA | NA | NA | 2 | 0.5 | 1 | 1 | NA | NA | NA | NA | NA | NA | 1 | 1 |
| 6_8587623 | 2 | 0 | 3 | 0 | NA | NA | 1 | 1 | 6 | 0.1667 | 2 | 0 | 4 | 0.25 | 2 | 0 | NA | NA | 8 | 0.125 |
| 6_8587641 | 2 | 0 | 4 | 0.5 | 1 | 0 | 1 | 0 | 8 | 0.25 | 3 | 0 | 3 | 0.3333 | 2 | 1 | NA | NA | 8 | 0.375 |
| 6_8587645 | 1 | 0 | 4 | 0.5 | 1 | 1 | 1 | 0 | 7 | 0.4286 | 2 | 0 | 3 | 0.3333 | 2 | 0.5 | NA | NA | 7 | 0.2857 |
| 6_8587679 | 1 | 1 | 2 | 0.5 | 1 | 1 | 1 | 1 | 5 | 0.8 | 2 | 0.5 | 3 | 0.6667 | 2 | 0.5 | NA | NA | 7 | 0.5714 |
| 6_8587686 | 2 | 0.5 | 4 | 0.25 | 1 | 1 | 1 | 0 | 8 | 0.375 | 1 | 1 | 3 | 1 | 2 | 1 | NA | NA | 6 | 1 |
| 6_8587691 | 2 | 0 | 4 | 0.25 | 1 | 0 | 1 | 0 | 8 | 0.125 | 2 | 0.5 | 2 | 0 | 2 | 0 | NA | NA | 6 | 0.1667 |
| 6_8587692 | 1 | 0 | 4 | 0.25 | NA | NA | 1 | 0 | 6 | 0.1667 | 2 | 0.5 | 2 | 1 | 2 | 0.5 | NA | NA | 6 | 0.6667 |
| 6_8587700 | 2 | 0.5 | 4 | 0.75 | 1 | 1 | 1 | 0 | 8 | 0.625 | 1 | 1 | 2 | 1 | 1 | 1 | NA | NA | 4 | 1 |
| 6_8587715 | 1 | 0 | 2 | 0.5 | 1 | 0 | NA | NA | 4 | 0.25 | 2 | 0.5 | 2 | 0.5 | 2 | 0.5 | NA | NA | 6 | 0.5 |
| 6_8587734 | 2 | 0 | 3 | 0.3333 | 1 | 0 | NA | NA | 6 | 0.1667 | NA | NA | 3 | 0.3333 | 2 | 0 | NA | NA | 5 | 0.2 |
| 6_8587801 | 2 | 0.5 | 2 | 0 | NA | NA | 1 | 0 | 5 | 0.2 | NA | NA | 3 | 0.3333 | NA | NA | NA | NA | 3 | 0.3333 |
| 6_8587802 | 2 | 0.5 | 2 | 0.5 | NA | NA | 1 | 0 | 5 | 0.4 | 1 | 1 | 3 | 0 | NA | NA | NA | NA | 4 | 0.25 |
| 6_8587804 | 2 | 1 | 2 | 1 | NA | NA | 1 | 1 | 5 | 1 | 1 | 1 | 2 | 1 | NA | NA | NA | NA | 3 | 1 |
| 6_8587813 | 2 | 0.5 | 2 | 0.5 | NA | NA | 1 | 1 | 5 | 0.6 | 1 | 1 | 3 | 0.6667 | NA | NA | NA | NA | 4 | 0.75 |
| 6_8587839 | 2 | 0.5 | 2 | 1 | NA | NA | NA | NA | 4 | 0.75 | 1 | 1 | 4 | 1 | NA | NA | NA | NA | 5 | 1 |
| 6_8588742 | 4 | 0.25 | 4 | 0.25 | 1 | 1 | 3 | 0.3333 | 12 | 0.3333 | 4 | 0.25 | 4 | 0 | 1 | 0 | 1 | 1 | 10 | 0.2 |
| 6_8588745 | 1 | 1 | 4 | 0.75 | NA | NA | 3 | 0 | 8 | 0.5 | 3 | 0.3333 | 4 | 0.25 | 1 | 0 | 1 | 1 | 9 | 0.3333 |
| 6_8588765 | 3 | 0 | 3 | 0.3333 | 1 | 0 | 2 | 0.5 | 9 | 0.2222 | 4 | 0.25 | 4 | 0 | 1 | 1 | 1 | 1 | 10 | 0.3 |
| 6_8588780 | 4 | 0.25 | 3 | 0.3333 | 1 | 0 | 2 | 0.5 | 10 | 0.3 | 4 | 0 | 2 | 0 | 1 | 1 | 1 | 1 | 8 | 0.25 |
| 6_8588789 | 4 | 0.25 | 3 | 0 | 1 | 0 | 3 | 0 | 11 | 0.0909 | 4 | 0.25 | 4 | 0.5 | 1 | 1 | NA | NA | 9 | 0.4444 |
| 6_8589214 | 1 | 0 | 2 | 0.5 | 3 | 0 | 1 | 0 | 7 | 0.1429 | 1 | 0 | 5 | 0.2 | 3 | 0 | 3 | 0 | 12 | 0.0833 |
| 6_8590231 | NA | NA | NA | NA | 2 | 1 | 1 | 0 | 3 | 0.6667 | 1 | 0 | 1 | 0 | NA | NA | 1 | 1 | 3 | 0.3333 |
| 6_8590308 | NA | NA | NA | NA | 2 | 0.5 | 1 | 1 | 3 | 0.6667 | 1 | 1 | NA | NA | NA | NA | 1 | 1 | 2 | 1 |
| 6_8590500 | NA | NA | 1 | 0 | NA | NA | 2 | 0.5 | 3 | 0.3333 | 1 | 0 | 4 | 0.5 | NA | NA | 1 | 0 | 6 | 0.3333 |
| 6_8590508 | NA | NA | 2 | 1 | NA | NA | 1 | 1 | 3 | 1 | 1 | 1 | 4 | 0.75 | 1 | 1 | 1 | 1 | 7 | 0.8571 |
| 6_8590576 | 8 | 0 | 4 | 0 | 7 | 0.2857 | 6 | 0.1667 | 25 | 0.12 | 4 | 0 | 15 | 0.0667 | 8 | 0.125 | 8 | 0 | 35 | 0.0571 |
| 6_8590721 | 7 | 0 | 6 | 0 | 12 | 0.0833 | 7 | 0.2857 | 32 | 0.0938 | 5 | 0.4 | 19 | 0.2105 | 7 | 0.2857 | 9 | 0.2222 | 40 | 0.25 |
| 6_8592032 | 2 | 0.5 | 3 | 0.3333 | 6 | 0.1667 | 4 | 0 | 15 | 0.2 | 3 | 0.3333 | 4 | 0 | 2 | 0.5 | 4 | 0.5 | 13 | 0.3077 |
| 6_8592445 | 1 | 0 | 4 | 0.25 | 4 | 0 | 6 | 0.3333 | 15 | 0.2 | 1 | 0 | 1 | 0 | 2 | 1 | 3 | 0.3333 | 7 | 0.4286 |
| 6_8592447 | 2 | 0.5 | 4 | 0.5 | 3 | 1 | 5 | 0.2 | 14 | 0.5 | NA | NA | NA | NA | 2 | 0 | 4 | 0.25 | 6 | 0.1667 |
| 6_8592448 | 2 | 0 | 4 | 0 | 4 | 0.25 | 7 | 0.5714 | 17 | 0.2941 | 2 | 0.5 | 1 | 0 | 2 | 1 | 2 | 1 | 7 | 0.7143 |
| 6_8592453 | 1 | 1 | 3 | 0 | 3 | 0.3333 | 5 | 0.2 | 12 | 0.25 | 4 | 0 | NA | NA | 4 | 0.25 | 4 | 0.25 | 12 | 0.1667 |
| 6_8592457 | 2 | 1 | 4 | 0.75 | 4 | 1 | 8 | 0.75 | 18 | 0.8333 | 3 | 1 | 2 | 1 | 4 | 1 | 4 | 1 | 13 | 1 |
| 6_8592459 | 2 | 0 | 4 | 0.25 | 5 | 0.4 | 8 | 0.25 | 19 | 0.2632 | 4 | 0 | 1 | 0 | 4 | 0.25 | 5 | 0 | 14 | 0.0714 |
| 6_8592462 | 2 | 0.5 | 4 | 0.5 | 5 | 0.8 | 7 | 0.8571 | 18 | 0.7222 | 4 | 0.5 | NA | NA | 3 | 0.3333 | 5 | 0.4 | 12 | 0.4167 |
| 6_8592464 | 2 | 1 | 4 | 0.75 | 4 | 1 | 7 | 1 | 17 | 0.9412 | 3 | 0.3333 | 1 | 1 | 4 | 0.75 | 3 | 1 | 11 | 0.7273 |
| 6_8592469 | 1 | 1 | 5 | 0.6 | 5 | 0.4 | 7 | 0.5714 | 18 | 0.5556 | 3 | 0.3333 | 1 | 0 | 4 | 0.5 | 5 | 0.6 | 13 | 0.4615 |
| 6_8609557 | 9 | 0.5556 | 9 | 0.3333 | 3 | 0.3333 | 5 | 0.8 | 26 | 0.5 | 5 | 0.6 | 4 | 0.75 | 5 | 0.4 | 4 | 0.75 | 18 | 0.6111 |
| 6_8609562 | 9 | 0.7778 | 10 | 0.9 | 3 | 0.6667 | 5 | 0.6 | 27 | 0.7778 | 6 | 0.5 | 4 | 1 | 4 | 1 | 4 | 1 | 18 | 0.8333 |
| 6_8609583 | 8 | 0.25 | 4 | 0 | 3 | 0 | 4 | 0 | 19 | 0.1053 | 6 | 0.1667 | 2 | 0.5 | 3 | 0.3333 | 2 | 0 | 13 | 0.2308 |
| 6_8609604 | 10 | 0.2 | 10 | 0.1 | 3 | 0 | 5 | 0 | 28 | 0.1071 | 7 | 0 | 4 | 0.5 | 5 | 0.2 | 3 | 0.3333 | 19 | 0.2105 |
| 6_8609626 | 11 | 0.0909 | 11 | 0.1818 | 3 | 0 | 5 | 0.2 | 30 | 0.1333 | 7 | 0 | 4 | 0.25 | 8 | 0 | 4 | 0.25 | 23 | 0.087 |
| 6_8609638 | 11 | 0.1818 | 7 | 0.2857 | 3 | 0.3333 | 5 | 0.6 | 26 | 0.3077 | 6 | 0 | 5 | 0 | 8 | 0.375 | 4 | 0.25 | 23 | 0.1739 |
| 6_8609684 | 14 | 0.0714 | 10 | 0.1 | 2 | 0 | 10 | 0.3 | 36 | 0.1389 | 6 | 0.1667 | 6 | 0 | 7 | 0 | 5 | 0.2 | 24 | 0.0833 |
| 6_8609697 | 11 | 0.3636 | 11 | 0.3636 | 3 | 0.3333 | 7 | 0.1429 | 32 | 0.3125 | 5 | 0.4 | 3 | 0.6667 | 7 | 0.5714 | 5 | 0.2 | 20 | 0.45 |
| 6_8609699 | 13 | 0.0769 | 10 | 0.1 | 1 | 0 | 7 | 0 | 31 | 0.0645 | 6 | 0.3333 | 6 | 0 | 8 | 0 | 6 | 0.1667 | 26 | 0.1154 |
| 6_8609715 | 11 | 0.1818 | 7 | 0 | 2 | 0 | 8 | 0.25 | 28 | 0.1429 | 5 | 0.2 | 4 | 0.75 | 6 | 0 | 4 | 0 | 19 | 0.2105 |
| 6_8609717 | 9 | 0.6667 | 6 | 0.3333 | 2 | 0 | 9 | 0.6667 | 26 | 0.5385 | 5 | 0.8 | 4 | 1 | 4 | 0.75 | 5 | 0.4 | 18 | 0.7222 |
| 6_8609719 | 11 | 0.2727 | 8 | 0.25 | 2 | 0 | 9 | 0.1111 | 30 | 0.2 | 6 | 0.1667 | 4 | 0 | 6 | 0.3333 | 6 | 0.1667 | 22 | 0.1818 |
| 6_8609721 | 12 | 0.8333 | 7 | 0.8571 | 3 | 1 | 7 | 0.7143 | 29 | 0.8276 | 5 | 1 | 3 | 0.6667 | 7 | 0.8571 | 5 | 0.6 | 20 | 0.8 |
| 6_8609726 | 9 | 1 | 8 | 1 | 3 | 1 | 8 | 0.625 | 28 | 0.8929 | 6 | 0.6667 | 4 | 1 | 6 | 0.8333 | 4 | 1 | 20 | 0.85 |
| 6_8609758 | 11 | 0.4545 | 7 | 0.5714 | 4 | 0.25 | 9 | 0.6667 | 31 | 0.5161 | 5 | 0.6 | 5 | 0.8 | 6 | 0.8333 | 6 | 0.1667 | 22 | 0.5909 |
| 6_8609776 | 10 | 0.3 | 9 | 0.1111 | 3 | 0 | 9 | 0.3333 | 31 | 0.2258 | 4 | 0.25 | 5 | 0 | 8 | 0.5 | 4 | 0.25 | 21 | 0.2857 |
| 6_8609810 | 16 | 0.125 | 12 | 0 | 5 | 0 | 12 | 0.25 | 45 | 0.1111 | 6 | 0 | 7 | 0.2857 | 8 | 0.25 | 6 | 0.3333 | 27 | 0.2222 |
| 6_8609832 | 15 | 0 | 14 | 0.1429 | 8 | 0.125 | 13 | 0.0769 | 50 | 0.08 | 8 | 0 | 10 | 0.3 | 8 | 0 | 7 | 0.2857 | 33 | 0.1515 |
| 6_8609838 | 17 | 0.2941 | 14 | 0.4286 | 9 | 0.4444 | 14 | 0.4286 | 54 | 0.3889 | 9 | 0.4444 | 11 | 0.5455 | 6 | 0.3333 | 6 | 0.3333 | 32 | 0.4375 |
| 6_8610070 | 22 | 0 | 21 | 0.0476 | 14 | 0.0714 | 24 | 0.1667 | 81 | 0.0741 | 12 | 0.1667 | 16 | 0 | 20 | 0.05 | 19 | 0.1053 | 67 | 0.0746 |

| | | | | | | | | | | | | | | | | | | | | |
|---|---|---|---|---|---|---|---|---|---|---|---|---|---|---|---|---|---|---|---|---|
| 6_86101258 | 37 | 0.0811 | 29 | 0.1034 | 24 | 0 | 30 | 0.0667 | 120 | 0.0667 | 18 | 0 | 24 | 0.0833 | 35 | 0.0857 | 31 | 0.1613 | 108 | 0.0926 |
| 6_86102679 | 2 | 0 | NA | NA | 1 | 0 | 3 | 0.3333 | 6 | 0.1667 | 1 | 0 | 1 | 1 | 1 | 0 | NA | NA | 3 | 0.3333 |
| 6_86249003 | 5 | 0.2 | 4 | 0.5 | 6 | 0.1667 | 6 | 0.5 | 21 | 0.3333 | 2 | 0.5 | 2 | 0.5 | 1 | 0 | 3 | 0 | 8 | 0.25 |
| 6_86249035 | 4 | 0.75 | 3 | 1 | 6 | 0.6667 | 4 | 1 | 17 | 0.8235 | 2 | 1 | 3 | 1 | 1 | 1 | 3 | 1 | 9 | 1 |
| 6_86249104 | 4 | 1 | 3 | 1 | 5 | 1 | 4 | 0.5 | 16 | 0.875 | 3 | 1 | 3 | 1 | 1 | 1 | 3 | 1 | 10 | 1 |
| 6_86249163 | 6 | 0.5 | 4 | 0.5 | 6 | 0.3333 | 5 | 0 | 21 | 0.3333 | 3 | 0.3333 | 3 | 0.3333 | 1 | 0 | 4 | 0 | 11 | 0.1818 |
| 6_86249177 | 6 | 0.5 | 4 | 0.25 | 5 | 0.6 | 6 | 0.3333 | 21 | 0.4286 | 3 | 0.3333 | 2 | 1 | NA | NA | 4 | 0.25 | 9 | 0.4444 |
| 6_86249186 | 5 | 0.8 | 4 | 0.75 | 5 | 0.2 | 4 | 0.5 | 18 | 0.5556 | 1 | 0 | NA | NA | 1 | 0 | 3 | 0.3333 | 5 | 0.2 |
| 6_86249192 | 3 | 0 | 1 | 0 | 4 | 0.25 | 4 | 0.75 | 12 | 0.3333 | 2 | 1 | NA | NA | NA | NA | 2 | 0 | 4 | 0.5 |
| 6_86249264 | 7 | 0.2857 | 4 | 0.25 | 4 | 0 | 6 | 0.5 | 21 | 0.2857 | 3 | 0.6667 | 3 | 0.3333 | 1 | 1 | 4 | 0 | 11 | 0.3636 |
| 6_86249272 | 7 | 1 | 3 | 1 | 4 | 0.75 | 6 | 1 | 20 | 0.95 | 3 | 1 | 4 | 1 | 1 | 1 | 3 | 0.6667 | 11 | 0.9091 |
| 6_86249313 | 6 | 0.1667 | 4 | 0 | 6 | 0.3333 | 6 | 0.6667 | 22 | 0.3182 | 2 | 0 | 3 | 0.3333 | 1 | 0 | 4 | 0.25 | 10 | 0.2 |
| 6_86249338 | 6 | 0.8333 | 4 | 1 | 5 | 0.8 | 6 | 1 | 21 | 0.9048 | 3 | 1 | 4 | 1 | 1 | 1 | 2 | 1 | 10 | 1 |
| 6_86250105 | 7 | 0 | 5 | 0.2 | 6 | 0.1667 | 8 | 0 | 26 | 0.0769 | 3 | 0 | 6 | 0.1667 | 1 | 0 | 5 | 0.2 | 15 | 0.1333 |
| 6_86250692 | 10 | 0.2 | 7 | 0.2857 | 5 | 0.4 | 7 | 0.4286 | 29 | 0.3103 | 4 | 0.25 | 3 | 0.3333 | 2 | 0.5 | 4 | 0 | 13 | 0.2308 |
| 6_86250889 | 10 | 0.2 | 6 | 0 | 6 | 0 | 7 | 0.1429 | 29 | 0.1034 | 4 | 0 | 6 | 0.1667 | 2 | 0 | 7 | 0.1429 | 19 | 0.1053 |
| 6_86345759 | 4 | 0.25 | 1 | 1 | 2 | 0.5 | 4 | 0.25 | 11 | 0.3636 | 3 | 0.3333 | 4 | 0.25 | 6 | 0.5 | 4 | 0.5 | 17 | 0.4118 |
| 6_86345776 | 3 | 0.3333 | 1 | 0 | 1 | 0 | 2 | 0 | 7 | 0.1429 | 3 | 0.3333 | 2 | 0.5 | 2 | 0 | 4 | 0 | 11 | 0.1818 |
| 6_86345917 | 4 | 0.5 | 1 | 0 | 3 | 0.3333 | 4 | 0.25 | 12 | 0.3333 | 3 | 0.3333 | 3 | 0 | 6 | 0.3333 | 4 | 0 | 16 | 0.1875 |
| 6_86345926 | 5 | 1 | 1 | 1 | 3 | 1 | 4 | 1 | 13 | 1 | 3 | 1 | 3 | 1 | 6 | 0.6667 | 4 | 0.75 | 16 | 0.8125 |
| 6_86346002 | 6 | 0.1667 | 1 | 0 | 3 | 0.3333 | 4 | 0 | 14 | 0.1429 | 3 | 0.3333 | 4 | 0 | 7 | 0.1429 | 4 | 0 | 18 | 0.1111 |
| 6_86346039 | 5 | 0.8 | 1 | 1 | 3 | 1 | 3 | 0.6667 | 12 | 0.8333 | 3 | 0.6667 | 3 | 1 | 5 | 0.6 | 3 | 1 | 14 | 0.7857 |
| 6_86346552 | 5 | 0.6 | 3 | 0.6667 | 2 | 1 | 4 | 0.25 | 14 | 0.5714 | 3 | 1 | 4 | 0 | 7 | 0.4286 | 5 | 0.4 | 19 | 0.4211 |
| 6_86346569 | 5 | 0 | 2 | 0.5 | 1 | 0 | 3 | 0 | 11 | 0.0909 | 2 | 0 | 4 | 0 | 8 | 0.125 | 4 | 0.25 | 18 | 0.1111 |
| 6_86346594 | 6 | 0.5 | 2 | 0 | 3 | 0.6667 | 4 | 0.5 | 15 | 0.4667 | 5 | 0.4 | 4 | 0.25 | 9 | 0.7778 | 5 | 0.8 | 23 | 0.6087 |
| 6_86346759 | 6 | 0.3333 | 3 | 0.6667 | 2 | 0.5 | 3 | 0 | 14 | 0.3571 | 6 | 0.5 | 2 | 0.5 | 9 | 0.3333 | 4 | 0.25 | 21 | 0.381 |
| 6_86363766 | 13 | 0.6923 | 8 | 0.625 | 5 | 0.4 | 10 | 1 | 36 | 0.7222 | 8 | 0.875 | 12 | 0.6667 | 5 | 0.4 | 6 | 0.8333 | 31 | 0.7097 |
| 6_86363921 | 11 | 0 | 10 | 0.1 | 7 | 0.1429 | 12 | 0.1667 | 40 | 0.1 | 8 | 0 | 12 | 0.4167 | 4 | 0.5 | 6 | 0 | 30 | 0.2333 |
| 6_86364274 | 12 | 0.1667 | 12 | 0 | 5 | 0 | 13 | 0.0769 | 42 | 0.0714 | 7 | 0.2857 | 15 | 0 | 5 | 0 | 6 | 0 | 33 | 0.0606 |
| 6_86364980 | 7 | 0.4286 | 9 | 0.1111 | 3 | 0 | 10 | 0.1 | 29 | 0.1724 | 5 | 0.2 | 10 | 0.1 | 4 | 0 | 4 | 0.25 | 23 | 0.1304 |
| 6_86365135 | 6 | 0.3333 | 7 | 0.2857 | 3 | 0 | 9 | 0.1111 | 25 | 0.2 | 3 | 0.3333 | 9 | 0.2222 | 4 | 0 | 3 | 0 | 19 | 0.1579 |
| 6_86365260 | 7 | 0.4286 | 7 | 0.5714 | 2 | 0.5 | 10 | 0.4 | 26 | 0.4615 | 3 | 0.6667 | 9 | 0.2222 | 2 | 0 | 6 | 0.5 | 20 | 0.35 |
| 6_86365269 | 7 | 0.2857 | 7 | 0.5714 | 3 | 0.3333 | 10 | 0.4 | 27 | 0.4074 | 2 | 0.5 | 11 | 0.4545 | 3 | 0.6667 | 7 | 0.1429 | 23 | 0.3913 |
| 6_86365423 | 7 | 0.2857 | 7 | 0.4286 | 3 | 0.3333 | 9 | 0.4444 | 26 | 0.3846 | 5 | 0.4 | 11 | 0.5455 | 1 | 1 | 6 | 0.1667 | 23 | 0.4348 |
| 6_86365509 | 8 | 0.25 | 6 | 0.3333 | 3 | 0 | 9 | 0.1111 | 26 | 0.1923 | 4 | 0 | 8 | 0.125 | 3 | 0.6667 | 7 | 0.1429 | 22 | 0.1818 |
| 6_86365629 | 7 | 0.7143 | 7 | 0.5714 | 3 | 0.6667 | 11 | 0.8182 | 28 | 0.7143 | 3 | 0.3333 | 10 | 0.6 | 2 | 0.5 | 7 | 0.7143 | 22 | 0.5909 |
| 6_86365654 | 5 | 0.8 | 6 | 0.8333 | 3 | 0.6667 | 7 | 0.7143 | 21 | 0.7619 | 2 | 1 | 8 | 1 | 3 | 1 | 5 | 0.6 | 18 | 0.8889 |
| 6_86365699 | 7 | 0.4286 | 5 | 1 | 1 | 0 | 10 | 0.1 | 23 | 0.3913 | 2 | 0.5 | 4 | 0 | 3 | 0.3333 | 5 | 0.2 | 14 | 0.2143 |
| 6_86365744 | 7 | 0.8571 | 7 | 1 | 1 | 1 | 10 | 0.9 | 25 | 0.92 | 4 | 1 | 9 | 0.8889 | 3 | 1 | 6 | 1 | 22 | 0.9545 |
| 6_86365965 | 4 | 0.25 | 6 | 0.1667 | 2 | 0.5 | 8 | 0.5 | 20 | 0.35 | 3 | 0.6667 | 8 | 0.375 | 1 | 0 | 2 | 0.5 | 14 | 0.4286 |
| 6_86365981 | 5 | 0.6 | 6 | 1 | 3 | 1 | 9 | 0.5556 | 23 | 0.7391 | 4 | 1 | 8 | 0.625 | 2 | 1 | 6 | 1 | 20 | 0.85 |
| 6_86366043 | 5 | 0.6 | 5 | 0.8 | 1 | 0 | 8 | 0.5 | 19 | 0.5789 | 4 | 0.75 | 6 | 0.5 | 2 | 0 | 7 | 0.8571 | 19 | 0.6316 |
| 6_86366071 | 3 | 1 | 5 | 0.8 | NA | NA | 8 | 0.25 | 16 | 0.5625 | 2 | 0.5 | 3 | 0.3333 | NA | NA | 4 | 0.25 | 9 | 0.3333 |
| 6_86366089 | 5 | 0.8 | 4 | 0.5 | 2 | 0.5 | 8 | 0.625 | 19 | 0.6316 | 4 | 0.5 | 7 | 0.5714 | 1 | 1 | 6 | 0.5 | 18 | 0.5556 |
| 6_86366160 | 5 | 0.8 | 4 | 0.75 | 3 | 0.6667 | 9 | 0.8889 | 21 | 0.8095 | 3 | 0.6667 | 6 | 0.5 | 1 | 1 | 2 | 1 | 12 | 0.6667 |
| 6_86366240 | 3 | 0.3333 | 5 | 0 | 2 | 0 | 7 | 0 | 17 | 0.0588 | 4 | 0.25 | 5 | 0 | 1 | 0 | 4 | 0 | 14 | 0.0714 |
| 6_86407253 | 2 | 0 | 3 | 0 | 3 | 0 | 9 | 0.1111 | 17 | 0.0588 | 1 | 1 | 4 | 0.25 | 4 | 0 | 5 | 0.2 | 14 | 0.2143 |
| 6_86407356 | 3 | 0.3333 | 3 | 0.3333 | 3 | 0 | 9 | 0 | 18 | 0.1111 | 1 | 0 | 3 | 0 | 4 | 0.25 | 5 | 0.6 | 13 | 0.3077 |
| 6_86407404 | 3 | 0 | 3 | 0 | 3 | 0 | 8 | 0.25 | 17 | 0.1176 | 1 | 0 | 5 | 0.4 | 4 | 0 | 5 | 0 | 15 | 0.1333 |
| 6_86407665 | 3 | 0 | 3 | 0.3333 | 4 | 0.5 | 8 | 0.125 | 18 | 0.2222 | 1 | 1 | 4 | 0.5 | 6 | 0.5 | 4 | 0 | 15 | 0.4 |
| 6_86407845 | 4 | 0 | 1 | 0 | 3 | 0.6667 | 7 | 0.1429 | 15 | 0.2 | 1 | 0 | 4 | 0 | 5 | 0.2 | 5 | 0.2 | 15 | 0.1333 |
| 6_86408018 | 4 | 0 | 3 | 0.3333 | 5 | 0.4 | 7 | 0.2857 | 19 | 0.2632 | 1 | 1 | 4 | 0.5 | 4 | 0.75 | 7 | 0.2857 | 16 | 0.5 |
| 6_86408092 | 3 | 0.6667 | 3 | 1 | 5 | 0.6 | 7 | 0.8571 | 18 | 0.7778 | 1 | 1 | 4 | 0.5 | 8 | 1 | 5 | 0.2 | 18 | 0.6667 |
| 6_86408132 | 3 | 0 | 3 | 0.3333 | 4 | 0.25 | 8 | 0 | 18 | 0.1111 | 1 | 0 | 4 | 0.25 | 8 | 0.125 | 6 | 0 | 19 | 0.1053 |
| 6_86408148 | 3 | 0 | 2 | 0 | 5 | 0 | 7 | 0.1429 | 17 | 0.0588 | 2 | 0.5 | 4 | 0.25 | 7 | 0 | 6 | 0 | 19 | 0.1053 |
| 6_86409434 | 3 | 0.3333 | 5 | 0.2 | 3 | 0.3333 | 6 | 0.3333 | 17 | 0.2941 | 2 | 0.5 | 1 | 0 | 7 | 0.7143 | 6 | 0.1667 | 16 | 0.4375 |
| 6_86409647 | 3 | 0 | 5 | 0.2 | 3 | 0 | 6 | 0.1667 | 17 | 0.1176 | 3 | 0 | 2 | 0 | 6 | 0 | 6 | 0.1667 | 17 | 0.0588 |
| 6_86415731 | 2 | 0.5 | NA | NA | 1 | 0 | NA | NA | 3 | 0.3333 | 1 | 1 | 2 | 0.5 | 1 | 0 | NA | NA | 4 | 0.5 |
| 6_86416210 | 2 | 0.5 | NA | NA | 1 | 0 | NA | NA | 3 | 0.3333 | 1 | 0 | 1 | 1 | NA | NA | NA | NA | 2 | 0.5 |
| 6_86416230 | 1 | 1 | NA | NA | 1 | 1 | NA | NA | 2 | 1 | 1 | 1 | NA | NA | NA | NA | NA | NA | 1 | 1 |
| 6_86416329 | 1 | 1 | NA | NA | 1 | 0 | NA | NA | 2 | 0.5 | 1 | 1 | NA | NA | NA | NA | NA | NA | 1 | 1 |
| 6_86422449 | 1 | 0 | 1 | 0 | 2 | 0 | 2 | 0.5 | 6 | 0.1667 | NA | NA | 3 | 0 | 1 | 0 | 1 | 1 | 5 | 0.2 |
| 6_86422474 | 1 | 1 | 1 | 0 | 2 | 0.5 | 2 | 0 | 6 | 0.3333 | NA | NA | 3 | 1 | 1 | 1 | 1 | 0 | 5 | 0.8 |
| 6_86422530 | 1 | 1 | 1 | 1 | 2 | 0.5 | 2 | 0.5 | 6 | 0.6667 | NA | NA | 4 | 0.5 | 1 | 1 | 1 | 1 | 6 | 0.6667 |
| 6_86422563 | 2 | 0 | 1 | 0 | 1 | 1 | 2 | 0 | 6 | 0.1667 | 1 | 1 | 3 | 0.3333 | 1 | 0 | 1 | 0 | 6 | 0.3333 |

| | | | | | | | | | | | | | | | | | | | | |
|---|---|---|---|---|---|---|---|---|---|---|---|---|---|---|---|---|---|---|---|---|
| 6_8642271 | 2 | 1 | 1 | 0 | 2 | 0.5 | 2 | 1 | 7 | 0.7143 | 1 | 0 | 4 | 0.75 | NA | NA | 1 | 1 | 6 | 0.6667 |
| 6_8642298 | 1 | 1 | 2 | 1 | 1 | 0 | 2 | 1 | 6 | 0.8333 | NA | NA | 1 | 1 | 1 | 0 | NA | NA | 2 | 0.5 |
| 6_8642322 | 2 | 1 | 2 | 1 | 2 | 1 | 3 | 1 | 9 | 1 | 1 | 1 | 4 | 1 | 1 | 1 | 1 | 1 | 7 | 1 |
| 6_8644685 | 2 | 0 | 2 | 0 | 3 | 0.3333 | 2 | 0 | 9 | 0.1111 | 1 | 1 | 4 | 0 | 1 | 0 | 1 | 1 | 7 | 0.2857 |
| 6_8644689 | 1 | 1 | NA | NA | 4 | 0.25 | 2 | 0.5 | 7 | 0.4286 | 1 | 0 | 2 | 1 | 1 | 0 | NA | NA | 4 | 0.5 |
| 6_8644698 | 1 | 1 | 1 | 1 | 4 | 1 | 4 | 1 | 10 | 1 | NA | NA | 4 | 1 | 1 | 1 | 1 | 1 | 6 | 1 |
| 6_8644708 | 2 | 0.5 | 2 | 1 | 3 | 0.3333 | 4 | 0.5 | 11 | 0.5455 | 1 | 1 | 5 | 0.4 | 1 | 0 | 1 | 1 | 8 | 0.5 |
| 6_8646038 | NA | NA | NA | NA | 1 | 1 | NA | NA | 1 | 1 | NA | NA | 1 | 1 | NA | NA | 2 | 0.5 | 3 | 0.6667 |
| 6_8646046 | NA | NA | NA | NA | 1 | 1 | NA | NA | 1 | 1 | NA | NA | 1 | 1 | NA | NA | 1 | 1 | 2 | 1 |
| 6_8650203 | 18 | 0.5556 | 30 | 0.5 | 24 | 0.5417 | 30 | 0.7667 | 102 | 0.598 | 25 | 0.68 | 22 | 0.6818 | 21 | 0.7619 | 25 | 0.64 | 93 | 0.6882 |
| 6_8650204 | 26 | 0.9615 | 28 | 1 | 27 | 0.963 | 34 | 1 | 115 | 0.9826 | 33 | 0.9697 | 26 | 0.9615 | 24 | 0.9583 | 22 | 0.9545 | 105 | 0.9619 |
| 6_8650211 | 34 | 0.2941 | 36 | 0.1944 | 35 | 0.2 | 36 | 0.25 | 141 | 0.234 | 35 | 0.3714 | 29 | 0.1724 | 33 | 0.3636 | 27 | 0.1481 | 124 | 0.2742 |
| 6_8650222 | 29 | 0.3793 | 34 | 0.2353 | 31 | 0.2581 | 41 | 0.2683 | 135 | 0.2815 | 37 | 0.2703 | 27 | 0.3704 | 31 | 0.2258 | 26 | 0.4615 | 121 | 0.3223 |
| 6_8650225 | 28 | 0.1429 | 32 | 0.0312 | 30 | 0.0667 | 33 | 0.0606 | 123 | 0.0732 | 39 | 0.0256 | 21 | 0.0952 | 29 | 0.1379 | 24 | 0.2083 | 113 | 0.1062 |
| 6_8650271 | 26 | 0.1154 | 30 | 0.0667 | 29 | 0.1724 | 32 | 0.0625 | 117 | 0.1026 | 30 | 0.2 | 21 | 0.1905 | 22 | 0.0909 | 20 | 0.05 | 93 | 0.1398 |
| 6_8650287 | 26 | 0.1923 | 23 | 0.2609 | 27 | 0.2593 | 31 | 0.2581 | 107 | 0.243 | 27 | 0.1852 | 20 | 0.2 | 21 | 0.2857 | 17 | 0.2941 | 85 | 0.2353 |
| 6_8650294 | 24 | 0.2917 | 19 | 0.4211 | 23 | 0.2174 | 25 | 0.32 | 91 | 0.3077 | 24 | 0.2917 | 15 | 0.4 | 19 | 0.2632 | 14 | 0.2857 | 72 | 0.3056 |
| 6_8662427 | 4 | 0 | 2 | 0 | 3 | 0 | 2 | 0.5 | 11 | 0.0909 | 2 | 0 | 5 | 0.2 | 2 | 0.5 | 1 | 0 | 10 | 0.2 |
| 6_8662449 | 3 | 0.6667 | 1 | 0 | 3 | 0.3333 | 2 | 0 | 9 | 0.3333 | 2 | 0 | 6 | 0 | 2 | 0 | 2 | 0.5 | 12 | 0.0833 |
| 6_8662494 | 2 | 0 | 1 | 0 | 2 | 0 | 1 | 1 | 6 | 0.1667 | 3 | 0 | 5 | 0.2 | 2 | 0 | 1 | 0 | 11 | 0.0909 |
| 6_8662499 | 2 | 0.5 | NA | NA | 2 | 0 | 1 | 1 | 5 | 0.4 | 2 | 1 | 4 | 0.75 | 2 | 1 | 1 | 1 | 9 | 0.8889 |
| 6_8662501 | 4 | 0.5 | 1 | 1 | 2 | 0.5 | 1 | 0 | 8 | 0.5 | 3 | 0 | 4 | 0.25 | 2 | 0.5 | 1 | 0 | 10 | 0.2 |
| 6_8662525 | 3 | 0 | 1 | 0 | 2 | 0.5 | 1 | 1 | 7 | 0.2857 | 3 | 0 | 5 | 0 | 1 | 0 | 1 | 1 | 10 | 0.1 |
| 6_8662527 | 3 | 0 | 1 | 0 | 1 | 1 | 1 | 0 | 6 | 0.1667 | 3 | 0.3333 | 5 | 0.4 | 1 | 0 | 1 | 1 | 10 | 0.4 |
| 6_8662531 | 4 | 0.25 | 1 | 0 | 2 | 0 | 1 | 1 | 8 | 0.25 | 2 | 0.5 | 3 | 0.6667 | 1 | 1 | 1 | 1 | 7 | 0.7143 |
| 6_8662562 | 4 | 0 | 1 | 0 | 2 | 0 | 2 | 0.5 | 9 | 0.1111 | 3 | 0.3333 | 4 | 0 | 1 | 0 | 1 | 0 | 9 | 0.1111 |
| 6_8662660 | 3 | 0.3333 | 1 | 0 | 1 | 0 | 2 | 0.5 | 7 | 0.2857 | 2 | 1 | 7 | 0.1429 | 1 | 0 | 1 | 1 | 11 | 0.3636 |
| 6_8662664 | 4 | 0.25 | NA | NA | 2 | 0 | 1 | 0 | 7 | 0.1429 | 1 | 1 | 4 | 0 | 1 | 0 | 1 | 0 | 7 | 0.1429 |
| 6_8662711 | 3 | 0.3333 | 1 | 1 | 3 | 0 | 2 | 0 | 9 | 0.2222 | 1 | 0 | 3 | 0.3333 | NA | NA | 1 | 0 | 5 | 0.2 |
| 6_8662722 | 3 | 0 | 1 | 0 | 1 | 0 | 2 | 0.5 | 7 | 0.1429 | 1 | 0 | 6 | 0.6667 | 1 | 0 | 1 | 1 | 9 | 0.5556 |
| 6_8662732 | 2 | 0.5 | NA | NA | 2 | 0.5 | 2 | 0.5 | 6 | 0.5 | 1 | 0 | 6 | 0.6667 | NA | NA | NA | NA | 7 | 0.5714 |
| 6_8662747 | 3 | 0.6667 | 1 | 1 | 3 | 1 | 1 | 1 | 8 | 0.875 | 1 | 1 | 4 | 1 | 1 | 1 | 1 | 1 | 7 | 1 |
| 6_8662748 | 3 | 0 | 1 | 1 | 2 | 0 | 1 | 0 | 7 | 0.1429 | 1 | 1 | 6 | 0 | 1 | 0 | 1 | 0 | 9 | 0.1111 |
| 6_8662767 | 4 | 0.75 | NA | NA | 3 | 1 | 1 | 0 | 8 | 0.75 | 1 | 1 | 6 | 0.8333 | 1 | 1 | 1 | 1 | 9 | 0.8889 |
| 6_8662781 | 4 | 0.25 | NA | NA | 3 | 0.3333 | 2 | 0 | 9 | 0.2222 | NA | NA | 6 | 0.3333 | 1 | 0 | 1 | 1 | 8 | 0.375 |
| 6_8662796 | 2 | 0 | NA | NA | 3 | 0.6667 | 1 | 0 | 6 | 0.3333 | NA | NA | 3 | 0.3333 | NA | NA | 1 | 0 | 4 | 0.25 |
| 6_8662799 | 3 | 0 | NA | NA | 3 | 0.3333 | 2 | 0 | 8 | 0.125 | NA | NA | 5 | 0.2 | 1 | 0 | 1 | 1 | 7 | 0.2857 |
| 6_8666501 | 1 | 1 | 1 | 1 | NA | NA | NA | NA | 2 | 1 | NA | NA | 1 | 1 | NA | NA | NA | NA | 1 | 1 |
| 6_8666508 | 1 | 1 | 1 | 1 | NA | NA | NA | NA | 2 | 1 | 1 | 1 | 1 | 1 | NA | NA | NA | NA | 2 | 1 |
| 6_8666536 | 1 | 1 | 1 | 1 | NA | NA | NA | NA | 2 | 1 | NA | NA | 1 | 1 | NA | NA | NA | NA | 1 | 1 |
| 6_8666562 | 2 | 0 | 1 | 1 | NA | NA | NA | NA | 3 | 0.3333 | 2 | 1 | 1 | 0 | NA | NA | NA | NA | 3 | 0.6667 |
| 6_8666576 | 1 | 1 | NA | NA | NA | NA | NA | NA | 1 | 1 | 2 | 0.5 | NA | NA | NA | NA | NA | NA | 2 | 0.5 |
| 6_8666905 | 3 | 0.3333 | 6 | 0 | 1 | 0 | 3 | 0 | 13 | 0.0769 | 1 | 0 | 8 | 0.125 | 1 | 1 | NA | NA | 10 | 0.2 |
| 6_8666926 | 5 | 0.4 | 7 | 0.1429 | 3 | 0.3333 | 4 | 0.25 | 19 | 0.2632 | 4 | 0.25 | 8 | 0.25 | 1 | 1 | 2 | 0.5 | 15 | 0.3333 |
| 6_8666981 | 8 | 0.25 | 6 | 0 | 2 | 0 | 4 | 0.25 | 20 | 0.15 | 4 | 0 | 6 | 0.1667 | 1 | 0 | 4 | 0 | 15 | 0.0667 |
| 6_8666982 | 4 | 1 | 6 | 1 | 2 | 1 | 3 | 1 | 15 | 1 | 4 | 0.75 | 4 | 1 | 1 | 1 | 1 | 1 | 10 | 0.9 |
| 6_8666991 | 7 | 0 | 7 | 0.1429 | 1 | 0 | 3 | 0.3333 | 18 | 0.1111 | 3 | 0.6667 | 5 | 0.6 | NA | NA | 4 | 0.25 | 12 | 0.5 |
| 6_8666994 | 7 | 0.4286 | 7 | 0.5714 | 2 | 0.5 | 4 | 1 | 20 | 0.6 | 4 | 0.75 | 6 | 0.5 | 1 | 1 | 3 | 0.3333 | 14 | 0.5714 |
| 6_8671435 | NA | NA | 2 | 0.5 | NA | NA | NA | NA | 2 | 0.5 | 1 | 1 | NA | NA | NA | NA | NA | NA | 1 | 1 |
| 6_8682562 | 1 | 1 | NA | NA | 2 | 1 | NA | NA | 3 | 1 | 1 | 1 | NA | NA | 1 | 1 | NA | NA | 2 | 1 |
| 6_8682598 | 2 | 1 | NA | NA | 2 | 1 | NA | NA | 4 | 1 | NA | NA | NA | NA | 1 | 1 | NA | NA | 1 | 1 |
| 6_8682602 | 2 | 0.5 | NA | NA | 1 | 0 | NA | NA | 3 | 0.3333 | 1 | 1 | NA | NA | 1 | 0 | NA | NA | 2 | 0.5 |
| 6_8682608 | 1 | 1 | NA | NA | 2 | 0.5 | NA | NA | 3 | 0.6667 | 1 | 1 | NA | NA | 1 | 1 | NA | NA | 2 | 1 |
| 6_8682621 | 2 | 0 | NA | NA | 2 | 0.5 | NA | NA | 4 | 0.25 | NA | NA | NA | NA | 1 | 1 | NA | NA | 1 | 1 |
| 6_8682698 | 1 | 1 | 1 | 1 | NA | NA | 2 | 0.5 | 4 | 0.75 | 1 | 0 | 1 | 1 | 2 | 1 | 1 | 0 | 5 | 0.6 |
| 6_8696459 | 4 | 0 | 5 | 0.4 | 5 | 0 | 5 | 0.8 | 19 | 0.3158 | 7 | 0.2857 | 6 | 0.1667 | 4 | 0.25 | 8 | 0.25 | 25 | 0.24 |
| 6_8696462 | 3 | 1 | 5 | 0.8 | 4 | 0.75 | 5 | 0.8 | 17 | 0.8235 | 8 | 0.625 | 7 | 1 | 4 | 1 | 8 | 0.875 | 27 | 0.8519 |
| 6_8696473 | 3 | 0 | 5 | 0 | 5 | 0.4 | 4 | 0.5 | 17 | 0.2353 | 8 | 0.625 | 7 | 0.2857 | 6 | 0.3333 | 5 | 0.6 | 26 | 0.4615 |
| 6_8696488 | 4 | 0.5 | 4 | 0.25 | 4 | 0.25 | 3 | 0.6667 | 15 | 0.4 | 5 | 0.2 | 3 | 0.3333 | 4 | 0 | 7 | 0.8571 | 19 | 0.4211 |
| 6_8696498 | 5 | 0.4 | 7 | 0 | 4 | 0.5 | 5 | 0 | 21 | 0.1905 | 8 | 0.375 | 7 | 0.1429 | 5 | 0 | 7 | 0 | 27 | 0.1481 |
| 6_8696509 | 2 | 0 | 3 | 0.3333 | 3 | 0 | 3 | 0.3333 | 11 | 0.1818 | 6 | 0.1667 | 6 | 0.3333 | 5 | 0 | 6 | 0 | 23 | 0.1304 |
| 6_8696512 | 4 | 0.75 | 3 | 0 | 4 | 0 | 3 | 0.3333 | 14 | 0.2857 | 7 | 0.2857 | 5 | 0.4 | 4 | 0 | 6 | 0.1667 | 22 | 0.2273 |
| 6_8696594 | 7 | 0.8571 | 9 | 0.7778 | 7 | 0.7143 | 8 | 0.875 | 31 | 0.8065 | 8 | 0.625 | 8 | 0.75 | 4 | 0.75 | 8 | 0.625 | 28 | 0.6786 |
| 6_8696596 | 6 | 0.8333 | 10 | 0.7 | 6 | 0.3333 | 6 | 1 | 28 | 0.7143 | 8 | 0.625 | 6 | 0.8333 | 4 | 0.75 | 6 | 0.5 | 24 | 0.6667 |
| 6_8696611 | 5 | 0.8 | 11 | 1 | 7 | 0.8571 | 9 | 0.8889 | 32 | 0.9062 | 9 | 0.8889 | 9 | 0.8889 | 3 | 1 | 6 | 0.8333 | 27 | 0.8889 |

| | | | | | | | | | | | | | | | | | | | | |
|---|---|---|---|---|---|---|---|---|---|---|---|---|---|---|---|---|---|---|---|---|
| 6_8696613 | 6 | 0.1667 | 12 | 0.4167 | 7 | 0.1429 | 7 | 0.2857 | 32 | 0.2812 | 8 | 0.5 | 8 | 0.375 | 3 | 0.3333 | 7 | 0.1429 | 26 | 0.3462 |
| 6_8696643 | 7 | 0 | 12 | 0 | 9 | 0 | 10 | 0.2 | 38 | 0.0526 | 9 | 0.1111 | 10 | 0.1 | 5 | 0 | 8 | 0.125 | 32 | 0.0938 |
| 6_8696657 | 5 | 0 | 14 | 0.2143 | 9 | 0 | 10 | 0 | 38 | 0.0789 | 9 | 0 | 10 | 0.1 | 5 | 0 | 8 | 0.125 | 32 | 0.0625 |
| 6_8696666 | 7 | 0.2857 | 13 | 0.0769 | 10 | 0.1 | 10 | 0 | 40 | 0.1 | 10 | 0.1 | 9 | 0.1111 | 6 | 0 | 7 | 0 | 32 | 0.0625 |
| 6_8696682 | 8 | 0 | 10 | 0 | 8 | 0 | 9 | 0.2222 | 35 | 0.0571 | 7 | 0 | 8 | 0.25 | 5 | 0 | 7 | 0.1429 | 27 | 0.1111 |
| 6_8696765 | 11 | 0 | 19 | 0.1053 | 14 | 0.2143 | 14 | 0.2143 | 58 | 0.1379 | 9 | 0 | 11 | 0.0909 | 10 | 0 | 15 | 0.2667 | 45 | 0.1111 |
| 6_8696784 | 12 | 0.0833 | 19 | 0.2105 | 18 | 0.3889 | 16 | 0.4375 | 65 | 0.2923 | 12 | 0.25 | 14 | 0.1429 | 10 | 0 | 14 | 0.3571 | 50 | 0.2 |
| 6_8696840 | 12 | 0.25 | 21 | 0.0952 | 18 | 0.3333 | 15 | 0.0667 | 66 | 0.1818 | 15 | 0.1333 | 16 | 0.3125 | 11 | 0.1818 | 16 | 0.125 | 58 | 0.1897 |
| 6_8697317 | NA | NA | NA | NA | NA | NA | 2 | 0.5 | 2 | 0.5 | NA | NA | 1 | 0 | 1 | 1 | NA | NA | 2 | 0.5 |
| 6_8697354 | NA | NA | NA | NA | NA | NA | 3 | 0.6667 | 3 | 0.6667 | NA | NA | 1 | 1 | 1 | 0 | NA | NA | 2 | 0.5 |
| 6_8697366 | NA | NA | NA | NA | NA | NA | 3 | 1 | 3 | 1 | NA | NA | NA | NA | 1 | 1 | NA | NA | 1 | 1 |
| 6_8697387 | NA | NA | NA | NA | 1 | 1 | 3 | 0.3333 | 4 | 0.5 | NA | NA | 1 | 1 | 1 | 1 | NA | NA | 2 | 1 |
| 6_8697421 | NA | NA | NA | NA | 1 | 1 | 3 | 0.6667 | 4 | 0.75 | NA | NA | 1 | 0 | 1 | 1 | NA | NA | 2 | 0.5 |
| 6_8697430 | NA | NA | NA | NA | 1 | 1 | 2 | 1 | 3 | 1 | NA | NA | 1 | 1 | 1 | 1 | NA | NA | 2 | 1 |
| 6_8697436 | NA | NA | NA | NA | 1 | 1 | 1 | 1 | 2 | 1 | NA | NA | 1 | 1 | 1 | 1 | NA | NA | 2 | 1 |
| 6_8697459 | NA | NA | NA | NA | NA | NA | 2 | 1 | 2 | 1 | NA | NA | 1 | 1 | NA | NA | NA | NA | 1 | 1 |
| 6_8697467 | NA | NA | NA | NA | 1 | 0 | 2 | 1 | 3 | 0.6667 | NA | NA | 1 | 0 | 1 | 1 | NA | NA | 2 | 0.5 |
| 6_8697476 | NA | NA | NA | NA | 1 | 0 | 3 | 0.6667 | 4 | 0.5 | NA | NA | 1 | 0 | 1 | 1 | NA | NA | 2 | 0.5 |
| 6_8697481 | NA | NA | NA | NA | 1 | 1 | 3 | 1 | 4 | 1 | NA | NA | 1 | 1 | 1 | 1 | NA | NA | 2 | 1 |
| 6_8697484 | NA | NA | NA | NA | NA | NA | 3 | 0.6667 | 3 | 0.6667 | NA | NA | 1 | 1 | 1 | 1 | NA | NA | 2 | 1 |
| 6_8697484 | NA | NA | NA | NA | NA | NA | 1 | 1 | 1 | 1 | NA | NA | 1 | 1 | NA | NA | NA | NA | 1 | 1 |
| 6_8697491 | NA | NA | NA | NA | 1 | 1 | 3 | 0.3333 | 4 | 0.5 | NA | NA | NA | NA | 1 | 1 | NA | NA | 1 | 1 |
| 6_8697512 | NA | NA | NA | NA | 1 | 1 | 3 | 0.3333 | 4 | 0.5 | 1 | 1 | 1 | 1 | 1 | 1 | NA | NA | 3 | 1 |
| 6_8697526 | NA | NA | 1 | 0 | 1 | 1 | 4 | 0 | 6 | 0.1667 | 1 | 0 | 1 | 0 | 1 | 1 | NA | NA | 3 | 0.3333 |
| 6_8697614 | NA | NA | 2 | 0.5 | 2 | 0 | 2 | 0.5 | 6 | 0.3333 | 2 | 0 | 1 | 0 | 1 | 1 | NA | NA | 4 | 0.25 |
| 6_8697630 | NA | NA | 2 | 0.5 | 2 | 0.5 | 2 | 0.5 | 6 | 0.5 | 2 | 0.5 | 1 | 1 | 1 | 0 | NA | NA | 4 | 0.5 |
| 6_8697646 | NA | NA | 2 | 0.5 | 1 | 1 | 2 | 0.5 | 5 | 0.6 | 2 | 0 | 1 | 0 | 2 | 0.5 | NA | NA | 5 | 0.2 |
| 6_8697651 | NA | NA | 1 | 0 | 2 | 0.5 | 1 | 1 | 4 | 0.5 | 2 | 0 | 1 | 1 | 2 | 0 | NA | NA | 5 | 0.2 |
| 6_8697677 | NA | NA | 2 | 1 | 2 | 0 | 3 | 0.3333 | 7 | 0.4286 | 2 | 0.5 | 1 | 1 | 2 | 0.5 | NA | NA | 5 | 0.6 |
| 6_8697712 | NA | NA | 2 | 0 | NA | NA | 3 | 0.6667 | 5 | 0.4 | 2 | 0 | 1 | 1 | 3 | 0.3333 | NA | NA | 6 | 0.3333 |
| 6_8697893 | NA | NA | 3 | 0 | 4 | 0.25 | 3 | 0.3333 | 10 | 0.2 | 2 | 0 | 1 | 1 | 4 | 0 | 2 | 0 | 9 | 0.1111 |
| 6_8697942 | NA | NA | 4 | 0.25 | 4 | 0.25 | 3 | 0 | 11 | 0.1818 | 3 | 0.3333 | NA | NA | 5 | 0.4 | NA | NA | 8 | 0.375 |
| 6_8697946 | NA | NA | 4 | 0.25 | 3 | 0 | 5 | 0.4 | 12 | 0.25 | 3 | 0.3333 | 1 | 0 | 2 | 0 | 2 | 0.5 | 8 | 0.25 |
| 6_8697983 | NA | NA | 4 | 0.5 | 5 | 0.2 | 5 | 0.2 | 14 | 0.2857 | 4 | 0 | 1 | 0 | 5 | 0.4 | 1 | 0 | 11 | 0.1818 |
| 6_8697989 | NA | NA | 4 | 0.5 | 5 | 0.4 | 4 | 0.5 | 13 | 0.4615 | 4 | 0 | 1 | 0 | 4 | 0.25 | 2 | 0.5 | 11 | 0.1818 |
| 6_8706866 | 7 | 0 | 5 | 0.2 | 7 | 0 | 7 | 0.1429 | 26 | 0.0769 | 5 | 0.2 | 5 | 0.2 | 12 | 0.25 | 5 | 0.6 | 27 | 0.2963 |
| 6_8743646 | NA | NA | 2 | 0.5 | 1 | 1 | NA | NA | 3 | 0.6667 | 1 | 1 | 1 | 1 | NA | NA | NA | NA | 2 | 1 |
| 6_8743661 | NA | NA | 2 | 0.5 | 1 | 0 | NA | NA | 3 | 0.3333 | 1 | 1 | 1 | 1 | NA | NA | 1 | 0 | 3 | 0.6667 |
| 6_8743664 | NA | NA | 2 | 1 | 1 | 0 | NA | NA | 3 | 0.6667 | 1 | 0 | 1 | 1 | NA | NA | NA | NA | 2 | 0.5 |
| 6_8743667 | NA | NA | 2 | 1 | 1 | 1 | NA | NA | 3 | 1 | 1 | 1 | 1 | 1 | NA | NA | NA | NA | 2 | 1 |
| 6_8743673 | NA | NA | 2 | 1 | 1 | 1 | NA | NA | 3 | 1 | 1 | 1 | NA | NA | NA | NA | 1 | 1 | 2 | 1 |
| 6_8743681 | NA | NA | 2 | 1 | 1 | 1 | NA | NA | 3 | 1 | 1 | 1 | NA | NA | NA | NA | NA | NA | 1 | 1 |
| 6_8743687 | NA | NA | 1 | 1 | 1 | 0 | NA | NA | 2 | 0.5 | NA | NA | 1 | 1 | NA | NA | NA | NA | 1 | 1 |
| 6_8744001 | NA | NA | 2 | 1 | NA | NA | NA | NA | 2 | 1 | NA | NA | 1 | 1 | NA | NA | 1 | 0 | 2 | 0.5 |
| 6_8744014 | NA | NA | 2 | 1 | 1 | 1 | NA | NA | 3 | 1 | NA | NA | 1 | 1 | NA | NA | 1 | 1 | 2 | 1 |
| 6_8744019 | NA | NA | 2 | 0.5 | 1 | 1 | NA | NA | 3 | 0.6667 | NA | NA | 1 | 1 | NA | NA | NA | NA | 1 | 1 |
| 6_8744032 | NA | NA | 2 | 1 | 1 | 1 | NA | NA | 3 | 1 | NA | NA | 1 | 1 | NA | NA | NA | NA | 1 | 1 |
| 6_8760696 | 1 | 1 | 3 | 0.3333 | NA | NA | 1 | 1 | 5 | 0.6 | NA | NA | 2 | 0.5 | NA | NA | NA | NA | 2 | 0.5 |
| 6_8762323 | NA | NA | 3 | 0 | NA | NA | 1 | 1 | 4 | 0.25 | NA | NA | 2 | 0.5 | NA | NA | NA | NA | 2 | 0.5 |
| 6_8770670 | 3 | 0 | 1 | 0 | 1 | 0 | 5 | 0.4 | 10 | 0.2 | 2 | 1 | 5 | 0.4 | 3 | 0.3333 | NA | NA | 10 | 0.5 |
| 6_8770675 | 3 | 0.3333 | 1 | 0 | 1 | 1 | 7 | 1 | 12 | 0.75 | 2 | 1 | 5 | 1 | 2 | 1 | NA | NA | 9 | 1 |
| 6_8770680 | 3 | 0.3333 | 2 | 0 | 1 | 0 | 6 | 0.1667 | 12 | 0.1667 | 2 | 0 | 6 | 0.1667 | 4 | 0 | NA | NA | 12 | 0.0833 |
| 6_8770685 | 2 | 0.5 | 2 | 0 | 1 | 0 | 8 | 0 | 13 | 0.0769 | 1 | 0 | 5 | 0.2 | 3 | 0 | NA | NA | 9 | 0.1111 |
| 6_8770712 | 3 | 0 | 3 | 0.3333 | 2 | 0 | 6 | 0.1667 | 14 | 0.1429 | 2 | 0 | 7 | 0.1429 | 4 | 0 | 1 | 0 | 14 | 0.0714 |
| 6_8770719 | 3 | 0 | 2 | 0 | 2 | 0 | 6 | 0.1667 | 13 | 0.0769 | 1 | 1 | 6 | 0.1667 | 3 | 0 | 1 | 0 | 11 | 0.1818 |
| 6_8770743 | 3 | 0.3333 | 3 | 0 | 2 | 0 | 6 | 0 | 14 | 0.0714 | 2 | 0.5 | 7 | 0.1429 | 3 | 0 | 1 | 0 | 13 | 0.1538 |
| 6_8770749 | 2 | 0 | 3 | 0 | 2 | 0 | 6 | 0.1667 | 13 | 0.0769 | 1 | 0 | 7 | 0.1429 | 3 | 0 | NA | NA | 11 | 0.0909 |
| 6_8770774 | 1 | 0 | 1 | 0 | 2 | 1 | 5 | 0.2 | 9 | 0.3333 | 1 | 0 | 5 | 0.4 | 1 | 1 | NA | NA | 7 | 0.4286 |
| 6_8770826 | 2 | 0.5 | 1 | 0 | 1 | 1 | 4 | 0.25 | 8 | 0.375 | 1 | 1 | 4 | 0.25 | NA | NA | NA | NA | 5 | 0.4 |
| 6_8770890 | 1 | 1 | NA | NA | 1 | 0 | NA | NA | 2 | 0.5 | 1 | 1 | NA | NA | NA | NA | NA | NA | 1 | 1 |
| 6_8773400 | NA | NA | NA | NA | NA | NA | 1 | 1 | 1 | 1 | NA | NA | NA | NA | 1 | 1 | 1 | 1 | 2 | 1 |
| 6_8773408 | NA | NA | NA | NA | NA | NA | 1 | 1 | 1 | 1 | NA | NA | NA | NA | 1 | 1 | NA | NA | 1 | 1 |
| 6_8776588 | 1 | 0 | NA | NA | 4 | 0.25 | 4 | 0 | 9 | 0.1111 | 2 | 0 | 1 | 0 | 2 | 0 | 1 | 1 | 6 | 0.1667 |
| 6_8776735 | 1 | 0 | NA | NA | 3 | 0.6667 | 2 | 1 | 6 | 0.6667 | 2 | 1 | 1 | 0 | 2 | 0 | 1 | 0 | 6 | 0.3333 |
| 6_8776742 | 1 | 0 | NA | NA | 3 | 0.3333 | 2 | 0.5 | 6 | 0.3333 | 2 | 0 | 1 | 1 | 1 | 0 | 1 | 0 | 5 | 0.2 |

| | | | | | | | | | | | | | | | | | | | | |
|---|---|---|---|---|---|---|---|---|---|---|---|---|---|---|---|---|---|---|---|---|
| 6_87768299 | 1 | 0 | NA | NA | 2 | 0.5 | 1 | 1 | 4 | 0.5 | NA | NA | 1 | 1 | 2 | 0 | NA | NA | 3 | 0.3333 |
| 6_87769135 | 1 | 1 | NA | NA | 1 | 1 | 1 | 1 | 3 | 1 | NA | NA | 1 | 1 | 2 | 0.5 | NA | NA | 3 | 0.6667 |
| 6_87769325 | 1 | 1 | NA | NA | 1 | 1 | 1 | 1 | 3 | 1 | NA | NA | NA | NA | 2 | 1 | NA | NA | 2 | 1 |
| 6_87769348 | NA | NA | NA | NA | 1 | 1 | 1 | 1 | 2 | 1 | NA | NA | NA | NA | 2 | 0.5 | NA | NA | 2 | 0.5 |
| 6_87769603 | NA | NA | NA | NA | 1 | 1 | 1 | 1 | 2 | 1 | NA | NA | NA | NA | 1 | 1 | NA | NA | 1 | 1 |
| 6_87796055 | 2 | 0 | 1 | 0 | 3 | 0.3333 | 5 | 0.2 | 11 | 0.1818 | 1 | 1 | 2 | 0 | 2 | 0.5 | 1 | 0 | 6 | 0.3333 |
| 6_87820403 | 53 | 0.1132 | 57 | 0.0877 | 44 | 0.0909 | 51 | 0.1176 | 205 | 0.1024 | 39 | 0.1795 | 56 | 0.0714 | 62 | 0.1613 | 51 | 0.0784 | 208 | 0.1202 |
| 6_87867598 | 2 | 0 | 5 | 0.4 | 6 | 0.1667 | 3 | 0 | 16 | 0.1875 | 1 | 0 | 6 | 0.3333 | 3 | 0.6667 | 5 | 0 | 15 | 0.2667 |
| 6_87867616 | 2 | 0 | 6 | 0 | 5 | 0.2 | 2 | 0.5 | 15 | 0.1333 | 3 | 0 | 5 | 0.4 | 2 | 0.5 | 4 | 0 | 14 | 0.2143 |
| 6_87879289 | 19 | 0.0526 | 9 | 0.2222 | 11 | 0 | 16 | 0.0625 | 55 | 0.0727 | 8 | 0.125 | 12 | 0.0833 | 13 | 0 | 22 | 0.2273 | 55 | 0.1273 |
| 6_87889208 | 22 | 0.0909 | 21 | 0.0476 | 22 | 0 | 27 | 0.1111 | 92 | 0.0652 | 12 | 0 | 22 | 0.0455 | 17 | 0.0588 | 35 | 0.0857 | 86 | 0.0581 |
| 6_87889501 | 23 | 0.3478 | 25 | 0.48 | 24 | 0.3333 | 25 | 0.2 | 97 | 0.3402 | 14 | 0.1429 | 25 | 0.36 | 21 | 0.4286 | 38 | 0.3158 | 98 | 0.3265 |
| 6_87889662 | 21 | 0.381 | 20 | 0.3 | 18 | 0.2222 | 19 | 0.3158 | 78 | 0.3077 | 10 | 0.3 | 19 | 0.4211 | 15 | 0.1333 | 30 | 0.3 | 74 | 0.2973 |
| 6_87889929 | 20 | 0.05 | 15 | 0 | 18 | 0.0556 | 21 | 0.0952 | 74 | 0.0541 | 10 | 0.2 | 17 | 0.1176 | 12 | 0.0833 | 27 | 0.1852 | 66 | 0.1515 |
| 6_87889989 | 20 | 0.15 | 18 | 0.1111 | 18 | 0.1111 | 20 | 0.2 | 76 | 0.1447 | 10 | 0.1 | 18 | 0.2222 | 14 | 0.0714 | 32 | 0.125 | 74 | 0.1351 |
| 6_87890368 | 23 | 0.1739 | 17 | 0.1176 | 23 | 0.1739 | 25 | 0.16 | 88 | 0.1591 | 10 | 0 | 16 | 0.0625 | 15 | 0.0667 | 32 | 0.1562 | 73 | 0.0959 |
| 6_87890484 | 28 | 0.0714 | 15 | 0 | 24 | 0.125 | 31 | 0.0645 | 98 | 0.0714 | 20 | 0.05 | 19 | 0.0526 | 21 | 0.0952 | 36 | 0.0278 | 96 | 0.0521 |
| 6_87913359 | 6 | 0.6667 | 2 | 0.5 | 4 | 0.5 | NA | NA | 12 | 0.5833 | 5 | 0.4 | 4 | 0.5 | 4 | 0.75 | 4 | 0.5 | 17 | 0.5294 |
| 6_87913419 | 4 | 0.25 | 2 | 0 | 4 | 0.25 | 1 | 0 | 11 | 0.1818 | 3 | 0.3333 | 3 | 0.3333 | 4 | 0.5 | 4 | 0.25 | 14 | 0.3571 |
| 6_87957905 | 15 | 0 | 15 | 0.2 | 13 | 0.4615 | 17 | 0.2353 | 60 | 0.2167 | 11 | 0.2727 | 15 | 0.2 | 15 | 0.2 | 13 | 0.1538 | 54 | 0.2037 |
| 6_87977058 | 44 | 0.25 | 51 | 0.1765 | 35 | 0.0857 | 33 | 0.2727 | 163 | 0.1963 | 39 | 0.359 | 37 | 0.2973 | 42 | 0.1667 | 29 | 0.3793 | 147 | 0.2925 |
| 6_87977126 | 49 | 0.2449 | 62 | 0.2581 | 41 | 0.2683 | 36 | 0.2778 | 188 | 0.2606 | 40 | 0.425 | 41 | 0.2683 | 52 | 0.2692 | 35 | 0.2857 | 168 | 0.3095 |
| 6_88182338 | 1 | 1 | 3 | 0.6667 | NA | NA | NA | NA | 4 | 0.75 | NA | NA | NA | NA | 1 | 1 | NA | NA | 1 | 1 |
| 6_88182503 | NA | NA | 2 | 1 | NA | NA | NA | NA | 2 | 1 | NA | NA | NA | NA | 1 | 1 | NA | NA | 1 | 1 |
| 6_88182765 | 1 | 1 | 3 | 0.6667 | NA | NA | NA | NA | 4 | 0.75 | NA | NA | NA | NA | 1 | 1 | NA | NA | 1 | 1 |
| 6_88182794 | 1 | 1 | 3 | 0 | NA | NA | NA | NA | 4 | 0.25 | NA | NA | NA | NA | 1 | 1 | NA | NA | 1 | 1 |
| 6_88234604 | NA | NA | 1 | 1 | NA | NA | NA | NA | 1 | 1 | NA | NA | 1 | 0 | 1 | 1 | 1 | 1 | 3 | 0.6667 |
| 6_88235033 | 1 | 0 | 5 | 0.2 | 1 | 0 | 2 | 0 | 9 | 0.1111 | 2 | 0.5 | 4 | 0.25 | 3 | 0.3333 | 2 | 0 | 11 | 0.2727 |
| 6_88235051 | 2 | 0 | 5 | 0 | 1 | 1 | 1 | 1 | 9 | 0.2222 | 4 | 0 | 4 | 0 | 3 | 0.3333 | 4 | 0 | 15 | 0.0667 |
| 6_88235178 | 2 | 0 | 5 | 0.2 | 1 | 0 | 1 | 0 | 9 | 0.1111 | 3 | 0 | 2 | 0 | 3 | 0.3333 | 4 | 0 | 12 | 0.0833 |
| 6_88235191 | 2 | 0.5 | 7 | 0.7143 | 1 | 0 | 2 | 0 | 12 | 0.5 | 3 | 1 | 5 | 0.2 | 3 | 0.6667 | 4 | 0.5 | 15 | 0.5333 |
| 6_88235248 | 2 | 0 | 6 | 0.1667 | 1 | 0 | 2 | 0 | 11 | 0.0909 | 3 | 0.3333 | 3 | 0 | 3 | 0 | 4 | 0 | 13 | 0.0769 |
| 6_88235299 | 1 | 0 | 3 | 0 | 1 | 1 | 1 | 0 | 6 | 0.1667 | 1 | 1 | 5 | 0 | 2 | 0 | 3 | 0 | 11 | 0.0909 |
| 6_88274629 | 2 | 0 | 5 | 0.6 | 1 | 1 | 2 | 0 | 10 | 0.4 | 3 | 0.3333 | 4 | 0.5 | 2 | 0 | 2 | 0 | 11 | 0.2727 |
| 6_88274635 | 1 | 0 | 5 | 0 | 1 | 0 | 2 | 0.5 | 9 | 0.1111 | 3 | 0 | 5 | 0 | 2 | 0 | 3 | 0.3333 | 13 | 0.0769 |
| 6_88274706 | 2 | 0.5 | 3 | 0.6667 | 1 | 0 | 2 | 0 | 8 | 0.375 | 3 | 0 | 4 | 0.5 | 1 | 0 | 3 | 0 | 11 | 0.1818 |
| 6_88274818 | 2 | 0.5 | 4 | 0.75 | 1 | 0 | 2 | 0 | 9 | 0.4444 | 3 | 0.6667 | 4 | 0 | 1 | 1 | 3 | 0 | 11 | 0.2727 |
| 6_88274978 | 1 | 0 | 3 | 1 | 1 | 0 | 1 | 1 | 6 | 0.6667 | 2 | 1 | 3 | 0.6667 | 1 | 1 | 3 | 0.6667 | 9 | 0.7778 |
| 6_88275024 | 2 | 0 | 4 | 0.25 | 1 | 0 | 2 | 0.5 | 9 | 0.2222 | 3 | 0.3333 | 4 | 0 | 1 | 0 | 3 | 0 | 11 | 0.0909 |
| 6_88480658 | 9 | 0.3333 | 14 | 0.3571 | 14 | 0.3571 | 3 | 0 | 40 | 0.325 | 8 | 0.25 | 9 | 0.3333 | 13 | 0.3077 | 9 | 0.3333 | 39 | 0.3077 |
| 6_88480724 | 8 | 0.125 | 11 | 0.0909 | 10 | 0 | 2 | 0 | 31 | 0.0645 | 7 | 0.1429 | 7 | 0.1429 | 9 | 0.1111 | 6 | 0 | 29 | 0.1034 |
| 6_88481993 | 5 | 0.6 | 11 | 0.3636 | 9 | 0.3333 | 3 | 0.3333 | 28 | 0.3929 | 3 | 1 | 5 | 0 | 7 | 0.5714 | 3 | 1 | 18 | 0.5556 |
| 6_88482045 | 3 | 0 | 9 | 0.1111 | 6 | 0.1667 | 3 | 0 | 21 | 0.0952 | 3 | 0.3333 | 6 | 0.1667 | 7 | 0 | 1 | 0 | 17 | 0.1176 |
| 6_88482071 | 5 | 0 | 8 | 0 | 7 | 0.2857 | 1 | 0 | 21 | 0.0952 | 3 | 0.3333 | 5 | 0.6 | 5 | 0.6 | 3 | 1 | 16 | 0.625 |
| 6_88631494 | 1 | 1 | NA | NA | NA | NA | NA | NA | 1 | 1 | NA | NA | NA | NA | 1 | 1 | NA | NA | 1 | 1 |
| 6_88631837 | 1 | 1 | NA | NA | NA | NA | NA | NA | 1 | 1 | NA | NA | NA | NA | 1 | 1 | NA | NA | 1 | 1 |
| 6_88631897 | 1 | 1 | NA | NA | NA | NA | NA | NA | 1 | 1 | NA | NA | NA | NA | 1 | 1 | NA | NA | 1 | 1 |
| 6_88694664 | 1 | 1 | NA | NA | NA | NA | NA | NA | 1 | 1 | NA | NA | 1 | 1 | NA | NA | 1 | 0 | 2 | 0.5 |
| 6_88695051 | 1 | 1 | NA | NA | NA | NA | NA | NA | 1 | 1 | NA | NA | 1 | 1 | NA | NA | 1 | 0 | 2 | 0.5 |
| 6_88695057 | 1 | 1 | NA | NA | NA | NA | NA | NA | 1 | 1 | NA | NA | 1 | 1 | NA | NA | 1 | 0 | 2 | 0.5 |
| 6_88695509 | 1 | 1 | NA | NA | NA | NA | NA | NA | 1 | 1 | NA | NA | 1 | 1 | NA | NA | 1 | 1 | 2 | 1 |
| 6_88803647 | 20 | 0.1 | 17 | 0.1176 | 7 | 0.1429 | 13 | 0 | 57 | 0.0877 | 14 | 0.0714 | 12 | 0.0833 | 18 | 0.1667 | 13 | 0.0769 | 57 | 0.1053 |
| 6_88804669 | 28 | 0.1071 | 24 | 0.2083 | 16 | 0.125 | 23 | 0.0435 | 91 | 0.1209 | 18 | 0.1111 | 22 | 0 | 34 | 0.0882 | 25 | 0.12 | 99 | 0.0808 |
| 6_88804867 | 26 | 0.0385 | 17 | 0.0588 | 15 | 0.0667 | 22 | 0.1364 | 80 | 0.075 | 20 | 0.1 | 22 | 0.0455 | 34 | 0.0588 | 22 | 0.1818 | 98 | 0.0918 |
| 6_88804994 | 20 | 0.55 | 23 | 0.6522 | 19 | 0.5789 | 25 | 0.6 | 87 | 0.5977 | 21 | 0.381 | 19 | 0.4737 | 33 | 0.5455 | 23 | 0.6957 | 96 | 0.5312 |
| 6_88813036 | 4 | 0.5 | 3 | 0 | 3 | 0.3333 | 1 | 0 | 11 | 0.2727 | 7 | 0.2857 | 2 | 0 | 4 | 0 | 4 | 0.25 | 17 | 0.1765 |
| 6_88813143 | 2 | 0.5 | 3 | 0.3333 | 2 | 0 | 1 | 1 | 8 | 0.375 | 4 | 0.5 | NA | NA | 5 | 0.2 | 4 | 0.5 | 13 | 0.3846 |
| 6_88813235 | 2 | 1 | 5 | 1 | 2 | 1 | 1 | 1 | 10 | 1 | 5 | 1 | 1 | 1 | 3 | 1 | 2 | 1 | 11 | 1 |
| 6_88813259 | 3 | 0.3333 | 5 | 0 | 2 | 0 | 1 | 0 | 11 | 0.0909 | 5 | 0 | 2 | 0.5 | 6 | 0.3333 | 3 | 0 | 16 | 0.1875 |
| 6_88813601 | 2 | 0 | 3 | 0.3333 | 3 | 0 | NA | NA | 8 | 0.125 | 5 | 0.2 | NA | NA | 5 | 0.2 | 3 | 0 | 13 | 0.1538 |
| 6_88819577 | 20 | 0.45 | 16 | 0.3125 | 17 | 0.6471 | 14 | 0.5714 | 67 | 0.4925 | 19 | 0.5789 | 17 | 0.5294 | 13 | 0.6154 | 16 | 0.5625 | 65 | 0.5692 |
| 6_88819762 | 19 | 0.3158 | 17 | 0.2941 | 18 | 0.2222 | 15 | 0.5333 | 69 | 0.3333 | 19 | 0.3684 | 17 | 0.3529 | 11 | 0.3636 | 14 | 0.3571 | 61 | 0.3607 |
| 6_88819783 | 13 | 0.5385 | 16 | 0.3125 | 15 | 0.5333 | 12 | 0.0833 | 56 | 0.375 | 19 | 0.2105 | 17 | 0.4706 | 14 | 0.5 | 17 | 0.4706 | 67 | 0.403 |
| 6_88826297 | 11 | 0.2727 | 16 | 0 | 11 | 0.1818 | 9 | 0 | 47 | 0.1064 | 12 | 0.1667 | 9 | 0 | 8 | 0 | 10 | 0.2 | 39 | 0.1026 |
| 6_88826313 | 12 | 0 | 13 | 0.1538 | 11 | 0.0909 | 11 | 0.1818 | 47 | 0.1064 | 13 | 0.2308 | 10 | 0 | 9 | 0.1111 | 9 | 0.1111 | 41 | 0.122 |

| | | | | | | | | | | | | | | | | | | | | |
|---|---|---|---|---|---|---|---|---|---|---|---|---|---|---|---|---|---|---|---|---|
| 6_8882635 | 11 | 0.1818 | 16 | 0.1875 | 11 | 0.1818 | 13 | 0.3077 | 51 | 0.2157 | 8 | 0.25 | 9 | 0.2222 | 7 | 0.4286 | 12 | 0 | 36 | 0.1944 |
| 6_8882642 | 11 | 0.1818 | 16 | 0.0625 | 11 | 0 | 13 | 0.0769 | 51 | 0.0784 | 12 | 0.0833 | 9 | 0.1111 | 9 | 0.1111 | 8 | 0 | 38 | 0.0789 |
| 6_8882652 | 10 | 0.5 | 10 | 0.6 | 8 | 0.125 | 12 | 0.6667 | 40 | 0.5 | 9 | 0.3333 | 8 | 0.5 | 7 | 0.1429 | 7 | 0.5714 | 31 | 0.3871 |
| 6_8884216 | 8 | 0.25 | 11 | 0.1818 | 6 | 0 | 5 | 0.4 | 30 | 0.2 | 7 | 0.1429 | 4 | 0.25 | 5 | 0.2 | 2 | 0 | 18 | 0.1667 |
| 6_8886081 | NA | NA | 1 | 0 | 1 | 1 | NA | NA | 2 | 0.5 | NA | NA | NA | NA | 1 | 1 | NA | NA | 1 | 1 |
| 6_8888882 | 7 | 0 | 6 | 0 | 3 | 0 | 5 | 0.4 | 21 | 0.0952 | 3 | 0 | 4 | 0 | 4 | 0 | 4 | 0.25 | 15 | 0.0667 |
| 6_8888896 | 4 | 1 | 4 | 0.5 | 4 | 0.75 | 6 | 0.6667 | 18 | 0.7222 | 4 | 0.5 | 3 | 1 | 3 | 1 | 3 | 0.6667 | 13 | 0.7692 |
| 6_8888897 | 4 | 0.25 | 4 | 0 | 3 | 0.6667 | 2 | 1 | 13 | 0.3846 | 5 | 0.4 | 4 | 0.25 | 5 | 0.8 | 2 | 0.5 | 16 | 0.5 |
| 6_8929371 | 1 | 0 | 4 | 0.25 | 4 | 0 | 9 | 0 | 18 | 0.0556 | 4 | 0 | 5 | 0.2 | 3 | 0.6667 | 6 | 0.1667 | 18 | 0.2222 |
| 6_8929376 | NA | NA | 5 | 0 | 5 | 0 | 8 | 0.125 | 18 | 0.0556 | 5 | 0 | 3 | 0 | 3 | 0 | 6 | 0.1667 | 17 | 0.0588 |
| 6_8929386 | 1 | 1 | 3 | 0 | 5 | 0 | 7 | 0 | 16 | 0.0625 | 4 | 0 | 4 | 0.25 | 2 | 0 | 4 | 0.75 | 14 | 0.2857 |
| 6_8929413 | 1 | 1 | 3 | 0.3333 | 2 | 0.5 | 4 | 0.25 | 10 | 0.4 | 3 | 0 | 4 | 0.5 | 2 | 0 | 3 | 0.3333 | 12 | 0.25 |
| 6_8929487 | 1 | 0 | 4 | 0 | 4 | 0 | 6 | 0.1667 | 15 | 0.0667 | 3 | 0 | 3 | 0 | 1 | 0 | 5 | 0.2 | 12 | 0.0833 |
| 6_8929505 | 1 | 1 | 4 | 0.25 | 4 | 0 | 5 | 0.2 | 14 | 0.2143 | 4 | 0 | 2 | 0.5 | 2 | 0 | 6 | 0 | 14 | 0.0714 |
| 6_8929521 | 1 | 0 | 5 | 0 | 4 | 0 | 5 | 0.2 | 15 | 0.0667 | 3 | 0 | 3 | 0 | 1 | 0 | 6 | 0.1667 | 13 | 0.0769 |
| 6_8929531 | NA | NA | 4 | 0.25 | 3 | 0.3333 | 5 | 0.2 | 12 | 0.25 | 4 | 0.25 | 3 | 0.6667 | NA | NA | 6 | 0 | 13 | 0.2308 |
| 6_8929536 | 1 | 0 | 4 | 0.75 | 3 | 1 | 5 | 0.6 | 13 | 0.6923 | 4 | 0.5 | 3 | 0.6667 | 1 | 0 | 6 | 0.5 | 14 | 0.5 |
| 6_8929547 | 1 | 0 | 5 | 0.2 | 3 | 0.6667 | 5 | 0.6 | 14 | 0.4286 | 3 | 0.6667 | 3 | 1 | 1 | 0 | 5 | 0 | 12 | 0.4167 |
| 6_8929572 | 1 | 1 | 4 | 1 | 4 | 0.75 | 4 | 1 | 13 | 0.9231 | 4 | 1 | 3 | 1 | 1 | 1 | 4 | 0.75 | 12 | 0.9167 |
| 6_8929573 | 1 | 1 | 4 | 0.25 | 4 | 0 | 5 | 0.2 | 14 | 0.2143 | 3 | 1 | 2 | 0 | 1 | 1 | 4 | 0.25 | 10 | 0.5 |
| 6_8929598 | 1 | 0 | 2 | 0 | 3 | 0.6667 | 5 | 0.6 | 11 | 0.4545 | 2 | 0.5 | 2 | 1 | 1 | 0 | 4 | 0.25 | 9 | 0.4444 |
| 6_8929600 | 1 | 0 | 4 | 0.75 | 3 | 0.6667 | 5 | 1 | 13 | 0.7692 | 3 | 0.6667 | 3 | 1 | 1 | 1 | 5 | 0.6 | 12 | 0.75 |
| 6_8929630 | 1 | 0 | 2 | 0 | 2 | 0 | 5 | 0.4 | 10 | 0.2 | 3 | 0.3333 | 2 | 0.5 | 1 | 0 | 5 | 0 | 11 | 0.1818 |
| 6_8929638 | NA | NA | 4 | 0.25 | 3 | 0.3333 | 4 | 0.25 | 11 | 0.2727 | 2 | 1 | 2 | 0 | 1 | 1 | 3 | 0.3333 | 8 | 0.5 |
| 6_9035886 | 1 | 0 | 2 | 0.5 | 4 | 0.5 | 1 | 1 | 8 | 0.5 | 2 | 1 | 2 | 0.5 | 2 | 0.5 | NA | NA | 6 | 0.6667 |
| 6_9035897 | NA | NA | 3 | 1 | 5 | 1 | 2 | 1 | 10 | 1 | 3 | 1 | 2 | 1 | 3 | 1 | NA | NA | 8 | 1 |
| 6_9035903 | 1 | 0 | 3 | 0.3333 | 4 | 0.25 | 1 | 0 | 9 | 0.2222 | 3 | 0.3333 | 2 | 0 | 2 | 0 | NA | NA | 7 | 0.1429 |
| 6_9035904 | 1 | 0 | 1 | 0 | 2 | 0.5 | 2 | 0 | 6 | 0.1667 | 3 | 0.6667 | 2 | 1 | 2 | 0.5 | NA | NA | 7 | 0.7143 |
| 6_9035905 | 1 | 1 | 3 | 0.3333 | 5 | 0.6 | 2 | 1 | 11 | 0.6364 | 3 | 0 | 3 | 1 | 3 | 0.6667 | NA | NA | 9 | 0.5556 |
| 6_9035910 | 1 | 1 | 3 | 1 | 1 | 1 | 2 | 1 | 7 | 1 | 3 | 1 | 3 | 1 | NA | NA | NA | NA | 6 | 1 |
| 6_9035912 | 1 | 0 | 4 | 0.75 | 4 | 1 | 2 | 1 | 11 | 0.8182 | 2 | 0 | 2 | 0.5 | 2 | 1 | NA | NA | 6 | 0.5 |
| 6_9035922 | 1 | 1 | 1 | 1 | 6 | 0.8333 | 2 | 0.5 | 10 | 0.8 | 2 | 1 | 3 | 1 | 4 | 0.75 | NA | NA | 9 | 0.8889 |
| 6_9035924 | 1 | 1 | 4 | 0.5 | 6 | 0.6667 | 2 | 1 | 13 | 0.6923 | 3 | 0.3333 | 3 | 0.3333 | 4 | 0.75 | NA | NA | 10 | 0.5 |
| 6_9035926 | NA | NA | 3 | 0 | 6 | 0.1667 | NA | NA | 9 | 0.1111 | 2 | 0 | 3 | 0.3333 | 2 | 0 | NA | NA | 7 | 0.1429 |
| 6_9035927 | 1 | 1 | 3 | 0.3333 | 6 | 0 | 2 | 0 | 12 | 0.1667 | 2 | 0 | 3 | 0 | 4 | 0.25 | NA | NA | 9 | 0.1111 |
| 6_9035936 | NA | NA | 2 | 0.5 | 6 | 0.8333 | 2 | 1 | 10 | 0.8 | 2 | 1 | 2 | 1 | 4 | 1 | NA | NA | 8 | 1 |
| 6_9035946 | 1 | 1 | 3 | 1 | 6 | 1 | 1 | 0 | 11 | 0.9091 | 3 | 0.6667 | 2 | 1 | 4 | 1 | NA | NA | 9 | 0.8889 |
| 6_9035961 | 1 | 1 | 2 | 1 | 4 | 1 | 1 | 0 | 8 | 0.875 | 1 | 1 | 2 | 1 | 3 | 1 | NA | NA | 6 | 1 |
| 6_9035964 | 1 | 1 | 2 | 1 | 3 | 0.3333 | 1 | 0 | 7 | 0.5714 | 1 | 0 | 1 | 0 | 3 | 0.6667 | NA | NA | 5 | 0.4 |
| 6_9035965 | NA | NA | 1 | 0 | 3 | 1 | NA | NA | 4 | 0.75 | NA | NA | 2 | 0.5 | 2 | 0.5 | NA | NA | 4 | 0.5 |
| 6_9035967 | 1 | 1 | 2 | 1 | 4 | 1 | 1 | 1 | 8 | 1 | 1 | 1 | 2 | 1 | 2 | 1 | NA | NA | 5 | 1 |
| 6_9035981 | 1 | 0 | 2 | 0 | 4 | 0.25 | 1 | 0 | 8 | 0.125 | 1 | 1 | 2 | 0.5 | 3 | 0 | NA | NA | 6 | 0.3333 |
| 6_9035991 | 1 | 0 | 3 | 0 | 4 | 0.25 | 1 | 0 | 9 | 0.1111 | 1 | 0 | 3 | 0.6667 | 3 | 0.3333 | NA | NA | 7 | 0.4286 |
| 6_9036001 | NA | NA | 2 | 0.5 | 4 | 0.5 | 1 | 1 | 7 | 0.5714 | 1 | 0 | 3 | 1 | 3 | 0.3333 | NA | NA | 7 | 0.5714 |
| 6_9037916 | NA | NA | 2 | 1 | 2 | 1 | 3 | 0.3333 | 7 | 0.7143 | 2 | 0.5 | 1 | 1 | 1 | 1 | NA | NA | 4 | 0.75 |
| 6_9037920 | NA | NA | NA | NA | 2 | 1 | 3 | 1 | 5 | 1 | 2 | 1 | 1 | 1 | 1 | 1 | NA | NA | 4 | 1 |
| 6_9044343 | 3 | 0 | 5 | 0 | 4 | 0 | 3 | 0.3333 | 15 | 0.0667 | 3 | 0 | 1 | 0 | 5 | 0.2 | 3 | 0 | 12 | 0.0833 |
| 6_9044359 | 4 | 0.5 | 6 | 0.5 | 5 | 0.6 | 5 | 0 | 20 | 0.4 | 3 | 0 | 1 | 1 | 5 | 0 | 5 | 0.2 | 14 | 0.1429 |
| 6_9044373 | 4 | 0 | 5 | 0.6 | 6 | 0.3333 | 4 | 0 | 19 | 0.2632 | 3 | 1 | 1 | 0 | 6 | 0.5 | 5 | 0.6 | 15 | 0.6 |
| 6_9044377 | 3 | 1 | 4 | 1 | 4 | 0.5 | 4 | 1 | 15 | 0.8667 | 3 | 0.6667 | 1 | 1 | 3 | 1 | 3 | 1 | 10 | 0.9 |
| 6_9044380 | 4 | 0.25 | 5 | 0.4 | 5 | 0.4 | 5 | 0.8 | 19 | 0.4737 | 3 | 1 | NA | NA | 4 | 0.5 | 5 | 1 | 12 | 0.8333 |
| 6_9044387 | 5 | 0.2 | 5 | 0.2 | 5 | 0.2 | 2 | 0.5 | 17 | 0.2353 | 3 | 0 | 1 | 0 | 4 | 0 | 5 | 0.2 | 13 | 0.0769 |
| 6_9044392 | 5 | 0.8 | 6 | 0.8333 | 6 | 0.6667 | 5 | 1 | 22 | 0.8182 | 4 | 1 | NA | NA | 6 | 1 | 5 | 0.8 | 15 | 0.9333 |
| 6_9044392 | 4 | 0 | 5 | 0 | 6 | 0 | 4 | 0.5 | 19 | 0.1053 | 2 | 0.5 | 1 | 1 | 5 | 0.2 | 4 | 0.25 | 12 | 0.3333 |
| 6_9044424 | 3 | 0 | 6 | 0.3333 | 7 | 0.5714 | 3 | 0 | 19 | 0.3158 | 3 | 0.3333 | 1 | 0 | 4 | 0.75 | 5 | 0.6 | 13 | 0.5385 |
| 6_9044435 | 5 | 0.2 | 6 | 0 | 6 | 0.1667 | 6 | 0.5 | 23 | 0.2174 | 4 | 0.5 | 1 | 0 | 4 | 0 | 6 | 0.3333 | 15 | 0.2667 |
| 6_9044437 | 5 | 0.6 | 6 | 0.8333 | 6 | 0.6667 | 4 | 1 | 21 | 0.7619 | 3 | 0.6667 | 2 | 0.5 | 4 | 1 | 5 | 0.6 | 14 | 0.7143 |
| 6_9045074 | 1 | 0 | NA | NA | 1 | 0 | 2 | 0.5 | 4 | 0.25 | NA | NA | NA | NA | NA | NA | 3 | 0.6667 | 3 | 0.6667 |
| 6_9045090 | 1 | 1 | NA | NA | 1 | 0 | 2 | 0 | 4 | 0.25 | NA | NA | NA | NA | NA | NA | 3 | 0.3333 | 3 | 0.3333 |
| 6_9045135 | 7 | 0 | 5 | 0 | 5 | 0.2 | 6 | 0.1667 | 23 | 0.087 | 3 | 0 | NA | NA | 5 | 0.2 | 8 | 0.125 | 16 | 0.125 |
| 6_9045171 | 9 | 0 | 5 | 0 | 5 | 0 | 10 | 0.2 | 29 | 0.069 | 5 | 0.2 | 1 | 1 | 6 | 0 | 10 | 0.2 | 22 | 0.1818 |
| 6_9045396 | NA | NA | NA | NA | 1 | 1 | 1 | 1 | 2 | 1 | NA | NA | 1 | 1 | NA | NA | 2 | 1 | 3 | 1 |
| 6_9045401 | NA | NA | NA | NA | 1 | 1 | 1 | 1 | 2 | 1 | NA | NA | 2 | 1 | NA | NA | 1 | 1 | 3 | 1 |
| 6_9045403 | NA | NA | NA | NA | 1 | 1 | 1 | 0 | 2 | 0.5 | NA | NA | 1 | 0 | NA | NA | 2 | 0.5 | 3 | 0.3333 |
| 6_9045404 | NA | NA | NA | NA | 1 | 1 | 1 | 1 | 2 | 1 | NA | NA | 2 | 0.5 | NA | NA | 2 | 1 | 4 | 0.75 |

| | | | | | | | | | | | | | | | | | | | | |
|---|---|---|---|---|---|---|---|---|---|---|---|---|---|---|---|---|---|---|---|---|
| 6_9045414 | NA | NA | NA | NA | 1 | 1 | 1 | 1 | 2 | 1 | NA | NA | 2 | 1 | NA | NA | 2 | 1 | 4 | 1 |
| 6_9045420 | NA | NA | NA | NA | NA | NA | 1 | 1 | 1 | 1 | NA | NA | 2 | 1 | NA | NA | 1 | 1 | 3 | 1 |
| 6_9045421 | NA | NA | NA | NA | 1 | 1 | 1 | 1 | 2 | 1 | NA | NA | 1 | 1 | NA | NA | 2 | 1 | 3 | 1 |
| 6_9045440 | NA | NA | NA | NA | 1 | 1 | 1 | 0 | 2 | 0.5 | NA | NA | 1 | 1 | NA | NA | 1 | 1 | 2 | 1 |
| 6_9045480 | NA | NA | NA | NA | 2 | 1 | 1 | 1 | 3 | 1 | NA | NA | 1 | 1 | NA | NA | 2 | 0.5 | 3 | 0.6667 |
| 6_9045493 | NA | NA | NA | NA | 1 | 0 | 1 | 1 | 2 | 0.5 | NA | NA | 2 | 0.5 | NA | NA | 2 | 0.5 | 4 | 0.5 |
| 6_9057551 | NA | NA | 1 | 0 | 2 | 0.5 | NA | NA | 3 | 0.3333 | NA | NA | NA | NA | 1 | 1 | 1 | 0 | 2 | 0.5 |
| 6_9057593 | 11 | 0.0909 | 6 | 0 | 5 | 0.4 | 5 | 0.4 | 27 | 0.1852 | 10 | 0.1 | 7 | 0.1429 | 4 | 0.25 | 9 | 0 | 30 | 0.1 |
| 6_9057598 | 10 | 0 | 6 | 0.1667 | 5 | 0.2 | 6 | 0 | 27 | 0.0741 | 9 | 0 | 7 | 0.1429 | 4 | 0 | 9 | 0.2222 | 29 | 0.1034 |
| 6_9062269 | 1 | 1 | NA | NA | 2 | 1 | NA | NA | 3 | 1 | NA | NA | NA | NA | 1 | 1 | 1 | 1 | 2 | 1 |
| 6_9063728 | 8 | 0.125 | 10 | 0.1 | 6 | 0.1667 | 10 | 0.2 | 34 | 0.1471 | 8 | 0.125 | 13 | 0.0769 | 16 | 0.0625 | 11 | 0 | 48 | 0.0625 |
| 6_9063745 | 21 | 0.381 | 17 | 0.6471 | 12 | 0.5833 | 18 | 0.5556 | 68 | 0.5294 | 26 | 0.5385 | 24 | 0.5833 | 27 | 0.7407 | 20 | 0.6 | 97 | 0.6186 |
| 6_9063747 | 21 | 0.5238 | 18 | 0.3333 | 15 | 0.4667 | 18 | 0.5 | 72 | 0.4583 | 25 | 0.52 | 24 | 0.7083 | 27 | 0.6667 | 21 | 0.381 | 97 | 0.5773 |
| 6_9063756 | 22 | 0.3636 | 18 | 0.3333 | 15 | 0.2667 | 18 | 0.4444 | 73 | 0.3562 | 27 | 0.3704 | 24 | 0.4167 | 27 | 0.2963 | 20 | 0.4 | 98 | 0.3673 |
| 6_9063758 | 20 | 0.95 | 19 | 1 | 16 | 0.9375 | 16 | 0.9375 | 71 | 0.9577 | 20 | 0.9 | 20 | 0.95 | 18 | 0.9444 | 21 | 0.9524 | 79 | 0.9367 |
| 6_9063765 | 24 | 0.125 | 20 | 0.2 | 17 | 0.1176 | 18 | 0.0556 | 79 | 0.1266 | 25 | 0.2 | 25 | 0.16 | 27 | 0.1852 | 20 | 0 | 97 | 0.1443 |
| 6_9063772 | 20 | 0.3 | 17 | 0.4706 | 12 | 0.4167 | 13 | 0.2308 | 62 | 0.3548 | 18 | 0.3333 | 17 | 0.6471 | 22 | 0.5455 | 14 | 0.5 | 71 | 0.507 |
| 6_9063786 | 23 | 0.3478 | 21 | 0.4286 | 14 | 0.6429 | 15 | 0.6 | 73 | 0.4795 | 25 | 0.68 | 24 | 0.4583 | 25 | 0.6 | 22 | 0.5455 | 96 | 0.5729 |
| 6_9063794 | 19 | 0.7895 | 17 | 0.8824 | 13 | 0.6923 | 15 | 0.8 | 64 | 0.7969 | 22 | 0.7273 | 18 | 0.6667 | 25 | 0.84 | 15 | 0.7333 | 80 | 0.75 |
| 6_9063795 | 20 | 0.1 | 20 | 0 | 14 | 0.2143 | 15 | 0.2 | 69 | 0.1159 | 23 | 0.1304 | 24 | 0.0833 | 26 | 0.1923 | 19 | 0.1579 | 92 | 0.1413 |
| 6_9063971 | 8 | 0.125 | 6 | 0 | 4 | 0 | 3 | 0.3333 | 21 | 0.0952 | 10 | 0.2 | 5 | 0.4 | 8 | 0 | 3 | 0 | 26 | 0.1538 |
| 6_9064165 | NA | NA | 1 | 0 | 1 | 1 | NA | NA | 2 | 0.5 | 1 | 1 | 1 | 1 | NA | NA | 2 | 1 | 4 | 1 |
| 6_9064215 | NA | NA | 1 | 0 | 1 | 1 | NA | NA | 2 | 0.5 | 2 | 1 | NA | NA | NA | NA | 2 | 0 | 4 | 0.5 |
| 6_9064229 | NA | NA | NA | NA | 1 | 1 | NA | NA | 1 | 1 | 1 | 1 | 1 | 1 | NA | NA | NA | NA | 2 | 1 |
| 6_9064253 | NA | NA | NA | NA | 1 | 1 | NA | NA | 1 | 1 | 2 | 1 | 1 | 1 | NA | NA | NA | NA | 3 | 1 |
| 6_9064297 | NA | NA | NA | NA | 1 | 1 | NA | NA | 1 | 1 | 1 | 1 | 1 | 1 | NA | NA | 1 | 1 | 3 | 1 |
| 6_9066663 | 10 | 0.1 | 6 | 0.1667 | 3 | 0 | 4 | 0.5 | 23 | 0.1739 | 8 | 0 | 13 | 0.0769 | 10 | 0.1 | 9 | 0.1111 | 40 | 0.075 |
| 6_9066667 | 10 | 0.3 | 5 | 0 | 3 | 0 | 4 | 0.25 | 22 | 0.1818 | 6 | 0.3333 | 15 | 0.1333 | 8 | 0.125 | 9 | 0.2222 | 38 | 0.1842 |
| 6_9066669 | 5 | 0.2 | 4 | 0 | 2 | 0 | 4 | 0.25 | 15 | 0.1333 | 5 | 0.4 | 10 | 0.2 | 9 | 0.1111 | 6 | 0 | 30 | 0.1667 |
| 6_9066672 | 5 | 0 | 3 | 0.3333 | 2 | 0.5 | 3 | 0 | 13 | 0.1538 | 4 | 0.25 | 12 | 0.3333 | 9 | 0.2222 | 7 | 0.1429 | 32 | 0.25 |
| 6_9066676 | 10 | 0.8 | 5 | 1 | 2 | 0.5 | 4 | 0.5 | 21 | 0.7619 | 6 | 0.6667 | 12 | 0.5833 | 9 | 0.8889 | 9 | 0.8889 | 36 | 0.75 |
| 6_9066684 | 7 | 0.1429 | 5 | 0 | 1 | 0 | 4 | 0 | 17 | 0.0588 | 6 | 0.1667 | 13 | 0.1538 | 9 | 0 | 10 | 0 | 38 | 0.0789 |
| 6_9066699 | 8 | 0.125 | 4 | 0 | 2 | 0 | 5 | 0.2 | 19 | 0.1053 | 5 | 0 | 13 | 0.1538 | 8 | 0 | 8 | 0.125 | 34 | 0.0882 |
| 6_9066699 | 6 | 0.8333 | 3 | 1 | 2 | 0 | 4 | 0.75 | 15 | 0.7333 | 6 | 0.6667 | 9 | 0.5556 | 6 | 0.6667 | 9 | 0.3333 | 30 | 0.5333 |
| 6_9066710 | 8 | 0.25 | 3 | 0.6667 | 1 | 0 | 4 | 0.25 | 16 | 0.3125 | 5 | 0.4 | 11 | 0.6364 | 7 | 0.7143 | 7 | 0.5714 | 30 | 0.6 |
| 6_9066712 | 7 | 0.2857 | 4 | 0 | 1 | 1 | 4 | 0 | 16 | 0.1875 | 6 | 0 | 11 | 0.2727 | 7 | 0.1429 | 9 | 0.6667 | 33 | 0.303 |
| 6_9066714 | 5 | 0.8 | 2 | 0.5 | 2 | 1 | 4 | 1 | 13 | 0.8462 | 5 | 1 | 9 | 1 | 5 | 0.8 | 9 | 1 | 28 | 0.9643 |
| 6_9066718 | 4 | 0.25 | 4 | 0 | 2 | 0 | 4 | 0.25 | 14 | 0.1429 | 4 | 0.25 | 11 | 0 | 7 | 0.2857 | 7 | 0.4286 | 29 | 0.2069 |
| 6_9066730 | 7 | 0.1429 | 3 | 0 | 1 | 1 | 2 | 0 | 13 | 0.1538 | 4 | 0.25 | 9 | 0.1111 | 8 | 0.375 | 7 | 0.1429 | 28 | 0.2143 |
| 6_9066745 | 6 | 0.1667 | 3 | 0 | 2 | 0 | 2 | 0.5 | 13 | 0.1538 | 4 | 0.25 | 8 | 0.125 | 7 | 0.1429 | 8 | 0.125 | 27 | 0.1481 |
| 6_9066748 | 5 | 0.4 | 2 | 0.5 | 2 | 0.5 | 2 | 0.5 | 11 | 0.4545 | 3 | 0.3333 | 9 | 0.4444 | 6 | 0.6667 | 7 | 0.5714 | 25 | 0.52 |
| 6_9066757 | 6 | 0.3333 | 3 | 0 | 1 | 1 | 2 | 0 | 12 | 0.25 | 3 | 0.3333 | 10 | 0.4 | 6 | 0.6667 | 8 | 0.5 | 27 | 0.4815 |
| 6_9066771 | 4 | 0.5 | 2 | 0 | 2 | 0.5 | 2 | 0 | 10 | 0.3 | 4 | 0.5 | 9 | 0.3333 | 8 | 0.125 | 7 | 0.4286 | 28 | 0.3214 |
| 6_9099016 | 2 | 1 | 3 | 0.6667 | NA | NA | 4 | 1 | 9 | 0.8889 | 2 | 1 | 4 | 0.5 | 4 | 1 | 1 | 1 | 11 | 0.8182 |
| 6_9099026 | 2 | 0.5 | 2 | 1 | NA | NA | 4 | 0.75 | 8 | 0.75 | 2 | 1 | 4 | 1 | 5 | 1 | 1 | 1 | 12 | 1 |
| 6_9099046 | 4 | 0.25 | 2 | 0.5 | NA | NA | 4 | 0.75 | 10 | 0.5 | 3 | 0.3333 | 4 | 0.5 | 7 | 0.7143 | 1 | 1 | 15 | 0.6 |
| 6_9099067 | 4 | 0.5 | 3 | 0 | NA | NA | 4 | 0 | 11 | 0.1818 | NA | NA | 3 | 0 | 5 | 0.2 | 1 | 0 | 9 | 0.1111 |
| 6_9099133 | 2 | 0.5 | 1 | 0 | NA | NA | 3 | 0 | 6 | 0.1667 | 1 | 0 | 1 | 1 | 3 | 0.3333 | NA | NA | 5 | 0.4 |
| 6_9115010 | 9 | 0.1111 | 19 | 0 | 16 | 0.0625 | 21 | 0.1429 | 65 | 0.0769 | 9 | 0.1111 | 20 | 0.05 | 8 | 0.125 | 14 | 0.0714 | 51 | 0.0784 |
| 6_9115020 | 13 | 0.1538 | 22 | 0.0455 | 16 | 0.25 | 23 | 0.0435 | 74 | 0.1081 | 14 | 0.0714 | 21 | 0.0952 | 8 | 0 | 17 | 0.3529 | 60 | 0.15 |
| 6_9115026 | 8 | 0.125 | 15 | 0.2 | 10 | 0.6 | 17 | 0.5882 | 50 | 0.4 | 10 | 0.7 | 15 | 0.4 | 8 | 0.5 | 11 | 0.5455 | 44 | 0.5227 |
| 6_9115132 | 15 | 0.4 | 28 | 0.5 | 26 | 0.3462 | 26 | 0.3462 | 95 | 0.4 | 26 | 0.4615 | 25 | 0.32 | 12 | 0.4167 | 19 | 0.4211 | 82 | 0.4024 |
| 6_9115151 | 20 | 0.05 | 29 | 0.069 | 25 | 0.16 | 27 | 0.1481 | 101 | 0.1089 | 31 | 0.129 | 26 | 0.2308 | 17 | 0 | 13 | 0 | 87 | 0.1149 |
| 6_9124876 | NA | NA | 1 | 1 | 2 | 1 | 2 | 0.5 | 5 | 0.8 | 2 | 0.5 | NA | NA | NA | NA | NA | NA | 2 | 0.5 |
| 6_9124894 | NA | NA | 1 | 0 | 2 | 0.5 | 2 | 0.5 | 5 | 0.4 | 2 | 0.5 | NA | NA | NA | NA | NA | NA | 2 | 0.5 |
| 6_9124894 | NA | NA | 1 | 0 | 2 | 0.5 | 2 | 0 | 5 | 0.2 | 2 | 0.5 | NA | NA | NA | NA | NA | NA | 2 | 0.5 |
| 6_9124909 | NA | NA | 1 | 0 | 2 | 0.5 | 2 | 0 | 5 | 0.2 | 2 | 1 | NA | NA | NA | NA | NA | NA | 2 | 1 |
| 6_9134114 | 3 | 0.3333 | NA | NA | 2 | 0 | 1 | 0 | 6 | 0.1667 | 1 | 0 | 6 | 0 | 1 | 0 | 3 | 0.3333 | 11 | 0.0909 |
| 6_9134201 | 2 | 1 | NA | NA | 2 | 0.5 | 1 | 1 | 5 | 0.8 | NA | NA | 4 | 1 | NA | NA | 1 | 1 | 5 | 1 |
| 6_9134240 | 3 | 1 | NA | NA | 2 | 1 | NA | NA | 5 | 1 | NA | NA | 2 | 1 | NA | NA | 1 | 1 | 3 | 1 |
| 6_9134241 | 3 | 0.6667 | NA | NA | 2 | 0.5 | NA | NA | 5 | 0.6 | NA | NA | 1 | 0 | NA | NA | 1 | 1 | 2 | 0.5 |
| 6_9134244 | 2 | 1 | NA | NA | 2 | 1 | 1 | 1 | 5 | 1 | NA | NA | 1 | 1 | NA | NA | 1 | 1 | 2 | 1 |
| 6_9134253 | 1 | 1 | NA | NA | 1 | 0 | NA | NA | 2 | 0.5 | 1 | 1 | 2 | 0 | NA | NA | NA | NA | 3 | 0.3333 |
| 6_9134269 | 1 | 0 | NA | NA | 1 | 1 | NA | NA | 2 | 0.5 | NA | NA | 1 | 0 | 1 | 1 | NA | NA | 2 | 0.5 |
| 6_9134303 | 1 | 1 | NA | NA | 1 | 0 | NA | NA | 2 | 0.5 | 1 | 1 | NA | NA | 1 | 1 | NA | NA | 2 | 1 |

| | | | | | | | | | | | | | | | | | | | | |
|---|---|---|---|---|---|---|---|---|---|---|---|---|---|---|---|---|---|---|---|---|
| 6_9134308 | 1 | 1 | NA | NA | NA | NA | NA | NA | 1 | 1 | 1 | 1 | NA | NA | 1 | 1 | NA | NA | 2 | 1 |
| 6_9144173 | 8 | 0 | 7 | 0.2857 | 7 | 0.1429 | 12 | 0.0833 | 34 | 0.1176 | 6 | 0.3333 | 13 | 0.2308 | 9 | 0.1111 | 6 | 0.1667 | 34 | 0.2059 |
| 6_9144189 | 7 | 0 | 6 | 0.1667 | 8 | 0.375 | 7 | 0.1429 | 28 | 0.1786 | 3 | 0.3333 | 10 | 0.2 | 6 | 0.1667 | 7 | 0.1429 | 26 | 0.1923 |
| 6_9144196 | 8 | 1 | 7 | 1 | 9 | 1 | 12 | 1 | 36 | 1 | 4 | 1 | 11 | 1 | 7 | 1 | 6 | 1 | 28 | 1 |
| 6_9144200 | 5 | 0.6 | 6 | 0.5 | 9 | 0.2222 | 9 | 0.2222 | 29 | 0.3448 | 5 | 0 | 11 | 0.4545 | 7 | 0.2857 | 5 | 0.4 | 28 | 0.3214 |
| 6_9144216 | 7 | 0.8571 | 3 | 0.6667 | 7 | 0.7143 | 10 | 1 | 27 | 0.8519 | 2 | 1 | 10 | 0.7 | 5 | 0.6 | 5 | 0.6 | 22 | 0.6818 |
| 6_9146381 | 6 | 0.3333 | 4 | 0.25 | 3 | 0.3333 | 5 | 0.6 | 18 | 0.3889 | 4 | 0.25 | 2 | 0.5 | 1 | 1 | 2 | 0.5 | 9 | 0.4444 |
| 6_9146390 | 5 | 0.6 | 5 | 0.6 | 2 | 1 | 6 | 0.8333 | 18 | 0.7222 | 4 | 0.75 | 2 | 0.5 | 2 | 1 | 2 | 1 | 10 | 0.8 |
| 6_9146415 | 6 | 0.3333 | 5 | 0.4 | 1 | 0 | 6 | 0.1667 | 18 | 0.2778 | 2 | 0.5 | 5 | 0.2 | 2 | 0.5 | 2 | 0 | 11 | 0.2727 |
| 6_9146419 | 4 | 0 | 4 | 0.5 | 2 | 0 | 5 | 0 | 15 | 0.1333 | 5 | 0.2 | 3 | 0 | 2 | 0 | 2 | 0.5 | 12 | 0.1667 |
| 6_9146423 | 4 | 1 | 2 | 1 | 2 | 1 | 5 | 0.8 | 13 | 0.9231 | 3 | 1 | 3 | 1 | 2 | 0.5 | 1 | 1 | 9 | 0.8889 |
| 6_9146433 | 4 | 1 | 6 | 0.6667 | 3 | 0.6667 | 6 | 0.8333 | 19 | 0.7895 | 5 | 1 | 5 | 0.8 | 2 | 1 | 1 | 1 | 13 | 0.9231 |
| 6_9146446 | 6 | 1 | 3 | 1 | 2 | 1 | 4 | 1 | 15 | 1 | 4 | 1 | 3 | 0.6667 | 2 | 1 | 2 | 1 | 11 | 0.9091 |
| 6_9146450 | 7 | 0.5714 | 4 | 0.75 | 2 | 0.5 | 6 | 0.3333 | 19 | 0.5263 | 4 | 0.75 | 4 | 0.75 | 2 | 1 | 2 | 1 | 12 | 0.8333 |
| 6_9146455 | 7 | 0.2857 | 4 | 0.25 | 2 | 0 | 5 | 0.4 | 18 | 0.2778 | 5 | 0.4 | 3 | 0 | 1 | 1 | 1 | 0 | 10 | 0.3 |
| 6_9146459 | 7 | 1 | 4 | 0.75 | 3 | 1 | 5 | 1 | 19 | 0.9474 | 4 | 0.75 | 4 | 1 | 1 | 1 | 2 | 1 | 11 | 0.9091 |
| 6_9146475 | 7 | 0.5714 | 5 | 1 | 2 | 1 | 5 | 0.8 | 19 | 0.7895 | 3 | 1 | 4 | 1 | 2 | 1 | 3 | 0.6667 | 12 | 0.9167 |
| 6_9146480 | 4 | 1 | 4 | 1 | 4 | 1 | 8 | 0.875 | 20 | 0.95 | 5 | 1 | 3 | 1 | 2 | 1 | 2 | 1 | 12 | 1 |
| 6_9146491 | 5 | 0.8 | 5 | 0.8 | 2 | 0.5 | 7 | 0.8571 | 19 | 0.7895 | 3 | 1 | 5 | 0.6 | 1 | 1 | 3 | 1 | 12 | 0.8333 |
| 6_9146670 | 4 | 0.25 | 1 | 0 | 2 | 0.5 | 3 | 0 | 10 | 0.2 | NA | NA | NA | NA | 1 | 1 | 3 | 0.6667 | 4 | 0.75 |
| 6_9146686 | 6 | 1 | 1 | 1 | 5 | 1 | 3 | 1 | 15 | 1 | NA | NA | 1 | 0 | 2 | 1 | 1 | 1 | 4 | 0.75 |
| 6_9146689 | 5 | 0.2 | NA | NA | 5 | 0.2 | 1 | 0 | 11 | 0.1818 | NA | NA | NA | NA | 2 | 0 | 2 | 1 | 4 | 0.5 |
| 6_9146698 | 5 | 0.4 | 1 | 1 | 2 | 0.5 | 2 | 0.5 | 10 | 0.5 | NA | NA | 1 | 0 | 2 | 0 | 3 | 0.3333 | 6 | 0.1667 |
| 6_9147640 | 3 | 0.6667 | NA | NA | NA | NA | NA | NA | 3 | 0.6667 | NA | NA | 1 | 1 | NA | NA | NA | NA | 1 | 1 |
| 6_9147648 | 3 | 1 | NA | NA | 1 | 1 | NA | NA | 4 | 1 | NA | NA | 1 | 1 | NA | NA | NA | NA | 1 | 1 |
| 6_9147669 | 1 | 0 | NA | NA | 2 | 1 | NA | NA | 3 | 0.6667 | NA | NA | 1 | 1 | NA | NA | NA | NA | 1 | 1 |
| 6_9149933 | 6 | 0.1667 | 4 | 0.25 | 6 | 0.5 | 5 | 0.2 | 21 | 0.2857 | 5 | 0.2 | 3 | 0.3333 | 4 | 0.25 | 5 | 0.4 | 17 | 0.2941 |
| 6_9149936 | 5 | 0.4 | 3 | 0.3333 | 8 | 0.375 | 5 | 0.4 | 21 | 0.381 | 4 | 0.25 | 3 | 0 | 4 | 0.5 | 3 | 0.6667 | 14 | 0.3571 |
| 6_9173228 | 5 | 0.4 | 3 | 0 | 1 | 0 | 4 | 0 | 13 | 0.1538 | 3 | 0.3333 | 3 | 0 | 5 | 0.2 | 3 | 0 | 14 | 0.1429 |
| 6_9173266 | 6 | 0 | 4 | 0 | 2 | 0 | 5 | 0.2 | 17 | 0.0588 | 4 | 0.25 | 3 | 0 | 7 | 0 | 6 | 0 | 20 | 0.05 |
| 6_9173888 | NA | NA | 1 | 0 | 1 | 0 | 1 | 1 | 3 | 0.3333 | 1 | 1 | NA | NA | 3 | 0.3333 | 4 | 0.5 | 8 | 0.5 |
| 6_9173897 | NA | NA | 2 | 1 | 1 | 1 | 1 | 1 | 4 | 1 | 1 | 1 | 1 | 1 | 4 | 0.75 | 3 | 1 | 9 | 0.8889 |
| 6_9173902 | NA | NA | 1 | 1 | 1 | 0 | NA | NA | 2 | 0.5 | 1 | 1 | 1 | 1 | 3 | 0.6667 | 2 | 1 | 7 | 0.8571 |
| 6_9173907 | NA | NA | 2 | 0.5 | 1 | 0 | 1 | 1 | 4 | 0.5 | 1 | 1 | 1 | 1 | 3 | 0 | 3 | 0.6667 | 8 | 0.5 |
| 6_9173910 | NA | NA | 1 | 0 | NA | NA | 1 | 1 | 2 | 0.5 | 1 | 0 | 1 | 1 | 4 | 0 | 1 | 0 | 7 | 0.1429 |
| 6_9173915 | NA | NA | 1 | 1 | 1 | 1 | NA | NA | 2 | 1 | 1 | 1 | 1 | 1 | 2 | 1 | 1 | 1 | 5 | 1 |
| 6_9173922 | NA | NA | 1 | 1 | NA | NA | 1 | 1 | 2 | 1 | 1 | 0 | 1 | 1 | 3 | 1 | 3 | 0.6667 | 8 | 0.75 |
| 6_9173940 | NA | NA | 2 | 0.5 | NA | NA | 1 | 1 | 3 | 0.6667 | 1 | 1 | NA | NA | 3 | 0.6667 | 1 | 0 | 5 | 0.6 |
| 6_9173941 | NA | NA | 1 | 1 | NA | NA | 1 | 1 | 2 | 1 | 1 | 1 | 1 | 1 | 3 | 0.6667 | 3 | 0.6667 | 8 | 0.75 |
| 6_9173948 | NA | NA | 2 | 0.5 | NA | NA | NA | NA | 2 | 0.5 | 1 | 0 | 1 | 1 | 3 | 0.3333 | 2 | 0.5 | 7 | 0.4286 |
| 6_9173955 | NA | NA | 2 | 1 | NA | NA | 1 | 1 | 3 | 1 | 1 | 1 | 1 | 1 | 4 | 1 | 2 | 1 | 8 | 1 |
| 6_9173967 | NA | NA | 2 | 1 | NA | NA | 1 | 1 | 3 | 1 | 1 | 1 | 1 | 1 | 4 | 0 | 3 | 0 | 9 | 0.2222 |
| 6_9173984 | NA | NA | 2 | 0 | NA | NA | 1 | 1 | 3 | 0.3333 | 1 | 1 | 1 | 1 | 4 | 0.25 | 3 | 0.3333 | 9 | 0.4444 |
| 6_9174016 | NA | NA | 2 | 1 | NA | NA | 1 | 1 | 3 | 1 | 1 | 1 | 1 | 1 | 3 | 1 | 3 | 1 | 8 | 1 |
| 6_9174047 | NA | NA | 1 | 0 | NA | NA | 1 | 1 | 2 | 0.5 | NA | NA | 1 | 0 | 1 | 0 | 3 | 0.3333 | 5 | 0.2 |
| 6_9187637 | 1 | 0 | 2 | 0 | NA | NA | 3 | 0.6667 | 6 | 0.3333 | 2 | 0 | 1 | 1 | 4 | 0.25 | 1 | 1 | 8 | 0.375 |
| 6_9187643 | 1 | 1 | 2 | 0.5 | NA | NA | 4 | 0.75 | 7 | 0.7143 | 3 | 0.6667 | 2 | 1 | 5 | 0.6 | 1 | 1 | 11 | 0.7273 |
| 6_9187661 | NA | NA | 1 | 0 | NA | NA | 5 | 0.2 | 6 | 0.1667 | 2 | 0 | 1 | 1 | 4 | 0 | 1 | 0 | 8 | 0.125 |
| 6_9205178 | NA | NA | NA | NA | NA | NA | 2 | 0.5 | 2 | 0.5 | NA | NA | 1 | 1 | NA | NA | NA | NA | 1 | 1 |
| 6_9205184 | NA | NA | NA | NA | NA | NA | 2 | 1 | 2 | 1 | NA | NA | 1 | 1 | NA | NA | NA | NA | 1 | 1 |
| 6_9205207 | NA | NA | NA | NA | NA | NA | 2 | 1 | 2 | 1 | NA | NA | 1 | 1 | NA | NA | NA | NA | 1 | 1 |
| 6_9205265 | NA | NA | NA | NA | NA | NA | 3 | 1 | 3 | 1 | NA | NA | 1 | 1 | NA | NA | NA | NA | 1 | 1 |
| 6_9205286 | NA | NA | NA | NA | NA | NA | 3 | 0.6667 | 3 | 0.6667 | NA | NA | 1 | 1 | NA | NA | NA | NA | 1 | 1 |
| 6_9205293 | NA | NA | NA | NA | NA | NA | 3 | 1 | 3 | 1 | NA | NA | 1 | 1 | NA | NA | NA | NA | 1 | 1 |
| 6_9214450 | 1 | 1 | 3 | 0.6667 | 1 | 0 | 1 | 0 | 6 | 0.5 | 3 | 0.3333 | 2 | 0.5 | NA | NA | 2 | 1 | 7 | 0.5714 |
| 6_9214454 | 1 | 1 | 2 | 0.5 | 1 | 1 | 3 | 0.6667 | 7 | 0.7143 | 3 | 0.3333 | 4 | 0.75 | 1 | 1 | 2 | 0.5 | 10 | 0.6 |
| 6_9214465 | 1 | 1 | 2 | 0.5 | NA | NA | 4 | 0.75 | 7 | 0.7143 | 3 | 0.6667 | 4 | 1 | NA | NA | 2 | 1 | 9 | 0.8889 |
| 6_9214476 | 1 | 1 | 3 | 0.6667 | 1 | 1 | 4 | 1 | 9 | 0.8889 | 2 | 0.5 | 5 | 0.6 | 1 | 1 | 2 | 1 | 10 | 0.7 |
| 6_9214881 | 7 | 0 | 6 | 0 | 5 | 0 | 6 | 0.3333 | 24 | 0.0833 | 10 | 0.2 | 10 | 0.1 | 6 | 0 | 8 | 0.125 | 34 | 0.1176 |
| 6_9215116 | 3 | 0.3333 | 3 | 0 | 1 | 0 | 3 | 0 | 10 | 0.1 | 3 | 0 | 5 | 0 | 1 | 0 | 5 | 0.4 | 14 | 0.1429 |
| 6_9215188 | 15 | 0.2667 | 14 | 0.5 | 6 | 0.5 | 9 | 0.6667 | 44 | 0.4545 | 7 | 0.5714 | 8 | 0.25 | 7 | 0.4286 | 10 | 0.5 | 32 | 0.4375 |
| 6_9215192 | 13 | 0.4615 | 12 | 0.6667 | 4 | 0 | 4 | 0.25 | 33 | 0.4545 | 6 | 0.1667 | 5 | 0.2 | 7 | 0.4286 | 8 | 0.5 | 26 | 0.3462 |
| 6_9215196 | 13 | 0.2308 | 13 | 0.2308 | 6 | 0.1667 | 8 | 0.125 | 40 | 0.2 | 8 | 0.125 | 8 | 0.375 | 8 | 0.25 | 9 | 0.2222 | 33 | 0.2424 |
| 6_9215197 | 15 | 0.6667 | 10 | 0.7 | 7 | 0.8571 | 6 | 0.5 | 38 | 0.6842 | 5 | 0.8 | 7 | 1 | 7 | 0.7143 | 8 | 0.75 | 27 | 0.8148 |
| 6_9216399 | 4 | 0.25 | 4 | 0.5 | 1 | 0 | 4 | 0.5 | 13 | 0.3846 | 3 | 0 | 4 | 0.25 | 5 | 0 | 4 | 0.25 | 16 | 0.125 |

| | | | | | | | | | | | | | | | | | | | | |
|---|---|---|---|---|---|---|---|---|---|---|---|---|---|---|---|---|---|---|---|---|
| 6_9216423 | 4 | 1 | 4 | 0.75 | 1 | 0 | 4 | 0.25 | 13 | 0.6154 | 5 | 0.4 | 4 | 0.5 | 6 | 0.5 | 3 | 0.3333 | 18 | 0.4444 |
| 6_9216461 | 4 | 0.25 | 6 | 0.1667 | 3 | 0.3333 | 4 | 0 | 17 | 0.1765 | 6 | 0 | 3 | 0 | 4 | 0.25 | 5 | 0 | 18 | 0.0556 |
| 6_9216597 | 3 | 0.3333 | 3 | 0.3333 | 1 | 1 | 3 | 0.6667 | 10 | 0.5 | 3 | 0 | 1 | 0 | 3 | 0 | 2 | 0.5 | 9 | 0.1111 |
| 6_9216606 | 3 | 0.6667 | 3 | 0.3333 | 1 | 0 | 3 | 0.3333 | 10 | 0.4 | 4 | 0.25 | 1 | 1 | 3 | 0 | 4 | 0.5 | 12 | 0.3333 |
| 6_9216616 | 3 | 0 | 2 | 0 | NA | NA | 2 | 0.5 | 7 | 0.1429 | 4 | 0.25 | 1 | 1 | 3 | 0 | 3 | 0.6667 | 11 | 0.3636 |
| 6_9216627 | 2 | 0 | 2 | 0.5 | NA | NA | 2 | 0.5 | 6 | 0.3333 | 2 | 1 | NA | NA | 3 | 0.6667 | 3 | 0.6667 | 8 | 0.75 |
| 6_9216629 | 3 | 0 | 2 | 0 | NA | NA | 2 | 0.5 | 7 | 0.1429 | 3 | 0 | NA | NA | 3 | 0 | 3 | 0.6667 | 9 | 0.2222 |
| 6_9216645 | 3 | 0.3333 | 2 | 0.5 | NA | NA | 2 | 1 | 7 | 0.5714 | 2 | 0.5 | NA | NA | 2 | 0 | 3 | 0.3333 | 7 | 0.2857 |
| 6_9216647 | 3 | 1 | 2 | 1 | NA | NA | 2 | 1 | 7 | 1 | 3 | 1 | NA | NA | 3 | 1 | 3 | 1 | 9 | 1 |
| 6_9216691 | 1 | 1 | 2 | 1 | NA | NA | 1 | 1 | 4 | 1 | 2 | 1 | NA | NA | 3 | 1 | 3 | 1 | 8 | 1 |
| 6_9216699 | NA | NA | 2 | 1 | NA | NA | NA | NA | 2 | 1 | NA | NA | NA | NA | 3 | 1 | 2 | 0.5 | 5 | 0.8 |
| 6_9216701 | 1 | 1 | 2 | 0.5 | NA | NA | 1 | 1 | 4 | 0.75 | 1 | 1 | NA | NA | 3 | 1 | 4 | 1 | 8 | 1 |
| 6_9216714 | NA | NA | 2 | 0 | NA | NA | 1 | 1 | 3 | 0.3333 | 1 | 1 | NA | NA | 3 | 0.3333 | 2 | 0.5 | 6 | 0.5 |
| 6_9234799 | 25 | 0.52 | 32 | 0.5938 | 17 | 0.5294 | 18 | 0.6667 | 92 | 0.5761 | 23 | 0.5652 | 23 | 0.6087 | 25 | 0.56 | 19 | 0.5263 | 90 | 0.5667 |
| 6_9234821 | 22 | 0.5455 | 28 | 0.6071 | 16 | 0.5 | 13 | 0.6923 | 79 | 0.5823 | 21 | 0.6667 | 22 | 0.5 | 22 | 0.4545 | 16 | 0.625 | 81 | 0.5556 |
| 6_9235151 | 7 | 0.1429 | 6 | 0.3333 | 8 | 0.125 | 7 | 0.2857 | 28 | 0.2143 | 7 | 0.4286 | 6 | 0.1667 | 5 | 0.4 | 3 | 0.3333 | 21 | 0.3333 |
| 6_9235231 | 5 | 0.2 | 6 | 0 | 8 | 0.125 | 5 | 0 | 24 | 0.0833 | 3 | 0 | 6 | 0 | 4 | 0.25 | 4 | 0 | 17 | 0.0588 |
| 6_9235244 | 4 | 0 | 5 | 0.2 | 6 | 0.5 | 4 | 0.25 | 19 | 0.2632 | 2 | 0 | 6 | 0.5 | 4 | 0 | 2 | 0.5 | 14 | 0.2857 |
| 6_9235274 | 5 | 0.6 | 5 | 0.6 | 5 | 0.6 | 2 | 1 | 17 | 0.6471 | 1 | 1 | 5 | 0.4 | 3 | 0.6667 | 2 | 0.5 | 11 | 0.5455 |
| 6_9235294 | 5 | 1 | 4 | 1 | 5 | 1 | 1 | 1 | 15 | 1 | 2 | 1 | 3 | 1 | 3 | 1 | NA | NA | 8 | 1 |
| 6_9235295 | 5 | 0.6 | 5 | 0.6 | 6 | 0.3333 | 2 | 1 | 18 | 0.5556 | 2 | 1 | 4 | 0.5 | 3 | 0.6667 | 2 | 1 | 11 | 0.7273 |
| 6_9235297 | 5 | 1 | 5 | 1 | 7 | 1 | 2 | 1 | 19 | 1 | 2 | 1 | 4 | 1 | 3 | 1 | 1 | 1 | 10 | 1 |
| 6_9235308 | 3 | 0 | 5 | 0 | 6 | 0.1667 | 2 | 0 | 16 | 0.0625 | 1 | 0 | 3 | 0.6667 | 3 | 0.3333 | 1 | 1 | 8 | 0.5 |
| 6_9235316 | 3 | 1 | 4 | 1 | 4 | 0.75 | 1 | 1 | 12 | 0.9167 | NA | NA | 4 | 0.75 | 3 | 1 | 1 | 1 | 8 | 0.875 |
| 6_9235320 | 5 | 0.8 | 5 | 0.6 | 5 | 0.4 | 2 | 0.5 | 17 | 0.5882 | 2 | 0.5 | 4 | 0.25 | 2 | 0.5 | 2 | 0.5 | 10 | 0.4 |
| 6_9235321 | 5 | 1 | 4 | 1 | 5 | 1 | 1 | 1 | 15 | 1 | 2 | 1 | 4 | 1 | 3 | 1 | 1 | 1 | 10 | 1 |
| 6_9235329 | 5 | 0.2 | 4 | 0.25 | 6 | 0 | 1 | 1 | 16 | 0.1875 | 2 | 0 | 4 | 0.25 | 2 | 0 | 2 | 0 | 10 | 0.1 |
| 6_9235329 | 5 | 1 | 3 | 1 | 5 | 1 | 1 | 1 | 14 | 1 | 2 | 1 | 4 | 1 | 2 | 1 | 2 | 1 | 10 | 1 |
| 6_9235344 | 2 | 1 | 3 | 0.6667 | 2 | 0 | 1 | 0 | 8 | 0.5 | 2 | 1 | 4 | 0.75 | 2 | 0.5 | NA | NA | 8 | 0.75 |
| 6_9235349 | 4 | 1 | 3 | 1 | 4 | 1 | 1 | 1 | 12 | 1 | 2 | 1 | 3 | 0.6667 | 1 | 1 | 2 | 1 | 8 | 0.875 |
| 6_9235356 | 4 | 1 | 2 | 0.5 | 5 | 1 | 1 | 1 | 12 | 0.9167 | 2 | 1 | 3 | 1 | 1 | 0 | 2 | 0.5 | 8 | 0.75 |
| 6_9235360 | 4 | 0.25 | 3 | 0 | 4 | 0 | 1 | 1 | 12 | 0.1667 | 2 | 0 | 3 | 0.3333 | 1 | 0 | 1 | 0 | 7 | 0.1429 |
| 6_9235372 | 4 | 1 | 3 | 1 | 4 | 1 | 1 | 1 | 12 | 1 | 2 | 1 | 3 | 1 | 1 | 1 | NA | NA | 6 | 1 |
| 6_9238798 | 3 | 1 | 3 | 1 | 4 | 1 | 1 | 1 | 11 | 1 | 1 | 1 | 2 | 0.5 | NA | NA | NA | NA | 3 | 0.6667 |
| 6_9238822 | 2 | 0.5 | 1 | 0 | 3 | 0 | 1 | 0 | 7 | 0.1429 | 2 | 0 | 2 | 0.5 | NA | NA | NA | NA | 4 | 0.25 |
| 6_9238829 | 2 | 0 | 1 | 0 | 2 | 0.5 | 1 | 0 | 6 | 0.1667 | 1 | 1 | 1 | 0 | NA | NA | NA | NA | 2 | 0.5 |
| 6_9238856 | 2 | 0 | 1 | 0 | 2 | 0.5 | 1 | 0 | 6 | 0.1667 | 1 | 0 | 2 | 0 | NA | NA | 2 | 0.5 | 5 | 0.2 |
| 6_9365281 | 5 | 0.8 | 4 | 0.5 | 2 | 0.5 | 2 | 0 | 13 | 0.5385 | NA | NA | 5 | 1 | 6 | 0.5 | 3 | 0.6667 | 14 | 0.7143 |
| 6_9365296 | 5 | 0.8 | 3 | 1 | NA | NA | 1 | 1 | 9 | 0.8889 | NA | NA | 5 | 0.4 | 4 | 0.75 | 3 | 0.6667 | 12 | 0.5833 |
| 6_9365325 | 6 | 0.3333 | 2 | 0.5 | 1 | 0 | 1 | 1 | 10 | 0.4 | NA | NA | 4 | 0.75 | 3 | 1 | 3 | 0.3333 | 10 | 0.7 |
| 6_9365345 | 5 | 0.2 | 2 | 0 | 1 | 1 | 2 | 1 | 10 | 0.4 | NA | NA | 5 | 0.4 | 4 | 0.75 | 3 | 1 | 12 | 0.6667 |
| 6_9365358 | 4 | 0.25 | 2 | 0 | NA | NA | 1 | 0 | 7 | 0.1429 | NA | NA | 3 | 0 | 3 | 0.3333 | 3 | 0.3333 | 9 | 0.2222 |
| 6_9365426 | 4 | 0.75 | 2 | 1 | 1 | 1 | 1 | 1 | 8 | 0.875 | NA | NA | 5 | 0.8 | 1 | 1 | 2 | 1 | 8 | 0.875 |
| 6_9365438 | 5 | 0.2 | 2 | 0.5 | 1 | 0 | 1 | 0 | 9 | 0.2222 | NA | NA | 4 | 0 | 1 | 1 | 1 | 0 | 6 | 0.1667 |
| 6_9365460 | 5 | 0.2 | 1 | 1 | 1 | 1 | NA | NA | 7 | 0.4286 | NA | NA | 6 | 0.5 | 1 | 1 | 1 | 1 | 8 | 0.625 |
| 6_9365476 | 5 | 0.8 | NA | NA | 1 | 1 | NA | NA | 6 | 0.8333 | NA | NA | 4 | 1 | NA | NA | 2 | 1 | 6 | 1 |
| 6_9365477 | 5 | 0.2 | 1 | 0 | 1 | 0 | 1 | 0 | 8 | 0.125 | NA | NA | 5 | 0.2 | 1 | 0 | 2 | 0.5 | 8 | 0.25 |
| 6_9365484 | 5 | 0.4 | 1 | 1 | 1 | 1 | 1 | 0 | 8 | 0.5 | NA | NA | 5 | 0.4 | 1 | 1 | 2 | 0.5 | 8 | 0.5 |
| 6_9365486 | 4 | 0.25 | NA | NA | 1 | 0 | 1 | 0 | 6 | 0.1667 | NA | NA | 4 | 0 | 1 | 1 | 2 | 0.5 | 7 | 0.2857 |
| 6_9365510 | 4 | 1 | 2 | 1 | 3 | 1 | 4 | 1 | 13 | 1 | NA | NA | 1 | 1 | 2 | 1 | 3 | 1 | 6 | 1 |
| 6_9365513 | 6 | 0.1667 | 2 | 0 | 3 | 0 | 4 | 0 | 15 | 0.0667 | NA | NA | 5 | 0.4 | 2 | 0.5 | 4 | 0.25 | 11 | 0.3636 |
| 6_9365522 | 4 | 1 | 1 | 1 | 3 | 1 | 4 | 0.5 | 12 | 0.8333 | NA | NA | 5 | 0.8 | 2 | 1 | 3 | 1 | 10 | 0.9 |
| 6_9365536 | 4 | 0.25 | 2 | 0 | 1 | 0 | 4 | 0 | 11 | 0.0909 | NA | NA | 5 | 0 | 1 | 1 | 4 | 0.25 | 10 | 0.2 |
| 6_9365537 | 4 | 0.75 | 2 | 0 | 2 | 0 | 4 | 0.75 | 12 | 0.5 | NA | NA | 3 | 0.6667 | 1 | 1 | 3 | 0.6667 | 7 | 0.7143 |
| 6_9365549 | 3 | 0.6667 | 2 | 1 | 3 | 0.3333 | 2 | 0.5 | 10 | 0.6 | NA | NA | 5 | 0.4 | 2 | 1 | 4 | 0.75 | 11 | 0.6364 |
| 6_9365568 | 2 | 0.5 | 1 | 0 | 3 | 0 | 1 | 1 | 7 | 0.2857 | NA | NA | 3 | 1 | 1 | 1 | 3 | 1 | 7 | 1 |
| 6_9365574 | 3 | 1 | NA | NA | 2 | 0.5 | 1 | 0 | 6 | 0.6667 | NA | NA | 4 | 1 | NA | NA | 3 | 1 | 7 | 1 |
| 6_9365576 | 3 | 0.6667 | 1 | 0 | 3 | 0 | NA | NA | 7 | 0.2857 | NA | NA | 1 | 0 | 2 | 0.5 | 3 | 0.6667 | 6 | 0.5 |
| 6_9365586 | 3 | 0 | 1 | 0 | 2 | 0 | 1 | 1 | 7 | 0.1429 | NA | NA | 2 | 0.5 | 1 | 1 | 4 | 0.25 | 7 | 0.4286 |
| 6_9365799 | 1 | 1 | 1 | 0 | NA | NA | NA | NA | 2 | 0.5 | NA | NA | 2 | 0.5 | NA | NA | 1 | 1 | 3 | 0.6667 |
| 6_9365801 | 1 | 1 | NA | NA | NA | NA | NA | NA | 1 | 1 | NA | NA | 2 | 0 | NA | NA | 1 | 1 | 3 | 0.3333 |
| 6_9365895 | 1 | 1 | 1 | 1 | 2 | 0 | NA | NA | 4 | 0.5 | NA | NA | 1 | 0 | NA | NA | 1 | 1 | 2 | 0.5 |
| 6_9365964 | 2 | 1 | 1 | 1 | NA | NA | NA | NA | 3 | 1 | NA | NA | 1 | 1 | NA | NA | NA | NA | 1 | 1 |
| 6_9365974 | 2 | 1 | 1 | 1 | NA | NA | NA | NA | 3 | 1 | NA | NA | 1 | 1 | NA | NA | NA | NA | 1 | 1 |
| 6_9366000 | 5 | 0.2 | 1 | 1 | 2 | 0 | 1 | 0 | 9 | 0.2222 | NA | NA | 3 | 0.3333 | 1 | 0 | 2 | 0 | 6 | 0.1667 |

| | | | | | | | | | | | | | | | | | | | | |
|---|---|---|---|---|---|---|---|---|---|---|---|---|---|---|---|---|---|---|---|---|
| 6_9372292 | 2 | 0.5 | NA | NA | NA | NA | NA | NA | 2 | 0.5 | NA | NA | NA | NA | NA | NA | 1 | 1 | 1 | 1 |
| 6_9372293 | 1 | 0 | NA | NA | NA | NA | 1 | 1 | 2 | 0.5 | NA | NA | NA | NA | NA | NA | 1 | 1 | 1 | 1 |
| 6_9458076 | 1 | 1 | 2 | 1 | NA | NA | 1 | 1 | 4 | 1 | 1 | 1 | 2 | 0.5 | NA | NA | 1 | 1 | 4 | 0.75 |
| 6_9458185 | 7 | 0.1429 | 2 | 0 | 5 | 0 | 1 | 0 | 15 | 0.0667 | 2 | 0 | 3 | 0 | 3 | 0 | 3 | 0.3333 | 11 | 0.0909 |
| 6_9458194 | 6 | 0.1667 | 2 | 0 | 3 | 0.6667 | NA | NA | 11 | 0.2727 | 2 | 0.5 | 3 | 0 | 4 | 0.25 | 2 | 0.5 | 11 | 0.2727 |
| 6_9458248 | 4 | 0.5 | 1 | 0 | 3 | 0.3333 | 1 | 0 | 9 | 0.3333 | 1 | 0 | 2 | 0 | 4 | 0.5 | 3 | 0.3333 | 10 | 0.3 |
| 6_9458257 | 4 | 0.75 | 2 | 0.5 | 3 | 0.6667 | 1 | 0 | 10 | 0.6 | 2 | 1 | 2 | 0.5 | 4 | 0.75 | 3 | 0.6667 | 11 | 0.7273 |
| 6_9458265 | 4 | 0.5 | 2 | 0 | 3 | 0 | 1 | 0 | 10 | 0.2 | 1 | 0 | 1 | 0 | 4 | 0.5 | 2 | 0.5 | 8 | 0.375 |
| 6_9458276 | 3 | 0.6667 | 2 | 0 | 3 | 0 | 1 | 0 | 9 | 0.2222 | 2 | 1 | 1 | 0 | 4 | 0.75 | 2 | 0 | 9 | 0.5556 |
| 6_9458282 | 4 | 0 | 2 | 0 | 4 | 0.75 | 1 | 0 | 11 | 0.2727 | 2 | 0.5 | 2 | 0.5 | 3 | 0 | 3 | 0.6667 | 10 | 0.4 |
| 6_9458297 | 2 | 1 | 2 | 0.5 | 3 | 0.6667 | 1 | 1 | 8 | 0.75 | 1 | 1 | 2 | 0.5 | 4 | 0.25 | 2 | 1 | 9 | 0.5556 |
| 6_9458318 | 2 | 0.5 | 1 | 0 | 1 | 1 | 1 | 0 | 5 | 0.4 | NA | NA | 1 | 1 | 4 | 0.25 | 1 | 1 | 6 | 0.5 |
| 6_9501671 | 2 | 0.5 | 2 | 1 | NA | NA | NA | NA | 4 | 0.75 | NA | NA | NA | NA | NA | NA | 1 | 1 | 1 | 1 |
| 6_9501680 | 2 | 0 | 2 | 0.5 | NA | NA | 1 | 0 | 5 | 0.2 | NA | NA | NA | NA | NA | NA | 1 | 1 | 1 | 1 |
| 6_9501717 | 1 | 0 | 2 | 0.5 | NA | NA | 1 | 0 | 4 | 0.25 | NA | NA | NA | NA | NA | NA | 1 | 1 | 1 | 1 |
| 6_9501739 | 2 | 1 | 2 | 0.5 | NA | NA | 1 | 0 | 5 | 0.6 | NA | NA | NA | NA | NA | NA | 1 | 1 | 1 | 1 |
| 6_9501776 | 2 | 1 | 1 | 0 | NA | NA | 1 | 1 | 4 | 0.75 | NA | NA | NA | NA | NA | NA | 1 | 1 | 1 | 1 |
| 6_9509933 | NA | NA | NA | NA | 2 | 1 | NA | NA | 2 | 1 | 1 | 0 | NA | NA | 1 | 1 | 1 | 1 | 3 | 0.6667 |
| 6_9510355 | NA | NA | NA | NA | 2 | 1 | NA | NA | 2 | 1 | 1 | 1 | NA | NA | NA | NA | 1 | 1 | 2 | 1 |
| 6_9510364 | NA | NA | NA | NA | 2 | 1 | NA | NA | 2 | 1 | 1 | 1 | NA | NA | 1 | 1 | 1 | 1 | 3 | 1 |
| 6_9510365 | NA | NA | NA | NA | 2 | 1 | NA | NA | 2 | 1 | 1 | 1 | NA | NA | 1 | 1 | 1 | 1 | 3 | 1 |
| 6_9510374 | NA | NA | NA | NA | 2 | 1 | NA | NA | 2 | 1 | 1 | 1 | NA | NA | 1 | 1 | 1 | 1 | 3 | 1 |
| 6_9510385 | NA | NA | NA | NA | 1 | 1 | NA | NA | 1 | 1 | 1 | 1 | NA | NA | 1 | 1 | 1 | 0 | 3 | 0.6667 |
| 6_9510388 | NA | NA | NA | NA | 2 | 0.5 | NA | NA | 2 | 0.5 | 1 | 1 | NA | NA | 1 | 1 | 1 | 1 | 3 | 1 |
| 6_9510404 | NA | NA | NA | NA | 2 | 1 | NA | NA | 2 | 1 | 1 | 1 | NA | NA | 1 | 1 | 1 | 1 | 3 | 1 |
| 6_9510426 | NA | NA | NA | NA | 3 | 0.3333 | NA | NA | 3 | 0.3333 | 1 | 1 | NA | NA | NA | NA | 1 | 0 | 2 | 0.5 |
| 6_9510545 | NA | NA | NA | NA | 2 | 0.5 | NA | NA | 2 | 0.5 | 1 | 1 | NA | NA | NA | NA | 1 | 0 | 2 | 0.5 |
| 6_9510550 | NA | NA | NA | NA | 3 | 0.6667 | NA | NA | 3 | 0.6667 | 2 | 0.5 | NA | NA | 1 | 0 | 1 | 0 | 4 | 0.25 |
| 6_9545191 | 4 | 0.75 | 2 | 0.5 | 2 | 0.5 | 2 | 0 | 10 | 0.5 | 1 | 0 | NA | NA | 5 | 0.4 | 1 | 0 | 7 | 0.2857 |
| 6_9545192 | 4 | 0 | 2 | 0 | 2 | 0.5 | 2 | 0 | 10 | 0.1 | 1 | 0 | 1 | 0 | 6 | 0.3333 | 1 | 0 | 9 | 0.2222 |
| 6_9609366 | NA | NA | NA | NA | 1 | 1 | NA | NA | 1 | 1 | NA | NA | NA | NA | 1 | 1 | NA | NA | 1 | 1 |
| 6_9609369 | NA | NA | NA | NA | 1 | 1 | NA | NA | 1 | 1 | NA | NA | NA | NA | 1 | 1 | NA | NA | 1 | 1 |
| 6_9663177 | 3 | 0 | 3 | 0.3333 | NA | NA | 1 | 0 | 7 | 0.1429 | 3 | 0.3333 | NA | NA | NA | NA | 1 | 1 | 4 | 0.5 |
| 6_9663191 | 3 | 0 | 4 | 0.25 | NA | NA | 1 | 0 | 8 | 0.125 | 3 | 0.3333 | NA | NA | NA | NA | 1 | 1 | 4 | 0.5 |
| 6_9663202 | 2 | 1 | 4 | 1 | NA | NA | 1 | 1 | 7 | 1 | 2 | 1 | NA | NA | NA | NA | 1 | 1 | 3 | 1 |
| 6_9663210 | 2 | 0 | NA | NA | 1 | 1 | NA | NA | 3 | 0.3333 | 1 | 1 | NA | NA | NA | NA | 1 | 1 | 2 | 1 |
| 6_9663253 | 1 | 0 | 5 | 0.2 | 2 | 0 | 1 | 0 | 9 | 0.1111 | 1 | 1 | NA | NA | NA | NA | 1 | 1 | 2 | 1 |
| 6_9680899 | NA | NA | NA | NA | NA | NA | 1 | 1 | 1 | 1 | NA | NA | 1 | 1 | NA | NA | NA | NA | 1 | 1 |
| 6_9680900 | NA | NA | NA | NA | NA | NA | 1 | 1 | 1 | 1 | NA | NA | 1 | 1 | NA | NA | NA | NA | 1 | 1 |
| 6_9680917 | NA | NA | NA | NA | NA | NA | 1 | 1 | 1 | 1 | NA | NA | 1 | 1 | NA | NA | NA | NA | 1 | 1 |
| 6_9680925 | NA | NA | NA | NA | NA | NA | 1 | 1 | 1 | 1 | NA | NA | 1 | 1 | NA | NA | NA | NA | 1 | 1 |
| 6_9680944 | NA | NA | NA | NA | NA | NA | 1 | 1 | 1 | 1 | NA | NA | 1 | 1 | NA | NA | NA | NA | 1 | 1 |
| 6_9708723 | 4 | 0 | 1 | 1 | 1 | 0 | 4 | 0.25 | 10 | 0.2 | NA | NA | 2 | 0.5 | 5 | 0.4 | 1 | 1 | 8 | 0.5 |
| 6_9708732 | 4 | 0.5 | 1 | 1 | NA | NA | 4 | 0.75 | 9 | 0.6667 | NA | NA | 1 | 0 | 4 | 0.25 | 2 | 0 | 7 | 0.1429 |
| 6_9708887 | 1 | 1 | NA | NA | 2 | 0.5 | 2 | 1 | 5 | 0.8 | NA | NA | 2 | 0 | 3 | 1 | 2 | 1 | 7 | 0.7143 |
| 6_9708897 | NA | NA | NA | NA | 2 | 1 | NA | NA | 2 | 1 | NA | NA | 1 | 0 | 3 | 1 | 2 | 0.5 | 6 | 0.6667 |
| 6_9713233 | 3 | 0.6667 | NA | NA | 1 | 1 | 2 | 0 | 6 | 0.5 | 3 | 0.6667 | 2 | 0 | NA | NA | NA | NA | 5 | 0.4 |
| 6_9715298 | 1 | 0 | NA | NA | 1 | 1 | NA | NA | 2 | 0.5 | 2 | 0.5 | 1 | 1 | NA | NA | NA | NA | 3 | 0.6667 |
| 6_9715310 | 1 | 0 | NA | NA | 1 | 1 | NA | NA | 2 | 0.5 | 2 | 1 | 1 | 1 | 1 | 1 | NA | NA | 4 | 1 |
| 6_9716151 | 8 | 0.375 | 6 | 0.5 | 2 | 1 | 4 | 0.25 | 20 | 0.45 | 2 | 0.5 | 12 | 0.5833 | 3 | 0.3333 | 9 | 0.2222 | 26 | 0.4231 |
| 6_9716155 | 13 | 0.3077 | 9 | 0.2222 | 4 | 0.25 | 5 | 0.4 | 31 | 0.2903 | 5 | 0 | 13 | 0.4615 | 4 | 0.5 | 12 | 0.6667 | 34 | 0.4706 |
| 6_9716177 | 15 | 0.2 | 8 | 0.125 | 5 | 0.2 | 5 | 0 | 33 | 0.1515 | 4 | 0.5 | 13 | 0.0769 | 5 | 0.2 | 11 | 0.1818 | 33 | 0.1818 |
| 6_9718786 | 24 | 0.0833 | 18 | 0.0556 | 13 | 0.0769 | 30 | 0.1 | 85 | 0.0824 | 18 | 0.1111 | 20 | 0.3 | 26 | 0.0769 | 21 | 0.0952 | 85 | 0.1412 |
| 6_9720959 | 30 | 0.0333 | 24 | 0.0833 | 11 | 0.1818 | 31 | 0.1935 | 96 | 0.1146 | 19 | 0.0526 | 21 | 0.0476 | 33 | 0.1515 | 25 | 0.08 | 98 | 0.0918 |
| 6_9720972 | 29 | 0.4483 | 25 | 0.6 | 13 | 0.6923 | 28 | 0.4643 | 95 | 0.5263 | 23 | 0.6087 | 25 | 0.72 | 34 | 0.5294 | 25 | 0.52 | 107 | 0.5888 |
| 6_9720973 | 30 | 0.1 | 25 | 0.12 | 14 | 0.1429 | 33 | 0.1818 | 102 | 0.1373 | 24 | 0.125 | 25 | 0.08 | 37 | 0.1081 | 26 | 0 | 112 | 0.0804 |
| 6_9720986 | 28 | 0.1429 | 22 | 0.1818 | 16 | 0.4375 | 33 | 0.303 | 99 | 0.2525 | 22 | 0.2727 | 24 | 0.1667 | 33 | 0.3333 | 24 | 0.25 | 103 | 0.2621 |
| 6_9720994 | 29 | 0.3103 | 23 | 0.3913 | 16 | 0.4375 | 30 | 0.5333 | 98 | 0.4184 | 26 | 0.5 | 25 | 0.48 | 36 | 0.4444 | 30 | 0.3333 | 117 | 0.4359 |
| 6_9726410 | 3 | 0.6667 | 2 | 0.5 | 1 | 1 | 2 | 0 | 8 | 0.5 | 1 | 0 | 2 | 0.5 | 1 | 0 | 1 | 1 | 5 | 0.4 |
| 6_9726412 | 3 | 0.3333 | 1 | 1 | 2 | 0.5 | 2 | 0.5 | 8 | 0.5 | 1 | 1 | 3 | 0 | 1 | 0 | 1 | 0 | 6 | 0.1667 |
| 6_9726422 | 2 | 0 | 2 | 0.5 | 1 | 1 | 2 | 0 | 7 | 0.2857 | 1 | 0 | 2 | 0.5 | 1 | 0 | 1 | 0 | 5 | 0.2 |
| 6_9726436 | 3 | 0 | 2 | 0.5 | 2 | 0 | 2 | 0 | 9 | 0.1111 | 1 | 0 | 3 | 0.6667 | 1 | 0 | 1 | 0 | 6 | 0.3333 |
| 6_9726439 | 2 | 0 | 2 | 0.5 | 2 | 0 | 2 | 0.5 | 8 | 0.25 | NA | NA | 3 | 0 | 1 | 1 | 1 | 1 | 5 | 0.4 |
| 6_9726447 | 2 | 0 | 2 | 0 | 2 | 0.5 | 2 | 0 | 8 | 0.125 | 1 | 0 | 3 | 0.6667 | 1 | 0 | 1 | 0 | 6 | 0.3333 |
| 6_9726448 | 2 | 0 | 2 | 0.5 | 2 | 0.5 | 2 | 0 | 8 | 0.25 | 1 | 0 | 3 | 0.3333 | 1 | 1 | 1 | 1 | 6 | 0.5 |

| | | | | | | | | | | | | | | | | | | | | |
|---|---|---|---|---|---|---|---|---|---|---|---|---|---|---|---|---|---|---|---|---|
| 6_9726457( | 3 | 0 | 1 | 0 | 2 | 1 | 2 | 0.5 | 8 | 0.375 | 1 | 1 | 3 | 0 | 1 | 0 | 1 | 0 | 6 | 0.1667 |
| 6_9726457! | NA | NA | 2 | 1 | 2 | 1 | 1 | 1 | 5 | 1 | NA | NA | 1 | 1 | 1 | 1 | 1 | 1 | 3 | 1 |
| 6_9726466 | 2 | 0 | 2 | 0 | 2 | 0.5 | 2 | 0.5 | 8 | 0.25 | 1 | 1 | 3 | 0.3333 | NA | NA | NA | NA | 4 | 0.5 |
| 6_9726482 | 3 | 0 | 2 | 0 | 2 | 0.5 | 2 | 0 | 9 | 0.1111 | 1 | 0 | 2 | 0.5 | 1 | 0 | NA | NA | 4 | 0.25 |
| 6_9726484 | 3 | 1 | 2 | 1 | 2 | 0.5 | 1 | 0 | 8 | 0.75 | 1 | 1 | 2 | 0.5 | 1 | 1 | NA | NA | 4 | 0.75 |
| 6_9726488 | 3 | 0 | 2 | 0.5 | 2 | 1 | 2 | 0.5 | 9 | 0.4444 | 1 | 1 | 2 | 1 | 1 | 1 | NA | NA | 4 | 1 |
| 6_9726510( | 3 | 0.6667 | 1 | 0 | 2 | 0.5 | 1 | 0 | 7 | 0.4286 | 1 | 1 | 1 | 0 | 1 | 0 | NA | NA | 3 | 0.3333 |
| 6_9726514 | 3 | 0 | 2 | 0 | 2 | 0 | 2 | 0.5 | 9 | 0.1111 | 1 | 0 | 1 | 1 | NA | NA | NA | NA | 2 | 0.5 |
| 6_9726516( | 3 | 1 | 2 | 1 | 2 | 1 | 2 | 1 | 9 | 1 | 1 | 1 | 1 | 1 | 1 | 1 | NA | NA | 3 | 1 |
| 6_9726522 | 2 | 0.5 | 2 | 0 | 2 | 0.5 | 2 | 0.5 | 8 | 0.375 | 1 | 0 | 1 | 1 | 1 | 1 | NA | NA | 3 | 0.6667 |
| 6_9726531( | 3 | 0.3333 | 2 | 0.5 | 2 | 0 | 2 | 0.5 | 9 | 0.3333 | 1 | 0 | 1 | 0 | 1 | 1 | NA | NA | 3 | 0.3333 |
| 6_9727266 | 2 | 0 | 1 | 1 | 1 | 0 | 2 | 0 | 6 | 0.1667 | NA | NA | NA | NA | 1 | 1 | NA | NA | 1 | 1 |
| 6_9727283 | 2 | 0 | NA | NA | 1 | 0 | 1 | 1 | 4 | 0.25 | 1 | 1 | NA | NA | 1 | 1 | NA | NA | 2 | 1 |
| 6_9759425 | NA | NA | 1 | 0 | NA | NA | 1 | 1 | 2 | 0.5 | 1 | 0 | NA | NA | 1 | 1 | NA | NA | 2 | 0.5 |
| 6_9763322 | NA | NA | 1 | 1 | NA | NA | NA | NA | 1 | 1 | NA | NA | NA | NA | NA | NA | 1 | 1 | 1 | 1 |
| 6_9799476 | NA | NA | 3 | 0.6667 | 1 | 1 | 1 | 1 | 5 | 0.8 | NA | NA | NA | NA | 1 | 1 | NA | NA | 1 | 1 |
| 6_9799485 | NA | NA | 4 | 0.75 | 1 | 1 | 1 | 1 | 6 | 0.8333 | NA | NA | NA | NA | 1 | 1 | NA | NA | 1 | 1 |
| 6_9799516( | NA | NA | 4 | 0.75 | 1 | 1 | 1 | 1 | 6 | 0.8333 | NA | NA | NA | NA | 1 | 1 | NA | NA | 1 | 1 |
| 6_9799519( | NA | NA | 4 | 0.75 | 1 | 1 | 1 | 1 | 6 | 0.8333 | NA | NA | NA | NA | 1 | 1 | NA | NA | 1 | 1 |
| 6_9799530 | NA | NA | 4 | 0.75 | 1 | 1 | 1 | 1 | 6 | 0.8333 | NA | NA | NA | NA | 1 | 1 | NA | NA | 1 | 1 |
| 6_9799537 | NA | NA | 4 | 1 | 1 | 1 | NA | NA | 5 | 1 | NA | NA | NA | NA | 1 | 1 | NA | NA | 1 | 1 |
| 6_9799574 | NA | NA | 4 | 0.5 | 1 | 0 | 1 | 0 | 6 | 0.3333 | NA | NA | NA | NA | 1 | 1 | NA | NA | 1 | 1 |
| 6_9799682 | NA | NA | 2 | 1 | 1 | 1 | 1 | 1 | 4 | 1 | NA | NA | NA | NA | 1 | 1 | NA | NA | 1 | 1 |
| 6_9799695 | NA | NA | 3 | 0.6667 | NA | NA | 1 | 0 | 4 | 0.5 | NA | NA | NA | NA | 1 | 1 | NA | NA | 1 | 1 |
| 6_9799791 | NA | NA | 4 | 0 | 2 | 0.5 | 2 | 0 | 8 | 0.125 | NA | NA | NA | NA | 1 | 1 | 1 | 0 | 2 | 0.5 |
| 6_9799800( | NA | NA | 5 | 0.2 | 2 | 1 | 2 | 0.5 | 9 | 0.4444 | NA | NA | NA | NA | 1 | 0 | 3 | 0.3333 | 4 | 0.25 |
| 6_9799813 | NA | NA | 5 | 0 | 2 | 1 | 2 | 0.5 | 9 | 0.3333 | NA | NA | NA | NA | 1 | 0 | 3 | 0.6667 | 4 | 0.5 |
| 6_9890262 | 7 | 0.4286 | 1 | 1 | 4 | 0.5 | 11 | 0.7273 | 23 | 0.6087 | 5 | 0.8 | 6 | 0.6667 | 6 | 0.6667 | 10 | 0.3 | 27 | 0.5556 |
| 6_9890555 | 6 | 0 | 4 | 0 | 2 | 0 | 7 | 0.1429 | 19 | 0.0526 | 4 | 0.25 | 5 | 0.2 | 8 | 0.125 | 6 | 0.1667 | 23 | 0.1739 |
| 6_9890663( | 7 | 0.4286 | 7 | 0.5714 | 6 | 0.3333 | 10 | 0.6 | 30 | 0.5 | 5 | 0.6 | 4 | 0.5 | 7 | 0.2857 | 6 | 0.8333 | 22 | 0.5455 |
| 6_9890667( | 3 | 0.6667 | 5 | 1 | 4 | 1 | 6 | 0.8333 | 18 | 0.8889 | 4 | 1 | 3 | 1 | 6 | 1 | 5 | 0.8 | 18 | 0.9444 |
| 6_9890668 | 4 | 1 | 2 | 1 | 3 | 0.6667 | 9 | 0.8889 | 18 | 0.8889 | 2 | 1 | 4 | 0.25 | 5 | 0.8 | 3 | 0.6667 | 14 | 0.6429 |
| 6_9890676 | 7 | 1 | 7 | 1 | 5 | 1 | 9 | 1 | 28 | 1 | 4 | 1 | 4 | 1 | 6 | 1 | 6 | 1 | 20 | 1 |
| 6_9890679 | 6 | 0 | 6 | 0.1667 | 5 | 0.2 | 8 | 0 | 25 | 0.08 | 5 | 0 | 3 | 0 | 4 | 0 | 4 | 0.5 | 16 | 0.125 |
| 6_9890685 | 7 | 0.4286 | 4 | 0.75 | 3 | 0 | 7 | 0.2857 | 21 | 0.381 | 2 | 0.5 | 2 | 1 | 2 | 0.5 | 3 | 0.3333 | 9 | 0.5556 |
| 6_9890705( | 4 | 0.5 | 6 | 0.1667 | 3 | 0 | 8 | 0.75 | 21 | 0.4286 | 4 | 0.5 | 1 | 0 | 4 | 0.5 | 4 | 0.25 | 13 | 0.3846 |
| 6_9890706 | 5 | 1 | 5 | 0.6 | 5 | 0.8 | 7 | 0.8571 | 22 | 0.8182 | 4 | 0.75 | 3 | 1 | 6 | 1 | 3 | 1 | 16 | 0.9375 |
| 6_9960233 | 1 | 1 | 5 | 0 | NA | NA | 4 | 0.5 | 10 | 0.3 | 1 | 1 | 2 | 0 | 1 | 0 | 4 | 0.5 | 8 | 0.375 |
| 6_9960287 | 1 | 0 | 3 | 0 | NA | NA | 4 | 0.25 | 8 | 0.125 | 2 | 1 | 1 | 1 | 1 | 1 | 2 | 0.5 | 6 | 0.8333 |
| 6_9960366 | 1 | 1 | 4 | 0.25 | NA | NA | 3 | 0.3333 | 8 | 0.375 | 1 | 0 | 2 | 0 | 1 | 0 | 3 | 0.6667 | 7 | 0.2857 |
| 6_9960382 | 1 | 1 | 4 | 0.75 | NA | NA | 2 | 0.5 | 7 | 0.7143 | 1 | 1 | 1 | 0 | 1 | 0 | 3 | 0.6667 | 6 | 0.5 |
| 6_9967033 | 4 | 1 | 3 | 0.6667 | 7 | 0.8571 | 7 | 1 | 21 | 0.9048 | 5 | 1 | 1 | 1 | 7 | 0.8571 | 2 | 1 | 15 | 0.9333 |
| 6_9967063( | 6 | 0.3333 | 6 | 0.5 | 6 | 0.1667 | 9 | 0.1111 | 27 | 0.2593 | 2 | 0.5 | 4 | 0.25 | 7 | 0.2857 | 4 | 0.5 | 17 | 0.3529 |
| 6_9967069 | 6 | 1 | 7 | 1 | 6 | 1 | 9 | 1 | 28 | 1 | 5 | 1 | 4 | 1 | 8 | 1 | 4 | 1 | 21 | 1 |
| 6_9967082 | 8 | 0.125 | 8 | 0.125 | 9 | 0.1111 | 9 | 0.2222 | 34 | 0.1471 | 5 | 0.2 | 4 | 0.25 | 9 | 0.1111 | 5 | 0.4 | 23 | 0.2174 |
| 6_9967087 | 8 | 0.75 | 8 | 0.75 | 8 | 0.875 | 9 | 0.3333 | 33 | 0.6667 | 6 | 0.6667 | 6 | 0.6667 | 8 | 1 | 5 | 1 | 25 | 0.84 |
| 6_9967089 | 7 | 0.8571 | 7 | 1 | 8 | 1 | 10 | 0.9 | 32 | 0.9375 | 5 | 1 | 5 | 0.8 | 7 | 0.8571 | 5 | 1 | 22 | 0.9091 |
| 6_9967109 | 10 | 0.2 | 10 | 0.2 | 9 | 0.3333 | 10 | 0.5 | 39 | 0.3077 | 6 | 0.3333 | 6 | 0 | 9 | 0.3333 | 6 | 0.5 | 27 | 0.2963 |
| 6_9967120( | 10 | 0 | 9 | 0 | 10 | 0.2 | 9 | 0.1111 | 38 | 0.0789 | 4 | 0.5 | 5 | 0.2 | 10 | 0.1 | 6 | 0.3333 | 25 | 0.24 |
| 6_9967123 | 5 | 0.4 | 7 | 0.5714 | 8 | 0.5 | 5 | 0.8 | 25 | 0.56 | 2 | 1 | 3 | 0.6667 | 6 | 0.5 | 2 | 1 | 13 | 0.6923 |
| 6_9967128( | 7 | 0 | 3 | 0 | 8 | 0 | 7 | 0.2857 | 25 | 0.08 | 4 | 0 | 3 | 0 | 7 | 0.1429 | 4 | 0.25 | 18 | 0.1111 |
| 6_9967130 | 11 | 0.2727 | 9 | 0 | 6 | 0 | 9 | 0.2222 | 35 | 0.1429 | 4 | 0 | 8 | 0.25 | 7 | 0.2857 | 7 | 0.1429 | 26 | 0.1923 |
| 6_9967145 | 12 | 0 | 12 | 0.1667 | 10 | 0.2 | 10 | 0.2 | 44 | 0.1364 | 6 | 0.3333 | 9 | 0.4444 | 14 | 0.3571 | 9 | 0.4444 | 38 | 0.3947 |
| 6_9967171 | 18 | 0.1111 | 12 | 0.0833 | 13 | 0 | 12 | 0 | 55 | 0.0545 | 8 | 0.125 | 11 | 0.1818 | 17 | 0.1176 | 13 | 0.0769 | 49 | 0.1224 |
| 6_9967189( | 15 | 0.6667 | 13 | 0.6154 | 12 | 0.6667 | 10 | 0.6 | 50 | 0.64 | 8 | 0.75 | 7 | 0.7143 | 17 | 0.5294 | 12 | 0.5 | 44 | 0.5909 |
| 6_9968860 | 1 | 0 | 2 | 0.5 | 1 | 0 | 3 | 0.3333 | 7 | 0.2857 | NA | NA | 1 | 0 | 6 | 0.1667 | 2 | 0 | 9 | 0.1111 |
| 6_9968888 | 1 | 0 | 2 | 0 | NA | NA | 5 | 0.2 | 8 | 0.125 | NA | NA | 1 | 0 | 4 | 0.25 | 1 | 0 | 6 | 0.1667 |
| 6_9968908 | 1 | 1 | 1 | 0 | NA | NA | 3 | 0.6667 | 5 | 0.6 | NA | NA | 1 | 1 | 5 | 0.6 | 2 | 0 | 8 | 0.5 |
| 6_9968919( | 1 | 0 | 1 | 1 | NA | NA | 4 | 0 | 6 | 0.1667 | NA | NA | 1 | 0 | 6 | 0.3333 | 2 | 0 | 9 | 0.2222 |
| 6_9968935 | 1 | 1 | 1 | 1 | 1 | 1 | 5 | 0.4 | 8 | 0.625 | NA | NA | 1 | 0 | 6 | 0.8333 | 2 | 1 | 9 | 0.7778 |
| 6_9985469 | 22 | 0.3636 | 23 | 0.5652 | 12 | 0.25 | 17 | 0.2353 | 74 | 0.3784 | 22 | 0.3182 | 18 | 0.2222 | 23 | 0.3478 | 15 | 0.4 | 78 | 0.3205 |
| 6_1002056 | 7 | 0.2857 | 3 | 0 | 3 | 0 | 3 | 0.3333 | 16 | 0.1875 | 3 | 0 | 5 | 0 | 4 | 0 | 4 | 0.25 | 16 | 0.0625 |
| 6_1002057 | 7 | 0 | 3 | 0.3333 | 4 | 0.25 | 3 | 0 | 17 | 0.1176 | 3 | 0 | 4 | 0 | 3 | 0 | 4 | 0.25 | 14 | 0.0714 |
| 6_1002058 | 4 | 0 | 3 | 0.3333 | 3 | 0 | 2 | 0 | 12 | 0.0833 | 1 | 0 | 5 | 0.2 | 2 | 0 | 3 | 0 | 11 | 0.0909 |
| 6_1002075 | 3 | 0 | 3 | 0.3333 | 2 | 0 | 1 | 0 | 9 | 0.1111 | NA | NA | 3 | 0.3333 | 2 | 0 | 2 | 0 | 7 | 0.1429 |

| | | | | | | | | | | | | | | | | | | | | |
|---|---|---|---|---|---|---|---|---|---|---|---|---|---|---|---|---|---|---|---|---|
| 6_10020839 | NA | NA | 1 | 1 | 2 | 0 | 2 | 0.5 | 5 | 0.4 | 2 | 0.5 | 3 | 0.3333 | 2 | 0.5 | 1 | 0 | 8 | 0.375 |
| 6_10020855 | 1 | 1 | 1 | 0 | 1 | 1 | 2 | 0 | 5 | 0.4 | 2 | 0.5 | 1 | 1 | 2 | 0.5 | 1 | 1 | 6 | 0.6667 |
| 6_10020911 | 1 | 1 | 1 | 1 | 1 | 1 | 1 | 1 | 4 | 1 | NA | NA | 1 | 1 | 2 | 0 | NA | NA | 3 | 0.3333 |
| 6_10020929 | 1 | 0 | 1 | 1 | 1 | 0 | 2 | 1 | 5 | 0.6 | 1 | 0 | 2 | 0.5 | 2 | 0.5 | 1 | 1 | 6 | 0.5 |
| 6_10020930 | 1 | 1 | 1 | 1 | 1 | 1 | 2 | 1 | 5 | 1 | 1 | 1 | 2 | 0.5 | 2 | 1 | 1 | 1 | 6 | 0.8333 |
| 6_10020933 | 1 | 0 | 1 | 0 | 1 | 0 | 2 | 0.5 | 5 | 0.2 | 1 | 0 | 1 | 1 | 2 | 0 | 1 | 0 | 5 | 0.2 |
| 6_10055022 | NA | NA | 1 | 1 | NA | NA | NA | NA | 1 | 1 | NA | NA | 1 | 1 | NA | NA | NA | NA | 1 | 1 |
| 6_10055055 | NA | NA | NA | NA | NA | NA | 1 | 1 | 1 | 1 | NA | NA | 1 | 1 | NA | NA | NA | NA | 1 | 1 |
| 6_10058220 | NA | NA | NA | NA | NA | NA | 1 | 1 | 1 | 1 | NA | NA | NA | NA | 1 | 1 | NA | NA | 1 | 1 |
| 6_10084285 | 4 | 0.25 | 7 | 0.4286 | 8 | 0.5 | 12 | 0.5833 | 31 | 0.4839 | 8 | 0.375 | 6 | 0.5 | 3 | 0 | 7 | 0.2857 | 24 | 0.3333 |
| 6_10084288 | 3 | 0.3333 | 6 | 0.1667 | 8 | 0.25 | 13 | 0.0769 | 30 | 0.1667 | 9 | 0.3333 | 7 | 0 | 4 | 0 | 9 | 0 | 29 | 0.1034 |
| 6_10084296 | 4 | 0 | 8 | 0 | 8 | 0.125 | 11 | 0.2727 | 31 | 0.129 | 7 | 0 | 6 | 0.3333 | 3 | 0 | 8 | 0 | 24 | 0.0833 |
| 6_10084330 | 6 | 1 | 10 | 0.6 | 8 | 0.75 | 11 | 0.9091 | 35 | 0.8 | 10 | 0.8 | 6 | 0.8333 | 5 | 0.6 | 9 | 0.7778 | 30 | 0.7667 |
| 6_10084335 | 5 | 0.8 | 9 | 0.5556 | 7 | 0.7143 | 13 | 0.8462 | 34 | 0.7353 | 8 | 0.625 | 7 | 0.7143 | 4 | 0.5 | 9 | 0.8889 | 28 | 0.7143 |
| 6_10084338 | 4 | 0.75 | 8 | 0.5 | 5 | 0.8 | 4 | 0.75 | 21 | 0.6667 | 5 | 0.4 | 8 | 0.625 | 3 | 0.6667 | 7 | 0.2857 | 23 | 0.4783 |
| 6_10084341 | 5 | 0 | 10 | 0 | 8 | 0 | 11 | 0.1818 | 34 | 0.0588 | 7 | 0.1429 | 7 | 0 | 6 | 0.1667 | 9 | 0.2222 | 29 | 0.1379 |
| 6_10084345 | 6 | 1 | 9 | 1 | 7 | 0.8571 | 11 | 1 | 33 | 0.9697 | 7 | 1 | 4 | 0.75 | 4 | 1 | 9 | 0.7778 | 24 | 0.875 |
| 6_10084353 | 7 | 0.1429 | 11 | 0.0909 | 8 | 0 | 12 | 0.0833 | 38 | 0.0789 | 10 | 0.2 | 10 | 0 | 6 | 0 | 11 | 0 | 37 | 0.0541 |
| 6_10084354 | 6 | 0 | 11 | 0.0909 | 8 | 0.25 | 10 | 0.2 | 35 | 0.1429 | 9 | 0.2222 | 10 | 0.1 | 7 | 0.2857 | 12 | 0.3333 | 38 | 0.2368 |
| 6_10084359 | 5 | 1 | 8 | 0.625 | 9 | 0.7778 | 12 | 0.8333 | 34 | 0.7941 | 11 | 0.8182 | 9 | 1 | 6 | 0.8333 | 10 | 1 | 36 | 0.9167 |
| 6_10084361 | 5 | 0.8 | 7 | 0.4286 | 7 | 0.4286 | 12 | 0.4167 | 31 | 0.4839 | 9 | 0.7778 | 6 | 1 | 7 | 0.5714 | 9 | 0.6667 | 31 | 0.7419 |
| 6_10084365 | 6 | 0 | 11 | 0.0909 | 9 | 0.1111 | 14 | 0.1429 | 40 | 0.1 | 10 | 0.2 | 9 | 0.1111 | 6 | 0 | 13 | 0.0769 | 38 | 0.1053 |
| 6_10084412 | 6 | 0 | 10 | 0.1 | 9 | 0 | 13 | 0.0769 | 38 | 0.0526 | 11 | 0.1818 | 11 | 0.1818 | 7 | 0.1429 | 15 | 0.1333 | 44 | 0.1591 |
| 6_10084419 | 9 | 0.1111 | 11 | 0.0909 | 9 | 0.1111 | 15 | 0.0667 | 44 | 0.0909 | 11 | 0.0909 | 13 | 0.1538 | 9 | 0 | 16 | 0.125 | 49 | 0.102 |
| 6_10084444 | 10 | 0.1 | 13 | 0 | 9 | 0 | 12 | 0.1667 | 44 | 0.0682 | 8 | 0.25 | 11 | 0 | 9 | 0.1111 | 14 | 0.1429 | 42 | 0.119 |
| 6_10084453 | 13 | 0.2308 | 9 | 0.2222 | 9 | 0.3333 | 16 | 0.3125 | 47 | 0.2766 | 11 | 0.4545 | 10 | 0.4 | 8 | 0.75 | 15 | 0.5333 | 44 | 0.5227 |
| 6_10084470 | 12 | 0 | 12 | 0.1667 | 11 | 0.0909 | 13 | 0.4615 | 48 | 0.1875 | 9 | 0.3333 | 9 | 0.1111 | 10 | 0 | 12 | 0.3333 | 40 | 0.2 |
| 6_10084504 | 5 | 0 | 7 | 0 | 4 | 0.5 | 10 | 0.1 | 26 | 0.1154 | 4 | 0 | 6 | 0 | 7 | 0 | 8 | 0.25 | 25 | 0.08 |
| 6_10084679 | 7 | 0 | 11 | 0.0909 | 2 | 0 | 10 | 0.1 | 30 | 0.0667 | 6 | 0 | 7 | 0 | 5 | 0 | 10 | 0.4 | 28 | 0.1429 |
| 6_10112660 | 4 | 0.25 | 2 | 0 | 4 | 0.25 | NA | NA | 10 | 0.2 | 1 | 0 | 1 | 1 | 1 | 0 | NA | NA | 3 | 0.3333 |
| 6_10112744 | 2 | 0.5 | 1 | 1 | 3 | 0.3333 | NA | NA | 6 | 0.5 | NA | NA | NA | NA | 1 | 1 | NA | NA | 1 | 1 |
| 6_10112748 | 4 | 0.5 | 1 | 1 | 3 | 0.3333 | NA | NA | 8 | 0.5 | NA | NA | 1 | 1 | 1 | 1 | NA | NA | 2 | 1 |
| 6_10112754 | 3 | 1 | 1 | 1 | 4 | 1 | NA | NA | 8 | 1 | NA | NA | 1 | 1 | 1 | 1 | NA | NA | 2 | 1 |
| 6_10112755 | 3 | 0 | 2 | 0 | 4 | 0.25 | NA | NA | 9 | 0.1111 | NA | NA | 1 | 0 | 1 | 1 | NA | NA | 2 | 0.5 |
| 6_10112761 | 4 | 0.25 | 1 | 1 | 4 | 1 | NA | NA | 9 | 0.6667 | NA | NA | 1 | 1 | 1 | 1 | NA | NA | 2 | 1 |
| 6_10112795 | 3 | 1 | 1 | 0 | 4 | 1 | NA | NA | 8 | 0.875 | NA | NA | NA | NA | 1 | 1 | NA | NA | 1 | 1 |
| 6_10112833 | 3 | 0.3333 | 1 | 0 | 4 | 0 | NA | NA | 8 | 0.125 | NA | NA | 1 | 0 | 1 | 1 | NA | NA | 2 | 0.5 |
| 6_10112872 | 3 | 0.6667 | 1 | 1 | 3 | 0 | NA | NA | 7 | 0.4286 | NA | NA | 1 | 0 | 1 | 1 | NA | NA | 2 | 0.5 |
| 6_10214091 | NA | NA | NA | NA | NA | NA | 4 | 0.25 | 4 | 0.25 | 3 | 0 | 1 | 0 | 3 | 0.3333 | 1 | 0 | 8 | 0.125 |
| 6_10214595 | NA | NA | NA | NA | NA | NA | 3 | 0.3333 | 3 | 0.3333 | 4 | 0.25 | 1 | 0 | 1 | 0 | NA | NA | 6 | 0.1667 |
| 6_10362640 | NA | NA | 2 | 0.5 | 1 | 0 | 3 | 0 | 6 | 0.1667 | 2 | 0 | 1 | 0 | 1 | 1 | 2 | 0 | 6 | 0.1667 |
| 6_10370615 | 1 | 0 | 4 | 0 | 2 | 0 | 4 | 0.25 | 11 | 0.0909 | 4 | 0 | 2 | 0 | 3 | 0 | 4 | 0.25 | 13 | 0.0769 |
| 6_10370629 | 1 | 0 | 2 | 0.5 | 1 | 0 | 3 | 0.3333 | 7 | 0.2857 | 4 | 0.5 | 3 | 0.3333 | 2 | 0.5 | 3 | 0.3333 | 12 | 0.4167 |
| 6_10370633 | 1 | 1 | 4 | 0.75 | 2 | 1 | 2 | 0.5 | 9 | 0.7778 | 4 | 1 | 3 | 1 | 3 | 1 | 4 | 1 | 14 | 1 |
| 6_10370640 | 1 | 1 | 3 | 1 | 2 | 1 | 4 | 1 | 10 | 1 | 4 | 0.75 | 2 | 1 | 3 | 1 | 3 | 1 | 12 | 0.9167 |
| 6_10370646 | 1 | 0 | 3 | 1 | 2 | 1 | 4 | 1 | 10 | 0.9 | 4 | 0.75 | 3 | 1 | 3 | 1 | 4 | 0.75 | 14 | 0.8571 |
| 6_10370651 | 1 | 0 | 3 | 0.6667 | 1 | 0 | 2 | 0 | 7 | 0.2857 | 3 | 0 | 2 | 0 | 1 | 0 | 3 | 0.3333 | 9 | 0.1111 |
| 6_10370654 | 1 | 1 | 3 | 0.6667 | 2 | 1 | 3 | 1 | 9 | 0.8889 | 4 | 0.75 | 3 | 1 | 3 | 1 | 3 | 1 | 13 | 0.9231 |
| 6_10370667 | 1 | 1 | 1 | 1 | 1 | 1 | 2 | 1 | 5 | 1 | 3 | 0.6667 | 2 | 1 | 3 | 1 | 4 | 1 | 12 | 0.9167 |
| 6_10370777 | 3 | 0.3333 | 6 | 0 | 5 | 0.2 | 5 | 0.2 | 19 | 0.1579 | 3 | 0 | 2 | 0 | 3 | 0 | 3 | 0.3333 | 11 | 0.0909 |
| 6_10370837 | 2 | 0.5 | 4 | 0.25 | 4 | 0.25 | 3 | 0 | 13 | 0.2308 | 5 | 0.4 | 2 | 0 | 3 | 0.3333 | 4 | 0 | 14 | 0.2143 |
| 6_10370860 | 3 | 0 | 7 | 0.1429 | 4 | 0 | 6 | 0.3333 | 20 | 0.15 | 4 | 0 | 2 | 0 | 3 | 0 | 5 | 0.2 | 14 | 0.0714 |
| 6_10370867 | 4 | 0.75 | 7 | 0.5714 | 4 | 0.75 | 5 | 0.4 | 20 | 0.6 | 4 | 0.75 | 2 | 0.5 | 3 | 0.6667 | 5 | 0.6 | 14 | 0.6429 |
| 6_10370908 | 3 | 0.3333 | 10 | 0.3 | 4 | 0 | 4 | 0 | 21 | 0.1905 | 5 | 0.2 | 3 | 0 | 4 | 0.25 | 8 | 0 | 20 | 0.1 |
| 6_10370951 | 6 | 0.5 | 18 | 0.3889 | 4 | 0.75 | 7 | 0.2857 | 35 | 0.4286 | 6 | 0.5 | 3 | 0.6667 | 4 | 0.25 | 11 | 0.2727 | 24 | 0.375 |
| 6_10370967 | 8 | 0.125 | 18 | 0.1111 | 4 | 0 | 8 | 0.125 | 38 | 0.1053 | 5 | 0 | 3 | 0.3333 | 5 | 0.4 | 12 | 0 | 25 | 0.12 |
| 6_10484028 | 2 | 0.5 | NA | NA | 4 | 0.25 | 3 | 0 | 9 | 0.2222 | 4 | 0.25 | 4 | 0.25 | 2 | 0 | 5 | 0 | 15 | 0.1333 |
| 6_10565071 | NA | NA | NA | NA | NA | NA | 1 | 1 | 1 | 1 | NA | NA | 1 | 1 | 1 | 0 | NA | NA | 2 | 0.5 |
| 6_10565137 | NA | NA | NA | NA | NA | NA | 1 | 1 | 1 | 1 | NA | NA | 2 | 0.5 | 1 | 0 | 2 | 0.5 | 5 | 0.4 |
| 6_10565149 | NA | NA | 1 | 0 | NA | NA | 1 | 1 | 2 | 0.5 | NA | NA | 2 | 0.5 | 1 | 0 | 2 | 0 | 5 | 0.2 |
| 6_10565159 | NA | NA | 1 | 0 | NA | NA | 1 | 1 | 2 | 0.5 | NA | NA | 1 | 1 | 1 | 1 | 2 | 0.5 | 4 | 0.75 |
| 6_10565175 | NA | NA | 1 | 0 | NA | NA | 1 | 1 | 2 | 0.5 | NA | NA | 3 | 0 | 1 | 0 | 2 | 0.5 | 6 | 0.1667 |
| 6_10667460 | 2 | 1 | 5 | 0.8 | 1 | 1 | 5 | 1 | 13 | 0.9231 | 1 | 1 | 1 | 1 | 2 | 0.5 | 3 | 0.6667 | 7 | 0.7143 |
| 6_10667503 | 2 | 0.5 | 5 | 0.2 | 1 | 0 | 5 | 0 | 13 | 0.1538 | 2 | 0 | 2 | 0.5 | 2 | 0 | 3 | 0 | 9 | 0.1111 |
| 6_10667616 | 2 | 0 | 9 | 0.1111 | 2 | 0 | 4 | 0 | 17 | 0.0588 | 1 | 0 | 2 | 0.5 | 1 | 0 | 3 | 0 | 7 | 0.1429 |

| | | | | | | | | | | | | | | | | | | | | |
|---|---|---|---|---|---|---|---|---|---|---|---|---|---|---|---|---|---|---|---|---|
| 6_1067558 | 4 | 0 | 6 | 0.1667 | 4 | 0.25 | 8 | 0 | 22 | 0.0909 | 4 | 0.25 | 2 | 0 | 3 | 0 | 5 | 0 | 14 | 0.0714 |
| 6_1067558 | 4 | 0 | 7 | 0.1429 | 3 | 0 | 6 | 0.5 | 20 | 0.2 | 5 | 0.6 | 1 | 0 | 3 | 0.6667 | 6 | 0.5 | 15 | 0.5333 |
| 6_1067562 | 2 | 1 | 8 | 0 | 2 | 0 | 4 | 0.25 | 16 | 0.1875 | 4 | 0.5 | 2 | 0 | 3 | 0 | 6 | 0.1667 | 15 | 0.2 |
| 6_1067563 | 3 | 0 | 5 | 0 | 3 | 0.3333 | 6 | 0 | 17 | 0.0588 | 2 | 0.5 | 1 | 0 | 3 | 0 | 5 | 0 | 11 | 0.0909 |
| 6_1067566 | 1 | 0 | 4 | 0.25 | 6 | 0.3333 | 3 | 0 | 14 | 0.2143 | NA | NA | 6 | 0.3333 | 3 | 0.6667 | 8 | 0.25 | 17 | 0.3529 |
| 6_1067570 | 1 | 1 | 5 | 0.2 | 4 | 0 | 2 | 0.5 | 12 | 0.25 | NA | NA | 5 | 0 | 3 | 0 | 7 | 0.1429 | 15 | 0.0667 |
| 6_1067570 | 1 | 0 | 5 | 0.4 | 5 | 0 | 2 | 0.5 | 13 | 0.2308 | NA | NA | 4 | 0.5 | 3 | 0.6667 | 6 | 0.3333 | 13 | 0.4615 |
| 6_1067575 | 9 | 0.1111 | 10 | 0.1 | 13 | 0.0769 | 4 | 0 | 36 | 0.0833 | 4 | 0 | 10 | 0.1 | 6 | 0.1667 | 14 | 0 | 34 | 0.0588 |
| 6_1067579 | 7 | 0.2857 | 9 | 0.8889 | 11 | 0.4545 | 4 | 0.75 | 31 | 0.5806 | 3 | 0.6667 | 8 | 0.375 | 4 | 0.5 | 11 | 0.6364 | 26 | 0.5385 |
| 6_1067579 | 7 | 0.2857 | 9 | 0.4444 | 8 | 0 | 4 | 0 | 28 | 0.2143 | 3 | 0 | 8 | 0.5 | 5 | 0.4 | 10 | 0.1 | 26 | 0.2692 |
| 6_1067587 | 3 | 0 | 8 | 0.125 | 4 | 0.25 | 6 | 0 | 21 | 0.0952 | 5 | 0.2 | 2 | 0.5 | 3 | 0.6667 | 4 | 0.25 | 14 | 0.3571 |
| 6_1067588 | 4 | 0 | 9 | 0.1111 | 6 | 0.3333 | 8 | 0 | 27 | 0.1111 | 5 | 0.2 | 3 | 0 | 4 | 0 | 6 | 0.1667 | 18 | 0.1111 |
| 6_1067588 | 4 | 0.25 | 9 | 0.2222 | 5 | 0.2 | 8 | 0 | 26 | 0.1538 | 5 | 0.2 | 4 | 0.25 | 4 | 0 | 6 | 0.1667 | 19 | 0.1579 |
| 6_1067588 | 4 | 0.25 | 8 | 0.25 | 4 | 0.25 | 6 | 0.1667 | 22 | 0.2273 | 4 | 0.25 | 3 | 0.3333 | 3 | 0 | 4 | 0 | 14 | 0.1429 |
| 6_1067590 | 1 | 0 | 9 | 0.3333 | 4 | 0 | 8 | 0.25 | 22 | 0.2273 | 3 | 0.6667 | 2 | 0 | 3 | 0 | 4 | 0.25 | 12 | 0.25 |
| 6_1067592 | 4 | 0 | 9 | 0 | 8 | 0.25 | 7 | 0.1429 | 28 | 0.1071 | 4 | 0 | 5 | 0 | 4 | 0.5 | 4 | 0 | 17 | 0.1176 |
| 6_1075439 | 5 | 0.2 | 3 | 0 | 2 | 0 | 4 | 0 | 14 | 0.0714 | 5 | 0.2 | 3 | 0.3333 | 4 | 0.5 | 5 | 0.6 | 17 | 0.4118 |
| 6_1075440 | 6 | 1 | 5 | 1 | 3 | 1 | 4 | 0.75 | 18 | 0.9444 | 3 | 1 | 2 | 0.5 | 3 | 1 | 4 | 1 | 12 | 0.9167 |
| 6_1075440 | 6 | 0.1667 | 5 | 0.2 | 4 | 0.5 | 4 | 0.25 | 19 | 0.2632 | 3 | 0 | 3 | 0 | 3 | 0.3333 | 5 | 0 | 14 | 0.0714 |
| 6_1075441 | 6 | 0.8333 | 5 | 0.8 | 4 | 0.75 | 4 | 1 | 19 | 0.8421 | 6 | 0.8333 | 3 | 0.6667 | 3 | 0.6667 | 5 | 1 | 17 | 0.8235 |
| 6_1075441 | 4 | 1 | 5 | 0.8 | 4 | 1 | 2 | 1 | 15 | 0.9333 | 5 | 1 | 2 | 1 | 2 | 1 | 5 | 1 | 14 | 1 |
| 6_1075442 | 6 | 0.1667 | 4 | 0.25 | 4 | 0.25 | 3 | 0 | 17 | 0.1765 | 6 | 0.3333 | 3 | 0.3333 | 2 | 0 | 4 | 0.25 | 15 | 0.2667 |
| 6_1075442 | 4 | 1 | 3 | 0.6667 | 4 | 1 | 2 | 1 | 13 | 0.9231 | 3 | 1 | 3 | 1 | 1 | 1 | 1 | 1 | 8 | 1 |
| 6_1075442 | 6 | 0.5 | 5 | 0.6 | 4 | 0.75 | 3 | 0 | 18 | 0.5 | 5 | 0.6 | 3 | 0.3333 | 3 | 1 | 5 | 0.2 | 16 | 0.5 |
| 6_1075444 | 6 | 0.3333 | 4 | 0.25 | 3 | 0 | 3 | 0.3333 | 16 | 0.25 | 6 | 0.1667 | 3 | 0.3333 | 4 | 0 | 5 | 0 | 18 | 0.1111 |
| 6_1075446 | 5 | 0 | 5 | 0.2 | 4 | 0.25 | 3 | 0 | 17 | 0.1176 | 7 | 0.1429 | 2 | 0 | 4 | 0 | 5 | 0.2 | 18 | 0.1111 |
| 6_1075448 | 7 | 0.4286 | 6 | 0.6667 | 4 | 0.5 | 5 | 0.6 | 22 | 0.5455 | 7 | 0.5714 | 3 | 0.3333 | 4 | 1 | 5 | 0.6 | 19 | 0.6316 |
| 6_1075448 | 7 | 0.5714 | 5 | 0 | 2 | 0.5 | 4 | 0.25 | 18 | 0.3333 | 5 | 0.8 | 3 | 0.6667 | 1 | 1 | 3 | 0 | 12 | 0.5833 |
| 6_1075449 | 5 | 1 | 6 | 0.8333 | 3 | 0.6667 | 4 | 1 | 18 | 0.8889 | 5 | 0.6 | 3 | 1 | 3 | 0.6667 | 5 | 0.6 | 16 | 0.6875 |
| 6_1075450 | 7 | 0.1429 | 6 | 0 | 5 | 0 | 4 | 0.5 | 22 | 0.1364 | 6 | 0.1667 | 3 | 0.3333 | 3 | 0 | 5 | 0.2 | 17 | 0.1765 |
| 6_1075450 | 5 | 0.6 | 5 | 0.4 | 5 | 0.4 | 4 | 1 | 19 | 0.5789 | 6 | 0.5 | 2 | 0 | 3 | 0.6667 | 3 | 0.6667 | 14 | 0.5 |
| 6_1075450 | 7 | 1 | 6 | 0.8333 | 4 | 0.5 | 5 | 0.6 | 22 | 0.7727 | 7 | 0.8571 | 3 | 1 | 3 | 0.3333 | 5 | 0.6 | 18 | 0.7222 |
| 6_1075451 | 6 | 0.5 | 6 | 0 | 3 | 0.3333 | 4 | 0.5 | 19 | 0.3158 | 5 | 0.2 | 2 | 0 | 3 | 0.6667 | 4 | 0.5 | 14 | 0.3571 |
| 6_1075451 | 5 | 0.2 | 6 | 0.3333 | 5 | 0.4 | 2 | 0 | 18 | 0.2778 | 5 | 0.4 | 3 | 0.3333 | 2 | 0 | 4 | 0.25 | 14 | 0.2857 |
| 6_1075452 | 7 | 0.1429 | 6 | 0.1667 | 3 | 0 | 4 | 0.5 | 20 | 0.2 | 3 | 0 | 3 | 0 | 1 | 0 | 4 | 0.25 | 11 | 0.0909 |
| 6_1075452 | 7 | 0.1429 | 6 | 0.1667 | 3 | 0 | 3 | 0 | 19 | 0.1053 | 4 | 0 | 3 | 0 | 3 | 0 | 5 | 0.2 | 15 | 0.0667 |
| 6_1075453 | 7 | 0.4286 | 7 | 0.1429 | 6 | 0.3333 | 4 | 0.25 | 24 | 0.2917 | 6 | 0.6667 | 3 | 0 | 3 | 0.3333 | 5 | 0.2 | 17 | 0.3529 |
| 6_1075454 | 8 | 0.875 | 8 | 1 | 4 | 0.25 | 3 | 0.3333 | 23 | 0.7391 | 6 | 0.3333 | 2 | 0.5 | 3 | 1 | 4 | 0.75 | 15 | 0.6 |
| 6_1075455 | 8 | 1 | 6 | 1 | 5 | 1 | 5 | 1 | 24 | 1 | 7 | 1 | 3 | 1 | NA | NA | 4 | 1 | 14 | 1 |
| 6_1075455 | 8 | 0.875 | 6 | 0.6667 | 4 | 0.75 | 4 | 1 | 22 | 0.8182 | 5 | 0.8 | 3 | 0.6667 | 2 | 0.5 | 3 | 0.6667 | 13 | 0.6923 |
| 6_1075456 | 7 | 0.1429 | 7 | 0 | 5 | 0 | 5 | 0.8 | 24 | 0.2083 | 6 | 0 | 2 | 0 | 3 | 0.3333 | 4 | 0 | 15 | 0.0667 |
| 6_1075456 | 7 | 0 | 8 | 0.25 | 6 | 0 | 3 | 0 | 24 | 0.0833 | 7 | 0.1429 | 4 | 0 | 3 | 0.6667 | 3 | 0.3333 | 17 | 0.2353 |
| 6_1075457 | 8 | 0 | 8 | 0.125 | 6 | 0 | 4 | 0.25 | 26 | 0.0769 | 6 | 0.3333 | 4 | 0 | 4 | 0.25 | 4 | 0.5 | 18 | 0.2778 |
| 6_1075458 | 8 | 0.125 | 8 | 0 | 6 | 0.3333 | 3 | 0 | 25 | 0.12 | 7 | 0.2857 | 4 | 0 | 3 | 0.3333 | 4 | 0.25 | 18 | 0.2222 |
| 6_1075458 | 7 | 0.1429 | 6 | 0.5 | 4 | 0.25 | 5 | 0.6 | 22 | 0.3636 | 7 | 0.5714 | 4 | 0.5 | 2 | 0 | 4 | 0.75 | 17 | 0.5294 |
| 6_1075460 | 7 | 0 | 5 | 0.4 | 6 | 0.1667 | 6 | 0.6667 | 24 | 0.2917 | 9 | 0.2222 | 4 | 0.5 | 4 | 0.5 | 4 | 0.25 | 21 | 0.3333 |
| 6_1075460 | 9 | 0.7778 | 7 | 1 | 6 | 0.6667 | 5 | 0.8 | 27 | 0.8148 | 9 | 0.6667 | 4 | 0.5 | 3 | 1 | 5 | 0.6 | 21 | 0.6667 |
| 6_1075462 | 8 | 0 | 6 | 0.3333 | 4 | 0.25 | 4 | 0 | 22 | 0.1364 | 9 | 0.1111 | 2 | 0 | 3 | 0 | 4 | 0 | 18 | 0.0556 |
| 6_1075462 | 11 | 0 | 6 | 0.1667 | 5 | 0.8 | 6 | 0.5 | 28 | 0.2857 | 8 | 0.125 | 4 | 0 | 1 | 0 | 3 | 0.3333 | 16 | 0.125 |
| 6_1075462 | 10 | 0.5 | 7 | 0.2857 | 6 | 0 | 7 | 0.1429 | 30 | 0.2667 | 10 | 0 | 4 | 0 | 4 | 0.25 | 4 | 0.5 | 22 | 0.1364 |
| 6_1075462 | 9 | 1 | 6 | 1 | 5 | 1 | 4 | 1 | 24 | 1 | 9 | 0.8889 | 4 | 1 | 4 | 0.75 | 4 | 1 | 21 | 0.9048 |
| 6_1075463 | 7 | 1 | 7 | 0.7143 | 5 | 0.8 | 2 | 1 | 21 | 0.8571 | 8 | 1 | 3 | 1 | 3 | 0.6667 | 4 | 0.75 | 18 | 0.8889 |
| 6_1075463 | 10 | 1 | 8 | 1 | 7 | 1 | 7 | 1 | 32 | 1 | 10 | 1 | 4 | 1 | 4 | 0.75 | 4 | 1 | 22 | 0.9545 |
| 6_1075464 | 11 | 1 | 8 | 0.5 | 7 | 0.8571 | 7 | 0.4286 | 33 | 0.7273 | 11 | 0.9091 | 4 | 0.75 | 4 | 0.75 | 4 | 1 | 23 | 0.8696 |
| 6_1075464 | 10 | 0.6 | 7 | 0.7143 | 5 | 0.8 | 6 | 0.8333 | 28 | 0.7143 | 10 | 0.8 | 4 | 0.75 | 5 | 0.8 | 5 | 0.8 | 24 | 0.7917 |
| 6_1075465 | 10 | 0 | 7 | 0 | 4 | 0.25 | 6 | 0.1667 | 27 | 0.0741 | 7 | 0.1429 | 4 | 0.25 | 5 | 0.4 | 3 | 0.3333 | 19 | 0.2632 |
| 6_1075468 | 14 | 0.8571 | 10 | 1 | 6 | 1 | 8 | 0.875 | 38 | 0.9211 | 12 | 0.8333 | 4 | 1 | 4 | 1 | 5 | 0.8 | 25 | 0.88 |
| 6_1075468 | 13 | 0.2308 | 9 | 0.2222 | 8 | 0.375 | 7 | 0.4286 | 37 | 0.2973 | 12 | 0.4167 | 4 | 0.25 | 4 | 0 | 5 | 0.4 | 25 | 0.32 |
| 6_1075468 | 15 | 0.2667 | 9 | 0.2222 | 8 | 0.375 | 8 | 0 | 40 | 0.225 | 14 | 0.0714 | 3 | 0 | 4 | 0 | 5 | 0.2 | 26 | 0.0769 |
| 6_1075469 | 11 | 0.1818 | 9 | 0.4444 | 6 | 0.6667 | 4 | 0.75 | 30 | 0.4333 | 11 | 0.1818 | 2 | 0 | 3 | 1 | 4 | 0 | 20 | 0.25 |
| 6_1080528 | 3 | 0.3333 | NA | NA | 3 | 0 | NA | NA | 6 | 0.1667 | NA | NA | NA | NA | NA | NA | 1 | 1 | 1 | 1 |
| 6_1080529 | 3 | 0.6667 | NA | NA | 3 | 0.6667 | NA | NA | 6 | 0.6667 | 1 | 1 | NA | NA | NA | NA | 1 | 1 | 2 | 1 |
| 6_1080529 | 4 | 0.75 | NA | NA | 3 | 1 | NA | NA | 7 | 0.8571 | 1 | 1 | NA | NA | NA | NA | 1 | 1 | 2 | 1 |
| 6_1080531 | 4 | 1 | NA | NA | 2 | 1 | NA | NA | 6 | 1 | 1 | 1 | 1 | 1 | NA | NA | 1 | 1 | 3 | 1 |
| 6_1080531 | 3 | 0 | NA | NA | 3 | 0.6667 | NA | NA | 6 | 0.3333 | 1 | 0 | NA | NA | NA | NA | 1 | 1 | 2 | 0.5 |

| | | | | | | | | | | | | | | | | | | | | |
|---|---|---|---|---|---|---|---|---|---|---|---|---|---|---|---|---|---|---|---|---|
| 6_10805322 | 4 | 1 | NA | NA | 2 | 1 | NA | NA | 6 | 1 | 1 | 1 | NA | NA | NA | NA | NA | NA | 1 | 1 |
| 6_10805328 | 4 | 1 | NA | NA | 1 | 0 | NA | NA | 5 | 0.8 | 1 | 1 | NA | NA | NA | NA | 1 | 1 | 2 | 1 |
| 6_10805334 | 4 | 0.75 | NA | NA | 2 | 1 | NA | NA | 6 | 0.8333 | 1 | 1 | 1 | 1 | NA | NA | 1 | 1 | 3 | 1 |
| 6_10805341 | 4 | 0.25 | NA | NA | 2 | 0.5 | NA | NA | 6 | 0.3333 | 1 | 1 | 1 | 0 | NA | NA | 1 | 0 | 3 | 0.3333 |
| 6_10805393 | 4 | 0.25 | NA | NA | 2 | 0.5 | 1 | 1 | 7 | 0.4286 | NA | NA | 1 | 0 | NA | NA | 1 | 1 | 2 | 0.5 |
| 6_10808502 | 3 | 0 | 2 | 0 | 3 | 0.3333 | 3 | 0 | 11 | 0.0909 | 2 | 0 | 5 | 0.2 | 2 | 0.5 | 2 | 0 | 11 | 0.1818 |
| 6_10808504 | 3 | 0 | 2 | 0.5 | 2 | 0.5 | 1 | 0 | 8 | 0.25 | 2 | 0 | 3 | 0.3333 | 2 | 0.5 | 2 | 0.5 | 9 | 0.3333 |
| 6_10808516 | 2 | 0 | 1 | 0 | 2 | 0.5 | 3 | 0 | 8 | 0.125 | 1 | 0 | 5 | 0.4 | 2 | 0 | 2 | 0.5 | 10 | 0.3 |
| 6_10808530 | NA | NA | 1 | 1 | 2 | 1 | 3 | 0.6667 | 6 | 0.8333 | 1 | 1 | 3 | 1 | 1 | 1 | 2 | 1 | 7 | 1 |
| 6_10808536 | 1 | 1 | 2 | 1 | 1 | 0 | 3 | 1 | 7 | 0.8571 | 1 | 1 | 3 | 1 | 2 | 1 | 2 | 0.5 | 8 | 0.875 |
| 6_10811731 | 28 | 0.0714 | 27 | 0.037 | 24 | 0.0833 | 27 | 0.037 | 106 | 0.0566 | 32 | 0.0312 | 27 | 0 | 32 | 0.1562 | 34 | 0.0882 | 125 | 0.072 |
| 6_10845825 | 19 | 0.2105 | 21 | 0.0952 | 14 | 0.2143 | 9 | 0.2222 | 63 | 0.1746 | 7 | 0.4286 | 22 | 0.1818 | 10 | 0.3 | 21 | 0.3333 | 60 | 0.2833 |
| 6_10845829 | 22 | 0.0909 | 16 | 0.1875 | 15 | 0.1333 | 12 | 0.25 | 65 | 0.1538 | 9 | 0.1111 | 23 | 0.087 | 14 | 0.3571 | 22 | 0.0909 | 68 | 0.1471 |
| 6_10852681 | 35 | 0.1714 | 28 | 0.1786 | 30 | 0.2 | 37 | 0.0811 | 130 | 0.1538 | 19 | 0.1579 | 39 | 0.1282 | 31 | 0.0645 | 40 | 0.1 | 129 | 0.1085 |
| 6_10852696 | 36 | 0.1667 | 30 | 0.1333 | 26 | 0.1538 | 29 | 0.1724 | 121 | 0.157 | 17 | 0.1176 | 35 | 0.1429 | 30 | 0.2 | 41 | 0.1951 | 123 | 0.1707 |
| 6_10852704 | 43 | 0.0465 | 30 | 0.1 | 31 | 0.1613 | 36 | 0.0278 | 140 | 0.0786 | 18 | 0.1111 | 40 | 0.1 | 34 | 0.0294 | 43 | 0.0698 | 135 | 0.0741 |
| 6_10864149 | 6 | 0.3333 | 7 | 0 | 5 | 0 | NA | NA | 18 | 0.1111 | 3 | 0 | 3 | 0 | 2 | 0 | 6 | 0.1667 | 14 | 0.0714 |
| 6_10864163 | 4 | 0.5 | 1 | 1 | 6 | 0.8333 | 2 | 1 | 13 | 0.7692 | 1 | 1 | 2 | 1 | 3 | 0.6667 | 5 | 1 | 11 | 0.9091 |
| 6_10864165 | 4 | 1 | 7 | 1 | 9 | 0.8889 | 2 | 1 | 22 | 0.9545 | 2 | 1 | 3 | 0.6667 | 4 | 1 | 6 | 1 | 15 | 0.9333 |
| 6_10864171 | 4 | 0.5 | 7 | 0.7143 | 6 | 0.6667 | 2 | 1 | 19 | 0.6842 | 3 | 0.3333 | 3 | 1 | 3 | 0.6667 | 5 | 0.2 | 14 | 0.5 |
| 6_10864173 | 3 | 1 | 5 | 0.8 | 2 | 1 | NA | NA | 10 | 0.9 | 2 | 0.5 | 2 | 1 | 3 | 1 | 3 | 1 | 10 | 0.9 |
| 6_10864177 | 4 | 1 | 7 | 0.7143 | 9 | 1 | 2 | 1 | 22 | 0.9091 | 2 | 0.5 | 2 | 1 | 5 | 0.8 | 5 | 1 | 14 | 0.8571 |
| 6_10864178 | 3 | 0 | 6 | 0.1667 | 5 | 0.2 | 1 | 0 | 15 | 0.1333 | 3 | 0 | 2 | 0.5 | 4 | 0 | 5 | 0 | 14 | 0.0714 |
| 6_10864181 | 5 | 1 | 6 | 0.8333 | 8 | 0.75 | 2 | 0.5 | 21 | 0.8095 | 3 | 0.3333 | 3 | 0.3333 | 5 | 0.6 | 6 | 0.8333 | 17 | 0.5882 |
| 6_10864215 | 5 | 0.4 | 7 | 0.5714 | 8 | 0.375 | 2 | 1 | 22 | 0.5 | 3 | 0.3333 | 2 | 0 | 4 | 0.25 | 8 | 0.25 | 17 | 0.2353 |
| 6_10864219 | 7 | 1 | 7 | 1 | 10 | 0.8 | 2 | 1 | 26 | 0.9231 | 3 | 1 | 3 | 1 | 5 | 1 | 8 | 0.875 | 19 | 0.9474 |
| 6_10864227 | 6 | 0.3333 | 6 | 0.1667 | 10 | 0.3 | 2 | 0 | 24 | 0.25 | 3 | 0 | 3 | 0 | 5 | 0 | 8 | 0.125 | 19 | 0.0526 |
| 6_10864228 | 4 | 0.25 | 5 | 0 | 8 | 0 | 1 | 0 | 18 | 0.0556 | 3 | 0 | 3 | 0.3333 | 6 | 0 | 8 | 0 | 20 | 0.05 |
| 6_10864228 | 7 | 1 | 5 | 1 | 10 | 0.9 | 2 | 0.5 | 24 | 0.9167 | 4 | 1 | 2 | 1 | 5 | 1 | 6 | 1 | 17 | 1 |
| 6_10864229 | 5 | 0.4 | 4 | 0 | 6 | 0.5 | NA | NA | 15 | 0.3333 | 2 | 0 | 2 | 0 | 4 | 0.25 | 4 | 0.25 | 12 | 0.1667 |
| 6_10864253 | 8 | 0 | 5 | 0.2 | 11 | 0.2727 | 3 | 0 | 27 | 0.1481 | 2 | 0.5 | 3 | 0.6667 | 6 | 0.6667 | 8 | 0.5 | 19 | 0.5789 |
| 6_10864254 | 7 | 1 | 7 | 0.7143 | 10 | 0.5 | 3 | 1 | 27 | 0.7407 | 3 | 1 | 3 | 1 | 7 | 1 | 8 | 0.625 | 21 | 0.8571 |
| 6_10864266 | 8 | 0.125 | 6 | 0.1667 | 9 | 0.1111 | 3 | 0 | 26 | 0.1154 | 4 | 0.25 | 3 | 0.3333 | 6 | 0.1667 | 8 | 0.25 | 21 | 0.2381 |
| 6_10864296 | 11 | 0.2727 | 7 | 0.1429 | 11 | 0 | 4 | 0 | 33 | 0.1212 | 3 | 0.3333 | 2 | 0.5 | 8 | 0.125 | 8 | 0.125 | 21 | 0.1905 |
| 6_10876024 | 9 | 0.1111 | 13 | 0.1538 | 11 | 0 | 15 | 0.2 | 48 | 0.125 | 9 | 0.1111 | 11 | 0 | 13 | 0 | 11 | 0.3636 | 44 | 0.1136 |
| 6_10879891 | 20 | 0.2 | 28 | 0.3571 | 23 | 0.1304 | 28 | 0.1786 | 99 | 0.2222 | 16 | 0.1875 | 18 | 0.1111 | 21 | 0.0952 | 18 | 0.3333 | 73 | 0.1781 |
| 6_10879948 | 25 | 0.08 | 30 | 0.1333 | 26 | 0.0769 | 32 | 0.0312 | 113 | 0.0796 | 18 | 0.0556 | 21 | 0.0952 | 26 | 0.0385 | 17 | 0.2941 | 82 | 0.1098 |
| 6_10879998 | 28 | 0 | 27 | 0.1481 | 25 | 0.08 | 24 | 0.0417 | 104 | 0.0673 | 15 | 0.0667 | 21 | 0.0476 | 25 | 0.08 | 17 | 0.1176 | 78 | 0.0769 |
| 6_10880056 | 10 | 0.1 | 13 | 0.2308 | 12 | 0.3333 | 8 | 0.375 | 43 | 0.2558 | 4 | 0.5 | 5 | 0.2 | 8 | 0.25 | 9 | 0 | 26 | 0.1923 |
| 6_10880060 | 11 | 0.3636 | 13 | 0.1538 | 15 | 0.2 | 10 | 0 | 49 | 0.1837 | 6 | 0.1667 | 7 | 0.1429 | 10 | 0 | 8 | 0.375 | 31 | 0.1613 |
| 6_10880197 | 2 | 1 | 1 | 0 | 4 | 0.75 | NA | NA | 7 | 0.7143 | NA | NA | 1 | 1 | NA | NA | NA | NA | 1 | 1 |
| 6_10883254 | NA | NA | 5 | 0.8 | 1 | 0 | 2 | 1 | 8 | 0.75 | NA | NA | NA | NA | NA | NA | 2 | 1 | 2 | 1 |
| 6_10883347 | NA | NA | 2 | 0 | 2 | 0 | 2 | 0.5 | 6 | 0.1667 | 1 | 0 | NA | NA | NA | NA | 1 | 1 | 2 | 0.5 |
| 6_11133522 | 2 | 0.5 | 1 | 1 | 2 | 0.5 | 2 | 0.5 | 7 | 0.5714 | 1 | 0 | 7 | 0.2857 | 1 | 0 | 4 | 1 | 13 | 0.4615 |
| 6_11133530 | 2 | 0 | 1 | 1 | 2 | 0 | 3 | 0.3333 | 8 | 0.25 | 1 | 0 | 6 | 0.1667 | 1 | 0 | 4 | 0 | 12 | 0.0833 |
| 6_11133531 | 1 | 1 | 1 | 0 | 2 | 1 | 3 | 1 | 7 | 0.8571 | 1 | 1 | 5 | 0.4 | 1 | 0 | 4 | 1 | 11 | 0.6364 |
| 6_11133547 | 2 | 0 | 1 | 0 | 2 | 0.5 | 3 | 0 | 8 | 0.125 | 1 | 1 | 7 | 0.2857 | 1 | 0 | 5 | 0.6 | 14 | 0.4286 |
| 6_11133549 | 1 | 0 | 1 | 0 | 2 | 0 | 2 | 0.5 | 6 | 0.1667 | 1 | 0 | 7 | 0.1429 | 1 | 0 | 4 | 0 | 13 | 0.0769 |
| 6_11133567 | 3 | 0.3333 | 1 | 0 | 2 | 0.5 | 2 | 1 | 8 | 0.5 | 1 | 0 | 6 | 0.5 | 1 | 1 | 6 | 0.5 | 14 | 0.5 |
| 6_11133578 | 3 | 0.6667 | 2 | 1 | 2 | 1 | 3 | 1 | 10 | 0.9 | 1 | 0 | 6 | 0.3333 | NA | NA | 6 | 0.6667 | 13 | 0.4615 |
| 6_11133580 | 3 | 0 | 2 | 0.5 | 1 | 1 | 2 | 0.5 | 8 | 0.375 | 1 | 0 | 4 | 0.75 | 1 | 1 | 6 | 0.3333 | 12 | 0.5 |
| 6_11133584 | 3 | 0.3333 | 2 | 0.5 | 2 | 0 | 3 | 0 | 10 | 0.2 | 1 | 0 | 6 | 0.1667 | 1 | 0 | 6 | 0.6667 | 14 | 0.3571 |
| 6_11133591 | 3 | 0.3333 | 1 | 0 | 2 | 0 | 4 | 0 | 10 | 0.1 | 1 | 0 | 5 | 0.2 | 1 | 1 | 6 | 0.5 | 13 | 0.3846 |
| 6_11147272 | 5 | 0 | 2 | 0 | 1 | 0 | 3 | 0.3333 | 11 | 0.0909 | 1 | 0 | 8 | 0.125 | NA | NA | 5 | 0.4 | 14 | 0.2143 |
| 6_11154309 | 4 | 1 | 2 | 0.5 | 3 | 0 | 3 | 0.6667 | 12 | 0.5833 | NA | NA | 8 | 0.75 | 2 | 0.5 | 5 | 0.8 | 15 | 0.7333 |
| 6_11154318 | 4 | 0.25 | 2 | 0 | 2 | 0 | 4 | 0.25 | 12 | 0.1667 | NA | NA | 7 | 0 | NA | NA | 4 | 0.25 | 11 | 0.0909 |
| 6_11154319 | 4 | 0.25 | 1 | 0 | 3 | 0.3333 | 4 | 0.5 | 12 | 0.3333 | NA | NA | 7 | 0.2857 | 2 | 0.5 | 5 | 0.4 | 14 | 0.3571 |
| 6_11154326 | 2 | 1 | 2 | 0.5 | 2 | 1 | 3 | 1 | 9 | 0.8889 | NA | NA | 6 | 1 | 1 | 1 | 4 | 0.75 | 11 | 0.9091 |
| 6_11154334 | 4 | 1 | 2 | 1 | 3 | 1 | 4 | 1 | 13 | 1 | 1 | 1 | 7 | 1 | 2 | 1 | 6 | 0.6667 | 16 | 0.875 |
| 6_11154342 | 4 | 0.75 | 2 | 0.5 | 5 | 0.4 | 4 | 1 | 15 | 0.6667 | 1 | 1 | 8 | 0.75 | 2 | 1 | 4 | 0.75 | 15 | 0.8 |
| 6_11154361 | 4 | 0 | 2 | 0.5 | 5 | 0.2 | 4 | 0.25 | 15 | 0.2 | 1 | 0 | 8 | 0.375 | 2 | 0 | 7 | 0.5714 | 18 | 0.3889 |
| 6_11154369 | 2 | 0 | 2 | 0.5 | 5 | 0 | 4 | 0.25 | 13 | 0.1538 | 1 | 1 | 6 | 0.5 | 1 | 0 | 6 | 0.1667 | 14 | 0.3571 |
| 6_11154398 | 4 | 0 | 2 | 0.5 | 6 | 0 | 5 | 0 | 17 | 0.0588 | 1 | 0 | 9 | 0.1111 | 2 | 0 | 8 | 0 | 20 | 0.05 |
| 6_11154400 | 3 | 0.3333 | 2 | 0.5 | 4 | 0.25 | 5 | 0.6 | 14 | 0.4286 | 1 | 1 | 9 | 0.4444 | 3 | 0 | 8 | 0.5 | 21 | 0.4286 |
| 6_11245235 | 2 | 1 | 3 | 1 | NA | NA | NA | NA | 5 | 1 | 1 | 1 | 1 | 1 | NA | NA | NA | NA | 2 | 1 |

| | | | | | | | | | | | | | | | | | | | | |
|---|---|---|---|---|---|---|---|---|---|---|---|---|---|---|---|---|---|---|---|---|
| 6_1124526 | 1 | 1 | 3 | 0.6667 | NA | NA | NA | NA | 4 | 0.75 | 1 | 0 | 1 | 1 | NA | NA | 1 | 1 | 3 | 0.6667 |
| 6_1124526 | 1 | 0 | 3 | 0.3333 | NA | NA | NA | NA | 4 | 0.25 | 1 | 1 | 1 | 1 | NA | NA | 1 | 0 | 3 | 0.6667 |
| 6_1124530 | 1 | 1 | 2 | 0 | NA | NA | NA | NA | 3 | 0.3333 | NA | NA | 1 | 1 | NA | NA | 1 | 0 | 2 | 0.5 |
| 6_1125429 | NA | NA | NA | NA | 1 | 1 | NA | NA | 1 | 1 | 1 | 1 | NA | NA | NA | NA | NA | NA | 1 | 1 |
| 6_1126951 | 19 | 0.2105 | 18 | 0.2778 | 18 | 0.1667 | 17 | 0.4118 | 72 | 0.2639 | 11 | 0.2727 | 15 | 0.2 | 19 | 0.2105 | 16 | 0.25 | 61 | 0.2295 |
| 6_1126952 | 21 | 0.2381 | 16 | 0.3125 | 22 | 0.2273 | 18 | 0.2222 | 77 | 0.2468 | 10 | 0.3 | 18 | 0.3333 | 20 | 0.2 | 19 | 0.3684 | 67 | 0.2985 |
| 6_1126953 | 16 | 0.3125 | 14 | 0.0714 | 20 | 0.2 | 15 | 0.4 | 65 | 0.2462 | 10 | 0.3 | 20 | 0.05 | 21 | 0.2381 | 18 | 0.0556 | 69 | 0.1449 |
| 6_1126954 | 16 | 0.0625 | 12 | 0.25 | 18 | 0.2222 | 14 | 0.1429 | 60 | 0.1667 | 9 | 0.2222 | 13 | 0.0769 | 17 | 0.1176 | 18 | 0.2222 | 57 | 0.1579 |
| 6_1126954 | 18 | 0.4444 | 15 | 0.4 | 22 | 0.5455 | 20 | 0.5 | 75 | 0.48 | 12 | 0.6667 | 17 | 0.5882 | 20 | 0.6 | 18 | 0.3333 | 67 | 0.5373 |
| 6_1126955 | 13 | 0.1538 | 18 | 0.2222 | 17 | 0.2941 | 9 | 0.1111 | 57 | 0.2105 | 8 | 0 | 13 | 0 | 15 | 0.2667 | 16 | 0.1875 | 52 | 0.1346 |
| 6_1126958 | 18 | 0.9444 | 14 | 0.9286 | 18 | 0.9444 | 16 | 0.8125 | 66 | 0.9091 | 11 | 0.9091 | 17 | 1 | 20 | 1 | 19 | 1 | 67 | 0.9851 |
| 6_1126959 | 17 | 0.5882 | 11 | 0.1818 | 13 | 0.6154 | 15 | 0.7333 | 56 | 0.5536 | 8 | 0.75 | 13 | 0.4615 | 18 | 0.5556 | 20 | 0.75 | 59 | 0.6271 |
| 6_1126959 | 16 | 0.75 | 12 | 0.9167 | 13 | 0.8462 | 15 | 0.9333 | 56 | 0.8571 | 11 | 0.9091 | 16 | 0.75 | 19 | 0.8421 | 18 | 0.8333 | 64 | 0.8281 |
| 6_1126960 | 11 | 1 | 10 | 0.9 | 12 | 1 | 7 | 0.8571 | 40 | 0.95 | 6 | 1 | 9 | 1 | 15 | 0.9333 | 10 | 1 | 40 | 0.975 |
| 6_1126960 | 11 | 0.5455 | 11 | 0.6364 | 12 | 0.5 | 14 | 0.6429 | 48 | 0.5833 | 9 | 0.4444 | 15 | 0.8 | 15 | 0.7333 | 11 | 0.6364 | 50 | 0.68 |
| 6_1126960 | 10 | 0.2 | 14 | 0.1429 | 16 | 0 | 13 | 0.1538 | 53 | 0.1132 | 10 | 0 | 14 | 0.0714 | 19 | 0 | 17 | 0.1176 | 60 | 0.05 |
| 6_1126961 | 15 | 0.7333 | 12 | 0.75 | 11 | 0.6364 | 13 | 0.5385 | 51 | 0.6667 | 7 | 0.4286 | 12 | 0.6667 | 13 | 0.9231 | 15 | 0.8 | 47 | 0.7447 |
| 6_1126962 | 14 | 0.7857 | 13 | 0.8462 | 15 | 0.7333 | 13 | 1 | 55 | 0.8364 | 9 | 1 | 14 | 0.6429 | 19 | 0.7895 | 16 | 0.8125 | 58 | 0.7931 |
| 6_1126964 | 12 | 0.5833 | 11 | 0.3636 | 13 | 0.4615 | 10 | 0.7 | 46 | 0.5217 | 10 | 0.4 | 12 | 0.5 | 18 | 0.5 | 14 | 0.4286 | 54 | 0.463 |
| 6_1126965 | 11 | 0.7273 | 12 | 0.6667 | 14 | 0.9286 | 10 | 0.7 | 47 | 0.766 | 11 | 0.9091 | 9 | 0.7778 | 13 | 0.7692 | 14 | 0.7857 | 47 | 0.8085 |
| 6_1126965 | 12 | 0.0833 | 12 | 0.25 | 10 | 0.3 | 10 | 0.3 | 44 | 0.2273 | 9 | 0.2222 | 11 | 0.1818 | 15 | 0.2667 | 12 | 0.25 | 47 | 0.234 |
| 6_1126973 | 6 | 0.3333 | 9 | 0 | 3 | 0 | 8 | 0.125 | 26 | 0.1154 | 2 | 0 | 6 | 0.1667 | 5 | 0.2 | 6 | 0.1667 | 19 | 0.1579 |
| 6_1126985 | 5 | 0.2 | 3 | 0 | 6 | 0.3333 | 6 | 0.3333 | 20 | 0.25 | 2 | 0 | 5 | 0.2 | 8 | 0.375 | 8 | 0 | 23 | 0.1739 |
| 6_1126987 | 4 | 0.25 | 4 | 0 | 6 | 0.3333 | 4 | 0.5 | 18 | 0.2778 | 2 | 0.5 | 5 | 0.6 | 7 | 0 | 7 | 0 | 21 | 0.1905 |
| 6_1126988 | 3 | 0.3333 | 4 | 0 | 4 | 0.5 | 5 | 0 | 16 | 0.1875 | 1 | 0 | 5 | 0.2 | 4 | 0 | 7 | 0.2857 | 17 | 0.1765 |
| 6_1126989 | 5 | 0.4 | 4 | 0 | 7 | 0.1429 | 5 | 0 | 21 | 0.1429 | 2 | 0 | 5 | 0.2 | 7 | 0.1429 | 5 | 0.2 | 19 | 0.1579 |
| 6_1126990 | 5 | 0.2 | 3 | 0 | 4 | 0 | 4 | 0.25 | 16 | 0.125 | 1 | 0 | 5 | 0.2 | 5 | 0.2 | 7 | 0.2857 | 18 | 0.2222 |
| 6_1126990 | 5 | 0 | 4 | 0 | 5 | 0.4 | 4 | 0.75 | 18 | 0.2778 | 2 | 0.5 | 5 | 0.4 | 8 | 0.375 | 7 | 0.1429 | 22 | 0.3182 |
| 6_1126991 | 5 | 0.6 | 3 | 0 | 3 | 0 | 4 | 0.5 | 15 | 0.3333 | 2 | 0.5 | 4 | 0.25 | 6 | 0.5 | 7 | 0.7143 | 19 | 0.5263 |
| 6_1126992 | 3 | 0.3333 | 2 | 0 | 4 | 0.5 | 3 | 0 | 12 | 0.25 | 2 | 0.5 | 5 | 0.2 | 5 | 0.2 | 7 | 0.4286 | 19 | 0.3158 |
| 6_1126994 | 4 | 1 | 3 | 1 | 5 | 1 | 4 | 0.75 | 16 | 0.9375 | 2 | 1 | 3 | 0.6667 | 6 | 1 | 6 | 0.8333 | 17 | 0.8824 |
| 6_1126995 | 3 | 0.3333 | 3 | 0 | 4 | 0.75 | 5 | 0.8 | 15 | 0.5333 | 2 | 0.5 | 3 | 0 | 6 | 0.5 | 6 | 0.5 | 17 | 0.4118 |
| 6_1126995 | 4 | 0 | 3 | 0.3333 | 4 | 0.5 | 5 | 0.8 | 16 | 0.4375 | 1 | 0 | 4 | 0.75 | 7 | 0.2857 | 4 | 0.5 | 16 | 0.4375 |
| 6_1126997 | 4 | 0 | 3 | 0 | 3 | 0.3333 | 5 | 0.8 | 15 | 0.3333 | 2 | 1 | 3 | 0.3333 | 6 | 0.5 | 5 | 0.8 | 16 | 0.625 |
| 6_1126997 | 3 | 1 | 3 | 0.6667 | 3 | 1 | 5 | 0.8 | 14 | 0.8571 | 2 | 1 | 2 | 1 | 6 | 0.8333 | 3 | 0.6667 | 13 | 0.8462 |
| 6_1126997 | 3 | 0.6667 | 3 | 0.3333 | 3 | 0 | 4 | 0.75 | 13 | 0.4615 | 2 | 1 | 3 | 0.6667 | 4 | 0.75 | 5 | 0 | 14 | 0.5 |
| 6_1127000 | 2 | 0 | 3 | 0 | 2 | 0 | 2 | 0.5 | 9 | 0.1111 | 2 | 0 | 1 | 0 | 3 | 0 | 3 | 0.3333 | 9 | 0.1111 |
| 6_1127001 | 2 | 0 | 2 | 0 | 2 | 0 | 3 | 0.3333 | 9 | 0.1111 | 1 | 0 | 2 | 0 | 3 | 0.3333 | 2 | 0 | 8 | 0.125 |
| 6_1127001 | 3 | 0 | 3 | 0 | 2 | 0 | 3 | 0.6667 | 11 | 0.1818 | 2 | 0.5 | 3 | 0 | 4 | 0.25 | 3 | 0.3333 | 12 | 0.25 |
| 6_1127044 | 2 | 0 | 2 | 0 | 2 | 0.5 | 3 | 0 | 9 | 0.1111 | 1 | 0 | 1 | 0 | 3 | 0 | 4 | 0.25 | 9 | 0.1111 |
| 6_1129235 | 2 | 1 | 3 | 0.6667 | 1 | 1 | 2 | 0.5 | 8 | 0.75 | NA | NA | NA | NA | NA | NA | 1 | 1 | 1 | 1 |
| 6_1129237 | 2 | 0 | 3 | 0 | 1 | 1 | 2 | 0.5 | 8 | 0.25 | 1 | 0 | NA | NA | NA | NA | 1 | 1 | 2 | 0.5 |
| 6_1130327 | 3 | 0 | 1 | 0 | 2 | 0.5 | 4 | 0 | 10 | 0.1 | 3 | 0.3333 | 1 | 0 | 3 | 0 | 2 | 0 | 9 | 0.1111 |
| 6_1130328 | 2 | 1 | 1 | 1 | 2 | 0.5 | 3 | 0.6667 | 8 | 0.75 | 2 | 1 | 1 | 1 | 3 | 1 | 3 | 1 | 9 | 1 |
| 6_1130329 | 3 | 0.3333 | 1 | 0 | 1 | 0 | 3 | 0 | 8 | 0.125 | 3 | 0.3333 | 1 | 0 | 2 | 0 | 3 | 0.6667 | 9 | 0.3333 |
| 6_1130329 | 4 | 0 | 1 | 0 | 2 | 0.5 | 3 | 0 | 10 | 0.1 | 3 | 0.3333 | 1 | 1 | 3 | 0 | 3 | 0 | 10 | 0.2 |
| 6_1130448 | 3 | 0.3333 | 1 | 0 | 1 | 0 | 2 | 0.5 | 7 | 0.2857 | 2 | 0.5 | 1 | 0 | 4 | 0.5 | 3 | 0.3333 | 10 | 0.4 |
| 6_1130449 | 4 | 0.25 | 3 | 0 | 4 | 0 | 4 | 0.25 | 15 | 0.1333 | 2 | 0 | 4 | 0.25 | 5 | 0 | 3 | 0 | 14 | 0.0714 |
| 6_1130483 | 2 | 0.5 | 5 | 0 | 2 | 0 | 1 | 0 | 10 | 0.1 | 1 | 1 | 1 | 0 | 3 | 0 | 3 | 0 | 8 | 0.125 |
| 6_1130830 | NA | NA | 1 | 0 | 2 | 0.5 | 4 | 0 | 7 | 0.1429 | 1 | 0 | 1 | 0 | 1 | 1 | NA | NA | 3 | 0.3333 |
| 6_1131207 | 1 | 1 | 2 | 1 | 2 | 0.5 | NA | NA | 5 | 0.8 | 1 | 1 | 2 | 1 | NA | NA | 3 | 0.6667 | 6 | 0.8333 |
| 6_1131208 | 1 | 1 | 2 | 1 | 1 | 1 | NA | NA | 4 | 1 | 1 | 1 | 3 | 1 | NA | NA | 4 | 1 | 8 | 1 |
| 6_1131209 | 1 | 0 | 2 | 0 | 2 | 0.5 | NA | NA | 5 | 0.2 | 1 | 0 | 3 | 0.3333 | NA | NA | 4 | 0.5 | 8 | 0.375 |
| 6_1131280 | 1 | 1 | 3 | 1 | 2 | 1 | NA | NA | 6 | 1 | 1 | 1 | 1 | 1 | 1 | 1 | 4 | 1 | 7 | 1 |
| 6_1131280 | NA | NA | 3 | 0.6667 | 3 | 0.6667 | NA | NA | 6 | 0.6667 | 2 | 0.5 | 3 | 0.3333 | 1 | 1 | 4 | 0.25 | 10 | 0.4 |
| 6_1131281 | 1 | 1 | 3 | 1 | 3 | 1 | NA | NA | 7 | 1 | 2 | 1 | 3 | 1 | 1 | 1 | 4 | 0.75 | 10 | 0.9 |
| 6_1131281 | 1 | 1 | 1 | 1 | 3 | 0.3333 | NA | NA | 5 | 0.6 | 1 | 1 | 3 | 1 | 1 | 1 | 4 | 0.5 | 9 | 0.7778 |
| 6_1131282 | NA | NA | 2 | 1 | 3 | 0 | NA | NA | 5 | 0.4 | 1 | 1 | 3 | 0.6667 | 1 | 0 | 4 | 0.5 | 9 | 0.5556 |
| 6_1131282 | 1 | 1 | 3 | 1 | 2 | 0.5 | NA | NA | 6 | 0.8333 | NA | NA | 2 | 0.5 | 1 | 1 | 3 | 1 | 6 | 0.8333 |
| 6_1131282 | 1 | 1 | 1 | 1 | 3 | 0.6667 | NA | NA | 5 | 0.8 | 1 | 1 | 3 | 0.3333 | 1 | 1 | 4 | 1 | 9 | 0.7778 |
| 6_1131282 | 1 | 1 | 3 | 1 | 3 | 0.6667 | NA | NA | 7 | 0.8571 | 1 | 1 | 3 | 0.6667 | 1 | 1 | 3 | 0.6667 | 8 | 0.75 |
| 6_1131284 | 1 | 1 | 2 | 1 | 2 | 1 | NA | NA | 5 | 1 | 1 | 1 | 3 | 1 | 1 | 1 | 4 | 0.5 | 9 | 0.7778 |
| 6_1131284 | 1 | 1 | 1 | 1 | 3 | 1 | NA | NA | 5 | 1 | 1 | 1 | 1 | 1 | 1 | 1 | 2 | 1 | 5 | 1 |
| 6_1131284 | NA | NA | 2 | 0.5 | 1 | 0 | NA | NA | 3 | 0.3333 | NA | NA | 2 | 0.5 | 1 | 1 | 3 | 0 | 6 | 0.3333 |
| 6_1131284 | NA | NA | 3 | 0.6667 | 3 | 1 | NA | NA | 6 | 0.8333 | NA | NA | 3 | 1 | 1 | 1 | 4 | 0.75 | 8 | 0.875 |

| | | | | | | | | | | | | | | | | | | | | |
|---|---|---|---|---|---|---|---|---|---|---|---|---|---|---|---|---|---|---|---|---|
| 6_1131285 | 1 | 1 | 3 | 0.6667 | 3 | 1 | 1 | 1 | 8 | 0.875 | 1 | 1 | 2 | 1 | 1 | 1 | 4 | 1 | 8 | 1 |
| 6_1131286 | 1 | 1 | 2 | 1 | 4 | 0.5 | 1 | 1 | 8 | 0.75 | 2 | 1 | 3 | 0.6667 | 1 | 1 | 4 | 0.5 | 10 | 0.7 |
| 6_1131452 | 11 | 0.2727 | 11 | 0.0909 | 6 | 0.3333 | 11 | 0.4545 | 39 | 0.2821 | 5 | 0.2 | 12 | 0.25 | 17 | 0.2353 | 6 | 0.1667 | 40 | 0.225 |
| 6_1131458 | 9 | 0 | 7 | 0 | 3 | 0 | 9 | 0.2222 | 28 | 0.0714 | 5 | 0 | 7 | 0 | 7 | 0.2857 | 4 | 0 | 23 | 0.087 |
| 6_1131479 | 2 | 1 | NA | NA | NA | NA | 2 | 1 | 4 | 1 | 1 | 0 | 1 | 1 | 2 | 0.5 | NA | NA | 4 | 0.5 |
| 6_1131479 | 2 | 0 | NA | NA | NA | NA | 2 | 1 | 4 | 0.5 | 1 | 0 | NA | NA | 1 | 1 | NA | NA | 2 | 0.5 |
| 6_1131480 | 2 | 0.5 | NA | NA | NA | NA | 2 | 1 | 4 | 0.75 | 1 | 0 | NA | NA | 2 | 0.5 | NA | NA | 3 | 0.3333 |
| 6_1131480 | 2 | 0.5 | NA | NA | NA | NA | 2 | 1 | 4 | 0.75 | 1 | 1 | 1 | 1 | 2 | 0.5 | NA | NA | 4 | 0.75 |
| 6_1131483 | 2 | 1 | NA | NA | NA | NA | 2 | 1 | 4 | 1 | 1 | 0 | 1 | 1 | 2 | 0.5 | NA | NA | 4 | 0.5 |
| 6_1131484 | 2 | 1 | NA | NA | NA | NA | 2 | 1 | 4 | 1 | 1 | 1 | 1 | 1 | 1 | 0 | NA | NA | 3 | 0.6667 |
| 6_1131484 | 2 | 0 | NA | NA | NA | NA | 2 | 1 | 4 | 0.5 | 1 | 0 | 1 | 1 | 1 | 1 | NA | NA | 3 | 0.6667 |
| 6_1131486 | 2 | 1 | NA | NA | NA | NA | 2 | 1 | 4 | 1 | 1 | 0 | NA | NA | 1 | 1 | NA | NA | 2 | 0.5 |
| 6_1131487 | 1 | 1 | NA | NA | NA | NA | 2 | 0.5 | 3 | 0.6667 | 1 | 0 | 1 | 0 | 1 | 1 | NA | NA | 3 | 0.3333 |
| 6_1131487 | 1 | 1 | NA | NA | NA | NA | 2 | 1 | 3 | 1 | NA | NA | 1 | 1 | 1 | 1 | NA | NA | 2 | 1 |
| 6_1131533 | NA | NA | NA | NA | NA | NA | 1 | 1 | 1 | 1 | NA | NA | 1 | 1 | NA | NA | NA | NA | 1 | 1 |
| 6_1131534 | 1 | 0 | NA | NA | NA | NA | 3 | 0.3333 | 4 | 0.25 | 1 | 0 | 2 | 0.5 | 1 | 0 | NA | NA | 4 | 0.25 |
| 6_1131535 | 1 | 1 | NA | NA | NA | NA | 3 | 1 | 4 | 1 | 1 | 1 | 2 | 0.5 | 1 | 1 | NA | NA | 4 | 0.75 |
| 6_1131535 | 1 | 0 | NA | NA | NA | NA | 3 | 0.6667 | 4 | 0.5 | 1 | 0 | 2 | 0.5 | 1 | 1 | NA | NA | 4 | 0.5 |
| 6_1132576 | NA | NA | 4 | 0.75 | 2 | 0 | NA | NA | 6 | 0.5 | 3 | 0.6667 | 4 | 0.25 | 3 | 0.3333 | NA | NA | 10 | 0.4 |
| 6_1132579 | NA | NA | 4 | 0 | 2 | 1 | NA | NA | 6 | 0.3333 | 3 | 0.3333 | 4 | 0.5 | 3 | 1 | NA | NA | 10 | 0.6 |
| 6_1132815 | 3 | 0.3333 | 1 | 0 | 3 | 0.3333 | 1 | 0 | 8 | 0.25 | NA | NA | 1 | 1 | 2 | 0.5 | 2 | 1 | 5 | 0.8 |
| 6_1132817 | 2 | 0 | 1 | 0 | 3 | 0.3333 | 1 | 0 | 7 | 0.1429 | 1 | 0 | 1 | 0 | 2 | 0.5 | 2 | 0 | 6 | 0.1667 |
| 6_1132872 | 1 | 1 | 2 | 1 | NA | NA | 1 | 1 | 4 | 1 | NA | NA | 1 | 0 | NA | NA | 1 | 1 | 2 | 0.5 |
| 6_1132917 | 1 | 1 | 2 | 1 | NA | NA | 1 | 1 | 4 | 1 | NA | NA | 1 | 1 | NA | NA | 1 | 1 | 2 | 1 |
| 6_1132919 | 1 | 1 | 2 | 0 | NA | NA | 1 | 0 | 4 | 0.25 | NA | NA | 1 | 1 | NA | NA | NA | NA | 1 | 1 |
| 6_1132921 | 1 | 1 | 1 | 0 | NA | NA | 1 | 0 | 3 | 0.3333 | NA | NA | 1 | 0 | 1 | 0 | 1 | 1 | 3 | 0.3333 |
| 6_1132961 | 1 | 0 | 3 | 0.3333 | 1 | 0 | 2 | 0 | 7 | 0.1429 | NA | NA | 2 | 0 | 2 | 0.5 | 1 | 0 | 5 | 0.2 |
| 6_1133011 | 2 | 0.5 | 3 | 0.6667 | 2 | 0 | 2 | 1 | 9 | 0.5556 | NA | NA | 1 | 1 | 1 | 1 | 2 | 1 | 4 | 1 |
| 6_1133012 | 4 | 0.75 | 3 | 0.6667 | 2 | 0.5 | 1 | 1 | 10 | 0.7 | NA | NA | 3 | 0.6667 | 3 | 1 | 2 | 1 | 8 | 0.875 |
| 6_1133436 | 1 | 0 | 1 | 0 | 5 | 0.2 | 2 | 0 | 9 | 0.1111 | 4 | 0 | 4 | 0.25 | 6 | 0 | 4 | 0 | 18 | 0.0556 |
| 6_1133437 | 1 | 1 | 1 | 1 | 5 | 0.4 | 2 | 1 | 9 | 0.6667 | 4 | 0.5 | 5 | 0.8 | 6 | 0.5 | 5 | 0.8 | 20 | 0.65 |
| 6_1133438 | 1 | 0 | 1 | 0 | 5 | 0.2 | 1 | 0 | 8 | 0.125 | 4 | 0 | 5 | 0.2 | 6 | 0.1667 | 3 | 0.6667 | 18 | 0.2222 |
| 6_1133517 | 1 | 0 | NA | NA | 4 | 0.5 | 2 | 1 | 7 | 0.5714 | 2 | 0.5 | 2 | 1 | 5 | 0 | 1 | 0 | 10 | 0.3 |
| 6_1133659 | 41 | 0.0732 | 28 | 0.1429 | 20 | 0.2 | 14 | 0.2857 | 103 | 0.1456 | 20 | 0 | 24 | 0.0833 | 21 | 0.0476 | 24 | 0.0833 | 89 | 0.0562 |
| 6_1133659 | 50 | 0.12 | 33 | 0.0606 | 22 | 0.2273 | 14 | 0 | 119 | 0.1092 | 25 | 0.2 | 35 | 0.1429 | 25 | 0.08 | 28 | 0.2143 | 113 | 0.1593 |
| 6_1133660 | 55 | 0.2364 | 34 | 0.2353 | 24 | 0.2083 | 19 | 0.0526 | 132 | 0.2045 | 22 | 0.1364 | 36 | 0.1389 | 25 | 0.04 | 29 | 0.2414 | 112 | 0.1429 |
| 6_1133662 | 55 | 0.0545 | 33 | 0.0606 | 26 | 0.0769 | 21 | 0 | 135 | 0.0519 | 29 | 0.0345 | 36 | 0.0833 | 30 | 0.1 | 23 | 0.1304 | 118 | 0.0847 |
| 6_1133662 | 61 | 0.1639 | 37 | 0.1892 | 29 | 0.1379 | 21 | 0.1429 | 148 | 0.1622 | 29 | 0.1379 | 37 | 0.2162 | 33 | 0.1515 | 29 | 0.3448 | 128 | 0.2109 |
| 6_1133665 | 49 | 0.102 | 28 | 0.1071 | 25 | 0.12 | 21 | 0.1905 | 123 | 0.122 | 24 | 0.0833 | 30 | 0.0333 | 30 | 0.1 | 25 | 0.24 | 109 | 0.1101 |
| 6_1133861 | NA | NA | 3 | 0.3333 | 1 | 1 | NA | NA | 4 | 0.5 | NA | NA | 1 | 1 | NA | NA | 1 | 1 | 2 | 1 |
| 6_1133861 | NA | NA | 3 | 0 | 1 | 1 | NA | NA | 4 | 0.25 | NA | NA | 1 | 0 | NA | NA | 1 | 1 | 2 | 0.5 |
| 6_1133899 | NA | NA | 2 | 0.5 | 1 | 1 | NA | NA | 3 | 0.6667 | NA | NA | 1 | 1 | NA | NA | 1 | 0 | 2 | 0.5 |
| 6_1133913 | NA | NA | 2 | 1 | 2 | 0.5 | NA | NA | 4 | 0.75 | NA | NA | 1 | 1 | NA | NA | NA | NA | 1 | 1 |
| 6_1134243 | 4 | 0 | 6 | 0 | 5 | 0.2 | 3 | 0.6667 | 18 | 0.1667 | 5 | 0.6 | 6 | 0.1667 | 4 | 0 | 7 | 0.2857 | 22 | 0.2727 |
| 6_1134246 | 2 | 1 | 6 | 0.8333 | 3 | 0.6667 | 1 | 1 | 12 | 0.8333 | 4 | 0.5 | 7 | 0.8571 | 2 | 1 | 5 | 0.8 | 18 | 0.7778 |
| 6_1134246 | 6 | 0.5 | 6 | 1 | 5 | 0.8 | 1 | 1 | 18 | 0.7778 | 6 | 0.8333 | 6 | 0.8333 | 3 | 1 | 6 | 1 | 21 | 0.9048 |
| 6_1134246 | 3 | 0.6667 | 5 | 0.6 | 2 | 0 | 4 | 0.5 | 14 | 0.5 | 4 | 0.5 | 6 | 0.5 | 4 | 0 | 6 | 0.6667 | 20 | 0.45 |
| 6_1134247 | 6 | 0.5 | 8 | 0.5 | 6 | 0.5 | 5 | 0.4 | 25 | 0.48 | 7 | 0.2857 | 7 | 0.4286 | 4 | 0.5 | 7 | 0.2857 | 25 | 0.36 |
| 6_1134248 | 4 | 0.25 | 8 | 0.125 | 5 | 0 | 5 | 0.2 | 22 | 0.1364 | 7 | 0.1429 | 6 | 0.1667 | 4 | 0.5 | 5 | 0 | 22 | 0.1818 |
| 6_1134249 | 6 | 0.6667 | 9 | 0.8889 | 5 | 0.8 | 5 | 0.8 | 25 | 0.8 | 6 | 0.6667 | 7 | 0.5714 | 4 | 0.75 | 8 | 0.375 | 25 | 0.56 |
| 6_1134249 | 7 | 0.2857 | 9 | 0.4444 | 6 | 0.5 | 6 | 0.1667 | 28 | 0.3571 | 9 | 0 | 7 | 0.5714 | 5 | 0.6 | 6 | 0.3333 | 27 | 0.3333 |
| 6_1134251 | 5 | 0 | 5 | 0 | 5 | 0.4 | 5 | 0 | 20 | 0.1 | 4 | 0 | 4 | 0 | 2 | 0 | 5 | 0.2 | 15 | 0.0667 |
| 6_1134600 | 1 | 0 | NA | NA | 1 | 1 | 1 | 0 | 3 | 0.3333 | NA | NA | 1 | 1 | 3 | 0.3333 | NA | NA | 4 | 0.5 |
| 6_1134600 | 1 | 1 | NA | NA | 1 | 0 | 1 | 0 | 3 | 0.3333 | NA | NA | 1 | 0 | 3 | 0.3333 | NA | NA | 4 | 0.25 |
| 6_1134600 | 1 | 1 | NA | NA | 1 | 0 | 1 | 0 | 3 | 0.3333 | NA | NA | 1 | 0 | 3 | 0.3333 | NA | NA | 4 | 0.25 |
| 6_1134699 | 5 | 0 | 11 | 0.1818 | 13 | 0.3846 | 12 | 0.3333 | 41 | 0.2683 | 11 | 0.1818 | 21 | 0.1905 | 11 | 0.1818 | 20 | 0.3 | 63 | 0.2222 |
| 6_1134699 | 3 | 0.6667 | 9 | 0 | 13 | 0.0769 | 8 | 0.125 | 33 | 0.1212 | 10 | 0.1 | 21 | 0.2857 | 10 | 0.3 | 16 | 0.125 | 57 | 0.2105 |
| 6_1134700 | 5 | 0.2 | 12 | 0.0833 | 14 | 0.0714 | 12 | 0.0833 | 43 | 0.093 | 13 | 0 | 24 | 0.125 | 9 | 0 | 24 | 0.125 | 70 | 0.0857 |
| 6_1134709 | 9 | 0.7778 | 9 | 0.7778 | 9 | 0.6667 | 5 | 0.2 | 32 | 0.6562 | 8 | 0.75 | 2 | 0 | 6 | 0.8333 | 7 | 0.5714 | 23 | 0.6522 |
| 6_1134709 | 8 | 0.125 | 6 | 0.6667 | 8 | 0.375 | 6 | 0.3333 | 28 | 0.3571 | 4 | 0.25 | 3 | 0.3333 | 5 | 0.2 | 6 | 0.5 | 18 | 0.3333 |
| 6_1134710 | 12 | 0.0833 | 6 | 0.3333 | 5 | 0 | 2 | 0 | 25 | 0.12 | 7 | 0.2857 | 2 | 0 | 3 | 0.3333 | 7 | 0.1429 | 19 | 0.2105 |
| 6_1134711 | 9 | 0.8889 | 7 | 1 | 6 | 1 | 5 | 0.6 | 27 | 0.8889 | 4 | 1 | 3 | 1 | 2 | 0.5 | 6 | 1 | 15 | 0.9333 |
| 6_1134711 | 13 | 0.3846 | 8 | 0.25 | 7 | 0.1429 | 4 | 0.25 | 32 | 0.2812 | 8 | 0.375 | 2 | 0.5 | 4 | 0.25 | 7 | 0.4286 | 21 | 0.381 |
| 6_1134712 | 13 | 0.8462 | 11 | 1 | 7 | 0.8571 | 5 | 1 | 36 | 0.9167 | 6 | 1 | 1 | 1 | 3 | 1 | 8 | 0.625 | 18 | 0.8333 |
| 6_1134713 | 8 | 1 | 4 | 1 | 4 | 1 | 2 | 1 | 18 | 1 | 6 | 1 | 2 | 1 | 4 | 1 | 5 | 0.8 | 17 | 0.9412 |

| | | | | | | | | | | | | | | | | | | | | |
|---|---|---|---|---|---|---|---|---|---|---|---|---|---|---|---|---|---|---|---|---|
| 6_1134714 | 12 | 0.0833 | 9 | 0.4444 | 6 | 0.1667 | 4 | 0.25 | 31 | 0.2258 | 8 | 0.25 | 3 | 0 | 4 | 0.25 | 7 | 0.2857 | 22 | 0.2273 |
| 6_1134714 | 11 | 1 | 6 | 0.8333 | 4 | 0.75 | 3 | 1 | 24 | 0.9167 | 6 | 0.5 | 2 | 1 | 2 | 1 | 8 | 1 | 18 | 0.8333 |
| 6_1134714 | 11 | 0.8182 | 11 | 0.6364 | 6 | 0.5 | 4 | 0.5 | 32 | 0.6562 | 6 | 0.6667 | 2 | 1 | 4 | 0.75 | 8 | 0.75 | 20 | 0.75 |
| 6_1134718 | 8 | 0.75 | 8 | 0.5 | 6 | 0.8333 | 4 | 1 | 26 | 0.7308 | 5 | 0.8 | 2 | 0.5 | 3 | 1 | 7 | 0.5714 | 17 | 0.7059 |
| 6_1134719 | 11 | 0.6364 | 10 | 0.4 | 6 | 0.5 | 5 | 0.2 | 32 | 0.4688 | 4 | 0.25 | 2 | 0.5 | 5 | 0.4 | 8 | 0.375 | 19 | 0.3684 |
| 6_1134719 | 6 | 0.3333 | 3 | 0.3333 | 3 | 0.3333 | 2 | 0.5 | 14 | 0.3571 | 3 | 0.6667 | 1 | 1 | 3 | 0.6667 | 4 | 0.5 | 11 | 0.6364 |
| 6_1134720 | 11 | 0.5455 | 10 | 0.5 | 6 | 0.6667 | 4 | 0.5 | 31 | 0.5484 | 6 | 0.6667 | 2 | 0.5 | 5 | 0.2 | 8 | 0.625 | 21 | 0.5238 |
| 6_1134721 | 9 | 0.1111 | 6 | 0 | 6 | 0.1667 | 3 | 0 | 24 | 0.0833 | 4 | 0 | 2 | 0 | 4 | 0 | 7 | 0.1429 | 17 | 0.0588 |
| 6_1134724 | 10 | 0.1 | 7 | 0.2857 | 5 | 0.8 | 3 | 0.3333 | 25 | 0.32 | 5 | 0.2 | 2 | 0.5 | 5 | 0.2 | 8 | 0.625 | 20 | 0.4 |
| 6_1134727 | 8 | 0.125 | 8 | 0.25 | 4 | 0.25 | 3 | 0 | 23 | 0.1739 | 4 | 0 | 1 | 0 | 5 | 0.2 | 7 | 0.1429 | 17 | 0.1176 |
| 6_1134743 | 7 | 0.1429 | 6 | 0.3333 | 3 | 0 | 1 | 0 | 17 | 0.1765 | 4 | 0.25 | 1 | 0 | 5 | 0 | 8 | 0 | 18 | 0.0556 |
| 6_1134745 | 6 | 0.1667 | 4 | 0 | 1 | 0 | 1 | 0 | 12 | 0.0833 | 4 | 0 | 1 | 0 | 3 | 0 | 5 | 0.2 | 13 | 0.0769 |
| 6_1134745 | 8 | 0.25 | 4 | 0 | 2 | 0.5 | 1 | 0 | 15 | 0.2 | 4 | 0.25 | 1 | 1 | 3 | 0 | 6 | 0.3333 | 14 | 0.2857 |
| 6_1134746 | 7 | 0.1429 | 6 | 0.1667 | 3 | 0 | NA | NA | 16 | 0.125 | 5 | 0 | 1 | 1 | 4 | 0 | 7 | 0 | 17 | 0.0588 |
| 6_1134746 | 8 | 0.625 | 6 | 0.5 | 2 | 0.5 | NA | NA | 16 | 0.5625 | 4 | 0.5 | 1 | 0 | 3 | 1 | 4 | 1 | 12 | 0.75 |
| 6_1134748 | 8 | 0.25 | 4 | 0.25 | 3 | 0 | 1 | 0 | 16 | 0.1875 | 4 | 0 | 1 | 0 | 3 | 0 | 8 | 0.125 | 16 | 0.0625 |
| 6_1134748 | 7 | 0.2857 | 3 | 0.6667 | 3 | 0.6667 | NA | NA | 13 | 0.4615 | 3 | 0 | 1 | 1 | 2 | 1 | 7 | 0.4286 | 13 | 0.4615 |
| 6_1134749 | 9 | 0.6667 | 6 | 0.8333 | 3 | 1 | 1 | 0 | 19 | 0.7368 | 3 | 0.6667 | 1 | 0 | 3 | 1 | 7 | 0.8571 | 14 | 0.7857 |
| 6_1134749 | 8 | 0.375 | 5 | 0 | 2 | 0.5 | 1 | 1 | 16 | 0.3125 | 2 | 0 | 1 | 0 | 3 | 0 | 6 | 0.1667 | 12 | 0.0833 |
| 6_1134750 | 7 | 0.1429 | 4 | 0 | 3 | 0 | 1 | 0 | 15 | 0.0667 | 3 | 0 | 1 | 0 | 3 | 0.3333 | 8 | 0.25 | 15 | 0.2 |
| 6_1134751 | 8 | 0.75 | 5 | 0.6 | 3 | 0.3333 | 1 | 0 | 17 | 0.5882 | 3 | 1 | 1 | 1 | 2 | 0.5 | 6 | 1 | 12 | 0.9167 |
| 6_1134928 | 18 | 0.3333 | 11 | 0.2727 | 11 | 0.0909 | 7 | 0.2857 | 47 | 0.2553 | 11 | 0.0909 | 6 | 0.1667 | 10 | 0 | 12 | 0.1667 | 39 | 0.1026 |
| 6_1135729 | 2 | 1 | 1 | 0 | NA | NA | 2 | 0.5 | 5 | 0.6 | 2 | 0.5 | NA | NA | NA | NA | NA | NA | 2 | 0.5 |
| 6_1135818 | 40 | 0.125 | 42 | 0.0952 | 20 | 0.05 | 24 | 0.0833 | 126 | 0.0952 | 37 | 0.1081 | 26 | 0.0769 | 32 | 0.0938 | 24 | 0.1667 | 119 | 0.1092 |
| 6_1135932 | 47 | 0.0638 | 44 | 0.0227 | 33 | 0 | 33 | 0.1212 | 157 | 0.051 | 43 | 0.0233 | 34 | 0.1176 | 35 | 0.0857 | 32 | 0.25 | 144 | 0.1111 |
| 6_1135933 | 40 | 0.1 | 37 | 0.1622 | 25 | 0.2 | 25 | 0.12 | 127 | 0.1417 | 37 | 0.0811 | 29 | 0.1724 | 32 | 0.1875 | 28 | 0.2143 | 126 | 0.1587 |
| 6_1135933 | 37 | 0.3243 | 35 | 0.2857 | 29 | 0.3103 | 24 | 0.25 | 125 | 0.296 | 41 | 0.2195 | 28 | 0.3214 | 32 | 0.3438 | 31 | 0.3548 | 132 | 0.303 |
| 6_1135933 | 46 | 0.2609 | 43 | 0.1395 | 34 | 0.0294 | 28 | 0.1786 | 151 | 0.1589 | 44 | 0.2273 | 37 | 0.2162 | 34 | 0.1765 | 30 | 0.1667 | 145 | 0.2 |
| 6_1135934 | 52 | 0.0769 | 46 | 0.0217 | 37 | 0.0811 | 33 | 0.1212 | 168 | 0.0714 | 48 | 0.0625 | 37 | 0.1622 | 40 | 0.05 | 33 | 0.1515 | 158 | 0.1013 |
| 6_1135935 | 41 | 0.4878 | 40 | 0.4 | 35 | 0.5429 | 30 | 0.4 | 146 | 0.4589 | 46 | 0.4348 | 36 | 0.5556 | 30 | 0.4667 | 34 | 0.5882 | 146 | 0.5068 |
| 6_1135936 | 19 | 0.4737 | 22 | 0.3182 | 20 | 0.4 | 13 | 0.3077 | 74 | 0.3784 | 29 | 0.2069 | 21 | 0.2857 | 23 | 0.3043 | 13 | 0.3846 | 86 | 0.2791 |
| 6_1135938 | 18 | 0.3333 | 17 | 0.1176 | 15 | 0.1333 | 14 | 0.0714 | 64 | 0.1719 | 21 | 0.0952 | 19 | 0.1053 | 23 | 0.0435 | 10 | 0.3 | 73 | 0.1096 |
| 6_1135938 | 20 | 0.65 | 24 | 0.3333 | 20 | 0.35 | 16 | 0.4375 | 80 | 0.4375 | 24 | 0.5833 | 19 | 0.6316 | 20 | 0.6 | 13 | 0.4615 | 76 | 0.5789 |
| 6_1136064 | 4 | 0.25 | 3 | 0.6667 | 7 | 0.1429 | 9 | 0.1111 | 23 | 0.2174 | 3 | 0 | 6 | 0.3333 | 7 | 0.1429 | 8 | 0.625 | 24 | 0.3333 |
| 6_1136065 | 5 | 0.4 | 2 | 1 | 7 | 0.7143 | 7 | 0.7143 | 21 | 0.6667 | 3 | 0.3333 | 6 | 0.6667 | 5 | 0.4 | 8 | 0.625 | 22 | 0.5455 |
| 6_1136068 | 5 | 0 | 4 | 0.5 | 6 | 0.3333 | 10 | 0.1 | 25 | 0.2 | 2 | 0 | 7 | 0.4286 | 7 | 0.2857 | 9 | 0 | 25 | 0.2 |
| 6_1136069 | 3 | 0.6667 | 2 | 0 | 3 | 0.6667 | 7 | 0.7143 | 15 | 0.6 | 1 | 1 | 6 | 0.5 | 4 | 0.25 | 6 | 0.5 | 17 | 0.4706 |
| 6_1136082 | 10 | 0 | 4 | 0.25 | 8 | 0.25 | 9 | 0 | 31 | 0.0968 | 7 | 0 | 8 | 0.25 | 9 | 0 | 12 | 0.0833 | 36 | 0.0833 |
| 6_1136707 | 2 | 0 | 1 | 0 | 4 | 0.25 | NA | NA | 7 | 0.1429 | 3 | 1 | 2 | 0 | 2 | 0 | 2 | 0 | 9 | 0.3333 |
| 6_1136707 | 2 | 0.5 | 1 | 1 | 4 | 0.75 | NA | NA | 7 | 0.7143 | 2 | 0.5 | 2 | 0.5 | 1 | 1 | 2 | 1 | 7 | 0.7143 |
| 6_1136711 | 2 | 0.5 | 3 | 1 | 5 | 0.8 | NA | NA | 10 | 0.8 | 3 | 0.6667 | 2 | 1 | 2 | 1 | 1 | 1 | 8 | 0.875 |
| 6_1136711 | 2 | 0 | 3 | 0.3333 | 3 | 0 | NA | NA | 8 | 0.125 | 3 | 0 | 2 | 0 | 3 | 0.3333 | 3 | 0.3333 | 11 | 0.1818 |
| 6_1136712 | 1 | 0 | 3 | 0 | 4 | 0.25 | NA | NA | 8 | 0.125 | 3 | 0.3333 | 3 | 0.6667 | 3 | 0 | 2 | 0.5 | 11 | 0.3636 |
| 6_1136712 | 1 | 1 | 2 | 1 | 5 | 0.8 | NA | NA | 8 | 0.875 | 2 | 1 | 2 | 1 | 3 | 1 | 2 | 0.5 | 9 | 0.8889 |
| 6_1136712 | 2 | 1 | 3 | 1 | 6 | 1 | NA | NA | 11 | 1 | 2 | 1 | 3 | 1 | 3 | 1 | 2 | 1 | 10 | 1 |
| 6_1136713 | 2 | 1 | 3 | 0.3333 | 6 | 0.6667 | NA | NA | 11 | 0.6364 | 3 | 0.6667 | 3 | 0.3333 | 2 | 1 | 2 | 0.5 | 10 | 0.6 |
| 6_1136713 | 1 | 0 | 3 | 0.3333 | 6 | 0.3333 | NA | NA | 10 | 0.3 | 4 | 0.5 | 3 | 0.6667 | 3 | 0.6667 | 3 | 0.3333 | 13 | 0.5385 |
| 6_1136811 | NA | NA | NA | NA | 1 | 1 | NA | NA | 1 | 1 | NA | NA | NA | NA | NA | NA | 1 | 1 | 1 | 1 |
| 6_1136815 | NA | NA | NA | NA | 1 | 1 | NA | NA | 1 | 1 | NA | NA | NA | NA | NA | NA | 1 | 1 | 1 | 1 |
| 6_1136924 | NA | NA | NA | NA | 1 | 1 | NA | NA | 1 | 1 | 1 | 0 | 1 | 1 | NA | NA | NA | NA | 2 | 0.5 |
| 6_1137051 | 20 | 0.25 | 8 | 0.125 | 9 | 0.1111 | 13 | 0.2308 | 50 | 0.2 | 6 | 0.1667 | 8 | 0.125 | 9 | 0.4444 | 5 | 0.2 | 28 | 0.25 |
| 6_1137209 | 11 | 0.1818 | 11 | 0 | 12 | 0.0833 | 11 | 0.0909 | 45 | 0.0889 | 4 | 0 | 7 | 0.1429 | 5 | 0.2 | 3 | 0 | 19 | 0.1053 |
| 6_1137215 | 7 | 0 | 9 | 0.5556 | 8 | 0.125 | 3 | 0.3333 | 27 | 0.2593 | 3 | 0 | 8 | 0 | 2 | 0.5 | 2 | 0 | 15 | 0.0667 |
| 6_1137215 | 11 | 0 | 13 | 0.0769 | 7 | 0 | 6 | 0.1667 | 37 | 0.0541 | 6 | 0 | 10 | 0.1 | 5 | 0.2 | 4 | 0.25 | 25 | 0.12 |
| 6_1137218 | 12 | 0.25 | 13 | 0 | 8 | 0 | 9 | 0.2222 | 42 | 0.119 | 7 | 0 | 8 | 0 | 5 | 0.4 | 6 | 0 | 26 | 0.0769 |
| 6_1137231 | 50 | 0.42 | 42 | 0.381 | 39 | 0.4872 | 38 | 0.4211 | 169 | 0.426 | 35 | 0.3714 | 38 | 0.4211 | 41 | 0.4146 | 38 | 0.3947 | 152 | 0.4013 |
| 6_1137232 | 43 | 0.1628 | 41 | 0.122 | 34 | 0.1176 | 38 | 0.1316 | 156 | 0.1346 | 36 | 0.1389 | 37 | 0.0541 | 44 | 0.2045 | 36 | 0.0556 | 153 | 0.1176 |
| 6_1137233 | 43 | 0.3256 | 43 | 0.3721 | 34 | 0.2059 | 34 | 0.2647 | 154 | 0.2987 | 33 | 0.3333 | 39 | 0.3077 | 39 | 0.2308 | 30 | 0.2667 | 141 | 0.2837 |
| 6_1137234 | 28 | 0.6071 | 33 | 0.6061 | 24 | 0.75 | 22 | 0.5 | 107 | 0.6168 | 20 | 0.45 | 26 | 0.5385 | 21 | 0.7619 | 25 | 0.68 | 92 | 0.6087 |
| 6_1137234 | 5 | 0.6 | 5 | 0.8 | 3 | 0.6667 | 5 | 0.6 | 18 | 0.6667 | 8 | 0.75 | 6 | 0.8333 | 5 | 0.8 | 3 | 0.6667 | 22 | 0.7727 |
| 6_1137235 | 42 | 0.5 | 42 | 0.3571 | 36 | 0.4722 | 37 | 0.3514 | 157 | 0.4204 | 33 | 0.3333 | 41 | 0.4878 | 40 | 0.625 | 31 | 0.3548 | 145 | 0.4621 |
| 6_1137236 | 30 | 0.4667 | 37 | 0.2973 | 27 | 0.4074 | 27 | 0.3333 | 121 | 0.3719 | 15 | 0.3333 | 34 | 0.4706 | 34 | 0.5 | 21 | 0.4286 | 104 | 0.4519 |
| 6_1137236 | 38 | 0.6579 | 40 | 0.6 | 33 | 0.7273 | 35 | 0.5429 | 146 | 0.6301 | 33 | 0.5455 | 38 | 0.6053 | 38 | 0.6316 | 29 | 0.8276 | 138 | 0.6449 |
| 6_1137237 | 39 | 0.1026 | 39 | 0.2308 | 30 | 0.1333 | 32 | 0.0625 | 140 | 0.1357 | 29 | 0.2069 | 35 | 0.1143 | 35 | 0.2857 | 29 | 0.2069 | 128 | 0.2031 |
| 6_1137237 | 34 | 0.3824 | 39 | 0.359 | 35 | 0.5429 | 33 | 0.303 | 141 | 0.3972 | 30 | 0.3667 | 37 | 0.3243 | 37 | 0.4595 | 28 | 0.5 | 132 | 0.4091 |

| | | | | | | | | | | | | | | | | | | | | |
|---|---|---|---|---|---|---|---|---|---|---|---|---|---|---|---|---|---|---|---|---|
| 6_1137237 | 30 | 0.6 | 30 | 0.4333 | 27 | 0.5185 | 26 | 0.4615 | 113 | 0.5044 | 26 | 0.5 | 33 | 0.5455 | 25 | 0.6 | 20 | 0.6 | 104 | 0.5577 |
| 6_1137240 | 34 | 0.1471 | 38 | 0.0789 | 30 | 0 | 37 | 0.1081 | 139 | 0.0863 | 27 | 0.037 | 42 | 0.0714 | 33 | 0.0909 | 25 | 0.12 | 127 | 0.0787 |
| 6_1137241 | 31 | 0.0645 | 39 | 0.0256 | 32 | 0.0312 | 33 | 0.2727 | 135 | 0.0963 | 31 | 0.0968 | 40 | 0.025 | 27 | 0.1111 | 23 | 0 | 121 | 0.0579 |
| 6_1137242 | 27 | 0.963 | 34 | 0.9412 | 30 | 0.8333 | 28 | 1 | 119 | 0.9328 | 28 | 0.9643 | 38 | 0.9737 | 27 | 1 | 23 | 0.9565 | 116 | 0.9741 |
| 6_1137243 | 29 | 0.6552 | 38 | 0.7895 | 32 | 0.8438 | 27 | 0.8148 | 126 | 0.7778 | 26 | 0.5385 | 38 | 0.7368 | 27 | 0.6667 | 25 | 0.8 | 116 | 0.6897 |
| 6_1137245 | 24 | 0.125 | 31 | 0.2903 | 28 | 0.1786 | 23 | 0.1304 | 106 | 0.1887 | 24 | 0.2083 | 33 | 0.303 | 26 | 0.1154 | 23 | 0.3913 | 106 | 0.2547 |
| 6_1137246 | 24 | 1 | 27 | 0.8519 | 26 | 1 | 25 | 1 | 102 | 0.9608 | 25 | 0.96 | 31 | 1 | 24 | 0.9583 | 23 | 1 | 103 | 0.9806 |
| 6_1137248 | 27 | 0.3704 | 27 | 0.2222 | 33 | 0.2727 | 26 | 0.3077 | 113 | 0.292 | 19 | 0.5263 | 32 | 0.25 | 29 | 0.3793 | 24 | 0.25 | 104 | 0.3365 |
| 6_1137248 | 30 | 0.6667 | 22 | 0.7273 | 28 | 0.6429 | 24 | 0.75 | 104 | 0.6923 | 22 | 0.7273 | 28 | 0.75 | 28 | 0.6071 | 23 | 0.6522 | 101 | 0.6832 |
| 6_1137249 | 24 | 0.625 | 19 | 0.5789 | 25 | 0.56 | 18 | 0.5556 | 86 | 0.5814 | 14 | 0.7857 | 17 | 0.4706 | 21 | 0.7143 | 22 | 0.7727 | 74 | 0.6892 |
| 6_1137249 | 22 | 0.4091 | 22 | 0.5 | 21 | 0.381 | 19 | 0.4211 | 84 | 0.4286 | 14 | 0.5 | 22 | 0.4091 | 22 | 0.3636 | 16 | 0.6875 | 74 | 0.473 |
| 6_1137250 | 28 | 0.8571 | 22 | 0.9545 | 29 | 0.8621 | 24 | 0.7917 | 103 | 0.8641 | 21 | 0.9524 | 31 | 0.8065 | 26 | 0.9615 | 23 | 0.913 | 101 | 0.901 |
| 6_1137250 | 26 | 0.2692 | 23 | 0.1304 | 27 | 0 | 20 | 0 | 96 | 0.1042 | 19 | 0.0526 | 25 | 0.04 | 24 | 0.125 | 23 | 0.087 | 91 | 0.0769 |
| 6_1137250 | 25 | 0.88 | 22 | 0.8636 | 31 | 0.9355 | 21 | 1 | 99 | 0.9192 | 21 | 1 | 26 | 0.8846 | 21 | 0.9048 | 23 | 0.913 | 91 | 0.9231 |
| 6_1137250 | 24 | 0.7083 | 19 | 0.8421 | 19 | 0.8947 | 16 | 0.75 | 78 | 0.7949 | 20 | 0.85 | 27 | 0.7407 | 24 | 0.75 | 22 | 0.7273 | 93 | 0.7634 |
| 6_1137251 | 23 | 1 | 21 | 0.9524 | 29 | 1 | 22 | 1 | 95 | 0.9895 | 18 | 1 | 32 | 0.9688 | 22 | 1 | 23 | 1 | 95 | 0.9895 |
| 6_1137251 | 24 | 1 | 23 | 0.913 | 26 | 0.9231 | 23 | 1 | 96 | 0.9583 | 19 | 1 | 32 | 1 | 26 | 1 | 25 | 1 | 102 | 1 |
| 6_1137251 | 22 | 0.4545 | 21 | 0.2857 | 21 | 0.4286 | 23 | 0.2609 | 87 | 0.3563 | 17 | 0.3529 | 25 | 0.4 | 23 | 0.4348 | 23 | 0.3043 | 88 | 0.375 |
| 6_1137252 | 19 | 0.2105 | 16 | 0.1875 | 19 | 0.3684 | 17 | 0.1765 | 71 | 0.2394 | 15 | 0.2 | 20 | 0.35 | 18 | 0.2222 | 16 | 0.1875 | 69 | 0.2464 |
| 6_1137253 | 9 | 1 | 12 | 1 | 16 | 0.9375 | 19 | 0.9474 | 56 | 0.9643 | 10 | 1 | 16 | 1 | 17 | 1 | 18 | 1 | 61 | 1 |
| 6_1137253 | 20 | 0.3 | 19 | 0.4737 | 22 | 0.1364 | 22 | 0.3636 | 83 | 0.3133 | 14 | 0.0714 | 25 | 0.2 | 22 | 0.2273 | 23 | 0.3043 | 84 | 0.2143 |
| 6_1137253 | 22 | 0.0455 | 18 | 0.0556 | 25 | 0.24 | 25 | 0.16 | 90 | 0.1333 | 17 | 0.0588 | 26 | 0.0769 | 24 | 0.0833 | 23 | 0 | 90 | 0.0556 |
| 6_1137253 | 21 | 0.0476 | 18 | 0.2222 | 23 | 0.1739 | 21 | 0.0476 | 83 | 0.1205 | 17 | 0.1765 | 26 | 0.0385 | 19 | 0.0526 | 22 | 0.0909 | 84 | 0.0833 |
| 6_1137254 | 15 | 0.9333 | 17 | 0.9412 | 20 | 1 | 21 | 0.9524 | 73 | 0.9589 | 15 | 1 | 26 | 1 | 17 | 0.9412 | 18 | 1 | 76 | 0.9868 |
| 6_1137256 | 18 | 0.1667 | 18 | 0.1111 | 22 | 0.2273 | 20 | 0.2 | 78 | 0.1795 | 15 | 0.2 | 26 | 0.0769 | 16 | 0.1875 | 16 | 0 | 73 | 0.1096 |
| 6_1137256 | 15 | 0.0667 | 15 | 0.0667 | 24 | 0.0417 | 20 | 0.05 | 74 | 0.0541 | 16 | 0.1875 | 25 | 0.12 | 20 | 0.15 | 18 | 0.1667 | 79 | 0.1519 |
| 6_1138193 | 5 | 0.4 | 5 | 0.6 | 8 | 0.25 | 5 | 0.4 | 23 | 0.3913 | 3 | 0.3333 | 3 | 0 | 5 | 0.2 | 3 | 0.3333 | 14 | 0.2143 |
| 6_1138194 | 5 | 0.8 | 4 | 1 | 7 | 0.7143 | 4 | 0.75 | 20 | 0.8 | 3 | 0 | 3 | 1 | 3 | 0.6667 | 3 | 0.3333 | 12 | 0.5 |
| 6_1138195 | 3 | 0.3333 | 4 | 0 | 6 | 0.1667 | 3 | 0 | 16 | 0.125 | 3 | 0.3333 | 2 | 0 | 3 | 0.6667 | 2 | 0 | 10 | 0.3 |
| 6_1142246 | 8 | 0.25 | 6 | 0.5 | 6 | 0 | 11 | 0 | 31 | 0.1613 | 9 | 0.3333 | 6 | 0.3333 | 7 | 0.5714 | 10 | 0.2 | 32 | 0.3438 |
| 6_1142248 | 9 | 0.7778 | 6 | 1 | 6 | 0.8333 | 11 | 0.8182 | 32 | 0.8438 | 9 | 1 | 6 | 1 | 9 | 0.8889 | 11 | 0.8182 | 35 | 0.9143 |
| 6_1142249 | 9 | 0.1111 | 6 | 0.3333 | 6 | 0 | 9 | 0.1111 | 30 | 0.1333 | 7 | 0.1429 | 7 | 0 | 11 | 0.0909 | 12 | 0.1667 | 37 | 0.1081 |
| 6_1142252 | 10 | 0.2 | 8 | 0.125 | 9 | 0.1111 | 12 | 0.1667 | 39 | 0.1538 | 9 | 0.2222 | 7 | 0.4286 | 11 | 0.2727 | 12 | 0.1667 | 39 | 0.2564 |
| 6_1142253 | 13 | 0.0769 | 10 | 0.1 | 9 | 0.1111 | 12 | 0 | 44 | 0.0682 | 9 | 0.2222 | 7 | 0.1429 | 13 | 0.1538 | 14 | 0.3571 | 43 | 0.2326 |
| 6_1142260 | 14 | 0.0714 | 11 | 0 | 11 | 0.2727 | 13 | 0.0769 | 49 | 0.102 | 8 | 0 | 8 | 0 | 13 | 0.0769 | 15 | 0.2 | 44 | 0.0909 |
| 6_1142264 | 13 | 0 | 20 | 0 | 11 | 0.0909 | 11 | 0.1818 | 55 | 0.0545 | 10 | 0.1 | 7 | 0 | 11 | 0.0909 | 14 | 0.1429 | 42 | 0.0952 |
| 6_1142924 | 21 | 0.5714 | 28 | 0.5357 | 28 | 0.3929 | 31 | 0.5161 | 108 | 0.5 | 16 | 0.4375 | 28 | 0.4643 | 33 | 0.3333 | 24 | 0.375 | 101 | 0.396 |
| 6_1142927 | 21 | 0.6667 | 37 | 0.4324 | 34 | 0.3235 | 36 | 0.5278 | 128 | 0.4688 | 23 | 0.3913 | 35 | 0.3429 | 37 | 0.4054 | 34 | 0.2353 | 129 | 0.3411 |
| 6_1142927 | 26 | 0.0769 | 39 | 0.1026 | 30 | 0.1333 | 41 | 0.0976 | 136 | 0.1029 | 27 | 0.1111 | 41 | 0.1463 | 38 | 0.1316 | 30 | 0.0667 | 136 | 0.1176 |
| 6_1142928 | 29 | 0.1034 | 47 | 0.0851 | 37 | 0.0811 | 45 | 0.0667 | 158 | 0.0823 | 28 | 0.1429 | 46 | 0 | 43 | 0.093 | 35 | 0 | 152 | 0.0526 |
| 6_1142930 | 27 | 0.2222 | 35 | 0.3714 | 32 | 0.25 | 34 | 0.1765 | 128 | 0.2578 | 20 | 0.15 | 35 | 0.1714 | 35 | 0.3143 | 30 | 0.3 | 120 | 0.2417 |
| 6_1142931 | 32 | 0.125 | 45 | 0.2444 | 38 | 0.1053 | 43 | 0.0698 | 158 | 0.1392 | 27 | 0.2593 | 48 | 0.0417 | 44 | 0.1591 | 40 | 0.15 | 159 | 0.1384 |
| 6_1142934 | 32 | 0.2188 | 51 | 0.1569 | 43 | 0.093 | 45 | 0.1333 | 171 | 0.1462 | 29 | 0.0345 | 47 | 0.0426 | 51 | 0.1373 | 42 | 0.1905 | 169 | 0.1065 |
| 6_1142935 | 33 | 0.0606 | 50 | 0.06 | 44 | 0.1364 | 50 | 0.1 | 177 | 0.0904 | 26 | 0.0769 | 52 | 0.0385 | 50 | 0.1 | 45 | 0.0667 | 173 | 0.0694 |
| 6_1142938 | 45 | 0.1111 | 54 | 0.037 | 47 | 0.1064 | 57 | 0.0526 | 203 | 0.0739 | 31 | 0.0645 | 57 | 0.0526 | 59 | 0.0508 | 48 | 0.0417 | 195 | 0.0513 |
| 6_1144567 | NA | NA | 1 | 0 | 2 | 0.5 | NA | NA | 3 | 0.3333 | NA | NA | NA | NA | NA | NA | 1 | 1 | 1 | 1 |
| 6_1144577 | NA | NA | 1 | 1 | 3 | 0 | NA | NA | 4 | 0.25 | 1 | 1 | NA | NA | NA | NA | 2 | 0 | 3 | 0.3333 |
| 6_1144578 | NA | NA | 1 | 0 | 2 | 0.5 | NA | NA | 3 | 0.3333 | 1 | 0 | NA | NA | NA | NA | 2 | 0.5 | 3 | 0.3333 |
| 6_1144579 | NA | NA | 1 | 0 | 2 | 0.5 | NA | NA | 3 | 0.3333 | NA | NA | NA | NA | NA | NA | 2 | 0.5 | 2 | 0.5 |
| 6_1144584 | NA | NA | 1 | 1 | 3 | 0.6667 | NA | NA | 4 | 0.75 | 2 | 1 | NA | NA | NA | NA | 1 | 1 | 3 | 1 |
| 6_1144584 | NA | NA | 1 | 0 | 3 | 0.3333 | NA | NA | 4 | 0.25 | 2 | 0.5 | NA | NA | NA | NA | 2 | 0.5 | 4 | 0.5 |
| 6_1144585 | NA | NA | 1 | 1 | 2 | 1 | NA | NA | 3 | 1 | 2 | 1 | NA | NA | NA | NA | 2 | 1 | 4 | 1 |
| 6_1144585 | NA | NA | NA | NA | 3 | 1 | NA | NA | 3 | 1 | NA | NA | NA | NA | NA | NA | 1 | 1 | 1 | 1 |
| 6_1144587 | NA | NA | 1 | 1 | 3 | 0.3333 | NA | NA | 4 | 0.5 | 2 | 0 | NA | NA | NA | NA | 2 | 0.5 | 4 | 0.25 |
| 6_1144589 | 1 | 1 | 1 | 1 | 3 | 1 | NA | NA | 5 | 1 | NA | NA | NA | NA | NA | NA | 2 | 1 | 2 | 1 |
| 6_1144589 | 1 | 1 | 2 | 0 | 3 | 0.6667 | NA | NA | 6 | 0.5 | 1 | 1 | NA | NA | NA | NA | 2 | 1 | 3 | 1 |
| 6_1144590 | 1 | 0 | 1 | 1 | 3 | 0.3333 | NA | NA | 5 | 0.4 | 2 | 1 | NA | NA | NA | NA | 2 | 0.5 | 4 | 0.75 |
| 6_1144590 | NA | NA | 2 | 1 | 3 | 1 | NA | NA | 5 | 1 | 2 | 1 | NA | NA | NA | NA | NA | NA | 2 | 1 |
| 6_1144591 | 1 | 1 | 1 | 1 | 3 | 1 | NA | NA | 5 | 1 | 2 | 1 | NA | NA | NA | NA | 1 | 1 | 3 | 1 |
| 6_1144591 | 1 | 1 | 2 | 1 | 3 | 1 | NA | NA | 6 | 1 | 1 | 1 | NA | NA | NA | NA | 2 | 1 | 3 | 1 |
| 6_1144591 | 1 | 0 | 2 | 0 | 1 | 1 | NA | NA | 4 | 0.25 | 1 | 0 | NA | NA | NA | NA | 2 | 1 | 3 | 0.6667 |
| 6_1148372 | 3 | 0.6667 | 5 | 0.6 | 3 | 0.6667 | 5 | 0.4 | 16 | 0.5625 | 1 | 0 | 1 | 0 | 4 | 0.5 | 3 | 0.3333 | 9 | 0.3333 |
| 6_1148379 | 2 | 1 | 2 | 0 | 1 | 0 | 4 | 0 | 9 | 0.2222 | NA | NA | 3 | 0 | 5 | 0 | 1 | 1 | 9 | 0.1111 |
| 6_1148381 | 1 | 0 | 2 | 0 | 2 | 0 | 5 | 0.2 | 10 | 0.1 | NA | NA | 3 | 0 | 5 | 0.2 | 1 | 0 | 9 | 0.1111 |
| 6_1148384 | NA | NA | 1 | 0 | 2 | 0.5 | 5 | 0.4 | 8 | 0.375 | NA | NA | 3 | 0.3333 | 5 | 0 | 1 | 1 | 9 | 0.2222 |

| | | | | | | | | | | | | | | | | | | | | |
|---|---|---|---|---|---|---|---|---|---|---|---|---|---|---|---|---|---|---|---|---|
| 6_11483849 | NA | NA | 1 | 1 | 1 | 0 | 5 | 0.6 | 7 | 0.5714 | NA | NA | 1 | 0 | 4 | 0.5 | 1 | 1 | 6 | 0.5 |
| 6_11483882 | NA | NA | 1 | 0 | 1 | 1 | 4 | 0.5 | 6 | 0.5 | NA | NA | 3 | 0 | 5 | 0.2 | 1 | 0 | 9 | 0.1111 |
| 6_11483898 | NA | NA | NA | NA | NA | NA | 4 | 0.75 | 4 | 0.75 | NA | NA | 3 | 0.3333 | 5 | 0.6 | 1 | 1 | 9 | 0.5556 |
| 6_11483907 | NA | NA | NA | NA | NA | NA | 2 | 0.5 | 2 | 0.5 | NA | NA | NA | NA | 1 | 1 | NA | NA | 1 | 1 |
| 6_11511204 | 15 | 0.0667 | 20 | 0.1 | 17 | 0 | 13 | 0.1538 | 65 | 0.0769 | 10 | 0.1 | 9 | 0 | 10 | 0.3 | 13 | 0.1538 | 42 | 0.1429 |
| 6_11522289 | 6 | 0.1667 | 4 | 0 | 7 | 0.1429 | 4 | 0 | 21 | 0.0952 | 2 | 0.5 | 3 | 0 | 4 | 0 | 4 | 0.25 | 13 | 0.1538 |
| 6_11522289 | 5 | 0.2 | 3 | 0 | 8 | 0 | 4 | 0 | 20 | 0.05 | 2 | 0 | 3 | 0 | 4 | 0 | 4 | 0.25 | 13 | 0.0769 |
| 6_11522324 | 5 | 0.2 | 5 | 0 | 8 | 0 | 4 | 0.25 | 22 | 0.0909 | 2 | 0 | 3 | 0 | 2 | 0.5 | 4 | 0 | 11 | 0.0909 |
| 6_11522338 | 5 | 0.4 | 5 | 0 | 6 | 0.3333 | 2 | 1 | 18 | 0.3333 | 2 | 0 | 3 | 0 | 3 | 1 | 4 | 0 | 12 | 0.25 |
| 6_11525141 | 21 | 0.0952 | 27 | 0.2593 | 20 | 0.2 | 23 | 0.3478 | 91 | 0.2308 | 12 | 0.0833 | 17 | 0.2941 | 26 | 0.2692 | 27 | 0.2963 | 82 | 0.2561 |
| 6_11525175 | 29 | 0.1034 | 31 | 0.0645 | 28 | 0.0714 | 29 | 0.0345 | 117 | 0.0684 | 11 | 0 | 25 | 0.08 | 27 | 0.1111 | 35 | 0.0571 | 98 | 0.0714 |
| 6_11525823 | 12 | 0.25 | 10 | 0.5 | 9 | 0.1111 | 8 | 0.375 | 39 | 0.3077 | 15 | 0.4667 | 11 | 0.4545 | 14 | 0.5714 | 10 | 0.5 | 50 | 0.5 |
| 6_11525918 | NA | NA | NA | NA | 1 | 1 | NA | NA | 1 | 1 | NA | NA | 2 | 0.5 | NA | NA | NA | NA | 2 | 0.5 |
| 6_11545003 | NA | NA | 1 | 1 | NA | NA | NA | NA | 1 | 1 | 1 | 1 | NA | NA | NA | NA | NA | NA | 1 | 1 |
| 6_11546714 | NA | NA | 1 | 1 | NA | NA | NA | NA | 1 | 1 | 1 | 1 | NA | NA | NA | NA | NA | NA | 1 | 1 |
| 6_11553675 | NA | NA | 4 | 0.25 | 1 | 0 | 5 | 0 | 10 | 0.1 | 1 | 1 | 3 | 0.3333 | 2 | 0 | 3 | 0.6667 | 9 | 0.4444 |
| 6_11553677 | NA | NA | 2 | 0 | NA | NA | 1 | 1 | 3 | 0.3333 | NA | NA | 2 | 0.5 | 1 | 1 | 1 | 0 | 4 | 0.5 |
| 6_11553681 | NA | NA | 3 | 0.6667 | 1 | 1 | 4 | 0.5 | 8 | 0.625 | NA | NA | 3 | 0.3333 | NA | NA | 3 | 0.3333 | 6 | 0.3333 |
| 6_11555499 | NA | NA | 2 | 0.5 | 1 | 0 | 8 | 0.25 | 11 | 0.2727 | 1 | 0 | 3 | 0.3333 | 1 | 1 | 3 | 0 | 8 | 0.25 |
| 6_11555510 | NA | NA | 5 | 1 | 1 | 1 | 8 | 0.875 | 14 | 0.9286 | 1 | 1 | 3 | 1 | 1 | 1 | 4 | 1 | 9 | 1 |
| 6_11555512 | NA | NA | 4 | 0.25 | NA | NA | 8 | 0 | 12 | 0.0833 | 1 | 0 | 3 | 0 | 2 | 0.5 | 3 | 0.3333 | 9 | 0.2222 |
| 6_11555519 | NA | NA | 4 | 1 | NA | NA | 5 | 0.4 | 9 | 0.6667 | NA | NA | 3 | 0 | 1 | 1 | 2 | 1 | 6 | 0.5 |
| 6_11555522 | NA | NA | 5 | 0.2 | NA | NA | 7 | 0.1429 | 12 | 0.1667 | 1 | 0 | 3 | 0 | 2 | 0 | 3 | 0.3333 | 9 | 0.1111 |
| 6_11555526 | NA | NA | 3 | 0.3333 | NA | NA | 7 | 0.1429 | 10 | 0.2 | 1 | 0 | 2 | 0 | 2 | 1 | 4 | 0.5 | 9 | 0.4444 |
| 6_11559494 | NA | NA | 5 | 0.4 | 3 | 0 | 4 | 0 | 12 | 0.1667 | 3 | 0.3333 | 5 | 0 | 4 | 0 | 4 | 0 | 16 | 0.0625 |
| 6_11559496 | 1 | 0 | 4 | 0.5 | 2 | 0 | 6 | 0.5 | 13 | 0.3846 | 2 | 0.5 | 5 | 0.2 | 2 | 1 | 4 | 0.25 | 13 | 0.3846 |
| 6_11559512 | 1 | 0 | 4 | 0 | 1 | 0 | 5 | 0.2 | 11 | 0.0909 | 4 | 0 | 5 | 0.2 | 2 | 0 | 2 | 0 | 13 | 0.0769 |
| 6_11559516 | 1 | 1 | 2 | 0 | 2 | 0.5 | 5 | 0.2 | 10 | 0.3 | 3 | 0 | 5 | 0.4 | 2 | 0 | 1 | 0 | 11 | 0.1818 |
| 6_11559529 | 2 | 0 | 4 | 0.75 | 2 | 0.5 | 4 | 0 | 12 | 0.3333 | 4 | 0.5 | 5 | 0.4 | 3 | 0.6667 | 2 | 0 | 14 | 0.4286 |
| 6_11559538 | 2 | 0.5 | 1 | 1 | 3 | 0 | 2 | 0.5 | 8 | 0.375 | 4 | 0.5 | 4 | 0.25 | 2 | 1 | 3 | 0.6667 | 13 | 0.5385 |
| 6_11559547 | 2 | 0 | 3 | 0 | 2 | 0.5 | 4 | 0.25 | 11 | 0.1818 | 2 | 0 | 3 | 1 | 2 | 0 | 2 | 0.5 | 9 | 0.4444 |
| 6_11559551 | 2 | 0 | 3 | 0 | 2 | 0 | 5 | 0.2 | 12 | 0.0833 | 4 | 0.25 | 5 | 0.4 | 4 | 0 | 3 | 0 | 16 | 0.1875 |
| 6_11559581 | 2 | 0.5 | 2 | 0 | 12 | 0.6667 | 8 | 0.5 | 24 | 0.5417 | 4 | 0.25 | 11 | 0.5455 | 4 | 0.25 | 2 | 0.5 | 21 | 0.4286 |
| 6_11559581 | 2 | 1 | 5 | 0.2 | 14 | 0.2143 | 9 | 0.3333 | 30 | 0.3 | 4 | 0.75 | 9 | 0.3333 | 5 | 0.4 | 4 | 0.75 | 22 | 0.5 |
| 6_11559582 | 3 | 1 | 5 | 0 | 13 | 0.6923 | 9 | 0.8889 | 30 | 0.6667 | 2 | 1 | 11 | 0.7273 | 3 | 0.6667 | 4 | 0.75 | 20 | 0.75 |
| 6_11559589 | 3 | 1 | 3 | 1 | 9 | 1 | 7 | 1 | 22 | 1 | 4 | 1 | 10 | 0.9 | 3 | 0.6667 | 2 | 1 | 19 | 0.8947 |
| 6_11559592 | 2 | 0 | 4 | 0.5 | 8 | 0.75 | 6 | 0.8333 | 20 | 0.65 | 3 | 1 | 10 | 0.7 | 3 | 0.3333 | 3 | 0.3333 | 19 | 0.6316 |
| 6_11559593 | 4 | 0.75 | 4 | 0.5 | 12 | 0.4167 | 10 | 0.6 | 30 | 0.5333 | 4 | 0.75 | 11 | 0.7273 | 4 | 1 | 3 | 1 | 22 | 0.8182 |
| 6_11559610 | 3 | 0 | 4 | 0 | 10 | 0.3 | 7 | 0.4286 | 24 | 0.25 | 3 | 0.3333 | 11 | 0.0909 | 5 | 0 | 4 | 0 | 23 | 0.087 |
| 6_11559611 | 4 | 0.75 | 4 | 0.5 | 8 | 0.875 | 7 | 0.7143 | 23 | 0.7391 | 4 | 0.5 | 11 | 0.7273 | 4 | 0.25 | 4 | 0.75 | 23 | 0.6087 |
| 6_11559627 | 3 | 0 | 4 | 0.5 | 10 | 0.2 | 6 | 0.3333 | 23 | 0.2609 | 4 | 0.25 | 10 | 0.2 | 5 | 0.4 | 4 | 0.25 | 23 | 0.2609 |
| 6_11559730 | NA | NA | NA | NA | NA | NA | 1 | 1 | 1 | 1 | 1 | 0 | NA | NA | 2 | 1 | NA | NA | 3 | 0.6667 |
| 6_11570614 | NA | NA | 1 | 1 | NA | NA | 1 | 1 | 2 | 1 | 1 | 1 | NA | NA | 2 | 0 | NA | NA | 3 | 0.3333 |
| 6_11570621 | NA | NA | NA | NA | NA | NA | 1 | 1 | 1 | 1 | NA | NA | NA | NA | 1 | 1 | NA | NA | 1 | 1 |
| 6_11576870 | NA | NA | 1 | 1 | NA | NA | 1 | 1 | 2 | 1 | 1 | 1 | NA | NA | NA | NA | NA | NA | 1 | 1 |
| 6_11576883 | NA | NA | 1 | 1 | NA | NA | 1 | 1 | 2 | 1 | 1 | 1 | NA | NA | NA | NA | NA | NA | 1 | 1 |
| 6_11576926 | NA | NA | 1 | 1 | NA | NA | 1 | 1 | 2 | 1 | 1 | 1 | NA | NA | NA | NA | NA | NA | 1 | 1 |
| 6_11576930 | NA | NA | 1 | 1 | NA | NA | 1 | 1 | 2 | 1 | 1 | 1 | NA | NA | NA | NA | NA | NA | 1 | 1 |
| 6_11576987 | NA | NA | 1 | 0 | NA | NA | 1 | 1 | 2 | 0.5 | 1 | 1 | NA | NA | NA | NA | NA | NA | 1 | 1 |
| 6_11578080 | NA | NA | 1 | 1 | NA | NA | 2 | 1 | 3 | 1 | 1 | 0 | NA | NA | 1 | 1 | 1 | 0 | 3 | 0.3333 |
| 6_11578771 | NA | NA | NA | NA | NA | NA | 2 | 0.5 | 2 | 0.5 | 1 | 0 | NA | NA | 2 | 1 | 1 | 0 | 4 | 0.5 |
| 6_11578799 | NA | NA | NA | NA | NA | NA | 2 | 1 | 2 | 1 | 1 | 1 | NA | NA | 1 | 1 | NA | NA | 2 | 1 |
| 6_11582609 | NA | NA | 1 | 1 | NA | NA | 1 | 1 | 2 | 1 | NA | NA | 1 | 1 | NA | NA | NA | NA | 1 | 1 |
| 6_11582637 | NA | NA | 1 | 1 | NA | NA | NA | NA | 1 | 1 | NA | NA | 1 | 1 | NA | NA | NA | NA | 1 | 1 |
| 6_11590322 | 4 | 0.25 | 1 | 0 | 3 | 0.3333 | 2 | 0 | 10 | 0.2 | 1 | 0 | 1 | 0 | 2 | 0 | 5 | 0.4 | 9 | 0.2222 |
| 6_11590333 | 4 | 0.5 | 1 | 0 | 3 | 0 | 2 | 0 | 10 | 0.2 | 3 | 0.6667 | 2 | 0.5 | 2 | 0 | 4 | 0.25 | 11 | 0.3636 |
| 6_11593181 | 7 | 0 | 10 | 0.5 | 7 | 0.1429 | 8 | 0.25 | 32 | 0.25 | 5 | 0 | 6 | 0 | 13 | 0.2308 | 6 | 0.6667 | 30 | 0.2333 |
| 6_11593193 | 4 | 0.25 | 9 | 0.2222 | 3 | 0 | 7 | 0.1429 | 23 | 0.1739 | 5 | 0.2 | 5 | 0.6 | 11 | 0.0909 | 6 | 0.3333 | 27 | 0.2593 |
| 6_11593200 | 7 | 0 | 11 | 0 | 7 | 0.1429 | 7 | 0.1429 | 32 | 0.0625 | 2 | 0 | 6 | 0.1667 | 11 | 0.0909 | 4 | 0.25 | 23 | 0.1304 |
| 6_11593202 | 1 | 0 | 5 | 0.2 | 2 | 0.5 | 3 | 0 | 11 | 0.1818 | 5 | 0 | 3 | 0 | 6 | 0.1667 | 5 | 0.2 | 19 | 0.1053 |
| 6_11593209 | 6 | 0 | 10 | 0.2 | 7 | 0.4286 | 6 | 0.1667 | 29 | 0.2069 | 4 | 0.5 | 7 | 0.4286 | 15 | 0.2 | 6 | 0.5 | 32 | 0.3438 |
| 6_11593215 | 5 | 0 | 5 | 0.4 | 5 | 0 | 5 | 0 | 20 | 0.1 | 2 | 0 | 5 | 0.4 | 7 | 0.1429 | 3 | 0 | 17 | 0.1765 |
| 6_11593234 | 5 | 0 | 6 | 0.3333 | 5 | 0.4 | 4 | 0.25 | 20 | 0.25 | 5 | 0 | 2 | 0 | 11 | 0.1818 | 5 | 0.6 | 23 | 0.2174 |
| 6_11593240 | 4 | 0.25 | 10 | 0.2 | 5 | 0 | 5 | 0.2 | 24 | 0.1667 | 3 | 0 | 7 | 0.1429 | 12 | 0.25 | 6 | 0.5 | 28 | 0.25 |
| 6_11593241 | 7 | 0.2857 | 10 | 0.2 | 5 | 0.4 | 7 | 0 | 29 | 0.2069 | 5 | 0.4 | 7 | 0 | 11 | 0.0909 | 6 | 0.1667 | 29 | 0.1379 |

| | | | | | | | | | | | | | | | | | | | | |
|---|---|---|---|---|---|---|---|---|---|---|---|---|---|---|---|---|---|---|---|---|
| 6_11593243 | 7 | 0.8571 | 10 | 0.9 | 6 | 0.8333 | 6 | 1 | 29 | 0.8966 | 5 | 1 | 6 | 1 | 11 | 0.8182 | 5 | 1 | 27 | 0.9259 |
| 6_11593256 | 5 | 0 | 9 | 0.5556 | 5 | 0.2 | 7 | 0.4286 | 26 | 0.3462 | 4 | 0.75 | 6 | 0.1667 | 12 | 0.1667 | 4 | 0.25 | 26 | 0.2692 |
| 6_11593260 | 6 | 0.5 | 11 | 0.6364 | 4 | 0.75 | 4 | 0.5 | 25 | 0.6 | 3 | 0.3333 | 4 | 0.75 | 13 | 0.5385 | 5 | 0.6 | 25 | 0.56 |
| 6_11597050 | NA | NA | 3 | 1 | 1 | 1 | 2 | 1 | 6 | 1 | 3 | 1 | NA | NA | 1 | 0 | 2 | 0.5 | 6 | 0.6667 |
| 6_11597082 | NA | NA | 1 | 1 | NA | NA | 1 | 1 | 2 | 1 | 3 | 1 | NA | NA | 1 | 0 | 2 | 1 | 6 | 0.8333 |
| 6_11597118 | NA | NA | 2 | 1 | 1 | 1 | 1 | 1 | 4 | 1 | 2 | 0.5 | NA | NA | 1 | 1 | 1 | 1 | 4 | 0.75 |
| 6_11599574 | 5 | 0 | 7 | 0.5714 | 17 | 0.1176 | 7 | 0.1429 | 36 | 0.1944 | 3 | 0.3333 | 10 | 0.1 | 7 | 0.1429 | 4 | 0.25 | 24 | 0.1667 |
| 6_11599581 | 5 | 0.2 | 4 | 0.25 | 15 | 0 | 5 | 0.2 | 29 | 0.1034 | 3 | 0.3333 | 8 | 0.25 | 6 | 0 | 4 | 0 | 21 | 0.1429 |
| 6_11599598 | 2 | 0 | 6 | 0 | 13 | 0.0769 | 5 | 0.4 | 26 | 0.1154 | 3 | 0.6667 | 7 | 0 | 6 | 0.1667 | 3 | 0.3333 | 19 | 0.2105 |
| 6_11599753 | 3 | 0 | 2 | 0 | 9 | 0 | 5 | 0.2 | 19 | 0.0526 | 2 | 0 | 5 | 0.2 | 4 | 0 | 4 | 0 | 15 | 0.0667 |
| 6_11599787 | 3 | 0.3333 | 2 | 0.5 | 7 | 0.1429 | 4 | 0.25 | 16 | 0.25 | 2 | 0 | 5 | 0 | 3 | 0 | 4 | 0.25 | 14 | 0.0714 |
| 6_11599842 | 7 | 0.5714 | 5 | 0 | 8 | 0.375 | 6 | 0.1667 | 26 | 0.3077 | 4 | 0.25 | 4 | 0.25 | 5 | 0.2 | 5 | 0.2 | 18 | 0.2222 |
| 6_11599846 | 5 | 0.4 | 3 | 0.3333 | 9 | 0.3333 | 9 | 0.3333 | 26 | 0.3462 | 5 | 0.2 | 5 | 0.2 | 3 | 0.3333 | 7 | 0.2857 | 20 | 0.25 |
| 6_11599850 | 6 | 0.3333 | 4 | 0 | 7 | 0 | 8 | 0.125 | 25 | 0.12 | 5 | 0 | 3 | 0 | 3 | 0.3333 | 6 | 0.1667 | 17 | 0.1176 |
| 6_11599852 | 8 | 0.25 | 4 | 0.5 | 9 | 0 | 7 | 0.5714 | 28 | 0.2857 | 5 | 0.2 | 4 | 0 | 4 | 0.5 | 7 | 0.1429 | 20 | 0.2 |
| 6_11599865 | 4 | 0.25 | 1 | 1 | 6 | 0.5 | 5 | 0.6 | 16 | 0.5 | 3 | 0.3333 | 3 | 0.6667 | 4 | 0.5 | 2 | 0.5 | 12 | 0.5 |
| 6_11599865 | 6 | 0.3333 | 3 | 0.6667 | 8 | 0.25 | 8 | 0.625 | 25 | 0.44 | 3 | 0.3333 | 6 | 0.3333 | 5 | 0.4 | 5 | 0.2 | 19 | 0.3158 |
| 6_11599870 | 6 | 0.1667 | 3 | 0 | 7 | 0.1429 | 7 | 0 | 23 | 0.087 | 3 | 0.3333 | 6 | 0 | 4 | 0 | 6 | 0 | 19 | 0.0526 |
| 6_11599871 | 7 | 0 | 4 | 0.25 | 6 | 0 | 8 | 0.125 | 25 | 0.08 | 4 | 0 | 6 | 0.1667 | 4 | 0 | 6 | 0.1667 | 20 | 0.1 |
| 6_11599873 | 7 | 0.8571 | 4 | 1 | 9 | 0.8889 | 9 | 1 | 29 | 0.931 | 5 | 1 | 6 | 1 | 5 | 1 | 6 | 0.6667 | 22 | 0.9091 |
| 6_11599878 | 4 | 1 | 2 | 1 | 5 | 0.8 | 5 | 1 | 16 | 0.9375 | 3 | 1 | 3 | 1 | 5 | 1 | 2 | 1 | 13 | 1 |
| 6_11599907 | 7 | 0.2857 | 2 | 0.5 | 9 | 0.3333 | 8 | 0.5 | 26 | 0.3846 | 3 | 0.3333 | 3 | 0 | 3 | 0 | 6 | 0.3333 | 15 | 0.2 |
| 6_11599916 | 7 | 0 | 2 | 0 | 9 | 0 | 9 | 0.2222 | 27 | 0.0741 | 3 | 0 | 5 | 0 | 4 | 0 | 6 | 0.3333 | 18 | 0.1111 |
| 6_11599924 | 7 | 0.2857 | 3 | 0 | 8 | 0.25 | 9 | 0.3333 | 27 | 0.2593 | 3 | 0 | 4 | 0.25 | 4 | 0 | 3 | 0 | 14 | 0.0714 |
| 6_11601987 | 7 | 0.5714 | 3 | 0.3333 | 8 | 0.625 | 7 | 0.2857 | 25 | 0.48 | 2 | 1 | 4 | 0.75 | 2 | 1 | 6 | 0.3333 | 14 | 0.6429 |
| 6_11601991 | 6 | 0.1667 | 3 | 0.6667 | 7 | 0.2857 | 7 | 0.7143 | 23 | 0.4348 | 2 | 0.5 | 3 | 0.6667 | 3 | 0.6667 | 6 | 0.3333 | 14 | 0.5 |
| 6_11601991 | 6 | 0 | 3 | 0.3333 | 8 | 0 | 7 | 0.1429 | 24 | 0.0833 | 2 | 0 | 4 | 0 | 3 | 0 | 6 | 0.1667 | 15 | 0.0667 |
| 6_11602011 | 6 | 0.8333 | 2 | 0 | 4 | 0.5 | 5 | 0.4 | 17 | 0.5294 | 1 | 0 | 3 | 0 | 2 | 0.5 | 6 | 0.1667 | 12 | 0.1667 |
| 6_11602015 | 4 | 0.75 | 2 | 0.5 | 6 | 0.8333 | 6 | 1 | 18 | 0.8333 | 2 | 1 | 2 | 0.5 | 2 | 1 | 4 | 1 | 10 | 0.9 |
| 6_11602017 | 7 | 1 | 2 | 1 | 5 | 1 | 6 | 1 | 20 | 1 | 2 | 1 | 3 | 1 | 3 | 1 | 5 | 1 | 13 | 1 |
| 6_11602019 | 7 | 0 | 2 | 0 | 6 | 0.3333 | 6 | 0 | 21 | 0.0952 | 2 | 0 | 3 | 0 | 3 | 0 | 5 | 0.4 | 13 | 0.1538 |
| 6_11602021 | 6 | 0 | 2 | 0.5 | 6 | 0.3333 | 6 | 0.3333 | 20 | 0.25 | 1 | 0 | 3 | 0 | 3 | 0.3333 | 5 | 0 | 12 | 0.0833 |
| 6_11602030 | 7 | 0.2857 | 1 | 1 | 5 | 0.8 | 4 | 0 | 17 | 0.4118 | 2 | 1 | 4 | 0.25 | 3 | 0.3333 | 4 | 0.5 | 13 | 0.4615 |
| 6_11602033 | 5 | 0.6 | 2 | 0 | 6 | 0.5 | 5 | 0.4 | 18 | 0.4444 | 2 | 0 | 3 | 0.3333 | 3 | 0.6667 | 4 | 0.75 | 12 | 0.5 |
| 6_11602037 | 7 | 0.4286 | 2 | 0.5 | 5 | 0 | 3 | 0.6667 | 17 | 0.3529 | 2 | 1 | 2 | 0 | 2 | 0.5 | 4 | 0.5 | 10 | 0.5 |
| 6_11603388 | 2 | 1 | NA | NA | NA | NA | 3 | 0.6667 | 5 | 0.8 | NA | NA | NA | NA | NA | NA | 1 | 1 | 1 | 1 |
| 6_11611093 | 3 | 0.6667 | 1 | 1 | 2 | 0.5 | NA | NA | 6 | 0.6667 | NA | NA | 3 | 0.3333 | 1 | 1 | 3 | 1 | 7 | 0.7143 |
| 6_11611104 | 1 | 1 | NA | NA | 2 | 0.5 | NA | NA | 3 | 0.6667 | NA | NA | 2 | 0.5 | 1 | 0 | 1 | 1 | 4 | 0.5 |
| 6_11611116 | 3 | 0.6667 | 1 | 0 | 2 | 1 | NA | NA | 6 | 0.6667 | NA | NA | 2 | 0.5 | 1 | 1 | 3 | 0.3333 | 6 | 0.5 |
| 6_11611121 | 3 | 0.3333 | 1 | 0 | 2 | 0 | NA | NA | 6 | 0.1667 | NA | NA | 3 | 0 | 1 | 0 | 3 | 0.3333 | 7 | 0.1429 |
| 6_11611125 | 3 | 0.3333 | 1 | 0 | 1 | 1 | NA | NA | 5 | 0.4 | NA | NA | 3 | 0.3333 | 1 | 1 | 1 | 0 | 5 | 0.4 |
| 6_11611127 | 3 | 0.6667 | NA | NA | 2 | 0.5 | NA | NA | 5 | 0.6 | NA | NA | 3 | 0.3333 | 1 | 1 | 2 | 0 | 6 | 0.3333 |
| 6_11615111 | 8 | 0.75 | 8 | 0.25 | 2 | 0 | 2 | 0 | 20 | 0.4 | 10 | 0.2 | 5 | 0.2 | 3 | 0 | 9 | 0.3333 | 27 | 0.2222 |
| 6_11615115 | 9 | 0.3333 | 9 | 0.4444 | 5 | 0.2 | 2 | 0.5 | 25 | 0.36 | 9 | 0.7778 | 6 | 0.5 | 3 | 1 | 8 | 0.625 | 26 | 0.6923 |
| 6_11617254 | 1 | 1 | NA | NA | NA | NA | NA | NA | 1 | 1 | NA | NA | NA | NA | NA | NA | 1 | 1 | 1 | 1 |
| 6_11617258 | 1 | 1 | NA | NA | NA | NA | NA | NA | 1 | 1 | NA | NA | NA | NA | NA | NA | 1 | 1 | 1 | 1 |
| 6_11623569 | 7 | 0.7143 | 9 | 0.5556 | 3 | 0 | 5 | 0.6 | 24 | 0.5417 | 9 | 0.6667 | 4 | 0.5 | 5 | 0.4 | 2 | 0.5 | 20 | 0.55 |
| 6_11623570 | 8 | 0.625 | 10 | 0.2 | 3 | 0 | 6 | 0.5 | 27 | 0.3704 | 9 | 0.2222 | 4 | 0 | 8 | 0.375 | 3 | 0 | 24 | 0.2083 |
| 6_11623578 | 4 | 0.25 | 6 | 0.5 | 1 | 1 | 4 | 0.5 | 15 | 0.4667 | 5 | 0.4 | 3 | 0.3333 | 6 | 0.1667 | 2 | 0 | 16 | 0.25 |
| 6_11623583 | 8 | 0.25 | 10 | 0 | 3 | 0.6667 | 6 | 0 | 27 | 0.1481 | 10 | 0.1 | 5 | 0 | 9 | 0.1111 | 3 | 0.3333 | 27 | 0.1111 |
| 6_11623588 | 8 | 0.375 | 8 | 0 | 2 | 0 | 6 | 0.5 | 24 | 0.25 | 11 | 0.0909 | 5 | 0.2 | 8 | 0 | 3 | 0 | 27 | 0.0741 |
| 6_11623591 | 9 | 0.4444 | 9 | 0.6667 | 1 | 0 | 4 | 0.5 | 23 | 0.5217 | 10 | 0.3 | 5 | 0.4 | 7 | 0.1429 | 3 | 0.3333 | 25 | 0.28 |
| 6_11625069 | 2 | 0.5 | NA | NA | NA | NA | NA | NA | 2 | 0.5 | NA | NA | 2 | 1 | 1 | 0 | 1 | 0 | 4 | 0.5 |
| 6_11625073 | 2 | 0.5 | NA | NA | NA | NA | NA | NA | 2 | 0.5 | NA | NA | 2 | 0.5 | 1 | 1 | 1 | 1 | 4 | 0.75 |
| 6_11629167 | 1 | 1 | NA | NA | NA | NA | NA | NA | 1 | 1 | NA | NA | NA | NA | 1 | 1 | NA | NA | 1 | 1 |
| 6_11638304 | 1 | 1 | 2 | 1 | 1 | 1 | 1 | 1 | 5 | 1 | 1 | 1 | 5 | 1 | 1 | 1 | 3 | 0.6667 | 10 | 0.9 |
| 6_11638309 | 1 | 1 | 3 | 0.6667 | 1 | 1 | 1 | 1 | 6 | 0.8333 | 1 | 1 | 4 | 1 | 1 | 1 | 3 | 1 | 9 | 1 |
| 6_11638313 | 1 | 1 | 3 | 1 | 1 | 1 | 1 | 1 | 6 | 1 | NA | NA | 5 | 0.8 | 2 | 1 | 2 | 1 | 9 | 0.8889 |
| 6_11638320 | 1 | 1 | 3 | 1 | 1 | 0 | NA | NA | 5 | 0.8 | 1 | 1 | 3 | 0.3333 | 2 | 1 | 1 | 1 | 7 | 0.7143 |
| 6_11660208 | 3 | 0.3333 | 7 | 0 | 4 | 0 | 4 | 0 | 18 | 0.0556 | 2 | 0 | 3 | 0 | 8 | 0.125 | 5 | 0 | 18 | 0.0556 |
| 6_11660208 | 4 | 0.5 | 6 | 0.6667 | 4 | 0.5 | 3 | 0.6667 | 17 | 0.5882 | 2 | 1 | 4 | 0.75 | 11 | 0.7273 | 5 | 0.8 | 22 | 0.7727 |
| 6_11660213 | 4 | 0 | 3 | 0 | 4 | 0.25 | 4 | 0 | 15 | 0.0667 | 1 | 0 | 3 | 0.3333 | 11 | 0.1818 | 4 | 0 | 19 | 0.1579 |
| 6_11660216 | 5 | 1 | 5 | 1 | 4 | 1 | 4 | 1 | 18 | 1 | 2 | 1 | 4 | 1 | 9 | 1 | 2 | 1 | 17 | 1 |
| 6_11660218 | 5 | 0 | 7 | 0.1429 | 2 | 0 | 4 | 0 | 18 | 0.0556 | 2 | 0 | 4 | 0 | 11 | 0.1818 | 5 | 0.2 | 22 | 0.1364 |
| 6_11660222 | 5 | 0.6 | 7 | 0.5714 | 2 | 0.5 | 3 | 0.6667 | 17 | 0.5882 | 2 | 1 | 3 | 1 | 11 | 0.9091 | 5 | 1 | 21 | 0.9524 |

| | | | | | | | | | | | | | | | | | | | | |
|---|---|---|---|---|---|---|---|---|---|---|---|---|---|---|---|---|---|---|---|---|
| 6_1166025 | 5 | 0 | 6 | 0.1667 | 3 | 0.3333 | 3 | 0.3333 | 17 | 0.1765 | 2 | 0.5 | 4 | 0.25 | 10 | 0.4 | 5 | 0 | 21 | 0.2857 |
| 6_1166028 | 4 | 0 | 5 | 0 | 4 | 0.5 | 3 | 0 | 16 | 0.125 | 2 | 0 | 3 | 0 | 8 | 0.25 | 5 | 0.2 | 18 | 0.1667 |
| 6_1166031 | 5 | 0 | 3 | 0 | 4 | 0 | 2 | 0.5 | 14 | 0.0714 | 2 | 0 | 4 | 0 | 7 | 0 | 4 | 0.25 | 17 | 0.0588 |
| 6_1166031 | 4 | 1 | 3 | 1 | 2 | 1 | 3 | 1 | 12 | 1 | 2 | 1 | 2 | 1 | 5 | 1 | 5 | 1 | 14 | 1 |
| 6_1166031 | 4 | 0.5 | 4 | 0 | 4 | 0.25 | 2 | 0 | 14 | 0.2143 | 2 | 0.5 | 3 | 0.3333 | 6 | 0.1667 | 4 | 0 | 15 | 0.2 |
| 6_1166031 | 3 | 0 | 4 | 0.25 | 4 | 0.25 | 3 | 0 | 14 | 0.1429 | 2 | 0 | 3 | 0.6667 | 7 | 0 | 5 | 0.2 | 17 | 0.1765 |
| 6_1166032 | 3 | 0.6667 | 4 | 0.75 | 4 | 0.5 | 3 | 0.6667 | 14 | 0.6429 | 2 | 0 | 3 | 0.6667 | 8 | 0.75 | 5 | 0.8 | 18 | 0.6667 |
| 6_1166035 | 4 | 0 | 4 | 0 | 3 | 0.3333 | 2 | 0 | 13 | 0.0769 | 1 | 0 | 3 | 0 | 7 | 0 | 5 | 0.2 | 16 | 0.0625 |
| 6_1166109 | 1 | 0 | NA | NA | 1 | 0 | 1 | 1 | 3 | 0.3333 | 1 | 1 | NA | NA | 1 | 1 | NA | NA | 2 | 1 |
| 6_1166109 | 1 | 1 | NA | NA | 1 | 0 | 1 | 0 | 3 | 0.3333 | 1 | 1 | NA | NA | 1 | 0 | 1 | 0 | 3 | 0.3333 |
| 6_1166133 | NA | NA | 1 | 1 | NA | NA | NA | NA | 1 | 1 | 1 | 0 | 1 | 0 | NA | NA | 1 | 1 | 3 | 0.3333 |
| 6_1166135 | NA | NA | 1 | 1 | NA | NA | NA | NA | 1 | 1 | 1 | 1 | NA | NA | NA | NA | NA | NA | 1 | 1 |
| 6_1166289 | 1 | 1 | NA | NA | NA | NA | 1 | 1 | 2 | 1 | NA | NA | NA | NA | NA | NA | 1 | 1 | 1 | 1 |
| 6_1166290 | 1 | 1 | NA | NA | NA | NA | 1 | 1 | 2 | 1 | NA | NA | NA | NA | 1 | 0 | 1 | 1 | 2 | 0.5 |
| 6_1166290 | 1 | 1 | NA | NA | NA | NA | 1 | 1 | 2 | 1 | NA | NA | NA | NA | 1 | 1 | NA | NA | 1 | 1 |
| 6_1166292 | 1 | 1 | NA | NA | NA | NA | NA | NA | 1 | 1 | NA | NA | NA | NA | 1 | 1 | 1 | 1 | 2 | 1 |
| 6_1166293 | 1 | 1 | NA | NA | NA | NA | 1 | 0 | 2 | 0.5 | NA | NA | NA | NA | 1 | 1 | 1 | 1 | 2 | 1 |
| 6_1171505 | NA | NA | 1 | 0 | 2 | 0.5 | NA | NA | 3 | 0.3333 | 1 | 1 | NA | NA | 1 | 0 | 2 | 0.5 | 4 | 0.5 |
| 6_1178203 | 11 | 0 | 12 | 0.25 | 3 | 0.3333 | 3 | 0 | 29 | 0.1379 | 3 | 0 | 9 | 0 | 7 | 0.2857 | 3 | 0 | 22 | 0.0909 |
| 6_1178223 | 13 | 0.6923 | 13 | 0.6154 | 4 | 1 | 4 | 1 | 34 | 0.7353 | 3 | 1 | 11 | 0.7273 | 8 | 0.625 | 4 | 0.75 | 26 | 0.7308 |
| 6_1178223 | 13 | 0.7692 | 7 | 0.7143 | 3 | 0.3333 | 4 | 0.75 | 27 | 0.7037 | 1 | 1 | 10 | 0.4 | 6 | 0.8333 | 4 | 0.75 | 21 | 0.619 |
| 6_1178226 | 14 | 0.5 | 13 | 0.3846 | 4 | 0.5 | 6 | 0.5 | 37 | 0.4595 | 3 | 0.3333 | 13 | 0.3077 | 9 | 0.4444 | 4 | 0 | 29 | 0.3103 |
| 6_1178228 | 13 | 0.2308 | 12 | 0.3333 | 4 | 0 | 6 | 0.5 | 35 | 0.2857 | 4 | 0.75 | 11 | 0.2727 | 9 | 0.3333 | 2 | 0 | 26 | 0.3462 |
| 6_1178228 | 11 | 0 | 13 | 0.1538 | 4 | 0 | 6 | 0 | 34 | 0.0588 | 4 | 0 | 11 | 0.0909 | 10 | 0.1 | 3 | 0 | 28 | 0.0714 |
| 6_1178228 | 13 | 0.6154 | 11 | 0.8182 | 3 | 0.6667 | 5 | 0.8 | 32 | 0.7188 | 1 | 1 | 11 | 0.7273 | 10 | 0.5 | 3 | 0.6667 | 25 | 0.64 |
| 6_1178400 | 1 | 0 | 1 | 1 | NA | NA | NA | NA | 2 | 0.5 | NA | NA | NA | NA | 1 | 1 | NA | NA | 1 | 1 |
| 6_1178439 | 1 | 1 | NA | NA | NA | NA | NA | NA | 1 | 1 | 1 | 1 | 1 | 1 | NA | NA | NA | NA | 2 | 1 |
| 6_1178473 | 4 | 0.5 | 4 | 0.5 | 1 | 1 | 4 | 0.5 | 13 | 0.5385 | 3 | 1 | 1 | 1 | 3 | 1 | 3 | 0.6667 | 10 | 0.9 |
| 6_1178474 | 4 | 0.5 | 4 | 0.75 | 1 | 1 | 5 | 0.2 | 14 | 0.5 | 3 | 0.3333 | 1 | 0 | 3 | 0 | 3 | 0.6667 | 10 | 0.3 |
| 6_1178474 | 4 | 1 | 4 | 0.75 | 1 | 1 | 4 | 1 | 13 | 0.9231 | 3 | 0.6667 | 1 | 0 | 3 | 1 | 1 | 1 | 8 | 0.75 |
| 6_1178476 | 4 | 1 | 5 | 1 | 1 | 1 | NA | NA | 10 | 1 | 1 | 1 | 1 | 1 | 3 | 1 | NA | NA | 5 | 1 |
| 6_1178478 | 4 | 1 | 5 | 1 | 1 | 1 | 6 | 0.8333 | 16 | 0.9375 | 3 | 1 | 2 | 1 | 3 | 1 | 2 | 1 | 10 | 1 |
| 6_1178478 | 5 | 0.2 | 5 | 0 | 1 | 0 | 6 | 0 | 17 | 0.0588 | 3 | 0.3333 | 2 | 0 | 3 | 0 | 2 | 0 | 10 | 0.1 |
| 6_1178478 | 5 | 0.8 | 5 | 0.4 | 1 | 1 | 5 | 0.2 | 16 | 0.5 | 3 | 0 | 2 | 0 | 3 | 0.6667 | 3 | 0.6667 | 11 | 0.3636 |
| 6_1178479 | 4 | 0.75 | 5 | 0.6 | NA | NA | 5 | 0.6 | 14 | 0.6429 | 2 | 0.5 | 2 | 1 | 4 | 0.75 | 1 | 1 | 9 | 0.7778 |
| 6_1178479 | 5 | 1 | 5 | 0.8 | 1 | 1 | 4 | 1 | 15 | 0.9333 | 2 | 1 | 1 | 0 | 4 | 1 | 2 | 1 | 9 | 0.8889 |
| 6_1178480 | 7 | 0 | 5 | 0.2 | 1 | 0 | 7 | 0 | 20 | 0.05 | 1 | 0 | 2 | 0 | 4 | 0.25 | 3 | 0 | 10 | 0.1 |
| 6_1178481 | 6 | 0.6667 | 5 | 0.8 | 1 | 1 | 6 | 0.3333 | 18 | 0.6111 | 3 | 1 | 2 | 0.5 | 3 | 1 | 3 | 0.3333 | 11 | 0.7273 |
| 6_1178481 | 7 | 0.2857 | 3 | 0 | 1 | 0 | 7 | 0.2857 | 18 | 0.2222 | 3 | 0.3333 | 2 | 0 | 2 | 0 | 3 | 0.3333 | 10 | 0.2 |
| 6_1178482 | 7 | 0 | 4 | 0.25 | 1 | 0 | 5 | 0 | 17 | 0.0588 | 2 | 0 | 2 | 0 | 4 | 0 | 3 | 0.3333 | 11 | 0.0909 |
| 6_1178483 | 7 | 0 | 5 | 0 | 1 | 1 | 5 | 0.2 | 18 | 0.1111 | 3 | 0 | 2 | 1 | 3 | 0 | 2 | 0 | 10 | 0.2 |
| 6_1178485 | 4 | 0 | 3 | 0.3333 | 2 | 0 | 5 | 0.2 | 14 | 0.1429 | 3 | 0.3333 | 2 | 0 | 3 | 0 | 2 | 0 | 10 | 0.1 |
| 6_1178488 | 7 | 0.1429 | 2 | 0 | 2 | 0 | 3 | 0 | 14 | 0.0714 | 2 | 0.5 | 2 | 0 | 5 | 0 | 2 | 0 | 11 | 0.0909 |
| 6_1178488 | 8 | 0.875 | 5 | 0.8 | 1 | 1 | 6 | 1 | 20 | 0.9 | 3 | 1 | 1 | 1 | 6 | 0.8333 | 4 | 1 | 14 | 0.9286 |
| 6_1178492 | 6 | 0.1667 | 4 | 0 | 2 | 0 | 5 | 0.2 | 17 | 0.1176 | 2 | 0.5 | 2 | 0 | 2 | 0 | 2 | 0 | 8 | 0.125 |
| 6_1178495 | 7 | 0.4286 | 4 | 0.25 | 2 | 0.5 | 5 | 0.2 | 18 | 0.3333 | 3 | 0.3333 | 4 | 0.25 | 5 | 0 | 1 | 0 | 13 | 0.1538 |
| 6_1178610 | 2 | 0.5 | NA | NA | 1 | 0 | NA | NA | 3 | 0.3333 | 1 | 1 | NA | NA | NA | NA | NA | NA | 1 | 1 |
| 6_1179008 | 12 | 0.25 | 12 | 0.3333 | 6 | 0.3333 | 8 | 0.625 | 38 | 0.3684 | 8 | 0.25 | 3 | 0.3333 | 5 | 0.4 | 9 | 0.4444 | 25 | 0.36 |
| 6_1179009 | 13 | 0.2308 | 11 | 0.0909 | 6 | 0.3333 | 8 | 0.125 | 38 | 0.1842 | 8 | 0.125 | 6 | 0.1667 | 5 | 0.2 | 9 | 0.1111 | 28 | 0.1429 |
| 6_1179009 | 11 | 0.0909 | 14 | 0.2143 | 6 | 0.1667 | 7 | 0 | 38 | 0.1316 | 7 | 0.1429 | 6 | 0 | 5 | 0 | 11 | 0.0909 | 29 | 0.069 |
| 6_1179010 | 12 | 0.4167 | 14 | 0.6429 | 7 | 0.4286 | 9 | 0.6667 | 42 | 0.5476 | 8 | 0.75 | 7 | 0.2857 | 5 | 0.2 | 8 | 0.375 | 28 | 0.4286 |
| 6_1179017 | 10 | 0.1 | 13 | 0.2308 | 9 | 0.3333 | 8 | 0.25 | 40 | 0.225 | 10 | 0 | 8 | 0.125 | 6 | 0.3333 | 11 | 0.2727 | 35 | 0.1714 |
| 6_1179017 | 9 | 0.3333 | 12 | 0.4167 | 9 | 0.3333 | 5 | 0.6 | 35 | 0.4 | 12 | 0.25 | 7 | 0.2857 | 7 | 0.5714 | 11 | 0.7273 | 37 | 0.4595 |
| 6_1180652 | 23 | 0.1739 | 18 | 0.0556 | 12 | 0.1667 | 9 | 0.1111 | 62 | 0.129 | 15 | 0.0667 | 17 | 0.0588 | 10 | 0.1 | 7 | 0 | 49 | 0.0612 |
| 6_1180657 | 1 | 1 | NA | NA | 1 | 0 | 1 | 0 | 3 | 0.3333 | 1 | 0 | 2 | 0.5 | 2 | 0 | 1 | 0 | 6 | 0.1667 |
| 6_1180661 | 19 | 0.0526 | 15 | 0.0667 | 9 | 0 | 9 | 0.1111 | 52 | 0.0577 | 15 | 0.0667 | 13 | 0.0769 | 10 | 0 | 7 | 0.1429 | 45 | 0.0667 |
| 6_1180668 | 22 | 0.0455 | 15 | 0 | 12 | 0.1667 | 12 | 0.1667 | 61 | 0.082 | 19 | 0 | 15 | 0.1333 | 13 | 0.1538 | 9 | 0 | 56 | 0.0714 |
| 6_1180676 | 33 | 0.1515 | 39 | 0.2308 | 21 | 0.0476 | 24 | 0.0833 | 117 | 0.1453 | 25 | 0.08 | 35 | 0.0571 | 21 | 0.0952 | 18 | 0.1667 | 99 | 0.0909 |
| 6_1180678 | 38 | 0.0789 | 41 | 0.0732 | 25 | 0.04 | 27 | 0.1111 | 131 | 0.0763 | 25 | 0.12 | 34 | 0.0588 | 26 | 0.1154 | 22 | 0.0455 | 107 | 0.0841 |
| 6_1180679 | 26 | 0.1923 | 34 | 0.2353 | 23 | 0.1739 | 24 | 0.2083 | 107 | 0.2056 | 21 | 0.2381 | 33 | 0.1515 | 21 | 0.3333 | 19 | 0.2632 | 94 | 0.234 |
| 6_1180859 | 2 | 0 | 3 | 0.6667 | NA | NA | NA | NA | 5 | 0.4 | 1 | 0 | 3 | 1 | 1 | 0 | 1 | 1 | 6 | 0.6667 |
| 6_1180859 | NA | NA | 1 | 1 | NA | NA | 1 | 1 | 2 | 1 | NA | NA | NA | NA | 2 | 1 | NA | NA | 2 | 1 |
| 6_1180860 | 2 | 0.5 | 3 | 0.6667 | 1 | 0 | 1 | 0 | 7 | 0.4286 | 1 | 0 | 3 | 1 | 3 | 0.6667 | 1 | 0 | 8 | 0.625 |
| 6_1180862 | 2 | 0.5 | 4 | 0.75 | 1 | 0 | 1 | 0 | 8 | 0.5 | 1 | 0 | 3 | 0.3333 | 3 | 0.3333 | 1 | 1 | 8 | 0.375 |
| 6_1180862 | 2 | 1 | 4 | 1 | 1 | 1 | 1 | 1 | 8 | 1 | 1 | 1 | 3 | 0.6667 | 3 | 1 | 1 | 0 | 8 | 0.75 |

| | | | | | | | | | | | | | | | | | | | | |
|---|---|---|---|---|---|---|---|---|---|---|---|---|---|---|---|---|---|---|---|---|
| 6_1181032 | 2 | 1 | 2 | 0 | NA | NA | NA | NA | 4 | 0.5 | NA | NA | 1 | 1 | 2 | 0.5 | NA | NA | 3 | 0.6667 |
| 6_1181057 | 2 | 1 | 2 | 1 | NA | NA | 1 | 0 | 5 | 0.8 | NA | NA | 1 | 1 | 1 | 0 | NA | NA | 2 | 0.5 |
| 6_1181059 | 2 | 0.5 | 1 | 0 | NA | NA | 1 | 0 | 4 | 0.25 | NA | NA | 1 | 1 | 3 | 0.3333 | NA | NA | 4 | 0.5 |
| 6_1181062 | 2 | 1 | 2 | 1 | NA | NA | NA | NA | 4 | 1 | NA | NA | 1 | 1 | 3 | 0.6667 | NA | NA | 4 | 0.75 |
| 6_1181063 | 1 | 1 | 1 | 1 | NA | NA | NA | NA | 2 | 1 | NA | NA | NA | NA | 3 | 0.3333 | NA | NA | 3 | 0.3333 |
| 6_1183604 | 5 | 0.8 | 4 | 0.25 | 2 | 1 | 1 | 0 | 12 | 0.5833 | 5 | 0.8 | 2 | 0 | 4 | 1 | 1 | 1 | 12 | 0.75 |
| 6_1183605 | 4 | 1 | 3 | 0.6667 | 1 | 1 | 3 | 1 | 11 | 0.9091 | 3 | 1 | 1 | 1 | 6 | 0.8333 | 1 | 1 | 11 | 0.9091 |
| 6_1183817 | 4 | 0.25 | 2 | 0 | 1 | 0 | 1 | 1 | 8 | 0.25 | 2 | 0.5 | NA | NA | 3 | 0.3333 | 1 | 1 | 6 | 0.5 |
| 6_1183817 | 2 | 0.5 | 2 | 0 | 1 | 0 | 1 | 1 | 6 | 0.3333 | 2 | 0 | NA | NA | 2 | 0.5 | 1 | 0 | 5 | 0.2 |
| 6_1184057 | 2 | 0.5 | 2 | 0 | 2 | 0 | NA | NA | 6 | 0.1667 | NA | NA | NA | NA | 3 | 0.6667 | NA | NA | 3 | 0.6667 |
| 6_1184059 | 2 | 0 | 2 | 0.5 | 3 | 0 | 1 | 0 | 8 | 0.125 | NA | NA | 1 | 1 | 3 | 0 | 1 | 0 | 5 | 0.2 |
| 6_1184059 | NA | NA | 2 | 1 | 2 | 1 | NA | NA | 4 | 1 | NA | NA | NA | NA | 2 | 1 | 1 | 1 | 3 | 1 |
| 6_1184060 | 2 | 1 | 1 | 1 | 2 | 0.5 | 1 | 1 | 6 | 0.8333 | NA | NA | NA | NA | 1 | 1 | NA | NA | 1 | 1 |
| 6_1184061 | 2 | 0.5 | 1 | 1 | 2 | 0 | NA | NA | 5 | 0.4 | NA | NA | 1 | 0 | 3 | 0.6667 | NA | NA | 4 | 0.5 |
| 6_1184061 | 1 | 0 | 2 | 0 | 2 | 0.5 | 1 | 0 | 6 | 0.1667 | NA | NA | 1 | 0 | 2 | 0.5 | 1 | 0 | 4 | 0.25 |
| 6_1184062 | 2 | 1 | 1 | 1 | 3 | 1 | 1 | 0 | 7 | 0.8571 | NA | NA | 1 | 1 | 3 | 1 | 1 | 1 | 5 | 1 |
| 6_1184063 | 2 | 0.5 | 2 | 0 | 2 | 0 | 1 | 1 | 7 | 0.2857 | NA | NA | 1 | 0 | 2 | 0.5 | 1 | 0 | 4 | 0.25 |
| 6_1184066 | 2 | 1 | 1 | 1 | 2 | 1 | 1 | 1 | 6 | 1 | NA | NA | 1 | 1 | 2 | 0.5 | 1 | 0 | 4 | 0.5 |
| 6_1184067 | 2 | 0 | 2 | 1 | 3 | 0 | 1 | 1 | 8 | 0.375 | NA | NA | 1 | 1 | 2 | 1 | 1 | 0 | 4 | 0.75 |
| 6_1184067 | 2 | 1 | 2 | 0 | 3 | 0.6667 | 1 | 0 | 8 | 0.5 | NA | NA | 1 | 1 | 3 | 0 | 1 | 1 | 5 | 0.4 |
| 6_1184068 | 2 | 0.5 | 2 | 1 | 3 | 1 | 1 | 1 | 8 | 0.875 | NA | NA | 1 | 1 | 3 | 1 | 1 | 1 | 5 | 1 |
| 6_1184068 | 2 | 1 | 1 | 0 | 2 | 1 | 1 | 0 | 6 | 0.6667 | NA | NA | 1 | 0 | 3 | 0.6667 | 1 | 1 | 5 | 0.6 |
| 6_1184068 | 1 | 1 | 1 | 1 | 2 | 1 | NA | NA | 4 | 1 | NA | NA | NA | NA | 2 | 1 | 1 | 1 | 3 | 1 |
| 6_1184070 | NA | NA | 2 | 1 | 2 | 0.5 | NA | NA | 4 | 0.75 | NA | NA | 1 | 1 | 2 | 0 | 1 | 1 | 4 | 0.5 |
| 6_1184071 | 2 | 1 | 2 | 1 | 3 | 1 | 1 | 1 | 8 | 1 | NA | NA | 1 | 1 | 2 | 1 | 1 | 1 | 4 | 1 |
| 6_1184071 | 2 | 0.5 | 2 | 0.5 | 2 | 0.5 | 1 | 1 | 7 | 0.5714 | NA | NA | 1 | 1 | 2 | 0 | 1 | 0 | 4 | 0.25 |
| 6_1184072 | 2 | 0.5 | 2 | 0 | 3 | 0.6667 | 1 | 1 | 8 | 0.5 | NA | NA | 1 | 1 | 1 | 1 | 1 | 0 | 3 | 0.6667 |
| 6_1184073 | 2 | 0.5 | 2 | 0.5 | 3 | 1 | 1 | 1 | 8 | 0.75 | NA | NA | 1 | 1 | 1 | 1 | 1 | 1 | 3 | 1 |
| 6_1184324 | NA | NA | 2 | 0.5 | NA | NA | NA | NA | 2 | 0.5 | NA | NA | NA | NA | 1 | 1 | NA | NA | 1 | 1 |
| 6_1184325 | 1 | 1 | 2 | 1 | NA | NA | NA | NA | 3 | 1 | NA | NA | NA | NA | 1 | 1 | NA | NA | 1 | 1 |
| 6_1184397 | 3 | 0 | 3 | 0.3333 | 1 | 0 | 3 | 0 | 10 | 0.1 | 1 | 0 | 3 | 0.3333 | 2 | 0 | 3 | 0.3333 | 9 | 0.2222 |
| 6_1184422 | 1 | 1 | 2 | 0.5 | NA | NA | 2 | 0.5 | 5 | 0.6 | NA | NA | 1 | 1 | 1 | 1 | NA | NA | 2 | 1 |
| 6_1184799 | NA | NA | NA | NA | 3 | 1 | NA | NA | 3 | 1 | NA | NA | 1 | 1 | 1 | 1 | 2 | 0.5 | 4 | 0.75 |
| 6_1184800 | NA | NA | NA | NA | 3 | 0.6667 | NA | NA | 3 | 0.6667 | NA | NA | 1 | 1 | 1 | 1 | 3 | 1 | 5 | 1 |
| 6_1185005 | 1 | 0 | NA | NA | 1 | 1 | NA | NA | 2 | 0.5 | NA | NA | NA | NA | NA | NA | 2 | 1 | 2 | 1 |
| 6_1185696 | 2 | 0 | 7 | 0.1429 | 1 | 0 | 4 | 0 | 14 | 0.0714 | 1 | 1 | 4 | 0 | 3 | 0 | 3 | 0 | 11 | 0.0909 |
| 6_1185698 | 1 | 1 | 5 | 0.4 | 1 | 1 | 2 | 0.5 | 9 | 0.5556 | 1 | 1 | 4 | 1 | 3 | 0.6667 | 3 | 0.6667 | 11 | 0.8182 |
| 6_1185706 | NA | NA | 3 | 0.3333 | 1 | 0 | 4 | 0.75 | 8 | 0.5 | 1 | 0 | 4 | 0.5 | 2 | 1 | 3 | 1 | 10 | 0.7 |
| 6_1185708 | NA | NA | 3 | 0.6667 | 1 | 0 | 2 | 1 | 6 | 0.6667 | 1 | 1 | 2 | 1 | 1 | 1 | 1 | 1 | 5 | 1 |
| 6_1185709 | NA | NA | 3 | 0.6667 | 1 | 1 | 3 | 0.6667 | 7 | 0.7143 | 1 | 0 | 1 | 0 | 2 | 0.5 | 2 | 1 | 6 | 0.5 |
| 6_1185709 | NA | NA | 4 | 0.75 | 1 | 1 | 3 | 0.6667 | 8 | 0.75 | 1 | 1 | 2 | 0.5 | 1 | 1 | 2 | 1 | 6 | 0.8333 |
| 6_1185709 | NA | NA | 3 | 1 | NA | NA | 1 | 1 | 4 | 1 | 1 | 0 | 2 | 1 | 1 | 1 | 2 | 1 | 6 | 0.8333 |
| 6_1185711 | NA | NA | 3 | 0.3333 | NA | NA | 3 | 0.3333 | 6 | 0.3333 | NA | NA | 2 | 0 | 2 | 0 | 2 | 0.5 | 6 | 0.1667 |
| 6_1185711 | NA | NA | 4 | 0 | 1 | 0 | 3 | 0.6667 | 8 | 0.25 | NA | NA | 2 | 0 | 1 | 0 | 2 | 1 | 5 | 0.4 |
| 6_1191918 | 3 | 0 | 5 | 0.6 | 1 | 0 | 2 | 0.5 | 11 | 0.3636 | NA | NA | NA | NA | NA | NA | 3 | 0.3333 | 3 | 0.3333 |
| 6_1191919 | 2 | 1 | 4 | 1 | 1 | 1 | 1 | 1 | 8 | 1 | NA | NA | NA | NA | NA | NA | 1 | 1 | 1 | 1 |
| 6_1191920 | 2 | 0.5 | 1 | 1 | 1 | 0 | 1 | 1 | 5 | 0.6 | NA | NA | NA | NA | NA | NA | 2 | 0.5 | 2 | 0.5 |
| 6_1191924 | 3 | 0.3333 | 5 | 0.4 | 2 | 0 | 2 | 0.5 | 12 | 0.3333 | NA | NA | NA | NA | 1 | 1 | 1 | 0 | 2 | 0.5 |
| 6_1191947 | 4 | 0.25 | 6 | 0 | 3 | 0 | 3 | 0 | 16 | 0.0625 | 1 | 0 | NA | NA | 1 | 1 | 2 | 0 | 4 | 0.25 |
| 6_1191992 | NA | NA | NA | NA | NA | NA | 1 | 1 | 1 | 1 | NA | NA | NA | NA | NA | NA | 1 | 1 | 1 | 1 |
| 6_1192921 | 10 | 0.1 | 7 | 0.1429 | 3 | 0 | 20 | 0.1 | 40 | 0.1 | 5 | 0.2 | 11 | 0 | 12 | 0.0833 | 4 | 0.25 | 32 | 0.0938 |
| 6_1192922 | 7 | 0.1429 | 11 | 0.0909 | 3 | 0 | 17 | 0.1176 | 38 | 0.1053 | 7 | 0.1429 | 8 | 0.125 | 16 | 0.375 | 4 | 0 | 35 | 0.2286 |
| 6_1192925 | 9 | 0.1111 | 13 | 0.0769 | 5 | 0 | 18 | 0.1111 | 45 | 0.0889 | 7 | 0 | 11 | 0 | 13 | 0.1538 | 5 | 0 | 36 | 0.0556 |
| 6_1192926 | 5 | 0.4 | 12 | 0.25 | 5 | 0.2 | 14 | 0.1429 | 36 | 0.2222 | 6 | 0.1667 | 10 | 0.1 | 8 | 0.125 | 6 | 0.1667 | 30 | 0.1333 |
| 6_1192927 | 6 | 0 | 10 | 0.4 | 5 | 0.6 | 15 | 0.2 | 36 | 0.2778 | 5 | 0.6 | 11 | 0.2727 | 10 | 0.3 | 6 | 0.5 | 32 | 0.375 |
| 6_1192928 | 5 | 0 | 11 | 0.3636 | 4 | 0.75 | 19 | 0.2105 | 39 | 0.2821 | 7 | 0.2857 | 11 | 0.3636 | 13 | 0.3077 | 5 | 0.4 | 36 | 0.3333 |
| 6_1192934 | 4 | 0 | 11 | 0.0909 | 4 | 0 | 11 | 0.2727 | 30 | 0.1333 | 6 | 0 | 9 | 0 | 9 | 0 | 6 | 0.3333 | 30 | 0.0667 |
| 6_1192936 | 3 | 0 | 10 | 0.2 | 4 | 0 | 10 | 0.3 | 27 | 0.1852 | 6 | 0 | 9 | 0.2222 | 10 | 0.1 | 6 | 0.3333 | 31 | 0.1613 |
| 6_1192936 | 4 | 0.25 | 7 | 0.2857 | 4 | 0 | 8 | 0.25 | 23 | 0.2174 | 6 | 0.6667 | 7 | 0.1429 | 9 | 0.1111 | 5 | 0.2 | 27 | 0.2593 |
| 6_1192937 | 4 | 0.5 | 10 | 1 | 4 | 0.25 | 8 | 0.5 | 26 | 0.6538 | 5 | 0.8 | 9 | 0.7778 | 10 | 0.9 | 6 | 0.8333 | 30 | 0.8333 |
| 6_1192937 | 4 | 0.5 | 10 | 0.3 | 4 | 0.25 | 7 | 0.4286 | 25 | 0.36 | 5 | 0.4 | 9 | 0.4444 | 10 | 0.5 | 4 | 0.25 | 28 | 0.4286 |
| 6_1192938 | 4 | 0.25 | 8 | 0.375 | 2 | 0 | 9 | 0 | 23 | 0.1739 | 4 | 0.5 | 8 | 0.25 | 7 | 0.1429 | 4 | 0.25 | 23 | 0.2609 |
| 6_1192939 | 4 | 1 | 8 | 1 | 2 | 1 | 9 | 0.8889 | 23 | 0.9565 | 5 | 1 | 10 | 1 | 10 | 0.8 | 6 | 1 | 31 | 0.9355 |
| 6_1192939 | 4 | 0 | 8 | 0.125 | 2 | 0 | 5 | 0 | 19 | 0.0526 | 4 | 0.25 | 9 | 0.1111 | 8 | 0.125 | 5 | 0.2 | 26 | 0.1538 |
| 6_1192939 | 3 | 1 | 6 | 1 | 3 | 1 | 8 | 0.875 | 20 | 0.95 | 5 | 1 | 8 | 1 | 8 | 1 | 4 | 0.75 | 25 | 0.96 |

| | | | | | | | | | | | | | | | | | | | | |
|---|---|---|---|---|---|---|---|---|---|---|---|---|---|---|---|---|---|---|---|---|
| 6_11929398 | 4 | 1 | 10 | 0.4 | 4 | 0.25 | 9 | 0.5556 | 27 | 0.5185 | 5 | 0.8 | 10 | 0.5 | 9 | 0.5556 | 4 | 0.75 | 28 | 0.6071 |
| 6_11929399 | 3 | 0 | 9 | 0.7778 | 4 | 0.25 | 9 | 0.3333 | 25 | 0.44 | 5 | 0.4 | 9 | 0.5556 | 10 | 0.4 | 5 | 0.4 | 29 | 0.4483 |
| 6_11929406 | 5 | 0.8 | 7 | 0.7143 | 3 | 1 | 8 | 0.875 | 23 | 0.8261 | 5 | 0.8 | 10 | 0.8 | 9 | 1 | 5 | 0.8 | 29 | 0.8621 |
| 6_11929433 | 4 | 0.5 | 9 | 0.5556 | 4 | 0.5 | 6 | 0.8333 | 23 | 0.6087 | 5 | 0.2 | 6 | 0.3333 | 8 | 0.5 | 5 | 0.6 | 24 | 0.4167 |
| 6_11929441 | 4 | 1 | 7 | 0.5714 | 4 | 1 | 7 | 0.8571 | 22 | 0.8182 | 5 | 0.8 | 4 | 0.25 | 6 | 0.6667 | 5 | 0.8 | 20 | 0.65 |
| 6_11929750 | 6 | 0.3333 | 5 | 0 | 4 | 0.25 | 7 | 0.2857 | 22 | 0.2273 | 4 | 0.5 | 7 | 0.1429 | 7 | 0.1429 | 5 | 0.4 | 23 | 0.2609 |
| 6_11929761 | 5 | 0 | 4 | 0 | 4 | 0 | 6 | 0.1667 | 19 | 0.0526 | 3 | 0 | 6 | 0 | 7 | 0.1429 | 4 | 0 | 20 | 0.05 |
| 6_11929763 | 5 | 0.4 | 5 | 0 | 3 | 0 | 6 | 0.3333 | 19 | 0.2105 | 4 | 0 | 7 | 0.4286 | 6 | 0 | 4 | 0.25 | 21 | 0.1905 |
| 6_11929764 | 3 | 0.3333 | 5 | 0 | 4 | 0 | 4 | 0 | 16 | 0.0625 | 1 | 0 | 4 | 0 | 3 | 0.3333 | 2 | 0 | 10 | 0.1 |
| 6_11929765 | 5 | 0.4 | 4 | 0 | 2 | 0.5 | 5 | 0.2 | 16 | 0.25 | 3 | 0 | 6 | 0 | 6 | 0 | 3 | 0.6667 | 18 | 0.1111 |
| 6_11929768 | 3 | 0.3333 | 4 | 0 | 3 | 0 | 6 | 0 | 16 | 0.0625 | 4 | 0 | 7 | 0 | 5 | 0.2 | 4 | 0 | 20 | 0.05 |
| 6_11929769 | 5 | 1 | 4 | 1 | 4 | 1 | 4 | 1 | 17 | 1 | 4 | 0.75 | 5 | 1 | 6 | 0.8333 | 1 | 1 | 16 | 0.875 |
| 6_11929774 | 4 | 1 | 5 | 0 | 4 | 0.25 | 4 | 0.5 | 17 | 0.4118 | 2 | 0.5 | 5 | 0.2 | 7 | 0.4286 | 4 | 0.5 | 18 | 0.3889 |
| 6_11929781 | 3 | 0.3333 | 4 | 0.25 | 3 | 0.3333 | 2 | 0 | 12 | 0.25 | 1 | 1 | 2 | 0 | 3 | 0.3333 | 1 | 0 | 7 | 0.2857 |
| 6_11929788 | 4 | 0.25 | 3 | 0.6667 | 3 | 0.6667 | 4 | 0.75 | 14 | 0.5714 | 3 | 0.6667 | 4 | 1 | 5 | 0.6 | 2 | 0.5 | 14 | 0.7143 |
| 6_11929790 | 5 | 0.2 | 3 | 0 | 3 | 0.3333 | 3 | 0.6667 | 14 | 0.2857 | 3 | 0.6667 | 4 | 0 | 2 | 0 | 4 | 0 | 13 | 0.1538 |
| 6_11933043 | 13 | 0.0769 | 11 | 0.0909 | 10 | 0.1 | 13 | 0.1538 | 47 | 0.1064 | 5 | 0 | 8 | 0.125 | 8 | 0.125 | 4 | 0.25 | 25 | 0.12 |
| 6_11933062 | 11 | 0.3636 | 10 | 0.4 | 10 | 0.4 | 9 | 0.3333 | 40 | 0.375 | 6 | 0.1667 | 7 | 0.5714 | 7 | 0.2857 | 5 | 0.8 | 25 | 0.44 |
| 6_11933068 | 9 | 0.4444 | 10 | 0.3 | 10 | 0.5 | 13 | 0.3846 | 42 | 0.4048 | 5 | 0.2 | 7 | 0.4286 | 8 | 0.5 | 3 | 0.3333 | 23 | 0.3913 |
| 6_11933080 | 10 | 0.1 | 7 | 0 | 11 | 0.1818 | 15 | 0.2 | 43 | 0.1395 | 6 | 0.3333 | 7 | 0.1429 | 8 | 0.125 | 4 | 0 | 25 | 0.16 |
| 6_11933090 | 9 | 0 | 8 | 0 | 7 | 0.2857 | 10 | 0.1 | 34 | 0.0882 | 4 | 0.25 | 8 | 0.25 | 6 | 0 | 3 | 0 | 21 | 0.1429 |
| 6_11933127 | 8 | 0.25 | 9 | 0.4444 | 9 | 0.2222 | 12 | 0.6667 | 38 | 0.4211 | 5 | 0.4 | 5 | 0.2 | 7 | 0.4286 | 3 | 0.6667 | 20 | 0.4 |
| 6_11933143 | 8 | 0.125 | 5 | 0 | 7 | 0.2857 | 7 | 0 | 27 | 0.1111 | 4 | 0 | 5 | 0.2 | 5 | 0 | 2 | 0 | 16 | 0.0625 |
| 6_11933194 | 6 | 0.3333 | 8 | 0.125 | 8 | 0.125 | 9 | 0.1111 | 31 | 0.1613 | 2 | 0.5 | 5 | 0 | 6 | 0.1667 | 2 | 0 | 15 | 0.1333 |
| 6_11933196 | 6 | 0 | 8 | 0.125 | 7 | 0.1429 | 7 | 0 | 28 | 0.0714 | 2 | 0 | 4 | 0.25 | 6 | 0.1667 | 3 | 0 | 15 | 0.1333 |
| 6_11933200 | 3 | 0.3333 | 7 | 0.5714 | 4 | 0 | 7 | 0.2857 | 21 | 0.3333 | 2 | 0.5 | 2 | 0 | 4 | 0.25 | 1 | 0 | 9 | 0.2222 |
| 6_11933217 | 5 | 0.2 | 4 | 0 | 4 | 0.5 | 5 | 0 | 18 | 0.1667 | 2 | 0.5 | 3 | 0 | 2 | 0 | 2 | 0 | 9 | 0.1111 |
| 6_11933259 | 4 | 0.25 | 4 | 0 | 5 | 0 | 5 | 0.2 | 18 | 0.1111 | 1 | 0 | 3 | 0.6667 | 4 | 0 | 3 | 0 | 11 | 0.1818 |
| 6_11933260 | 8 | 0.5 | 5 | 0.6 | 5 | 0.6 | 6 | 0.6667 | 24 | 0.5833 | 1 | 1 | 4 | 0.25 | 5 | 0.4 | 3 | 1 | 13 | 0.5385 |
| 6_11933265 | 4 | 1 | 3 | 1 | 1 | 1 | 3 | 0.6667 | 11 | 0.9091 | 1 | 1 | 2 | 1 | 4 | 1 | 2 | 1 | 9 | 1 |
| 6_11933266 | 5 | 0.2 | 3 | 0 | 4 | 0 | 4 | 0.5 | 16 | 0.1875 | NA | NA | 2 | 0.5 | 4 | 0.5 | 2 | 0 | 8 | 0.375 |
| 6_11933274 | 7 | 1 | 5 | 1 | 6 | 1 | 6 | 0.8333 | 24 | 0.9583 | 1 | 1 | 3 | 1 | 5 | 1 | 3 | 1 | 12 | 1 |
| 6_11934724 | 6 | 0.1667 | 2 | 0 | 3 | 0 | 4 | 0.25 | 15 | 0.1333 | 1 | 0 | 2 | 0 | 3 | 0.6667 | 2 | 0 | 8 | 0.25 |
| 6_11945745 | 1 | 1 | 3 | 0.3333 | 2 | 1 | 3 | 1 | 9 | 0.7778 | 3 | 0.6667 | 3 | 0.3333 | 1 | 1 | 2 | 1 | 9 | 0.6667 |
| 6_11945746 | 2 | 0.5 | 5 | 0 | 1 | 0 | 3 | 0 | 11 | 0.0909 | 2 | 0 | 3 | 0 | 1 | 1 | 3 | 0 | 9 | 0.1111 |
| 6_11945759 | 2 | 0 | 4 | 0.25 | 2 | 0 | 4 | 0.25 | 12 | 0.1667 | 3 | 0 | 4 | 0.25 | 2 | 0 | 3 | 0 | 12 | 0.0833 |
| 6_11945768 | 2 | 0 | 6 | 0.5 | 1 | 0 | 4 | 0.25 | 13 | 0.3077 | 3 | 0 | 2 | 0 | 2 | 0 | 4 | 0.25 | 11 | 0.0909 |
| 6_11945772 | 2 | 1 | 5 | 1 | 1 | 0 | 4 | 0.75 | 12 | 0.8333 | 2 | 1 | 3 | 1 | 2 | 0.5 | 4 | 1 | 11 | 0.9091 |
| 6_11945783 | 2 | 1 | 6 | 0.8333 | 1 | 1 | 4 | 1 | 13 | 0.9231 | 2 | 1 | 2 | 1 | 2 | 1 | 3 | 1 | 9 | 1 |
| 6_11945785 | 3 | 1 | 7 | 1 | 2 | 1 | 3 | 1 | 15 | 1 | 2 | 0.5 | 4 | 0.75 | NA | NA | 5 | 0.8 | 11 | 0.7273 |
| 6_11945798 | 2 | 1 | 6 | 1 | NA | NA | 5 | 1 | 13 | 1 | 3 | 1 | 1 | 1 | 2 | 1 | 3 | 1 | 9 | 1 |
| 6_11945800 | 2 | 1 | 7 | 1 | 2 | 1 | 2 | 0.5 | 13 | 0.9231 | 1 | 1 | 3 | 1 | 1 | 1 | 4 | 1 | 9 | 1 |
| 6_11945806 | 2 | 0.5 | 7 | 0.5714 | NA | NA | 3 | 0.3333 | 12 | 0.5 | 2 | 0.5 | 3 | 0.3333 | 1 | 1 | 5 | 0.4 | 11 | 0.4545 |
| 6_11945824 | 5 | 0.8 | 7 | 0.5714 | 2 | 1 | 5 | 0.8 | 19 | 0.7368 | 2 | 0 | 3 | 1 | 3 | 1 | 5 | 0.8 | 13 | 0.7692 |
| 6_11945828 | 4 | 1 | 6 | 1 | 3 | 0.6667 | 5 | 0.8 | 18 | 0.8889 | 3 | 1 | 3 | 1 | 2 | 1 | 5 | 0.8 | 13 | 0.9231 |
| 6_11945846 | 4 | 0 | 4 | 0 | 2 | 0.5 | 5 | 0 | 15 | 0.0667 | 3 | 0 | 3 | 0 | 2 | 0 | 4 | 0.25 | 12 | 0.0833 |
| 6_11945942 | 8 | 0.125 | 11 | 0.0909 | 5 | 0 | 8 | 0 | 32 | 0.0625 | 4 | 0.25 | 3 | 0 | 4 | 0 | 5 | 0 | 16 | 0.0625 |
| 6_11945944 | 8 | 0.25 | 11 | 0.0909 | 5 | 0.2 | 8 | 0.375 | 32 | 0.2188 | 4 | 0 | 3 | 0.3333 | 5 | 0 | 6 | 0.1667 | 18 | 0.1111 |
| 6_11945979 | 1 | 1 | 2 | 0 | 1 | 0 | 1 | 0 | 5 | 0.2 | NA | NA | 1 | 1 | 1 | 0 | 1 | 0 | 3 | 0.3333 |
| 6_11954862 | NA | NA | 4 | 0.25 | 2 | 0 | 2 | 0 | 8 | 0.125 | 1 | 0 | 1 | 0 | 2 | 0.5 | 4 | 0.25 | 8 | 0.25 |
| 6_11954908 | 1 | 0 | 4 | 0.5 | 5 | 0.4 | 3 | 0.3333 | 13 | 0.3846 | 2 | 0.5 | 1 | 1 | NA | NA | 2 | 0.5 | 5 | 0.6 |
| 6_11954909 | 1 | 1 | 4 | 0.5 | 4 | 0.75 | 2 | 0.5 | 11 | 0.6364 | 2 | 0 | 1 | 0 | 1 | 1 | 2 | 0.5 | 6 | 0.3333 |
| 6_11954920 | 1 | 0 | 4 | 0.5 | 3 | 0.3333 | 3 | 1 | 11 | 0.5455 | 2 | 1 | 1 | 1 | NA | NA | 2 | 0.5 | 5 | 0.8 |
| 6_11954921 | 1 | 1 | 4 | 0.75 | 2 | 1 | 2 | 1 | 9 | 0.8889 | 1 | 1 | 1 | 1 | NA | NA | 1 | 1 | 3 | 1 |
| 6_11954936 | 1 | 0 | 4 | 0.25 | 4 | 0.25 | 3 | 0 | 12 | 0.1667 | 2 | 0.5 | 1 | 0 | 1 | 0 | 3 | 0 | 7 | 0.1429 |
| 6_11954945 | 1 | 1 | 4 | 1 | 4 | 1 | 1 | 1 | 10 | 1 | 2 | 1 | 1 | 1 | 1 | 1 | 3 | 1 | 7 | 1 |
| 6_11954947 | 1 | 1 | 3 | 1 | 4 | 1 | 1 | 1 | 9 | 1 | 1 | 1 | 1 | 1 | 1 | 1 | 3 | 1 | 6 | 1 |
| 6_11954949 | 1 | 1 | 3 | 1 | 3 | 1 | 2 | 1 | 9 | 1 | 2 | 1 | 1 | 1 | 1 | 1 | 3 | 1 | 7 | 1 |
| 6_11954967 | 1 | 0 | 4 | 0.75 | 4 | 0.75 | 1 | 0 | 10 | 0.6 | 1 | 0 | 1 | 1 | 1 | 1 | 3 | 0.6667 | 6 | 0.6667 |
| 6_11954983 | 1 | 1 | 3 | 1 | 2 | 1 | 2 | 1 | 8 | 1 | 1 | 1 | 1 | 1 | 1 | 1 | 2 | 1 | 5 | 1 |
| 6_11954985 | 1 | 0 | 3 | 0 | 3 | 0 | 2 | 1 | 9 | 0.2222 | 2 | 0 | 1 | 1 | 1 | 1 | 3 | 0 | 7 | 0.2857 |
| 6_11954988 | 1 | 1 | 3 | 0.6667 | 2 | 0.5 | 2 | 1 | 8 | 0.75 | 2 | 0.5 | 1 | 1 | 1 | 0 | 3 | 0.3333 | 7 | 0.4286 |
| 6_11954995 | 1 | 0 | 3 | 0 | 3 | 0 | 2 | 0.5 | 9 | 0.1111 | 2 | 0 | 1 | 1 | 1 | 0 | 3 | 0 | 7 | 0.1429 |
| 6_11955174 | NA | NA | 2 | 0.5 | NA | NA | 1 | 1 | 3 | 0.6667 | 1 | 1 | NA | NA | 1 | 1 | 3 | 0.3333 | 5 | 0.6 |
| 6_11955199 | NA | NA | 2 | 0.5 | NA | NA | 1 | 1 | 3 | 0.6667 | 1 | 0 | NA | NA | 1 | 1 | 2 | 1 | 4 | 0.75 |

| | | | | | | | | | | | | | | | | | | | | |
|---|---|---|---|---|---|---|---|---|---|---|---|---|---|---|---|---|---|---|---|---|
| 6_1195522 | NA | NA | 2 | 1 | NA | NA | NA | NA | 2 | 1 | 1 | 1 | NA | NA | NA | NA | 2 | 0.5 | 3 | 0.6667 |
| 6_1195525 | NA | NA | 2 | 1 | NA | NA | NA | NA | 2 | 1 | NA | NA | NA | NA | NA | NA | 2 | 1 | 2 | 1 |
| 6_1197739 | NA | NA | NA | NA | NA | NA | 1 | 1 | 1 | 1 | NA | NA | NA | NA | NA | NA | 1 | 1 | 1 | 1 |
| 6_1198957 | NA | NA | 1 | 1 | NA | NA | 1 | 0 | 2 | 0.5 | 2 | 0.5 | NA | NA | 2 | 0.5 | NA | NA | 4 | 0.5 |
| 6_1198957 | 1 | 1 | 1 | 1 | NA | NA | 1 | 1 | 3 | 1 | 2 | 0 | NA | NA | 4 | 1 | 2 | 0.5 | 8 | 0.625 |
| 6_1198959 | 1 | 0 | 1 | 1 | NA | NA | 2 | 0 | 4 | 0.25 | 2 | 0 | NA | NA | 4 | 0.25 | 1 | 1 | 7 | 0.2857 |
| 6_1198995 | 2 | 0 | 1 | 1 | NA | NA | 2 | 1 | 5 | 0.6 | 2 | 1 | NA | NA | 4 | 1 | 4 | 0.75 | 10 | 0.9 |
| 6_1198995 | 2 | 0 | 1 | 0 | NA | NA | 2 | 0.5 | 5 | 0.2 | 2 | 0 | NA | NA | 2 | 1 | 2 | 0 | 6 | 0.3333 |
| 6_1198995 | 3 | 0.6667 | 1 | 0 | NA | NA | 2 | 0.5 | 6 | 0.5 | 2 | 1 | NA | NA | 4 | 0.75 | 4 | 1 | 10 | 0.9 |
| 6_1198996 | 2 | 0.5 | NA | NA | NA | NA | 2 | 0 | 4 | 0.25 | 2 | 0.5 | NA | NA | 3 | 0.6667 | 1 | 1 | 6 | 0.6667 |
| 6_1198997 | 2 | 0 | 1 | 0 | NA | NA | 1 | 1 | 4 | 0.25 | 2 | 0 | NA | NA | 4 | 0.25 | 4 | 0.25 | 10 | 0.2 |
| 6_1199011 | 14 | 0 | 22 | 0.2273 | 18 | 0.0556 | 13 | 0.1538 | 67 | 0.1194 | 10 | 0.1 | 14 | 0.2143 | 11 | 0.1818 | 13 | 0.1538 | 48 | 0.1667 |
| 6_1199012 | 14 | 0.0714 | 21 | 0.2381 | 19 | 0.1579 | 13 | 0.2308 | 67 | 0.1791 | 11 | 0 | 15 | 0.4 | 10 | 0.2 | 14 | 0 | 50 | 0.16 |
| 6_1199024 | 10 | 0.2 | 13 | 0.2308 | 12 | 0.0833 | 9 | 0 | 44 | 0.1364 | 9 | 0.3333 | 12 | 0.25 | 9 | 0.3333 | 6 | 0 | 36 | 0.25 |
| 6_1199027 | 8 | 0 | 16 | 0.125 | 11 | 0.0909 | 11 | 0.0909 | 46 | 0.087 | 10 | 0 | 11 | 0.1818 | 7 | 0 | 7 | 0 | 35 | 0.0571 |
| 6_1199030 | 8 | 0 | 14 | 0.1429 | 9 | 0 | 10 | 0.2 | 41 | 0.0976 | 9 | 0.1111 | 14 | 0.2143 | 4 | 0.5 | 8 | 0.25 | 35 | 0.2286 |
| 6_1199033 | 4 | 0.25 | 8 | 0 | 5 | 0.2 | 7 | 0 | 24 | 0.0833 | 4 | 0.5 | 9 | 0.3333 | 4 | 0.25 | 6 | 0.3333 | 23 | 0.3478 |
| 6_1199034 | 4 | 0 | 12 | 0.25 | 5 | 0 | 9 | 0 | 30 | 0.1 | 8 | 0 | 13 | 0.2308 | 6 | 0.1667 | 7 | 0 | 34 | 0.1176 |
| 6_1199146 | 4 | 0.25 | 7 | 0.8571 | 4 | 0.75 | 3 | 0.6667 | 18 | 0.6667 | 6 | 0.8333 | 9 | 0.5556 | 1 | 0 | 5 | 0.6 | 21 | 0.619 |
| 6_1199146 | 2 | 0.5 | 6 | 0.1667 | 4 | 0.25 | 2 | 0 | 14 | 0.2143 | 5 | 0.4 | 7 | 0.2857 | 2 | 0.5 | 3 | 0 | 17 | 0.2941 |
| 6_1199252 | 3 | 0 | 5 | 0 | 3 | 0 | 4 | 0.5 | 15 | 0.1333 | 4 | 0.25 | 6 | 0.1667 | 2 | 0.5 | 1 | 0 | 13 | 0.2308 |
| 6_1199252 | 3 | 0 | 5 | 1 | 3 | 1 | 4 | 0.25 | 15 | 0.6 | 5 | 0.2 | 6 | 0.8333 | 1 | 1 | 1 | 1 | 13 | 0.6154 |
| 6_1199261 | 1 | 0 | 4 | 0.25 | 1 | 0 | 3 | 0 | 9 | 0.1111 | 5 | 0 | 3 | 0.6667 | 1 | 0 | NA | NA | 9 | 0.2222 |
| 6_1199261 | 2 | 0 | 5 | 0.2 | 1 | 0 | 2 | 0 | 10 | 0.1 | 5 | 0 | 5 | 0.2 | 1 | 1 | NA | NA | 11 | 0.1818 |
| 6_1199262 | 2 | 0.5 | 4 | 0 | 1 | 1 | 1 | 0 | 8 | 0.25 | 4 | 0 | 5 | 0.2 | 1 | 0 | NA | NA | 10 | 0.1 |
| 6_1199262 | 2 | 0 | 5 | 0.2 | 1 | 0 | 2 | 0 | 10 | 0.1 | 4 | 0 | 5 | 0.4 | 1 | 0 | NA | NA | 10 | 0.2 |
| 6_1199262 | 2 | 0 | 5 | 0.2 | 1 | 0 | 2 | 0 | 10 | 0.1 | 4 | 0 | 5 | 0.2 | 1 | 0 | NA | NA | 10 | 0.1 |
| 6_1199262 | 1 | 0 | 5 | 0.6 | 1 | 0 | 2 | 0.5 | 9 | 0.4444 | 3 | 0.3333 | 5 | 0.4 | 1 | 1 | NA | NA | 9 | 0.4444 |
| 6_1199397 | 2 | 1 | NA | NA | NA | NA | NA | NA | 2 | 1 | 1 | 1 | 1 | 1 | NA | NA | NA | NA | 2 | 1 |
| 6_1199397 | 2 | 0.5 | NA | NA | NA | NA | NA | NA | 2 | 0.5 | NA | NA | 1 | 1 | NA | NA | NA | NA | 1 | 1 |
| 6_1199400 | 2 | 0.5 | NA | NA | NA | NA | NA | NA | 2 | 0.5 | 1 | 0 | 1 | 1 | NA | NA | NA | NA | 2 | 0.5 |
| 6_1199403 | 2 | 1 | NA | NA | NA | NA | NA | NA | 2 | 1 | 1 | 0 | 1 | 1 | NA | NA | NA | NA | 2 | 0.5 |
| 6_1200147 | 1 | 1 | NA | NA | NA | NA | 1 | 0 | 2 | 0.5 | 1 | 0 | 2 | 0.5 | NA | NA | NA | NA | 3 | 0.3333 |
| 6_1200148 | 1 | 1 | NA | NA | NA | NA | 1 | 1 | 2 | 1 | 1 | 1 | 2 | 1 | NA | NA | NA | NA | 3 | 1 |
| 6_1200152 | 1 | 0 | NA | NA | NA | NA | 1 | 1 | 2 | 0.5 | 1 | 1 | 1 | 0 | NA | NA | NA | NA | 2 | 0.5 |
| 6_1204039 | 2 | 0.5 | 1 | 1 | 1 | 1 | NA | NA | 4 | 0.75 | NA | NA | 2 | 1 | 1 | 1 | 2 | 1 | 5 | 1 |
| 6_1204040 | 2 | 1 | 1 | 1 | 1 | 1 | NA | NA | 4 | 1 | NA | NA | 2 | 1 | 1 | 1 | 2 | 1 | 5 | 1 |
| 6_1204042 | 1 | 0 | 1 | 0 | 1 | 1 | NA | NA | 3 | 0.3333 | NA | NA | 2 | 0.5 | 1 | 1 | 1 | 1 | 4 | 0.75 |
| 6_1204088 | 2 | 0 | 3 | 0.3333 | 1 | 0 | NA | NA | 6 | 0.1667 | NA | NA | 2 | 0.5 | 2 | 0 | 2 | 0.5 | 6 | 0.3333 |
| 6_1204089 | 2 | 0.5 | 3 | 1 | 1 | 1 | NA | NA | 6 | 0.8333 | NA | NA | 3 | 1 | 1 | 1 | 3 | 0.6667 | 7 | 0.8571 |
| 6_1204160 | 2 | 1 | 4 | 1 | 1 | 1 | 1 | 0 | 8 | 0.875 | NA | NA | 3 | 1 | 2 | 0.5 | 2 | 1 | 7 | 0.8571 |
| 6_1204161 | 3 | 0.6667 | 4 | 0.25 | 2 | 0.5 | 3 | 0.3333 | 12 | 0.4167 | NA | NA | 3 | 0.6667 | 3 | 0.6667 | 3 | 0.6667 | 9 | 0.6667 |
| 6_1204161 | 2 | 1 | 4 | 0.75 | 1 | 1 | 1 | 1 | 8 | 0.875 | NA | NA | 3 | 1 | 3 | 0.6667 | 3 | 0.3333 | 9 | 0.6667 |
| 6_1204162 | 3 | 0.3333 | 3 | 0.3333 | 2 | 1 | 2 | 1 | 10 | 0.6 | NA | NA | 3 | 0.6667 | 3 | 0.6667 | 3 | 0.3333 | 9 | 0.5556 |
| 6_1204162 | 3 | 1 | 3 | 1 | 2 | 1 | 2 | 1 | 10 | 1 | NA | NA | 3 | 1 | 3 | 1 | 2 | 1 | 8 | 1 |
| 6_1204164 | 2 | 0.5 | 2 | 0.5 | 2 | 0.5 | 3 | 0 | 9 | 0.3333 | NA | NA | 2 | 0.5 | 3 | 0.6667 | 3 | 0.6667 | 8 | 0.625 |
| 6_1204179 | 2 | 0 | 4 | 0 | 2 | 1 | 3 | 0 | 11 | 0.1818 | 1 | 1 | 4 | 0.25 | 2 | 0 | 2 | 0 | 9 | 0.2222 |
| 6_1204204 | NA | NA | NA | NA | NA | NA | 1 | 1 | 1 | 1 | NA | NA | 2 | 0.5 | NA | NA | 1 | 1 | 3 | 0.6667 |
| 6_1204660 | 2 | 1 | 8 | 0.625 | 4 | 0.5 | 6 | 0.6667 | 20 | 0.65 | NA | NA | 8 | 0.875 | 8 | 0.5 | 8 | 0.375 | 24 | 0.5833 |
| 6_1204662 | 6 | 0.1667 | 8 | 0.125 | 4 | 0.25 | 7 | 0.1429 | 25 | 0.16 | NA | NA | 7 | 0.4286 | 6 | 0.1667 | 6 | 0.1667 | 19 | 0.2632 |
| 6_1204664 | 7 | 0.4286 | 7 | 0 | 4 | 0.25 | 4 | 0 | 22 | 0.1818 | NA | NA | 6 | 0 | 7 | 0 | 6 | 0.1667 | 19 | 0.0526 |
| 6_1204666 | 4 | 0 | 7 | 0.2857 | 4 | 0 | 5 | 0 | 20 | 0.1 | NA | NA | 6 | 0 | 5 | 0 | 6 | 0.1667 | 17 | 0.0588 |
| 6_1204667 | 7 | 0.5714 | 7 | 0.2857 | 4 | 0.25 | 5 | 0.6 | 23 | 0.4348 | NA | NA | 5 | 0.2 | 3 | 0.3333 | 5 | 0.2 | 13 | 0.2308 |
| 6_1204667 | 7 | 0.2857 | 7 | 0.1429 | 4 | 0.25 | 5 | 0.6 | 23 | 0.3043 | NA | NA | 7 | 0.2857 | 6 | 0 | 6 | 0.3333 | 19 | 0.2105 |
| 6_1204667 | 6 | 0.5 | 7 | 0.8571 | 4 | 0.75 | 7 | 0.8571 | 24 | 0.75 | NA | NA | 7 | 1 | 6 | 0.3333 | 6 | 0.5 | 19 | 0.6316 |
| 6_1204669 | 4 | 0.75 | 4 | 1 | 4 | 1 | 6 | 1 | 18 | 0.9444 | NA | NA | 5 | 1 | 3 | 0.6667 | 3 | 1 | 11 | 0.9091 |
| 6_1204669 | 5 | 0.6 | 6 | 1 | 4 | 0.75 | 5 | 0.8 | 20 | 0.8 | NA | NA | 5 | 0.6 | 3 | 0.3333 | 5 | 1 | 13 | 0.6923 |
| 6_1204670 | 2 | 1 | 3 | 1 | 4 | 1 | 1 | 1 | 10 | 1 | NA | NA | 3 | 1 | 2 | 1 | 2 | 1 | 7 | 1 |
| 6_1204670 | 1 | 0 | 4 | 0.5 | 1 | 0 | 4 | 0.5 | 10 | 0.4 | NA | NA | 2 | 0 | 1 | 1 | 3 | 0.6667 | 6 | 0.5 |
| 6_1204670 | 3 | 0.3333 | 6 | 0.5 | 2 | 0.5 | 5 | 0.4 | 16 | 0.4375 | NA | NA | 6 | 0.6667 | 5 | 0.2 | 4 | 0.5 | 15 | 0.4667 |
| 6_1204671 | 4 | 0.25 | 5 | 0 | 3 | 0 | NA | NA | 12 | 0.0833 | NA | NA | 3 | 0 | 4 | 0.5 | 3 | 0.3333 | 10 | 0.3 |
| 6_1204671 | 5 | 0.4 | 5 | 0.6 | 3 | 0.3333 | 4 | 0.75 | 17 | 0.5294 | NA | NA | 5 | 0.2 | 5 | 0.6 | 5 | 0.6 | 15 | 0.4667 |
| 6_1204674 | 2 | 0 | 4 | 0.5 | 2 | 0 | 4 | 0.5 | 12 | 0.3333 | NA | NA | 6 | 0.1667 | 3 | 0.6667 | 3 | 0.3333 | 12 | 0.3333 |
| 6_1204685 | 1 | 1 | 1 | 1 | NA | NA | 2 | 1 | 4 | 1 | NA | NA | 3 | 1 | NA | NA | 5 | 1 | 8 | 1 |
| 6_1204688 | 1 | 1 | 1 | 1 | NA | NA | 3 | 1 | 5 | 1 | NA | NA | 4 | 1 | 3 | 1 | 2 | 1 | 9 | 1 |

| | | | | | | | | | | | | | | | | | | | | |
|---|---|---|---|---|---|---|---|---|---|---|---|---|---|---|---|---|---|---|---|---|
| 6_1204691 | 1 | 1 | 1 | 1 | NA | NA | 2 | 1 | 4 | 1 | NA | NA | 3 | 1 | 2 | 1 | 2 | 1 | 7 | 1 |
| 6_1204693 | NA | NA | 1 | 1 | NA | NA | 1 | 1 | 2 | 1 | NA | NA | 1 | 1 | 1 | 1 | NA | NA | 2 | 1 |
| 6_1204695 | NA | NA | 1 | 1 | NA | NA | 2 | 1 | 3 | 1 | NA | NA | 2 | 1 | 1 | 1 | 1 | 1 | 4 | 1 |
| 6_1204695 | NA | NA | 1 | 1 | NA | NA | 2 | 1 | 3 | 1 | NA | NA | 2 | 1 | 2 | 1 | 1 | 1 | 5 | 1 |
| 6_1204697 | NA | NA | 1 | 1 | NA | NA | 2 | 1 | 3 | 1 | NA | NA | 2 | 1 | NA | NA | 1 | 1 | 3 | 1 |
| 6_1204700 | NA | NA | 1 | 0 | NA | NA | 2 | 0.5 | 3 | 0.3333 | NA | NA | 1 | 0 | 2 | 0.5 | 2 | 0 | 5 | 0.2 |
| 6_1204700 | NA | NA | 1 | 0 | NA | NA | 2 | 0.5 | 3 | 0.3333 | NA | NA | 1 | 0 | 2 | 1 | 1 | 0 | 4 | 0.5 |
| 6_1204703 | NA | NA | 1 | 1 | NA | NA | 2 | 0.5 | 3 | 0.6667 | NA | NA | NA | NA | 2 | 1 | 1 | 0 | 3 | 0.6667 |
| 6_1204833 | NA | NA | NA | NA | 1 | 1 | NA | NA | 1 | 1 | 1 | 1 | NA | NA | NA | NA | NA | NA | 1 | 1 |
| 6_1204865 | 2 | 0 | 2 | 0 | 3 | 0 | 1 | 1 | 8 | 0.125 | 2 | 0 | 2 | 0 | 2 | 0 | 3 | 0.3333 | 9 | 0.1111 |
| 6_1204866 | 2 | 1 | 3 | 0.3333 | 3 | 1 | 1 | 1 | 9 | 0.7778 | 2 | 1 | 3 | 1 | 2 | 1 | 3 | 1 | 10 | 1 |
| 6_1204869 | 1 | 0 | 3 | 0.3333 | 3 | 0 | 1 | 0 | 8 | 0.125 | 2 | 0.5 | 3 | 0.6667 | 1 | 0 | 3 | 0.3333 | 9 | 0.4444 |
| 6_1204869 | 2 | 1 | 1 | 1 | 2 | 0.5 | 1 | 1 | 6 | 0.8333 | 1 | 1 | 2 | 1 | 2 | 0.5 | 1 | 1 | 6 | 0.8333 |
| 6_1204872 | 2 | 0 | 3 | 0.6667 | 3 | 0.6667 | NA | NA | 8 | 0.5 | 2 | 0.5 | 3 | 1 | 2 | 1 | 3 | 0.3333 | 10 | 0.7 |
| 6_1204872 | 1 | 0 | 3 | 0.3333 | 3 | 0.6667 | 1 | 1 | 8 | 0.5 | 2 | 0.5 | 3 | 0.3333 | 2 | 1 | 3 | 1 | 10 | 0.7 |
| 6_1207725 | 78 | 0.1795 | 84 | 0.2262 | 70 | 0.1571 | 59 | 0.2373 | 291 | 0.1993 | 66 | 0.1818 | 65 | 0.2923 | 71 | 0.1408 | 71 | 0.3521 | 273 | 0.2418 |
| 6_1207726 | 27 | 0.3333 | 30 | 0.4 | 29 | 0.2759 | 29 | 0.5517 | 115 | 0.3913 | 23 | 0.4783 | 18 | 0.4444 | 21 | 0.4286 | 26 | 0.2692 | 88 | 0.3977 |
| 6_1208637 | 1 | 1 | NA | NA | 2 | 0 | 2 | 0 | 5 | 0.2 | NA | NA | 1 | 0 | 3 | 0.3333 | 4 | 0.5 | 8 | 0.375 |
| 6_1208637 | 1 | 0 | NA | NA | 2 | 1 | 2 | 0 | 5 | 0.4 | NA | NA | 1 | 0 | 3 | 0 | 3 | 0.3333 | 7 | 0.1429 |
| 6_1208638 | 1 | 1 | NA | NA | 2 | 0.5 | 3 | 1 | 6 | 0.8333 | NA | NA | 1 | 0 | 3 | 1 | 4 | 0.75 | 8 | 0.75 |
| 6_1208641 | 1 | 0 | 1 | 1 | 3 | 0.6667 | 3 | 0 | 8 | 0.375 | NA | NA | 1 | 0 | 3 | 0.6667 | 4 | 0 | 8 | 0.25 |
| 6_1208643 | 2 | 0 | 2 | 0 | 3 | 0 | 2 | 0.5 | 9 | 0.1111 | 2 | 0 | 1 | 0 | 3 | 0 | 4 | 0.25 | 10 | 0.1 |
| 6_1208643 | 2 | 0 | 2 | 0.5 | 3 | 0 | 3 | 0 | 10 | 0.1 | 2 | 0.5 | 1 | 0 | 2 | 0.5 | 4 | 0.5 | 9 | 0.4444 |
| 6_1208643 | 2 | 0.5 | 2 | 1 | 3 | 0.3333 | 3 | 0.6667 | 10 | 0.6 | 2 | 0 | 1 | 0 | 4 | 0.5 | 3 | 0.3333 | 10 | 0.3 |
| 6_1208644 | 2 | 0.5 | 1 | 1 | 2 | 0 | 2 | 0.5 | 7 | 0.4286 | 2 | 0 | NA | NA | 4 | 1 | 3 | 0.6667 | 9 | 0.6667 |
| 6_1208644 | 2 | 1 | 2 | 0.5 | 2 | 0.5 | 2 | 0.5 | 8 | 0.625 | 1 | 1 | 1 | 1 | 3 | 1 | 3 | 0.3333 | 8 | 0.75 |
| 6_1208644 | 2 | 1 | 2 | 1 | 3 | 1 | 3 | 1 | 10 | 1 | 1 | 1 | 1 | 1 | 4 | 1 | 4 | 1 | 10 | 1 |
| 6_1208644 | 2 | 0.5 | 2 | 0.5 | 3 | 1 | 3 | 0.6667 | 10 | 0.7 | NA | NA | 1 | 1 | 4 | 1 | 2 | 0.5 | 7 | 0.8571 |
| 6_1208644 | 2 | 0.5 | 2 | 1 | 3 | 0.6667 | 2 | 1 | 9 | 0.7778 | 2 | 0 | 1 | 0 | 4 | 0.5 | 4 | 1 | 11 | 0.5455 |
| 6_1208654 | 2 | 0.5 | 2 | 0.5 | 3 | 0 | 3 | 0.3333 | 10 | 0.3 | 1 | 0 | 1 | 1 | 2 | 0.5 | 6 | 0.1667 | 10 | 0.3 |
| 6_1208656 | 2 | 0.5 | 2 | 0.5 | 2 | 0 | 3 | 0 | 9 | 0.2222 | 2 | 0.5 | 2 | 0 | 4 | 0.25 | 5 | 0.2 | 13 | 0.2308 |
| 6_1208656 | 2 | 0.5 | 1 | 0 | 3 | 0 | 3 | 0 | 9 | 0.1111 | 1 | 0 | 2 | 0.5 | 4 | 0.25 | 6 | 0 | 13 | 0.1538 |
| 6_1208657 | 2 | 0 | 2 | 0.5 | 3 | 0.3333 | 3 | 0 | 10 | 0.2 | 1 | 0 | 1 | 0 | 4 | 0.5 | 6 | 0 | 12 | 0.1667 |
| 6_1208657 | 2 | 1 | 1 | 0 | 3 | 0 | 2 | 0 | 8 | 0.25 | 2 | 0 | 2 | 0.5 | 4 | 0.25 | 6 | 0.1667 | 14 | 0.2143 |
| 6_1208658 | 1 | 0 | 2 | 0.5 | 3 | 0 | 3 | 0 | 9 | 0.1111 | 1 | 0 | 3 | 0 | 3 | 0.3333 | 6 | 0 | 13 | 0.0769 |
| 6_1208671 | 2 | 0 | NA | NA | 1 | 1 | 2 | 0 | 5 | 0.2 | 1 | 0 | 3 | 0.3333 | 1 | 0 | 2 | 1 | 7 | 0.4286 |
| 6_1208676 | 2 | 0.5 | NA | NA | 1 | 1 | 2 | 0 | 5 | 0.4 | 1 | 0 | 2 | 0 | 1 | 1 | 3 | 0 | 7 | 0.1429 |
| 6_1208676 | 2 | 0 | NA | NA | 1 | 0 | 2 | 0.5 | 5 | 0.2 | 1 | 0 | 2 | 0 | 1 | 1 | 3 | 0 | 7 | 0.1429 |
| 6_1208702 | NA | NA | NA | NA | NA | NA | 1 | 1 | 1 | 1 | 1 | 1 | NA | NA | NA | NA | NA | NA | 1 | 1 |
| 6_1208703 | 1 | 1 | NA | NA | NA | NA | 2 | 0.5 | 3 | 0.6667 | NA | NA | NA | NA | NA | NA | 2 | 1 | 2 | 1 |
| 6_1208703 | 2 | 1 | 2 | 0.5 | NA | NA | 2 | 1 | 6 | 0.8333 | 3 | 0.6667 | NA | NA | NA | NA | 2 | 0.5 | 5 | 0.6 |
| 6_1208705 | 1 | 1 | 1 | 1 | NA | NA | 2 | 0.5 | 4 | 0.75 | 3 | 0.3333 | NA | NA | NA | NA | 2 | 0 | 5 | 0.2 |
| 6_1208705 | 2 | 0.5 | 1 | 1 | NA | NA | 2 | 0.5 | 5 | 0.6 | 3 | 0.3333 | NA | NA | NA | NA | 2 | 0.5 | 5 | 0.4 |
| 6_1208706 | 2 | 1 | 1 | 1 | NA | NA | 1 | 1 | 4 | 1 | 2 | 1 | NA | NA | NA | NA | 2 | 1 | 4 | 1 |
| 6_1208707 | 2 | 1 | 1 | 0 | NA | NA | 1 | 0 | 4 | 0.5 | 3 | 0 | NA | NA | NA | NA | 1 | 1 | 4 | 0.25 |
| 6_1208708 | 2 | 0.5 | 1 | 1 | NA | NA | 2 | 0.5 | 5 | 0.6 | 2 | 0 | NA | NA | NA | NA | 2 | 1 | 4 | 0.5 |
| 6_1208708 | 1 | 1 | NA | NA | NA | NA | 1 | 1 | 2 | 1 | 2 | 0 | NA | NA | NA | NA | 1 | 1 | 3 | 0.3333 |
| 6_1208709 | 2 | 1 | 1 | 1 | NA | NA | 2 | 1 | 5 | 1 | 2 | 1 | NA | NA | NA | NA | 1 | 1 | 3 | 1 |
| 6_1208711 | 2 | 0.5 | 1 | 1 | NA | NA | 1 | 0 | 4 | 0.5 | 3 | 1 | NA | NA | NA | NA | 2 | 1 | 5 | 1 |
| 6_1209088 | 3 | 0.6667 | 9 | 0.6667 | 6 | 0.6667 | 8 | 0.75 | 26 | 0.6923 | 3 | 1 | 5 | 0.8 | 7 | 0.7143 | 5 | 0.8 | 20 | 0.8 |
| 6_1209088 | 4 | 0.25 | 11 | 0.5455 | 6 | 0.5 | 7 | 0.8571 | 28 | 0.5714 | 2 | 1 | 6 | 0.5 | 8 | 0.625 | 5 | 0.4 | 21 | 0.5714 |
| 6_1209089 | 2 | 0.5 | 8 | 0.375 | 5 | 0.2 | 9 | 0.4444 | 24 | 0.375 | 2 | 0 | 5 | 0.4 | 8 | 0 | 6 | 0.5 | 21 | 0.2381 |
| 6_1209089 | 3 | 1 | 11 | 0.8182 | 5 | 0.6 | 8 | 0.75 | 27 | 0.7778 | 2 | 0.5 | 4 | 0.75 | 9 | 0.8889 | 5 | 0.8 | 20 | 0.8 |
| 6_1209090 | 3 | 0.6667 | 8 | 1 | 3 | 1 | 7 | 1 | 21 | 0.9524 | 1 | 1 | 2 | 1 | 8 | 1 | 3 | 1 | 14 | 1 |
| 6_1209090 | 3 | 0 | 12 | 0.25 | 7 | 0.1429 | 8 | 0.25 | 30 | 0.2 | 2 | 0 | 5 | 0.2 | 8 | 0 | 4 | 0.25 | 19 | 0.1053 |
| 6_1209093 | 2 | 1 | 12 | 0.9167 | 6 | 1 | 7 | 0.8571 | 27 | 0.9259 | 3 | 1 | 5 | 1 | 7 | 1 | 5 | 0.8 | 20 | 0.95 |
| 6_1209094 | 2 | 0.5 | 5 | 0 | 5 | 0 | 5 | 0.2 | 17 | 0.1176 | 2 | 0.5 | 3 | 0.3333 | 7 | 0 | 5 | 0 | 17 | 0.1176 |
| 6_1209096 | 3 | 0 | 11 | 0.0909 | 5 | 0 | 5 | 0.2 | 24 | 0.0833 | 3 | 0 | 4 | 0.25 | 8 | 0 | 5 | 0 | 20 | 0.05 |
| 6_1209096 | 2 | 0.5 | 10 | 0.2 | 4 | 0.25 | 5 | 0.4 | 21 | 0.2857 | 2 | 0 | 4 | 0.25 | 8 | 0.125 | 5 | 0.2 | 19 | 0.1579 |
| 6_1209096 | NA | NA | 8 | 0 | 4 | 0.75 | 2 | 0 | 14 | 0.2143 | 1 | 1 | 4 | 0.5 | 6 | 0.5 | 4 | 0 | 15 | 0.4 |
| 6_1209096 | 3 | 0.6667 | 7 | 0.8571 | 5 | 0.8 | 4 | 0.5 | 19 | 0.7368 | 3 | 0.6667 | 4 | 1 | 6 | 1 | 4 | 1 | 17 | 0.9412 |
| 6_1209096 | 2 | 0 | 8 | 0.125 | 3 | 0.3333 | 4 | 0.25 | 17 | 0.1765 | 1 | 0 | 3 | 0.3333 | 5 | 0 | 3 | 0 | 12 | 0.0833 |
| 6_1209097 | 2 | 0.5 | 8 | 1 | 3 | 1 | 3 | 1 | 16 | 0.9375 | 1 | 1 | 4 | 1 | 7 | 0.8571 | 4 | 1 | 16 | 0.9375 |
| 6_1209099 | 2 | 1 | 10 | 0.5 | 3 | 0.6667 | 3 | 0.3333 | 18 | 0.5556 | 3 | 0 | 5 | 1 | 4 | 0.5 | 3 | 0.6667 | 15 | 0.6 |
| 6_1209100 | 2 | 0.5 | 8 | 0.5 | 3 | 0.6667 | 2 | 1 | 15 | 0.6 | 2 | 1 | 4 | 0.5 | 4 | 1 | 3 | 1 | 13 | 0.8462 |

| | | | | | | | | | | | | | | | | | | | | |
|---|---|---|---|---|---|---|---|---|---|---|---|---|---|---|---|---|---|---|---|---|
| 6_1209101 | 1 | 0 | 10 | 0.2 | 3 | 0 | 3 | 0 | 17 | 0.1176 | 2 | 0 | 5 | 0.2 | 6 | 0.1667 | 5 | 0 | 18 | 0.1111 |
| 6_1209102 | 1 | 1 | 9 | 0.2222 | 1 | 0 | 2 | 0 | 13 | 0.2308 | 1 | 0 | 4 | 0.25 | 3 | 0.3333 | 5 | 0.2 | 13 | 0.2308 |
| 6_1209102 | 2 | 0 | 9 | 0.1111 | 4 | 0 | 3 | 0 | 18 | 0.0556 | 2 | 0.5 | 3 | 0 | 6 | 0 | 5 | 0 | 16 | 0.0625 |
| 6_1209102 | 1 | 1 | 9 | 0.1111 | 4 | 0.25 | 2 | 0 | 16 | 0.1875 | 2 | 0 | 3 | 0.3333 | 6 | 0.3333 | 4 | 0.5 | 15 | 0.3333 |
| 6_1209103 | 2 | 0.5 | 9 | 0.5556 | 4 | 0.25 | 3 | 0.3333 | 18 | 0.4444 | 3 | 0.6667 | 4 | 0.25 | 6 | 0.5 | 5 | 0.8 | 18 | 0.5556 |
| 6_1209104 | 2 | 1 | 6 | 1 | 3 | 1 | 3 | 1 | 14 | 1 | 2 | 1 | 3 | 1 | 6 | 0.8333 | 5 | 1 | 16 | 0.9375 |
| 6_1209107 | 2 | 1 | 7 | 0.8571 | 3 | 1 | 2 | 0.5 | 14 | 0.8571 | 2 | 1 | 1 | 1 | 6 | 1 | 1 | 0 | 10 | 0.9 |
| 6_1209107 | 2 | 0 | 9 | 0.3333 | 3 | 1 | 3 | 0 | 17 | 0.3529 | 1 | 0 | 4 | 0.25 | 6 | 0.1667 | 4 | 0.75 | 15 | 0.3333 |
| 6_1209108 | 2 | 0.5 | 8 | 0.625 | 4 | 0.75 | 3 | 0.6667 | 17 | 0.6471 | 1 | 0 | 3 | 0.6667 | 6 | 0.6667 | 3 | 1 | 13 | 0.6923 |
| 6_1209109 | 3 | 1 | 5 | 1 | 3 | 1 | 2 | 0.5 | 13 | 0.9231 | 1 | 1 | 3 | 1 | 6 | 1 | 5 | 0.8 | 15 | 0.9333 |
| 6_1209109 | 3 | 0.3333 | 7 | 0 | 4 | 0 | 1 | 0 | 15 | 0.0667 | 2 | 0 | 2 | 0 | 6 | 0.1667 | 5 | 0 | 15 | 0.0667 |
| 6_1209110 | 2 | 0.5 | 7 | 0.8571 | 3 | 1 | 2 | 1 | 14 | 0.8571 | 2 | 1 | 2 | 1 | 4 | 1 | 4 | 1 | 12 | 1 |
| 6_1209111 | 1 | 0 | 6 | 1 | 3 | 0.6667 | 2 | 0.5 | 12 | 0.75 | 2 | 1 | 3 | 1 | 3 | 0.6667 | 3 | 0.6667 | 11 | 0.8182 |
| 6_1209114 | 2 | 0 | 5 | 0.8 | 3 | 0.6667 | 2 | 0.5 | 12 | 0.5833 | 2 | 0 | 2 | 0 | 6 | 0.6667 | 3 | 0 | 13 | 0.3077 |
| 6_1209115 | 1 | 0 | 8 | 0.25 | 3 | 0 | 2 | 0 | 14 | 0.1429 | 2 | 0.5 | 2 | 0 | 6 | 0.1667 | 3 | 0.3333 | 13 | 0.2308 |
| 6_1209116 | 2 | 0.5 | 7 | 1 | 3 | 1 | 2 | 1 | 14 | 0.9286 | 2 | 1 | 1 | 1 | 5 | 1 | 4 | 1 | 12 | 1 |
| 6_1209116 | 2 | 0.5 | 8 | 0.25 | 3 | 0.3333 | 2 | 0 | 15 | 0.2667 | 2 | 0.5 | 2 | 0.5 | 5 | 0.2 | 4 | 0 | 13 | 0.2308 |
| 6_1209116 | 2 | 0 | 5 | 0.2 | 3 | 0 | 2 | 0 | 12 | 0.0833 | 1 | 0 | 2 | 0 | 4 | 0.25 | 4 | 0 | 11 | 0.0909 |
| 6_1209129 | NA | NA | 2 | 0.5 | NA | NA | NA | NA | 2 | 0.5 | NA | NA | NA | NA | 1 | 1 | NA | NA | 1 | 1 |
| 6_1209359 | 1 | 0 | 4 | 0.25 | NA | NA | NA | NA | 5 | 0.2 | 1 | 0 | NA | NA | 3 | 0.3333 | 2 | 0.5 | 6 | 0.3333 |
| 6_1209360 | 1 | 0 | 2 | 0.5 | NA | NA | NA | NA | 3 | 0.3333 | 1 | 1 | NA | NA | 3 | 0.3333 | 2 | 1 | 6 | 0.6667 |
| 6_1209361 | 1 | 0 | 4 | 0.5 | NA | NA | NA | NA | 5 | 0.4 | 1 | 1 | 1 | 0 | 1 | 0 | 2 | 0 | 5 | 0.2 |
| 6_1209362 | 1 | 1 | 4 | 0.5 | NA | NA | NA | NA | 5 | 0.6 | NA | NA | 1 | 1 | 2 | 0.5 | 2 | 0 | 5 | 0.4 |
| 6_1209366 | 1 | 1 | 2 | 0.5 | NA | NA | NA | NA | 3 | 0.6667 | 1 | 1 | 1 | 0 | 2 | 0.5 | 1 | 1 | 5 | 0.6 |
| 6_1211705 | 1 | 0 | 1 | 0 | 2 | 0.5 | 1 | 1 | 5 | 0.4 | 1 | 1 | 2 | 0.5 | NA | NA | 1 | 1 | 4 | 0.75 |
| 6_1211707 | 2 | 0.5 | 1 | 0 | 2 | 0.5 | 1 | 1 | 6 | 0.5 | 1 | 0 | 1 | 0 | NA | NA | 2 | 1 | 4 | 0.5 |
| 6_1211707 | 2 | 0.5 | NA | NA | 2 | 0 | 1 | 1 | 5 | 0.4 | 1 | 0 | 1 | 0 | NA | NA | 2 | 0.5 | 4 | 0.25 |
| 6_1211725 | 2 | 0.5 | 2 | 0 | 3 | 0 | 2 | 0 | 9 | 0.1111 | 2 | 0 | 4 | 0.5 | 1 | 0 | 2 | 0 | 9 | 0.2222 |
| 6_1211726 | 2 | 0.5 | 2 | 0.5 | 3 | 0.6667 | 2 | 0 | 9 | 0.4444 | 2 | 1 | 4 | 0.25 | 1 | 0 | 2 | 1 | 9 | 0.5556 |
| 6_1211727 | 2 | 0 | 1 | 1 | 2 | 0 | 2 | 0 | 7 | 0.1429 | 1 | 0 | 4 | 0 | 1 | 1 | 2 | 0 | 8 | 0.125 |
| 6_1211729 | 1 | 0 | 2 | 0.5 | 3 | 0 | 2 | 0.5 | 8 | 0.25 | 2 | 0 | 3 | 0.3333 | 1 | 0 | 1 | 1 | 7 | 0.2857 |
| 6_1211999 | NA | NA | 4 | 0.75 | 2 | 0.5 | NA | NA | 6 | 0.6667 | 1 | 1 | 2 | 0.5 | 1 | 0 | NA | NA | 4 | 0.5 |
| 6_1212000 | NA | NA | 6 | 1 | 3 | 0.6667 | NA | NA | 9 | 0.8889 | 1 | 1 | 2 | 1 | 1 | 1 | NA | NA | 4 | 1 |
| 6_1212029 | 1 | 0 | 5 | 0.6 | 3 | 1 | NA | NA | 9 | 0.6667 | 2 | 1 | 2 | 1 | 3 | 0.6667 | NA | NA | 7 | 0.8571 |
| 6_1213144 | 2 | 0.5 | 4 | 0.25 | 4 | 0 | 4 | 0 | 14 | 0.1429 | NA | NA | NA | NA | 1 | 1 | 1 | 0 | 2 | 0.5 |
| 6_1213425 | 1 | 1 | NA | NA | NA | NA | 1 | 1 | 2 | 1 | 1 | 0 | 1 | 1 | NA | NA | NA | NA | 2 | 0.5 |
| 6_1213425 | 1 | 0 | NA | NA | 1 | 1 | 2 | 0 | 4 | 0.25 | 1 | 1 | 1 | 0 | NA | NA | NA | NA | 2 | 0.5 |
| 6_1213507 | 9 | 0 | 11 | 0.0909 | 6 | 0.1667 | 10 | 0.1 | 36 | 0.0833 | 7 | 0.1429 | 9 | 0 | 11 | 0.0909 | 9 | 0 | 36 | 0.0556 |
| 6_1213517 | 5 | 0.2 | 7 | 0.1429 | 4 | 0.5 | 6 | 0 | 22 | 0.1818 | 4 | 0.25 | 7 | 0 | 9 | 0.1111 | 3 | 0 | 23 | 0.087 |
| 6_1213519 | 6 | 0 | 9 | 0.2222 | 5 | 0 | 7 | 0 | 27 | 0.0741 | 5 | 0.2 | 9 | 0 | 10 | 0.1 | 5 | 0 | 29 | 0.069 |
| 6_1213523 | 4 | 0.5 | 8 | 0.5 | 1 | 0 | 6 | 0.3333 | 19 | 0.4211 | 6 | 0.5 | 7 | 0.2857 | 8 | 0.5 | 4 | 0.25 | 25 | 0.4 |
| 6_1213527 | 5 | 0.4 | 4 | 0 | 3 | 0.3333 | 5 | 0.6 | 17 | 0.3529 | 4 | 0.5 | 9 | 0.4444 | 8 | 0.625 | 6 | 0.5 | 27 | 0.5185 |
| 6_1213528 | 4 | 0.5 | 5 | 1 | 2 | 0 | 3 | 0.6667 | 14 | 0.6429 | 2 | 0.5 | 7 | 0.5714 | 4 | 0.75 | 4 | 1 | 17 | 0.7059 |
| 6_1213531 | 5 | 0.6 | 4 | 0.75 | 2 | 0 | 5 | 0.4 | 16 | 0.5 | 4 | 0.25 | 7 | 0.1429 | 4 | 0.25 | 3 | 0.3333 | 18 | 0.2222 |
| 6_1213532 | 5 | 0.2 | 2 | 0 | 2 | 0.5 | 4 | 0.25 | 13 | 0.2308 | 1 | 0 | 6 | 0.5 | 4 | 0.5 | 3 | 0.3333 | 14 | 0.4286 |
| 6_1213898 | 2 | 0 | 1 | 1 | 1 | 0 | 1 | 0 | 5 | 0.2 | NA | NA | 1 | 1 | 2 | 0.5 | 1 | 0 | 4 | 0.5 |
| 6_1213898 | 2 | 0 | 1 | 1 | 1 | 1 | 1 | 0 | 5 | 0.4 | NA | NA | 1 | 0 | 2 | 0.5 | NA | NA | 3 | 0.3333 |
| 6_1213902 | 2 | 1 | NA | NA | 1 | 0 | 1 | 1 | 4 | 0.75 | 1 | 1 | NA | NA | 1 | 1 | NA | NA | 2 | 1 |
| 6_1214501 | 2 | 0 | NA | NA | 1 | 1 | 1 | 0 | 4 | 0.25 | NA | NA | NA | NA | 3 | 0.6667 | NA | NA | 3 | 0.6667 |
| 6_1222904 | NA | NA | 1 | 1 | 1 | 1 | 1 | 0 | 3 | 0.6667 | 1 | 1 | NA | NA | 1 | 0 | 2 | 0 | 4 | 0.25 |
| 6_1222906 | NA | NA | 1 | 0 | 1 | 1 | 2 | 0 | 4 | 0.25 | 1 | 1 | NA | NA | 1 | 0 | 2 | 0 | 4 | 0.25 |
| 6_1222906 | NA | NA | 1 | 1 | 1 | 0 | NA | NA | 2 | 0.5 | 1 | 1 | NA | NA | 1 | 0 | 2 | 0 | 4 | 0.25 |
| 6_1222907 | NA | NA | 1 | 1 | 1 | 0 | 2 | 0.5 | 4 | 0.5 | 1 | 1 | NA | NA | 1 | 0 | 1 | 0 | 3 | 0.3333 |
| 6_1222936 | 17 | 0.0588 | 9 | 0 | 10 | 0 | 10 | 0.2 | 46 | 0.0652 | 6 | 0 | 12 | 0 | 16 | 0.1875 | 15 | 0.2 | 49 | 0.1224 |
| 6_1222939 | 17 | 0.1765 | 8 | 0.25 | 11 | 0.1818 | 13 | 0.0769 | 49 | 0.1633 | 5 | 0.4 | 14 | 0 | 11 | 0.0909 | 15 | 0.2667 | 45 | 0.1556 |
| 6_1222947 | 3 | 0 | 5 | 0.2 | 1 | 0 | 3 | 0 | 12 | 0.0833 | 3 | 0.3333 | 1 | 0 | 2 | 0 | 2 | 0 | 8 | 0.125 |
| 6_1222952 | 4 | 0.5 | 5 | 0.2 | 1 | 1 | 2 | 1 | 12 | 0.5 | 3 | 0 | NA | NA | 2 | 1 | 2 | 0.5 | 7 | 0.4286 |
| 6_1222952 | 4 | 0 | 5 | 0 | 1 | 1 | 2 | 0 | 12 | 0.0833 | 3 | 0 | 1 | 0 | 2 | 0.5 | 2 | 0 | 8 | 0.125 |
| 6_1222953 | 5 | 0.2 | 6 | 0.1667 | 2 | 0.5 | 2 | 0 | 15 | 0.2 | 6 | 0 | 2 | 0 | 4 | 0.25 | 8 | 0 | 20 | 0.05 |
| 6_1222953 | 5 | 0.4 | 4 | 0.25 | 2 | 0.5 | 3 | 0.3333 | 14 | 0.3571 | 5 | 0.4 | 3 | 0.6667 | 5 | 0.2 | 7 | 0 | 20 | 0.25 |
| 6_1222953 | 5 | 0.2 | 5 | 0.6 | 2 | 0.5 | 3 | 0.3333 | 15 | 0.4 | 6 | 0.1667 | 4 | 0.5 | 4 | 0.25 | 8 | 0.125 | 22 | 0.2273 |
| 6_1222954 | 4 | 0.5 | 4 | 0 | 3 | 0 | 3 | 0 | 14 | 0.1429 | 4 | 0.75 | 3 | 0 | 5 | 0.4 | 8 | 0.375 | 20 | 0.4 |
| 6_1222954 | 5 | 0.2 | 6 | 0.1667 | 3 | 0 | 3 | 0 | 17 | 0.1176 | 5 | 0.2 | 4 | 0 | 6 | 0.1667 | 8 | 0.125 | 23 | 0.1304 |
| 6_1222955 | 4 | 0.25 | 5 | 0.4 | 1 | 0 | 3 | 0 | 13 | 0.2308 | 4 | 0.5 | 2 | 0 | 3 | 0.3333 | 4 | 0.25 | 13 | 0.3077 |
| 6_1222955 | 5 | 0.4 | 5 | 0.4 | 3 | 0.3333 | 2 | 0.5 | 15 | 0.4 | 6 | 0.5 | 3 | 0.3333 | 5 | 0.6 | 5 | 0.4 | 19 | 0.4737 |

| | | | | | | | | | | | | | | | | | | | | |
|---|---|---|---|---|---|---|---|---|---|---|---|---|---|---|---|---|---|---|---|---|
| 6_1223004( | 2 | 0 | 4 | 0.25 | 2 | 0 | 2 | 0 | 10 | 0.1 | 2 | 0 | 2 | 0 | 2 | 0 | 5 | 0.2 | 11 | 0.0909 |
| 6_1223004' | 4 | 0 | 4 | 0 | 1 | 1 | 3 | 0 | 12 | 0.0833 | 3 | 0.3333 | 3 | 0 | 3 | 0.3333 | 5 | 0 | 14 | 0.1429 |
| 6_1223004! | 3 | 0.6667 | 4 | 0.5 | 2 | 1 | 3 | 0.3333 | 12 | 0.5833 | 3 | 0.6667 | 2 | 0 | 4 | 0.25 | 5 | 0.2 | 14 | 0.2857 |
| 6_1223025! | 4 | 0.5 | 3 | 0.6667 | 2 | 0 | 2 | 0 | 11 | 0.3636 | 1 | 0 | NA | NA | 4 | 0.5 | 2 | 0.5 | 7 | 0.4286 |
| 6_1223026: | 2 | 0 | 3 | 0.3333 | 1 | 0 | 2 | 0.5 | 8 | 0.25 | 3 | 0 | NA | NA | 3 | 0.6667 | 2 | 0 | 8 | 0.25 |
| 6_1223027! | 2 | 0 | 1 | 1 | 1 | 0 | 1 | 0 | 5 | 0.2 | 3 | 0.3333 | NA | NA | 3 | 0 | 1 | 0 | 7 | 0.1429 |
| 6_1224302! | 1 | 1 | NA | NA | NA | NA | 1 | 1 | 2 | 1 | NA | NA | NA | NA | 2 | 1 | NA | NA | 2 | 1 |
| 6_1224304! | 1 | 1 | NA | NA | NA | NA | NA | NA | 1 | 1 | NA | NA | NA | NA | 2 | 1 | NA | NA | 2 | 1 |
| 6_1224305' | 1 | 1 | NA | NA | NA | NA | NA | NA | 1 | 1 | NA | NA | NA | NA | 2 | 1 | NA | NA | 2 | 1 |
| 6_1224315( | 2 | 0.5 | NA | NA | 1 | 1 | 1 | 1 | 4 | 0.75 | NA | NA | 1 | 1 | 1 | 0 | NA | NA | 2 | 0.5 |
| 6_1224315! | 4 | 1 | NA | NA | 1 | 1 | NA | NA | 5 | 1 | NA | NA | 1 | 1 | 2 | 1 | NA | NA | 3 | 1 |
| 6_1224323: | 3 | 0 | 1 | 0 | 3 | 0.3333 | 1 | 0 | 8 | 0.125 | NA | NA | 2 | 0 | 3 | 0.3333 | 1 | 0 | 6 | 0.1667 |
| 6_1224324: | 4 | 0 | 1 | 1 | 4 | 0 | 2 | 0 | 11 | 0.0909 | 1 | 0 | 2 | 0 | 3 | 0.3333 | NA | NA | 6 | 0.1667 |
| 6_1224326( | 2 | 0 | 1 | 1 | 4 | 1 | 2 | 0 | 9 | 0.5556 | 1 | 0 | 3 | 1 | 3 | 0.3333 | 1 | 1 | 8 | 0.625 |
| 6_1224331' | 3 | 0.3333 | NA | NA | 4 | 1 | 2 | 0 | 9 | 0.5556 | 1 | 1 | 1 | 0 | 3 | 1 | NA | NA | 5 | 0.8 |
| 6_1224331! | 3 | 0.6667 | NA | NA | 3 | 0.6667 | 1 | 0 | 7 | 0.5714 | 1 | 1 | 3 | 0.3333 | 2 | 0 | 1 | 0 | 7 | 0.2857 |
| 6_1224333: | 3 | 0.6667 | NA | NA | 4 | 0.5 | 2 | 0 | 9 | 0.4444 | 1 | 1 | 3 | 0.6667 | 2 | 0 | 1 | 1 | 7 | 0.5714 |
| 6_1224334' | 3 | 0.6667 | NA | NA | 4 | 1 | 2 | 0 | 9 | 0.6667 | 1 | 1 | 3 | 0.6667 | 1 | 0 | NA | NA | 5 | 0.6 |
| 6_1227049! | 22 | 0.0909 | 20 | 0 | 18 | 0.2222 | 21 | 0 | 81 | 0.0741 | 13 | 0.3077 | 14 | 0.0714 | 23 | 0.2174 | 17 | 0.1765 | 67 | 0.194 |
| 6_1227049! | 3 | 0 | 1 | 0 | 2 | 0.5 | 1 | 0 | 7 | 0.1429 | NA | NA | 1 | 0 | 3 | 0.3333 | NA | NA | 4 | 0.25 |
| 6_1227051: | 20 | 0.25 | 22 | 0.3182 | 18 | 0.1667 | 22 | 0.1818 | 82 | 0.2317 | 11 | 0.0909 | 15 | 0.0667 | 26 | 0.3077 | 16 | 0.375 | 68 | 0.2353 |
| 6_1227066: | 15 | 0.1333 | 12 | 0.1667 | 12 | 0.0833 | 14 | 0.0714 | 53 | 0.1132 | 6 | 0.1667 | 11 | 0.0909 | 12 | 0.0833 | 13 | 0.0769 | 42 | 0.0952 |
| 6_1227067( | 11 | 0.1818 | 11 | 0.0909 | 10 | 0.2 | 12 | 0 | 44 | 0.1136 | 5 | 0 | 9 | 0.1111 | 10 | 0.1 | 10 | 0.1 | 34 | 0.0882 |
| 6_1227068( | 14 | 1 | 12 | 1 | 10 | 1 | 15 | 0.9333 | 51 | 0.9804 | 6 | 1 | 9 | 1 | 13 | 1 | 11 | 0.9091 | 39 | 0.9744 |
| 6_1227069! | 8 | 0.875 | 9 | 0.4444 | 8 | 0.5 | 13 | 0.6923 | 38 | 0.6316 | 5 | 0.8 | 9 | 0.6667 | 8 | 0.625 | 8 | 0.75 | 30 | 0.7 |
| 6_1227070( | 11 | 0.5455 | 9 | 0.7778 | 11 | 0.7273 | 12 | 0.8333 | 43 | 0.7209 | 6 | 0.6667 | 10 | 0.6 | 8 | 0.5 | 10 | 0.8 | 34 | 0.6471 |
| 6_1227084! | 11 | 0 | 8 | 0.25 | 8 | 0.125 | 13 | 0 | 40 | 0.075 | 6 | 0.1667 | 9 | 0 | 8 | 0.125 | 9 | 0.1111 | 32 | 0.0938 |
| 6_1227168' | 5 | 0.2 | 2 | 0 | 1 | 0 | 7 | 0.1429 | 15 | 0.1333 | 3 | 0.3333 | 3 | 0.3333 | 3 | 0 | 4 | 0.25 | 13 | 0.2308 |
| 6_1227973 | 2 | 1 | NA | NA | NA | NA | 1 | 1 | 3 | 1 | NA | NA | NA | NA | NA | NA | 1 | 1 | 1 | 1 |
| 6_1227974' | 1 | 1 | NA | NA | NA | NA | 1 | 1 | 2 | 1 | NA | NA | NA | NA | NA | NA | 1 | 1 | 1 | 1 |
| 6_1228051( | 2 | 0.5 | 1 | 0 | NA | NA | 1 | 0 | 4 | 0.25 | NA | NA | 1 | 0 | NA | NA | 2 | 0.5 | 3 | 0.3333 |
| 6_1228197! | 2 | 0.5 | 2 | 1 | 1 | 1 | 1 | 0 | 6 | 0.6667 | 1 | 0 | 1 | 0 | NA | NA | 2 | 0.5 | 4 | 0.25 |
| 6_1228644: | 23 | 0.087 | 21 | 0.0476 | 10 | 0.1 | 21 | 0.0476 | 75 | 0.0667 | 16 | 0.125 | 14 | 0.1429 | 26 | 0.2308 | 19 | 0.2105 | 75 | 0.1867 |
| 6_1228645: | 20 | 0.1 | 22 | 0 | 10 | 0.1 | 19 | 0.1579 | 71 | 0.0845 | 16 | 0.0625 | 15 | 0.0667 | 24 | 0.0833 | 18 | 0 | 73 | 0.0548 |
| 6_1228647 | 23 | 0.1304 | 24 | 0.0417 | 13 | 0.1538 | 25 | 0.04 | 85 | 0.0824 | 18 | 0 | 16 | 0.125 | 27 | 0.1111 | 22 | 0.0909 | 83 | 0.0843 |
| 6_1228656 | 26 | 0.1154 | 29 | 0.1034 | 17 | 0 | 28 | 0.1429 | 100 | 0.1 | 23 | 0.1304 | 18 | 0 | 30 | 0.1667 | 26 | 0.0769 | 97 | 0.1031 |
| 6_1228657: | 21 | 0.1905 | 30 | 0.1333 | 15 | 0 | 26 | 0.1154 | 92 | 0.1196 | 22 | 0.1364 | 19 | 0.1053 | 28 | 0.1786 | 25 | 0.08 | 94 | 0.1277 |
| 6_1242893: | NA | NA | 1 | 1 | NA | NA | 2 | 0.5 | 3 | 0.6667 | NA | NA | 1 | 1 | NA | NA | 1 | 1 | 2 | 1 |
| 6_1242906( | NA | NA | 1 | 0 | NA | NA | 1 | 1 | 2 | 0.5 | NA | NA | 1 | 1 | NA | NA | NA | NA | 1 | 1 |
| 6_1242908 | NA | NA | NA | NA | 1 | 0 | 3 | 0.3333 | 4 | 0.25 | 1 | 1 | 1 | 1 | 3 | 0.3333 | 1 | 1 | 6 | 0.6667 |
| 6_1242909: | NA | NA | 1 | 0 | 1 | 1 | 3 | 0.3333 | 5 | 0.4 | 1 | 1 | 1 | 1 | 3 | 0.6667 | 1 | 1 | 6 | 0.8333 |
| 6_1242929 | 2 | 1 | 1 | 1 | NA | NA | 2 | 0.5 | 5 | 0.8 | NA | NA | 2 | 0.5 | 2 | 0 | 1 | 1 | 5 | 0.4 |
| 6_1242930! | 2 | 0.5 | NA | NA | NA | NA | NA | NA | 2 | 0.5 | NA | NA | 1 | 0 | NA | NA | 1 | 1 | 2 | 0.5 |
| 6_1242938: | 1 | 0 | 1 | 1 | NA | NA | 2 | 0.5 | 4 | 0.5 | NA | NA | 2 | 0.5 | 2 | 1 | NA | NA | 4 | 0.75 |
| 6_1242938! | NA | NA | 1 | 0 | NA | NA | 2 | 0.5 | 3 | 0.3333 | NA | NA | 1 | 1 | NA | NA | NA | NA | 1 | 1 |
| 6_1242938( | 1 | 1 | NA | NA | NA | NA | 1 | 1 | 2 | 1 | NA | NA | 1 | 1 | 2 | 1 | NA | NA | 3 | 1 |
| 6_1242940 | 1 | 1 | NA | NA | NA | NA | 1 | 1 | 2 | 1 | NA | NA | 1 | 1 | 1 | 1 | NA | NA | 2 | 1 |
| 6_1242941' | 1 | 0 | 1 | 1 | NA | NA | 2 | 0 | 4 | 0.25 | NA | NA | 1 | 1 | 2 | 1 | NA | NA | 3 | 1 |
| 6_1242941 | NA | NA | 1 | 1 | NA | NA | 1 | 1 | 2 | 1 | NA | NA | NA | NA | 1 | 1 | NA | NA | 1 | 1 |
| 6_1242942( | 1 | 1 | 1 | 1 | NA | NA | 2 | 1 | 4 | 1 | NA | NA | NA | NA | 2 | 1 | NA | NA | 2 | 1 |
| 6_1242943( | 1 | 0 | 1 | 1 | NA | NA | 2 | 1 | 4 | 0.75 | NA | NA | 1 | 1 | 2 | 0.5 | 1 | 1 | 4 | 0.75 |
| 6_1242944! | 1 | 1 | 1 | 1 | NA | NA | 2 | 1 | 4 | 1 | NA | NA | 1 | 1 | 2 | 1 | 1 | 1 | 4 | 1 |
| 6_1242946: | 1 | 1 | 1 | 1 | NA | NA | 1 | 1 | 3 | 1 | NA | NA | 1 | 1 | 2 | 1 | 1 | 1 | 4 | 1 |
| 6_1242947( | NA | NA | 1 | 0 | NA | NA | 2 | 1 | 3 | 0.6667 | NA | NA | NA | NA | 2 | 0.5 | 1 | 1 | 3 | 0.6667 |
| 6_1242947( | 1 | 1 | 1 | 1 | NA | NA | 2 | 1 | 4 | 1 | NA | NA | NA | NA | 1 | 1 | 1 | 1 | 2 | 1 |
| 6_1242948 | 1 | 0 | 1 | 1 | NA | NA | NA | NA | 2 | 0.5 | NA | NA | 1 | 0 | 2 | 0 | 1 | 1 | 4 | 0.25 |
| 6_1242970! | 1 | 1 | NA | NA | NA | NA | NA | NA | 1 | 1 | NA | NA | NA | NA | 1 | 1 | NA | NA | 1 | 1 |
| 6_1242973' | 1 | 1 | NA | NA | NA | NA | NA | NA | 1 | 1 | NA | NA | NA | NA | 1 | 1 | NA | NA | 1 | 1 |
| 6_1242974: | 2 | 1 | NA | NA | NA | NA | NA | NA | 2 | 1 | NA | NA | NA | NA | 1 | 1 | NA | NA | 1 | 1 |
| 6_1243739! | 5 | 0 | 10 | 0.2 | 7 | 0.1429 | 5 | 0.2 | 27 | 0.1481 | 5 | 0 | NA | NA | 6 | 0.5 | 2 | 0 | 13 | 0.2308 |
| 6_1243741 | 7 | 0.1429 | 7 | 0.1429 | 7 | 0.2857 | 6 | 0.5 | 27 | 0.2593 | 4 | 0.25 | NA | NA | 5 | 0.6 | 3 | 0.3333 | 12 | 0.4167 |
| 6_1243743( | 6 | 0.1667 | 6 | 0.1667 | 6 | 0.1667 | 5 | 0 | 23 | 0.1304 | 4 | 0.5 | NA | NA | 6 | 0.1667 | 2 | 0 | 12 | 0.25 |
| 6_1243749: | 3 | 0 | 6 | 0.1667 | 7 | 0 | 5 | 0.2 | 21 | 0.0952 | 1 | 1 | NA | NA | 2 | 0 | 2 | 0 | 5 | 0.2 |
| 6_1244084( | 36 | 0.9722 | 31 | 0.9677 | 26 | 0.9615 | 19 | 0.9474 | 112 | 0.9643 | 21 | 0.9048 | 25 | 1 | 20 | 0.9 | 19 | 0.9474 | 85 | 0.9412 |
| 6_1244084( | NA | NA | 2 | 1 | NA | NA | NA | NA | 2 | 1 | 1 | 1 | 1 | 1 | 1 | 1 | 1 | 0 | 4 | 0.75 |

| | | | | | | | | | | | | | | | | | | | | |
|---|---|---|---|---|---|---|---|---|---|---|---|---|---|---|---|---|---|---|---|---|
| 6_12441582 | 7 | 0.2857 | 10 | 0.2 | 9 | 0 | 4 | 0.25 | 30 | 0.1667 | 6 | 0 | 7 | 0 | 10 | 0.1 | 6 | 0.3333 | 29 | 0.1034 |
| 6_12443858 | 11 | 0 | 8 | 0 | 7 | 0.2857 | 2 | 0 | 28 | 0.0714 | 7 | 0.2857 | 10 | 0.1 | 7 | 0 | 6 | 0.6667 | 30 | 0.2333 |
| 6_12444158 | 11 | 0.3636 | 3 | 0.3333 | 10 | 0.3 | 3 | 0 | 27 | 0.2963 | 6 | 0.6667 | 11 | 0.3636 | 8 | 0.125 | 6 | 0 | 31 | 0.2903 |
| 6_12468141 | 12 | 0.0833 | 13 | 0.1538 | 2 | 0.5 | 4 | 0.25 | 31 | 0.1613 | 8 | 0.375 | 10 | 0 | 9 | 0.1111 | 5 | 0 | 32 | 0.125 |
| 6_12468147 | 14 | 0.8571 | 11 | 0.9091 | 2 | 1 | 5 | 1 | 32 | 0.9062 | 7 | 1 | 13 | 0.8462 | 8 | 0.625 | 7 | 0.8571 | 35 | 0.8286 |
| 6_12468244 | 4 | 0 | 2 | 0 | 1 | 1 | 1 | 0 | 8 | 0.125 | 1 | 0 | 3 | 0.3333 | 2 | 0 | 1 | 0 | 7 | 0.1429 |
| 6_12468246 | 4 | 0.5 | 2 | 0.5 | 1 | 0 | 1 | 0 | 8 | 0.375 | 1 | 1 | 4 | 0.5 | 2 | 0 | 1 | 0 | 8 | 0.375 |
| 6_12468884 | 6 | 0 | 8 | 0 | NA | NA | 3 | 0.3333 | 17 | 0.0588 | 3 | 0 | 6 | 0.1667 | 3 | 0 | 3 | 0 | 15 | 0.0667 |
| 6_12468941 | 2 | 0 | 9 | 0.1111 | 8 | 0.125 | 6 | 0.1667 | 25 | 0.12 | 10 | 0.2 | 9 | 0.3333 | 13 | 0.1538 | 9 | 0.1111 | 41 | 0.1951 |
| 6_12469370 | 7 | 0.8571 | 17 | 0.8235 | 12 | 0.8333 | 11 | 0.8182 | 47 | 0.8298 | 13 | 0.5385 | 16 | 0.875 | 18 | 0.7778 | 12 | 0.8333 | 59 | 0.7627 |
| 6_12469374 | 6 | 0.1667 | 17 | 0.0588 | 18 | 0.1111 | 11 | 0.2727 | 52 | 0.1346 | 15 | 0.1333 | 14 | 0.0714 | 17 | 0.0588 | 15 | 0.0667 | 61 | 0.082 |
| 6_12469380 | 7 | 0 | 13 | 0.2308 | 13 | 0.1538 | 10 | 0 | 43 | 0.1163 | 15 | 0.3333 | 16 | 0.1875 | 18 | 0.2778 | 14 | 0.1429 | 63 | 0.2381 |
| 6_12472216 | 12 | 0.6667 | 15 | 0.7333 | 9 | 0.6667 | 17 | 0.4706 | 53 | 0.6226 | 7 | 0.5714 | 14 | 0.3571 | 14 | 0.5714 | 16 | 0.875 | 51 | 0.6078 |
| 6_12472322 | 9 | 0.3333 | 13 | 0.3077 | 9 | 0.2222 | 15 | 0.1333 | 46 | 0.2391 | 5 | 0 | 12 | 0.25 | 11 | 0.0909 | 12 | 0.3333 | 40 | 0.2 |
| 6_12472332 | 4 | 0 | 8 | 0.125 | 7 | 0 | 9 | 0.1111 | 28 | 0.0714 | 2 | 0 | 10 | 0.1 | 8 | 0.125 | 10 | 0 | 30 | 0.0667 |
| 6_12472420 | 2 | 1 | 8 | 1 | 5 | 0.6 | 7 | 0.5714 | 22 | 0.7727 | 4 | 1 | 7 | 0.8571 | 6 | 0.6667 | 4 | 1 | 21 | 0.8571 |
| 6_12472468 | 4 | 0.25 | 8 | 0.125 | 5 | 0.2 | 6 | 0.1667 | 23 | 0.1739 | 2 | 0 | 6 | 0 | 6 | 0 | 6 | 0.1667 | 20 | 0.05 |
| 6_12472479 | 4 | 0.75 | 8 | 0.625 | 4 | 0.75 | 5 | 0.4 | 21 | 0.619 | 3 | 0.3333 | 5 | 1 | 6 | 1 | 5 | 1 | 19 | 0.8947 |
| 6_12472480 | 3 | 0.6667 | 7 | 0.1429 | 4 | 0 | 5 | 0.2 | 19 | 0.2105 | 2 | 0 | 5 | 0.4 | 6 | 0 | 5 | 0.2 | 18 | 0.1667 |
| 6_12473710 | 26 | 0.2308 | 27 | 0.1852 | 22 | 0.2727 | 21 | 0.2381 | 96 | 0.2292 | 30 | 0.3667 | 25 | 0.24 | 44 | 0.2955 | 23 | 0.2609 | 122 | 0.2951 |
| 6_12473720 | 25 | 0.64 | 25 | 0.72 | 20 | 0.65 | 17 | 0.8235 | 87 | 0.7011 | 24 | 0.7917 | 30 | 0.8333 | 35 | 0.8286 | 21 | 0.6667 | 110 | 0.7909 |
| 6_12473730 | 24 | 0.9167 | 22 | 1 | 21 | 0.9048 | 19 | 0.8421 | 86 | 0.9186 | 28 | 0.9643 | 26 | 0.8846 | 41 | 0.9268 | 22 | 0.8636 | 117 | 0.9145 |
| 6_12473751 | 62 | 0.2903 | 82 | 0.3171 | 74 | 0.3378 | 65 | 0.2154 | 283 | 0.2933 | 43 | 0.2558 | 76 | 0.3158 | 85 | 0.3647 | 63 | 0.3175 | 267 | 0.3221 |
| 6_12473764 | 104 | 0.0288 | 126 | 0.0635 | 103 | 0.0777 | 119 | 0.0336 | 452 | 0.0509 | 69 | 0.1159 | 133 | 0.0602 | 124 | 0.0484 | 107 | 0.0374 | 433 | 0.06 |
| 6_12473784 | 80 | 0.0875 | 89 | 0.1798 | 72 | 0.0972 | 87 | 0.0575 | 328 | 0.1067 | 59 | 0.1186 | 99 | 0.1313 | 102 | 0.098 | 80 | 0.1875 | 340 | 0.1324 |
| 6_12473785 | 80 | 0.075 | 100 | 0.1 | 80 | 0.0875 | 87 | 0.0345 | 347 | 0.0749 | 62 | 0.129 | 115 | 0.087 | 107 | 0.0467 | 89 | 0.0449 | 373 | 0.0724 |
| 6_12473838 | 1 | 1 | NA | NA | NA | NA | NA | NA | 1 | 1 | NA | NA | 2 | 1 | NA | NA | 2 | 0 | 4 | 0.5 |
| 6_12478633 | NA | NA | NA | NA | 1 | 1 | NA | NA | 1 | 1 | 1 | 1 | NA | NA | 1 | 1 | NA | NA | 2 | 1 |
| 6_12478673 | NA | NA | NA | NA | 1 | 1 | NA | NA | 1 | 1 | 1 | 1 | NA | NA | 1 | 0 | NA | NA | 2 | 0.5 |
| 6_12478676 | NA | NA | NA | NA | 1 | 1 | NA | NA | 1 | 1 | 1 | 1 | NA | NA | 2 | 0.5 | NA | NA | 3 | 0.6667 |
| 6_12478679 | NA | NA | NA | NA | 1 | 1 | NA | NA | 1 | 1 | 1 | 1 | NA | NA | 1 | 1 | NA | NA | 2 | 1 |
| 6_12478798 | 87 | 0.0575 | 123 | 0.0976 | 113 | 0.0531 | 88 | 0.1023 | 411 | 0.0779 | 84 | 0.0595 | 103 | 0.068 | 89 | 0.0787 | 95 | 0.0632 | 371 | 0.0674 |
| 6_12478819 | 89 | 0.2472 | 120 | 0.35 | 97 | 0.3711 | 95 | 0.3684 | 401 | 0.3367 | 82 | 0.2683 | 99 | 0.4242 | 88 | 0.3523 | 96 | 0.4688 | 365 | 0.3836 |
| 6_12478996 | NA | NA | 1 | 0 | 1 | 1 | 1 | 0 | 3 | 0.3333 | 2 | 0.5 | 1 | 0 | 1 | 0 | NA | NA | 4 | 0.25 |
| 6_12479229 | 12 | 0.3333 | 29 | 0.1724 | 19 | 0.2105 | 15 | 0.1333 | 75 | 0.2 | 22 | 0.1818 | 18 | 0.2778 | 29 | 0.1724 | 13 | 0.1538 | 82 | 0.1951 |
| 6_12479234 | 9 | 0.1111 | 18 | 0 | 18 | 0 | 12 | 0.25 | 57 | 0.0702 | 15 | 0.1333 | 18 | 0.0556 | 19 | 0.0526 | 11 | 0.1818 | 63 | 0.0952 |
| 6_12479235 | 12 | 0.5 | 21 | 0.5714 | 15 | 0.4667 | 13 | 0.6923 | 61 | 0.5574 | 18 | 0.3889 | 17 | 0.5882 | 23 | 0.4783 | 9 | 0.6667 | 67 | 0.5075 |
| 6_12479245 | 10 | 0.3 | 23 | 0.1739 | 24 | 0.2083 | 13 | 0.4615 | 70 | 0.2571 | 22 | 0.3636 | 19 | 0.3158 | 32 | 0.3125 | 12 | 0.4167 | 85 | 0.3412 |
| 6_12479249 | 8 | 0 | 23 | 0.0435 | 19 | 0.1579 | 12 | 0 | 62 | 0.0645 | 19 | 0.0526 | 15 | 0 | 27 | 0.0741 | 11 | 0.1818 | 72 | 0.0694 |
| 6_12481819 | NA | NA | 4 | 0.25 | 2 | 0 | 1 | 0 | 7 | 0.1429 | 1 | 1 | 3 | 0.3333 | 2 | 0 | 3 | 0.3333 | 9 | 0.3333 |
| 6_12481840 | NA | NA | 4 | 0 | 1 | 0 | 1 | 1 | 6 | 0.1667 | 1 | 0 | 5 | 0 | 4 | 0.25 | NA | NA | 10 | 0.1 |
| 6_12481840 | NA | NA | 3 | 0.6667 | 1 | 1 | 1 | 0 | 5 | 0.6 | 1 | 1 | 5 | 0.6 | 3 | 0.3333 | 2 | 1 | 11 | 0.6364 |
| 6_12481848 | NA | NA | 4 | 0.75 | NA | NA | 1 | 1 | 5 | 0.8 | 1 | 0 | 5 | 0.4 | 3 | 0 | 1 | 0 | 10 | 0.2 |
| 6_12481850 | NA | NA | 4 | 1 | 1 | 0 | 1 | 1 | 6 | 0.8333 | 1 | 1 | 4 | 0.75 | 2 | 1 | 1 | 1 | 8 | 0.875 |
| 6_12481852 | NA | NA | 3 | 1 | 1 | 0 | 1 | 1 | 5 | 0.8 | 1 | 1 | 5 | 0.6 | 2 | 0.5 | 1 | 1 | 9 | 0.6667 |
| 6_12481859 | NA | NA | 3 | 0.6667 | 1 | 0 | 1 | 1 | 5 | 0.6 | 1 | 0 | 5 | 0.6 | 3 | 0.3333 | 1 | 0 | 10 | 0.4 |
| 6_12481864 | NA | NA | 3 | 0.3333 | NA | NA | 1 | 0 | 4 | 0.25 | 1 | 0 | 5 | 0.6 | 1 | 0 | 1 | 1 | 8 | 0.5 |
| 6_12483169 | NA | NA | 1 | 1 | NA | NA | NA | NA | 1 | 1 | 1 | 1 | 2 | 0.5 | 1 | 1 | 1 | 0 | 5 | 0.6 |
| 6_12483575 | 11 | 1 | 29 | 0.9655 | 30 | 0.9667 | 8 | 1 | 78 | 0.9744 | 19 | 1 | 11 | 1 | 17 | 1 | 18 | 0.9444 | 65 | 0.9846 |
| 6_12483581 | 18 | 0.0556 | 37 | 0.1081 | 31 | 0.0645 | 10 | 0.2 | 96 | 0.0938 | 19 | 0.2105 | 14 | 0.1429 | 18 | 0 | 18 | 0.0556 | 69 | 0.1014 |
| 6_12483583 | 14 | 0.2143 | 35 | 0.4 | 31 | 0.4839 | 9 | 0.6667 | 89 | 0.427 | 17 | 0.4118 | 14 | 0.3571 | 20 | 0.35 | 19 | 0.3158 | 70 | 0.3571 |
| 6_12483586 | 14 | 0.0714 | 35 | 0 | 33 | 0.0909 | 7 | 0.1429 | 89 | 0.0562 | 19 | 0 | 10 | 0.3 | 21 | 0.0476 | 19 | 0 | 69 | 0.058 |
| 6_12483602 | 15 | 0.2667 | 31 | 0.2581 | 30 | 0.5 | 9 | 0.1111 | 85 | 0.3294 | 17 | 0.2353 | 11 | 0.1818 | 21 | 0.381 | 18 | 0.2778 | 67 | 0.2836 |
| 6_12483602 | 13 | 0.8462 | 26 | 0.6923 | 22 | 0.8636 | 10 | 0.8 | 71 | 0.7887 | 15 | 0.8667 | 12 | 0.9167 | 19 | 0.6842 | 12 | 0.6667 | 58 | 0.7759 |
| 6_12483668 | 1 | 0 | 2 | 0.5 | 3 | 0 | NA | NA | 6 | 0.1667 | NA | NA | 4 | 0.25 | 1 | 0 | 3 | 0.3333 | 8 | 0.25 |
| 6_12483671 | 1 | 0 | 4 | 0.25 | 8 | 0.25 | 2 | 0.5 | 15 | 0.2667 | 1 | 0 | 5 | 0 | 2 | 0 | 3 | 0.3333 | 11 | 0.0909 |
| 6_12483803 | 1 | 1 | NA | NA | 2 | 1 | NA | NA | 3 | 1 | 1 | 1 | NA | NA | NA | NA | NA | NA | 1 | 1 |
| 6_12483803 | 5 | 1 | 14 | 0.8571 | 11 | 0.7273 | 3 | 1 | 33 | 0.8485 | 8 | 0.875 | 6 | 0.8333 | 7 | 1 | 7 | 0.7143 | 28 | 0.8571 |
| 6_12483839 | 7 | 0.4286 | 17 | 0.4118 | 13 | 0.6154 | 2 | 0 | 39 | 0.4615 | 7 | 0.4286 | 4 | 0.75 | 8 | 0.5 | 6 | 0.8333 | 25 | 0.6 |
| 6_12483840 | 8 | 0.375 | 20 | 0.6 | 14 | 0.5 | 3 | 0.6667 | 45 | 0.5333 | 6 | 0.5 | 6 | 0.1667 | 7 | 0.1429 | 6 | 0.5 | 25 | 0.32 |
| 6_12483874 | 5 | 1 | 16 | 1 | 11 | 1 | 3 | 1 | 35 | 1 | 7 | 1 | 7 | 1 | 5 | 1 | 5 | 1 | 24 | 1 |
| 6_12483878 | 4 | 0.75 | 16 | 0.9375 | 10 | 0.9 | 2 | 1 | 32 | 0.9062 | 6 | 1 | 7 | 0.8571 | 5 | 0.8 | 5 | 0.8 | 23 | 0.8696 |
| 6_12483887 | 5 | 0.8 | 18 | 0.8333 | 11 | 0.7273 | 3 | 1 | 37 | 0.8108 | 7 | 1 | 6 | 0.8333 | 5 | 1 | 4 | 1 | 22 | 0.9545 |
| 6_12483890 | 5 | 0.2 | 13 | 0.1538 | 9 | 0.1111 | 2 | 0 | 29 | 0.1379 | 5 | 0.4 | 5 | 0.2 | 4 | 0.25 | 3 | 1 | 17 | 0.4118 |
| 6_12484282 | 1 | 0 | NA | NA | 1 | 0 | 5 | 0.4 | 7 | 0.2857 | NA | NA | 4 | 0 | 1 | 1 | 2 | 0.5 | 7 | 0.2857 |

| | | | | | | | | | | | | | | | | | | | | |
|---|---|---|---|---|---|---|---|---|---|---|---|---|---|---|---|---|---|---|---|---|
| 6_1248428 | 1 | 1 | 1 | 1 | 1 | 1 | 4 | 0.75 | 7 | 0.8571 | NA | NA | 6 | 1 | 3 | 1 | 3 | 1 | 12 | 1 |
| 6_1248429 | 1 | 0 | 1 | 0 | NA | NA | 5 | 0.2 | 7 | 0.1429 | NA | NA | 6 | 0.1667 | 3 | 0.3333 | 1 | 0 | 10 | 0.2 |
| 6_1248429 | 1 | 1 | 1 | 1 | 1 | 1 | 4 | 0.75 | 7 | 0.8571 | NA | NA | 6 | 1 | 3 | 0.6667 | 2 | 1 | 11 | 0.9091 |
| 6_1248446 | 1 | 0 | 1 | 0 | 1 | 1 | 3 | 0 | 6 | 0.1667 | NA | NA | 4 | 0.25 | 2 | 0 | 2 | 1 | 8 | 0.375 |
| 6_1248941 | 1 | 1 | NA | NA | NA | NA | 1 | 1 | 2 | 1 | 3 | 1 | NA | NA | 1 | 1 | 1 | 1 | 5 | 1 |
| 6_1249126 | 101 | 0.0594 | 96 | 0.0833 | 78 | 0.1154 | 64 | 0.0625 | 339 | 0.0796 | 65 | 0.0923 | 109 | 0.0734 | 66 | 0.0606 | 65 | 0.0769 | 305 | 0.0754 |
| 6_1249129 | 102 | 0.1961 | 104 | 0.2404 | 76 | 0.25 | 73 | 0.1096 | 355 | 0.2028 | 73 | 0.3014 | 116 | 0.2155 | 76 | 0.2237 | 71 | 0.2113 | 336 | 0.2351 |
| 6_1249354 | 3 | 0 | 1 | 0 | 4 | 0 | 6 | 0.6667 | 14 | 0.2857 | 2 | 0.5 | 1 | 0 | 2 | 0 | 1 | 0 | 6 | 0.1667 |
| 6_1249354 | 3 | 0 | 1 | 0 | 4 | 0.25 | 7 | 0 | 15 | 0.0667 | 2 | 0.5 | 1 | 0 | 2 | 0 | 1 | 0 | 6 | 0.1667 |
| 6_1249356 | 3 | 0.3333 | 1 | 0 | 3 | 0 | 6 | 0 | 13 | 0.0769 | 1 | 1 | 2 | 0 | 2 | 0 | 1 | 0 | 6 | 0.1667 |
| 6_1249360 | 5 | 0 | 18 | 0.1111 | 18 | 0.2222 | 16 | 0.25 | 57 | 0.1754 | 12 | 0.1667 | 20 | 0.25 | 9 | 0.4444 | 14 | 0.3571 | 55 | 0.2909 |
| 6_1249365 | 2 | 0 | 18 | 0.2778 | 22 | 0.3636 | 15 | 0.2 | 57 | 0.2807 | 14 | 0.3571 | 21 | 0.1905 | 11 | 0.2727 | 13 | 0.4615 | 59 | 0.3051 |
| 6_1249365 | 3 | 0 | 16 | 0.0625 | 22 | 0.0909 | 15 | 0.0667 | 56 | 0.0714 | 14 | 0.2143 | 23 | 0.0435 | 10 | 0 | 13 | 0.1538 | 60 | 0.1 |
| 6_1249765 | 12 | 0.4167 | 13 | 0.0769 | 15 | 0.4 | 13 | 0.4615 | 53 | 0.3396 | 14 | 0.2857 | 17 | 0.4706 | 21 | 0.4762 | 7 | 0.5714 | 59 | 0.4407 |
| 6_1249785 | 20 | 0 | 27 | 0.1111 | 16 | 0.0625 | 22 | 0.0909 | 85 | 0.0706 | 21 | 0.0952 | 21 | 0.0476 | 25 | 0.04 | 14 | 0.2143 | 81 | 0.0864 |
| 6_1250010 | 2 | 0 | 1 | 1 | 1 | 0 | 1 | 0 | 5 | 0.2 | 1 | 0 | 2 | 0 | 2 | 0 | 3 | 0.3333 | 8 | 0.125 |
| 6_1250135 | 1 | 0 | 5 | 0.4 | 1 | 1 | 1 | 0 | 8 | 0.375 | 2 | 0 | 3 | 0 | 4 | 0.75 | 5 | 0.6 | 14 | 0.4286 |
| 6_1250135 | 1 | 0 | 4 | 0.75 | 1 | 0 | 2 | 0.5 | 8 | 0.5 | 2 | 0.5 | 2 | 0 | 4 | 0.25 | 4 | 0.5 | 12 | 0.3333 |
| 6_1250136 | 1 | 0 | 5 | 0.2 | 1 | 1 | 2 | 0 | 9 | 0.2222 | 2 | 0.5 | 3 | 0 | 6 | 0.3333 | 5 | 0.4 | 16 | 0.3125 |
| 6_1250139 | 1 | 0 | 5 | 0.2 | 2 | 0.5 | 2 | 0.5 | 10 | 0.3 | 2 | 1 | 3 | 0 | 6 | 0.5 | 3 | 0.3333 | 14 | 0.4286 |
| 6_1250140 | 1 | 0 | 6 | 0.1667 | 2 | 0 | 2 | 0.5 | 11 | 0.1818 | 2 | 1 | 2 | 0 | 6 | 0.3333 | 3 | 0 | 13 | 0.3077 |
| 6_1250141 | 1 | 0 | 4 | 0.25 | 2 | 0.5 | 2 | 0 | 9 | 0.2222 | 2 | 0.5 | 3 | 0.3333 | 5 | 0.4 | 4 | 0.5 | 14 | 0.4286 |
| 6_1250144 | 1 | 0 | 5 | 0.8 | 3 | 1 | 2 | 0.5 | 11 | 0.7273 | 2 | 1 | 1 | 0 | 5 | 0.8 | 5 | 0.6 | 13 | 0.6923 |
| 6_1250144 | 2 | 1 | 3 | 0.6667 | 2 | 0.5 | NA | NA | 7 | 0.7143 | 2 | 0.5 | 3 | 1 | 2 | 1 | 6 | 0.5 | 13 | 0.6923 |
| 6_1250144 | 3 | 0.3333 | 6 | 0 | 3 | 0 | 2 | 0.5 | 14 | 0.1429 | 2 | 0 | 4 | 0.25 | 6 | 0.1667 | 6 | 0.1667 | 18 | 0.1667 |
| 6_1250254 | 5 | 0 | 7 | 0.2857 | 3 | 0 | 4 | 0 | 19 | 0.1053 | 5 | 0.2 | 9 | 0.2222 | 5 | 0 | 4 | 0 | 23 | 0.1304 |
| 6_1250254 | 3 | 0 | 7 | 0.4286 | 4 | 0.25 | 4 | 0 | 18 | 0.2222 | 7 | 0 | 6 | 0.5 | 3 | 0 | 6 | 0.1667 | 22 | 0.1818 |
| 6_1250256 | 6 | 0.5 | 5 | 0.2 | 4 | 0.5 | 4 | 0.5 | 19 | 0.4211 | 7 | 0.4286 | 10 | 0.5 | 4 | 0.75 | 5 | 0 | 26 | 0.4231 |
| 6_1250259 | 4 | 0.25 | 8 | 0.375 | 2 | 0.5 | 3 | 0.6667 | 17 | 0.4118 | 7 | 0.2857 | 6 | 0.3333 | 5 | 0 | 3 | 0.6667 | 21 | 0.2857 |
| 6_1250260 | 4 | 0 | 8 | 0.125 | 1 | 0 | 2 | 0 | 15 | 0.0667 | 7 | 0.1429 | 7 | 0 | 4 | 0 | 4 | 0.25 | 22 | 0.0909 |
| 6_1250261 | 3 | 0.3333 | 8 | 0.375 | 3 | 0.6667 | 2 | 1 | 16 | 0.5 | 7 | 0.5714 | 7 | 0.1429 | 5 | 0.2 | 4 | 0.25 | 23 | 0.3043 |
| 6_1250308 | 1 | 1 | 1 | 1 | NA | NA | NA | NA | 2 | 1 | NA | NA | 1 | 1 | 1 | 1 | NA | NA | 2 | 1 |
| 6_1250309 | 1 | 0 | 1 | 1 | NA | NA | NA | NA | 2 | 0.5 | NA | NA | 1 | 1 | NA | NA | NA | NA | 1 | 1 |
| 6_1251070 | 14 | 0.2857 | 19 | 0.3684 | 20 | 0.25 | 15 | 0.2 | 68 | 0.2794 | 5 | 0.4 | 15 | 0.2667 | 14 | 0.2857 | 10 | 0.3 | 44 | 0.2955 |
| 6_1251071 | 18 | 0.1667 | 20 | 0.1 | 20 | 0.1 | 16 | 0.0625 | 74 | 0.1081 | 6 | 0 | 14 | 0.1429 | 14 | 0.0714 | 11 | 0.1818 | 45 | 0.1111 |
| 6_1251072 | 16 | 0.4375 | 23 | 0.3043 | 19 | 0.1579 | 13 | 0.3846 | 71 | 0.3099 | 6 | 0.3333 | 17 | 0.4118 | 14 | 0.2857 | 13 | 0.3846 | 50 | 0.36 |
| 6_1251072 | 14 | 0.7143 | 17 | 0.6471 | 17 | 0.7059 | 12 | 0.25 | 60 | 0.6 | 6 | 1 | 15 | 0.6667 | 13 | 0.4615 | 10 | 0.5 | 44 | 0.6136 |
| 6_1251075 | 17 | 0.2353 | 19 | 0.2105 | 19 | 0.3158 | 15 | 0.1333 | 70 | 0.2286 | 8 | 0.5 | 16 | 0.3125 | 13 | 0.2308 | 9 | 0.1111 | 46 | 0.2826 |
| 6_1251141 | 2 | 0.5 | 2 | 0 | 2 | 0 | NA | NA | 6 | 0.1667 | 1 | 0 | 1 | 1 | 3 | 0 | 2 | 0 | 7 | 0.1429 |
| 6_1251212 | 3 | 0 | 3 | 0 | 2 | 0 | 3 | 0.3333 | 11 | 0.0909 | 1 | 0 | 8 | 0 | 3 | 0.3333 | 2 | 0.5 | 14 | 0.1429 |
| 6_1251213 | 4 | 0.75 | 3 | 1 | 2 | 0.5 | 3 | 1 | 12 | 0.8333 | 1 | 1 | 7 | 0.5714 | 4 | 0.75 | 2 | 1 | 14 | 0.7143 |
| 6_1251213 | 4 | 1 | 3 | 1 | 2 | 1 | 4 | 1 | 13 | 1 | 1 | 1 | 8 | 0.875 | 3 | 0.6667 | 2 | 1 | 14 | 0.8571 |
| 6_1251215 | 4 | 0 | 3 | 0.3333 | 2 | 0.5 | 4 | 0.25 | 13 | 0.2308 | 1 | 1 | 7 | 0.4286 | 4 | 0.75 | 2 | 0.5 | 14 | 0.5714 |
| 6_1251216 | 4 | 1 | 3 | 1 | 3 | 1 | 4 | 1 | 14 | 1 | 1 | 1 | 7 | 0.8571 | 4 | 1 | 2 | 1 | 14 | 0.9286 |
| 6_1251216 | 4 | 0.5 | 3 | 0.3333 | 3 | 0 | 4 | 0 | 14 | 0.2143 | 1 | 0 | 8 | 0 | 4 | 0 | 2 | 0.5 | 15 | 0.0667 |
| 6_1251216 | 2 | 0 | 2 | 0 | 3 | 0.3333 | 4 | 0.25 | 11 | 0.1818 | NA | NA | 7 | 0.1429 | 4 | 0 | NA | NA | 11 | 0.0909 |
| 6_1251216 | 2 | 0.5 | 3 | 0.3333 | 2 | 0.5 | 3 | 0 | 10 | 0.3 | 1 | 1 | 3 | 0.6667 | 2 | 0 | 1 | 1 | 7 | 0.5714 |
| 6_1251227 | 4 | 0.75 | 3 | 0 | 1 | 0 | 4 | 0.5 | 12 | 0.4167 | 3 | 0.3333 | 5 | 0.6 | 1 | 1 | 1 | 0 | 10 | 0.5 |
| 6_1251228 | 4 | 0.5 | 4 | 0.75 | 1 | 1 | 5 | 0.8 | 14 | 0.7143 | 3 | 0.6667 | 5 | 0.6 | 1 | 1 | 1 | 1 | 10 | 0.7 |
| 6_1251377 | 6 | 0.6667 | 7 | 0.7143 | 5 | 1 | 6 | 0.5 | 24 | 0.7083 | 5 | 0.6 | 7 | 0.4286 | 2 | 0 | 4 | 0.25 | 18 | 0.3889 |
| 6_1251378 | 6 | 0.3333 | 7 | 0.2857 | 6 | 0.1667 | 6 | 0.5 | 25 | 0.32 | 7 | 0.1429 | 7 | 0.5714 | 2 | 0 | 4 | 0 | 20 | 0.25 |
| 6_1251378 | 6 | 0.3333 | 9 | 0.2222 | 5 | 0 | 5 | 0 | 25 | 0.16 | 7 | 0.2857 | 7 | 0.2857 | 2 | 0 | 4 | 0 | 20 | 0.2 |
| 6_1251383 | 6 | 0.5 | 7 | 0.2857 | 4 | 0.5 | 3 | 0.6667 | 20 | 0.45 | 6 | 0.1667 | 7 | 0.2857 | 2 | 0 | 1 | 1 | 16 | 0.25 |
| 6_1251390 | 4 | 0 | 2 | 0 | 3 | 0 | 2 | 0.5 | 11 | 0.0909 | 4 | 0 | 5 | 0.2 | 4 | 0 | 6 | 0 | 19 | 0.0526 |
| 6_1251393 | 2 | 0 | 1 | 1 | NA | NA | 1 | 0 | 4 | 0.25 | NA | NA | 5 | 0 | 2 | 0 | 1 | 1 | 8 | 0.125 |
| 6_1251691 | 1 | 0 | NA | NA | 1 | 1 | 3 | 0.3333 | 5 | 0.4 | NA | NA | 2 | 0.5 | 1 | 1 | 1 | 0 | 4 | 0.5 |
| 6_1251702 | 19 | 0.1053 | 14 | 0 | 16 | 0.0625 | 17 | 0.0588 | 66 | 0.0606 | 8 | 0 | 16 | 0.0625 | 11 | 0.0909 | 13 | 0.0769 | 48 | 0.0625 |
| 6_1251703 | 22 | 0.1818 | 16 | 0.4375 | 19 | 0.2632 | 16 | 0.1875 | 73 | 0.2603 | 12 | 0 | 18 | 0.1667 | 13 | 0.0769 | 14 | 0.3571 | 57 | 0.1579 |
| 6_1251703 | 22 | 0 | 16 | 0 | 19 | 0.2105 | 17 | 0.1176 | 74 | 0.0811 | 11 | 0.0909 | 15 | 0.2 | 15 | 0.2667 | 15 | 0.2667 | 56 | 0.2143 |
| 6_1251704 | 18 | 1 | 15 | 0.9333 | 17 | 0.9412 | 16 | 0.9375 | 66 | 0.9545 | 13 | 1 | 14 | 1 | 14 | 1 | 13 | 1 | 54 | 1 |
| 6_1251707 | 1 | 0 | 2 | 0.5 | 3 | 0.6667 | 2 | 0.5 | 8 | 0.5 | 1 | 1 | NA | NA | NA | NA | 1 | 1 | 2 | 1 |
| 6_1251967 | NA | NA | 1 | 1 | NA | NA | NA | NA | 1 | 1 | NA | NA | 1 | 1 | NA | NA | NA | NA | 1 | 1 |
| 6_1251989 | 14 | 0.0714 | 12 | 0.25 | 17 | 0.1765 | 10 | 0.1 | 53 | 0.1509 | 7 | 0.1429 | 12 | 0.0833 | 9 | 0 | 9 | 0.1111 | 37 | 0.0811 |
| 6_1252014 | 1 | 1 | NA | NA | NA | NA | NA | NA | 1 | 1 | 1 | 1 | NA | NA | NA | NA | NA | NA | 1 | 1 |
| 6_1252530 | NA | NA | NA | NA | 1 | 1 | NA | NA | 1 | 1 | NA | NA | NA | NA | 1 | 1 | NA | NA | 1 | 1 |

| | | | | | | | | | | | | | | | | | | | | |
|---|---|---|---|---|---|---|---|---|---|---|---|---|---|---|---|---|---|---|---|---|
| 6_1252535 | NA | NA | NA | NA | 1 | 1 | NA | NA | 1 | 1 | NA | NA | NA | NA | 1 | 1 | NA | NA | 1 | 1 |
| 6_1252542 | NA | NA | NA | NA | 1 | 1 | NA | NA | 1 | 1 | NA | NA | NA | NA | 1 | 1 | NA | NA | 1 | 1 |
| 6_1252546 | NA | NA | NA | NA | 1 | 1 | NA | NA | 1 | 1 | NA | NA | NA | NA | 1 | 1 | NA | NA | 1 | 1 |
| 6_1252548 | NA | NA | NA | NA | 1 | 1 | NA | NA | 1 | 1 | NA | NA | NA | NA | 1 | 1 | NA | NA | 1 | 1 |
| 6_1252837 | 4 | 0.5 | 2 | 0.5 | 2 | 0 | 2 | 1 | 10 | 0.5 | 4 | 0.5 | NA | NA | 1 | 1 | 1 | 1 | 6 | 0.6667 |
| 6_1252837 | 4 | 0 | 2 | 1 | 1 | 0 | 3 | 0 | 10 | 0.2 | 4 | 0.25 | NA | NA | 1 | 0 | 1 | 0 | 6 | 0.1667 |
| 6_1252838 | 4 | 0.5 | 2 | 0 | 1 | 0 | 3 | 0 | 10 | 0.2 | 4 | 0.25 | NA | NA | 2 | 0 | 1 | 0 | 7 | 0.1429 |
| 6_1252839 | 4 | 0.75 | 2 | 0.5 | 1 | 0 | 2 | 1 | 9 | 0.6667 | 3 | 0.6667 | NA | NA | 1 | 1 | 1 | 0 | 5 | 0.6 |
| 6_1252839 | 4 | 1 | 1 | 1 | 1 | 1 | 3 | 1 | 9 | 1 | 4 | 1 | NA | NA | 2 | 0.5 | 1 | 1 | 7 | 0.8571 |
| 6_1252840 | 3 | 1 | 2 | 0.5 | 1 | 1 | 1 | 0 | 7 | 0.7143 | 4 | 0.75 | NA | NA | 2 | 1 | 1 | 1 | 7 | 0.8571 |
| 6_1252840 | 3 | 1 | 1 | 0 | 1 | 1 | 3 | 0.6667 | 8 | 0.75 | 4 | 0.5 | NA | NA | 2 | 0.5 | 1 | 1 | 7 | 0.5714 |
| 6_1252840 | 2 | 1 | 1 | 1 | 1 | 1 | 1 | 1 | 5 | 1 | 4 | 0.75 | NA | NA | NA | NA | NA | NA | 4 | 0.75 |
| 6_1252841 | 3 | 0.3333 | 2 | 0.5 | 1 | 1 | 3 | 0.3333 | 9 | 0.4444 | 4 | 0.25 | NA | NA | 2 | 0.5 | 1 | 0 | 7 | 0.2857 |
| 6_1252842 | 3 | 0.3333 | 2 | 1 | 1 | 0 | 3 | 0.6667 | 9 | 0.5556 | 3 | 0.6667 | NA | NA | 2 | 0.5 | NA | NA | 5 | 0.6 |
| 6_1253389 | 1 | 1 | 4 | 0.75 | 1 | 1 | 1 | 1 | 7 | 0.8571 | 4 | 0.75 | 4 | 1 | 3 | 1 | 3 | 1 | 14 | 0.9286 |
| 6_1253389 | 2 | 0.5 | 4 | 0.25 | NA | NA | NA | NA | 6 | 0.3333 | 3 | 1 | 4 | 0.75 | 3 | 0.6667 | 3 | 0.3333 | 13 | 0.6923 |
| 6_1253390 | 2 | 1 | 5 | 0.6 | NA | NA | 1 | 0 | 8 | 0.625 | 3 | 0.6667 | 4 | 0.75 | 3 | 1 | 4 | 0.75 | 14 | 0.7857 |
| 6_1253391 | 2 | 0.5 | 5 | 1 | 1 | 1 | 1 | 1 | 9 | 0.8889 | 3 | 1 | 3 | 1 | 3 | 1 | 4 | 1 | 13 | 1 |
| 6_1253392 | 3 | 0.3333 | 6 | 0.1667 | 1 | 0 | 1 | 0 | 11 | 0.1818 | 4 | 0.5 | 3 | 1 | 3 | 0.3333 | 4 | 0.25 | 14 | 0.5 |
| 6_1253393 | 1 | 0 | 3 | 0.3333 | NA | NA | 1 | 0 | 5 | 0.2 | 5 | 1 | 1 | 0 | 3 | 0.6667 | 3 | 0.6667 | 12 | 0.75 |
| 6_1260168 | 2 | 0.5 | NA | NA | NA | NA | 4 | 0 | 6 | 0.1667 | NA | NA | 4 | 0.5 | 6 | 0.1667 | 1 | 0 | 11 | 0.2727 |
| 6_1265095 | 7 | 0 | 4 | 0.25 | 5 | 0 | 6 | 0.1667 | 22 | 0.0909 | NA | NA | 6 | 0.1667 | 7 | 0.1429 | 3 | 0.3333 | 16 | 0.1875 |
| 6_1265096 | 6 | 0 | 2 | 0 | 5 | 0 | 6 | 0.3333 | 19 | 0.1053 | NA | NA | 4 | 0 | 9 | 0 | 3 | 0.3333 | 16 | 0.0625 |
| 6_1265097 | 9 | 0.5556 | 5 | 0.4 | 5 | 0.4 | 7 | 0.5714 | 26 | 0.5 | NA | NA | 10 | 0.6 | 9 | 0.3333 | 3 | 0 | 22 | 0.4091 |
| 6_1265097 | 8 | 0 | 5 | 0.2 | 5 | 0.2 | 5 | 0 | 23 | 0.087 | NA | NA | 9 | 0.1111 | 8 | 0 | 2 | 0 | 19 | 0.0526 |
| 6_1265099 | 5 | 1 | 3 | 1 | 4 | 1 | 9 | 0.8889 | 21 | 0.9524 | NA | NA | 9 | 0.8889 | 8 | 0.625 | 1 | 1 | 18 | 0.7778 |
| 6_1265103 | 6 | 0.5 | 4 | 0.5 | 4 | 0.25 | 8 | 0.375 | 22 | 0.4091 | NA | NA | 7 | 0.5714 | 10 | 0.7 | 3 | 0 | 20 | 0.55 |
| 6_1265103 | 7 | 1 | 5 | 1 | 5 | 1 | 6 | 1 | 23 | 1 | NA | NA | 10 | 1 | 11 | 1 | 2 | 1 | 23 | 1 |
| 6_1265103 | 7 | 0 | 5 | 0.4 | 4 | 0 | 7 | 0 | 23 | 0.087 | NA | NA | 10 | 0.2 | 10 | 0.1 | 3 | 0 | 23 | 0.1304 |
| 6_1265104 | 6 | 0.5 | 4 | 0.25 | 4 | 0 | 6 | 0.5 | 20 | 0.35 | NA | NA | 8 | 0.5 | 9 | 0.6667 | 2 | 0 | 19 | 0.5263 |
| 6_1265104 | 7 | 0.1429 | 4 | 0.5 | 4 | 0.5 | 8 | 0.5 | 23 | 0.3913 | NA | NA | 9 | 0.3333 | 11 | 0.0909 | 3 | 0.3333 | 23 | 0.2174 |
| 6_1265104 | 4 | 0.5 | 5 | 1 | 4 | 0.5 | 7 | 0.5714 | 20 | 0.65 | NA | NA | 9 | 0.7778 | 8 | 0.75 | 3 | 0.6667 | 20 | 0.75 |
| 6_1265105 | 6 | 0.1667 | 5 | 0.8 | 4 | 0.25 | 7 | 0.2857 | 22 | 0.3636 | NA | NA | 7 | 0.4286 | 9 | 0.2222 | 2 | 0 | 18 | 0.2778 |
| 6_1265107 | 5 | 1 | 3 | 1 | 5 | 1 | 7 | 1 | 20 | 1 | NA | NA | 9 | 0.8889 | 9 | 0.7778 | 2 | 1 | 20 | 0.85 |
| 6_1265109 | 6 | 0.8333 | 3 | 0.6667 | 5 | 1 | 6 | 1 | 20 | 0.9 | NA | NA | 9 | 1 | 6 | 0.8333 | 1 | 1 | 16 | 0.9375 |
| 6_1265110 | 4 | 1 | 1 | 1 | 3 | 1 | 4 | 0.75 | 12 | 0.9167 | NA | NA | 8 | 1 | 5 | 0.8 | 1 | 1 | 14 | 0.9286 |
| 6_1265111 | 4 | 1 | 2 | 1 | 4 | 1 | 6 | 1 | 16 | 1 | NA | NA | 9 | 0.7778 | 6 | 1 | 1 | 1 | 16 | 0.875 |
| 6_1265113 | 4 | 1 | 3 | 0.6667 | 3 | 0.3333 | 5 | 0.6 | 15 | 0.6667 | NA | NA | 7 | 0.5714 | 6 | 0.6667 | 1 | 1 | 14 | 0.6429 |
| 6_1265118 | 4 | 0.5 | 3 | 0.6667 | 2 | 0.5 | 4 | 1 | 13 | 0.6923 | NA | NA | 7 | 0.5714 | 5 | 0.4 | 1 | 1 | 13 | 0.5385 |
| 6_1265120 | 2 | 1 | 1 | 0 | 1 | 1 | 4 | 0.75 | 8 | 0.75 | NA | NA | 6 | 0.3333 | 5 | 0.6 | 1 | 1 | 12 | 0.5 |
| 6_1265263 | 2 | 0 | 5 | 0 | 4 | 0.5 | 1 | 0 | 12 | 0.1667 | 1 | 1 | NA | NA | 1 | 0 | 2 | 0 | 4 | 0.25 |
| 6_1265265 | 3 | 0.3333 | 4 | 0 | 2 | 0 | NA | NA | 9 | 0.1111 | 1 | 1 | NA | NA | 1 | 0 | 2 | 0.5 | 4 | 0.5 |
| 6_1265265 | 3 | 1 | 2 | 1 | 4 | 0.5 | 1 | 1 | 10 | 0.8 | 1 | 1 | NA | NA | NA | NA | 1 | 1 | 2 | 1 |
| 6_1265266 | 3 | 0.6667 | 4 | 1 | 4 | 1 | 1 | 1 | 12 | 0.9167 | 1 | 1 | NA | NA | 1 | 0 | 1 | 0 | 3 | 0.3333 |
| 6_1265267 | 3 | 0 | 5 | 0 | 3 | 0.3333 | 1 | 0 | 12 | 0.0833 | 1 | 1 | NA | NA | 1 | 0 | 1 | 0 | 3 | 0.3333 |
| 6_1265268 | 2 | 0 | 4 | 0.25 | 2 | 0.5 | NA | NA | 8 | 0.25 | NA | NA | NA | NA | 1 | 0 | 1 | 1 | 2 | 0.5 |
| 6_1265268 | 3 | 0.6667 | 5 | 1 | 4 | 1 | 1 | 1 | 13 | 0.9231 | 1 | 1 | NA | NA | 1 | 1 | 1 | 1 | 3 | 1 |
| 6_1265269 | 3 | 0 | 5 | 0.4 | 4 | 0.25 | 1 | 0 | 13 | 0.2308 | 1 | 1 | NA | NA | 1 | 0 | 1 | 0 | 3 | 0.3333 |
| 6_1265270 | 1 | 0 | 5 | 0.8 | 4 | 0.75 | NA | NA | 10 | 0.7 | 1 | 1 | NA | NA | 1 | 1 | 1 | 0 | 3 | 0.6667 |
| 6_1266136 | 28 | 0.0714 | 27 | 0.1481 | 31 | 0.1613 | 38 | 0.0789 | 124 | 0.1129 | 26 | 0.0769 | 18 | 0.1111 | 21 | 0.0476 | 27 | 0.1481 | 92 | 0.0978 |
| 6_1266137 | 26 | 0.1154 | 28 | 0.0357 | 29 | 0.069 | 33 | 0.0606 | 116 | 0.069 | 27 | 0 | 18 | 0.1667 | 21 | 0.0476 | 25 | 0.04 | 91 | 0.0549 |
| 6_1266148 | 11 | 0 | 22 | 0.0909 | 23 | 0.0435 | 21 | 0.0476 | 77 | 0.0519 | 14 | 0 | 12 | 0 | 10 | 0.1 | 19 | 0.1579 | 55 | 0.0727 |
| 6_1266149 | 8 | 0 | 19 | 0 | 22 | 0.1364 | 24 | 0.0417 | 73 | 0.0548 | 16 | 0.25 | 15 | 0.0667 | 8 | 0.125 | 20 | 0.1 | 59 | 0.1356 |
| 6_1266157 | 6 | 0.3333 | 9 | 0.4444 | 13 | 0.3077 | 17 | 0.3529 | 45 | 0.3556 | 9 | 0.4444 | 14 | 0.3571 | 8 | 0.125 | 10 | 0.2 | 41 | 0.2927 |
| 6_1266158 | 8 | 0.125 | 12 | 0.5833 | 14 | 0.5 | 18 | 0.7222 | 52 | 0.5385 | 10 | 0.7 | 13 | 0.6923 | 6 | 0.6667 | 12 | 0.5 | 41 | 0.6341 |
| 6_1266158 | 9 | 0.8889 | 12 | 0.3333 | 14 | 0.2143 | 21 | 0.2857 | 56 | 0.375 | 13 | 0.3846 | 14 | 0.2857 | 7 | 0.5714 | 14 | 0.4286 | 48 | 0.3958 |
| 6_1266159 | 8 | 0.875 | 10 | 0.8 | 7 | 1 | 14 | 0.9286 | 39 | 0.8974 | 8 | 0.75 | 8 | 1 | 8 | 1 | 11 | 1 | 35 | 0.9429 |
| 6_1266159 | 8 | 0.625 | 15 | 0.2667 | 15 | 0.3333 | 19 | 0.4211 | 57 | 0.386 | 12 | 0.4167 | 14 | 0.3571 | 8 | 0.625 | 15 | 0.4667 | 49 | 0.449 |
| 6_1266159 | 9 | 0.6667 | 11 | 0.4545 | 11 | 0.6364 | 18 | 0.5556 | 49 | 0.5714 | 12 | 0.5 | 15 | 0.5333 | 8 | 0.5 | 14 | 0.7143 | 49 | 0.5714 |
| 6_1266159 | 5 | 1 | 10 | 1 | 12 | 1 | 18 | 0.9444 | 45 | 0.9778 | 13 | 1 | 14 | 0.8571 | 7 | 1 | 14 | 1 | 48 | 0.9583 |
| 6_1266160 | 7 | 0.1429 | 11 | 0.2727 | 11 | 0.2727 | 15 | 0.1333 | 44 | 0.2045 | 10 | 0.2 | 12 | 0.5 | 7 | 0.1429 | 12 | 0.4167 | 41 | 0.3415 |
| 6_1266160 | 7 | 0.4286 | 13 | 0.1538 | 13 | 0.0769 | 15 | 0.2 | 48 | 0.1875 | 10 | 0 | 13 | 0.1538 | 8 | 0 | 14 | 0.3571 | 45 | 0.1556 |
| 6_1266161 | 7 | 0.5714 | 13 | 0.6154 | 13 | 0.7692 | 12 | 0.75 | 45 | 0.6889 | 11 | 0.7273 | 10 | 0.6 | 5 | 1 | 11 | 0.9091 | 37 | 0.7838 |
| 6_1266161 | 8 | 1 | 14 | 1 | 13 | 1 | 16 | 0.9375 | 51 | 0.9804 | 13 | 1 | 13 | 1 | 8 | 1 | 15 | 1 | 49 | 1 |
| 6_1266162 | 8 | 0.5 | 13 | 0.3077 | 13 | 0.6154 | 16 | 0.3125 | 50 | 0.42 | 13 | 0.5385 | 13 | 0.5385 | 8 | 0.375 | 13 | 0.7692 | 47 | 0.5745 |

| | | | | | | | | | | | | | | | | | | | | |
|---|---|---|---|---|---|---|---|---|---|---|---|---|---|---|---|---|---|---|---|---|
| 6_1266162 | 8 | 0.375 | 13 | 0.0769 | 15 | 0.3333 | 13 | 0.4615 | 49 | 0.3061 | 12 | 0.25 | 11 | 0.4545 | 8 | 0 | 14 | 0.2143 | 45 | 0.2444 |
| 6_1266162 | 5 | 0.2 | 12 | 0.25 | 15 | 0.2 | 13 | 0.1538 | 45 | 0.2 | 11 | 0.2727 | 14 | 0.2143 | 7 | 0.2857 | 12 | 0.25 | 44 | 0.25 |
| 6_1266165 | 7 | 0.1429 | 11 | 0.5455 | 13 | 0.3846 | 14 | 0.5 | 45 | 0.4222 | 11 | 0.3636 | 11 | 0.4545 | 7 | 0.7143 | 12 | 0.3333 | 41 | 0.439 |
| 6_1266166 | 7 | 0.7143 | 10 | 0.3 | 12 | 0.3333 | 12 | 0.5 | 41 | 0.439 | 9 | 0.2222 | 10 | 0.4 | 7 | 0.5714 | 13 | 0.5385 | 39 | 0.4359 |
| 6_1266166 | 7 | 1 | 10 | 1 | 14 | 1 | 12 | 1 | 43 | 1 | 8 | 1 | 12 | 1 | 7 | 0.8571 | 12 | 1 | 39 | 0.9744 |
| 6_1266167 | 6 | 1 | 11 | 0.8182 | 12 | 1 | 12 | 1 | 41 | 0.9512 | 7 | 1 | 10 | 0.9 | 6 | 0.8333 | 10 | 0.7 | 33 | 0.8485 |
| 6_1266168 | 2 | 0.5 | 9 | 0.7778 | 11 | 0.7273 | 12 | 0.75 | 34 | 0.7353 | 7 | 0.7143 | 10 | 0.7 | 4 | 1 | 11 | 0.6364 | 32 | 0.7188 |
| 6_1266168 | 5 | 0.6 | 8 | 0.5 | 11 | 0.5455 | 11 | 0.5455 | 35 | 0.5429 | 8 | 0.75 | 9 | 0.5556 | 6 | 0.6667 | 10 | 0.6 | 33 | 0.6364 |
| 6_1266168 | 4 | 0.25 | 10 | 0.1 | 12 | 0.1667 | 11 | 0.5455 | 37 | 0.2703 | 8 | 0.125 | 8 | 0.375 | 4 | 0.5 | 8 | 0.25 | 28 | 0.2857 |
| 6_1266168 | 2 | 1 | 7 | 0.8571 | 12 | 0.9167 | 7 | 1 | 28 | 0.9286 | 8 | 0.875 | 6 | 1 | 5 | 1 | 8 | 1 | 27 | 0.963 |
| 6_1266168 | 5 | 0 | 11 | 0.3636 | 14 | 0.3571 | 12 | 0.4167 | 42 | 0.3333 | 11 | 0.4545 | 11 | 0.2727 | 7 | 0.5714 | 12 | 0.5833 | 41 | 0.4634 |
| 6_1266171 | 5 | 0.8 | 10 | 0.3 | 11 | 0.2727 | 7 | 0.1429 | 33 | 0.3333 | 8 | 0.125 | 7 | 0.1429 | 5 | 0.6 | 10 | 0.3 | 30 | 0.2667 |
| 6_1266182 | 2 | 1 | 3 | 0.3333 | 3 | 0.6667 | 3 | 0.6667 | 11 | 0.6364 | 3 | 0 | 4 | 0.25 | 2 | 0.5 | 3 | 0 | 12 | 0.1667 |
| 6_1266183 | 2 | 0 | 5 | 0 | 3 | 0.3333 | 5 | 0 | 15 | 0.0667 | 5 | 0.2 | 3 | 0 | 4 | 0 | 5 | 0 | 17 | 0.0588 |
| 6_1266188 | 3 | 0.3333 | 6 | 0 | 3 | 0.3333 | 6 | 0.3333 | 18 | 0.2222 | 6 | 0.5 | 5 | 0.2 | 6 | 0.1667 | 4 | 0.25 | 21 | 0.2857 |
| 6_1266188 | 2 | 0 | 6 | 0.1667 | 5 | 0.2 | 4 | 0 | 17 | 0.1176 | 4 | 0.25 | 5 | 0 | 6 | 0 | 4 | 0 | 19 | 0.0526 |
| 6_1266188 | 3 | 0.6667 | 4 | 1 | 5 | 1 | 5 | 1 | 17 | 0.9412 | 4 | 1 | 4 | 0.5 | 5 | 1 | 4 | 0.5 | 17 | 0.7647 |
| 6_1266189 | 2 | 0.5 | 3 | 0 | 6 | 0.1667 | 3 | 0.3333 | 14 | 0.2143 | 3 | 0.6667 | 3 | 0 | 5 | 0.4 | 3 | 0.6667 | 14 | 0.4286 |
| 6_1266190 | 3 | 0 | 6 | 0 | 4 | 0.25 | 2 | 0 | 15 | 0.0667 | 6 | 0 | 5 | 0.2 | 5 | 0 | 3 | 0 | 19 | 0.0526 |
| 6_1266190 | 2 | 0.5 | 5 | 0.6 | 5 | 0.2 | 5 | 0.2 | 17 | 0.3529 | 2 | 1 | 4 | 0.25 | 6 | 0.6667 | 5 | 0.6 | 17 | 0.5882 |
| 6_1266191 | 1 | 0 | 6 | 0 | 2 | 0 | 5 | 0.4 | 14 | 0.1429 | 4 | 0.25 | 4 | 0 | 4 | 0 | 5 | 0.2 | 17 | 0.1176 |
| 6_1266191 | 3 | 0 | 6 | 0.1667 | 2 | 0 | 4 | 0.25 | 15 | 0.1333 | 6 | 0.1667 | 3 | 0 | 4 | 0 | 5 | 0.4 | 18 | 0.1667 |
| 6_1266194 | 3 | 0 | 3 | 0 | 4 | 0.5 | 3 | 0 | 13 | 0.1538 | 5 | 0 | 3 | 0.3333 | 4 | 0.25 | 5 | 0.2 | 17 | 0.1765 |
| 6_1266195 | NA | NA | 3 | 1 | 2 | 1 | 2 | 1 | 7 | 1 | 4 | 1 | 4 | 1 | 3 | 0.6667 | 4 | 0.75 | 15 | 0.8667 |
| 6_1266195 | 3 | 0.6667 | 4 | 0.25 | 3 | 0 | 2 | 0.5 | 12 | 0.3333 | 3 | 0.3333 | 3 | 0 | 3 | 0.6667 | 3 | 0.6667 | 12 | 0.4167 |
| 6_1266195 | 3 | 1 | 4 | 0.75 | 3 | 1 | 2 | 1 | 12 | 0.9167 | 4 | 1 | 4 | 1 | 3 | 0.6667 | 4 | 1 | 15 | 0.9333 |
| 6_1266195 | 3 | 0.3333 | 4 | 0 | 3 | 0.3333 | 2 | 0 | 12 | 0.1667 | 4 | 0.25 | 4 | 0.5 | 3 | 0.3333 | 5 | 0.2 | 16 | 0.3125 |
| 6_1266196 | 3 | 0.3333 | 4 | 0.25 | 3 | 0.3333 | 2 | 0.5 | 12 | 0.3333 | 4 | 0.75 | 3 | 0.3333 | 3 | 0 | 5 | 0.4 | 15 | 0.4 |
| 6_1266197 | 3 | 0.6667 | 4 | 0.5 | 3 | 0.6667 | 2 | 1 | 12 | 0.6667 | 4 | 0.75 | 4 | 1 | 4 | 1 | 4 | 1 | 16 | 0.9375 |
| 6_1266197 | 3 | 0 | 4 | 0.5 | 1 | 1 | 2 | 0 | 10 | 0.3 | 4 | 0 | 4 | 0.5 | 2 | 0 | 4 | 0 | 14 | 0.1429 |
| 6_1266198 | 2 | 1 | 4 | 0.25 | 3 | 0 | 2 | 0.5 | 11 | 0.3636 | 4 | 0 | 3 | 0.3333 | 4 | 0 | 5 | 0.8 | 16 | 0.3125 |
| 6_1266199 | 2 | 0.5 | 3 | 1 | 3 | 0.6667 | 1 | 0 | 9 | 0.6667 | 2 | 0.5 | 2 | 1 | 2 | 1 | 4 | 0.75 | 10 | 0.8 |
| 6_1266201 | 2 | 1 | 2 | 0.5 | 2 | 0.5 | 2 | 0.5 | 8 | 0.625 | 4 | 0.75 | 4 | 1 | 4 | 0.5 | 4 | 0.75 | 16 | 0.75 |
| 6_1266204 | 2 | 1 | 2 | 0.5 | 2 | 0.5 | 2 | 1 | 8 | 0.75 | 4 | 0.5 | 4 | 0.5 | 1 | 0 | 5 | 0.4 | 14 | 0.4286 |
| 6_1266204 | 2 | 0 | 2 | 0.5 | 3 | 0 | 2 | 0 | 9 | 0.1111 | 3 | 0 | 4 | 0 | 1 | 0 | 4 | 0.25 | 12 | 0.0833 |
| 6_1266204 | 2 | 0 | 2 | 0 | 2 | 0.5 | 2 | 0 | 8 | 0.125 | 4 | 0 | 4 | 0.25 | 1 | 0 | 5 | 0.2 | 14 | 0.1429 |
| 6_1266217 | NA | NA | 1 | 1 | 1 | 0 | NA | NA | 2 | 0.5 | 1 | 1 | 1 | 0 | 1 | 0 | NA | NA | 3 | 0.3333 |
| 6_1266853 | 8 | 0.25 | 13 | 0.0769 | 11 | 0.0909 | 9 | 0 | 41 | 0.0976 | 9 | 0 | 6 | 0 | 8 | 0.125 | 7 | 0.2857 | 30 | 0.1 |
| 6_1266855 | 9 | 0.6667 | 11 | 0.7273 | 13 | 0.7692 | 10 | 0.6 | 43 | 0.6977 | 9 | 0.8889 | 7 | 0.5714 | 4 | 0.5 | 5 | 0.8 | 25 | 0.72 |
| 6_1266855 | 8 | 0.125 | 8 | 0.125 | 7 | 0 | 10 | 0.2 | 33 | 0.1212 | 7 | 0.1429 | 6 | 0 | 5 | 0.2 | 4 | 0.25 | 22 | 0.1364 |
| 6_1266856 | 4 | 0 | 13 | 0.2308 | 13 | 0.0769 | 10 | 0.1 | 40 | 0.125 | 10 | 0.2 | 7 | 0 | 8 | 0.125 | 5 | 0 | 30 | 0.1 |
| 6_1267122 | 5 | 0 | 10 | 0.5 | 7 | 0.1429 | 7 | 0.1429 | 29 | 0.2414 | 6 | 0.1667 | 5 | 0.2 | 3 | 0.3333 | 6 | 0.1667 | 20 | 0.2 |
| 6_1267122 | 6 | 0.5 | 13 | 0.4615 | 8 | 0.375 | 8 | 0.375 | 35 | 0.4286 | 9 | 0.7778 | 4 | 0 | 3 | 1 | 3 | 0.6667 | 19 | 0.6316 |
| 6_1267123 | 4 | 0.25 | 11 | 0.0909 | 7 | 0.2857 | 5 | 0 | 27 | 0.1481 | 4 | 0.5 | 2 | 0.5 | 4 | 0.5 | 4 | 0.25 | 14 | 0.4286 |
| 6_1267123 | 6 | 0.1667 | 11 | 0.2727 | 10 | 0.2 | 7 | 0.1429 | 34 | 0.2059 | 6 | 0.3333 | 4 | 0 | 4 | 0.5 | 2 | 0.5 | 16 | 0.3125 |
| 6_1267123 | 6 | 0.1667 | 11 | 0.1818 | 8 | 0.25 | 8 | 0 | 33 | 0.1515 | 9 | 0.3333 | 3 | 0 | 4 | 0 | 3 | 0.3333 | 19 | 0.2105 |
| 6_1267124 | 6 | 0.5 | 9 | 0.7778 | 7 | 0.2857 | 8 | 0.5 | 30 | 0.5333 | 8 | 0.625 | 3 | 0.3333 | 4 | 0.75 | 4 | 0.25 | 19 | 0.5263 |
| 6_1267124 | 5 | 0.8 | 9 | 0.6667 | 6 | 0.5 | 8 | 0.5 | 28 | 0.6071 | 6 | 0.5 | 3 | 0.3333 | 5 | 0.8 | 2 | 1 | 16 | 0.625 |
| 6_1267124 | 5 | 0.2 | 9 | 0.3333 | 5 | 0.2 | 7 | 0 | 26 | 0.1923 | 3 | 0.3333 | 3 | 0 | 5 | 0.4 | 4 | 0 | 15 | 0.2 |
| 6_1267125 | 5 | 0 | 10 | 0.2 | 3 | 0 | 5 | 0 | 23 | 0.087 | 5 | 0 | 3 | 0.3333 | 4 | 0 | 3 | 0.3333 | 15 | 0.1333 |
| 6_1267126 | 5 | 0.8 | 9 | 0.4444 | 4 | 0.75 | 5 | 0.6 | 23 | 0.6087 | 6 | 0.8333 | 2 | 1 | 4 | 0.5 | 3 | 0.6667 | 15 | 0.7333 |
| 6_1267126 | 4 | 0.25 | 8 | 0.25 | 4 | 0.5 | 5 | 0.2 | 21 | 0.2857 | 5 | 0.4 | 3 | 0.3333 | 4 | 0.5 | 3 | 0.3333 | 15 | 0.4 |
| 6_1267128 | 4 | 1 | 8 | 0.75 | 4 | 0.5 | 4 | 1 | 20 | 0.8 | 4 | 0.75 | 3 | 0.6667 | 4 | 0.75 | 2 | 0 | 13 | 0.6154 |
| 6_1267128 | 4 | 1 | 8 | 1 | 3 | 0.6667 | 2 | 1 | 17 | 0.9412 | 3 | 0.6667 | 1 | 1 | 2 | 1 | 1 | 1 | 7 | 0.8571 |
| 6_1267128 | 3 | 0.6667 | 7 | 0.4286 | 3 | 1 | 3 | 0.6667 | 16 | 0.625 | 3 | 0.3333 | 3 | 0 | 3 | 0.6667 | 3 | 0.6667 | 12 | 0.4167 |
| 6_1267129 | 5 | 0.2 | 10 | 0.2 | 4 | 0 | 4 | 0 | 23 | 0.1304 | 4 | 0.25 | 3 | 0 | 3 | 0 | 2 | 0 | 12 | 0.0833 |
| 6_1267130 | 4 | 1 | 9 | 0.5556 | 3 | 0 | 3 | 0 | 19 | 0.4737 | 3 | 0.6667 | 2 | 0.5 | 3 | 0.6667 | 3 | 0.6667 | 11 | 0.6364 |
| 6_1267130 | 5 | 0.8 | 6 | 0.8333 | 1 | 1 | 3 | 0.6667 | 15 | 0.8 | 3 | 0.6667 | NA | NA | 1 | 1 | 2 | 1 | 6 | 0.8333 |
| 6_1267131 | 5 | 0.2 | 7 | 0.1429 | 3 | 0 | 4 | 0.5 | 19 | 0.2105 | 4 | 0.5 | 2 | 0 | 2 | 0 | 3 | 0.6667 | 11 | 0.3636 |
| 6_1267134 | 3 | 0.3333 | 3 | 0.3333 | 3 | 0 | 3 | 0.3333 | 12 | 0.25 | 3 | 0.3333 | 1 | 0 | 2 | 0.5 | 3 | 0.3333 | 9 | 0.3333 |
| 6_1267138 | 3 | 0.3333 | 5 | 0.4 | 2 | 0 | 3 | 0.3333 | 13 | 0.3077 | 3 | 0 | 2 | 0.5 | 2 | 0.5 | 3 | 0 | 10 | 0.2 |
| 6_1267138 | 3 | 1 | 6 | 1 | 3 | 0.6667 | 3 | 1 | 15 | 0.9333 | 3 | 0.6667 | 2 | 1 | 2 | 1 | 3 | 0.6667 | 10 | 0.8 |
| 6_1267139 | 2 | 0.5 | 6 | 0.5 | 3 | 0.3333 | 3 | 0 | 14 | 0.3571 | 2 | 0 | 2 | 0.5 | 2 | 1 | 2 | 0.5 | 8 | 0.5 |
| 6_1267143 | 2 | 1 | 7 | 1 | NA | NA | 1 | 1 | 10 | 1 | 3 | 1 | NA | NA | 1 | 1 | 3 | 0.6667 | 7 | 0.8571 |
| 6_1267144 | 1 | 1 | 6 | 1 | 2 | 1 | 2 | 1 | 11 | 1 | 3 | 1 | 2 | 0.5 | 2 | 0.5 | 1 | 1 | 8 | 0.75 |

| | | | | | | | | | | | | | | | | | | | | |
|---|---|---|---|---|---|---|---|---|---|---|---|---|---|---|---|---|---|---|---|---|
| 6_1267144 | 2 | 0.5 | 8 | 0.75 | 2 | 0 | 2 | 0 | 14 | 0.5 | 2 | 0.5 | 2 | 0.5 | 1 | 0 | 3 | 0.3333 | 8 | 0.375 |
| 6_1267144 | 2 | 1 | 8 | 0.875 | 2 | 1 | 2 | 1 | 14 | 0.9286 | 3 | 1 | 2 | 1 | 1 | 1 | 2 | 1 | 8 | 1 |
| 6_1267144 | 2 | 0 | 7 | 0.2857 | 2 | 0 | 1 | 0 | 12 | 0.1667 | 3 | 0.3333 | 1 | 0 | 2 | 0 | 3 | 0 | 9 | 0.1111 |
| 6_1267145 | 2 | 1 | 7 | 0.7143 | 2 | 0.5 | 2 | 0.5 | 13 | 0.6923 | 2 | 0.5 | 1 | 0 | 1 | 1 | 2 | 0.5 | 6 | 0.5 |
| 6_1267146 | 2 | 0 | 6 | 0.3333 | 1 | 0 | 2 | 0.5 | 11 | 0.2727 | 2 | 0 | 2 | 0.5 | 1 | 0 | 2 | 0 | 7 | 0.1429 |
| 6_1267147 | 2 | 0.5 | 7 | 1 | 1 | 1 | 1 | 1 | 11 | 0.9091 | 2 | 1 | 2 | 0.5 | 1 | 1 | 1 | 1 | 6 | 0.8333 |
| 6_1267147 | 2 | 1 | 4 | 1 | NA | NA | 1 | 1 | 7 | 1 | 1 | 1 | 1 | 1 | 1 | 0 | 2 | 1 | 5 | 0.8 |
| 6_1267149 | 2 | 0.5 | 8 | 0.125 | 1 | 0 | 2 | 0 | 13 | 0.1538 | 1 | 0 | 2 | 0.5 | 1 | 0 | 2 | 0 | 6 | 0.1667 |
| 6_1267150 | 2 | 0.5 | 5 | 0.8 | 1 | 1 | NA | NA | 8 | 0.75 | 1 | 1 | 2 | 1 | 1 | 1 | 2 | 1 | 6 | 1 |
| 6_1267150 | 2 | 0 | 6 | 0.6667 | 1 | 0 | 1 | 1 | 10 | 0.5 | 2 | 1 | 2 | 0 | 1 | 1 | 1 | 0 | 6 | 0.5 |
| 6_1267150 | 2 | 1 | 6 | 0.6667 | 1 | 0 | 2 | 0.5 | 11 | 0.6364 | 2 | 0.5 | 2 | 0 | 2 | 0.5 | 2 | 0.5 | 8 | 0.375 |
| 6_1267151 | 2 | 1 | 5 | 1 | 1 | 1 | 2 | 1 | 10 | 1 | 2 | 1 | 2 | 1 | 2 | 1 | 2 | 1 | 8 | 1 |
| 6_1267151 | 2 | 1 | 6 | 1 | 1 | 1 | NA | NA | 9 | 1 | 2 | 1 | 2 | 1 | 1 | 1 | 2 | 0.5 | 7 | 0.8571 |
| 6_1267151 | 2 | 1 | 6 | 0.5 | 1 | 0 | 2 | 0 | 11 | 0.4545 | 2 | 0.5 | 1 | 1 | 2 | 0 | 1 | 0 | 6 | 0.3333 |
| 6_1267152 | 2 | 0 | 7 | 0.1429 | 1 | 0 | 2 | 0.5 | 12 | 0.1667 | 2 | 0.5 | 1 | 0 | 1 | 1 | 2 | 1 | 6 | 0.6667 |
| 6_1267153 | 2 | 0.5 | 5 | 0.6 | 1 | 0 | 2 | 0 | 10 | 0.4 | 2 | 0 | 1 | 0 | 1 | 1 | 2 | 0.5 | 6 | 0.3333 |
| 6_1267154 | 1 | 1 | 3 | 0.3333 | 1 | 1 | 1 | 1 | 6 | 0.6667 | 1 | 1 | 1 | 1 | NA | NA | 2 | 0.5 | 4 | 0.75 |
| 6_1267154 | 2 | 1 | 7 | 1 | 1 | 1 | 1 | 0 | 11 | 0.9091 | 2 | 0.5 | 1 | 1 | 1 | 0 | 2 | 1 | 6 | 0.6667 |
| 6_1267156 | 2 | 0.5 | 5 | 0.6 | 1 | 1 | 2 | 0.5 | 10 | 0.6 | 2 | 1 | 1 | 1 | NA | NA | 2 | 0 | 5 | 0.6 |
| 6_1267156 | 2 | 0.5 | 4 | 0.25 | 1 | 0 | 1 | 0 | 8 | 0.25 | 2 | 0.5 | NA | NA | 1 | 0 | NA | NA | 3 | 0.3333 |
| 6_1267159 | 2 | 0.5 | 4 | 0.25 | 1 | 1 | 1 | 1 | 8 | 0.5 | 2 | 0.5 | 1 | 1 | 1 | 1 | 2 | 1 | 6 | 0.8333 |
| 6_1267159 | 2 | 1 | 4 | 0.75 | NA | NA | NA | NA | 6 | 0.8333 | 1 | 1 | 1 | 0 | 1 | 1 | 2 | 1 | 5 | 0.8 |
| 6_1267161 | 2 | 0.5 | 3 | 0.3333 | 1 | 1 | NA | NA | 6 | 0.5 | 2 | 1 | NA | NA | 1 | 0 | 2 | 1 | 5 | 0.8 |
| 6_1267165 | 2 | 1 | 4 | 1 | 1 | 1 | NA | NA | 7 | 1 | 1 | 1 | NA | NA | 1 | 1 | 1 | 1 | 3 | 1 |
| 6_1267173 | NA | NA | NA | NA | 2 | 1 | NA | NA | 2 | 1 | 1 | 1 | 1 | 1 | NA | NA | NA | NA | 2 | 1 |
| 6_1269011 | NA | NA | NA | NA | 2 | 0.5 | NA | NA | 2 | 0.5 | NA | NA | NA | NA | NA | NA | 1 | 1 | 1 | 1 |
| 6_1269521 | 5 | 0.4 | 4 | 0 | 4 | 0 | 5 | 0 | 18 | 0.1111 | 4 | 0.25 | 2 | 0 | 4 | 0.25 | 6 | 0.1667 | 16 | 0.1875 |
| 6_1269522 | 7 | 0.5714 | 3 | 0.3333 | 5 | 0 | 7 | 0.4286 | 22 | 0.3636 | 4 | 0.25 | 3 | 0.6667 | 5 | 0.2 | 5 | 0.2 | 17 | 0.2941 |
| 6_1269522 | 7 | 0.5714 | 4 | 1 | 5 | 0.8 | 6 | 0.5 | 22 | 0.6818 | 4 | 0.75 | 2 | 1 | 5 | 0.8 | 5 | 0.6 | 16 | 0.75 |
| 6_1269522 | 5 | 0 | 4 | 0 | 4 | 0 | 6 | 0.1667 | 19 | 0.0526 | 3 | 0 | 3 | 0 | 4 | 0.25 | 6 | 0.1667 | 16 | 0.125 |
| 6_1269524 | 4 | 0.75 | 3 | 1 | 4 | 0.75 | 6 | 0.6667 | 17 | 0.7647 | 2 | 1 | 3 | 0.6667 | 5 | 1 | 6 | 0.6667 | 16 | 0.8125 |
| 6_1269557 | NA | NA | NA | NA | 1 | 1 | NA | NA | 1 | 1 | NA | NA | NA | NA | 1 | 1 | NA | NA | 1 | 1 |
| 6_1270631 | 3 | 0 | 3 | 0 | 5 | 0.2 | 1 | 0 | 12 | 0.0833 | 1 | 1 | 4 | 0 | 2 | 0.5 | 1 | 0 | 8 | 0.25 |
| 6_1270646 | 1 | 1 | NA | NA | 2 | 0.5 | 3 | 0.6667 | 6 | 0.6667 | NA | NA | 2 | 1 | 2 | 1 | NA | NA | 4 | 1 |
| 6_1270647 | 1 | 1 | NA | NA | 2 | 1 | 2 | 1 | 5 | 1 | 1 | 1 | 2 | 1 | 2 | 1 | NA | NA | 5 | 1 |
| 6_1270648 | 1 | 1 | NA | NA | 3 | 0.3333 | 4 | 0.5 | 8 | 0.5 | 1 | 1 | 2 | 0 | 2 | 1 | NA | NA | 5 | 0.6 |
| 6_1270648 | 2 | 0 | NA | NA | 3 | 0.3333 | 4 | 0 | 9 | 0.1111 | 1 | 0 | 3 | 0 | 3 | 0.3333 | NA | NA | 7 | 0.1429 |
| 6_1270649 | 1 | 1 | NA | NA | 1 | 1 | 4 | 0.5 | 6 | 0.6667 | 1 | 1 | 2 | 1 | 1 | 1 | NA | NA | 4 | 1 |
| 6_1270649 | 2 | 0.5 | NA | NA | 1 | 1 | 4 | 1 | 7 | 0.8571 | 1 | 1 | 3 | 0.6667 | 3 | 1 | NA | NA | 7 | 0.8571 |
| 6_1270649 | 2 | 0 | NA | NA | 2 | 0.5 | 4 | 0.75 | 8 | 0.5 | 1 | 1 | 3 | 0.3333 | 3 | 0.6667 | NA | NA | 7 | 0.5714 |
| 6_1270650 | 1 | 0 | NA | NA | 1 | 0 | 2 | 0.5 | 4 | 0.25 | 1 | 0 | 3 | 0.3333 | 2 | 0 | NA | NA | 6 | 0.1667 |
| 6_1271021 | 4 | 0 | 4 | 0.25 | 8 | 0 | 1 | 0 | 17 | 0.0588 | 5 | 0 | 4 | 0 | 1 | 1 | 3 | 0 | 13 | 0.0769 |
| 6_1271068 | 2 | 0.5 | 1 | 0 | NA | NA | NA | NA | 3 | 0.3333 | NA | NA | 2 | 0.5 | NA | NA | 2 | 0.5 | 4 | 0.5 |
| 6_1271886 | NA | NA | NA | NA | 2 | 0.5 | NA | NA | 2 | 0.5 | NA | NA | 1 | 1 | NA | NA | NA | NA | 1 | 1 |
| 6_1271886 | NA | NA | NA | NA | 2 | 1 | NA | NA | 2 | 1 | NA | NA | 1 | 1 | NA | NA | NA | NA | 1 | 1 |
| 6_1271888 | NA | NA | NA | NA | 2 | 0.5 | NA | NA | 2 | 0.5 | NA | NA | 1 | 1 | NA | NA | NA | NA | 1 | 1 |
| 6_1271890 | NA | NA | NA | NA | 2 | 1 | NA | NA | 2 | 1 | NA | NA | 1 | 1 | NA | NA | NA | NA | 1 | 1 |
| 6_1271890 | NA | NA | NA | NA | 2 | 1 | NA | NA | 2 | 1 | NA | NA | 1 | 1 | NA | NA | NA | NA | 1 | 1 |
| 6_1274685 | NA | NA | 4 | 0.75 | NA | NA | 2 | 0.5 | 6 | 0.6667 | NA | NA | NA | NA | 4 | 0.75 | NA | NA | 4 | 0.75 |
| 6_1274685 | NA | NA | 2 | 0.5 | NA | NA | 3 | 0.3333 | 5 | 0.4 | NA | NA | NA | NA | 2 | 1 | NA | NA | 2 | 1 |
| 6_1274686 | NA | NA | 2 | 1 | NA | NA | 2 | 1 | 4 | 1 | NA | NA | NA | NA | 2 | 1 | NA | NA | 2 | 1 |
| 6_1274690 | NA | NA | 2 | 0.5 | NA | NA | 3 | 0.6667 | 5 | 0.6 | NA | NA | NA | NA | 2 | 0 | 1 | 1 | 3 | 0.3333 |
| 6_1274819 | NA | NA | 1 | 1 | 1 | 0 | NA | NA | 2 | 0.5 | NA | NA | NA | NA | 1 | 1 | NA | NA | 1 | 1 |
| 6_1274819 | NA | NA | 1 | 1 | 1 | 0 | NA | NA | 2 | 0.5 | NA | NA | NA | NA | 1 | 1 | NA | NA | 1 | 1 |
| 6_1274820 | NA | NA | 1 | 1 | 1 | 0 | NA | NA | 2 | 0.5 | NA | NA | NA | NA | 1 | 1 | NA | NA | 1 | 1 |
| 6_1274821 | NA | NA | 1 | 1 | 1 | 0 | NA | NA | 2 | 0.5 | NA | NA | NA | NA | 1 | 1 | NA | NA | 1 | 1 |
| 6_1274822 | NA | NA | 1 | 1 | 1 | 0 | NA | NA | 2 | 0.5 | NA | NA | NA | NA | 1 | 1 | NA | NA | 1 | 1 |
| 6_1274825 | NA | NA | 1 | 1 | 1 | 0 | NA | NA | 2 | 0.5 | NA | NA | NA | NA | 1 | 1 | NA | NA | 1 | 1 |
| 6_1276660 | 14 | 0.0714 | 13 | 0.2308 | 9 | 0.3333 | 12 | 0.1667 | 48 | 0.1875 | 7 | 0.1429 | 11 | 0.1818 | 10 | 0.1 | 10 | 0 | 38 | 0.1053 |
| 6_1276664 | 8 | 0.625 | 10 | 0.5 | 7 | 0.2857 | 8 | 0.5 | 33 | 0.4848 | 7 | 0.7143 | 9 | 0.6667 | 6 | 0 | 10 | 0.5 | 32 | 0.5 |
| 6_1276666 | 8 | 0.375 | 11 | 0.4545 | 3 | 0.3333 | 9 | 0.2222 | 31 | 0.3548 | 5 | 0 | 7 | 0.7143 | 4 | 0.25 | 10 | 0.3 | 26 | 0.3462 |
| 6_1276667 | 9 | 0.3333 | 13 | 0.3846 | 5 | 0 | 12 | 0.0833 | 39 | 0.2308 | 8 | 0.25 | 8 | 0.125 | 8 | 0.125 | 10 | 0.2 | 34 | 0.1765 |
| 6_1276668 | 10 | 0.1 | 13 | 0.1538 | 8 | 0.125 | 13 | 0 | 44 | 0.0909 | 9 | 0.1111 | 10 | 0.2 | 7 | 0.2857 | 13 | 0.3077 | 39 | 0.2308 |
| 6_1276668 | 12 | 0.25 | 14 | 0.3571 | 8 | 0.25 | 11 | 0.3636 | 45 | 0.3111 | 10 | 0.2 | 11 | 0.2727 | 7 | 0.2857 | 15 | 0.4667 | 43 | 0.3256 |
| 6_1277458 | 6 | 0.6667 | 7 | 0.5714 | 3 | 0.6667 | 6 | 0.5 | 22 | 0.5909 | 6 | 0.1667 | 8 | 0.5 | 5 | 0.4 | 8 | 0.25 | 27 | 0.3333 |

| | | | | | | | | | | | | | | | | | | | | |
|---|---|---|---|---|---|---|---|---|---|---|---|---|---|---|---|---|---|---|---|---|
| 6_1277459 | 4 | 0.25 | 5 | 0.6 | 3 | 0 | 5 | 0.4 | 17 | 0.3529 | 3 | 0 | 7 | 0.1429 | 4 | 0 | 7 | 0.1429 | 21 | 0.0952 |
| 6_1277460 | 4 | 0 | 6 | 0.1667 | 2 | 0 | 3 | 0.3333 | 15 | 0.1333 | 3 | 0 | 6 | 0.1667 | 3 | 0 | 7 | 0.1429 | 19 | 0.1053 |
| 6_1277513 | NA | NA | NA | NA | NA | NA | 1 | 1 | 1 | 1 | NA | NA | 1 | 0 | 2 | 0.5 | NA | NA | 3 | 0.3333 |
| 6_1277514 | 1 | 1 | NA | NA | NA | NA | 1 | 1 | 2 | 1 | NA | NA | NA | NA | NA | NA | 1 | 1 | 1 | 1 |
| 6_1277517 | 1 | 0 | NA | NA | NA | NA | 1 | 1 | 2 | 0.5 | NA | NA | 1 | 1 | 2 | 1 | 1 | 1 | 4 | 1 |
| 6_1277518 | NA | NA | NA | NA | NA | NA | 1 | 1 | 1 | 1 | NA | NA | 1 | 1 | 2 | 1 | 1 | 1 | 4 | 1 |
| 6_1277520 | 1 | 1 | NA | NA | NA | NA | 1 | 1 | 2 | 1 | NA | NA | NA | NA | NA | NA | 2 | 1 | 2 | 1 |
| 6_1277524 | 1 | 0 | NA | NA | NA | NA | 1 | 1 | 2 | 0.5 | NA | NA | NA | NA | NA | NA | 1 | 1 | 1 | 1 |
| 6_1279387 | 5 | 0 | 4 | 0 | 4 | 0.5 | 2 | 0 | 15 | 0.1333 | 1 | 0 | NA | NA | 5 | 0.2 | 2 | 0 | 8 | 0.125 |
| 6_1279406 | 2 | 0.5 | 1 | 0 | 2 | 0.5 | 1 | 1 | 6 | 0.5 | NA | NA | 1 | 1 | 3 | 0 | 1 | 1 | 5 | 0.4 |
| 6_1279410 | 2 | 1 | 1 | 0 | 1 | 1 | 1 | 0 | 5 | 0.6 | NA | NA | 1 | 0 | 2 | 1 | 1 | 1 | 4 | 0.75 |
| 6_1279411 | 2 | 1 | 1 | 0 | 1 | 0 | 1 | 1 | 5 | 0.6 | NA | NA | NA | NA | 2 | 0 | 1 | 1 | 3 | 0.3333 |
| 6_1279412 | 2 | 1 | NA | NA | 1 | 0 | 1 | 1 | 4 | 0.75 | NA | NA | 1 | 1 | 1 | 1 | NA | NA | 2 | 1 |
| 6_1280113 | 2 | 1 | NA | NA | 1 | 1 | NA | NA | 3 | 1 | NA | NA | 1 | 1 | NA | NA | NA | NA | 1 | 1 |
| 6_1281467 | 4 | 0 | NA | NA | 5 | 0.2 | 1 | 0 | 10 | 0.1 | 1 | 1 | 1 | 1 | 3 | 0 | 2 | 0.5 | 7 | 0.4286 |
| 6_1281468 | 4 | 0 | NA | NA | 5 | 0.2 | 1 | 0 | 10 | 0.1 | 1 | 1 | 1 | 0 | 3 | 0.3333 | 2 | 0 | 7 | 0.2857 |
| 6_1281469 | 3 | 1 | NA | NA | 3 | 1 | 1 | 1 | 7 | 1 | 1 | 1 | 1 | 1 | NA | NA | 1 | 1 | 3 | 1 |
| 6_1281471 | 3 | 0.6667 | NA | NA | 4 | 1 | 1 | 1 | 8 | 0.875 | 1 | 1 | NA | NA | 2 | 0.5 | 1 | 1 | 4 | 0.75 |
| 6_1281472 | 3 | 0.6667 | NA | NA | 4 | 0 | 1 | 1 | 8 | 0.375 | NA | NA | 1 | 0 | 2 | 0.5 | 2 | 0.5 | 5 | 0.4 |
| 6_1281473 | 2 | 0 | NA | NA | 4 | 0.25 | 1 | 0 | 7 | 0.1429 | NA | NA | 1 | 0 | 2 | 1 | 1 | 0 | 4 | 0.5 |
| 6_1281483 | 3 | 0.3333 | NA | NA | 2 | 0 | 1 | 1 | 6 | 0.3333 | NA | NA | NA | NA | 2 | 0.5 | 2 | 0 | 4 | 0.25 |
| 6_1281488 | 3 | 1 | NA | NA | 2 | 1 | 1 | 1 | 6 | 1 | NA | NA | NA | NA | 2 | 1 | 2 | 1 | 4 | 1 |
| 6_1281488 | 2 | 1 | NA | NA | 2 | 1 | NA | NA | 4 | 1 | NA | NA | NA | NA | 2 | 0.5 | 1 | 1 | 3 | 0.6667 |
| 6_1281489 | 2 | 0 | NA | NA | 1 | 1 | 1 | 0 | 4 | 0.25 | NA | NA | NA | NA | 2 | 0.5 | 2 | 0 | 4 | 0.25 |
| 6_1281489 | 3 | 1 | NA | NA | 2 | 1 | NA | NA | 5 | 1 | NA | NA | NA | NA | 2 | 1 | 2 | 1 | 4 | 1 |
| 6_1281490 | 3 | 0.6667 | NA | NA | 1 | 0 | 1 | 1 | 5 | 0.6 | NA | NA | NA | NA | 2 | 1 | 2 | 1 | 4 | 1 |
| 6_1281490 | 3 | 1 | NA | NA | 1 | 1 | 1 | 1 | 5 | 1 | NA | NA | NA | NA | 2 | 1 | 1 | 1 | 3 | 1 |
| 6_1281491 | 3 | 1 | NA | NA | 1 | 1 | 1 | 1 | 5 | 1 | NA | NA | NA | NA | 2 | 0 | 2 | 0.5 | 4 | 0.25 |
| 6_1281598 | 2 | 0.5 | NA | NA | 1 | 1 | 1 | 1 | 4 | 0.75 | NA | NA | NA | NA | 2 | 0.5 | NA | NA | 2 | 0.5 |
| 6_1281600 | 2 | 0 | NA | NA | 1 | 1 | 1 | 0 | 4 | 0.25 | NA | NA | NA | NA | 2 | 0.5 | NA | NA | 2 | 0.5 |
| 6_1281603 | 2 | 1 | NA | NA | 1 | 0 | NA | NA | 3 | 0.6667 | NA | NA | NA | NA | 1 | 1 | NA | NA | 1 | 1 |
| 6_1282982 | 3 | 0.6667 | 3 | 0.6667 | 2 | 1 | 2 | 1 | 10 | 0.8 | 5 | 0.8 | 3 | 0.3333 | 1 | 0 | 1 | 1 | 10 | 0.6 |
| 6_1282982 | 3 | 0 | 3 | 0 | 3 | 0 | 2 | 0.5 | 11 | 0.0909 | 6 | 0.1667 | 3 | 0 | NA | NA | 2 | 0 | 11 | 0.0909 |
| 6_1282985 | 3 | 0.6667 | 2 | 1 | 1 | 1 | 1 | 1 | 7 | 0.8571 | 5 | 1 | 2 | 0.5 | 1 | 1 | 2 | 0.5 | 10 | 0.8 |
| 6_1282986 | 2 | 0 | 3 | 0 | 3 | 0.3333 | 3 | 1 | 11 | 0.3636 | 6 | 0.1667 | 3 | 0 | 1 | 0 | 2 | 0.5 | 12 | 0.1667 |
| 6_1282988 | 1 | 0 | 2 | 0 | 3 | 0 | 3 | 0.3333 | 9 | 0.1111 | 5 | 0.4 | 3 | 0 | 1 | 0 | 2 | 0 | 11 | 0.1818 |
| 6_1282989 | 1 | 0 | 2 | 0.5 | 2 | 0 | 2 | 0.5 | 7 | 0.2857 | 4 | 0 | 3 | 0.3333 | 1 | 0 | 2 | 0.5 | 10 | 0.2 |
| 6_1282990 | 1 | 0 | 2 | 0.5 | 2 | 0 | 2 | 0.5 | 7 | 0.2857 | 5 | 0.4 | 3 | 0.3333 | 1 | 0 | 1 | 1 | 10 | 0.4 |
| 6_1282996 | 1 | 0 | 2 | 0 | 1 | 0 | 2 | 1 | 6 | 0.3333 | 5 | 0.4 | 2 | 0.5 | NA | NA | NA | NA | 7 | 0.4286 |
| 6_1283001 | NA | NA | 1 | 1 | 1 | 0 | 1 | 0 | 3 | 0.3333 | 1 | 0 | 1 | 1 | 1 | 0 | NA | NA | 3 | 0.3333 |
| 6_1283344 | 4 | 0 | 4 | 0.25 | 5 | 0.2 | 3 | 0 | 16 | 0.125 | 5 | 0.2 | 3 | 0 | 3 | 0 | NA | NA | 11 | 0.0909 |
| 6_1283344 | 4 | 0.25 | 3 | 0.3333 | 3 | 0 | 2 | 1 | 12 | 0.3333 | 5 | 0.2 | 2 | 0 | 4 | 0 | NA | NA | 11 | 0.0909 |
| 6_1283344 | 3 | 1 | 5 | 1 | 5 | 0.8 | 3 | 1 | 16 | 0.9375 | 4 | 1 | 2 | 1 | 3 | 1 | NA | NA | 9 | 1 |
| 6_1283345 | 3 | 0.3333 | 4 | 0.25 | 4 | 0.25 | 1 | 0 | 12 | 0.25 | 3 | 0.3333 | 2 | 0.5 | 1 | 0 | NA | NA | 6 | 0.3333 |
| 6_1283346 | 4 | 0.5 | 6 | 0 | 5 | 0.2 | 3 | 0 | 18 | 0.1667 | 4 | 0.25 | 3 | 0 | 3 | 0.3333 | NA | NA | 10 | 0.2 |
| 6_1283347 | 5 | 0.4 | 5 | 0.6 | 5 | 0.4 | 2 | 0.5 | 17 | 0.4706 | 5 | 0.6 | 3 | 0.3333 | 3 | 0.6667 | NA | NA | 11 | 0.5455 |
| 6_1283347 | 5 | 0.6 | 5 | 0.2 | 4 | 0 | 3 | 0.3333 | 17 | 0.2941 | 5 | 0.4 | 2 | 0.5 | 3 | 0.6667 | NA | NA | 10 | 0.5 |
| 6_1283348 | 5 | 0.4 | 5 | 0.4 | 5 | 0 | 2 | 0 | 17 | 0.2353 | 3 | 0.3333 | 3 | 0 | 3 | 0.3333 | NA | NA | 9 | 0.2222 |
| 6_1283349 | 4 | 1 | 2 | 1 | 5 | 1 | 2 | 0.5 | 13 | 0.9231 | 3 | 0.6667 | 3 | 1 | 3 | 1 | NA | NA | 9 | 0.8889 |
| 6_1283349 | 5 | 0.8 | 4 | 0.25 | 5 | 0.4 | 2 | 1 | 16 | 0.5625 | 3 | 0.6667 | 3 | 0 | 3 | 0.6667 | NA | NA | 9 | 0.4444 |
| 6_1283852 | 3 | 0.3333 | NA | NA | 1 | 0 | 2 | 0 | 6 | 0.1667 | 2 | 0 | 5 | 0 | 3 | 0.3333 | 1 | 0 | 11 | 0.0909 |
| 6_1283853 | 3 | 0.3333 | NA | NA | 1 | 0 | 2 | 0 | 6 | 0.1667 | 1 | 0 | 4 | 0.25 | 3 | 0 | 1 | 0 | 9 | 0.1111 |
| 6_1283869 | 4 | 0.75 | 4 | 0.75 | 1 | 1 | 2 | 1 | 11 | 0.8182 | 1 | 1 | 4 | 1 | 5 | 1 | 2 | 0 | 12 | 0.8333 |
| 6_1283871 | 3 | 0.3333 | 4 | 0 | NA | NA | NA | NA | 7 | 0.1429 | 1 | 0 | 2 | 1 | 2 | 1 | 2 | 0.5 | 7 | 0.7143 |
| 6_1283872 | 3 | 0 | 4 | 0.25 | 2 | 0.5 | 2 | 0.5 | 11 | 0.2727 | 1 | 0 | 4 | 0 | 2 | 0 | 1 | 1 | 8 | 0.125 |
| 6_1283873 | 4 | 0.25 | 3 | 0 | 2 | 0 | 2 | 0.5 | 11 | 0.1818 | 1 | 0 | 3 | 0.3333 | 3 | 0.3333 | 2 | 0 | 9 | 0.2222 |
| 6_1284076 | 15 | 0.1333 | 15 | 0.3333 | 16 | 0.1875 | 13 | 0.1538 | 59 | 0.2034 | 11 | 0.1818 | 14 | 0 | 15 | 0.1333 | 14 | 0.2857 | 54 | 0.1481 |
| 6_1286236 | 2 | 0 | NA | NA | 4 | 0.25 | 4 | 0.25 | 10 | 0.2 | 4 | 0 | 2 | 0 | 4 | 0 | 2 | 0.5 | 12 | 0.0833 |
| 6_1286237 | 2 | 0 | NA | NA | 3 | 0.3333 | 4 | 0 | 9 | 0.1111 | 4 | 0.25 | 2 | 0 | 5 | 0 | 2 | 0 | 13 | 0.0769 |
| 6_1288263 | NA | NA | NA | NA | 1 | 1 | NA | NA | 1 | 1 | NA | NA | NA | NA | 1 | 1 | NA | NA | 1 | 1 |
| 6_1288267 | NA | NA | NA | NA | 1 | 1 | NA | NA | 1 | 1 | NA | NA | NA | NA | 1 | 1 | NA | NA | 1 | 1 |
| 6_1289830 | 1 | 1 | NA | NA | NA | NA | NA | NA | 1 | 1 | 1 | 0 | NA | NA | 1 | 1 | NA | NA | 2 | 0.5 |
| 6_1294398 | NA | NA | NA | NA | NA | NA | 1 | 1 | 1 | 1 | 1 | 1 | NA | NA | NA | NA | 1 | 1 | 2 | 1 |
| 6_1294400 | NA | NA | NA | NA | NA | NA | 1 | 1 | 1 | 1 | NA | NA | NA | NA | NA | NA | 1 | 1 | 1 | 1 |
| 6_1295091 | NA | NA | 1 | 0 | 1 | 1 | NA | NA | 2 | 0.5 | NA | NA | 1 | 0 | 1 | 1 | NA | NA | 2 | 0.5 |

| | | | | | | | | | | | | | | | | | | | | |
|---|---|---|---|---|---|---|---|---|---|---|---|---|---|---|---|---|---|---|---|---|
| 6_12950927 | NA | NA | 1 | 1 | 1 | 1 | NA | NA | 2 | 1 | NA | NA | 1 | 1 | 1 | 1 | NA | NA | 2 | 1 |
| 6_12950964 | NA | NA | 1 | 0 | 1 | 1 | NA | NA | 2 | 0.5 | NA | NA | 1 | 0 | 1 | 1 | NA | NA | 2 | 0.5 |
| 6_12951032 | 31 | 0.1613 | 23 | 0.1739 | 23 | 0.087 | 30 | 0.1333 | 107 | 0.1402 | 24 | 0.0417 | 28 | 0.1071 | 17 | 0.0588 | 18 | 0.2222 | 87 | 0.1034 |
| 6_12951816 | 48 | 0.8542 | 50 | 0.86 | 43 | 0.9535 | 44 | 0.8409 | 185 | 0.8757 | 42 | 0.9524 | 52 | 0.8654 | 36 | 0.7778 | 31 | 0.9032 | 161 | 0.8758 |
| 6_12951819 | 52 | 0.2308 | 52 | 0.25 | 46 | 0.3261 | 39 | 0.5128 | 189 | 0.3175 | 37 | 0.3243 | 52 | 0.3462 | 37 | 0.2703 | 31 | 0.2903 | 157 | 0.3121 |
| 6_12951833 | 80 | 0.225 | 74 | 0.3784 | 64 | 0.2812 | 60 | 0.4167 | 278 | 0.3201 | 53 | 0.3962 | 73 | 0.3425 | 56 | 0.3571 | 47 | 0.3404 | 229 | 0.3581 |
| 6_12951883 | 90 | 0.2667 | 88 | 0.2841 | 77 | 0.3247 | 67 | 0.3881 | 322 | 0.3106 | 60 | 0.3167 | 80 | 0.3625 | 67 | 0.3134 | 60 | 0.3667 | 267 | 0.3408 |
| 6_12951911 | 77 | 0.2597 | 87 | 0.2989 | 82 | 0.2317 | 64 | 0.3594 | 310 | 0.2839 | 65 | 0.2 | 80 | 0.325 | 76 | 0.3158 | 61 | 0.3115 | 282 | 0.2908 |
| 6_13134225 | 1 | 0 | 6 | 0.5 | 5 | 0.6 | 7 | 0.5714 | 19 | 0.5263 | 5 | 1 | 6 | 0.5 | 7 | 0.7143 | 5 | 0.8 | 23 | 0.7391 |
| 6_13308233 | 18 | 0.1667 | 18 | 0 | 17 | 0.0588 | 16 | 0 | 69 | 0.058 | 10 | 0 | 15 | 0.0667 | 14 | 0.0714 | 14 | 0.1429 | 53 | 0.0755 |
| 6_13308251 | 21 | 0.5238 | 17 | 0.5882 | 18 | 0.3333 | 14 | 0.6429 | 70 | 0.5143 | 10 | 0.4 | 9 | 0.4444 | 13 | 0.3846 | 14 | 0.5 | 46 | 0.4348 |
| 6_13308261 | 22 | 0.1364 | 20 | 0.2 | 16 | 0.0625 | 16 | 0.0625 | 74 | 0.1216 | 12 | 0.1667 | 14 | 0 | 10 | 0.2 | 16 | 0.0625 | 52 | 0.0962 |
| 6_13308290 | 20 | 0.05 | 20 | 0.1 | 20 | 0.3 | 17 | 0.2353 | 77 | 0.1688 | 12 | 0.0833 | 17 | 0.0588 | 14 | 0 | 17 | 0.1765 | 60 | 0.0833 |
| 6_13308292 | 15 | 0.3333 | 21 | 0.1429 | 19 | 0.2105 | 15 | 0.2667 | 70 | 0.2286 | 12 | 0.1667 | 15 | 0.4 | 13 | 0.2308 | 18 | 0.1667 | 58 | 0.2414 |
| 6_13308295 | 20 | 1 | 24 | 1 | 17 | 1 | 16 | 1 | 77 | 1 | 12 | 1 | 17 | 1 | 14 | 1 | 18 | 1 | 61 | 1 |
| 6_13308296 | 21 | 0.5238 | 20 | 0.35 | 19 | 0.1579 | 17 | 0.4706 | 77 | 0.3766 | 13 | 0.3846 | 14 | 0.3571 | 11 | 0.6364 | 19 | 0.4211 | 57 | 0.4386 |
| 6_13308300 | 14 | 0.9286 | 21 | 0.5714 | 16 | 0.5 | 12 | 0.6667 | 63 | 0.6508 | 14 | 0.8571 | 15 | 0.6667 | 13 | 0.4615 | 16 | 0.8125 | 58 | 0.7069 |
| 6_13308306 | 20 | 0.15 | 24 | 0.25 | 20 | 0.35 | 19 | 0.3158 | 83 | 0.2651 | 13 | 0.0769 | 17 | 0.3529 | 14 | 0.0714 | 20 | 0.3 | 64 | 0.2188 |
| 6_13308312 | 19 | 1 | 23 | 1 | 17 | 1 | 17 | 1 | 76 | 1 | 13 | 0.9231 | 16 | 1 | 14 | 1 | 19 | 1 | 62 | 0.9839 |
| 6_13308315 | NA | NA | 1 | 1 | NA | NA | NA | NA | 1 | 1 | 1 | 1 | NA | NA | NA | NA | NA | NA | 1 | 1 |
| 6_13308319 | 20 | 0.95 | 20 | 0.9 | 18 | 0.9444 | 15 | 0.8667 | 73 | 0.9178 | 12 | 1 | 14 | 0.7857 | 11 | 0.8182 | 19 | 0.7895 | 56 | 0.8393 |
| 6_13308322 | 25 | 0.32 | 23 | 0.3043 | 20 | 0.3 | 18 | 0.4444 | 86 | 0.3372 | 15 | 0.3333 | 16 | 0.3125 | 15 | 0.3333 | 21 | 0.1429 | 67 | 0.2687 |
| 6_13308330 | 17 | 0.7059 | 18 | 0.3889 | 14 | 0.7143 | 12 | 0.6667 | 61 | 0.6066 | 12 | 0.5833 | 14 | 0.5714 | 10 | 0.4 | 16 | 0.5 | 52 | 0.5192 |
| 6_13308333 | 26 | 0.1923 | 21 | 0.1905 | 14 | 0.2143 | 20 | 0.1 | 81 | 0.1728 | 14 | 0.1429 | 17 | 0.0588 | 14 | 0.0714 | 22 | 0.0909 | 67 | 0.0896 |
| 6_13308335 | 25 | 0.48 | 22 | 0.4545 | 19 | 0.5263 | 22 | 0.4091 | 88 | 0.4659 | 15 | 0.5333 | 19 | 0.5263 | 15 | 0.4 | 20 | 0.65 | 69 | 0.5362 |
| 6_13308335 | 26 | 0.1923 | 21 | 0.3333 | 19 | 0.2632 | 21 | 0.2857 | 87 | 0.2644 | 13 | 0 | 16 | 0.25 | 15 | 0.0667 | 19 | 0.0526 | 63 | 0.0952 |
| 6_13308341 | 22 | 0.3182 | 23 | 0.2609 | 17 | 0.3529 | 14 | 0.4286 | 76 | 0.3289 | 14 | 0.4286 | 12 | 0.3333 | 15 | 0.2667 | 16 | 0.5 | 57 | 0.386 |
| 6_13308344 | 27 | 0.963 | 22 | 1 | 21 | 1 | 20 | 1 | 90 | 0.9889 | 15 | 0.9333 | 14 | 1 | 14 | 1 | 20 | 1 | 63 | 0.9841 |
| 6_13308345 | 22 | 0.1364 | 20 | 0.3 | 19 | 0.2632 | 18 | 0.4444 | 79 | 0.2785 | 12 | 0.1667 | 13 | 0.1538 | 12 | 0.25 | 13 | 0.3846 | 50 | 0.24 |
| 6_13308352 | 25 | 0.88 | 21 | 0.9524 | 16 | 0.9375 | 19 | 1 | 81 | 0.9383 | 14 | 1 | 15 | 0.9333 | 19 | 0.7895 | 20 | 1 | 68 | 0.9265 |
| 6_13308385 | 24 | 0.0417 | 23 | 0.1739 | 12 | 0.3333 | 15 | 0 | 74 | 0.1216 | 11 | 0.1818 | 13 | 0 | 9 | 0 | 22 | 0.2727 | 55 | 0.1455 |
| 6_13308388 | 26 | 0.2308 | 25 | 0.16 | 14 | 0.2857 | 17 | 0.3529 | 82 | 0.2439 | 11 | 0 | 16 | 0.3125 | 11 | 0 | 22 | 0.1818 | 60 | 0.15 |
| 6_13308460 | 20 | 0.7 | 25 | 0.68 | 18 | 0.6667 | 24 | 0.4167 | 87 | 0.6092 | 16 | 0.625 | 16 | 0.6875 | 10 | 0.2 | 23 | 0.4783 | 65 | 0.5231 |
| 6_13308462 | 19 | 0.6316 | 27 | 0.4074 | 22 | 0.4545 | 28 | 0.6429 | 96 | 0.5312 | 16 | 0.3125 | 18 | 0.3889 | 15 | 0.5333 | 23 | 0.3913 | 72 | 0.4028 |
| 6_13308783 | NA | NA | NA | NA | 2 | 1 | 1 | 0 | 3 | 0.6667 | NA | NA | NA | NA | 2 | 0.5 | NA | NA | 2 | 0.5 |
| 6_13424344 | 1 | 1 | 1 | 1 | NA | NA | NA | NA | 2 | 1 | 1 | 0 | 1 | 1 | 2 | 1 | NA | NA | 4 | 0.75 |
| 6_13424346 | 1 | 1 | 1 | 1 | NA | NA | NA | NA | 2 | 1 | 1 | 0 | 1 | 1 | 2 | 0.5 | NA | NA | 4 | 0.5 |
| 6_13424353 | 1 | 1 | 1 | 1 | NA | NA | NA | NA | 2 | 1 | 1 | 1 | NA | NA | 2 | 0.5 | NA | NA | 3 | 0.6667 |
| 6_13424676 | 1 | 0 | 1 | 1 | 1 | 1 | NA | NA | 3 | 0.6667 | NA | NA | 2 | 1 | 4 | 0.75 | NA | NA | 6 | 0.8333 |
| 6_13424691 | 1 | 0 | 2 | 0.5 | 1 | 0 | NA | NA | 4 | 0.25 | NA | NA | 2 | 0 | 4 | 0.5 | NA | NA | 6 | 0.3333 |
| 6_13424698 | 1 | 0 | 2 | 1 | 2 | 0 | NA | NA | 5 | 0.4 | NA | NA | 1 | 1 | 4 | 0 | NA | NA | 5 | 0.2 |
| 6_13442442 | 1 | 0 | 2 | 0.5 | 2 | 0 | 5 | 0 | 10 | 0.1 | NA | NA | 3 | 0 | 6 | 0.1667 | 3 | 0 | 12 | 0.0833 |
| 6_13442705 | 1 | 0 | NA | NA | NA | NA | 3 | 0.3333 | 4 | 0.25 | NA | NA | 1 | 1 | NA | NA | 1 | 0 | 2 | 0.5 |
| 6_13442724 | NA | NA | NA | NA | NA | NA | 4 | 0.25 | 4 | 0.25 | NA | NA | 1 | 1 | NA | NA | 1 | 0 | 2 | 0.5 |
| 6_13442725 | 1 | 1 | NA | NA | NA | NA | 4 | 0.25 | 5 | 0.4 | NA | NA | 1 | 0 | NA | NA | 1 | 1 | 2 | 0.5 |
| 6_13442745 | 1 | 0 | NA | NA | NA | NA | 2 | 0.5 | 3 | 0.3333 | NA | NA | 1 | 1 | NA | NA | 1 | 1 | 2 | 1 |
| 6_13442775 | 1 | 1 | NA | NA | 1 | 1 | 2 | 1 | 4 | 1 | NA | NA | 1 | 1 | NA | NA | 1 | 1 | 2 | 1 |
| 6_13451804 | 2 | 0.5 | NA | NA | 1 | 1 | 2 | 0 | 5 | 0.4 | 1 | 1 | NA | NA | 4 | 0 | 1 | 0 | 6 | 0.1667 |
| 6_13451851 | 1 | 1 | NA | NA | 1 | 1 | 1 | 0 | 3 | 0.6667 | 1 | 1 | NA | NA | 3 | 1 | 1 | 0 | 5 | 0.8 |
| 6_13451873 | 1 | 1 | NA | NA | 1 | 1 | NA | NA | 2 | 1 | 1 | 1 | NA | NA | 3 | 1 | 1 | 1 | 5 | 1 |
| 6_13451881 | 1 | 1 | NA | NA | 1 | 1 | NA | NA | 2 | 1 | NA | NA | NA | NA | 2 | 0.5 | 1 | 1 | 3 | 0.6667 |
| 6_13451883 | 1 | 1 | NA | NA | 1 | 0 | NA | NA | 2 | 0.5 | NA | NA | NA | NA | 2 | 1 | 1 | 1 | 3 | 1 |
| 6_13451889 | 1 | 1 | NA | NA | 1 | 1 | NA | NA | 2 | 1 | 1 | 1 | NA | NA | 3 | 0.6667 | 1 | 1 | 5 | 0.8 |
| 6_13451897 | 1 | 0 | NA | NA | 1 | 1 | NA | NA | 2 | 0.5 | 1 | 1 | NA | NA | 3 | 0.6667 | 1 | 0 | 5 | 0.6 |
| 6_13458684 | 2 | 1 | 1 | 1 | NA | NA | NA | NA | 3 | 1 | 1 | 1 | 2 | 1 | NA | NA | NA | NA | 3 | 1 |
| 6_13458689 | 1 | 1 | 1 | 1 | NA | NA | NA | NA | 2 | 1 | 1 | 1 | 2 | 1 | NA | NA | NA | NA | 3 | 1 |
| 6_13458706 | 2 | 1 | 1 | 1 | NA | NA | NA | NA | 3 | 1 | 1 | 1 | 1 | 1 | NA | NA | NA | NA | 2 | 1 |
| 6_13458715 | 2 | 0 | 1 | 1 | NA | NA | NA | NA | 3 | 0.3333 | 1 | 1 | 1 | 1 | NA | NA | NA | NA | 2 | 1 |
| 6_13458716 | 2 | 0.5 | 1 | 1 | NA | NA | NA | NA | 3 | 0.6667 | 1 | 1 | NA | NA | NA | NA | NA | NA | 1 | 1 |
| 6_13458727 | 2 | 1 | 1 | 1 | NA | NA | NA | NA | 3 | 1 | 1 | 1 | 2 | 1 | NA | NA | NA | NA | 3 | 1 |
| 6_13458741 | 2 | 1 | 1 | 1 | NA | NA | NA | NA | 3 | 1 | NA | NA | 1 | 1 | NA | NA | NA | NA | 1 | 1 |
| 6_13458752 | 2 | 0.5 | 1 | 0 | NA | NA | NA | NA | 3 | 0.3333 | 1 | 0 | 2 | 1 | NA | NA | NA | NA | 3 | 0.6667 |
| 6_13468731 | 6 | 1 | 1 | 1 | 3 | 1 | 9 | 1 | 19 | 1 | 6 | 1 | 9 | 0.8889 | 1 | 1 | 5 | 1 | 21 | 0.9524 |
| 6_13468735 | 6 | 1 | 3 | 1 | 4 | 1 | 8 | 1 | 21 | 1 | 7 | 1 | 9 | 1 | 3 | 1 | 4 | 1 | 23 | 1 |
| 6_13468738 | 6 | 1 | 3 | 1 | 3 | 0.6667 | 6 | 1 | 18 | 0.9444 | 6 | 1 | 5 | 1 | 3 | 1 | 5 | 1 | 19 | 1 |

| | | | | | | | | | | | | | | | | | | | | |
|---|---|---|---|---|---|---|---|---|---|---|---|---|---|---|---|---|---|---|---|---|
| 6_1346874 | 6 | 0.6667 | 2 | 0.5 | 4 | 0.75 | 9 | 0.7778 | 21 | 0.7143 | 6 | 0.6667 | 8 | 0.625 | 2 | 0.5 | 5 | 1 | 21 | 0.7143 |
| 6_1346874 | 5 | 1 | 3 | 1 | 4 | 1 | 7 | 0.8571 | 19 | 0.9474 | 6 | 0.8333 | 8 | 0.875 | 3 | 1 | 5 | 1 | 22 | 0.9091 |
| 6_1346875 | 7 | 0.2857 | 3 | 0.3333 | 4 | 0.75 | 9 | 0.3333 | 23 | 0.3913 | 7 | 0.1429 | 9 | 0.3333 | 3 | 0.6667 | 6 | 0.5 | 25 | 0.36 |
| 6_1346875 | 5 | 0.2 | 2 | 0.5 | 3 | 0.3333 | 8 | 0.75 | 18 | 0.5 | 6 | 0.6667 | 7 | 0.8571 | 4 | 0.75 | 6 | 0.6667 | 23 | 0.7391 |
| 6_1346876 | 7 | 0.8571 | 1 | 1 | 3 | 0.3333 | 9 | 0.6667 | 20 | 0.7 | 6 | 0.1667 | 9 | 0.5556 | 4 | 0.75 | 5 | 0.6 | 24 | 0.5 |
| 6_1346878 | 5 | 0.6 | 3 | 1 | 5 | 0.8 | 9 | 0.6667 | 22 | 0.7273 | 7 | 1 | 9 | 1 | 3 | 0.6667 | 6 | 0.1667 | 25 | 0.76 |
| 6_1346878 | 6 | 0.3333 | 3 | 0 | 5 | 0 | 9 | 0 | 23 | 0.087 | 7 | 0.2857 | 8 | 0 | 4 | 0.25 | 8 | 0 | 27 | 0.1111 |
| 6_1346878 | 6 | 0.6667 | 2 | 0.5 | 4 | 0.75 | 7 | 0.8571 | 19 | 0.7368 | 5 | 0.4 | 9 | 0.5556 | 1 | 1 | 6 | 0.6667 | 21 | 0.5714 |
| 6_1346879 | 7 | 1 | 3 | 1 | 5 | 1 | 10 | 0.9 | 25 | 0.96 | 7 | 1 | 10 | 0.9 | 4 | 1 | 7 | 0.8571 | 28 | 0.9286 |
| 6_1346926 | NA | NA | 1 | 1 | 2 | 1 | NA | NA | 3 | 1 | NA | NA | NA | NA | 3 | 1 | 1 | 1 | 4 | 1 |
| 6_1346928 | NA | NA | 1 | 1 | 2 | 1 | NA | NA | 3 | 1 | NA | NA | NA | NA | 1 | 1 | NA | NA | 1 | 1 |
| 6_1346928 | 1 | 1 | 1 | 1 | NA | NA | NA | NA | 2 | 1 | NA | NA | NA | NA | 3 | 0.3333 | 1 | 0 | 4 | 0.25 |
| 6_1346953 | 1 | 1 | 2 | 1 | NA | NA | NA | NA | 3 | 1 | NA | NA | NA | NA | 1 | 1 | NA | NA | 1 | 1 |
| 6_1348330 | 3 | 1 | 2 | 1 | 2 | 1 | 6 | 1 | 13 | 1 | 1 | 1 | NA | NA | 1 | 1 | 6 | 1 | 8 | 1 |
| 6_1348330 | 3 | 0.6667 | 2 | 0.5 | 2 | 0.5 | 7 | 0.1429 | 14 | 0.3571 | 1 | 1 | NA | NA | 1 | 0 | 6 | 0.1667 | 8 | 0.25 |
| 6_1348331 | 1 | 0 | 1 | 1 | 2 | 1 | 6 | 0.5 | 10 | 0.6 | 1 | 1 | NA | NA | 1 | 1 | 6 | 0.5 | 8 | 0.625 |
| 6_1348333 | 3 | 0.3333 | 3 | 0.6667 | 2 | 0 | 8 | 0.375 | 16 | 0.375 | 1 | 0 | NA | NA | 2 | 1 | 6 | 0.5 | 9 | 0.5556 |
| 6_1348344 | 3 | 0 | 4 | 0.5 | 2 | 0 | 7 | 0.2857 | 16 | 0.25 | 2 | 0 | 1 | 0 | 3 | 0.3333 | 5 | 0.6 | 11 | 0.3636 |
| 6_1348346 | 3 | 0.3333 | 5 | 0.2 | 3 | 0 | 5 | 0.2 | 16 | 0.1875 | 2 | 0.5 | NA | NA | 2 | 0 | 6 | 0.1667 | 10 | 0.2 |
| 6_1348350 | 3 | 0 | 5 | 0.6 | 2 | 0.5 | 6 | 0.5 | 16 | 0.4375 | 3 | 0.3333 | 2 | 0 | 3 | 0 | 5 | 0.2 | 13 | 0.1538 |
| 6_1348355 | 4 | 0.25 | 5 | 0.4 | 2 | 0 | 6 | 0.3333 | 17 | 0.2941 | 3 | 0 | 3 | 0.3333 | 3 | 0 | 4 | 0 | 13 | 0.0769 |
| 6_1348356 | 4 | 0 | 3 | 0.3333 | 4 | 0 | 6 | 0 | 17 | 0.0588 | 3 | 0.3333 | 4 | 0 | 4 | 0 | 5 | 0 | 16 | 0.0625 |
| 6_1348461 | 2 | 0 | 2 | 0.5 | NA | NA | 5 | 0.2 | 9 | 0.2222 | 1 | 0 | 3 | 0.3333 | 2 | 0 | NA | NA | 6 | 0.1667 |
| 6_1348462 | 2 | 0 | 3 | 1 | NA | NA | 3 | 0.3333 | 8 | 0.5 | NA | NA | 3 | 0.6667 | 2 | 1 | NA | NA | 5 | 0.8 |
| 6_1348462 | 2 | 1 | 2 | 1 | NA | NA | 5 | 1 | 9 | 1 | 1 | 1 | NA | NA | 1 | 1 | NA | NA | 2 | 1 |
| 6_1348462 | 2 | 1 | 3 | 0.6667 | NA | NA | 5 | 1 | 10 | 0.9 | 1 | 1 | 3 | 1 | 2 | 0.5 | NA | NA | 6 | 0.8333 |
| 6_1348463 | 2 | 1 | 3 | 1 | NA | NA | 6 | 0.8333 | 11 | 0.9091 | 1 | 0 | 3 | 0.6667 | 2 | 0.5 | NA | NA | 6 | 0.5 |
| 6_1348463 | 2 | 1 | 3 | 1 | NA | NA | 5 | 1 | 10 | 1 | 1 | 1 | 2 | 1 | 2 | 1 | NA | NA | 5 | 1 |
| 6_1348466 | 1 | 1 | NA | NA | NA | NA | 2 | 0 | 3 | 0.3333 | 1 | 0 | 2 | 1 | NA | NA | NA | NA | 3 | 0.6667 |
| 6_1348467 | 2 | 0 | 2 | 0.5 | NA | NA | 4 | 0 | 8 | 0.125 | 1 | 0 | 2 | 0 | 1 | 1 | NA | NA | 4 | 0.25 |
| 6_1348470 | 1 | 1 | 2 | 0.5 | NA | NA | 5 | 0.2 | 8 | 0.375 | 1 | 0 | 2 | 0.5 | 1 | 0 | NA | NA | 4 | 0.25 |
| 6_1348471 | 1 | 1 | 1 | 1 | NA | NA | 3 | 1 | 5 | 1 | 1 | 0 | 1 | 1 | 1 | 1 | NA | NA | 3 | 0.6667 |
| 6_1348472 | 2 | 0.5 | 2 | 0.5 | NA | NA | 3 | 0.3333 | 7 | 0.4286 | NA | NA | 1 | 1 | 1 | 1 | NA | NA | 2 | 1 |
| 6_1348977 | 2 | 0 | 1 | 1 | NA | NA | 1 | 0 | 4 | 0.25 | 2 | 0.5 | 1 | 0 | 2 | 0 | 1 | 1 | 6 | 0.3333 |
| 6_1348980 | 2 | 1 | 2 | 0.5 | NA | NA | 1 | 0 | 5 | 0.6 | 2 | 0.5 | 1 | 0 | 2 | 0 | 1 | 0 | 6 | 0.1667 |
| 6_1349013 | 3 | 0.6667 | 4 | 0.5 | NA | NA | 2 | 0.5 | 9 | 0.5556 | 2 | 1 | 4 | 0.75 | 3 | 0.6667 | 1 | 1 | 10 | 0.8 |
| 6_1349013 | 3 | 0.3333 | 4 | 0 | NA | NA | 2 | 0 | 9 | 0.1111 | 2 | 0 | 4 | 0.25 | 3 | 0 | 1 | 0 | 10 | 0.1 |
| 6_1349609 | 1 | 1 | NA | NA | 1 | 0 | NA | NA | 2 | 0.5 | 1 | 0 | 1 | 1 | NA | NA | 1 | 1 | 3 | 0.6667 |
| 6_1349611 | 1 | 1 | NA | NA | 1 | 1 | NA | NA | 2 | 1 | NA | NA | 1 | 1 | NA | NA | 1 | 1 | 2 | 1 |
| 6_1349611 | 1 | 1 | NA | NA | NA | NA | NA | NA | 1 | 1 | NA | NA | 1 | 1 | NA | NA | 1 | 1 | 2 | 1 |
| 6_1349613 | 1 | 1 | NA | NA | 1 | 1 | NA | NA | 2 | 1 | 1 | 1 | 1 | 1 | NA | NA | 1 | 1 | 3 | 1 |
| 6_1349614 | 1 | 1 | NA | NA | 1 | 1 | NA | NA | 2 | 1 | NA | NA | 1 | 1 | NA | NA | 1 | 1 | 2 | 1 |
| 6_1349617 | 1 | 0 | NA | NA | 1 | 1 | NA | NA | 2 | 0.5 | 1 | 1 | 1 | 0 | NA | NA | NA | NA | 2 | 0.5 |
| 6_1349619 | 1 | 1 | NA | NA | 1 | 0 | NA | NA | 2 | 0.5 | 1 | 0 | 1 | 1 | NA | NA | 1 | 0 | 3 | 0.3333 |
| 6_1350003 | 8 | 0.25 | 4 | 0.5 | 6 | 0.5 | 4 | 0.25 | 22 | 0.3636 | 3 | 0.6667 | 7 | 0.4286 | 7 | 0.4286 | 10 | 0.5 | 27 | 0.4815 |
| 6_1350004 | 8 | 0.625 | 4 | 0.75 | 7 | 0.8571 | 3 | 0.6667 | 22 | 0.7273 | 2 | 1 | 6 | 0.8333 | 10 | 0.7 | 9 | 0.4444 | 27 | 0.6667 |
| 6_1350005 | 6 | 1 | 3 | 1 | 5 | 0.6 | 4 | 1 | 18 | 0.8889 | 3 | 1 | 7 | 0.7143 | 9 | 0.7778 | 10 | 0.9 | 29 | 0.8276 |
| 6_1350006 | 7 | 0.4286 | 4 | 0.25 | 5 | 0.4 | 3 | 1 | 19 | 0.4737 | 4 | 0.25 | 6 | 0.6667 | 11 | 0.3636 | 9 | 0.5556 | 30 | 0.4667 |
| 6_1351146 | 5 | 0.2 | 10 | 0.1 | 6 | 0 | 4 | 0.25 | 25 | 0.12 | 5 | 0.2 | 8 | 0.125 | 6 | 0 | NA | NA | 19 | 0.1053 |
| 6_1351148 | 3 | 0.6667 | 7 | 0.8571 | 2 | 1 | 5 | 1 | 17 | 0.8824 | 1 | 1 | 4 | 0.75 | 5 | 0.8 | 1 | 1 | 11 | 0.8182 |
| 6_1352103 | 9 | 0 | 17 | 0 | 14 | 0.2143 | 10 | 0.2 | 50 | 0.1 | 14 | 0 | 14 | 0.2143 | 21 | 0 | 16 | 0.25 | 65 | 0.1077 |
| 6_1352110 | 7 | 0.8571 | 12 | 0.5 | 11 | 0.6364 | 10 | 0.6 | 40 | 0.625 | 13 | 0.8462 | 15 | 0.8 | 16 | 0.75 | 12 | 0.8333 | 56 | 0.8036 |
| 6_1352110 | 9 | 0.1111 | 16 | 0.0625 | 16 | 0.125 | 11 | 0.1818 | 52 | 0.1154 | 14 | 0.1429 | 16 | 0.125 | 19 | 0.0526 | 15 | 0.0667 | 64 | 0.0938 |
| 6_1352111 | 9 | 0.4444 | 12 | 0.8333 | 15 | 0.7333 | 10 | 0.6 | 46 | 0.6739 | 14 | 0.5714 | 16 | 0.5 | 20 | 0.7 | 16 | 0.5 | 66 | 0.5758 |
| 6_1352212 | NA | NA | 2 | 1 | NA | NA | NA | NA | 2 | 1 | NA | NA | 1 | 1 | NA | NA | NA | NA | 1 | 1 |
| 6_1352212 | NA | NA | 2 | 0.5 | NA | NA | NA | NA | 2 | 0.5 | NA | NA | 1 | 1 | NA | NA | NA | NA | 1 | 1 |
| 6_1352477 | NA | NA | 3 | 1 | NA | NA | NA | NA | 3 | 1 | NA | NA | NA | NA | NA | NA | 1 | 1 | 1 | 1 |
| 6_1356904 | 16 | 0.4375 | 22 | 0.5909 | 13 | 0.2308 | 7 | 0.5714 | 58 | 0.4655 | 16 | 0.375 | 12 | 0.5 | 11 | 0.7273 | 13 | 0.6154 | 52 | 0.5385 |
| 6_1356905 | 14 | 0.1429 | 15 | 0.0667 | 11 | 0.1818 | 5 | 0.2 | 45 | 0.1333 | 12 | 0.0833 | 11 | 0 | 9 | 0 | 14 | 0.2143 | 46 | 0.087 |
| 6_1356908 | 9 | 0.1111 | 9 | 0.4444 | 9 | 0.4444 | 4 | 1 | 31 | 0.4194 | 9 | 0.3333 | 9 | 0.3333 | 5 | 0.2 | 9 | 0 | 32 | 0.2188 |
| 6_1364658 | NA | NA | 1 | 1 | 1 | 1 | NA | NA | 2 | 1 | NA | NA | NA | NA | 1 | 1 | 1 | 1 | 2 | 1 |
| 6_1364660 | NA | NA | 1 | 1 | 1 | 1 | NA | NA | 2 | 1 | NA | NA | NA | NA | 2 | 0.5 | 1 | 1 | 3 | 0.6667 |
| 6_1365376 | NA | NA | NA | NA | 1 | 1 | NA | NA | 1 | 1 | NA | NA | 2 | 0.5 | NA | NA | NA | NA | 2 | 0.5 |
| 6_1365381 | NA | NA | NA | NA | 1 | 1 | NA | NA | 1 | 1 | 1 | 0 | 2 | 0 | NA | NA | 1 | 1 | 4 | 0.25 |
| 6_1365641 | NA | NA | NA | NA | 2 | 0.5 | NA | NA | 2 | 0.5 | 1 | 0 | 3 | 0.3333 | 1 | 0 | 1 | 0 | 6 | 0.1667 |

| | | | | | | | | | | | | | | | | | | | | |
|---|---|---|---|---|---|---|---|---|---|---|---|---|---|---|---|---|---|---|---|---|
| 6_13656429 | NA | NA | NA | NA | 2 | 0.5 | NA | NA | 2 | 0.5 | NA | NA | 3 | 1 | 1 | 1 | 1 | 1 | 5 | 1 |
| 6_13656445 | NA | NA | NA | NA | 2 | 0.5 | NA | NA | 2 | 0.5 | 1 | 0 | 3 | 0.3333 | NA | NA | 1 | 1 | 5 | 0.4 |
| 6_13656454 | NA | NA | NA | NA | 2 | 1 | NA | NA | 2 | 1 | 1 | 1 | 3 | 1 | 1 | 1 | 1 | 1 | 6 | 1 |
| 6_13656457 | NA | NA | NA | NA | 2 | 1 | NA | NA | 2 | 1 | 1 | 1 | 3 | 0.6667 | 1 | 1 | NA | NA | 5 | 0.8 |
| 6_13658363 | 3 | 0.3333 | 4 | 0.25 | 2 | 0 | 2 | 0 | 11 | 0.1818 | 2 | 0.5 | 3 | 0.3333 | 2 | 0 | 1 | 0 | 8 | 0.25 |
| 6_13658381 | 3 | 0.6667 | 4 | 0.75 | 4 | 1 | 3 | 1 | 14 | 0.8571 | 2 | 1 | 3 | 1 | 2 | 1 | 1 | 1 | 8 | 1 |
| 6_13658384 | 3 | 0.6667 | 4 | 0 | 2 | 0 | 3 | 0 | 12 | 0.1667 | 3 | 0 | 3 | 0.6667 | 2 | 0.5 | 2 | 0 | 10 | 0.3 |
| 6_13658388 | 2 | 1 | 3 | 0.6667 | 4 | 0.5 | 2 | 0.5 | 11 | 0.6364 | 1 | 1 | 2 | 1 | NA | NA | NA | NA | 3 | 1 |
| 6_13658389 | 3 | 0.3333 | 4 | 0 | 4 | 0.25 | 3 | 0 | 14 | 0.1429 | 3 | 0 | 4 | 0.75 | 2 | 0 | 2 | 0 | 11 | 0.2727 |
| 6_13658400 | 3 | 0.3333 | 4 | 0.75 | 4 | 0.5 | 2 | 0.5 | 13 | 0.5385 | 4 | 0.5 | 4 | 1 | 1 | 1 | 1 | 1 | 10 | 0.8 |
| 6_13658400 | 2 | 1 | 2 | 1 | 4 | 1 | 2 | 1 | 10 | 1 | 3 | 1 | 3 | 1 | 1 | 1 | 1 | 0 | 8 | 0.875 |
| 6_13658407 | 1 | 1 | 3 | 0.6667 | 2 | 0.5 | 1 | 0 | 7 | 0.5714 | 2 | 0 | 2 | 0.5 | NA | NA | NA | NA | 4 | 0.25 |
| 6_13658508 | 2 | 0 | 2 | 0.5 | 1 | 0 | 4 | 0 | 9 | 0.1111 | 4 | 0.5 | 3 | 0.3333 | 2 | 0.5 | 2 | 0.5 | 11 | 0.4545 |
| 6_13678366 | NA | NA | 1 | 1 | NA | NA | NA | NA | 1 | 1 | NA | NA | 1 | 1 | NA | NA | NA | NA | 1 | 1 |
| 6_13678375 | NA | NA | 1 | 1 | NA | NA | NA | NA | 1 | 1 | NA | NA | 1 | 1 | NA | NA | NA | NA | 1 | 1 |
| 6_13678377 | NA | NA | 2 | 1 | NA | NA | NA | NA | 2 | 1 | NA | NA | 2 | 1 | NA | NA | NA | NA | 2 | 1 |
| 6_13678383 | NA | NA | 2 | 1 | NA | NA | NA | NA | 2 | 1 | NA | NA | 1 | 1 | NA | NA | NA | NA | 1 | 1 |
| 6_13678392 | NA | NA | 2 | 1 | NA | NA | NA | NA | 2 | 1 | NA | NA | 1 | 1 | NA | NA | NA | NA | 1 | 1 |
| 6_13678395 | NA | NA | 2 | 1 | NA | NA | NA | NA | 2 | 1 | NA | NA | 1 | 1 | NA | NA | NA | NA | 1 | 1 |
| 6_13678582 | 1 | 1 | 2 | 1 | NA | NA | 1 | 1 | 4 | 1 | 1 | 1 | NA | NA | 3 | 0.3333 | 1 | 1 | 5 | 0.6 |
| 6_13678590 | 1 | 0 | 2 | 0.5 | NA | NA | 1 | 0 | 4 | 0.25 | 1 | 0 | NA | NA | 3 | 0.3333 | 1 | 0 | 5 | 0.2 |
| 6_13678595 | 1 | 0 | 2 | 0.5 | NA | NA | 2 | 0.5 | 5 | 0.4 | 1 | 1 | NA | NA | 3 | 0.6667 | 1 | 1 | 5 | 0.8 |
| 6_13678604 | 1 | 0 | 2 | 0.5 | NA | NA | 2 | 0 | 5 | 0.2 | 1 | 0 | NA | NA | 2 | 0.5 | 1 | 0 | 4 | 0.25 |
| 6_13678700 | 1 | 0 | 3 | 0.3333 | 1 | 0 | 3 | 0 | 8 | 0.125 | NA | NA | NA | NA | 3 | 0 | 2 | 0.5 | 5 | 0.2 |
| 6_13678704 | 1 | 0 | 3 | 0 | 1 | 1 | 3 | 0 | 8 | 0.125 | NA | NA | NA | NA | 3 | 0 | 3 | 0.6667 | 6 | 0.3333 |
| 6_13679067 | 16 | 0.0625 | 16 | 0.375 | 7 | 0.1429 | 6 | 0.3333 | 45 | 0.2222 | 13 | 0.2308 | 15 | 0.2667 | 17 | 0.2353 | 15 | 0.2 | 60 | 0.2333 |
| 6_13679085 | 16 | 0 | 14 | 0.2143 | 8 | 0 | 5 | 0.2 | 43 | 0.093 | 12 | 0 | 18 | 0.1111 | 15 | 0.1333 | 15 | 0.1333 | 60 | 0.1 |
| 6_13679091 | 13 | 0.0769 | 11 | 0.0909 | 5 | 0 | 5 | 0.4 | 34 | 0.1176 | 7 | 0.1429 | 12 | 0.0833 | 11 | 0.3636 | 9 | 0.1111 | 39 | 0.1795 |
| 6_13679103 | 15 | 0.1333 | 14 | 0.2143 | 8 | 0.125 | 6 | 0 | 43 | 0.1395 | 9 | 0.1111 | 17 | 0.0588 | 14 | 0.0714 | 13 | 0.3077 | 53 | 0.1321 |
| 6_13679115 | 14 | 0.2857 | 11 | 0.2727 | 7 | 0.1429 | 4 | 0 | 36 | 0.2222 | 11 | 0 | 13 | 0 | 12 | 0.25 | 13 | 0 | 49 | 0.0612 |
| 6_13679118 | 16 | 0.375 | 12 | 0.5 | 9 | 0.5556 | 6 | 0.5 | 43 | 0.4651 | 12 | 0.5 | 17 | 0.4118 | 11 | 0.3636 | 12 | 0.3333 | 52 | 0.4038 |
| 6_13679125 | 13 | 0.2308 | 14 | 0.3571 | 7 | 0 | 3 | 0 | 37 | 0.2162 | 11 | 0.2727 | 15 | 0.1333 | 11 | 0.2727 | 9 | 0.1111 | 46 | 0.1957 |
| 6_13679132 | 16 | 0.0625 | 11 | 0.2727 | 7 | 0 | 6 | 0.1667 | 40 | 0.125 | 9 | 0.2222 | 18 | 0.1667 | 12 | 0.0833 | 8 | 0 | 47 | 0.1277 |
| 6_13680726 | 2 | 0 | 3 | 0.3333 | 2 | 0 | 1 | 0 | 8 | 0.125 | NA | NA | 1 | 0 | 2 | 0 | 1 | 1 | 4 | 0.25 |
| 6_13680747 | 2 | 1 | 3 | 0.6667 | 1 | 1 | 2 | 0.5 | 8 | 0.75 | NA | NA | 1 | 1 | 1 | 1 | NA | NA | 2 | 1 |
| 6_13680772 | 3 | 1 | 5 | 1 | 2 | 1 | 4 | 1 | 14 | 1 | NA | NA | NA | NA | 1 | 1 | 2 | 1 | 3 | 1 |
| 6_13680773 | 2 | 1 | 4 | 1 | 2 | 1 | 4 | 1 | 12 | 1 | NA | NA | 1 | 1 | 1 | 1 | 2 | 1 | 4 | 1 |
| 6_13680775 | 2 | 0.5 | 4 | 0.25 | 2 | 0.5 | 4 | 0.5 | 12 | 0.4167 | NA | NA | 1 | 0 | 1 | 0 | 2 | 0.5 | 4 | 0.25 |
| 6_13680780 | 3 | 0.3333 | 5 | 0 | 2 | 0.5 | 4 | 0 | 14 | 0.1429 | NA | NA | 1 | 1 | 1 | 0 | 2 | 0 | 4 | 0.25 |
| 6_13680781 | 3 | 0 | 5 | 0 | 2 | 0.5 | 4 | 0.5 | 14 | 0.2143 | NA | NA | 1 | 1 | 1 | 1 | 2 | 0.5 | 4 | 0.75 |
| 6_13680782 | 1 | 1 | 5 | 0.6 | 2 | 0.5 | 4 | 0.25 | 12 | 0.5 | NA | NA | 1 | 0 | 1 | 1 | 2 | 0 | 4 | 0.25 |
| 6_13680783 | 2 | 1 | 5 | 1 | 1 | 1 | 3 | 1 | 11 | 1 | NA | NA | 1 | 1 | 1 | 1 | 2 | 1 | 4 | 1 |
| 6_13680788 | 3 | 0.3333 | 5 | 0.6 | 2 | 0 | 4 | 0.75 | 14 | 0.5 | NA | NA | 1 | 0 | 1 | 1 | 1 | 0 | 3 | 0.3333 |
| 6_13680795 | 3 | 0.6667 | 5 | 0 | 2 | 1 | 4 | 0.25 | 14 | 0.3571 | NA | NA | 1 | 1 | 1 | 0 | 1 | 0 | 3 | 0.3333 |
| 6_13680819 | 2 | 0.5 | 4 | 0.75 | 2 | 1 | 3 | 1 | 11 | 0.8182 | NA | NA | 1 | 1 | 1 | 1 | NA | NA | 2 | 1 |
| 6_13680831 | 3 | 0.3333 | 5 | 0.4 | 2 | 1 | 3 | 1 | 13 | 0.6154 | NA | NA | 1 | 1 | 1 | 1 | 1 | 1 | 3 | 1 |
| 6_13680840 | 3 | 1 | 5 | 1 | 1 | 1 | 3 | 1 | 12 | 1 | NA | NA | 1 | 1 | NA | NA | 1 | 1 | 2 | 1 |
| 6_13680847 | 3 | 1 | 4 | 0.5 | 1 | 0 | 4 | 0.5 | 12 | 0.5833 | NA | NA | 1 | 0 | 1 | 1 | 1 | 1 | 3 | 0.6667 |
| 6_13680860 | 3 | 0 | 5 | 0.2 | 1 | 0 | 2 | 0.5 | 11 | 0.1818 | NA | NA | 1 | 0 | 1 | 1 | 1 | 0 | 3 | 0.3333 |
| 6_13680863 | 2 | 0.5 | 5 | 0.8 | 1 | 1 | 1 | 0 | 9 | 0.6667 | NA | NA | 1 | 1 | 1 | 1 | 1 | 1 | 3 | 1 |
| 6_13680867 | 3 | 0.3333 | 5 | 0 | 1 | 0 | 1 | 0 | 10 | 0.1 | NA | NA | 1 | 1 | 1 | 0 | 1 | 0 | 3 | 0.3333 |
| 6_13680867 | 3 | 0.6667 | 5 | 0 | 1 | 1 | 1 | 0 | 10 | 0.3 | NA | NA | 1 | 1 | 1 | 0 | 1 | 0 | 3 | 0.3333 |
| 6_13680879 | 2 | 0.5 | 5 | 0.4 | 1 | 0 | 2 | 1 | 10 | 0.5 | NA | NA | 1 | 1 | 1 | 1 | 1 | 0 | 3 | 0.6667 |
| 6_13690092 | 3 | 0.6667 | 5 | 0 | NA | NA | 1 | 0 | 9 | 0.2222 | 1 | 1 | 4 | 0.5 | 2 | 0.5 | 6 | 0.3333 | 13 | 0.4615 |
| 6_13703257 | 5 | 0.2 | 5 | 0 | 3 | 0 | 1 | 0 | 14 | 0.0714 | NA | NA | 3 | 0 | 5 | 0.2 | 4 | 0 | 12 | 0.0833 |
| 6_13703273 | 4 | 0 | 5 | 0.2 | 5 | 0.2 | 2 | 0 | 16 | 0.125 | NA | NA | 3 | 0.6667 | 5 | 0.2 | 3 | 0 | 11 | 0.2727 |
| 6_13703294 | 3 | 0 | 5 | 0 | 3 | 0.3333 | 2 | 0 | 13 | 0.0769 | NA | NA | 3 | 0.3333 | 4 | 0.25 | 4 | 0 | 11 | 0.1818 |
| 6_13703331 | 2 | 0.5 | 2 | 1 | 3 | 0 | 2 | 0 | 9 | 0.3333 | NA | NA | 4 | 0.25 | 4 | 0.5 | 4 | 0.75 | 12 | 0.5 |
| 6_13703336 | 3 | 0.3333 | 3 | 0.3333 | 2 | 1 | 2 | 0.5 | 10 | 0.5 | NA | NA | 3 | 0 | 4 | 0.25 | 3 | 0 | 10 | 0.1 |
| 6_13703339 | 2 | 1 | 2 | 1 | 3 | 0.6667 | 2 | 0.5 | 9 | 0.7778 | NA | NA | 4 | 0.75 | 4 | 1 | 3 | 1 | 11 | 0.9091 |
| 6_13722961 | 1 | 0 | 1 | 0 | NA | NA | 2 | 0.5 | 4 | 0.25 | 2 | 0 | NA | NA | 2 | 0.5 | NA | NA | 4 | 0.25 |
| 6_13735725 | 1 | 0 | 1 | 0 | NA | NA | 3 | 0.3333 | 5 | 0.2 | 2 | 0 | 1 | 0 | 2 | 0.5 | NA | NA | 5 | 0.2 |
| 6_13735740 | 2 | 0 | 1 | 0 | NA | NA | 3 | 0.3333 | 6 | 0.1667 | 2 | 0.5 | 1 | 0 | 3 | 0.6667 | NA | NA | 6 | 0.5 |
| 6_13735744 | 2 | 1 | 1 | 0 | NA | NA | 3 | 0.3333 | 6 | 0.5 | 1 | 0 | 1 | 1 | 2 | 0.5 | NA | NA | 4 | 0.5 |
| 6_13735747 | 2 | 0.5 | 1 | 1 | NA | NA | 3 | 0.6667 | 6 | 0.6667 | 3 | 1 | 1 | 1 | 3 | 0.6667 | NA | NA | 7 | 0.8571 |

| | | | | | | | | | | | | | | | | | | | | |
|---|---|---|---|---|---|---|---|---|---|---|---|---|---|---|---|---|---|---|---|---|
| 6_13743919 | 3 | 0.6667 | 2 | 0.5 | NA | NA | 4 | 0.75 | 9 | 0.6667 | 2 | 1 | 2 | 0.5 | 4 | 0 | 4 | 0.75 | 12 | 0.5 |
| 6_13743928 | 4 | 1 | 3 | 1 | 1 | 1 | 8 | 1 | 16 | 1 | 4 | 1 | 4 | 1 | 6 | 1 | 6 | 1 | 20 | 1 |
| 6_13743958 | 4 | 0.25 | 3 | 0.6667 | 1 | 0 | 10 | 0 | 18 | 0.1667 | 3 | 0 | 4 | 0 | 6 | 0.3333 | 7 | 0.1429 | 20 | 0.15 |
| 6_13743974 | 3 | 0 | 3 | 0.3333 | 1 | 1 | 9 | 0.2222 | 16 | 0.25 | 3 | 0.6667 | 5 | 0.2 | 6 | 0.1667 | 7 | 0.2857 | 21 | 0.2857 |
| 6_13743975 | 5 | 0.2 | 2 | 0.5 | 1 | 0 | 10 | 0.2 | 18 | 0.2222 | 4 | 0.25 | 6 | 0 | 6 | 0 | 7 | 0.1429 | 23 | 0.087 |
| 6_13743987 | 5 | 0 | 4 | 0 | 1 | 1 | 11 | 0.0909 | 21 | 0.0952 | 4 | 0 | 7 | 0 | 7 | 0.2857 | 8 | 0 | 26 | 0.0769 |
| 6_13743987 | 6 | 0.3333 | 4 | 0.25 | 1 | 1 | 10 | 0.6 | 21 | 0.4762 | 4 | 0.75 | 7 | 0.2857 | 6 | 0.3333 | 8 | 0.5 | 25 | 0.44 |
| 6_13744003 | 5 | 0.4 | 5 | 0 | 2 | 0 | 9 | 0.1111 | 21 | 0.1429 | 3 | 0 | 6 | 0.3333 | 6 | 0 | 8 | 0 | 23 | 0.087 |
| 6_13744004 | 5 | 0.6 | 5 | 0.4 | 2 | 0.5 | 12 | 0.5833 | 24 | 0.5417 | 4 | 0.25 | 6 | 0.6667 | 6 | 0.5 | 8 | 0.75 | 24 | 0.5833 |
| 6_13744006 | 5 | 0.4 | 3 | 0.6667 | 2 | 0 | 10 | 0.3 | 20 | 0.35 | 2 | 0 | 7 | 0.1429 | 6 | 0.3333 | 7 | 0.2857 | 22 | 0.2273 |
| 6_13744007 | 5 | 0.8 | 5 | 0.4 | 2 | 0.5 | 11 | 0.1818 | 23 | 0.3913 | 3 | 0.6667 | 6 | 0.5 | 6 | 0.6667 | 8 | 0.125 | 23 | 0.4348 |
| 6_13745529 | 2 | 0 | 3 | 0 | 4 | 0.25 | 2 | 0 | 11 | 0.0909 | 4 | 0 | 4 | 0 | 4 | 0.25 | 5 | 0 | 17 | 0.0588 |
| 6_13745596 | 4 | 0.5 | 5 | 0.6 | 3 | 0.6667 | 3 | 0.6667 | 15 | 0.6 | 4 | 0.5 | 4 | 1 | 3 | 0.6667 | 2 | 0.5 | 13 | 0.6923 |
| 6_13745600 | 4 | 0.5 | 4 | 0.25 | 2 | 0.5 | 2 | 0.5 | 12 | 0.4167 | 4 | 0.25 | 5 | 0.6 | 4 | 0.25 | NA | NA | 13 | 0.3846 |
| 6_13772764 | 21 | 0.6667 | 19 | 0.6842 | 9 | 0.8889 | 22 | 0.6364 | 71 | 0.6901 | 10 | 0.6 | 23 | 0.7826 | 19 | 0.8947 | 24 | 0.8333 | 76 | 0.8026 |
| 6_13772774 | 21 | 0.3333 | 17 | 0.2353 | 8 | 0.25 | 22 | 0.3182 | 68 | 0.2941 | 13 | 0.3846 | 18 | 0.2778 | 14 | 0.0714 | 19 | 0.1579 | 64 | 0.2188 |
| 6_13772793 | 25 | 0.36 | 17 | 0.1765 | 12 | 0.1667 | 26 | 0.2308 | 80 | 0.25 | 11 | 0.3636 | 25 | 0.2 | 21 | 0.1905 | 23 | 0.3043 | 80 | 0.25 |
| 6_13772803 | 25 | 0.56 | 20 | 0.55 | 12 | 0.75 | 22 | 0.7727 | 79 | 0.6456 | 12 | 0.9167 | 25 | 0.6 | 18 | 0.7778 | 27 | 0.7037 | 82 | 0.7195 |
| 6_13772819 | 20 | 0.2 | 12 | 0.0833 | 8 | 0.125 | 17 | 0.0588 | 57 | 0.1228 | 9 | 0.1111 | 20 | 0.15 | 13 | 0 | 13 | 0.1538 | 55 | 0.1091 |
| 6_13833999 | 13 | 0.0769 | 34 | 0.0588 | 23 | 0.0435 | 17 | 0.0588 | 87 | 0.0575 | 25 | 0.08 | 14 | 0.1429 | 9 | 0.1111 | 20 | 0.2 | 68 | 0.1324 |
| 6_13834019 | 11 | 1 | 33 | 0.9394 | 22 | 0.9545 | 15 | 0.8667 | 81 | 0.9383 | 22 | 0.9091 | 14 | 0.8571 | 11 | 1 | 15 | 0.8667 | 62 | 0.9032 |
| 6_13834153 | 27 | 0.1481 | 40 | 0.075 | 28 | 0.0357 | 22 | 0.0909 | 117 | 0.0855 | 34 | 0.1471 | 18 | 0.0556 | 13 | 0.2308 | 21 | 0.0952 | 86 | 0.1279 |
| 6_13834183 | 15 | 0.4667 | 37 | 0.4054 | 22 | 0.5909 | 15 | 0.4 | 89 | 0.4607 | 21 | 0.4762 | 12 | 0.5 | 10 | 0.6 | 14 | 0.4286 | 57 | 0.4912 |
| 6_13834220 | 17 | 0.2353 | 29 | 0.2069 | 20 | 0.3 | 15 | 0.2667 | 81 | 0.2469 | 25 | 0.12 | 15 | 0.1333 | 9 | 0.2222 | 16 | 0.0625 | 65 | 0.1231 |
| 6_13834265 | 16 | 0.1875 | 23 | 0.1304 | 13 | 0.2308 | 15 | 0.2 | 67 | 0.1791 | 24 | 0.2083 | 9 | 0.2222 | 8 | 0.25 | 17 | 0.2353 | 58 | 0.2241 |
| 6_13834279 | 19 | 0.6316 | 31 | 0.8065 | 24 | 0.8333 | 17 | 0.8824 | 91 | 0.7912 | 30 | 0.8333 | 15 | 0.8667 | 10 | 0.7 | 18 | 0.7222 | 73 | 0.7945 |
| 6_13834285 | 19 | 0.0526 | 32 | 0 | 23 | 0.0435 | 16 | 0.25 | 90 | 0.0667 | 28 | 0 | 13 | 0.1538 | 11 | 0.0909 | 17 | 0.0588 | 69 | 0.058 |
| 6_13834301 | 17 | 0.1176 | 25 | 0.04 | 17 | 0.0588 | 12 | 0.0833 | 71 | 0.0704 | 19 | 0.1053 | 11 | 0.0909 | 7 | 0.1429 | 14 | 0.1429 | 51 | 0.1176 |
| 6_13839961 | 9 | 0.1111 | 13 | 0.0769 | 7 | 0.1429 | 8 | 0 | 37 | 0.0811 | 9 | 0.2222 | 9 | 0 | 5 | 0 | 8 | 0 | 31 | 0.0645 |
| 6_13840282 | NA | NA | 1 | 0 | 3 | 0.6667 | 1 | 0 | 5 | 0.4 | 1 | 1 | NA | NA | 1 | 0 | 1 | 0 | 3 | 0.3333 |
| 6_14036935 | NA | NA | NA | NA | 1 | 1 | NA | NA | 1 | 1 | NA | NA | NA | NA | 1 | 1 | NA | NA | 1 | 1 |
| 6_14036943 | NA | NA | NA | NA | 2 | 0.5 | NA | NA | 2 | 0.5 | NA | NA | 1 | 1 | NA | NA | NA | NA | 1 | 1 |
| 6_14048962 | 1 | 0 | 2 | 0.5 | 1 | 0 | 1 | 0 | 5 | 0.2 | 2 | 0 | 1 | 0 | NA | NA | 4 | 0.5 | 7 | 0.2857 |
| 6_14048965 | 1 | 1 | 2 | 0.5 | 2 | 0 | 2 | 1 | 7 | 0.5714 | 2 | 0.5 | 1 | 0 | NA | NA | 2 | 0 | 5 | 0.2 |
| 6_14049656 | 1 | 0 | 2 | 0.5 | 2 | 1 | 2 | 0 | 7 | 0.4286 | 3 | 0 | 1 | 0 | NA | NA | 5 | 0.2 | 9 | 0.1111 |
| 6_14054231 | 8 | 0 | 6 | 0 | 9 | 0.1111 | 5 | 0.2 | 28 | 0.0714 | 7 | 0.1429 | 8 | 0 | 9 | 0.2222 | 10 | 0.4 | 34 | 0.2059 |
| 6_14056987 | 1 | 0 | 3 | 0 | 1 | 0 | 2 | 0.5 | 7 | 0.1429 | NA | NA | NA | NA | 1 | 0 | 5 | 0.4 | 6 | 0.3333 |
| 6_14059666 | 6 | 0.5 | 3 | 0.3333 | 3 | 0.3333 | 6 | 0.5 | 18 | 0.4444 | 1 | 1 | NA | NA | 1 | 1 | 7 | 0.7143 | 9 | 0.7778 |
| 6_14059675 | 6 | 0.1667 | 3 | 0 | 5 | 0 | 6 | 0 | 20 | 0.05 | 1 | 0 | NA | NA | 2 | 0 | 7 | 0.1429 | 10 | 0.1 |
| 6_14059704 | 7 | 0.1429 | 5 | 0 | 6 | 0.1667 | 7 | 0.2857 | 25 | 0.16 | 1 | 0 | NA | NA | 3 | 0 | 10 | 0.1 | 14 | 0.0714 |
| 6_14059731 | 5 | 0.4 | 5 | 0 | 4 | 0 | 8 | 0 | 22 | 0.0909 | 1 | 0 | 1 | 0 | 4 | 0 | 9 | 0.1111 | 15 | 0.0667 |
| 6_14059776 | 9 | 0.4444 | 8 | 0.25 | 7 | 0.1429 | 9 | 0.3333 | 33 | 0.303 | 2 | 0.5 | 2 | 0.5 | 4 | 0.25 | 13 | 0.5385 | 21 | 0.4762 |
| 6_14059894 | 6 | 0.3333 | 3 | 0.6667 | 8 | 0.125 | 6 | 0.3333 | 23 | 0.3043 | NA | NA | 4 | 0.25 | 5 | 0 | 4 | 0.25 | 13 | 0.1538 |
| 6_14059924 | 7 | 0.4286 | 3 | 0.3333 | 9 | 0.3333 | 8 | 0 | 27 | 0.2593 | NA | NA | 5 | 0.2 | 6 | 0 | 4 | 0 | 15 | 0.0667 |
| 6_14059950 | 6 | 0.3333 | 3 | 0.6667 | 7 | 0.4286 | 10 | 0.4 | 26 | 0.4231 | 2 | 0.5 | 6 | 0.3333 | 5 | 0.8 | 5 | 0.4 | 18 | 0.5 |
| 6_14059957 | 2 | 0 | 1 | 0 | 1 | 1 | 5 | 0.2 | 9 | 0.2222 | 3 | 0 | 3 | 0 | 4 | 0.25 | 2 | 0 | 12 | 0.0833 |
| 6_14059991 | 2 | 0.5 | 2 | 0.5 | NA | NA | 5 | 0.6 | 9 | 0.5556 | 2 | 0 | 2 | 0.5 | 2 | 0.5 | 2 | 0 | 8 | 0.25 |
| 6_14059996 | 2 | 0 | 2 | 0 | 2 | 0 | 5 | 0.4 | 11 | 0.1818 | 3 | 0 | 3 | 0.3333 | 4 | 0.5 | 3 | 0.3333 | 13 | 0.3077 |
| 6_14060008 | 2 | 0.5 | 2 | 0.5 | 3 | 1 | 4 | 0.25 | 11 | 0.5455 | 2 | 0 | 3 | 0.3333 | 4 | 0.75 | 3 | 0.6667 | 12 | 0.5 |
| 6_14060057 | 3 | 0.3333 | 1 | 0 | 2 | 0 | 6 | 0.5 | 12 | 0.3333 | 4 | 0.25 | 2 | 0.5 | 4 | 0.75 | 2 | 0.5 | 12 | 0.5 |
| 6_14060078 | 2 | 1 | 1 | 0 | 1 | 0 | 6 | 0.6667 | 10 | 0.6 | 3 | 0.3333 | 2 | 0 | 6 | 0.1667 | 1 | 1 | 12 | 0.25 |
| 6_14147035 | 2 | 0 | 6 | 0.1667 | 6 | 0 | 8 | 0.25 | 22 | 0.1364 | 4 | 0.25 | 5 | 0 | 3 | 0 | 2 | 0 | 14 | 0.0714 |
| 6_14175167 | 1 | 0 | 7 | 0.2857 | 2 | 0 | 2 | 0 | 12 | 0.1667 | 1 | 0 | 4 | 0.25 | 5 | 0.2 | 3 | 0 | 13 | 0.1538 |
| 6_14175217 | 1 | 0 | 5 | 0 | 1 | 1 | 4 | 0 | 11 | 0.0909 | 2 | 0 | 5 | 0.2 | 5 | 0 | 4 | 0 | 16 | 0.0625 |
| 6_14175260 | NA | NA | 6 | 0.8333 | 1 | 0 | 4 | 0.75 | 11 | 0.7273 | 2 | 1 | 4 | 0.5 | 6 | 0.8333 | 4 | 1 | 16 | 0.8125 |
| 6_14175280 | 1 | 0 | 5 | 0 | 3 | 0.3333 | 4 | 0 | 13 | 0.0769 | 1 | 0 | 4 | 0 | 6 | 0.1667 | 5 | 0 | 16 | 0.0625 |
| 6_14230284 | NA | NA | 3 | 1 | 1 | 1 | 1 | 1 | 5 | 1 | NA | NA | NA | NA | 1 | 1 | NA | NA | 1 | 1 |
| 6_14230341 | NA | NA | 2 | 0.5 | 1 | 0 | 1 | 1 | 4 | 0.5 | NA | NA | NA | NA | 1 | 1 | NA | NA | 1 | 1 |
| 6_14230359 | NA | NA | 3 | 1 | 1 | 0 | 1 | 1 | 5 | 0.8 | NA | NA | NA | NA | 1 | 1 | NA | NA | 1 | 1 |
| 6_14230363 | NA | NA | 3 | 1 | 1 | 1 | 1 | 1 | 5 | 1 | NA | NA | NA | NA | 1 | 1 | NA | NA | 1 | 1 |
| 6_14230364 | NA | NA | 2 | 1 | 1 | 1 | NA | NA | 3 | 1 | NA | NA | NA | NA | 1 | 1 | NA | NA | 1 | 1 |
| 6_14230427 | NA | NA | 3 | 1 | NA | NA | 1 | 1 | 4 | 1 | NA | NA | NA | NA | 1 | 1 | NA | NA | 1 | 1 |
| 6_14230491 | NA | NA | 3 | 1 | NA | NA | NA | NA | 3 | 1 | NA | NA | NA | NA | 1 | 1 | NA | NA | 1 | 1 |
| 6_14230499 | NA | NA | 3 | 1 | NA | NA | 1 | 1 | 4 | 1 | NA | NA | NA | NA | 1 | 1 | NA | NA | 1 | 1 |
| 6_14230503 | NA | NA | 3 | 0.3333 | NA | NA | 1 | 1 | 4 | 0.5 | NA | NA | NA | NA | 1 | 1 | NA | NA | 1 | 1 |

| | | | | | | | | | | | | | | | | | | | | |
|---|---|---|---|---|---|---|---|---|---|---|---|---|---|---|---|---|---|---|---|---|
| 6_1423052 | NA | NA | 3 | 0.3333 | NA | NA | 1 | 1 | 4 | 0.5 | NA | NA | NA | NA | 1 | 1 | NA | NA | 1 | 1 |
| 6_1423076 | NA | NA | 1 | 1 | 1 | 1 | 2 | 0.5 | 4 | 0.75 | NA | NA | NA | NA | 4 | 0.75 | NA | NA | 4 | 0.75 |
| 6_1423076 | NA | NA | NA | NA | 1 | 1 | 2 | 0.5 | 3 | 0.6667 | NA | NA | NA | NA | 2 | 1 | NA | NA | 2 | 1 |
| 6_1423078 | NA | NA | 1 | 1 | NA | NA | NA | NA | 1 | 1 | NA | NA | NA | NA | 2 | 1 | NA | NA | 2 | 1 |
| 6_1423079 | NA | NA | 1 | 1 | NA | NA | NA | NA | 1 | 1 | NA | NA | NA | NA | 2 | 1 | NA | NA | 2 | 1 |
| 6_1423082 | NA | NA | 1 | 1 | NA | NA | NA | NA | 1 | 1 | NA | NA | NA | NA | 2 | 1 | NA | NA | 2 | 1 |
| 6_1423473 | 3 | 0 | 6 | 0.3333 | 3 | 0.3333 | 5 | 0.4 | 17 | 0.2941 | 2 | 0 | 5 | 0 | 4 | 0.25 | 5 | 0.4 | 16 | 0.1875 |
| 6_1423473 | 3 | 0.6667 | 5 | 0 | 2 | 0 | 4 | 0 | 14 | 0.1429 | 2 | 0 | 5 | 0 | 5 | 0.2 | 4 | 0.25 | 16 | 0.125 |
| 6_1423473 | 3 | 0.6667 | 5 | 0.6 | 2 | 0.5 | 4 | 1 | 14 | 0.7143 | 2 | 0.5 | 6 | 0.5 | 5 | 0.8 | 3 | 0.3333 | 16 | 0.5625 |
| 6_1423475 | 3 | 0.6667 | 5 | 0.6 | 4 | 1 | 4 | 0.75 | 16 | 0.75 | 3 | 0.3333 | 4 | 0.75 | 3 | 0.6667 | 4 | 1 | 14 | 0.7143 |
| 6_1423475 | 3 | 0 | 7 | 0.2857 | 4 | 0.5 | 5 | 0 | 19 | 0.2105 | 3 | 0.3333 | 4 | 0 | 5 | 0 | 5 | 0.4 | 17 | 0.1765 |
| 6_1423476 | 3 | 0 | 7 | 0 | 3 | 0.3333 | 4 | 0 | 17 | 0.0588 | 2 | 0.5 | 5 | 0 | 3 | 0 | 4 | 0 | 14 | 0.0714 |
| 6_1423478 | 3 | 0 | 7 | 0.1429 | 4 | 0.25 | 4 | 0 | 18 | 0.1111 | 3 | 0 | 8 | 0.25 | 4 | 0.5 | 5 | 0.2 | 20 | 0.25 |
| 6_1423478 | 2 | 0.5 | 5 | 0.4 | 4 | 0.5 | 5 | 0.2 | 16 | 0.375 | 1 | 1 | 7 | 0.2857 | 4 | 0 | 4 | 0.5 | 16 | 0.3125 |
| 6_1423479 | 3 | 0 | 6 | 0.1667 | 3 | 0 | 3 | 0.3333 | 15 | 0.1333 | 1 | 0 | 7 | 0.2857 | 4 | 0 | 3 | 0.3333 | 15 | 0.2 |
| 6_1423481 | 3 | 0.3333 | 6 | 0.3333 | 5 | 0.6 | 5 | 0.8 | 19 | 0.5263 | 1 | 1 | 7 | 0.8571 | 4 | 0.5 | 4 | 0.75 | 16 | 0.75 |
| 6_1423482 | 2 | 0 | 6 | 0.3333 | 6 | 0.1667 | 5 | 0.4 | 19 | 0.2632 | 1 | 0 | 5 | 0.4 | 4 | 0.25 | 4 | 0.75 | 14 | 0.4286 |
| 6_1423486 | 4 | 0.25 | 5 | 0.4 | 6 | 0 | 5 | 0 | 20 | 0.15 | 2 | 0 | 7 | 0 | 4 | 0.25 | 4 | 0 | 17 | 0.0588 |
| 6_1423487 | 5 | 0.4 | 3 | 0 | 2 | 0 | 3 | 0.3333 | 13 | 0.2308 | 2 | 0 | 7 | 0.1429 | 5 | 0.2 | 3 | 0.6667 | 17 | 0.2353 |
| 6_1423489 | 3 | 0.3333 | 3 | 0 | 6 | 0.3333 | 6 | 0.3333 | 18 | 0.2778 | 1 | 0 | 7 | 0 | 5 | 0 | 2 | 0.5 | 15 | 0.0667 |
| 6_1423491 | 5 | 0 | 6 | 0.3333 | 5 | 0 | 6 | 0 | 22 | 0.0909 | NA | NA | 6 | 0 | 5 | 0 | 3 | 0.3333 | 14 | 0.0714 |
| 6_1423509 | NA | NA | 2 | 0.5 | 1 | 0 | NA | NA | 3 | 0.3333 | NA | NA | 1 | 0 | 1 | 1 | NA | NA | 2 | 0.5 |
| 6_1423510 | NA | NA | 2 | 1 | 2 | 0.5 | NA | NA | 4 | 0.75 | NA | NA | 1 | 0 | 1 | 1 | NA | NA | 2 | 0.5 |
| 6_1423645 | NA | NA | NA | NA | 1 | 1 | NA | NA | 1 | 1 | NA | NA | NA | NA | NA | NA | 1 | 1 | 1 | 1 |
| 6_1424361 | 148 | 0.1014 | 142 | 0.1056 | 127 | 0.1417 | 139 | 0.1727 | 556 | 0.1295 | 99 | 0.1414 | 138 | 0.1304 | 139 | 0.1511 | 113 | 0.0973 | 489 | 0.1309 |
| 6_1424361 | 137 | 0.1314 | 132 | 0.1061 | 118 | 0.178 | 126 | 0.119 | 513 | 0.1326 | 93 | 0.1183 | 132 | 0.1364 | 131 | 0.1603 | 106 | 0.1887 | 462 | 0.1515 |
| 6_1425112 | 1 | 1 | 1 | 1 | 2 | 1 | NA | NA | 4 | 1 | 1 | 1 | NA | NA | NA | NA | NA | NA | 1 | 1 |
| 6_1425113 | 1 | 0 | 1 | 1 | 3 | 0.6667 | NA | NA | 5 | 0.6 | 1 | 1 | NA | NA | 1 | 1 | NA | NA | 2 | 1 |
| 6_1425114 | 1 | 1 | 1 | 1 | 1 | 1 | NA | NA | 3 | 1 | 1 | 1 | NA | NA | 1 | 1 | NA | NA | 2 | 1 |
| 6_1425115 | NA | NA | NA | NA | 3 | 1 | NA | NA | 3 | 1 | 1 | 1 | NA | NA | 1 | 1 | NA | NA | 2 | 1 |
| 6_1425116 | 1 | 0 | 1 | 1 | 3 | 1 | NA | NA | 5 | 0.8 | 1 | 0 | NA | NA | 1 | 1 | NA | NA | 2 | 0.5 |
| 6_1425116 | 1 | 0 | 1 | 1 | 2 | 0.5 | NA | NA | 4 | 0.5 | 1 | 1 | NA | NA | 1 | 0 | NA | NA | 2 | 0.5 |
| 6_1427178 | 11 | 0 | 14 | 0.2143 | 11 | 0.0909 | 12 | 0.0833 | 48 | 0.1042 | 9 | 0.2222 | 9 | 0.1111 | 12 | 0.0833 | 16 | 0.0625 | 46 | 0.1087 |
| 6_1427693 | NA | NA | 5 | 0 | 2 | 0 | 2 | 0.5 | 9 | 0.1111 | 1 | 0 | 1 | 1 | 1 | 0 | 1 | 0 | 4 | 0.25 |
| 6_1427724 | NA | NA | 2 | 0.5 | NA | NA | NA | NA | 2 | 0.5 | NA | NA | NA | NA | 1 | 1 | NA | NA | 1 | 1 |
| 6_1427727 | NA | NA | 2 | 0 | 1 | 1 | 1 | 0 | 4 | 0.25 | 2 | 0.5 | NA | NA | 1 | 0 | NA | NA | 3 | 0.3333 |
| 6_1427740 | NA | NA | 2 | 0.5 | 1 | 0 | 1 | 1 | 4 | 0.5 | 1 | 0 | NA | NA | 1 | 1 | NA | NA | 2 | 0.5 |
| 6_1427740 | NA | NA | 2 | 0.5 | 1 | 0 | 1 | 1 | 4 | 0.5 | 2 | 0.5 | NA | NA | 1 | 0 | NA | NA | 3 | 0.3333 |
| 6_1427740 | NA | NA | 2 | 1 | 1 | 1 | 1 | 0 | 4 | 0.75 | 2 | 1 | NA | NA | 1 | 1 | NA | NA | 3 | 1 |
| 6_1427741 | NA | NA | 1 | 0 | 1 | 1 | 1 | 1 | 3 | 0.6667 | 2 | 1 | NA | NA | 1 | 1 | NA | NA | 3 | 1 |
| 6_1427742 | NA | NA | 1 | 1 | 1 | 1 | NA | NA | 2 | 1 | NA | NA | NA | NA | 1 | 1 | NA | NA | 1 | 1 |
| 6_1427743 | NA | NA | 1 | 1 | 1 | 1 | 1 | 1 | 3 | 1 | 1 | 1 | NA | NA | 1 | 1 | NA | NA | 2 | 1 |
| 6_1427744 | NA | NA | NA | NA | 1 | 1 | NA | NA | 1 | 1 | 1 | 1 | NA | NA | 1 | 0 | NA | NA | 2 | 0.5 |
| 6_1427744 | NA | NA | 1 | 1 | 1 | 0 | 1 | 1 | 3 | 0.6667 | 2 | 1 | NA | NA | NA | NA | NA | NA | 2 | 1 |
| 6_1427745 | NA | NA | 1 | 1 | 1 | 1 | 1 | 1 | 3 | 1 | 2 | 1 | NA | NA | 1 | 1 | NA | NA | 3 | 1 |
| 6_1427747 | NA | NA | 1 | 1 | 1 | 0 | 1 | 1 | 3 | 0.6667 | 2 | 1 | NA | NA | NA | NA | NA | NA | 2 | 1 |
| 6_1429570 | 5 | 0.6 | 7 | 0 | 3 | 0.6667 | 1 | 0 | 16 | 0.3125 | 2 | 0.5 | 8 | 0.25 | 5 | 0 | 3 | 0 | 18 | 0.1667 |
| 6_1429571 | 4 | 0.25 | 5 | 0.2 | 3 | 0 | 2 | 0 | 14 | 0.1429 | 2 | 0 | 4 | 0.25 | 5 | 0.2 | 3 | 0 | 14 | 0.1429 |
| 6_1429572 | 4 | 0.25 | 6 | 0 | 3 | 0 | 3 | 0 | 16 | 0.0625 | 2 | 0 | 7 | 0 | 4 | 0.25 | 3 | 0.3333 | 16 | 0.125 |
| 6_1431265 | 3 | 0 | 7 | 0 | 6 | 0.3333 | 4 | 0.5 | 20 | 0.2 | 2 | 0 | 4 | 0.25 | 6 | 0.1667 | 4 | 0 | 16 | 0.125 |
| 6_1431510 | 11 | 0.6364 | 15 | 0.5333 | 12 | 0.5 | 16 | 0.625 | 54 | 0.5741 | 7 | 0.8571 | 12 | 0.6667 | 17 | 0.6471 | 13 | 0.6923 | 49 | 0.6939 |
| 6_1431512 | 12 | 0.4167 | 18 | 0.2778 | 13 | 0.2308 | 20 | 0.4 | 63 | 0.3333 | 8 | 0.375 | 14 | 0.3571 | 18 | 0.4444 | 17 | 0.3529 | 57 | 0.386 |
| 6_1431513 | 13 | 0.2308 | 16 | 0.375 | 14 | 0.5 | 20 | 0.2 | 63 | 0.3175 | 9 | 0.3333 | 15 | 0.1333 | 17 | 0.5294 | 18 | 0.3333 | 59 | 0.339 |
| 6_1431531 | 17 | 0.2353 | 27 | 0.0741 | 20 | 0.15 | 28 | 0.2143 | 92 | 0.163 | 22 | 0.1364 | 22 | 0.1364 | 23 | 0.1739 | 24 | 0.125 | 91 | 0.1429 |
| 6_1437745 | 2 | 0 | 5 | 0 | 6 | 0.1667 | 1 | 0 | 14 | 0.0714 | 3 | 0 | 2 | 0 | 1 | 0 | 1 | 1 | 7 | 0.1429 |
| 6_1437758 | 1 | 0 | 2 | 0 | 4 | 0 | 1 | 1 | 8 | 0.125 | 1 | 0 | 2 | 0 | 1 | 0 | 1 | 1 | 5 | 0.2 |
| 6_1449450 | 9 | 0.3333 | 19 | 0.4211 | 12 | 0.5 | 8 | 0.375 | 48 | 0.4167 | 10 | 0.5 | 6 | 0.5 | 11 | 0.3636 | 12 | 0.25 | 39 | 0.3846 |
| 6_1451604 | NA | NA | 3 | 0.3333 | NA | NA | NA | NA | 3 | 0.3333 | NA | NA | 1 | 1 | NA | NA | NA | NA | 1 | 1 |
| 6_1451607 | NA | NA | 2 | 1 | NA | NA | NA | NA | 2 | 1 | NA | NA | 2 | 0.5 | 1 | 0 | NA | NA | 3 | 0.3333 |
| 6_1451610 | 3 | 0.3333 | 3 | 0.3333 | 2 | 1 | 2 | 0.5 | 10 | 0.5 | NA | NA | 2 | 0.5 | 3 | 0 | 2 | 0.5 | 7 | 0.2857 |
| 6_1451611 | 3 | 0.3333 | 4 | 0 | 2 | 0.5 | 2 | 0 | 11 | 0.1818 | NA | NA | 2 | 1 | 4 | 0 | 1 | 0 | 7 | 0.2857 |
| 6_1451619 | 3 | 0.3333 | 4 | 0 | 3 | 0 | 2 | 0.5 | 12 | 0.1667 | NA | NA | 3 | 0 | 4 | 0 | 3 | 0.3333 | 10 | 0.1 |
| 6_1451644 | 3 | 0 | 4 | 0 | 10 | 0.4 | 4 | 0 | 21 | 0.1905 | 2 | 0 | NA | NA | 3 | 0 | 5 | 0.2 | 10 | 0.1 |
| 6_1451649 | 5 | 0 | 11 | 0.2727 | 9 | 0.3333 | 9 | 0.2222 | 34 | 0.2353 | 4 | 0 | 5 | 0.6 | 7 | 0.1429 | 9 | 0.1111 | 25 | 0.2 |
| 6_1451649 | 5 | 0 | 11 | 0.0909 | 7 | 0.2857 | 4 | 0.75 | 27 | 0.2222 | 2 | 0.5 | 4 | 0.25 | 3 | 0 | 5 | 0.2 | 14 | 0.2143 |

| | | | | | | | | | | | | | | | | | | | | |
|---|---|---|---|---|---|---|---|---|---|---|---|---|---|---|---|---|---|---|---|---|
| 6_14516583 | 11 | 0.1818 | 13 | 0.1538 | 11 | 0.0909 | 10 | 0.1 | 45 | 0.1333 | 6 | 0.3333 | 6 | 0 | 8 | 0.25 | 8 | 0.125 | 28 | 0.1786 |
| 6_14516586 | 12 | 0.0833 | 13 | 0.1538 | 12 | 0.0833 | 10 | 0 | 47 | 0.0851 | 6 | 0.5 | 7 | 0 | 7 | 0.2857 | 8 | 0 | 28 | 0.1786 |
| 6_14516590 | 12 | 0.3333 | 13 | 0.1538 | 12 | 0.1667 | 11 | 0 | 48 | 0.1667 | 6 | 0.8333 | 7 | 0.2857 | 8 | 0.25 | 7 | 0.2857 | 28 | 0.3929 |
| 6_14516592 | 8 | 0.125 | 8 | 0.625 | 11 | 0.6364 | 7 | 0.5714 | 34 | 0.5 | 1 | 1 | 6 | 0.5 | 3 | 0.3333 | 5 | 0.6 | 15 | 0.5333 |
| 6_14516602 | 5 | 0.2 | 7 | 0.2857 | 4 | 0.25 | 8 | 0.25 | 24 | 0.25 | 4 | 0 | 3 | 0 | 3 | 0.6667 | 6 | 0.1667 | 16 | 0.1875 |
| 6_14516608 | 11 | 0.2727 | 13 | 0.1538 | 12 | 0.25 | 11 | 0.1818 | 47 | 0.2128 | 6 | 0.1667 | 7 | 0.1429 | 6 | 0.1667 | 7 | 0.1429 | 26 | 0.1538 |
| 6_14516610 | 9 | 0.7778 | 11 | 0.7273 | 10 | 0.7 | 8 | 0.75 | 38 | 0.7368 | 5 | 0.6 | 6 | 0.8333 | 6 | 0.3333 | 6 | 1 | 23 | 0.6957 |
| 6_14516612 | 11 | 1 | 10 | 0.8 | 11 | 1 | 11 | 0.8182 | 43 | 0.907 | 6 | 1 | 6 | 0.8333 | 7 | 0.7143 | 6 | 0.8333 | 25 | 0.84 |
| 6_14516617 | 9 | 0.2222 | 11 | 0.2727 | 12 | 0.3333 | 11 | 0.3636 | 43 | 0.3023 | 5 | 0.6 | 6 | 0 | 7 | 0.5714 | 6 | 0.5 | 24 | 0.4167 |
| 6_14516621 | 10 | 0.1 | 9 | 0 | 10 | 0.2 | 10 | 0.2 | 39 | 0.1282 | 6 | 0.1667 | 6 | 0.1667 | 7 | 0.1429 | 5 | 0.2 | 24 | 0.1667 |
| 6_14516635 | 9 | 0 | 11 | 0.0909 | 12 | 0.0833 | 10 | 0.1 | 42 | 0.0714 | 4 | 0 | 7 | 0.1429 | 7 | 0.1429 | 7 | 0.2857 | 25 | 0.16 |
| 6_14519718 | 1 | 0 | NA | NA | NA | NA | 2 | 1 | 3 | 0.6667 | 1 | 1 | NA | NA | NA | NA | 2 | 0.5 | 3 | 0.6667 |
| 6_14519719 | 1 | 0 | NA | NA | NA | NA | 1 | 1 | 2 | 0.5 | 1 | 0 | NA | NA | NA | NA | 2 | 1 | 3 | 0.6667 |
| 6_14519735 | 1 | 1 | NA | NA | NA | NA | 1 | 1 | 2 | 1 | NA | NA | 1 | 1 | NA | NA | 2 | 0 | 3 | 0.3333 |
| 6_14572375 | NA | NA | NA | NA | NA | NA | 1 | 1 | 1 | 1 | NA | NA | NA | NA | 1 | 1 | NA | NA | 1 | 1 |
| 6_14572388 | NA | NA | NA | NA | NA | NA | 1 | 1 | 1 | 1 | NA | NA | NA | NA | 1 | 1 | NA | NA | 1 | 1 |
| 6_14572458 | NA | NA | NA | NA | NA | NA | 1 | 1 | 1 | 1 | NA | NA | NA | NA | 1 | 1 | NA | NA | 1 | 1 |
| 6_14572492 | NA | NA | NA | NA | NA | NA | 1 | 1 | 1 | 1 | NA | NA | NA | NA | 1 | 1 | NA | NA | 1 | 1 |
| 6_14572512 | NA | NA | NA | NA | NA | NA | 1 | 1 | 1 | 1 | NA | NA | NA | NA | 1 | 1 | NA | NA | 1 | 1 |
| 6_14572533 | NA | NA | NA | NA | NA | NA | 1 | 1 | 1 | 1 | NA | NA | NA | NA | 1 | 1 | NA | NA | 1 | 1 |
| 6_14576097 | NA | NA | 1 | 1 | NA | NA | NA | NA | 1 | 1 | NA | NA | NA | NA | 1 | 1 | 1 | 1 | 2 | 1 |
| 6_14576109 | NA | NA | 1 | 1 | NA | NA | NA | NA | 1 | 1 | NA | NA | NA | NA | 1 | 0 | 1 | 1 | 2 | 0.5 |
| 6_14576112 | NA | NA | 1 | 1 | NA | NA | NA | NA | 1 | 1 | NA | NA | NA | NA | 1 | 1 | 1 | 1 | 2 | 1 |
| 6_14576143 | NA | NA | 1 | 1 | NA | NA | NA | NA | 1 | 1 | NA | NA | NA | NA | 1 | 1 | 1 | 1 | 2 | 1 |
| 6_14576295 | NA | NA | NA | NA | 1 | 1 | NA | NA | 1 | 1 | NA | NA | NA | NA | NA | NA | 1 | 1 | 1 | 1 |
| 6_14576321 | NA | NA | 1 | 0 | 1 | 1 | NA | NA | 2 | 0.5 | NA | NA | NA | NA | 1 | 1 | 1 | 0 | 2 | 0.5 |
| 6_14576332 | NA | NA | 1 | 0 | 1 | 1 | NA | NA | 2 | 0.5 | NA | NA | NA | NA | 1 | 1 | 1 | 1 | 2 | 1 |
| 6_14576494 | NA | NA | 2 | 0.5 | 1 | 0 | 2 | 1 | 5 | 0.6 | 1 | 0 | NA | NA | 1 | 1 | 3 | 0.3333 | 5 | 0.4 |
| 6_14576582 | 1 | 1 | 4 | 0.5 | 3 | 0 | 4 | 0.5 | 12 | 0.4167 | 2 | 0 | NA | NA | 3 | 0.3333 | 3 | 0.3333 | 8 | 0.25 |
| 6_14576596 | 1 | 1 | 4 | 0.25 | 3 | 0.6667 | 4 | 0.5 | 12 | 0.5 | 2 | 0.5 | NA | NA | 3 | 0.6667 | 3 | 0.3333 | 8 | 0.5 |
| 6_14576604 | 1 | 1 | 3 | 1 | 3 | 0.6667 | 3 | 0.6667 | 10 | 0.8 | 2 | 0.5 | NA | NA | 3 | 1 | 2 | 0.5 | 7 | 0.7143 |
| 6_14576643 | 1 | 1 | 3 | 0 | 3 | 0 | 4 | 0.5 | 11 | 0.2727 | 2 | 0.5 | NA | NA | 3 | 0.6667 | 2 | 0 | 7 | 0.4286 |
| 6_14580720 | 10 | 0.2 | 7 | 0.5714 | 8 | 0.25 | 7 | 0.2857 | 32 | 0.3125 | 8 | 0.5 | 8 | 0.25 | 5 | 0 | 5 | 0.2 | 26 | 0.2692 |
| 6_14580773 | 7 | 0 | 2 | 0.5 | 4 | 0.75 | 3 | 0.3333 | 16 | 0.3125 | 5 | 0 | 7 | 0.1429 | 5 | 0.2 | 6 | 0.1667 | 23 | 0.1304 |
| 6_14580827 | 6 | 0.5 | 4 | 0.25 | 2 | 0.5 | 1 | 0 | 13 | 0.3846 | 3 | 0 | 4 | 0.25 | 5 | 0 | 3 | 0 | 15 | 0.0667 |
| 6_14580845 | 6 | 0.3333 | 2 | 0.5 | 4 | 0.5 | 3 | 0.3333 | 15 | 0.4 | 3 | 0.3333 | 3 | 0.6667 | 5 | 0.2 | 4 | 0 | 15 | 0.2667 |
| 6_14580857 | 5 | 0 | 4 | 0 | 4 | 0.25 | 3 | 0 | 16 | 0.0625 | 2 | 0.5 | 4 | 0 | 5 | 0.2 | 4 | 0 | 15 | 0.1333 |
| 6_14580863 | 6 | 0.3333 | 4 | 0 | 2 | 0 | 4 | 0.25 | 16 | 0.1875 | 4 | 0.25 | 5 | 0.2 | 4 | 0 | 4 | 0.75 | 17 | 0.2941 |
| 6_14580886 | 4 | 1 | 3 | 1 | 2 | 1 | 3 | 0.6667 | 12 | 0.9167 | 4 | 1 | 3 | 1 | 3 | 1 | 4 | 1 | 14 | 1 |
| 6_14580888 | 4 | 0.25 | 3 | 0 | 2 | 0 | 3 | 0 | 12 | 0.0833 | 4 | 0.25 | 4 | 0 | 4 | 0 | 4 | 0 | 16 | 0.0625 |
| 6_14580907 | 4 | 1 | 3 | 1 | 2 | 1 | 2 | 1 | 11 | 1 | 3 | 1 | 3 | 1 | 1 | 1 | 3 | 1 | 10 | 1 |
| 6_14580914 | 3 | 0.6667 | 3 | 0.6667 | 2 | 0.5 | 3 | 0.3333 | 11 | 0.5455 | 3 | 0 | 2 | 0.5 | 3 | 1 | 3 | 0.6667 | 11 | 0.5455 |
| 6_14580917 | 4 | 1 | 2 | 1 | 2 | 1 | 2 | 1 | 10 | 1 | 3 | 1 | 3 | 1 | 2 | 1 | 3 | 1 | 11 | 1 |
| 6_14580935 | 3 | 0.3333 | 2 | 0.5 | 1 | 0 | 2 | 0 | 8 | 0.25 | 3 | 0 | 2 | 0 | 3 | 0.3333 | 3 | 0.6667 | 11 | 0.2727 |
| 6_14581078 | NA | NA | 3 | 0 | 1 | 1 | 1 | 0 | 5 | 0.2 | 1 | 0 | 1 | 0 | 2 | 0 | 2 | 1 | 6 | 0.3333 |
| 6_14600985 | NA | NA | 4 | 0.25 | 1 | 0 | NA | NA | 5 | 0.2 | 2 | 0.5 | NA | NA | 2 | 0.5 | 1 | 0 | 5 | 0.4 |
| 6_14600995 | NA | NA | 4 | 0.5 | NA | NA | NA | NA | 4 | 0.5 | 2 | 0.5 | NA | NA | 3 | 0.3333 | 1 | 0 | 6 | 0.3333 |
| 6_14600998 | NA | NA | 3 | 0.3333 | 1 | 0 | NA | NA | 4 | 0.25 | 2 | 0 | NA | NA | 3 | 0.3333 | 1 | 0 | 6 | 0.1667 |
| 6_14601004 | NA | NA | 4 | 0.5 | 1 | 0 | NA | NA | 5 | 0.4 | 1 | 1 | NA | NA | 3 | 0.3333 | 1 | 1 | 5 | 0.6 |
| 6_14601008 | NA | NA | 3 | 0.6667 | 1 | 0 | NA | NA | 4 | 0.5 | 2 | 0.5 | NA | NA | 1 | 0 | 1 | 0 | 4 | 0.25 |
| 6_14601015 | NA | NA | 4 | 0.25 | 1 | 0 | NA | NA | 5 | 0.2 | 2 | 0 | NA | NA | 2 | 0.5 | 1 | 0 | 5 | 0.2 |
| 6_14601025 | NA | NA | 3 | 0.6667 | 1 | 0 | NA | NA | 4 | 0.5 | NA | NA | NA | NA | 2 | 0.5 | 1 | 1 | 3 | 0.6667 |
| 6_14601028 | NA | NA | 4 | 0.25 | 1 | 0 | NA | NA | 5 | 0.2 | 2 | 0.5 | NA | NA | 2 | 0 | 1 | 0 | 5 | 0.2 |
| 6_14601033 | NA | NA | 4 | 0.5 | 1 | 0 | NA | NA | 5 | 0.4 | 2 | 0.5 | NA | NA | 1 | 1 | 1 | 1 | 4 | 0.75 |
| 6_14601041 | NA | NA | 4 | 0.25 | 1 | 0 | NA | NA | 5 | 0.2 | 2 | 0 | NA | NA | 1 | 1 | 1 | 0 | 4 | 0.25 |
| 6_14601056 | NA | NA | 4 | 0.25 | 1 | 0 | NA | NA | 5 | 0.2 | 2 | 1 | NA | NA | 2 | 0.5 | 1 | 1 | 5 | 0.8 |
| 6_14601062 | NA | NA | 4 | 0.25 | 1 | 0 | NA | NA | 5 | 0.2 | 2 | 0.5 | NA | NA | 2 | 0 | 1 | 0 | 5 | 0.2 |
| 6_14601079 | NA | NA | 4 | 0.25 | 1 | 0 | NA | NA | 5 | 0.2 | 2 | 0.5 | NA | NA | NA | NA | 1 | 0 | 3 | 0.3333 |
| 6_14601243 | NA | NA | 3 | 0.3333 | NA | NA | NA | NA | 3 | 0.3333 | 1 | 1 | NA | NA | 1 | 0 | NA | NA | 2 | 0.5 |
| 6_14601248 | NA | NA | 3 | 1 | NA | NA | NA | NA | 3 | 1 | 1 | 1 | NA | NA | 1 | 0 | NA | NA | 2 | 0.5 |
| 6_14601251 | NA | NA | 3 | 1 | NA | NA | NA | NA | 3 | 1 | 1 | 1 | NA | NA | 1 | 1 | NA | NA | 2 | 1 |
| 6_14601263 | NA | NA | 1 | 1 | NA | NA | NA | NA | 1 | 1 | 1 | 1 | NA | NA | NA | NA | NA | NA | 1 | 1 |
| 6_14601271 | 2 | 0 | 2 | 0.5 | NA | NA | NA | NA | 4 | 0.25 | 1 | 1 | NA | NA | 1 | 0 | 1 | 1 | 3 | 0.6667 |
| 6_14601280 | 1 | 0 | 3 | 0.6667 | NA | NA | NA | NA | 4 | 0.5 | NA | NA | NA | NA | 1 | 1 | 1 | 1 | 2 | 1 |
| 6_14601292 | 2 | 1 | 1 | 1 | NA | NA | NA | NA | 3 | 1 | 1 | 1 | NA | NA | 1 | 1 | NA | NA | 2 | 1 |

| | | | | | | | | | | | | | | | | | | | | |
|---|---|---|---|---|---|---|---|---|---|---|---|---|---|---|---|---|---|---|---|---|
| 6_1460130 | 2 | 0.5 | 3 | 0.6667 | NA | NA | NA | NA | 5 | 0.6 | 1 | 0 | NA | NA | 1 | 0 | 1 | 1 | 3 | 0.3333 |
| 6_1464513 | 5 | 0.2 | 3 | 0.3333 | 5 | 0.6 | 1 | 1 | 14 | 0.4286 | 3 | 0.6667 | 3 | 1 | 6 | 0.5 | 4 | 0.75 | 16 | 0.6875 |
| 6_1464988 | 12 | 0.75 | 15 | 0.7333 | 9 | 0.6667 | 10 | 0.8 | 46 | 0.7391 | 4 | 0.5 | 17 | 0.7647 | 12 | 0.6667 | 12 | 0.75 | 45 | 0.7111 |
| 6_1464989 | 13 | 0.2308 | 16 | 0.0625 | 11 | 0.2727 | 9 | 0.3333 | 49 | 0.2041 | 3 | 0 | 20 | 0.05 | 10 | 0.1 | 12 | 0.0833 | 45 | 0.0667 |
| 6_1464989 | 11 | 0.3636 | 11 | 0.1818 | 8 | 0.25 | 8 | 0 | 38 | 0.2105 | 3 | 0 | 18 | 0.1111 | 10 | 0.2 | 13 | 0 | 44 | 0.0909 |
| 6_1464990 | 12 | 0.8333 | 16 | 1 | 11 | 1 | 9 | 0.6667 | 48 | 0.8958 | 4 | 1 | 21 | 0.9048 | 12 | 1 | 14 | 0.8571 | 51 | 0.9216 |
| 6_1464990 | 11 | 0.3636 | 15 | 0.4 | 9 | 0.1111 | 10 | 0.2 | 45 | 0.2889 | 4 | 0.25 | 19 | 0.4211 | 11 | 0.3636 | 13 | 0.1538 | 47 | 0.3191 |
| 6_1465001 | 16 | 0.125 | 22 | 0.0909 | 13 | 0.2308 | 12 | 0.25 | 63 | 0.1587 | 8 | 0.375 | 20 | 0.25 | 16 | 0.0625 | 16 | 0.1875 | 60 | 0.2 |
| 6_1465049 | 7 | 0 | 3 | 0.3333 | 2 | 0 | NA | NA | 12 | 0.0833 | 2 | 0 | 4 | 0 | 1 | 0 | 3 | 0.3333 | 10 | 0.1 |
| 6_1465053 | 8 | 0.25 | 6 | 0 | 3 | 0 | NA | NA | 17 | 0.1176 | 6 | 0 | 5 | 0.4 | 3 | 0 | 7 | 0 | 21 | 0.0952 |
| 6_1465518 | 3 | 0.6667 | 3 | 0.6667 | 6 | 0.8333 | 5 | 0.8 | 17 | 0.7647 | 1 | 1 | 7 | 0.8571 | 3 | 1 | 2 | 1 | 13 | 0.9231 |
| 6_1465518 | 4 | 0.5 | 7 | 0.2857 | 8 | 0.25 | 8 | 0.125 | 27 | 0.2593 | 2 | 0.5 | 6 | 0.3333 | 2 | 0 | 3 | 0.3333 | 13 | 0.3077 |
| 6_1465656 | NA | NA | 1 | 1 | NA | NA | NA | NA | 1 | 1 | 1 | 1 | 1 | 1 | NA | NA | NA | NA | 2 | 1 |
| 6_1465657 | NA | NA | 1 | 1 | NA | NA | NA | NA | 1 | 1 | 1 | 1 | 1 | 1 | NA | NA | NA | NA | 2 | 1 |
| 6_1465657 | NA | NA | 1 | 1 | NA | NA | NA | NA | 1 | 1 | 1 | 1 | 1 | 1 | NA | NA | NA | NA | 2 | 1 |
| 6_1465658 | NA | NA | 1 | 1 | NA | NA | NA | NA | 1 | 1 | 1 | 1 | 1 | 1 | NA | NA | NA | NA | 2 | 1 |
| 6_1465659 | NA | NA | 1 | 1 | NA | NA | NA | NA | 1 | 1 | 1 | 1 | 1 | 1 | NA | NA | NA | NA | 2 | 1 |
| 6_1466771 | 2 | 0.5 | 1 | 1 | 2 | 0.5 | 1 | 0 | 6 | 0.5 | 1 | 1 | NA | NA | 3 | 0.6667 | 1 | 1 | 5 | 0.8 |
| 6_1466772 | 3 | 0.3333 | 1 | 1 | 1 | 0 | 1 | 1 | 6 | 0.5 | 1 | 0 | NA | NA | 5 | 0.4 | 1 | 1 | 7 | 0.4286 |
| 6_1466773 | 3 | 0.3333 | 1 | 1 | 2 | 1 | 1 | 0 | 7 | 0.5714 | 2 | 0.5 | NA | NA | 5 | 0.8 | 1 | 0 | 8 | 0.625 |
| 6_1466774 | 3 | 0.3333 | 2 | 0.5 | 2 | 0.5 | NA | NA | 7 | 0.4286 | 2 | 0 | NA | NA | 3 | 0.3333 | NA | NA | 5 | 0.2 |
| 6_1466774 | 3 | 0 | 1 | 0 | 1 | 0 | 1 | 1 | 6 | 0.1667 | 2 | 0 | NA | NA | 3 | 0.3333 | 1 | 0 | 6 | 0.1667 |
| 6_1466774 | 2 | 0.5 | 1 | 1 | 1 | 1 | 1 | 0 | 5 | 0.6 | 2 | 0.5 | NA | NA | 5 | 0.8 | 1 | 1 | 8 | 0.75 |
| 6_1466775 | 2 | 1 | 1 | 1 | 2 | 0.5 | 1 | 0 | 6 | 0.6667 | 1 | 1 | NA | NA | 3 | 1 | 1 | 1 | 5 | 1 |
| 6_1466776 | 3 | 0.6667 | 1 | 0 | 2 | 0.5 | NA | NA | 6 | 0.5 | 3 | 0.6667 | NA | NA | 6 | 0.6667 | 1 | 1 | 10 | 0.7 |
| 6_1466776 | 3 | 1 | 1 | 1 | 2 | 1 | 2 | 1 | 8 | 1 | 3 | 1 | NA | NA | 6 | 1 | 2 | 1 | 11 | 1 |
| 6_1466777 | 2 | 0.5 | 1 | 0 | 1 | 1 | 2 | 0.5 | 6 | 0.5 | 3 | 0 | NA | NA | 6 | 0 | 2 | 1 | 11 | 0.1818 |
| 6_1466777 | 3 | 0.6667 | 1 | 1 | 2 | 0 | 2 | 0 | 8 | 0.375 | 3 | 0 | NA | NA | 6 | 0.3333 | 2 | 0.5 | 11 | 0.2727 |
| 6_1466778 | 3 | 0.6667 | 2 | 0.5 | 2 | 0.5 | 1 | 0 | 8 | 0.5 | 2 | 0.5 | NA | NA | 6 | 0.3333 | 2 | 0.5 | 10 | 0.4 |
| 6_1466779 | 2 | 1 | NA | NA | 2 | 1 | 2 | 0.5 | 6 | 0.8333 | 3 | 1 | NA | NA | 3 | 1 | 2 | 0.5 | 8 | 0.875 |
| 6_1466780 | 1 | 1 | NA | NA | NA | NA | NA | NA | 1 | 1 | 1 | 0 | NA | NA | 4 | 1 | 1 | 1 | 6 | 0.8333 |
| 6_1469320 | 5 | 0.6 | 3 | 0.3333 | 2 | 1 | 3 | 0.6667 | 13 | 0.6154 | 1 | 1 | 1 | 1 | 2 | 1 | 2 | 0.5 | 6 | 0.8333 |
| 6_1469320 | 3 | 0.3333 | 4 | 0.75 | 1 | 1 | 3 | 1 | 11 | 0.7273 | 1 | 1 | 1 | 0 | 2 | 0.5 | 3 | 1 | 7 | 0.7143 |
| 6_1469321 | 5 | 0.6 | 4 | 0 | 3 | 0 | 3 | 0.3333 | 15 | 0.2667 | 1 | 1 | NA | NA | 1 | 0 | 3 | 0.3333 | 5 | 0.4 |
| 6_1469325 | 4 | 0.25 | 5 | 0 | 3 | 0 | 2 | 0 | 14 | 0.0714 | 1 | 1 | 1 | 0 | 2 | 0.5 | 4 | 0 | 8 | 0.25 |
| 6_1469325 | 5 | 0.4 | 5 | 0.8 | 4 | 1 | 3 | 1 | 17 | 0.7647 | NA | NA | 2 | 1 | 2 | 0.5 | 4 | 0.5 | 8 | 0.625 |
| 6_1469325 | 4 | 0 | 5 | 0 | 4 | 0.25 | 3 | 0 | 16 | 0.0625 | 1 | 1 | 2 | 0 | 2 | 0.5 | 3 | 0 | 8 | 0.25 |
| 6_1469326 | 5 | 0.2 | 4 | 0 | 4 | 0 | 2 | 0 | 15 | 0.0667 | 1 | 0 | 2 | 0 | 2 | 0.5 | 4 | 0 | 9 | 0.1111 |
| 6_1469328 | 4 | 0.25 | 5 | 0.2 | 5 | 0.4 | 3 | 0.3333 | 17 | 0.2941 | 1 | 0 | 2 | 0.5 | 2 | 0.5 | 4 | 0 | 9 | 0.2222 |
| 6_1469330 | 4 | 0 | 5 | 0 | 5 | 0.2 | 2 | 0 | 16 | 0.0625 | 2 | 0 | 2 | 0.5 | 2 | 0 | 3 | 0 | 9 | 0.1111 |
| 6_1469333 | 6 | 0.1667 | 7 | 0 | 6 | 0 | 4 | 0.25 | 23 | 0.087 | 1 | 0 | 2 | 0 | 2 | 0 | 3 | 0.3333 | 8 | 0.125 |
| 6_1469334 | 5 | 0.4 | 6 | 0.1667 | 6 | 0 | 3 | 0 | 20 | 0.15 | 2 | 0.5 | 2 | 0 | 2 | 0 | 2 | 0.5 | 8 | 0.25 |
| 6_1469723 | 2 | 1 | 4 | 1 | 4 | 1 | 2 | 1 | 12 | 1 | 7 | 1 | 4 | 1 | 4 | 0.75 | 7 | 0.8571 | 22 | 0.9091 |
| 6_1469724 | NA | NA | NA | NA | NA | NA | 2 | 0.5 | 2 | 0.5 | 1 | 1 | 1 | 0 | NA | NA | 2 | 0 | 4 | 0.25 |
| 6_1469746 | 1 | 1 | NA | NA | NA | NA | 1 | 1 | 2 | 1 | 1 | 1 | NA | NA | NA | NA | NA | NA | 1 | 1 |
| 6_1469934 | 2 | 1 | 1 | 1 | NA | NA | NA | NA | 3 | 1 | NA | NA | NA | NA | NA | NA | 1 | 1 | 1 | 1 |
| 6_1469937 | 2 | 0.5 | 1 | 1 | 1 | 0 | 1 | 0 | 5 | 0.4 | 1 | 1 | NA | NA | NA | NA | 1 | 0 | 2 | 0.5 |
| 6_1470093 | 1 | 1 | 1 | 1 | 1 | 1 | 1 | 1 | 4 | 1 | 1 | 1 | NA | NA | NA | NA | 2 | 0.5 | 3 | 0.6667 |
| 6_1470093 | 1 | 1 | 1 | 1 | 1 | 0 | 1 | 1 | 4 | 0.75 | 1 | 0 | NA | NA | NA | NA | 3 | 0.3333 | 4 | 0.25 |
| 6_1470094 | 1 | 1 | 1 | 0 | 1 | 1 | 1 | 1 | 4 | 0.75 | 1 | 1 | NA | NA | NA | NA | 3 | 1 | 4 | 1 |
| 6_1470095 | 2 | 1 | 1 | 1 | 1 | 1 | 3 | 1 | 7 | 1 | 2 | 1 | NA | NA | NA | NA | 3 | 1 | 5 | 1 |
| 6_1470096 | 2 | 1 | 1 | 0 | 1 | 1 | 2 | 1 | 6 | 0.8333 | 1 | 1 | NA | NA | NA | NA | 3 | 0.6667 | 4 | 0.75 |
| 6_1470096 | 1 | 1 | 1 | 1 | NA | NA | 2 | 1 | 4 | 1 | 1 | 1 | NA | NA | NA | NA | 2 | 0.5 | 3 | 0.6667 |
| 6_1470097 | 2 | 1 | 1 | 1 | 1 | 1 | 2 | 0.5 | 6 | 0.8333 | 2 | 1 | NA | NA | NA | NA | 4 | 0.5 | 6 | 0.6667 |
| 6_1470097 | 1 | 1 | NA | NA | 1 | 0 | 2 | 0.5 | 4 | 0.5 | 2 | 0.5 | NA | NA | NA | NA | 4 | 0.75 | 6 | 0.6667 |
| 6_1470097 | 2 | 0.5 | NA | NA | NA | NA | 1 | 1 | 3 | 0.6667 | 2 | 1 | NA | NA | NA | NA | 3 | 0.3333 | 5 | 0.6 |
| 6_1470099 | 2 | 0 | NA | NA | 1 | 1 | 1 | 0 | 4 | 0.25 | 1 | 1 | NA | NA | NA | NA | 4 | 0.5 | 5 | 0.6 |
| 6_1470100 | 1 | 0 | NA | NA | 1 | 1 | 1 | 0 | 3 | 0.3333 | 2 | 0 | NA | NA | NA | NA | 4 | 0.25 | 6 | 0.1667 |
| 6_1470101 | NA | NA | NA | NA | 1 | 1 | 1 | 1 | 2 | 1 | 2 | 0.5 | NA | NA | NA | NA | 4 | 0.75 | 6 | 0.6667 |
| 6_1470109 | 1 | 1 | NA | NA | 1 | 1 | 1 | 1 | 3 | 1 | 2 | 0.5 | NA | NA | NA | NA | 5 | 0.6 | 7 | 0.5714 |
| 6_1470109 | 1 | 1 | NA | NA | 1 | 0 | 2 | 0.5 | 4 | 0.5 | 2 | 0 | NA | NA | NA | NA | 5 | 0.2 | 7 | 0.1429 |
| 6_1470111 | 1 | 1 | NA | NA | 1 | 0 | NA | NA | 2 | 0.5 | 1 | 0 | NA | NA | NA | NA | 5 | 0.2 | 6 | 0.1667 |
| 6_1470111 | 1 | 1 | NA | NA | 1 | 1 | 1 | 1 | 3 | 1 | 2 | 0.5 | NA | NA | NA | NA | 5 | 0 | 7 | 0.1429 |
| 6_1470119 | 1 | 0 | NA | NA | 1 | 0 | 1 | 1 | 3 | 0.3333 | 1 | 0 | NA | NA | NA | NA | 4 | 0.25 | 5 | 0.2 |
| 6_1470120 | 2 | 0 | NA | NA | 1 | 0 | 1 | 1 | 4 | 0.25 | 2 | 0.5 | NA | NA | NA | NA | 3 | 0 | 5 | 0.2 |

| | | | | | | | | | | | | | | | | | | | | |
|---|---|---|---|---|---|---|---|---|---|---|---|---|---|---|---|---|---|---|---|---|
| 6_14701364 | 4 | 0 | 2 | 0.5 | 2 | 0 | 1 | 0 | 9 | 0.1111 | 3 | 0 | NA | NA | NA | NA | 5 | 0.2 | 8 | 0.125 |
| 6_14701366 | 4 | 0.5 | 2 | 0.5 | 3 | 0 | 4 | 0.25 | 13 | 0.3077 | 3 | 0 | NA | NA | NA | NA | 6 | 0.1667 | 9 | 0.1111 |
| 6_14701372 | 5 | 0.2 | 2 | 0.5 | 2 | 0 | 3 | 0.6667 | 12 | 0.3333 | 3 | 0 | NA | NA | NA | NA | 5 | 0.4 | 8 | 0.25 |
| 6_14701381 | 4 | 0.5 | 1 | 0 | 2 | 0 | 2 | 0.5 | 9 | 0.3333 | 3 | 0 | NA | NA | NA | NA | 6 | 0.6667 | 9 | 0.4444 |
| 6_14753317 | 3 | 0 | 5 | 0.2 | 3 | 0 | 6 | 0 | 17 | 0.0588 | 2 | 0.5 | 5 | 0.2 | 1 | 0 | 4 | 0 | 12 | 0.1667 |
| 6_14753318 | 2 | 0 | 5 | 0 | 3 | 0 | 6 | 0.1667 | 16 | 0.0625 | 2 | 0.5 | 4 | 0.5 | 2 | 0.5 | 4 | 0.25 | 12 | 0.4167 |
| 6_14753334 | 2 | 1 | 5 | 0.8 | 3 | 1 | 6 | 0.5 | 16 | 0.75 | 2 | 1 | 6 | 0.8333 | 2 | 1 | 4 | 0.75 | 14 | 0.8571 |
| 6_14753336 | 3 | 0.3333 | 5 | 0.2 | 3 | 0.3333 | 6 | 0.5 | 17 | 0.3529 | 2 | 1 | 5 | 0.6 | 2 | 0.5 | 4 | 0.5 | 13 | 0.6154 |
| 6_14753381 | 3 | 0 | 6 | 0 | 3 | 0.3333 | 3 | 0 | 15 | 0.0667 | 2 | 0 | 4 | 0.25 | 4 | 0 | 4 | 0 | 14 | 0.0714 |
| 6_14753401 | 3 | 0 | 7 | 0.1429 | 4 | 0.25 | 7 | 0.1429 | 21 | 0.1429 | 3 | 0 | 6 | 0.1667 | 4 | 0.25 | 5 | 0 | 18 | 0.1111 |
| 6_14808222 | 4 | 0 | 3 | 0.6667 | 5 | 0 | 4 | 0 | 16 | 0.125 | 3 | 0.3333 | 2 | 0 | 2 | 0 | NA | NA | 7 | 0.1429 |
| 6_14808234 | 4 | 0 | 3 | 0 | 3 | 0 | 3 | 0.3333 | 13 | 0.0769 | 2 | 0.5 | 1 | 1 | 3 | 0.3333 | NA | NA | 6 | 0.5 |
| 6_14808248 | 4 | 0.25 | 3 | 0.3333 | 5 | 0 | 4 | 0.25 | 16 | 0.1875 | 3 | 0.3333 | 1 | 0 | 3 | 0 | NA | NA | 7 | 0.1429 |
| 6_14808259 | 4 | 0.5 | 3 | 0 | 4 | 0 | 4 | 0.25 | 15 | 0.2 | 3 | 0 | NA | NA | 2 | 0.5 | NA | NA | 5 | 0.2 |
| 6_14808270 | 5 | 0.4 | 4 | 0 | 6 | 0.3333 | 4 | 0.25 | 19 | 0.2632 | 4 | 0 | 1 | 1 | 2 | 0 | 2 | 0 | 9 | 0.1111 |
| 6_14808274 | 3 | 0.6667 | 1 | 1 | 4 | 0.25 | 3 | 0.3333 | 11 | 0.4545 | 1 | 1 | NA | NA | 2 | 0 | 2 | 0.5 | 5 | 0.4 |
| 6_14808274 | 1 | 1 | 1 | 1 | 1 | 1 | NA | NA | 3 | 1 | 1 | 1 | 1 | 1 | NA | NA | NA | NA | 2 | 1 |
| 6_14808274 | NA | NA | 2 | 0.5 | NA | NA | NA | NA | 2 | 0.5 | 1 | 1 | NA | NA | NA | NA | NA | NA | 1 | 1 |
| 6_14808275 | 4 | 0.25 | 4 | 0.25 | 5 | 0.4 | 4 | 0.25 | 17 | 0.2941 | 4 | 0.5 | 1 | 1 | 3 | 0.3333 | 2 | 0 | 10 | 0.4 |
| 6_14808289 | 5 | 0.6 | 8 | 0.375 | 7 | 0.2857 | 8 | 0.375 | 28 | 0.3929 | 4 | 0.75 | 2 | 0.5 | 6 | 0.3333 | 2 | 0.5 | 14 | 0.5 |
| 6_14808302 | 2 | 0.5 | 5 | 0.4 | 6 | 0.8333 | 7 | 0.4286 | 20 | 0.55 | 2 | 0.5 | 2 | 1 | 5 | 0.6 | 2 | 1 | 11 | 0.7273 |
| 6_14808307 | 4 | 0 | 8 | 0.125 | 7 | 0.1429 | 7 | 0 | 26 | 0.0769 | 4 | 0 | 2 | 0 | 6 | 0.1667 | 2 | 0.5 | 14 | 0.1429 |
| 6_14808314 | 5 | 0.4 | 8 | 0.5 | 7 | 0.1429 | 7 | 0.4286 | 27 | 0.3704 | 4 | 0 | 2 | 0.5 | 6 | 0.5 | 2 | 0.5 | 14 | 0.3571 |
| 6_14808315 | 4 | 0.5 | 6 | 0.6667 | 6 | 0.8333 | 4 | 0.75 | 20 | 0.7 | 3 | 1 | 1 | 0 | 6 | 0.6667 | 2 | 0.5 | 12 | 0.6667 |
| 6_14808318 | 5 | 0 | 8 | 0 | 7 | 0.2857 | 4 | 0 | 24 | 0.0833 | 4 | 0 | 2 | 0 | 6 | 0.1667 | 2 | 0 | 14 | 0.0714 |
| 6_14813897 | 1 | 1 | 2 | 0.5 | 1 | 0 | 2 | 0 | 6 | 0.3333 | 1 | 0 | 1 | 0 | 1 | 1 | 1 | 1 | 4 | 0.5 |
| 6_14813900 | 1 | 0 | 2 | 0 | 1 | 1 | 3 | 0.6667 | 7 | 0.4286 | 1 | 0 | 1 | 0 | 1 | 0 | 1 | 1 | 4 | 0.25 |
| 6_14813915 | 1 | 1 | 2 | 0.5 | 1 | 0 | 2 | 0.5 | 6 | 0.5 | 1 | 1 | 1 | 1 | NA | NA | 1 | 1 | 3 | 1 |
| 6_14813918 | 1 | 1 | 1 | 1 | 1 | 1 | 2 | 0 | 5 | 0.6 | 1 | 1 | NA | NA | NA | NA | 1 | 0 | 2 | 0.5 |
| 6_14813926 | 1 | 1 | 2 | 0.5 | 1 | 1 | 2 | 1 | 6 | 0.8333 | 1 | 0 | 1 | 1 | NA | NA | 1 | 0 | 3 | 0.3333 |
| 6_14813928 | NA | NA | NA | NA | 1 | 1 | 2 | 1 | 3 | 1 | 1 | 0 | NA | NA | NA | NA | 1 | 1 | 2 | 0.5 |
| 6_14825696 | 3 | 0 | 3 | 0 | 3 | 0.3333 | 4 | 0 | 13 | 0.0769 | 7 | 0 | 5 | 0 | 5 | 0.4 | 4 | 0 | 21 | 0.0952 |
| 6_14825700 | 3 | 1 | 3 | 1 | 2 | 1 | 3 | 1 | 11 | 1 | 5 | 1 | 6 | 1 | 4 | 0.75 | 3 | 1 | 18 | 0.9444 |
| 6_14825701 | 3 | 1 | 4 | 1 | 3 | 1 | 4 | 1 | 14 | 1 | 6 | 1 | 5 | 1 | 5 | 1 | 4 | 1 | 20 | 1 |
| 6_14867277 | 3 | 0 | 1 | 1 | 5 | 0.4 | 2 | 0 | 11 | 0.2727 | 6 | 0 | 2 | 0.5 | 4 | 0 | 6 | 0.1667 | 18 | 0.1111 |
| 6_14867284 | 2 | 0 | 3 | 0 | 7 | 0.2857 | 3 | 0.6667 | 15 | 0.2667 | 6 | 0 | 2 | 0.5 | 5 | 0.2 | 5 | 0.6 | 18 | 0.2778 |
| 6_14867347 | 2 | 0 | 4 | 0 | 6 | 0 | 3 | 0.3333 | 15 | 0.0667 | 4 | 0 | 2 | 0 | 4 | 0.25 | 7 | 0.1429 | 17 | 0.1176 |
| 6_14867363 | 1 | 0 | 3 | 0.3333 | 5 | 0 | 3 | 0.6667 | 12 | 0.25 | 4 | 0 | 2 | 0.5 | 4 | 0.5 | 7 | 0.4286 | 17 | 0.3529 |
| 6_14867401 | 1 | 0 | 4 | 0.25 | 5 | 0.4 | 3 | 0 | 13 | 0.2308 | 4 | 0.25 | 2 | 0.5 | 4 | 0.5 | 5 | 0.2 | 15 | 0.3333 |
| 6_14868427 | 1 | 0 | 2 | 0 | 4 | 0.25 | 3 | 0 | 10 | 0.1 | 2 | 0 | 1 | 0 | NA | NA | 4 | 0.25 | 7 | 0.1429 |
| 6_14882430 | NA | NA | 1 | 1 | NA | NA | NA | NA | 1 | 1 | 1 | 1 | NA | NA | NA | NA | NA | NA | 1 | 1 |
| 6_14889118 | 2 | 0 | 5 | 0.4 | 2 | 0 | 3 | 0 | 12 | 0.1667 | 1 | 0 | 4 | 0 | 2 | 0 | 6 | 0.3333 | 13 | 0.1538 |
| 6_14889211 | 2 | 0 | 5 | 0 | 2 | 0 | 3 | 0.3333 | 12 | 0.0833 | 1 | 1 | 3 | 0 | 3 | 0.3333 | 5 | 0 | 12 | 0.1667 |
| 6_14889306 | 1 | 0 | 3 | 0.3333 | 1 | 0 | 2 | 0 | 7 | 0.1429 | 1 | 0 | 1 | 0 | NA | NA | 4 | 0.5 | 6 | 0.3333 |
| 6_14889435 | 1 | 0 | 3 | 0 | NA | NA | 2 | 0.5 | 6 | 0.1667 | NA | NA | 2 | 0.5 | NA | NA | 2 | 0 | 4 | 0.25 |
| 6_14889502 | 1 | 0 | 2 | 0 | NA | NA | 2 | 0.5 | 5 | 0.2 | 1 | 1 | 1 | 0 | NA | NA | 2 | 0 | 4 | 0.25 |
| 6_14889507 | 1 | 0 | 2 | 0.5 | NA | NA | 1 | 1 | 4 | 0.5 | NA | NA | 1 | 1 | NA | NA | 1 | 1 | 2 | 1 |
| 6_14889513 | 1 | 0 | 3 | 0.6667 | NA | NA | 2 | 0.5 | 6 | 0.5 | 1 | 0 | 1 | 0 | NA | NA | 2 | 1 | 4 | 0.5 |
| 6_14889514 | 1 | 1 | 3 | 0.6667 | NA | NA | 2 | 0.5 | 6 | 0.6667 | 1 | 1 | 1 | 0 | NA | NA | 2 | 0.5 | 4 | 0.5 |
| 6_14889518 | 1 | 1 | 1 | 0 | NA | NA | 2 | 0 | 4 | 0.25 | 1 | 1 | 1 | 0 | NA | NA | 2 | 0 | 4 | 0.25 |
| 6_14889529 | 1 | 1 | 3 | 0.6667 | NA | NA | 2 | 1 | 6 | 0.8333 | 1 | 0 | 1 | 0 | NA | NA | 2 | 1 | 4 | 0.5 |
| 6_14894599 | NA | NA | NA | NA | NA | NA | 2 | 1 | 2 | 1 | 1 | 0 | NA | NA | NA | NA | 1 | 1 | 2 | 0.5 |
| 6_14894612 | NA | NA | NA | NA | NA | NA | 2 | 0.5 | 2 | 0.5 | 1 | 1 | NA | NA | NA | NA | 1 | 1 | 2 | 1 |
| 6_14894622 | NA | NA | NA | NA | NA | NA | 2 | 1 | 2 | 1 | 1 | 1 | NA | NA | NA | NA | 1 | 1 | 2 | 1 |
| 6_14896966 | NA | NA | NA | NA | NA | NA | 2 | 0.5 | 2 | 0.5 | 1 | 1 | NA | NA | NA | NA | 1 | 1 | 2 | 1 |
| 6_14896998 | NA | NA | NA | NA | NA | NA | 2 | 1 | 2 | 1 | 1 | 1 | NA | NA | NA | NA | 1 | 1 | 2 | 1 |
| 6_14900392 | NA | NA | 1 | 1 | NA | NA | NA | NA | 1 | 1 | NA | NA | 1 | 1 | NA | NA | NA | NA | 1 | 1 |
| 6_14923073 | 3 | 0.3333 | 1 | 1 | NA | NA | 1 | 0 | 5 | 0.4 | 1 | 1 | NA | NA | NA | NA | NA | NA | 1 | 1 |
| 6_14923083 | 1 | 0 | 1 | 1 | NA | NA | 1 | 1 | 3 | 0.6667 | 1 | 1 | NA | NA | NA | NA | NA | NA | 1 | 1 |
| 6_14923639 | 3 | 1 | 2 | 0 | NA | NA | 1 | 1 | 6 | 0.6667 | 1 | 1 | NA | NA | NA | NA | NA | NA | 1 | 1 |
| 6_14946308 | 2 | 0 | 6 | 0.3333 | 4 | 0.25 | 2 | 0 | 14 | 0.2143 | 1 | 0 | 1 | 0 | 3 | 0 | 3 | 0.3333 | 8 | 0.125 |
| 6_14946314 | 2 | 0 | 6 | 0 | 4 | 0.25 | 3 | 0 | 15 | 0.0667 | 1 | 0 | 2 | 0.5 | 4 | 0.25 | 3 | 0 | 10 | 0.2 |
| 7_3231587 | 1 | 0 | 6 | 0 | 4 | 0.5 | 4 | 0.25 | 15 | 0.2 | 2 | 0 | 4 | 0.25 | 3 | 0 | 3 | 0 | 12 | 0.0833 |
| 7_3231599 | 2 | 0.5 | 6 | 0 | 3 | 0 | 3 | 0 | 14 | 0.0714 | 2 | 0.5 | 4 | 0 | 4 | 0 | 3 | 0 | 13 | 0.0769 |
| 7_3345402 | 1 | 1 | 1 | 1 | 1 | 1 | 2 | 1 | 5 | 1 | 2 | 0.5 | NA | NA | 3 | 0.6667 | 2 | 0.5 | 7 | 0.5714 |

| | | | | | | | | | | | | | | | | | | | | |
|---|---|---|---|---|---|---|---|---|---|---|---|---|---|---|---|---|---|---|---|---|
| 7_3345418 | NA | NA | 2 | 0.5 | 3 | 0 | 4 | 0.25 | 9 | 0.2222 | 4 | 0.25 | NA | NA | 3 | 0.3333 | 2 | 0.5 | 9 | 0.3333 |
| 7_3345488 | 1 | 1 | 2 | 0.5 | 3 | 1 | 4 | 1 | 10 | 0.9 | 4 | 1 | 1 | 1 | 2 | 1 | 2 | 1 | 9 | 1 |
| 7_3345625 | 1 | 1 | 2 | 0 | 3 | 0.6667 | 4 | 0.75 | 10 | 0.6 | 4 | 0.5 | 1 | 1 | 3 | 0.6667 | 1 | 0 | 9 | 0.5556 |
| 7_3345693 | 2 | 1 | 3 | 1 | 3 | 1 | 4 | 1 | 12 | 1 | 4 | 1 | 1 | 1 | 2 | 1 | 2 | 1 | 9 | 1 |
| 7_3345762 | 2 | 0.5 | 3 | 0.3333 | 3 | 0 | 4 | 0.5 | 12 | 0.3333 | 5 | 0.4 | NA | NA | 4 | 0.5 | 1 | 0 | 10 | 0.4 |
| 7_3346105 | 3 | 0.3333 | 5 | 0.4 | 2 | 0 | 4 | 0.75 | 14 | 0.4286 | 4 | 0.75 | NA | NA | 2 | 1 | 2 | 0 | 8 | 0.625 |
| 7_3378792 | 29 | 0.1724 | 30 | 0.1333 | 14 | 0.0714 | 16 | 0.125 | 89 | 0.1348 | 29 | 0 | 17 | 0.1765 | 16 | 0 | 18 | 0.1111 | 80 | 0.0625 |
| 7_3379530 | 31 | 0.129 | 37 | 0.1351 | 23 | 0.0435 | 18 | 0.1111 | 109 | 0.1101 | 40 | 0.175 | 16 | 0.25 | 28 | 0.0714 | 29 | 0.1034 | 113 | 0.1416 |
| 7_3385192 | 7 | 0.1429 | 6 | 0.1667 | 9 | 0.1111 | 4 | 0.25 | 26 | 0.1538 | 4 | 0 | 8 | 0.125 | 6 | 0.1667 | 6 | 0.1667 | 24 | 0.125 |
| 7_3415433 | 8 | 0 | 11 | 0.0909 | 13 | 0.1538 | 9 | 0.2222 | 41 | 0.122 | 9 | 0.2222 | 12 | 0.0833 | 8 | 0 | 8 | 0.375 | 37 | 0.1622 |
| 7_3415541 | 7 | 0.5714 | 7 | 0.2857 | 12 | 0.5 | 6 | 0.3333 | 32 | 0.4375 | 5 | 0.4 | 9 | 0.1111 | 11 | 0.6364 | 7 | 0.4286 | 32 | 0.4062 |
| 7_3415975 | 10 | 0 | 17 | 0.2353 | 13 | 0.1538 | 10 | 0.1 | 50 | 0.14 | 12 | 0.4167 | 14 | 0 | 9 | 0.4444 | 11 | 0.1818 | 46 | 0.2391 |
| 7_3416394 | 6 | 0 | 16 | 0.1875 | 13 | 0.1538 | 9 | 0.1111 | 44 | 0.1364 | 8 | 0.125 | 7 | 0 | 8 | 0.5 | 10 | 0.3 | 33 | 0.2424 |
| 7_3442740 | 1 | 0 | 3 | 0.3333 | 3 | 0 | 4 | 0.5 | 11 | 0.2727 | 5 | 0.8 | NA | NA | 3 | 0.3333 | 5 | 0.8 | 13 | 0.6923 |
| 7_3442865 | 3 | 0.3333 | 3 | 0 | 2 | 0.5 | 3 | 0.3333 | 11 | 0.2727 | 6 | 0.3333 | NA | NA | 2 | 0 | 4 | 0.5 | 12 | 0.3333 |
| 7_3442929 | 2 | 0.5 | 1 | 0 | 2 | 0.5 | 4 | 0.25 | 9 | 0.3333 | 6 | 0.3333 | NA | NA | 3 | 0.3333 | 4 | 0.5 | 13 | 0.3846 |
| 7_3442970 | 3 | 0.3333 | 2 | 0.5 | 3 | 0.3333 | 4 | 0.75 | 12 | 0.5 | 7 | 0 | NA | NA | 2 | 1 | 4 | 0.75 | 13 | 0.3846 |
| 7_3443220 | 2 | 1 | NA | NA | 1 | 1 | NA | NA | 3 | 1 | NA | NA | NA | NA | 1 | 1 | 4 | 1 | 5 | 1 |
| 7_3443311 | 2 | 0.5 | NA | NA | 1 | 1 | 1 | 1 | 4 | 0.75 | NA | NA | NA | NA | NA | NA | 4 | 1 | 4 | 1 |
| 7_3463568 | 2 | 1 | 3 | 0.6667 | 3 | 0.6667 | 3 | 1 | 11 | 0.8182 | 6 | 0.8333 | NA | NA | 2 | 0.5 | 2 | 1 | 10 | 0.8 |
| 7_3463757 | 2 | 1 | 4 | 1 | 4 | 1 | 4 | 0.75 | 14 | 0.9286 | 8 | 0.75 | 1 | 1 | 2 | 0.5 | 2 | 1 | 13 | 0.7692 |
| 7_3463910 | 2 | 0.5 | 4 | 0.75 | 3 | 0.6667 | 4 | 0 | 13 | 0.4615 | 6 | 0.1667 | NA | NA | 3 | 0 | 2 | 1 | 11 | 0.2727 |
| 7_3464056 | 2 | 0 | 6 | 0.5 | 5 | 0.2 | 4 | 0 | 17 | 0.2353 | 7 | 0.4286 | 1 | 0 | 3 | 0.3333 | 3 | 0.3333 | 14 | 0.3571 |
| 7_3464239 | 2 | 0.5 | 5 | 0.6 | 5 | 1 | 5 | 0.8 | 17 | 0.7647 | 7 | 0.8571 | 2 | 1 | 2 | 1 | 3 | 0.6667 | 14 | 0.8571 |
| 7_3464323 | 2 | 0.5 | 4 | 0.75 | 4 | 1 | 5 | 0.6 | 15 | 0.7333 | 6 | 0.6667 | 2 | 1 | 2 | 0.5 | 1 | 1 | 11 | 0.7273 |
| 7_3464753 | 8 | 0.125 | 7 | 0.1429 | 8 | 0 | 5 | 0.2 | 28 | 0.1071 | 11 | 0.0909 | 5 | 0 | 13 | 0.0769 | 6 | 0.1667 | 35 | 0.0857 |
| 7_3467176 | 4 | 0 | 3 | 0.3333 | 1 | 0 | 1 | 0 | 9 | 0.1111 | 5 | 0 | 2 | 0 | 1 | 1 | NA | NA | 8 | 0.125 |
| 7_3468426 | 1 | 0 | 1 | 0 | 3 | 0.6667 | 4 | 0 | 9 | 0.2222 | 1 | 1 | NA | NA | NA | NA | NA | NA | 1 | 1 |
| 7_3470447 | 1 | 0 | 4 | 0.25 | 5 | 0.2 | 2 | 0 | 12 | 0.1667 | 1 | 1 | NA | NA | 3 | 0 | 1 | 0 | 5 | 0.2 |
| 7_3470831 | 2 | 0 | 6 | 0.3333 | 6 | 0.5 | 4 | 0 | 18 | 0.2778 | 2 | 1 | NA | NA | 4 | 0.25 | 1 | 1 | 7 | 0.5714 |
| 7_3470878 | 2 | 0 | 7 | 0.1429 | 5 | 0 | 4 | 0 | 18 | 0.0556 | 2 | 0.5 | NA | NA | 4 | 0 | 1 | 1 | 7 | 0.2857 |
| 7_3470920 | 2 | 0.5 | 6 | 0.5 | 5 | 0.6 | 3 | 0.3333 | 16 | 0.5 | 2 | 0 | NA | NA | 4 | 0.25 | 2 | 0 | 8 | 0.125 |
| 7_3471005 | 1 | 1 | 7 | 0.8571 | 7 | 1 | 4 | 0.75 | 19 | 0.8947 | 2 | 0.5 | 1 | 1 | 3 | 0.6667 | 2 | 1 | 8 | 0.75 |
| 7_3613687 | NA | NA | NA | NA | 1 | 1 | NA | NA | 1 | 1 | NA | NA | NA | NA | 1 | 1 | 1 | 1 | 2 | 1 |
| 7_3613893 | NA | NA | NA | NA | 1 | 1 | NA | NA | 1 | 1 | NA | NA | NA | NA | 1 | 1 | 1 | 0 | 2 | 0.5 |
| 7_3614202 | NA | NA | NA | NA | 1 | 1 | NA | NA | 1 | 1 | NA | NA | NA | NA | 1 | 0 | 1 | 1 | 2 | 0.5 |
| 7_3620851 | NA | NA | NA | NA | 2 | 0.5 | NA | NA | 2 | 0.5 | 1 | 1 | 1 | 1 | 2 | 0.5 | 1 | 1 | 5 | 0.8 |
| 7_3620865 | NA | NA | NA | NA | 3 | 0.3333 | NA | NA | 3 | 0.3333 | 1 | 0 | 1 | 0 | 3 | 0 | 1 | 1 | 6 | 0.1667 |
| 7_3621113 | NA | NA | NA | NA | 3 | 0.3333 | 1 | 0 | 4 | 0.25 | 2 | 0 | 1 | 1 | 3 | 0 | 1 | 0 | 7 | 0.1429 |
| 7_3633731 | 2 | 0.5 | 6 | 0.3333 | NA | NA | 3 | 0 | 11 | 0.2727 | 3 | 0 | 1 | 0 | 5 | 0 | 3 | 0.6667 | 12 | 0.1667 |
| 7_3633821 | NA | NA | NA | NA | NA | NA | 1 | 1 | 1 | 1 | 1 | 1 | NA | NA | NA | NA | 1 | 1 | 2 | 1 |
| 7_3643958 | 2 | 0.5 | 6 | 0.3333 | 4 | 1 | 6 | 0.5 | 18 | 0.5556 | 2 | 0.5 | 4 | 0.5 | 6 | 0.6667 | 1 | 0 | 13 | 0.5385 |
| 7_3643967 | 3 | 1 | 6 | 1 | 2 | 1 | 5 | 0.8 | 16 | 0.9375 | 2 | 1 | 4 | 0.75 | 6 | 1 | 1 | 1 | 13 | 0.9231 |
| 7_3643985 | 3 | 0 | 7 | 0 | 3 | 0 | 5 | 0.4 | 18 | 0.1111 | 2 | 0.5 | 3 | 0 | 6 | 0 | 2 | 0.5 | 13 | 0.1538 |
| 7_3644081 | 2 | 0 | 7 | 0.2857 | 4 | 0.25 | 6 | 0.5 | 19 | 0.3158 | 2 | 0 | 3 | 0.6667 | 6 | 0.3333 | 2 | 0.5 | 13 | 0.3846 |
| 7_3644228 | NA | NA | 3 | 0.6667 | NA | NA | NA | NA | 3 | 0.6667 | NA | NA | 1 | 1 | NA | NA | NA | NA | 1 | 1 |
| 7_3663388 | NA | NA | 2 | 0.5 | NA | NA | NA | NA | 2 | 0.5 | NA | NA | 1 | 0 | NA | NA | 3 | 0.3333 | 4 | 0.25 |
| 7_3664038 | 1 | 0 | 6 | 0.1667 | 1 | 1 | 5 | 0 | 13 | 0.1538 | 4 | 0.25 | 3 | 0 | 2 | 0 | 2 | 0 | 11 | 0.0909 |
| 7_3665920 | 1 | 1 | 3 | 0.3333 | NA | NA | 1 | 0 | 5 | 0.4 | NA | NA | NA | NA | NA | NA | 2 | 0.5 | 2 | 0.5 |
| 7_3668817 | NA | NA | NA | NA | 2 | 0.5 | NA | NA | 2 | 0.5 | NA | NA | NA | NA | 1 | 1 | 1 | 0 | 2 | 0.5 |
| 7_3668881 | 1 | 1 | NA | NA | 2 | 0.5 | NA | NA | 3 | 0.6667 | NA | NA | NA | NA | 1 | 1 | 1 | 0 | 2 | 0.5 |
| 7_3681055 | 14 | 0.2143 | 17 | 0.3529 | 10 | 0.3 | 15 | 0.2 | 56 | 0.2679 | 14 | 0.3571 | 17 | 0.5294 | 22 | 0.2273 | 9 | 0.2222 | 62 | 0.3387 |
| 7_3681087 | 14 | 0 | 20 | 0.15 | 12 | 0.0833 | 19 | 0.0526 | 65 | 0.0769 | 10 | 0.1 | 18 | 0.1111 | 18 | 0.0556 | 10 | 0 | 56 | 0.0714 |
| 7_3681115 | 16 | 0 | 18 | 0.1111 | 13 | 0.0769 | 17 | 0.0588 | 64 | 0.0625 | 12 | 0 | 19 | 0.1053 | 19 | 0.1053 | 12 | 0.0833 | 62 | 0.0806 |
| 7_3681404 | 13 | 0.0769 | 17 | 0.2353 | 11 | 0 | 17 | 0.1176 | 58 | 0.1207 | 12 | 0.1667 | 13 | 0.1538 | 17 | 0.1176 | 12 | 0.0833 | 54 | 0.1296 |
| 7_3681693 | 13 | 0.6923 | 15 | 0.8667 | 7 | 0.8571 | 12 | 0.5833 | 47 | 0.7447 | 8 | 0.625 | 14 | 0.8571 | 21 | 0.7619 | 11 | 0.7273 | 54 | 0.7593 |
| 7_3686916 | 14 | 0 | 15 | 0.2667 | 8 | 0 | 13 | 0 | 50 | 0.08 | 9 | 0 | 12 | 0 | 18 | 0.1667 | 8 | 0.125 | 47 | 0.0851 |
| 7_3687252 | 6 | 0.8333 | 9 | 0.6667 | 2 | 1 | 9 | 0.6667 | 26 | 0.7308 | 2 | 1 | 9 | 0.8889 | 13 | 1 | 6 | 1 | 30 | 0.9667 |
| 7_3687836 | 5 | 0.2 | 5 | 0 | 2 | 0 | 8 | 0.25 | 20 | 0.15 | 3 | 0.3333 | 7 | 0.1429 | 10 | 0.1 | 4 | 0 | 24 | 0.125 |
| 7_3696599 | NA | NA | NA | NA | 1 | 1 | NA | NA | 1 | 1 | NA | NA | NA | NA | 1 | 1 | NA | NA | 1 | 1 |
| 7_3703722 | 17 | 0.2941 | 19 | 0.0526 | 19 | 0.2632 | 9 | 0.4444 | 64 | 0.2344 | 17 | 0.1765 | 7 | 0.2857 | 19 | 0.1579 | 14 | 0.2143 | 57 | 0.193 |
| 7_3703827 | 17 | 0.2941 | 16 | 0.1875 | 16 | 0.25 | 10 | 0.2 | 59 | 0.2373 | 16 | 0.1875 | 7 | 0.1429 | 15 | 0.0667 | 13 | 0.3846 | 51 | 0.1961 |
| 7_3707655 | 96 | 0.0729 | 98 | 0.0918 | 88 | 0.0682 | 95 | 0.1158 | 377 | 0.0875 | 70 | 0.0714 | 85 | 0.1294 | 87 | 0.069 | 71 | 0.1127 | 313 | 0.0958 |
| 7_3709843 | 97 | 0.1856 | 111 | 0.1081 | 88 | 0.1364 | 96 | 0.1354 | 392 | 0.1403 | 70 | 0.1429 | 82 | 0.122 | 94 | 0.1489 | 70 | 0.1 | 316 | 0.1297 |
| 7_4012201 | NA | NA | 1 | 0 | 2 | 1 | NA | NA | 3 | 0.6667 | NA | NA | NA | NA | 1 | 1 | 2 | 1 | 3 | 1 |

| | | | | | | | | | | | | | | | | | | | | |
|---|---|---|---|---|---|---|---|---|---|---|---|---|---|---|---|---|---|---|---|---|
| 7_4012210 | 1 | 0 | 1 | 1 | 3 | 1 | NA | NA | 5 | 0.8 | NA | NA | 1 | 1 | 1 | 1 | 2 | 1 | 4 | 1 |
| 7_4012330 | 1 | 0 | 1 | 0 | 3 | 0.3333 | NA | NA | 5 | 0.2 | NA | NA | 1 | 1 | 1 | 1 | NA | NA | 2 | 1 |
| 7_4012336 | 1 | 1 | 1 | 0 | 2 | 0 | NA | NA | 4 | 0.25 | NA | NA | 1 | 0 | 1 | 0 | 2 | 0.5 | 4 | 0.25 |
| 7_4012487 | 1 | 1 | 1 | 0 | 2 | 0 | NA | NA | 4 | 0.25 | NA | NA | 1 | 0 | 1 | 1 | 2 | 0 | 4 | 0.25 |
| 7_4012696 | 1 | 0 | 1 | 0 | 2 | 1 | NA | NA | 4 | 0.5 | NA | NA | 1 | 0 | 1 | 1 | 2 | 0.5 | 4 | 0.5 |
| 7_4073658 | 1 | 1 | NA | NA | NA | NA | NA | NA | 1 | 1 | NA | NA | NA | NA | 1 | 1 | NA | NA | 1 | 1 |
| 7_4073773 | 1 | 1 | NA | NA | NA | NA | NA | NA | 1 | 1 | NA | NA | NA | NA | 1 | 1 | NA | NA | 1 | 1 |
| 7_4078110 | NA | NA | NA | NA | 1 | 0 | 1 | 1 | 2 | 0.5 | 1 | 1 | NA | NA | NA | NA | NA | NA | 1 | 1 |
| 7_4078199 | NA | NA | NA | NA | 1 | 0 | 1 | 1 | 2 | 0.5 | 1 | 1 | NA | NA | NA | NA | NA | NA | 1 | 1 |
| 7_4078272 | NA | NA | NA | NA | 1 | 0 | 1 | 1 | 2 | 0.5 | 1 | 1 | NA | NA | NA | NA | NA | NA | 1 | 1 |
| 7_4136943 | 51 | 0.098 | 51 | 0.0392 | 43 | 0.1628 | 56 | 0.125 | 201 | 0.1045 | 39 | 0.1538 | 59 | 0.0508 | 60 | 0.1 | 43 | 0.1163 | 201 | 0.0995 |
| 7_4137042 | 52 | 0.1538 | 54 | 0.0741 | 40 | 0.075 | 61 | 0.1475 | 207 | 0.1159 | 36 | 0.1667 | 57 | 0.1579 | 56 | 0.1429 | 41 | 0.122 | 190 | 0.1474 |
| 7_4137305 | 41 | 0.0732 | 34 | 0.0882 | 37 | 0.2162 | 45 | 0.1333 | 157 | 0.1274 | 35 | 0.0571 | 45 | 0.1111 | 45 | 0.1333 | 30 | 0.1667 | 155 | 0.1161 |
| 7_4138019 | 41 | 0.0976 | 45 | 0.1333 | 41 | 0.1463 | 51 | 0.1373 | 178 | 0.1292 | 34 | 0.1176 | 55 | 0.0727 | 34 | 0.1176 | 31 | 0.1613 | 154 | 0.1104 |
| 7_4138280 | 52 | 0.25 | 54 | 0.2037 | 50 | 0.3 | 62 | 0.2419 | 218 | 0.2477 | 44 | 0.3409 | 59 | 0.3051 | 43 | 0.3256 | 36 | 0.25 | 182 | 0.3077 |
| 7_4138364 | 14 | 0.0714 | 15 | 0.0667 | 18 | 0.1111 | 15 | 0.0667 | 62 | 0.0806 | 15 | 0.3333 | 9 | 0.2222 | 9 | 0.1111 | 7 | 0 | 40 | 0.2 |
| 7_4138476 | 10 | 0.3 | 16 | 0.3125 | 15 | 0.5333 | 13 | 0.3077 | 54 | 0.3704 | 8 | 0.25 | 10 | 0.3 | 7 | 0.2857 | 9 | 0.4444 | 34 | 0.3235 |
| 7_4138534 | 12 | 0 | 16 | 0 | 16 | 0.0625 | 16 | 0.125 | 60 | 0.05 | 12 | 0.25 | 9 | 0.2222 | 9 | 0.2222 | 8 | 0.125 | 38 | 0.2105 |
| 7_4148175 | 2 | 0.5 | 6 | 0 | 2 | 0.5 | 3 | 0 | 13 | 0.1538 | 4 | 0 | 8 | 0.125 | 3 | 0 | 4 | 0 | 19 | 0.0526 |
| 7_4148421 | 3 | 0.3333 | 7 | 0.7143 | 4 | 0.5 | 3 | 0.3333 | 17 | 0.5294 | 2 | 1 | 5 | 1 | 2 | 0.5 | 4 | 0.25 | 13 | 0.6923 |
| 7_4148909 | 4 | 0.25 | 5 | 0.2 | 4 | 0 | 1 | 0 | 14 | 0.1429 | 4 | 0.25 | 8 | 0.125 | 4 | 0.25 | 4 | 0.25 | 20 | 0.2 |
| 7_4149274 | NA | NA | 4 | 0 | 4 | 0.25 | 1 | 0 | 9 | 0.1111 | 2 | 0 | 8 | 0.125 | 2 | 0 | 2 | 0.5 | 14 | 0.1429 |
| 7_4150065 | 1 | 0 | 4 | 0.25 | 4 | 0.25 | 2 | 0.5 | 11 | 0.2727 | 4 | 0 | 7 | 0.1429 | 1 | 0 | 5 | 0 | 17 | 0.0588 |
| 7_4150173 | 4 | 0.25 | 10 | 0.5 | 4 | 1 | 2 | 0 | 20 | 0.5 | 4 | 0.75 | 9 | 0.2222 | 4 | 0.25 | 4 | 0.75 | 21 | 0.4286 |
| 7_4150539 | 5 | 0.8 | 10 | 0.9 | 3 | 1 | 3 | 1 | 21 | 0.9048 | 6 | 1 | 9 | 0.8889 | 7 | 1 | 4 | 1 | 26 | 0.9615 |
| 7_4150597 | 3 | 1 | 9 | 0.8889 | 6 | 1 | 4 | 0.75 | 22 | 0.9091 | 6 | 1 | 11 | 0.9091 | 7 | 1 | 4 | 1 | 28 | 0.9643 |
| 7_4150682 | 3 | 0 | 10 | 0.2 | 5 | 0 | 4 | 0 | 22 | 0.0909 | 7 | 0 | 11 | 0.0909 | 6 | 0 | 5 | 0.2 | 29 | 0.069 |
| 7_4151037 | 4 | 0.75 | 10 | 0.5 | 6 | 0.5 | 5 | 0.4 | 25 | 0.52 | 7 | 0.4286 | 11 | 0.5455 | 9 | 0.5556 | 7 | 0.4286 | 34 | 0.5 |
| 7_4151394 | 3 | 0.3333 | 2 | 0 | 1 | 0 | NA | NA | 6 | 0.1667 | 1 | 1 | NA | NA | 1 | 0 | NA | NA | 2 | 0.5 |
| 7_4151515 | 2 | 1 | NA | NA | 1 | 0 | NA | NA | 3 | 0.6667 | 2 | 1 | NA | NA | 1 | 1 | NA | NA | 3 | 1 |
| 7_4151929 | 1 | 1 | NA | NA | 1 | 1 | NA | NA | 2 | 1 | 1 | 1 | NA | NA | 1 | 1 | NA | NA | 2 | 1 |
| 7_4152770 | NA | NA | 1 | 1 | NA | NA | NA | NA | 1 | 1 | NA | NA | NA | NA | 1 | 1 | NA | NA | 1 | 1 |
| 7_4153269 | NA | NA | 1 | 1 | NA | NA | NA | NA | 1 | 1 | NA | NA | NA | NA | 1 | 1 | NA | NA | 1 | 1 |
| 7_4428803 | 6 | 0.3333 | 2 | 1 | 4 | 0.5 | 3 | 0 | 15 | 0.4 | 2 | 0.5 | 2 | 0 | 7 | 0.7143 | 3 | 0.3333 | 14 | 0.5 |
| 7_4429312 | 7 | 0 | 2 | 0 | 5 | 0.4 | 3 | 0.3333 | 17 | 0.1765 | 2 | 0 | 3 | 0.6667 | 5 | 0 | 3 | 0 | 13 | 0.1538 |
| 7_4429377 | 4 | 0.25 | 2 | 0 | 2 | 0.5 | 3 | 0.3333 | 11 | 0.2727 | 2 | 0 | 3 | 0.6667 | 4 | 0 | 2 | 0 | 11 | 0.1818 |
| 7_4429567 | 7 | 0 | 1 | 0 | 3 | 0.3333 | 4 | 0 | 15 | 0.0667 | 1 | 0 | 2 | 0 | 3 | 0 | 3 | 0.6667 | 9 | 0.2222 |
| 7_4430399 | 4 | 0 | NA | NA | 4 | 0.25 | 4 | 0.5 | 12 | 0.25 | 1 | 0 | 4 | 0 | 4 | 0.75 | 2 | 0 | 11 | 0.2727 |
| 7_4430418 | 4 | 0.5 | 1 | 1 | 3 | 0.6667 | 4 | 0.5 | 12 | 0.5833 | 1 | 0 | 4 | 0.5 | 4 | 0 | 4 | 0.5 | 13 | 0.3077 |
| 7_4430459 | 3 | 0.6667 | NA | NA | 1 | 1 | 4 | 1 | 8 | 0.875 | 1 | 1 | 3 | 0.3333 | 4 | 0.75 | 4 | 0.75 | 12 | 0.6667 |
| 7_4430522 | 3 | 0.6667 | NA | NA | 3 | 0.6667 | 3 | 0.6667 | 9 | 0.6667 | 1 | 1 | 4 | 0.75 | 4 | 0.25 | 4 | 0.75 | 13 | 0.6154 |
| 7_4430547 | 4 | 0.5 | 1 | 0 | 4 | 0 | 3 | 0 | 12 | 0.1667 | 1 | 0 | 4 | 0.5 | 4 | 0 | 4 | 0.5 | 13 | 0.3077 |
| 7_4430575 | 4 | 0.5 | 1 | 0 | 4 | 1 | 3 | 0.3333 | 12 | 0.5833 | 1 | 1 | 4 | 0.25 | 4 | 0.75 | 4 | 1 | 13 | 0.6923 |
| 7_4430638 | 2 | 0 | 1 | 0 | 4 | 0.75 | 2 | 1 | 9 | 0.5556 | 1 | 1 | 2 | 0.5 | 4 | 0 | 4 | 0.25 | 11 | 0.2727 |
| 7_4430692 | 3 | 0 | NA | NA | 3 | 0.6667 | 1 | 1 | 7 | 0.4286 | NA | NA | 4 | 0.75 | 4 | 1 | 4 | 0.5 | 12 | 0.75 |
| 7_4430725 | 4 | 0 | 1 | 0 | 3 | 0 | 3 | 0.3333 | 11 | 0.0909 | 1 | 0 | 3 | 0 | 4 | 0 | 4 | 0.25 | 12 | 0.0833 |
| 7_4482094 | NA | NA | 2 | 0 | 1 | 0 | 2 | 0.5 | 5 | 0.2 | NA | NA | 1 | 1 | NA | NA | NA | NA | 1 | 1 |
| 7_4482145 | NA | NA | NA | NA | 1 | 0 | 2 | 0.5 | 3 | 0.3333 | NA | NA | 1 | 1 | NA | NA | NA | NA | 1 | 1 |
| 7_4482156 | NA | NA | 1 | 0 | 1 | 1 | 2 | 0.5 | 4 | 0.5 | NA | NA | 1 | 1 | NA | NA | NA | NA | 1 | 1 |
| 7_4484794 | 16 | 0.0625 | 21 | 0.1429 | 14 | 0.1429 | 15 | 0.1333 | 66 | 0.1212 | 13 | 0.0769 | 20 | 0.1 | 13 | 0.0769 | 7 | 0.1429 | 53 | 0.0943 |
| 7_4484838 | 11 | 0.0909 | 16 | 0.0625 | 11 | 0.0909 | 12 | 0.25 | 50 | 0.12 | 9 | 0.1111 | 14 | 0 | 9 | 0 | 7 | 0.1429 | 39 | 0.0513 |
| 7_4484901 | 14 | 0.0714 | 22 | 0.0455 | 14 | 0.1429 | 10 | 0.2 | 60 | 0.1 | 12 | 0.0833 | 20 | 0.3 | 11 | 0 | 5 | 0.2 | 48 | 0.1667 |
| 7_4484948 | 13 | 0.1538 | 21 | 0.0476 | 15 | 0 | 12 | 0.1667 | 61 | 0.082 | 12 | 0.0833 | 20 | 0.1 | 11 | 0 | 6 | 0 | 49 | 0.0612 |
| 7_4485071 | 14 | 0.4286 | 18 | 0.1111 | 14 | 0.0714 | 10 | 0.1 | 56 | 0.1786 | 10 | 0.1 | 17 | 0.1176 | 12 | 0 | 5 | 0 | 44 | 0.0682 |
| 7_4504452 | NA | NA | NA | NA | 1 | 1 | NA | NA | 1 | 1 | NA | NA | 1 | 1 | NA | NA | NA | NA | 1 | 1 |
| 7_4504793 | NA | NA | NA | NA | 1 | 1 | NA | NA | 1 | 1 | NA | NA | 1 | 1 | NA | NA | NA | NA | 1 | 1 |
| 7_4631331 | 1 | 1 | NA | NA | 1 | 1 | NA | NA | 2 | 1 | NA | NA | 1 | 1 | 1 | 0 | NA | NA | 2 | 0.5 |
| 7_4631485 | NA | NA | NA | NA | 1 | 1 | NA | NA | 1 | 1 | NA | NA | 1 | 1 | 1 | 1 | NA | NA | 2 | 1 |
| 7_4631575 | NA | NA | NA | NA | 1 | 1 | NA | NA | 1 | 1 | NA | NA | 1 | 1 | 1 | 0 | NA | NA | 2 | 0.5 |
| 7_4718996 | 1 | 0 | 8 | 0.125 | 6 | 0 | 1 | 0 | 16 | 0.0625 | 3 | 0 | 5 | 0 | 5 | 0.2 | 1 | 0 | 14 | 0.0714 |
| 7_4749655 | 1 | 1 | 3 | 1 | 2 | 1 | 1 | 1 | 7 | 1 | 1 | 1 | 1 | 1 | NA | NA | 2 | 1 | 4 | 1 |
| 7_4749704 | 1 | 0 | 3 | 0 | 2 | 0.5 | 1 | 0 | 7 | 0.1429 | 2 | 0 | NA | NA | 2 | 0 | 2 | 0.5 | 6 | 0.1667 |
| 7_4749868 | 1 | 1 | 2 | 0 | 1 | 1 | NA | NA | 4 | 0.5 | 1 | 0 | 1 | 0 | 2 | 0.5 | 2 | 0.5 | 6 | 0.3333 |
| 7_4750106 | 1 | 0 | 3 | 0.3333 | 2 | 0.5 | NA | NA | 6 | 0.3333 | 3 | 0.6667 | 1 | 0 | NA | NA | 2 | 0.5 | 6 | 0.5 |
| 7_4787949 | NA | NA | NA | NA | 1 | 1 | NA | NA | 1 | 1 | 1 | 1 | NA | NA | NA | NA | NA | NA | 1 | 1 |

| | | | | | | | | | | | | | | | | | | | | |
|---|---|---|---|---|---|---|---|---|---|---|---|---|---|---|---|---|---|---|---|---|
| 7_4829944 | 8 | 0.125 | 10 | 0.3 | 8 | 0 | 11 | 0.1818 | 37 | 0.1622 | 5 | 0 | 12 | 0.3333 | 9 | 0.1111 | 8 | 0.375 | 34 | 0.2353 |
| 7_4830058 | 6 | 0.3333 | 6 | 0.1667 | 8 | 0.125 | 7 | 0.1429 | 27 | 0.1852 | 3 | 0 | 7 | 0.1429 | 5 | 0.4 | 5 | 0.4 | 20 | 0.25 |
| 7_4830745 | 6 | 0.1667 | 10 | 0.1 | 8 | 0 | 8 | 0.5 | 32 | 0.1875 | 5 | 0 | 10 | 0 | 8 | 0 | 7 | 0.2857 | 30 | 0.0667 |
| 7_4831229 | 5 | 0.6 | 9 | 0.4444 | 7 | 0.2857 | 7 | 0.5714 | 28 | 0.4643 | 3 | 0 | 7 | 0.4286 | 5 | 0.6 | 7 | 0.4286 | 22 | 0.4091 |
| 7_4831280 | 4 | 1 | 9 | 0.7778 | 6 | 1 | 8 | 1 | 27 | 0.9259 | 2 | 1 | 7 | 0.8571 | 6 | 1 | 6 | 0.8333 | 21 | 0.9048 |
| 7_4831391 | 5 | 0.4 | 5 | 0 | 6 | 0.1667 | 7 | 0 | 23 | 0.1304 | 2 | 0 | 7 | 0.1429 | 6 | 0.3333 | 7 | 0.1429 | 22 | 0.1818 |
| 7_4831511 | 9 | 0.7778 | 12 | 0.75 | 7 | 1 | 9 | 0.7778 | 37 | 0.8108 | 4 | 1 | 6 | 1 | 4 | 1 | 7 | 0.7143 | 21 | 0.9048 |
| 7_4831573 | 9 | 0.8889 | 14 | 0.9286 | 3 | 1 | 10 | 1 | 36 | 0.9444 | 5 | 1 | 8 | 0.875 | 6 | 0.8333 | 7 | 1 | 26 | 0.9231 |
| 7_4831753 | 10 | 0.3 | 9 | 0.1111 | 7 | 0.2857 | 8 | 0 | 34 | 0.1765 | 6 | 0 | 8 | 0.25 | 6 | 0.1667 | 6 | 0.1667 | 26 | 0.1538 |
| 7_4831813 | 8 | 1 | 9 | 0.8889 | 7 | 0.8571 | 7 | 1 | 31 | 0.9355 | 4 | 1 | 7 | 1 | 6 | 1 | 5 | 1 | 22 | 1 |
| 7_4831865 | 9 | 0.8889 | 6 | 1 | 6 | 0.8333 | 8 | 1 | 29 | 0.931 | 7 | 1 | 8 | 1 | 7 | 1 | 5 | 0.8 | 27 | 0.963 |
| 7_4831931 | 10 | 0.9 | 8 | 1 | 8 | 1 | 7 | 1 | 33 | 0.9697 | 7 | 1 | 7 | 1 | 7 | 0.8571 | 6 | 1 | 27 | 0.963 |
| 7_4831966 | 8 | 0 | 6 | 0.3333 | 7 | 0 | 7 | 0.1429 | 28 | 0.1071 | 7 | 0.1429 | 7 | 0.2857 | 5 | 0 | 4 | 0.5 | 23 | 0.2174 |
| 7_4831995 | 9 | 1 | 8 | 0.875 | 7 | 0.8571 | 8 | 0.75 | 32 | 0.875 | 6 | 0.6667 | 8 | 0.75 | 5 | 0.6 | 5 | 1 | 24 | 0.75 |
| 7_4832009 | 5 | 0.6 | 7 | 0.5714 | 6 | 0.8333 | 7 | 0.7143 | 25 | 0.68 | 4 | 1 | 6 | 0.6667 | 4 | 0.75 | 5 | 0.4 | 19 | 0.6842 |
| 7_4832122 | 8 | 0.25 | 7 | 0.4286 | 8 | 0.375 | 8 | 0.375 | 31 | 0.3548 | 5 | 0.4 | 8 | 0.125 | 6 | 0.1667 | 5 | 0 | 24 | 0.1667 |
| 7_4832143 | 9 | 0 | 6 | 0 | 4 | 0.25 | 8 | 0.25 | 27 | 0.1111 | 3 | 0.3333 | 8 | 0.125 | 4 | 0 | 4 | 0.25 | 19 | 0.1579 |
| 7_4832162 | 9 | 0.4444 | 7 | 0.7143 | 7 | 0.4286 | 8 | 0.25 | 31 | 0.4516 | 6 | 0.5 | 8 | 0.625 | 6 | 0.5 | 6 | 0.6667 | 26 | 0.5769 |
| 7_4832194 | 6 | 0.1667 | 7 | 0.2857 | 4 | 0 | 5 | 0 | 22 | 0.1364 | 6 | 0 | 8 | 0 | 2 | 0 | 3 | 0.3333 | 19 | 0.0526 |
| 7_4832252 | 8 | 0.375 | 2 | 1 | 5 | 0.6 | 5 | 0.8 | 20 | 0.6 | 4 | 0.75 | 5 | 0.8 | 4 | 0.75 | 4 | 1 | 17 | 0.8235 |
| 7_4832348 | 9 | 0.7778 | 5 | 1 | 5 | 1 | 7 | 1 | 26 | 0.9231 | 5 | 0.8 | 7 | 1 | 6 | 0.8333 | 3 | 1 | 21 | 0.9048 |
| 7_4832449 | 9 | 0.1111 | 5 | 0 | 5 | 0.2 | 5 | 0.2 | 24 | 0.125 | 4 | 0 | 7 | 0 | 4 | 0.5 | 4 | 0.75 | 19 | 0.2632 |
| 7_4832520 | 9 | 0.4444 | 6 | 0.6667 | 6 | 0.3333 | 6 | 0.5 | 27 | 0.4815 | 5 | 0.6 | 7 | 0.5714 | 6 | 0.5 | 3 | 1 | 21 | 0.619 |
| 7_4832537 | 8 | 0 | 8 | 0.125 | 6 | 0.1667 | 7 | 0.1429 | 29 | 0.1034 | 5 | 0 | 8 | 0 | 6 | 0.1667 | 5 | 0.2 | 24 | 0.0833 |
| 7_4832642 | 8 | 0.875 | 6 | 0.8333 | 6 | 0.8333 | 4 | 1 | 24 | 0.875 | 3 | 0.6667 | 8 | 1 | 2 | 1 | 3 | 1 | 16 | 0.9375 |
| 7_4833015 | 4 | 0 | 5 | 0 | 5 | 0.2 | 5 | 0.4 | 19 | 0.1579 | 3 | 0 | 5 | 0.2 | 5 | 0 | 3 | 0.3333 | 16 | 0.125 |
| 7_4848184 | 1 | 1 | NA | NA | NA | NA | NA | NA | 1 | 1 | 1 | 1 | NA | NA | NA | NA | NA | NA | 1 | 1 |
| 7_4848339 | 1 | 1 | NA | NA | NA | NA | NA | NA | 1 | 1 | 1 | 1 | NA | NA | NA | NA | NA | NA | 1 | 1 |
| 7_4898504 | NA | NA | 1 | 0 | 2 | 0.5 | NA | NA | 3 | 0.3333 | 3 | 1 | NA | NA | 1 | 0 | NA | NA | 4 | 0.75 |
| 7_4921976 | 1 | 0 | NA | NA | 1 | 1 | NA | NA | 2 | 0.5 | NA | NA | 1 | 1 | 1 | 0 | 1 | 1 | 3 | 0.6667 |
| 7_4922249 | 1 | 0 | NA | NA | 2 | 1 | NA | NA | 3 | 0.6667 | NA | NA | 1 | 0 | 1 | 1 | 1 | 0 | 3 | 0.3333 |
| 7_4922898 | 1 | 1 | 1 | 1 | 2 | 1 | NA | NA | 4 | 1 | NA | NA | 1 | 1 | 1 | 1 | 2 | 0.5 | 4 | 0.75 |
| 7_4923042 | 1 | 0 | 1 | 0 | 2 | 0.5 | NA | NA | 4 | 0.25 | NA | NA | 1 | 0 | 1 | 1 | 2 | 1 | 4 | 0.75 |
| 7_4923179 | 1 | 1 | 1 | 1 | 2 | 1 | NA | NA | 4 | 1 | NA | NA | 1 | 0 | 1 | 1 | 2 | 0.5 | 4 | 0.5 |
| 7_4923904 | 1 | 1 | 1 | 1 | 2 | 1 | 1 | 0 | 5 | 0.8 | 1 | 1 | 1 | 1 | 1 | 1 | 2 | 1 | 5 | 1 |
| 7_4924062 | NA | NA | 1 | 1 | 2 | 1 | NA | NA | 3 | 1 | 2 | 0 | 1 | 1 | 1 | 1 | 2 | 0.5 | 6 | 0.5 |
| 7_4924492 | 1 | 0 | 3 | 0.3333 | 2 | 0 | 1 | 0 | 7 | 0.1429 | 2 | 0 | NA | NA | 1 | 1 | 2 | 0 | 5 | 0.2 |
| 7_4924629 | 1 | 1 | 2 | 1 | 3 | 1 | 1 | 1 | 7 | 1 | 2 | 0.5 | 1 | 1 | 1 | 1 | 1 | 1 | 5 | 0.8 |
| 7_4924635 | NA | NA | 2 | 0.5 | 2 | 0.5 | 1 | 0 | 5 | 0.4 | 2 | 0 | 1 | 0 | 1 | 1 | 2 | 0 | 6 | 0.1667 |
| 7_4924976 | 1 | 0 | 3 | 0.3333 | 3 | 0.3333 | NA | NA | 7 | 0.2857 | 2 | 0.5 | 2 | 0.5 | 1 | 0 | 2 | 0.5 | 7 | 0.4286 |
| 7_4926991 | 14 | 0.3571 | 12 | 0.25 | 8 | 0.375 | 18 | 0.0556 | 52 | 0.2308 | 12 | 0.4167 | 16 | 0.1875 | 14 | 0 | 11 | 0.3636 | 53 | 0.2264 |
| 7_4927443 | 18 | 0.2222 | 17 | 0 | 13 | 0.0769 | 21 | 0 | 69 | 0.0725 | 13 | 0.2308 | 27 | 0.1481 | 17 | 0.0588 | 16 | 0.25 | 73 | 0.1644 |
| 7_4927588 | 17 | 0.8235 | 17 | 0.8824 | 16 | 0.875 | 20 | 0.6 | 70 | 0.7857 | 14 | 0.9286 | 28 | 0.8929 | 18 | 0.8333 | 17 | 0.7647 | 77 | 0.8571 |
| 7_4927625 | 18 | 0.6667 | 19 | 0.8947 | 16 | 0.75 | 21 | 0.8095 | 74 | 0.7838 | 14 | 0.8571 | 28 | 0.7143 | 16 | 0.6875 | 17 | 0.8235 | 75 | 0.76 |
| 7_4995931 | 2 | 0.5 | 5 | 0 | 2 | 1 | NA | NA | 9 | 0.3333 | 1 | 1 | 2 | 0 | 4 | 1 | 1 | 0 | 8 | 0.625 |
| 7_4995950 | 2 | 1 | 5 | 0.6 | 3 | 1 | 2 | 0.5 | 12 | 0.75 | 2 | 1 | 3 | 1 | 2 | 1 | NA | NA | 7 | 1 |
| 7_4995957 | 3 | 0.3333 | 7 | 0.2857 | 2 | 0 | 3 | 0.3333 | 15 | 0.2667 | 3 | 0 | 3 | 0.3333 | 6 | 0.3333 | 2 | 0 | 14 | 0.2143 |
| 7_4995996 | 3 | 1 | 8 | 0.75 | 3 | 1 | 2 | 1 | 16 | 0.875 | 3 | 0.6667 | 3 | 1 | 5 | 1 | 1 | 1 | 12 | 0.9167 |
| 7_4996018 | 4 | 1 | 4 | 0.75 | 3 | 1 | 3 | 1 | 14 | 0.9286 | 3 | 1 | 2 | 1 | 7 | 1 | 2 | 1 | 14 | 1 |
| 7_4996066 | 4 | 0.5 | 9 | 0.6667 | 3 | 1 | 2 | 0.5 | 18 | 0.6667 | 3 | 0 | 3 | 0.6667 | 6 | 0.3333 | 2 | 0 | 14 | 0.2857 |
| 7_4996264 | 3 | 1 | 6 | 1 | 2 | 1 | 2 | 1 | 13 | 1 | 3 | 1 | 2 | 1 | 5 | 1 | 1 | 1 | 11 | 1 |
| 7_4996888 | 3 | 1 | 3 | 1 | 1 | 1 | 3 | 1 | 10 | 1 | 1 | 1 | 1 | 1 | 2 | 1 | 1 | 0 | 5 | 0.8 |
| 7_4997014 | 1 | 0 | 3 | 0.6667 | 1 | 0 | 3 | 0 | 8 | 0.25 | 1 | 1 | 1 | 0 | 3 | 0 | 1 | 1 | 6 | 0.3333 |
| 7_4997133 | 3 | 0 | 2 | 0 | 1 | 0 | 3 | 0.3333 | 9 | 0.1111 | 1 | 0 | 2 | 0 | 3 | 0.3333 | 1 | 1 | 7 | 0.2857 |
| 7_4997383 | 1 | 0 | 3 | 0.6667 | 2 | 1 | 3 | 1 | 9 | 0.7778 | 1 | 1 | 1 | 1 | 2 | 0.5 | 1 | 1 | 5 | 0.8 |
| 7_5010081 | 5 | 0.4 | 3 | 0 | 7 | 0.1429 | 4 | 0 | 19 | 0.1579 | 4 | 0 | 3 | 0.3333 | 3 | 0 | 3 | 0 | 13 | 0.0769 |
| 7_5010184 | 4 | 0 | 2 | 0 | 6 | 0.1667 | 3 | 0 | 15 | 0.0667 | 3 | 0 | 1 | 0 | 2 | 0 | 3 | 0.3333 | 9 | 0.1111 |
| 7_5010414 | 5 | 0.6 | 3 | 0 | 5 | 0.4 | 5 | 0 | 18 | 0.2778 | 4 | 0.25 | 4 | 0.5 | 4 | 0 | 3 | 0 | 15 | 0.2 |
| 7_5010651 | 5 | 0 | 2 | 0.5 | 6 | 0 | 4 | 0 | 17 | 0.0588 | 4 | 0.25 | 4 | 0 | 3 | 0 | 3 | 0 | 14 | 0.0714 |
| 7_5010793 | 4 | 1 | 3 | 0.6667 | 7 | 0.7143 | 4 | 1 | 18 | 0.8333 | 2 | 0 | 4 | 1 | 3 | 0.6667 | 3 | 1 | 12 | 0.75 |
| 7_5010849 | 4 | 1 | 3 | 1 | 7 | 0.8571 | 3 | 1 | 17 | 0.9412 | 3 | 1 | 4 | 1 | 4 | 0.75 | 3 | 1 | 14 | 0.9286 |
| 7_5010888 | 3 | 0.6667 | 2 | 0.5 | 4 | 0.25 | 4 | 0.25 | 13 | 0.3846 | 3 | 0.3333 | 3 | 0.3333 | 3 | 0.3333 | 2 | 0.5 | 11 | 0.3636 |
| 7_5010967 | 4 | 0.25 | 3 | 0 | 2 | 0 | 3 | 0 | 12 | 0.0833 | 2 | 0 | 2 | 0 | 3 | 0.3333 | 2 | 0 | 9 | 0.1111 |
| 7_5010975 | 4 | 0.5 | 3 | 0.3333 | 6 | 0.1667 | 4 | 0 | 17 | 0.2353 | 4 | 0.25 | 4 | 0 | 4 | 0 | 2 | 0 | 14 | 0.0714 |
| 7_5011004 | 4 | 0.25 | 2 | 0 | 6 | 0.5 | 5 | 0.2 | 17 | 0.2941 | 4 | 0 | 4 | 0.25 | 3 | 0.3333 | 3 | 0.3333 | 14 | 0.2143 |

| | | | | | | | | | | | | | | | | | | | | |
|---|---|---|---|---|---|---|---|---|---|---|---|---|---|---|---|---|---|---|---|---|
| 7_5011018 | 4 | 1 | 1 | 1 | 5 | 1 | 4 | 1 | 14 | 1 | 3 | 1 | 3 | 1 | 4 | 1 | 2 | 1 | 12 | 1 |
| 7_5011296 | 4 | 0 | 3 | 0 | 3 | 0 | 5 | 0.2 | 15 | 0.0667 | 2 | 0 | 4 | 0.5 | 4 | 0.25 | 3 | 0 | 13 | 0.2308 |
| 7_5011493 | 4 | 0.75 | 3 | 0.3333 | 3 | 0.3333 | 4 | 0.5 | 14 | 0.5 | 2 | 0.5 | 4 | 0.75 | 4 | 0.75 | 2 | 0.5 | 12 | 0.6667 |
| 7_5011519 | 4 | 0 | 3 | 0 | 3 | 0.3333 | 4 | 0 | 14 | 0.0714 | 3 | 0 | 4 | 0.25 | 3 | 0 | 3 | 0 | 13 | 0.0769 |
| 7_5011649 | 2 | 0 | 2 | 0 | 3 | 0 | 1 | 1 | 8 | 0.125 | 3 | 0.3333 | 3 | 0 | 2 | 0 | 2 | 0 | 10 | 0.1 |
| 7_5011876 | 2 | 0 | NA | NA | 3 | 0.3333 | 2 | 0.5 | 7 | 0.2857 | 3 | 0 | 2 | 0 | 1 | 0 | 3 | 0.6667 | 9 | 0.2222 |
| 7_5011950 | 1 | 0 | 1 | 1 | 3 | 1 | 2 | 1 | 7 | 0.8571 | 3 | 0.6667 | 3 | 1 | NA | NA | 1 | 1 | 7 | 0.8571 |
| 7_5020606 | NA | NA | 5 | 1 | 1 | 1 | 1 | 1 | 7 | 1 | 1 | 1 | 3 | 1 | 4 | 1 | 1 | 1 | 9 | 1 |
| 7_5020685 | NA | NA | 5 | 0 | 1 | 1 | 1 | 1 | 7 | 0.2857 | 1 | 1 | 2 | 0 | 3 | 0.6667 | 1 | 1 | 7 | 0.5714 |
| 7_5021170 | NA | NA | 3 | 0.3333 | 1 | 0 | 1 | 0 | 5 | 0.2 | NA | NA | 4 | 0.25 | 4 | 0 | 1 | 1 | 9 | 0.2222 |
| 7_5021435 | NA | NA | 3 | 1 | 1 | 1 | 1 | 1 | 5 | 1 | NA | NA | NA | NA | 4 | 1 | 1 | 1 | 5 | 1 |
| 7_5032466 | 3 | 0.3333 | 2 | 0.5 | NA | NA | 1 | 0 | 6 | 0.3333 | NA | NA | 1 | 1 | 1 | 0 | NA | NA | 2 | 0.5 |
| 7_5032532 | 3 | 1 | 2 | 1 | 1 | 1 | 1 | 0 | 7 | 0.8571 | NA | NA | 1 | 1 | 1 | 1 | NA | NA | 2 | 1 |
| 7_5032996 | 2 | 0.5 | 1 | 0 | 1 | 0 | NA | NA | 4 | 0.25 | NA | NA | NA | NA | NA | NA | 1 | 1 | 1 | 1 |
| 7_5033316 | 3 | 1 | 3 | 1 | 1 | 1 | 1 | 1 | 8 | 1 | NA | NA | 1 | 1 | 1 | 1 | 1 | 0 | 3 | 0.6667 |
| 7_5033497 | 3 | 0.6667 | 3 | 1 | 1 | 1 | 1 | 0 | 8 | 0.75 | NA | NA | 1 | 1 | 1 | 0 | 1 | 0 | 3 | 0.3333 |
| 7_5033946 | 3 | 0.3333 | 3 | 0.3333 | 2 | 1 | 1 | 0 | 9 | 0.4444 | 1 | 1 | 2 | 0 | 1 | 1 | NA | NA | 4 | 0.5 |
| 7_5034019 | 2 | 1 | 2 | 0.5 | 2 | 1 | 1 | 0 | 7 | 0.7143 | 1 | 1 | 2 | 1 | 1 | 0 | 1 | 0 | 5 | 0.6 |
| 7_5034133 | 3 | 0.3333 | 3 | 0.3333 | 1 | 0 | 1 | 0 | 8 | 0.25 | 1 | 0 | 2 | 0.5 | 1 | 0 | 1 | 1 | 5 | 0.4 |
| 7_5034259 | 3 | 0 | 4 | 0 | 2 | 0 | 1 | 1 | 10 | 0.1 | 1 | 0 | 2 | 0 | 1 | 0 | 1 | 1 | 5 | 0.2 |
| 7_5034266 | 3 | 1 | 4 | 1 | 2 | 1 | 1 | 1 | 10 | 1 | 1 | 1 | 2 | 1 | 1 | 1 | 1 | 1 | 5 | 1 |
| 7_5034366 | 3 | 0.6667 | 2 | 0.5 | 2 | 1 | NA | NA | 7 | 0.7143 | 2 | 0 | 3 | 0.6667 | 1 | 0 | 1 | 1 | 7 | 0.4286 |
| 7_5034438 | 2 | 1 | 3 | 1 | 1 | 1 | 1 | 1 | 7 | 1 | 2 | 1 | 3 | 0.6667 | 1 | 1 | 1 | 1 | 7 | 0.8571 |
| 7_5034468 | 3 | 0.6667 | 4 | 0.5 | 3 | 1 | 1 | 1 | 11 | 0.7273 | 2 | 0.5 | 3 | 1 | 1 | 0 | 1 | 0 | 7 | 0.5714 |
| 7_5034831 | 4 | 0.25 | 5 | 0.2 | 2 | 0 | 1 | 0 | 12 | 0.1667 | 2 | 0 | 2 | 0.5 | 2 | 0 | 1 | 0 | 7 | 0.1429 |
| 7_5035239 | 5 | 1 | 4 | 1 | 4 | 1 | NA | NA | 13 | 1 | 2 | 1 | 3 | 1 | 1 | 1 | NA | NA | 6 | 1 |
| 7_5035267 | 5 | 1 | 4 | 0.5 | 4 | 0.75 | 1 | 1 | 14 | 0.7857 | 2 | 1 | 4 | 0.5 | 2 | 1 | 1 | 0 | 9 | 0.6667 |
| 7_5035305 | 4 | 1 | 3 | 1 | 2 | 1 | 1 | 1 | 10 | 1 | 2 | 1 | 3 | 1 | 2 | 0.5 | 1 | 1 | 8 | 0.875 |
| 7_5035348 | 4 | 0.5 | 5 | 0.2 | 4 | 0.5 | 1 | 1 | 14 | 0.4286 | 2 | 1 | 3 | 0.6667 | 1 | 1 | 2 | 0.5 | 8 | 0.75 |
| 7_5035408 | 2 | 0 | 4 | 0.25 | 4 | 0 | 1 | 0 | 11 | 0.0909 | 2 | 0.5 | 2 | 0 | 2 | 0.5 | 1 | 0 | 7 | 0.2857 |
| 7_5035415 | 2 | 1 | 5 | 1 | 4 | 1 | 1 | 1 | 12 | 1 | 2 | 1 | 3 | 0.6667 | 1 | 1 | 2 | 1 | 8 | 0.875 |
| 7_5035490 | 3 | 1 | 5 | 1 | 4 | 1 | 1 | 1 | 13 | 1 | 2 | 1 | 2 | 1 | 2 | 1 | 1 | 1 | 7 | 1 |
| 7_5035514 | 3 | 0.3333 | 5 | 0.2 | 2 | 0 | 1 | 0 | 11 | 0.1818 | 2 | 0.5 | 2 | 0.5 | 1 | 0 | 2 | 0 | 7 | 0.2857 |
| 7_5035552 | 1 | 1 | 6 | 0.6667 | 4 | 0.5 | 1 | 1 | 12 | 0.6667 | 2 | 1 | 3 | 0.6667 | 3 | 1 | 2 | 1 | 10 | 0.9 |
| 7_5035559 | 5 | 1 | 7 | 1 | 3 | 1 | 1 | 1 | 16 | 1 | 2 | 1 | 3 | 1 | 3 | 1 | 2 | 0.5 | 10 | 0.9 |
| 7_5035656 | 5 | 0.2 | 5 | 0 | 3 | 0.3333 | 1 | 0 | 14 | 0.1429 | 1 | 0 | 3 | 0 | 2 | 0.5 | 2 | 0.5 | 8 | 0.25 |
| 7_5035829 | 1 | 1 | NA | NA | NA | NA | NA | NA | 1 | 1 | NA | NA | NA | NA | 1 | 0 | 1 | 1 | 2 | 0.5 |
| 7_5036051 | 1 | 1 | NA | NA | NA | NA | NA | NA | 1 | 1 | NA | NA | NA | NA | 1 | 1 | 1 | 0 | 2 | 0.5 |
| 7_5038617 | 2 | 0.5 | 1 | 1 | NA | NA | NA | NA | 3 | 0.6667 | NA | NA | 2 | 1 | NA | NA | 1 | 1 | 3 | 1 |
| 7_5038650 | 1 | 0 | 1 | 1 | NA | NA | NA | NA | 2 | 0.5 | NA | NA | 2 | 0 | NA | NA | 1 | 1 | 3 | 0.3333 |
| 7_5038729 | 2 | 0.5 | 1 | 1 | NA | NA | NA | NA | 3 | 0.6667 | NA | NA | 2 | 0.5 | NA | NA | NA | NA | 2 | 0.5 |
| 7_5039048 | 2 | 0.5 | 1 | 0 | NA | NA | NA | NA | 3 | 0.3333 | NA | NA | 2 | 0.5 | NA | NA | NA | NA | 2 | 0.5 |
| 7_5055874 | 6 | 0.3333 | 2 | 0 | 6 | 0.3333 | 3 | 0.3333 | 17 | 0.2941 | 5 | 0.2 | 5 | 0.6 | 2 | 0 | 5 | 0 | 17 | 0.2353 |
| 7_5055891 | 3 | 0.3333 | 1 | 0 | 3 | 0 | 1 | 0 | 8 | 0.125 | 4 | 0.25 | 3 | 0 | 1 | 0 | 2 | 0 | 10 | 0.1 |
| 7_5056162 | 4 | 0.25 | 2 | 0.5 | 6 | 0.5 | 3 | 0.3333 | 15 | 0.4 | 5 | 0.6 | 4 | 1 | 2 | 0.5 | 3 | 0.3333 | 14 | 0.6429 |
| 7_5056492 | 3 | 0 | 3 | 0 | 9 | 0 | 3 | 0.3333 | 18 | 0.0556 | 6 | 0 | 8 | 0.25 | 3 | 0 | 5 | 0 | 22 | 0.0909 |
| 7_5056532 | 5 | 0.6 | 4 | 0.75 | 9 | 0.4444 | 3 | 0 | 21 | 0.4762 | 3 | 0.3333 | 8 | 0.5 | 2 | 0 | 5 | 0.6 | 18 | 0.4444 |
| 7_5060243 | 1 | 0 | 1 | 1 | 1 | 0 | 1 | 0 | 4 | 0.25 | 1 | 0 | 2 | 0.5 | 4 | 0 | 3 | 0 | 10 | 0.1 |
| 7_5060462 | 1 | 1 | 4 | 0 | 2 | 0 | 2 | 0 | 9 | 0.1111 | 2 | 0 | 5 | 0 | 9 | 0.1111 | 4 | 0 | 20 | 0.05 |
| 7_5060855 | 8 | 0.25 | 4 | 0 | 7 | 0 | 5 | 0 | 24 | 0.0833 | 4 | 0 | 9 | 0.1111 | 11 | 0.0909 | 10 | 0.1 | 34 | 0.0882 |
| 7_5061069 | 8 | 0.125 | 5 | 0 | 6 | 0 | 6 | 0.1667 | 25 | 0.08 | 4 | 0 | 10 | 0.1 | 13 | 0.1538 | 10 | 0.1 | 37 | 0.1081 |
| 7_5061135 | 8 | 0.125 | 3 | 0 | 5 | 0 | 4 | 0 | 20 | 0.05 | 2 | 0 | 8 | 0.25 | 8 | 0 | 10 | 0.1 | 28 | 0.1071 |
| 7_5063305 | 8 | 0.125 | 5 | 0.4 | 4 | 0 | 6 | 0 | 23 | 0.1304 | 4 | 0 | 8 | 0.125 | 10 | 0.2 | 10 | 0.1 | 32 | 0.125 |
| 7_5063451 | 9 | 0 | 5 | 0.4 | 5 | 0 | 6 | 0 | 25 | 0.08 | 4 | 0.25 | 4 | 0.25 | 8 | 0.25 | 11 | 0.2727 | 27 | 0.2593 |
| 7_5063731 | 10 | 0.1 | 6 | 0 | 9 | 0.2222 | 8 | 0.125 | 33 | 0.1212 | 4 | 0 | 9 | 0.1111 | 14 | 0.0714 | 12 | 0.0833 | 39 | 0.0769 |
| 7_5063765 | 7 | 0.1429 | 4 | 0.75 | 5 | 0 | 6 | 0.1667 | 22 | 0.2273 | 2 | 0 | 9 | 0.2222 | 8 | 0.125 | 8 | 0.375 | 27 | 0.2222 |
| 7_5082305 | 11 | 0.2727 | 15 | 0.2 | 12 | 0.1667 | 9 | 0.5556 | 47 | 0.2766 | 7 | 0.4286 | 14 | 0.4286 | 11 | 0.3636 | 10 | 0.6 | 42 | 0.4524 |
| 7_5093012 | 25 | 0.12 | 42 | 0.0238 | 38 | 0.0789 | 23 | 0.0435 | 128 | 0.0625 | 23 | 0.1739 | 34 | 0.0588 | 27 | 0.0741 | 12 | 0.0833 | 96 | 0.0938 |
| 7_5100964 | 24 | 0.2083 | 47 | 0.4468 | 45 | 0.4444 | 22 | 0.4091 | 138 | 0.3986 | 22 | 0.5 | 38 | 0.3421 | 37 | 0.3243 | 19 | 0.4737 | 116 | 0.3879 |
| 7_6134847 | 1 | 1 | 2 | 0.5 | 4 | 0.75 | 5 | 1 | 12 | 0.8333 | 2 | 0 | 2 | 1 | 1 | 0 | 2 | 0.5 | 7 | 0.4286 |
| 7_6134927 | 1 | 1 | 1 | 1 | 4 | 0.75 | 4 | 0.75 | 10 | 0.8 | 2 | 0 | 2 | 0.5 | 1 | 0 | 2 | 1 | 7 | 0.4286 |
| 7_6135445 | 1 | 1 | 2 | 0.5 | 2 | 0 | 3 | 0.3333 | 8 | 0.375 | 2 | 1 | 2 | 0.5 | 1 | 0 | NA | NA | 5 | 0.6 |
| 7_6187044 | 4 | 0.25 | 6 | 0 | 5 | 0.2 | 5 | 0.4 | 20 | 0.2 | 3 | 0 | 6 | 0.3333 | 3 | 0.6667 | 3 | 0.3333 | 15 | 0.3333 |
| 7_6192767 | 5 | 0.2 | 7 | 0.1429 | 6 | 0.3333 | 6 | 0.1667 | 24 | 0.2083 | 4 | 0.25 | 8 | 0.125 | 4 | 0 | 4 | 0 | 20 | 0.1 |
| 7_6192923 | 6 | 1 | 5 | 1 | 7 | 1 | 5 | 1 | 23 | 1 | 4 | 1 | 8 | 1 | 5 | 1 | 4 | 1 | 21 | 1 |

| | | | | | | | | | | | | | | | | | | | | |
|---|---|---|---|---|---|---|---|---|---|---|---|---|---|---|---|---|---|---|---|---|
| 7_6193116 | 7 | 0.2857 | 10 | 0 | 8 | 0 | 7 | 0 | 32 | 0.0625 | 5 | 0 | 9 | 0.1111 | 5 | 0 | 4 | 0.25 | 23 | 0.087 |
| 7_6193162 | 6 | 0.3333 | 9 | 0.6667 | 8 | 0.5 | 8 | 0.5 | 31 | 0.5161 | 5 | 0.4 | 10 | 0.3 | 6 | 0.3333 | 4 | 0.25 | 25 | 0.32 |
| 7_6193174 | 6 | 0.1667 | 9 | 0.2222 | 7 | 0.1429 | 4 | 1 | 26 | 0.3077 | 5 | 0 | 10 | 0.2 | 6 | 0.1667 | 3 | 1 | 24 | 0.25 |
| 7_6193259 | 6 | 0.6667 | 6 | 0.1667 | 5 | 0.6 | 7 | 0.2857 | 24 | 0.4167 | 3 | 0 | 7 | 0.4286 | 4 | 0.5 | 4 | 0.5 | 18 | 0.3889 |
| 7_6193293 | 7 | 0 | 6 | 0.3333 | 7 | 0.1429 | 8 | 0 | 28 | 0.1071 | 5 | 0.2 | 8 | 0 | 3 | 0 | 4 | 0 | 20 | 0.05 |
| 7_6193401 | 4 | 0.5 | 9 | 0.3333 | 7 | 0.2857 | 8 | 0.375 | 28 | 0.3571 | 5 | 0.4 | 9 | 0.3333 | 6 | 0.3333 | 4 | 0.75 | 24 | 0.4167 |
| 7_6193405 | 3 | 0 | 8 | 0.125 | 6 | 0.1667 | 8 | 0.125 | 25 | 0.12 | 5 | 0.2 | 9 | 0 | 5 | 0 | 4 | 0.25 | 23 | 0.087 |
| 7_6193503 | 1 | 0 | NA | NA | 2 | 0.5 | 1 | 1 | 4 | 0.5 | 1 | 0 | 1 | 0 | NA | NA | 1 | 1 | 3 | 0.3333 |
| 7_6196452 | NA | NA | 3 | 0.3333 | NA | NA | NA | NA | 3 | 0.3333 | NA | NA | NA | NA | NA | NA | 2 | 1 | 2 | 1 |
| 7_6196762 | NA | NA | 4 | 0.75 | NA | NA | 1 | 1 | 5 | 0.8 | NA | NA | NA | NA | 1 | 1 | 1 | 1 | 2 | 1 |
| 7_6307824 | NA | NA | 1 | 1 | 4 | 0.75 | 2 | 0.5 | 7 | 0.7143 | NA | NA | 1 | 1 | 1 | 0 | NA | NA | 2 | 0.5 |
| 7_6307953 | NA | NA | 1 | 1 | 4 | 0.75 | 3 | 1 | 8 | 0.875 | NA | NA | 2 | 1 | 2 | 1 | NA | NA | 4 | 1 |
| 7_6307973 | NA | NA | 1 | 0 | 3 | 0.3333 | 2 | 0.5 | 6 | 0.3333 | NA | NA | 2 | 0.5 | 2 | 1 | NA | NA | 4 | 0.75 |
| 7_6308427 | NA | NA | 1 | 1 | 4 | 0.75 | 3 | 0.6667 | 8 | 0.75 | NA | NA | 2 | 1 | 1 | 1 | NA | NA | 3 | 1 |
| 7_6308569 | 1 | 0 | 1 | 0 | 3 | 0.3333 | 2 | 0.5 | 7 | 0.2857 | NA | NA | 2 | 0 | 2 | 0.5 | NA | NA | 4 | 0.25 |
| 7_6308817 | 1 | 1 | NA | NA | 2 | 1 | 1 | 1 | 4 | 1 | NA | NA | 2 | 0.5 | 1 | 0 | NA | NA | 3 | 0.3333 |
| 7_6308989 | 1 | 1 | 1 | 1 | 5 | 0.4 | 2 | 0.5 | 9 | 0.5556 | NA | NA | 1 | 1 | 2 | 0 | NA | NA | 3 | 0.3333 |
| 7_6309205 | 1 | 0 | 1 | 1 | 5 | 1 | 3 | 0.6667 | 10 | 0.8 | NA | NA | 2 | 1 | 2 | 1 | NA | NA | 4 | 1 |
| 7_6309259 | 1 | 0 | 1 | 1 | 5 | 0.4 | 3 | 0 | 10 | 0.3 | NA | NA | 2 | 1 | 2 | 1 | NA | NA | 4 | 1 |
| 7_6309310 | 1 | 0 | 1 | 0 | 5 | 0.4 | 2 | 0.5 | 9 | 0.3333 | NA | NA | 1 | 1 | 2 | 0.5 | NA | NA | 3 | 0.6667 |
| 7_6309324 | NA | NA | 1 | 0 | 4 | 0.5 | 2 | 0 | 7 | 0.2857 | NA | NA | 1 | 0 | 2 | 0.5 | NA | NA | 3 | 0.3333 |
| 7_6309795 | 1 | 0 | 2 | 0 | 4 | 0.25 | 3 | 0 | 10 | 0.1 | 1 | 0 | 2 | 0 | 1 | 1 | NA | NA | 4 | 0.25 |
| 7_6310337 | 1 | 0 | 2 | 1 | 4 | 0.75 | 2 | 0.5 | 9 | 0.6667 | NA | NA | 2 | 0.5 | 1 | 1 | 1 | 1 | 4 | 0.75 |
| 7_6310362 | 1 | 0 | 2 | 0 | 4 | 0.5 | 3 | 0 | 10 | 0.2 | NA | NA | 2 | 0.5 | 2 | 0.5 | 1 | 0 | 5 | 0.4 |
| 7_6310434 | 1 | 0 | 2 | 1 | 4 | 0.75 | 1 | 1 | 8 | 0.75 | NA | NA | 1 | 1 | 2 | 1 | 1 | 1 | 4 | 1 |
| 7_6310453 | 1 | 1 | 2 | 1 | 4 | 1 | 2 | 1 | 9 | 1 | NA | NA | 2 | 1 | 2 | 1 | 1 | 1 | 5 | 1 |
| 7_6310534 | 1 | 1 | 2 | 0 | 4 | 0 | 2 | 0 | 9 | 0.1111 | 1 | 0 | 2 | 0.5 | 2 | 0.5 | 1 | 1 | 6 | 0.5 |
| 7_6356544 | NA | NA | 1 | 0 | NA | NA | 1 | 1 | 2 | 0.5 | NA | NA | 2 | 0.5 | NA | NA | NA | NA | 2 | 0.5 |
| 7_6356572 | NA | NA | 1 | 0 | NA | NA | 1 | 1 | 2 | 0.5 | NA | NA | 2 | 0.5 | NA | NA | NA | NA | 2 | 0.5 |
| 7_6356943 | 1 | 1 | 1 | 1 | NA | NA | 1 | 1 | 3 | 1 | NA | NA | 2 | 1 | NA | NA | NA | NA | 2 | 1 |
| 7_6357158 | 2 | 1 | 1 | 1 | 1 | 1 | NA | NA | 4 | 1 | NA | NA | 1 | 1 | NA | NA | NA | NA | 1 | 1 |
| 7_6357303 | 2 | 0.5 | 1 | 0 | 1 | 1 | NA | NA | 4 | 0.5 | NA | NA | 2 | 0.5 | NA | NA | NA | NA | 2 | 0.5 |
| 7_6357577 | 2 | 1 | 1 | 1 | 1 | 1 | NA | NA | 4 | 1 | NA | NA | 2 | 0.5 | NA | NA | NA | NA | 2 | 0.5 |
| 7_6357593 | 2 | 0 | 1 | 1 | NA | NA | 1 | 0 | 4 | 0.25 | NA | NA | 1 | 1 | NA | NA | NA | NA | 1 | 1 |
| 7_6357603 | 2 | 1 | 1 | 0 | 1 | 0 | 1 | 0 | 5 | 0.4 | NA | NA | 2 | 1 | NA | NA | NA | NA | 2 | 1 |
| 7_6357720 | 2 | 1 | 1 | 1 | NA | NA | NA | NA | 3 | 1 | NA | NA | 2 | 1 | NA | NA | NA | NA | 2 | 1 |
| 7_6382484 | NA | NA | NA | NA | 2 | 1 | NA | NA | 2 | 1 | NA | NA | NA | NA | NA | NA | 1 | 1 | 1 | 1 |
| 7_6382530 | NA | NA | NA | NA | 2 | 0.5 | 1 | 0 | 3 | 0.3333 | NA | NA | NA | NA | 1 | 1 | 1 | 1 | 2 | 1 |
| 7_6382806 | NA | NA | NA | NA | 1 | 1 | NA | NA | 1 | 1 | NA | NA | NA | NA | NA | NA | 1 | 1 | 1 | 1 |
| 7_6396518 | NA | NA | NA | NA | 1 | 1 | NA | NA | 1 | 1 | 1 | 0 | 1 | 1 | 1 | 1 | NA | NA | 3 | 0.6667 |
| 7_6396624 | NA | NA | NA | NA | 1 | 1 | NA | NA | 1 | 1 | 1 | 1 | 1 | 1 | 1 | 0 | NA | NA | 3 | 0.6667 |
| 7_6396691 | NA | NA | NA | NA | 1 | 1 | NA | NA | 1 | 1 | 1 | 1 | 1 | 1 | NA | NA | NA | NA | 2 | 1 |
| 7_6396719 | NA | NA | NA | NA | 1 | 1 | NA | NA | 1 | 1 | NA | NA | 1 | 1 | 1 | 0 | NA | NA | 2 | 0.5 |
| 7_6397433 | NA | NA | NA | NA | 1 | 1 | NA | NA | 1 | 1 | 2 | 0.5 | 1 | 1 | 1 | 1 | NA | NA | 4 | 0.75 |
| 7_6397600 | NA | NA | NA | NA | 1 | 1 | NA | NA | 1 | 1 | 2 | 0.5 | 1 | 0 | 1 | 0 | NA | NA | 4 | 0.25 |
| 7_6397831 | NA | NA | NA | NA | 1 | 1 | NA | NA | 1 | 1 | 2 | 0.5 | 1 | 1 | NA | NA | NA | NA | 3 | 0.6667 |
| 7_6397910 | 1 | 0 | NA | NA | 1 | 1 | NA | NA | 2 | 0.5 | 2 | 1 | 2 | 1 | 1 | 0 | NA | NA | 5 | 0.8 |
| 7_6427807 | 5 | 0 | NA | NA | 6 | 0.3333 | 1 | 0 | 12 | 0.1667 | NA | NA | 1 | 1 | 2 | 0 | NA | NA | 3 | 0.3333 |
| 7_6675294 | NA | NA | NA | NA | NA | NA | 1 | 1 | 1 | 1 | NA | NA | NA | NA | 1 | 1 | NA | NA | 1 | 1 |
| 7_6675628 | NA | NA | NA | NA | NA | NA | 1 | 1 | 1 | 1 | NA | NA | NA | NA | 1 | 1 | NA | NA | 1 | 1 |
| 7_6675985 | NA | NA | NA | NA | NA | NA | 1 | 1 | 1 | 1 | NA | NA | NA | NA | 1 | 1 | NA | NA | 1 | 1 |
| 7_6676078 | NA | NA | NA | NA | NA | NA | 1 | 1 | 1 | 1 | NA | NA | NA | NA | 1 | 1 | NA | NA | 1 | 1 |
| 7_6679169 | 2 | 0.5 | NA | NA | NA | NA | 1 | 1 | 3 | 0.6667 | NA | NA | NA | NA | 1 | 1 | NA | NA | 1 | 1 |
| 7_6679198 | 1 | 1 | NA | NA | NA | NA | 1 | 0 | 2 | 0.5 | NA | NA | NA | NA | 1 | 1 | NA | NA | 1 | 1 |
| 7_6679306 | 2 | 1 | NA | NA | NA | NA | NA | NA | 2 | 1 | NA | NA | NA | NA | 1 | 1 | NA | NA | 1 | 1 |
| 7_6679321 | 2 | 1 | NA | NA | NA | NA | 1 | 1 | 3 | 1 | NA | NA | NA | NA | 1 | 1 | NA | NA | 1 | 1 |
| 7_6679374 | 2 | 1 | NA | NA | NA | NA | 1 | 1 | 3 | 1 | NA | NA | NA | NA | 1 | 1 | NA | NA | 1 | 1 |
| 7_6707020 | 26 | 0.1154 | 23 | 0.3043 | 13 | 0.0769 | 27 | 0.1481 | 89 | 0.1685 | 18 | 0.1111 | 23 | 0.2174 | 33 | 0.4545 | 30 | 0.1333 | 104 | 0.25 |
| 7_6707323 | 16 | 0.125 | 18 | 0.2222 | 11 | 0.1818 | 25 | 0.2 | 70 | 0.1857 | 14 | 0.2857 | 15 | 0.2667 | 23 | 0.3043 | 24 | 0.2917 | 76 | 0.2895 |
| 7_6707951 | 16 | 0.0625 | 15 | 0 | 9 | 0.1111 | 16 | 0.125 | 56 | 0.0714 | 12 | 0.25 | 13 | 0.1538 | 22 | 0.0909 | 19 | 0.1579 | 66 | 0.1515 |
| 7_6709766 | 7 | 0.7143 | 8 | 0.25 | 6 | 0.3333 | 12 | 0.25 | 33 | 0.3636 | 5 | 0.2 | 12 | 0.0833 | 13 | 0.0769 | 14 | 0.5 | 44 | 0.2273 |
| 7_6710194 | 4 | 0.25 | 7 | 0.1429 | 4 | 0.75 | 12 | 0.1667 | 27 | 0.2593 | 4 | 0.25 | 12 | 0.3333 | 10 | 0.1 | 11 | 0.3636 | 37 | 0.2703 |
| 7_6710247 | 3 | 0 | 5 | 0.2 | 5 | 0.4 | 11 | 0 | 24 | 0.125 | 5 | 0.2 | 11 | 0 | 8 | 0 | 9 | 0.3333 | 33 | 0.1212 |
| 7_6710279 | 4 | 0.25 | 6 | 0.1667 | 4 | 0.25 | 12 | 0 | 26 | 0.1154 | 4 | 0 | 9 | 0 | 9 | 0.1111 | 7 | 0.2857 | 29 | 0.1034 |
| 7_6710309 | 4 | 1 | 5 | 1 | 5 | 1 | 9 | 0.8889 | 23 | 0.9565 | 4 | 1 | 7 | 1 | 9 | 1 | 10 | 0.9 | 30 | 0.9667 |

| | | | | | | | | | | | | | | | | | | | | |
|---|---|---|---|---|---|---|---|---|---|---|---|---|---|---|---|---|---|---|---|---|
| 7_6710324 | 3 | 1 | 7 | 0.8571 | 5 | 1 | 13 | 0.9231 | 28 | 0.9286 | 5 | 1 | 11 | 0.9091 | 10 | 0.9 | 10 | 0.9 | 36 | 0.9167 |
| 7_6710340 | 2 | 1 | 6 | 0.8333 | 4 | 1 | 6 | 1 | 18 | 0.9444 | 2 | 0.5 | 7 | 0.8571 | 8 | 1 | 5 | 1 | 22 | 0.9091 |
| 7_6710359 | 4 | 1 | 6 | 0.8333 | 5 | 1 | 10 | 1 | 25 | 0.96 | 4 | 1 | 6 | 0.8333 | 8 | 1 | 7 | 0.5714 | 25 | 0.84 |
| 7_6710373 | 3 | 0.6667 | 5 | 1 | 4 | 1 | 11 | 0.7273 | 23 | 0.8261 | 3 | 0.6667 | 9 | 1 | 5 | 0.6 | 6 | 0.3333 | 23 | 0.6957 |
| 7_6710451 | 4 | 0 | 7 | 0 | 5 | 0.4 | 12 | 0.1667 | 28 | 0.1429 | 5 | 0 | 11 | 0.0909 | 9 | 0.1111 | 9 | 0 | 34 | 0.0588 |
| 7_6711507 | 2 | 0.5 | 7 | 0.2857 | 2 | 0 | 8 | 0.125 | 19 | 0.2105 | 3 | 0.3333 | 7 | 0 | 7 | 0.1429 | 6 | 0 | 23 | 0.087 |
| 7_6711534 | 3 | 0.3333 | 7 | 0 | 3 | 0 | 6 | 0.1667 | 19 | 0.1053 | 3 | 0 | 8 | 0.125 | 8 | 0.125 | 6 | 0.3333 | 25 | 0.16 |
| 7_6711576 | 2 | 0 | 7 | 0 | 4 | 0 | 7 | 0.1429 | 20 | 0.05 | 4 | 0.25 | 6 | 0.3333 | 7 | 0.1429 | 5 | 0.2 | 22 | 0.2273 |
| 7_6711786 | 3 | 0.3333 | 7 | 0.1429 | 4 | 0 | 8 | 0.125 | 22 | 0.1364 | 4 | 0.25 | 8 | 0.125 | 8 | 0.125 | 6 | 0 | 26 | 0.1154 |
| 7_6711937 | 2 | 1 | 5 | 0.4 | 4 | 0.75 | 8 | 0.75 | 19 | 0.6842 | 4 | 1 | 6 | 0.8333 | 6 | 0.8333 | 6 | 0.8333 | 22 | 0.8636 |
| 7_6712382 | 3 | 0 | 3 | 0 | 4 | 0 | 7 | 0.1429 | 17 | 0.0588 | 3 | 0 | 8 | 0.125 | 4 | 0 | 5 | 0.2 | 20 | 0.1 |
| 7_6712415 | 2 | 1 | 5 | 0.6 | 4 | 0.75 | 6 | 0.8333 | 17 | 0.7647 | 3 | 0.3333 | 5 | 0.8 | 4 | 0.75 | 5 | 1 | 17 | 0.7647 |
| 7_6713068 | 2 | 0.5 | 2 | 1 | NA | NA | 4 | 0.75 | 8 | 0.75 | 1 | 1 | 5 | 1 | 4 | 0.5 | 3 | 0.6667 | 13 | 0.7692 |
| 7_6713184 | 2 | 0.5 | 3 | 0.3333 | 1 | 1 | 1 | 1 | 7 | 0.5714 | 2 | 0 | 2 | 1 | 4 | 0.75 | 3 | 0.6667 | 11 | 0.6364 |
| 7_6713671 | 2 | 0 | 3 | 0 | 1 | 1 | 3 | 0 | 9 | 0.1111 | 2 | 0 | 2 | 0.5 | 4 | 0 | 2 | 0 | 10 | 0.1 |
| 7_6714169 | 2 | 0.5 | 2 | 0 | NA | NA | 3 | 0 | 7 | 0.1429 | NA | NA | 2 | 0.5 | 4 | 0 | 2 | 0 | 8 | 0.125 |
| 7_6714541 | NA | NA | 1 | 1 | NA | NA | 2 | 0.5 | 3 | 0.6667 | NA | NA | 3 | 0.6667 | 3 | 1 | 2 | 1 | 8 | 0.875 |
| 7_6733863 | 2 | 0.5 | 2 | 0 | NA | NA | 1 | 0 | 5 | 0.2 | NA | NA | 2 | 1 | 1 | 1 | NA | NA | 3 | 1 |
| 7_6734189 | 3 | 0.6667 | 2 | 0 | NA | NA | 2 | 0 | 7 | 0.2857 | 1 | 0 | 3 | 0.3333 | 1 | 0 | NA | NA | 5 | 0.2 |
| 7_6734239 | 2 | 1 | 1 | 0 | NA | NA | 2 | 0.5 | 5 | 0.6 | NA | NA | 3 | 0.3333 | NA | NA | NA | NA | 3 | 0.3333 |
| 7_6964371 | 3 | 1 | 1 | 1 | NA | NA | 1 | 1 | 5 | 1 | NA | NA | 1 | 1 | 1 | 1 | NA | NA | 2 | 1 |
| 7_6964727 | 3 | 0.6667 | 1 | 0 | NA | NA | 1 | 1 | 5 | 0.6 | NA | NA | 2 | 0.5 | 1 | 0 | NA | NA | 3 | 0.3333 |
| 7_6964956 | 3 | 1 | 1 | 1 | NA | NA | 1 | 1 | 5 | 1 | 1 | 1 | 2 | 1 | 1 | 1 | NA | NA | 4 | 1 |
| 7_6965105 | 1 | 1 | 1 | 1 | NA | NA | NA | NA | 2 | 1 | 1 | 1 | 1 | 1 | NA | NA | 1 | 1 | 3 | 1 |
| 7_6965189 | 3 | 1 | 1 | 0 | NA | NA | NA | NA | 4 | 0.75 | 1 | 1 | 2 | 1 | 1 | 0 | NA | NA | 4 | 0.75 |
| 7_6965277 | 2 | 0.5 | 1 | 0 | NA | NA | 1 | 1 | 4 | 0.5 | 1 | 1 | 1 | 0 | 1 | 0 | 1 | 1 | 4 | 0.5 |
| 7_6965325 | 3 | 0.3333 | 1 | 0 | NA | NA | 1 | 0 | 5 | 0.2 | 1 | 0 | 2 | 0.5 | 1 | 0 | 1 | 0 | 5 | 0.2 |
| 7_6965530 | 3 | 0.3333 | 1 | 0 | NA | NA | 1 | 0 | 5 | 0.2 | 1 | 1 | 2 | 0.5 | 1 | 0 | 1 | 0 | 5 | 0.4 |
| 7_6965702 | 3 | 0.6667 | 2 | 0 | NA | NA | 1 | 1 | 6 | 0.5 | 1 | 1 | 2 | 0.5 | 1 | 0 | 2 | 0.5 | 6 | 0.5 |
| 7_6966197 | 3 | 1 | 2 | 0 | 1 | 0 | 1 | 1 | 7 | 0.5714 | 1 | 1 | 1 | 1 | 2 | 0.5 | 3 | 1 | 7 | 0.8571 |
| 7_6966347 | 2 | 0.5 | 2 | 0 | 1 | 0 | 1 | 1 | 6 | 0.3333 | 1 | 0 | 1 | 1 | 2 | 0.5 | 2 | 0 | 6 | 0.3333 |
| 7_6966521 | 2 | 1 | 1 | 0 | 1 | 0 | 1 | 1 | 5 | 0.6 | 1 | 1 | 2 | 1 | 1 | 1 | 1 | 0 | 5 | 0.8 |
| 7_6966651 | 1 | 1 | 1 | 1 | 2 | 1 | 1 | 1 | 5 | 1 | NA | NA | 1 | 0 | 2 | 1 | 3 | 0.6667 | 6 | 0.6667 |
| 7_6966710 | 3 | 0.6667 | 2 | 0.5 | 3 | 1 | 1 | 1 | 9 | 0.7778 | 1 | 1 | 2 | 1 | 2 | 1 | 3 | 0.6667 | 8 | 0.875 |
| 7_6966835 | 3 | 0.6667 | 2 | 0.5 | 3 | 0.3333 | NA | NA | 8 | 0.5 | 1 | 1 | 2 | 1 | 2 | 1 | 3 | 1 | 8 | 1 |
| 7_6966853 | 3 | 0.6667 | 2 | 0.5 | 4 | 0.5 | 1 | 1 | 10 | 0.6 | 1 | 1 | 2 | 1 | 2 | 1 | 3 | 0.6667 | 8 | 0.875 |
| 7_6966871 | 2 | 1 | 2 | 1 | 4 | 0.75 | 1 | 1 | 9 | 0.8889 | 1 | 1 | 1 | 1 | 1 | 1 | 2 | 1 | 5 | 1 |
| 7_6966881 | 2 | 0 | 1 | 0 | 4 | 0.5 | 1 | 1 | 8 | 0.375 | 1 | 1 | 1 | 1 | 1 | 0 | 2 | 1 | 5 | 0.8 |
| 7_6966992 | 3 | 0.3333 | 2 | 0 | 4 | 0 | 1 | 0 | 10 | 0.1 | 1 | 0 | 2 | 0 | 3 | 0 | 3 | 0.3333 | 9 | 0.1111 |
| 7_6967208 | 3 | 0 | 2 | 0 | 4 | 0.25 | 1 | 0 | 10 | 0.1 | 1 | 0 | 2 | 0 | 3 | 0.3333 | 5 | 0 | 11 | 0.0909 |
| 7_6967232 | 3 | 0.3333 | 2 | 0 | 4 | 0.25 | 1 | 1 | 10 | 0.3 | 1 | 0 | 2 | 0.5 | 3 | 0.3333 | 4 | 0.25 | 10 | 0.3 |
| 7_6967707 | 4 | 0.25 | 2 | 0 | 3 | 0.3333 | 3 | 0.3333 | 12 | 0.25 | 1 | 0 | 3 | 0.6667 | 3 | 0.3333 | 3 | 0 | 10 | 0.3 |
| 7_7117821 | 3 | 0 | 6 | 0.3333 | 1 | 1 | 1 | 1 | 11 | 0.3636 | 1 | 1 | NA | NA | 1 | 0 | 5 | 0.2 | 7 | 0.2857 |
| 7_7117840 | 3 | 0 | 5 | 0.2 | 1 | 0 | 1 | 0 | 10 | 0.1 | 2 | 0.5 | NA | NA | 1 | 1 | 7 | 0.2857 | 10 | 0.4 |
| 7_7117892 | 3 | 1 | 5 | 1 | 1 | 1 | 1 | 1 | 10 | 1 | 1 | 1 | 1 | 1 | 1 | 1 | 5 | 1 | 8 | 1 |
| 7_7117903 | 3 | 0.6667 | 4 | 0 | 1 | 0 | 1 | 1 | 9 | 0.3333 | 1 | 0 | 1 | 1 | 1 | 0 | 7 | 0.4286 | 10 | 0.4 |
| 7_7117944 | 3 | 0.6667 | 5 | 0.2 | 1 | 0 | 1 | 0 | 10 | 0.3 | 2 | 0 | NA | NA | 1 | 0 | 6 | 0.3333 | 9 | 0.2222 |
| 7_7117980 | 3 | 0 | 5 | 0.2 | 1 | 0 | 1 | 0 | 10 | 0.1 | 2 | 0 | 1 | 0 | 1 | 1 | 7 | 0 | 11 | 0.0909 |
| 7_7118093 | 3 | 0.6667 | 5 | 1 | NA | NA | 1 | 1 | 9 | 0.8889 | 1 | 1 | 1 | 1 | 1 | 1 | 7 | 1 | 10 | 1 |
| 7_7118135 | 3 | 1 | 5 | 1 | NA | NA | 1 | 1 | 9 | 1 | 1 | 1 | 1 | 1 | 1 | 1 | 7 | 1 | 10 | 1 |
| 7_7118242 | 2 | 0 | 4 | 0.5 | 1 | 0 | NA | NA | 7 | 0.2857 | NA | NA | 1 | 1 | 1 | 1 | 3 | 0 | 5 | 0.4 |
| 7_7118400 | 2 | 0.5 | 2 | 0 | NA | NA | NA | NA | 4 | 0.25 | 1 | 0 | NA | NA | NA | NA | 6 | 0.5 | 7 | 0.4286 |
| 7_7118721 | 3 | 1 | 4 | 1 | 1 | 1 | 1 | 1 | 9 | 1 | 1 | 1 | NA | NA | NA | NA | 7 | 0.8571 | 8 | 0.875 |
| 7_7118953 | 3 | 0.3333 | 4 | 0.25 | 1 | 1 | 1 | 0 | 9 | 0.3333 | NA | NA | NA | NA | NA | NA | 7 | 0.4286 | 7 | 0.4286 |
| 7_7119020 | 2 | 0 | 3 | 0.6667 | 1 | 0 | 1 | 0 | 7 | 0.2857 | NA | NA | NA | NA | NA | NA | 7 | 0.2857 | 7 | 0.2857 |
| 7_7185604 | NA | NA | NA | NA | NA | NA | 1 | 1 | 1 | 1 | NA | NA | 1 | 1 | 1 | 0 | NA | NA | 2 | 0.5 |
| 7_7185771 | NA | NA | NA | NA | NA | NA | 1 | 1 | 1 | 1 | NA | NA | 1 | 1 | 1 | 0 | NA | NA | 2 | 0.5 |
| 7_7205315 | 1 | 1 | 2 | 1 | 1 | 1 | NA | NA | 4 | 1 | NA | NA | 1 | 1 | NA | NA | NA | NA | 1 | 1 |
| 7_7205424 | 1 | 1 | 2 | 0.5 | NA | NA | NA | NA | 3 | 0.6667 | NA | NA | 1 | 0 | NA | NA | 2 | 0.5 | 3 | 0.3333 |
| 7_7205451 | NA | NA | 2 | 0.5 | 1 | 1 | NA | NA | 3 | 0.6667 | NA | NA | 1 | 0 | NA | NA | 2 | 0.5 | 3 | 0.3333 |
| 7_7205571 | 1 | 0 | 1 | 0 | 1 | 1 | NA | NA | 3 | 0.3333 | NA | NA | 1 | 1 | NA | NA | 2 | 0 | 3 | 0.3333 |
| 7_7205677 | 1 | 1 | 2 | 0 | 1 | 0 | NA | NA | 4 | 0.25 | NA | NA | 1 | 0 | NA | NA | 1 | 1 | 2 | 0.5 |
| 7_7205759 | 1 | 1 | 2 | 0.5 | 1 | 1 | NA | NA | 4 | 0.75 | NA | NA | 1 | 0 | NA | NA | 2 | 1 | 3 | 0.6667 |
| 7_7205779 | 1 | 1 | 2 | 0.5 | 1 | 1 | NA | NA | 4 | 0.75 | NA | NA | 1 | 0 | NA | NA | 2 | 1 | 3 | 0.6667 |
| 7_7206137 | 1 | 1 | 2 | 0.5 | 1 | 0 | 1 | 0 | 5 | 0.4 | NA | NA | 1 | 0 | NA | NA | 2 | 0.5 | 3 | 0.3333 |

| | | | | | | | | | | | | | | | | | | | | |
|---|---|---|---|---|---|---|---|---|---|---|---|---|---|---|---|---|---|---|---|---|
| 7_7206151 | 1 | 1 | 2 | 1 | 1 | 1 | 1 | 1 | 5 | 1 | NA | NA | 1 | 1 | NA | NA | 2 | 1 | 3 | 1 |
| 7_7206364 | 1 | 1 | 2 | 1 | 1 | 0 | 1 | 0 | 5 | 0.6 | NA | NA | 1 | 0 | NA | NA | 2 | 0.5 | 3 | 0.3333 |
| 7_7206491 | NA | NA | 2 | 1 | NA | NA | 1 | 0 | 3 | 0.6667 | NA | NA | 1 | 0 | NA | NA | 1 | 1 | 2 | 0.5 |
| 7_7206531 | 1 | 1 | 2 | 0 | NA | NA | 1 | 0 | 4 | 0.25 | NA | NA | 1 | 0 | NA | NA | 1 | 1 | 2 | 0.5 |
| 7_7207100 | 1 | 1 | 1 | 1 | NA | NA | 1 | 1 | 3 | 1 | NA | NA | 1 | 1 | NA | NA | 1 | 1 | 2 | 1 |
| 7_7207232 | 1 | 1 | 1 | 1 | NA | NA | 1 | 1 | 3 | 1 | NA | NA | 1 | 1 | NA | NA | 1 | 1 | 2 | 1 |
| 7_7207251 | 1 | 0 | 1 | 0 | NA | NA | 1 | 1 | 3 | 0.3333 | NA | NA | 1 | 0 | NA | NA | 1 | 1 | 2 | 0.5 |
| 7_7285691 | 18 | 0.0556 | 11 | 0.4545 | 21 | 0.0952 | 13 | 0.2308 | 63 | 0.1746 | 8 | 0.125 | 12 | 0.1667 | 19 | 0.1579 | 21 | 0.0952 | 60 | 0.1333 |
| 7_7286010 | 15 | 0.0667 | 10 | 0 | 16 | 0.0625 | 13 | 0.1538 | 54 | 0.0741 | 7 | 0.1429 | 7 | 0 | 11 | 0 | 15 | 0.1333 | 40 | 0.075 |
| 7_7286038 | 19 | 0.3158 | 7 | 0.8571 | 16 | 0.25 | 12 | 0.4167 | 54 | 0.3889 | 7 | 0.1429 | 9 | 0.4444 | 14 | 0.4286 | 17 | 0.4706 | 47 | 0.4043 |
| 7_7286405 | 21 | 0.2381 | 8 | 0.25 | 17 | 0.2353 | 14 | 0 | 60 | 0.1833 | 6 | 0.3333 | 13 | 0.3846 | 11 | 0.4545 | 20 | 0.45 | 50 | 0.42 |
| 7_7286426 | 19 | 0.3158 | 8 | 0.625 | 15 | 0.2 | 13 | 0.3077 | 55 | 0.3273 | 6 | 0.1667 | 12 | 0.3333 | 8 | 0.5 | 19 | 0.4737 | 45 | 0.4 |
| 7_7286447 | 21 | 0.9048 | 6 | 0.8333 | 16 | 0.875 | 13 | 0.7692 | 56 | 0.8571 | 6 | 1 | 10 | 1 | 11 | 0.8182 | 17 | 0.9412 | 44 | 0.9318 |
| 7_7286466 | 22 | 0.1364 | 9 | 0 | 18 | 0.0556 | 13 | 0 | 62 | 0.0645 | 5 | 0 | 13 | 0.3077 | 10 | 0.1 | 20 | 0 | 48 | 0.1042 |
| 7_7286473 | 18 | 0.8889 | 6 | 1 | 14 | 0.9286 | 13 | 0.9231 | 51 | 0.9216 | 3 | 0.6667 | 12 | 0.8333 | 9 | 0.8889 | 18 | 0.8333 | 42 | 0.8333 |
| 7_7286486 | 17 | 0.5882 | 5 | 0.4 | 13 | 0.4615 | 12 | 0.4167 | 47 | 0.4894 | 5 | 0.6 | 11 | 0.6364 | 8 | 0.75 | 15 | 0.8 | 39 | 0.7179 |
| 7_7286514 | 21 | 0.1905 | 9 | 0.1111 | 18 | 0.4444 | 13 | 0.3077 | 61 | 0.2787 | 5 | 0 | 12 | 0.3333 | 9 | 0.2222 | 21 | 0.2857 | 47 | 0.2553 |
| 7_7286773 | 19 | 0.1579 | 7 | 0.7143 | 16 | 0.25 | 10 | 0.3 | 52 | 0.2885 | 5 | 0.4 | 12 | 0.25 | 8 | 0.625 | 18 | 0.2222 | 43 | 0.3256 |
| 7_7286858 | 18 | 0.2222 | 5 | 0.4 | 16 | 0.1875 | 9 | 0.1111 | 48 | 0.2083 | 4 | 0 | 12 | 0.1667 | 7 | 0.1429 | 16 | 0.1875 | 39 | 0.1538 |
| 7_7286908 | 12 | 0.3333 | 1 | 0 | 10 | 0.2 | 7 | 0 | 30 | 0.2 | 2 | 0.5 | 8 | 0.125 | 4 | 0.5 | 8 | 0.375 | 22 | 0.3182 |
| 7_7286975 | 12 | 0.0833 | 2 | 0.5 | 11 | 0.0909 | 6 | 0.1667 | 31 | 0.129 | 2 | 0 | 10 | 0.1 | 5 | 0.4 | 11 | 0.1818 | 28 | 0.1786 |
| 7_7287001 | 13 | 0.1538 | 2 | 0 | 10 | 0.3 | 9 | 0.1111 | 34 | 0.1765 | 3 | 0.3333 | 10 | 0.3 | 5 | 0.6 | 12 | 0.0833 | 30 | 0.2667 |
| 7_7287093 | 9 | 0.1111 | 2 | 1 | 6 | 0.6667 | 4 | 0 | 21 | 0.3333 | 2 | 0.5 | 5 | 0.2 | 2 | 0.5 | 8 | 0.5 | 17 | 0.4118 |
| 7_7287208 | 10 | 0.3 | 2 | 0 | 11 | 0.5455 | 7 | 0.2857 | 30 | 0.3667 | 2 | 0.5 | 7 | 0.2857 | 7 | 0.5714 | 11 | 0.4545 | 27 | 0.4444 |
| 7_7292222 | 6 | 0 | 1 | 0 | 6 | 0.1667 | 4 | 0.25 | 17 | 0.1176 | NA | NA | 5 | 0 | 2 | 0.5 | 8 | 0.125 | 15 | 0.1333 |
| 7_7292226 | 3 | 0.3333 | 1 | 0 | 7 | 0.2857 | 3 | 0 | 14 | 0.2143 | NA | NA | 4 | 0.5 | 2 | 1 | 6 | 0.1667 | 12 | 0.4167 |
| 7_7294806 | 3 | 1 | 1 | 1 | 2 | 1 | 3 | 1 | 9 | 1 | NA | NA | 4 | 0.75 | 2 | 1 | 4 | 0.75 | 10 | 0.8 |
| 7_7299823 | 3 | 0.3333 | NA | NA | 1 | 0 | 3 | 0.3333 | 7 | 0.2857 | NA | NA | 3 | 0.3333 | NA | NA | 4 | 0 | 7 | 0.1429 |
| 7_1022285 | 1 | 0 | 1 | 1 | 1 | 0 | 1 | 1 | 4 | 0.5 | 1 | 0 | 2 | 0 | 2 | 0.5 | 1 | 0 | 6 | 0.1667 |
| 7_1022291 | 1 | 0 | 1 | 1 | 1 | 0 | 1 | 0 | 4 | 0.25 | 1 | 0 | 2 | 0 | 2 | 0.5 | 1 | 0 | 6 | 0.1667 |
| 7_1022325 | 1 | 0 | 1 | 0 | 1 | 1 | 1 | 1 | 4 | 0.5 | 1 | 0 | 2 | 0 | 2 | 0 | 1 | 1 | 6 | 0.1667 |
| 7_1022331 | 1 | 0 | 1 | 0 | 1 | 1 | 1 | 0 | 4 | 0.25 | 1 | 0 | 2 | 0 | 2 | 0.5 | 1 | 0 | 6 | 0.1667 |
| 7_1022481 | 1 | 0 | NA | NA | 1 | 1 | NA | NA | 2 | 0.5 | NA | NA | 2 | 0.5 | 1 | 0 | 1 | 0 | 4 | 0.25 |
| 7_1214711 | 1 | 1 | NA | NA | NA | NA | 2 | 0.5 | 3 | 0.6667 | 1 | 0 | 3 | 0 | 2 | 0.5 | NA | NA | 6 | 0.1667 |
| 7_1214805 | 2 | 1 | NA | NA | NA | NA | 2 | 0 | 4 | 0.5 | NA | NA | 3 | 0.6667 | 1 | 0 | NA | NA | 4 | 0.5 |
| 7_1214836 | 1 | 1 | NA | NA | NA | NA | 2 | 0 | 3 | 0.3333 | NA | NA | 3 | 0.3333 | 2 | 0.5 | NA | NA | 5 | 0.4 |
| 7_1214894 | 1 | 0 | NA | NA | NA | NA | 2 | 0.5 | 3 | 0.3333 | NA | NA | 3 | 0.6667 | 1 | 0 | NA | NA | 4 | 0.5 |
| 7_1214908 | 1 | 0 | NA | NA | NA | NA | 2 | 0.5 | 3 | 0.3333 | NA | NA | 2 | 0.5 | 1 | 0 | NA | NA | 3 | 0.3333 |
| 7_1214909 | 1 | 0 | NA | NA | NA | NA | 2 | 0.5 | 3 | 0.3333 | NA | NA | 2 | 0.5 | 1 | 0 | NA | NA | 3 | 0.6667 |
| 7_1214939 | 1 | 1 | NA | NA | NA | NA | 2 | 1 | 3 | 1 | NA | NA | 3 | 1 | 1 | 1 | NA | NA | 4 | 1 |
| 7_1214947 | 1 | 1 | NA | NA | NA | NA | 2 | 0.5 | 3 | 0.6667 | NA | NA | 3 | 0.6667 | 1 | 0 | NA | NA | 4 | 0.5 |
| 7_1214958 | NA | NA | NA | NA | NA | NA | 2 | 1 | 2 | 1 | NA | NA | 2 | 1 | 1 | 1 | NA | NA | 3 | 1 |
| 7_1214973 | NA | NA | NA | NA | NA | NA | 1 | 1 | 1 | 1 | NA | NA | 3 | 0.6667 | 1 | 0 | NA | NA | 4 | 0.5 |
| 7_1215001 | 1 | 1 | NA | NA | NA | NA | NA | NA | 1 | 1 | NA | NA | 2 | 1 | 1 | 1 | NA | NA | 3 | 1 |
| 7_1215019 | 1 | 1 | NA | NA | NA | NA | 1 | 0 | 2 | 0.5 | NA | NA | 2 | 0.5 | 1 | 1 | NA | NA | 3 | 0.6667 |
| 7_1215035 | 1 | 1 | NA | NA | NA | NA | 1 | 1 | 2 | 1 | NA | NA | 1 | 1 | 1 | 0 | NA | NA | 2 | 0.5 |
| 7_1215048 | 1 | 1 | NA | NA | NA | NA | 1 | 0 | 2 | 0.5 | NA | NA | 2 | 1 | NA | NA | NA | NA | 2 | 1 |
| 7_1215050 | 1 | 1 | NA | NA | NA | NA | 1 | 0 | 2 | 0.5 | NA | NA | 1 | 0 | 1 | 1 | NA | NA | 2 | 0.5 |
| 7_1215097 | 1 | 1 | NA | NA | NA | NA | NA | NA | 1 | 1 | NA | NA | 2 | 0.5 | 1 | 0 | NA | NA | 3 | 0.3333 |
| 7_1222652 | NA | NA | 1 | 1 | NA | NA | NA | NA | 1 | 1 | NA | NA | NA | NA | NA | NA | 2 | 1 | 2 | 1 |
| 7_1222723 | NA | NA | 1 | 1 | NA | NA | NA | NA | 1 | 1 | NA | NA | NA | NA | NA | NA | 2 | 1 | 2 | 1 |
| 7_1222739 | NA | NA | 1 | 1 | NA | NA | NA | NA | 1 | 1 | NA | NA | NA | NA | NA | NA | 2 | 1 | 2 | 1 |
| 7_1222757 | NA | NA | 1 | 1 | NA | NA | NA | NA | 1 | 1 | NA | NA | NA | NA | NA | NA | 1 | 1 | 1 | 1 |
| 7_1222782 | NA | NA | 1 | 1 | NA | NA | NA | NA | 1 | 1 | NA | NA | NA | NA | NA | NA | 2 | 1 | 2 | 1 |
| 7_1222828 | 1 | 1 | 1 | 1 | NA | NA | NA | NA | 2 | 1 | NA | NA | NA | NA | NA | NA | 2 | 1 | 2 | 1 |
| 7_1250281 | 1 | 1 | 1 | 1 | 2 | 1 | 2 | 0.5 | 6 | 0.8333 | NA | NA | 2 | 0.5 | 1 | 0 | NA | NA | 3 | 0.3333 |
| 7_1250312 | 3 | 0.6667 | NA | NA | 3 | 0 | 2 | 0 | 8 | 0.25 | NA | NA | 2 | 0.5 | 1 | 0 | NA | NA | 3 | 0.3333 |
| 7_1250328 | 3 | 0.3333 | 1 | 1 | 1 | 1 | 2 | 1 | 7 | 0.7143 | NA | NA | NA | NA | 2 | 0.5 | NA | NA | 2 | 0.5 |
| 7_1250329 | 2 | 0.5 | 1 | 1 | 3 | 1 | 2 | 0.5 | 8 | 0.75 | NA | NA | 2 | 1 | 2 | 1 | NA | NA | 4 | 1 |
| 7_1250342 | 3 | 0.6667 | 1 | 1 | 3 | 0.3333 | 1 | 1 | 8 | 0.625 | NA | NA | 1 | 1 | 2 | 1 | NA | NA | 3 | 1 |
| 7_1250824 | 1 | 1 | NA | NA | 1 | 1 | 1 | 1 | 3 | 1 | NA | NA | 1 | 1 | 2 | 0.5 | NA | NA | 3 | 0.6667 |
| 7_1253937 | 1 | 1 | 1 | 0 | 4 | 0.5 | 4 | 0.75 | 10 | 0.6 | 3 | 0.6667 | 1 | 1 | 2 | 0.5 | 1 | 1 | 7 | 0.7143 |
| 7_1254149 | NA | NA | NA | NA | 2 | 0.5 | 1 | 0 | 3 | 0.3333 | NA | NA | NA | NA | NA | NA | 1 | 1 | 1 | 1 |
| 7_1254335 | NA | NA | 1 | 1 | 2 | 0 | 1 | 0 | 4 | 0.25 | NA | NA | NA | NA | 1 | 1 | 2 | 0 | 3 | 0.3333 |
| 7_1254365 | NA | NA | 1 | 0 | 2 | 0 | 1 | 1 | 4 | 0.25 | NA | NA | NA | NA | 1 | 0 | 1 | 1 | 2 | 0.5 |

| | | | | | | | | | | | | | | | | | | | | |
|---|---|---|---|---|---|---|---|---|---|---|---|---|---|---|---|---|---|---|---|---|
| 7_1254368 | NA | NA | 1 | 1 | 2 | 1 | 1 | 1 | 4 | 1 | NA | NA | NA | NA | 1 | 1 | 1 | 1 | 2 | 1 |
| 7_1254379 | NA | NA | NA | NA | 2 | 1 | 1 | 1 | 3 | 1 | NA | NA | NA | NA | 1 | 1 | 1 | 1 | 2 | 1 |
| 7_1254390 | NA | NA | NA | NA | 2 | 0.5 | NA | NA | 2 | 0.5 | NA | NA | NA | NA | 1 | 1 | 1 | 0 | 2 | 0.5 |
| 7_1254395 | NA | NA | 1 | 1 | 2 | 1 | NA | NA | 3 | 1 | NA | NA | NA | NA | NA | NA | 1 | 1 | 1 | 1 |
| 7_1254407 | NA | NA | 1 | 1 | 2 | 0.5 | 1 | 1 | 4 | 0.75 | NA | NA | NA | NA | 1 | 1 | 1 | 1 | 2 | 1 |
| 7_1254409 | NA | NA | 1 | 0 | 2 | 0.5 | 1 | 1 | 4 | 0.5 | NA | NA | NA | NA | 1 | 1 | 1 | 1 | 2 | 1 |
| 7_1254425 | NA | NA | 1 | 1 | 2 | 1 | NA | NA | 3 | 1 | NA | NA | NA | NA | 1 | 1 | NA | NA | 1 | 1 |
| 7_1254426 | NA | NA | NA | NA | 1 | 1 | 1 | 1 | 2 | 1 | NA | NA | NA | NA | 1 | 1 | NA | NA | 1 | 1 |
| 7_1254449 | NA | NA | 1 | 1 | 2 | 1 | 1 | 0 | 4 | 0.75 | NA | NA | NA | NA | 1 | 1 | NA | NA | 1 | 1 |
| 7_1254465 | NA | NA | 1 | 1 | 1 | 1 | 1 | 1 | 3 | 1 | NA | NA | NA | NA | 1 | 1 | NA | NA | 1 | 1 |
| 7_1254485 | NA | NA | 1 | 1 | 1 | 0 | NA | NA | 2 | 0.5 | NA | NA | NA | NA | 1 | 1 | NA | NA | 1 | 1 |
| 7_1254491 | NA | NA | 1 | 1 | 2 | 1 | NA | NA | 3 | 1 | NA | NA | NA | NA | 1 | 1 | NA | NA | 1 | 1 |
| 7_1254503 | NA | NA | 1 | 1 | 2 | 0.5 | NA | NA | 3 | 0.6667 | NA | NA | NA | NA | 1 | 1 | NA | NA | 1 | 1 |
| 7_1254504 | NA | NA | 1 | 1 | 1 | 1 | NA | NA | 2 | 1 | NA | NA | NA | NA | 1 | 1 | NA | NA | 1 | 1 |
| 7_1254528 | NA | NA | 1 | 1 | NA | NA | NA | NA | 1 | 1 | NA | NA | NA | NA | 1 | 1 | NA | NA | 1 | 1 |
| 7_1254546 | NA | NA | 1 | 1 | NA | NA | NA | NA | 1 | 1 | NA | NA | NA | NA | 1 | 1 | NA | NA | 1 | 1 |
| 7_1258271 | NA | NA | NA | NA | NA | NA | 2 | 0.5 | 2 | 0.5 | 2 | 1 | 1 | 1 | NA | NA | NA | NA | 3 | 1 |
| 7_1258289 | NA | NA | 1 | 1 | 1 | 1 | 2 | 1 | 4 | 1 | 2 | 1 | 2 | 1 | 1 | 1 | NA | NA | 5 | 1 |
| 7_1258417 | 2 | 0 | 4 | 0.25 | 1 | 1 | 2 | 0 | 9 | 0.2222 | 2 | 0.5 | 4 | 0.25 | 2 | 0 | NA | NA | 8 | 0.25 |
| 7_1258431 | 2 | 0 | 4 | 0.25 | 1 | 1 | 1 | 0 | 8 | 0.25 | 2 | 0.5 | 2 | 0.5 | 1 | 0 | NA | NA | 5 | 0.4 |
| 7_1262409 | NA | NA | 1 | 0 | 2 | 0.5 | NA | NA | 3 | 0.3333 | NA | NA | 1 | 0 | 2 | 0.5 | 1 | 0 | 4 | 0.25 |
| 7_1262415 | NA | NA | 1 | 1 | 1 | 1 | NA | NA | 2 | 1 | NA | NA | 1 | 1 | 2 | 1 | 1 | 1 | 4 | 1 |
| 7_1262423 | NA | NA | 1 | 1 | 2 | 0 | NA | NA | 3 | 0.3333 | NA | NA | 1 | 0 | 2 | 0.5 | 1 | 0 | 4 | 0.25 |
| 7_1262428 | NA | NA | 1 | 1 | 2 | 1 | NA | NA | 3 | 1 | NA | NA | 1 | 1 | 2 | 0.5 | 1 | 1 | 4 | 0.75 |
| 7_1262436 | NA | NA | 1 | 1 | 2 | 0.5 | NA | NA | 3 | 0.6667 | NA | NA | 1 | 1 | 1 | 1 | 1 | 0 | 3 | 0.6667 |
| 7_1262465 | NA | NA | 2 | 1 | 2 | 1 | NA | NA | 4 | 1 | NA | NA | 1 | 1 | 2 | 1 | 1 | 1 | 4 | 1 |
| 7_1262470 | NA | NA | 1 | 1 | NA | NA | NA | NA | 1 | 1 | NA | NA | 1 | 1 | 2 | 0 | 1 | 0 | 4 | 0.25 |
| 7_1262476 | NA | NA | 2 | 0.5 | 1 | 0 | NA | NA | 3 | 0.3333 | NA | NA | 1 | 1 | 2 | 0.5 | 1 | 0 | 4 | 0.5 |
| 7_1262479 | NA | NA | 2 | 1 | 2 | 1 | NA | NA | 4 | 1 | NA | NA | 1 | 1 | 2 | 1 | 1 | 1 | 4 | 1 |
| 7_1262495 | NA | NA | 2 | 1 | 2 | 1 | NA | NA | 4 | 1 | NA | NA | NA | NA | 2 | 1 | 1 | 1 | 3 | 1 |
| 7_1262500 | NA | NA | 2 | 1 | 2 | 1 | NA | NA | 4 | 1 | NA | NA | 1 | 1 | 2 | 1 | 1 | 1 | 4 | 1 |
| 7_1262516 | NA | NA | 2 | 0.5 | 2 | 1 | NA | NA | 4 | 0.75 | NA | NA | 1 | 0 | 2 | 0.5 | 1 | 0 | 4 | 0.25 |
| 7_1262517 | NA | NA | NA | NA | 2 | 0.5 | NA | NA | 2 | 0.5 | NA | NA | 1 | 1 | 2 | 0.5 | 1 | 1 | 4 | 0.75 |
| 7_1262522 | NA | NA | 1 | 1 | 3 | 0 | NA | NA | 4 | 0.25 | NA | NA | 1 | 1 | 2 | 0.5 | 1 | 1 | 4 | 0.75 |
| 7_1262523 | NA | NA | 2 | 0 | 2 | 0.5 | NA | NA | 4 | 0.25 | NA | NA | 2 | 0.5 | 2 | 0.5 | 1 | 0 | 5 | 0.4 |
| 7_1262544 | 1 | 0 | 1 | 1 | 2 | 0 | NA | NA | 4 | 0.25 | NA | NA | 2 | 0 | 2 | 0.5 | NA | NA | 4 | 0.25 |
| 7_1262604 | 1 | 0 | 1 | 1 | 1 | 1 | NA | NA | 3 | 0.6667 | NA | NA | 2 | 0.5 | 1 | 1 | NA | NA | 3 | 0.6667 |
| 7_1262608 | 1 | 1 | 1 | 1 | 1 | 1 | NA | NA | 3 | 1 | NA | NA | 3 | 1 | 1 | 1 | NA | NA | 4 | 1 |
| 7_1262644 | 1 | 1 | 1 | 1 | NA | NA | NA | NA | 2 | 1 | NA | NA | 3 | 0.6667 | 1 | 1 | NA | NA | 4 | 0.75 |
| 7_1264052 | NA | NA | NA | NA | 2 | 0.5 | 1 | 0 | 3 | 0.3333 | NA | NA | NA | NA | 2 | 1 | NA | NA | 2 | 1 |
| 7_1264101 | NA | NA | NA | NA | 2 | 1 | NA | NA | 2 | 1 | NA | NA | NA | NA | 2 | 1 | NA | NA | 2 | 1 |
| 7_1264108 | NA | NA | 1 | 1 | 2 | 1 | 1 | 1 | 4 | 1 | NA | NA | NA | NA | 2 | 1 | NA | NA | 2 | 1 |
| 7_1264116 | NA | NA | 1 | 1 | 2 | 1 | 1 | 1 | 4 | 1 | NA | NA | NA | NA | 2 | 1 | NA | NA | 2 | 1 |
| 7_1264137 | NA | NA | 1 | 1 | 2 | 1 | NA | NA | 3 | 1 | NA | NA | NA | NA | 2 | 1 | NA | NA | 2 | 1 |
| 7_1264146 | NA | NA | NA | NA | 2 | 1 | 1 | 0 | 3 | 0.6667 | NA | NA | NA | NA | 2 | 0.5 | NA | NA | 2 | 0.5 |
| 7_1270572 | NA | NA | 3 | 1 | NA | NA | NA | NA | 3 | 1 | NA | NA | NA | NA | 1 | 1 | NA | NA | 1 | 1 |
| 7_1270574 | NA | NA | 3 | 1 | 1 | 1 | NA | NA | 4 | 1 | NA | NA | NA | NA | 1 | 1 | NA | NA | 1 | 1 |
| 7_1270579 | NA | NA | 2 | 1 | NA | NA | NA | NA | 2 | 1 | NA | NA | NA | NA | 1 | 1 | NA | NA | 1 | 1 |
| 7_1271333 | NA | NA | NA | NA | NA | NA | 1 | 1 | 1 | 1 | NA | NA | NA | NA | 1 | 1 | NA | NA | 1 | 1 |
| 7_1271369 | NA | NA | NA | NA | NA | NA | 2 | 0.5 | 2 | 0.5 | NA | NA | NA | NA | 1 | 1 | NA | NA | 1 | 1 |
| 7_1271578 | NA | NA | NA | NA | NA | NA | 3 | 0.6667 | 3 | 0.6667 | NA | NA | 1 | 1 | NA | NA | NA | NA | 1 | 1 |
| 7_1271612 | NA | NA | NA | NA | NA | NA | 2 | 0.5 | 2 | 0.5 | NA | NA | 1 | 1 | NA | NA | NA | NA | 1 | 1 |
| 7_1271622 | NA | NA | NA | NA | NA | NA | 4 | 1 | 4 | 1 | NA | NA | 1 | 1 | NA | NA | NA | NA | 1 | 1 |
| 7_1271631 | NA | NA | NA | NA | NA | NA | 3 | 1 | 3 | 1 | NA | NA | 1 | 1 | NA | NA | NA | NA | 1 | 1 |
| 7_1271654 | NA | NA | NA | NA | 1 | 1 | 2 | 1 | 3 | 1 | NA | NA | 1 | 1 | NA | NA | NA | NA | 1 | 1 |
| 7_1271740 | NA | NA | NA | NA | 1 | 0 | 2 | 0.5 | 3 | 0.3333 | NA | NA | 1 | 1 | NA | NA | NA | NA | 1 | 1 |
| 7_1273960 | 7 | 0.8571 | 3 | 1 | 4 | 1 | 7 | 0.4286 | 21 | 0.7619 | 3 | 0.6667 | 5 | 1 | 6 | 0.8333 | 5 | 1 | 19 | 0.8947 |
| 7_1273964 | 7 | 0.1429 | 3 | 0 | 5 | 0.2 | 8 | 0 | 23 | 0.087 | 3 | 0 | 5 | 0 | 6 | 0.3333 | 6 | 0 | 20 | 0.1 |
| 7_1273977 | 8 | 0.25 | 3 | 0.3333 | 4 | 0 | 7 | 0.2857 | 22 | 0.2273 | 3 | 0 | 6 | 0.1667 | 6 | 0 | 5 | 0 | 20 | 0.05 |
| 7_1273979 | 8 | 0 | 3 | 0 | 4 | 0 | 7 | 0.2857 | 22 | 0.0909 | 3 | 0 | 5 | 0 | 5 | 0 | 5 | 0.2 | 18 | 0.0556 |
| 7_1273983 | 8 | 0.375 | 3 | 0.3333 | 4 | 0.5 | 7 | 0.7143 | 22 | 0.5 | 2 | 0 | 4 | 0.5 | 7 | 0.2857 | 5 | 0.6 | 18 | 0.3889 |
| 7_1273986 | 8 | 0.875 | 3 | 0.6667 | 5 | 0.8 | 6 | 0.6667 | 22 | 0.7727 | 3 | 0.6667 | 6 | 0.8333 | 5 | 0.4 | 5 | 0.8 | 19 | 0.6842 |
| 7_1273990 | 8 | 0.25 | 3 | 0.3333 | 4 | 0 | 6 | 0.3333 | 21 | 0.2381 | 3 | 0 | 6 | 0.3333 | 7 | 0 | 6 | 0.1667 | 22 | 0.1364 |
| 7_1273991 | 8 | 0.5 | 3 | 0.6667 | 4 | 0.5 | 7 | 0.7143 | 22 | 0.5909 | 3 | 0.3333 | 6 | 0.1667 | 7 | 0.2857 | 5 | 0.4 | 21 | 0.2857 |
| 7_1273992 | 8 | 0.875 | 3 | 1 | 4 | 1 | 7 | 1 | 22 | 0.9545 | NA | NA | 5 | 1 | 7 | 1 | 4 | 1 | 16 | 1 |

| | | | | | | | | | | | | | | | | | | | | |
|---|---|---|---|---|---|---|---|---|---|---|---|---|---|---|---|---|---|---|---|---|
| 7_12739939 | 7 | 0.1429 | 3 | 0 | 2 | 0 | 6 | 0.1667 | 18 | 0.1111 | 2 | 0.5 | 5 | 0 | 6 | 0 | 6 | 0 | 19 | 0.0526 |
| 7_12739962 | 6 | 0.8333 | 3 | 0.6667 | 4 | 1 | 7 | 0.7143 | 20 | 0.8 | 3 | 0 | 5 | 1 | 7 | 1 | 5 | 0.8 | 20 | 0.8 |
| 7_12740004 | 6 | 1 | 3 | 1 | 2 | 1 | 6 | 0.8333 | 17 | 0.9412 | 3 | 0.6667 | 5 | 1 | 6 | 1 | 4 | 1 | 18 | 0.9444 |
| 7_12740010 | 6 | 1 | 2 | 1 | 3 | 1 | 5 | 1 | 16 | 1 | 2 | 1 | 6 | 0.8333 | 6 | 1 | 4 | 1 | 18 | 0.9444 |
| 7_12741278 | 7 | 0.1429 | 2 | 0 | 4 | 0 | 6 | 0.3333 | 19 | 0.1579 | 3 | 0.3333 | 5 | 0.2 | 6 | 0 | 4 | 0.25 | 18 | 0.1667 |
| 7_12741422 | 7 | 0.7143 | 3 | 1 | 3 | 1 | 5 | 0.6 | 18 | 0.7778 | 1 | 1 | 4 | 1 | 5 | 0.6 | 3 | 1 | 13 | 0.8462 |
| 7_12741499 | 5 | 0.2 | 1 | 0 | 2 | 0.5 | 5 | 0 | 13 | 0.1538 | 1 | 0 | 4 | 0.5 | 5 | 0.2 | 4 | 0 | 14 | 0.2143 |
| 7_12742022 | 6 | 0 | 2 | 0 | 3 | 0.3333 | 6 | 0 | 17 | 0.0588 | 2 | 0 | 4 | 0.25 | 2 | 0 | 3 | 0 | 11 | 0.0909 |
| 7_12742074 | 3 | 1 | 2 | 1 | 2 | 1 | 6 | 0.5 | 13 | 0.7692 | 2 | 1 | 3 | 1 | 1 | 0 | 2 | 0.5 | 8 | 0.75 |
| 7_12742216 | 4 | 0.25 | 3 | 0.3333 | 3 | 0.6667 | 5 | 0.2 | 15 | 0.3333 | 1 | 0 | 4 | 0.25 | 3 | 0 | 1 | 0 | 9 | 0.1111 |
| 7_12765947 | 16 | 0.1875 | 14 | 0 | 7 | 0.1429 | 15 | 0.1333 | 52 | 0.1154 | 13 | 0 | 12 | 0.1667 | 28 | 0.2143 | 14 | 0.0714 | 67 | 0.1343 |
| 7_12766006 | 22 | 0.0455 | 22 | 0.0455 | 13 | 0.1538 | 16 | 0.1875 | 73 | 0.0959 | 20 | 0.05 | 15 | 0.1333 | 28 | 0.0357 | 20 | 0.1 | 83 | 0.0723 |
| 7_12766643 | 21 | 0.1905 | 21 | 0 | 13 | 0 | 16 | 0 | 71 | 0.0563 | 15 | 0.0667 | 15 | 0.1333 | 26 | 0.0385 | 13 | 0 | 69 | 0.058 |
| 7_12777092 | NA | NA | 3 | 0.6667 | 2 | 0.5 | 2 | 0 | 7 | 0.4286 | 2 | 0.5 | NA | NA | 2 | 0 | 1 | 0 | 5 | 0.2 |
| 7_12777171 | 1 | 0 | 3 | 0.6667 | 1 | 0 | 2 | 0 | 7 | 0.2857 | 2 | 0 | NA | NA | 2 | 0.5 | 1 | 0 | 5 | 0.2 |
| 7_12777262 | 1 | 1 | 3 | 1 | 1 | 1 | 2 | 1 | 7 | 1 | 2 | 1 | NA | NA | 2 | 1 | 1 | 1 | 5 | 1 |
| 7_12777548 | 1 | 0 | 2 | 1 | NA | NA | 2 | 0.5 | 5 | 0.6 | 1 | 1 | NA | NA | 2 | 1 | 1 | 1 | 4 | 1 |
| 7_12777871 | 1 | 1 | 3 | 0.3333 | NA | NA | 1 | 0 | 5 | 0.4 | 1 | 1 | NA | NA | 1 | 1 | NA | NA | 2 | 1 |
| 7_12778352 | 1 | 1 | 3 | 1 | NA | NA | 1 | 1 | 5 | 1 | 1 | 1 | NA | NA | 1 | 1 | 1 | 1 | 3 | 1 |
| 7_12778453 | 1 | 1 | 3 | 0.3333 | NA | NA | 1 | 0 | 5 | 0.4 | 1 | 0 | NA | NA | 1 | 0 | 1 | 1 | 3 | 0.3333 |
| 7_12778530 | 1 | 1 | 2 | 1 | NA | NA | 1 | 1 | 4 | 1 | 1 | 1 | NA | NA | 1 | 1 | 1 | 1 | 3 | 1 |
| 7_12778609 | 1 | 0 | 2 | 1 | NA | NA | 1 | 0 | 4 | 0.5 | 1 | 1 | NA | NA | NA | NA | 1 | 1 | 2 | 1 |
| 7_12786865 | NA | NA | 2 | 1 | NA | NA | 1 | 1 | 3 | 1 | NA | NA | NA | NA | 1 | 1 | NA | NA | 1 | 1 |
| 7_12993364 | 1 | 1 | NA | NA | NA | NA | NA | NA | 1 | 1 | NA | NA | 1 | 1 | 1 | 1 | NA | NA | 2 | 1 |
| 7_12993561 | 1 | 1 | NA | NA | NA | NA | NA | NA | 1 | 1 | NA | NA | NA | NA | 1 | 1 | NA | NA | 1 | 1 |
| 7_12993645 | 1 | 1 | NA | NA | NA | NA | NA | NA | 1 | 1 | NA | NA | NA | NA | 1 | 1 | NA | NA | 1 | 1 |
| 7_14297125 | NA | NA | NA | NA | 1 | 1 | NA | NA | 1 | 1 | NA | NA | NA | NA | 1 | 1 | NA | NA | 1 | 1 |
| 7_15651412 | 231 | 0.0563 | 238 | 0.0966 | 217 | 0.0829 | 213 | 0.0469 | 899 | 0.0712 | 241 | 0.1037 | 206 | 0.068 | 241 | 0.083 | 193 | 0.0777 | 881 | 0.084 |
| 7_15651565 | NA | NA | 5 | 0 | 3 | 0.3333 | 3 | 0 | 11 | 0.0909 | 1 | 0 | 3 | 0 | 2 | 0 | 5 | 0.2 | 11 | 0.0909 |
| 7_15656539 | NA | NA | NA | NA | NA | NA | 1 | 1 | 1 | 1 | NA | NA | NA | NA | 1 | 1 | NA | NA | 1 | 1 |
| 7_15683172 | 2 | 0.5 | NA | NA | NA | NA | 1 | 1 | 3 | 0.6667 | NA | NA | NA | NA | NA | NA | 2 | 1 | 2 | 1 |
| 7_15683773 | NA | NA | 1 | 1 | NA | NA | 1 | 1 | 2 | 1 | NA | NA | NA | NA | NA | NA | 2 | 0.5 | 2 | 0.5 |
| 7_15683971 | 1 | 1 | 2 | 0.5 | NA | NA | NA | NA | 3 | 0.6667 | NA | NA | NA | NA | NA | NA | 1 | 1 | 1 | 1 |
| 7_15683997 | 2 | 0.5 | 2 | 0.5 | NA | NA | NA | NA | 4 | 0.5 | NA | NA | NA | NA | NA | NA | 2 | 0.5 | 2 | 0.5 |
| 7_15684028 | 2 | 1 | 2 | 1 | NA | NA | 1 | 1 | 5 | 1 | NA | NA | NA | NA | NA | NA | 2 | 1 | 2 | 1 |
| 7_15684452 | 2 | 0.5 | 2 | 1 | NA | NA | NA | NA | 4 | 0.75 | NA | NA | NA | NA | NA | NA | 2 | 1 | 2 | 1 |
| 7_15697886 | 2 | 0.5 | 1 | 1 | NA | NA | NA | NA | 3 | 0.6667 | NA | NA | NA | NA | NA | NA | 2 | 0.5 | 2 | 0.5 |
| 7_15705265 | NA | NA | NA | NA | 2 | 1 | 1 | 1 | 3 | 1 | 3 | 1 | 2 | 0.5 | 1 | 1 | NA | NA | 6 | 0.8333 |
| 7_15705420 | NA | NA | NA | NA | 2 | 1 | 1 | 1 | 3 | 1 | 2 | 1 | 2 | 1 | 1 | 0 | NA | NA | 5 | 0.8 |
| 7_15705710 | NA | NA | NA | NA | 1 | 1 | NA | NA | 1 | 1 | 2 | 1 | 2 | 1 | NA | NA | NA | NA | 4 | 1 |
| 7_15705837 | NA | NA | NA | NA | 2 | 1 | 1 | 1 | 3 | 1 | 3 | 1 | 2 | 1 | 1 | 1 | NA | NA | 6 | 1 |
| 7_15706041 | NA | NA | NA | NA | 2 | 1 | NA | NA | 2 | 1 | 3 | 0.3333 | 2 | 0 | 1 | 0 | NA | NA | 6 | 0.1667 |
| 7_15706084 | NA | NA | NA | NA | 2 | 1 | NA | NA | 2 | 1 | 2 | 1 | 1 | 1 | NA | NA | NA | NA | 3 | 1 |
| 7_15706298 | NA | NA | NA | NA | 2 | 0.5 | NA | NA | 2 | 0.5 | 2 | 0.5 | 2 | 1 | 1 | 0 | NA | NA | 5 | 0.6 |
| 7_15706629 | NA | NA | NA | NA | 2 | 0.5 | NA | NA | 2 | 0.5 | 2 | 0 | 2 | 0.5 | 1 | 0 | NA | NA | 5 | 0.2 |
| 7_15721992 | NA | NA | NA | NA | 1 | 1 | NA | NA | 1 | 1 | 1 | 1 | NA | NA | NA | NA | NA | NA | 1 | 1 |
| 7_15722321 | NA | NA | NA | NA | 1 | 1 | NA | NA | 1 | 1 | 1 | 1 | NA | NA | NA | NA | NA | NA | 1 | 1 |
| 7_15722897 | NA | NA | NA | NA | 1 | 1 | NA | NA | 1 | 1 | 1 | 1 | NA | NA | NA | NA | NA | NA | 1 | 1 |
| 7_15723059 | NA | NA | NA | NA | 1 | 1 | NA | NA | 1 | 1 | 1 | 1 | NA | NA | NA | NA | NA | NA | 1 | 1 |
| 7_15782569 | NA | NA | 1 | 1 | NA | NA | NA | NA | 1 | 1 | 2 | 0.5 | 3 | 1 | 1 | 0 | NA | NA | 6 | 0.6667 |
| 7_15782589 | NA | NA | 2 | 1 | NA | NA | NA | NA | 2 | 1 | 2 | 0.5 | 4 | 0.5 | 1 | 1 | NA | NA | 7 | 0.5714 |
| 7_15782632 | NA | NA | 2 | 0.5 | NA | NA | NA | NA | 2 | 0.5 | 2 | 1 | 4 | 0.75 | NA | NA | NA | NA | 6 | 0.8333 |
| 7_15782695 | NA | NA | 2 | 1 | NA | NA | NA | NA | 2 | 1 | 2 | 1 | 4 | 1 | 2 | 1 | NA | NA | 8 | 1 |
| 7_15782879 | NA | NA | 2 | 1 | NA | NA | NA | NA | 2 | 1 | 1 | 1 | 4 | 0.75 | 2 | 1 | NA | NA | 7 | 0.8571 |
| 7_15850700 | 41 | 0.4878 | 42 | 0.381 | 39 | 0.3333 | 44 | 0.2955 | 166 | 0.3735 | 32 | 0.3438 | 36 | 0.1944 | 32 | 0.4688 | 30 | 0.4 | 130 | 0.3462 |
| 7_15850718 | 39 | 0.1795 | 45 | 0.3333 | 41 | 0.2439 | 48 | 0.3333 | 173 | 0.2775 | 32 | 0.2188 | 41 | 0.2195 | 35 | 0.1429 | 34 | 0.3529 | 142 | 0.2324 |
| 7_15850828 | 34 | 0.0882 | 42 | 0.0952 | 41 | 0.0732 | 43 | 0.093 | 160 | 0.0875 | 30 | 0.1 | 34 | 0.0882 | 32 | 0.0938 | 34 | 0.1765 | 130 | 0.1154 |
| 7_15850845 | 42 | 0.2381 | 51 | 0.3137 | 44 | 0.25 | 44 | 0.2045 | 181 | 0.2541 | 34 | 0.2059 | 41 | 0.2195 | 34 | 0.2059 | 37 | 0.2162 | 146 | 0.2123 |
| 7_15851027 | 38 | 0.8158 | 50 | 0.88 | 44 | 0.7273 | 44 | 0.7727 | 176 | 0.8011 | 37 | 0.6757 | 42 | 0.8571 | 36 | 0.6389 | 36 | 0.7778 | 151 | 0.7417 |
| 7_15851098 | 7 | 0.8571 | 7 | 0.2857 | 7 | 0.2857 | 7 | 0.5714 | 28 | 0.5 | 4 | 0 | 10 | 0.5 | 3 | 0 | 5 | 0.2 | 22 | 0.2727 |
| 7_15851173 | 7 | 1 | 7 | 0.7143 | 7 | 1 | 7 | 0.4286 | 28 | 0.7857 | 4 | 0.75 | 10 | 0.9 | 2 | 0.5 | 3 | 0.3333 | 19 | 0.7368 |
| 7_15851259 | 7 | 0 | 10 | 0.2 | 8 | 0.125 | 8 | 0 | 33 | 0.0909 | 4 | 0.25 | 10 | 0.2 | 3 | 0 | 7 | 0.2857 | 24 | 0.2083 |
| 7_15851532 | 9 | 0 | 6 | 0 | 5 | 0.2 | 6 | 0.1667 | 26 | 0.0769 | 5 | 0 | 10 | 0 | 3 | 0 | 7 | 0.2857 | 25 | 0.08 |
| 7_15851623 | 8 | 0.125 | 10 | 0.4 | 8 | 0.25 | 7 | 0.2857 | 33 | 0.2727 | 5 | 0.2 | 14 | 0.4286 | 3 | 0 | 8 | 0.5 | 30 | 0.3667 |
| 7_15851647 | 10 | 0 | 11 | 0.1818 | 8 | 0.375 | 8 | 0.125 | 37 | 0.1622 | 5 | 0.4 | 14 | 0.5 | 3 | 0.3333 | 9 | 0.1111 | 31 | 0.3548 |

| | | | | | | | | | | | | | | | | | | | | |
|---|---|---|---|---|---|---|---|---|---|---|---|---|---|---|---|---|---|---|---|---|
| 7_15851681 | 10 | 0.1 | 11 | 0.0909 | 7 | 0 | 8 | 0.125 | 36 | 0.0833 | 4 | 0.25 | 13 | 0.0769 | 2 | 0.5 | 8 | 0.25 | 27 | 0.1852 |
| 7_15851701 | 7 | 0.1429 | 7 | 0 | 5 | 0.4 | 6 | 0.5 | 25 | 0.24 | 3 | 0 | 9 | 0.2222 | 1 | 0 | 5 | 0.2 | 18 | 0.1667 |
| 7_15851781 | 9 | 0.5556 | 11 | 0.7273 | 7 | 0.4286 | 9 | 0.5556 | 36 | 0.5833 | 5 | 0.6 | 13 | 0.7692 | 3 | 1 | 7 | 0.4286 | 28 | 0.6786 |
| 7_15851806 | 10 | 0 | 10 | 0 | 7 | 0.2857 | 9 | 0.1111 | 36 | 0.0833 | 5 | 0 | 12 | 0.25 | 3 | 0 | 7 | 0 | 27 | 0.1111 |
| 7_15861256 | 2 | 0.5 | 4 | 0.25 | 2 | 1 | 4 | 0.25 | 12 | 0.4167 | 1 | 0 | 5 | 0.4 | 5 | 0.2 | 2 | 0.5 | 13 | 0.3077 |
| 7_15867873 | 4 | 0 | 3 | 0.3333 | 5 | 0 | 4 | 0 | 16 | 0.0625 | 5 | 0 | 1 | 0 | 7 | 0.1429 | 1 | 0 | 14 | 0.0714 |
| 7_15867954 | 5 | 0.2 | 3 | 0 | 6 | 0.6667 | 3 | 0 | 17 | 0.2941 | 5 | 0.4 | 3 | 0.6667 | 7 | 0 | 1 | 0 | 16 | 0.25 |
| 7_15868205 | 6 | 1 | 3 | 1 | 6 | 0.8333 | 6 | 1 | 21 | 0.9524 | 6 | 0.8333 | 3 | 1 | 7 | 0.7143 | 1 | 1 | 17 | 0.8235 |
| 7_15874674 | 5 | 0.6 | 3 | 0.3333 | 5 | 0.2 | 6 | 0 | 19 | 0.2632 | 5 | 0.4 | 3 | 0.3333 | 9 | 0.6667 | 2 | 0.5 | 19 | 0.5263 |
| 7_15874715 | 7 | 0.8571 | 4 | 0.75 | 7 | 0.8571 | 7 | 0.5714 | 25 | 0.76 | 7 | 0.7143 | 4 | 0.75 | 10 | 0.7 | 2 | 1 | 23 | 0.7391 |
| 7_15874742 | 6 | 0.3333 | 4 | 0 | 6 | 0.3333 | 7 | 0.4286 | 23 | 0.3043 | 6 | 0.5 | 4 | 0.5 | 10 | 0.6 | 2 | 0 | 22 | 0.5 |
| 7_15874837 | 6 | 0.1667 | 6 | 0 | 7 | 0.2857 | 3 | 0 | 22 | 0.1364 | 6 | 0.1667 | 4 | 0 | 9 | 0.1111 | 1 | 0 | 20 | 0.1 |
| 7_15874976 | 6 | 0 | 4 | 0 | 6 | 0.3333 | 6 | 0 | 22 | 0.0909 | 6 | 0.5 | 6 | 0.1667 | 9 | 0.3333 | 2 | 0 | 23 | 0.3043 |
| 7_15875042 | 6 | 0 | 6 | 0.1667 | 7 | 0.2857 | 9 | 0 | 28 | 0.1071 | 4 | 0.25 | 5 | 0 | 7 | 0.1429 | NA | NA | 16 | 0.125 |
| 7_15878966 | 7 | 0 | 8 | 0.125 | 8 | 0 | 10 | 0.2 | 33 | 0.0909 | 7 | 0.1429 | 6 | 0.1667 | 9 | 0.2222 | 3 | 0 | 25 | 0.16 |
| 7_15879101 | 5 | 0.2 | 8 | 0.125 | 7 | 0.2857 | 9 | 0.2222 | 29 | 0.2069 | 8 | 0.25 | 8 | 0.25 | 10 | 0.1 | 3 | 0 | 29 | 0.1724 |
| 7_15891303 | 10 | 0.5 | 12 | 0.5833 | 9 | 0.5556 | 11 | 0.6364 | 42 | 0.5714 | 8 | 0.75 | 12 | 0.5 | 12 | 0.4167 | 4 | 0 | 36 | 0.4722 |
| 7_15893009 | 14 | 0.1429 | 15 | 0.0667 | 10 | 0.2 | 11 | 0.0909 | 50 | 0.12 | 8 | 0.125 | 12 | 0.3333 | 13 | 0.2308 | 6 | 0.1667 | 39 | 0.2308 |
| 7_15893134 | 17 | 0.4118 | 14 | 0.5 | 12 | 0.0833 | 11 | 0.3636 | 54 | 0.3519 | 8 | 0.375 | 13 | 0.5385 | 12 | 0.4167 | 7 | 0.4286 | 40 | 0.45 |
| 7_15893156 | 13 | 0.3077 | 13 | 0.5385 | 11 | 0.5455 | 12 | 0.6667 | 49 | 0.5102 | 7 | 0.5714 | 13 | 0.4615 | 14 | 0.4286 | 6 | 0.1667 | 40 | 0.425 |
| 7_15893178 | 14 | 0 | 13 | 0.1538 | 14 | 0.0714 | 9 | 0.1111 | 50 | 0.08 | 7 | 0.1429 | 12 | 0.3333 | 15 | 0 | 5 | 0.2 | 39 | 0.1538 |
| 7_15893401 | 14 | 0.4286 | 9 | 0.4444 | 8 | 0.25 | 7 | 0.2857 | 38 | 0.3684 | 6 | 0.1667 | 10 | 0.3 | 12 | 0.5 | 4 | 0.25 | 32 | 0.3438 |
| 7_15893828 | 18 | 0.1111 | 14 | 0.0714 | 15 | 0.0667 | 11 | 0.0909 | 58 | 0.0862 | 10 | 0 | 16 | 0.3125 | 14 | 0.4286 | 9 | 0 | 49 | 0.2245 |
| 7_15893854 | 18 | 0.3333 | 20 | 0.3 | 16 | 0.25 | 13 | 0.3846 | 67 | 0.3134 | 11 | 0.1818 | 19 | 0.3158 | 19 | 0.2632 | 11 | 0.1818 | 60 | 0.25 |
| 7_15893889 | 19 | 0.1579 | 19 | 0.3158 | 13 | 0.2308 | 12 | 0.1667 | 63 | 0.2222 | 9 | 0.3333 | 14 | 0.2857 | 15 | 0.1333 | 9 | 0.1111 | 47 | 0.2128 |
| 7_15893895 | 18 | 0.1667 | 15 | 0.0667 | 15 | 0.2 | 11 | 0 | 59 | 0.1186 | 13 | 0.0769 | 17 | 0.0588 | 16 | 0.0625 | 11 | 0.1818 | 57 | 0.0877 |
| 7_15894037 | 19 | 0.1053 | 20 | 0.25 | 18 | 0.2778 | 15 | 0.1333 | 72 | 0.1944 | 16 | 0.1875 | 19 | 0.1579 | 19 | 0.2105 | 11 | 0.2727 | 65 | 0.2 |
| 7_15894915 | 5 | 0.6 | 5 | 0.8 | 2 | 0.5 | 2 | 1 | 14 | 0.7143 | 4 | 0.75 | 5 | 0.8 | 6 | 0.5 | 2 | 1 | 17 | 0.7059 |
| 7_15920054 | 5 | 0 | 6 | 0 | 8 | 0.125 | 6 | 0.1667 | 25 | 0.08 | 1 | 0 | 5 | 0 | 8 | 0.125 | 5 | 0.2 | 19 | 0.1053 |
| 7_15920276 | 5 | 0 | 6 | 0.5 | 9 | 0.4444 | 5 | 0.4 | 25 | 0.36 | 1 | 0 | 5 | 0.2 | 9 | 0.2222 | 3 | 0.3333 | 18 | 0.2222 |
| 7_16006105 | 6 | 0.3333 | 9 | 0.1111 | 11 | 0 | 5 | 0 | 31 | 0.0968 | 6 | 0.1667 | 7 | 0 | 8 | 0 | 6 | 0.1667 | 27 | 0.0741 |
| 7_16006117 | 4 | 0.5 | 9 | 0.4444 | 10 | 0.2 | 4 | 0.25 | 27 | 0.3333 | 6 | 0 | 5 | 0.4 | 7 | 0.5714 | 5 | 0.2 | 23 | 0.3043 |
| 7_16006210 | 3 | 0 | 7 | 0.2857 | 7 | 0.2857 | 4 | 0.75 | 21 | 0.3333 | 4 | 0.5 | 3 | 0.3333 | 6 | 0.1667 | 6 | 0.6667 | 19 | 0.4211 |
| 7_16006268 | 5 | 0.8 | 11 | 0.6364 | 10 | 0.5 | 4 | 0.75 | 30 | 0.6333 | 6 | 1 | 3 | 1 | 6 | 0.8333 | 5 | 0.6 | 20 | 0.85 |
| 7_16006301 | 5 | 0 | 10 | 0.1 | 11 | 0.1818 | 4 | 0.25 | 30 | 0.1333 | 5 | 0.2 | 4 | 0.5 | 8 | 0.25 | 5 | 0.4 | 22 | 0.3182 |
| 7_16006426 | 3 | 0.3333 | 9 | 0.1111 | 9 | 0.3333 | 2 | 0.5 | 23 | 0.2609 | 4 | 0 | 4 | 0.5 | 8 | 0.125 | 4 | 0 | 20 | 0.15 |
| 7_16006445 | 3 | 0.3333 | 10 | 0.2 | 11 | 0 | 2 | 0 | 26 | 0.1154 | 6 | 0 | 3 | 0 | 8 | 0.25 | 3 | 0.3333 | 20 | 0.15 |
| 7_16006671 | 3 | 0.6667 | 10 | 0.5 | 10 | 0.7 | 1 | 1 | 24 | 0.625 | 4 | 0.75 | 2 | 0.5 | 7 | 0.4286 | 2 | 1 | 15 | 0.6 |
| 7_16006693 | 3 | 0.6667 | 8 | 0.875 | 11 | 0.8182 | 2 | 1 | 24 | 0.8333 | 2 | 1 | 3 | 0.6667 | 9 | 0.8889 | 2 | 1 | 16 | 0.875 |
| 7_16006901 | 3 | 0.3333 | 8 | 0.75 | 9 | 0.3333 | 1 | 0 | 21 | 0.4762 | 3 | 0.6667 | 3 | 1 | 7 | 0.7143 | 3 | 0.3333 | 16 | 0.6875 |
| 7_16007959 | NA | NA | NA | NA | 1 | 1 | NA | NA | 1 | 1 | NA | NA | NA | NA | 1 | 1 | NA | NA | 1 | 1 |
| 7_16009649 | 1 | 1 | NA | NA | 2 | 1 | 2 | 1 | 5 | 1 | 2 | 0.5 | 2 | 0.5 | 3 | 0.6667 | 1 | 1 | 8 | 0.625 |
| 7_16009715 | 1 | 0 | NA | NA | 3 | 0 | 2 | 0.5 | 6 | 0.1667 | 1 | 0 | 4 | 0.25 | 2 | 0.5 | NA | NA | 7 | 0.2857 |
| 7_16051108 | 1 | 0 | NA | NA | NA | NA | 1 | 1 | 2 | 0.5 | 2 | 0 | NA | NA | 1 | 1 | 2 | 1 | 5 | 0.6 |
| 7_16051242 | 2 | 0 | NA | NA | NA | NA | 3 | 0.6667 | 5 | 0.4 | 2 | 1 | NA | NA | 1 | 0 | 2 | 1 | 5 | 0.8 |
| 7_16051270 | 1 | 1 | NA | NA | NA | NA | 3 | 1 | 4 | 1 | 2 | 1 | NA | NA | 1 | 1 | 2 | 1 | 5 | 1 |
| 7_16051352 | NA | NA | NA | NA | NA | NA | 1 | 1 | 1 | 1 | 1 | 1 | NA | NA | NA | NA | 2 | 1 | 3 | 1 |
| 7_16051402 | 2 | 1 | NA | NA | NA | NA | 3 | 1 | 5 | 1 | 2 | 1 | NA | NA | NA | NA | 1 | 1 | 3 | 1 |
| 7_16051762 | 2 | 0.5 | NA | NA | NA | NA | NA | NA | 2 | 0.5 | 2 | 0 | 1 | 0 | 1 | 0 | 2 | 0.5 | 6 | 0.1667 |
| 7_16061398 | 10 | 0.4 | 10 | 0.3 | 9 | 0.5556 | 9 | 0.7778 | 38 | 0.5 | 10 | 0.3 | 13 | 0.3846 | 9 | 0.6667 | 15 | 0.4667 | 47 | 0.4468 |
| 7_16061462 | 9 | 0.7778 | 10 | 0.2 | 7 | 0.4286 | 9 | 0.3333 | 35 | 0.4286 | 8 | 0.625 | 10 | 0.6 | 6 | 0.3333 | 12 | 0.5 | 36 | 0.5278 |
| 7_16061519 | 8 | 0.875 | 8 | 0.875 | 7 | 0.7143 | 9 | 1 | 32 | 0.875 | 7 | 0.7143 | 9 | 0.7778 | 8 | 1 | 13 | 0.7692 | 37 | 0.8108 |
| 7_16061585 | 6 | 0.3333 | 8 | 0.125 | 7 | 0 | 7 | 0.4286 | 28 | 0.2143 | 8 | 0.25 | 11 | 0.7273 | 7 | 0.4286 | 12 | 0.5 | 38 | 0.5 |
| 7_16061653 | 10 | 0.3 | 10 | 0.5 | 9 | 0.4444 | 8 | 0.25 | 37 | 0.3784 | 8 | 0.75 | 12 | 0.5 | 8 | 0.25 | 13 | 0.5385 | 41 | 0.5122 |
| 7_16168173 | 1 | 1 | 3 | 1 | 2 | 1 | 3 | 0.6667 | 9 | 0.8889 | NA | NA | NA | NA | 1 | 1 | NA | NA | 1 | 1 |
| 7_16168198 | 1 | 1 | 4 | 0.5 | 3 | 1 | 1 | 0 | 9 | 0.6667 | 2 | 1 | 1 | 1 | 1 | 1 | 1 | 1 | 5 | 1 |
| 7_16168298 | NA | NA | 4 | 0.75 | 3 | 0.6667 | 1 | 1 | 8 | 0.75 | 2 | 0.5 | NA | NA | 1 | 1 | NA | NA | 3 | 0.6667 |
| 7_16168354 | 1 | 1 | 4 | 0.75 | 3 | 0 | 3 | 1 | 11 | 0.6364 | 1 | 1 | 1 | 1 | NA | NA | 1 | 1 | 3 | 1 |
| 7_16168366 | 1 | 1 | 1 | 1 | 2 | 0 | 3 | 0 | 7 | 0.2857 | 2 | 0.5 | 1 | 0 | 1 | 1 | 1 | 1 | 5 | 0.6 |
| 7_16168379 | 1 | 0 | 4 | 0.75 | 3 | 0 | 3 | 0.3333 | 11 | 0.3636 | 2 | 0.5 | 1 | 0 | 1 | 0 | 1 | 0 | 5 | 0.2 |
| 7_16168489 | 1 | 0 | 3 | 0.3333 | 3 | 0.3333 | 3 | 0 | 10 | 0.2 | 2 | 0.5 | 1 | 0 | 1 | 1 | 1 | 1 | 5 | 0.6 |
| 7_16168660 | 1 | 1 | 4 | 0.5 | 3 | 0.3333 | 2 | 0 | 10 | 0.4 | 2 | 1 | 1 | 0 | 1 | 1 | 1 | 0 | 5 | 0.6 |
| 7_16168667 | 1 | 0 | 4 | 0.75 | 3 | 1 | 2 | 1 | 10 | 0.8 | 2 | 0.5 | NA | NA | 1 | 0 | 1 | 1 | 4 | 0.5 |
| 7_16168972 | 1 | 0 | 7 | 0.1429 | 3 | 0 | 3 | 0 | 14 | 0.0714 | 2 | 0.5 | 1 | 0 | 1 | 0 | 2 | 0 | 6 | 0.1667 |
| 7_16168980 | 1 | 1 | 5 | 1 | 3 | 1 | 3 | 1 | 12 | 1 | 2 | 1 | 1 | 1 | 1 | 1 | 2 | 1 | 6 | 1 |

| | | | | | | | | | | | | | | | | | | | | |
|---|---|---|---|---|---|---|---|---|---|---|---|---|---|---|---|---|---|---|---|---|
| 7_16169001 | 1 | 0 | 7 | 0.4286 | 3 | 0 | 2 | 0.5 | 13 | 0.3077 | 2 | 0 | 1 | 0 | NA | NA | 2 | 0.5 | 5 | 0.2 |
| 7_16169124 | 1 | 1 | 5 | 0.6 | 1 | 1 | 3 | 0.6667 | 10 | 0.7 | 2 | 0 | 1 | 0 | 1 | 0 | 2 | 0.5 | 6 | 0.1667 |
| 7_16169161 | 1 | 1 | 5 | 1 | 2 | 1 | 2 | 1 | 10 | 1 | 2 | 1 | NA | NA | 1 | 1 | 2 | 0.5 | 5 | 0.8 |
| 7_16169295 | 2 | 1 | 6 | 1 | 3 | 1 | 4 | 1 | 15 | 1 | 2 | 1 | 1 | 1 | 1 | 1 | 2 | 1 | 6 | 1 |
| 7_16169717 | 1 | 1 | 8 | 0.5 | 2 | 0.5 | 5 | 0.8 | 16 | 0.625 | 1 | 0 | 1 | 1 | 1 | 1 | 1 | 1 | 4 | 0.75 |
| 7_16169728 | 1 | 1 | 4 | 1 | 2 | 0 | 3 | 0.3333 | 10 | 0.6 | 2 | 1 | NA | NA | 1 | 1 | 1 | 1 | 4 | 1 |
| 7_16170058 | 3 | 0.6667 | 7 | 0.2857 | 3 | 0.6667 | 4 | 1 | 17 | 0.5882 | 1 | 1 | 1 | 0 | 1 | 1 | 2 | 1 | 5 | 0.8 |
| 7_16170073 | 2 | 0.5 | 7 | 0 | 3 | 0.3333 | 4 | 0.5 | 16 | 0.25 | 2 | 0 | 1 | 0 | 1 | 0 | 2 | 0.5 | 6 | 0.1667 |
| 7_16170105 | 2 | 0.5 | 5 | 1 | 2 | 0 | 4 | 0.5 | 13 | 0.6154 | 2 | 0.5 | 1 | 0 | 1 | 0 | 1 | 1 | 5 | 0.4 |
| 7_16170576 | 4 | 0 | 8 | 0.125 | 2 | 0 | 5 | 0 | 19 | 0.0526 | 2 | 0.5 | NA | NA | 1 | 0 | 2 | 0 | 5 | 0.2 |
| 7_16171307 | 2 | 0 | 8 | 0.25 | 3 | 0.6667 | 3 | 0 | 16 | 0.25 | 1 | 0 | 1 | 0 | 3 | 0 | 2 | 0.5 | 7 | 0.1429 |
| 7_16171337 | 4 | 0.75 | 10 | 0.5 | 3 | 0.6667 | 6 | 0.1667 | 23 | 0.4783 | 1 | 0 | 1 | 0 | 2 | 0 | 1 | 1 | 5 | 0.2 |
| 7_16189267 | 20 | 0.2 | 17 | 0.2941 | 14 | 0.3571 | 12 | 0.1667 | 63 | 0.254 | 12 | 0.1667 | 14 | 0.5 | 12 | 0.0833 | 10 | 0.1 | 48 | 0.2292 |
| 7_16189369 | 16 | 0.125 | 12 | 0 | 12 | 0.3333 | 10 | 0 | 50 | 0.12 | 11 | 0.0909 | 10 | 0.3 | 8 | 0 | 8 | 0.25 | 37 | 0.1622 |
| 7_16189392 | 19 | 0.2632 | 16 | 0.25 | 12 | 0.3333 | 11 | 0.2727 | 58 | 0.2759 | 15 | 0.0667 | 12 | 0.4167 | 11 | 0.0909 | 10 | 0.2 | 48 | 0.1875 |
| 7_16189423 | NA | NA | 1 | 0 | NA | NA | 1 | 1 | 2 | 0.5 | NA | NA | 1 | 1 | NA | NA | NA | NA | 1 | 1 |
| 7_16228481 | 3 | 0 | 3 | 0 | 5 | 0.4 | 2 | 0 | 13 | 0.1538 | 6 | 0.1667 | 4 | 0 | 7 | 0.1429 | 6 | 0 | 23 | 0.087 |
| 7_16228724 | 2 | 0 | 4 | 0 | 3 | 0 | 2 | 0.5 | 11 | 0.0909 | 5 | 0 | 1 | 1 | 6 | 0 | 5 | 0 | 17 | 0.0588 |
| 7_16230303 | 4 | 0 | 9 | 0.2222 | 11 | 0 | 9 | 0.1111 | 33 | 0.0909 | 9 | 0 | 9 | 0.2222 | 12 | 0.0833 | 11 | 0.1818 | 41 | 0.122 |
| 7_16230538 | 3 | 0 | 9 | 0.1111 | 12 | 0.1667 | 9 | 0.1111 | 33 | 0.1212 | 12 | 0 | 10 | 0 | 13 | 0.1538 | 12 | 0.1667 | 47 | 0.0851 |
| 7_16230822 | 1 | 0 | 8 | 0.5 | 9 | 0.5556 | 9 | 0.4444 | 27 | 0.4815 | 12 | 0.5 | 8 | 0.5 | 7 | 0.7143 | 12 | 0.3333 | 39 | 0.4872 |
| 7_16230930 | 3 | 0 | 9 | 0.1111 | 10 | 0.2 | 8 | 0.25 | 30 | 0.1667 | 11 | 0.1818 | 8 | 0.5 | 11 | 0.2727 | 10 | 0.2 | 40 | 0.275 |
| 7_16230963 | 2 | 0.5 | 6 | 0 | 7 | 0.2857 | 8 | 0 | 23 | 0.1304 | 9 | 0.1111 | 7 | 0.2857 | 11 | 0 | 10 | 0 | 37 | 0.0811 |
| 7_16230992 | 3 | 0 | 9 | 0.3333 | 10 | 0.4 | 9 | 0.3333 | 31 | 0.3226 | 12 | 0.4167 | 9 | 0.6667 | 10 | 0.2 | 8 | 0.25 | 39 | 0.3846 |
| 7_16231085 | 2 | 0.5 | 6 | 0.5 | 7 | 0.8571 | 8 | 0.625 | 23 | 0.6522 | 10 | 0.5 | 8 | 0.5 | 9 | 0.5556 | 10 | 0.4 | 37 | 0.4865 |
| 7_16231117 | 1 | 0 | 8 | 0.25 | 10 | 0.3 | 8 | 0 | 27 | 0.1852 | 10 | 0.2 | 6 | 0.3333 | 10 | 0.2 | 9 | 0 | 35 | 0.1714 |
| 7_16231179 | 2 | 1 | 7 | 0.8571 | 9 | 0.8889 | 8 | 0.75 | 26 | 0.8462 | 9 | 1 | 6 | 1 | 9 | 0.7778 | 9 | 0.7778 | 33 | 0.8788 |
| 7_16231243 | 1 | 1 | 7 | 0.7143 | 7 | 0.4286 | 8 | 0.625 | 23 | 0.6087 | 9 | 0.7778 | 7 | 1 | 7 | 0.8571 | 10 | 0.5 | 33 | 0.7576 |
| 7_16231295 | 1 | 0 | 7 | 0.4286 | 10 | 0.4 | 8 | 0.25 | 26 | 0.3462 | 9 | 0.3333 | 7 | 0.2857 | 9 | 0.2222 | 10 | 0.3 | 35 | 0.2857 |
| 7_16231336 | 1 | 0 | 5 | 1 | 5 | 1 | 5 | 1 | 16 | 0.9375 | 6 | 1 | 1 | 1 | 5 | 0.8 | 5 | 1 | 17 | 0.9412 |
| 7_16231341 | 1 | 1 | 6 | 1 | 6 | 1 | 7 | 1 | 20 | 1 | 7 | 1 | 5 | 0.8 | 8 | 1 | 7 | 1 | 27 | 0.963 |
| 7_16231463 | 1 | 1 | 4 | 1 | 5 | 1 | 5 | 1 | 15 | 1 | 3 | 1 | 7 | 1 | 5 | 1 | 5 | 1 | 20 | 1 |
| 7_16231513 | 1 | 0 | 2 | 0 | 9 | 0.3333 | 7 | 0.4286 | 19 | 0.3158 | 4 | 0.25 | 6 | 0.6667 | 9 | 0.2222 | 8 | 0.25 | 27 | 0.3333 |
| 7_16231590 | NA | NA | 3 | 0 | 6 | 0.1667 | 4 | 1 | 13 | 0.3846 | 4 | 0.25 | 3 | 1 | 5 | 0.6 | 6 | 0.5 | 18 | 0.5556 |
| 7_16295462 | 1 | 1 | 5 | 0.8 | 9 | 0.5556 | 5 | 0.6 | 20 | 0.65 | 4 | 0.75 | 5 | 0.8 | 6 | 0.5 | 8 | 0.625 | 23 | 0.6522 |
| 7_16295489 | NA | NA | 5 | 0 | 9 | 0.1111 | 5 | 0 | 19 | 0.0526 | 3 | 0.3333 | 7 | 0 | 5 | 0 | 6 | 0.3333 | 21 | 0.1429 |
| 7_16295512 | 1 | 0 | 5 | 0 | 8 | 0.25 | 4 | 0.5 | 18 | 0.2222 | 4 | 0.25 | 7 | 0.2857 | 6 | 0.1667 | 7 | 0.2857 | 24 | 0.25 |
| 7_16295586 | 1 | 1 | 5 | 0.2 | 8 | 0.25 | 4 | 0.25 | 18 | 0.2778 | 4 | 0 | 7 | 0.2857 | 4 | 0 | 6 | 0.3333 | 21 | 0.1905 |
| 7_16295648 | 1 | 1 | 5 | 1 | 8 | 1 | 5 | 1 | 19 | 1 | 4 | 1 | 4 | 0.75 | 5 | 0.8 | 6 | 0.8333 | 19 | 0.8421 |
| 7_16295660 | 1 | 1 | 4 | 0 | 6 | 0.3333 | 1 | 0 | 12 | 0.25 | 2 | 0.5 | 7 | 0 | 6 | 0.1667 | 5 | 0 | 20 | 0.1 |
| 7_16295702 | 1 | 0 | 5 | 0.2 | 7 | 0.1429 | 5 | 0 | 18 | 0.1111 | 2 | 0 | 5 | 0.2 | 2 | 0.5 | 4 | 0.25 | 13 | 0.2308 |
| 7_16295866 | NA | NA | 4 | 0.25 | 8 | 0.5 | 3 | 0.3333 | 15 | 0.4 | 2 | 0 | 5 | 0.2 | 3 | 0.3333 | 6 | 0.3333 | 16 | 0.25 |
| 7_16295960 | 1 | 0 | 4 | 0.25 | 8 | 0 | 4 | 0.25 | 17 | 0.1176 | 2 | 0 | 5 | 0.2 | 3 | 0 | 6 | 0 | 16 | 0.0625 |
| 7_16295965 | NA | NA | 4 | 0.25 | 8 | 0 | 3 | 0 | 15 | 0.0667 | 2 | 0 | 5 | 0 | 4 | 0.25 | 5 | 0.2 | 16 | 0.125 |
| 7_16296016 | NA | NA | 4 | 0.5 | 8 | 0.5 | 3 | 0.6667 | 15 | 0.5333 | 2 | 0 | 5 | 0.8 | 4 | 0.75 | 6 | 0.3333 | 17 | 0.5294 |
| 7_16296113 | 1 | 0 | 5 | 0.4 | 8 | 0.625 | 3 | 0.6667 | 17 | 0.5294 | 2 | 0 | 6 | 0.8333 | 5 | 0.4 | 5 | 0.4 | 18 | 0.5 |
| 7_16296227 | 1 | 1 | 4 | 0 | 5 | 0 | 3 | 0.6667 | 13 | 0.2308 | 2 | 0.5 | 6 | 0.3333 | 4 | 0 | 3 | 0.3333 | 15 | 0.2667 |
| 7_16296274 | 1 | 0 | 3 | 0 | 6 | 0.3333 | 4 | 0 | 14 | 0.1429 | 2 | 0.5 | 6 | 0.3333 | 5 | 0 | 2 | 0 | 15 | 0.2 |
| 7_16296368 | 1 | 0 | 3 | 0 | 6 | 0.5 | 4 | 0.25 | 14 | 0.2857 | 2 | 0.5 | 6 | 0.1667 | 5 | 0.2 | 2 | 0.5 | 15 | 0.2667 |
| 7_16296374 | 1 | 0 | 3 | 0 | 7 | 0.1429 | 4 | 0.25 | 15 | 0.1333 | 2 | 0 | 6 | 0 | 5 | 0.2 | 3 | 0 | 16 | 0.0625 |
| 7_16296583 | 1 | 1 | 2 | 0.5 | 4 | 0.75 | 3 | 1 | 10 | 0.8 | 1 | 1 | 6 | 0.8333 | 4 | 0.5 | 2 | 1 | 13 | 0.7692 |
| 7_16296595 | NA | NA | 2 | 0.5 | 6 | 0.5 | 2 | 0.5 | 10 | 0.5 | 2 | 0 | 4 | 0 | 5 | 0.4 | 2 | 0 | 13 | 0.1538 |
| 7_16296722 | 1 | 0 | 3 | 0.3333 | 6 | 0.6667 | 3 | 0.6667 | 13 | 0.5385 | 2 | 1 | 6 | 0.5 | 5 | 0.4 | 2 | 0.5 | 15 | 0.5333 |
| 7_16296785 | 1 | 0 | 2 | 0 | 5 | 0.4 | 2 | 0.5 | 10 | 0.3 | 2 | 0 | 6 | 0.1667 | 4 | 0.5 | 1 | 0 | 13 | 0.2308 |
| 7_16296793 | 1 | 0 | 3 | 0.3333 | 5 | 0.6 | 3 | 0.6667 | 12 | 0.5 | 2 | 0.5 | 5 | 0.4 | 5 | 0.2 | 1 | 0 | 13 | 0.3077 |
| 7_16296927 | 1 | 0 | 3 | 0.3333 | 6 | 0.1667 | 3 | 0 | 13 | 0.1538 | 2 | 1 | 6 | 0.5 | 5 | 0.2 | 2 | 0.5 | 15 | 0.4667 |
| 7_16296941 | 1 | 0 | 3 | 0 | 5 | 0.2 | 3 | 0 | 12 | 0.0833 | 2 | 0 | 6 | 0.1667 | 4 | 0.25 | 1 | 0 | 13 | 0.1538 |
| 7_16297118 | 1 | 1 | 3 | 0.6667 | 6 | 0.6667 | 3 | 0.6667 | 13 | 0.6923 | 1 | 1 | 6 | 0.8333 | 4 | 0.5 | 2 | 1 | 13 | 0.7692 |
| 7_16297301 | 1 | 0 | 3 | 0 | 5 | 0.8 | 2 | 0.5 | 11 | 0.4545 | 2 | 0.5 | 6 | 0.6667 | 4 | 0.25 | 2 | 0.5 | 14 | 0.5 |
| 7_16297351 | 1 | 1 | 2 | 0.5 | 3 | 0 | 1 | 1 | 7 | 0.4286 | 2 | 1 | 5 | 0.2 | 5 | 0.4 | 2 | 0.5 | 14 | 0.4286 |
| 7_16297640 | 1 | 0 | 1 | 0 | 4 | 0.5 | 1 | 1 | 7 | 0.4286 | 2 | 0.5 | 5 | 0.4 | 4 | 0.75 | 2 | 0 | 13 | 0.4615 |
| 7_16298144 | 1 | 0 | 1 | 1 | 3 | 0.3333 | 1 | 1 | 6 | 0.5 | 2 | 0.5 | 6 | 0.5 | 3 | 0 | 2 | 0.5 | 13 | 0.3846 |
| 7_16298192 | NA | NA | 1 | 1 | 3 | 0 | 1 | 0 | 5 | 0.2 | 1 | 0 | 4 | 0.25 | 2 | 0 | 2 | 0 | 9 | 0.1111 |
| 7_16298240 | NA | NA | 1 | 0 | 3 | 1 | 1 | 1 | 5 | 0.8 | 1 | 1 | 6 | 0 | 3 | 0.3333 | 2 | 0 | 12 | 0.1667 |
| 7_16298354 | NA | NA | 1 | 1 | 4 | 1 | NA | NA | 5 | 1 | 1 | 1 | 5 | 1 | 3 | 1 | 2 | 1 | 11 | 1 |

| | | | | | | | | | | | | | | | | | | | | |
|---|---|---|---|---|---|---|---|---|---|---|---|---|---|---|---|---|---|---|---|---|
| 7_1629838 | NA | NA | 1 | 0 | 3 | 0.3333 | NA | NA | 4 | 0.25 | 2 | 0.5 | 6 | 0.1667 | 3 | 0 | 2 | 0 | 13 | 0.1538 |
| 7_1629854 | NA | NA | 1 | 0 | 3 | 0.3333 | NA | NA | 4 | 0.25 | 1 | 0 | 6 | 0.1667 | 3 | 0.3333 | NA | NA | 10 | 0.2 |
| 7_1629863 | NA | NA | 1 | 1 | 2 | 0 | NA | NA | 3 | 0.3333 | 1 | 0 | 6 | 0.3333 | 3 | 0.6667 | NA | NA | 10 | 0.4 |
| 7_1629864 | NA | NA | 1 | 0 | 2 | 0.5 | NA | NA | 3 | 0.3333 | NA | NA | 6 | 0.5 | 3 | 1 | NA | NA | 9 | 0.6667 |
| 7_1629875 | NA | NA | 1 | 1 | NA | NA | NA | NA | 1 | 1 | NA | NA | 4 | 0.25 | 3 | 0.3333 | NA | NA | 7 | 0.2857 |
| 7_1629877 | NA | NA | 1 | 1 | NA | NA | NA | NA | 1 | 1 | 1 | 0 | 4 | 0 | 3 | 0.6667 | NA | NA | 8 | 0.25 |
| 7_1629887 | NA | NA | 1 | 1 | NA | NA | NA | NA | 1 | 1 | 1 | 1 | 5 | 1 | 3 | 0.6667 | NA | NA | 9 | 0.8889 |
| 7_1629911 | NA | NA | 1 | 1 | NA | NA | NA | NA | 1 | 1 | 1 | 0 | 3 | 0.6667 | 3 | 0.3333 | NA | NA | 7 | 0.4286 |
| 7_1653167 | 1 | 1 | 1 | 1 | NA | NA | NA | NA | 2 | 1 | NA | NA | 1 | 1 | 1 | 1 | NA | NA | 2 | 1 |
| 7_1653173 | 1 | 1 | NA | NA | NA | NA | NA | NA | 1 | 1 | NA | NA | 1 | 1 | NA | NA | NA | NA | 1 | 1 |
| 7_1654341 | 7 | 0.5714 | 6 | 0.6667 | 5 | 0.2 | 8 | 0.375 | 26 | 0.4615 | 9 | 0.5556 | 4 | 0.5 | 6 | 0.1667 | 3 | 0.3333 | 22 | 0.4091 |
| 7_1654346 | 6 | 0.1667 | 3 | 0 | 4 | 0.25 | 6 | 0 | 19 | 0.1053 | 5 | 0.2 | 3 | 1 | 5 | 0.2 | 2 | 0 | 15 | 0.3333 |
| 7_1654367 | 7 | 0 | 4 | 0 | 2 | 0 | 5 | 0.4 | 18 | 0.1111 | 2 | 0 | 1 | 1 | 4 | 0 | 2 | 0 | 9 | 0.1111 |
| 7_1654368 | 5 | 0.8 | 3 | 1 | 3 | 1 | 5 | 1 | 16 | 0.9375 | 6 | 1 | 2 | 1 | 3 | 1 | 2 | 1 | 13 | 1 |
| 7_1654378 | 5 | 0.6 | 4 | 1 | 3 | 0.3333 | 5 | 0.6 | 17 | 0.6471 | 5 | 0.4 | 1 | 1 | 4 | 0.5 | 2 | 0.5 | 12 | 0.5 |
| 7_1654398 | 7 | 0.7143 | 6 | 0.6667 | 2 | 0.5 | 7 | 0.2857 | 22 | 0.5455 | 5 | 0.6 | 3 | 1 | 5 | 0.6 | 2 | 1 | 15 | 0.7333 |
| 7_1654403 | 2 | 0 | 4 | 0.25 | 3 | 0.3333 | 4 | 0.25 | 13 | 0.2308 | 3 | 0 | 2 | 0 | 4 | 0.5 | 2 | 0 | 11 | 0.1818 |
| 7_1654416 | 7 | 0.8571 | 5 | 1 | 2 | 1 | 6 | 1 | 20 | 0.95 | 1 | 0 | 2 | 1 | 4 | 1 | 2 | 1 | 9 | 0.8889 |
| 7_1654420 | 6 | 0.8333 | 5 | 1 | 3 | 1 | 7 | 0.8571 | 21 | 0.9048 | 1 | 1 | 3 | 1 | 4 | 1 | 2 | 1 | 10 | 1 |
| 7_1654423 | 5 | 0.4 | 3 | 1 | 2 | 0.5 | 3 | 1 | 13 | 0.6923 | NA | NA | 2 | 1 | 4 | 0.25 | 1 | 0 | 7 | 0.4286 |
| 7_1654434 | 3 | 0 | 4 | 0.75 | 3 | 0.3333 | 6 | 0 | 16 | 0.25 | 1 | 0 | 2 | 0 | 3 | 0.3333 | 1 | 0 | 7 | 0.1429 |
| 7_1654438 | 5 | 0 | 4 | 0.25 | 3 | 0 | 6 | 0 | 18 | 0.0556 | 1 | 0 | 3 | 0 | 4 | 0.5 | 2 | 0.5 | 10 | 0.3 |
| 7_1654442 | 6 | 0.1667 | 5 | 0.2 | 3 | 0 | 5 | 0 | 19 | 0.1053 | 1 | 0 | 3 | 0.3333 | 5 | 0 | 2 | 0 | 11 | 0.0909 |
| 7_1654456 | 6 | 0.6667 | 4 | 0.75 | 3 | 1 | 3 | 0.6667 | 16 | 0.75 | 1 | 1 | 2 | 0.5 | 5 | 0.8 | 2 | 1 | 10 | 0.8 |
| 7_1654457 | 5 | 1 | 3 | 1 | 3 | 1 | 3 | 1 | 14 | 1 | NA | NA | 3 | 0.6667 | 2 | 1 | NA | NA | 5 | 0.8 |
| 7_1654461 | 6 | 1 | 4 | 1 | 3 | 1 | 3 | 1 | 16 | 1 | NA | NA | 3 | 0.6667 | 5 | 1 | 1 | 0 | 9 | 0.7778 |
| 7_1654478 | 5 | 1 | 5 | 0.8 | 3 | 0.6667 | 4 | 1 | 17 | 0.8824 | 1 | 1 | 3 | 1 | 4 | 1 | NA | NA | 8 | 1 |
| 7_1654488 | 5 | 0.4 | 4 | 0.25 | 3 | 0.3333 | 4 | 0.75 | 16 | 0.4375 | 1 | 0 | 2 | 0 | 3 | 0.3333 | NA | NA | 6 | 0.1667 |
| 7_1654492 | 4 | 0.25 | 5 | 0.4 | 3 | 0.3333 | 4 | 1 | 16 | 0.5 | 1 | 0 | 1 | 1 | 4 | 0.5 | NA | NA | 6 | 0.5 |
| 7_1654515 | 3 | 1 | 4 | 1 | 2 | 1 | 1 | 1 | 10 | 1 | NA | NA | NA | NA | 2 | 1 | NA | NA | 2 | 1 |
| 7_1654532 | 2 | 1 | 5 | 0.8 | 3 | 1 | 1 | 1 | 11 | 0.9091 | 1 | 1 | 2 | 1 | 2 | 1 | NA | NA | 5 | 1 |
| 7_1654553 | 2 | 0.5 | 4 | 0.5 | 2 | 0.5 | 2 | 0.5 | 10 | 0.5 | 1 | 0 | NA | NA | 3 | 1 | NA | NA | 4 | 0.75 |
| 7_1654564 | 2 | 0.5 | 5 | 0.2 | 3 | 0 | 3 | 0.3333 | 13 | 0.2308 | 1 | 1 | 2 | 1 | 3 | 0 | NA | NA | 6 | 0.5 |
| 7_1654573 | 3 | 1 | 5 | 1 | 3 | 1 | 3 | 1 | 14 | 1 | NA | NA | 2 | 1 | 3 | 1 | NA | NA | 5 | 1 |
| 7_1654576 | 2 | 0 | 4 | 0.25 | 3 | 0.3333 | 1 | 0 | 10 | 0.2 | NA | NA | 2 | 0 | 1 | 1 | NA | NA | 3 | 0.3333 |
| 7_1660413 | NA | NA | NA | NA | NA | NA | 1 | 1 | 1 | 1 | NA | NA | 2 | 1 | 1 | 1 | NA | NA | 3 | 1 |
| 7_1660819 | NA | NA | 1 | 1 | NA | NA | NA | NA | 1 | 1 | 1 | 1 | NA | NA | NA | NA | NA | NA | 1 | 1 |
| 7_1660851 | NA | NA | 1 | 1 | NA | NA | NA | NA | 1 | 1 | 1 | 1 | NA | NA | NA | NA | NA | NA | 1 | 1 |
| 7_1661973 | 9 | 0.1111 | 29 | 0.069 | 20 | 0.1 | 11 | 0 | 69 | 0.0725 | 26 | 0.0385 | 18 | 0.1111 | 18 | 0.1111 | 12 | 0 | 74 | 0.0676 |
| 7_1662036 | 11 | 0.6364 | 32 | 0.375 | 21 | 0.3333 | 12 | 0.4167 | 76 | 0.4079 | 30 | 0.4333 | 17 | 0.5294 | 16 | 0.4375 | 16 | 0.625 | 79 | 0.4937 |
| 7_1662043 | 12 | 0 | 39 | 0.0769 | 23 | 0.1304 | 16 | 0.0625 | 90 | 0.0778 | 36 | 0.1667 | 20 | 0.2 | 17 | 0.0588 | 18 | 0.0556 | 91 | 0.1319 |
| 7_1662046 | 12 | 0 | 38 | 0.1316 | 23 | 0.0435 | 17 | 0.0588 | 90 | 0.0778 | 34 | 0.0588 | 22 | 0.0455 | 22 | 0.0909 | 19 | 0.1053 | 97 | 0.0722 |
| 7_1662082 | 18 | 0.2778 | 43 | 0.186 | 25 | 0.28 | 17 | 0.2353 | 103 | 0.233 | 39 | 0.359 | 22 | 0.3182 | 24 | 0.2083 | 18 | 0.3333 | 103 | 0.3107 |
| 7_1662093 | 20 | 1 | 37 | 0.973 | 28 | 0.9286 | 18 | 0.9444 | 103 | 0.9612 | 38 | 1 | 21 | 0.9524 | 25 | 1 | 19 | 0.9474 | 103 | 0.9806 |
| 7_1662095 | 14 | 0.7857 | 28 | 0.75 | 19 | 0.8421 | 12 | 0.5833 | 73 | 0.7534 | 29 | 0.8276 | 14 | 0.7857 | 20 | 0.8 | 15 | 0.7333 | 78 | 0.7949 |
| 7_1664945 | 111 | 0.1351 | 223 | 0.1883 | 175 | 0.1829 | 103 | 0.165 | 612 | 0.1732 | 170 | 0.2059 | 176 | 0.1591 | 159 | 0.2013 | 109 | 0.156 | 614 | 0.1824 |
| 7_1664956 | 87 | 0.023 | 176 | 0.0568 | 137 | 0.0584 | 82 | 0.0976 | 482 | 0.0581 | 147 | 0.0408 | 151 | 0.0662 | 128 | 0.0547 | 98 | 0.0408 | 524 | 0.0515 |
| 7_1665076 | 70 | 0.2 | 103 | 0.233 | 98 | 0.2449 | 59 | 0.1864 | 330 | 0.2212 | 115 | 0.2174 | 100 | 0.24 | 94 | 0.2021 | 69 | 0.2319 | 378 | 0.2222 |
| 7_1667875 | 5 | 0.2 | 1 | 0 | 4 | 0.25 | 9 | 0.3333 | 19 | 0.2632 | NA | NA | 3 | 0 | 7 | 0.2857 | 8 | 0 | 18 | 0.1111 |
| 7_1667890 | 4 | 0.75 | 1 | 1 | 1 | 1 | 5 | 1 | 11 | 0.9091 | NA | NA | 3 | 0.6667 | 6 | 0.8333 | 7 | 0.7143 | 16 | 0.75 |
| 7_1667914 | 8 | 0.125 | 1 | 0 | 4 | 0 | 8 | 0.125 | 21 | 0.0952 | NA | NA | 3 | 0 | 5 | 0 | 12 | 0.0833 | 20 | 0.05 |
| 7_1667917 | 8 | 0.75 | 1 | 1 | 5 | 0.8 | 10 | 0.8 | 24 | 0.7917 | NA | NA | 3 | 0.6667 | 7 | 0.8571 | 13 | 0.8462 | 23 | 0.8261 |
| 7_1667981 | 9 | 0 | 2 | 0 | 5 | 0 | 12 | 0.1667 | 28 | 0.0714 | 2 | 0 | 8 | 0 | 10 | 0.1 | 14 | 0.0714 | 34 | 0.0588 |
| 7_1667982 | 9 | 0.1111 | 2 | 0 | 4 | 0.25 | 14 | 0.3571 | 29 | 0.2414 | 2 | 0.5 | 7 | 0 | 9 | 0.1111 | 14 | 0.1429 | 32 | 0.125 |
| 7_1667992 | 10 | 0 | 2 | 0 | 5 | 0.2 | 14 | 0.2857 | 31 | 0.1613 | 2 | 1 | 7 | 0 | 9 | 0 | 14 | 0.0714 | 32 | 0.0938 |
| 7_1668009 | 10 | 0.1 | 4 | 0 | 7 | 0.1429 | 13 | 0.0769 | 34 | 0.0882 | 1 | 0 | 11 | 0.0909 | 14 | 0.1429 | 16 | 0.0625 | 42 | 0.0952 |
| 7_1668014 | 11 | 0 | 5 | 0 | 6 | 0 | 16 | 0.125 | 38 | 0.0526 | 4 | 0.25 | 7 | 0.2857 | 14 | 0.1429 | 15 | 0.1333 | 40 | 0.175 |
| 7_1668017 | 11 | 0.3636 | 3 | 0.3333 | 7 | 0.4286 | 15 | 0.2667 | 36 | 0.3333 | 1 | 1 | 10 | 0.1 | 8 | 0 | 14 | 0.5 | 33 | 0.2727 |
| 7_1668021 | 11 | 0.3636 | 5 | 0.2 | 7 | 0.5714 | 13 | 0.3846 | 36 | 0.3889 | 4 | 1 | 11 | 0.6364 | 14 | 0.6429 | 16 | 0.3125 | 45 | 0.5556 |
| 7_1668034 | 8 | 0 | 5 | 0.2 | 5 | 0.4 | 13 | 0.3846 | 31 | 0.2581 | 4 | 0 | 11 | 0.1818 | 16 | 0.4375 | 13 | 0.3846 | 44 | 0.3182 |
| 7_1668038 | 8 | 0.375 | 3 | 0.3333 | 6 | 0.1667 | 10 | 0.4 | 27 | 0.3333 | 5 | 0.6 | 10 | 0.2 | 14 | 0.2857 | 14 | 0.7143 | 43 | 0.4419 |
| 7_1668063 | 14 | 0.5 | 6 | 0.5 | 6 | 0.6667 | 20 | 0.6 | 46 | 0.5652 | 8 | 0.625 | 17 | 0.7059 | 23 | 0.6087 | 17 | 0.7647 | 65 | 0.6769 |
| 7_1668084 | 14 | 0.6429 | 10 | 0.5 | 10 | 0.4 | 22 | 0.5 | 56 | 0.5179 | 9 | 0.8889 | 13 | 0.7692 | 21 | 0.619 | 19 | 0.4737 | 62 | 0.6452 |
| 7_1668092 | 13 | 0.1538 | 9 | 0.2222 | 13 | 0.5385 | 23 | 0.3478 | 58 | 0.3276 | 7 | 0.4286 | 13 | 0.5385 | 17 | 0.5882 | 18 | 0.3889 | 55 | 0.4909 |
| 7_1668104 | 16 | 0.9375 | 9 | 1 | 15 | 0.9333 | 22 | 1 | 62 | 0.9677 | 7 | 0.8571 | 15 | 0.8667 | 25 | 1 | 23 | 1 | 70 | 0.9571 |

| | | | | | | | | | | | | | | | | | | | | |
|---|---|---|---|---|---|---|---|---|---|---|---|---|---|---|---|---|---|---|---|---|
| 7_1668118 | 10 | 0.5 | 6 | 0.8333 | 7 | 0.4286 | 15 | 0.6667 | 38 | 0.6053 | 9 | 0.5556 | 8 | 1 | 15 | 0.8 | 18 | 0.6667 | 50 | 0.74 |
| 7_1668124 | 19 | 0.4211 | 10 | 0.6 | 17 | 0.5882 | 29 | 0.3103 | 75 | 0.44 | 13 | 0.4615 | 21 | 0.2381 | 26 | 0.5769 | 25 | 0.28 | 85 | 0.3882 |
| 7_1668124 | 17 | 0.1176 | 10 | 0.1 | 16 | 0.25 | 28 | 0.0714 | 71 | 0.1268 | 13 | 0.3077 | 22 | 0.1818 | 23 | 0.087 | 25 | 0.12 | 83 | 0.1566 |
| 7_1668129 | 21 | 0.0476 | 8 | 0.375 | 17 | 0.1765 | 31 | 0.2581 | 77 | 0.1948 | 12 | 0.1667 | 18 | 0.2778 | 25 | 0.2 | 24 | 0.0833 | 79 | 0.1772 |
| 7_1668141 | 17 | 0.5294 | 10 | 0.8 | 14 | 0.5714 | 20 | 0.35 | 61 | 0.5246 | 7 | 0.8571 | 11 | 0.6364 | 17 | 0.6471 | 13 | 0.4615 | 48 | 0.625 |
| 7_1668164 | 25 | 0.2 | 15 | 0 | 23 | 0.1304 | 37 | 0.0541 | 100 | 0.1 | 17 | 0.0588 | 27 | 0.037 | 39 | 0.1282 | 30 | 0.1333 | 113 | 0.0973 |
| 7_1668246 | 38 | 0.1579 | 33 | 0.2121 | 37 | 0.2162 | 49 | 0.2857 | 157 | 0.2229 | 24 | 0.2917 | 43 | 0.3023 | 63 | 0.2222 | 39 | 0.2308 | 169 | 0.2544 |
| 7_1668247 | 35 | 0.3429 | 30 | 0.1333 | 31 | 0.2581 | 43 | 0.3721 | 139 | 0.2878 | 20 | 0.3 | 42 | 0.2143 | 58 | 0.3276 | 35 | 0.3143 | 155 | 0.2903 |
| 7_1668254 | 38 | 0.3158 | 34 | 0.4412 | 39 | 0.3846 | 49 | 0.4082 | 160 | 0.3875 | 25 | 0.4 | 47 | 0.4255 | 71 | 0.3239 | 43 | 0.5349 | 186 | 0.4086 |
| 7_1668255 | 34 | 0.2647 | 28 | 0.4286 | 33 | 0.3636 | 45 | 0.3556 | 140 | 0.35 | 22 | 0.4091 | 33 | 0.4242 | 61 | 0.3607 | 36 | 0.4167 | 152 | 0.3947 |
| 7_1668261 | 37 | 0.4865 | 29 | 0.6207 | 30 | 0.5667 | 42 | 0.5238 | 138 | 0.5435 | 22 | 0.5909 | 36 | 0.5833 | 58 | 0.6034 | 36 | 0.5556 | 152 | 0.5855 |
| 7_1668262 | 32 | 0.9375 | 28 | 0.9286 | 30 | 0.9667 | 44 | 0.9091 | 134 | 0.9328 | 23 | 0.9565 | 35 | 0.9429 | 62 | 0.9355 | 38 | 0.9474 | 158 | 0.943 |
| 7_1669420 | 4 | 1 | 5 | 0.8 | 3 | 0.3333 | 7 | 0.5714 | 19 | 0.6842 | 1 | 1 | 5 | 0.4 | 4 | 0.5 | 4 | 0.75 | 14 | 0.5714 |
| 7_1669458 | 1 | 0 | 4 | 0.75 | 4 | 0.5 | 10 | 0.6 | 19 | 0.5789 | 3 | 1 | 6 | 0.5 | 5 | 1 | 6 | 1 | 20 | 0.85 |
| 7_1669481 | 4 | 0.5 | 8 | 0.125 | 6 | 0.1667 | 11 | 0.1818 | 29 | 0.2069 | 4 | 0 | 7 | 0.1429 | 4 | 0.25 | 10 | 0.3 | 25 | 0.2 |
| 7_1669505 | 7 | 0.1429 | 10 | 0 | 6 | 0 | 13 | 0.0769 | 36 | 0.0556 | 3 | 0 | 12 | 0.25 | 6 | 0 | 11 | 0.1818 | 32 | 0.1562 |
| 7_1669602 | 6 | 0.6667 | 11 | 0.5455 | 6 | 0.5 | 14 | 0.4286 | 37 | 0.5135 | 5 | 0.4 | 13 | 0.5385 | 11 | 0.8182 | 12 | 0.6667 | 41 | 0.6341 |
| 7_1669605 | 8 | 0.75 | 7 | 0.7143 | 6 | 0.8333 | 14 | 0.6429 | 35 | 0.7143 | 5 | 0.8 | 14 | 0.6429 | 7 | 0.8571 | 12 | 1 | 38 | 0.8158 |
| 7_1669606 | 8 | 0.75 | 10 | 0.8 | 4 | 0.75 | 17 | 0.3529 | 39 | 0.5897 | 6 | 0.5 | 12 | 0.8333 | 6 | 0.5 | 10 | 0.8 | 34 | 0.7059 |
| 7_1669621 | 8 | 0.125 | 10 | 0.2 | 7 | 0.2857 | 15 | 0 | 40 | 0.125 | 4 | 0.25 | 13 | 0.3077 | 9 | 0.2222 | 12 | 0.0833 | 38 | 0.2105 |
| 7_1669638 | 11 | 0.1818 | 12 | 0.25 | 6 | 0.1667 | 15 | 0.2 | 44 | 0.2045 | 5 | 0.2 | 10 | 0.2 | 9 | 0.3333 | 15 | 0.2 | 39 | 0.2308 |
| 7_1669663 | 12 | 0.4167 | 13 | 0.1538 | 6 | 0 | 17 | 0.0588 | 48 | 0.1667 | 6 | 0.1667 | 11 | 0.0909 | 11 | 0.2727 | 15 | 0.2667 | 43 | 0.2093 |
| 7_1669665 | 10 | 0.3 | 12 | 0.0833 | 4 | 0 | 15 | 0.0667 | 41 | 0.122 | 5 | 0 | 10 | 0.2 | 8 | 0 | 10 | 0.2 | 33 | 0.1212 |
| 7_1669675 | 10 | 0.1 | 9 | 0.1111 | 6 | 0.3333 | 13 | 0.2308 | 38 | 0.1842 | 4 | 0 | 9 | 0.2222 | 9 | 0.2222 | 9 | 0.2222 | 31 | 0.1935 |
| 7_1669814 | 16 | 0.3125 | 14 | 0.2143 | 8 | 0.5 | 15 | 0.1333 | 53 | 0.2642 | 5 | 0.2 | 8 | 0.375 | 10 | 0.5 | 18 | 0.2222 | 41 | 0.3171 |
| 7_1669830 | 23 | 0.2609 | 17 | 0.1765 | 13 | 0.2308 | 22 | 0.2273 | 75 | 0.2267 | 7 | 0.1429 | 9 | 0.3333 | 15 | 0.2 | 19 | 0.2105 | 50 | 0.22 |
| 7_1669838 | 19 | 0.2632 | 15 | 0.1333 | 10 | 0.3 | 17 | 0.2941 | 61 | 0.2459 | 8 | 0 | 10 | 0.4 | 10 | 0.3 | 22 | 0.4091 | 50 | 0.32 |
| 7_1669844 | 23 | 0.087 | 16 | 0.0625 | 10 | 0.1 | 21 | 0.1429 | 70 | 0.1 | 9 | 0.2222 | 13 | 0.0769 | 16 | 0.0625 | 18 | 0.2778 | 56 | 0.1607 |
| 7_1671835 | NA | NA | NA | NA | NA | NA | 2 | 0.5 | 2 | 0.5 | NA | NA | 1 | 1 | NA | NA | 1 | 1 | 2 | 1 |
| 7_1671925 | 1 | 1 | 3 | 0.6667 | NA | NA | 7 | 0.4286 | 11 | 0.5455 | NA | NA | 1 | 0 | NA | NA | 2 | 0.5 | 3 | 0.3333 |
| 7_1671933 | 2 | 0 | 1 | 0 | NA | NA | 7 | 0.7143 | 10 | 0.5 | NA | NA | NA | NA | NA | NA | 2 | 1 | 2 | 1 |
| 7_1671935 | 2 | 0.5 | 2 | 1 | NA | NA | 5 | 1 | 9 | 0.8889 | NA | NA | 1 | 1 | NA | NA | 1 | 1 | 2 | 1 |
| 7_1671942 | 2 | 0.5 | 3 | 1 | NA | NA | 4 | 1 | 9 | 0.8889 | NA | NA | 1 | 1 | NA | NA | 2 | 0.5 | 3 | 0.6667 |
| 7_1671943 | NA | NA | 3 | 0.3333 | NA | NA | 7 | 0.7143 | 10 | 0.6 | NA | NA | 1 | 1 | NA | NA | 2 | 1 | 3 | 1 |
| 7_1671945 | 1 | 0 | 3 | 0.3333 | NA | NA | 7 | 0.4286 | 11 | 0.3636 | NA | NA | 1 | 0 | NA | NA | 2 | 0.5 | 3 | 0.3333 |
| 7_1671951 | 2 | 0 | 2 | 1 | NA | NA | 7 | 0.8571 | 11 | 0.7273 | NA | NA | 1 | 1 | NA | NA | 1 | 1 | 2 | 1 |
| 7_1671952 | 1 | 0 | 3 | 0.3333 | NA | NA | 7 | 0.2857 | 11 | 0.2727 | NA | NA | 1 | 1 | NA | NA | 1 | 0 | 2 | 0.5 |
| 7_1671954 | 1 | 0 | 3 | 0.6667 | NA | NA | 6 | 0.3333 | 10 | 0.4 | NA | NA | 1 | 0 | NA | NA | 2 | 0.5 | 3 | 0.3333 |
| 7_1671958 | 1 | 1 | 3 | 1 | NA | NA | 7 | 1 | 11 | 1 | NA | NA | NA | NA | NA | NA | 2 | 1 | 2 | 1 |
| 7_1672003 | 2 | 0.5 | 2 | 0.5 | NA | NA | 4 | 0 | 8 | 0.25 | NA | NA | 2 | 0.5 | NA | NA | 2 | 0 | 4 | 0.25 |
| 7_1672063 | 2 | 0 | 4 | 0 | NA | NA | 7 | 0.1429 | 13 | 0.0769 | 1 | 1 | 3 | 0 | NA | NA | 3 | 0 | 7 | 0.1429 |
| 7_1672168 | 10 | 0.1 | 9 | 0.4444 | 1 | 0 | 13 | 0.0769 | 33 | 0.1818 | 6 | 0.3333 | 13 | 0.1538 | 3 | 0 | 6 | 0.1667 | 28 | 0.1786 |
| 7_1672185 | 13 | 0.2308 | 9 | 0.3333 | 2 | 0.5 | 14 | 0.2143 | 38 | 0.2632 | 6 | 0.1667 | 13 | 0.0769 | 4 | 0.25 | 7 | 0.2857 | 30 | 0.1667 |
| 7_1672188 | 13 | 0.0769 | 10 | 0.1 | 2 | 0 | 13 | 0.1538 | 38 | 0.1053 | 8 | 0.125 | 14 | 0 | 5 | 0 | 8 | 0.125 | 35 | 0.0571 |
| 7_1672189 | 7 | 0 | 8 | 0.375 | NA | NA | 9 | 0.5556 | 24 | 0.3333 | 6 | 0.3333 | 9 | 0.4444 | 3 | 0.3333 | 2 | 0 | 20 | 0.35 |
| 7_1673106 | NA | NA | 1 | 1 | NA | NA | NA | NA | 1 | 1 | NA | NA | 1 | 1 | 1 | 0 | NA | NA | 2 | 0.5 |
| 7_1676558 | 4 | 0 | 2 | 0 | 1 | 1 | 3 | 0 | 10 | 0.1 | 1 | 0 | 3 | 0.3333 | NA | NA | 1 | 1 | 5 | 0.4 |
| 7_1676955 | 4 | 0.25 | NA | NA | NA | NA | 1 | 1 | 5 | 0.4 | NA | NA | 2 | 0.5 | NA | NA | NA | NA | 2 | 0.5 |
| 7_1676975 | 3 | 0 | NA | NA | NA | NA | 1 | 1 | 4 | 0.25 | NA | NA | 3 | 0.3333 | NA | NA | NA | NA | 3 | 0.3333 |
| 7_1862427 | 3 | 0 | 17 | 0.1765 | 8 | 0.125 | 8 | 0 | 36 | 0.1111 | 12 | 0.0833 | 4 | 0 | 5 | 0 | 8 | 0.125 | 29 | 0.069 |
| 7_1869149 | 3 | 0 | 9 | 0 | 8 | 0.25 | 7 | 0.1429 | 27 | 0.1111 | 6 | 0.1667 | 7 | 0.1429 | 3 | 0 | 8 | 0.125 | 24 | 0.125 |
| 7_1869231 | 2 | 1 | 14 | 0.7857 | 8 | 0.875 | 8 | 0.75 | 32 | 0.8125 | 7 | 0.7143 | 9 | 0.6667 | 3 | 1 | 7 | 0.8571 | 26 | 0.7692 |
| 7_1869242 | 2 | 0 | 12 | 0.1667 | 5 | 0.4 | 8 | 0.125 | 27 | 0.1852 | 7 | 0.1429 | 8 | 0 | 3 | 0.3333 | 7 | 0.1429 | 25 | 0.12 |
| 7_1869261 | 4 | 0.5 | 15 | 0.1333 | 10 | 0 | 9 | 0.4444 | 38 | 0.2105 | 6 | 0.1667 | 11 | 0 | 5 | 0.4 | 7 | 0.1429 | 29 | 0.1379 |
| 7_1869263 | 4 | 0.25 | 16 | 0.25 | 10 | 0.2 | 10 | 0.1 | 40 | 0.2 | 6 | 0.1667 | 10 | 0.2 | 5 | 0.6 | 8 | 0.25 | 29 | 0.2759 |
| 7_1869373 | 3 | 0 | 2 | 0 | 4 | 0 | 3 | 0.3333 | 12 | 0.0833 | 5 | 0 | 5 | 0 | 4 | 0.25 | 3 | 0 | 17 | 0.0588 |
| 7_1869581 | 8 | 0 | 10 | 0.1 | 7 | 0.1429 | 5 | 0 | 30 | 0.0667 | 7 | 0 | 6 | 0.3333 | 5 | 0.2 | 3 | 0 | 21 | 0.1429 |
| 7_1869610 | 8 | 0.125 | 9 | 0.1111 | 6 | 0.1667 | 4 | 0.25 | 27 | 0.1481 | 7 | 0.1429 | 5 | 0 | 4 | 0 | 2 | 0 | 18 | 0.0556 |
| 7_1869617 | 8 | 0 | 11 | 0.2727 | 7 | 0 | 5 | 0 | 31 | 0.0968 | 6 | 0.3333 | 7 | 0 | 4 | 0.25 | 5 | 0.2 | 22 | 0.1818 |
| 7_1869637 | 8 | 0.75 | 13 | 0.8462 | 7 | 0.7143 | 5 | 0.8 | 33 | 0.7879 | 7 | 1 | 6 | 0.8333 | 3 | 0.6667 | 3 | 1 | 19 | 0.8947 |
| 7_1869652 | 9 | 0.5556 | 13 | 0.4615 | 8 | 0.25 | 6 | 0.3333 | 36 | 0.4167 | 6 | 0.3333 | 8 | 0.375 | 4 | 0.5 | 5 | 0.2 | 23 | 0.3478 |
| 7_1869665 | 9 | 1 | 12 | 1 | 7 | 1 | 7 | 1 | 35 | 1 | 8 | 1 | 8 | 1 | 5 | 1 | 5 | 1 | 26 | 1 |
| 7_1869676 | 7 | 0.2857 | 9 | 0.4444 | 6 | 0.5 | 3 | 1 | 25 | 0.48 | 5 | 0.4 | 3 | 0.3333 | 3 | 0.3333 | 5 | 0.4 | 16 | 0.375 |
| 7_1869684 | 5 | 1 | 12 | 0.9167 | 6 | 0.8333 | 2 | 1 | 25 | 0.92 | 4 | 1 | 7 | 1 | 3 | 1 | 4 | 0.75 | 18 | 0.9444 |
| 7_1869692 | 9 | 0.4444 | 15 | 0.8667 | 8 | 0.5 | 8 | 0.625 | 40 | 0.65 | 6 | 0.5 | 9 | 0.3333 | 5 | 0.4 | 8 | 0.25 | 28 | 0.3571 |

| | | | | | | | | | | | | | | | | | | | | |
|---|---|---|---|---|---|---|---|---|---|---|---|---|---|---|---|---|---|---|---|---|
| 7_1869703 | 10 | 0.3 | 14 | 0.0714 | 7 | 0 | 7 | 0 | 38 | 0.1053 | 9 | 0.1111 | 8 | 0.125 | 5 | 0 | 8 | 0 | 30 | 0.0667 |
| 7_1869709 | 11 | 0 | 15 | 0.4 | 8 | 0.25 | 8 | 0.5 | 42 | 0.2857 | 8 | 0.125 | 10 | 0.2 | 7 | 0.4286 | 7 | 0.2857 | 32 | 0.25 |
| 7_1869713 | 11 | 0.3636 | 15 | 0.5333 | 8 | 0.625 | 7 | 0.2857 | 41 | 0.4634 | 9 | 0.4444 | 9 | 0.5556 | 7 | 0.7143 | 8 | 0.25 | 33 | 0.4848 |
| 7_1869723 | 7 | 0.1429 | 12 | 0.4167 | 7 | 0.2857 | 8 | 0.25 | 34 | 0.2941 | 7 | 0.2857 | 10 | 0.4 | 5 | 0 | 11 | 0.3636 | 33 | 0.303 |
| 7_1869730 | 11 | 0.2727 | 13 | 0.4615 | 7 | 0.1429 | 9 | 0.1111 | 40 | 0.275 | 8 | 0.375 | 11 | 0.2727 | 6 | 0.3333 | 11 | 0.2727 | 36 | 0.3056 |
| 7_1869732 | 7 | 0.8571 | 8 | 0.75 | 6 | 0.6667 | 6 | 1 | 27 | 0.8148 | 6 | 1 | 8 | 0.75 | 5 | 0.6 | 8 | 0.875 | 27 | 0.8148 |
| 7_1869759 | 5 | 0.2 | 10 | 0 | 7 | 0 | 5 | 0.2 | 27 | 0.0741 | 6 | 0.3333 | 6 | 0.3333 | 5 | 0.4 | 7 | 0.1429 | 24 | 0.2917 |
| 7_1869785 | 9 | 0.2222 | 11 | 0 | 6 | 0.3333 | 7 | 0 | 33 | 0.1212 | 6 | 0 | 8 | 0 | 6 | 0.1667 | 8 | 0.125 | 28 | 0.0714 |
| 7_1869786 | 9 | 0.2222 | 9 | 0.4444 | 7 | 0.1429 | 6 | 0.1667 | 31 | 0.2581 | 7 | 0.4286 | 9 | 0.4444 | 5 | 0.6 | 10 | 0.5 | 31 | 0.4839 |
| 7_1869796 | 10 | 0 | 15 | 0 | 10 | 0.1 | 10 | 0.3 | 45 | 0.0889 | 9 | 0 | 10 | 0.1 | 6 | 0.1667 | 11 | 0.0909 | 36 | 0.0833 |
| 7_1869808 | 12 | 0.0833 | 13 | 0.0769 | 10 | 0.1 | 9 | 0.1111 | 44 | 0.0909 | 8 | 0.25 | 13 | 0 | 7 | 0.2857 | 10 | 0.2 | 38 | 0.1579 |
| 7_1871134 | NA | NA | 1 | 1 | NA | NA | NA | NA | 1 | 1 | NA | NA | 2 | 1 | NA | NA | NA | NA | 2 | 1 |
| 7_1871149 | NA | NA | 2 | 1 | NA | NA | NA | NA | 2 | 1 | NA | NA | 2 | 1 | NA | NA | NA | NA | 2 | 1 |
| 7_1872591 | 3 | 0.3333 | 3 | 0.3333 | 4 | 0 | 3 | 0 | 13 | 0.1538 | 3 | 0.6667 | 9 | 0 | NA | NA | 2 | 0 | 14 | 0.1429 |
| 7_1872594 | 6 | 0.1667 | 3 | 0 | 3 | 0 | 4 | 0 | 16 | 0.0625 | 5 | 0.2 | 7 | 0 | NA | NA | 2 | 0.5 | 14 | 0.1429 |
| 7_1872621 | 6 | 0.6667 | 3 | 0.3333 | 4 | 0.25 | 6 | 0.1667 | 19 | 0.3684 | 5 | 0.4 | 9 | 0.4444 | NA | NA | 3 | 0 | 17 | 0.3529 |
| 7_1872638 | 6 | 0.8333 | 5 | 0.4 | 3 | 0.3333 | 7 | 0.4286 | 21 | 0.5238 | 4 | 0.75 | 10 | 0.3 | 1 | 1 | 4 | 0.5 | 19 | 0.4737 |
| 7_1872640 | 6 | 0.5 | 4 | 0.5 | 2 | 0.5 | 2 | 0.5 | 14 | 0.5 | 3 | 0.3333 | 9 | 0.7778 | 1 | 0 | 3 | 0.6667 | 16 | 0.625 |
| 7_1873466 | 6 | 0.1667 | 3 | 0.6667 | 5 | 0.6 | 9 | 0.4444 | 23 | 0.4348 | 5 | 0.8 | 9 | 0.6667 | 1 | 0 | 3 | 0.3333 | 18 | 0.6111 |
| 7_1873469 | 7 | 0.4286 | 5 | 0.2 | 8 | 0.375 | 10 | 0.3 | 30 | 0.3333 | 5 | 0.2 | 11 | 0.3636 | 3 | 0 | 4 | 0 | 23 | 0.2174 |
| 7_1873473 | 9 | 0.5556 | 6 | 0.1667 | 7 | 0.5714 | 11 | 0.5455 | 33 | 0.4848 | 4 | 0.5 | 11 | 0.5455 | 3 | 0.3333 | 4 | 0.5 | 22 | 0.5 |
| 7_1873526 | 12 | 0.0833 | 5 | 0 | 8 | 0 | 12 | 0.0833 | 37 | 0.0541 | 8 | 0.125 | 12 | 0.1667 | 2 | 0 | 4 | 0 | 26 | 0.1154 |
| 7_1873553 | 10 | 0.2 | 8 | 0 | 9 | 0 | 11 | 0.0909 | 38 | 0.0789 | 7 | 0.2857 | 14 | 0.2143 | 5 | 0 | 3 | 0 | 29 | 0.1724 |
| 7_1873555 | 10 | 0.3 | 6 | 0 | 8 | 0.375 | 9 | 0 | 33 | 0.1818 | 7 | 0.2857 | 11 | 0.1818 | 4 | 0.25 | 3 | 0.3333 | 25 | 0.24 |
| 7_1873844 | 5 | 0.6 | 4 | 0.5 | 4 | 0.75 | 4 | 1 | 17 | 0.7059 | 1 | 1 | 8 | 0.625 | 2 | 0 | 4 | 0.5 | 15 | 0.5333 |
| 7_1873848 | 8 | 0 | 3 | 0.3333 | 5 | 0.2 | 4 | 0.25 | 20 | 0.15 | NA | NA | 9 | 0.2222 | 3 | 0 | 2 | 0 | 14 | 0.1429 |
| 7_1873859 | 9 | 0.8889 | 3 | 0.6667 | 3 | 1 | 2 | 1 | 17 | 0.8824 | 1 | 1 | 9 | 0.8889 | 2 | 0 | 1 | 1 | 13 | 0.7692 |
| 7_1873866 | 12 | 0.25 | 5 | 0.2 | 5 | 0 | 5 | 0.4 | 27 | 0.2222 | 1 | 0 | 11 | 0.0909 | 3 | 0 | 4 | 0.25 | 19 | 0.1053 |
| 7_1873869 | 11 | 0.3636 | 4 | 0.5 | 6 | 0.5 | 5 | 0.4 | 26 | 0.4231 | NA | NA | 13 | 0.6154 | 3 | 0 | 5 | 0.6 | 21 | 0.5238 |
| 7_1873897 | 13 | 0.8462 | 2 | 1 | 6 | 1 | 3 | 1 | 24 | 0.9167 | 1 | 1 | 14 | 1 | 4 | 0.75 | 3 | 0.3333 | 22 | 0.8636 |
| 7_1873899 | 10 | 0.9 | 4 | 0.75 | 5 | 1 | 5 | 1 | 24 | 0.9167 | 1 | 1 | 11 | 0.8182 | 3 | 1 | 5 | 0.8 | 20 | 0.85 |
| 7_1873910 | 11 | 1 | 3 | 1 | 5 | 1 | 5 | 1 | 24 | 1 | 1 | 1 | 12 | 1 | 4 | 1 | 5 | 1 | 22 | 1 |
| 7_1873942 | 5 | 0.4 | 2 | 0 | 3 | 0.3333 | 3 | 0 | 13 | 0.2308 | NA | NA | 12 | 0.0833 | NA | NA | 3 | 0 | 15 | 0.0667 |
| 7_1873960 | 12 | 0.0833 | 6 | 0.1667 | 6 | 0.1667 | 5 | 0.4 | 29 | 0.1724 | 1 | 0 | 18 | 0.1667 | 5 | 0 | 8 | 0.375 | 32 | 0.1875 |
| 7_1873967 | 11 | 1 | 4 | 1 | 4 | 1 | 4 | 1 | 23 | 1 | 1 | 1 | 14 | 1 | 1 | 1 | 8 | 1 | 24 | 1 |
| 7_1873993 | 16 | 0.375 | 5 | 0 | 4 | 0.25 | 9 | 0.1111 | 34 | 0.2353 | 1 | 1 | 20 | 0.5 | 6 | 0.5 | 8 | 0.75 | 35 | 0.5714 |
| 7_1874010 | 17 | 0.2353 | 6 | 0 | 6 | 0.3333 | 10 | 0.2 | 39 | 0.2051 | 2 | 0 | 16 | 0.25 | 6 | 0.3333 | 7 | 0.2857 | 31 | 0.2581 |
| 7_1874042 | 12 | 0.3333 | 7 | 0.4286 | 7 | 0 | 11 | 0.5455 | 37 | 0.3514 | 4 | 0.5 | 17 | 0.4118 | 6 | 0.5 | 5 | 0.2 | 32 | 0.4062 |
| 7_1874049 | 11 | 0.1818 | 5 | 0 | 7 | 0.2857 | 6 | 0.1667 | 29 | 0.1724 | 1 | 0 | 13 | 0.0769 | 4 | 0.25 | 6 | 0.1667 | 24 | 0.125 |
| 7_1874057 | 4 | 0.75 | 2 | 1 | 3 | 1 | 2 | 1 | 11 | 0.9091 | NA | NA | 2 | 1 | 2 | 1 | 3 | 1 | 7 | 1 |
| 7_1874059 | 3 | 0.6667 | 2 | 0 | 4 | 0.75 | 2 | 1 | 11 | 0.6364 | NA | NA | 2 | 0.5 | 3 | 0.3333 | 3 | 1 | 8 | 0.625 |
| 7_1878519 | 3 | 0.3333 | NA | NA | NA | NA | 1 | 0 | 4 | 0.25 | NA | NA | NA | NA | 3 | 0.3333 | 3 | 0.6667 | 6 | 0.5 |
| 7_1878819 | 9 | 0.1111 | 12 | 0.1667 | 6 | 0 | 5 | 0 | 32 | 0.0938 | 5 | 0.2 | 6 | 0 | 13 | 0.0769 | 4 | 0 | 28 | 0.0714 |
| 7_1881436 | 12 | 0.1667 | 16 | 0.125 | 7 | 0 | 9 | 0.1111 | 44 | 0.1136 | 6 | 0 | 15 | 0.2 | 11 | 0 | 11 | 0.0909 | 43 | 0.093 |
| 7_1881438 | 12 | 0.0833 | 15 | 0.2667 | 8 | 0.125 | 9 | 0.2222 | 44 | 0.1818 | 7 | 0.1429 | 15 | 0.0667 | 11 | 0.1818 | 8 | 0.125 | 41 | 0.122 |
| 7_1881463 | 11 | 0.6364 | 18 | 0.3889 | 8 | 0.5 | 10 | 0.5 | 47 | 0.4894 | 8 | 0.625 | 15 | 0.5333 | 15 | 0.4667 | 15 | 0.5333 | 53 | 0.5283 |
| 7_1881465 | 14 | 0.7857 | 20 | 0.7 | 10 | 0.6 | 10 | 0.6 | 54 | 0.6852 | 7 | 0.4286 | 14 | 0.4286 | 13 | 0.8462 | 14 | 0.7857 | 48 | 0.6458 |
| 7_1881476 | 11 | 0.4545 | 13 | 0.5385 | 10 | 0.3 | 11 | 0.3636 | 45 | 0.4222 | 7 | 0.1429 | 16 | 0.4375 | 13 | 0.3846 | 12 | 0.3333 | 48 | 0.3542 |
| 7_1881492 | 14 | 0 | 17 | 0.1765 | 9 | 0 | 9 | 0.3333 | 49 | 0.1224 | 8 | 0.375 | 12 | 0.0833 | 12 | 0.25 | 16 | 0.25 | 48 | 0.2292 |
| 7_1881494 | 14 | 0.0714 | 19 | 0.2105 | 8 | 0.25 | 12 | 0.0833 | 53 | 0.1509 | 8 | 0 | 18 | 0.1111 | 15 | 0.2 | 10 | 0.3 | 51 | 0.1569 |
| 7_1881500 | 16 | 0.125 | 21 | 0.1429 | 9 | 0.4444 | 9 | 0.3333 | 55 | 0.2182 | 11 | 0.0909 | 17 | 0.0588 | 16 | 0.125 | 15 | 0.0667 | 59 | 0.0847 |
| 7_1881504 | 16 | 0.4375 | 19 | 0.4211 | 11 | 0.0909 | 13 | 0.3846 | 59 | 0.3559 | 9 | 0.3333 | 17 | 0.2941 | 17 | 0.4706 | 16 | 0.375 | 59 | 0.3729 |
| 7_1881632 | 16 | 0.0625 | 21 | 0.0952 | 8 | 0 | 14 | 0.3571 | 59 | 0.1356 | 9 | 0.1111 | 20 | 0.3 | 17 | 0.0588 | 14 | 0 | 60 | 0.1333 |
| 7_1881634 | 14 | 0.8571 | 23 | 0.7391 | 11 | 1 | 15 | 0.7333 | 63 | 0.8095 | 13 | 0.8462 | 18 | 0.8889 | 17 | 0.7059 | 17 | 0.8235 | 65 | 0.8154 |
| 7_1881651 | 16 | 0.25 | 18 | 0.3333 | 11 | 0.3636 | 15 | 0.2 | 60 | 0.2833 | 6 | 0.8333 | 15 | 0.2667 | 17 | 0.2353 | 16 | 0.375 | 54 | 0.3519 |
| 7_1881653 | 16 | 0.3125 | 22 | 0.1818 | 11 | 0.1818 | 13 | 0.0769 | 62 | 0.1935 | 12 | 0.25 | 17 | 0.0588 | 17 | 0.0588 | 18 | 0 | 64 | 0.0781 |
| 7_1881667 | 13 | 0.0769 | 21 | 0 | 10 | 0.2 | 12 | 0.1667 | 56 | 0.0893 | 8 | 0.125 | 18 | 0.1111 | 13 | 0 | 14 | 0.1429 | 53 | 0.0943 |
| 7_1882703 | 3 | 0 | 4 | 0.25 | 3 | 0.3333 | 5 | 0 | 15 | 0.1333 | 2 | 0.5 | 3 | 0 | 2 | 0 | 1 | 0 | 8 | 0.125 |
| 7_1882762 | 4 | 0.5 | 6 | 0.5 | 3 | 0.3333 | 6 | 0.3333 | 19 | 0.4211 | 2 | 0 | 5 | 0.2 | 2 | 0 | 2 | 0.5 | 11 | 0.1818 |
| 7_1882765 | 4 | 0 | 4 | 0.25 | 2 | 0 | 6 | 0.1667 | 16 | 0.125 | 2 | 0 | 4 | 0 | 2 | 0.5 | 2 | 0 | 10 | 0.1 |
| 7_1883158 | NA | NA | NA | NA | 1 | 1 | NA | NA | 1 | 1 | 1 | 1 | NA | NA | NA | NA | NA | NA | 1 | 1 |
| 7_1886951 | 2 | 0 | 1 | 1 | 1 | 1 | NA | NA | 4 | 0.5 | NA | NA | NA | NA | 1 | 1 | NA | NA | 1 | 1 |
| 7_1886964 | 2 | 1 | 1 | 0 | 1 | 0 | NA | NA | 4 | 0.5 | NA | NA | NA | NA | 1 | 1 | 1 | 0 | 2 | 0.5 |
| 7_1886992 | 2 | 0.5 | 1 | 0 | 1 | 0 | NA | NA | 4 | 0.25 | NA | NA | NA | NA | 2 | 1 | 1 | 0 | 3 | 0.6667 |
| 7_1887005 | 2 | 1 | 1 | 1 | 1 | 1 | NA | NA | 4 | 1 | NA | NA | NA | NA | 1 | 1 | 1 | 1 | 2 | 1 |

| | | | | | | | | | | | | | | | | | | | | |
|---|---|---|---|---|---|---|---|---|---|---|---|---|---|---|---|---|---|---|---|---|
| 7_1887009 | 1 | 1 | 1 | 1 | 1 | 1 | NA | NA | 3 | 1 | NA | NA | NA | NA | 2 | 1 | NA | NA | 2 | 1 |
| 7_1887026 | 2 | 0.5 | NA | NA | 1 | 1 | NA | NA | 3 | 0.6667 | NA | NA | NA | NA | 2 | 0 | 1 | 1 | 3 | 0.3333 |
| 7_1887027 | 2 | 1 | 1 | 1 | 1 | 0 | NA | NA | 4 | 0.75 | NA | NA | NA | NA | 2 | 1 | NA | NA | 2 | 1 |
| 7_1887041 | 1 | 1 | 1 | 1 | 1 | 1 | NA | NA | 3 | 1 | NA | NA | NA | NA | 2 | 1 | NA | NA | 2 | 1 |
| 7_1887045 | 2 | 1 | 1 | 1 | 1 | 1 | NA | NA | 4 | 1 | NA | NA | NA | NA | 2 | 1 | NA | NA | 2 | 1 |
| 7_1887055 | 2 | 1 | 1 | 1 | 1 | 1 | NA | NA | 4 | 1 | NA | NA | NA | NA | 2 | 1 | NA | NA | 2 | 1 |
| 7_1887070 | 2 | 1 | 1 | 1 | 1 | 1 | NA | NA | 4 | 1 | NA | NA | NA | NA | 2 | 0.5 | 1 | 1 | 3 | 0.6667 |
| 7_1887092 | 2 | 1 | 1 | 1 | 1 | 1 | NA | NA | 4 | 1 | NA | NA | NA | NA | 2 | 1 | 1 | 1 | 3 | 1 |
| 7_1887094 | 2 | 0.5 | 1 | 1 | 1 | 0 | NA | NA | 4 | 0.5 | NA | NA | NA | NA | 2 | 1 | 1 | 1 | 3 | 1 |
| 7_1887121 | 2 | 0.5 | 2 | 0 | 2 | 1 | NA | NA | 6 | 0.5 | NA | NA | 1 | 1 | 2 | 0 | 1 | 0 | 4 | 0.25 |
| 7_1887126 | 2 | 1 | 2 | 1 | 2 | 1 | NA | NA | 6 | 1 | NA | NA | 1 | 1 | 2 | 1 | 1 | 1 | 4 | 1 |
| 7_1887132 | 1 | 1 | 2 | 1 | 1 | 1 | NA | NA | 4 | 1 | NA | NA | 1 | 1 | 2 | 1 | 1 | 1 | 4 | 1 |
| 7_1887132 | 1 | 1 | 1 | 0 | 1 | 0 | NA | NA | 3 | 0.3333 | 1 | 1 | NA | NA | 1 | 0 | 1 | 0 | 3 | 0.3333 |
| 7_1887140 | 2 | 0.5 | 1 | 0 | 2 | 1 | NA | NA | 5 | 0.6 | NA | NA | 1 | 1 | 3 | 1 | 2 | 1 | 6 | 1 |
| 7_1887150 | 2 | 0.5 | 1 | 0 | 1 | 0 | NA | NA | 4 | 0.25 | NA | NA | NA | NA | 1 | 0 | 2 | 0.5 | 3 | 0.3333 |
| 7_1887182 | 1 | 0 | 2 | 1 | 1 | 1 | NA | NA | 4 | 0.75 | NA | NA | NA | NA | 3 | 0.6667 | 1 | 1 | 4 | 0.75 |
| 7_1887201 | 1 | 1 | 2 | 0.5 | NA | NA | NA | NA | 3 | 0.6667 | 1 | 1 | NA | NA | 2 | 0.5 | 1 | 0 | 4 | 0.5 |
| 7_1887421 | NA | NA | NA | NA | 1 | 1 | 1 | 0 | 2 | 0.5 | NA | NA | NA | NA | NA | NA | 2 | 1 | 2 | 1 |
| 7_1887426 | 1 | 0 | 1 | 0 | 2 | 1 | 3 | 0.3333 | 7 | 0.4286 | 2 | 0.5 | 3 | 0 | NA | NA | 3 | 0.6667 | 8 | 0.375 |
| 7_1887432 | 2 | 0 | 1 | 0 | 2 | 0 | 3 | 0.3333 | 8 | 0.125 | 2 | 0 | 2 | 0 | NA | NA | 2 | 0.5 | 6 | 0.1667 |
| 7_1887435 | 1 | 0 | 1 | 0 | 2 | 1 | 3 | 1 | 7 | 0.7143 | 1 | 1 | 2 | 1 | NA | NA | 3 | 0.3333 | 6 | 0.6667 |
| 7_1887441 | 3 | 0 | 1 | 0 | 2 | 0.5 | 3 | 0 | 9 | 0.1111 | 2 | 0 | 2 | 0.5 | NA | NA | 3 | 0 | 7 | 0.1429 |
| 7_1887444 | 2 | 0.5 | 1 | 0 | 2 | 0.5 | 3 | 0 | 8 | 0.25 | 1 | 1 | 3 | 0.3333 | NA | NA | 2 | 0 | 6 | 0.3333 |
| 7_1887445 | 2 | 0.5 | 1 | 1 | 1 | 1 | 2 | 1 | 6 | 0.8333 | 2 | 1 | 1 | 1 | NA | NA | NA | NA | 3 | 1 |
| 7_1887454 | 2 | 0.5 | 1 | 0 | 2 | 0 | 2 | 1 | 7 | 0.4286 | 2 | 0 | 3 | 1 | NA | NA | 3 | 0.6667 | 8 | 0.625 |
| 7_1887461 | 3 | 1 | NA | NA | 1 | 1 | 2 | 1 | 6 | 1 | 2 | 1 | 1 | 1 | NA | NA | 1 | 0 | 4 | 0.75 |
| 7_1887472 | 3 | 1 | 1 | 0 | NA | NA | 3 | 0.6667 | 7 | 0.7143 | 2 | 1 | 2 | 0.5 | NA | NA | 3 | 0.6667 | 7 | 0.7143 |
| 7_1887482 | 3 | 0 | NA | NA | 1 | 1 | 3 | 0 | 7 | 0.1429 | 2 | 0 | 4 | 0 | NA | NA | 3 | 0.3333 | 9 | 0.1111 |
| 7_1887485 | 3 | 1 | 1 | 0 | 2 | 0.5 | 3 | 0.6667 | 9 | 0.6667 | 2 | 0.5 | 4 | 1 | NA | NA | 3 | 0.3333 | 9 | 0.6667 |
| 7_1887493 | 4 | 0.5 | 1 | 0 | 1 | 0 | 3 | 1 | 9 | 0.5556 | 2 | 0.5 | 3 | 0 | NA | NA | 3 | 0 | 8 | 0.125 |
| 7_1887502 | 4 | 0.5 | NA | NA | 2 | 0 | 2 | 0.5 | 8 | 0.375 | 1 | 0 | 2 | 0 | NA | NA | 2 | 0.5 | 5 | 0.2 |
| 7_1887503 | 4 | 1 | 1 | 1 | 2 | 1 | 2 | 1 | 9 | 1 | 1 | 1 | 3 | 1 | NA | NA | 3 | 1 | 7 | 1 |
| 7_1887508 | 3 | 1 | NA | NA | 2 | 1 | 3 | 0.3333 | 8 | 0.75 | 2 | 1 | 2 | 0.5 | NA | NA | 3 | 0.6667 | 7 | 0.7143 |
| 7_1888644 | 32 | 0 | 27 | 0.1111 | 20 | 0.15 | 21 | 0 | 100 | 0.06 | 17 | 0.0588 | 24 | 0.125 | 22 | 0 | 21 | 0.0952 | 84 | 0.0714 |
| 7_1888678 | NA | NA | 1 | 1 | NA | NA | NA | NA | 1 | 1 | 1 | 1 | NA | NA | 2 | 0.5 | NA | NA | 3 | 0.6667 |
| 7_1889102 | NA | NA | NA | NA | NA | NA | 1 | 1 | 1 | 1 | 1 | 1 | NA | NA | NA | NA | NA | NA | 1 | 1 |
| 7_1889105 | NA | NA | NA | NA | NA | NA | 1 | 1 | 1 | 1 | 1 | 1 | NA | NA | NA | NA | NA | NA | 1 | 1 |
| 7_1895625 | 2 | 1 | NA | NA | NA | NA | 2 | 1 | 4 | 1 | NA | NA | NA | NA | 1 | 1 | NA | NA | 1 | 1 |
| 7_1895642 | 2 | 0.5 | NA | NA | NA | NA | 2 | 1 | 4 | 0.75 | NA | NA | NA | NA | 1 | 1 | NA | NA | 1 | 1 |
| 7_1895685 | 2 | 1 | NA | NA | NA | NA | 2 | 1 | 4 | 1 | NA | NA | NA | NA | 1 | 1 | NA | NA | 1 | 1 |
| 7_1895713 | 2 | 1 | 1 | 0 | NA | NA | NA | NA | 3 | 0.6667 | NA | NA | NA | NA | 1 | 1 | NA | NA | 1 | 1 |
| 7_1897885 | 4 | 0 | 2 | 0 | 1 | 0 | 2 | 0.5 | 9 | 0.1111 | 5 | 0.2 | 4 | 0.25 | 4 | 0.5 | 1 | 0 | 14 | 0.2857 |
| 7_1897887 | 3 | 0 | 2 | 0 | 1 | 1 | 1 | 0 | 7 | 0.1429 | 3 | 0 | 4 | 0 | 2 | 0.5 | NA | NA | 9 | 0.1111 |
| 7_1897909 | 4 | 1 | 4 | 1 | 1 | 1 | 2 | 1 | 11 | 1 | 5 | 1 | 6 | 1 | 5 | 1 | 3 | 1 | 19 | 1 |
| 7_1897910 | 4 | 0.25 | 4 | 0.25 | 1 | 0 | 2 | 0.5 | 11 | 0.2727 | 5 | 0 | 5 | 0.2 | 5 | 0 | 3 | 0 | 18 | 0.0556 |
| 7_1897917 | 4 | 0 | 4 | 0.25 | 1 | 0 | 2 | 0 | 11 | 0.0909 | 5 | 0.2 | 4 | 0 | 5 | 0.2 | 3 | 0 | 17 | 0.1176 |
| 7_1897918 | 3 | 1 | 4 | 1 | 1 | 1 | 2 | 1 | 10 | 1 | 6 | 1 | 6 | 1 | 5 | 1 | 2 | 1 | 19 | 1 |
| 7_1897919 | 3 | 0 | 4 | 0.5 | 1 | 0 | 2 | 0 | 10 | 0.2 | 4 | 0.25 | 7 | 0 | 4 | 0 | 3 | 0 | 18 | 0.0556 |
| 7_1897922 | 4 | 0.25 | 4 | 0 | NA | NA | 2 | 1 | 10 | 0.3 | 3 | 0.6667 | 5 | 0.2 | 2 | 1 | 2 | 0.5 | 12 | 0.5 |
| 7_1897928 | 4 | 0 | 4 | 0.5 | 2 | 0.5 | 2 | 0.5 | 12 | 0.3333 | 5 | 0.4 | 6 | 0.1667 | 6 | 0.5 | 3 | 0 | 20 | 0.3 |
| 7_1897931 | 3 | 1 | 4 | 1 | 2 | 1 | 2 | 1 | 11 | 1 | 5 | 1 | 8 | 1 | 6 | 1 | 3 | 1 | 22 | 1 |
| 7_1897933 | 4 | 0.75 | NA | NA | 1 | 1 | 2 | 0.5 | 7 | 0.7143 | 6 | 0.3333 | 7 | 0.7143 | 5 | 0.8 | 3 | 0.3333 | 21 | 0.5714 |
| 7_1897939 | 4 | 0.25 | 6 | 0.3333 | 2 | 0.5 | 2 | 0 | 14 | 0.2857 | 5 | 0 | 6 | 0.3333 | 5 | 0.6 | 4 | 0.5 | 20 | 0.35 |
| 7_1898040 | 6 | 0.3333 | 8 | 0.25 | 2 | 0 | 6 | 0.3333 | 22 | 0.2727 | 4 | 0.5 | 8 | 0.375 | 8 | 0.375 | 3 | 0.3333 | 23 | 0.3913 |
| 7_1898052 | 5 | 0 | 7 | 0 | 1 | 1 | 7 | 0.2857 | 20 | 0.15 | 6 | 0 | 9 | 0 | 7 | 0.2857 | 5 | 0 | 27 | 0.0741 |
| 7_1898083 | 3 | 0 | 6 | 0 | 2 | 0.5 | 6 | 0 | 17 | 0.0588 | 5 | 0.2 | 9 | 0.2222 | 7 | 0 | 4 | 0 | 25 | 0.12 |
| 7_1898162 | 6 | 0.3333 | 10 | 0.8 | 4 | 0.5 | 7 | 0.5714 | 27 | 0.5926 | 8 | 0.25 | 9 | 0.7778 | 8 | 0.5 | 4 | 0.75 | 29 | 0.5517 |
| 7_1898177 | 5 | 0.4 | 8 | 0.125 | 5 | 0.2 | 7 | 0.1429 | 25 | 0.2 | 9 | 0.2222 | 11 | 0.0909 | 8 | 0 | 5 | 0.2 | 33 | 0.1212 |
| 7_1899137 | NA | NA | 3 | 0.6667 | 3 | 0 | 1 | 0 | 7 | 0.2857 | 2 | 0 | 2 | 0.5 | 1 | 1 | NA | NA | 5 | 0.4 |
| 7_1902069 | 11 | 0.0909 | 8 | 0.125 | 9 | 0.1111 | 14 | 0 | 42 | 0.0714 | 13 | 0.0769 | 13 | 0 | 7 | 0.1429 | 6 | 0.1667 | 39 | 0.0769 |
| 7_1902076 | 11 | 1 | 9 | 1 | 10 | 0.5 | 15 | 0.7333 | 45 | 0.8 | 15 | 0.8 | 14 | 0.8571 | 13 | 0.9231 | 11 | 0.5455 | 53 | 0.7925 |
| 7_1902081 | 10 | 0.1 | 12 | 0.0833 | 9 | 0.1111 | 14 | 0.0714 | 45 | 0.0889 | 15 | 0.1333 | 13 | 0 | 10 | 0 | 10 | 0.1 | 48 | 0.0625 |
| 7_1902082 | 10 | 0.3 | 13 | 0.2308 | 4 | 0 | 13 | 0.2308 | 40 | 0.225 | 15 | 0.2667 | 13 | 0.2308 | 11 | 0.2727 | 8 | 0.25 | 47 | 0.2553 |
| 7_1902094 | 10 | 0.5 | 9 | 0.4444 | 7 | 0.2857 | 10 | 0.2 | 36 | 0.3611 | 13 | 0.3846 | 11 | 0.1818 | 9 | 0.4444 | 10 | 0.6 | 43 | 0.3953 |
| 7_1902118 | 10 | 0.3 | 14 | 0.3571 | 12 | 0.0833 | 14 | 0.2857 | 50 | 0.26 | 17 | 0.2353 | 16 | 0.0625 | 15 | 0.4667 | 11 | 0.3636 | 59 | 0.2712 |

| | | | | | | | | | | | | | | | | | | | | |
|---|---|---|---|---|---|---|---|---|---|---|---|---|---|---|---|---|---|---|---|---|
| 7_1902130 | 14 | 0.0714 | 15 | 0 | 10 | 0 | 15 | 0.1333 | 54 | 0.0556 | 18 | 0.1111 | 13 | 0.1538 | 12 | 0.1667 | 8 | 0.125 | 51 | 0.1373 |
| 7_1902985 | 20 | 0.45 | 21 | 0.5238 | 17 | 0.5294 | 19 | 0.4211 | 77 | 0.4805 | 19 | 0.4737 | 17 | 0.4118 | 24 | 0.3333 | 17 | 0.4118 | 77 | 0.4026 |
| 7_1902991 | 18 | 0.8333 | 23 | 0.6957 | 20 | 0.9 | 23 | 0.8696 | 84 | 0.8214 | 27 | 0.7778 | 20 | 0.75 | 24 | 0.9167 | 17 | 0.7059 | 88 | 0.7955 |
| 7_1903695 | 2 | 0 | 2 | 0.5 | 2 | 0.5 | 1 | 1 | 7 | 0.4286 | 1 | 0 | 3 | 0.6667 | 4 | 0.25 | 2 | 0 | 10 | 0.3 |
| 7_1903851 | 1 | 0 | 2 | 0.5 | 1 | 0 | 1 | 0 | 5 | 0.2 | 2 | 0 | 2 | 0.5 | 2 | 0 | 2 | 0.5 | 8 | 0.25 |
| 7_1903854 | 1 | 1 | NA | NA | 1 | 1 | NA | NA | 2 | 1 | 2 | 1 | 2 | 1 | 2 | 1 | 1 | 1 | 7 | 1 |
| 7_1903861 | 1 | 0 | 1 | 0 | 1 | 0 | 1 | 1 | 4 | 0.25 | 2 | 0.5 | 1 | 1 | 2 | 0.5 | 2 | 0.5 | 7 | 0.5714 |
| 7_1903862 | 1 | 0 | 1 | 0 | 1 | 1 | 1 | 1 | 4 | 0.5 | 2 | 1 | 2 | 0 | 2 | 0 | 1 | 1 | 7 | 0.4286 |
| 7_1903866 | 1 | 1 | 1 | 0 | 1 | 0 | 1 | 0 | 4 | 0.25 | 2 | 0 | 1 | 0 | 2 | 0.5 | 1 | 1 | 6 | 0.3333 |
| 7_1903866 | 1 | 0 | 1 | 0 | 1 | 0 | 1 | 1 | 4 | 0.25 | 2 | 0 | 1 | 1 | 2 | 1 | 2 | 0.5 | 7 | 0.5714 |
| 7_1903875 | 1 | 0 | 1 | 0 | 1 | 0 | 1 | 1 | 4 | 0.25 | 1 | 0 | 2 | 0.5 | 2 | 0 | 1 | 0 | 6 | 0.1667 |
| 7_1903876 | 1 | 1 | 1 | 1 | 1 | 1 | 1 | 1 | 4 | 1 | 2 | 1 | 1 | 1 | 1 | 1 | 2 | 0.5 | 6 | 0.8333 |
| 7_1903879 | 1 | 1 | 1 | 1 | NA | NA | 1 | 0 | 3 | 0.6667 | 2 | 1 | 2 | 1 | NA | NA | 2 | 0.5 | 6 | 0.8333 |
| 7_1904351 | 1 | 1 | 1 | 1 | 1 | 0 | NA | NA | 3 | 0.6667 | 2 | 0.5 | NA | NA | 1 | 0 | NA | NA | 3 | 0.3333 |
| 7_1904358 | 1 | 1 | 1 | 1 | NA | NA | NA | NA | 2 | 1 | 2 | 1 | NA | NA | 1 | 0 | 1 | 1 | 4 | 0.75 |
| 7_1904375 | 1 | 1 | 1 | 1 | NA | NA | NA | NA | 2 | 1 | 2 | 1 | NA | NA | 1 | 0 | 1 | 0 | 4 | 0.5 |
| 7_1909008 | 2 | 0 | 4 | 0.25 | NA | NA | 1 | 0 | 7 | 0.1429 | 1 | 0 | 2 | 0.5 | 1 | 0 | 1 | 0 | 5 | 0.2 |
| 7_1909018 | 3 | 0.3333 | 1 | 1 | NA | NA | NA | NA | 4 | 0.5 | NA | NA | 1 | 0 | NA | NA | 1 | 1 | 2 | 0.5 |
| 7_1909019 | 3 | 1 | 2 | 1 | NA | NA | 1 | 1 | 6 | 1 | NA | NA | 1 | 1 | NA | NA | 1 | 1 | 2 | 1 |
| 7_1909042 | 2 | 1 | 3 | 0.3333 | NA | NA | 1 | 1 | 6 | 0.6667 | 1 | 1 | 2 | 0.5 | NA | NA | 1 | 1 | 4 | 0.75 |
| 7_1909049 | 4 | 1 | 7 | 0.7143 | NA | NA | 1 | 1 | 12 | 0.8333 | 3 | 0.3333 | 3 | 0.3333 | 2 | 1 | 1 | 0 | 9 | 0.4444 |
| 7_1909052 | 4 | 1 | 7 | 0.5714 | NA | NA | 2 | 0 | 13 | 0.6154 | 2 | 1 | 4 | 0.25 | 1 | 1 | NA | NA | 7 | 0.5714 |
| 7_1909057 | 4 | 0 | 5 | 0 | NA | NA | 2 | 0.5 | 11 | 0.0909 | 3 | 0 | 4 | 0.25 | 1 | 0 | 1 | 1 | 9 | 0.2222 |
| 7_1909059 | 4 | 0.5 | 6 | 0.3333 | NA | NA | 2 | 0 | 12 | 0.3333 | 2 | 0.5 | 4 | 0.25 | 2 | 0.5 | 1 | 0 | 9 | 0.3333 |
| 7_1909064 | 4 | 0.25 | 5 | 0 | NA | NA | 1 | 0 | 10 | 0.1 | 3 | 0.3333 | 4 | 0 | 2 | 0 | 1 | 1 | 10 | 0.2 |
| 7_1909083 | 1 | 0 | 6 | 0.1667 | NA | NA | 1 | 1 | 8 | 0.25 | 3 | 0.3333 | 4 | 0 | 1 | 1 | 1 | 0 | 9 | 0.2222 |
| 7_1909096 | 3 | 0.3333 | 6 | 0 | NA | NA | 1 | 0 | 10 | 0.1 | 2 | 0 | 4 | 0.25 | 2 | 0 | 1 | 0 | 9 | 0.1111 |
| 7_1909107 | 2 | 0.5 | 4 | 0 | NA | NA | 1 | 0 | 7 | 0.1429 | 1 | 0 | 3 | 0.3333 | 1 | 0 | 1 | 0 | 6 | 0.1667 |
| 7_1909113 | 1 | 0 | 3 | 0.6667 | NA | NA | 1 | 0 | 5 | 0.4 | 2 | 0 | 4 | 0 | 2 | 0 | 1 | 1 | 9 | 0.1111 |
| 7_1909124 | 2 | 0.5 | 4 | 0.5 | NA | NA | 1 | 0 | 7 | 0.4286 | 1 | 0 | 4 | 0.25 | 1 | 0 | NA | NA | 6 | 0.1667 |
| 7_1909129 | 2 | 0 | 5 | 0 | NA | NA | 1 | 1 | 8 | 0.125 | 1 | 0 | 4 | 0.25 | 1 | 0 | NA | NA | 6 | 0.1667 |
| 7_1910700 | NA | NA | NA | NA | 1 | 1 | NA | NA | 1 | 1 | 2 | 0.5 | NA | NA | NA | NA | NA | NA | 2 | 0.5 |
| 7_1912880 | 1 | 0 | 1 | 0 | 2 | 0.5 | NA | NA | 4 | 0.25 | NA | NA | 1 | 0 | 2 | 0.5 | NA | NA | 3 | 0.3333 |
| 7_1912885 | 3 | 0.3333 | 1 | 0 | 1 | 0 | NA | NA | 5 | 0.2 | 2 | 0 | 2 | 0.5 | 2 | 0 | 1 | 0 | 7 | 0.1429 |
| 7_1912889 | 2 | 0 | 1 | 1 | 1 | 1 | NA | NA | 4 | 0.5 | 2 | 0.5 | 2 | 0 | 1 | 1 | 1 | 1 | 6 | 0.5 |
| 7_1912891 | 2 | 1 | 1 | 0 | 2 | 0 | NA | NA | 5 | 0.4 | 2 | 0.5 | 2 | 0 | 2 | 0 | 1 | 0 | 7 | 0.1429 |
| 7_1912893 | 2 | 0 | 1 | 0 | 2 | 0.5 | NA | NA | 5 | 0.2 | 1 | 0 | 1 | 0 | 2 | 0 | 2 | 0.5 | 6 | 0.1667 |
| 7_1912896 | 3 | 0.6667 | NA | NA | 2 | 0.5 | NA | NA | 5 | 0.6 | 1 | 1 | 2 | 1 | 2 | 1 | 1 | 1 | 6 | 1 |
| 7_1912907 | 3 | 0.3333 | 1 | 1 | 1 | 1 | NA | NA | 5 | 0.6 | 2 | 0.5 | 2 | 0.5 | 1 | 0 | 2 | 0.5 | 7 | 0.4286 |
| 7_1912909 | 1 | 1 | 1 | 1 | 2 | 1 | NA | NA | 4 | 1 | 2 | 0.5 | 2 | 1 | 2 | 1 | 1 | 1 | 7 | 0.8571 |
| 7_1912912 | 2 | 0 | 1 | 0 | 1 | 1 | NA | NA | 4 | 0.25 | 2 | 0.5 | 2 | 0.5 | 1 | 1 | 1 | 0 | 6 | 0.5 |
| 7_1912914 | 3 | 0.6667 | NA | NA | 1 | 1 | NA | NA | 4 | 0.75 | 2 | 0.5 | 1 | 1 | 2 | 0.5 | 1 | 1 | 6 | 0.6667 |
| 7_1912922 | 2 | 0 | 1 | 0 | 1 | 1 | NA | NA | 4 | 0.25 | 1 | 0 | 2 | 0 | 1 | 1 | 1 | 0 | 5 | 0.2 |
| 7_1915964 | 9 | 0.4444 | 6 | 0.3333 | 7 | 0.2857 | 5 | 0.2 | 27 | 0.3333 | 8 | 0.25 | 6 | 0.5 | 4 | 0.25 | 4 | 0.5 | 22 | 0.3636 |
| 7_1915965 | 12 | 0.25 | 5 | 0 | 8 | 0.25 | 7 | 0.1429 | 32 | 0.1875 | 9 | 0.1111 | 9 | 0 | 6 | 0 | 9 | 0.1111 | 33 | 0.0606 |
| 7_1915972 | 13 | 0.4615 | 7 | 0.5714 | 7 | 0.5714 | 7 | 0.5714 | 34 | 0.5294 | 9 | 0.3333 | 10 | 0.3 | 7 | 0.2857 | 9 | 0.3333 | 35 | 0.3143 |
| 7_1915983 | 13 | 0.0769 | 7 | 0.2857 | 8 | 0.125 | 6 | 0.3333 | 34 | 0.1765 | 9 | 0.3333 | 9 | 0.2222 | 7 | 0.1429 | 9 | 0.4444 | 34 | 0.2941 |
| 7_1915984 | 8 | 1 | 4 | 0.75 | 3 | 1 | 5 | 0.4 | 20 | 0.8 | 6 | 0.5 | 6 | 0.6667 | 6 | 0.6667 | 7 | 0.8571 | 25 | 0.68 |
| 7_1915985 | 10 | 0.7 | 4 | 0.75 | 8 | 0.75 | 7 | 0.4286 | 29 | 0.6552 | 7 | 0.4286 | 4 | 0.75 | 6 | 0.8333 | 8 | 0.5 | 25 | 0.6 |
| 7_1915988 | 13 | 0.2308 | 6 | 0.1667 | 7 | 0.1429 | 7 | 0.2857 | 33 | 0.2121 | 10 | 0.2 | 9 | 0.2222 | 8 | 0.125 | 8 | 0.25 | 35 | 0.2 |
| 7_1915990 | 11 | 0.3636 | 7 | 0.4286 | 6 | 0.1667 | 7 | 0.1429 | 31 | 0.2903 | 7 | 0.1429 | 6 | 0.5 | 6 | 0.8333 | 6 | 0.1667 | 25 | 0.4 |
| 7_1916000 | 10 | 0.3 | 6 | 0.3333 | 6 | 0.5 | 6 | 0.6667 | 28 | 0.4286 | 7 | 0.4286 | 8 | 0.75 | 8 | 0.625 | 6 | 0.5 | 29 | 0.5862 |
| 7_1924036 | NA | NA | 2 | 0 | 2 | 1 | 2 | 1 | 6 | 0.6667 | 3 | 1 | 1 | 1 | NA | NA | 2 | 1 | 6 | 1 |
| 7_1924050 | 1 | 0 | 1 | 0 | 2 | 0 | 2 | 0.5 | 6 | 0.1667 | 3 | 0 | 1 | 0 | NA | NA | 3 | 0.3333 | 7 | 0.1429 |
| 7_1924087 | 1 | 1 | 2 | 1 | 2 | 1 | 2 | 1 | 7 | 1 | 4 | 0.75 | NA | NA | NA | NA | 3 | 1 | 7 | 0.8571 |
| 7_1924089 | 1 | 0 | 1 | 1 | 2 | 1 | 2 | 0.5 | 6 | 0.6667 | 4 | 0.75 | 1 | 1 | NA | NA | 2 | 0.5 | 7 | 0.7143 |
| 7_1924091 | 1 | 1 | 2 | 0.5 | 2 | 0.5 | 3 | 0.3333 | 8 | 0.5 | 3 | 1 | 1 | 1 | NA | NA | 2 | 0.5 | 6 | 0.8333 |
| 7_1924095 | 1 | 1 | 2 | 1 | 2 | 1 | 3 | 0.6667 | 8 | 0.875 | 2 | 1 | 1 | 1 | NA | NA | 2 | 0.5 | 5 | 0.8 |
| 7_1924095 | 1 | 0 | 2 | 0 | 2 | 0 | 3 | 0.3333 | 8 | 0.125 | 3 | 0.6667 | 1 | 0 | NA | NA | 2 | 0.5 | 6 | 0.5 |
| 7_1924099 | 1 | 1 | 1 | 1 | 2 | 0.5 | 3 | 1 | 7 | 0.8571 | 4 | 1 | 1 | 1 | NA | NA | 2 | 1 | 7 | 1 |
| 7_1924105 | 1 | 1 | 1 | 1 | 2 | 1 | 3 | 1 | 7 | 1 | 3 | 1 | 1 | 1 | NA | NA | 1 | 1 | 5 | 1 |
| 7_1924127 | 1 | 0 | NA | NA | 1 | 1 | 3 | 0.3333 | 5 | 0.4 | 3 | 0.6667 | 1 | 1 | NA | NA | 2 | 1 | 6 | 0.8333 |
| 7_1924142 | NA | NA | 1 | 1 | 1 | 1 | 1 | 0 | 3 | 0.6667 | 2 | 0.5 | NA | NA | NA | NA | 2 | 0 | 4 | 0.25 |
| 7_1926487 | 22 | 0.1818 | 18 | 0.1111 | 17 | 0.1176 | 10 | 0.3 | 67 | 0.1642 | 18 | 0.1667 | 24 | 0.0417 | 21 | 0 | 12 | 0.25 | 75 | 0.0933 |
| 7_1926493 | 18 | 0.2222 | 13 | 0 | 8 | 0.125 | 7 | 0.1429 | 46 | 0.1304 | 13 | 0.2308 | 22 | 0.1364 | 15 | 0.0667 | 12 | 0.1667 | 62 | 0.1452 |

| | | | | | | | | | | | | | | | | | | | | |
|---|---|---|---|---|---|---|---|---|---|---|---|---|---|---|---|---|---|---|---|---|
| 7_1926500 | 22 | 0.0909 | 19 | 0.1579 | 14 | 0.2143 | 10 | 0.3 | 65 | 0.1692 | 20 | 0.25 | 27 | 0.1852 | 22 | 0.3636 | 14 | 0.1429 | 83 | 0.241 |
| 7_1926519 | 21 | 1 | 15 | 1 | 16 | 0.9375 | 10 | 1 | 62 | 0.9839 | 16 | 1 | 25 | 1 | 17 | 0.9412 | 13 | 0.9231 | 71 | 0.9718 |
| 7_1926586 | 19 | 0.1579 | 16 | 0.1875 | 17 | 0 | 8 | 0 | 60 | 0.1 | 16 | 0 | 24 | 0.125 | 19 | 0.0526 | 9 | 0 | 68 | 0.0588 |
| 7_1926587 | 18 | 0.1667 | 14 | 0.1429 | 15 | 0.0667 | 9 | 0.1111 | 56 | 0.125 | 14 | 0.2857 | 24 | 0.2083 | 17 | 0.2353 | 8 | 0.25 | 63 | 0.2381 |
| 7_1926609 | 14 | 0.1429 | 12 | 0.4167 | 14 | 0.2143 | 8 | 0.375 | 48 | 0.2708 | 14 | 0.3571 | 19 | 0.1579 | 14 | 0.2857 | 9 | 0.2222 | 56 | 0.25 |
| 7_1926617 | 17 | 0 | 11 | 0 | 17 | 0.1176 | 7 | 0.2857 | 52 | 0.0769 | 14 | 0.0714 | 21 | 0.1429 | 16 | 0.1875 | 10 | 0.1 | 61 | 0.1311 |
| 7_1926620 | 13 | 0.3846 | 9 | 0.2222 | 10 | 0.3 | 8 | 0.375 | 40 | 0.325 | 11 | 0.1818 | 16 | 0.5 | 18 | 0.5 | 9 | 0.3333 | 54 | 0.4074 |
| 7_1926621 | 15 | 0.1333 | 13 | 0.1538 | 17 | 0.6471 | 7 | 0.1429 | 52 | 0.3077 | 11 | 0.0909 | 22 | 0.2273 | 15 | 0.4667 | 10 | 0.2 | 58 | 0.2586 |
| 7_1926622 | 16 | 0.1875 | 12 | 0.1667 | 18 | 0.3333 | 8 | 0.25 | 54 | 0.2407 | 12 | 0.0833 | 22 | 0.1364 | 16 | 0.25 | 9 | 0.5556 | 59 | 0.2203 |
| 7_1926623 | 16 | 0 | 9 | 0.1111 | 16 | 0.125 | 5 | 0 | 46 | 0.0652 | 14 | 0.2143 | 23 | 0.1739 | 17 | 0.2353 | 11 | 0.0909 | 65 | 0.1846 |
| 7_1926837 | 13 | 0 | 7 | 0.1429 | 13 | 0.0769 | 7 | 0 | 40 | 0.05 | 9 | 0 | 19 | 0.0526 | 13 | 0.2308 | 8 | 0.125 | 49 | 0.102 |
| 7_1929895 | 9 | 0.5556 | 9 | 0.5556 | 11 | 0.3636 | 10 | 0.7 | 39 | 0.5385 | 5 | 0.4 | 9 | 0.3333 | 6 | 0.5 | 3 | 0.3333 | 23 | 0.3913 |
| 7_1929899 | 7 | 0.1429 | 5 | 0.2 | 4 | 0 | 5 | 0 | 21 | 0.0952 | 3 | 0 | 4 | 0.5 | 5 | 0.2 | 4 | 0.25 | 16 | 0.25 |
| 7_1929906 | 10 | 0.8 | 6 | 0.8333 | 10 | 0.9 | 8 | 0.25 | 34 | 0.7059 | 4 | 0.5 | 8 | 0.625 | 6 | 1 | 4 | 1 | 22 | 0.7727 |
| 7_1929907 | 8 | 0.625 | 9 | 0.5556 | 9 | 0.6667 | 7 | 0.5714 | 33 | 0.6061 | 3 | 0.6667 | 8 | 0.75 | 4 | 0.25 | 4 | 0.25 | 19 | 0.5263 |
| 7_1929909 | 10 | 0.4 | 9 | 0.4444 | 11 | 0.4545 | 9 | 0.3333 | 39 | 0.4103 | 4 | 0.75 | 8 | 0.625 | 6 | 0.5 | 4 | 0.5 | 22 | 0.5909 |
| 7_1929911 | 10 | 0.9 | 9 | 1 | 10 | 1 | 8 | 1 | 37 | 0.973 | 3 | 1 | 8 | 0.875 | 5 | 1 | 4 | 1 | 20 | 0.95 |
| 7_1929914 | 8 | 0.625 | 8 | 0.5 | 9 | 0.5556 | 8 | 0.25 | 33 | 0.4848 | 3 | 0.6667 | 8 | 0.125 | 6 | 0.5 | 3 | 0.3333 | 20 | 0.35 |
| 7_1929915 | 7 | 0.4286 | 7 | 0.4286 | 7 | 0.4286 | 7 | 0.4286 | 28 | 0.4286 | 2 | 0 | 8 | 0.25 | 4 | 0 | 3 | 0.6667 | 17 | 0.2353 |
| 7_1929920 | 8 | 0.875 | 7 | 1 | 5 | 1 | 4 | 1 | 24 | 0.9583 | 2 | 1 | 4 | 1 | 4 | 1 | 3 | 1 | 13 | 1 |
| 7_1929923 | 9 | 0.4444 | 9 | 0.5556 | 10 | 0.6 | 9 | 0.7778 | 37 | 0.5946 | 3 | 0.6667 | 8 | 0.75 | 5 | 0.4 | 4 | 0.5 | 20 | 0.6 |
| 7_1929927 | 6 | 0.3333 | 7 | 0.8571 | 8 | 0.625 | 9 | 0.4444 | 30 | 0.5667 | 1 | 1 | 4 | 0.5 | 2 | 0 | 2 | 0.5 | 9 | 0.4444 |
| 7_1929943 | 2 | 0.5 | 6 | 0 | 7 | 0 | 9 | 0.1111 | 24 | 0.0833 | 2 | 0 | 7 | 0.2857 | 3 | 0 | 3 | 0 | 15 | 0.1333 |
| 7_1934026 | 1 | 0 | 1 | 1 | 1 | 0 | 1 | 0 | 4 | 0.25 | 1 | 1 | 1 | 1 | NA | NA | 1 | 1 | 3 | 1 |
| 7_1934060 | 1 | 1 | NA | NA | 1 | 0 | 2 | 1 | 4 | 0.75 | NA | NA | 1 | 1 | NA | NA | NA | NA | 1 | 1 |
| 7_1936598 | 2 | 0.5 | 2 | 0 | NA | NA | 1 | 0 | 5 | 0.2 | NA | NA | NA | NA | 2 | 0.5 | NA | NA | 2 | 0.5 |
| 7_1936752 | 22 | 0.7273 | 23 | 0.3478 | 20 | 0.4 | 15 | 0.5333 | 80 | 0.5 | 21 | 0.5714 | 23 | 0.5217 | 21 | 0.6667 | 13 | 0.6154 | 78 | 0.5897 |
| 7_1936758 | 26 | 0.3077 | 30 | 0.3667 | 25 | 0.36 | 22 | 0.6364 | 103 | 0.4078 | 25 | 0.48 | 30 | 0.3333 | 27 | 0.2963 | 15 | 0.3333 | 97 | 0.3608 |
| 7_1936763 | 29 | 0.1724 | 32 | 0.1562 | 29 | 0.1034 | 28 | 0.1786 | 118 | 0.1525 | 27 | 0.2593 | 35 | 0.1143 | 28 | 0.1429 | 18 | 0.2222 | 108 | 0.1759 |
| 7_1943047 | 458 | 0.0808 | 398 | 0.0955 | 397 | 0.0756 | 417 | 0.0935 | 1670 | 0.0862 | 477 | 0.0692 | 404 | 0.0941 | 415 | 0.0795 | 383 | 0.0757 | 1679 | 0.0792 |
| 7_1943047 | 378 | 0.1032 | 323 | 0.1053 | 334 | 0.0659 | 349 | 0.0688 | 1384 | 0.086 | 393 | 0.0662 | 347 | 0.0865 | 347 | 0.098 | 329 | 0.0881 | 1416 | 0.084 |
| 7_1943057 | 402 | 0.0896 | 321 | 0.0498 | 332 | 0.0542 | 366 | 0.0738 | 1421 | 0.0683 | 404 | 0.0545 | 350 | 0.0686 | 338 | 0.0799 | 326 | 0.0583 | 1418 | 0.0649 |
| 7_1943101 | 397 | 0.1713 | 343 | 0.1866 | 367 | 0.139 | 355 | 0.1634 | 1462 | 0.1648 | 409 | 0.1443 | 364 | 0.1676 | 372 | 0.1344 | 330 | 0.1485 | 1475 | 0.1485 |
| 7_1943527 | 8 | 0 | 8 | 0.375 | 8 | 0 | NA | NA | 24 | 0.125 | 4 | 0.25 | 3 | 0.3333 | 3 | 0.3333 | 1 | 1 | 11 | 0.3636 |
| 7_1943529 | 13 | 0.1538 | 15 | 0.6 | 14 | 0.5 | 6 | 0.6667 | 48 | 0.4583 | 9 | 0.4444 | 10 | 0.6 | 14 | 0.5 | 10 | 0.7 | 43 | 0.5581 |
| 7_1943540 | 11 | 0.5455 | 18 | 0.3889 | 15 | 0.4 | 6 | 0.5 | 50 | 0.44 | 11 | 0.4545 | 10 | 0.6 | 14 | 0.4286 | 9 | 0.4444 | 44 | 0.4773 |
| 7_1943551 | 10 | 0.2 | 13 | 0.1538 | 11 | 0.0909 | 3 | 0.3333 | 37 | 0.1622 | 8 | 0 | 8 | 0.375 | 10 | 0.4 | 6 | 0.3333 | 32 | 0.2812 |
| 7_1945816 | 1 | 1 | NA | NA | NA | NA | 2 | 1 | 3 | 1 | 1 | 1 | NA | NA | NA | NA | NA | NA | 1 | 1 |
| 7_1945822 | 1 | 1 | NA | NA | NA | NA | 2 | 0.5 | 3 | 0.6667 | 1 | 1 | NA | NA | NA | NA | NA | NA | 1 | 1 |
| 7_1945824 | 1 | 0 | NA | NA | NA | NA | 2 | 0.5 | 3 | 0.3333 | 1 | 1 | NA | NA | NA | NA | NA | NA | 1 | 1 |
| 7_1945830 | 1 | 1 | NA | NA | NA | NA | 1 | 1 | 2 | 1 | 1 | 1 | NA | NA | NA | NA | NA | NA | 1 | 1 |
| 7_1945831 | 1 | 0 | NA | NA | NA | NA | 2 | 1 | 3 | 0.6667 | 1 | 1 | NA | NA | NA | NA | NA | NA | 1 | 1 |
| 7_1945838 | 1 | 0 | NA | NA | NA | NA | 2 | 1 | 3 | 0.6667 | 1 | 1 | NA | NA | NA | NA | NA | NA | 1 | 1 |
| 7_1949050 | 4 | 0.5 | 4 | 0.5 | NA | NA | 2 | 0 | 10 | 0.4 | NA | NA | 2 | 0 | 3 | 0.3333 | 4 | 0 | 9 | 0.1111 |
| 7_1963889 | 2 | 0 | 1 | 0 | 2 | 0.5 | 1 | 1 | 6 | 0.3333 | NA | NA | 2 | 1 | 2 | 0 | 2 | 0 | 6 | 0.3333 |
| 7_1963896 | 2 | 0.5 | 1 | 1 | 1 | 1 | NA | NA | 4 | 0.75 | NA | NA | 2 | 0.5 | 2 | 1 | 1 | 1 | 5 | 0.8 |
| 7_1965081 | 2 | 0.5 | NA | NA | 1 | 0 | 1 | 0 | 4 | 0.25 | NA | NA | 1 | 0 | 1 | 0 | 2 | 0.5 | 4 | 0.25 |
| 7_1965257 | 1 | 0 | NA | NA | 2 | 1 | NA | NA | 3 | 0.6667 | NA | NA | 1 | 0 | 2 | 0.5 | 2 | 0.5 | 5 | 0.4 |
| 7_1965273 | 2 | 0 | NA | NA | 2 | 0.5 | NA | NA | 4 | 0.25 | NA | NA | 1 | 0 | 2 | 0.5 | 2 | 0.5 | 5 | 0.4 |
| 7_2380396 | 4 | 0 | 2 | 0.5 | 4 | 0.25 | 2 | 0.5 | 12 | 0.25 | 2 | 1 | 2 | 1 | 3 | 0.3333 | 4 | 0.75 | 11 | 0.7273 |
| 7_2380413 | 2 | 0 | 1 | 0 | 1 | 0 | 2 | 0.5 | 6 | 0.1667 | 2 | 0 | 2 | 0.5 | 1 | 0 | 3 | 0.3333 | 8 | 0.25 |
| 7_2380430 | 3 | 1 | 2 | 1 | 4 | 1 | 4 | 1 | 13 | 1 | 2 | 1 | 3 | 1 | 2 | 1 | 3 | 1 | 10 | 1 |
| 7_2380437 | 3 | 0 | 1 | 0 | 4 | 0.25 | 4 | 0 | 12 | 0.0833 | 2 | 0.5 | 3 | 0.3333 | 1 | 0 | 3 | 0 | 9 | 0.2222 |
| 7_2380476 | 3 | 1 | 1 | 1 | 4 | 1 | 4 | 0.75 | 12 | 0.9167 | 1 | 1 | 3 | 1 | 1 | 1 | 4 | 1 | 9 | 1 |
| 7_2380483 | 3 | 0 | 1 | 0 | 6 | 0.1667 | 4 | 0 | 14 | 0.0714 | 2 | 0 | 5 | 0.2 | 2 | 0.5 | 4 | 0 | 13 | 0.1538 |
| 7_2380484 | 4 | 0 | 2 | 0 | 5 | 0 | 4 | 0.25 | 15 | 0.0667 | 1 | 1 | 6 | 0 | 3 | 0 | 4 | 0 | 14 | 0.0714 |
| 7_2380499 | 4 | 0.25 | 1 | 0 | 4 | 0 | 4 | 0 | 13 | 0.0769 | 2 | 1 | 6 | 0.1667 | 3 | 0.6667 | 4 | 0.5 | 15 | 0.4667 |
| 7_2380501 | 3 | 0.6667 | 1 | 1 | 5 | 0.8 | 4 | 0.75 | 13 | 0.7692 | 1 | 1 | 6 | 0.3333 | 2 | 1 | 2 | 1 | 11 | 0.6364 |
| 7_2380518 | 4 | 0.25 | 1 | 0 | 6 | 0 | 4 | 0.25 | 15 | 0.1333 | 2 | 1 | 7 | 0 | 2 | 0 | 2 | 0.5 | 13 | 0.2308 |
| 7_2380535 | 5 | 0.6 | 2 | 0.5 | 6 | 0.5 | 3 | 1 | 16 | 0.625 | 2 | 1 | 9 | 0.5556 | 3 | 0.3333 | 3 | 1 | 17 | 0.6471 |
| 7_2380543 | 4 | 0.25 | 2 | 0 | 5 | 0.4 | 2 | 0.5 | 13 | 0.3077 | 1 | 0 | 6 | 0.3333 | 1 | 0 | 2 | 0.5 | 10 | 0.3 |
| 7_2380557 | 6 | 0.5 | 2 | 0 | 6 | 0.3333 | 5 | 0.6 | 19 | 0.4211 | 2 | 0 | 9 | 0.2222 | 6 | 0.5 | 4 | 0.5 | 21 | 0.3333 |
| 7_2380558 | 6 | 1 | 2 | 1 | 7 | 1 | 4 | 1 | 19 | 1 | 2 | 1 | 7 | 1 | 5 | 1 | 3 | 1 | 17 | 1 |
| 7_2380563 | 6 | 0.5 | 1 | 0 | 7 | 0.1429 | 4 | 0.25 | 18 | 0.2778 | 3 | 0.6667 | 8 | 0.5 | 6 | 0.5 | 4 | 0.5 | 21 | 0.5238 |
| 7_2380566 | 7 | 0.2857 | 1 | 1 | 6 | 0.6667 | 3 | 0.3333 | 17 | 0.4706 | 3 | 0.3333 | 9 | 0.7778 | 3 | 0 | 4 | 0.25 | 19 | 0.4737 |

| | | | | | | | | | | | | | | | | | | | | |
|---|---|---|---|---|---|---|---|---|---|---|---|---|---|---|---|---|---|---|---|---|
| 7_2380574 | 5 | 0.8 | 2 | 1 | 4 | 0.25 | 3 | 0.6667 | 14 | 0.6429 | 3 | 1 | 5 | 0.6 | 1 | 0 | 2 | 0.5 | 11 | 0.6364 |
| 7_2380578 | 6 | 1 | 2 | 1 | 7 | 0.8571 | 4 | 1 | 19 | 0.9474 | 4 | 1 | 6 | 1 | 4 | 1 | 2 | 1 | 16 | 1 |
| 7_2380583 | 6 | 0 | 2 | 0.5 | 7 | 0 | 5 | 0.2 | 20 | 0.1 | 2 | 0 | 8 | 0.125 | 4 | 0 | 3 | 0 | 17 | 0.0588 |
| 7_2380589 | 6 | 0.8333 | 2 | 0.5 | 9 | 0.6667 | 5 | 0.4 | 22 | 0.6364 | 3 | 0.6667 | 9 | 0.6667 | 3 | 0.3333 | 2 | 0 | 17 | 0.5294 |
| 7_2380605 | 5 | 0.6 | 1 | 0 | 5 | 0 | 5 | 0 | 16 | 0.1875 | 3 | 0 | 7 | 0.2857 | 2 | 0 | 4 | 0.25 | 16 | 0.1875 |
| 7_2380644 | 6 | 0.6667 | 4 | 0.5 | 9 | 0.5556 | 6 | 0.5 | 25 | 0.56 | 3 | 0.6667 | 11 | 0.9091 | 6 | 0.5 | 4 | 1 | 24 | 0.7917 |
| 7_2380653 | 2 | 0 | 2 | 0 | 9 | 0.2222 | 6 | 0 | 19 | 0.1053 | 3 | 0 | 8 | 0 | 5 | 0 | 4 | 0.25 | 20 | 0.05 |
| 7_2380662 | 6 | 0.1667 | 2 | 1 | 11 | 0.1818 | 7 | 0.1429 | 26 | 0.2308 | 3 | 0.3333 | 10 | 0.1 | 7 | 0.2857 | 5 | 0.4 | 25 | 0.24 |
| 7_2380679 | 7 | 0.7143 | 3 | 0.6667 | 8 | 0.625 | 8 | 0.125 | 26 | 0.5 | 3 | 0.6667 | 11 | 0.4545 | 5 | 0.6 | 3 | 0.6667 | 22 | 0.5455 |
| 7_2380680 | 5 | 0 | 2 | 0 | 6 | 0.1667 | 4 | 0 | 17 | 0.0588 | 2 | 0.5 | 11 | 0.3636 | 3 | 0.3333 | 4 | 0.5 | 20 | 0.4 |
| 7_2380680 | 6 | 0.3333 | 3 | 0.6667 | 11 | 0.5455 | 6 | 0.3333 | 26 | 0.4615 | 3 | 0.6667 | 12 | 0.4167 | 8 | 0.5 | 3 | 0.3333 | 26 | 0.4615 |
| 7_2380682 | 7 | 0.1429 | 3 | 0 | 11 | 0.0909 | 7 | 0.1429 | 28 | 0.1071 | 3 | 0.3333 | 12 | 0.1667 | 8 | 0 | 4 | 0 | 27 | 0.1111 |
| 7_2380697 | 7 | 0.1429 | 2 | 0 | 10 | 0.4 | 8 | 0 | 27 | 0.1852 | 3 | 0.3333 | 11 | 0.3636 | 10 | 0.3 | 4 | 0 | 28 | 0.2857 |
| 7_2382540 | 1 | 1 | NA | NA | NA | NA | NA | NA | 1 | 1 | NA | NA | 1 | 1 | NA | NA | NA | NA | 1 | 1 |
| 7_2382547 | 2 | 0.5 | NA | NA | 1 | 0 | NA | NA | 3 | 0.3333 | NA | NA | 1 | 1 | NA | NA | NA | NA | 1 | 1 |
| 7_2382563 | 2 | 1 | NA | NA | NA | NA | NA | NA | 2 | 1 | NA | NA | 1 | 1 | NA | NA | NA | NA | 1 | 1 |
| 7_2382603 | 2 | 0.5 | NA | NA | 1 | 1 | NA | NA | 3 | 0.6667 | NA | NA | 1 | 1 | NA | NA | NA | NA | 1 | 1 |
| 7_2382682 | 2 | 0.5 | NA | NA | 1 | 0 | NA | NA | 3 | 0.3333 | NA | NA | 1 | 1 | NA | NA | NA | NA | 1 | 1 |
| 7_2382698 | 2 | 0.5 | NA | NA | 1 | 0 | NA | NA | 3 | 0.3333 | NA | NA | 1 | 0 | 1 | 1 | NA | NA | 2 | 0.5 |
| 7_2382704 | 1 | 1 | NA | NA | 1 | 1 | NA | NA | 2 | 1 | NA | NA | NA | NA | 1 | 1 | NA | NA | 1 | 1 |
| 7_2383296 | NA | NA | NA | NA | NA | NA | 1 | 1 | 1 | 1 | NA | NA | 1 | 1 | NA | NA | NA | NA | 1 | 1 |
| 7_2383308 | NA | NA | NA | NA | NA | NA | 1 | 1 | 1 | 1 | NA | NA | 1 | 1 | NA | NA | NA | NA | 1 | 1 |
| 7_2384004 | 1 | 1 | 1 | 1 | 1 | 1 | 1 | 1 | 4 | 1 | NA | NA | 3 | 0.6667 | 1 | 1 | NA | NA | 4 | 0.75 |
| 7_2384008 | 2 | 1 | 1 | 1 | 1 | 1 | NA | NA | 4 | 1 | NA | NA | 3 | 1 | 1 | 1 | NA | NA | 4 | 1 |
| 7_2384018 | 2 | 0.5 | NA | NA | 1 | 1 | 1 | 1 | 4 | 0.75 | NA | NA | 3 | 0.3333 | 1 | 0 | NA | NA | 4 | 0.25 |
| 7_2384020 | 2 | 0 | 1 | 1 | NA | NA | 1 | 1 | 4 | 0.5 | NA | NA | 3 | 1 | 1 | 1 | NA | NA | 4 | 1 |
| 7_2384032 | 2 | 1 | 1 | 0 | 1 | 1 | 1 | 0 | 5 | 0.6 | NA | NA | 3 | 0.3333 | 1 | 0 | NA | NA | 4 | 0.25 |
| 7_2384048 | 2 | 0 | 1 | 1 | 1 | 1 | 1 | 0 | 5 | 0.4 | NA | NA | 2 | 0.5 | 1 | 0 | NA | NA | 3 | 0.3333 |
| 7_2384077 | 2 | 0.5 | 1 | 0 | 1 | 1 | NA | NA | 4 | 0.5 | NA | NA | 3 | 0.3333 | NA | NA | 1 | 1 | 4 | 0.5 |
| 7_2384118 | 3 | 0.6667 | 1 | 1 | 2 | 1 | 1 | 1 | 7 | 0.8571 | NA | NA | 3 | 1 | 1 | 1 | 1 | 1 | 5 | 1 |
| 7_2384136 | 2 | 0 | 1 | 0 | 2 | 1 | 1 | 0 | 6 | 0.3333 | NA | NA | 4 | 0.5 | 1 | 1 | 1 | 0 | 6 | 0.5 |
| 7_2384148 | 3 | 0.6667 | 1 | 1 | 2 | 1 | 1 | 1 | 7 | 0.8571 | NA | NA | 4 | 1 | 1 | 1 | 1 | 1 | 6 | 1 |
| 7_2384174 | 3 | 0.3333 | 1 | 1 | 2 | 0 | 2 | 0.5 | 8 | 0.375 | NA | NA | 4 | 0.25 | 1 | 1 | 1 | 1 | 6 | 0.5 |
| 7_2384186 | 2 | 0.5 | NA | NA | 1 | 0 | 2 | 0.5 | 5 | 0.4 | NA | NA | 4 | 1 | NA | NA | 2 | 1 | 6 | 1 |
| 7_2384188 | 1 | 0 | NA | NA | 2 | 0.5 | 2 | 0.5 | 5 | 0.4 | 1 | 1 | 4 | 0.75 | NA | NA | NA | NA | 5 | 0.8 |
| 7_2384215 | 4 | 0.25 | 1 | 1 | 1 | 1 | 2 | 0 | 8 | 0.375 | 1 | 1 | 3 | 0.6667 | 1 | 0 | 2 | 0.5 | 7 | 0.5714 |
| 7_2384222 | 4 | 0.75 | NA | NA | 2 | 0.5 | 2 | 0.5 | 8 | 0.625 | 1 | 0 | 4 | 0.25 | 1 | 1 | 2 | 0 | 8 | 0.25 |
| 7_2384228 | 4 | 0 | 1 | 0 | 2 | 0 | 2 | 0.5 | 9 | 0.1111 | 1 | 0 | 4 | 0.25 | 1 | 0 | 2 | 0.5 | 8 | 0.25 |
| 7_2384234 | 4 | 0 | 1 | 0 | 2 | 0 | 2 | 0.5 | 9 | 0.1111 | 1 | 1 | 4 | 0 | 1 | 0 | 2 | 0.5 | 8 | 0.25 |
| 7_2384247 | 4 | 0.25 | 1 | 1 | 2 | 0.5 | 2 | 0 | 9 | 0.3333 | 1 | 1 | 4 | 0.75 | 1 | 0 | 2 | 0.5 | 8 | 0.625 |
| 7_2384253 | 4 | 0.25 | 1 | 1 | 2 | 1 | 1 | 1 | 8 | 0.625 | 2 | 1 | 4 | 0.75 | 2 | 0.5 | 2 | 0.5 | 10 | 0.7 |
| 7_2389410 | 1 | 1 | NA | NA | NA | NA | NA | NA | 1 | 1 | 1 | 1 | NA | NA | NA | NA | 1 | 0 | 2 | 0.5 |
| 7_2389727 | NA | NA | NA | NA | 1 | 1 | NA | NA | 1 | 1 | 1 | 0 | 1 | 0 | NA | NA | 1 | 1 | 3 | 0.3333 |
| 7_2389784 | 1 | 1 | NA | NA | 1 | 1 | NA | NA | 2 | 1 | 1 | 1 | NA | NA | NA | NA | 1 | 1 | 2 | 1 |
| 7_2389788 | 1 | 1 | NA | NA | 1 | 1 | NA | NA | 2 | 1 | 1 | 1 | NA | NA | NA | NA | 1 | 1 | 2 | 1 |
| 7_2389805 | 1 | 1 | NA | NA | 1 | 0 | NA | NA | 2 | 0.5 | 1 | 1 | NA | NA | NA | NA | 1 | 0 | 2 | 0.5 |
| 7_2389844 | 1 | 1 | NA | NA | NA | NA | NA | NA | 1 | 1 | 1 | 0 | NA | NA | 1 | 1 | NA | NA | 2 | 0.5 |
| 7_2389869 | 1 | 1 | NA | NA | NA | NA | NA | NA | 1 | 1 | 1 | 0 | NA | NA | NA | NA | 1 | 1 | 2 | 0.5 |
| 7_2389910 | 1 | 1 | NA | NA | NA | NA | NA | NA | 1 | 1 | 1 | 0 | NA | NA | NA | NA | 1 | 1 | 2 | 0.5 |
| 7_2392501 | 2 | 0.5 | 2 | 0.5 | NA | NA | NA | NA | 4 | 0.5 | 1 | 0 | 2 | 0.5 | NA | NA | 2 | 0 | 5 | 0.2 |
| 7_2392506 | 2 | 0.5 | 2 | 0.5 | NA | NA | NA | NA | 4 | 0.5 | 1 | 0 | 2 | 0.5 | NA | NA | 1 | 0 | 4 | 0.25 |
| 7_2392515 | 2 | 1 | NA | NA | NA | NA | NA | NA | 2 | 1 | NA | NA | 2 | 0.5 | NA | NA | 1 | 1 | 3 | 0.6667 |
| 7_2392529 | 2 | 0.5 | 2 | 0 | NA | NA | NA | NA | 4 | 0.25 | 1 | 0 | 1 | 1 | NA | NA | 1 | 0 | 3 | 0.3333 |
| 7_2392568 | 2 | 0.5 | 1 | 1 | NA | NA | NA | NA | 3 | 0.6667 | 1 | 0 | 1 | 1 | NA | NA | NA | NA | 2 | 0.5 |
| 7_2392574 | 2 | 0.5 | 1 | 0 | NA | NA | NA | NA | 3 | 0.3333 | NA | NA | NA | NA | NA | NA | 1 | 1 | 1 | 1 |
| 7_2392581 | 2 | 1 | 2 | 1 | NA | NA | NA | NA | 4 | 1 | 1 | 1 | 1 | 1 | NA | NA | 1 | 0 | 3 | 0.6667 |
| 7_2392704 | 2 | 1 | 2 | 0 | NA | NA | NA | NA | 4 | 0.5 | 1 | 1 | 1 | 1 | NA | NA | 1 | 0 | 3 | 0.6667 |
| 7_2392759 | 1 | 1 | 1 | 1 | NA | NA | NA | NA | 2 | 1 | 1 | 1 | 1 | 1 | NA | NA | 1 | 1 | 3 | 1 |
| 7_2392774 | 1 | 0 | 2 | 0.5 | NA | NA | NA | NA | 3 | 0.3333 | 1 | 1 | 1 | 0 | NA | NA | 1 | 1 | 3 | 0.6667 |
| 7_2392787 | 1 | 1 | 2 | 1 | NA | NA | NA | NA | 3 | 1 | 1 | 1 | 1 | 1 | NA | NA | 1 | 1 | 3 | 1 |
| 7_2392878 | 1 | 1 | 1 | 1 | NA | NA | NA | NA | 2 | 1 | 1 | 1 | 1 | 1 | NA | NA | 1 | 1 | 3 | 1 |
| 7_2392893 | 1 | 1 | 1 | 1 | NA | NA | NA | NA | 2 | 1 | 1 | 1 | 1 | 1 | NA | NA | 1 | 1 | 3 | 1 |
| 7_2392904 | 1 | 1 | 1 | 1 | NA | NA | NA | NA | 2 | 1 | 1 | 1 | 1 | 1 | NA | NA | 1 | 1 | 3 | 1 |
| 7_2395987 | NA | NA | 1 | 1 | 1 | 1 | NA | NA | 2 | 1 | NA | NA | 2 | 0.5 | NA | NA | NA | NA | 2 | 0.5 |
| 7_2395990 | NA | NA | 1 | 0 | 3 | 0.6667 | NA | NA | 4 | 0.5 | NA | NA | 2 | 0.5 | NA | NA | NA | NA | 2 | 0.5 |

| | | | | | | | | | | | | | | | | | | | | |
|---|---|---|---|---|---|---|---|---|---|---|---|---|---|---|---|---|---|---|---|---|
| 7_2396019 | NA | NA | 1 | 1 | 2 | 0.5 | NA | NA | 3 | 0.6667 | NA | NA | 2 | 0.5 | NA | NA | NA | NA | 2 | 0.5 |
| 7_2396030 | NA | NA | 1 | 0 | 3 | 0.3333 | NA | NA | 4 | 0.25 | NA | NA | 2 | 0.5 | 1 | 0 | NA | NA | 3 | 0.3333 |
| 7_2396045 | NA | NA | 1 | 0 | 3 | 0 | 1 | 1 | 5 | 0.2 | NA | NA | 2 | 0.5 | 1 | 0 | NA | NA | 3 | 0.3333 |
| 7_2396051 | NA | NA | 1 | 1 | 3 | 0.6667 | NA | NA | 4 | 0.75 | NA | NA | 2 | 1 | 1 | 1 | NA | NA | 3 | 1 |
| 7_2396099 | NA | NA | 1 | 1 | 2 | 0.5 | 1 | 1 | 4 | 0.75 | NA | NA | 2 | 1 | 1 | 1 | NA | NA | 3 | 1 |
| 7_2396105 | NA | NA | 2 | 0 | 3 | 0.6667 | 1 | 0 | 6 | 0.3333 | NA | NA | 2 | 1 | 1 | 1 | NA | NA | 3 | 1 |
| 7_2396120 | NA | NA | 1 | 1 | 1 | 1 | 1 | 1 | 3 | 1 | NA | NA | 1 | 1 | 1 | 1 | NA | NA | 2 | 1 |
| 7_2396137 | NA | NA | 1 | 1 | 3 | 0.6667 | 1 | 1 | 5 | 0.8 | NA | NA | 2 | 0 | 1 | 1 | NA | NA | 3 | 0.3333 |
| 7_2396144 | NA | NA | 2 | 0.5 | 2 | 1 | 1 | 0 | 5 | 0.6 | NA | NA | 2 | 1 | 1 | 1 | NA | NA | 3 | 1 |
| 7_2396157 | NA | NA | 1 | 1 | 2 | 0.5 | NA | NA | 3 | 0.6667 | NA | NA | 2 | 0.5 | 1 | 1 | NA | NA | 3 | 0.6667 |
| 7_2396161 | NA | NA | 2 | 1 | 3 | 1 | 1 | 1 | 6 | 1 | NA | NA | 2 | 1 | 1 | 1 | NA | NA | 3 | 1 |
| 7_2396165 | NA | NA | 2 | 0.5 | 3 | 0.3333 | 1 | 0 | 6 | 0.3333 | NA | NA | 2 | 0 | 1 | 1 | NA | NA | 3 | 0.3333 |
| 7_2396169 | NA | NA | 3 | 0.3333 | 1 | 1 | 1 | 1 | 5 | 0.6 | NA | NA | 2 | 1 | 1 | 0 | 1 | 0 | 4 | 0.5 |
| 7_2396180 | NA | NA | NA | NA | 2 | 1 | NA | NA | 2 | 1 | NA | NA | 2 | 1 | 1 | 1 | NA | NA | 3 | 1 |
| 7_2396182 | NA | NA | NA | NA | 2 | 1 | NA | NA | 2 | 1 | NA | NA | 2 | 1 | 1 | 1 | NA | NA | 3 | 1 |
| 7_2396375 | NA | NA | NA | NA | 1 | 1 | NA | NA | 1 | 1 | NA | NA | 1 | 1 | NA | NA | NA | NA | 1 | 1 |
| 7_2397513 | 1 | 1 | NA | NA | 1 | 1 | 1 | 1 | 3 | 1 | 1 | 1 | NA | NA | 1 | 1 | NA | NA | 2 | 1 |
| 7_2397539 | 1 | 1 | NA | NA | 1 | 0 | 1 | 0 | 3 | 0.3333 | 2 | 0.5 | NA | NA | 2 | 0 | NA | NA | 4 | 0.25 |
| 7_2397595 | 3 | 1 | 1 | 1 | 1 | 1 | 2 | 0 | 7 | 0.7143 | 2 | 1 | 1 | 1 | 2 | 1 | NA | NA | 5 | 1 |
| 7_2399087 | NA | NA | 2 | 1 | NA | NA | NA | NA | 2 | 1 | 2 | 0.5 | 1 | 1 | 2 | 1 | 3 | 1 | 8 | 0.875 |
| 7_2399136 | NA | NA | 1 | 1 | NA | NA | NA | NA | 1 | 1 | 1 | 1 | 1 | 1 | 2 | 1 | 1 | 1 | 5 | 1 |
| 7_2399173 | NA | NA | 1 | 1 | NA | NA | NA | NA | 1 | 1 | 1 | 1 | 1 | 1 | 1 | 1 | 1 | 1 | 4 | 1 |
| 7_2400137 | 1 | 0 | 2 | 0.5 | 1 | 1 | 1 | 1 | 5 | 0.6 | 1 | 1 | NA | NA | NA | NA | NA | NA | 1 | 1 |
| 7_2400155 | 1 | 1 | 2 | 0.5 | 1 | 0 | 1 | 0 | 5 | 0.4 | 1 | 1 | NA | NA | NA | NA | NA | NA | 1 | 1 |
| 7_2400162 | 1 | 1 | 2 | 1 | 1 | 1 | 1 | 1 | 5 | 1 | 1 | 1 | NA | NA | NA | NA | NA | NA | 1 | 1 |
| 7_2400165 | 1 | 1 | 1 | 1 | NA | NA | NA | NA | 2 | 1 | 1 | 1 | NA | NA | NA | NA | NA | NA | 1 | 1 |
| 7_2400182 | NA | NA | 2 | 0.5 | 1 | 1 | 1 | 0 | 4 | 0.5 | 1 | 1 | NA | NA | NA | NA | NA | NA | 1 | 1 |
| 7_2400189 | NA | NA | 2 | 0.5 | 1 | 1 | 1 | 1 | 4 | 0.75 | 1 | 1 | NA | NA | NA | NA | NA | NA | 1 | 1 |
| 7_2412186 | 1 | 1 | 2 | 0.5 | 5 | 0.2 | 6 | 0.1667 | 14 | 0.2857 | 3 | 0.3333 | 4 | 0.5 | NA | NA | 3 | 0.3333 | 10 | 0.4 |
| 7_2412205 | 1 | 1 | 3 | 0 | 5 | 0 | 7 | 0 | 16 | 0.0625 | 3 | 0.3333 | 4 | 0 | NA | NA | 3 | 0 | 10 | 0.1 |
| 7_2412211 | 3 | 0 | 3 | 0 | 5 | 0.2 | 6 | 0 | 17 | 0.0588 | 3 | 0.3333 | 2 | 0 | 1 | 0 | 3 | 0 | 9 | 0.1111 |
| 7_2417486 | NA | NA | NA | NA | NA | NA | 1 | 1 | 1 | 1 | NA | NA | NA | NA | NA | NA | 1 | 1 | 1 | 1 |
| 7_2420308 | 7 | 0 | 5 | 0.2 | NA | NA | 5 | 0 | 17 | 0.0588 | 6 | 0.1667 | 5 | 0.4 | 5 | 0.4 | 5 | 0.2 | 21 | 0.2857 |
| 7_2420321 | 10 | 0.3 | 8 | 0 | 5 | 0.2 | 9 | 0.1111 | 32 | 0.1562 | 8 | 0 | 5 | 0.2 | 6 | 0.3333 | 7 | 0.1429 | 26 | 0.1538 |
| 7_2421489 | 9 | 0.1111 | 7 | 0.2857 | 7 | 0.4286 | 12 | 0.3333 | 35 | 0.2857 | 6 | 0.3333 | 18 | 0.4444 | 13 | 0.3077 | 5 | 0.6 | 42 | 0.4048 |
| 7_2423093 | 1 | 1 | 2 | 0 | NA | NA | NA | NA | 3 | 0.3333 | 1 | 1 | NA | NA | NA | NA | 3 | 0.6667 | 4 | 0.75 |
| 7_2423121 | 1 | 1 | 2 | 0 | NA | NA | NA | NA | 3 | 0.3333 | 1 | 0 | 1 | 0 | 1 | 0 | 1 | 1 | 4 | 0.25 |
| 7_2423283 | 1 | 0 | 1 | 1 | NA | NA | NA | NA | 2 | 0.5 | 2 | 1 | 1 | 1 | NA | NA | 3 | 1 | 6 | 1 |
| 7_2423298 | 1 | 0 | 2 | 0.5 | NA | NA | NA | NA | 3 | 0.3333 | 2 | 1 | 1 | 0 | 1 | 1 | 3 | 0.6667 | 7 | 0.7143 |
| 7_2423321 | 2 | 1 | 2 | 1 | NA | NA | NA | NA | 4 | 1 | 1 | 1 | NA | NA | 2 | 1 | 3 | 1 | 6 | 1 |
| 7_2423326 | 2 | 0.5 | 2 | 1 | NA | NA | NA | NA | 4 | 0.75 | 2 | 1 | 1 | 1 | 2 | 1 | 3 | 1 | 8 | 1 |
| 7_2423357 | 1 | 1 | NA | NA | NA | NA | NA | NA | 1 | 1 | 2 | 1 | NA | NA | 1 | 1 | 3 | 1 | 6 | 1 |
| 7_2423400 | 1 | 0 | 2 | 0.5 | 1 | 0 | NA | NA | 4 | 0.25 | 2 | 0 | NA | NA | 2 | 1 | 3 | 0.3333 | 7 | 0.4286 |
| 7_2423414 | 1 | 1 | 2 | 1 | NA | NA | NA | NA | 3 | 1 | 2 | 1 | NA | NA | 4 | 1 | 4 | 0.75 | 10 | 0.9 |
| 7_2423475 | 1 | 0 | 3 | 0.3333 | 1 | 0 | NA | NA | 5 | 0.2 | 1 | 0 | NA | NA | NA | NA | 5 | 0.2 | 6 | 0.1667 |
| 7_2423515 | 2 | 0 | 4 | 0.25 | 3 | 0.3333 | 2 | 0.5 | 11 | 0.2727 | 3 | 0 | NA | NA | 4 | 0.25 | 5 | 0.2 | 12 | 0.1667 |
| 7_2423561 | 1 | 1 | 4 | 0 | 3 | 0 | 2 | 0.5 | 10 | 0.2 | 3 | 0 | 1 | 0 | 3 | 0 | 7 | 0.1429 | 14 | 0.0714 |
| 7_2423779 | 5 | 0.6 | 6 | 0.5 | 10 | 0.2 | 10 | 0.5 | 31 | 0.4194 | 7 | 0.4286 | 5 | 0.8 | 10 | 0.4 | 8 | 0.375 | 30 | 0.4667 |
| 7_2426731 | 2 | 0.5 | 3 | 0 | 2 | 0.5 | 1 | 0 | 8 | 0.25 | 4 | 0.25 | 3 | 0.6667 | 2 | 0 | 1 | 1 | 10 | 0.4 |
| 7_2428433 | NA | NA | 1 | 0 | 2 | 0.5 | 1 | 1 | 4 | 0.5 | 2 | 0.5 | NA | NA | NA | NA | NA | NA | 2 | 0.5 |
| 7_2428478 | NA | NA | 1 | 1 | NA | NA | NA | NA | 1 | 1 | 2 | 0.5 | NA | NA | NA | NA | NA | NA | 2 | 0.5 |
| 7_2428479 | NA | NA | 1 | 1 | NA | NA | 1 | 1 | 2 | 1 | 2 | 1 | NA | NA | NA | NA | NA | NA | 2 | 1 |
| 7_2428484 | NA | NA | NA | NA | NA | NA | 1 | 1 | 1 | 1 | 2 | 1 | NA | NA | NA | NA | NA | NA | 2 | 1 |
| 7_2428487 | 1 | 0 | 3 | 0 | 2 | 0 | 4 | 0.25 | 10 | 0.1 | 3 | 0.3333 | 2 | 0 | NA | NA | 1 | 0 | 6 | 0.1667 |
| 7_2428525 | NA | NA | 2 | 0 | 2 | 0 | 4 | 0.25 | 8 | 0.125 | 1 | 0 | 2 | 0.5 | NA | NA | NA | NA | 3 | 0.3333 |
| 7_2456202 | 20 | 0.3 | 33 | 0.5152 | 28 | 0.5714 | 36 | 0.4722 | 117 | 0.4786 | 27 | 0.5185 | 38 | 0.3421 | 27 | 0.4444 | 24 | 0.125 | 116 | 0.3621 |
| 7_2462598 | 3 | 0 | 5 | 0.2 | 1 | 0 | NA | NA | 9 | 0.1111 | 3 | 0.3333 | 3 | 0 | 2 | 0 | 7 | 0 | 15 | 0.0667 |
| 7_2470944 | 24 | 0.5833 | 22 | 0.5455 | 21 | 0.7143 | 15 | 0.7333 | 82 | 0.6341 | 20 | 0.7 | 25 | 0.68 | 28 | 0.7857 | 20 | 0.75 | 93 | 0.7312 |
| 7_2470948 | 30 | 0.2 | 31 | 0.1935 | 27 | 0.3333 | 30 | 0.2333 | 118 | 0.2373 | 29 | 0.1724 | 35 | 0.2571 | 40 | 0.225 | 35 | 0.2286 | 139 | 0.223 |
| 7_2470950 | 23 | 0.4348 | 29 | 0.7931 | 21 | 0.619 | 21 | 0.3333 | 94 | 0.5638 | 20 | 0.6 | 31 | 0.6129 | 29 | 0.6552 | 31 | 0.6129 | 111 | 0.6216 |
| 7_2470956 | 26 | 0.8846 | 34 | 0.9412 | 26 | 0.9231 | 27 | 0.8889 | 113 | 0.9115 | 24 | 0.9167 | 33 | 0.8788 | 40 | 0.975 | 28 | 1 | 125 | 0.944 |
| 7_2470958 | 29 | 0.8966 | 33 | 0.8182 | 23 | 0.7826 | 29 | 0.7931 | 114 | 0.8246 | 26 | 0.8462 | 34 | 0.7353 | 38 | 0.7895 | 33 | 0.9091 | 131 | 0.8168 |
| 7_2470958 | 29 | 0.2069 | 32 | 0.2812 | 26 | 0.2692 | 27 | 0.2593 | 114 | 0.2544 | 31 | 0.2258 | 34 | 0.1471 | 41 | 0.0976 | 32 | 0.2812 | 138 | 0.1812 |
| 7_2470965 | 22 | 0.8182 | 20 | 0.75 | 20 | 0.85 | 25 | 0.84 | 87 | 0.8161 | 22 | 1 | 22 | 0.7727 | 33 | 0.9091 | 23 | 0.7391 | 100 | 0.86 |
| 7_2470973 | 28 | 0.25 | 31 | 0.3226 | 25 | 0.24 | 30 | 0.2667 | 114 | 0.2719 | 23 | 0.1304 | 32 | 0.2812 | 35 | 0.3429 | 32 | 0.375 | 122 | 0.2951 |

| | | | | | | | | | | | | | | | | | | | | |
|---|---|---|---|---|---|---|---|---|---|---|---|---|---|---|---|---|---|---|---|---|
| 7_2470976 | 23 | 0.1304 | 30 | 0.2 | 17 | 0.2353 | 22 | 0.1364 | 92 | 0.1739 | 17 | 0.3529 | 28 | 0.1071 | 34 | 0.1765 | 28 | 0.1071 | 107 | 0.1682 |
| 7_2470999 | 19 | 0.1053 | 21 | 0.0952 | 17 | 0.1176 | 18 | 0.0556 | 75 | 0.0933 | 18 | 0 | 22 | 0.1364 | 21 | 0.0476 | 24 | 0.125 | 85 | 0.0824 |
| 7_2476005 | 13 | 0.0769 | 12 | 0.0833 | 9 | 0.2222 | 14 | 0.1429 | 48 | 0.125 | 9 | 0 | 15 | 0.2 | 17 | 0.2353 | 12 | 0.0833 | 53 | 0.1509 |
| 7_2477203 | 4 | 0.25 | 4 | 0 | 6 | 0.3333 | 2 | 0.5 | 16 | 0.25 | 4 | 0 | 12 | 0.1667 | 7 | 0 | 6 | 0.3333 | 29 | 0.1379 |
| 7_2477205 | 3 | 0.3333 | 4 | 0 | 5 | 0.4 | 2 | 1 | 14 | 0.3571 | 5 | 0 | 10 | 0.2 | 6 | 0.1667 | 5 | 0.2 | 26 | 0.1538 |
| 7_2477908 | 1 | 1 | NA | NA | NA | NA | 2 | 0 | 3 | 0.3333 | 2 | 0.5 | 1 | 1 | NA | NA | 1 | 1 | 4 | 0.75 |
| 7_2477936 | 3 | 1 | NA | NA | 1 | 1 | 3 | 1 | 7 | 1 | 1 | 1 | 2 | 1 | 1 | 1 | 1 | 1 | 5 | 1 |
| 7_2477951 | 3 | 0.6667 | NA | NA | 1 | 1 | 3 | 0.3333 | 7 | 0.5714 | NA | NA | 1 | 1 | 2 | 1 | NA | NA | 3 | 1 |
| 7_2477991 | 1 | 1 | NA | NA | 1 | 1 | 2 | 0.5 | 4 | 0.75 | 2 | 1 | 2 | 1 | NA | NA | 1 | 1 | 5 | 1 |
| 7_2478004 | 3 | 0.3333 | NA | NA | 1 | 1 | 3 | 0 | 7 | 0.2857 | 2 | 0 | 2 | 0.5 | 2 | 0.5 | 1 | 0 | 7 | 0.2857 |
| 7_2478005 | 4 | 0 | NA | NA | 1 | 0 | 4 | 0.25 | 9 | 0.1111 | 2 | 0 | 2 | 0 | 2 | 0.5 | 1 | 0 | 7 | 0.1429 |
| 7_2478009 | 3 | 0.6667 | NA | NA | NA | NA | 4 | 1 | 7 | 0.8571 | 1 | 1 | 2 | 0.5 | 2 | 1 | 1 | 1 | 6 | 0.8333 |
| 7_2478031 | 5 | 0.4 | 2 | 0.5 | 1 | 0 | 4 | 0.5 | 12 | 0.4167 | 1 | 0 | 3 | 0.3333 | 3 | 0 | 1 | 0 | 8 | 0.125 |
| 7_2478056 | 3 | 1 | 2 | 1 | 1 | 1 | 4 | 0.75 | 10 | 0.9 | 1 | 1 | 4 | 0.5 | 3 | 1 | 2 | 0.5 | 10 | 0.7 |
| 7_2478065 | 5 | 1 | 2 | 0.5 | 1 | 1 | 4 | 1 | 12 | 0.9167 | 2 | 0.5 | 4 | 1 | 3 | 1 | 2 | 1 | 11 | 0.9091 |
| 7_2478099 | 6 | 0.8333 | 2 | 0 | 1 | 0 | 4 | 0.25 | 13 | 0.4615 | 1 | 0 | 6 | 0.3333 | 4 | 0.25 | 3 | 0.6667 | 14 | 0.3571 |
| 7_2478130 | 4 | 0.5 | 2 | 1 | 1 | 0 | 5 | 1 | 12 | 0.75 | 2 | 1 | 5 | 0.6 | 3 | 0.6667 | 4 | 1 | 14 | 0.7857 |
| 7_2478133 | 5 | 0 | 1 | 1 | 1 | 0 | 4 | 0.25 | 11 | 0.1818 | 1 | 0 | 7 | 0.2857 | 2 | 0 | 4 | 0.5 | 14 | 0.2857 |
| 7_2478146 | 5 | 0.6 | 1 | 1 | NA | NA | 4 | 0.25 | 10 | 0.5 | 2 | 0.5 | 6 | 0.5 | 4 | 0.5 | 4 | 1 | 16 | 0.625 |
| 7_2478159 | 5 | 1 | 1 | 1 | 2 | 1 | 4 | 1 | 12 | 1 | 1 | 0 | 5 | 1 | 4 | 0.75 | 3 | 0.6667 | 13 | 0.7692 |
| 7_2478189 | 4 | 0.75 | 1 | 1 | 2 | 1 | 5 | 1 | 12 | 0.9167 | 2 | 0.5 | 5 | 1 | 3 | 1 | 2 | 1 | 12 | 0.9167 |
| 7_2478190 | 6 | 1 | 2 | 1 | 1 | 1 | 5 | 1 | 14 | 1 | 1 | 1 | 2 | 1 | 1 | 1 | 3 | 1 | 7 | 1 |
| 7_2489946 | NA | NA | 2 | 0.5 | NA | NA | 1 | 0 | 3 | 0.3333 | 1 | 1 | NA | NA | 2 | 0.5 | 3 | 0.3333 | 6 | 0.5 |
| 7_2489985 | 1 | 0 | 2 | 0.5 | NA | NA | NA | NA | 3 | 0.3333 | 2 | 0.5 | NA | NA | 2 | 0.5 | 3 | 0.3333 | 7 | 0.4286 |
| 7_2490230 | 1 | 1 | 4 | 0.5 | 1 | 0 | 1 | 0 | 7 | 0.4286 | 1 | 0 | 1 | 1 | 4 | 1 | 4 | 0 | 10 | 0.5 |
| 7_2490237 | 3 | 0 | 4 | 0.25 | 1 | 0 | 4 | 0.5 | 12 | 0.25 | 2 | 0 | 1 | 0 | 5 | 0.2 | 4 | 0.5 | 12 | 0.25 |
| 7_2490243 | 2 | 1 | 4 | 0.75 | 1 | 0 | 3 | 0.6667 | 10 | 0.7 | 2 | 0 | 1 | 1 | 5 | 0.8 | 2 | 0 | 10 | 0.5 |
| 7_2490246 | 1 | 1 | 2 | 1 | NA | NA | 4 | 1 | 7 | 1 | 1 | 1 | NA | NA | 3 | 1 | 3 | 1 | 7 | 1 |
| 7_2490246 | 3 | 0.6667 | 4 | 0 | 1 | 0 | 5 | 0 | 13 | 0.1538 | 2 | 0 | NA | NA | 5 | 0.2 | 4 | 0.25 | 11 | 0.1818 |
| 7_2490288 | 2 | 0 | 2 | 0.5 | 1 | 0 | 4 | 0 | 9 | 0.1111 | 3 | 0 | 1 | 0 | 5 | 0.2 | 4 | 0 | 13 | 0.0769 |
| 7_2490294 | 2 | 1 | 2 | 1 | 1 | 1 | 3 | 0.6667 | 8 | 0.875 | 3 | 1 | 1 | 1 | 4 | 0.75 | 3 | 1 | 11 | 0.9091 |
| 7_2490310 | 2 | 1 | 2 | 0.5 | 1 | 0 | 2 | 1 | 7 | 0.7143 | 1 | 1 | NA | NA | 2 | 0.5 | 4 | 0.5 | 7 | 0.5714 |
| 7_2490321 | 2 | 1 | 3 | 1 | 1 | 0 | 3 | 0.6667 | 9 | 0.7778 | 2 | 0.5 | 1 | 1 | 4 | 0.5 | 3 | 0.6667 | 10 | 0.6 |
| 7_2490324 | 2 | 0.5 | 3 | 0 | 1 | 0 | 4 | 0.5 | 10 | 0.3 | 2 | 0 | 1 | 1 | 4 | 0.25 | 4 | 0.25 | 11 | 0.2727 |
| 7_2492485 | NA | NA | NA | NA | NA | NA | 1 | 1 | 1 | 1 | NA | NA | NA | NA | 1 | 1 | NA | NA | 1 | 1 |
| 7_2492535 | NA | NA | NA | NA | NA | NA | 1 | 1 | 1 | 1 | NA | NA | NA | NA | 2 | 1 | NA | NA | 2 | 1 |
| 7_2492562 | NA | NA | NA | NA | 1 | 0 | 1 | 1 | 2 | 0.5 | NA | NA | NA | NA | 2 | 0.5 | NA | NA | 2 | 0.5 |
| 7_2492569 | NA | NA | NA | NA | 1 | 1 | 1 | 1 | 2 | 1 | NA | NA | NA | NA | 1 | 1 | NA | NA | 1 | 1 |
| 7_2492574 | NA | NA | NA | NA | 1 | 1 | 1 | 1 | 2 | 1 | NA | NA | NA | NA | 2 | 1 | NA | NA | 2 | 1 |
| 7_2492591 | NA | NA | NA | NA | 1 | 1 | 1 | 1 | 2 | 1 | NA | NA | NA | NA | 2 | 0.5 | NA | NA | 2 | 0.5 |
| 7_2492595 | NA | NA | NA | NA | 1 | 1 | 1 | 1 | 2 | 1 | NA | NA | NA | NA | 2 | 1 | NA | NA | 2 | 1 |
| 7_2492618 | NA | NA | NA | NA | 1 | 0 | 1 | 1 | 2 | 0.5 | NA | NA | NA | NA | 2 | 0.5 | NA | NA | 2 | 0.5 |
| 7_2492655 | NA | NA | NA | NA | 1 | 1 | 1 | 0 | 2 | 0.5 | NA | NA | NA | NA | 2 | 0.5 | 2 | 0.5 | 4 | 0.5 |
| 7_2492928 | NA | NA | NA | NA | 1 | 0 | 2 | 0.5 | 3 | 0.3333 | NA | NA | 1 | 1 | 1 | 0 | NA | NA | 2 | 0.5 |
| 7_2494243 | 2 | 0 | NA | NA | 1 | 0 | 3 | 0.3333 | 6 | 0.1667 | 2 | 0.5 | 1 | 0 | NA | NA | 1 | 0 | 4 | 0.25 |
| 7_2494379 | 6 | 0.3333 | 1 | 0 | 5 | 0.2 | 4 | 0.25 | 16 | 0.25 | 5 | 0 | 2 | 0 | 1 | 1 | 3 | 0 | 11 | 0.0909 |
| 7_2494385 | 5 | 0.8 | 2 | 0.5 | 3 | 0.6667 | 4 | 1 | 14 | 0.7857 | 6 | 0.6667 | NA | NA | 1 | 1 | 4 | 0.75 | 11 | 0.7273 |
| 7_2494388 | 5 | 0 | 2 | 0 | 3 | 0.3333 | 5 | 0.2 | 15 | 0.1333 | 6 | 0 | 2 | 0 | 1 | 0 | 4 | 0.25 | 13 | 0.0769 |
| 7_2494395 | 4 | 0.25 | 3 | 0 | 4 | 0.25 | 5 | 0 | 16 | 0.125 | 4 | 0.25 | 3 | 0 | 1 | 0 | 5 | 0.2 | 13 | 0.1538 |
| 7_2494400 | 5 | 0.2 | 2 | 0 | 4 | 0.75 | 5 | 0.2 | 16 | 0.3125 | 6 | 0 | 2 | 0 | 1 | 0 | 4 | 0.25 | 13 | 0.0769 |
| 7_2494410 | 4 | 1 | 2 | 1 | 2 | 0.5 | 3 | 1 | 11 | 0.9091 | 6 | 0.5 | 2 | 1 | 1 | 1 | 3 | 1 | 12 | 0.75 |
| 7_2494440 | 4 | 0.25 | 2 | 0 | 2 | 0.5 | 4 | 0.75 | 12 | 0.4167 | 6 | 0.6667 | 3 | 0.6667 | 1 | 0 | 3 | 0.6667 | 13 | 0.6154 |
| 7_2494476 | 5 | 0.8 | 2 | 0.5 | 2 | 1 | 4 | 1 | 13 | 0.8462 | 7 | 0.8571 | 2 | 1 | 1 | 1 | 3 | 1 | 13 | 0.9231 |
| 7_2497097 | 1 | 1 | NA | NA | NA | NA | NA | NA | 1 | 1 | 1 | 1 | NA | NA | NA | NA | NA | NA | 1 | 1 |
| 7_2497122 | 1 | 1 | NA | NA | NA | NA | NA | NA | 1 | 1 | 1 | 1 | NA | NA | NA | NA | NA | NA | 1 | 1 |
| 7_2497144 | 1 | 1 | NA | NA | NA | NA | NA | NA | 1 | 1 | 1 | 1 | NA | NA | NA | NA | NA | NA | 1 | 1 |
| 7_2497189 | 1 | 1 | NA | NA | NA | NA | NA | NA | 1 | 1 | 1 | 1 | NA | NA | NA | NA | NA | NA | 1 | 1 |
| 7_2497241 | 1 | 1 | NA | NA | NA | NA | NA | NA | 1 | 1 | 1 | 1 | NA | NA | NA | NA | NA | NA | 1 | 1 |
| 7_2497243 | 1 | 1 | NA | NA | NA | NA | NA | NA | 1 | 1 | NA | NA | NA | NA | NA | NA | 1 | 1 | 1 | 1 |
| 7_2497250 | 1 | 1 | NA | NA | NA | NA | NA | NA | 1 | 1 | 1 | 1 | NA | NA | NA | NA | 1 | 1 | 2 | 1 |
| 7_2497251 | 1 | 1 | NA | NA | NA | NA | NA | NA | 1 | 1 | 1 | 1 | NA | NA | NA | NA | 1 | 1 | 2 | 1 |
| 7_2497263 | 1 | 1 | NA | NA | NA | NA | NA | NA | 1 | 1 | 1 | 1 | NA | NA | NA | NA | 1 | 1 | 2 | 1 |
| 7_2497268 | 1 | 1 | NA | NA | NA | NA | NA | NA | 1 | 1 | 1 | 0 | NA | NA | NA | NA | 1 | 1 | 2 | 0.5 |
| 7_2498769 | 2 | 0 | 3 | 0 | 1 | 1 | 3 | 0 | 9 | 0.1111 | 5 | 0.2 | 2 | 0 | 4 | 0 | 4 | 0.25 | 15 | 0.1333 |
| 7_2498783 | 2 | 1 | 3 | 1 | 2 | 0 | 3 | 0 | 10 | 0.5 | 5 | 0.4 | 2 | 0.5 | 6 | 0.5 | 3 | 0 | 16 | 0.375 |

| | | | | | | | | | | | | | | | | | | | | |
|---|---|---|---|---|---|---|---|---|---|---|---|---|---|---|---|---|---|---|---|---|
| 7_2498791 | NA | NA | 3 | 1 | 2 | 0 | 3 | 0.6667 | 8 | 0.625 | 6 | 0.5 | 2 | 0 | 3 | 0.6667 | 4 | 0.25 | 15 | 0.4 |
| 7_2498813 | 2 | 0.5 | 3 | 0.3333 | 1 | 0 | 2 | 0 | 8 | 0.25 | 7 | 0.2857 | 2 | 0 | 5 | 0 | 3 | 0 | 17 | 0.1176 |
| 7_2498815 | 1 | 1 | 3 | 1 | 2 | 1 | 3 | 1 | 9 | 1 | 7 | 0.7143 | 3 | 1 | 5 | 1 | 3 | 1 | 18 | 0.8889 |
| 7_2498826 | 2 | 0 | 3 | 0 | 2 | 0 | 2 | 0.5 | 9 | 0.1111 | 6 | 0.1667 | 3 | 0.3333 | 6 | 0 | 4 | 0 | 19 | 0.1053 |
| 7_2499282 | 7 | 0.4286 | 11 | 0.6364 | 8 | 1 | 6 | 0.5 | 32 | 0.6562 | 10 | 0.7 | 10 | 0.7 | 14 | 0.3571 | 6 | 0.5 | 40 | 0.55 |
| 7_2499301 | 5 | 0.2 | 9 | 0.2222 | 10 | 0.2 | 6 | 0 | 30 | 0.1667 | 10 | 0.4 | 11 | 0.2727 | 10 | 0.4 | 6 | 0.1667 | 37 | 0.3243 |
| 7_2499335 | 4 | 0 | 9 | 0.1111 | 11 | 0.0909 | 8 | 0 | 32 | 0.0625 | 9 | 0 | 6 | 0 | 13 | 0.1538 | 6 | 0 | 34 | 0.0588 |
| 7_2501531 | 1 | 1 | 1 | 1 | 1 | 1 | 3 | 0.3333 | 6 | 0.6667 | 1 | 1 | 1 | 0 | 3 | 0.6667 | 2 | 1 | 7 | 0.7143 |
| 7_2501537 | 1 | 0 | 1 | 0 | 1 | 1 | 3 | 0 | 6 | 0.1667 | 1 | 0 | 2 | 0 | 3 | 0 | 2 | 1 | 8 | 0.25 |
| 7_2501542 | 1 | 0 | 1 | 0 | 2 | 0 | 3 | 0.3333 | 7 | 0.1429 | 1 | 0 | 2 | 0 | 3 | 0.3333 | 2 | 0.5 | 8 | 0.25 |
| 7_2501551 | 1 | 1 | 2 | 0.5 | 2 | 1 | 3 | 0.3333 | 8 | 0.625 | 1 | 1 | 2 | 0 | 3 | 0.6667 | 2 | 0.5 | 8 | 0.5 |
| 7_2501679 | 1 | 1 | NA | NA | NA | NA | NA | NA | 1 | 1 | 1 | 1 | NA | NA | NA | NA | NA | NA | 1 | 1 |
| 7_2501681 | 1 | 1 | NA | NA | NA | NA | NA | NA | 1 | 1 | 1 | 1 | NA | NA | NA | NA | NA | NA | 1 | 1 |
| 7_2506294 | 11 | 0.9091 | 6 | 0.8333 | 2 | 1 | 3 | 0.6667 | 22 | 0.8636 | 6 | 1 | 1 | 1 | 6 | 0.8333 | 3 | 1 | 16 | 0.9375 |
| 7_2506297 | 11 | 0.5455 | 6 | 1 | 3 | 0.3333 | 3 | 0.6667 | 23 | 0.6522 | 6 | 0.8333 | 3 | 0.6667 | 6 | 0.6667 | 3 | 0.6667 | 18 | 0.7222 |
| 7_2506300 | 6 | 0.8333 | 6 | 0.5 | 1 | 1 | 3 | 1 | 16 | 0.75 | 3 | 0.3333 | 1 | 1 | 5 | 0.6 | 3 | 0.6667 | 12 | 0.5833 |
| 7_2506312 | 11 | 0.7273 | 6 | 0.8333 | 3 | 0.6667 | 1 | 1 | 21 | 0.7619 | 7 | 0.4286 | 3 | 0.3333 | 6 | 0.6667 | 3 | 1 | 19 | 0.5789 |
| 7_2506313 | 12 | 0.9167 | 6 | 1 | 3 | 1 | 4 | 1 | 25 | 0.96 | 6 | 0.8333 | 2 | 1 | 6 | 1 | 3 | 1 | 17 | 0.9412 |
| 7_2506322 | 9 | 0.1111 | 5 | 0 | 3 | 0 | 4 | 0.25 | 21 | 0.0952 | 5 | 0.2 | 2 | 0 | 5 | 0.2 | 2 | 0.5 | 14 | 0.2143 |
| 7_2506325 | 11 | 0.9091 | 6 | 0.6667 | 2 | 0.5 | 3 | 1 | 22 | 0.8182 | 6 | 0.8333 | 4 | 0.75 | 6 | 1 | 2 | 1 | 18 | 0.8889 |
| 7_2506337 | 9 | 0.6667 | 5 | 0.6 | 3 | 1 | 4 | 0.75 | 21 | 0.7143 | 5 | 0.4 | 2 | 0.5 | 6 | 0.6667 | 3 | 1 | 16 | 0.625 |
| 7_2506341 | 10 | 0.2 | 6 | 0 | 2 | 0.5 | 4 | 0.25 | 22 | 0.1818 | 7 | 0.4286 | 4 | 0.25 | 6 | 0.3333 | 2 | 0.5 | 19 | 0.3684 |
| 7_2506350 | 12 | 0.5833 | 5 | 0.2 | 4 | 0 | 2 | 0.5 | 23 | 0.3913 | 6 | 0.8333 | 3 | 0 | 5 | 0 | 2 | 1 | 16 | 0.4375 |
| 7_2506359 | 12 | 0.25 | 6 | 0 | 3 | 0.3333 | 5 | 0 | 26 | 0.1538 | 7 | 0.2857 | 3 | 0.3333 | 6 | 0 | 3 | 0 | 19 | 0.1579 |
| 7_2506371 | 11 | 0.6364 | 6 | 0.6667 | 5 | 0.8 | 3 | 0.6667 | 25 | 0.68 | 6 | 0.6667 | 2 | 1 | 7 | 0.7143 | 3 | 0.6667 | 18 | 0.7222 |
| 7_2506377 | 11 | 0.6364 | 4 | 0.75 | 5 | 1 | 3 | 0.6667 | 23 | 0.7391 | 5 | 0.8 | 4 | 0.5 | 8 | 0.75 | 2 | 1 | 19 | 0.7368 |
| 7_2506394 | 10 | 0.6 | 6 | 0.5 | 4 | 0.5 | 3 | 0.3333 | 23 | 0.5217 | 6 | 0.6667 | 3 | 0 | 6 | 0.3333 | 3 | 1 | 18 | 0.5 |
| 7_2506397 | 11 | 0.6364 | 7 | 0.4286 | 4 | 0.75 | 4 | 0.75 | 26 | 0.6154 | 9 | 0.5556 | 2 | 0.5 | 9 | 0.7778 | 4 | 0.75 | 24 | 0.6667 |
| 7_2506399 | 12 | 0.4167 | 7 | 0.4286 | 5 | 0.4 | 5 | 0.2 | 29 | 0.3793 | 9 | 0.4444 | 3 | 0 | 7 | 0.2857 | 4 | 0.75 | 23 | 0.3913 |
| 7_2506407 | 8 | 0.125 | 4 | 0 | 5 | 0.4 | 4 | 0.25 | 21 | 0.1905 | 8 | 0.125 | 2 | 0 | 8 | 0.125 | 3 | 0 | 21 | 0.0952 |
| 7_2506408 | 14 | 0.6429 | 6 | 0.5 | 4 | 0.75 | 5 | 0.8 | 29 | 0.6552 | 9 | 0.6667 | 2 | 1 | 8 | 0.75 | 2 | 1 | 21 | 0.7619 |
| 7_2506438 | 17 | 0.0588 | 7 | 0.2857 | 4 | 0 | 5 | 0 | 33 | 0.0909 | 11 | 0.1818 | 3 | 0 | 8 | 0.25 | 6 | 0 | 28 | 0.1429 |
| 7_2506452 | 12 | 0.0833 | 4 | 0 | 6 | 0.5 | 5 | 0.2 | 27 | 0.1852 | 9 | 0.3333 | 4 | 0 | 9 | 0.1111 | 7 | 0.1429 | 29 | 0.1724 |
| 7_2506486 | 15 | 0.0667 | 6 | 0 | 5 | 0.2 | 8 | 0 | 34 | 0.0588 | 10 | 0 | 4 | 0.25 | 7 | 0 | 9 | 0.1111 | 30 | 0.0667 |
| 7_2506506 | 16 | 0.1875 | 8 | 0.5 | 7 | 0.5714 | 6 | 0.3333 | 37 | 0.3514 | 14 | 0.3571 | 3 | 0 | 10 | 0.2 | 10 | 0.2 | 37 | 0.2432 |
| 7_2506510 | 18 | 0.5556 | 7 | 0.7143 | 6 | 0.6667 | 8 | 0.625 | 39 | 0.6154 | 16 | 0.5 | 5 | 0.4 | 11 | 0.3636 | 8 | 0.875 | 40 | 0.525 |
| 7_2506515 | 19 | 0.1579 | 7 | 0.4286 | 6 | 0.1667 | 8 | 0.125 | 40 | 0.2 | 17 | 0.1765 | 5 | 0 | 13 | 0.1538 | 11 | 0.3636 | 46 | 0.1957 |
| 7_2507939 | NA | NA | NA | NA | 2 | 0 | 1 | 1 | 3 | 0.3333 | NA | NA | 1 | 1 | NA | NA | NA | NA | 1 | 1 |
| 7_2508027 | NA | NA | 1 | 1 | 3 | 0.3333 | 1 | 1 | 5 | 0.6 | NA | NA | 1 | 1 | NA | NA | NA | NA | 1 | 1 |
| 7_2516168 | NA | NA | NA | NA | NA | NA | 2 | 0.5 | 2 | 0.5 | NA | NA | NA | NA | 1 | 1 | NA | NA | 1 | 1 |
| 7_2516241 | NA | NA | NA | NA | NA | NA | 2 | 1 | 2 | 1 | NA | NA | NA | NA | 1 | 1 | NA | NA | 1 | 1 |
| 7_2516261 | NA | NA | NA | NA | NA | NA | 2 | 1 | 2 | 1 | NA | NA | NA | NA | 1 | 1 | NA | NA | 1 | 1 |
| 7_2526846 | 1 | 1 | 2 | 0.5 | NA | NA | NA | NA | 3 | 0.6667 | NA | NA | NA | NA | NA | NA | 1 | 1 | 1 | 1 |
| 7_2526863 | 1 | 0 | 2 | 0.5 | NA | NA | NA | NA | 3 | 0.3333 | NA | NA | NA | NA | NA | NA | 1 | 1 | 1 | 1 |
| 7_2532443 | 11 | 0.2727 | 12 | 0.3333 | 6 | 0 | 11 | 0.1818 | 40 | 0.225 | 5 | 0.2 | 12 | 0.0833 | 10 | 0.3 | 6 | 0 | 33 | 0.1515 |
| 7_2532444 | 12 | 0.75 | 10 | 0.7 | 6 | 0.8333 | 9 | 0.7778 | 37 | 0.7568 | 5 | 0.8 | 12 | 0.4167 | 7 | 0.8571 | 8 | 0.75 | 32 | 0.6562 |
| 7_2532451 | 11 | 0 | 11 | 0.1818 | 5 | 0.2 | 12 | 0 | 39 | 0.0769 | 6 | 0 | 13 | 0.0769 | 11 | 0.0909 | 7 | 0 | 37 | 0.0541 |
| 7_2532456 | 11 | 0.9091 | 12 | 0.75 | 5 | 0.6 | 9 | 1 | 37 | 0.8378 | 5 | 0.8 | 12 | 0.75 | 10 | 0.9 | 8 | 0.875 | 35 | 0.8286 |
| 7_2532460 | 10 | 0.6 | 13 | 0.8462 | 5 | 0.6 | 11 | 0.8182 | 39 | 0.7436 | 5 | 0.8 | 13 | 0.6154 | 11 | 0.6364 | 9 | 0.6667 | 38 | 0.6579 |
| 7_2532461 | 12 | 0.1667 | 13 | 0.0769 | 6 | 0.1667 | 12 | 0.25 | 43 | 0.1628 | 4 | 0.25 | 13 | 0.1538 | 11 | 0 | 8 | 0.25 | 36 | 0.1389 |
| 7_2532477 | 13 | 0.8462 | 14 | 0.7143 | 5 | 0.6 | 12 | 0.75 | 44 | 0.75 | 6 | 0.8333 | 13 | 0.6923 | 12 | 0.6667 | 9 | 0.7778 | 40 | 0.725 |
| 7_2532478 | 12 | 0.75 | 12 | 0.6667 | 4 | 0.75 | 10 | 1 | 38 | 0.7895 | 5 | 0.8 | 11 | 0.6364 | 11 | 0.2727 | 9 | 0.7778 | 36 | 0.5833 |
| 7_2532958 | 13 | 0.3077 | 7 | 0 | 9 | 0.2222 | 2 | 0 | 31 | 0.1935 | 7 | 0.4286 | 7 | 0.1429 | 8 | 0.125 | 3 | 0 | 25 | 0.2 |
| 7_2532972 | 12 | 0 | 7 | 0.1429 | 6 | 0.1667 | 1 | 1 | 26 | 0.1154 | 5 | 0 | 6 | 0.3333 | 7 | 0.1429 | 2 | 0 | 20 | 0.15 |
| 7_2536268 | NA | NA | 1 | 1 | NA | NA | NA | NA | 1 | 1 | NA | NA | NA | NA | NA | NA | 1 | 1 | 1 | 1 |
| 7_2536364 | 1 | 1 | 1 | 1 | NA | NA | NA | NA | 2 | 1 | NA | NA | 1 | 1 | 1 | 0 | NA | NA | 2 | 0.5 |
| 7_2536494 | 4 | 0 | 2 | 0.5 | 1 | 0 | 3 | 0 | 10 | 0.1 | 4 | 0.5 | 5 | 0.2 | 3 | 0.3333 | 2 | 0.5 | 14 | 0.3571 |
| 7_2536736 | 4 | 0 | 3 | 0.3333 | NA | NA | 3 | 0 | 10 | 0.1 | 6 | 0.1667 | 6 | 0.1667 | 3 | 0 | 3 | 0 | 18 | 0.1111 |
| 7_2538551 | 2 | 0.5 | 4 | 0 | 3 | 0 | 4 | 0.25 | 13 | 0.1538 | 2 | 0 | 6 | 0.5 | 4 | 0.75 | 3 | 0 | 15 | 0.4 |
| 7_2538572 | 5 | 1 | 8 | 0.5 | 4 | 1 | 6 | 0.8333 | 23 | 0.7826 | 4 | 1 | 6 | 1 | 5 | 0.6 | 6 | 0.8333 | 21 | 0.8571 |
| 7_2538579 | 5 | 1 | 7 | 0.7143 | 2 | 1 | 5 | 0.8 | 19 | 0.8421 | 6 | 0.6667 | 4 | 0.75 | 5 | 1 | 7 | 0.8571 | 22 | 0.8182 |
| 7_2538580 | 5 | 0.6 | 8 | 0.5 | 4 | 0.75 | 7 | 0.7143 | 24 | 0.625 | 6 | 0.1667 | 5 | 0.2 | 6 | 0.5 | 7 | 0.2857 | 24 | 0.2917 |
| 7_2538584 | 4 | 0.5 | 6 | 0.1667 | 3 | 0 | 7 | 0.4286 | 20 | 0.3 | 6 | 1 | 4 | 1 | 5 | 1 | 7 | 0.8571 | 22 | 0.9545 |
| 7_2538586 | 4 | 0 | 7 | 0.2857 | 2 | 0 | 5 | 0.2 | 18 | 0.1667 | 4 | 0 | 4 | 0 | 4 | 0.25 | 6 | 0.1667 | 18 | 0.1111 |
| 7_2538588 | 4 | 0.75 | 7 | 0.4286 | 5 | 0.8 | 6 | 0.8333 | 22 | 0.6818 | 5 | 0.6 | 5 | 0.4 | 3 | 1 | 3 | 0 | 16 | 0.5 |

| | | | | | | | | | | | | | | | | | | | | |
|---|---|---|---|---|---|---|---|---|---|---|---|---|---|---|---|---|---|---|---|---|
| 7_2538594 | 4 | 0.75 | 7 | 0.4286 | 4 | 0.75 | 6 | 0.6667 | 21 | 0.619 | 6 | 0.5 | 5 | 0.6 | 4 | 0.5 | 7 | 0.2857 | 22 | 0.4545 |
| 7_2538674 | NA | NA | 3 | 0.3333 | NA | NA | NA | NA | 3 | 0.3333 | 1 | 1 | 1 | 0 | NA | NA | 1 | 1 | 3 | 0.6667 |
| 7_2538679 | NA | NA | 3 | 0.3333 | NA | NA | NA | NA | 3 | 0.3333 | 1 | 0 | 1 | 1 | NA | NA | 1 | 1 | 3 | 0.6667 |
| 7_2538690 | NA | NA | 3 | 0.3333 | NA | NA | NA | NA | 3 | 0.3333 | 1 | 0 | 1 | 1 | NA | NA | NA | NA | 2 | 0.5 |
| 7_2538698 | NA | NA | 3 | 0.6667 | NA | NA | NA | NA | 3 | 0.6667 | 1 | 0 | 1 | 0 | NA | NA | 1 | 1 | 3 | 0.3333 |
| 7_2538701 | NA | NA | 3 | 0.6667 | NA | NA | NA | NA | 3 | 0.6667 | 1 | 0 | 1 | 1 | NA | NA | 1 | 0 | 3 | 0.3333 |
| 7_2538704 | NA | NA | 2 | 0.5 | NA | NA | NA | NA | 2 | 0.5 | 1 | 1 | 1 | 0 | NA | NA | NA | NA | 2 | 0.5 |
| 7_2538717 | NA | NA | 3 | 1 | NA | NA | NA | NA | 3 | 1 | 1 | 1 | NA | NA | NA | NA | 1 | 1 | 2 | 1 |
| 7_2538738 | NA | NA | 3 | 0.3333 | NA | NA | NA | NA | 3 | 0.3333 | 1 | 0 | 1 | 0 | NA | NA | 1 | 1 | 3 | 0.3333 |
| 7_2540444 | 1 | 1 | 4 | 0 | NA | NA | 1 | 0 | 6 | 0.1667 | 2 | 0 | 1 | 0 | NA | NA | 1 | 1 | 4 | 0.25 |
| 7_2541063 | 3 | 0.3333 | 5 | 0.2 | 6 | 0 | 3 | 0 | 17 | 0.1176 | 3 | 0 | 3 | 0.3333 | 2 | 0 | 4 | 0.25 | 12 | 0.1667 |
| 7_2541070 | 5 | 0.8 | 4 | 1 | 4 | 0.75 | 4 | 1 | 17 | 0.8824 | 4 | 0.25 | 4 | 0.75 | 5 | 0.6 | 6 | 0.6667 | 19 | 0.5789 |
| 7_2541075 | 5 | 0 | 4 | 0 | 5 | 0.2 | 3 | 0.3333 | 17 | 0.1176 | 5 | 0.2 | 5 | 0.4 | 4 | 0 | 6 | 0 | 20 | 0.15 |
| 7_2541079 | 4 | 0.25 | 4 | 0.25 | 4 | 0 | 4 | 0 | 16 | 0.125 | 4 | 0.5 | 3 | 0 | 3 | 0 | 6 | 0.3333 | 16 | 0.25 |
| 7_2541083 | 4 | 0.25 | 3 | 0 | 3 | 0 | 1 | 1 | 11 | 0.1818 | 1 | 0 | 3 | 0 | 5 | 0.2 | 4 | 0 | 13 | 0.0769 |
| 7_2541084 | 4 | 0 | 4 | 0.5 | 7 | 0.7143 | 3 | 1 | 18 | 0.5556 | 4 | 0.5 | 4 | 0.5 | 4 | 0 | 7 | 0.2857 | 19 | 0.3158 |
| 7_2541093 | 5 | 0.8 | 4 | 0.75 | 7 | 0.2857 | 3 | 0.3333 | 19 | 0.5263 | 4 | 0.25 | 5 | 0.8 | 4 | 0.75 | 6 | 0.8333 | 19 | 0.6842 |
| 7_2541094 | 5 | 0 | 4 | 0 | 7 | 0.1429 | 2 | 0 | 18 | 0.0556 | 4 | 0.25 | 5 | 0.2 | 5 | 0 | 7 | 0 | 21 | 0.0952 |
| 7_2541096 | 5 | 0.2 | 4 | 0 | 6 | 0 | 4 | 0 | 19 | 0.0526 | 4 | 0.25 | 5 | 0 | 5 | 0 | 6 | 0 | 20 | 0.05 |
| 7_2541100 | 3 | 0.3333 | 3 | 0 | 4 | 0.25 | 3 | 0.3333 | 13 | 0.2308 | 4 | 0 | 5 | 0.2 | 5 | 0 | 6 | 0 | 20 | 0.05 |
| 7_2541110 | 4 | 0.5 | 3 | 0 | 3 | 0 | 2 | 0.5 | 12 | 0.25 | 3 | 0.3333 | 4 | 0.25 | 5 | 0.2 | 6 | 0.1667 | 18 | 0.2222 |
| 7_2541236 | 5 | 0.2 | 3 | 0.3333 | 5 | 0.2 | 4 | 0 | 17 | 0.1765 | 3 | 0.3333 | 5 | 0 | 3 | 0.6667 | 5 | 0.4 | 16 | 0.3125 |
| 7_2541242 | 4 | 0.25 | 3 | 0.6667 | 5 | 0.2 | 3 | 0.3333 | 15 | 0.3333 | 3 | 0.6667 | 3 | 0.3333 | 3 | 0 | 5 | 0.2 | 14 | 0.2857 |
| 7_2541247 | 5 | 0 | 3 | 0 | 6 | 0.1667 | 4 | 0 | 18 | 0.0556 | 3 | 0 | 4 | 0 | 3 | 0 | 4 | 0.25 | 14 | 0.0714 |
| 7_2541250 | 3 | 0 | 2 | 0 | 4 | 0.25 | 2 | 0 | 11 | 0.0909 | 2 | 0.5 | 3 | 0.3333 | 3 | 0.3333 | 5 | 0.2 | 13 | 0.3077 |
| 7_2541437 | 2 | 0 | 2 | 0 | 4 | 0.25 | 3 | 0 | 11 | 0.0909 | 3 | 0 | 3 | 0 | 2 | 0.5 | 2 | 0 | 10 | 0.1 |
| 7_2545377 | 3 | 0.3333 | 3 | 0.3333 | 5 | 0.8 | 6 | 0.5 | 17 | 0.5294 | 3 | 1 | 8 | 0.625 | 5 | 0.4 | 4 | 1 | 20 | 0.7 |
| 7_2545386 | 3 | 0 | 2 | 0 | 4 | 0.25 | 5 | 0 | 14 | 0.0714 | 2 | 0 | 8 | 0.125 | 4 | 0.25 | 3 | 0.6667 | 17 | 0.2353 |
| 7_2545399 | 2 | 0 | 1 | 1 | 3 | 0.6667 | 2 | 1 | 8 | 0.625 | 1 | 0 | 6 | 0.6667 | 2 | 0.5 | 4 | 0.5 | 13 | 0.5385 |
| 7_2545675 | 1 | 0 | 4 | 0.25 | 2 | 0 | 1 | 0 | 8 | 0.125 | 3 | 0.6667 | 2 | 0 | NA | NA | 2 | 0.5 | 7 | 0.4286 |
| 7_2545715 | 8 | 0.125 | 7 | 0.2857 | 5 | 0.2 | 4 | 0 | 24 | 0.1667 | 6 | 0 | 3 | 0 | NA | NA | 3 | 0.3333 | 12 | 0.0833 |
| 7_2545805 | 6 | 0.3333 | 7 | 0.2857 | 5 | 0 | 5 | 0.4 | 23 | 0.2609 | 6 | 0.1667 | 4 | 0.25 | 1 | 0 | 2 | 0.5 | 13 | 0.2308 |
| 7_2545806 | 4 | 0.75 | 5 | 0 | 4 | 0.75 | 4 | 0.5 | 17 | 0.4706 | 5 | 0.6 | 4 | 0 | 1 | 0 | 2 | 0 | 12 | 0.25 |
| 7_2545808 | 6 | 0 | 7 | 0.2857 | 5 | 0.2 | 5 | 0.4 | 23 | 0.2174 | 5 | 0 | 3 | 0.3333 | 1 | 0 | 1 | 0 | 10 | 0.1 |
| 7_2545814 | 6 | 0.3333 | 6 | 0.1667 | 5 | 0.4 | 4 | 0.25 | 21 | 0.2857 | 5 | 0.4 | 4 | 0.25 | 1 | 0 | 2 | 0 | 12 | 0.25 |
| 7_2545841 | 5 | 0 | 6 | 0 | 5 | 0 | 4 | 0.25 | 20 | 0.05 | 5 | 0 | 2 | 0 | 1 | 0 | 1 | 1 | 9 | 0.1111 |
| 7_2550216 | 4 | 0.75 | NA | NA | 1 | 1 | NA | NA | 5 | 0.8 | 1 | 0 | NA | NA | 1 | 1 | NA | NA | 2 | 0.5 |
| 7_2550253 | 4 | 0.75 | NA | NA | 1 | 1 | NA | NA | 5 | 0.8 | 1 | 1 | NA | NA | 1 | 1 | NA | NA | 2 | 1 |
| 7_2645668 | 3 | 0.3333 | NA | NA | NA | NA | 1 | 0 | 4 | 0.25 | NA | NA | NA | NA | 1 | 1 | NA | NA | 1 | 1 |
| 7_2645692 | NA | NA | NA | NA | NA | NA | 1 | 1 | 1 | 1 | NA | NA | NA | NA | 1 | 1 | NA | NA | 1 | 1 |
| 7_2685817 | 11 | 0.5455 | 10 | 0.5 | 19 | 0.5789 | 14 | 0.7143 | 54 | 0.5926 | 5 | 0.8 | 6 | 0.6667 | 11 | 0.8182 | 15 | 0.6667 | 37 | 0.7297 |
| 7_2685836 | 9 | 0.1111 | 12 | 0.0833 | 18 | 0.1667 | 15 | 0.2 | 54 | 0.1481 | 4 | 0 | 7 | 0.2857 | 13 | 0.2308 | 15 | 0.1333 | 39 | 0.1795 |
| 7_2685843 | 10 | 0 | 13 | 0.0769 | 20 | 0.05 | 12 | 0.0833 | 55 | 0.0545 | 5 | 0 | 7 | 0.1429 | 16 | 0.0625 | 14 | 0.0714 | 42 | 0.0714 |
| 7_2686417 | 8 | 0.125 | 9 | 0.2222 | 10 | 0 | 7 | 0 | 34 | 0.0882 | 4 | 0 | 5 | 0.2 | 9 | 0.1111 | 13 | 0.0769 | 31 | 0.0968 |
| 7_2686430 | 4 | 0.5 | 5 | 0.4 | 17 | 0.7059 | 6 | 0.5 | 32 | 0.5938 | 4 | 0.25 | 4 | 1 | 3 | 0.3333 | 12 | 0.75 | 23 | 0.6522 |
| 7_2686434 | 7 | 1 | 4 | 1 | 11 | 1 | 5 | 1 | 27 | 1 | 4 | 1 | 2 | 1 | 5 | 1 | 10 | 1 | 21 | 1 |
| 7_2686446 | 6 | 0 | 4 | 0.25 | 14 | 0.1429 | 6 | 0 | 30 | 0.1 | 3 | 0.3333 | 3 | 0 | 6 | 0.1667 | 12 | 0 | 24 | 0.0833 |
| 7_2686474 | 4 | 0.25 | 5 | 0.2 | 9 | 0.6667 | 4 | 0.25 | 22 | 0.4091 | 3 | 0.6667 | 1 | 1 | 5 | 0.2 | 10 | 0.3 | 19 | 0.3684 |
| 7_2686491 | 3 | 0.3333 | 2 | 0.5 | 7 | 0.5714 | 3 | 0.6667 | 15 | 0.5333 | 3 | 0.6667 | NA | NA | 3 | 0.6667 | 9 | 0.5556 | 15 | 0.6 |
| 7_2686798 | 9 | 0 | 6 | 0.1667 | 11 | 0.0909 | 14 | 0 | 40 | 0.05 | 14 | 0.0714 | 7 | 0.4286 | 14 | 0 | 12 | 0 | 47 | 0.0851 |
| 7_2690340 | 1 | 0 | 1 | 1 | NA | NA | 1 | 0 | 3 | 0.3333 | NA | NA | 2 | 0 | 3 | 0.6667 | 2 | 1 | 7 | 0.5714 |
| 7_2690343 | NA | NA | 2 | 0.5 | 1 | 1 | 1 | 0 | 4 | 0.5 | 2 | 1 | 3 | 0.6667 | 3 | 1 | 3 | 0.3333 | 11 | 0.7273 |
| 7_2690353 | 1 | 0 | 2 | 0 | 1 | 1 | NA | NA | 4 | 0.25 | 2 | 0 | 3 | 0.3333 | 3 | 0.3333 | 2 | 0 | 10 | 0.2 |
| 7_2690354 | 1 | 1 | 2 | 1 | 1 | 1 | NA | NA | 4 | 1 | 2 | 0 | 2 | 0.5 | 3 | 0.3333 | 3 | 0.3333 | 10 | 0.3 |
| 7_2690359 | 1 | 0 | 1 | 1 | NA | NA | NA | NA | 2 | 0.5 | 1 | 1 | 2 | 0.5 | 3 | 0.3333 | 1 | 1 | 7 | 0.5714 |
| 7_2690451 | 1 | 1 | NA | NA | NA | NA | 1 | 0 | 2 | 0.5 | NA | NA | 2 | 0.5 | 1 | 0 | 2 | 0 | 5 | 0.2 |
| 7_2690672 | 1 | 1 | 1 | 1 | 2 | 1 | NA | NA | 4 | 1 | 3 | 1 | NA | NA | 1 | 1 | NA | NA | 4 | 1 |
| 7_2690687 | 1 | 1 | 1 | 1 | 2 | 1 | NA | NA | 4 | 1 | 2 | 1 | NA | NA | NA | NA | 1 | 1 | 3 | 1 |
| 7_2690688 | 1 | 1 | NA | NA | NA | NA | NA | NA | 1 | 1 | 1 | 1 | NA | NA | NA | NA | 2 | 1 | 3 | 1 |
| 7_2690690 | 1 | 1 | 1 | 1 | 1 | 1 | NA | NA | 3 | 1 | 2 | 1 | NA | NA | NA | NA | 2 | 1 | 4 | 1 |
| 7_2690706 | 1 | 0 | 1 | 1 | NA | NA | NA | NA | 2 | 0.5 | NA | NA | 3 | 0.6667 | 2 | 0.5 | NA | NA | 5 | 0.6 |
| 7_2690710 | NA | NA | 1 | 0 | 2 | 1 | NA | NA | 3 | 0.6667 | 3 | 0.3333 | NA | NA | 1 | 1 | 2 | 0.5 | 6 | 0.5 |
| 7_2690720 | NA | NA | 1 | 1 | 1 | 0 | NA | NA | 2 | 0.5 | 1 | 1 | NA | NA | 1 | 1 | 2 | 0.5 | 4 | 0.75 |
| 7_2690760 | 1 | 0 | 2 | 0.5 | NA | NA | NA | NA | 3 | 0.3333 | NA | NA | 3 | 0 | 2 | 1 | NA | NA | 5 | 0.4 |
| 7_2690770 | NA | NA | 1 | 1 | NA | NA | NA | NA | 1 | 1 | 3 | 1 | NA | NA | 1 | 1 | 2 | 1 | 6 | 1 |

| | | | | | | | | | | | | | | | | | | | | |
|---|---|---|---|---|---|---|---|---|---|---|---|---|---|---|---|---|---|---|---|---|
| 7_2692692! | NA | NA | NA | NA | 1 | 1 | NA | NA | 1 | 1 | NA | NA | NA | NA | NA | NA | 1 | 1 | 1 | 1 |
| 7_2692716: | NA | NA | NA | NA | 1 | 1 | NA | NA | 1 | 1 | NA | NA | NA | NA | NA | NA | 1 | 1 | 1 | 1 |
| 7_2695673! | NA | NA | 3 | 0.3333 | 3 | 1 | NA | NA | 6 | 0.6667 | 3 | 0 | 6 | 0.5 | 2 | 0.5 | 1 | 1 | 12 | 0.4167 |
| 7_2695695 | NA | NA | 6 | 0.1667 | 3 | 0.6667 | 2 | 0 | 11 | 0.2727 | 4 | 0 | 6 | 0.1667 | 2 | 0 | 1 | 1 | 13 | 0.1538 |
| 7_2698041( | 6 | 0 | 7 | 0.2857 | 5 | 0.4 | 3 | 0 | 21 | 0.1905 | 3 | 0.3333 | 8 | 0.125 | 7 | 0.2857 | 2 | 0.5 | 20 | 0.25 |
| 7_2698072! | 9 | 0.1111 | 8 | 0 | 6 | 0.1667 | 3 | 0 | 26 | 0.0769 | 6 | 0 | 7 | 0.2857 | 11 | 0.2727 | 4 | 0.5 | 28 | 0.25 |
| 7_2698080: | 8 | 0.25 | 11 | 0.1818 | 5 | 0.2 | 4 | 0.25 | 28 | 0.2143 | 6 | 0 | 12 | 0.0833 | 14 | 0.2143 | 5 | 0 | 37 | 0.1081 |
| 7_2698082! | 9 | 1 | 9 | 0.6667 | 5 | 1 | 3 | 1 | 26 | 0.8846 | 6 | 0.3333 | 9 | 0.7778 | 7 | 0.7143 | 7 | 0.5714 | 29 | 0.6207 |
| 7_2698103! | 9 | 0.4444 | 11 | 0.2727 | 6 | 0.1667 | 3 | 0.3333 | 29 | 0.3103 | 7 | 0.1429 | 14 | 0.3571 | 13 | 0.3077 | 7 | 0.1429 | 41 | 0.2683 |
| 7_2698107( | 6 | 0.1667 | 10 | 0.5 | 7 | 0.5714 | 3 | 0 | 26 | 0.3846 | 5 | 0.4 | 10 | 0.7 | 6 | 0.5 | 5 | 0.4 | 26 | 0.5385 |
| 7_2700613( | 5 | 0 | 8 | 0.125 | 3 | 0.3333 | 6 | 0.3333 | 22 | 0.1818 | 3 | 0 | 2 | 0.5 | 1 | 0 | 1 | 0 | 7 | 0.1429 |
| 7_2704169! | 2 | 0.5 | 3 | 0 | 5 | 0 | 5 | 0.2 | 15 | 0.1333 | 4 | 0 | 2 | 0.5 | 1 | 0 | 6 | 0.1667 | 13 | 0.1538 |
| 7_2704170! | 1 | 1 | NA | NA | 2 | 1 | 2 | 0 | 5 | 0.6 | 1 | 1 | 2 | 0.5 | 1 | 1 | 2 | 1 | 6 | 0.8333 |
| 7_2705604 | 9 | 0 | 15 | 0 | 8 | 0.125 | 5 | 0.4 | 37 | 0.0811 | 7 | 0 | 15 | 0 | 16 | 0.125 | 7 | 0.2857 | 45 | 0.0889 |
| 7_2705631! | 6 | 0.1667 | 13 | 0.4615 | 5 | 0.4 | 5 | 0.8 | 29 | 0.4483 | 5 | 0.4 | 6 | 0.5 | 12 | 0.6667 | 3 | 0.6667 | 26 | 0.5769 |
| 7_2705645! | 7 | 0 | 13 | 0.1538 | 6 | 0.5 | 4 | 0 | 30 | 0.1667 | 4 | 0.25 | 3 | 0.3333 | 6 | 0.3333 | 4 | 0.75 | 17 | 0.4118 |
| 7_2705657! | 6 | 1 | 17 | 0.8824 | 10 | 1 | 4 | 0.75 | 37 | 0.9189 | 5 | 0.8 | 11 | 0.9091 | 14 | 0.8571 | 5 | 1 | 35 | 0.8857 |
| 7_2705661: | 9 | 0 | 19 | 0.1579 | 11 | 0.4545 | 6 | 0.1667 | 45 | 0.2 | 7 | 0 | 10 | 0.3 | 13 | 0.3077 | 5 | 0.4 | 35 | 0.2571 |
| 7_2705664' | 8 | 0.125 | 17 | 0.1176 | 10 | 0.1 | 3 | 0.3333 | 38 | 0.1316 | 7 | 0.4286 | 10 | 0.1 | 13 | 0.3846 | 4 | 0 | 34 | 0.2647 |
| 7_2705664! | 9 | 0.6667 | 19 | 0.5789 | 11 | 0.6364 | 6 | 0.6667 | 45 | 0.6222 | 5 | 0.6 | 11 | 0.4545 | 13 | 0.5385 | 4 | 0.5 | 33 | 0.5152 |
| 7_2706379 | 4 | 0.5 | 8 | 0.25 | 2 | 0 | 3 | 0.3333 | 17 | 0.2941 | 2 | 1 | 4 | 0.5 | 6 | 0 | 1 | 0 | 13 | 0.3077 |
| 7_2706380! | 4 | 0.5 | 6 | 0.3333 | 3 | 0.3333 | 2 | 0.5 | 15 | 0.4 | 3 | 0 | 4 | 0 | 6 | 0.3333 | 2 | 0 | 15 | 0.1333 |
| 7_2706381 | 2 | 0 | 3 | 0 | 5 | 0.2 | 2 | 0 | 12 | 0.0833 | 2 | 0 | 4 | 0.5 | 4 | 0.5 | 2 | 0.5 | 12 | 0.4167 |
| 7_2706386( | 4 | 0.25 | 9 | 0.1111 | 6 | 0.3333 | 3 | 0 | 22 | 0.1818 | 3 | 0 | 4 | 0.75 | 6 | 0.1667 | 3 | 0 | 16 | 0.25 |
| 7_2706393: | 5 | 0.6 | 10 | 0.5 | 6 | 0.5 | 3 | 0.6667 | 24 | 0.5417 | 3 | 0.3333 | 5 | 0.8 | 6 | 1 | 3 | 1 | 17 | 0.8235 |
| 7_2706395! | 5 | 1 | 9 | 0.6667 | 6 | 0.8333 | 2 | 1 | 22 | 0.8182 | 2 | 1 | 3 | 0.3333 | 6 | 0.8333 | 3 | 1 | 14 | 0.7857 |
| 7_2706404! | 5 | 0.4 | 8 | 0.25 | 6 | 0.3333 | 3 | 0 | 22 | 0.2727 | 2 | 1 | 5 | 0.4 | 7 | 0.2857 | 3 | 0.6667 | 17 | 0.4706 |
| 7_2706410 | 4 | 0.5 | 9 | 0.1111 | 6 | 0 | 3 | 0 | 22 | 0.1364 | 3 | 0 | 5 | 0.2 | 5 | 0 | 3 | 0.3333 | 16 | 0.125 |
| 7_2706428' | 3 | 0.3333 | 2 | 0 | 2 | 0 | NA | NA | 7 | 0.1429 | 1 | 1 | NA | NA | 1 | 0 | 1 | 1 | 3 | 0.6667 |
| 7_2706882' | 2 | 0 | 3 | 0.3333 | 4 | 0.5 | 2 | 0 | 11 | 0.2727 | 1 | 0 | 3 | 0.3333 | 4 | 0.25 | 1 | 1 | 9 | 0.3333 |
| 7_2716857( | 11 | 0.0909 | 17 | 0.1765 | 13 | 0.1538 | 13 | 0.0769 | 54 | 0.1296 | 14 | 0.1429 | 14 | 0.2143 | 14 | 0.1429 | 9 | 0.1111 | 51 | 0.1569 |
| 7_2716895( | 2 | 0 | 7 | 0.2857 | 3 | 0 | 2 | 0 | 14 | 0.1429 | 3 | 0.3333 | 3 | 0 | 2 | 0.5 | 2 | 0 | 10 | 0.2 |
| 7_2716911 | 3 | 0 | 5 | 0 | 3 | 0.3333 | 2 | 0 | 13 | 0.0769 | 5 | 0 | 2 | 0 | 2 | 0 | 1 | 1 | 10 | 0.1 |
| 7_2717562! | NA | NA | NA | NA | 1 | 1 | 1 | 0 | 2 | 0.5 | NA | NA | NA | NA | NA | NA | 1 | 1 | 1 | 1 |
| 7_2717600' | NA | NA | NA | NA | 1 | 0 | 1 | 1 | 2 | 0.5 | NA | NA | NA | NA | NA | NA | 1 | 1 | 1 | 1 |
| 7_2717602( | NA | NA | NA | NA | 1 | 1 | 1 | 1 | 2 | 1 | NA | NA | NA | NA | NA | NA | 1 | 1 | 1 | 1 |
| 7_2717625 | NA | NA | NA | NA | 1 | 1 | 1 | 1 | 2 | 1 | NA | NA | NA | NA | NA | NA | 1 | 1 | 1 | 1 |
| 7_2717629 | NA | NA | NA | NA | 1 | 1 | 1 | 1 | 2 | 1 | NA | NA | NA | NA | NA | NA | 1 | 1 | 1 | 1 |
| 7_2717638( | NA | NA | NA | NA | 1 | 0 | 1 | 1 | 2 | 0.5 | NA | NA | NA | NA | NA | NA | 1 | 1 | 1 | 1 |
| 7_2717652( | NA | NA | NA | NA | 1 | 1 | 1 | 1 | 2 | 1 | NA | NA | NA | NA | NA | NA | 1 | 1 | 1 | 1 |
| 7_2717658 | NA | NA | NA | NA | 1 | 1 | 1 | 1 | 2 | 1 | NA | NA | NA | NA | NA | NA | 1 | 1 | 1 | 1 |
| 7_2718874! | 1 | 0 | 1 | 1 | 1 | 0 | NA | NA | 3 | 0.3333 | 3 | 0.3333 | 3 | 0.3333 | 1 | 0 | 2 | 0.5 | 9 | 0.3333 |
| 7_2718906( | 3 | 0.6667 | 1 | 0 | 1 | 0 | NA | NA | 5 | 0.4 | 3 | 0.3333 | 4 | 0.5 | 2 | 1 | 2 | 0 | 11 | 0.4545 |
| 7_2718914 | 3 | 1 | 1 | 1 | 1 | 1 | 1 | 1 | 6 | 1 | 4 | 1 | 4 | 1 | 2 | 1 | 2 | 1 | 12 | 1 |
| 7_2718918 | 3 | 0.6667 | 1 | 1 | 1 | 1 | 1 | 1 | 6 | 0.8333 | 4 | 1 | 5 | 0.6 | 2 | 1 | 2 | 1 | 13 | 0.8462 |
| 7_2718920 | 3 | 0 | 1 | 0 | 1 | 1 | 2 | 0 | 7 | 0.1429 | 4 | 0.25 | 5 | 0.4 | 2 | 0 | 2 | 0 | 13 | 0.2308 |
| 7_2718923! | 3 | 1 | NA | NA | NA | NA | 1 | 1 | 4 | 1 | 3 | 0.6667 | 4 | 1 | 2 | 1 | 2 | 1 | 11 | 0.9091 |
| 7_2718932 | 3 | 0.6667 | 1 | 1 | 1 | 1 | 2 | 0.5 | 7 | 0.7143 | 4 | 1 | 4 | 1 | 2 | 0.5 | 2 | 0.5 | 12 | 0.8333 |
| 7_2718940( | 3 | 0.6667 | 1 | 1 | 1 | 0 | 1 | 0 | 6 | 0.5 | 4 | 0.5 | 5 | 0.4 | 2 | 1 | 2 | 0 | 13 | 0.4615 |
| 7_2718941! | 3 | 0.6667 | 1 | 1 | 1 | 1 | 1 | 0 | 6 | 0.6667 | 4 | 1 | 5 | 0.8 | 1 | 1 | 2 | 1 | 12 | 0.9167 |
| 7_2718951 | 3 | 1 | NA | NA | 1 | 1 | 1 | 1 | 5 | 1 | 4 | 1 | 5 | 1 | 2 | 1 | 3 | 1 | 14 | 1 |
| 7_2718953 | 3 | 0 | 2 | 0.5 | 1 | 0 | 2 | 0 | 8 | 0.125 | 4 | 0 | 4 | 0.5 | 2 | 0 | 3 | 0.3333 | 13 | 0.2308 |
| 7_2718961' | 3 | 0.3333 | 2 | 0.5 | NA | NA | 2 | 0.5 | 7 | 0.4286 | 3 | 0 | 3 | 0.3333 | 2 | 1 | 3 | 0.3333 | 11 | 0.3636 |
| 7_2718969 | 3 | 0 | 2 | 0.5 | 1 | 0 | 2 | 0.5 | 8 | 0.25 | 4 | 0.75 | 5 | 0 | 2 | 1 | 3 | 0.3333 | 14 | 0.4286 |
| 7_2722251! | NA | NA | 1 | 1 | NA | NA | NA | NA | 1 | 1 | 1 | 0 | NA | NA | 1 | 1 | 2 | 0 | 4 | 0.25 |
| 7_2722254 | NA | NA | 1 | 1 | NA | NA | NA | NA | 1 | 1 | 1 | 0 | NA | NA | 1 | 1 | 2 | 1 | 4 | 0.75 |
| 7_2722295 | NA | NA | 1 | 1 | NA | NA | NA | NA | 1 | 1 | NA | NA | NA | NA | 1 | 0 | 2 | 1 | 3 | 0.6667 |
| 7_2722298( | NA | NA | 1 | 1 | NA | NA | NA | NA | 1 | 1 | NA | NA | NA | NA | 1 | 1 | 2 | 1 | 3 | 1 |
| 7_2722317 | NA | NA | 1 | 1 | NA | NA | NA | NA | 1 | 1 | NA | NA | NA | NA | 1 | 1 | 2 | 0 | 3 | 0.3333 |
| 7_2722323( | NA | NA | 1 | 1 | NA | NA | NA | NA | 1 | 1 | NA | NA | NA | NA | 1 | 1 | 1 | 1 | 2 | 1 |
| 7_2722344 | NA | NA | 1 | 1 | NA | NA | NA | NA | 1 | 1 | NA | NA | NA | NA | 1 | 1 | NA | NA | 1 | 1 |
| 7_2722348 | NA | NA | 1 | 1 | NA | NA | NA | NA | 1 | 1 | NA | NA | NA | NA | 1 | 1 | 2 | 1 | 3 | 1 |
| 7_2722437' | NA | NA | 1 | 1 | NA | NA | NA | NA | 1 | 1 | 1 | 1 | NA | NA | 1 | 0 | 2 | 0 | 4 | 0.25 |
| 7_2722771' | 5 | 0 | 1 | 0 | 2 | 0 | 4 | 0.25 | 12 | 0.0833 | 1 | 0 | 2 | 0 | 7 | 0.1429 | 3 | 0 | 13 | 0.0769 |
| 7_2722778 | 4 | 0.5 | 3 | 0.6667 | 2 | 0 | 5 | 0.4 | 14 | 0.4286 | 3 | 0.6667 | 3 | 0.6667 | 9 | 0.3333 | 3 | 0.3333 | 18 | 0.4444 |

| | | | | | | | | | | | | | | | | | | | | |
|---|---|---|---|---|---|---|---|---|---|---|---|---|---|---|---|---|---|---|---|---|
| 7_2722833 | 6 | 0.1667 | 2 | 0 | 3 | 0 | 4 | 0.25 | 15 | 0.1333 | 1 | 1 | 4 | 0 | 5 | 0.2 | 4 | 0 | 14 | 0.1429 |
| 7_2722856 | 6 | 1 | 2 | 1 | 5 | 1 | 4 | 1 | 17 | 1 | 4 | 1 | 5 | 1 | 7 | 1 | 2 | 1 | 18 | 1 |
| 7_2722859 | 3 | 1 | 2 | 1 | 3 | 1 | 4 | 1 | 12 | 1 | 3 | 0.6667 | 4 | 1 | 6 | 0.8333 | 2 | 1 | 15 | 0.8667 |
| 7_2722866 | 6 | 0.8333 | 2 | 1 | 4 | 1 | 4 | 0.5 | 16 | 0.8125 | 4 | 1 | 4 | 0.75 | 6 | 0.8333 | 3 | 0.6667 | 17 | 0.8235 |
| 7_2722878 | 6 | 0.1667 | 3 | 0 | 4 | 0.25 | 4 | 0.5 | 17 | 0.2353 | 3 | 0 | 5 | 1 | 7 | 0.4286 | 3 | 0 | 18 | 0.4444 |
| 7_2722885 | 6 | 0.1667 | 2 | 0 | 3 | 0.3333 | 3 | 0.6667 | 14 | 0.2857 | 2 | 0 | 4 | 0.25 | 6 | 0.3333 | 3 | 0 | 15 | 0.2 |
| 7_2722890 | 6 | 0.5 | 2 | 1 | 4 | 0.75 | 3 | 1 | 15 | 0.7333 | 4 | 0.75 | 5 | 0.6 | 7 | 0.7143 | 3 | 0.6667 | 19 | 0.6842 |
| 7_2722891 | 5 | 0 | 2 | 0 | 4 | 0.25 | 3 | 0 | 14 | 0.0714 | 4 | 0 | 5 | 0.2 | 6 | 0 | 3 | 0 | 18 | 0.0556 |
| 7_2722895 | 5 | 0.8 | 1 | 1 | 4 | 0.75 | 3 | 1 | 13 | 0.8462 | 4 | 0.75 | 5 | 1 | 6 | 1 | 3 | 0.3333 | 18 | 0.8333 |
| 7_2722916 | 5 | 0 | 2 | 0 | 4 | 0.5 | 1 | 0 | 12 | 0.1667 | 2 | 0 | 5 | 0.2 | 5 | 0.2 | 3 | 0 | 15 | 0.1333 |
| 7_2723014 | 4 | 0 | 2 | 0 | 3 | 0 | 5 | 0.2 | 14 | 0.0714 | 1 | 0 | 5 | 0 | 5 | 0.4 | 5 | 0 | 16 | 0.125 |
| 7_2723167 | 57 | 0.5263 | 56 | 0.4286 | 49 | 0.449 | 41 | 0.4634 | 203 | 0.468 | 56 | 0.625 | 55 | 0.4182 | 49 | 0.5918 | 34 | 0.4118 | 194 | 0.5206 |
| 7_2723170 | 72 | 0.25 | 69 | 0.2319 | 58 | 0.2759 | 49 | 0.2857 | 248 | 0.2581 | 58 | 0.2759 | 60 | 0.25 | 61 | 0.2623 | 46 | 0.1957 | 225 | 0.2489 |
| 7_2723179 | 80 | 0.4125 | 71 | 0.2535 | 67 | 0.3881 | 52 | 0.25 | 270 | 0.3333 | 65 | 0.4923 | 62 | 0.3871 | 57 | 0.5088 | 46 | 0.3261 | 230 | 0.4348 |
| 7_2279823 | 3 | 0.3333 | 1 | 0 | NA | NA | 1 | 1 | 5 | 0.4 | 1 | 0 | 5 | 0.4 | 2 | 0 | 5 | 0.8 | 13 | 0.4615 |
| 7_2728005 | 3 | 0.6667 | NA | NA | NA | NA | 3 | 0 | 6 | 0.3333 | 2 | 0.5 | 5 | 0.2 | 3 | 0.6667 | 5 | 0.8 | 15 | 0.5333 |
| 7_2728009 | 3 | 0.3333 | 1 | 0 | NA | NA | 2 | 0 | 6 | 0.1667 | 2 | 0 | 5 | 0.2 | 3 | 0 | 4 | 0.25 | 14 | 0.1429 |
| 7_2728015 | 1 | 1 | 1 | 1 | NA | NA | 2 | 1 | 4 | 1 | 1 | 1 | 3 | 1 | 1 | 1 | NA | NA | 5 | 1 |
| 7_2728021 | 1 | 1 | 1 | 1 | NA | NA | 2 | 0.5 | 4 | 0.75 | 1 | 1 | 3 | 0.3333 | 1 | 0 | 2 | 1 | 7 | 0.5714 |
| 7_2728024 | 2 | 0.5 | 1 | 1 | NA | NA | 2 | 0 | 5 | 0.4 | NA | NA | 2 | 0.5 | 1 | 0 | 2 | 0.5 | 5 | 0.4 |
| 7_2728025 | 2 | 1 | 1 | 1 | NA | NA | 2 | 1 | 5 | 1 | NA | NA | 3 | 1 | NA | NA | 2 | 0.5 | 5 | 0.8 |
| 7_2728047 | 2 | 0.5 | 1 | 0 | NA | NA | 2 | 0 | 5 | 0.2 | 1 | 0 | 1 | 0 | NA | NA | 1 | 1 | 3 | 0.3333 |
| 7_2728086 | 2 | 1 | 3 | 0 | NA | NA | 3 | 0.3333 | 8 | 0.375 | 1 | 1 | 8 | 0.375 | 2 | 0 | 3 | 0.6667 | 14 | 0.4286 |
| 7_2728090 | 3 | 0.3333 | 2 | 0 | 1 | 0 | 3 | 0 | 9 | 0.1111 | 2 | 0 | 6 | 0 | 3 | 0 | 3 | 0.6667 | 14 | 0.1429 |
| 7_2728104 | 4 | 0 | 3 | 0 | 1 | 0 | 2 | 0.5 | 10 | 0.1 | 1 | 1 | 7 | 0 | 3 | 0 | 4 | 0 | 15 | 0.0667 |
| 7_2728104 | 4 | 0 | 2 | 0.5 | 1 | 0 | 3 | 0.6667 | 10 | 0.3 | 1 | 1 | 8 | 0.375 | 3 | 0 | 4 | 0 | 16 | 0.25 |
| 7_2728113 | 4 | 0 | 2 | 0 | NA | NA | 3 | 0.6667 | 9 | 0.2222 | 1 | 0 | 7 | 0 | 2 | 0 | 3 | 0.3333 | 13 | 0.0769 |
| 7_2733572 | 4 | 0 | 7 | 0.2857 | 3 | 0.3333 | 3 | 0.3333 | 17 | 0.2353 | 3 | 0.3333 | 7 | 0.2857 | 8 | 0.375 | 4 | 0.25 | 22 | 0.3182 |
| 7_2733758 | 11 | 0.1818 | 10 | 0.1 | 10 | 0.1 | 7 | 0 | 38 | 0.1053 | 5 | 0.2 | 12 | 0.3333 | 11 | 0.0909 | 7 | 0 | 35 | 0.1714 |
| 7_2733763 | 8 | 0.375 | 11 | 0.5455 | 12 | 0.4167 | 5 | 0.4 | 36 | 0.4444 | 4 | 0.5 | 9 | 0.5556 | 9 | 0.3333 | 10 | 0.3 | 32 | 0.4062 |
| 7_2733771 | 8 | 0.125 | 11 | 0.1818 | 13 | 0.3077 | 7 | 0.1429 | 39 | 0.2051 | 1 | 0 | 14 | 0.1429 | 11 | 0.3636 | 11 | 0.2727 | 37 | 0.2432 |
| 7_2733778 | 11 | 0.3636 | 12 | 0.1667 | 13 | 0.1538 | 4 | 0.5 | 40 | 0.25 | 2 | 0.5 | 11 | 0.2727 | 9 | 0 | 10 | 0.2 | 32 | 0.1875 |
| 7_2733789 | 12 | 0 | 13 | 0.0769 | 14 | 0.2143 | 7 | 0.1429 | 46 | 0.1087 | 4 | 0.25 | 13 | 0.0769 | 12 | 0 | 12 | 0.25 | 41 | 0.122 |
| 7_2733801 | 11 | 0.0909 | 13 | 0.0769 | 14 | 0.0714 | 6 | 0.3333 | 44 | 0.1136 | 5 | 0 | 13 | 0.1538 | 11 | 0 | 8 | 0 | 37 | 0.0541 |
| 7_2733830 | 9 | 0.1111 | 13 | 0.1538 | 13 | 0 | 4 | 0 | 39 | 0.0769 | 3 | 0.3333 | 13 | 0 | 8 | 0.125 | 8 | 0 | 32 | 0.0625 |
| 7_2733832 | 8 | 0.125 | 11 | 0.6364 | 17 | 0.4706 | 8 | 0.25 | 44 | 0.4091 | 5 | 0.8 | 13 | 0.4615 | 9 | 0.5556 | 13 | 0.5385 | 40 | 0.55 |
| 7_2735612 | 17 | 0.8235 | 22 | 0.7727 | 24 | 0.75 | 7 | 0.7143 | 70 | 0.7714 | 8 | 1 | 11 | 0.7273 | 16 | 0.875 | 8 | 0.75 | 43 | 0.8372 |
| 7_2735615 | 13 | 0.9231 | 17 | 0.9412 | 24 | 0.875 | 8 | 1 | 62 | 0.9194 | 8 | 0.625 | 14 | 0.8571 | 18 | 1 | 8 | 1 | 48 | 0.8958 |
| 7_2735615 | 11 | 0.5455 | 20 | 0.65 | 19 | 0.6316 | 10 | 0.7 | 60 | 0.6333 | 6 | 0.6667 | 13 | 0.6923 | 12 | 0.6667 | 5 | 0.8 | 36 | 0.6944 |
| 7_2735623 | 13 | 0.3846 | 21 | 0.4286 | 25 | 0.6 | 10 | 0.4 | 69 | 0.4783 | 10 | 0.6 | 12 | 0.5833 | 19 | 0.5263 | 8 | 0.25 | 49 | 0.5102 |
| 7_2735625 | 14 | 0.0714 | 20 | 0.1 | 20 | 0.1 | 10 | 0 | 64 | 0.0781 | 8 | 0.125 | 9 | 0 | 13 | 0.0769 | 6 | 0.1667 | 36 | 0.0833 |
| 7_2735639 | 8 | 0.25 | 16 | 0.25 | 13 | 0.3077 | 7 | 0.1429 | 44 | 0.25 | 7 | 0.1429 | 10 | 0.1 | 14 | 0.1429 | 5 | 0.4 | 36 | 0.1667 |
| 7_2735658 | 1 | 1 | NA | NA | NA | NA | NA | NA | 1 | 1 | NA | NA | NA | NA | 1 | 1 | 1 | 1 | 2 | 1 |
| 7_2735659 | 1 | 1 | NA | NA | NA | NA | NA | NA | 1 | 1 | NA | NA | NA | NA | 1 | 0 | 1 | 1 | 2 | 0.5 |
| 7_2735679 | 1 | 1 | NA | NA | 1 | 1 | NA | NA | 2 | 1 | NA | NA | NA | NA | 1 | 1 | 1 | 1 | 2 | 1 |
| 7_2735681 | 1 | 1 | NA | NA | 1 | 1 | NA | NA | 2 | 1 | NA | NA | NA | NA | 1 | 1 | 1 | 1 | 2 | 1 |
| 7_2735724 | 1 | 1 | NA | NA | 1 | 1 | NA | NA | 2 | 1 | NA | NA | NA | NA | NA | NA | 1 | 1 | 1 | 1 |
| 7_2735739 | 13 | 0.2308 | 19 | 0.0526 | 25 | 0.04 | 11 | 0.0909 | 68 | 0.0882 | 10 | 0.1 | 14 | 0 | 17 | 0.0588 | 6 | 0.3333 | 47 | 0.0851 |
| 7_2735750 | 13 | 0 | 19 | 0.0526 | 18 | 0.1111 | 10 | 0.2 | 60 | 0.0833 | 7 | 0 | 12 | 0.0833 | 14 | 0.0714 | 5 | 0.2 | 38 | 0.0789 |
| 7_2735753 | 10 | 0.2 | 18 | 0.1111 | 20 | 0.35 | 5 | 0.2 | 53 | 0.2264 | 7 | 0.2857 | 10 | 0.3 | 15 | 0.3333 | 5 | 0.4 | 37 | 0.3243 |
| 7_2735768 | 10 | 0.9 | 13 | 0.7692 | 21 | 0.9524 | 9 | 0.8889 | 53 | 0.8868 | 7 | 1 | 12 | 0.8333 | 15 | 0.9333 | 6 | 0.8333 | 40 | 0.9 |
| 7_2735780 | 14 | 0.2143 | 17 | 0.1765 | 20 | 0.25 | 9 | 0 | 60 | 0.1833 | 7 | 0.1429 | 11 | 0.3636 | 15 | 0.2667 | 5 | 0 | 38 | 0.2368 |
| 7_2736279 | 13 | 0 | 12 | 0.0833 | 12 | 0.1667 | 8 | 0.125 | 45 | 0.0889 | 7 | 0 | 12 | 0 | 9 | 0.2222 | 1 | 0 | 29 | 0.069 |
| 7_2736368 | 4 | 0.5 | 8 | 0.875 | 5 | 0.6 | 2 | 1 | 19 | 0.7368 | 3 | 0.6667 | 2 | 1 | 2 | 0 | 7 | 0.7143 | 14 | 0.6429 |
| 7_2736540 | 2 | 0 | 1 | 0 | 3 | 0.3333 | 1 | 0 | 7 | 0.1429 | 1 | 1 | 1 | 0 | 2 | 0 | 2 | 0 | 6 | 0.1667 |
| 7_2736641 | 1 | 1 | NA | NA | 1 | 0 | NA | NA | 2 | 0.5 | 1 | 1 | NA | NA | 1 | 0 | 1 | 0 | 3 | 0.3333 |
| 7_2737282 | 10 | 0.9 | 15 | 1 | 14 | 0.8571 | 4 | 1 | 43 | 0.9302 | 9 | 0.8889 | 10 | 0.9 | 4 | 1 | 7 | 1 | 30 | 0.9333 |
| 7_2737303 | 12 | 0.5 | 13 | 0.3846 | 14 | 0.2143 | 5 | 0.2 | 44 | 0.3409 | 10 | 0.3 | 12 | 0.1667 | 5 | 0.2 | 6 | 0.5 | 33 | 0.2727 |
| 7_2737304 | 13 | 0.0769 | 14 | 0.2143 | 14 | 0.2857 | 7 | 0.2857 | 48 | 0.2083 | 9 | 0.3333 | 12 | 0.5 | 8 | 0.375 | 8 | 0.125 | 37 | 0.3514 |
| 7_2737306 | 12 | 0.5833 | 13 | 0.4615 | 15 | 0.5333 | 7 | 0.5714 | 47 | 0.5319 | 8 | 0.25 | 12 | 0.5 | 8 | 0.75 | 7 | 0.8571 | 35 | 0.5714 |
| 7_2737327 | 10 | 0.5 | 9 | 0.8889 | 14 | 0.2857 | 4 | 0.25 | 37 | 0.4865 | 8 | 0.5 | 10 | 0.6 | 8 | 0.375 | 7 | 0.4286 | 33 | 0.4848 |
| 7_2737330 | 10 | 0.9 | 11 | 0.9091 | 14 | 0.9286 | 6 | 0.8333 | 41 | 0.9024 | 8 | 1 | 12 | 1 | 8 | 1 | 7 | 1 | 35 | 1 |
| 7_2737354 | 9 | 0.6667 | 14 | 0.9286 | 16 | 0.9375 | 5 | 0.8 | 44 | 0.8636 | 8 | 0.875 | 12 | 0.9167 | 8 | 1 | 5 | 0.4 | 33 | 0.8485 |
| 7_2737371 | 9 | 0 | 8 | 0.25 | 10 | 0 | 4 | 0 | 31 | 0.0645 | 5 | 0.2 | 9 | 0.1111 | 8 | 0 | 5 | 0 | 27 | 0.0741 |
| 7_2740195 | NA | NA | 1 | 0 | 1 | 1 | NA | NA | 2 | 0.5 | NA | NA | NA | NA | NA | NA | 1 | 1 | 1 | 1 |

| | | | | | | | | | | | | | | | | | | | | |
|---|---|---|---|---|---|---|---|---|---|---|---|---|---|---|---|---|---|---|---|---|
| 7_2740247 | NA | NA | NA | NA | 1 | 1 | NA | NA | 1 | 1 | 1 | 1 | NA | NA | NA | NA | 1 | 0 | 2 | 0.5 |
| 7_2740290 | NA | NA | NA | NA | 1 | 1 | NA | NA | 1 | 1 | 1 | 1 | NA | NA | NA | NA | 1 | 0 | 2 | 0.5 |
| 7_2740311 | NA | NA | NA | NA | 1 | 1 | NA | NA | 1 | 1 | 1 | 0 | NA | NA | NA | NA | 1 | 1 | 2 | 0.5 |
| 7_2740322 | NA | NA | NA | NA | 1 | 1 | NA | NA | 1 | 1 | NA | NA | NA | NA | NA | NA | 1 | 1 | 1 | 1 |
| 7_2740334 | NA | NA | NA | NA | 1 | 1 | NA | NA | 1 | 1 | 1 | 1 | NA | NA | NA | NA | 1 | 0 | 2 | 0.5 |
| 7_2740353 | NA | NA | NA | NA | 1 | 1 | NA | NA | 1 | 1 | NA | NA | NA | NA | 1 | 1 | 1 | 1 | 2 | 1 |
| 7_2740359 | NA | NA | 1 | 1 | 1 | 1 | NA | NA | 2 | 1 | 1 | 1 | NA | NA | 1 | 1 | 1 | 1 | 3 | 1 |
| 7_2740469 | NA | NA | 1 | 0 | 1 | 1 | NA | NA | 2 | 0.5 | NA | NA | NA | NA | 1 | 0 | 2 | 1 | 3 | 0.6667 |
| 7_2740486 | 1 | 0 | 1 | 0 | 1 | 1 | NA | NA | 3 | 0.3333 | 2 | 0.5 | NA | NA | 1 | 1 | 2 | 0.5 | 5 | 0.6 |
| 7_2744768 | 3 | 0.3333 | 2 | 0 | 2 | 0 | 1 | 1 | 8 | 0.25 | 1 | 0 | 2 | 0 | 2 | 0.5 | 2 | 0 | 7 | 0.1429 |
| 7_2744782 | 5 | 0.6 | 2 | 1 | 3 | 0.3333 | 1 | 0 | 11 | 0.5455 | 1 | 0 | 4 | 0.25 | 3 | 0.3333 | 3 | 1 | 11 | 0.4545 |
| 7_2744784 | 4 | 0 | 3 | 0 | 3 | 0.3333 | 1 | 0 | 11 | 0.0909 | 2 | 0 | 4 | 0 | 4 | 0.25 | 3 | 0 | 13 | 0.0769 |
| 7_2744786 | 5 | 1 | 2 | 1 | 3 | 1 | NA | NA | 10 | 1 | 2 | 1 | 4 | 1 | 4 | 1 | 2 | 1 | 12 | 1 |
| 7_2744794 | 5 | 1 | 3 | 1 | 3 | 0.6667 | 1 | 1 | 12 | 0.9167 | 1 | 1 | 4 | 0.75 | 4 | 1 | 3 | 1 | 12 | 0.9167 |
| 7_2744801 | 5 | 0.8 | 3 | 1 | 3 | 0.6667 | 1 | 1 | 12 | 0.8333 | 2 | 0.5 | 4 | 1 | 4 | 0.5 | 3 | 0.3333 | 13 | 0.6154 |
| 7_2744816 | 5 | 1 | 3 | 1 | 3 | 1 | 1 | 1 | 12 | 1 | 2 | 1 | 4 | 1 | 4 | 1 | 3 | 1 | 13 | 1 |
| 7_2744826 | 5 | 0.8 | 2 | 0.5 | 3 | 0.6667 | NA | NA | 10 | 0.7 | 2 | 0.5 | 4 | 0.75 | 3 | 0.6667 | NA | NA | 9 | 0.6667 |
| 7_2744833 | 6 | 0.1667 | 2 | 0 | 3 | 0.6667 | 1 | 0 | 12 | 0.25 | 2 | 0 | 5 | 0.8 | 4 | 0.5 | 3 | 0.3333 | 14 | 0.5 |
| 7_2744857 | 5 | 0.2 | 2 | 0 | 4 | 0 | NA | NA | 11 | 0.0909 | 2 | 0 | 4 | 0 | 4 | 0.25 | 2 | 0 | 12 | 0.0833 |
| 7_2744870 | 6 | 1 | 3 | 1 | 4 | 1 | 1 | 1 | 14 | 1 | 3 | 1 | 5 | 1 | 3 | 1 | 3 | 1 | 14 | 1 |
| 7_2744872 | 5 | 0.4 | 3 | 0 | 4 | 0 | 1 | 0 | 13 | 0.1538 | 3 | 0.3333 | 4 | 0.5 | 4 | 0 | 3 | 0 | 14 | 0.2143 |
| 7_2744893 | 6 | 0 | 2 | 0.5 | 3 | 0 | NA | NA | 11 | 0.0909 | 1 | 0 | 5 | 0 | 4 | 0.25 | 2 | 0 | 12 | 0.0833 |
| 7_2744902 | 5 | 0.6 | 3 | 0.3333 | 3 | 0.3333 | 1 | 1 | 12 | 0.5 | 3 | 0 | 5 | 0.8 | 4 | 0 | 3 | 0.3333 | 15 | 0.3333 |
| 7_2744903 | 5 | 1 | 2 | 1 | 4 | 1 | 1 | 1 | 12 | 1 | 2 | 1 | 3 | 1 | 4 | 1 | 4 | 1 | 13 | 1 |
| 7_2744914 | 6 | 0.1667 | 3 | 0 | 3 | 0.3333 | 1 | 0 | 13 | 0.1538 | 3 | 0.3333 | 5 | 0.4 | 4 | 0.5 | 4 | 0.75 | 16 | 0.5 |
| 7_2744933 | 5 | 1 | 2 | 1 | 4 | 1 | 2 | 1 | 13 | 1 | 2 | 1 | 5 | 1 | 3 | 1 | 4 | 1 | 14 | 1 |
| 7_2744935 | 5 | 0 | 3 | 0.3333 | 3 | 0.3333 | 2 | 0 | 13 | 0.1538 | 2 | 0 | 4 | 0.25 | 3 | 0 | 4 | 1 | 13 | 0.3846 |
| 7_2744949 | 6 | 0.1667 | 3 | 0 | 4 | 0 | 2 | 0 | 15 | 0.0667 | 4 | 0 | 5 | 0 | 4 | 0 | 4 | 0.25 | 17 | 0.0588 |
| 7_2744971 | 6 | 0 | 3 | 0.3333 | 4 | 0.5 | 3 | 0 | 16 | 0.1875 | 4 | 0 | 4 | 0.25 | 6 | 0 | 5 | 0.2 | 19 | 0.1053 |
| 7_2744976 | 5 | 0.2 | 3 | 0 | 3 | 0 | 3 | 0 | 14 | 0.0714 | 4 | 0 | 5 | 0 | 6 | 0 | 5 | 0.4 | 20 | 0.1 |
| 7_2744978 | 5 | 0.8 | 3 | 0.3333 | 4 | 0.25 | 3 | 0 | 15 | 0.4 | 4 | 0.25 | 5 | 0.6 | 6 | 0.1667 | 5 | 0 | 20 | 0.25 |
| 7_2744981 | 3 | 0.3333 | 2 | 0.5 | 1 | 1 | 1 | 1 | 7 | 0.5714 | 1 | 1 | 2 | 0 | 4 | 0.5 | 4 | 0.25 | 11 | 0.3636 |
| 7_2744991 | 7 | 1 | 2 | 1 | 3 | 1 | 3 | 1 | 15 | 1 | 4 | 1 | 5 | 0.6 | 7 | 0.8571 | 5 | 0.8 | 21 | 0.8095 |
| 7_2744993 | 5 | 0.2 | 3 | 0 | 3 | 0.3333 | 2 | 0.5 | 13 | 0.2308 | 4 | 0.25 | 4 | 0 | 5 | 0 | 4 | 0.25 | 17 | 0.1176 |
| 7_2744995 | 8 | 0.125 | 3 | 0 | 4 | 0.25 | 2 | 0.5 | 17 | 0.1765 | 4 | 0.25 | 5 | 0.4 | 4 | 0.25 | 5 | 0.4 | 18 | 0.3333 |
| 7_2744996 | 6 | 1 | 1 | 1 | 3 | 1 | 3 | 1 | 13 | 1 | 2 | 1 | 2 | 1 | 5 | 1 | 3 | 1 | 12 | 1 |
| 7_2744998 | 9 | 0.6667 | 3 | 1 | 4 | 1 | 3 | 0.6667 | 19 | 0.7895 | 4 | 0.75 | 5 | 0.6 | 7 | 0.8571 | 5 | 1 | 21 | 0.8095 |
| 7_2745010 | 8 | 0.5 | 3 | 0 | 4 | 0.5 | 3 | 0 | 18 | 0.3333 | 3 | 0.3333 | 4 | 0.75 | 5 | 0.4 | 5 | 1 | 17 | 0.6471 |
| 7_2745020 | 9 | 0.2222 | 3 | 0 | 4 | 0 | 3 | 0 | 19 | 0.1053 | 3 | 0 | 4 | 0.25 | 7 | 0.2857 | 5 | 0 | 19 | 0.1579 |
| 7_2745022 | 8 | 0.875 | 3 | 0 | 3 | 0.3333 | 3 | 0.3333 | 17 | 0.5294 | 4 | 0.5 | 4 | 0.75 | 7 | 0.4286 | 5 | 0.4 | 20 | 0.5 |
| 7_2751869 | NA | NA | 1 | 1 | 1 | 1 | NA | NA | 2 | 1 | NA | NA | NA | NA | 1 | 1 | NA | NA | 1 | 1 |
| 7_2752175 | 1 | 1 | 2 | 0.5 | 4 | 0 | NA | NA | 7 | 0.2857 | NA | NA | 1 | 0 | 6 | 0.3333 | NA | NA | 7 | 0.2857 |
| 7_2755290 | 1 | 1 | NA | NA | NA | NA | NA | NA | 1 | 1 | NA | NA | NA | NA | 1 | 1 | 1 | 1 | 2 | 1 |
| 7_2755360 | 1 | 1 | NA | NA | NA | NA | NA | NA | 1 | 1 | NA | NA | NA | NA | 1 | 1 | NA | NA | 1 | 1 |
| 7_2755515 | 1 | 0 | 1 | 1 | NA | NA | NA | NA | 2 | 0.5 | NA | NA | NA | NA | 1 | 0 | 2 | 1 | 3 | 0.6667 |
| 7_2755516 | 2 | 1 | 1 | 1 | NA | NA | NA | NA | 3 | 1 | NA | NA | NA | NA | 2 | 0.5 | 1 | 0 | 3 | 0.3333 |
| 7_2755532 | 1 | 1 | NA | NA | NA | NA | NA | NA | 1 | 1 | NA | NA | NA | NA | 2 | 1 | 1 | 0 | 3 | 0.6667 |
| 7_2755554 | 2 | 0.5 | 1 | 0 | NA | NA | 1 | 1 | 4 | 0.5 | NA | NA | NA | NA | 2 | 1 | 2 | 0.5 | 4 | 0.75 |
| 7_2755560 | 2 | 0.5 | 1 | 0 | NA | NA | 1 | 1 | 4 | 0.5 | NA | NA | NA | NA | 2 | 0.5 | 1 | 1 | 3 | 0.6667 |
| 7_2755575 | 2 | 1 | 1 | 1 | NA | NA | NA | NA | 3 | 1 | NA | NA | 1 | 1 | 2 | 1 | 1 | 1 | 4 | 1 |
| 7_2755598 | 2 | 1 | 1 | 0 | NA | NA | NA | NA | 3 | 0.6667 | NA | NA | 1 | 1 | 2 | 0.5 | 1 | 1 | 4 | 0.75 |
| 7_2758189 | 1 | 1 | 1 | 0 | 1 | 0 | 3 | 0.6667 | 6 | 0.5 | 1 | 1 | 1 | 1 | NA | NA | NA | NA | 2 | 1 |
| 7_2758203 | NA | NA | 1 | 0 | 2 | 0.5 | 3 | 0 | 6 | 0.1667 | 1 | 1 | NA | NA | 2 | 0 | NA | NA | 3 | 0.3333 |
| 7_2760487 | 1 | 1 | 1 | 0 | NA | NA | NA | NA | 2 | 0.5 | NA | NA | NA | NA | 1 | 0 | 2 | 0.5 | 3 | 0.3333 |
| 7_2760503 | 1 | 1 | 1 | 0 | NA | NA | NA | NA | 2 | 0.5 | NA | NA | NA | NA | 1 | 1 | 1 | 0 | 2 | 0.5 |
| 7_2760509 | 1 | 1 | 1 | 0 | NA | NA | NA | NA | 2 | 0.5 | NA | NA | NA | NA | 1 | 1 | 1 | 0 | 2 | 0.5 |
| 7_2760520 | 1 | 1 | 1 | 0 | NA | NA | NA | NA | 2 | 0.5 | NA | NA | NA | NA | 1 | 1 | 1 | 1 | 2 | 1 |
| 7_2760635 | 1 | 1 | 1 | 0 | NA | NA | NA | NA | 2 | 0.5 | NA | NA | NA | NA | 1 | 1 | 1 | 1 | 2 | 1 |
| 7_2760645 | 1 | 1 | 1 | 1 | NA | NA | NA | NA | 2 | 1 | NA | NA | NA | NA | 1 | 1 | 1 | 0 | 2 | 0.5 |
| 7_2760688 | 1 | 1 | NA | NA | NA | NA | NA | NA | 1 | 1 | NA | NA | NA | NA | 1 | 1 | NA | NA | 1 | 1 |
| 7_2760837 | 1 | 0 | 1 | 1 | NA | NA | NA | NA | 2 | 0.5 | 1 | 1 | NA | NA | 3 | 0 | 1 | 1 | 5 | 0.4 |
| 7_2760933 | NA | NA | 1 | 1 | NA | NA | NA | NA | 1 | 1 | 1 | 0 | NA | NA | 3 | 0.6667 | 1 | 0 | 5 | 0.4 |
| 7_2760947 | NA | NA | 1 | 1 | NA | NA | NA | NA | 1 | 1 | 1 | 0 | NA | NA | 3 | 1 | 1 | 1 | 5 | 0.8 |
| 7_2760957 | NA | NA | 1 | 1 | NA | NA | NA | NA | 1 | 1 | NA | NA | NA | NA | 3 | 1 | 1 | 1 | 4 | 1 |
| 7_2760961 | NA | NA | 1 | 1 | NA | NA | NA | NA | 1 | 1 | NA | NA | NA | NA | 3 | 0.6667 | NA | NA | 3 | 0.6667 |

| | | | | | | | | | | | | | | | | | | | | |
|---|---|---|---|---|---|---|---|---|---|---|---|---|---|---|---|---|---|---|---|---|
| 7_2766079 | 1 | 0 | NA | NA | NA | NA | 2 | 1 | 3 | 0.6667 | 1 | 0 | 2 | 0.5 | 1 | 0 | 1 | 1 | 5 | 0.4 |
| 7_2766144 | 1 | 1 | 1 | 0 | NA | NA | 2 | 1 | 4 | 0.75 | NA | NA | 2 | 1 | 1 | 1 | 1 | 1 | 4 | 1 |
| 7_2766146 | 1 | 0 | 1 | 0 | NA | NA | 2 | 0.5 | 4 | 0.25 | 1 | 0 | 1 | 0 | 2 | 1 | 1 | 1 | 5 | 0.6 |
| 7_2766148 | 1 | 1 | 1 | 0 | NA | NA | 2 | 1 | 4 | 0.75 | 1 | 1 | 1 | 1 | 2 | 0.5 | NA | NA | 4 | 0.75 |
| 7_2766149 | 1 | 1 | 1 | 0 | NA | NA | 2 | 0 | 4 | 0.25 | 1 | 0 | 2 | 0.5 | 2 | 0.5 | 1 | 1 | 6 | 0.5 |
| 7_2766149 | 1 | 1 | 1 | 0 | NA | NA | 2 | 0 | 4 | 0.25 | 1 | 1 | 2 | 0.5 | 2 | 0.5 | 1 | 1 | 6 | 0.6667 |
| 7_2766161 | 1 | 1 | 1 | 0 | NA | NA | 2 | 0 | 4 | 0.25 | 1 | 0 | 2 | 0 | 2 | 0.5 | 1 | 0 | 6 | 0.1667 |
| 7_2766168 | 1 | 1 | 1 | 0 | NA | NA | 2 | 1 | 4 | 0.75 | 1 | 1 | 2 | 1 | 2 | 1 | 1 | 1 | 6 | 1 |
| 7_2766174 | 1 | 1 | 1 | 0 | NA | NA | 2 | 1 | 4 | 0.75 | 1 | 0 | 2 | 1 | 2 | 1 | 1 | 1 | 6 | 0.8333 |
| 7_2766197 | 1 | 1 | 1 | 0 | NA | NA | 2 | 0.5 | 4 | 0.5 | 1 | 0 | 2 | 0 | 2 | 0.5 | 1 | 1 | 6 | 0.3333 |
| 7_2766203 | 1 | 1 | 1 | 0 | NA | NA | 2 | 0 | 4 | 0.25 | 1 | 1 | 2 | 0 | 1 | 0 | 1 | 0 | 5 | 0.2 |
| 7_2766223 | 1 | 0 | 1 | 0 | NA | NA | 2 | 0.5 | 4 | 0.25 | 1 | 1 | 2 | 0 | 1 | 1 | 1 | 0 | 5 | 0.4 |
| 7_2766225 | 1 | 1 | 1 | 0 | NA | NA | 2 | 1 | 4 | 0.75 | 1 | 1 | 2 | 0.5 | 1 | 1 | 1 | 1 | 5 | 0.8 |
| 7_2766228 | NA | NA | 1 | 1 | NA | NA | 2 | 0.5 | 3 | 0.6667 | 1 | 0 | 2 | 1 | 1 | 0 | 1 | 0 | 5 | 0.4 |
| 7_2766261 | 1 | 1 | 1 | 0 | NA | NA | 2 | 0 | 4 | 0.25 | 1 | 0 | 1 | 1 | 1 | 1 | 1 | 0 | 4 | 0.5 |
| 7_2766265 | 1 | 1 | 1 | 1 | NA | NA | 2 | 1 | 4 | 1 | 1 | 1 | 1 | 1 | 1 | 1 | NA | NA | 3 | 1 |
| 7_2766270 | 1 | 1 | 1 | 0 | NA | NA | 2 | 0 | 4 | 0.25 | 1 | 1 | 1 | 0 | 1 | 0 | 1 | 0 | 4 | 0.25 |
| 7_2766275 | 1 | 1 | 1 | 0 | NA | NA | 2 | 0 | 4 | 0.25 | 1 | 0 | 1 | 1 | NA | NA | NA | NA | 2 | 0.5 |
| 7_2766289 | 1 | 1 | NA | NA | NA | NA | 1 | 1 | 2 | 1 | 1 | 1 | 1 | 1 | NA | NA | NA | NA | 2 | 1 |
| 7_2766308 | 1 | 1 | 1 | 0 | NA | NA | 2 | 0.5 | 4 | 0.5 | 1 | 1 | 1 | 1 | 1 | 1 | NA | NA | 3 | 1 |
| 7_2766320 | 1 | 0 | 1 | 0 | NA | NA | 2 | 1 | 4 | 0.5 | 1 | 1 | NA | NA | 1 | 1 | 1 | 1 | 3 | 1 |
| 7_2766348 | 1 | 0 | 1 | 0 | NA | NA | 2 | 0.5 | 4 | 0.25 | 1 | 0 | 1 | 0 | 1 | 1 | 1 | 0 | 4 | 0.25 |
| 7_2766361 | 1 | 1 | 1 | 1 | NA | NA | 1 | 1 | 3 | 1 | 1 | 1 | 1 | 1 | NA | NA | 1 | 1 | 3 | 1 |
| 7_2766362 | 1 | 1 | 1 | 1 | NA | NA | 1 | 0 | 3 | 0.6667 | 1 | 1 | 1 | 1 | 1 | 1 | 1 | 1 | 4 | 1 |
| 7_2766407 | 1 | 1 | 1 | 0 | NA | NA | 1 | 1 | 3 | 0.6667 | 1 | 1 | 1 | 1 | NA | NA | 1 | 1 | 3 | 1 |
| 7_2766414 | 1 | 1 | 1 | 1 | NA | NA | 1 | 1 | 3 | 1 | 1 | 1 | 1 | 1 | NA | NA | NA | NA | 2 | 1 |
| 7_2766418 | 1 | 1 | 1 | 1 | NA | NA | 1 | 1 | 3 | 1 | 1 | 1 | 1 | 1 | NA | NA | NA | NA | 2 | 1 |
| 7_2766420 | 1 | 1 | 1 | 0 | NA | NA | 1 | 0 | 3 | 0.3333 | 1 | 0 | 1 | 1 | NA | NA | NA | NA | 2 | 0.5 |
| 7_2768793 | NA | NA | NA | NA | 1 | 1 | 1 | 1 | 2 | 1 | NA | NA | NA | NA | 1 | 1 | 1 | 1 | 2 | 1 |
| 7_2768795 | NA | NA | NA | NA | 1 | 1 | 1 | 1 | 2 | 1 | NA | NA | NA | NA | 1 | 1 | 1 | 1 | 2 | 1 |
| 7_2768796 | NA | NA | 1 | 1 | 1 | 1 | 1 | 1 | 3 | 1 | NA | NA | NA | NA | 1 | 0 | 1 | 1 | 2 | 0.5 |
| 7_2768823 | NA | NA | 1 | 0 | 1 | 1 | 1 | 1 | 3 | 0.6667 | NA | NA | 1 | 1 | 1 | 1 | NA | NA | 2 | 1 |
| 7_2768839 | NA | NA | 1 | 0 | 1 | 1 | 1 | 0 | 3 | 0.3333 | NA | NA | 1 | 1 | 1 | 0 | NA | NA | 2 | 0.5 |
| 7_2768856 | NA | NA | 1 | 1 | 1 | 1 | 1 | 1 | 3 | 1 | NA | NA | 1 | 1 | 1 | 1 | NA | NA | 2 | 1 |
| 7_2768873 | NA | NA | 1 | 1 | 1 | 0 | 1 | 1 | 3 | 0.6667 | NA | NA | 1 | 1 | 1 | 0 | NA | NA | 2 | 0.5 |
| 7_2768890 | NA | NA | 1 | 0 | 1 | 1 | 1 | 1 | 3 | 0.6667 | NA | NA | 1 | 1 | 1 | 1 | NA | NA | 2 | 1 |
| 7_2768929 | NA | NA | 1 | 0 | 1 | 1 | NA | NA | 2 | 0.5 | NA | NA | 1 | 1 | 1 | 1 | NA | NA | 2 | 1 |
| 7_2768970 | NA | NA | 1 | 1 | 1 | 0 | NA | NA | 2 | 0.5 | NA | NA | 1 | 1 | 1 | 1 | NA | NA | 2 | 1 |
| 7_2768979 | NA | NA | 1 | 1 | 1 | 1 | NA | NA | 2 | 1 | NA | NA | 1 | 1 | 1 | 1 | NA | NA | 2 | 1 |
| 7_2768984 | NA | NA | 1 | 1 | 1 | 1 | NA | NA | 2 | 1 | NA | NA | NA | NA | 1 | 1 | NA | NA | 1 | 1 |
| 7_2786678 | 1 | 1 | 1 | 1 | NA | NA | NA | NA | 2 | 1 | NA | NA | NA | NA | 1 | 1 | NA | NA | 1 | 1 |
| 7_2786680 | 1 | 0 | 1 | 1 | NA | NA | NA | NA | 2 | 0.5 | NA | NA | NA | NA | 1 | 1 | NA | NA | 1 | 1 |
| 7_2786681 | 1 | 1 | NA | NA | NA | NA | NA | NA | 1 | 1 | NA | NA | NA | NA | 1 | 1 | NA | NA | 1 | 1 |
| 7_2786691 | 1 | 1 | 1 | 0 | NA | NA | NA | NA | 2 | 0.5 | NA | NA | NA | NA | 1 | 1 | NA | NA | 1 | 1 |
| 7_2788607 | 1 | 1 | 3 | 0.3333 | 2 | 0 | NA | NA | 6 | 0.3333 | 1 | 1 | 4 | 0 | 1 | 0 | NA | NA | 6 | 0.1667 |
| 7_2788617 | 2 | 0 | 5 | 0.4 | 3 | 0 | NA | NA | 10 | 0.2 | 1 | 0 | 5 | 0.2 | 1 | 0 | NA | NA | 7 | 0.1429 |
| 7_2788623 | 2 | 1 | 5 | 0.6 | 3 | 0.6667 | NA | NA | 10 | 0.7 | 1 | 0 | 6 | 0.8333 | 2 | 1 | NA | NA | 9 | 0.7778 |
| 7_2788643 | 2 | 1 | 5 | 0.2 | 2 | 0.5 | NA | NA | 9 | 0.4444 | 1 | 0 | 5 | 0 | 1 | 1 | NA | NA | 7 | 0.1429 |
| 7_2788643 | 1 | 1 | 1 | 1 | 1 | 1 | NA | NA | 3 | 1 | 1 | 1 | 4 | 1 | 2 | 1 | NA | NA | 7 | 1 |
| 7_2788647 | 1 | 0 | 5 | 0.2 | 2 | 0 | NA | NA | 8 | 0.125 | 1 | 0 | 4 | 0.25 | 1 | 1 | NA | NA | 6 | 0.3333 |
| 7_2788653 | 1 | 0 | 4 | 0.5 | 2 | 0.5 | NA | NA | 7 | 0.4286 | 1 | 1 | 5 | 0.6 | 2 | 1 | NA | NA | 8 | 0.75 |
| 7_2788654 | 2 | 0.5 | 6 | 1 | 2 | 0.5 | NA | NA | 10 | 0.8 | NA | NA | 4 | 0.5 | 2 | 0.5 | NA | NA | 6 | 0.5 |
| 7_2788659 | 1 | 1 | 4 | 0.25 | 2 | 0.5 | NA | NA | 7 | 0.4286 | 1 | 0 | 4 | 0.25 | 1 | 1 | NA | NA | 6 | 0.3333 |
| 7_2788763 | NA | NA | 1 | 0 | 1 | 1 | NA | NA | 2 | 0.5 | NA | NA | NA | NA | 1 | 1 | NA | NA | 1 | 1 |
| 7_2789613 | 3 | 0.6667 | 2 | 0 | 1 | 0 | 3 | 0 | 9 | 0.2222 | 2 | 0 | 1 | 0 | 4 | 0 | 5 | 0.4 | 12 | 0.1667 |
| 7_2789661 | 1 | 0 | 3 | 0 | 1 | 0 | 3 | 0.6667 | 8 | 0.25 | NA | NA | 2 | 1 | 4 | 0.75 | 4 | 0 | 10 | 0.5 |
| 7_2796786 | 7 | 0 | 8 | 0.25 | 5 | 0 | 10 | 0 | 30 | 0.0667 | 6 | 0 | 8 | 0 | 12 | 0.1667 | 7 | 0 | 33 | 0.0606 |
| 7_2796799 | 11 | 0.7273 | 8 | 0.875 | 6 | 0.6667 | 11 | 0.6364 | 36 | 0.7222 | 13 | 0.6923 | 6 | 0.8333 | 9 | 0.6667 | 9 | 0.6667 | 37 | 0.7027 |
| 7_2796802 | 10 | 0.4 | 7 | 0.5714 | 2 | 0 | 8 | 0.25 | 27 | 0.3704 | 10 | 0.4 | 6 | 0.5 | 10 | 0.5 | 6 | 0.1667 | 32 | 0.4062 |
| 7_2796805 | 12 | 0.6667 | 7 | 1 | 6 | 0.8333 | 10 | 0.8 | 35 | 0.8 | 11 | 0.5455 | 8 | 1 | 7 | 0.7143 | 8 | 0.625 | 34 | 0.7059 |
| 7_2796809 | 11 | 0.9091 | 9 | 1 | 4 | 0.75 | 11 | 0.8182 | 35 | 0.8857 | 13 | 0.9231 | 8 | 0.75 | 13 | 0.9231 | 9 | 1 | 43 | 0.907 |
| 7_2796825 | 13 | 0.5385 | 11 | 0.5455 | 5 | 0.6 | 13 | 0.3846 | 42 | 0.5 | 14 | 0.4286 | 10 | 0.5 | 12 | 0.5 | 9 | 0.4444 | 45 | 0.4667 |
| 7_2796835 | 12 | 0.8333 | 10 | 0.9 | 7 | 0.8571 | 11 | 0.9091 | 40 | 0.875 | 13 | 0.6154 | 10 | 0.9 | 11 | 0.6364 | 10 | 0.7 | 44 | 0.7045 |
| 7_2796836 | 12 | 1 | 11 | 1 | 7 | 1 | 11 | 1 | 41 | 1 | 14 | 1 | 11 | 1 | 12 | 1 | 11 | 1 | 48 | 1 |
| 7_2796844 | 10 | 0.8 | 11 | 0.3636 | 7 | 0.5714 | 9 | 0.3333 | 37 | 0.5135 | 14 | 0.2143 | 7 | 0.4286 | 9 | 0.5556 | 9 | 0.3333 | 39 | 0.359 |

| | | | | | | | | | | | | | | | | | | | | |
|---|---|---|---|---|---|---|---|---|---|---|---|---|---|---|---|---|---|---|---|---|
| 7_27968528 | 9 | 0.2222 | 11 | 0.2727 | 6 | 0 | 11 | 0.1818 | 37 | 0.1892 | 19 | 0.1579 | 8 | 0.125 | 13 | 0.3846 | 12 | 0.0833 | 52 | 0.1923 |
| 7_2797140: | NA | NA | 1 | 1 | NA | NA | NA | NA | 1 | 1 | NA | NA | 1 | 0 | NA | NA | 1 | 1 | 2 | 0.5 |
| 7_27977594 | 3 | 1 | 3 | 1 | 1 | 1 | 4 | 1 | 11 | 1 | 6 | 1 | 3 | 1 | 2 | 1 | 1 | 1 | 12 | 1 |
| 7_27977705 | 2 | 0.5 | 5 | 0 | 2 | 0 | 4 | 0 | 13 | 0.0769 | 6 | 0.3333 | 3 | 0.3333 | 3 | 0 | 2 | 0.5 | 14 | 0.2857 |
| 7_27977710 | 1 | 1 | 3 | 1 | 2 | 1 | 4 | 1 | 10 | 1 | 4 | 0.75 | 3 | 1 | 3 | 1 | 2 | 1 | 12 | 0.9167 |
| 7_27977767 | 4 | 1 | 4 | 1 | 2 | 1 | 4 | 1 | 14 | 1 | 6 | 1 | 4 | 1 | 3 | 1 | 2 | 1 | 15 | 1 |
| 7_27977824 | 4 | 0 | 5 | 0.2 | 1 | 0 | 4 | 0.25 | 14 | 0.1429 | 6 | 0 | 4 | 0 | 3 | 0.6667 | 2 | 0 | 15 | 0.1333 |
| 7_27977971 | 4 | 1 | 5 | 1 | 2 | 1 | 4 | 1 | 15 | 1 | 6 | 1 | 4 | 1 | 3 | 1 | 2 | 1 | 15 | 1 |
| 7_27977980 | 3 | 0 | 3 | 0 | 2 | 0.5 | 4 | 0.25 | 12 | 0.1667 | 6 | 0.1667 | 3 | 0 | 2 | 0 | 2 | 0.5 | 13 | 0.1538 |
| 7_27977994 | 2 | 1 | 2 | 1 | 2 | 1 | 3 | 1 | 9 | 1 | 5 | 0.8 | 3 | 1 | 2 | 1 | 1 | 1 | 11 | 0.9091 |
| 7_27978102 | 4 | 0.25 | 5 | 0.2 | 2 | 0 | 4 | 0.25 | 15 | 0.2 | 5 | 1 | 4 | 0.5 | 2 | 0 | 2 | 0 | 13 | 0.5385 |
| 7_27978133 | 4 | 0 | 3 | 0.3333 | 2 | 0 | 4 | 0.25 | 13 | 0.1538 | 6 | 0.3333 | 3 | 0.3333 | 2 | 0.5 | 2 | 0 | 13 | 0.3077 |
| 7_27978207 | 3 | 1 | 5 | 1 | 2 | 1 | 3 | 1 | 13 | 1 | 4 | 1 | 3 | 1 | 3 | 1 | 2 | 1 | 12 | 1 |
| 7_27978350 | 4 | 0.25 | 4 | 0 | 3 | 0 | 3 | 0 | 14 | 0.0714 | 6 | 0 | 5 | 0.2 | 3 | 0 | 2 | 0 | 16 | 0.0625 |
| 7_27978406 | 2 | 0 | 4 | 0 | 3 | 0.6667 | 3 | 0 | 12 | 0.1667 | 4 | 0 | 5 | 0.2 | 3 | 0.3333 | 2 | 0 | 14 | 0.1429 |
| 7_27978413 | 3 | 0.6667 | 4 | 0.25 | 3 | 0 | 3 | 0.3333 | 13 | 0.3077 | 4 | 0.75 | 3 | 0.3333 | 3 | 0 | 2 | 0 | 12 | 0.3333 |
| 7_27978535 | 3 | 0.6667 | 4 | 0.25 | 3 | 1 | 2 | 1 | 12 | 0.6667 | 4 | 0.75 | 3 | 1 | 3 | 1 | 2 | 1 | 12 | 0.9167 |
| 7_27978581 | 3 | 0.3333 | 5 | 0.4 | 3 | 0.3333 | 2 | 0 | 13 | 0.3077 | 5 | 0 | 5 | 0 | 3 | 0 | 2 | 0.5 | 15 | 0.0667 |
| 7_27978695 | 3 | 0 | 4 | 0.25 | 3 | 0 | 2 | 0 | 12 | 0.0833 | 5 | 0.2 | 5 | 0 | 3 | 0 | 2 | 0 | 15 | 0.0667 |
| 7_27979425 | NA | NA | NA | NA | NA | NA | 2 | 0.5 | 2 | 0.5 | NA | NA | 1 | 0 | 1 | 0 | 1 | 1 | 3 | 0.3333 |
| 7_27979589 | NA | NA | 1 | 1 | NA | NA | 1 | 1 | 2 | 1 | NA | NA | 2 | 0 | 1 | 0 | 3 | 0.6667 | 6 | 0.3333 |
| 7_27979651 | NA | NA | NA | NA | NA | NA | 2 | 0.5 | 2 | 0.5 | 1 | 0 | 2 | 0 | 1 | 0 | 2 | 0.5 | 6 | 0.1667 |
| 7_27980053 | 1 | 0 | 1 | 0 | NA | NA | 3 | 0.3333 | 5 | 0.2 | NA | NA | 2 | 0 | 3 | 0 | 1 | 1 | 6 | 0.1667 |
| 7_27980493 | NA | NA | 1 | 0 | NA | NA | 2 | 0.5 | 3 | 0.3333 | NA | NA | 3 | 0.3333 | 2 | 0 | 2 | 0 | 7 | 0.1429 |
| 7_27980666 | 1 | 0 | 2 | 0 | NA | NA | 3 | 0.3333 | 6 | 0.1667 | 1 | 1 | 4 | 0 | 3 | 0 | 2 | 0 | 10 | 0.1 |
| 7_27980804 | 1 | 0 | 2 | 0 | NA | NA | 3 | 0.3333 | 6 | 0.1667 | 1 | 1 | 4 | 0 | 3 | 0 | 3 | 0.6667 | 11 | 0.2727 |
| 7_28005010 | 4 | 0.5 | 2 | 1 | 8 | 0.5 | 2 | 0.5 | 16 | 0.5625 | 5 | 0.2 | 3 | 0.3333 | 3 | 0.3333 | 3 | 0.6667 | 14 | 0.3571 |
| 7_28005113 | 4 | 0.25 | 3 | 0 | 7 | 0.1429 | 2 | 0 | 16 | 0.125 | 4 | 0 | 3 | 0.3333 | 2 | 0 | 3 | 0 | 12 | 0.0833 |
| 7_28005180 | 3 | 1 | 2 | 0.5 | 8 | 0.625 | 2 | 0.5 | 15 | 0.6667 | 2 | 0.5 | 2 | 0.5 | 2 | 0.5 | 2 | 1 | 8 | 0.625 |
| 7_28005201 | 3 | 1 | 3 | 0.3333 | 8 | 0.5 | 1 | 1 | 15 | 0.6 | 5 | 0.8 | 2 | 1 | 3 | 0.6667 | 3 | 1 | 13 | 0.8462 |
| 7_28005280 | 1 | 0 | 6 | 0.3333 | 9 | 0.2222 | 3 | 0.3333 | 19 | 0.2632 | 6 | 0 | 3 | 0.6667 | 4 | 0.25 | 6 | 0.3333 | 19 | 0.2632 |
| 7_28005334 | 2 | 1 | 6 | 0.3333 | 11 | 0.7273 | 3 | 0.3333 | 22 | 0.5909 | 8 | 0.25 | 4 | 0.5 | 5 | 0.8 | 6 | 0.6667 | 23 | 0.5217 |
| 7_28059179 | NA | NA | 1 | 1 | NA | NA | 2 | 1 | 3 | 1 | 1 | 1 | NA | NA | NA | NA | NA | NA | 1 | 1 |
| 7_28059204 | NA | NA | 1 | 1 | NA | NA | NA | NA | 1 | 1 | 1 | 1 | NA | NA | NA | NA | NA | NA | 1 | 1 |
| 7_28076669 | 1 | 1 | 1 | 1 | NA | NA | 1 | 1 | 3 | 1 | NA | NA | 3 | 0.3333 | 2 | 0.5 | NA | NA | 5 | 0.4 |
| 7_28076910 | 2 | 0.5 | 1 | 0 | NA | NA | NA | NA | 3 | 0.3333 | NA | NA | 1 | 0 | 3 | 0.3333 | NA | NA | 4 | 0.25 |
| 7_28076929 | 2 | 1 | 1 | 1 | NA | NA | 1 | 1 | 4 | 1 | NA | NA | 1 | 0 | 3 | 0.3333 | NA | NA | 4 | 0.25 |
| 7_28077236 | 3 | 0 | 1 | 1 | NA | NA | 1 | 0 | 5 | 0.2 | NA | NA | 1 | 0 | 3 | 0.3333 | NA | NA | 4 | 0.25 |
| 7_28077455 | 2 | 1 | NA | NA | NA | NA | 1 | 1 | 3 | 1 | NA | NA | 1 | 1 | 2 | 0.5 | NA | NA | 3 | 0.6667 |
| 7_28077532 | 2 | 0.5 | 1 | 0 | NA | NA | 1 | 1 | 4 | 0.5 | NA | NA | NA | NA | 3 | 1 | NA | NA | 3 | 1 |
| 7_28077636 | 2 | 0.5 | 1 | 1 | NA | NA | 1 | 1 | 4 | 0.75 | NA | NA | 1 | 1 | 2 | 1 | NA | NA | 3 | 1 |
| 7_28077836 | 2 | 0.5 | NA | NA | NA | NA | 1 | 0 | 3 | 0.3333 | NA | NA | NA | NA | 3 | 0.3333 | NA | NA | 3 | 0.3333 |
| 7_28077856 | 2 | 0.5 | 1 | 1 | NA | NA | 1 | 0 | 4 | 0.5 | NA | NA | NA | NA | 3 | 0.3333 | NA | NA | 3 | 0.3333 |
| 7_28086245 | 6 | 0.5 | 7 | 0.4286 | 5 | 0.4 | 5 | 0.8 | 23 | 0.5217 | 10 | 0.2 | 8 | 0.375 | 4 | 0.5 | 5 | 0 | 27 | 0.2593 |
| 7_28086271 | 9 | 1 | 7 | 0.7143 | 6 | 0.8333 | 6 | 1 | 28 | 0.8929 | 8 | 0.625 | 8 | 1 | 6 | 0.8333 | 5 | 1 | 27 | 0.8519 |
| 7_28086292 | 9 | 0.5556 | 9 | 0.5556 | 6 | 0.6667 | 7 | 0.4286 | 31 | 0.5484 | 9 | 0.7778 | 8 | 0.75 | 4 | 0.75 | 6 | 0.1667 | 27 | 0.6296 |
| 7_28092339 | NA | NA | NA | NA | NA | NA | 1 | 1 | 1 | 1 | NA | NA | NA | NA | NA | NA | 1 | 1 | 1 | 1 |
| 7_28098461 | 13 | 0.0769 | 11 | 0.0909 | 8 | 0.25 | 9 | 0.2222 | 41 | 0.1463 | 10 | 0.1 | 12 | 0.0833 | 10 | 0 | 5 | 0.2 | 37 | 0.0811 |
| 7_28098476 | 6 | 0.6667 | 10 | 0.3 | 7 | 0.5714 | 7 | 0.7143 | 30 | 0.5333 | 7 | 0.5714 | 9 | 0.5556 | 11 | 0.7273 | 4 | 0.25 | 31 | 0.5806 |
| 7_28098503 | 12 | 0.0833 | 10 | 0.2 | 9 | 0 | 7 | 0 | 38 | 0.0789 | 9 | 0.2222 | 10 | 0.1 | 8 | 0.125 | 4 | 0.25 | 31 | 0.1613 |
| 7_28098509 | 11 | 0 | 10 | 0 | 10 | 0.1 | 7 | 0.2857 | 38 | 0.0789 | 9 | 0 | 9 | 0.3333 | 11 | 0 | 6 | 0 | 35 | 0.0857 |
| 7_28098515 | 10 | 0.4 | 10 | 0.3 | 8 | 0.125 | 6 | 0.1667 | 34 | 0.2647 | 8 | 0.375 | 9 | 0.1111 | 8 | 0 | 6 | 0.3333 | 31 | 0.1935 |
| 7_28098665 | 11 | 0.3636 | 13 | 0.2308 | 12 | 0.25 | 7 | 0.4286 | 43 | 0.3023 | 9 | 0.2222 | 12 | 0.3333 | 9 | 0.2222 | 5 | 0.2 | 35 | 0.2571 |
| 7_28099849 | 5 | 0 | 9 | 0 | 2 | 0 | 9 | 0.2222 | 25 | 0.08 | 7 | 0.2857 | 8 | 0 | 6 | 0 | 1 | 1 | 22 | 0.1364 |
| 7_28100781 | 7 | 0 | 4 | 0.25 | 1 | 0 | 7 | 0 | 19 | 0.0526 | 6 | 0.1667 | 6 | 0 | 4 | 0 | 1 | 0 | 17 | 0.0588 |
| 7_28113367 | 6 | 0.6667 | 2 | 1 | 1 | 1 | 6 | 0.8333 | 15 | 0.8 | 2 | 1 | 5 | 0.6 | 1 | 1 | 2 | 1 | 10 | 0.8 |
| 7_28113389 | 6 | 0 | 2 | 0 | 1 | 0 | 6 | 0.1667 | 15 | 0.0667 | 3 | 0 | 6 | 0 | 3 | 0.3333 | 2 | 0 | 14 | 0.0714 |
| 7_28113433 | 7 | 0.2857 | 2 | 0 | 1 | 0 | 7 | 0 | 17 | 0.1176 | 2 | 0 | 5 | 0.4 | 2 | 0 | 2 | 0 | 11 | 0.1818 |
| 7_28113548 | 7 | 0.1429 | 1 | 0 | NA | NA | 6 | 0.1667 | 14 | 0.1429 | 3 | 0 | 6 | 0.6667 | NA | NA | 1 | 1 | 10 | 0.5 |
| 7_28113655 | 7 | 0.2857 | 1 | 0 | NA | NA | 7 | 0.2857 | 15 | 0.2667 | 3 | 0.3333 | 4 | 0.25 | 2 | 0 | 2 | 0.5 | 11 | 0.2727 |
| 7_28158579 | NA | NA | 4 | 0.25 | 3 | 0 | NA | NA | 7 | 0.1429 | 2 | 0.5 | 1 | 0 | 1 | 0 | NA | NA | 4 | 0.25 |
| 7_28158642 | NA | NA | 3 | 0.6667 | 3 | 1 | NA | NA | 6 | 0.8333 | 1 | 1 | NA | NA | 1 | 1 | NA | NA | 2 | 1 |
| 7_28158663 | NA | NA | 3 | 0.3333 | 3 | 1 | NA | NA | 6 | 0.6667 | 2 | 0.5 | 1 | 0 | 2 | 1 | NA | NA | 5 | 0.6 |
| 7_28158729 | NA | NA | 3 | 0.6667 | 2 | 1 | NA | NA | 5 | 0.8 | 2 | 1 | 1 | 1 | 2 | 1 | NA | NA | 5 | 1 |
| 7_28158769 | NA | NA | 2 | 1 | 3 | 1 | NA | NA | 5 | 1 | 1 | 1 | NA | NA | 1 | 1 | NA | NA | 2 | 1 |

| | | | | | | | | | | | | | | | | | | | | |
|---|---|---|---|---|---|---|---|---|---|---|---|---|---|---|---|---|---|---|---|---|
| 7_28158859 | NA | NA | 3 | 0.3333 | 1 | 1 | NA | NA | 4 | 0.5 | 2 | 0.5 | NA | NA | 1 | 1 | NA | NA | 3 | 0.6667 |
| 7_28159110 | NA | NA | 4 | 0.25 | 1 | 0 | NA | NA | 5 | 0.2 | 2 | 0.5 | 1 | 0 | 2 | 0.5 | NA | NA | 5 | 0.4 |
| 7_28159121 | NA | NA | 4 | 1 | 1 | 1 | NA | NA | 5 | 1 | 1 | 1 | 1 | 1 | 2 | 1 | NA | NA | 4 | 1 |
| 7_28159173 | NA | NA | 3 | 1 | 3 | 1 | NA | NA | 6 | 1 | 2 | 1 | 1 | 1 | 2 | 1 | NA | NA | 5 | 1 |
| 7_28159378 | 1 | 1 | 3 | 1 | 3 | 1 | NA | NA | 7 | 1 | 1 | 1 | 1 | 0 | 2 | 0.5 | NA | NA | 4 | 0.5 |
| 7_28166304 | 1 | 0 | 6 | 0.8333 | 4 | 1 | 1 | 0 | 12 | 0.75 | 6 | 1 | 1 | 1 | 1 | 1 | NA | NA | 8 | 1 |
| 7_28166366 | 1 | 0 | 5 | 0 | 4 | 0.25 | 1 | 0 | 11 | 0.0909 | 5 | 0.2 | NA | NA | 2 | 0 | NA | NA | 7 | 0.1429 |
| 7_28166412 | 1 | 0 | 5 | 0.2 | 4 | 0 | 1 | 0 | 11 | 0.0909 | 6 | 0.1667 | 1 | 0 | 2 | 0 | NA | NA | 9 | 0.1111 |
| 7_28166564 | 1 | 1 | 8 | 0.75 | 4 | 1 | 1 | 0 | 14 | 0.7857 | 3 | 1 | 1 | 1 | 2 | 0.5 | NA | NA | 6 | 0.8333 |
| 7_28166718 | 1 | 1 | 7 | 1 | 4 | 1 | 1 | 1 | 13 | 1 | 7 | 1 | NA | NA | 1 | 1 | NA | NA | 8 | 1 |
| 7_28166768 | 1 | 0 | 6 | 1 | 3 | 0.6667 | 1 | 1 | 11 | 0.8182 | 6 | 0.3333 | 1 | 1 | 1 | 1 | 1 | 1 | 9 | 0.5556 |
| 7_28166835 | 1 | 0 | 6 | 0.1667 | 3 | 0.6667 | 1 | 0 | 11 | 0.2727 | 8 | 0.375 | 1 | 1 | 2 | 0 | 1 | 0 | 12 | 0.3333 |
| 7_28167090 | 1 | 0 | 7 | 0.5714 | 4 | 0.25 | 2 | 0.5 | 14 | 0.4286 | 5 | 0.2 | NA | NA | 1 | 0 | NA | NA | 6 | 0.1667 |
| 7_28167163 | 2 | 0.5 | 10 | 0.1 | 6 | 0 | 2 | 0 | 20 | 0.1 | 8 | 0.25 | NA | NA | 2 | 0 | 1 | 0 | 11 | 0.1818 |
| 7_28167188 | 2 | 1 | 5 | 0.2 | 3 | 0 | 1 | 1 | 11 | 0.3636 | 6 | 0.6667 | 1 | 0 | 2 | 0.5 | 1 | 1 | 10 | 0.6 |
| 7_28167232 | 2 | 1 | 9 | 0.6667 | 6 | 1 | 2 | 1 | 19 | 0.8421 | 7 | 0.8571 | 1 | 1 | 2 | 1 | NA | NA | 10 | 0.9 |
| 7_28167536 | 2 | 0 | 10 | 0 | 6 | 0.3333 | 3 | 0 | 21 | 0.0952 | 9 | 0.1111 | 1 | 0 | 2 | 0 | 2 | 0 | 14 | 0.0714 |
| 7_28258782 | NA | NA | 3 | 0.3333 | NA | NA | 1 | 0 | 4 | 0.25 | NA | NA | 2 | 0.5 | NA | NA | 1 | 0 | 3 | 0.3333 |
| 7_28258891 | NA | NA | 1 | 1 | NA | NA | NA | NA | 1 | 1 | NA | NA | 2 | 0.5 | NA | NA | 1 | 0 | 3 | 0.3333 |
| 7_28258931 | 1 | 1 | 3 | 1 | 1 | 1 | 1 | 1 | 6 | 1 | NA | NA | 2 | 1 | NA | NA | 1 | 1 | 3 | 1 |
| 7_28397832 | NA | NA | 3 | 0 | 1 | 1 | 1 | 1 | 5 | 0.4 | 2 | 0.5 | 1 | 0 | NA | NA | 1 | 1 | 4 | 0.5 |
| 7_28398299 | NA | NA | 2 | 0.5 | 1 | 0 | 1 | 0 | 4 | 0.25 | 2 | 0 | 1 | 0 | NA | NA | 1 | 1 | 4 | 0.25 |
| 7_28437355 | 1 | 0 | NA | NA | 1 | 1 | 1 | 0 | 3 | 0.3333 | NA | NA | NA | NA | 2 | 0.5 | NA | NA | 2 | 0.5 |
| 7_28437361 | 1 | 1 | NA | NA | 1 | 0 | 1 | 1 | 3 | 0.6667 | NA | NA | NA | NA | 1 | 1 | NA | NA | 1 | 1 |
| 7_28439201 | 14 | 0.7143 | 13 | 0.6154 | 13 | 0.5385 | 13 | 0.6154 | 53 | 0.6226 | 10 | 0.7 | 10 | 0.7 | 11 | 0.6364 | 12 | 0.6667 | 43 | 0.6744 |
| 7_28439294 | 12 | 0.5 | 13 | 0.6154 | 8 | 0.375 | 14 | 0.3571 | 47 | 0.4681 | 8 | 0.5 | 7 | 0.2857 | 13 | 0.4615 | 13 | 0.4615 | 41 | 0.439 |
| 7_28439322 | 15 | 0.3333 | 13 | 0.0769 | 13 | 0.0769 | 12 | 0.0833 | 53 | 0.1509 | 10 | 0.2 | 10 | 0.3 | 12 | 0.1667 | 15 | 0.2 | 47 | 0.2128 |
| 7_28439348 | 15 | 0.1333 | 12 | 0 | 11 | 0 | 15 | 0.0667 | 53 | 0.0566 | 11 | 0.0909 | 8 | 0.125 | 14 | 0.1429 | 15 | 0.1333 | 48 | 0.125 |
| 7_28439364 | 15 | 0.0667 | 16 | 0.0625 | 11 | 0.2727 | 14 | 0.2143 | 56 | 0.1429 | 11 | 0.1818 | 11 | 0.1818 | 13 | 0 | 16 | 0.0625 | 51 | 0.098 |
| 7_28453050 | 1 | 1 | 2 | 0 | 1 | 1 | 1 | 0 | 5 | 0.4 | 1 | 0 | NA | NA | 1 | 1 | 2 | 0.5 | 4 | 0.5 |
| 7_28453104 | 1 | 1 | 2 | 0 | 1 | 1 | 3 | 0.6667 | 7 | 0.5714 | NA | NA | NA | NA | NA | NA | 2 | 0.5 | 2 | 0.5 |
| 7_28487892 | 14 | 0.5 | 22 | 0.5 | 25 | 0.64 | 23 | 0.4348 | 84 | 0.5238 | 11 | 0.7273 | 25 | 0.44 | 22 | 0.5 | 11 | 0.5455 | 69 | 0.5217 |
| 7_28516185 | NA | NA | 2 | 0.5 | NA | NA | 3 | 0.3333 | 5 | 0.4 | 3 | 0.3333 | 3 | 0.3333 | 3 | 0 | 3 | 0.6667 | 12 | 0.3333 |
| 7_28516597 | 1 | 0 | 2 | 0 | NA | NA | 5 | 0.2 | 8 | 0.125 | 3 | 0.3333 | 3 | 0 | 2 | 0 | 3 | 0.3333 | 11 | 0.1818 |
| 7_28516691 | 1 | 0 | 2 | 0 | NA | NA | 5 | 0.2 | 8 | 0.125 | 3 | 0.3333 | 4 | 0.5 | 3 | 0 | 3 | 0.3333 | 13 | 0.3077 |
| 7_28516738 | 1 | 0 | 2 | 0 | NA | NA | 4 | 0.25 | 7 | 0.1429 | 3 | 0.3333 | 4 | 0 | 3 | 0 | 3 | 0 | 13 | 0.0769 |
| 7_28516870 | 1 | 0 | 3 | 0 | 1 | 0 | 4 | 0.25 | 9 | 0.1111 | 3 | 0.6667 | 4 | 0 | 3 | 0 | 3 | 0.3333 | 13 | 0.2308 |
| 7_28521523 | 32 | 0.3438 | 25 | 0.4 | 29 | 0.4483 | 29 | 0.2414 | 115 | 0.3565 | 23 | 0.2609 | 31 | 0.5484 | 18 | 0.1667 | 25 | 0.32 | 97 | 0.3505 |
| 7_28530870 | 11 | 0 | 4 | 0.25 | 9 | 0 | 6 | 0.3333 | 30 | 0.1 | 5 | 0.2 | 6 | 0.1667 | 7 | 0 | 4 | 0.25 | 22 | 0.1364 |
| 7_28531502 | 16 | 0 | 11 | 0 | 11 | 0.1818 | 7 | 0.1429 | 45 | 0.0667 | 12 | 0.1667 | 13 | 0.3846 | 8 | 0 | 9 | 0 | 42 | 0.1667 |
| 7_28531545 | 13 | 0.0769 | 10 | 0 | 11 | 0.0909 | 7 | 0.1429 | 41 | 0.0732 | 9 | 0 | 14 | 0.1429 | 7 | 0.1429 | 9 | 0 | 39 | 0.0769 |
| 7_28531561 | 15 | 0.0667 | 11 | 0 | 11 | 0.4545 | 5 | 0.4 | 42 | 0.1905 | 12 | 0.1667 | 11 | 0.1818 | 6 | 0 | 7 | 0 | 36 | 0.1111 |
| 7_28593131 | 3 | 0 | 13 | 0.1538 | 14 | 0.0714 | 12 | 0 | 42 | 0.0714 | 12 | 0.25 | 14 | 0.0714 | 8 | 0 | 12 | 0 | 46 | 0.087 |
| 7_28593219 | 6 | 0.1667 | 16 | 0.125 | 15 | 0 | 10 | 0 | 47 | 0.0638 | 12 | 0.1667 | 13 | 0.0769 | 9 | 0.1111 | 12 | 0.0833 | 46 | 0.1087 |
| 7_28593582 | 4 | 0.25 | 9 | 0 | 14 | 0.1429 | 11 | 0 | 38 | 0.0789 | 8 | 0 | 13 | 0.1538 | 8 | 0.125 | 8 | 0.125 | 37 | 0.1081 |
| 7_28593629 | 5 | 0.2 | 10 | 0.5 | 12 | 0.0833 | 10 | 0.3 | 37 | 0.2703 | 11 | 0.5455 | 14 | 0.5714 | 7 | 0.4286 | 10 | 0.2 | 42 | 0.4524 |
| 7_28856900 | 1 | 1 | NA | NA | 2 | 1 | 1 | 1 | 4 | 1 | 2 | 1 | NA | NA | 1 | 1 | 3 | 1 | 6 | 1 |
| 7_28857006 | NA | NA | NA | NA | 1 | 1 | NA | NA | 1 | 1 | 3 | 0.3333 | NA | NA | 3 | 0.3333 | 1 | 0 | 7 | 0.2857 |
| 7_28857035 | 1 | 0 | NA | NA | 2 | 0.5 | NA | NA | 3 | 0.3333 | 3 | 0 | NA | NA | 4 | 0.25 | 1 | 1 | 8 | 0.25 |
| 7_28858504 | 2 | 0 | 2 | 0.5 | 2 | 0.5 | 2 | 0 | 8 | 0.25 | 1 | 0 | 2 | 0.5 | 1 | 0 | 2 | 1 | 6 | 0.5 |
| 7_28859822 | 1 | 0 | 1 | 0 | 1 | 1 | 2 | 0.5 | 5 | 0.4 | 1 | 0 | 3 | 0.3333 | 1 | 0 | NA | NA | 5 | 0.2 |
| 7_28860368 | NA | NA | 1 | 0 | 1 | 0 | 3 | 0.3333 | 5 | 0.2 | 1 | 0 | 2 | 0.5 | NA | NA | NA | NA | 3 | 0.3333 |
| 7_28860395 | NA | NA | NA | NA | 1 | 1 | 2 | 0 | 3 | 0.3333 | 1 | 0 | 2 | 0.5 | 1 | 0 | NA | NA | 4 | 0.25 |
| 7_28860601 | NA | NA | 1 | 0 | 1 | 0 | 3 | 0.3333 | 5 | 0.2 | 1 | 0 | 1 | 1 | 1 | 0 | NA | NA | 3 | 0.3333 |
| 7_28860648 | NA | NA | 1 | 1 | 1 | 0 | 3 | 0.3333 | 5 | 0.4 | 1 | 0 | 3 | 0.3333 | 1 | 0 | NA | NA | 5 | 0.2 |
| 7_28860793 | NA | NA | 1 | 1 | 1 | 1 | 3 | 1 | 5 | 1 | 1 | 1 | 2 | 1 | NA | NA | NA | NA | 3 | 1 |
| 7_28860800 | NA | NA | 1 | 0 | 1 | 1 | 3 | 0 | 5 | 0.2 | 1 | 1 | 3 | 0 | NA | NA | NA | NA | 4 | 0.25 |
| 7_28860846 | NA | NA | NA | NA | 1 | 1 | 3 | 1 | 4 | 1 | 1 | 1 | 3 | 0.3333 | NA | NA | NA | NA | 4 | 0.5 |
| 7_28860876 | NA | NA | NA | NA | 1 | 0 | 3 | 1 | 4 | 0.75 | 1 | 1 | 3 | 0.3333 | 1 | 0 | NA | NA | 5 | 0.4 |
| 7_28860894 | 1 | 0 | 1 | 1 | 1 | 0 | 3 | 0.3333 | 6 | 0.3333 | 1 | 1 | 3 | 0 | 1 | 0 | NA | NA | 5 | 0.2 |
| 7_28861129 | 1 | 1 | 1 | 1 | NA | NA | 3 | 0.6667 | 5 | 0.8 | NA | NA | 2 | 1 | NA | NA | NA | NA | 2 | 1 |
| 7_28861454 | NA | NA | NA | NA | NA | NA | 2 | 0.5 | 2 | 0.5 | NA | NA | 2 | 0.5 | NA | NA | NA | NA | 2 | 0.5 |
| 7_28861522 | NA | NA | NA | NA | NA | NA | 2 | 0.5 | 2 | 0.5 | 1 | 0 | 1 | 1 | NA | NA | NA | NA | 2 | 0.5 |
| 7_28873762 | 16 | 0 | 26 | 0.1154 | 18 | 0.0556 | 10 | 0.1 | 70 | 0.0714 | 15 | 0.1333 | 18 | 0.1111 | 18 | 0.1667 | 10 | 0.4 | 61 | 0.1803 |
| 7_28873806 | 11 | 0.5455 | 27 | 0.4074 | 16 | 0.375 | 10 | 0.5 | 64 | 0.4375 | 13 | 0.6154 | 16 | 0.5625 | 19 | 0.5789 | 10 | 0.6 | 58 | 0.5862 |

| | | | | | | | | | | | | | | | | | | | | |
|---|---|---|---|---|---|---|---|---|---|---|---|---|---|---|---|---|---|---|---|---|
| 7_2898127 | 1 | 0 | NA | NA | 4 | 0.25 | 4 | 0 | 9 | 0.1111 | NA | NA | 1 | 0 | NA | NA | 1 | 1 | 2 | 0.5 |
| 7_2901652 | 8 | 0.25 | 12 | 0 | 7 | 0.2857 | 12 | 0.0833 | 39 | 0.1282 | 15 | 0.1333 | 9 | 0.2222 | 5 | 0.2 | 8 | 0.125 | 37 | 0.1622 |
| 7_2901653 | 6 | 0.1667 | 13 | 0 | 6 | 0.3333 | 12 | 0 | 37 | 0.0811 | 14 | 0.0714 | 10 | 0.2 | 7 | 0 | 6 | 0 | 37 | 0.0811 |
| 7_2901668 | 6 | 0.5 | 10 | 0.1 | 6 | 0.3333 | 8 | 0.125 | 30 | 0.2333 | 16 | 0.5 | 10 | 0 | 5 | 0.2 | 5 | 0.2 | 36 | 0.2778 |
| 7_2902056 | 5 | 0 | 5 | 0.2 | 3 | 0 | NA | NA | 13 | 0.0769 | 8 | 0.125 | 4 | 0 | 15 | 0.2 | 3 | 0.3333 | 30 | 0.1667 |
| 7_2902107 | 6 | 0 | 4 | 0.25 | 1 | 0 | 1 | 0 | 12 | 0.0833 | 10 | 0.1 | 4 | 0 | 19 | 0.2105 | 2 | 0 | 35 | 0.1429 |
| 7_2902122 | 4 | 0 | 3 | 0.3333 | 2 | 0 | 1 | 0 | 10 | 0.1 | 8 | 0.25 | 3 | 0.3333 | 18 | 0.1667 | 3 | 0 | 32 | 0.1875 |
| 7_2902124 | 3 | 0 | 4 | 0.25 | 2 | 0 | NA | NA | 9 | 0.1111 | 7 | 0 | 3 | 0 | 15 | 0.1333 | 2 | 0 | 27 | 0.0741 |
| 7_2902145 | 5 | 0 | 4 | 0 | 2 | 0.5 | 1 | 1 | 12 | 0.1667 | 7 | 0.1429 | 2 | 0 | 14 | 0.2857 | 3 | 0.6667 | 26 | 0.2692 |
| 7_2907055 | 3 | 0.6667 | 2 | 0 | NA | NA | NA | NA | 5 | 0.4 | 4 | 0.5 | NA | NA | 5 | 0.6 | 2 | 1 | 11 | 0.6364 |
| 7_2907698 | 2 | 0.5 | 2 | 0 | NA | NA | NA | NA | 4 | 0.25 | 5 | 0 | 1 | 0 | 4 | 0.25 | 2 | 0 | 12 | 0.0833 |
| 7_2907702 | 3 | 0 | 3 | 0.3333 | NA | NA | NA | NA | 6 | 0.1667 | 4 | 0.25 | 1 | 0 | 6 | 0.1667 | 1 | 0 | 12 | 0.1667 |
| 7_2907703 | 3 | 0.3333 | 3 | 0.3333 | NA | NA | NA | NA | 6 | 0.3333 | 4 | 0.25 | 1 | 1 | 6 | 0.1667 | 1 | 1 | 12 | 0.3333 |
| 7_2907712 | 2 | 0.5 | 1 | 1 | NA | NA | NA | NA | 3 | 0.6667 | 5 | 0.2 | 1 | 1 | 5 | 0.6 | 1 | 1 | 12 | 0.5 |
| 7_2907715 | 3 | 0.3333 | 3 | 0.3333 | NA | NA | NA | NA | 6 | 0.3333 | 4 | 0.75 | 1 | 1 | 5 | 1 | 2 | 1 | 12 | 0.9167 |
| 7_2907719 | 3 | 0.3333 | 2 | 0 | NA | NA | NA | NA | 5 | 0.2 | 2 | 0 | 1 | 1 | 6 | 0 | 2 | 0 | 11 | 0.0909 |
| 7_2907729 | 3 | 1 | 3 | 1 | NA | NA | NA | NA | 6 | 1 | 3 | 1 | 1 | 0 | 5 | 0.8 | NA | NA | 9 | 0.7778 |
| 7_2907730 | 3 | 0 | 3 | 0.3333 | NA | NA | NA | NA | 6 | 0.1667 | 3 | 0.6667 | 1 | 1 | 6 | 0.5 | 1 | 1 | 11 | 0.6364 |
| 7_2907740 | 3 | 0.6667 | 2 | 1 | NA | NA | NA | NA | 5 | 0.8 | 4 | 1 | 1 | 0 | 5 | 0.8 | 2 | 1 | 12 | 0.8333 |
| 7_2909876 | 3 | 0.3333 | 1 | 0 | NA | NA | NA | NA | 4 | 0.25 | 3 | 1 | NA | NA | 4 | 0.25 | 2 | 0 | 9 | 0.4444 |
| 7_2909904 | 3 | 0.3333 | 1 | 0 | NA | NA | NA | NA | 4 | 0.25 | 4 | 0.5 | NA | NA | 4 | 0.25 | 1 | 0 | 9 | 0.3333 |
| 7_2909907 | 2 | 0.5 | 1 | 0 | NA | NA | NA | NA | 3 | 0.3333 | 4 | 0.75 | NA | NA | 3 | 0.6667 | NA | NA | 7 | 0.7143 |
| 7_2909910 | 3 | 0.3333 | 1 | 0 | NA | NA | NA | NA | 4 | 0.25 | 4 | 0 | NA | NA | 3 | 0.6667 | 1 | 0 | 8 | 0.25 |
| 7_2909932 | 2 | 0.5 | 1 | 0 | NA | NA | NA | NA | 3 | 0.3333 | 4 | 0.75 | NA | NA | 4 | 1 | 1 | 1 | 9 | 0.8889 |
| 7_2909940 | 2 | 0.5 | 1 | 1 | NA | NA | NA | NA | 3 | 0.6667 | 4 | 0.5 | NA | NA | 4 | 0.5 | 1 | 0 | 9 | 0.4444 |
| 7_2909991 | 2 | 0.5 | 1 | 0 | NA | NA | NA | NA | 3 | 0.3333 | 4 | 0.5 | NA | NA | 1 | 1 | NA | NA | 5 | 0.6 |
| 7_2910017 | 2 | 0.5 | 1 | 0 | NA | NA | NA | NA | 3 | 0.3333 | 3 | 0.3333 | NA | NA | 1 | 0 | NA | NA | 4 | 0.25 |
| 7_2910027 | 2 | 0.5 | 1 | 1 | NA | NA | NA | NA | 3 | 0.6667 | 2 | 0.5 | NA | NA | 1 | 1 | NA | NA | 3 | 0.6667 |
| 7_2910034 | 2 | 1 | 1 | 0 | NA | NA | NA | NA | 3 | 0.6667 | 3 | 0.3333 | NA | NA | 1 | 0 | NA | NA | 4 | 0.25 |
| 7_2910036 | 2 | 0 | 1 | 1 | NA | NA | NA | NA | 3 | 0.3333 | 3 | 1 | NA | NA | NA | NA | NA | NA | 3 | 1 |
| 7_2945987 | NA | NA | 1 | 1 | 1 | 0 | NA | NA | 2 | 0.5 | NA | NA | NA | NA | 1 | 1 | NA | NA | 1 | 1 |
| 7_2946002 | NA | NA | 1 | 1 | 1 | 1 | NA | NA | 2 | 1 | NA | NA | NA | NA | 1 | 1 | NA | NA | 1 | 1 |
| 7_2946003 | NA | NA | 1 | 1 | 1 | 1 | NA | NA | 2 | 1 | NA | NA | NA | NA | 1 | 1 | NA | NA | 1 | 1 |
| 7_2946027 | NA | NA | NA | NA | 1 | 1 | NA | NA | 1 | 1 | NA | NA | NA | NA | 1 | 1 | NA | NA | 1 | 1 |
| 7_2946138 | NA | NA | NA | NA | 2 | 0 | 1 | 1 | 3 | 0.3333 | NA | NA | NA | NA | 1 | 1 | NA | NA | 1 | 1 |
| 7_2946142 | NA | NA | NA | NA | 2 | 0.5 | 1 | 1 | 3 | 0.6667 | NA | NA | NA | NA | 1 | 1 | NA | NA | 1 | 1 |
| 7_2946152 | NA | NA | NA | NA | 2 | 0.5 | NA | NA | 2 | 0.5 | NA | NA | NA | NA | 1 | 1 | NA | NA | 1 | 1 |
| 7_2946153 | NA | NA | NA | NA | 2 | 0 | 1 | 1 | 3 | 0.3333 | NA | NA | NA | NA | 1 | 1 | NA | NA | 1 | 1 |
| 7_2946156 | NA | NA | NA | NA | 2 | 0.5 | 1 | 0 | 3 | 0.3333 | NA | NA | NA | NA | 1 | 1 | NA | NA | 1 | 1 |
| 7_2946175 | NA | NA | NA | NA | 2 | 0 | 1 | 1 | 3 | 0.3333 | NA | NA | NA | NA | 1 | 1 | NA | NA | 1 | 1 |
| 7_2946180 | NA | NA | NA | NA | 1 | 1 | 1 | 0 | 2 | 0.5 | NA | NA | NA | NA | 1 | 1 | NA | NA | 1 | 1 |
| 7_2946272 | NA | NA | NA | NA | 2 | 0.5 | NA | NA | 2 | 0.5 | NA | NA | NA | NA | 1 | 1 | NA | NA | 1 | 1 |
| 7_2946299 | NA | NA | NA | NA | 2 | 0.5 | NA | NA | 2 | 0.5 | NA | NA | NA | NA | 1 | 1 | NA | NA | 1 | 1 |
| 7_2947669 | 2 | 0 | 1 | 0 | NA | NA | 2 | 1 | 5 | 0.4 | NA | NA | NA | NA | NA | NA | 1 | 1 | 1 | 1 |
| 7_2947676 | 2 | 0.5 | 1 | 0 | NA | NA | 2 | 0.5 | 5 | 0.4 | NA | NA | NA | NA | NA | NA | 1 | 1 | 1 | 1 |
| 7_2947699 | 3 | 0.6667 | 1 | 0 | NA | NA | 1 | 1 | 5 | 0.6 | NA | NA | NA | NA | NA | NA | 1 | 1 | 1 | 1 |
| 7_2947706 | 2 | 1 | 1 | 1 | NA | NA | NA | NA | 3 | 1 | NA | NA | NA | NA | NA | NA | 1 | 1 | 1 | 1 |
| 7_2947710 | 2 | 1 | 2 | 1 | NA | NA | 2 | 1 | 6 | 1 | NA | NA | 1 | 1 | NA | NA | 1 | 1 | 2 | 1 |
| 7_2947715 | 3 | 1 | 3 | 1 | NA | NA | 1 | 1 | 7 | 1 | NA | NA | 1 | 1 | NA | NA | 1 | 1 | 2 | 1 |
| 7_2947725 | 3 | 0 | 3 | 0.3333 | NA | NA | 1 | 0 | 7 | 0.1429 | NA | NA | 1 | 0 | NA | NA | 1 | 1 | 2 | 0.5 |
| 7_2947726 | 2 | 0 | NA | NA | NA | NA | 2 | 0.5 | 4 | 0.25 | NA | NA | 1 | 1 | NA | NA | 1 | 1 | 2 | 1 |
| 7_2947767 | 3 | 1 | 2 | 1 | NA | NA | NA | NA | 5 | 1 | NA | NA | NA | NA | NA | NA | 1 | 1 | 1 | 1 |
| 7_2947783 | 3 | 1 | 2 | 1 | NA | NA | 2 | 1 | 7 | 1 | NA | NA | NA | NA | NA | NA | 1 | 1 | 1 | 1 |
| 7_2947787 | 3 | 1 | 1 | 1 | NA | NA | 2 | 1 | 6 | 1 | NA | NA | NA | NA | NA | NA | 1 | 1 | 1 | 1 |
| 7_2947789 | 3 | 0.3333 | 2 | 0.5 | NA | NA | 2 | 1 | 7 | 0.5714 | NA | NA | NA | NA | NA | NA | 1 | 1 | 1 | 1 |
| 7_2947823 | 1 | 0 | 1 | 1 | NA | NA | 1 | 0 | 3 | 0.3333 | NA | NA | NA | NA | NA | NA | 2 | 0.5 | 2 | 0.5 |
| 7_2947923 | NA | NA | 1 | 0 | 1 | 0 | 3 | 0.3333 | 5 | 0.2 | NA | NA | NA | NA | NA | NA | 2 | 0.5 | 2 | 0.5 |
| 7_2947939 | 1 | 0 | 1 | 1 | 1 | 0 | 3 | 0 | 6 | 0.1667 | NA | NA | NA | NA | NA | NA | 2 | 0.5 | 2 | 0.5 |
| 7_2948353 | 5 | 0.2 | 2 | 0 | 1 | 0 | 1 | 0 | 9 | 0.1111 | 1 | 1 | NA | NA | NA | NA | NA | NA | 1 | 1 |
| 7_2948379 | 6 | 0.3333 | 3 | 0.3333 | 1 | 0 | 1 | 0 | 11 | 0.2727 | 1 | 1 | NA | NA | NA | NA | NA | NA | 1 | 1 |
| 7_2948411 | 6 | 0 | 3 | 0.3333 | 2 | 0 | 1 | 0 | 12 | 0.0833 | 1 | 1 | NA | NA | 1 | 0 | NA | NA | 2 | 0.5 |
| 7_2949191 | 6 | 0.6667 | 1 | 1 | 2 | 1 | 3 | 1 | 12 | 0.8333 | 1 | 1 | NA | NA | 1 | 1 | 1 | 1 | 3 | 1 |
| 7_2949194 | 1 | 1 | 1 | 1 | 2 | 1 | 1 | 1 | 5 | 1 | 1 | 1 | NA | NA | 1 | 1 | 1 | 1 | 3 | 1 |
| 7_2949227 | 6 | 0.3333 | 1 | 0 | 2 | 0 | 3 | 0.6667 | 12 | 0.3333 | 1 | 1 | NA | NA | 2 | 1 | 1 | 0 | 4 | 0.75 |
| 7_2949235 | 5 | 0.2 | 1 | 0 | 3 | 0 | 3 | 0 | 12 | 0.0833 | 1 | 0 | NA | NA | 2 | 1 | NA | NA | 3 | 0.6667 |

| | | | | | | | | | | | | | | | | | | | | |
|---|---|---|---|---|---|---|---|---|---|---|---|---|---|---|---|---|---|---|---|---|
| 7_29492545 | 5 | 0.2 | 1 | 0 | 1 | 1 | 2 | 0.5 | 9 | 0.3333 | 1 | 0 | NA | NA | 2 | 0.5 | 1 | 1 | 4 | 0.5 |
| 7_29492823 | 4 | 1 | NA | NA | 3 | 0.6667 | 2 | 0.5 | 9 | 0.7778 | 2 | 1 | 1 | 1 | 2 | 1 | NA | NA | 5 | 1 |
| 7_29492867 | 5 | 0.6 | 1 | 0 | 2 | 0 | 2 | 0.5 | 10 | 0.4 | 1 | 0 | NA | NA | 2 | 1 | 1 | 1 | 4 | 0.75 |
| 7_29492927 | 6 | 0.5 | 1 | 0 | 2 | 0.5 | 3 | 0.3333 | 12 | 0.4167 | 2 | 1 | 1 | 1 | 1 | 1 | 1 | 0 | 5 | 0.8 |
| 7_29492935 | 5 | 0.4 | 1 | 1 | 2 | 0.5 | 3 | 1 | 11 | 0.6364 | 2 | 0.5 | NA | NA | 2 | 0.5 | 1 | 0 | 5 | 0.4 |
| 7_29493154 | 6 | 1 | NA | NA | 3 | 1 | 3 | 1 | 12 | 1 | 2 | 1 | 1 | 1 | 2 | 1 | 1 | 1 | 6 | 1 |
| 7_29493295 | 6 | 0.3333 | 1 | 0 | 1 | 0 | 2 | 0 | 10 | 0.2 | 2 | 0 | 1 | 1 | 2 | 1 | NA | NA | 5 | 0.6 |
| 7_29493439 | 1 | 1 | NA | NA | 1 | 1 | NA | NA | 2 | 1 | 1 | 0 | NA | NA | 1 | 1 | NA | NA | 2 | 0.5 |
| 7_29493726 | 1 | 0 | NA | NA | 2 | 1 | NA | NA | 3 | 0.6667 | 1 | 0 | NA | NA | 1 | 1 | NA | NA | 2 | 0.5 |
| 7_29527757 | NA | NA | NA | NA | NA | NA | 1 | 1 | 1 | 1 | NA | NA | NA | NA | NA | NA | 1 | 1 | 1 | 1 |
| 7_29528133 | NA | NA | 2 | 1 | NA | NA | 2 | 1 | 4 | 1 | NA | NA | NA | NA | 1 | 1 | 1 | 1 | 2 | 1 |
| 7_29528755 | NA | NA | 3 | 1 | NA | NA | 3 | 1 | 6 | 1 | NA | NA | NA | NA | 1 | 1 | 1 | 0 | 2 | 0.5 |
| 7_29528958 | NA | NA | 4 | 0 | NA | NA | 3 | 0.3333 | 7 | 0.1429 | NA | NA | NA | NA | 2 | 0 | 1 | 1 | 3 | 0.3333 |
| 7_29529071 | NA | NA | 4 | 0.25 | NA | NA | 3 | 0.3333 | 7 | 0.2857 | NA | NA | NA | NA | 1 | 0 | 1 | 1 | 2 | 0.5 |
| 7_29529123 | NA | NA | 4 | 0.5 | NA | NA | 1 | 1 | 5 | 0.6 | NA | NA | NA | NA | 2 | 0 | 1 | 1 | 3 | 0.3333 |
| 7_29529175 | NA | NA | 4 | 1 | 1 | 1 | 3 | 1 | 8 | 1 | NA | NA | NA | NA | 1 | 1 | 1 | 1 | 2 | 1 |
| 7_29529278 | NA | NA | 4 | 0.75 | 1 | 1 | 2 | 1 | 7 | 0.8571 | NA | NA | NA | NA | 1 | 0 | 1 | 1 | 2 | 0.5 |
| 7_29529296 | NA | NA | 4 | 0.75 | 1 | 0 | 3 | 0.3333 | 8 | 0.5 | NA | NA | NA | NA | 2 | 1 | 1 | 1 | 3 | 1 |
| 7_29529878 | 1 | 0 | 4 | 0.25 | 1 | 0 | 3 | 0 | 9 | 0.1111 | NA | NA | 2 | 0.5 | 1 | 0 | 1 | 1 | 4 | 0.5 |
| 7_29529902 | 1 | 0 | 4 | 0.25 | 1 | 1 | 3 | 0.6667 | 9 | 0.4444 | NA | NA | 2 | 0.5 | 2 | 1 | 1 | 1 | 5 | 0.8 |
| 7_29529931 | 1 | 1 | 4 | 0.75 | 1 | 0 | 2 | 0.5 | 8 | 0.625 | NA | NA | 2 | 0 | 2 | 0 | 2 | 1 | 6 | 0.3333 |
| 7_29529960 | NA | NA | 3 | 1 | 1 | 1 | 2 | 1 | 6 | 1 | NA | NA | 2 | 1 | 2 | 1 | 1 | 1 | 5 | 1 |
| 7_29529982 | 1 | 0 | 4 | 0.5 | 1 | 0 | 2 | 1 | 8 | 0.5 | NA | NA | 2 | 0.5 | 2 | 0.5 | 1 | 0 | 5 | 0.4 |
| 7_29530007 | 1 | 1 | 2 | 1 | 1 | 0 | 3 | 0.6667 | 7 | 0.7143 | NA | NA | NA | NA | 2 | 1 | 1 | 0 | 3 | 0.6667 |
| 7_29530031 | 1 | 1 | 4 | 1 | 1 | 1 | 3 | 1 | 9 | 1 | NA | NA | 1 | 1 | 2 | 1 | 2 | 1 | 5 | 1 |
| 7_29530134 | 1 | 1 | 3 | 1 | 1 | 1 | 3 | 1 | 8 | 1 | NA | NA | 1 | 1 | 1 | 1 | 1 | 1 | 3 | 1 |
| 7_29530278 | 1 | 0 | 3 | 0.3333 | 1 | 0 | 3 | 0.3333 | 8 | 0.25 | NA | NA | 2 | 0.5 | 2 | 0 | 1 | 0 | 5 | 0.2 |
| 7_29543643 | 4 | 0.5 | 1 | 0 | 2 | 0.5 | 1 | 0 | 8 | 0.375 | 1 | 0 | NA | NA | 3 | 0.3333 | 1 | 1 | 5 | 0.4 |
| 7_29543679 | 3 | 0.6667 | 1 | 1 | 1 | 1 | NA | NA | 5 | 0.8 | 1 | 0 | NA | NA | 2 | 1 | NA | NA | 3 | 0.6667 |
| 7_29544011 | 3 | 1 | 2 | 0.5 | 2 | 1 | 1 | 1 | 8 | 0.875 | 1 | 1 | NA | NA | 3 | 1 | NA | NA | 4 | 1 |
| 7_29544356 | 2 | 1 | 2 | 1 | 2 | 1 | NA | NA | 6 | 1 | 1 | 1 | NA | NA | 1 | 1 | 1 | 1 | 3 | 1 |
| 7_29544677 | 3 | 0 | 2 | 0.5 | 2 | 0 | 1 | 0 | 8 | 0.125 | 2 | 0 | NA | NA | 2 | 0.5 | 1 | 0 | 5 | 0.2 |
| 7_29544716 | 4 | 1 | 2 | 1 | 2 | 1 | NA | NA | 8 | 1 | 1 | 1 | NA | NA | 2 | 0.5 | 1 | 0 | 4 | 0.5 |
| 7_29544950 | 4 | 0.5 | 2 | 0.5 | 1 | 1 | NA | NA | 7 | 0.5714 | 2 | 1 | NA | NA | NA | NA | 1 | 0 | 3 | 0.6667 |
| 7_29545151 | 4 | 0.75 | 2 | 0.5 | 2 | 1 | 1 | 1 | 9 | 0.7778 | 1 | 1 | NA | NA | 1 | 1 | 1 | 1 | 3 | 1 |
| 7_29545301 | 2 | 0 | 1 | 0 | 1 | 1 | 1 | 1 | 5 | 0.4 | 1 | 1 | NA | NA | 2 | 1 | 1 | 1 | 4 | 1 |
| 7_29545316 | 4 | 0 | 1 | 0 | 2 | 0.5 | 1 | 0 | 8 | 0.125 | 2 | 0 | NA | NA | 2 | 0.5 | 1 | 1 | 5 | 0.4 |
| 7_29545426 | 4 | 1 | 1 | 1 | 1 | 1 | NA | NA | 6 | 1 | 1 | 1 | NA | NA | 1 | 1 | 1 | 1 | 3 | 1 |
| 7_29573682 | 2 | 0.5 | 1 | 0 | 1 | 1 | 2 | 1 | 6 | 0.6667 | 1 | 0 | NA | NA | NA | NA | 1 | 1 | 2 | 0.5 |
| 7_29573692 | 2 | 1 | 1 | 1 | 1 | 0 | 2 | 1 | 6 | 0.8333 | 1 | 0 | 1 | 1 | NA | NA | 1 | 1 | 3 | 0.6667 |
| 7_29573706 | 2 | 1 | 1 | 0 | 1 | 1 | 2 | 0.5 | 6 | 0.6667 | 1 | 1 | 1 | 1 | NA | NA | 1 | 1 | 3 | 1 |
| 7_29574012 | 3 | 1 | 1 | 1 | 1 | 1 | 2 | 0 | 7 | 0.7143 | 1 | 1 | 2 | 0.5 | 1 | 1 | 1 | 1 | 5 | 0.8 |
| 7_29574239 | 3 | 1 | 1 | 0 | 1 | 1 | 1 | 1 | 6 | 0.8333 | 1 | 1 | 2 | 1 | 1 | 1 | NA | NA | 4 | 1 |
| 7_29574264 | 3 | 0.6667 | 1 | 1 | 1 | 1 | 2 | 1 | 7 | 0.8571 | 1 | 0 | 1 | 1 | 1 | 1 | 1 | 1 | 4 | 0.75 |
| 7_29574646 | 4 | 0.25 | 1 | 0 | NA | NA | 1 | 0 | 6 | 0.1667 | 2 | 0 | 2 | 1 | 1 | 0 | 1 | 0 | 6 | 0.3333 |
| 7_29574834 | 4 | 1 | 1 | 1 | NA | NA | 1 | 0 | 6 | 0.8333 | 2 | 1 | 2 | 1 | 2 | 1 | 1 | 1 | 7 | 1 |
| 7_29575020 | 4 | 1 | 1 | 1 | 1 | 1 | 2 | 1 | 8 | 1 | 2 | 1 | 2 | 1 | 2 | 1 | 1 | 1 | 7 | 1 |
| 7_29575189 | 3 | 1 | NA | NA | 1 | 1 | 2 | 1 | 6 | 1 | 1 | 1 | 2 | 1 | 1 | 1 | 1 | 1 | 5 | 1 |
| 7_29575206 | 4 | 0.25 | 1 | 1 | 1 | 1 | 2 | 0 | 8 | 0.375 | 2 | 0.5 | 2 | 0 | 1 | 0 | 1 | 0 | 6 | 0.1667 |
| 7_29575374 | 4 | 0.25 | NA | NA | 1 | 1 | 2 | 0 | 7 | 0.2857 | 2 | 0.5 | 2 | 0 | 1 | 0 | 1 | 0 | 6 | 0.1667 |
| 7_29575466 | 4 | 1 | 1 | 1 | NA | NA | 3 | 1 | 8 | 1 | 1 | 1 | 2 | 1 | 2 | 1 | 1 | 1 | 6 | 1 |
| 7_29575487 | 4 | 1 | 1 | 1 | 1 | 1 | 3 | 1 | 9 | 1 | 2 | 1 | 2 | 1 | 2 | 0.5 | 1 | 0 | 7 | 0.7143 |
| 7_29575601 | 4 | 1 | 1 | 1 | 1 | 1 | 2 | 1 | 8 | 1 | 2 | 1 | 2 | 1 | 1 | 1 | 1 | 1 | 6 | 1 |
| 7_29576252 | 3 | 0 | 2 | 0 | 1 | 0 | 2 | 0.5 | 8 | 0.125 | 1 | 1 | 2 | 0 | 2 | 0 | 1 | 0 | 6 | 0.1667 |
| 7_29592812 | NA | NA | NA | NA | 2 | 0.5 | NA | NA | 2 | 0.5 | NA | NA | 1 | 0 | 1 | 1 | 1 | 1 | 3 | 0.6667 |
| 7_29592823 | 1 | 1 | NA | NA | 2 | 0 | 1 | 0 | 4 | 0.25 | NA | NA | 1 | 1 | 1 | 0 | 1 | 0 | 3 | 0.3333 |
| 7_29592863 | 1 | 1 | NA | NA | 2 | 1 | NA | NA | 3 | 1 | NA | NA | 1 | 1 | 1 | 0 | 1 | 1 | 3 | 0.6667 |
| 7_29593028 | 1 | 1 | NA | NA | 2 | 0.5 | 1 | 0 | 4 | 0.5 | 3 | 0.3333 | NA | NA | 2 | 0.5 | NA | NA | 5 | 0.4 |
| 7_29593097 | 1 | 1 | NA | NA | 2 | 0.5 | NA | NA | 3 | 0.6667 | 1 | 1 | NA | NA | 1 | 1 | NA | NA | 2 | 1 |
| 7_29593140 | 1 | 1 | 1 | 0 | 2 | 1 | 1 | 0 | 5 | 0.6 | 2 | 0.5 | 1 | 1 | 2 | 1 | NA | NA | 5 | 0.8 |
| 7_29593270 | 1 | 1 | 1 | 1 | 3 | 1 | 2 | 1 | 7 | 1 | 2 | 1 | 1 | 1 | 2 | 1 | 1 | 1 | 6 | 1 |
| 7_29593346 | 1 | 1 | 1 | 1 | 3 | 1 | 2 | 1 | 7 | 1 | 1 | 1 | 2 | 1 | 2 | 1 | 2 | 1 | 7 | 1 |
| 7_29593455 | 1 | 1 | 1 | 1 | 3 | 1 | 3 | 1 | 8 | 1 | 2 | 0.5 | 1 | 1 | 1 | 1 | 1 | 1 | 5 | 0.8 |
| 7_29593486 | NA | NA | 1 | 1 | 3 | 1 | 3 | 0.6667 | 7 | 0.8571 | 1 | 1 | 2 | 1 | 1 | 1 | 1 | 1 | 5 | 1 |
| 7_29593497 | 1 | 1 | 1 | 1 | 3 | 1 | 2 | 0.5 | 7 | 0.8571 | 2 | 1 | 2 | 1 | 2 | 1 | 2 | 1 | 8 | 1 |

| | | | | | | | | | | | | | | | | | | | | |
|---|---|---|---|---|---|---|---|---|---|---|---|---|---|---|---|---|---|---|---|---|
| 7_2959359 | 1 | 0 | 1 | 1 | 2 | 0.5 | 3 | 0 | 7 | 0.2857 | 2 | 0.5 | 2 | 0 | 2 | 0 | 2 | 0 | 8 | 0.125 |
| 7_2959361 | NA | NA | 1 | 1 | 2 | 0 | 2 | 0 | 5 | 0.2 | 1 | 0 | 2 | 0 | 1 | 0 | 2 | 0.5 | 6 | 0.1667 |
| 7_2959416 | 1 | 0 | 1 | 0 | 1 | 0 | 3 | 1 | 6 | 0.5 | 1 | 1 | 1 | 0 | 1 | 1 | 2 | 0.5 | 5 | 0.6 |
| 7_2959417 | 1 | 1 | 1 | 1 | 1 | 1 | 1 | 0 | 4 | 0.75 | 1 | 1 | 2 | 0 | 1 | 0 | 2 | 0.5 | 6 | 0.3333 |
| 7_2959441 | NA | NA | NA | NA | NA | NA | 1 | 1 | 1 | 1 | NA | NA | NA | NA | 1 | 1 | NA | NA | 1 | 1 |
| 7_2959466 | 1 | 1 | 1 | 0 | 1 | 1 | 3 | 1 | 6 | 0.8333 | NA | NA | 2 | 0.5 | 1 | 1 | 2 | 1 | 5 | 0.8 |
| 7_2959499 | NA | NA | 1 | 0 | NA | NA | 2 | 0.5 | 3 | 0.3333 | NA | NA | NA | NA | NA | NA | 2 | 0.5 | 2 | 0.5 |
| 7_2959525 | 1 | 1 | 1 | 1 | NA | NA | 3 | 0.3333 | 5 | 0.6 | NA | NA | 2 | 1 | NA | NA | 2 | 0 | 4 | 0.5 |
| 7_2959551 | 1 | 1 | 1 | 0 | 1 | 0 | 3 | 1 | 6 | 0.6667 | NA | NA | 2 | 0.5 | 1 | 1 | 2 | 1 | 5 | 0.8 |
| 7_2959559 | 1 | 1 | 1 | 0 | 1 | 0 | 3 | 1 | 6 | 0.6667 | NA | NA | 2 | 0 | 1 | 1 | 1 | 0 | 4 | 0.25 |
| 7_2959562 | 1 | 1 | 1 | 1 | NA | NA | 3 | 1 | 5 | 1 | NA | NA | 2 | 1 | NA | NA | 2 | 1 | 4 | 1 |
| 7_2959566 | 1 | 0 | 1 | 0 | 1 | 1 | 3 | 0.3333 | 6 | 0.3333 | NA | NA | 2 | 0 | 1 | 0 | 2 | 0.5 | 5 | 0.2 |
| 7_2959570 | 1 | 1 | 1 | 1 | NA | NA | 2 | 1 | 4 | 1 | NA | NA | 2 | 1 | 1 | 0 | 2 | 0 | 5 | 0.4 |
| 7_2961412 | 1 | 1 | 1 | 1 | NA | NA | 2 | 1 | 4 | 1 | NA | NA | NA | NA | 1 | 1 | NA | NA | 1 | 1 |
| 7_2961433 | 1 | 1 | 1 | 1 | 1 | 1 | 1 | 1 | 4 | 1 | NA | NA | NA | NA | 1 | 1 | NA | NA | 1 | 1 |
| 7_2961523 | 1 | 1 | NA | NA | 1 | 0 | 1 | 0 | 3 | 0.3333 | NA | NA | NA | NA | 1 | 1 | NA | NA | 1 | 1 |
| 7_2961565 | 2 | 0 | 1 | 0 | 1 | 0 | 2 | 0.5 | 6 | 0.1667 | 1 | 0 | 1 | 1 | 1 | 1 | NA | NA | 3 | 0.6667 |
| 7_2963292 | 3 | 0.3333 | 3 | 0.3333 | 2 | 0 | 3 | 0.3333 | 11 | 0.2727 | NA | NA | 1 | 1 | NA | NA | 2 | 0 | 3 | 0.3333 |
| 7_2963295 | 2 | 0.5 | 3 | 0.3333 | 2 | 0 | 1 | 0 | 8 | 0.25 | NA | NA | 2 | 0.5 | NA | NA | 2 | 0 | 4 | 0.25 |
| 7_2963313 | 2 | 1 | 3 | 1 | 2 | 1 | 2 | 1 | 9 | 1 | NA | NA | 1 | 1 | NA | NA | 2 | 0.5 | 3 | 0.6667 |
| 7_2963335 | 3 | 1 | 2 | 1 | 1 | 1 | 3 | 1 | 9 | 1 | NA | NA | 1 | 1 | NA | NA | 1 | 1 | 2 | 1 |
| 7_2963337 | 2 | 0.5 | 1 | 1 | 1 | 0 | 2 | 1 | 6 | 0.6667 | NA | NA | NA | NA | NA | NA | 2 | 1 | 2 | 1 |
| 7_2963343 | 1 | 1 | 2 | 0.5 | NA | NA | 2 | 1 | 5 | 0.8 | NA | NA | 1 | 1 | NA | NA | 1 | 1 | 2 | 1 |
| 7_2963350 | 3 | 1 | 1 | 1 | NA | NA | 3 | 1 | 7 | 1 | NA | NA | 2 | 1 | NA | NA | 2 | 1 | 4 | 1 |
| 7_2963352 | 2 | 1 | 2 | 0 | NA | NA | 3 | 0.3333 | 7 | 0.4286 | NA | NA | 2 | 0.5 | NA | NA | 2 | 0.5 | 4 | 0.5 |
| 7_2963354 | 2 | 0.5 | 2 | 0 | NA | NA | 2 | 0.5 | 6 | 0.3333 | NA | NA | 1 | 0 | NA | NA | 2 | 0.5 | 3 | 0.3333 |
| 7_2963358 | 2 | 1 | 2 | 1 | NA | NA | 3 | 0.3333 | 7 | 0.7143 | NA | NA | 1 | 0 | NA | NA | 2 | 0.5 | 3 | 0.3333 |
| 7_2963361 | 1 | 0 | 2 | 0 | NA | NA | 2 | 1 | 5 | 0.4 | NA | NA | 1 | 0 | NA | NA | 2 | 0.5 | 3 | 0.3333 |
| 7_2963363 | 2 | 1 | 1 | 1 | NA | NA | 3 | 1 | 6 | 1 | NA | NA | 1 | 1 | NA | NA | 1 | 1 | 2 | 1 |
| 7_2963498 | 1 | 0 | NA | NA | NA | NA | 2 | 0.5 | 3 | 0.3333 | NA | NA | 1 | 0 | NA | NA | 2 | 0.5 | 3 | 0.3333 |
| 7_2963511 | 3 | 0.3333 | 1 | 0 | 1 | 0 | 2 | 0.5 | 7 | 0.2857 | NA | NA | 1 | 0 | NA | NA | 2 | 0.5 | 3 | 0.3333 |
| 7_2963512 | 1 | 1 | 1 | 1 | 1 | 0 | 2 | 0.5 | 5 | 0.6 | NA | NA | 1 | 1 | NA | NA | 2 | 0.5 | 3 | 0.6667 |
| 7_2963523 | 2 | 0.5 | 1 | 1 | NA | NA | 1 | 1 | 4 | 0.75 | NA | NA | NA | NA | NA | NA | 2 | 1 | 2 | 1 |
| 7_2963534 | NA | NA | 1 | 1 | NA | NA | 2 | 0.5 | 3 | 0.6667 | NA | NA | 1 | 1 | NA | NA | 2 | 0 | 3 | 0.3333 |
| 7_2966464 | NA | NA | 1 | 1 | 1 | 1 | 1 | 0 | 3 | 0.6667 | NA | NA | 2 | 0.5 | 2 | 0.5 | 2 | 0.5 | 6 | 0.5 |
| 7_2966473 | NA | NA | 1 | 1 | 1 | 1 | 1 | 1 | 3 | 1 | NA | NA | 2 | 1 | 2 | 0.5 | 2 | 0.5 | 6 | 0.6667 |
| 7_2966486 | NA | NA | 1 | 1 | 1 | 0 | 1 | 0 | 3 | 0.3333 | NA | NA | 2 | 0 | 2 | 0.5 | 2 | 0 | 6 | 0.1667 |
| 7_2966493 | NA | NA | 1 | 1 | 1 | 0 | 1 | 0 | 3 | 0.3333 | NA | NA | 2 | 0 | 2 | 0.5 | 2 | 0 | 6 | 0.1667 |
| 7_2966496 | NA | NA | 1 | 1 | NA | NA | 1 | 1 | 2 | 1 | NA | NA | 1 | 0 | 2 | 0.5 | 2 | 0.5 | 5 | 0.4 |
| 7_2966500 | NA | NA | 1 | 1 | 1 | 1 | 1 | 1 | 3 | 1 | NA | NA | 2 | 1 | 2 | 1 | 2 | 1 | 6 | 1 |
| 7_2966505 | NA | NA | 1 | 1 | 1 | 1 | 1 | 1 | 3 | 1 | 1 | 0 | 2 | 0.5 | NA | NA | 2 | 0.5 | 5 | 0.4 |
| 7_2966510 | NA | NA | 1 | 1 | 1 | 1 | 1 | 1 | 3 | 1 | 1 | 1 | 1 | 1 | 2 | 1 | 2 | 1 | 6 | 1 |
| 7_2966511 | NA | NA | 1 | 1 | 1 | 0 | NA | NA | 2 | 0.5 | 1 | 1 | 2 | 0 | 1 | 0 | 2 | 0 | 6 | 0.1667 |
| 7_2972732 | NA | NA | 1 | 0 | 1 | 1 | NA | NA | 2 | 0.5 | 2 | 0.5 | NA | NA | NA | NA | NA | NA | 2 | 0.5 |
| 7_2973636 | 1 | 0 | 2 | 0 | 2 | 0.5 | NA | NA | 5 | 0.2 | 2 | 0.5 | NA | NA | NA | NA | 4 | 0 | 6 | 0.1667 |
| 7_2973723 | 1 | 0 | 4 | 1 | 1 | 0 | 1 | 1 | 7 | 0.7143 | 2 | 0.5 | NA | NA | NA | NA | 2 | 0.5 | 4 | 0.5 |
| 7_2973739 | NA | NA | 4 | 1 | 1 | 1 | 1 | 1 | 6 | 1 | 3 | 1 | NA | NA | NA | NA | 1 | 1 | 4 | 1 |
| 7_2973761 | NA | NA | 4 | 0.75 | NA | NA | 1 | 1 | 5 | 0.8 | 3 | 1 | NA | NA | NA | NA | 1 | 1 | 4 | 1 |
| 7_2973762 | NA | NA | 4 | 1 | 1 | 1 | 1 | 1 | 6 | 1 | 3 | 1 | NA | NA | NA | NA | 1 | 1 | 4 | 1 |
| 7_2973789 | NA | NA | 4 | 0.5 | NA | NA | 1 | 0 | 5 | 0.4 | 3 | 0.6667 | NA | NA | NA | NA | 1 | 0 | 4 | 0.5 |
| 7_2973804 | NA | NA | 4 | 0.5 | 1 | 1 | 1 | 0 | 6 | 0.5 | 3 | 1 | NA | NA | NA | NA | 1 | 1 | 4 | 1 |
| 7_2973815 | NA | NA | 4 | 0.25 | NA | NA | 1 | 1 | 5 | 0.4 | 3 | 0.6667 | NA | NA | NA | NA | 1 | 1 | 4 | 0.75 |
| 7_2973816 | NA | NA | 4 | 0.75 | 1 | 0 | NA | NA | 5 | 0.6 | 3 | 0.3333 | NA | NA | NA | NA | 1 | 1 | 4 | 0.5 |
| 7_2973835 | NA | NA | 4 | 0.25 | NA | NA | 1 | 1 | 5 | 0.4 | 2 | 0.5 | NA | NA | NA | NA | NA | NA | 2 | 0.5 |
| 7_2973838 | NA | NA | 3 | 1 | NA | NA | 1 | 1 | 4 | 1 | 2 | 1 | NA | NA | NA | NA | 1 | 1 | 3 | 1 |
| 7_2973867 | NA | NA | 2 | 0 | NA | NA | 1 | 1 | 3 | 0.3333 | 1 | 1 | NA | NA | NA | NA | 1 | 1 | 2 | 1 |
| 7_2973869 | NA | NA | 3 | 1 | 1 | 1 | 1 | 1 | 5 | 1 | 1 | 0 | NA | NA | NA | NA | 1 | 1 | 2 | 0.5 |
| 7_2977682 | 2 | 0 | NA | NA | NA | NA | 2 | 0.5 | 4 | 0.25 | NA | NA | 2 | 0 | 1 | 1 | 3 | 0.3333 | 6 | 0.3333 |
| 7_2977689 | 2 | 0 | NA | NA | NA | NA | 2 | 0.5 | 4 | 0.25 | NA | NA | 2 | 1 | 1 | 0 | 2 | 0.5 | 5 | 0.6 |
| 7_2977697 | 2 | 1 | NA | NA | NA | NA | 2 | 1 | 4 | 1 | NA | NA | 2 | 1 | 1 | 1 | 4 | 1 | 7 | 1 |
| 7_2977698 | 2 | 1 | NA | NA | NA | NA | 2 | 1 | 4 | 1 | NA | NA | 2 | 1 | 1 | 0 | 3 | 0.6667 | 6 | 0.6667 |
| 7_2977706 | 2 | 0.5 | NA | NA | NA | NA | 2 | 0.5 | 4 | 0.5 | NA | NA | 2 | 0 | 1 | 1 | 4 | 0.25 | 7 | 0.2857 |
| 7_2977750 | 1 | 1 | NA | NA | NA | NA | 2 | 1 | 3 | 1 | NA | NA | 2 | 1 | 1 | 1 | 4 | 1 | 7 | 1 |
| 7_2977766 | 2 | 1 | NA | NA | NA | NA | 2 | 0.5 | 4 | 0.75 | NA | NA | 2 | 1 | 1 | 0 | 4 | 1 | 7 | 0.8571 |
| 7_2977785 | 2 | 0 | NA | NA | NA | NA | 2 | 0.5 | 4 | 0.25 | NA | NA | 1 | 1 | 1 | 0 | 4 | 0 | 6 | 0.1667 |

| | | | | | | | | | | | | | | | | | | | | |
|---|---|---|---|---|---|---|---|---|---|---|---|---|---|---|---|---|---|---|---|---|
| 7_2977786! | 2 | 0.5 | NA | NA | NA | NA | 2 | 0 | 4 | 0.25 | NA | NA | 1 | 0 | 1 | 0 | 4 | 0.25 | 6 | 0.1667 |
| 7_2977821! | 1 | 1 | NA | NA | NA | NA | NA | NA | 1 | 1 | NA | NA | NA | NA | NA | NA | 1 | 1 | 1 | 1 |
| 7_2980373( | NA | NA | 6 | 0.6667 | 2 | 0.5 | 1 | 0 | 9 | 0.5556 | 2 | 0.5 | 1 | 1 | 3 | 1 | 5 | 1 | 11 | 0.9091 |
| 7_2980376( | NA | NA | 6 | 0 | 2 | 0.5 | 1 | 0 | 9 | 0.1111 | 1 | 0 | 1 | 0 | 2 | 0 | 4 | 0.25 | 8 | 0.125 |
| 7_2980379' | NA | NA | 6 | 0.1667 | 2 | 1 | NA | NA | 8 | 0.375 | 2 | 0.5 | 1 | 1 | 3 | 1 | 5 | 0.8 | 11 | 0.8182 |
| 7_2980384. | NA | NA | 6 | 1 | 2 | 0.5 | 1 | 1 | 9 | 0.8889 | 3 | 1 | 1 | 1 | 2 | 1 | 5 | 1 | 11 | 1 |
| 7_2980385: | NA | NA | 4 | 0.5 | 2 | 1 | 1 | 1 | 7 | 0.7143 | 3 | 0.6667 | 1 | 0 | 3 | 0.3333 | 4 | 0.25 | 11 | 0.3636 |
| 7_2980388! | NA | NA | 3 | 0.6667 | 2 | 1 | 1 | 1 | 6 | 0.8333 | 3 | 1 | 1 | 1 | 3 | 0.6667 | 4 | 0.75 | 11 | 0.8182 |
| 7_2980391. | NA | NA | 6 | 1 | 2 | 1 | 1 | 1 | 9 | 1 | 3 | 1 | 1 | 1 | 3 | 1 | 5 | 1 | 12 | 1 |
| 7_2980399! | NA | NA | 6 | 1 | 2 | 1 | 1 | 1 | 9 | 1 | 2 | 1 | 1 | 1 | 3 | 1 | 4 | 1 | 10 | 1 |
| 7_2980422' | NA | NA | 5 | 0 | 1 | 0 | 1 | 1 | 7 | 0.1429 | 3 | 0 | 1 | 1 | 3 | 0 | 3 | 0 | 10 | 0.1 |
| 7_2980430( | NA | NA | 6 | 0.1667 | 2 | 0 | 1 | 0 | 9 | 0.1111 | 3 | 0 | NA | NA | 2 | 0.5 | 4 | 0 | 9 | 0.1111 |
| 7_2980448( | NA | NA | 7 | 0.2857 | 3 | 0 | 1 | 0 | 11 | 0.1818 | 3 | 0 | NA | NA | 3 | 0 | 5 | 0.2 | 11 | 0.0909 |
| 7_2980454! | NA | NA | 6 | 0.3333 | 4 | 0.25 | NA | NA | 10 | 0.3 | 3 | 0.6667 | 1 | 0 | 3 | 0.3333 | 5 | 0.2 | 12 | 0.3333 |
| 7_2980479. | NA | NA | 7 | 0.1429 | 4 | 0 | 1 | 0 | 12 | 0.0833 | 3 | 0 | 2 | 0 | 3 | 0 | 5 | 0.2 | 13 | 0.0769 |
| 7_2980487( | NA | NA | 6 | 1 | 4 | 1 | 1 | 1 | 11 | 1 | 3 | 0.6667 | 1 | 1 | 3 | 1 | 5 | 0.8 | 12 | 0.8333 |
| 7_2980495( | NA | NA | 4 | 0.25 | 3 | 0 | 1 | 0 | 8 | 0.125 | 3 | 0.3333 | 1 | 0 | 4 | 0 | 4 | 0.5 | 12 | 0.25 |
| 7_2980496! | NA | NA | 7 | 0 | 4 | 0.25 | 1 | 0 | 12 | 0.0833 | 3 | 0 | 2 | 0 | 4 | 0 | 5 | 0.2 | 14 | 0.0714 |
| 7_2980535: | NA | NA | 8 | 0.375 | 3 | 0 | 1 | 0 | 12 | 0.25 | 3 | 0.3333 | 2 | 0 | 6 | 0.5 | 4 | 0.25 | 15 | 0.3333 |
| 7_2980596( | NA | NA | 9 | 0.1111 | 4 | 0 | 3 | 0 | 16 | 0.0625 | 3 | 0 | 2 | 0 | 6 | 0.1667 | 6 | 0 | 17 | 0.0588 |
| 7_2980599‹ | NA | NA | 9 | 0 | 3 | 0.3333 | 2 | 0 | 14 | 0.0714 | 3 | 0.3333 | 2 | 0 | 6 | 0 | 5 | 0 | 16 | 0.0625 |
| 7_2980701' | 4 | 0 | 10 | 0.1 | 7 | 0.2857 | 3 | 0 | 24 | 0.125 | 3 | 0.3333 | 4 | 0.25 | 3 | 0 | 5 | 0.2 | 15 | 0.2 |
| 7_2983319! | 2 | 0.5 | NA | NA | NA | NA | NA | NA | 2 | 0.5 | NA | NA | NA | NA | NA | NA | 1 | 1 | 1 | 1 |
| 7_2983350‹ | 2 | 1 | NA | NA | NA | NA | NA | NA | 2 | 1 | NA | NA | NA | NA | NA | NA | 1 | 1 | 1 | 1 |
| 7_2983372: | 2 | 1 | 1 | 1 | NA | NA | NA | NA | 3 | 1 | NA | NA | NA | NA | NA | NA | 1 | 1 | 1 | 1 |
| 7_2983384( | 2 | 0.5 | NA | NA | NA | NA | NA | NA | 2 | 0.5 | NA | NA | NA | NA | NA | NA | 1 | 1 | 1 | 1 |
| 7_2983416: | 2 | 0.5 | NA | NA | NA | NA | NA | NA | 2 | 0.5 | NA | NA | NA | NA | NA | NA | 1 | 1 | 1 | 1 |
| 7_2983880! | 2 | 0.5 | 1 | 0 | 2 | 0 | 2 | 0.5 | 7 | 0.2857 | NA | NA | 2 | 0.5 | 1 | 0 | 1 | 0 | 4 | 0.25 |
| 7_2983973( | 2 | 1 | 2 | 1 | 5 | 1 | 2 | 0.5 | 11 | 0.9091 | NA | NA | 1 | 0 | 1 | 1 | 3 | 1 | 5 | 0.8 |
| 7_2983974: | 2 | 0 | 2 | 0.5 | 4 | 0.5 | 2 | 0 | 10 | 0.3 | NA | NA | 2 | 0 | 1 | 1 | 3 | 0 | 6 | 0.1667 |
| 7_2983983! | 2 | 0 | 2 | 0.5 | 6 | 0.5 | 2 | 0.5 | 12 | 0.4167 | NA | NA | 3 | 0.3333 | 1 | 0 | 3 | 0.6667 | 7 | 0.4286 |
| 7_2983994( | 2 | 0 | 3 | 0 | 6 | 0.1667 | 1 | 0 | 12 | 0.0833 | NA | NA | 3 | 0 | 1 | 0 | 4 | 0.25 | 8 | 0.125 |
| 7_2985996! | 4 | 0 | 8 | 0 | 5 | 0.6 | 4 | 0.5 | 21 | 0.2381 | 1 | 0 | 3 | 0 | 5 | 0.4 | 5 | 0 | 14 | 0.1429 |
| 7_2986018' | 4 | 0.5 | 6 | 0.5 | 4 | 0.25 | 2 | 0.5 | 16 | 0.4375 | 1 | 1 | 3 | 0.3333 | 4 | 0.75 | 4 | 0.5 | 12 | 0.5833 |
| 7_2986024‹ | 1 | 0 | 4 | 0.25 | 2 | 1 | 1 | 0 | 8 | 0.375 | 1 | 0 | 2 | 0 | 2 | 0.5 | 2 | 0 | 7 | 0.1429 |
| 7_2986037' | 1 | 1 | 1 | 1 | 1 | 1 | 1 | 1 | 4 | 1 | 1 | 1 | 1 | 1 | 3 | 0.6667 | 2 | 0.5 | 7 | 0.7143 |
| 7_2986042! | 2 | 0.5 | 2 | 1 | 2 | 1 | 1 | 1 | 7 | 0.8571 | NA | NA | 2 | 1 | 3 | 1 | 2 | 1 | 7 | 1 |
| 7_2986048: | 2 | 1 | 3 | 1 | 2 | 1 | 1 | 1 | 8 | 1 | NA | NA | 2 | 1 | 3 | 1 | 2 | 1 | 7 | 1 |
| 7_2986052! | 1 | 0 | 2 | 0.5 | NA | NA | 1 | 1 | 4 | 0.5 | NA | NA | 1 | 0 | 2 | 0.5 | 2 | 0.5 | 5 | 0.4 |
| 7_2986065‹ | NA | NA | 2 | 1 | NA | NA | 1 | 1 | 3 | 1 | NA | NA | 1 | 1 | 2 | 1 | 1 | 1 | 4 | 1 |
| 7_2986080( | 1 | 1 | 2 | 1 | 2 | 0.5 | 1 | 1 | 6 | 0.8333 | NA | NA | 1 | 1 | 2 | 1 | 2 | 1 | 5 | 1 |
| 7_2986082. | 1 | 1 | 2 | 0 | 2 | 0.5 | 1 | 1 | 6 | 0.5 | NA | NA | 1 | 0 | 2 | 0.5 | 1 | 1 | 4 | 0.5 |
| 7_2986084‹ | 1 | 1 | 2 | 0.5 | 1 | 0 | 1 | 1 | 5 | 0.6 | NA | NA | 1 | 1 | NA | NA | 1 | 0 | 2 | 0.5 |
| 7_2986094. | 1 | 1 | 2 | 1 | 2 | 1 | NA | NA | 5 | 1 | NA | NA | 1 | 1 | 2 | 1 | 2 | 1 | 5 | 1 |
| 7_2986101( | NA | NA | 2 | 0 | 2 | 1 | 1 | 0 | 5 | 0.4 | NA | NA | 1 | 1 | 1 | 1 | 2 | 0.5 | 4 | 0.75 |
| 7_2986106: | 1 | 0 | 2 | 0 | 2 | 0.5 | 1 | 0 | 6 | 0.1667 | NA | NA | 1 | 0 | 2 | 0 | 2 | 0.5 | 5 | 0.2 |
| 7_2986168( | NA | NA | 1 | 1 | 1 | 0 | NA | NA | 2 | 0.5 | NA | NA | 1 | 0 | 2 | 0.5 | 2 | 0.5 | 5 | 0.4 |
| 7_2986193( | NA | NA | 1 | 1 | 1 | 0 | NA | NA | 2 | 0.5 | NA | NA | 1 | 1 | 2 | 1 | NA | NA | 3 | 1 |
| 7_2986209( | NA | NA | 2 | 1 | 2 | 1 | NA | NA | 4 | 1 | NA | NA | 1 | 1 | 1 | 1 | 1 | 1 | 3 | 1 |
| 7_2986214‹ | NA | NA | 2 | 1 | 1 | 0 | NA | NA | 3 | 0.6667 | NA | NA | 1 | 1 | NA | NA | 1 | 1 | 2 | 1 |
| 7_2987731: | NA | NA | NA | NA | NA | NA | 3 | 0.3333 | 3 | 0.3333 | NA | NA | 2 | 1 | 1 | 1 | NA | NA | 3 | 1 |
| 7_2987733( | NA | NA | NA | NA | NA | NA | 4 | 0.25 | 4 | 0.25 | NA | NA | 2 | 0.5 | 1 | 0 | NA | NA | 3 | 0.3333 |
| 7_2987753‹ | NA | NA | NA | NA | NA | NA | 5 | 0.2 | 5 | 0.2 | NA | NA | 2 | 1 | 1 | 1 | NA | NA | 3 | 1 |
| 7_2987755: | NA | NA | NA | NA | NA | NA | 3 | 0.6667 | 3 | 0.6667 | NA | NA | 2 | 0.5 | 1 | 1 | NA | NA | 3 | 0.6667 |
| 7_2987758( | NA | NA | NA | NA | NA | NA | 3 | 1 | 3 | 1 | NA | NA | 2 | 1 | 1 | 1 | NA | NA | 3 | 1 |
| 7_2987759! | NA | NA | NA | NA | NA | NA | 5 | 1 | 5 | 1 | NA | NA | 2 | 0.5 | 1 | 1 | NA | NA | 3 | 0.6667 |
| 7_2987775! | NA | NA | NA | NA | NA | NA | 5 | 1 | 5 | 1 | NA | NA | 2 | 1 | 1 | 1 | NA | NA | 3 | 1 |
| 7_2987783( | NA | NA | NA | NA | NA | NA | 3 | 1 | 3 | 1 | NA | NA | 2 | 1 | 1 | 1 | NA | NA | 3 | 1 |
| 7_2987794' | NA | NA | NA | NA | NA | NA | 4 | 0.5 | 4 | 0.5 | NA | NA | 2 | 0.5 | NA | NA | NA | NA | 2 | 0.5 |
| 7_2987799: | NA | NA | NA | NA | NA | NA | 2 | 0.5 | 2 | 0.5 | NA | NA | 2 | 0 | 1 | 1 | NA | NA | 3 | 0.3333 |
| 7_2987801: | NA | NA | NA | NA | NA | NA | 3 | 1 | 3 | 1 | NA | NA | 1 | 1 | 1 | 1 | NA | NA | 2 | 1 |
| 7_2987840‹ | NA | NA | NA | NA | NA | NA | 3 | 1 | 3 | 1 | NA | NA | 3 | 1 | 1 | 1 | NA | NA | 4 | 1 |
| 7_2987975! | 1 | 0 | 1 | 0 | NA | NA | 4 | 0.75 | 6 | 0.5 | NA | NA | 3 | 0 | 1 | 0 | 1 | 1 | 5 | 0.2 |
| 7_2988172' | 1 | 0 | 1 | 0 | 2 | 0.5 | 5 | 0 | 9 | 0.1111 | NA | NA | 6 | 0 | 3 | 0 | 2 | 0.5 | 11 | 0.0909 |
| 7_2988640( | 6 | 0 | 10 | 0.1 | 14 | 0 | 11 | 0.1818 | 41 | 0.0732 | 11 | 0 | 8 | 0.125 | 13 | 0.3846 | 6 | 0 | 38 | 0.1579 |

| | | | | | | | | | | | | | | | | | | | | |
|---|---|---|---|---|---|---|---|---|---|---|---|---|---|---|---|---|---|---|---|---|
| 7_2988652 | 18 | 0.1111 | 24 | 0.125 | 21 | 0 | 17 | 0.1176 | 80 | 0.0875 | 14 | 0.1429 | 19 | 0.1053 | 22 | 0 | 14 | 0.0714 | 69 | 0.0725 |
| 7_2993123 | 1 | 1 | 3 | 1 | 1 | 0 | 1 | 0 | 6 | 0.6667 | 2 | 0.5 | 1 | 1 | 1 | 1 | 3 | 0 | 7 | 0.4286 |
| 7_2993151 | 1 | 1 | 2 | 1 | 1 | 0 | 1 | 1 | 5 | 0.8 | 1 | 1 | 1 | 1 | 1 | 1 | 2 | 0 | 5 | 0.6 |
| 7_3000740 | 40 | 0.05 | 48 | 0.125 | 53 | 0.0755 | 45 | 0.0444 | 186 | 0.0753 | 39 | 0.1026 | 49 | 0.0816 | 39 | 0.0513 | 44 | 0.0909 | 171 | 0.0819 |
| 7_3000762 | 43 | 0.1628 | 51 | 0.1569 | 55 | 0.1636 | 43 | 0.1395 | 192 | 0.1562 | 48 | 0.1458 | 45 | 0.1111 | 43 | 0.1395 | 39 | 0.1282 | 175 | 0.1314 |
| 7_3000771 | 42 | 0.0714 | 56 | 0.125 | 55 | 0.0909 | 41 | 0.0732 | 194 | 0.0928 | 50 | 0.16 | 41 | 0.0488 | 41 | 0.0488 | 36 | 0.0833 | 168 | 0.0893 |
| 7_3001362 | 3 | 0 | 6 | 0 | 11 | 0 | 8 | 0.25 | 28 | 0.0714 | 8 | 0.125 | 7 | 0 | 8 | 0.125 | 4 | 0.25 | 27 | 0.1111 |
| 7_3001363 | 3 | 0 | 5 | 0.2 | 12 | 0.0833 | 6 | 0 | 26 | 0.0769 | 8 | 0 | 4 | 0 | 10 | 0.1 | 5 | 0.2 | 27 | 0.0741 |
| 7_3002093 | 6 | 0.8333 | 2 | 1 | 3 | 1 | 4 | 1 | 15 | 0.9333 | 6 | 1 | 5 | 0.6 | 4 | 0.75 | 3 | 1 | 18 | 0.8333 |
| 7_3002099 | 6 | 0.3333 | 1 | 0 | 2 | 0 | 4 | 0.25 | 13 | 0.2308 | 6 | 0 | 4 | 0.25 | 4 | 0.25 | 3 | 0.3333 | 17 | 0.1765 |
| 7_3002100 | 5 | 0.2 | 1 | 0 | 3 | 0 | 4 | 0.25 | 13 | 0.1538 | 5 | 0 | 5 | 0.2 | 4 | 0.25 | 2 | 0 | 16 | 0.125 |
| 7_3002101 | 5 | 0.2 | 1 | 1 | 2 | 0 | 3 | 0.6667 | 11 | 0.3636 | 6 | 0 | 5 | 0.2 | 4 | 0 | 3 | 0 | 18 | 0.0556 |
| 7_3002105 | 5 | 0.8 | 1 | 1 | 2 | 0.5 | 3 | 1 | 11 | 0.8182 | 4 | 0.75 | 5 | 0.8 | 3 | 1 | 2 | 1 | 14 | 0.8571 |
| 7_3002111 | 4 | 0.5 | 1 | 0 | 2 | 0.5 | 2 | 0 | 9 | 0.3333 | 5 | 0 | 4 | 0 | 3 | 0.3333 | 2 | 0.5 | 14 | 0.1429 |
| 7_3002112 | 5 | 0 | 1 | 1 | 3 | 0 | 3 | 0.3333 | 12 | 0.1667 | 3 | 0 | 5 | 0.2 | 4 | 0.25 | 3 | 0 | 15 | 0.1333 |
| 7_3002122 | 5 | 0 | 1 | 0 | 3 | 0.6667 | 4 | 0 | 13 | 0.1538 | 5 | 0.2 | 5 | 0 | 4 | 0 | 3 | 0 | 17 | 0.0588 |
| 7_3002123 | 6 | 1 | 1 | 1 | 1 | 1 | 3 | 0.6667 | 11 | 0.9091 | 5 | 0.8 | 4 | 1 | 4 | 1 | 2 | 0.5 | 15 | 0.8667 |
| 7_3002127 | 6 | 0.1667 | 1 | 0 | 3 | 0 | 3 | 0.3333 | 13 | 0.1538 | 5 | 0 | 5 | 0 | 4 | 0.75 | 3 | 0.3333 | 17 | 0.2353 |
| 7_3002134 | 5 | 0.2 | 1 | 0 | 2 | 0.5 | 4 | 0.5 | 12 | 0.3333 | 5 | 0.4 | 4 | 0.25 | 4 | 0.25 | 3 | 0.6667 | 16 | 0.375 |
| 7_3002139 | 5 | 0.2 | 2 | 0 | 1 | 0 | 4 | 0 | 12 | 0.0833 | 4 | 0 | 4 | 0 | 3 | 0.3333 | 3 | 0 | 14 | 0.0714 |
| 7_3002139 | 5 | 0.2 | 1 | 0 | 3 | 0 | 4 | 0 | 13 | 0.0769 | 5 | 0 | 5 | 0 | 4 | 0.25 | 3 | 0 | 17 | 0.0588 |
| 7_3005497 | 4 | 1 | 7 | 1 | 3 | 1 | 5 | 0.8 | 19 | 0.9474 | 4 | 1 | 5 | 0.8 | 3 | 0.6667 | 6 | 1 | 18 | 0.8889 |
| 7_3005500 | 5 | 0.6 | 6 | 0.8333 | 4 | 0.75 | 6 | 1 | 21 | 0.8095 | 4 | 1 | 6 | 1 | 2 | 1 | 5 | 1 | 17 | 1 |
| 7_3005501 | 6 | 0.1667 | 8 | 0.375 | 5 | 0.2 | 6 | 0 | 25 | 0.2 | 6 | 0 | 6 | 0.1667 | 3 | 0.6667 | 6 | 0.1667 | 21 | 0.1905 |
| 7_3005502 | 6 | 0 | 7 | 0.5714 | 3 | 0 | 6 | 0.1667 | 22 | 0.2273 | 5 | 0.4 | 5 | 0 | 3 | 0.3333 | 3 | 0.3333 | 16 | 0.25 |
| 7_3005506 | 6 | 0.6667 | 7 | 0.4286 | 3 | 0.6667 | 7 | 0.4286 | 23 | 0.5217 | 6 | 0.5 | 6 | 0.8333 | 2 | 0.5 | 6 | 0.6667 | 20 | 0.65 |
| 7_3005507 | 3 | 0.6667 | 4 | 0.5 | 4 | 0.25 | 5 | 0.6 | 16 | 0.5 | 6 | 0.6667 | 5 | 0.8 | 2 | 0.5 | 6 | 1 | 19 | 0.7895 |
| 7_3005510 | 4 | 1 | 6 | 1 | 3 | 1 | 6 | 1 | 19 | 1 | 6 | 1 | 3 | 1 | 3 | 1 | 6 | 1 | 18 | 1 |
| 7_3005518 | 5 | 0 | 6 | 0.3333 | 4 | 0 | 6 | 0.3333 | 21 | 0.1905 | 5 | 0 | 6 | 0.1667 | 3 | 0.3333 | 6 | 0 | 20 | 0.1 |
| 7_3005543 | 4 | 0.5 | 4 | 0.5 | 4 | 0.75 | 5 | 0.8 | 17 | 0.6471 | 3 | 1 | 6 | 0.5 | 2 | 0 | 5 | 0.8 | 16 | 0.625 |
| 7_3005551 | 4 | 0.25 | 4 | 0 | 4 | 0.25 | 4 | 0 | 16 | 0.125 | 5 | 0 | 5 | 0.2 | 2 | 0 | 4 | 0 | 16 | 0.0625 |
| 7_3005892 | 2 | 1 | 7 | 1 | 3 | 1 | 4 | 1 | 16 | 1 | 3 | 1 | 4 | 1 | 3 | 1 | 4 | 1 | 14 | 1 |
| 7_3005895 | 3 | 0.3333 | 7 | 0.2857 | 3 | 0.3333 | 7 | 0.2857 | 20 | 0.3 | 5 | 0 | 5 | 0 | 3 | 0.3333 | 5 | 0.2 | 18 | 0.1111 |
| 7_3005908 | 3 | 0.3333 | 7 | 0.4286 | 3 | 0.3333 | 5 | 0.2 | 18 | 0.3333 | 5 | 0.2 | 4 | 0.5 | 3 | 0.6667 | 5 | 0.8 | 17 | 0.5294 |
| 7_3005917 | 1 | 1 | 6 | 0.8333 | 3 | 0.3333 | 7 | 0.8571 | 17 | 0.7647 | 5 | 1 | 3 | 1 | 3 | 0.6667 | 4 | 0.5 | 15 | 0.8 |
| 7_3005925 | 3 | 0 | 7 | 0.4286 | 2 | 0 | 6 | 0.3333 | 18 | 0.2778 | 4 | 0.25 | 2 | 0 | 2 | 0.5 | 4 | 0.25 | 12 | 0.25 |
| 7_3005959 | 3 | 0.6667 | 6 | 0.8333 | 2 | 0.5 | 7 | 1 | 18 | 0.8333 | 5 | 1 | 3 | 1 | 3 | 1 | 4 | 0.75 | 15 | 0.9333 |
| 7_3005972 | 2 | 0 | 6 | 0 | 2 | 0.5 | 3 | 0 | 13 | 0.0769 | 2 | 0 | 1 | 0 | 3 | 0.3333 | 2 | 0 | 8 | 0.125 |
| 7_3005974 | 2 | 1 | 5 | 0.4 | 2 | 1 | 4 | 1 | 13 | 0.7692 | 5 | 0.8 | 2 | 1 | 3 | 0.3333 | 4 | 1 | 14 | 0.7857 |
| 7_3005987 | 1 | 1 | 4 | 1 | 1 | 1 | 1 | 1 | 7 | 1 | 2 | 1 | 2 | 0.5 | 2 | 1 | 2 | 1 | 8 | 0.875 |
| 7_3006145 | 1 | 0 | 3 | 0 | 2 | 0.5 | 1 | 0 | 7 | 0.1429 | 4 | 0.25 | 3 | 0 | 3 | 0 | 4 | 0 | 14 | 0.0714 |
| 7_3006147 | 1 | 0 | 3 | 0.3333 | 2 | 0 | 3 | 0.6667 | 9 | 0.3333 | 3 | 0.3333 | 3 | 0.3333 | 2 | 0 | 4 | 0 | 12 | 0.1667 |
| 7_3006156 | 1 | 0 | 3 | 0.3333 | 2 | 1 | 3 | 0 | 9 | 0.3333 | 5 | 0.4 | 1 | 0 | 2 | 1 | 4 | 1 | 12 | 0.6667 |
| 7_3006167 | 1 | 1 | 3 | 1 | 1 | 1 | 2 | 1 | 7 | 1 | 5 | 0.6 | 3 | 0.6667 | 2 | 1 | 2 | 1 | 12 | 0.75 |
| 7_3006173 | 1 | 0 | 3 | 0.3333 | 2 | 0 | 3 | 0 | 9 | 0.1111 | 5 | 0 | 3 | 0.3333 | 3 | 0.3333 | 4 | 0 | 15 | 0.1333 |
| 7_3006180 | NA | NA | 2 | 1 | 1 | 1 | 1 | 1 | 4 | 1 | 4 | 0.5 | 3 | 0.6667 | 2 | 0.5 | 3 | 1 | 12 | 0.6667 |
| 7_3006184 | 1 | 0 | 2 | 0 | 2 | 0 | 3 | 0.3333 | 8 | 0.125 | 3 | 0 | 3 | 0.3333 | 2 | 0 | 4 | 0 | 12 | 0.0833 |
| 7_3006186 | 1 | 0 | 2 | 0.5 | 2 | 0.5 | 3 | 0.3333 | 8 | 0.375 | 4 | 0.5 | 3 | 0.6667 | 2 | 0 | 4 | 0.25 | 13 | 0.3846 |
| 7_3006191 | 1 | 0 | 2 | 1 | 2 | 0.5 | 3 | 1 | 8 | 0.75 | 3 | 0.6667 | 3 | 0.3333 | 3 | 0.6667 | 2 | 0 | 11 | 0.4545 |
| 7_3006192 | 1 | 1 | 2 | 1 | 2 | 1 | 3 | 1 | 8 | 1 | 4 | 0.5 | 3 | 1 | 2 | 0.5 | 3 | 1 | 12 | 0.75 |
| 7_3006207 | 1 | 1 | 2 | 0.5 | 2 | 1 | 3 | 0 | 8 | 0.5 | 3 | 0.3333 | 2 | 0 | 2 | 0.5 | 3 | 0.3333 | 10 | 0.3 |
| 7_3006208 | 1 | 1 | 1 | 1 | 2 | 0 | 2 | 0.5 | 6 | 0.5 | 4 | 0.25 | 3 | 0.3333 | 2 | 0 | 4 | 0.5 | 13 | 0.3077 |
| 7_3013454 | 192 | 0.0469 | 198 | 0.0202 | 219 | 0.0685 | 188 | 0.0851 | 797 | 0.0552 | 183 | 0.0656 | 206 | 0.0291 | 212 | 0.0802 | 175 | 0.0629 | 776 | 0.0593 |
| 7_3014701 | 2 | 0.5 | 1 | 0 | 2 | 1 | 1 | 0 | 6 | 0.5 | 1 | 0 | 3 | 0.3333 | NA | NA | 2 | 1 | 6 | 0.5 |
| 7_3014727 | 1 | 1 | NA | NA | 2 | 0 | 1 | 1 | 4 | 0.5 | 1 | 1 | 3 | 0.6667 | NA | NA | 1 | 1 | 5 | 0.8 |
| 7_3014729 | 2 | 0.5 | NA | NA | 2 | 0 | 1 | 0 | 5 | 0.2 | 1 | 1 | 3 | 0.3333 | NA | NA | 2 | 0.5 | 6 | 0.5 |
| 7_3014732 | 2 | 0 | NA | NA | 2 | 1 | 1 | 0 | 5 | 0.4 | 1 | 1 | 3 | 0.3333 | NA | NA | 2 | 0.5 | 6 | 0.5 |
| 7_3014739 | 2 | 0.5 | NA | NA | 2 | 0 | 1 | 0 | 5 | 0.2 | 1 | 0 | 3 | 0.3333 | NA | NA | 2 | 1 | 6 | 0.5 |
| 7_3014743 | 1 | 0 | NA | NA | 2 | 0.5 | 1 | 0 | 4 | 0.25 | 1 | 0 | 3 | 0.3333 | NA | NA | NA | NA | 4 | 0.25 |
| 7_3014743 | 2 | 1 | NA | NA | 1 | 0 | 1 | 0 | 4 | 0.5 | 1 | 0 | 3 | 0.6667 | NA | NA | 1 | 0 | 5 | 0.4 |
| 7_3014755 | 2 | 0.5 | NA | NA | 2 | 0 | 1 | 0 | 5 | 0.2 | 3 | 0 | 2 | 0.5 | NA | NA | 2 | 0 | 7 | 0.1429 |
| 7_3014760 | 1 | 0 | NA | NA | 2 | 0.5 | 1 | 0 | 4 | 0.25 | 2 | 0 | 3 | 0.6667 | NA | NA | 2 | 0 | 7 | 0.2857 |
| 7_3014763 | 2 | 0.5 | NA | NA | 1 | 1 | 1 | 0 | 4 | 0.5 | 2 | 0.5 | 3 | 0.3333 | NA | NA | 2 | 0.5 | 7 | 0.4286 |
| 7_3014771 | 2 | 0.5 | NA | NA | 2 | 0 | 1 | 1 | 5 | 0.4 | 2 | 0 | 3 | 0.3333 | NA | NA | 1 | 1 | 6 | 0.3333 |
| 7_3014773 | 2 | 1 | NA | NA | NA | NA | 1 | 0 | 3 | 0.6667 | 3 | 0.3333 | 2 | 0.5 | NA | NA | 1 | 1 | 6 | 0.5 |

| | | | | | | | | | | | | | | | | | | | | |
|---|---|---|---|---|---|---|---|---|---|---|---|---|---|---|---|---|---|---|---|---|
| 7_30221819 | 10 | 0.3 | 10 | 0.2 | 19 | 0.3158 | 17 | 0.1765 | 56 | 0.25 | 11 | 0.2727 | 22 | 0.5 | 21 | 0.3333 | 15 | 0.4 | 69 | 0.3913 |
| 7_30221995 | 12 | 0.0833 | 15 | 0.4667 | 20 | 0.2 | 14 | 0.5 | 61 | 0.3115 | 10 | 0.3 | 24 | 0.1667 | 21 | 0.2381 | 15 | 0.2 | 70 | 0.2143 |
| 7_30222202 | 7 | 0.1429 | 14 | 0 | 20 | 0.1 | 13 | 0.2308 | 54 | 0.1111 | 10 | 0.1 | 19 | 0.1579 | 19 | 0.0526 | 10 | 0.1 | 58 | 0.1034 |
| 7_30222662 | 8 | 0.125 | 12 | 0.0833 | 23 | 0.2609 | 12 | 0.0833 | 55 | 0.1636 | 11 | 0 | 21 | 0.1429 | 17 | 0.1765 | 11 | 0.0909 | 60 | 0.1167 |
| 7_30223035 | 6 | 0.1667 | 10 | 0.5 | 17 | 0.4118 | 8 | 0.375 | 41 | 0.3902 | 7 | 0.2857 | 10 | 0.2 | 17 | 0.0588 | 8 | 0.25 | 42 | 0.1667 |
| 7_30223581 | 4 | 0.5 | 10 | 0.2 | 14 | 0.3571 | 6 | 0.3333 | 34 | 0.3235 | 9 | 0.2222 | 14 | 0.3571 | 15 | 0.2667 | 10 | 0.2 | 48 | 0.2708 |
| 7_30223613 | 6 | 1 | 10 | 0.8 | 18 | 0.5556 | 8 | 0.75 | 42 | 0.7143 | 10 | 0.7 | 16 | 0.8125 | 14 | 0.7143 | 9 | 0.7778 | 49 | 0.7551 |
| 7_30223701 | 5 | 0 | 9 | 0 | 15 | 0.0667 | 9 | 0.3333 | 38 | 0.1053 | 10 | 0 | 16 | 0.0625 | 17 | 0.1176 | 10 | 0 | 53 | 0.0566 |
| 7_30223746 | 5 | 0.4 | 11 | 0.0909 | 17 | 0.2353 | 9 | 0.4444 | 42 | 0.2619 | 10 | 0.2 | 17 | 0.1176 | 13 | 0.1538 | 8 | 0.25 | 48 | 0.1667 |
| 7_30223751 | 5 | 0.2 | 9 | 0.5556 | 18 | 0.3889 | 9 | 0.4444 | 41 | 0.4146 | 11 | 0.3636 | 16 | 0.4375 | 16 | 0.5 | 10 | 0.5 | 53 | 0.4528 |
| 7_30223760 | 4 | 0.75 | 10 | 0.7 | 17 | 0.7059 | 9 | 0.8889 | 40 | 0.75 | 10 | 0.8 | 14 | 0.7857 | 17 | 0.6471 | 10 | 0.8 | 51 | 0.7451 |
| 7_30238755 | 1 | 1 | NA | NA | NA | NA | NA | NA | 1 | 1 | NA | NA | NA | NA | 1 | 1 | NA | NA | 1 | 1 |
| 7_30254664 | NA | NA | 1 | 1 | 1 | 1 | 4 | 0.25 | 6 | 0.5 | 2 | 0.5 | NA | NA | NA | NA | 1 | 1 | 3 | 0.6667 |
| 7_30261303 | 12 | 0.0833 | 12 | 0.1667 | 14 | 0.0714 | 9 | 0.1111 | 47 | 0.1064 | 13 | 0 | 15 | 0.1333 | 16 | 0.25 | 12 | 0.1667 | 56 | 0.1429 |
| 7_30263809 | NA | NA | 3 | 0.6667 | NA | NA | NA | NA | 3 | 0.6667 | NA | NA | NA | NA | 1 | 1 | 1 | 0 | 2 | 0.5 |
| 7_30265568 | NA | NA | 1 | 1 | 1 | 1 | NA | NA | 2 | 1 | NA | NA | NA | NA | 1 | 0 | 1 | 1 | 2 | 0.5 |
| 7_30274018 | 1 | 1 | NA | NA | 2 | 1 | 1 | 0 | 4 | 0.75 | 2 | 0.5 | NA | NA | 2 | 0.5 | NA | NA | 4 | 0.5 |
| 7_30274314 | 2 | 1 | NA | NA | 2 | 1 | 1 | 1 | 5 | 1 | 2 | 1 | 2 | 1 | 2 | 1 | NA | NA | 6 | 1 |
| 7_30274356 | 1 | 1 | NA | NA | 2 | 0 | 1 | 0 | 4 | 0.25 | 2 | 1 | NA | NA | 1 | 1 | NA | NA | 3 | 1 |
| 7_30274650 | 2 | 0 | NA | NA | 2 | 0.5 | 1 | 0 | 5 | 0.2 | 2 | 0.5 | 2 | 1 | 2 | 0.5 | NA | NA | 6 | 0.6667 |
| 7_30274717 | 2 | 1 | 1 | 0 | 2 | 1 | 1 | 1 | 6 | 0.8333 | 2 | 0.5 | 1 | 0 | 1 | 1 | NA | NA | 4 | 0.5 |
| 7_30274740 | 1 | 1 | 1 | 0 | 2 | 1 | 1 | 0 | 5 | 0.6 | 1 | 1 | 2 | 1 | 1 | 1 | NA | NA | 4 | 1 |
| 7_30274760 | 2 | 1 | 1 | 0 | 2 | 0 | 1 | 1 | 6 | 0.5 | 2 | 0 | 1 | 0 | 1 | 1 | NA | NA | 4 | 0.25 |
| 7_30274788 | 2 | 1 | 1 | 0 | 2 | 0 | 1 | 0 | 6 | 0.3333 | 2 | 0 | 2 | 1 | 1 | 0 | NA | NA | 5 | 0.4 |
| 7_30427262 | 18 | 0.0556 | 26 | 0 | 15 | 0.0667 | 16 | 0.125 | 75 | 0.0533 | 19 | 0.0526 | 20 | 0.05 | 22 | 0.1818 | 16 | 0.0625 | 77 | 0.0909 |
| 7_30599011 | 16 | 0.125 | 25 | 0.16 | 33 | 0.0606 | 20 | 0.05 | 94 | 0.0957 | 13 | 0.0769 | 27 | 0.037 | 12 | 0.1667 | 14 | 0.0714 | 66 | 0.0758 |
| 7_30610945 | 8 | 0.125 | 17 | 0.2941 | 12 | 0.3333 | 7 | 0.1429 | 44 | 0.25 | 6 | 0.1667 | 12 | 0.3333 | 6 | 0 | 7 | 0.1429 | 31 | 0.1935 |
| 7_30645622 | 9 | 0.1111 | 17 | 0.1176 | 12 | 0.0833 | 16 | 0 | 54 | 0.0741 | 13 | 0.0769 | 18 | 0.0556 | 18 | 0.1111 | 16 | 0.1875 | 65 | 0.1077 |
| 7_30645876 | 13 | 0.0769 | 17 | 0 | 13 | 0.1538 | 17 | 0.1176 | 60 | 0.0833 | 14 | 0.1429 | 19 | 0.1053 | 22 | 0.0455 | 17 | 0.1765 | 72 | 0.1111 |
| 7_30646198 | 11 | 0.1818 | 11 | 0.3636 | 10 | 0.4 | 15 | 0.2667 | 47 | 0.2979 | 12 | 0.3333 | 19 | 0.1579 | 16 | 0.375 | 16 | 0.25 | 63 | 0.2698 |
| 7_30657589 | 3 | 0.3333 | 2 | 0 | 1 | 0 | NA | NA | 6 | 0.1667 | 3 | 0 | NA | NA | 2 | 0.5 | 1 | 0 | 6 | 0.1667 |
| 7_30657709 | 3 | 1 | 2 | 1 | NA | NA | NA | NA | 5 | 1 | 4 | 1 | NA | NA | 2 | 1 | 2 | 0.5 | 8 | 0.875 |
| 7_30658252 | 1 | 1 | NA | NA | NA | NA | NA | NA | 1 | 1 | 1 | 1 | 1 | 0 | NA | NA | NA | NA | 2 | 0.5 |
| 7_30658469 | 1 | 1 | NA | NA | NA | NA | NA | NA | 1 | 1 | 1 | 1 | 1 | 1 | NA | NA | NA | NA | 2 | 1 |
| 7_30658483 | 1 | 1 | NA | NA | NA | NA | NA | NA | 1 | 1 | 1 | 1 | 1 | 0 | NA | NA | NA | NA | 2 | 0.5 |
| 7_30658739 | 1 | 1 | NA | NA | NA | NA | NA | NA | 1 | 1 | 1 | 1 | NA | NA | NA | NA | NA | NA | 1 | 1 |
| 7_30658808 | 1 | 1 | NA | NA | NA | NA | NA | NA | 1 | 1 | 1 | 1 | 1 | 0 | NA | NA | NA | NA | 2 | 0.5 |
| 7_30659085 | 1 | 1 | NA | NA | NA | NA | NA | NA | 1 | 1 | 1 | 1 | 1 | 1 | NA | NA | NA | NA | 2 | 1 |
| 7_30659641 | 1 | 1 | NA | NA | NA | NA | NA | NA | 1 | 1 | 1 | 1 | NA | NA | NA | NA | NA | NA | 1 | 1 |
| 7_30671388 | 2 | 0.5 | 1 | 0 | 1 | 0 | NA | NA | 4 | 0.25 | 4 | 0.25 | NA | NA | 2 | 0 | NA | NA | 6 | 0.1667 |
| 7_30740317 | 1 | 1 | NA | NA | 1 | 1 | NA | NA | 2 | 1 | NA | NA | NA | NA | NA | NA | 2 | 0.5 | 2 | 0.5 |
| 7_30740474 | 1 | 1 | NA | NA | 1 | 1 | NA | NA | 2 | 1 | NA | NA | NA | NA | NA | NA | 1 | 1 | 1 | 1 |
| 7_30742103 | 6 | 0 | 12 | 0.3333 | 11 | 0.1818 | 10 | 0.4 | 39 | 0.2564 | 16 | 0.3125 | 7 | 0.1429 | 7 | 0.4286 | 10 | 0.3 | 40 | 0.3 |
| 7_30767266 | 2 | 1 | 3 | 0.6667 | 1 | 0 | 2 | 1 | 8 | 0.75 | 2 | 0 | 2 | 0.5 | 2 | 0.5 | 3 | 0.6667 | 9 | 0.4444 |
| 7_30768628 | 3 | 0.6667 | 4 | 0.5 | 3 | 0.3333 | 4 | 0.25 | 14 | 0.4286 | 1 | 1 | 3 | 0.3333 | 2 | 1 | 5 | 0.6 | 11 | 0.6364 |
| 7_30768720 | 3 | 0 | 4 | 0.25 | 2 | 0 | 4 | 0.5 | 13 | 0.2308 | 3 | 0 | 3 | 0.3333 | 2 | 0 | 5 | 0 | 13 | 0.0769 |
| 7_30768859 | 2 | 1 | 3 | 0 | 2 | 0 | 4 | 0.75 | 11 | 0.4545 | 4 | 0.75 | 3 | 0.3333 | 2 | 0 | 4 | 0 | 13 | 0.3077 |
| 7_30768964 | 3 | 0.3333 | 4 | 0.5 | 2 | 0 | 3 | 0.3333 | 12 | 0.3333 | 5 | 0 | 3 | 0.3333 | 2 | 0 | 5 | 0.2 | 15 | 0.1333 |
| 7_30769257 | 1 | 0 | 4 | 0.5 | 2 | 0 | 3 | 0 | 10 | 0.2 | 3 | 0.3333 | 3 | 0 | 2 | 0 | 5 | 0 | 13 | 0.0769 |
| 7_30769350 | 1 | 0 | 3 | 1 | 2 | 0.5 | 4 | 0.5 | 10 | 0.6 | 4 | 0.5 | 4 | 0.25 | 2 | 0.5 | 3 | 0.3333 | 13 | 0.3846 |
| 7_30769549 | 3 | 0.3333 | 7 | 0.1429 | 2 | 0 | 3 | 0 | 15 | 0.1333 | 5 | 0 | 4 | 0.25 | 3 | 0.3333 | 6 | 0.1667 | 18 | 0.1667 |
| 7_30769988 | 3 | 0.6667 | 3 | 0.6667 | 1 | 1 | 3 | 0.6667 | 10 | 0.7 | 5 | 0.4 | 2 | 1 | 4 | 0.5 | 5 | 0.6 | 16 | 0.5625 |
| 7_30770233 | 4 | 0.25 | 2 | 0 | 2 | 0.5 | 4 | 0.5 | 12 | 0.3333 | 4 | 0.25 | 4 | 0.25 | 5 | 0.2 | 4 | 0 | 17 | 0.1765 |
| 7_30816100 | 51 | 0.098 | 76 | 0.1053 | 69 | 0.029 | 29 | 0.0345 | 225 | 0.0711 | 55 | 0.0545 | 46 | 0.1522 | 62 | 0.0806 | 35 | 0 | 198 | 0.0758 |
| 7_30816155 | 42 | 0.2143 | 70 | 0.0857 | 53 | 0.1132 | 26 | 0.1923 | 191 | 0.1361 | 38 | 0.2105 | 40 | 0.175 | 55 | 0.1455 | 35 | 0.1429 | 168 | 0.1667 |
| 7_30816394 | 52 | 0.0962 | 75 | 0.0933 | 61 | 0.0328 | 22 | 0.0455 | 210 | 0.0714 | 50 | 0.1 | 41 | 0.0488 | 52 | 0.0577 | 40 | 0.05 | 183 | 0.0656 |
| 7_30816595 | 13 | 0.4615 | 26 | 0.3462 | 11 | 0.3636 | 6 | 0.5 | 56 | 0.3929 | 8 | 0.125 | 9 | 0.5556 | 16 | 0.25 | 10 | 0.5 | 43 | 0.3488 |
| 7_30816635 | 10 | 0 | 20 | 0.1 | 14 | 0.0714 | 6 | 0.1667 | 50 | 0.08 | 8 | 0.125 | 10 | 0.1 | 16 | 0.0625 | 10 | 0 | 44 | 0.0682 |
| 7_30829770 | 11 | 0.2727 | 10 | 0.3 | 2 | 0 | 12 | 0.25 | 35 | 0.2571 | 5 | 0 | 15 | 0.2667 | 13 | 0.3846 | 10 | 0.1 | 43 | 0.2326 |
| 7_30829874 | 8 | 0.375 | 8 | 0.25 | 3 | 0 | 11 | 0.3636 | 30 | 0.3 | 5 | 0.4 | 14 | 0.0714 | 12 | 0.5 | 9 | 0.3333 | 40 | 0.3 |
| 7_30829882 | 11 | 0.4545 | 9 | 0.3333 | 4 | 0 | 12 | 0.1667 | 36 | 0.2778 | 3 | 0.6667 | 15 | 0.4 | 14 | 0.4286 | 9 | 0.3333 | 41 | 0.4146 |
| 7_30829914 | 10 | 0.1 | 6 | 0 | 4 | 0 | 11 | 0.2727 | 31 | 0.129 | 4 | 0 | 15 | 0.2 | 12 | 0.25 | 9 | 0.2222 | 40 | 0.2 |
| 7_30829951 | 8 | 0.125 | 8 | 0.125 | 2 | 0 | 10 | 0 | 28 | 0.0714 | 3 | 0.3333 | 12 | 0.0833 | 14 | 0 | 9 | 0.1111 | 38 | 0.0789 |
| 7_33809105 | NA | NA | 3 | 0.3333 | 1 | 0 | 2 | 0 | 6 | 0.1667 | 1 | 1 | 1 | 0 | NA | NA | 1 | 0 | 3 | 0.3333 |
| 7_33809387 | 1 | 1 | 2 | 1 | 2 | 1 | 2 | 1 | 7 | 1 | 1 | 1 | 1 | 1 | 1 | 1 | 1 | 1 | 4 | 1 |

| | | | | | | | | | | | | | | | | | | | | |
|---|---|---|---|---|---|---|---|---|---|---|---|---|---|---|---|---|---|---|---|---|
| 7_3384061 | 18 | 0.2222 | 16 | 0.5625 | 12 | 0.25 | 13 | 0.2308 | 59 | 0.322 | 11 | 0.1818 | 14 | 0.2143 | 15 | 0.4667 | 7 | 0.4286 | 47 | 0.3191 |
| 7_3384062 | 17 | 0.2941 | 18 | 0.2222 | 12 | 0.5 | 13 | 0.4615 | 60 | 0.35 | 11 | 0.2727 | 11 | 0.1818 | 14 | 0.4286 | 4 | 0 | 40 | 0.275 |
| 7_3389236 | 3 | 0.3333 | 2 | 0.5 | 1 | 1 | 1 | 0 | 7 | 0.4286 | 1 | 0 | 2 | 0.5 | 1 | 0 | 3 | 0 | 7 | 0.1429 |
| 7_3389239 | 2 | 1 | 1 | 0 | 1 | 1 | 1 | 1 | 5 | 0.8 | 1 | 1 | 2 | 0.5 | 1 | 1 | 2 | 0 | 6 | 0.5 |
| 7_3390170 | 89 | 0.2584 | 97 | 0.1959 | 91 | 0.1758 | 74 | 0.2432 | 351 | 0.2165 | 73 | 0.2329 | 74 | 0.2432 | 71 | 0.2113 | 73 | 0.1781 | 291 | 0.2165 |
| 7_3390173 | 72 | 0.2083 | 83 | 0.253 | 68 | 0.2353 | 61 | 0.2459 | 284 | 0.2359 | 68 | 0.2059 | 69 | 0.2464 | 60 | 0.2833 | 55 | 0.2545 | 252 | 0.246 |
| 7_3390188 | 95 | 0.2947 | 98 | 0.3163 | 91 | 0.3297 | 81 | 0.2963 | 365 | 0.3096 | 70 | 0.3286 | 74 | 0.2973 | 70 | 0.2429 | 73 | 0.2877 | 287 | 0.2892 |
| 7_3390935 | NA | NA | 1 | 1 | NA | NA | 3 | 0.3333 | 4 | 0.5 | 2 | 0.5 | 1 | 0 | NA | NA | NA | NA | 3 | 0.3333 |
| 7_3393827 | 3 | 0.3333 | 2 | 0 | 3 | 0.3333 | 1 | 0 | 9 | 0.2222 | 1 | 0 | 3 | 0.3333 | 3 | 0 | 1 | 0 | 8 | 0.125 |
| 7_3393849 | 2 | 0 | 3 | 0 | 3 | 0.6667 | NA | NA | 8 | 0.25 | 1 | 0 | 3 | 0 | 3 | 0 | 1 | 1 | 8 | 0.125 |
| 7_3394447 | 2 | 0 | 2 | 0 | 2 | 0.5 | 1 | 0 | 7 | 0.1429 | 1 | 0 | 4 | 0.5 | 2 | 0.5 | 1 | 1 | 8 | 0.5 |
| 7_3394449 | 2 | 0.5 | 1 | 0 | 2 | 0.5 | 1 | 1 | 6 | 0.5 | NA | NA | 2 | 0 | 2 | 1 | 1 | 1 | 5 | 0.6 |
| 7_3394451 | 2 | 1 | 2 | 1 | 1 | 0 | 1 | 1 | 6 | 0.8333 | 1 | 1 | 4 | 0.75 | 1 | 1 | 1 | 0 | 7 | 0.7143 |
| 7_3394455 | 2 | 0.5 | 2 | 0 | 1 | 0 | 1 | 0 | 6 | 0.1667 | 1 | 0 | 4 | 0.25 | 2 | 0 | NA | NA | 7 | 0.1429 |
| 7_3394467 | 2 | 0.5 | 2 | 0 | 2 | 0 | 1 | 0 | 7 | 0.1429 | 1 | 1 | 4 | 0.25 | 1 | 0 | NA | NA | 6 | 0.3333 |
| 7_3394486 | 1 | 0 | 2 | 0 | 2 | 0.5 | 1 | 0 | 6 | 0.1667 | 1 | 0 | 3 | 0.3333 | 1 | 0 | NA | NA | 5 | 0.2 |
| 7_3394487 | 2 | 0 | 2 | 0.5 | 2 | 0 | 1 | 0 | 7 | 0.1429 | 1 | 0 | 4 | 0.25 | 1 | 0 | 1 | 0 | 7 | 0.1429 |
| 7_3394499 | 1 | 1 | 1 | 1 | 2 | 0.5 | 1 | 0 | 5 | 0.6 | 1 | 1 | 4 | 1 | 1 | 1 | 1 | 1 | 7 | 1 |
| 7_3394503 | 1 | 0 | 2 | 0.5 | 2 | 0.5 | 1 | 0 | 6 | 0.3333 | 1 | 0 | 2 | 0.5 | NA | NA | 1 | 1 | 4 | 0.5 |
| 7_3394511 | 1 | 1 | 2 | 0.5 | 2 | 0 | NA | NA | 5 | 0.4 | 1 | 1 | 4 | 0.75 | NA | NA | 1 | 0 | 6 | 0.6667 |
| 7_3394513 | 1 | 1 | 2 | 1 | 2 | 0 | 1 | 0 | 6 | 0.5 | 1 | 1 | 2 | 1 | NA | NA | 1 | 0 | 4 | 0.75 |
| 7_3394529 | 1 | 0 | 1 | 0 | 2 | 0.5 | NA | NA | 4 | 0.25 | NA | NA | 4 | 0.25 | NA | NA | 1 | 0 | 5 | 0.2 |
| 7_3404448 | 12 | 0.25 | 12 | 0.3333 | 7 | 0.4286 | 7 | 0.2857 | 38 | 0.3158 | 5 | 0.6 | 8 | 0.25 | 10 | 0.5 | 10 | 0.5 | 33 | 0.4545 |
| 7_3404455 | 9 | 0 | 9 | 0 | 6 | 0.1667 | 8 | 0.125 | 32 | 0.0625 | 4 | 0.25 | 8 | 0.125 | 8 | 0.25 | 10 | 0.2 | 30 | 0.2 |
| 7_3404509 | 12 | 0.5 | 11 | 0.0909 | 5 | 0.2 | 9 | 0.4444 | 37 | 0.3243 | 5 | 0.4 | 8 | 0.375 | 10 | 0.3 | 9 | 0.6667 | 32 | 0.4375 |
| 7_3433860 | 2 | 0.5 | 1 | 0 | 2 | 1 | NA | NA | 5 | 0.6 | NA | NA | NA | NA | 1 | 1 | 2 | 1 | 3 | 1 |
| 7_3433872 | 2 | 0.5 | 2 | 0.5 | 2 | 0.5 | NA | NA | 6 | 0.5 | NA | NA | NA | NA | 2 | 0 | 3 | 1 | 5 | 0.6 |
| 7_3433899 | 2 | 0 | 1 | 1 | 1 | 0 | NA | NA | 4 | 0.25 | NA | NA | NA | NA | 3 | 0.3333 | 2 | 0.5 | 5 | 0.4 |
| 7_3433908 | 1 | 0 | 2 | 0.5 | 2 | 0.5 | NA | NA | 5 | 0.4 | NA | NA | NA | NA | 1 | 1 | 1 | 0 | 2 | 0.5 |
| 7_3433909 | 1 | 0 | 1 | 0 | 2 | 1 | NA | NA | 4 | 0.5 | NA | NA | NA | NA | 2 | 0.5 | 2 | 0.5 | 4 | 0.5 |
| 7_3433918 | 1 | 1 | 2 | 1 | 2 | 1 | NA | NA | 5 | 1 | NA | NA | NA | NA | 3 | 0.6667 | 1 | 1 | 4 | 0.75 |
| 7_3433930 | 2 | 0.5 | 2 | 0 | 2 | 0 | NA | NA | 6 | 0.1667 | NA | NA | NA | NA | 3 | 0.3333 | 2 | 0 | 5 | 0.2 |
| 7_3433934 | 1 | 1 | 1 | 0 | 2 | 1 | NA | NA | 4 | 0.75 | NA | NA | NA | NA | 3 | 0.3333 | 2 | 0 | 5 | 0.2 |
| 7_3433937 | 2 | 0.5 | 2 | 0 | 1 | 0 | NA | NA | 5 | 0.2 | NA | NA | NA | NA | 2 | 0 | 2 | 0.5 | 4 | 0.25 |
| 7_3433952 | 1 | 0 | 2 | 0 | 2 | 0.5 | NA | NA | 5 | 0.2 | NA | NA | NA | NA | 2 | 0.5 | 2 | 0 | 4 | 0.25 |
| 7_3433953 | 2 | 0.5 | 2 | 1 | 2 | 1 | NA | NA | 6 | 0.8333 | NA | NA | NA | NA | 2 | 1 | 2 | 1 | 4 | 1 |
| 7_3433960 | NA | NA | 1 | 1 | 1 | 1 | NA | NA | 2 | 1 | NA | NA | NA | NA | 2 | 1 | 1 | 1 | 3 | 1 |
| 7_3433966 | 2 | 1 | 1 | 1 | 2 | 1 | NA | NA | 5 | 1 | NA | NA | NA | NA | 3 | 0.6667 | 2 | 1 | 5 | 0.8 |
| 7_3433972 | 2 | 1 | 1 | 1 | 1 | 1 | NA | NA | 4 | 1 | NA | NA | NA | NA | 3 | 1 | 1 | 1 | 4 | 1 |
| 7_3433981 | 1 | 1 | 1 | 1 | 1 | 1 | NA | NA | 3 | 1 | NA | NA | NA | NA | 2 | 0.5 | 2 | 0.5 | 4 | 0.5 |
| 7_3474372 | 3 | 0 | 7 | 0.1429 | 2 | 0 | 4 | 0.25 | 16 | 0.125 | 4 | 0.25 | 4 | 0.5 | 4 | 0.75 | 1 | 0 | 13 | 0.4615 |
| 7_3474380 | 4 | 0.25 | 6 | 0.5 | 2 | 0.5 | 5 | 0.4 | 17 | 0.4118 | 2 | 0.5 | 2 | 0 | 4 | 0.25 | 1 | 0 | 9 | 0.2222 |
| 7_3474381 | 3 | 0.6667 | 4 | 0.25 | 2 | 0.5 | 4 | 0.5 | 13 | 0.4615 | 4 | 0.5 | 3 | 0.6667 | 3 | 0.3333 | 1 | 0 | 11 | 0.4545 |
| 7_3474399 | 5 | 0.8 | 7 | 0.7143 | 3 | 0.3333 | 4 | 0.75 | 19 | 0.6842 | 4 | 0.5 | 4 | 0.25 | 3 | 0.3333 | 1 | 1 | 12 | 0.4167 |
| 7_3474412 | 4 | 0.25 | 2 | 0 | 2 | 0 | 3 | 0.3333 | 11 | 0.1818 | 1 | 0 | 2 | 0 | 2 | 0.5 | NA | NA | 5 | 0.2 |
| 7_3474729 | 2 | 0 | 4 | 0.25 | 3 | 0 | 4 | 0.25 | 13 | 0.1538 | 2 | 0 | 5 | 0.2 | 3 | 0.6667 | 6 | 0.1667 | 16 | 0.25 |
| 7_3474760 | 1 | 1 | 3 | 0 | 1 | 0 | 1 | 0 | 6 | 0.1667 | 1 | 0 | 4 | 0.25 | NA | NA | 4 | 0.25 | 9 | 0.2222 |
| 7_3474952 | NA | NA | 2 | 1 | NA | NA | 2 | 1 | 4 | 1 | 2 | 0.5 | 2 | 1 | NA | NA | 2 | 0.5 | 6 | 0.6667 |
| 7_3474958 | NA | NA | 4 | 1 | NA | NA | 2 | 1 | 6 | 1 | 2 | 1 | 2 | 1 | NA | NA | 2 | 1 | 6 | 1 |
| 7_3474968 | NA | NA | 3 | 0.3333 | NA | NA | 1 | 1 | 4 | 0.5 | NA | NA | 2 | 1 | NA | NA | NA | NA | 2 | 1 |
| 7_3474973 | NA | NA | 3 | 0.3333 | NA | NA | 2 | 0 | 5 | 0.2 | 2 | 0.5 | 2 | 1 | NA | NA | 1 | 1 | 5 | 0.8 |
| 7_3474977 | NA | NA | 3 | 0.3333 | NA | NA | 2 | 1 | 5 | 0.6 | 1 | 1 | 1 | 0 | NA | NA | NA | NA | 2 | 0.5 |
| 7_3474978 | NA | NA | 3 | 1 | NA | NA | 2 | 1 | 5 | 1 | NA | NA | 2 | 1 | NA | NA | 1 | 0 | 3 | 0.6667 |
| 7_3475004 | NA | NA | 3 | 0.3333 | NA | NA | 2 | 0.5 | 5 | 0.4 | 1 | 1 | 1 | 1 | NA | NA | 1 | 0 | 3 | 0.6667 |
| 7_3481980 | NA | NA | 1 | 1 | 4 | 0.25 | NA | NA | 5 | 0.4 | 2 | 0.5 | NA | NA | 2 | 1 | NA | NA | 4 | 0.75 |
| 7_3481984 | NA | NA | NA | NA | 3 | 1 | NA | NA | 3 | 1 | 3 | 1 | 1 | 1 | NA | NA | 2 | 1 | 6 | 1 |
| 7_3481985 | NA | NA | NA | NA | 3 | 1 | NA | NA | 3 | 1 | 2 | 1 | 1 | 1 | 2 | 1 | 1 | 1 | 6 | 1 |
| 7_3481994 | NA | NA | 1 | 0 | 4 | 0.25 | NA | NA | 5 | 0.2 | 2 | 0 | 1 | 0 | 2 | 1 | 2 | 0.5 | 7 | 0.4286 |
| 7_3481996 | NA | NA | 1 | 0 | 4 | 0.25 | NA | NA | 5 | 0.2 | 3 | 0.3333 | 1 | 0 | 2 | 0 | 2 | 0 | 8 | 0.125 |
| 7_3482002 | NA | NA | 1 | 1 | 4 | 1 | NA | NA | 5 | 1 | 4 | 1 | 1 | 0 | 1 | 1 | 2 | 1 | 8 | 0.875 |
| 7_3482009 | NA | NA | 1 | 1 | 3 | 1 | NA | NA | 4 | 1 | 3 | 1 | 1 | 1 | 2 | 1 | 3 | 1 | 9 | 1 |
| 7_3482015 | NA | NA | 1 | 1 | 4 | 1 | NA | NA | 5 | 1 | 4 | 1 | 1 | 1 | 2 | 1 | 3 | 1 | 10 | 1 |
| 7_3482023 | NA | NA | 1 | 1 | 3 | 0.3333 | NA | NA | 4 | 0.5 | 2 | 0.5 | 1 | 1 | 2 | 0.5 | 2 | 0.5 | 7 | 0.5714 |
| 7_3482043 | NA | NA | 1 | 0 | 4 | 0.25 | NA | NA | 5 | 0.2 | 3 | 0.3333 | 1 | 0 | 1 | 0 | 1 | 0 | 6 | 0.1667 |
| 7_3482075 | NA | NA | 1 | 1 | 4 | 0.5 | NA | NA | 5 | 0.6 | 4 | 0 | 1 | 0 | 2 | 0.5 | 3 | 0.3333 | 10 | 0.2 |

| | | | | | | | | | | | | | | | | | | | | |
|---|---|---|---|---|---|---|---|---|---|---|---|---|---|---|---|---|---|---|---|---|
| 7_3482078 | NA | NA | 2 | 1 | 4 | 0.25 | NA | NA | 6 | 0.5 | 4 | 0.75 | 1 | 0 | 1 | 0 | 3 | 0.6667 | 9 | 0.5556 |
| 7_3482096 | NA | NA | 1 | 0 | 4 | 0.25 | NA | NA | 5 | 0.2 | 3 | 0.3333 | 1 | 1 | 1 | 1 | 2 | 0.5 | 7 | 0.5714 |
| 7_3490028 | 2 | 0.5 | 2 | 0 | 5 | 0.2 | NA | NA | 9 | 0.2222 | 4 | 0.25 | 5 | 0 | 6 | 0.3333 | 5 | 0 | 20 | 0.15 |
| 7_3490054 | 3 | 1 | 9 | 0.6667 | 5 | 0.6 | 3 | 0 | 20 | 0.6 | 3 | 0.6667 | 3 | 0.6667 | 10 | 0.6 | 1 | 0 | 17 | 0.5882 |
| 7_3490080 | 3 | 1 | 7 | 0.8571 | 4 | 0.75 | 2 | 0.5 | 16 | 0.8125 | 3 | 0.6667 | 5 | 0.4 | 9 | 0.7778 | 1 | 0 | 18 | 0.6111 |
| 7_3490110 | 3 | 1 | 5 | 0.6 | 2 | 0.5 | 1 | 1 | 11 | 0.7273 | NA | NA | 2 | 1 | 5 | 1 | 1 | 0 | 8 | 0.875 |
| 7_3490148 | 8 | 0.375 | 5 | 0.4 | 5 | 0.2 | 5 | 0.6 | 23 | 0.3913 | 3 | 0.6667 | 3 | 0 | 7 | 0.5714 | 3 | 0.6667 | 16 | 0.5 |
| 7_3490149 | 5 | 0.6 | 5 | 0.8 | 4 | 1 | 4 | 0.75 | 18 | 0.7778 | 3 | 1 | 4 | 0.5 | 6 | 1 | 2 | 0.5 | 15 | 0.8 |
| 7_3490163 | 4 | 0.5 | 5 | 0.6 | 3 | 0.6667 | 3 | 0.6667 | 15 | 0.6 | 3 | 0.6667 | 3 | 0 | 7 | 0.7143 | 3 | 1 | 16 | 0.625 |
| 7_3490167 | 6 | 0.6667 | 5 | 1 | 4 | 1 | 6 | 0.6667 | 21 | 0.8095 | 2 | 1 | 2 | 0.5 | 5 | 0.8 | 3 | 1 | 12 | 0.8333 |
| 7_3490173 | 6 | 0.8333 | 5 | 1 | 4 | 0.75 | 4 | 0.5 | 19 | 0.7895 | 1 | 1 | 3 | 0.6667 | 6 | 0.6667 | 3 | 0.3333 | 13 | 0.6154 |
| 7_3490185 | 2 | 0 | 5 | 0.8 | 3 | 1 | 3 | 0.6667 | 13 | 0.6923 | 2 | 1 | 3 | 0.6667 | 6 | 0.6667 | 3 | 1 | 14 | 0.7857 |
| 7_3490192 | 5 | 0.4 | 4 | 0.5 | 2 | 0 | 2 | 0 | 13 | 0.3077 | NA | NA | 3 | 0.3333 | 6 | 0.3333 | 3 | 1 | 12 | 0.5 |
| 7_3490272 | 1 | 0 | 3 | 0 | 4 | 0.5 | NA | NA | 8 | 0.25 | NA | NA | 1 | 0 | 3 | 0 | 2 | 0.5 | 6 | 0.1667 |
| 7_3490275 | 3 | 1 | 5 | 0.6 | 5 | 0.8 | 3 | 1 | 16 | 0.8125 | NA | NA | 2 | 1 | 5 | 0.8 | 2 | 1 | 9 | 0.8889 |
| 7_3490284 | 2 | 0.5 | 5 | 1 | 3 | 1 | 2 | 1 | 12 | 0.9167 | NA | NA | 1 | 1 | 3 | 1 | 3 | 1 | 7 | 1 |
| 7_3490289 | 3 | 0 | 5 | 0.4 | 5 | 0.2 | 3 | 0 | 16 | 0.1875 | NA | NA | 1 | 0 | 4 | 0.25 | 3 | 0.3333 | 8 | 0.25 |
| 7_3490293 | 3 | 1 | 3 | 1 | 5 | 1 | 3 | 0.3333 | 14 | 0.8571 | 1 | 0 | 2 | 0 | 4 | 0.5 | 2 | 1 | 9 | 0.4444 |
| 7_3490302 | 2 | 0.5 | 4 | 0.75 | 5 | 0.8 | 4 | 1 | 15 | 0.8 | 1 | 1 | 2 | 0 | 5 | 0.6 | 3 | 0.3333 | 11 | 0.4545 |
| 7_3490332 | 3 | 1 | 4 | 0.75 | 5 | 0.4 | 2 | 0 | 14 | 0.5714 | NA | NA | 1 | 0 | 5 | 0.6 | 2 | 1 | 8 | 0.625 |
| 7_3490336 | 2 | 0.5 | 3 | 0 | 5 | 0.4 | 3 | 0 | 13 | 0.2308 | 1 | 1 | 2 | 0.5 | 3 | 0 | 2 | 0.5 | 8 | 0.375 |
| 7_3490341 | 3 | 0.3333 | 3 | 0 | 5 | 0.2 | 4 | 0.25 | 15 | 0.2 | 1 | 0 | 2 | 0 | 5 | 0.4 | 2 | 0 | 10 | 0.2 |
| 7_3490342 | 2 | 0 | 2 | 0 | 5 | 0.2 | 3 | 0.3333 | 12 | 0.1667 | NA | NA | 2 | 0 | 5 | 0.2 | 1 | 0 | 8 | 0.125 |
| 7_3490344 | 2 | 0 | 3 | 0.3333 | 4 | 0.5 | 3 | 0.6667 | 12 | 0.4167 | NA | NA | 2 | 1 | 5 | 0.6 | 2 | 1 | 9 | 0.7778 |
| 7_3490347 | 3 | 0.3333 | 4 | 0.25 | 5 | 0.4 | 1 | 1 | 13 | 0.3846 | 1 | 1 | 2 | 0.5 | 5 | 0.2 | 2 | 0 | 10 | 0.3 |
| 7_3490356 | 2 | 1 | 4 | 1 | 5 | 0.8 | 3 | 0.6667 | 14 | 0.8571 | 1 | 1 | 2 | 1 | 5 | 1 | 2 | 1 | 10 | 1 |
| 7_3509021 | 2 | 0 | 1 | 0 | 2 | 0.5 | 2 | 0 | 7 | 0.1429 | NA | NA | 1 | 0 | NA | NA | 2 | 0.5 | 3 | 0.3333 |
| 7_3509032 | 1 | 1 | 1 | 0 | 1 | 0 | 2 | 0.5 | 5 | 0.4 | NA | NA | NA | NA | NA | NA | 2 | 0.5 | 2 | 0.5 |
| 7_3509050 | 1 | 1 | 2 | 0.5 | 2 | 1 | 1 | 1 | 6 | 0.8333 | NA | NA | 1 | 1 | NA | NA | 2 | 1 | 3 | 1 |
| 7_3509053 | 2 | 1 | 2 | 1 | 2 | 0.5 | 1 | 1 | 7 | 0.8571 | NA | NA | 1 | 1 | NA | NA | 2 | 1 | 3 | 1 |
| 7_3509057 | 1 | 0 | 2 | 0.5 | 2 | 0.5 | 2 | 1 | 7 | 0.5714 | NA | NA | 1 | 1 | NA | NA | 2 | 0.5 | 3 | 0.6667 |
| 7_3509117 | 2 | 0 | 2 | 0.5 | 2 | 0 | 2 | 0 | 8 | 0.125 | NA | NA | 1 | 0 | NA | NA | 2 | 0.5 | 3 | 0.3333 |
| 7_3509148 | 2 | 1 | 3 | 0 | 1 | 0 | 1 | 0 | 7 | 0.2857 | NA | NA | 1 | 0 | NA | NA | 2 | 0.5 | 3 | 0.3333 |
| 7_3509197 | 1 | 1 | 1 | 0 | 1 | 0 | 2 | 1 | 5 | 0.6 | 1 | 1 | NA | NA | 2 | 1 | 2 | 1 | 5 | 1 |
| 7_3509202 | 1 | 1 | 1 | 1 | 1 | 1 | 2 | 1 | 5 | 1 | 1 | 1 | NA | NA | 2 | 1 | 2 | 1 | 5 | 1 |
| 7_3509215 | 1 | 1 | NA | NA | 1 | 1 | 1 | 1 | 3 | 1 | NA | NA | NA | NA | 1 | 1 | 1 | 1 | 2 | 1 |
| 7_3509216 | 1 | 0 | 1 | 0 | 1 | 1 | 2 | 0.5 | 5 | 0.4 | 1 | 1 | NA | NA | 2 | 0.5 | 2 | 0 | 5 | 0.4 |
| 7_3509223 | 1 | 0 | 1 | 0 | 1 | 0 | 2 | 0.5 | 5 | 0.2 | NA | NA | NA | NA | 2 | 0.5 | 2 | 0 | 4 | 0.25 |
| 7_3509580 | NA | NA | 1 | 1 | 1 | 0 | 2 | 0.5 | 4 | 0.5 | 1 | 0 | NA | NA | 2 | 0 | 2 | 0.5 | 5 | 0.2 |
| 7_3513787 | 3 | 0.3333 | 5 | 0.8 | NA | NA | NA | NA | 8 | 0.625 | 1 | 1 | NA | NA | NA | NA | 1 | 0 | 2 | 0.5 |
| 7_3524680 | 8 | 0.375 | 15 | 0.1333 | 12 | 0 | 13 | 0.0769 | 48 | 0.125 | 11 | 0.0909 | 12 | 0.0833 | 16 | 0.125 | 12 | 0 | 51 | 0.0784 |
| 7_3524694 | 6 | 0 | 15 | 0.3333 | 10 | 0.1 | 11 | 0.1818 | 42 | 0.1905 | 11 | 0.2727 | 11 | 0.2727 | 14 | 0.2857 | 11 | 0.2727 | 47 | 0.2766 |
| 7_3524710 | 7 | 0 | 13 | 0 | 8 | 0.125 | 11 | 0.0909 | 39 | 0.0513 | 9 | 0.1111 | 12 | 0.3333 | 11 | 0 | 7 | 0.1429 | 39 | 0.1538 |
| 7_3524719 | 11 | 0 | 15 | 0.2 | 11 | 0.2727 | 13 | 0.2308 | 50 | 0.18 | 14 | 0.0714 | 14 | 0.3571 | 18 | 0.2222 | 13 | 0.0769 | 59 | 0.1864 |
| 7_3524729 | 8 | 0.125 | 17 | 0.3529 | 10 | 0.2 | 12 | 0.25 | 47 | 0.2553 | 15 | 0.2 | 17 | 0.2941 | 14 | 0.2857 | 13 | 0.3077 | 59 | 0.2712 |
| 7_3524737 | 10 | 0.1 | 15 | 0.0667 | 10 | 0.2 | 14 | 0.1429 | 49 | 0.1224 | 14 | 0.0714 | 17 | 0.0588 | 16 | 0.0625 | 12 | 0 | 59 | 0.0508 |
| 7_3524748 | 9 | 0.3333 | 16 | 0.1875 | 12 | 0.1667 | 14 | 0.1429 | 51 | 0.1961 | 15 | 0.2667 | 16 | 0.0625 | 19 | 0 | 11 | 0 | 61 | 0.082 |
| 7_3533762 | 2 | 0.5 | 4 | 0.25 | 2 | 0 | 2 | 0 | 10 | 0.2 | 5 | 0.2 | 1 | 0 | 2 | 0 | NA | NA | 8 | 0.125 |
| 7_3533772 | 2 | 0.5 | 5 | 0.4 | 1 | 1 | 2 | 0 | 10 | 0.4 | 5 | 0.4 | 1 | 0 | 1 | 0 | NA | NA | 7 | 0.2857 |
| 7_3533774 | 1 | 0 | 4 | 0 | 3 | 0.6667 | 2 | 1 | 10 | 0.4 | 4 | 0.25 | 1 | 1 | 2 | 0.5 | NA | NA | 7 | 0.4286 |
| 7_3533777 | 1 | 1 | 2 | 0.5 | 3 | 0.3333 | 1 | 1 | 7 | 0.5714 | 2 | 0.5 | NA | NA | NA | NA | NA | NA | 2 | 0.5 |
| 7_3533778 | 2 | 1 | 4 | 1 | 3 | 1 | 2 | 1 | 11 | 1 | 5 | 0.8 | 1 | 1 | 2 | 1 | NA | NA | 8 | 0.875 |
| 7_3533784 | 2 | 0 | 4 | 0.75 | 1 | 0 | 2 | 1 | 9 | 0.5556 | 5 | 0.2 | NA | NA | 2 | 1 | NA | NA | 7 | 0.4286 |
| 7_3533789 | 1 | 0 | 4 | 0.75 | 3 | 1 | 2 | 1 | 10 | 0.8 | 4 | 0.75 | 1 | 1 | 1 | 0 | NA | NA | 6 | 0.6667 |
| 7_3533807 | 2 | 0 | 5 | 0.2 | 3 | 0 | 2 | 0.5 | 12 | 0.1667 | 5 | 0.2 | 1 | 0 | 1 | 0 | NA | NA | 7 | 0.1429 |
| 7_3533809 | 1 | 0 | 4 | 0.75 | 3 | 0.6667 | 1 | 0 | 9 | 0.5556 | 4 | 0 | 1 | 1 | 2 | 0.5 | NA | NA | 7 | 0.2857 |
| 7_3533813 | 2 | 0.5 | 5 | 0.4 | 3 | 1 | 2 | 0.5 | 12 | 0.5833 | 5 | 1 | 1 | 1 | 2 | 1 | NA | NA | 8 | 1 |
| 7_3533841 | 2 | 0 | 5 | 0.2 | 3 | 0 | 3 | 0 | 13 | 0.0769 | 5 | 0 | 1 | 1 | 2 | 0 | NA | NA | 8 | 0.125 |
| 7_3538571 | 4 | 0 | 4 | 0.25 | 3 | 0 | 3 | 0 | 14 | 0.0714 | 2 | 0.5 | 4 | 0 | 4 | 0 | 3 | 0 | 13 | 0.0769 |
| 7_3538579 | 4 | 0.25 | 4 | 0 | 3 | 0.3333 | 3 | 0.3333 | 14 | 0.2143 | 2 | 0.5 | 4 | 0.25 | 3 | 0.3333 | 5 | 0 | 14 | 0.2143 |
| 7_3538582 | 4 | 0.25 | 4 | 0.75 | 3 | 0 | 3 | 0 | 14 | 0.2857 | NA | NA | 4 | 0 | 3 | 0.3333 | 4 | 0.25 | 11 | 0.1818 |
| 7_3538846 | 2 | 0 | 2 | 0 | 4 | 0.25 | 1 | 0 | 9 | 0.1111 | 1 | 1 | 2 | 0 | 2 | 0 | 3 | 0 | 8 | 0.125 |
| 7_3538999 | 6 | 0.5 | 2 | 0.5 | 2 | 1 | 4 | 0.25 | 14 | 0.5 | 2 | 0.5 | 8 | 0.375 | 9 | 0.4444 | 6 | 0.5 | 25 | 0.44 |
| 7_3539014 | 2 | 1 | 1 | 1 | 2 | 1 | 3 | 1 | 8 | 1 | 2 | 1 | 6 | 0.6667 | 7 | 1 | 5 | 0.4 | 20 | 0.75 |
| 7_3539018 | 5 | 0.6 | 1 | 0 | 2 | 0 | 4 | 0 | 12 | 0.25 | 2 | 1 | 7 | 0.4286 | 6 | 0.6667 | 6 | 0.1667 | 21 | 0.4762 |

| | | | | | | | | | | | | | | | | | | | | |
|---|---|---|---|---|---|---|---|---|---|---|---|---|---|---|---|---|---|---|---|---|
| 7_3539024 | 5 | 0 | 2 | 0 | 2 | 0 | 5 | 0.2 | 14 | 0.0714 | 2 | 1 | 7 | 0.4286 | 9 | 0.1111 | 6 | 0.1667 | 24 | 0.2917 |
| 7_3544922 | 2 | 0 | NA | NA | NA | NA | 3 | 0.3333 | 5 | 0.2 | 1 | 0 | 2 | 0.5 | 4 | 0 | 3 | 0 | 10 | 0.1 |
| 7_3547186 | NA | NA | 1 | 1 | 2 | 1 | 1 | 0 | 4 | 0.75 | NA | NA | 3 | 0.3333 | 2 | 0.5 | 2 | 1 | 7 | 0.5714 |
| 7_3547321 | 3 | 0 | NA | NA | 1 | 0 | 1 | 1 | 5 | 0.2 | 1 | 1 | 1 | 1 | 1 | 0 | NA | NA | 3 | 0.6667 |
| 7_3547454 | 3 | 0.3333 | NA | NA | NA | NA | NA | NA | 3 | 0.3333 | 1 | 1 | NA | NA | NA | NA | NA | NA | 1 | 1 |
| 7_3547472 | 3 | 0.3333 | NA | NA | NA | NA | NA | NA | 3 | 0.3333 | 1 | 1 | NA | NA | NA | NA | NA | NA | 1 | 1 |
| 7_3547535 | 1 | 1 | NA | NA | 1 | 1 | NA | NA | 2 | 1 | 1 | 1 | NA | NA | NA | NA | NA | NA | 1 | 1 |
| 7_3547545 | 1 | 1 | NA | NA | NA | NA | NA | NA | 1 | 1 | 1 | 1 | NA | NA | NA | NA | NA | NA | 1 | 1 |
| 7_3549302 | 1 | 0 | NA | NA | 1 | 1 | NA | NA | 2 | 0.5 | 1 | 1 | NA | NA | NA | NA | NA | NA | 1 | 1 |
| 7_3646819 | 1 | 0 | 1 | 1 | NA | NA | NA | NA | 2 | 0.5 | NA | NA | NA | NA | 1 | 1 | NA | NA | 1 | 1 |
| 7_3646827 | 1 | 1 | 1 | 0 | NA | NA | NA | NA | 2 | 0.5 | NA | NA | NA | NA | 1 | 1 | NA | NA | 1 | 1 |
| 7_3646866 | 1 | 1 | 1 | 0 | NA | NA | NA | NA | 2 | 0.5 | NA | NA | NA | NA | 1 | 1 | NA | NA | 1 | 1 |
| 7_3646956 | 2 | 1 | 2 | 1 | NA | NA | 1 | 1 | 5 | 1 | NA | NA | NA | NA | 1 | 1 | NA | NA | 1 | 1 |
| 7_3646984 | 2 | 0.5 | 1 | 0 | NA | NA | 1 | 0 | 4 | 0.25 | NA | NA | NA | NA | 1 | 1 | NA | NA | 1 | 1 |
| 7_3647014 | 2 | 0 | 2 | 0.5 | NA | NA | 1 | 0 | 5 | 0.2 | NA | NA | NA | NA | 1 | 1 | NA | NA | 1 | 1 |
| 7_3647038 | 2 | 1 | 1 | 1 | NA | NA | NA | NA | 3 | 1 | NA | NA | NA | NA | 1 | 1 | NA | NA | 1 | 1 |
| 7_3647076 | 2 | 0.5 | 2 | 0 | NA | NA | 1 | 0 | 5 | 0.2 | NA | NA | NA | NA | 1 | 1 | NA | NA | 1 | 1 |
| 7_3647089 | 1 | 1 | 1 | 0 | NA | NA | NA | NA | 2 | 0.5 | NA | NA | NA | NA | 1 | 1 | NA | NA | 1 | 1 |
| 7_3647097 | 2 | 1 | 1 | 1 | NA | NA | 1 | 1 | 4 | 1 | NA | NA | NA | NA | 1 | 1 | NA | NA | 1 | 1 |
| 7_3647102 | 2 | 1 | 2 | 1 | NA | NA | 1 | 1 | 5 | 1 | NA | NA | NA | NA | 1 | 1 | NA | NA | 1 | 1 |
| 7_3647109 | 1 | 1 | NA | NA | NA | NA | NA | NA | 1 | 1 | NA | NA | NA | NA | 1 | 1 | NA | NA | 1 | 1 |
| 7_3647112 | 2 | 0.5 | 1 | 1 | NA | NA | 1 | 1 | 4 | 0.75 | NA | NA | NA | NA | 1 | 1 | NA | NA | 1 | 1 |
| 7_3647114 | 2 | 0.5 | 2 | 0.5 | NA | NA | 1 | 1 | 5 | 0.6 | NA | NA | NA | NA | 1 | 1 | NA | NA | 1 | 1 |
| 7_3647115 | 2 | 1 | 2 | 1 | NA | NA | 1 | 1 | 5 | 1 | NA | NA | NA | NA | 1 | 1 | NA | NA | 1 | 1 |
| 7_3647118 | 1 | 0 | 2 | 0.5 | NA | NA | NA | NA | 3 | 0.3333 | NA | NA | NA | NA | 1 | 1 | NA | NA | 1 | 1 |
| 7_3647128 | 2 | 1 | 2 | 0.5 | NA | NA | 1 | 1 | 5 | 0.8 | NA | NA | NA | NA | 1 | 1 | NA | NA | 1 | 1 |
| 7_3647137 | 2 | 1 | 2 | 1 | NA | NA | 1 | 1 | 5 | 1 | NA | NA | NA | NA | 1 | 1 | NA | NA | 1 | 1 |
| 7_3647139 | 1 | 1 | 2 | 1 | NA | NA | 1 | 0 | 4 | 0.75 | NA | NA | NA | NA | 1 | 1 | NA | NA | 1 | 1 |
| 7_3647148 | 2 | 1 | 2 | 1 | NA | NA | 1 | 0 | 5 | 0.8 | NA | NA | NA | NA | 1 | 1 | NA | NA | 1 | 1 |
| 7_3647159 | 2 | 0 | 3 | 0.3333 | NA | NA | 1 | 0 | 6 | 0.1667 | NA | NA | NA | NA | 1 | 1 | NA | NA | 1 | 1 |
| 7_3717178 | 2 | 0.5 | 8 | 0.875 | 6 | 0.5 | 2 | 1 | 18 | 0.7222 | 2 | 0.5 | 2 | 0.5 | 4 | 1 | 3 | 0.3333 | 11 | 0.6364 |
| 7_3717188 | 1 | 0 | 7 | 0.1429 | 5 | 0 | 2 | 0 | 15 | 0.0667 | 1 | 0 | 3 | 0 | 3 | 0.3333 | 3 | 0 | 10 | 0.1 |
| 7_3717908 | NA | NA | 2 | 1 | 1 | 1 | NA | NA | 3 | 1 | NA | NA | NA | NA | NA | NA | 2 | 0.5 | 2 | 0.5 |
| 7_3717909 | NA | NA | 1 | 0 | 1 | 1 | NA | NA | 2 | 0.5 | NA | NA | NA | NA | NA | NA | 3 | 0.3333 | 3 | 0.3333 |
| 7_3717929 | NA | NA | 2 | 1 | 1 | 1 | NA | NA | 3 | 1 | NA | NA | NA | NA | NA | NA | 2 | 0.5 | 2 | 0.5 |
| 7_3717934 | NA | NA | 3 | 1 | 1 | 1 | NA | NA | 4 | 1 | NA | NA | NA | NA | NA | NA | 3 | 1 | 3 | 1 |
| 7_3717983 | NA | NA | 3 | 0.3333 | 1 | 0 | NA | NA | 4 | 0.25 | NA | NA | NA | NA | NA | NA | 3 | 0.6667 | 3 | 0.6667 |
| 7_3718002 | NA | NA | 3 | 0.6667 | NA | NA | NA | NA | 3 | 0.6667 | NA | NA | NA | NA | NA | NA | 3 | 0.6667 | 3 | 0.6667 |
| 7_3726756 | NA | NA | 3 | 0.6667 | 1 | 0 | NA | NA | 4 | 0.5 | NA | NA | NA | NA | NA | NA | 2 | 0.5 | 2 | 0.5 |
| 7_3726763 | NA | NA | 2 | 0.5 | NA | NA | NA | NA | 2 | 0.5 | NA | NA | NA | NA | NA | NA | 2 | 0.5 | 2 | 0.5 |
| 7_3726786 | NA | NA | 3 | 1 | NA | NA | NA | NA | 3 | 1 | NA | NA | NA | NA | NA | NA | 2 | 1 | 2 | 1 |
| 7_3726830 | NA | NA | 2 | 1 | NA | NA | NA | NA | 2 | 1 | NA | NA | NA | NA | NA | NA | 1 | 1 | 1 | 1 |
| 7_3765948 | 1 | 0 | 9 | 0 | 9 | 0.1111 | 8 | 0.25 | 27 | 0.1111 | 1 | 0 | 7 | 0.1429 | 8 | 0 | 2 | 0 | 18 | 0.0556 |
| 7_3766061 | 1 | 0 | 9 | 0.2222 | 9 | 0.5556 | 8 | 0.25 | 27 | 0.3333 | 2 | 0.5 | 4 | 0 | 7 | 0 | 4 | 0 | 17 | 0.0588 |
| 7_3766062 | 1 | 0 | 9 | 0.3333 | 9 | 0.7778 | 6 | 0.1667 | 25 | 0.44 | 2 | 0 | 4 | 0.25 | 7 | 0.2857 | 4 | 0.25 | 17 | 0.2353 |
| 7_3766076 | 1 | 0 | 7 | 0.1429 | 9 | 0.1111 | 8 | 0 | 25 | 0.08 | 2 | 0.5 | 2 | 0.5 | 6 | 0 | 4 | 0 | 14 | 0.1429 |
| 7_3766090 | NA | NA | 8 | 0.875 | 9 | 0.6667 | 6 | 0.6667 | 23 | 0.7391 | 2 | 0.5 | NA | NA | 7 | 0.4286 | 4 | 0.25 | 13 | 0.3846 |
| 7_3766111 | NA | NA | 8 | 0.75 | 7 | 0.8571 | 7 | 0.5714 | 22 | 0.7273 | 1 | 1 | NA | NA | 5 | 0.8 | 4 | 0.5 | 10 | 0.7 |
| 7_3766485 | 1 | 0 | 7 | 0.2857 | 7 | 0.4286 | 7 | 0.2857 | 22 | 0.3182 | 2 | 0.5 | NA | NA | 7 | 0.4286 | 3 | 0.3333 | 12 | 0.4167 |
| 7_3766680 | NA | NA | 6 | 0.1667 | 7 | 0 | 7 | 0 | 20 | 0.05 | 2 | 0 | NA | NA | 7 | 0 | 3 | 0.3333 | 12 | 0.0833 |
| 7_3771867 | NA | NA | 6 | 1 | 6 | 1 | 6 | 0.8333 | 18 | 0.9444 | 1 | 1 | NA | NA | 5 | 1 | 3 | 0.3333 | 9 | 0.7778 |
| 7_3771869 | NA | NA | 4 | 0.25 | 5 | 0.6 | 4 | 0.5 | 13 | 0.4615 | 1 | 0 | NA | NA | 4 | 0.25 | 2 | 0.5 | 7 | 0.2857 |
| 7_3779747 | 1 | 0 | 5 | 0.4 | 3 | 0.3333 | 1 | 0 | 10 | 0.3 | 7 | 0.1429 | 1 | 0 | 1 | 0 | 8 | 0.375 | 17 | 0.2353 |
| 7_3779766 | 1 | 1 | 6 | 1 | 4 | 1 | NA | NA | 11 | 1 | 6 | 1 | 2 | 1 | 1 | 1 | 6 | 0.8333 | 15 | 0.9333 |
| 7_3779772 | 1 | 0 | 2 | 0 | 1 | 1 | 1 | 0 | 5 | 0.2 | 5 | 0 | 1 | 1 | 1 | 0 | 3 | 0 | 10 | 0.1 |
| 7_3779775 | NA | NA | 5 | 0.2 | 4 | 0 | 2 | 0 | 11 | 0.0909 | 5 | 0.2 | 2 | 0 | 1 | 0 | 8 | 0.125 | 16 | 0.125 |
| 7_3789417 | NA | NA | 2 | 0 | 1 | 0 | 5 | 0.2 | 8 | 0.125 | 1 | 1 | NA | NA | 1 | 1 | 1 | 1 | 3 | 1 |
| 7_3789426 | NA | NA | 2 | 0 | 1 | 0 | 5 | 0.2 | 8 | 0.125 | 2 | 1 | NA | NA | 2 | 0.5 | 2 | 0 | 6 | 0.5 |
| 7_3789426 | NA | NA | 2 | 1 | 1 | 1 | 4 | 1 | 7 | 1 | 3 | 1 | NA | NA | 2 | 1 | 2 | 1 | 7 | 1 |
| 7_3789427 | NA | NA | 2 | 0.5 | 1 | 0 | 3 | 0.6667 | 6 | 0.5 | 1 | 1 | NA | NA | 2 | 0.5 | NA | NA | 3 | 0.6667 |
| 7_3789429 | NA | NA | 2 | 0 | 1 | 0 | 4 | 0.25 | 7 | 0.1429 | 2 | 0 | NA | NA | 1 | 0 | 2 | 0.5 | 5 | 0.2 |
| 7_3789439 | NA | NA | 1 | 1 | 1 | 1 | 3 | 1 | 5 | 1 | 2 | 0.5 | NA | NA | 1 | 1 | NA | NA | 3 | 0.6667 |
| 7_3789447 | 1 | 0 | 3 | 0.3333 | 1 | 0 | 4 | 0.25 | 9 | 0.2222 | 3 | 0 | NA | NA | 2 | 0 | 2 | 1 | 7 | 0.2857 |
| 7_3789454 | NA | NA | 1 | 1 | NA | NA | 4 | 0.5 | 5 | 0.6 | 2 | 1 | NA | NA | 1 | 0 | 2 | 0 | 5 | 0.4 |
| 7_3789463 | NA | NA | 2 | 1 | 2 | 0.5 | 5 | 0.8 | 9 | 0.7778 | 3 | 0.3333 | NA | NA | 2 | 0.5 | 2 | 0.5 | 7 | 0.4286 |

| | | | | | | | | | | | | | | | | | | | | |
|---|---|---|---|---|---|---|---|---|---|---|---|---|---|---|---|---|---|---|---|---|
| 7_3789476 | 1 | 0 | 3 | 0 | 2 | 0 | 4 | 0.75 | 10 | 0.3 | 3 | 0 | NA | NA | 2 | 1 | 2 | 0 | 7 | 0.2857 |
| 7_3789484 | NA | NA | 3 | 0 | 2 | 0.5 | 5 | 0.2 | 10 | 0.2 | 2 | 0 | NA | NA | 2 | 0.5 | NA | NA | 4 | 0.25 |
| 7_3789503 | 1 | 1 | 2 | 0.5 | 3 | 0 | 3 | 0.3333 | 9 | 0.3333 | 2 | 0 | NA | NA | 2 | 0.5 | 1 | 0 | 5 | 0.2 |
| 7_3789519 | NA | NA | 4 | 0 | 2 | 0.5 | 6 | 0.1667 | 12 | 0.1667 | 3 | 0.3333 | 1 | 0 | 2 | 0.5 | 1 | 0 | 7 | 0.2857 |
| 7_3791686 | 1 | 1 | NA | NA | NA | NA | NA | NA | 1 | 1 | NA | NA | 1 | 1 | 1 | 1 | NA | NA | 2 | 1 |
| 7_3791689 | 1 | 1 | NA | NA | NA | NA | NA | NA | 1 | 1 | NA | NA | 1 | 0 | 1 | 1 | NA | NA | 2 | 0.5 |
| 7_3791696 | 1 | 1 | NA | NA | NA | NA | NA | NA | 1 | 1 | NA | NA | 1 | 0 | 1 | 1 | NA | NA | 2 | 0.5 |
| 7_3791707 | 1 | 1 | NA | NA | NA | NA | NA | NA | 1 | 1 | NA | NA | 1 | 0 | 1 | 1 | NA | NA | 2 | 0.5 |
| 7_3792016 | 1 | 1 | 3 | 0 | 3 | 0.3333 | 3 | 0 | 10 | 0.2 | 3 | 0.6667 | NA | NA | NA | NA | NA | NA | 3 | 0.6667 |
| 7_3792051 | NA | NA | 1 | 1 | 2 | 0.5 | 3 | 1 | 6 | 0.8333 | 1 | 1 | NA | NA | NA | NA | NA | NA | 1 | 1 |
| 7_3792061 | 1 | 1 | NA | NA | 1 | 1 | 1 | 1 | 3 | 1 | 1 | 1 | 1 | 0 | 1 | 1 | NA | NA | 3 | 0.6667 |
| 7_3792065 | NA | NA | 1 | 0 | 1 | 0 | 2 | 0.5 | 4 | 0.25 | 1 | 0 | 1 | 1 | NA | NA | NA | NA | 2 | 0.5 |
| 7_3792078 | 1 | 1 | 1 | 1 | 2 | 1 | 2 | 1 | 6 | 1 | 1 | 1 | 1 | 1 | 1 | 1 | NA | NA | 3 | 1 |
| 7_3792081 | 1 | 1 | NA | NA | 2 | 1 | 2 | 1 | 5 | 1 | 1 | 1 | 1 | 1 | 1 | 1 | NA | NA | 3 | 1 |
| 7_3792110 | 1 | 1 | 1 | 1 | 2 | 1 | 1 | 1 | 5 | 1 | 1 | 1 | 1 | 1 | 1 | 1 | NA | NA | 3 | 1 |
| 7_3792111 | 1 | 1 | 1 | 1 | 2 | 1 | 3 | 1 | 7 | 1 | 1 | 1 | 1 | 1 | 1 | 1 | NA | NA | 3 | 1 |
| 7_3796229 | 3 | 0.3333 | 5 | 0.2 | 2 | 0 | 6 | 0 | 16 | 0.125 | 2 | 0 | 1 | 0 | NA | NA | 3 | 0.3333 | 6 | 0.1667 |
| 7_3796255 | 4 | 0.5 | 7 | 0.1429 | 3 | 0.6667 | 5 | 0.4 | 19 | 0.3684 | 2 | 1 | 2 | 0.5 | 1 | 1 | 3 | 0.3333 | 8 | 0.625 |
| 7_3796259 | 2 | 0.5 | 6 | 0.3333 | 3 | 0 | 7 | 0.2857 | 18 | 0.2778 | 2 | 0 | 1 | 1 | 1 | 0 | 3 | 0.3333 | 7 | 0.2857 |
| 7_3912326 | 1 | 1 | 1 | 1 | NA | NA | NA | NA | 2 | 1 | NA | NA | 1 | 1 | 1 | 1 | NA | NA | 2 | 1 |
| 7_3912395 | 1 | 1 | 1 | 1 | NA | NA | NA | NA | 2 | 1 | NA | NA | NA | NA | 1 | 1 | 1 | 1 | 2 | 1 |
| 7_3912414 | 2 | 0.5 | 1 | 0 | NA | NA | NA | NA | 3 | 0.3333 | 1 | 0 | NA | NA | 1 | 1 | 1 | 1 | 3 | 0.6667 |
| 7_3912681 | 1 | 0 | 2 | 0.5 | NA | NA | NA | NA | 3 | 0.3333 | 1 | 0 | 1 | 1 | NA | NA | NA | NA | 2 | 0.5 |
| 7_3921647 | NA | NA | 2 | 0.5 | 1 | 0 | NA | NA | 3 | 0.3333 | 1 | 1 | NA | NA | NA | NA | NA | NA | 1 | 1 |
| 7_3921656 | NA | NA | 2 | 1 | 1 | 0 | NA | NA | 3 | 0.6667 | 1 | 0 | NA | NA | NA | NA | 1 | 1 | 2 | 0.5 |
| 7_3921660 | NA | NA | 1 | 1 | 1 | 1 | NA | NA | 2 | 1 | 1 | 1 | NA | NA | NA | NA | 2 | 1 | 3 | 1 |
| 7_3921672 | NA | NA | 1 | 1 | NA | NA | NA | NA | 1 | 1 | 1 | 1 | NA | NA | NA | NA | 2 | 0 | 3 | 0.3333 |
| 7_3921673 | NA | NA | 2 | 1 | 1 | 1 | NA | NA | 3 | 1 | 1 | 1 | NA | NA | NA | NA | NA | NA | 1 | 1 |
| 7_3921681 | NA | NA | 2 | 0.5 | 1 | 1 | NA | NA | 3 | 0.6667 | 1 | 0 | NA | NA | NA | NA | 2 | 0.5 | 3 | 0.3333 |
| 7_3921685 | NA | NA | 2 | 1 | 1 | 1 | NA | NA | 3 | 1 | NA | NA | NA | NA | NA | NA | 1 | 1 | 1 | 1 |
| 7_3921699 | NA | NA | 2 | 1 | 1 | 1 | NA | NA | 3 | 1 | 1 | 1 | NA | NA | NA | NA | 1 | 0 | 2 | 0.5 |
| 7_3921767 | NA | NA | 2 | 0.5 | 1 | 0 | NA | NA | 3 | 0.3333 | 1 | 0 | NA | NA | NA | NA | 1 | 1 | 2 | 0.5 |
| 7_4050010 | NA | NA | NA | NA | NA | NA | 1 | 1 | 1 | 1 | 1 | 1 | NA | NA | NA | NA | 2 | 0 | 3 | 0.3333 |
| 7_4054848 | 3 | 0 | 11 | 0.3636 | 4 | 0 | 3 | 0 | 21 | 0.1905 | 5 | 0.2 | 2 | 0 | 3 | 0.3333 | 3 | 0.6667 | 13 | 0.3077 |
| 7_4054852 | 3 | 1 | 10 | 0.4 | 5 | 0.2 | 3 | 0.6667 | 21 | 0.4762 | 4 | 0.25 | 3 | 0 | 3 | 0 | 3 | 0.6667 | 13 | 0.2308 |
| 7_4055194 | 6 | 0.3333 | 14 | 0.0714 | 6 | 0.5 | 7 | 0.4286 | 33 | 0.2727 | 11 | 0.2727 | 4 | 0.5 | 7 | 0.4286 | 4 | 0.25 | 26 | 0.3462 |
| 7_4057891 | 6 | 0.5 | 20 | 0.1 | 12 | 0.0833 | 9 | 0.1111 | 47 | 0.1489 | 9 | 0.2222 | 5 | 0.2 | 11 | 0 | 6 | 0 | 31 | 0.0968 |
| 7_4057893 | 6 | 0.5 | 17 | 0.7059 | 11 | 0.8182 | 6 | 0.5 | 40 | 0.675 | 10 | 0.8 | 5 | 0.6 | 9 | 0.8889 | 6 | 0.8333 | 30 | 0.8 |
| 7_4057894 | 6 | 1 | 16 | 0.875 | 9 | 0.7778 | 7 | 0.8571 | 38 | 0.8684 | 8 | 0.875 | 6 | 0.8333 | 7 | 1 | 5 | 1 | 26 | 0.9231 |
| 7_4057901 | 6 | 0 | 20 | 0.45 | 12 | 0.75 | 8 | 0.25 | 46 | 0.4348 | 12 | 0.4167 | 6 | 0.1667 | 11 | 0.6364 | 6 | 0.3333 | 35 | 0.4286 |
| 7_4057914 | 5 | 0.8 | 15 | 0.8667 | 9 | 1 | 5 | 1 | 34 | 0.9118 | 10 | 0.8 | 3 | 0.3333 | 6 | 1 | 3 | 1 | 22 | 0.8182 |
| 7_4102270 | NA | NA | 2 | 0 | 3 | 0.3333 | 2 | 0 | 7 | 0.1429 | 1 | 0 | 1 | 0 | 1 | 0 | 2 | 0.5 | 5 | 0.2 |
| 7_4102291 | NA | NA | 2 | 0 | NA | NA | 2 | 0.5 | 4 | 0.25 | NA | NA | 1 | 1 | NA | NA | 1 | 0 | 2 | 0.5 |
| 7_4102292 | NA | NA | 2 | 0.5 | 2 | 0.5 | 2 | 0.5 | 6 | 0.5 | 1 | 0 | NA | NA | 1 | 0 | 2 | 1 | 4 | 0.5 |
| 7_4102300 | NA | NA | 1 | 1 | 2 | 1 | 1 | 0 | 4 | 0.75 | 1 | 1 | 1 | 1 | 1 | 1 | 2 | 0.5 | 5 | 0.8 |
| 7_4102301 | NA | NA | 2 | 1 | 2 | 0.5 | NA | NA | 4 | 0.75 | 1 | 1 | 1 | 1 | 1 | 1 | 2 | 1 | 5 | 1 |
| 7_4102304 | NA | NA | 1 | 1 | 2 | 0.5 | 1 | 0 | 4 | 0.5 | 1 | 1 | 1 | 0 | 1 | 1 | 2 | 0.5 | 5 | 0.6 |
| 7_4102305 | NA | NA | 2 | 0 | 2 | 0.5 | 2 | 0 | 6 | 0.1667 | 1 | 1 | 1 | 0 | NA | NA | 2 | 0 | 4 | 0.25 |
| 7_4102323 | NA | NA | 2 | 0 | 1 | 1 | 1 | 1 | 4 | 0.5 | 1 | 1 | 1 | 1 | 1 | 1 | 2 | 0.5 | 5 | 0.8 |
| 7_4102328 | NA | NA | 1 | 1 | 2 | 0.5 | 2 | 1 | 5 | 0.8 | NA | NA | 1 | 0 | 1 | 1 | 1 | 0 | 3 | 0.3333 |
| 7_4102421 | NA | NA | 2 | 0.5 | 1 | 0 | 2 | 0.5 | 5 | 0.4 | 1 | 0 | 1 | 1 | 1 | 1 | 2 | 0 | 5 | 0.4 |
| 7_4102446 | NA | NA | 1 | 1 | 1 | 1 | 2 | 1 | 4 | 1 | 1 | 1 | NA | NA | NA | NA | 2 | 0.5 | 3 | 0.6667 |
| 7_4112887 | NA | NA | 1 | 1 | NA | NA | NA | NA | 1 | 1 | NA | NA | NA | NA | 1 | 1 | 3 | 1 | 4 | 1 |
| 7_4112891 | NA | NA | 1 | 1 | NA | NA | NA | NA | 1 | 1 | 1 | 1 | NA | NA | 1 | 1 | 2 | 1 | 4 | 1 |
| 7_4112899 | NA | NA | 1 | 1 | NA | NA | NA | NA | 1 | 1 | 1 | 1 | NA | NA | NA | NA | 2 | 1 | 3 | 1 |
| 7_4112906 | NA | NA | 1 | 1 | NA | NA | NA | NA | 1 | 1 | 1 | 0 | NA | NA | 1 | 1 | 2 | 1 | 4 | 0.75 |
| 7_4112923 | NA | NA | 1 | 1 | NA | NA | NA | NA | 1 | 1 | 1 | 1 | NA | NA | 1 | 1 | 1 | 0 | 3 | 0.6667 |
| 7_4126134 | NA | NA | 1 | 1 | NA | NA | 1 | 0 | 2 | 0.5 | NA | NA | 2 | 0.5 | 1 | 0 | 1 | 0 | 4 | 0.25 |
| 7_4127472 | 1 | 1 | NA | NA | NA | NA | 1 | 1 | 2 | 1 | NA | NA | 2 | 0.5 | 1 | 0 | 1 | 0 | 4 | 0.25 |
| 7_4127482 | 1 | 1 | 1 | 1 | NA | NA | 1 | 1 | 3 | 1 | NA | NA | 2 | 1 | 1 | 1 | 1 | 1 | 4 | 1 |
| 7_4127502 | 1 | 1 | 1 | 1 | NA | NA | 1 | 0 | 3 | 0.6667 | NA | NA | 2 | 1 | 1 | 1 | 1 | 0 | 4 | 0.75 |
| 7_4127534 | 1 | 1 | 1 | 0 | NA | NA | 1 | 0 | 3 | 0.3333 | NA | NA | 2 | 0.5 | 1 | 0 | 1 | 0 | 4 | 0.25 |
| 7_4127540 | 1 | 0 | 1 | 1 | NA | NA | 1 | 0 | 3 | 0.3333 | NA | NA | 2 | 0.5 | 1 | 0 | 1 | 0 | 4 | 0.25 |
| 7_4127587 | 1 | 0 | 1 | 1 | NA | NA | NA | NA | 2 | 0.5 | NA | NA | 4 | 0 | 1 | 1 | 1 | 1 | 6 | 0.3333 |
| 7_4127662 | 1 | 0 | 5 | 0.4 | NA | NA | 2 | 0 | 8 | 0.25 | NA | NA | 4 | 0 | 1 | 1 | 1 | 0 | 6 | 0.1667 |

| | | | | | | | | | | | | | | | | | | | | |
|---|---|---|---|---|---|---|---|---|---|---|---|---|---|---|---|---|---|---|---|---|
| 7_4127677 | 1 | 1 | 5 | 1 | NA | NA | 2 | 1 | 8 | 1 | NA | NA | 4 | 0.75 | 1 | 0 | 1 | 1 | 6 | 0.6667 |
| 7_4127685 | NA | NA | 4 | 0 | NA | NA | 2 | 0.5 | 6 | 0.1667 | NA | NA | 4 | 0.25 | 1 | 0 | 1 | 0 | 6 | 0.1667 |
| 7_4127688 | 1 | 0 | 3 | 0.3333 | NA | NA | 2 | 0.5 | 6 | 0.3333 | NA | NA | 3 | 0.6667 | 1 | 0 | 2 | 0 | 6 | 0.3333 |
| 7_4127700 | 1 | 1 | 5 | 1 | NA | NA | 2 | 1 | 8 | 1 | NA | NA | 4 | 1 | NA | NA | 1 | 1 | 5 | 1 |
| 7_4127719 | 1 | 1 | 4 | 0 | NA | NA | 2 | 0 | 7 | 0.1429 | NA | NA | 3 | 0.3333 | 1 | 0 | 1 | 0 | 5 | 0.2 |
| 7_4127727 | 1 | 1 | 4 | 0.25 | NA | NA | 1 | 0 | 6 | 0.3333 | NA | NA | 4 | 0.5 | NA | NA | 3 | 0.3333 | 7 | 0.4286 |
| 7_4127734 | NA | NA | 5 | 1 | NA | NA | 3 | 0.6667 | 8 | 0.875 | NA | NA | 4 | 1 | NA | NA | 3 | 1 | 7 | 1 |
| 7_4127749 | 1 | 1 | 4 | 0 | NA | NA | 3 | 0 | 8 | 0.125 | NA | NA | 5 | 0.4 | 1 | 0 | 3 | 0 | 9 | 0.2222 |
| 7_4127772 | 3 | 0 | 4 | 0.25 | NA | NA | 3 | 0 | 10 | 0.1 | NA | NA | 6 | 0 | 1 | 0 | 2 | 0.5 | 9 | 0.1111 |
| 7_4129763 | 2 | 0 | 3 | 0.6667 | 4 | 0.5 | 4 | 0.25 | 13 | 0.3846 | 2 | 0.5 | 1 | 0 | 7 | 0.7143 | 6 | 0.5 | 16 | 0.5625 |
| 7_4129776 | 2 | 1 | 3 | 1 | 3 | 1 | 3 | 0.6667 | 11 | 0.9091 | NA | NA | 1 | 1 | 5 | 1 | 7 | 0.7143 | 13 | 0.8462 |
| 7_4129777 | 2 | 0.5 | 3 | 0.6667 | 4 | 0 | 2 | 0.5 | 11 | 0.3636 | 1 | 1 | 1 | 0 | 5 | 0.6 | 7 | 0.4286 | 14 | 0.5 |
| 7_4129789 | 1 | 0 | 3 | 0 | 4 | 0.25 | 3 | 0 | 11 | 0.0909 | 1 | 0 | 1 | 0 | 6 | 0.1667 | 7 | 0 | 15 | 0.0667 |
| 7_4129792 | NA | NA | NA | NA | 4 | 0 | 3 | 0.3333 | 7 | 0.1429 | 2 | 0.5 | 1 | 0 | 7 | 0.1429 | 4 | 0 | 14 | 0.1429 |
| 7_4129793 | 1 | 1 | 2 | 1 | 4 | 1 | 2 | 0.5 | 9 | 0.8889 | 1 | 1 | 1 | 1 | 6 | 1 | 5 | 1 | 13 | 1 |
| 7_4129797 | 1 | 0 | 3 | 0.6667 | 4 | 0.5 | 3 | 0.3333 | 11 | 0.4545 | 2 | 0 | 1 | 0 | 6 | 0.3333 | 7 | 0.1429 | 16 | 0.1875 |
| 7_4129799 | 1 | 0 | 2 | 1 | 4 | 1 | 2 | 1 | 9 | 0.8889 | NA | NA | 1 | 1 | 5 | 0.8 | 5 | 1 | 11 | 0.9091 |
| 7_4129864 | 1 | 0 | 3 | 0 | 4 | 0.25 | 2 | 0 | 10 | 0.1 | 1 | 1 | 2 | 0 | 3 | 0 | 7 | 0.1429 | 13 | 0.1538 |
| 7_4129872 | 1 | 1 | 3 | 0 | 4 | 0 | 3 | 0 | 11 | 0.0909 | 1 | 1 | 2 | 0 | 7 | 0.1429 | 7 | 0.2857 | 17 | 0.2353 |
| 7_4130050 | 3 | 0.3333 | 2 | 0.5 | 3 | 0.3333 | 6 | 0.1667 | 14 | 0.2857 | 1 | 1 | 2 | 0.5 | 6 | 0.1667 | 8 | 0.25 | 17 | 0.2941 |
| 7_4212296 | 1 | 0 | 3 | 0 | NA | NA | 3 | 0.3333 | 7 | 0.1429 | 2 | 0 | 1 | 0 | 1 | 0 | 2 | 0.5 | 6 | 0.1667 |
| 7_4212301 | 1 | 0 | 3 | 0.3333 | NA | NA | 3 | 0.3333 | 7 | 0.2857 | 2 | 0.5 | 1 | 1 | 2 | 0.5 | 2 | 0 | 7 | 0.4286 |
| 7_4212312 | 1 | 0 | 3 | 0.3333 | NA | NA | 3 | 0.6667 | 7 | 0.4286 | 2 | 0.5 | 1 | 0 | 2 | 0.5 | 1 | 1 | 6 | 0.5 |
| 7_4212320 | 1 | 0 | 3 | 0.6667 | NA | NA | 3 | 0.3333 | 7 | 0.4286 | 1 | 0 | 1 | 0 | 2 | 0.5 | 2 | 0.5 | 6 | 0.3333 |
| 7_4212331 | NA | NA | 4 | 1 | NA | NA | 3 | 1 | 7 | 1 | 1 | 1 | 1 | 1 | 2 | 0 | NA | NA | 4 | 0.5 |
| 7_4212343 | 1 | 0 | 4 | 0.25 | NA | NA | 3 | 0.6667 | 8 | 0.375 | 2 | 0 | 1 | 0 | 2 | 0 | 2 | 0.5 | 7 | 0.1429 |
| 7_4212345 | 1 | 0 | 4 | 0 | NA | NA | 3 | 0.3333 | 8 | 0.125 | 1 | 0 | 1 | 0 | 2 | 0.5 | 1 | 0 | 5 | 0.2 |
| 7_4212365 | 1 | 0 | 4 | 0.25 | NA | NA | 3 | 0.3333 | 8 | 0.25 | 2 | 0.5 | 1 | 0 | 2 | 0 | 2 | 0 | 7 | 0.1429 |
| 7_4212366 | 1 | 0 | 3 | 0.6667 | NA | NA | 3 | 0 | 7 | 0.2857 | 2 | 0 | 1 | 0 | 2 | 0.5 | 2 | 0 | 7 | 0.1429 |
| 7_4212385 | 1 | 0 | 6 | 0.3333 | NA | NA | 2 | 0 | 9 | 0.2222 | 2 | 0 | 1 | 0 | 2 | 0 | 3 | 0.3333 | 8 | 0.125 |
| 7_4212388 | NA | NA | 5 | 0.6 | NA | NA | NA | NA | 5 | 0.6 | 2 | 1 | 1 | 0 | 2 | 0.5 | 2 | 1 | 7 | 0.7143 |
| 7_4212424 | 1 | 1 | 5 | 0.2 | NA | NA | 3 | 0.6667 | 9 | 0.4444 | 2 | 0 | 1 | 0 | 2 | 0.5 | 2 | 1 | 7 | 0.4286 |
| 7_4212426 | NA | NA | 6 | 0.5 | NA | NA | 1 | 0 | 7 | 0.4286 | 2 | 1 | NA | NA | 1 | 1 | 3 | 0.3333 | 6 | 0.6667 |
| 7_4212445 | NA | NA | 6 | 1 | NA | NA | 3 | 1 | 9 | 1 | 2 | 1 | NA | NA | 2 | 1 | 3 | 1 | 7 | 1 |
| 7_4212449 | NA | NA | 6 | 0.3333 | NA | NA | 3 | 0 | 9 | 0.2222 | 2 | 0 | 1 | 0 | 2 | 0.5 | 3 | 0 | 8 | 0.125 |
| 7_4212465 | NA | NA | 5 | 0.2 | NA | NA | 3 | 0.3333 | 8 | 0.25 | 1 | 0 | 1 | 1 | 2 | 0 | 3 | 0 | 7 | 0.1429 |
| 7_4212477 | NA | NA | 5 | 0.2 | NA | NA | 3 | 0 | 8 | 0.125 | 1 | 1 | 1 | 0 | 2 | 0 | 3 | 0 | 7 | 0.1429 |
| 7_4212491 | NA | NA | 5 | 0.8 | NA | NA | 2 | 1 | 7 | 0.8571 | NA | NA | 1 | 1 | 1 | 0 | 3 | 0.6667 | 5 | 0.6 |
| 7_4212494 | NA | NA | 5 | 0.8 | NA | NA | 3 | 0.6667 | 8 | 0.75 | 1 | 1 | 1 | 1 | 1 | 1 | 3 | 0.3333 | 6 | 0.6667 |
| 7_4212524 | NA | NA | 4 | 0 | NA | NA | 3 | 0.3333 | 7 | 0.1429 | 1 | 1 | 1 | 0 | 1 | 1 | 3 | 0 | 6 | 0.3333 |
| 7_4212535 | NA | NA | 3 | 0.3333 | NA | NA | 3 | 0.3333 | 6 | 0.3333 | 1 | 0 | 1 | 0 | 1 | 1 | 2 | 0.5 | 5 | 0.4 |
| 7_4212546 | NA | NA | 4 | 0.75 | NA | NA | 2 | 1 | 6 | 0.8333 | 1 | 1 | NA | NA | 1 | 1 | 1 | 0 | 3 | 0.6667 |
| 7_4212552 | NA | NA | 3 | 0.6667 | NA | NA | 3 | 0.6667 | 6 | 0.6667 | 1 | 1 | NA | NA | 1 | 0 | 2 | 0 | 4 | 0.25 |
| 7_4212585 | NA | NA | 3 | 1 | NA | NA | 3 | 1 | 6 | 1 | 1 | 1 | NA | NA | 1 | 1 | 1 | 0 | 3 | 0.6667 |
| 7_4212596 | NA | NA | 4 | 0.75 | NA | NA | 2 | 0.5 | 6 | 0.6667 | 1 | 1 | NA | NA | 1 | 1 | 2 | 0.5 | 4 | 0.75 |
| 7_4212606 | NA | NA | 4 | 1 | NA | NA | 3 | 0.6667 | 7 | 0.8571 | 1 | 0 | NA | NA | 1 | 1 | 2 | 0.5 | 4 | 0.5 |
| 7_4231021 | NA | NA | 2 | 1 | 2 | 1 | 2 | 1 | 6 | 1 | NA | NA | 1 | 0 | 2 | 0 | 2 | 1 | 5 | 0.4 |
| 7_4231031 | NA | NA | 2 | 0 | 2 | 0.5 | 1 | 1 | 5 | 0.4 | NA | NA | 1 | 1 | 1 | 0 | 1 | 0 | 3 | 0.3333 |
| 7_4231046 | NA | NA | 2 | 1 | 1 | 1 | 2 | 1 | 5 | 1 | NA | NA | 1 | 1 | 2 | 1 | 2 | 1 | 5 | 1 |
| 7_4231056 | NA | NA | 2 | 0 | 1 | 1 | 2 | 1 | 5 | 0.6 | NA | NA | 1 | 0 | 2 | 0.5 | 2 | 0 | 5 | 0.2 |
| 7_4231058 | NA | NA | 2 | 0 | 1 | 0 | 2 | 0.5 | 5 | 0.2 | NA | NA | 1 | 1 | 2 | 0 | 1 | 0 | 4 | 0.25 |
| 7_4231096 | NA | NA | 1 | 1 | 1 | 0 | 1 | 1 | 3 | 0.6667 | NA | NA | NA | NA | 2 | 1 | 1 | 1 | 3 | 1 |
| 7_4231140 | NA | NA | 1 | 1 | 1 | 1 | 1 | 0 | 3 | 0.6667 | NA | NA | NA | NA | 2 | 0.5 | 2 | 0.5 | 4 | 0.5 |
| 7_4231182 | NA | NA | NA | NA | 1 | 1 | NA | NA | 1 | 1 | NA | NA | NA | NA | 2 | 1 | 2 | 1 | 4 | 1 |
| 7_4231207 | NA | NA | NA | NA | 1 | 1 | 1 | 0 | 2 | 0.5 | NA | NA | NA | NA | 2 | 0.5 | 2 | 0 | 4 | 0.25 |
| 7_4231214 | NA | NA | NA | NA | 1 | 1 | 1 | 1 | 2 | 1 | NA | NA | NA | NA | 2 | 0 | 2 | 0.5 | 4 | 0.25 |
| 7_4231224 | NA | NA | NA | NA | 1 | 0 | 1 | 1 | 2 | 0.5 | NA | NA | NA | NA | 1 | 1 | 2 | 0.5 | 3 | 0.6667 |
| 7_4294629 | 5 | 0 | 5 | 0.2 | 5 | 0 | 4 | 0.25 | 19 | 0.1053 | 3 | 0.3333 | 3 | 0.3333 | NA | NA | 1 | 0 | 7 | 0.2857 |
| 7_4294709 | 4 | 0.5 | 2 | 0.5 | 5 | 0.8 | 5 | 0.2 | 16 | 0.5 | 1 | 0 | 2 | 0.5 | NA | NA | 1 | 1 | 4 | 0.5 |
| 7_4294714 | 5 | 0.6 | 3 | 1 | 5 | 1 | 5 | 1 | 18 | 0.8889 | NA | NA | 2 | 0.5 | NA | NA | 1 | 1 | 3 | 0.6667 |
| 7_4294716 | 4 | 0.25 | 3 | 0.3333 | 4 | 0.25 | 4 | 0.5 | 15 | 0.3333 | NA | NA | 2 | 0.5 | NA | NA | NA | NA | 2 | 0.5 |
| 7_4294717 | 4 | 0 | 2 | 0 | 1 | 0 | 5 | 0.6 | 12 | 0.25 | 1 | 0 | 2 | 1 | NA | NA | 1 | 1 | 4 | 0.75 |
| 7_4294728 | 2 | 1 | 1 | 1 | 5 | 1 | 3 | 0.6667 | 11 | 0.9091 | 1 | 1 | NA | NA | NA | NA | NA | NA | 1 | 1 |
| 7_4294747 | 5 | 0.6 | 2 | 1 | 4 | 1 | 5 | 1 | 16 | 0.875 | 1 | 1 | NA | NA | NA | NA | 1 | 1 | 2 | 1 |
| 7_4294755 | 5 | 1 | 3 | 0.6667 | 4 | 0.25 | 5 | 0.6 | 17 | 0.6471 | 1 | 0 | 2 | 0 | NA | NA | 1 | 1 | 4 | 0.25 |

| | | | | | | | | | | | | | | | | | | | | |
|---|---|---|---|---|---|---|---|---|---|---|---|---|---|---|---|---|---|---|---|---|
| 7_42947592 | 2 | 1 | 2 | 1 | 3 | 1 | 4 | 1 | 11 | 1 | 1 | 1 | 1 | 0 | NA | NA | 1 | 1 | 3 | 0.6667 |
| 7_42947791 | 5 | 1 | 3 | 1 | 4 | 1 | 5 | 1 | 17 | 1 | NA | NA | 1 | 1 | NA | NA | 1 | 1 | 2 | 1 |
| 7_42948141 | 4 | 0.5 | 2 | 0.5 | 3 | 0.3333 | 5 | 0 | 14 | 0.2857 | NA | NA | 1 | 1 | NA | NA | NA | NA | 1 | 1 |
| 7_42948376 | 5 | 0.6 | 2 | 0.5 | 2 | 1 | 3 | 1 | 12 | 0.75 | NA | NA | 1 | 1 | NA | NA | NA | NA | 1 | 1 |
| 7_42948628 | 4 | 0.25 | 3 | 0.3333 | 3 | 0 | 5 | 1 | 15 | 0.4667 | NA | NA | 1 | 1 | NA | NA | NA | NA | 1 | 1 |
| 7_42948715 | 3 | 0.3333 | 2 | 1 | 3 | 0 | 4 | 0.25 | 12 | 0.3333 | NA | NA | 1 | 1 | NA | NA | NA | NA | 1 | 1 |
| 7_43122505 | 2 | 0 | 6 | 0.1667 | 1 | 0 | 5 | 0.2 | 14 | 0.1429 | 3 | 0.3333 | 1 | 0 | 4 | 0 | 6 | 0.3333 | 14 | 0.2143 |
| 7_43178130 | 1 | 1 | NA | NA | NA | NA | 1 | 1 | 2 | 1 | NA | NA | NA | NA | 3 | 0.6667 | NA | NA | 3 | 0.6667 |
| 7_43178152 | 1 | 1 | NA | NA | NA | NA | 1 | 1 | 2 | 1 | NA | NA | NA | NA | 2 | 1 | NA | NA | 2 | 1 |
| 7_43178371 | 1 | 1 | NA | NA | NA | NA | 1 | 1 | 2 | 1 | NA | NA | NA | NA | 2 | 0.5 | NA | NA | 2 | 0.5 |
| 7_43178457 | 1 | 1 | NA | NA | NA | NA | 1 | 1 | 2 | 1 | NA | NA | NA | NA | 1 | 1 | NA | NA | 1 | 1 |
| 7_43222128 | 2 | 1 | NA | NA | 1 | 0 | NA | NA | 3 | 0.6667 | 3 | 0.6667 | NA | NA | NA | NA | NA | NA | 3 | 0.6667 |
| 7_43222232 | 2 | 1 | NA | NA | 1 | 1 | NA | NA | 3 | 1 | 3 | 1 | NA | NA | NA | NA | NA | NA | 3 | 1 |
| 7_43222805 | 2 | 1 | NA | NA | 1 | 1 | NA | NA | 3 | 1 | 2 | 1 | NA | NA | NA | NA | NA | NA | 2 | 1 |
| 7_43222936 | 1 | 1 | NA | NA | 1 | 0 | NA | NA | 2 | 0.5 | 3 | 0.3333 | NA | NA | NA | NA | NA | NA | 3 | 0.3333 |
| 7_43223190 | 2 | 1 | NA | NA | 1 | 0 | NA | NA | 3 | 0.6667 | 2 | 0.5 | NA | NA | NA | NA | NA | NA | 2 | 0.5 |
| 7_43223611 | 2 | 0.5 | NA | NA | 1 | 0 | NA | NA | 3 | 0.3333 | 3 | 0.3333 | NA | NA | NA | NA | NA | NA | 3 | 0.3333 |
| 7_43223935 | 2 | 1 | NA | NA | 1 | 1 | 1 | 1 | 4 | 1 | 3 | 1 | NA | NA | NA | NA | NA | NA | 3 | 1 |
| 7_43223981 | 1 | 1 | NA | NA | 1 | 1 | NA | NA | 2 | 1 | 3 | 1 | NA | NA | NA | NA | NA | NA | 3 | 1 |
| 7_43224132 | 2 | 0.5 | NA | NA | 2 | 1 | 1 | 1 | 5 | 0.8 | 3 | 1 | NA | NA | NA | NA | NA | NA | 3 | 1 |
| 7_43224283 | 2 | 1 | NA | NA | 2 | 0.5 | 1 | 0 | 5 | 0.6 | 2 | 0.5 | NA | NA | NA | NA | NA | NA | 2 | 0.5 |
| 7_43224367 | 2 | 1 | NA | NA | 2 | 1 | 1 | 1 | 5 | 1 | 3 | 1 | NA | NA | NA | NA | NA | NA | 3 | 1 |
| 7_43224509 | 2 | 0.5 | NA | NA | 2 | 1 | 1 | 1 | 5 | 0.8 | 3 | 1 | NA | NA | NA | NA | NA | NA | 3 | 1 |
| 7_43238794 | 3 | 0.6667 | 3 | 1 | NA | NA | NA | NA | 6 | 0.8333 | NA | NA | 2 | 1 | 2 | 0 | 1 | 0 | 5 | 0.4 |
| 7_43238825 | 3 | 0.3333 | 3 | 0.6667 | NA | NA | NA | NA | 6 | 0.5 | NA | NA | 2 | 1 | 2 | 0.5 | 1 | 0 | 5 | 0.6 |
| 7_43238888 | 3 | 0 | 2 | 0.5 | NA | NA | NA | NA | 5 | 0.2 | NA | NA | 2 | 0 | 2 | 0.5 | 1 | 0 | 5 | 0.2 |
| 7_43238907 | 1 | 1 | 3 | 0.6667 | NA | NA | NA | NA | 4 | 0.75 | NA | NA | 2 | 1 | 2 | 1 | 1 | 1 | 5 | 1 |
| 7_43238988 | 3 | 0.3333 | 3 | 0.6667 | 1 | 1 | NA | NA | 7 | 0.5714 | NA | NA | 1 | 1 | 1 | 1 | 1 | 0 | 3 | 0.6667 |
| 7_43239260 | 3 | 0 | 3 | 0.3333 | 2 | 0.5 | NA | NA | 8 | 0.25 | NA | NA | 2 | 0.5 | 2 | 0 | 1 | 0 | 5 | 0.2 |
| 7_43239314 | 3 | 0 | 2 | 0.5 | 2 | 0 | NA | NA | 7 | 0.1429 | NA | NA | 2 | 0.5 | 2 | 0 | 1 | 0 | 5 | 0.2 |
| 7_43239329 | 3 | 0 | 3 | 0.3333 | 2 | 0 | NA | NA | 8 | 0.125 | NA | NA | 2 | 0.5 | 2 | 0 | 1 | 0 | 5 | 0.2 |
| 7_43239363 | 3 | 1 | 3 | 0.6667 | 2 | 0.5 | NA | NA | 8 | 0.75 | NA | NA | 2 | 0 | 1 | 1 | 1 | 0 | 4 | 0.25 |
| 7_43239531 | 4 | 1 | 3 | 0.6667 | 1 | 1 | NA | NA | 8 | 0.875 | NA | NA | 2 | 0.5 | 2 | 1 | 1 | 1 | 5 | 0.8 |
| 7_43239615 | 2 | 1 | 2 | 1 | 1 | 1 | NA | NA | 5 | 1 | NA | NA | 2 | 1 | 2 | 1 | 1 | 1 | 5 | 1 |
| 7_43239655 | 3 | 0.6667 | 3 | 0 | 2 | 1 | NA | NA | 8 | 0.5 | NA | NA | 2 | 0.5 | 1 | 0 | 1 | 0 | 4 | 0.25 |
| 7_43239699 | 4 | 0.75 | 3 | 1 | 2 | 0.5 | NA | NA | 9 | 0.7778 | NA | NA | 2 | 1 | 1 | 1 | 1 | 1 | 4 | 1 |
| 7_43239760 | 4 | 1 | 2 | 1 | 1 | 1 | NA | NA | 7 | 1 | NA | NA | 2 | 1 | 2 | 0.5 | 1 | 1 | 5 | 0.8 |
| 7_43239867 | 3 | 0.3333 | 3 | 0.3333 | 1 | 0 | NA | NA | 7 | 0.2857 | NA | NA | 1 | 0 | 2 | 0.5 | 1 | 1 | 4 | 0.5 |
| 7_43239951 | 3 | 1 | 2 | 1 | 2 | 1 | NA | NA | 7 | 1 | NA | NA | 2 | 1 | 2 | 1 | 1 | 1 | 5 | 1 |
| 7_43240102 | 4 | 0.5 | 2 | 0.5 | 2 | 0.5 | NA | NA | 8 | 0.5 | NA | NA | 2 | 0.5 | 2 | 1 | 1 | 1 | 5 | 0.8 |
| 7_43240143 | 4 | 0.25 | 2 | 0.5 | 2 | 1 | NA | NA | 8 | 0.5 | NA | NA | 2 | 0.5 | 1 | 0 | 1 | 0 | 4 | 0.25 |
| 7_43240203 | 2 | 1 | 3 | 0.6667 | 2 | 0.5 | NA | NA | 7 | 0.7143 | NA | NA | 2 | 1 | 1 | 1 | NA | NA | 3 | 1 |
| 7_43240371 | 4 | 1 | 4 | 1 | 3 | 1 | NA | NA | 11 | 1 | NA | NA | 2 | 1 | 1 | 1 | 1 | 1 | 4 | 1 |
| 7_43240495 | 4 | 1 | 3 | 0.6667 | 1 | 1 | NA | NA | 8 | 0.875 | NA | NA | 2 | 1 | 2 | 1 | 1 | 1 | 5 | 1 |
| 7_43240574 | 4 | 0.5 | 4 | 0.25 | 3 | 0.3333 | NA | NA | 11 | 0.3636 | NA | NA | 2 | 0.5 | 2 | 0.5 | 1 | 1 | 5 | 0.6 |
| 7_43240598 | 2 | 0 | 3 | 0.3333 | 2 | 0 | NA | NA | 7 | 0.1429 | NA | NA | 1 | 1 | 1 | 0 | NA | NA | 2 | 0.5 |
| 7_43240758 | 4 | 0 | 5 | 0.2 | 3 | 1 | NA | NA | 12 | 0.3333 | NA | NA | 2 | 1 | 2 | 0.5 | 1 | 1 | 5 | 0.8 |
| 7_43240944 | 4 | 0 | 4 | 0 | 4 | 0.5 | NA | NA | 12 | 0.1667 | NA | NA | 2 | 0.5 | 2 | 0 | 2 | 0 | 6 | 0.1667 |
| 7_43240989 | 4 | 0.25 | 4 | 0.5 | 3 | 0.3333 | NA | NA | 11 | 0.3636 | NA | NA | 2 | 0 | 1 | 1 | 1 | 0 | 4 | 0.25 |
| 7_43241042 | 1 | 0 | 5 | 0.2 | 3 | 0 | NA | NA | 9 | 0.1111 | NA | NA | 2 | 0 | 2 | 0.5 | 2 | 0 | 6 | 0.1667 |
| 7_43241184 | 3 | 1 | 5 | 1 | 5 | 1 | NA | NA | 13 | 1 | NA | NA | 2 | 1 | 2 | 1 | 2 | 1 | 6 | 1 |
| 7_43324750 | 1 | 1 | NA | NA | NA | NA | NA | NA | 1 | 1 | NA | NA | NA | NA | 1 | 1 | NA | NA | 1 | 1 |
| 7_43326091 | 1 | 1 | 1 | 0 | NA | NA | NA | NA | 2 | 0.5 | NA | NA | 1 | 1 | 1 | 1 | 1 | 0 | 3 | 0.6667 |
| 7_43326130 | 1 | 1 | NA | NA | NA | NA | NA | NA | 1 | 1 | NA | NA | 1 | 0 | 1 | 1 | 1 | 1 | 3 | 0.6667 |
| 7_43326380 | 1 | 0 | 1 | 0 | NA | NA | 1 | 1 | 3 | 0.3333 | NA | NA | 1 | 0 | 1 | 1 | 1 | 0 | 3 | 0.3333 |
| 7_43326526 | 2 | 1 | 1 | 1 | NA | NA | 1 | 1 | 4 | 1 | NA | NA | 1 | 1 | 1 | 1 | 1 | 1 | 3 | 1 |
| 7_43326550 | 1 | 1 | 1 | 1 | NA | NA | NA | NA | 2 | 1 | NA | NA | NA | NA | 1 | 0 | 1 | 1 | 2 | 0.5 |
| 7_43326727 | 1 | 1 | 1 | 1 | NA | NA | NA | NA | 2 | 1 | NA | NA | 1 | 1 | 1 | 1 | 1 | 1 | 3 | 1 |
| 7_43327086 | 1 | 1 | 1 | 0 | NA | NA | 1 | 0 | 3 | 0.3333 | NA | NA | 2 | 0 | 1 | 1 | NA | NA | 3 | 0.3333 |
| 7_43327117 | 2 | 0.5 | 1 | 0 | NA | NA | NA | NA | 3 | 0.3333 | NA | NA | NA | NA | 1 | 0 | 1 | 1 | 2 | 0.5 |
| 7_43327301 | 1 | 0 | 1 | 1 | NA | NA | NA | NA | 2 | 0.5 | NA | NA | 1 | 0 | 1 | 0 | 1 | 1 | 3 | 0.3333 |
| 7_43900085 | 3 | 1 | 4 | 1 | 4 | 0.75 | 7 | 0.8571 | 18 | 0.8889 | 2 | 1 | 4 | 0.75 | 5 | 0.8 | 5 | 1 | 16 | 0.875 |
| 7_43900140 | 4 | 0.25 | 4 | 0 | 4 | 0 | 7 | 0 | 19 | 0.0526 | 3 | 0 | 5 | 0.2 | 5 | 0 | 5 | 0 | 18 | 0.0556 |
| 7_43900352 | 4 | 0.25 | 5 | 0 | 6 | 0.1667 | 10 | 0 | 25 | 0.08 | 4 | 0.25 | 5 | 0.2 | 4 | 0 | 9 | 0 | 22 | 0.0909 |
| 7_43900442 | 3 | 0.3333 | 4 | 0 | 5 | 0.2 | 7 | 0.1429 | 19 | 0.1579 | 4 | 0.25 | 4 | 0 | 5 | 0.2 | 6 | 0 | 19 | 0.1053 |

| | | | | | | | | | | | | | | | | | | | | |
|---|---|---|---|---|---|---|---|---|---|---|---|---|---|---|---|---|---|---|---|---|
| 7_4390052 | 4 | 0.25 | 4 | 0.5 | 6 | 0.8333 | 10 | 0.6 | 24 | 0.5833 | 3 | 0.3333 | 4 | 0.75 | 4 | 0.75 | 8 | 0.625 | 19 | 0.6316 |
| 7_4390066 | 2 | 1 | 6 | 0.8333 | 5 | 0.6 | 8 | 0.75 | 21 | 0.7619 | 3 | 0.3333 | 4 | 1 | 6 | 0.8333 | 7 | 0.7143 | 20 | 0.75 |
| 7_4390074 | 3 | 1 | 4 | 1 | 7 | 0.8571 | 9 | 0.8889 | 23 | 0.913 | 4 | 1 | 3 | 1 | 5 | 1 | 10 | 1 | 22 | 1 |
| 7_4390080 | 3 | 0.3333 | 2 | 1 | 6 | 0.3333 | 11 | 0 | 22 | 0.2273 | 2 | 0 | 6 | 0 | 5 | 0.2 | 9 | 0.2222 | 22 | 0.1364 |
| 7_4390082 | 2 | 1 | 6 | 0.8333 | 3 | 0.6667 | 5 | 0.8 | 16 | 0.8125 | 5 | 0.8 | 3 | 1 | 7 | 0.5714 | 8 | 0.875 | 23 | 0.7826 |
| 7_4390095 | 5 | 1 | 5 | 1 | 5 | 0.8 | 11 | 1 | 26 | 0.9615 | 5 | 1 | 5 | 1 | 8 | 1 | 11 | 0.9091 | 29 | 0.9655 |
| 7_4390097 | 5 | 0.6 | 5 | 0.6 | 6 | 1 | 10 | 0.8 | 26 | 0.7692 | 4 | 0.5 | 6 | 0.6667 | 6 | 0.8333 | 10 | 0.7 | 26 | 0.6923 |
| 7_4390107 | 6 | 1 | 6 | 0.8333 | 8 | 0.75 | 10 | 0.9 | 30 | 0.8667 | 4 | 0.75 | 7 | 0.8571 | 7 | 1 | 10 | 0.9 | 28 | 0.8929 |
| 7_4390129 | 8 | 0.25 | 7 | 0 | 7 | 0 | 12 | 0.25 | 34 | 0.1471 | 6 | 0.3333 | 6 | 0 | 9 | 0.2222 | 10 | 0 | 31 | 0.129 |
| 7_4395405 | NA | NA | 1 | 1 | 3 | 0.3333 | NA | NA | 4 | 0.5 | NA | NA | NA | NA | 1 | 1 | NA | NA | 1 | 1 |
| 7_4400315 | 3 | 1 | 8 | 0.875 | 2 | 1 | 2 | 1 | 15 | 0.9333 | 1 | 1 | NA | NA | 1 | 1 | 1 | 1 | 3 | 1 |
| 7_4400333 | 3 | 0.3333 | 8 | 0.875 | 1 | 1 | 2 | 1 | 14 | 0.7857 | 2 | 1 | NA | NA | 1 | 0 | 1 | 1 | 4 | 0.75 |
| 7_4400347 | 4 | 0.75 | 11 | 0.9091 | 1 | 1 | 2 | 0.5 | 18 | 0.8333 | 1 | 0 | NA | NA | 1 | 0 | 1 | 1 | 3 | 0.3333 |
| 7_4400352 | 3 | 1 | 14 | 1 | 3 | 1 | 2 | 0.5 | 22 | 0.9545 | 3 | 1 | NA | NA | 2 | 1 | 1 | 1 | 6 | 1 |
| 7_4400355 | 1 | 1 | 9 | 0.6667 | 3 | 0.6667 | 2 | 0.5 | 15 | 0.6667 | 1 | 0 | NA | NA | 2 | 1 | 1 | 0 | 4 | 0.5 |
| 7_4400359 | 4 | 1 | 13 | 1 | 1 | 1 | 2 | 1 | 20 | 1 | 3 | 0.6667 | NA | NA | 2 | 1 | 1 | 1 | 6 | 0.8333 |
| 7_4400360 | 4 | 0.25 | 11 | 0.1818 | 3 | 0.6667 | 2 | 0 | 20 | 0.25 | 3 | 0.3333 | NA | NA | 1 | 0 | NA | NA | 4 | 0.25 |
| 7_4400362 | 3 | 1 | 10 | 0.5 | 1 | 1 | 1 | 1 | 15 | 0.6667 | 1 | 1 | NA | NA | 1 | 0 | 1 | 1 | 3 | 0.6667 |
| 7_4400371 | 4 | 1 | 10 | 1 | 1 | 1 | 2 | 1 | 17 | 1 | 1 | 1 | NA | NA | 2 | 1 | 1 | 1 | 4 | 1 |
| 7_4400391 | 2 | 0.5 | 10 | 0.4 | 2 | 0.5 | 1 | 0 | 15 | 0.4 | 2 | 0.5 | NA | NA | 2 | 0.5 | 1 | 0 | 5 | 0.4 |
| 7_4400610 | 2 | 1 | 12 | 0.5 | 1 | 1 | 2 | 1 | 17 | 0.6471 | 2 | 0.5 | 4 | 0.5 | 1 | 1 | 2 | 0.5 | 9 | 0.5556 |
| 7_4400635 | 3 | 0.6667 | 11 | 0.5455 | 3 | 0.3333 | 3 | 0.3333 | 20 | 0.5 | 3 | 0.6667 | 4 | 0.5 | 2 | 0.5 | 2 | 1 | 11 | 0.6364 |
| 7_4400653 | 2 | 0.5 | 14 | 0.4286 | 2 | 0.5 | 3 | 0.6667 | 21 | 0.4762 | 2 | 0.5 | 4 | 0.75 | 2 | 0.5 | 2 | 1 | 10 | 0.7 |
| 7_4400657 | 4 | 0.75 | 16 | 0.875 | 2 | 1 | 3 | 1 | 25 | 0.88 | 2 | 1 | 4 | 1 | 3 | 0.6667 | 2 | 0.5 | 11 | 0.8182 |
| 7_4400663 | 2 | 0 | 10 | 0.6 | 2 | 1 | 3 | 1 | 17 | 0.6471 | 3 | 0.6667 | 3 | 0.6667 | 3 | 0.3333 | 2 | 0 | 11 | 0.4545 |
| 7_4400668 | 3 | 1 | 16 | 1 | 2 | 1 | 4 | 1 | 25 | 1 | 3 | 1 | 4 | 1 | 4 | 1 | 2 | 1 | 13 | 1 |
| 7_4400669 | 4 | 1 | 12 | 0.8333 | 3 | 1 | 3 | 1 | 22 | 0.9091 | 5 | 0.8 | 2 | 1 | 4 | 1 | 2 | 1 | 13 | 0.9231 |
| 7_4400672 | 2 | 1 | 6 | 1 | 2 | 1 | 3 | 1 | 13 | 1 | 2 | 1 | 2 | 1 | 2 | 1 | 2 | 1 | 8 | 1 |
| 7_4400679 | 4 | 0.75 | 15 | 1 | 3 | 1 | 4 | 1 | 26 | 0.9615 | 5 | 1 | 5 | 1 | 4 | 1 | 2 | 1 | 16 | 1 |
| 7_4400683 | 2 | 0.5 | 14 | 0.0714 | 1 | 0 | 1 | 0 | 18 | 0.1111 | 1 | 0 | 5 | 0.2 | 4 | 0.25 | 2 | 0 | 12 | 0.1667 |
| 7_4400687 | 2 | 1 | 16 | 0.9375 | 2 | 1 | 3 | 1 | 23 | 0.9565 | 3 | 1 | 5 | 0.8 | 4 | 1 | 2 | 1 | 14 | 0.9286 |
| 7_4400688 | 3 | 0.3333 | 14 | 0.5714 | 3 | 0.6667 | 2 | 0.5 | 22 | 0.5455 | 5 | 0.8 | 5 | 0.6 | 3 | 0.3333 | 2 | 0 | 15 | 0.5333 |
| 7_4400691 | 3 | 0.3333 | 17 | 0.4118 | 2 | 0.5 | 2 | 0 | 24 | 0.375 | 5 | 0.4 | 6 | 0.3333 | 3 | 1 | 2 | 0 | 16 | 0.4375 |
| 7_4400693 | 4 | 0.5 | 16 | 0.0625 | 3 | 0 | 4 | 0 | 27 | 0.1111 | 6 | 0 | 6 | 0.1667 | 3 | 0 | 3 | 0 | 18 | 0.0556 |
| 7_4400694 | 3 | 0 | 12 | 0.1667 | 2 | 0.5 | 2 | 0.5 | 19 | 0.2105 | 2 | 0 | 3 | 0.3333 | 4 | 0 | 2 | 0 | 11 | 0.0909 |
| 7_4400698 | 2 | 1 | 16 | 0.375 | 1 | 1 | 3 | 1 | 22 | 0.5455 | 3 | 1 | 4 | 1 | 2 | 1 | 2 | 0.5 | 11 | 0.9091 |
| 7_4400702 | 4 | 0.25 | 17 | 0 | 3 | 0.3333 | 3 | 0.3333 | 27 | 0.1111 | 5 | 0.2 | 6 | 0.1667 | 4 | 0 | 2 | 0 | 17 | 0.1176 |
| 7_4400930 | 19 | 0.2105 | 33 | 0.1212 | 13 | 0.2308 | 7 | 0 | 72 | 0.1528 | 20 | 0.2 | 18 | 0.3889 | 9 | 0.3333 | 18 | 0.1667 | 65 | 0.2615 |
| 7_4404002 | 6 | 0 | 2 | 0 | 7 | 0.4286 | 2 | 0.5 | 17 | 0.2353 | 3 | 0 | 4 | 0.25 | 5 | 0.2 | 4 | 0 | 16 | 0.125 |
| 7_4404006 | 5 | 0.2 | 2 | 0.5 | 2 | 0 | 3 | 1 | 12 | 0.4167 | 2 | 1 | 2 | 1 | 6 | 0.8333 | 3 | 0 | 13 | 0.6923 |
| 7_4404025 | 2 | 0 | 2 | 0 | 6 | 0.1667 | 2 | 0.5 | 12 | 0.1667 | 2 | 0.5 | 5 | 0.2 | 2 | 0.5 | 3 | 0 | 12 | 0.25 |
| 7_4404026 | 4 | 0.75 | 2 | 1 | 4 | 1 | 1 | 1 | 11 | 0.9091 | 4 | 1 | 4 | 1 | 2 | 0.5 | 2 | 0.5 | 12 | 0.8333 |
| 7_4404198 | 7 | 0.1429 | 3 | 0 | 6 | 0.1667 | 2 | 0.5 | 18 | 0.1667 | 3 | 0.3333 | 8 | 0.25 | 7 | 0.2857 | 4 | 0.75 | 22 | 0.3636 |
| 7_4404212 | 8 | 0 | 1 | 1 | 10 | 0.1 | 3 | 0 | 22 | 0.0909 | 3 | 0.6667 | 6 | 0.5 | 6 | 0.1667 | 3 | 0 | 18 | 0.3333 |
| 7_4404493 | 9 | 0.2222 | 3 | 0 | 6 | 0 | 6 | 0.3333 | 24 | 0.1667 | 2 | 0.5 | 5 | 0 | 8 | 0 | 4 | 0.25 | 19 | 0.1053 |
| 7_4409150 | 1 | 0 | 4 | 0 | 2 | 0.5 | 2 | 0 | 9 | 0.1111 | 2 | 0.5 | 1 | 0 | 1 | 0 | NA | NA | 4 | 0.25 |
| 7_4411055 | 9 | 0.2222 | 19 | 0.3158 | 10 | 0.3 | 10 | 0.5 | 48 | 0.3333 | 9 | 0.5556 | 7 | 0.2857 | 20 | 0.25 | 10 | 0.6 | 46 | 0.3913 |
| 7_4411075 | 11 | 0 | 25 | 0.2 | 11 | 0.2727 | 11 | 0.0909 | 58 | 0.1552 | 13 | 0.0769 | 13 | 0.1538 | 23 | 0.1739 | 15 | 0 | 64 | 0.1094 |
| 7_4411079 | 11 | 1 | 26 | 0.9615 | 13 | 0.9231 | 10 | 0.9 | 60 | 0.95 | 12 | 1 | 12 | 1 | 22 | 0.9091 | 14 | 0.9286 | 60 | 0.95 |
| 7_4411086 | 10 | 0.6 | 23 | 0.6087 | 12 | 0.75 | 8 | 0.625 | 53 | 0.6415 | 13 | 0.6154 | 13 | 0.7692 | 21 | 0.7143 | 13 | 0.6154 | 60 | 0.6833 |
| 7_4411106 | 10 | 0 | 28 | 0.0357 | 12 | 0.1667 | 10 | 0.1 | 60 | 0.0667 | 8 | 0.125 | 11 | 0.0909 | 16 | 0 | 13 | 0.0769 | 48 | 0.0625 |
| 7_4411108 | 5 | 0.8 | 18 | 1 | 13 | 0.8462 | 10 | 1 | 46 | 0.9348 | 8 | 1 | 9 | 1 | 15 | 0.9333 | 6 | 0.8333 | 38 | 0.9474 |
| 7_4411112 | 10 | 0.1 | 28 | 0.4643 | 14 | 0.2143 | 12 | 0.5 | 64 | 0.3594 | 12 | 0.3333 | 12 | 0.0833 | 23 | 0.2174 | 15 | 0.1333 | 62 | 0.1935 |
| 7_4411119 | 11 | 0.1818 | 30 | 0.1 | 17 | 0.0588 | 13 | 0.1538 | 71 | 0.1127 | 15 | 0.2 | 13 | 0.1538 | 25 | 0.16 | 17 | 0.0588 | 70 | 0.1429 |
| 7_4411125 | 9 | 0.4444 | 27 | 0.7037 | 16 | 0.6875 | 13 | 0.6154 | 65 | 0.6462 | 11 | 0.5455 | 9 | 0.5556 | 24 | 0.4167 | 14 | 0.3571 | 58 | 0.4483 |
| 7_4411142 | 15 | 1 | 30 | 1 | 21 | 0.8571 | 15 | 0.8667 | 81 | 0.9383 | 13 | 0.6923 | 16 | 1 | 25 | 0.88 | 18 | 0.8333 | 72 | 0.8611 |
| 7_4411145 | 15 | 0.0667 | 32 | 0.1562 | 19 | 0.1579 | 13 | 0.0769 | 79 | 0.1266 | 13 | 0.1538 | 15 | 0.0667 | 25 | 0.2 | 18 | 0.0556 | 71 | 0.1268 |
| 7_4411145 | 6 | 0.3333 | 28 | 0.5357 | 16 | 0.375 | 11 | 0.8182 | 61 | 0.5246 | 13 | 0.6923 | 12 | 0.5 | 22 | 0.5 | 12 | 0.5833 | 59 | 0.5593 |
| 7_4411150 | 14 | 0.5714 | 33 | 0.697 | 21 | 0.6667 | 13 | 0.6154 | 81 | 0.6543 | 14 | 0.7857 | 14 | 0.8571 | 21 | 0.6667 | 17 | 0.6471 | 66 | 0.7273 |
| 7_4411182 | 19 | 0.8421 | 34 | 0.9118 | 22 | 0.8182 | 20 | 0.9 | 95 | 0.8737 | 19 | 0.8947 | 21 | 0.9524 | 35 | 0.9429 | 23 | 0.9565 | 98 | 0.9388 |
| 7_4411206 | 15 | 0.1333 | 22 | 0.1364 | 15 | 0.2 | 11 | 0.1818 | 63 | 0.1587 | 12 | 0.1667 | 11 | 0 | 20 | 0.3 | 14 | 0.0714 | 57 | 0.1579 |
| 7_4411220 | 22 | 0.6818 | 23 | 0.6957 | 16 | 0.625 | 11 | 0.8182 | 72 | 0.6944 | 17 | 0.7059 | 15 | 0.8 | 28 | 0.75 | 17 | 0.6471 | 77 | 0.7273 |
| 7_4411224 | 29 | 1 | 38 | 0.8684 | 24 | 0.9583 | 23 | 0.8696 | 114 | 0.9211 | 26 | 0.8077 | 25 | 0.96 | 40 | 0.95 | 25 | 0.96 | 116 | 0.9224 |
| 7_4411235 | 25 | 0.92 | 29 | 0.931 | 19 | 0.9474 | 17 | 0.7647 | 90 | 0.9 | 21 | 0.9524 | 20 | 0.8 | 25 | 0.96 | 19 | 0.9474 | 85 | 0.9176 |
| 7_4411236 | 26 | 0.1538 | 29 | 0.2414 | 22 | 0.4091 | 21 | 0.1429 | 98 | 0.2347 | 26 | 0.2692 | 24 | 0.25 | 39 | 0.3846 | 21 | 0.3333 | 110 | 0.3182 |

| | | | | | | | | | | | | | | | | | | | | |
|---|---|---|---|---|---|---|---|---|---|---|---|---|---|---|---|---|---|---|---|---|
| 7_44112587 | 35 | 0.5714 | 43 | 0.3488 | 29 | 0.4483 | 26 | 0.5385 | 133 | 0.4662 | 32 | 0.3438 | 30 | 0.4333 | 41 | 0.6585 | 30 | 0.4333 | 133 | 0.4812 |
| 7_44112682 | 35 | 1 | 41 | 0.878 | 26 | 0.9615 | 24 | 1 | 126 | 0.9524 | 28 | 0.9643 | 31 | 0.9355 | 38 | 0.9211 | 26 | 0.9231 | 123 | 0.935 |
| 7_44118684 | NA | NA | 1 | 1 | 2 | 0 | 1 | 0 | 4 | 0.25 | 1 | 0 | 1 | 1 | NA | NA | 1 | 0 | 3 | 0.3333 |
| 7_44120191 | 2 | 0.5 | 2 | 0 | NA | NA | NA | NA | 4 | 0.25 | 2 | 0 | NA | NA | 2 | 0.5 | NA | NA | 4 | 0.25 |
| 7_44120213 | 2 | 1 | 1 | 1 | NA | NA | NA | NA | 3 | 1 | 2 | 0.5 | NA | NA | 2 | 0 | NA | NA | 4 | 0.25 |
| 7_44120500 | 2 | 0 | 2 | 0 | 1 | 1 | NA | NA | 5 | 0.2 | 3 | 0 | NA | NA | 2 | 0 | 1 | 1 | 6 | 0.1667 |
| 7_44120683 | 5 | 0.2 | 4 | 0.25 | 6 | 0.3333 | 5 | 0 | 20 | 0.2 | 4 | 0 | 6 | 0 | 4 | 0 | 12 | 0.1667 | 26 | 0.0769 |
| 7_44140004 | 31 | 0.0645 | 38 | 0 | 48 | 0.1042 | 42 | 0.0238 | 159 | 0.0503 | 23 | 0.087 | 41 | 0.0488 | 39 | 0.0256 | 33 | 0.1515 | 136 | 0.0735 |
| 7_44146193 | 1 | 1 | NA | NA | NA | NA | NA | NA | 1 | 1 | NA | NA | 1 | 1 | 1 | 1 | NA | NA | 2 | 1 |
| 7_44150433 | NA | NA | NA | NA | 1 | 0 | 1 | 1 | 2 | 0.5 | NA | NA | NA | NA | 1 | 1 | NA | NA | 1 | 1 |
| 7_44199331 | 8 | 0.5 | 16 | 0.125 | 10 | 0.3 | 5 | 0.4 | 39 | 0.2821 | 8 | 0.25 | 9 | 0.2222 | 11 | 0.2727 | 5 | 0.2 | 33 | 0.2424 |
| 7_44199361 | 9 | 0.4444 | 16 | 0.4375 | 10 | 0.4 | 7 | 0.5714 | 42 | 0.4524 | 7 | 0.7143 | 9 | 0.4444 | 9 | 0.7778 | 6 | 0.3333 | 31 | 0.5806 |
| 7_44200204 | 8 | 0.125 | 17 | 0.1176 | 8 | 0 | 5 | 0 | 38 | 0.0789 | 6 | 0 | 8 | 0.125 | 11 | 0.0909 | 3 | 0 | 28 | 0.0714 |
| 7_44200821 | 6 | 0.1667 | 15 | 0.2667 | 7 | 0 | 7 | 0.1429 | 35 | 0.1714 | 5 | 0.2 | 8 | 0.25 | 10 | 0.4 | 3 | 0.3333 | 26 | 0.3077 |
| 7_44202135 | 2 | 0 | 8 | 0 | 3 | 0.3333 | 4 | 0 | 17 | 0.0588 | NA | NA | 4 | 0 | 4 | 0.25 | 2 | 0 | 10 | 0.1 |
| 7_44233693 | NA | NA | NA | NA | NA | NA | 1 | 1 | 1 | 1 | 1 | 0 | NA | NA | NA | NA | 1 | 1 | 2 | 0.5 |
| 7_44233755 | NA | NA | NA | NA | NA | NA | 1 | 1 | 1 | 1 | 1 | 0 | NA | NA | NA | NA | 1 | 1 | 2 | 0.5 |
| 7_44233768 | NA | NA | NA | NA | NA | NA | 1 | 1 | 1 | 1 | 1 | 1 | NA | NA | NA | NA | 1 | 1 | 2 | 1 |
| 7_44240649 | 2 | 0 | 2 | 0.5 | NA | NA | NA | NA | 4 | 0.25 | NA | NA | 1 | 0 | 4 | 0.5 | 1 | 1 | 6 | 0.5 |
| 7_44240914 | 2 | 0.5 | 2 | 1 | NA | NA | NA | NA | 4 | 0.75 | NA | NA | 1 | 0 | 4 | 1 | 2 | 1 | 7 | 0.8571 |
| 7_44240932 | 2 | 0 | 1 | 1 | NA | NA | 1 | 0 | 4 | 0.25 | NA | NA | 1 | 0 | 4 | 0.5 | 1 | 0 | 6 | 0.3333 |
| 7_44240981 | 2 | 0 | 1 | 0 | NA | NA | 1 | 1 | 4 | 0.25 | NA | NA | 1 | 0 | 3 | 0.3333 | NA | NA | 4 | 0.25 |
| 7_44245180 | 2 | 1 | 2 | 1 | NA | NA | NA | NA | 4 | 1 | NA | NA | 1 | 0 | 1 | 1 | 3 | 0.3333 | 5 | 0.4 |
| 7_44245237 | 2 | 1 | 2 | 1 | NA | NA | 1 | 0 | 5 | 0.8 | NA | NA | 2 | 0.5 | 3 | 0 | 3 | 0.3333 | 8 | 0.25 |
| 7_44245562 | 5 | 0.6 | 2 | 0.5 | NA | NA | 1 | 1 | 8 | 0.625 | NA | NA | 2 | 1 | 3 | 0.3333 | 4 | 0.5 | 9 | 0.5556 |
| 7_44419031 | 1 | 1 | NA | NA | NA | NA | 6 | 0 | 7 | 0.1429 | 6 | 0 | 2 | 0.5 | 4 | 0 | 2 | 0 | 14 | 0.0714 |
| 7_44462494 | 2 | 0 | 6 | 0.1667 | 3 | 0 | 4 | 0 | 15 | 0.0667 | 2 | 0 | 5 | 0 | 3 | 0.3333 | 2 | 0 | 12 | 0.0833 |
| 7_44462644 | 3 | 0.3333 | 8 | 0.125 | 5 | 0.2 | 4 | 0.5 | 20 | 0.25 | 2 | 0 | 7 | 0.4286 | 3 | 0 | 2 | 0.5 | 14 | 0.2857 |
| 7_44462921 | 2 | 0.5 | 7 | 0.1429 | 3 | 0 | 4 | 0 | 16 | 0.125 | 1 | 0 | 5 | 0.2 | 3 | 0 | 2 | 0 | 11 | 0.0909 |
| 7_44464514 | 2 | 0.5 | 7 | 0.1429 | 3 | 0 | 3 | 0.3333 | 15 | 0.2 | 2 | 0 | 5 | 0.2 | 3 | 0 | 1 | 0 | 11 | 0.0909 |
| 7_44464567 | 2 | 0.5 | 6 | 0 | 1 | 0 | NA | NA | 9 | 0.1111 | 2 | 0.5 | 3 | 0.3333 | 2 | 0 | 1 | 1 | 8 | 0.375 |
| 7_44478044 | 1 | 0 | 1 | 0 | 3 | 0.6667 | 2 | 0.5 | 7 | 0.4286 | NA | NA | 2 | 0 | 2 | 1 | 2 | 0.5 | 6 | 0.5 |
| 7_44478053 | 2 | 1 | 2 | 1 | 4 | 1 | 1 | 1 | 9 | 1 | NA | NA | NA | NA | 2 | 1 | 1 | 1 | 3 | 1 |
| 7_44478128 | NA | NA | 2 | 1 | 4 | 0.75 | 2 | 1 | 8 | 0.875 | NA | NA | 2 | 0.5 | 1 | 0 | NA | NA | 3 | 0.3333 |
| 7_44478392 | 1 | 0 | 2 | 0 | 4 | 0.75 | 2 | 1 | 9 | 0.5556 | NA | NA | 2 | 0.5 | 2 | 0.5 | 1 | 0 | 5 | 0.4 |
| 7_44478581 | 2 | 0 | 3 | 0.3333 | 2 | 0 | 2 | 0 | 9 | 0.1111 | NA | NA | 3 | 0 | 2 | 0.5 | 3 | 0 | 8 | 0.125 |
| 7_44478749 | 2 | 0.5 | 4 | 0.75 | 3 | 1 | 2 | 1 | 11 | 0.8182 | 1 | 1 | 3 | 0.6667 | 2 | 1 | 2 | 1 | 8 | 0.875 |
| 7_44478962 | 1 | 1 | 4 | 1 | 5 | 1 | 2 | 1 | 12 | 1 | 1 | 1 | 3 | 0.6667 | 3 | 1 | 2 | 1 | 9 | 0.8889 |
| 7_44479083 | 2 | 0 | 4 | 0.25 | 5 | 0.4 | 3 | 0.3333 | 14 | 0.2857 | 1 | 0 | 5 | 0.2 | 3 | 0 | 3 | 0 | 12 | 0.0833 |
| 7_44479179 | 2 | 0.5 | 6 | 0.8333 | 6 | 1 | 3 | 1 | 17 | 0.8824 | 1 | 1 | 4 | 0.75 | 3 | 0.6667 | 3 | 1 | 11 | 0.8182 |
| 7_44479314 | 1 | 0 | 6 | 0 | 6 | 0 | 3 | 0.6667 | 16 | 0.125 | NA | NA | 5 | 0.2 | 3 | 0 | 2 | 0 | 10 | 0.1 |
| 7_44479347 | 2 | 0 | 5 | 0.2 | 6 | 0.1667 | 3 | 0.6667 | 16 | 0.25 | 1 | 0 | 5 | 0.2 | 4 | 0.5 | 2 | 0.5 | 12 | 0.3333 |
| 7_44479374 | 1 | 0 | 5 | 0.2 | 5 | 0 | 3 | 0 | 14 | 0.0714 | 1 | 0 | 2 | 0.5 | 3 | 0.3333 | 3 | 0 | 9 | 0.2222 |
| 7_44479446 | 2 | 1 | 5 | 1 | 4 | 1 | 3 | 1 | 14 | 1 | 1 | 1 | 3 | 1 | 3 | 1 | 3 | 1 | 10 | 1 |
| 7_44479464 | 1 | 0 | 4 | 0.25 | 5 | 0.6 | 2 | 1 | 12 | 0.5 | 2 | 1 | 4 | 0.5 | 4 | 0.75 | 2 | 1 | 12 | 0.75 |
| 7_44479534 | 1 | 0 | 4 | 0.25 | 2 | 0.5 | 2 | 0.5 | 9 | 0.3333 | NA | NA | 2 | 0 | 2 | 0.5 | 1 | 0 | 5 | 0.2 |
| 7_44479563 | 1 | 0 | 5 | 0.8 | 3 | 1 | 3 | 0 | 12 | 0.5833 | 2 | 1 | 4 | 0.75 | 3 | 0 | 1 | 1 | 10 | 0.6 |
| 7_44479724 | 2 | 0.5 | 6 | 0 | 4 | 0 | 3 | 0 | 15 | 0.0667 | 2 | 0 | 5 | 0 | 4 | 0 | 3 | 0.3333 | 14 | 0.0714 |
| 7_44479744 | 1 | 1 | 6 | 1 | 5 | 1 | 2 | 1 | 14 | 1 | 2 | 1 | 3 | 1 | 3 | 0.6667 | 3 | 1 | 11 | 0.9091 |
| 7_44479825 | 1 | 0 | 3 | 0.6667 | 4 | 1 | 2 | 1 | 10 | 0.8 | NA | NA | 5 | 0.6 | 3 | 1 | 1 | 0 | 9 | 0.6667 |
| 7_44479884 | 2 | 1 | 7 | 1 | 6 | 1 | 3 | 1 | 18 | 1 | 1 | 1 | 6 | 1 | 2 | 1 | 2 | 1 | 11 | 1 |
| 7_44479901 | 2 | 0 | 8 | 0 | 5 | 0 | 3 | 0.3333 | 18 | 0.0556 | 1 | 0 | 6 | 0 | 4 | 0 | 2 | 0.5 | 13 | 0.0769 |
| 7_44480036 | 2 | 1 | 5 | 1 | 5 | 0.8 | 2 | 1 | 14 | 0.9286 | 2 | 1 | 2 | 1 | 3 | 1 | 1 | 1 | 8 | 1 |
| 7_44480040 | 3 | 0.3333 | 8 | 0.5 | 5 | 0.2 | 2 | 0 | 18 | 0.3333 | 1 | 0 | 4 | 0.75 | 4 | 0.5 | 3 | 0.6667 | 12 | 0.5833 |
| 7_44490435 | NA | NA | 1 | 0 | 1 | 1 | 1 | 1 | 3 | 0.6667 | NA | NA | 2 | 1 | 2 | 1 | 1 | 0 | 5 | 0.8 |
| 7_44490456 | 4 | 0 | 5 | 0.6 | 3 | 0 | 8 | 0.125 | 20 | 0.2 | 3 | 0.3333 | 7 | 0.2857 | 6 | 0.3333 | 2 | 1 | 18 | 0.3889 |
| 7_44490565 | 6 | 0 | 6 | 0.1667 | 7 | 0.1429 | 8 | 0 | 27 | 0.0741 | 4 | 0 | 10 | 0.3 | 11 | 0.1818 | 6 | 0.5 | 31 | 0.2581 |
| 7_44494036 | NA | NA | 1 | 1 | NA | NA | NA | NA | 1 | 1 | NA | NA | NA | NA | 1 | 1 | NA | NA | 1 | 1 |
| 7_44499530 | NA | NA | 3 | 0.6667 | 3 | 0.3333 | NA | NA | 6 | 0.5 | NA | NA | NA | NA | 1 | 1 | NA | NA | 1 | 1 |
| 7_44502025 | 5 | 0 | 8 | 0.125 | 4 | 0 | 3 | 0.3333 | 20 | 0.1 | 6 | 0.3333 | 5 | 0.2 | 8 | 0.125 | 6 | 0 | 25 | 0.16 |
| 7_44504164 | 8 | 0.125 | 6 | 0.1667 | 7 | 0 | 1 | 0 | 22 | 0.0909 | 7 | 0 | 8 | 0 | 9 | 0.1111 | 6 | 0.1667 | 30 | 0.0667 |
| 7_44504215 | 3 | 0.6667 | 7 | 0.2857 | 6 | 0.5 | 2 | 0.5 | 18 | 0.4444 | 5 | 0.6 | 6 | 0.5 | 5 | 0.2 | 8 | 0.75 | 24 | 0.5417 |
| 7_44504584 | 8 | 0.125 | 9 | 0.1111 | 7 | 0 | 3 | 0 | 27 | 0.0741 | 11 | 0.0909 | 5 | 0 | 8 | 0 | 7 | 0.1429 | 31 | 0.0645 |
| 7_44513223 | NA | NA | NA | NA | NA | NA | 2 | 1 | 2 | 1 | NA | NA | 1 | 1 | NA | NA | NA | NA | 1 | 1 |
| 7_44513301 | NA | NA | NA | NA | NA | NA | 1 | 1 | 1 | 1 | NA | NA | 1 | 1 | NA | NA | NA | NA | 1 | 1 |

| | | | | | | | | | | | | | | | | | | | | |
|---|---|---|---|---|---|---|---|---|---|---|---|---|---|---|---|---|---|---|---|---|
| 7_4451334 | NA | NA | NA | NA | NA | NA | 2 | 0.5 | 2 | 0.5 | NA | NA | 1 | 1 | NA | NA | NA | NA | 1 | 1 |
| 7_4451334 | NA | NA | NA | NA | NA | NA | 2 | 0.5 | 2 | 0.5 | NA | NA | 1 | 0 | NA | NA | NA | NA | 1 | 1 |
| 7_4451352 | NA | NA | NA | NA | NA | NA | 2 | 1 | 2 | 1 | NA | NA | 1 | 1 | NA | NA | NA | NA | 1 | 1 |
| 7_4451360 | NA | NA | NA | NA | NA | NA | 2 | 1 | 2 | 1 | NA | NA | 1 | 1 | NA | NA | NA | NA | 1 | 1 |
| 7_4451386 | NA | NA | NA | NA | NA | NA | 1 | 1 | 1 | 1 | NA | NA | 1 | 1 | NA | NA | NA | NA | 1 | 1 |
| 7_4451582 | NA | NA | 3 | 0.3333 | 1 | 0 | NA | NA | 4 | 0.25 | 1 | 0 | 1 | 0 | 2 | 0 | 2 | 0.5 | 6 | 0.1667 |
| 7_4452626 | 1 | 0 | 1 | 1 | 2 | 1 | 4 | 0 | 8 | 0.375 | 2 | 0.5 | 1 | 0 | 3 | 0.6667 | 3 | 0 | 9 | 0.3333 |
| 7_4452646 | 2 | 0.5 | 1 | 0 | NA | NA | NA | NA | 3 | 0.3333 | 1 | 0 | 3 | 1 | NA | NA | 4 | 0 | 8 | 0.375 |
| 7_4452649 | 3 | 0 | 1 | 1 | 3 | 0 | 3 | 0 | 10 | 0.1 | 2 | 0 | 2 | 0.5 | 3 | 0.3333 | 4 | 0 | 11 | 0.1818 |
| 7_4452676 | 2 | 1 | 1 | 1 | 3 | 0.6667 | 2 | 1 | 8 | 0.875 | 1 | 1 | 3 | 1 | 2 | 1 | 3 | 1 | 9 | 1 |
| 7_4452676 | 4 | 1 | 1 | 1 | 2 | 0.5 | 2 | 0 | 9 | 0.6667 | 1 | 0 | 3 | 0.3333 | 2 | 0 | 4 | 0.25 | 10 | 0.2 |
| 7_4452680 | 2 | 1 | 1 | 0 | 2 | 0 | 2 | 0.5 | 7 | 0.4286 | 1 | 0 | 2 | 0.5 | 1 | 1 | 3 | 0.6667 | 7 | 0.5714 |
| 7_4452682 | 3 | 0 | 1 | 0 | 1 | 0 | 3 | 0.3333 | 8 | 0.125 | 1 | 1 | 3 | 0 | 2 | 0 | 3 | 0 | 9 | 0.1111 |
| 7_4452686 | 2 | 1 | 1 | 0 | 2 | 0.5 | 3 | 0 | 8 | 0.375 | 1 | 0 | 3 | 0.6667 | 2 | 0 | 4 | 0 | 10 | 0.2 |
| 7_4452689 | 3 | 0.3333 | 2 | 1 | 2 | 0 | 3 | 0 | 10 | 0.3 | 2 | 0.5 | 2 | 0.5 | 2 | 0 | 4 | 0 | 10 | 0.2 |
| 7_4452690 | 1 | 1 | 1 | 0 | 2 | 0.5 | 3 | 0.6667 | 7 | 0.5714 | 1 | 0 | 3 | 1 | NA | NA | 4 | 0.75 | 8 | 0.75 |
| 7_4452695 | 3 | 1 | 1 | 1 | 3 | 1 | 3 | 0.6667 | 10 | 0.9 | 2 | 1 | 3 | 0.6667 | 2 | 1 | 3 | 1 | 10 | 0.9 |
| 7_4452697 | 3 | 0.6667 | 2 | 0.5 | 2 | 0.5 | 3 | 0.3333 | 10 | 0.5 | 2 | 0.5 | 3 | 0.3333 | 2 | 0 | 4 | 0.25 | 11 | 0.2727 |
| 7_4452699 | 3 | 1 | 1 | 1 | 3 | 1 | 3 | 0.6667 | 10 | 0.9 | 1 | 1 | 2 | 0.5 | 1 | 0 | 4 | 0.5 | 8 | 0.5 |
| 7_4452709 | 3 | 0.3333 | 2 | 0 | 3 | 0 | 3 | 0 | 11 | 0.0909 | 2 | 0 | 2 | 0 | 2 | 0 | 4 | 0.25 | 10 | 0.1 |
| 7_4452721 | 2 | 0.5 | 1 | 0 | 3 | 1 | 3 | 1 | 9 | 0.7778 | 2 | 1 | 3 | 1 | 2 | 0.5 | 4 | 0.75 | 11 | 0.8182 |
| 7_4452732 | 3 | 0.3333 | 2 | 0.5 | 2 | 0 | 2 | 0 | 9 | 0.2222 | 2 | 0 | 3 | 0.3333 | 2 | 0.5 | 4 | 0.25 | 11 | 0.2727 |
| 7_4452749 | 3 | 0.6667 | 1 | 0 | 3 | 0.3333 | 2 | 0 | 9 | 0.3333 | 1 | 0 | 2 | 0.5 | 2 | 0 | 3 | 0 | 8 | 0.125 |
| 7_4452765 | 3 | 0 | 1 | 0 | 3 | 0.3333 | 2 | 0 | 9 | 0.1111 | NA | NA | 3 | 0 | 2 | 0.5 | 4 | 0.25 | 9 | 0.2222 |
| 7_4452885 | 10 | 0 | 9 | 0.1111 | 7 | 0 | 11 | 0.0909 | 37 | 0.0541 | 9 | 0.1111 | 8 | 0.125 | 10 | 0 | 6 | 0 | 33 | 0.0606 |
| 7_4455009 | 11 | 0.0909 | 24 | 0.0417 | 14 | 0.0714 | 12 | 0.0833 | 61 | 0.0656 | 23 | 0.0435 | 15 | 0 | 25 | 0.12 | 9 | 0 | 72 | 0.0556 |
| 7_4455015 | 24 | 0.0833 | 51 | 0.0784 | 31 | 0.0968 | 22 | 0.0909 | 128 | 0.0859 | 29 | 0.2414 | 29 | 0.069 | 37 | 0.0811 | 16 | 0.125 | 111 | 0.1261 |
| 7_4460896 | 7 | 0.2857 | 10 | 0.2 | 11 | 0.3636 | 4 | 0 | 32 | 0.25 | 6 | 0.6667 | 18 | 0.3889 | 15 | 0.4 | 6 | 0 | 45 | 0.3778 |
| 7_4462413 | 3 | 0.3333 | 3 | 0.3333 | 4 | 0.25 | 2 | 0 | 12 | 0.25 | 2 | 0 | 4 | 0.25 | 7 | 0 | NA | NA | 13 | 0.0769 |
| 7_4462609 | 2 | 0 | 1 | 0 | 3 | 0 | 1 | 1 | 7 | 0.1429 | 1 | 0 | NA | NA | 1 | 1 | NA | NA | 2 | 0.5 |
| 7_4464944 | 1 | 0 | 5 | 0.2 | NA | NA | 2 | 0 | 8 | 0.125 | 1 | 0 | 3 | 0 | 2 | 0.5 | 3 | 0 | 9 | 0.1111 |
| 7_4465022 | 2 | 1 | 4 | 0 | NA | NA | 3 | 0.3333 | 9 | 0.3333 | 2 | 0 | 4 | 0 | 2 | 0 | 3 | 0.6667 | 11 | 0.1818 |
| 7_4465166 | 1 | 1 | 5 | 0.2 | NA | NA | 2 | 0.5 | 8 | 0.375 | 2 | 0 | 3 | 0.6667 | 1 | 0 | 2 | 0 | 8 | 0.25 |
| 7_4465172 | 2 | 1 | 4 | 0 | 2 | 0 | 2 | 0 | 10 | 0.2 | 2 | 0 | 5 | 0.4 | 3 | 0.3333 | 4 | 0.25 | 14 | 0.2857 |
| 7_4465173 | 1 | 0 | 5 | 0.4 | 2 | 0.5 | 3 | 0 | 11 | 0.2727 | 2 | 0.5 | 5 | 0.2 | 2 | 0 | 4 | 0 | 13 | 0.1538 |
| 7_4465174 | 1 | 1 | 5 | 0.2 | 1 | 1 | 2 | 1 | 9 | 0.5556 | 2 | 0.5 | 3 | 0 | 1 | 0 | 4 | 0.5 | 10 | 0.3 |
| 7_4465256 | 17 | 0.2353 | 13 | 0.3077 | 13 | 0.3846 | 18 | 0.4444 | 61 | 0.3443 | 11 | 0.4545 | 10 | 0.2 | 16 | 0.5 | 22 | 0.5455 | 59 | 0.4576 |
| 7_4465575 | 13 | 0.8462 | 16 | 0.5625 | 10 | 0.7 | 17 | 0.8235 | 56 | 0.7321 | 6 | 1 | 6 | 0.8333 | 15 | 0.6 | 19 | 0.7895 | 46 | 0.7609 |
| 7_4465584 | 15 | 0.1333 | 15 | 0 | 12 | 0 | 20 | 0.2 | 62 | 0.0968 | 8 | 0.125 | 7 | 0.1429 | 15 | 0.0667 | 22 | 0.2273 | 52 | 0.1538 |
| 7_4465599 | 11 | 0.3636 | 13 | 0.4615 | 9 | 0.5556 | 20 | 0.55 | 53 | 0.4906 | 9 | 0.7778 | 7 | 0.5714 | 13 | 0.5385 | 19 | 0.3684 | 48 | 0.5208 |
| 7_4465605 | 12 | 0.1667 | 13 | 0.3077 | 7 | 0.2857 | 16 | 0.375 | 48 | 0.2917 | 9 | 0.2222 | 7 | 0.4286 | 12 | 0.3333 | 17 | 0.2941 | 45 | 0.3111 |
| 7_4465606 | 12 | 0.25 | 11 | 0.0909 | 7 | 0.4286 | 18 | 0.2778 | 48 | 0.25 | 9 | 0.1111 | 7 | 0.4286 | 13 | 0.3846 | 15 | 0.4 | 44 | 0.3409 |
| 7_4465616 | 12 | 0.25 | 9 | 0.1111 | 6 | 0 | 15 | 0.3333 | 42 | 0.2143 | 8 | 0.25 | 7 | 0.4286 | 10 | 0.6 | 14 | 0.1429 | 39 | 0.3333 |
| 7_4465654 | 7 | 0.1429 | 8 | 0.125 | 8 | 0 | 9 | 0 | 32 | 0.0625 | 5 | 0.2 | 5 | 0.2 | 6 | 0 | 9 | 0 | 25 | 0.08 |
| 7_4465663 | 11 | 0.7273 | 10 | 0.4 | 7 | 0.5714 | 13 | 0.7692 | 41 | 0.6341 | 7 | 0.7143 | 4 | 0.25 | 11 | 0.4545 | 14 | 0.7143 | 36 | 0.5833 |
| 7_4465676 | 12 | 0.0833 | 8 | 0.125 | 8 | 0.125 | 15 | 0.2 | 43 | 0.1395 | 5 | 0.4 | 5 | 0.2 | 9 | 0.3333 | 15 | 0 | 34 | 0.1765 |
| 7_4465680 | 10 | 0.2 | 7 | 0.2857 | 6 | 0.3333 | 16 | 0.4375 | 39 | 0.3333 | 6 | 0.8333 | 5 | 0.6 | 8 | 0.625 | 13 | 0.6154 | 32 | 0.6562 |
| 7_4465689 | 6 | 0.1667 | 5 | 0.4 | 7 | 0 | 7 | 0.1429 | 25 | 0.16 | 5 | 0 | 3 | 0 | 8 | 0.125 | 9 | 0.4444 | 25 | 0.2 |
| 7_4465693 | 12 | 0.0833 | 6 | 0.1667 | 7 | 0 | 13 | 0 | 38 | 0.0526 | 6 | 0.1667 | 3 | 0.6667 | 7 | 0.1429 | 13 | 0.1538 | 29 | 0.2069 |
| 7_4465701 | 9 | 0.5556 | 7 | 0.7143 | 7 | 0.4286 | 9 | 0.4444 | 32 | 0.5312 | 6 | 1 | 4 | 0.5 | 6 | 0.8333 | 11 | 0.5455 | 27 | 0.7037 |
| 7_4465745 | 6 | 0 | 5 | 0.2 | 5 | 0.4 | 11 | 0.4545 | 27 | 0.2963 | 5 | 0.2 | 4 | 0.25 | 6 | 0 | 12 | 0.1667 | 27 | 0.1481 |
| 7_4465751 | 5 | 0.6 | 4 | 0.5 | 5 | 0.2 | 9 | 0.4444 | 23 | 0.4348 | 5 | 0.4 | 4 | 0.25 | 6 | 0 | 12 | 0.25 | 27 | 0.2222 |
| 7_4465754 | 6 | 0.5 | 4 | 0.75 | 5 | 0.8 | 10 | 0.8 | 25 | 0.72 | 4 | 0.75 | 4 | 0.5 | 4 | 0.75 | 9 | 0.7778 | 21 | 0.7143 |
| 7_4465756 | 8 | 0.125 | 4 | 0.25 | 6 | 0.1667 | 11 | 0.1818 | 29 | 0.1724 | 5 | 0.2 | 4 | 0.25 | 6 | 0.5 | 12 | 0.3333 | 27 | 0.3333 |
| 7_4465783 | 7 | 0.8571 | 3 | 0.6667 | 4 | 1 | 11 | 0.9091 | 25 | 0.88 | 4 | 0.5 | 4 | 0.75 | 4 | 1 | 9 | 0.6667 | 21 | 0.7143 |
| 7_4465788 | 7 | 0.4286 | 4 | 0.5 | 5 | 0.8 | 9 | 0.2222 | 25 | 0.44 | 5 | 0.4 | 3 | 0.6667 | 5 | 0.4 | 7 | 0.1429 | 20 | 0.35 |
| 7_4465796 | 8 | 0.125 | 3 | 0 | 5 | 0.4 | 10 | 0.1 | 26 | 0.1538 | 4 | 0.5 | 3 | 0.3333 | 6 | 0.5 | 12 | 0.0833 | 25 | 0.28 |
| 7_4466499 | 1 | 0 | NA | NA | NA | NA | 3 | 0.3333 | 4 | 0.25 | NA | NA | NA | NA | 2 | 0.5 | 4 | 1 | 6 | 0.8333 |
| 7_4466502 | 1 | 0 | 1 | 0 | NA | NA | 3 | 1 | 5 | 0.6 | NA | NA | NA | NA | 2 | 0.5 | 4 | 0.75 | 6 | 0.6667 |
| 7_4467782 | NA | NA | 7 | 0.1429 | 8 | 0 | 3 | 0 | 18 | 0.0556 | 4 | 0 | 6 | 0 | 6 | 0.1667 | 4 | 0 | 20 | 0.05 |
| 7_4467785 | NA | NA | 6 | 0.1667 | 8 | 0 | 2 | 0 | 16 | 0.0625 | 3 | 0.3333 | 5 | 0 | 5 | 0 | 4 | 0 | 17 | 0.0588 |
| 7_4467786 | NA | NA | 4 | 0.5 | 7 | 0.7143 | 2 | 1 | 13 | 0.6923 | 4 | 1 | 4 | 1 | 5 | 0.8 | 4 | 0.75 | 17 | 0.8824 |
| 7_4468334 | NA | NA | 5 | 0.2 | 6 | 0.1667 | 2 | 0.5 | 13 | 0.2308 | 3 | 0.6667 | 4 | 0 | 3 | 0.6667 | 3 | 0.3333 | 13 | 0.3846 |
| 7_4468336 | NA | NA | 5 | 0.4 | 5 | 0.8 | 2 | 1 | 12 | 0.6667 | 2 | 0.5 | 4 | 1 | 3 | 0 | 3 | 1 | 12 | 0.6667 |
| 7_4469293 | NA | NA | 3 | 0 | 4 | 0.25 | 1 | 0 | 8 | 0.125 | 2 | 0 | 6 | 0.1667 | 3 | 0.3333 | 3 | 0 | 14 | 0.1429 |

| | | | | | | | | | | | | | | | | | | | | |
|---|---|---|---|---|---|---|---|---|---|---|---|---|---|---|---|---|---|---|---|---|
| 7_4469531 | NA | NA | 2 | 0 | 3 | 0 | 2 | 0.5 | 7 | 0.1429 | 2 | 0 | 4 | 0.25 | 3 | 0 | 3 | 0 | 12 | 0.0833 |
| 7_4469563 | NA | NA | 1 | 1 | 3 | 0 | 1 | 1 | 5 | 0.4 | 1 | 0 | 2 | 0.5 | 2 | 0 | 3 | 0 | 8 | 0.125 |
| 7_4469572 | NA | NA | NA | NA | 3 | 0.6667 | 1 | 0 | 4 | 0.5 | 1 | 1 | 3 | 0.6667 | 2 | 0.5 | 3 | 0.3333 | 9 | 0.5556 |
| 7_4469634 | NA | NA | 1 | 0 | 3 | 0 | 2 | 0.5 | 6 | 0.1667 | 1 | 1 | 3 | 0.3333 | 1 | 0 | 2 | 0.5 | 7 | 0.4286 |
| 7_4469651 | NA | NA | NA | NA | 2 | 1 | 2 | 0.5 | 4 | 0.75 | 1 | 1 | 3 | 1 | 1 | 1 | 2 | 1 | 7 | 1 |
| 7_4469661 | NA | NA | 1 | 1 | 2 | 0.5 | 2 | 0.5 | 5 | 0.6 | 1 | 0 | 3 | 0.3333 | 1 | 1 | 1 | 0 | 6 | 0.3333 |
| 7_4469692 | NA | NA | 1 | 1 | 3 | 0 | 2 | 0 | 6 | 0.1667 | 1 | 1 | 3 | 0.6667 | 1 | 1 | 2 | 0.5 | 7 | 0.7143 |
| 7_4469693 | NA | NA | 1 | 0 | 3 | 0.3333 | 2 | 0 | 6 | 0.1667 | 1 | 1 | 2 | 0.5 | 1 | 0 | 2 | 0 | 6 | 0.3333 |
| 7_4469695 | NA | NA | 1 | 1 | 1 | 1 | 1 | 1 | 3 | 1 | 1 | 0 | 3 | 1 | 1 | 0 | 2 | 1 | 7 | 0.7143 |
| 7_4469700 | NA | NA | 1 | 1 | 2 | 0.5 | 1 | 0 | 4 | 0.5 | 1 | 1 | 2 | 0 | NA | NA | 2 | 0 | 5 | 0.2 |
| 7_4469717 | NA | NA | 1 | 0 | 2 | 0.5 | 1 | 0 | 4 | 0.25 | 1 | 0 | 1 | 1 | 1 | 1 | 2 | 1 | 5 | 0.8 |
| 7_4469728 | NA | NA | 1 | 1 | 2 | 0.5 | 1 | 1 | 4 | 0.75 | 1 | 1 | 1 | 1 | NA | NA | 1 | 1 | 3 | 1 |
| 7_4469770 | NA | NA | 1 | 0 | 2 | 0.5 | 1 | 1 | 4 | 0.5 | 1 | 1 | NA | NA | 1 | 1 | 2 | 0 | 4 | 0.5 |
| 7_4469774 | NA | NA | NA | NA | 2 | 0.5 | NA | NA | 2 | 0.5 | 1 | 1 | NA | NA | NA | NA | 2 | 0.5 | 3 | 0.6667 |
| 7_4469780 | NA | NA | 1 | 1 | 2 | 0.5 | 1 | 1 | 4 | 0.75 | 1 | 0 | NA | NA | 1 | 0 | 2 | 0.5 | 4 | 0.25 |
| 7_4469790 | NA | NA | 1 | 1 | 2 | 1 | 1 | 1 | 4 | 1 | 1 | 1 | NA | NA | 1 | 1 | 2 | 1 | 4 | 1 |
| 7_4469796 | NA | NA | 1 | 1 | 2 | 0.5 | 1 | 1 | 4 | 0.75 | 1 | 1 | NA | NA | 1 | 1 | 2 | 1 | 4 | 1 |
| 7_4469852 | NA | NA | 1 | 0 | 2 | 0.5 | 1 | 1 | 4 | 0.5 | 1 | 1 | NA | NA | 1 | 1 | 1 | 1 | 3 | 1 |
| 7_4469873 | NA | NA | NA | NA | 2 | 0.5 | 1 | 0 | 3 | 0.3333 | 1 | 1 | NA | NA | 1 | 1 | 2 | 0.5 | 4 | 0.75 |
| 7_4469906 | NA | NA | NA | NA | 2 | 1 | 1 | 0 | 3 | 0.6667 | 1 | 0 | NA | NA | 1 | 0 | 2 | 0.5 | 4 | 0.25 |
| 7_4472698 | 6 | 0.5 | 3 | 0.3333 | 2 | 0.5 | 5 | 0.2 | 16 | 0.375 | 6 | 0.1667 | 3 | 0 | 6 | 0 | 9 | 0.4444 | 24 | 0.2083 |
| 7_4472707 | 4 | 0.5 | 4 | 0.25 | 2 | 0 | 4 | 0.25 | 14 | 0.2857 | 5 | 0 | 3 | 0 | 7 | 0.2857 | 6 | 0.1667 | 21 | 0.1429 |
| 7_4472754 | 6 | 0.1667 | 4 | 0 | 1 | 0 | 6 | 0.3333 | 17 | 0.1765 | 4 | 0.25 | 3 | 0 | 7 | 0.1429 | 7 | 0.1429 | 21 | 0.1429 |
| 7_4472758 | 5 | 0 | 5 | 0.4 | 1 | 0 | 4 | 0 | 15 | 0.1333 | 6 | 0.1667 | 4 | 0 | 5 | 0.4 | 7 | 0 | 22 | 0.1364 |
| 7_4474458 | 4 | 0 | 5 | 0.2 | 2 | 0.5 | 2 | 0 | 13 | 0.1538 | 1 | 0 | 4 | 0 | 8 | 0.125 | 3 | 0 | 16 | 0.0625 |
| 7_4475148 | 3 | 0.3333 | 4 | 0.5 | 2 | 0 | 1 | 0 | 10 | 0.3 | 1 | 1 | 5 | 0.6 | 8 | 0.375 | 1 | 0 | 15 | 0.4667 |
| 7_4475157 | 3 | 0 | 6 | 0.1667 | 2 | 0.5 | 3 | 0.3333 | 14 | 0.2143 | 1 | 0 | 3 | 0 | 7 | 0.2857 | 3 | 0 | 14 | 0.1429 |
| 7_4475160 | 3 | 0 | 6 | 0.1667 | 2 | 0 | 3 | 0 | 14 | 0.0714 | 1 | 0 | 5 | 0 | 5 | 0.2 | 3 | 0 | 14 | 0.0714 |
| 7_4475261 | 1 | 0 | 2 | 0.5 | 2 | 0 | 2 | 0 | 7 | 0.1429 | 1 | 0 | 4 | 0 | 4 | 0 | 1 | 1 | 10 | 0.1 |
| 7_4477185 | 109 | 0.1376 | 86 | 0.0814 | 91 | 0.1099 | 89 | 0.191 | 375 | 0.1307 | 73 | 0.1233 | 70 | 0.1286 | 89 | 0.1573 | 62 | 0.1774 | 294 | 0.1463 |
| 7_4477525 | 10 | 0.1 | 7 | 0.1429 | 9 | 0 | 12 | 0.25 | 38 | 0.1316 | 11 | 0 | 7 | 0.1429 | 8 | 0.125 | 10 | 0.1 | 36 | 0.0833 |
| 7_4477535 | 158 | 0.1582 | 167 | 0.1497 | 119 | 0.1765 | 203 | 0.1872 | 647 | 0.1685 | 184 | 0.212 | 165 | 0.1152 | 168 | 0.1667 | 146 | 0.2534 | 663 | 0.1855 |
| 7_4478076 | 1 | 0 | NA | NA | 1 | 1 | NA | NA | 2 | 0.5 | 1 | 1 | NA | NA | NA | NA | NA | NA | 1 | 1 |
| 7_4479127 | 2 | 0.5 | NA | NA | 1 | 0 | NA | NA | 3 | 0.3333 | 1 | 0 | NA | NA | 1 | 1 | 2 | 0 | 4 | 0.25 |
| 7_4479127 | 1 | 1 | NA | NA | 1 | 0 | NA | NA | 2 | 0.5 | 1 | 0 | NA | NA | 1 | 1 | 1 | 0 | 3 | 0.3333 |
| 7_4479136 | 1 | 1 | NA | NA | 2 | 0 | NA | NA | 3 | 0.3333 | 1 | 0 | NA | NA | 1 | 1 | 2 | 1 | 4 | 0.75 |
| 7_4480377 | NA | NA | 2 | 0.5 | 2 | 0 | NA | NA | 4 | 0.25 | NA | NA | 2 | 0 | 1 | 1 | NA | NA | 3 | 0.3333 |
| 7_4481823 | 31 | 0.0323 | 60 | 0.0667 | 29 | 0.1034 | 13 | 0.0769 | 133 | 0.0677 | 56 | 0.0357 | 23 | 0 | 16 | 0.125 | 38 | 0.0789 | 133 | 0.0526 |
| 7_4481926 | 38 | 0.1842 | 71 | 0.0986 | 42 | 0.1429 | 17 | 0.1176 | 168 | 0.131 | 67 | 0.0896 | 27 | 0.1111 | 17 | 0 | 44 | 0.0909 | 155 | 0.0839 |
| 7_4482446 | 57 | 0.614 | 92 | 0.6304 | 57 | 0.5614 | 22 | 0.3636 | 228 | 0.5833 | 78 | 0.5256 | 31 | 0.6452 | 21 | 0.6667 | 52 | 0.6731 | 182 | 0.6044 |
| 7_4482446 | 2 | 0 | NA | NA | 1 | 0 | 3 | 0.3333 | 6 | 0.1667 | 2 | 0.5 | 3 | 0 | NA | NA | 2 | 0.5 | 7 | 0.2857 |
| 7_4482450 | 57 | 0.7018 | 106 | 0.7075 | 64 | 0.6406 | 22 | 0.8182 | 249 | 0.6988 | 81 | 0.5926 | 30 | 0.4667 | 22 | 0.5909 | 53 | 0.717 | 186 | 0.6075 |
| 7_4482476 | 74 | 0.3514 | 129 | 0.2868 | 85 | 0.2941 | 28 | 0.3214 | 316 | 0.307 | 101 | 0.3366 | 48 | 0.2708 | 25 | 0.24 | 68 | 0.3382 | 242 | 0.314 |
| 7_4482496 | 75 | 0.2533 | 137 | 0.3942 | 90 | 0.3778 | 35 | 0.6 | 337 | 0.3798 | 127 | 0.4331 | 47 | 0.4255 | 29 | 0.4483 | 90 | 0.3889 | 293 | 0.4198 |
| 7_4488316 | 2 | 1 | 1 | 1 | 1 | 0 | NA | NA | 4 | 0.75 | NA | NA | NA | NA | 1 | 1 | NA | NA | 1 | 1 |
| 7_4495295 | 1 | 0 | 1 | 0 | 1 | 1 | NA | NA | 3 | 0.3333 | NA | NA | 1 | 0 | NA | NA | 1 | 1 | 2 | 0.5 |
| 7_4498931 | 3 | 0.6667 | 15 | 0.4667 | 6 | 0.6667 | 1 | 0 | 25 | 0.52 | 4 | 0.25 | 5 | 0.8 | 8 | 0.875 | 8 | 0.625 | 25 | 0.68 |
| 7_4498934 | 6 | 0.3333 | 23 | 0.6087 | 8 | 0.75 | 4 | 0.75 | 41 | 0.6098 | 6 | 0.3333 | 6 | 0.6667 | 9 | 0.4444 | 11 | 0.5455 | 32 | 0.5 |
| 7_4498939 | 4 | 0 | 23 | 0.1304 | 10 | 0 | 4 | 0.25 | 41 | 0.0976 | 7 | 0 | 8 | 0.375 | 10 | 0 | 11 | 0.1818 | 36 | 0.1389 |
| 7_4501102 | 1 | 1 | 4 | 0 | 2 | 0 | NA | NA | 7 | 0.1429 | 2 | 0 | NA | NA | 2 | 0 | 4 | 0.25 | 8 | 0.125 |
| 7_4501845 | 3 | 0.3333 | 1 | 0 | 3 | 0 | 2 | 0 | 9 | 0.1111 | 1 | 0 | 2 | 0 | 1 | 1 | NA | NA | 4 | 0.25 |
| 7_4502508 | NA | NA | 1 | 1 | 1 | 0 | NA | NA | 2 | 0.5 | 1 | 1 | NA | NA | 1 | 0 | NA | NA | 2 | 0.5 |
| 7_4502600 | 17 | 0.6471 | 21 | 0.7619 | 16 | 0.5625 | 16 | 0.75 | 70 | 0.6857 | 12 | 0.3333 | 19 | 0.6316 | 15 | 0.5333 | 13 | 0.7692 | 59 | 0.5763 |
| 7_4502613 | 11 | 0.4545 | 17 | 0.5882 | 13 | 0.6923 | 12 | 0.5833 | 53 | 0.5849 | 11 | 0.6364 | 16 | 0.5 | 9 | 0.7778 | 8 | 0.75 | 44 | 0.6364 |
| 7_4502623 | 11 | 0.2727 | 15 | 0.5333 | 9 | 0.3333 | 11 | 0.1818 | 46 | 0.3478 | 9 | 0.2222 | 12 | 0.1667 | 13 | 0.1538 | 10 | 0.4 | 44 | 0.2273 |
| 7_4502637 | 13 | 0.2308 | 18 | 0.2778 | 10 | 0.2 | 11 | 0.1818 | 52 | 0.2308 | 10 | 0.1 | 14 | 0.1429 | 12 | 0.0833 | 10 | 0.2 | 46 | 0.1304 |
| 7_4502651 | 12 | 0 | 10 | 0.3 | 6 | 0 | 9 | 0.4444 | 37 | 0.1892 | 10 | 0.1 | 12 | 0.3333 | 7 | 0.1429 | 6 | 0.3333 | 35 | 0.2286 |
| 7_4502652 | 12 | 0 | 11 | 0.1818 | 8 | 0.25 | 11 | 0.0909 | 42 | 0.119 | 10 | 0.4 | 12 | 0.0833 | 7 | 0.2857 | 7 | 0.1429 | 36 | 0.2222 |
| 7_4507141 | 1 | 0 | 3 | 0.3333 | 5 | 0 | 7 | 0 | 16 | 0.0625 | 5 | 0 | 3 | 0.3333 | 4 | 0 | 8 | 0 | 20 | 0.05 |
| 7_4510471 | 2 | 1 | 2 | 0.5 | 4 | 0.5 | 3 | 1 | 11 | 0.7273 | 1 | 1 | NA | NA | NA | NA | 4 | 0.5 | 5 | 0.6 |
| 7_4510488 | 2 | 0 | 3 | 0.6667 | 3 | 0 | 3 | 0.6667 | 11 | 0.3636 | 1 | 1 | NA | NA | NA | NA | 3 | 0.3333 | 4 | 0.5 |
| 7_4510493 | 3 | 0.3333 | 4 | 0 | 4 | 0 | 4 | 0.25 | 15 | 0.1333 | 1 | 1 | NA | NA | NA | NA | 2 | 0 | 3 | 0.3333 |
| 7_4510504 | 2 | 0.5 | 4 | 0 | 5 | 0 | 4 | 0.25 | 15 | 0.1333 | 1 | 1 | NA | NA | NA | NA | 4 | 0 | 5 | 0.2 |
| 7_4510519 | 2 | 0 | 3 | 0 | 5 | 0.2 | 3 | 0 | 13 | 0.0769 | 1 | 0 | NA | NA | NA | NA | 4 | 0.25 | 5 | 0.2 |
| 7_4510524 | 3 | 0.3333 | 4 | 0 | 5 | 0 | 5 | 0.4 | 17 | 0.1765 | 1 | 1 | NA | NA | NA | NA | 4 | 0.25 | 5 | 0.4 |

| | | | | | | | | | | | | | | | | | | | | |
|---|---|---|---|---|---|---|---|---|---|---|---|---|---|---|---|---|---|---|---|---|
| 7_4510556 | 2 | 0.5 | 4 | 0.5 | 2 | 0.5 | 5 | 0.4 | 13 | 0.4615 | NA | NA | NA | NA | NA | NA | 2 | 0.5 | 2 | 0.5 |
| 7_4510564 | 3 | 0 | 3 | 0 | 3 | 0.3333 | 5 | 0 | 14 | 0.0714 | 1 | 1 | NA | NA | NA | NA | 3 | 0 | 4 | 0.25 |
| 7_4511126 | 5 | 1 | 5 | 0.2 | 4 | 0.25 | 1 | 0 | 15 | 0.4667 | 3 | 0.6667 | 4 | 0.75 | 2 | 1 | 3 | 0.6667 | 12 | 0.75 |
| 7_4512463 | NA | NA | 2 | 1 | 1 | 0 | NA | NA | 3 | 0.6667 | NA | NA | 2 | 0.5 | NA | NA | NA | NA | 2 | 0.5 |
| 7_4514277 | 5 | 0.4 | 9 | 0.1111 | 6 | 0.3333 | 4 | 0 | 24 | 0.2083 | 7 | 0.2857 | 10 | 0.2 | 9 | 0.2222 | 5 | 0.4 | 31 | 0.2581 |
| 7_4514289 | 5 | 0.2 | 9 | 0.2222 | 7 | 0.2857 | 4 | 0.5 | 25 | 0.28 | 8 | 0.5 | 9 | 0.3333 | 8 | 0.375 | 4 | 0.25 | 29 | 0.3793 |
| 7_4517343 | 1 | 0 | 1 | 0 | 2 | 0 | 3 | 0.3333 | 7 | 0.1429 | 2 | 0 | NA | NA | 1 | 0 | 1 | 1 | 4 | 0.25 |
| 7_4517346 | 1 | 0 | 1 | 1 | 2 | 0.5 | 3 | 0 | 7 | 0.2857 | 2 | 0.5 | NA | NA | 3 | 0 | 1 | 1 | 6 | 0.3333 |
| 7_4517398 | 1 | 0 | 1 | 0 | 2 | 0.5 | 3 | 0.3333 | 7 | 0.2857 | 1 | 0 | NA | NA | 2 | 1 | 1 | 1 | 4 | 0.75 |
| 7_4517445 | 1 | 0 | 1 | 1 | 2 | 0 | 2 | 0 | 6 | 0.1667 | 1 | 1 | NA | NA | 3 | 0.3333 | NA | NA | 4 | 0.5 |
| 7_4517448 | 1 | 0 | 1 | 0 | 2 | 1 | 3 | 0.6667 | 7 | 0.5714 | 1 | 1 | NA | NA | 3 | 0 | 1 | 1 | 5 | 0.4 |
| 7_4517455 | 1 | 0 | 1 | 0 | 2 | 0 | 3 | 0.3333 | 7 | 0.1429 | 1 | 0 | NA | NA | 3 | 0 | 1 | 1 | 5 | 0.2 |
| 7_4523884 | NA | NA | 1 | 0 | 5 | 0.6 | 4 | 0.25 | 10 | 0.4 | NA | NA | 4 | 0 | 1 | 1 | NA | NA | 5 | 0.2 |
| 7_4523893 | NA | NA | 1 | 0 | 6 | 0.1667 | 4 | 0.5 | 11 | 0.2727 | NA | NA | 2 | 0 | 1 | 1 | NA | NA | 3 | 0.3333 |
| 7_4528546 | NA | NA | NA | NA | 1 | 1 | 1 | 1 | 2 | 1 | 1 | 1 | 3 | 1 | 1 | 0 | NA | NA | 5 | 0.8 |
| 7_4528571 | 1 | 1 | NA | NA | 2 | 1 | 1 | 1 | 4 | 1 | 1 | 1 | 2 | 1 | NA | NA | NA | NA | 3 | 1 |
| 7_4528591 | 1 | 1 | NA | NA | NA | NA | 1 | 1 | 2 | 1 | 2 | 1 | 1 | 1 | NA | NA | NA | NA | 3 | 1 |
| 7_4528593 | 1 | 1 | NA | NA | 2 | 1 | 1 | 1 | 4 | 1 | 2 | 1 | 1 | 1 | NA | NA | NA | NA | 3 | 1 |
| 7_4528625 | 1 | 1 | 1 | 1 | 1 | 1 | 1 | 1 | 4 | 1 | 1 | 1 | NA | NA | NA | NA | NA | NA | 1 | 1 |
| 7_4528845 | 1 | 0 | 1 | 1 | NA | NA | NA | NA | 2 | 0.5 | 1 | 0 | NA | NA | 1 | 0 | 1 | 1 | 3 | 0.3333 |
| 7_4534381 | NA | NA | 1 | 1 | NA | NA | NA | NA | 1 | 1 | 2 | 0.5 | NA | NA | NA | NA | 1 | 1 | 3 | 0.6667 |
| 7_4534391 | NA | NA | 1 | 1 | NA | NA | NA | NA | 1 | 1 | 1 | 1 | NA | NA | NA | NA | 1 | 1 | 2 | 1 |
| 7_4534946 | 7 | 0 | 11 | 0 | 4 | 0.25 | 13 | 0.0769 | 35 | 0.0571 | 6 | 0.1667 | 10 | 0 | 13 | 0.1538 | 7 | 0 | 36 | 0.0833 |
| 7_4535228 | 19 | 0 | 27 | 0.1481 | 18 | 0 | 32 | 0.1562 | 96 | 0.0938 | 16 | 0.125 | 29 | 0.069 | 32 | 0.1875 | 15 | 0.1333 | 92 | 0.1304 |
| 7_4535402 | 17 | 0.2353 | 32 | 0.2812 | 25 | 0.4 | 45 | 0.2444 | 119 | 0.2857 | 23 | 0.1304 | 44 | 0.2045 | 48 | 0.3125 | 22 | 0.2727 | 137 | 0.2409 |
| 7_4535471 | 14 | 0.0714 | 9 | 0 | 6 | 0.3333 | 7 | 0.1429 | 36 | 0.1111 | 2 | 0 | 14 | 0.2143 | 7 | 0 | 5 | 0 | 28 | 0.1071 |
| 7_4535478 | 18 | 0.0556 | 10 | 0.1 | 9 | 0.1111 | 10 | 0.1 | 47 | 0.0851 | 3 | 0 | 17 | 0.0588 | 10 | 0.1 | 10 | 0.1 | 40 | 0.075 |
| 7_4535483 | 20 | 0.2 | 9 | 0.3333 | 13 | 0.2308 | 11 | 0.7273 | 53 | 0.3396 | 5 | 0.2 | 16 | 0.3125 | 10 | 0.2 | 9 | 0.5556 | 40 | 0.325 |
| 7_4535486 | 20 | 0.15 | 10 | 0.2 | 11 | 0.2727 | 14 | 0.1429 | 55 | 0.1818 | 5 | 0.2 | 15 | 0.0667 | 13 | 0.2308 | 10 | 0.3 | 43 | 0.186 |
| 7_4535486 | 19 | 0.0526 | 10 | 0.2 | 11 | 0.0909 | 13 | 0.0769 | 53 | 0.0943 | 5 | 0.2 | 19 | 0.1053 | 12 | 0.25 | 13 | 0.0769 | 49 | 0.1429 |
| 7_4535494 | 21 | 0.0952 | 10 | 0 | 14 | 0.2143 | 11 | 0.2727 | 56 | 0.1429 | 4 | 0.25 | 18 | 0.2222 | 12 | 0.1667 | 12 | 0.3333 | 46 | 0.2391 |
| 7_4535670 | 3 | 0.3333 | 1 | 0 | NA | NA | 2 | 0.5 | 6 | 0.3333 | 1 | 0 | 1 | 1 | 1 | 0 | NA | NA | 3 | 0.3333 |
| 7_4535671 | 4 | 0 | 1 | 0 | NA | NA | 2 | 0.5 | 7 | 0.1429 | 1 | 0 | 2 | 1 | NA | NA | NA | NA | 3 | 0.6667 |
| 7_4535690 | 5 | 0.2 | 1 | 0 | NA | NA | 2 | 0.5 | 8 | 0.25 | 1 | 0 | 2 | 1 | 1 | 0 | NA | NA | 4 | 0.5 |
| 7_4535929 | 30 | 0.2667 | 17 | 0.5294 | 21 | 0.381 | 28 | 0.5357 | 96 | 0.4167 | 14 | 0.3571 | 34 | 0.2941 | 18 | 0.5 | 22 | 0.3636 | 88 | 0.3636 |
| 7_4535934 | 31 | 0.2903 | 13 | 0.3846 | 19 | 0.5263 | 24 | 0.1667 | 87 | 0.3218 | 15 | 0.2667 | 36 | 0.1944 | 16 | 0.4375 | 21 | 0.3333 | 88 | 0.2841 |
| 7_4536005 | 3 | 0.3333 | 4 | 0.5 | 3 | 1 | NA | NA | 10 | 0.6 | 3 | 0.3333 | 1 | 1 | 1 | 1 | 5 | 0.6 | 10 | 0.6 |
| 7_4536033 | 2 | 0 | 5 | 0 | 6 | 0.1667 | NA | NA | 13 | 0.0769 | 4 | 0 | 2 | 0 | 4 | 0 | 6 | 0.1667 | 16 | 0.0625 |
| 7_4536052 | 3 | 0.6667 | 4 | 0.5 | 6 | 0.8333 | NA | NA | 13 | 0.6923 | 3 | 0.6667 | 1 | 0 | 3 | 1 | 5 | 1 | 12 | 0.8333 |
| 7_4536056 | 2 | 1 | 4 | 0.5 | 6 | 0.6667 | NA | NA | 12 | 0.6667 | 3 | 0.3333 | NA | NA | 3 | 0.3333 | 4 | 0.5 | 10 | 0.4 |
| 7_4536074 | 2 | 0.5 | 5 | 0.2 | 3 | 0.3333 | NA | NA | 10 | 0.3 | 2 | 0.5 | 1 | 0 | 3 | 0.3333 | 4 | 1 | 10 | 0.6 |
| 7_4536074 | 2 | 0 | 3 | 0.3333 | 4 | 0.25 | NA | NA | 9 | 0.2222 | 3 | 0 | 1 | 0 | 4 | 0.5 | 3 | 0 | 11 | 0.1818 |
| 7_4536090 | 2 | 0.5 | 5 | 0.2 | 4 | 0.25 | NA | NA | 11 | 0.2727 | 3 | 0 | 1 | 1 | 3 | 0 | 6 | 0 | 13 | 0.0769 |
| 7_4536107 | 1 | 1 | 5 | 1 | 1 | 1 | NA | NA | 7 | 1 | 3 | 1 | 1 | 1 | 3 | 1 | 2 | 1 | 9 | 1 |
| 7_4536115 | 2 | 0 | 3 | 0.3333 | 1 | 0 | NA | NA | 6 | 0.1667 | 2 | 0 | 1 | 0 | 3 | 0 | 4 | 0.25 | 10 | 0.1 |
| 7_4536279 | 1 | 1 | 3 | 0 | 2 | 0 | NA | NA | 6 | 0.1667 | 3 | 0 | NA | NA | 2 | 0.5 | 3 | 0 | 8 | 0.125 |
| 7_4536286 | 1 | 1 | 4 | 0 | 2 | 0 | NA | NA | 7 | 0.1429 | 3 | 0.3333 | 1 | 0 | 3 | 0 | 4 | 0 | 11 | 0.0909 |
| 7_4536310 | NA | NA | 5 | 0.8 | 2 | 1 | NA | NA | 7 | 0.8571 | 1 | 0 | 1 | 0 | 2 | 0.5 | 2 | 0.5 | 6 | 0.3333 |
| 7_4536447 | NA | NA | 2 | 1 | 1 | 0 | NA | NA | 3 | 0.6667 | NA | NA | NA | NA | 2 | 0.5 | 1 | 1 | 3 | 0.6667 |
| 7_4539173 | 2 | 0 | 1 | 0 | 2 | 0.5 | NA | NA | 5 | 0.2 | NA | NA | 1 | 1 | 1 | 0 | NA | NA | 2 | 0.5 |
| 7_4544273 | 8 | 0.125 | 18 | 0.1111 | 13 | 0.1538 | 10 | 0.2 | 49 | 0.1429 | 8 | 0 | 7 | 0 | 12 | 0.0833 | 7 | 0.2857 | 34 | 0.0882 |
| 7_4544311 | 11 | 0.1818 | 16 | 0.0625 | 13 | 0 | 12 | 0 | 52 | 0.0577 | 8 | 0.125 | 8 | 0.125 | 10 | 0 | 8 | 0 | 34 | 0.0588 |
| 7_4544334 | 11 | 0.2727 | 17 | 0.4118 | 14 | 0.1429 | 12 | 0.3333 | 54 | 0.2963 | 11 | 0.0909 | 5 | 0.2 | 10 | 0.2 | 9 | 0.2222 | 35 | 0.1714 |
| 7_4544358 | 12 | 0 | 19 | 0.1579 | 14 | 0.1429 | 13 | 0.0769 | 58 | 0.1034 | 15 | 0.0667 | 6 | 0.3333 | 14 | 0.1429 | 9 | 0.1111 | 44 | 0.1364 |
| 7_4544376 | 13 | 0.2308 | 16 | 0.3125 | 15 | 0.0667 | 14 | 0.1429 | 58 | 0.1897 | 13 | 0.0769 | 7 | 0.2857 | 14 | 0.1429 | 8 | 0.375 | 42 | 0.1905 |
| 7_4544400 | 12 | 0.1667 | 18 | 0.1667 | 13 | 0.1538 | 12 | 0 | 55 | 0.1273 | 14 | 0.0714 | 10 | 0 | 16 | 0.125 | 10 | 0.1 | 50 | 0.08 |
| 7_4544401 | 13 | 0.4615 | 20 | 0.55 | 14 | 0.4286 | 14 | 0.2857 | 61 | 0.4426 | 14 | 0.3571 | 6 | 0.1667 | 13 | 0.2308 | 6 | 0.3333 | 39 | 0.2821 |
| 7_4544404 | 15 | 0.4 | 16 | 0.375 | 14 | 0.4286 | 12 | 0.3333 | 57 | 0.386 | 11 | 0.2727 | 10 | 0.1 | 12 | 0.4167 | 10 | 0.5 | 43 | 0.3256 |
| 7_4544407 | 6 | 0 | 18 | 0.3333 | 12 | 0.3333 | 11 | 0.1818 | 47 | 0.2553 | 9 | 0.3333 | 7 | 0.2857 | 16 | 0.0625 | 7 | 0.1429 | 39 | 0.1795 |
| 7_4544410 | 12 | 0.5 | 20 | 0.5 | 13 | 0.2308 | 14 | 0.2857 | 59 | 0.3898 | 15 | 0.0667 | 7 | 0.1429 | 12 | 0.1667 | 8 | 0.5 | 42 | 0.1905 |
| 7_4544420 | 14 | 0.0714 | 16 | 0.125 | 14 | 0.2143 | 14 | 0.2143 | 58 | 0.1552 | 14 | 0.1429 | 9 | 0.3333 | 22 | 0 | 9 | 0 | 54 | 0.0926 |
| 7_4544438 | 14 | 0.0714 | 22 | 0.2727 | 14 | 0.1429 | 15 | 0.2 | 65 | 0.1846 | 16 | 0.3125 | 9 | 0.4444 | 22 | 0.2273 | 9 | 0.2222 | 56 | 0.2857 |
| 7_4544440 | 13 | 0.8462 | 17 | 0.8824 | 15 | 0.6667 | 16 | 0.75 | 61 | 0.7869 | 15 | 0.8667 | 10 | 0.7 | 20 | 0.65 | 11 | 1 | 56 | 0.7857 |
| 7_4544444 | 10 | 0.4 | 15 | 0.1333 | 14 | 0.4286 | 11 | 0.1818 | 50 | 0.28 | 12 | 0.1667 | 10 | 0.4 | 21 | 0.2381 | 9 | 0.2222 | 52 | 0.25 |
| 7_4544456 | 14 | 0 | 23 | 0.1304 | 15 | 0.0667 | 15 | 0.3333 | 67 | 0.1343 | 17 | 0.1765 | 11 | 0.0909 | 21 | 0.0952 | 10 | 0.1 | 59 | 0.1186 |

| | | | | | | | | | | | | | | | | | | | | |
|---|---|---|---|---|---|---|---|---|---|---|---|---|---|---|---|---|---|---|---|---|
| 7_45444572 | 15 | 0.1333 | 25 | 0.24 | 14 | 0.2857 | 17 | 0.1765 | 71 | 0.2113 | 17 | 0.3529 | 11 | 0.0909 | 21 | 0.2381 | 11 | 0.1818 | 60 | 0.2333 |
| 7_45444580 | 14 | 0.7857 | 23 | 0.8696 | 14 | 0.6429 | 15 | 0.8 | 66 | 0.7879 | 15 | 0.8667 | 13 | 1 | 22 | 0.8182 | 7 | 0.7143 | 57 | 0.8596 |
| 7_45444641 | 13 | 1 | 21 | 1 | 15 | 1 | 16 | 1 | 65 | 1 | 18 | 1 | 12 | 1 | 18 | 1 | 11 | 0.9091 | 59 | 0.9831 |
| 7_45444947 | 13 | 0.1538 | 23 | 0.087 | 13 | 0.0769 | 13 | 0.0769 | 62 | 0.0968 | 14 | 0.2857 | 9 | 0 | 19 | 0.0526 | 7 | 0 | 49 | 0.102 |
| 7_45453068 | 21 | 0.0476 | 27 | 0.037 | 15 | 0 | 24 | 0.125 | 87 | 0.0575 | 13 | 0.1538 | 14 | 0.0714 | 23 | 0 | 8 | 0 | 58 | 0.0517 |
| 7_45453163 | 16 | 0.0625 | 22 | 0.0455 | 15 | 0 | 19 | 0.1053 | 72 | 0.0556 | 17 | 0.0588 | 17 | 0 | 15 | 0.0667 | 11 | 0.0909 | 60 | 0.05 |
| 7_45453327 | 13 | 0.1538 | 26 | 0.0769 | 13 | 0.1538 | 15 | 0.0667 | 67 | 0.1045 | 10 | 0.2 | 17 | 0.0588 | 23 | 0.1739 | 7 | 0.1429 | 57 | 0.1404 |
| 7_45453345 | 24 | 0.5833 | 34 | 0.6471 | 16 | 0.75 | 21 | 0.6667 | 95 | 0.6526 | 22 | 0.6364 | 20 | 0.6 | 24 | 0.5833 | 14 | 0.5714 | 80 | 0.6 |
| 7_45453350 | 16 | 0.25 | 24 | 0.5833 | 8 | 0.625 | 12 | 0.5833 | 60 | 0.5 | 10 | 0.6 | 13 | 0.6923 | 15 | 0.3333 | 9 | 0.6667 | 47 | 0.5532 |
| 7_45453360 | 19 | 0.8947 | 21 | 0.9524 | 13 | 0.7692 | 19 | 0.9474 | 72 | 0.9028 | 12 | 0.75 | 9 | 0.7778 | 21 | 0.8571 | 12 | 0.5833 | 54 | 0.7593 |
| 7_45453545 | 18 | 0.0556 | 27 | 0.0741 | 13 | 0.0769 | 20 | 0.15 | 78 | 0.0897 | 20 | 0.05 | 15 | 0 | 22 | 0.0455 | 11 | 0.1818 | 68 | 0.0588 |
| 7_45466104 | NA | NA | NA | NA | 2 | 0.5 | NA | NA | 2 | 0.5 | NA | NA | NA | NA | NA | NA | 1 | 1 | 1 | 1 |
| 7_45466421 | NA | NA | NA | NA | 2 | 0.5 | NA | NA | 2 | 0.5 | NA | NA | NA | NA | NA | NA | 1 | 1 | 1 | 1 |
| 7_45466467 | NA | NA | NA | NA | 2 | 0.5 | NA | NA | 2 | 0.5 | NA | NA | NA | NA | NA | NA | 1 | 1 | 1 | 1 |
| 7_45466796 | NA | NA | NA | NA | 2 | 1 | NA | NA | 2 | 1 | NA | NA | NA | NA | NA | NA | 1 | 1 | 1 | 1 |
| 7_45467172 | NA | NA | NA | NA | 2 | 1 | NA | NA | 2 | 1 | NA | NA | NA | NA | NA | NA | 1 | 1 | 1 | 1 |
| 7_45474678 | 2 | 0 | 2 | 0 | 1 | 0 | 4 | 1 | 9 | 0.4444 | 2 | 0 | 5 | 0.2 | 2 | 1 | NA | NA | 9 | 0.3333 |
| 7_45474865 | 2 | 0.5 | 2 | 1 | 1 | 1 | 2 | 0.5 | 7 | 0.7143 | 2 | 0.5 | 3 | 1 | 2 | 1 | NA | NA | 7 | 0.8571 |
| 7_45474896 | 2 | 0.5 | 1 | 1 | 1 | 1 | 2 | 0.5 | 6 | 0.6667 | 3 | 0.6667 | 4 | 1 | 2 | 0.5 | NA | NA | 9 | 0.7778 |
| 7_45474910 | 2 | 1 | 2 | 1 | 1 | 1 | 3 | 0.6667 | 8 | 0.875 | 3 | 0.6667 | 4 | 1 | 2 | 1 | NA | NA | 9 | 0.8889 |
| 7_45474999 | 2 | 1 | 2 | 1 | 1 | 1 | 2 | 0.5 | 7 | 0.8571 | 2 | 1 | 5 | 0.8 | 1 | 1 | NA | NA | 8 | 0.875 |
| 7_45475061 | 2 | 1 | 2 | 1 | 1 | 1 | 3 | 1 | 8 | 1 | 2 | 1 | 5 | 1 | 2 | 1 | 1 | 1 | 10 | 1 |
| 7_45475118 | 2 | 0.5 | 1 | 1 | NA | NA | 1 | 1 | 4 | 0.75 | 2 | 1 | 3 | 0.6667 | 1 | 0 | NA | NA | 6 | 0.6667 |
| 7_45475158 | 2 | 0.5 | 2 | 1 | 1 | 1 | 2 | 1 | 7 | 0.8571 | 1 | 1 | 5 | 1 | 2 | 0 | NA | NA | 8 | 0.75 |
| 7_45475176 | 2 | 1 | 2 | 0.5 | 1 | 0 | 3 | 0.6667 | 8 | 0.625 | 2 | 1 | 5 | 0.6 | 2 | 1 | NA | NA | 9 | 0.7778 |
| 7_45475298 | 1 | 1 | 1 | 1 | NA | NA | 1 | 0 | 3 | 0.6667 | 3 | 0 | 2 | 1 | 2 | 0 | NA | NA | 7 | 0.2857 |
| 7_45475351 | 1 | 0 | 2 | 0.5 | 1 | 0 | 2 | 0 | 6 | 0.1667 | 3 | 0.3333 | 4 | 0 | 2 | 0 | NA | NA | 9 | 0.1111 |
| 7_45475355 | NA | NA | 2 | 0.5 | NA | NA | 3 | 0.3333 | 5 | 0.4 | 3 | 1 | 3 | 0.6667 | 1 | 1 | NA | NA | 7 | 0.8571 |
| 7_45475396 | 2 | 0.5 | 2 | 0 | 1 | 1 | 3 | 0 | 8 | 0.25 | 2 | 0.5 | 5 | 0 | 2 | 0 | NA | NA | 9 | 0.1111 |
| 7_45475447 | 2 | 0.5 | 2 | 0 | 1 | 1 | 2 | 0.5 | 7 | 0.4286 | 3 | 0.6667 | 5 | 0.4 | 2 | 0.5 | NA | NA | 10 | 0.5 |
| 7_45481648 | NA | NA | 1 | 0 | NA | NA | 2 | 0.5 | 3 | 0.3333 | 1 | 1 | NA | NA | NA | NA | 2 | 0 | 3 | 0.3333 |
| 7_45481781 | NA | NA | 1 | 0 | NA | NA | 2 | 1 | 3 | 0.6667 | 1 | 1 | NA | NA | 1 | 0 | 2 | 0.5 | 4 | 0.5 |
| 7_45482019 | NA | NA | 1 | 1 | NA | NA | 2 | 1 | 3 | 1 | NA | NA | NA | NA | 1 | 1 | 2 | 0.5 | 3 | 0.6667 |
| 7_45482129 | NA | NA | 1 | 1 | NA | NA | NA | NA | 1 | 1 | NA | NA | NA | NA | 1 | 0 | 2 | 1 | 3 | 0.6667 |
| 7_45482709 | NA | NA | NA | NA | NA | NA | 1 | 1 | 1 | 1 | NA | NA | NA | NA | NA | NA | 2 | 1 | 2 | 1 |
| 7_45482889 | NA | NA | NA | NA | NA | NA | 1 | 1 | 1 | 1 | NA | NA | NA | NA | NA | NA | 2 | 0.5 | 2 | 0.5 |
| 7_45484039 | 4 | 0.75 | 1 | 0 | NA | NA | 2 | 0.5 | 7 | 0.5714 | 1 | 1 | NA | NA | 1 | 1 | NA | NA | 2 | 1 |
| 7_45484257 | 4 | 0.5 | 1 | 0 | NA | NA | 2 | 1 | 7 | 0.5714 | 2 | 0 | 1 | 0 | 1 | 1 | NA | NA | 4 | 0.25 |
| 7_45484272 | 4 | 0.25 | NA | NA | NA | NA | 2 | 0.5 | 6 | 0.3333 | 2 | 0 | 1 | 1 | 1 | 0 | NA | NA | 4 | 0.25 |
| 7_45484487 | 2 | 1 | NA | NA | NA | NA | 1 | 0 | 3 | 0.6667 | 1 | 1 | NA | NA | NA | NA | NA | NA | 1 | 1 |
| 7_45484581 | 4 | 0.75 | NA | NA | NA | NA | 2 | 0.5 | 6 | 0.6667 | 2 | 0.5 | 1 | 0 | 1 | 0 | NA | NA | 4 | 0.25 |
| 7_45532519 | 15 | 0.0667 | 18 | 0.0556 | 11 | 0 | 8 | 0.125 | 52 | 0.0577 | 10 | 0.1 | 12 | 0 | 18 | 0 | 12 | 0.25 | 52 | 0.0769 |
| 7_45532608 | 16 | 0 | 20 | 0.2 | 10 | 0.1 | 10 | 0.1 | 56 | 0.1071 | 11 | 0.0909 | 13 | 0 | 20 | 0.1 | 18 | 0.1111 | 62 | 0.0806 |
| 7_45532692 | 17 | 0.0588 | 20 | 0.05 | 11 | 0.0909 | 9 | 0.1111 | 57 | 0.0702 | 9 | 0.2222 | 12 | 0 | 18 | 0 | 17 | 0.1765 | 56 | 0.0893 |
| 7_45532729 | 15 | 0.4 | 18 | 0.4444 | 12 | 0.3333 | 9 | 0.3333 | 54 | 0.3889 | 8 | 0.625 | 9 | 0.3333 | 14 | 0.5 | 16 | 0.3125 | 47 | 0.4255 |
| 7_45565911 | 2 | 1 | 1 | 0 | 3 | 0.3333 | NA | NA | 6 | 0.5 | 2 | 1 | 1 | 0 | 1 | 0 | 4 | 0.5 | 8 | 0.5 |
| 7_45565956 | 2 | 1 | 1 | 1 | 5 | 1 | NA | NA | 8 | 1 | 2 | 0.5 | 3 | 0.6667 | 2 | 1 | 4 | 0.75 | 11 | 0.7273 |
| 7_45566148 | 2 | 1 | 1 | 0 | 5 | 0.6 | NA | NA | 8 | 0.625 | 2 | 1 | 2 | 0.5 | 2 | 0.5 | 6 | 0.8333 | 12 | 0.75 |
| 7_45566353 | 2 | 0.5 | 1 | 0 | 6 | 0.1667 | 1 | 0 | 10 | 0.2 | 2 | 0 | 2 | 0 | 2 | 0 | 7 | 0.2857 | 13 | 0.1538 |
| 7_45566504 | 1 | 0 | 2 | 0.5 | 6 | 0.3333 | 2 | 0 | 11 | 0.2727 | 2 | 0.5 | 3 | 0 | 2 | 0.5 | 7 | 0.1429 | 14 | 0.2143 |
| 7_45566558 | 2 | 0.5 | 1 | 0 | 3 | 0 | 2 | 0.5 | 8 | 0.25 | 2 | 0.5 | 1 | 0 | 2 | 0 | 7 | 0 | 12 | 0.0833 |
| 7_45733218 | 13 | 0.1538 | 15 | 0.1333 | 11 | 0.3636 | 23 | 0.3478 | 62 | 0.2581 | 12 | 0.3333 | 13 | 0.3077 | 13 | 0.2308 | 13 | 0.5385 | 51 | 0.3529 |
| 7_45733332 | 18 | 0.8333 | 13 | 0.7692 | 14 | 0.7857 | 22 | 0.8636 | 67 | 0.8209 | 15 | 0.6 | 13 | 0.8462 | 12 | 0.8333 | 13 | 0.8462 | 53 | 0.7736 |
| 7_45733353 | 11 | 0.2727 | 12 | 0 | 8 | 0 | 22 | 0.3182 | 53 | 0.1887 | 13 | 0.1538 | 8 | 0.125 | 8 | 0 | 10 | 0.2 | 39 | 0.1282 |
| 7_45733437 | 15 | 0.3333 | 17 | 0.2941 | 15 | 0.4 | 24 | 0.1667 | 71 | 0.2817 | 15 | 0.1333 | 15 | 0.0667 | 14 | 0.4286 | 12 | 0.25 | 56 | 0.2143 |
| 7_45733511 | 20 | 0.95 | 15 | 0.7333 | 13 | 1 | 28 | 0.8571 | 76 | 0.8816 | 14 | 0.9286 | 17 | 1 | 15 | 0.9333 | 14 | 0.9286 | 60 | 0.95 |
| 7_45733699 | NA | NA | NA | NA | NA | NA | 1 | 1 | 1 | 1 | NA | NA | NA | NA | 1 | 1 | NA | NA | 1 | 1 |
| 7_45737073 | NA | NA | 1 | 1 | NA | NA | NA | NA | 1 | 1 | NA | NA | NA | NA | 1 | 1 | 1 | 1 | 2 | 1 |
| 7_45737544 | NA | NA | NA | NA | 1 | 1 | NA | NA | 1 | 1 | NA | NA | NA | NA | NA | NA | 1 | 1 | 1 | 1 |
| 7_45746812 | 2 | 0 | 4 | 0.25 | 2 | 0.5 | 3 | 0 | 11 | 0.1818 | 4 | 0.25 | 3 | 0.3333 | 4 | 0 | 5 | 0.4 | 16 | 0.25 |
| 7_45747077 | NA | NA | 4 | 0.25 | 2 | 0.5 | 3 | 0.3333 | 9 | 0.3333 | 2 | 0.5 | 4 | 0.25 | 3 | 0.6667 | 2 | 1 | 11 | 0.5455 |
| 7_45747153 | 1 | 1 | 4 | 0.75 | 1 | 1 | 3 | 0.3333 | 9 | 0.6667 | 3 | 0 | 3 | 0.6667 | 3 | 0.3333 | 4 | 1 | 13 | 0.5385 |
| 7_45747611 | 1 | 0 | 3 | 0 | 1 | 0 | 3 | 0.3333 | 8 | 0.125 | 1 | 0 | 5 | 0.4 | 3 | 0.6667 | 1 | 0 | 10 | 0.4 |
| 7_45747780 | 1 | 0 | 3 | 0 | 1 | 1 | 2 | 0 | 7 | 0.1429 | 1 | 1 | 5 | 0 | 3 | 0 | 1 | 0 | 10 | 0.1 |
| 7_45747898 | NA | NA | 3 | 0.6667 | NA | NA | 1 | 1 | 4 | 0.75 | NA | NA | 4 | 1 | 2 | 1 | 1 | 1 | 7 | 1 |

| | | | | | | | | | | | | | | | | | | | | |
|---|---|---|---|---|---|---|---|---|---|---|---|---|---|---|---|---|---|---|---|---|
| 7_4574803( | NA | NA | 2 | 1 | NA | NA | 2 | 1 | 4 | 1 | 1 | 1 | 4 | 0.75 | 2 | 1 | 1 | 1 | 8 | 0.875 |
| 7_4574826 | NA | NA | 1 | 1 | NA | NA | 2 | 1 | 3 | 1 | 1 | 1 | 3 | 1 | 1 | 1 | 1 | 0 | 6 | 0.8333 |
| 7_4574833( | NA | NA | 1 | 1 | NA | NA | 2 | 1 | 3 | 1 | 1 | 1 | 3 | 1 | 1 | 1 | 1 | 1 | 6 | 1 |
| 7_4574844! | NA | NA | NA | NA | NA | NA | 2 | 0.5 | 2 | 0.5 | 1 | 1 | 3 | 0.3333 | 1 | 0 | 1 | 0 | 6 | 0.3333 |
| 7_4574848: | NA | NA | 1 | 1 | NA | NA | 2 | 1 | 3 | 1 | 1 | 1 | 3 | 0 | 1 | 1 | NA | NA | 5 | 0.4 |
| 7_4574871 | NA | NA | 1 | 0 | NA | NA | 2 | 0.5 | 3 | 0.3333 | NA | NA | 3 | 0.3333 | 1 | 1 | NA | NA | 4 | 0.5 |
| 7_4574912( | NA | NA | NA | NA | NA | NA | 2 | 0.5 | 2 | 0.5 | NA | NA | 3 | 0.3333 | 1 | 0 | NA | NA | 4 | 0.25 |
| 7_4574917! | NA | NA | NA | NA | NA | NA | 1 | 1 | 1 | 1 | NA | NA | 2 | 1 | NA | NA | NA | NA | 2 | 1 |
| 7_4575404( | 5 | 0.2 | 5 | 0 | 5 | 0 | 1 | 0 | 16 | 0.0625 | 2 | 0 | 1 | 0 | 1 | 0 | 1 | 1 | 5 | 0.2 |
| 7_4575751 | 3 | 0.3333 | 4 | 0.25 | 3 | 0 | NA | NA | 10 | 0.2 | 2 | 0.5 | 1 | 0 | 1 | 0 | 1 | 0 | 5 | 0.2 |
| 7_4576639: | 4 | 0.25 | 1 | 0 | 3 | 0 | NA | NA | 8 | 0.125 | 2 | 0.5 | 1 | 0 | 1 | 0 | 1 | 0 | 5 | 0.2 |
| 7_4576664! | 4 | 0 | 2 | 0 | 3 | 0.3333 | NA | NA | 9 | 0.1111 | 1 | 1 | 1 | 0 | 1 | 0 | NA | NA | 3 | 0.3333 |
| 7_4604687( | 1 | 1 | NA | NA | 1 | 0 | NA | NA | 2 | 0.5 | 2 | 0.5 | NA | NA | NA | NA | 1 | 0 | 3 | 0.3333 |
| 7_4604730 | 1 | 1 | NA | NA | 2 | 0.5 | 1 | 1 | 4 | 0.75 | 1 | 1 | 1 | 0 | 2 | 0.5 | NA | NA | 4 | 0.5 |
| 7_4604746( | 1 | 1 | NA | NA | 1 | 0 | 1 | 1 | 3 | 0.6667 | 2 | 0 | 2 | 0.5 | 2 | 0.5 | NA | NA | 6 | 0.3333 |
| 7_4604764( | 1 | 1 | NA | NA | 1 | 1 | NA | NA | 2 | 1 | 4 | 0.25 | 2 | 0 | 2 | 0 | 1 | 1 | 9 | 0.2222 |
| 7_4607702( | 1 | 1 | NA | NA | 1 | 0 | 2 | 1 | 4 | 0.75 | 3 | 0.6667 | 1 | 1 | 3 | 0.6667 | 1 | 1 | 8 | 0.75 |
| 7_4607704 | 1 | 1 | NA | NA | 2 | 0.5 | 2 | 1 | 5 | 0.8 | 5 | 1 | 1 | 1 | 3 | 1 | NA | NA | 9 | 1 |
| 7_4607733! | NA | NA | 1 | 0 | 1 | 1 | 1 | 1 | 3 | 0.6667 | 6 | 0.6667 | 2 | 1 | 4 | 0.75 | 1 | 1 | 13 | 0.7692 |
| 7_4607735( | 1 | 0 | 1 | 1 | 2 | 0 | 2 | 0 | 6 | 0.1667 | 6 | 0.1667 | 2 | 1 | 3 | 0.3333 | 1 | 0 | 12 | 0.3333 |
| 7_4607741: | NA | NA | NA | NA | 2 | 1 | 4 | 0.5 | 6 | 0.6667 | 5 | 0.6 | 2 | 0.5 | 4 | 0.5 | 2 | 0.5 | 13 | 0.5385 |
| 7_4607750! | 1 | 1 | 1 | 1 | 1 | 1 | 4 | 0.5 | 7 | 0.7143 | 5 | 0.8 | 2 | 0.5 | 3 | 0.6667 | NA | NA | 10 | 0.7 |
| 7_4607755: | 1 | 1 | 1 | 0 | 2 | 0 | 4 | 0.5 | 8 | 0.375 | 6 | 0.1667 | 2 | 0 | 2 | 0.5 | 2 | 0 | 12 | 0.1667 |
| 7_4607762! | 2 | 0.5 | 1 | 1 | NA | NA | 2 | 0 | 5 | 0.4 | 4 | 0.5 | 2 | 0 | 2 | 0.5 | 2 | 0.5 | 10 | 0.4 |
| 7_4607765( | 2 | 0.5 | 1 | 0 | 2 | 0.5 | 4 | 0.25 | 9 | 0.3333 | 5 | 0.2 | 2 | 0 | 3 | 0 | 2 | 0.5 | 12 | 0.1667 |
| 7_4607798: | 1 | 1 | NA | NA | NA | NA | 3 | 0.6667 | 4 | 0.75 | 2 | 1 | NA | NA | NA | NA | 1 | 1 | 3 | 1 |
| 7_4607840: | 1 | 1 | NA | NA | NA | NA | 3 | 0.3333 | 4 | 0.5 | 3 | 0.3333 | 1 | 0 | NA | NA | 1 | 0 | 5 | 0.2 |
| 7_4607841 | 1 | 0 | NA | NA | NA | NA | 4 | 0.25 | 5 | 0.2 | 3 | 0.3333 | 1 | 1 | NA | NA | 1 | 0 | 5 | 0.4 |
| 7_4607875! | 2 | 0.5 | NA | NA | 1 | 1 | 4 | 0.75 | 7 | 0.7143 | 2 | 1 | 1 | 1 | 1 | 1 | NA | NA | 4 | 1 |
| 7_4608219( | 2 | 0.5 | 3 | 0.6667 | 2 | 0.5 | 9 | 0.1111 | 16 | 0.3125 | 9 | 0.1111 | 2 | 0 | 2 | 0 | 3 | 0 | 16 | 0.0625 |
| 7_4608231( | 5 | 0.6 | 3 | 0.3333 | 3 | 0.6667 | 12 | 0.4167 | 23 | 0.4783 | 8 | 0.375 | NA | NA | 2 | 0 | 1 | 1 | 11 | 0.3636 |
| 7_4608269( | 5 | 0 | 3 | 0 | 3 | 0.3333 | 13 | 0.2308 | 24 | 0.1667 | 9 | 0.1111 | 2 | 0 | 2 | 0 | 2 | 0.5 | 15 | 0.1333 |
| 7_4608675! | 2 | 0.5 | 1 | 0 | 1 | 1 | 1 | 1 | 5 | 0.6 | 2 | 0.5 | 3 | 0.6667 | 5 | 0.6 | NA | NA | 10 | 0.6 |
| 7_4608678( | 3 | 1 | 1 | 0 | 2 | 1 | 2 | 0.5 | 8 | 0.75 | 4 | 0.75 | 5 | 0.8 | 5 | 0.8 | 1 | 1 | 15 | 0.8 |
| 7_4608683 | 3 | 0.3333 | 1 | 1 | 2 | 1 | 1 | 1 | 7 | 0.7143 | 4 | 0.5 | 5 | 0.6 | 3 | 0.6667 | 1 | 1 | 13 | 0.6154 |
| 7_4608708! | 2 | 0 | 4 | 0.75 | 2 | 0.5 | 3 | 0 | 11 | 0.3636 | 4 | 0.25 | 5 | 0.6 | 5 | 0.4 | 3 | 0.6667 | 17 | 0.4706 |
| 7_4608718( | 3 | 0.3333 | 4 | 0.25 | 2 | 0 | 4 | 0 | 13 | 0.1538 | 4 | 0.25 | 7 | 0 | 3 | 0.3333 | 2 | 0 | 16 | 0.125 |
| 7_4608738 | 4 | 0.5 | 4 | 0.75 | 5 | 0.8 | 4 | 1 | 17 | 0.7647 | 6 | 0.8333 | 6 | 0.8333 | 8 | 0.625 | 5 | 0.6 | 25 | 0.72 |
| 7_4608748! | 2 | 0.5 | 6 | 0.5 | 6 | 0 | 5 | 0.2 | 19 | 0.2632 | 6 | 0.5 | 5 | 0 | 7 | 0.1429 | 4 | 0.25 | 22 | 0.2273 |
| 7_4608767: | 5 | 0.2 | 6 | 0.3333 | 7 | 0.5714 | 5 | 0.4 | 23 | 0.3913 | 6 | 0 | 8 | 0.375 | 5 | 0 | 5 | 0.2 | 24 | 0.1667 |
| 7_4608790( | 2 | 0 | NA | NA | 3 | 0.3333 | NA | NA | 5 | 0.2 | 2 | 0.5 | 1 | 0 | 4 | 0 | 1 | 0 | 8 | 0.125 |
| 7_4609143 | 1 | 1 | NA | NA | NA | NA | NA | NA | 1 | 1 | 1 | 1 | NA | NA | 1 | 0 | NA | NA | 2 | 0.5 |
| 7_4609269 | 5 | 0.2 | 1 | 1 | 3 | 0.3333 | 1 | 0 | 10 | 0.3 | NA | NA | 1 | 1 | 2 | 0.5 | 3 | 0.6667 | 6 | 0.6667 |
| 7_4609270 | 5 | 0 | NA | NA | 2 | 0 | 1 | 1 | 8 | 0.125 | 1 | 0 | 1 | 0 | 2 | 0 | 3 | 0.3333 | 7 | 0.1429 |
| 7_4609270 | 1 | 0 | 1 | 0 | 3 | 1 | 1 | 0 | 6 | 0.5 | 1 | 1 | 1 | 1 | 2 | 0 | 2 | 0.5 | 6 | 0.5 |
| 7_4609273 | 5 | 0.2 | NA | NA | 2 | 0 | 1 | 0 | 8 | 0.125 | 1 | 0 | 1 | 0 | 1 | 1 | 1 | 1 | 4 | 0.5 |
| 7_4616139! | NA | NA | 1 | 1 | NA | NA | 1 | 0 | 2 | 0.5 | NA | NA | NA | NA | 1 | 0 | 1 | 1 | 2 | 0.5 |
| 7_4616169: | NA | NA | NA | NA | NA | NA | 1 | 1 | 1 | 1 | NA | NA | NA | NA | 1 | 1 | 1 | 1 | 2 | 1 |
| 7_4616203 | NA | NA | 1 | 0 | NA | NA | 1 | 1 | 2 | 0.5 | NA | NA | NA | NA | 1 | 1 | NA | NA | 1 | 1 |
| 7_4628516 | NA | NA | 1 | 1 | NA | NA | NA | NA | 1 | 1 | NA | NA | NA | NA | NA | NA | 1 | 1 | 1 | 1 |
| 7_4633681 | NA | NA | NA | NA | NA | NA | 1 | 1 | 1 | 1 | NA | NA | NA | NA | NA | NA | 1 | 1 | 1 | 1 |
| 7_4633750: | NA | NA | NA | NA | NA | NA | 1 | 1 | 1 | 1 | NA | NA | NA | NA | NA | NA | 1 | 1 | 1 | 1 |
| 7_4633757( | NA | NA | NA | NA | NA | NA | 1 | 1 | 1 | 1 | NA | NA | NA | NA | NA | NA | 1 | 1 | 1 | 1 |
| 7_4633770: | NA | NA | NA | NA | NA | NA | 1 | 1 | 1 | 1 | NA | NA | NA | NA | NA | NA | 1 | 1 | 1 | 1 |
| 7_4633869! | NA | NA | NA | NA | NA | NA | 1 | 1 | 1 | 1 | 1 | 1 | NA | NA | NA | NA | 1 | 1 | 2 | 1 |
| 7_4633875 | NA | NA | 1 | 0 | NA | NA | 1 | 1 | 2 | 0.5 | 2 | 0.5 | NA | NA | NA | NA | 3 | 0.3333 | 5 | 0.4 |
| 7_4641044: | 1 | 0 | 2 | 0 | 4 | 0.25 | 1 | 0 | 8 | 0.125 | NA | NA | 2 | 0 | NA | NA | 1 | 1 | 3 | 0.3333 |
| 7_4642111 | 1 | 0 | 1 | 1 | 1 | 1 | NA | NA | 3 | 0.6667 | NA | NA | 1 | 1 | NA | NA | 1 | 0 | 2 | 0.5 |
| 7_4642123( | 1 | 1 | 1 | 1 | 1 | 1 | NA | NA | 3 | 1 | NA | NA | 1 | 1 | NA | NA | 1 | 1 | 2 | 1 |
| 7_4642150 | 1 | 0 | 2 | 0.5 | NA | NA | NA | NA | 3 | 0.3333 | NA | NA | 1 | 0 | NA | NA | 1 | 1 | 2 | 0.5 |
| 7_4647237 | 6 | 0 | 4 | 0.5 | 4 | 0 | 5 | 0.4 | 19 | 0.2105 | 3 | 0 | 9 | 0.1111 | 3 | 0.3333 | 4 | 0 | 19 | 0.1053 |
| 7_4647249: | 10 | 0.3 | 4 | 0 | 4 | 0.25 | 5 | 0.2 | 23 | 0.2174 | 4 | 0.25 | 9 | 0.1111 | 4 | 0 | 4 | 0 | 21 | 0.0952 |
| 7_4647249! | 7 | 0 | 4 | 0.25 | 4 | 0 | 5 | 0 | 20 | 0.05 | 4 | 0.25 | 8 | 0 | 3 | 0 | 3 | 0 | 18 | 0.0556 |
| 7_4647252( | 6 | 0.1667 | 2 | 0 | 4 | 0.5 | 3 | 0.3333 | 15 | 0.2667 | 3 | 0.3333 | 5 | 0.6 | 4 | 0.25 | 3 | 0.6667 | 15 | 0.4667 |
| 7_4647261( | 8 | 0.5 | 5 | 0.6 | 4 | 0.75 | 6 | 0.6667 | 23 | 0.6087 | 5 | 0.2 | 10 | 0.6 | 6 | 0.8333 | 4 | 0.75 | 25 | 0.6 |

| | | | | | | | | | | | | | | | | | | | | |
|---|---|---|---|---|---|---|---|---|---|---|---|---|---|---|---|---|---|---|---|---|
| 7_4657612 | 1 | 1 | NA | NA | NA | NA | 1 | 0 | 2 | 0.5 | 1 | 1 | NA | NA | 1 | 0 | NA | NA | 2 | 0.5 |
| 7_4664027 | 3 | 0 | 4 | 0.5 | 2 | 0.5 | 2 | 0.5 | 11 | 0.3636 | 1 | 0 | 1 | 1 | 2 | 0 | NA | NA | 4 | 0.25 |
| 7_4664029 | 2 | 0 | 4 | 0.25 | 2 | 0 | 2 | 0 | 10 | 0.1 | 1 | 0 | 1 | 0 | 2 | 0.5 | NA | NA | 4 | 0.25 |
| 7_4664062 | 4 | 0.25 | 4 | 0.25 | 2 | 0 | 4 | 0.75 | 14 | 0.3571 | 2 | 0 | 2 | 0 | 1 | 1 | 2 | 0 | 7 | 0.1429 |
| 7_4664072 | 3 | 0.3333 | 4 | 0 | 2 | 0 | 2 | 0 | 11 | 0.0909 | 2 | 0.5 | 2 | 0 | 1 | 0 | 2 | 0 | 7 | 0.1429 |
| 7_4664079 | 4 | 0 | 4 | 0 | 2 | 0.5 | 4 | 0 | 14 | 0.0714 | 1 | 0 | 2 | 0 | 1 | 1 | 2 | 0 | 6 | 0.1667 |
| 7_4664183 | 2 | 0.5 | 4 | 0.25 | NA | NA | 4 | 0.5 | 10 | 0.4 | NA | NA | 2 | 0.5 | 1 | 0 | 2 | 0 | 5 | 0.2 |
| 7_4664187 | 1 | 1 | 2 | 0.5 | NA | NA | 4 | 0.5 | 7 | 0.5714 | NA | NA | 2 | 0.5 | 1 | 0 | 2 | 0 | 5 | 0.2 |
| 7_4664194 | 2 | 0 | 4 | 0 | NA | NA | 4 | 0.25 | 10 | 0.1 | NA | NA | 1 | 0 | NA | NA | 2 | 0.5 | 3 | 0.3333 |
| 7_4664311 | 1 | 1 | 2 | 0 | NA | NA | 3 | 0 | 6 | 0.1667 | NA | NA | 1 | 0 | NA | NA | 1 | 1 | 2 | 0.5 |
| 7_4664741 | 1 | 1 | 2 | 0.5 | NA | NA | 2 | 1 | 5 | 0.8 | NA | NA | NA | NA | NA | NA | 2 | 1 | 2 | 1 |
| 7_4664748 | NA | NA | NA | NA | NA | NA | 1 | 1 | 1 | 1 | NA | NA | NA | NA | NA | NA | 2 | 1 | 2 | 1 |
| 7_4664770 | 2 | 0.5 | 2 | 0.5 | NA | NA | 1 | 1 | 5 | 0.6 | NA | NA | NA | NA | NA | NA | 2 | 0.5 | 2 | 0.5 |
| 7_4664793 | 1 | 1 | 2 | 0.5 | NA | NA | 1 | 0 | 4 | 0.5 | NA | NA | NA | NA | NA | NA | 2 | 1 | 2 | 1 |
| 7_4664795 | 1 | 0 | 2 | 0.5 | NA | NA | 2 | 1 | 5 | 0.6 | NA | NA | NA | NA | NA | NA | 1 | 1 | 1 | 1 |
| 7_4665864 | NA | NA | NA | NA | 1 | 1 | 2 | 0.5 | 3 | 0.6667 | NA | NA | NA | NA | 1 | 0 | 1 | 1 | 2 | 0.5 |
| 7_4665869 | NA | NA | NA | NA | 1 | 1 | 1 | 1 | 2 | 1 | NA | NA | NA | NA | 1 | 1 | 1 | 0 | 2 | 0.5 |
| 7_4665883 | NA | NA | NA | NA | 1 | 1 | NA | NA | 1 | 1 | NA | NA | NA | NA | NA | NA | 1 | 1 | 1 | 1 |
| 7_4670358 | NA | NA | NA | NA | 2 | 0.5 | 1 | 0 | 3 | 0.3333 | NA | NA | 3 | 0.3333 | 3 | 0 | 1 | 0 | 7 | 0.1429 |
| 7_4670360 | NA | NA | NA | NA | 3 | 0.6667 | 1 | 1 | 4 | 0.75 | NA | NA | 3 | 1 | 4 | 1 | 3 | 0.6667 | 10 | 0.9 |
| 7_4670364 | NA | NA | NA | NA | 2 | 1 | 1 | 1 | 3 | 1 | NA | NA | 1 | 1 | 3 | 1 | 2 | 1 | 6 | 1 |
| 7_4670382 | NA | NA | NA | NA | 3 | 0.6667 | 1 | 1 | 4 | 0.75 | NA | NA | 3 | 0.6667 | 3 | 0.3333 | 2 | 0.5 | 8 | 0.5 |
| 7_4670389 | NA | NA | NA | NA | 3 | 0.3333 | 1 | 0 | 4 | 0.25 | NA | NA | 2 | 0 | 4 | 0.25 | 2 | 0 | 8 | 0.125 |
| 7_4670411 | NA | NA | 1 | 0 | 2 | 0 | 1 | 1 | 4 | 0.25 | NA | NA | 3 | 0 | 3 | 0.3333 | 3 | 0 | 9 | 0.1111 |
| 7_4670429 | NA | NA | 1 | 0 | 3 | 0 | 1 | 1 | 5 | 0.2 | NA | NA | 2 | 0.5 | 4 | 0 | 3 | 0 | 9 | 0.1111 |
| 7_4672782 | 36 | 0.1389 | 31 | 0.0323 | 35 | 0.0571 | 49 | 0.0816 | 151 | 0.0795 | 17 | 0.0588 | 47 | 0.1064 | 50 | 0.08 | 42 | 0.1667 | 156 | 0.109 |
| 7_4678318 | 2 | 0.5 | NA | NA | 1 | 0 | NA | NA | 3 | 0.3333 | NA | NA | 1 | 1 | 2 | 0 | NA | NA | 3 | 0.3333 |
| 7_4847669 | 4 | 0 | 6 | 0.1667 | 8 | 0.125 | 4 | 0.25 | 22 | 0.1364 | 4 | 0 | 6 | 0 | 4 | 0.25 | 2 | 0 | 16 | 0.0625 |
| 7_4847678 | 4 | 0.5 | 6 | 0.5 | 8 | 0.5 | 4 | 0.75 | 22 | 0.5455 | 4 | 0.5 | 6 | 0.3333 | 4 | 0.25 | 3 | 0.6667 | 17 | 0.4118 |
| 7_4847688 | 4 | 0.25 | 7 | 0.1429 | 6 | 0.3333 | 4 | 0.25 | 21 | 0.2381 | 3 | 0.3333 | 7 | 0.2857 | 4 | 0 | 3 | 0.6667 | 17 | 0.2941 |
| 7_4847694 | 3 | 0 | 6 | 0 | 8 | 0.125 | 3 | 0.3333 | 20 | 0.1 | 4 | 0 | 7 | 0.1429 | 3 | 0 | 3 | 0 | 17 | 0.0588 |
| 7_4847697 | 4 | 0.25 | 6 | 0.1667 | 6 | 0.1667 | 4 | 0.25 | 20 | 0.2 | 2 | 0 | 6 | 0.3333 | 3 | 0.3333 | 2 | 0.5 | 13 | 0.3077 |
| 7_4847699 | 3 | 0 | 7 | 0 | 5 | 0.2 | 4 | 0 | 19 | 0.0526 | 4 | 0 | 7 | 0.1429 | 4 | 0 | 1 | 0 | 16 | 0.0625 |
| 7_4925674 | NA | NA | 2 | 0.5 | 2 | 0.5 | NA | NA | 4 | 0.5 | 1 | 1 | NA | NA | 1 | 0 | 1 | 0 | 3 | 0.3333 |
| 7_4925697 | NA | NA | 2 | 0.5 | 2 | 1 | NA | NA | 4 | 0.75 | 1 | 1 | NA | NA | 1 | 1 | 1 | 1 | 3 | 1 |
| 7_4925834 | 1 | 0 | 3 | 0 | 4 | 0 | 2 | 0.5 | 10 | 0.1 | 4 | 0.5 | NA | NA | 2 | 0.5 | 1 | 0 | 7 | 0.4286 |
| 7_4925910 | 1 | 0 | 6 | 0 | 4 | 0.25 | 3 | 0 | 14 | 0.0714 | 3 | 0 | NA | NA | 5 | 0.2 | 1 | 0 | 9 | 0.1111 |
| 7_4925911 | 1 | 0 | 6 | 0.1667 | 4 | 0 | 3 | 0 | 14 | 0.0714 | 3 | 0.3333 | NA | NA | 5 | 0.2 | 1 | 0 | 9 | 0.2222 |
| 7_4942321 | 3 | 0 | 3 | 0.3333 | 2 | 1 | NA | NA | 8 | 0.375 | NA | NA | NA | NA | NA | NA | 2 | 0.5 | 2 | 0.5 |
| 7_4942328 | 2 | 0.5 | 2 | 0.5 | 2 | 1 | NA | NA | 6 | 0.6667 | NA | NA | NA | NA | NA | NA | 2 | 1 | 2 | 1 |
| 7_4950719 | NA | NA | 3 | 0.3333 | 2 | 0 | 1 | 0 | 6 | 0.1667 | 2 | 0 | 1 | 0 | 1 | 1 | NA | NA | 4 | 0.25 |
| 7_4950722 | NA | NA | 5 | 0.2 | 2 | 0 | 1 | 0 | 8 | 0.125 | 1 | 0 | 1 | 0 | 1 | 0 | 2 | 0.5 | 5 | 0.2 |
| 7_4950732 | NA | NA | 5 | 0.6 | 3 | 0.6667 | 1 | 1 | 9 | 0.6667 | 1 | 1 | 2 | 0 | 2 | 0.5 | 1 | 1 | 6 | 0.5 |
| 7_4950749 | NA | NA | 6 | 0.1667 | 3 | 0 | 1 | 0 | 10 | 0.1 | 2 | 0 | 3 | 0.3333 | 3 | 0 | 1 | 1 | 9 | 0.2222 |
| 7_4950784 | NA | NA | 4 | 0.25 | 1 | 0 | 1 | 0 | 6 | 0.1667 | 3 | 0 | 3 | 0 | 3 | 0 | 3 | 0.3333 | 12 | 0.0833 |
| 7_5051271 | 7 | 0.4286 | 8 | 1 | 13 | 0.6154 | 6 | 0.6667 | 34 | 0.6765 | 9 | 0.4444 | 6 | 0.5 | 2 | 0.5 | 3 | 1 | 20 | 0.55 |
| 7_5051273 | 6 | 0.5 | 8 | 0.25 | 14 | 0.4286 | 6 | 0.3333 | 34 | 0.3824 | 9 | 0.3333 | 6 | 0.3333 | 2 | 0.5 | 4 | 0.75 | 21 | 0.4286 |
| 7_5051302 | 10 | 0 | 10 | 0.5 | 16 | 0.1875 | 9 | 0.4444 | 45 | 0.2667 | 10 | 0.2 | 6 | 0.1667 | 2 | 0.5 | 4 | 0.25 | 22 | 0.2273 |
| 7_5118833 | NA | NA | NA | NA | 1 | 1 | NA | NA | 1 | 1 | NA | NA | 3 | 0.6667 | NA | NA | NA | NA | 3 | 0.6667 |
| 7_5131908 | NA | NA | 2 | 1 | NA | NA | NA | NA | 2 | 1 | NA | NA | 1 | 1 | NA | NA | NA | NA | 1 | 1 |
| 7_5151086 | NA | NA | 3 | 1 | NA | NA | NA | NA | 3 | 1 | NA | NA | 3 | 0.3333 | NA | NA | 1 | 1 | 4 | 0.5 |
| 7_5151094 | NA | NA | 1 | 1 | NA | NA | NA | NA | 1 | 1 | NA | NA | 2 | 1 | NA | NA | 1 | 1 | 3 | 1 |
| 7_5151101 | 1 | 1 | 2 | 1 | NA | NA | NA | NA | 3 | 1 | NA | NA | 2 | 1 | NA | NA | 1 | 0 | 3 | 0.6667 |
| 7_5151115 | 1 | 1 | 3 | 1 | NA | NA | NA | NA | 4 | 1 | NA | NA | 3 | 1 | NA | NA | 1 | 1 | 4 | 1 |
| 7_5151123 | 1 | 1 | 3 | 1 | NA | NA | NA | NA | 4 | 1 | NA | NA | 3 | 1 | NA | NA | 1 | 1 | 4 | 1 |
| 7_5151137 | 1 | 1 | 3 | 0.3333 | NA | NA | NA | NA | 4 | 0.5 | NA | NA | 3 | 0.6667 | NA | NA | 1 | 0 | 4 | 0.5 |
| 7_5151147 | NA | NA | 2 | 1 | NA | NA | NA | NA | 2 | 1 | NA | NA | 3 | 1 | NA | NA | 1 | 0 | 4 | 0.75 |
| 7_5151214 | NA | NA | NA | NA | 2 | 1 | NA | NA | 2 | 1 | 1 | 1 | NA | NA | NA | NA | NA | NA | 1 | 1 |
| 7_5164871 | 3 | 0 | 2 | 0.5 | 4 | 0 | 9 | 0 | 18 | 0.0556 | 4 | 0.5 | 6 | 0 | 6 | 0 | 6 | 0 | 22 | 0.0909 |
| 7_5164941 | 5 | 0 | 2 | 0.5 | 6 | 0 | 8 | 0.125 | 21 | 0.0952 | 3 | 0.3333 | 8 | 0.25 | 5 | 0.2 | 3 | 0 | 19 | 0.2105 |
| 7_5406166 | 4 | 0 | 3 | 0.3333 | 2 | 0 | 6 | 0 | 15 | 0.0667 | 4 | 0.25 | 4 | 0 | 4 | 0 | 7 | 0.1429 | 19 | 0.1053 |
| 7_5406176 | 9 | 0.3333 | 7 | 0.5714 | 2 | 1 | 6 | 0.3333 | 24 | 0.4583 | 5 | 0.8 | 4 | 0.75 | 4 | 0.75 | 6 | 0.5 | 19 | 0.6842 |
| 7_5406186 | 6 | 0.3333 | 5 | 0.6 | 3 | 0.3333 | 7 | 0.4286 | 21 | 0.4286 | 4 | 0.25 | 4 | 0.25 | 3 | 0 | 7 | 0.1429 | 18 | 0.1667 |
| 7_5406208 | 6 | 0 | 2 | 0 | 2 | 0 | 7 | 0.1429 | 17 | 0.0588 | 3 | 0 | 1 | 0 | 1 | 0 | 4 | 0.25 | 9 | 0.1111 |
| 7_5491521 | 14 | 0.1429 | 17 | 0.1765 | 18 | 0.1667 | 5 | 0 | 54 | 0.1481 | 9 | 0.2222 | 11 | 0.1818 | 9 | 0.1111 | 12 | 0.25 | 41 | 0.1951 |

| | | | | | | | | | | | | | | | | | | | | |
|---|---|---|---|---|---|---|---|---|---|---|---|---|---|---|---|---|---|---|---|---|
| 7_5491522 | 13 | 0.0769 | 18 | 0.1667 | 18 | 0 | 4 | 0 | 53 | 0.0755 | 10 | 0.1 | 10 | 0 | 9 | 0.1111 | 12 | 0.1667 | 41 | 0.0976 |
| 7_5491523 | 11 | 0.6364 | 13 | 0.6154 | 18 | 0.7778 | 5 | 1 | 47 | 0.7234 | 9 | 0.8889 | 9 | 0.5556 | 9 | 0.5556 | 10 | 0.7 | 37 | 0.6757 |
| 7_5491527 | 11 | 0 | 17 | 0.2353 | 21 | 0.0476 | 6 | 0 | 55 | 0.0909 | 10 | 0.1 | 12 | 0.1667 | 8 | 0.25 | 12 | 0.1667 | 42 | 0.1667 |
| 7_5491771 | 21 | 0.1429 | 27 | 0.3333 | 24 | 0.125 | 15 | 0.1333 | 87 | 0.1954 | 16 | 0.125 | 16 | 0.25 | 13 | 0.3077 | 15 | 0.2 | 60 | 0.2167 |
| 7_5491781 | 18 | 0.0556 | 31 | 0.1935 | 25 | 0.08 | 14 | 0.1429 | 88 | 0.125 | 16 | 0.25 | 15 | 0.0667 | 12 | 0.1667 | 16 | 0.25 | 59 | 0.1864 |
| 7_5491835 | 10 | 0.1 | 17 | 0.2941 | 14 | 0.0714 | 13 | 0 | 54 | 0.1296 | 6 | 0 | 10 | 0.2 | 9 | 0.3333 | 9 | 0.2222 | 34 | 0.2059 |
| 7_5541835 | NA | NA | 3 | 0.3333 | 3 | 0 | 1 | 1 | 7 | 0.2857 | 1 | 0 | 1 | 0 | 3 | 0.6667 | NA | NA | 5 | 0.4 |
| 7_5558256 | 8 | 0 | 5 | 0.2 | 3 | 0 | 8 | 0.625 | 24 | 0.25 | 6 | 0.1667 | 6 | 0 | 3 | 0 | 6 | 0.1667 | 21 | 0.0952 |
| 7_5558266 | 10 | 0.4 | 4 | 0.5 | 1 | 0 | 10 | 0.3 | 25 | 0.36 | 5 | 0.2 | 6 | 0.3333 | 3 | 0.6667 | 6 | 0.8333 | 20 | 0.5 |
| 7_5558290 | 9 | 1 | 4 | 0.75 | 3 | 1 | 9 | 0.8889 | 25 | 0.92 | 5 | 1 | 6 | 1 | 3 | 1 | 5 | 1 | 19 | 1 |
| 7_5558296 | 9 | 0.4444 | 3 | 0.3333 | 2 | 0.5 | 9 | 0.1111 | 23 | 0.3043 | 6 | 0.1667 | 6 | 0.3333 | 3 | 0.3333 | 5 | 0 | 20 | 0.2 |
| 7_5558299 | 8 | 0.125 | 5 | 0.4 | 4 | 0.25 | 7 | 0.2857 | 24 | 0.25 | 6 | 0.1667 | 5 | 0.4 | 3 | 0 | 7 | 0.4286 | 21 | 0.2857 |
| 7_5558302 | 8 | 1 | 4 | 1 | 4 | 1 | 8 | 1 | 24 | 1 | 6 | 1 | 6 | 1 | 2 | 1 | 7 | 1 | 21 | 1 |
| 7_5558321 | 9 | 0.1111 | 3 | 0 | 3 | 0 | 8 | 0.375 | 23 | 0.1739 | 6 | 0.3333 | 6 | 0.1667 | 2 | 0.5 | 7 | 0 | 21 | 0.1905 |
| 7_5558322 | 10 | 0.9 | 3 | 1 | 3 | 0.3333 | 8 | 0.75 | 24 | 0.7917 | 5 | 0.6 | 4 | 0.75 | 2 | 0.5 | 6 | 1 | 17 | 0.7647 |
| 7_5558323 | 8 | 0.125 | 3 | 0 | 3 | 0.3333 | 8 | 0.125 | 22 | 0.1364 | 6 | 0 | 6 | 0.1667 | 2 | 0 | 7 | 0.1429 | 21 | 0.0952 |
| 7_5562390 | 7 | 0.5714 | 9 | 0.4444 | 7 | 0.5714 | 8 | 0.5 | 31 | 0.5161 | 8 | 0.25 | 10 | 0.4 | 10 | 0.6 | 6 | 0.6667 | 34 | 0.4706 |
| 7_5562610 | NA | NA | 5 | 0 | 5 | 0.2 | 3 | 0 | 13 | 0.0769 | 2 | 0.5 | 2 | 0 | 7 | 0 | 3 | 0 | 14 | 0.0714 |
| 7_5562820 | 1 | 0 | 6 | 0.1667 | 5 | 0.4 | 2 | 0 | 14 | 0.2143 | 1 | 0 | 1 | 1 | 4 | 0 | 1 | 1 | 7 | 0.2857 |
| 7_5562840 | 5 | 0 | 14 | 0.0714 | 9 | 0.1111 | 9 | 0 | 37 | 0.0541 | 3 | 0 | 2 | 0 | 7 | 0.1429 | 5 | 0 | 17 | 0.0588 |
| 7_5562902 | 5 | 1 | 9 | 0.8889 | 3 | 1 | 7 | 0.8571 | 24 | 0.9167 | 2 | 0.5 | NA | NA | 5 | 1 | 4 | 0.5 | 11 | 0.7273 |
| 7_5562903 | 2 | 1 | 11 | 0.5455 | 5 | 0.6 | 5 | 0.4 | 23 | 0.5652 | 3 | 0.6667 | NA | NA | 6 | 0.8333 | 3 | 1 | 12 | 0.8333 |
| 7_5562906 | 4 | 1 | 13 | 1 | 5 | 0.8 | 7 | 1 | 29 | 0.9655 | 3 | 1 | NA | NA | 7 | 0.8571 | 5 | 1 | 15 | 0.9333 |
| 7_5562911 | 4 | 0.25 | 8 | 0.125 | 2 | 0 | 3 | 0.3333 | 17 | 0.1765 | 3 | 0.3333 | NA | NA | 4 | 0 | 2 | 0.5 | 9 | 0.2222 |
| 7_5562914 | 3 | 1 | 10 | 0.9 | 4 | 1 | 6 | 1 | 23 | 0.9565 | 3 | 1 | NA | NA | 4 | 1 | 4 | 1 | 11 | 1 |
| 7_5562916 | 3 | 0.6667 | 8 | 0.5 | 2 | 0 | 7 | 0.1429 | 20 | 0.35 | 2 | 0 | NA | NA | 4 | 0 | 1 | 1 | 7 | 0.1429 |
| 7_5562926 | 3 | 1 | 9 | 1 | 4 | 0.75 | 6 | 1 | 22 | 0.9545 | 3 | 1 | NA | NA | 3 | 1 | 4 | 1 | 10 | 1 |
| 7_5562948 | 4 | 0.75 | 9 | 0.5556 | 5 | 0.2 | 7 | 0.4286 | 25 | 0.48 | 3 | 0.3333 | NA | NA | 3 | 0.3333 | 4 | 0.75 | 10 | 0.5 |
| 7_5585654 | 2 | 0.5 | 4 | 0 | 1 | 0 | 2 | 0 | 9 | 0.1111 | 2 | 0 | 4 | 0 | 5 | 0.2 | 2 | 0 | 13 | 0.0769 |
| 7_5588075 | 7 | 0.4286 | 11 | 0.6364 | 14 | 0.4286 | 8 | 0.875 | 40 | 0.575 | 8 | 0.375 | 6 | 0.6667 | 9 | 0.6667 | 8 | 0.875 | 31 | 0.6452 |
| 7_5588075 | 6 | 0.8333 | 10 | 0.5 | 13 | 0.2308 | 3 | 1 | 32 | 0.5 | 8 | 0.5 | 8 | 0.5 | 11 | 0.4545 | 9 | 0.2222 | 36 | 0.4167 |
| 7_5588081 | 8 | 1 | 13 | 1 | 14 | 0.9286 | 9 | 0.7778 | 44 | 0.9318 | 8 | 0.875 | 10 | 0.9 | 12 | 0.8333 | 10 | 0.7 | 40 | 0.825 |
| 7_5588084 | 8 | 0.5 | 14 | 0.2857 | 15 | 0.2667 | 8 | 0.375 | 45 | 0.3333 | 9 | 0.1111 | 10 | 0.5 | 11 | 0.0909 | 10 | 0.4 | 40 | 0.275 |
| 7_5588086 | 8 | 0.375 | 13 | 0.2308 | 16 | 0.0625 | 8 | 0.25 | 45 | 0.2 | 10 | 0.2 | 9 | 0.2222 | 11 | 0.2727 | 11 | 0.0909 | 41 | 0.1951 |
| 7_5588087 | 6 | 0.1667 | 16 | 0.6875 | 15 | 0.6667 | 6 | 0.8333 | 43 | 0.6279 | 11 | 0.7273 | 10 | 0.7 | 9 | 0.6667 | 10 | 0.8 | 40 | 0.725 |
| 7_5588100 | 8 | 0.875 | 14 | 0.7857 | 16 | 0.75 | 8 | 0.75 | 46 | 0.7826 | 11 | 0.8182 | 8 | 0.875 | 9 | 0.7778 | 10 | 0.9 | 38 | 0.8421 |
| 7_5588101 | 6 | 0.3333 | 14 | 0.3571 | 14 | 0.2143 | 7 | 0.1429 | 41 | 0.2683 | 7 | 0 | 6 | 0 | 11 | 0.2727 | 6 | 0.1667 | 30 | 0.1333 |
| 7_5588101 | 8 | 0.25 | 13 | 0.2308 | 17 | 0.1765 | 9 | 0.1111 | 47 | 0.1915 | 8 | 0.25 | 8 | 0.125 | 9 | 0.2222 | 7 | 0.2857 | 32 | 0.2188 |
| 7_5588102 | 7 | 0.8571 | 15 | 0.9333 | 16 | 0.9375 | 9 | 0.8889 | 47 | 0.9149 | 11 | 0.9091 | 9 | 0.8889 | 9 | 0.7778 | 8 | 0.875 | 37 | 0.8649 |
| 7_5588103 | 7 | 0.1429 | 16 | 0.0625 | 16 | 0.125 | 7 | 0.4286 | 46 | 0.1522 | 9 | 0.2222 | 9 | 0 | 12 | 0.1667 | 10 | 0.1 | 40 | 0.125 |
| 7_5588106 | 4 | 0.5 | 11 | 0.5455 | 14 | 0.2857 | 6 | 0.3333 | 35 | 0.4 | 7 | 0.2857 | 7 | 0.5714 | 10 | 0.3 | 7 | 0.5714 | 31 | 0.4194 |
| 7_5588111 | 9 | 0.3333 | 16 | 0.375 | 19 | 0.0526 | 8 | 0 | 52 | 0.1923 | 11 | 0 | 10 | 0.2 | 12 | 0.0833 | 11 | 0.2727 | 44 | 0.1364 |
| 7_5588114 | 7 | 0.7143 | 16 | 0.6875 | 19 | 0.6842 | 10 | 0.6 | 52 | 0.6731 | 11 | 0.5455 | 10 | 0.8 | 13 | 0.5385 | 10 | 0.6 | 44 | 0.6136 |
| 7_5636655 | 2 | 0.5 | 4 | 0.25 | 2 | 0 | 1 | 0 | 9 | 0.2222 | 2 | 0 | 1 | 0 | 4 | 0 | 5 | 0.2 | 12 | 0.0833 |
| 7_5636662 | 4 | 0.25 | 3 | 0.3333 | 2 | 0 | 2 | 0 | 11 | 0.1818 | 2 | 0 | NA | NA | 3 | 0 | 6 | 0.1667 | 11 | 0.0909 |
| 7_5637406 | 1 | 1 | NA | NA | NA | NA | 1 | 0 | 2 | 0.5 | NA | NA | 1 | 0 | NA | NA | 3 | 0.3333 | 4 | 0.25 |
| 7_5637424 | 1 | 1 | NA | NA | NA | NA | 1 | 1 | 2 | 1 | NA | NA | 1 | 1 | NA | NA | 3 | 0.6667 | 4 | 0.75 |
| 7_5637432 | NA | NA | NA | NA | NA | NA | 1 | 1 | 1 | 1 | NA | NA | 1 | 0 | NA | NA | 4 | 0.75 | 5 | 0.6 |
| 7_5657924 | NA | NA | NA | NA | NA | NA | 1 | 1 | 1 | 1 | NA | NA | NA | NA | 1 | 1 | NA | NA | 1 | 1 |
| 7_5657925 | NA | NA | NA | NA | NA | NA | 1 | 1 | 1 | 1 | NA | NA | NA | NA | 1 | 1 | NA | NA | 1 | 1 |
| 7_5657960 | NA | NA | NA | NA | NA | NA | 1 | 1 | 1 | 1 | NA | NA | NA | NA | 1 | 1 | NA | NA | 1 | 1 |
| 7_5682847 | NA | NA | 2 | 0 | NA | NA | 1 | 1 | 3 | 0.3333 | 1 | 1 | NA | NA | NA | NA | NA | NA | 1 | 1 |
| 7_5682942 | NA | NA | 1 | 0 | NA | NA | 1 | 1 | 2 | 0.5 | 1 | 1 | NA | NA | NA | NA | NA | NA | 1 | 1 |
| 7_5705765 | 4 | 0 | 8 | 0.125 | 3 | 0 | 3 | 0 | 18 | 0.0556 | 3 | 0 | 4 | 0 | 5 | 0.2 | 3 | 0 | 15 | 0.0667 |
| 7_5705784 | 3 | 0.3333 | 6 | 0.5 | 3 | 0.3333 | 3 | 0.3333 | 15 | 0.4 | 3 | 0.3333 | 3 | 0 | 4 | 0.25 | 2 | 0 | 12 | 0.1667 |
| 7_5705791 | 4 | 0.25 | 5 | 0.4 | 3 | 0 | 3 | 0 | 15 | 0.2 | 3 | 0 | 5 | 0.2 | 2 | 0.5 | 1 | 0 | 11 | 0.1818 |
| 7_5705792 | 4 | 0.25 | 7 | 0.1429 | 3 | 0 | 3 | 0 | 17 | 0.1176 | 3 | 0 | 5 | 0.2 | 2 | 0 | 2 | 0 | 12 | 0.0833 |
| 7_5705800 | 3 | 0 | 4 | 0.5 | 2 | 0 | 3 | 0 | 12 | 0.1667 | 2 | 0.5 | 3 | 0.3333 | 1 | 1 | 2 | 0 | 8 | 0.375 |
| 7_5705807 | 4 | 0.25 | 5 | 0 | 3 | 0 | 3 | 0 | 15 | 0.0667 | 1 | 0 | 5 | 0.2 | 1 | 0 | 1 | 0 | 8 | 0.125 |
| 7_5705813 | 2 | 0 | 6 | 0.3333 | 3 | 0 | 3 | 0 | 14 | 0.1429 | 2 | 0 | 5 | 0.4 | 2 | 0.5 | NA | NA | 9 | 0.3333 |
| 7_5705814 | 2 | 0 | 6 | 0.1667 | 3 | 0 | 3 | 0.3333 | 14 | 0.1429 | 2 | 0 | 3 | 0 | 2 | 0.5 | 2 | 0.5 | 9 | 0.2222 |
| 7_5705829 | 3 | 1 | 6 | 1 | 2 | 1 | 3 | 1 | 14 | 1 | 3 | 1 | 5 | 1 | 2 | 1 | 2 | 1 | 12 | 1 |
| 7_5705832 | 2 | 1 | 6 | 0.3333 | 2 | 0 | 2 | 0.5 | 12 | 0.4167 | 2 | 0 | 5 | 0.4 | 2 | 0.5 | 2 | 0.5 | 11 | 0.3636 |
| 7_5705849 | 3 | 0 | 6 | 0.5 | 2 | 0 | 3 | 0.3333 | 14 | 0.2857 | 3 | 0.3333 | 5 | 0 | 2 | 0 | 2 | 0.5 | 12 | 0.1667 |
| 7_5705851 | 2 | 0.5 | 6 | 0 | 2 | 0 | 3 | 0 | 13 | 0.0769 | 3 | 0 | 5 | 0.2 | 2 | 0.5 | 2 | 0 | 12 | 0.1667 |

| | | | | | | | | | | | | | | | | | | | | |
|---|---|---|---|---|---|---|---|---|---|---|---|---|---|---|---|---|---|---|---|---|
| 7_5705864 | 2 | 0.5 | 6 | 0.3333 | 2 | 0.5 | 3 | 0 | 13 | 0.3077 | 1 | 0 | 5 | 0 | 2 | 0.5 | 2 | 0 | 10 | 0.1 |
| 7_5705873 | NA | NA | NA | NA | NA | NA | 1 | 1 | 1 | 1 | 1 | 1 | NA | NA | NA | NA | NA | NA | 1 | 1 |
| 7_5706346 | 2 | 1 | 6 | 0.6667 | 2 | 0.5 | 1 | 1 | 11 | 0.7273 | 3 | 0.6667 | 4 | 1 | 2 | 0 | 2 | 0.5 | 11 | 0.6364 |
| 7_5746607 | 4 | 0.25 | 8 | 0.25 | 8 | 0.375 | 16 | 0.125 | 36 | 0.2222 | 6 | 0.5 | 7 | 0.4286 | 12 | 0.0833 | 10 | 0.1 | 35 | 0.2286 |
| 7_5747427 | 4 | 0 | 7 | 0.1429 | 9 | 0 | 17 | 0.0588 | 37 | 0.0541 | 6 | 0.1667 | 8 | 0 | 14 | 0.1429 | 14 | 0.1429 | 42 | 0.119 |
| 7_5747429 | 5 | 0 | 6 | 0.1667 | 8 | 0.375 | 12 | 0.0833 | 31 | 0.1613 | 6 | 0.1667 | 5 | 0.2 | 10 | 0.1 | 12 | 0.0833 | 33 | 0.1212 |
| 7_5747433 | 4 | 0.5 | 6 | 0 | 11 | 0 | 15 | 0.2 | 36 | 0.1389 | 6 | 0 | 8 | 0.125 | 15 | 0.0667 | 15 | 0.0667 | 44 | 0.0682 |
| 7_5747443 | 5 | 0.8 | 7 | 0.7143 | 10 | 0.6 | 13 | 0.5385 | 35 | 0.6286 | 6 | 0.6667 | 7 | 1 | 12 | 0.5833 | 12 | 0.6667 | 37 | 0.7027 |
| 7_5747451 | 5 | 0.8 | 6 | 1 | 11 | 1 | 16 | 0.9375 | 38 | 0.9474 | 5 | 1 | 8 | 0.875 | 14 | 0.7857 | 16 | 0.875 | 43 | 0.8605 |
| 7_5747453 | 5 | 1 | 6 | 0.6667 | 10 | 1 | 17 | 0.7647 | 38 | 0.8421 | 5 | 0.8 | 9 | 0.7778 | 16 | 0.75 | 14 | 0.8571 | 44 | 0.7955 |
| 7_5747466 | 5 | 0 | 8 | 0.25 | 10 | 0.2 | 15 | 0.1333 | 38 | 0.1579 | 5 | 0.4 | 11 | 0.1818 | 14 | 0.2857 | 12 | 0.3333 | 42 | 0.2857 |
| 7_5747479 | 4 | 0.75 | 7 | 1 | 12 | 0.4167 | 17 | 0.4706 | 40 | 0.575 | 5 | 0.8 | 10 | 0.8 | 15 | 0.4667 | 16 | 0.625 | 46 | 0.6304 |
| 7_5747586 | 9 | 0.4444 | 14 | 0.4286 | 16 | 0.3125 | 19 | 0.4737 | 58 | 0.4138 | 8 | 0.625 | 16 | 0.1875 | 16 | 0.25 | 17 | 0.5294 | 57 | 0.3684 |
| 7_5747619 | 4 | 0 | 5 | 0.2 | 10 | 0 | 12 | 0.0833 | 31 | 0.0645 | 4 | 0 | 10 | 0.1 | 9 | 0.2222 | 9 | 0.1111 | 32 | 0.125 |
| 7_5747620 | 9 | 0 | 18 | 0.2222 | 16 | 0.3125 | 19 | 0.2105 | 62 | 0.2097 | 8 | 0 | 20 | 0.35 | 18 | 0.0556 | 20 | 0.05 | 66 | 0.1364 |
| 7_5747752 | 16 | 0.125 | 25 | 0.08 | 17 | 0.1176 | 27 | 0.0741 | 85 | 0.0941 | 11 | 0.2727 | 22 | 0.2727 | 24 | 0.0833 | 20 | 0.05 | 77 | 0.1558 |
| 7_5747814 | 20 | 0.15 | 27 | 0.3704 | 19 | 0.2632 | 26 | 0.3462 | 92 | 0.2935 | 17 | 0.2941 | 24 | 0.1667 | 27 | 0.2963 | 23 | 0.1739 | 91 | 0.2308 |
| 7_5747853 | 14 | 0.1429 | 24 | 0.125 | 17 | 0.1176 | 22 | 0.1818 | 77 | 0.1429 | 16 | 0.25 | 25 | 0.2 | 19 | 0.2105 | 19 | 0.1579 | 79 | 0.2025 |
| 7_5747855 | NA | NA | 1 | 0 | 1 | 1 | 2 | 0 | 4 | 0.25 | 3 | 0.6667 | 1 | 1 | 1 | 0 | 1 | 0 | 6 | 0.5 |
| 7_5847797 | 2 | 1 | 2 | 0 | 1 | 1 | 3 | 0.3333 | 8 | 0.5 | NA | NA | NA | NA | 1 | 1 | NA | NA | 1 | 1 |
| 7_5847799 | 2 | 0 | 2 | 0.5 | 1 | 0 | 3 | 0.6667 | 8 | 0.375 | 1 | 0 | 2 | 0.5 | 1 | 1 | NA | NA | 4 | 0.5 |
| 7_5847803 | 2 | 0.5 | 2 | 0 | 1 | 0 | 3 | 0.3333 | 8 | 0.25 | 1 | 0 | 2 | 0.5 | 1 | 1 | NA | NA | 4 | 0.5 |
| 7_5847804 | 3 | 0.6667 | 2 | 0 | 1 | 0 | 3 | 1 | 9 | 0.5556 | 1 | 1 | 2 | 0.5 | 1 | 1 | NA | NA | 4 | 0.75 |
| 7_5847805 | 3 | 0.3333 | 2 | 0.5 | 1 | 1 | 3 | 0.6667 | 9 | 0.5556 | 1 | 1 | 1 | 1 | 1 | 1 | NA | NA | 3 | 1 |
| 7_5847817 | 3 | 0 | 2 | 1 | 1 | 1 | 3 | 1 | 9 | 0.6667 | 1 | 1 | 2 | 1 | 1 | 1 | 1 | 1 | 5 | 1 |
| 7_5847825 | 2 | 0.5 | 1 | 1 | NA | NA | 3 | 0 | 6 | 0.3333 | 1 | 1 | 2 | 0.5 | 1 | 1 | NA | NA | 4 | 0.75 |
| 7_5847827 | 3 | 0.3333 | 2 | 0.5 | 1 | 1 | 3 | 1 | 9 | 0.6667 | 1 | 1 | 2 | 1 | 1 | 1 | 1 | 1 | 5 | 1 |
| 7_5847844 | 3 | 0 | 2 | 0 | 1 | 1 | 3 | 0 | 9 | 0.1111 | 1 | 0 | 2 | 0 | 1 | 0 | 1 | 1 | 5 | 0.2 |
| 7_5847846 | 2 | 0 | 1 | 0 | 1 | 1 | 3 | 0.3333 | 7 | 0.2857 | 1 | 0 | 3 | 1 | 1 | 0 | NA | NA | 5 | 0.6 |
| 7_5887774 | NA | NA | NA | NA | 1 | 1 | 1 | 1 | 2 | 1 | NA | NA | NA | NA | 1 | 0 | 3 | 0.3333 | 4 | 0.25 |
| 7_5887781 | NA | NA | NA | NA | 1 | 1 | 1 | 1 | 2 | 1 | NA | NA | NA | NA | 1 | 1 | 2 | 0.5 | 3 | 0.6667 |
| 7_5887786 | NA | NA | NA | NA | 1 | 1 | 1 | 1 | 2 | 1 | NA | NA | NA | NA | 1 | 1 | 2 | 0.5 | 3 | 0.6667 |
| 7_5887795 | NA | NA | NA | NA | 1 | 1 | 1 | 1 | 2 | 1 | NA | NA | NA | NA | 1 | 1 | 2 | 0.5 | 3 | 0.6667 |
| 7_5887798 | NA | NA | NA | NA | 1 | 1 | NA | NA | 1 | 1 | NA | NA | NA | NA | 1 | 1 | 2 | 0.5 | 3 | 0.6667 |
| 7_5887805 | NA | NA | NA | NA | NA | NA | 1 | 1 | 1 | 1 | NA | NA | NA | NA | NA | NA | 2 | 1 | 2 | 1 |
| 7_5887818 | NA | NA | NA | NA | 1 | 1 | 1 | 1 | 2 | 1 | NA | NA | NA | NA | 1 | 0 | 2 | 0.5 | 3 | 0.3333 |
| 7_5887822 | NA | NA | NA | NA | 1 | 0 | 1 | 1 | 2 | 0.5 | NA | NA | NA | NA | 1 | 0 | 2 | 0.5 | 3 | 0.3333 |
| 7_5887827 | NA | NA | NA | NA | 1 | 1 | 1 | 1 | 2 | 1 | NA | NA | NA | NA | 1 | 1 | 2 | 1 | 3 | 1 |
| 7_5887852 | NA | NA | NA | NA | 1 | 1 | 1 | 1 | 2 | 1 | NA | NA | NA | NA | 1 | 0 | 1 | 1 | 2 | 0.5 |
| 7_5887879 | 4 | 0.75 | 2 | 0 | NA | NA | 1 | 1 | 7 | 0.5714 | NA | NA | 3 | 0.3333 | 2 | 0 | 1 | 0 | 6 | 0.1667 |
| 7_5887901 | 3 | 0 | 2 | 0 | 2 | 0 | 3 | 0.6667 | 10 | 0.2 | NA | NA | 2 | 0 | 3 | 0.3333 | 2 | 1 | 7 | 0.4286 |
| 7_5892584 | 3 | 1 | 1 | 1 | 2 | 1 | NA | NA | 6 | 1 | 1 | 1 | 2 | 1 | 3 | 1 | 3 | 1 | 9 | 1 |
| 7_5892590 | 3 | 1 | 2 | 1 | 2 | 0.5 | NA | NA | 7 | 0.8571 | 1 | 0 | 3 | 1 | 2 | 0.5 | 3 | 0.3333 | 9 | 0.5556 |
| 7_5892595 | 3 | 0 | 2 | 0.5 | 2 | 0 | NA | NA | 7 | 0.1429 | 1 | 1 | 3 | 0 | 2 | 0 | 2 | 0 | 8 | 0.125 |
| 7_5892611 | 3 | 0.3333 | 2 | 0.5 | NA | NA | 1 | 0 | 6 | 0.3333 | 1 | 0 | 3 | 0.3333 | 3 | 0.6667 | 3 | 0 | 10 | 0.3 |
| 7_5892631 | 2 | 1 | 2 | 0.5 | 2 | 1 | 1 | 0 | 7 | 0.7143 | 1 | 1 | 3 | 1 | 3 | 0.6667 | 3 | 0.6667 | 10 | 0.8 |
| 7_5892643 | 4 | 0.75 | 2 | 0.5 | 2 | 1 | NA | NA | 8 | 0.75 | 1 | 0 | 3 | 0.3333 | 3 | 1 | 2 | 0 | 9 | 0.4444 |
| 7_5892645 | 3 | 1 | 1 | 1 | 1 | 1 | 1 | 1 | 6 | 1 | 1 | 1 | 2 | 1 | 1 | 0 | 2 | 0.5 | 6 | 0.6667 |
| 7_5892650 | 3 | 0 | 2 | 0.5 | 2 | 0 | 1 | 0 | 8 | 0.125 | 1 | 0 | 3 | 0.3333 | 3 | 0 | 2 | 0.5 | 9 | 0.2222 |
| 7_5892652 | 4 | 0.75 | 2 | 1 | 3 | 0.6667 | 1 | 1 | 10 | 0.8 | 1 | 1 | 3 | 1 | 3 | 1 | 2 | 1 | 9 | 1 |
| 7_5892663 | 2 | 0.5 | 2 | 0 | 3 | 0 | 1 | 0 | 8 | 0.125 | 1 | 0 | 4 | 0 | 2 | 0.5 | 3 | 0 | 10 | 0.1 |
| 7_5892669 | 4 | 0.25 | 2 | 0 | 3 | 0 | 1 | 0 | 10 | 0.1 | 1 | 0 | 4 | 0.25 | 3 | 0 | 3 | 0 | 11 | 0.0909 |
| 7_5892671 | 3 | 0.3333 | 2 | 0 | 3 | 0.3333 | 1 | 0 | 9 | 0.2222 | 1 | 0 | 4 | 0 | 3 | 0.3333 | 2 | 0 | 10 | 0.1 |
| 7_5895462 | 6 | 0.1667 | 2 | 0 | 5 | 0 | 2 | 0 | 15 | 0.0667 | 2 | 0 | 3 | 0.3333 | 3 | 0 | 4 | 0 | 12 | 0.0833 |
| 7_5895482 | 5 | 0.4 | 2 | 0.5 | 5 | 0.2 | 3 | 0 | 15 | 0.2667 | 2 | 0 | 1 | 0 | 3 | 0.3333 | 3 | 0.6667 | 9 | 0.3333 |
| 7_5895514 | 7 | 0.2857 | 2 | 1 | 4 | 0 | 3 | 0.3333 | 16 | 0.3125 | 2 | 0 | 2 | 0 | 4 | 0 | 5 | 0.2 | 13 | 0.0769 |
| 7_5931754 | NA | NA | 1 | 1 | NA | NA | NA | NA | 1 | 1 | 2 | 1 | 1 | 1 | NA | NA | NA | NA | 3 | 1 |
| 7_5958883 | 2 | 1 | 4 | 0.5 | 1 | 1 | NA | NA | 7 | 0.7143 | 1 | 1 | 1 | 0 | 1 | 1 | 1 | 1 | 4 | 0.75 |
| 7_5959013 | 1 | 0 | 3 | 0.3333 | 1 | 0 | NA | NA | 5 | 0.2 | 1 | 1 | 1 | 0 | 1 | 0 | 1 | 0 | 4 | 0.25 |
| 7_5959052 | NA | NA | 4 | 0.75 | 1 | 0 | 1 | 0 | 6 | 0.5 | NA | NA | NA | NA | 1 | 1 | 1 | 1 | 2 | 1 |
| 7_5962017 | 1 | 0 | 4 | 0.5 | 2 | 1 | NA | NA | 7 | 0.5714 | NA | NA | NA | NA | 1 | 1 | 1 | 1 | 2 | 1 |
| 7_5962019 | 1 | 1 | 4 | 0.5 | 2 | 0.5 | NA | NA | 7 | 0.5714 | NA | NA | NA | NA | 1 | 1 | 1 | 1 | 2 | 1 |
| 7_5962021 | 1 | 0 | 3 | 0.3333 | 2 | 1 | NA | NA | 6 | 0.5 | NA | NA | NA | NA | 1 | 1 | 1 | 0 | 2 | 0.5 |
| 7_5962317 | 1 | 1 | 4 | 0.5 | 2 | 1 | NA | NA | 7 | 0.7143 | NA | NA | NA | NA | 1 | 1 | 1 | 0 | 2 | 0.5 |
| 7_5962325 | 1 | 0 | 4 | 0.25 | 1 | 1 | NA | NA | 6 | 0.3333 | NA | NA | NA | NA | 1 | 0 | 1 | 1 | 2 | 0.5 |

| | | | | | | | | | | | | | | | | | | | | |
|---|---|---|---|---|---|---|---|---|---|---|---|---|---|---|---|---|---|---|---|---|
| 7_59623323 | 1 | 1 | 2 | 0 | 1 | 1 | NA | NA | 4 | 0.5 | NA | NA | NA | NA | 1 | 1 | 1 | 1 | 2 | 1 |
| 7_59623651 | 1 | 1 | 2 | 0.5 | 1 | 0 | NA | NA | 4 | 0.5 | NA | NA | NA | NA | 1 | 1 | 1 | 0 | 2 | 0.5 |
| 7_59806330 | NA | NA | NA | NA | 1 | 1 | 1 | 0 | 2 | 0.5 | NA | NA | NA | NA | 1 | 1 | 1 | 1 | 2 | 1 |
| 7_60565603 | 1 | 1 | NA | NA | NA | NA | 1 | 0 | 2 | 0.5 | NA | NA | NA | NA | NA | NA | 1 | 1 | 1 | 1 |
| 7_60847959 | 1 | 0 | NA | NA | NA | NA | 1 | 1 | 2 | 0.5 | NA | NA | NA | NA | NA | NA | 1 | 1 | 1 | 1 |
| 7_60933012 | 1 | 0 | NA | NA | 2 | 0.5 | 4 | 1 | 7 | 0.7143 | 2 | 0.5 | 2 | 1 | 1 | 1 | 4 | 0.5 | 9 | 0.6667 |
| 7_61172321 | NA | NA | NA | NA | 2 | 0.5 | NA | NA | 2 | 0.5 | NA | NA | NA | NA | NA | NA | 1 | 1 | 1 | 1 |
| 7_61177344 | 10 | 0.2 | 13 | 0.1538 | 9 | 0.3333 | 18 | 0.2222 | 50 | 0.22 | 10 | 0.1 | 15 | 0.2667 | 10 | 0.6 | 12 | 0.4167 | 47 | 0.3404 |
| 7_61177567 | 9 | 0.2222 | 11 | 0 | 8 | 0.375 | 16 | 0.125 | 44 | 0.1591 | 8 | 0.125 | 11 | 0.1818 | 9 | 0.3333 | 10 | 0.1 | 38 | 0.1842 |
| 7_61178128 | 6 | 0.1667 | 7 | 0 | 3 | 0 | 12 | 0.0833 | 28 | 0.0714 | 5 | 0 | 10 | 0.1 | 10 | 0 | 9 | 0.1111 | 34 | 0.0588 |
| 7_61180274 | 5 | 0.4 | 3 | 0.6667 | 2 | 0.5 | 7 | 0.7143 | 17 | 0.5882 | 3 | 0.3333 | 4 | 0.25 | 7 | 0.4286 | 4 | 0.5 | 18 | 0.3889 |
| 7_61180384 | 4 | 0.25 | 6 | 0 | 2 | 0 | 8 | 0 | 20 | 0.05 | 4 | 0 | 7 | 0.1429 | 8 | 0.125 | 3 | 0 | 22 | 0.0909 |
| 7_61184915 | 6 | 0.3333 | 6 | 0 | 2 | 0 | 8 | 0.125 | 22 | 0.1364 | 4 | 0 | 7 | 0 | 5 | 0 | 4 | 0.25 | 20 | 0.05 |
| 7_61355719 | NA | NA | 1 | 1 | NA | NA | NA | NA | 1 | 1 | NA | NA | NA | NA | 2 | 1 | NA | NA | 2 | 1 |
| 7_61401333 | 1 | 1 | NA | NA | 3 | 0.6667 | 2 | 0.5 | 6 | 0.6667 | NA | NA | 2 | 0.5 | 1 | 0 | 1 | 1 | 4 | 0.5 |
| 7_61402759 | NA | NA | NA | NA | 2 | 1 | 1 | 0 | 3 | 0.6667 | NA | NA | 2 | 0.5 | 1 | 1 | 1 | 1 | 4 | 0.75 |
| 7_61402807 | NA | NA | NA | NA | 1 | 1 | 1 | 0 | 2 | 0.5 | NA | NA | 2 | 0 | 1 | 1 | 1 | 0 | 4 | 0.25 |
| 7_61403172 | NA | NA | NA | NA | 1 | 1 | 1 | 0 | 2 | 0.5 | NA | NA | 1 | 1 | NA | NA | NA | NA | 1 | 1 |
| 7_61403313 | NA | NA | NA | NA | 2 | 0.5 | NA | NA | 2 | 0.5 | NA | NA | 1 | 1 | 1 | 1 | 1 | 0 | 3 | 0.6667 |
| 7_61403477 | NA | NA | NA | NA | 2 | 0.5 | NA | NA | 2 | 0.5 | NA | NA | NA | NA | 1 | 0 | 1 | 1 | 2 | 0.5 |
| 7_61999129 | 24 | 1 | 21 | 0.9524 | 18 | 1 | 30 | 0.9667 | 93 | 0.9785 | 10 | 0.8 | 19 | 0.8947 | 25 | 1 | 15 | 0.7333 | 69 | 0.8841 |
| 7_61999442 | 24 | 0.7083 | 23 | 0.6957 | 19 | 0.7368 | 35 | 0.7429 | 101 | 0.7228 | 15 | 0.8667 | 17 | 0.7647 | 24 | 0.5 | 21 | 0.6667 | 77 | 0.6753 |
| 7_61999487 | 28 | 0.8571 | 24 | 0.9167 | 22 | 0.7727 | 37 | 0.8108 | 111 | 0.8378 | 16 | 0.6875 | 24 | 0.8333 | 30 | 0.9333 | 23 | 0.8696 | 93 | 0.8495 |
| 7_61999578 | 25 | 0.96 | 24 | 0.9167 | 18 | 1 | 36 | 0.9444 | 103 | 0.9515 | 15 | 0.9333 | 21 | 0.9524 | 27 | 0.963 | 21 | 0.9048 | 84 | 0.9405 |
| 7_61999588 | 25 | 0.24 | 21 | 0.1905 | 18 | 0.1667 | 26 | 0.2308 | 90 | 0.2111 | 15 | 0.2 | 21 | 0.2381 | 27 | 0.2593 | 17 | 0.2353 | 80 | 0.2375 |
| 7_61999593 | 26 | 0.8846 | 22 | 0.9091 | 21 | 0.9524 | 32 | 1 | 101 | 0.9406 | 16 | 0.9375 | 22 | 0.9091 | 27 | 0.9259 | 22 | 0.9091 | 87 | 0.9195 |
| 7_62067755 | 1 | 1 | NA | NA | NA | NA | 2 | 1 | 3 | 1 | 1 | 1 | NA | NA | NA | NA | 1 | 0 | 2 | 0.5 |
| 7_62068084 | 1 | 1 | NA | NA | NA | NA | 1 | 1 | 2 | 1 | 1 | 1 | NA | NA | NA | NA | 1 | 0 | 2 | 0.5 |
| 7_62068759 | 1 | 1 | NA | NA | NA | NA | 2 | 1 | 3 | 1 | 1 | 1 | NA | NA | NA | NA | 1 | 0 | 2 | 0.5 |
| 7_62069112 | 1 | 1 | NA | NA | NA | NA | 2 | 0.5 | 3 | 0.6667 | 1 | 1 | NA | NA | NA | NA | 1 | 0 | 2 | 0.5 |
| 7_62069245 | NA | NA | NA | NA | NA | NA | 2 | 0.5 | 2 | 0.5 | 1 | 1 | NA | NA | NA | NA | 1 | 0 | 2 | 0.5 |
| 7_62069813 | NA | NA | NA | NA | NA | NA | 2 | 0.5 | 2 | 0.5 | 1 | 1 | NA | NA | NA | NA | NA | NA | 1 | 1 |
| 7_62745760 | 4 | 0.5 | 1 | 0 | 3 | 0.3333 | 3 | 0.6667 | 11 | 0.4545 | 2 | 0 | 3 | 0.3333 | 3 | 0.3333 | 2 | 1 | 10 | 0.4 |
| 7_62748884 | NA | NA | 1 | 1 | NA | NA | 2 | 0.5 | 3 | 0.6667 | NA | NA | NA | NA | 2 | 1 | NA | NA | 2 | 1 |
| 7_62748982 | NA | NA | 1 | 1 | NA | NA | 1 | 1 | 2 | 1 | NA | NA | NA | NA | 2 | 1 | NA | NA | 2 | 1 |
| 7_62749078 | NA | NA | 1 | 1 | NA | NA | 1 | 1 | 2 | 1 | 1 | 1 | NA | NA | 2 | 0.5 | NA | NA | 3 | 0.6667 |
| 7_62749276 | NA | NA | 1 | 0 | NA | NA | 2 | 1 | 3 | 0.6667 | NA | NA | NA | NA | 2 | 1 | NA | NA | 2 | 1 |
| 7_63404449 | 3 | 0 | 6 | 0 | 5 | 0 | 3 | 0.3333 | 17 | 0.0588 | NA | NA | 3 | 0.3333 | 4 | 0.25 | 5 | 0 | 12 | 0.1667 |
| 7_63407131 | 3 | 0 | 3 | 0.3333 | 5 | 0.6 | 2 | 0.5 | 13 | 0.3846 | NA | NA | 5 | 0.2 | 7 | 0.2857 | 4 | 0.75 | 16 | 0.375 |
| 7_63407137 | 3 | 0 | 5 | 0.4 | 5 | 0 | 3 | 0.3333 | 16 | 0.1875 | NA | NA | 5 | 0.2 | 6 | 0 | 4 | 0 | 15 | 0.0667 |
| 7_63407191 | 3 | 0.3333 | 6 | 0.5 | 3 | 0.3333 | 4 | 0.5 | 16 | 0.4375 | NA | NA | 5 | 0.2 | 5 | 0.2 | 4 | 0.5 | 14 | 0.2857 |
| 7_63407218 | 3 | 0 | 5 | 0.4 | 5 | 0.2 | 3 | 0.3333 | 16 | 0.25 | NA | NA | 5 | 0.2 | 7 | 0.1429 | 3 | 0 | 15 | 0.1333 |
| 7_63407239 | 2 | 0 | 4 | 0.5 | 3 | 0 | 4 | 0.5 | 13 | 0.3077 | NA | NA | 3 | 0.3333 | 7 | 0.1429 | 4 | 0 | 14 | 0.1429 |
| 7_63407299 | 3 | 0 | 3 | 0 | 3 | 0 | 4 | 0.25 | 13 | 0.0769 | NA | NA | 3 | 0 | 6 | 0.5 | 3 | 0 | 12 | 0.25 |
| 7_63407485 | 2 | 0.5 | 6 | 0 | 6 | 0.3333 | 3 | 0 | 17 | 0.1765 | NA | NA | 6 | 0.6667 | 6 | 0 | 4 | 0.25 | 16 | 0.3125 |
| 7_63407682 | 3 | 0 | 7 | 0 | 6 | 0.1667 | 4 | 0 | 20 | 0.05 | NA | NA | 5 | 0 | 9 | 0 | 4 | 0.25 | 18 | 0.0556 |
| 7_63407717 | 1 | 1 | 3 | 0.6667 | NA | NA | 1 | 1 | 5 | 0.8 | NA | NA | 3 | 1 | 5 | 0.8 | 2 | 1 | 10 | 0.9 |
| 7_63407898 | 3 | 1 | 6 | 1 | 5 | 1 | 4 | 1 | 18 | 1 | NA | NA | 6 | 1 | 8 | 1 | 4 | 1 | 18 | 1 |
| 7_63407993 | 1 | 0 | 6 | 0.5 | 6 | 0.8333 | 2 | 1 | 15 | 0.6667 | NA | NA | 6 | 0.8333 | 8 | 0.875 | 2 | 0.5 | 16 | 0.8125 |
| 7_63408124 | 2 | 0 | 4 | 0.25 | 5 | 0.2 | 3 | 0.3333 | 14 | 0.2143 | NA | NA | 3 | 0.3333 | 7 | 0.5714 | 4 | 0.25 | 14 | 0.4286 |
| 7_63408224 | 3 | 0.3333 | 7 | 0 | 6 | 0.3333 | 4 | 0.25 | 20 | 0.2 | NA | NA | 6 | 0.6667 | 9 | 0.1111 | 5 | 0.2 | 20 | 0.3 |
| 7_63408246 | 2 | 0 | 6 | 0.3333 | 5 | 0 | 3 | 0.3333 | 16 | 0.1875 | NA | NA | 6 | 0.1667 | 7 | 0.1429 | 5 | 0.2 | 18 | 0.1667 |
| 7_63408379 | 3 | 0.6667 | 4 | 0.75 | 5 | 1 | 1 | 0 | 13 | 0.7692 | NA | NA | 4 | 0.75 | 2 | 0.5 | 2 | 1 | 8 | 0.75 |
| 7_63408405 | 2 | 0 | 8 | 0.125 | 6 | 0.5 | 4 | 0 | 20 | 0.2 | NA | NA | 6 | 0.1667 | 10 | 0.1 | 4 | 0.5 | 20 | 0.2 |
| 7_63408490 | 3 | 1 | 8 | 0.875 | 6 | 1 | 4 | 0.75 | 21 | 0.9048 | NA | NA | 6 | 1 | 10 | 1 | 4 | 1 | 20 | 1 |
| 7_63408605 | 3 | 0.6667 | 8 | 0.25 | 6 | 0.3333 | 4 | 0.25 | 21 | 0.3333 | NA | NA | 4 | 0 | 8 | 0.125 | 4 | 0 | 16 | 0.0625 |
| 7_63408703 | 2 | 0 | 6 | 0.5 | 7 | 0.1429 | 5 | 0.2 | 20 | 0.25 | NA | NA | 6 | 0.1667 | 9 | 0.1111 | 4 | 0 | 19 | 0.1053 |
| 7_63408818 | NA | NA | 2 | 1 | NA | NA | NA | NA | 2 | 1 | NA | NA | 3 | 0.3333 | NA | NA | NA | NA | 3 | 0.3333 |
| 7_63409207 | NA | NA | 1 | 1 | 2 | 0 | NA | NA | 3 | 0.3333 | NA | NA | 3 | 0.6667 | 2 | 0.5 | NA | NA | 5 | 0.6 |
| 7_63409281 | NA | NA | 2 | 0.5 | 1 | 1 | NA | NA | 3 | 0.6667 | NA | NA | 3 | 0.3333 | 2 | 0 | NA | NA | 5 | 0.2 |
| 7_63409753 | NA | NA | NA | NA | 2 | 0.5 | NA | NA | 2 | 0.5 | NA | NA | 2 | 0.5 | 3 | 0.6667 | NA | NA | 5 | 0.6 |
| 7_63409802 | NA | NA | 2 | 0.5 | 2 | 1 | NA | NA | 4 | 0.75 | NA | NA | 3 | 0 | 3 | 0.3333 | NA | NA | 6 | 0.1667 |
| 7_63536373 | 18 | 0.2222 | 23 | 0.3043 | 13 | 0.3846 | 14 | 0.4286 | 68 | 0.3235 | 14 | 0.3571 | 9 | 0.4444 | 8 | 0.125 | 19 | 0.3684 | 50 | 0.34 |
| 7_63536463 | 12 | 0.4167 | 15 | 0.2 | 12 | 0.25 | 10 | 0.2 | 49 | 0.2653 | 4 | 0.5 | 3 | 0 | 8 | 0.5 | 14 | 0.1429 | 29 | 0.2759 |
| 7_63536492 | 16 | 0.1875 | 24 | 0.0833 | 15 | 0.0667 | 14 | 0 | 69 | 0.087 | 13 | 0.0769 | 14 | 0 | 11 | 0.0909 | 18 | 0.0556 | 56 | 0.0536 |

| | | | | | | | | | | | | | | | | | | | | |
|---|---|---|---|---|---|---|---|---|---|---|---|---|---|---|---|---|---|---|---|---|
| 7_6353650 | 15 | 0.2 | 15 | 0.1333 | 9 | 0.1111 | 11 | 0.1818 | 50 | 0.16 | 9 | 0.1111 | 11 | 0.1818 | 6 | 0.3333 | 14 | 0.1429 | 40 | 0.175 |
| 7_6353660 | 15 | 0.6 | 20 | 0.55 | 10 | 0.5 | 13 | 0.5385 | 58 | 0.5517 | 10 | 0.7 | 11 | 0.6364 | 9 | 0.5556 | 18 | 0.5 | 48 | 0.5833 |
| 7_6353764 | 11 | 0 | 10 | 0.1 | 4 | 0.25 | 9 | 0.1111 | 34 | 0.0882 | 10 | 0 | 10 | 0 | 6 | 0.1667 | 16 | 0.1875 | 42 | 0.0952 |
| 7_6353772 | 13 | 0.7692 | 10 | 0.6 | 5 | 0.4 | 10 | 0.7 | 38 | 0.6579 | 8 | 0.5 | 8 | 0.75 | 6 | 0.8333 | 14 | 0.6429 | 36 | 0.6667 |
| 7_6353781 | 12 | 0.1667 | 10 | 0 | 5 | 0 | 8 | 0 | 35 | 0.0571 | 8 | 0.125 | 7 | 0 | 7 | 0.1429 | 15 | 0 | 37 | 0.0541 |
| 7_6353783 | 11 | 0 | 10 | 0.1 | 5 | 0 | 11 | 0.1818 | 37 | 0.0811 | 10 | 0 | 9 | 0 | 7 | 0 | 14 | 0.2143 | 40 | 0.075 |
| 7_6353794 | 8 | 0.75 | 7 | 0.2857 | 5 | 0 | 7 | 0.4286 | 27 | 0.4074 | 9 | 0.4444 | 8 | 0.5 | 6 | 0.5 | 13 | 0.1538 | 36 | 0.3611 |
| 7_6353801 | 10 | 0.3 | 11 | 0.1818 | 5 | 0 | 9 | 0.2222 | 35 | 0.2 | 9 | 0.4444 | 7 | 0.4286 | 6 | 0.1667 | 15 | 0.2667 | 37 | 0.3243 |
| 7_6353809 | 8 | 0.25 | 6 | 0.5 | 3 | 0 | 6 | 0.3333 | 23 | 0.3043 | 7 | 0.4286 | 6 | 0.1667 | 5 | 0.2 | 12 | 0.4167 | 30 | 0.3333 |
| 7_6353827 | 8 | 0 | 9 | 0 | 2 | 0.5 | 6 | 0.1667 | 25 | 0.08 | 7 | 0.1429 | 5 | 0 | 7 | 0.1429 | 12 | 0 | 31 | 0.0645 |
| 7_6353893 | 8 | 0.375 | 7 | 0.5714 | 3 | 0.6667 | 3 | 0.3333 | 21 | 0.4762 | 7 | 0.5714 | 4 | 0.5 | 6 | 0.6667 | 7 | 0.7143 | 24 | 0.625 |
| 7_6353955 | 3 | 0 | 6 | 0.5 | 3 | 0 | 3 | 0 | 15 | 0.2 | 5 | 0 | 3 | 0 | 4 | 0.25 | 4 | 0.25 | 16 | 0.125 |
| 7_6353973 | 5 | 0 | 4 | 0.25 | 3 | 0 | 2 | 0 | 14 | 0.0714 | 2 | 0 | 1 | 0 | 5 | 0.2 | 2 | 0 | 10 | 0.1 |
| 7_6353981 | 5 | 0.4 | 5 | 0.2 | 3 | 0.3333 | 3 | 0 | 16 | 0.25 | 2 | 0.5 | 2 | 0.5 | 4 | 0.5 | 2 | 0 | 10 | 0.4 |
| 7_6353984 | 1 | 1 | 4 | 1 | 2 | 0 | 2 | 0.5 | 9 | 0.6667 | NA | NA | 2 | 0.5 | 2 | 0 | NA | NA | 4 | 0.25 |
| 7_6354002 | 2 | 0.5 | 2 | 0 | 3 | 0 | 1 | 0 | 8 | 0.125 | NA | NA | NA | NA | 3 | 0.3333 | 2 | 0 | 5 | 0.2 |
| 7_6354016 | 2 | 1 | 3 | 1 | 1 | 1 | 1 | 1 | 7 | 1 | 2 | 0.5 | NA | NA | 4 | 0.75 | 2 | 0.5 | 8 | 0.625 |
| 7_6354018 | 3 | 1 | 2 | 1 | 3 | 1 | 2 | 1 | 10 | 1 | 2 | 1 | NA | NA | 4 | 1 | 2 | 1 | 8 | 1 |
| 7_6354019 | 1 | 1 | 3 | 0.6667 | 3 | 1 | 1 | 1 | 8 | 0.875 | 2 | 1 | NA | NA | 2 | 1 | 1 | 1 | 5 | 1 |
| 7_6354021 | 2 | 0 | 2 | 0 | 3 | 0.3333 | 2 | 0.5 | 9 | 0.2222 | 2 | 0 | NA | NA | 4 | 0.25 | 2 | 0.5 | 8 | 0.25 |
| 7_6354023 | 3 | 1 | 3 | 0.6667 | 2 | 1 | 1 | 1 | 9 | 0.8889 | 1 | 1 | NA | NA | 3 | 1 | 2 | 1 | 6 | 1 |
| 7_6354023 | 3 | 0.3333 | 3 | 0 | 3 | 0 | 2 | 0.5 | 11 | 0.1818 | 2 | 0 | NA | NA | 4 | 0.25 | 2 | 0 | 8 | 0.125 |
| 7_6354031 | 2 | 1 | 2 | 1 | 2 | 1 | 1 | 1 | 7 | 1 | 2 | 1 | NA | NA | 4 | 1 | 1 | 1 | 7 | 1 |
| 7_6354038 | 1 | 1 | 1 | 0 | 3 | 1 | 1 | 1 | 6 | 0.8333 | 1 | 0 | NA | NA | 2 | 0.5 | 2 | 1 | 5 | 0.6 |
| 7_6354038 | 3 | 1 | 2 | 1 | 2 | 1 | 1 | 1 | 8 | 1 | 1 | 0 | NA | NA | 4 | 0.75 | 2 | 1 | 7 | 0.7143 |
| 7_6354045 | 2 | 1 | 2 | 1 | 3 | 1 | 1 | 0 | 8 | 0.875 | 2 | 1 | NA | NA | 4 | 1 | 2 | 1 | 8 | 1 |
| 7_6354048 | 1 | 1 | 2 | 1 | 3 | 0.6667 | 1 | 1 | 7 | 0.8571 | 1 | 1 | NA | NA | 4 | 1 | 2 | 1 | 7 | 1 |
| 7_6354051 | 3 | 1 | 2 | 1 | 3 | 0.6667 | 1 | 1 | 9 | 0.8889 | 2 | 1 | NA | NA | 2 | 1 | 2 | 0.5 | 6 | 0.8333 |
| 7_6354069 | 3 | 0.6667 | 2 | 0 | NA | NA | 1 | 1 | 6 | 0.5 | 1 | 0 | NA | NA | 1 | 0 | 2 | 0.5 | 4 | 0.25 |
| 7_6354070 | 2 | 1 | 2 | 1 | 2 | 1 | 1 | 1 | 7 | 1 | 1 | 1 | NA | NA | 2 | 1 | 2 | 1 | 5 | 1 |
| 7_6354072 | 2 | 1 | 2 | 0.5 | 2 | 1 | 1 | 1 | 7 | 0.8571 | 1 | 1 | NA | NA | 3 | 0.6667 | 2 | 0.5 | 6 | 0.6667 |
| 7_6354082 | 3 | 1 | 2 | 0 | 3 | 0.6667 | 1 | 1 | 9 | 0.6667 | 1 | 1 | NA | NA | 3 | 0.6667 | 2 | 0.5 | 6 | 0.6667 |
| 7_6354088 | 3 | 1 | 2 | 1 | 3 | 0.3333 | 1 | 1 | 9 | 0.7778 | 1 | 0 | NA | NA | 3 | 0.6667 | 2 | 0.5 | 6 | 0.5 |
| 7_6354103 | 3 | 0 | 2 | 0.5 | 2 | 0.5 | 1 | 0 | 8 | 0.25 | 1 | 0 | NA | NA | 2 | 0.5 | 1 | 0 | 4 | 0.25 |
| 7_6354119 | 2 | 1 | 2 | 1 | 2 | 0.5 | 1 | 1 | 7 | 0.8571 | 1 | 1 | NA | NA | 2 | 1 | 1 | 1 | 4 | 1 |
| 7_6354124 | 2 | 0 | 1 | 0 | 2 | 0.5 | NA | NA | 5 | 0.2 | 1 | 1 | NA | NA | 1 | 0 | NA | NA | 2 | 0.5 |
| 7_6354969 | NA | NA | NA | NA | NA | NA | 2 | 0.5 | 2 | 0.5 | NA | NA | NA | NA | 1 | 1 | NA | NA | 1 | 1 |
| 7_6354984 | 1 | 1 | NA | NA | NA | NA | 2 | 0.5 | 3 | 0.6667 | NA | NA | NA | NA | 1 | 1 | NA | NA | 1 | 1 |
| 7_6358820 | 1 | 0 | NA | NA | 2 | 0.5 | NA | NA | 3 | 0.3333 | 1 | 1 | NA | NA | 2 | 0.5 | NA | NA | 3 | 0.6667 |
| 7_6398742 | 2 | 1 | 1 | 1 | 2 | 1 | 3 | 1 | 8 | 1 | 3 | 0.6667 | 1 | 1 | NA | NA | NA | NA | 4 | 0.75 |
| 7_6398751 | 2 | 1 | 1 | 1 | 2 | 1 | 3 | 0.3333 | 8 | 0.75 | 2 | 1 | 1 | 1 | NA | NA | NA | NA | 3 | 1 |
| 7_6398758 | 2 | 0.5 | 1 | 0 | 2 | 0.5 | 3 | 1 | 8 | 0.625 | 3 | 0.6667 | 1 | 1 | NA | NA | NA | NA | 4 | 0.75 |
| 7_6398763 | 2 | 1 | 1 | 0 | 2 | 0 | 3 | 1 | 8 | 0.625 | 2 | 0.5 | 1 | 1 | NA | NA | NA | NA | 3 | 0.6667 |
| 7_6398777 | 2 | 0.5 | 1 | 0 | 1 | 0 | 3 | 0.3333 | 7 | 0.2857 | 3 | 0.3333 | 1 | 0 | NA | NA | NA | NA | 4 | 0.25 |
| 7_6398779 | NA | NA | 1 | 0 | 2 | 1 | 3 | 0.6667 | 6 | 0.6667 | 2 | 1 | 1 | 1 | NA | NA | NA | NA | 3 | 1 |
| 7_6398781 | 2 | 1 | 1 | 0 | 1 | 1 | 3 | 0.6667 | 7 | 0.7143 | 3 | 1 | 1 | 1 | NA | NA | NA | NA | 4 | 1 |
| 7_6398783 | 1 | 1 | 1 | 1 | 1 | 1 | 3 | 1 | 6 | 1 | 1 | 1 | NA | NA | NA | NA | NA | NA | 1 | 1 |
| 7_6398794 | 2 | 0.5 | NA | NA | 1 | 0 | 3 | 0 | 6 | 0.1667 | 3 | 0.3333 | 1 | 0 | NA | NA | NA | NA | 4 | 0.25 |
| 7_6398873 | 3 | 0.3333 | NA | NA | 3 | 0.3333 | 3 | 0.6667 | 9 | 0.4444 | 3 | 0.3333 | 1 | 0 | NA | NA | NA | NA | 4 | 0.25 |
| 7_6398896 | 1 | 1 | NA | NA | 3 | 0.3333 | 2 | 0 | 6 | 0.3333 | 3 | 0.3333 | 1 | 1 | NA | NA | NA | NA | 4 | 0.5 |
| 7_6398947 | 3 | 0.3333 | NA | NA | 3 | 0.6667 | 3 | 0.6667 | 9 | 0.5556 | 3 | 0.3333 | 1 | 0 | NA | NA | NA | NA | 4 | 0.25 |
| 7_6398966 | 2 | 0.5 | NA | NA | 3 | 0.3333 | 3 | 0.6667 | 8 | 0.5 | 2 | 0.5 | 2 | 1 | NA | NA | NA | NA | 4 | 0.75 |
| 7_6398969 | 3 | 0.3333 | NA | NA | 3 | 0 | 3 | 0 | 9 | 0.1111 | 3 | 0.3333 | 2 | 0 | NA | NA | NA | NA | 5 | 0.2 |
| 7_6398981 | 2 | 0.5 | NA | NA | 2 | 0 | 3 | 0 | 7 | 0.1429 | 3 | 0.3333 | 2 | 1 | NA | NA | NA | NA | 5 | 0.6 |
| 7_6398984 | 2 | 0.5 | NA | NA | 1 | 1 | 3 | 0.6667 | 6 | 0.6667 | 3 | 0.6667 | 2 | 1 | NA | NA | NA | NA | 5 | 0.8 |
| 7_6398987 | 3 | 0.3333 | NA | NA | 2 | 0.5 | 3 | 0.3333 | 8 | 0.375 | 3 | 0 | 2 | 0.5 | NA | NA | NA | NA | 5 | 0.2 |
| 7_6398990 | 3 | 0 | NA | NA | 3 | 0.3333 | 3 | 0 | 9 | 0.1111 | 4 | 0.25 | 2 | 0 | NA | NA | NA | NA | 6 | 0.1667 |
| 7_6398992 | 3 | 1 | NA | NA | 3 | 1 | 4 | 1 | 10 | 1 | 4 | 0.75 | 2 | 1 | NA | NA | NA | NA | 6 | 0.8333 |
| 7_6398994 | 2 | 0 | NA | NA | 3 | 0.3333 | 3 | 1 | 8 | 0.5 | 3 | 1 | 2 | 0.5 | NA | NA | NA | NA | 5 | 0.8 |
| 7_6399001 | 3 | 0.3333 | NA | NA | 4 | 0.25 | 3 | 0 | 10 | 0.2 | 4 | 0.5 | 2 | 1 | NA | NA | NA | NA | 6 | 0.6667 |
| 7_6399012 | 2 | 0.5 | NA | NA | 4 | 0.5 | 3 | 0.3333 | 9 | 0.4444 | 2 | 0.5 | 2 | 0.5 | NA | NA | NA | NA | 4 | 0.5 |
| 7_6399030 | NA | NA | NA | NA | 1 | 1 | 2 | 1 | 3 | 1 | NA | NA | 1 | 1 | NA | NA | NA | NA | 1 | 1 |
| 7_6399596 | NA | NA | NA | NA | NA | NA | 1 | 1 | 1 | 1 | NA | NA | 1 | 0 | 1 | 1 | NA | NA | 2 | 0.5 |
| 7_6399600 | NA | NA | NA | NA | NA | NA | 1 | 1 | 1 | 1 | NA | NA | NA | NA | 1 | 1 | NA | NA | 1 | 1 |
| 7_6403873 | 2 | 0.5 | 2 | 0 | 4 | 0 | 1 | 0 | 9 | 0.1111 | 1 | 0 | 2 | 0.5 | 4 | 0 | 1 | 0 | 8 | 0.125 |

| | | | | | | | | | | | | | | | | | | | | |
|---|---|---|---|---|---|---|---|---|---|---|---|---|---|---|---|---|---|---|---|---|
| 7_64039348 | 2 | 0 | 1 | 0 | 4 | 0.25 | 1 | 0 | 8 | 0.125 | 1 | 1 | 2 | 0 | 4 | 0 | NA | NA | 7 | 0.1429 |
| 7_64039445 | 2 | 0 | 1 | 1 | 4 | 0 | 1 | 0 | 8 | 0.125 | 1 | 0 | 2 | 0 | 4 | 0.75 | NA | NA | 7 | 0.4286 |
| 7_64039714 | 2 | 0 | NA | NA | 4 | 0.25 | 1 | 0 | 7 | 0.1429 | 1 | 0 | 1 | 1 | 2 | 0.5 | 1 | 0 | 5 | 0.4 |
| 7_64039763 | 1 | 0 | NA | NA | 2 | 0 | 1 | 1 | 4 | 0.25 | 1 | 0 | 2 | 0.5 | 1 | 1 | 1 | 0 | 5 | 0.4 |
| 7_64039774 | 1 | 1 | NA | NA | 4 | 0 | 1 | 1 | 6 | 0.3333 | 1 | 1 | 2 | 0.5 | 4 | 0.5 | 1 | 0 | 8 | 0.5 |
| 7_64048611 | 1 | 1 | NA | NA | NA | NA | NA | NA | 1 | 1 | NA | NA | 1 | 0 | NA | NA | 1 | 1 | 2 | 0.5 |
| 7_64050119 | 2 | 0 | 3 | 0.3333 | 6 | 0 | 3 | 0 | 14 | 0.0714 | 1 | 0 | 6 | 0.1667 | 2 | 0 | 3 | 0 | 12 | 0.0833 |
| 7_64050167 | 2 | 0 | 4 | 0.75 | 6 | 0.3333 | 3 | 0.6667 | 15 | 0.4667 | 1 | 0 | 7 | 0.2857 | 2 | 0 | 3 | 0.3333 | 13 | 0.2308 |
| 7_64061698 | 3 | 0.6667 | 8 | 0.375 | 5 | 0.8 | 9 | 0.6667 | 25 | 0.6 | 3 | 0.6667 | 8 | 0.625 | 4 | 1 | 3 | 1 | 18 | 0.7778 |
| 7_64061720 | 3 | 0 | 8 | 0.125 | 8 | 0.125 | 9 | 0.1111 | 28 | 0.1071 | 4 | 0 | 11 | 0.0909 | 5 | 0.2 | 6 | 0.1667 | 26 | 0.1154 |
| 7_64345023 | NA | NA | 2 | 0 | 1 | 0 | 3 | 0.3333 | 6 | 0.1667 | 3 | 0.3333 | 1 | 0 | 2 | 0 | 6 | 0.1667 | 12 | 0.1667 |
| 7_64345065 | NA | NA | 2 | 0 | 1 | 1 | 2 | 0.5 | 5 | 0.4 | 2 | 0.5 | 1 | 0 | 2 | 0.5 | 7 | 0.5714 | 12 | 0.5 |
| 7_64345155 | NA | NA | 3 | 1 | NA | NA | 3 | 0.6667 | 6 | 0.8333 | 2 | 0 | 1 | 0 | 2 | 0.5 | 7 | 0.4286 | 12 | 0.3333 |
| 7_64345230 | NA | NA | 3 | 0.3333 | 1 | 1 | 2 | 0.5 | 6 | 0.5 | 3 | 0 | 1 | 0 | 2 | 0 | 7 | 0.2857 | 13 | 0.1538 |
| 7_64345239 | NA | NA | 3 | 0.3333 | 1 | 0 | 2 | 0.5 | 6 | 0.3333 | 3 | 0.6667 | 1 | 1 | 2 | 0.5 | 5 | 0.4 | 11 | 0.5455 |
| 7_64345395 | 1 | 1 | 6 | 0.6667 | 1 | 0 | 3 | 0.3333 | 11 | 0.5455 | 4 | 0.5 | 1 | 0 | 3 | 0 | 6 | 0.5 | 14 | 0.3571 |
| 7_64345633 | 1 | 0 | 7 | 0.2857 | 1 | 0 | 3 | 0.6667 | 12 | 0.3333 | 4 | 0.25 | 2 | 0.5 | 4 | 0 | 7 | 0.4286 | 17 | 0.2941 |
| 7_64402963 | 7 | 0.4286 | 10 | 0.3 | 8 | 0.125 | 6 | 0.3333 | 31 | 0.2903 | 9 | 0.2222 | 5 | 0.2 | 12 | 0.25 | 9 | 0 | 35 | 0.1714 |
| 7_64403008 | 7 | 0.4286 | 9 | 0.2222 | 6 | 0 | 7 | 0.4286 | 29 | 0.2759 | 6 | 0.6667 | 5 | 0.4 | 9 | 0.4444 | 7 | 0.2857 | 27 | 0.4444 |
| 7_64403528 | 9 | 0.3333 | 8 | 0.125 | 8 | 0 | 11 | 0.0909 | 36 | 0.1389 | 9 | 0.2222 | 6 | 0 | 16 | 0.25 | 9 | 0.3333 | 40 | 0.225 |
| 7_64403660 | 6 | 0.8333 | 8 | 1 | 6 | 0.5 | 10 | 0.7 | 30 | 0.7667 | 6 | 0.6667 | 4 | 0.5 | 10 | 1 | 4 | 0.75 | 24 | 0.7917 |
| 7_64403739 | 8 | 0 | 9 | 0 | 7 | 0 | 10 | 0.2 | 34 | 0.0588 | 5 | 0 | 4 | 0 | 10 | 0.2 | 5 | 0 | 24 | 0.0833 |
| 7_64403812 | 5 | 0.2 | 8 | 0.125 | 7 | 0.1429 | 8 | 0.375 | 28 | 0.2143 | 6 | 0.6667 | 2 | 0 | 9 | 0.5556 | 5 | 0.6 | 22 | 0.5455 |
| 7_64404074 | 6 | 0.3333 | 4 | 0.25 | 5 | 0 | 7 | 0.1429 | 22 | 0.1818 | 5 | 0.4 | 4 | 0.25 | 10 | 0.6 | 3 | 0.6667 | 22 | 0.5 |
| 7_64406105 | 4 | 1 | 4 | 0.75 | 1 | 1 | 6 | 0.8333 | 15 | 0.8667 | 4 | 1 | 5 | 0.8 | 10 | 0.8 | 1 | 1 | 20 | 0.85 |
| 7_64406309 | 4 | 0.25 | 4 | 0 | 4 | 0 | 6 | 0 | 18 | 0.0556 | 4 | 0 | 6 | 0 | 7 | 0.1429 | 1 | 0 | 18 | 0.0556 |
| 7_64406996 | 4 | 0 | 2 | 0.5 | 3 | 0 | 4 | 0 | 13 | 0.0769 | 4 | 0 | 6 | 0.1667 | 7 | 0 | 2 | 0 | 19 | 0.0526 |
| 7_64407774 | 4 | 0 | NA | NA | 3 | 0.3333 | 2 | 0 | 9 | 0.1111 | NA | NA | 2 | 0 | 6 | 0 | 2 | 0.5 | 10 | 0.1 |
| 7_64407842 | 4 | 0 | NA | NA | 2 | 0 | 2 | 0.5 | 8 | 0.125 | 4 | 0.25 | 3 | 0.6667 | 5 | 0.2 | 1 | 0 | 13 | 0.3077 |
| 7_64408053 | 5 | 0 | NA | NA | 1 | 0 | 1 | 1 | 7 | 0.1429 | 4 | 0.5 | 5 | 0.2 | 6 | 0.1667 | NA | NA | 15 | 0.2667 |
| 7_64408510 | 4 | 0.25 | 1 | 0 | 4 | 0.25 | 4 | 0.75 | 13 | 0.3846 | 5 | 0.6 | 3 | 0.6667 | 5 | 0.6 | NA | NA | 13 | 0.6154 |
| 7_64408518 | 3 | 0.3333 | 1 | 0 | 4 | 0 | 3 | 0.3333 | 11 | 0.1818 | 5 | 0.4 | 2 | 0 | 4 | 0.25 | NA | NA | 11 | 0.2727 |
| 7_64408529 | 4 | 0.75 | 1 | 1 | 3 | 0.6667 | 4 | 0.75 | 12 | 0.75 | 5 | 0.6 | 3 | 0.6667 | 5 | 1 | NA | NA | 13 | 0.7692 |
| 7_64408536 | 3 | 0.3333 | NA | NA | 4 | 0 | 4 | 0 | 11 | 0.0909 | 5 | 0.4 | 2 | 1 | 5 | 0.2 | NA | NA | 12 | 0.4167 |
| 7_64408569 | 4 | 0.5 | 1 | 1 | 3 | 0.3333 | 3 | 0.6667 | 11 | 0.5455 | 4 | 0.5 | 3 | 1 | 2 | 1 | NA | NA | 9 | 0.7778 |
| 7_64408635 | 4 | 0.5 | NA | NA | 3 | 0 | 4 | 0.5 | 11 | 0.3636 | 4 | 0.5 | 2 | 0 | 4 | 0.25 | NA | NA | 10 | 0.3 |
| 7_64408658 | 4 | 1 | 1 | 1 | 4 | 1 | 5 | 1 | 14 | 1 | 4 | 1 | 3 | 1 | 5 | 0.8 | NA | NA | 12 | 0.9167 |
| 7_64408695 | 4 | 0.75 | 1 | 1 | 3 | 1 | 5 | 1 | 13 | 0.9231 | 3 | 1 | 3 | 1 | 5 | 1 | NA | NA | 11 | 1 |
| 7_64408739 | 3 | 0.3333 | 1 | 0 | 3 | 0 | 3 | 0 | 10 | 0.1 | 3 | 0.3333 | 3 | 0.3333 | 5 | 0.2 | NA | NA | 11 | 0.2727 |
| 7_64408821 | 3 | 0.6667 | 1 | 1 | 3 | 1 | 5 | 1 | 12 | 0.9167 | 4 | 1 | 1 | 1 | 3 | 1 | NA | NA | 8 | 1 |
| 7_64408855 | 3 | 0 | 1 | 0 | 4 | 0 | 3 | 0.3333 | 11 | 0.0909 | 4 | 0.25 | 2 | 0.5 | 3 | 0 | NA | NA | 9 | 0.2222 |
| 7_64408971 | 3 | 0.6667 | 1 | 0 | 3 | 0.3333 | 2 | 1 | 9 | 0.5556 | 3 | 0.3333 | NA | NA | 2 | 0.5 | NA | NA | 5 | 0.4 |
| 7_64409063 | 2 | 0.5 | 1 | 1 | 3 | 0.3333 | 4 | 0.25 | 10 | 0.4 | 4 | 0.75 | 2 | 0.5 | 2 | 1 | NA | NA | 8 | 0.75 |
| 7_64409075 | 3 | 0 | NA | NA | 3 | 0 | 4 | 0.5 | 10 | 0.2 | 4 | 0.25 | 2 | 0.5 | 2 | 0.5 | NA | NA | 8 | 0.375 |
| 7_64521778 | 4 | 0.75 | 3 | 1 | 4 | 0.25 | 8 | 0.75 | 19 | 0.6842 | 5 | 0.6 | 1 | 1 | 3 | 0.6667 | 4 | 0.5 | 13 | 0.6154 |
| 7_64521802 | 5 | 1 | 3 | 1 | 4 | 1 | 12 | 1 | 24 | 1 | 5 | 1 | 2 | 1 | 5 | 1 | 7 | 0.8571 | 19 | 0.9474 |
| 7_64521826 | 3 | 1 | 2 | 1 | 4 | 0.75 | 9 | 0.8889 | 18 | 0.8889 | 5 | 1 | 2 | 1 | 5 | 1 | 6 | 1 | 18 | 1 |
| 7_64521915 | 4 | 1 | 2 | 1 | 4 | 1 | 7 | 1 | 17 | 1 | 3 | 1 | 1 | 1 | 4 | 1 | 5 | 1 | 13 | 1 |
| 7_64521924 | 2 | 0 | 3 | 0.3333 | 5 | 0.2 | 9 | 0.4444 | 19 | 0.3158 | 4 | 0.25 | 2 | 0 | 4 | 0.25 | 4 | 0.25 | 14 | 0.2143 |
| 7_64521985 | 4 | 0.25 | 2 | 0 | 2 | 0 | 9 | 0.1111 | 17 | 0.1176 | 5 | 0 | 2 | 0 | 4 | 0.25 | 6 | 0 | 17 | 0.0588 |
| 7_64522011 | 4 | 1 | 2 | 1 | 4 | 1 | 8 | 1 | 18 | 1 | 4 | 1 | NA | NA | 3 | 1 | 7 | 1 | 14 | 1 |
| 7_64522068 | 2 | 0 | 1 | 0 | 4 | 0 | 8 | 0.375 | 15 | 0.2 | 4 | 0 | 1 | 1 | 5 | 0.2 | 6 | 0.1667 | 16 | 0.1875 |
| 7_64522082 | 3 | 0.3333 | 2 | 0.5 | 4 | 0 | 10 | 0.3 | 19 | 0.2632 | 4 | 0.25 | 2 | 0.5 | 5 | 0.6 | 7 | 0.2857 | 18 | 0.3889 |
| 7_64522109 | 1 | 1 | 2 | 0 | 2 | 1 | 8 | 0.5 | 13 | 0.5385 | 3 | 0.3333 | 2 | 0 | 4 | 0 | 5 | 0.2 | 14 | 0.1429 |
| 7_64522146 | 3 | 1 | 2 | 0.5 | 3 | 0.6667 | 9 | 0.6667 | 17 | 0.7059 | 3 | 1 | 1 | 1 | 5 | 0.8 | 6 | 1 | 15 | 0.9333 |
| 7_64522159 | 2 | 0 | 2 | 0.5 | NA | NA | 7 | 0.1429 | 11 | 0.1818 | 2 | 0 | 2 | 0 | 5 | 0.6 | 6 | 0 | 15 | 0.2 |
| 7_64522583 | 3 | 0.3333 | 2 | 0 | 5 | 0 | 8 | 0.125 | 18 | 0.1111 | 3 | 0 | 2 | 0 | 4 | 0.25 | 3 | 0 | 12 | 0.0833 |
| 7_64522608 | 1 | 0 | 1 | 1 | 3 | 0 | 4 | 0.25 | 9 | 0.2222 | 2 | 0 | 1 | 0 | 4 | 0 | 4 | 0.25 | 11 | 0.0909 |
| 7_64946066 | 12 | 0 | 9 | 0.1111 | 5 | 0.2 | 11 | 0 | 37 | 0.0541 | 12 | 0 | 7 | 0.1429 | 10 | 0.2 | 11 | 0.0909 | 40 | 0.1 |
| 7_64946271 | 11 | 0.0909 | 8 | 0 | 3 | 0 | 12 | 0.0833 | 34 | 0.0588 | 9 | 0 | 6 | 0.1667 | 7 | 0.1429 | 8 | 0 | 30 | 0.0667 |
| 7_64946543 | 12 | 0.4167 | 10 | 0.2 | 2 | 1 | 11 | 0.1818 | 35 | 0.3143 | 7 | 0 | 5 | 0.4 | 10 | 0.2 | 11 | 0.2727 | 33 | 0.2121 |
| 7_64946559 | 12 | 0.1667 | 10 | 0.1 | 2 | 0 | 7 | 0.1429 | 31 | 0.129 | 8 | 0.125 | 6 | 0.1667 | 9 | 0.1111 | 12 | 0.25 | 35 | 0.1714 |
| 7_64947108 | 3 | 0.3333 | 5 | 0 | NA | NA | 2 | 0 | 10 | 0.1 | 3 | 0 | 4 | 0.5 | 4 | 0.25 | 3 | 0.3333 | 14 | 0.2857 |
| 7_64947185 | 6 | 0.1667 | 4 | 0.25 | NA | NA | 4 | 0.25 | 14 | 0.2143 | 2 | 0 | 4 | 0.25 | 3 | 0 | 6 | 0.6667 | 15 | 0.3333 |
| 7_64947274 | 5 | 0 | 5 | 0 | NA | NA | 4 | 0.25 | 14 | 0.0714 | 3 | 0.3333 | 3 | 0 | 4 | 0.25 | 6 | 0.3333 | 16 | 0.25 |

| | | | | | | | | | | | | | | | | | | | | |
|---|---|---|---|---|---|---|---|---|---|---|---|---|---|---|---|---|---|---|---|---|
| 7_6494729 | 2 | 0.5 | 5 | 0.8 | NA | NA | 3 | 0 | 10 | 0.5 | 2 | 0.5 | 3 | 0.6667 | 3 | 0.6667 | 6 | 1 | 14 | 0.7857 |
| 7_6494730 | 6 | 0 | 5 | 0.2 | NA | NA | 3 | 0.3333 | 14 | 0.1429 | 2 | 0.5 | 4 | 0.5 | 4 | 0 | 5 | 0 | 15 | 0.2 |
| 7_6495256 | 4 | 0.25 | 5 | 0.2 | NA | NA | 3 | 0 | 12 | 0.1667 | 2 | 1 | 3 | 0 | 4 | 0.25 | 3 | 0 | 12 | 0.25 |
| 7_6495262 | 2 | 0 | 3 | 0.3333 | NA | NA | 2 | 0.5 | 7 | 0.2857 | 1 | 1 | 4 | 0.25 | 3 | 0.6667 | 4 | 0.75 | 12 | 0.5833 |
| 7_6495267 | 4 | 0.25 | 5 | 0.4 | NA | NA | 3 | 0.6667 | 12 | 0.4167 | 1 | 1 | 3 | 0 | 4 | 0.5 | 5 | 0.6 | 13 | 0.4615 |
| 7_6495279 | 4 | 0.5 | 5 | 0.6 | NA | NA | 3 | 0.6667 | 12 | 0.5833 | 2 | 0 | 3 | 0 | 4 | 0.75 | 5 | 0.4 | 14 | 0.3571 |
| 7_6495285 | 4 | 0.25 | 4 | 0 | NA | NA | 3 | 0.3333 | 11 | 0.1818 | 2 | 0 | 3 | 0.3333 | 3 | 0.3333 | 5 | 0.4 | 13 | 0.3077 |
| 7_6495293 | 4 | 0 | 5 | 0 | NA | NA | 2 | 0.5 | 11 | 0.0909 | NA | NA | 3 | 0 | 4 | 0.5 | 4 | 0 | 11 | 0.1818 |
| 7_6496283 | NA | NA | 2 | 0.5 | NA | NA | 1 | 0 | 3 | 0.3333 | NA | NA | 1 | 0 | 3 | 0.3333 | 4 | 0.25 | 8 | 0.25 |
| 7_6496301 | NA | NA | 4 | 0 | NA | NA | 1 | 1 | 5 | 0.2 | NA | NA | 1 | 0 | 3 | 0 | 4 | 0.25 | 8 | 0.125 |
| 7_6496305 | NA | NA | 3 | 0.3333 | NA | NA | 1 | 0 | 4 | 0.25 | NA | NA | 1 | 0 | 3 | 0 | 4 | 0.5 | 8 | 0.25 |
| 7_6496459 | NA | NA | 2 | 0 | NA | NA | 1 | 1 | 3 | 0.3333 | NA | NA | 1 | 0 | 3 | 0 | 4 | 0.25 | 8 | 0.125 |
| 7_6534110 | 4 | 0 | 4 | 0 | 5 | 0.2 | 6 | 0 | 19 | 0.0526 | 5 | 0 | 3 | 0 | 7 | 0.1429 | 3 | 0 | 18 | 0.0556 |
| 7_6534169 | 7 | 0 | 6 | 0.1667 | 6 | 0.1667 | 6 | 0 | 25 | 0.08 | 4 | 0.5 | 3 | 0 | 7 | 0.2857 | 4 | 0 | 18 | 0.2222 |
| 7_6569895 | 2 | 0.5 | NA | NA | 1 | 0 | 3 | 0 | 6 | 0.1667 | 1 | 0 | NA | NA | 1 | 0 | 1 | 1 | 3 | 0.3333 |
| 7_6569897 | NA | NA | NA | NA | 2 | 1 | 2 | 1 | 4 | 1 | NA | NA | 1 | 1 | 1 | 1 | NA | NA | 2 | 1 |
| 7_6569900 | 3 | 0.6667 | NA | NA | 2 | 0.5 | 3 | 1 | 8 | 0.75 | 1 | 1 | 1 | 1 | 1 | 1 | 1 | 1 | 4 | 1 |
| 7_6569903 | 3 | 0.6667 | NA | NA | 1 | 0 | 2 | 0 | 6 | 0.3333 | 1 | 0 | 1 | 1 | 1 | 0 | 1 | 1 | 4 | 0.5 |
| 7_6569906 | 3 | 0 | 1 | 0 | 2 | 0.5 | 2 | 0.5 | 8 | 0.25 | 1 | 0 | 1 | 0 | 1 | 0 | 1 | 1 | 4 | 0.25 |
| 7_6569908 | 3 | 0 | 1 | 0 | 2 | 0.5 | 3 | 0 | 9 | 0.1111 | 1 | 0 | 1 | 0 | 1 | 1 | 1 | 0 | 4 | 0.25 |
| 7_6569918 | 3 | 0.3333 | 1 | 0 | 2 | 1 | 3 | 0.3333 | 9 | 0.4444 | 1 | 0 | 1 | 1 | 1 | 1 | 1 | 1 | 4 | 0.75 |
| 7_6569926 | 3 | 1 | 1 | 1 | 2 | 1 | 3 | 0.6667 | 9 | 0.8889 | NA | NA | 1 | 1 | 1 | 0 | 1 | 1 | 3 | 0.6667 |
| 7_6569927 | 3 | 0.3333 | 1 | 1 | 1 | 1 | 3 | 1 | 8 | 0.75 | NA | NA | 1 | 1 | NA | NA | 1 | 1 | 2 | 1 |
| 7_6569956 | 2 | 0.5 | 1 | 0 | 2 | 1 | 3 | 0.6667 | 8 | 0.625 | NA | NA | 1 | 1 | 1 | 1 | 1 | 0 | 3 | 0.6667 |
| 7_6569968 | 3 | 1 | 1 | 1 | 2 | 1 | 3 | 1 | 9 | 1 | NA | NA | NA | NA | 1 | 1 | 1 | 1 | 2 | 1 |
| 7_6569995 | 2 | 0.5 | 2 | 0.5 | 1 | 0 | 3 | 0 | 8 | 0.25 | 1 | 0 | 1 | 1 | 1 | 0 | 1 | 0 | 4 | 0.25 |
| 7_6572421 | 4 | 0.25 | 3 | 1 | 1 | 1 | 4 | 0.5 | 12 | 0.5833 | 2 | 0 | 2 | 0.5 | 1 | 1 | 2 | 0.5 | 7 | 0.4286 |
| 7_6573868 | 7 | 0.1429 | 9 | 0.1111 | 5 | 0 | 7 | 0.1429 | 28 | 0.1071 | 4 | 0 | 6 | 0.1667 | 8 | 0.125 | 5 | 0.2 | 23 | 0.1304 |
| 7_6582095 | NA | NA | 1 | 1 | NA | NA | NA | NA | 1 | 1 | 2 | 1 | 1 | 1 | 1 | 1 | NA | NA | 4 | 1 |
| 7_6582126 | NA | NA | 1 | 1 | NA | NA | NA | NA | 1 | 1 | 3 | 1 | 1 | 1 | 2 | 1 | NA | NA | 6 | 1 |
| 7_6582134 | NA | NA | 1 | 1 | NA | NA | NA | NA | 1 | 1 | 3 | 0.3333 | 1 | 0 | 1 | 0 | NA | NA | 5 | 0.2 |
| 7_6582142 | NA | NA | 1 | 1 | NA | NA | NA | NA | 1 | 1 | 3 | 0.6667 | NA | NA | 1 | 1 | NA | NA | 4 | 0.75 |
| 7_6582166 | NA | NA | 1 | 1 | NA | NA | NA | NA | 1 | 1 | 3 | 1 | 1 | 1 | 2 | 1 | NA | NA | 6 | 1 |
| 7_6582179 | NA | NA | 1 | 1 | NA | NA | NA | NA | 1 | 1 | 3 | 0.3333 | 1 | 0 | 3 | 0 | NA | NA | 7 | 0.1429 |
| 7_6582183 | NA | NA | 1 | 1 | NA | NA | NA | NA | 1 | 1 | 3 | 1 | 1 | 1 | 2 | 1 | NA | NA | 6 | 1 |
| 7_6582192 | NA | NA | 1 | 1 | 1 | 1 | NA | NA | 2 | 1 | 2 | 0.5 | 2 | 0.5 | 3 | 0.3333 | NA | NA | 7 | 0.4286 |
| 7_6582196 | NA | NA | 1 | 0 | 1 | 1 | NA | NA | 2 | 0.5 | 2 | 0.5 | 2 | 0.5 | 1 | 1 | NA | NA | 5 | 0.6 |
| 7_6582203 | NA | NA | 1 | 1 | 1 | 0 | NA | NA | 2 | 0.5 | 3 | 0 | 2 | 0.5 | 2 | 0.5 | NA | NA | 7 | 0.2857 |
| 7_6582229 | NA | NA | 1 | 1 | 1 | 1 | NA | NA | 2 | 1 | 3 | 1 | 2 | 1 | 2 | 0.5 | NA | NA | 7 | 0.8571 |
| 7_6582241 | 1 | 0 | 1 | 0 | 1 | 1 | NA | NA | 3 | 0.3333 | 3 | 0 | 4 | 0.25 | 2 | 0 | NA | NA | 9 | 0.1111 |
| 7_6582285 | 2 | 0.5 | 1 | 1 | 2 | 0.5 | NA | NA | 5 | 0.6 | 3 | 1 | 4 | 0.5 | 4 | 0.75 | NA | NA | 11 | 0.7273 |
| 7_6582288 | 1 | 0 | 1 | 0 | 1 | 1 | NA | NA | 3 | 0.3333 | 2 | 0 | 5 | 0.4 | 3 | 0.3333 | NA | NA | 10 | 0.3 |
| 7_6590893 | NA | NA | NA | NA | NA | NA | 2 | 0.5 | 2 | 0.5 | 1 | 1 | NA | NA | 1 | 0 | NA | NA | 2 | 0.5 |
| 7_6590918 | NA | NA | NA | NA | NA | NA | 2 | 0.5 | 2 | 0.5 | 1 | 1 | NA | NA | 1 | 0 | NA | NA | 2 | 0.5 |
| 7_6591220 | NA | NA | NA | NA | NA | NA | 2 | 1 | 2 | 1 | NA | NA | NA | NA | 1 | 1 | NA | NA | 1 | 1 |
| 7_6629486 | 1 | 0 | 2 | 0 | 1 | 1 | 3 | 0 | 7 | 0.1429 | NA | NA | 3 | 0.3333 | 1 | 0 | 3 | 0 | 7 | 0.1429 |
| 7_6629751 | 1 | 0 | 1 | 0 | NA | NA | 1 | 1 | 3 | 0.3333 | NA | NA | 1 | 1 | 1 | 0 | 2 | 0.5 | 4 | 0.5 |
| 7_6629752 | 1 | 1 | 1 | 1 | NA | NA | 1 | 1 | 3 | 1 | NA | NA | 1 | 1 | 1 | 1 | 3 | 1 | 5 | 1 |
| 7_6629756 | 1 | 1 | 1 | 0 | NA | NA | 1 | 1 | 3 | 0.6667 | NA | NA | NA | NA | 1 | 1 | 3 | 1 | 4 | 1 |
| 7_6630947 | 1 | 1 | NA | NA | NA | NA | NA | NA | 1 | 1 | NA | NA | NA | NA | NA | NA | 2 | 1 | 2 | 1 |
| 7_6630948 | NA | NA | NA | NA | NA | NA | 1 | 1 | 1 | 1 | NA | NA | NA | NA | NA | NA | 2 | 1 | 2 | 1 |
| 7_6630959 | 1 | 1 | NA | NA | NA | NA | NA | NA | 1 | 1 | NA | NA | NA | NA | NA | NA | 1 | 1 | 1 | 1 |
| 7_6630982 | 1 | 1 | NA | NA | NA | NA | 1 | 1 | 2 | 1 | NA | NA | NA | NA | NA | NA | 1 | 1 | 1 | 1 |
| 7_6630986 | 1 | 1 | NA | NA | NA | NA | 1 | 1 | 2 | 1 | NA | NA | NA | NA | NA | NA | 1 | 1 | 1 | 1 |
| 7_6630989 | 1 | 1 | NA | NA | NA | NA | 1 | 1 | 2 | 1 | NA | NA | NA | NA | NA | NA | 1 | 1 | 1 | 1 |
| 7_6635828 | NA | NA | 1 | 1 | 1 | 0 | NA | NA | 2 | 0.5 | NA | NA | NA | NA | NA | NA | 2 | 0.5 | 2 | 0.5 |
| 7_6636026 | NA | NA | NA | NA | 1 | 1 | NA | NA | 1 | 1 | NA | NA | NA | NA | NA | NA | 2 | 0.5 | 2 | 0.5 |
| 7_6636370 | NA | NA | NA | NA | 1 | 1 | NA | NA | 1 | 1 | NA | NA | 1 | 1 | NA | NA | 2 | 1 | 3 | 1 |
| 7_6636374 | NA | NA | NA | NA | 1 | 1 | NA | NA | 1 | 1 | NA | NA | 1 | 1 | NA | NA | 1 | 0 | 2 | 0.5 |
| 7_6636375 | NA | NA | NA | NA | 1 | 1 | NA | NA | 1 | 1 | NA | NA | 1 | 1 | NA | NA | 2 | 0.5 | 3 | 0.6667 |
| 7_6636390 | NA | NA | NA | NA | 1 | 1 | NA | NA | 1 | 1 | NA | NA | 1 | 1 | NA | NA | 2 | 1 | 3 | 1 |
| 7_6636414 | NA | NA | NA | NA | 1 | 1 | NA | NA | 1 | 1 | NA | NA | 1 | 1 | NA | NA | 1 | 1 | 2 | 1 |
| 7_6636424 | NA | NA | NA | NA | 1 | 1 | NA | NA | 1 | 1 | NA | NA | 1 | 1 | NA | NA | 2 | 1 | 3 | 1 |
| 7_6636448 | NA | NA | NA | NA | 1 | 1 | NA | NA | 1 | 1 | NA | NA | NA | NA | NA | NA | 1 | 1 | 1 | 1 |
| 7_6648962 | 1 | 1 | NA | NA | NA | NA | NA | NA | 1 | 1 | 1 | 1 | NA | NA | NA | NA | NA | NA | 1 | 1 |

| | | | | | | | | | | | | | | | | | | | | |
|---|---|---|---|---|---|---|---|---|---|---|---|---|---|---|---|---|---|---|---|---|
| 7_66873467 | 3 | 0.3333 | 2 | 0 | 3 | 0.3333 | 5 | 0.2 | 13 | 0.2308 | 1 | 0 | 1 | 1 | 4 | 0.25 | 1 | 0 | 7 | 0.2857 |
| 7_66873476 | 3 | 0 | 3 | 0 | 4 | 0 | 3 | 0.3333 | 13 | 0.0769 | 1 | 0 | 1 | 0 | 4 | 0.25 | 2 | 0 | 8 | 0.125 |
| 7_66874854 | 1 | 0 | NA | NA | NA | NA | 2 | 0.5 | 3 | 0.3333 | NA | NA | NA | NA | NA | NA | 1 | 1 | 1 | 1 |
| 7_66875725 | 1 | 1 | 2 | 1 | 1 | 1 | 3 | 1 | 7 | 1 | NA | NA | NA | NA | NA | NA | 1 | 1 | 1 | 1 |
| 7_66875832 | 1 | 1 | 2 | 0 | 2 | 1 | 2 | 0.5 | 7 | 0.5714 | NA | NA | NA | NA | NA | NA | 1 | 1 | 1 | 1 |
| 7_66875856 | 1 | 1 | 1 | 1 | 2 | 1 | 2 | 1 | 6 | 1 | NA | NA | NA | NA | NA | NA | 1 | 1 | 1 | 1 |
| 7_66876183 | NA | NA | 2 | 1 | 1 | 0 | 2 | 0 | 5 | 0.4 | NA | NA | NA | NA | NA | NA | 1 | 1 | 1 | 1 |
| 7_66876246 | 1 | 1 | 2 | 0.5 | 3 | 1 | 4 | 0.75 | 10 | 0.8 | NA | NA | NA | NA | NA | NA | 1 | 1 | 1 | 1 |
| 7_66876312 | 1 | 0 | 2 | 1 | 3 | 1 | 3 | 1 | 9 | 0.8889 | NA | NA | NA | NA | NA | NA | 1 | 1 | 1 | 1 |
| 7_66876405 | 1 | 1 | 2 | 1 | 3 | 1 | 3 | 0.6667 | 9 | 0.8889 | NA | NA | NA | NA | NA | NA | 1 | 1 | 1 | 1 |
| 7_66876560 | 1 | 0 | 2 | 1 | 3 | 0.3333 | 4 | 0.5 | 10 | 0.5 | NA | NA | NA | NA | NA | NA | 1 | 1 | 1 | 1 |
| 7_66876682 | 1 | 0 | 2 | 0.5 | 2 | 0 | 4 | 0.25 | 9 | 0.2222 | NA | NA | NA | NA | NA | NA | 1 | 1 | 1 | 1 |
| 7_66876820 | 1 | 0 | NA | NA | 3 | 0.6667 | 4 | 1 | 8 | 0.75 | NA | NA | NA | NA | 1 | 1 | NA | NA | 1 | 1 |
| 7_66876852 | 1 | 1 | 2 | 1 | 2 | 1 | 3 | 1 | 8 | 1 | NA | NA | NA | NA | 1 | 1 | 1 | 1 | 2 | 1 |
| 7_66877922 | 4 | 0 | 1 | 0 | 3 | 0.3333 | 3 | 0 | 11 | 0.0909 | 2 | 0.5 | NA | NA | NA | NA | 2 | 0 | 4 | 0.25 |
| 7_66881113 | 16 | 0.25 | 22 | 0.2727 | 20 | 0.35 | 25 | 0.36 | 83 | 0.3133 | 8 | 0.125 | 11 | 0.2727 | 20 | 0.2 | 19 | 0.1053 | 58 | 0.1724 |
| 7_66881310 | 13 | 0.0769 | 22 | 0.0909 | 20 | 0.1 | 32 | 0.0938 | 87 | 0.092 | 7 | 0.1429 | 11 | 0.0909 | 23 | 0.1304 | 23 | 0.0435 | 64 | 0.0938 |
| 7_66881608 | 12 | 0.1667 | 23 | 0.0435 | 18 | 0.2222 | 28 | 0.2143 | 81 | 0.1605 | 4 | 0 | 10 | 0.2 | 20 | 0.4 | 23 | 0.1304 | 57 | 0.2281 |
| 7_66881751 | 14 | 0 | 19 | 0 | 15 | 0.2667 | 27 | 0.1111 | 75 | 0.0933 | 5 | 0.2 | 9 | 0 | 18 | 0.2222 | 19 | 0.1053 | 51 | 0.1373 |
| 7_66881776 | 14 | 0.0714 | 23 | 0.0435 | 18 | 0.0556 | 28 | 0.0714 | 83 | 0.0602 | 5 | 0 | 10 | 0.1 | 20 | 0.15 | 20 | 0.05 | 55 | 0.0909 |
| 7_66881906 | 14 | 0.2857 | 21 | 0.0476 | 17 | 0.1765 | 26 | 0.1154 | 78 | 0.141 | 5 | 0 | 7 | 0 | 19 | 0.2105 | 20 | 0.05 | 51 | 0.098 |
| 7_66881918 | 14 | 0.0714 | 24 | 0.125 | 16 | 0.0625 | 26 | 0.0385 | 80 | 0.075 | 5 | 0.2 | 8 | 0.125 | 16 | 0.125 | 20 | 0.05 | 49 | 0.102 |
| 7_66883564 | 6 | 0.1667 | 12 | 0 | 7 | 0 | 15 | 0.1333 | 40 | 0.075 | 5 | 0 | 6 | 0.1667 | 11 | 0.0909 | 13 | 0.0769 | 35 | 0.0857 |
| 7_66884329 | 4 | 0.25 | 9 | 0 | 8 | 0 | 13 | 0.3077 | 34 | 0.1471 | 5 | 0.4 | 2 | 0 | 8 | 0.125 | 10 | 0.1 | 25 | 0.16 |
| 7_66884541 | 3 | 0.6667 | 11 | 0.0909 | 7 | 0.2857 | 13 | 0.3077 | 34 | 0.2647 | 4 | 0.25 | 5 | 0.4 | 9 | 0 | 11 | 0.2727 | 29 | 0.2069 |
| 7_66884574 | 5 | 0.2 | 8 | 0.25 | 5 | 0 | 11 | 0.1818 | 29 | 0.1724 | 4 | 0.25 | 5 | 0.2 | 8 | 0.125 | 10 | 0.3 | 27 | 0.2222 |
| 7_66884632 | 5 | 1 | 12 | 1 | 6 | 1 | 12 | 1 | 35 | 1 | 5 | 0.8 | 4 | 1 | 9 | 1 | 10 | 1 | 28 | 0.9643 |
| 7_66884668 | 4 | 0.5 | 11 | 0.6364 | 6 | 0.3333 | 12 | 0.5833 | 33 | 0.5455 | 5 | 0.2 | 5 | 0.6 | 6 | 0.8333 | 9 | 0.4444 | 25 | 0.52 |
| 7_66884773 | 4 | 1 | 11 | 1 | 6 | 0.8333 | 12 | 0.9167 | 33 | 0.9394 | 4 | 0.75 | 3 | 0.3333 | 5 | 0.8 | 8 | 0.875 | 20 | 0.75 |
| 7_67163334 | 2 | 0.5 | 3 | 0 | 1 | 1 | 3 | 0 | 9 | 0.2222 | 5 | 0.2 | 1 | 0 | 2 | 0.5 | 2 | 0.5 | 10 | 0.3 |
| 7_67163596 | NA | NA | 3 | 0.3333 | 1 | 1 | 3 | 0.3333 | 7 | 0.4286 | 5 | 0.2 | 2 | 0.5 | 3 | 0 | 3 | 0 | 13 | 0.1538 |
| 7_67163727 | NA | NA | 4 | 0 | 1 | 0 | 4 | 0.25 | 9 | 0.1111 | 4 | 0 | 2 | 0.5 | 1 | 0 | 2 | 0 | 9 | 0.1111 |
| 7_67163778 | NA | NA | 3 | 0 | 1 | 0 | 4 | 0.25 | 8 | 0.125 | 4 | 0.25 | 2 | 0.5 | 2 | 0.5 | 2 | 0 | 10 | 0.3 |
| 7_67164481 | NA | NA | 3 | 0.3333 | NA | NA | 4 | 0.75 | 7 | 0.5714 | 4 | 0.5 | 1 | 1 | 2 | 0 | 2 | 0 | 9 | 0.3333 |
| 7_67164510 | NA | NA | 2 | 0.5 | 1 | 1 | 4 | 1 | 7 | 0.8571 | 4 | 0.25 | NA | NA | 3 | 0.6667 | 1 | 0 | 8 | 0.375 |
| 7_67164568 | NA | NA | 3 | 0.3333 | NA | NA | 4 | 0.25 | 7 | 0.2857 | 4 | 0.5 | 1 | 1 | 4 | 0 | 2 | 0.5 | 11 | 0.3636 |
| 7_67164705 | NA | NA | 3 | 1 | 1 | 1 | 5 | 1 | 9 | 1 | 5 | 1 | 1 | 1 | 4 | 1 | 2 | 1 | 12 | 1 |
| 7_67164753 | NA | NA | 4 | 0 | 1 | 0 | 6 | 0.1667 | 11 | 0.0909 | 4 | 0.25 | 1 | 0 | 7 | 0 | 1 | 0 | 13 | 0.0769 |
| 7_67164874 | NA | NA | 4 | 1 | 1 | 1 | 5 | 1 | 10 | 1 | 3 | 1 | 1 | 1 | 6 | 0.5 | 2 | 1 | 12 | 0.75 |
| 7_67297358 | 2 | 1 | 1 | 0 | NA | NA | NA | NA | 3 | 0.6667 | NA | NA | NA | NA | NA | NA | 1 | 1 | 1 | 1 |
| 7_67375886 | 2 | 1 | NA | NA | NA | NA | NA | NA | 2 | 1 | NA | NA | NA | NA | NA | NA | 1 | 1 | 1 | 1 |
| 7_67376118 | 2 | 1 | 1 | 0 | NA | NA | NA | NA | 3 | 0.6667 | NA | NA | NA | NA | NA | NA | 1 | 1 | 1 | 1 |
| 7_67376285 | 2 | 0.5 | 1 | 0 | NA | NA | NA | NA | 3 | 0.3333 | NA | NA | NA | NA | NA | NA | 1 | 1 | 1 | 1 |
| 7_67380174 | 2 | 0.5 | NA | NA | 1 | 1 | 1 | 0 | 4 | 0.5 | 2 | 0.5 | 2 | 0.5 | 1 | 0 | NA | NA | 5 | 0.4 |
| 7_67380267 | 2 | 0.5 | NA | NA | 1 | 0 | 1 | 0 | 4 | 0.25 | 1 | 0 | 3 | 0.3333 | 1 | 1 | NA | NA | 5 | 0.4 |
| 7_67380471 | 5 | 0.2 | NA | NA | 2 | 0.5 | 3 | 0.3333 | 10 | 0.3 | 4 | 0.5 | 2 | 0.5 | 1 | 1 | 1 | 0 | 8 | 0.5 |
| 7_67380917 | 3 | 0.3333 | NA | NA | 2 | 0 | 2 | 0 | 7 | 0.1429 | 2 | 0.5 | 1 | 0 | 1 | 0 | 1 | 1 | 5 | 0.4 |
| 7_67381072 | 4 | 1 | NA | NA | 3 | 0 | 1 | 1 | 8 | 0.625 | 2 | 1 | 1 | 1 | 1 | 1 | NA | NA | 4 | 1 |
| 7_67381517 | 4 | 0.25 | NA | NA | 3 | 0 | 1 | 0 | 8 | 0.125 | 3 | 0.3333 | 1 | 0 | 1 | 0 | NA | NA | 5 | 0.2 |
| 7_67381547 | 3 | 1 | NA | NA | 2 | 1 | 1 | 1 | 6 | 1 | 3 | 0.6667 | 1 | 1 | 1 | 1 | NA | NA | 5 | 0.8 |
| 7_67381618 | 3 | 0.3333 | NA | NA | 3 | 0.6667 | 1 | 0 | 7 | 0.4286 | 3 | 0.3333 | 1 | 0 | 1 | 1 | NA | NA | 5 | 0.4 |
| 7_67381820 | 4 | 0.5 | NA | NA | 3 | 0.3333 | 1 | 1 | 8 | 0.5 | 3 | 0.3333 | 1 | 0 | 1 | 1 | NA | NA | 5 | 0.4 |
| 7_67384252 | 1 | 0 | NA | NA | 2 | 1 | 1 | 0 | 4 | 0.5 | 1 | 1 | NA | NA | 1 | 1 | NA | NA | 2 | 1 |
| 7_67384617 | 1 | 1 | NA | NA | 1 | 1 | 1 | 1 | 3 | 1 | 1 | 1 | NA | NA | 1 | 1 | NA | NA | 2 | 1 |
| 7_67384639 | 1 | 1 | NA | NA | 2 | 1 | NA | NA | 3 | 1 | NA | NA | NA | NA | 1 | 1 | NA | NA | 1 | 1 |
| 7_67384669 | NA | NA | NA | NA | 2 | 1 | 1 | 1 | 3 | 1 | 1 | 1 | NA | NA | 1 | 1 | NA | NA | 2 | 1 |
| 7_67385370 | NA | NA | NA | NA | 1 | 0 | 1 | 1 | 2 | 0.5 | 1 | 0 | NA | NA | 1 | 1 | NA | NA | 2 | 0.5 |
| 7_67385507 | 1 | 1 | NA | NA | 1 | 1 | 1 | 1 | 3 | 1 | NA | NA | NA | NA | 1 | 1 | NA | NA | 1 | 1 |
| 7_67386181 | NA | NA | NA | NA | 1 | 1 | NA | NA | 1 | 1 | NA | NA | NA | NA | 1 | 1 | NA | NA | 1 | 1 |
| 7_67876002 | NA | NA | NA | NA | NA | NA | 1 | 1 | 1 | 1 | NA | NA | NA | NA | 1 | 0 | 1 | 1 | 2 | 0.5 |
| 7_67876011 | NA | NA | NA | NA | NA | NA | 1 | 1 | 1 | 1 | NA | NA | NA | NA | 1 | 0 | 1 | 1 | 2 | 0.5 |
| 7_67876333 | NA | NA | NA | NA | 2 | 0.5 | 1 | 0 | 3 | 0.3333 | NA | NA | NA | NA | 2 | 1 | 1 | 1 | 3 | 1 |
| 7_67877069 | NA | NA | NA | NA | 1 | 0 | 1 | 1 | 2 | 0.5 | NA | NA | NA | NA | 2 | 0.5 | NA | NA | 2 | 0.5 |
| 7_67877246 | NA | NA | NA | NA | 1 | 1 | 1 | 0 | 2 | 0.5 | NA | NA | NA | NA | 2 | 0.5 | NA | NA | 2 | 0.5 |
| 7_67880519 | 3 | 0 | 2 | 0 | 4 | 0 | 3 | 0.3333 | 12 | 0.0833 | 1 | 1 | NA | NA | 5 | 0 | 2 | 0 | 8 | 0.125 |

| | | | | | | | | | | | | | | | | | | | | |
|---|---|---|---|---|---|---|---|---|---|---|---|---|---|---|---|---|---|---|---|---|
| 7_6788061 | 2 | 0.5 | 2 | 0.5 | 5 | 0.6 | 2 | 0 | 11 | 0.4545 | 3 | 1 | NA | NA | 5 | 0.8 | 3 | 0.3333 | 11 | 0.7273 |
| 7_6788066 | 3 | 0.3333 | 3 | 0.3333 | 3 | 0.6667 | 3 | 0.6667 | 12 | 0.5 | 3 | 1 | NA | NA | 6 | 0.6667 | 2 | 0.5 | 11 | 0.7273 |
| 7_6788102 | 3 | 0 | 4 | 0 | 5 | 0 | 4 | 0.25 | 16 | 0.0625 | 3 | 0.3333 | NA | NA | 6 | 0 | 3 | 0 | 12 | 0.0833 |
| 7_6788125 | 3 | 0 | 4 | 0 | 5 | 0.2 | 4 | 0 | 16 | 0.0625 | 3 | 0.6667 | 1 | 0 | 6 | 0 | 3 | 0 | 13 | 0.1538 |
| 7_6788127 | 3 | 0 | 4 | 0.25 | 4 | 0 | 2 | 1 | 13 | 0.2308 | 3 | 0.3333 | 1 | 1 | 6 | 0.3333 | 3 | 0 | 13 | 0.3077 |
| 7_6788130 | 3 | 0 | 3 | 0.3333 | 5 | 0.2 | 4 | 0 | 15 | 0.1333 | 1 | 0 | 1 | 0 | 6 | 0 | 3 | 0.3333 | 11 | 0.0909 |
| 7_6788133 | 3 | 0 | 4 | 0 | 5 | 0.2 | 4 | 0 | 16 | 0.0625 | 1 | 0 | 1 | 1 | 5 | 0 | 3 | 0 | 10 | 0.1 |
| 7_6788134 | 3 | 0 | 3 | 0.3333 | 4 | 0.5 | 4 | 0 | 14 | 0.2143 | 3 | 0.6667 | 1 | 1 | 3 | 0.6667 | 3 | 0 | 10 | 0.5 |
| 7_6788140 | 3 | 0.3333 | 2 | 0.5 | 5 | 1 | 2 | 0.5 | 12 | 0.6667 | 3 | 0.6667 | 1 | 1 | 3 | 0.6667 | 3 | 0.3333 | 10 | 0.6 |
| 7_6788152 | 3 | 0.3333 | 3 | 0.3333 | 5 | 0.6 | 4 | 0.5 | 15 | 0.4667 | 2 | 0.5 | 1 | 1 | 6 | 0.6667 | 3 | 0.3333 | 12 | 0.5833 |
| 7_6788153 | 3 | 0 | 4 | 0.75 | 5 | 1 | 4 | 1 | 16 | 0.75 | NA | NA | 1 | 1 | 6 | 0.8333 | 3 | 1 | 10 | 0.9 |
| 7_6788160 | 2 | 0 | 4 | 0.5 | 5 | 0.4 | 3 | 0.3333 | 14 | 0.3571 | 4 | 0.5 | 1 | 0 | 5 | 0.2 | 3 | 0.3333 | 13 | 0.3077 |
| 7_6788166 | 3 | 0.3333 | 4 | 0.5 | 4 | 0.25 | 4 | 0.75 | 15 | 0.4667 | 3 | 0.6667 | 2 | 1 | 5 | 0.6 | 3 | 0.3333 | 13 | 0.6154 |
| 7_6788235 | 4 | 0.25 | 4 | 0.25 | 4 | 0 | 4 | 0.5 | 16 | 0.25 | 4 | 0 | 1 | 0 | 4 | 0.25 | 3 | 0.3333 | 12 | 0.1667 |
| 7_6788239 | 3 | 0 | 5 | 0.4 | 4 | 0.25 | 5 | 0.2 | 17 | 0.2353 | 4 | 0 | 1 | 0 | 8 | 0.25 | 3 | 0 | 16 | 0.125 |
| 7_6788252 | 5 | 0.4 | 6 | 0 | 6 | 0.3333 | 5 | 0.2 | 22 | 0.2273 | 5 | 0.2 | 1 | 0 | 9 | 0 | 3 | 0.3333 | 18 | 0.1111 |
| 7_6788279 | 3 | 0 | 4 | 0 | 4 | 0 | 6 | 0.1667 | 17 | 0.0588 | 6 | 0.1667 | NA | NA | 6 | 0 | 2 | 0 | 14 | 0.0714 |
| 7_6839140 | 2 | 1 | 1 | 0 | NA | NA | 1 | 1 | 4 | 0.75 | 1 | 0 | NA | NA | 1 | 1 | 2 | 0.5 | 4 | 0.5 |
| 7_6839153 | 1 | 1 | 1 | 0 | NA | NA | NA | NA | 2 | 0.5 | 1 | 1 | NA | NA | 1 | 1 | 1 | 0 | 3 | 0.6667 |
| 7_7000361 | NA | NA | 2 | 0.5 | 1 | 0 | 1 | 0 | 4 | 0.25 | NA | NA | NA | NA | 2 | 1 | NA | NA | 2 | 1 |
| 7_7000422 | NA | NA | 1 | 1 | 2 | 1 | 1 | 1 | 4 | 1 | NA | NA | NA | NA | 1 | 1 | NA | NA | 1 | 1 |
| 7_7000426 | NA | NA | 2 | 1 | 2 | 1 | 2 | 1 | 6 | 1 | NA | NA | NA | NA | 2 | 0.5 | NA | NA | 2 | 0.5 |
| 7_7000435 | NA | NA | NA | NA | 2 | 1 | 1 | 1 | 3 | 1 | NA | NA | NA | NA | 2 | 0.5 | NA | NA | 2 | 0.5 |
| 7_7000761 | NA | NA | NA | NA | 1 | 1 | 2 | 0 | 3 | 0.3333 | NA | NA | NA | NA | 1 | 1 | NA | NA | 1 | 1 |
| 7_7000783 | NA | NA | NA | NA | 2 | 0.5 | 1 | 1 | 3 | 0.6667 | NA | NA | NA | NA | 3 | 0.3333 | NA | NA | 3 | 0.3333 |
| 7_7000784 | NA | NA | 1 | 0 | 2 | 1 | 1 | 1 | 4 | 0.75 | NA | NA | NA | NA | 3 | 0.6667 | NA | NA | 3 | 0.6667 |
| 7_7000971 | NA | NA | 1 | 0 | 2 | 0 | 2 | 0.5 | 5 | 0.2 | NA | NA | NA | NA | 2 | 0.5 | NA | NA | 2 | 0.5 |
| 7_7001068 | NA | NA | NA | NA | 2 | 1 | 2 | 1 | 4 | 1 | NA | NA | NA | NA | 1 | 1 | NA | NA | 1 | 1 |
| 7_7001069 | NA | NA | NA | NA | 1 | 0 | 2 | 0.5 | 3 | 0.3333 | NA | NA | NA | NA | 1 | 1 | NA | NA | 1 | 1 |
| 7_7232429 | 1 | 1 | NA | NA | NA | NA | 1 | 1 | 2 | 1 | NA | NA | 1 | 1 | 1 | 0 | NA | NA | 2 | 0.5 |
| 7_7232430 | 1 | 1 | 1 | 1 | NA | NA | NA | NA | 2 | 1 | NA | NA | 1 | 1 | 1 | 1 | NA | NA | 2 | 1 |
| 7_7232448 | NA | NA | NA | NA | NA | NA | 1 | 1 | 1 | 1 | 1 | 1 | 1 | 1 | NA | NA | NA | NA | 2 | 1 |
| 7_7296580 | NA | NA | 3 | 1 | NA | NA | 1 | 0 | 4 | 0.75 | NA | NA | 1 | 1 | NA | NA | NA | NA | 1 | 1 |
| 7_7296584 | NA | NA | 3 | 1 | NA | NA | 1 | 1 | 4 | 1 | NA | NA | 1 | 1 | NA | NA | NA | NA | 1 | 1 |
| 7_7305911 | 5 | 0.2 | 3 | 0 | NA | NA | 1 | 0 | 9 | 0.1111 | 1 | 0 | 5 | 0 | 8 | 0 | 1 | 1 | 15 | 0.0667 |
| 7_7305917 | 5 | 0.8 | 4 | 0.75 | NA | NA | 1 | 1 | 10 | 0.8 | 2 | 1 | 4 | 0.5 | 9 | 0.8889 | 1 | 1 | 16 | 0.8125 |
| 7_7305935 | 5 | 0.4 | 4 | 0.25 | NA | NA | 1 | 0 | 10 | 0.3 | 1 | 0 | 5 | 0 | 7 | 0.1429 | 1 | 0 | 14 | 0.0714 |
| 7_7305936 | 5 | 1 | 4 | 0.5 | NA | NA | 1 | 0 | 10 | 0.7 | 2 | 0.5 | 5 | 0.8 | 9 | 0.8889 | NA | NA | 16 | 0.8125 |
| 7_7306710 | 7 | 0 | 4 | 0.25 | NA | NA | 1 | 0 | 12 | 0.0833 | 3 | 0.3333 | 5 | 0 | 9 | 0.1111 | 1 | 0 | 18 | 0.1111 |
| 7_7306713 | 4 | 0.25 | 1 | 1 | NA | NA | NA | NA | 5 | 0.4 | 2 | 0 | 3 | 1 | 7 | 0.4286 | NA | NA | 12 | 0.5 |
| 7_7306717 | 5 | 0.2 | 4 | 0 | NA | NA | NA | NA | 9 | 0.1111 | 1 | 0 | 2 | 0 | 8 | 0.125 | 1 | 0 | 12 | 0.0833 |
| 7_7306733 | 8 | 0.75 | 4 | 0.25 | NA | NA | NA | NA | 12 | 0.5833 | 3 | 0.6667 | 4 | 0.5 | 10 | 0.6 | 1 | 1 | 18 | 0.6111 |
| 7_7306734 | 10 | 0 | 5 | 0.2 | NA | NA | NA | NA | 15 | 0.0667 | 3 | 0 | 5 | 0.4 | 9 | 0 | 1 | 0 | 18 | 0.1111 |
| 7_7306747 | 9 | 0.5556 | 5 | 0.4 | NA | NA | NA | NA | 14 | 0.5 | 3 | 1 | 5 | 0.2 | 8 | 0.375 | 1 | 0 | 17 | 0.4118 |
| 7_7306748 | 9 | 0.3333 | 4 | 0 | NA | NA | 1 | 0 | 14 | 0.2143 | 3 | 0.3333 | 6 | 0.1667 | 8 | 0.5 | NA | NA | 17 | 0.3529 |
| 7_7306757 | 10 | 0.7 | 4 | 1 | NA | NA | 1 | 0 | 15 | 0.7333 | 3 | 1 | 5 | 1 | 10 | 0.6 | 1 | 1 | 19 | 0.7895 |
| 7_7306763 | 11 | 0.8182 | 7 | 1 | NA | NA | 1 | 0 | 19 | 0.8421 | 2 | 0.5 | 6 | 1 | 9 | 0.5556 | 1 | 1 | 18 | 0.7222 |
| 7_7306766 | 6 | 0.8333 | 6 | 0.8333 | NA | NA | 1 | 1 | 13 | 0.8462 | 1 | 1 | 5 | 1 | 9 | 1 | 1 | 1 | 16 | 1 |
| 7_7306778 | 8 | 0.375 | 8 | 0.5 | 1 | 1 | 2 | 0 | 19 | 0.4211 | 3 | 1 | 6 | 0.3333 | 9 | 0.1111 | 1 | 1 | 19 | 0.3684 |
| 7_7306792 | 10 | 0 | 9 | 0.2222 | 1 | 0 | 1 | 0 | 21 | 0.0952 | 3 | 0 | 6 | 0 | 10 | 0.1 | 1 | 0 | 20 | 0.05 |
| 7_7306804 | 8 | 0.5 | 7 | 0.2857 | 1 | 1 | 1 | 0 | 17 | 0.4118 | 2 | 0.5 | 4 | 0.75 | 9 | 0.3333 | 1 | 0 | 16 | 0.4375 |
| 7_7306810 | 11 | 0.3636 | 10 | 0.1 | 1 | 0 | NA | NA | 22 | 0.2273 | 2 | 0 | 6 | 0 | 11 | 0.1818 | NA | NA | 19 | 0.1053 |
| 7_7306822 | 9 | 0.4444 | 10 | 0.5 | 1 | 1 | 3 | 0.3333 | 23 | 0.4783 | 3 | 0.6667 | 6 | 0.6667 | 12 | 0.4167 | 1 | 0 | 22 | 0.5 |
| 7_7306856 | 10 | 0.3 | 10 | 0.6 | 1 | 0 | 3 | 0 | 24 | 0.375 | 2 | 0.5 | 6 | 0.1667 | 12 | 0.25 | 2 | 0.5 | 22 | 0.2727 |
| 7_7306896 | 7 | 0 | 7 | 0.1429 | 1 | 0 | 1 | 1 | 16 | 0.125 | 4 | 0 | 2 | 0 | 10 | 0.1 | 2 | 0 | 18 | 0.0556 |
| 7_7306902 | 11 | 0.1818 | 10 | 0.2 | NA | NA | 4 | 0 | 25 | 0.16 | 4 | 0 | 6 | 0.1667 | 7 | 0.1429 | 2 | 0 | 19 | 0.1053 |
| 7_7306932 | 16 | 0.25 | 11 | 0.2727 | 2 | 0 | 2 | 0 | 31 | 0.2258 | 6 | 0.1667 | 5 | 0.8 | 9 | 0.2222 | 3 | 0.3333 | 23 | 0.3478 |
| 7_7307227 | 1 | 1 | 1 | 1 | 1 | 1 | NA | NA | 3 | 1 | NA | NA | NA | NA | NA | NA | 1 | 1 | 1 | 1 |
| 7_7307288 | 1 | 1 | 2 | 0 | 1 | 0 | NA | NA | 4 | 0.25 | NA | NA | NA | NA | 1 | 1 | NA | NA | 1 | 1 |
| 7_7307311 | 1 | 1 | 2 | 1 | 1 | 1 | NA | NA | 4 | 1 | NA | NA | NA | NA | 1 | 1 | 1 | 1 | 2 | 1 |
| 7_7307314 | 1 | 0 | 2 | 1 | 1 | 1 | NA | NA | 4 | 0.75 | NA | NA | NA | NA | 1 | 1 | 1 | 0 | 2 | 0.5 |
| 7_7307318 | 1 | 1 | 2 | 0.5 | 1 | 1 | NA | NA | 4 | 0.75 | NA | NA | NA | NA | 1 | 1 | 1 | 1 | 2 | 1 |
| 7_7307348 | 1 | 1 | 1 | 1 | 1 | 1 | NA | NA | 3 | 1 | NA | NA | NA | NA | 1 | 1 | 1 | 1 | 2 | 1 |
| 7_7307656 | 2 | 0.5 | 1 | 1 | 2 | 0.5 | 4 | 1 | 9 | 0.7778 | 6 | 0.6667 | 2 | 0.5 | 3 | 1 | NA | NA | 11 | 0.7273 |
| 7_7307673 | 2 | 0 | 2 | 0 | 2 | 0 | 4 | 0.25 | 10 | 0.1 | 4 | 0 | 2 | 0.5 | 3 | 0 | 2 | 0 | 11 | 0.0909 |

| | | | | | | | | | | | | | | | | | | | | |
|---|---|---|---|---|---|---|---|---|---|---|---|---|---|---|---|---|---|---|---|---|
| 7_7307703 | 2 | 0 | 1 | 0 | 2 | 0.5 | 4 | 0 | 9 | 0.1111 | 5 | 0.6 | NA | NA | 4 | 0.25 | 2 | 0 | 11 | 0.3636 |
| 7_7307730 | 2 | 0 | 1 | 0 | 1 | 0 | 4 | 0.25 | 8 | 0.125 | 6 | 0 | 1 | 1 | 4 | 0 | 2 | 0 | 13 | 0.0769 |
| 7_7307760 | 1 | 0 | 1 | 0 | 1 | 1 | 3 | 0.3333 | 6 | 0.3333 | 4 | 0.5 | 1 | 0 | 4 | 0 | 2 | 0 | 11 | 0.1818 |
| 7_7307793 | 1 | 0 | NA | NA | 1 | 1 | NA | NA | 2 | 0.5 | 3 | 0.3333 | 1 | 1 | NA | NA | 1 | 1 | 5 | 0.6 |
| 7_7319393 | 6 | 0.8333 | 6 | 0.8333 | 5 | 0.6 | 3 | 1 | 20 | 0.8 | 6 | 1 | 8 | 0.75 | 10 | 0.7 | 5 | 1 | 29 | 0.8276 |
| 7_7319395 | 6 | 0.8333 | 6 | 0.8333 | 6 | 1 | 3 | 1 | 21 | 0.9048 | 5 | 1 | 9 | 0.8889 | 7 | 1 | 4 | 1 | 25 | 0.96 |
| 7_7319410 | 6 | 0.3333 | 6 | 0.6667 | 6 | 0.6667 | 3 | 1 | 21 | 0.619 | 6 | 0.8333 | 7 | 0.5714 | 8 | 0.25 | 5 | 0.6 | 26 | 0.5385 |
| 7_7338391 | 1 | 1 | NA | NA | NA | NA | 1 | 1 | 2 | 1 | NA | NA | 1 | 0 | NA | NA | 1 | 1 | 2 | 0.5 |
| 7_7342476 | 1 | 1 | 1 | 1 | NA | NA | NA | NA | 2 | 1 | 2 | 1 | 1 | 1 | 1 | 0 | 2 | 1 | 6 | 0.8333 |
| 7_7342535 | 2 | 0 | 5 | 0.2 | 1 | 0 | NA | NA | 8 | 0.125 | 2 | 0 | 2 | 0 | 6 | 0.1667 | 3 | 0 | 13 | 0.0769 |
| 7_7342561 | 3 | 1 | 5 | 1 | 1 | 1 | NA | NA | 9 | 1 | 3 | 1 | 2 | 1 | 6 | 0.6667 | 3 | 0.6667 | 14 | 0.7857 |
| 7_7342563 | 3 | 0 | 4 | 0.25 | 1 | 0 | NA | NA | 8 | 0.125 | 3 | 0 | 2 | 0 | 6 | 0.1667 | 3 | 0.3333 | 14 | 0.1429 |
| 7_7342574 | 2 | 0 | 4 | 0.25 | 1 | 0 | NA | NA | 7 | 0.1429 | 4 | 0.75 | 1 | 1 | 5 | 0.4 | 3 | 0.3333 | 13 | 0.5385 |
| 7_7342575 | 3 | 0.6667 | 4 | 0.25 | 1 | 1 | NA | NA | 8 | 0.5 | 3 | 0.6667 | 2 | 1 | 6 | 0.5 | 3 | 0.6667 | 14 | 0.6429 |
| 7_7342579 | 3 | 0 | 3 | 1 | 1 | 1 | NA | NA | 7 | 0.5714 | 5 | 0.4 | 2 | 1 | 5 | 0 | 2 | 0 | 14 | 0.2857 |
| 7_7342579 | 3 | 0 | 4 | 0.25 | NA | NA | NA | NA | 7 | 0.1429 | 5 | 0.6 | 1 | 0 | 7 | 0.2857 | 3 | 0 | 16 | 0.3125 |
| 7_7342580 | 2 | 1 | 3 | 1 | 1 | 1 | NA | NA | 6 | 1 | 5 | 1 | 2 | 1 | 6 | 0.8333 | 2 | 0.5 | 15 | 0.8667 |
| 7_7342580 | 3 | 0 | 4 | 0 | 1 | 0 | 1 | 1 | 9 | 0.1111 | 5 | 0 | 2 | 0 | 7 | 0.2857 | 2 | 0 | 16 | 0.125 |
| 7_7342590 | 3 | 1 | 3 | 0.6667 | 1 | 1 | 1 | 1 | 8 | 0.875 | 2 | 1 | 1 | 1 | 5 | 1 | 3 | 1 | 11 | 1 |
| 7_7342600 | 3 | 1 | 5 | 0.6 | 1 | 1 | 1 | 1 | 10 | 0.8 | 6 | 1 | 2 | 0.5 | 7 | 0.8571 | 3 | 1 | 18 | 0.8889 |
| 7_7342612 | 2 | 0 | 4 | 0.25 | 1 | 0 | 1 | 0 | 8 | 0.125 | 4 | 0.25 | 1 | 0 | 6 | 0.1667 | 3 | 0 | 14 | 0.1429 |
| 7_7342622 | 3 | 0 | 5 | 0.2 | 1 | 0 | 1 | 0 | 10 | 0.1 | 6 | 0.3333 | 2 | 0 | 6 | 0.6667 | 3 | 0 | 17 | 0.3529 |
| 7_7342623 | 3 | 0.3333 | 4 | 0.25 | 1 | 1 | 1 | 0 | 9 | 0.3333 | 6 | 0.5 | 1 | 0 | 4 | 0.25 | 3 | 0.3333 | 14 | 0.3571 |
| 7_7342635 | 2 | 1 | 5 | 0.4 | 1 | 0 | 1 | 0 | 9 | 0.4444 | 3 | 0 | 2 | 0 | 4 | 0.75 | 1 | 1 | 10 | 0.4 |
| 7_7342636 | 3 | 0 | 4 | 0.5 | 1 | 1 | 1 | 1 | 9 | 0.4444 | 7 | 0.2857 | 2 | 0.5 | 7 | 0.1429 | 3 | 0.3333 | 19 | 0.2632 |
| 7_7342638 | 2 | 0.5 | 4 | 1 | NA | NA | 1 | 1 | 7 | 0.8571 | 6 | 1 | 2 | 0.5 | 5 | 1 | 2 | 1 | 15 | 0.9333 |
| 7_7342645 | 3 | 0 | 5 | 0.2 | 1 | 0 | 1 | 0 | 10 | 0.1 | 7 | 0.4286 | 1 | 0 | 7 | 0.2857 | 2 | 0 | 17 | 0.2941 |
| 7_7342647 | 3 | 0.6667 | 6 | 0.1667 | 1 | 0 | 1 | 0 | 11 | 0.2727 | 5 | 0.6 | 2 | 0 | 6 | 0.5 | 2 | 0 | 15 | 0.4 |
| 7_7342652 | 2 | 0.5 | 3 | 0 | 1 | 0 | 1 | 1 | 7 | 0.2857 | 7 | 0.1429 | 2 | 0 | 5 | 0 | NA | NA | 14 | 0.0714 |
| 7_7342656 | 3 | 0 | 6 | 0.3333 | 1 | 0 | 2 | 0 | 12 | 0.1667 | 9 | 0.2222 | 1 | 1 | 5 | 0 | 3 | 0.3333 | 18 | 0.2222 |
| 7_7342658 | 3 | 1 | 5 | 0.8 | 1 | 1 | 2 | 0.5 | 11 | 0.8182 | 9 | 0.8889 | 2 | 0.5 | 7 | 0.8571 | 3 | 1 | 21 | 0.8571 |
| 7_7342660 | 1 | 1 | 3 | 0.3333 | 1 | 0 | 2 | 0.5 | 7 | 0.4286 | 9 | 0.1111 | NA | NA | 7 | 0.2857 | 3 | 0.6667 | 19 | 0.2632 |
| 7_7392530 | 5 | 0.6 | 4 | 0.5 | 2 | 0.5 | NA | NA | 11 | 0.5455 | 2 | 0.5 | 3 | 0.3333 | 2 | 0 | 4 | 0.5 | 11 | 0.3636 |
| 7_7392550 | 3 | 0.6667 | 5 | 0.6 | 1 | 0 | NA | NA | 9 | 0.5556 | 2 | 0.5 | 2 | 0.5 | 2 | 0 | 3 | 0.3333 | 9 | 0.3333 |
| 7_7392557 | 2 | 1 | 5 | 0.6 | 1 | 1 | NA | NA | 8 | 0.75 | 1 | 0 | 2 | 0.5 | 1 | 1 | 4 | 1 | 8 | 0.75 |
| 7_7392577 | 4 | 0 | 3 | 0.3333 | 1 | 0 | NA | NA | 8 | 0.125 | 1 | 1 | 2 | 0 | 2 | 0 | 4 | 0 | 9 | 0.1111 |
| 7_7393334 | NA | NA | 1 | 1 | 1 | 1 | NA | NA | 2 | 1 | NA | NA | NA | NA | 1 | 0 | 2 | 1 | 3 | 0.6667 |
| 7_7393339 | 1 | 0 | 1 | 1 | 1 | 0 | NA | NA | 3 | 0.3333 | NA | NA | NA | NA | NA | NA | 2 | 1 | 2 | 1 |
| 7_7393366 | 1 | 1 | 1 | 1 | 1 | 1 | NA | NA | 3 | 1 | NA | NA | NA | NA | 2 | 1 | 1 | 1 | 3 | 1 |
| 7_7393369 | 1 | 0 | 1 | 0 | 1 | 1 | NA | NA | 3 | 0.3333 | NA | NA | 1 | 0 | 1 | 1 | NA | NA | 2 | 0.5 |
| 7_7393379 | 1 | 1 | 1 | 1 | 1 | 1 | NA | NA | 3 | 1 | NA | NA | 1 | 1 | 1 | 1 | 1 | 1 | 3 | 1 |
| 7_7393401 | NA | NA | 1 | 1 | 1 | 1 | NA | NA | 2 | 1 | NA | NA | 1 | 1 | 2 | 0.5 | 1 | 1 | 4 | 0.75 |
| 7_7393410 | NA | NA | NA | NA | 1 | 1 | NA | NA | 1 | 1 | NA | NA | 1 | 1 | 1 | 0 | 1 | 1 | 3 | 0.6667 |
| 7_7393410 | NA | NA | 1 | 0 | 1 | 1 | NA | NA | 2 | 0.5 | NA | NA | 1 | 1 | NA | NA | 1 | 1 | 2 | 1 |
| 7_7393412 | NA | NA | 1 | 1 | 1 | 1 | NA | NA | 2 | 1 | NA | NA | 1 | 1 | 1 | 1 | NA | NA | 2 | 1 |
| 7_7393421 | 3 | 1 | 2 | 0 | 1 | 1 | NA | NA | 6 | 0.6667 | 2 | 0.5 | 1 | 0 | 3 | 1 | 3 | 0.6667 | 9 | 0.6667 |
| 7_7415394 | 1 | 0 | 2 | 0.5 | 1 | 0 | NA | NA | 4 | 0.25 | 1 | 0 | 3 | 0 | 1 | 1 | 1 | 0 | 6 | 0.1667 |
| 7_7415405 | 1 | 1 | 1 | 1 | 1 | 0 | NA | NA | 3 | 0.6667 | 1 | 0 | 3 | 0 | 1 | 0 | 1 | 1 | 6 | 0.1667 |
| 7_7415406 | NA | NA | 2 | 0.5 | 1 | 0 | NA | NA | 3 | 0.3333 | NA | NA | NA | NA | 1 | 0 | 1 | 1 | 2 | 0.5 |
| 7_7415408 | 1 | 0 | 2 | 0.5 | 1 | 1 | NA | NA | 4 | 0.5 | 1 | 0 | 3 | 0.3333 | 1 | 0 | 1 | 1 | 6 | 0.3333 |
| 7_7476477 | 25 | 0.12 | 47 | 0.0638 | 35 | 0.0857 | 20 | 0.3 | 127 | 0.1181 | 23 | 0.1304 | 18 | 0.0556 | 13 | 0.0769 | 27 | 0.1481 | 81 | 0.1111 |
| 7_7476480 | 25 | 0.12 | 47 | 0.234 | 30 | 0.0333 | 21 | 0.1905 | 123 | 0.1545 | 22 | 0.1364 | 18 | 0.2222 | 13 | 0.1538 | 29 | 0.4138 | 82 | 0.2561 |
| 7_7476483 | 19 | 0.2632 | 42 | 0.0238 | 29 | 0.069 | 18 | 0.2222 | 108 | 0.1111 | 21 | 0.0952 | 15 | 0.0667 | 10 | 0 | 24 | 0.0833 | 70 | 0.0714 |
| 7_7476585 | 19 | 0.0526 | 30 | 0.0667 | 23 | 0.1304 | 15 | 0.1333 | 87 | 0.092 | 18 | 0.1111 | 16 | 0.1875 | 11 | 0.0909 | 19 | 0 | 64 | 0.0938 |
| 7_7476585 | 18 | 0.0556 | 29 | 0.0345 | 22 | 0 | 13 | 0.3077 | 82 | 0.0732 | 18 | 0.1111 | 17 | 0 | 10 | 0.3 | 22 | 0 | 67 | 0.0746 |
| 7_7476646 | 15 | 0.3333 | 26 | 0.1538 | 21 | 0.1429 | 9 | 0.2222 | 71 | 0.1972 | 15 | 0.1333 | 10 | 0.2 | 10 | 0.4 | 16 | 0.4375 | 51 | 0.2941 |
| 7_7476656 | 16 | 0.1875 | 28 | 0.25 | 20 | 0.35 | 10 | 0.1 | 74 | 0.2432 | 15 | 0.2 | 13 | 0.0769 | 10 | 0.1 | 20 | 0.3 | 58 | 0.1897 |
| 7_7476671 | 12 | 0.5 | 23 | 0.4348 | 17 | 0.4706 | 9 | 0.5556 | 61 | 0.4754 | 12 | 0.5 | 11 | 0.4545 | 8 | 0.125 | 17 | 0.7059 | 48 | 0.5 |
| 7_7476674 | 15 | 0.7333 | 27 | 0.5556 | 15 | 0.5333 | 6 | 0.5 | 63 | 0.5873 | 14 | 0.4286 | 13 | 0.4615 | 9 | 0.3333 | 21 | 0.2857 | 57 | 0.3684 |
| 7_7476676 | 17 | 0.8235 | 28 | 0.9286 | 14 | 0.8571 | 9 | 0.8889 | 68 | 0.8824 | 13 | 0.8462 | 12 | 1 | 10 | 0.8 | 19 | 0.8947 | 54 | 0.8889 |
| 7_7476683 | 11 | 0.2727 | 20 | 0 | 14 | 0 | 5 | 0.2 | 50 | 0.08 | 9 | 0.1111 | 10 | 0.1 | 4 | 0 | 17 | 0 | 40 | 0.05 |
| 7_7476699 | 13 | 0.1538 | 20 | 0.3 | 13 | 0.2308 | 9 | 0.4444 | 55 | 0.2727 | 12 | 0.5 | 13 | 0.3077 | 6 | 0 | 18 | 0.2222 | 49 | 0.2857 |
| 7_7476700 | 11 | 0.9091 | 19 | 0.8947 | 12 | 1 | 8 | 1 | 50 | 0.94 | 10 | 0.8 | 13 | 0.9231 | 7 | 0.7143 | 17 | 0.8824 | 47 | 0.8511 |
| 7_7476702 | 12 | 0.1667 | 20 | 0.1 | 14 | 0.1429 | 8 | 0.125 | 54 | 0.1296 | 9 | 0.1111 | 13 | 0 | 6 | 0.1667 | 16 | 0.0625 | 44 | 0.0682 |
| 7_7476708 | 15 | 0.2 | 22 | 0.3182 | 14 | 0.2857 | 9 | 0.5556 | 60 | 0.3167 | 11 | 0.2727 | 13 | 0.3077 | 9 | 0.2222 | 19 | 0.2632 | 52 | 0.2692 |

| | | | | | | | | | | | | | | | | | | | | |
|---|---|---|---|---|---|---|---|---|---|---|---|---|---|---|---|---|---|---|---|---|
| 7_7476720 | 14 | 0 | 23 | 0.0435 | 15 | 0.1333 | 8 | 0 | 60 | 0.05 | 11 | 0.0909 | 11 | 0.1818 | 9 | 0.1111 | 18 | 0.0556 | 49 | 0.102 |
| 7_7476722 | 14 | 0.9286 | 22 | 1 | 15 | 1 | 9 | 1 | 60 | 0.9833 | 11 | 1 | 10 | 1 | 9 | 1 | 19 | 0.8947 | 49 | 0.9592 |
| 7_7476724 | 10 | 0 | 17 | 0.1765 | 12 | 0.1667 | 8 | 0.125 | 47 | 0.1277 | 9 | 0.1111 | 8 | 0.125 | 6 | 0.1667 | 17 | 0.0588 | 40 | 0.1 |
| 7_7476730 | 12 | 0.75 | 18 | 0.5 | 13 | 0.3077 | 8 | 0.625 | 51 | 0.5294 | 9 | 0.3333 | 10 | 0.8 | 7 | 0.7143 | 17 | 0.5882 | 43 | 0.6047 |
| 7_7476732 | 14 | 0.3571 | 20 | 0.2 | 13 | 0.4615 | 8 | 0.125 | 55 | 0.2909 | 7 | 0.2857 | 10 | 0.2 | 7 | 0 | 18 | 0.4444 | 42 | 0.2857 |
| 7_7476736 | 13 | 0.7692 | 20 | 0.65 | 14 | 0.7143 | 7 | 0.8571 | 54 | 0.7222 | 10 | 0.9 | 10 | 0.9 | 9 | 0.5556 | 17 | 0.8824 | 46 | 0.8261 |
| 7_7476747 | 13 | 0 | 20 | 0.2 | 13 | 0.1538 | 8 | 0.125 | 54 | 0.1296 | 11 | 0.0909 | 9 | 0.2222 | 7 | 0 | 17 | 0.1765 | 44 | 0.1364 |
| 7_7476749 | 14 | 0.8571 | 19 | 1 | 13 | 0.9231 | 8 | 1 | 54 | 0.9444 | 10 | 1 | 10 | 1 | 7 | 0.5714 | 18 | 0.9444 | 45 | 0.9111 |
| 7_7477388 | 12 | 0.25 | 18 | 0.1111 | 13 | 0.1538 | 8 | 0 | 51 | 0.1373 | 10 | 0.3 | 9 | 0.3333 | 7 | 0 | 15 | 0 | 41 | 0.1463 |
| 7_7485587 | 7 | 0 | 10 | 0.2 | 6 | 0 | 5 | 0 | 28 | 0.0714 | 5 | 0 | 1 | 0 | 4 | 0.5 | 13 | 0.1538 | 23 | 0.1739 |
| 7_7485596 | 7 | 1 | 9 | 0.3333 | 6 | 0.5 | 5 | 0.8 | 27 | 0.6296 | 5 | 0.6 | 2 | 1 | 6 | 0.6667 | 11 | 0.6364 | 24 | 0.6667 |
| 7_7485605 | 6 | 0.1667 | 10 | 0.3 | 5 | 0.4 | 5 | 0.4 | 26 | 0.3077 | 6 | 0.1667 | 2 | 0.5 | 4 | 0 | 13 | 0.3846 | 25 | 0.28 |
| 7_7485626 | 6 | 0.6667 | 7 | 1 | 4 | 0.75 | 5 | 1 | 22 | 0.8636 | 6 | 1 | 2 | 1 | 3 | 0.3333 | 10 | 0.6 | 21 | 0.7143 |
| 7_7485630 | 6 | 0 | 8 | 0.125 | 4 | 0.5 | 5 | 0 | 23 | 0.1304 | 6 | 0.1667 | 1 | 0 | 3 | 0.3333 | 13 | 0 | 23 | 0.087 |
| 7_7485635 | 7 | 0.4286 | 4 | 0.25 | 2 | 0.5 | 3 | 0.3333 | 16 | 0.375 | 3 | 0.3333 | NA | NA | 2 | 0.5 | 9 | 0.5556 | 14 | 0.5 |
| 7_7485653 | 6 | 0 | 8 | 0.25 | 3 | 0 | 5 | 0.2 | 22 | 0.1364 | 5 | 0 | 1 | 0 | 3 | 0 | 13 | 0.1538 | 22 | 0.0909 |
| 7_7485658 | 6 | 0.6667 | 8 | 0.125 | 2 | 0.5 | 5 | 0 | 21 | 0.2857 | 4 | 0.25 | NA | NA | 3 | 0.3333 | 10 | 0.1 | 17 | 0.1765 |
| 7_7539903 | 1 | 0 | 1 | 1 | 2 | 1 | 1 | 1 | 5 | 0.8 | NA | NA | 1 | 0 | NA | NA | 1 | 1 | 2 | 0.5 |
| 7_7539904 | 1 | 1 | NA | NA | 2 | 0.5 | 1 | 1 | 4 | 0.75 | NA | NA | 1 | 1 | NA | NA | 1 | 0 | 2 | 0.5 |
| 7_7540325 | 1 | 0 | 2 | 0.5 | 4 | 0.25 | 2 | 0.5 | 9 | 0.3333 | NA | NA | 2 | 0 | 2 | 1 | 2 | 0.5 | 6 | 0.5 |
| 7_7551625 | 1 | 1 | NA | NA | NA | NA | NA | NA | 1 | 1 | NA | NA | NA | NA | 1 | 1 | NA | NA | 1 | 1 |
| 7_7551636 | 1 | 1 | NA | NA | NA | NA | NA | NA | 1 | 1 | NA | NA | NA | NA | 1 | 1 | 1 | 1 | 2 | 1 |
| 7_7551651 | 1 | 1 | NA | NA | NA | NA | NA | NA | 1 | 1 | NA | NA | NA | NA | 1 | 1 | 1 | 0 | 2 | 0.5 |
| 7_7551666 | 1 | 1 | NA | NA | NA | NA | NA | NA | 1 | 1 | NA | NA | NA | NA | 1 | 0 | 1 | 1 | 2 | 0.5 |
| 7_7551676 | 1 | 1 | NA | NA | NA | NA | NA | NA | 1 | 1 | 1 | 1 | NA | NA | 1 | 1 | 1 | 1 | 3 | 1 |
| 7_7551679 | 1 | 1 | NA | NA | NA | NA | NA | NA | 1 | 1 | 1 | 1 | NA | NA | 1 | 1 | NA | NA | 2 | 1 |
| 7_7552273 | 1 | 1 | 1 | 1 | NA | NA | NA | NA | 2 | 1 | 1 | 1 | NA | NA | NA | NA | 1 | 1 | 2 | 1 |
| 7_7552278 | NA | NA | 1 | 1 | NA | NA | NA | NA | 1 | 1 | 1 | 1 | NA | NA | 1 | 0 | 1 | 0 | 3 | 0.3333 |
| 7_7552298 | NA | NA | 1 | 1 | NA | NA | NA | NA | 1 | 1 | 1 | 1 | NA | NA | NA | NA | 1 | 1 | 2 | 1 |
| 7_7552314 | 1 | 1 | 1 | 0 | 1 | 1 | NA | NA | 3 | 0.6667 | 1 | 1 | NA | NA | 1 | 1 | 1 | 1 | 3 | 1 |
| 7_7552321 | 1 | 1 | 1 | 1 | 1 | 1 | NA | NA | 3 | 1 | 1 | 0 | NA | NA | 1 | 1 | 1 | 1 | 3 | 0.6667 |
| 7_7552326 | 1 | 0 | 1 | 0 | 1 | 1 | NA | NA | 3 | 0.3333 | 1 | 0 | NA | NA | 1 | 1 | 1 | 1 | 3 | 0.6667 |
| 7_7552343 | 1 | 0 | NA | NA | 1 | 1 | NA | NA | 2 | 0.5 | 1 | 0 | NA | NA | 1 | 1 | 1 | 0 | 3 | 0.3333 |
| 7_7552366 | 1 | 0 | 1 | 0 | 1 | 1 | NA | NA | 3 | 0.3333 | 1 | 0 | NA | NA | 1 | 0 | 1 | 1 | 3 | 0.3333 |
| 7_7552372 | 1 | 1 | 1 | 1 | 1 | 1 | NA | NA | 3 | 1 | 1 | 1 | NA | NA | 1 | 1 | 1 | 1 | 3 | 1 |
| 7_7552376 | 1 | 1 | 1 | 1 | 1 | 1 | NA | NA | 3 | 1 | 1 | 1 | NA | NA | 2 | 1 | 1 | 1 | 4 | 1 |
| 7_7552380 | NA | NA | 1 | 0 | 1 | 1 | NA | NA | 2 | 0.5 | 1 | 0 | NA | NA | 1 | 1 | 1 | 1 | 3 | 0.6667 |
| 7_7552387 | NA | NA | NA | NA | 1 | 1 | NA | NA | 1 | 1 | 1 | 1 | NA | NA | 2 | 0.5 | 1 | 1 | 4 | 0.75 |
| 7_7782670 | 3 | 0 | 4 | 0.5 | 3 | 0.3333 | 9 | 0.4444 | 19 | 0.3684 | 2 | 0.5 | 5 | 0.2 | 6 | 0.6667 | 7 | 0.4286 | 20 | 0.45 |
| 7_7782689 | 2 | 0 | 4 | 0.25 | 2 | 0 | 9 | 0.3333 | 17 | 0.2353 | 1 | 1 | 5 | 0 | 5 | 0.6 | 5 | 0 | 16 | 0.25 |
| 7_7782726 | 2 | 0.5 | 4 | 0.75 | 1 | 0 | 7 | 0.8571 | 14 | 0.7143 | 1 | 0 | 5 | 0.6 | 5 | 1 | 5 | 0.8 | 16 | 0.75 |
| 7_7782736 | NA | NA | 4 | 0.5 | 1 | 1 | 4 | 0.5 | 9 | 0.5556 | 1 | 1 | NA | NA | 3 | 0 | 1 | 1 | 5 | 0.4 |
| 7_7783550 | 1 | 1 | 1 | 0 | NA | NA | 2 | 0 | 4 | 0.25 | NA | NA | 1 | 1 | NA | NA | NA | NA | 1 | 1 |
| 7_7783887 | 2 | 0.5 | NA | NA | NA | NA | NA | NA | 2 | 0.5 | NA | NA | 1 | 0 | NA | NA | 1 | 1 | 2 | 0.5 |
| 7_7784029 | 2 | 1 | NA | NA | NA | NA | NA | NA | 2 | 1 | NA | NA | NA | NA | NA | NA | 1 | 1 | 1 | 1 |
| 7_7784042 | 2 | 0.5 | NA | NA | NA | NA | NA | NA | 2 | 0.5 | NA | NA | NA | NA | NA | NA | 1 | 1 | 1 | 1 |
| 7_7784047 | 2 | 0.5 | NA | NA | NA | NA | NA | NA | 2 | 0.5 | NA | NA | NA | NA | NA | NA | 1 | 1 | 1 | 1 |
| 7_7784067 | 2 | 0.5 | NA | NA | NA | NA | NA | NA | 2 | 0.5 | NA | NA | NA | NA | NA | NA | 1 | 1 | 1 | 1 |
| 7_7784071 | 2 | 1 | NA | NA | NA | NA | NA | NA | 2 | 1 | NA | NA | NA | NA | NA | NA | 1 | 1 | 1 | 1 |
| 7_7784133 | 2 | 1 | NA | NA | NA | NA | NA | NA | 2 | 1 | NA | NA | NA | NA | NA | NA | 1 | 1 | 1 | 1 |
| 7_7795140 | 5 | 1 | 2 | 0.5 | 5 | 0.8 | 4 | 0.75 | 16 | 0.8125 | NA | NA | 2 | 0.5 | 3 | 1 | 1 | 0 | 6 | 0.6667 |
| 7_7795144 | 5 | 0.4 | 4 | 0.5 | 4 | 0 | 4 | 0 | 17 | 0.2353 | NA | NA | 4 | 0 | 3 | 0.3333 | NA | NA | 7 | 0.1429 |
| 7_7795145 | 4 | 0.5 | 3 | 0.6667 | 5 | 0.8 | 3 | 0.6667 | 15 | 0.6667 | NA | NA | 3 | 0.6667 | 3 | 0.6667 | 1 | 1 | 7 | 0.7143 |
| 7_7795152 | 5 | 0.6 | 4 | 0.5 | 5 | 0.2 | 4 | 0.5 | 18 | 0.4444 | NA | NA | 4 | 0.5 | 4 | 0.25 | 1 | 0 | 9 | 0.3333 |
| 7_7795153 | 5 | 1 | 4 | 1 | 4 | 1 | 4 | 1 | 17 | 1 | NA | NA | 4 | 1 | 3 | 1 | 1 | 1 | 8 | 1 |
| 7_7795154 | 5 | 0.4 | 4 | 0.5 | 5 | 0.6 | 3 | 0.3333 | 17 | 0.4706 | NA | NA | 2 | 0.5 | 2 | 0.5 | 1 | 0 | 5 | 0.4 |
| 7_7795171 | 5 | 1 | 4 | 0.75 | 5 | 0.8 | 4 | 1 | 18 | 0.8889 | NA | NA | 3 | 1 | 4 | 0.5 | 1 | 1 | 8 | 0.75 |
| 7_7795179 | 3 | 0 | 3 | 0 | 4 | 0.25 | 3 | 0 | 13 | 0.0769 | NA | NA | 4 | 0 | 3 | 0.3333 | 1 | 1 | 8 | 0.25 |
| 7_7795193 | 4 | 0 | 3 | 0 | 4 | 0.25 | 3 | 0 | 14 | 0.0714 | NA | NA | 3 | 0.3333 | 4 | 0 | 1 | 0 | 8 | 0.125 |
| 7_7795224 | 2 | 1 | 3 | 0.6667 | 4 | 1 | 1 | 1 | 10 | 0.9 | NA | NA | 1 | 1 | 2 | 0.5 | 1 | 1 | 4 | 0.75 |
| 7_7795235 | 1 | 1 | 3 | 1 | 3 | 0.6667 | 1 | 0 | 8 | 0.75 | NA | NA | 1 | 1 | 2 | 0.5 | 1 | 1 | 4 | 0.75 |
| 7_7795245 | 2 | 0.5 | 4 | 0.5 | 4 | 0.5 | 3 | 0.6667 | 13 | 0.5385 | NA | NA | 1 | 0 | 3 | 0 | 1 | 1 | 5 | 0.2 |
| 7_7822741 | 1 | 0 | 3 | 0.3333 | 1 | 0 | NA | NA | 5 | 0.2 | NA | NA | 1 | 1 | 2 | 0 | NA | NA | 3 | 0.3333 |
| 7_7822746 | 1 | 1 | 2 | 0.5 | 1 | 1 | 1 | 1 | 5 | 0.8 | NA | NA | 1 | 0 | 2 | 1 | NA | NA | 3 | 0.6667 |
| 7_7822863 | 1 | 1 | NA | NA | NA | NA | 1 | 1 | 2 | 1 | NA | NA | NA | NA | NA | NA | 1 | 1 | 1 | 1 |

| | | | | | | | | | | | | | | | | | | | | |
|---|---|---|---|---|---|---|---|---|---|---|---|---|---|---|---|---|---|---|---|---|
| 7_7822877 | 1 | 1 | NA | NA | NA | NA | 2 | 1 | 3 | 1 | NA | NA | NA | NA | NA | NA | 1 | 1 | 1 | 1 |
| 7_7822907 | 1 | 1 | NA | NA | NA | NA | 1 | 1 | 2 | 1 | NA | NA | NA | NA | NA | NA | 1 | 1 | 1 | 1 |
| 7_7842502 | 4 | 1 | 2 | 1 | 1 | 0 | NA | NA | 7 | 0.8571 | 1 | 0 | 1 | 0 | 1 | 0 | 1 | 1 | 4 | 0.25 |
| 7_7842514 | 9 | 1 | 4 | 1 | 4 | 0.75 | 5 | 1 | 22 | 0.9545 | 5 | 1 | 4 | 1 | 4 | 0.5 | 4 | 0.75 | 17 | 0.8235 |
| 7_7842515 | 11 | 0.0909 | 5 | 0.2 | 4 | 0 | 7 | 0 | 27 | 0.0741 | 5 | 0.2 | 6 | 0.1667 | 4 | 0 | 6 | 0.3333 | 21 | 0.1905 |
| 7_7842516 | 12 | 0.5833 | 4 | 0 | 4 | 0.25 | 6 | 0.3333 | 26 | 0.3846 | 4 | 0.75 | 5 | 0.2 | 3 | 0 | 6 | 0.3333 | 18 | 0.3333 |
| 7_7842517 | 10 | 0.4 | 4 | 0.25 | 2 | 0.5 | 6 | 0.1667 | 22 | 0.3182 | 4 | 0 | 5 | 0.2 | 4 | 0.5 | 3 | 0.6667 | 16 | 0.3125 |
| 7_7842522 | 12 | 1 | 4 | 1 | 4 | 1 | 7 | 1 | 27 | 1 | 4 | 1 | 6 | 1 | 3 | 1 | 5 | 0.8 | 18 | 0.9444 |
| 7_7842523 | 11 | 1 | 4 | 0.75 | 4 | 0.75 | 7 | 1 | 26 | 0.9231 | 4 | 1 | 4 | 0.75 | 3 | 0.6667 | 6 | 1 | 17 | 0.8824 |
| 7_7842533 | 11 | 0.9091 | 5 | 0.8 | 4 | 0.75 | 6 | 0.5 | 26 | 0.7692 | 4 | 0.25 | 6 | 1 | 4 | 0.25 | 6 | 0.5 | 20 | 0.55 |
| 7_7844240 | 6 | 0.1667 | 7 | 0.1429 | 5 | 1 | 5 | 0.6 | 23 | 0.4348 | 6 | 0.1667 | 4 | 0.75 | 5 | 0.2 | 9 | 0.6667 | 24 | 0.4583 |
| 7_7844266 | 4 | 1 | 3 | 0.6667 | 4 | 1 | 4 | 1 | 15 | 0.9333 | 2 | 0.5 | 2 | 0.5 | 5 | 0.6 | 7 | 0.7143 | 16 | 0.625 |
| 7_7847751 | NA | NA | 1 | 1 | NA | NA | NA | NA | 1 | 1 | 1 | 1 | NA | NA | 1 | 0 | 1 | 1 | 3 | 0.6667 |
| 7_7847764 | NA | NA | 1 | 1 | NA | NA | NA | NA | 1 | 1 | 1 | 0 | NA | NA | 1 | 1 | 1 | 1 | 3 | 0.6667 |
| 7_7847772 | NA | NA | 1 | 1 | NA | NA | NA | NA | 1 | 1 | 1 | 0 | NA | NA | 1 | 0 | 1 | 1 | 3 | 0.3333 |
| 7_7849313 | NA | NA | 2 | 1 | NA | NA | NA | NA | 2 | 1 | 1 | 1 | NA | NA | 1 | 1 | 1 | 1 | 3 | 1 |
| 7_7849343 | NA | NA | 2 | 1 | NA | NA | NA | NA | 2 | 1 | 1 | 1 | NA | NA | 1 | 1 | NA | NA | 2 | 1 |
| 7_7855237 | NA | NA | 1 | 1 | NA | NA | NA | NA | 1 | 1 | 1 | 1 | NA | NA | NA | NA | 2 | 1 | 3 | 1 |
| 7_7855250 | NA | NA | 1 | 1 | NA | NA | NA | NA | 1 | 1 | 1 | 1 | NA | NA | NA | NA | NA | NA | 1 | 1 |
| 7_7855711 | NA | NA | 3 | 0.3333 | NA | NA | NA | NA | 3 | 0.3333 | 3 | 0.6667 | 1 | 0 | 1 | 0 | 1 | 0 | 6 | 0.3333 |
| 7_7855722 | NA | NA | 3 | 0 | 1 | 1 | NA | NA | 4 | 0.25 | 3 | 0.3333 | 2 | 0 | NA | NA | 3 | 0.3333 | 8 | 0.25 |
| 7_7855737 | 1 | 1 | 3 | 0 | 2 | 0 | NA | NA | 6 | 0.1667 | 3 | 0 | 3 | 0 | 1 | 1 | 3 | 0 | 10 | 0.1 |
| 7_7855804 | 1 | 0 | 2 | 0 | 2 | 1 | 1 | 0 | 6 | 0.3333 | 3 | 0 | 3 | 0 | 2 | 0 | 3 | 0.6667 | 11 | 0.1818 |
| 7_7874826 | 1 | 1 | NA | NA | NA | NA | NA | NA | 1 | 1 | NA | NA | NA | NA | 1 | 1 | 1 | 0 | 2 | 0.5 |
| 7_7874844 | 1 | 1 | NA | NA | NA | NA | NA | NA | 1 | 1 | NA | NA | NA | NA | 1 | 1 | 1 | 1 | 2 | 1 |
| 7_7874890 | 1 | 1 | NA | NA | NA | NA | NA | NA | 1 | 1 | NA | NA | NA | NA | 1 | 1 | 1 | 1 | 2 | 1 |
| 7_7874904 | 1 | 1 | NA | NA | NA | NA | NA | NA | 1 | 1 | NA | NA | NA | NA | 1 | 1 | 1 | 1 | 2 | 1 |
| 7_7874918 | 1 | 1 | NA | NA | NA | NA | NA | NA | 1 | 1 | NA | NA | NA | NA | 1 | 1 | 1 | 1 | 2 | 1 |
| 7_7874929 | 1 | 1 | NA | NA | NA | NA | NA | NA | 1 | 1 | NA | NA | NA | NA | 1 | 0 | 1 | 1 | 2 | 0.5 |
| 7_7874992 | 2 | 1 | NA | NA | NA | NA | NA | NA | 2 | 1 | NA | NA | NA | NA | NA | NA | 1 | 1 | 1 | 1 |
| 7_7874996 | 1 | 1 | NA | NA | NA | NA | NA | NA | 1 | 1 | NA | NA | NA | NA | 1 | 1 | 1 | 1 | 2 | 1 |
| 7_7875001 | 2 | 0.5 | NA | NA | NA | NA | NA | NA | 2 | 0.5 | NA | NA | NA | NA | 1 | 1 | 1 | 1 | 2 | 1 |
| 7_7875028 | 2 | 0.5 | NA | NA | NA | NA | NA | NA | 2 | 0.5 | NA | NA | NA | NA | 1 | 0 | 1 | 1 | 2 | 0.5 |
| 7_7875037 | 2 | 1 | NA | NA | NA | NA | NA | NA | 2 | 1 | NA | NA | NA | NA | NA | NA | 1 | 1 | 1 | 1 |
| 7_7875050 | 2 | 0.5 | NA | NA | NA | NA | NA | NA | 2 | 0.5 | NA | NA | NA | NA | 1 | 0 | 1 | 1 | 2 | 0.5 |
| 7_7875073 | 2 | 1 | NA | NA | NA | NA | NA | NA | 2 | 1 | NA | NA | NA | NA | 1 | 0 | 1 | 1 | 2 | 0.5 |
| 7_7875084 | 2 | 1 | NA | NA | NA | NA | NA | NA | 2 | 1 | NA | NA | NA | NA | 1 | 0 | 1 | 1 | 2 | 0.5 |
| 7_7876428 | 2 | 0.5 | NA | NA | NA | NA | NA | NA | 2 | 0.5 | NA | NA | NA | NA | 1 | 0 | 1 | 1 | 2 | 0.5 |
| 7_7878363 | 6 | 0 | 21 | 0.1905 | 28 | 0.1786 | 9 | 0.1111 | 64 | 0.1562 | 23 | 0.1304 | 27 | 0 | 22 | 0.0909 | 12 | 0.1667 | 84 | 0.0833 |
| 7_7878381 | 5 | 0.2 | 21 | 0.2381 | 25 | 0.44 | 8 | 0.25 | 59 | 0.322 | 22 | 0.3636 | 23 | 0.2174 | 25 | 0.36 | 14 | 0.1429 | 84 | 0.2857 |
| 7_7878391 | 4 | 0.25 | 17 | 0.1176 | 22 | 0.1364 | 6 | 0.1667 | 49 | 0.1429 | 17 | 0.1176 | 20 | 0.1 | 18 | 0.0556 | 6 | 0.1667 | 61 | 0.0984 |
| 7_7900991 | 4 | 0.75 | 1 | 1 | 2 | 0.5 | 3 | 0.3333 | 10 | 0.6 | 3 | 1 | 3 | 0.6667 | 2 | 1 | 2 | 1 | 10 | 0.9 |
| 7_7901021 | 4 | 0.75 | 2 | 0 | 2 | 0.5 | 5 | 0.6 | 13 | 0.5385 | 4 | 0.25 | 4 | 0.5 | 1 | 0 | 2 | 1 | 11 | 0.4545 |
| 7_7901083 | 2 | 0 | 3 | 0 | 3 | 0 | 6 | 0.1667 | 14 | 0.0714 | 2 | 0.5 | 5 | 0 | 1 | 0 | 1 | 0 | 9 | 0.1111 |
| 7_7901085 | 3 | 0.3333 | 2 | 0 | 3 | 0 | 6 | 0.1667 | 14 | 0.1429 | 4 | 0.25 | 4 | 0 | 1 | 0 | 3 | 0 | 12 | 0.0833 |
| 7_7901123 | 3 | 0.6667 | 3 | 0.3333 | 2 | 1 | 6 | 0.6667 | 14 | 0.6429 | 3 | 0.6667 | 5 | 0.2 | NA | NA | 4 | 0.5 | 12 | 0.4167 |
| 7_7901129 | 3 | 0 | 4 | 0 | 3 | 0 | 7 | 0.1429 | 17 | 0.0588 | 5 | 0 | 5 | 0 | 1 | 0 | 5 | 0.2 | 16 | 0.0625 |
| 7_7901139 | 4 | 0 | 5 | 0 | 2 | 0.5 | 4 | 0 | 15 | 0.0667 | 4 | 0 | 2 | 0.5 | 1 | 0 | 1 | 0 | 8 | 0.125 |
| 7_7901150 | 5 | 0.4 | 5 | 0.6 | 2 | 0.5 | 7 | 0.4286 | 19 | 0.4737 | 4 | 1 | 5 | 0.6 | 1 | 1 | 5 | 1 | 15 | 0.8667 |
| 7_7901249 | 4 | 0 | 6 | 0.1667 | 4 | 0.25 | 8 | 0.125 | 22 | 0.1364 | 6 | 0.3333 | 8 | 0 | 3 | 0 | 7 | 0 | 24 | 0.0833 |
| 7_7901503 | 16 | 0.5 | 16 | 0.5625 | 13 | 0.3846 | 18 | 0.3889 | 63 | 0.4603 | 12 | 0.3333 | 19 | 0.4737 | 6 | 0.3333 | 16 | 0.5 | 53 | 0.434 |
| 7_7902506 | NA | NA | 4 | 0.25 | 7 | 0 | 2 | 0 | 13 | 0.0769 | 9 | 0.1111 | 5 | 0 | 2 | 0 | 4 | 0 | 20 | 0.05 |
| 7_7902539 | NA | NA | 3 | 0 | 7 | 0.2857 | 3 | 0 | 13 | 0.1538 | 7 | 0 | 5 | 0 | 2 | 0 | 4 | 0.25 | 18 | 0.0556 |
| 7_7909644 | NA | NA | NA | NA | 4 | 0.25 | NA | NA | 4 | 0.25 | NA | NA | NA | NA | 3 | 0.3333 | 1 | 0 | 4 | 0.25 |
| 7_7909865 | 1 | 1 | NA | NA | 2 | 0.5 | NA | NA | 3 | 0.6667 | NA | NA | NA | NA | 1 | 0 | 1 | 1 | 2 | 0.5 |
| 7_7909870 | 2 | 0 | 3 | 0.3333 | 3 | 0 | NA | NA | 8 | 0.125 | 2 | 0 | 1 | 0 | 3 | 0.3333 | 1 | 1 | 7 | 0.2857 |
| 7_7909878 | 2 | 0 | 3 | 0 | 3 | 0.6667 | NA | NA | 8 | 0.25 | 2 | 0 | 1 | 1 | 3 | 0 | 1 | 0 | 7 | 0.1429 |
| 7_7909888 | 2 | 0 | 3 | 0.3333 | 1 | 0 | NA | NA | 6 | 0.1667 | 1 | 0 | 1 | 0 | 3 | 0.3333 | 1 | 0 | 6 | 0.1667 |
| 7_7909913 | 1 | 0 | 4 | 0.5 | 2 | 0 | 2 | 0 | 9 | 0.2222 | 4 | 0 | 3 | 0 | 7 | 0.2857 | 4 | 0 | 18 | 0.1111 |
| 7_7909926 | 2 | 0 | 6 | 0.5 | 3 | 0 | 2 | 0 | 13 | 0.2308 | 5 | 0 | 2 | 0 | 8 | 0.125 | 2 | 0.5 | 17 | 0.1176 |
| 7_7909927 | 1 | 0 | 5 | 0.6 | 3 | 0.3333 | 1 | 1 | 10 | 0.5 | 3 | 0.3333 | 2 | 0 | 6 | 0.3333 | 2 | 0 | 13 | 0.2308 |
| 7_7909931 | 2 | 0 | 4 | 0.25 | 2 | 0 | 1 | 0 | 9 | 0.1111 | 3 | 0 | 3 | 0.3333 | 9 | 0.3333 | 3 | 0 | 18 | 0.2222 |
| 7_7909945 | 3 | 0 | 5 | 0.4 | 4 | 0 | 5 | 0 | 17 | 0.1176 | 3 | 0 | 5 | 0.4 | 8 | 0.125 | 6 | 0 | 22 | 0.1364 |
| 7_7909953 | 2 | 1 | 5 | 0.4 | 3 | 0.3333 | 4 | 0.25 | 14 | 0.4286 | 3 | 0.3333 | 5 | 0 | 9 | 0.3333 | 6 | 0.1667 | 23 | 0.2174 |
| 7_7909957 | NA | NA | NA | NA | 1 | 1 | NA | NA | 1 | 1 | NA | NA | NA | NA | NA | NA | 1 | 1 | 1 | 1 |

| | | | | | | | | | | | | | | | | | | | | |
|---|---|---|---|---|---|---|---|---|---|---|---|---|---|---|---|---|---|---|---|---|
| 7_7909971 | 1 | 0 | 7 | 0.1429 | 4 | 0.25 | 4 | 0 | 16 | 0.125 | 4 | 0.25 | 4 | 0 | 10 | 0.2 | 5 | 0.2 | 23 | 0.1739 |
| 7_7909975 | 1 | 1 | 5 | 0.2 | 3 | 0 | 3 | 0.3333 | 12 | 0.25 | 1 | 0 | 5 | 0 | 6 | 0 | 5 | 0.2 | 17 | 0.0588 |
| 7_7910028 | NA | NA | 6 | 0.1667 | 2 | 0 | 3 | 0.6667 | 11 | 0.2727 | 4 | 0.25 | 4 | 0 | 4 | 0.25 | 4 | 0 | 16 | 0.125 |
| 7_7911448 | 1 | 1 | 2 | 0.5 | NA | NA | NA | NA | 3 | 0.6667 | NA | NA | NA | NA | 1 | 1 | 2 | 0.5 | 3 | 0.6667 |
| 7_7918087 | 1 | 1 | 2 | 0 | 2 | 0.5 | 2 | 0 | 7 | 0.2857 | NA | NA | 1 | 0 | 1 | 0 | 2 | 0.5 | 4 | 0.25 |
| 7_7918091 | NA | NA | 2 | 0 | 2 | 0.5 | 2 | 0 | 6 | 0.1667 | NA | NA | 2 | 0.5 | 1 | 1 | 2 | 1 | 5 | 0.8 |
| 7_7918104 | 5 | 0 | 4 | 0 | 5 | 0.2 | 5 | 0 | 19 | 0.0526 | NA | NA | 7 | 0 | 2 | 0 | 4 | 0.25 | 13 | 0.0769 |
| 7_7918112 | 7 | 0.8571 | 3 | 0.3333 | 5 | 1 | 5 | 1 | 20 | 0.85 | NA | NA | 6 | 1 | 2 | 1 | 3 | 1 | 11 | 1 |
| 7_7918126 | 7 | 0.4286 | 4 | 0.75 | 3 | 1 | 5 | 0.6 | 19 | 0.6316 | NA | NA | 5 | 0.6 | 2 | 1 | 4 | 0.5 | 11 | 0.6364 |
| 7_7918129 | 8 | 0.875 | 4 | 1 | 5 | 1 | 5 | 1 | 22 | 0.9545 | NA | NA | 7 | 0.8571 | 2 | 1 | 4 | 0.75 | 13 | 0.8462 |
| 7_7918132 | 7 | 0.2857 | 4 | 0 | 4 | 0.25 | 5 | 0 | 20 | 0.15 | NA | NA | 8 | 0.375 | 1 | 1 | 3 | 0.6667 | 12 | 0.5 |
| 7_7918136 | 6 | 0.6667 | 4 | 0.75 | 4 | 0.5 | 5 | 0.4 | 19 | 0.5789 | NA | NA | 7 | 0.7143 | 2 | 0.5 | 4 | 0.75 | 13 | 0.6923 |
| 7_7918176 | 10 | 0.3 | 5 | 0.2 | 5 | 0.4 | 5 | 0.6 | 25 | 0.36 | 1 | 1 | 8 | 0.125 | 2 | 0 | 5 | 0.4 | 16 | 0.25 |
| 7_7918183 | 9 | 0.2222 | 6 | 0 | 5 | 0.2 | 5 | 0.2 | 25 | 0.16 | 1 | 0 | 8 | 0.125 | 2 | 0 | 3 | 0.3333 | 14 | 0.1429 |
| 7_7918186 | 8 | 0.125 | 7 | 0 | 5 | 0 | 5 | 0.4 | 25 | 0.12 | 1 | 0 | 9 | 0 | 2 | 0 | 6 | 0.1667 | 18 | 0.0556 |
| 7_7918202 | 10 | 0.2 | 6 | 0.1667 | 4 | 0.25 | 5 | 0.2 | 25 | 0.2 | 1 | 0 | 9 | 0.4444 | 3 | 0.3333 | 6 | 0.3333 | 19 | 0.3684 |
| 7_7918219 | 11 | 0.7273 | 7 | 0.4286 | 6 | 0.5 | 5 | 0.4 | 29 | 0.5517 | 1 | 0 | 9 | 0.3333 | 3 | 0.6667 | 4 | 1 | 17 | 0.5294 |
| 7_7918225 | 10 | 0.5 | 8 | 0.25 | 5 | 0.4 | 4 | 0.25 | 27 | 0.3704 | 1 | 1 | 9 | 0.4444 | 3 | 0.3333 | 5 | 0.6 | 18 | 0.5 |
| 7_7918310 | 1 | 1 | NA | NA | NA | NA | 1 | 1 | 2 | 1 | 1 | 1 | 1 | 1 | 1 | 1 | NA | NA | 3 | 1 |
| 7_7918397 | 1 | 0 | NA | NA | 1 | 1 | 1 | 0 | 3 | 0.3333 | NA | NA | 1 | 1 | 1 | 0 | NA | NA | 2 | 0.5 |
| 7_7920507 | NA | NA | NA | NA | 1 | 1 | 1 | 0 | 2 | 0.5 | 4 | 0 | 1 | 1 | 1 | 0 | 1 | 1 | 7 | 0.2857 |
| 7_7920524 | NA | NA | NA | NA | 1 | 0 | 1 | 1 | 2 | 0.5 | 4 | 0.5 | 1 | 0 | 1 | 0 | 1 | 1 | 7 | 0.4286 |
| 7_7920527 | NA | NA | NA | NA | 1 | 1 | 1 | 0 | 2 | 0.5 | 4 | 0.25 | 1 | 1 | 1 | 0 | 1 | 0 | 7 | 0.2857 |
| 7_7920544 | NA | NA | NA | NA | 2 | 1 | 1 | 1 | 3 | 1 | 4 | 0.25 | 1 | 1 | NA | NA | 1 | 0 | 6 | 0.3333 |
| 7_7920606 | 3 | 0.6667 | NA | NA | 2 | 1 | 1 | 1 | 6 | 0.8333 | 4 | 0.75 | 1 | 1 | 1 | 1 | 1 | 1 | 7 | 0.8571 |
| 7_7920611 | 3 | 0.3333 | NA | NA | 1 | 0 | 1 | 1 | 5 | 0.4 | 3 | 0.6667 | 1 | 0 | 1 | 1 | 1 | 0 | 6 | 0.5 |
| 7_7920646 | 2 | 0 | NA | NA | 2 | 0.5 | 1 | 0 | 5 | 0.2 | 4 | 0.25 | 1 | 0 | 1 | 0 | 1 | 1 | 7 | 0.2857 |
| 7_7920688 | 3 | 0.6667 | NA | NA | 2 | 0.5 | NA | NA | 5 | 0.6 | 3 | 0.3333 | 1 | 1 | NA | NA | 1 | 0 | 5 | 0.4 |
| 7_7920697 | 3 | 0.6667 | NA | NA | 2 | 1 | NA | NA | 5 | 0.8 | 2 | 0 | 1 | 1 | NA | NA | 1 | 1 | 4 | 0.5 |
| 7_7920711 | 2 | 1 | NA | NA | 3 | 1 | NA | NA | 5 | 1 | 2 | 1 | 1 | 1 | NA | NA | 1 | 1 | 4 | 1 |
| 7_7920726 | 4 | 0.5 | NA | NA | 3 | 1 | NA | NA | 7 | 0.7143 | 3 | 0.3333 | 1 | 0 | NA | NA | 1 | 0 | 5 | 0.2 |
| 7_7920730 | 4 | 0.25 | NA | NA | 3 | 0.3333 | NA | NA | 7 | 0.2857 | 3 | 0 | 1 | 0 | NA | NA | 1 | 1 | 5 | 0.2 |
| 7_7920734 | 3 | 0.6667 | NA | NA | 1 | 1 | NA | NA | 4 | 0.75 | 3 | 1 | 1 | 1 | NA | NA | 2 | 1 | 6 | 1 |
| 7_7920741 | 4 | 0 | NA | NA | 3 | 0.3333 | NA | NA | 7 | 0.1429 | 3 | 0.3333 | 1 | 0 | NA | NA | 2 | 0.5 | 6 | 0.3333 |
| 7_7920755 | 2 | 1 | NA | NA | 1 | 1 | NA | NA | 3 | 1 | 3 | 1 | 1 | 1 | NA | NA | 1 | 1 | 5 | 1 |
| 7_7920756 | 4 | 0 | NA | NA | 3 | 0.3333 | NA | NA | 7 | 0.1429 | 3 | 0 | 1 | 1 | NA | NA | 1 | 0 | 5 | 0.2 |
| 7_7920766 | 3 | 0 | NA | NA | 3 | 0.3333 | NA | NA | 6 | 0.1667 | 3 | 0.3333 | 1 | 0 | NA | NA | 2 | 0.5 | 6 | 0.3333 |
| 7_7920817 | 1 | 1 | NA | NA | 4 | 0.25 | NA | NA | 5 | 0.4 | 1 | 0 | 1 | 0 | NA | NA | 2 | 0.5 | 4 | 0.25 |
| 7_7920885 | 2 | 0.5 | NA | NA | 3 | 0.3333 | 1 | 0 | 6 | 0.3333 | 2 | 0 | 3 | 0.3333 | NA | NA | 3 | 0 | 8 | 0.125 |
| 7_7920889 | 3 | 0.3333 | NA | NA | 3 | 0 | 1 | 1 | 7 | 0.2857 | 3 | 0 | 4 | 0.25 | NA | NA | 3 | 0 | 10 | 0.1 |
| 7_7920904 | 3 | 0.6667 | NA | NA | 2 | 1 | NA | NA | 5 | 0.8 | 1 | 0 | 3 | 1 | NA | NA | 2 | 0.5 | 6 | 0.6667 |
| 7_7920921 | 1 | 1 | NA | NA | 2 | 0.5 | 1 | 1 | 4 | 0.75 | 1 | 0 | 2 | 0.5 | NA | NA | NA | NA | 3 | 0.3333 |
| 7_7920930 | 2 | 1 | NA | NA | 2 | 1 | 1 | 1 | 5 | 1 | 1 | 0 | 4 | 0.75 | NA | NA | 3 | 0.6667 | 8 | 0.625 |
| 7_7920961 | 2 | 0.5 | 1 | 0 | 1 | 0 | NA | NA | 4 | 0.25 | 1 | 0 | 4 | 0.25 | NA | NA | 2 | 0 | 7 | 0.1429 |
| 7_7920968 | 2 | 1 | NA | NA | 3 | 1 | NA | NA | 5 | 1 | 1 | 0 | 1 | 1 | NA | NA | 2 | 0.5 | 4 | 0.5 |
| 7_7921035 | 1 | 0 | NA | NA | 1 | 1 | 2 | 0 | 4 | 0.25 | NA | NA | 1 | 1 | 1 | 0 | 1 | 0 | 3 | 0.3333 |
| 7_7921160 | 1 | 0 | NA | NA | 2 | 0 | 2 | 0.5 | 5 | 0.2 | 2 | 0 | 2 | 0.5 | 2 | 0 | 1 | 0 | 7 | 0.1429 |
| 7_7921204 | 1 | 0 | NA | NA | 2 | 0.5 | 3 | 0 | 6 | 0.1667 | 3 | 0 | 3 | 0.3333 | 2 | 0 | 1 | 0 | 9 | 0.1111 |
| 7_7921240 | 1 | 0 | NA | NA | 2 | 0.5 | 3 | 0 | 6 | 0.1667 | 5 | 0 | 3 | 0 | 1 | 0 | 1 | 1 | 10 | 0.1 |
| 7_7921246 | 1 | 0 | NA | NA | 2 | 1 | 3 | 0.6667 | 6 | 0.6667 | 4 | 1 | 3 | 1 | 2 | 1 | 1 | 1 | 10 | 1 |
| 7_7924321 | 2 | 1 | NA | NA | NA | NA | NA | NA | 2 | 1 | 1 | 0 | NA | NA | NA | NA | 1 | 1 | 2 | 0.5 |
| 7_7938581 | 1 | 0 | 2 | 1 | 1 | 0 | 1 | 0 | 5 | 0.4 | NA | NA | NA | NA | 1 | 1 | 2 | 1 | 3 | 1 |
| 7_7938586 | 1 | 0 | 3 | 1 | 1 | 1 | NA | NA | 5 | 0.8 | NA | NA | NA | NA | NA | NA | 1 | 1 | 1 | 1 |
| 7_7938643 | 1 | 0 | 3 | 0.3333 | 1 | 1 | 1 | 0 | 6 | 0.3333 | 1 | 1 | 1 | 0 | NA | NA | 2 | 0 | 4 | 0.25 |
| 7_7938716 | 1 | 0 | 2 | 0.5 | 1 | 1 | 1 | 0 | 5 | 0.4 | 1 | 1 | 1 | 1 | NA | NA | 2 | 1 | 4 | 1 |
| 7_7938731 | 1 | 1 | 2 | 1 | 1 | 1 | 1 | 1 | 5 | 1 | 1 | 1 | 1 | 0 | NA | NA | 2 | 1 | 4 | 0.75 |
| 7_7938734 | 1 | 1 | 2 | 0.5 | 1 | 1 | 1 | 1 | 5 | 0.8 | NA | NA | 1 | 0 | NA | NA | 2 | 1 | 3 | 0.6667 |
| 7_7938739 | 1 | 0 | 2 | 1 | 1 | 1 | 1 | 1 | 5 | 0.8 | NA | NA | 1 | 0 | NA | NA | 2 | 1 | 3 | 0.6667 |
| 7_7938753 | 1 | 1 | 1 | 1 | 1 | 0 | 1 | 0 | 4 | 0.5 | 1 | 0 | 1 | 0 | NA | NA | 2 | 0.5 | 4 | 0.25 |
| 7_7952513 | 2 | 0 | 4 | 0 | 7 | 0.1429 | 3 | 0.3333 | 16 | 0.125 | 1 | 0 | 5 | 0.2 | 6 | 0.1667 | NA | NA | 12 | 0.1667 |
| 7_7952517 | 2 | 0 | 6 | 0.1667 | 3 | 0 | 3 | 0 | 14 | 0.0714 | 1 | 1 | 4 | 0.5 | 3 | 0 | NA | NA | 8 | 0.375 |
| 7_7952526 | 2 | 0 | 5 | 0.2 | 6 | 0 | 3 | 0.3333 | 16 | 0.125 | 1 | 0 | 6 | 0.3333 | 6 | 0 | 1 | 0 | 14 | 0.1429 |
| 7_7952549 | 2 | 0 | 6 | 0 | 5 | 0 | 3 | 0.3333 | 16 | 0.0625 | 1 | 0 | 5 | 0.2 | 5 | 0 | NA | NA | 11 | 0.0909 |
| 7_7952563 | 2 | 0 | 6 | 0 | 7 | 0 | 3 | 0.3333 | 18 | 0.0556 | 1 | 0 | 6 | 0 | 6 | 0 | 1 | 1 | 14 | 0.0714 |
| 7_7952594 | 1 | 1 | 5 | 0 | 5 | 0 | 3 | 0.3333 | 14 | 0.1429 | 1 | 0 | 4 | 0.25 | 4 | 0 | NA | NA | 9 | 0.1111 |

| | | | | | | | | | | | | | | | | | | | | |
|---|---|---|---|---|---|---|---|---|---|---|---|---|---|---|---|---|---|---|---|---|
| 7_7952752( | 5 | 0.2 | 4 | 0.25 | 4 | 0 | 2 | 0 | 15 | 0.1333 | 1 | 0 | 4 | 0 | 5 | 0 | 3 | 0.3333 | 13 | 0.0769 |
| 7_7952861! | 2 | 0 | 1 | 1 | 3 | 0.3333 | 1 | 0 | 7 | 0.2857 | NA | NA | 4 | 0.75 | 4 | 0.25 | 2 | 1 | 10 | 0.6 |
| 7_7952902 | 2 | 0 | 3 | 0 | 5 | 0.4 | 2 | 0 | 12 | 0.1667 | NA | NA | 4 | 0.25 | 5 | 0 | 1 | 0 | 10 | 0.1 |
| 7_7952931( | 3 | 0.3333 | 3 | 0 | 5 | 0 | 1 | 0 | 12 | 0.0833 | NA | NA | 3 | 0.3333 | 5 | 0 | 1 | 0 | 9 | 0.1111 |
| 7_7952957! | 6 | 0 | 2 | 0 | 8 | 0.125 | 2 | 0 | 18 | 0.0556 | 3 | 0.3333 | 7 | 0.4286 | 6 | 0.3333 | 2 | 1 | 18 | 0.4444 |
| 7_7952993 | 3 | 0 | 2 | 0 | 5 | 0.2 | 1 | 0 | 11 | 0.0909 | 1 | 0 | 7 | 0.2857 | 5 | 0 | 2 | 0 | 15 | 0.1333 |
| 7_7952994 | 4 | 0 | 2 | 0 | 3 | 0.3333 | 1 | 0 | 10 | 0.1 | 2 | 0 | 4 | 0 | 4 | 0.25 | 1 | 0 | 11 | 0.0909 |
| 7_7952994 | 4 | 1 | 1 | 1 | 6 | 1 | 1 | 1 | 12 | 1 | 2 | 1 | 7 | 0.8571 | 6 | 0.8333 | 2 | 1 | 17 | 0.8824 |
| 7_7952998! | 4 | 1 | 2 | 1 | 6 | 1 | 1 | 1 | 13 | 1 | 2 | 1 | 7 | 1 | 5 | 1 | 2 | 1 | 16 | 1 |
| 7_7953005 | 4 | 0.25 | 1 | 1 | 4 | 0.5 | 1 | 0 | 10 | 0.4 | 2 | 0 | 7 | 0.5714 | 5 | 0.4 | 1 | 0 | 15 | 0.4 |
| 7_7953006! | 4 | 0 | 2 | 0 | 5 | 0.2 | 1 | 1 | 12 | 0.1667 | 1 | 1 | 7 | 0.1429 | 6 | 0.3333 | 2 | 0 | 16 | 0.25 |
| 7_7953021 | 1 | 0 | 2 | 0 | 3 | 0.6667 | NA | NA | 6 | 0.3333 | 2 | 0.5 | 6 | 0.5 | 6 | 0.5 | 1 | 1 | 15 | 0.5333 |
| 7_7953022 | 4 | 0.5 | 1 | 0 | 5 | 0 | 1 | 1 | 11 | 0.2727 | 1 | 0 | 5 | 0.2 | 5 | 0.2 | 2 | 0.5 | 13 | 0.2308 |
| 7_7957249 | 4 | 0.25 | NA | NA | 3 | 0 | 2 | 0 | 9 | 0.1111 | 1 | 0 | 1 | 0 | 2 | 0.5 | 2 | 0.5 | 6 | 0.3333 |
| 7_7957257 | 4 | 0 | NA | NA | 2 | 0 | 4 | 0.5 | 10 | 0.2 | 1 | 0 | 2 | 0 | 3 | 0.3333 | 2 | 0 | 8 | 0.125 |
| 7_7957633 | 2 | 0.5 | 2 | 0 | 4 | 0.5 | NA | NA | 8 | 0.375 | NA | NA | 3 | 0.3333 | 1 | 0 | NA | NA | 4 | 0.25 |
| 7_7974211! | 2 | 0 | 2 | 0 | 3 | 0.3333 | NA | NA | 7 | 0.1429 | 1 | 0 | 2 | 0 | 2 | 0 | 2 | 0.5 | 7 | 0.1429 |
| 7_7987461 | 1 | 1 | 1 | 0 | NA | NA | 2 | 0.5 | 4 | 0.5 | NA | NA | NA | NA | 2 | 1 | 3 | 1 | 5 | 1 |
| 7_7987470 | NA | NA | 1 | 0 | NA | NA | 2 | 0.5 | 3 | 0.3333 | NA | NA | NA | NA | 2 | 1 | 2 | 0.5 | 4 | 0.75 |
| 7_7987480 | 1 | 0 | 1 | 1 | NA | NA | 3 | 0.3333 | 5 | 0.4 | NA | NA | NA | NA | 2 | 0.5 | 3 | 0.3333 | 5 | 0.4 |
| 7_7987500 | 1 | 1 | NA | NA | NA | NA | 4 | 0.75 | 5 | 0.8 | NA | NA | NA | NA | NA | NA | 3 | 0.3333 | 3 | 0.3333 |
| 7_7987507( | 1 | 1 | 1 | 1 | NA | NA | 4 | 1 | 6 | 1 | NA | NA | NA | NA | 2 | 0.5 | 3 | 1 | 5 | 0.8 |
| 7_7987509( | NA | NA | 1 | 1 | NA | NA | 5 | 0.6 | 6 | 0.6667 | NA | NA | NA | NA | 2 | 1 | 2 | 0 | 4 | 0.5 |
| 7_7987529! | 2 | 0.5 | 1 | 0 | NA | NA | 5 | 0.4 | 8 | 0.375 | NA | NA | NA | NA | 1 | 1 | 3 | 0 | 4 | 0.25 |
| 7_7987541( | 2 | 1 | 1 | 1 | NA | NA | 5 | 1 | 8 | 1 | NA | NA | NA | NA | 2 | 1 | 3 | 1 | 5 | 1 |
| 7_7987547 | NA | NA | 1 | 1 | NA | NA | 4 | 1 | 5 | 1 | NA | NA | NA | NA | 1 | 1 | 3 | 1 | 4 | 1 |
| 7_7987556 | 2 | 1 | 1 | 1 | NA | NA | 4 | 0.75 | 7 | 0.8571 | NA | NA | NA | NA | 2 | 1 | 4 | 0.75 | 6 | 0.8333 |
| 7_7987559! | 1 | 1 | 1 | 1 | NA | NA | 4 | 0.5 | 6 | 0.6667 | NA | NA | NA | NA | 2 | 0.5 | 4 | 1 | 6 | 0.8333 |
| 7_7987566 | 2 | 1 | NA | NA | NA | NA | 4 | 1 | 6 | 1 | NA | NA | NA | NA | 2 | 1 | 4 | 1 | 6 | 1 |
| 7_7987573! | 2 | 0 | 1 | 0 | NA | NA | 5 | 0.2 | 8 | 0.125 | NA | NA | NA | NA | 2 | 0.5 | 4 | 0.5 | 6 | 0.5 |
| 7_7987576( | 2 | 0.5 | 1 | 1 | NA | NA | 4 | 0.5 | 7 | 0.5714 | NA | NA | NA | NA | 1 | 1 | 3 | 1 | 4 | 1 |
| 7_7987586 | 2 | 1 | NA | NA | NA | NA | 3 | 0.3333 | 5 | 0.6 | 1 | 0 | NA | NA | NA | NA | 4 | 0.5 | 5 | 0.4 |
| 7_7987589 | 2 | 0 | 1 | 1 | NA | NA | 5 | 0.6 | 8 | 0.5 | NA | NA | NA | NA | 2 | 0.5 | 4 | 0.75 | 6 | 0.6667 |
| 7_7987598( | 2 | 0.5 | 1 | 0 | NA | NA | 4 | 0.5 | 7 | 0.4286 | 1 | 0 | NA | NA | 2 | 0.5 | 3 | 0.6667 | 6 | 0.5 |
| 7_7987606 | 2 | 1 | 1 | 0 | NA | NA | 4 | 0.5 | 7 | 0.5714 | 1 | 1 | NA | NA | 1 | 1 | 3 | 0.6667 | 5 | 0.8 |
| 7_7988805 | NA | NA | 2 | 1 | NA | NA | NA | NA | 2 | 1 | 1 | 1 | 1 | 1 | NA | NA | 2 | 1 | 4 | 1 |
| 7_7988812! | NA | NA | 2 | 1 | NA | NA | NA | NA | 2 | 1 | 1 | 0 | 1 | 1 | NA | NA | 2 | 1 | 4 | 0.75 |
| 7_7988818( | NA | NA | 1 | 1 | NA | NA | NA | NA | 1 | 1 | 1 | 1 | NA | NA | NA | NA | 2 | 0.5 | 3 | 0.6667 |
| 7_7988861 | NA | NA | 2 | 1 | NA | NA | NA | NA | 2 | 1 | 1 | 1 | 1 | 1 | 1 | 0 | 1 | 0 | 4 | 0.5 |
| 7_7988880( | NA | NA | 2 | 1 | NA | NA | NA | NA | 2 | 1 | 2 | 1 | 1 | 0 | 1 | 0 | 2 | 1 | 6 | 0.6667 |
| 7_7988881 | NA | NA | 2 | 0.5 | NA | NA | NA | NA | 2 | 0.5 | 2 | 1 | 1 | 1 | 1 | 1 | 2 | 0.5 | 6 | 0.8333 |
| 7_7988901( | NA | NA | 2 | 1 | NA | NA | NA | NA | 2 | 1 | 2 | 1 | 1 | 1 | 1 | 1 | 2 | 1 | 6 | 1 |
| 7_7988915 | NA | NA | 1 | 1 | NA | NA | NA | NA | 1 | 1 | 2 | 1 | 1 | 0 | 1 | 1 | NA | NA | 4 | 0.75 |
| 7_7988924! | NA | NA | 2 | 1 | NA | NA | NA | NA | 2 | 1 | 1 | 1 | 1 | 1 | 1 | 1 | 2 | 1 | 5 | 1 |
| 7_7991146 | 2 | 0 | 3 | 0.3333 | 4 | 0 | 4 | 0 | 13 | 0.0769 | 2 | 0 | NA | NA | NA | NA | 1 | 1 | 3 | 0.3333 |
| 7_7991424! | 2 | 0.5 | 4 | 1 | 3 | 0.6667 | 5 | 1 | 14 | 0.8571 | 6 | 0.8333 | 3 | 0.6667 | 3 | 1 | 2 | 1 | 14 | 0.8571 |
| 7_7991456! | 6 | 0.6667 | 5 | 0.4 | 4 | 0 | 5 | 0.6 | 20 | 0.45 | 6 | 0.5 | 2 | 0.5 | 4 | 0.5 | 2 | 0.5 | 14 | 0.5 |
| 7_7991469 | 5 | 0.2 | 6 | 0.3333 | 6 | 0.1667 | 6 | 0 | 23 | 0.1739 | 7 | 0.1429 | 3 | 0 | 3 | 0 | 3 | 0 | 16 | 0.0625 |
| 7_7991471 | 6 | 0.3333 | 6 | 0.1667 | 6 | 0.1667 | 8 | 0 | 26 | 0.1538 | 6 | 0.5 | 3 | 0 | 3 | 0 | 3 | 0.6667 | 15 | 0.3333 |
| 7_7991476 | 6 | 0 | 6 | 0 | 5 | 0 | 8 | 0.25 | 25 | 0.08 | 8 | 0.125 | 2 | 0.5 | 3 | 0.3333 | 3 | 0 | 16 | 0.1875 |
| 7_7991477 | 7 | 1 | 5 | 1 | 6 | 0.6667 | 6 | 0.6667 | 24 | 0.8333 | 8 | 1 | 2 | 1 | 2 | 0.5 | 2 | 1 | 14 | 0.9286 |
| 7_7991478( | 7 | 0.4286 | 5 | 0.6 | 6 | 0.3333 | 8 | 0.625 | 26 | 0.5 | 6 | 0.6667 | 3 | 0.6667 | 4 | 0.5 | 3 | 0.3333 | 16 | 0.5625 |
| 7_7991487 | 7 | 0.4286 | 6 | 0.5 | 6 | 0.1667 | 8 | 0.125 | 27 | 0.2963 | 8 | 0.125 | 4 | 0.25 | 4 | 0.25 | 2 | 0 | 18 | 0.1667 |
| 7_7991505 | 5 | 0.6 | 4 | 0.5 | 2 | 0 | 7 | 0.5714 | 18 | 0.5 | 6 | 0.5 | 5 | 0.8 | 5 | 0.8 | 2 | 1 | 18 | 0.7222 |
| 7_7994378 | NA | NA | NA | NA | NA | NA | 1 | 1 | 1 | 1 | NA | NA | NA | NA | 2 | 0.5 | NA | NA | 2 | 0.5 |
| 7_7994384 | NA | NA | NA | NA | NA | NA | 1 | 1 | 1 | 1 | NA | NA | NA | NA | 2 | 0.5 | NA | NA | 2 | 0.5 |
| 7_7994424 | NA | NA | NA | NA | NA | NA | 1 | 1 | 1 | 1 | NA | NA | NA | NA | 1 | 1 | NA | NA | 1 | 1 |
| 7_7994467 | NA | NA | NA | NA | NA | NA | 1 | 1 | 1 | 1 | NA | NA | NA | NA | 2 | 0.5 | NA | NA | 2 | 0.5 |
| 7_7996882 | NA | NA | NA | NA | 1 | 0 | 2 | 1 | 3 | 0.6667 | NA | NA | NA | NA | 1 | 1 | NA | NA | 1 | 1 |
| 7_7996889 | NA | NA | NA | NA | 1 | 1 | 2 | 0.5 | 3 | 0.6667 | NA | NA | NA | NA | 1 | 1 | NA | NA | 1 | 1 |
| 7_7997504! | 2 | 0.5 | 5 | 0.2 | 2 | 0 | 1 | 0 | 10 | 0.2 | 3 | 0 | 4 | 0.5 | 2 | 0.5 | 2 | 0 | 11 | 0.2727 |
| 7_7997512 | 4 | 0.75 | 7 | 0.8571 | 2 | 1 | 3 | 0 | 16 | 0.6875 | 4 | 0 | 4 | 0.5 | 3 | 1 | 4 | 0.75 | 15 | 0.5333 |
| 7_7997524! | 5 | 1 | 7 | 0.7143 | 2 | 1 | 3 | 0.6667 | 17 | 0.8235 | 3 | 0 | 4 | 0.75 | 3 | 1 | 4 | 0.5 | 14 | 0.5714 |
| 7_7997539 | 4 | 0.5 | 7 | 0.1429 | 1 | 0 | 3 | 0 | 15 | 0.2 | 4 | 0 | 3 | 0.6667 | 3 | 0.6667 | 4 | 0 | 14 | 0.2857 |
| 7_7998159! | 2 | 0 | 4 | 0 | NA | NA | 1 | 1 | 7 | 0.1429 | 3 | 0 | 1 | 1 | NA | NA | 3 | 0 | 7 | 0.1429 |

| | | | | | | | | | | | | | | | | | | | | |
|---|---|---|---|---|---|---|---|---|---|---|---|---|---|---|---|---|---|---|---|---|
| 7_7999417 | NA | NA | NA | NA | 1 | 0 | 1 | 1 | 2 | 0.5 | 1 | 0 | NA | NA | 1 | 1 | 1 | 0 | 3 | 0.3333 |
| 7_7999463 | NA | NA | NA | NA | 1 | 0 | 1 | 1 | 2 | 0.5 | 1 | 0 | NA | NA | 1 | 1 | 1 | 1 | 3 | 0.6667 |
| 7_7999546 | NA | NA | 1 | 1 | 1 | 1 | 1 | 0 | 3 | 0.6667 | 2 | 0.5 | 2 | 0.5 | 1 | 1 | 2 | 0.5 | 7 | 0.5714 |
| 7_7999578 | NA | NA | 1 | 0 | 1 | 1 | 1 | 1 | 3 | 0.6667 | 1 | 0 | 3 | 0 | 1 | 1 | 2 | 1 | 7 | 0.4286 |
| 7_7999907 | 17 | 0 | 12 | 0.0833 | 17 | 0.1176 | 15 | 0.2667 | 61 | 0.1148 | 13 | 0.0769 | 18 | 0.0556 | 15 | 0.0667 | 19 | 0.0526 | 65 | 0.0615 |
| 7_7999944 | 11 | 0 | 9 | 0 | 13 | 0.0769 | 12 | 0.1667 | 45 | 0.0667 | 11 | 0 | 14 | 0 | 17 | 0.1176 | 16 | 0.125 | 58 | 0.069 |
| 7_7999956 | 11 | 0.3636 | 11 | 0.2727 | 14 | 0.3571 | 13 | 0.2308 | 49 | 0.3061 | 10 | 0.4 | 14 | 0.2143 | 14 | 0.2143 | 17 | 0.1765 | 55 | 0.2364 |
| 7_7999964 | 12 | 0 | 15 | 0.2667 | 14 | 0.1429 | 14 | 0.1429 | 55 | 0.1455 | 12 | 0.0833 | 14 | 0.0714 | 15 | 0.1333 | 18 | 0.0556 | 59 | 0.0847 |
| 7_7999977 | 11 | 0.1818 | 13 | 0.4615 | 15 | 0.4667 | 11 | 0 | 50 | 0.3 | 9 | 0.1111 | 13 | 0.2308 | 14 | 0.3571 | 17 | 0.1765 | 53 | 0.2264 |
| 7_7999992 | 11 | 0.7273 | 12 | 1 | 14 | 1 | 12 | 0.9167 | 49 | 0.9184 | 9 | 0.8889 | 12 | 0.9167 | 10 | 0.8 | 18 | 0.8333 | 49 | 0.8571 |
| 7_8000006 | 6 | 0.1667 | 8 | 0.75 | 12 | 0.75 | 10 | 0.6 | 36 | 0.6111 | 7 | 0.8571 | 7 | 0.5714 | 9 | 0.5556 | 16 | 0.5625 | 39 | 0.6154 |
| 7_8000007 | 8 | 1 | 8 | 1 | 13 | 0.7692 | 10 | 1 | 39 | 0.9231 | 6 | 1 | 9 | 0.6667 | 10 | 0.9 | 7 | 0.8571 | 32 | 0.8438 |
| 7_8000027 | 8 | 0.25 | 13 | 0.5385 | 15 | 0.5333 | 10 | 0.3 | 46 | 0.4348 | 9 | 0.4444 | 11 | 0.4545 | 9 | 0.5556 | 19 | 0.6316 | 48 | 0.5417 |
| 7_8000030 | 8 | 0.5 | 11 | 0.6364 | 11 | 0.4545 | 12 | 0.5 | 42 | 0.5238 | 6 | 0.5 | 9 | 0.1111 | 9 | 0.2222 | 18 | 0.4444 | 42 | 0.3333 |
| 7_8000037 | 9 | 0.1111 | 12 | 0.1667 | 18 | 0.2222 | 13 | 0.0769 | 52 | 0.1538 | 9 | 0.1111 | 10 | 0.1 | 12 | 0.0833 | 18 | 0.2778 | 49 | 0.1633 |
| 7_8000038 | 8 | 0 | 13 | 0.0769 | 15 | 0.0667 | 12 | 0.0833 | 48 | 0.0625 | 8 | 0.125 | 10 | 0.3 | 12 | 0 | 15 | 0.0667 | 45 | 0.1111 |
| 7_8000047 | 10 | 0.3 | 14 | 0.0714 | 18 | 0.0556 | 14 | 0 | 56 | 0.0893 | 9 | 0.2222 | 10 | 0.1 | 11 | 0.1818 | 17 | 0.2353 | 47 | 0.1915 |
| 7_8000050 | 9 | 0.2222 | 13 | 0.5385 | 14 | 0.2143 | 14 | 0.1429 | 50 | 0.28 | 9 | 0.2222 | 8 | 0.25 | 12 | 0.1667 | 17 | 0.2353 | 46 | 0.2174 |
| 7_8000080 | 8 | 0.125 | 11 | 0.1818 | 11 | 0.1818 | 10 | 0.2 | 40 | 0.175 | 7 | 0 | 8 | 0 | 10 | 0.1 | 17 | 0.1176 | 42 | 0.0714 |
| 7_8000082 | 8 | 0.125 | 11 | 0.0909 | 12 | 0.1667 | 12 | 0.0833 | 43 | 0.1163 | 8 | 0.5 | 8 | 0 | 11 | 0 | 13 | 0.0769 | 40 | 0.125 |
| 7_8000124 | 5 | 0.8 | 5 | 0.2 | 7 | 0.1429 | 9 | 0 | 26 | 0.2308 | 8 | 0 | 5 | 0 | 9 | 0.1111 | 13 | 0.1538 | 35 | 0.0857 |
| 7_8000128 | 5 | 0.8 | 5 | 0.2 | 6 | 0.5 | 7 | 0.5714 | 23 | 0.5217 | 3 | 0.3333 | 5 | 0.2 | 3 | 0.6667 | 10 | 0.5 | 21 | 0.4286 |
| 7_8000132 | 7 | 1 | 8 | 0.75 | 10 | 0.9 | 9 | 1 | 34 | 0.9118 | 9 | 0.6667 | 6 | 0.8333 | 10 | 0.8 | 18 | 0.5 | 43 | 0.6512 |
| 7_8000133 | 5 | 0.2 | 6 | 0 | 10 | 0.1 | 8 | 0 | 29 | 0.069 | 8 | 0 | 6 | 0.1667 | 9 | 0.1111 | 16 | 0.0625 | 39 | 0.0769 |
| 7_8000137 | 4 | 0.25 | 4 | 0.5 | 4 | 0 | 7 | 0.1429 | 19 | 0.2105 | 5 | 0.6 | 3 | 0.6667 | 5 | 0.2 | 9 | 0.3333 | 22 | 0.4091 |
| 7_8000138 | 6 | 0.3333 | 6 | 0.5 | 6 | 0 | 7 | 0 | 25 | 0.2 | 9 | 0 | 4 | 0.75 | 6 | 0.3333 | 14 | 0.1429 | 33 | 0.2121 |
| 7_8000140 | 7 | 0.8571 | 6 | 0.6667 | 7 | 0.8571 | 8 | 0.875 | 28 | 0.8214 | 8 | 1 | 5 | 0.8 | 8 | 0.75 | 17 | 0.8235 | 38 | 0.8421 |
| 7_8000149 | 4 | 0 | 5 | 0.2 | 6 | 0 | 9 | 0.2222 | 24 | 0.125 | 9 | 0.1111 | 4 | 0 | 8 | 0.125 | 14 | 0.0714 | 35 | 0.0857 |
| 7_8000153 | 6 | 0.3333 | 6 | 0.1667 | 7 | 0 | 7 | 0.1429 | 26 | 0.1538 | 6 | 0.1667 | 4 | 0 | 7 | 0.2857 | 15 | 0.2 | 32 | 0.1875 |
| 7_8003921 | 3 | 0.3333 | 6 | 0.1667 | 3 | 0.3333 | 4 | 0.25 | 16 | 0.25 | 4 | 0.25 | 7 | 0 | 2 | 0 | 3 | 0 | 16 | 0.0625 |
| 7_8003997 | 4 | 0.25 | 4 | 0.5 | 2 | 0.5 | 1 | 1 | 11 | 0.4545 | 5 | 0.6 | 4 | 0 | 1 | 0 | 3 | 0 | 13 | 0.2308 |
| 7_8004020 | 4 | 1 | 5 | 1 | 1 | 1 | 2 | 1 | 12 | 1 | 3 | 1 | 3 | 1 | NA | NA | 2 | 1 | 8 | 1 |
| 7_8004033 | 4 | 1 | 4 | 1 | 2 | 1 | 2 | 1 | 12 | 1 | 5 | 1 | 4 | 1 | 1 | 1 | 2 | 1 | 12 | 1 |
| 7_8004036 | 4 | 1 | 3 | 1 | 2 | 1 | 2 | 1 | 11 | 1 | 4 | 1 | 4 | 1 | 1 | 1 | 3 | 1 | 12 | 1 |
| 7_8004047 | 3 | 0 | 5 | 0.2 | 2 | 0.5 | 2 | 0 | 12 | 0.1667 | 5 | 0.2 | 4 | 0 | 1 | 0 | 3 | 0.6667 | 13 | 0.2308 |
| 7_8004048 | 2 | 1 | 4 | 0.25 | 2 | 0 | 1 | 0 | 9 | 0.3333 | 4 | 0.5 | 4 | 0.75 | 1 | 1 | 3 | 0.6667 | 12 | 0.6667 |
| 7_8004051 | 1 | 1 | 5 | 0.8 | 2 | 1 | 2 | 1 | 10 | 0.9 | 3 | 1 | 4 | 1 | 1 | 0 | 3 | 1 | 11 | 0.9091 |
| 7_8004058 | 3 | 0.6667 | 3 | 0.3333 | 2 | 0.5 | 1 | 1 | 9 | 0.5556 | 5 | 0.8 | 2 | 0.5 | NA | NA | 3 | 0.3333 | 10 | 0.6 |
| 7_8004083 | 3 | 0.6667 | 5 | 0.6 | 2 | 1 | 2 | 0.5 | 12 | 0.6667 | 4 | 0.5 | 3 | 0.6667 | 1 | 1 | 2 | 0.5 | 10 | 0.6 |
| 7_8004086 | 2 | 1 | 5 | 0.8 | 2 | 1 | 1 | 1 | 10 | 0.9 | 3 | 0.6667 | 3 | 1 | 1 | 1 | 3 | 0.6667 | 10 | 0.8 |
| 7_8023646 | 1 | 0 | 2 | 0.5 | 3 | 0.3333 | 1 | 1 | 7 | 0.4286 | 4 | 0.25 | 2 | 0.5 | 3 | 0.3333 | 2 | 0 | 11 | 0.2727 |
| 7_8023701 | 1 | 0 | 2 | 0 | 3 | 0.3333 | 1 | 0 | 7 | 0.1429 | 3 | 0.6667 | 2 | 0 | 3 | 0.3333 | 1 | 0 | 9 | 0.3333 |
| 7_8023789 | 1 | 1 | 1 | 0 | 2 | 0 | NA | NA | 4 | 0.25 | NA | NA | 1 | 1 | 2 | 1 | 2 | 1 | 5 | 1 |
| 7_8023861 | NA | NA | 1 | 1 | 1 | 1 | NA | NA | 2 | 1 | NA | NA | 1 | 1 | 2 | 1 | 1 | 1 | 4 | 1 |
| 7_8023873 | NA | NA | 1 | 1 | 1 | 0 | NA | NA | 2 | 0.5 | NA | NA | 1 | 1 | 2 | 1 | 2 | 1 | 5 | 1 |
| 7_8023929 | NA | NA | 1 | 0 | 1 | 1 | NA | NA | 2 | 0.5 | NA | NA | 1 | 1 | 2 | 0.5 | 2 | 0 | 5 | 0.4 |
| 7_8023947 | NA | NA | 1 | 1 | 1 | 0 | NA | NA | 2 | 0.5 | NA | NA | NA | NA | NA | NA | 1 | 1 | 1 | 1 |
| 7_8023956 | NA | NA | 1 | 1 | 1 | 0 | NA | NA | 2 | 0.5 | NA | NA | 1 | 1 | 1 | 1 | 2 | 1 | 4 | 1 |
| 7_8036140 | NA | NA | 2 | 1 | 1 | 0 | 1 | 1 | 4 | 0.75 | NA | NA | 2 | 0 | 3 | 0.3333 | 2 | 0 | 7 | 0.1429 |
| 7_8036174 | NA | NA | 2 | 0.5 | NA | NA | 1 | 0 | 3 | 0.3333 | 1 | 0 | 2 | 0.5 | 2 | 0 | 2 | 0 | 7 | 0.1429 |
| 7_8036287 | NA | NA | 2 | 1 | NA | NA | 2 | 0 | 4 | 0.5 | 1 | 0 | 2 | 0 | 1 | 0 | 1 | 1 | 5 | 0.2 |
| 7_8036289 | NA | NA | 3 | 0.6667 | NA | NA | 2 | 0.5 | 5 | 0.6 | 1 | 0 | 2 | 0 | 1 | 1 | 1 | 0 | 5 | 0.2 |
| 7_8036295 | NA | NA | 2 | 1 | NA | NA | 2 | 1 | 4 | 1 | 1 | 1 | 2 | 1 | 1 | 1 | 1 | 1 | 5 | 1 |
| 7_8036302 | NA | NA | 2 | 1 | NA | NA | 2 | 1 | 4 | 1 | 1 | 1 | 2 | 0.5 | 1 | 1 | 1 | 1 | 5 | 0.8 |
| 7_8036348 | NA | NA | 2 | 1 | NA | NA | 2 | 0.5 | 4 | 0.75 | NA | NA | 2 | 1 | 1 | 1 | 1 | 0 | 4 | 0.75 |
| 7_8052480 | NA | NA | 1 | 1 | NA | NA | NA | NA | 1 | 1 | 1 | 0 | 1 | 1 | 1 | 1 | NA | NA | 3 | 0.6667 |
| 7_8052481 | NA | NA | NA | NA | NA | NA | 1 | 1 | 1 | 1 | 1 | 1 | 1 | 1 | 1 | 1 | NA | NA | 3 | 1 |
| 7_8052482 | NA | NA | 1 | 1 | NA | NA | NA | NA | 1 | 1 | NA | NA | 1 | 1 | 1 | 1 | NA | NA | 2 | 1 |
| 7_8052496 | NA | NA | 1 | 1 | NA | NA | 1 | 0 | 2 | 0.5 | 1 | 0 | 1 | 1 | 2 | 0.5 | NA | NA | 4 | 0.5 |
| 7_8052507 | NA | NA | 1 | 1 | NA | NA | 1 | 1 | 2 | 1 | NA | NA | 1 | 1 | 2 | 1 | NA | NA | 3 | 1 |
| 7_8052512 | NA | NA | 1 | 0 | NA | NA | 1 | 1 | 2 | 0.5 | 1 | 1 | 1 | 0 | 1 | 0 | NA | NA | 3 | 0.3333 |
| 7_8052555 | NA | NA | 1 | 0 | NA | NA | 1 | 1 | 2 | 0.5 | 1 | 0 | 1 | 1 | 2 | 0.5 | NA | NA | 4 | 0.5 |
| 7_8052617 | NA | NA | 2 | 0 | NA | NA | 1 | 1 | 3 | 0.3333 | 1 | 0 | 2 | 0 | 2 | 0.5 | NA | NA | 5 | 0.2 |
| 7_8054095 | 1 | 1 | NA | NA | 1 | 1 | NA | NA | 2 | 1 | NA | NA | NA | NA | 3 | 1 | 1 | 1 | 4 | 1 |
| 7_8054096 | 3 | 1 | NA | NA | 1 | 1 | NA | NA | 4 | 1 | 1 | 1 | NA | NA | 4 | 0.75 | 2 | 1 | 7 | 0.8571 |

| | | | | | | | | | | | | | | | | | | | | |
|---|---|---|---|---|---|---|---|---|---|---|---|---|---|---|---|---|---|---|---|---|
| 7_80541067 | 7 | 0.5714 | 1 | 0 | 3 | 0.6667 | 2 | 0.5 | 13 | 0.5385 | 3 | 0 | NA | NA | 6 | 0.6667 | 3 | 0.6667 | 12 | 0.5 |
| 7_80541218 | 4 | 0.25 | 1 | 1 | 1 | 0 | 2 | 1 | 8 | 0.5 | 2 | 0.5 | NA | NA | 3 | 0 | 2 | 0.5 | 7 | 0.2857 |
| 7_80541352 | 7 | 0.1429 | 1 | 0 | 4 | 0 | 3 | 0 | 15 | 0.0667 | 3 | 0.3333 | NA | NA | 7 | 0 | 4 | 0 | 14 | 0.0714 |
| 7_80541363 | 6 | 0.8333 | 1 | 1 | 4 | 1 | 3 | 1 | 14 | 0.9286 | 4 | 1 | NA | NA | 5 | 1 | 4 | 1 | 13 | 1 |
| 7_80541450 | 5 | 0.6 | 1 | 1 | 4 | 0.5 | 3 | 0.6667 | 13 | 0.6154 | 4 | 0.75 | NA | NA | 5 | 0.8 | 1 | 1 | 10 | 0.8 |
| 7_80541493 | 4 | 0.75 | 1 | 1 | 3 | 1 | 3 | 0.6667 | 11 | 0.8182 | 3 | 0.3333 | NA | NA | 4 | 0.75 | 2 | 0.5 | 9 | 0.5556 |
| 7_80541543 | 6 | 0.5 | NA | NA | 3 | 1 | 3 | 0.3333 | 12 | 0.5833 | 5 | 0.2 | NA | NA | 6 | 0.1667 | 3 | 0.3333 | 14 | 0.2143 |
| 7_80541646 | 6 | 0.5 | 1 | 1 | 2 | 0.5 | 2 | 1 | 11 | 0.6364 | 5 | 0.2 | NA | NA | 4 | 0.5 | 3 | 0.3333 | 12 | 0.3333 |
| 7_80541886 | 1 | 0 | 1 | 1 | NA | NA | NA | NA | 2 | 0.5 | NA | NA | NA | NA | NA | NA | 1 | 1 | 1 | 1 |
| 7_80542967 | 7 | 0.1429 | NA | NA | 2 | 0.5 | 3 | 0.3333 | 12 | 0.25 | 5 | 0 | NA | NA | 3 | 0 | 3 | 0.3333 | 11 | 0.0909 |
| 7_80565402 | 4 | 0.25 | 3 | 0.3333 | 3 | 0.3333 | 5 | 0.8 | 15 | 0.4667 | 3 | 0.6667 | 3 | 0 | 6 | 0.3333 | 7 | 0.8571 | 19 | 0.5263 |
| 7_80565509 | 3 | 0.3333 | 4 | 0 | 3 | 0 | 5 | 0.2 | 15 | 0.1333 | 2 | 0.5 | 3 | 0 | 6 | 0.1667 | 7 | 0 | 18 | 0.1111 |
| 7_80642616 | NA | NA | NA | NA | NA | NA | 2 | 1 | 2 | 1 | NA | NA | NA | NA | NA | NA | 2 | 0.5 | 2 | 0.5 |
| 7_80642662 | 1 | 1 | NA | NA | NA | NA | 2 | 0 | 3 | 0.3333 | NA | NA | NA | NA | NA | NA | 2 | 0.5 | 2 | 0.5 |
| 7_80642710 | 1 | 1 | NA | NA | NA | NA | 1 | 1 | 2 | 1 | NA | NA | NA | NA | NA | NA | 2 | 0.5 | 2 | 0.5 |
| 7_80673560 | 1 | 1 | 2 | 0.5 | NA | NA | 2 | 1 | 5 | 0.8 | NA | NA | NA | NA | 2 | 1 | NA | NA | 2 | 1 |
| 7_80673610 | 1 | 1 | 2 | 0 | NA | NA | 1 | 0 | 4 | 0.25 | NA | NA | 1 | 1 | 2 | 0 | NA | NA | 3 | 0.3333 |
| 7_80674410 | 2 | 0.5 | 2 | 0.5 | NA | NA | 2 | 0.5 | 6 | 0.5 | NA | NA | 1 | 1 | 1 | 0 | NA | NA | 2 | 0.5 |
| 7_80674563 | 1 | 1 | 2 | 0.5 | NA | NA | 2 | 1 | 5 | 0.8 | NA | NA | 1 | 1 | 1 | 0 | NA | NA | 2 | 0.5 |
| 7_80674595 | 1 | 1 | 2 | 0 | NA | NA | 2 | 0 | 5 | 0.2 | NA | NA | 1 | 0 | 2 | 0.5 | NA | NA | 3 | 0.3333 |
| 7_80674960 | 2 | 0.5 | 2 | 0.5 | NA | NA | 2 | 0.5 | 6 | 0.5 | NA | NA | 1 | 1 | 2 | 1 | NA | NA | 3 | 1 |
| 7_80675100 | 2 | 1 | 1 | 1 | NA | NA | 3 | 0.6667 | 6 | 0.8333 | NA | NA | 1 | 1 | 2 | 1 | NA | NA | 3 | 1 |
| 7_80687229 | 1 | 0 | 2 | 0 | 1 | 1 | 3 | 0 | 7 | 0.1429 | NA | NA | 1 | 0 | 2 | 0 | 1 | 1 | 4 | 0.25 |
| 7_80691223 | 1 | 0 | 2 | 0.5 | 1 | 1 | 2 | 1 | 6 | 0.6667 | 1 | 1 | 3 | 0.6667 | 2 | 0 | NA | NA | 6 | 0.5 |
| 7_80691230 | 2 | 0.5 | 2 | 0 | 1 | 0 | 3 | 0.3333 | 8 | 0.25 | 1 | 0 | 5 | 0.4 | 3 | 0.3333 | 1 | 1 | 10 | 0.4 |
| 7_80691268 | 2 | 0.5 | 2 | 1 | 1 | 1 | 3 | 0.6667 | 8 | 0.75 | 1 | 1 | 5 | 0.8 | 3 | 1 | 1 | 1 | 10 | 0.9 |
| 7_80691382 | 2 | 0.5 | 1 | 0 | 1 | 0 | 3 | 0 | 7 | 0.1429 | 2 | 0.5 | 4 | 0 | 3 | 0 | 1 | 0 | 10 | 0.1 |
| 7_80691464 | 1 | 1 | 2 | 1 | 1 | 1 | 1 | 1 | 5 | 1 | 2 | 1 | 5 | 1 | 2 | 1 | 1 | 1 | 10 | 1 |
| 7_80691535 | 2 | 0.5 | 2 | 0 | 1 | 1 | 3 | 0 | 8 | 0.25 | 2 | 0 | 5 | 0.2 | 3 | 0.3333 | 1 | 1 | 11 | 0.2727 |
| 7_80691577 | 2 | 0 | 2 | 0 | 1 | 1 | 3 | 0.3333 | 8 | 0.25 | 2 | 0.5 | 6 | 0.1667 | 3 | 0.6667 | 1 | 1 | 12 | 0.4167 |
| 7_80691666 | 2 | 1 | 2 | 1 | 1 | 1 | 3 | 0.6667 | 8 | 0.875 | 2 | 0.5 | 6 | 0.1667 | 2 | 0.5 | 1 | 1 | 11 | 0.3636 |
| 7_80692145 | NA | NA | 3 | 0.3333 | 1 | 0 | 2 | 0.5 | 6 | 0.3333 | 1 | 1 | 1 | 1 | 3 | 0.6667 | NA | NA | 5 | 0.8 |
| 7_80983476 | 1 | 1 | 1 | 1 | 1 | 1 | 2 | 1 | 5 | 1 | NA | NA | 1 | 1 | 2 | 0.5 | 1 | 1 | 4 | 0.75 |
| 7_80997222 | 4 | 0 | 4 | 0 | 7 | 0 | 7 | 0.2857 | 22 | 0.0909 | 2 | 0.5 | 2 | 0 | 1 | 0 | 1 | 0 | 6 | 0.1667 |
| 7_80998093 | NA | NA | 3 | 0.3333 | 6 | 0 | 4 | 0.25 | 13 | 0.1538 | 1 | 1 | 2 | 0 | 1 | 0 | NA | NA | 4 | 0.25 |
| 7_81105317 | NA | NA | NA | NA | 2 | 0.5 | 1 | 1 | 3 | 0.6667 | NA | NA | NA | NA | 1 | 0 | 1 | 1 | 2 | 0.5 |
| 7_81105522 | NA | NA | NA | NA | 2 | 0.5 | 1 | 0 | 3 | 0.3333 | NA | NA | NA | NA | 1 | 1 | 1 | 0 | 2 | 0.5 |
| 7_81110399 | 17 | 0.0588 | 17 | 0.0588 | 18 | 0.0556 | 8 | 0 | 60 | 0.05 | 8 | 0 | 12 | 0.25 | 16 | 0 | 11 | 0 | 47 | 0.0638 |
| 7_81181052 | 2 | 0.5 | NA | NA | 1 | 1 | 1 | 0 | 4 | 0.5 | NA | NA | 1 | 1 | 2 | 0.5 | 1 | 1 | 4 | 0.75 |
| 7_81221455 | 1 | 0 | 1 | 0 | 1 | 1 | NA | NA | 3 | 0.3333 | NA | NA | 1 | 0 | NA | NA | 1 | 1 | 2 | 0.5 |
| 7_81243790 | 4 | 0.25 | 3 | 0 | 1 | 0 | NA | NA | 8 | 0.125 | 2 | 1 | 3 | 0 | NA | NA | 5 | 0 | 10 | 0.2 |
| 7_81245890 | 4 | 0.75 | 2 | 1 | NA | NA | NA | NA | 6 | 0.8333 | 2 | 1 | 3 | 0.6667 | 1 | 1 | 3 | 1 | 9 | 0.8889 |
| 7_81246059 | 3 | 0.3333 | 2 | 0.5 | 1 | 0 | NA | NA | 6 | 0.3333 | 1 | 0 | 1 | 1 | 1 | 1 | 4 | 0.75 | 7 | 0.7143 |
| 7_81246067 | 5 | 0.2 | 2 | 0 | 1 | 1 | NA | NA | 8 | 0.25 | 1 | 0 | 2 | 0 | 1 | 0 | 3 | 0.6667 | 7 | 0.2857 |
| 7_81246072 | 5 | 0.6 | 3 | 0.3333 | 1 | 0 | NA | NA | 9 | 0.4444 | 2 | 0.5 | 3 | 0.6667 | 1 | 1 | 4 | 0.5 | 10 | 0.6 |
| 7_81246185 | 4 | 0 | 2 | 0.5 | NA | NA | NA | NA | 6 | 0.1667 | 2 | 0 | 2 | 0 | 1 | 1 | 4 | 0 | 9 | 0.1111 |
| 7_81250449 | 1 | 0 | 4 | 0.25 | 3 | 0.3333 | 2 | 0 | 10 | 0.2 | 7 | 0.1429 | 3 | 0 | NA | NA | 3 | 0.3333 | 13 | 0.1538 |
| 7_81255423 | NA | NA | 2 | 0 | 4 | 0.25 | NA | NA | 6 | 0.1667 | 4 | 0 | 2 | 0 | NA | NA | 3 | 0.3333 | 9 | 0.1111 |
| 7_81255580 | NA | NA | 3 | 0 | 4 | 0.5 | NA | NA | 7 | 0.2857 | 3 | 0.3333 | 2 | 0 | NA | NA | 3 | 0.6667 | 8 | 0.375 |
| 7_81259835 | 1 | 1 | 4 | 0.75 | 1 | 0 | 1 | 0 | 7 | 0.5714 | 4 | 0.25 | NA | NA | NA | NA | NA | NA | 4 | 0.25 |
| 7_81418306 | 18 | 0.1667 | 14 | 0.0714 | 16 | 0 | 19 | 0.0526 | 67 | 0.0746 | 10 | 0 | 13 | 0.0769 | 14 | 0.0714 | 10 | 0.1 | 47 | 0.0638 |
| 7_81418334 | 17 | 0.4118 | 18 | 0.1667 | 17 | 0.2941 | 22 | 0.3182 | 74 | 0.2973 | 13 | 0.2308 | 12 | 0.3333 | 18 | 0.0556 | 11 | 0.1818 | 54 | 0.1852 |
| 7_81418436 | 19 | 0.0526 | 20 | 0.15 | 17 | 0 | 21 | 0.0952 | 77 | 0.0779 | 13 | 0.2308 | 13 | 0 | 16 | 0.0625 | 11 | 0.1818 | 53 | 0.1132 |
| 7_81418569 | 14 | 0.1429 | 16 | 0.1875 | 11 | 0 | 14 | 0.2857 | 55 | 0.1636 | 9 | 0.2222 | 9 | 0.2222 | 14 | 0.1429 | 10 | 0.1 | 42 | 0.1667 |
| 7_81418685 | 19 | 0.1053 | 20 | 0.3 | 16 | 0.4375 | 20 | 0.2 | 75 | 0.2533 | 12 | 0.3333 | 13 | 0.3077 | 18 | 0.3333 | 11 | 0.3636 | 54 | 0.3333 |
| 7_81430143 | 3 | 0.3333 | 2 | 0 | NA | NA | 2 | 0.5 | 7 | 0.2857 | 1 | 0 | NA | NA | 2 | 0.5 | 1 | 0 | 4 | 0.25 |
| 7_81430183 | 3 | 0.3333 | 2 | 0.5 | NA | NA | 2 | 0.5 | 7 | 0.4286 | 1 | 0 | NA | NA | 2 | 0.5 | 2 | 0.5 | 5 | 0.4 |
| 7_81432067 | NA | NA | 2 | 0 | NA | NA | 2 | 0.5 | 4 | 0.25 | 2 | 0.5 | 1 | 0 | NA | NA | 1 | 0 | 4 | 0.25 |
| 7_81442022 | 24 | 0.2917 | 27 | 0.4444 | 26 | 0.2692 | 16 | 0.25 | 93 | 0.3226 | 10 | 0.4 | 13 | 0.1538 | 25 | 0.28 | 20 | 0.25 | 68 | 0.2647 |
| 7_81442126 | 24 | 0.0417 | 26 | 0.0769 | 21 | 0.1429 | 15 | 0.0667 | 86 | 0.0814 | 13 | 0.0769 | 13 | 0.0769 | 23 | 0.087 | 19 | 0.1053 | 68 | 0.0882 |
| 7_81442419 | 20 | 0.15 | 25 | 0.04 | 20 | 0 | 14 | 0 | 79 | 0.0506 | 11 | 0.0909 | 13 | 0.0769 | 22 | 0.0455 | 15 | 0.0667 | 61 | 0.0656 |
| 7_81442652 | 17 | 0.4118 | 21 | 0.2381 | 18 | 0.2778 | 12 | 0.25 | 68 | 0.2941 | 10 | 0.2 | 12 | 0.4167 | 21 | 0.0952 | 14 | 0.3571 | 57 | 0.2456 |
| 7_81442917 | 15 | 0.0667 | 22 | 0.1818 | 15 | 0.1333 | 13 | 0.2308 | 65 | 0.1538 | 8 | 0.25 | 14 | 0 | 19 | 0 | 14 | 0.2857 | 55 | 0.1091 |
| 7_81443157 | 15 | 0.2 | 20 | 0.2 | 17 | 0.0588 | 12 | 0.1667 | 64 | 0.1562 | 10 | 0.1 | 12 | 0.1667 | 16 | 0.125 | 12 | 0.0833 | 50 | 0.12 |
| 7_81443346 | 12 | 0.3333 | 21 | 0.4762 | 15 | 0.4 | 14 | 0.6429 | 62 | 0.4677 | 12 | 0.75 | 10 | 0.7 | 16 | 0.5625 | 15 | 0.6 | 53 | 0.6415 |

| | | | | | | | | | | | | | | | | | | | | |
|---|---|---|---|---|---|---|---|---|---|---|---|---|---|---|---|---|---|---|---|---|
| 7_81443405 | 14 | 0.0714 | 22 | 0.0909 | 15 | 0.2 | 13 | 0.0769 | 64 | 0.1094 | 12 | 0.25 | 12 | 0 | 16 | 0.125 | 14 | 0.2857 | 54 | 0.1667 |
| 7_81443445 | 14 | 0.0714 | 22 | 0.1364 | 17 | 0.1765 | 13 | 0 | 66 | 0.1061 | 12 | 0.0833 | 12 | 0.1667 | 17 | 0.0588 | 13 | 0.2308 | 54 | 0.1296 |
| 7_81443989 | 11 | 0.1818 | 23 | 0.2609 | 16 | 0.0625 | 11 | 0.0909 | 61 | 0.1639 | 10 | 0.3 | 11 | 0.0909 | 16 | 0.1875 | 10 | 0.3 | 47 | 0.2128 |
| 7_81531278 | 1 | 0 | 7 | 0 | 8 | 0.25 | 10 | 0 | 26 | 0.0769 | 3 | 0.3333 | 7 | 0 | 4 | 0 | 4 | 0 | 18 | 0.0556 |
| 7_81531332 | 2 | 0 | 7 | 0.1429 | 10 | 0.2 | 9 | 0.1111 | 28 | 0.1429 | 3 | 0.6667 | 8 | 0 | 4 | 0 | 5 | 0.4 | 20 | 0.2 |
| 7_81531366 | 2 | 0 | 5 | 0 | 7 | 0.4286 | 7 | 0.4286 | 21 | 0.2857 | 3 | 0.3333 | 4 | 0.5 | 4 | 0 | 4 | 0 | 15 | 0.2 |
| 7_81531452 | 2 | 0 | 5 | 0.4 | 7 | 0.5714 | 6 | 0.1667 | 20 | 0.35 | 2 | 0.5 | 7 | 0.8571 | 4 | 0.25 | 3 | 0 | 16 | 0.5 |
| 7_81531463 | 1 | 0 | 5 | 0 | 8 | 0 | 6 | 0.1667 | 20 | 0.05 | 3 | 0 | 7 | 0.1429 | 3 | 0 | 5 | 0 | 18 | 0.0556 |
| 7_81531475 | 2 | 0 | 6 | 0 | 9 | 0.4444 | 9 | 0 | 26 | 0.1538 | 3 | 0 | 8 | 0.125 | 3 | 0 | 4 | 0.5 | 18 | 0.1667 |
| 7_81531646 | 1 | 0 | 5 | 0 | 8 | 0.125 | 6 | 0.1667 | 20 | 0.1 | 1 | 0 | 7 | 0.2857 | 3 | 0 | 3 | 0 | 14 | 0.1429 |
| 7_81533021 | NA | NA | 5 | 0 | 7 | 0.1429 | 5 | 0 | 17 | 0.0588 | 1 | 1 | 6 | 0 | 5 | 0 | 4 | 0 | 16 | 0.0625 |
| 7_81533128 | 1 | 0 | 3 | 0 | 3 | 0.3333 | 4 | 0.25 | 11 | 0.1818 | 5 | 0.2 | 1 | 1 | NA | NA | 1 | 0 | 7 | 0.2857 |
| 7_81533194 | 1 | 1 | 3 | 1 | 3 | 1 | 5 | 0.8 | 12 | 0.9167 | 3 | 1 | 1 | 1 | NA | NA | NA | NA | 4 | 1 |
| 7_81534305 | 8 | 0.25 | 5 | 0.4 | 12 | 0 | 8 | 0.25 | 33 | 0.1818 | 2 | 0 | 4 | 0 | 9 | 0.2222 | 4 | 0 | 19 | 0.1053 |
| 7_81534417 | 9 | 0.1111 | 7 | 0.2857 | 11 | 0.2727 | 8 | 0.375 | 35 | 0.2571 | 2 | 0 | 5 | 0.2 | 10 | 0.2 | 5 | 0.2 | 22 | 0.1818 |
| 7_81535050 | 7 | 0 | 9 | 0 | 10 | 0.3 | 6 | 0.1667 | 32 | 0.125 | 2 | 0 | 6 | 0.1667 | 10 | 0.3 | 5 | 0 | 23 | 0.1739 |
| 7_81535076 | 7 | 0 | 9 | 0 | 10 | 0.2 | 6 | 0.1667 | 32 | 0.0938 | 2 | 0 | 6 | 0.1667 | 9 | 0.1111 | 4 | 0 | 21 | 0.0952 |
| 7_81535183 | 7 | 0.4286 | 9 | 0.5556 | 9 | 0.6667 | 4 | 0.25 | 29 | 0.5172 | 2 | 0 | 7 | 0.2857 | 10 | 0.4 | 4 | 0.5 | 23 | 0.3478 |
| 7_81535629 | 6 | 0 | 8 | 0.25 | 8 | 0.25 | 6 | 0.5 | 28 | 0.25 | NA | NA | 5 | 0 | 10 | 0.2 | 5 | 0 | 20 | 0.1 |
| 7_81535652 | 5 | 0.6 | 7 | 0.8571 | 10 | 0.8 | 6 | 1 | 28 | 0.8214 | NA | NA | 5 | 0.4 | 10 | 0.6 | 4 | 1 | 19 | 0.6316 |
| 7_81535738 | 6 | 0 | 6 | 0.1667 | 9 | 0.2222 | 5 | 0 | 26 | 0.1154 | NA | NA | 5 | 0 | 10 | 0.3 | 3 | 0 | 18 | 0.1667 |
| 7_81956088 | 8 | 0.125 | 11 | 0.2727 | 12 | 0.25 | 13 | 0.0769 | 44 | 0.1818 | 5 | 0 | 14 | 0 | 13 | 0.3077 | 6 | 0 | 38 | 0.1053 |
| 7_81956121 | 7 | 0.1429 | 8 | 0 | 8 | 0.125 | 11 | 0.2727 | 34 | 0.1471 | 5 | 0.2 | 9 | 0.2222 | 8 | 0.25 | 5 | 0 | 27 | 0.1852 |
| 7_81956217 | 9 | 0.6667 | 14 | 0.6429 | 11 | 0.5455 | 11 | 0.7273 | 45 | 0.6444 | 6 | 0.3333 | 16 | 0.5 | 12 | 0.3333 | 6 | 0.6667 | 40 | 0.45 |
| 7_81956230 | 7 | 0.2857 | 10 | 0.3 | 6 | 0.5 | 7 | 0.7143 | 30 | 0.4333 | 3 | 0 | 13 | 0.2308 | 11 | 0.4545 | 6 | 0.6667 | 33 | 0.3636 |
| 7_82295495 | NA | NA | 1 | 1 | 2 | 0.5 | 1 | 0 | 4 | 0.5 | 1 | 1 | 1 | 0 | 1 | 0 | 1 | 1 | 4 | 0.5 |
| 7_82295602 | 1 | 0 | 1 | 1 | 2 | 0 | 1 | 0 | 5 | 0.2 | 1 | 1 | 1 | 1 | 1 | 0 | NA | NA | 3 | 0.6667 |
| 7_82295658 | 1 | 0 | 1 | 0 | 1 | 1 | 1 | 0 | 4 | 0.25 | 1 | 1 | 1 | 0 | 2 | 0 | NA | NA | 4 | 0.25 |
| 7_82411853 | NA | NA | 3 | 1 | NA | NA | 2 | 0 | 5 | 0.6 | 1 | 0 | 1 | 1 | NA | NA | NA | NA | 2 | 0.5 |
| 7_82412065 | NA | NA | 3 | 0.6667 | NA | NA | 1 | 1 | 4 | 0.75 | 1 | 1 | 1 | 1 | 2 | 1 | NA | NA | 4 | 1 |
| 7_82412146 | NA | NA | 1 | 1 | NA | NA | 2 | 0 | 3 | 0.3333 | 1 | 1 | 1 | 0 | 2 | 0.5 | NA | NA | 4 | 0.5 |
| 7_82412203 | NA | NA | 4 | 1 | NA | NA | 2 | 1 | 6 | 1 | 1 | 1 | 1 | 1 | 3 | 1 | NA | NA | 5 | 1 |
| 7_82412324 | NA | NA | 3 | 0 | NA | NA | 2 | 0.5 | 5 | 0.2 | NA | NA | 1 | 0 | 1 | 0 | 1 | 1 | 3 | 0.3333 |
| 7_82412473 | NA | NA | 4 | 0.25 | NA | NA | 3 | 0 | 7 | 0.1429 | 1 | 0 | 1 | 1 | 2 | 0 | 1 | 0 | 5 | 0.2 |
| 7_82412488 | NA | NA | 3 | 0.3333 | NA | NA | 3 | 0.6667 | 6 | 0.5 | 1 | 1 | 1 | 1 | 2 | 1 | 1 | 1 | 5 | 1 |
| 7_82412498 | NA | NA | 4 | 0.75 | NA | NA | 2 | 1 | 6 | 0.8333 | 1 | 1 | 1 | 1 | 2 | 0.5 | 1 | 0 | 5 | 0.6 |
| 7_82516958 | 1 | 1 | 1 | 1 | 2 | 0 | 1 | 1 | 5 | 0.6 | 1 | 1 | 1 | 0 | NA | NA | 2 | 0 | 4 | 0.25 |
| 7_82517014 | 1 | 1 | 2 | 0 | 2 | 0.5 | 1 | 0 | 6 | 0.3333 | 2 | 0 | 1 | 0 | NA | NA | 2 | 0.5 | 5 | 0.2 |
| 7_82517019 | 1 | 1 | 1 | 1 | NA | NA | 1 | 0 | 3 | 0.6667 | 2 | 0.5 | NA | NA | NA | NA | 2 | 0 | 4 | 0.25 |
| 7_82518184 | 1 | 1 | 2 | 1 | 2 | 1 | 2 | 1 | 7 | 1 | 3 | 0.6667 | 2 | 1 | 1 | 0 | 3 | 1 | 9 | 0.7778 |
| 7_82518188 | 1 | 0 | 2 | 0.5 | 2 | 0 | 2 | 0 | 7 | 0.1429 | 3 | 0 | 2 | 0.5 | 1 | 0 | 2 | 0 | 8 | 0.125 |
| 7_82518275 | 1 | 1 | 2 | 0 | 2 | 0.5 | 3 | 0 | 8 | 0.25 | 3 | 0 | 2 | 0.5 | 1 | 0 | 2 | 0 | 8 | 0.125 |
| 7_82518600 | NA | NA | 2 | 1 | 2 | 1 | 2 | 1 | 6 | 1 | 2 | 0.5 | 2 | 1 | 1 | 1 | 1 | 1 | 6 | 0.8333 |
| 7_82518620 | 1 | 0 | 2 | 0.5 | 1 | 0 | 3 | 0 | 7 | 0.1429 | 2 | 0 | 1 | 1 | NA | NA | 1 | 0 | 4 | 0.25 |
| 7_82518657 | 2 | 1 | 2 | 1 | 2 | 1 | 2 | 1 | 8 | 1 | 4 | 1 | 1 | 1 | 1 | 1 | 4 | 1 | 10 | 1 |
| 7_82518702 | 2 | 0.5 | 2 | 0 | 2 | 0 | 2 | 0.5 | 8 | 0.25 | 3 | 0.3333 | 1 | 0 | NA | NA | 1 | 0 | 5 | 0.2 |
| 7_82518821 | 2 | 1 | 2 | 0.5 | 2 | 1 | 2 | 0.5 | 8 | 0.75 | 3 | 0.6667 | 3 | 0.6667 | NA | NA | 4 | 0.75 | 10 | 0.7 |
| 7_82519173 | 2 | 1 | 3 | 1 | 2 | 1 | 2 | 1 | 9 | 1 | 4 | 1 | 3 | 1 | 1 | 1 | 3 | 1 | 11 | 1 |
| 7_82519609 | 1 | 1 | 4 | 1 | 4 | 1 | 2 | 1 | 11 | 1 | 3 | 1 | 2 | 0.5 | NA | NA | 1 | 1 | 6 | 0.8333 |
| 7_82519785 | 1 | 1 | 3 | 1 | 4 | 1 | 2 | 1 | 10 | 1 | 3 | 0.6667 | 1 | 1 | 1 | 1 | 3 | 0.6667 | 8 | 0.75 |
| 7_82519831 | 1 | 1 | 3 | 0.6667 | 4 | 1 | 3 | 1 | 11 | 0.9091 | 4 | 1 | 2 | 1 | 1 | 1 | 2 | 1 | 9 | 1 |
| 7_82519942 | 2 | 0 | 4 | 0.25 | 4 | 0 | 3 | 0 | 13 | 0.0769 | 4 | 0 | 3 | 0.3333 | 1 | 0 | 5 | 0.2 | 13 | 0.1538 |
| 7_82519967 | 2 | 0 | 3 | 0 | 4 | 0.25 | 2 | 0 | 11 | 0.0909 | 3 | 0 | 3 | 0 | 1 | 0 | 4 | 0.25 | 11 | 0.0909 |
| 7_82520127 | 2 | 0 | 3 | 0 | 3 | 0 | 3 | 0.3333 | 11 | 0.0909 | 4 | 0.25 | 3 | 0 | 1 | 0 | 5 | 0.2 | 13 | 0.1538 |
| 7_83273407 | NA | NA | NA | NA | 1 | 1 | 1 | 1 | 2 | 1 | NA | NA | NA | NA | NA | NA | 1 | 1 | 1 | 1 |
| 7_83273425 | NA | NA | NA | NA | 1 | 0 | 1 | 1 | 2 | 0.5 | NA | NA | 1 | 1 | NA | NA | 1 | 1 | 2 | 1 |
| 7_83273487 | NA | NA | NA | NA | 1 | 1 | 1 | 1 | 2 | 1 | NA | NA | 1 | 1 | NA | NA | 1 | 0 | 2 | 0.5 |
| 7_83273504 | NA | NA | NA | NA | NA | NA | 1 | 1 | 1 | 1 | NA | NA | 1 | 1 | NA | NA | 1 | 1 | 2 | 1 |
| 7_83273569 | NA | NA | NA | NA | 1 | 1 | 1 | 1 | 2 | 1 | NA | NA | 1 | 1 | NA | NA | 1 | 1 | 2 | 1 |
| 7_83289704 | 3 | 0.3333 | 3 | 0 | NA | NA | 6 | 0.3333 | 12 | 0.25 | 1 | 0 | 5 | 0.2 | 2 | 0.5 | 4 | 0.25 | 12 | 0.25 |
| 7_83289818 | 2 | 0 | 3 | 0.3333 | NA | NA | 6 | 0.1667 | 11 | 0.1818 | 1 | 0 | 5 | 0.6 | 2 | 0.5 | 4 | 0.25 | 12 | 0.4167 |
| 7_83289842 | 3 | 0.6667 | 3 | 1 | NA | NA | 6 | 0.5 | 12 | 0.6667 | 1 | 1 | 4 | 0.75 | 2 | 1 | 4 | 0.5 | 11 | 0.7273 |
| 7_83290240 | 3 | 0.6667 | 3 | 0.6667 | NA | NA | 4 | 0.75 | 10 | 0.7 | 1 | 0 | 5 | 0.4 | 4 | 0.25 | 4 | 0 | 14 | 0.2143 |
| 7_83292962 | 1 | 1 | 1 | 0 | 1 | 1 | NA | NA | 3 | 0.6667 | NA | NA | 1 | 1 | NA | NA | NA | NA | 1 | 1 |
| 7_83301407 | 1 | 1 | 2 | 1 | NA | NA | NA | NA | 3 | 1 | NA | NA | 1 | 1 | NA | NA | NA | NA | 1 | 1 |

| | | | | | | | | | | | | | | | | | | | | |
|---|---|---|---|---|---|---|---|---|---|---|---|---|---|---|---|---|---|---|---|---|
| 7_8330165 | 1 | 0 | 2 | 1 | NA | NA | NA | NA | 3 | 0.6667 | NA | NA | 1 | 1 | NA | NA | NA | NA | 1 | 1 |
| 7_8340678 | 5 | 0.2 | 2 | 0 | 6 | 0 | 3 | 0.3333 | 16 | 0.125 | 3 | 0 | 2 | 0.5 | 2 | 0 | 1 | 1 | 8 | 0.25 |
| 7_8348388 | NA | NA | NA | NA | NA | NA | 1 | 1 | 1 | 1 | NA | NA | 1 | 1 | 1 | 1 | NA | NA | 2 | 1 |
| 7_8348389 | NA | NA | NA | NA | 1 | 1 | 1 | 0 | 2 | 0.5 | NA | NA | 1 | 0 | 1 | 1 | NA | NA | 2 | 0.5 |
| 7_8348390 | NA | NA | NA | NA | 1 | 1 | 1 | 0 | 2 | 0.5 | NA | NA | 1 | 1 | 2 | 0.5 | NA | NA | 3 | 0.6667 |
| 7_8348423 | NA | NA | NA | NA | 1 | 0 | 1 | 1 | 2 | 0.5 | NA | NA | 1 | 0 | 3 | 0.3333 | NA | NA | 4 | 0.25 |
| 7_8348643 | NA | NA | NA | NA | 1 | 1 | 1 | 1 | 2 | 1 | NA | NA | 1 | 1 | 3 | 0.3333 | NA | NA | 4 | 0.5 |
| 7_8348660 | 1 | 1 | NA | NA | 1 | 1 | 1 | 1 | 3 | 1 | NA | NA | NA | NA | 2 | 0.5 | NA | NA | 2 | 0.5 |
| 7_8348690 | 1 | 1 | NA | NA | 1 | 1 | 1 | 1 | 3 | 1 | NA | NA | 1 | 1 | 3 | 1 | NA | NA | 4 | 1 |
| 7_8348707 | NA | NA | NA | NA | 1 | 1 | 1 | 1 | 2 | 1 | NA | NA | 1 | 1 | 3 | 1 | NA | NA | 4 | 1 |
| 7_8348708 | 1 | 1 | NA | NA | 1 | 1 | 1 | 1 | 3 | 1 | NA | NA | 1 | 1 | 3 | 1 | NA | NA | 4 | 1 |
| 7_8348712 | 1 | 1 | NA | NA | 1 | 0 | 1 | 0 | 3 | 0.3333 | NA | NA | 1 | 0 | 1 | 1 | NA | NA | 2 | 0.5 |
| 7_8348720 | 1 | 0 | NA | NA | 1 | 1 | 1 | 0 | 3 | 0.3333 | NA | NA | 2 | 0.5 | 3 | 0.3333 | NA | NA | 5 | 0.4 |
| 7_8348738 | 1 | 1 | NA | NA | 1 | 0 | 1 | 0 | 3 | 0.3333 | NA | NA | 2 | 1 | 3 | 0.6667 | NA | NA | 5 | 0.8 |
| 7_8353112 | 4 | 0.5 | 5 | 0.4 | 1 | 0 | 10 | 0.3 | 20 | 0.35 | 1 | 0 | 10 | 0.7 | 12 | 0.6667 | 6 | 0.5 | 29 | 0.6207 |
| 7_8353127 | 7 | 0.1429 | 5 | 0 | 3 | 0.3333 | 9 | 0 | 24 | 0.0833 | 3 | 0.3333 | 9 | 0 | 12 | 0.25 | 7 | 0.1429 | 31 | 0.1613 |
| 7_8353137 | 5 | 1 | 4 | 1 | 2 | 1 | 8 | 0.875 | 19 | 0.9474 | 2 | 1 | 9 | 1 | 9 | 1 | 4 | 1 | 24 | 1 |
| 7_8353144 | 6 | 0.3333 | 5 | 0.2 | 2 | 0.5 | 7 | 0.1429 | 20 | 0.25 | 1 | 0 | 6 | 0.3333 | 11 | 0.3636 | 5 | 0.2 | 23 | 0.3043 |
| 7_8353146 | 6 | 1 | 5 | 1 | 1 | 1 | 7 | 1 | 19 | 1 | 1 | 1 | 7 | 1 | 9 | 1 | 4 | 0.75 | 21 | 0.9524 |
| 7_8353179 | 3 | 1 | 4 | 0.75 | NA | NA | 3 | 0.6667 | 10 | 0.8 | 1 | 1 | 5 | 0.2 | 4 | 1 | 3 | 0.6667 | 13 | 0.6154 |
| 7_8353181 | 5 | 0.6 | 6 | 1 | 2 | 0.5 | 8 | 0.5 | 21 | 0.6667 | 1 | 0 | 3 | 0.3333 | 10 | 0.8 | 6 | 0.6667 | 20 | 0.65 |
| 7_8353190 | 5 | 0.8 | 5 | 1 | 2 | 1 | 8 | 0.875 | 20 | 0.9 | 1 | 1 | 4 | 1 | 9 | 0.8889 | 6 | 0.8333 | 20 | 0.9 |
| 7_8353194 | 5 | 0.4 | 5 | 0.2 | 2 | 0 | 6 | 0 | 18 | 0.1667 | NA | NA | 5 | 0 | 8 | 0.125 | 6 | 0.1667 | 19 | 0.1053 |
| 7_8353198 | 5 | 1 | 5 | 0.8 | 2 | 1 | 9 | 0.6667 | 21 | 0.8095 | 1 | 0 | 7 | 0.2857 | 11 | 0.9091 | 6 | 0.5 | 25 | 0.6 |
| 7_8353208 | 4 | 1 | 5 | 1 | 1 | 1 | 7 | 0.7143 | 17 | 0.8824 | 1 | 1 | 6 | 1 | 9 | 1 | 4 | 1 | 20 | 1 |
| 7_8353210 | 4 | 0.75 | 3 | 0 | 1 | 1 | 6 | 0.1667 | 14 | 0.3571 | 1 | 1 | 3 | 1 | 9 | 0.1111 | 4 | 0.25 | 17 | 0.3529 |
| 7_8353219 | 4 | 0.25 | 6 | 0.3333 | 2 | 0 | 6 | 0 | 18 | 0.1667 | 1 | 0 | 5 | 0.4 | 8 | 0.375 | 6 | 0 | 20 | 0.25 |
| 7_8353267 | 3 | 0.3333 | 6 | 0 | 2 | 0.5 | 5 | 0.4 | 16 | 0.25 | NA | NA | 5 | 0.4 | 7 | 0.2857 | 6 | 0.5 | 18 | 0.3889 |
| 7_8353270 | 3 | 0.6667 | 5 | 1 | 1 | 1 | 5 | 0.8 | 14 | 0.8571 | NA | NA | 5 | 1 | 6 | 0.8333 | 4 | 0.75 | 15 | 0.8667 |
| 7_8353279 | 4 | 0 | 6 | 0.1667 | 2 | 0 | 5 | 0.6 | 17 | 0.2353 | NA | NA | 6 | 0 | 6 | 0.3333 | 6 | 0.1667 | 18 | 0.1667 |
| 7_8354131 | NA | NA | 3 | 0.6667 | 1 | 1 | NA | NA | 4 | 0.75 | NA | NA | 1 | 0 | 1 | 1 | NA | NA | 2 | 0.5 |
| 7_8354142 | 2 | 0.5 | 4 | 0.25 | 1 | 0 | NA | NA | 7 | 0.2857 | NA | NA | 2 | 0.5 | 1 | 1 | NA | NA | 3 | 0.6667 |
| 7_8354163 | 8 | 0.125 | 12 | 0.0833 | 7 | 0 | 4 | 0 | 31 | 0.0645 | 4 | 0 | 2 | 0 | 5 | 0 | 3 | 0.3333 | 14 | 0.0714 |
| 7_8354193 | NA | NA | NA | NA | 1 | 1 | NA | NA | 1 | 1 | NA | NA | NA | NA | 1 | 1 | NA | NA | 1 | 1 |
| 7_8354702 | 10 | 0.8 | 10 | 1 | 7 | 1 | 4 | 0.75 | 31 | 0.9032 | 6 | 1 | 3 | 1 | 8 | 1 | 4 | 0.75 | 21 | 0.9524 |
| 7_8354707 | 8 | 0.75 | 8 | 0.75 | 6 | 0.8333 | 5 | 0.4 | 27 | 0.7037 | 5 | 1 | 3 | 1 | 8 | 1 | 3 | 0.3333 | 19 | 0.8947 |
| 7_8354724 | 11 | 0.4545 | 12 | 0.5 | 6 | 0 | 5 | 0.4 | 34 | 0.3824 | 6 | 0.3333 | 3 | 0 | 9 | 0.2222 | 4 | 0 | 22 | 0.1818 |
| 7_8354726 | 11 | 0.5455 | 12 | 0.75 | 7 | 0.4286 | 5 | 0.4 | 35 | 0.5714 | 6 | 0.5 | 3 | 0.6667 | 8 | 0.625 | 4 | 0.25 | 21 | 0.5238 |
| 7_8354728 | 10 | 0.4 | 9 | 0.2222 | 6 | 0.3333 | 4 | 0.25 | 29 | 0.3103 | 6 | 0 | 3 | 0.3333 | 8 | 0.125 | 5 | 0 | 22 | 0.0909 |
| 7_8354730 | 10 | 0.7 | 11 | 0.4545 | 7 | 0.5714 | 4 | 0.75 | 32 | 0.5938 | 5 | 0.4 | 3 | 0.3333 | 8 | 0.375 | 5 | 0.4 | 21 | 0.381 |
| 7_8354741 | 10 | 0.9 | 11 | 1 | 7 | 1 | 4 | 1 | 32 | 0.9688 | 6 | 1 | 2 | 1 | 9 | 1 | 5 | 0.8 | 22 | 0.9545 |
| 7_8354744 | 12 | 0.6667 | 11 | 0.5455 | 7 | 0.2857 | 4 | 0.25 | 34 | 0.5 | 6 | 0.8333 | 3 | 0.6667 | 9 | 0.5556 | 5 | 0.8 | 23 | 0.6957 |
| 7_8354745 | 13 | 0.2308 | 12 | 0.1667 | 5 | 0 | 5 | 0 | 35 | 0.1429 | 6 | 0.3333 | 3 | 0 | 9 | 0.2222 | 5 | 0 | 23 | 0.1739 |
| 7_8354760 | 14 | 0.2143 | 12 | 0.25 | 6 | 0 | 5 | 0.2 | 37 | 0.1892 | 6 | 0.1667 | 3 | 0 | 9 | 0.3333 | 6 | 0.3333 | 24 | 0.25 |
| 7_8354764 | 10 | 0.1 | 10 | 0.2 | 8 | 0.125 | 3 | 0.3333 | 31 | 0.1613 | 4 | 0 | 3 | 0.3333 | 6 | 0.3333 | 4 | 0 | 17 | 0.1765 |
| 7_8354772 | 11 | 0.4545 | 13 | 0.1538 | 6 | 0 | 6 | 0.1667 | 36 | 0.2222 | 3 | 0 | 2 | 0 | 8 | 0.125 | 6 | 0 | 19 | 0.0526 |
| 7_8354776 | 11 | 0.1818 | 14 | 0.0714 | 8 | 0.25 | 4 | 0.25 | 37 | 0.1622 | 5 | 0.4 | 2 | 0 | 10 | 0.1 | 2 | 0 | 19 | 0.1579 |
| 7_8354780 | 9 | 1 | 14 | 0.7143 | 5 | 0.8 | 4 | 0.75 | 32 | 0.8125 | 3 | 0.3333 | 1 | 1 | 5 | 0.6 | 6 | 0.5 | 15 | 0.5333 |
| 7_8354785 | 12 | 0.0833 | 14 | 0 | 8 | 0.125 | 4 | 0 | 38 | 0.0526 | 6 | 0 | 3 | 0 | 10 | 0.2 | 4 | 0 | 23 | 0.087 |
| 7_8354800 | 14 | 0.4286 | 13 | 0.3077 | 9 | 0.1111 | 6 | 0.5 | 42 | 0.3333 | 6 | 0.5 | 3 | 0.6667 | 11 | 0.5455 | 5 | 0.8 | 25 | 0.6 |
| 7_8354803 | 12 | 0.5833 | 13 | 0.9231 | 7 | 0.7143 | 5 | 0.8 | 37 | 0.7568 | 6 | 0.6667 | 3 | 0.3333 | 10 | 0.7 | 7 | 0.5714 | 26 | 0.6154 |
| 7_8354808 | 13 | 0 | 14 | 0 | 9 | 0.1111 | 4 | 0.25 | 40 | 0.05 | 5 | 0 | 3 | 0 | 11 | 0.1818 | 6 | 0 | 25 | 0.08 |
| 7_8354816 | 9 | 0.7778 | 10 | 1 | 6 | 0.8333 | 2 | 1 | 27 | 0.8889 | 3 | 1 | 3 | 1 | 7 | 1 | 4 | 0.5 | 17 | 0.8824 |
| 7_8354819 | 12 | 0.9167 | 14 | 0.9286 | 6 | 1 | 4 | 1 | 36 | 0.9444 | 5 | 1 | 3 | 1 | 10 | 0.9 | 6 | 0.8333 | 24 | 0.9167 |
| 7_8375902 | 7 | 0.1429 | 9 | 0.1111 | 2 | 0 | 12 | 0 | 30 | 0.0667 | 7 | 0 | 4 | 0 | 10 | 0.2 | 13 | 0.0769 | 34 | 0.0882 |
| 7_8375928 | 6 | 0.3333 | 10 | 0.1 | 1 | 0 | 11 | 0.2727 | 28 | 0.2143 | 8 | 0.375 | 3 | 0.3333 | 7 | 0.2857 | 11 | 0.2727 | 29 | 0.3103 |
| 7_8375934 | 6 | 0 | 11 | 0.1818 | 1 | 0 | 11 | 0.0909 | 29 | 0.1034 | 8 | 0.25 | 3 | 0.3333 | 10 | 0.2 | 9 | 0 | 30 | 0.1667 |
| 7_8375936 | 5 | 0.2 | 9 | 0.4444 | 1 | 0 | 11 | 0.3636 | 26 | 0.3462 | 6 | 0.5 | 3 | 0.6667 | 10 | 0.2 | 9 | 0.6667 | 28 | 0.4643 |
| 7_8375953 | 6 | 0.5 | 10 | 0.4 | 1 | 1 | 11 | 0 | 28 | 0.2857 | 8 | 0.25 | 3 | 0.6667 | 7 | 0.2857 | 7 | 0.2857 | 25 | 0.32 |
| 7_8375955 | 4 | 1 | 8 | 0.375 | 1 | 1 | 9 | 0.7778 | 22 | 0.6818 | 7 | 0.8571 | 2 | 1 | 7 | 0.7143 | 8 | 1 | 24 | 0.875 |
| 7_8375972 | 4 | 0.25 | 10 | 0.6 | 1 | 1 | 6 | 0.3333 | 21 | 0.4762 | 6 | 0.6667 | 2 | 1 | 9 | 0.5556 | 6 | 0.8333 | 23 | 0.6957 |
| 7_8375982 | 5 | 0 | 9 | 0.2222 | 1 | 0 | 7 | 0.1429 | 22 | 0.1364 | 5 | 0 | 3 | 0.3333 | 9 | 0 | 6 | 0.1667 | 23 | 0.087 |
| 7_8376087 | NA | NA | 1 | 1 | NA | NA | NA | NA | 1 | 1 | NA | NA | NA | NA | 1 | 1 | NA | NA | 1 | 1 |
| 7_8380064 | 4 | 0.5 | 6 | 0 | NA | NA | 7 | 0.2857 | 17 | 0.2353 | 4 | 0.25 | 3 | 0.3333 | 5 | 0.2 | 6 | 0.3333 | 18 | 0.2778 |
| 7_8380077 | 3 | 0 | 6 | 0 | NA | NA | 6 | 0.1667 | 15 | 0.0667 | 4 | 0 | 2 | 0 | 5 | 0.2 | 5 | 0 | 16 | 0.0625 |

| | | | | | | | | | | | | | | | | | | | | |
|---|---|---|---|---|---|---|---|---|---|---|---|---|---|---|---|---|---|---|---|---|
| 7_8389562 | 21 | 0.0476 | 21 | 0.0476 | 20 | 0.1 | 16 | 0.0625 | 78 | 0.0641 | 18 | 0.0556 | 18 | 0 | 19 | 0.1053 | 17 | 0.1176 | 72 | 0.0694 |
| 7_8389690 | 18 | 0 | 13 | 0.2308 | 13 | 0 | 15 | 0.2 | 59 | 0.1017 | 13 | 0 | 14 | 0.1429 | 16 | 0.3125 | 11 | 0.0909 | 54 | 0.1481 |
| 7_8389899 | 4 | 0.75 | 7 | 0.5714 | 5 | 0.4 | 7 | 0.5714 | 23 | 0.5652 | 4 | 0.75 | 4 | 0.75 | 9 | 0.6667 | 3 | 0.3333 | 20 | 0.65 |
| 7_8389912 | 3 | 0.6667 | 5 | 0.2 | 4 | 0.25 | 5 | 0 | 17 | 0.2353 | 3 | 0.3333 | 2 | 0 | 8 | 0.25 | 3 | 0.3333 | 16 | 0.25 |
| 7_8389930 | 4 | 0 | 6 | 0 | 4 | 0.25 | 6 | 0.5 | 20 | 0.2 | 3 | 0.3333 | 1 | 0 | 7 | 0.5714 | 4 | 0.25 | 15 | 0.4 |
| 7_8389932 | 4 | 0 | 6 | 0 | 4 | 0.25 | 5 | 0.2 | 19 | 0.1053 | 3 | 0.3333 | 2 | 0 | 8 | 0 | 4 | 0 | 17 | 0.0588 |
| 7_8422615 | NA | NA | 2 | 1 | NA | NA | 3 | 0.3333 | 5 | 0.6 | 2 | 0 | 1 | 1 | NA | NA | 1 | 1 | 4 | 0.5 |
| 7_8422624 | 1 | 1 | 3 | 1 | NA | NA | 2 | 0 | 6 | 0.6667 | 2 | 0.5 | 1 | 1 | NA | NA | 1 | 1 | 4 | 0.75 |
| 7_8422632 | 1 | 0 | 3 | 0.6667 | NA | NA | 3 | 1 | 7 | 0.7143 | 2 | 0 | 1 | 1 | NA | NA | 1 | 1 | 4 | 0.5 |
| 7_8422637 | 1 | 1 | 3 | 0.3333 | NA | NA | 3 | 0.3333 | 7 | 0.4286 | 2 | 0 | 1 | 0 | NA | NA | 1 | 1 | 4 | 0.25 |
| 7_8422642 | 1 | 0 | 2 | 1 | NA | NA | 3 | 1 | 6 | 0.8333 | 2 | 1 | NA | NA | NA | NA | 1 | 1 | 3 | 1 |
| 7_8422655 | NA | NA | 3 | 0.6667 | NA | NA | 3 | 0 | 6 | 0.3333 | 2 | 0 | 1 | 1 | NA | NA | 1 | 0 | 4 | 0.25 |
| 7_8422660 | 1 | 1 | 3 | 0.3333 | NA | NA | 3 | 1 | 7 | 0.7143 | 2 | 1 | NA | NA | NA | NA | 1 | 0 | 3 | 0.6667 |
| 7_8422665 | 1 | 1 | 2 | 0 | NA | NA | 3 | 0.3333 | 6 | 0.3333 | 2 | 1 | NA | NA | NA | NA | 1 | 0 | 3 | 0.6667 |
| 7_8422670 | 1 | 0 | 3 | 0.3333 | NA | NA | 3 | 0.3333 | 7 | 0.2857 | 2 | 0.5 | 1 | 0 | NA | NA | 1 | 1 | 4 | 0.5 |
| 7_8422673 | 1 | 1 | 2 | 0 | NA | NA | NA | NA | 3 | 0.3333 | 1 | 1 | 1 | 1 | NA | NA | 1 | 0 | 3 | 0.6667 |
| 7_8423009 | 1 | 1 | NA | NA | NA | NA | NA | NA | 1 | 1 | 1 | 1 | NA | NA | NA | NA | NA | NA | 1 | 1 |
| 7_8423246 | 1 | 1 | 1 | 1 | 1 | 1 | NA | NA | 3 | 1 | 1 | 1 | NA | NA | NA | NA | NA | NA | 1 | 1 |
| 7_8426485 | 7 | 0 | 7 | 0.1429 | 5 | 0.2 | 10 | 0.2 | 29 | 0.1379 | 5 | 0.2 | 2 | 0 | 7 | 0 | 4 | 0.25 | 18 | 0.1111 |
| 7_8426486 | 8 | 0 | 8 | 0.375 | 5 | 0.2 | 12 | 0.25 | 33 | 0.2121 | 5 | 0.2 | 5 | 0.4 | 8 | 0.5 | 7 | 0 | 25 | 0.28 |
| 7_8426497 | 9 | 0.2222 | 7 | 0.4286 | 4 | 0.5 | 10 | 0.4 | 30 | 0.3667 | 3 | 0 | 5 | 0.6 | 8 | 0.25 | 4 | 0.75 | 20 | 0.4 |
| 7_8426502 | 9 | 0 | 8 | 0 | 6 | 0.1667 | 11 | 0.2727 | 34 | 0.1176 | 5 | 0 | 5 | 0 | 7 | 0.4286 | 6 | 0 | 23 | 0.1304 |
| 7_8636883 | 1 | 1 | NA | NA | NA | NA | 1 | 0 | 2 | 0.5 | NA | NA | NA | NA | 1 | 1 | 1 | 0 | 2 | 0.5 |
| 7_8636888 | 1 | 0 | NA | NA | NA | NA | 1 | 1 | 2 | 0.25 | NA | NA | NA | NA | 1 | 1 | 1 | 1 | 2 | 1 |
| 7_8767580 | 2 | 0 | 3 | 0 | 4 | 0.5 | 3 | 0.3333 | 12 | 0.25 | 1 | 0 | 2 | 0 | 5 | 0.2 | 7 | 0 | 15 | 0.0667 |
| 7_8772352 | 3 | 0.6667 | 6 | 0.6667 | 4 | 0.75 | 5 | 0.8 | 18 | 0.7222 | 2 | 1 | 3 | 0.6667 | 6 | 1 | 10 | 0.7 | 21 | 0.8095 |
| 7_8772355 | 3 | 0 | 4 | 0 | 3 | 0 | 6 | 0.5 | 16 | 0.1875 | 1 | 0 | 2 | 1 | 4 | 0.75 | 9 | 0.1111 | 16 | 0.375 |
| 7_8772387 | 3 | 0.3333 | 4 | 0.5 | 4 | 0.75 | 7 | 0.4286 | 18 | 0.5 | 1 | 1 | 3 | 0.6667 | 6 | 0.6667 | 11 | 0.8182 | 21 | 0.7619 |
| 7_8772398 | 3 | 0 | 6 | 0.1667 | 4 | 0.25 | 8 | 0.25 | 21 | 0.1905 | 2 | 0 | 4 | 0.5 | 7 | 0 | 12 | 0 | 25 | 0.08 |
| 7_8777995 | 5 | 1 | 6 | 1 | 8 | 1 | 8 | 0.875 | 27 | 0.963 | 3 | 1 | 8 | 1 | 12 | 1 | 14 | 1 | 37 | 1 |
| 7_8777995 | 3 | 1 | 7 | 0.8571 | 7 | 1 | 8 | 1 | 25 | 0.96 | 3 | 1 | 7 | 1 | 11 | 0.9091 | 13 | 0.9231 | 34 | 0.9412 |
| 7_8777998 | 6 | 0.3333 | 7 | 0.4286 | 9 | 0.2222 | 11 | 0.1818 | 33 | 0.2727 | 4 | 0 | 8 | 0.125 | 11 | 0.1818 | 15 | 0.2 | 38 | 0.1579 |
| 7_8778003 | 4 | 0.75 | 7 | 0.4286 | 8 | 0.375 | 10 | 0.6 | 29 | 0.5172 | 4 | 0.75 | 5 | 0.4 | 12 | 0.9167 | 16 | 0.6875 | 37 | 0.7297 |
| 7_8778010 | 5 | 0.8 | 2 | 0.5 | 6 | 0.3333 | 5 | 0.6 | 18 | 0.5556 | 2 | 0.5 | 7 | 0.8571 | 8 | 0.75 | 10 | 0.7 | 27 | 0.7407 |
| 7_8778011 | 7 | 0.4286 | 8 | 0.625 | 9 | 0.3333 | 11 | 0.0909 | 35 | 0.3429 | 4 | 0.25 | 9 | 0.1111 | 17 | 0.1176 | 19 | 0.2105 | 49 | 0.1633 |
| 7_8778016 | 6 | 0.3333 | 6 | 0.3333 | 6 | 0.6667 | 11 | 0.2727 | 29 | 0.3793 | 3 | 0.6667 | 9 | 0.6667 | 13 | 0.3846 | 16 | 0.625 | 41 | 0.561 |
| 7_8778020 | 9 | 0.2222 | 7 | 0.4286 | 9 | 0.5556 | 11 | 0.5455 | 36 | 0.4444 | 4 | 0.25 | 11 | 0.2727 | 19 | 0.5789 | 19 | 0.4737 | 53 | 0.4528 |
| 7_8778029 | 9 | 0 | 8 | 0 | 8 | 0 | 9 | 0.4444 | 34 | 0.1176 | 3 | 0 | 9 | 0.1111 | 19 | 0.0526 | 18 | 0.0556 | 49 | 0.0612 |
| 7_8778031 | 9 | 1 | 6 | 0.8333 | 9 | 0.4444 | 10 | 0.8 | 34 | 0.7647 | 4 | 1 | 11 | 0.7273 | 16 | 0.875 | 18 | 0.7778 | 49 | 0.8163 |
| 7_8778034 | 11 | 0 | 8 | 0 | 9 | 0 | 11 | 0.1818 | 39 | 0.0513 | 4 | 0 | 12 | 0.0833 | 17 | 0.1176 | 17 | 0 | 50 | 0.06 |
| 7_8778041 | 3 | 0 | 4 | 0.5 | 5 | 0.4 | 1 | 1 | 13 | 0.3846 | 1 | 1 | 2 | 0 | 3 | 0.6667 | 5 | 0.4 | 11 | 0.4545 |
| 7_8778044 | 11 | 1 | 7 | 1 | 9 | 1 | 11 | 1 | 38 | 1 | 4 | 1 | 10 | 0.9 | 19 | 1 | 20 | 0.95 | 53 | 0.9623 |
| 7_8778050 | 10 | 0 | 7 | 0.2857 | 8 | 0.375 | 11 | 0.2727 | 36 | 0.2222 | 4 | 0.25 | 8 | 0.25 | 15 | 0.2 | 14 | 0.1429 | 41 | 0.1951 |
| 7_8778052 | 11 | 0 | 8 | 0.25 | 5 | 0 | 8 | 0.125 | 32 | 0.0938 | 2 | 0 | 9 | 0.1111 | 14 | 0 | 11 | 0.0909 | 36 | 0.0556 |
| 7_8778052 | 7 | 0.2857 | 5 | 0.2 | 4 | 0.25 | 7 | 0.4286 | 23 | 0.3043 | 3 | 0.6667 | 8 | 0.5 | 13 | 0.6923 | 13 | 0.6154 | 37 | 0.6216 |
| 7_8778054 | 9 | 0.8889 | 7 | 0.7143 | 8 | 0.375 | 9 | 0.7778 | 33 | 0.697 | 4 | 0.5 | 9 | 1 | 17 | 0.8235 | 18 | 0.8889 | 48 | 0.8542 |
| 7_8778072 | 10 | 0.4 | 7 | 0 | 7 | 0 | 11 | 0.1818 | 35 | 0.1714 | 4 | 0.25 | 10 | 0.2 | 16 | 0.3125 | 18 | 0.2222 | 48 | 0.25 |
| 7_8778074 | 12 | 0.8333 | 8 | 1 | 10 | 0.9 | 13 | 0.9231 | 43 | 0.907 | 4 | 0.75 | 11 | 0.9091 | 15 | 0.9333 | 20 | 1 | 50 | 0.94 |
| 7_8778082 | 7 | 1 | 8 | 0.75 | 13 | 1 | 13 | 0.9231 | 41 | 0.9268 | 5 | 1 | 11 | 0.9091 | 19 | 1 | 18 | 1 | 53 | 0.9811 |
| 7_8778092 | 7 | 0.4286 | 7 | 0.4286 | 8 | 0.625 | 10 | 0.6 | 32 | 0.5312 | 4 | 0.75 | 11 | 0.2727 | 20 | 0.5 | 17 | 0.4118 | 52 | 0.4423 |
| 7_8778122 | 11 | 0.7273 | 10 | 0.2 | 11 | 0.3636 | 15 | 0.2 | 47 | 0.3617 | 6 | 0 | 14 | 0.4286 | 22 | 0.2727 | 19 | 0.3158 | 61 | 0.2951 |
| 7_8778134 | 13 | 0.1538 | 10 | 0.1 | 12 | 0.0833 | 11 | 0 | 46 | 0.087 | 6 | 0.1667 | 14 | 0.2143 | 23 | 0.1739 | 23 | 0.1818 | 66 | 0.1818 |
| 7_8778280 | 20 | 0.1 | 15 | 0 | 16 | 0.0625 | 23 | 0.087 | 74 | 0.0676 | 7 | 0 | 21 | 0 | 30 | 0.1 | 23 | 0.1304 | 81 | 0.0741 |
| 7_8778326 | 19 | 0.2632 | 16 | 0 | 19 | 0.1053 | 26 | 0.1923 | 80 | 0.15 | 6 | 0.1667 | 23 | 0.1304 | 34 | 0.1471 | 26 | 0.0385 | 89 | 0.1124 |
| 7_8778382 | 22 | 0.1364 | 19 | 0.3158 | 19 | 0.3158 | 28 | 0.2143 | 88 | 0.2386 | 9 | 0.3333 | 24 | 0.375 | 41 | 0.2683 | 31 | 0.2903 | 105 | 0.3048 |
| 7_8793810 | 1 | 1 | NA | NA | NA | NA | NA | NA | 1 | 1 | NA | NA | NA | NA | NA | NA | 1 | 1 | 1 | 1 |
| 7_8793884 | 1 | 1 | NA | NA | NA | NA | NA | NA | 1 | 1 | NA | NA | NA | NA | NA | NA | 1 | 1 | 1 | 1 |
| 7_8795882 | 1 | 1 | 3 | 0.6667 | 2 | 1 | NA | NA | 6 | 0.8333 | 2 | 1 | NA | NA | NA | NA | NA | NA | 2 | 1 |
| 7_8795899 | NA | NA | 1 | 1 | 2 | 0.5 | NA | NA | 3 | 0.6667 | 1 | 1 | NA | NA | NA | NA | NA | NA | 1 | 1 |
| 7_8795905 | 1 | 0 | 4 | 0.5 | 3 | 0.3333 | NA | NA | 8 | 0.375 | 2 | 0.5 | NA | NA | NA | NA | NA | NA | 2 | 0.5 |
| 7_8878947 | 9 | 0.1111 | 8 | 0.125 | 7 | 0.1429 | 5 | 0 | 29 | 0.1034 | 5 | 0 | 5 | 0.4 | 6 | 0.6667 | 6 | 0.3333 | 22 | 0.3636 |
| 7_8878953 | 9 | 0.1111 | 9 | 0.2222 | 8 | 0.125 | 4 | 0 | 30 | 0.1333 | 5 | 0 | 7 | 0 | 5 | 0.6 | 5 | 0 | 22 | 0.1364 |
| 7_8878957 | 10 | 0 | 9 | 0.2222 | 8 | 0.125 | 6 | 0.1667 | 33 | 0.1212 | 4 | 0 | 6 | 0 | 8 | 0.25 | 6 | 0.1667 | 24 | 0.125 |
| 7_8879102 | 10 | 0.7 | 12 | 0.8333 | 8 | 0.5 | 8 | 0.625 | 38 | 0.6842 | 3 | 1 | 9 | 0.5556 | 8 | 0.75 | 7 | 0.8571 | 27 | 0.7407 |
| 7_8879104 | 7 | 0.1429 | 10 | 0.2 | 6 | 0 | 6 | 0.1667 | 29 | 0.1379 | 3 | 0 | 8 | 0.125 | 8 | 0.25 | 6 | 0.1667 | 25 | 0.16 |

| | | | | | | | | | | | | | | | | | | | | |
|---|---|---|---|---|---|---|---|---|---|---|---|---|---|---|---|---|---|---|---|---|
| 7_88791056 | 9 | 0.3333 | 9 | 0.5556 | 7 | 0.5714 | 8 | 0.625 | 33 | 0.5152 | 3 | 1 | 9 | 0.6667 | 7 | 0.4286 | 3 | 1 | 22 | 0.6818 |
| 7_89056385 | 2 | 0 | 1 | 1 | 1 | 1 | NA | NA | 4 | 0.5 | 1 | 1 | NA | NA | NA | NA | NA | NA | 1 | 1 |
| 7_89056550 | 2 | 1 | 2 | 1 | 1 | 1 | NA | NA | 5 | 1 | 1 | 1 | 1 | 0 | NA | NA | 2 | 1 | 4 | 0.75 |
| 7_89056842 | 2 | 1 | 2 | 1 | 1 | 1 | NA | NA | 5 | 1 | 1 | 1 | NA | NA | NA | NA | 2 | 1 | 3 | 1 |
| 7_89056950 | 2 | 1 | 2 | 1 | 1 | 1 | NA | NA | 5 | 1 | 1 | 1 | 1 | 1 | NA | NA | 2 | 1 | 4 | 1 |
| 7_89057052 | 2 | 1 | 2 | 1 | 1 | 1 | NA | NA | 5 | 1 | NA | NA | 1 | 1 | NA | NA | 2 | 0.5 | 3 | 0.6667 |
| 7_89057070 | 2 | 1 | NA | NA | 1 | 1 | NA | NA | 3 | 1 | 1 | 1 | 1 | 1 | NA | NA | 1 | 1 | 3 | 1 |
| 7_89057074 | 1 | 0 | 1 | 1 | 1 | 1 | NA | NA | 3 | 0.6667 | 1 | 1 | NA | NA | NA | NA | 1 | 1 | 2 | 1 |
| 7_89057096 | 2 | 0.5 | 1 | 1 | NA | NA | NA | NA | 3 | 0.6667 | 1 | 0 | 1 | 0 | NA | NA | 1 | 1 | 3 | 0.3333 |
| 7_89057227 | 2 | 0.5 | 2 | 1 | 1 | 1 | NA | NA | 5 | 0.8 | 1 | 1 | 1 | 0 | NA | NA | 2 | 0 | 4 | 0.25 |
| 7_89057233 | 1 | 1 | 2 | 0 | NA | NA | NA | NA | 3 | 0.3333 | 1 | 1 | NA | NA | NA | NA | 2 | 0.5 | 3 | 0.6667 |
| 7_89057336 | 2 | 1 | 2 | 0.5 | 1 | 0 | NA | NA | 5 | 0.6 | 1 | 1 | 1 | 0 | NA | NA | 1 | 0 | 3 | 0.3333 |
| 7_89057366 | 2 | 1 | 2 | 1 | 1 | 0 | NA | NA | 5 | 0.8 | 1 | 1 | 1 | 1 | NA | NA | 2 | 1 | 4 | 1 |
| 7_89057452 | 2 | 1 | 2 | 0 | 1 | 0 | NA | NA | 5 | 0.4 | 1 | 0 | 1 | 1 | NA | NA | 2 | 0 | 4 | 0.25 |
| 7_89057557 | 2 | 1 | 2 | 1 | 1 | 1 | NA | NA | 5 | 1 | 1 | 1 | 1 | 1 | NA | NA | 2 | 0.5 | 4 | 0.75 |
| 7_89057570 | 2 | 0 | 2 | 0.5 | 1 | 0 | NA | NA | 5 | 0.2 | 1 | 1 | 1 | 0 | NA | NA | 2 | 0 | 4 | 0.25 |
| 7_89057722 | 2 | 0 | 2 | 0.5 | 1 | 1 | NA | NA | 5 | 0.4 | 1 | 0 | NA | NA | NA | NA | 2 | 0.5 | 3 | 0.3333 |
| 7_89058458 | 2 | 0 | 1 | 1 | 1 | 0 | NA | NA | 4 | 0.25 | 1 | 1 | 1 | 0 | NA | NA | 2 | 0.5 | 4 | 0.5 |
| 7_89058513 | 2 | 0 | 2 | 0.5 | 1 | 0 | NA | NA | 5 | 0.2 | 1 | 0 | 1 | 0 | NA | NA | 2 | 0.5 | 4 | 0.25 |
| 7_89059672 | 2 | 0 | 2 | 0.5 | 2 | 0 | 1 | 0 | 7 | 0.1429 | 1 | 0 | 2 | 0 | NA | NA | 2 | 0.5 | 5 | 0.2 |
| 7_89061985 | 3 | 0 | 3 | 0.3333 | 3 | 0 | 1 | 1 | 10 | 0.2 | 2 | 0 | 3 | 0.6667 | NA | NA | 4 | 0 | 9 | 0.2222 |
| 7_89157349 | NA | NA | 2 | 0.5 | NA | NA | NA | NA | 2 | 0.5 | 2 | 0.5 | NA | NA | NA | NA | NA | NA | 2 | 0.5 |
| 7_89157474 | NA | NA | 2 | 0.5 | NA | NA | NA | NA | 2 | 0.5 | 2 | 0.5 | NA | NA | NA | NA | NA | NA | 2 | 0.5 |
| 7_89158382 | 5 | 0.2 | 12 | 0.3333 | 7 | 0.1429 | 6 | 0.6667 | 30 | 0.3333 | 13 | 0.1538 | 3 | 0.3333 | 1 | 1 | 10 | 0.3 | 27 | 0.2593 |
| 7_89158713 | 6 | 1 | 12 | 0.9167 | 6 | 0.8333 | 5 | 1 | 29 | 0.931 | 11 | 0.8182 | 2 | 1 | 1 | 1 | 9 | 0.6667 | 23 | 0.7826 |
| 7_89158768 | 5 | 0.2 | 11 | 0.1818 | 3 | 0.3333 | 3 | 0.3333 | 22 | 0.2273 | 8 | 0.375 | 2 | 0 | 1 | 1 | 8 | 0.25 | 19 | 0.3158 |
| 7_89158848 | 5 | 0.6 | 11 | 0.2727 | 4 | 0.75 | 4 | 0.25 | 24 | 0.4167 | 9 | 0.3333 | 2 | 1 | NA | NA | 7 | 0.2857 | 18 | 0.3889 |
| 7_89159031 | 6 | 1 | 10 | 0.7 | 4 | 0.75 | 3 | 1 | 23 | 0.8261 | 11 | 0.8182 | 2 | 1 | 1 | 1 | 6 | 1 | 20 | 0.9 |
| 7_89159090 | 5 | 0.2 | 13 | 0.3077 | 3 | 0 | 3 | 0 | 24 | 0.2083 | 11 | 0.0909 | 2 | 0 | 1 | 0 | 6 | 0.3333 | 20 | 0.15 |
| 7_89159170 | 6 | 0 | 13 | 0.0769 | 3 | 0.3333 | 5 | 0 | 27 | 0.0741 | 10 | 0.1 | 2 | 0 | 1 | 0 | 6 | 0 | 19 | 0.0526 |
| 7_89159223 | 5 | 0.6 | 12 | 0.1667 | 3 | 0 | 5 | 0.2 | 25 | 0.24 | 11 | 0.3636 | 2 | 0.5 | 1 | 0 | 6 | 0.1667 | 20 | 0.3 |
| 7_89159297 | 5 | 1 | 7 | 1 | 2 | 1 | 4 | 1 | 18 | 1 | 7 | 1 | 2 | 1 | 1 | 1 | 6 | 1 | 16 | 1 |
| 7_89159318 | 5 | 0.2 | 9 | 0.6667 | 3 | 0.3333 | 5 | 0.2 | 22 | 0.4091 | 11 | 0.5455 | 2 | 0.5 | 1 | 0 | 6 | 0.5 | 20 | 0.5 |
| 7_89159454 | 4 | 0 | 11 | 0.3636 | 3 | 0 | 5 | 0.6 | 23 | 0.3043 | 8 | 0 | 2 | 0 | 1 | 0 | 5 | 0.2 | 16 | 0.0625 |
| 7_89159504 | 5 | 0.6 | 7 | 0.4286 | 2 | 0.5 | 3 | 0 | 17 | 0.4118 | 7 | 0.2857 | 2 | 0.5 | 1 | 1 | 4 | 0.5 | 14 | 0.4286 |
| 7_89159512 | 4 | 1 | 11 | 1 | 2 | 1 | 5 | 0.8 | 22 | 0.9545 | 8 | 1 | 2 | 1 | 1 | 1 | 5 | 1 | 16 | 1 |
| 7_89159643 | 5 | 1 | 10 | 1 | 3 | 1 | 3 | 1 | 21 | 1 | 8 | 1 | 2 | 1 | 1 | 1 | 5 | 1 | 16 | 1 |
| 7_89159721 | 5 | 1 | 10 | 0.9 | 3 | 1 | 4 | 0.5 | 22 | 0.8636 | 10 | 0.9 | 2 | 1 | 1 | 1 | 5 | 1 | 18 | 0.9444 |
| 7_89159776 | 5 | 1 | 11 | 1 | 3 | 1 | 5 | 1 | 24 | 1 | 10 | 1 | 2 | 1 | 1 | 1 | 5 | 1 | 18 | 1 |
| 7_89159806 | 3 | 0 | 10 | 0.4 | 3 | 0.3333 | 2 | 0 | 18 | 0.2778 | 9 | 0 | 1 | 0 | 1 | 0 | 5 | 0.2 | 16 | 0.0625 |
| 7_89159854 | 3 | 0.3333 | 5 | 0 | 3 | 0 | 2 | 0 | 13 | 0.0769 | 8 | 0.125 | 2 | 0 | 1 | 0 | 4 | 0.25 | 15 | 0.1333 |
| 7_89159873 | 5 | 0.8 | 9 | 0.5556 | 1 | 0 | 4 | 0.25 | 19 | 0.5263 | 8 | 0.5 | 1 | 1 | 1 | 0 | 4 | 0.75 | 14 | 0.5714 |
| 7_89159916 | 4 | 0 | 11 | 0.3636 | 3 | 0.3333 | 4 | 0.25 | 22 | 0.2727 | 9 | 0.1111 | 1 | 0 | 1 | 0 | 5 | 0 | 16 | 0.0625 |
| 7_89467898 | 2 | 1 | NA | NA | 6 | 0.8333 | 2 | 1 | 10 | 0.9 | 1 | 1 | 3 | 0.6667 | 1 | 1 | 1 | 1 | 6 | 0.8333 |
| 7_89467971 | 1 | 1 | 1 | 0 | 6 | 0.5 | 2 | 1 | 10 | 0.6 | 1 | 1 | 2 | 1 | 1 | 1 | NA | NA | 4 | 1 |
| 7_89467992 | 2 | 1 | NA | NA | 5 | 0.8 | 2 | 1 | 9 | 0.8889 | NA | NA | 2 | 1 | 1 | 1 | 1 | 1 | 4 | 1 |
| 7_89468103 | 2 | 0 | 1 | 0 | 6 | 0.3333 | 2 | 0.5 | 11 | 0.2727 | 1 | 0 | 3 | 0.3333 | 1 | 1 | NA | NA | 5 | 0.4 |
| 7_89473673 | 1 | 0 | 3 | 0 | 5 | 0.4 | 3 | 0 | 12 | 0.1667 | 1 | 1 | NA | NA | 2 | 0 | NA | NA | 3 | 0.3333 |
| 7_89473851 | 1 | 0 | 1 | 0 | 4 | 0.5 | 3 | 0.6667 | 9 | 0.4444 | 1 | 0 | NA | NA | NA | NA | 1 | 1 | 2 | 0.5 |
| 7_89473906 | 1 | 0 | 3 | 0.3333 | 4 | 0.25 | 3 | 0 | 11 | 0.1818 | 1 | 0 | NA | NA | 2 | 0.5 | 1 | 0 | 4 | 0.25 |
| 7_89474421 | 1 | 0 | 4 | 0 | 5 | 0.4 | 4 | 0 | 14 | 0.1429 | NA | NA | NA | NA | 2 | 0 | 1 | 1 | 3 | 0.3333 |
| 7_89491032 | NA | NA | NA | NA | NA | NA | 1 | 1 | 1 | 1 | NA | NA | 1 | 1 | 1 | 1 | NA | NA | 2 | 1 |
| 7_89491213 | NA | NA | NA | NA | NA | NA | 1 | 1 | 1 | 1 | NA | NA | 1 | 1 | 1 | 1 | NA | NA | 2 | 1 |
| 7_89491343 | NA | NA | NA | NA | NA | NA | 1 | 1 | 1 | 1 | NA | NA | 1 | 1 | 1 | 1 | NA | NA | 2 | 1 |
| 7_89501496 | NA | NA | 1 | 1 | NA | NA | 3 | 0.3333 | 4 | 0.5 | 1 | 1 | 2 | 0 | 1 | 0 | 1 | 1 | 5 | 0.4 |
| 7_89565567 | 1 | 0 | 1 | 1 | 1 | 0 | 1 | 0 | 4 | 0.25 | 1 | 0 | 1 | 0 | 1 | 0 | 1 | 1 | 4 | 0.25 |
| 7_89568038 | 9 | 0.1111 | 10 | 0.2 | 13 | 0.1538 | 7 | 0 | 39 | 0.1282 | 7 | 0 | 11 | 0 | 7 | 0.1429 | 9 | 0.1111 | 34 | 0.0588 |
| 7_89568065 | 11 | 0 | 9 | 0.1111 | 13 | 0.0769 | 9 | 0.1111 | 42 | 0.0714 | 6 | 0 | 12 | 0.1667 | 9 | 0.1111 | 10 | 0.1 | 37 | 0.1081 |
| 7_89635541 | NA | NA | NA | NA | 1 | 1 | 1 | 0 | 2 | 0.5 | 1 | 1 | 1 | 0 | 1 | 0 | NA | NA | 3 | 0.3333 |
| 7_89635671 | NA | NA | NA | NA | 1 | 1 | 1 | 0 | 2 | 0.5 | 1 | 1 | 1 | 1 | 1 | 0 | NA | NA | 3 | 0.6667 |
| 7_89856882 | 16 | 0.0625 | 17 | 0.1176 | 14 | 0.0714 | 7 | 0 | 54 | 0.0741 | 7 | 0.1429 | 13 | 0.0769 | 11 | 0 | 8 | 0.125 | 39 | 0.0769 |
| 7_89856952 | 15 | 0.5333 | 17 | 0.3529 | 14 | 0.2857 | 5 | 0.2 | 51 | 0.3725 | 7 | 0.5714 | 11 | 0.5455 | 12 | 0.75 | 7 | 0.2857 | 37 | 0.5676 |
| 7_89857014 | 16 | 0.25 | 16 | 0.1875 | 14 | 0.3571 | 8 | 0.375 | 54 | 0.2778 | 9 | 0.2222 | 11 | 0.3636 | 12 | 0.1667 | 6 | 0.1667 | 38 | 0.2368 |
| 7_89857036 | 14 | 0.1429 | 18 | 0.1111 | 12 | 0.0833 | 8 | 0.25 | 52 | 0.1346 | 9 | 0.1111 | 14 | 0 | 11 | 0.3636 | 7 | 0 | 41 | 0.122 |
| 7_89858349 | 5 | 0.2 | 2 | 0 | 1 | 0 | NA | NA | 8 | 0.125 | NA | NA | 2 | 0 | 2 | 0 | 2 | 0.5 | 6 | 0.1667 |

| | | | | | | | | | | | | | | | | | | | | |
|---|---|---|---|---|---|---|---|---|---|---|---|---|---|---|---|---|---|---|---|---|
| 7_90058806 | 1 | 0 | 2 | 1 | 1 | 1 | 3 | 0.6667 | 7 | 0.7143 | 1 | 1 | 1 | 1 | 1 | 1 | 1 | 0 | 4 | 0.75 |
| 7_90058827 | 1 | 0 | 1 | 1 | 1 | 1 | 2 | 1 | 5 | 0.8 | 1 | 0 | 1 | 1 | NA | NA | NA | NA | 2 | 0.5 |
| 7_90058856 | 1 | 1 | 1 | 1 | NA | NA | 3 | 0.6667 | 5 | 0.8 | 1 | 1 | NA | NA | NA | NA | 1 | 0 | 2 | 0.5 |
| 7_90059000 | 2 | 1 | 2 | 0 | 1 | 0 | 3 | 1 | 8 | 0.625 | 1 | 1 | 1 | 1 | 1 | 0 | 1 | 0 | 4 | 0.5 |
| 7_90059146 | 2 | 0.5 | 1 | 0 | 1 | 0 | 3 | 1 | 7 | 0.5714 | NA | NA | NA | NA | NA | NA | 1 | 1 | 1 | 1 |
| 7_90059175 | 2 | 0 | 2 | 0 | 1 | 0 | 3 | 0.3333 | 8 | 0.125 | 1 | 1 | 1 | 1 | 1 | 0 | NA | NA | 3 | 0.6667 |
| 7_90059235 | 2 | 0.5 | 2 | 0 | NA | NA | 3 | 0.3333 | 7 | 0.2857 | 1 | 0 | 1 | 0 | 3 | 0.3333 | 1 | 0 | 6 | 0.1667 |
| 7_90059263 | 2 | 0.5 | 2 | 1 | 1 | 1 | 2 | 0.5 | 7 | 0.7143 | 1 | 1 | 1 | 0 | 3 | 1 | NA | NA | 5 | 0.8 |
| 7_90059267 | 2 | 1 | 1 | 1 | 1 | 0 | 2 | 0 | 6 | 0.5 | 1 | 1 | 1 | 0 | 2 | 0 | 1 | 0 | 5 | 0.2 |
| 7_90059314 | 2 | 1 | 2 | 0.5 | 1 | 1 | 3 | 0.6667 | 8 | 0.75 | 1 | 0 | 1 | 1 | 3 | 1 | 1 | 1 | 6 | 0.8333 |
| 7_90059348 | 2 | 1 | 2 | 1 | 1 | 1 | 3 | 1 | 8 | 1 | 1 | 1 | 1 | 1 | 3 | 1 | 1 | 1 | 6 | 1 |
| 7_90092122 | 2 | 0 | 1 | 1 | NA | NA | 5 | 0.4 | 8 | 0.375 | NA | NA | 3 | 0.6667 | 4 | 0.75 | 1 | 0 | 8 | 0.625 |
| 7_90092539 | 2 | 1 | 1 | 1 | 1 | 1 | 4 | 0.25 | 8 | 0.625 | 1 | 1 | 3 | 1 | 5 | 0.8 | 3 | 1 | 12 | 0.9167 |
| 7_90092590 | 2 | 1 | 1 | 1 | 1 | 1 | 5 | 0.4 | 9 | 0.6667 | 1 | 0 | 2 | 0.5 | 2 | 0 | 1 | 1 | 6 | 0.3333 |
| 7_90092948 | 4 | 1 | 2 | 0.5 | 2 | 1 | 5 | 0.8 | 13 | 0.8462 | 1 | 0 | 3 | 0.6667 | 6 | 0.6667 | 2 | 0.5 | 12 | 0.5833 |
| 7_90093057 | 3 | 0 | 3 | 0.3333 | 2 | 0 | 6 | 0 | 14 | 0.0714 | 1 | 0 | 3 | 0 | 3 | 0 | 4 | 0.25 | 11 | 0.0909 |
| 7_90093062 | 3 | 0.3333 | 3 | 0.6667 | NA | NA | 7 | 0.5714 | 13 | 0.5385 | 1 | 1 | 2 | 1 | 3 | 1 | 4 | 1 | 10 | 1 |
| 7_90093098 | 4 | 1 | 3 | 0 | 2 | 0 | 6 | 0 | 15 | 0.2667 | 1 | 0 | 3 | 0.6667 | 5 | 0.2 | 2 | 0.5 | 11 | 0.3636 |
| 7_90093127 | 3 | 0.6667 | 3 | 0.6667 | 1 | 1 | 6 | 0.6667 | 13 | 0.6923 | 1 | 0 | 2 | 0 | 5 | 0.4 | 4 | 0.5 | 12 | 0.3333 |
| 7_90093229 | 4 | 1 | 4 | 1 | 1 | 0 | 5 | 0.4 | 14 | 0.7143 | 1 | 1 | 5 | 0.4 | 5 | 0.6 | 3 | 0.6667 | 14 | 0.5714 |
| 7_90093373 | 2 | 0.5 | 3 | 0.6667 | 1 | 1 | 5 | 0.6 | 11 | 0.6364 | 2 | 1 | 3 | 0.6667 | 6 | 0.6667 | 3 | 0 | 14 | 0.5714 |
| 7_90094139 | 3 | 0.3333 | 4 | 0 | 2 | 0 | 3 | 0 | 12 | 0.0833 | 2 | 0 | 4 | 0 | 5 | 0.2 | 3 | 0.3333 | 14 | 0.1429 |
| 7_90095014 | 4 | 0.5 | 6 | 0 | 4 | 0.5 | 6 | 0.6667 | 20 | 0.4 | 3 | 0 | 6 | 0.3333 | 9 | 0.3333 | 4 | 0.25 | 22 | 0.2727 |
| 7_90095220 | 2 | 0.5 | 6 | 0.1667 | 2 | 0 | 5 | 0 | 15 | 0.1333 | 2 | 0 | 4 | 0.25 | 8 | 0 | 3 | 0 | 17 | 0.0588 |
| 7_90095397 | 1 | 0 | 4 | 0 | 2 | 0.5 | 3 | 0.3333 | 10 | 0.2 | 1 | 0 | 5 | 0.2 | 3 | 0 | 2 | 0 | 11 | 0.0909 |
| 7_90096099 | NA | NA | 2 | 0.5 | 1 | 0 | 1 | 0 | 4 | 0.25 | 1 | 0 | 2 | 0 | 2 | 0 | 1 | 1 | 6 | 0.1667 |
| 7_90096458 | 2 | 1 | 3 | 0.6667 | 1 | 1 | 3 | 0 | 9 | 0.5556 | 2 | 1 | 1 | 1 | 3 | 1 | NA | NA | 6 | 1 |
| 7_90096518 | 2 | 1 | 3 | 0 | 1 | 0 | 3 | 0 | 9 | 0.2222 | 2 | 0.5 | 2 | 0 | 2 | 0 | 3 | 0 | 9 | 0.1111 |
| 7_90096613 | 2 | 1 | 3 | 0.3333 | 1 | 0 | 3 | 1 | 9 | 0.6667 | 2 | 1 | 2 | 0 | 2 | 1 | 3 | 1 | 9 | 0.7778 |
| 7_90096633 | 2 | 0.5 | 3 | 0.3333 | 1 | 0 | 3 | 0.6667 | 9 | 0.4444 | 2 | 0 | 2 | 0 | 3 | 0.3333 | 3 | 0.6667 | 10 | 0.3 |
| 7_90096783 | 2 | 1 | 3 | 1 | NA | NA | 3 | 0.6667 | 8 | 0.875 | 2 | 1 | 2 | 0 | 3 | 1 | 3 | 1 | 10 | 0.8 |
| 7_90096800 | 2 | 0 | 4 | 0.25 | 1 | 0 | 3 | 0 | 10 | 0.1 | 2 | 0 | 2 | 0 | 3 | 0 | 4 | 0.25 | 11 | 0.0909 |
| 7_90100040 | 11 | 0.0909 | 11 | 0 | 5 | 0 | 13 | 0.1538 | 40 | 0.075 | 6 | 0.1667 | 10 | 0.2 | 10 | 0.1 | 5 | 0 | 31 | 0.129 |
| 7_90116714 | 1 | 0 | 1 | 1 | 3 | 0.3333 | 2 | 0.5 | 7 | 0.4286 | 2 | 0 | NA | NA | 1 | 0 | 2 | 0.5 | 5 | 0.2 |
| 7_90116991 | 7 | 0.4286 | 4 | 0.5 | 3 | 0.3333 | 6 | 0.6667 | 20 | 0.5 | 3 | 0.3333 | 8 | 0.375 | 3 | 0.6667 | 3 | 0.6667 | 17 | 0.4706 |
| 7_90117010 | 5 | 0.4 | 3 | 0.6667 | 3 | 1 | 6 | 0.8333 | 17 | 0.7059 | 1 | 0 | 7 | 0.8571 | 2 | 0.5 | NA | NA | 10 | 0.7 |
| 7_90117024 | 6 | 0.8333 | 5 | 0.8 | 5 | 1 | 7 | 0.8571 | 23 | 0.8696 | 2 | 1 | 7 | 0.7143 | 3 | 0.6667 | NA | NA | 12 | 0.75 |
| 7_90117043 | 8 | 0 | 5 | 0.2 | 6 | 0 | 8 | 0.125 | 27 | 0.0741 | 3 | 0.6667 | 8 | 0.25 | 3 | 0.3333 | 3 | 0 | 17 | 0.2941 |
| 7_90117166 | 6 | 0.3333 | 5 | 0.8 | 6 | 0.8333 | 7 | 0.5714 | 24 | 0.625 | 3 | 0.6667 | 7 | 0.2857 | 3 | 1 | 2 | 0.5 | 15 | 0.5333 |
| 7_90117379 | 4 | 0.25 | 4 | 0.25 | 4 | 0.5 | 6 | 0.3333 | 18 | 0.3333 | 2 | 0 | 6 | 0.3333 | 3 | 0.3333 | 2 | 0 | 13 | 0.2308 |
| 7_90117387 | 6 | 0.8333 | 5 | 0.8 | 5 | 1 | 7 | 0.8571 | 23 | 0.8696 | 1 | 0 | 7 | 0.8571 | 3 | 0.3333 | 2 | 1 | 13 | 0.6923 |
| 7_90117818 | 4 | 0 | 3 | 0.3333 | 4 | 0.25 | 7 | 0.1429 | 18 | 0.1667 | 3 | 0.6667 | 6 | 0 | 2 | 0.5 | 2 | 0.5 | 13 | 0.3077 |
| 7_90117824 | 4 | 0 | 3 | 0.3333 | 4 | 0.25 | 7 | 0.1429 | 18 | 0.1667 | 3 | 0.3333 | 4 | 0.25 | 2 | 0 | 1 | 0 | 10 | 0.2 |
| 7_90118073 | 6 | 0.6667 | 5 | 0 | 4 | 0.75 | 7 | 0.2857 | 22 | 0.4091 | 3 | 0.6667 | 5 | 0.2 | 1 | 1 | 2 | 1 | 11 | 0.5455 |
| 7_90118095 | 6 | 0.6667 | 5 | 0.8 | 4 | 0.75 | 7 | 0.8571 | 22 | 0.7727 | 3 | 1 | 4 | 0.75 | 1 | 0 | 3 | 1 | 11 | 0.8182 |
| 7_90118125 | 6 | 1 | 5 | 0.8 | 3 | 1 | 7 | 1 | 21 | 0.9524 | 3 | 1 | 5 | 1 | 1 | 1 | 1 | 1 | 10 | 1 |
| 7_90118141 | 3 | 0.6667 | 2 | 1 | 3 | 1 | 5 | 1 | 13 | 0.9231 | 3 | 0.6667 | 2 | 1 | 1 | 1 | 1 | 1 | 7 | 0.8571 |
| 7_90118170 | 5 | 1 | 5 | 0.6 | 2 | 0.5 | 6 | 0.6667 | 18 | 0.7222 | 2 | 1 | 2 | 1 | NA | NA | 1 | 1 | 5 | 1 |
| 7_90118179 | 4 | 0.5 | 4 | 0.5 | 3 | 0.6667 | 5 | 0.2 | 16 | 0.4375 | 3 | 0.6667 | 4 | 0.25 | 1 | 0 | 1 | 0 | 9 | 0.3333 |
| 7_90118305 | 6 | 0 | 4 | 0 | 1 | 0 | 6 | 0.1667 | 17 | 0.0588 | 3 | 0 | 4 | 0.25 | 1 | 0 | 2 | 0 | 10 | 0.1 |
| 7_90126039 | NA | NA | NA | NA | NA | NA | 1 | 1 | 1 | 1 | 1 | 0 | 1 | 1 | NA | NA | 1 | 1 | 3 | 0.6667 |
| 7_90126097 | NA | NA | NA | NA | NA | NA | 1 | 1 | 1 | 1 | 1 | 1 | 1 | 1 | NA | NA | 1 | 1 | 3 | 1 |
| 7_92093789 | 7 | 0 | 14 | 0.2857 | 10 | 0.1 | 14 | 0.2143 | 45 | 0.1778 | 4 | 0.75 | 12 | 0.25 | 8 | 0.125 | 21 | 0.1905 | 45 | 0.2444 |
| 7_92093967 | 5 | 0.4 | 16 | 0.3125 | 6 | 0.1667 | 13 | 0.3077 | 40 | 0.3 | 4 | 0 | 14 | 0.0714 | 11 | 0.0909 | 20 | 0.25 | 49 | 0.1429 |
| 7_92095408 | 6 | 0 | 21 | 0.1429 | 11 | 0 | 18 | 0 | 56 | 0.0536 | 7 | 0.2857 | 19 | 0 | 24 | 0 | 24 | 0.0833 | 74 | 0.0541 |
| 7_92097511 | 14 | 0.1429 | 38 | 0.3421 | 22 | 0.3182 | 28 | 0.3214 | 102 | 0.3039 | 16 | 0.375 | 29 | 0.2759 | 40 | 0.325 | 37 | 0.2973 | 122 | 0.3115 |
| 7_92097589 | 12 | 0 | 37 | 0.0811 | 22 | 0 | 29 | 0.1379 | 100 | 0.07 | 15 | 0.3333 | 29 | 0.069 | 36 | 0.0833 | 35 | 0.0857 | 115 | 0.113 |
| 7_92210741 | 1 | 0 | 1 | 0 | 3 | 0 | 3 | 0.3333 | 8 | 0.125 | 1 | 0 | NA | NA | 2 | 0.5 | 1 | 0 | 4 | 0.25 |
| 7_92231157 | 4 | 0.5 | 2 | 0.5 | 5 | 0 | 4 | 0.25 | 15 | 0.2667 | 1 | 1 | NA | NA | 2 | 0 | 1 | 0 | 4 | 0.25 |
| 7_92233560 | 1 | 1 | 1 | 0 | 2 | 1 | 1 | 1 | 5 | 0.8 | NA | NA | 3 | 0.3333 | NA | NA | NA | NA | 3 | 0.3333 |
| 7_92285646 | NA | NA | NA | NA | NA | NA | 1 | 1 | 1 | 1 | NA | NA | 2 | 1 | NA | NA | NA | NA | 2 | 1 |
| 7_92293169 | 1 | 0 | 2 | 0.5 | NA | NA | 5 | 0.6 | 8 | 0.5 | 1 | 0 | 5 | 0 | 2 | 0.5 | 1 | 1 | 9 | 0.2222 |
| 7_92293271 | 1 | 1 | 2 | 0.5 | NA | NA | 3 | 0 | 6 | 0.3333 | 1 | 1 | 5 | 0.4 | 1 | 1 | NA | NA | 7 | 0.5714 |
| 7_92293340 | 1 | 0 | 2 | 0 | NA | NA | 3 | 0.6667 | 6 | 0.3333 | 1 | 1 | 6 | 0.1667 | 2 | 0.5 | NA | NA | 9 | 0.3333 |
| 7_92306615 | NA | NA | 1 | 1 | NA | NA | 3 | 0.3333 | 4 | 0.5 | 2 | 1 | 3 | 1 | 2 | 0.5 | 3 | 1 | 10 | 0.9 |

| | | | | | | | | | | | | | | | | | | | | |
|---|---|---|---|---|---|---|---|---|---|---|---|---|---|---|---|---|---|---|---|---|
| 7_92306669 | NA | NA | 1 | 1 | NA | NA | 3 | 1 | 4 | 1 | 1 | 0 | 3 | 1 | 3 | 1 | 3 | 1 | 10 | 0.9 |
| 7_92306684 | NA | NA | 1 | 1 | NA | NA | 3 | 1 | 4 | 1 | 1 | 0 | 3 | 0.6667 | 3 | 1 | 3 | 0.3333 | 10 | 0.6 |
| 7_92306714 | NA | NA | 1 | 0 | NA | NA | 3 | 0.3333 | 4 | 0.25 | 1 | 1 | 3 | 0.6667 | 3 | 0.6667 | 3 | 1 | 10 | 0.8 |
| 7_92306830 | NA | NA | 1 | 1 | NA | NA | 3 | 0.6667 | 4 | 0.75 | 1 | 0 | 3 | 1 | 3 | 1 | 3 | 1 | 10 | 0.9 |
| 7_92306930 | NA | NA | 1 | 0 | NA | NA | 2 | 0.5 | 3 | 0.3333 | 1 | 1 | 3 | 0.3333 | 3 | 1 | 3 | 0.6667 | 10 | 0.7 |
| 7_92306939 | NA | NA | 1 | 1 | NA | NA | 3 | 0.6667 | 4 | 0.75 | 1 | 0 | 3 | 0 | 3 | 0.6667 | 3 | 0.3333 | 10 | 0.3 |
| 7_92306971 | NA | NA | 1 | 0 | NA | NA | 3 | 0.6667 | 4 | 0.5 | NA | NA | 3 | 1 | 1 | 1 | 1 | 1 | 5 | 1 |
| 7_92307044 | NA | NA | NA | NA | NA | NA | 3 | 0.3333 | 3 | 0.3333 | 1 | 1 | 3 | 0.6667 | 2 | 0.5 | 1 | 1 | 7 | 0.7143 |
| 7_92307821 | NA | NA | NA | NA | NA | NA | 2 | 0.5 | 2 | 0.5 | 1 | 1 | 3 | 1 | 2 | 1 | 1 | 1 | 7 | 1 |
| 7_92307905 | NA | NA | NA | NA | NA | NA | 2 | 0.5 | 2 | 0.5 | NA | NA | 3 | 0.6667 | 2 | 1 | 1 | 1 | 6 | 0.8333 |
| 7_92310409 | NA | NA | NA | NA | NA | NA | 2 | 0.5 | 2 | 0.5 | NA | NA | 2 | 1 | 1 | 0 | NA | NA | 3 | 0.6667 |
| 7_92310836 | NA | NA | NA | NA | NA | NA | 1 | 1 | 1 | 1 | NA | NA | 3 | 1 | 1 | 0 | NA | NA | 4 | 0.75 |
| 7_92484969 | 2 | 0.5 | NA | NA | 6 | 0.3333 | 3 | 0.6667 | 11 | 0.4545 | 1 | 0 | 1 | 1 | 3 | 0.6667 | 5 | 0.2 | 10 | 0.4 |
| 7_92485018 | 2 | 0 | NA | NA | 4 | 0.25 | 3 | 0 | 9 | 0.1111 | 1 | 0 | 1 | 0 | 3 | 0 | 5 | 0.4 | 10 | 0.2 |
| 7_92485077 | 2 | 1 | NA | NA | 4 | 0.25 | 2 | 0 | 8 | 0.375 | 1 | 0 | 1 | 1 | 2 | 0.5 | 4 | 0.25 | 8 | 0.375 |
| 7_92485127 | 2 | 1 | NA | NA | 6 | 0.3333 | 1 | 1 | 9 | 0.5556 | 1 | 0 | 1 | 0 | 3 | 0.6667 | 4 | 0.5 | 9 | 0.4444 |
| 7_92485224 | 2 | 0.5 | NA | NA | 5 | 0.4 | 4 | 0.25 | 11 | 0.3636 | 1 | 0 | 1 | 0 | 3 | 0.3333 | 4 | 0.5 | 9 | 0.3333 |
| 7_92485244 | 1 | 1 | NA | NA | 6 | 0.8333 | 2 | 1 | 9 | 0.8889 | 1 | 1 | 1 | 1 | 3 | 1 | 6 | 1 | 11 | 1 |
| 7_92485252 | 2 | 0.5 | NA | NA | 7 | 0.2857 | 4 | 0.25 | 13 | 0.3077 | 1 | 0 | 1 | 0 | 3 | 0.3333 | 6 | 0 | 11 | 0.0909 |
| 7_92485322 | 2 | 0 | NA | NA | 6 | 0.3333 | 4 | 0 | 12 | 0.1667 | 1 | 0 | 1 | 0 | 3 | 0.6667 | 6 | 0.3333 | 11 | 0.3636 |
| 7_92485693 | 2 | 0.5 | NA | NA | 6 | 0.1667 | 4 | 0 | 12 | 0.1667 | 1 | 0 | 1 | 0 | 3 | 0 | 5 | 0.4 | 10 | 0.2 |
| 7_92486116 | 1 | 0 | 1 | 0 | 2 | 1 | 1 | 0 | 5 | 0.4 | NA | NA | NA | NA | 2 | 0.5 | 4 | 0.75 | 6 | 0.6667 |
| 7_92486123 | 1 | 0 | 1 | 0 | 1 | 1 | 1 | 1 | 4 | 0.5 | NA | NA | NA | NA | 2 | 0 | 3 | 0.6667 | 5 | 0.4 |
| 7_92486767 | 1 | 1 | 1 | 0 | 1 | 0 | 1 | 0 | 4 | 0.25 | NA | NA | 1 | 1 | NA | NA | 2 | 0 | 3 | 0.3333 |
| 7_92582569 | NA | NA | 2 | 0 | 2 | 0.5 | 2 | 0.5 | 6 | 0.3333 | NA | NA | 1 | 1 | NA | NA | 2 | 0.5 | 3 | 0.6667 |
| 7_92582623 | NA | NA | 2 | 0 | 2 | 0.5 | 2 | 1 | 6 | 0.5 | NA | NA | NA | NA | NA | NA | 2 | 0.5 | 2 | 0.5 |
| 7_92582859 | NA | NA | 2 | 0 | 1 | 1 | 2 | 1 | 5 | 0.6 | NA | NA | 1 | 1 | NA | NA | 2 | 1 | 3 | 1 |
| 7_92583011 | NA | NA | 3 | 1 | 1 | 0 | 1 | 0 | 5 | 0.6 | NA | NA | 1 | 1 | NA | NA | 2 | 1 | 3 | 1 |
| 7_92729440 | 4 | 0.25 | 1 | 1 | 4 | 0.75 | 2 | 0.5 | 11 | 0.5455 | 1 | 1 | 3 | 0.3333 | 7 | 0.4286 | 6 | 0.1667 | 17 | 0.3529 |
| 7_92729473 | 5 | 0.6 | NA | NA | 2 | 1 | 1 | 0 | 8 | 0.625 | NA | NA | NA | NA | 5 | 0.2 | 2 | 0.5 | 7 | 0.2857 |
| 7_92729556 | 5 | 0.2 | 1 | 1 | 4 | 0.5 | 4 | 0.5 | 14 | 0.4286 | 1 | 0 | 2 | 1 | 7 | 0.4286 | 6 | 0.3333 | 16 | 0.4375 |
| 7_92729977 | 9 | 0 | 2 | 0 | 5 | 0.2 | 4 | 0 | 20 | 0.05 | 2 | 0 | 3 | 0 | 7 | 0 | 6 | 0.3333 | 18 | 0.1111 |
| 7_92730028 | 8 | 0.125 | 1 | 0 | 5 | 0.2 | 2 | 0 | 16 | 0.125 | 2 | 0 | 3 | 0 | 6 | 0.1667 | 5 | 0.2 | 16 | 0.125 |
| 7_92730070 | 9 | 0.4444 | 1 | 0 | 4 | 0.25 | 4 | 0.25 | 18 | 0.3333 | 1 | 1 | 3 | 0.6667 | 7 | 0.4286 | 5 | 0.4 | 16 | 0.5 |
| 7_92730289 | 3 | 1 | 1 | 1 | 2 | 1 | 2 | 1 | 8 | 1 | 2 | 1 | 1 | 1 | 4 | 1 | 4 | 1 | 11 | 1 |
| 7_92730428 | 8 | 0 | 2 | 0 | 5 | 0.2 | 4 | 0 | 19 | 0.0526 | 2 | 0 | 3 | 0 | 8 | 0 | 6 | 0.1667 | 19 | 0.0526 |
| 7_92730478 | 7 | 0.5714 | 2 | 0 | 4 | 1 | 1 | 1 | 14 | 0.6429 | 2 | 0 | 2 | 1 | 7 | 0 | 3 | 0.3333 | 14 | 0.2143 |
| 7_92730482 | 5 | 0.8 | NA | NA | 4 | 0.5 | 3 | 0.3333 | 12 | 0.5833 | 2 | 1 | 2 | 0.5 | 4 | 0.75 | 4 | 0.75 | 12 | 0.75 |
| 7_92730492 | 9 | 0 | 2 | 1 | 5 | 0.6 | 4 | 0.75 | 20 | 0.4 | 2 | 0 | 3 | 0.3333 | 7 | 0.4286 | 6 | 0.8333 | 18 | 0.5 |
| 7_92730511 | 5 | 0.6 | 1 | 1 | 2 | 0.5 | 2 | 0.5 | 10 | 0.6 | 2 | 0.5 | 3 | 0.3333 | 6 | 0 | 4 | 0.75 | 15 | 0.3333 |
| 7_92730601 | 7 | 0.1429 | 2 | 0 | 5 | 0.2 | 3 | 0 | 17 | 0.1176 | 3 | 0 | 3 | 0 | 7 | 0 | 5 | 0.2 | 18 | 0.0556 |
| 7_92730613 | 7 | 0.1429 | 2 | 0 | 5 | 0.4 | 4 | 0 | 18 | 0.1667 | 2 | 0 | 3 | 0 | 7 | 0.2857 | 5 | 0 | 17 | 0.1176 |
| 7_92730629 | 8 | 0.125 | 2 | 0 | 2 | 0 | 2 | 0 | 14 | 0.0714 | 3 | 0 | 2 | 0 | 8 | 0.25 | 3 | 0.3333 | 16 | 0.1875 |
| 7_92730743 | 7 | 0.7143 | 2 | 1 | 4 | 0.75 | 2 | 1 | 15 | 0.8 | 3 | 0.6667 | 2 | 1 | 6 | 0.8333 | 4 | 1 | 15 | 0.8667 |
| 7_92730767 | 8 | 0.125 | 1 | 1 | 3 | 0 | 2 | 0 | 14 | 0.1429 | 1 | 0 | 2 | 0.5 | 7 | 0.2857 | 5 | 0.4 | 15 | 0.3333 |
| 7_96492566 | 1 | 0 | 1 | 1 | 4 | 0.25 | 1 | 0 | 7 | 0.2857 | NA | NA | 2 | 0 | 2 | 1 | 2 | 1 | 6 | 0.6667 |
| 7_96544054 | NA | NA | 1 | 1 | NA | NA | NA | NA | 1 | 1 | NA | NA | NA | NA | 3 | 0.6667 | 1 | 0 | 4 | 0.5 |
| 7_96544133 | NA | NA | 1 | 1 | NA | NA | NA | NA | 1 | 1 | 1 | 1 | NA | NA | 3 | 0.6667 | 1 | 0 | 5 | 0.6 |
| 7_96544565 | NA | NA | 1 | 1 | 1 | 0 | NA | NA | 2 | 0.5 | 1 | 1 | NA | NA | 3 | 1 | 2 | 0 | 6 | 0.6667 |
| 7_96544618 | NA | NA | 1 | 0 | 1 | 1 | 1 | 0 | 3 | 0.3333 | 1 | 0 | NA | NA | 3 | 0.6667 | 2 | 1 | 6 | 0.6667 |
| 7_96544676 | NA | NA | 1 | 0 | 1 | 1 | 1 | 0 | 3 | 0.3333 | 1 | 0 | NA | NA | 3 | 0.6667 | 1 | 0 | 5 | 0.4 |
| 7_96545095 | NA | NA | 1 | 1 | 1 | 1 | NA | NA | 2 | 1 | 1 | 0 | NA | NA | 4 | 0.75 | 1 | 1 | 6 | 0.6667 |
| 7_96545183 | NA | NA | NA | NA | 1 | 0 | 1 | 1 | 2 | 0.5 | 1 | 1 | NA | NA | 4 | 0.75 | 2 | 1 | 7 | 0.8571 |
| 7_96545238 | NA | NA | 1 | 1 | 1 | 1 | 1 | 0 | 3 | 0.6667 | 1 | 1 | NA | NA | 4 | 1 | 2 | 1 | 7 | 1 |
| 7_96545351 | NA | NA | 1 | 1 | 1 | 0 | 1 | 0 | 3 | 0.3333 | 1 | 1 | NA | NA | 4 | 0.25 | 3 | 0 | 8 | 0.25 |
| 7_96545368 | NA | NA | 1 | 1 | 1 | 1 | 1 | 1 | 3 | 1 | NA | NA | NA | NA | 1 | 1 | 3 | 1 | 4 | 1 |
| 7_96555284 | NA | NA | 1 | 1 | 1 | 0 | 1 | 0 | 3 | 0.3333 | NA | NA | NA | NA | 4 | 0.75 | 4 | 0.75 | 8 | 0.75 |
| 7_96555305 | NA | NA | 1 | 0 | 1 | 0 | 2 | 0.5 | 4 | 0.25 | 1 | 0 | NA | NA | 2 | 0 | 4 | 0.25 | 7 | 0.1429 |
| 7_96555332 | NA | NA | 1 | 1 | 1 | 0 | 1 | 0 | 3 | 0.3333 | 1 | 1 | NA | NA | 4 | 0 | 3 | 0 | 8 | 0.125 |
| 7_96555357 | NA | NA | 1 | 0 | 2 | 0.5 | 2 | 0.5 | 5 | 0.4 | 1 | 1 | NA | NA | 5 | 1 | 4 | 0.75 | 10 | 0.9 |
| 7_96555466 | NA | NA | 1 | 0 | 1 | 1 | 2 | 0.5 | 4 | 0.5 | 2 | 0.5 | NA | NA | 4 | 1 | 4 | 0.75 | 10 | 0.8 |
| 7_96555529 | NA | NA | 1 | 0 | 2 | 0.5 | 2 | 0.5 | 5 | 0.4 | 2 | 1 | NA | NA | 5 | 0.8 | 4 | 0.25 | 11 | 0.6364 |
| 7_96555615 | NA | NA | 1 | 1 | 1 | 1 | 1 | 1 | 3 | 1 | 2 | 1 | NA | NA | 3 | 1 | 4 | 1 | 9 | 1 |
| 7_96555649 | 1 | 1 | 1 | 1 | 1 | 1 | 2 | 1 | 5 | 1 | 2 | 1 | NA | NA | 4 | 1 | 4 | 1 | 10 | 1 |
| 7_96555677 | NA | NA | 1 | 1 | 1 | 1 | 1 | 0 | 3 | 0.6667 | 2 | 0.5 | NA | NA | 4 | 0.75 | 1 | 0 | 7 | 0.5714 |

| | | | | | | | | | | | | | | | | | | | | |
|---|---|---|---|---|---|---|---|---|---|---|---|---|---|---|---|---|---|---|---|---|
| 7_9655573( | NA | NA | 1 | 1 | 1 | 1 | 2 | 1 | 4 | 1 | 2 | 0.5 | NA | NA | 5 | 0.8 | 4 | 0.75 | 11 | 0.7273 |
| 7_9655577 | 1 | 0 | 1 | 0 | 2 | 0 | 2 | 0.5 | 6 | 0.1667 | 2 | 0 | NA | NA | 3 | 0 | 3 | 0.3333 | 8 | 0.125 |
| 7_9655581 | 1 | 1 | 1 | 1 | 1 | 1 | 2 | 0.5 | 5 | 0.8 | 2 | 1 | NA | NA | 5 | 1 | 4 | 0.75 | 11 | 0.9091 |
| 7_9655583 | 1 | 1 | 1 | 1 | 1 | 0 | 2 | 0 | 5 | 0.4 | 1 | 1 | NA | NA | 4 | 0.5 | 4 | 0.5 | 9 | 0.5556 |
| 7_9655586 | 1 | 1 | 1 | 1 | 2 | 1 | 3 | 0.6667 | 7 | 0.8571 | 1 | 1 | NA | NA | 5 | 0.8 | 4 | 0.75 | 10 | 0.8 |
| 7_9655592( | NA | NA | 1 | 1 | 2 | 0 | 2 | 0 | 5 | 0.2 | 2 | 1 | 1 | 1 | 5 | 0.2 | 4 | 0.5 | 12 | 0.5 |
| 7_9655595 | 1 | 0 | 1 | 1 | 1 | 0 | 2 | 0 | 5 | 0.2 | 3 | 0.3333 | 1 | 0 | 5 | 0 | 5 | 0.2 | 14 | 0.1429 |
| 7_9655663 | 3 | 0 | 2 | 1 | 3 | 0 | 2 | 0 | 10 | 0.2 | 3 | 0 | 1 | 0 | 5 | 0.2 | 3 | 0 | 12 | 0.0833 |
| 7_9655732( | 5 | 0.4 | 4 | 0 | 4 | 0 | 3 | 0.3333 | 16 | 0.1875 | 4 | 0.75 | 2 | 0 | 7 | 0.1429 | 6 | 0.6667 | 19 | 0.4211 |
| 7_9655746 | 3 | 0.3333 | 4 | 0 | 4 | 0 | 2 | 0 | 13 | 0.0769 | 5 | 0.6 | 2 | 0 | 9 | 0.2222 | 6 | 0.3333 | 22 | 0.3182 |
| 7_9655754( | 3 | 0 | 3 | 0 | 3 | 0.3333 | 3 | 0.6667 | 12 | 0.25 | 2 | 1 | 2 | 0 | 9 | 0.3333 | 5 | 0.8 | 18 | 0.5 |
| 7_9655943( | 7 | 0.2857 | 9 | 0.4444 | 6 | 0.1667 | 8 | 0.375 | 30 | 0.3333 | 6 | 0 | 6 | 0.1667 | 11 | 0.0909 | 3 | 0.3333 | 26 | 0.1154 |
| 7_9655952( | 5 | 0.4 | 7 | 0.2857 | 7 | 0.2857 | 8 | 0.375 | 27 | 0.3333 | 8 | 0.25 | 7 | 0.1429 | 13 | 0.3846 | 3 | 0.6667 | 31 | 0.3226 |
| 7_9655957 | 7 | 0.5714 | 10 | 0.1 | 7 | 0 | 11 | 0.3636 | 35 | 0.2571 | 8 | 0.5 | 8 | 0.25 | 15 | 0.2667 | 7 | 0.5714 | 38 | 0.3684 |
| 7_9655959 | 9 | 0.1111 | 10 | 0.2 | 8 | 0.125 | 11 | 0 | 38 | 0.1053 | 7 | 0 | 8 | 0 | 15 | 0 | 8 | 0.375 | 38 | 0.0789 |
| 7_9655980 | 9 | 0 | 9 | 0.2222 | 7 | 0.2857 | 10 | 0.2 | 35 | 0.1714 | 6 | 0 | 9 | 0 | 15 | 0.1333 | 7 | 0.1429 | 37 | 0.0811 |
| 7_9655999 | 9 | 1 | 9 | 1 | 8 | 1 | 12 | 0.75 | 38 | 0.9211 | 8 | 0.875 | 7 | 0.8571 | 15 | 0.8 | 7 | 0.7143 | 37 | 0.8108 |
| 7_9656006 | 10 | 0.8 | 11 | 0.6364 | 9 | 0.7778 | 10 | 0.7 | 40 | 0.725 | 8 | 0.75 | 9 | 0.4444 | 15 | 0.5333 | 7 | 0.8571 | 39 | 0.6154 |
| 7_9656010 | 7 | 0.2857 | 12 | 0.4167 | 10 | 0.6 | 11 | 0.1818 | 40 | 0.375 | 7 | 0.4286 | 7 | 0.5714 | 14 | 0.6429 | 8 | 0.5 | 36 | 0.5556 |
| 7_9656016 | 10 | 0.3 | 13 | 0.0769 | 8 | 0 | 10 | 0 | 41 | 0.0976 | 8 | 0 | 7 | 0.4286 | 15 | 0.0667 | 8 | 0.375 | 38 | 0.1842 |
| 7_9671192 | 3 | 1 | 5 | 0.6 | 3 | 1 | 1 | 1 | 12 | 0.8333 | 2 | 0.5 | 2 | 1 | NA | NA | 3 | 1 | 7 | 0.8571 |
| 7_9671198 | 3 | 1 | 5 | 0.4 | 3 | 0 | 1 | 1 | 12 | 0.5 | 2 | 0 | 2 | 1 | NA | NA | 2 | 1 | 6 | 0.6667 |
| 7_9671210 | 3 | 0.6667 | 3 | 1 | 1 | 1 | NA | NA | 7 | 0.8571 | 1 | 0 | 1 | 1 | NA | NA | 1 | 1 | 3 | 0.6667 |
| 7_9671214 | 3 | 0.6667 | 4 | 1 | 3 | 0.6667 | 1 | 1 | 11 | 0.8182 | 2 | 1 | 2 | 1 | NA | NA | 3 | 0.6667 | 7 | 0.8571 |
| 7_9671216 | 2 | 0.5 | 6 | 0.3333 | 3 | 0 | 1 | 1 | 12 | 0.3333 | 1 | 0 | 1 | 0 | NA | NA | 3 | 0.3333 | 5 | 0.2 |
| 7_9671228 | 3 | 1 | 5 | 0.8 | 3 | 0 | 1 | 0 | 12 | 0.5833 | 2 | 0 | 2 | 0.5 | NA | NA | 3 | 0.3333 | 7 | 0.2857 |
| 7_9694819( | NA | NA | NA | NA | 1 | 0 | 1 | 1 | 2 | 0.5 | NA | NA | NA | NA | 1 | 1 | 1 | 0 | 2 | 0.5 |
| 7_9694826 | NA | NA | NA | NA | 1 | 1 | 1 | 0 | 2 | 0.5 | NA | NA | NA | NA | 1 | 0 | 1 | 1 | 2 | 0.5 |
| 7_9694833 | NA | NA | NA | NA | 1 | 0 | 1 | 1 | 2 | 0.5 | NA | NA | NA | NA | 1 | 0 | 1 | 1 | 2 | 0.5 |
| 7_9695417 | NA | NA | 1 | 0 | 2 | 0.5 | 1 | 1 | 4 | 0.5 | NA | NA | NA | NA | NA | NA | 1 | 1 | 1 | 1 |
| 7_9695436 | NA | NA | 1 | 1 | 1 | 0 | 1 | 1 | 3 | 0.6667 | NA | NA | NA | NA | 1 | 1 | 1 | 1 | 2 | 1 |
| 7_9695443 | NA | NA | 1 | 1 | 2 | 0 | 1 | 1 | 4 | 0.5 | NA | NA | NA | NA | 1 | 0 | 1 | 1 | 2 | 0.5 |
| 7_9695452 | NA | NA | 1 | 1 | 1 | 1 | 1 | 1 | 3 | 1 | NA | NA | NA | NA | 1 | 1 | NA | NA | 1 | 1 |
| 7_9695486( | NA | NA | 2 | 1 | 1 | 1 | 1 | 1 | 4 | 1 | NA | NA | NA | NA | 1 | 1 | 1 | 1 | 2 | 1 |
| 7_9695494( | NA | NA | 2 | 0 | 1 | 0 | 1 | 1 | 4 | 0.25 | NA | NA | NA | NA | 1 | 0 | 1 | 1 | 2 | 0.5 |
| 7_9695645 | 1 | 1 | 3 | 0 | 1 | 0 | 1 | 1 | 6 | 0.3333 | NA | NA | 3 | 0 | 1 | 0 | 1 | 1 | 5 | 0.2 |
| 7_9695868 | 5 | 0.2 | 1 | 0 | 1 | 0 | 3 | 0 | 10 | 0.1 | 3 | 0 | 1 | 0 | 3 | 0.3333 | NA | NA | 7 | 0.1429 |
| 7_9695872 | 4 | 0.25 | 1 | 0 | 1 | 0 | 3 | 0.3333 | 9 | 0.2222 | 3 | 1 | 1 | 0 | 3 | 0.6667 | NA | NA | 7 | 0.7143 |
| 7_9695955 | 1 | 0 | 1 | 0 | 1 | 1 | 2 | 0.5 | 5 | 0.4 | 1 | 1 | 1 | 1 | 2 | 0.5 | NA | NA | 4 | 0.75 |
| 7_9696060 | 4 | 0 | NA | NA | NA | NA | 3 | 0.3333 | 7 | 0.1429 | 1 | 0 | 1 | 0 | 3 | 0.3333 | NA | NA | 5 | 0.2 |
| 7_9696070 | 5 | 0.8 | NA | NA | 1 | 1 | 2 | 1 | 8 | 0.875 | 1 | 1 | NA | NA | 3 | 1 | NA | NA | 4 | 1 |
| 7_9696078 | 5 | 0.4 | NA | NA | 1 | 0 | 3 | 0.3333 | 9 | 0.3333 | 1 | 0 | NA | NA | 3 | 0.3333 | NA | NA | 4 | 0.25 |
| 7_9696111 | 3 | 0 | NA | NA | NA | NA | 3 | 0.3333 | 6 | 0.1667 | 1 | 0 | NA | NA | 3 | 0.3333 | NA | NA | 4 | 0.25 |
| 7_9696116 | 2 | 1 | NA | NA | NA | NA | 3 | 1 | 5 | 1 | 1 | 1 | NA | NA | 3 | 1 | NA | NA | 4 | 1 |
| 7_9696145 | 3 | 0.3333 | NA | NA | NA | NA | 2 | 0 | 5 | 0.2 | NA | NA | NA | NA | 1 | 1 | NA | NA | 1 | 1 |
| 7_9699706 | 1 | 0 | NA | NA | 2 | 0.5 | 2 | 0.5 | 5 | 0.4 | 1 | 1 | 1 | 0 | NA | NA | NA | NA | 2 | 0.5 |
| 7_9699708 | 1 | 1 | 1 | 0 | 2 | 1 | 2 | 1 | 6 | 0.8333 | NA | NA | 1 | 1 | NA | NA | NA | NA | 1 | 1 |
| 7_9699726 | 2 | 1 | 1 | 1 | 1 | 1 | 2 | 1 | 6 | 1 | 1 | 1 | 1 | 1 | NA | NA | NA | NA | 2 | 1 |
| 7_9699740 | 2 | 0.5 | 1 | 0 | 1 | 1 | 2 | 0.5 | 6 | 0.5 | 1 | 0 | 1 | 1 | NA | NA | NA | NA | 2 | 0.5 |
| 7_9699743 | 1 | 1 | 1 | 1 | 2 | 0 | 2 | 1 | 6 | 0.6667 | 1 | 1 | 1 | 1 | NA | NA | NA | NA | 2 | 1 |
| 7_9699779 | 3 | 1 | 1 | 1 | 2 | 1 | 2 | 1 | 8 | 1 | 1 | 1 | 1 | 1 | NA | NA | 1 | 1 | 3 | 1 |
| 7_9699785 | 3 | 0.3333 | NA | NA | 3 | 0.6667 | 2 | 0 | 8 | 0.375 | 1 | 1 | 1 | 1 | NA | NA | 1 | 0 | 3 | 0.6667 |
| 7_9699813 | 4 | 1 | 1 | 1 | 3 | 1 | 2 | 1 | 10 | 1 | 1 | 1 | 1 | 1 | NA | NA | 1 | 1 | 3 | 1 |
| 7_9699841 | 5 | 0.2 | 2 | 0.5 | 4 | 0 | 3 | 0 | 14 | 0.1429 | 1 | 0 | 1 | 0 | NA | NA | 1 | 1 | 3 | 0.3333 |
| 7_9699844 | 4 | 1 | 2 | 1 | 3 | 1 | 2 | 1 | 11 | 1 | 1 | 1 | 1 | 1 | NA | NA | 1 | 1 | 3 | 1 |
| 7_9699861 | 5 | 0.6 | NA | NA | 3 | 0.6667 | 3 | 0.6667 | 11 | 0.6364 | 1 | 1 | 1 | 1 | NA | NA | 1 | 0 | 3 | 0.6667 |
| 7_9699891 | 3 | 0.3333 | 2 | 0.5 | 4 | 0.25 | 2 | 0 | 11 | 0.2727 | 1 | 1 | NA | NA | 1 | 0 | 1 | 0 | 3 | 0.3333 |
| 7_9702910 | 1 | 0 | 2 | 0.5 | 3 | 0 | 2 | 0 | 8 | 0.125 | 1 | 0 | NA | NA | 1 | 0 | 3 | 0.3333 | 5 | 0.2 |
| 7_9704093 | 2 | 0.5 | 2 | 0.5 | 4 | 0.75 | 2 | 0.5 | 10 | 0.6 | NA | NA | 1 | 1 | 2 | 0.5 | 1 | 0 | 4 | 0.5 |
| 7_9704096( | 2 | 0 | 2 | 0 | 3 | 0.6667 | 2 | 0 | 9 | 0.2222 | 1 | 1 | 1 | 1 | 2 | 0.5 | 1 | 0 | 5 | 0.6 |
| 7_9704102 | 2 | 1 | NA | NA | 1 | 1 | NA | NA | 3 | 1 | 1 | 1 | 1 | 0 | 1 | 0 | 2 | 0.5 | 5 | 0.4 |
| 7_9704107( | 1 | 1 | 1 | 1 | 3 | 1 | NA | NA | 5 | 1 | 1 | 1 | 1 | 1 | 1 | 1 | 2 | 0.5 | 5 | 0.8 |
| 7_9704117( | 2 | 0 | 1 | 1 | 2 | 0 | NA | NA | 5 | 0.2 | NA | NA | 1 | 1 | 1 | 0 | 1 | 0 | 3 | 0.3333 |
| 7_9704121 | 2 | 0.5 | 1 | 0 | 4 | 0.25 | NA | NA | 7 | 0.2857 | 1 | 0 | NA | NA | 1 | 0 | 2 | 0.5 | 4 | 0.25 |
| 7_9704952 | 51 | 0.0588 | 51 | 0.098 | 44 | 0.1364 | 42 | 0.1667 | 188 | 0.1117 | 34 | 0.0588 | 51 | 0.1765 | 37 | 0.1622 | 40 | 0.125 | 162 | 0.1358 |

| | | | | | | | | | | | | | | | | | | | | |
|---|---|---|---|---|---|---|---|---|---|---|---|---|---|---|---|---|---|---|---|---|
| 7_9706259 | 4 | 0.75 | 2 | 1 | 4 | 0.75 | 2 | 0 | 12 | 0.6667 | 4 | 0.75 | 2 | 0.5 | 4 | 0.75 | 2 | 1 | 12 | 0.75 |
| 7_9706637 | 4 | 0.25 | 2 | 0 | 3 | 0 | 2 | 0 | 11 | 0.0909 | 3 | 0.3333 | 2 | 0 | 3 | 0 | NA | NA | 8 | 0.125 |
| 7_9706649 | 4 | 0.5 | 1 | 0 | 2 | 0 | 1 | 1 | 8 | 0.375 | 2 | 0.5 | NA | NA | 4 | 0.25 | NA | NA | 6 | 0.3333 |
| 7_9719048 | 4 | 0 | 4 | 0 | 2 | 0 | 6 | 0.1667 | 16 | 0.0625 | 3 | 0 | 2 | 0 | 5 | 0 | 3 | 0.3333 | 13 | 0.0769 |
| 7_9719970 | 2 | 0.5 | 7 | 0.5714 | 3 | 0 | 4 | 0.25 | 16 | 0.375 | 4 | 0.25 | 1 | 1 | 8 | 0.875 | 7 | 0.4286 | 20 | 0.6 |
| 7_9719972 | 3 | 0.6667 | 7 | 0.2857 | 2 | 0.5 | 5 | 0.6 | 17 | 0.4706 | 3 | 0.6667 | 3 | 1 | 9 | 0.5556 | 8 | 0.875 | 23 | 0.7391 |
| 7_9731048 | 2 | 0 | 2 | 0.5 | 1 | 0 | NA | NA | 5 | 0.2 | NA | NA | 1 | 1 | NA | NA | NA | NA | 1 | 1 |
| 7_9731055 | 1 | 1 | 2 | 0 | 1 | 0 | NA | NA | 4 | 0.25 | 1 | 0 | 2 | 0.5 | NA | NA | 1 | 0 | 4 | 0.25 |
| 7_9731089 | 2 | 1 | 2 | 1 | 1 | 1 | NA | NA | 5 | 1 | 1 | 0 | 2 | 0.5 | NA | NA | 1 | 1 | 4 | 0.5 |
| 7_9731133 | 1 | 1 | 2 | 0.5 | 1 | 1 | NA | NA | 4 | 0.75 | 1 | 1 | 1 | 1 | NA | NA | 1 | 0 | 3 | 0.6667 |
| 7_9731153 | 2 | 0.5 | 2 | 0 | 1 | 0 | NA | NA | 5 | 0.2 | NA | NA | 2 | 0 | NA | NA | 2 | 0.5 | 4 | 0.25 |
| 7_9731154 | 2 | 0 | 2 | 0.5 | 1 | 0 | NA | NA | 5 | 0.2 | 1 | 1 | 1 | 0 | NA | NA | 1 | 0 | 3 | 0.3333 |
| 7_9733004 | 3 | 0.3333 | 4 | 0 | 2 | 0 | NA | NA | 9 | 0.1111 | 2 | 0 | 2 | 0 | 3 | 0 | 4 | 0.25 | 11 | 0.0909 |
| 7_9733492 | 3 | 0 | 4 | 0.5 | 3 | 0.3333 | NA | NA | 10 | 0.3 | 2 | 0 | 2 | 0 | 3 | 0 | 4 | 0.5 | 11 | 0.1818 |
| 7_9733495 | 3 | 1 | 2 | 1 | 1 | 1 | NA | NA | 6 | 1 | 2 | 1 | 3 | 1 | 3 | 1 | 4 | 1 | 12 | 1 |
| 7_9733496 | 3 | 1 | 4 | 0.25 | 3 | 0.3333 | NA | NA | 10 | 0.5 | 2 | 0 | 3 | 0 | 3 | 0.6667 | 4 | 0.5 | 12 | 0.3333 |
| 7_9733498 | 2 | 1 | 4 | 1 | 3 | 1 | NA | NA | 9 | 1 | 1 | 1 | 2 | 0.5 | 2 | 1 | 4 | 1 | 9 | 0.8889 |
| 7_9733501 | 3 | 1 | 3 | 0.6667 | 3 | 1 | NA | NA | 9 | 0.8889 | 1 | 1 | 2 | 1 | 2 | 1 | 4 | 1 | 9 | 1 |
| 7_9733505 | 3 | 1 | 3 | 1 | 3 | 1 | NA | NA | 9 | 1 | 1 | 1 | 2 | 1 | 3 | 1 | 4 | 1 | 10 | 1 |
| 7_9733509 | 2 | 1 | 3 | 0.3333 | 3 | 0.6667 | NA | NA | 8 | 0.625 | 2 | 0 | 2 | 0.5 | 2 | 1 | 4 | 1 | 10 | 0.7 |
| 7_9733510 | 2 | 1 | NA | NA | 2 | 0.5 | NA | NA | 4 | 0.75 | 1 | 1 | 1 | 0 | 1 | 1 | 2 | 1 | 5 | 0.8 |
| 7_9733515 | 3 | 0.6667 | 4 | 0.75 | 2 | 1 | NA | NA | 9 | 0.7778 | 2 | 1 | 2 | 1 | 2 | 1 | 3 | 0.6667 | 9 | 0.8889 |
| 7_9733518 | 2 | 1 | 2 | 1 | 2 | 0.5 | NA | NA | 6 | 0.8333 | 1 | 1 | 2 | 1 | 3 | 0.6667 | 1 | 1 | 7 | 0.8571 |
| 7_9733519 | 3 | 0.6667 | 4 | 0.75 | 3 | 1 | NA | NA | 10 | 0.8 | 1 | 0 | 3 | 1 | 3 | 1 | 3 | 0.6667 | 10 | 0.8 |
| 7_9733535 | 2 | 1 | 3 | 0.6667 | 3 | 1 | NA | NA | 8 | 0.875 | 2 | 1 | 3 | 0.6667 | 1 | 1 | 4 | 0.75 | 10 | 0.8 |
| 7_9733548 | 2 | 0.5 | 4 | 0 | 3 | 0 | NA | NA | 9 | 0.1111 | 2 | 0 | 3 | 0 | 3 | 0 | 3 | 0.3333 | 11 | 0.0909 |
| 7_9733565 | 2 | 0 | 2 | 0.5 | 2 | 0.5 | NA | NA | 6 | 0.3333 | NA | NA | NA | NA | 2 | 1 | 2 | 0.5 | 4 | 0.75 |
| 7_9733868 | 5 | 0.2 | 1 | 0 | NA | NA | 2 | 0 | 8 | 0.125 | NA | NA | 2 | 0.5 | 1 | 0 | 1 | 0 | 4 | 0.25 |
| 7_9736756 | 17 | 0.5882 | 9 | 0.5556 | 10 | 0.4 | 12 | 0.5833 | 48 | 0.5417 | 12 | 0.9167 | 10 | 0.6 | 9 | 0.8889 | 13 | 0.5385 | 44 | 0.7273 |
| 7_9736759 | 17 | 0.5882 | 9 | 0 | 8 | 0 | 12 | 0.0833 | 46 | 0.2391 | 12 | 0.25 | 11 | 0.1818 | 11 | 0.2727 | 12 | 0.4167 | 46 | 0.2826 |
| 7_9736778 | 16 | 0.0625 | 9 | 0.3333 | 8 | 0 | 13 | 0.2308 | 46 | 0.1522 | 14 | 0.0714 | 10 | 0.2 | 9 | 0.1111 | 13 | 0.3077 | 46 | 0.1739 |
| 7_9737570 | 1 | 1 | 2 | 1 | 3 | 0.6667 | 1 | 1 | 7 | 0.8571 | 1 | 1 | 1 | 1 | 1 | 0 | 4 | 0.75 | 7 | 0.7143 |
| 7_9737581 | 1 | 0 | 1 | 1 | NA | NA | 2 | 1 | 4 | 0.75 | NA | NA | 1 | 1 | 1 | 1 | 2 | 0.5 | 4 | 0.75 |
| 7_9738696 | NA | NA | 2 | 1 | 3 | 1 | 2 | 1 | 7 | 1 | NA | NA | 1 | 1 | NA | NA | 4 | 0.75 | 5 | 0.8 |
| 7_9738698 | 1 | 1 | 2 | 1 | 3 | 0.6667 | 2 | 0.5 | 8 | 0.75 | 1 | 0 | 1 | 0 | NA | NA | 4 | 0.5 | 6 | 0.3333 |
| 7_9756043 | 26 | 0.1923 | 36 | 0.1389 | 45 | 0.1556 | 17 | 0.1176 | 124 | 0.1532 | 17 | 0.0588 | 27 | 0.1481 | 19 | 0.1053 | 22 | 0.0455 | 85 | 0.0941 |
| 7_9756045 | 26 | 0.3077 | 31 | 0.1935 | 36 | 0.2222 | 10 | 0.5 | 103 | 0.2621 | 20 | 0.1 | 27 | 0.1852 | 19 | 0.3684 | 21 | 0.1905 | 87 | 0.2069 |
| 7_9756051 | 27 | 0.3704 | 34 | 0.2941 | 40 | 0.375 | 16 | 0.4375 | 117 | 0.359 | 21 | 0.4286 | 28 | 0.3214 | 19 | 0.4211 | 21 | 0.1905 | 89 | 0.3371 |
| 7_9756053 | 25 | 0.12 | 36 | 0.1667 | 36 | 0.1389 | 17 | 0.2353 | 114 | 0.1579 | 19 | 0.2105 | 27 | 0.1852 | 19 | 0.1579 | 21 | 0.2857 | 86 | 0.2093 |
| 7_9756063 | 21 | 0.0952 | 33 | 0.1515 | 35 | 0.1429 | 12 | 0 | 101 | 0.1188 | 17 | 0.1765 | 27 | 0.1481 | 17 | 0.2353 | 19 | 0.2105 | 80 | 0.1875 |
| 7_9756072 | 27 | 0.037 | 39 | 0.1026 | 36 | 0.0556 | 16 | 0.125 | 118 | 0.0763 | 16 | 0.125 | 28 | 0.1071 | 18 | 0.2222 | 15 | 0 | 77 | 0.1169 |
| 7_9756125 | 32 | 0.2188 | 38 | 0.2368 | 40 | 0.125 | 15 | 0.2 | 125 | 0.192 | 22 | 0.0909 | 30 | 0.1 | 21 | 0.1429 | 22 | 0.2273 | 95 | 0.1368 |
| 7_9756137 | 28 | 0.1429 | 38 | 0.0526 | 35 | 0.0857 | 14 | 0 | 115 | 0.0783 | 23 | 0.1304 | 29 | 0.069 | 16 | 0.125 | 24 | 0.0417 | 92 | 0.087 |
| 7_9777097 | 1 | 0 | 3 | 0 | 2 | 0 | 2 | 0.5 | 8 | 0.125 | 1 | 0 | NA | NA | 3 | 0 | 4 | 0.25 | 8 | 0.125 |
| 7_9777098 | 2 | 0 | 3 | 0 | 2 | 0 | 4 | 0.5 | 11 | 0.1818 | 2 | 0 | 1 | 0 | 3 | 0 | 4 | 0.25 | 10 | 0.1 |
| 7_9777101 | 2 | 0 | 5 | 0 | 3 | 0.3333 | 5 | 0.2 | 15 | 0.1333 | 3 | 0 | 1 | 0 | 4 | 0 | 4 | 0.25 | 12 | 0.0833 |
| 7_9777136 | 2 | 0 | 4 | 0 | 2 | 0 | 4 | 0.5 | 12 | 0.1667 | 3 | 0.3333 | 1 | 0 | 4 | 0.5 | 3 | 0 | 11 | 0.2727 |
| 7_9777162 | 1 | 0 | 3 | 0.6667 | 2 | 0.5 | 4 | 0.5 | 10 | 0.5 | 2 | 0.5 | 1 | 0 | 4 | 0.5 | 3 | 1 | 10 | 0.6 |
| 7_9777176 | 2 | 0 | 5 | 0.2 | 2 | 0.5 | 2 | 0.5 | 11 | 0.2727 | 3 | 0.3333 | NA | NA | 3 | 0 | 3 | 0.6667 | 9 | 0.3333 |
| 7_9777207 | 1 | 0 | 4 | 0 | 2 | 0.5 | 2 | 0.5 | 9 | 0.2222 | 2 | 0 | 1 | 1 | 4 | 0 | 3 | 0.3333 | 10 | 0.2 |
| 7_9777214 | 1 | 0 | 3 | 0.3333 | 2 | 0.5 | 3 | 1 | 9 | 0.5556 | 2 | 0.5 | NA | NA | 4 | 0.75 | 3 | 0.6667 | 9 | 0.6667 |
| 7_9777216 | 1 | 0 | 4 | 0.25 | 1 | 0 | 3 | 0.3333 | 9 | 0.2222 | 2 | 0.5 | 1 | 0 | 4 | 0.25 | 2 | 0 | 9 | 0.2222 |
| 7_9777274 | 1 | 0 | 3 | 0 | 2 | 0 | 3 | 0.3333 | 9 | 0.1111 | 2 | 0 | 1 | 0 | 2 | 0 | 3 | 0.3333 | 8 | 0.125 |
| 7_9777299 | 1 | 0 | 3 | 0 | 2 | 0 | 2 | 0.5 | 8 | 0.125 | 2 | 0 | 1 | 0 | 5 | 0.2 | 1 | 1 | 9 | 0.2222 |
| 7_9777301 | 1 | 0 | 3 | 0 | 2 | 0.5 | 2 | 0 | 8 | 0.125 | 2 | 0 | 1 | 0 | 4 | 0 | 1 | 1 | 8 | 0.125 |
| 7_9786584 | 2 | 0 | 1 | 1 | NA | NA | 1 | 1 | 4 | 0.5 | 3 | 0.3333 | NA | NA | 4 | 0.25 | NA | NA | 7 | 0.2857 |
| 7_9786592 | 2 | 0 | 1 | 0 | NA | NA | 1 | 1 | 4 | 0.25 | 2 | 0.5 | NA | NA | 4 | 0.5 | NA | NA | 6 | 0.5 |
| 7_9786596 | 3 | 0.6667 | 1 | 0 | NA | NA | NA | NA | 4 | 0.5 | 2 | 0.5 | NA | NA | 4 | 0.75 | NA | NA | 6 | 0.6667 |
| 7_9795860 | 3 | 0.3333 | NA | NA | NA | NA | NA | NA | 3 | 0.3333 | 1 | 1 | NA | NA | 2 | 0 | NA | NA | 3 | 0.3333 |
| 7_9823828 | 1 | 1 | 4 | 0 | 1 | 0 | 1 | 0 | 7 | 0.1429 | 1 | 1 | 3 | 0.3333 | 1 | 0 | 2 | 0 | 7 | 0.2857 |
| 7_9823832 | 1 | 1 | 2 | 0.5 | NA | NA | 1 | 0 | 4 | 0.5 | 1 | 1 | 2 | 0 | 1 | 1 | 2 | 0 | 6 | 0.3333 |
| 7_9823870 | 2 | 0 | 3 | 0.6667 | NA | NA | 2 | 0 | 7 | 0.2857 | 1 | 0 | 3 | 0 | 1 | 0 | 3 | 0.3333 | 8 | 0.125 |
| 7_9823871 | 2 | 0 | 8 | 0.125 | 4 | 0 | 6 | 0 | 20 | 0.05 | 3 | 0.3333 | 5 | 0 | 5 | 0.2 | 6 | 0.1667 | 19 | 0.1579 |
| 7_9823901 | 2 | 1 | 5 | 0.6 | 5 | 0.8 | 7 | 0.8571 | 19 | 0.7895 | 2 | 0.5 | 3 | 0.6667 | 5 | 0.6 | 9 | 0.4444 | 19 | 0.5263 |
| 7_9823902 | 2 | 1 | 4 | 1 | 4 | 1 | 5 | 1 | 15 | 1 | 2 | 1 | 3 | 1 | 4 | 1 | 7 | 0.8571 | 16 | 0.9375 |

| | | | | | | | | | | | | | | | | | | | | |
|---|---|---|---|---|---|---|---|---|---|---|---|---|---|---|---|---|---|---|---|---|
| 7_98239118 | 3 | 1 | 3 | 1 | 5 | 0.8 | 4 | 0.75 | 15 | 0.8667 | 2 | 0.5 | 1 | 1 | 4 | 0.75 | 8 | 0.875 | 15 | 0.8 |
| 7_98239363 | 3 | 0 | 5 | 0 | 5 | 0.2 | 6 | 0.1667 | 19 | 0.1053 | 1 | 0 | 4 | 0 | 4 | 0 | 8 | 0.25 | 17 | 0.1176 |
| 7_98239502 | 2 | 1 | 4 | 0.75 | 4 | 1 | 6 | 0.6667 | 16 | 0.8125 | 1 | 1 | 3 | 1 | 2 | 0.5 | 7 | 0.8571 | 13 | 0.8462 |
| 7_98239617 | 3 | 1 | 5 | 0.8 | 5 | 1 | 6 | 0.8333 | 19 | 0.8947 | 1 | 1 | 3 | 0.6667 | 2 | 1 | 7 | 1 | 13 | 0.9231 |
| 7_98239733 | 2 | 0.5 | 3 | 0.3333 | 5 | 0.4 | 6 | 0.5 | 16 | 0.4375 | 1 | 0 | 4 | 0.5 | 1 | 0 | 6 | 0.3333 | 12 | 0.3333 |
| 7_98239759 | 1 | 1 | 4 | 1 | 5 | 1 | 6 | 1 | 16 | 1 | 1 | 1 | 4 | 1 | 1 | 1 | 6 | 1 | 12 | 1 |
| 7_98239840 | 1 | 1 | 4 | 0.25 | 5 | 0 | 6 | 0.3333 | 16 | 0.25 | 1 | 0 | 4 | 0 | 1 | 0 | 7 | 0.5714 | 13 | 0.3077 |
| 7_98239938 | 1 | 1 | 2 | 1 | 5 | 0.8 | 6 | 0.6667 | 14 | 0.7857 | 1 | 1 | 4 | 1 | 1 | 0 | 6 | 0.8333 | 12 | 0.8333 |
| 7_98239958 | 1 | 0 | 3 | 0 | 5 | 0.2 | 6 | 0 | 15 | 0.0667 | 1 | 0 | 4 | 0 | 1 | 0 | 6 | 0.1667 | 12 | 0.0833 |
| 7_98239964 | 1 | 1 | 3 | 0.6667 | 5 | 0.4 | 5 | 0 | 14 | 0.3571 | NA | NA | 3 | 0.6667 | 1 | 0 | 6 | 0.6667 | 10 | 0.6 |
| 7_98242799 | NA | NA | 2 | 0 | 3 | 0.3333 | 2 | 0.5 | 7 | 0.2857 | 1 | 0 | 2 | 0 | NA | NA | 5 | 0.2 | 8 | 0.125 |
| 7_98242848 | 1 | 0 | 3 | 0.3333 | 4 | 0.25 | 4 | 0 | 12 | 0.1667 | 1 | 0 | 2 | 0 | NA | NA | 6 | 0.3333 | 9 | 0.2222 |
| 7_98244105 | NA | NA | 1 | 0 | 2 | 0.5 | 4 | 0.25 | 7 | 0.2857 | NA | NA | 3 | 0.6667 | NA | NA | 2 | 0.5 | 5 | 0.6 |
| 7_98244143 | NA | NA | 2 | 0 | 5 | 0.4 | 4 | 0.75 | 11 | 0.4545 | 1 | 1 | 4 | 0.25 | NA | NA | 4 | 0 | 9 | 0.2222 |
| 7_98244220 | NA | NA | 1 | 1 | 3 | 0.6667 | 5 | 0.6 | 9 | 0.6667 | 1 | 1 | 3 | 0.6667 | NA | NA | 4 | 1 | 8 | 0.875 |
| 7_98244259 | NA | NA | 2 | 0.5 | 2 | 0.5 | 5 | 0.4 | 9 | 0.4444 | 1 | 0 | 3 | 0.3333 | NA | NA | 3 | 0 | 7 | 0.1429 |
| 7_98244345 | NA | NA | 1 | 1 | 5 | 0.4 | 5 | 0.6 | 11 | 0.5455 | 1 | 1 | 4 | 0.5 | NA | NA | 4 | 0.5 | 9 | 0.5556 |
| 7_98244443 | NA | NA | 1 | 1 | 4 | 0.75 | 5 | 0.2 | 10 | 0.5 | 1 | 1 | 4 | 0.5 | NA | NA | 3 | 0.3333 | 8 | 0.5 |
| 7_98244471 | NA | NA | 1 | 1 | 5 | 0.2 | 5 | 0 | 11 | 0.1818 | 1 | 0 | 3 | 0 | NA | NA | 4 | 0.25 | 8 | 0.125 |
| 7_98306483 | NA | NA | NA | NA | 1 | 1 | NA | NA | 1 | 1 | 1 | 1 | NA | NA | NA | NA | NA | NA | 1 | 1 |
| 7_98306877 | NA | NA | NA | NA | 1 | 1 | NA | NA | 1 | 1 | 1 | 1 | NA | NA | NA | NA | NA | NA | 1 | 1 |
| 7_98307017 | NA | NA | NA | NA | 1 | 1 | NA | NA | 1 | 1 | 1 | 1 | NA | NA | NA | NA | NA | NA | 1 | 1 |
| 7_98347398 | 1 | 1 | NA | NA | NA | NA | NA | NA | 1 | 1 | NA | NA | NA | NA | 1 | 1 | NA | NA | 1 | 1 |
| 7_98347498 | 1 | 1 | NA | NA | NA | NA | NA | NA | 1 | 1 | NA | NA | NA | NA | NA | NA | 1 | 1 | 1 | 1 |
| 7_98347583 | 1 | 1 | NA | NA | NA | NA | NA | NA | 1 | 1 | NA | NA | NA | NA | 1 | 0 | 1 | 1 | 2 | 0.5 |
| 7_98347755 | 1 | 1 | NA | NA | NA | NA | NA | NA | 1 | 1 | NA | NA | NA | NA | 1 | 1 | 1 | 1 | 2 | 1 |
| 7_98364360 | 4 | 0 | 4 | 0.25 | 5 | 0 | 4 | 0 | 17 | 0.0588 | 1 | 0 | 4 | 0 | 4 | 0.25 | 4 | 0.75 | 13 | 0.3077 |
| 7_98364417 | 6 | 0.1667 | 7 | 0 | 2 | 0.5 | 3 | 0 | 18 | 0.1111 | 1 | 0 | 4 | 0.25 | 3 | 0 | 3 | 0 | 11 | 0.0909 |
| 7_98364569 | 9 | 0.6667 | 6 | 0.5 | 5 | 0.4 | 6 | 0.1667 | 26 | 0.4615 | 2 | 0 | 5 | 0.2 | 4 | 0.5 | 3 | 0.6667 | 14 | 0.3571 |
| 7_98364588 | 10 | 1 | 7 | 0.8571 | 4 | 1 | 5 | 1 | 26 | 0.9615 | 2 | 1 | 5 | 0.6 | 4 | 1 | 5 | 1 | 16 | 0.875 |
| 7_98364627 | 9 | 0.8889 | 7 | 0.8571 | 3 | 1 | 5 | 0.8 | 24 | 0.875 | 2 | 1 | 3 | 0.6667 | 5 | 1 | 6 | 0.8333 | 16 | 0.875 |
| 7_98364764 | 9 | 1 | 6 | 1 | 3 | 1 | 4 | 1 | 22 | 1 | 3 | 1 | 3 | 1 | 1 | 1 | 3 | 1 | 10 | 1 |
| 7_98364859 | 10 | 1 | 6 | 1 | 3 | 1 | 6 | 0.8333 | 25 | 0.96 | 3 | 1 | 4 | 1 | 5 | 0.8 | 4 | 1 | 16 | 0.9375 |
| 7_98365021 | 9 | 0.7778 | 6 | 0.6667 | 3 | 1 | 4 | 1 | 22 | 0.8182 | 3 | 0.6667 | 4 | 0.5 | 5 | 0.8 | 4 | 0.5 | 16 | 0.625 |
| 7_98365075 | 10 | 0.9 | 6 | 1 | 4 | 1 | 4 | 1 | 24 | 0.9583 | 3 | 1 | 3 | 0.6667 | 3 | 1 | 5 | 1 | 14 | 0.9286 |
| 7_98365103 | 7 | 0.8571 | 4 | 1 | 3 | 1 | 4 | 0.25 | 18 | 0.7778 | 2 | 0.5 | 4 | 0.75 | 3 | 1 | 4 | 0.75 | 13 | 0.7692 |
| 7_98365185 | 10 | 0.4 | 7 | 0.4286 | 5 | 0.2 | 7 | 0.2857 | 29 | 0.3448 | 4 | 0.5 | 5 | 0.4 | 4 | 0 | 4 | 0 | 17 | 0.2353 |
| 7_98365206 | 7 | 0 | 9 | 0.4444 | 1 | 0 | 5 | 0 | 22 | 0.1818 | 3 | 0 | 5 | 0.2 | 3 | 0.3333 | 4 | 0 | 15 | 0.1333 |
| 7_98365251 | 9 | 0.8889 | 7 | 0.8571 | 4 | 1 | 5 | 1 | 25 | 0.92 | 4 | 1 | 4 | 1 | 4 | 1 | 5 | 1 | 17 | 1 |
| 7_98365330 | 10 | 0.8 | 9 | 0.3333 | 5 | 0.2 | 6 | 0.1667 | 30 | 0.4333 | 4 | 0.75 | 3 | 1 | 3 | 0.3333 | 6 | 0.5 | 16 | 0.625 |
| 7_98365367 | 10 | 0.1 | 9 | 0.2222 | 5 | 0 | 8 | 0.125 | 32 | 0.125 | 5 | 0 | 5 | 0.2 | 5 | 0 | 5 | 0.2 | 20 | 0.1 |
| 7_98365459 | 9 | 0.1111 | 9 | 0.1111 | 5 | 0.2 | 6 | 0.1667 | 29 | 0.1379 | 4 | 0.25 | 4 | 0 | 4 | 0.25 | 6 | 0.1667 | 18 | 0.1667 |
| 7_98365464 | 7 | 0.4286 | 9 | 0.5556 | 2 | 0 | 7 | 0.4286 | 25 | 0.44 | 3 | 0.3333 | 5 | 0.2 | 4 | 0.5 | 5 | 0.2 | 17 | 0.2941 |
| 7_98365551 | 11 | 0.8182 | 9 | 0.8889 | 5 | 1 | 4 | 0.75 | 29 | 0.8621 | 6 | 0.8333 | 4 | 0.75 | 4 | 0.75 | 3 | 1 | 17 | 0.8235 |
| 7_98365591 | 10 | 0.8 | 10 | 0.9 | 4 | 1 | 7 | 1 | 31 | 0.9032 | 6 | 1 | 5 | 0.6 | 5 | 1 | 6 | 1 | 22 | 0.9091 |
| 7_98365612 | 9 | 0.3333 | 9 | 0.5556 | 4 | 0.5 | 4 | 0.25 | 26 | 0.4231 | 4 | 0 | 3 | 0.3333 | 3 | 0.3333 | 4 | 0.5 | 14 | 0.2857 |
| 7_98365716 | 10 | 0 | 10 | 0.1 | 5 | 0.4 | 7 | 0.1429 | 32 | 0.125 | 6 | 0 | 5 | 0.4 | 5 | 0.2 | 6 | 0.1667 | 22 | 0.1818 |
| 7_98365953 | 9 | 0.3333 | 11 | 0.6364 | 6 | 0.6667 | 6 | 0.8333 | 32 | 0.5938 | 7 | 0.7143 | 3 | 0 | 4 | 0.75 | 4 | 0.75 | 18 | 0.6111 |
| 7_98365965 | 10 | 0.1 | 12 | 0.1667 | 6 | 0.1667 | 7 | 0.2857 | 35 | 0.1714 | 6 | 0 | 4 | 0.5 | 7 | 0.2857 | 6 | 0 | 23 | 0.1739 |
| 7_98365995 | 10 | 0 | 10 | 0 | 5 | 0.2 | 6 | 0.1667 | 31 | 0.0645 | 8 | 0.125 | 4 | 0.25 | 7 | 0 | 6 | 0 | 25 | 0.08 |
| 7_98366014 | 11 | 0.0909 | 11 | 0.2727 | 6 | 0.1667 | 7 | 0.1429 | 35 | 0.1714 | 7 | 0.2857 | 5 | 0.4 | 7 | 0.2857 | 6 | 0 | 25 | 0.24 |
| 7_98366037 | 11 | 1 | 11 | 0.9091 | 6 | 1 | 8 | 1 | 36 | 0.9722 | 8 | 1 | 5 | 1 | 7 | 1 | 5 | 1 | 25 | 1 |
| 7_98366050 | 8 | 0.875 | 11 | 0.7273 | 5 | 0.8 | 5 | 0.8 | 29 | 0.7931 | 7 | 0.8571 | 5 | 0.6 | 7 | 0.7143 | 4 | 1 | 23 | 0.7826 |
| 7_98366093 | 9 | 0 | 13 | 0.0769 | 4 | 0.25 | 8 | 0 | 34 | 0.0588 | 8 | 0.125 | 4 | 0 | 6 | 0.1667 | 6 | 0.1667 | 24 | 0.125 |
| 7_98366186 | 12 | 0.3333 | 13 | 0.8462 | 5 | 0.8 | 9 | 0.4444 | 39 | 0.5897 | 8 | 0.875 | 5 | 0.6 | 7 | 0.7143 | 6 | 0.3333 | 26 | 0.6538 |
| 7_98366198 | 12 | 0.0833 | 12 | 0.1667 | 6 | 0 | 9 | 0 | 39 | 0.0769 | 8 | 0 | 5 | 0.2 | 6 | 0.1667 | 6 | 0 | 25 | 0.08 |
| 7_98366260 | 10 | 0.4 | 6 | 0.5 | 4 | 0.5 | 8 | 0.375 | 28 | 0.4286 | 7 | 0.5714 | 5 | 0.6 | 4 | 0.5 | 7 | 0.4286 | 23 | 0.5217 |
| 7_98366404 | 10 | 0.2 | 12 | 0.5833 | 6 | 0.1667 | 6 | 0 | 34 | 0.2941 | 6 | 0.3333 | 2 | 0 | 5 | 0.2 | 8 | 0.125 | 21 | 0.1905 |
| 7_98366677 | 10 | 0.1 | 10 | 0.3 | 3 | 0 | 3 | 0 | 26 | 0.1538 | 4 | 0.25 | 1 | 1 | 5 | 0.4 | 6 | 0.1667 | 16 | 0.3125 |
| 7_98367107 | 10 | 0.2 | 11 | 0.4545 | 5 | 0.4 | 4 | 0.25 | 30 | 0.3333 | 5 | 0.2 | 1 | 1 | 5 | 0.4 | 6 | 0.5 | 17 | 0.4118 |
| 7_98536936 | 4 | 0.75 | 1 | 1 | 3 | 0.6667 | 2 | 1 | 10 | 0.8 | 1 | 0 | 1 | 0 | 4 | 0.5 | 1 | 1 | 7 | 0.4286 |
| 7_98536945 | 4 | 0.25 | 2 | 0.5 | 3 | 0.6667 | 2 | 1 | 11 | 0.5455 | NA | NA | 1 | 1 | 4 | 0.75 | 1 | 1 | 6 | 0.8333 |
| 7_98536954 | 3 | 1 | 2 | 1 | 2 | 1 | 1 | 1 | 8 | 1 | NA | NA | 2 | 1 | 2 | 1 | 1 | 1 | 5 | 1 |
| 7_98537224 | 4 | 0.75 | 2 | 1 | 3 | 0.6667 | 2 | 0.5 | 11 | 0.7273 | NA | NA | 2 | 0.5 | 4 | 0.25 | 1 | 0 | 7 | 0.2857 |
| 7_98537458 | 2 | 1 | 2 | 1 | 2 | 1 | 2 | 1 | 8 | 1 | 1 | 1 | 2 | 1 | 4 | 1 | 1 | 1 | 8 | 1 |

| | | | | | | | | | | | | | | | | | | | | |
|---|---|---|---|---|---|---|---|---|---|---|---|---|---|---|---|---|---|---|---|---|
| 7_98537530 | 2 | 0.5 | 2 | 1 | 3 | 1 | 2 | 1 | 9 | 0.8889 | 1 | 1 | 2 | 1 | 3 | 1 | 1 | 1 | 7 | 1 |
| 7_98537542 | 1 | 0 | 1 | 0 | 3 | 0.3333 | 1 | 0 | 6 | 0.1667 | 1 | 0 | 1 | 1 | 2 | 0 | 1 | 0 | 5 | 0.2 |
| 7_98537581 | 1 | 1 | 2 | 0 | 3 | 0.6667 | 2 | 0.5 | 8 | 0.5 | 1 | 0 | 2 | 1 | 2 | 0.5 | NA | NA | 5 | 0.6 |
| 7_98537658 | 1 | 1 | 2 | 0.5 | 3 | 1 | 2 | 0.5 | 8 | 0.75 | 1 | 1 | 2 | 0.5 | 3 | 0.6667 | 1 | 1 | 7 | 0.7143 |
| 7_98803144 | 13 | 0.2308 | 20 | 0.45 | 11 | 0.3636 | 19 | 0.2105 | 63 | 0.3175 | 13 | 0.3846 | 27 | 0.3704 | 20 | 0.2 | 15 | 0.2667 | 75 | 0.3067 |
| 7_98803306 | 7 | 0.5714 | 6 | 0.5 | 5 | 0.4 | 7 | 0 | 25 | 0.36 | 7 | 0.5714 | 11 | 0.5455 | 7 | 0.5714 | 3 | 1 | 28 | 0.6071 |
| 7_98803757 | 11 | 0.1818 | 13 | 0.0769 | 10 | 0.1 | 12 | 0.1667 | 46 | 0.1304 | 6 | 0.1667 | 17 | 0.2941 | 17 | 0.1765 | 10 | 0.1 | 50 | 0.2 |
| 7_98803776 | 14 | 0.5714 | 17 | 0.4706 | 10 | 0.4 | 14 | 0.5714 | 55 | 0.5091 | 9 | 0.7778 | 26 | 0.6154 | 19 | 0.6316 | 14 | 0.3571 | 68 | 0.5882 |
| 7_98916968 | 1 | 0 | 4 | 0.5 | 2 | 1 | 1 | 1 | 8 | 0.625 | 2 | 0 | 2 | 0.5 | 3 | 0.3333 | 2 | 0.5 | 9 | 0.3333 |
| 7_98917053 | 1 | 0 | 3 | 0 | 1 | 1 | NA | NA | 5 | 0.2 | 1 | 0 | 1 | 1 | 2 | 1 | 1 | 1 | 5 | 0.8 |
| 7_98917062 | 1 | 1 | 3 | 1 | 2 | 1 | 2 | 1 | 8 | 1 | 2 | 1 | 2 | 1 | 4 | 0.75 | 2 | 1 | 10 | 0.9 |
| 7_98917296 | 1 | 1 | 5 | 0.4 | 2 | 0.5 | 3 | 0.3333 | 11 | 0.4545 | 3 | 0.3333 | 2 | 0 | 4 | 0.5 | 3 | 0 | 12 | 0.25 |
| 7_98917499 | 1 | 0 | 7 | 0.5714 | 3 | 0.3333 | 4 | 0.5 | 15 | 0.4667 | 2 | 0.5 | 1 | 0 | 6 | 0.6667 | 2 | 1 | 11 | 0.6364 |
| 7_98917575 | 1 | 0 | 8 | 0.25 | 3 | 0.3333 | 4 | 0.5 | 16 | 0.3125 | 4 | 0.25 | 2 | 0.5 | 5 | 0.2 | 3 | 0.6667 | 14 | 0.3571 |
| 7_98917593 | 1 | 1 | 8 | 0.25 | 2 | 0 | 4 | 0.25 | 15 | 0.2667 | 3 | 0 | 3 | 0 | 8 | 0.25 | 3 | 0 | 17 | 0.1176 |
| 7_98917698 | 1 | 1 | 8 | 0.75 | 3 | 0.6667 | 3 | 1 | 15 | 0.8 | 4 | 0.5 | 3 | 0.6667 | 6 | 0.6667 | 3 | 1 | 16 | 0.6875 |
| 7_98917863 | 1 | 0 | 7 | 0.2857 | 3 | 0.3333 | 5 | 0 | 16 | 0.1875 | 5 | 0 | 4 | 0 | 8 | 0.125 | 3 | 0 | 20 | 0.05 |
| 7_98917964 | 1 | 1 | 9 | 0.2222 | 4 | 0.25 | 4 | 0.25 | 18 | 0.2778 | 5 | 0 | 4 | 0 | 8 | 0.25 | 4 | 0.25 | 21 | 0.1429 |
| 7_98918102 | 1 | 0 | 10 | 0.1 | 4 | 0 | 5 | 0.4 | 20 | 0.15 | 7 | 0 | 3 | 0 | 9 | 0.1111 | 5 | 0.2 | 24 | 0.0833 |
| 7_98918224 | 1 | 0 | 9 | 0 | 4 | 0.25 | 6 | 0 | 20 | 0.05 | 7 | 0.1429 | 2 | 0.5 | 8 | 0.125 | 7 | 0 | 24 | 0.125 |
| 7_98918398 | 1 | 0 | 7 | 0.1429 | 4 | 0.25 | 5 | 0.2 | 17 | 0.1765 | 6 | 0.1667 | 3 | 0 | 5 | 0.4 | 6 | 0.1667 | 20 | 0.2 |
| 7_98966858 | 2 | 0.5 | 12 | 0.0833 | 5 | 0 | 5 | 0 | 24 | 0.0833 | 5 | 0.2 | 3 | 0 | 5 | 0 | 6 | 0 | 19 | 0.0526 |
| 7_98969652 | 2 | 0.5 | NA | NA | NA | NA | NA | NA | 2 | 0.5 | NA | NA | NA | NA | NA | NA | 1 | 1 | 1 | 1 |
| 7_98985333 | 6 | 1 | 15 | 0.7333 | 7 | 0.8571 | 9 | 1 | 37 | 0.8649 | 6 | 1 | 5 | 1 | 6 | 1 | 13 | 0.9231 | 30 | 0.9667 |
| 7_98985482 | 6 | 0.5 | 13 | 0.3077 | 7 | 0.1429 | 9 | 0.3333 | 35 | 0.3143 | 5 | 0 | 6 | 0.1667 | 5 | 0.2 | 14 | 0.2143 | 30 | 0.1667 |
| 7_98985846 | 10 | 0.7 | 17 | 0.7647 | 12 | 0.6667 | 11 | 0.7273 | 50 | 0.72 | 5 | 0.4 | 6 | 0.8333 | 7 | 0.7143 | 13 | 0.5385 | 31 | 0.6129 |
| 7_98985944 | 9 | 0.1111 | 14 | 0.2143 | 13 | 0.0769 | 11 | 0 | 47 | 0.1064 | 10 | 0.2 | 6 | 0.3333 | 5 | 0 | 14 | 0.0714 | 35 | 0.1429 |
| 7_98986108 | 10 | 0.2 | 17 | 0.1765 | 14 | 0.1429 | 12 | 0.5 | 53 | 0.2453 | 12 | 0.4167 | 8 | 0.25 | 7 | 0.2857 | 14 | 0.1429 | 41 | 0.2683 |
| 7_98994767 | 5 | 0 | 3 | 0.3333 | 3 | 0 | 3 | 0.3333 | 14 | 0.1429 | 3 | 0.3333 | 2 | 0 | 1 | 0 | 1 | 0 | 7 | 0.1429 |
| 7_98995140 | 6 | 0.1667 | 2 | 0 | 5 | 0 | 3 | 0 | 16 | 0.0625 | 3 | 0 | 1 | 0 | 3 | 0 | 3 | 0.3333 | 10 | 0.1 |
| 7_98995505 | 5 | 0 | 2 | 0 | 3 | 0 | 2 | 0.5 | 12 | 0.0833 | 3 | 0 | 1 | 0 | 1 | 0 | 2 | 1 | 7 | 0.2857 |
| 7_98995550 | 5 | 0.2 | 2 | 0 | 4 | 0.5 | 2 | 0.5 | 13 | 0.3077 | 3 | 0.3333 | 1 | 0 | 2 | 0 | 2 | 0 | 8 | 0.125 |
| 7_98998435 | 3 | 0.6667 | 2 | 0 | 3 | 0.6667 | NA | NA | 8 | 0.5 | NA | NA | NA | NA | NA | NA | 1 | 1 | 1 | 1 |
| 7_99030976 | NA | NA | 1 | 0 | 4 | 0.5 | 1 | 0 | 6 | 0.3333 | NA | NA | 2 | 0 | 3 | 0.3333 | 1 | 0 | 6 | 0.1667 |
| 7_99031133 | NA | NA | 1 | 0 | 3 | 0.3333 | 1 | 0 | 5 | 0.2 | NA | NA | 1 | 0 | 3 | 1 | NA | NA | 4 | 0.75 |
| 7_99031220 | NA | NA | 1 | 0 | 4 | 0.5 | 1 | 0 | 6 | 0.3333 | NA | NA | 2 | 0 | 2 | 0.5 | NA | NA | 4 | 0.25 |
| 7_99109521 | 4 | 0 | 3 | 0.3333 | 6 | 0 | 3 | 0.3333 | 16 | 0.125 | 1 | 0 | 8 | 0.125 | 7 | 0.2857 | 3 | 0 | 19 | 0.1579 |
| 7_99109527 | 3 | 0.6667 | 3 | 0.6667 | 5 | 0.4 | 4 | 1 | 15 | 0.6667 | 1 | 1 | 7 | 0.5714 | 6 | 0.8333 | 3 | 0 | 17 | 0.5882 |
| 7_99109739 | 3 | 0.3333 | 3 | 0.6667 | 5 | 0.2 | 4 | 0 | 15 | 0.2667 | 1 | 0 | 9 | 0.2222 | 6 | 0.5 | 4 | 0.25 | 20 | 0.3 |
| 7_99109944 | 5 | 0.2 | 3 | 0.3333 | 7 | 0.1429 | 4 | 0 | 19 | 0.1579 | 1 | 1 | 8 | 0.125 | 7 | 0.2857 | 3 | 0.3333 | 19 | 0.2632 |
| 7_99110023 | 5 | 0.2 | 3 | 0.3333 | 8 | 0 | 4 | 0 | 20 | 0.1 | 1 | 0 | 8 | 0.25 | 7 | 0 | 3 | 0.3333 | 19 | 0.1579 |
| 7_99110039 | 3 | 0.3333 | 3 | 1 | 7 | 1 | 2 | 1 | 15 | 0.8667 | NA | NA | 7 | 1 | 6 | 1 | 2 | 1 | 15 | 1 |
| 7_99110170 | 4 | 0.5 | 2 | 0 | 8 | 0.875 | 4 | 0.25 | 18 | 0.5556 | 1 | 0 | 6 | 0 | 8 | 0.625 | 4 | 0.75 | 19 | 0.4211 |
| 7_99127272 | 4 | 0 | 1 | 0 | 6 | 0.1667 | 2 | 0 | 13 | 0.0769 | 2 | 0 | 2 | 0 | 2 | 0.5 | NA | NA | 6 | 0.1667 |
| 7_99127864 | 3 | 0 | 1 | 0 | 3 | 0.3333 | 1 | 0 | 8 | 0.125 | 2 | 0 | 2 | 0.5 | 3 | 0 | NA | NA | 7 | 0.1429 |
| 7_99127894 | 3 | 0 | 1 | 1 | 3 | 0 | NA | NA | 7 | 0.1429 | 2 | 0 | 2 | 0 | 3 | 0.3333 | NA | NA | 7 | 0.1429 |
| 7_99127955 | 2 | 0 | 1 | 0 | 1 | 0 | 1 | 1 | 5 | 0.2 | 1 | 0 | 2 | 0 | 2 | 0.5 | NA | NA | 5 | 0.2 |
| 7_99128117 | 87 | 0.0345 | 100 | 0.11 | 97 | 0.1031 | 93 | 0.086 | 377 | 0.0849 | 110 | 0.1 | 131 | 0.1298 | 129 | 0.062 | 75 | 0.1067 | 445 | 0.0989 |
| 7_99250328 | 6 | 0 | NA | NA | 2 | 0 | 3 | 0.3333 | 11 | 0.0909 | 4 | 0.5 | 2 | 0 | 2 | 0 | 2 | 0 | 10 | 0.2 |
| 7_99250417 | 5 | 0 | NA | NA | 2 | 0.5 | 2 | 0.5 | 9 | 0.2222 | 2 | 0 | 3 | 0.3333 | 3 | 0.3333 | 1 | 1 | 9 | 0.3333 |
| 7_99250855 | 9 | 0.1111 | 2 | 0.5 | 6 | 0 | 3 | 0 | 20 | 0.1 | 4 | 0.5 | 4 | 0 | 8 | 0.25 | 5 | 0.4 | 21 | 0.2857 |
| 7_99250913 | 9 | 0.1111 | 1 | 0 | 6 | 0.1667 | 2 | 0 | 18 | 0.1111 | 4 | 0 | 3 | 0 | 6 | 0 | 6 | 0.1667 | 19 | 0.0526 |
| 7_99250999 | 8 | 0.25 | 3 | 0 | 5 | 0.4 | 3 | 0 | 19 | 0.2105 | 3 | 0.3333 | 4 | 0 | 5 | 0.2 | 8 | 0.125 | 20 | 0.15 |
| 7_99253264 | 11 | 0.1818 | 5 | 0.2 | 10 | 0.1 | 7 | 0.2857 | 33 | 0.1818 | 4 | 0.25 | 7 | 0.1429 | 13 | 0.0769 | 10 | 0.2 | 34 | 0.1471 |
| 7_99253293 | 8 | 0.5 | 3 | 0.6667 | 9 | 0.4444 | 7 | 0.8571 | 27 | 0.5926 | 4 | 0.25 | 5 | 0.2 | 11 | 0.7273 | 9 | 0.7778 | 29 | 0.5862 |
| 7_99253382 | 12 | 0 | 6 | 0.1667 | 9 | 0.1111 | 6 | 0.3333 | 33 | 0.1212 | 4 | 0.25 | 7 | 0 | 13 | 0.1538 | 10 | 0.1 | 34 | 0.1176 |
| 7_99255236 | 22 | 0.1364 | 13 | 0.1538 | 19 | 0.1579 | 10 | 0.2 | 64 | 0.1562 | 9 | 0 | 12 | 0.0833 | 25 | 0.12 | 14 | 0.2857 | 60 | 0.1333 |
| 7_99255977 | 2 | 0 | 6 | 0.1667 | NA | NA | 2 | 0 | 10 | 0.1 | 3 | 0 | 4 | 0.25 | 2 | 0 | 1 | 0 | 10 | 0.1 |
| 7_99273610 | 3 | 0.3333 | NA | NA | 2 | 0.5 | 5 | 0.8 | 10 | 0.6 | NA | NA | 1 | 1 | NA | NA | NA | NA | 1 | 1 |
| 7_99274193 | 3 | 0.3333 | NA | NA | NA | NA | 3 | 0.3333 | 6 | 0.3333 | 1 | 1 | 1 | 0 | NA | NA | NA | NA | 2 | 0.5 |
| 7_99274294 | 3 | 0.3333 | NA | NA | 1 | 0 | 3 | 0 | 7 | 0.1429 | 1 | 1 | 1 | 0 | NA | NA | NA | NA | 2 | 0.5 |
| 7_99274580 | 3 | 0 | NA | NA | 1 | 0 | 3 | 0.6667 | 7 | 0.2857 | 1 | 1 | 2 | 1 | NA | NA | NA | NA | 3 | 1 |
| 7_99274659 | 2 | 1 | NA | NA | NA | NA | 2 | 1 | 4 | 1 | 1 | 1 | 2 | 1 | NA | NA | NA | NA | 3 | 1 |
| 7_99275035 | 3 | 0.6667 | NA | NA | NA | NA | 1 | 1 | 4 | 0.75 | 1 | 0 | 2 | 0.5 | NA | NA | NA | NA | 3 | 0.3333 |
| 7_99276018 | 2 | 0 | NA | NA | NA | NA | 1 | 1 | 3 | 0.3333 | 1 | 1 | NA | NA | NA | NA | NA | NA | 1 | 1 |

| | | | | | | | | | | | | | | | | | | | | |
|---|---|---|---|---|---|---|---|---|---|---|---|---|---|---|---|---|---|---|---|---|
| 7_9927740 | NA | NA | 2 | 0 | NA | NA | 1 | 1 | 3 | 0.3333 | NA | NA | NA | NA | 1 | 1 | 1 | 1 | 2 | 1 |
| 7_9927742 | NA | NA | 2 | 1 | NA | NA | 1 | 1 | 3 | 1 | NA | NA | NA | NA | 1 | 1 | 1 | 1 | 2 | 1 |
| 7_9927751 | NA | NA | 2 | 0.5 | 1 | 0 | 1 | 1 | 4 | 0.5 | NA | NA | NA | NA | 1 | 0 | 1 | 1 | 2 | 0.5 |
| 7_9927758 | NA | NA | 1 | 1 | 1 | 0 | NA | NA | 2 | 0.5 | NA | NA | NA | NA | 1 | 0 | 1 | 1 | 2 | 0.5 |
| 7_9927784 | NA | NA | 3 | 0.6667 | 2 | 0.5 | 1 | 0 | 6 | 0.5 | NA | NA | NA | NA | 1 | 0 | 1 | 1 | 2 | 0.5 |
| 7_9927789 | NA | NA | 2 | 1 | 2 | 0.5 | 1 | 1 | 5 | 0.8 | NA | NA | NA | NA | 1 | 1 | 1 | 1 | 2 | 1 |
| 7_9927814 | NA | NA | 3 | 1 | 1 | 0 | NA | NA | 4 | 0.75 | 1 | 1 | NA | NA | 1 | 1 | NA | NA | 2 | 1 |
| 7_9927819 | NA | NA | 4 | 0.25 | 1 | 0 | 1 | 0 | 6 | 0.1667 | 1 | 0 | NA | NA | 1 | 1 | 1 | 1 | 3 | 0.6667 |
| 7_9930788 | 1 | 1 | 5 | 0.4 | NA | NA | 2 | 1 | 8 | 0.625 | 1 | 1 | 4 | 0.75 | NA | NA | NA | NA | 5 | 0.8 |
| 7_9930801 | 1 | 0 | 8 | 0.375 | NA | NA | 1 | 0 | 10 | 0.3 | 1 | 0 | 3 | 0 | NA | NA | 1 | 1 | 5 | 0.2 |
| 7_9930840 | 1 | 1 | 8 | 0.5 | NA | NA | 2 | 0.5 | 11 | 0.5455 | 1 | 1 | 2 | 0 | NA | NA | 1 | 0 | 4 | 0.25 |
| 7_9930853 | 1 | 0 | 8 | 0.125 | NA | NA | NA | NA | 9 | 0.1111 | NA | NA | 2 | 0 | NA | NA | 1 | 1 | 3 | 0.3333 |
| 7_9930888 | 1 | 0 | 7 | 0.2857 | NA | NA | NA | NA | 8 | 0.25 | 1 | 1 | 1 | 0 | NA | NA | 1 | 0 | 3 | 0.3333 |
| 7_9930895 | 1 | 0 | 7 | 0.1429 | NA | NA | NA | NA | 8 | 0.125 | 1 | 0 | 1 | 1 | NA | NA | 1 | 1 | 3 | 0.6667 |
| 7_9930906 | NA | NA | 5 | 0.2 | NA | NA | NA | NA | 5 | 0.2 | 1 | 1 | 1 | 0 | NA | NA | 1 | 0 | 3 | 0.3333 |
| 7_9930907 | 1 | 1 | 8 | 1 | NA | NA | NA | NA | 9 | 1 | NA | NA | 1 | 1 | NA | NA | 1 | 1 | 2 | 1 |
| 7_9948895 | 19 | 0.1053 | 27 | 0.1481 | 16 | 0.125 | 21 | 0.0952 | 83 | 0.1205 | 13 | 0.1538 | 9 | 0.1111 | 16 | 0.125 | 12 | 0.1667 | 50 | 0.14 |
| 7_9948898 | 15 | 0.2667 | 22 | 0.3182 | 15 | 0.3333 | 19 | 0.2105 | 71 | 0.2817 | 11 | 0.4545 | 10 | 0.5 | 16 | 0.4375 | 10 | 0.2 | 47 | 0.4043 |
| 7_9956688 | NA | NA | NA | NA | 2 | 1 | NA | NA | 2 | 1 | NA | NA | NA | NA | NA | NA | 1 | 1 | 1 | 1 |
| 7_9956695 | NA | NA | NA | NA | 2 | 0.5 | NA | NA | 2 | 0.5 | NA | NA | NA | NA | NA | NA | 1 | 1 | 1 | 1 |
| 7_9956947 | 2 | 0 | 5 | 0.2 | 10 | 0.2 | 3 | 0.3333 | 20 | 0.2 | 5 | 0 | 7 | 0 | 8 | 0.125 | 5 | 0.4 | 25 | 0.12 |
| 7_9956963 | 3 | 0 | 3 | 0 | 10 | 0 | 3 | 0.3333 | 19 | 0.0526 | 4 | 0 | 6 | 0 | 7 | 0.1429 | 4 | 0.25 | 21 | 0.0952 |
| 7_9956963 | 3 | 0.3333 | 4 | 0.25 | 10 | 0.3 | 3 | 0.6667 | 20 | 0.35 | 4 | 0.25 | 7 | 0.5714 | 8 | 0.25 | 5 | 0.4 | 24 | 0.375 |
| 7_9956994 | 2 | 0.5 | 2 | 0 | 7 | 0.4286 | 3 | 0.3333 | 14 | 0.3571 | 4 | 0 | 8 | 0.25 | 6 | 0 | 5 | 0.2 | 23 | 0.1304 |
| 7_9957011 | 1 | 0 | 3 | 0.3333 | 6 | 0.1667 | 4 | 0.25 | 14 | 0.2143 | 2 | 0.5 | 7 | 0.1429 | 5 | 0 | 4 | 0 | 18 | 0.1111 |
| 7_9957015 | 2 | 0.5 | 3 | 0 | 6 | 0 | 4 | 0 | 15 | 0.0667 | 2 | 0 | 8 | 0.125 | 5 | 0 | 3 | 0 | 18 | 0.0556 |
| 7_9957023 | 1 | 0 | 3 | 0.3333 | 5 | 0 | 2 | 0 | 11 | 0.0909 | 2 | 0 | 8 | 0.125 | 4 | 0 | 2 | 0 | 16 | 0.0625 |
| 7_9957042 | 2 | 0 | 3 | 0 | 5 | 0.2 | 4 | 0 | 14 | 0.0714 | 1 | 0 | 8 | 0.125 | 4 | 0 | 3 | 0 | 16 | 0.0625 |
| 7_9957304 | 1 | 1 | 2 | 0 | 1 | 0 | 2 | 0.5 | 6 | 0.3333 | 1 | 0 | 2 | 0 | 2 | 0 | 2 | 0.5 | 7 | 0.1429 |
| 7_9957328 | NA | NA | 2 | 0 | 1 | 1 | 2 | 0 | 5 | 0.2 | 1 | 1 | 4 | 0.25 | 2 | 0 | 2 | 0 | 9 | 0.2222 |
| 7_9957334 | NA | NA | 2 | 0 | 1 | 1 | 1 | 1 | 4 | 0.5 | 1 | 1 | 4 | 0.5 | 2 | 0 | 1 | 0 | 8 | 0.375 |
| 7_9957346 | 1 | 1 | 2 | 1 | 1 | 1 | 1 | 1 | 5 | 1 | 1 | 1 | 3 | 1 | 2 | 1 | 1 | 1 | 7 | 1 |
| 7_9957353 | NA | NA | 1 | 1 | NA | NA | 1 | 1 | 2 | 1 | 1 | 1 | 4 | 0.75 | 1 | 1 | 1 | 1 | 7 | 0.8571 |
| 7_9957359 | NA | NA | 1 | 0 | 1 | 1 | 2 | 0 | 4 | 0.25 | NA | NA | 3 | 0.3333 | 1 | 0 | 1 | 0 | 5 | 0.2 |
| 7_9957365 | NA | NA | 2 | 0.5 | 1 | 0 | 2 | 0 | 5 | 0.2 | 1 | 0 | 4 | 0.25 | 1 | 0 | 1 | 1 | 7 | 0.2857 |
| 7_9957366 | NA | NA | 2 | 0.5 | 1 | 0 | 2 | 0 | 5 | 0.2 | 1 | 1 | 4 | 0 | 1 | 1 | 1 | 1 | 7 | 0.4286 |
| 7_9957371 | NA | NA | 1 | 0 | 1 | 1 | 2 | 0 | 4 | 0.25 | 1 | 1 | 3 | 0 | 1 | 0 | 1 | 0 | 6 | 0.1667 |
| 7_9957372 | 1 | 0 | 2 | 0 | NA | NA | 2 | 0.5 | 5 | 0.2 | 1 | 0 | 3 | 0.6667 | 1 | 0 | 1 | 0 | 6 | 0.3333 |
| 7_9957397 | 1 | 0 | 2 | 1 | 1 | 1 | 2 | 0.5 | 6 | 0.6667 | NA | NA | 4 | 1 | 1 | 1 | 1 | 1 | 6 | 1 |
| 7_9957408 | 1 | 0 | 2 | 0.5 | 1 | 0 | 2 | 0.5 | 6 | 0.3333 | NA | NA | 4 | 0 | NA | NA | 1 | 1 | 5 | 0.2 |
| 7_9957432 | 1 | 0 | 1 | 1 | 1 | 0 | 2 | 0.5 | 5 | 0.4 | NA | NA | 4 | 0.25 | NA | NA | 1 | 1 | 5 | 0.4 |
| 7_9957458 | 1 | 1 | 1 | 1 | 1 | 1 | 2 | 1 | 5 | 1 | NA | NA | 3 | 1 | 1 | 1 | 1 | 0 | 5 | 0.8 |
| 7_9957463 | 1 | 1 | 1 | 1 | 1 | 0 | 1 | 0 | 4 | 0.5 | NA | NA | 3 | 0.3333 | 1 | 0 | 1 | 0 | 5 | 0.2 |
| 7_9957467 | 1 | 1 | 1 | 1 | 1 | 1 | 2 | 1 | 5 | 1 | NA | NA | 3 | 0.6667 | 1 | 1 | 1 | 1 | 5 | 0.8 |
| 7_9957472 | NA | NA | 1 | 1 | NA | NA | 1 | 1 | 2 | 1 | NA | NA | 1 | 1 | NA | NA | NA | NA | 1 | 1 |
| 7_9957485 | 1 | 1 | 1 | 1 | 1 | 0 | 2 | 1 | 5 | 0.8 | NA | NA | 2 | 1 | NA | NA | 1 | 1 | 3 | 1 |
| 7_9957492 | 1 | 1 | 1 | 0 | 1 | 1 | 2 | 0.5 | 5 | 0.6 | NA | NA | 2 | 0.5 | 1 | 0 | 1 | 0 | 4 | 0.25 |
| 7_9957494 | NA | NA | 1 | 0 | 1 | 0 | 1 | 1 | 3 | 0.3333 | NA | NA | 2 | 0.5 | 1 | 0 | 1 | 0 | 4 | 0.25 |
| 7_9957508 | 1 | 1 | 1 | 1 | 1 | 1 | 2 | 1 | 5 | 1 | NA | NA | 2 | 1 | 1 | 1 | 1 | 1 | 4 | 1 |
| 7_9957544 | 1 | 1 | 1 | 1 | 1 | 1 | 2 | 1 | 5 | 1 | NA | NA | 1 | 1 | 1 | 1 | 1 | 1 | 3 | 1 |
| 7_9973148 | 4 | 0.75 | 5 | 0.6 | 4 | 0.5 | 3 | 0.3333 | 16 | 0.5625 | 1 | 1 | 3 | 0.3333 | 1 | 1 | 5 | 0.4 | 10 | 0.5 |
| 7_9973157 | 4 | 0.5 | 3 | 0 | 3 | 0 | 3 | 0.6667 | 13 | 0.3077 | 1 | 0 | 3 | 0 | 1 | 0 | 5 | 0.2 | 10 | 0.1 |
| 7_9973267 | 3 | 1 | 2 | 0.5 | 3 | 1 | 2 | 1 | 10 | 0.9 | NA | NA | 3 | 1 | 1 | 1 | 6 | 0.6667 | 10 | 0.8 |
| 7_9973273 | 4 | 0.25 | 2 | 1 | 4 | 0.25 | 2 | 0 | 12 | 0.3333 | NA | NA | 3 | 0.3333 | 1 | 0 | 5 | 0.4 | 9 | 0.3333 |
| 7_9973280 | 4 | 1 | 2 | 1 | 4 | 0.75 | 1 | 1 | 11 | 0.9091 | NA | NA | 4 | 0.25 | 1 | 1 | 6 | 0.8333 | 11 | 0.6364 |
| 7_9973284 | 4 | 1 | 2 | 1 | 4 | 1 | 2 | 1 | 12 | 1 | NA | NA | 4 | 0.5 | 1 | 1 | 5 | 1 | 10 | 0.8 |
| 7_9980947 | 1 | 1 | NA | NA | 1 | 1 | 1 | 1 | 3 | 1 | NA | NA | NA | NA | NA | NA | 1 | 1 | 1 | 1 |
| 7_9980951 | 1 | 0 | NA | NA | 1 | 0 | 1 | 1 | 3 | 0.3333 | NA | NA | NA | NA | NA | NA | 3 | 0.3333 | 3 | 0.3333 |
| 7_9980956 | 1 | 1 | NA | NA | 1 | 1 | 1 | 1 | 3 | 1 | NA | NA | NA | NA | NA | NA | 3 | 0.6667 | 3 | 0.6667 |
| 7_9980961 | 1 | 1 | NA | NA | 1 | 1 | 1 | 1 | 3 | 1 | NA | NA | NA | NA | NA | NA | 3 | 1 | 3 | 1 |
| 7_9992369 | 4 | 0.25 | 4 | 0.25 | 1 | 0 | 6 | 0.1667 | 15 | 0.2 | 4 | 0.25 | 2 | 0 | 5 | 0 | 4 | 0.25 | 15 | 0.1333 |
| 7_9992450 | 6 | 0.1667 | 5 | 0 | NA | NA | 7 | 0 | 18 | 0.0556 | 5 | 0.2 | 3 | 0.3333 | 8 | 0.25 | 4 | 0 | 20 | 0.2 |
| 7_9992464 | 7 | 0.4286 | 5 | 0.4 | 2 | 0.5 | 6 | 0 | 20 | 0.3 | 4 | 0.25 | 2 | 0 | 7 | 0.1429 | 4 | 0 | 17 | 0.1176 |
| 7_9992746 | 14 | 0.4286 | 24 | 0.25 | 11 | 0.3636 | 15 | 0.1333 | 64 | 0.2812 | 14 | 0.2143 | 9 | 0.4444 | 18 | 0.3889 | 13 | 0.1538 | 54 | 0.2963 |
| 7_9996830 | NA | NA | NA | NA | 2 | 1 | NA | NA | 2 | 1 | NA | NA | NA | NA | NA | NA | 1 | 1 | 1 | 1 |

| | | | | | | | | | | | | | | | | | | | | |
|---|---|---|---|---|---|---|---|---|---|---|---|---|---|---|---|---|---|---|---|---|
| 7_9996838 | NA | NA | NA | NA | 2 | 0.5 | NA | NA | 2 | 0.5 | NA | NA | NA | NA | NA | NA | 2 | 1 | 2 | 1 |
| 7_9997608 | 23 | 0.0435 | 34 | 0.1176 | 28 | 0 | 29 | 0.0345 | 114 | 0.0526 | 22 | 0 | 19 | 0.2632 | 29 | 0.069 | 18 | 0.2222 | 88 | 0.125 |
| 7_9997620 | 17 | 0.1765 | 37 | 0.1081 | 27 | 0.0741 | 27 | 0.0741 | 108 | 0.1019 | 19 | 0.0526 | 20 | 0.1 | 30 | 0.1667 | 14 | 0.1429 | 83 | 0.1205 |
| 7_1000814 | 2 | 1 | NA | NA | NA | NA | NA | NA | 2 | 1 | 1 | 1 | 1 | 1 | 1 | 1 | 2 | 1 | 5 | 1 |
| 7_1000815 | 2 | 1 | NA | NA | NA | NA | NA | NA | 2 | 1 | NA | NA | 1 | 1 | 1 | 1 | 2 | 1 | 4 | 1 |
| 7_1000850 | 2 | 0.5 | NA | NA | NA | NA | 1 | 1 | 3 | 0.6667 | 1 | 1 | 2 | 1 | 2 | 0.5 | 1 | 1 | 6 | 0.8333 |
| 7_1001041 | 2 | 0 | 1 | 0 | 1 | 0 | 1 | 1 | 5 | 0.2 | 1 | 0 | 3 | 0.3333 | 5 | 0.2 | 3 | 0 | 12 | 0.1667 |
| 7_1001041 | 2 | 0.5 | 1 | 0 | 1 | 0 | 1 | 0 | 5 | 0.2 | 1 | 0 | 3 | 0.3333 | 5 | 0.2 | 4 | 0.5 | 13 | 0.3077 |
| 7_1001041 | 2 | 0.5 | 1 | 0 | 1 | 1 | 1 | 0 | 5 | 0.4 | 1 | 0 | 3 | 0 | 4 | 0.5 | 4 | 0 | 12 | 0.1667 |
| 7_1001043 | 2 | 0 | 1 | 1 | 2 | 0 | 2 | 0 | 7 | 0.1429 | 2 | 0.5 | 4 | 0 | 6 | 0.1667 | 4 | 0 | 16 | 0.125 |
| 7_1001045 | 2 | 0.5 | 1 | 1 | 2 | 0 | 1 | 1 | 6 | 0.5 | 2 | 0.5 | 4 | 0.25 | 5 | 0.4 | 4 | 0.75 | 15 | 0.4667 |
| 7_1001186 | 2 | 0 | 1 | 1 | 2 | 0 | 2 | 0 | 7 | 0.1429 | 2 | 0 | 4 | 0.25 | 5 | 0.2 | 4 | 0.25 | 15 | 0.2 |
| 7_1001187 | 2 | 1 | 1 | 0 | 2 | 1 | 1 | 1 | 6 | 0.8333 | 1 | 1 | 4 | 1 | 5 | 1 | 3 | 1 | 13 | 1 |
| 7_1001187 | 2 | 0 | 1 | 0 | 2 | 0.5 | 2 | 0.5 | 7 | 0.2857 | 2 | 0.5 | 4 | 0.25 | 6 | 0.6667 | 4 | 0 | 16 | 0.375 |
| 7_1001188 | 2 | 0.5 | 1 | 0 | 2 | 0 | 3 | 0 | 8 | 0.125 | 2 | 0.5 | 5 | 0 | 6 | 0 | 4 | 0.25 | 17 | 0.1176 |
| 7_1001189 | 2 | 0 | 2 | 0.5 | 2 | 0.5 | 2 | 0.5 | 8 | 0.375 | 2 | 1 | 5 | 0.8 | 6 | 0 | 4 | 0.5 | 17 | 0.4706 |
| 7_1001189 | 1 | 1 | 2 | 0 | 2 | 0 | 3 | 0 | 8 | 0.125 | 2 | 0 | 4 | 0 | 6 | 0.1667 | 4 | 0 | 16 | 0.0625 |
| 7_1001189 | 2 | 0 | 2 | 0 | 1 | 0 | 3 | 0.3333 | 8 | 0.125 | 2 | 0 | 5 | 0 | 5 | 0.6 | 4 | 0.5 | 16 | 0.3125 |
| 7_1001189 | 1 | 0 | 2 | 0.5 | NA | NA | 2 | 0.5 | 5 | 0.4 | 2 | 0 | NA | NA | 4 | 0.5 | NA | NA | 6 | 0.3333 |
| 7_1001190 | 2 | 0.5 | 2 | 0 | 2 | 1 | 3 | 0.3333 | 9 | 0.4444 | 2 | 0 | 5 | 0.4 | 5 | 0.4 | 3 | 0.3333 | 15 | 0.3333 |
| 7_1001884 | 2 | 1 | NA | NA | 1 | 0 | 2 | 0 | 5 | 0.4 | 1 | 1 | NA | NA | NA | NA | 1 | 0 | 2 | 0.5 |
| 7_1001886 | 2 | 0.5 | NA | NA | 1 | 0 | 2 | 0 | 5 | 0.2 | 1 | 1 | NA | NA | NA | NA | 2 | 0 | 3 | 0.3333 |
| 7_1001886 | 1 | 1 | NA | NA | NA | NA | 2 | 1 | 3 | 1 | NA | NA | NA | NA | NA | NA | 2 | 1 | 2 | 1 |
| 7_1001888 | 3 | 0.6667 | NA | NA | 1 | 1 | 1 | 1 | 5 | 0.8 | 1 | 1 | NA | NA | NA | NA | 2 | 1 | 3 | 1 |
| 7_1001889 | 3 | 1 | NA | NA | 1 | 1 | 2 | 1 | 6 | 1 | 1 | 1 | NA | NA | NA | NA | 2 | 0 | 3 | 0.3333 |
| 7_1001889 | 3 | 0.3333 | NA | NA | 1 | 0 | 2 | 1 | 6 | 0.5 | 1 | 0 | NA | NA | NA | NA | 2 | 0.5 | 3 | 0.3333 |
| 7_1001952 | 1 | 0 | 2 | 0.5 | 1 | 1 | 1 | 0 | 5 | 0.4 | 1 | 0 | NA | NA | 2 | 0.5 | NA | NA | 3 | 0.3333 |
| 7_1001963 | 1 | 0 | 2 | 1 | 1 | 1 | 1 | 1 | 5 | 0.8 | 1 | 1 | NA | NA | 1 | 1 | NA | NA | 2 | 1 |
| 7_1001963 | 1 | 0 | 2 | 0.5 | 1 | 1 | 1 | 1 | 5 | 0.6 | 1 | 1 | NA | NA | NA | NA | NA | NA | 1 | 1 |
| 7_1001983 | 9 | 0.1111 | 11 | 0.0909 | 9 | 0 | 13 | 0.1538 | 42 | 0.0952 | 7 | 0.1429 | 9 | 0 | 15 | 0.2667 | 11 | 0.0909 | 42 | 0.1429 |
| 7_1001985 | 9 | 0 | 10 | 0.2 | 10 | 0.1 | 11 | 0.2727 | 40 | 0.15 | 6 | 0.1667 | 9 | 0.1111 | 14 | 0.2143 | 11 | 0.1818 | 40 | 0.175 |
| 7_1002571 | 2 | 0 | 3 | 0.3333 | 1 | 0 | 3 | 0 | 9 | 0.1111 | 1 | 1 | 1 | 0 | 3 | 0 | 1 | 0 | 6 | 0.1667 |
| 7_1002573 | 28 | 0.1429 | 47 | 0.1489 | 39 | 0.1538 | 27 | 0.0741 | 141 | 0.1348 | 34 | 0.1176 | 35 | 0.1429 | 28 | 0.1786 | 38 | 0.2105 | 135 | 0.163 |
| 7_1002885 | 34 | 0.2059 | 49 | 0.2449 | 43 | 0.3488 | 40 | 0.175 | 166 | 0.247 | 41 | 0.2439 | 38 | 0.2632 | 42 | 0.3095 | 45 | 0.2889 | 166 | 0.2771 |
| 7_1002940 | 31 | 0.0968 | 44 | 0.1591 | 43 | 0.0698 | 32 | 0.125 | 150 | 0.1133 | 46 | 0.0435 | 35 | 0.0857 | 30 | 0.1667 | 31 | 0.129 | 142 | 0.0986 |
| 7_1004859 | 3 | 0 | 5 | 0 | 4 | 0 | 8 | 0.125 | 20 | 0.05 | 3 | 0 | 2 | 0 | 4 | 0 | 3 | 0.3333 | 12 | 0.0833 |
| 7_1004861 | 3 | 0.3333 | 5 | 0.2 | 4 | 0.25 | 6 | 0.5 | 18 | 0.3333 | 3 | 1 | 3 | 0.3333 | 3 | 0.3333 | 3 | 0.6667 | 12 | 0.5833 |
| 7_1004863 | 2 | 0.5 | 4 | 0.25 | 4 | 0.25 | 8 | 0.5 | 18 | 0.3889 | 3 | 0.3333 | 3 | 0.6667 | 2 | 0.5 | 3 | 0.6667 | 11 | 0.5455 |
| 7_1004863 | 3 | 0.6667 | 6 | 0.3333 | 5 | 0.4 | 9 | 0.3333 | 23 | 0.3913 | 2 | 0.5 | 5 | 0.2 | 2 | 0.5 | 3 | 1 | 12 | 0.5 |
| 7_1004864 | 3 | 0.3333 | 6 | 0.1667 | 5 | 0.2 | 8 | 0.25 | 22 | 0.2273 | 3 | 0.3333 | 6 | 0.1667 | 6 | 0.3333 | 4 | 0.75 | 19 | 0.3684 |
| 7_1004865 | 1 | 0 | 5 | 0.4 | 6 | 1 | 8 | 0.5 | 20 | 0.6 | 2 | 1 | 4 | 1 | 5 | 1 | 2 | 1 | 13 | 1 |
| 7_1004867 | 2 | 0.5 | 6 | 0.3333 | 5 | 0.4 | 9 | 0.3333 | 22 | 0.3636 | 2 | 0 | 7 | 0.4286 | 3 | 0.3333 | 4 | 0.25 | 16 | 0.3125 |
| 7_1004867 | 3 | 1 | 5 | 1 | 4 | 1 | 7 | 1 | 19 | 1 | 3 | 0.6667 | 7 | 0.8571 | 6 | 0.8333 | 4 | 1 | 20 | 0.85 |
| 7_1004868 | 3 | 0 | 6 | 0 | 6 | 0.3333 | 11 | 0.1818 | 26 | 0.1538 | 3 | 0.3333 | 7 | 0 | 6 | 0 | 3 | 0 | 19 | 0.0526 |
| 7_1004869 | 5 | 1 | 6 | 0.8333 | 7 | 0.7143 | 10 | 0.9 | 28 | 0.8571 | 3 | 1 | 6 | 0.8333 | 4 | 1 | 4 | 1 | 17 | 0.9412 |
| 7_1004869 | 2 | 0 | 6 | 0.3333 | 5 | 0 | 9 | 0.3333 | 22 | 0.2273 | 3 | 0.3333 | 7 | 0.1429 | 5 | 0 | 3 | 0 | 18 | 0.1111 |
| 7_1004869 | 5 | 1 | 5 | 0.8 | 6 | 0.8333 | 8 | 1 | 24 | 0.9167 | 3 | 1 | 7 | 1 | 5 | 0.8 | 4 | 1 | 19 | 0.9474 |
| 7_1004870 | 5 | 0.6 | 6 | 0.3333 | 5 | 0.2 | 6 | 0.1667 | 22 | 0.3182 | 2 | 0 | 5 | 0.4 | 6 | 0.3333 | 2 | 0.5 | 15 | 0.3333 |
| 7_1004871 | 6 | 0 | 6 | 0 | 8 | 0.25 | 10 | 0.3 | 30 | 0.1667 | 4 | 0.25 | 5 | 0.2 | 7 | 0 | 4 | 0 | 20 | 0.1 |
| 7_1004883 | 8 | 0.125 | 14 | 0.3571 | 11 | 0.4545 | 14 | 0.2143 | 47 | 0.2979 | 6 | 0.5 | 9 | 0.1111 | 10 | 0.4 | 5 | 0.4 | 30 | 0.3333 |
| 7_1004885 | 4 | 0.25 | 13 | 0.2308 | 14 | 0.3571 | 15 | 0.3333 | 46 | 0.3043 | 4 | 0 | 13 | 0.0769 | 12 | 0.25 | 8 | 0.125 | 37 | 0.1351 |
| 7_1004886 | 9 | 0.5556 | 14 | 0.5 | 14 | 0.3571 | 16 | 0.5 | 53 | 0.4717 | 6 | 0.3333 | 10 | 0.6 | 13 | 0.4615 | 7 | 0.2857 | 36 | 0.4444 |
| 7_1004886 | 7 | 0.2857 | 16 | 0.0625 | 14 | 0.4286 | 18 | 0.1111 | 55 | 0.2 | 6 | 0.3333 | 12 | 0.25 | 15 | 0.4 | 8 | 0.125 | 41 | 0.2927 |
| 7_1004887 | 6 | 0.3333 | 17 | 0.1765 | 13 | 0.1538 | 15 | 0.1333 | 51 | 0.1765 | 4 | 0.5 | 9 | 0.2222 | 14 | 0.0714 | 6 | 0 | 33 | 0.1515 |
| 7_1004888 | 8 | 0.125 | 15 | 0.0667 | 14 | 0.0714 | 18 | 0 | 55 | 0.0545 | 5 | 0.2 | 13 | 0 | 12 | 0 | 8 | 0.125 | 38 | 0.0526 |
| 7_1004889 | 9 | 0.2222 | 17 | 0.3529 | 15 | 0.4667 | 13 | 0.1538 | 54 | 0.3148 | 5 | 0.4 | 15 | 0.2 | 13 | 0.2308 | 9 | 0.5556 | 42 | 0.3095 |
| 7_1004889 | 5 | 0.6 | 18 | 0.6667 | 16 | 0.75 | 17 | 0.7647 | 56 | 0.7143 | 5 | 0.6 | 12 | 0.8333 | 12 | 0.5833 | 7 | 0.8571 | 36 | 0.7222 |
| 7_1004889 | 6 | 0.3333 | 18 | 0.5556 | 12 | 0.5833 | 16 | 0.6875 | 52 | 0.5769 | 3 | 0.6667 | 14 | 0.3571 | 13 | 0.3846 | 9 | 0.4444 | 39 | 0.4103 |
| 7_1004890 | 7 | 0.4286 | 18 | 0 | 11 | 0.1818 | 11 | 0.6364 | 47 | 0.2553 | 5 | 0.4 | 7 | 0.2857 | 9 | 0.2222 | 3 | 0 | 24 | 0.25 |
| 7_1004891 | 8 | 1 | 13 | 0.8462 | 12 | 1 | 18 | 1 | 51 | 0.9608 | 4 | 1 | 12 | 1 | 15 | 1 | 8 | 1 | 39 | 1 |
| 7_1004891 | 9 | 0.1111 | 18 | 0.1111 | 14 | 0.0714 | 18 | 0.2222 | 59 | 0.1356 | 6 | 0 | 16 | 0.4375 | 14 | 0.2857 | 9 | 0.1111 | 45 | 0.2667 |
| 7_1004896 | 11 | 0 | 20 | 0.05 | 13 | 0.1538 | 20 | 0.1 | 64 | 0.0781 | 7 | 0.1429 | 16 | 0.375 | 16 | 0.0625 | 7 | 0.1429 | 46 | 0.1957 |
| 7_1004896 | 10 | 0 | 19 | 0.1053 | 13 | 0.0769 | 17 | 0.1176 | 59 | 0.0847 | 7 | 0.2857 | 12 | 0.1667 | 13 | 0.1538 | 7 | 0.1429 | 39 | 0.1795 |
| 7_1004958 | 1 | 1 | 1 | 1 | 2 | 1 | NA | NA | 4 | 1 | 2 | 0 | 3 | 0.6667 | 4 | 0.5 | 1 | 0 | 10 | 0.4 |
| 7_1005197 | NA | NA | 2 | 0.5 | 1 | 0 | 1 | 0 | 4 | 0.25 | 2 | 0.5 | 1 | 0 | NA | NA | 2 | 0 | 5 | 0.2 |

| | | | | | | | | | | | | | | | | | | | | |
|---|---|---|---|---|---|---|---|---|---|---|---|---|---|---|---|---|---|---|---|---|
| 7_1005207 | NA | NA | 2 | 0.5 | NA | NA | NA | NA | 2 | 0.5 | 2 | 0 | 1 | 0 | NA | NA | 2 | 0.5 | 5 | 0.2 |
| 7_1005223 | 10 | 0.1 | 4 | 0 | 5 | 0.4 | 14 | 0 | 33 | 0.0909 | 4 | 0 | 17 | 0.1176 | 12 | 0 | 5 | 0.2 | 38 | 0.0789 |
| 7_1005223 | 11 | 0.4545 | 5 | 0.8 | 5 | 0 | 16 | 0.4375 | 37 | 0.4324 | 9 | 0.5556 | 13 | 0.5385 | 11 | 0.4545 | 7 | 0.5714 | 40 | 0.525 |
| 7_1005223 | 11 | 1 | 6 | 1 | 6 | 1 | 19 | 1 | 42 | 1 | 11 | 1 | 18 | 0.8889 | 17 | 0.8824 | 6 | 1 | 52 | 0.9231 |
| 7_1005224 | 9 | 0.3333 | 3 | 0 | 4 | 0 | 12 | 0.1667 | 28 | 0.1786 | 9 | 0.1111 | 15 | 0.0667 | 11 | 0.0909 | 4 | 0.25 | 39 | 0.1026 |
| 7_1005224 | 12 | 1 | 7 | 1 | 6 | 1 | 16 | 1 | 41 | 1 | 10 | 1 | 21 | 0.9524 | 17 | 1 | 6 | 1 | 54 | 0.9815 |
| 7_1005225 | 11 | 0.0909 | 5 | 0 | 4 | 0.25 | 17 | 0.2941 | 37 | 0.1892 | 9 | 0.2222 | 19 | 0.3684 | 15 | 0.2667 | 6 | 0.1667 | 49 | 0.2857 |
| 7_1005225 | 11 | 0.5455 | 5 | 0.4 | 4 | 0.75 | 16 | 0.75 | 36 | 0.6389 | 7 | 0.7143 | 21 | 0.7143 | 16 | 0.5 | 6 | 0.5 | 50 | 0.62 |
| 7_1005226 | 11 | 0.2727 | 5 | 0 | 5 | 0.8 | 15 | 0.5333 | 36 | 0.4167 | 8 | 0.25 | 21 | 0.4762 | 16 | 0.5 | 4 | 0.5 | 49 | 0.449 |
| 7_1005227 | 9 | 0.1111 | 4 | 0.25 | 5 | 0.2 | 14 | 0.2143 | 32 | 0.1875 | 7 | 0.1429 | 19 | 0.1053 | 14 | 0.1429 | 3 | 0 | 43 | 0.1163 |
| 7_1005869 | 1 | 1 | NA | NA | 1 | 1 | NA | NA | 2 | 1 | 1 | 0 | NA | NA | NA | NA | 1 | 1 | 2 | 0.5 |
| 7_1005871 | NA | NA | NA | NA | 1 | 1 | NA | NA | 1 | 1 | 1 | 1 | NA | NA | NA | NA | 1 | 1 | 2 | 1 |
| 7_1005874 | 1 | 1 | NA | NA | 1 | 1 | NA | NA | 2 | 1 | NA | NA | NA | NA | NA | NA | 1 | 1 | 1 | 1 |
| 7_1009261 | 2 | 0 | 1 | 0 | 2 | 0 | 4 | 0.25 | 9 | 0.1111 | NA | NA | 5 | 0.2 | 3 | 0.3333 | NA | NA | 8 | 0.25 |
| 7_1009318 | 2 | 0.5 | 2 | 1 | 2 | 0 | 9 | 0.4444 | 15 | 0.4667 | 2 | 0.5 | 5 | 0.4 | 5 | 0.8 | 2 | 1 | 14 | 0.6429 |
| 7_1009318 | 2 | 0.5 | 3 | 0.6667 | 1 | 1 | 8 | 0.625 | 14 | 0.6429 | 2 | 1 | 3 | 1 | 7 | 0.7143 | 2 | 1 | 14 | 0.8571 |
| 7_1009320 | 3 | 0.3333 | 3 | 0.3333 | 1 | 0 | 7 | 0.2857 | 14 | 0.2857 | 2 | 0.5 | 3 | 0.3333 | 7 | 0.2857 | 2 | 0.5 | 14 | 0.3571 |
| 7_1009320 | 2 | 0 | 2 | 0.5 | 1 | 0 | 8 | 0.5 | 13 | 0.3846 | 1 | 0 | 4 | 0.5 | 6 | 0.5 | NA | NA | 11 | 0.4545 |
| 7_1009321 | 2 | 0.5 | 3 | 0.3333 | 2 | 0.5 | 8 | 0.25 | 15 | 0.3333 | 3 | 0.6667 | 3 | 0 | 6 | 0.3333 | 2 | 0.5 | 14 | 0.3571 |
| 7_1009322 | 3 | 0.6667 | 3 | 0.3333 | 1 | 1 | 9 | 0.6667 | 16 | 0.625 | 3 | 0.6667 | 6 | 0.3333 | 7 | 0.5714 | 2 | 1 | 18 | 0.5556 |
| 7_1009327 | 3 | 0.3333 | 4 | 0.25 | 2 | 0.5 | 10 | 0.2 | 19 | 0.2632 | 3 | 0.6667 | 6 | 0.6667 | 7 | 0 | 2 | 0 | 18 | 0.3333 |
| 7_1009330 | 2 | 0 | 4 | 0 | 2 | 0.5 | 9 | 0.1111 | 17 | 0.1176 | 3 | 0 | 6 | 0 | 6 | 0.1667 | 1 | 0 | 16 | 0.0625 |
| 7_1009389 | 3 | 0.3333 | 2 | 0.5 | NA | NA | 2 | 0 | 7 | 0.2857 | 2 | 0.5 | 1 | 0 | 1 | 0 | 1 | 1 | 5 | 0.4 |
| 7_1009390 | 3 | 0.3333 | 2 | 1 | NA | NA | 1 | 0 | 6 | 0.5 | 2 | 0.5 | 1 | 1 | 1 | 0 | 2 | 0.5 | 6 | 0.5 |
| 7_1009410 | 2 | 0 | 2 | 0.5 | NA | NA | 1 | 0 | 5 | 0.2 | NA | NA | 1 | 0 | 1 | 1 | 1 | 1 | 3 | 0.6667 |
| 7_1009709 | 15 | 0.0667 | 14 | 0.0714 | 19 | 0.1053 | 19 | 0.0526 | 67 | 0.0746 | 9 | 0.2222 | 11 | 0.0909 | 17 | 0.0588 | 9 | 0.1111 | 46 | 0.1087 |
| 7_1009711 | 16 | 0.0625 | 15 | 0.2 | 25 | 0.12 | 18 | 0.1667 | 74 | 0.1351 | 12 | 0.1667 | 15 | 0 | 20 | 0.25 | 13 | 0.1538 | 60 | 0.15 |
| 7_1009952 | 21 | 0.0476 | 22 | 0.0455 | 30 | 0.1 | 20 | 0.1 | 93 | 0.0753 | 18 | 0.1667 | 25 | 0.08 | 26 | 0.0769 | 13 | 0.0769 | 82 | 0.0976 |
| 7_1009953 | 17 | 0.1176 | 22 | 0.2273 | 27 | 0.2222 | 17 | 0.2353 | 83 | 0.2048 | 13 | 0.5385 | 22 | 0.2727 | 23 | 0.2174 | 12 | 0.5 | 70 | 0.3429 |
| 7_1009953 | 18 | 0.1667 | 21 | 0.1429 | 31 | 0.2581 | 18 | 0.3889 | 88 | 0.2386 | 17 | 0.3529 | 23 | 0.1304 | 27 | 0.2222 | 9 | 0.1111 | 76 | 0.2105 |
| 7_1009953 | 21 | 0 | 20 | 0.1 | 28 | 0.2143 | 18 | 0.1111 | 87 | 0.1149 | 16 | 0.1875 | 21 | 0.0476 | 26 | 0.1154 | 14 | 0.0714 | 77 | 0.1039 |
| 7_1009954 | 17 | 0.4118 | 18 | 0.7222 | 26 | 0.6154 | 20 | 0.6 | 81 | 0.5926 | 16 | 0.5625 | 24 | 0.5833 | 28 | 0.6786 | 15 | 0.6667 | 83 | 0.6265 |
| 7_1009955 | 10 | 0 | 11 | 0.1818 | 22 | 0.1364 | 15 | 0.0667 | 58 | 0.1034 | 14 | 0.1429 | 14 | 0.2857 | 13 | 0.2308 | 10 | 0.2 | 51 | 0.2157 |
| 7_1009956 | 3 | 1 | 6 | 1 | 13 | 0.8462 | 7 | 0.4286 | 29 | 0.7931 | 4 | 1 | 9 | 0.8889 | 9 | 1 | 4 | 1 | 26 | 0.9615 |
| 7_1009957 | 1 | 0 | 3 | 0.3333 | 5 | 0 | 2 | 0.5 | 11 | 0.1818 | 3 | 0 | 4 | 0.25 | 3 | 0 | 1 | 0 | 11 | 0.0909 |
| 7_1010608 | 2 | 1 | 1 | 1 | NA | NA | 1 | 1 | 4 | 1 | NA | NA | 1 | 1 | 1 | 1 | NA | NA | 2 | 1 |
| 7_1010609 | 2 | 0.5 | 1 | 1 | NA | NA | 1 | 1 | 4 | 0.75 | NA | NA | 2 | 0.5 | 1 | 0 | NA | NA | 3 | 0.3333 |
| 7_1010710 | 7 | 0.1429 | 4 | 0 | 6 | 0.1667 | 11 | 0.4545 | 28 | 0.25 | 3 | 0 | 3 | 0 | 15 | 0.1333 | 4 | 0.25 | 25 | 0.12 |
| 7_1011483 | 1 | 1 | NA | NA | NA | NA | NA | NA | 1 | 1 | NA | NA | 1 | 1 | 2 | 0 | 1 | 0 | 4 | 0.25 |
| 7_1011494 | 1 | 0 | 1 | 0 | 1 | 1 | NA | NA | 3 | 0.3333 | NA | NA | 2 | 1 | 2 | 0 | 2 | 0.5 | 6 | 0.5 |
| 7_1011608 | 48 | 0.1667 | 42 | 0.119 | 38 | 0.0789 | 43 | 0.0698 | 171 | 0.1111 | 29 | 0.2414 | 53 | 0.0377 | 67 | 0.0896 | 41 | 0.0732 | 190 | 0.0947 |
| 7_1011609 | 44 | 0.1364 | 35 | 0.2286 | 32 | 0.0625 | 38 | 0.1053 | 149 | 0.1342 | 32 | 0.0938 | 45 | 0.1556 | 55 | 0.1636 | 37 | 0.1351 | 169 | 0.142 |
| 7_1011610 | 54 | 0.8519 | 41 | 0.8293 | 36 | 0.8889 | 41 | 0.7805 | 172 | 0.8372 | 32 | 0.9062 | 57 | 0.8246 | 62 | 0.8387 | 42 | 0.8333 | 193 | 0.8446 |
| 7_1011611 | 52 | 0.1154 | 40 | 0.3 | 38 | 0.2105 | 39 | 0.2564 | 169 | 0.213 | 31 | 0.1935 | 53 | 0.2642 | 53 | 0.2642 | 37 | 0.1892 | 174 | 0.2356 |
| 7_1011611 | 48 | 0.1875 | 40 | 0.175 | 39 | 0.2308 | 39 | 0.2051 | 166 | 0.1988 | 31 | 0.1613 | 54 | 0.2037 | 63 | 0.2063 | 35 | 0.2 | 183 | 0.1967 |
| 7_1011612 | 55 | 0.1636 | 42 | 0.1429 | 40 | 0.175 | 43 | 0.093 | 180 | 0.1444 | 34 | 0.1765 | 52 | 0.1538 | 71 | 0.1127 | 39 | 0.1795 | 196 | 0.148 |
| 7_1011613 | 54 | 0.5556 | 45 | 0.6 | 46 | 0.4783 | 45 | 0.5778 | 190 | 0.5526 | 32 | 0.5312 | 60 | 0.5167 | 71 | 0.6479 | 41 | 0.6585 | 204 | 0.5931 |
| 7_1011613 | 62 | 0.371 | 47 | 0.5319 | 47 | 0.4468 | 46 | 0.3913 | 202 | 0.4307 | 34 | 0.4118 | 60 | 0.4333 | 74 | 0.473 | 44 | 0.3864 | 212 | 0.434 |
| 7_1011618 | 53 | 0.1132 | 40 | 0.075 | 45 | 0.1556 | 39 | 0.1026 | 177 | 0.113 | 33 | 0.1515 | 54 | 0.0926 | 68 | 0.1618 | 44 | 0.1591 | 199 | 0.1407 |
| 7_1011619 | 66 | 0.1818 | 47 | 0.1064 | 53 | 0.1887 | 49 | 0.0816 | 215 | 0.1442 | 37 | 0.1622 | 66 | 0.2121 | 71 | 0.1972 | 42 | 0.119 | 216 | 0.1806 |
| 7_1014367 | 2 | 0 | 4 | 0 | 11 | 0.1818 | 6 | 0 | 23 | 0.087 | 1 | 0 | 7 | 0 | 7 | 0.1429 | 5 | 0 | 20 | 0.05 |
| 7_1014368 | 2 | 0 | 4 | 0.25 | 8 | 0 | 5 | 0 | 19 | 0.0526 | 1 | 0 | 6 | 0 | 7 | 0.2857 | 4 | 0 | 18 | 0.1111 |
| 7_1014368 | 2 | 0.5 | 5 | 0.6 | 12 | 0.5 | 6 | 0.1667 | 25 | 0.44 | 1 | 0 | 7 | 0.4286 | 7 | 0.5714 | 3 | 0.6667 | 18 | 0.5 |
| 7_1014369 | 2 | 0.5 | 5 | 0.8 | 10 | 0.3 | 7 | 0.2857 | 24 | 0.4167 | 1 | 0 | 6 | 0.3333 | 7 | 0.5714 | 5 | 0.2 | 19 | 0.3684 |
| 7_1014369 | 2 | 0 | 3 | 0.3333 | 7 | 0.1429 | 4 | 0.5 | 16 | 0.25 | 1 | 0 | 6 | 0.1667 | 4 | 0.25 | 4 | 0.25 | 15 | 0.2 |
| 7_1014372 | NA | NA | NA | NA | 1 | 1 | 1 | 1 | 2 | 1 | NA | NA | 1 | 1 | NA | NA | NA | NA | 1 | 1 |
| 7_1014426 | NA | NA | 1 | 0 | 1 | 1 | 2 | 0.5 | 4 | 0.5 | 1 | 0 | 2 | 0.5 | 3 | 0.6667 | NA | NA | 6 | 0.5 |
| 7_1014434 | NA | NA | 1 | 1 | 2 | 1 | 2 | 1 | 5 | 1 | 2 | 0.5 | 2 | 1 | 3 | 0.3333 | 1 | 1 | 8 | 0.625 |
| 7_1014435 | NA | NA | 1 | 0 | 2 | 0.5 | 1 | 0 | 4 | 0.25 | 2 | 0.5 | 1 | 0 | 3 | 0 | 1 | 1 | 7 | 0.2857 |
| 7_1014439 | 2 | 0 | 2 | 0.5 | 4 | 0.25 | 3 | 0.6667 | 11 | 0.3636 | 2 | 0 | 2 | 0 | 3 | 0.6667 | 1 | 1 | 8 | 0.375 |
| 7_1014439 | 2 | 0 | 3 | 0.3333 | 5 | 0.8 | 1 | 0 | 11 | 0.4545 | 2 | 0.5 | 2 | 1 | 3 | 1 | 1 | 1 | 8 | 0.875 |
| 7_1014440 | 1 | 0 | 3 | 0.6667 | 4 | 1 | 3 | 1 | 11 | 0.8182 | 1 | 1 | 2 | 1 | 3 | 1 | 1 | 1 | 7 | 1 |
| 7_1014442 | 1 | 0 | 3 | 0 | 5 | 0.8 | 3 | 1 | 12 | 0.5833 | 2 | 0.5 | 2 | 0.5 | 2 | 0.5 | 1 | 1 | 7 | 0.5714 |
| 7_1014719 | 2 | 0 | 2 | 0.5 | 5 | 0.4 | 1 | 0 | 10 | 0.3 | 3 | 0.6667 | 2 | 0.5 | 4 | 0.25 | 4 | 0.5 | 13 | 0.4615 |
| 7_1014720 | 2 | 1 | 2 | 0 | 4 | 1 | 1 | 0 | 9 | 0.6667 | 3 | 0 | 2 | 0.5 | 3 | 0.3333 | 3 | 0 | 11 | 0.1818 |

| | | | | | | | | | | | | | | | | | | | | |
|---|---|---|---|---|---|---|---|---|---|---|---|---|---|---|---|---|---|---|---|---|
| 7_1014723 | 1 | 1 | NA | NA | 3 | 0.6667 | 1 | 1 | 5 | 0.8 | 2 | 0.5 | 1 | 1 | 2 | 0.5 | 2 | 1 | 7 | 0.7143 |
| 7_1014731 | NA | NA | 1 | 0 | 1 | 0 | 1 | 1 | 3 | 0.3333 | 2 | 0.5 | NA | NA | 1 | 0 | 2 | 0.5 | 5 | 0.4 |
| 7_1014734 | NA | NA | 2 | 0.5 | 2 | 0 | 1 | 0 | 5 | 0.2 | 2 | 0.5 | NA | NA | 2 | 0 | 2 | 0.5 | 6 | 0.3333 |
| 7_1015482 | 3 | 0 | 2 | 0 | NA | NA | 3 | 0.3333 | 8 | 0.125 | 1 | 1 | 6 | 0 | 2 | 0 | NA | NA | 9 | 0.1111 |
| 7_1015485 | 4 | 0 | 3 | 0 | 1 | 0 | 2 | 0.5 | 10 | 0.1 | 1 | 0 | 5 | 0.2 | 2 | 0 | NA | NA | 8 | 0.125 |
| 7_1015485 | 5 | 0.2 | 3 | 0 | 1 | 1 | 3 | 0 | 12 | 0.1667 | 1 | 1 | 6 | 0 | 2 | 0 | NA | NA | 9 | 0.1111 |
| 7_1015487 | 3 | 0.6667 | 3 | 0 | 1 | 1 | 4 | 0 | 11 | 0.2727 | 1 | 1 | 6 | 0.3333 | 2 | 0.5 | NA | NA | 9 | 0.4444 |
| 7_1015488 | NA | NA | NA | NA | 2 | 1 | 1 | 1 | 3 | 1 | NA | NA | NA | NA | 1 | 1 | NA | NA | 1 | 1 |
| 7_1015488 | 1 | 1 | NA | NA | 2 | 0.5 | 1 | 0 | 4 | 0.5 | NA | NA | 1 | 0 | 1 | 1 | 1 | 1 | 3 | 0.6667 |
| 7_1015501 | 1 | 0 | NA | NA | 1 | 1 | 2 | 0 | 4 | 0.25 | 1 | 0 | 2 | 0.5 | 1 | 1 | 1 | 1 | 5 | 0.6 |
| 7_1015605 | 11 | 0.2727 | 13 | 0.1538 | 17 | 0 | 15 | 0.2 | 56 | 0.1429 | 20 | 0 | 11 | 0.2727 | 20 | 0.2 | 11 | 0.0909 | 62 | 0.129 |
| 7_1015607 | 14 | 0.9286 | 15 | 1 | 16 | 0.8125 | 18 | 1 | 63 | 0.9365 | 23 | 0.913 | 14 | 0.8571 | 20 | 0.85 | 14 | 0.8571 | 71 | 0.8732 |
| 7_1015608 | 14 | 0.0714 | 9 | 0 | 12 | 0.1667 | 18 | 0 | 53 | 0.0566 | 26 | 0.0385 | 17 | 0.0588 | 18 | 0.0556 | 11 | 0.0909 | 72 | 0.0556 |
| 7_1015625 | NA | NA | NA | NA | 1 | 1 | NA | NA | 1 | 1 | NA | NA | NA | NA | 1 | 1 | NA | NA | 1 | 1 |
| 7_1016481 | 1 | 1 | 1 | 0 | NA | NA | 1 | 1 | 3 | 0.6667 | NA | NA | 3 | 0.3333 | 3 | 0.3333 | 2 | 0.5 | 8 | 0.375 |
| 7_1016485 | 1 | 0 | 1 | 0 | NA | NA | 2 | 0.5 | 4 | 0.25 | NA | NA | 3 | 0 | 4 | 0.5 | 2 | 0 | 9 | 0.2222 |
| 7_1016494 | 2 | 0.5 | 2 | 0.5 | NA | NA | NA | NA | 4 | 0.5 | NA | NA | 1 | 1 | 3 | 0.6667 | 1 | 1 | 5 | 0.8 |
| 7_1016494 | 3 | 0.3333 | 1 | 0 | NA | NA | 2 | 0.5 | 6 | 0.3333 | NA | NA | 3 | 0.3333 | 5 | 0.8 | 2 | 0.5 | 10 | 0.6 |
| 7_1016496 | 3 | 0.3333 | 2 | 0 | NA | NA | 2 | 0 | 7 | 0.1429 | NA | NA | 3 | 0 | 5 | 0.6 | 2 | 0.5 | 10 | 0.4 |
| 7_1016501 | 4 | 0.25 | 2 | 1 | 1 | 0 | 3 | 0.3333 | 10 | 0.4 | NA | NA | 3 | 0.3333 | 5 | 0.4 | 1 | 1 | 9 | 0.4444 |
| 7_1016506 | 3 | 0.6667 | 2 | 1 | 2 | 1 | 3 | 0 | 10 | 0.6 | 2 | 0.5 | 2 | 0.5 | 5 | 0.8 | 2 | 0.5 | 11 | 0.6364 |
| 7_1016506 | 3 | 0 | 1 | 0 | 1 | 0 | 3 | 0.3333 | 8 | 0.125 | 2 | 0.5 | 4 | 0.25 | 4 | 0.75 | 2 | 0 | 12 | 0.4167 |
| 7_1016636 | 9 | 0.6667 | 9 | 0.6667 | 5 | 1 | 10 | 0.7 | 33 | 0.7273 | 4 | 0.5 | 11 | 0.9091 | 5 | 0.6 | 4 | 1 | 24 | 0.7917 |
| 7_1016636 | 9 | 0.6667 | 9 | 0.2222 | 10 | 0.6 | 13 | 0.5385 | 41 | 0.5122 | 3 | 1 | 12 | 0.8333 | 5 | 0.8 | 4 | 0.25 | 24 | 0.75 |
| 7_1016638 | 9 | 0.1111 | 10 | 0.1 | 11 | 0.1818 | 13 | 0.1538 | 43 | 0.1395 | 3 | 0 | 13 | 0.0769 | 6 | 0 | 4 | 0.25 | 26 | 0.0769 |
| 7_1016690 | 2 | 0 | 3 | 0 | 8 | 0.125 | 5 | 0.2 | 18 | 0.1111 | NA | NA | 6 | 0 | 2 | 0.5 | 3 | 0 | 11 | 0.0909 |
| 7_1016690 | 1 | 0 | 2 | 0 | 5 | 0.2 | 5 | 0 | 13 | 0.0769 | NA | NA | 5 | 0.2 | 2 | 0.5 | 4 | 0 | 11 | 0.1818 |
| 7_1016694 | 1 | 0 | 3 | 0.3333 | 1 | 0 | 2 | 0 | 7 | 0.1429 | NA | NA | 4 | 0 | 3 | 0.3333 | 1 | 0 | 8 | 0.125 |
| 7_1016694 | 2 | 1 | 4 | 1 | 6 | 0.6667 | 4 | 1 | 16 | 0.875 | NA | NA | 5 | 1 | 5 | 0.8 | 4 | 1 | 14 | 0.9286 |
| 7_1016695 | 2 | 0.5 | 4 | 0.5 | 7 | 0.7143 | 6 | 0.8333 | 19 | 0.6842 | NA | NA | 6 | 0.6667 | 5 | 0.6 | 3 | 0.3333 | 14 | 0.5714 |
| 7_1016695 | 2 | 1 | 5 | 0.4 | 6 | 0.8333 | 5 | 1 | 18 | 0.7778 | NA | NA | 3 | 1 | 4 | 0.75 | 3 | 0.3333 | 10 | 0.7 |
| 7_1017314 | 2 | 0.5 | NA | NA | 1 | 0 | 1 | 1 | 4 | 0.5 | 1 | 0 | NA | NA | 4 | 0.25 | 4 | 0.25 | 9 | 0.2222 |
| 7_1017315 | 2 | 1 | NA | NA | 1 | 1 | 1 | 0 | 4 | 0.75 | 1 | 1 | NA | NA | 3 | 1 | 2 | 0.5 | 6 | 0.8333 |
| 7_1017316 | 2 | 0 | NA | NA | 1 | 1 | 1 | 0 | 4 | 0.25 | 1 | 0 | NA | NA | 4 | 1 | 4 | 0.75 | 9 | 0.7778 |
| 7_1017317 | 3 | 0.3333 | NA | NA | 1 | 0 | 1 | 0 | 5 | 0.2 | 1 | 0 | NA | NA | 2 | 0.5 | 3 | 0.6667 | 6 | 0.5 |
| 7_1017687 | 7 | 0.4286 | 2 | 0.5 | 5 | 0.4 | 3 | 0.3333 | 17 | 0.4118 | 2 | 0 | 6 | 0.1667 | 6 | 0 | 6 | 0.3333 | 20 | 0.15 |
| 7_1017688 | 6 | 0.5 | 1 | 0 | 4 | 0.5 | 3 | 0 | 14 | 0.3571 | 1 | 1 | 4 | 0.25 | 4 | 1 | 4 | 0.5 | 13 | 0.6154 |
| 7_1017690 | 3 | 0.6667 | 2 | 1 | 2 | 0.5 | 2 | 0.5 | 9 | 0.6667 | 1 | 1 | 4 | 1 | 7 | 0.8571 | 5 | 1 | 17 | 0.9412 |
| 7_1017691 | 5 | 0.8 | 2 | 1 | 2 | 1 | 3 | 0.6667 | 12 | 0.8333 | 1 | 1 | 5 | 0.6 | 6 | 0.8333 | 5 | 0.8 | 17 | 0.7647 |
| 7_1017692 | 5 | 0 | 2 | 0 | 2 | 0.5 | 3 | 0 | 12 | 0.0833 | 1 | 0 | 4 | 0 | 6 | 0.1667 | 4 | 0 | 15 | 0.0667 |
| 7_1017692 | 5 | 0.6 | 1 | 1 | 2 | 0.5 | 2 | 1 | 10 | 0.7 | 2 | 1 | 5 | 0.6 | 7 | 0.5714 | 4 | 0.25 | 18 | 0.5556 |
| 7_1017692 | 4 | 0.25 | 2 | 0.5 | 2 | 0 | 3 | 0.6667 | 11 | 0.3636 | 2 | 0 | 5 | 0.4 | 7 | 0.2857 | 2 | 0 | 16 | 0.25 |
| 7_1017692 | 4 | 0 | 2 | 0 | 2 | 0 | 3 | 0.3333 | 11 | 0.0909 | 1 | 0 | 5 | 0 | 7 | 0.1429 | 4 | 0 | 17 | 0.0588 |
| 7_1017692 | 4 | 1 | 2 | 1 | 2 | 1 | 3 | 1 | 11 | 1 | 2 | 1 | 5 | 1 | 7 | 0.7143 | 4 | 1 | 18 | 0.8889 |
| 7_1017696 | 3 | 0.6667 | 1 | 0 | 2 | 1 | 3 | 0.6667 | 9 | 0.6667 | 1 | 1 | 1 | 1 | 3 | 0.6667 | 2 | 1 | 7 | 0.8571 |
| 7_1017696 | 2 | 0.5 | 1 | 0 | 1 | 0 | 1 | 0 | 5 | 0.2 | NA | NA | 2 | 0 | 2 | 0.5 | 1 | 0 | 5 | 0.2 |
| 7_1017697 | 3 | 0 | 1 | 1 | 2 | 0 | 2 | 0 | 8 | 0.125 | 2 | 0 | 4 | 0 | 7 | 0.4286 | 3 | 0 | 16 | 0.1875 |
| 7_1017697 | 2 | 0.5 | 1 | 0 | 1 | 1 | 1 | 0 | 5 | 0.4 | 1 | 0 | 3 | 0.3333 | 6 | 0.1667 | 3 | 0 | 13 | 0.1538 |
| 7_1017743 | 2 | 0 | 1 | 1 | 1 | 0 | 3 | 0 | 7 | 0.1429 | 2 | 0 | 2 | 0 | 6 | 0.1667 | 2 | 0 | 12 | 0.0833 |
| 7_1017838 | 1 | 1 | 1 | 0 | 1 | 0 | 2 | 0.5 | 5 | 0.4 | 1 | 1 | 3 | 0.6667 | 6 | 0.3333 | 2 | 0 | 12 | 0.4167 |
| 7_1017838 | 2 | 1 | 1 | 1 | 1 | 1 | 1 | 1 | 5 | 1 | NA | NA | 3 | 0.6667 | 4 | 0.75 | 2 | 1 | 9 | 0.7778 |
| 7_1017839 | 2 | 1 | 1 | 0 | 1 | 1 | 1 | 1 | 5 | 0.8 | NA | NA | 3 | 1 | 3 | 1 | 2 | 1 | 8 | 1 |
| 7_1017840 | 1 | 0 | NA | NA | 1 | 1 | 1 | 0 | 3 | 0.3333 | NA | NA | 1 | 0 | 3 | 0 | 3 | 0.6667 | 7 | 0.2857 |
| 7_1017841 | 2 | 0.5 | 1 | 0 | 1 | 0 | 1 | 0 | 5 | 0.2 | NA | NA | 3 | 0.3333 | 5 | 0 | 3 | 0 | 11 | 0.0909 |
| 7_1017947 | 2 | 0 | 1 | 0 | 1 | 1 | 1 | 0 | 5 | 0.2 | 1 | 0 | 2 | 0 | 4 | 0.5 | 3 | 0.3333 | 10 | 0.3 |
| 7_1018855 | 3 | 0 | 2 | 0 | 1 | 1 | 1 | 1 | 7 | 0.2857 | NA | NA | 2 | 1 | 3 | 0.6667 | 1 | 1 | 6 | 0.8333 |
| 7_1018858 | 2 | 0.5 | NA | NA | NA | NA | 1 | 0 | 3 | 0.3333 | NA | NA | 1 | 0 | 1 | 1 | NA | NA | 2 | 0.5 |
| 7_1018858 | 2 | 1 | NA | NA | NA | NA | NA | NA | 2 | 1 | NA | NA | 1 | 1 | NA | NA | NA | NA | 1 | 1 |
| 7_1018860 | 2 | 0.5 | NA | NA | NA | NA | 1 | 1 | 3 | 0.6667 | NA | NA | NA | NA | 1 | 1 | NA | NA | 1 | 1 |
| 7_1018862 | 2 | 0 | NA | NA | NA | NA | 1 | 1 | 3 | 0.3333 | NA | NA | 1 | 0 | 2 | 0.5 | NA | NA | 3 | 0.3333 |
| 7_1018866 | 6 | 0.3333 | 3 | 0.6667 | 1 | 0 | 1 | 1 | 11 | 0.4545 | 1 | 0 | 2 | 0.5 | 5 | 0.2 | 1 | 1 | 9 | 0.3333 |
| 7_1018869 | 4 | 1 | 3 | 0.3333 | 1 | 1 | 2 | 1 | 10 | 0.8 | 1 | 1 | 2 | 0.5 | 5 | 0.6 | 1 | 1 | 9 | 0.6667 |
| 7_1018871 | 6 | 0 | 3 | 0 | 1 | 1 | 3 | 0 | 13 | 0.0769 | NA | NA | 3 | 0 | 4 | 0.25 | 1 | 0 | 8 | 0.125 |
| 7_1018875 | 7 | 0.1429 | 2 | 0 | NA | NA | 2 | 0 | 11 | 0.0909 | 1 | 0 | 3 | 0.3333 | 6 | 0.1667 | NA | NA | 10 | 0.2 |
| 7_1018883 | 7 | 0.5714 | 7 | 0.2857 | 2 | 0 | 5 | 0.2 | 21 | 0.3333 | 8 | 0 | 7 | 0 | 6 | 0.1667 | 1 | 1 | 22 | 0.0909 |

| | | | | | | | | | | | | | | | | | | | | |
|---|---|---|---|---|---|---|---|---|---|---|---|---|---|---|---|---|---|---|---|---|
| 7_10188840 | 7 | 0.4286 | 7 | 0.4286 | 4 | 0.5 | 8 | 0.625 | 26 | 0.5 | 10 | 0.2 | 8 | 0.25 | 4 | 0.25 | 2 | 0 | 24 | 0.2083 |
| 7_10188841 | 9 | 0.8889 | 8 | 0.625 | 4 | 1 | 10 | 0.8 | 31 | 0.8065 | 12 | 0.5833 | 11 | 0.9091 | 7 | 0.7143 | 3 | 1 | 33 | 0.7576 |
| 7_10188851 | 8 | 0.875 | 7 | 0.8571 | 3 | 0.3333 | 7 | 0.5714 | 25 | 0.72 | 10 | 0.8 | 9 | 0.8889 | 7 | 1 | 3 | 0.6667 | 29 | 0.8621 |
| 7_10188876 | 8 | 0.125 | 8 | 0 | 3 | 0 | 7 | 0.1429 | 26 | 0.0769 | 10 | 0.1 | 9 | 0.1111 | 7 | 0 | 3 | 0 | 29 | 0.069 |
| 7_10188878 | 8 | 1 | 8 | 1 | 3 | 1 | 6 | 1 | 25 | 1 | 9 | 1 | 8 | 0.875 | 6 | 0.8333 | 3 | 1 | 26 | 0.9231 |
| 7_10188887 | 8 | 0.125 | 8 | 0.375 | 3 | 0.3333 | 7 | 0 | 26 | 0.1923 | 10 | 0 | 9 | 0 | 7 | 0.1429 | 2 | 1 | 28 | 0.1071 |
| 7_10188909 | 6 | 0.3333 | 7 | 0.4286 | 2 | 0.5 | 7 | 0.5714 | 22 | 0.4545 | 10 | 0.5 | 8 | 0.75 | 6 | 0.5 | 3 | 0.6667 | 27 | 0.5926 |
| 7_10188928 | 7 | 1 | 5 | 1 | 3 | 1 | 5 | 1 | 20 | 1 | 7 | 1 | 7 | 1 | 5 | 0.6 | 2 | 1 | 21 | 0.9048 |
| 7_10188929 | 7 | 0.4286 | 7 | 0 | 3 | 0.3333 | 6 | 0.1667 | 23 | 0.2174 | 10 | 0.2 | 5 | 0 | 6 | 0 | 2 | 0.5 | 23 | 0.1304 |
| 7_10188929 | 6 | 0.5 | 5 | 0.2 | 2 | 0.5 | 4 | 0.5 | 17 | 0.4118 | 8 | 0.25 | 4 | 0.5 | 3 | 0.3333 | 2 | 1 | 17 | 0.4118 |
| 7_10188933 | 6 | 0.1667 | 6 | 0 | 2 | 0 | 5 | 0 | 19 | 0.0526 | 8 | 0.125 | 6 | 0.1667 | 5 | 0.2 | 3 | 0 | 22 | 0.1364 |
| 7_10188946 | 6 | 0 | 7 | 0.1429 | 3 | 0.3333 | 7 | 0.1429 | 23 | 0.1304 | 7 | 0.1429 | 8 | 0 | 6 | 0 | 3 | 0.3333 | 24 | 0.0833 |
| 7_10191749 | 1 | 0 | 4 | 0 | 1 | 0 | 2 | 1 | 8 | 0.25 | NA | NA | 1 | 1 | 1 | 0 | 1 | 0 | 3 | 0.3333 |
| 7_10191753 | 2 | 0 | 4 | 0 | 1 | 0 | 3 | 0.3333 | 10 | 0.1 | NA | NA | 3 | 0.3333 | 4 | 0 | 2 | 0 | 9 | 0.1111 |
| 7_10191760 | 3 | 0.6667 | 4 | 1 | 1 | 0 | 2 | 1 | 10 | 0.8 | NA | NA | 3 | 0.3333 | 2 | 0 | 3 | 0.3333 | 8 | 0.25 |
| 7_10208480 | 6 | 0.1667 | 4 | 0.25 | 3 | 0.3333 | 5 | 0 | 18 | 0.1667 | 1 | 0 | 8 | 0 | 12 | 0.1667 | 5 | 0.2 | 26 | 0.1154 |
| 7_10211751 | 2 | 0 | 5 | 0.2 | NA | NA | 2 | 0.5 | 9 | 0.2222 | 2 | 0.5 | 1 | 0 | 4 | 0.5 | NA | NA | 7 | 0.4286 |
| 7_10211807 | 2 | 0 | 8 | 0 | 1 | 1 | 2 | 0 | 13 | 0.0769 | 2 | 0 | 1 | 0 | 2 | 0.5 | 1 | 0 | 6 | 0.1667 |
| 7_10405883 | 1 | 1 | NA | NA | 1 | 0 | 1 | 1 | 3 | 0.6667 | NA | NA | NA | NA | NA | NA | 1 | 1 | 1 | 1 |
| 7_10405968 | NA | NA | NA | NA | 1 | 0 | 1 | 1 | 2 | 0.5 | NA | NA | 1 | 0 | 1 | 1 | NA | NA | 2 | 0.5 |
| 7_10412088 | NA | NA | 1 | 1 | NA | NA | NA | NA | 1 | 1 | NA | NA | NA | NA | 1 | 1 | NA | NA | 1 | 1 |
| 7_10502055 | 2 | 0.5 | 4 | 0.5 | 2 | 0.5 | 2 | 0.5 | 10 | 0.5 | NA | NA | 3 | 0.6667 | 5 | 0.8 | 2 | 0.5 | 10 | 0.7 |
| 7_10502057 | 1 | 0 | 6 | 0.3333 | 3 | 0.3333 | 3 | 0.3333 | 13 | 0.3077 | 2 | 0.5 | 5 | 0.8 | 7 | 0.8571 | 2 | 0.5 | 16 | 0.75 |
| 7_10502062 | 2 | 0.5 | 5 | 0.6 | 3 | 1 | 1 | 1 | 11 | 0.7273 | 1 | 1 | 4 | 0.75 | 7 | 0.8571 | 2 | 0.5 | 14 | 0.7857 |
| 7_10502209 | 1 | 0 | 5 | 0.2 | 1 | 0 | 2 | 0 | 9 | 0.1111 | NA | NA | 4 | 0.25 | 3 | 0 | 2 | 0 | 9 | 0.1111 |
| 7_10502785 | 1 | 0 | 4 | 0.25 | 3 | 0.3333 | 4 | 0.25 | 12 | 0.25 | 4 | 0.25 | 3 | 0 | 5 | 0.6 | 4 | 0 | 16 | 0.25 |
| 7_10502788 | 1 | 1 | 3 | 1 | 3 | 1 | 5 | 1 | 12 | 1 | 6 | 1 | 3 | 0.3333 | 5 | 1 | 4 | 0.75 | 18 | 0.8333 |
| 7_10502792 | 1 | 0 | 4 | 0.25 | 3 | 0 | 5 | 0.4 | 13 | 0.2308 | 4 | 0.75 | 3 | 0.3333 | 4 | 0.25 | 5 | 0.2 | 16 | 0.375 |
| 7_10502796 | 1 | 1 | 3 | 1 | 2 | 1 | 2 | 1 | 8 | 1 | 3 | 1 | 1 | 1 | 5 | 1 | 2 | 1 | 11 | 1 |
| 7_10502822 | 1 | 0 | 1 | 1 | NA | NA | 4 | 1 | 6 | 0.8333 | 1 | 1 | 2 | 1 | 2 | 1 | 4 | 1 | 9 | 1 |
| 7_10502822 | 1 | 0 | 4 | 1 | 2 | 1 | 3 | 0.6667 | 10 | 0.8 | 2 | 1 | 1 | 1 | 2 | 1 | 4 | 0.5 | 9 | 0.7778 |
| 7_10502823 | 1 | 0 | 4 | 0.5 | 2 | 0 | 4 | 0.25 | 11 | 0.2727 | 4 | 0 | 1 | 0 | 2 | 0 | 3 | 0.3333 | 10 | 0.1 |
| 7_10503319 | 1 | 0 | 3 | 0 | 2 | 0 | 2 | 0.5 | 8 | 0.125 | 2 | 0 | NA | NA | 2 | 0 | 4 | 0.25 | 8 | 0.125 |
| 7_10503320 | 1 | 0 | 3 | 0.6667 | 1 | 1 | 4 | 0.5 | 9 | 0.5556 | 5 | 0 | NA | NA | 3 | 0.3333 | 4 | 0.5 | 12 | 0.25 |
| 7_10503340 | 1 | 0 | 3 | 0.3333 | 2 | 0.5 | 3 | 0.3333 | 9 | 0.3333 | 4 | 0.5 | NA | NA | 2 | 1 | 4 | 0.75 | 10 | 0.7 |
| 7_10503344 | 1 | 1 | 3 | 1 | 2 | 0.5 | 3 | 1 | 9 | 0.8889 | 3 | 0.6667 | NA | NA | 2 | 0.5 | 4 | 1 | 9 | 0.7778 |
| 7_10503884 | 1 | 1 | 2 | 0.5 | NA | NA | 3 | 0.6667 | 6 | 0.6667 | 1 | 0 | NA | NA | 1 | 1 | 3 | 0.6667 | 5 | 0.6 |
| 7_10503885 | 1 | 0 | 2 | 1 | 1 | 1 | 1 | 1 | 5 | 0.8 | 1 | 1 | NA | NA | NA | NA | 4 | 0.25 | 5 | 0.4 |
| 7_10503935 | 1 | 1 | 2 | 1 | 1 | 1 | 2 | 0.5 | 6 | 0.8333 | NA | NA | NA | NA | 1 | 0 | 3 | 1 | 4 | 0.75 |
| 7_10503954 | 1 | 0 | 2 | 0 | 1 | 0 | 3 | 0.3333 | 7 | 0.1429 | 1 | 0 | NA | NA | 1 | 0 | 2 | 0.5 | 4 | 0.25 |
| 7_10508449 | 4 | 0.25 | 4 | 0.5 | 1 | 1 | NA | NA | 9 | 0.4444 | 2 | 0.5 | 1 | 1 | NA | NA | NA | NA | 3 | 0.6667 |
| 7_10508462 | 4 | 0.25 | 4 | 1 | 1 | 0 | NA | NA | 9 | 0.5556 | 2 | 0.5 | 1 | 1 | NA | NA | NA | NA | 3 | 0.6667 |
| 7_10508515 | 5 | 0 | 4 | 0.25 | 1 | 0 | NA | NA | 10 | 0.1 | 3 | 0 | 2 | 0.5 | NA | NA | NA | NA | 5 | 0.2 |
| 7_10508521 | 5 | 0.2 | 4 | 0.25 | 1 | 0 | NA | NA | 10 | 0.2 | 2 | 0.5 | 1 | 0 | NA | NA | NA | NA | 3 | 0.3333 |
| 7_10508523 | 4 | 0.5 | 4 | 0.25 | NA | NA | NA | NA | 8 | 0.375 | 3 | 0.3333 | 1 | 0 | NA | NA | NA | NA | 4 | 0.25 |
| 7_10512988 | 4 | 0 | 6 | 0.1667 | 8 | 0 | 2 | 0 | 20 | 0.05 | 5 | 0.2 | 5 | 0 | 4 | 0.25 | 3 | 0.3333 | 17 | 0.1765 |
| 7_10513009 | 4 | 0 | 6 | 0 | 8 | 0.125 | 3 | 0.3333 | 21 | 0.0952 | 5 | 0.2 | 5 | 0 | 4 | 0 | 3 | 0 | 17 | 0.0588 |
| 7_10513010 | 3 | 0 | 5 | 0 | 4 | 0.25 | 2 | 0 | 14 | 0.0714 | 3 | 0 | 4 | 0.5 | 3 | 0 | 1 | 0 | 11 | 0.1818 |
| 7_10520379 | 9 | 0.1111 | 10 | 0.2 | 6 | 0.1667 | 6 | 0.1667 | 31 | 0.1613 | 4 | 0 | 6 | 0.1667 | 15 | 0.3333 | 8 | 0.125 | 33 | 0.2121 |
| 7_10520383 | 8 | 0.25 | 8 | 0.125 | 7 | 0.5714 | 7 | 0.4286 | 30 | 0.3333 | 3 | 0.3333 | 8 | 0.25 | 11 | 0.2727 | 6 | 0.3333 | 28 | 0.2857 |
| 7_10520455 | 12 | 0.5833 | 12 | 0.4167 | 9 | 0.6667 | 10 | 0.6 | 43 | 0.5581 | 3 | 0.3333 | 8 | 0.75 | 14 | 0.6429 | 10 | 0.4 | 35 | 0.5714 |
| 7_10520461 | 15 | 0.9333 | 13 | 1 | 9 | 1 | 11 | 0.8182 | 48 | 0.9375 | 4 | 1 | 10 | 0.8 | 18 | 0.9444 | 9 | 0.8889 | 41 | 0.9024 |
| 7_10520462 | 13 | 0.6923 | 11 | 0.5455 | 8 | 0.5 | 7 | 0.4286 | 39 | 0.5641 | 3 | 0.6667 | 8 | 0.5 | 18 | 0.6111 | 10 | 0.3 | 39 | 0.5128 |
| 7_10520474 | 13 | 0.8462 | 10 | 1 | 8 | 0.875 | 8 | 0.875 | 39 | 0.8974 | 5 | 0.6 | 9 | 0.6667 | 17 | 0.7059 | 12 | 0.9167 | 43 | 0.7442 |
| 7_10520490 | 6 | 0.8333 | 2 | 1 | 2 | 0.5 | 2 | 0 | 12 | 0.6667 | 2 | 1 | 1 | 0 | 8 | 0.875 | 4 | 1 | 15 | 0.8667 |
| 7_10520501 | 16 | 0 | 16 | 0.0625 | 9 | 0.1111 | 14 | 0.0714 | 55 | 0.0545 | 6 | 0 | 12 | 0.1667 | 20 | 0.05 | 13 | 0 | 51 | 0.0588 |
| 7_10520698 | 18 | 0.5 | 18 | 0.6667 | 13 | 0.9231 | 18 | 0.5556 | 67 | 0.6418 | 12 | 0.3333 | 13 | 0.7692 | 19 | 0.5789 | 17 | 0.4118 | 61 | 0.5246 |
| 7_10520725 | 17 | 0.1176 | 25 | 0.44 | 10 | 0.4 | 18 | 0.2778 | 70 | 0.3143 | 12 | 0.4167 | 11 | 0.0909 | 22 | 0.2273 | 16 | 0.1875 | 61 | 0.2295 |
| 7_10520752 | 22 | 0.6818 | 26 | 0.6923 | 13 | 0.7692 | 21 | 0.619 | 82 | 0.6829 | 14 | 0.6429 | 17 | 0.8235 | 25 | 0.68 | 17 | 0.7059 | 73 | 0.7123 |
| 7_10520758 | 16 | 0.0625 | 23 | 0.0435 | 12 | 0.0833 | 21 | 0.0952 | 72 | 0.0694 | 13 | 0.0769 | 15 | 0.0667 | 24 | 0.0833 | 16 | 0.0625 | 68 | 0.0735 |
| 7_10520789 | 43 | 0.5116 | 52 | 0.4231 | 36 | 0.3611 | 41 | 0.3659 | 172 | 0.4186 | 45 | 0.4222 | 38 | 0.5 | 37 | 0.5405 | 27 | 0.7407 | 147 | 0.5306 |
| 7_10522289 | 22 | 0.2727 | 27 | 0.0741 | 24 | 0.0833 | 27 | 0.1481 | 100 | 0.14 | 30 | 0.1 | 21 | 0.2381 | 23 | 0.2174 | 14 | 0.2143 | 88 | 0.1818 |
| 7_10522294 | 22 | 0.7727 | 23 | 0.7391 | 21 | 0.619 | 22 | 0.7727 | 88 | 0.7273 | 24 | 0.5 | 18 | 0.8889 | 23 | 0.7391 | 11 | 0.6364 | 76 | 0.6842 |
| 7_10522309 | 20 | 0.7 | 22 | 0.6364 | 21 | 0.6667 | 17 | 0.3529 | 80 | 0.6 | 21 | 0.6667 | 13 | 0.6154 | 20 | 0.8 | 10 | 0.9 | 64 | 0.7344 |
| 7_10522309 | 1 | 1 | 2 | 0.5 | 1 | 1 | 1 | 0 | 5 | 0.6 | 2 | 0.5 | 1 | 0 | NA | NA | 1 | 0 | 4 | 0.25 |

| | | | | | | | | | | | | | | | | | | | | |
|---|---|---|---|---|---|---|---|---|---|---|---|---|---|---|---|---|---|---|---|---|
| 7_10522317 | 21 | 0.1905 | 24 | 0.0833 | 23 | 0.3043 | 21 | 0.2381 | 89 | 0.2022 | 20 | 0 | 14 | 0.2143 | 23 | 0.087 | 11 | 0.0909 | 68 | 0.0882 |
| 7_10522333 | 16 | 0.25 | 20 | 0 | 19 | 0.1579 | 18 | 0.1111 | 73 | 0.1233 | 17 | 0.1176 | 11 | 0.0909 | 20 | 0.2 | 9 | 0.2222 | 57 | 0.1579 |
| 7_10522334 | 16 | 0.125 | 18 | 0.1111 | 20 | 0.15 | 18 | 0 | 72 | 0.0972 | 18 | 0.1111 | 13 | 0.2308 | 20 | 0 | 11 | 0.2727 | 62 | 0.129 |
| 7_10522338 | 16 | 0 | 17 | 0.1176 | 14 | 0 | 13 | 0.0769 | 60 | 0.05 | 13 | 0.2308 | 13 | 0 | 16 | 0.125 | 8 | 0.125 | 50 | 0.12 |
| 7_10522346 | 16 | 0.625 | 18 | 0.7778 | 23 | 0.6522 | 17 | 0.6471 | 74 | 0.6757 | 19 | 0.6842 | 14 | 0.7143 | 21 | 0.6667 | 8 | 0.875 | 62 | 0.7097 |
| 7_10526006 | 5 | 0.6 | 13 | 0.3077 | 7 | 0 | 9 | 0.3333 | 34 | 0.2941 | 6 | 0.1667 | 8 | 0.25 | 10 | 0.3 | 9 | 0.1111 | 33 | 0.2121 |
| 7_10526007 | 7 | 0 | 14 | 0.1429 | 15 | 0.1333 | 13 | 0 | 49 | 0.0816 | 9 | 0 | 10 | 0 | 10 | 0.1 | 9 | 0.1111 | 38 | 0.0526 |
| 7_10526021 | 6 | 0.5 | 13 | 0.3846 | 15 | 0.5333 | 13 | 0.4615 | 47 | 0.4681 | 9 | 0.3333 | 13 | 0.1538 | 6 | 0.3333 | 12 | 0.5 | 40 | 0.325 |
| 7_10526022 | 7 | 1 | 15 | 0.8 | 15 | 0.8 | 11 | 0.9091 | 48 | 0.8542 | 11 | 0.9091 | 13 | 0.8462 | 9 | 0.8889 | 11 | 0.7273 | 44 | 0.8409 |
| 7_10526716 | 3 | 0.6667 | 11 | 0.4545 | 5 | 0.4 | 5 | 0.8 | 24 | 0.5417 | 7 | 0.5714 | 5 | 0.6 | 6 | 0.3333 | 6 | 0.8333 | 24 | 0.5833 |
| 7_10526723 | 3 | 0.3333 | 11 | 0.0909 | 5 | 0.6 | 7 | 0.2857 | 26 | 0.2692 | 10 | 0.5 | 3 | 0 | 6 | 0 | 3 | 0.3333 | 22 | 0.2727 |
| 7_10526733 | 4 | 0.5 | 11 | 0.1818 | 6 | 0.1667 | 5 | 0.2 | 26 | 0.2308 | 6 | 0 | 4 | 0 | 5 | 0 | 5 | 0.6 | 20 | 0.15 |
| 7_10526740 | 2 | 0 | 8 | 0.125 | 6 | 0.3333 | 4 | 0.25 | 20 | 0.2 | 5 | 0 | 5 | 0.2 | 6 | 0.1667 | 3 | 0.3333 | 19 | 0.1579 |
| 7_10526748 | 4 | 0.5 | 11 | 0.3636 | 5 | 0.8 | 3 | 0.3333 | 23 | 0.4783 | 7 | 0.5714 | 6 | 1 | 7 | 0.2857 | 4 | 0.5 | 24 | 0.5833 |
| 7_10526759 | 4 | 0.5 | 8 | 0.375 | 4 | 0.25 | 4 | 0 | 20 | 0.3 | 8 | 0.125 | 6 | 0.5 | 8 | 0 | 2 | 0 | 24 | 0.1667 |
| 7_10528413 | 4 | 0.25 | 3 | 0.3333 | 1 | 1 | 4 | 0.25 | 12 | 0.3333 | 6 | 0.1667 | 1 | 0 | 3 | 0 | 1 | 0 | 11 | 0.0909 |
| 7_10528497 | 5 | 0.2 | 3 | 0 | NA | NA | 4 | 0 | 12 | 0.0833 | 3 | 0 | 2 | 0.5 | 3 | 0.3333 | 3 | 0.3333 | 11 | 0.2727 |
| 7_10528497 | 9 | 0 | 8 | 0 | 9 | 0.1111 | 13 | 0.1538 | 39 | 0.0769 | 11 | 0.0909 | 10 | 0.1 | 11 | 0 | 7 | 0 | 39 | 0.0513 |
| 7_10528532 | 7 | 0 | 7 | 0.1429 | 6 | 0 | 13 | 0.2308 | 33 | 0.1212 | 13 | 0.2308 | 12 | 0.1667 | 9 | 0.2222 | 8 | 0 | 42 | 0.1667 |
| 7_10528537 | 7 | 0.1429 | 9 | 0.3333 | 8 | 0.125 | 11 | 0.5455 | 35 | 0.3143 | 11 | 0.3636 | 10 | 0.2 | 11 | 0.4545 | 8 | 0.5 | 40 | 0.375 |
| 7_10528541 | 9 | 0.8889 | 8 | 0.75 | 8 | 0.875 | 13 | 0.9231 | 38 | 0.8684 | 7 | 0.8571 | 10 | 0.9 | 8 | 0.75 | 6 | 0.6667 | 31 | 0.8065 |
| 7_10528567 | 1 | 1 | 1 | 1 | 1 | 1 | NA | NA | 3 | 1 | NA | NA | 1 | 1 | 2 | 0.5 | NA | NA | 3 | 0.6667 |
| 7_10528612 | 9 | 0 | 9 | 0.1111 | 8 | 0 | 11 | 0.0909 | 37 | 0.0541 | 13 | 0.1538 | 13 | 0.1538 | 9 | 0.1111 | 6 | 0 | 41 | 0.122 |
| 7_10529039 | 10 | 0.6 | 10 | 0.2 | 5 | 0 | 9 | 0.1111 | 34 | 0.2647 | 9 | 0.6667 | 4 | 0.25 | 9 | 0.3333 | 7 | 0.4286 | 29 | 0.4483 |
| 7_10529046 | 9 | 0.7778 | 8 | 0.75 | 4 | 0.75 | 4 | 1 | 25 | 0.8 | 6 | 0.6667 | 4 | 0.75 | 9 | 0.8889 | 4 | 0.75 | 23 | 0.7826 |
| 7_10529104 | NA | NA | 4 | 0 | 2 | 0.5 | 3 | 0 | 9 | 0.1111 | 1 | 0 | NA | NA | 5 | 0.2 | 2 | 0 | 8 | 0.125 |
| 7_10530820 | NA | NA | NA | NA | NA | NA | 3 | 0.6667 | 3 | 0.6667 | 1 | 0 | NA | NA | NA | NA | 1 | 1 | 2 | 0.5 |
| 7_10538182 | 2 | 0.5 | 7 | 0.1429 | 2 | 0 | 2 | 0 | 13 | 0.1538 | 2 | 0.5 | 4 | 0 | 2 | 0 | 3 | 0 | 11 | 0.0909 |
| 7_10538194 | 2 | 0.5 | 8 | 0.875 | 1 | 1 | 3 | 1 | 14 | 0.8571 | 2 | 1 | 5 | 0.6 | 2 | 1 | 4 | 0.75 | 13 | 0.7692 |
| 7_10538211 | NA | NA | 8 | 0.625 | 2 | 1 | 1 | 1 | 11 | 0.7273 | NA | NA | 4 | 0.5 | 1 | 0 | 2 | 0.5 | 7 | 0.4286 |
| 7_10538217 | 1 | 1 | 8 | 0.875 | 2 | 1 | 2 | 1 | 13 | 0.9231 | 1 | 0 | 5 | 0.8 | 2 | 1 | 4 | 1 | 12 | 0.8333 |
| 7_10538218 | 1 | 1 | 8 | 0.75 | 1 | 0 | 2 | 1 | 12 | 0.75 | 1 | 1 | 3 | 1 | 2 | 0 | 3 | 0 | 9 | 0.4444 |
| 7_10538219 | 1 | 0 | 7 | 0.5714 | 2 | 0.5 | 2 | 0 | 12 | 0.4167 | 1 | 1 | 5 | 0 | 2 | 0 | 4 | 0 | 12 | 0.0833 |
| 7_10538222 | 1 | 0 | 7 | 0.2857 | 2 | 0 | 2 | 1 | 12 | 0.3333 | 1 | 0 | 4 | 0.75 | 2 | 0 | 4 | 0.5 | 11 | 0.4545 |
| 7_10538228 | 1 | 1 | 8 | 0 | 2 | 0 | 2 | 0 | 13 | 0.0769 | 1 | 0 | 5 | 0.4 | 2 | 0 | 4 | 0 | 12 | 0.1667 |
| 7_10538335 | 1 | 1 | 4 | 0.75 | 1 | 1 | 1 | 1 | 7 | 0.8571 | NA | NA | 1 | 1 | 1 | 1 | 2 | 1 | 4 | 1 |
| 7_10538404 | 1 | 0 | 7 | 0.1429 | 2 | 0 | 2 | 0 | 12 | 0.0833 | 1 | 0 | 1 | 1 | 1 | 0 | 2 | 0.5 | 5 | 0.4 |
| 7_10539194 | 24 | 0.0417 | 19 | 0.1579 | 14 | 0.0714 | 17 | 0.1176 | 74 | 0.0946 | 8 | 0.375 | 19 | 0.1053 | 12 | 0.0833 | 18 | 0.1111 | 57 | 0.1404 |
| 7_10539199 | 21 | 0.1429 | 19 | 0.1579 | 13 | 0.1538 | 16 | 0.1875 | 69 | 0.1594 | 7 | 0.1429 | 19 | 0.2105 | 13 | 0.0769 | 18 | 0.1667 | 57 | 0.1579 |
| 7_10539211 | 13 | 0.1538 | 13 | 0 | 10 | 0 | 11 | 0.0909 | 47 | 0.0638 | 14 | 0 | 16 | 0.125 | 11 | 0.0909 | 7 | 0.1429 | 48 | 0.0833 |
| 7_10539307 | 15 | 0.2 | 11 | 0 | 12 | 0.3333 | 12 | 0.1667 | 50 | 0.18 | 8 | 0 | 16 | 0.125 | 13 | 0.1538 | 6 | 0.1667 | 43 | 0.1163 |
| 7_10539428 | 5 | 0 | 4 | 0.25 | 3 | 0 | NA | NA | 12 | 0.0833 | 2 | 0.5 | 1 | 0 | 3 | 0 | 2 | 0.5 | 8 | 0.25 |
| 7_10539542 | 15 | 0.0667 | 13 | 0 | 8 | 0 | 4 | 0.25 | 40 | 0.05 | 8 | 0.125 | 8 | 0 | 14 | 0.0714 | 6 | 0 | 36 | 0.0556 |
| 7_10539551 | 9 | 0.2222 | 9 | 0 | 6 | 0 | 3 | 0 | 27 | 0.0741 | 4 | 0 | 6 | 0 | 7 | 0.4286 | 5 | 0 | 22 | 0.1364 |
| 7_10539565 | 14 | 0 | 12 | 0.1667 | 8 | 0.125 | 4 | 0.25 | 38 | 0.1053 | 8 | 0.125 | 8 | 0 | 13 | 0.0769 | 4 | 0 | 33 | 0.0606 |
| 7_10539569 | 13 | 0.4615 | 12 | 0.4167 | 8 | 0.375 | 4 | 1 | 37 | 0.4865 | 8 | 0.5 | 8 | 0.5 | 13 | 0.6154 | 4 | 0.5 | 33 | 0.5455 |
| 7_10539574 | 10 | 0 | 11 | 0.0909 | 7 | 0.2857 | 3 | 0.6667 | 31 | 0.1613 | 6 | 0.3333 | 8 | 0.125 | 12 | 0.1667 | 3 | 0 | 29 | 0.1724 |
| 7_10539581 | 12 | 0.6667 | 11 | 0.5455 | 7 | 0.5714 | 4 | 1 | 34 | 0.6471 | 8 | 0.625 | 7 | 1 | 10 | 0.8 | 2 | 1 | 27 | 0.8148 |
| 7_10539615 | 11 | 0 | 9 | 0.2222 | 7 | 0.1429 | 4 | 0 | 31 | 0.0968 | 5 | 0 | 7 | 0 | 13 | 0.1538 | 3 | 0 | 28 | 0.0714 |
| 7_10539941 | 6 | 0.1667 | 11 | 0.0909 | 7 | 0.1429 | 4 | 0.5 | 28 | 0.1786 | 5 | 0 | 5 | 0 | 7 | 0.1429 | 2 | 0 | 19 | 0.0526 |
| 7_10540221 | NA | NA | NA | NA | 1 | 0 | 2 | 0.5 | 3 | 0.3333 | 1 | 0 | 2 | 0.5 | 4 | 0 | NA | NA | 7 | 0.1429 |
| 7_10540234 | NA | NA | NA | NA | 1 | 0 | 2 | 1 | 3 | 0.6667 | 1 | 1 | 2 | 0.5 | 4 | 1 | NA | NA | 7 | 0.8571 |
| 7_10540247 | NA | NA | NA | NA | 1 | 1 | 2 | 0.5 | 3 | 0.6667 | 1 | 0 | 2 | 1 | 4 | 0 | NA | NA | 7 | 0.2857 |
| 7_10540251 | NA | NA | NA | NA | 1 | 0 | 2 | 1 | 3 | 0.6667 | NA | NA | 1 | 0 | 3 | 0.6667 | NA | NA | 4 | 0.5 |
| 7_10540252 | NA | NA | NA | NA | 1 | 0 | 2 | 0.5 | 3 | 0.3333 | 1 | 0 | 1 | 1 | 4 | 0 | NA | NA | 6 | 0.1667 |
| 7_10540256 | NA | NA | NA | NA | 1 | 1 | NA | NA | 1 | 1 | 1 | 0 | NA | NA | 2 | 1 | NA | NA | 3 | 0.6667 |
| 7_10540257 | NA | NA | NA | NA | 1 | 1 | 1 | 1 | 2 | 1 | 1 | 1 | 1 | 0 | 4 | 0.75 | NA | NA | 6 | 0.6667 |
| 7_10540263 | NA | NA | NA | NA | NA | NA | 2 | 0.5 | 2 | 0.5 | NA | NA | 2 | 1 | 3 | 0 | NA | NA | 5 | 0.4 |
| 7_10540307 | NA | NA | NA | NA | 1 | 1 | 1 | 1 | 2 | 1 | 1 | 1 | 1 | 1 | 1 | 1 | NA | NA | 3 | 1 |
| 7_10540316 | NA | NA | NA | NA | 1 | 1 | 1 | 1 | 2 | 1 | 1 | 0 | NA | NA | 3 | 1 | NA | NA | 4 | 0.75 |
| 7_10540336 | NA | NA | NA | NA | 1 | 0 | 2 | 1 | 3 | 0.6667 | 1 | 0 | 1 | 1 | NA | NA | NA | NA | 2 | 0.5 |
| 7_10540354 | NA | NA | NA | NA | 1 | 1 | 1 | 1 | 2 | 1 | NA | NA | 1 | 1 | 1 | 1 | NA | NA | 2 | 1 |
| 7_10540380 | NA | NA | NA | NA | 1 | 1 | 1 | 1 | 2 | 1 | NA | NA | 1 | 1 | 2 | 1 | NA | NA | 3 | 1 |
| 7_10540394 | NA | NA | NA | NA | 1 | 1 | 1 | 1 | 2 | 1 | 1 | 1 | 1 | 1 | 2 | 1 | NA | NA | 4 | 1 |
| 7_10540434 | NA | NA | NA | NA | 1 | 1 | 1 | 1 | 2 | 1 | 1 | 1 | 1 | 0 | 2 | 0.5 | NA | NA | 4 | 0.5 |

| | | | | | | | | | | | | | | | | | | | | |
|---|---|---|---|---|---|---|---|---|---|---|---|---|---|---|---|---|---|---|---|---|
| 7_1054043 | NA | NA | NA | NA | 1 | 1 | 1 | 1 | 2 | 1 | 1 | 1 | 1 | 1 | 1 | 1 | NA | NA | 3 | 1 |
| 7_1054044 | NA | NA | NA | NA | 1 | 1 | 1 | 0 | 2 | 0.5 | NA | NA | 1 | 1 | 2 | 0.5 | NA | NA | 3 | 0.6667 |
| 7_1054045 | NA | NA | NA | NA | NA | NA | 1 | 1 | 1 | 1 | 1 | 1 | 1 | 1 | 2 | 1 | NA | NA | 4 | 1 |
| 7_1054045 | NA | NA | NA | NA | 1 | 1 | 1 | 1 | 2 | 1 | 1 | 1 | 1 | 1 | 2 | 1 | NA | NA | 4 | 1 |
| 7_1054047 | NA | NA | NA | NA | 1 | 1 | 1 | 0 | 2 | 0.5 | 1 | 1 | 1 | 1 | 1 | 0 | NA | NA | 3 | 0.6667 |
| 7_1054047 | NA | NA | NA | NA | 1 | 1 | NA | NA | 1 | 1 | NA | NA | 1 | 1 | 2 | 1 | NA | NA | 3 | 1 |
| 7_1054048 | NA | NA | NA | NA | 1 | 1 | 1 | 1 | 2 | 1 | 1 | 0 | 1 | 1 | 2 | 1 | NA | NA | 4 | 0.75 |
| 7_1054048 | NA | NA | NA | NA | 1 | 1 | 1 | 1 | 2 | 1 | 1 | 1 | 1 | 1 | 2 | 1 | NA | NA | 4 | 1 |
| 7_1054050 | NA | NA | NA | NA | 1 | 1 | 1 | 1 | 2 | 1 | 1 | 1 | 1 | 1 | 2 | 0.5 | NA | NA | 4 | 0.75 |
| 7_1054587 | 2 | 0.5 | 3 | 0 | NA | NA | 2 | 0 | 7 | 0.1429 | 2 | 0 | 1 | 0 | 1 | 0 | 2 | 0.5 | 6 | 0.1667 |
| 7_1054588 | 3 | 0.3333 | 4 | 0.25 | 3 | 0 | 1 | 0 | 11 | 0.1818 | 3 | 0.3333 | 2 | 0 | 2 | 0 | 5 | 0.2 | 12 | 0.1667 |
| 7_1054588 | 2 | 0.5 | 2 | 0.5 | 2 | 1 | 1 | 0 | 7 | 0.5714 | 3 | 0.6667 | 1 | 1 | 1 | 0 | 5 | 0.4 | 10 | 0.5 |
| 7_1054589 | 6 | 0.1667 | 7 | 0 | 10 | 0.1 | 3 | 0.3333 | 26 | 0.1154 | 10 | 0.1 | 7 | 0 | 9 | 0 | 11 | 0.1818 | 37 | 0.0811 |
| 7_1054590 | 5 | 0.4 | 6 | 0.8333 | 7 | 1 | 6 | 0.8333 | 24 | 0.7917 | 9 | 0.7778 | 3 | 1 | 5 | 1 | 10 | 0.8 | 27 | 0.8519 |
| 7_1054591 | 8 | 0 | 9 | 0 | 9 | 0.1111 | 6 | 0.1667 | 32 | 0.0625 | 10 | 0.1 | 6 | 0 | 8 | 0.125 | 10 | 0.2 | 34 | 0.1176 |
| 7_1054591 | 6 | 0.8333 | 9 | 0.3333 | 9 | 0.4444 | 4 | 0.25 | 28 | 0.4643 | 9 | 0.3333 | 6 | 0.3333 | 7 | 0.2857 | 9 | 0.5556 | 31 | 0.3871 |
| 7_1054591 | 8 | 0.125 | 9 | 0.1111 | 8 | 0 | 6 | 0.1667 | 31 | 0.0968 | 9 | 0.2222 | 6 | 0 | 7 | 0.1429 | 10 | 0.1 | 32 | 0.125 |
| 7_1054592 | 2 | 0 | 6 | 0 | 6 | 0 | 4 | 0.25 | 18 | 0.0556 | 9 | 0.3333 | 5 | 0 | 5 | 0 | 9 | 0.1111 | 28 | 0.1429 |
| 7_1071895 | 2 | 0 | NA | NA | 4 | 0.25 | NA | NA | 6 | 0.1667 | 2 | 1 | 2 | 0.5 | 2 | 1 | 1 | 1 | 7 | 0.8571 |
| 7_1071896 | 2 | 1 | NA | NA | 4 | 1 | NA | NA | 6 | 1 | 1 | 1 | 1 | 1 | 2 | 1 | 1 | 1 | 5 | 1 |
| 7_1071896 | 2 | 0.5 | NA | NA | 5 | 0.6 | NA | NA | 7 | 0.5714 | 2 | 0 | 2 | 1 | 2 | 0.5 | NA | NA | 6 | 0.5 |
| 7_1071897 | 2 | 1 | NA | NA | 5 | 1 | NA | NA | 7 | 1 | 1 | 1 | 2 | 1 | 2 | 1 | 1 | 1 | 6 | 1 |
| 7_1071897 | 2 | 1 | NA | NA | 4 | 0.75 | NA | NA | 6 | 0.8333 | 2 | 1 | 2 | 1 | 2 | 0.5 | 1 | 0 | 7 | 0.7143 |
| 7_1071898 | 1 | 0 | NA | NA | 3 | 0.3333 | NA | NA | 4 | 0.25 | 2 | 0.5 | 1 | 0 | 2 | 0.5 | 1 | 0 | 6 | 0.3333 |
| 7_1071898 | 1 | 1 | NA | NA | 4 | 1 | NA | NA | 5 | 1 | 1 | 0 | 2 | 1 | 1 | 1 | 1 | 1 | 5 | 0.8 |
| 7_1071901 | 2 | 1 | 1 | 0 | 4 | 0.5 | 1 | 1 | 8 | 0.625 | 1 | 1 | 2 | 1 | 2 | 0.5 | NA | NA | 5 | 0.8 |
| 7_1071901 | 2 | 0.5 | 1 | 0 | 5 | 0.6 | 1 | 0 | 9 | 0.4444 | 2 | 0.5 | 2 | 0 | 2 | 1 | NA | NA | 6 | 0.5 |
| 7_1071901 | 2 | 1 | NA | NA | 5 | 0.8 | 1 | 1 | 8 | 0.875 | 2 | 1 | 2 | 1 | 2 | 1 | NA | NA | 6 | 1 |
| 7_1071905 | 1 | 0 | 1 | 1 | 3 | 0 | 1 | 0 | 6 | 0.1667 | 2 | 0.5 | 1 | 0 | 1 | 0 | 1 | 0 | 5 | 0.2 |
| 7_1071905 | 1 | 1 | 1 | 0 | 2 | 0.5 | 1 | 1 | 5 | 0.6 | 2 | 0.5 | 1 | 0 | 1 | 1 | NA | NA | 4 | 0.5 |
| 7_1085241 | NA | NA | 1 | 1 | NA | NA | NA | NA | 1 | 1 | NA | NA | 1 | 1 | NA | NA | NA | NA | 1 | 1 |
| 7_1085243 | NA | NA | 1 | 1 | NA | NA | NA | NA | 1 | 1 | NA | NA | 1 | 1 | NA | NA | NA | NA | 1 | 1 |
| 7_1085249 | NA | NA | 1 | 1 | NA | NA | NA | NA | 1 | 1 | NA | NA | 1 | 1 | NA | NA | NA | NA | 1 | 1 |
| 7_1085250 | NA | NA | 1 | 1 | NA | NA | NA | NA | 1 | 1 | NA | NA | 1 | 1 | NA | NA | NA | NA | 1 | 1 |
| 7_1085408 | 2 | 1 | 2 | 0.5 | 2 | 0.5 | NA | NA | 6 | 0.6667 | 1 | 1 | 2 | 1 | 2 | 1 | 1 | 0 | 6 | 0.8333 |
| 7_1086098 | NA | NA | 1 | 1 | NA | NA | NA | NA | 1 | 1 | NA | NA | 1 | 1 | NA | NA | NA | NA | 1 | 1 |
| 7_1086098 | NA | NA | 1 | 1 | NA | NA | NA | NA | 1 | 1 | NA | NA | 1 | 0 | NA | NA | 1 | 1 | 2 | 0.5 |
| 7_1086293 | 3 | 0.3333 | 1 | 1 | NA | NA | 1 | 0 | 5 | 0.4 | 2 | 0 | 3 | 0.3333 | NA | NA | 2 | 1 | 7 | 0.4286 |
| 7_1086295 | 3 | 0.3333 | 3 | 0.3333 | NA | NA | 1 | 0 | 7 | 0.2857 | 3 | 0.3333 | 3 | 0 | 1 | 0 | 2 | 0.5 | 9 | 0.2222 |
| 7_1086295 | 4 | 1 | 3 | 0.6667 | NA | NA | 1 | 1 | 8 | 0.875 | 3 | 0.6667 | 2 | 1 | NA | NA | 2 | 1 | 7 | 0.8571 |
| 7_1086296 | 3 | 0.3333 | 3 | 0.3333 | NA | NA | 1 | 0 | 7 | 0.2857 | 2 | 0 | 3 | 0.3333 | 1 | 0 | 1 | 1 | 7 | 0.2857 |
| 7_1086298 | 4 | 0.75 | 3 | 0.3333 | NA | NA | 1 | 1 | 8 | 0.625 | 3 | 0.3333 | 2 | 1 | 1 | 1 | 2 | 1 | 8 | 0.75 |
| 7_1086307 | 4 | 0.5 | 3 | 0.6667 | NA | NA | 3 | 0 | 10 | 0.4 | 2 | 0 | 6 | 0.1667 | 2 | 0 | 2 | 0 | 12 | 0.0833 |
| 7_1086307 | 4 | 0.25 | 3 | 0 | NA | NA | 3 | 0 | 10 | 0.1 | 2 | 0.5 | 6 | 0 | 2 | 0 | 2 | 0.5 | 12 | 0.1667 |
| 7_1086314 | 4 | 0.5 | 4 | 0.25 | 1 | 1 | 5 | 0.6 | 14 | 0.5 | 6 | 0.8333 | 6 | 0.3333 | 3 | 0 | 3 | 0.6667 | 18 | 0.5 |
| 7_1086315 | 6 | 0.1667 | 5 | 0 | NA | NA | 5 | 0 | 16 | 0.0625 | 6 | 0 | 5 | 0.2 | 3 | 0 | 3 | 0.6667 | 17 | 0.1765 |
| 7_1086317 | 5 | 0.2 | 7 | 0.2857 | 3 | 0.3333 | 6 | 0 | 21 | 0.1905 | 6 | 0.3333 | 6 | 0.5 | 3 | 0 | 3 | 0.3333 | 18 | 0.3333 |
| 7_1086321 | 5 | 0.6 | 7 | 0.5714 | 4 | 0.75 | 6 | 0.8333 | 22 | 0.6818 | 5 | 0.8 | 8 | 0.75 | 3 | 0.3333 | 3 | 0.6667 | 19 | 0.6842 |
| 7_1086322 | 6 | 0.1667 | 5 | 0.4 | 5 | 0 | 6 | 0.5 | 22 | 0.2727 | 6 | 0.5 | 5 | 0.8 | 3 | 0.6667 | 4 | 0.5 | 18 | 0.6111 |
| 7_1086326 | 7 | 0.1429 | 7 | 0.1429 | 6 | 0 | 8 | 0 | 28 | 0.0714 | 6 | 0.1667 | 7 | 0.1429 | 4 | 0 | 3 | 0.3333 | 20 | 0.15 |
| 7_1086331 | 7 | 0 | 9 | 0.1111 | 8 | 0.25 | 11 | 0 | 35 | 0.0857 | 7 | 0.1429 | 8 | 0 | 3 | 0.3333 | 3 | 0 | 21 | 0.0952 |
| 7_1086336 | 5 | 0.4 | 2 | 0 | 3 | 0.3333 | 3 | 0.3333 | 13 | 0.3077 | NA | NA | 2 | 0 | 2 | 1 | 1 | 0 | 5 | 0.4 |
| 7_1086337 | 3 | 0.6667 | 2 | 0.5 | 2 | 1 | 2 | 1 | 9 | 0.7778 | NA | NA | 1 | 1 | 2 | 0.5 | 1 | 1 | 4 | 0.75 |
| 7_1086342 | 2 | 0.5 | 2 | 0.5 | 2 | 0 | 1 | 0 | 7 | 0.2857 | NA | NA | 2 | 0.5 | 1 | 0 | 1 | 0 | 4 | 0.25 |
| 7_1086342 | 3 | 0.3333 | 2 | 1 | 3 | 0.3333 | 2 | 0 | 10 | 0.4 | NA | NA | 3 | 0.6667 | 1 | 0 | 1 | 0 | 5 | 0.4 |
| 7_1086351 | 4 | 0.25 | 5 | 0.2 | 3 | 0 | 5 | 0.2 | 17 | 0.1765 | NA | NA | 4 | 0.5 | 2 | 1 | 1 | 1 | 7 | 0.7143 |
| 7_1086357 | 1 | 1 | 4 | 0 | 2 | 0 | 5 | 0.2 | 12 | 0.1667 | NA | NA | 3 | 0.3333 | 1 | 0 | NA | NA | 4 | 0.25 |
| 7_1086364 | 5 | 0.6 | 2 | 0 | 2 | 0.5 | 7 | 0.1429 | 16 | 0.3125 | NA | NA | 4 | 0.5 | 2 | 0 | 1 | 1 | 7 | 0.4286 |
| 7_1086368 | 6 | 0.5 | 2 | 0 | 3 | 0.6667 | 6 | 0.1667 | 17 | 0.3529 | NA | NA | 4 | 0.75 | 2 | 0.5 | 1 | 1 | 7 | 0.7143 |
| 7_1086369 | 6 | 0.3333 | 2 | 0.5 | 1 | 0 | 4 | 0.25 | 13 | 0.3077 | NA | NA | 4 | 1 | 1 | 0 | NA | NA | 5 | 0.8 |
| 7_1086369 | 6 | 0.5 | 2 | 1 | 1 | 1 | 4 | 0.25 | 13 | 0.5385 | NA | NA | 4 | 0 | 2 | 1 | 1 | 1 | 7 | 0.4286 |
| 7_1086371 | 6 | 0.3333 | 1 | 0 | 2 | 0.5 | 6 | 0.1667 | 15 | 0.2667 | NA | NA | 4 | 0.5 | 2 | 0.5 | 1 | 0 | 7 | 0.4286 |
| 7_1086372 | 6 | 0.1667 | 2 | 0 | 2 | 0 | 5 | 0.2 | 15 | 0.1333 | NA | NA | 4 | 0.5 | 2 | 0.5 | NA | NA | 6 | 0.5 |
| 7_1086373 | 6 | 0.8333 | 2 | 0 | 2 | 0.5 | 7 | 0.8571 | 17 | 0.7059 | 1 | 0 | 4 | 1 | 2 | 1 | 1 | 0 | 8 | 0.75 |
| 7_1086374 | 4 | 0.5 | 2 | 0 | 1 | 0 | 6 | 0.6667 | 13 | 0.4615 | NA | NA | 4 | 0 | 1 | 1 | 1 | 1 | 6 | 0.3333 |

| | | | | | | | | | | | | | | | | | | | | |
|---|---|---|---|---|---|---|---|---|---|---|---|---|---|---|---|---|---|---|---|---|
| 7_1086377 | 3 | 1 | 2 | 1 | NA | NA | 4 | 1 | 9 | 1 | NA | NA | 4 | 1 | 1 | 1 | 1 | 1 | 6 | 1 |
| 7_1086379 | 3 | 0.6667 | 2 | 0 | 1 | 1 | 4 | 0 | 10 | 0.3 | 1 | 1 | 4 | 0.5 | 1 | 1 | 1 | 0 | 7 | 0.5714 |
| 7_1086380 | 3 | 0 | 2 | 0 | 1 | 0 | 5 | 0.2 | 11 | 0.0909 | NA | NA | 4 | 0.25 | 1 | 0 | 1 | 0 | 6 | 0.1667 |
| 7_1086468 | 1 | 0 | 2 | 0.5 | NA | NA | NA | NA | 3 | 0.3333 | 2 | 1 | 1 | 1 | NA | NA | NA | NA | 3 | 1 |
| 7_1086471 | 4 | 0 | 4 | 0.25 | 1 | 0 | 6 | 0 | 15 | 0.0667 | 3 | 0 | 5 | 0.2 | 1 | 0 | 1 | 0 | 10 | 0.1 |
| 7_1093110 | 1 | 0 | 1 | 1 | 1 | 1 | 1 | 1 | 4 | 0.75 | 1 | 1 | 1 | 0 | 1 | 1 | NA | NA | 3 | 0.6667 |
| 7_1093112 | 1 | 0 | 1 | 1 | 1 | 1 | NA | NA | 3 | 0.6667 | 1 | 1 | 1 | 0 | NA | NA | NA | NA | 2 | 0.5 |
| 7_1093351 | 12 | 0.5833 | 16 | 0.6875 | 24 | 0.5417 | 14 | 0.5714 | 66 | 0.5909 | 11 | 0.6364 | 18 | 0.7222 | 11 | 0.3636 | 26 | 0.6154 | 66 | 0.6061 |
| 7_1093351 | 13 | 0.4615 | 14 | 0.5 | 21 | 0.4762 | 14 | 0.2143 | 62 | 0.4194 | 10 | 0.4 | 18 | 0.5556 | 12 | 0.25 | 21 | 0.3333 | 61 | 0.3934 |
| 7_1093352 | 16 | 0.1875 | 16 | 0.25 | 23 | 0.1739 | 15 | 0.1333 | 70 | 0.1857 | 13 | 0.3077 | 15 | 0.5333 | 14 | 0.2857 | 27 | 0.4074 | 69 | 0.3913 |
| 7_1093352 | 16 | 0.5 | 18 | 0.6667 | 23 | 0.4783 | 16 | 0.5 | 73 | 0.5342 | 15 | 0.5333 | 17 | 0.7647 | 15 | 0.4667 | 29 | 0.6552 | 76 | 0.6184 |
| 7_1093353 | 13 | 0.0769 | 15 | 0.0667 | 24 | 0.125 | 14 | 0.2143 | 66 | 0.1212 | 15 | 0.2 | 17 | 0.3529 | 14 | 0.1429 | 26 | 0.0769 | 72 | 0.1806 |
| 7_1093355 | 15 | 0.0667 | 15 | 0.1333 | 20 | 0.2 | 15 | 0.3333 | 65 | 0.1846 | 14 | 0.2143 | 17 | 0.2353 | 13 | 0 | 26 | 0.2308 | 70 | 0.1857 |
| 7_1093356 | 13 | 0.1538 | 15 | 0.2 | 19 | 0.1053 | 15 | 0.2 | 62 | 0.1613 | 13 | 0.1538 | 16 | 0.125 | 12 | 0.0833 | 28 | 0.1071 | 69 | 0.1159 |
| 7_1093358 | 11 | 0.1818 | 16 | 0.3125 | 18 | 0.2778 | 18 | 0.5 | 63 | 0.3333 | 14 | 0.3571 | 15 | 0.2667 | 14 | 0.2143 | 25 | 0.24 | 68 | 0.2647 |
| 7_1093358 | 14 | 0.7143 | 18 | 0.9444 | 19 | 0.8947 | 13 | 0.6923 | 64 | 0.8281 | 14 | 0.7143 | 13 | 0.9231 | 15 | 0.8 | 23 | 0.913 | 65 | 0.8462 |
| 7_1093359 | 13 | 0.2308 | 19 | 0.0526 | 21 | 0.0952 | 17 | 0 | 70 | 0.0857 | 13 | 0.1538 | 17 | 0.1765 | 16 | 0 | 25 | 0.08 | 71 | 0.0986 |
| 7_1093360 | 13 | 0 | 17 | 0 | 22 | 0.1818 | 16 | 0.125 | 68 | 0.0882 | 13 | 0 | 18 | 0.1111 | 14 | 0.0714 | 22 | 0.0909 | 67 | 0.0746 |
| 7_1093360 | 11 | 0.7273 | 15 | 0.7333 | 18 | 0.6111 | 17 | 0.4706 | 61 | 0.623 | 12 | 0.3333 | 14 | 0.5714 | 14 | 0.7143 | 21 | 0.7143 | 61 | 0.6066 |
| 7_1093519 | 1 | 1 | NA | NA | 1 | 0 | 1 | 0 | 3 | 0.3333 | NA | NA | 2 | 1 | NA | NA | 1 | 1 | 3 | 1 |
| 7_1093520 | 1 | 0 | NA | NA | 1 | 0 | 1 | 1 | 3 | 0.3333 | NA | NA | 1 | 1 | NA | NA | 1 | 1 | 2 | 1 |
| 7_1093520 | NA | NA | NA | NA | NA | NA | 1 | 1 | 1 | 1 | NA | NA | 2 | 0.5 | NA | NA | 1 | 1 | 3 | 0.6667 |
| 7_1093521 | 1 | 1 | NA | NA | 2 | 0 | 1 | 1 | 4 | 0.5 | NA | NA | 1 | 0 | NA | NA | 1 | 1 | 2 | 0.5 |
| 7_1093598 | 1 | 1 | 7 | 0 | 4 | 0 | 1 | 0 | 13 | 0.0769 | 6 | 0 | 3 | 0 | 3 | 0 | 7 | 0.1429 | 19 | 0.0526 |
| 7_1093602 | 1 | 0 | 7 | 0 | 4 | 0.25 | 2 | 0 | 14 | 0.0714 | 4 | 0 | 2 | 0.5 | 1 | 0 | 5 | 0 | 12 | 0.0833 |
| 7_1093603 | 2 | 0.5 | 7 | 0.2857 | 5 | 0 | 2 | 0 | 16 | 0.1875 | 6 | 0.1667 | 1 | 0 | 3 | 0 | 2 | 0 | 12 | 0.0833 |
| 7_1093603 | 2 | 0.5 | 6 | 0.3333 | 3 | 0.3333 | 1 | 0 | 12 | 0.3333 | 5 | 0.2 | 2 | 0 | 3 | 0.6667 | 5 | 0.6 | 15 | 0.4 |
| 7_1094932 | 23 | 0.1739 | 32 | 0.0938 | 20 | 0.1 | 24 | 0.1667 | 99 | 0.1313 | 22 | 0.1364 | 31 | 0.0968 | 38 | 0.0526 | 19 | 0 | 110 | 0.0727 |
| 7_1094936 | 18 | 0.1111 | 25 | 0.2 | 12 | 0.0833 | 16 | 0.125 | 71 | 0.1408 | 15 | 0.2 | 23 | 0.2174 | 28 | 0.1429 | 14 | 0.0714 | 80 | 0.1625 |
| 7_1094937 | 20 | 0.1 | 27 | 0.1481 | 11 | 0.2727 | 20 | 0.05 | 78 | 0.1282 | 16 | 0.0625 | 28 | 0 | 30 | 0.2333 | 19 | 0.1053 | 93 | 0.1075 |
| 7_1094937 | 18 | 0.4444 | 27 | 0.2963 | 12 | 0.3333 | 19 | 0.4211 | 76 | 0.3684 | 16 | 0.1875 | 26 | 0.3846 | 30 | 0.2333 | 19 | 0.5789 | 91 | 0.3407 |
| 7_1094938 | 19 | 0.1579 | 29 | 0.2069 | 13 | 0.0769 | 21 | 0.2381 | 82 | 0.1829 | 17 | 0.1176 | 27 | 0.2222 | 35 | 0.2 | 17 | 0.2941 | 96 | 0.2083 |
| 7_1094938 | 17 | 0.7059 | 27 | 0.7037 | 12 | 0.25 | 12 | 0.75 | 68 | 0.6324 | 15 | 0.7333 | 21 | 0.8095 | 23 | 0.6522 | 16 | 0.5625 | 75 | 0.6933 |
| 7_1094939 | 19 | 0.1579 | 30 | 0.1 | 12 | 0.1667 | 14 | 0.2143 | 75 | 0.1467 | 16 | 0.0625 | 27 | 0.0741 | 29 | 0.069 | 12 | 0 | 84 | 0.0595 |
| 7_1095206 | 6 | 0.5 | 12 | 0.1667 | 6 | 0.1667 | 11 | 0.1818 | 35 | 0.2286 | 3 | 0 | 11 | 0.1818 | 13 | 0.5385 | 6 | 0.3333 | 33 | 0.3333 |
| 7_1095328 | 5 | 0.6 | 8 | 0.875 | 3 | 0.6667 | 10 | 0.8 | 26 | 0.7692 | 3 | 1 | 4 | 0.75 | 6 | 0.6667 | 3 | 0.6667 | 16 | 0.75 |
| 7_1095329 | 5 | 0.2 | 8 | 0.375 | 5 | 0.4 | 10 | 0.3 | 28 | 0.3214 | 3 | 0 | 8 | 0.25 | 7 | 0.4286 | 5 | 0.6 | 23 | 0.3478 |
| 7_1095331 | 6 | 1 | 7 | 0.8571 | 4 | 1 | 8 | 0.75 | 25 | 0.88 | 3 | 0.3333 | 7 | 0.7143 | 7 | 0.5714 | 5 | 0.6 | 22 | 0.5909 |
| 7_1095332 | 5 | 0.2 | 6 | 0.1667 | 3 | 0 | 8 | 0.25 | 22 | 0.1818 | 3 | 0 | 9 | 0.2222 | 7 | 0.4286 | 5 | 0.4 | 24 | 0.2917 |
| 7_1095333 | 4 | 0 | 7 | 0 | 3 | 0 | 6 | 0.1667 | 20 | 0.05 | 3 | 0 | 9 | 0.1111 | 5 | 0.2 | 5 | 0 | 22 | 0.0909 |
| 7_1095333 | 3 | 0.6667 | 5 | 0.8 | 3 | 1 | 8 | 0.625 | 19 | 0.7368 | 2 | 0.5 | 7 | 0.7143 | 6 | 0.6667 | 4 | 1 | 19 | 0.7368 |
| 7_1095720 | 1 | 1 | 2 | 0.5 | 1 | 0 | NA | NA | 4 | 0.5 | 1 | 1 | 3 | 0.6667 | 1 | 0 | 1 | 0 | 6 | 0.5 |
| 7_1095722 | 2 | 0 | 2 | 1 | 1 | 0 | NA | NA | 5 | 0.4 | 2 | 1 | 3 | 0.6667 | 1 | 0 | 2 | 0.5 | 8 | 0.625 |
| 7_1095726 | 2 | 0 | 3 | 0.3333 | 1 | 1 | NA | NA | 6 | 0.3333 | 3 | 0.6667 | 2 | 0 | 1 | 0 | 2 | 0 | 8 | 0.25 |
| 7_1095727 | 6 | 0.6667 | 4 | 0.25 | 6 | 0.6667 | 1 | 0 | 17 | 0.5294 | 9 | 0.7778 | 2 | 0.5 | 3 | 0.6667 | 6 | 0.5 | 20 | 0.65 |
| 7_1095728 | 5 | 0.2 | 4 | 0 | 4 | 1 | NA | NA | 13 | 0.3846 | 9 | 0.1111 | 2 | 0 | 4 | 0.25 | 4 | 0.25 | 19 | 0.1579 |
| 7_1095728 | 5 | 0.4 | 4 | 0.25 | 5 | 0.6 | 1 | 0 | 15 | 0.4 | 9 | 0.3333 | 3 | 0.6667 | 4 | 0.25 | 6 | 0.1667 | 22 | 0.3182 |
| 7_1095735 | 5 | 0 | 4 | 0 | 8 | 0.125 | 1 | 0 | 18 | 0.0556 | 7 | 0.2857 | 3 | 0.6667 | 2 | 0 | 6 | 0 | 18 | 0.2222 |
| 7_1095736 | 2 | 0.5 | 5 | 0.4 | 6 | 0 | 2 | 0 | 15 | 0.2 | 7 | 0.4286 | 4 | 0.5 | 3 | 0 | 5 | 0.4 | 19 | 0.3684 |
| 7_1095736 | 3 | 0 | 5 | 0.2 | 7 | 0.2857 | 2 | 0 | 17 | 0.1765 | 9 | 0.3333 | 5 | 0.4 | 3 | 0 | 5 | 0.4 | 22 | 0.3182 |
| 7_1095738 | 3 | 0 | 4 | 0 | 3 | 0.3333 | 1 | 0 | 11 | 0.0909 | 5 | 0 | 4 | 0.25 | 2 | 0 | 5 | 0.2 | 16 | 0.125 |
| 7_1095738 | 4 | 1 | 5 | 1 | 6 | 1 | 1 | 1 | 16 | 1 | 7 | 0.7143 | 5 | 0.8 | 3 | 1 | 4 | 0.75 | 19 | 0.7895 |
| 7_1095738 | 3 | 0 | 6 | 0.1667 | 6 | 0.5 | NA | NA | 15 | 0.2667 | 4 | 0 | 3 | 0.3333 | 3 | 0.3333 | 4 | 0.75 | 14 | 0.3571 |
| 7_1095739 | 3 | 0.3333 | 6 | 0.8333 | 8 | 0.25 | 1 | 1 | 18 | 0.5 | 7 | 0.2857 | 5 | 0.4 | 3 | 0.3333 | 5 | 0.2 | 20 | 0.3 |
| 7_1096544 | 5 | 0.4 | 7 | 0 | 6 | 0 | 4 | 0.25 | 22 | 0.1364 | NA | NA | 2 | 0.5 | 3 | 0 | 5 | 0.6 | 10 | 0.4 |
| 7_1096544 | 7 | 0.8571 | 8 | 0.625 | 3 | 0.3333 | 6 | 0.6667 | 24 | 0.6667 | 2 | 1 | 2 | 1 | 3 | 1 | 7 | 0.7143 | 14 | 0.8571 |
| 7_1096544 | 10 | 0.4 | 9 | 0.5556 | 6 | 0.3333 | 7 | 0.4286 | 32 | 0.4375 | 2 | 0.5 | 3 | 0 | 2 | 1 | 8 | 0.375 | 15 | 0.4 |
| 7_1096545 | 9 | 0.2222 | 10 | 0.4 | 6 | 0.3333 | 6 | 0.1667 | 31 | 0.2903 | 5 | 0.4 | 7 | 0.4286 | 8 | 0.75 | 12 | 0.6667 | 32 | 0.5938 |
| 7_1096546 | 10 | 0.5 | 12 | 0.25 | 7 | 0.2857 | 11 | 0.7273 | 40 | 0.45 | 6 | 0.5 | 9 | 0.6667 | 8 | 0.375 | 12 | 0.3333 | 35 | 0.4571 |
| 7_1096548 | 9 | 0.3333 | 9 | 0 | 7 | 0.1429 | 11 | 0.0909 | 36 | 0.1389 | 6 | 0.1667 | 12 | 0 | 8 | 0.25 | 15 | 0.1333 | 41 | 0.122 |
| 7_1096551 | 12 | 0.1667 | 14 | 0 | 8 | 0.125 | 11 | 0.0909 | 45 | 0.0889 | 8 | 0.125 | 13 | 0.0769 | 8 | 0.125 | 12 | 0.1667 | 41 | 0.122 |
| 7_1096936 | 1 | 1 | 1 | 1 | 1 | 1 | 2 | 1 | 5 | 1 | 1 | 1 | NA | NA | 1 | 1 | NA | NA | 2 | 1 |
| 7_1096937 | 2 | 1 | 1 | 1 | 1 | 1 | 2 | 1 | 6 | 1 | 1 | 1 | NA | NA | 2 | 1 | NA | NA | 3 | 1 |
| 7_1096939 | 2 | 0 | 1 | 0 | 2 | 0 | 2 | 0.5 | 7 | 0.1429 | 1 | 0 | 1 | 0 | 2 | 0.5 | 1 | 0 | 5 | 0.2 |
| 7_1096944 | 2 | 0 | 2 | 0 | NA | NA | 2 | 0.5 | 6 | 0.1667 | 1 | 0 | 1 | 0 | 2 | 0 | 1 | 1 | 5 | 0.2 |

| | | | | | | | | | | | | | | | | | | | | |
|---|---|---|---|---|---|---|---|---|---|---|---|---|---|---|---|---|---|---|---|---|
| 7_1096945 | 1 | 0 | 1 | 0 | 2 | 0 | 2 | 0.5 | 6 | 0.1667 | 1 | 0 | 1 | 0 | 3 | 0.3333 | 1 | 1 | 6 | 0.3333 |
| 7_1097413 | 2 | 1 | NA | NA | NA | NA | NA | NA | 2 | 1 | 3 | 1 | 2 | 0.5 | 4 | 1 | 1 | 1 | 10 | 0.9 |
| 7_1097414 | 3 | 0.3333 | NA | NA | NA | NA | 2 | 0.5 | 5 | 0.4 | 3 | 0.6667 | 2 | 0 | 4 | 0.25 | 1 | 0 | 10 | 0.3 |
| 7_1097414 | 3 | 0 | NA | NA | NA | NA | 2 | 0.5 | 5 | 0.2 | 3 | 0.3333 | 2 | 0 | 4 | 0 | 1 | 0 | 10 | 0.1 |
| 7_1097416 | 4 | 0.75 | NA | NA | NA | NA | 2 | 0.5 | 6 | 0.6667 | 3 | 1 | 2 | 1 | 3 | 0.3333 | 1 | 1 | 9 | 0.7778 |
| 7_1097417 | 4 | 0.75 | NA | NA | NA | NA | 2 | 0.5 | 6 | 0.6667 | 3 | 1 | 2 | 1 | 4 | 0.25 | 1 | 1 | 10 | 0.7 |
| 7_1097420 | NA | NA | NA | NA | NA | NA | 2 | 0.5 | 2 | 0.5 | 1 | 1 | 2 | 0 | 4 | 0.5 | NA | NA | 7 | 0.4286 |
| 7_1098809 | NA | NA | NA | NA | NA | NA | 1 | 1 | 1 | 1 | NA | NA | 1 | 0 | NA | NA | 3 | 0.6667 | 4 | 0.5 |
| 7_1098811 | NA | NA | NA | NA | NA | NA | 1 | 1 | 1 | 1 | NA | NA | 2 | 1 | NA | NA | 3 | 0.6667 | 5 | 0.8 |
| 7_1099075 | 7 | 0.2857 | 6 | 0 | 5 | 0.2 | 7 | 0.5714 | 25 | 0.28 | 4 | 0 | 2 | 0 | 4 | 0.25 | 3 | 0.3333 | 13 | 0.1538 |
| 7_1099078 | 4 | 1 | 5 | 0.4 | 5 | 1 | 7 | 0.8571 | 21 | 0.8095 | 2 | 0.5 | 1 | 1 | 7 | 0.8571 | 2 | 1 | 12 | 0.8333 |
| 7_1099079 | 5 | 0.2 | 6 | 0.5 | 4 | 0 | 7 | 0.2857 | 22 | 0.2727 | 3 | 0 | 3 | 0.3333 | 7 | 0.4286 | 2 | 0.5 | 15 | 0.3333 |
| 7_1099079 | 4 | 0.75 | 6 | 0.3333 | 3 | 0.3333 | 6 | 0.5 | 19 | 0.4737 | 2 | 0.5 | 2 | 0.5 | 6 | 0.1667 | 2 | 0.5 | 12 | 0.3333 |
| 7_1099079 | 4 | 0.5 | 4 | 0.25 | 4 | 0.25 | 5 | 0.6 | 17 | 0.4118 | 3 | 0.6667 | 2 | 0.5 | 7 | 0.2857 | 2 | 0 | 14 | 0.3571 |
| 7_1099079 | 4 | 0.75 | 2 | 0 | 3 | 0.3333 | 4 | 0.25 | 13 | 0.3846 | 3 | 0.6667 | 3 | 0 | 7 | 0.4286 | 1 | 1 | 14 | 0.4286 |
| 7_1099079 | 5 | 0.4 | 4 | 0.75 | 5 | 0.8 | 5 | 0.6 | 19 | 0.6316 | 3 | 0.6667 | 3 | 0.6667 | 7 | 0.5714 | 2 | 0.5 | 15 | 0.6 |
| 7_1099079 | 5 | 0.6 | 4 | 0.75 | 5 | 0.4 | 4 | 0.25 | 18 | 0.5 | 3 | 0.3333 | 2 | 0.5 | 7 | 0 | 1 | 1 | 13 | 0.2308 |
| 7_1099079 | 5 | 0.6 | 5 | 0.8 | 4 | 1 | 6 | 0.8333 | 20 | 0.8 | 2 | 0 | 2 | 0 | 7 | 0.5714 | 2 | 0 | 13 | 0.3077 |
| 7_1099080 | 5 | 0.4 | 4 | 0.5 | 5 | 0.8 | 6 | 0.3333 | 20 | 0.5 | 3 | 0.6667 | 2 | 0 | 7 | 0.1429 | 2 | 0 | 14 | 0.2143 |
| 7_1099080 | 5 | 0 | 4 | 0 | 5 | 0.2 | 6 | 0 | 20 | 0.05 | 3 | 0 | 2 | 0.5 | 6 | 0.1667 | 2 | 0 | 13 | 0.1538 |
| 7_1099082 | 3 | 0.3333 | 6 | 0.1667 | 4 | 0.5 | 4 | 0.25 | 17 | 0.2941 | 3 | 0 | 2 | 0 | 5 | 0.2 | 2 | 0.5 | 12 | 0.1667 |
| 7_1099083 | 5 | 0.4 | 5 | 0.6 | 2 | 1 | 6 | 0.3333 | 18 | 0.5 | 3 | 0.6667 | NA | NA | 6 | 0.8333 | 2 | 0.5 | 11 | 0.7273 |
| 7_1099083 | 4 | 0.75 | 4 | 0.5 | 3 | 1 | 6 | 0.6667 | 17 | 0.7059 | 2 | 0.5 | 2 | 0.5 | 6 | 1 | 2 | 1 | 12 | 0.8333 |
| 7_1099084 | 5 | 0.8 | 5 | 0.6 | 4 | 0.75 | 5 | 0.8 | 19 | 0.7368 | 2 | 1 | 2 | 0 | 6 | 0.8333 | 2 | 1 | 12 | 0.75 |
| 7_1099085 | 4 | 0.75 | 4 | 0.5 | 4 | 0.75 | 4 | 1 | 16 | 0.75 | 3 | 0.3333 | 1 | 0 | 5 | 0.4 | 2 | 1 | 11 | 0.4545 |
| 7_1099086 | 5 | 0.6 | 4 | 0 | 4 | 0.75 | 4 | 0.25 | 17 | 0.4118 | 3 | 0 | 1 | 0 | 5 | 0.2 | 2 | 0 | 11 | 0.0909 |
| 7_1104102 | 1 | 0 | 3 | 0.3333 | 1 | 0 | NA | NA | 5 | 0.2 | 2 | 0 | 4 | 0.75 | 1 | 0 | 1 | 0 | 8 | 0.375 |
| 7_1104103 | 1 | 1 | 3 | 0 | 2 | 0.5 | 1 | 1 | 7 | 0.4286 | 2 | 0 | 4 | 0.25 | 3 | 0.3333 | 1 | 0 | 10 | 0.2 |
| 7_1104104 | 1 | 0 | 3 | 0.3333 | NA | NA | 1 | 1 | 5 | 0.4 | 1 | 1 | 4 | 0.5 | 3 | 0.3333 | 1 | 0 | 9 | 0.4444 |
| 7_1104104 | 2 | 0.5 | 3 | 0.3333 | 2 | 0.5 | 1 | 0 | 8 | 0.375 | 2 | 0 | 2 | 0 | 2 | 1 | 1 | 1 | 7 | 0.4286 |
| 7_1104109 | 3 | 0 | 3 | 0.3333 | 3 | 0.3333 | 2 | 0 | 11 | 0.1818 | 2 | 0 | 3 | 0.6667 | 5 | 0 | 1 | 0 | 11 | 0.1818 |
| 7_1104153 | 1 | 0 | 8 | 0.375 | 2 | 0 | 4 | 0.75 | 15 | 0.4 | 4 | 0 | 1 | 1 | NA | NA | 4 | 0.5 | 9 | 0.3333 |
| 7_1104153 | 1 | 0 | 9 | 0 | 1 | 0 | 4 | 0.25 | 15 | 0.0667 | 4 | 0.5 | 1 | 0 | 2 | 0 | 3 | 0 | 10 | 0.2 |
| 7_1104153 | 1 | 0 | 10 | 0 | 1 | 1 | 4 | 0.5 | 16 | 0.1875 | 3 | 0 | 1 | 0 | 2 | 0 | 4 | 0.25 | 10 | 0.1 |
| 7_1104158 | 1 | 0 | 5 | 0.4 | 2 | 0 | 5 | 0.2 | 13 | 0.2308 | 4 | 0.25 | 1 | 0 | 2 | 0 | 1 | 0 | 8 | 0.125 |
| 7_1106280 | NA | NA | NA | NA | NA | NA | 1 | 1 | 1 | 1 | 1 | 1 | NA | NA | NA | NA | NA | NA | 1 | 1 |
| 7_1106329 | 2 | 0.5 | 3 | 0.6667 | 2 | 0 | 4 | 0 | 11 | 0.2727 | 1 | 1 | 1 | 0 | 5 | 0.4 | NA | NA | 7 | 0.4286 |
| 7_1106547 | 6 | 0.5 | 4 | 0.75 | 3 | 0 | 9 | 0.1111 | 22 | 0.3182 | 5 | 0.6 | 7 | 0.2857 | 10 | 0.1 | 7 | 0.2857 | 29 | 0.2759 |
| 7_1106548 | 3 | 0.3333 | 5 | 0.4 | 3 | 0 | 6 | 0.5 | 17 | 0.3529 | 5 | 0.8 | 6 | 0.5 | 6 | 0.3333 | 5 | 0.4 | 22 | 0.5 |
| 7_1106549 | 8 | 0.125 | 5 | 0 | 3 | 0 | 9 | 0.2222 | 25 | 0.12 | 5 | 0.2 | 7 | 0 | 9 | 0.1111 | 6 | 0 | 27 | 0.0741 |
| 7_1106550 | 7 | 0.8571 | 6 | 0.8333 | 1 | 1 | 7 | 1 | 21 | 0.9048 | 3 | 1 | 4 | 1 | 8 | 0.875 | 6 | 0.8333 | 21 | 0.9048 |
| 7_1106550 | 8 | 0.25 | 6 | 0 | 3 | 0 | 10 | 0.1 | 27 | 0.1111 | 3 | 0 | 7 | 0.2857 | 9 | 0.1111 | 7 | 0 | 26 | 0.1154 |
| 7_1106551 | 6 | 0.6667 | 6 | 0.5 | 3 | 0.6667 | 11 | 0.6364 | 26 | 0.6154 | 6 | 0.6667 | 6 | 0.8333 | 9 | 0.5556 | 6 | 0.8333 | 27 | 0.7037 |
| 7_1106552 | 8 | 0.125 | 7 | 0 | 2 | 0 | 10 | 0.1 | 27 | 0.0741 | 7 | 0.1429 | 7 | 0.2857 | 11 | 0.2727 | 7 | 0.2857 | 32 | 0.25 |
| 7_1106554 | 9 | 0 | 7 | 0.1429 | 3 | 0.3333 | 12 | 0.25 | 31 | 0.1613 | 7 | 0.2857 | 7 | 0.1429 | 7 | 0.2857 | 7 | 0.2857 | 28 | 0.25 |
| 7_1106554 | 8 | 0.5 | 5 | 0 | 2 | 0 | 11 | 0.3636 | 26 | 0.3077 | 4 | 0 | 7 | 0.1429 | 7 | 0.2857 | 4 | 0.75 | 22 | 0.2727 |
| 7_1106716 | 55 | 0.0909 | 71 | 0.1127 | 74 | 0.0541 | 68 | 0.1029 | 268 | 0.0896 | 43 | 0.0465 | 68 | 0.1029 | 56 | 0.0536 | 56 | 0.0893 | 223 | 0.0762 |
| 7_1106737 | 48 | 0.0833 | 52 | 0.0769 | 56 | 0.1429 | 64 | 0.1562 | 220 | 0.1182 | 37 | 0.1351 | 54 | 0.1481 | 48 | 0.1458 | 48 | 0.0833 | 187 | 0.1283 |
| 7_1110711 | 3 | 0.6667 | 7 | 0.5714 | 6 | 0.6667 | 4 | 0.5 | 20 | 0.6 | 4 | 0.75 | 7 | 0.7143 | 13 | 0.7692 | 5 | 0.2 | 29 | 0.6552 |
| 7_1110711 | 6 | 0.1667 | 6 | 0.1667 | 8 | 0 | 4 | 0 | 24 | 0.0833 | 5 | 0 | 6 | 0.1667 | 11 | 0 | 6 | 0.1667 | 28 | 0.0714 |
| 7_1113787 | 4 | 0.25 | 3 | 0.3333 | 4 | 0 | 1 | 0 | 12 | 0.1667 | 4 | 0.25 | 1 | 1 | 5 | 0.4 | 1 | 0 | 11 | 0.3636 |
| 7_1113787 | 3 | 0.3333 | 2 | 0.5 | 4 | 0 | 1 | 0 | 10 | 0.2 | 3 | 0.3333 | 1 | 0 | 5 | 0.2 | 1 | 0 | 10 | 0.2 |
| 7_1113788 | 3 | 0 | 2 | 0.5 | 3 | 0 | 1 | 0 | 9 | 0.1111 | 2 | 0 | 1 | 1 | 2 | 0 | 1 | 0 | 6 | 0.1667 |
| 7_1113788 | 2 | 0.5 | 1 | 1 | 2 | 1 | 1 | 0 | 6 | 0.6667 | 3 | 1 | 1 | 0 | 4 | 0.5 | NA | NA | 8 | 0.625 |
| 7_1116920 | 4 | 0.25 | 3 | 0 | 3 | 0 | 4 | 0 | 14 | 0.0714 | 1 | 0 | 4 | 0 | 4 | 0 | 3 | 0.3333 | 12 | 0.0833 |
| 7_1116920 | 6 | 0.6667 | 4 | 0.25 | 5 | 0.4 | 4 | 0.25 | 19 | 0.4211 | 1 | 0 | 4 | 0.5 | 6 | 0.5 | 3 | 0 | 14 | 0.3571 |
| 7_1116921 | 9 | 0.4444 | 4 | 0.75 | 5 | 0.6 | 5 | 0.2 | 23 | 0.4783 | 1 | 1 | 3 | 0.3333 | 7 | 0.4286 | 4 | 0.25 | 15 | 0.4 |
| 7_1116921 | 6 | 0.1667 | 2 | 0 | 5 | 0.4 | 5 | 0.6 | 18 | 0.3333 | NA | NA | 4 | 0.5 | 6 | 0.3333 | 3 | 0.6667 | 13 | 0.4615 |
| 7_1116922 | 9 | 1 | 4 | 0.5 | 5 | 0.8 | 4 | 0.75 | 22 | 0.8182 | NA | NA | 4 | 1 | 6 | 1 | 3 | 1 | 13 | 1 |
| 7_1116923 | 9 | 0.6667 | 3 | 1 | 5 | 0.8 | 5 | 0.4 | 22 | 0.6818 | 1 | 1 | 3 | 0.3333 | 4 | 0.75 | 4 | 0.75 | 12 | 0.6667 |
| 7_1116923 | 8 | 0.25 | 3 | 0.6667 | 6 | 0.1667 | 4 | 0.25 | 21 | 0.2857 | 1 | 0 | 3 | 0 | 5 | 0.4 | 4 | 0 | 13 | 0.1538 |
| 7_1116923 | 6 | 0.1667 | 2 | 0 | 3 | 0.3333 | 4 | 0.5 | 15 | 0.2667 | 1 | 1 | 3 | 0 | 4 | 0 | 4 | 0 | 12 | 0.0833 |
| 7_1116923 | 7 | 0.4286 | 4 | 0.25 | 5 | 0.4 | 5 | 0.4 | 21 | 0.381 | 1 | 0 | 4 | 0.5 | 7 | 0.7143 | 2 | 0.5 | 14 | 0.5714 |
| 7_1116924 | 7 | 0 | 3 | 0.3333 | 6 | 0 | 5 | 0.2 | 21 | 0.0952 | 2 | 0 | 4 | 0 | 7 | 0.1429 | 4 | 0 | 17 | 0.0588 |
| 7_1116924 | 7 | 0.5714 | 4 | 0.75 | 6 | 1 | 5 | 1 | 22 | 0.8182 | 2 | 0.5 | 4 | 1 | 7 | 1 | 4 | 1 | 17 | 0.9412 |

| | | | | | | | | | | | | | | | | | | | | |
|---|---|---|---|---|---|---|---|---|---|---|---|---|---|---|---|---|---|---|---|---|
| 7_1116924 | 7 | 0.2857 | 3 | 0 | 3 | 0 | 5 | 0 | 18 | 0.1111 | 1 | 0 | 4 | 0 | 7 | 0.2857 | 4 | 0 | 16 | 0.125 |
| 7_1116925 | 9 | 0.1111 | 5 | 0.2 | 5 | 0 | 5 | 0 | 24 | 0.0833 | 2 | 0 | 3 | 0.3333 | 5 | 0.2 | 3 | 0.3333 | 13 | 0.2308 |
| 7_1116925 | 7 | 0.7143 | 3 | 0.3333 | 7 | 0.4286 | 5 | 0.2 | 22 | 0.4545 | 3 | 0.6667 | 3 | 0.6667 | 6 | 1 | 3 | 1 | 15 | 0.8667 |
| 7_1116926 | 10 | 0.2 | 4 | 0 | 8 | 0.25 | 5 | 0 | 27 | 0.1481 | 3 | 0.3333 | 4 | 0.5 | 7 | 0.2857 | 3 | 0 | 17 | 0.2941 |
| 7_1117093 | 10 | 0.2 | 4 | 0.5 | 6 | 0.3333 | 7 | 0.4286 | 27 | 0.3333 | 3 | 0.3333 | 4 | 0.25 | 5 | 0.4 | 5 | 0.2 | 17 | 0.2941 |
| 7_1117093 | 10 | 0.4 | 5 | 0.2 | 8 | 0.375 | 7 | 0.2857 | 30 | 0.3333 | 2 | 0 | 6 | 0.3333 | 8 | 0 | 4 | 0 | 20 | 0.1 |
| 7_1117094 | 11 | 1 | 6 | 0.6667 | 8 | 0.875 | 7 | 0.8571 | 32 | 0.875 | 2 | 0.5 | 6 | 0.8333 | 8 | 0.875 | 5 | 0.8 | 21 | 0.8095 |
| 7_1117094 | 8 | 0.875 | 5 | 0.8 | 9 | 0.7778 | 7 | 1 | 29 | 0.8621 | 3 | 1 | 5 | 0.8 | 9 | 0.6667 | 5 | 0.8 | 22 | 0.7727 |
| 7_1117094 | 8 | 0 | 3 | 0.3333 | 7 | 0.1429 | 6 | 0.1667 | 24 | 0.125 | 1 | 1 | 7 | 0 | 7 | 0.1429 | 4 | 0.25 | 19 | 0.1579 |
| 7_1117096 | 11 | 0.2727 | 7 | 0.2857 | 9 | 0.3333 | 6 | 0 | 33 | 0.2424 | 3 | 0 | 6 | 0 | 10 | 0.1 | 5 | 0.2 | 24 | 0.0833 |
| 7_1117162 | 15 | 0.0667 | 16 | 0.3125 | 10 | 0.3 | 9 | 0 | 50 | 0.18 | 11 | 0.0909 | 9 | 0 | 10 | 0 | 16 | 0.1875 | 46 | 0.087 |
| 7_1117168 | 9 | 0.1111 | 15 | 0 | 7 | 0 | 7 | 0.1429 | 38 | 0.0526 | 9 | 0.2222 | 5 | 0 | 8 | 0.125 | 12 | 0 | 34 | 0.0882 |
| 7_1117170 | 12 | 0.5833 | 15 | 0.3333 | 8 | 0.875 | 8 | 0.375 | 43 | 0.5116 | 8 | 0.5 | 8 | 0.5 | 9 | 0.6667 | 16 | 0.5 | 41 | 0.5366 |
| 7_1117172 | 11 | 0.2727 | 15 | 0.2667 | 6 | 0 | 7 | 0 | 39 | 0.1795 | 8 | 0 | 6 | 0 | 7 | 0.1429 | 8 | 0.25 | 29 | 0.1034 |
| 7_1118705 | 5 | 0.2 | 9 | 0 | 2 | 0 | 2 | 0 | 18 | 0.0556 | 4 | 0 | 5 | 0.2 | 2 | 0 | 6 | 0 | 17 | 0.0588 |
| 7_1118705 | 4 | 0 | 9 | 0.1111 | 2 | 0 | 2 | 0 | 17 | 0.0588 | 4 | 0 | 5 | 0.2 | 2 | 0 | 5 | 0.2 | 16 | 0.125 |
| 7_1120121 | NA | NA | NA | NA | NA | NA | 1 | 1 | 1 | 1 | 1 | 1 | NA | NA | NA | NA | NA | NA | 1 | 1 |
| 7_1121879 | 2 | 0.5 | 1 | 0 | 1 | 0 | 5 | 0.8 | 9 | 0.5556 | NA | NA | 3 | 0.6667 | 3 | 0.6667 | NA | NA | 6 | 0.6667 |
| 7_1121880 | 1 | 0 | 2 | 0.5 | 1 | 0 | 4 | 0 | 8 | 0.125 | 1 | 0 | 4 | 0.5 | 3 | 0.3333 | 2 | 0.5 | 10 | 0.4 |
| 7_1125036 | NA | NA | 1 | 1 | 2 | 1 | 1 | 1 | 4 | 1 | 1 | 1 | NA | NA | 1 | 1 | NA | NA | 2 | 1 |
| 7_1125038 | NA | NA | 3 | 1 | 1 | 0 | 1 | 1 | 5 | 0.8 | NA | NA | NA | NA | 1 | 1 | 1 | 0 | 2 | 0.5 |
| 7_1125039 | NA | NA | 3 | 0.3333 | 2 | 0 | 1 | 0 | 6 | 0.1667 | 1 | 0 | NA | NA | 1 | 1 | 1 | 0 | 3 | 0.3333 |
| 7_1125040 | NA | NA | 3 | 0.6667 | 2 | 1 | 2 | 1 | 7 | 0.8571 | 1 | 1 | NA | NA | 1 | 1 | 1 | 1 | 3 | 1 |
| 7_1125049 | 1 | 0 | 3 | 0 | 2 | 0.5 | 1 | 1 | 7 | 0.2857 | 1 | 1 | NA | NA | 1 | 0 | 1 | 1 | 3 | 0.6667 |
| 7_1125546 | NA | NA | 1 | 1 | NA | NA | 1 | 1 | 2 | 1 | NA | NA | 1 | 1 | NA | NA | NA | NA | 1 | 1 |
| 7_1125546 | NA | NA | 1 | 1 | NA | NA | 1 | 1 | 2 | 1 | NA | NA | 1 | 1 | NA | NA | NA | NA | 1 | 1 |
| 7_1125546 | NA | NA | 1 | 1 | NA | NA | 1 | 1 | 2 | 1 | NA | NA | 1 | 1 | NA | NA | NA | NA | 1 | 1 |
| 7_1125548 | NA | NA | 1 | 1 | NA | NA | 1 | 1 | 2 | 1 | NA | NA | 1 | 1 | NA | NA | NA | NA | 1 | 1 |
| 7_1125552 | NA | NA | 1 | 1 | NA | NA | 1 | 1 | 2 | 1 | NA | NA | 1 | 1 | NA | NA | NA | NA | 1 | 1 |
| 7_1125553 | NA | NA | 1 | 1 | NA | NA | 1 | 1 | 2 | 1 | NA | NA | 1 | 1 | NA | NA | NA | NA | 1 | 1 |
| 7_1129127 | 2 | 0.5 | 5 | 0.2 | 2 | 0 | 6 | 0.3333 | 15 | 0.2667 | 3 | 0 | 5 | 0.2 | 3 | 0 | 4 | 0 | 15 | 0.0667 |
| 7_1129128 | 3 | 0 | 7 | 0.1429 | 4 | 0.5 | 6 | 0.5 | 20 | 0.3 | 4 | 0.5 | 5 | 0.2 | 6 | 0.3333 | 4 | 0.75 | 19 | 0.4211 |
| 7_1129130 | 3 | 0 | 6 | 0.1667 | 4 | 0.25 | 6 | 0.1667 | 19 | 0.1579 | 3 | 0 | 5 | 0.4 | 5 | 0 | 4 | 0.5 | 17 | 0.2353 |
| 7_1129155 | 6 | 0 | 8 | 0.125 | 8 | 0.25 | 4 | 0.25 | 26 | 0.1538 | 12 | 0.3333 | 5 | 0 | 9 | 0.4444 | 6 | 0.5 | 32 | 0.3438 |
| 7_1129157 | 7 | 0.2857 | 7 | 0.1429 | 6 | 0.3333 | 3 | 0.3333 | 23 | 0.2609 | 12 | 0.1667 | 5 | 0.2 | 9 | 0.1111 | 7 | 0.4286 | 33 | 0.2121 |
| 7_1129157 | 7 | 0.5714 | 8 | 0.375 | 7 | 0.2857 | 3 | 0.3333 | 25 | 0.4 | 12 | 0.25 | 5 | 0.2 | 9 | 0.3333 | 6 | 0.5 | 32 | 0.3125 |
| 7_1129158 | 4 | 1 | 8 | 0.625 | 6 | 0.6667 | NA | NA | 18 | 0.7222 | 10 | 0.7 | 5 | 1 | 7 | 0.8571 | 6 | 0.8333 | 28 | 0.8214 |
| 7_1129158 | 6 | 0 | 8 | 0.125 | 6 | 0.1667 | 3 | 0 | 23 | 0.087 | 12 | 0 | 4 | 0.25 | 8 | 0 | 6 | 0.1667 | 30 | 0.0667 |
| 7_1131675 | 5 | 0.4 | 5 | 0.2 | 6 | 0 | 8 | 0.125 | 24 | 0.1667 | 1 | 0 | 4 | 0 | 8 | 0.125 | 8 | 0.125 | 21 | 0.0952 |
| 7_1131680 | 6 | 0.5 | 5 | 0 | 7 | 0.1429 | 7 | 0.2857 | 25 | 0.24 | 1 | 0 | 4 | 0.5 | 8 | 0.25 | 10 | 0.2 | 23 | 0.2609 |
| 7_1131681 | 7 | 0.4286 | 4 | 0 | 5 | 0.2 | 8 | 0.125 | 24 | 0.2083 | 1 | 0 | 5 | 0 | 7 | 0 | 10 | 0.2 | 23 | 0.087 |
| 7_1131682 | 6 | 0.3333 | 4 | 0 | 6 | 0.1667 | 5 | 0 | 21 | 0.1429 | 1 | 0 | 2 | 0.5 | 5 | 0 | 7 | 0.2857 | 15 | 0.2 |
| 7_1131683 | 6 | 0.1667 | 7 | 0.5714 | 6 | 0.1667 | 8 | 0.125 | 27 | 0.2593 | NA | NA | 3 | 0.3333 | 6 | 0.5 | 8 | 0.125 | 17 | 0.2941 |
| 7_1131702 | 6 | 0 | 4 | 0 | 5 | 0.4 | 6 | 0 | 21 | 0.0952 | NA | NA | 3 | 0 | 5 | 0 | 9 | 0.1111 | 17 | 0.0588 |
| 7_1136358 | 2 | 0 | 10 | 0.3 | 6 | 0.3333 | 1 | 0 | 19 | 0.2632 | 4 | 0.25 | 3 | 0 | 5 | 0 | 2 | 0 | 14 | 0.0714 |
| 7_1136359 | 2 | 0 | 11 | 0.5455 | 6 | 0.1667 | 1 | 0 | 20 | 0.35 | 5 | 0.8 | 4 | 0 | 5 | 0.2 | 6 | 0.8333 | 20 | 0.5 |
| 7_1136390 | 3 | 0 | 9 | 0.1111 | 9 | 0.2222 | 1 | 0 | 22 | 0.1364 | 5 | 0 | 3 | 0.3333 | 6 | 0 | 5 | 0.4 | 19 | 0.1579 |
| 7_1136391 | 3 | 0.6667 | 10 | 0.2 | 7 | 0 | 1 | 0 | 21 | 0.1905 | 5 | 0.2 | 4 | 0.25 | 6 | 0.1667 | 6 | 0 | 21 | 0.1429 |
| 7_1136393 | 3 | 0.6667 | 11 | 0.4545 | 6 | 0.5 | 2 | 0 | 22 | 0.4545 | 5 | 0.6 | 4 | 0.25 | 8 | 0.5 | 6 | 0.8333 | 23 | 0.5652 |
| 7_1136394 | 2 | 0 | 8 | 0 | 5 | 0.2 | 2 | 0 | 17 | 0.0588 | 5 | 0 | 3 | 0.3333 | 4 | 0 | 7 | 0.1429 | 19 | 0.1053 |
| 7_1136395 | 3 | 0 | 12 | 0.25 | 7 | 0.1429 | 2 | 0 | 24 | 0.1667 | 5 | 0.6 | 4 | 0.25 | 8 | 0.375 | 7 | 0 | 24 | 0.2917 |
| 7_1136402 | 3 | 0.6667 | 9 | 0.4444 | 8 | 0.125 | 1 | 0 | 21 | 0.3333 | 2 | 1 | 2 | 0.5 | 5 | 0.2 | 2 | 0 | 11 | 0.3636 |
| 7_1136410 | 1 | 0 | 2 | 1 | 3 | 0.3333 | 2 | 0 | 8 | 0.375 | 2 | 0 | 1 | 1 | 2 | 0 | 5 | 0.4 | 10 | 0.3 |
| 7_1136410 | 1 | 1 | 2 | 0.5 | 3 | 0 | 2 | 0.5 | 8 | 0.375 | 2 | 0.5 | 1 | 1 | 2 | 0 | 5 | 0.2 | 10 | 0.3 |
| 7_1136412 | 2 | 1 | 1 | 1 | 1 | 1 | NA | NA | 4 | 1 | 1 | 0 | NA | NA | 2 | 0.5 | 4 | 0 | 7 | 0.1429 |
| 7_1136415 | 2 | 0.5 | 2 | 1 | 1 | 0 | 1 | 1 | 6 | 0.6667 | 2 | 1 | NA | NA | 4 | 0.5 | 4 | 0.5 | 10 | 0.6 |
| 7_1136468 | NA | NA | 1 | 0 | 2 | 0.5 | 2 | 0.5 | 5 | 0.4 | 1 | 0 | 2 | 0.5 | 2 | 0.5 | 2 | 0.5 | 7 | 0.4286 |
| 7_1136472 | NA | NA | 1 | 1 | 2 | 0 | 2 | 0.5 | 5 | 0.4 | 1 | 0 | 1 | 1 | 2 | 1 | 3 | 0 | 7 | 0.4286 |
| 7_1138150 | 11 | 0 | 15 | 0.1333 | 6 | 0.1667 | 9 | 0.2222 | 41 | 0.122 | 1 | 0 | 6 | 0.5 | 12 | 0 | 6 | 0 | 25 | 0.12 |
| 7_1138641 | 18 | 0 | 25 | 0.08 | 19 | 0 | 20 | 0.15 | 82 | 0.061 | 14 | 0.2143 | 19 | 0.1053 | 18 | 0.1111 | 20 | 0.1 | 71 | 0.1268 |
| 7_1141354 | NA | NA | NA | NA | 2 | 1 | NA | NA | 2 | 1 | 1 | 1 | NA | NA | 2 | 1 | NA | NA | 3 | 1 |
| 7_1141356 | NA | NA | NA | NA | NA | NA | 1 | 1 | 1 | 1 | 1 | 1 | NA | NA | 1 | 1 | NA | NA | 2 | 1 |
| 7_1141356 | NA | NA | NA | NA | 1 | 0 | 2 | 0.5 | 3 | 0.3333 | 1 | 0 | 1 | 1 | 1 | 1 | NA | NA | 3 | 0.6667 |
| 7_1141357 | NA | NA | NA | NA | 2 | 0 | 2 | 0.5 | 4 | 0.25 | 1 | 1 | 1 | 0 | 1 | 0 | NA | NA | 3 | 0.3333 |
| 7_1141358 | NA | NA | NA | NA | 2 | 0.5 | 2 | 0 | 4 | 0.25 | 1 | 0 | 1 | 0 | 2 | 0.5 | NA | NA | 4 | 0.25 |

| | | | | | | | | | | | | | | | | | | | | |
|---|---|---|---|---|---|---|---|---|---|---|---|---|---|---|---|---|---|---|---|---|
| 7_11413603 | NA | NA | NA | NA | 2 | 0.5 | 2 | 0.5 | 4 | 0.5 | NA | NA | 1 | 1 | 1 | 1 | NA | NA | 2 | 1 |
| 7_11413603 | NA | NA | NA | NA | 2 | 0 | 2 | 0.5 | 4 | 0.25 | 1 | 0 | 1 | 0 | 2 | 0.5 | NA | NA | 4 | 0.25 |
| 7_11414081 | NA | NA | NA | NA | NA | NA | 1 | 1 | 1 | 1 | NA | NA | NA | NA | 1 | 1 | NA | NA | 1 | 1 |
| 7_11414115 | NA | NA | NA | NA | NA | NA | 1 | 1 | 1 | 1 | NA | NA | NA | NA | 1 | 1 | NA | NA | 1 | 1 |
| 7_11414122 | NA | NA | NA | NA | NA | NA | 1 | 1 | 1 | 1 | NA | NA | NA | NA | 1 | 1 | NA | NA | 1 | 1 |
| 7_11414137 | NA | NA | NA | NA | NA | NA | 1 | 1 | 1 | 1 | NA | NA | NA | NA | 1 | 1 | NA | NA | 1 | 1 |
| 7_11414166 | NA | NA | NA | NA | NA | NA | 1 | 1 | 1 | 1 | NA | NA | NA | NA | 1 | 1 | NA | NA | 1 | 1 |
| 7_11414212 | NA | NA | NA | NA | NA | NA | 1 | 1 | 1 | 1 | NA | NA | NA | NA | 1 | 1 | NA | NA | 1 | 1 |
| 7_11414224 | NA | NA | NA | NA | NA | NA | 1 | 1 | 1 | 1 | NA | NA | NA | NA | 1 | 1 | NA | NA | 1 | 1 |
| 7_11507040 | NA | NA | 2 | 1 | NA | NA | 3 | 0.6667 | 5 | 0.8 | 1 | 1 | NA | NA | NA | NA | 3 | 0.6667 | 4 | 0.75 |
| 7_11507044 | NA | NA | 2 | 1 | NA | NA | 2 | 1 | 4 | 1 | NA | NA | NA | NA | 1 | 1 | 3 | 1 | 4 | 1 |
| 7_11507048 | NA | NA | 1 | 1 | NA | NA | 3 | 0.6667 | 4 | 0.75 | NA | NA | NA | NA | 1 | 1 | 3 | 1 | 4 | 1 |
| 7_11507058 | NA | NA | 2 | 1 | NA | NA | 3 | 1 | 5 | 1 | 1 | 1 | NA | NA | 1 | 1 | 4 | 1 | 6 | 1 |
| 7_11507089 | NA | NA | 2 | 0.5 | NA | NA | 4 | 0.75 | 6 | 0.6667 | 1 | 1 | NA | NA | NA | NA | 3 | 0.6667 | 4 | 0.75 |
| 7_11507102 | NA | NA | 2 | 0.5 | NA | NA | 4 | 1 | 6 | 0.8333 | 2 | 0 | NA | NA | NA | NA | 2 | 1 | 4 | 0.5 |
| 7_11507374 | NA | NA | NA | NA | NA | NA | 3 | 0.3333 | 3 | 0.3333 | 1 | 1 | NA | NA | NA | NA | 3 | 0.6667 | 4 | 0.75 |
| 7_11507569 | NA | NA | 1 | 1 | NA | NA | 2 | 0.5 | 3 | 0.6667 | 1 | 0 | NA | NA | NA | NA | 1 | 1 | 2 | 0.5 |
| 7_11507590 | NA | NA | 1 | 1 | NA | NA | 2 | 1 | 3 | 1 | 1 | 1 | NA | NA | NA | NA | 1 | 1 | 2 | 1 |
| 7_11507597 | NA | NA | 1 | 0 | NA | NA | 2 | 0.5 | 3 | 0.3333 | 1 | 0 | NA | NA | NA | NA | 1 | 1 | 2 | 0.5 |
| 7_11507600 | NA | NA | 1 | 0 | NA | NA | 2 | 0.5 | 3 | 0.3333 | 1 | 0 | NA | NA | NA | NA | 1 | 1 | 2 | 0.5 |
| 7_11507612 | NA | NA | NA | NA | NA | NA | 2 | 1 | 2 | 1 | 1 | 1 | NA | NA | NA | NA | 1 | 1 | 2 | 1 |
| 7_11507612 | NA | NA | 1 | 1 | NA | NA | 2 | 0.5 | 3 | 0.6667 | NA | NA | NA | NA | NA | NA | 1 | 1 | 1 | 1 |
| 7_11507619 | NA | NA | 1 | 1 | NA | NA | 2 | 1 | 3 | 1 | 1 | 1 | NA | NA | NA | NA | 1 | 1 | 2 | 1 |
| 7_11563880 | NA | NA | 1 | 1 | NA | NA | NA | NA | 1 | 1 | NA | NA | NA | NA | 1 | 1 | NA | NA | 1 | 1 |
| 7_11570373 | 20 | 0.35 | 21 | 0.4286 | 13 | 0.4615 | 17 | 0.1765 | 71 | 0.3521 | 11 | 0.5455 | 14 | 0.4286 | 17 | 0.3529 | 17 | 0.2941 | 59 | 0.3898 |
| 7_11570419 | 25 | 0.08 | 29 | 0.069 | 17 | 0.0588 | 20 | 0.05 | 91 | 0.0659 | 17 | 0.1176 | 20 | 0.05 | 18 | 0.1667 | 18 | 0.1111 | 73 | 0.1096 |
| 7_11572297 | 2 | 0.5 | 1 | 1 | NA | NA | 2 | 0 | 5 | 0.4 | NA | NA | 1 | 0 | 2 | 0 | 2 | 0.5 | 5 | 0.2 |
| 7_11572314 | 1 | 0 | 1 | 0 | NA | NA | 2 | 0.5 | 4 | 0.25 | NA | NA | 2 | 1 | 2 | 0.5 | 1 | 1 | 5 | 0.8 |
| 7_11572319 | 2 | 0.5 | 1 | 1 | NA | NA | 1 | 1 | 4 | 0.75 | NA | NA | 1 | 0 | 2 | 0.5 | 2 | 1 | 5 | 0.6 |
| 7_11572322 | 2 | 1 | 1 | 1 | NA | NA | 2 | 0 | 5 | 0.6 | 1 | 1 | NA | NA | 2 | 1 | 2 | 1 | 5 | 1 |
| 7_11572373 | 2 | 1 | 1 | 1 | NA | NA | 2 | 1 | 5 | 1 | NA | NA | NA | NA | 1 | 1 | 2 | 1 | 3 | 1 |
| 7_11573403 | NA | NA | 1 | 1 | NA | NA | 1 | 0 | 2 | 0.5 | 1 | 0 | NA | NA | 1 | 1 | 1 | 1 | 3 | 0.6667 |
| 7_11576336 | NA | NA | 1 | 1 | NA | NA | 2 | 0.5 | 3 | 0.6667 | 1 | 1 | NA | NA | 1 | 1 | NA | NA | 2 | 1 |
| 7_11576338 | NA | NA | 1 | 1 | NA | NA | 2 | 0 | 3 | 0.3333 | 1 | 1 | NA | NA | 1 | 0 | NA | NA | 2 | 0.5 |
| 7_11576353 | NA | NA | NA | NA | NA | NA | 2 | 0.5 | 2 | 0.5 | 1 | 1 | NA | NA | 1 | 1 | NA | NA | 2 | 1 |
| 7_11576361 | NA | NA | 1 | 1 | NA | NA | 2 | 0.5 | 3 | 0.6667 | 1 | 0 | NA | NA | 1 | 1 | NA | NA | 2 | 0.5 |
| 7_11593747 | 2 | 0 | 1 | 0 | NA | NA | 2 | 0.5 | 5 | 0.2 | 2 | 0.5 | 3 | 0.3333 | 2 | 0 | 2 | 0 | 9 | 0.2222 |
| 7_11593904 | NA | NA | 2 | 0.5 | NA | NA | NA | NA | 2 | 0.5 | 2 | 0 | 2 | 0 | 1 | 1 | 1 | 1 | 6 | 0.3333 |
| 7_11593916 | NA | NA | 3 | 0.6667 | NA | NA | NA | NA | 3 | 0.6667 | 2 | 0.5 | 1 | 0 | 1 | 0 | 1 | 1 | 5 | 0.4 |
| 7_11593920 | NA | NA | 3 | 1 | NA | NA | NA | NA | 3 | 1 | 2 | 1 | 2 | 1 | NA | NA | 1 | 1 | 5 | 1 |
| 7_11593931 | NA | NA | 3 | 0.3333 | NA | NA | NA | NA | 3 | 0.3333 | 2 | 0.5 | 2 | 1 | NA | NA | 1 | 1 | 5 | 0.8 |
| 7_11601690 | NA | NA | 1 | 1 | NA | NA | NA | NA | 1 | 1 | 1 | 0 | 1 | 1 | NA | NA | NA | NA | 2 | 0.5 |
| 7_11601702 | NA | NA | 1 | 1 | NA | NA | NA | NA | 1 | 1 | 1 | 1 | 1 | 1 | NA | NA | NA | NA | 2 | 1 |
| 7_11601718 | NA | NA | 1 | 1 | NA | NA | NA | NA | 1 | 1 | 1 | 1 | 1 | 1 | NA | NA | NA | NA | 2 | 1 |
| 7_11604329 | NA | NA | NA | NA | 2 | 0.5 | 1 | 1 | 3 | 0.6667 | 1 | 0 | 2 | 1 | NA | NA | NA | NA | 3 | 0.6667 |
| 7_11604334 | NA | NA | NA | NA | 2 | 1 | NA | NA | 2 | 1 | 1 | 0 | 2 | 1 | NA | NA | 1 | 0 | 4 | 0.5 |
| 7_11613980 | 1 | 0 | 4 | 1 | NA | NA | 2 | 1 | 7 | 0.8571 | NA | NA | 1 | 1 | 2 | 1 | NA | NA | 3 | 1 |
| 7_11614325 | NA | NA | 1 | 1 | NA | NA | NA | NA | 1 | 1 | NA | NA | 1 | 1 | NA | NA | NA | NA | 1 | 1 |
| 7_11642367 | 5 | 0.4 | 1 | 0 | 3 | 0 | 2 | 0.5 | 11 | 0.2727 | 2 | 0.5 | 2 | 0.5 | 4 | 0 | NA | NA | 8 | 0.25 |
| 7_11642367 | 7 | 0.1429 | 1 | 1 | 3 | 0 | 3 | 0 | 14 | 0.1429 | 1 | 0 | 1 | 0 | 5 | 0.4 | 1 | 0 | 8 | 0.25 |
| 7_11725587 | 1 | 0 | 2 | 1 | NA | NA | NA | NA | 3 | 0.6667 | 1 | 0 | NA | NA | NA | NA | 2 | 1 | 3 | 0.6667 |
| 7_11726668 | 1 | 1 | NA | NA | NA | NA | NA | NA | 1 | 1 | 1 | 0 | 1 | 1 | 1 | 1 | 2 | 1 | 5 | 0.8 |
| 7_11726670 | 1 | 0 | 2 | 0.5 | NA | NA | NA | NA | 3 | 0.3333 | 2 | 0 | 1 | 1 | 1 | 1 | 2 | 1 | 6 | 0.6667 |
| 7_11726670 | 1 | 1 | 1 | 1 | NA | NA | NA | NA | 2 | 1 | 2 | 1 | 1 | 1 | 1 | 1 | 2 | 1 | 6 | 1 |
| 7_11726671 | 1 | 1 | 2 | 0.5 | NA | NA | NA | NA | 3 | 0.6667 | 2 | 1 | 1 | 1 | 1 | 0 | 2 | 1 | 6 | 0.8333 |
| 7_11726686 | 1 | 1 | 2 | 0.5 | NA | NA | NA | NA | 3 | 0.6667 | 4 | 0.75 | NA | NA | 1 | 0 | 2 | 0.5 | 7 | 0.5714 |
| 7_11726692 | 1 | 1 | 2 | 1 | NA | NA | NA | NA | 3 | 1 | 2 | 0 | 1 | 0 | 2 | 0 | 2 | 0.5 | 7 | 0.1429 |
| 7_11726694 | 1 | 1 | 1 | 1 | NA | NA | NA | NA | 2 | 1 | 3 | 0.3333 | NA | NA | 1 | 1 | 1 | 0 | 5 | 0.4 |
| 7_11726714 | 1 | 1 | 2 | 0 | NA | NA | NA | NA | 3 | 0.3333 | 5 | 0.4 | 1 | 1 | 1 | 0 | 2 | 0.5 | 9 | 0.4444 |
| 7_11726758 | 1 | 0 | 3 | 0.3333 | NA | NA | NA | NA | 4 | 0.25 | 4 | 0.5 | 1 | 0 | 2 | 0 | 2 | 0.5 | 9 | 0.3333 |
| 7_11726786 | 1 | 0 | 2 | 1 | NA | NA | NA | NA | 3 | 0.6667 | 5 | 0.8 | 1 | 1 | 2 | 1 | 2 | 1 | 10 | 0.9 |
| 7_11726898 | 1 | 0 | 2 | 0.5 | NA | NA | 1 | 0 | 4 | 0.25 | 3 | 0 | 1 | 0 | 2 | 0.5 | NA | NA | 6 | 0.1667 |
| 7_11726906 | 1 | 0 | 4 | 0.25 | NA | NA | 1 | 0 | 6 | 0.1667 | 2 | 0 | 1 | 0 | 2 | 0.5 | 2 | 0 | 7 | 0.1429 |
| 7_11770733 | NA | NA | 1 | 0 | 1 | 0 | 3 | 0.3333 | 5 | 0.2 | NA | NA | 3 | 0 | NA | NA | 2 | 0.5 | 5 | 0.2 |
| 7_11770746 | NA | NA | 1 | 0 | 1 | 1 | 3 | 0.3333 | 5 | 0.4 | NA | NA | 3 | 0.3333 | NA | NA | 2 | 0 | 5 | 0.2 |

| | | | | | | | | | | | | | | | | | | | | |
|---|---|---|---|---|---|---|---|---|---|---|---|---|---|---|---|---|---|---|---|---|
| 7_11770829 | 29 | 0.3793 | 33 | 0.4242 | 27 | 0.2963 | 39 | 0.3846 | 128 | 0.375 | 33 | 0.4545 | 48 | 0.4583 | 41 | 0.3902 | 24 | 0.375 | 146 | 0.4247 |
| 7_11771845 | 44 | 0.0455 | 41 | 0.0732 | 28 | 0.0357 | 34 | 0.0882 | 147 | 0.0612 | 24 | 0.0417 | 33 | 0.1212 | 28 | 0 | 48 | 0.2083 | 133 | 0.1128 |
| 7_11771928 | 35 | 0.1429 | 38 | 0.1316 | 25 | 0.12 | 33 | 0.0909 | 131 | 0.1221 | 27 | 0.0741 | 31 | 0.129 | 29 | 0.069 | 45 | 0.0444 | 132 | 0.0758 |
| 7_11773382 | 3 | 0.6667 | 5 | 0.2 | 3 | 0.6667 | 4 | 0.25 | 15 | 0.4 | 1 | 1 | 2 | 0.5 | 2 | 0 | NA | NA | 5 | 0.4 |
| 7_11773388 | 3 | 0.6667 | 1 | 1 | 1 | 0 | 3 | 0.3333 | 8 | 0.5 | 1 | 1 | 2 | 0.5 | 4 | 0.25 | NA | NA | 7 | 0.4286 |
| 7_11773391 | 3 | 0.3333 | 5 | 0.6 | 2 | 0 | 4 | 0 | 14 | 0.2857 | 1 | 0 | 2 | 0 | 4 | 1 | NA | NA | 7 | 0.5714 |
| 7_11773403 | 1 | 1 | 3 | 1 | 2 | 1 | 3 | 1 | 9 | 1 | 1 | 1 | 3 | 0.6667 | 4 | 1 | NA | NA | 8 | 0.875 |
| 7_11773406 | 2 | 1 | 5 | 1 | 3 | 0.6667 | 1 | 1 | 11 | 0.9091 | 1 | 1 | 2 | 0.5 | 4 | 0.75 | NA | NA | 7 | 0.7143 |
| 7_11773417 | 2 | 1 | 3 | 1 | 3 | 0.3333 | 1 | 1 | 9 | 0.7778 | 1 | 1 | 2 | 0 | 3 | 0.6667 | NA | NA | 6 | 0.5 |
| 7_11773419 | 2 | 0 | 3 | 0.6667 | 3 | 0.3333 | 3 | 0.6667 | 11 | 0.4545 | 1 | 0 | 2 | 0.5 | 3 | 0.6667 | NA | NA | 6 | 0.5 |
| 7_11773422 | 2 | 1 | 5 | 1 | 1 | 1 | 2 | 1 | 10 | 1 | 1 | 1 | 2 | 1 | 5 | 1 | NA | NA | 8 | 1 |
| 7_11773430 | 2 | 1 | 3 | 1 | 2 | 1 | 2 | 1 | 9 | 1 | NA | NA | 3 | 1 | 4 | 1 | NA | NA | 7 | 1 |
| 7_11773434 | 1 | 0 | 3 | 0.6667 | 3 | 0.6667 | 3 | 0.6667 | 10 | 0.6 | NA | NA | 3 | 0.3333 | 5 | 0.4 | NA | NA | 8 | 0.375 |
| 7_11773442 | 2 | 1 | 4 | 0.5 | 3 | 1 | 1 | 1 | 10 | 0.8 | 1 | 0 | 3 | 0.6667 | 4 | 0.5 | NA | NA | 8 | 0.5 |
| 7_11773443 | 2 | 0 | 3 | 0.6667 | 2 | 0 | 3 | 0.3333 | 10 | 0.3 | 1 | 0 | 2 | 0.5 | 3 | 0.6667 | NA | NA | 6 | 0.5 |
| 7_11773449 | 2 | 0 | 2 | 0 | 3 | 0 | 3 | 0.3333 | 10 | 0.1 | 1 | 1 | 1 | 1 | 5 | 0 | NA | NA | 7 | 0.2857 |
| 7_11773459 | 2 | 0.5 | 4 | 0.5 | 3 | 0 | 3 | 0 | 12 | 0.25 | 1 | 1 | 3 | 0 | 5 | 0.6 | NA | NA | 9 | 0.4444 |
| 7_11773475 | 2 | 0 | 3 | 0 | 3 | 0 | 2 | 0.5 | 10 | 0.1 | 1 | 0 | 2 | 0 | 3 | 0.3333 | NA | NA | 6 | 0.1667 |
| 7_11773477 | 2 | 0 | 5 | 0.6 | 2 | 1 | 2 | 0 | 11 | 0.4545 | 1 | 0 | 2 | 0.5 | 5 | 0.8 | NA | NA | 8 | 0.625 |
| 7_11773971 | 2 | 0.5 | 3 | 0 | 1 | 0 | 2 | 0 | 8 | 0.125 | NA | NA | 3 | 0 | 3 | 0.3333 | NA | NA | 6 | 0.1667 |
| 7_11774478 | 1 | 0 | 3 | 0.6667 | NA | NA | 1 | 0 | 5 | 0.4 | NA | NA | 2 | 0.5 | 3 | 1 | NA | NA | 5 | 0.8 |
| 7_11774484 | NA | NA | 2 | 0.5 | NA | NA | 1 | 1 | 3 | 0.6667 | NA | NA | 2 | 0.5 | 3 | 0.6667 | NA | NA | 5 | 0.6 |
| 7_11774486 | 1 | 0 | 3 | 0.3333 | NA | NA | 1 | 1 | 5 | 0.4 | NA | NA | 2 | 0 | 3 | 0.6667 | NA | NA | 5 | 0.4 |
| 7_11774498 | 1 | 0 | 3 | 0 | NA | NA | 1 | 1 | 5 | 0.2 | NA | NA | 2 | 1 | 3 | 0 | NA | NA | 5 | 0.4 |
| 7_11798045 | 8 | 0.125 | 3 | 0 | 3 | 0.6667 | NA | NA | 14 | 0.2143 | 10 | 0.1 | 2 | 0 | NA | NA | 3 | 0.3333 | 15 | 0.1333 |
| 7_11798083 | 10 | 0.5 | 7 | 0.7143 | 6 | 0.6667 | NA | NA | 23 | 0.6087 | 8 | 0.625 | 2 | 1 | 1 | 1 | 7 | 0.7143 | 18 | 0.7222 |
| 7_11798088 | 12 | 0.1667 | 7 | 0.2857 | 6 | 0 | NA | NA | 25 | 0.16 | 9 | 0 | 2 | 0.5 | 1 | 0 | 8 | 0 | 20 | 0.05 |
| 7_11798114 | 6 | 0.3333 | 5 | 0.2 | 4 | 0 | NA | NA | 15 | 0.2 | 7 | 0 | 1 | 1 | 1 | 0 | 4 | 0.25 | 13 | 0.1538 |
| 7_11803310 | 10 | 0.8 | 5 | 1 | 5 | 1 | NA | NA | 20 | 0.9 | 3 | 0.6667 | 1 | 1 | NA | NA | 7 | 1 | 11 | 0.9091 |
| 7_11803316 | 8 | 0.125 | 3 | 0 | 4 | 0 | NA | NA | 15 | 0.0667 | 4 | 0 | 1 | 0 | 1 | 0 | 3 | 0.3333 | 9 | 0.1111 |
| 7_11803319 | 9 | 0.6667 | NA | NA | 4 | 0.25 | NA | NA | 13 | 0.5385 | 3 | 0.3333 | 1 | 1 | 1 | 1 | 6 | 0.8333 | 11 | 0.7273 |
| 7_11803328 | 9 | 0.1111 | 3 | 0 | 5 | 0 | NA | NA | 17 | 0.0588 | 3 | 0 | 1 | 0 | 1 | 0 | 7 | 0.1429 | 12 | 0.0833 |
| 7_11803333 | 10 | 0.2 | 3 | 0 | 5 | 0.6 | NA | NA | 18 | 0.2778 | 3 | 0.6667 | 1 | 0 | 1 | 0 | 6 | 0.3333 | 11 | 0.3636 |
| 7_11803337 | 9 | 0.5556 | 4 | 0.25 | 5 | 0.2 | NA | NA | 18 | 0.3889 | 4 | 0.5 | 1 | 0 | 1 | 0 | 6 | 0.3333 | 12 | 0.3333 |
| 7_11803348 | 9 | 0.4444 | 3 | 0.3333 | 4 | 0.5 | NA | NA | 16 | 0.4375 | 4 | 0 | 1 | 1 | 1 | 1 | 7 | 0.7143 | 13 | 0.5385 |
| 7_11803349 | 8 | 1 | 3 | 0.3333 | 3 | 1 | NA | NA | 14 | 0.8571 | 3 | 0.3333 | 1 | 0 | 1 | 1 | 5 | 1 | 10 | 0.7 |
| 7_11813728 | 1 | 0 | 2 | 0.5 | 2 | 1 | 1 | 0 | 6 | 0.5 | 1 | 1 | NA | NA | NA | NA | NA | NA | 1 | 1 |
| 7_11828781 | 36 | 0.1111 | 54 | 0.0741 | 45 | 0.0889 | 45 | 0.0444 | 180 | 0.0778 | 39 | 0.0256 | 43 | 0.0465 | 34 | 0.0882 | 40 | 0.1 | 156 | 0.0641 |
| 7_11828790 | 33 | 0.1515 | 53 | 0.2264 | 45 | 0.2222 | 43 | 0.2558 | 174 | 0.2184 | 35 | 0.1429 | 42 | 0.2381 | 33 | 0.2121 | 39 | 0.2051 | 149 | 0.2013 |
| 7_11828813 | 39 | 0.0513 | 55 | 0.1091 | 45 | 0.1111 | 47 | 0.0851 | 186 | 0.0914 | 34 | 0.1176 | 40 | 0.1 | 31 | 0.0645 | 40 | 0.05 | 145 | 0.0828 |
| 7_11828816 | 31 | 0.8065 | 45 | 0.8 | 34 | 0.7059 | 34 | 0.6765 | 144 | 0.75 | 27 | 0.8148 | 37 | 0.7027 | 25 | 0.8 | 29 | 0.8276 | 118 | 0.7797 |
| 7_11828817 | 38 | 0.7105 | 48 | 0.6667 | 42 | 0.6667 | 43 | 0.6744 | 171 | 0.6784 | 32 | 0.625 | 40 | 0.725 | 31 | 0.6452 | 37 | 0.6216 | 140 | 0.6571 |
| 7_11828824 | 37 | 0.2703 | 52 | 0.1538 | 37 | 0.1351 | 42 | 0.1905 | 168 | 0.1845 | 31 | 0.0645 | 39 | 0.1795 | 32 | 0.1875 | 35 | 0.2857 | 137 | 0.1825 |
| 7_11828832 | 36 | 0.6667 | 48 | 0.7083 | 34 | 0.5588 | 40 | 0.55 | 158 | 0.6266 | 25 | 0.6 | 41 | 0.7317 | 26 | 0.6538 | 34 | 0.7647 | 126 | 0.6984 |
| 7_11829088 | 29 | 0.069 | 39 | 0.1026 | 28 | 0.1071 | 33 | 0.0909 | 129 | 0.093 | 21 | 0.0952 | 32 | 0.0625 | 24 | 0.0417 | 29 | 0.069 | 106 | 0.066 |
| 7_11832080 | NA | NA | NA | NA | 1 | 1 | 1 | 0 | 2 | 0.5 | 1 | 1 | NA | NA | NA | NA | 1 | 0 | 2 | 0.5 |
| 7_11832185 | NA | NA | 1 | 0 | 2 | 0 | 2 | 0.5 | 5 | 0.2 | 3 | 0.3333 | 1 | 0 | NA | NA | 1 | 0 | 5 | 0.2 |
| 7_11832190 | NA | NA | 1 | 1 | NA | NA | 2 | 0.5 | 3 | 0.6667 | 3 | 0.3333 | 1 | 0 | NA | NA | 1 | 0 | 5 | 0.2 |
| 7_11832203 | NA | NA | NA | NA | 2 | 0.5 | 3 | 0.3333 | 5 | 0.4 | 2 | 1 | NA | NA | NA | NA | 2 | 0 | 4 | 0.5 |
| 7_11832210 | NA | NA | 3 | 0.3333 | 2 | 0 | 2 | 0 | 7 | 0.1429 | 3 | 0.3333 | 1 | 0 | NA | NA | 1 | 0 | 5 | 0.2 |
| 7_11833463 | NA | NA | 2 | 1 | 5 | 0.8 | 3 | 0 | 10 | 0.6 | 4 | 0.5 | 1 | 0 | 1 | 1 | 1 | 1 | 7 | 0.5714 |
| 7_11833469 | NA | NA | 4 | 0.25 | 5 | 0.6 | 3 | 0 | 12 | 0.3333 | 3 | 0.6667 | 2 | 0.5 | NA | NA | 1 | 0 | 6 | 0.5 |
| 7_11833573 | NA | NA | 7 | 0.1429 | 5 | 0 | 7 | 0 | 19 | 0.0526 | 5 | 0.2 | 3 | 0 | 2 | 0 | 3 | 0 | 13 | 0.0769 |
| 7_11833587 | 2 | 0 | 6 | 0 | 6 | 0.1667 | 7 | 0.2857 | 21 | 0.1429 | 5 | 0.4 | 3 | 0 | 2 | 0 | 3 | 0.3333 | 13 | 0.2308 |
| 7_11843991 | 11 | 0 | 5 | 0.4 | 6 | 0 | 6 | 0 | 28 | 0.0714 | 2 | 0 | 4 | 0.25 | 1 | 0 | 7 | 0.1429 | 14 | 0.1429 |
| 7_11844052 | 12 | 0.0833 | 6 | 0 | 7 | 0.1429 | 6 | 0 | 31 | 0.0645 | 4 | 0 | 4 | 0 | 1 | 0 | 8 | 0.125 | 17 | 0.0588 |
| 7_11844199 | NA | NA | 3 | 0 | NA | NA | 3 | 0.6667 | 6 | 0.3333 | 2 | 0 | 1 | 0 | 3 | 0.3333 | 3 | 0 | 9 | 0.1111 |
| 7_11844238 | NA | NA | 3 | 0.3333 | NA | NA | 3 | 0 | 6 | 0.1667 | 1 | 0 | 1 | 1 | 4 | 0 | 2 | 0 | 8 | 0.125 |
| 7_11844240 | NA | NA | 3 | 0 | NA | NA | 2 | 0.5 | 5 | 0.2 | 1 | 1 | NA | NA | 3 | 0 | 3 | 0 | 7 | 0.1429 |
| 7_11844241 | NA | NA | 3 | 1 | NA | NA | 2 | 1 | 5 | 1 | NA | NA | NA | NA | 3 | 1 | 1 | 1 | 4 | 1 |
| 7_11844242 | NA | NA | 3 | 0.3333 | NA | NA | 3 | 0.3333 | 6 | 0.3333 | 1 | 1 | 1 | 1 | 1 | 0 | 1 | 1 | 4 | 0.75 |
| 7_11844251 | NA | NA | 2 | 0.5 | NA | NA | 3 | 1 | 5 | 0.8 | 1 | 0 | 1 | 1 | 3 | 1 | 3 | 0.3333 | 8 | 0.625 |
| 7_11844265 | NA | NA | 2 | 0 | NA | NA | 2 | 1 | 4 | 0.5 | 1 | 1 | 1 | 1 | 3 | 0.3333 | 2 | 0 | 7 | 0.4286 |
| 7_11844330 | NA | NA | 1 | 0 | NA | NA | 3 | 0.6667 | 4 | 0.5 | 1 | 1 | NA | NA | 2 | 0.5 | 1 | 1 | 4 | 0.75 |
| 7_11844343 | NA | NA | 1 | 1 | NA | NA | 3 | 0.6667 | 4 | 0.75 | 1 | 1 | NA | NA | 3 | 0.6667 | 2 | 1 | 6 | 0.8333 |

| | | | | | | | | | | | | | | | | | | | | |
|---|---|---|---|---|---|---|---|---|---|---|---|---|---|---|---|---|---|---|---|---|
| 7_1184435( | NA | NA | 1 | 0 | NA | NA | 2 | 1 | 3 | 0.6667 | 1 | 0 | NA | NA | 3 | 0 | 2 | 0.5 | 6 | 0.1667 |
| 7_1184436 | NA | NA | NA | NA | NA | NA | 3 | 1 | 3 | 1 | NA | NA | NA | NA | 1 | 1 | 2 | 1 | 3 | 1 |
| 7_1184437( | NA | NA | NA | NA | NA | NA | 2 | 1 | 2 | 1 | NA | NA | NA | NA | 3 | 1 | 1 | 1 | 4 | 1 |
| 7_1184437! | NA | NA | NA | NA | NA | NA | 3 | 0.6667 | 3 | 0.6667 | 1 | 0 | NA | NA | 3 | 0.6667 | 2 | 0.5 | 6 | 0.5 |
| 7_1184439! | NA | NA | NA | NA | NA | NA | 3 | 0.3333 | 3 | 0.3333 | NA | NA | 1 | 0 | 3 | 0.3333 | 1 | 1 | 5 | 0.4 |
| 7_1184440 | NA | NA | NA | NA | NA | NA | 3 | 1 | 3 | 1 | NA | NA | 1 | 0 | 2 | 0.5 | 2 | 0.5 | 5 | 0.4 |
| 7_1184441' | NA | NA | NA | NA | NA | NA | 3 | 0.6667 | 3 | 0.6667 | NA | NA | 1 | 1 | 3 | 1 | 2 | 1 | 6 | 1 |
| 7_1184441! | NA | NA | NA | NA | NA | NA | 3 | 0.3333 | 3 | 0.3333 | NA | NA | NA | NA | 2 | 1 | 1 | 1 | 3 | 1 |
| 7_1184442( | NA | NA | NA | NA | NA | NA | 3 | 0.3333 | 3 | 0.3333 | NA | NA | 1 | 1 | 3 | 0.3333 | 1 | 1 | 5 | 0.6 |
| 7_1184443( | NA | NA | NA | NA | NA | NA | 3 | 1 | 3 | 1 | NA | NA | 1 | 1 | 3 | 1 | 2 | 1 | 6 | 1 |
| 7_1184444 | NA | NA | NA | NA | NA | NA | 2 | 1 | 2 | 1 | NA | NA | 1 | 1 | 2 | 1 | 2 | 1 | 5 | 1 |
| 7_1184444( | NA | NA | NA | NA | NA | NA | 2 | 1 | 2 | 1 | NA | NA | 1 | 1 | 3 | 0.6667 | 2 | 1 | 6 | 0.8333 |
| 7_1184445 | NA | NA | NA | NA | NA | NA | 2 | 0.5 | 2 | 0.5 | NA | NA | 1 | 0 | 2 | 0.5 | 2 | 0 | 5 | 0.2 |
| 7_1184448' | 2 | 0 | 1 | 1 | NA | NA | 2 | 0.5 | 5 | 0.4 | NA | NA | 4 | 0 | 3 | 0 | 2 | 0.5 | 9 | 0.1111 |
| 7_1184448 | 1 | 1 | NA | NA | NA | NA | 1 | 0 | 2 | 0.5 | NA | NA | 2 | 0.5 | NA | NA | 1 | 1 | 3 | 0.6667 |
| 7_1184449' | 1 | 0 | 1 | 1 | NA | NA | 1 | 0 | 3 | 0.3333 | NA | NA | 3 | 0.6667 | 3 | 1 | 1 | 1 | 7 | 0.8571 |
| 7_1184451 | 2 | 1 | 3 | 0.6667 | NA | NA | 1 | 1 | 6 | 0.8333 | 1 | 1 | 4 | 1 | 3 | 1 | 1 | 1 | 9 | 1 |
| 7_1184524! | 2 | 1 | NA | NA | NA | NA | 1 | 0 | 3 | 0.6667 | NA | NA | 2 | 0.5 | 2 | 0.5 | 1 | 1 | 5 | 0.6 |
| 7_1185716( | 9 | 0.4444 | 17 | 0.1176 | 7 | 0.2857 | 15 | 0.2 | 48 | 0.2292 | 10 | 0.1 | 14 | 0.1429 | 11 | 0.4545 | 7 | 0.2857 | 42 | 0.2381 |
| 7_1185716 | 8 | 0.25 | 12 | 0.3333 | 7 | 0.2857 | 14 | 0.2143 | 41 | 0.2683 | 9 | 0.2222 | 10 | 0.3 | 10 | 0.2 | 7 | 0 | 36 | 0.1944 |
| 7_1185717( | 5 | 0.4 | 16 | 0.5 | 7 | 0.1429 | 14 | 0.2143 | 42 | 0.3333 | 8 | 0 | 13 | 0.3077 | 10 | 0.6 | 4 | 0.75 | 35 | 0.3714 |
| 7_1185718 | 7 | 0.8571 | 13 | 0.9231 | 5 | 1 | 16 | 1 | 41 | 0.9512 | 11 | 0.6364 | 13 | 0.9231 | 8 | 0.75 | 6 | 1 | 38 | 0.8158 |
| 7_1185719 | 7 | 0.4286 | 16 | 0.1875 | 7 | 0.2857 | 12 | 0.0833 | 42 | 0.2143 | 11 | 0.1818 | 12 | 0.3333 | 10 | 0.3 | 7 | 0.2857 | 40 | 0.275 |
| 7_1185720( | 7 | 0.7143 | 15 | 0.7333 | 8 | 0.625 | 11 | 0.8182 | 41 | 0.7317 | 8 | 0.75 | 11 | 0.6364 | 9 | 0.8889 | 7 | 0.8571 | 35 | 0.7714 |
| 7_1185730 | 7 | 1 | 15 | 0.9333 | 10 | 0.9 | 20 | 0.8 | 52 | 0.8846 | 14 | 0.7857 | 10 | 1 | 17 | 0.8824 | 13 | 0.7692 | 54 | 0.8519 |
| 7_1185737! | 7 | 0.7143 | 12 | 0.75 | 10 | 0.6 | 16 | 0.6875 | 45 | 0.6889 | 14 | 0.8571 | 8 | 0.875 | 14 | 0.7143 | 14 | 0.5714 | 50 | 0.74 |
| 7_1185738( | 6 | 0.6667 | 11 | 0.3636 | 10 | 0.6 | 17 | 0.2941 | 44 | 0.4318 | 14 | 0.4286 | 9 | 0.5556 | 16 | 0.4375 | 12 | 0.4167 | 51 | 0.451 |
| 7_1185739 | 3 | 0.6667 | 10 | 0.7 | 8 | 0.875 | 15 | 0.6667 | 36 | 0.7222 | 10 | 0.7 | 7 | 0.7143 | 13 | 0.9231 | 10 | 0.8 | 40 | 0.8 |
| 7_1185740 | 4 | 0 | 11 | 0.1818 | 8 | 0.125 | 16 | 0.0625 | 39 | 0.1026 | 12 | 0.0833 | 7 | 0.4286 | 14 | 0.2857 | 9 | 0.1111 | 42 | 0.2143 |
| 7_1185740( | 6 | 0.1667 | 13 | 0.2308 | 6 | 0.1667 | 14 | 0.1429 | 39 | 0.1795 | 13 | 0.1538 | 5 | 0.2 | 10 | 0 | 9 | 0.3333 | 37 | 0.1622 |
| 7_1185740 | 7 | 0 | 12 | 0.0833 | 8 | 0.25 | 16 | 0.125 | 43 | 0.1163 | 14 | 0.0714 | 7 | 0.1429 | 13 | 0.0769 | 11 | 0 | 45 | 0.0667 |
| 7_1185741' | 5 | 0.2 | 11 | 0.6364 | 10 | 0.5 | 14 | 0.2857 | 40 | 0.425 | 11 | 0.1818 | 6 | 0.3333 | 10 | 0.1 | 9 | 0.3333 | 36 | 0.2222 |
| 7_1185741 | 4 | 0.25 | 12 | 0.1667 | 10 | 0.2 | 16 | 0 | 42 | 0.119 | 10 | 0.1 | 5 | 0.2 | 10 | 0 | 11 | 0 | 36 | 0.0556 |
| 7_1185742' | 1 | 1 | 6 | 0.5 | 7 | 0.7143 | 13 | 0.5385 | 27 | 0.5926 | 10 | 0.9 | 6 | 0.6667 | 12 | 0.9167 | 9 | 0.7778 | 37 | 0.8378 |
| 7_1185742 | 5 | 0.2 | 11 | 0 | 9 | 0.2222 | 16 | 0.0625 | 41 | 0.0976 | 14 | 0.0714 | 8 | 0.125 | 13 | 0.0769 | 10 | 0.1 | 45 | 0.0889 |
| 7_1185742 | 6 | 0.1667 | 9 | 0.2222 | 10 | 0 | 17 | 0.2353 | 42 | 0.1667 | 14 | 0.0714 | 6 | 0.3333 | 13 | 0.2308 | 10 | 0.3 | 43 | 0.2093 |
| 7_1185828 | 6 | 0 | 7 | 0.2857 | 6 | 0 | 10 | 0 | 29 | 0.069 | 11 | 0.0909 | 6 | 0 | 9 | 0.1111 | 3 | 0.6667 | 29 | 0.1379 |
| 7_1185833( | 8 | 0 | 9 | 0.1111 | 8 | 0.125 | 11 | 0.0909 | 36 | 0.0833 | 10 | 0.1 | 7 | 0 | 10 | 0.1 | 5 | 0 | 32 | 0.0625 |
| 7_1185833' | 7 | 0.8571 | 10 | 0.5 | 8 | 0.875 | 12 | 1 | 37 | 0.8108 | 9 | 1 | 6 | 0.8333 | 10 | 0.9 | 5 | 0.6 | 30 | 0.8667 |
| 7_1185835' | 7 | 0.2857 | 7 | 0.1429 | 5 | 0 | 10 | 0 | 29 | 0.1034 | 8 | 0.125 | 5 | 0 | 8 | 0.25 | 3 | 0.6667 | 24 | 0.2083 |
| 7_1185835( | 1 | 1 | 2 | 1 | 3 | 1 | 6 | 0.6667 | 12 | 0.8333 | 6 | 0.6667 | 3 | 0.6667 | 6 | 0.8333 | 4 | 1 | 19 | 0.7895 |
| 7_1185836' | 6 | 0.8333 | 7 | 0.7143 | 5 | 0.2 | 11 | 0.6364 | 29 | 0.6207 | 7 | 0.7143 | 4 | 1 | 6 | 0.5 | 3 | 0.3333 | 20 | 0.65 |
| 7_1185836 | 6 | 0.3333 | 5 | 0.4 | 5 | 0 | 9 | 0.1111 | 25 | 0.2 | 6 | 0.5 | 3 | 0.3333 | 7 | 0.4286 | 5 | 0 | 21 | 0.3333 |
| 7_1185837 | 5 | 0 | 8 | 0.5 | 7 | 0.1429 | 11 | 0.4545 | 31 | 0.3226 | 7 | 0.4286 | 3 | 0.6667 | 8 | 0.5 | 4 | 0.5 | 22 | 0.5 |
| 7_1185837! | 6 | 0.6667 | 7 | 0.4286 | 5 | 0.6 | 8 | 0.625 | 26 | 0.5769 | 8 | 0.75 | 3 | 1 | 7 | 0.7143 | 4 | 0.5 | 22 | 0.7273 |
| 7_1187402 | NA | NA | NA | NA | 1 | 1 | NA | NA | 1 | 1 | 1 | 0 | 1 | 1 | NA | NA | NA | NA | 2 | 0.5 |
| 7_1187404' | NA | NA | NA | NA | 1 | 1 | NA | NA | 1 | 1 | 1 | 0 | 1 | 1 | NA | NA | NA | NA | 2 | 0.5 |
| 7_1187405 | NA | NA | NA | NA | 1 | 1 | NA | NA | 1 | 1 | 1 | 1 | NA | NA | NA | NA | NA | NA | 1 | 1 |
| 7_1193143 | 10 | 0.2 | 7 | 0.1429 | 6 | 0 | 7 | 0 | 30 | 0.1 | 10 | 0.1 | 8 | 0 | 6 | 0.1667 | 6 | 0 | 30 | 0.0667 |
| 7_1193146 | 10 | 0.2 | 6 | 0.5 | 4 | 0.25 | 8 | 0.25 | 28 | 0.2857 | 9 | 0.2222 | 8 | 0.125 | 6 | 0.3333 | 7 | 0.4286 | 30 | 0.2667 |
| 7_1193154' | 8 | 0 | 4 | 0 | 2 | 0 | 5 | 0.2 | 19 | 0.0526 | 7 | 0 | 4 | 0 | 5 | 0.2 | 3 | 0 | 19 | 0.0526 |
| 7_1193156' | 5 | 0.6 | 2 | 0.5 | 2 | 0.5 | 2 | 0 | 11 | 0.4545 | 4 | 0 | 4 | 0 | 4 | 0.5 | 2 | 1 | 14 | 0.2857 |
| 7_1193156 | 5 | 0.2 | 3 | 0.3333 | NA | NA | 1 | 0 | 9 | 0.2222 | 3 | 0.3333 | 3 | 0 | 4 | 0.5 | 1 | 1 | 11 | 0.3636 |
| 7_1193156 | 6 | 0.3333 | 3 | 0.6667 | NA | NA | 2 | 1 | 11 | 0.5455 | 4 | 0.5 | 4 | 0.5 | 4 | 0.25 | 2 | 1 | 14 | 0.5 |
| 7_1193160 | 7 | 0.1429 | 3 | 0.3333 | 2 | 0 | 3 | 0 | 15 | 0.1333 | 4 | 0.25 | 3 | 0.3333 | 5 | 0.4 | 1 | 0 | 13 | 0.3077 |
| 7_1193161 | 6 | 1 | 2 | 0.5 | 2 | 1 | 4 | 1 | 14 | 0.9286 | 4 | 0.75 | 3 | 1 | 4 | 0.75 | 1 | 1 | 12 | 0.8333 |
| 7_1193173' | 7 | 0.7143 | 1 | 0 | 2 | 0.5 | 2 | 0.5 | 12 | 0.5833 | 4 | 0.5 | 2 | 0 | 4 | 0.75 | 1 | 1 | 11 | 0.5455 |
| 7_1193173 | 6 | 0.1667 | 1 | 0 | 2 | 0 | 2 | 0 | 11 | 0.0909 | 5 | 0 | 3 | 0 | 5 | 0.2 | 1 | 0 | 14 | 0.0714 |
| 7_1193173! | 7 | 0.1429 | 1 | 0 | 2 | 0 | 3 | 0 | 13 | 0.0769 | 3 | 0 | 3 | 0 | 4 | 0 | 1 | 1 | 11 | 0.0909 |
| 7_1193394( | 3 | 0.3333 | 2 | 0 | 1 | 0 | 7 | 0.1429 | 13 | 0.1538 | 5 | 0.2 | 7 | 0.2857 | 5 | 0 | 6 | 0.3333 | 23 | 0.2174 |
| 7_1193395' | 3 | 0.3333 | 3 | 0.3333 | 2 | 0 | 6 | 0 | 14 | 0.1429 | 5 | 0.2 | 6 | 0.3333 | 5 | 0 | 8 | 0 | 24 | 0.125 |
| 7_1193396 | 2 | 1 | 4 | 0.25 | 1 | 1 | 4 | 0.25 | 11 | 0.4545 | 4 | 0.25 | 5 | 0.2 | 4 | 0.75 | 6 | 0.3333 | 19 | 0.3684 |
| 7_1193396 | 2 | 0 | 3 | 0.3333 | 1 | 1 | 4 | 0.25 | 10 | 0.3 | 4 | 0.25 | 7 | 0.1429 | 5 | 0.2 | 5 | 0 | 21 | 0.1429 |
| 7_1193397( | 3 | 0.3333 | 6 | 0 | 3 | 0 | 7 | 0 | 19 | 0.0526 | 4 | 0 | 6 | 0.1667 | 4 | 0 | 8 | 0.125 | 22 | 0.0909 |
| 7_1193398' | 3 | 0.3333 | 6 | 0.1667 | 2 | 0.5 | 5 | 0.2 | 16 | 0.25 | 5 | 0.2 | 7 | 0.1429 | 5 | 0 | 8 | 0.25 | 25 | 0.16 |

| | | | | | | | | | | | | | | | | | | | | |
|---|---|---|---|---|---|---|---|---|---|---|---|---|---|---|---|---|---|---|---|---|
| 7_11934034 | 2 | 1 | 6 | 1 | 4 | 1 | 5 | 0.6 | 17 | 0.8824 | 3 | 1 | 5 | 1 | 8 | 0.875 | 10 | 0.8 | 26 | 0.8846 |
| 7_11934038 | 2 | 0 | 5 | 0 | 3 | 0.3333 | 4 | 0.5 | 14 | 0.2143 | 5 | 0.4 | 5 | 0.6 | 6 | 0 | 9 | 0.2222 | 25 | 0.28 |
| 7_11934051 | 3 | 0 | 6 | 0.1667 | 6 | 0.1667 | 6 | 0 | 21 | 0.0952 | 5 | 0 | 5 | 0.2 | 7 | 0 | 11 | 0.2727 | 28 | 0.1429 |
| 7_11934077 | 3 | 0.3333 | 6 | 0 | 6 | 0.1667 | 7 | 0.1429 | 22 | 0.1364 | 2 | 0.5 | 9 | 0.3333 | 7 | 0.1429 | 8 | 0.5 | 26 | 0.3462 |
| 7_11934095 | 2 | 1 | 2 | 0.5 | 2 | 1 | 1 | 1 | 7 | 0.8571 | 2 | 0.5 | 2 | 0.5 | 2 | 0 | 5 | 0.2 | 11 | 0.2727 |
| 7_11934132 | 10 | 0 | 11 | 0.1818 | 8 | 0 | 9 | 0.1111 | 38 | 0.0789 | 9 | 0.1111 | 14 | 0.2857 | 10 | 0 | 8 | 0 | 41 | 0.122 |
| 7_11938311 | 3 | 0 | NA | NA | 5 | 0.6 | 2 | 0.5 | 10 | 0.4 | NA | NA | NA | NA | NA | NA | 3 | 0.6667 | 3 | 0.6667 |
| 7_11938412 | 2 | 0 | NA | NA | 4 | 0.25 | 2 | 0 | 8 | 0.125 | NA | NA | NA | NA | NA | NA | 4 | 0.25 | 4 | 0.25 |
| 7_11938420 | NA | NA | NA | NA | 4 | 0.75 | 1 | 1 | 5 | 0.8 | NA | NA | 1 | 0 | NA | NA | 1 | 1 | 2 | 0.5 |
| 7_11938429 | 3 | 1 | NA | NA | 3 | 0.6667 | 2 | 1 | 8 | 0.875 | NA | NA | NA | NA | NA | NA | 1 | 1 | 1 | 1 |
| 7_11938439 | 3 | 1 | NA | NA | 5 | 1 | 2 | 1 | 10 | 1 | NA | NA | 1 | 0 | NA | NA | 4 | 1 | 5 | 0.8 |
| 7_11938441 | 3 | 0.3333 | NA | NA | 4 | 0 | 1 | 1 | 8 | 0.25 | NA | NA | 1 | 0 | NA | NA | 2 | 0.5 | 3 | 0.3333 |
| 7_11938442 | 2 | 1 | NA | NA | 3 | 1 | NA | NA | 5 | 1 | NA | NA | 1 | 1 | NA | NA | 2 | 0.5 | 3 | 0.6667 |
| 7_11938443 | 2 | 0.5 | NA | NA | 5 | 0.4 | 2 | 1 | 9 | 0.5556 | NA | NA | NA | NA | NA | NA | 3 | 0.6667 | 3 | 0.6667 |
| 7_11938445 | 3 | 0 | NA | NA | 5 | 0 | 2 | 1 | 10 | 0.2 | NA | NA | 1 | 0 | NA | NA | 4 | 0.25 | 5 | 0.2 |
| 7_11938449 | 2 | 0.5 | NA | NA | 5 | 0 | 1 | 0 | 8 | 0.125 | NA | NA | 1 | 0 | NA | NA | 4 | 0.25 | 5 | 0.2 |
| 7_11938456 | 3 | 0.6667 | NA | NA | 4 | 1 | 1 | 1 | 8 | 0.875 | NA | NA | 1 | 1 | NA | NA | 3 | 1 | 4 | 1 |
| 7_11938458 | 3 | 0.3333 | NA | NA | 3 | 0.6667 | 2 | 0.5 | 8 | 0.5 | NA | NA | NA | NA | NA | NA | 3 | 0.3333 | 3 | 0.3333 |
| 7_11938458 | 2 | 0.5 | NA | NA | 2 | 0 | 2 | 0.5 | 6 | 0.3333 | NA | NA | NA | NA | NA | NA | 2 | 0.5 | 2 | 0.5 |
| 7_11938462 | 3 | 0 | NA | NA | 4 | 0.5 | 2 | 0.5 | 9 | 0.3333 | NA | NA | NA | NA | NA | NA | 2 | 1 | 2 | 1 |
| 7_11938474 | 3 | 0.6667 | NA | NA | 4 | 0 | 2 | 1 | 9 | 0.4444 | NA | NA | 1 | 0 | NA | NA | 4 | 0.75 | 5 | 0.6 |
| 7_11938481 | 3 | 0.3333 | NA | NA | 2 | 0 | 2 | 0 | 7 | 0.1429 | NA | NA | 1 | 0 | NA | NA | 3 | 0.3333 | 4 | 0.25 |
| 7_11938523 | 1 | 1 | NA | NA | 3 | 1 | 2 | 1 | 6 | 1 | NA | NA | 1 | 1 | NA | NA | 3 | 0.6667 | 4 | 0.75 |
| 7_11938530 | 1 | 1 | NA | NA | 3 | 0.6667 | 2 | 1 | 6 | 0.8333 | NA | NA | 1 | 1 | NA | NA | 4 | 1 | 5 | 1 |
| 7_11938548 | 2 | 0 | NA | NA | 3 | 0.3333 | 2 | 0 | 7 | 0.1429 | NA | NA | NA | NA | NA | NA | 4 | 0.25 | 4 | 0.25 |
| 7_11945673 | 1 | 1 | 3 | 1 | 1 | 1 | 2 | 1 | 7 | 1 | 2 | 1 | NA | NA | 1 | 1 | 1 | 1 | 4 | 1 |
| 7_11945680 | 1 | 0 | 3 | 0.6667 | 1 | 1 | 1 | 0 | 6 | 0.5 | 2 | 0.5 | NA | NA | NA | NA | 1 | 1 | 3 | 0.6667 |
| 7_11945698 | NA | NA | 3 | 0.6667 | 1 | 1 | NA | NA | 4 | 0.75 | 1 | 0 | NA | NA | 1 | 1 | NA | NA | 2 | 0.5 |
| 7_11945698 | NA | NA | 3 | 1 | 1 | 1 | 1 | 0 | 5 | 0.8 | 2 | 1 | NA | NA | NA | NA | 1 | 1 | 3 | 1 |
| 7_11945704 | NA | NA | 3 | 1 | 1 | 1 | 1 | 1 | 5 | 1 | 2 | 1 | NA | NA | 1 | 1 | 1 | 1 | 4 | 1 |
| 7_11945706 | NA | NA | 1 | 1 | 1 | 1 | NA | NA | 2 | 1 | 1 | 1 | NA | NA | NA | NA | NA | NA | 1 | 1 |
| 7_11945713 | NA | NA | 3 | 1 | 1 | 1 | NA | NA | 4 | 1 | 2 | 0.5 | NA | NA | 1 | 1 | NA | NA | 3 | 0.6667 |
| 7_11945875 | NA | NA | 3 | 0.3333 | 1 | 0 | NA | NA | 4 | 0.25 | 2 | 0.5 | NA | NA | 1 | 1 | 1 | 1 | 4 | 0.75 |
| 7_11945886 | NA | NA | 3 | 1 | 1 | 1 | NA | NA | 4 | 1 | 2 | 0.5 | NA | NA | NA | NA | 1 | 1 | 3 | 0.6667 |
| 7_11951548 | 1 | 1 | 3 | 0.6667 | 4 | 1 | NA | NA | 8 | 0.875 | 3 | 1 | 2 | 1 | 2 | 1 | 2 | 1 | 9 | 1 |
| 7_11951555 | 1 | 1 | 2 | 0.5 | 4 | 0 | NA | NA | 7 | 0.2857 | 3 | 0 | 1 | 0 | 2 | 0 | 2 | 0.5 | 8 | 0.125 |
| 7_11951556 | 1 | 0 | 1 | 0 | 4 | 0.25 | NA | NA | 6 | 0.1667 | 2 | 0 | 2 | 0.5 | 2 | 0 | 1 | 0 | 7 | 0.1429 |
| 7_11951566 | 1 | 1 | 2 | 1 | 4 | 0.75 | NA | NA | 7 | 0.8571 | 2 | 0.5 | 2 | 1 | 2 | 1 | 1 | 0 | 7 | 0.7143 |
| 7_11951569 | 1 | 1 | 3 | 1 | 3 | 1 | NA | NA | 7 | 1 | 3 | 1 | 2 | 1 | 2 | 1 | 1 | 1 | 8 | 1 |
| 7_11951588 | 1 | 0 | 3 | 0 | 4 | 0.25 | NA | NA | 8 | 0.125 | 3 | 0 | 2 | 0 | 3 | 0.3333 | 2 | 0 | 10 | 0.1 |
| 7_11951596 | 1 | 0 | 3 | 0 | 3 | 0.6667 | NA | NA | 7 | 0.2857 | 3 | 0.3333 | 2 | 0 | 4 | 0 | 2 | 0.5 | 11 | 0.1818 |
| 7_11955090 | 1 | 1 | NA | NA | NA | NA | NA | NA | 1 | 1 | NA | NA | NA | NA | NA | NA | 1 | 1 | 1 | 1 |
| 7_11955110 | 1 | 1 | NA | NA | NA | NA | NA | NA | 1 | 1 | NA | NA | NA | NA | NA | NA | 1 | 1 | 1 | 1 |
| 7_11955120 | 1 | 1 | NA | NA | NA | NA | NA | NA | 1 | 1 | NA | NA | NA | NA | NA | NA | 1 | 1 | 1 | 1 |
| 7_11971956 | 1 | 0 | 2 | 0.5 | 1 | 0 | 1 | 1 | 5 | 0.4 | NA | NA | 1 | 0 | 1 | 1 | 1 | 0 | 3 | 0.3333 |
| 7_12027423 | NA | NA | NA | NA | 1 | 1 | NA | NA | 1 | 1 | NA | NA | 1 | 1 | NA | NA | NA | NA | 1 | 1 |
| 7_12027427 | NA | NA | NA | NA | 1 | 1 | NA | NA | 1 | 1 | NA | NA | 1 | 1 | 1 | 0 | NA | NA | 2 | 0.5 |
| 7_12032547 | 1 | 0 | NA | NA | 4 | 0.75 | 1 | 1 | 6 | 0.6667 | 2 | 1 | 1 | 1 | 4 | 1 | NA | NA | 7 | 1 |
| 7_12032578 | NA | NA | NA | NA | 2 | 0 | 2 | 0.5 | 4 | 0.25 | 3 | 0.3333 | 1 | 0 | 2 | 0 | 1 | 0 | 7 | 0.1429 |
| 7_12032984 | 1 | 0 | 1 | 0 | 2 | 0.5 | 1 | 0 | 5 | 0.2 | NA | NA | 1 | 1 | 3 | 0.6667 | NA | NA | 4 | 0.75 |
| 7_12033020 | 2 | 0 | 1 | 0 | 2 | 0.5 | NA | NA | 5 | 0.2 | NA | NA | 1 | 0 | 3 | 0 | 1 | 1 | 5 | 0.2 |
| 7_12033155 | 4 | 0 | 8 | 0 | 2 | 0 | 5 | 0.2 | 19 | 0.0526 | NA | NA | 3 | 0.3333 | 5 | 0.2 | 4 | 0 | 12 | 0.1667 |
| 7_12033174 | 6 | 0.1667 | 7 | 0 | 3 | 0.3333 | 6 | 0.5 | 22 | 0.2273 | NA | NA | 2 | 0.5 | 7 | 0.2857 | 5 | 0 | 14 | 0.2143 |
| 7_12033183 | 6 | 0 | 6 | 0.1667 | 3 | 0.3333 | 5 | 0 | 20 | 0.1 | NA | NA | 3 | 0 | 5 | 0.2 | 5 | 0 | 13 | 0.0769 |
| 7_12033184 | 5 | 1 | 8 | 1 | 2 | 1 | 3 | 1 | 18 | 1 | NA | NA | 2 | 1 | 6 | 1 | 5 | 0.8 | 13 | 0.9231 |
| 7_12033233 | 9 | 0.2222 | 9 | 0.3333 | 3 | 0.3333 | 6 | 0.5 | 27 | 0.3333 | 1 | 0 | 2 | 0 | 7 | 0.4286 | 4 | 1 | 14 | 0.5 |
| 7_12033237 | 9 | 0 | 9 | 0.1111 | 3 | 0 | 7 | 0.1429 | 28 | 0.0714 | 1 | 0 | 4 | 0.25 | 6 | 0.1667 | 4 | 0 | 15 | 0.1333 |
| 7_12033260 | 12 | 0.1667 | 10 | 0 | 3 | 0 | 8 | 0 | 33 | 0.0606 | 1 | 0 | 3 | 0 | 7 | 0.2857 | 6 | 0 | 17 | 0.1176 |
| 7_12033304 | 12 | 0 | 10 | 0.1 | 3 | 0 | 7 | 0.2857 | 32 | 0.0938 | 1 | 0 | 6 | 0.1667 | 9 | 0.1111 | 6 | 0 | 22 | 0.0909 |
| 7_12050266 | 6 | 0.5 | 3 | 1 | 3 | 0.3333 | 1 | 0 | 13 | 0.5385 | NA | NA | 1 | 1 | 3 | 0 | NA | NA | 4 | 0.25 |
| 7_12050295 | 6 | 0.1667 | 3 | 1 | 5 | 0.4 | 1 | 0 | 15 | 0.4 | 1 | 0 | 3 | 0 | 3 | 0.3333 | NA | NA | 7 | 0.1429 |
| 7_12050500 | 1 | 1 | 2 | 0.5 | 1 | 0 | NA | NA | 4 | 0.5 | NA | NA | NA | NA | 1 | 1 | NA | NA | 1 | 1 |
| 7_12053973 | 1 | 1 | 2 | 0.5 | 2 | 1 | 4 | 0.75 | 9 | 0.7778 | 1 | 1 | 4 | 0.75 | 1 | 1 | 1 | 1 | 7 | 0.8571 |
| 7_12053981 | 1 | 0 | 3 | 0.3333 | 1 | 0 | 3 | 0.6667 | 8 | 0.375 | 1 | 1 | 2 | 0 | NA | NA | 1 | 1 | 4 | 0.5 |
| 7_12053982 | 1 | 1 | 3 | 1 | 1 | 1 | 4 | 0.75 | 9 | 0.8889 | 1 | 1 | 4 | 0.75 | 2 | 1 | 1 | 1 | 8 | 0.875 |

| | | | | | | | | | | | | | | | | | | | | |
|---|---|---|---|---|---|---|---|---|---|---|---|---|---|---|---|---|---|---|---|---|
| 7_12053986 | 1 | 0 | 3 | 0.3333 | 1 | 1 | 3 | 0 | 8 | 0.25 | NA | NA | 3 | 1 | 2 | 1 | 1 | 1 | 6 | 1 |
| 7_12053989 | NA | NA | 3 | 0.3333 | 1 | 1 | 4 | 0.75 | 8 | 0.625 | 1 | 0 | 4 | 0.75 | 3 | 0.6667 | 1 | 1 | 9 | 0.6667 |
| 7_12053992 | 1 | 0 | 4 | 0.25 | 2 | 0 | 6 | 0 | 13 | 0.0769 | NA | NA | 4 | 0 | 1 | 1 | 3 | 0.3333 | 8 | 0.25 |
| 7_12054470 | 1 | 1 | 4 | 0.5 | 1 | 1 | 6 | 0.8333 | 12 | 0.75 | 2 | 1 | 4 | 1 | 3 | 0.6667 | 3 | 0.6667 | 12 | 0.8333 |
| 7_12054474 | 1 | 1 | 3 | 0.6667 | 1 | 1 | 5 | 0.8 | 10 | 0.8 | 2 | 0 | 4 | 1 | 2 | 0.5 | 2 | 1 | 10 | 0.7 |
| 7_12054476 | 2 | 0 | 5 | 0 | 1 | 1 | 6 | 0.3333 | 14 | 0.2143 | 2 | 0 | 4 | 0 | 2 | 0.5 | 3 | 0.3333 | 11 | 0.1818 |
| 7_12054482 | 2 | 0 | 5 | 0.4 | 2 | 0 | 7 | 0.1429 | 16 | 0.1875 | 2 | 0 | 5 | 0 | 2 | 0 | 3 | 0.3333 | 12 | 0.0833 |
| 7_12054489 | 2 | 0.5 | 5 | 1 | 2 | 1 | 7 | 0.8571 | 16 | 0.875 | 3 | 1 | 5 | 1 | 3 | 1 | 1 | 1 | 12 | 1 |
| 7_12054495 | 2 | 0 | 5 | 0.6 | 2 | 0.5 | 7 | 0.4286 | 16 | 0.4375 | 4 | 0.25 | 4 | 0 | 3 | 0.3333 | 3 | 0 | 14 | 0.1429 |
| 7_12054520 | NA | NA | 4 | 0.75 | 1 | 0 | 3 | 0.3333 | 8 | 0.5 | 3 | 0 | 4 | 0.5 | 2 | 0 | 2 | 0 | 11 | 0.1818 |
| 7_12055735 | 1 | 1 | 3 | 0.6667 | 1 | 1 | NA | NA | 5 | 0.8 | 5 | 0.8 | 2 | 0 | NA | NA | NA | NA | 7 | 0.5714 |
| 7_12055736 | 1 | 0 | 3 | 0.6667 | 1 | 0 | NA | NA | 5 | 0.4 | 5 | 0.4 | 2 | 0 | NA | NA | NA | NA | 7 | 0.2857 |
| 7_12055741 | 1 | 0 | 3 | 0.6667 | 1 | 1 | NA | NA | 5 | 0.6 | 4 | 0.75 | 2 | 0 | NA | NA | NA | NA | 6 | 0.5 |
| 7_12055747 | 1 | 1 | 3 | 0.6667 | 1 | 1 | NA | NA | 5 | 0.8 | 5 | 1 | 2 | 1 | NA | NA | NA | NA | 7 | 1 |
| 7_12055754 | 1 | 1 | 1 | 1 | 1 | 1 | NA | NA | 3 | 1 | 4 | 0.75 | 2 | 0.5 | NA | NA | NA | NA | 6 | 0.6667 |
| 7_12055755 | 1 | 0 | 1 | 0 | 1 | 1 | NA | NA | 3 | 0.3333 | 4 | 0.25 | 2 | 0 | NA | NA | NA | NA | 6 | 0.1667 |
| 7_12063403 | 13 | 0.1538 | 13 | 0.0769 | 14 | 0.2143 | 6 | 0 | 46 | 0.1304 | 8 | 0.125 | 17 | 0.1765 | 15 | 0.2 | 11 | 0.0909 | 51 | 0.1569 |
| 7_12065136 | 15 | 0.2 | 15 | 0.2 | 16 | 0.0625 | 9 | 0 | 55 | 0.1273 | 14 | 0.3571 | 16 | 0.1875 | 19 | 0.1579 | 14 | 0.0714 | 63 | 0.1905 |
| 7_12065138 | 13 | 0.1538 | 16 | 0.25 | 16 | 0 | 9 | 0.1111 | 54 | 0.1296 | 14 | 0 | 17 | 0.1765 | 16 | 0.1875 | 14 | 0 | 61 | 0.0984 |
| 7_12067547 | 16 | 0.375 | 15 | 0.3333 | 16 | 0.5 | 10 | 0.4 | 57 | 0.4035 | 12 | 0.4167 | 13 | 0.5385 | 19 | 0.2105 | 12 | 0.5 | 56 | 0.3929 |
| 7_12067551 | 19 | 0.3158 | 18 | 0.0556 | 18 | 0.2222 | 11 | 0.0909 | 66 | 0.1818 | 13 | 0.2308 | 18 | 0.2222 | 17 | 0.2353 | 17 | 0.2353 | 65 | 0.2308 |
| 7_12067554 | 19 | 0.2105 | 19 | 0 | 19 | 0.0526 | 12 | 0 | 69 | 0.0725 | 13 | 0.0769 | 20 | 0 | 21 | 0 | 15 | 0.2 | 69 | 0.058 |
| 7_12067564 | 19 | 0.4211 | 18 | 0.2778 | 17 | 0.1765 | 12 | 0.25 | 66 | 0.2879 | 12 | 0 | 20 | 0.25 | 20 | 0.35 | 16 | 0.375 | 68 | 0.2647 |
| 7_12067565 | 19 | 0.8421 | 17 | 0.8235 | 16 | 0.875 | 10 | 0.7 | 62 | 0.8226 | 11 | 1 | 17 | 0.8824 | 18 | 0.8889 | 16 | 0.875 | 62 | 0.9032 |
| 7_12067585 | 22 | 0.3182 | 16 | 0.4375 | 18 | 0.3889 | 12 | 0.0833 | 68 | 0.3235 | 14 | 0.3571 | 20 | 0.15 | 19 | 0.2105 | 17 | 0.2941 | 70 | 0.2429 |
| 7_12067592 | 18 | 0.1111 | 15 | 0.0667 | 18 | 0.0556 | 13 | 0 | 64 | 0.0625 | 13 | 0.1538 | 20 | 0.2 | 20 | 0.05 | 17 | 0.3529 | 70 | 0.1857 |
| 7_12099185 | 5 | 0.2 | 1 | 0 | 6 | 0 | 2 | 0 | 14 | 0.0714 | 1 | 1 | 2 | 0 | NA | NA | 2 | 0.5 | 5 | 0.4 |
| 7_12099233 | 5 | 0.2 | 4 | 0 | 5 | 0.4 | 2 | 0 | 16 | 0.1875 | 3 | 0.3333 | 3 | 0 | 1 | 0 | 4 | 0 | 11 | 0.0909 |
| 7_12099477 | NA | NA | 1 | 1 | NA | NA | NA | NA | 1 | 1 | 1 | 1 | NA | NA | NA | NA | NA | NA | 1 | 1 |
| 7_12099495 | NA | NA | 1 | 1 | NA | NA | NA | NA | 1 | 1 | 1 | 1 | NA | NA | NA | NA | NA | NA | 1 | 1 |
| 7_12099534 | NA | NA | 1 | 1 | 1 | 0 | NA | NA | 2 | 0.5 | 1 | 1 | NA | NA | NA | NA | NA | NA | 1 | 1 |
| 7_12099545 | NA | NA | 1 | 1 | 1 | 1 | NA | NA | 2 | 1 | 1 | 1 | NA | NA | NA | NA | NA | NA | 1 | 1 |
| 7_12099548 | NA | NA | 1 | 1 | 1 | 1 | NA | NA | 2 | 1 | 1 | 1 | NA | NA | NA | NA | NA | NA | 1 | 1 |
| 7_12109978 | 6 | 0 | 5 | 0.2 | 5 | 0 | 3 | 0 | 19 | 0.0526 | 4 | 0 | 3 | 0.3333 | 1 | 0 | 5 | 0.2 | 13 | 0.1538 |
| 7_12109979 | 6 | 0.1667 | 5 | 0 | 5 | 0.2 | 3 | 0 | 19 | 0.1053 | 4 | 0.25 | 3 | 0 | 1 | 0 | 5 | 0.2 | 13 | 0.1538 |
| 7_12109980 | 6 | 0.6667 | 5 | 0.4 | 4 | 0.5 | 3 | 0.3333 | 18 | 0.5 | 3 | 0.3333 | 3 | 0 | 1 | 0 | 5 | 0.4 | 12 | 0.25 |
| 7_12109982 | 6 | 0.3333 | 5 | 0.4 | 5 | 0 | 3 | 0.3333 | 19 | 0.2632 | 4 | 0.5 | 1 | 1 | 1 | 0 | 3 | 0.3333 | 9 | 0.4444 |
| 7_12110002 | 7 | 0.8571 | 5 | 1 | 5 | 1 | 2 | 1 | 19 | 0.9474 | 5 | 1 | 3 | 1 | 1 | 0 | 5 | 1 | 14 | 0.9286 |
| 7_12110015 | 5 | 0.4 | 5 | 1 | 5 | 0.8 | 2 | 1 | 17 | 0.7647 | 6 | 0.6667 | 3 | 0 | NA | NA | 7 | 0.2857 | 16 | 0.375 |
| 7_12110018 | 8 | 0.125 | 4 | 0 | 5 | 0 | 3 | 0 | 20 | 0.05 | 5 | 0 | 3 | 0.3333 | 2 | 0 | 8 | 0 | 18 | 0.0556 |
| 7_12110020 | 8 | 0.875 | 4 | 1 | 5 | 1 | 3 | 1 | 20 | 0.95 | 5 | 1 | 3 | 0.6667 | 2 | 1 | 6 | 0.8333 | 16 | 0.875 |
| 7_12110023 | 6 | 1 | 4 | 1 | 2 | 1 | 2 | 1 | 14 | 1 | 5 | 1 | 3 | 1 | 1 | 1 | 5 | 1 | 14 | 1 |
| 7_12110024 | 8 | 0.875 | 5 | 1 | 5 | 0.8 | 3 | 1 | 21 | 0.9048 | 6 | 1 | 3 | 1 | 2 | 1 | 5 | 1 | 16 | 1 |
| 7_12110035 | 6 | 0 | 4 | 0 | 5 | 0 | 3 | 0.3333 | 18 | 0.0556 | 5 | 0 | 3 | 0.3333 | 2 | 0 | 7 | 0 | 17 | 0.0588 |
| 7_12110038 | 7 | 0.7143 | 5 | 0.6 | 3 | 1 | 2 | 1 | 17 | 0.7647 | 6 | 0.5 | 2 | 1 | 2 | 0.5 | 6 | 0.6667 | 16 | 0.625 |
| 7_12110040 | 9 | 1 | 5 | 1 | 5 | 1 | 3 | 1 | 22 | 1 | 5 | 1 | 3 | 1 | 2 | 1 | 7 | 1 | 17 | 1 |
| 7_12110057 | 8 | 0.875 | 6 | 0.5 | 3 | 1 | 2 | 1 | 19 | 0.7895 | 5 | 0.6 | 3 | 0.6667 | 1 | 1 | 5 | 1 | 14 | 0.7857 |
| 7_12110076 | 8 | 1 | 7 | 1 | 8 | 0.875 | 4 | 1 | 27 | 0.963 | 6 | 1 | 3 | 1 | 2 | 1 | 9 | 1 | 20 | 1 |
| 7_12110079 | 8 | 0.5 | 7 | 0.7143 | 4 | 0.5 | 2 | 0.5 | 21 | 0.5714 | 5 | 0.8 | 3 | 0.6667 | 2 | 0.5 | 8 | 0.875 | 18 | 0.7778 |
| 7_12110085 | 9 | 0.6667 | 7 | 0.8571 | 6 | 1 | 4 | 0.75 | 26 | 0.8077 | 6 | 0.5 | 3 | 0.3333 | 2 | 0.5 | 6 | 0.8333 | 17 | 0.5882 |
| 7_12110087 | 10 | 0.5 | 10 | 0.6 | 9 | 0.2222 | 5 | 0.8 | 34 | 0.5 | 8 | 0.625 | 3 | 0.6667 | 2 | 1 | 7 | 0.8571 | 20 | 0.75 |
| 7_12124185 | 6 | 0.1667 | 5 | 0.4 | 6 | 0.3333 | 6 | 0.3333 | 23 | 0.3043 | 5 | 0 | 6 | 0.3333 | 5 | 0 | 7 | 0.1429 | 23 | 0.1304 |
| 7_12124266 | 3 | 0.3333 | 4 | 0 | 6 | 0 | 4 | 0 | 17 | 0.0588 | 4 | 0 | 5 | 0.2 | 4 | 0 | 5 | 0 | 18 | 0.0556 |
| 7_12124275 | 4 | 0.75 | 5 | 0.2 | 5 | 0 | 4 | 0 | 18 | 0.2222 | 4 | 0.25 | 5 | 0.2 | 4 | 0 | 6 | 0.3333 | 19 | 0.2105 |
| 7_12124278 | 3 | 0.3333 | 6 | 0 | 6 | 0 | 5 | 0.4 | 20 | 0.15 | 4 | 0 | 5 | 0.2 | 3 | 0.3333 | 6 | 0.1667 | 18 | 0.1667 |
| 7_12124284 | 3 | 0.6667 | 6 | 0.5 | 5 | 0.2 | 3 | 0 | 17 | 0.3529 | 4 | 0.25 | 4 | 0.25 | 2 | 0.5 | 6 | 0.3333 | 16 | 0.3125 |
| 7_12124323 | 4 | 0 | 4 | 0.25 | 4 | 0 | 3 | 0.3333 | 15 | 0.1333 | 4 | 0 | 1 | 0 | 4 | 0.25 | 5 | 0 | 14 | 0.0714 |
| 7_12124399 | 3 | 0 | 2 | 0 | 3 | 0 | 2 | 0.5 | 10 | 0.1 | 2 | 0.5 | 1 | 0 | 3 | 0 | 5 | 0 | 11 | 0.0909 |
| 7_12124423 | 2 | 0.5 | 3 | 0.6667 | 3 | 0.6667 | 2 | 1 | 10 | 0.7 | 2 | 0.5 | 1 | 0 | 2 | 1 | 5 | 0.8 | 10 | 0.7 |
| 7_12124692 | 3 | 0 | 3 | 0.3333 | 3 | 0.3333 | 2 | 0 | 11 | 0.1818 | NA | NA | 3 | 0.3333 | 2 | 0 | 4 | 0.5 | 9 | 0.3333 |
| 7_12124709 | 2 | 0.5 | 3 | 1 | 3 | 1 | 3 | 0.6667 | 11 | 0.8182 | NA | NA | 1 | 0 | 3 | 0.6667 | 4 | 0.5 | 8 | 0.5 |
| 7_12124710 | 3 | 0 | 3 | 0 | 3 | 0.3333 | 3 | 0 | 12 | 0.0833 | NA | NA | 1 | 0 | 3 | 0.6667 | 4 | 0.5 | 8 | 0.5 |
| 7_12124740 | 3 | 1 | 2 | 0 | 2 | 1 | 3 | 0 | 10 | 0.5 | NA | NA | 2 | 1 | 2 | 0.5 | 3 | 0.6667 | 7 | 0.7143 |
| 7_12124743 | 4 | 0.25 | 2 | 0 | 2 | 0 | 3 | 0.3333 | 11 | 0.1818 | NA | NA | 2 | 0 | 2 | 0 | 4 | 0.5 | 8 | 0.25 |
| 7_12124756 | 3 | 0 | 3 | 0.6667 | 2 | 0 | 1 | 0 | 9 | 0.2222 | NA | NA | 1 | 0 | 1 | 0 | 4 | 0.5 | 6 | 0.3333 |

| | | | | | | | | | | | | | | | | | | | | |
|---|---|---|---|---|---|---|---|---|---|---|---|---|---|---|---|---|---|---|---|---|
| 7_12124757 | 4 | 1 | 3 | 1 | 2 | 1 | 3 | 1 | 12 | 1 | NA | NA | 1 | 1 | 2 | 1 | 4 | 1 | 7 | 1 |
| 7_12124758 | 4 | 0.5 | 2 | 0.5 | 1 | 1 | 2 | 0.5 | 9 | 0.5556 | NA | NA | 2 | 0 | 1 | 0 | 2 | 0.5 | 5 | 0.2 |
| 7_12124764 | 4 | 0.25 | 1 | 0 | 2 | 0.5 | 3 | 0.6667 | 10 | 0.4 | NA | NA | 1 | 0 | 1 | 0 | 3 | 0.3333 | 5 | 0.2 |
| 7_12124766 | 3 | 1 | 3 | 0.3333 | 2 | 1 | 3 | 0.6667 | 11 | 0.7273 | NA | NA | 2 | 0.5 | 1 | 0 | 4 | 0.75 | 7 | 0.5714 |
| 7_12124775 | 4 | 0.25 | 3 | 0.3333 | 2 | 0.5 | 3 | 0 | 12 | 0.25 | NA | NA | 2 | 1 | 2 | 0.5 | 4 | 0.5 | 8 | 0.625 |
| 7_12124778 | 4 | 1 | 3 | 1 | 2 | 1 | 3 | 1 | 12 | 1 | NA | NA | 2 | 1 | 2 | 1 | 4 | 0.75 | 8 | 0.875 |
| 7_12124780 | 2 | 1 | 2 | 1 | 1 | 1 | 2 | 1 | 7 | 1 | NA | NA | 2 | 0 | 2 | 0.5 | 3 | 1 | 7 | 0.5714 |
| 7_12124789 | 2 | 0.5 | 3 | 0.3333 | 1 | 0 | 3 | 0.3333 | 9 | 0.3333 | NA | NA | 1 | 0 | 2 | 0.5 | 2 | 0.5 | 5 | 0.4 |
| 7_12124798 | 4 | 0.25 | 3 | 0.3333 | 1 | 0 | 3 | 0 | 11 | 0.1818 | NA | NA | 2 | 0 | 1 | 0 | 4 | 0.25 | 7 | 0.1429 |
| 7_12124802 | 4 | 1 | 2 | 0.5 | 1 | 1 | 3 | 1 | 10 | 0.9 | NA | NA | 2 | 1 | 2 | 1 | 3 | 1 | 7 | 1 |
| 7_12124848 | 2 | 0.5 | 2 | 1 | NA | NA | 3 | 0.6667 | 7 | 0.7143 | NA | NA | 1 | 1 | 2 | 1 | 2 | 0.5 | 5 | 0.8 |
| 7_12124857 | 2 | 0 | 3 | 0.3333 | NA | NA | 3 | 0.3333 | 8 | 0.25 | NA | NA | 2 | 0 | 2 | 0 | 2 | 0.5 | 6 | 0.1667 |
| 7_12124858 | 2 | 1 | 2 | 1 | NA | NA | 2 | 1 | 6 | 1 | NA | NA | 1 | 0 | 2 | 1 | 2 | 0.5 | 5 | 0.6 |
| 7_12124860 | 2 | 0.5 | 2 | 1 | NA | NA | 3 | 1 | 7 | 0.8571 | NA | NA | 2 | 1 | 2 | 0.5 | 1 | 1 | 5 | 0.8 |
| 7_12124868 | 1 | 1 | 2 | 1 | NA | NA | 1 | 1 | 4 | 1 | NA | NA | 2 | 0.5 | NA | NA | NA | NA | 2 | 0.5 |
| 7_12124880 | 1 | 1 | 2 | 1 | NA | NA | 3 | 1 | 6 | 1 | NA | NA | 1 | 0 | 2 | 1 | 2 | 1 | 5 | 0.8 |
| 7_12152219 | 9 | 0.3333 | 6 | 0 | 7 | 0.1429 | 11 | 0.0909 | 33 | 0.1515 | 5 | 0.2 | 9 | 0.4444 | 8 | 0.25 | 4 | 0.5 | 26 | 0.3462 |
| 7_12152219 | 7 | 0.1429 | 5 | 0 | 5 | 0.4 | 11 | 0.0909 | 28 | 0.1429 | 3 | 0 | 7 | 0.4286 | 6 | 0 | 3 | 0.3333 | 19 | 0.2105 |
| 7_12152248 | 9 | 0.2222 | 5 | 0.4 | 7 | 0 | 10 | 0.1 | 31 | 0.1613 | 5 | 0 | 8 | 0.25 | 7 | 0 | 4 | 0 | 24 | 0.0833 |
| 7_12152248 | 10 | 0.7 | 6 | 1 | 7 | 0.5714 | 9 | 0.7778 | 32 | 0.75 | 5 | 1 | 7 | 1 | 7 | 0.7143 | 3 | 0.6667 | 22 | 0.8636 |
| 7_12158076 | 5 | 0.2 | 2 | 1 | 2 | 0 | 5 | 0.4 | 14 | 0.3571 | NA | NA | 4 | 0.25 | 2 | 0.5 | 2 | 0 | 8 | 0.25 |
| 7_12158080 | 5 | 0.2 | 2 | 0.5 | 3 | 0 | 7 | 0.2857 | 17 | 0.2353 | 1 | 1 | 5 | 0 | 1 | 0 | 2 | 0 | 9 | 0.1111 |
| 7_12158885 | 8 | 0.5 | 1 | 0 | 4 | 0.75 | 4 | 0.5 | 17 | 0.5294 | NA | NA | 1 | 1 | 9 | 0.6667 | 4 | 0.5 | 14 | 0.6429 |
| 7_12163380 | NA | NA | NA | NA | 1 | 1 | NA | NA | 1 | 1 | NA | NA | 3 | 0.3333 | NA | NA | NA | NA | 3 | 0.3333 |
| 7_12163805 | NA | NA | NA | NA | 1 | 1 | 1 | 1 | 2 | 1 | NA | NA | 1 | 1 | NA | NA | 1 | 1 | 2 | 1 |
| 7_12163809 | NA | NA | NA | NA | 1 | 1 | 1 | 0 | 2 | 0.5 | NA | NA | 1 | 0 | NA | NA | 1 | 1 | 2 | 0.5 |
| 7_12163820 | NA | NA | NA | NA | 1 | 0 | 1 | 1 | 2 | 0.5 | NA | NA | 1 | 1 | NA | NA | 1 | 1 | 2 | 1 |
| 7_12164741 | NA | NA | NA | NA | 1 | 1 | 1 | 1 | 2 | 1 | NA | NA | 1 | 1 | NA | NA | NA | NA | 1 | 1 |
| 7_12166804 | 3 | 0 | 6 | 0 | 3 | 0.3333 | 2 | 0.5 | 14 | 0.1429 | NA | NA | 4 | 0.5 | 4 | 0 | 3 | 0.6667 | 11 | 0.3636 |
| 7_12167775 | 4 | 0.75 | 7 | 0.5714 | 2 | 0.5 | 4 | 0.25 | 17 | 0.5294 | 1 | 0 | 6 | 0.3333 | 4 | 0.5 | 3 | 0.6667 | 14 | 0.4286 |
| 7_12167781 | 3 | 0 | 7 | 0.2857 | 3 | 0 | 3 | 0 | 16 | 0.125 | 1 | 0 | 5 | 0.2 | 3 | 0.6667 | 4 | 0.25 | 13 | 0.3077 |
| 7_12167782 | 4 | 0.25 | 6 | 0.6667 | 4 | 0.25 | 2 | 1 | 16 | 0.5 | 1 | 1 | 6 | 0.8333 | 3 | 0.6667 | 4 | 0.75 | 14 | 0.7857 |
| 7_12167787 | 4 | 1 | 7 | 1 | 4 | 1 | 3 | 0.6667 | 18 | 0.9444 | 1 | 1 | 6 | 1 | 4 | 1 | 4 | 1 | 15 | 1 |
| 7_12167792 | 4 | 0.5 | 6 | 0.1667 | 3 | 0 | 3 | 0 | 16 | 0.1875 | 1 | 0 | 6 | 0 | 4 | 0.25 | 3 | 0 | 14 | 0.0714 |
| 7_12167797 | 4 | 0.25 | 6 | 0.5 | 3 | 0 | 3 | 0.6667 | 16 | 0.375 | 1 | 0 | 5 | 0.2 | 4 | 0.5 | 4 | 0.25 | 14 | 0.2857 |
| 7_12167905 | 3 | 0 | 8 | 0.125 | 4 | 0 | 5 | 0 | 20 | 0.05 | 1 | 0 | 6 | 0 | 4 | 0.25 | 6 | 0 | 17 | 0.0588 |
| 7_12167922 | 2 | 0 | 5 | 0.2 | 4 | 0 | 3 | 0 | 14 | 0.0714 | 1 | 0 | 6 | 0.1667 | 5 | 0.4 | 2 | 0 | 14 | 0.2143 |
| 7_12174740 | 13 | 1 | 6 | 0.5 | 7 | 0.5714 | 7 | 0.8571 | 33 | 0.7879 | 7 | 0.7143 | 10 | 0.8 | 7 | 0.7143 | 14 | 1 | 38 | 0.8421 |
| 7_12174742 | 13 | 0.0769 | 10 | 0.2 | 10 | 0.1 | 15 | 0 | 48 | 0.0833 | 10 | 0 | 13 | 0.1538 | 10 | 0.1 | 16 | 0.0625 | 49 | 0.0816 |
| 7_12174744 | 13 | 0.3846 | 10 | 0 | 11 | 0.1818 | 15 | 0.1333 | 49 | 0.1837 | 10 | 0.1 | 13 | 0.2308 | 10 | 0.2 | 16 | 0.0625 | 49 | 0.1429 |
| 7_12174760 | 17 | 0.2353 | 15 | 0.2667 | 11 | 0.2727 | 14 | 0.0714 | 57 | 0.2105 | 9 | 0.5556 | 15 | 0.4 | 9 | 0.6667 | 16 | 0.25 | 49 | 0.4286 |
| 7_12174769 | 15 | 0.0667 | 14 | 0.1429 | 14 | 0.2143 | 12 | 0 | 55 | 0.1091 | 8 | 0 | 15 | 0.2667 | 10 | 0 | 20 | 0 | 53 | 0.0755 |
| 7_12174771 | 16 | 0.5 | 15 | 0.2667 | 14 | 0.7143 | 14 | 0.5 | 59 | 0.4915 | 10 | 0.4 | 16 | 0.5625 | 10 | 0.5 | 20 | 0.5 | 56 | 0.5 |
| 7_12174778 | 16 | 0.3125 | 12 | 0.0833 | 15 | 0.2667 | 12 | 0.25 | 55 | 0.2364 | 9 | 0.3333 | 13 | 0.4615 | 10 | 0.4 | 18 | 0.3333 | 50 | 0.38 |
| 7_12174797 | 15 | 0.4 | 12 | 0.3333 | 14 | 0.5 | 14 | 0.2857 | 55 | 0.3818 | 9 | 0.2222 | 15 | 0.4667 | 8 | 0.5 | 19 | 0.5263 | 51 | 0.451 |
| 7_12174805 | 14 | 0.1429 | 14 | 0.1429 | 15 | 0 | 13 | 0.1538 | 56 | 0.1071 | 6 | 0 | 15 | 0.1333 | 11 | 0 | 20 | 0.2 | 52 | 0.1154 |
| 7_12222700 | 11 | 0.2727 | 7 | 0.1429 | 7 | 0.1429 | 12 | 0.3333 | 37 | 0.2432 | 5 | 0 | 13 | 0.1538 | 11 | 0.1818 | 9 | 0.2222 | 38 | 0.1579 |
| 7_12222802 | 18 | 0.1111 | 15 | 0.1333 | 13 | 0.0769 | 18 | 0 | 64 | 0.0781 | 8 | 0.125 | 12 | 0 | 19 | 0 | 12 | 0.1667 | 51 | 0.0588 |
| 7_12223280 | 55 | 0.1273 | 49 | 0.1429 | 55 | 0.1091 | 52 | 0.1154 | 211 | 0.1232 | 33 | 0.0303 | 45 | 0.1778 | 53 | 0.1509 | 45 | 0.1333 | 176 | 0.1307 |
| 7_12223302 | 63 | 0.127 | 48 | 0.0833 | 51 | 0.3529 | 49 | 0.0816 | 211 | 0.1611 | 36 | 0.1667 | 40 | 0.075 | 52 | 0.1538 | 50 | 0.22 | 178 | 0.1573 |
| 7_12223304 | 69 | 0.0725 | 56 | 0.0893 | 64 | 0.0312 | 55 | 0.0364 | 244 | 0.0574 | 40 | 0.075 | 51 | 0.098 | 65 | 0.0923 | 56 | 0.0714 | 212 | 0.0849 |
| 7_12223305 | 68 | 0.3235 | 51 | 0.4118 | 65 | 0.3692 | 59 | 0.2881 | 243 | 0.3457 | 39 | 0.4103 | 53 | 0.3962 | 67 | 0.3134 | 59 | 0.4237 | 218 | 0.3807 |
| 7_12223306 | 61 | 0.1639 | 55 | 0.0909 | 64 | 0.2031 | 55 | 0.2364 | 235 | 0.1745 | 39 | 0.2564 | 44 | 0.0909 | 56 | 0.125 | 57 | 0.2632 | 196 | 0.1837 |
| 7_12227080 | 6 | 0.1667 | 11 | 0.4545 | 8 | 0.5 | 6 | 0.5 | 31 | 0.4194 | 5 | 0 | 9 | 0.4444 | 8 | 0.25 | 12 | 0.3333 | 34 | 0.2941 |
| 7_12227081 | 6 | 0.1667 | 10 | 0.1 | 9 | 0.3333 | 8 | 0.125 | 33 | 0.1818 | 4 | 0 | 8 | 0.25 | 9 | 0 | 11 | 0.0909 | 32 | 0.0938 |
| 7_12227106 | 8 | 0 | 16 | 0.0625 | 11 | 0.1818 | 7 | 0 | 42 | 0.0714 | 4 | 0 | 9 | 0.1111 | 9 | 0.1111 | 13 | 0 | 35 | 0.0571 |
| 7_12227108 | 8 | 0.75 | 15 | 0.7333 | 11 | 1 | 7 | 1 | 41 | 0.8537 | 5 | 0.8 | 11 | 1 | 10 | 1 | 14 | 0.9286 | 40 | 0.95 |
| 7_12227112 | 8 | 0.125 | 17 | 0.4706 | 11 | 0.4545 | 9 | 0.5556 | 45 | 0.4222 | 4 | 0.75 | 11 | 0.6364 | 10 | 0.4 | 14 | 0.7857 | 39 | 0.641 |
| 7_12236762 | 9 | 0.7778 | 22 | 0.7727 | 10 | 0.4 | 10 | 0.8 | 51 | 0.7059 | 7 | 0.5714 | 14 | 0.6429 | 14 | 0.7143 | 17 | 0.8824 | 52 | 0.7308 |
| 7_12236785 | 11 | 0.1818 | 24 | 0.0833 | 12 | 0 | 10 | 0 | 57 | 0.0702 | 8 | 0 | 13 | 0.0769 | 15 | 0.1333 | 16 | 0 | 52 | 0.0577 |
| 7_12236793 | 11 | 0.3636 | 24 | 0.0417 | 11 | 0 | 10 | 0 | 56 | 0.0893 | 7 | 0 | 12 | 0 | 15 | 0.2667 | 17 | 0 | 51 | 0.0784 |
| 7_12236795 | 9 | 0 | 24 | 0.3333 | 13 | 0.2308 | 9 | 0.3333 | 55 | 0.2545 | 8 | 0.25 | 14 | 0.3571 | 15 | 0.3333 | 16 | 0.3125 | 53 | 0.3208 |
| 7_12236797 | 9 | 0.3333 | 25 | 0.52 | 13 | 0.4615 | 10 | 0.1 | 57 | 0.4035 | 8 | 0.25 | 11 | 0.6364 | 15 | 0.5333 | 17 | 0.4118 | 51 | 0.4706 |
| 7_12236819 | 11 | 0.9091 | 24 | 0.9583 | 15 | 1 | 9 | 0.8889 | 59 | 0.9492 | 8 | 0.875 | 12 | 1 | 15 | 1 | 17 | 1 | 52 | 0.9808 |
| 7_12236834 | 10 | 0.1 | 23 | 0.087 | 14 | 0.0714 | 8 | 0 | 55 | 0.0727 | 8 | 0 | 10 | 0.1 | 14 | 0.0714 | 8 | 0.25 | 40 | 0.1 |

| | | | | | | | | | | | | | | | | | | | | |
|---|---|---|---|---|---|---|---|---|---|---|---|---|---|---|---|---|---|---|---|---|
| 7_12236837 | 11 | 0.8182 | 26 | 0.8077 | 15 | 0.9333 | 12 | 0.8333 | 64 | 0.8438 | 10 | 1 | 14 | 0.9286 | 16 | 0.875 | 18 | 0.9444 | 58 | 0.931 |
| 7_12236846 | 10 | 0.1 | 22 | 0 | 13 | 0.1538 | 12 | 0.1667 | 57 | 0.0877 | 6 | 0 | 13 | 0 | 12 | 0.0833 | 12 | 0.25 | 43 | 0.093 |
| 7_12256970 | 3 | 1 | 6 | 1 | 5 | 1 | 5 | 1 | 19 | 1 | 1 | 1 | 5 | 1 | 4 | 0.75 | 1 | 1 | 11 | 0.9091 |
| 7_12256977 | 3 | 1 | 8 | 1 | 6 | 1 | 4 | 0.75 | 21 | 0.9524 | 1 | 1 | 6 | 1 | 8 | 1 | 2 | 1 | 17 | 1 |
| 7_12256980 | 1 | 1 | 6 | 1 | 3 | 1 | 6 | 0.8333 | 16 | 0.9375 | 2 | 1 | 3 | 1 | 4 | 0.75 | 2 | 1 | 11 | 0.9091 |
| 7_12256982 | 3 | 1 | 10 | 1 | 6 | 1 | 7 | 1 | 26 | 1 | 2 | 1 | 7 | 1 | 7 | 1 | 1 | 1 | 17 | 1 |
| 7_12256984 | 3 | 1 | 9 | 1 | 5 | 1 | 5 | 0.8 | 22 | 0.9545 | 2 | 1 | 7 | 1 | 7 | 1 | 1 | 1 | 17 | 1 |
| 7_12256985 | 3 | 0.3333 | 9 | 0.5556 | 4 | 0.5 | 7 | 0.5714 | 23 | 0.5217 | 1 | 1 | 6 | 0.8333 | 7 | 0.7143 | 2 | 1 | 16 | 0.8125 |
| 7_12256990 | 3 | 0 | 9 | 0.1111 | 6 | 0.5 | 7 | 0.1429 | 25 | 0.2 | 2 | 0.5 | 5 | 0.2 | 8 | 0.25 | 2 | 1 | 17 | 0.3529 |
| 7_12256991 | 3 | 0 | 9 | 0.1111 | 6 | 0.1667 | 7 | 0.1429 | 25 | 0.12 | 2 | 0.5 | 6 | 0.5 | 8 | 0 | 2 | 0 | 18 | 0.2222 |
| 7_12257002 | 3 | 0.6667 | 4 | 0.25 | 2 | 0.5 | 6 | 0.3333 | 15 | 0.4 | 2 | 1 | 5 | 0.4 | 7 | 0.5714 | 2 | 1 | 16 | 0.625 |
| 7_12257839 | 3 | 0 | 6 | 0 | 5 | 0.2 | 1 | 0 | 15 | 0.0667 | 2 | 0 | 1 | 0 | 5 | 0.2 | 2 | 0 | 10 | 0.1 |
| 7_12257845 | 3 | 0 | 5 | 0 | 5 | 0.2 | 2 | 0 | 15 | 0.0667 | 2 | 0 | 1 | 0 | 5 | 0.2 | 2 | 0 | 10 | 0.1 |
| 7_12259815 | 2 | 0 | 2 | 0 | 1 | 1 | 1 | 0 | 6 | 0.1667 | NA | NA | 5 | 0.4 | 2 | 0.5 | 1 | 0 | 8 | 0.375 |
| 7_12259826 | 2 | 0.5 | 2 | 0.5 | 2 | 1 | 2 | 0.5 | 8 | 0.625 | 1 | 1 | 5 | 0.6 | 2 | 0.5 | 1 | 0 | 9 | 0.5556 |
| 7_12259828 | 2 | 1 | 2 | 0.5 | 2 | 1 | 3 | 1 | 9 | 0.8889 | 1 | 0 | 4 | 1 | 2 | 1 | 1 | 1 | 8 | 0.875 |
| 7_12259841 | 2 | 1 | 2 | 1 | 2 | 1 | 3 | 0.3333 | 9 | 0.7778 | 1 | 1 | 4 | 0.25 | 2 | 0 | 1 | 0 | 8 | 0.25 |
| 7_12259860 | 2 | 0 | 3 | 0.3333 | 2 | 0 | 3 | 0 | 10 | 0.1 | 1 | 0 | 5 | 0.2 | 4 | 0 | 1 | 0 | 11 | 0.0909 |
| 7_12259878 | 3 | 0 | 2 | 0 | 2 | 0.5 | 3 | 0.3333 | 10 | 0.2 | 1 | 1 | 5 | 0 | 4 | 0 | 1 | 0 | 11 | 0.0909 |
| 7_12259881 | 3 | 0.3333 | 1 | 0 | 2 | 0 | 2 | 0 | 8 | 0.125 | NA | NA | 4 | 0 | 4 | 0 | 1 | 1 | 9 | 0.1111 |
| 7_12259890 | 1 | 0 | 2 | 1 | 2 | 0.5 | 3 | 0.6667 | 8 | 0.625 | 1 | 1 | 2 | 0.5 | 3 | 0.6667 | 1 | 1 | 7 | 0.7143 |
| 7_12259903 | 2 | 0.5 | 2 | 1 | 2 | 1 | 2 | 1 | 8 | 0.875 | 1 | 0 | 5 | 0.4 | 3 | 1 | 1 | 0 | 10 | 0.5 |
| 7_12259933 | 2 | 0.5 | 2 | 0 | 2 | 0 | 3 | 0 | 9 | 0.1111 | 1 | 0 | 5 | 0.4 | 4 | 0 | 1 | 0 | 11 | 0.1818 |
| 7_12259986 | 2 | 0 | 2 | 0.5 | 1 | 0 | 3 | 0.3333 | 8 | 0.25 | 1 | 0 | 6 | 0.1667 | 3 | 0 | 1 | 0 | 11 | 0.0909 |
| 7_12259997 | 2 | 0.5 | 1 | 0 | 2 | 0.5 | 4 | 0 | 9 | 0.2222 | 1 | 0 | 6 | 0.3333 | 3 | 0 | NA | NA | 10 | 0.2 |
| 7_12260029 | 2 | 0.5 | 2 | 0 | 2 | 0 | 3 | 0 | 9 | 0.1111 | NA | NA | 6 | 0.1667 | 1 | 0 | 1 | 0 | 8 | 0.125 |
| 7_12260031 | 4 | 1 | 2 | 1 | 3 | 0.6667 | 4 | 1 | 13 | 0.9231 | 1 | 1 | 6 | 0.8333 | 3 | 0.6667 | 1 | 1 | 11 | 0.8182 |
| 7_12260032 | 3 | 0.3333 | 1 | 1 | 3 | 0.3333 | 3 | 0.6667 | 10 | 0.5 | 1 | 0 | 5 | 0.8 | 4 | 0.75 | 1 | 1 | 11 | 0.7273 |
| 7_12260035 | 4 | 0.25 | 2 | 0 | 3 | 0.3333 | 4 | 0.75 | 13 | 0.3846 | 1 | 0 | 6 | 0.5 | 4 | 0 | 1 | 0 | 12 | 0.25 |
| 7_12260039 | 5 | 0.4 | 2 | 0.5 | 3 | 0.3333 | 4 | 0 | 14 | 0.2857 | 1 | 1 | 4 | 0.75 | 4 | 0.25 | 1 | 0 | 10 | 0.5 |
| 7_12260041 | 5 | 0 | 3 | 0 | 3 | 0 | 4 | 0.25 | 15 | 0.0667 | 1 | 0 | 6 | 0 | 4 | 0.25 | 1 | 0 | 12 | 0.0833 |
| 7_12260042 | 6 | 0 | 2 | 1 | 3 | 0.3333 | 4 | 0.25 | 15 | 0.2667 | 1 | 0 | 5 | 0.6 | 2 | 0 | 1 | 0 | 9 | 0.3333 |
| 7_12260049 | 6 | 0.1667 | 2 | 0 | 3 | 0 | 4 | 0.25 | 15 | 0.1333 | 1 | 0 | 5 | 0 | 3 | 0.6667 | 1 | 1 | 10 | 0.3 |
| 7_12260061 | 6 | 0.3333 | 2 | 0.5 | 3 | 0.6667 | 4 | 0.75 | 15 | 0.5333 | 1 | 1 | 6 | 0.3333 | 3 | 0.3333 | 1 | 0 | 11 | 0.3636 |
| 7_12260068 | 3 | 0.6667 | 2 | 0.5 | 2 | 0.5 | 3 | 0.3333 | 10 | 0.5 | 1 | 1 | 6 | 1 | 5 | 0.8 | 1 | 0 | 13 | 0.8462 |
| 7_12260098 | 7 | 0.7143 | 2 | 1 | 3 | 1 | 6 | 0.5 | 18 | 0.7222 | 1 | 0 | 7 | 0.8571 | 6 | 0.5 | 1 | 0 | 15 | 0.6 |
| 7_12260102 | 5 | 0.4 | 1 | 0 | 3 | 0 | 5 | 0.4 | 14 | 0.2857 | 1 | 0 | 6 | 0.1667 | 6 | 0.1667 | 1 | 0 | 14 | 0.1429 |
| 7_12260138 | 6 | 0.6667 | 3 | 0.6667 | 2 | 0.5 | 5 | 0.6 | 16 | 0.625 | 1 | 0 | 6 | 0.6667 | 4 | 0.5 | 2 | 0.5 | 13 | 0.5385 |
| 7_12260153 | 5 | 1 | 4 | 1 | 3 | 1 | 5 | 0.8 | 17 | 0.9412 | 1 | 1 | 5 | 0.8 | 5 | 0.8 | 2 | 1 | 13 | 0.8462 |
| 7_12260156 | 6 | 0.1667 | 3 | 0.3333 | 2 | 0 | 3 | 0.3333 | 14 | 0.2143 | 1 | 0 | 7 | 0 | 4 | 0 | 2 | 0.5 | 14 | 0.0714 |
| 7_12260171 | 5 | 0 | 4 | 0 | 2 | 0.5 | 3 | 0.3333 | 14 | 0.1429 | 1 | 0 | 7 | 0 | 7 | 0.1429 | 1 | 0 | 16 | 0.0625 |
| 7_12266560 | NA | NA | 1 | 0 | NA | NA | 1 | 1 | 2 | 0.5 | NA | NA | 1 | 1 | NA | NA | NA | NA | 1 | 1 |
| 7_12277002 | 1 | 0 | NA | NA | NA | NA | 2 | 0.5 | 3 | 0.3333 | 1 | 1 | NA | NA | NA | NA | NA | NA | 1 | 1 |
| 7_12277044 | 1 | 1 | NA | NA | NA | NA | 1 | 1 | 2 | 1 | 1 | 1 | NA | NA | NA | NA | NA | NA | 1 | 1 |
| 7_12277069 | 1 | 0 | NA | NA | NA | NA | 1 | 1 | 2 | 0.5 | 1 | 1 | NA | NA | NA | NA | NA | NA | 1 | 1 |
| 7_12277083 | NA | NA | NA | NA | NA | NA | 2 | 1 | 2 | 1 | 1 | 1 | NA | NA | NA | NA | NA | NA | 1 | 1 |
| 7_12277108 | 2 | 0 | NA | NA | NA | NA | 2 | 0.5 | 4 | 0.25 | 2 | 1 | NA | NA | NA | NA | NA | NA | 2 | 1 |
| 7_12279197 | 3 | 0.6667 | 3 | 0.6667 | 3 | 1 | 4 | 0.5 | 13 | 0.6923 | 3 | 0.6667 | 2 | 1 | 1 | 1 | 1 | 1 | 7 | 0.8571 |
| 7_12279279 | 4 | 0 | 3 | 0 | 6 | 0 | 6 | 0.1667 | 19 | 0.0526 | 4 | 0 | 2 | 0.5 | 3 | 0 | NA | NA | 9 | 0.1111 |
| 7_12279337 | 4 | 0.25 | 4 | 0 | 7 | 0.1429 | 8 | 0 | 23 | 0.087 | 3 | 0 | 4 | 0 | 4 | 0 | 2 | 0.5 | 13 | 0.0769 |
| 7_12279358 | 4 | 0 | 5 | 0 | 7 | 0 | 9 | 0.2222 | 25 | 0.08 | 4 | 0.25 | 4 | 0 | 4 | 0 | 2 | 0 | 14 | 0.0714 |
| 7_12279391 | 3 | 0.6667 | 4 | 0.25 | 3 | 0 | 4 | 0.25 | 14 | 0.2857 | 1 | 1 | 4 | 0 | 1 | 0 | NA | NA | 6 | 0.1667 |
| 7_12279392 | 4 | 0.5 | 2 | 0.5 | 5 | 0.2 | 4 | 0.75 | 15 | 0.4667 | 2 | 0.5 | 4 | 0.25 | 1 | 0 | NA | NA | 7 | 0.2857 |
| 7_12279395 | 2 | 0 | 2 | 0.5 | 5 | 0.2 | 2 | 0 | 11 | 0.1818 | 2 | 0.5 | 4 | 0 | 1 | 0 | NA | NA | 7 | 0.1429 |
| 7_12279396 | 3 | 0.6667 | 4 | 0.5 | 5 | 0.4 | 4 | 0.75 | 16 | 0.5625 | 1 | 1 | 4 | 0.75 | 1 | 0 | NA | NA | 6 | 0.6667 |
| 7_12279407 | 3 | 0.3333 | 3 | 0 | 4 | 0.25 | 4 | 0 | 14 | 0.1429 | 2 | 0.5 | 4 | 0 | 1 | 0 | NA | NA | 7 | 0.1429 |
| 7_12279409 | 4 | 0.25 | 4 | 0 | 4 | 0.25 | 3 | 0.3333 | 15 | 0.2 | 2 | 0.5 | 4 | 0.25 | 1 | 0 | NA | NA | 7 | 0.2857 |
| 7_12279416 | 3 | 1 | 4 | 1 | 4 | 1 | 3 | 0.6667 | 14 | 0.9286 | 2 | 0.5 | 3 | 1 | 1 | 1 | NA | NA | 6 | 0.8333 |
| 7_12279427 | 4 | 0.75 | 3 | 0.3333 | 5 | 0.4 | 2 | 0.5 | 14 | 0.5 | 2 | 0.5 | 4 | 0.25 | 1 | 0 | NA | NA | 7 | 0.2857 |
| 7_12279438 | 4 | 0 | 4 | 0.25 | 5 | 0.8 | 4 | 0.25 | 17 | 0.3529 | 3 | 0.3333 | 4 | 0.25 | 1 | 0 | NA | NA | 8 | 0.25 |
| 7_12287876 | NA | NA | 1 | 1 | NA | NA | NA | NA | 1 | 1 | NA | NA | 3 | 0.3333 | 2 | 1 | 2 | 0.5 | 7 | 0.5714 |
| 7_12287880 | NA | NA | 1 | 1 | NA | NA | NA | NA | 1 | 1 | NA | NA | 3 | 1 | 3 | 1 | 2 | 1 | 8 | 1 |
| 7_12287887 | NA | NA | 1 | 1 | NA | NA | NA | NA | 1 | 1 | NA | NA | 4 | 0.5 | 2 | 0.5 | 2 | 0 | 8 | 0.375 |
| 7_12287896 | NA | NA | 1 | 1 | NA | NA | NA | NA | 1 | 1 | NA | NA | 4 | 1 | 3 | 0.6667 | 2 | 1 | 9 | 0.8889 |
| 7_12287908 | NA | NA | 1 | 1 | NA | NA | NA | NA | 1 | 1 | NA | NA | 4 | 1 | 1 | 1 | 2 | 1 | 7 | 1 |

| | | | | | | | | | | | | | | | | | | | | |
|---|---|---|---|---|---|---|---|---|---|---|---|---|---|---|---|---|---|---|---|---|
| 7_1229706 | NA | NA | 1 | 1 | NA | NA | NA | NA | 1 | 1 | 1 | 1 | NA | NA | NA | NA | NA | NA | 1 | 1 |
| 7_1229711 | NA | NA | 1 | 0 | NA | NA | 1 | 1 | 2 | 0.5 | 1 | 1 | NA | NA | NA | NA | NA | NA | 1 | 1 |
| 7_1229712 | NA | NA | 1 | 1 | NA | NA | 1 | 1 | 2 | 1 | 1 | 1 | NA | NA | NA | NA | NA | NA | 1 | 1 |
| 7_1229715 | NA | NA | 1 | 0 | NA | NA | 1 | 1 | 2 | 0.5 | 1 | 1 | NA | NA | NA | NA | NA | NA | 1 | 1 |
| 7_1229720 | NA | NA | NA | NA | NA | NA | 1 | 1 | 1 | 1 | 1 | 1 | NA | NA | NA | NA | NA | NA | 1 | 1 |
| 7_1229731 | NA | NA | 1 | 1 | NA | NA | 1 | 1 | 2 | 1 | 1 | 1 | NA | NA | NA | NA | NA | NA | 1 | 1 |
| 7_1229738 | NA | NA | 1 | 1 | NA | NA | 1 | 1 | 2 | 1 | 1 | 1 | NA | NA | NA | NA | NA | NA | 1 | 1 |
| 7_1229738 | NA | NA | 1 | 1 | NA | NA | 1 | 1 | 2 | 1 | 1 | 1 | NA | NA | NA | NA | NA | NA | 1 | 1 |
| 7_1229738 | NA | NA | 1 | 1 | NA | NA | 1 | 1 | 2 | 1 | 1 | 1 | NA | NA | NA | NA | NA | NA | 1 | 1 |
| 7_1229738 | NA | NA | 1 | 1 | NA | NA | 1 | 1 | 2 | 1 | 1 | 1 | NA | NA | NA | NA | NA | NA | 1 | 1 |
| 7_1229741 | NA | NA | 1 | 0 | NA | NA | 1 | 1 | 2 | 0.5 | 1 | 1 | NA | NA | NA | NA | NA | NA | 1 | 1 |
| 7_1230748 | NA | NA | 1 | 0 | NA | NA | 5 | 0.2 | 6 | 0.1667 | 1 | 0 | 1 | 0 | 1 | 0 | 2 | 0.5 | 5 | 0.2 |
| 7_1230749 | NA | NA | 1 | 0 | NA | NA | 5 | 1 | 6 | 0.8333 | 1 | 0 | 1 | 0 | 1 | 0 | 2 | 0.5 | 5 | 0.2 |
| 7_1230750 | NA | NA | 1 | 1 | NA | NA | 4 | 0.25 | 5 | 0.4 | NA | NA | 2 | 0 | 1 | 0 | 2 | 0.5 | 5 | 0.2 |
| 7_1230752 | NA | NA | 1 | 0 | NA | NA | 3 | 0.3333 | 4 | 0.25 | 1 | 0 | 2 | 0.5 | 1 | 0 | 2 | 0.5 | 6 | 0.3333 |
| 7_1230754 | NA | NA | 1 | 0 | NA | NA | 2 | 0.5 | 3 | 0.3333 | NA | NA | 1 | 0 | 1 | 0 | 2 | 0.5 | 4 | 0.25 |
| 7_1230755 | NA | NA | 1 | 1 | NA | NA | 2 | 0.5 | 3 | 0.6667 | 1 | 0 | 2 | 1 | 1 | 0 | 2 | 1 | 6 | 0.6667 |
| 7_1230758 | NA | NA | NA | NA | NA | NA | 1 | 1 | 1 | 1 | 1 | 0 | NA | NA | 1 | 1 | NA | NA | 2 | 0.5 |
| 7_1230759 | NA | NA | 1 | 1 | NA | NA | 2 | 0.5 | 3 | 0.6667 | 1 | 0 | 1 | 0 | 1 | 1 | 2 | 0.5 | 5 | 0.4 |
| 7_1230759 | NA | NA | 1 | 1 | NA | NA | 2 | 0.5 | 3 | 0.6667 | 1 | 0 | 2 | 0 | 1 | 1 | 2 | 0.5 | 6 | 0.3333 |
| 7_1230765 | NA | NA | 1 | 1 | NA | NA | 2 | 0 | 3 | 0.3333 | 1 | 0 | 2 | 0 | 1 | 1 | 2 | 0 | 6 | 0.1667 |
| 7_1230772 | NA | NA | 1 | 1 | NA | NA | 2 | 0.5 | 3 | 0.6667 | 1 | 0 | 2 | 0.5 | NA | NA | 2 | 1 | 5 | 0.6 |
| 7_1230773 | NA | NA | 1 | 1 | NA | NA | 1 | 0 | 2 | 0.5 | NA | NA | 2 | 0.5 | NA | NA | 2 | 1 | 4 | 0.75 |
| 7_1230783 | NA | NA | 1 | 1 | NA | NA | 1 | 1 | 2 | 1 | 1 | 0 | 2 | 0.5 | NA | NA | 1 | 0 | 4 | 0.25 |
| 7_1230786 | NA | NA | 1 | 1 | NA | NA | 1 | 1 | 2 | 1 | NA | NA | 1 | 1 | NA | NA | NA | NA | 1 | 1 |
| 7_1230786 | NA | NA | 1 | 1 | NA | NA | 1 | 1 | 2 | 1 | NA | NA | 2 | 0.5 | NA | NA | NA | NA | 2 | 0.5 |
| 7_1230787 | NA | NA | 1 | 1 | NA | NA | 1 | 1 | 2 | 1 | NA | NA | 2 | 1 | NA | NA | NA | NA | 2 | 1 |
| 7_1230787 | NA | NA | NA | NA | NA | NA | 1 | 1 | 1 | 1 | NA | NA | 2 | 1 | NA | NA | NA | NA | 2 | 1 |
| 7_1230790 | NA | NA | 1 | 1 | NA | NA | NA | NA | 1 | 1 | NA | NA | 2 | 1 | NA | NA | NA | NA | 2 | 1 |
| 7_1230795 | NA | NA | 1 | 1 | NA | NA | NA | NA | 1 | 1 | NA | NA | 2 | 0.5 | 1 | 1 | NA | NA | 3 | 0.6667 |
| 7_1230795 | NA | NA | 1 | 1 | NA | NA | NA | NA | 1 | 1 | NA | NA | 1 | 1 | NA | NA | NA | NA | 1 | 1 |
| 7_1230797 | NA | NA | 1 | 1 | NA | NA | NA | NA | 1 | 1 | NA | NA | 1 | 1 | 1 | 1 | NA | NA | 2 | 1 |
| 7_1230798 | NA | NA | 1 | 1 | NA | NA | NA | NA | 1 | 1 | NA | NA | 2 | 1 | 1 | 0 | NA | NA | 3 | 0.6667 |
| 7_1230799 | NA | NA | 1 | 1 | NA | NA | NA | NA | 1 | 1 | NA | NA | 2 | 0.5 | 1 | 1 | NA | NA | 3 | 0.6667 |
| 7_1230890 | NA | NA | 1 | 1 | NA | NA | NA | NA | 1 | 1 | NA | NA | 1 | 1 | 1 | 1 | NA | NA | 2 | 1 |
| 7_1230890 | NA | NA | 2 | 0.5 | NA | NA | NA | NA | 2 | 0.5 | NA | NA | 1 | 1 | 1 | 0 | NA | NA | 2 | 0.5 |
| 7_1230892 | NA | NA | 2 | 1 | NA | NA | NA | NA | 2 | 1 | NA | NA | NA | NA | 1 | 1 | NA | NA | 1 | 1 |
| 7_1230989 | 1 | 1 | NA | NA | NA | NA | NA | NA | 1 | 1 | NA | NA | NA | NA | 1 | 1 | NA | NA | 1 | 1 |
| 7_1230991 | 1 | 1 | 1 | 1 | NA | NA | NA | NA | 2 | 1 | NA | NA | NA | NA | 1 | 1 | NA | NA | 1 | 1 |
| 7_1239961 | NA | NA | 2 | 0.5 | NA | NA | 2 | 0 | 4 | 0.25 | NA | NA | NA | NA | 3 | 0.3333 | 1 | 0 | 4 | 0.25 |
| 7_1239962 | 1 | 1 | 2 | 1 | NA | NA | 3 | 1 | 6 | 1 | NA | NA | NA | NA | 3 | 1 | 1 | 1 | 4 | 1 |
| 7_1239962 | 1 | 1 | 2 | 0 | NA | NA | 1 | 1 | 4 | 0.5 | NA | NA | NA | NA | 3 | 1 | 1 | 1 | 4 | 1 |
| 7_1239962 | 1 | 1 | 2 | 1 | NA | NA | 2 | 0.5 | 5 | 0.8 | NA | NA | NA | NA | 3 | 1 | 2 | 1 | 5 | 1 |
| 7_1239963 | NA | NA | 2 | 1 | NA | NA | 2 | 0.5 | 4 | 0.75 | NA | NA | NA | NA | 3 | 0.6667 | 2 | 1 | 5 | 0.8 |
| 7_1239964 | NA | NA | 2 | 0.5 | NA | NA | 2 | 1 | 4 | 0.75 | NA | NA | NA | NA | 4 | 0.75 | 2 | 1 | 6 | 0.8333 |
| 7_1239964 | 1 | 1 | 2 | 0.5 | NA | NA | 3 | 1 | 6 | 0.8333 | NA | NA | NA | NA | 4 | 1 | 1 | 1 | 5 | 1 |
| 7_1239965 | 1 | 1 | 2 | 1 | NA | NA | 3 | 1 | 6 | 1 | NA | NA | NA | NA | 3 | 0.6667 | 2 | 1 | 5 | 0.8 |
| 7_1239965 | 1 | 1 | 2 | 0.5 | NA | NA | 3 | 0.6667 | 6 | 0.6667 | NA | NA | NA | NA | 4 | 0.75 | 2 | 0.5 | 6 | 0.6667 |
| 7_1239966 | NA | NA | 2 | 0.5 | NA | NA | 2 | 0.5 | 4 | 0.5 | NA | NA | NA | NA | 3 | 1 | 1 | 1 | 4 | 1 |
| 7_1239979 | 1 | 1 | 4 | 0.75 | 1 | 1 | 5 | 0.6 | 11 | 0.7273 | 1 | 0 | 1 | 1 | 5 | 0.2 | 3 | 1 | 10 | 0.5 |
| 7_1247866 | 1 | 1 | NA | NA | NA | NA | 1 | 1 | 2 | 1 | NA | NA | NA | NA | 3 | 1 | NA | NA | 3 | 1 |
| 7_1247866 | 1 | 1 | NA | NA | NA | NA | 1 | 0 | 2 | 0.5 | NA | NA | NA | NA | 3 | 0 | 1 | 1 | 4 | 0.25 |
| 7_1247868 | 1 | 1 | NA | NA | NA | NA | NA | NA | 1 | 1 | NA | NA | NA | NA | 3 | 1 | 1 | 1 | 4 | 1 |
| 7_1247871 | 1 | 1 | NA | NA | NA | NA | 1 | 1 | 2 | 1 | NA | NA | NA | NA | 3 | 1 | NA | NA | 3 | 1 |
| 7_1247873 | 1 | 1 | NA | NA | NA | NA | 2 | 1 | 3 | 1 | NA | NA | NA | NA | 3 | 0.6667 | NA | NA | 3 | 0.6667 |
| 7_1247873 | NA | NA | NA | NA | NA | NA | 2 | 1 | 2 | 1 | NA | NA | NA | NA | 3 | 0.6667 | 1 | 1 | 4 | 0.75 |
| 7_1247874 | 1 | 1 | NA | NA | NA | NA | 2 | 0.5 | 3 | 0.6667 | NA | NA | NA | NA | 3 | 0.3333 | NA | NA | 3 | 0.3333 |
| 7_1247874 | NA | NA | NA | NA | NA | NA | 1 | 1 | 1 | 1 | NA | NA | NA | NA | 3 | 0.6667 | NA | NA | 3 | 0.6667 |
| 7_1251750 | NA | NA | NA | NA | 1 | 1 | NA | NA | 1 | 1 | 1 | 1 | NA | NA | NA | NA | NA | NA | 1 | 1 |
| 7_1251753 | NA | NA | NA | NA | 1 | 1 | NA | NA | 1 | 1 | 1 | 1 | NA | NA | 1 | 1 | NA | NA | 2 | 1 |
| 7_1251756 | NA | NA | NA | NA | 1 | 1 | NA | NA | 1 | 1 | 1 | 1 | NA | NA | 1 | 1 | NA | NA | 2 | 1 |
| 7_1251757 | NA | NA | NA | NA | 1 | 1 | NA | NA | 1 | 1 | 1 | 1 | NA | NA | 1 | 1 | NA | NA | 2 | 1 |
| 7_1251757 | NA | NA | NA | NA | 1 | 1 | NA | NA | 1 | 1 | 1 | 1 | NA | NA | NA | NA | NA | NA | 1 | 1 |
| 7_1252402 | 8 | 0.125 | 13 | 0.3077 | 13 | 0.3846 | 18 | 0.1111 | 52 | 0.2308 | 9 | 0.5556 | 15 | 0.4 | 10 | 0.1 | 11 | 0.6364 | 45 | 0.4222 |
| 7_1252403 | 10 | 0.2 | 18 | 0.1111 | 13 | 0.1538 | 22 | 0.1818 | 63 | 0.1587 | 14 | 0.2857 | 13 | 0.3077 | 13 | 0.1538 | 11 | 0 | 51 | 0.1961 |

| | | | | | | | | | | | | | | | | | | | | |
|---|---|---|---|---|---|---|---|---|---|---|---|---|---|---|---|---|---|---|---|---|
| 7_12524046 | 7 | 0.1429 | 18 | 0.1667 | 14 | 0.0714 | 21 | 0.3333 | 60 | 0.2 | 12 | 0.25 | 14 | 0.2857 | 11 | 0.2727 | 11 | 0.3636 | 48 | 0.2917 |
| 7_12524062 | 9 | 0.7778 | 17 | 0.7059 | 16 | 0.8125 | 22 | 0.6364 | 64 | 0.7188 | 13 | 0.6154 | 13 | 0.9231 | 11 | 0.8182 | 11 | 0.9091 | 48 | 0.8125 |
| 7_12524074 | 9 | 0.6667 | 14 | 0.7143 | 12 | 0.4167 | 13 | 0.7692 | 48 | 0.6458 | 12 | 0.5833 | 10 | 0.6 | 10 | 0.7 | 8 | 0.625 | 40 | 0.625 |
| 7_12524087 | 6 | 0 | 20 | 0.1 | 11 | 0.0909 | 19 | 0.1053 | 56 | 0.0893 | 11 | 0 | 9 | 0 | 9 | 0.1111 | 10 | 0.1 | 39 | 0.0513 |
| 7_12524329 | 6 | 0.3333 | 16 | 0.5625 | 6 | 0.3333 | 18 | 0.2778 | 46 | 0.3913 | 10 | 0.5 | 7 | 0.2857 | 8 | 0.5 | 8 | 0.5 | 33 | 0.4545 |
| 7_12524469 | 2 | 0.5 | 5 | 0 | 3 | 0 | 8 | 0 | 18 | 0.0556 | 5 | 0.2 | 5 | 0 | 5 | 0 | 5 | 0 | 20 | 0.05 |
| 7_12524662 | 4 | 0.5 | 12 | 0.25 | 5 | 0.4 | 14 | 0.0714 | 35 | 0.2286 | 5 | 0.2 | 5 | 0.2 | 8 | 0.375 | 6 | 0.1667 | 24 | 0.25 |
| 7_12524664 | 2 | 0 | 4 | 0 | 2 | 0 | 10 | 0.1 | 18 | 0.0556 | 3 | 0.3333 | 4 | 0.5 | 5 | 0 | 7 | 0 | 19 | 0.1579 |
| 7_12524668 | 4 | 0 | 9 | 0 | 6 | 0 | 14 | 0.1429 | 33 | 0.0606 | 5 | 0 | 5 | 0.2 | 6 | 0 | 7 | 0.1429 | 23 | 0.087 |
| 7_12547401 | NA | NA | NA | NA | NA | NA | 1 | 1 | 1 | 1 | 1 | 1 | NA | NA | NA | NA | NA | NA | 1 | 1 |
| 7_12547784 | 7 | 0.2857 | 7 | 0.4286 | 9 | 0 | 8 | 0.5 | 31 | 0.2903 | 5 | 0.4 | 7 | 0.2857 | 14 | 0.4286 | 9 | 0.4444 | 35 | 0.4 |
| 7_12547800 | 6 | 0.6667 | 6 | 0.3333 | 10 | 0.3 | 8 | 0.25 | 30 | 0.3667 | 5 | 0.2 | 5 | 0 | 12 | 0.4167 | 7 | 0.7143 | 29 | 0.3793 |
| 7_12547814 | 7 | 0.1429 | 5 | 0 | 7 | 0.1429 | 7 | 0 | 26 | 0.0769 | 2 | 0 | 2 | 0 | 11 | 0.0909 | 5 | 0 | 20 | 0.05 |
| 7_12547838 | 7 | 0.2857 | 8 | 0.125 | 6 | 0 | 5 | 0.4 | 26 | 0.1923 | 4 | 0.5 | 5 | 0 | 12 | 0.25 | 6 | 0 | 27 | 0.1852 |
| 7_12547845 | 7 | 0.2857 | 7 | 0.1429 | 5 | 0.2 | 4 | 0.25 | 23 | 0.2174 | 4 | 0 | 4 | 0 | 14 | 0.1429 | 7 | 0 | 29 | 0.069 |
| 7_12547864 | 7 | 0.5714 | 5 | 0.8 | 5 | 0.8 | 4 | 0.75 | 21 | 0.7143 | 3 | 0.6667 | 5 | 1 | 12 | 0.75 | 6 | 0.6667 | 26 | 0.7692 |
| 7_12547870 | 7 | 0.7143 | 5 | 0.6 | 4 | 0.5 | 4 | 0.5 | 20 | 0.6 | 3 | 1 | 3 | 0 | 11 | 0.7273 | 6 | 0.6667 | 23 | 0.6522 |
| 7_12548011 | 4 | 0.25 | 3 | 1 | 2 | 1 | 1 | 1 | 10 | 0.7 | 2 | 1 | 3 | 1 | 7 | 0.8571 | 2 | 1 | 14 | 0.9286 |
| 7_12548013 | 4 | 1 | 2 | 1 | 1 | 1 | 1 | 1 | 8 | 1 | 2 | 1 | 2 | 1 | 6 | 0.6667 | 1 | 1 | 11 | 0.8182 |
| 7_12548019 | 3 | 0.6667 | 2 | 1 | 2 | 0 | 1 | 1 | 8 | 0.625 | 2 | 1 | 3 | 0.6667 | 5 | 0.6 | 2 | 1 | 12 | 0.75 |
| 7_12548023 | 4 | 0.75 | 3 | 1 | 2 | 1 | 1 | 0 | 10 | 0.8 | 1 | 1 | 2 | 1 | 5 | 0.8 | 2 | 1 | 10 | 0.9 |
| 7_12569203 | NA | NA | 1 | 1 | NA | NA | 1 | 0 | 2 | 0.5 | NA | NA | 1 | 1 | NA | NA | NA | NA | 1 | 1 |
| 7_12569226 | NA | NA | NA | NA | NA | NA | 1 | 1 | 1 | 1 | NA | NA | 1 | 1 | NA | NA | NA | NA | 1 | 1 |
| 7_12569405 | NA | NA | NA | NA | 1 | 1 | 1 | 1 | 2 | 1 | 1 | 1 | NA | NA | NA | NA | NA | NA | 1 | 1 |
| 7_12569587 | NA | NA | NA | NA | 1 | 1 | 1 | 1 | 2 | 1 | 1 | 1 | NA | NA | NA | NA | NA | NA | 1 | 1 |
| 7_12569589 | NA | NA | NA | NA | 1 | 0 | 1 | 1 | 2 | 0.5 | 1 | 1 | NA | NA | NA | NA | NA | NA | 1 | 1 |
| 7_12569598 | NA | NA | NA | NA | 1 | 1 | 1 | 1 | 2 | 1 | 1 | 1 | NA | NA | NA | NA | NA | NA | 1 | 1 |
| 7_12569603 | NA | NA | NA | NA | 1 | 1 | NA | NA | 1 | 1 | 1 | 1 | NA | NA | NA | NA | NA | NA | 1 | 1 |
| 7_12569613 | NA | NA | NA | NA | 1 | 1 | 1 | 1 | 2 | 1 | 1 | 1 | NA | NA | NA | NA | NA | NA | 1 | 1 |
| 7_12569694 | NA | NA | NA | NA | 1 | 1 | 1 | 0 | 2 | 0.5 | 1 | 1 | NA | NA | NA | NA | NA | NA | 1 | 1 |
| 7_12569977 | NA | NA | NA | NA | NA | NA | 2 | 1 | 2 | 1 | 1 | 1 | NA | NA | 2 | 1 | NA | NA | 3 | 1 |
| 7_12569992 | 1 | 1 | NA | NA | NA | NA | 2 | 0.5 | 3 | 0.6667 | 1 | 1 | NA | NA | 1 | 1 | NA | NA | 2 | 1 |
| 7_12570058 | 1 | 0 | NA | NA | NA | NA | 1 | 1 | 2 | 0.5 | 2 | 1 | NA | NA | 1 | 0 | NA | NA | 3 | 0.6667 |
| 7_12570060 | 1 | 1 | NA | NA | NA | NA | 2 | 1 | 3 | 1 | 2 | 0.5 | NA | NA | 1 | 1 | NA | NA | 3 | 0.6667 |
| 7_12570076 | 1 | 1 | NA | NA | NA | NA | 1 | 0 | 2 | 0.5 | 2 | 0.5 | NA | NA | 2 | 1 | NA | NA | 4 | 0.75 |
| 7_12570082 | NA | NA | NA | NA | NA | NA | 1 | 1 | 1 | 1 | 1 | 1 | NA | NA | 2 | 0 | NA | NA | 3 | 0.3333 |
| 7_12570089 | NA | NA | NA | NA | NA | NA | 2 | 1 | 2 | 1 | 1 | 1 | NA | NA | 1 | 0 | NA | NA | 2 | 0.5 |
| 7_12570092 | 14 | 0.5714 | 7 | 0.7143 | 7 | 0.2857 | 15 | 0.6 | 43 | 0.5581 | 9 | 0.3333 | 10 | 0.3 | 10 | 0.6 | 8 | 0.375 | 37 | 0.4054 |
| 7_12570097 | 15 | 0.8 | 8 | 0.5 | 8 | 0.375 | 15 | 0.6667 | 46 | 0.6304 | 8 | 0.75 | 12 | 0.6667 | 10 | 0.7 | 9 | 0.4444 | 39 | 0.641 |
| 7_12570104 | 13 | 0.9231 | 5 | 0.8 | 8 | 0.75 | 15 | 1 | 41 | 0.9024 | 6 | 1 | 10 | 0.7 | 9 | 1 | 9 | 0.6667 | 34 | 0.8235 |
| 7_12570113 | 13 | 0.6154 | 8 | 0.875 | 7 | 0.7143 | 17 | 0.7647 | 45 | 0.7333 | 8 | 0.625 | 12 | 0.9167 | 11 | 0.8182 | 9 | 0.5556 | 40 | 0.75 |
| 7_12570119 | 13 | 0.6154 | 7 | 0.1429 | 8 | 0.375 | 17 | 0.4118 | 45 | 0.4222 | 8 | 0.5 | 10 | 0.3 | 10 | 0.5 | 7 | 0.1429 | 35 | 0.3714 |
| 7_12570122 | 13 | 0.6154 | 5 | 1 | 7 | 0.7143 | 17 | 0.7647 | 42 | 0.7381 | 5 | 1 | 10 | 0.8 | 11 | 0.8182 | 9 | 0.8889 | 35 | 0.8571 |
| 7_12570126 | 11 | 0.4545 | 5 | 0.4 | 8 | 0.625 | 15 | 0.5333 | 39 | 0.5128 | 7 | 0.7143 | 9 | 0.4444 | 10 | 0.8 | 9 | 0.4444 | 35 | 0.6 |
| 7_12575212 | 5 | 0.6 | 2 | 0 | 1 | 1 | 3 | 1 | 11 | 0.6364 | 2 | 1 | 3 | 1 | 1 | 1 | 1 | 0 | 7 | 0.8571 |
| 7_12575214 | 6 | 0.8333 | 2 | 1 | 1 | 1 | 3 | 0.6667 | 12 | 0.8333 | 3 | 0.6667 | 2 | 1 | NA | NA | 1 | 1 | 6 | 0.8333 |
| 7_12575220 | 5 | 1 | 1 | 1 | NA | NA | 2 | 1 | 8 | 1 | 3 | 1 | 3 | 1 | 1 | 0 | 1 | 1 | 8 | 0.875 |
| 7_12587206 | 1 | 0 | 1 | 1 | 1 | 0 | 2 | 0.5 | 5 | 0.4 | 5 | 0.4 | 1 | 0 | 2 | 0.5 | 3 | 0.3333 | 11 | 0.3636 |
| 7_12587212 | 1 | 0 | 1 | 0 | NA | NA | 2 | 0.5 | 4 | 0.25 | 5 | 0 | 1 | 0 | 2 | 0 | 3 | 0.3333 | 11 | 0.0909 |
| 7_12589105 | 3 | 1 | 3 | 0.6667 | NA | NA | 1 | 1 | 7 | 0.8571 | 7 | 0.8571 | NA | NA | 2 | 1 | 4 | 1 | 13 | 0.9231 |
| 7_12589125 | 3 | 1 | 3 | 0.6667 | 1 | 1 | 2 | 1 | 9 | 0.8889 | 7 | 1 | 2 | 0.5 | 3 | 0.6667 | 4 | 1 | 16 | 0.875 |
| 7_12589157 | 2 | 0 | 3 | 1 | 2 | 0 | 3 | 0.6667 | 10 | 0.5 | 6 | 0 | 2 | 0.5 | 2 | 1 | 4 | 0.5 | 14 | 0.3571 |
| 7_12589175 | 3 | 0 | 4 | 0.5 | 2 | 0.5 | 4 | 0 | 13 | 0.2308 | 9 | 0.1111 | 2 | 0 | 3 | 0.3333 | 5 | 0.2 | 19 | 0.1579 |
| 7_12589176 | 2 | 0.5 | 4 | 0.75 | 2 | 1 | 4 | 0.75 | 12 | 0.75 | 8 | 0.875 | 2 | 1 | 2 | 0.5 | 4 | 1 | 16 | 0.875 |
| 7_12589187 | 2 | 0 | 3 | 0 | 2 | 0 | 4 | 0.25 | 11 | 0.0909 | 7 | 0.4286 | NA | NA | 4 | 0 | 3 | 0 | 14 | 0.2143 |
| 7_12589206 | 3 | 0 | 5 | 0 | 2 | 0 | 4 | 0.25 | 14 | 0.0714 | 8 | 0.125 | 1 | 1 | 5 | 0 | 5 | 0 | 19 | 0.1053 |
| 7_12589219 | 4 | 0 | 5 | 0 | 2 | 0.5 | 4 | 0.5 | 15 | 0.2 | 6 | 0.1667 | 2 | 0 | 4 | 0 | 7 | 0.1429 | 19 | 0.1053 |
| 7_12589233 | 3 | 0.3333 | 5 | 0.6 | 2 | 0 | 4 | 0.75 | 14 | 0.5 | 6 | 0.1667 | 2 | 0 | 3 | 0 | 6 | 0.1667 | 17 | 0.1176 |
| 7_12589279 | 4 | 0 | 6 | 0.1667 | 2 | 0 | 5 | 0 | 17 | 0.0588 | 8 | 0.25 | 3 | 0 | 6 | 0 | 8 | 0.125 | 25 | 0.12 |
| 7_12589281 | 5 | 0.4 | 6 | 0.3333 | 3 | 0 | 7 | 0.1429 | 21 | 0.2381 | 11 | 0.1818 | 3 | 0 | 10 | 0.2 | 7 | 0.1429 | 31 | 0.1613 |
| 7_12589347 | 8 | 0.25 | 5 | 0.2 | 4 | 0.25 | 8 | 0.25 | 25 | 0.24 | 12 | 0 | 5 | 0.2 | 11 | 0.0909 | 10 | 0 | 38 | 0.0526 |
| 7_12589348 | 8 | 0.125 | 6 | 0.1667 | 3 | 0.3333 | 7 | 0 | 24 | 0.125 | 11 | 0.2727 | 5 | 0.2 | 11 | 0 | 10 | 0.1 | 37 | 0.1351 |
| 7_12589395 | 7 | 0.1429 | 9 | 0.1111 | 4 | 0.5 | 9 | 0.2222 | 29 | 0.2069 | 12 | 0.1667 | 8 | 0.25 | 12 | 0.3333 | 13 | 0.2308 | 45 | 0.2444 |
| 7_12604456 | NA | NA | 1 | 0 | 2 | 1 | NA | NA | 3 | 0.6667 | NA | NA | 5 | 0.4 | 1 | 0 | 1 | 1 | 7 | 0.4286 |
| 7_12604463 | 1 | 1 | 2 | 1 | 2 | 0 | NA | NA | 5 | 0.6 | NA | NA | 3 | 0.6667 | 1 | 1 | 1 | 1 | 5 | 0.8 |

| | | | | | | | | | | | | | | | | | | | | |
|---|---|---|---|---|---|---|---|---|---|---|---|---|---|---|---|---|---|---|---|---|
| 7_12604465 | 1 | 0 | 2 | 0 | 3 | 0.3333 | NA | NA | 6 | 0.1667 | NA | NA | 4 | 0.25 | NA | NA | 1 | 0 | 5 | 0.2 |
| 7_12604496 | 1 | 1 | 2 | 1 | 3 | 0.3333 | NA | NA | 6 | 0.6667 | NA | NA | 5 | 0.6 | 1 | 0 | 1 | 1 | 7 | 0.5714 |
| 7_12606635 | 6 | 0.1667 | 5 | 0 | 5 | 0.2 | 12 | 0.1667 | 28 | 0.1429 | 3 | 0 | 13 | 0.1538 | 8 | 0 | 7 | 0.1429 | 31 | 0.0968 |
| 7_12606652 | 7 | 0.1429 | 4 | 0 | 7 | 0.2857 | 12 | 0.0833 | 30 | 0.1333 | 5 | 0.2 | 13 | 0 | 8 | 0.25 | 7 | 0.2857 | 33 | 0.1515 |
| 7_12606667 | 5 | 0 | 5 | 0.2 | 5 | 0.4 | 9 | 0.1111 | 24 | 0.1667 | 4 | 0.75 | 13 | 0.2308 | 8 | 0 | 7 | 0.1429 | 32 | 0.2188 |
| 7_12606689 | 3 | 0.3333 | 2 | 0.5 | 1 | 0 | 5 | 0.4 | 11 | 0.3636 | 2 | 0 | 8 | 0.25 | 3 | 0 | 4 | 0.25 | 17 | 0.1765 |
| 7_12607151 | 2 | 0.5 | 2 | 1 | 4 | 1 | 7 | 0.8571 | 15 | 0.8667 | 1 | 1 | 5 | 1 | 5 | 0.8 | 4 | 0.75 | 15 | 0.8667 |
| 7_12607156 | 2 | 1 | 3 | 1 | 4 | 1 | 5 | 1 | 14 | 1 | 2 | 1 | 7 | 0.8571 | 4 | 1 | 5 | 0.8 | 18 | 0.8889 |
| 7_12607165 | 2 | 0.5 | 3 | 0 | 3 | 0 | 3 | 0.3333 | 11 | 0.1818 | NA | NA | 4 | 0.25 | 3 | 0.3333 | 5 | 0 | 12 | 0.1667 |
| 7_12607179 | 2 | 0.5 | 3 | 0.3333 | 2 | 0.5 | 5 | 0.2 | 12 | 0.3333 | 2 | 0 | 4 | 0.5 | 5 | 0.4 | 3 | 0.3333 | 14 | 0.3571 |
| 7_12608967 | 13 | 0.6923 | 13 | 0.9231 | 7 | 0.7143 | 12 | 0.6667 | 45 | 0.7556 | 10 | 0.9 | 16 | 0.5625 | 6 | 0.8333 | 11 | 0.7273 | 43 | 0.7209 |
| 7_12608979 | 10 | 0.2 | 13 | 0.1538 | 9 | 0 | 12 | 0 | 44 | 0.0909 | 11 | 0.0909 | 16 | 0.0625 | 5 | 0.2 | 7 | 0.1429 | 39 | 0.1026 |
| 7_12608983 | 7 | 0.8571 | 11 | 0.6364 | 6 | 0.6667 | 6 | 0.5 | 30 | 0.6667 | 9 | 0.8889 | 11 | 0.6364 | 3 | 1 | 8 | 0.75 | 31 | 0.7742 |
| 7_12609095 | 10 | 0.1 | 25 | 0 | 14 | 0.2143 | 16 | 0 | 65 | 0.0615 | 9 | 0 | 14 | 0.2857 | 17 | 0.2353 | 11 | 0.1818 | 51 | 0.1961 |
| 7_12614596 | 3 | 0.3333 | 5 | 0 | 2 | 0 | 3 | 0 | 13 | 0.0769 | 5 | 0 | 6 | 0.1667 | 3 | 0 | 4 | 0 | 18 | 0.0556 |
| 7_12614600 | 4 | 0.25 | 6 | 0.1667 | 3 | 0.3333 | 2 | 0.5 | 15 | 0.2667 | 6 | 0.1667 | 5 | 0 | 6 | 0.3333 | 3 | 0.6667 | 20 | 0.25 |
| 7_12614603 | 4 | 0 | 7 | 0.4286 | 4 | 0.25 | 5 | 0.6 | 20 | 0.35 | 6 | 0.1667 | 7 | 0.1429 | 7 | 0 | 3 | 0.6667 | 23 | 0.1739 |
| 7_12622248 | 3 | 0 | NA | NA | 1 | 1 | 1 | 0 | 5 | 0.2 | 4 | 0.5 | 2 | 0.5 | 1 | 0 | 2 | 0.5 | 9 | 0.4444 |
| 7_12627208 | 7 | 0.1429 | 8 | 0.25 | 6 | 0.5 | 4 | 0.25 | 25 | 0.28 | 3 | 0.3333 | NA | NA | 5 | 0.2 | 4 | 0.25 | 12 | 0.25 |
| 7_12627211 | 8 | 0 | 8 | 0 | 6 | 0.1667 | 5 | 0.2 | 27 | 0.0741 | 3 | 0.3333 | 2 | 0 | 4 | 0 | 4 | 0 | 13 | 0.0769 |
| 7_12627219 | 9 | 0.4444 | 7 | 0.1429 | 4 | 0.5 | 5 | 0 | 25 | 0.28 | 3 | 0.6667 | 2 | 0 | 5 | 0.2 | 3 | 0.6667 | 13 | 0.3846 |
| 7_12629165 | 1 | 1 | 1 | 0 | 1 | 1 | NA | NA | 3 | 0.6667 | NA | NA | 1 | 1 | 1 | 1 | NA | NA | 2 | 1 |
| 7_12629584 | 13 | 0.2308 | 15 | 0.2 | 13 | 0.3846 | 13 | 0.3077 | 54 | 0.2778 | 18 | 0.2778 | 14 | 0.1429 | 13 | 0.3077 | 9 | 0.3333 | 54 | 0.2593 |
| 7_12637928 | 32 | 0.125 | 42 | 0.119 | 34 | 0.0294 | 32 | 0.0938 | 140 | 0.0929 | 27 | 0.0741 | 30 | 0.1 | 26 | 0.1923 | 38 | 0.1579 | 121 | 0.1322 |
| 7_12637937 | 27 | 0.0741 | 30 | 0.2 | 29 | 0.1724 | 28 | 0.0714 | 114 | 0.1316 | 27 | 0.1481 | 32 | 0.1562 | 28 | 0.1071 | 36 | 0.0833 | 123 | 0.122 |
| 7_12637943 | 35 | 0.2571 | 33 | 0.3636 | 32 | 0.3438 | 28 | 0.2143 | 128 | 0.2969 | 30 | 0.4 | 31 | 0.2581 | 31 | 0.3226 | 38 | 0.3421 | 130 | 0.3308 |
| 7_12638522 | 7 | 0.7143 | 3 | 1 | 2 | 0 | 2 | 0.5 | 14 | 0.6429 | 6 | 0.8333 | NA | NA | 4 | 0.75 | 3 | 1 | 13 | 0.8462 |
| 7_12638529 | 14 | 0.0714 | 9 | 0.1111 | 2 | 0 | 7 | 0.1429 | 32 | 0.0938 | 10 | 0 | 6 | 0 | 6 | 0.1667 | 8 | 0.125 | 30 | 0.0667 |
| 7_12638530 | 12 | 0.1667 | 6 | 0.1667 | 3 | 0 | 6 | 0.1667 | 27 | 0.1481 | 12 | 0.25 | 4 | 0.25 | 4 | 0 | 7 | 0.1429 | 27 | 0.1852 |
| 7_12642866 | 3 | 0.3333 | 12 | 0.25 | 9 | 0.2222 | 8 | 0.125 | 32 | 0.2188 | 7 | 0.4286 | 8 | 0.375 | 6 | 0.1667 | 6 | 0 | 27 | 0.2593 |
| 7_12642880 | 4 | 0 | 15 | 0.1333 | 9 | 0.1111 | 10 | 0 | 38 | 0.0789 | 8 | 0.25 | 10 | 0.1 | 7 | 0 | 8 | 0.125 | 33 | 0.1212 |
| 7_12642894 | 2 | 0 | 10 | 0.1 | 8 | 0.125 | 5 | 0 | 25 | 0.08 | 4 | 0 | 5 | 0.2 | 4 | 0 | 7 | 0.1429 | 20 | 0.1 |
| 7_12642896 | 5 | 0 | 8 | 0.125 | 6 | 0.1667 | 9 | 0.1111 | 28 | 0.1071 | 8 | 0.125 | 7 | 0 | 7 | 0 | 8 | 0.125 | 30 | 0.0667 |
| 7_12642898 | 4 | 0 | 15 | 0.0667 | 10 | 0.1 | 10 | 0.1 | 39 | 0.0769 | 9 | 0.2222 | 10 | 0.1 | 7 | 0.1429 | 7 | 0 | 33 | 0.1212 |
| 7_12645093 | 2 | 1 | 1 | 1 | 2 | 0 | 1 | 0 | 6 | 0.5 | 1 | 1 | 1 | 0 | 1 | 0 | 1 | 0 | 4 | 0.25 |
| 7_12645094 | 2 | 0 | 5 | 0.2 | NA | NA | NA | NA | 7 | 0.1429 | 4 | 0 | 2 | 0 | NA | NA | 1 | 1 | 7 | 0.1429 |
| 7_12645122 | 2 | 1 | 1 | 1 | 3 | 0.3333 | 1 | 0 | 7 | 0.5714 | NA | NA | 2 | 1 | 2 | 1 | 1 | 1 | 5 | 1 |
| 7_12645133 | 2 | 0 | 1 | 1 | 3 | 0.3333 | 1 | 0 | 7 | 0.2857 | 1 | 0 | 3 | 0.6667 | 2 | 0.5 | 1 | 1 | 7 | 0.5714 |
| 7_12645136 | 4 | 0.75 | 10 | 0.4 | 1 | 0 | NA | NA | 15 | 0.4667 | 5 | 0.4 | 2 | 0 | 1 | 0 | 1 | 0 | 9 | 0.2222 |
| 7_12645168 | 11 | 0.0909 | 7 | 0 | 7 | 0.1429 | 4 | 0 | 29 | 0.069 | 8 | 0.125 | 12 | 0.0833 | 5 | 0 | 6 | 0 | 31 | 0.0645 |
| 7_12645174 | 8 | 0.125 | 6 | 0.1667 | 4 | 0.25 | 2 | 0 | 20 | 0.15 | 7 | 0 | 10 | 0 | 3 | 0.3333 | 5 | 0.2 | 25 | 0.08 |
| 7_12645179 | 4 | 0.25 | 10 | 0.2 | 2 | 0.5 | NA | NA | 16 | 0.25 | 7 | 0.1429 | 2 | 0 | 1 | 0 | 1 | 0 | 11 | 0.0909 |
| 7_12646100 | 5 | 0 | 2 | 0.5 | 5 | 0 | 3 | 0 | 15 | 0.0667 | 4 | 0 | 5 | 0 | 5 | 0.2 | 4 | 0.5 | 18 | 0.1667 |
| 7_12646346 | NA | NA | NA | NA | NA | NA | 1 | 1 | 1 | 1 | NA | NA | 1 | 0 | NA | NA | 2 | 1 | 3 | 0.6667 |
| 7_12646408 | 2 | 0 | 2 | 0.5 | 1 | 0 | 3 | 0.3333 | 8 | 0.25 | 1 | 0 | 1 | 0 | 3 | 0.3333 | 2 | 0 | 7 | 0.1429 |
| 7_12646414 | 2 | 0.5 | 2 | 0.5 | 1 | 0 | 1 | 0 | 6 | 0.3333 | 1 | 0 | 4 | 0.25 | 3 | 0.3333 | 3 | 0.3333 | 11 | 0.2727 |
| 7_12646424 | 3 | 0.6667 | 1 | 1 | 1 | 1 | 2 | 0.5 | 7 | 0.7143 | 1 | 1 | 4 | 1 | 2 | 1 | 3 | 1 | 10 | 1 |
| 7_12646426 | 3 | 0.6667 | 2 | 0.5 | 1 | 0 | 3 | 0.6667 | 9 | 0.5556 | 1 | 0 | 4 | 0.5 | 2 | 0.5 | 3 | 0.3333 | 10 | 0.4 |
| 7_12646496 | 3 | 0.3333 | 6 | 0.1667 | 6 | 0 | 10 | 0.2 | 25 | 0.16 | 8 | 0 | 5 | 0 | 3 | 0 | 11 | 0.2727 | 27 | 0.1111 |
| 7_12646519 | 3 | 0 | 6 | 0.5 | 4 | 0.5 | 7 | 0 | 20 | 0.25 | 8 | 0.625 | 4 | 0 | 3 | 0 | 9 | 0.3333 | 24 | 0.3333 |
| 7_12646530 | 4 | 0.25 | 4 | 0 | 5 | 0.2 | 6 | 0.3333 | 19 | 0.2105 | 7 | 0 | 5 | 0.4 | 4 | 0 | 5 | 0 | 21 | 0.0952 |
| 7_12646577 | 3 | 0.3333 | 2 | 0 | 4 | 0 | 4 | 0 | 13 | 0.0769 | 6 | 0 | 3 | 0 | 4 | 0.25 | 7 | 0 | 20 | 0.05 |
| 7_12646598 | 3 | 0 | 2 | 0 | 2 | 0 | 4 | 0.25 | 11 | 0.0909 | 7 | 0.2857 | 2 | 0.5 | 2 | 0 | 5 | 0.2 | 16 | 0.25 |
| 7_12646997 | 1 | 1 | 3 | 0.3333 | 6 | 0.6667 | 1 | 1 | 11 | 0.6364 | 5 | 0.6 | 2 | 0.5 | 2 | 0.5 | NA | NA | 9 | 0.5556 |
| 7_12647039 | 1 | 1 | 3 | 0.3333 | 1 | 0 | NA | NA | 5 | 0.4 | 2 | 0.5 | NA | NA | 2 | 0.5 | NA | NA | 4 | 0.5 |
| 7_12647052 | 1 | 1 | 3 | 0 | 2 | 0 | NA | NA | 6 | 0.1667 | 2 | 0.5 | 1 | 0 | 2 | 0 | NA | NA | 5 | 0.2 |
| 7_12647078 | 2 | 0 | 5 | 0.4 | 2 | 0.5 | 2 | 0 | 11 | 0.2727 | 9 | 0.2222 | 3 | 0.3333 | 3 | 0.3333 | 5 | 0.4 | 20 | 0.3 |
| 7_12647081 | 2 | 0 | 5 | 0.4 | 2 | 0.5 | 2 | 0 | 11 | 0.2727 | 10 | 0.1 | 3 | 0 | 2 | 0.5 | 6 | 0 | 21 | 0.0952 |
| 7_12648542 | 3 | 0 | 2 | 0 | 5 | 0 | 2 | 0.5 | 12 | 0.0833 | 4 | 0 | NA | NA | 7 | 0 | 4 | 0.25 | 15 | 0.0667 |
| 7_12654428 | 13 | 0.5385 | 25 | 0.68 | 13 | 0.5385 | 22 | 0.5455 | 73 | 0.589 | 14 | 0.7857 | 18 | 0.6667 | 14 | 0.5 | 12 | 0.4167 | 58 | 0.6034 |
| 7_12654460 | 18 | 0.3889 | 33 | 0.4545 | 14 | 0.3571 | 26 | 0.3077 | 91 | 0.3846 | 18 | 0.3333 | 20 | 0.25 | 21 | 0.4286 | 17 | 0.2353 | 76 | 0.3158 |
| 7_12654522 | 5 | 0 | 3 | 0.3333 | 1 | 0 | 2 | 0 | 11 | 0.0909 | 7 | 0.4286 | NA | NA | 2 | 0 | 3 | 0.3333 | 12 | 0.3333 |
| 7_12654769 | 5 | 0.4 | 3 | 0 | 2 | 0 | 3 | 0.6667 | 13 | 0.3077 | 4 | 0.25 | 1 | 0 | 1 | 0 | 4 | 0.25 | 10 | 0.2 |
| 7_12654770 | 6 | 0 | 4 | 0.75 | 2 | 0 | 3 | 0 | 15 | 0.2 | 4 | 0.5 | 1 | 0 | 1 | 0 | 5 | 0.4 | 11 | 0.3636 |
| 7_12654984 | 4 | 0 | 7 | 0.1429 | 4 | 0 | 8 | 0.125 | 23 | 0.087 | 4 | 0 | 8 | 0.125 | 8 | 0 | 8 | 0.25 | 28 | 0.1071 |

| | | | | | | | | | | | | | | | | | | | | |
|---|---|---|---|---|---|---|---|---|---|---|---|---|---|---|---|---|---|---|---|---|
| 7_12655003 | 6 | 0.1667 | 10 | 0.2 | 7 | 0 | 10 | 0 | 33 | 0.0909 | 4 | 0.25 | 12 | 0 | 8 | 0 | 10 | 0.2 | 34 | 0.0882 |
| 7_12656950 | 11 | 0.2727 | 17 | 0.5882 | 19 | 0.3684 | 24 | 0.3333 | 71 | 0.3944 | 22 | 0.3182 | 17 | 0.2941 | 11 | 0.1818 | 16 | 0.25 | 66 | 0.2727 |
| 7_12656957 | 23 | 0.3478 | 23 | 0.3478 | 21 | 0.4762 | 25 | 0.32 | 92 | 0.3696 | 22 | 0.2727 | 23 | 0.6087 | 12 | 0.5 | 17 | 0.4118 | 74 | 0.4459 |
| 7_12657904 | 19 | 0.2105 | 32 | 0.125 | 24 | 0.0833 | 13 | 0 | 88 | 0.1136 | 20 | 0.05 | 16 | 0.125 | 19 | 0.0526 | 19 | 0.0526 | 74 | 0.0676 |
| 7_12657917 | 20 | 0.6 | 34 | 0.5882 | 20 | 0.45 | 14 | 0.4286 | 88 | 0.5341 | 21 | 0.5238 | 16 | 0.625 | 21 | 0.619 | 17 | 0.7059 | 75 | 0.6133 |
| 7_12657918 | 14 | 0.1429 | 28 | 0.1429 | 20 | 0 | 12 | 0.0833 | 74 | 0.0946 | 19 | 0.1579 | 14 | 0.2143 | 20 | 0.05 | 18 | 0.1667 | 71 | 0.1408 |
| 7_12657920 | 20 | 0.05 | 31 | 0.0645 | 21 | 0.1429 | 14 | 0 | 86 | 0.0698 | 23 | 0.087 | 13 | 0.1538 | 21 | 0.1429 | 19 | 0.0526 | 76 | 0.1053 |
| 7_12657922 | 12 | 0.5 | 27 | 0.5926 | 19 | 0.5789 | 10 | 0.2 | 68 | 0.5147 | 16 | 0.5625 | 14 | 0.6429 | 18 | 0.5 | 15 | 0.3333 | 63 | 0.5079 |
| 7_12661436 | 14 | 0.2857 | 13 | 0.1538 | 10 | 0.3 | 10 | 0.2 | 47 | 0.234 | 2 | 0.5 | 8 | 0.125 | 8 | 0.25 | 10 | 0.4 | 28 | 0.2857 |
| 7_12661441 | 9 | 0.1111 | 12 | 0.5833 | 11 | 0.4545 | 10 | 0.4 | 42 | 0.4048 | 2 | 0 | 6 | 0.3333 | 6 | 0 | 8 | 0.5 | 22 | 0.2727 |
| 7_12661444 | 12 | 0.4167 | 7 | 0.5714 | 6 | 0.1667 | 6 | 0.1667 | 31 | 0.3548 | 2 | 0.5 | 5 | 0.4 | 4 | 0.25 | 8 | 0.25 | 19 | 0.3158 |
| 7_12661540 | 10 | 0.1 | 13 | 0.0769 | 9 | 0.1111 | 8 | 0.125 | 40 | 0.1 | 1 | 0 | 9 | 0.1111 | 7 | 0 | 10 | 0.1 | 27 | 0.0741 |
| 7_12661574 | 13 | 0.0769 | 9 | 0 | 8 | 0.375 | 9 | 0.3333 | 39 | 0.1795 | 1 | 1 | 5 | 0.4 | 5 | 0.4 | 10 | 0 | 21 | 0.2381 |
| 7_12661599 | 10 | 0.5 | 6 | 0 | 7 | 0.2857 | 7 | 0.5714 | 30 | 0.3667 | 1 | 0 | 4 | 0.5 | 3 | 0.6667 | 6 | 0.3333 | 14 | 0.4286 |
| 7_12661620 | 10 | 0.3 | 7 | 0 | 5 | 0 | 6 | 0.3333 | 28 | 0.1786 | 2 | 0 | 9 | 0.2222 | 5 | 0 | 5 | 0.2 | 21 | 0.1429 |
| 7_12661700 | 31 | 0.0968 | 24 | 0.0417 | 18 | 0 | 31 | 0.0645 | 104 | 0.0577 | 13 | 0.1538 | 23 | 0.0435 | 24 | 0.0417 | 19 | 0.1053 | 79 | 0.0759 |
| 7_12661910 | 16 | 0.1875 | 12 | 0.25 | 11 | 0 | 15 | 0.2 | 54 | 0.1667 | 9 | 0.1111 | 16 | 0.1875 | 17 | 0.3529 | 17 | 0.1176 | 59 | 0.2034 |
| 7_12661928 | 16 | 0.1875 | 11 | 0.0909 | 13 | 0.1538 | 11 | 0.6364 | 51 | 0.2549 | 7 | 0.4286 | 15 | 0.3333 | 14 | 0.2857 | 16 | 0.0625 | 52 | 0.25 |
| 7_12662193 | 20 | 0.05 | 18 | 0.1667 | 13 | 0.1538 | 9 | 0.1111 | 60 | 0.1167 | 15 | 0.1333 | 15 | 0.1333 | 18 | 0.2222 | 17 | 0.1176 | 65 | 0.1538 |
| 7_12662195 | 12 | 0.4167 | 11 | 0.3636 | 14 | 0.5714 | 6 | 0.5 | 43 | 0.4651 | 11 | 0.4545 | 13 | 0.4615 | 13 | 0.6154 | 11 | 0.4545 | 48 | 0.5 |
| 7_12662424 | 6 | 0.6667 | 2 | 0.5 | 1 | 1 | 4 | 1 | 13 | 0.7692 | 1 | 1 | NA | NA | 1 | 1 | 1 | 1 | 3 | 1 |
| 7_12662456 | 18 | 0.6111 | 9 | 0.6667 | 11 | 0.6364 | 7 | 1 | 45 | 0.6889 | 11 | 0.7273 | 7 | 1 | 7 | 0.8571 | 10 | 0.6 | 35 | 0.7714 |
| 7_12662515 | 9 | 0.1111 | 8 | 0.25 | 9 | 0.3333 | 5 | 0.2 | 31 | 0.2258 | 8 | 0.25 | 4 | 0.25 | 5 | 0.4 | 6 | 0.1667 | 23 | 0.2609 |
| 7_12662526 | 9 | 0.8889 | 7 | 0.7143 | 9 | 0.6667 | 5 | 0.6 | 30 | 0.7333 | 6 | 0.3333 | 4 | 0.5 | 5 | 0.2 | 7 | 0.8571 | 22 | 0.5 |
| 7_12662550 | 5 | 0 | 7 | 0.1429 | 6 | 0.1667 | 2 | 0 | 20 | 0.1 | 7 | 0.1429 | 3 | 0.3333 | 3 | 0.3333 | 5 | 0.2 | 18 | 0.2222 |
| 7_12662553 | 4 | 0.75 | 5 | 0.4 | 5 | 1 | 1 | 1 | 15 | 0.7333 | 2 | 0 | 1 | 1 | 4 | 1 | 6 | 0.8333 | 13 | 0.7692 |
| 7_12670724 | NA | NA | 1 | 1 | NA | NA | NA | NA | 1 | 1 | 1 | 1 | NA | NA | NA | NA | NA | NA | 1 | 1 |
| 7_12679103 | 1 | 1 | NA | NA | NA | NA | NA | NA | 1 | 1 | NA | NA | 1 | 1 | 1 | 0 | NA | NA | 2 | 0.5 |
| 7_12679137 | 4 | 0.5 | 10 | 0.2 | 8 | 0.125 | 8 | 0.125 | 30 | 0.2 | 8 | 0.375 | 10 | 0.3 | 10 | 0 | 4 | 0 | 32 | 0.1875 |
| 7_12679253 | 1 | 0 | 6 | 0.1667 | 2 | 1 | 3 | 0 | 12 | 0.25 | 4 | 0.25 | 5 | 0.2 | 5 | 0.4 | 2 | 0.5 | 16 | 0.3125 |
| 7_12679257 | 2 | 0.5 | 4 | 0 | 2 | 0.5 | 3 | 0 | 11 | 0.1818 | 6 | 0.1667 | 4 | 0.25 | 5 | 0.2 | 2 | 0 | 17 | 0.1765 |
| 7_12679258 | 2 | 0 | 3 | 0 | 2 | 0 | 4 | 0.25 | 11 | 0.0909 | 6 | 0.1667 | 4 | 0 | 5 | 0 | 2 | 0.5 | 17 | 0.1176 |
| 7_12679268 | 2 | 0 | 4 | 0.25 | 1 | 0 | 2 | 0 | 9 | 0.1111 | 6 | 0.3333 | 5 | 0 | 5 | 0 | 2 | 0.5 | 18 | 0.1667 |
| 7_12679277 | 2 | 0.5 | 5 | 0.2 | 2 | 0.5 | 3 | 0.3333 | 12 | 0.3333 | 6 | 0.3333 | 2 | 1 | 5 | 0.2 | 2 | 1 | 15 | 0.4667 |
| 7_12679286 | 6 | 0.6667 | 5 | 0.2 | 5 | 0.4 | 4 | 0 | 20 | 0.35 | 7 | 0.1429 | 2 | 0.5 | 8 | 0.125 | 3 | 0.3333 | 20 | 0.2 |
| 7_12679294 | 4 | 0.25 | 3 | 0.3333 | 3 | 0.6667 | 2 | 0.5 | 12 | 0.4167 | 8 | 0.375 | 2 | 0.5 | 8 | 0.375 | 3 | 1 | 21 | 0.4762 |
| 7_12679343 | 3 | 0 | 4 | 0.25 | NA | NA | 2 | 0 | 9 | 0.1111 | 5 | 0.4 | 3 | 0.3333 | 9 | 0.4444 | 2 | 0 | 19 | 0.3684 |
| 7_12679361 | 6 | 0.5 | 4 | 0.25 | 2 | 0 | 1 | 1 | 13 | 0.3846 | 6 | 0.3333 | 3 | 0.6667 | 8 | 0.25 | 2 | 1 | 19 | 0.4211 |
| 7_12679368 | 4 | 0.5 | 3 | 1 | 2 | 1 | 1 | 0 | 10 | 0.7 | 5 | 1 | 2 | 1 | 7 | 0.7143 | 1 | 0 | 15 | 0.8 |
| 7_12679370 | 6 | 0.5 | 4 | 0.5 | 1 | 1 | 1 | 0 | 12 | 0.5 | 6 | 0.3333 | 3 | 0.6667 | 7 | 0.5714 | 1 | 1 | 17 | 0.5294 |
| 7_12679756 | 3 | 0.6667 | 8 | 0.5 | 6 | 0.6667 | 10 | 0.4 | 27 | 0.5185 | 8 | 0.5 | 12 | 0.9167 | 6 | 0.5 | 6 | 0.3333 | 32 | 0.625 |
| 7_12679758 | 1 | 1 | 8 | 0.125 | 3 | 0.3333 | 7 | 0.2857 | 19 | 0.2632 | 6 | 0.3333 | 9 | 0.4444 | 6 | 0.5 | 4 | 0.25 | 25 | 0.4 |
| 7_12679761 | 3 | 0 | 8 | 0.125 | 6 | 0 | 9 | 0.1111 | 26 | 0.0769 | 8 | 0.25 | 12 | 0.1667 | 6 | 0 | 6 | 0 | 32 | 0.125 |
| 7_12679774 | 3 | 0 | 9 | 0.2222 | 6 | 0.1667 | 10 | 0 | 28 | 0.1071 | 8 | 0.125 | 8 | 0.125 | 6 | 0.3333 | 4 | 0 | 26 | 0.1538 |
| 7_12679778 | 3 | 0.6667 | 8 | 0.25 | 5 | 0.8 | 7 | 0.8571 | 23 | 0.6087 | 5 | 0.6 | 8 | 0.25 | 5 | 0 | 3 | 0 | 21 | 0.2381 |
| 7_12679783 | 3 | 0 | 7 | 0.1429 | 4 | 0.25 | 7 | 0.2857 | 21 | 0.1905 | 5 | 0 | 8 | 0.125 | 3 | 0.3333 | 5 | 0 | 21 | 0.0952 |
| 7_12679808 | 1 | 0 | 6 | 0.8333 | 6 | 0.6667 | 8 | 0.75 | 21 | 0.7143 | 5 | 0.6 | 9 | 0.8889 | 6 | 0.8333 | 4 | 0.75 | 24 | 0.7917 |
| 7_12679848 | 2 | 0 | 6 | 0.3333 | 6 | 0.1667 | 5 | 0.2 | 19 | 0.2105 | 5 | 0.2 | 7 | 0.1429 | 6 | 0.1667 | 2 | 0 | 20 | 0.15 |
| 7_12680670 | NA | NA | 4 | 0 | 3 | 0 | 4 | 0.25 | 11 | 0.0909 | 4 | 0 | 4 | 0 | 2 | 0.5 | NA | NA | 10 | 0.1 |
| 7_12680676 | NA | NA | 4 | 1 | 4 | 0.75 | 5 | 1 | 13 | 0.9231 | 4 | 0.75 | 4 | 1 | 2 | 1 | 1 | 0 | 11 | 0.8182 |
| 7_12680681 | NA | NA | 3 | 0.6667 | 2 | 0.5 | 5 | 0.4 | 10 | 0.5 | 2 | 0.5 | 4 | 0.75 | 2 | 1 | 1 | 0 | 9 | 0.6667 |
| 7_12680685 | NA | NA | 3 | 0.3333 | 1 | 1 | 4 | 0.5 | 8 | 0.5 | 4 | 0 | 3 | 0.3333 | 2 | 0 | 1 | 0 | 10 | 0.1 |
| 7_12680692 | NA | NA | 4 | 1 | 2 | 1 | 5 | 0.6 | 11 | 0.8182 | 3 | 0.3333 | 5 | 0.8 | 2 | 0.5 | 1 | 1 | 11 | 0.6364 |
| 7_12680701 | NA | NA | 3 | 0.6667 | 3 | 0.3333 | 3 | 0.6667 | 9 | 0.5556 | 3 | 0 | 5 | 0.6 | 2 | 1 | 1 | 1 | 11 | 0.5455 |
| 7_12680703 | NA | NA | 3 | 0.6667 | 2 | 1 | 4 | 0.5 | 9 | 0.6667 | 3 | 1 | 5 | 0.6 | 2 | 1 | 1 | 0 | 11 | 0.7273 |
| 7_12680711 | NA | NA | 3 | 0 | 3 | 0 | 4 | 1 | 10 | 0.4 | 2 | 0.5 | 5 | 0.4 | 2 | 0.5 | NA | NA | 9 | 0.4444 |
| 7_12680713 | NA | NA | 3 | 0 | 3 | 0 | 5 | 0.2 | 11 | 0.0909 | 3 | 0.3333 | 5 | 0.2 | 2 | 0 | 1 | 0 | 11 | 0.1818 |
| 7_12680713 | NA | NA | 3 | 0.6667 | 3 | 0.3333 | 5 | 0.4 | 11 | 0.4545 | 3 | 0.3333 | 5 | 0.2 | 2 | 0 | NA | NA | 10 | 0.2 |
| 7_12680718 | NA | NA | 2 | 0 | 1 | 0 | 5 | 0.2 | 8 | 0.125 | 3 | 0 | 4 | 0.5 | 1 | 0 | 1 | 0 | 9 | 0.2222 |
| 7_12680718 | NA | NA | 2 | 0.5 | 2 | 0 | 3 | 0.6667 | 7 | 0.4286 | 2 | 0 | 3 | 0.3333 | NA | NA | 1 | 0 | 6 | 0.1667 |
| 7_12683459 | NA | NA | NA | NA | NA | NA | 1 | 1 | 1 | 1 | NA | NA | NA | NA | 1 | 1 | 2 | 1 | 3 | 1 |
| 7_12683481 | NA | NA | NA | NA | NA | NA | 1 | 1 | 1 | 1 | NA | NA | NA | NA | 1 | 1 | 2 | 0.5 | 3 | 0.6667 |
| 7_12683505 | NA | NA | NA | NA | NA | NA | 1 | 1 | 1 | 1 | NA | NA | NA | NA | 1 | 1 | 2 | 0.5 | 3 | 0.6667 |
| 7_12683508 | NA | NA | NA | NA | NA | NA | 1 | 1 | 1 | 1 | NA | NA | NA | NA | 1 | 0 | 2 | 0.5 | 3 | 0.3333 |
| 7_12683517 | NA | NA | NA | NA | NA | NA | 2 | 0.5 | 2 | 0.5 | 1 | 0 | NA | NA | 1 | 0 | 2 | 0.5 | 4 | 0.25 |

| | | | | | | | | | | | | | | | | | | | | |
|---|---|---|---|---|---|---|---|---|---|---|---|---|---|---|---|---|---|---|---|---|
| 7_1268355 | NA | NA | NA | NA | 1 | 1 | 2 | 1 | 3 | 1 | 1 | 1 | NA | NA | 2 | 1 | 2 | 1 | 5 | 1 |
| 7_1268358 | NA | NA | NA | NA | 1 | 1 | 2 | 1 | 3 | 1 | 2 | 0.5 | NA | NA | 2 | 1 | 1 | 1 | 5 | 0.8 |
| 7_1268362 | NA | NA | NA | NA | 1 | 1 | 2 | 0.5 | 3 | 0.6667 | NA | NA | NA | NA | 3 | 1 | 3 | 0.3333 | 6 | 0.6667 |
| 7_1268369 | 2 | 0 | 1 | 0 | 1 | 1 | 4 | 0 | 8 | 0.125 | 4 | 0.25 | 6 | 0.1667 | 3 | 0 | 5 | 0.2 | 18 | 0.1667 |
| 7_1268370 | 2 | 0 | 1 | 0 | 1 | 1 | 4 | 0 | 8 | 0.125 | 4 | 0 | 6 | 0 | 2 | 0 | 6 | 0.3333 | 18 | 0.1111 |
| 7_1268370 | NA | NA | 1 | 1 | NA | NA | 4 | 0 | 5 | 0.2 | 4 | 0.75 | 8 | 0.375 | 1 | 0 | 6 | 0.6667 | 19 | 0.5263 |
| 7_1268372 | 1 | 1 | NA | NA | NA | NA | 1 | 1 | 2 | 1 | 1 | 1 | NA | NA | 1 | 0 | 2 | 1 | 4 | 0.75 |
| 7_1268530 | 4 | 0 | 5 | 0.2 | 3 | 0.3333 | 9 | 0.3333 | 21 | 0.2381 | 5 | 0.4 | 7 | 0.5714 | 6 | 0.3333 | 9 | 0.3333 | 27 | 0.4074 |
| 7_1268532 | 5 | 0 | 6 | 0.1667 | 4 | 0.25 | 10 | 0.1 | 25 | 0.12 | 7 | 0 | 9 | 0.1111 | 6 | 0 | 12 | 0.0833 | 34 | 0.0588 |
| 7_1268533 | 10 | 1 | 8 | 0.875 | 5 | 1 | 10 | 1 | 33 | 0.9697 | 10 | 1 | 10 | 0.8 | 7 | 0.8571 | 13 | 1 | 40 | 0.925 |
| 7_1268537 | 11 | 0.2727 | 11 | 0.3636 | 5 | 0.2 | 9 | 0.5556 | 36 | 0.3611 | 9 | 0.3333 | 7 | 0.1429 | 6 | 0.5 | 13 | 0.4615 | 35 | 0.3714 |
| 7_1268538 | 10 | 0.9 | 11 | 0.9091 | 6 | 1 | 9 | 1 | 36 | 0.9444 | 9 | 1 | 7 | 1 | 6 | 1 | 12 | 1 | 34 | 1 |
| 7_1268538 | 9 | 0.8889 | 9 | 0.8889 | 5 | 0.8 | 5 | 1 | 28 | 0.8929 | 9 | 0.8889 | 5 | 0.8 | 7 | 0.7143 | 12 | 0.8333 | 33 | 0.8182 |
| 7_1268562 | 5 | 0 | NA | NA | 5 | 0.2 | 4 | 0 | 14 | 0.0714 | 2 | 0 | 4 | 0 | 1 | 0 | 5 | 0.2 | 12 | 0.0833 |
| 7_1268710 | 7 | 0.7143 | 5 | 0.6 | 5 | 0.2 | 5 | 0.8 | 22 | 0.5909 | 5 | 0 | 4 | 0.5 | 9 | 0.6667 | 7 | 0.8571 | 25 | 0.56 |
| 7_1268711 | 12 | 0.0833 | 6 | 0.1667 | 6 | 0.3333 | 8 | 0.25 | 32 | 0.1875 | 6 | 0 | 5 | 0.2 | 9 | 0 | 8 | 0.25 | 28 | 0.1071 |
| 7_1268712 | 8 | 0 | 7 | 0.2857 | 4 | 0 | 6 | 0.1667 | 25 | 0.12 | 3 | 0 | 4 | 0.25 | 7 | 0 | 7 | 0.1429 | 21 | 0.0952 |
| 7_1268713 | 12 | 0.3333 | 8 | 0.5 | 4 | 0.75 | 6 | 0.6667 | 30 | 0.5 | 7 | 0.1429 | 5 | 0.2 | 7 | 0.2857 | 8 | 0.25 | 27 | 0.2222 |
| 7_1268713 | 10 | 0.3 | 8 | 0.625 | 6 | 0.3333 | 5 | 0.8 | 29 | 0.4828 | 5 | 0.2 | 5 | 0.4 | 9 | 0.5556 | 7 | 0.5714 | 26 | 0.4615 |
| 7_1268716 | 10 | 0.5 | 6 | 0.5 | 6 | 0.5 | 6 | 0.8333 | 28 | 0.5714 | 5 | 0.8 | 4 | 0.75 | 7 | 0.4286 | 6 | 0.5 | 22 | 0.5909 |
| 7_1268716 | 9 | 1 | 5 | 1 | 5 | 1 | 5 | 1 | 24 | 1 | 6 | 1 | 5 | 1 | 9 | 1 | 7 | 1 | 27 | 1 |
| 7_1268717 | 10 | 0.9 | 7 | 1 | 5 | 0.6 | 6 | 1 | 28 | 0.8929 | 4 | 1 | 4 | 1 | 8 | 0.875 | 7 | 1 | 23 | 0.9565 |
| 7_1268718 | 9 | 0.1111 | 7 | 0.1429 | 5 | 0 | 4 | 0 | 25 | 0.08 | 3 | 0.6667 | 4 | 0.25 | 7 | 0 | 5 | 0.2 | 19 | 0.2105 |
| 7_1268721 | 10 | 0.4 | 6 | 0.6667 | 5 | 0.2 | 5 | 0.6 | 26 | 0.4615 | 5 | 0.2 | 4 | 0.75 | 6 | 0.6667 | 7 | 0.5714 | 22 | 0.5455 |
| 7_1268721 | 10 | 0.9 | 6 | 1 | 4 | 1 | 4 | 1 | 24 | 0.9583 | 6 | 1 | 3 | 1 | 9 | 0.8889 | 7 | 1 | 25 | 0.96 |
| 7_1268721 | 8 | 1 | 6 | 1 | 5 | 1 | 4 | 1 | 23 | 1 | 5 | 1 | 4 | 1 | 9 | 1 | 8 | 1 | 26 | 1 |
| 7_1268723 | 6 | 0 | 6 | 0.5 | 5 | 0.2 | 3 | 0 | 20 | 0.2 | 5 | 0.2 | 3 | 0 | 9 | 0.3333 | 7 | 0.1429 | 24 | 0.2083 |
| 7_1268724 | 8 | 0.125 | 5 | 0.2 | 5 | 0.4 | 4 | 0 | 22 | 0.1818 | 5 | 0.4 | 2 | 0 | 8 | 0.375 | 7 | 0.1429 | 22 | 0.2727 |
| 7_1268724 | 7 | 0.4286 | 5 | 0 | 5 | 0.2 | 4 | 0.5 | 21 | 0.2857 | 5 | 0.4 | 2 | 1 | 8 | 0.375 | 6 | 0.3333 | 21 | 0.4286 |
| 7_1268727 | 6 | 0.8333 | 5 | 0.8 | 5 | 0.6 | 4 | 0.25 | 20 | 0.65 | 4 | 1 | 2 | 0.5 | 6 | 0.5 | 8 | 0.75 | 20 | 0.7 |
| 7_1268728 | 7 | 0.4286 | 3 | 0.3333 | 3 | 0.3333 | 4 | 0.25 | 17 | 0.3529 | 4 | 0.25 | 2 | 0 | 6 | 0.6667 | 6 | 0.3333 | 18 | 0.3889 |
| 7_1268729 | 5 | 0.2 | 4 | 0.25 | 4 | 0 | 2 | 0 | 15 | 0.1333 | 3 | 0.3333 | 2 | 0 | 5 | 0 | 6 | 0 | 16 | 0.0625 |
| 7_1269815 | NA | NA | 2 | 1 | NA | NA | NA | NA | 2 | 1 | 1 | 1 | NA | NA | NA | NA | NA | NA | 1 | 1 |
| 7_1270030 | 4 | 0.25 | 1 | 0 | NA | NA | 1 | 0 | 6 | 0.1667 | 1 | 1 | 1 | 0 | NA | NA | 1 | 0 | 3 | 0.3333 |
| 7_1270033 | 4 | 1 | 1 | 0 | NA | NA | NA | NA | 5 | 0.8 | NA | NA | 1 | 1 | NA | NA | 1 | 0 | 2 | 0.5 |
| 7_1270037 | 3 | 0.3333 | 1 | 0 | NA | NA | 1 | 0 | 5 | 0.2 | NA | NA | 1 | 0 | NA | NA | 2 | 0.5 | 3 | 0.3333 |
| 7_1270038 | 4 | 0.5 | 1 | 1 | NA | NA | 1 | 0 | 6 | 0.5 | NA | NA | 1 | 1 | NA | NA | 1 | 1 | 2 | 1 |
| 7_1270039 | 4 | 1 | 1 | 1 | NA | NA | 1 | 0 | 6 | 0.8333 | NA | NA | 1 | 1 | NA | NA | 2 | 1 | 3 | 1 |
| 7_1270040 | 4 | 0.75 | 1 | 1 | NA | NA | NA | NA | 5 | 0.8 | NA | NA | 1 | 0 | NA | NA | 1 | 1 | 2 | 0.5 |
| 7_1270043 | 3 | 0 | 1 | 1 | NA | NA | 1 | 1 | 5 | 0.4 | NA | NA | 1 | 1 | NA | NA | 2 | 0.5 | 3 | 0.6667 |
| 7_1270044 | 2 | 1 | 1 | 1 | NA | NA | 1 | 1 | 4 | 1 | NA | NA | 1 | 1 | NA | NA | 2 | 1 | 3 | 1 |
| 7_1270045 | 4 | 0.5 | 1 | 1 | NA | NA | 1 | 0 | 6 | 0.5 | NA | NA | 1 | 1 | NA | NA | 2 | 1 | 3 | 1 |
| 7_1270050 | NA | NA | NA | NA | NA | NA | 1 | 1 | 1 | 1 | NA | NA | NA | NA | NA | NA | 1 | 1 | 1 | 1 |
| 7_1270052 | NA | NA | NA | NA | NA | NA | 1 | 1 | 1 | 1 | NA | NA | NA | NA | NA | NA | 1 | 1 | 1 | 1 |
| 7_1270053 | NA | NA | NA | NA | NA | NA | 1 | 1 | 1 | 1 | NA | NA | NA | NA | NA | NA | 1 | 1 | 1 | 1 |
| 7_1270182 | 2 | 0.5 | 6 | 0.5 | 1 | 0 | 4 | 0 | 13 | 0.3077 | 3 | 0 | 3 | 0 | 5 | 0.4 | NA | NA | 11 | 0.1818 |
| 7_1270182 | 1 | 1 | 5 | 1 | 1 | 1 | 3 | 1 | 10 | 1 | 3 | 1 | 3 | 1 | 5 | 1 | NA | NA | 11 | 1 |
| 7_1270182 | 1 | 1 | 5 | 0.6 | NA | NA | 3 | 0 | 9 | 0.4444 | 3 | 0.6667 | 2 | 0.5 | 4 | 0 | NA | NA | 9 | 0.3333 |
| 7_1270183 | 2 | 1 | 6 | 0.6667 | 1 | 1 | 3 | 0.3333 | 12 | 0.6667 | 2 | 0.5 | 3 | 1 | 4 | 0.5 | NA | NA | 9 | 0.6667 |
| 7_1270185 | 2 | 1 | 6 | 0.8333 | NA | NA | 2 | 1 | 10 | 0.9 | 3 | 1 | 3 | 1 | 5 | 1 | NA | NA | 11 | 1 |
| 7_1270187 | 1 | 1 | 4 | 1 | NA | NA | 4 | 0.5 | 9 | 0.7778 | 2 | 1 | 2 | 1 | 3 | 1 | NA | NA | 7 | 1 |
| 7_1270433 | 1 | 1 | 1 | 1 | NA | NA | NA | NA | 2 | 1 | NA | NA | 2 | 1 | NA | NA | NA | NA | 2 | 1 |
| 7_1270434 | 1 | 1 | NA | NA | NA | NA | 1 | 1 | 2 | 1 | NA | NA | 2 | 1 | NA | NA | NA | NA | 2 | 1 |
| 7_1270435 | NA | NA | NA | NA | NA | NA | 1 | 1 | 1 | 1 | NA | NA | 1 | 1 | NA | NA | NA | NA | 1 | 1 |
| 7_1270436 | 1 | 1 | NA | NA | NA | NA | 1 | 1 | 2 | 1 | NA | NA | 1 | 1 | NA | NA | NA | NA | 1 | 1 |
| 7_1270437 | 1 | 1 | NA | NA | NA | NA | 1 | 1 | 2 | 1 | NA | NA | 2 | 1 | NA | NA | NA | NA | 2 | 1 |
| 7_1270439 | NA | NA | NA | NA | NA | NA | 1 | 1 | 1 | 1 | NA | NA | 2 | 1 | NA | NA | NA | NA | 2 | 1 |
| 7_1270478 | 1 | 1 | NA | NA | NA | NA | 1 | 0 | 2 | 0.5 | NA | NA | 2 | 0.5 | NA | NA | NA | NA | 2 | 0.5 |
| 7_1270620 | NA | NA | NA | NA | 1 | 1 | NA | NA | 1 | 1 | 1 | 1 | 3 | 0.6667 | 2 | 1 | NA | NA | 6 | 0.8333 |
| 7_1270623 | NA | NA | NA | NA | 1 | 1 | NA | NA | 1 | 1 | 2 | 0 | 3 | 0.6667 | 2 | 1 | NA | NA | 7 | 0.5714 |
| 7_1270630 | NA | NA | NA | NA | 1 | 1 | NA | NA | 1 | 1 | NA | NA | 1 | 1 | NA | NA | NA | NA | 1 | 1 |
| 7_1270658 | NA | NA | 1 | 0 | 1 | 0 | 2 | 0.5 | 4 | 0.25 | 1 | 0 | NA | NA | NA | NA | 4 | 0.25 | 5 | 0.2 |
| 7_1270658 | NA | NA | 1 | 0 | 2 | 0 | 2 | 0.5 | 5 | 0.2 | 1 | 0 | NA | NA | NA | NA | 3 | 0.3333 | 4 | 0.25 |
| 7_1270661 | NA | NA | 2 | 0.5 | 2 | 0 | 2 | 0.5 | 6 | 0.3333 | 1 | 0 | NA | NA | NA | NA | 4 | 0.5 | 5 | 0.4 |
| 7_1270710 | NA | NA | NA | NA | 1 | 1 | NA | NA | 1 | 1 | NA | NA | 1 | 0 | NA | NA | 1 | 1 | 2 | 0.5 |

| | | | | | | | | | | | | | | | | | | | | |
|---|---|---|---|---|---|---|---|---|---|---|---|---|---|---|---|---|---|---|---|---|
| 7_1270754 | 3 | 0 | 1 | 0 | 1 | 0 | 3 | 0.3333 | 8 | 0.125 | 2 | 0 | 2 | 0.5 | 1 | 0 | 3 | 0 | 8 | 0.125 |
| 7_1270755 | 3 | 0.3333 | 1 | 0 | 1 | 0 | 2 | 0 | 7 | 0.1429 | 1 | 0 | 2 | 0.5 | 1 | 0 | 2 | 0 | 6 | 0.1667 |
| 7_1270756 | 3 | 0 | 1 | 0 | 1 | 0 | 2 | 0.5 | 7 | 0.1429 | 1 | 0 | 2 | 0.5 | 1 | 0 | 3 | 0 | 7 | 0.1429 |
| 7_1270892 | 3 | 1 | 1 | 1 | 1 | 1 | 3 | 0.6667 | 8 | 0.875 | NA | NA | 1 | 0 | NA | NA | 1 | 1 | 2 | 0.5 |
| 7_1270892 | 6 | 0.8333 | 1 | 0 | 2 | 0.5 | 4 | 0.5 | 13 | 0.6154 | NA | NA | 3 | 1 | NA | NA | 1 | 1 | 4 | 1 |
| 7_1270895 | 4 | 0.25 | NA | NA | 3 | 0.3333 | 8 | 0 | 15 | 0.1333 | NA | NA | 2 | 0.5 | NA | NA | NA | NA | 2 | 0.5 |
| 7_1270896 | 5 | 1 | 1 | 1 | 3 | 0.3333 | 10 | 0.8 | 19 | 0.7895 | 1 | 1 | 5 | 0.4 | NA | NA | 1 | 1 | 7 | 0.5714 |
| 7_1271207 | NA | NA | 1 | 1 | 1 | 0 | 1 | 0 | 3 | 0.3333 | NA | NA | NA | NA | NA | NA | 1 | 1 | 1 | 1 |
| 7_1271407 | NA | NA | NA | NA | 2 | 0.5 | 1 | 1 | 3 | 0.6667 | NA | NA | 1 | 1 | 2 | 0 | NA | NA | 3 | 0.3333 |
| 7_1271408 | 1 | 1 | NA | NA | 3 | 0.6667 | 2 | 0.5 | 6 | 0.6667 | NA | NA | NA | NA | 2 | 0 | 2 | 1 | 4 | 0.5 |
| 7_1271410 | 2 | 0 | NA | NA | 3 | 0.3333 | 1 | 0 | 6 | 0.1667 | 1 | 1 | 1 | 1 | 2 | 0 | NA | NA | 4 | 0.5 |
| 7_1271410 | 3 | 0.6667 | NA | NA | 3 | 0.3333 | 2 | 0.5 | 8 | 0.5 | 1 | 1 | 1 | 1 | 4 | 0.25 | 2 | 0 | 8 | 0.375 |
| 7_1271411 | 3 | 0.3333 | NA | NA | 2 | 1 | 2 | 1 | 7 | 0.7143 | 1 | 0 | 1 | 0 | 4 | 0.75 | 2 | 0 | 8 | 0.375 |
| 7_1271412 | 3 | 0.3333 | NA | NA | 3 | 0.3333 | 2 | 1 | 8 | 0.5 | 1 | 1 | 2 | 0.5 | 4 | 1 | 1 | 0 | 8 | 0.75 |
| 7_1271413 | 3 | 0 | NA | NA | 3 | 0 | 2 | 0.5 | 8 | 0.125 | 1 | 0 | 2 | 0.5 | 2 | 0.5 | 2 | 0.5 | 7 | 0.4286 |
| 7_1271413 | 2 | 1 | NA | NA | 3 | 0.6667 | 2 | 1 | 7 | 0.8571 | 1 | 0 | 2 | 1 | 3 | 1 | 2 | 0.5 | 8 | 0.75 |
| 7_1271413 | 2 | 1 | NA | NA | 2 | 0 | NA | NA | 4 | 0.5 | 1 | 1 | 2 | 0.5 | 3 | 0.6667 | 1 | 1 | 7 | 0.7143 |
| 7_1271414 | 2 | 0.5 | NA | NA | 3 | 0.6667 | 2 | 0.5 | 7 | 0.5714 | 1 | 0 | 2 | 0 | 3 | 0.3333 | 2 | 0 | 8 | 0.125 |
| 7_1271415 | 2 | 0.5 | NA | NA | 3 | 1 | 1 | 1 | 6 | 0.8333 | 1 | 0 | 1 | 1 | 2 | 0.5 | 1 | 0 | 5 | 0.4 |
| 7_1271416 | 3 | 0.6667 | NA | NA | 3 | 1 | 2 | 0.5 | 8 | 0.75 | 1 | 1 | 2 | 1 | 4 | 0.75 | 2 | 0 | 9 | 0.6667 |
| 7_1271417 | 2 | 0 | NA | NA | 2 | 0 | 2 | 0.5 | 6 | 0.1667 | 1 | 0 | 2 | 0.5 | 3 | 0 | 2 | 0 | 8 | 0.125 |
| 7_1271417 | 3 | 0.6667 | NA | NA | 3 | 0.6667 | 2 | 0.5 | 8 | 0.625 | 1 | 0 | 1 | 0 | 4 | 0.5 | 2 | 0 | 8 | 0.25 |
| 7_1271483 | 1 | 0 | NA | NA | 2 | 0.5 | 1 | 0 | 4 | 0.25 | 1 | 1 | 1 | 0 | 4 | 0 | 1 | 0 | 7 | 0.1429 |
| 7_1271484 | 2 | 0 | 1 | 0 | 3 | 0.3333 | 2 | 0.5 | 8 | 0.25 | 1 | 0 | 1 | 1 | 4 | 0 | 2 | 1 | 8 | 0.375 |
| 7_1271572 | 4 | 0.5 | 3 | 0.3333 | 4 | 0.5 | 3 | 0 | 14 | 0.3571 | 2 | 0.5 | 5 | 0.2 | 6 | 0.3333 | 2 | 0 | 15 | 0.2667 |
| 7_1271573 | 4 | 0.75 | 4 | 0.75 | 5 | 0.8 | 3 | 0.3333 | 16 | 0.6875 | 2 | 1 | 4 | 1 | 6 | 0.6667 | 3 | 1 | 15 | 0.8667 |
| 7_1271579 | 5 | 1 | 4 | 0.75 | 4 | 1 | 3 | 0.3333 | 16 | 0.8125 | 2 | 1 | 5 | 1 | 6 | 1 | 4 | 1 | 17 | 1 |
| 7_1271580 | 5 | 0.4 | 3 | 0.3333 | 4 | 0.25 | 4 | 0.75 | 16 | 0.4375 | 2 | 0 | 6 | 0 | 5 | 0.4 | 4 | 0.25 | 17 | 0.1765 |
| 7_1271581 | 3 | 0 | 4 | 0.25 | 5 | 0.2 | 3 | 0.3333 | 15 | 0.2 | 2 | 0.5 | 5 | 0.4 | 6 | 0.1667 | 4 | 0.5 | 17 | 0.3529 |
| 7_1271581 | 5 | 0 | 3 | 0 | 5 | 0 | 4 | 0.5 | 17 | 0.1176 | 2 | 0 | 6 | 0 | 6 | 0 | 4 | 0.25 | 18 | 0.0556 |
| 7_1271582 | 5 | 0.6 | 4 | 0.5 | 5 | 0.6 | 4 | 0.5 | 18 | 0.5556 | 2 | 0.5 | 6 | 0.6667 | 5 | 0.8 | 4 | 0.25 | 17 | 0.5882 |
| 7_1271582 | 4 | 0.75 | 3 | 0.6667 | 5 | 1 | 4 | 0.25 | 16 | 0.6875 | 2 | 0 | 5 | 0.6 | 5 | 0.8 | 3 | 0.6667 | 15 | 0.6 |
| 7_1271583 | 3 | 1 | 3 | 1 | 4 | 1 | 3 | 0.6667 | 13 | 0.9231 | 2 | 1 | 3 | 0 | 3 | 0.6667 | 3 | 0.6667 | 11 | 0.5455 |
| 7_1271584 | 5 | 0.4 | 4 | 0.25 | 6 | 0.3333 | 5 | 0 | 20 | 0.25 | 2 | 0 | 5 | 0.2 | 6 | 0.5 | 4 | 0.25 | 17 | 0.2941 |
| 7_1271584 | 5 | 0.2 | 4 | 0.25 | 4 | 0 | 5 | 0.4 | 18 | 0.2222 | 2 | 0 | 6 | 0.1667 | 6 | 0.3333 | 2 | 0 | 16 | 0.1875 |
| 7_1271585 | 3 | 0 | 3 | 0.3333 | 4 | 0.5 | 3 | 1 | 13 | 0.4615 | 2 | 0 | 2 | 0 | 5 | 0.4 | 3 | 0.3333 | 12 | 0.25 |
| 7_1271585 | 5 | 0.4 | 4 | 0.5 | 6 | 0.1667 | 4 | 0.75 | 19 | 0.4211 | 2 | 0 | 3 | 1 | 7 | 0.1429 | 4 | 0.25 | 16 | 0.3125 |
| 7_1271587 | 4 | 0.75 | 4 | 0.75 | 5 | 0.8 | 4 | 1 | 17 | 0.8235 | 1 | 1 | 5 | 0.6 | 7 | 0.5714 | 2 | 1 | 15 | 0.6667 |
| 7_1271587 | 4 | 0 | 4 | 0.25 | 5 | 0 | 5 | 0.2 | 18 | 0.1111 | 2 | 0 | 4 | 0 | 7 | 0.1429 | 4 | 0 | 17 | 0.0588 |
| 7_1271588 | 4 | 0 | 4 | 0 | 4 | 0 | 5 | 0.2 | 17 | 0.0588 | 1 | 1 | 5 | 0.4 | 8 | 0.125 | 1 | 0 | 15 | 0.2667 |
| 7_1271589 | 1 | 1 | 3 | 1 | 3 | 1 | 4 | 0.75 | 11 | 0.9091 | 1 | 1 | 2 | 1 | 5 | 1 | 3 | 0.6667 | 11 | 0.9091 |
| 7_1271589 | 4 | 0 | 3 | 0 | 6 | 0.1667 | 5 | 0 | 18 | 0.0556 | 1 | 0 | 6 | 0 | 6 | 0.6667 | 4 | 0 | 17 | 0.2353 |
| 7_1271590 | 5 | 0.4 | 3 | 0.6667 | 6 | 0.3333 | 6 | 0.1667 | 20 | 0.35 | 2 | 0 | 7 | 0.1429 | 8 | 0.25 | 4 | 0 | 21 | 0.1429 |
| 7_1271590 | 5 | 0 | 4 | 0 | 5 | 0.6 | 6 | 0 | 20 | 0.15 | 2 | 0.5 | 7 | 0 | 8 | 0 | 3 | 0 | 20 | 0.05 |
| 7_1271590 | 5 | 0 | 4 | 0 | 5 | 0 | 5 | 0.2 | 19 | 0.0526 | 2 | 0 | 6 | 0.1667 | 6 | 0 | 4 | 0 | 18 | 0.0556 |
| 7_1271591 | 5 | 0.8 | 2 | 1 | 4 | 0.75 | 4 | 0.75 | 15 | 0.8 | NA | NA | 6 | 1 | 6 | 0.8333 | 3 | 1 | 15 | 0.9333 |
| 7_1271591 | 4 | 0.25 | 4 | 0 | 4 | 0 | 5 | 0 | 17 | 0.0588 | 1 | 0 | 3 | 0.3333 | 8 | 0 | 4 | 0.5 | 16 | 0.1875 |
| 7_1271593 | 5 | 0.8 | 4 | 1 | 7 | 0.5714 | 7 | 0.4286 | 23 | 0.6522 | 3 | 0 | 4 | 0.5 | 10 | 0.3 | 4 | 1 | 21 | 0.4286 |
| 7_1271593 | 5 | 0.8 | 4 | 1 | 7 | 1 | 5 | 1 | 21 | 0.9524 | 3 | 1 | 7 | 1 | 9 | 1 | 4 | 1 | 23 | 1 |
| 7_1271595 | 4 | 0.75 | 3 | 1 | 7 | 1 | 5 | 0.8 | 19 | 0.8947 | 2 | 1 | 6 | 1 | 8 | 0.875 | 3 | 1 | 19 | 0.9474 |
| 7_1271595 | 3 | 0.3333 | 4 | 0 | 8 | 0 | 6 | 0.3333 | 21 | 0.1429 | 3 | 0.3333 | 7 | 0 | 9 | 0.3333 | 4 | 0.25 | 23 | 0.2174 |
| 7_1271596 | 5 | 1 | 3 | 0.6667 | 6 | 0.8333 | 7 | 0.8571 | 21 | 0.8571 | 3 | 1 | 7 | 0.8571 | 9 | 0.8889 | 3 | 1 | 22 | 0.9091 |
| 7_1271598 | 3 | 0.3333 | 4 | 0.25 | 9 | 0.4444 | 6 | 0.1667 | 22 | 0.3182 | 2 | 0.5 | 6 | 0.1667 | 10 | 0.2 | 2 | 0.5 | 20 | 0.25 |
| 7_1271600 | 5 | 0.2 | 5 | 0 | 8 | 0.125 | 6 | 0.3333 | 24 | 0.1667 | 5 | 0.4 | 11 | 0.0909 | 11 | 0.0909 | 3 | 0 | 30 | 0.1333 |
| 7_1271640 | NA | NA | 1 | 0 | 1 | 1 | 2 | 0.5 | 4 | 0.5 | 2 | 0.5 | NA | NA | NA | NA | 1 | 1 | 3 | 0.6667 |
| 7_1271819 | 2 | 1 | 8 | 0.25 | 6 | 0 | 9 | 0.1111 | 25 | 0.2 | 2 | 0.5 | 12 | 0.1667 | 6 | 0.1667 | 5 | 0.2 | 25 | 0.2 |
| 7_1271821 | 4 | 0.5 | 5 | 0.2 | 6 | 0.3333 | 10 | 0.2 | 25 | 0.28 | 3 | 0.3333 | 11 | 0 | 9 | 0 | 7 | 0.1429 | 30 | 0.0667 |
| 7_1271821 | 4 | 0 | 8 | 0 | 7 | 0.1429 | 11 | 0.0909 | 30 | 0.0667 | 3 | 0.3333 | 13 | 0.0769 | 9 | 0 | 7 | 0.1429 | 32 | 0.0938 |
| 7_1271822 | 4 | 0.75 | 9 | 0.8889 | 4 | 0.75 | 11 | 0.7273 | 28 | 0.7857 | 2 | 1 | 13 | 0.8462 | 8 | 0.625 | 7 | 0.5714 | 30 | 0.7333 |
| 7_1271822 | 4 | 0.75 | 8 | 0.75 | 7 | 0.5714 | 12 | 0.75 | 31 | 0.7097 | 3 | 0.6667 | 13 | 0.8462 | 10 | 0.6 | 7 | 0.8571 | 33 | 0.7576 |
| 7_1271822 | 4 | 0.75 | 9 | 0.8889 | 6 | 0.8333 | 11 | 0.9091 | 30 | 0.8667 | 3 | 1 | 12 | 0.8333 | 11 | 0.8182 | 7 | 0.7143 | 33 | 0.8182 |
| 7_1271823 | 2 | 1 | 9 | 0.7778 | 7 | 0.8571 | 10 | 0.9 | 28 | 0.8571 | 4 | 0.75 | 10 | 0.8 | 6 | 0.8333 | 5 | 0.6 | 25 | 0.76 |
| 7_1271955 | 2 | 0.5 | 1 | 0 | NA | NA | NA | NA | 3 | 0.3333 | 3 | 0 | 2 | 0 | 3 | 0 | 2 | 0.5 | 10 | 0.1 |
| 7_1272018 | 8 | 0.25 | 2 | 1 | 3 | 0 | 2 | 0 | 15 | 0.2667 | 3 | 0.3333 | 3 | 0.6667 | 8 | 0.125 | 4 | 0.25 | 18 | 0.2778 |
| 7_1272018 | 7 | 1 | 3 | 1 | 3 | 1 | 2 | 1 | 15 | 1 | 3 | 1 | 3 | 1 | 7 | 1 | 5 | 1 | 18 | 1 |

| | | | | | | | | | | | | | | | | | | | | |
|---|---|---|---|---|---|---|---|---|---|---|---|---|---|---|---|---|---|---|---|---|
| 7_1272019 | 7 | 0.4286 | 3 | 1 | 2 | 0 | 1 | 1 | 13 | 0.5385 | 3 | 0.3333 | 2 | 0.5 | 6 | 0.3333 | 4 | 0.25 | 15 | 0.3333 |
| 7_1272021 | 7 | 0.1429 | 4 | 0.5 | 3 | 0 | 2 | 0 | 16 | 0.1875 | 3 | 0 | 3 | 0.3333 | 8 | 0.5 | 5 | 0.2 | 19 | 0.3158 |
| 7_1272023 | 5 | 0 | 4 | 0.25 | 1 | 0 | 2 | 0 | 12 | 0.0833 | 2 | 0 | 5 | 0 | 7 | 0.1429 | 2 | 0 | 16 | 0.0625 |
| 7_1272023 | 9 | 0 | 5 | 0.2 | 4 | 0.25 | 2 | 0 | 20 | 0.1 | 3 | 0 | 3 | 0 | 9 | 0.1111 | 5 | 0 | 20 | 0.05 |
| 7_1272026 | 8 | 0.625 | 6 | 0.6667 | 2 | 0.5 | 3 | 1 | 19 | 0.6842 | 3 | 0.6667 | 3 | 0.6667 | 7 | 0.7143 | NA | NA | 13 | 0.6923 |
| 7_1272064 | 3 | 0.6667 | 7 | 0 | 1 | 0 | NA | NA | 11 | 0.1818 | 5 | 0.2 | 5 | 0 | 1 | 0 | 3 | 0.3333 | 14 | 0.1429 |
| 7_1272064 | 3 | 0.3333 | 7 | 0 | 1 | 0 | NA | NA | 11 | 0.0909 | 4 | 0.25 | 4 | 0 | 1 | 0 | 3 | 0 | 12 | 0.0833 |
| 7_1272067 | 2 | 0 | 7 | 0.2857 | 1 | 0 | NA | NA | 10 | 0.2 | 4 | 0 | 5 | 0 | 2 | 0 | 3 | 0.3333 | 14 | 0.0714 |
| 7_1272067 | 2 | 0.5 | 6 | 0.1667 | 1 | 1 | NA | NA | 9 | 0.3333 | 3 | 0 | 5 | 0.2 | 2 | 0.5 | 2 | 0.5 | 12 | 0.25 |
| 7_1272068 | 1 | 0 | 6 | 0.3333 | 1 | 0 | NA | NA | 8 | 0.25 | 3 | 0.3333 | 4 | 0.25 | 2 | 0 | 2 | 0 | 11 | 0.1818 |
| 7_1272072 | 1 | 0 | 6 | 0.6667 | 1 | 0 | NA | NA | 8 | 0.5 | 2 | 0 | 3 | 0.3333 | 1 | 0 | 2 | 0 | 8 | 0.125 |
| 7_1272073 | 1 | 0 | 6 | 0.5 | 1 | 0 | NA | NA | 8 | 0.375 | 2 | 0 | 4 | 0.5 | 1 | 0 | 2 | 0 | 9 | 0.2222 |
| 7_1272073 | 1 | 1 | 6 | 0.1667 | 1 | 0 | NA | NA | 8 | 0.25 | 2 | 0 | 4 | 0 | 1 | 1 | 2 | 0 | 9 | 0.1111 |
| 7_1272074 | 1 | 1 | 6 | 0.6667 | 1 | 1 | NA | NA | 8 | 0.75 | 2 | 0 | 4 | 1 | NA | NA | 2 | 0 | 8 | 0.5 |
| 7_1272074 | 1 | 0 | 4 | 0 | 1 | 1 | NA | NA | 6 | 0.1667 | 2 | 0.5 | 3 | 0.6667 | 1 | 0 | 2 | 0.5 | 8 | 0.5 |
| 7_1272075 | NA | NA | 4 | 0.5 | NA | NA | NA | NA | 4 | 0.5 | 2 | 0.5 | 2 | 0.5 | 1 | 1 | 1 | 1 | 6 | 0.6667 |
| 7_1272075 | NA | NA | 5 | 1 | NA | NA | NA | NA | 5 | 1 | 2 | 0.5 | 3 | 1 | NA | NA | 2 | 1 | 7 | 0.8571 |
| 7_1272075 | 1 | 0 | 6 | 0.6667 | 1 | 0 | NA | NA | 8 | 0.5 | 1 | 1 | 3 | 0.6667 | NA | NA | 1 | 0 | 5 | 0.6 |
| 7_1272076 | 2 | 1 | 4 | 0.75 | 1 | 1 | NA | NA | 7 | 0.8571 | NA | NA | 2 | 1 | 1 | 1 | 1 | 1 | 4 | 1 |
| 7_1272076 | 2 | 0.5 | 5 | 0.6 | 1 | 1 | NA | NA | 8 | 0.625 | 2 | 1 | NA | NA | 1 | 1 | 2 | 1 | 5 | 1 |
| 7_1272076 | 2 | 0.5 | 4 | 1 | 1 | 1 | NA | NA | 7 | 0.8571 | 2 | 0.5 | 3 | 0.3333 | 1 | 1 | 2 | 1 | 8 | 0.625 |
| 7_1272077 | 1 | 0 | 5 | 0.8 | 1 | 1 | NA | NA | 7 | 0.7143 | 1 | 1 | 3 | 0.6667 | NA | NA | 1 | 1 | 5 | 0.8 |
| 7_1272078 | NA | NA | 3 | 0.6667 | 1 | 0 | NA | NA | 4 | 0.5 | NA | NA | NA | NA | NA | NA | 1 | 1 | 1 | 1 |
| 7_1272078 | 2 | 1 | 3 | 1 | 1 | 1 | NA | NA | 6 | 1 | 1 | 1 | 1 | 1 | NA | NA | 1 | 1 | 3 | 1 |
| 7_1272078 | 1 | 0 | 6 | 0 | 1 | 1 | NA | NA | 8 | 0.125 | 2 | 0.5 | 3 | 0.3333 | 1 | 1 | 2 | 0 | 8 | 0.375 |
| 7_1272079 | 1 | 1 | 4 | 0.25 | 1 | 0 | NA | NA | 6 | 0.3333 | 2 | 0.5 | 2 | 0.5 | 1 | 0 | 1 | 0 | 6 | 0.3333 |
| 7_1272079 | 2 | 0 | 6 | 0.1667 | 1 | 0 | NA | NA | 9 | 0.1111 | 1 | 1 | 2 | 0 | NA | NA | 2 | 0 | 5 | 0.2 |
| 7_1272079 | 2 | 0.5 | 6 | 0.3333 | 1 | 1 | NA | NA | 9 | 0.4444 | 2 | 0 | 3 | 0.6667 | NA | NA | 2 | 0.5 | 7 | 0.4286 |
| 7_1272084 | 1 | 1 | 3 | 0 | 1 | 0 | 1 | 0 | 6 | 0.1667 | 1 | 1 | 2 | 0 | NA | NA | NA | NA | 3 | 0.3333 |
| 7_1272085 | 1 | 0 | 4 | 0.25 | 1 | 0 | 1 | 0 | 7 | 0.1429 | 1 | 1 | 2 | 0 | NA | NA | 1 | 0 | 4 | 0.25 |
| 7_1272091 | NA | NA | 3 | 0.6667 | NA | NA | 1 | 0 | 4 | 0.5 | 1 | 1 | 2 | 0.5 | 1 | 0 | 1 | 1 | 5 | 0.6 |
| 7_1272097 | NA | NA | 1 | 1 | NA | NA | 1 | 1 | 2 | 1 | 1 | 1 | 2 | 1 | NA | NA | NA | NA | 3 | 1 |
| 7_1272098 | NA | NA | 2 | 1 | NA | NA | NA | NA | 2 | 1 | 1 | 1 | 1 | 1 | 1 | 1 | NA | NA | 3 | 1 |
| 7_1272104 | NA | NA | 1 | 1 | NA | NA | 1 | 0 | 2 | 0.5 | 1 | 1 | NA | NA | NA | NA | NA | NA | 1 | 1 |
| 7_1272115 | NA | NA | NA | NA | NA | NA | 1 | 1 | 1 | 1 | NA | NA | 1 | 1 | NA | NA | NA | NA | 1 | 1 |
| 7_1272467 | NA | NA | NA | NA | 1 | 1 | NA | NA | 1 | 1 | NA | NA | 1 | 1 | NA | NA | NA | NA | 1 | 1 |
| 7_1273513 | 2 | 0.5 | 1 | 0 | 1 | 0 | 1 | 0 | 5 | 0.2 | NA | NA | 2 | 0 | 1 | 1 | NA | NA | 3 | 0.3333 |
| 7_1273514 | 2 | 0 | 1 | 1 | 1 | 0 | 1 | 1 | 5 | 0.4 | NA | NA | 3 | 0.3333 | 1 | 1 | NA | NA | 4 | 0.5 |
| 7_1273670 | 2 | 1 | 1 | 1 | 1 | 1 | 1 | 1 | 5 | 1 | NA | NA | 2 | 1 | 1 | 1 | NA | NA | 3 | 1 |
| 7_1273671 | 2 | 0.5 | 1 | 1 | 1 | 0 | 1 | 1 | 5 | 0.6 | NA | NA | 3 | 1 | 1 | 1 | NA | NA | 4 | 1 |
| 7_1273671 | 3 | 0.3333 | 1 | 0 | 3 | 0.3333 | 1 | 0 | 8 | 0.25 | NA | NA | 3 | 0.6667 | 1 | 0 | NA | NA | 4 | 0.5 |
| 7_1273672 | 3 | 0 | NA | NA | 3 | 0.6667 | 1 | 0 | 7 | 0.2857 | NA | NA | 4 | 0.75 | 1 | 1 | NA | NA | 5 | 0.8 |
| 7_1273672 | 3 | 1 | 1 | 1 | 3 | 0.6667 | NA | NA | 7 | 0.8571 | NA | NA | 4 | 1 | 1 | 1 | 1 | 1 | 6 | 1 |
| 7_1273676 | 3 | 0 | 3 | 0 | 1 | 1 | 2 | 0 | 9 | 0.1111 | 1 | 0 | 4 | 0.25 | NA | NA | NA | NA | 5 | 0.2 |
| 7_1273678 | 2 | 0 | 4 | 0.5 | 3 | 0 | 4 | 0.5 | 13 | 0.3077 | 1 | 0 | 4 | 0.25 | 1 | 0 | 2 | 0 | 8 | 0.125 |
| 7_1273679 | 2 | 1 | 3 | 0.6667 | 3 | 0.3333 | 5 | 0.8 | 13 | 0.6923 | 2 | 1 | 3 | 0.3333 | 1 | 1 | 2 | 1 | 8 | 0.75 |
| 7_1273679 | 3 | 1 | 4 | 0 | 4 | 0 | 4 | 0.5 | 15 | 0.3333 | 2 | 0.5 | 3 | 0.3333 | 2 | 1 | 2 | 1 | 9 | 0.6667 |
| 7_1273681 | 1 | 0 | 4 | 0.25 | 3 | 0 | 3 | 0 | 11 | 0.0909 | 2 | 0 | 3 | 0.3333 | NA | NA | 1 | 1 | 6 | 0.3333 |
| 7_1273682 | NA | NA | 3 | 1 | 3 | 0.3333 | 2 | 0 | 8 | 0.5 | 1 | 0 | 4 | 0.25 | 1 | 1 | 1 | 0 | 7 | 0.2857 |
| 7_1273682 | NA | NA | 4 | 0 | 4 | 0 | 5 | 0.2 | 13 | 0.0769 | 3 | 0 | 4 | 0.25 | 1 | 1 | 3 | 0.6667 | 11 | 0.3636 |
| 7_1273692 | NA | NA | 1 | 0 | 1 | 0 | 1 | 1 | 3 | 0.3333 | 1 | 0 | 1 | 0 | 3 | 0 | 2 | 0.5 | 7 | 0.1429 |
| 7_1273730 | NA | NA | 1 | 1 | 1 | 1 | 1 | 1 | 3 | 1 | 2 | 1 | 1 | 1 | 3 | 1 | 2 | 1 | 8 | 1 |
| 7_1273732 | 1 | 1 | NA | NA | 1 | 1 | 1 | 1 | 3 | 1 | NA | NA | 1 | 1 | 3 | 0.6667 | 2 | 0 | 6 | 0.5 |
| 7_1273732 | 1 | 1 | 1 | 1 | 1 | 1 | 1 | 1 | 4 | 1 | 2 | 1 | 1 | 1 | 1 | 1 | 3 | 1 | 7 | 1 |
| 7_1273733 | 1 | 1 | 1 | 0 | 1 | 0 | 1 | 0 | 4 | 0.25 | 3 | 0 | 1 | 0 | 2 | 0.5 | NA | NA | 6 | 0.1667 |
| 7_1273733 | NA | NA | 1 | 1 | NA | NA | 1 | 1 | 2 | 1 | 3 | 1 | NA | NA | 3 | 1 | 3 | 0.6667 | 9 | 0.8889 |
| 7_1273735 | NA | NA | 1 | 1 | 1 | 1 | NA | NA | 2 | 1 | 3 | 0.6667 | 1 | 1 | 3 | 0.6667 | 3 | 0.6667 | 10 | 0.7 |
| 7_1273738 | 2 | 0.5 | 2 | 1 | 2 | 0.5 | 1 | 0 | 7 | 0.5714 | 3 | 0.6667 | 1 | 1 | 3 | 0.3333 | 4 | 0.25 | 11 | 0.4545 |
| 7_1273738 | 1 | 1 | 2 | 1 | 2 | 0.5 | 2 | 0.5 | 7 | 0.7143 | 3 | 0.3333 | 1 | 1 | 3 | 0.6667 | 4 | 0.5 | 11 | 0.5455 |
| 7_1273738 | 2 | 0.5 | 2 | 0.5 | 2 | 0.5 | 2 | 0.5 | 8 | 0.5 | 3 | 0 | 1 | 0 | 2 | 0 | 4 | 0.5 | 10 | 0.2 |
| 7_1273739 | 2 | 0 | 2 | 0.5 | 2 | 0.5 | 3 | 0.3333 | 9 | 0.3333 | 3 | 0 | 1 | 1 | 1 | 0 | 4 | 0.25 | 9 | 0.2222 |
| 7_1273748 | 1 | 1 | NA | NA | NA | NA | NA | NA | 1 | 1 | NA | NA | 2 | 0.5 | NA | NA | NA | NA | 2 | 0.5 |
| 7_1273748 | 1 | 1 | NA | NA | NA | NA | NA | NA | 1 | 1 | 1 | 1 | 2 | 0.5 | NA | NA | NA | NA | 3 | 0.6667 |
| 7_1273748 | 1 | 1 | NA | NA | NA | NA | NA | NA | 1 | 1 | 1 | 1 | 1 | 1 | NA | NA | NA | NA | 2 | 1 |
| 7_1273749 | 1 | 1 | NA | NA | NA | NA | NA | NA | 1 | 1 | 1 | 1 | 2 | 1 | NA | NA | NA | NA | 3 | 1 |

| | | | | | | | | | | | | | | | | | | | | |
|---|---|---|---|---|---|---|---|---|---|---|---|---|---|---|---|---|---|---|---|---|
| 7_1273750( | 1 | 1 | NA | NA | NA | NA | NA | NA | 1 | 1 | 1 | 1 | 2 | 0 | NA | NA | NA | NA | 3 | 0.3333 |
| 7_1273833! | NA | NA | 2 | 0 | 2 | 0.5 | NA | NA | 4 | 0.25 | 1 | 0 | NA | NA | 2 | 0.5 | 1 | 0 | 4 | 0.25 |
| 7_1273843: | NA | NA | 1 | 0 | 2 | 0 | 1 | 1 | 4 | 0.25 | 1 | 1 | NA | NA | 2 | 0.5 | 1 | 0 | 4 | 0.5 |
| 7_1273844: | NA | NA | 2 | 0.5 | 1 | 1 | NA | NA | 3 | 0.6667 | 1 | 1 | NA | NA | 1 | 0 | 1 | 0 | 3 | 0.3333 |
| 7_1273845( | 1 | 0 | 1 | 0 | 2 | 1 | NA | NA | 4 | 0.5 | NA | NA | NA | NA | 2 | 1 | 2 | 0 | 4 | 0.5 |
| 7_1273846' | 1 | 0 | 2 | 0 | 2 | 0.5 | NA | NA | 5 | 0.2 | 1 | 0 | 1 | 1 | 1 | 0 | 2 | 0 | 5 | 0.2 |
| 7_1273853: | 1 | 1 | 2 | 0.5 | 2 | 0.5 | NA | NA | 5 | 0.6 | 1 | 1 | 1 | 0 | 2 | 0.5 | 3 | 0.3333 | 7 | 0.4286 |
| 7_1273854: | NA | NA | 1 | 1 | 2 | 0 | NA | NA | 3 | 0.3333 | 1 | 0 | 1 | 1 | NA | NA | 3 | 0.3333 | 5 | 0.4 |
| 7_1273949! | 1 | 0 | 5 | 0 | 5 | 0.6 | NA | NA | 11 | 0.2727 | 2 | 0.5 | 3 | 0 | 3 | 0 | 1 | 0 | 9 | 0.1111 |
| 7_1273950' | 2 | 0.5 | 5 | 0 | 4 | 0 | 1 | 0 | 12 | 0.0833 | 2 | 0.5 | 3 | 0.3333 | 3 | 0.6667 | 3 | 0.6667 | 11 | 0.5455 |
| 7_1273950! | 2 | 0 | 5 | 0.4 | 4 | 0 | NA | NA | 11 | 0.1818 | 2 | 0.5 | 3 | 0.6667 | 2 | 0 | 3 | 0 | 10 | 0.3 |
| 7_1273952( | 2 | 0.5 | 3 | 0.3333 | 4 | 0.75 | NA | NA | 9 | 0.5556 | 3 | 0.6667 | 2 | 0.5 | 3 | 0 | 3 | 0 | 11 | 0.2727 |
| 7_1273952' | 2 | 0.5 | 5 | 0 | 5 | 0.2 | NA | NA | 12 | 0.1667 | 2 | 0 | 3 | 0 | 3 | 0.3333 | 3 | 0.3333 | 11 | 0.1818 |
| 7_1273953( | 1 | 0 | 5 | 0.2 | 4 | 0.5 | 1 | 0 | 11 | 0.2727 | 2 | 0.5 | 3 | 0 | 2 | 1 | 4 | 0.5 | 11 | 0.4545 |
| 7_1273957! | 2 | 0.5 | 5 | 0.8 | 5 | 0.4 | NA | NA | 12 | 0.5833 | 3 | 0.3333 | 3 | 1 | 5 | 0.8 | 3 | 1 | 14 | 0.7857 |
| 7_1273957: | 3 | 0 | 5 | 0 | 5 | 0.2 | 1 | 0 | 14 | 0.0714 | 3 | 0 | 5 | 0 | 5 | 0 | 4 | 0.25 | 17 | 0.0588 |
| 7_1273958: | 3 | 0.3333 | 4 | 0.25 | 3 | 0.3333 | NA | NA | 10 | 0.3 | 3 | 0.6667 | 4 | 0.5 | 5 | 0.6 | 4 | 0 | 16 | 0.4375 |
| 7_1273965: | 2 | 0.5 | 6 | 0 | 5 | 0 | 1 | 0 | 14 | 0.0714 | 4 | 0 | 5 | 0.2 | 4 | 0 | 3 | 0.3333 | 16 | 0.125 |
| 7_1273986( | 5 | 0 | 5 | 0.2 | 7 | 0 | 1 | 0 | 18 | 0.0556 | 4 | 0.25 | 6 | 0 | 5 | 0 | 3 | 0 | 18 | 0.0556 |
| 7_1273989! | 5 | 0 | 5 | 0 | 5 | 0.2 | 1 | 0 | 16 | 0.0625 | 2 | 0 | 5 | 0.2 | 3 | 0 | 3 | 0.3333 | 13 | 0.1538 |
| 7_1273990! | 4 | 0 | 7 | 0 | 5 | 0.2 | 3 | 0 | 19 | 0.0526 | 3 | 0.3333 | 7 | 0 | 4 | 0 | 3 | 0.3333 | 17 | 0.1176 |
| 7_1274020: | NA | NA | NA | NA | 2 | 1 | 1 | 1 | 3 | 1 | NA | NA | 3 | 0.3333 | 2 | 0.5 | 1 | 0 | 6 | 0.3333 |
| 7_1274023: | NA | NA | NA | NA | 2 | 0.5 | 1 | 1 | 3 | 0.6667 | NA | NA | 2 | 0.5 | 2 | 1 | 1 | 1 | 5 | 0.8 |
| 7_1274024: | NA | NA | NA | NA | 2 | 0.5 | 1 | 0 | 3 | 0.3333 | NA | NA | 3 | 0.3333 | 1 | 0 | 1 | 0 | 5 | 0.2 |
| 7_1274024! | 1 | 0 | NA | NA | 1 | 1 | 1 | 0 | 3 | 0.3333 | NA | NA | 2 | 0 | 2 | 0.5 | 1 | 0 | 5 | 0.2 |
| 7_1274025( | 1 | 1 | NA | NA | 2 | 0 | 2 | 0 | 5 | 0.2 | NA | NA | 3 | 0.6667 | 2 | 0.5 | 1 | 0 | 6 | 0.5 |
| 7_1274025: | 1 | 0 | NA | NA | 2 | 0.5 | 1 | 1 | 4 | 0.5 | NA | NA | 2 | 1 | 2 | 0 | 1 | 0 | 5 | 0.4 |
| 7_1274026: | 1 | 0 | NA | NA | 2 | 0.5 | NA | NA | 3 | 0.3333 | NA | NA | 2 | 1 | 1 | 0 | 1 | 0 | 4 | 0.5 |
| 7_1274026! | 1 | 1 | NA | NA | 2 | 0.5 | 2 | 0.5 | 5 | 0.6 | NA | NA | 3 | 0 | 2 | 1 | 1 | 1 | 6 | 0.5 |
| 7_1274410' | 3 | 0.6667 | 1 | 1 | 4 | 0.5 | 2 | 0.5 | 10 | 0.6 | 1 | 0 | NA | NA | 1 | 1 | NA | NA | 2 | 0.5 |
| 7_1274602( | NA | NA | 1 | 0 | 2 | 1 | 2 | 0 | 5 | 0.4 | NA | NA | 4 | 0.5 | NA | NA | 1 | 0 | 5 | 0.4 |
| 7_1274649' | NA | NA | NA | NA | 1 | 1 | 2 | 0.5 | 3 | 0.6667 | 1 | 1 | NA | NA | NA | NA | NA | NA | 1 | 1 |
| 7_1274651! | 1 | 0 | 1 | 0 | 2 | 0.5 | 3 | 0 | 7 | 0.1429 | 2 | 0 | NA | NA | 3 | 0.3333 | 4 | 0.5 | 9 | 0.3333 |
| 7_1274655! | NA | NA | 1 | 0 | 2 | 0.5 | 2 | 0 | 5 | 0.2 | 2 | 0 | NA | NA | 2 | 0.5 | 1 | 0 | 5 | 0.2 |
| 7_1274656' | NA | NA | 1 | 1 | 1 | 1 | 1 | 1 | 3 | 1 | 1 | 1 | NA | NA | 1 | 1 | 3 | 0.3333 | 5 | 0.6 |
| 7_1274656! | NA | NA | 1 | 0 | 2 | 0.5 | 3 | 0.3333 | 6 | 0.3333 | NA | NA | NA | NA | 1 | 0 | 4 | 0.5 | 5 | 0.4 |
| 7_1274657: | NA | NA | 1 | 0 | 2 | 0.5 | 3 | 0 | 6 | 0.1667 | 1 | 1 | NA | NA | 1 | 0 | 4 | 0.25 | 6 | 0.3333 |
| 7_1274657· | NA | NA | 1 | 0 | 1 | 1 | 2 | 1 | 4 | 0.75 | NA | NA | NA | NA | 1 | 0 | 2 | 0.5 | 3 | 0.3333 |
| 7_1274658! | NA | NA | 1 | 1 | 1 | 1 | 1 | 0 | 3 | 0.6667 | 1 | 1 | NA | NA | 1 | 1 | 4 | 0.75 | 6 | 0.8333 |
| 7_1274659: | NA | NA | 1 | 0 | 1 | 1 | 1 | 0 | 3 | 0.3333 | 1 | 1 | NA | NA | 1 | 0 | 4 | 0 | 6 | 0.1667 |
| 7_1274661( | NA | NA | 1 | 0 | 2 | 0.5 | 3 | 0.3333 | 6 | 0.3333 | 1 | 0 | NA | NA | 1 | 0 | 3 | 0.3333 | 5 | 0.2 |
| 7_1274661: | NA | NA | 1 | 1 | 2 | 1 | 3 | 1 | 6 | 1 | 1 | 1 | NA | NA | 1 | 1 | 4 | 1 | 6 | 1 |
| 7_1274661! | NA | NA | 1 | 1 | 2 | 1 | 2 | 1 | 5 | 1 | 1 | 1 | NA | NA | NA | NA | 4 | 1 | 5 | 1 |
| 7_1274662: | NA | NA | 1 | 1 | 2 | 1 | 3 | 1 | 6 | 1 | 1 | 1 | NA | NA | 1 | 1 | 3 | 1 | 5 | 1 |
| 7_1274664: | NA | NA | 1 | 1 | 2 | 0 | 1 | 0 | 4 | 0.25 | 1 | 1 | NA | NA | 1 | 0 | 2 | 0 | 4 | 0.25 |
| 7_1274669· | NA | NA | NA | NA | 2 | 0 | 3 | 0.6667 | 5 | 0.4 | 1 | 0 | NA | NA | 1 | 1 | 3 | 0.3333 | 5 | 0.4 |
| 7_1274674: | NA | NA | NA | NA | 2 | 1 | 2 | 1 | 4 | 1 | 1 | 1 | NA | NA | 1 | 0 | 3 | 0 | 5 | 0.2 |
| 7_1274674! | NA | NA | NA | NA | 1 | 1 | 2 | 1 | 3 | 1 | 1 | 0 | NA | NA | 1 | 0 | 3 | 1 | 5 | 0.6 |
| 7_1274796! | NA | NA | 1 | 0 | 1 | 1 | 1 | 0 | 3 | 0.3333 | NA | NA | NA | NA | 1 | 1 | NA | NA | 1 | 1 |
| 7_1274799( | NA | NA | 1 | 1 | 1 | 1 | 1 | 1 | 3 | 1 | NA | NA | NA | NA | 1 | 1 | NA | NA | 1 | 1 |
| 7_1274800! | NA | NA | 1 | 0 | 1 | 1 | NA | NA | 2 | 0.5 | NA | NA | NA | NA | 1 | 1 | NA | NA | 1 | 1 |
| 7_1274801: | NA | NA | 1 | 0 | 1 | 1 | 1 | 0 | 3 | 0.3333 | NA | NA | NA | NA | 1 | 1 | NA | NA | 1 | 1 |
| 7_1274804! | NA | NA | 1 | 1 | 1 | 1 | NA | NA | 2 | 1 | NA | NA | NA | NA | 1 | 1 | NA | NA | 1 | 1 |
| 7_1274807: | NA | NA | 1 | 1 | 1 | 0 | 1 | 1 | 3 | 0.6667 | NA | NA | 1 | 0 | 1 | 1 | NA | NA | 2 | 0.5 |
| 7_1274813( | NA | NA | 1 | 1 | 1 | 1 | 1 | 0 | 3 | 0.6667 | 2 | 0.5 | 1 | 0 | NA | NA | NA | NA | 3 | 0.3333 |
| 7_1274814( | NA | NA | 1 | 0 | 1 | 1 | 1 | 0 | 3 | 0.3333 | 2 | 1 | 1 | 0 | NA | NA | NA | NA | 3 | 0.6667 |
| 7_1274815! | NA | NA | 1 | 1 | 1 | 0 | NA | NA | 2 | 0.5 | 1 | 1 | 1 | 1 | NA | NA | NA | NA | 2 | 1 |
| 7_1274816( | NA | NA | 1 | 1 | 1 | 0 | 1 | 0 | 3 | 0.3333 | 1 | 1 | 1 | 0 | NA | NA | NA | NA | 2 | 0.5 |
| 7_1274816' | NA | NA | NA | NA | 1 | 0 | 1 | 1 | 2 | 0.5 | 1 | 0 | 1 | 1 | NA | NA | NA | NA | 2 | 0.5 |
| 7_1274816· | NA | NA | 1 | 1 | NA | NA | 1 | 1 | 2 | 1 | 2 | 1 | 1 | 1 | NA | NA | NA | NA | 3 | 1 |
| 7_1274817: | NA | NA | 1 | 1 | 1 | 1 | 1 | 1 | 3 | 1 | 1 | 0 | 1 | 0 | 1 | 1 | NA | NA | 3 | 0.3333 |
| 7_1274817! | NA | NA | 1 | 1 | 1 | 1 | 1 | 0 | 3 | 0.6667 | 2 | 0.5 | 1 | 1 | 1 | 0 | NA | NA | 4 | 0.5 |
| 7_1274818! | NA | NA | 1 | 0 | 1 | 0 | 1 | 1 | 3 | 0.3333 | 2 | 0.5 | 1 | 0 | 1 | 0 | NA | NA | 4 | 0.25 |
| 7_1274819( | NA | NA | 1 | 1 | 1 | 0 | 1 | 0 | 3 | 0.3333 | 2 | 0.5 | 1 | 0 | 1 | 1 | NA | NA | 4 | 0.5 |
| 7_1274819' | NA | NA | 1 | 1 | 1 | 0 | 1 | 1 | 3 | 0.6667 | 1 | 1 | 1 | 0 | 1 | 1 | NA | NA | 3 | 0.6667 |

| | | | | | | | | | | | | | | | | | | | | |
|---|---|---|---|---|---|---|---|---|---|---|---|---|---|---|---|---|---|---|---|---|
| 7_1274821 | NA | NA | 1 | 0 | 1 | 0 | 1 | 1 | 3 | 0.3333 | 3 | 1 | 2 | 0 | NA | NA | NA | NA | 5 | 0.6 |
| 7_1274823 | NA | NA | 1 | 1 | 1 | 1 | 1 | 1 | 3 | 1 | 1 | 1 | 2 | 1 | 1 | 1 | NA | NA | 4 | 1 |
| 7_1274825 | 1 | 1 | 1 | 1 | 1 | 1 | 1 | 1 | 4 | 1 | 3 | 0.3333 | 3 | 0 | 1 | 0 | NA | NA | 7 | 0.1429 |
| 7_1274828 | 1 | 1 | 1 | 1 | 2 | 1 | 1 | 1 | 5 | 1 | 2 | 1 | 2 | 1 | 2 | 1 | NA | NA | 6 | 1 |
| 7_1274828 | 1 | 0 | 1 | 0 | 2 | 0.5 | 1 | 0 | 5 | 0.2 | 2 | 0 | 3 | 0.3333 | 2 | 0 | NA | NA | 7 | 0.1429 |
| 7_1274829 | 1 | 1 | 1 | 0 | 2 | 0.5 | 1 | 1 | 5 | 0.6 | 2 | 0.5 | 3 | 0.6667 | 1 | 0 | NA | NA | 6 | 0.5 |
| 7_1274847 | NA | NA | 1 | 0 | 3 | 0.3333 | NA | NA | 4 | 0.25 | 1 | 0 | 3 | 0.3333 | 2 | 1 | NA | NA | 6 | 0.5 |
| 7_1274847 | 1 | 1 | NA | NA | 2 | 0 | 1 | 0 | 4 | 0.25 | 1 | 0 | 3 | 0.3333 | 2 | 0 | NA | NA | 6 | 0.1667 |
| 7_1274857 | NA | NA | 1 | 1 | NA | NA | 1 | 0 | 2 | 0.5 | NA | NA | 2 | 0 | 2 | 0.5 | NA | NA | 4 | 0.25 |
| 7_1275041 | 5 | 0.2 | 5 | 0.2 | 4 | 0.5 | 3 | 0 | 17 | 0.2353 | 2 | 0 | 4 | 0 | 1 | 1 | 3 | 0 | 10 | 0.1 |
| 7_1275075 | 5 | 0.2 | 8 | 0.375 | 7 | 0.1429 | 3 | 0 | 23 | 0.2174 | 3 | 0 | 4 | 0 | 2 | 0 | 7 | 0.1429 | 16 | 0.0625 |
| 7_1275075 | 7 | 0.4286 | 9 | 0.3333 | 5 | 0.4 | 4 | 0.25 | 25 | 0.36 | 4 | 0.25 | 4 | 0 | 1 | 0 | 6 | 0.3333 | 15 | 0.2 |
| 7_1275075 | 9 | 0.2222 | 10 | 0 | 7 | 0.2857 | 4 | 0 | 30 | 0.1333 | 5 | 0 | 5 | 0 | 2 | 0.5 | 6 | 0.1667 | 18 | 0.1111 |
| 7_1275076 | 8 | 0.75 | 7 | 0.5714 | 6 | 0.8333 | 2 | 0.5 | 23 | 0.6957 | 2 | 0.5 | 4 | 0.75 | 1 | 1 | 7 | 1 | 14 | 0.8571 |
| 7_1275077 | 6 | 1 | 7 | 0.4286 | 4 | 0.75 | 3 | 0.3333 | 20 | 0.65 | 5 | 0.6 | 4 | 0.75 | 3 | 0.3333 | 6 | 0.5 | 18 | 0.5556 |
| 7_1275146 | 2 | 0.5 | 2 | 0 | 4 | 0 | 6 | 0 | 14 | 0.0714 | 1 | 0 | NA | NA | 1 | 0 | 4 | 0.25 | 6 | 0.1667 |
| 7_1275247 | 3 | 0 | 2 | 0 | 5 | 0.2 | 4 | 0.5 | 14 | 0.2143 | 4 | 0 | 1 | 0 | 3 | 0.3333 | 4 | 0.5 | 12 | 0.25 |
| 7_1275318 | NA | NA | 1 | 1 | 2 | 1 | 2 | 1 | 5 | 1 | NA | NA | 3 | 1 | NA | NA | 3 | 0.6667 | 6 | 0.8333 |
| 7_1275320 | NA | NA | 2 | 0 | 1 | 1 | 2 | 0.5 | 5 | 0.4 | NA | NA | 3 | 0 | NA | NA | 3 | 0.3333 | 6 | 0.1667 |
| 7_1275733 | NA | NA | 3 | 0 | 1 | 1 | 1 | 0 | 5 | 0.2 | 1 | 0 | 1 | 0 | 1 | 1 | NA | NA | 3 | 0.3333 |
| 7_1277166 | 1 | 0 | 1 | 0 | 2 | 0.5 | 1 | 0 | 5 | 0.2 | 1 | 0 | 2 | 0 | 1 | 1 | NA | NA | 4 | 0.25 |
| 7_1278046 | NA | NA | 2 | 0.5 | 1 | 0 | NA | NA | 3 | 0.3333 | 2 | 1 | NA | NA | NA | NA | NA | NA | 2 | 1 |
| 7_1278339 | 1 | 0 | 1 | 1 | NA | NA | 1 | 1 | 3 | 0.6667 | 1 | 1 | NA | NA | NA | NA | 1 | 0 | 2 | 0.5 |
| 7_1278343 | 1 | 1 | 1 | 1 | NA | NA | NA | NA | 2 | 1 | NA | NA | NA | NA | NA | NA | 1 | 1 | 1 | 1 |
| 7_1278345 | 1 | 1 | 1 | 1 | NA | NA | 1 | 1 | 3 | 1 | NA | NA | NA | NA | NA | NA | 1 | 1 | 1 | 1 |
| 7_1278347 | 1 | 1 | 1 | 1 | NA | NA | 1 | 1 | 3 | 1 | NA | NA | NA | NA | NA | NA | 1 | 1 | 1 | 1 |
| 7_1278348 | 1 | 0 | 1 | 1 | NA | NA | 1 | 0 | 3 | 0.3333 | NA | NA | NA | NA | NA | NA | 1 | 1 | 1 | 1 |
| 7_1278354 | NA | NA | NA | NA | NA | NA | 1 | 1 | 1 | 1 | NA | NA | NA | NA | NA | NA | 1 | 1 | 1 | 1 |
| 7_1278357 | NA | NA | NA | NA | NA | NA | 1 | 1 | 1 | 1 | NA | NA | NA | NA | NA | NA | 1 | 1 | 1 | 1 |
| 7_1278359 | NA | NA | NA | NA | NA | NA | 1 | 1 | 1 | 1 | NA | NA | NA | NA | NA | NA | 1 | 1 | 1 | 1 |
| 7_1278399 | 1 | 1 | 1 | 0 | NA | NA | NA | NA | 2 | 0.5 | 1 | 1 | 2 | 1 | 1 | 0 | NA | NA | 4 | 0.75 |
| 7_1278438 | 2 | 1 | 2 | 1 | 1 | 1 | 1 | 1 | 6 | 1 | 1 | 1 | 2 | 1 | 2 | 1 | 2 | 1 | 7 | 1 |
| 7_1278438 | 1 | 1 | 2 | 0 | 1 | 0 | 1 | 0 | 5 | 0.2 | NA | NA | 2 | 0.5 | 3 | 0.6667 | 1 | 0 | 6 | 0.5 |
| 7_1278440 | 3 | 0 | 2 | 0 | 1 | 0 | 1 | 1 | 7 | 0.1429 | 1 | 1 | 2 | 0 | 3 | 0.3333 | 2 | 0 | 8 | 0.25 |
| 7_1278441 | 3 | 0.3333 | 2 | 0 | 1 | 0 | 1 | 0 | 7 | 0.1429 | 1 | 1 | 2 | 0.5 | 2 | 0 | 2 | 0 | 7 | 0.2857 |
| 7_1278442 | 1 | 0 | 2 | 0.5 | 1 | 0 | NA | NA | 4 | 0.25 | NA | NA | 2 | 0.5 | 2 | 0.5 | 1 | 1 | 5 | 0.6 |
| 7_1278525 | 2 | 1 | 1 | 1 | NA | NA | 1 | 1 | 4 | 1 | 2 | 1 | NA | NA | 2 | 1 | NA | NA | 4 | 1 |
| 7_1278707 | 2 | 0 | 1 | 0 | NA | NA | 2 | 0.5 | 5 | 0.2 | 1 | 1 | 4 | 0.25 | 1 | 0 | 1 | 0 | 7 | 0.2857 |
| 7_1278709 | 2 | 0.5 | NA | NA | 1 | 0 | 1 | 0 | 4 | 0.25 | 1 | 1 | 4 | 0.75 | 1 | 1 | 1 | 0 | 7 | 0.7143 |
| 7_1278711 | 1 | 1 | NA | NA | 1 | 1 | 1 | 1 | 3 | 1 | 1 | 1 | 3 | 1 | 1 | 1 | 1 | 0 | 6 | 0.8333 |
| 7_1278711 | 1 | 1 | NA | NA | 1 | 1 | 1 | 1 | 3 | 1 | 1 | 1 | 2 | 1 | 1 | 1 | NA | NA | 4 | 1 |
| 7_1278712 | 2 | 0.5 | NA | NA | 1 | 0 | 1 | 1 | 4 | 0.5 | 1 | 0 | 2 | 0 | 1 | 1 | NA | NA | 4 | 0.25 |
| 7_1278715 | 2 | 1 | NA | NA | NA | NA | 1 | 1 | 3 | 1 | 1 | 1 | 2 | 1 | NA | NA | NA | NA | 3 | 1 |
| 7_1278715 | 1 | 1 | NA | NA | NA | NA | NA | NA | 1 | 1 | 1 | 1 | NA | NA | NA | NA | NA | NA | 1 | 1 |
| 7_1279755 | 11 | 0.3636 | 11 | 0.5455 | 11 | 0.1818 | 12 | 0.4167 | 45 | 0.3778 | 5 | 0.4 | 5 | 0.6 | 12 | 0.4167 | 4 | 0.5 | 26 | 0.4615 |
| 7_1279755 | 12 | 0.5833 | 10 | 0.4 | 12 | 0.6667 | 10 | 0.8 | 44 | 0.6136 | 4 | 0 | 8 | 0.625 | 12 | 0.6667 | 3 | 0.3333 | 27 | 0.5185 |
| 7_1279755 | 12 | 0.5 | 11 | 0.3636 | 12 | 0.25 | 12 | 0.5833 | 47 | 0.4255 | 4 | 0.75 | 6 | 0.1667 | 12 | 0.3333 | 2 | 0.5 | 24 | 0.375 |
| 7_1280417 | 4 | 0.5 | 8 | 0.125 | 3 | 0.3333 | 7 | 0.1429 | 22 | 0.2273 | 4 | 0.25 | 3 | 0.3333 | 10 | 0.4 | 6 | 0.1667 | 23 | 0.3043 |
| 7_1280417 | 4 | 0.25 | 8 | 0.625 | 3 | 0.3333 | 7 | 0 | 22 | 0.3182 | 3 | 0 | 3 | 0.6667 | 7 | 0.2857 | 9 | 0.2222 | 22 | 0.2727 |
| 7_1280433 | 4 | 0 | 6 | 0 | NA | NA | 6 | 0.1667 | 16 | 0.0625 | 5 | 0.2 | 2 | 0 | 6 | 0.1667 | 6 | 0 | 19 | 0.1053 |
| 7_1280433 | 4 | 0 | 5 | 0.2 | NA | NA | 5 | 0.2 | 14 | 0.1429 | 5 | 0.2 | 2 | 0 | 5 | 0 | 5 | 0.4 | 17 | 0.1765 |
| 7_1280436 | 4 | 0 | 4 | 0.25 | NA | NA | 4 | 0.5 | 12 | 0.25 | 4 | 0.25 | 1 | 0 | 5 | 0 | 5 | 0.4 | 15 | 0.2 |
| 7_1280436 | NA | NA | NA | NA | NA | NA | 2 | 0.5 | 2 | 0.5 | NA | NA | NA | NA | 1 | 1 | NA | NA | 1 | 1 |
| 7_1280436 | 4 | 0.25 | 5 | 0.4 | NA | NA | 6 | 0.1667 | 15 | 0.2667 | 5 | 0.2 | 2 | 0 | 6 | 0.3333 | 5 | 0.6 | 18 | 0.3333 |
| 7_1280436 | 4 | 0.75 | 5 | 0.6 | NA | NA | 6 | 0.5 | 15 | 0.6 | 4 | 0.25 | 2 | 0.5 | 6 | 0.1667 | 6 | 0.5 | 18 | 0.3333 |
| 7_1280436 | 2 | 1 | 3 | 0.6667 | NA | NA | 3 | 0.6667 | 8 | 0.75 | 2 | 0 | 1 | 1 | 5 | 0.2 | 4 | 0.75 | 12 | 0.4167 |
| 7_1280438 | 4 | 0 | 5 | 0.2 | NA | NA | 5 | 0.2 | 14 | 0.1429 | 4 | 0 | 2 | 0.5 | 6 | 0 | 4 | 0 | 16 | 0.0625 |
| 7_1280438 | 1 | 1 | 3 | 1 | NA | NA | 3 | 0.6667 | 7 | 0.8571 | 3 | 0.6667 | NA | NA | 3 | 1 | 2 | 0.5 | 8 | 0.75 |
| 7_1280439 | 4 | 0.75 | 4 | 0.75 | NA | NA | 5 | 0.8 | 13 | 0.7692 | 4 | 0.25 | 2 | 1 | 5 | 0.4 | 5 | 0.2 | 16 | 0.375 |
| 7_1280439 | 4 | 0.5 | 3 | 0.3333 | 2 | 0 | 7 | 0.4286 | 16 | 0.375 | 4 | 0.5 | NA | NA | 4 | 0.5 | 7 | 0.4286 | 15 | 0.4667 |
| 7_1280440 | 6 | 0.6667 | 10 | 0.6 | 5 | 0.6 | 10 | 0.7 | 31 | 0.6452 | 6 | 0.3333 | 3 | 0.3333 | 7 | 0.7143 | 9 | 0.5556 | 25 | 0.52 |
| 7_1280440 | 5 | 0.2 | 3 | 0 | 3 | 0 | 10 | 0.2 | 21 | 0.1429 | 3 | 0 | 3 | 0.3333 | 6 | 0.1667 | 7 | 0 | 19 | 0.1053 |
| 7_1280440 | 8 | 0.625 | 10 | 0.1 | 6 | 0.3333 | 12 | 0.25 | 36 | 0.3056 | 6 | 0.3333 | 5 | 0.4 | 7 | 0.2857 | 11 | 0.2727 | 29 | 0.3103 |
| 7_1280441 | 8 | 1 | 10 | 0.7 | 6 | 0.5 | 14 | 0.7857 | 38 | 0.7632 | 6 | 0.8333 | 5 | 0.8 | 8 | 1 | 10 | 0.7 | 29 | 0.8276 |
| 7_1280441 | 8 | 0.5 | 10 | 0.3 | 5 | 0.4 | 14 | 0.4286 | 37 | 0.4054 | 6 | 0.5 | 3 | 0 | 7 | 0.5714 | 13 | 0.0769 | 29 | 0.2759 |

| | | | | | | | | | | | | | | | | | | | | |
|---|---|---|---|---|---|---|---|---|---|---|---|---|---|---|---|---|---|---|---|---|
| 7_12804425 | 8 | 0.75 | 9 | 1 | 6 | 0.6667 | 10 | 0.8 | 33 | 0.8182 | 5 | 0.6 | 5 | 1 | 7 | 0.8571 | 10 | 0.9 | 27 | 0.8519 |
| 7_12804561 | 9 | 0.1111 | 7 | 0 | 4 | 0 | 13 | 0.3846 | 33 | 0.1818 | 6 | 0.1667 | 4 | 0.25 | 4 | 0 | 7 | 0 | 21 | 0.0952 |
| 7_12814761 | 1 | 0 | 4 | 0.25 | NA | NA | 2 | 0 | 7 | 0.1429 | NA | NA | 1 | 0 | NA | NA | 2 | 0.5 | 3 | 0.3333 |
| 7_12814774 | 1 | 0 | 4 | 0.5 | NA | NA | 2 | 0.5 | 7 | 0.4286 | NA | NA | 1 | 1 | NA | NA | 1 | 1 | 2 | 1 |
| 7_12814788 | 1 | 0 | 4 | 0.25 | NA | NA | 2 | 0 | 7 | 0.1429 | NA | NA | 1 | 1 | NA | NA | 1 | 1 | 2 | 1 |
| 7_12814792 | 1 | 1 | 3 | 0 | NA | NA | 2 | 0.5 | 6 | 0.3333 | NA | NA | NA | NA | NA | NA | 1 | 1 | 1 | 1 |
| 7_12814803 | NA | NA | 4 | 0.5 | NA | NA | 1 | 1 | 5 | 0.6 | NA | NA | NA | NA | NA | NA | 1 | 1 | 1 | 1 |
| 7_12814813 | 1 | 0 | 3 | 0.3333 | NA | NA | 2 | 0.5 | 6 | 0.3333 | 1 | 0 | 1 | 0 | NA | NA | 1 | 1 | 3 | 0.3333 |
| 7_12814814 | 1 | 1 | 4 | 1 | NA | NA | 2 | 1 | 7 | 1 | 1 | 1 | 1 | 1 | NA | NA | 1 | 1 | 3 | 1 |
| 7_12814817 | NA | NA | 3 | 0.3333 | NA | NA | 1 | 0 | 4 | 0.25 | 1 | 1 | NA | NA | NA | NA | 1 | 0 | 2 | 0.5 |
| 7_12814822 | 1 | 0 | 4 | 0.5 | NA | NA | 2 | 1 | 7 | 0.5714 | 1 | 0 | 1 | 0 | NA | NA | 1 | 1 | 3 | 0.3333 |
| 7_12814847 | 1 | 0 | 4 | 0 | NA | NA | 2 | 1 | 7 | 0.2857 | 1 | 1 | NA | NA | NA | NA | 1 | 1 | 2 | 1 |
| 7_12829606 | 7 | 0.1429 | 4 | 0.5 | 9 | 0.3333 | 12 | 0 | 32 | 0.1875 | 4 | 0.25 | 4 | 0 | 12 | 0.0833 | 6 | 0.1667 | 26 | 0.1154 |
| 7_12829609 | 8 | 0.125 | 4 | 0.25 | 10 | 0.1 | 13 | 0.4615 | 35 | 0.2571 | 4 | 0.5 | 3 | 0 | 10 | 0.3 | 3 | 0 | 20 | 0.25 |
| 7_12829610 | 5 | 0.2 | 3 | 0.6667 | 8 | 0.125 | 12 | 0.0833 | 28 | 0.1786 | 2 | 0 | 3 | 0 | 11 | 0.2727 | 5 | 0.2 | 21 | 0.1905 |
| 7_12829611 | 1 | 1 | NA | NA | NA | NA | NA | NA | 1 | 1 | NA | NA | NA | NA | NA | NA | 1 | 1 | 1 | 1 |
| 7_12829614 | 8 | 0.125 | 4 | 0.5 | 10 | 0.3 | 12 | 0.0833 | 34 | 0.2059 | 4 | 0.25 | 4 | 0.25 | 13 | 0.1538 | 5 | 0.2 | 26 | 0.1923 |
| 7_12829618 | 1 | 1 | NA | NA | NA | NA | NA | NA | 1 | 1 | NA | NA | NA | NA | NA | NA | 1 | 1 | 1 | 1 |
| 7_12829620 | 5 | 0.8 | 5 | 0.8 | 10 | 1 | 12 | 0.8333 | 32 | 0.875 | 2 | 1 | 4 | 1 | 12 | 0.75 | 5 | 1 | 23 | 0.8696 |
| 7_12829643 | 9 | 0.1111 | 4 | 0 | 10 | 0.2 | 13 | 0.1538 | 36 | 0.1389 | 3 | 0 | 5 | 0.2 | 13 | 0.2308 | 7 | 0.1429 | 28 | 0.1786 |
| 7_12829650 | 8 | 1 | 4 | 1 | 8 | 1 | 12 | 0.9167 | 32 | 0.9688 | 3 | 1 | 6 | 1 | 11 | 1 | 8 | 1 | 28 | 1 |
| 7_12829655 | 7 | 0 | 4 | 0 | 10 | 0.1 | 8 | 0.125 | 29 | 0.069 | 3 | 0.3333 | 6 | 0.3333 | 13 | 0.2308 | 6 | 0 | 28 | 0.2143 |
| 7_12829662 | 7 | 0.2857 | 5 | 0.4 | 9 | 0.3333 | 12 | 0.25 | 33 | 0.303 | 5 | 0 | 5 | 0.4 | 12 | 0.1667 | 8 | 0.25 | 30 | 0.2 |
| 7_12829665 | 1 | 1 | NA | NA | NA | NA | NA | NA | 1 | 1 | NA | NA | NA | NA | NA | NA | 1 | 1 | 1 | 1 |
| 7_12829666 | 9 | 0.8889 | 5 | 0.8 | 10 | 0.6 | 10 | 0.4 | 34 | 0.6471 | 5 | 0.2 | 6 | 0.5 | 13 | 0.6154 | 8 | 0.375 | 32 | 0.4688 |
| 7_12829674 | 8 | 0.25 | 4 | 0.25 | 9 | 0.2222 | 13 | 0.0769 | 34 | 0.1765 | 5 | 0.6 | 5 | 0.2 | 13 | 0.1538 | 7 | 0 | 30 | 0.2 |
| 7_12829679 | 7 | 0.4286 | 5 | 0.4 | 9 | 0.5556 | 13 | 0.3077 | 34 | 0.4118 | 4 | 0.25 | 6 | 0.6667 | 13 | 0.4615 | 7 | 0.2857 | 30 | 0.4333 |
| 7_12829691 | 8 | 0.375 | 6 | 0.1667 | 8 | 0.125 | 13 | 0.1538 | 35 | 0.2 | 5 | 0 | 4 | 0.25 | 14 | 0.0714 | 8 | 0.25 | 31 | 0.129 |
| 7_12829700 | 7 | 0.8571 | 6 | 1 | 7 | 1 | 13 | 0.9231 | 33 | 0.9394 | 6 | 1 | 5 | 1 | 10 | 0.9 | 5 | 0.8 | 26 | 0.9231 |
| 7_12829722 | 9 | 0 | 6 | 0 | 11 | 0.0909 | 15 | 0.1333 | 41 | 0.0732 | 6 | 0 | 5 | 0.6 | 17 | 0.1176 | 9 | 0 | 37 | 0.1351 |
| 7_12829739 | 2 | 0 | NA | NA | 1 | 1 | NA | NA | 3 | 0.3333 | NA | NA | NA | NA | 2 | 1 | 1 | 1 | 3 | 1 |
| 7_12829940 | 3 | 0 | NA | NA | 1 | 1 | NA | NA | 4 | 0.25 | 1 | 0 | NA | NA | 5 | 0.4 | 3 | 0 | 9 | 0.2222 |
| 7_12829954 | 6 | 0.5 | 2 | 0.5 | 2 | 1 | 1 | 1 | 11 | 0.6364 | 2 | 1 | 2 | 0.5 | 4 | 1 | 2 | 0.5 | 10 | 0.8 |
| 7_12829965 | 6 | 0.1667 | 3 | 0.3333 | 2 | 0.5 | 1 | 0 | 12 | 0.25 | 1 | 0 | 2 | 0.5 | 9 | 0.8889 | 3 | 0.6667 | 15 | 0.7333 |
| 7_12829970 | 5 | 0 | 3 | 0 | 2 | 0.5 | 1 | 0 | 11 | 0.0909 | 2 | 0 | 2 | 0.5 | 10 | 0 | 3 | 0 | 17 | 0.0588 |
| 7_12830006 | 6 | 0.5 | 4 | 0.5 | 4 | 0.75 | 1 | 0 | 15 | 0.5333 | NA | NA | 3 | 0 | 8 | 0.875 | 3 | 0.6667 | 14 | 0.6429 |
| 7_12835191 | NA | NA | 1 | 0 | 1 | 1 | 1 | 1 | 3 | 0.6667 | 3 | 0.6667 | 1 | 1 | 1 | 1 | NA | NA | 5 | 0.8 |
| 7_12835194 | 1 | 0 | 3 | 0.3333 | NA | NA | 4 | 0.25 | 8 | 0.25 | 3 | 0.3333 | NA | NA | 3 | 0.6667 | 1 | 0 | 7 | 0.4286 |
| 7_12835215 | 3 | 0.6667 | 5 | 0.4 | 1 | 1 | 3 | 0.6667 | 12 | 0.5833 | 3 | 0.3333 | 3 | 0.3333 | 5 | 0.4 | 1 | 1 | 12 | 0.4167 |
| 7_12838525 | 7 | 0.4286 | 6 | 0.6667 | 2 | 0.5 | 5 | 0.2 | 20 | 0.45 | 6 | 0.1667 | 7 | 0.2857 | 6 | 0.3333 | 2 | 0.5 | 21 | 0.2857 |
| 7_12838569 | 2 | 0.5 | 3 | 0 | 1 | 0 | 3 | 0 | 9 | 0.1111 | 2 | 0 | 1 | 0 | 4 | 0 | 2 | 1 | 9 | 0.2222 |
| 7_12838603 | 2 | 0.5 | 3 | 0 | 1 | 0 | 3 | 0 | 9 | 0.1111 | 2 | 0.5 | 1 | 0 | 4 | 0 | 3 | 0 | 10 | 0.1 |
| 7_12838604 | 1 | 0 | 3 | 0.3333 | 1 | 0 | 3 | 0 | 8 | 0.125 | 2 | 0 | 1 | 0 | 3 | 0.3333 | 3 | 0.3333 | 9 | 0.2222 |
| 7_12838638 | 4 | 0 | 2 | 0 | 1 | 0 | 3 | 0.3333 | 10 | 0.1 | 2 | 0.5 | 1 | 0 | 4 | 0 | 2 | 0 | 9 | 0.1111 |
| 7_12899280 | NA | NA | 2 | 1 | NA | NA | 1 | 1 | 3 | 1 | 4 | 0.75 | NA | NA | 2 | 1 | NA | NA | 6 | 0.8333 |
| 7_12899297 | 1 | 1 | 2 | 0.5 | NA | NA | 1 | 1 | 4 | 0.75 | 5 | 0.4 | NA | NA | 2 | 1 | NA | NA | 7 | 0.5714 |
| 7_12899298 | 1 | 0 | 2 | 0.5 | NA | NA | 1 | 0 | 4 | 0.25 | 5 | 0.2 | NA | NA | 2 | 0.5 | NA | NA | 7 | 0.2857 |
| 7_12900272 | 1 | 1 | NA | NA | NA | NA | NA | NA | 1 | 1 | 1 | 1 | NA | NA | NA | NA | NA | NA | 1 | 1 |
| 7_12900275 | 1 | 1 | NA | NA | NA | NA | NA | NA | 1 | 1 | 1 | 1 | NA | NA | NA | NA | NA | NA | 1 | 1 |
| 7_12900288 | 1 | 1 | NA | NA | NA | NA | NA | NA | 1 | 1 | 1 | 1 | NA | NA | NA | NA | NA | NA | 1 | 1 |
| 7_12923248 | 5 | 0 | 2 | 0 | 2 | 0 | 3 | 0.3333 | 12 | 0.0833 | 1 | 0 | 1 | 0 | 4 | 0.25 | 1 | 0 | 7 | 0.1429 |
| 7_12923713 | 6 | 0 | 2 | 0 | 5 | 0.2 | 3 | 0 | 16 | 0.0625 | 3 | 0 | 2 | 0 | 5 | 0.2 | 3 | 0.3333 | 13 | 0.1538 |
| 7_12923722 | 6 | 0.1667 | 2 | 0 | 5 | 0.2 | 4 | 0.25 | 17 | 0.1765 | 2 | 0 | 1 | 0 | 5 | 0 | 1 | 1 | 9 | 0.1111 |
| 7_12976445 | 1 | 1 | 4 | 0 | 6 | 0 | 4 | 0.25 | 15 | 0.1333 | 9 | 0.1111 | 5 | 0 | 3 | 0 | 2 | 0 | 19 | 0.0526 |
| 7_12976451 | 1 | 0 | 5 | 0.8 | 6 | 0.5 | 4 | 0.25 | 16 | 0.5 | 10 | 0.6 | 5 | 0.6 | 2 | 0.5 | 2 | 1 | 19 | 0.6316 |
| 7_12976470 | 1 | 1 | 5 | 0.8 | 6 | 0.5 | 2 | 0.5 | 14 | 0.6429 | 8 | 0.125 | 5 | 0.4 | 3 | 0.3333 | 2 | 0.5 | 18 | 0.2778 |
| 7_12976477 | 1 | 0 | 4 | 0.25 | 4 | 0 | 3 | 0 | 12 | 0.0833 | 10 | 0.1 | 4 | 0 | 3 | 0 | 2 | 0 | 19 | 0.0526 |
| 7_12976496 | NA | NA | 3 | 0.6667 | 5 | 0.6 | 3 | 0.6667 | 11 | 0.6364 | 8 | 0.375 | 4 | 0.75 | 3 | 0.6667 | 2 | 0.5 | 17 | 0.5294 |
| 7_12976498 | 1 | 0 | 5 | 0.2 | 6 | 0.1667 | 3 | 0.3333 | 15 | 0.2 | 9 | 0.3333 | 3 | 0.3333 | 3 | 0 | 2 | 0 | 17 | 0.2353 |
| 7_12976540 | 2 | 1 | 4 | 1 | 6 | 1 | 2 | 0.5 | 14 | 0.9286 | 6 | 1 | 3 | 0.6667 | 2 | 1 | 2 | 1 | 13 | 0.9231 |
| 7_12976544 | 2 | 0 | 5 | 0 | 5 | 0.2 | 4 | 0 | 16 | 0.0625 | 8 | 0.125 | 4 | 0 | 1 | 0 | 4 | 0 | 17 | 0.0588 |
| 7_12976547 | 2 | 1 | 5 | 1 | 6 | 0.6667 | 4 | 1 | 17 | 0.8824 | 9 | 0.8889 | 4 | 1 | 3 | 0.6667 | 3 | 0.6667 | 19 | 0.8421 |
| 7_12976549 | 2 | 1 | 4 | 0.75 | 6 | 0.6667 | 4 | 1 | 16 | 0.8125 | 7 | 0.5714 | 4 | 1 | 3 | 1 | 4 | 1 | 18 | 0.8333 |
| 7_12976560 | 2 | 0 | 5 | 0.2 | 5 | 0.2 | 3 | 0.3333 | 15 | 0.2 | 9 | 0.1111 | 3 | 0.3333 | 3 | 0 | 4 | 0 | 19 | 0.1053 |
| 7_12976564 | 2 | 0.5 | 5 | 0.2 | 5 | 0.6 | 2 | 0.5 | 14 | 0.4286 | 9 | 0.2222 | 4 | 0.5 | 3 | 0.3333 | 4 | 0.5 | 20 | 0.35 |

| | | | | | | | | | | | | | | | | | | | | |
|---|---|---|---|---|---|---|---|---|---|---|---|---|---|---|---|---|---|---|---|---|
| 7_1297656 | 2 | 0 | 5 | 0.4 | 5 | 0 | 2 | 0.5 | 14 | 0.2143 | 8 | 0.125 | 3 | 0 | 3 | 0 | 4 | 0 | 18 | 0.0556 |
| 7_1297657 | 2 | 0.5 | 5 | 0.8 | 5 | 0.6 | 3 | 0.6667 | 15 | 0.6667 | 7 | 0.7143 | 4 | 0.5 | 3 | 0.6667 | 4 | 1 | 18 | 0.7222 |
| 7_1297658 | 1 | 0 | 2 | 0 | 2 | 0.5 | 2 | 0 | 7 | 0.1429 | 5 | 0.2 | 1 | 0 | 3 | 0.6667 | 2 | 0 | 11 | 0.2727 |
| 7_1299937 | 4 | 0 | 3 | 0.3333 | 5 | 0 | 4 | 0 | 16 | 0.0625 | 6 | 0 | 1 | 0 | 5 | 0 | 5 | 0.2 | 17 | 0.0588 |
| 7_1299959 | 1 | 0 | 1 | 0 | 1 | 1 | 2 | 0.5 | 5 | 0.4 | 3 | 0.3333 | 1 | 1 | 2 | 0 | 2 | 0.5 | 8 | 0.375 |
| 7_1299960 | 1 | 0 | NA | NA | 1 | 1 | 2 | 0.5 | 4 | 0.5 | 3 | 0.3333 | 1 | 0 | 2 | 0 | 1 | 1 | 7 | 0.2857 |
| 7_1299963 | 1 | 0 | 1 | 1 | 1 | 0 | 1 | 1 | 4 | 0.5 | 3 | 0 | 2 | 0 | 2 | 0.5 | 2 | 0 | 9 | 0.1111 |
| 7_1303305 | 1 | 1 | 2 | 1 | 3 | 1 | 2 | 0.5 | 8 | 0.875 | NA | NA | 1 | 1 | 1 | 1 | 1 | 0 | 3 | 0.6667 |
| 7_1303309 | 3 | 0.3333 | 3 | 1 | 3 | 0 | 2 | 0.5 | 11 | 0.4545 | NA | NA | 2 | 0.5 | 1 | 0 | 1 | 1 | 4 | 0.5 |
| 7_1303310 | 3 | 1 | 2 | 0.5 | 1 | 1 | 1 | 1 | 7 | 0.8571 | NA | NA | 2 | 1 | 2 | 1 | 1 | 1 | 5 | 1 |
| 7_1303310 | 1 | 1 | 2 | 0.5 | 4 | 0.5 | 2 | 1 | 9 | 0.6667 | NA | NA | 2 | 1 | 1 | 1 | 2 | 0.5 | 5 | 0.8 |
| 7_1303310 | 2 | 0 | 2 | 0 | 4 | 0.5 | 1 | 1 | 9 | 0.3333 | NA | NA | 2 | 0.5 | 2 | 0 | 2 | 0 | 6 | 0.1667 |
| 7_1303310 | 3 | 0.6667 | 3 | 0.3333 | 4 | 0.5 | 1 | 0 | 11 | 0.4545 | NA | NA | 2 | 0.5 | 2 | 0.5 | 2 | 0 | 6 | 0.3333 |
| 7_1303312 | 2 | 0.5 | 3 | 0.3333 | 3 | 0 | 2 | 0 | 10 | 0.2 | NA | NA | 2 | 0.5 | 2 | 0 | 2 | 0 | 6 | 0.1667 |
| 7_1303313 | 3 | 0 | 3 | 0 | 5 | 0.4 | 2 | 0 | 13 | 0.1538 | NA | NA | 2 | 0.5 | 2 | 0 | 1 | 0 | 5 | 0.2 |
| 7_1303660 | 4 | 0.25 | 3 | 0 | 8 | 0 | 2 | 0 | 17 | 0.0588 | 2 | 0 | 6 | 0.3333 | 6 | 0 | 3 | 0 | 17 | 0.1176 |
| 7_1303660 | 2 | 0 | 8 | 0.375 | 9 | 0.3333 | 3 | 0 | 22 | 0.2727 | 2 | 0 | 9 | 0.1111 | 6 | 0.1667 | 2 | 0 | 19 | 0.1053 |
| 7_1303662 | 4 | 0.5 | 10 | 0.5 | 12 | 0.5833 | 4 | 0.25 | 30 | 0.5 | 2 | 0 | 9 | 0.4444 | 7 | 0.4286 | 4 | 0.75 | 22 | 0.4545 |
| 7_1303663 | 4 | 0.75 | 8 | 0.875 | 10 | 0.7 | 4 | 0.75 | 26 | 0.7692 | 2 | 0.5 | 7 | 0.5714 | 5 | 0.8 | 3 | 0.3333 | 17 | 0.5882 |
| 7_1303663 | 5 | 0.2 | 8 | 0 | 10 | 0.2 | 4 | 0.25 | 27 | 0.1481 | 1 | 0 | 8 | 0 | 6 | 0 | 2 | 0.5 | 17 | 0.0588 |
| 7_1303663 | 4 | 0.75 | 10 | 0.8 | 11 | 0.7273 | 5 | 0.8 | 30 | 0.7667 | 3 | 0.6667 | 9 | 0.6667 | 6 | 0.8333 | 4 | 0.5 | 22 | 0.6818 |
| 7_1304634 | NA | NA | NA | NA | 1 | 1 | NA | NA | 1 | 1 | 1 | 1 | NA | NA | NA | NA | NA | NA | 1 | 1 |
| 7_1304639 | NA | NA | NA | NA | 1 | 1 | NA | NA | 1 | 1 | 1 | 1 | NA | NA | NA | NA | NA | NA | 1 | 1 |
| 7_1304642 | NA | NA | NA | NA | 1 | 1 | NA | NA | 1 | 1 | 1 | 1 | NA | NA | NA | NA | NA | NA | 1 | 1 |
| 7_1304643 | NA | NA | NA | NA | 1 | 1 | NA | NA | 1 | 1 | 1 | 1 | NA | NA | NA | NA | NA | NA | 1 | 1 |
| 7_1305127 | 7 | 0 | 12 | 0.1667 | 6 | 0.1667 | 7 | 0 | 32 | 0.0938 | 4 | 0 | 3 | 0 | 7 | 0 | 4 | 0.5 | 18 | 0.1111 |
| 7_1305127 | 6 | 0.3333 | 10 | 0.3 | 7 | 0.1429 | 6 | 0.3333 | 29 | 0.2759 | 5 | 0.6 | 4 | 0.25 | 7 | 0.4286 | 3 | 0.3333 | 19 | 0.4211 |
| 7_1305127 | 6 | 1 | 10 | 0.9 | 5 | 1 | 6 | 0.8333 | 27 | 0.9259 | 5 | 0.8 | 3 | 1 | 6 | 0.8333 | 3 | 1 | 17 | 0.8824 |
| 7_1305127 | 5 | 0 | 7 | 0.1429 | 8 | 0 | 6 | 0.1667 | 26 | 0.0769 | 4 | 0.25 | 3 | 0 | 7 | 0 | 4 | 0.25 | 18 | 0.1111 |
| 7_1305127 | 7 | 0.1429 | 12 | 0 | 8 | 0 | 7 | 0.2857 | 34 | 0.0882 | 5 | 0 | 4 | 0.25 | 9 | 0.2222 | 4 | 0 | 22 | 0.1364 |
| 7_1305128 | 7 | 0.7143 | 13 | 1 | 6 | 0.8333 | 6 | 0.8333 | 32 | 0.875 | 5 | 0.8 | 4 | 1 | 8 | 1 | 4 | 1 | 21 | 0.9524 |
| 7_1305129 | 5 | 0.6 | 14 | 0.8571 | 7 | 1 | 5 | 0.8 | 31 | 0.8387 | 4 | 1 | 3 | 1 | 11 | 0.9091 | 4 | 1 | 22 | 0.9545 |
| 7_1305130 | 8 | 0.625 | 15 | 0.3333 | 8 | 0.5 | 7 | 0 | 38 | 0.3684 | 4 | 0.25 | 3 | 0.3333 | 10 | 0.2 | 4 | 0 | 21 | 0.1905 |
| 7_1305133 | 8 | 0.5 | 14 | 0.5 | 9 | 0.2222 | 6 | 0.5 | 37 | 0.4324 | 4 | 0.25 | 2 | 1 | 9 | 0.2222 | 4 | 0.75 | 19 | 0.4211 |
| 7_1305145 | 7 | 0.2857 | 17 | 0.0588 | 14 | 0.1429 | 9 | 0.1111 | 47 | 0.1277 | 5 | 0 | 7 | 0.2857 | 15 | 0.2667 | 6 | 0.1667 | 33 | 0.2121 |
| 7_1305147 | 12 | 0.25 | 17 | 0.3529 | 12 | 0.3333 | 10 | 0.2 | 51 | 0.2941 | 8 | 0 | 8 | 0.375 | 14 | 0.4286 | 7 | 0.1429 | 37 | 0.2703 |
| 7_1305150 | 5 | 0 | 9 | 0.2222 | 9 | 0.2222 | 6 | 0 | 29 | 0.1379 | 1 | 0 | 5 | 0 | 6 | 0.1667 | 4 | 0.75 | 16 | 0.25 |
| 7_1305381 | 24 | 0.3333 | 22 | 0.5 | 22 | 0.4545 | 34 | 0.3235 | 102 | 0.3922 | 32 | 0.5 | 23 | 0.5217 | 19 | 0.5789 | 27 | 0.4815 | 101 | 0.5149 |
| 7_1305637 | 26 | 0.3077 | 29 | 0.1379 | 27 | 0.1481 | 36 | 0.1667 | 118 | 0.1864 | 38 | 0.2632 | 26 | 0.1538 | 28 | 0.1786 | 22 | 0.2273 | 114 | 0.2105 |
| 7_1305811 | 27 | 0.0741 | 27 | 0.0741 | 29 | 0.0345 | 39 | 0.1538 | 122 | 0.0902 | 44 | 0.1136 | 24 | 0.0417 | 28 | 0.0714 | 24 | 0 | 120 | 0.0667 |
| 7_1305867 | 26 | 0.1538 | 26 | 0.1923 | 30 | 0.1333 | 37 | 0.1351 | 119 | 0.1513 | 36 | 0.25 | 21 | 0.0952 | 29 | 0.2069 | 26 | 0.0769 | 112 | 0.1696 |
| 7_1305868 | 34 | 0.1765 | 37 | 0.0811 | 34 | 0 | 43 | 0.0233 | 148 | 0.0676 | 46 | 0.1304 | 30 | 0.0667 | 36 | 0.0556 | 35 | 0.0857 | 147 | 0.0884 |
| 7_1305868 | 29 | 0.4483 | 32 | 0.5 | 31 | 0.5484 | 38 | 0.4737 | 130 | 0.4923 | 42 | 0.4048 | 27 | 0.4444 | 33 | 0.5758 | 33 | 0.4545 | 135 | 0.4667 |
| 7_1305869 | 30 | 0.7 | 35 | 0.7714 | 32 | 0.75 | 41 | 0.7073 | 138 | 0.7319 | 45 | 0.7556 | 25 | 0.76 | 31 | 0.6774 | 32 | 0.75 | 133 | 0.7368 |
| 7_1305869 | 35 | 0.2857 | 41 | 0.2439 | 34 | 0.2353 | 38 | 0.1579 | 148 | 0.2297 | 49 | 0.2245 | 29 | 0.2069 | 36 | 0.3611 | 34 | 0.2647 | 148 | 0.2635 |
| 7_1305869 | 34 | 0.5294 | 39 | 0.641 | 37 | 0.5405 | 42 | 0.4762 | 152 | 0.5461 | 48 | 0.3958 | 30 | 0.5 | 37 | 0.4595 | 36 | 0.5278 | 151 | 0.4636 |
| 7_1305873 | 28 | 0.2857 | 34 | 0.1471 | 31 | 0.129 | 39 | 0.2564 | 132 | 0.2045 | 43 | 0.2093 | 31 | 0.1935 | 40 | 0.2 | 32 | 0.125 | 146 | 0.1849 |
| 7_1309450 | 3 | 0.3333 | 2 | 0 | NA | NA | 2 | 0 | 7 | 0.1429 | 4 | 0.5 | 4 | 0.25 | 4 | 0.5 | 1 | 0 | 13 | 0.3846 |
| 7_1309451 | 4 | 0.5 | 1 | 0 | 1 | 0 | 1 | 0 | 7 | 0.2857 | 6 | 0.1667 | 3 | 0 | 4 | 0.25 | 1 | 0 | 14 | 0.1429 |
| 7_1309453 | 4 | 1 | 1 | 1 | 1 | 1 | 2 | 1 | 8 | 1 | 4 | 1 | 3 | 1 | 5 | 1 | 1 | 1 | 13 | 1 |
| 7_1309453 | 4 | 0.5 | 1 | 1 | 1 | 1 | 2 | 1 | 8 | 0.75 | 4 | 0.75 | 2 | 0.5 | 5 | 0.8 | 2 | 0.5 | 13 | 0.6923 |
| 7_1309453 | 4 | 1 | 1 | 1 | 1 | 1 | 2 | 1 | 8 | 1 | 3 | 1 | 2 | 1 | 4 | 1 | 2 | 1 | 11 | 1 |
| 7_1309454 | 3 | 0.6667 | 1 | 1 | 1 | 1 | 1 | 1 | 6 | 0.8333 | 4 | 0.75 | 2 | 0 | 5 | 0.4 | 2 | 0.5 | 13 | 0.4615 |
| 7_1309454 | 4 | 0.25 | 1 | 0 | 1 | 1 | 2 | 1 | 8 | 0.5 | 3 | 0.3333 | 1 | 0 | 2 | 0.5 | 2 | 0.5 | 8 | 0.375 |
| 7_1309524 | 4 | 0.75 | 1 | 0 | 1 | 0 | 2 | 0 | 8 | 0.375 | 4 | 0.5 | 1 | 1 | 3 | 0.3333 | 1 | 0 | 9 | 0.4444 |
| 7_1309525 | 4 | 0 | 1 | 0 | 1 | 0 | 2 | 0.5 | 8 | 0.125 | 5 | 0.2 | 2 | 0 | 4 | 0 | 1 | 0 | 12 | 0.0833 |
| 7_1309526 | 2 | 1 | NA | NA | 1 | 1 | 1 | 1 | 4 | 1 | 2 | 1 | 2 | 0.5 | 1 | 1 | 1 | 1 | 6 | 0.8333 |
| 7_1309635 | 1 | 1 | 1 | 0 | 1 | 0 | 2 | 0 | 5 | 0.2 | 4 | 0.25 | 1 | 0 | 2 | 0 | NA | NA | 7 | 0.1429 |
| 7_1309753 | 2 | 0 | 1 | 0 | 3 | 0.3333 | 1 | 0 | 7 | 0.1429 | 1 | 0 | 2 | 0.5 | NA | NA | 1 | 0 | 4 | 0.25 |
| 7_1309880 | 2 | 0 | 2 | 0 | NA | NA | 1 | 1 | 5 | 0.2 | 2 | 0 | NA | NA | 1 | 0 | 1 | 1 | 4 | 0.25 |
| 7_1309911 | 5 | 0 | 5 | 0 | 7 | 0.2857 | 4 | 0 | 21 | 0.0952 | 3 | 0 | 1 | 0 | 6 | 0.1667 | 3 | 0.3333 | 13 | 0.1538 |
| 7_1309913 | 3 | 0.6667 | 5 | 0 | 2 | 1 | 3 | 0.6667 | 13 | 0.4615 | 3 | 0.3333 | NA | NA | 3 | 0.3333 | NA | NA | 6 | 0.3333 |
| 7_1309913 | 2 | 0 | NA | NA | 2 | 0 | 1 | 1 | 5 | 0.2 | 1 | 0 | 1 | 0 | 3 | 0.6667 | 3 | 0.3333 | 8 | 0.375 |
| 7_1309914 | 4 | 0 | 3 | 0 | 6 | 0 | 3 | 0.3333 | 16 | 0.0625 | 3 | 0 | 1 | 0 | 6 | 0.1667 | 3 | 0 | 13 | 0.0769 |
| 7_1309916 | 5 | 0 | 4 | 0.25 | 5 | 0 | 4 | 0 | 18 | 0.0556 | 4 | 0.25 | 1 | 0 | 5 | 0.4 | 3 | 0 | 13 | 0.2308 |

| | | | | | | | | | | | | | | | | | | | | |
|---|---|---|---|---|---|---|---|---|---|---|---|---|---|---|---|---|---|---|---|---|
| 7_13099172 | 4 | 0.25 | 4 | 0 | 5 | 0.2 | 4 | 0 | 17 | 0.1176 | 4 | 0.25 | 1 | 1 | 6 | 0.1667 | 2 | 0 | 13 | 0.2308 |
| 7_13099190 | 5 | 0.4 | 4 | 0 | 5 | 0.2 | 3 | 0 | 17 | 0.1765 | 4 | 0 | 1 | 0 | 6 | 0.1667 | 2 | 0 | 13 | 0.0769 |
| 7_13099336 | 1 | 0 | 3 | 0 | 4 | 0.5 | 2 | 0.5 | 10 | 0.3 | 3 | 0 | NA | NA | 5 | 0.4 | 2 | 0.5 | 10 | 0.3 |
| 7_13099341 | 3 | 0.6667 | 2 | 1 | 6 | 0.8333 | 2 | 0.5 | 13 | 0.7692 | 2 | 1 | NA | NA | 4 | 0.25 | 2 | 0.5 | 8 | 0.5 |
| 7_13099350 | 3 | 0 | 3 | 0 | 5 | 0.2 | 2 | 0 | 13 | 0.0769 | 4 | 0 | NA | NA | 5 | 0.2 | 2 | 0 | 11 | 0.0909 |
| 7_13099365 | NA | NA | 1 | 1 | NA | NA | 1 | 0 | 2 | 0.5 | 1 | 1 | NA | NA | NA | NA | NA | NA | 1 | 1 |
| 7_13099365 | 3 | 0.3333 | 2 | 0.5 | 4 | 0 | 2 | 0.5 | 11 | 0.2727 | 3 | 0 | NA | NA | 4 | 0.75 | 2 | 0 | 9 | 0.3333 |
| 7_13099381 | 1 | 0 | 2 | 0 | 6 | 0.1667 | 1 | 0 | 10 | 0.1 | 4 | 0.25 | NA | NA | 4 | 0.5 | 2 | 0 | 10 | 0.3 |
| 7_13099384 | 2 | 1 | 3 | 1 | 6 | 0.8333 | 2 | 0 | 13 | 0.7692 | 3 | 1 | NA | NA | 3 | 0.6667 | 2 | 0.5 | 8 | 0.75 |
| 7_13099389 | 2 | 1 | 1 | 1 | 5 | 1 | 2 | 1 | 10 | 1 | 3 | 1 | NA | NA | 3 | 1 | 1 | 1 | 7 | 1 |
| 7_13099390 | 2 | 1 | 1 | 1 | 6 | 0.8333 | 2 | 0.5 | 11 | 0.8182 | 4 | 0.5 | NA | NA | 4 | 1 | 2 | 1 | 10 | 0.8 |
| 7_13099395 | 2 | 0.5 | 2 | 0.5 | 4 | 0.5 | 2 | 0.5 | 10 | 0.5 | 3 | 0.3333 | NA | NA | 4 | 0.25 | 2 | 1 | 9 | 0.4444 |
| 7_13099395 | 2 | 1 | 2 | 1 | 5 | 1 | 1 | 1 | 10 | 1 | 4 | 0.75 | NA | NA | 3 | 1 | 2 | 0.5 | 9 | 0.7778 |
| 7_13099399 | 2 | 1 | 2 | 1 | 6 | 0.8333 | NA | NA | 10 | 0.9 | 3 | 0.3333 | NA | NA | 4 | 0.75 | 2 | 1 | 9 | 0.6667 |
| 7_13099400 | 2 | 0 | 3 | 0.3333 | 5 | 0.2 | 2 | 0.5 | 12 | 0.25 | 3 | 0 | NA | NA | 4 | 0.25 | 2 | 0 | 9 | 0.1111 |
| 7_13099428 | NA | NA | 3 | 0.3333 | 1 | 1 | 1 | 0 | 5 | 0.4 | 2 | 0.5 | NA | NA | NA | NA | 1 | 0 | 3 | 0.3333 |
| 7_13099435 | NA | NA | 4 | 0.5 | 1 | 1 | 1 | 0 | 6 | 0.5 | 3 | 0.3333 | NA | NA | NA | NA | 1 | 1 | 4 | 0.5 |
| 7_13099438 | NA | NA | 4 | 0.25 | 1 | 1 | 1 | 0 | 6 | 0.3333 | 3 | 0.3333 | NA | NA | NA | NA | 1 | 1 | 4 | 0.5 |
| 7_13104603 | 5 | 0.4 | 1 | 0 | 4 | 0.75 | 7 | 0.2857 | 17 | 0.4118 | 6 | 0.1667 | 4 | 0.5 | 5 | 0.4 | 3 | 0.6667 | 18 | 0.3889 |
| 7_13104613 | 4 | 0.75 | 1 | 1 | 4 | 0.5 | 6 | 0.5 | 15 | 0.6 | 4 | 0.75 | 4 | 1 | 6 | 0.3333 | 3 | 0.6667 | 17 | 0.6471 |
| 7_13104659 | 7 | 0.1429 | 3 | 0 | 7 | 0 | 4 | 0.25 | 21 | 0.0952 | 4 | 0 | 5 | 0.2 | 6 | 0 | 4 | 0.25 | 19 | 0.1053 |
| 7_13116262 | 9 | 0.3333 | 4 | 0.25 | 6 | 0.1667 | 9 | 0.3333 | 28 | 0.2857 | 6 | 0.3333 | 8 | 0.25 | 6 | 0.1667 | 2 | 1 | 22 | 0.3182 |
| 7_13117269 | 15 | 0.3333 | 13 | 0.3077 | 12 | 0.25 | 12 | 0.4167 | 52 | 0.3269 | 10 | 0.2 | 13 | 0.3077 | 10 | 0.3 | 4 | 0.25 | 37 | 0.2703 |
| 7_13117327 | 6 | 0.3333 | 9 | 0.4444 | 5 | 0.4 | 7 | 0.1429 | 27 | 0.3333 | 2 | 0 | 8 | 0.625 | 8 | 0.25 | 3 | 0 | 21 | 0.3333 |
| 7_13117840 | NA | NA | 1 | 1 | NA | NA | NA | NA | 1 | 1 | NA | NA | NA | NA | NA | NA | 1 | 1 | 1 | 1 |
| 7_13156829 | 1 | 0 | NA | NA | 1 | 0 | 1 | 1 | 3 | 0.3333 | 2 | 0 | 2 | 0.5 | 1 | 1 | NA | NA | 5 | 0.4 |
| 7_13156836 | 1 | 1 | NA | NA | 1 | 0 | 1 | 0 | 3 | 0.3333 | 3 | 0.6667 | 2 | 0 | 1 | 1 | NA | NA | 6 | 0.5 |
| 7_13156837 | 1 | 1 | NA | NA | NA | NA | 1 | 1 | 2 | 1 | 3 | 0.3333 | 2 | 0 | 1 | 0 | NA | NA | 6 | 0.1667 |
| 7_13156866 | 2 | 1 | 2 | 1 | NA | NA | 2 | 1 | 6 | 1 | 3 | 1 | 1 | 1 | 1 | 1 | NA | NA | 5 | 1 |
| 7_13156888 | 2 | 0 | 2 | 0.5 | 1 | 1 | 2 | 1 | 7 | 0.5714 | 4 | 0.25 | 2 | 0.5 | 2 | 0 | NA | NA | 8 | 0.25 |
| 7_13157608 | 3 | 0.3333 | NA | NA | 1 | 0 | 3 | 0 | 7 | 0.1429 | 3 | 0.3333 | 3 | 0 | 4 | 0 | 1 | 0 | 11 | 0.0909 |
| 7_13158582 | 2 | 0 | 2 | 0 | 1 | 0 | 3 | 0.3333 | 8 | 0.125 | 3 | 0 | 3 | 0.6667 | 4 | 0 | 1 | 0 | 11 | 0.1818 |
| 7_13158600 | 2 | 1 | 2 | 1 | 2 | 0.5 | 3 | 0.3333 | 9 | 0.6667 | 3 | 0.6667 | 3 | 0.3333 | 3 | 0.6667 | 1 | 0 | 10 | 0.5 |
| 7_13158602 | 2 | 1 | 2 | 0.5 | 2 | 1 | 3 | 0.3333 | 9 | 0.6667 | 1 | 0 | 3 | 0.6667 | 4 | 0.5 | NA | NA | 8 | 0.5 |
| 7_13158604 | 2 | 1 | 3 | 0.3333 | 2 | 1 | 3 | 0.6667 | 10 | 0.7 | 3 | 0.6667 | 4 | 0.75 | 4 | 0.5 | NA | NA | 11 | 0.6364 |
| 7_13158610 | 2 | 1 | 3 | 1 | 2 | 1 | 4 | 1 | 11 | 1 | 4 | 1 | 4 | 1 | 4 | 0.75 | 1 | 1 | 13 | 0.9231 |
| 7_13158612 | 1 | 0 | 3 | 0.6667 | 1 | 1 | 3 | 0.3333 | 8 | 0.5 | 4 | 0.75 | 3 | 0.6667 | 4 | 0.75 | 1 | 1 | 12 | 0.75 |
| 7_13158612 | 2 | 0.5 | 2 | 0 | 2 | 0 | 3 | 0 | 9 | 0.1111 | 4 | 0.5 | 2 | 0 | 2 | 0 | 1 | 0 | 9 | 0.2222 |
| 7_13158621 | 3 | 1 | 3 | 1 | 2 | 1 | 4 | 1 | 12 | 1 | 2 | 1 | 2 | 1 | 3 | 1 | 1 | 1 | 8 | 1 |
| 7_13158629 | 2 | 0.5 | 3 | 0 | 1 | 0 | 4 | 0 | 10 | 0.1 | 1 | 0 | 3 | 0.3333 | 2 | 0 | 1 | 0 | 7 | 0.1429 |
| 7_13158634 | 3 | 0.3333 | 4 | 0.5 | 2 | 0 | 4 | 0.5 | 13 | 0.3846 | 2 | 0.5 | 4 | 0.25 | 4 | 0.5 | 1 | 0 | 11 | 0.3636 |
| 7_13158642 | 2 | 1 | 4 | 0.75 | 1 | 1 | 5 | 0.8 | 12 | 0.8333 | 4 | 1 | 4 | 0.75 | 3 | 1 | 1 | 1 | 12 | 0.9167 |
| 7_13158664 | 2 | 0.5 | 3 | 0.6667 | 1 | 0 | 6 | 1 | 12 | 0.75 | 4 | 0 | 3 | 0.6667 | 5 | 0.6 | 1 | 0 | 13 | 0.3846 |
| 7_13158914 | 5 | 0 | 5 | 0.2 | 1 | 0 | 6 | 0.1667 | 17 | 0.1176 | 4 | 0.5 | 7 | 0 | 3 | 0 | 1 | 0 | 15 | 0.1333 |
| 7_13159140 | 5 | 0.2 | 3 | 0 | 1 | 1 | 2 | 0 | 11 | 0.1818 | 3 | 0 | 4 | 0 | 6 | 0.1667 | 2 | 0 | 15 | 0.0667 |
| 7_13159256 | 6 | 0 | 8 | 0.125 | 4 | 0.5 | 4 | 0.25 | 22 | 0.1818 | 5 | 0.4 | 5 | 0.4 | 10 | 0.3 | 6 | 0.5 | 26 | 0.3846 |
| 7_13159289 | 5 | 0 | 8 | 0.25 | 5 | 0.2 | 4 | 0.25 | 22 | 0.1818 | 4 | 0 | 5 | 0 | 8 | 0.375 | 7 | 0.2857 | 24 | 0.2083 |
| 7_13159295 | 5 | 0.4 | 9 | 0.4444 | 4 | 0.75 | 3 | 0.3333 | 21 | 0.4762 | 5 | 0.4 | 6 | 0.8333 | 11 | 0.7273 | 8 | 0.625 | 30 | 0.6667 |
| 7_13159553 | 9 | 0.4444 | 22 | 0.6818 | 14 | 0.7143 | 9 | 0.7778 | 54 | 0.6667 | 7 | 0.5714 | 12 | 0.8333 | 16 | 0.8125 | 15 | 0.7333 | 50 | 0.76 |
| 7_13159594 | 14 | 0.7857 | 20 | 0.65 | 11 | 0.8182 | 7 | 0.8571 | 52 | 0.75 | 10 | 0.8 | 11 | 0.6364 | 14 | 0.5714 | 13 | 0.7692 | 48 | 0.6875 |
| 7_13183767 | 12 | 0 | 7 | 0.1429 | 9 | 0.1111 | 15 | 0.0667 | 43 | 0.0698 | 10 | 0.3 | 17 | 0.1765 | 15 | 0.0667 | 5 | 0 | 47 | 0.1489 |
| 7_13183784 | 12 | 0.0833 | 6 | 0 | 7 | 0 | 14 | 0.0714 | 39 | 0.0513 | 9 | 0 | 18 | 0.1111 | 16 | 0.0625 | 5 | 0.4 | 48 | 0.1042 |
| 7_13183804 | 11 | 0.2727 | 7 | 0.1429 | 7 | 0.2857 | 12 | 0.25 | 37 | 0.2432 | 9 | 0.2222 | 15 | 0.2 | 15 | 0.1333 | 6 | 0.3333 | 45 | 0.2 |
| 7_13183809 | 11 | 0.2727 | 7 | 0.1429 | 7 | 0.2857 | 13 | 0.1538 | 38 | 0.2105 | 9 | 0.1111 | 15 | 0.2667 | 14 | 0.1429 | 4 | 0 | 42 | 0.1667 |
| 7_13183844 | 9 | 0.8889 | 5 | 1 | 3 | 1 | 9 | 0.8889 | 26 | 0.9231 | 6 | 1 | 14 | 0.9286 | 16 | 0.9375 | 5 | 1 | 41 | 0.9512 |
| 7_13183850 | 7 | 1 | 6 | 1 | 7 | 1 | 10 | 0.9 | 30 | 0.9667 | 7 | 0.8571 | 12 | 1 | 15 | 1 | 5 | 1 | 39 | 0.9744 |
| 7_13183854 | 10 | 1 | 5 | 1 | 6 | 1 | 9 | 1 | 30 | 1 | 7 | 1 | 16 | 1 | 15 | 1 | 6 | 1 | 44 | 1 |
| 7_13183877 | 8 | 0.875 | 3 | 0.6667 | 6 | 0.3333 | 9 | 0.6667 | 26 | 0.6538 | 7 | 0.5714 | 17 | 0.8235 | 15 | 0.8 | 5 | 1 | 44 | 0.7955 |
| 7_13183880 | 7 | 0.1429 | 1 | 0 | 5 | 0.2 | 9 | 0.2222 | 22 | 0.1818 | 7 | 0.1429 | 11 | 0.2727 | 11 | 0.2727 | 5 | 0.2 | 34 | 0.2353 |
| 7_13183882 | 7 | 0 | 3 | 0.6667 | 6 | 0.3333 | 8 | 0.375 | 24 | 0.2917 | 8 | 0.25 | 14 | 0.2143 | 11 | 0.3636 | 4 | 1 | 37 | 0.3514 |
| 7_13183890 | 3 | 0.6667 | 3 | 1 | 4 | 0.75 | 6 | 0.5 | 16 | 0.6875 | 4 | 0.75 | 12 | 0.8333 | 9 | 0.7778 | 5 | 0.6 | 30 | 0.7667 |
| 7_13183895 | 4 | 0 | 2 | 1 | 4 | 0 | 4 | 0.25 | 14 | 0.2143 | 6 | 0.5 | 9 | 0.4444 | 7 | 0.4286 | 3 | 0.3333 | 25 | 0.44 |
| 7_13183896 | 5 | 0 | 4 | 0.5 | 6 | 0 | 7 | 0 | 22 | 0.0909 | 8 | 0 | 15 | 0.1333 | 9 | 0.1111 | 5 | 0.2 | 37 | 0.1081 |
| 7_13183917 | 5 | 0 | 4 | 0.5 | 6 | 0.3333 | 4 | 0.25 | 19 | 0.2632 | 5 | 0.4 | 9 | 0.2222 | 5 | 0.2 | 2 | 0.5 | 21 | 0.2857 |
| 7_13183918 | 6 | 0 | 2 | 0 | 6 | 0.1667 | 5 | 0.2 | 19 | 0.1053 | 8 | 0.125 | 13 | 0.2308 | 10 | 0 | 2 | 0 | 33 | 0.1212 |

| | | | | | | | | | | | | | | | | | | | | |
|---|---|---|---|---|---|---|---|---|---|---|---|---|---|---|---|---|---|---|---|---|
| 7_1318391 | 5 | 0.2 | 4 | 0 | 5 | 0.2 | 5 | 0 | 19 | 0.1053 | 7 | 0 | 11 | 0.0909 | 8 | 0.125 | 1 | 0 | 27 | 0.0741 |
| 7_1318392 | 6 | 1 | 5 | 1 | 5 | 1 | 6 | 1 | 22 | 1 | 6 | 1 | 15 | 0.9333 | 11 | 1 | 3 | 1 | 35 | 0.9714 |
| 7_1318392 | 6 | 0.5 | 2 | 0.5 | 3 | 0.6667 | 7 | 0.5714 | 18 | 0.5556 | 7 | 0.5714 | 11 | 0.4545 | 8 | 0.625 | 3 | 0.6667 | 29 | 0.5517 |
| 7_1318393 | 6 | 0.5 | 4 | 1 | 3 | 0.6667 | 7 | 0.7143 | 20 | 0.7 | 7 | 0.5714 | 12 | 0.8333 | 9 | 0.7778 | 2 | 1 | 30 | 0.7667 |
| 7_1318394 | 6 | 0.3333 | 4 | 0 | 6 | 0 | 5 | 0.2 | 21 | 0.1429 | 7 | 0.1429 | 11 | 0.0909 | 10 | 0.1 | 3 | 0.3333 | 31 | 0.129 |
| 7_1318395 | 7 | 1 | 2 | 1 | 4 | 0.5 | 6 | 1 | 19 | 0.8947 | 4 | 1 | 10 | 1 | 8 | 0.875 | 3 | 0.6667 | 25 | 0.92 |
| 7_1318396 | 7 | 0.8571 | 1 | 1 | 3 | 1 | 6 | 1 | 17 | 0.9412 | 5 | 1 | 10 | 1 | 9 | 1 | 2 | 1 | 26 | 1 |
| 7_1318396 | 7 | 0.2857 | 3 | 0.3333 | 5 | 0 | 6 | 0.5 | 21 | 0.2857 | 7 | 0.1429 | 14 | 0.1429 | 8 | 0 | 3 | 0.3333 | 32 | 0.125 |
| 7_1318398 | 6 | 1 | 3 | 1 | 6 | 0.8333 | 5 | 1 | 20 | 0.95 | 6 | 1 | 11 | 0.7273 | 8 | 0.75 | 1 | 1 | 26 | 0.8077 |
| 7_1318399 | 3 | 1 | 2 | 1 | 5 | 1 | 6 | 0.8333 | 16 | 0.9375 | 7 | 1 | 12 | 0.9167 | 6 | 1 | 1 | 1 | 26 | 0.9615 |
| 7_1318399 | 4 | 0.75 | 2 | 1 | 5 | 1 | 4 | 0.5 | 15 | 0.8 | 5 | 0.2 | 11 | 0.4545 | 6 | 0.3333 | 1 | 0 | 23 | 0.3478 |
| 7_1318399 | 5 | 1 | 2 | 0.5 | 5 | 0.8 | 6 | 0.6667 | 18 | 0.7778 | 7 | 1 | 11 | 0.8182 | 8 | 0.875 | 1 | 1 | 27 | 0.8889 |
| 7_1318684 | NA | NA | NA | NA | 4 | 0.25 | 2 | 0 | 6 | 0.1667 | 3 | 0.3333 | 9 | 0 | 3 | 0 | 1 | 0 | 16 | 0.0625 |
| 7_1318717 | 3 | 0 | 1 | 0 | 3 | 0 | 2 | 1 | 9 | 0.2222 | 2 | 1 | 9 | 0 | 4 | 0 | 1 | 0 | 16 | 0.125 |
| 7_1318718 | 3 | 1 | 2 | 1 | 3 | 1 | 2 | 1 | 10 | 1 | 1 | 1 | 9 | 0.8889 | 4 | 0.75 | 1 | 0 | 15 | 0.8 |
| 7_1318719 | 3 | 1 | NA | NA | 2 | 0.5 | 2 | 1 | 7 | 0.8571 | 2 | 1 | 6 | 0.6667 | 3 | 0.6667 | 1 | 1 | 12 | 0.75 |
| 7_1318720 | 2 | 0.5 | 2 | 0 | 3 | 0.3333 | 2 | 0.5 | 9 | 0.3333 | 3 | 0.6667 | 9 | 0.1111 | 4 | 0.25 | 1 | 0 | 17 | 0.2353 |
| 7_1318720 | 1 | 0 | 1 | 1 | 3 | 0.6667 | NA | NA | 5 | 0.6 | 2 | 1 | 6 | 0.8333 | 3 | 0.3333 | 1 | 1 | 12 | 0.75 |
| 7_1318720 | 2 | 1 | 2 | 0.5 | 3 | 1 | 2 | 1 | 9 | 0.8889 | 3 | 1 | 8 | 0.875 | 3 | 1 | NA | NA | 14 | 0.9286 |
| 7_1318721 | 3 | 0.3333 | 1 | 0 | 2 | 0 | 1 | 0 | 7 | 0.1429 | 2 | 0.5 | 7 | 0.2857 | 4 | 0.75 | 1 | 0 | 14 | 0.4286 |
| 7_1318721 | NA | NA | 2 | 0.5 | 3 | 0.6667 | 2 | 0.5 | 7 | 0.5714 | 3 | 0.3333 | 6 | 0.3333 | 4 | 0.75 | 1 | 0 | 14 | 0.4286 |
| 7_1318723 | 2 | 0.5 | 2 | 1 | 3 | 1 | 2 | 1 | 9 | 0.8889 | 2 | 1 | 5 | 1 | 4 | 1 | NA | NA | 11 | 1 |
| 7_1318723 | 3 | 0.3333 | 2 | 0 | 3 | 0.3333 | 2 | 0.5 | 10 | 0.3 | 2 | 1 | 6 | 0.6667 | 4 | 0.75 | 1 | 0 | 13 | 0.6923 |
| 7_1318724 | 3 | 0.6667 | 2 | 1 | 2 | 1 | 2 | 1 | 9 | 0.8889 | NA | NA | 6 | 1 | 3 | 0.6667 | NA | NA | 9 | 0.8889 |
| 7_1318724 | 3 | 0.6667 | 2 | 0.5 | 3 | 1 | 1 | 0 | 9 | 0.6667 | 1 | 1 | 6 | 0.8333 | 4 | 0.75 | 1 | 1 | 12 | 0.8333 |
| 7_1318725 | 3 | 0 | 1 | 1 | 3 | 0.3333 | 1 | 0 | 8 | 0.25 | 1 | 0 | 6 | 0.1667 | 3 | 0 | 1 | 0 | 11 | 0.0909 |
| 7_1319169 | NA | NA | NA | NA | 1 | 1 | NA | NA | 1 | 1 | NA | NA | NA | NA | 1 | 1 | NA | NA | 1 | 1 |
| 7_1321595 | 6 | 0 | 12 | 0.1667 | 13 | 0 | 9 | 0 | 40 | 0.05 | 17 | 0.1176 | 9 | 0 | 12 | 0 | 14 | 0.0714 | 52 | 0.0577 |
| 7_1321596 | 10 | 0.2 | 14 | 0 | 11 | 0.1818 | 14 | 0 | 49 | 0.0816 | 16 | 0.125 | 10 | 0.1 | 15 | 0.2 | 18 | 0.2222 | 59 | 0.1695 |
| 7_1321596 | 10 | 0.7 | 11 | 0.2727 | 14 | 0.4286 | 12 | 0.3333 | 47 | 0.4255 | 16 | 0.4375 | 10 | 0.3 | 14 | 0.1429 | 13 | 0.3846 | 53 | 0.3208 |
| 7_1321597 | 8 | 0.625 | 11 | 0.3636 | 12 | 0.3333 | 13 | 0.3846 | 44 | 0.4091 | 17 | 0.4118 | 11 | 0.2727 | 13 | 0.3846 | 16 | 0.5 | 57 | 0.4035 |
| 7_1321598 | 10 | 0.3 | 9 | 0.1111 | 12 | 0.0833 | 13 | 0.1538 | 44 | 0.1591 | 15 | 0.2 | 9 | 0.1111 | 15 | 0.2 | 15 | 0.2 | 54 | 0.1852 |
| 7_1321598 | 8 | 0.75 | 8 | 0.625 | 10 | 0.8 | 11 | 0.4545 | 37 | 0.6486 | 14 | 0.4286 | 8 | 0.375 | 11 | 0.7273 | 8 | 0.5 | 41 | 0.5122 |
| 7_1321599 | 9 | 0.1111 | 9 | 0 | 12 | 0.4167 | 10 | 0.2 | 40 | 0.2 | 14 | 0.0714 | 8 | 0.125 | 14 | 0.0714 | 17 | 0.0588 | 53 | 0.0755 |
| 7_1323075 | 3 | 0.3333 | 2 | 0 | 1 | 0 | 2 | 0 | 8 | 0.125 | 4 | 0 | 3 | 0.3333 | 3 | 0.6667 | 3 | 0.3333 | 13 | 0.3077 |
| 7_1323079 | 4 | 0.5 | 2 | 0 | 2 | 0.5 | 2 | 0.5 | 10 | 0.4 | 4 | 0 | 3 | 0 | 4 | 0.25 | 3 | 0 | 14 | 0.0714 |
| 7_1323080 | 4 | 0 | 2 | 0 | 2 | 0.5 | 1 | 0 | 9 | 0.1111 | 4 | 0 | 2 | 0 | 5 | 0 | 3 | 0.3333 | 14 | 0.0714 |
| 7_1323080 | 5 | 0.2 | 2 | 0 | 2 | 0 | 2 | 0 | 11 | 0.0909 | 4 | 0 | 3 | 0 | 5 | 0.2 | 3 | 0 | 15 | 0.0667 |
| 7_1323173 | 1 | 1 | 2 | 1 | NA | NA | 1 | 1 | 4 | 1 | 1 | 1 | NA | NA | NA | NA | 1 | 1 | 2 | 1 |
| 7_1323175 | 1 | 1 | 1 | 0 | NA | NA | 1 | 1 | 3 | 0.6667 | 1 | 1 | NA | NA | NA | NA | 1 | 1 | 2 | 1 |
| 7_1323175 | 1 | 1 | 2 | 0.5 | NA | NA | 1 | 0 | 4 | 0.5 | 1 | 0 | NA | NA | NA | NA | 1 | 1 | 2 | 0.5 |
| 7_1323175 | 1 | 1 | 1 | 1 | NA | NA | 1 | 1 | 3 | 1 | 1 | 1 | NA | NA | NA | NA | NA | NA | 1 | 1 |
| 7_1323458 | NA | NA | 1 | 1 | NA | NA | NA | NA | 1 | 1 | NA | NA | 1 | 1 | 1 | 1 | NA | NA | 2 | 1 |
| 7_1323458 | NA | NA | 1 | 1 | NA | NA | NA | NA | 1 | 1 | NA | NA | 1 | 1 | 1 | 1 | NA | NA | 2 | 1 |
| 7_1323459 | NA | NA | 1 | 1 | NA | NA | NA | NA | 1 | 1 | NA | NA | 1 | 1 | NA | NA | NA | NA | 1 | 1 |
| 7_1323459 | NA | NA | 1 | 1 | NA | NA | NA | NA | 1 | 1 | NA | NA | 1 | 1 | 1 | 0 | NA | NA | 2 | 0.5 |
| 7_1323461 | NA | NA | 1 | 1 | NA | NA | NA | NA | 1 | 1 | NA | NA | 1 | 1 | 1 | 1 | NA | NA | 2 | 1 |
| 7_1323462 | NA | NA | 1 | 1 | NA | NA | NA | NA | 1 | 1 | NA | NA | 1 | 1 | 1 | 1 | NA | NA | 2 | 1 |
| 7_1323464 | NA | NA | 1 | 1 | NA | NA | NA | NA | 1 | 1 | NA | NA | 1 | 1 | 1 | 1 | NA | NA | 2 | 1 |
| 7_1323465 | NA | NA | 1 | 1 | NA | NA | NA | NA | 1 | 1 | NA | NA | 1 | 1 | 1 | 1 | NA | NA | 2 | 1 |
| 7_1323467 | NA | NA | 1 | 1 | NA | NA | NA | NA | 1 | 1 | NA | NA | NA | NA | 1 | 1 | NA | NA | 1 | 1 |
| 7_1323614 | 1 | 1 | 3 | 1 | NA | NA | 1 | 1 | 5 | 1 | 1 | 1 | NA | NA | NA | NA | NA | NA | 1 | 1 |
| 7_1323616 | NA | NA | 2 | 1 | NA | NA | 1 | 0 | 3 | 0.6667 | 1 | 1 | NA | NA | NA | NA | NA | NA | 1 | 1 |
| 7_1323617 | 1 | 1 | 1 | 1 | NA | NA | 1 | 1 | 3 | 1 | 1 | 1 | NA | NA | NA | NA | NA | NA | 1 | 1 |
| 7_1324253 | NA | NA | NA | NA | 3 | 0.3333 | NA | NA | 3 | 0.3333 | 1 | 0 | NA | NA | 1 | 1 | NA | NA | 2 | 0.5 |
| 7_1324255 | NA | NA | NA | NA | 2 | 0.5 | NA | NA | 2 | 0.5 | 1 | 1 | NA | NA | 1 | 0 | NA | NA | 2 | 0.5 |
| 7_1324268 | NA | NA | NA | NA | 1 | 1 | NA | NA | 1 | 1 | 1 | 1 | NA | NA | NA | NA | NA | NA | 1 | 1 |
| 7_1324293 | 3 | 0 | 7 | 0.2857 | 3 | 0.6667 | 3 | 0 | 16 | 0.25 | 3 | 0.6667 | 5 | 0.4 | 4 | 0.75 | 3 | 0.6667 | 15 | 0.6 |
| 7_1324294 | 3 | 0 | 9 | 0.1111 | 4 | 0 | 3 | 0 | 19 | 0.0526 | 3 | 0.3333 | 4 | 0 | 3 | 0 | 4 | 0 | 14 | 0.0714 |
| 7_1324295 | 2 | 0 | 6 | 0.3333 | 4 | 0.25 | 4 | 0.5 | 16 | 0.3125 | 1 | 0 | 5 | 0.4 | 3 | 0.6667 | 2 | 0 | 11 | 0.3636 |
| 7_1324295 | 2 | 0 | 7 | 0.5714 | 3 | 0.3333 | 4 | 0.25 | 16 | 0.375 | 3 | 0.3333 | 4 | 0.5 | 2 | 0 | 4 | 1 | 13 | 0.5385 |
| 7_1324296 | 2 | 0 | 8 | 0 | 3 | 0.3333 | 4 | 0 | 17 | 0.0588 | 3 | 0.3333 | 5 | 0 | 3 | 0.3333 | 3 | 0 | 14 | 0.1429 |
| 7_1324332 | 2 | 0 | 5 | 0.2 | 2 | 1 | 2 | 0.5 | 11 | 0.3636 | 3 | 0.6667 | 4 | 0.25 | 1 | 0 | 3 | 0 | 11 | 0.2727 |
| 7_1324850 | 1 | 1 | 4 | 1 | 1 | 1 | 6 | 0.8333 | 12 | 0.9167 | 2 | 0.5 | NA | NA | NA | NA | 2 | 1 | 4 | 0.75 |
| 7_1324850 | 2 | 0.5 | 1 | 1 | 1 | 0 | 6 | 0.1667 | 10 | 0.3 | NA | NA | 1 | 1 | NA | NA | 2 | 0 | 3 | 0.3333 |

| | | | | | | | | | | | | | | | | | | | | |
|---|---|---|---|---|---|---|---|---|---|---|---|---|---|---|---|---|---|---|---|---|
| 7_13248510 | 2 | 0.5 | 4 | 0 | 1 | 0 | 6 | 0 | 13 | 0.0769 | 2 | 0.5 | 1 | 1 | NA | NA | 2 | 1 | 5 | 0.8 |
| 7_13248521 | 3 | 0.3333 | 2 | 0.5 | 1 | 1 | 6 | 0.8333 | 12 | 0.6667 | 1 | 1 | 1 | 1 | NA | NA | 2 | 1 | 4 | 1 |
| 7_13248524 | 3 | 0.6667 | 3 | 0.3333 | 1 | 1 | 4 | 1 | 11 | 0.7273 | 2 | 1 | NA | NA | NA | NA | 1 | 0 | 3 | 0.6667 |
| 7_13248529 | 3 | 1 | 4 | 1 | 1 | 1 | 5 | 1 | 13 | 1 | 2 | 1 | 1 | 0 | NA | NA | 1 | 1 | 4 | 0.75 |
| 7_13248536 | 2 | 0 | 3 | 0.3333 | 1 | 1 | 6 | 0.3333 | 12 | 0.3333 | 2 | 0 | 1 | 0 | NA | NA | 2 | 0.5 | 5 | 0.2 |
| 7_13248554 | 3 | 0.3333 | 5 | 0.8 | 1 | 0 | 6 | 0.5 | 15 | 0.5333 | 2 | 0.5 | 1 | 1 | NA | NA | 1 | 1 | 4 | 0.75 |
| 7_13248565 | 4 | 1 | 5 | 0.8 | 1 | 0 | 6 | 0.6667 | 16 | 0.75 | 2 | 1 | 1 | 1 | NA | NA | 2 | 1 | 5 | 1 |
| 7_13248573 | 3 | 0 | 4 | 0 | 1 | 0 | 3 | 0.3333 | 11 | 0.0909 | 2 | 0 | 1 | 1 | NA | NA | 2 | 0 | 5 | 0.2 |
| 7_13248639 | 6 | 0.6667 | 3 | 0.6667 | 1 | 0 | 5 | 0.6 | 15 | 0.6 | 2 | 1 | 1 | 1 | NA | NA | 1 | 0 | 4 | 0.75 |
| 7_13248650 | 6 | 0 | 5 | 0.2 | 2 | 0 | 6 | 0 | 19 | 0.0526 | 3 | 0.3333 | 2 | 0 | NA | NA | 3 | 0 | 8 | 0.125 |
| 7_13249131 | NA | NA | NA | NA | 1 | 1 | NA | NA | 1 | 1 | NA | NA | 1 | 1 | NA | NA | NA | NA | 1 | 1 |
| 7_13255155 | NA | NA | 2 | 0.5 | 3 | 0.6667 | 1 | 0 | 6 | 0.5 | 1 | 0 | 2 | 0.5 | 1 | 1 | NA | NA | 4 | 0.5 |
| 7_13255164 | NA | NA | 2 | 0.5 | 2 | 1 | 2 | 1 | 6 | 0.8333 | 1 | 1 | 2 | 1 | 1 | 1 | NA | NA | 4 | 1 |
| 7_13255179 | NA | NA | 3 | 1 | 3 | 0.3333 | 1 | 1 | 7 | 0.7143 | 2 | 1 | 2 | 0.5 | 1 | 0 | NA | NA | 5 | 0.6 |
| 7_13255181 | NA | NA | 2 | 1 | 3 | 1 | 2 | 1 | 7 | 1 | 2 | 1 | 3 | 1 | 1 | 1 | NA | NA | 6 | 1 |
| 7_13255183 | NA | NA | 3 | 0.6667 | 3 | 0 | 1 | 0 | 7 | 0.2857 | 1 | 1 | 2 | 0 | 1 | 1 | NA | NA | 4 | 0.5 |
| 7_13255191 | NA | NA | 3 | 0.6667 | 3 | 1 | 2 | 1 | 8 | 0.875 | 1 | 1 | 3 | 0.6667 | 1 | 1 | NA | NA | 5 | 0.8 |
| 7_13255201 | NA | NA | 3 | 1 | 2 | 1 | 2 | 0.5 | 7 | 0.8571 | 2 | 1 | 3 | 1 | 1 | 1 | NA | NA | 6 | 1 |
| 7_13255209 | NA | NA | 3 | 1 | 2 | 1 | 2 | 1 | 7 | 1 | 2 | 1 | 2 | 1 | 1 | 1 | NA | NA | 5 | 1 |
| 7_13255213 | NA | NA | 3 | 0 | 3 | 0.3333 | 1 | 1 | 7 | 0.2857 | 2 | 0.5 | 2 | 0.5 | 1 | 0 | NA | NA | 5 | 0.4 |
| 7_13258455 | 1 | 0 | 4 | 0.25 | 4 | 0 | 1 | 0 | 10 | 0.1 | 2 | 0.5 | 7 | 0 | 1 | 0 | 1 | 0 | 11 | 0.0909 |
| 7_13258550 | 2 | 1 | 5 | 0.8 | 4 | 0.75 | 2 | 1 | 13 | 0.8462 | 2 | 1 | 6 | 0.6667 | 1 | 1 | 1 | 1 | 10 | 0.8 |
| 7_13258551 | 1 | 1 | 5 | 0.4 | 5 | 0.2 | 4 | 0.25 | 15 | 0.3333 | 2 | 0 | 8 | 0.25 | 2 | 0.5 | 1 | 0 | 13 | 0.2308 |
| 7_13258561 | 3 | 0 | 4 | 0.5 | 5 | 0.2 | 3 | 0 | 15 | 0.2 | 2 | 0 | 7 | 0 | 2 | 0 | 1 | 1 | 12 | 0.0833 |
| 7_13258570 | 2 | 0 | 4 | 0 | 5 | 0.2 | 4 | 0.25 | 15 | 0.1333 | 1 | 0 | 7 | 0.1429 | 2 | 0 | 1 | 0 | 11 | 0.0909 |
| 7_13258584 | 4 | 0.75 | 6 | 0.8333 | 4 | 0.75 | 3 | 0.6667 | 17 | 0.7647 | 2 | 0.5 | 8 | 1 | 2 | 0 | 1 | 1 | 13 | 0.7692 |
| 7_13258586 | 4 | 1 | 7 | 1 | 5 | 1 | 5 | 1 | 21 | 1 | 2 | 1 | 8 | 1 | 2 | 1 | 1 | 1 | 13 | 1 |
| 7_13258587 | 3 | 0.6667 | 6 | 0.1667 | 3 | 0.6667 | 5 | 0.6 | 17 | 0.4706 | 2 | 0.5 | 7 | 0.7143 | 2 | 0 | 1 | 1 | 12 | 0.5833 |
| 7_13258596 | 3 | 0 | 5 | 0.2 | 5 | 0 | 2 | 0.5 | 15 | 0.1333 | 1 | 0 | 3 | 0.6667 | 2 | 1 | 1 | 1 | 7 | 0.7143 |
| 7_13258596 | 3 | 0.6667 | 7 | 0.4286 | 3 | 0.6667 | 2 | 0.5 | 15 | 0.5333 | 2 | 1 | 3 | 0.6667 | 2 | 1 | NA | NA | 7 | 0.8571 |
| 7_13258633 | 6 | 0 | 6 | 0 | 4 | 0 | 4 | 0.25 | 20 | 0.05 | 1 | 0 | 7 | 0.1429 | 2 | 0 | 1 | 0 | 11 | 0.0909 |
| 7_13258660 | 4 | 1 | 3 | 0.3333 | 3 | 0.6667 | 6 | 0.5 | 16 | 0.625 | 2 | 0.5 | 7 | 0.2857 | 3 | 0 | 2 | 0.5 | 14 | 0.2857 |
| 7_13258664 | 1 | 0 | 1 | 1 | NA | NA | 1 | 0 | 3 | 0.3333 | 1 | 1 | 1 | 0 | 1 | 0 | NA | NA | 3 | 0.3333 |
| 7_13258775 | 4 | 0 | 5 | 0.2 | 1 | 0 | 2 | 0.5 | 12 | 0.1667 | 1 | 0 | 3 | 0.6667 | NA | NA | 1 | 1 | 5 | 0.6 |
| 7_13258795 | 5 | 0.2 | 4 | 1 | 1 | 0 | 4 | 0.5 | 14 | 0.5 | 2 | 0.5 | 4 | 0.5 | NA | NA | 1 | 1 | 7 | 0.5714 |
| 7_13258795 | 3 | 0 | 3 | 0.3333 | 1 | 0 | 4 | 0 | 11 | 0.0909 | 2 | 0.5 | 4 | 0 | NA | NA | NA | NA | 6 | 0.1667 |
| 7_13258798 | 4 | 0.25 | 5 | 0.2 | 1 | 1 | 4 | 0.25 | 14 | 0.2857 | 2 | 0.5 | 2 | 0.5 | NA | NA | 1 | 0 | 5 | 0.4 |
| 7_13258807 | 5 | 0.6 | 6 | 0.8333 | 1 | 0 | 5 | 0.6 | 17 | 0.6471 | 2 | 0.5 | 4 | 0.25 | NA | NA | 1 | 1 | 7 | 0.4286 |
| 7_13258910 | 3 | 0.6667 | NA | NA | 1 | 1 | NA | NA | 4 | 0.75 | 1 | 1 | 1 | 0 | 1 | 0 | 1 | 0 | 4 | 0.25 |
| 7_13258932 | 1 | 1 | NA | NA | 1 | 1 | NA | NA | 2 | 1 | 1 | 0 | 1 | 0 | 1 | 1 | NA | NA | 3 | 0.3333 |
| 7_13258946 | 4 | 0.75 | NA | NA | 1 | 1 | NA | NA | 5 | 0.8 | 1 | 1 | 1 | 1 | 1 | 0 | 2 | 1 | 5 | 0.8 |
| 7_13258981 | 2 | 0.5 | NA | NA | 1 | 0 | NA | NA | 3 | 0.3333 | 2 | 1 | 4 | 0.75 | 1 | 0 | 1 | 1 | 8 | 0.75 |
| 7_13258982 | 3 | 0.3333 | NA | NA | 1 | 1 | NA | NA | 4 | 0.5 | 3 | 0 | 4 | 0.25 | 1 | 0 | 2 | 0.5 | 10 | 0.2 |
| 7_13258985 | 2 | 1 | NA | NA | 1 | 1 | NA | NA | 3 | 1 | 3 | 0.6667 | 1 | 0 | 1 | 0 | 1 | 1 | 6 | 0.5 |
| 7_13258986 | 4 | 0.25 | NA | NA | 1 | 0 | NA | NA | 5 | 0.2 | 3 | 0 | 3 | 0.3333 | 1 | 0 | 2 | 0 | 9 | 0.1111 |
| 7_13258992 | 3 | 0.3333 | NA | NA | 1 | 0 | NA | NA | 4 | 0.25 | 3 | 0.3333 | 3 | 0.3333 | 1 | 0 | 2 | 0 | 9 | 0.2222 |
| 7_13327328 | 5 | 0.8 | 2 | 1 | 1 | 0 | 2 | 1 | 10 | 0.8 | NA | NA | 2 | 0 | 4 | 1 | 1 | 1 | 7 | 0.7143 |
| 7_13327334 | 5 | 0.4 | 2 | 0.5 | 1 | 0 | 2 | 1 | 10 | 0.5 | NA | NA | 2 | 0.5 | 4 | 0.25 | 1 | 0 | 7 | 0.2857 |
| 7_13327336 | 4 | 0.25 | 2 | 0 | 1 | 1 | 2 | 0.5 | 9 | 0.3333 | NA | NA | 2 | 0.5 | 3 | 0 | 1 | 0 | 6 | 0.1667 |
| 7_13327354 | 5 | 0.2 | 2 | 0 | 1 | 0 | 2 | 0 | 10 | 0.1 | NA | NA | 2 | 0 | 4 | 0 | 1 | 1 | 7 | 0.1429 |
| 7_13328833 | 2 | 0 | 6 | 0 | 6 | 0.1667 | 1 | 0 | 15 | 0.0667 | 2 | 0 | 1 | 0 | 3 | 0 | 1 | 1 | 7 | 0.1429 |
| 7_13328834 | 1 | 0 | 6 | 0 | 6 | 0.1667 | 1 | 0 | 14 | 0.0714 | 2 | 0 | 1 | 0 | 3 | 0.3333 | NA | NA | 6 | 0.1667 |
| 7_13328845 | 1 | 1 | 6 | 0.5 | 6 | 0.8333 | 1 | 1 | 14 | 0.7143 | 2 | 1 | NA | NA | 4 | 0.75 | 1 | 0 | 7 | 0.7143 |
| 7_13328849 | 1 | 1 | 5 | 0.6 | 6 | 0.3333 | 2 | 1 | 14 | 0.5714 | 2 | 1 | 1 | 0 | 3 | 0.3333 | 1 | 1 | 7 | 0.5714 |
| 7_13328867 | 1 | 0 | 5 | 0.6 | 7 | 0.2857 | 2 | 0.5 | 15 | 0.4 | 1 | 0 | 1 | 0 | 3 | 0.3333 | 1 | 0 | 6 | 0.1667 |
| 7_13329171 | 1 | 0 | 1 | 0 | 1 | 1 | NA | NA | 3 | 0.3333 | 1 | 0 | NA | NA | NA | NA | 1 | 1 | 2 | 0.5 |
| 7_13329200 | 1 | 0 | 1 | 1 | 2 | 0 | NA | NA | 4 | 0.25 | 1 | 0 | NA | NA | NA | NA | 1 | 1 | 2 | 0.5 |
| 7_13332253 | 6 | 0.1667 | 8 | 0.5 | 7 | 0.4286 | 9 | 0.4444 | 30 | 0.4 | 6 | 0.1667 | 10 | 0.2 | 13 | 0.1538 | 4 | 0.5 | 33 | 0.2121 |
| 7_13332270 | 5 | 0.6 | 6 | 0.3333 | 2 | 0 | 6 | 0.6667 | 19 | 0.4737 | 7 | 0.2857 | 6 | 0.3333 | 8 | 0.25 | 3 | 0.6667 | 24 | 0.3333 |
| 7_13332288 | 4 | 0 | 5 | 0.4 | 2 | 0 | 6 | 0.5 | 17 | 0.2941 | 5 | 0 | 5 | 0.6 | 10 | 0.3 | 3 | 0.6667 | 23 | 0.3478 |
| 7_13332289 | 1 | 0 | 3 | 0.3333 | 2 | 0 | 5 | 0.2 | 11 | 0.1818 | 3 | 0 | 3 | 0 | 7 | 0.2857 | 1 | 1 | 14 | 0.2143 |
| 7_13332293 | 4 | 0.5 | 5 | 0.6 | 2 | 1 | 6 | 0.3333 | 17 | 0.5294 | 5 | 0.6 | 5 | 0.4 | 11 | 0.3636 | 3 | 0.6667 | 24 | 0.4583 |
| 7_13332306 | 4 | 0 | 5 | 0.2 | 2 | 0 | 7 | 0 | 18 | 0.0556 | 5 | 0 | 4 | 0 | 11 | 0.1818 | 3 | 0 | 23 | 0.087 |
| 7_13333573 | 2 | 0.5 | NA | NA | NA | NA | 3 | 0 | 5 | 0.2 | 1 | 0 | 1 | 0 | 6 | 0.3333 | 1 | 1 | 9 | 0.3333 |
| 7_13333576 | 2 | 0.5 | 1 | 0 | 1 | 0 | 5 | 0 | 9 | 0.1111 | 3 | 0.3333 | 2 | 0 | 7 | 0 | 3 | 0 | 15 | 0.0667 |

| | | | | | | | | | | | | | | | | | | | | |
|---|---|---|---|---|---|---|---|---|---|---|---|---|---|---|---|---|---|---|---|---|
| 7_1333444 | 2 | 0.5 | 1 | 0 | 1 | 0 | 6 | 0.5 | 10 | 0.4 | 1 | 0 | 2 | 0.5 | 5 | 0.8 | 2 | 0 | 10 | 0.5 |
| 7_1333611 | 2 | 1 | 1 | 1 | 1 | 1 | 5 | 0.6 | 9 | 0.7778 | 2 | 0 | 1 | 0 | 4 | 0.25 | 1 | 1 | 8 | 0.25 |
| 7_1333612 | 2 | 1 | NA | NA | 1 | 1 | 3 | 0.6667 | 6 | 0.8333 | 2 | 0 | 1 | 0 | 3 | 0.6667 | 2 | 1 | 8 | 0.5 |
| 7_1333614 | 2 | 1 | 1 | 1 | 1 | 1 | 5 | 0.8 | 9 | 0.8889 | 2 | 1 | 1 | 0 | 5 | 0.8 | 2 | 1 | 10 | 0.8 |
| 7_1333614 | 1 | 1 | 1 | 0 | 1 | 1 | 6 | 0.8333 | 9 | 0.7778 | 2 | 1 | 1 | 1 | 5 | 1 | 1 | 1 | 9 | 1 |
| 7_1333615 | NA | NA | 1 | 0 | NA | NA | 5 | 0.2 | 6 | 0.1667 | 1 | 0 | 1 | 0 | 5 | 0 | 2 | 0.5 | 9 | 0.1111 |
| 7_1334825 | NA | NA | 1 | 0 | 2 | 1 | NA | NA | 3 | 0.6667 | NA | NA | NA | NA | NA | NA | 1 | 1 | 1 | 1 |
| 7_1338676 | 1 | 1 | 2 | 0.5 | 3 | 1 | 1 | 1 | 7 | 0.8571 | 1 | 1 | 3 | 0.3333 | 3 | 0.3333 | 1 | 1 | 8 | 0.5 |
| 7_1338678 | 1 | 1 | 3 | 1 | 4 | 0.75 | 1 | 0 | 9 | 0.7778 | 1 | 1 | 2 | 1 | 4 | 0.5 | 1 | 1 | 8 | 0.75 |
| 7_1338681 | NA | NA | 2 | 0.5 | 2 | 0.5 | 1 | 0 | 5 | 0.4 | 1 | 0 | 2 | 0.5 | 4 | 0.5 | NA | NA | 7 | 0.4286 |
| 7_1338681 | NA | NA | 2 | 0 | 2 | 1 | NA | NA | 4 | 0.5 | 1 | 0 | 2 | 0.5 | 4 | 0.25 | NA | NA | 7 | 0.2857 |
| 7_1338682 | NA | NA | 2 | 0.5 | 2 | 0 | 1 | 0 | 5 | 0.2 | 1 | 0 | 1 | 0 | 3 | 0.3333 | NA | NA | 5 | 0.2 |
| 7_1339505 | NA | NA | 2 | 1 | 2 | 1 | 1 | 1 | 5 | 1 | 1 | 0 | NA | NA | 2 | 1 | NA | NA | 3 | 0.6667 |
| 7_1339507 | NA | NA | 2 | 1 | 1 | 1 | NA | NA | 3 | 1 | 1 | 1 | NA | NA | 1 | 1 | NA | NA | 2 | 1 |
| 7_1339508 | NA | NA | 1 | 1 | 1 | 0 | NA | NA | 2 | 0.5 | NA | NA | NA | NA | 2 | 0.5 | NA | NA | 2 | 0.5 |
| 7_1339510 | NA | NA | 1 | 0 | 2 | 0.5 | NA | NA | 3 | 0.3333 | 1 | 1 | NA | NA | 2 | 0 | NA | NA | 3 | 0.3333 |
| 7_1339512 | NA | NA | NA | NA | 2 | 1 | NA | NA | 2 | 1 | 1 | 1 | NA | NA | 2 | 1 | NA | NA | 3 | 1 |
| 7_1339513 | NA | NA | NA | NA | 2 | 1 | NA | NA | 2 | 1 | 1 | 0 | NA | NA | 1 | 1 | NA | NA | 2 | 0.5 |
| 7_1344837 | 2 | 0 | NA | NA | 3 | 0 | 3 | 0.3333 | 8 | 0.125 | 1 | 0 | NA | NA | 1 | 1 | 1 | 0 | 3 | 0.3333 |
| 7_1344839 | 2 | 0 | NA | NA | 2 | 0 | 4 | 0.5 | 8 | 0.25 | 1 | 0 | 1 | 0 | 2 | 0 | 1 | 1 | 5 | 0.2 |
| 7_1344839 | 1 | 1 | NA | NA | 5 | 0.4 | 3 | 0.6667 | 9 | 0.5556 | 1 | 1 | NA | NA | 2 | 1 | 1 | 1 | 4 | 1 |
| 7_1344845 | 2 | 0 | NA | NA | 2 | 0.5 | 3 | 0.3333 | 7 | 0.2857 | NA | NA | 1 | 1 | 1 | 0 | 1 | 1 | 3 | 0.6667 |
| 7_1344855 | 1 | 1 | NA | NA | 1 | 1 | 2 | 0 | 4 | 0.5 | NA | NA | NA | NA | 2 | 1 | NA | NA | 2 | 1 |
| 7_1344857 | 1 | 1 | NA | NA | 1 | 1 | 2 | 1 | 4 | 1 | NA | NA | NA | NA | 1 | 1 | NA | NA | 1 | 1 |
| 7_1344858 | 1 | 1 | NA | NA | 1 | 0 | 2 | 0 | 4 | 0.25 | NA | NA | NA | NA | 1 | 1 | NA | NA | 1 | 1 |
| 7_1347742 | 2 | 0.5 | 3 | 0 | 3 | 0 | 3 | 0 | 11 | 0.0909 | NA | NA | 2 | 0.5 | 1 | 1 | 2 | 0 | 5 | 0.4 |
| 7_1347744 | 1 | 0 | 3 | 0 | 2 | 0.5 | 3 | 0.3333 | 9 | 0.2222 | 1 | 0 | 1 | 0 | 1 | 1 | 3 | 0.3333 | 6 | 0.3333 |
| 7_1352866 | 2 | 0 | 2 | 0 | 2 | 0 | 2 | 0.5 | 8 | 0.125 | NA | NA | 2 | 0.5 | 1 | 0 | 3 | 0.6667 | 6 | 0.5 |
| 7_1352894 | NA | NA | 1 | 0 | 1 | 1 | NA | NA | 2 | 0.5 | NA | NA | 2 | 1 | NA | NA | NA | NA | 2 | 1 |
| 7_1352894 | NA | NA | 1 | 1 | NA | NA | NA | NA | 1 | 1 | NA | NA | 2 | 1 | NA | NA | NA | NA | 2 | 1 |
| 7_1357695 | 1 | 1 | NA | NA | NA | NA | NA | NA | 1 | 1 | NA | NA | NA | NA | 1 | 1 | NA | NA | 1 | 1 |
| 7_1357696 | 1 | 1 | NA | NA | NA | NA | NA | NA | 1 | 1 | NA | NA | NA | NA | 1 | 1 | 1 | 1 | 2 | 1 |
| 7_1357699 | 1 | 1 | NA | NA | NA | NA | NA | NA | 1 | 1 | NA | NA | NA | NA | 1 | 1 | 1 | 1 | 2 | 1 |
| 7_1357700 | 1 | 1 | NA | NA | NA | NA | NA | NA | 1 | 1 | NA | NA | NA | NA | 1 | 1 | 1 | 1 | 2 | 1 |
| 7_1357701 | 1 | 1 | NA | NA | NA | NA | NA | NA | 1 | 1 | NA | NA | NA | NA | 1 | 1 | NA | NA | 1 | 1 |
| 7_1357705 | 1 | 1 | NA | NA | NA | NA | NA | NA | 1 | 1 | NA | NA | NA | NA | 1 | 1 | NA | NA | 1 | 1 |
| 7_1357708 | 1 | 1 | NA | NA | NA | NA | NA | NA | 1 | 1 | NA | NA | NA | NA | 1 | 1 | 1 | 0 | 2 | 0.5 |
| 7_1369774 | 1 | 1 | NA | NA | NA | NA | 2 | 0.5 | 3 | 0.6667 | NA | NA | NA | NA | NA | NA | 3 | 0.3333 | 3 | 0.3333 |
| 7_1369774 | 1 | 1 | NA | NA | NA | NA | 2 | 1 | 3 | 1 | NA | NA | 1 | 1 | NA | NA | 3 | 0.3333 | 4 | 0.5 |
| 7_1369776 | 1 | 0 | NA | NA | NA | NA | 2 | 0.5 | 3 | 0.3333 | NA | NA | 1 | 1 | NA | NA | 3 | 0.3333 | 4 | 0.5 |
| 7_1370697 | 19 | 0 | 19 | 0.3158 | 14 | 0.2143 | 19 | 0.1579 | 71 | 0.169 | 15 | 0.2667 | 19 | 0.1053 | 16 | 0.0625 | 16 | 0.0625 | 66 | 0.1212 |
| 7_1370697 | 18 | 0.1111 | 18 | 0 | 13 | 0.0769 | 14 | 0.0714 | 63 | 0.0635 | 12 | 0.0833 | 18 | 0.1111 | 15 | 0 | 13 | 0 | 58 | 0.0517 |
| 7_1370697 | 19 | 0.1579 | 19 | 0.1053 | 11 | 0.1818 | 16 | 0.125 | 65 | 0.1385 | 12 | 0.0833 | 18 | 0.0556 | 17 | 0.0588 | 13 | 0 | 60 | 0.05 |
| 7_1378109 | 3 | 0.3333 | 1 | 1 | 1 | 0 | NA | NA | 5 | 0.4 | NA | NA | 1 | 1 | 3 | 0 | NA | NA | 4 | 0.25 |
| 7_1378109 | 3 | 0 | 1 | 1 | 1 | 0 | NA | NA | 5 | 0.2 | 1 | 1 | 3 | 0 | 2 | 0 | NA | NA | 6 | 0.1667 |
| 7_1378109 | 3 | 0.6667 | 1 | 0 | 1 | 0 | NA | NA | 5 | 0.4 | NA | NA | 3 | 1 | 3 | 0.6667 | NA | NA | 6 | 0.8333 |
| 7_1378110 | 3 | 0.3333 | 1 | 0 | 1 | 1 | NA | NA | 5 | 0.4 | 1 | 1 | 3 | 0.3333 | 2 | 0.5 | NA | NA | 6 | 0.5 |
| 7_1378113 | 2 | 0.5 | NA | NA | 1 | 1 | NA | NA | 3 | 0.6667 | NA | NA | 3 | 0.3333 | 3 | 0 | NA | NA | 6 | 0.1667 |
| 7_1378114 | 3 | 0.3333 | 1 | 0 | 1 | 0 | NA | NA | 5 | 0.2 | 1 | 0 | 2 | 0 | 3 | 0.3333 | NA | NA | 6 | 0.1667 |
| 7_1384376 | 1 | 0 | 7 | 0.2857 | 3 | 0 | 4 | 0 | 15 | 0.1333 | 7 | 0.1429 | 3 | 0.3333 | 4 | 0.5 | 2 | 0.5 | 16 | 0.3125 |
| 7_1384376 | 5 | 0.2 | 8 | 0.125 | 5 | 0.2 | 7 | 0.2857 | 25 | 0.2 | 8 | 0.625 | 9 | 0.4444 | 7 | 0.4286 | 7 | 0.1429 | 31 | 0.4194 |
| 7_1384376 | 4 | 1 | 8 | 0.5 | 4 | 0.75 | 7 | 0.8571 | 23 | 0.7391 | 8 | 0.5 | 8 | 0.75 | 5 | 0.8 | 8 | 0.625 | 29 | 0.6552 |
| 7_1384377 | 5 | 0.8 | 10 | 0.8 | 6 | 0.8333 | 7 | 1 | 28 | 0.8571 | 7 | 1 | 9 | 0.8889 | 7 | 1 | 7 | 1 | 30 | 0.9667 |
| 7_1384377 | 5 | 0.4 | 8 | 0.5 | 5 | 0 | 6 | 0.5 | 24 | 0.375 | 7 | 0.5714 | 9 | 0.3333 | 7 | 0.4286 | 6 | 0.5 | 29 | 0.4483 |
| 7_1384379 | 5 | 0 | 10 | 0.2 | 6 | 0.5 | 7 | 0 | 28 | 0.1786 | 8 | 0.125 | 9 | 0.2222 | 8 | 0 | 7 | 0.1429 | 32 | 0.125 |
| 7_1384380 | 5 | 0.4 | 10 | 0.6 | 6 | 0.3333 | 6 | 0.8333 | 27 | 0.5556 | 8 | 0.25 | 9 | 0.3333 | 7 | 0.5714 | 6 | 0.6667 | 30 | 0.4333 |
| 7_1384381 | 4 | 1 | 8 | 0.875 | 4 | 0.75 | 3 | 1 | 19 | 0.8947 | 7 | 1 | 6 | 1 | 5 | 1 | 4 | 1 | 22 | 1 |
| 7_1384383 | 2 | 0 | 6 | 0.1667 | 6 | 0 | 3 | 0 | 17 | 0.0588 | 5 | 0.2 | 8 | 0.375 | 4 | 0.5 | 5 | 0.2 | 22 | 0.3182 |
| 7_1384384 | 2 | 1 | 6 | 1 | 6 | 1 | 5 | 1 | 19 | 1 | 5 | 1 | 7 | 0.7143 | 5 | 1 | 5 | 1 | 22 | 0.9091 |
| 7_1384385 | 3 | 0 | 6 | 0.1667 | 4 | 0.5 | 5 | 0.2 | 18 | 0.2222 | 6 | 0.3333 | 7 | 0.4286 | 4 | 0 | 5 | 0.4 | 22 | 0.3182 |
| 7_1384386 | 4 | 0.5 | 8 | 0.375 | 5 | 0.4 | 4 | 0.25 | 21 | 0.381 | 5 | 0 | 7 | 0.7143 | 4 | 0.5 | 6 | 0.1667 | 22 | 0.3636 |
| 7_1384386 | 3 | 0.3333 | 6 | 0.3333 | 5 | 0.6 | 5 | 0.6 | 19 | 0.4737 | 5 | 0.8 | 7 | 0.7143 | 6 | 0.6667 | 7 | 0.8571 | 25 | 0.76 |
| 7_1384476 | 2 | 0 | 3 | 0.3333 | 3 | 0 | 1 | 0 | 9 | 0.1111 | 3 | 0.3333 | 4 | 0 | 1 | 0 | 1 | 0 | 9 | 0.1111 |
| 7_1384478 | 1 | 0 | 2 | 1 | 2 | 0.5 | 1 | 1 | 6 | 0.6667 | 3 | 1 | 4 | 1 | 1 | 1 | NA | NA | 8 | 1 |
| 7_1384478 | 2 | 0 | 3 | 0.6667 | 3 | 0 | 1 | 1 | 9 | 0.3333 | 3 | 0.6667 | 3 | 0.3333 | 1 | 0 | 1 | 0 | 8 | 0.375 |

| | | | | | | | | | | | | | | | | | | | | |
|---|---|---|---|---|---|---|---|---|---|---|---|---|---|---|---|---|---|---|---|---|
| 7_13847829 | 12 | 0.4167 | 7 | 0.2857 | 8 | 0.25 | 9 | 0.3333 | 36 | 0.3333 | 8 | 0.25 | 7 | 0.4286 | 11 | 0.4545 | 8 | 0.5 | 34 | 0.4118 |
| 7_13848410 | 10 | 0.8 | 7 | 0.8571 | 9 | 0.4444 | 10 | 0.3 | 36 | 0.5833 | 6 | 0.6667 | 5 | 1 | 12 | 0.6667 | 6 | 0.8333 | 29 | 0.7586 |
| 7_13848411 | 11 | 0.6364 | 9 | 0.3333 | 11 | 0.5455 | 12 | 0.4167 | 43 | 0.4884 | 8 | 0.375 | 9 | 0.4444 | 13 | 0.6923 | 8 | 0.5 | 38 | 0.5263 |
| 7_13848418 | 12 | 0.4167 | 8 | 0.375 | 9 | 0.1111 | 8 | 0.125 | 37 | 0.2703 | 7 | 0 | 8 | 0.5 | 14 | 0.5 | 8 | 0.375 | 37 | 0.3784 |
| 7_13848420 | 14 | 0.4286 | 8 | 0.625 | 9 | 0.6667 | 10 | 0.8 | 41 | 0.6098 | 8 | 0.625 | 10 | 0.4 | 13 | 0.7692 | 9 | 0.8889 | 40 | 0.675 |
| 7_13848422 | 9 | 0.2222 | 8 | 0.625 | 9 | 0.1111 | 7 | 0.4286 | 33 | 0.3333 | 5 | 0.2 | 7 | 0.7143 | 10 | 0.4 | 7 | 0.2857 | 29 | 0.4138 |
| 7_13848422 | 9 | 0.3333 | 8 | 0.375 | 8 | 0.125 | 10 | 0.8 | 35 | 0.4286 | 5 | 0.2 | 6 | 0.3333 | 10 | 0.4 | 6 | 0.6667 | 27 | 0.4074 |
| 7_13848423 | 11 | 0.4545 | 8 | 0.5 | 9 | 0.1111 | 11 | 0 | 39 | 0.2564 | 9 | 0.1111 | 10 | 0.2 | 12 | 0.3333 | 10 | 0.7 | 41 | 0.3415 |
| 7_13848442 | 14 | 1 | 9 | 1 | 11 | 1 | 11 | 0.9091 | 45 | 0.9778 | 8 | 0.875 | 10 | 1 | 11 | 1 | 9 | 1 | 38 | 0.9737 |
| 7_13848449 | 10 | 0.4 | 8 | 0 | 8 | 0.25 | 10 | 0.4 | 36 | 0.2778 | 9 | 0.2222 | 7 | 0.2857 | 13 | 0.3846 | 9 | 0.3333 | 38 | 0.3158 |
| 7_13848450 | 5 | 0.4 | 2 | 0 | 5 | 0.2 | 8 | 0.375 | 20 | 0.3 | 5 | 0 | 4 | 0.5 | 7 | 0.5714 | 5 | 0.2 | 21 | 0.3333 |
| 7_13848459 | 4 | 0.75 | 6 | 0.3333 | 7 | 0.7143 | 1 | 1 | 18 | 0.6111 | 4 | 0.25 | 2 | 1 | 5 | 0.2 | 5 | 0.2 | 16 | 0.3125 |
| 7_13849113 | 3 | 0.6667 | 4 | 1 | 2 | 0.5 | 2 | 1 | 11 | 0.8182 | 1 | 0 | 1 | 1 | 4 | 1 | 3 | 1 | 9 | 0.8889 |
| 7_13849113 | 5 | 1 | 4 | 0.5 | 2 | 1 | 3 | 1 | 14 | 0.8571 | 1 | 1 | 1 | 1 | 4 | 0.75 | 3 | 1 | 9 | 0.8889 |
| 7_13849118 | 3 | 0.3333 | 4 | 1 | 1 | 1 | 3 | 1 | 11 | 0.8182 | 1 | 0 | 1 | 1 | 3 | 1 | 2 | 1 | 7 | 0.8571 |
| 7_13849119 | 4 | 1 | 4 | 1 | 2 | 1 | 3 | 0.6667 | 13 | 0.9231 | 1 | 1 | 1 | 1 | 5 | 1 | 3 | 1 | 10 | 1 |
| 7_13849126 | 3 | 1 | 4 | 0.75 | 1 | 0 | 3 | 1 | 11 | 0.8182 | 1 | 0 | NA | NA | 2 | 1 | 2 | 1 | 5 | 0.8 |
| 7_13849156 | 4 | 1 | 4 | 1 | 2 | 1 | 2 | 1 | 12 | 1 | 1 | 1 | 1 | 1 | 5 | 1 | 2 | 1 | 9 | 1 |
| 7_13849168 | 3 | 0.6667 | 3 | 1 | 2 | 1 | 2 | 1 | 10 | 0.9 | 1 | 0 | 1 | 1 | 5 | 0.8 | 2 | 0.5 | 9 | 0.6667 |
| 7_13849196 | 2 | 1 | 2 | 1 | 2 | 1 | 2 | 0.5 | 8 | 0.875 | 1 | 1 | 1 | 1 | 4 | 0.75 | 2 | 1 | 8 | 0.875 |
| 7_13849209 | 3 | 0.3333 | 2 | 0.5 | 1 | 0 | 2 | 1 | 8 | 0.5 | NA | NA | 1 | 1 | 4 | 0.5 | 2 | 1 | 7 | 0.7143 |
| 7_13849210 | 4 | 0.25 | 1 | 1 | 2 | 0.5 | 1 | 0 | 8 | 0.375 | NA | NA | NA | NA | 4 | 0.5 | 2 | 0.5 | 6 | 0.5 |
| 7_13849211 | 3 | 0 | 2 | 0.5 | 2 | 0 | 2 | 0 | 9 | 0.1111 | NA | NA | 1 | 0 | 3 | 0.3333 | 2 | 0 | 6 | 0.1667 |
| 7_13849292 | 24 | 0.2083 | 20 | 0.2 | 21 | 0.0952 | 24 | 0.1667 | 89 | 0.1685 | 20 | 0.35 | 28 | 0.3214 | 24 | 0.0833 | 18 | 0.1111 | 90 | 0.2222 |
| 7_13849348 | 22 | 0.0909 | 18 | 0.0556 | 18 | 0.1667 | 23 | 0.087 | 81 | 0.0988 | 17 | 0.1176 | 25 | 0.04 | 22 | 0.0909 | 17 | 0 | 81 | 0.0617 |
| 7_13864540 | NA | NA | 1 | 1 | 1 | 1 | NA | NA | 2 | 1 | 1 | 1 | NA | NA | 2 | 1 | 2 | 1 | 5 | 1 |
| 7_13864541 | NA | NA | 1 | 1 | NA | NA | NA | NA | 1 | 1 | NA | NA | NA | NA | 1 | 1 | 1 | 1 | 2 | 1 |
| 7_13864545 | NA | NA | NA | NA | 1 | 1 | NA | NA | 1 | 1 | 1 | 1 | NA | NA | 2 | 0.5 | 2 | 0 | 5 | 0.4 |
| 7_13864556 | NA | NA | NA | NA | 1 | 1 | NA | NA | 1 | 1 | NA | NA | NA | NA | 2 | 0.5 | 1 | 0 | 3 | 0.3333 |
| 7_13864577 | NA | NA | NA | NA | 1 | 1 | NA | NA | 1 | 1 | 1 | 0 | NA | NA | 3 | 0.6667 | 1 | 0 | 5 | 0.4 |
| 7_13866238 | NA | NA | 3 | 0 | 1 | 1 | 1 | 1 | 5 | 0.4 | 1 | 0 | 1 | 0 | 2 | 0 | 5 | 0.2 | 9 | 0.1111 |
| 7_13866262 | NA | NA | 3 | 0.3333 | 2 | 0.5 | 2 | 0 | 7 | 0.2857 | 1 | 0 | 1 | 0 | 4 | 0.25 | 5 | 0.2 | 11 | 0.1818 |
| 7_13866286 | NA | NA | 4 | 0.25 | 3 | 0.6667 | 1 | 0 | 8 | 0.375 | 1 | 0 | NA | NA | 2 | 0 | 4 | 0.25 | 7 | 0.1429 |
| 7_13866298 | NA | NA | 4 | 0 | 2 | 0.5 | 2 | 0.5 | 8 | 0.25 | 1 | 0 | 1 | 0 | 3 | 0.3333 | 3 | 0.3333 | 8 | 0.25 |
| 7_13866313 | NA | NA | 4 | 0.25 | 2 | 1 | 2 | 0.5 | 8 | 0.5 | 1 | 0 | 1 | 1 | 3 | 0.3333 | 5 | 0.4 | 10 | 0.4 |
| 7_13866334 | NA | NA | 4 | 0.75 | 3 | 1 | 2 | 1 | 9 | 0.8889 | 1 | 0 | 1 | 0 | 4 | 1 | 5 | 0.8 | 11 | 0.7273 |
| 7_13866336 | NA | NA | 4 | 0.25 | 3 | 0.3333 | 2 | 0 | 9 | 0.2222 | 1 | 0 | 1 | 0 | 4 | 0.25 | 4 | 0.25 | 10 | 0.2 |
| 7_13868091 | 1 | 1 | 4 | 0 | 9 | 0 | 5 | 0 | 19 | 0.0526 | 3 | 0.3333 | 2 | 0 | 3 | 0.3333 | 2 | 0.5 | 10 | 0.3 |
| 7_13868355 | 8 | 0.125 | 6 | 0.1667 | 8 | 0.125 | 3 | 0.3333 | 25 | 0.16 | 5 | 0 | 6 | 0.3333 | 5 | 0 | 4 | 0 | 20 | 0.1 |
| 7_13868362 | 8 | 0.125 | 5 | 0.2 | 10 | 0.1 | 5 | 0 | 28 | 0.1071 | 7 | 0.1429 | 7 | 0.2857 | 5 | 0 | 4 | 0.25 | 23 | 0.1739 |
| 7_13868381 | 7 | 0.2857 | 6 | 0 | 6 | 0.1667 | 4 | 0.25 | 23 | 0.1739 | 6 | 0.1667 | 6 | 0.3333 | 5 | 0 | 1 | 0 | 18 | 0.1667 |
| 7_13868408 | 4 | 0 | 5 | 0.2 | 4 | 0 | 2 | 0 | 15 | 0.0667 | 4 | 0 | 4 | 0.25 | 3 | 0.3333 | 1 | 0 | 12 | 0.1667 |
| 7_13868412 | 5 | 0.2 | 6 | 0.3333 | 5 | 0 | 3 | 0.3333 | 19 | 0.2105 | 6 | 0.1667 | 5 | 0.2 | 3 | 0 | 1 | 0 | 15 | 0.1333 |
| 7_13883491 | 4 | 0 | 1 | 0 | 2 | 0.5 | NA | NA | 7 | 0.1429 | NA | NA | 1 | 1 | 2 | 0 | NA | NA | 3 | 0.3333 |
| 7_13883544 | 3 | 1 | 1 | 1 | 2 | 1 | NA | NA | 6 | 1 | NA | NA | 1 | 1 | 1 | 0 | NA | NA | 2 | 0.5 |
| 7_13883553 | 4 | 0 | 1 | 0 | 2 | 0.5 | NA | NA | 7 | 0.1429 | NA | NA | 1 | 0 | 2 | 1 | NA | NA | 3 | 0.6667 |
| 7_13883572 | 4 | 1 | 1 | 1 | 2 | 1 | NA | NA | 7 | 1 | NA | NA | 1 | 1 | 2 | 1 | NA | NA | 3 | 1 |
| 7_13883581 | 3 | 1 | 1 | 1 | 2 | 1 | NA | NA | 6 | 1 | NA | NA | NA | NA | 2 | 0.5 | NA | NA | 2 | 0.5 |
| 7_13883586 | 4 | 1 | 2 | 0.5 | 3 | 0.6667 | NA | NA | 9 | 0.7778 | NA | NA | 1 | 1 | 2 | 1 | NA | NA | 3 | 1 |
| 7_13883606 | 4 | 0.75 | 2 | 1 | 3 | 0.6667 | 1 | 1 | 10 | 0.8 | NA | NA | 1 | 1 | 2 | 1 | NA | NA | 3 | 1 |
| 7_13883621 | 2 | 1 | 1 | 1 | NA | NA | 1 | 0 | 4 | 0.75 | NA | NA | NA | NA | 1 | 1 | NA | NA | 1 | 1 |
| 7_13883624 | 2 | 1 | NA | NA | 2 | 1 | 2 | 1 | 6 | 1 | NA | NA | NA | NA | 1 | 1 | NA | NA | 1 | 1 |
| 7_13884670 | 4 | 0.25 | 2 | 0 | 3 | 0.3333 | 2 | 0.5 | 11 | 0.2727 | 1 | 0 | 5 | 0 | 3 | 0.3333 | NA | NA | 9 | 0.1111 |
| 7_13884697 | 4 | 0.25 | 2 | 0 | 4 | 0.5 | 2 | 0.5 | 12 | 0.3333 | 1 | 0 | 4 | 0.5 | 3 | 0.3333 | NA | NA | 8 | 0.375 |
| 7_13915776 | NA | NA | NA | NA | NA | NA | 1 | 1 | 1 | 1 | NA | NA | 1 | 1 | NA | NA | NA | NA | 1 | 1 |
| 7_13915845 | 10 | 0.1 | 6 | 0 | 8 | 0.25 | 7 | 0.1429 | 31 | 0.129 | 3 | 0 | 9 | 0.2222 | 7 | 0.2857 | 3 | 0.3333 | 22 | 0.2273 |
| 7_13915846 | 8 | 0.25 | 4 | 0 | 8 | 0.25 | 7 | 0 | 27 | 0.1481 | 3 | 0 | 9 | 0 | 9 | 0.2222 | 4 | 0 | 25 | 0.08 |
| 7_13915850 | 8 | 0.125 | 5 | 0.2 | 6 | 0.3333 | 8 | 0.125 | 27 | 0.1852 | 4 | 0 | 9 | 0.1111 | 7 | 0 | 4 | 0.25 | 24 | 0.0833 |
| 7_13915851 | 9 | 0.1111 | 3 | 0 | 7 | 0.1429 | 7 | 0.1429 | 26 | 0.1154 | 3 | 0 | 9 | 0.2222 | 8 | 0 | 4 | 0.25 | 24 | 0.125 |
| 7_13915853 | 10 | 0.1 | 7 | 0.1429 | 7 | 0.2857 | 10 | 0.2 | 34 | 0.1765 | 4 | 0.25 | 9 | 0 | 10 | 0 | 5 | 0.2 | 28 | 0.0714 |
| 7_13915869 | 10 | 0.2 | 7 | 0.1429 | 7 | 0.2857 | 10 | 0.2 | 34 | 0.2059 | 3 | 0.3333 | 9 | 0.2222 | 9 | 0.1111 | 7 | 0.1429 | 28 | 0.1786 |
| 7_13915901 | 11 | 0.4545 | 9 | 0.1111 | 8 | 0.625 | 13 | 0.2308 | 41 | 0.3415 | 4 | 0 | 9 | 0.1111 | 11 | 0.0909 | 5 | 0.4 | 29 | 0.1379 |
| 7_13915924 | 10 | 0.5 | 7 | 0.2857 | 6 | 0.8333 | 10 | 0.2 | 33 | 0.4242 | 8 | 0.25 | 7 | 0.2857 | 11 | 0.2727 | 6 | 0.5 | 32 | 0.3125 |
| 7_13915933 | 7 | 0.1429 | 9 | 0.1111 | 5 | 0.6 | 8 | 0.125 | 29 | 0.2069 | 3 | 0 | 6 | 0 | 10 | 0 | 5 | 0.4 | 24 | 0.0833 |
| 7_13915933 | NA | NA | 2 | 0.5 | 1 | 0 | 2 | 1 | 5 | 0.6 | 1 | 1 | NA | NA | 2 | 0.5 | 1 | 0 | 4 | 0.5 |

| | | | | | | | | | | | | | | | | | | | | |
|---|---|---|---|---|---|---|---|---|---|---|---|---|---|---|---|---|---|---|---|---|
| 7_1391594 | 9 | 0.1111 | 11 | 0.0909 | 7 | 0.1429 | 11 | 0.2727 | 38 | 0.1579 | 7 | 0.2857 | 9 | 0.1111 | 11 | 0.2727 | 6 | 0.3333 | 33 | 0.2424 |
| 7_1391609 | NA | NA | 2 | 0.5 | 2 | 1 | 1 | 1 | 5 | 0.8 | 2 | 1 | NA | NA | 2 | 0.5 | 1 | 1 | 5 | 0.8 |
| 7_1391610 | NA | NA | 1 | 1 | 2 | 0 | 2 | 0.5 | 5 | 0.4 | 2 | 0.5 | NA | NA | 2 | 0.5 | 1 | 0 | 5 | 0.4 |
| 7_1391610 | NA | NA | 2 | 0 | 1 | 1 | 2 | 0.5 | 5 | 0.4 | 2 | 1 | NA | NA | 2 | 0 | 1 | 1 | 5 | 0.6 |
| 7_1391613 | 1 | 0 | 1 | 0 | 5 | 0.4 | 4 | 0.75 | 11 | 0.4545 | 3 | 0.6667 | NA | NA | 3 | 0.3333 | 2 | 0.5 | 8 | 0.5 |
| 7_1391613 | 1 | 0 | 1 | 1 | 1 | 1 | 3 | 1 | 6 | 0.8333 | 1 | 1 | NA | NA | 2 | 0.5 | 1 | 1 | 4 | 0.75 |
| 7_1394551 | 1 | 1 | 2 | 1 | 1 | 1 | NA | NA | 4 | 1 | 1 | 1 | NA | NA | NA | NA | 1 | 1 | 2 | 1 |
| 7_1394552 | 1 | 1 | 2 | 1 | NA | NA | NA | NA | 3 | 1 | NA | NA | NA | NA | NA | NA | 2 | 0.5 | 2 | 0.5 |
| 7_1394552 | 1 | 1 | 2 | 1 | 2 | 1 | NA | NA | 5 | 1 | NA | NA | NA | NA | 1 | 1 | 2 | 1 | 3 | 1 |
| 7_1394554 | 1 | 1 | 2 | 1 | 2 | 0.5 | NA | NA | 5 | 0.8 | 2 | 1 | NA | NA | 2 | 1 | 3 | 1 | 7 | 1 |
| 7_1394556 | NA | NA | 2 | 1 | 2 | 1 | 1 | 0 | 5 | 0.8 | NA | NA | NA | NA | 2 | 1 | NA | NA | 2 | 1 |
| 7_1394559 | NA | NA | 2 | 0.5 | 2 | 0 | 1 | 0 | 5 | 0.2 | 3 | 0.3333 | NA | NA | 1 | 0 | 3 | 0.3333 | 7 | 0.2857 |
| 7_1394559 | 1 | 1 | 1 | 1 | 2 | 1 | 1 | 1 | 5 | 1 | 1 | 1 | NA | NA | 2 | 1 | 2 | 1 | 5 | 1 |
| 7_1394560 | 1 | 1 | 2 | 1 | 2 | 1 | 2 | 1 | 7 | 1 | 3 | 1 | NA | NA | 2 | 1 | 4 | 1 | 9 | 1 |
| 7_1394560 | 1 | 0 | 1 | 0 | 2 | 0.5 | 1 | 1 | 5 | 0.4 | 1 | 1 | NA | NA | 2 | 0 | 2 | 0 | 5 | 0.2 |
| 7_1394560 | 2 | 1 | 2 | 1 | 2 | 0.5 | 2 | 1 | 8 | 0.875 | 2 | 1 | NA | NA | 1 | 1 | 4 | 0.5 | 7 | 0.7143 |
| 7_1394560 | 1 | 0 | 2 | 0.5 | 2 | 1 | 2 | 0.5 | 7 | 0.5714 | 3 | 0.3333 | NA | NA | 2 | 0 | 4 | 0.25 | 9 | 0.2222 |
| 7_1394561 | 1 | 1 | 2 | 1 | 2 | 1 | 2 | 1 | 7 | 1 | 2 | 1 | NA | NA | 2 | 1 | 4 | 1 | 8 | 1 |
| 7_1394562 | 1 | 1 | 3 | 0.6667 | 1 | 0 | 2 | 0 | 7 | 0.4286 | 2 | 0.5 | NA | NA | 1 | 0 | 3 | 0.3333 | 6 | 0.3333 |
| 7_1394562 | 1 | 1 | 1 | 1 | 2 | 0.5 | 2 | 0.5 | 6 | 0.6667 | 1 | 1 | NA | NA | 1 | 1 | 4 | 0.5 | 6 | 0.6667 |
| 7_1394562 | 1 | 1 | 3 | 1 | 2 | 1 | 2 | 1 | 8 | 1 | 4 | 0.5 | NA | NA | 2 | 1 | 4 | 1 | 10 | 0.8 |
| 7_1394563 | NA | NA | 4 | 0 | 2 | 0.5 | 2 | 0 | 8 | 0.125 | 4 | 0 | NA | NA | 2 | 0.5 | 4 | 0.5 | 10 | 0.3 |
| 7_1394564 | 2 | 0 | 4 | 0.5 | 1 | 1 | 2 | 0 | 9 | 0.3333 | 4 | 0.25 | NA | NA | 2 | 0 | 4 | 0.75 | 10 | 0.4 |
| 7_1394567 | 1 | 1 | 4 | 0.5 | 3 | 0.6667 | 2 | 0.5 | 10 | 0.6 | 4 | 0 | NA | NA | 2 | 0 | 4 | 1 | 10 | 0.4 |
| 7_1394570 | 1 | 0 | 2 | 0 | 3 | 0.3333 | 3 | 0.3333 | 9 | 0.2222 | 5 | 0.2 | NA | NA | 2 | 0 | 4 | 0 | 11 | 0.0909 |
| 7_1394576 | 1 | 0 | 3 | 0.3333 | 4 | 0.25 | 2 | 0 | 10 | 0.2 | 1 | 1 | 1 | 0 | NA | NA | 3 | 0 | 5 | 0.2 |
| 7_1394576 | 1 | 1 | 4 | 0.5 | 4 | 0.5 | 2 | 0.5 | 11 | 0.5455 | 4 | 1 | 1 | 0 | 2 | 1 | 4 | 1 | 11 | 0.9091 |
| 7_1394904 | 1 | 0 | 1 | 1 | NA | NA | NA | NA | 2 | 0.5 | NA | NA | 1 | 1 | 1 | 1 | 1 | 1 | 3 | 1 |
| 7_1395014 | 3 | 0.3333 | 3 | 0 | 1 | 0 | NA | NA | 7 | 0.1429 | NA | NA | 2 | 0 | 1 | 0 | 1 | 1 | 4 | 0.25 |
| 7_1395038 | 3 | 0 | 2 | 0.5 | 1 | 0 | NA | NA | 6 | 0.1667 | 1 | 0 | 1 | 0 | 1 | 0 | 1 | 1 | 4 | 0.25 |
| 7_1395199 | 8 | 0.25 | 8 | 0.375 | 4 | 0.5 | 1 | 0 | 21 | 0.3333 | 1 | 0 | 3 | 0.3333 | 1 | 0 | 1 | 1 | 6 | 0.3333 |
| 7_1395199 | 7 | 0 | 7 | 0.1429 | 5 | 0 | NA | NA | 19 | 0.0526 | 1 | 0 | 3 | 0.3333 | 2 | 0 | NA | NA | 6 | 0.1667 |
| 7_1395200 | 6 | 0 | 7 | 0 | 4 | 0.25 | 1 | 0 | 18 | 0.0556 | 2 | 0.5 | 2 | 0 | 2 | 0 | NA | NA | 6 | 0.1667 |
| 7_1395206 | 6 | 0.5 | 10 | 0.3 | 5 | 0.2 | 1 | 1 | 22 | 0.3636 | 4 | 0.25 | 5 | 0.2 | 1 | 0 | 2 | 0 | 12 | 0.1667 |
| 7_1395208 | 9 | 0.3333 | 10 | 0.3 | 4 | 0.25 | 1 | 1 | 24 | 0.3333 | 5 | 0.2 | 5 | 0.2 | 1 | 0 | 2 | 0.5 | 13 | 0.2308 |
| 7_1395208 | 9 | 0.3333 | 10 | 0.1 | 5 | 0.4 | 1 | 1 | 25 | 0.28 | 5 | 0.2 | 4 | 0 | 1 | 0 | 2 | 0 | 12 | 0.0833 |
| 7_1395209 | 8 | 0.625 | 9 | 0.3333 | 3 | 0.6667 | 1 | 1 | 21 | 0.5238 | 4 | 0.5 | 5 | 0.4 | NA | NA | 2 | 1 | 11 | 0.5455 |
| 7_1395209 | 8 | 0.5 | 8 | 0.125 | 3 | 0.6667 | NA | NA | 19 | 0.3684 | 5 | 0.6 | 4 | 0.25 | 1 | 0 | 2 | 1 | 12 | 0.5 |
| 7_1395210 | 7 | 0.1429 | 10 | 0.1 | 5 | 0.6 | 1 | 0 | 23 | 0.2174 | 5 | 0.2 | 5 | 0 | 1 | 0 | 1 | 1 | 12 | 0.1667 |
| 7_1395211 | 8 | 1 | 5 | 0.8 | 5 | 1 | 1 | 1 | 19 | 0.9474 | 3 | 1 | 2 | 1 | 1 | 0 | 2 | 1 | 8 | 0.875 |
| 7_1395212 | 7 | 0.7143 | 6 | 0.5 | 4 | 0.75 | 1 | 1 | 18 | 0.6667 | 4 | 0.25 | 6 | 0.3333 | 2 | 1 | 2 | 1 | 14 | 0.5 |
| 7_1395212 | 9 | 0.5556 | 8 | 0.375 | 5 | 0.6 | 1 | 0 | 23 | 0.4783 | 5 | 0.2 | 6 | 0.5 | 2 | 0.5 | 3 | 1 | 16 | 0.5 |
| 7_1395754 | 1 | 1 | 2 | 1 | 1 | 1 | NA | NA | 4 | 1 | NA | NA | NA | NA | NA | NA | 1 | 1 | 1 | 1 |
| 7_1395848 | 2 | 0 | 2 | 0 | 5 | 0.4 | 2 | 0.5 | 11 | 0.2727 | 3 | 0 | 5 | 0 | 2 | 0 | 3 | 0.3333 | 13 | 0.0769 |
| 7_1395851 | NA | NA | NA | NA | 1 | 1 | 1 | 0 | 2 | 0.5 | 1 | 0 | 1 | 1 | NA | NA | NA | NA | 2 | 0.5 |
| 7_1396171 | 9 | 0 | 2 | 0 | 4 | 0 | 5 | 0.2 | 20 | 0.05 | 2 | 0 | 1 | 0 | 3 | 0.3333 | 1 | 0 | 7 | 0.1429 |
| 7_1396202 | 9 | 0.7778 | 2 | 1 | 3 | 1 | 6 | 1 | 20 | 0.9 | 2 | 1 | 3 | 0.3333 | 4 | 1 | 1 | 0 | 10 | 0.7 |
| 7_1396203 | 9 | 0.5556 | 3 | 0.3333 | 4 | 1 | 5 | 0.8 | 21 | 0.6667 | 3 | 0.6667 | 2 | 1 | 4 | 1 | 2 | 1 | 11 | 0.9091 |
| 7_1396203 | 7 | 0.1429 | 3 | 0.6667 | 7 | 0.4286 | 5 | 0.6 | 22 | 0.4091 | 3 | 0.6667 | 3 | 0 | 4 | 0.25 | 3 | 0 | 13 | 0.2308 |
| 7_1396204 | 6 | 0.3333 | 3 | 0 | 4 | 0.5 | 5 | 0 | 18 | 0.2222 | 3 | 0 | 2 | 0 | 3 | 0.3333 | 3 | 0.6667 | 11 | 0.2727 |
| 7_1396204 | 8 | 0.125 | 3 | 0.6667 | 4 | 0.5 | 3 | 0.3333 | 18 | 0.3333 | 2 | 0.5 | 4 | 0.75 | 3 | 0.3333 | 2 | 1 | 11 | 0.6364 |
| 7_1396499 | 6 | 0.1667 | 5 | 0.2 | 8 | 0.125 | 9 | 0.1111 | 28 | 0.1429 | 10 | 0.1 | 7 | 0 | 7 | 0.2857 | 10 | 0.2 | 34 | 0.1471 |
| 7_1396502 | 41 | 0.2927 | 40 | 0.375 | 50 | 0.44 | 53 | 0.2453 | 184 | 0.337 | 49 | 0.4694 | 50 | 0.42 | 57 | 0.3158 | 41 | 0.439 | 197 | 0.4061 |
| 7_1396503 | 40 | 0.075 | 38 | 0.1842 | 50 | 0.1 | 49 | 0.1224 | 177 | 0.1186 | 50 | 0.12 | 54 | 0.1481 | 54 | 0.1296 | 48 | 0.1042 | 206 | 0.1262 |
| 7_1396794 | 11 | 0.7273 | 13 | 0.5385 | 8 | 0.75 | 8 | 0.625 | 40 | 0.65 | 8 | 0.625 | 12 | 0.6667 | 5 | 0.8 | 9 | 0.5556 | 34 | 0.6471 |
| 7_1396795 | 8 | 0.25 | 5 | 0.2 | 2 | 0 | 7 | 0 | 22 | 0.1364 | 4 | 0 | 5 | 0.2 | 2 | 0.5 | 6 | 0 | 17 | 0.1176 |
| 7_1396795 | 10 | 0 | 13 | 0.0769 | 7 | 0.1429 | 9 | 0 | 39 | 0.0513 | 5 | 0 | 12 | 0.0833 | 5 | 0 | 8 | 0.375 | 30 | 0.1333 |
| 7_1396799 | 3 | 0.6667 | 2 | 1 | NA | NA | 1 | 1 | 6 | 0.8333 | 2 | 0.5 | 1 | 0 | NA | NA | 1 | 1 | 4 | 0.5 |
| 7_1396799 | 3 | 0 | 2 | 0.5 | NA | NA | 1 | 0 | 6 | 0.1667 | 2 | 0.5 | 1 | 0 | NA | NA | 1 | 1 | 4 | 0.5 |
| 7_1396862 | 4 | 0 | 4 | 0 | 13 | 0.0769 | 12 | 0.0833 | 33 | 0.0606 | 12 | 0.0833 | 8 | 0.125 | 7 | 0.2857 | 8 | 0.125 | 35 | 0.1429 |
| 7_1397064 | NA | NA | NA | NA | NA | NA | 1 | 1 | 1 | 1 | NA | NA | NA | NA | NA | NA | 2 | 0.5 | 2 | 0.5 |
| 7_1397066 | 1 | 0 | NA | NA | NA | NA | 1 | 1 | 2 | 0.5 | NA | NA | NA | NA | NA | NA | 2 | 0.5 | 2 | 0.5 |
| 7_1397302 | 2 | 0 | 1 | 0 | NA | NA | 5 | 0.2 | 8 | 0.125 | 7 | 0 | 4 | 0.25 | 4 | 0 | 5 | 0 | 20 | 0.05 |
| 7_1397302 | 3 | 1 | 1 | 0 | NA | NA | 4 | 1 | 8 | 0.875 | 6 | 0.6667 | 4 | 1 | 3 | 0.6667 | 3 | 0.6667 | 16 | 0.75 |
| 7_1397308 | 11 | 0.2727 | 6 | 0.6667 | 7 | 0.4286 | 6 | 0.3333 | 30 | 0.4 | 6 | 0.5 | 9 | 0.2222 | 5 | 0.4 | 14 | 0.4286 | 34 | 0.3824 |

| | | | | | | | | | | | | | | | | | | | | |
|---|---|---|---|---|---|---|---|---|---|---|---|---|---|---|---|---|---|---|---|---|
| 7_13973082 | 9 | 0.8889 | 6 | 1 | 8 | 0.75 | 5 | 0.6 | 28 | 0.8214 | 4 | 0.75 | 8 | 0.875 | 4 | 1 | 9 | 0.8889 | 25 | 0.88 |
| 7_13973093 | 10 | 0.8 | 11 | 1 | 9 | 1 | 11 | 0.9091 | 41 | 0.9268 | 3 | 0.6667 | 10 | 0.9 | 5 | 1 | 13 | 0.7692 | 31 | 0.8387 |
| 7_13973103 | 8 | 0.25 | 7 | 0.7143 | 10 | 0.3 | 10 | 0.5 | 35 | 0.4286 | 2 | 1 | 11 | 0.5455 | 5 | 0.6 | 17 | 0.6471 | 35 | 0.6286 |
| 7_13973123 | 7 | 0.5714 | 7 | 0.4286 | 10 | 0.6 | 9 | 0.4444 | 33 | 0.5152 | 1 | 1 | 12 | 0.5833 | 5 | 0.8 | 15 | 0.8 | 33 | 0.7273 |
| 7_13973124 | 6 | 0 | 5 | 0 | 8 | 0.25 | 6 | 0 | 25 | 0.08 | 1 | 0 | 8 | 0.375 | 4 | 0.25 | 12 | 0 | 25 | 0.16 |
| 7_13973138 | 7 | 0.4286 | 6 | 0.3333 | 10 | 0.5 | 11 | 0.3636 | 34 | 0.4118 | 2 | 1 | 11 | 0.5455 | 5 | 0.6 | 11 | 0.6364 | 29 | 0.6207 |
| 7_13973141 | 7 | 0.2857 | 4 | 0.25 | 8 | 0 | 11 | 0.0909 | 30 | 0.1333 | 1 | 0 | 7 | 0.1429 | 4 | 0.5 | 10 | 0.2 | 22 | 0.2273 |
| 7_13973152 | 5 | 0 | 5 | 0.2 | 7 | 0.2857 | 11 | 0.0909 | 28 | 0.1429 | 1 | 0 | 10 | 0.1 | 4 | 0.25 | 12 | 0.25 | 27 | 0.1852 |
| 7_13973158 | 5 | 0.2 | 4 | 0.25 | 8 | 0.125 | 8 | 0.25 | 25 | 0.2 | 1 | 0 | 10 | 0 | 4 | 0 | 15 | 0.2 | 30 | 0.1 |
| 7_13973159 | 3 | 0.6667 | 4 | 1 | 7 | 0.7143 | 9 | 0.6667 | 23 | 0.7391 | 2 | 1 | 11 | 0.7273 | 5 | 0.8 | 12 | 0.9167 | 30 | 0.8333 |
| 7_13973218 | 3 | 0 | 2 | 0 | 6 | 0 | 5 | 0.2 | 16 | 0.0625 | 2 | 0 | 7 | 0 | 1 | 0 | 10 | 0.1 | 20 | 0.05 |
| 7_13973249 | 3 | 0.6667 | 4 | 0 | 8 | 0 | 9 | 0.1111 | 24 | 0.125 | NA | NA | 8 | 0.375 | 2 | 0 | 12 | 0.1667 | 22 | 0.2273 |
| 7_13973250 | 4 | 0.25 | 4 | 0 | 6 | 0.1667 | 7 | 0 | 21 | 0.0952 | 1 | 0 | 9 | 0.2222 | 1 | 0 | 11 | 0.2727 | 22 | 0.2273 |
| 7_13973267 | 3 | 0 | 1 | 0 | 7 | 0.1429 | 8 | 0 | 19 | 0.0526 | 1 | 0 | 8 | 0.125 | 2 | 0 | 11 | 0.1818 | 22 | 0.1364 |
| 7_13973270 | 3 | 1 | 2 | 1 | 6 | 0.8333 | 8 | 0.5 | 19 | 0.7368 | 1 | 0 | 8 | 0.75 | 2 | 0 | 12 | 0.4167 | 23 | 0.4783 |
| 7_13973273 | 3 | 0.3333 | 1 | 0 | 6 | 0.1667 | 8 | 0.25 | 18 | 0.2222 | 1 | 0 | 8 | 0.125 | 2 | 0.5 | 11 | 0.2727 | 22 | 0.2273 |
| 7_13973280 | 2 | 0.5 | 2 | 0.5 | 4 | 0.5 | 4 | 0.25 | 12 | 0.4167 | 1 | 0 | 8 | 0.5 | 1 | 0 | 11 | 0.2727 | 21 | 0.3333 |
| 7_13973280 | 3 | 0.3333 | 2 | 0 | 5 | 0.2 | 8 | 0.125 | 18 | 0.1667 | 1 | 1 | 9 | 0.3333 | 2 | 0.5 | 12 | 0.4167 | 24 | 0.4167 |
| 7_13973285 | 1 | 0 | 2 | 0.5 | 3 | 0.6667 | 4 | 0.25 | 10 | 0.4 | NA | NA | 5 | 0.2 | 1 | 0 | 7 | 0.7143 | 13 | 0.4615 |
| 7_13973286 | 3 | 0.3333 | 2 | 0.5 | 4 | 0 | 7 | 0 | 16 | 0.125 | 1 | 0 | 9 | 0.1111 | 2 | 0 | 12 | 0.1667 | 24 | 0.125 |
| 7_13973290 | 3 | 0 | 2 | 0.5 | 5 | 0 | 7 | 0.1429 | 17 | 0.1176 | 1 | 0 | 8 | 0.125 | 2 | 0 | 12 | 0.1667 | 23 | 0.1304 |
| 7_13973314 | 2 | 1 | 1 | 1 | 4 | 0.5 | 5 | 1 | 12 | 0.8333 | 1 | 1 | 5 | 0.4 | 1 | 1 | 7 | 0.8571 | 14 | 0.7143 |
| 7_13973321 | 3 | 0.6667 | 1 | 1 | 2 | 0.5 | 6 | 0.3333 | 12 | 0.5 | 1 | 0 | 6 | 0.1667 | 1 | 1 | 9 | 0.4444 | 17 | 0.3529 |
| 7_13973356 | 1 | 0 | 1 | 0 | 3 | 0 | 4 | 0.25 | 9 | 0.1111 | 1 | 0 | 4 | 0.25 | 1 | 0 | 5 | 0.4 | 11 | 0.2727 |
| 7_13973356 | 2 | 0 | 1 | 0 | 1 | 0 | 6 | 0.1667 | 10 | 0.1 | NA | NA | 6 | 0.3333 | 2 | 0 | 7 | 0.1429 | 15 | 0.2 |
| 7_14038979 | 3 | 1 | 2 | 1 | 1 | 1 | 1 | 1 | 7 | 1 | 2 | 1 | 2 | 1 | 6 | 1 | NA | NA | 10 | 1 |
| 7_14039102 | 1 | 1 | NA | NA | NA | NA | NA | NA | 1 | 1 | 1 | 0 | 2 | 0.5 | 3 | 0.6667 | NA | NA | 6 | 0.5 |
| 7_14039114 | 2 | 0 | 2 | 0 | NA | NA | 1 | 1 | 5 | 0.2 | 1 | 1 | 2 | 0.5 | 2 | 1 | NA | NA | 5 | 0.8 |
| 7_14039117 | 2 | 1 | 2 | 1 | NA | NA | 1 | 1 | 5 | 1 | 1 | 1 | 1 | 1 | 3 | 1 | NA | NA | 5 | 1 |
| 7_14039120 | 2 | 1 | 1 | 1 | NA | NA | 1 | 1 | 4 | 1 | NA | NA | 2 | 1 | 1 | 1 | NA | NA | 3 | 1 |
| 7_14039147 | 1 | 0 | 2 | 1 | NA | NA | 1 | 1 | 4 | 0.75 | 1 | 1 | 1 | 1 | 3 | 1 | NA | NA | 5 | 1 |
| 7_14039154 | 1 | 0 | 2 | 1 | NA | NA | 1 | 0 | 4 | 0.5 | 1 | 0 | 1 | 0 | 3 | 0.6667 | NA | NA | 5 | 0.4 |
| 7_14039199 | NA | NA | 2 | 1 | NA | NA | 1 | 1 | 3 | 1 | 1 | 1 | NA | NA | 3 | 1 | NA | NA | 4 | 1 |
| 7_14039206 | 1 | 1 | 1 | 0 | NA | NA | 1 | 1 | 3 | 0.6667 | NA | NA | 1 | 1 | 3 | 0.6667 | NA | NA | 4 | 0.75 |
| 7_14039231 | 1 | 1 | 2 | 0 | NA | NA | 1 | 1 | 4 | 0.5 | 1 | 0 | 1 | 0 | 2 | 0.5 | NA | NA | 4 | 0.25 |
| 7_14039256 | 1 | 0 | 2 | 0.5 | NA | NA | 1 | 0 | 4 | 0.25 | 1 | 1 | 1 | 1 | 1 | 1 | NA | NA | 3 | 1 |
| 7_14039276 | 1 | 0 | 2 | 0.5 | NA | NA | 1 | 0 | 4 | 0.25 | 1 | 1 | 1 | 1 | 1 | 0 | NA | NA | 3 | 0.6667 |
| 7_14039280 | NA | NA | 1 | 1 | NA | NA | 1 | 1 | 2 | 1 | 1 | 1 | NA | NA | 1 | 1 | NA | NA | 2 | 1 |
| 7_14039280 | 1 | 1 | 2 | 1 | NA | NA | 1 | 1 | 4 | 1 | 1 | 1 | NA | NA | 1 | 1 | NA | NA | 2 | 1 |
| 7_14039290 | 1 | 0 | 2 | 0.5 | NA | NA | 1 | 0 | 4 | 0.25 | 1 | 0 | 1 | 0 | 1 | 1 | NA | NA | 3 | 0.3333 |
| 7_14039295 | 1 | 0 | 2 | 0.5 | NA | NA | 1 | 0 | 4 | 0.25 | 1 | 1 | 1 | 0 | 1 | 1 | NA | NA | 3 | 0.6667 |
| 7_14039299 | 1 | 1 | 2 | 0 | NA | NA | 1 | 1 | 4 | 0.5 | 1 | 1 | 1 | 1 | 1 | 1 | NA | NA | 3 | 1 |
| 7_14039304 | NA | NA | 1 | 1 | NA | NA | 1 | 0 | 2 | 0.5 | 1 | 1 | 1 | 0 | 1 | 0 | NA | NA | 3 | 0.3333 |
| 7_14043345 | 7 | 0.7143 | 5 | 0.2 | 3 | 0 | 5 | 0.6 | 20 | 0.45 | 4 | 0.5 | 4 | 0 | 5 | 0.2 | 1 | 0 | 14 | 0.2143 |
| 7_14043353 | 7 | 0.1429 | 5 | 0.2 | 1 | 0 | 5 | 0.2 | 18 | 0.1667 | 4 | 0.25 | 4 | 0 | 4 | 0 | 1 | 0 | 13 | 0.0769 |
| 7_14043366 | 6 | 0 | 5 | 0 | 3 | 0 | 5 | 0.2 | 19 | 0.0526 | 5 | 0 | 4 | 0.25 | 3 | 0.3333 | NA | NA | 12 | 0.1667 |
| 7_14043367 | 5 | 0.4 | 4 | 0.25 | 3 | 0.3333 | 5 | 0.6 | 17 | 0.4118 | 3 | 0.6667 | 4 | 0 | 4 | 0.25 | 1 | 0 | 12 | 0.25 |
| 7_14043378 | 5 | 1 | 4 | 1 | 1 | 1 | 5 | 0.8 | 15 | 0.9333 | 5 | 1 | 1 | 1 | 3 | 1 | 1 | 1 | 10 | 1 |
| 7_14043380 | 3 | 1 | 3 | 1 | 2 | 1 | 4 | 1 | 12 | 1 | 4 | 1 | 3 | 1 | 4 | 1 | 1 | 1 | 12 | 1 |
| 7_14043386 | 6 | 0.8333 | 4 | 1 | 3 | 1 | 5 | 0.8 | 18 | 0.8889 | 5 | 0.6 | 3 | 1 | 4 | 0.75 | 1 | 1 | 13 | 0.7692 |
| 7_14043394 | 6 | 0.3333 | 4 | 0.75 | 3 | 0 | 4 | 0.75 | 17 | 0.4706 | 5 | 0.2 | 2 | 0.5 | 4 | 0.5 | NA | NA | 11 | 0.3636 |
| 7_14043413 | 4 | 0.75 | 4 | 1 | 1 | 1 | 5 | 1 | 14 | 0.9286 | 4 | 1 | 3 | 1 | 3 | 1 | 1 | 1 | 11 | 1 |
| 7_14043430 | 5 | 1 | 4 | 1 | 3 | 1 | 5 | 1 | 17 | 1 | 5 | 1 | 3 | 1 | 3 | 1 | 1 | 1 | 12 | 1 |
| 7_14043438 | 5 | 0.2 | 3 | 0 | 3 | 0 | 5 | 0.4 | 16 | 0.1875 | 4 | 0 | 4 | 0 | 3 | 0 | 1 | 1 | 12 | 0.0833 |
| 7_14043453 | 5 | 0 | 4 | 0 | 2 | 0 | 6 | 0.1667 | 17 | 0.0588 | 4 | 0 | 4 | 0.25 | 2 | 0.5 | 1 | 0 | 11 | 0.1818 |
| 7_14043650 | NA | NA | NA | NA | NA | NA | 1 | 1 | 1 | 1 | 1 | 1 | NA | NA | NA | NA | 1 | 1 | 2 | 1 |
| 7_14043715 | NA | NA | NA | NA | 1 | 0 | 1 | 1 | 2 | 0.5 | 1 | 1 | NA | NA | NA | NA | 1 | 1 | 2 | 1 |
| 7_14043796 | 4 | 0 | 3 | 0 | 3 | 0 | 4 | 0.25 | 14 | 0.0714 | 3 | 0 | 3 | 0 | 4 | 0.25 | 4 | 0 | 14 | 0.0714 |
| 7_14043800 | 4 | 0.5 | 3 | 0.3333 | 3 | 1 | 4 | 0.75 | 14 | 0.6429 | 3 | 0.3333 | 3 | 0.3333 | 4 | 0.5 | 5 | 0.6 | 15 | 0.4667 |
| 7_14043820 | 4 | 1 | 4 | 1 | 2 | 1 | 3 | 0.3333 | 13 | 0.8462 | 3 | 1 | 3 | 1 | 2 | 1 | 5 | 1 | 13 | 1 |
| 7_14043825 | 4 | 0 | 3 | 0.3333 | 2 | 0 | 4 | 0 | 13 | 0.0769 | 3 | 0 | 3 | 0 | 3 | 0.3333 | 5 | 0 | 14 | 0.0714 |
| 7_14043835 | 4 | 0 | 1 | 0 | 3 | 0 | 4 | 0.25 | 12 | 0.0833 | 3 | 0.3333 | 1 | 0 | 3 | 0 | 5 | 0 | 12 | 0.0833 |
| 7_14044237 | 4 | 0.5 | 5 | 0 | 5 | 0.6 | 6 | 0.1667 | 20 | 0.3 | 3 | 0.3333 | 4 | 0.5 | 5 | 0 | 4 | 0.25 | 16 | 0.25 |
| 7_14044250 | 6 | 1 | 5 | 0.8 | 5 | 0.8 | 6 | 0.8333 | 22 | 0.8636 | 2 | 1 | 4 | 1 | 4 | 0.75 | 5 | 1 | 15 | 0.9333 |
| 7_14044262 | 7 | 0.8571 | 4 | 0.75 | 5 | 0.8 | 4 | 0.75 | 20 | 0.8 | 2 | 1 | 2 | 0.5 | 3 | 0.6667 | 2 | 0.5 | 9 | 0.6667 |

| | | | | | | | | | | | | | | | | | | | | |
|---|---|---|---|---|---|---|---|---|---|---|---|---|---|---|---|---|---|---|---|---|
| 7_1404430( | 10 | 0.2 | 9 | 0.2222 | 5 | 0.2 | 5 | 0.4 | 29 | 0.2414 | 7 | 0.1429 | 3 | 0 | 6 | 0 | 4 | 0.25 | 20 | 0.1 |
| 7_1404718( | NA | NA | NA | NA | NA | NA | 1 | 1 | 1 | 1 | 3 | 0.3333 | NA | NA | 3 | 0.3333 | NA | NA | 6 | 0.3333 |
| 7_1404718( | NA | NA | NA | NA | NA | NA | 2 | 0.5 | 2 | 0.5 | 4 | 0 | NA | NA | 3 | 0.6667 | NA | NA | 7 | 0.2857 |
| 7_1404718! | NA | NA | NA | NA | NA | NA | 1 | 1 | 1 | 1 | 3 | 0.3333 | NA | NA | 2 | 0.5 | NA | NA | 5 | 0.4 |
| 7_1404720 | NA | NA | NA | NA | NA | NA | 2 | 1 | 2 | 1 | 7 | 0.4286 | NA | NA | 6 | 0.6667 | NA | NA | 13 | 0.5385 |
| 7_1404783: | 12 | 0.4167 | 13 | 0.6154 | 13 | 0.3077 | 14 | 0.5 | 52 | 0.4615 | 10 | 0.3 | 10 | 0.3 | 10 | 0.2 | 17 | 0.5882 | 47 | 0.383 |
| 7_1404783 | 14 | 0.7857 | 12 | 0.4167 | 8 | 0.625 | 12 | 0.5833 | 46 | 0.6087 | 9 | 0.7778 | 10 | 0.8 | 9 | 0.2222 | 12 | 0.8333 | 40 | 0.675 |
| 7_1406479 | 2 | 0 | 3 | 0.6667 | 4 | 0.25 | 4 | 0 | 13 | 0.2308 | 3 | 0 | 1 | 1 | 1 | 0 | 3 | 0 | 8 | 0.125 |
| 7_1406480 | 1 | 0 | 2 | 1 | 4 | 0.5 | 3 | 1 | 10 | 0.7 | 2 | 1 | 1 | 1 | 1 | 1 | 2 | 1 | 6 | 1 |
| 7_1406480( | 2 | 0 | 4 | 0.75 | 3 | 0.3333 | 2 | 1 | 11 | 0.5455 | 3 | 0.6667 | NA | NA | 1 | 1 | 2 | 0.5 | 6 | 0.6667 |
| 7_1406480! | 2 | 0 | 2 | 0 | 4 | 0.75 | 4 | 0.25 | 12 | 0.3333 | 4 | 0.5 | 1 | 1 | NA | NA | 3 | 0.6667 | 8 | 0.625 |
| 7_1406481 | 2 | 0.5 | 3 | 0 | 4 | 0.75 | 3 | 0.6667 | 12 | 0.5 | 4 | 0.75 | 1 | 1 | 1 | 1 | 3 | 0.3333 | 9 | 0.6667 |
| 7_1406482: | 1 | 0 | 4 | 0.75 | 4 | 0.25 | 2 | 0.5 | 11 | 0.4545 | 3 | 0 | 1 | 1 | 1 | 1 | 3 | 0.3333 | 8 | 0.375 |
| 7_1406482 | 2 | 0 | 4 | 0.75 | 3 | 0 | 4 | 0.5 | 13 | 0.3846 | 3 | 0.6667 | 1 | 1 | 1 | 0 | 2 | 0.5 | 7 | 0.5714 |
| 7_1406482! | 2 | 0 | 3 | 1 | 4 | 0.75 | 4 | 0.25 | 13 | 0.5385 | 3 | 0.6667 | NA | NA | 1 | 1 | 4 | 0.75 | 8 | 0.75 |
| 7_1406483 | 2 | 1 | 4 | 1 | 2 | 1 | 3 | 1 | 11 | 1 | 3 | 1 | 1 | 1 | 1 | 1 | 3 | 1 | 8 | 1 |
| 7_1406483 | 2 | 0.5 | 4 | 0.25 | 3 | 0.6667 | 2 | 0 | 11 | 0.3636 | 3 | 0 | NA | NA | NA | NA | 4 | 0.25 | 7 | 0.1429 |
| 7_1406483 | 2 | 0 | 4 | 0.25 | NA | NA | 2 | 1 | 8 | 0.375 | 1 | 0 | NA | NA | 1 | 1 | 3 | 0.3333 | 5 | 0.4 |
| 7_1406485 | 2 | 1 | 7 | 1 | 3 | 1 | 5 | 0.8 | 17 | 0.9412 | 4 | 0.75 | NA | NA | NA | NA | 4 | 0.5 | 8 | 0.625 |
| 7_1406485( | 3 | 0 | 6 | 0 | 4 | 0.25 | 4 | 0.25 | 17 | 0.1176 | 4 | 0.25 | 1 | 0 | 1 | 0 | 4 | 0.25 | 10 | 0.2 |
| 7_1406487: | 2 | 0.5 | 1 | 1 | 1 | 1 | 1 | 0 | 5 | 0.6 | NA | NA | NA | NA | NA | NA | 1 | 1 | 1 | 1 |
| 7_1406488: | 1 | 0 | 1 | 1 | NA | NA | NA | NA | 2 | 0.5 | NA | NA | NA | NA | 1 | 1 | 1 | 1 | 2 | 1 |
| 7_1406489 | 1 | 1 | 1 | 1 | NA | NA | NA | NA | 2 | 1 | 1 | 1 | NA | NA | 1 | 1 | 2 | 0.5 | 4 | 0.75 |
| 7_1406507 | 6 | 0.3333 | 8 | 0 | 5 | 0.2 | 3 | 0 | 22 | 0.1364 | 6 | 0 | 3 | 0.6667 | 2 | 0 | 2 | 0 | 13 | 0.1538 |
| 7_1406521 | 10 | 0.4 | 9 | 0.4444 | 4 | 0.25 | 4 | 0.5 | 27 | 0.4074 | 7 | 0.4286 | 2 | 0.5 | 5 | 0.6 | 2 | 0 | 16 | 0.4375 |
| 7_1407327 | 3 | 0 | 3 | 0.3333 | 4 | 0 | 4 | 0.25 | 14 | 0.1429 | 2 | 0 | 4 | 0 | 3 | 0 | 3 | 0.3333 | 12 | 0.0833 |
| 7_1407329 | 5 | 0.2 | 3 | 0 | 4 | 0.25 | 5 | 0.2 | 17 | 0.1765 | 3 | 0 | 4 | 0.25 | 3 | 0 | 6 | 0.1667 | 16 | 0.125 |
| 7_1407348 | 5 | 0.4 | 1 | 1 | 1 | 0 | 1 | 0 | 8 | 0.375 | 1 | 1 | 3 | 0.3333 | 3 | 0 | 5 | 0.2 | 12 | 0.25 |
| 7_1407348 | 6 | 0.6667 | 4 | 0.75 | 3 | 0 | 4 | 0.5 | 17 | 0.5294 | 3 | 0.6667 | 6 | 0.3333 | 4 | 0.25 | 6 | 0.1667 | 19 | 0.3158 |
| 7_1407352 | 7 | 0.1429 | 3 | 0.3333 | 4 | 0 | 4 | 0 | 18 | 0.1111 | 3 | 0.3333 | 6 | 0 | 4 | 0 | 7 | 0 | 20 | 0.05 |
| 7_1407353 | 6 | 0 | 4 | 0.5 | 2 | 0 | 5 | 0.6 | 17 | 0.2941 | 3 | 0 | 5 | 0.4 | 5 | 0.4 | 6 | 0.1667 | 19 | 0.2632 |
| 7_1407353 | 5 | 0.2 | 4 | 0.5 | 3 | 0.3333 | 5 | 0.2 | 17 | 0.2941 | 3 | 0 | 6 | 0.3333 | 3 | 0 | 6 | 0.3333 | 18 | 0.2222 |
| 7_1407354 | 6 | 0 | 4 | 0 | 4 | 0.5 | 2 | 0 | 16 | 0.125 | 3 | 0.3333 | 4 | 0 | 3 | 0 | 3 | 0 | 13 | 0.0769 |
| 7_1407359 | 8 | 0.875 | 4 | 0.75 | 7 | 0.4286 | 5 | 0.2 | 24 | 0.5833 | 3 | 0 | 7 | 0.7143 | 4 | 0.5 | 6 | 0.5 | 20 | 0.5 |
| 7_1407359 | 8 | 0 | 3 | 0.6667 | 6 | 0.1667 | 5 | 0.6 | 22 | 0.2727 | 3 | 0.6667 | 6 | 0.3333 | 4 | 0.25 | 6 | 0.1667 | 19 | 0.3158 |
| 7_1407709 | 24 | 0.9167 | 31 | 0.9032 | 27 | 0.8519 | 28 | 0.8929 | 110 | 0.8909 | 24 | 0.875 | 29 | 0.9655 | 32 | 0.9688 | 28 | 1 | 113 | 0.9558 |
| 7_1407709 | 27 | 0.7778 | 31 | 0.8065 | 32 | 0.8125 | 34 | 0.9412 | 124 | 0.8387 | 23 | 0.8696 | 34 | 0.8235 | 23 | 0.8261 | 31 | 0.8387 | 111 | 0.8378 |
| 7_1407709 | 30 | 0.6667 | 34 | 0.5882 | 28 | 0.6786 | 30 | 0.5667 | 122 | 0.623 | 22 | 0.6364 | 42 | 0.6667 | 33 | 0.7879 | 32 | 0.7188 | 129 | 0.7054 |
| 7_1407709 | 20 | 0.05 | 30 | 0.0667 | 21 | 0.2857 | 20 | 0.05 | 91 | 0.1099 | 17 | 0.1176 | 30 | 0.3 | 27 | 0.1481 | 22 | 0.1818 | 96 | 0.1979 |
| 7_1407712 | 8 | 0.375 | 13 | 0.7692 | 9 | 0.4444 | 14 | 0.4286 | 44 | 0.5227 | 7 | 0.7143 | 17 | 0.5882 | 7 | 0.2857 | 15 | 0.4 | 46 | 0.5 |
| 7_1407712 | 9 | 0.2222 | 15 | 0.2 | 11 | 0 | 14 | 0.0714 | 49 | 0.1224 | 9 | 0.2222 | 16 | 0.0625 | 11 | 0 | 16 | 0 | 52 | 0.0577 |
| 7_1407984 | NA | NA | NA | NA | NA | NA | 1 | 1 | 1 | 1 | NA | NA | NA | NA | NA | NA | 1 | 1 | 1 | 1 |
| 7_1408381 | 1 | 1 | 2 | 0.5 | 2 | 0 | NA | NA | 5 | 0.4 | 1 | 0 | 3 | 0.6667 | 2 | 1 | NA | NA | 6 | 0.6667 |
| 7_1408382 | 1 | 1 | 2 | 0 | 2 | 0 | NA | NA | 5 | 0.2 | 1 | 1 | 3 | 0.6667 | 1 | 1 | NA | NA | 5 | 0.8 |
| 7_1408384 | 1 | 0 | 2 | 0.5 | 2 | 0.5 | NA | NA | 5 | 0.4 | 1 | 0 | 3 | 0.3333 | 2 | 0 | NA | NA | 6 | 0.1667 |
| 7_1408388 | 1 | 1 | 1 | 1 | 1 | 1 | NA | NA | 3 | 1 | 1 | 1 | 2 | 1 | 2 | 0.5 | 1 | 0 | 6 | 0.6667 |
| 7_1408389 | 1 | 1 | 2 | 0.5 | 2 | 0.5 | NA | NA | 5 | 0.6 | 1 | 0 | 3 | 0.6667 | 1 | 0 | 1 | 1 | 6 | 0.5 |
| 7_1408389 | 1 | 1 | 2 | 1 | 3 | 0 | NA | NA | 6 | 0.5 | 2 | 0 | 3 | 0.6667 | 1 | 0 | 1 | 0 | 7 | 0.2857 |
| 7_1408389 | 1 | 0 | 2 | 1 | 3 | 1 | NA | NA | 6 | 0.8333 | 2 | 1 | 3 | 1 | 1 | 1 | 1 | 1 | 7 | 1 |
| 7_1408391 | 1 | 0 | 3 | 0.3333 | 2 | 0 | NA | NA | 6 | 0.1667 | 1 | 1 | 3 | 0.3333 | 1 | 0 | 1 | 1 | 6 | 0.5 |
| 7_1408392 | 1 | 1 | 2 | 0.5 | 2 | 1 | NA | NA | 5 | 0.8 | 2 | 0.5 | 3 | 0.6667 | 1 | 0 | 1 | 1 | 7 | 0.5714 |
| 7_1408397 | 2 | 0 | 3 | 0 | 3 | 0.3333 | NA | NA | 8 | 0.125 | 2 | 0.5 | 4 | 0 | 2 | 0 | 1 | 0 | 9 | 0.1111 |
| 7_1408401 | 3 | 0.3333 | 3 | 0 | 2 | 0 | 1 | 0 | 9 | 0.1111 | 2 | 1 | 5 | 0 | 2 | 0 | 1 | 1 | 10 | 0.3 |
| 7_1408401 | 3 | 1 | 3 | 1 | NA | NA | 1 | 1 | 7 | 1 | 2 | 1 | 5 | 1 | 2 | 1 | 1 | 1 | 10 | 1 |
| 7_1408402 | 3 | 0.3333 | 3 | 0.3333 | 3 | 0.3333 | 1 | 1 | 10 | 0.4 | 2 | 0.5 | 5 | 0.4 | 2 | 0 | 1 | 0 | 10 | 0.3 |
| 7_1408402 | 3 | 0.3333 | 2 | 1 | 2 | 1 | 1 | 1 | 8 | 0.75 | 2 | 0.5 | 5 | 0.6 | 3 | 0 | 1 | 0 | 11 | 0.3636 |
| 7_1408404 | 2 | 1 | 3 | 0.3333 | 4 | 0.5 | 2 | 0 | 11 | 0.4545 | 2 | 0.5 | 3 | 0.6667 | 1 | 1 | 1 | 1 | 7 | 0.7143 |
| 7_1408423 | 4 | 0 | 4 | 0 | 6 | 0.1667 | 3 | 0 | 17 | 0.0588 | 4 | 0.25 | 5 | 0 | 5 | 0.4 | 3 | 0 | 17 | 0.1765 |
| 7_1408425 | 3 | 1 | 3 | 0.3333 | 5 | 0.8 | 2 | 0.5 | 13 | 0.6923 | 2 | 0.5 | 6 | 0.6667 | 5 | 0.6 | 3 | 1 | 16 | 0.6875 |
| 7_1408772 | 8 | 0.125 | 7 | 0.1429 | 3 | 0.3333 | 4 | 0.25 | 22 | 0.1818 | 4 | 0.25 | 8 | 0.125 | 4 | 0 | 4 | 0.5 | 20 | 0.2 |
| 7_1408773 | 6 | 0.1667 | 8 | 0.125 | 2 | 0 | 4 | 0 | 20 | 0.1 | 3 | 0 | 7 | 0.1429 | 4 | 0.25 | 3 | 0 | 17 | 0.1176 |
| 7_1408773 | 5 | 0.8 | 5 | 0.8 | 1 | 1 | 3 | 1 | 14 | 0.8571 | 3 | 0.6667 | 7 | 0.8571 | 3 | 1 | 4 | 0.75 | 17 | 0.8235 |
| 7_1409132 | 1 | 0 | NA | NA | 1 | 1 | NA | NA | 2 | 0.5 | 1 | 1 | NA | NA | NA | NA | NA | NA | 1 | 1 |
| 7_1409161 | 2 | 0 | 2 | 0 | 1 | 0 | 3 | 0.3333 | 8 | 0.125 | 2 | 1 | 3 | 0 | NA | NA | 1 | 0 | 6 | 0.3333 |
| 7_1409163 | 2 | 0.5 | 2 | 0.5 | 1 | 1 | 3 | 0.3333 | 8 | 0.5 | 2 | 0.5 | 5 | 0.4 | NA | NA | 2 | 0.5 | 9 | 0.4444 |

| | | | | | | | | | | | | | | | | | | | | |
|---|---|---|---|---|---|---|---|---|---|---|---|---|---|---|---|---|---|---|---|---|
| 7_1409424 | 1 | 0 | 2 | 0 | NA | NA | 1 | 1 | 4 | 0.25 | NA | NA | NA | NA | NA | NA | 1 | 1 | 1 | 1 |
| 7_1409879 | 1 | 1 | NA | NA | NA | NA | NA | NA | 1 | 1 | NA | NA | NA | NA | 1 | 1 | NA | NA | 1 | 1 |
| 7_1409881 | 4 | 0.25 | 2 | 0 | 3 | 0 | 1 | 0 | 10 | 0.1 | 3 | 0 | 2 | 0 | 6 | 0.1667 | 1 | 0 | 12 | 0.0833 |
| 7_1409884 | 8 | 0.5 | 6 | 0.3333 | 4 | 0.25 | 4 | 0.25 | 22 | 0.3636 | 3 | 0.6667 | 9 | 0.3333 | 7 | 0.5714 | 5 | 0.6 | 24 | 0.5 |
| 7_1409886 | 8 | 0.125 | 8 | 0.125 | 3 | 0 | 6 | 0 | 25 | 0.08 | 3 | 0 | 5 | 0 | 7 | 0.4286 | 4 | 0 | 19 | 0.1579 |
| 7_1409887 | 4 | 0.75 | 7 | 0.8571 | 2 | 0.5 | 5 | 0.6 | 18 | 0.7222 | 3 | 0.6667 | 6 | 0.8333 | 6 | 0.8333 | 4 | 0.75 | 19 | 0.7895 |
| 7_1409887 | 5 | 0 | 7 | 0.1429 | 3 | 0.3333 | 5 | 0.2 | 20 | 0.15 | 2 | 0 | 6 | 0.1667 | 6 | 0 | 4 | 0 | 18 | 0.0556 |
| 7_1409888 | 5 | 0.2 | 6 | 0.1667 | 2 | 0 | 5 | 0.4 | 18 | 0.2222 | 3 | 0 | 6 | 0.1667 | 7 | 0.1429 | 4 | 0 | 20 | 0.1 |
| 7_1410063 | 45 | 0.2667 | 36 | 0.3333 | 39 | 0.359 | 29 | 0.2759 | 149 | 0.3087 | 36 | 0.3611 | 27 | 0.4815 | 38 | 0.3421 | 38 | 0.3684 | 139 | 0.3813 |
| 7_1410064 | 39 | 0.5385 | 38 | 0.5 | 49 | 0.4898 | 34 | 0.5294 | 160 | 0.5125 | 37 | 0.4865 | 28 | 0.5357 | 39 | 0.5385 | 35 | 0.6286 | 139 | 0.5468 |
| 7_1410064 | 25 | 0.16 | 25 | 0.04 | 32 | 0.0938 | 22 | 0.1818 | 104 | 0.1154 | 25 | 0.16 | 19 | 0.2105 | 29 | 0.1724 | 30 | 0.1 | 103 | 0.1553 |
| 7_1410069 | 28 | 0.1071 | 26 | 0.1154 | 27 | 0 | 18 | 0.2778 | 99 | 0.1111 | 23 | 0.2609 | 17 | 0.0588 | 28 | 0.1429 | 23 | 0.1739 | 91 | 0.1648 |
| 7_1410070 | 35 | 0.1714 | 30 | 0.2667 | 29 | 0.1724 | 27 | 0.2963 | 121 | 0.2231 | 32 | 0.1562 | 19 | 0.3158 | 31 | 0.2903 | 27 | 0.1852 | 109 | 0.2294 |
| 7_1410073 | 35 | 0.1143 | 29 | 0.2069 | 33 | 0.0909 | 27 | 0.2593 | 124 | 0.1613 | 32 | 0.2188 | 17 | 0.0588 | 34 | 0.1471 | 27 | 0.0741 | 110 | 0.1364 |
| 7_1410074 | 29 | 1 | 27 | 1 | 27 | 1 | 24 | 1 | 107 | 1 | 27 | 1 | 17 | 1 | 25 | 0.96 | 24 | 1 | 93 | 0.9892 |
| 7_1410076 | 22 | 0.0455 | 23 | 0 | 28 | 0 | 21 | 0.1905 | 94 | 0.0532 | 18 | 0 | 12 | 0 | 25 | 0.16 | 21 | 0.1429 | 76 | 0.0921 |
| 7_1410099 | 32 | 0.0625 | 33 | 0.0909 | 24 | 0.125 | 25 | 0.08 | 114 | 0.0877 | 24 | 0.0417 | 28 | 0.1429 | 17 | 0 | 25 | 0.04 | 94 | 0.0638 |
| 7_1410101 | 28 | 0.1071 | 31 | 0.1935 | 20 | 0.2 | 25 | 0.2 | 104 | 0.1731 | 22 | 0.2273 | 24 | 0.375 | 18 | 0.2222 | 27 | 0.2222 | 91 | 0.2637 |
| 7_1410112 | 16 | 0 | 18 | 0.1111 | 12 | 0 | 13 | 0.0769 | 59 | 0.0508 | 16 | 0.0625 | 17 | 0.0588 | 10 | 0 | 12 | 0.0833 | 55 | 0.0545 |
| 7_1410184 | 1 | 1 | NA | NA | NA | NA | NA | NA | 1 | 1 | NA | NA | NA | NA | 2 | 0.5 | NA | NA | 2 | 0.5 |
| 7_1410397 | 7 | 0.7143 | 5 | 0.8 | 10 | 0.4 | NA | NA | 22 | 0.5909 | 5 | 0.8 | 4 | 1 | 2 | 0.5 | 6 | 0.3333 | 17 | 0.6471 |
| 7_1410397 | 8 | 0.375 | 7 | 0.1429 | 10 | 0.4 | NA | NA | 25 | 0.32 | 6 | 0.3333 | 5 | 0.2 | 2 | 0 | 7 | 0.2857 | 20 | 0.25 |
| 7_1410398 | 7 | 0.5714 | 7 | 0.2857 | 9 | 0.2222 | 1 | 0 | 24 | 0.3333 | 6 | 0.3333 | 5 | 0.4 | 1 | 0 | 5 | 0.8 | 17 | 0.4706 |
| 7_1410507 | 4 | 0 | 1 | 0 | 1 | 1 | 2 | 0 | 8 | 0.125 | 4 | 0.25 | 3 | 0.3333 | 2 | 0 | 1 | 0 | 10 | 0.2 |
| 7_1410507 | 4 | 0.25 | 1 | 1 | 1 | 1 | 3 | 0 | 9 | 0.3333 | 4 | 0.5 | 5 | 0.4 | NA | NA | 1 | 0 | 10 | 0.4 |
| 7_1410510 | 9 | 0.1111 | 3 | 0 | 3 | 0 | 2 | 0.5 | 17 | 0.1176 | 3 | 0 | 6 | 0.1667 | 1 | 1 | 2 | 0 | 12 | 0.1667 |
| 7_1410530 | 14 | 0.5 | 9 | 1 | 13 | 0.7692 | 14 | 0.5714 | 50 | 0.68 | 8 | 0.875 | 8 | 0.875 | 3 | 0.6667 | 11 | 0.6364 | 30 | 0.7667 |
| 7_1410531 | 11 | 0.4545 | 8 | 0.375 | 12 | 0.0833 | 12 | 0.5 | 43 | 0.3488 | 7 | 0.5714 | 6 | 0.5 | 4 | 0.5 | 9 | 0.2222 | 26 | 0.4231 |
| 7_1410729 | 6 | 0.1667 | 1 | 0 | 7 | 0.5714 | 1 | 1 | 15 | 0.4 | 2 | 0 | 8 | 0.5 | 4 | 0.75 | 2 | 0 | 16 | 0.4375 |
| 7_1410730 | 5 | 0 | 3 | 0 | 6 | 0 | 1 | 1 | 15 | 0.0667 | 3 | 0 | 6 | 0.3333 | 3 | 1 | 3 | 0.3333 | 15 | 0.4 |
| 7_1410730 | 6 | 0.6667 | 3 | 0 | 7 | 0.7143 | 1 | 1 | 17 | 0.5882 | 3 | 0.3333 | 8 | 0.375 | 3 | 0 | 3 | 0.6667 | 17 | 0.3529 |
| 7_1410732 | 1 | 1 | NA | NA | 1 | 0 | 1 | 0 | 3 | 0.3333 | NA | NA | NA | NA | 1 | 1 | NA | NA | 1 | 1 |
| 7_1410733 | 1 | 1 | NA | NA | 1 | 0 | NA | NA | 2 | 0.5 | NA | NA | NA | NA | 1 | 1 | NA | NA | 1 | 1 |
| 7_1410748 | 1 | 1 | 2 | 0 | 1 | 1 | NA | NA | 4 | 0.5 | NA | NA | NA | NA | 2 | 0.5 | NA | NA | 2 | 0.5 |
| 7_1410753 | 5 | 0.6 | 3 | 0 | 3 | 0.3333 | 3 | 0 | 14 | 0.2857 | 1 | 0 | 2 | 0.5 | 2 | 0.5 | 6 | 0 | 11 | 0.1818 |
| 7_1410753 | 4 | 0.75 | 4 | 0.75 | 2 | 1 | 3 | 1 | 13 | 0.8462 | NA | NA | 2 | 0.5 | 2 | 1 | 5 | 0.8 | 9 | 0.7778 |
| 7_1410754 | 6 | 0.1667 | 4 | 0.25 | 3 | 0 | 3 | 0 | 16 | 0.125 | 1 | 0 | 1 | 0 | 2 | 0.5 | 3 | 0.3333 | 7 | 0.2857 |
| 7_1410754 | 5 | 1 | 3 | 1 | 3 | 0.6667 | 4 | 0.75 | 15 | 0.8667 | 1 | 1 | 1 | 0 | 1 | 1 | 5 | 0.4 | 8 | 0.5 |
| 7_1410756 | 6 | 1 | 4 | 1 | 3 | 1 | 4 | 1 | 17 | 1 | 1 | 1 | 5 | 0.6 | 3 | 1 | 7 | 0.8571 | 16 | 0.8125 |
| 7_1410756 | 6 | 0.3333 | 4 | 0.25 | 2 | 0.5 | 3 | 0 | 15 | 0.2667 | 1 | 0 | 4 | 0.25 | 3 | 0.3333 | 6 | 0.1667 | 14 | 0.2143 |
| 7_1410757 | 7 | 0.8571 | 4 | 0.75 | 4 | 0.5 | 4 | 1 | 19 | 0.7895 | NA | NA | 4 | 0.5 | 2 | 0.5 | 7 | 0.7143 | 13 | 0.6154 |
| 7_1410757 | 7 | 0.5714 | 3 | 0.6667 | 3 | 0.6667 | 4 | 0 | 17 | 0.4706 | 1 | 0 | 4 | 0.75 | 2 | 0 | 7 | 0.4286 | 14 | 0.4286 |
| 7_1410758 | 7 | 0 | 4 | 0.25 | 2 | 0 | 4 | 0 | 17 | 0.0588 | NA | NA | 4 | 0 | 3 | 0 | 7 | 0.1429 | 14 | 0.0714 |
| 7_1410759 | 7 | 0.5714 | 8 | 0.5 | 5 | 0.4 | 6 | 0.6667 | 26 | 0.5385 | 2 | 1 | 5 | 1 | 3 | 1 | 11 | 0.4545 | 21 | 0.7143 |
| 7_1410759 | 6 | 0.8333 | 6 | 0.6667 | 4 | 0.75 | 6 | 0.5 | 22 | 0.6818 | 2 | 0.5 | 3 | 0.6667 | 3 | 1 | 11 | 0.9091 | 19 | 0.8421 |
| 7_1410759 | 5 | 0.8 | 7 | 0.4286 | 6 | 0.5 | 5 | 0.8 | 23 | 0.6087 | 2 | 0.5 | 4 | 0.25 | 3 | 0.6667 | 11 | 0.2727 | 20 | 0.35 |
| 7_1412114 | 23 | 0.1739 | 16 | 0.25 | 20 | 0.2 | 24 | 0.2083 | 83 | 0.2048 | 13 | 0.2308 | 16 | 0.0625 | 14 | 0 | 25 | 0.08 | 68 | 0.0882 |
| 7_1412122 | 20 | 0.3 | 18 | 0.3889 | 20 | 0.25 | 26 | 0.4615 | 84 | 0.3571 | 11 | 0.4545 | 17 | 0.3529 | 13 | 0.3077 | 23 | 0.3913 | 64 | 0.375 |
| 7_1414335 | NA | NA | 1 | 1 | NA | NA | NA | NA | 1 | 1 | 2 | 0.5 | 1 | 0 | 1 | 1 | NA | NA | 4 | 0.5 |
| 7_1414354 | 19 | 0 | 15 | 0.1333 | 9 | 0.2222 | 21 | 0.0952 | 64 | 0.0938 | 7 | 0 | 17 | 0.0588 | 17 | 0.1765 | 15 | 0 | 56 | 0.0714 |
| 7_1414356 | 15 | 0.2 | 10 | 0 | 7 | 0.5714 | 9 | 0.1111 | 41 | 0.1951 | 5 | 0 | 13 | 0.0769 | 8 | 0.5 | 9 | 0.1111 | 35 | 0.1714 |
| 7_1414368 | 9 | 0.2222 | 11 | 0.0909 | 7 | 0.2857 | 13 | 0.1538 | 40 | 0.175 | 6 | 0.3333 | 11 | 0.1818 | 12 | 0 | 11 | 0.0909 | 40 | 0.125 |
| 7_1414368 | 10 | 0.1 | 9 | 0.5556 | 6 | 0.1667 | 11 | 0.1818 | 36 | 0.25 | 6 | 0.3333 | 10 | 0.1 | 12 | 0.25 | 12 | 0.5 | 40 | 0.3 |
| 7_1414376 | 6 | 0.1667 | 5 | 0 | 7 | 0 | 9 | 0.4444 | 27 | 0.1852 | 2 | 0 | 9 | 0.1111 | 9 | 0 | 12 | 0.0833 | 32 | 0.0625 |
| 7_1414379 | 6 | 1 | 4 | 0.5 | 8 | 0.375 | 9 | 0.4444 | 27 | 0.5556 | 3 | 1 | 7 | 0.4286 | 5 | 0.8 | 12 | 0.75 | 27 | 0.7037 |
| 7_1414380 | 4 | 0 | 4 | 0.25 | 8 | 0.25 | 10 | 0.1 | 26 | 0.1538 | 3 | 0 | 9 | 0.2222 | 6 | 0 | 11 | 0.1818 | 29 | 0.1379 |
| 7_1414380 | 4 | 1 | 3 | 0.6667 | 6 | 0.6667 | 7 | 1 | 20 | 0.85 | 3 | 1 | 5 | 1 | 8 | 1 | 10 | 1 | 26 | 1 |
| 7_1414381 | 5 | 0.6 | 3 | 0.6667 | 8 | 0.5 | 11 | 0.6364 | 27 | 0.5926 | 2 | 1 | 8 | 0.875 | 6 | 0.5 | 11 | 0.7273 | 27 | 0.7407 |
| 7_1414382 | 4 | 0.5 | 4 | 0.75 | 7 | 0 | 9 | 0.3333 | 24 | 0.3333 | 1 | 1 | 8 | 0.25 | 7 | 0.2857 | 10 | 0.4 | 26 | 0.3462 |
| 7_1415468 | NA | NA | NA | NA | 1 | 1 | NA | NA | 1 | 1 | NA | NA | NA | NA | NA | NA | 1 | 1 | 1 | 1 |
| 7_1415558 | 1 | 0 | 2 | 0.5 | 2 | 0 | 4 | 0 | 9 | 0.1111 | 5 | 0 | 4 | 0.25 | 2 | 0 | 2 | 0 | 13 | 0.0769 |
| 7_1415559 | NA | NA | 1 | 1 | 1 | 1 | 4 | 0.25 | 6 | 0.5 | 5 | 0.6 | 4 | 0 | 4 | 0.25 | 5 | 0.2 | 18 | 0.2778 |
| 7_1415560 | 1 | 1 | 2 | 0.5 | 3 | 0 | 3 | 0.6667 | 9 | 0.4444 | 6 | 0.6667 | 4 | 1 | 4 | 0.25 | 5 | 0.4 | 19 | 0.5789 |
| 7_1415560 | 1 | 1 | 1 | 0 | 3 | 0 | 1 | 0 | 6 | 0.1667 | 5 | 0.2 | 4 | 0 | 4 | 0 | 5 | 0.2 | 18 | 0.1111 |
| 7_1415562 | 5 | 0.4 | 4 | 0.5 | 5 | 0.4 | 9 | 0.5556 | 23 | 0.4783 | 8 | 0.75 | 8 | 0.5 | 10 | 0.6 | 3 | 0.6667 | 29 | 0.6207 |

| | | | | | | | | | | | | | | | | | | | | |
|---|---|---|---|---|---|---|---|---|---|---|---|---|---|---|---|---|---|---|---|---|
| 7_1415563 | 4 | 0 | 5 | 0.4 | 4 | 0 | 9 | 0.1111 | 22 | 0.1364 | 8 | 0.25 | 8 | 0.125 | 5 | 0.4 | 8 | 0.125 | 29 | 0.2069 |
| 7_1415564 | 5 | 0 | 7 | 0 | 4 | 0.25 | 11 | 0.1818 | 27 | 0.1111 | 11 | 0 | 10 | 0.1 | 11 | 0.1818 | 8 | 0 | 40 | 0.075 |
| 7_1415564 | 6 | 0.6667 | 7 | 0.2857 | 5 | 0.4 | 10 | 0.5 | 28 | 0.4643 | 11 | 0.2727 | 10 | 0.8 | 12 | 0.25 | 10 | 0.5 | 43 | 0.4419 |
| 7_1415565 | 4 | 1 | 5 | 1 | 4 | 1 | 10 | 1 | 23 | 1 | 6 | 1 | 7 | 0.8571 | 11 | 0.9091 | 5 | 1 | 29 | 0.931 |
| 7_1415566 | 5 | 0.6 | 8 | 0.375 | 6 | 0.1667 | 12 | 0.4167 | 31 | 0.3871 | 10 | 0.5 | 13 | 0.3077 | 10 | 0.7 | 9 | 0.3333 | 42 | 0.4524 |
| 7_1415569 | 4 | 0.5 | 5 | 0.2 | 5 | 0 | 5 | 0.2 | 19 | 0.2105 | 6 | 0 | 9 | 0.2222 | 11 | 0.1818 | 6 | 0.1667 | 32 | 0.1562 |
| 7_1415569 | 6 | 0.1667 | 9 | 0 | 8 | 0.125 | 13 | 0.0769 | 36 | 0.0833 | 12 | 0.1667 | 13 | 0.0769 | 12 | 0.1667 | 11 | 0.0909 | 48 | 0.125 |
| 7_1415570 | 5 | 0 | 6 | 0.1667 | 4 | 0 | 10 | 0.1 | 25 | 0.08 | 10 | 0 | 14 | 0.0714 | 7 | 0 | 4 | 0.25 | 35 | 0.0571 |
| 7_1415578 | 10 | 0.3 | 13 | 0.4615 | 8 | 0.5 | 12 | 0.4167 | 43 | 0.4186 | 11 | 0.2727 | 15 | 0.4667 | 13 | 0.3846 | 8 | 0.5 | 47 | 0.4043 |
| 7_1415579 | 10 | 0.4 | 12 | 0.3333 | 6 | 0.1667 | 14 | 0.4286 | 42 | 0.3571 | 11 | 0.3636 | 17 | 0.1765 | 12 | 0.1667 | 8 | 0.5 | 48 | 0.2708 |
| 7_1415579 | 8 | 0 | 10 | 0 | 8 | 0 | 13 | 0.1538 | 39 | 0.0513 | 16 | 0 | 16 | 0.0625 | 12 | 0.1667 | 9 | 0.1111 | 53 | 0.0755 |
| 7_1415623 | 6 | 0.6667 | 5 | 0.4 | 4 | 0.5 | 5 | 0.2 | 20 | 0.45 | 3 | 0.6667 | 5 | 0.8 | 5 | 0.2 | 4 | 0.5 | 17 | 0.5294 |
| 7_1415627 | 6 | 1 | 5 | 0.8 | 4 | 0.75 | 3 | 1 | 18 | 0.8889 | 3 | 1 | 5 | 1 | 5 | 0.8 | 4 | 1 | 17 | 0.9412 |
| 7_1415629 | 6 | 0.3333 | 6 | 0.6667 | 3 | 0 | 4 | 0.25 | 19 | 0.3684 | 4 | 0.5 | 6 | 0.3333 | 4 | 0.5 | 4 | 0.75 | 18 | 0.5 |
| 7_1415630 | 6 | 0.1667 | 6 | 0 | 4 | 0 | 4 | 0 | 20 | 0.05 | 3 | 0 | 4 | 0.25 | 5 | 0.2 | 4 | 0 | 16 | 0.125 |
| 7_1415631 | 5 | 0 | 4 | 0 | 4 | 0.25 | 3 | 0 | 16 | 0.0625 | 3 | 0 | 6 | 0 | 5 | 0.2 | 2 | 0 | 16 | 0.0625 |
| 7_1415631 | 7 | 1 | 5 | 1 | 4 | 1 | 3 | 1 | 19 | 1 | 4 | 1 | 6 | 1 | 3 | 1 | 4 | 0.5 | 17 | 0.8824 |
| 7_1416139 | 1 | 1 | NA | NA | NA | NA | NA | NA | 1 | 1 | NA | NA | 1 | 1 | NA | NA | 2 | 0 | 3 | 0.3333 |
| 7_1416140 | NA | NA | 1 | 1 | NA | NA | NA | NA | 1 | 1 | NA | NA | 1 | 1 | NA | NA | 1 | 0 | 2 | 0.5 |
| 7_1416142 | NA | NA | 1 | 1 | NA | NA | NA | NA | 1 | 1 | NA | NA | 1 | 1 | NA | NA | 2 | 1 | 3 | 1 |
| 7_1416144 | NA | NA | 1 | 1 | NA | NA | NA | NA | 1 | 1 | NA | NA | 1 | 1 | NA | NA | 2 | 1 | 3 | 1 |
| 7_1416333 | 19 | 0.1053 | 22 | 0.0909 | 14 | 0 | 24 | 0.0833 | 79 | 0.0759 | 16 | 0.0625 | 24 | 0.2083 | 25 | 0.04 | 18 | 0 | 83 | 0.0843 |
| 7_1416337 | 17 | 0.0588 | 21 | 0 | 17 | 0.0588 | 22 | 0.0909 | 77 | 0.0519 | 10 | 0 | 24 | 0.0417 | 24 | 0.0417 | 16 | 0.125 | 74 | 0.0541 |
| 7_1416353 | 8 | 0.125 | 10 | 0 | 9 | 0 | 13 | 0.1538 | 40 | 0.075 | 8 | 0.125 | 8 | 0 | 11 | 0 | 7 | 0.1429 | 34 | 0.0588 |
| 7_1416357 | 5 | 0.4 | 6 | 0 | 8 | 0.25 | 11 | 0 | 30 | 0.1333 | 9 | 0.1111 | 3 | 0.3333 | 10 | 0.2 | 7 | 0.1429 | 29 | 0.1724 |
| 7_1416357 | 6 | 0.3333 | 7 | 0 | 7 | 0.4286 | 11 | 0.1818 | 31 | 0.2258 | 9 | 0.2222 | 5 | 0.4 | 10 | 0.1 | 7 | 0.1429 | 31 | 0.1935 |
| 7_1416359 | 4 | 0.5 | 6 | 0.1667 | 6 | 0.3333 | 9 | 0.1111 | 25 | 0.24 | 6 | 0.3333 | 5 | 0.2 | 7 | 0 | 5 | 0.2 | 23 | 0.1739 |
| 7_1416360 | 4 | 1 | 4 | 1 | 6 | 1 | 5 | 1 | 19 | 1 | 6 | 1 | 2 | 1 | 9 | 1 | 5 | 1 | 22 | 1 |
| 7_1416361 | 3 | 1 | 3 | 0 | 6 | 0.8333 | 10 | 0.7 | 22 | 0.6818 | 6 | 0.3333 | 4 | 0.75 | 7 | 0.5714 | 5 | 0.6 | 22 | 0.5455 |
| 7_1416361 | 5 | 0 | 4 | 0 | 6 | 0.3333 | 8 | 0.25 | 23 | 0.1739 | 7 | 0.1429 | 5 | 0.2 | 8 | 0 | 6 | 0.1667 | 26 | 0.1154 |
| 7_1416361 | 3 | 1 | 4 | 1 | 5 | 1 | 7 | 0.8571 | 19 | 0.9474 | 7 | 1 | 4 | 1 | 7 | 1 | 4 | 1 | 22 | 1 |
| 7_1416362 | 5 | 1 | 4 | 1 | 6 | 1 | 8 | 1 | 23 | 1 | 6 | 1 | 5 | 1 | 6 | 0.8333 | 5 | 1 | 22 | 0.9545 |
| 7_1416363 | 2 | 0.5 | 1 | 1 | 4 | 0.25 | 3 | 0.6667 | 10 | 0.5 | 2 | 0.5 | 3 | 0.6667 | 5 | 1 | 4 | 0.5 | 14 | 0.7143 |
| 7_1416363 | 5 | 0.8 | 3 | 0.6667 | 6 | 0.6667 | 6 | 0.8333 | 20 | 0.75 | 4 | 0.75 | 5 | 0.8 | 6 | 0.8333 | 4 | 0.75 | 19 | 0.7895 |
| 7_1416364 | 4 | 0.5 | 4 | 0.5 | 3 | 0 | 7 | 0.1429 | 18 | 0.2778 | 5 | 0.6 | 4 | 0 | 7 | 0.2857 | 4 | 0.75 | 20 | 0.4 |
| 7_1416366 | 5 | 0.6 | 2 | 0 | 6 | 0.3333 | 6 | 0.3333 | 19 | 0.3684 | 5 | 0.2 | 5 | 0.2 | 8 | 0.125 | 2 | 0 | 20 | 0.15 |
| 7_1416366 | 4 | 1 | 2 | 1 | 6 | 1 | 7 | 1 | 19 | 1 | 7 | 0.8571 | 5 | 1 | 6 | 1 | 1 | 1 | 19 | 0.9474 |
| 7_1416366 | 5 | 0.8 | 2 | 1 | 4 | 1 | 7 | 0.8571 | 18 | 0.8889 | 6 | 1 | 4 | 0.75 | 5 | 0.8 | 3 | 1 | 18 | 0.8889 |
| 7_1419092 | 1 | 1 | 2 | 0 | 2 | 0 | 3 | 0 | 8 | 0.125 | NA | NA | 2 | 0 | 1 | 0 | 3 | 0.3333 | 6 | 0.1667 |
| 7_1419096 | 2 | 1 | 2 | 0.5 | 3 | 1 | 3 | 0.6667 | 10 | 0.8 | NA | NA | 2 | 0.5 | 1 | 1 | 4 | 1 | 7 | 0.8571 |
| 7_1419298 | 52 | 0.3077 | 56 | 0.3571 | 34 | 0.3235 | 55 | 0.3455 | 197 | 0.335 | 59 | 0.3729 | 34 | 0.3529 | 32 | 0.3125 | 37 | 0.3514 | 162 | 0.3519 |
| 7_1419298 | 70 | 0.0571 | 63 | 0.1111 | 53 | 0.0566 | 60 | 0.0833 | 246 | 0.0772 | 74 | 0.1216 | 44 | 0.1364 | 42 | 0.119 | 40 | 0.05 | 200 | 0.11 |
| 7_1419301 | 55 | 0.2182 | 57 | 0.1404 | 44 | 0.2955 | 56 | 0.1786 | 212 | 0.2028 | 60 | 0.25 | 38 | 0.2105 | 44 | 0.1591 | 43 | 0.2558 | 185 | 0.2216 |
| 7_1419304 | 53 | 0.0943 | 54 | 0.0556 | 39 | 0.0513 | 45 | 0.0889 | 191 | 0.0733 | 57 | 0.0351 | 34 | 0.0588 | 41 | 0.0976 | 38 | 0.0789 | 170 | 0.0647 |
| 7_1420482 | 2 | 0 | 1 | 1 | NA | NA | 1 | 0 | 4 | 0.25 | 2 | 1 | 1 | 0 | NA | NA | 1 | 1 | 4 | 0.75 |
| 7_1425873 | NA | NA | 1 | 1 | 1 | 1 | 2 | 0.5 | 4 | 0.75 | 2 | 0.5 | 1 | 0 | NA | NA | NA | NA | 3 | 0.3333 |
| 7_1426233 | NA | NA | 1 | 0 | 1 | 0 | 3 | 0.3333 | 5 | 0.2 | 2 | 0 | 4 | 0.25 | 2 | 0.5 | 3 | 0.3333 | 11 | 0.2727 |
| 7_1426234 | NA | NA | 2 | 0 | 1 | 1 | 3 | 0.6667 | 6 | 0.5 | 1 | 0 | 4 | 0.75 | 2 | 0.5 | 4 | 0.5 | 11 | 0.5455 |
| 7_1426236 | NA | NA | 1 | 1 | NA | NA | 1 | 0 | 2 | 0.5 | NA | NA | 2 | 0.5 | 1 | 0 | 3 | 0.3333 | 6 | 0.3333 |
| 7_1426236 | NA | NA | NA | NA | NA | NA | 1 | 1 | 1 | 1 | NA | NA | 2 | 1 | 1 | 1 | 2 | 1 | 5 | 1 |
| 7_1426237 | 3 | 1 | 5 | 0.6 | 3 | 0.6667 | 7 | 0.8571 | 18 | 0.7778 | 4 | 0.25 | 4 | 0.75 | 5 | 0.8 | 7 | 0.5714 | 20 | 0.6 |
| 7_1426237 | 3 | 0.3333 | 5 | 0.2 | 3 | 0.3333 | 7 | 0.1429 | 18 | 0.2222 | 4 | 0.75 | 4 | 0.75 | 5 | 0.4 | 8 | 0.375 | 21 | 0.5238 |
| 7_1426238 | 3 | 0.3333 | 5 | 0.2 | 3 | 0.6667 | 8 | 0.25 | 19 | 0.3158 | 4 | 0.5 | 4 | 0.75 | 5 | 0.2 | 9 | 0.1111 | 22 | 0.3182 |
| 7_1426238 | NA | NA | 2 | 0 | 3 | 0.3333 | 4 | 0 | 9 | 0.1111 | 3 | 0.6667 | 2 | 0.5 | 4 | 0.25 | 9 | 0.2222 | 18 | 0.3333 |
| 7_1426240 | 3 | 0.3333 | 5 | 0.4 | 3 | 0.3333 | 6 | 0.1667 | 17 | 0.2941 | 2 | 0 | 4 | 0.75 | 3 | 0.3333 | 7 | 0.5714 | 16 | 0.5 |
| 7_1426240 | 3 | 0 | 5 | 0.2 | 2 | 0 | 7 | 0 | 17 | 0.0588 | 3 | 0 | 2 | 0 | 5 | 0 | 10 | 0.2 | 20 | 0.1 |
| 7_1426240 | 3 | 0 | 6 | 0 | 3 | 0.6667 | 8 | 0.125 | 20 | 0.15 | 2 | 0.5 | 4 | 0.25 | 4 | 0 | 10 | 0.1 | 20 | 0.15 |
| 7_1426241 | 3 | 0.3333 | 5 | 0 | 2 | 0.5 | 5 | 0.2 | 15 | 0.2 | 1 | 0 | 4 | 0.25 | 4 | 0.5 | 7 | 0.4286 | 16 | 0.375 |
| 7_1426241 | 3 | 0 | 4 | 0 | 3 | 0 | 7 | 0.1429 | 17 | 0.0588 | 2 | 0 | 4 | 0.5 | 4 | 0 | 7 | 0 | 17 | 0.1176 |
| 7_1426241 | 3 | 0.3333 | 5 | 0.2 | 3 | 0 | 8 | 0.25 | 19 | 0.2105 | 3 | 0.3333 | 2 | 0.5 | 4 | 0.5 | 10 | 0.4 | 19 | 0.4211 |
| 7_1426242 | 2 | 1 | 5 | 0.2 | 2 | 0.5 | 7 | 0.7143 | 16 | 0.5625 | 3 | 0.3333 | 2 | 1 | 5 | 1 | 10 | 0.7 | 20 | 0.75 |
| 7_1426243 | 3 | 0 | 3 | 0.3333 | 3 | 0 | 8 | 0 | 17 | 0.0588 | 2 | 0 | 3 | 0 | 3 | 0.3333 | 8 | 0.125 | 16 | 0.125 |
| 7_1426244 | 4 | 0 | 6 | 0 | 3 | 0 | 9 | 0.2222 | 22 | 0.0909 | 4 | 0 | 5 | 0.4 | 6 | 0.1667 | 9 | 0 | 24 | 0.125 |
| 7_1426244 | 2 | 0.5 | 3 | 0.6667 | NA | NA | 6 | 0.3333 | 11 | 0.4545 | 1 | 0 | 4 | 0.25 | 2 | 0 | 5 | 0 | 12 | 0.0833 |
| 7_1426246 | 3 | 0.6667 | 7 | 0.1429 | 3 | 0 | 8 | 0.375 | 21 | 0.2857 | 4 | 0.5 | 6 | 0.5 | 6 | 0.1667 | 8 | 0.375 | 24 | 0.375 |

| | | | | | | | | | | | | | | | | | | | | |
|---|---|---|---|---|---|---|---|---|---|---|---|---|---|---|---|---|---|---|---|---|
| 7_1426247 | 5 | 1 | 8 | 0.875 | 3 | 1 | 9 | 0.7778 | 25 | 0.88 | 5 | 1 | 7 | 1 | 7 | 0.8571 | 10 | 1 | 29 | 0.9655 |
| 7_1430677 | 13 | 0.0769 | 9 | 0.2222 | 14 | 0.0714 | 8 | 0 | 44 | 0.0909 | 14 | 0.0714 | 8 | 0.5 | 13 | 0.1538 | 10 | 0.1 | 45 | 0.1778 |
| 7_1430677 | 7 | 0.4286 | 6 | 0.3333 | 8 | 0.375 | 6 | 0.6667 | 27 | 0.4444 | 10 | 0.7 | 9 | 0.5556 | 10 | 0.4 | 8 | 0.5 | 37 | 0.5405 |
| 7_1430680 | 6 | 0 | 5 | 0.2 | 8 | 0.25 | 6 | 0 | 25 | 0.12 | 10 | 0 | 6 | 0.3333 | 9 | 0.2222 | 8 | 0.125 | 33 | 0.1515 |
| 7_1430680 | 9 | 0.2222 | 7 | 0.1429 | 8 | 0 | 4 | 0 | 28 | 0.1071 | 8 | 0 | 8 | 0 | 10 | 0.2 | 7 | 0.1429 | 33 | 0.0909 |
| 7_1430681 | 10 | 0.2 | 9 | 0.4444 | 10 | 0.1 | 7 | 0.1429 | 36 | 0.2222 | 12 | 0.25 | 10 | 0.3 | 11 | 0.5455 | 9 | 0.1111 | 42 | 0.3095 |
| 7_1430682 | 6 | 0.1667 | 5 | 0 | 10 | 0.1 | 6 | 0 | 27 | 0.0741 | 12 | 0.0833 | 8 | 0.25 | 10 | 0.1 | 9 | 0.1111 | 39 | 0.1282 |
| 7_1431114 | 4 | 0.5 | 4 | 0.5 | 10 | 0.5 | 2 | 0.5 | 20 | 0.5 | 2 | 0 | 4 | 0.75 | 6 | 0.5 | 5 | 0.8 | 17 | 0.5882 |
| 7_1431115 | 5 | 0.6 | 4 | 1 | 10 | 0.5 | 3 | 0.3333 | 22 | 0.5909 | 3 | 0.3333 | 5 | 0.8 | 5 | 0.8 | 4 | 0.5 | 17 | 0.6471 |
| 7_1431116 | 6 | 0 | 4 | 0.25 | 10 | 0.1 | 3 | 0 | 23 | 0.087 | 3 | 0 | 4 | 0.25 | 5 | 0.4 | 5 | 0 | 17 | 0.1765 |
| 7_1432430 | NA | NA | 1 | 1 | 2 | 0.5 | 2 | 1 | 5 | 0.8 | NA | NA | 1 | 0 | NA | NA | 1 | 1 | 2 | 0.5 |
| 7_1432430 | NA | NA | NA | NA | 2 | 1 | 1 | 0 | 3 | 0.6667 | NA | NA | 1 | 0 | NA | NA | 1 | 1 | 2 | 0.5 |
| 7_1432430 | NA | NA | 1 | 0 | 2 | 1 | 2 | 1 | 5 | 0.8 | NA | NA | 1 | 0 | NA | NA | 1 | 1 | 2 | 0.5 |
| 7_1432431 | NA | NA | 1 | 0 | 2 | 0 | 2 | 1 | 5 | 0.4 | NA | NA | 1 | 1 | NA | NA | 1 | 1 | 2 | 1 |
| 7_1432433 | NA | NA | 1 | 0 | 1 | 0 | 2 | 0.5 | 4 | 0.25 | NA | NA | 1 | 1 | NA | NA | 1 | 0 | 2 | 0.5 |
| 7_1432434 | NA | NA | 1 | 1 | 1 | 0 | 2 | 0.5 | 4 | 0.5 | NA | NA | 1 | 1 | NA | NA | NA | NA | 1 | 1 |
| 7_1433322 | 1 | 1 | 1 | 0 | 1 | 1 | 2 | 0.5 | 5 | 0.6 | NA | NA | 1 | 0 | NA | NA | 1 | 1 | 2 | 0.5 |
| 7_1433329 | NA | NA | NA | NA | NA | NA | 2 | 0.5 | 2 | 0.5 | NA | NA | NA | NA | NA | NA | 1 | 1 | 1 | 1 |
| 7_1433329 | 1 | 1 | 1 | 0 | 1 | 0 | 2 | 0 | 5 | 0.2 | NA | NA | 1 | 0 | NA | NA | 2 | 0.5 | 3 | 0.3333 |
| 7_1433329 | 1 | 1 | NA | NA | 1 | 0 | 2 | 1 | 4 | 0.75 | NA | NA | 1 | 1 | NA | NA | 2 | 1 | 3 | 1 |
| 7_1433329 | 1 | 1 | 1 | 0 | NA | NA | 2 | 0.5 | 4 | 0.5 | NA | NA | 1 | 0 | NA | NA | 1 | 1 | 2 | 0.5 |
| 7_1433331 | 1 | 1 | 1 | 1 | NA | NA | 1 | 0 | 3 | 0.6667 | NA | NA | 1 | 1 | NA | NA | 2 | 1 | 3 | 1 |
| 7_1433915 | NA | NA | NA | NA | NA | NA | 5 | 0.2 | 5 | 0.2 | 4 | 0.25 | 1 | 0 | 9 | 0 | 3 | 0 | 17 | 0.0588 |
| 7_1434009 | 1 | 1 | 6 | 0.6667 | 2 | 0.5 | 5 | 0.6 | 14 | 0.6429 | 5 | 0.6 | 4 | 0.75 | 7 | 0.8571 | 4 | 0.5 | 20 | 0.7 |
| 7_1434009 | 2 | 1 | 5 | 0.4 | 2 | 0.5 | 6 | 0.5 | 15 | 0.5333 | 6 | 0.3333 | 4 | 0.75 | 8 | 0.375 | 6 | 0.1667 | 24 | 0.375 |
| 7_1434010 | 1 | 0 | 4 | 0.5 | 2 | 1 | 4 | 0 | 11 | 0.3636 | 6 | 0 | 4 | 0.25 | 4 | 0.5 | 5 | 0.4 | 19 | 0.2632 |
| 7_1434011 | 2 | 0.5 | 5 | 1 | 2 | 0.5 | 5 | 1 | 14 | 0.8571 | 5 | 0.6 | 2 | 1 | 6 | 0.1667 | 5 | 0.4 | 18 | 0.4444 |
| 7_1434011 | 1 | 1 | 4 | 0.5 | 2 | 1 | 2 | 0 | 9 | 0.5556 | 7 | 0 | 4 | 0.25 | 4 | 0 | 4 | 0 | 19 | 0.0526 |
| 7_1434013 | 2 | 0 | 6 | 0.1667 | 5 | 0 | 4 | 0.25 | 17 | 0.1176 | 8 | 0.375 | 4 | 0 | 8 | 0.125 | 6 | 0.1667 | 26 | 0.1923 |
| 7_1434018 | 3 | 0.6667 | 6 | 0.8333 | 7 | 0.4286 | 9 | 0.4444 | 25 | 0.56 | 9 | 0.4444 | 4 | 0.25 | 13 | 0.3846 | 5 | 0 | 31 | 0.3226 |
| 7_1434019 | 3 | 0 | 5 | 0 | 7 | 0.2857 | 10 | 0 | 25 | 0.08 | 10 | 0.3 | 5 | 0.4 | 12 | 0.1667 | 5 | 0 | 32 | 0.2188 |
| 7_1434019 | 2 | 0 | 6 | 0.1667 | 7 | 0.1429 | 10 | 0.1 | 25 | 0.12 | 10 | 0.3 | 5 | 0 | 14 | 0.2857 | 6 | 0.1667 | 35 | 0.2286 |
| 7_1434020 | 3 | 0.3333 | 6 | 0.3333 | 8 | 0.25 | 9 | 0.3333 | 26 | 0.3077 | 10 | 0.1 | 4 | 0 | 13 | 0.3077 | 5 | 0.4 | 32 | 0.2188 |
| 7_1439632 | 1 | 0 | 1 | 0 | 3 | 0.3333 | 3 | 0 | 8 | 0.125 | 1 | 1 | 2 | 0 | 3 | 0 | NA | NA | 6 | 0.1667 |
| 7_1439633 | 1 | 1 | 2 | 1 | 3 | 0.3333 | 1 | 1 | 7 | 0.7143 | 1 | 1 | 2 | 0.5 | 3 | 0.6667 | NA | NA | 6 | 0.6667 |
| 7_1439634 | 1 | 0 | 2 | 0 | 3 | 0 | 3 | 0.3333 | 9 | 0.1111 | 1 | 0 | 2 | 0.5 | 4 | 0 | NA | NA | 7 | 0.1429 |
| 7_1439634 | 1 | 0 | 2 | 0.5 | 3 | 0.6667 | 3 | 0.3333 | 9 | 0.4444 | 1 | 1 | 2 | 0.5 | 3 | 0.6667 | NA | NA | 6 | 0.6667 |
| 7_1439635 | 1 | 1 | 2 | 0 | 3 | 0.6667 | 4 | 0.5 | 10 | 0.5 | 1 | 0 | 2 | 0 | 3 | 0.6667 | NA | NA | 6 | 0.3333 |
| 7_1439635 | 1 | 1 | 1 | 1 | 1 | 1 | 3 | 0.6667 | 6 | 0.8333 | 1 | 1 | 2 | 0.5 | 3 | 0.3333 | NA | NA | 6 | 0.5 |
| 7_1439641 | 1 | 0 | 2 | 0.5 | 3 | 0 | 4 | 0 | 10 | 0.1 | 3 | 0 | 2 | 0.5 | 4 | 0 | 2 | 0 | 11 | 0.0909 |
| 7_1439644 | 2 | 0 | 2 | 0 | 3 | 0.3333 | 5 | 0 | 12 | 0.0833 | 2 | 0 | 2 | 0.5 | 4 | 0 | 2 | 0 | 10 | 0.1 |
| 7_1439645 | 3 | 0.6667 | 2 | 0.5 | 3 | 0 | 5 | 0.4 | 13 | 0.3846 | 3 | 0.3333 | NA | NA | 3 | 0.6667 | 2 | 0.5 | 8 | 0.5 |
| 7_1439646 | 2 | 0.5 | 1 | 0 | 1 | 0 | 6 | 0.3333 | 10 | 0.3 | 3 | 0 | 2 | 0.5 | 5 | 0.2 | 2 | 0 | 12 | 0.1667 |
| 7_1439648 | 3 | 1 | 3 | 1 | 2 | 1 | 4 | 1 | 12 | 1 | 4 | 1 | 2 | 1 | 5 | 1 | 2 | 1 | 13 | 1 |
| 7_1439648 | 3 | 1 | 3 | 1 | 3 | 1 | 5 | 0.8 | 14 | 0.9286 | 2 | 1 | 2 | 1 | 5 | 1 | 2 | 1 | 11 | 1 |
| 7_1439650 | 5 | 0.8 | 2 | 0.5 | 3 | 0.6667 | 5 | 0.8 | 15 | 0.7333 | 4 | 0.25 | 2 | 0.5 | 6 | 0.5 | 1 | 1 | 13 | 0.4615 |
| 7_1439651 | 4 | 0.75 | 1 | 0 | 2 | 0.5 | 3 | 0 | 10 | 0.4 | 3 | 0.3333 | 1 | 1 | 5 | 0 | 2 | 0 | 11 | 0.1818 |
| 7_1439651 | 5 | 0.2 | 3 | 0 | 3 | 0 | 6 | 0 | 17 | 0.0588 | 3 | 0 | 2 | 0.5 | 6 | 0 | 2 | 0 | 13 | 0.0769 |
| 7_1439652 | 5 | 0.8 | 3 | 1 | 3 | 0.6667 | 6 | 1 | 17 | 0.8824 | 4 | 1 | 2 | 1 | 5 | 0.6 | 2 | 0.5 | 13 | 0.7692 |
| 7_1439653 | 5 | 0.2 | 3 | 0 | 2 | 0 | 6 | 0.1667 | 16 | 0.125 | 4 | 0.25 | 3 | 0.3333 | 6 | 0.3333 | 3 | 0 | 16 | 0.25 |
| 7_1439654 | 5 | 1 | 3 | 0.6667 | 3 | 0.6667 | 6 | 0.5 | 17 | 0.7059 | 3 | 0.6667 | 3 | 1 | 6 | 1 | 3 | 1 | 15 | 0.9333 |
| 7_1439744 | 4 | 0.25 | 3 | 0.3333 | 3 | 0.3333 | 7 | 0.1429 | 17 | 0.2353 | 4 | 0.5 | 3 | 0.3333 | 8 | 0.625 | 3 | 0.6667 | 18 | 0.5556 |
| 7_1439745 | 4 | 0 | 3 | 0.6667 | 3 | 0.3333 | 8 | 0.375 | 18 | 0.3333 | 3 | 0.3333 | 3 | 0 | 7 | 0.2857 | 3 | 0.3333 | 16 | 0.25 |
| 7_1439745 | 4 | 0.25 | 3 | 0 | 3 | 0 | 7 | 0.1429 | 17 | 0.1176 | 4 | 0 | 4 | 0 | 5 | 0.2 | 3 | 0 | 16 | 0.0625 |
| 7_1439747 | 1 | 1 | 3 | 0.3333 | 3 | 0.3333 | 6 | 1 | 13 | 0.6923 | 5 | 0.4 | 5 | 0.8 | 5 | 0.8 | 2 | 0.5 | 17 | 0.6471 |
| 7_1439747 | 5 | 1 | 3 | 1 | 3 | 1 | 7 | 1 | 18 | 1 | 4 | 0.75 | 5 | 0.8 | 9 | 1 | 3 | 1 | 21 | 0.9048 |
| 7_1439747 | 5 | 1 | 2 | 1 | 2 | 1 | 7 | 0.8571 | 16 | 0.9375 | 4 | 1 | 5 | 1 | 8 | 1 | 4 | 1 | 21 | 1 |
| 7_1439747 | 5 | 0.6 | 2 | 0.5 | 3 | 1 | 8 | 0.5 | 18 | 0.6111 | 5 | 0.6 | 4 | 0.25 | 7 | 0.1429 | 3 | 0.6667 | 19 | 0.3684 |
| 7_1439748 | 6 | 0.6667 | 3 | 0.6667 | 3 | 0.6667 | 7 | 0.2857 | 19 | 0.5263 | 4 | 0 | 5 | 0.6 | 7 | 0.5714 | 4 | 0.25 | 20 | 0.4 |
| 7_1439748 | 3 | 0.3333 | 1 | 0 | 3 | 0.3333 | 6 | 0.6667 | 13 | 0.4615 | 2 | 0.5 | 4 | 0.75 | 7 | 0.4286 | 4 | 0.5 | 17 | 0.5294 |
| 7_1439748 | 3 | 1 | 2 | 1 | 2 | 1 | 5 | 1 | 12 | 1 | 1 | 1 | 4 | 1 | 5 | 0.8 | 3 | 1 | 13 | 0.9231 |
| 7_1439748 | 5 | 1 | 3 | 1 | 4 | 1 | 7 | 1 | 19 | 1 | 3 | 1 | 5 | 1 | 9 | 0.6667 | 4 | 1 | 21 | 0.8571 |
| 7_1439750 | 5 | 0.2 | 1 | 0 | 3 | 0 | 5 | 0.4 | 14 | 0.2143 | 3 | 0.3333 | 1 | 1 | 6 | 0 | 3 | 0.6667 | 13 | 0.3077 |
| 7_1439753 | 7 | 0.1429 | 4 | 0.25 | 5 | 0.2 | 9 | 0.3333 | 25 | 0.24 | 4 | 0.25 | 5 | 0.4 | 10 | 0.1 | 4 | 0.5 | 23 | 0.2609 |
| 7_1439753 | 7 | 0 | 4 | 0 | 4 | 0.25 | 9 | 0.2222 | 24 | 0.125 | 5 | 0.4 | 5 | 0 | 8 | 0.375 | 3 | 0.6667 | 21 | 0.3333 |

| | | | | | | | | | | | | | | | | | | | | |
|---|---|---|---|---|---|---|---|---|---|---|---|---|---|---|---|---|---|---|---|---|
| 7_1439754 | 7 | 0.2857 | 4 | 0 | 5 | 0.2 | 8 | 0.25 | 24 | 0.2083 | 5 | 0.2 | 5 | 0 | 8 | 0.125 | 3 | 0 | 21 | 0.0952 |
| 7_1439756 | 8 | 0.125 | 4 | 0 | 5 | 0 | 8 | 0.125 | 25 | 0.08 | 5 | 0.2 | 5 | 0 | 8 | 0.25 | 4 | 0.25 | 22 | 0.1818 |
| 7_1439757 | 9 | 0.1111 | 3 | 0 | 6 | 0.1667 | 8 | 0.25 | 26 | 0.1538 | 4 | 0.25 | 4 | 0.25 | 10 | 0.1 | 4 | 0 | 22 | 0.1364 |
| 7_1439757 | 8 | 0 | 3 | 0 | 6 | 0 | 8 | 0.375 | 25 | 0.12 | 5 | 0 | 5 | 0.2 | 10 | 0.1 | 4 | 0 | 24 | 0.0833 |
| 7_1439759 | 8 | 0.25 | 5 | 0.2 | 7 | 0.2857 | 9 | 0.2222 | 29 | 0.2414 | 4 | 0.25 | 5 | 0.4 | 11 | 0.2727 | 4 | 0.25 | 24 | 0.2917 |
| 7_1439760 | 8 | 0.25 | 4 | 0 | 1 | 0 | 9 | 0.2222 | 22 | 0.1818 | 4 | 0.25 | 3 | 0 | 7 | 0 | 5 | 0 | 19 | 0.0526 |
| 7_1439765 | 6 | 0.1667 | 4 | 0 | 4 | 0 | 8 | 0.25 | 22 | 0.1364 | 6 | 0 | 6 | 0 | 8 | 0.125 | 5 | 0.4 | 25 | 0.12 |
| 7_1439773 | 9 | 0.1111 | 8 | 0 | 6 | 0.1667 | 10 | 0 | 33 | 0.0606 | 5 | 0 | 10 | 0.1 | 14 | 0.0714 | 4 | 0 | 33 | 0.0606 |
| 7_1439774 | 15 | 0.2 | 6 | 0.1667 | 7 | 0 | 9 | 0 | 37 | 0.1081 | 8 | 0.125 | 7 | 0 | 12 | 0.0833 | 5 | 0.2 | 32 | 0.0938 |
| 7_1439777 | 14 | 0 | 7 | 0 | 9 | 0.1111 | 10 | 0.1 | 40 | 0.05 | 9 | 0 | 9 | 0 | 13 | 0.3077 | 7 | 0.1429 | 38 | 0.1316 |
| 7_1439785 | 3 | 0.6667 | 2 | 0.5 | 5 | 0.8 | 1 | 0 | 11 | 0.6364 | NA | NA | NA | NA | 2 | 1 | 1 | 1 | 3 | 1 |
| 7_1439787 | 4 | 1 | 1 | 1 | 4 | 0.5 | 2 | 0.5 | 11 | 0.7273 | 2 | 1 | NA | NA | 2 | 1 | 1 | 1 | 5 | 1 |
| 7_1439790 | 4 | 1 | 2 | 0.5 | 4 | 0.75 | 1 | 0 | 11 | 0.7273 | 1 | 0 | 1 | 1 | 2 | 1 | 1 | 1 | 5 | 0.8 |
| 7_1439790 | 4 | 0.25 | 2 | 0 | 5 | 0.4 | 2 | 0 | 13 | 0.2308 | 2 | 0.5 | 1 | 0 | 2 | 0 | 1 | 0 | 6 | 0.1667 |
| 7_1439793 | 4 | 0.75 | 1 | 0 | 5 | 0 | 2 | 0.5 | 12 | 0.3333 | 2 | 0 | 1 | 0 | 2 | 0.5 | 1 | 0 | 6 | 0.1667 |
| 7_1439794 | 3 | 0 | 2 | 0 | 4 | 0.25 | 3 | 0 | 12 | 0.0833 | 2 | 0 | 2 | 0 | 1 | 1 | 2 | 0 | 7 | 0.1429 |
| 7_1439795 | 4 | 0.25 | 2 | 0.5 | 2 | 0 | 3 | 0.6667 | 11 | 0.3636 | 2 | 0 | NA | NA | 1 | 1 | 1 | 0 | 4 | 0.25 |
| 7_1439796 | 3 | 1 | 2 | 0 | 4 | 0.5 | 1 | 1 | 10 | 0.6 | 2 | 1 | 1 | 1 | NA | NA | NA | NA | 3 | 1 |
| 7_1439796 | 5 | 0.2 | 3 | 0 | 4 | 0 | 3 | 0.3333 | 15 | 0.1333 | 2 | 0.5 | 2 | 0 | 1 | 0 | 2 | 0 | 7 | 0.1429 |
| 7_1439905 | 7 | 0.1429 | 5 | 0.2 | 7 | 0 | 13 | 0.1538 | 32 | 0.125 | 7 | 0.4286 | 6 | 0 | 7 | 0 | 12 | 0.1667 | 32 | 0.1562 |
| 7_1440308 | 3 | 0 | 2 | 0.5 | 3 | 0 | 7 | 0 | 15 | 0.0667 | 3 | 0 | 2 | 0 | 2 | 0 | 2 | 0.5 | 9 | 0.1111 |
| 7_1440308 | 5 | 0.2 | 3 | 0.3333 | 3 | 0.3333 | 4 | 0 | 15 | 0.2 | 4 | 0 | 3 | 0.3333 | 1 | 1 | 4 | 0 | 12 | 0.1667 |
| 7_1440310 | 4 | 0.25 | 2 | 0.5 | 4 | 0 | 4 | 0.25 | 14 | 0.2143 | 4 | 0 | 1 | 0 | 1 | 0 | 4 | 0.5 | 10 | 0.2 |
| 7_1440311 | 5 | 0.4 | 3 | 0.6667 | 4 | 0.25 | 6 | 0.1667 | 18 | 0.3333 | 4 | 0.25 | 3 | 0 | 2 | 0 | 4 | 0.25 | 13 | 0.1538 |
| 7_1440311 | 4 | 0 | 2 | 0.5 | 4 | 0 | 4 | 0.25 | 14 | 0.1429 | 4 | 0.5 | 3 | 0 | 2 | 0.5 | 3 | 0 | 12 | 0.25 |
| 7_1440311 | 4 | 0 | 3 | 0.3333 | 4 | 0 | 6 | 0.1667 | 17 | 0.1176 | 4 | 0 | 2 | 0.5 | 2 | 0.5 | 4 | 0.25 | 12 | 0.25 |
| 7_1440315 | 3 | 0.6667 | 2 | 0.5 | 4 | 0 | 6 | 0 | 15 | 0.2 | 4 | 0 | 3 | 0.3333 | 2 | 0.5 | 2 | 0.5 | 11 | 0.2727 |
| 7_1440349 | NA | NA | 1 | 0 | 3 | 0 | 3 | 0.3333 | 7 | 0.1429 | 3 | 0.3333 | 2 | 0 | 2 | 0 | 2 | 0 | 9 | 0.1111 |
| 7_1441330 | NA | NA | NA | NA | NA | NA | 1 | 1 | 1 | 1 | 1 | 1 | 1 | 1 | NA | NA | NA | NA | 2 | 1 |
| 7_1441337 | NA | NA | 1 | 1 | NA | NA | 2 | 1 | 3 | 1 | 1 | 1 | NA | NA | NA | NA | NA | NA | 1 | 1 |
| 7_1441338 | NA | NA | 1 | 1 | NA | NA | 2 | 1 | 3 | 1 | 1 | 1 | NA | NA | NA | NA | NA | NA | 1 | 1 |
| 7_1441338 | NA | NA | 1 | 1 | NA | NA | 2 | 1 | 3 | 1 | 1 | 1 | NA | NA | NA | NA | NA | NA | 1 | 1 |
| 7_1441339 | NA | NA | 1 | 1 | NA | NA | 2 | 0.5 | 3 | 0.6667 | 1 | 1 | NA | NA | NA | NA | NA | NA | 1 | 1 |
| 7_1441341 | NA | NA | 1 | 1 | NA | NA | 2 | 0.5 | 3 | 0.6667 | 1 | 1 | NA | NA | NA | NA | NA | NA | 1 | 1 |
| 7_1441342 | NA | NA | 1 | 1 | NA | NA | 2 | 1 | 3 | 1 | 1 | 1 | NA | NA | NA | NA | NA | NA | 1 | 1 |
| 7_1444735 | 1 | 0 | 1 | 1 | 1 | 0 | 2 | 0 | 5 | 0.2 | 1 | 0 | 2 | 0.5 | NA | NA | 1 | 0 | 4 | 0.25 |
| 7_1444828 | NA | NA | NA | NA | NA | NA | 1 | 1 | 1 | 1 | NA | NA | 2 | 1 | NA | NA | 1 | 1 | 3 | 1 |
| 7_1444832 | NA | NA | NA | NA | NA | NA | 1 | 1 | 1 | 1 | NA | NA | 2 | 1 | NA | NA | 1 | 1 | 3 | 1 |
| 7_1444840 | 6 | 0 | 4 | 0.25 | 6 | 0 | 4 | 0.25 | 20 | 0.1 | 6 | 0 | 3 | 0.3333 | 8 | 0.125 | 5 | 0 | 22 | 0.0909 |
| 7_1444842 | 6 | 0 | 2 | 0 | 5 | 0.2 | 5 | 0 | 18 | 0.0556 | 5 | 0.2 | 3 | 0.3333 | 5 | 0 | 4 | 0 | 17 | 0.1176 |
| 7_1444847 | 6 | 0.1667 | 4 | 0 | 4 | 0 | 4 | 0.25 | 18 | 0.1111 | 5 | 0 | 3 | 0 | 4 | 0 | 4 | 0.25 | 16 | 0.0625 |
| 7_1444863 | 2 | 0.5 | 2 | 0 | 1 | 0 | 3 | 0.3333 | 8 | 0.25 | 2 | 1 | NA | NA | 1 | 0 | 1 | 1 | 4 | 0.75 |
| 8_3201068 | 8 | 0.25 | 6 | 0 | 4 | 0.25 | 3 | 0 | 21 | 0.1429 | 6 | 0.5 | 1 | 1 | 1 | 0 | 1 | 0 | 9 | 0.4444 |
| 8_3201571 | 3 | 0 | 5 | 0.2 | 4 | 0 | 2 | 0 | 14 | 0.0714 | 4 | 0.25 | NA | NA | 2 | 0.5 | 1 | 0 | 7 | 0.2857 |
| 8_3201666 | 5 | 0.2 | 5 | 0.2 | 3 | 0 | 2 | 0 | 15 | 0.1333 | 4 | 0 | NA | NA | 1 | 1 | NA | NA | 5 | 0.2 |
| 8_3202437 | 4 | 0 | 5 | 0 | 4 | 0.75 | 1 | 0 | 14 | 0.2143 | 2 | 0.5 | NA | NA | NA | NA | 1 | 0 | 3 | 0.3333 |
| 8_3203555 | 3 | 0.3333 | 4 | 0.5 | 6 | 0.8333 | 3 | 1 | 16 | 0.6875 | 4 | 0.25 | 2 | 1 | 4 | 1 | 2 | 0 | 12 | 0.5833 |
| 8_3203639 | 3 | 0.3333 | 4 | 0.25 | 3 | 0.3333 | 5 | 0.4 | 15 | 0.3333 | 4 | 0.5 | 2 | 0 | 3 | 0.3333 | 3 | 0.6667 | 12 | 0.4167 |
| 8_3203663 | 5 | 0.6 | 3 | 0.3333 | 5 | 1 | 2 | 0.5 | 15 | 0.6667 | 5 | 0.6 | 2 | 0.5 | 4 | 0.75 | 3 | 0.6667 | 14 | 0.6429 |
| 8_3203752 | 5 | 0.6 | 3 | 1 | 2 | 1 | 3 | 1 | 13 | 0.8462 | 3 | 1 | 2 | 1 | 4 | 1 | 2 | 0.5 | 11 | 0.9091 |
| 8_3203813 | 5 | 0.6 | 4 | 0.75 | 5 | 0.8 | 3 | 0.3333 | 17 | 0.6471 | 3 | 0.3333 | 2 | 1 | 4 | 0.75 | 3 | 0.6667 | 12 | 0.6667 |
| 8_3203831 | 5 | 0.2 | 3 | 0.3333 | 2 | 0.5 | 4 | 0.25 | 14 | 0.2857 | 4 | 0.25 | 1 | 0 | 1 | 0 | 3 | 1 | 9 | 0.4444 |
| 8_3203852 | 6 | 1 | 4 | 1 | 5 | 1 | 4 | 1 | 19 | 1 | 5 | 0.8 | 2 | 1 | 4 | 1 | 3 | 0.6667 | 14 | 0.8571 |
| 8_3203881 | 6 | 0.8333 | 4 | 1 | 5 | 0.8 | 4 | 0.75 | 19 | 0.8421 | 5 | 0.8 | 2 | 1 | 4 | 0.75 | 3 | 0.6667 | 14 | 0.7857 |
| 8_3203948 | 5 | 0.6 | 4 | 0.75 | 3 | 0.6667 | 5 | 0.6 | 17 | 0.6471 | 5 | 0.6 | 2 | 1 | 4 | 0.75 | 2 | 1 | 13 | 0.7692 |
| 8_3204064 | 5 | 0.8 | 4 | 0.25 | 4 | 0.25 | 4 | 0 | 17 | 0.3529 | 4 | 0.25 | 1 | 1 | 3 | 0.3333 | 2 | 0.5 | 10 | 0.4 |
| 8_3204073 | 3 | 1 | 3 | 1 | 5 | 1 | 2 | 1 | 13 | 1 | 5 | 1 | 2 | 1 | 4 | 1 | 3 | 1 | 14 | 1 |
| 8_3204105 | 5 | 0.2 | 3 | 0 | 4 | 0 | 4 | 0 | 16 | 0.0625 | 4 | 0 | 2 | 0 | 3 | 0.3333 | 3 | 0 | 12 | 0.0833 |
| 8_3204253 | 3 | 0.3333 | 4 | 0 | 4 | 0.5 | 4 | 0.25 | 15 | 0.2667 | 5 | 0.4 | 2 | 1 | 2 | 0 | 2 | 0.5 | 11 | 0.4545 |
| 8_3204318 | 4 | 1 | 4 | 0.25 | 5 | 0.2 | 5 | 0.2 | 18 | 0.3889 | 4 | 0.5 | 2 | 1 | 3 | 0.6667 | 3 | 0 | 12 | 0.5 |
| 8_3204748 | 4 | 0.25 | 1 | 1 | 3 | 1 | 2 | 0.5 | 10 | 0.6 | 2 | 0 | 1 | 0 | 3 | 0.3333 | 2 | 0 | 8 | 0.125 |
| 8_3204828 | 3 | 0.3333 | 2 | 0 | 5 | 0.8 | 3 | 0.6667 | 13 | 0.5385 | 2 | 0 | 1 | 1 | 3 | 1 | 2 | 1 | 8 | 0.75 |
| 8_3205576 | 1 | 1 | 1 | 1 | NA | NA | 1 | 1 | 3 | 1 | 2 | 1 | NA | NA | NA | NA | 1 | 1 | 3 | 1 |
| 8_3205622 | 1 | 0 | 1 | 0 | NA | NA | 1 | 1 | 3 | 0.3333 | 2 | 0 | NA | NA | NA | NA | 1 | 1 | 3 | 0.3333 |
| 8_3205758 | 1 | 1 | 1 | 0 | NA | NA | 1 | 1 | 3 | 0.6667 | 2 | 1 | NA | NA | NA | NA | 1 | 1 | 3 | 1 |

| | | | | | | | | | | | | | | | | | | | | |
|---|---|---|---|---|---|---|---|---|---|---|---|---|---|---|---|---|---|---|---|---|
| 8_3502045 | 2 | 0 | 3 | 0.3333 | 2 | 0 | NA | NA | 7 | 0.1429 | NA | NA | 3 | 0 | 1 | 0 | 3 | 0.3333 | 7 | 0.1429 |
| 8_3502197 | 3 | 0 | 3 | 0.3333 | 2 | 0 | NA | NA | 8 | 0.125 | NA | NA | 2 | 0 | 1 | 0 | 2 | 0.5 | 5 | 0.2 |
| 8_3504607 | 6 | 0.5 | 2 | 0.5 | 4 | 0 | 4 | 0.5 | 16 | 0.375 | 4 | 0.25 | 5 | 0.8 | 1 | 1 | 2 | 1 | 12 | 0.6667 |
| 8_3504976 | 6 | 0.5 | 4 | 0 | 3 | 0.3333 | 2 | 0 | 15 | 0.2667 | 6 | 0.1667 | 4 | 0 | 1 | 0 | 2 | 0.5 | 13 | 0.1538 |
| 8_3505109 | 6 | 0.5 | 5 | 0.2 | 3 | 0.6667 | 3 | 0.3333 | 17 | 0.4118 | 5 | 0.6 | 4 | 0.25 | 1 | 0 | 2 | 0.5 | 12 | 0.4167 |
| 8_3505144 | 6 | 1 | 3 | 0.6667 | 4 | 0.75 | 3 | 1 | 16 | 0.875 | 6 | 0.8333 | 4 | 0.75 | NA | NA | 2 | 1 | 12 | 0.8333 |
| 8_3505393 | 7 | 0.2857 | 4 | 0.5 | 4 | 0 | 2 | 0.5 | 17 | 0.2941 | 5 | 0.4 | 5 | 0 | 2 | 0.5 | 1 | 0 | 13 | 0.2308 |
| 8_3505416 | 7 | 0.1429 | 5 | 0 | 4 | 0.25 | 2 | 0.5 | 18 | 0.1667 | 5 | 0 | 5 | 0 | 2 | 0.5 | 1 | 0 | 13 | 0.0769 |
| 8_3506374 | 7 | 0 | 5 | 0.2 | 7 | 0.2857 | 7 | 0.2857 | 26 | 0.1923 | 5 | 0.4 | 6 | 0 | 2 | 0 | 1 | 0 | 14 | 0.1429 |
| 8_3545481 | 1 | 1 | 2 | 0.5 | 4 | 0 | 3 | 0.3333 | 10 | 0.3 | 1 | 0 | 2 | 0 | 2 | 0.5 | 1 | 0 | 6 | 0.1667 |
| 8_3545508 | 1 | 0 | 1 | 0 | 4 | 0.5 | 2 | 0.5 | 8 | 0.375 | 1 | 0 | 2 | 0.5 | 2 | 0 | NA | NA | 5 | 0.2 |
| 8_3565071 | 9 | 0.4444 | 7 | 0.1429 | 3 | 0.6667 | 10 | 0.1 | 29 | 0.2759 | 2 | 0 | 6 | 0.5 | 8 | 0.375 | 6 | 0.3333 | 22 | 0.3636 |
| 8_3565178 | 11 | 0.8182 | 6 | 1 | 4 | 1 | 11 | 0.9091 | 32 | 0.9062 | 3 | 1 | 6 | 1 | 11 | 0.8182 | 7 | 1 | 27 | 0.9259 |
| 8_3594167 | 3 | 0.6667 | 7 | 0.4286 | 1 | 1 | 3 | 0.3333 | 14 | 0.5 | 3 | 0.3333 | 2 | 0.5 | NA | NA | 1 | 1 | 6 | 0.5 |
| 8_3654491 | 2 | 1 | 3 | 1 | 1 | 1 | 1 | 1 | 7 | 1 | 1 | 1 | NA | NA | NA | NA | NA | NA | 1 | 1 |
| 8_3654571 | 1 | 0 | 4 | 0.25 | 1 | 0 | 1 | 1 | 7 | 0.2857 | 1 | 1 | NA | NA | NA | NA | NA | NA | 1 | 1 |
| 8_3654820 | 2 | 0.5 | 2 | 0 | 1 | 0 | 1 | 0 | 6 | 0.1667 | 1 | 1 | NA | NA | NA | NA | NA | NA | 1 | 1 |
| 8_3654870 | 2 | 0.5 | 3 | 0.6667 | 1 | 1 | NA | NA | 6 | 0.6667 | 1 | 1 | NA | NA | NA | NA | NA | NA | 1 | 1 |
| 8_3658843 | 6 | 0.1667 | 4 | 0 | 3 | 0.3333 | 3 | 0 | 16 | 0.125 | 2 | 0 | 2 | 0.5 | 1 | 0 | 2 | 0.5 | 7 | 0.2857 |
| 8_3663602 | 3 | 0 | 5 | 0.2 | NA | NA | 4 | 0 | 12 | 0.0833 | 4 | 0 | 4 | 0 | 3 | 0.3333 | 1 | 0 | 12 | 0.0833 |
| 8_3673435 | 1 | 0 | 1 | 0 | 2 | 0.5 | NA | NA | 4 | 0.25 | 2 | 0.5 | 3 | 0 | NA | NA | 2 | 0.5 | 7 | 0.2857 |
| 8_3674073 | 1 | 1 | 1 | 0 | 2 | 0 | NA | NA | 4 | 0.25 | 2 | 0 | 3 | 0.3333 | NA | NA | 2 | 0 | 7 | 0.1429 |
| 8_3721276 | 2 | 1 | 3 | 0.3333 | 2 | 0.5 | 1 | 0 | 8 | 0.5 | 2 | 0 | 3 | 0 | 1 | 1 | 2 | 1 | 8 | 0.375 |
| 8_3721342 | 2 | 1 | 3 | 0 | 2 | 0.5 | 1 | 0 | 8 | 0.375 | 2 | 0.5 | 2 | 0 | 2 | 0.5 | 1 | 0 | 7 | 0.2857 |
| 8_3729016 | 6 | 0.1667 | 6 | 0.1667 | 4 | 0.25 | 8 | 0.125 | 24 | 0.1667 | 3 | 0 | 6 | 0.1667 | 8 | 0.125 | 5 | 0 | 22 | 0.0909 |
| 8_3729261 | 7 | 1 | 8 | 0.375 | 7 | 0.5714 | 7 | 1 | 29 | 0.7241 | 4 | 0.75 | 8 | 0.625 | 9 | 0.2222 | 5 | 0.4 | 26 | 0.4615 |
| 8_3729331 | 6 | 0.5 | 6 | 0.3333 | 6 | 0.5 | 7 | 0.2857 | 25 | 0.4 | 5 | 0.2 | 8 | 0.125 | 9 | 0.2222 | 4 | 0.25 | 26 | 0.1923 |
| 8_3729436 | 5 | 1 | 3 | 1 | 3 | 1 | 7 | 1 | 18 | 1 | 3 | 1 | 6 | 1 | 9 | 1 | 4 | 1 | 22 | 1 |
| 8_3729525 | 7 | 0.7143 | 6 | 0.6667 | 5 | 0.2 | 8 | 0.875 | 26 | 0.6538 | 4 | 0.25 | 8 | 0.5 | 12 | 0.4167 | 4 | 0.5 | 28 | 0.4286 |
| 8_3730112 | 5 | 0.4 | 3 | 0 | 7 | 0 | 5 | 0.2 | 20 | 0.15 | 3 | 0 | 6 | 0.1667 | 7 | 0.2857 | 2 | 0 | 18 | 0.1667 |
| 8_3730257 | 8 | 1 | 3 | 1 | 7 | 1 | 4 | 1 | 22 | 1 | 4 | 1 | 5 | 0.8 | 7 | 1 | 2 | 0.5 | 18 | 0.8889 |
| 8_3730359 | 7 | 0 | 3 | 0.3333 | 7 | 0.1429 | 4 | 0 | 21 | 0.0952 | 3 | 0 | 5 | 0.2 | 7 | 0 | 2 | 0 | 17 | 0.0588 |
| 8_3730404 | 6 | 0.8333 | 3 | 0.6667 | 6 | 0.8333 | 3 | 0.6667 | 18 | 0.7778 | 3 | 0.6667 | 5 | 0.8 | 6 | 1 | 2 | 0 | 16 | 0.75 |
| 8_3730438 | 7 | 0 | 3 | 0 | 7 | 0 | 2 | 0.5 | 19 | 0.0526 | 3 | 0 | 4 | 0 | 6 | 0.1667 | 2 | 0 | 15 | 0.0667 |
| 8_3730446 | 9 | 0 | 3 | 0.3333 | 7 | 0.1429 | 4 | 0 | 23 | 0.087 | 2 | 0.5 | 5 | 0.2 | 5 | 0 | 2 | 0 | 14 | 0.1429 |
| 8_3730473 | 6 | 0.8333 | 2 | 0.5 | 6 | 0.5 | 3 | 0.6667 | 17 | 0.6471 | 3 | 0.3333 | 5 | 0.6 | 4 | 0.75 | 2 | 0.5 | 14 | 0.5714 |
| 8_3736259 | NA | NA | NA | NA | 1 | 1 | 1 | 0 | 2 | 0.5 | 1 | 1 | NA | NA | NA | NA | NA | NA | 1 | 1 |
| 8_3736305 | NA | NA | NA | NA | 1 | 1 | 1 | 1 | 2 | 1 | 1 | 1 | NA | NA | NA | NA | NA | NA | 1 | 1 |
| 8_3736343 | NA | NA | NA | NA | 1 | 1 | NA | NA | 1 | 1 | 1 | 1 | NA | NA | NA | NA | NA | NA | 1 | 1 |
| 8_3736386 | NA | NA | NA | NA | 1 | 1 | 1 | 1 | 2 | 1 | 1 | 1 | NA | NA | NA | NA | NA | NA | 1 | 1 |
| 8_4256710 | 4 | 0.25 | 9 | 0.3333 | 2 | 0.5 | 4 | 0.75 | 19 | 0.4211 | 3 | 0.6667 | 4 | 0.5 | 5 | 0.4 | 5 | 0.4 | 17 | 0.4706 |
| 8_4256744 | 3 | 0.3333 | 9 | 0.7778 | 3 | 0.3333 | 4 | 0.25 | 19 | 0.5263 | 3 | 1 | 1 | 0 | 5 | 0.8 | 3 | 0.6667 | 12 | 0.75 |
| 8_4256776 | 2 | 0.5 | 7 | 0.7143 | 2 | 0.5 | 5 | 0.8 | 16 | 0.6875 | 3 | 0.6667 | 3 | 1 | 6 | 0.8333 | 3 | 0.3333 | 15 | 0.7333 |
| 8_4257066 | 2 | 1 | 8 | 0.5 | 4 | 0.5 | 4 | 0.75 | 18 | 0.6111 | 1 | 0 | 5 | 0.2 | 6 | 0.5 | 6 | 0.5 | 18 | 0.3889 |
| 8_4257095 | 4 | 1 | 10 | 0.8 | 5 | 0.6 | 5 | 0.6 | 24 | 0.75 | 3 | 1 | 6 | 0.5 | 8 | 1 | 6 | 0.8333 | 23 | 0.8261 |
| 8_4257145 | 2 | 0 | 9 | 0.1111 | 4 | 0.25 | 5 | 0.2 | 20 | 0.15 | 3 | 0 | 3 | 0 | 6 | 0 | 5 | 0.2 | 17 | 0.0588 |
| 8_4257177 | 3 | 0 | 9 | 0.2222 | 5 | 0.2 | 5 | 0 | 22 | 0.1364 | 3 | 0 | 5 | 0 | 8 | 0.25 | 6 | 0.1667 | 22 | 0.1364 |
| 8_4257209 | 5 | 0.6 | 11 | 0.8182 | 5 | 1 | 5 | 1 | 26 | 0.8462 | 2 | 1 | 5 | 0.2 | 6 | 0.8333 | 5 | 1 | 18 | 0.7222 |
| 8_4257230 | 4 | 0.25 | 7 | 0 | 4 | 0 | 5 | 0 | 20 | 0.05 | 4 | 0 | 4 | 0.25 | 7 | 0.1429 | 5 | 0 | 20 | 0.1 |
| 8_4257243 | 5 | 0 | 9 | 0.4444 | 4 | 0.25 | 5 | 0.2 | 23 | 0.2609 | 3 | 0.3333 | 5 | 0 | 7 | 0.7143 | 6 | 0.1667 | 21 | 0.3333 |
| 8_4257282 | 5 | 0 | 7 | 0.2857 | 5 | 0 | 3 | 0 | 20 | 0.1 | 4 | 0 | 5 | 0 | 6 | 0 | 5 | 0.2 | 20 | 0.05 |
| 8_4257299 | 5 | 0.6 | 11 | 0.8182 | 4 | 0.5 | 4 | 0.75 | 24 | 0.7083 | 3 | 0.6667 | 4 | 0.5 | 6 | 1 | 7 | 0.8571 | 20 | 0.8 |
| 8_4257372 | 1 | 0 | 7 | 0 | 4 | 0.25 | 5 | 0 | 17 | 0.0588 | 2 | 0 | 6 | 0 | 4 | 0 | 7 | 0.2857 | 19 | 0.1053 |
| 8_4257589 | 4 | 0 | 8 | 0.125 | 7 | 0.4286 | 5 | 0.2 | 24 | 0.2083 | 4 | 0.25 | 7 | 0 | 7 | 0.1429 | 6 | 0 | 24 | 0.0833 |
| 8_4259758 | 1 | 1 | 3 | 0 | 1 | 0 | 2 | 0 | 7 | 0.1429 | 1 | 0 | 3 | 0 | 2 | 0 | 3 | 0.3333 | 9 | 0.1111 |
| 8_4259878 | 2 | 0 | 2 | 0.5 | NA | NA | 2 | 1 | 6 | 0.5 | 1 | 0 | 2 | 0.5 | 3 | 0.3333 | 2 | 1 | 8 | 0.5 |
| 8_4259978 | 2 | 1 | 3 | 0.3333 | NA | NA | 2 | 1 | 7 | 0.7143 | NA | NA | 2 | 0.5 | 3 | 0.3333 | 3 | 0.6667 | 8 | 0.5 |
| 8_4260011 | 1 | 1 | 2 | 1 | NA | NA | 2 | 1 | 5 | 1 | NA | NA | 1 | 1 | 1 | 1 | 2 | 1 | 4 | 1 |
| 8_4260076 | 2 | 1 | 3 | 1 | NA | NA | 2 | 1 | 7 | 1 | NA | NA | 2 | 1 | 3 | 1 | 3 | 1 | 8 | 1 |
| 8_4260121 | 2 | 0.5 | 2 | 0.5 | NA | NA | 2 | 0.5 | 6 | 0.5 | 1 | 0 | 3 | 0.6667 | 2 | 0 | 2 | 0.5 | 8 | 0.375 |
| 8_4260129 | 2 | 0.5 | 1 | 0 | NA | NA | 3 | 0.6667 | 6 | 0.5 | NA | NA | 3 | 0.6667 | 2 | 1 | 3 | 0.6667 | 8 | 0.75 |
| 8_4260204 | 2 | 0.5 | 2 | 0 | NA | NA | 1 | 0 | 5 | 0.2 | NA | NA | 3 | 0 | 3 | 0.3333 | 1 | 0 | 7 | 0.1429 |
| 8_4260530 | 2 | 0.5 | 2 | 1 | NA | NA | 1 | 0 | 5 | 0.6 | NA | NA | 3 | 0.3333 | 2 | 0.5 | 3 | 0.3333 | 8 | 0.375 |
| 8_4260548 | 1 | 1 | 2 | 1 | NA | NA | 1 | 1 | 4 | 1 | NA | NA | 3 | 0.6667 | 2 | 1 | 2 | 1 | 7 | 0.8571 |
| 8_4260770 | 1 | 1 | 1 | 1 | NA | NA | 1 | 1 | 3 | 1 | NA | NA | NA | NA | 2 | 1 | 1 | 1 | 3 | 1 |

| | | | | | | | | | | | | | | | | | | | | |
|---|---|---|---|---|---|---|---|---|---|---|---|---|---|---|---|---|---|---|---|---|
| 8_4295785 | 3 | 0 | 4 | 0 | 5 | 0 | 2 | 0.5 | 14 | 0.0714 | 3 | 0.3333 | 4 | 0 | 6 | 0 | 1 | 0 | 14 | 0.0714 |
| 8_4295945 | 3 | 0.6667 | 3 | 0.6667 | 4 | 0.75 | 2 | 1 | 12 | 0.75 | 2 | 0.5 | 4 | 1 | 6 | 0.8333 | NA | NA | 12 | 0.8333 |
| 8_4295952 | 1 | 0 | 3 | 0.6667 | 3 | 0 | 1 | 0 | 8 | 0.25 | 2 | 1 | 2 | 0.5 | 4 | 0.5 | 1 | 1 | 9 | 0.6667 |
| 8_4296005 | 2 | 0.5 | 3 | 0.3333 | 5 | 0.4 | 1 | 0 | 11 | 0.3636 | 3 | 0.6667 | 4 | 0.75 | 5 | 0.2 | 1 | 1 | 13 | 0.5385 |
| 8_4296082 | 2 | 0 | 2 | 0 | 4 | 0.25 | 2 | 0 | 10 | 0.1 | 3 | 0.6667 | 4 | 0 | 6 | 0 | 1 | 0 | 14 | 0.1429 |
| 8_4296116 | 3 | 0.6667 | 3 | 0.3333 | 4 | 0 | 2 | 0.5 | 12 | 0.3333 | 3 | 0.3333 | 4 | 0 | 6 | 0.3333 | 1 | 0 | 14 | 0.2143 |
| 8_4296636 | 3 | 0.3333 | 3 | 0.3333 | 5 | 0.8 | 4 | 0 | 15 | 0.4 | 3 | 0 | 5 | 0.2 | 5 | 0.4 | 2 | 0 | 15 | 0.2 |
| 8_4297086 | 1 | 0 | 3 | 0 | 5 | 0.2 | 3 | 0.3333 | 12 | 0.1667 | 2 | 0 | 4 | 0 | 4 | 0.25 | 4 | 0 | 14 | 0.0714 |
| 8_4305302 | 6 | 0.3333 | 4 | 0.5 | 3 | 0 | 6 | 0.1667 | 19 | 0.2632 | 5 | 0.2 | NA | NA | 6 | 0.3333 | 4 | 0.25 | 15 | 0.2667 |
| 8_4305524 | 4 | 0.25 | 4 | 0 | 3 | 0 | 6 | 0 | 17 | 0.0588 | 5 | 0.2 | NA | NA | 5 | 0.4 | 4 | 0.25 | 14 | 0.2857 |
| 8_4305633 | 5 | 0.6 | 5 | 0.4 | 3 | 0 | 5 | 0.4 | 18 | 0.3889 | 7 | 0.4286 | NA | NA | 6 | 0.6667 | 4 | 0.5 | 17 | 0.5294 |
| 8_4305648 | 6 | 0.6667 | 5 | 0.6 | 3 | 0.6667 | 5 | 0.6 | 19 | 0.6316 | 5 | 0.8 | NA | NA | 7 | 0.5714 | 4 | 0.5 | 16 | 0.625 |
| 8_4305873 | 6 | 0.1667 | 5 | 0.2 | 3 | 0 | 7 | 0 | 21 | 0.0952 | 7 | 0 | NA | NA | 6 | 0.3333 | 5 | 0.6 | 18 | 0.2778 |
| 8_4305885 | 4 | 1 | 5 | 1 | 3 | 1 | 5 | 1 | 17 | 1 | 8 | 1 | NA | NA | 7 | 1 | 4 | 1 | 19 | 1 |
| 8_4306151 | 6 | 0.5 | 6 | 0.8333 | 3 | 0.6667 | 7 | 0.2857 | 22 | 0.5455 | 8 | 0.5 | 1 | 1 | 5 | 0.8 | 7 | 0.5714 | 21 | 0.619 |
| 8_4307910 | 99 | 0.1616 | 90 | 0.0444 | 79 | 0.0759 | 91 | 0.1319 | 359 | 0.1058 | 69 | 0.058 | 100 | 0.15 | 90 | 0.0889 | 64 | 0.0781 | 323 | 0.0991 |
| 8_4308051 | 99 | 0.3939 | 84 | 0.2262 | 74 | 0.2162 | 90 | 0.3889 | 347 | 0.3141 | 62 | 0.2258 | 94 | 0.2766 | 81 | 0.284 | 60 | 0.4167 | 297 | 0.2963 |
| 8_4308115 | 99 | 0.0505 | 85 | 0.1176 | 73 | 0.0959 | 93 | 0.086 | 350 | 0.0857 | 64 | 0.0469 | 96 | 0.0417 | 88 | 0.0568 | 67 | 0.0597 | 315 | 0.0508 |
| 8_4308319 | 78 | 0.9359 | 67 | 0.9254 | 61 | 0.9836 | 68 | 0.9265 | 274 | 0.9416 | 47 | 0.9149 | 77 | 0.9351 | 71 | 0.9296 | 53 | 1 | 248 | 0.9435 |
| 8_4308434 | 72 | 0.1667 | 71 | 0.1972 | 65 | 0.0615 | 71 | 0.1972 | 279 | 0.1577 | 49 | 0.1429 | 79 | 0.1519 | 62 | 0.1613 | 54 | 0.1667 | 244 | 0.1557 |
| 8_4309903 | 6 | 0.1667 | 2 | 0.5 | 9 | 0.3333 | 9 | 0.2222 | 26 | 0.2692 | 7 | 0 | 1 | 1 | 5 | 0.4 | 8 | 0.375 | 21 | 0.2857 |
| 8_4309964 | 8 | 0.375 | 2 | 0.5 | 5 | 0.8 | 9 | 0.7778 | 24 | 0.625 | 9 | 0.2222 | 1 | 1 | 6 | 0.6667 | 7 | 0.4286 | 23 | 0.4348 |
| 8_4310000 | 6 | 0.3333 | 2 | 0 | 2 | 0.5 | 10 | 0.3 | 20 | 0.3 | 8 | 0.25 | 1 | 0 | 7 | 0 | 5 | 0.2 | 21 | 0.1429 |
| 8_4310076 | 8 | 0.625 | 3 | 0.6667 | 7 | 0.2857 | 10 | 0.6 | 28 | 0.5357 | 8 | 0.5 | 2 | 0.5 | 7 | 0.4286 | 8 | 0.625 | 25 | 0.52 |
| 8_4310090 | 6 | 0.1667 | 2 | 0.5 | 5 | 0.2 | 8 | 0.25 | 21 | 0.2381 | 8 | 0.25 | 2 | 0 | 6 | 0.3333 | 8 | 0.25 | 24 | 0.25 |
| 8_4310170 | 6 | 0 | NA | NA | 5 | 0.2 | 6 | 0 | 17 | 0.0588 | 7 | 0 | 1 | 0 | 4 | 0 | 5 | 0.2 | 17 | 0.0588 |
| 8_4335694 | 6 | 0.3333 | 8 | 0.25 | 5 | 0.2 | 10 | 0.2 | 29 | 0.2414 | 8 | 0.25 | 9 | 0.5556 | 7 | 0.2857 | 6 | 0.3333 | 30 | 0.3667 |
| 8_4338548 | 6 | 0.1667 | 3 | 0.3333 | 6 | 0.3333 | 9 | 0 | 24 | 0.1667 | 2 | 0 | 7 | 0.2857 | 7 | 0.4286 | 3 | 0.3333 | 19 | 0.3158 |
| 8_4338910 | 5 | 0.2 | 3 | 0 | 5 | 0.2 | 9 | 0 | 22 | 0.0909 | 3 | 0 | 7 | 0.1429 | 7 | 0.2857 | 4 | 0.25 | 21 | 0.1905 |
| 8_4339206 | 6 | 0.5 | NA | NA | 7 | 0.1429 | 8 | 0.25 | 21 | 0.2857 | 4 | 0 | 6 | 0.1667 | 5 | 0.2 | 2 | 0 | 17 | 0.1176 |
| 8_4339242 | 6 | 0.6667 | NA | NA | 5 | 0.6 | 6 | 0.5 | 17 | 0.5882 | 4 | 0.75 | 5 | 0.8 | 5 | 0.4 | 3 | 1 | 17 | 0.7059 |
| 8_4339267 | 5 | 0.2 | NA | NA | 6 | 0.5 | 7 | 0.1429 | 18 | 0.2778 | 3 | 0 | 6 | 0.5 | 6 | 0.6667 | 3 | 0.3333 | 18 | 0.4444 |
| 8_4339305 | 5 | 0.4 | NA | NA | 7 | 0 | 7 | 0.2857 | 19 | 0.2105 | 3 | 0 | 6 | 0.6667 | 4 | 0 | 3 | 0.3333 | 16 | 0.3125 |
| 8_4339600 | 3 | 0 | NA | NA | 4 | 0 | 7 | 0.1429 | 14 | 0.0714 | 2 | 0 | 2 | 0 | 4 | 0 | 3 | 0.3333 | 11 | 0.0909 |
| 8_4339646 | 4 | 0.5 | NA | NA | 5 | 0.2 | 7 | 0.4286 | 16 | 0.375 | 3 | 0.6667 | 4 | 0.25 | 3 | 0.6667 | 3 | 0.3333 | 13 | 0.4615 |
| 8_4339700 | 5 | 0.2 | NA | NA | 5 | 0 | 7 | 0.4286 | 17 | 0.2353 | 3 | 0 | 4 | 0.5 | 3 | 0.3333 | 2 | 0.5 | 12 | 0.3333 |
| 8_4339710 | 5 | 0 | NA | NA | 5 | 0.2 | 7 | 0.1429 | 17 | 0.1176 | 3 | 0.3333 | 4 | 0 | 4 | 0 | 2 | 0 | 13 | 0.0769 |
| 8_4409599 | 1 | 1 | 1 | 0 | NA | NA | 2 | 1 | 4 | 0.75 | 1 | 0 | NA | NA | 1 | 1 | 2 | 0.5 | 4 | 0.5 |
| 8_4573791 | 2 | 1 | NA | NA | 1 | 1 | 3 | 0.6667 | 6 | 0.8333 | NA | NA | 2 | 0 | 1 | 1 | 2 | 1 | 5 | 0.6 |
| 8_4573888 | 2 | 0 | 1 | 1 | 2 | 1 | 3 | 1 | 8 | 0.75 | 1 | 0 | 2 | 0.5 | 2 | 0 | 3 | 0.6667 | 8 | 0.375 |
| 8_4573947 | 2 | 0 | 1 | 0 | 1 | 0 | 3 | 0.6667 | 7 | 0.2857 | NA | NA | 3 | 0 | 2 | 0 | 2 | 0.5 | 7 | 0.1429 |
| 8_4573964 | 1 | 1 | 1 | 1 | 2 | 1 | 2 | 1 | 6 | 1 | 1 | 1 | 2 | 0.5 | 2 | 1 | 3 | 1 | 8 | 0.875 |
| 8_4574032 | 2 | 0 | 1 | 1 | 2 | 1 | 3 | 1 | 8 | 0.75 | 1 | 1 | 3 | 0 | 1 | 0 | 3 | 0.6667 | 8 | 0.375 |
| 8_4574588 | 2 | 0 | 1 | 0 | 2 | 0.5 | 4 | 0.25 | 9 | 0.2222 | 1 | 0 | 1 | 0 | NA | NA | 2 | 0.5 | 4 | 0.25 |
| 8_4575406 | 1 | 1 | 1 | 0 | NA | NA | 3 | 0 | 5 | 0.2 | 3 | 0.3333 | NA | NA | NA | NA | NA | NA | 3 | 0.3333 |
| 8_4678402 | NA | NA | 1 | 1 | 1 | 0 | NA | NA | 2 | 0.5 | NA | NA | NA | NA | 5 | 0.4 | 1 | 1 | 6 | 0.5 |
| 8_4678430 | NA | NA | 1 | 1 | 1 | 1 | 1 | 1 | 3 | 1 | 1 | 1 | NA | NA | 4 | 1 | 1 | 1 | 6 | 1 |
| 8_4678514 | NA | NA | 1 | 1 | 1 | 1 | 1 | 0 | 3 | 0.6667 | 1 | 1 | NA | NA | 4 | 1 | 1 | 1 | 6 | 1 |
| 8_4679186 | NA | NA | 1 | 0 | 1 | 1 | NA | NA | 2 | 0.5 | 1 | 1 | NA | NA | 5 | 0.2 | 1 | 0 | 7 | 0.2857 |
| 8_5691118 | 1 | 0 | 1 | 0 | 1 | 1 | 1 | 0 | 4 | 0.25 | 3 | 0.3333 | 2 | 0.5 | 1 | 1 | 1 | 0 | 7 | 0.4286 |
| 8_5691150 | 1 | 0 | NA | NA | 2 | 0.5 | 1 | 0 | 4 | 0.25 | 4 | 0.5 | 3 | 0.3333 | 3 | 0.3333 | 1 | 1 | 11 | 0.4545 |
| 8_8668106 | 9 | 0.2222 | 5 | 0 | 11 | 0.0909 | 9 | 0 | 34 | 0.0882 | 3 | 0.3333 | 11 | 0.1818 | 19 | 0.2105 | 10 | 0.2 | 43 | 0.2093 |
| 8_8669846 | 7 | 0 | 3 | 0 | 5 | 0.2 | 5 | 0 | 20 | 0.05 | 2 | 0 | 4 | 0 | 9 | 0.1111 | 3 | 0 | 18 | 0.0556 |
| 8_8670074 | 7 | 0.2857 | 2 | 0 | 6 | 0.3333 | 4 | 0.25 | 19 | 0.2632 | 2 | 0.5 | 4 | 0 | 8 | 0.125 | 5 | 0.2 | 19 | 0.1579 |
| 8_8670212 | 9 | 0.6667 | 2 | 0.5 | 5 | 1 | 4 | 0.75 | 20 | 0.75 | 2 | 1 | 6 | 1 | 8 | 0.875 | 4 | 1 | 20 | 0.95 |
| 8_8670225 | 7 | 0.8571 | 2 | 1 | 4 | 0.75 | 2 | 1 | 15 | 0.8667 | 1 | 1 | 6 | 1 | 9 | 0.6667 | 3 | 1 | 19 | 0.8421 |
| 8_8670318 | 6 | 0.6667 | 2 | 0.5 | 4 | 0.25 | 4 | 0.25 | 16 | 0.4375 | 2 | 0.5 | 5 | 0.6 | 10 | 0.6 | 4 | 1 | 21 | 0.6667 |
| 8_8670369 | 9 | 0.3333 | NA | NA | 4 | 0 | 4 | 0.25 | 17 | 0.2353 | 2 | 0.5 | 5 | 0 | 7 | 0 | 4 | 0 | 18 | 0.0556 |
| 8_8670389 | 6 | 1 | NA | NA | 3 | 1 | 3 | 1 | 12 | 1 | 1 | 1 | 4 | 1 | 5 | 1 | 2 | 0.5 | 12 | 0.9167 |
| 8_8670412 | 9 | 0.5556 | 2 | 1 | 4 | 0.5 | 4 | 0.25 | 19 | 0.5263 | 1 | 1 | 6 | 0.1667 | 7 | 0.8571 | 2 | 0.5 | 16 | 0.5625 |
| 8_8670440 | 8 | 0.25 | 2 | 0 | 4 | 0.25 | 3 | 0 | 17 | 0.1765 | 2 | 0.5 | 5 | 0 | 7 | 0.4286 | 2 | 0 | 16 | 0.25 |
| 8_8670476 | 6 | 1 | 2 | 1 | 4 | 1 | 2 | 1 | 14 | 1 | 2 | 1 | 6 | 1 | 8 | 1 | 2 | 1 | 18 | 1 |
| 8_8670531 | 7 | 1 | 2 | 1 | 4 | 1 | 2 | 1 | 15 | 1 | 2 | 1 | 5 | 1 | 9 | 1 | 3 | 1 | 19 | 1 |
| 8_8670560 | 6 | 1 | 2 | 1 | 2 | 1 | 2 | 1 | 12 | 1 | 2 | 1 | 6 | 0.8333 | 5 | 1 | 3 | 1 | 16 | 0.9375 |
| 8_8670661 | 7 | 1 | 1 | 1 | 2 | 1 | 1 | 1 | 11 | 1 | 2 | 1 | 5 | 1 | 7 | 1 | 3 | 1 | 17 | 1 |

| | | | | | | | | | | | | | | | | | | | | |
|---|---|---|---|---|---|---|---|---|---|---|---|---|---|---|---|---|---|---|---|---|
| 8_8670707 | 7 | 1 | 2 | 1 | 3 | 1 | NA | NA | 12 | 1 | 2 | 1 | 5 | 1 | 6 | 1 | 2 | 1 | 15 | 1 |
| 8_8689407 | 4 | 0.75 | 2 | 0.5 | 3 | 0.3333 | 1 | 1 | 10 | 0.6 | 1 | 0 | 2 | 0.5 | 5 | 0.4 | 1 | 1 | 9 | 0.4444 |
| 8_8689415 | 5 | 1 | 2 | 1 | 2 | 1 | NA | NA | 9 | 1 | 1 | 1 | 3 | 1 | 4 | 0.75 | 1 | 1 | 9 | 0.8889 |
| 8_8710542 | 5 | 0.4 | 1 | 0 | 3 | 0.3333 | NA | NA | 9 | 0.3333 | NA | NA | 2 | 0 | 4 | 0.75 | 1 | 0 | 7 | 0.4286 |
| 8_8710564 | 3 | 1 | 2 | 1 | 2 | 1 | NA | NA | 7 | 1 | NA | NA | 3 | 1 | 3 | 1 | 1 | 1 | 7 | 1 |
| 8_8710589 | 4 | 0.75 | 2 | 1 | 3 | 1 | NA | NA | 9 | 0.8889 | NA | NA | 3 | 1 | 5 | 0.8 | 1 | 1 | 9 | 0.8889 |
| 8_8710632 | 5 | 0.2 | 1 | 1 | 2 | 0 | NA | NA | 8 | 0.25 | NA | NA | 2 | 0.5 | 2 | 1 | 1 | 0 | 5 | 0.6 |
| 8_8711049 | 3 | 0.3333 | NA | NA | 2 | 0.5 | NA | NA | 5 | 0.4 | NA | NA | 1 | 1 | 2 | 0.5 | NA | NA | 3 | 0.6667 |
| 8_8711110 | 4 | 0.5 | NA | NA | 3 | 0 | NA | NA | 7 | 0.2857 | NA | NA | 3 | 0 | 3 | 0.6667 | 1 | 0 | 7 | 0.2857 |
| 8_8715641 | 2 | 0 | 4 | 0 | 2 | 0 | 3 | 0.3333 | 11 | 0.0909 | NA | NA | NA | NA | 4 | 0 | 2 | 0.5 | 6 | 0.1667 |
| 8_8716663 | 28 | 0.7143 | 24 | 0.8333 | 11 | 0.5455 | 17 | 0.6471 | 80 | 0.7125 | 10 | 0.9 | 19 | 0.8421 | 31 | 0.6452 | 14 | 0.8571 | 74 | 0.7703 |
| 8_8716788 | 27 | 0.037 | 25 | 0.2 | 11 | 0.3636 | 17 | 0.1765 | 80 | 0.1625 | 10 | 0.1 | 20 | 0.25 | 27 | 0.1481 | 13 | 0.2308 | 70 | 0.1857 |
| 8_8716793 | 21 | 0.0952 | 22 | 0.0455 | 11 | 0 | 17 | 0.0588 | 71 | 0.0563 | 11 | 0 | 18 | 0 | 27 | 0.1481 | 14 | 0.1429 | 70 | 0.0857 |
| 8_8716913 | 27 | 0.1852 | 24 | 0.2083 | 11 | 0.3636 | 14 | 0 | 76 | 0.1842 | 9 | 0.2222 | 15 | 0.1333 | 27 | 0.037 | 10 | 0.1 | 61 | 0.0984 |
| 8_8716948 | 25 | 0.4 | 23 | 0.3913 | 13 | 0.3077 | 16 | 0.375 | 77 | 0.3766 | 10 | 0.6 | 17 | 0.2353 | 28 | 0.1786 | 10 | 0.1 | 65 | 0.2462 |
| 8_8717212 | 14 | 0.1429 | 13 | 0.1538 | 4 | 0.25 | 6 | 0.1667 | 37 | 0.1622 | 8 | 0.125 | 6 | 0.3333 | 17 | 0.3529 | 8 | 0.5 | 39 | 0.3333 |
| 8_8717213 | 1 | 0 | 3 | 0.3333 | 2 | 0.5 | 3 | 0.3333 | 9 | 0.3333 | NA | NA | 3 | 0.6667 | 3 | 0 | 1 | 0 | 7 | 0.2857 |
| 8_8717237 | 15 | 1 | 19 | 0.8947 | 4 | 1 | 9 | 1 | 47 | 0.9574 | 7 | 1 | 11 | 0.9091 | 18 | 0.8889 | 7 | 0.8571 | 43 | 0.907 |
| 8_8717244 | 17 | 0.4118 | 20 | 0.35 | 9 | 0.3333 | 13 | 0.1538 | 59 | 0.322 | 9 | 0.2222 | 10 | 0.4 | 22 | 0.2273 | 11 | 0.2727 | 52 | 0.2692 |
| 8_8717263 | 19 | 0.0526 | 21 | 0.0952 | 10 | 0 | 11 | 0.0909 | 61 | 0.0656 | 10 | 0 | 11 | 0.0909 | 26 | 0.0769 | 10 | 0.1 | 57 | 0.0702 |
| 8_8717282 | 18 | 0.9444 | 19 | 0.9474 | 10 | 0.7 | 12 | 0.9167 | 59 | 0.8983 | 8 | 0.75 | 11 | 0.9091 | 21 | 0.8095 | 10 | 0.9 | 50 | 0.84 |
| 8_8732274 | 1 | 0 | 8 | 0.125 | 2 | 1 | 2 | 0 | 13 | 0.2308 | 4 | 0 | 3 | 0 | 7 | 0 | 3 | 0.6667 | 17 | 0.1176 |
| 8_8732433 | 1 | 0 | 5 | 0.2 | 2 | 0 | NA | NA | 8 | 0.125 | 2 | 0 | 2 | 0 | 6 | 0.1667 | 4 | 0.5 | 14 | 0.2143 |
| 8_8733216 | 1 | 0 | 7 | 0.1429 | 2 | 0.5 | NA | NA | 10 | 0.2 | 2 | 0.5 | 2 | 0 | 4 | 0 | 3 | 0.6667 | 11 | 0.2727 |
| 8_8733474 | NA | NA | 6 | 0 | 3 | 0.3333 | NA | NA | 9 | 0.1111 | 2 | 0.5 | 2 | 0 | 5 | 0 | 3 | 0.3333 | 12 | 0.1667 |
| 8_8733494 | NA | NA | 8 | 0 | 2 | 0.5 | NA | NA | 10 | 0.1 | 2 | 0 | 1 | 0 | 5 | 0.2 | 3 | 0.3333 | 11 | 0.1818 |
| 8_8733583 | NA | NA | 7 | 0 | 3 | 0.3333 | NA | NA | 10 | 0.1 | 2 | 0 | 2 | 0 | 4 | 0.25 | 3 | 0 | 11 | 0.0909 |
| 8_8733719 | NA | NA | 5 | 0 | 2 | 0.5 | NA | NA | 7 | 0.1429 | 2 | 0 | 2 | 0 | 3 | 0 | 3 | 0.3333 | 10 | 0.1 |
| 8_8740487 | 10 | 0.1 | 11 | 0.0909 | 7 | 0.2857 | 1 | 1 | 29 | 0.1724 | NA | NA | 1 | 0 | 6 | 0.5 | 3 | 0 | 10 | 0.3 |
| 8_9256307 | 7 | 0.1429 | 15 | 0.1333 | 5 | 0 | 5 | 0 | 32 | 0.0938 | 5 | 0.2 | 1 | 0 | 4 | 0 | 6 | 0.1667 | 16 | 0.125 |
| 8_9256727 | 7 | 0 | 13 | 0 | 2 | 0 | 4 | 0.5 | 26 | 0.0769 | 4 | 0.25 | 4 | 0 | 5 | 0 | 4 | 0.25 | 17 | 0.1176 |
| 8_9256746 | 6 | 0 | 14 | 0.1429 | 5 | 0 | 4 | 0 | 29 | 0.069 | 4 | 0.25 | 3 | 0 | 6 | 0 | 6 | 0 | 19 | 0.0526 |
| 8_9257512 | 5 | 1 | 11 | 0.8182 | 5 | 1 | 3 | 1 | 24 | 0.9167 | 3 | 1 | 2 | 1 | 3 | 1 | 4 | 0.75 | 12 | 0.9167 |
| 8_9257552 | 5 | 1 | 10 | 0.9 | 4 | 1 | 4 | 1 | 23 | 0.9565 | 4 | 1 | 1 | 1 | 2 | 1 | 2 | 1 | 9 | 1 |
| 8_9257609 | 5 | 1 | 10 | 1 | 4 | 1 | 3 | 1 | 22 | 1 | 4 | 1 | 2 | 1 | 3 | 1 | 4 | 1 | 13 | 1 |
| 8_9257639 | 4 | 0.75 | 8 | 0.5 | 3 | 0.6667 | 2 | 0.5 | 17 | 0.5882 | 1 | 1 | 1 | 0 | 2 | 1 | 3 | 1 | 7 | 0.8571 |
| 8_9257674 | 6 | 0 | 7 | 0.1429 | 3 | 0 | 4 | 0 | 20 | 0.05 | 4 | 0 | 2 | 0 | 3 | 0 | 4 | 0.25 | 13 | 0.0769 |
| 8_9257701 | 6 | 0.8333 | 7 | 0.5714 | 3 | 0.6667 | 2 | 1 | 18 | 0.7222 | 4 | 0.75 | 1 | 1 | 3 | 1 | 4 | 0.5 | 12 | 0.75 |
| 8_9257747 | 4 | 1 | 6 | 1 | 2 | 1 | 3 | 1 | 15 | 1 | 3 | 1 | 2 | 1 | 3 | 1 | 4 | 1 | 12 | 1 |
| 8_9257765 | 6 | 0.3333 | 9 | 0.3333 | 3 | 0 | 4 | 0.25 | 22 | 0.2727 | 4 | 0 | 1 | 1 | 2 | 0 | 4 | 0.75 | 11 | 0.3636 |
| 8_9257797 | 5 | 1 | 2 | 1 | 2 | 1 | 3 | 1 | 12 | 1 | 1 | 1 | 1 | 1 | 1 | 1 | NA | NA | 3 | 1 |
| 8_9257807 | 6 | 0.1667 | 8 | 0.125 | 1 | 0 | 3 | 0 | 18 | 0.1111 | 4 | 0.25 | 1 | 0 | 3 | 0.3333 | 2 | 0 | 10 | 0.2 |
| 8_9257887 | 4 | 1 | 5 | 0.8 | 2 | 1 | 3 | 1 | 14 | 0.9286 | 3 | 1 | 2 | 1 | 1 | 1 | 3 | 1 | 9 | 1 |
| 8_9257916 | 5 | 0 | 8 | 0.125 | 2 | 0 | 4 | 0 | 19 | 0.0526 | 4 | 0.25 | 1 | 0 | 2 | 0.5 | 3 | 0 | 10 | 0.2 |
| 8_9257930 | 6 | 0.3333 | 8 | 0.375 | 3 | 0 | 4 | 0.25 | 21 | 0.2857 | 4 | 0 | 2 | 0 | 3 | 0.3333 | 2 | 0 | 11 | 0.0909 |
| 8_9820411 | NA | NA | NA | NA | 1 | 1 | NA | NA | 1 | 1 | 1 | 1 | NA | NA | NA | NA | NA | NA | 1 | 1 |
| 8_10020274 | 1 | 0 | 2 | 0 | 2 | 0 | 2 | 0.5 | 7 | 0.1429 | 2 | 0 | NA | NA | 1 | 1 | 2 | 0.5 | 5 | 0.4 |
| 8_10021158 | 1 | 1 | 1 | 0 | 2 | 0 | 3 | 0.3333 | 7 | 0.2857 | 2 | 0 | NA | NA | NA | NA | 2 | 0.5 | 4 | 0.25 |
| 8_10021296 | 1 | 1 | 1 | 0 | 2 | 0.5 | 3 | 0.6667 | 7 | 0.5714 | 1 | 1 | NA | NA | NA | NA | 2 | 1 | 3 | 1 |
| 8_10021317 | NA | NA | 1 | 1 | 2 | 1 | 3 | 1 | 6 | 1 | 2 | 1 | NA | NA | NA | NA | 2 | 1 | 4 | 1 |
| 8_10021362 | 1 | 0 | 1 | 0 | 1 | 0 | 2 | 1 | 5 | 0.4 | 2 | 0 | NA | NA | NA | NA | 1 | 1 | 3 | 0.3333 |
| 8_10021386 | 1 | 0 | 1 | 0 | 2 | 0.5 | 2 | 0 | 6 | 0.1667 | 2 | 1 | NA | NA | NA | NA | 1 | 0 | 3 | 0.6667 |
| 8_10021503 | 1 | 1 | 1 | 0 | 1 | 0 | 3 | 0.6667 | 6 | 0.5 | 2 | 0 | NA | NA | NA | NA | 2 | 0.5 | 4 | 0.25 |
| 8_10021585 | 1 | 0 | 1 | 1 | 2 | 0 | 3 | 0 | 7 | 0.1429 | 1 | 0 | NA | NA | NA | NA | 2 | 0.5 | 3 | 0.3333 |
| 8_10021647 | 1 | 1 | 1 | 0 | 2 | 0.5 | 3 | 0.6667 | 7 | 0.5714 | 1 | 0 | NA | NA | NA | NA | 1 | 1 | 2 | 0.5 |
| 8_10021740 | 1 | 0 | 1 | 1 | 2 | 0 | 3 | 0 | 7 | 0.1429 | 1 | 0 | NA | NA | NA | NA | 2 | 0.5 | 3 | 0.3333 |
| 8_10021853 | 1 | 1 | 1 | 1 | 2 | 1 | 2 | 1 | 6 | 1 | 1 | 1 | NA | NA | NA | NA | 2 | 1 | 3 | 1 |
| 8_10021913 | 1 | 0 | 1 | 1 | 1 | 1 | 3 | 0 | 6 | 0.3333 | 2 | 0.5 | NA | NA | NA | NA | 1 | 1 | 3 | 0.6667 |
| 8_10022121 | 1 | 0 | 1 | 1 | 2 | 1 | 3 | 1 | 7 | 0.8571 | 2 | 1 | NA | NA | NA | NA | NA | NA | 2 | 1 |
| 8_10022237 | 1 | 1 | 1 | 1 | NA | NA | 3 | 1 | 5 | 1 | 2 | 1 | NA | NA | NA | NA | 1 | 1 | 3 | 1 |
| 8_10022408 | 1 | 1 | 1 | 1 | 2 | 1 | 3 | 1 | 7 | 1 | 2 | 1 | NA | NA | NA | NA | 1 | 1 | 3 | 1 |
| 8_10022638 | 1 | 1 | 1 | 1 | 2 | 0.5 | 2 | 0 | 6 | 0.5 | 2 | 1 | NA | NA | NA | NA | NA | NA | 2 | 1 |
| 8_10022695 | 1 | 1 | 1 | 0 | 2 | 0.5 | 2 | 0.5 | 6 | 0.5 | 1 | 1 | NA | NA | NA | NA | 1 | 0 | 2 | 0.5 |
| 8_10022994 | 1 | 0 | NA | NA | 1 | 1 | 2 | 0.5 | 4 | 0.5 | 1 | 1 | NA | NA | NA | NA | 1 | 0 | 2 | 0.5 |
| 8_10023069 | 1 | 0 | 1 | 1 | 1 | 0 | 2 | 0 | 5 | 0.2 | 1 | 1 | NA | NA | NA | NA | 1 | 0 | 2 | 0.5 |

| | | | | | | | | | | | | | | | | | | | | |
|---|---|---|---|---|---|---|---|---|---|---|---|---|---|---|---|---|---|---|---|---|
| 8_1002320 | 1 | 1 | 1 | 1 | 1 | 1 | 1 | 1 | 4 | 1 | NA | NA | NA | NA | NA | NA | 1 | 1 | 1 | 1 |
| 8_1002328 | 1 | 0 | 1 | 0 | 1 | 1 | 1 | 0 | 4 | 0.25 | 1 | 0 | NA | NA | NA | NA | 1 | 1 | 2 | 0.5 |
| 8_1002332 | 1 | 1 | 1 | 1 | 1 | 1 | 1 | 1 | 4 | 1 | 1 | 1 | NA | NA | NA | NA | 1 | 1 | 2 | 1 |
| 8_1002340 | NA | NA | 1 | 1 | 1 | 1 | 1 | 1 | 3 | 1 | 1 | 1 | NA | NA | NA | NA | 1 | 0 | 2 | 0.5 |
| 8_1002348 | 1 | 0 | 1 | 0 | 1 | 0 | 1 | 1 | 4 | 0.25 | 1 | 1 | NA | NA | NA | NA | 1 | 0 | 2 | 0.5 |
| 8_1002359 | 1 | 1 | 1 | 1 | 1 | 1 | NA | NA | 3 | 1 | 1 | 1 | NA | NA | NA | NA | 1 | 1 | 2 | 1 |
| 8_1003751 | 4 | 1 | 6 | 1 | 1 | 1 | NA | NA | 11 | 1 | 1 | 1 | 4 | 1 | 1 | 1 | 4 | 1 | 10 | 1 |
| 8_1003761 | 4 | 0 | 6 | 0.6667 | 1 | 0 | 1 | 1 | 12 | 0.4167 | 2 | 0 | 3 | 0.3333 | 1 | 0 | 4 | 0.5 | 10 | 0.3 |
| 8_1003773 | 4 | 0.5 | 7 | 0.5714 | 1 | 1 | 2 | 0 | 14 | 0.5 | 2 | 0.5 | 4 | 0.5 | 1 | 1 | 5 | 0.8 | 12 | 0.6667 |
| 8_1003782 | 5 | 0 | 7 | 0.2857 | 1 | 0 | 2 | 1 | 15 | 0.2667 | 2 | 0.5 | 4 | 0 | 1 | 0 | 4 | 0 | 11 | 0.0909 |
| 8_1003792 | 4 | 0.75 | 5 | 1 | 1 | 1 | 2 | 1 | 12 | 0.9167 | 2 | 1 | 3 | 0.6667 | 1 | 1 | 5 | 0.8 | 11 | 0.8182 |
| 8_1003795 | 6 | 0.5 | 6 | 0.3333 | 1 | 0 | 2 | 0 | 15 | 0.3333 | 2 | 0.5 | 3 | 0 | 1 | 1 | 6 | 0.3333 | 12 | 0.3333 |
| 8_1003805 | 5 | 0.2 | 8 | 0.25 | 1 | 1 | 2 | 1 | 16 | 0.375 | 2 | 0.5 | 4 | 0 | 1 | 0 | 5 | 0.4 | 12 | 0.25 |
| 8_1003826 | 6 | 1 | 6 | 1 | 1 | 1 | 2 | 1 | 15 | 1 | 2 | 1 | 5 | 1 | 1 | 1 | 5 | 1 | 13 | 1 |
| 8_1003834 | 3 | 1 | 6 | 0.8333 | 1 | 0 | 1 | 0 | 11 | 0.7273 | NA | NA | 3 | 1 | NA | NA | 3 | 1 | 6 | 1 |
| 8_1003850 | 5 | 1 | 8 | 1 | 1 | 1 | 2 | 1 | 16 | 1 | 2 | 1 | 5 | 1 | NA | NA | 6 | 1 | 13 | 1 |
| 8_1003855 | 4 | 0.25 | 8 | 0.5 | 1 | 0 | 2 | 1 | 15 | 0.4667 | 1 | 1 | 5 | 0.4 | 1 | 0 | 4 | 0.5 | 11 | 0.4545 |
| 8_1003857 | 4 | 1 | 7 | 0.8571 | 1 | 1 | 2 | 1 | 14 | 0.9286 | 2 | 1 | 4 | 0.75 | 1 | 1 | 4 | 0.75 | 11 | 0.8182 |
| 8_1003860 | 3 | 0.3333 | 6 | 0.8333 | 1 | 0 | 2 | 0.5 | 12 | 0.5833 | 2 | 0.5 | 3 | 0.3333 | 1 | 1 | 5 | 0.6 | 11 | 0.5455 |
| 8_1003863 | 5 | 0 | 6 | 0.5 | 1 | 0 | 1 | 0 | 13 | 0.2308 | 2 | 0 | 4 | 0 | 1 | 0 | 5 | 0.4 | 12 | 0.1667 |
| 8_1003877 | 4 | 1 | 7 | 0.5714 | 2 | 1 | 2 | 1 | 15 | 0.8 | 2 | 1 | 4 | 1 | 1 | 1 | 5 | 0.8 | 12 | 0.9167 |
| 8_1003885 | 6 | 0 | 8 | 0 | 2 | 0.5 | 2 | 0 | 18 | 0.0556 | 2 | 0.5 | 5 | 0 | 2 | 0 | 6 | 0 | 15 | 0.0667 |
| 8_1003936 | 5 | 0.6 | 7 | 0.4286 | 2 | 0 | 2 | 0 | 16 | 0.375 | 1 | 0 | 3 | 0.6667 | 2 | 0 | 6 | 0.3333 | 12 | 0.3333 |
| 8_1004026 | 4 | 1 | 4 | 1 | 3 | 0.6667 | 1 | 0 | 12 | 0.8333 | 1 | 0 | 2 | 1 | 1 | 0 | 3 | 1 | 7 | 0.7143 |
| 8_1004093 | 4 | 0 | 5 | 0.2 | 2 | 0 | 1 | 1 | 12 | 0.1667 | 1 | 0 | 2 | 0 | 2 | 0 | 3 | 0.3333 | 8 | 0.125 |
| 8_1004165 | 5 | 0.4 | 6 | 0 | 3 | 0 | 3 | 0 | 17 | 0.1176 | 1 | 0 | 2 | 0 | 2 | 0.5 | 3 | 0 | 8 | 0.125 |
| 8_1029941 | 1 | 1 | NA | NA | 1 | 0 | NA | NA | 2 | 0.5 | 1 | 1 | 1 | 0 | NA | NA | 1 | 0 | 3 | 0.3333 |
| 8_1061958 | 3 | 1 | 5 | 1 | 5 | 1 | 3 | 1 | 16 | 1 | 2 | 1 | 3 | 1 | 5 | 1 | 2 | 1 | 12 | 1 |
| 8_1061961 | 2 | 0 | 3 | 0 | 3 | 0 | 3 | 0.6667 | 11 | 0.1818 | 1 | 0 | 1 | 0 | 5 | 0.2 | 2 | 0 | 9 | 0.1111 |
| 8_1061986 | 4 | 1 | 5 | 0.8 | 5 | 1 | 4 | 1 | 18 | 0.9444 | 2 | 1 | 3 | 1 | 5 | 0.6 | 2 | 1 | 12 | 0.8333 |
| 8_1061992 | 3 | 0 | 5 | 0 | 3 | 0 | 5 | 0.6 | 16 | 0.1875 | 1 | 0 | 3 | 0 | 5 | 0.2 | 4 | 0.25 | 13 | 0.1538 |
| 8_1062004 | 5 | 0 | 5 | 0 | 4 | 0.5 | 5 | 0.6 | 19 | 0.2632 | 2 | 0 | 3 | 0.3333 | 4 | 0.25 | 4 | 0.5 | 13 | 0.3077 |
| 8_1062006 | 4 | 0.25 | 5 | 0.2 | 5 | 0 | 5 | 0 | 19 | 0.1053 | 2 | 0 | 3 | 0.3333 | 5 | 0 | 4 | 0 | 14 | 0.0714 |
| 8_1062031 | 3 | 0 | 4 | 0.25 | 1 | 0 | 4 | 0 | 12 | 0.0833 | 2 | 0.5 | 2 | 0.5 | 4 | 0 | 3 | 0.3333 | 11 | 0.2727 |
| 8_1062046 | 4 | 0.5 | 3 | 0 | 3 | 0.3333 | 5 | 0.4 | 15 | 0.3333 | 2 | 0 | 4 | 0 | 4 | 0 | 3 | 0.3333 | 13 | 0.0769 |
| 8_1062047 | 4 | 0.25 | 4 | 0 | 3 | 0 | 5 | 0 | 16 | 0.0625 | 1 | 0 | 3 | 0 | 5 | 0.2 | 3 | 0.3333 | 12 | 0.1667 |
| 8_1068348 | 2 | 0 | 3 | 0.3333 | 3 | 0.3333 | 3 | 0 | 11 | 0.1818 | 2 | 1 | 5 | 0.4 | 6 | 0 | 3 | 0.3333 | 16 | 0.3125 |
| 8_1068363 | 5 | 0.2 | 5 | 0.4 | 2 | 0 | 5 | 0.8 | 17 | 0.4118 | 2 | 0.5 | 4 | 0 | 7 | 0.1429 | 4 | 0.25 | 17 | 0.1765 |
| 8_1068366 | 5 | 1 | 6 | 0.6667 | 5 | 1 | 5 | 1 | 21 | 0.9048 | 2 | 1 | 7 | 1 | 7 | 1 | 5 | 1 | 21 | 1 |
| 8_1068369 | 5 | 0.6 | 5 | 0.6 | 5 | 0.8 | 3 | 1 | 18 | 0.7222 | 2 | 0 | 7 | 0.8571 | 5 | 0.6 | 4 | 0.75 | 18 | 0.6667 |
| 8_1068376 | 5 | 0.6 | 4 | 0.75 | 4 | 0.75 | 3 | 1 | 16 | 0.75 | 2 | 1 | 7 | 0.8571 | 7 | 0.8571 | 5 | 0.8 | 21 | 0.8571 |
| 8_1068377 | 3 | 0 | 5 | 0 | 4 | 0 | 3 | 0.3333 | 15 | 0.0667 | 2 | 0 | 7 | 0.1429 | 7 | 0 | 4 | 0 | 20 | 0.05 |
| 8_1068403 | 8 | 0 | 9 | 0.1111 | 7 | 0 | 6 | 0.1667 | 30 | 0.0667 | 4 | 0.25 | 8 | 0 | 9 | 0 | 6 | 0.1667 | 27 | 0.0741 |
| 8_1068446 | 8 | 0.25 | 7 | 0.1429 | 5 | 0.6 | 7 | 0.7143 | 27 | 0.4074 | 4 | 0.75 | 9 | 0.1111 | 6 | 0.1667 | 6 | 0.5 | 25 | 0.32 |
| 8_1068450 | 8 | 0 | 5 | 0.2 | 4 | 0.25 | 6 | 0 | 23 | 0.087 | 4 | 0.25 | 7 | 0.2857 | 4 | 0.25 | 6 | 0 | 21 | 0.1905 |
| 8_1068468 | 8 | 0.25 | 6 | 0.3333 | 5 | 0.2 | 6 | 0.3333 | 25 | 0.28 | 4 | 0.5 | 8 | 0.25 | 8 | 0.125 | 7 | 0.4286 | 27 | 0.2963 |
| 8_1103485 | 12 | 0.25 | 11 | 0.3636 | 4 | 0.25 | 8 | 0.25 | 35 | 0.2857 | 9 | 0.4444 | 18 | 0.1667 | 8 | 0.375 | 9 | 0.2222 | 44 | 0.2727 |
| 8_1103495 | 12 | 0.0833 | 10 | 0.1 | 6 | 0.1667 | 9 | 0 | 37 | 0.0811 | 8 | 0.125 | 13 | 0 | 8 | 0.125 | 6 | 0 | 35 | 0.0571 |
| 8_1103505 | 12 | 0.0833 | 7 | 0 | 6 | 0 | 6 | 0.1667 | 31 | 0.0645 | 8 | 0 | 9 | 0.1111 | 8 | 0.125 | 8 | 0.125 | 33 | 0.0909 |
| 8_1103516 | 8 | 0.375 | 11 | 0.0909 | 6 | 0.1667 | 8 | 0.25 | 33 | 0.2121 | 8 | 0.125 | 13 | 0.0769 | 8 | 0.25 | 7 | 0.1429 | 36 | 0.1389 |
| 8_1103519 | 9 | 0.8889 | 11 | 0.8182 | 5 | 0.8 | 6 | 0.6667 | 31 | 0.8065 | 9 | 0.7778 | 12 | 0.8333 | 7 | 0.8571 | 5 | 0.8 | 33 | 0.8182 |
| 8_1103527 | 9 | 0.8889 | 10 | 1 | 5 | 1 | 6 | 1 | 30 | 0.9667 | 9 | 1 | 13 | 1 | 9 | 1 | 6 | 1 | 37 | 1 |
| 8_1103527 | 9 | 0.8889 | 7 | 0.7143 | 6 | 0.8333 | 8 | 0.75 | 30 | 0.8 | 7 | 0.7143 | 11 | 0.8182 | 7 | 0.8571 | 6 | 1 | 31 | 0.8387 |
| 8_1103586 | 6 | 0.6667 | 7 | 0.5714 | 5 | 0.6 | 5 | 0.6 | 23 | 0.6087 | 6 | 1 | 9 | 0.7778 | 6 | 0.6667 | 6 | 0.5 | 27 | 0.7407 |
| 8_1103591 | 6 | 0.1667 | 7 | 0.5714 | 5 | 0.4 | 5 | 0.4 | 23 | 0.3913 | 6 | 0.6667 | 10 | 0.3 | 5 | 0.4 | 6 | 0.1667 | 27 | 0.3704 |
| 8_1103595 | 1 | 1 | 3 | 0 | NA | NA | NA | NA | 4 | 0.25 | 2 | 0 | 2 | 0 | 2 | 0.5 | 1 | 0 | 7 | 0.1429 |
| 8_1103608 | 5 | 0.2 | 6 | 0.1667 | 3 | 0.3333 | 5 | 0 | 19 | 0.1579 | 6 | 0.5 | 9 | 0 | 6 | 0 | 6 | 0.1667 | 27 | 0.1481 |
| 8_1103611 | 2 | 0 | 6 | 0.1667 | 2 | 0.5 | 2 | 0 | 12 | 0.1667 | 4 | 0 | 7 | 0.2857 | 5 | 0.2 | 3 | 0.3333 | 19 | 0.2105 |
| 8_1103614 | 5 | 0 | 5 | 0.6 | 3 | 0.6667 | 4 | 0.5 | 17 | 0.4118 | 5 | 1 | 10 | 0.4 | 6 | 0.6667 | 4 | 0.75 | 25 | 0.64 |
| 8_1103622 | 5 | 0.2 | 5 | 0.2 | 2 | 0.5 | 2 | 0 | 14 | 0.2143 | 5 | 0.4 | 11 | 0.1818 | 5 | 0.4 | 5 | 0 | 26 | 0.2308 |
| 8_1103632 | NA | NA | 7 | 1 | 1 | 1 | 3 | 1 | 11 | 1 | 4 | 1 | 10 | 1 | 6 | 0.8333 | 4 | 1 | 24 | 0.9583 |
| 8_1103635 | 4 | 1 | 7 | 1 | 3 | 1 | 3 | 1 | 17 | 1 | 4 | 1 | 10 | 0.9 | 6 | 1 | 4 | 1 | 24 | 0.9583 |
| 8_1103638 | 3 | 1 | 7 | 0.8571 | 2 | 1 | 4 | 1 | 16 | 0.9375 | 5 | 1 | 11 | 1 | 6 | 1 | 3 | 1 | 25 | 1 |
| 8_1103643 | 5 | 1 | 6 | 1 | 3 | 1 | 4 | 1 | 18 | 1 | 4 | 1 | 10 | 1 | 6 | 1 | 3 | 1 | 23 | 1 |
| 8_1103646 | 4 | 0 | 6 | 0 | 2 | 0 | 4 | 0.25 | 16 | 0.0625 | 3 | 0.3333 | 9 | 0.1111 | 5 | 0 | 2 | 0 | 19 | 0.1053 |

| | | | | | | | | | | | | | | | | | | | | |
|---|---|---|---|---|---|---|---|---|---|---|---|---|---|---|---|---|---|---|---|---|
| 8_11036547 | 3 | 0 | 4 | 0 | NA | NA | 4 | 0.5 | 11 | 0.1818 | 2 | 0 | 6 | 0.1667 | 3 | 0 | 2 | 0 | 13 | 0.0769 |
| 8_11036744 | 2 | 1 | 5 | 1 | 1 | 1 | NA | NA | 8 | 1 | 2 | 1 | 10 | 1 | 4 | 1 | 3 | 1 | 19 | 1 |
| 8_11036828 | 1 | 1 | 6 | 0.3333 | 2 | 0 | 2 | 0.5 | 11 | 0.3636 | NA | NA | 4 | 0 | 3 | 0.3333 | 1 | 1 | 8 | 0.25 |
| 8_11036850 | 2 | 1 | 6 | 0.8333 | 3 | 0.3333 | 3 | 1 | 14 | 0.7857 | 3 | 1 | 8 | 0.875 | 6 | 1 | 2 | 0.5 | 19 | 0.8947 |
| 8_11036942 | 2 | 1 | 5 | 0.8 | 1 | 1 | 3 | 0.6667 | 11 | 0.8182 | 4 | 0.75 | 5 | 0.8 | 6 | 0.3333 | 3 | 0.6667 | 18 | 0.6111 |
| 8_11036947 | 2 | 0.5 | 2 | 0.5 | 1 | 0 | 3 | 0.3333 | 8 | 0.375 | 1 | 0 | 4 | 0.5 | 5 | 0.2 | 1 | 1 | 11 | 0.3636 |
| 8_11036967 | 2 | 0.5 | 6 | 0 | 1 | 0 | 3 | 0 | 12 | 0.0833 | 4 | 0 | 6 | 0.1667 | 6 | 0.3333 | 3 | 0 | 19 | 0.1579 |
| 8_11036973 | 2 | 0 | 4 | 0.25 | 1 | 0 | 2 | 0 | 9 | 0.1111 | 4 | 0 | 6 | 0 | 5 | 0.2 | 2 | 0 | 17 | 0.0588 |
| 8_11054637 | 2 | 0.5 | 6 | 0.5 | 1 | 0 | 4 | 0.25 | 13 | 0.3846 | 4 | 0.5 | 8 | 0.625 | 4 | 0.5 | 3 | 0.3333 | 19 | 0.5263 |
| 8_11054737 | 2 | 0 | 6 | 0.3333 | 1 | 1 | 4 | 0.75 | 13 | 0.4615 | 3 | 0 | 8 | 0.25 | 3 | 0.6667 | 3 | 0.3333 | 17 | 0.2941 |
| 8_11054855 | 2 | 0.5 | 4 | 0 | 1 | 0 | 3 | 0.6667 | 10 | 0.3 | 4 | 0 | 6 | 0.1667 | 2 | 0.5 | 3 | 0.3333 | 15 | 0.2 |
| 8_11055013 | 2 | 1 | 5 | 0.6 | 1 | 1 | 3 | 1 | 11 | 0.8182 | 3 | 0.6667 | 8 | 0.875 | 3 | 0.6667 | 3 | 0.3333 | 17 | 0.7059 |
| 8_11055048 | NA | NA | 4 | 0.25 | NA | NA | 2 | 0 | 6 | 0.1667 | 3 | 0 | 5 | 0.2 | 2 | 0 | 3 | 0.3333 | 13 | 0.1538 |
| 8_11055252 | 1 | 1 | 5 | 0.8 | NA | NA | 2 | 0.5 | 8 | 0.75 | 2 | 1 | 6 | 1 | 2 | 1 | 1 | 1 | 11 | 1 |
| 8_11055465 | 2 | 0 | 4 | 0.25 | 1 | 0 | 2 | 0 | 9 | 0.1111 | 3 | 1 | 5 | 0.2 | 2 | 0 | NA | NA | 10 | 0.4 |
| 8_11055510 | 1 | 0 | 5 | 1 | NA | NA | 2 | 1 | 8 | 0.875 | 2 | 1 | 4 | 1 | 2 | 1 | 1 | 1 | 9 | 1 |
| 8_11055516 | 1 | 0 | 3 | 0 | 1 | 0 | 2 | 1 | 7 | 0.2857 | 2 | 1 | 5 | 0.4 | 2 | 0 | 1 | 0 | 10 | 0.4 |
| 8_11055525 | 1 | 0 | 4 | 0.25 | NA | NA | 1 | 0 | 6 | 0.1667 | 1 | 0 | 3 | 0.3333 | 2 | 0.5 | 1 | 0 | 7 | 0.2857 |
| 8_11055687 | NA | NA | 2 | 1 | 1 | 1 | 3 | 1 | 6 | 1 | 1 | 1 | 4 | 0.75 | 1 | 1 | 1 | 1 | 7 | 0.8571 |
| 8_11055764 | 1 | 1 | 5 | 0.6 | 1 | 1 | 3 | 0.3333 | 10 | 0.6 | 2 | 1 | 6 | 0.8333 | 1 | 1 | 1 | 1 | 10 | 0.9 |
| 8_11055944 | 1 | 1 | 4 | 1 | 1 | 1 | 1 | 0 | 7 | 0.8571 | 1 | 1 | 5 | 1 | 1 | 1 | 1 | 1 | 8 | 1 |
| 8_11056110 | 1 | 0 | 3 | 0.3333 | 1 | 0 | NA | NA | 5 | 0.2 | 2 | 0 | 5 | 0.4 | NA | NA | 1 | 0 | 8 | 0.25 |
| 8_11056120 | 1 | 0 | 4 | 0.25 | 1 | 0 | NA | NA | 6 | 0.1667 | 2 | 0 | 5 | 0.2 | 1 | 1 | 1 | 0 | 9 | 0.2222 |
| 8_11056161 | 1 | 0 | 4 | 0.25 | NA | NA | NA | NA | 5 | 0.2 | 2 | 0.5 | 4 | 0 | 1 | 0 | NA | NA | 7 | 0.1429 |
| 8_11056239 | NA | NA | 4 | 0 | 1 | 1 | NA | NA | 5 | 0.2 | 2 | 0 | 5 | 0.2 | 1 | 1 | 1 | 0 | 9 | 0.2222 |
| 8_11056300 | 1 | 0 | 4 | 0.5 | 1 | 1 | NA | NA | 6 | 0.5 | 1 | 0 | 5 | 0.4 | 1 | 1 | 1 | 0 | 8 | 0.375 |
| 8_11056551 | NA | NA | 2 | 0.5 | NA | NA | NA | NA | 2 | 0.5 | 1 | 1 | 5 | 1 | 1 | 1 | NA | NA | 7 | 1 |
| 8_11056720 | NA | NA | 2 | 1 | NA | NA | NA | NA | 2 | 1 | 2 | 1 | 4 | 1 | 1 | 1 | NA | NA | 7 | 1 |
| 8_11056746 | NA | NA | 2 | 0.5 | NA | NA | NA | NA | 2 | 0.5 | 2 | 0.5 | 4 | 0 | 1 | 1 | NA | NA | 7 | 0.2857 |
| 8_11056829 | NA | NA | 2 | 1 | NA | NA | NA | NA | 2 | 1 | 1 | 1 | 4 | 1 | 1 | 1 | NA | NA | 6 | 1 |
| 8_11057191 | NA | NA | 1 | 1 | NA | NA | NA | NA | 1 | 1 | 1 | 1 | 1 | 1 | 1 | 1 | NA | NA | 3 | 1 |
| 8_11057709 | NA | NA | 1 | 1 | NA | NA | NA | NA | 1 | 1 | 2 | 0 | 2 | 1 | NA | NA | NA | NA | 4 | 0.5 |
| 8_11057822 | NA | NA | 2 | 0.5 | NA | NA | NA | NA | 2 | 0.5 | 2 | 1 | 3 | 0.6667 | NA | NA | NA | NA | 5 | 0.8 |
| 8_11058446 | NA | NA | 2 | 0.5 | NA | NA | NA | NA | 2 | 0.5 | 1 | 0 | 3 | 0.3333 | NA | NA | NA | NA | 4 | 0.25 |
| 8_11058663 | NA | NA | 2 | 0.5 | NA | NA | NA | NA | 2 | 0.5 | 1 | 1 | 3 | 0 | NA | NA | NA | NA | 4 | 0.25 |
| 8_11249269 | 1 | 1 | 2 | 0 | 1 | 0 | NA | NA | 4 | 0.25 | 1 | 0 | 4 | 0.25 | 1 | 0 | NA | NA | 6 | 0.1667 |
| 8_11249323 | 1 | 0 | 2 | 0.5 | 1 | 0 | 2 | 0 | 6 | 0.1667 | 2 | 0 | 4 | 0.25 | 1 | 1 | NA | NA | 7 | 0.2857 |
| 8_11498123 | 1 | 1 | 2 | 0 | 1 | 0 | 3 | 0.3333 | 7 | 0.2857 | 5 | 0.2 | 1 | 0 | 3 | 0 | 1 | 0 | 10 | 0.1 |
| 8_11498225 | 1 | 0 | 3 | 0.6667 | 1 | 0 | 3 | 0 | 8 | 0.25 | 5 | 0.2 | 2 | 0 | 3 | 0 | 1 | 0 | 11 | 0.0909 |
| 8_11498434 | NA | NA | 3 | 0.3333 | NA | NA | 3 | 0 | 6 | 0.1667 | 5 | 0.8 | 3 | 1 | 4 | 0 | 2 | 0.5 | 14 | 0.5714 |
| 8_11498735 | NA | NA | 2 | 0.5 | 1 | 0 | 4 | 0.5 | 7 | 0.4286 | 6 | 0.3333 | 1 | 1 | 3 | 0 | 3 | 0.3333 | 13 | 0.3077 |
| 8_11559308 | NA | NA | NA | NA | NA | NA | 1 | 1 | 1 | 1 | NA | NA | NA | NA | NA | NA | 1 | 1 | 1 | 1 |
| 8_11559563 | NA | NA | NA | NA | NA | NA | 1 | 1 | 1 | 1 | NA | NA | NA | NA | NA | NA | 1 | 1 | 1 | 1 |
| 8_11563069 | 13 | 0.2308 | 8 | 0.125 | 7 | 0.1429 | 6 | 0.3333 | 34 | 0.2059 | 8 | 0.125 | 5 | 0.2 | 7 | 0.1429 | 6 | 0.5 | 26 | 0.2308 |
| 8_11572047 | NA | NA | NA | NA | 1 | 1 | NA | NA | 1 | 1 | NA | NA | NA | NA | NA | NA | 1 | 1 | 1 | 1 |
| 8_11611601 | 3 | 0.6667 | 2 | 0.5 | 1 | 1 | NA | NA | 6 | 0.6667 | NA | NA | NA | NA | 2 | 0.5 | 3 | 0.6667 | 5 | 0.6 |
| 8_11611634 | 5 | 0.6 | 7 | 0.1429 | 3 | 0 | 2 | 0 | 17 | 0.2353 | NA | NA | 2 | 0 | 3 | 0.3333 | 3 | 0 | 8 | 0.125 |
| 8_11612019 | 6 | 0.1667 | 7 | 0 | 3 | 0.3333 | 2 | 0 | 18 | 0.1111 | 1 | 1 | 2 | 0.5 | 1 | 0 | 3 | 0 | 7 | 0.2857 |
| 8_11612075 | 6 | 1 | 7 | 1 | 2 | 1 | 1 | 1 | 16 | 1 | 1 | 1 | 3 | 1 | 1 | 1 | 3 | 1 | 8 | 1 |
| 8_11612129 | 6 | 1 | 9 | 1 | 3 | 1 | 3 | 1 | 21 | 1 | 1 | 1 | 4 | 1 | 3 | 1 | 4 | 1 | 12 | 1 |
| 8_11612149 | 5 | 0.4 | 10 | 0.5 | 2 | 0.5 | 2 | 0.5 | 19 | 0.4737 | 1 | 0 | 5 | 0.8 | 2 | 0.5 | 2 | 0 | 10 | 0.5 |
| 8_11612294 | 6 | 0.3333 | 9 | 0.3333 | 4 | 0.75 | 3 | 1 | 22 | 0.5 | 1 | 0 | 5 | 1 | 1 | 0 | 4 | 0.5 | 11 | 0.6364 |
| 8_11612345 | 5 | 0.2 | 10 | 0 | 3 | 0.3333 | 2 | 0 | 20 | 0.1 | 1 | 0 | 4 | 0 | 3 | 0 | 4 | 0.25 | 12 | 0.0833 |
| 8_11612365 | 4 | 0.25 | 5 | 0.6 | 3 | 0.6667 | 2 | 1 | 14 | 0.5714 | 1 | 1 | 3 | 1 | 3 | 0.6667 | 4 | 0.75 | 11 | 0.8182 |
| 8_11612389 | 7 | 0.1429 | 8 | 0.125 | 3 | 0 | 3 | 0.6667 | 21 | 0.1905 | 1 | 0 | 4 | 0.25 | 3 | 0 | 4 | 0.5 | 12 | 0.25 |
| 8_11612651 | 7 | 0.4286 | 10 | 0 | 4 | 0.25 | 3 | 0 | 24 | 0.1667 | 1 | 1 | 6 | 0 | 4 | 0 | 4 | 0 | 15 | 0.0667 |
| 8_11613236 | 4 | 0.25 | 9 | 0.3333 | 5 | 0 | 5 | 0.2 | 23 | 0.2174 | 2 | 0 | 3 | 0 | 5 | 0.4 | 2 | 0 | 12 | 0.1667 |
| 8_11661915 | 3 | 0 | 13 | 0.0769 | 8 | 0.125 | 6 | 0 | 30 | 0.0667 | 5 | 0 | 2 | 0 | 4 | 0 | 9 | 0.1111 | 20 | 0.05 |
| 8_11661974 | 5 | 0 | 12 | 0.1667 | 7 | 0 | 7 | 0 | 31 | 0.0645 | 4 | 0.25 | 2 | 0 | 4 | 0 | 8 | 0.25 | 18 | 0.1667 |
| 8_11662060 | 5 | 0 | 12 | 0.1667 | 7 | 0 | 7 | 0 | 31 | 0.0645 | 5 | 0 | 2 | 0 | 4 | 0 | 9 | 0.2222 | 20 | 0.1 |
| 8_11662153 | 3 | 0.6667 | 7 | 0.2857 | 3 | 0.6667 | 4 | 0.25 | 17 | 0.4118 | 4 | 0.75 | 1 | 1 | 2 | 0 | 4 | 0.5 | 11 | 0.5455 |
| 8_11662224 | 4 | 0.25 | 12 | 0.6667 | 2 | 0 | 4 | 0.25 | 22 | 0.4545 | 3 | 1 | 2 | 0.5 | 4 | 0 | 8 | 0.375 | 17 | 0.4118 |
| 8_11662260 | 5 | 0 | 7 | 0.2857 | 4 | 0 | 5 | 0 | 21 | 0.0952 | 2 | 0 | 1 | 0 | 4 | 0 | 7 | 0.4286 | 14 | 0.2143 |
| 8_11662351 | 5 | 0.2 | 11 | 0.1818 | 5 | 0.2 | 6 | 0 | 27 | 0.1481 | 5 | 0 | 2 | 0.5 | 4 | 0.25 | 10 | 0.2 | 21 | 0.1905 |
| 8_11662435 | 4 | 0 | 11 | 0.1818 | 3 | 0.6667 | 3 | 0 | 21 | 0.1905 | 4 | 0.25 | 2 | 0 | 4 | 0 | 7 | 0 | 17 | 0.0588 |

| | | | | | | | | | | | | | | | | | | | | |
|---|---|---|---|---|---|---|---|---|---|---|---|---|---|---|---|---|---|---|---|---|
| 8_1166260 | 3 | 0.3333 | 10 | 0.6 | 4 | 0.75 | 6 | 0.1667 | 23 | 0.4783 | 5 | 0.4 | 2 | 0.5 | 4 | 0.25 | 9 | 0.6667 | 20 | 0.5 |
| 8_1166262 | 5 | 0.2 | 9 | 0.5556 | 3 | 0.6667 | 5 | 0.4 | 22 | 0.4545 | 5 | 0.6 | 2 | 0.5 | 4 | 0.75 | 9 | 0.7778 | 20 | 0.7 |
| 8_1166264 | 3 | 0 | 10 | 0.4 | 4 | 0.25 | 5 | 0.6 | 22 | 0.3636 | 4 | 0.25 | 2 | 0 | 3 | 0 | 9 | 0.3333 | 18 | 0.2222 |
| 8_1166273 | 5 | 0.4 | 10 | 0.6 | 4 | 0.75 | 7 | 0.8571 | 26 | 0.6538 | 5 | 0.4 | 2 | 1 | 4 | 0.25 | 10 | 0.8 | 21 | 0.619 |
| 8_1166275 | 5 | 1 | 12 | 0.25 | 3 | 1 | 5 | 0.2 | 25 | 0.48 | 4 | 0.25 | 2 | 0.5 | 4 | 0.25 | 8 | 0.625 | 18 | 0.4444 |
| 8_1166279 | 4 | 0.25 | 9 | 0.3333 | 4 | 0.5 | 6 | 0.1667 | 23 | 0.3043 | 3 | 0 | 1 | 0 | 3 | 0 | 7 | 0.1429 | 14 | 0.0714 |
| 8_1166288 | 3 | 0 | 8 | 0.25 | 2 | 0 | 6 | 0.3333 | 19 | 0.2105 | 5 | 0.2 | 2 | 0.5 | 4 | 0.25 | 8 | 0.5 | 19 | 0.3684 |
| 8_1166288 | 4 | 0 | 7 | 0 | 4 | 0 | 5 | 0.2 | 20 | 0.05 | 4 | 0.25 | 2 | 0.5 | 2 | 0 | 8 | 0.5 | 16 | 0.375 |
| 8_1166524 | 1 | 0 | 2 | 0 | 1 | 1 | 2 | 0 | 6 | 0.1667 | 1 | 0 | NA | NA | 3 | 0 | 5 | 0.4 | 9 | 0.2222 |
| 8_1166552 | NA | NA | 2 | 0 | NA | NA | 2 | 0.5 | 4 | 0.25 | 1 | 0 | NA | NA | 3 | 0.3333 | 5 | 0 | 9 | 0.1111 |
| 8_1166573 | NA | NA | 2 | 0.5 | NA | NA | 1 | 0 | 3 | 0.3333 | 1 | 1 | NA | NA | 3 | 0.6667 | 4 | 0.5 | 8 | 0.625 |
| 8_1166580 | NA | NA | NA | NA | 1 | 1 | NA | NA | 1 | 1 | 1 | 0 | NA | NA | 2 | 0.5 | 2 | 0.5 | 5 | 0.4 |
| 8_1166580 | NA | NA | 1 | 0 | 1 | 1 | 2 | 0 | 4 | 0.25 | 1 | 1 | NA | NA | 3 | 0.6667 | 4 | 0.25 | 8 | 0.5 |
| 8_1168663 | NA | NA | 1 | 1 | NA | NA | NA | NA | 1 | 1 | NA | NA | 1 | 1 | NA | NA | NA | NA | 1 | 1 |
| 8_1187462 | 3 | 0 | 8 | 0.25 | 1 | 0 | 4 | 0 | 16 | 0.125 | 7 | 0 | 2 | 0 | 7 | 0.1429 | 5 | 0.4 | 21 | 0.1429 |
| 8_1187463 | 3 | 0.3333 | 7 | 0 | 1 | 0 | 3 | 0.3333 | 14 | 0.1429 | 5 | 0.2 | 2 | 0.5 | 6 | 0.1667 | 3 | 0.3333 | 16 | 0.25 |
| 8_1187464 | 3 | 0.6667 | 6 | 0.5 | 1 | 1 | 3 | 0.3333 | 13 | 0.5385 | 4 | 0.75 | 2 | 1 | 5 | 0.6 | 4 | 0.75 | 15 | 0.7333 |
| 8_1187471 | 3 | 0 | 4 | 0.5 | 1 | 1 | 2 | 0 | 10 | 0.3 | 4 | 0.5 | 2 | 0.5 | 3 | 0 | 5 | 0.2 | 14 | 0.2857 |
| 8_1187471 | 4 | 0 | 5 | 0.2 | NA | NA | 1 | 0 | 10 | 0.1 | 2 | 0 | 2 | 0 | 5 | 0.2 | 4 | 0 | 13 | 0.0769 |
| 8_1187488 | 5 | 0.4 | 5 | 0.2 | 1 | 0 | 2 | 0.5 | 13 | 0.3077 | 4 | 0 | 3 | 0.6667 | 6 | 0.3333 | 5 | 0.4 | 18 | 0.3333 |
| 8_1187501 | 5 | 0.8 | 6 | 0.3333 | 3 | 1 | 1 | 1 | 15 | 0.6667 | 5 | 0 | 3 | 0.6667 | 6 | 0.5 | 6 | 0.1667 | 20 | 0.3 |
| 8_1187542 | 5 | 0 | 4 | 0 | 4 | 0.25 | 3 | 0.3333 | 16 | 0.125 | 6 | 0 | 4 | 0.5 | 8 | 0.125 | 7 | 0.1429 | 25 | 0.16 |
| 8_1187668 | 5 | 0 | 7 | 0.1429 | 7 | 0.1429 | 3 | 0 | 22 | 0.0909 | 8 | 0 | 7 | 0.5714 | 9 | 0.2222 | 6 | 0.1667 | 30 | 0.2333 |
| 8_1188302 | 3 | 0 | 9 | 0 | 1 | 0 | 8 | 0.25 | 21 | 0.0952 | 2 | 0 | 1 | 1 | 3 | 0 | 3 | 0 | 9 | 0.1111 |
| 8_1188313 | 2 | 0 | 2 | 0 | NA | NA | 2 | 1 | 6 | 0.3333 | 1 | 0 | NA | NA | 3 | 0.3333 | 2 | 0 | 6 | 0.1667 |
| 8_1188316 | 2 | 0 | 8 | 0.625 | 1 | 1 | 8 | 0.25 | 19 | 0.4211 | 2 | 0.5 | 1 | 1 | 4 | 0.25 | 2 | 0 | 9 | 0.3333 |
| 8_1188317 | 3 | 0.3333 | 8 | 0.875 | 1 | 1 | 8 | 0.75 | 20 | 0.75 | 2 | 1 | NA | NA | 4 | 0.25 | 3 | 1 | 9 | 0.6667 |
| 8_1188325 | 2 | 0.5 | 9 | 0.4444 | 1 | 1 | 7 | 0.5714 | 19 | 0.5263 | 2 | 1 | NA | NA | 4 | 0.25 | 3 | 1 | 9 | 0.6667 |
| 8_1188330 | 1 | 1 | 9 | 0.6667 | 1 | 1 | 7 | 0.7143 | 18 | 0.7222 | 1 | 1 | 2 | 1 | 2 | 1 | 2 | 1 | 7 | 1 |
| 8_1188332 | 2 | 1 | 9 | 0.5556 | 1 | 1 | 8 | 0.625 | 20 | 0.65 | 2 | 1 | 2 | 0.5 | 4 | 0.75 | 4 | 0.75 | 12 | 0.75 |
| 8_1188340 | 3 | 0.3333 | 10 | 0.2 | 1 | 1 | 7 | 0.4286 | 21 | 0.3333 | 2 | 0 | 2 | 0 | 4 | 0.75 | 4 | 0.5 | 12 | 0.4167 |
| 8_1188461 | 3 | 0 | 13 | 0.1538 | 4 | 0 | 7 | 0.1429 | 27 | 0.1111 | 4 | 0.25 | 2 | 0 | 3 | 0 | 5 | 0 | 14 | 0.0714 |
| 8_1188466 | 3 | 0 | 13 | 0.8462 | 5 | 1 | 10 | 0.6 | 31 | 0.7097 | 3 | 0.6667 | 2 | 1 | 4 | 0.75 | 4 | 1 | 13 | 0.8462 |
| 8_1188474 | 3 | 0 | 10 | 0.2 | 3 | 0 | 9 | 0 | 25 | 0.08 | 3 | 0.3333 | 2 | 0 | 3 | 0 | 3 | 0.3333 | 11 | 0.1818 |
| 8_1188505 | 7 | 0 | 14 | 0.1429 | 4 | 0 | 12 | 0 | 37 | 0.0541 | 5 | 0 | 3 | 0 | 6 | 0.1667 | 5 | 0 | 19 | 0.0526 |
| 8_1188512 | 5 | 0.2 | 11 | 0.0909 | 5 | 0.2 | 10 | 0 | 31 | 0.0968 | 4 | 0 | 2 | 0.5 | 6 | 0.5 | 4 | 0.25 | 16 | 0.3125 |
| 8_1244612 | 3 | 0 | 1 | 0 | 3 | 0.3333 | 4 | 0.25 | 11 | 0.1818 | NA | NA | 3 | 0.6667 | 2 | 0 | 7 | 0.2857 | 12 | 0.3333 |
| 8_1244614 | 3 | 0.3333 | NA | NA | 3 | 0.6667 | 6 | 0.3333 | 12 | 0.4167 | NA | NA | 4 | 0.75 | 2 | 0 | 6 | 0.6667 | 12 | 0.5833 |
| 8_1244642 | 2 | 0.5 | 1 | 1 | 4 | 0.5 | 6 | 0.5 | 13 | 0.5385 | NA | NA | 3 | 0.3333 | 1 | 0 | 7 | 0.7143 | 11 | 0.5455 |
| 8_1244655 | 2 | 0 | 1 | 1 | 3 | 0.3333 | 4 | 0.25 | 10 | 0.3 | NA | NA | 3 | 0.6667 | 1 | 0 | 6 | 0 | 10 | 0.2 |
| 8_1244667 | 3 | 0.6667 | 2 | 0.5 | 6 | 1 | 6 | 0.8333 | 17 | 0.8235 | 1 | 0 | 5 | 0.8 | 2 | 1 | 8 | 0.875 | 16 | 0.8125 |
| 8_1244675 | 3 | 1 | 1 | 1 | 5 | 1 | 5 | 1 | 14 | 1 | NA | NA | 6 | 1 | 2 | 1 | 7 | 1 | 15 | 1 |
| 8_1244724 | 5 | 0.6 | 2 | 0 | 2 | 0.5 | 6 | 0.1667 | 15 | 0.3333 | 1 | 0 | 5 | 0.8 | 1 | 1 | 3 | 0.3333 | 10 | 0.6 |
| 8_1244737 | 5 | 0.2 | 5 | 0.4 | 8 | 0.625 | 9 | 0.3333 | 27 | 0.4074 | 2 | 0.5 | 5 | 1 | 3 | 0.6667 | 9 | 0.5556 | 19 | 0.6842 |
| 8_1244755 | 7 | 0.7143 | 5 | 0.8 | 6 | 0.8333 | 10 | 0.9 | 28 | 0.8214 | 2 | 0.5 | 6 | 0.8333 | 5 | 1 | 8 | 0.875 | 21 | 0.8571 |
| 8_1244768 | 6 | 0.1667 | 4 | 0 | 8 | 0.125 | 9 | 0.1111 | 27 | 0.1111 | 1 | 0 | 7 | 0.2857 | 5 | 0 | 9 | 0.2222 | 22 | 0.1818 |
| 8_1244771 | 6 | 0.6667 | 3 | 1 | 9 | 0.8889 | 11 | 0.6364 | 29 | 0.7586 | 2 | 1 | 6 | 1 | 5 | 0.6 | 9 | 0.8889 | 22 | 0.8636 |
| 8_1244773 | 7 | 0.4286 | 3 | 0.3333 | 8 | 0.25 | 9 | 0.3333 | 27 | 0.3333 | 1 | 0 | 6 | 0.3333 | 4 | 0.25 | 8 | 0.25 | 19 | 0.2632 |
| 8_1244774 | 4 | 1 | 1 | 1 | 9 | 1 | 3 | 1 | 17 | 1 | NA | NA | 4 | 1 | 3 | 1 | 4 | 1 | 11 | 1 |
| 8_1244775 | 5 | 1 | 1 | 1 | 7 | 1 | 6 | 1 | 19 | 1 | 2 | 1 | 5 | 1 | 5 | 1 | 7 | 1 | 19 | 1 |
| 8_1244777 | 8 | 0.5 | 5 | 0.6 | 8 | 0.25 | 10 | 0.4 | 31 | 0.4194 | 2 | 0.5 | 7 | 0.4286 | 5 | 0.4 | 10 | 0.5 | 24 | 0.4583 |
| 8_1244783 | 10 | 0.5 | 4 | 0.75 | 9 | 0.5556 | 11 | 0.1818 | 34 | 0.4412 | 1 | 0 | 7 | 0.1429 | 5 | 0.6 | 10 | 0.6 | 23 | 0.4348 |
| 8_1244784 | 9 | 0.2222 | 4 | 0 | 8 | 0.375 | 10 | 0.4 | 31 | 0.2903 | 3 | 0.6667 | 7 | 0.2857 | 5 | 0.6 | 11 | 0.3636 | 26 | 0.4231 |
| 8_1244787 | 5 | 0.8 | 3 | 1 | 6 | 0.6667 | 8 | 0.75 | 22 | 0.7727 | 2 | 0.5 | 8 | 0.75 | 5 | 0.4 | 9 | 0.7778 | 24 | 0.6667 |
| 8_1244789 | 6 | 0.3333 | 2 | 0 | 5 | 0.4 | 4 | 0.25 | 17 | 0.2941 | NA | NA | NA | NA | 2 | 0.5 | 5 | 0.2 | 7 | 0.2857 |
| 8_1244794 | 8 | 0.5 | 4 | 0.5 | 7 | 0.4286 | 8 | 0.375 | 27 | 0.4444 | 2 | 0.5 | 8 | 0.5 | 5 | 0.4 | 11 | 0.2727 | 26 | 0.3846 |
| 8_1244799 | 9 | 0.1111 | 3 | 0 | 6 | 0.5 | 9 | 0 | 27 | 0.1481 | 2 | 0.5 | 8 | 0.5 | 5 | 0 | 11 | 0.1818 | 26 | 0.2692 |
| 8_1244845 | 8 | 0.125 | 3 | 0.3333 | 8 | 0.25 | 7 | 0.1429 | 26 | 0.1923 | 2 | 0 | 5 | 0.2 | 4 | 0 | 8 | 0 | 19 | 0.0526 |
| 8_1244934 | 11 | 0.3636 | 5 | 0.2 | 5 | 0.2 | 6 | 0.3333 | 27 | 0.2963 | 2 | 0 | 6 | 0.3333 | 1 | 0 | 4 | 0.75 | 13 | 0.3846 |
| 8_1248104 | 1 | 0 | NA | NA | NA | NA | 2 | 0.5 | 3 | 0.3333 | 3 | 0.3333 | 1 | 0 | NA | NA | NA | NA | 4 | 0.25 |
| 8_1248110 | NA | NA | NA | NA | NA | NA | 1 | 1 | 1 | 1 | 3 | 0.6667 | NA | NA | NA | NA | NA | NA | 3 | 0.6667 |
| 8_1248111 | 1 | 1 | NA | NA | NA | NA | 2 | 1 | 3 | 1 | 3 | 0.3333 | 1 | 0 | NA | NA | NA | NA | 4 | 0.25 |
| 8_1248169 | 2 | 0.5 | NA | NA | 1 | 0 | 2 | 1 | 5 | 0.6 | 4 | 0.5 | 1 | 0 | 1 | 0 | 2 | 0 | 8 | 0.25 |
| 8_1248235 | NA | NA | NA | NA | 1 | 1 | 1 | 0 | 2 | 0.5 | 1 | 0 | NA | NA | 3 | 0.3333 | 2 | 0.5 | 6 | 0.3333 |
| 8_1248259 | NA | NA | NA | NA | 1 | 0 | 2 | 1 | 3 | 0.6667 | 1 | 1 | NA | NA | 2 | 0.5 | 2 | 0.5 | 5 | 0.6 |

| | | | | | | | | | | | | | | | | | | | | |
|---|---|---|---|---|---|---|---|---|---|---|---|---|---|---|---|---|---|---|---|---|
| 8_1248267( | NA | NA | NA | NA | 1 | 1 | 2 | 1 | 3 | 1 | 2 | 1 | NA | NA | 3 | 1 | 1 | 1 | 6 | 1 |
| 8_1248268! | NA | NA | NA | NA | 1 | 1 | 2 | 1 | 3 | 1 | 1 | 1 | NA | NA | 2 | 0.5 | 1 | 1 | 4 | 0.75 |
| 8_1248270! | NA | NA | NA | NA | NA | NA | 2 | 0.5 | 2 | 0.5 | 1 | 1 | NA | NA | 2 | 1 | 2 | 1 | 5 | 1 |
| 8_1248280! | NA | NA | NA | NA | 1 | 1 | 2 | 0 | 3 | 0.3333 | 2 | 1 | NA | NA | 3 | 0.6667 | 1 | 0 | 6 | 0.6667 |
| 8_1248284! | NA | NA | NA | NA | 1 | 1 | 3 | 0.6667 | 4 | 0.75 | 1 | 1 | NA | NA | 3 | 1 | 2 | 1 | 6 | 1 |
| 8_1248294! | NA | NA | NA | NA | 1 | 0 | 3 | 0.6667 | 4 | 0.5 | 2 | 1 | NA | NA | 2 | 1 | 1 | 0 | 5 | 0.8 |
| 8_1248300! | NA | NA | 1 | 1 | NA | NA | 3 | 0.6667 | 4 | 0.75 | 2 | 0.5 | NA | NA | 2 | 0 | 1 | 1 | 5 | 0.4 |
| 8_1248301 | NA | NA | 1 | 0 | 2 | 0 | 3 | 0.3333 | 6 | 0.1667 | 1 | 0 | NA | NA | 1 | 1 | 1 | 0 | 3 | 0.3333 |
| 8_1248312 | NA | NA | 1 | 1 | 2 | 1 | 3 | 1 | 6 | 1 | 2 | 1 | NA | NA | 2 | 1 | 1 | 1 | 5 | 1 |
| 8_1248318 | NA | NA | 1 | 1 | 2 | 0 | 3 | 0.3333 | 6 | 0.3333 | 2 | 0.5 | NA | NA | 3 | 0 | 1 | 0 | 6 | 0.1667 |
| 8_1248333 | NA | NA | 1 | 1 | 1 | 1 | 4 | 1 | 6 | 1 | 2 | 1 | NA | NA | 2 | 1 | 1 | 1 | 5 | 1 |
| 8_1248373( | NA | NA | 1 | 1 | 3 | 0 | 4 | 0.25 | 8 | 0.25 | 3 | 0.6667 | NA | NA | 5 | 0.2 | NA | NA | 8 | 0.375 |
| 8_1248455 | 1 | 0 | 1 | 0 | 4 | 0.25 | 4 | 0 | 10 | 0.1 | 3 | 0 | NA | NA | 9 | 0.1111 | 1 | 0 | 13 | 0.0769 |
| 8_1248505 | 1 | 0 | 1 | 0 | 5 | 0.6 | 3 | 0.6667 | 10 | 0.5 | 3 | 0.6667 | 1 | 0 | 8 | 0.625 | 1 | 1 | 13 | 0.6154 |
| 8_1248556 | 1 | 0 | 2 | 0 | 6 | 0.1667 | 4 | 0 | 13 | 0.0769 | 3 | 0 | 1 | 0 | 8 | 0.125 | 1 | 0 | 13 | 0.0769 |
| 8_1248577 | 1 | 0 | 2 | 0.5 | 6 | 0.1667 | 7 | 0.1429 | 16 | 0.1875 | 3 | 0 | 4 | 0.25 | 7 | 0 | 1 | 1 | 15 | 0.1333 |
| 8_1248579 | 1 | 0 | 2 | 0.5 | 6 | 0.5 | 7 | 0.8571 | 16 | 0.625 | 3 | 1 | 4 | 0.75 | 9 | 0.4444 | 1 | 0 | 17 | 0.5882 |
| 8_1248581 | 1 | 1 | 1 | 1 | 4 | 0.75 | 6 | 1 | 12 | 0.9167 | 3 | 0.3333 | 4 | 0.75 | 8 | 0.75 | 1 | 1 | 16 | 0.6875 |
| 8_1248594 | 1 | 1 | 2 | 0.5 | 6 | 1 | 7 | 0.8571 | 16 | 0.875 | 2 | 0 | 4 | 0.75 | 9 | 0.6667 | 1 | 1 | 16 | 0.625 |
| 8_1248601 | 1 | 0 | 2 | 0 | 5 | 0 | 7 | 0.2857 | 15 | 0.1333 | 3 | 0 | 5 | 0.2 | 9 | 0.3333 | 1 | 0 | 18 | 0.2222 |
| 8_1248608 | 1 | 0 | 2 | 0 | 6 | 0.1667 | 7 | 0.1429 | 16 | 0.125 | 3 | 0 | 4 | 0 | 9 | 0.1111 | 1 | 0 | 17 | 0.0588 |
| 8_1248618 | 1 | 0 | 2 | 0 | 6 | 0.1667 | 7 | 0.2857 | 16 | 0.1875 | 3 | 0.3333 | 5 | 0 | 9 | 0.1111 | 1 | 0 | 18 | 0.1111 |
| 8_1266487 | 3 | 0 | 5 | 0.4 | 5 | 0.6 | 3 | 0.3333 | 16 | 0.375 | 1 | 0 | 2 | 0 | 1 | 1 | 4 | 0.25 | 8 | 0.25 |
| 8_1266491 | 2 | 0.5 | 6 | 0 | 5 | 0.4 | 4 | 0 | 17 | 0.1765 | 2 | 0.5 | 2 | 0 | 2 | 0.5 | 4 | 0.25 | 10 | 0.3 |
| 8_1266493 | 3 | 0.3333 | 5 | 0 | 4 | 0 | 3 | 0 | 15 | 0.0667 | 2 | 0.5 | 2 | 0 | 2 | 0 | 3 | 0 | 9 | 0.1111 |
| 8_1266500 | 1 | 0 | 5 | 0.8 | 3 | 0.3333 | 3 | 0.3333 | 12 | 0.5 | 2 | 0.5 | 1 | 1 | 2 | 1 | 3 | 0.6667 | 8 | 0.75 |
| 8_1266521( | 2 | 0 | 4 | 0 | 4 | 0.5 | NA | NA | 10 | 0.2 | 1 | 0 | 2 | 0.5 | 2 | 0 | 2 | 0 | 7 | 0.1429 |
| 8_1266522( | 3 | 0 | 6 | 0.3333 | 3 | 0 | 3 | 0 | 15 | 0.1333 | 2 | 0 | 1 | 0 | 2 | 0.5 | 4 | 0.25 | 9 | 0.2222 |
| 8_1291499 | NA | NA | NA | NA | 1 | 1 | 2 | 0 | 3 | 0.3333 | 1 | 0 | 3 | 0 | 2 | 0.5 | 1 | 0 | 7 | 0.1429 |
| 8_1291515 | NA | NA | NA | NA | 1 | 1 | 1 | 1 | 2 | 1 | 3 | 0.6667 | 2 | 1 | 2 | 1 | 1 | 0 | 8 | 0.75 |
| 8_1291536( | 1 | 1 | NA | NA | 2 | 1 | 1 | 1 | 4 | 1 | 2 | 1 | 3 | 1 | 2 | 1 | 1 | 1 | 8 | 1 |
| 8_1291561 | 1 | 1 | NA | NA | 3 | 1 | 2 | 1 | 6 | 1 | 4 | 0.75 | 3 | 1 | 2 | 1 | 2 | 1 | 11 | 0.9091 |
| 8_1291586 | 1 | 0 | 1 | 0 | 3 | 0.3333 | 2 | 0 | 7 | 0.1429 | 3 | 0.3333 | 2 | 0.5 | 1 | 0 | 1 | 1 | 7 | 0.4286 |
| 8_1291605 | NA | NA | 1 | 1 | 4 | 0.75 | 1 | 1 | 6 | 0.8333 | 3 | 0.3333 | 2 | 1 | 2 | 0.5 | 2 | 1 | 9 | 0.6667 |
| 8_1291607 | 1 | 1 | 1 | 1 | 6 | 1 | 2 | 1 | 10 | 1 | 4 | 1 | 3 | 1 | 2 | 1 | 3 | 1 | 12 | 1 |
| 8_1291608 | 1 | 0 | 1 | 0 | 5 | 1 | 2 | 0 | 9 | 0.5556 | 4 | 1 | 4 | 0.75 | 2 | 1 | 3 | 0.6667 | 13 | 0.8462 |
| 8_1291615 | 1 | 0 | 1 | 0 | 6 | 0.3333 | 2 | 0 | 10 | 0.2 | 4 | 0.25 | 3 | 0 | 3 | 0 | 3 | 0 | 13 | 0.0769 |
| 8_1291635 | 1 | 0 | 1 | 0 | 6 | 0.1667 | 2 | 0 | 10 | 0.1 | 4 | 0.25 | 2 | 1 | 3 | 0 | 3 | 1 | 12 | 0.5 |
| 8_1291693 | 2 | 0.5 | 2 | 0 | 7 | 0.5714 | 3 | 0 | 14 | 0.3571 | 6 | 0.5 | 4 | 1 | 5 | 0.4 | 6 | 0.6667 | 21 | 0.619 |
| 8_1291735 | 4 | 0 | 5 | 0 | 6 | 0 | 3 | 0.3333 | 18 | 0.0556 | 5 | 0.2 | 5 | 0 | 4 | 0 | 6 | 0 | 20 | 0.05 |
| 8_1291771 | 4 | 0 | 4 | 0.25 | 4 | 0 | 3 | 0 | 15 | 0.0667 | 7 | 0.2857 | 5 | 0 | 7 | 0.1429 | 6 | 0.1667 | 25 | 0.16 |
| 8_1291812 | 4 | 0 | 8 | 0 | 7 | 0.2857 | 5 | 0.2 | 24 | 0.125 | 8 | 0.375 | 5 | 0 | 7 | 0.1429 | 9 | 0.1111 | 29 | 0.1724 |
| 8_1291813 | 5 | 0.4 | 8 | 0.5 | 7 | 0.4286 | 5 | 0.8 | 25 | 0.52 | 8 | 0.75 | 5 | 0.8 | 6 | 0.6667 | 9 | 0.5556 | 28 | 0.6786 |
| 8_1291867 | 1 | 0 | 6 | 0 | 6 | 0.6667 | 3 | 0 | 16 | 0.25 | 6 | 0 | 4 | 0 | 7 | 0.2857 | 9 | 0.2222 | 26 | 0.1538 |
| 8_1303450 | 2 | 0.5 | 1 | 1 | NA | NA | 1 | 0 | 4 | 0.5 | NA | NA | NA | NA | 2 | 1 | 1 | 1 | 3 | 1 |
| 8_1303478 | 1 | 1 | 1 | 1 | NA | NA | NA | NA | 2 | 1 | NA | NA | NA | NA | 1 | 1 | 1 | 0 | 2 | 0.5 |
| 8_1306893 | 8 | 0 | 10 | 0.1 | 3 | 0.3333 | 7 | 0.1429 | 28 | 0.1071 | 4 | 0 | 5 | 0.4 | 10 | 0 | 12 | 0.1667 | 31 | 0.129 |
| 8_1306921 | 12 | 0.9167 | 15 | 0.6 | 6 | 0.6667 | 10 | 0.9 | 43 | 0.7674 | 7 | 1 | 9 | 0.7778 | 13 | 0.9231 | 15 | 0.8667 | 44 | 0.8864 |
| 8_1306923 | 14 | 0.0714 | 17 | 0 | 6 | 0.3333 | 10 | 0 | 47 | 0.0638 | 8 | 0 | 10 | 0 | 16 | 0.125 | 15 | 0.3333 | 49 | 0.1429 |
| 8_1306929 | 12 | 0.75 | 16 | 0.625 | 5 | 0.6 | 9 | 0.8889 | 42 | 0.7143 | 8 | 0.75 | 8 | 0.875 | 16 | 0.75 | 12 | 0.6667 | 44 | 0.75 |
| 8_1306930 | 13 | 0.8462 | 15 | 0.7333 | 4 | 0.75 | 11 | 0.7273 | 43 | 0.7674 | 8 | 1 | 7 | 0.8571 | 13 | 0.7692 | 10 | 0.9 | 38 | 0.8684 |
| 8_1306945 | 12 | 0.75 | 12 | 0.3333 | 6 | 0.1667 | 7 | 0.4286 | 37 | 0.4595 | 7 | 0.1429 | 9 | 0.3333 | 12 | 0.4167 | 12 | 0.3333 | 40 | 0.325 |
| 8_1306949 | 13 | 0.8462 | 15 | 0.8667 | 9 | 0.8889 | 11 | 0.7273 | 48 | 0.8333 | 8 | 0.875 | 9 | 0.8889 | 18 | 0.8333 | 15 | 0.8667 | 50 | 0.86 |
| 8_1306960 | 13 | 0 | 17 | 0.1765 | 8 | 0.125 | 12 | 0.1667 | 50 | 0.12 | 9 | 0.1111 | 8 | 0 | 16 | 0 | 15 | 0.2 | 48 | 0.0833 |
| 8_1306991 | 15 | 0 | 19 | 0.0526 | 9 | 0.1111 | 12 | 0.0833 | 55 | 0.0545 | 10 | 0.1 | 13 | 0.1538 | 18 | 0.1111 | 14 | 0 | 55 | 0.0909 |
| 8_1307016 | 17 | 0.1765 | 17 | 0 | 8 | 0.125 | 11 | 0 | 53 | 0.0755 | 12 | 0.1667 | 12 | 0.0833 | 19 | 0.1053 | 15 | 0 | 58 | 0.0862 |
| 8_1312281 | NA | NA | 3 | 0.6667 | 1 | 1 | 3 | 0 | 7 | 0.4286 | 2 | 0.5 | 5 | 0.4 | 3 | 0 | 2 | 0.5 | 12 | 0.3333 |
| 8_1312301 | NA | NA | 2 | 0.5 | 1 | 1 | 3 | 0.3333 | 6 | 0.5 | 2 | 0 | 4 | 0 | 5 | 0 | 2 | 0.5 | 13 | 0.0769 |
| 8_1312324( | NA | NA | 2 | 0 | 1 | 1 | NA | NA | 3 | 0.3333 | 1 | 1 | 4 | 0.5 | 4 | 0 | 1 | 0 | 10 | 0.3 |
| 8_1312332 | NA | NA | NA | NA | NA | NA | 1 | 1 | 1 | 1 | 1 | 1 | 4 | 1 | 4 | 1 | 2 | 0.5 | 11 | 0.9091 |
| 8_1312336 | NA | NA | NA | NA | 1 | 1 | 2 | 1 | 3 | 1 | 1 | 0 | 3 | 0.6667 | 2 | 0 | 2 | 0 | 8 | 0.25 |
| 8_1312351 | NA | NA | NA | NA | 1 | 0 | 2 | 0.5 | 3 | 0.3333 | 1 | 0 | 4 | 0.5 | 4 | 0.5 | 2 | 0 | 11 | 0.3636 |
| 8_1312355 | NA | NA | NA | NA | 1 | 1 | 1 | 1 | 2 | 1 | NA | NA | 4 | 1 | 3 | 0.3333 | 2 | 1 | 9 | 0.7778 |
| 8_1312359 | NA | NA | NA | NA | NA | NA | 1 | 1 | 1 | 1 | 1 | 1 | 3 | 1 | 3 | 1 | 2 | 1 | 9 | 1 |
| 8_1312389( | NA | NA | NA | NA | NA | NA | 1 | 1 | 1 | 1 | 1 | 1 | 4 | 1 | 3 | 1 | 1 | 1 | 9 | 1 |

| | | | | | | | | | | | | | | | | | | | | |
|---|---|---|---|---|---|---|---|---|---|---|---|---|---|---|---|---|---|---|---|---|
| 8_1312477 | NA | NA | 1 | 1 | 1 | 0 | NA | NA | 2 | 0.5 | 1 | 1 | 2 | 0.5 | 2 | 1 | NA | NA | 5 | 0.8 |
| 8_1312804 | 3 | 0 | 1 | 1 | 3 | 0.6667 | 2 | 0.5 | 9 | 0.4444 | 2 | 0.5 | 2 | 1 | 3 | 1 | 3 | 0.6667 | 10 | 0.8 |
| 8_1312823 | 3 | 0 | 1 | 0 | 5 | 0 | 8 | 0.125 | 17 | 0.0588 | 2 | 0.5 | 2 | 0 | 5 | 0 | 4 | 0 | 13 | 0.0769 |
| 8_1312825 | 2 | 0 | 1 | 0 | 5 | 0 | 8 | 0.125 | 16 | 0.0625 | 2 | 1 | 2 | 0.5 | 4 | 0.25 | 4 | 0 | 12 | 0.3333 |
| 8_1315504 | NA | NA | NA | NA | 1 | 1 | NA | NA | 1 | 1 | NA | NA | NA | NA | 1 | 0 | 2 | 0.5 | 3 | 0.3333 |
| 8_1315518 | NA | NA | NA | NA | 1 | 1 | NA | NA | 1 | 1 | NA | NA | NA | NA | NA | NA | 2 | 0.5 | 2 | 0.5 |
| 8_1315524 | NA | NA | NA | NA | 1 | 1 | NA | NA | 1 | 1 | NA | NA | NA | NA | 1 | 0 | 2 | 0.5 | 3 | 0.3333 |
| 8_1318371 | 5 | 0.2 | 6 | 0 | 6 | 0 | 2 | 0 | 19 | 0.0526 | NA | NA | 4 | 0.25 | 3 | 0 | 1 | 0 | 8 | 0.125 |
| 8_1319656 | 8 | 0.25 | 7 | 0 | 11 | 0 | 5 | 0.2 | 31 | 0.0968 | 1 | 0 | 4 | 0 | 4 | 0 | 2 | 0.5 | 11 | 0.0909 |
| 8_1319657 | 7 | 0.5714 | 7 | 0.8571 | 8 | 0.625 | 6 | 0.8333 | 28 | 0.7143 | 1 | 1 | 5 | 0.8 | 4 | 0.75 | 2 | 1 | 12 | 0.8333 |
| 8_1319659 | 5 | 0.2 | 7 | 0.2857 | 7 | 0.4286 | 5 | 0.4 | 24 | 0.3333 | 1 | 1 | 4 | 0.5 | 3 | 0.6667 | 2 | 0 | 10 | 0.5 |
| 8_1319663 | 6 | 0 | 7 | 0.2857 | 10 | 0.2 | 6 | 0.5 | 29 | 0.2414 | 1 | 0 | 3 | 0.3333 | 2 | 1 | 2 | 0 | 8 | 0.375 |
| 8_1319694 | 8 | 0.75 | 8 | 0.375 | 11 | 0.4545 | 8 | 0.375 | 35 | 0.4857 | 2 | 1 | 4 | 0.25 | 3 | 0.3333 | 3 | 0.6667 | 12 | 0.5 |
| 8_1319698 | 8 | 0.375 | 7 | 0.4286 | 9 | 0.4444 | 8 | 0.125 | 32 | 0.3438 | NA | NA | 5 | 0.6 | 4 | 0.5 | 3 | 0.3333 | 12 | 0.5 |
| 8_1319703 | 8 | 0.25 | 7 | 0.4286 | 11 | 0.4545 | 8 | 0.25 | 34 | 0.3529 | 2 | 0 | 4 | 0.5 | 4 | 0 | 3 | 0 | 13 | 0.1538 |
| 8_1319710 | 8 | 0.375 | 8 | 0.25 | 12 | 0.4167 | 6 | 0.1667 | 34 | 0.3235 | 2 | 0.5 | 5 | 0.4 | 4 | 0.25 | 3 | 0.3333 | 14 | 0.3571 |
| 8_1319716 | 8 | 0.375 | 8 | 0 | 9 | 0 | 8 | 0.125 | 33 | 0.1212 | 3 | 0.6667 | 5 | 0 | 3 | 0 | 3 | 0 | 14 | 0.1429 |
| 8_1322465 | 56 | 0.125 | 48 | 0.0208 | 48 | 0.0417 | 61 | 0.082 | 213 | 0.0704 | 56 | 0.1786 | 53 | 0.0566 | 38 | 0.0526 | 32 | 0.0625 | 179 | 0.095 |
| 8_1322501 | 43 | 0.186 | 43 | 0.3256 | 29 | 0.2069 | 43 | 0.2093 | 158 | 0.2342 | 35 | 0.3714 | 32 | 0.2188 | 24 | 0.4167 | 17 | 0.2353 | 108 | 0.3148 |
| 8_1330637 | 5 | 0.8 | 7 | 0.5714 | 6 | 0.6667 | 7 | 0.5714 | 25 | 0.64 | 3 | 0.6667 | 7 | 0.5714 | 7 | 0.4286 | 1 | 1 | 18 | 0.5556 |
| 8_1330647 | 3 | 0.3333 | 7 | 0.5714 | 6 | 0.3333 | 7 | 0.7143 | 23 | 0.5217 | NA | NA | 7 | 0.7143 | 6 | 0.6667 | 1 | 1 | 14 | 0.7143 |
| 8_1330647 | 3 | 0 | 6 | 0.3333 | 6 | 0.1667 | 7 | 0.1429 | 22 | 0.1818 | 1 | 0 | 7 | 0.2857 | 6 | 0.3333 | 1 | 0 | 15 | 0.2667 |
| 8_1330650 | 5 | 0.8 | 6 | 0.6667 | 6 | 0.6667 | 7 | 0.8571 | 24 | 0.75 | 3 | 0.6667 | 9 | 0.5556 | 8 | 0.75 | 2 | 0.5 | 22 | 0.6364 |
| 8_1330651 | 5 | 0.8 | 7 | 0.4286 | 7 | 0.5714 | 7 | 0.8571 | 26 | 0.6538 | 2 | 0.5 | 9 | 0.4444 | 6 | 0.5 | 2 | 0.5 | 19 | 0.4737 |
| 8_1330659 | 2 | 1 | 5 | 1 | 5 | 0.8 | 6 | 0.8333 | 18 | 0.8889 | 2 | 1 | 7 | 1 | 5 | 1 | 1 | 1 | 15 | 1 |
| 8_1330663 | 5 | 1 | 6 | 1 | 4 | 1 | 7 | 1 | 22 | 1 | 3 | 1 | 8 | 1 | 7 | 1 | 2 | 1 | 20 | 1 |
| 8_1330675 | 4 | 1 | 7 | 1 | 7 | 1 | 7 | 1 | 25 | 1 | 3 | 1 | 6 | 1 | 6 | 1 | 2 | 1 | 17 | 1 |
| 8_1330676 | 4 | 1 | 7 | 0.8571 | 7 | 0.8571 | 6 | 0.8333 | 24 | 0.875 | 3 | 0.6667 | 9 | 0.8889 | 7 | 0.8571 | 2 | 0.5 | 21 | 0.8095 |
| 8_1330683 | 3 | 1 | 6 | 0.8333 | 6 | 0.8333 | 7 | 1 | 22 | 0.9091 | 3 | 0.6667 | 7 | 0.7143 | 5 | 0.8 | 2 | 1 | 17 | 0.7647 |
| 8_1330687 | 2 | 1 | 5 | 0.2 | 6 | 0.3333 | 5 | 0.2 | 18 | 0.3333 | 3 | 0 | 7 | 0.1429 | 5 | 0.6 | 1 | 0 | 16 | 0.25 |
| 8_1330718 | 4 | 0.25 | 6 | 0 | 3 | 0 | 5 | 0 | 18 | 0.0556 | 1 | 0 | 7 | 0.2857 | 3 | 0.3333 | 2 | 0.5 | 13 | 0.3077 |
| 8_1330736 | 4 | 1 | 6 | 1 | 4 | 1 | 6 | 1 | 20 | 1 | 1 | 1 | 6 | 1 | 2 | 1 | 2 | 1 | 11 | 1 |
| 8_1330749 | 4 | 0 | 4 | 0 | 5 | 0.2 | 6 | 0 | 19 | 0.0526 | 2 | 0 | 7 | 0 | 4 | 0.25 | 1 | 0 | 14 | 0.0714 |
| 8_1330758 | 5 | 0 | 5 | 0.4 | 5 | 0.2 | 6 | 0.3333 | 21 | 0.2381 | 3 | 0 | 7 | 0.1429 | 3 | 0 | 2 | 0.5 | 15 | 0.1333 |
| 8_1330767 | 4 | 0.75 | 6 | 0.3333 | 4 | 0.5 | 4 | 0.25 | 18 | 0.4444 | 1 | 0 | 9 | 0.3333 | 4 | 0.5 | 2 | 0.5 | 16 | 0.375 |
| 8_1330771 | 5 | 0.2 | 3 | 0 | 5 | 0.4 | 5 | 0.8 | 18 | 0.3889 | 2 | 0.5 | 8 | 0.125 | 3 | 0.6667 | 2 | 1 | 15 | 0.4 |
| 8_1330772 | 3 | 1 | 5 | 0.6 | 4 | 0.75 | 6 | 0.5 | 18 | 0.6667 | 2 | 0.5 | 9 | 0.5556 | 2 | 0.5 | 2 | 0.5 | 15 | 0.5333 |
| 8_1333668 | NA | NA | NA | NA | 1 | 1 | 1 | 1 | 2 | 1 | NA | NA | NA | NA | 1 | 1 | 1 | 0 | 2 | 0.5 |
| 8_1333683 | NA | NA | NA | NA | NA | NA | 1 | 1 | 1 | 1 | NA | NA | NA | NA | 1 | 1 | NA | NA | 1 | 1 |
| 8_1333688 | NA | NA | NA | NA | NA | NA | 1 | 1 | 1 | 1 | NA | NA | NA | NA | 1 | 1 | NA | NA | 1 | 1 |
| 8_1334196 | 3 | 0 | 3 | 0 | NA | NA | 2 | 0.5 | 8 | 0.125 | 2 | 0.5 | 1 | 0 | 2 | 0 | 4 | 0 | 9 | 0.1111 |
| 8_1337102 | 6 | 0.5 | 7 | 0.4286 | 4 | 0.75 | 3 | 1 | 20 | 0.6 | 6 | 0.6667 | 3 | 1 | 4 | 0.75 | 7 | 0.5714 | 20 | 0.7 |
| 8_1342735 | 5 | 0.6 | 3 | 0.3333 | 7 | 0.8571 | 7 | 0.2857 | 22 | 0.5455 | 5 | 1 | 5 | 0.6 | 5 | 0.4 | 6 | 0.5 | 21 | 0.619 |
| 8_1342763 | 10 | 0.4 | 3 | 0.3333 | 9 | 0.1111 | 7 | 0.1429 | 29 | 0.2414 | 7 | 0.2857 | 9 | 0.2222 | 5 | 0.4 | 6 | 0 | 27 | 0.2222 |
| 8_1342765 | 7 | 0.1429 | 3 | 0 | 7 | 0.2857 | 6 | 0.5 | 23 | 0.2609 | 6 | 0.3333 | 8 | 0.125 | 6 | 0 | 4 | 0.25 | 24 | 0.1667 |
| 8_1342776 | 11 | 0.0909 | 3 | 0 | 6 | 0 | 9 | 0.2222 | 29 | 0.1034 | 6 | 0.1667 | 7 | 0 | 6 | 0.1667 | 8 | 0 | 27 | 0.0741 |
| 8_1342783 | 9 | 0.4444 | 4 | 0.25 | 8 | 0.75 | 8 | 0.5 | 29 | 0.5172 | 6 | 0.8333 | 8 | 0.875 | 4 | 0.5 | 8 | 0.625 | 26 | 0.7308 |
| 8_1342786 | 9 | 0.2222 | 3 | 0.3333 | 9 | 0.3333 | 7 | 0 | 28 | 0.2143 | 6 | 0 | 7 | 0.1429 | 4 | 0.25 | 8 | 0 | 25 | 0.08 |
| 8_1351569 | 7 | 0.5714 | 13 | 0.6154 | 8 | 0.25 | 7 | 0.7143 | 35 | 0.5429 | 7 | 0.7143 | 6 | 0.6667 | 11 | 0.3636 | 2 | 0 | 26 | 0.5 |
| 8_1361744 | 7 | 0.1429 | 2 | 0 | 5 | 0 | 6 | 0 | 20 | 0.05 | 7 | 0.1429 | 2 | 0 | 6 | 0 | 5 | 0 | 20 | 0.05 |
| 8_1361754 | 7 | 0.2857 | 2 | 0 | 8 | 0.125 | 4 | 0 | 21 | 0.1429 | 6 | 0.1667 | 1 | 0 | 5 | 0 | 3 | 0.3333 | 15 | 0.1333 |
| 8_1361783 | 7 | 0.2857 | 2 | 0.5 | 5 | 0.2 | 3 | 0.3333 | 17 | 0.2941 | 6 | 0.6667 | 1 | 0 | 5 | 0 | 1 | 0 | 13 | 0.3077 |
| 8_1361807 | 7 | 0 | 2 | 0 | 6 | 0.1667 | 4 | 0 | 19 | 0.0526 | 6 | 0.1667 | 1 | 0 | 5 | 0 | 2 | 0 | 14 | 0.0714 |
| 8_1361838 | 6 | 0.8333 | 2 | 1 | 6 | 0.8333 | 4 | 1 | 18 | 0.8889 | 5 | 1 | 1 | 1 | 5 | 0.8 | 1 | 1 | 12 | 0.9167 |
| 8_1361852 | 4 | 1 | 2 | 0.5 | 4 | 0.75 | 3 | 1 | 13 | 0.8462 | 4 | 0.5 | 1 | 1 | 3 | 0.6667 | NA | NA | 8 | 0.625 |
| 8_1383146 | 6 | 0.5 | 8 | 0.5 | 8 | 0.25 | 10 | 0.3 | 32 | 0.375 | 5 | 0.2 | 8 | 0.25 | 7 | 0.5714 | 7 | 0.2857 | 27 | 0.3333 |
| 8_1383150 | 5 | 0.2 | 6 | 0.1667 | 7 | 0.2857 | 9 | 0.1111 | 27 | 0.1852 | 6 | 0.5 | 8 | 0.25 | 7 | 0.1429 | 6 | 0.3333 | 27 | 0.2963 |
| 8_1383156 | 5 | 0.8 | 8 | 1 | 8 | 0.75 | 8 | 0.875 | 29 | 0.8621 | 6 | 0.8333 | 7 | 0.7143 | 7 | 0.7143 | 7 | 0.8571 | 27 | 0.7778 |
| 8_1383179 | 6 | 0.1667 | 8 | 0 | 8 | 0 | 10 | 0.2 | 32 | 0.0938 | 6 | 0.1667 | 6 | 0 | 7 | 0 | 7 | 0.2857 | 26 | 0.1154 |
| 8_1384835 | 11 | 0.1818 | 5 | 0.4 | 14 | 0.2857 | 7 | 0 | 37 | 0.2162 | 3 | 0 | 5 | 0.2 | 6 | 0.1667 | 3 | 0.3333 | 17 | 0.1765 |
| 8_1384856 | 12 | 0.25 | 5 | 0.4 | 10 | 0.2 | 8 | 0 | 35 | 0.2 | 3 | 0.3333 | 5 | 0 | 6 | 0.1667 | 4 | 0 | 18 | 0.1111 |
| 8_1384868 | 12 | 0.3333 | 5 | 0.2 | 12 | 0 | 7 | 0 | 36 | 0.1389 | 5 | 0.4 | 5 | 0.2 | 7 | 0.2857 | 4 | 0 | 21 | 0.2381 |
| 8_1392857 | NA | NA | 3 | 0.3333 | NA | NA | 3 | 0 | 6 | 0.1667 | NA | NA | 1 | 0 | 2 | 0 | 3 | 0.6667 | 6 | 0.3333 |
| 8_1392858 | NA | NA | 3 | 0.3333 | NA | NA | 3 | 0 | 6 | 0.1667 | NA | NA | 2 | 0 | 2 | 0.5 | 3 | 0 | 7 | 0.1429 |
| 8_1392863 | NA | NA | 4 | 1 | NA | NA | 3 | 0.6667 | 7 | 0.8571 | NA | NA | 2 | 1 | 2 | 0.5 | 1 | 0 | 5 | 0.6 |

| | | | | | | | | | | | | | | | | | | | | |
|---|---|---|---|---|---|---|---|---|---|---|---|---|---|---|---|---|---|---|---|---|
| 8_1392867 | NA | NA | 4 | 0 | NA | NA | 3 | 0.6667 | 7 | 0.2857 | NA | NA | 2 | 1 | 2 | 0.5 | 3 | 0.6667 | 7 | 0.7143 |
| 8_1392880 | NA | NA | 4 | 0.5 | NA | NA | 4 | 0.25 | 8 | 0.375 | NA | NA | 2 | 0.5 | 2 | 0.5 | 3 | 0 | 7 | 0.2857 |
| 8_1392885 | NA | NA | 4 | 0.25 | NA | NA | 4 | 0.75 | 8 | 0.5 | NA | NA | 2 | 0.5 | 2 | 0.5 | 3 | 0.3333 | 7 | 0.4286 |
| 8_1392949 | NA | NA | 3 | 0.3333 | 1 | 1 | 3 | 0.6667 | 7 | 0.5714 | 2 | 0.5 | 1 | 1 | 1 | 1 | 2 | 0.5 | 6 | 0.6667 |
| 8_1392963 | 2 | 1 | 5 | 0.8 | 1 | 1 | 2 | 1 | 10 | 0.9 | 1 | 1 | 2 | 1 | 1 | 1 | 3 | 1 | 7 | 1 |
| 8_1392967 | 2 | 0.5 | 3 | 0.3333 | 1 | 1 | 3 | 1 | 9 | 0.6667 | 1 | 1 | 1 | 1 | 1 | 0 | 2 | 0 | 5 | 0.4 |
| 8_1392969 | 2 | 0 | 5 | 0.2 | NA | NA | 4 | 0 | 11 | 0.0909 | 1 | 0 | 2 | 0 | 2 | 0 | 2 | 0.5 | 7 | 0.1429 |
| 8_1392974 | 2 | 1 | 4 | 0.75 | 1 | 0 | 4 | 0.75 | 11 | 0.7273 | 3 | 0.6667 | 1 | 1 | 2 | 1 | 3 | 0.3333 | 9 | 0.6667 |
| 8_1392978 | 1 | 1 | 5 | 0.4 | NA | NA | 4 | 0 | 10 | 0.3 | 3 | 0 | 2 | 0 | 2 | 0 | 1 | 1 | 8 | 0.125 |
| 8_1392981 | NA | NA | 5 | 0.6 | 1 | 0 | 4 | 0.25 | 10 | 0.4 | 2 | 0 | 1 | 0 | 2 | 0.5 | 1 | 0 | 6 | 0.1667 |
| 8_1393007 | 2 | 0.5 | 4 | 0.5 | 2 | 0 | 4 | 0.25 | 12 | 0.3333 | 2 | 0 | 2 | 0.5 | 1 | 0 | 2 | 0 | 7 | 0.1429 |
| 8_1393015 | 2 | 1 | 5 | 1 | 2 | 1 | 4 | 1 | 13 | 1 | 3 | 1 | 1 | 1 | 2 | 1 | 1 | 1 | 7 | 1 |
| 8_1393016 | 1 | 1 | 3 | 0.3333 | NA | NA | 4 | 0 | 8 | 0.25 | NA | NA | 1 | 0 | 1 | 1 | 1 | 0 | 3 | 0.3333 |
| 8_1393026 | 2 | 0 | 4 | 0.5 | 1 | 0 | 3 | 0 | 10 | 0.2 | 4 | 0.25 | 2 | 0.5 | 1 | 0 | 2 | 0 | 9 | 0.2222 |
| 8_1393029 | 2 | 1 | 5 | 0.8 | 2 | 1 | 3 | 0.6667 | 12 | 0.8333 | 4 | 1 | 2 | 1 | 2 | 1 | 3 | 1 | 11 | 1 |
| 8_1393032 | 2 | 1 | 5 | 1 | 2 | 0.5 | 4 | 1 | 13 | 0.9231 | 4 | 0.75 | 2 | 1 | 2 | 0.5 | 3 | 1 | 11 | 0.8182 |
| 8_1393062 | 1 | 0 | 4 | 0.25 | 2 | 0 | 5 | 0.2 | 12 | 0.1667 | 5 | 0 | 1 | 1 | 3 | 0 | 2 | 0 | 11 | 0.0909 |
| 8_1393100 | 2 | 0.5 | 2 | 0.5 | 1 | 1 | 6 | 0.6667 | 11 | 0.6364 | 4 | 0.75 | 1 | 1 | 1 | 1 | 2 | 0.5 | 8 | 0.75 |
| 8_1393109 | 2 | 0 | 4 | 0 | 2 | 0 | 5 | 0.2 | 13 | 0.0769 | 5 | 0.2 | 1 | 0 | 3 | 0 | 2 | 0.5 | 11 | 0.1818 |
| 8_1393136 | 2 | 0 | 4 | 0.25 | 4 | 0.5 | 4 | 0 | 14 | 0.2143 | 5 | 0.6 | 1 | 1 | 1 | 0 | 1 | 0 | 8 | 0.5 |
| 8_1393152 | 2 | 0 | 4 | 0.5 | 3 | 0 | 6 | 0.1667 | 15 | 0.2 | 5 | 0.2 | 1 | 1 | 3 | 0.3333 | 3 | 0 | 12 | 0.25 |
| 8_1393486 | 2 | 0 | 14 | 0.3571 | 2 | 1 | 5 | 0.2 | 23 | 0.3478 | 7 | 0.4286 | 8 | 0 | 4 | 0.75 | 6 | 0.3333 | 25 | 0.32 |
| 8_1393505 | 4 | 0.5 | 14 | 0.5 | 3 | 0.3333 | 4 | 0.5 | 25 | 0.48 | 7 | 0.4286 | 7 | 0.5714 | 4 | 0.5 | 4 | 0.75 | 22 | 0.5455 |
| 8_1398956 | 10 | 0.1 | 8 | 0 | 12 | 0.0833 | 8 | 0 | 38 | 0.0526 | 7 | 0 | 10 | 0.2 | 11 | 0.0909 | 12 | 0.0833 | 40 | 0.1 |
| 8_1398973 | 10 | 0.1 | 7 | 0.2857 | 13 | 0.0769 | 9 | 0 | 39 | 0.1026 | 8 | 0 | 12 | 0.0833 | 10 | 0.1 | 10 | 0 | 40 | 0.05 |
| 8_1398975 | 10 | 0.1 | 7 | 0.1429 | 12 | 0.25 | 9 | 0.3333 | 38 | 0.2105 | 6 | 0 | 10 | 0.1 | 10 | 0.5 | 12 | 0.4167 | 38 | 0.2895 |
| 8_1398988 | 9 | 0.1111 | 8 | 0 | 12 | 0.0833 | 4 | 0.25 | 33 | 0.0909 | 6 | 0.1667 | 11 | 0.1818 | 11 | 0 | 10 | 0 | 38 | 0.0789 |
| 8_1398997 | 8 | 0.25 | 5 | 0.4 | 9 | 0.3333 | 4 | 1 | 26 | 0.4231 | 6 | 0.6667 | 9 | 0.4444 | 7 | 0.2857 | 9 | 0.5556 | 31 | 0.4839 |
| 8_1399002 | 10 | 0.2 | 7 | 0.4286 | 10 | 0.3 | 8 | 0.625 | 35 | 0.3714 | 5 | 0.6 | 9 | 0.7778 | 8 | 0.625 | 9 | 0.5556 | 31 | 0.6452 |
| 8_1399019 | 8 | 0.25 | 9 | 0.7778 | 12 | 0.5 | 9 | 0.7778 | 38 | 0.5789 | 9 | 0.5556 | 12 | 0.3333 | 10 | 0.3 | 10 | 0.4 | 41 | 0.3902 |
| 8_1399027 | 14 | 0.2143 | 10 | 0.2 | 13 | 0 | 8 | 0.25 | 45 | 0.1556 | 7 | 0 | 11 | 0.1818 | 12 | 0.0833 | 11 | 0 | 41 | 0.0732 |
| 8_1400211 | 5 | 0.6 | 1 | 1 | 8 | 0.5 | 6 | 1 | 20 | 0.7 | NA | NA | 4 | 1 | 5 | 0.6 | 3 | 0.6667 | 12 | 0.75 |
| 8_1400222 | 4 | 0.5 | 1 | 0 | 6 | 0 | 4 | 0.25 | 15 | 0.2 | NA | NA | 3 | 0 | 4 | 0.25 | 3 | 0 | 10 | 0.1 |
| 8_1400355 | 4 | 0.5 | NA | NA | 5 | 0.6 | 5 | 0.4 | 14 | 0.5 | NA | NA | 2 | 0 | 6 | 0.6667 | 2 | 0 | 10 | 0.4 |
| 8_1400378 | 3 | 0 | NA | NA | 6 | 0.1667 | 4 | 0.5 | 13 | 0.2308 | NA | NA | 1 | 0 | 5 | 0.4 | 2 | 1 | 8 | 0.5 |
| 8_1400380 | 4 | 1 | NA | NA | 5 | 0.6 | 5 | 0.6 | 14 | 0.7143 | NA | NA | 2 | 0.5 | 5 | 0.8 | 2 | 1 | 9 | 0.7778 |
| 8_1400404 | 4 | 0 | NA | NA | 6 | 0.1667 | 4 | 0 | 14 | 0.0714 | NA | NA | 2 | 0 | 4 | 0 | 2 | 0.5 | 8 | 0.125 |
| 8_1407785 | NA | NA | 1 | 1 | 1 | 0 | 3 | 0.6667 | 5 | 0.6 | NA | NA | NA | NA | 2 | 1 | NA | NA | 2 | 1 |
| 8_1407787 | NA | NA | 1 | 1 | 1 | 0 | 3 | 0 | 5 | 0.2 | NA | NA | NA | NA | 2 | 0.5 | NA | NA | 2 | 0.5 |
| 8_1407794 | NA | NA | 1 | 1 | 1 | 1 | 3 | 0 | 5 | 0.4 | NA | NA | NA | NA | 1 | 1 | NA | NA | 1 | 1 |
| 8_1414086 | 2 | 0.5 | 3 | 0.3333 | 3 | 0 | 2 | 0.5 | 10 | 0.3 | NA | NA | NA | NA | 1 | 0 | 2 | 0.5 | 3 | 0.3333 |
| 8_1414088 | 2 | 0 | 3 | 0 | 3 | 0.6667 | 2 | 0 | 10 | 0.2 | 1 | 0 | NA | NA | 1 | 1 | 2 | 0 | 4 | 0.25 |
| 8_1414096 | 3 | 0.6667 | 4 | 0.25 | 3 | 0.6667 | 2 | 1 | 12 | 0.5833 | 2 | 0 | NA | NA | 1 | 0 | 3 | 0.3333 | 6 | 0.1667 |
| 8_1414096 | 2 | 0.5 | 4 | 0.75 | 4 | 0.75 | 1 | 1 | 11 | 0.7273 | 1 | 0 | NA | NA | 1 | 0 | 3 | 1 | 5 | 0.6 |
| 8_1414108 | 2 | 0 | 4 | 0.5 | 4 | 0.25 | 1 | 1 | 11 | 0.3636 | 1 | 0 | NA | NA | 1 | 1 | 3 | 0 | 5 | 0.2 |
| 8_1414115 | 3 | 1 | 4 | 1 | 3 | 1 | 2 | 1 | 12 | 1 | 1 | 1 | NA | NA | 1 | 1 | 3 | 1 | 5 | 1 |
| 8_1414116 | 1 | 0 | 3 | 0 | 3 | 0.3333 | 1 | 0 | 8 | 0.125 | 1 | 1 | NA | NA | NA | NA | 3 | 0.3333 | 4 | 0.5 |
| 8_1414118 | 2 | 0.5 | 2 | 0 | 2 | 0.5 | 2 | 0.5 | 8 | 0.375 | 1 | 0 | NA | NA | 1 | 0 | 2 | 0.5 | 4 | 0.25 |
| 8_1414122 | 3 | 1 | 2 | 1 | 4 | 1 | NA | NA | 9 | 1 | 1 | 1 | NA | NA | NA | NA | 3 | 1 | 4 | 1 |
| 8_1414195 | NA | NA | 2 | 0.5 | 2 | 0 | NA | NA | 4 | 0.25 | NA | NA | NA | NA | NA | NA | 1 | 1 | 1 | 1 |
| 8_1414201 | NA | NA | 2 | 1 | 2 | 1 | NA | NA | 4 | 1 | 1 | 1 | NA | NA | NA | NA | 2 | 0.5 | 3 | 0.6667 |
| 8_1414205 | NA | NA | 2 | 0.5 | 1 | 1 | NA | NA | 3 | 0.6667 | NA | NA | NA | NA | NA | NA | 1 | 1 | 1 | 1 |
| 8_1414247 | NA | NA | 2 | 0.5 | 2 | 0.5 | NA | NA | 4 | 0.5 | NA | NA | NA | NA | NA | NA | 2 | 1 | 2 | 1 |
| 8_1414253 | NA | NA | 2 | 1 | 2 | 1 | NA | NA | 4 | 1 | 1 | 1 | NA | NA | NA | NA | 2 | 1 | 3 | 1 |
| 8_1414259 | NA | NA | 1 | 1 | NA | NA | NA | NA | 1 | 1 | NA | NA | NA | NA | NA | NA | 1 | 1 | 1 | 1 |
| 8_1414287 | NA | NA | 1 | 1 | 1 | 1 | NA | NA | 2 | 1 | NA | NA | NA | NA | NA | NA | 1 | 1 | 1 | 1 |
| 8_1414299 | NA | NA | 1 | 1 | 1 | 1 | NA | NA | 2 | 1 | NA | NA | NA | NA | NA | NA | 1 | 1 | 1 | 1 |
| 8_1477707 | 2 | 0.5 | NA | NA | 1 | 0 | 1 | 0 | 4 | 0.25 | 2 | 0 | 1 | 0 | 2 | 0 | 2 | 0.5 | 7 | 0.1429 |
| 8_1477716 | NA | NA | NA | NA | 1 | 1 | 1 | 0 | 2 | 0.5 | 1 | 0 | NA | NA | 1 | 1 | NA | NA | 2 | 0.5 |
| 8_1477753 | 2 | 0 | NA | NA | 1 | 1 | 1 | 1 | 4 | 0.5 | 4 | 0 | 2 | 0.5 | 2 | 0 | 2 | 0 | 10 | 0.1 |
| 8_1477767 | 2 | 0.5 | NA | NA | 1 | 1 | 1 | 1 | 4 | 0.75 | 3 | 0.3333 | 3 | 0.6667 | 2 | 0.5 | 2 | 1 | 10 | 0.6 |
| 8_1477769 | 1 | 0 | NA | NA | 1 | 1 | NA | NA | 2 | 0.5 | 2 | 0.5 | 2 | 0.5 | 1 | 0 | 2 | 1 | 7 | 0.5714 |
| 8_1477773 | 2 | 1 | NA | NA | 1 | 1 | 1 | 1 | 4 | 1 | 3 | 1 | 3 | 1 | NA | NA | 2 | 0.5 | 8 | 0.875 |
| 8_1477777 | 2 | 0 | NA | NA | 1 | 0 | 1 | 1 | 4 | 0.25 | 3 | 0.6667 | 2 | 1 | 2 | 0 | 1 | 1 | 8 | 0.625 |
| 8_1477779 | 2 | 1 | NA | NA | 1 | 1 | 1 | 1 | 4 | 1 | 3 | 0.6667 | 3 | 1 | 2 | 0.5 | 2 | 0.5 | 10 | 0.7 |

| | | | | | | | | | | | | | | | | | | | | |
|---|---|---|---|---|---|---|---|---|---|---|---|---|---|---|---|---|---|---|---|---|
| 8_14777853 | 2 | 0 | NA | NA | 1 | 1 | 1 | 1 | 4 | 0.5 | 4 | 0 | 3 | 0.3333 | 2 | 0 | 2 | 0 | 11 | 0.0909 |
| 8_14872529 | 4 | 0.75 | 1 | 1 | 1 | 1 | 2 | 1 | 8 | 0.875 | 5 | 0.8 | 3 | 0.6667 | 5 | 0.8 | 2 | 1 | 15 | 0.8 |
| 8_14881402 | 4 | 0.75 | 1 | 1 | 1 | 1 | 2 | 1 | 8 | 0.875 | 5 | 0.6 | 3 | 1 | 5 | 1 | 4 | 1 | 17 | 0.8824 |
| 8_14881439 | 3 | 0.3333 | 1 | 0 | 1 | 0 | 2 | 0 | 7 | 0.1429 | 4 | 0.25 | 4 | 0.75 | 3 | 0 | 4 | 0.25 | 15 | 0.3333 |
| 8_14881447 | 4 | 0.25 | 1 | 0 | 1 | 0 | 2 | 0 | 8 | 0.125 | 5 | 0 | 3 | 0.3333 | 6 | 0.1667 | 4 | 0.25 | 18 | 0.1667 |
| 8_14881457 | 4 | 0.25 | NA | NA | 1 | 0 | 2 | 0.5 | 7 | 0.2857 | 4 | 0.75 | 4 | 0.5 | 5 | 0.8 | 2 | 1 | 15 | 0.7333 |
| 8_14881716 | NA | NA | 1 | 1 | NA | NA | NA | NA | 1 | 1 | 1 | 1 | NA | NA | NA | NA | NA | NA | 1 | 1 |
| 8_14881871 | NA | NA | 1 | 1 | NA | NA | NA | NA | 1 | 1 | 1 | 1 | NA | NA | NA | NA | NA | NA | 1 | 1 |
| 8_14881880 | NA | NA | 1 | 1 | NA | NA | NA | NA | 1 | 1 | 1 | 1 | NA | NA | NA | NA | NA | NA | 1 | 1 |
| 8_14881997 | NA | NA | 1 | 1 | NA | NA | NA | NA | 1 | 1 | 1 | 1 | NA | NA | NA | NA | NA | NA | 1 | 1 |
| 8_14882056 | NA | NA | 1 | 1 | NA | NA | NA | NA | 1 | 1 | 1 | 1 | NA | NA | NA | NA | NA | NA | 1 | 1 |
| 8_14882092 | NA | NA | 1 | 1 | NA | NA | NA | NA | 1 | 1 | 1 | 1 | NA | NA | NA | NA | NA | NA | 1 | 1 |
| 8_14888211 | NA | NA | 2 | 1 | NA | NA | NA | NA | 2 | 1 | NA | NA | NA | NA | 1 | 1 | NA | NA | 1 | 1 |
| 8_14888236 | NA | NA | 2 | 1 | NA | NA | NA | NA | 2 | 1 | NA | NA | NA | NA | 1 | 1 | NA | NA | 1 | 1 |
| 8_14896569 | 4 | 0.25 | 6 | 0.5 | 1 | 1 | 2 | 0 | 13 | 0.3846 | 3 | 0.3333 | 5 | 0.2 | 8 | 0.625 | 5 | 0.4 | 21 | 0.4286 |
| 8_14896644 | 3 | 0.6667 | 5 | 0.8 | NA | NA | 1 | 0 | 9 | 0.6667 | 3 | 1 | 2 | 0.5 | 7 | 0.5714 | 5 | 0.6 | 17 | 0.6471 |
| 8_14896711 | 4 | 1 | 5 | 0.6 | NA | NA | 2 | 1 | 11 | 0.8182 | 5 | 0.8 | 5 | 0.8 | 10 | 0.8 | 7 | 0.4286 | 27 | 0.7037 |
| 8_14896839 | 5 | 1 | 5 | 1 | 1 | 1 | 2 | 1 | 13 | 1 | 4 | 1 | 4 | 1 | 8 | 1 | 6 | 0.6667 | 22 | 0.9091 |
| 8_14896889 | 5 | 0.4 | 6 | 0.6667 | 1 | 1 | 2 | 1 | 14 | 0.6429 | 3 | 0.3333 | 5 | 0.4 | 8 | 0.5 | 7 | 0.4286 | 23 | 0.4348 |
| 8_14896899 | 5 | 0.6 | 6 | 0.6667 | 1 | 0 | 2 | 1 | 14 | 0.6429 | 4 | 0.75 | 5 | 1 | 10 | 1 | 7 | 0.4286 | 26 | 0.8077 |
| 8_14896934 | 5 | 0.4 | 4 | 0 | 1 | 0 | 2 | 0 | 12 | 0.1667 | 5 | 0.6 | 4 | 0 | 11 | 0.1818 | 7 | 0 | 27 | 0.1852 |
| 8_14896982 | 5 | 1 | 6 | 1 | 2 | 1 | 2 | 1 | 15 | 1 | 5 | 1 | 5 | 1 | 11 | 0.9091 | 7 | 0.8571 | 28 | 0.9286 |
| 8_14897107 | 5 | 0 | 5 | 0.6 | 2 | 0.5 | 3 | 0 | 15 | 0.2667 | 6 | 0.3333 | 5 | 0.2 | 11 | 0.3636 | 7 | 0.1429 | 29 | 0.2759 |
| 8_14897424 | 6 | 0.1667 | 5 | 0.2 | 2 | 0 | 3 | 0 | 16 | 0.125 | 4 | 0 | 5 | 0.2 | 10 | 0.1 | 6 | 0.1667 | 25 | 0.12 |
| 8_14897517 | 6 | 0.8333 | 5 | 0.8 | 2 | 1 | 3 | 1 | 16 | 0.875 | 6 | 1 | 4 | 1 | 10 | 0.9 | 7 | 0.8571 | 27 | 0.9259 |
| 8_14897547 | 6 | 0.3333 | 6 | 0.1667 | 2 | 0 | 3 | 0 | 17 | 0.1765 | 8 | 0.125 | 4 | 0.25 | 10 | 0.5 | 7 | 0.2857 | 29 | 0.3103 |
| 8_14898280 | 2 | 0.5 | 3 | 0.3333 | 2 | 0 | 4 | 0 | 11 | 0.1818 | 5 | 0.2 | 1 | 0 | 6 | 0.1667 | 6 | 0 | 18 | 0.1111 |
| 8_14898940 | 1 | 0 | 4 | 0.25 | 1 | 0 | 3 | 0 | 9 | 0.1111 | 1 | 0 | 2 | 0.5 | 3 | 0 | 3 | 0 | 9 | 0.1111 |
| 8_14899590 | 1 | 0 | 4 | 0.5 | 1 | 0 | 3 | 0.6667 | 9 | 0.4444 | 1 | 0 | 1 | 0 | 2 | 0.5 | 4 | 0.75 | 8 | 0.5 |
| 8_14899628 | 1 | 0 | 3 | 0.6667 | NA | NA | 2 | 0.5 | 6 | 0.5 | 1 | 0 | 1 | 1 | 4 | 1 | 3 | 0.6667 | 9 | 0.7778 |
| 8_14899948 | 3 | 0 | 4 | 0 | 1 | 0 | 3 | 0.3333 | 11 | 0.0909 | 1 | 0 | 2 | 0.5 | 4 | 0.25 | 4 | 0 | 11 | 0.1818 |
| 8_14900298 | 3 | 0 | 4 | 0 | 2 | 0 | 3 | 0.3333 | 12 | 0.0833 | 2 | 0.5 | 2 | 0 | 4 | 0 | 6 | 0 | 14 | 0.0714 |
| 8_14900470 | 3 | 0 | 4 | 0.25 | 2 | 0 | 3 | 0.3333 | 12 | 0.1667 | 2 | 0 | 2 | 0 | 3 | 0 | 5 | 0.2 | 12 | 0.0833 |
| 8_14900512 | 4 | 0 | 4 | 0.25 | 2 | 0 | 3 | 0 | 13 | 0.0769 | 1 | 1 | 2 | 0 | 3 | 0 | 6 | 0.1667 | 12 | 0.1667 |
| 8_14900517 | 4 | 0.75 | 4 | 0.75 | 1 | 1 | 2 | 0.5 | 11 | 0.7273 | 1 | 1 | 2 | 0.5 | 3 | 0.3333 | 5 | 0.6 | 11 | 0.5455 |
| 8_14900679 | 3 | 0 | 3 | 0.3333 | 4 | 0.25 | 3 | 0 | 13 | 0.1538 | 3 | 0.3333 | 2 | 0 | 4 | 0 | 5 | 0 | 14 | 0.0714 |
| 8_14900696 | 3 | 0.3333 | 5 | 0.2 | 3 | 0.3333 | 3 | 0.3333 | 14 | 0.2857 | 3 | 0.6667 | 2 | 0.5 | 4 | 0.75 | 5 | 0.6 | 14 | 0.6429 |
| 8_14900730 | 4 | 0.75 | 4 | 1 | 3 | 0 | 3 | 0.6667 | 14 | 0.6429 | 3 | 0.6667 | 2 | 0.5 | 4 | 0.5 | 6 | 0.3333 | 15 | 0.4667 |
| 8_14900837 | 3 | 0 | 5 | 0 | 4 | 0 | 3 | 0.3333 | 15 | 0.0667 | 2 | 0 | 2 | 0 | 4 | 0.5 | 7 | 0 | 15 | 0.1333 |
| 8_14901174 | 4 | 1 | 4 | 1 | 4 | 0.75 | 5 | 1 | 17 | 0.9412 | 2 | 1 | 2 | 1 | 3 | 1 | 7 | 0.8571 | 14 | 0.9286 |
| 8_14901241 | 4 | 0 | 5 | 0.2 | 4 | 0 | 6 | 0 | 19 | 0.0526 | 2 | 0 | 2 | 0 | 4 | 0.5 | 7 | 0.2857 | 15 | 0.2667 |
| 8_14901458 | 4 | 0 | 5 | 0.4 | 4 | 0.5 | 6 | 0.5 | 19 | 0.3684 | 3 | 0.6667 | 2 | 0 | 4 | 0 | 7 | 0.4286 | 16 | 0.3125 |
| 8_14901804 | 2 | 0 | 4 | 0 | 3 | 0.3333 | 5 | 0.2 | 14 | 0.1429 | 3 | 0 | 4 | 0 | 3 | 0.6667 | 6 | 0 | 16 | 0.125 |
| 8_14901941 | 3 | 1 | 4 | 0.5 | 3 | 0.6667 | 4 | 0.5 | 14 | 0.6429 | 2 | 0.5 | 2 | 0.5 | 2 | 1 | 3 | 0.3333 | 9 | 0.5556 |
| 8_14901946 | 2 | 0.5 | 4 | 0.5 | 2 | 0.5 | 6 | 0.3333 | 14 | 0.4286 | 3 | 0.6667 | 2 | 1 | 4 | 1 | 6 | 0.5 | 15 | 0.7333 |
| 8_14902030 | 3 | 1 | 5 | 1 | 3 | 0.6667 | 4 | 1 | 15 | 0.9333 | 1 | 1 | 1 | 1 | 3 | 0.6667 | 4 | 0.75 | 9 | 0.7778 |
| 8_14902066 | 4 | 0.75 | 6 | 0.8333 | 4 | 0.5 | 6 | 0.5 | 20 | 0.65 | 3 | 0.6667 | 4 | 0.5 | 4 | 1 | 7 | 0.4286 | 18 | 0.6111 |
| 8_14902281 | 4 | 0 | 6 | 0 | 5 | 0.2 | 5 | 0.2 | 20 | 0.1 | 3 | 0.3333 | 4 | 0.25 | 5 | 0 | 5 | 0 | 17 | 0.1176 |
| 8_14902403 | 4 | 0 | 6 | 0 | 5 | 0.2 | 6 | 0.1667 | 21 | 0.0952 | 3 | 0 | 4 | 0 | 5 | 0.2 | 6 | 0 | 18 | 0.0556 |
| 8_14902449 | 4 | 0.25 | 5 | 0 | 5 | 0 | 5 | 0 | 19 | 0.0526 | 3 | 0.3333 | 5 | 0 | 5 | 0.2 | 6 | 0 | 19 | 0.1053 |
| 8_14902875 | 4 | 0.25 | 6 | 0.1667 | 6 | 0.1667 | 7 | 0 | 23 | 0.1304 | 4 | 0 | 5 | 0.4 | 5 | 0.4 | 6 | 0.3333 | 20 | 0.3 |
| 8_14903062 | 4 | 0 | 8 | 0.125 | 6 | 0.1667 | 7 | 0 | 25 | 0.08 | 4 | 0.25 | 6 | 0.1667 | 7 | 0 | 6 | 0 | 23 | 0.087 |
| 8_14944737 | 1 | 1 | 4 | 0.75 | 4 | 0.5 | 1 | 0 | 10 | 0.6 | NA | NA | 2 | 0.5 | 1 | 0 | NA | NA | 3 | 0.3333 |
| 8_14944989 | 4 | 1 | 4 | 1 | 6 | 1 | 1 | 0 | 15 | 0.9333 | NA | NA | 3 | 1 | 1 | 1 | NA | NA | 4 | 1 |
| 8_14945082 | 5 | 0.6 | 5 | 1 | 5 | 0.8 | NA | NA | 15 | 0.8 | NA | NA | 3 | 1 | 1 | 1 | NA | NA | 4 | 1 |
| 8_14946735 | 3 | 0.6667 | 5 | 0.6 | 3 | 0.6667 | NA | NA | 11 | 0.6364 | NA | NA | 2 | 1 | 1 | 1 | 1 | 1 | 4 | 1 |
| 8_14946778 | 4 | 1 | 4 | 0.75 | 4 | 1 | NA | NA | 12 | 0.9167 | NA | NA | 2 | 1 | NA | NA | 1 | 1 | 3 | 1 |
| 8_14946819 | 5 | 1 | 5 | 1 | 5 | 1 | 1 | 1 | 16 | 1 | NA | NA | 3 | 0.6667 | NA | NA | 1 | 1 | 4 | 0.75 |
| 8_14946871 | 4 | 0.25 | 5 | 0.6 | 6 | 0.5 | 1 | 0 | 16 | 0.4375 | NA | NA | 3 | 0.3333 | 1 | 1 | 1 | 1 | 5 | 0.6 |
| 8_14946880 | 4 | 0.25 | 4 | 0.25 | 6 | 0.3333 | 1 | 0 | 15 | 0.2667 | NA | NA | 3 | 0 | 1 | 1 | 1 | 0 | 5 | 0.2 |
| 8_14946914 | 4 | 0 | 5 | 0.6 | 5 | 0.4 | 1 | 0 | 15 | 0.3333 | NA | NA | 3 | 0 | 1 | 1 | 1 | 0 | 5 | 0.2 |
| 8_14946930 | 5 | 1 | 5 | 1 | 4 | 1 | 1 | 1 | 15 | 1 | 1 | 1 | 3 | 1 | 1 | 1 | 1 | 1 | 6 | 1 |
| 8_14946951 | 4 | 1 | 6 | 0.8333 | 6 | 1 | 1 | 1 | 17 | 0.9412 | 1 | 1 | 3 | 1 | 1 | 0 | 1 | 1 | 6 | 0.8333 |
| 8_14946983 | 5 | 0.6 | 5 | 0.4 | 5 | 0.8 | 1 | 0 | 16 | 0.5625 | 1 | 0 | 3 | 0.6667 | 1 | 0 | NA | NA | 5 | 0.4 |
| 8_14947067 | 5 | 0.4 | 3 | 0.6667 | 3 | 0.3333 | 1 | 1 | 12 | 0.5 | 1 | 1 | 2 | 1 | NA | NA | 1 | 0 | 4 | 0.75 |

| | | | | | | | | | | | | | | | | | | | | |
|---|---|---|---|---|---|---|---|---|---|---|---|---|---|---|---|---|---|---|---|---|
| 8_1494720 | 3 | 0.6667 | 5 | 1 | 5 | 1 | NA | NA | 13 | 0.9231 | 1 | 1 | 1 | 1 | NA | NA | 1 | 1 | 3 | 1 |
| 8_1494736 | 5 | 0.8 | 6 | 0.6667 | 6 | 0.8333 | 1 | 1 | 18 | 0.7778 | 1 | 1 | 3 | 0.3333 | 2 | 1 | 1 | 1 | 7 | 0.7143 |
| 8_1494960 | 1 | 0 | 2 | 0 | 1 | 0 | 2 | 0.5 | 6 | 0.1667 | NA | NA | 2 | 0.5 | NA | NA | NA | NA | 2 | 0.5 |
| 8_1501114 | 1 | 0 | 4 | 0.25 | 1 | 0 | 2 | 0 | 8 | 0.125 | 3 | 0 | 4 | 0 | 2 | 0.5 | NA | NA | 9 | 0.1111 |
| 8_1503191 | 2 | 0.5 | NA | NA | NA | NA | NA | NA | 2 | 0.5 | NA | NA | 1 | 0 | 2 | 0.5 | NA | NA | 3 | 0.3333 |
| 8_1504939 | 3 | 0 | 5 | 0.4 | 3 | 0.3333 | 2 | 0 | 13 | 0.2308 | 3 | 0 | 3 | 0.3333 | 6 | 0 | NA | NA | 12 | 0.0833 |
| 8_1504940 | 4 | 0.25 | 10 | 0.1 | 2 | 0 | 2 | 0 | 18 | 0.1111 | 3 | 0 | 3 | 0.3333 | 6 | 0.3333 | NA | NA | 12 | 0.25 |
| 8_1504945 | 4 | 0.75 | 10 | 0.6 | 3 | 0.6667 | 3 | 0.3333 | 20 | 0.6 | 4 | 0.5 | 4 | 0.5 | 7 | 0.2857 | NA | NA | 15 | 0.4 |
| 8_1504952 | 4 | 0.5 | 8 | 0 | 4 | 0 | 3 | 0 | 19 | 0.1053 | 5 | 0 | 5 | 0.2 | 7 | 0.1429 | 1 | 0 | 18 | 0.1111 |
| 8_1504958 | 4 | 0.25 | 9 | 0.4444 | 3 | 0 | 1 | 0 | 17 | 0.2941 | 4 | 0.5 | 5 | 0 | 8 | 0.25 | 2 | 0.5 | 19 | 0.2632 |
| 8_1504962 | 4 | 0.75 | 10 | 0.6 | 2 | 0 | 2 | 0.5 | 18 | 0.5556 | 5 | 0.8 | 4 | 0.5 | 7 | 0.7143 | 2 | 1 | 18 | 0.7222 |
| 8_1504964 | 4 | 0.25 | 11 | 0 | 4 | 0 | 3 | 0.3333 | 22 | 0.0909 | 5 | 0.2 | 5 | 0 | 8 | 0 | 2 | 0 | 20 | 0.05 |
| 8_1504965 | 4 | 1 | 10 | 0.8 | 4 | 1 | 3 | 1 | 21 | 0.9048 | 5 | 1 | 5 | 0.6 | 8 | 0.875 | 2 | 0.5 | 20 | 0.8 |
| 8_1504978 | 3 | 0.6667 | 10 | 0.7 | 4 | 0.5 | 2 | 1 | 19 | 0.6842 | 6 | 0.6667 | 3 | 0.6667 | 8 | 0.75 | 2 | 1 | 19 | 0.7368 |
| 8_1504987 | 2 | 1 | 9 | 0.8889 | 4 | 0.25 | 3 | 1 | 18 | 0.7778 | 4 | 1 | 3 | 1 | 7 | 0.7143 | 1 | 1 | 15 | 0.8667 |
| 8_1504992 | 3 | 0.3333 | 11 | 0.4545 | 4 | 0.25 | 3 | 1 | 21 | 0.4762 | 5 | 0.2 | 5 | 0.6 | 8 | 0.375 | 2 | 0 | 20 | 0.35 |
| 8_1505005 | 3 | 0 | 4 | 0 | 3 | 0.3333 | 3 | 0 | 13 | 0.0769 | 2 | 0 | 3 | 0 | 4 | 0.25 | 1 | 0 | 10 | 0.1 |
| 8_1505036 | 4 | 0.5 | 11 | 0.2727 | 3 | 0 | 3 | 0.3333 | 21 | 0.2857 | 6 | 0.1667 | 6 | 0.1667 | 10 | 0.3 | 3 | 0 | 25 | 0.2 |
| 8_1505056 | 4 | 0.5 | 8 | 0.25 | 3 | 0 | 2 | 0 | 17 | 0.2353 | 7 | 0.2857 | 5 | 0 | 7 | 0 | 3 | 0 | 22 | 0.0909 |
| 8_1505079 | 3 | 0.3333 | 9 | 0.3333 | 5 | 0.2 | 4 | 0.25 | 21 | 0.2857 | 7 | 0 | 6 | 0.3333 | 7 | 0.1429 | 3 | 0 | 23 | 0.1304 |
| 8_1505083 | 4 | 0.5 | 10 | 0.4 | 5 | 0.2 | 3 | 0.3333 | 22 | 0.3636 | 6 | 0.3333 | 6 | 0.3333 | 8 | 0.25 | 3 | 0.6667 | 23 | 0.3478 |
| 8_1505096 | 3 | 0.3333 | 11 | 0.7273 | 5 | 0.2 | 4 | 0.5 | 23 | 0.5217 | 7 | 0.2857 | 6 | 0.6667 | 8 | 0.625 | 3 | 0 | 24 | 0.4583 |
| 8_1505106 | 3 | 0.3333 | 7 | 0 | 1 | 0 | 3 | 0 | 14 | 0.0714 | 4 | 0.25 | 6 | 0.1667 | 4 | 0 | 2 | 0 | 16 | 0.125 |
| 8_1507696 | 1 | 1 | 1 | 1 | NA | NA | NA | NA | 2 | 1 | NA | NA | NA | NA | NA | NA | 1 | 1 | 1 | 1 |
| 8_1507732 | 2 | 1 | 2 | 0 | NA | NA | NA | NA | 4 | 0.5 | 1 | 1 | NA | NA | NA | NA | NA | NA | 1 | 1 |
| 8_1507745 | 2 | 1 | 2 | 1 | NA | NA | NA | NA | 4 | 1 | 1 | 1 | NA | NA | NA | NA | NA | NA | 1 | 1 |
| 8_1507776 | 2 | 0.5 | 2 | 0.5 | NA | NA | NA | NA | 4 | 0.5 | 1 | 0 | NA | NA | NA | NA | 1 | 1 | 2 | 0.5 |
| 8_1507812 | 1 | 1 | 2 | 1 | NA | NA | 1 | 1 | 4 | 1 | 1 | 1 | NA | NA | NA | NA | 1 | 0 | 2 | 0.5 |
| 8_1507832 | 2 | 1 | 2 | 0.5 | NA | NA | NA | NA | 4 | 0.75 | 1 | 1 | NA | NA | NA | NA | 1 | 1 | 2 | 1 |
| 8_1507893 | 4 | 0.5 | 2 | 0 | NA | NA | 1 | 1 | 7 | 0.4286 | 1 | 1 | NA | NA | NA | NA | 1 | 0 | 2 | 0.5 |
| 8_1507913 | 4 | 1 | 2 | 0.5 | 2 | 1 | 3 | 1 | 11 | 0.9091 | 1 | 1 | NA | NA | NA | NA | 2 | 0.5 | 3 | 0.6667 |
| 8_1507915 | 4 | 0.5 | 3 | 0.6667 | 2 | 1 | 3 | 0.3333 | 12 | 0.5833 | 1 | 0 | NA | NA | NA | NA | 2 | 1 | 3 | 0.6667 |
| 8_1507918 | 4 | 1 | 3 | 1 | 2 | 1 | 4 | 0.75 | 13 | 0.9231 | 1 | 1 | NA | NA | 1 | 1 | 1 | 1 | 3 | 1 |
| 8_1507927 | 4 | 1 | 3 | 1 | 1 | 1 | 4 | 1 | 12 | 1 | 1 | 1 | NA | NA | 1 | 1 | 2 | 1 | 4 | 1 |
| 8_1507930 | NA | NA | 2 | 1 | 1 | 1 | 3 | 1 | 6 | 1 | 1 | 0 | NA | NA | NA | NA | 2 | 0.5 | 3 | 0.3333 |
| 8_1507950 | 4 | 0.5 | 3 | 0.6667 | 3 | 0.6667 | 3 | 0.6667 | 13 | 0.6154 | 1 | 0 | 1 | 0 | 1 | 0 | 2 | 0.5 | 5 | 0.2 |
| 8_1507967 | 4 | 1 | 3 | 0.6667 | 4 | 0.75 | 5 | 1 | 16 | 0.875 | NA | NA | 1 | 0 | 1 | 1 | 1 | 0 | 3 | 0.3333 |
| 8_1507972 | 5 | 0.2 | 3 | 0.3333 | 4 | 0.75 | 7 | 0.5714 | 19 | 0.4737 | NA | NA | 2 | 0.5 | 1 | 0 | 2 | 0.5 | 5 | 0.4 |
| 8_1508000 | 5 | 0 | 2 | 0 | 4 | 0 | 7 | 0.1429 | 18 | 0.0556 | NA | NA | 2 | 0.5 | 1 | 0 | 2 | 0 | 5 | 0.2 |
| 8_1508097 | 6 | 0 | 6 | 0 | 4 | 0.25 | 15 | 0.2 | 31 | 0.129 | 4 | 0.5 | 4 | 0 | 6 | 0.3333 | 8 | 0 | 22 | 0.1818 |
| 8_1508219 | 12 | 0 | 7 | 0.1429 | 5 | 0.2 | 16 | 0.0625 | 40 | 0.075 | 5 | 0.2 | 3 | 0.3333 | 10 | 0 | 10 | 0.1 | 28 | 0.1071 |
| 8_1508224 | 14 | 0.2143 | 8 | 0.125 | 3 | 0.3333 | 18 | 0.1111 | 43 | 0.1628 | 8 | 0.25 | 4 | 0 | 10 | 0.1 | 13 | 0.2308 | 35 | 0.1714 |
| 8_1508288 | 17 | 0.1765 | 8 | 0 | 6 | 0 | 23 | 0.2174 | 54 | 0.1481 | 9 | 0.3333 | 9 | 0.1111 | 13 | 0.3077 | 10 | 0.6 | 41 | 0.3415 |
| 8_1508298 | 20 | 0.45 | 11 | 0.6364 | 8 | 0.625 | 21 | 0.2857 | 60 | 0.45 | 11 | 0.5455 | 10 | 0.4 | 16 | 0.3125 | 17 | 0.4118 | 54 | 0.4074 |
| 8_1508312 | 20 | 0 | 13 | 0.2308 | 8 | 0.125 | 23 | 0.087 | 64 | 0.0938 | 9 | 0.1111 | 10 | 0.1 | 15 | 0.0667 | 16 | 0 | 50 | 0.06 |
| 8_1508313 | 18 | 0.1111 | 11 | 0.0909 | 6 | 0.1667 | 20 | 0.05 | 55 | 0.0909 | 9 | 0 | 7 | 0 | 13 | 0.1538 | 11 | 0.0909 | 40 | 0.075 |
| 8_1594389 | 2 | 0.5 | 4 | 0.25 | 1 | 0 | 1 | 0 | 8 | 0.25 | 1 | 0 | 5 | 0.8 | 5 | 0.2 | 3 | 0 | 14 | 0.3571 |
| 8_1594401 | 1 | 0 | 4 | 0.25 | NA | NA | 2 | 0 | 7 | 0.1429 | 1 | 0 | 5 | 0.4 | 4 | 0 | 3 | 0.6667 | 13 | 0.3077 |
| 8_1594402 | 1 | 0 | 5 | 0 | 1 | 1 | 2 | 0 | 9 | 0.1111 | 1 | 0 | 3 | 0 | 5 | 0 | 3 | 0.3333 | 12 | 0.0833 |
| 8_1594464 | 1 | 0 | 3 | 0.3333 | 1 | 0 | 2 | 0 | 7 | 0.1429 | 1 | 0 | 2 | 0.5 | 5 | 0.2 | 3 | 0.3333 | 11 | 0.2727 |
| 8_1594503 | NA | NA | 3 | 0.3333 | 2 | 0.5 | 2 | 0 | 7 | 0.2857 | 1 | 0 | 4 | 0.75 | 1 | 0 | 2 | 0.5 | 8 | 0.5 |
| 8_1594523 | NA | NA | 2 | 0.5 | 1 | 0 | NA | NA | 3 | 0.3333 | 1 | 0 | 3 | 0 | 2 | 0 | 1 | 1 | 7 | 0.1429 |
| 8_1594525 | NA | NA | 3 | 1 | 1 | 1 | 2 | 1 | 6 | 1 | NA | NA | 4 | 1 | 2 | 1 | 2 | 0.5 | 8 | 0.875 |
| 8_1594540 | NA | NA | 2 | 1 | NA | NA | 1 | 1 | 3 | 1 | 1 | 1 | 4 | 0.75 | 3 | 1 | 1 | 1 | 9 | 0.8889 |
| 8_1594542 | NA | NA | 3 | 0.3333 | 1 | 1 | 2 | 1 | 6 | 0.6667 | 1 | 1 | 4 | 0.75 | 4 | 0.5 | 2 | 0.5 | 11 | 0.6364 |
| 8_1594543 | NA | NA | 3 | 1 | 1 | 1 | 2 | 1 | 6 | 1 | 1 | 1 | 4 | 0.75 | 2 | 1 | 2 | 1 | 9 | 0.8889 |
| 8_1594567 | NA | NA | 3 | 1 | 1 | 1 | 2 | 0.5 | 6 | 0.8333 | 1 | 1 | 2 | 1 | 4 | 1 | 2 | 1 | 9 | 1 |
| 8_1594569 | NA | NA | 3 | 1 | 1 | 1 | 2 | 1 | 6 | 1 | 1 | 1 | 1 | 1 | 4 | 0.75 | 2 | 1 | 8 | 0.875 |
| 8_1594578 | NA | NA | 3 | 1 | 1 | 1 | 1 | 0 | 5 | 0.8 | NA | NA | 1 | 0 | 2 | 1 | 2 | 1 | 5 | 0.8 |
| 8_1594582 | NA | NA | 3 | 0.6667 | 1 | 1 | 1 | 0 | 5 | 0.6 | NA | NA | 1 | 0 | 4 | 0.75 | 2 | 0.5 | 7 | 0.5714 |
| 8_1868192 | 3 | 0.6667 | 1 | 1 | NA | NA | NA | NA | 4 | 0.75 | 1 | 1 | NA | NA | 1 | 1 | 3 | 0.6667 | 5 | 0.8 |
| 8_1868203 | 3 | 0.6667 | 1 | 0 | NA | NA | NA | NA | 4 | 0.5 | NA | NA | NA | NA | 1 | 0 | 3 | 0.6667 | 4 | 0.5 |
| 8_1868207 | 3 | 1 | 1 | 1 | NA | NA | NA | NA | 4 | 1 | 1 | 1 | NA | NA | 1 | 1 | 3 | 0.6667 | 5 | 0.8 |
| 8_1868211 | 3 | 0.3333 | 1 | 1 | NA | NA | NA | NA | 4 | 0.5 | 1 | 1 | NA | NA | 1 | 0 | 2 | 0.5 | 4 | 0.5 |
| 8_1868226 | 2 | 1 | 1 | 1 | NA | NA | NA | NA | 3 | 1 | 1 | 1 | NA | NA | 1 | 1 | 3 | 1 | 5 | 1 |

| | | | | | | | | | | | | | | | | | | | | |
|---|---|---|---|---|---|---|---|---|---|---|---|---|---|---|---|---|---|---|---|---|
| 8_1868249 | 2 | 1 | 1 | 1 | NA | NA | NA | NA | 3 | 1 | 1 | 1 | NA | NA | 1 | 1 | 2 | 0.5 | 4 | 0.75 |
| 8_1893184 | 11 | 0.0909 | 12 | 0.3333 | 9 | 0 | 11 | 0 | 43 | 0.1163 | 8 | 0 | 11 | 0 | 10 | 0.3 | 14 | 0.0714 | 43 | 0.093 |
| 8_1893187 | 9 | 0.1111 | 12 | 0.0833 | 11 | 0.0909 | 11 | 0.1818 | 43 | 0.1163 | 5 | 0 | 8 | 0 | 8 | 0.25 | 14 | 0 | 35 | 0.0571 |
| 8_1893191 | 10 | 0.5 | 13 | 0.5385 | 10 | 0.1 | 11 | 0.4545 | 44 | 0.4091 | 6 | 0.5 | 9 | 0.3333 | 10 | 0.5 | 14 | 0.3571 | 39 | 0.4103 |
| 8_1893211 | 5 | 0 | 7 | 0.2857 | 5 | 0 | 9 | 0 | 26 | 0.0769 | 6 | 0 | 5 | 0 | 8 | 0.25 | 10 | 0.1 | 29 | 0.1034 |
| 8_1893412 | 1 | 1 | 1 | 0 | 2 | 0.5 | 3 | 0 | 7 | 0.2857 | 2 | 0.5 | 1 | 0 | 1 | 0 | NA | NA | 4 | 0.25 |
| 8_2001715 | 1 | 0 | NA | NA | NA | NA | 1 | 1 | 2 | 0.5 | NA | NA | 1 | 1 | NA | NA | 2 | 0.5 | 3 | 0.6667 |
| 8_2001716 | 1 | 1 | NA | NA | NA | NA | 1 | 1 | 2 | 1 | NA | NA | 1 | 1 | NA | NA | 2 | 1 | 3 | 1 |
| 8_2001722 | NA | NA | NA | NA | NA | NA | 1 | 1 | 1 | 1 | NA | NA | 1 | 1 | NA | NA | 2 | 1 | 3 | 1 |
| 8_2001738 | NA | NA | NA | NA | NA | NA | 1 | 1 | 1 | 1 | NA | NA | 1 | 1 | NA | NA | 1 | 1 | 2 | 1 |
| 8_2001755 | NA | NA | NA | NA | NA | NA | 1 | 1 | 1 | 1 | NA | NA | 1 | 0 | NA | NA | 2 | 1 | 3 | 0.6667 |
| 8_2003080 | 1 | 1 | 1 | 1 | 2 | 1 | 3 | 0.3333 | 7 | 0.7143 | 2 | 0 | 1 | 1 | 1 | 0 | 5 | 0.4 | 9 | 0.3333 |
| 8_2003597 | 1 | 0 | 1 | 0 | 3 | 0.3333 | 6 | 0 | 11 | 0.0909 | 2 | 0 | NA | NA | 1 | 1 | 5 | 0 | 8 | 0.125 |
| 8_2003598 | NA | NA | 1 | 0 | 2 | 0.5 | 5 | 0.2 | 8 | 0.25 | 2 | 0 | 2 | 0.5 | 1 | 0 | 4 | 0 | 9 | 0.1111 |
| 8_2003603 | 1 | 0 | 1 | 0 | 3 | 0.3333 | 5 | 0 | 10 | 0.1 | 2 | 0 | 3 | 0.3333 | 1 | 0 | 4 | 0 | 10 | 0.1 |
| 8_2004188 | 1 | 0 | NA | NA | 1 | 0 | 5 | 0.4 | 7 | 0.2857 | 2 | 0.5 | 4 | 0.75 | NA | NA | 4 | 0.75 | 10 | 0.7 |
| 8_2004189 | 1 | 0 | 1 | 0 | 2 | 1 | 6 | 0.5 | 10 | 0.5 | 1 | 1 | 3 | 0 | NA | NA | 4 | 1 | 8 | 0.625 |
| 8_2004190 | 2 | 0 | 1 | 0 | 2 | 1 | 6 | 0.6667 | 11 | 0.5455 | 2 | 0.5 | 4 | 0.25 | NA | NA | 4 | 1 | 10 | 0.6 |
| 8_2004191 | 1 | 0 | 1 | 0 | 1 | 1 | 6 | 0.6667 | 9 | 0.5556 | 2 | 0 | 4 | 0 | NA | NA | 4 | 1 | 10 | 0.4 |
| 8_2004213 | 1 | 0 | 1 | 0 | 2 | 0.5 | 6 | 0.6667 | 10 | 0.5 | 2 | 0.5 | 4 | 0 | NA | NA | 5 | 0.8 | 11 | 0.4545 |
| 8_2004220 | 1 | 0 | NA | NA | NA | NA | 5 | 0.2 | 6 | 0.1667 | 2 | 0 | 3 | 0 | NA | NA | 4 | 0.5 | 9 | 0.2222 |
| 8_2004263 | NA | NA | NA | NA | 2 | 0 | 5 | 0.6 | 7 | 0.4286 | 2 | 0 | 4 | 0.25 | NA | NA | 5 | 0.2 | 11 | 0.1818 |
| 8_2004348 | NA | NA | 2 | 0 | 2 | 0.5 | 6 | 0 | 10 | 0.1 | 2 | 0 | 4 | 0.25 | NA | NA | 4 | 0 | 10 | 0.1 |
| 8_2004418 | NA | NA | 2 | 0 | 2 | 0 | 6 | 0.1667 | 10 | 0.1 | 2 | 0 | 2 | 0.5 | 3 | 0 | 4 | 0 | 11 | 0.0909 |
| 8_2004544 | NA | NA | 5 | 0.4 | 1 | 0 | 2 | 0.5 | 8 | 0.375 | 2 | 0.5 | 2 | 1 | NA | NA | 2 | 1 | 6 | 0.8333 |
| 8_2004586 | 1 | 1 | 2 | 1 | NA | NA | 1 | 0 | 4 | 0.75 | 1 | 1 | NA | NA | 1 | 1 | 1 | 1 | 3 | 1 |
| 8_2004588 | 2 | 0.5 | 1 | 1 | 1 | 0 | 1 | 0 | 5 | 0.4 | 1 | 1 | NA | NA | 1 | 1 | 1 | 1 | 3 | 1 |
| 8_2004617 | 1 | 0 | 1 | 1 | 1 | 0 | 1 | 0 | 4 | 0.25 | NA | NA | NA | NA | 1 | 1 | 1 | 0 | 2 | 0.5 |
| 8_2005852 | NA | NA | 2 | 0.5 | NA | NA | 1 | 0 | 3 | 0.3333 | 2 | 0.5 | NA | NA | NA | NA | NA | NA | 2 | 0.5 |
| 8_2057680 | NA | NA | NA | NA | NA | NA | 1 | 1 | 1 | 1 | NA | NA | NA | NA | NA | NA | 1 | 1 | 1 | 1 |
| 8_2071731 | NA | NA | 1 | 0 | 2 | 0.5 | 4 | 0.25 | 7 | 0.2857 | 1 | 0 | NA | NA | 1 | 1 | 2 | 0.5 | 4 | 0.5 |
| 8_2072318 | NA | NA | 1 | 0 | 3 | 0.6667 | 2 | 0 | 6 | 0.3333 | 1 | 0 | NA | NA | 3 | 0.3333 | 3 | 0 | 7 | 0.1429 |
| 8_2072322 | NA | NA | 2 | 1 | 3 | 0 | 3 | 0.3333 | 8 | 0.375 | 1 | 0 | NA | NA | 3 | 0.3333 | 3 | 0 | 7 | 0.1429 |
| 8_2072323 | NA | NA | 1 | 1 | 4 | 0.5 | 3 | 0.6667 | 8 | 0.625 | 2 | 0 | NA | NA | 3 | 1 | 3 | 0.6667 | 8 | 0.625 |
| 8_2072324 | NA | NA | 2 | 1 | 4 | 0.5 | 2 | 1 | 8 | 0.75 | 2 | 1 | NA | NA | 2 | 1 | 3 | 0.6667 | 7 | 0.8571 |
| 8_2072325 | NA | NA | 3 | 0 | 4 | 0.25 | 2 | 1 | 9 | 0.3333 | 2 | 0 | 1 | 0 | 3 | 0 | 3 | 1 | 9 | 0.3333 |
| 8_2072347 | NA | NA | 3 | 0.6667 | 5 | 0.6 | 4 | 0.5 | 12 | 0.5833 | 2 | 1 | 1 | 0 | 3 | 0.3333 | 3 | 0.3333 | 9 | 0.4444 |
| 8_2072455 | 2 | 0.5 | 5 | 0.2 | 6 | 0.1667 | 4 | 0 | 17 | 0.1765 | 2 | 0.5 | 1 | 0 | 5 | 0.2 | 3 | 0 | 11 | 0.1818 |
| 8_2072653 | 1 | 0 | 4 | 0.25 | 5 | 0 | 3 | 0 | 13 | 0.0769 | NA | NA | NA | NA | 2 | 0 | 3 | 0.3333 | 5 | 0.2 |
| 8_2072679 | 1 | 1 | 2 | 1 | 2 | 1 | 1 | 0 | 6 | 0.8333 | NA | NA | NA | NA | 3 | 0.3333 | 3 | 0.6667 | 6 | 0.5 |
| 8_2072681 | 1 | 0 | 3 | 0.3333 | 1 | 1 | 1 | 0 | 6 | 0.3333 | NA | NA | 1 | 0 | 3 | 0 | 3 | 0.6667 | 7 | 0.2857 |
| 8_2072721 | NA | NA | 1 | 1 | NA | NA | 1 | 1 | 2 | 1 | NA | NA | NA | NA | 1 | 1 | NA | NA | 1 | 1 |
| 8_2105231 | NA | NA | NA | NA | 1 | 1 | NA | NA | 1 | 1 | NA | NA | NA | NA | 1 | 1 | NA | NA | 1 | 1 |
| 8_2105324 | NA | NA | NA | NA | 1 | 1 | NA | NA | 1 | 1 | NA | NA | NA | NA | NA | NA | 1 | 1 | 1 | 1 |
| 8_2112234 | 3 | 1 | NA | NA | 2 | 1 | 2 | 0.5 | 7 | 0.8571 | 1 | 1 | 1 | 1 | 1 | 1 | 1 | 1 | 4 | 1 |
| 8_2112235 | 3 | 0.3333 | NA | NA | 2 | 0.5 | 2 | 0 | 7 | 0.2857 | 1 | 0 | 3 | 0 | 1 | 0 | 1 | 1 | 6 | 0.1667 |
| 8_2112278 | NA | NA | NA | NA | NA | NA | 1 | 1 | 1 | 1 | NA | NA | NA | NA | 1 | 1 | NA | NA | 1 | 1 |
| 8_2113345 | 1 | 1 | 1 | 0 | NA | NA | NA | NA | 2 | 0.5 | 1 | 1 | 1 | 0 | 1 | 1 | NA | NA | 3 | 0.6667 |
| 8_2113360 | 1 | 1 | 1 | 1 | NA | NA | NA | NA | 2 | 1 | NA | NA | 1 | 1 | 1 | 1 | NA | NA | 2 | 1 |
| 8_2113423 | 1 | 0 | 2 | 0.5 | NA | NA | NA | NA | 3 | 0.3333 | 1 | 0 | 1 | 1 | 1 | 0 | NA | NA | 3 | 0.3333 |
| 8_2113472 | 1 | 0 | 2 | 0.5 | NA | NA | NA | NA | 3 | 0.3333 | NA | NA | 2 | 0.5 | 1 | 0 | NA | NA | 3 | 0.3333 |
| 8_2113500 | 1 | 1 | 2 | 0 | NA | NA | NA | NA | 3 | 0.3333 | 1 | 1 | 2 | 0 | 1 | 0 | NA | NA | 4 | 0.25 |
| 8_2255549 | 1 | 0 | 2 | 1 | 1 | 1 | 2 | 0.5 | 6 | 0.6667 | NA | NA | 2 | 0.5 | NA | NA | 1 | 1 | 3 | 0.6667 |
| 8_2255552 | NA | NA | 2 | 1 | 1 | 1 | 2 | 1 | 5 | 1 | NA | NA | 2 | 1 | NA | NA | 1 | 0 | 3 | 0.6667 |
| 8_2255559 | 1 | 0 | 2 | 0 | 1 | 0 | 2 | 1 | 6 | 0.3333 | NA | NA | 2 | 0.5 | NA | NA | 1 | 1 | 3 | 0.6667 |
| 8_2255560 | 1 | 0 | 2 | 0.5 | 1 | 1 | NA | NA | 4 | 0.5 | NA | NA | 2 | 1 | NA | NA | 1 | 1 | 3 | 1 |
| 8_2255567 | 1 | 1 | 2 | 0.5 | 1 | 0 | 2 | 0.5 | 6 | 0.5 | NA | NA | 2 | 0.5 | NA | NA | 1 | 1 | 3 | 0.6667 |
| 8_2255575 | 1 | 1 | 2 | 0 | NA | NA | 2 | 0.5 | 5 | 0.4 | NA | NA | 2 | 0.5 | NA | NA | 1 | 1 | 3 | 0.6667 |
| 8_2255796 | NA | NA | 2 | 0.5 | NA | NA | 3 | 0.6667 | 5 | 0.6 | NA | NA | 2 | 0.5 | NA | NA | 1 | 0 | 3 | 0.3333 |
| 8_2255798 | NA | NA | 2 | 0.5 | NA | NA | 3 | 0.3333 | 5 | 0.4 | NA | NA | 2 | 1 | NA | NA | 1 | 0 | 3 | 0.6667 |
| 8_2255802 | NA | NA | 2 | 0.5 | NA | NA | 3 | 0.6667 | 5 | 0.6 | NA | NA | 3 | 0.3333 | NA | NA | 1 | 1 | 4 | 0.5 |
| 8_2255806 | 1 | 1 | 2 | 0.5 | 1 | 0 | 3 | 0.3333 | 7 | 0.4286 | NA | NA | 3 | 0.6667 | NA | NA | 1 | 1 | 4 | 0.75 |
| 8_2255936 | 2 | 0.5 | 3 | 0 | 1 | 1 | 2 | 0.5 | 8 | 0.375 | NA | NA | 3 | 0 | 1 | 1 | 1 | 1 | 5 | 0.4 |
| 8_2256024 | 2 | 0 | 3 | 0 | NA | NA | 2 | 0.5 | 7 | 0.1429 | NA | NA | 4 | 0.25 | 1 | 0 | NA | NA | 5 | 0.2 |
| 8_2256083 | 1 | 0 | 3 | 0 | NA | NA | 5 | 0.2 | 9 | 0.1111 | 1 | 0 | 4 | 0.25 | 1 | 0 | NA | NA | 6 | 0.1667 |

| | | | | | | | | | | | | | | | | | | | | |
|---|---|---|---|---|---|---|---|---|---|---|---|---|---|---|---|---|---|---|---|---|
| 8_2256090 | 1 | 1 | 3 | 0 | 1 | 1 | 5 | 0.4 | 10 | 0.4 | 1 | 1 | 3 | 0.3333 | 2 | 0.5 | NA | NA | 6 | 0.5 |
| 8_2256104 | 2 | 0 | 4 | 0 | 1 | 0 | 5 | 0.2 | 12 | 0.0833 | 1 | 1 | 3 | 0 | 2 | 0.5 | NA | NA | 6 | 0.3333 |
| 8_2256146 | 3 | 0 | 5 | 0 | 2 | 0 | 4 | 0.25 | 14 | 0.0714 | NA | NA | 6 | 0.1667 | 2 | 0.5 | NA | NA | 8 | 0.25 |
| 8_2273316 | 1 | 1 | NA | NA | 1 | 1 | NA | NA | 2 | 1 | NA | NA | 2 | 0.5 | NA | NA | NA | NA | 2 | 0.5 |
| 8_2273320 | 1 | 0 | NA | NA | 1 | 1 | NA | NA | 2 | 0.5 | NA | NA | 2 | 0.5 | NA | NA | NA | NA | 2 | 0.5 |
| 8_2273327 | NA | NA | NA | NA | 1 | 1 | NA | NA | 1 | 1 | NA | NA | 2 | 0.5 | NA | NA | 1 | 1 | 3 | 0.6667 |
| 8_2273336 | 1 | 1 | NA | NA | 1 | 1 | NA | NA | 2 | 1 | NA | NA | 2 | 1 | NA | NA | NA | NA | 2 | 1 |
| 8_2273343 | 1 | 1 | NA | NA | 1 | 0 | NA | NA | 2 | 0.5 | NA | NA | 1 | 1 | NA | NA | NA | NA | 1 | 1 |
| 8_2273350 | 1 | 1 | NA | NA | NA | NA | NA | NA | 1 | 1 | NA | NA | 3 | 0.3333 | NA | NA | NA | NA | 3 | 0.3333 |
| 8_2273353 | 1 | 1 | NA | NA | 1 | 1 | NA | NA | 2 | 1 | NA | NA | 1 | 1 | NA | NA | NA | NA | 1 | 1 |
| 8_2273362 | 1 | 1 | NA | NA | 1 | 1 | NA | NA | 2 | 1 | NA | NA | 3 | 0.6667 | NA | NA | NA | NA | 3 | 0.6667 |
| 8_2273365 | 1 | 1 | NA | NA | 1 | 1 | NA | NA | 2 | 1 | NA | NA | 3 | 1 | NA | NA | 1 | 1 | 4 | 1 |
| 8_2273381 | 1 | 1 | NA | NA | NA | NA | NA | NA | 1 | 1 | NA | NA | 2 | 1 | NA | NA | 1 | 1 | 3 | 1 |
| 8_2274861 | 1 | 1 | NA | NA | 1 | 1 | NA | NA | 2 | 1 | NA | NA | 3 | 0.6667 | NA | NA | NA | NA | 3 | 0.6667 |
| 8_2274880 | 1 | 1 | NA | NA | 1 | 1 | NA | NA | 2 | 1 | NA | NA | 3 | 1 | NA | NA | NA | NA | 3 | 1 |
| 8_2274908 | 1 | 1 | NA | NA | 1 | 1 | NA | NA | 2 | 1 | NA | NA | 3 | 1 | NA | NA | NA | NA | 3 | 1 |
| 8_2274934 | 1 | 0 | NA | NA | 1 | 1 | NA | NA | 2 | 0.5 | NA | NA | 3 | 0.6667 | NA | NA | NA | NA | 3 | 0.6667 |
| 8_2274941 | 1 | 1 | NA | NA | 1 | 0 | NA | NA | 2 | 0.5 | NA | NA | 2 | 0.5 | NA | NA | NA | NA | 2 | 0.5 |
| 8_2274961 | 1 | 1 | NA | NA | 1 | 0 | NA | NA | 2 | 0.5 | NA | NA | 3 | 0.6667 | NA | NA | NA | NA | 3 | 0.6667 |
| 8_2275016 | NA | NA | NA | NA | 1 | 1 | NA | NA | 1 | 1 | NA | NA | 1 | 1 | NA | NA | NA | NA | 1 | 1 |
| 8_2275035 | NA | NA | NA | NA | 1 | 1 | NA | NA | 1 | 1 | NA | NA | 1 | 1 | NA | NA | NA | NA | 1 | 1 |
| 8_2286574 | 1 | 0 | 5 | 0.2 | 2 | 0.5 | 1 | 0 | 9 | 0.2222 | 3 | 0.3333 | 2 | 0 | NA | NA | 3 | 0.3333 | 8 | 0.25 |
| 8_2286750 | 2 | 0 | 4 | 0.5 | 3 | 0.6667 | NA | NA | 9 | 0.4444 | NA | NA | 2 | 0.5 | 1 | 1 | 2 | 1 | 5 | 0.8 |
| 8_2286758 | 2 | 0.5 | 4 | 0 | 3 | 0.3333 | NA | NA | 9 | 0.2222 | NA | NA | 2 | 0 | NA | NA | 2 | 0.5 | 4 | 0.25 |
| 8_2286776 | 2 | 0.5 | 3 | 0.3333 | 2 | 0.5 | NA | NA | 7 | 0.4286 | NA | NA | 2 | 0.5 | NA | NA | 1 | 0 | 3 | 0.3333 |
| 8_2286777 | 2 | 0.5 | 3 | 1 | 2 | 0.5 | NA | NA | 7 | 0.7143 | NA | NA | 2 | 0 | NA | NA | 2 | 1 | 4 | 0.5 |
| 8_2286779 | 2 | 0.5 | 3 | 0 | 1 | 1 | NA | NA | 6 | 0.3333 | NA | NA | 1 | 1 | NA | NA | 2 | 0 | 3 | 0.3333 |
| 8_2286783 | 2 | 1 | 3 | 1 | 2 | 1 | NA | NA | 7 | 1 | NA | NA | 1 | 1 | NA | NA | 2 | 1 | 3 | 1 |
| 8_2286797 | 2 | 0.5 | 3 | 1 | 2 | 0.5 | NA | NA | 7 | 0.7143 | NA | NA | NA | NA | NA | NA | 1 | 1 | 1 | 1 |
| 8_2290488 | 1 | 0 | 6 | 0 | 2 | 1 | 3 | 0.3333 | 12 | 0.25 | 3 | 0 | 1 | 0 | 6 | 0.3333 | 4 | 0.25 | 14 | 0.2143 |
| 8_2290489 | 2 | 0 | 6 | 0.3333 | 2 | 0 | 4 | 0 | 14 | 0.1429 | 3 | 0 | 4 | 0.25 | 7 | 0 | 4 | 0 | 18 | 0.0556 |
| 8_2295288 | 7 | 0.5714 | 4 | 0.5 | 3 | 0.6667 | 6 | 0.5 | 20 | 0.55 | 3 | 0.3333 | 5 | 0.4 | 3 | 0.6667 | 6 | 0.5 | 17 | 0.4706 |
| 8_2295289 | 6 | 0.3333 | 5 | 0.2 | 3 | 0 | 6 | 0.5 | 20 | 0.3 | 4 | 0 | 7 | 0.4286 | 2 | 0.5 | 6 | 0.6667 | 19 | 0.4211 |
| 8_2295295 | 6 | 0.1667 | 5 | 0.2 | 4 | 0.25 | 5 | 0.4 | 20 | 0.25 | 4 | 0.25 | 7 | 0.1429 | 3 | 0.3333 | 7 | 0.2857 | 21 | 0.2381 |
| 8_2295314 | 7 | 1 | 5 | 1 | 4 | 1 | 5 | 1 | 21 | 1 | 5 | 1 | 4 | 1 | 5 | 1 | 9 | 0.7778 | 23 | 0.913 |
| 8_2295315 | 5 | 1 | 5 | 0.4 | 3 | 0.6667 | 5 | 0.4 | 18 | 0.6111 | 4 | 0.75 | 5 | 0.8 | 4 | 0.75 | 9 | 0.7778 | 22 | 0.7727 |
| 8_2295320 | 7 | 0.8571 | 5 | 1 | 4 | 0.75 | 6 | 0.8333 | 22 | 0.8636 | 4 | 1 | 6 | 0.6667 | 5 | 0.8 | 10 | 0.8 | 25 | 0.8 |
| 8_2295331 | 6 | 0 | 7 | 0 | 4 | 0.25 | 5 | 0.2 | 22 | 0.0909 | 5 | 0.2 | 4 | 0 | 5 | 0.4 | 10 | 0.1 | 24 | 0.1667 |
| 8_2296652 | 5 | 0 | 4 | 0 | 4 | 0.25 | 5 | 0 | 18 | 0.0556 | 4 | 0 | 4 | 0 | 2 | 0 | 6 | 0.1667 | 16 | 0.0625 |
| 8_2297295 | NA | NA | 2 | 1 | NA | NA | NA | NA | 2 | 1 | NA | NA | 1 | 0 | 1 | 1 | 1 | 1 | 3 | 0.6667 |
| 8_2297297 | NA | NA | 2 | 0.5 | NA | NA | NA | NA | 2 | 0.5 | NA | NA | 1 | 0 | 1 | 1 | 1 | 0 | 3 | 0.3333 |
| 8_2297300 | NA | NA | 2 | 0.5 | NA | NA | NA | NA | 2 | 0.5 | NA | NA | 1 | 0 | 1 | 1 | 1 | 1 | 3 | 0.6667 |
| 8_2297330 | NA | NA | 2 | 1 | NA | NA | NA | NA | 2 | 1 | NA | NA | 1 | 0 | NA | NA | 1 | 1 | 2 | 0.5 |
| 8_2302509 | 3 | 0 | 1 | 0 | 2 | 0 | 3 | 0.3333 | 9 | 0.1111 | NA | NA | 1 | 0 | 1 | 0 | 4 | 0.5 | 6 | 0.3333 |
| 8_2302511 | 2 | 0.5 | 1 | 0 | 1 | 1 | 2 | 1 | 6 | 0.6667 | NA | NA | 2 | 1 | 1 | 1 | 3 | 1 | 6 | 1 |
| 8_2302521 | 3 | 0.3333 | 1 | 0 | 1 | 1 | 3 | 0.3333 | 8 | 0.375 | NA | NA | 2 | 1 | NA | NA | 3 | 1 | 5 | 1 |
| 8_2302527 | 3 | 0.3333 | 1 | 0 | 2 | 0.5 | 3 | 0 | 9 | 0.2222 | NA | NA | 1 | 1 | 1 | 1 | 4 | 0.75 | 6 | 0.8333 |
| 8_2302543 | 4 | 0.25 | 1 | 0 | 1 | 0 | 3 | 0 | 9 | 0.1111 | NA | NA | 2 | 0 | 1 | 1 | 4 | 0 | 7 | 0.1429 |
| 8_2302546 | 3 | 0.3333 | 1 | 0 | 2 | 0 | 3 | 0 | 9 | 0.1111 | NA | NA | NA | NA | 1 | 0 | 3 | 0.3333 | 4 | 0.25 |
| 8_2305101 | 4 | 1 | 4 | 0.5 | 4 | 0.75 | 3 | 1 | 15 | 0.8 | 2 | 1 | 2 | 0.5 | 2 | 1 | 5 | 0.8 | 11 | 0.8182 |
| 8_2305103 | 6 | 0.1667 | 4 | 0.5 | 4 | 1 | 3 | 1 | 17 | 0.5882 | 1 | 0 | 2 | 0.5 | 2 | 0.5 | 5 | 0.6 | 10 | 0.5 |
| 8_2305118 | 4 | 0.5 | 4 | 0 | 3 | 0.3333 | 4 | 0 | 15 | 0.2 | 2 | 0 | 1 | 0 | 4 | 0 | 6 | 0.1667 | 13 | 0.0769 |
| 8_2305120 | 6 | 0.5 | 4 | 0.25 | 3 | 0.6667 | 4 | 0.25 | 17 | 0.4118 | 1 | 0 | 2 | 1 | 3 | 0 | 7 | 0.7143 | 13 | 0.5385 |
| 8_2305140 | 6 | 0.1667 | 3 | 0 | 5 | 0.4 | 3 | 0 | 17 | 0.1765 | 1 | 1 | 2 | 0 | 3 | 0 | 5 | 0 | 11 | 0.0909 |
| 8_2305831 | 4 | 0.25 | 5 | 0.4 | 5 | 0.2 | 4 | 0.25 | 18 | 0.2778 | 2 | 0 | 1 | 0 | 4 | 0 | 7 | 0.1429 | 14 | 0.0714 |
| 8_2305833 | 3 | 0 | 3 | 0 | 3 | 0.3333 | 4 | 0.25 | 13 | 0.1538 | 3 | 0 | 2 | 0 | 4 | 0 | 5 | 0.2 | 14 | 0.0714 |
| 8_2305835 | 5 | 0.2 | 4 | 0 | 4 | 0 | 4 | 0.25 | 17 | 0.1176 | 3 | 0 | 1 | 1 | 3 | 0.3333 | 6 | 0 | 13 | 0.1538 |
| 8_2305837 | 5 | 0.4 | 5 | 0 | 2 | 0 | 4 | 0.25 | 16 | 0.1875 | 2 | 0.5 | 2 | 0.5 | 5 | 0.4 | 3 | 0.3333 | 12 | 0.4167 |
| 8_2305874 | 8 | 0.125 | 5 | 0.2 | 5 | 0 | 4 | 0 | 22 | 0.0909 | 3 | 0 | 3 | 0 | 6 | 0.1667 | 7 | 0 | 19 | 0.0526 |
| 8_2305887 | 9 | 0.2222 | 6 | 0.1667 | 4 | 0.25 | 5 | 0.6 | 24 | 0.2917 | 5 | 0.4 | 2 | 1 | 5 | 0.2 | 6 | 0.3333 | 18 | 0.3889 |
| 8_2305889 | 8 | 0.125 | 4 | 0.25 | 4 | 0 | 3 | 0 | 19 | 0.1053 | 5 | 0.4 | 3 | 0 | 6 | 0.1667 | 7 | 0.1429 | 21 | 0.1905 |
| 8_2305938 | 10 | 0.4 | 6 | 0 | 5 | 0.6 | 6 | 1 | 27 | 0.4815 | 5 | 0.2 | 3 | 0.3333 | 6 | 0.1667 | 9 | 0.3333 | 23 | 0.2609 |
| 8_2305942 | 10 | 0 | 5 | 0 | 5 | 0.2 | 5 | 0.2 | 25 | 0.08 | 5 | 0 | 3 | 0 | 5 | 0.2 | 7 | 0 | 20 | 0.05 |
| 8_2306742 | 22 | 0.4545 | 28 | 0.4286 | 22 | 0.4091 | 23 | 0.4348 | 95 | 0.4316 | 20 | 0.55 | 31 | 0.5161 | 37 | 0.5135 | 21 | 0.3333 | 109 | 0.4862 |
| 8_2306753 | 20 | 0.3 | 26 | 0.1923 | 20 | 0.1 | 17 | 0.0588 | 83 | 0.1687 | 22 | 0.0909 | 25 | 0.2 | 32 | 0.2188 | 23 | 0.2609 | 102 | 0.1961 |

| | | | | | | | | | | | | | | | | | | | | |
|---|---|---|---|---|---|---|---|---|---|---|---|---|---|---|---|---|---|---|---|---|
| 8_2312216 | 2 | 0 | 3 | 0 | 1 | 0 | 2 | 0.5 | 8 | 0.125 | NA | NA | 2 | 0.5 | 5 | 0.4 | 1 | 0 | 8 | 0.375 |
| 8_2314976 | 5 | 0.2 | 7 | 0 | 1 | 0 | 3 | 0 | 16 | 0.0625 | NA | NA | 2 | 0.5 | 2 | 0 | 1 | 0 | 5 | 0.2 |
| 8_2314988 | 5 | 0.4 | 6 | 0.5 | 1 | 0 | 2 | 0.5 | 14 | 0.4286 | NA | NA | 3 | 0 | 2 | 0 | 2 | 1 | 7 | 0.2857 |
| 8_2314994 | 6 | 0.1667 | 6 | 0.3333 | 1 | 0 | 2 | 0.5 | 15 | 0.2667 | NA | NA | 4 | 0.25 | 1 | 0 | 2 | 0 | 7 | 0.1429 |
| 8_2315005 | 6 | 0 | 6 | 0.3333 | 1 | 0 | 2 | 0 | 15 | 0.1333 | NA | NA | 3 | 0.3333 | 2 | 0 | 2 | 0 | 7 | 0.1429 |
| 8_2315013 | NA | NA | 1 | 1 | NA | NA | NA | NA | 1 | 1 | NA | NA | NA | NA | 1 | 1 | 1 | 1 | 2 | 1 |
| 8_2315034 | 5 | 0.2 | 7 | 0.1429 | 1 | 0 | 1 | 0 | 14 | 0.1429 | NA | NA | 3 | 0 | 2 | 0.5 | 2 | 0.5 | 7 | 0.2857 |
| 8_2315041 | 5 | 0.2 | 6 | 0.5 | 1 | 0 | 2 | 0.5 | 14 | 0.3571 | NA | NA | 4 | 0.25 | 2 | 0.5 | 2 | 0.5 | 8 | 0.375 |
| 8_2315046 | 5 | 0 | 6 | 0 | 1 | 0 | 2 | 0.5 | 14 | 0.0714 | NA | NA | 3 | 0.3333 | 2 | 0 | 2 | 0.5 | 7 | 0.2857 |
| 8_2315110 | NA | NA | NA | NA | 1 | 1 | 1 | 1 | 2 | 1 | NA | NA | 2 | 0.5 | NA | NA | 1 | 1 | 3 | 0.6667 |
| 8_2315132 | NA | NA | NA | NA | 1 | 1 | 1 | 1 | 2 | 1 | NA | NA | 1 | 1 | NA | NA | 1 | 1 | 2 | 1 |
| 8_2327209 | 3 | 0.3333 | 5 | 0 | 3 | 0.3333 | 4 | 0 | 15 | 0.1333 | 3 | 0 | 7 | 0.1429 | 8 | 0.25 | 2 | 0 | 20 | 0.15 |
| 8_2327787 | 12 | 0.1667 | 10 | 0.2 | 11 | 0 | 13 | 0.1538 | 46 | 0.1304 | 10 | 0.2 | 12 | 0 | 9 | 0.1111 | 7 | 0.4286 | 38 | 0.1579 |
| 8_2327828 | 6 | 0.1667 | 6 | 0.3333 | 6 | 0 | 9 | 0.4444 | 27 | 0.2593 | 7 | 0.1429 | 11 | 0.0909 | 8 | 0 | 7 | 0 | 33 | 0.0606 |
| 8_2327835 | 4 | 0.25 | 6 | 0.6667 | 7 | 0.2857 | 8 | 0.375 | 25 | 0.4 | 1 | 0 | 5 | 0.4 | 4 | 0.25 | 5 | 0.4 | 15 | 0.3333 |
| 8_2327841 | 5 | 0.2 | 8 | 0.375 | 8 | 0.125 | 10 | 0.2 | 31 | 0.2258 | 7 | 0 | 8 | 0.25 | 7 | 0 | 6 | 0.1667 | 28 | 0.1071 |
| 8_2327851 | 5 | 0.2 | 8 | 0.5 | 10 | 0.2 | 11 | 0.5455 | 34 | 0.3824 | 8 | 0.375 | 7 | 0.5714 | 7 | 0.2857 | 7 | 0.7143 | 29 | 0.4828 |
| 8_2327861 | 5 | 0 | 6 | 0.3333 | 10 | 0 | 10 | 0.1 | 31 | 0.0968 | 7 | 0 | 11 | 0.2727 | 7 | 0.1429 | 6 | 0.1667 | 31 | 0.1613 |
| 8_2327867 | 5 | 0.8 | 8 | 0.75 | 11 | 0.6364 | 8 | 0.875 | 32 | 0.75 | 7 | 0.5714 | 11 | 0.6364 | 8 | 0.5 | 6 | 0.6667 | 32 | 0.5938 |
| 8_2327882 | 2 | 0.5 | 7 | 0.7143 | 9 | 0.5556 | 8 | 0.875 | 26 | 0.6923 | 7 | 0.2857 | 7 | 0.4286 | 6 | 0.6667 | 7 | 0.7143 | 27 | 0.5185 |
| 8_2327892 | 2 | 1 | 5 | 0.6 | 9 | 0.7778 | 7 | 1 | 23 | 0.8261 | 4 | 0.5 | 6 | 0.8333 | 7 | 0.5714 | 3 | 1 | 20 | 0.7 |
| 8_2327912 | 2 | 0 | 6 | 0.1667 | 11 | 0.0909 | 9 | 0 | 28 | 0.0714 | 4 | 0.25 | 8 | 0.125 | 6 | 0.3333 | 5 | 0.2 | 23 | 0.2174 |
| 8_2327916 | 2 | 0.5 | 6 | 0.8333 | 9 | 0.5556 | 7 | 0.8571 | 24 | 0.7083 | 6 | 1 | 7 | 0.5714 | 3 | 0.3333 | 3 | 1 | 19 | 0.7368 |
| 8_2327933 | 3 | 0.6667 | 6 | 0.8333 | 10 | 1 | 8 | 0.875 | 27 | 0.8889 | 5 | 1 | 6 | 1 | 5 | 0.8 | 4 | 0.75 | 20 | 0.9 |
| 8_2327946 | 2 | 0 | 4 | 0.25 | 3 | 0 | 6 | 0.1667 | 15 | 0.1333 | 5 | 0 | 5 | 0 | 3 | 0 | 5 | 0.4 | 18 | 0.1111 |
| 8_2328254 | NA | NA | NA | NA | 1 | 1 | NA | NA | 1 | 1 | NA | NA | NA | NA | 1 | 1 | NA | NA | 1 | 1 |
| 8_2329402 | NA | NA | 3 | 0 | 3 | 0.3333 | 7 | 0.1429 | 13 | 0.1538 | 2 | 0 | 5 | 0 | 2 | 0 | 3 | 0.3333 | 12 | 0.0833 |
| 8_2342225 | NA | NA | 3 | 1 | NA | NA | NA | NA | 3 | 1 | NA | NA | NA | NA | 1 | 1 | NA | NA | 1 | 1 |
| 8_2342829 | NA | NA | 4 | 1 | NA | NA | NA | NA | 4 | 1 | NA | NA | 1 | 1 | 1 | 1 | 1 | 0 | 3 | 0.6667 |
| 8_2342870 | NA | NA | 4 | 0.75 | NA | NA | NA | NA | 4 | 0.75 | NA | NA | 1 | 1 | 1 | 1 | 1 | 1 | 3 | 1 |
| 8_2342896 | NA | NA | 4 | 0.5 | NA | NA | NA | NA | 4 | 0.5 | NA | NA | 1 | 1 | 1 | 1 | 1 | 0 | 3 | 0.6667 |
| 8_2342943 | NA | NA | 2 | 0.5 | NA | NA | NA | NA | 2 | 0.5 | NA | NA | 1 | 1 | 1 | 1 | 2 | 0.5 | 4 | 0.75 |
| 8_2342949 | NA | NA | 3 | 0.3333 | NA | NA | NA | NA | 3 | 0.3333 | NA | NA | 1 | 0 | 1 | 1 | 2 | 0.5 | 4 | 0.5 |
| 8_2342993 | NA | NA | 4 | 1 | 1 | 1 | NA | NA | 5 | 1 | NA | NA | NA | NA | 1 | 1 | 2 | 1 | 3 | 1 |
| 8_2342997 | NA | NA | 4 | 0.25 | 1 | 1 | NA | NA | 5 | 0.4 | NA | NA | 1 | 1 | 1 | 0 | 2 | 0 | 4 | 0.25 |
| 8_2343001 | NA | NA | 4 | 1 | NA | NA | NA | NA | 4 | 1 | NA | NA | 1 | 1 | 1 | 0 | 2 | 1 | 4 | 0.75 |
| 8_2343005 | NA | NA | 4 | 0.25 | 1 | 0 | NA | NA | 5 | 0.2 | NA | NA | 1 | 1 | 1 | 0 | 2 | 0 | 4 | 0.25 |
| 8_2343010 | NA | NA | 3 | 0.6667 | NA | NA | NA | NA | 3 | 0.6667 | NA | NA | 1 | 1 | NA | NA | 2 | 0.5 | 3 | 0.6667 |
| 8_2343033 | NA | NA | 4 | 1 | 2 | 1 | NA | NA | 6 | 1 | NA | NA | 2 | 1 | 1 | 1 | 1 | 1 | 4 | 1 |
| 8_2343039 | NA | NA | 4 | 1 | NA | NA | NA | NA | 4 | 1 | NA | NA | 2 | 1 | 1 | 1 | 2 | 1 | 5 | 1 |
| 8_2343041 | NA | NA | 2 | 1 | NA | NA | NA | NA | 2 | 1 | NA | NA | 2 | 1 | 1 | 1 | 1 | 1 | 4 | 1 |
| 8_2343042 | NA | NA | 4 | 0 | 2 | 0.5 | NA | NA | 6 | 0.1667 | NA | NA | 1 | 1 | 1 | 1 | 2 | 0.5 | 4 | 0.75 |
| 8_2343044 | 1 | 0 | 3 | 0.6667 | 2 | 0 | NA | NA | 6 | 0.3333 | NA | NA | 2 | 0.5 | 1 | 1 | 2 | 1 | 5 | 0.8 |
| 8_2343047 | 1 | 0 | 4 | 0.25 | 1 | 0 | NA | NA | 6 | 0.1667 | NA | NA | 2 | 1 | 1 | 0 | 2 | 0.5 | 5 | 0.6 |
| 8_2343056 | 1 | 1 | 3 | 0.6667 | 2 | 0 | NA | NA | 6 | 0.5 | NA | NA | 1 | 1 | 1 | 1 | 2 | 1 | 4 | 1 |
| 8_2343064 | 1 | 1 | 4 | 1 | 1 | 1 | NA | NA | 6 | 1 | NA | NA | 2 | 1 | 1 | 1 | 2 | 1 | 5 | 1 |
| 8_2343066 | 1 | 0 | 3 | 0.3333 | NA | NA | NA | NA | 4 | 0.25 | NA | NA | 2 | 0 | 1 | 1 | 2 | 0 | 5 | 0.2 |
| 8_2343070 | 2 | 1 | 4 | 1 | 2 | 1 | NA | NA | 8 | 1 | NA | NA | 2 | 1 | 1 | 1 | 1 | 1 | 4 | 1 |
| 8_2343074 | 2 | 1 | 4 | 1 | 2 | 1 | NA | NA | 8 | 1 | NA | NA | 2 | 1 | 1 | 1 | 2 | 1 | 5 | 1 |
| 8_2343099 | 2 | 0.5 | 5 | 0.4 | 1 | 0 | NA | NA | 8 | 0.375 | NA | NA | 1 | 0 | 1 | 0 | 2 | 0.5 | 4 | 0.25 |
| 8_2343152 | 4 | 0.25 | 3 | 0 | 1 | 0 | NA | NA | 8 | 0.125 | NA | NA | 2 | 0 | 1 | 0 | 2 | 0.5 | 5 | 0.2 |
| 8_2343268 | 8 | 0.125 | 10 | 0.1 | 5 | 0 | 5 | 0.2 | 28 | 0.1071 | NA | NA | 5 | 0.2 | 6 | 0 | 4 | 0.25 | 15 | 0.1333 |
| 8_2362232 | 2 | 0.5 | NA | NA | 1 | 0 | 1 | 0 | 4 | 0.25 | 2 | 0.5 | 1 | 0 | 2 | 0 | 1 | 0 | 6 | 0.1667 |
| 8_2362235 | 2 | 1 | NA | NA | 3 | 1 | 1 | 1 | 6 | 1 | 2 | 1 | 1 | 1 | 1 | 1 | 1 | 1 | 5 | 1 |
| 8_2362239 | 3 | 0 | NA | NA | 3 | 0.3333 | 1 | 0 | 7 | 0.1429 | 2 | 0 | 1 | 0 | 2 | 0.5 | NA | NA | 5 | 0.2 |
| 8_2362242 | 3 | 0 | NA | NA | 3 | 0.6667 | 1 | 0 | 7 | 0.2857 | 2 | 0 | 1 | 0 | 2 | 0 | 1 | 1 | 6 | 0.1667 |
| 8_2363836 | 2 | 0 | 1 | 0 | 4 | 0.25 | 1 | 0 | 8 | 0.125 | 4 | 0 | 1 | 0 | 2 | 0.5 | 1 | 0 | 8 | 0.125 |
| 8_2363852 | 5 | 0.2 | 1 | 0 | 3 | 0 | 1 | 0 | 10 | 0.1 | 4 | 0.25 | 1 | 1 | 2 | 0.5 | 1 | 0 | 8 | 0.375 |
| 8_2363870 | 6 | 0.6667 | 1 | 0 | 3 | 0.3333 | 1 | 1 | 11 | 0.5455 | 2 | 0 | 1 | 0 | 3 | 0.3333 | 1 | 0 | 7 | 0.1429 |
| 8_2363884 | 7 | 0.5714 | 1 | 1 | 4 | 0.5 | 1 | 0 | 13 | 0.5385 | 2 | 0.5 | 2 | 1 | 4 | 0.5 | 1 | 0 | 9 | 0.5556 |
| 8_2363894 | 6 | 0 | 1 | 0 | 6 | 0 | 1 | 1 | 14 | 0.0714 | 4 | 0 | 3 | 0.3333 | 3 | 0 | 1 | 0 | 11 | 0.0909 |
| 8_2363935 | 7 | 0.5714 | 2 | 0.5 | 5 | 0.6 | NA | NA | 14 | 0.5714 | 3 | 1 | 3 | 0.3333 | 5 | 0.4 | 1 | 1 | 12 | 0.5833 |
| 8_2363963 | 8 | 0.5 | 3 | 0.3333 | 4 | 0.25 | NA | NA | 15 | 0.4 | 3 | 0.3333 | 3 | 0.3333 | 4 | 0.25 | 1 | 0 | 11 | 0.2727 |
| 8_2363966 | 8 | 0.375 | 3 | 0.3333 | 5 | 0.4 | NA | NA | 16 | 0.375 | 4 | 0 | 2 | 1 | 3 | 0.3333 | 1 | 0 | 10 | 0.3 |
| 8_2363980 | 8 | 0.625 | 2 | 1 | 4 | 0.5 | NA | NA | 14 | 0.6429 | 4 | 0.75 | 3 | 1 | 4 | 0.75 | 2 | 0.5 | 13 | 0.7692 |

| | | | | | | | | | | | | | | | | | | | | |
|---|---|---|---|---|---|---|---|---|---|---|---|---|---|---|---|---|---|---|---|---|
| 8_2363988 | 7 | 0.2857 | 3 | 0.6667 | 6 | 0.5 | NA | NA | 16 | 0.4375 | 4 | 0.25 | 3 | 0 | 5 | 0.4 | 2 | 0 | 14 | 0.2143 |
| 8_2364038 | 7 | 0.1429 | 2 | 0 | 4 | 0 | NA | NA | 13 | 0.0769 | 3 | 0 | 3 | 0.3333 | 2 | 0 | 2 | 0 | 10 | 0.1 |
| 8_2366316 | 3 | 0.3333 | 1 | 0 | 3 | 0.3333 | 2 | 0.5 | 9 | 0.3333 | NA | NA | 1 | 1 | 1 | 1 | NA | NA | 2 | 1 |
| 8_2366316 | 4 | 0.25 | 1 | 0 | 4 | 0 | 2 | 0.5 | 11 | 0.1818 | NA | NA | 1 | 1 | 1 | 0 | 1 | 1 | 3 | 0.6667 |
| 8_2366384 | 17 | 0.4706 | 15 | 0.3333 | 11 | 0.0909 | 11 | 0.3636 | 54 | 0.3333 | 9 | 0.1111 | 12 | 0.3333 | 11 | 0.4545 | 13 | 0.1538 | 45 | 0.2667 |
| 8_2366398 | 15 | 0.2667 | 9 | 0.5556 | 10 | 0.4 | 11 | 0.3636 | 45 | 0.3778 | 8 | 0.125 | 10 | 0.3 | 11 | 0.2727 | 14 | 0.3571 | 43 | 0.2791 |
| 8_2366413 | 14 | 0.9286 | 11 | 1 | 10 | 0.9 | 11 | 0.9091 | 46 | 0.9348 | 8 | 0.875 | 11 | 0.7273 | 10 | 1 | 9 | 1 | 38 | 0.8947 |
| 8_2366419 | 14 | 0.7143 | 8 | 0.375 | 8 | 0.375 | 9 | 0.3333 | 39 | 0.4872 | 6 | 0.6667 | 9 | 0.3333 | 10 | 0.7 | 12 | 0.4167 | 37 | 0.5135 |
| 8_2366434 | 17 | 0.8824 | 10 | 0.9 | 11 | 0.6364 | 9 | 0.8889 | 47 | 0.8298 | 8 | 0.75 | 11 | 0.9091 | 9 | 0.7778 | 13 | 0.7692 | 41 | 0.8049 |
| 8_2366441 | 16 | 0.4375 | 11 | 0.2727 | 11 | 0.7273 | 11 | 0.3636 | 49 | 0.449 | 8 | 0.25 | 11 | 0.3636 | 8 | 0.75 | 11 | 0.6364 | 38 | 0.5 |
| 8_2366442 | 13 | 0.9231 | 5 | 0.8 | 9 | 0.6667 | 6 | 0.8333 | 33 | 0.8182 | 7 | 0.5714 | 9 | 0.5556 | 9 | 0.7778 | 12 | 0.9167 | 37 | 0.7297 |
| 8_2366444 | 16 | 0.9375 | 9 | 0.7778 | 10 | 0.7 | 12 | 0.5 | 47 | 0.7447 | 7 | 0.7143 | 11 | 0.8182 | 10 | 0.8 | 13 | 0.6154 | 41 | 0.7317 |
| 8_2368095 | 8 | 0.75 | 2 | 0.5 | 4 | 0.5 | 6 | 0.8333 | 20 | 0.7 | 9 | 0.6667 | 4 | 0.5 | 4 | 0.25 | 7 | 0.5714 | 24 | 0.5417 |
| 8_2368099 | 8 | 0.875 | 3 | 1 | 5 | 0.8 | 6 | 1 | 22 | 0.9091 | 8 | 0.875 | 5 | 1 | 3 | 1 | 10 | 0.7 | 26 | 0.8462 |
| 8_2368102 | 8 | 1 | 3 | 1 | 5 | 1 | 6 | 0.8333 | 22 | 0.9545 | 7 | 0.7143 | 2 | 1 | 4 | 1 | 8 | 1 | 21 | 0.9048 |
| 8_2368115 | 7 | 0.8571 | 3 | 0.3333 | 5 | 0.6 | 6 | 0.6667 | 21 | 0.6667 | 7 | 0.7143 | 4 | 0.5 | 4 | 1 | 8 | 0.875 | 23 | 0.7826 |
| 8_2368116 | 8 | 0.125 | 3 | 0.6667 | 5 | 0.4 | 6 | 0.5 | 22 | 0.3636 | 6 | 0.8333 | 3 | 1 | 4 | 0.25 | 7 | 0.2857 | 20 | 0.55 |
| 8_2368120 | 7 | 0.1429 | 2 | 0.5 | 4 | 0 | 4 | 0.25 | 17 | 0.1765 | 6 | 0 | 3 | 0 | 4 | 0 | 7 | 0.2857 | 20 | 0.1 |
| 8_2368127 | 8 | 0.25 | 3 | 0.3333 | 5 | 0.6 | 5 | 0.2 | 21 | 0.3333 | 7 | 0.2857 | 4 | 0.75 | 4 | 0.25 | 7 | 0.4286 | 22 | 0.4091 |
| 8_2368130 | 6 | 0.3333 | 1 | 0 | 4 | 0.25 | 4 | 0.75 | 15 | 0.4 | 6 | 0.3333 | 4 | 0.25 | 4 | 0.25 | 6 | 0.3333 | 20 | 0.3 |
| 8_2368144 | 7 | 0.7143 | 2 | 0.5 | 3 | 0.6667 | 3 | 1 | 15 | 0.7333 | 6 | 1 | 2 | 1 | 3 | 1 | 4 | 0.5 | 15 | 0.8667 |
| 8_2368146 | 6 | 1 | 2 | 1 | 4 | 1 | 3 | 0.6667 | 15 | 0.9333 | 7 | 0.7143 | NA | NA | 4 | 1 | 5 | 0.8 | 16 | 0.8125 |
| 8_2368147 | 8 | 0.25 | 2 | 0.5 | 4 | 0 | 3 | 0.3333 | 17 | 0.2353 | 7 | 0 | 3 | 0 | 4 | 0.5 | 5 | 0.4 | 19 | 0.2105 |
| 8_2368160 | 5 | 0.2 | 1 | 0 | 2 | 0.5 | 3 | 0.3333 | 11 | 0.2727 | 4 | 0 | 1 | 1 | 4 | 0.75 | 3 | 0 | 12 | 0.3333 |
| 8_2371665 | 2 | 0 | 3 | 0 | 2 | 0.5 | 1 | 0 | 8 | 0.125 | 1 | 0 | NA | NA | 1 | 0 | 3 | 0.3333 | 5 | 0.2 |
| 8_2371676 | 2 | 1 | 3 | 1 | 2 | 1 | 1 | 1 | 8 | 1 | 1 | 1 | NA | NA | 1 | 1 | 3 | 1 | 5 | 1 |
| 8_2371705 | 1 | 0 | 4 | 0.5 | 2 | 0 | 6 | 0.5 | 13 | 0.3846 | 2 | 1 | 3 | 0 | 3 | 1 | 6 | 0.5 | 14 | 0.5714 |
| 8_2371707 | 1 | 0 | 4 | 0.5 | 3 | 0 | 5 | 0.2 | 13 | 0.2308 | 2 | 0.5 | 2 | 0.5 | 3 | 0 | 4 | 0.25 | 11 | 0.2727 |
| 8_2373248 | 19 | 0.4211 | 15 | 0.4 | 22 | 0.5455 | 22 | 0.4545 | 78 | 0.4615 | 12 | 0.5833 | 13 | 0.7692 | 13 | 0.3077 | 13 | 0.4615 | 51 | 0.5294 |
| 8_2393651 | 1 | 1 | NA | NA | NA | NA | NA | NA | 1 | 1 | NA | NA | NA | NA | 2 | 0.5 | NA | NA | 2 | 0.5 |
| 8_2393665 | 1 | 1 | NA | NA | NA | NA | 1 | 0 | 2 | 0.5 | NA | NA | 1 | 1 | 1 | 0 | NA | NA | 2 | 0.5 |
| 8_2393689 | 1 | 1 | NA | NA | NA | NA | 1 | 0 | 2 | 0.5 | 1 | 1 | 1 | 1 | 3 | 0.3333 | NA | NA | 5 | 0.6 |
| 8_2393944 | 3 | 0.3333 | 2 | 0 | 1 | 0 | 1 | 0 | 7 | 0.1429 | 1 | 1 | NA | NA | 4 | 0.5 | 2 | 0.5 | 7 | 0.5714 |
| 8_2454987 | NA | NA | 1 | 1 | NA | NA | NA | NA | 1 | 1 | NA | NA | NA | NA | 1 | 1 | NA | NA | 1 | 1 |
| 8_2455036 | 3 | 0.6667 | 1 | 0 | NA | NA | 2 | 0.5 | 6 | 0.5 | 2 | 0.5 | NA | NA | 1 | 1 | 2 | 0 | 5 | 0.4 |
| 8_2455045 | 2 | 0.5 | 2 | 1 | 1 | 1 | 1 | 0 | 6 | 0.6667 | 2 | 1 | NA | NA | 2 | 1 | 2 | 1 | 6 | 1 |
| 8_2455072 | 4 | 1 | 2 | 1 | 1 | 1 | 2 | 1 | 9 | 1 | 2 | 1 | NA | NA | 1 | 0 | 2 | 0 | 5 | 0.4 |
| 8_2455073 | 3 | 0 | 2 | 0.5 | 1 | 0 | 2 | 0 | 8 | 0.125 | 1 | 0 | NA | NA | 2 | 0.5 | 2 | 0.5 | 5 | 0.4 |
| 8_2455075 | 4 | 0.5 | 2 | 0.5 | 1 | 1 | 1 | 0 | 8 | 0.5 | 2 | 0.5 | NA | NA | 2 | 1 | 2 | 0.5 | 6 | 0.6667 |
| 8_2455076 | 2 | 1 | 2 | 0.5 | NA | NA | 2 | 1 | 6 | 0.8333 | 2 | 1 | NA | NA | 1 | 1 | 2 | 1 | 5 | 1 |
| 8_2455087 | 4 | 0.25 | 2 | 0 | 1 | 0 | 2 | 0 | 9 | 0.1111 | 2 | 0 | NA | NA | 2 | 0.5 | 2 | 0.5 | 6 | 0.3333 |
| 8_2455088 | 2 | 0.5 | 2 | 1 | 1 | 1 | 1 | 0 | 6 | 0.6667 | 2 | 0.5 | NA | NA | 2 | 1 | 1 | 1 | 5 | 0.8 |
| 8_2455096 | 4 | 0.5 | 2 | 1 | 1 | 0 | 2 | 0.5 | 9 | 0.5556 | 2 | 1 | NA | NA | 2 | 1 | 2 | 1 | 6 | 1 |
| 8_2455100 | 3 | 0.3333 | 2 | 0.5 | 1 | 1 | 2 | 0.5 | 8 | 0.5 | 1 | 1 | NA | NA | 2 | 1 | 2 | 0.5 | 5 | 0.8 |
| 8_2455156 | 3 | 0 | 1 | 0 | 1 | 0 | 2 | 0.5 | 7 | 0.1429 | 2 | 0 | NA | NA | 2 | 0 | 3 | 0.6667 | 7 | 0.2857 |
| 8_2455196 | 2 | 0.5 | 2 | 0 | 2 | 0 | 1 | 0 | 7 | 0.1429 | 2 | 0 | NA | NA | 1 | 0 | 3 | 0.3333 | 6 | 0.1667 |
| 8_2455259 | 4 | 0.25 | 3 | 0 | 2 | 0 | 2 | 0 | 11 | 0.0909 | 3 | 0 | NA | NA | 2 | 0.5 | 2 | 0.5 | 7 | 0.2857 |
| 8_2455259 | 4 | 0.25 | 3 | 0 | 2 | 0 | 2 | 0 | 11 | 0.0909 | 3 | 0.3333 | NA | NA | 2 | 0 | 2 | 0 | 7 | 0.1429 |
| 8_2455274 | 4 | 0 | 3 | 0 | 2 | 0.5 | 1 | 0 | 10 | 0.1 | 4 | 0.25 | NA | NA | 1 | 0 | 2 | 0 | 7 | 0.1429 |
| 8_2455284 | 3 | 0.3333 | 2 | 0 | 2 | 0 | 2 | 0 | 9 | 0.1111 | 3 | 0 | NA | NA | 2 | 0 | 3 | 0.3333 | 8 | 0.125 |
| 8_2507449 | NA | NA | 2 | 1 | NA | NA | 1 | 1 | 3 | 1 | 1 | 1 | 1 | 1 | 3 | 1 | 1 | 1 | 6 | 1 |
| 8_2544021 | 3 | 0.3333 | 7 | 0.1429 | 3 | 0.3333 | 5 | 0 | 18 | 0.1667 | 4 | 0.25 | 6 | 0 | 6 | 0 | 5 | 0.2 | 21 | 0.0952 |
| 8_2544044 | 2 | 0.5 | 5 | 0.6 | 2 | 1 | 5 | 0.4 | 14 | 0.5714 | 5 | 0.4 | 4 | 0.5 | 3 | 0.3333 | 2 | 0.5 | 14 | 0.4286 |
| 8_2544057 | 2 | 0 | 7 | 0.1429 | 2 | 0 | 5 | 0 | 16 | 0.0625 | 6 | 0 | 5 | 0.2 | 4 | 0 | 4 | 0 | 19 | 0.0526 |
| 8_2544061 | 1 | 1 | 5 | 0 | 2 | 0 | 5 | 0 | 13 | 0.0769 | 6 | 0.1667 | 5 | 0 | 4 | 0.25 | 4 | 0.25 | 19 | 0.1579 |
| 8_2544089 | 2 | 0 | 7 | 0.1429 | 2 | 0.5 | 5 | 0 | 16 | 0.125 | 6 | 0.1667 | 4 | 0 | 4 | 0 | 4 | 0.25 | 18 | 0.1111 |
| 8_2551270 | 11 | 0.9091 | 11 | 1 | 17 | 1 | 17 | 0.9412 | 56 | 0.9643 | 11 | 1 | 11 | 1 | 19 | 1 | 7 | 1 | 48 | 1 |
| 8_2551286 | 6 | 0.6667 | 9 | 0.6667 | 13 | 0.7692 | 12 | 0.6667 | 40 | 0.7 | 10 | 0.5 | 8 | 0.75 | 13 | 0.6923 | 1 | 1 | 32 | 0.6562 |
| 8_2551290 | 13 | 0.4615 | 17 | 0.5294 | 15 | 0.3333 | 18 | 0.2222 | 63 | 0.381 | 16 | 0.1875 | 15 | 0.1333 | 21 | 0.381 | 10 | 0.6 | 62 | 0.3065 |
| 8_2551302 | 13 | 0.2308 | 17 | 0 | 15 | 0.2667 | 18 | 0.2778 | 63 | 0.1905 | 15 | 0.2 | 15 | 0.0667 | 18 | 0.1111 | 7 | 0.2857 | 55 | 0.1455 |
| 8_2551309 | 15 | 0.5333 | 19 | 0.3684 | 16 | 0.25 | 20 | 0.4 | 70 | 0.3857 | 15 | 0.6 | 15 | 0.3333 | 18 | 0.2778 | 12 | 0.5 | 60 | 0.4167 |
| 8_2551314 | 14 | 0 | 20 | 0.05 | 19 | 0.1053 | 20 | 0.05 | 73 | 0.0548 | 18 | 0 | 17 | 0 | 23 | 0.1739 | 9 | 0.2222 | 67 | 0.0896 |
| 8_2551320 | 11 | 0.5455 | 16 | 0.5625 | 13 | 0.6923 | 17 | 0.7059 | 57 | 0.6316 | 14 | 0.4286 | 13 | 0.7692 | 16 | 0.875 | 10 | 0.6 | 53 | 0.6792 |
| 8_2551323 | 15 | 0.2667 | 18 | 0.2222 | 14 | 0.4286 | 17 | 0.3529 | 64 | 0.3125 | 17 | 0.4118 | 16 | 0.375 | 22 | 0.3636 | 8 | 0.25 | 63 | 0.3651 |
| 8_2552962 | NA | NA | 2 | 0 | 2 | 1 | NA | NA | 4 | 0.5 | 2 | 0.5 | NA | NA | 1 | 0 | NA | NA | 3 | 0.3333 |

| | | | | | | | | | | | | | | | | | | | | |
|---|---|---|---|---|---|---|---|---|---|---|---|---|---|---|---|---|---|---|---|---|
| 8_2552965 | NA | NA | 1 | 0 | 2 | 0 | 1 | 1 | 4 | 0.25 | 3 | 0.3333 | NA | NA | 1 | 0 | 1 | 0 | 5 | 0.2 |
| 8_2553065 | 1 | 0 | 2 | 0.5 | NA | NA | NA | NA | 3 | 0.3333 | 2 | 0.5 | NA | NA | 1 | 0 | NA | NA | 3 | 0.3333 |
| 8_2553127 | NA | NA | 4 | 0 | 1 | 0 | 3 | 0.3333 | 8 | 0.125 | 5 | 0.2 | 1 | 0 | 1 | 0 | 2 | 0.5 | 9 | 0.2222 |
| 8_2553262 | 1 | 0 | 3 | 0.3333 | NA | NA | 2 | 0.5 | 6 | 0.3333 | 3 | 0.3333 | NA | NA | 2 | 0 | 2 | 0 | 7 | 0.1429 |
| 8_2553265 | NA | NA | 2 | 0.5 | NA | NA | 2 | 0 | 4 | 0.25 | 2 | 0.5 | 2 | 0.5 | 2 | 0.5 | 2 | 0 | 8 | 0.375 |
| 8_2553284 | NA | NA | 3 | 1 | NA | NA | 2 | 1 | 5 | 1 | 3 | 1 | 1 | 1 | 2 | 1 | 2 | 1 | 8 | 1 |
| 8_2553289 | NA | NA | 2 | 0.5 | NA | NA | 2 | 0.5 | 4 | 0.5 | 3 | 0.6667 | 1 | 0 | 2 | 0.5 | NA | NA | 6 | 0.5 |
| 8_2559167 | 1 | 1 | 2 | 0.5 | NA | NA | NA | NA | 3 | 0.6667 | 1 | 1 | 1 | 1 | NA | NA | NA | NA | 2 | 1 |
| 8_2559180 | 1 | 0 | 2 | 1 | NA | NA | NA | NA | 3 | 0.6667 | 1 | 1 | 1 | 0 | NA | NA | NA | NA | 2 | 0.5 |
| 8_2559196 | 1 | 1 | 1 | 1 | NA | NA | NA | NA | 2 | 1 | 1 | 1 | 1 | 1 | NA | NA | NA | NA | 2 | 1 |
| 8_2559202 | 1 | 1 | 2 | 0 | NA | NA | NA | NA | 3 | 0.3333 | 1 | 1 | 1 | 0 | NA | NA | NA | NA | 2 | 0.5 |
| 8_2564549 | 24 | 0.0417 | 10 | 0.1 | 20 | 0.2 | 23 | 0.087 | 77 | 0.1039 | 14 | 0.2857 | 17 | 0.0588 | 25 | 0.04 | 21 | 0.0952 | 77 | 0.1039 |
| 8_2564819 | 12 | 0 | 2 | 0 | 11 | 0.0909 | 13 | 0.0769 | 38 | 0.0526 | 5 | 0.2 | 8 | 0 | 12 | 0 | 11 | 0.2727 | 36 | 0.1111 |
| 8_2564891 | 7 | 0.2857 | 1 | 1 | 5 | 0.6 | 7 | 0.5714 | 20 | 0.5 | 2 | 0.5 | 4 | 0 | 6 | 0.1667 | 10 | 0.3 | 22 | 0.2273 |
| 8_2564924 | 4 | 0.5 | 2 | 0 | 9 | 0.4444 | 8 | 0.375 | 23 | 0.3913 | 3 | 0.3333 | 6 | 0.5 | 8 | 0.375 | 9 | 0.5556 | 26 | 0.4615 |
| 8_2564942 | 4 | 0.25 | 1 | 0 | 8 | 0.25 | 5 | 0.2 | 18 | 0.2222 | 2 | 0 | 5 | 0.2 | 9 | 0.2222 | 7 | 0.2857 | 23 | 0.2174 |
| 8_2564954 | 4 | 0 | 1 | 0 | 10 | 0.1 | 6 | 0.1667 | 21 | 0.0952 | 2 | 0 | 6 | 0 | 9 | 0.1111 | 7 | 0.1429 | 24 | 0.0833 |
| 8_2567043 | 1 | 1 | NA | NA | 1 | 1 | NA | NA | 2 | 1 | NA | NA | 2 | 1 | 1 | 1 | NA | NA | 3 | 1 |
| 8_2567162 | 1 | 1 | NA | NA | 1 | 0 | NA | NA | 2 | 0.5 | 2 | 0.5 | 2 | 0.5 | 2 | 0.5 | NA | NA | 6 | 0.5 |
| 8_2567182 | 1 | 1 | NA | NA | 1 | 0 | NA | NA | 2 | 0.5 | 1 | 1 | 1 | 1 | 1 | 1 | NA | NA | 3 | 1 |
| 8_2567187 | 2 | 0.5 | NA | NA | 2 | 0.5 | 1 | 0 | 5 | 0.4 | 2 | 0 | 3 | 1 | 3 | 0.3333 | 3 | 0.6667 | 11 | 0.5455 |
| 8_2567195 | 2 | 0 | NA | NA | 2 | 1 | 1 | 0 | 5 | 0.4 | 1 | 0 | 1 | 1 | 3 | 0.6667 | 2 | 1 | 7 | 0.7143 |
| 8_2567196 | 2 | 0 | NA | NA | 3 | 0 | 2 | 1 | 7 | 0.2857 | 2 | 0 | 2 | 0 | 3 | 0.3333 | 3 | 0.3333 | 10 | 0.2 |
| 8_2567202 | 1 | 1 | NA | NA | 2 | 0 | 2 | 0.5 | 5 | 0.4 | 1 | 0 | 1 | 0 | 3 | 0.3333 | 2 | 0.5 | 7 | 0.2857 |
| 8_2567213 | 2 | 0 | NA | NA | 3 | 0.3333 | 2 | 0 | 7 | 0.1429 | 2 | 0 | 1 | 0 | 4 | 0.25 | 3 | 0.3333 | 10 | 0.2 |
| 8_2567219 | 2 | 0.5 | NA | NA | 3 | 0.3333 | 2 | 0.5 | 7 | 0.4286 | 2 | 0 | NA | NA | 1 | 0 | 2 | 1 | 5 | 0.4 |
| 8_2567232 | 1 | 1 | NA | NA | 2 | 1 | NA | NA | 3 | 1 | NA | NA | 1 | 1 | 3 | 0.6667 | 2 | 0.5 | 6 | 0.6667 |
| 8_2567257 | NA | NA | NA | NA | 2 | 1 | 1 | 1 | 3 | 1 | 1 | 1 | 1 | 0 | 3 | 1 | 2 | 0.5 | 7 | 0.7143 |
| 8_2567258 | NA | NA | NA | NA | 2 | 0.5 | 1 | 0 | 3 | 0.3333 | 2 | 0.5 | NA | NA | 3 | 0 | 2 | 1 | 7 | 0.4286 |
| 8_2567263 | NA | NA | NA | NA | 1 | 1 | NA | NA | 1 | 1 | 1 | 1 | 1 | 1 | 1 | 1 | 1 | 1 | 4 | 1 |
| 8_2567272 | NA | NA | NA | NA | 2 | 0.5 | 1 | 1 | 3 | 0.6667 | 2 | 0.5 | NA | NA | 3 | 0.3333 | 1 | 0 | 6 | 0.3333 |
| 8_2606384 | 10 | 0.3 | 7 | 0.1429 | 1 | 0 | 5 | 0.2 | 23 | 0.2174 | 3 | 0.3333 | 4 | 0.25 | 7 | 0.5714 | 6 | 0.1667 | 20 | 0.35 |
| 8_2606392 | 11 | 0.9091 | 8 | 0.875 | 1 | 1 | 4 | 0.75 | 24 | 0.875 | 4 | 0.5 | 4 | 0.75 | 7 | 0.8571 | 4 | 1 | 19 | 0.7895 |
| 8_2606401 | 9 | 0.7778 | 7 | 0.5714 | 1 | 1 | 5 | 0.8 | 22 | 0.7273 | 3 | 0.6667 | 6 | 0.8333 | 7 | 1 | 5 | 1 | 21 | 0.9048 |
| 8_2606406 | 11 | 0.3636 | 11 | 0.5455 | 1 | 1 | 5 | 0.4 | 28 | 0.4643 | 4 | 0.25 | 5 | 0.2 | 8 | 0 | 6 | 0.1667 | 23 | 0.1304 |
| 8_2606420 | 9 | 0.3333 | 8 | 0.25 | 1 | 0 | 4 | 0.25 | 22 | 0.2727 | 4 | 0 | 5 | 0.4 | 8 | 0.25 | 5 | 0.2 | 22 | 0.2273 |
| 8_2606446 | 10 | 0.9 | 12 | 0.8333 | 1 | 1 | 4 | 0.75 | 27 | 0.8519 | 6 | 0.6667 | 6 | 1 | 8 | 0.875 | 6 | 1 | 26 | 0.8846 |
| 8_2606450 | 10 | 0 | 12 | 0.25 | 1 | 0 | 5 | 0.2 | 28 | 0.1429 | 7 | 0.1429 | 6 | 0.3333 | 8 | 0.125 | 7 | 0.2857 | 28 | 0.2143 |
| 8_2606466 | 10 | 0.1 | 9 | 0.1111 | NA | NA | 6 | 0 | 25 | 0.08 | 6 | 0 | 6 | 0 | 8 | 0.125 | 7 | 0.1429 | 27 | 0.0741 |
| 8_2606868 | 4 | 0.75 | 1 | 1 | 2 | 0.5 | NA | NA | 7 | 0.7143 | 1 | 1 | 1 | 1 | NA | NA | 2 | 1 | 4 | 1 |
| 8_2608384 | 2 | 0.5 | 1 | 0 | NA | NA | NA | NA | 3 | 0.3333 | NA | NA | NA | NA | NA | NA | 2 | 1 | 2 | 1 |
| 8_2608400 | 2 | 0.5 | 1 | 1 | NA | NA | NA | NA | 3 | 0.6667 | NA | NA | NA | NA | NA | NA | 1 | 1 | 1 | 1 |
| 8_2608401 | 1 | 1 | 1 | 0 | NA | NA | NA | NA | 2 | 0.5 | NA | NA | NA | NA | 1 | 0 | 2 | 0.5 | 3 | 0.3333 |
| 8_2613075 | 1 | 0 | 4 | 0.25 | NA | NA | NA | NA | 5 | 0.2 | 4 | 0.25 | 1 | 0 | NA | NA | 1 | 0 | 6 | 0.1667 |
| 8_2613088 | 1 | 0 | 5 | 0.2 | 1 | 1 | NA | NA | 7 | 0.2857 | 3 | 0.6667 | 1 | 0 | NA | NA | 2 | 0.5 | 6 | 0.5 |
| 8_2613095 | 1 | 1 | 5 | 0.6 | 1 | 0 | NA | NA | 7 | 0.5714 | 4 | 0.25 | 1 | 1 | NA | NA | 2 | 0 | 7 | 0.2857 |
| 8_2613096 | 1 | 1 | 6 | 0.5 | 1 | 0 | NA | NA | 8 | 0.5 | 4 | 0.5 | 1 | 1 | NA | NA | 1 | 1 | 6 | 0.6667 |
| 8_2613109 | 1 | 0 | 5 | 0.6 | NA | NA | NA | NA | 6 | 0.5 | 4 | 1 | 1 | 1 | NA | NA | 1 | 1 | 6 | 1 |
| 8_2613114 | NA | NA | 5 | 0 | 1 | 1 | NA | NA | 6 | 0.1667 | 5 | 0 | 1 | 1 | NA | NA | 1 | 0 | 7 | 0.1429 |
| 8_2615279 | 1 | 0 | 6 | 0.1667 | 1 | 0 | NA | NA | 8 | 0.125 | 5 | 0.6 | 1 | 0 | NA | NA | 2 | 1 | 8 | 0.625 |
| 8_2615298 | 1 | 0 | 7 | 0.1429 | 1 | 0 | NA | NA | 9 | 0.1111 | 4 | 0.25 | 1 | 0 | NA | NA | 1 | 0 | 6 | 0.1667 |
| 8_2615298 | 1 | 0 | 7 | 0.5714 | 1 | 0 | NA | NA | 9 | 0.4444 | 5 | 0.2 | 1 | 0 | NA | NA | 2 | 0.5 | 8 | 0.25 |
| 8_2616679 | 2 | 1 | 6 | 1 | NA | NA | NA | NA | 8 | 1 | 4 | 0.75 | 2 | 1 | 1 | 1 | 2 | 1 | 9 | 0.8889 |
| 8_2616695 | 1 | 0 | 6 | 0.5 | NA | NA | NA | NA | 7 | 0.4286 | 3 | 0.3333 | 1 | 1 | NA | NA | 2 | 0.5 | 6 | 0.5 |
| 8_2616707 | 2 | 0 | 7 | 0.2857 | NA | NA | NA | NA | 9 | 0.2222 | 3 | 0 | 1 | 0 | 1 | 0 | 3 | 0.3333 | 8 | 0.125 |
| 8_2616712 | 2 | 1 | 6 | 1 | NA | NA | NA | NA | 8 | 1 | 4 | 1 | 2 | 1 | 2 | 1 | 3 | 1 | 11 | 1 |
| 8_2616718 | 2 | 0.5 | 6 | 0.3333 | NA | NA | NA | NA | 8 | 0.375 | 4 | 0.5 | 1 | 0 | 2 | 0.5 | 3 | 0.6667 | 10 | 0.5 |
| 8_2616730 | 2 | 0 | 6 | 0.3333 | 1 | 1 | NA | NA | 9 | 0.3333 | 5 | 0 | 1 | 0 | NA | NA | 3 | 0.3333 | 9 | 0.1111 |
| 8_2616771 | 2 | 0.5 | 8 | 1 | 1 | 0 | 2 | 1 | 13 | 0.8462 | 4 | 1 | 1 | 1 | 2 | 1 | 4 | 1 | 11 | 1 |
| 8_2616779 | 2 | 0.5 | 5 | 0.8 | 1 | 0 | 2 | 0.5 | 10 | 0.6 | 4 | 0.25 | 1 | 1 | 1 | 0 | 3 | 0 | 9 | 0.2222 |
| 8_2616780 | 3 | 0 | 8 | 0.125 | 2 | 0 | 2 | 0 | 15 | 0.0667 | 2 | 0 | 2 | 0 | 2 | 0 | 4 | 0.25 | 10 | 0.1 |
| 8_2620655 | 1 | 1 | 2 | 0 | 1 | 0 | 1 | 0 | 5 | 0.2 | 1 | 0 | 4 | 0.25 | 1 | 0 | 1 | 0 | 7 | 0.1429 |
| 8_2620751 | 1 | 0 | 2 | 0.5 | 1 | 0 | 1 | 0 | 5 | 0.2 | 1 | 1 | 3 | 0.3333 | 1 | 0 | 2 | 0 | 7 | 0.2857 |
| 8_2620785 | 2 | 0.5 | 3 | 0.6667 | NA | NA | 1 | 0 | 6 | 0.5 | NA | NA | 5 | 0.4 | 1 | 0 | 2 | 0 | 8 | 0.25 |
| 8_2620803 | 1 | 0 | 1 | 0 | 1 | 1 | 1 | 0 | 4 | 0.25 | 1 | 0 | 5 | 0.4 | 2 | 0.5 | 2 | 0.5 | 10 | 0.4 |

| | | | | | | | | | | | | | | | | | | | | |
|---|---|---|---|---|---|---|---|---|---|---|---|---|---|---|---|---|---|---|---|---|
| 8_2620859 | 2 | 0.5 | 2 | 0.5 | 1 | 1 | 1 | 0 | 6 | 0.5 | 1 | 0 | 6 | 0.8333 | 3 | 0.6667 | 2 | 0.5 | 12 | 0.6667 |
| 8_2621417 | 5 | 0 | 6 | 0.1667 | 1 | 0 | 3 | 0 | 15 | 0.0667 | 3 | 0.3333 | 4 | 0.25 | 2 | 0 | 2 | 0 | 11 | 0.1818 |
| 8_2621431 | 5 | 0.4 | 5 | 0.6 | NA | NA | 2 | 0.5 | 12 | 0.5 | 1 | 0 | 3 | 0.6667 | 2 | 1 | 2 | 0 | 8 | 0.5 |
| 8_2621437 | 6 | 0.3333 | 4 | 0 | 1 | 0 | 3 | 0.3333 | 14 | 0.2143 | 3 | 0.3333 | 3 | 0.3333 | 2 | 0.5 | 2 | 0 | 10 | 0.3 |
| 8_2621447 | 4 | 1 | 6 | 0.5 | NA | NA | 3 | 0.6667 | 13 | 0.6923 | 3 | 0.6667 | 4 | 0.75 | 2 | 1 | 2 | 0 | 11 | 0.6364 |
| 8_2621563 | 10 | 0.2 | 8 | 0 | 5 | 0 | 3 | 0 | 26 | 0.0769 | 5 | 0.6 | 9 | 0.1111 | 5 | 0 | 3 | 0 | 22 | 0.1818 |
| 8_2621655 | 3 | 0 | 6 | 0 | 3 | 0 | 3 | 0.3333 | 15 | 0.0667 | 3 | 0 | 10 | 0.1 | 4 | 0.25 | 4 | 0 | 21 | 0.0952 |
| 8_2621676 | 2 | 0.5 | 9 | 0.4444 | 2 | 1 | 2 | 0 | 15 | 0.4667 | 2 | 0.5 | 10 | 0.6 | 2 | 0.5 | 5 | 0.6 | 19 | 0.5789 |
| 8_2621681 | 2 | 0.5 | 7 | 0.1429 | 2 | 0.5 | 2 | 0.5 | 13 | 0.3077 | 2 | 0 | 9 | 0.3333 | 2 | 0 | 5 | 0.2 | 18 | 0.2222 |
| 8_2621691 | NA | NA | 8 | 0.375 | 1 | 1 | 1 | 0 | 10 | 0.4 | 1 | 0 | 8 | 0 | 2 | 0.5 | 3 | 0 | 14 | 0.0714 |
| 8_2621726 | 1 | 0 | 6 | 0.5 | 2 | 0.5 | 1 | 0 | 10 | 0.4 | 1 | 0 | 9 | 0.1111 | 1 | 0 | 2 | 0 | 13 | 0.0769 |
| 8_2621737 | 1 | 0 | 6 | 0.1667 | 2 | 0 | 1 | 0 | 10 | 0.1 | 2 | 0 | 9 | 0.2222 | 1 | 0 | 2 | 1 | 14 | 0.2857 |
| 8_2621738 | 1 | 0 | 7 | 0.5714 | 2 | 1 | 1 | 1 | 11 | 0.6364 | 2 | 0.5 | 8 | 0.5 | NA | NA | 2 | 0.5 | 12 | 0.5 |
| 8_2621745 | 1 | 0 | 3 | 0.3333 | 2 | 0.5 | 1 | 0 | 7 | 0.2857 | NA | NA | 5 | 0.4 | 1 | 0 | 1 | 0 | 7 | 0.2857 |
| 8_2621747 | 1 | 0 | 3 | 0.3333 | 1 | 0 | 1 | 0 | 6 | 0.1667 | NA | NA | 5 | 0.2 | 1 | 1 | 1 | 0 | 7 | 0.2857 |
| 8_2621758 | 1 | 0 | 3 | 0.6667 | 2 | 1 | 1 | 0 | 7 | 0.5714 | NA | NA | 3 | 0.6667 | NA | NA | 1 | 1 | 4 | 0.75 |
| 8_2621759 | NA | NA | 3 | 0.6667 | 1 | 0 | NA | NA | 4 | 0.5 | NA | NA | 4 | 0.25 | NA | NA | 1 | 0 | 5 | 0.2 |
| 8_2621760 | 1 | 0 | 3 | 0.3333 | 2 | 0 | 1 | 0 | 7 | 0.1429 | NA | NA | 5 | 0.2 | 1 | 0 | 1 | 0 | 7 | 0.1429 |
| 8_2621761 | 1 | 0 | 3 | 0.6667 | 1 | 1 | 1 | 0 | 6 | 0.5 | NA | NA | 4 | 0.5 | NA | NA | 1 | 0 | 5 | 0.4 |
| 8_2621768 | 1 | 0 | 6 | 0.1667 | 1 | 0 | NA | NA | 8 | 0.125 | NA | NA | 5 | 0.2 | 1 | 0 | 1 | 0 | 7 | 0.1429 |
| 8_2625491 | 10 | 0 | 9 | 0.3333 | 8 | 0.25 | 12 | 0 | 39 | 0.1282 | 10 | 0.2 | 9 | 0.2222 | 11 | 0 | 6 | 0.1667 | 36 | 0.1389 |
| 8_2625721 | 3 | 0.6667 | 3 | 0 | 3 | 0.3333 | 5 | 0 | 14 | 0.2143 | 4 | 0.25 | 3 | 0.3333 | 7 | 0.2857 | 2 | 0 | 16 | 0.25 |
| 8_2625750 | 7 | 0.5714 | 3 | 1 | 6 | 0.3333 | 9 | 0.3333 | 25 | 0.48 | 1 | 1 | 5 | 0.2 | 9 | 0.3333 | 7 | 0.5714 | 22 | 0.4091 |
| 8_2625751 | 6 | 0.3333 | 3 | 0 | 5 | 0.4 | 8 | 0.125 | 22 | 0.2273 | 2 | 0.5 | 5 | 0.2 | 8 | 0.125 | 7 | 0.2857 | 22 | 0.2273 |
| 8_2625754 | 7 | 0.1429 | 2 | 0 | 6 | 0 | 6 | 0.1667 | 21 | 0.0952 | 2 | 1 | 5 | 0 | 8 | 0.25 | 5 | 0 | 20 | 0.2 |
| 8_2625757 | 6 | 1 | 2 | 1 | 5 | 1 | 9 | 0.8889 | 22 | 0.9545 | 2 | 1 | 4 | 1 | 7 | 1 | 6 | 1 | 19 | 1 |
| 8_2625763 | 6 | 1 | 2 | 1 | 5 | 0.8 | 9 | 1 | 22 | 0.9545 | 2 | 1 | 5 | 1 | 9 | 0.8889 | 6 | 1 | 22 | 0.9545 |
| 8_2625763 | NA | NA | NA | NA | 1 | 1 | NA | NA | 1 | 1 | NA | NA | NA | NA | NA | NA | 1 | 1 | 1 | 1 |
| 8_2625773 | 7 | 0 | 3 | 0 | 5 | 0 | 7 | 0.2857 | 22 | 0.0909 | 2 | 0 | 4 | 0 | 7 | 0.1429 | 5 | 0.2 | 18 | 0.1111 |
| 8_2625774 | 5 | 0.6 | 2 | 0 | 5 | 0.6 | 8 | 0.375 | 20 | 0.45 | NA | NA | 4 | 0.5 | 6 | 0.3333 | 4 | 0.5 | 14 | 0.4286 |
| 8_2625780 | 4 | 0.75 | 3 | 0.3333 | 6 | 0.5 | 6 | 0.3333 | 19 | 0.4737 | 1 | 1 | 4 | 0.5 | 6 | 0.3333 | 5 | 0.4 | 16 | 0.4375 |
| 8_2625782 | 7 | 0.2857 | 3 | 0 | 3 | 0.3333 | 6 | 0.5 | 19 | 0.3158 | 2 | 0 | 5 | 0.2 | 9 | 0.1111 | 7 | 0.2857 | 23 | 0.1739 |
| 8_2625788 | 1 | 1 | NA | NA | NA | NA | 1 | 1 | 2 | 1 | NA | NA | NA | NA | 1 | 1 | NA | NA | 1 | 1 |
| 8_2625790 | 6 | 1 | 2 | 1 | 6 | 1 | 7 | 1 | 21 | 1 | 2 | 1 | 2 | 1 | 7 | 1 | 5 | 1 | 16 | 1 |
| 8_2625792 | 7 | 0.8571 | 2 | 0.5 | 6 | 1 | 8 | 1 | 23 | 0.913 | 2 | 1 | 5 | 1 | 9 | 0.7778 | 4 | 0.25 | 20 | 0.75 |
| 8_2625794 | 5 | 0.4 | 1 | 0 | 4 | 0 | 1 | 0 | 11 | 0.1818 | 2 | 0.5 | 4 | 0 | 7 | 0.2857 | 3 | 0.3333 | 16 | 0.25 |
| 8_2625795 | 7 | 1 | 3 | 1 | 6 | 1 | 9 | 1 | 25 | 1 | 2 | 0.5 | 3 | 1 | 7 | 1 | 6 | 0.8333 | 18 | 0.8889 |
| 8_2625803 | 6 | 0.8333 | 3 | 1 | 5 | 1 | 7 | 0.7143 | 21 | 0.8571 | 2 | 1 | 5 | 0.8 | 8 | 0.75 | 7 | 0.7143 | 22 | 0.7727 |
| 8_2625804 | 6 | 0.3333 | 3 | 0 | 5 | 0.4 | 7 | 0.2857 | 21 | 0.2857 | NA | NA | 5 | 0.6 | 7 | 0.5714 | 7 | 0.4286 | 19 | 0.5263 |
| 8_2625806 | 4 | 1 | 2 | 1 | 4 | 1 | 7 | 1 | 17 | 1 | 1 | 1 | 3 | 0.6667 | 5 | 0.8 | 6 | 0.6667 | 15 | 0.7333 |
| 8_2625806 | 5 | 0.8 | 3 | 1 | 4 | 1 | 8 | 0.875 | 20 | 0.9 | 2 | 1 | 4 | 1 | 8 | 1 | 3 | 0.3333 | 17 | 0.8824 |
| 8_2625811 | 7 | 0.4286 | 2 | 0.5 | 5 | 0.6 | 9 | 0.7778 | 23 | 0.6087 | 1 | 0 | 5 | 0.6 | 9 | 0.7778 | 6 | 0.5 | 21 | 0.619 |
| 8_2625831 | NA | NA | NA | NA | NA | NA | 1 | 1 | 1 | 1 | NA | NA | NA | NA | 1 | 1 | NA | NA | 1 | 1 |
| 8_2625854 | NA | NA | NA | NA | NA | NA | 1 | 1 | 1 | 1 | NA | NA | NA | NA | 1 | 1 | NA | NA | 1 | 1 |
| 8_2625870 | NA | NA | NA | NA | NA | NA | 1 | 1 | 1 | 1 | NA | NA | NA | NA | 1 | 1 | NA | NA | 1 | 1 |
| 8_2627570 | 3 | 0 | 6 | 0 | 2 | 0 | 2 | 0.5 | 13 | 0.0769 | 3 | 0 | 7 | 0.1429 | 1 | 0 | NA | NA | 11 | 0.0909 |
| 8_2629210 | 5 | 0 | 6 | 0.3333 | 4 | 0 | 2 | 0 | 17 | 0.1176 | 5 | 0 | 9 | 0.1111 | 1 | 0 | 2 | 0 | 17 | 0.0588 |
| 8_2629214 | 5 | 0.6 | 6 | 1 | 6 | 1 | 2 | 0.5 | 19 | 0.8421 | 5 | 1 | 7 | 1 | NA | NA | 2 | 0.5 | 14 | 0.9286 |
| 8_2629237 | 3 | 0.3333 | 5 | 0.8 | 5 | 0.8 | NA | NA | 13 | 0.6923 | 5 | 0.4 | 5 | 0.6 | NA | NA | 2 | 0.5 | 12 | 0.5 |
| 8_2629246 | 6 | 0 | 6 | 0.1667 | 7 | 0.2857 | 2 | 0.5 | 21 | 0.1905 | 5 | 0 | 9 | 0.1111 | 2 | 0 | 2 | 0.5 | 18 | 0.1111 |
| 8_2629255 | 2 | 0 | 4 | 0.25 | 3 | 0 | 2 | 0.5 | 11 | 0.1818 | 4 | 0 | 6 | 0.3333 | NA | NA | 2 | 0 | 12 | 0.1667 |
| 8_2642779 | NA | NA | 1 | 0 | NA | NA | 1 | 1 | 2 | 0.5 | NA | NA | NA | NA | 1 | 1 | NA | NA | 1 | 1 |
| 8_2643462 | NA | NA | 5 | 0.2 | 2 | 1 | 1 | 0 | 8 | 0.375 | 1 | 1 | 3 | 0 | 2 | 1 | 2 | 0.5 | 8 | 0.5 |
| 8_2643467 | 3 | 1 | 6 | 0.6667 | 2 | 1 | 4 | 0.75 | 15 | 0.8 | 2 | 1 | 5 | 1 | 6 | 0.5 | 6 | 0.8333 | 19 | 0.7895 |
| 8_2643469 | 5 | 0.2 | 9 | 0.2222 | 3 | 0 | 11 | 0 | 28 | 0.1071 | 7 | 0.2857 | 9 | 0.2222 | 11 | 0 | 10 | 0.3 | 37 | 0.1892 |
| 8_2643474 | 2 | 0 | 10 | 0.5 | 2 | 0.5 | 10 | 0.3 | 24 | 0.375 | 5 | 0.4 | 6 | 0.1667 | 7 | 0 | 4 | 0.25 | 22 | 0.1818 |
| 8_2643475 | 4 | 0 | 6 | 0 | 2 | 0 | 6 | 0.1667 | 18 | 0.0556 | 5 | 0 | 5 | 0 | 5 | 0.2 | 2 | 0 | 17 | 0.0588 |
| 8_2643477 | 6 | 0.3333 | 9 | 0 | 4 | 0.25 | 11 | 0.0909 | 30 | 0.1333 | 5 | 0 | 8 | 0 | 10 | 0.1 | 11 | 0.0909 | 34 | 0.0588 |
| 8_2643482 | 4 | 0.25 | 10 | 0.6 | 2 | 0.5 | 2 | 0.5 | 18 | 0.5 | 3 | 0.3333 | 6 | 0.6667 | 3 | 0.3333 | 5 | 1 | 17 | 0.6471 |
| 8_2643486 | 7 | 0.8571 | 11 | 0.8182 | 4 | 0.75 | 12 | 1 | 34 | 0.8824 | 7 | 0.8571 | 9 | 0.7778 | 9 | 0.5556 | 11 | 0.9091 | 36 | 0.7778 |
| 8_2643489 | 2 | 1 | 8 | 1 | 3 | 1 | 9 | 0.8889 | 22 | 0.9545 | 6 | 1 | 5 | 0.8 | 7 | 1 | 6 | 1 | 24 | 0.9583 |
| 8_2643494 | 5 | 0.2 | 8 | 0.375 | 4 | 0.5 | 10 | 0.6 | 27 | 0.4444 | 5 | 0.2 | 8 | 0.375 | 8 | 0.375 | 7 | 0.4286 | 28 | 0.3571 |
| 8_2643501 | 5 | 0.2 | 9 | 0.4444 | 2 | 1 | 10 | 0.4 | 26 | 0.4231 | 7 | 0.2857 | 8 | 0.375 | 10 | 0.3 | 8 | 0.375 | 33 | 0.3333 |
| 8_2643504 | 6 | 0 | 10 | 0.1 | 4 | 0.25 | 11 | 0.1818 | 31 | 0.129 | 7 | 0.1429 | 9 | 0.1111 | 8 | 0 | 8 | 0.25 | 32 | 0.125 |
| 8_2647080 | NA | NA | 1 | 1 | NA | NA | NA | NA | 1 | 1 | NA | NA | NA | NA | NA | NA | 1 | 1 | 1 | 1 |

| | | | | | | | | | | | | | | | | | | | | |
|---|---|---|---|---|---|---|---|---|---|---|---|---|---|---|---|---|---|---|---|---|
| 8_2647087 | NA | NA | 1 | 1 | NA | NA | NA | NA | 1 | 1 | 1 | 1 | NA | NA | NA | NA | 1 | 0 | 2 | 0.5 |
| 8_2647160 | 2 | 0.5 | 2 | 0 | 5 | 0 | 4 | 0.25 | 13 | 0.1538 | 6 | 0 | 3 | 0.3333 | 5 | 0 | 3 | 0 | 17 | 0.0588 |
| 8_2647169 | 1 | 1 | 2 | 0 | 4 | 0 | 5 | 0 | 12 | 0.0833 | 6 | 0.3333 | 3 | 0.3333 | 6 | 0.1667 | NA | NA | 15 | 0.2667 |
| 8_2647172 | 3 | 1 | 2 | 1 | 6 | 0 | 5 | 0.4 | 16 | 0.4375 | 6 | 0.6667 | 3 | 1 | 6 | 0.6667 | 3 | 1 | 18 | 0.7778 |
| 8_2647183 | 2 | 0.5 | NA | NA | 2 | 0 | 4 | 0.25 | 8 | 0.25 | 3 | 0.3333 | 1 | 0 | 6 | 0.1667 | NA | NA | 10 | 0.2 |
| 8_2647188 | 3 | 1 | 2 | 1 | 5 | 0.8 | 4 | 0.5 | 14 | 0.7857 | 5 | 1 | 2 | 0 | 6 | 0.6667 | 2 | 1 | 15 | 0.7333 |
| 8_2647195 | 3 | 0.3333 | 2 | 1 | 6 | 0.1667 | 4 | 0 | 15 | 0.2667 | 5 | 0.2 | 3 | 0 | 6 | 0.1667 | 2 | 0 | 16 | 0.125 |
| 8_2647230 | 3 | 0 | 2 | 0 | 6 | 0 | 4 | 0.5 | 15 | 0.1333 | 5 | 0.4 | 4 | 0 | 6 | 0.1667 | 1 | 0 | 16 | 0.1875 |
| 8_2647248 | 1 | 0 | 2 | 0 | 1 | 0 | 4 | 0.25 | 8 | 0.125 | 3 | 0.3333 | 4 | 0 | 3 | 0 | 2 | 0 | 12 | 0.0833 |
| 8_2647253 | 3 | 1 | 2 | 1 | 6 | 0.8333 | 5 | 1 | 16 | 0.9375 | 5 | 1 | 5 | 1 | 3 | 1 | 2 | 1 | 15 | 1 |
| 8_2647259 | 3 | 1 | 2 | 1 | 5 | 0.8 | 5 | 0.8 | 15 | 0.8667 | 6 | 0.8333 | 5 | 0.8 | 3 | 1 | 1 | 1 | 15 | 0.8667 |
| 8_2647260 | 2 | 0 | 2 | 0 | 6 | 0.3333 | 4 | 0.25 | 14 | 0.2143 | 5 | 0.2 | 5 | 0.4 | 3 | 0.6667 | 1 | 1 | 14 | 0.4286 |
| 8_2647262 | 2 | 0 | 2 | 0 | 6 | 0.1667 | 5 | 0 | 15 | 0.0667 | 6 | 0 | 5 | 0.2 | 3 | 0 | 1 | 0 | 15 | 0.0667 |
| 8_2647280 | 3 | 0 | 2 | 0.5 | 6 | 0.1667 | 6 | 0.1667 | 17 | 0.1765 | 5 | 0.4 | 5 | 0 | 2 | 0.5 | 2 | 0 | 14 | 0.2143 |
| 8_2647293 | 3 | 0 | 2 | 0.5 | 4 | 0.5 | 3 | 0.3333 | 12 | 0.3333 | 5 | 0.2 | 5 | 0.6 | 3 | 0.6667 | 2 | 0.5 | 15 | 0.4667 |
| 8_2647299 | 3 | 0.3333 | 2 | 0 | 6 | 0.1667 | 5 | 0.2 | 16 | 0.1875 | 5 | 0.2 | 5 | 0 | 3 | 0.3333 | 2 | 0.5 | 15 | 0.2 |
| 8_2647324 | 3 | 0.6667 | 1 | 0 | 6 | 0.8333 | 5 | 1 | 15 | 0.8 | 4 | 1 | 5 | 1 | 3 | 1 | 2 | 1 | 14 | 1 |
| 8_2647325 | 3 | 0 | 1 | 1 | 5 | 0 | 5 | 0.2 | 14 | 0.1429 | 4 | 0 | 3 | 0.3333 | 3 | 0.3333 | 2 | 0 | 12 | 0.1667 |
| 8_2647332 | 3 | 0 | 1 | 0 | 4 | 0.5 | 4 | 0 | 12 | 0.1667 | 3 | 0 | 4 | 0 | 3 | 0.3333 | 1 | 1 | 11 | 0.1818 |
| 8_2647334 | 3 | 0 | NA | NA | 4 | 0.75 | 5 | 0.4 | 12 | 0.4167 | 3 | 0.6667 | 5 | 0.4 | 3 | 0.6667 | 2 | 0.5 | 13 | 0.5385 |
| 8_2647338 | 2 | 1 | 1 | 0 | 3 | 0.6667 | 3 | 0.3333 | 9 | 0.5556 | 2 | 1 | 3 | 0.6667 | 2 | 0.5 | 1 | 1 | 8 | 0.75 |
| 8_2647343 | 1 | 1 | NA | NA | NA | NA | 1 | 0 | 2 | 0.5 | 2 | 0.5 | NA | NA | 1 | 0 | 2 | 1 | 5 | 0.6 |
| 8_2647346 | 3 | 0.3333 | 1 | 0 | 5 | 0 | 5 | 0 | 14 | 0.0714 | 3 | 0 | 5 | 0.4 | 3 | 0 | 2 | 0 | 13 | 0.1538 |
| 8_2647360 | 2 | 1 | 1 | 0 | 3 | 0.3333 | 5 | 0.2 | 11 | 0.3636 | 2 | 0.5 | 5 | 0.6 | 3 | 0.3333 | 2 | 0 | 12 | 0.4167 |
| 8_2647640 | 3 | 0.3333 | 1 | 1 | 3 | 0 | 3 | 0 | 10 | 0.2 | 2 | 0.5 | 5 | 0.2 | 1 | 0 | 2 | 0 | 10 | 0.2 |
| 8_2648934 | 8 | 0 | 11 | 0.1818 | 6 | 0 | 11 | 0 | 36 | 0.0556 | 9 | 0 | 13 | 0.1538 | 9 | 0.2222 | 9 | 0.1111 | 40 | 0.125 |
| 8_2648935 | 7 | 0.4286 | 9 | 0.4444 | 8 | 0.5 | 9 | 0.3333 | 33 | 0.4242 | 7 | 0.4286 | 13 | 0.3846 | 8 | 0.375 | 8 | 0.375 | 36 | 0.3889 |
| 8_2648956 | 7 | 0 | 11 | 0.0909 | 6 | 0 | 9 | 0.2222 | 33 | 0.0909 | 8 | 0.125 | 11 | 0.0909 | 9 | 0 | 9 | 0 | 37 | 0.0541 |
| 8_2650543 | 4 | 0.25 | 5 | 0.2 | 2 | 0 | 6 | 0.1667 | 17 | 0.1765 | 4 | 0.25 | 8 | 0.125 | 8 | 0.125 | 4 | 0.25 | 24 | 0.1667 |
| 8_2663325 | 1 | 0 | NA | NA | 2 | 1 | 1 | 0 | 4 | 0.5 | 1 | 0 | 2 | 0.5 | 6 | 0.3333 | 1 | 1 | 10 | 0.4 |
| 8_2663333 | 1 | 1 | 1 | 0 | 1 | 0 | 1 | 0 | 4 | 0.25 | 1 | 0 | 2 | 0.5 | 6 | 0.3333 | 1 | 0 | 10 | 0.3 |
| 8_2663355 | 1 | 0 | 1 | 1 | 2 | 0.5 | 1 | 1 | 5 | 0.6 | 2 | 0.5 | 2 | 0 | 6 | 0 | 1 | 1 | 11 | 0.1818 |
| 8_2664915 | NA | NA | 1 | 1 | NA | NA | NA | NA | 1 | 1 | NA | NA | NA | NA | NA | NA | 1 | 1 | 1 | 1 |
| 8_2665279 | 1 | 0 | 4 | 0.25 | 1 | 1 | 1 | 1 | 7 | 0.4286 | 1 | 0 | 1 | 1 | 5 | 0.6 | 3 | 0.3333 | 10 | 0.5 |
| 8_2665285 | 2 | 0.5 | 5 | 0.4 | 1 | 1 | 1 | 1 | 9 | 0.5556 | NA | NA | 1 | 0 | 5 | 0.8 | 2 | 0.5 | 8 | 0.625 |
| 8_2665285 | 2 | 0.5 | 4 | 0.25 | 1 | 0 | 1 | 1 | 8 | 0.375 | 1 | 0 | 1 | 1 | 4 | 0.5 | 3 | 0 | 9 | 0.3333 |
| 8_2665331 | 1 | 0 | 6 | 0.1667 | 1 | 0 | 1 | 0 | 9 | 0.1111 | 1 | 1 | 1 | 0 | 5 | 0.4 | 2 | 0.5 | 9 | 0.4444 |
| 8_2665367 | 3 | 0 | 6 | 0.1667 | 2 | 0.5 | 1 | 0 | 12 | 0.1667 | 1 | 0 | 1 | 0 | 7 | 0.1429 | 2 | 0 | 11 | 0.0909 |
| 8_2746880 | 4 | 0.25 | 2 | 0.5 | 1 | 0 | 3 | 0.6667 | 10 | 0.4 | 4 | 0.5 | NA | NA | 3 | 0 | 2 | 0.5 | 9 | 0.3333 |
| 8_2746886 | 5 | 0.2 | 2 | 0 | 1 | 0 | 3 | 0 | 11 | 0.0909 | 6 | 0.1667 | 3 | 0 | 3 | 0 | 3 | 0.3333 | 15 | 0.1333 |
| 8_2746904 | 3 | 0.3333 | 1 | 1 | NA | NA | 2 | 0.5 | 6 | 0.5 | 3 | 0.3333 | 3 | 0.3333 | 3 | 1 | 1 | 1 | 10 | 0.6 |
| 8_2746907 | 3 | 0 | 2 | 0 | 1 | 0 | 3 | 0.3333 | 9 | 0.1111 | 4 | 0.5 | 2 | 0 | 2 | 0.5 | 3 | 0.6667 | 11 | 0.4545 |
| 8_2746908 | 3 | 0.3333 | 3 | 0.3333 | NA | NA | 4 | 0.5 | 10 | 0.4 | 7 | 0.5714 | 2 | 0.5 | 1 | 0 | 3 | 0.3333 | 13 | 0.4615 |
| 8_2746914 | 4 | 0.5 | 2 | 1 | 1 | 1 | 4 | 1 | 11 | 0.8182 | 7 | 0.7143 | 3 | 0.6667 | 3 | 1 | 3 | 1 | 16 | 0.8125 |
| 8_2746970 | 5 | 0.4 | 5 | 0 | 2 | 0.5 | 2 | 0 | 14 | 0.2143 | 4 | 0.5 | 4 | 0 | 3 | 0.3333 | 3 | 0.3333 | 14 | 0.2857 |
| 8_2746979 | 5 | 1 | 6 | 1 | 4 | 1 | 4 | 1 | 19 | 1 | 7 | 1 | 5 | 1 | 1 | 1 | 3 | 1 | 16 | 1 |
| 8_2746982 | 6 | 0.6667 | 5 | 0.8 | 4 | 1 | 3 | 0.6667 | 18 | 0.7778 | 7 | 1 | 5 | 0.6 | 3 | 0.6667 | 4 | 0.75 | 19 | 0.7895 |
| 8_2747005 | 3 | 1 | 6 | 0.6667 | 5 | 0.8 | 2 | 1 | 16 | 0.8125 | 4 | 0.75 | 3 | 1 | 3 | 1 | 3 | 1 | 13 | 0.9231 |
| 8_2747008 | 5 | 0.4 | 6 | 1 | 2 | 1 | 2 | 1 | 15 | 0.8 | 5 | 0.8 | 4 | 1 | NA | NA | 5 | 1 | 14 | 0.9286 |
| 8_2747008 | 7 | 1 | 5 | 1 | 4 | 1 | 4 | 1 | 20 | 1 | 7 | 1 | 7 | 0.8571 | 4 | 1 | 6 | 1 | 24 | 0.9583 |
| 8_2747025 | 5 | 1 | 7 | 1 | 5 | 1 | 4 | 1 | 21 | 1 | 6 | 1 | 7 | 1 | 2 | 1 | 5 | 1 | 20 | 1 |
| 8_2747029 | 6 | 0.1667 | 7 | 0.1429 | 4 | 0.25 | 2 | 0 | 19 | 0.1579 | 6 | 0.5 | 7 | 0.2857 | 3 | 0.3333 | 5 | 0 | 21 | 0.2857 |
| 8_2747056 | 6 | 0.3333 | 11 | 0.0909 | 4 | 0 | 4 | 0 | 25 | 0.12 | 5 | 0.2 | 8 | 0 | 3 | 0.3333 | 6 | 0 | 22 | 0.0909 |
| 8_2747093 | 7 | 0.1429 | 9 | 0.2222 | 4 | 0 | 3 | 0 | 23 | 0.1304 | 7 | 0 | 9 | 0.1111 | 7 | 0.1429 | 7 | 0.1429 | 30 | 0.1 |
| 8_2747132 | 8 | 0.125 | 12 | 0.0833 | 7 | 0 | 5 | 0 | 32 | 0.0625 | 9 | 0.1111 | 10 | 0.1 | 9 | 0.2222 | 8 | 0 | 36 | 0.1111 |
| 8_2752654 | 4 | 0 | 2 | 0.5 | 4 | 0 | 2 | 0 | 12 | 0.0833 | 6 | 0.3333 | 4 | 0.25 | 5 | 0 | 4 | 0.25 | 19 | 0.2105 |
| 8_2752666 | 4 | 0.75 | NA | NA | 3 | 0.3333 | 1 | 1 | 8 | 0.625 | 4 | 0.75 | 4 | 1 | 4 | 1 | 5 | 1 | 17 | 0.9412 |
| 8_2752666 | 3 | 0.3333 | 2 | 0.5 | 4 | 0 | 2 | 0 | 11 | 0.1818 | 5 | 0.2 | 2 | 0 | 5 | 0 | 3 | 0.3333 | 15 | 0.1333 |
| 8_2752676 | 4 | 1 | 2 | 1 | 4 | 1 | 2 | 0.5 | 12 | 0.9167 | 5 | 1 | 3 | 1 | 5 | 0.6 | 3 | 1 | 16 | 0.875 |
| 8_2752680 | 3 | 1 | 1 | 1 | 3 | 0.6667 | 2 | 1 | 9 | 0.8889 | 5 | 0.8 | 4 | 1 | 7 | 0.8571 | 4 | 1 | 20 | 0.9 |
| 8_2752712 | 3 | 0.3333 | NA | NA | 4 | 0.25 | 2 | 0 | 9 | 0.2222 | 5 | 0 | 4 | 0 | 5 | 0.2 | 3 | 0 | 17 | 0.0588 |
| 8_2752716 | 4 | 0.75 | 2 | 0 | 5 | 0.2 | 2 | 0 | 13 | 0.3077 | 4 | 0.25 | 3 | 0.3333 | 7 | 0.2857 | 4 | 0.75 | 18 | 0.3889 |
| 8_2752721 | 4 | 0.75 | 2 | 0.5 | 5 | 0.4 | 2 | 0 | 13 | 0.4615 | 6 | 0.3333 | 5 | 0.4 | 8 | 0.375 | 4 | 0.25 | 23 | 0.3478 |
| 8_2752814 | 3 | 0.3333 | 3 | 0 | 4 | 0 | 4 | 0 | 14 | 0.0714 | 5 | 0 | 7 | 0.1429 | 7 | 0.2857 | 7 | 0.1429 | 26 | 0.1538 |
| 8_2752907 | 6 | 0 | 5 | 0.4 | 4 | 0 | 5 | 0 | 20 | 0.1 | 7 | 0.1429 | 8 | 0 | 8 | 0 | 8 | 0.125 | 31 | 0.0645 |

| | | | | | | | | | | | | | | | | | | | | |
|---|---|---|---|---|---|---|---|---|---|---|---|---|---|---|---|---|---|---|---|---|
| 8_2753961( | NA | NA | NA | NA | 1 | 0 | 1 | 1 | 2 | 0.5 | NA | NA | 1 | 1 | NA | NA | NA | NA | 1 | 1 |
| 8_2754139: | 4 | 0.25 | 6 | 0.1667 | 5 | 0.2 | 4 | 0.25 | 19 | 0.2105 | NA | NA | 4 | 0 | 9 | 0.3333 | 3 | 0 | 16 | 0.1875 |
| 8_2754336( | 4 | 0 | 6 | 0.1667 | 6 | 0.3333 | 5 | 0 | 21 | 0.1429 | 4 | 0 | 3 | 0.3333 | 9 | 0.1111 | 6 | 0 | 22 | 0.0909 |
| 8_2754343( | 7 | 0.1429 | 8 | 0.5 | 6 | 0.1667 | 7 | 0.1429 | 28 | 0.25 | 4 | 0.25 | 4 | 0.25 | 10 | 0 | 7 | 0.2857 | 25 | 0.16 |
| 8_2754360: | 8 | 0.25 | 9 | 0.1111 | 7 | 0.4286 | 7 | 0.1429 | 31 | 0.2258 | 4 | 0.5 | 3 | 0.3333 | 10 | 0.7 | 7 | 0.5714 | 24 | 0.5833 |
| 8_2754362 | 9 | 0.1111 | 9 | 0.1111 | 7 | 0.4286 | 6 | 0 | 31 | 0.1613 | 3 | 0.3333 | 4 | 0.25 | 8 | 0.125 | 6 | 0.1667 | 21 | 0.1905 |
| 8_2754365( | 9 | 0.2222 | 9 | 0.1111 | 7 | 0.1429 | 7 | 0 | 32 | 0.125 | 4 | 0.5 | 4 | 0.25 | 10 | 0.4 | 7 | 0 | 25 | 0.28 |
| 8_2754371 | 8 | 0.625 | 9 | 0.7778 | 5 | 1 | 6 | 0.6667 | 28 | 0.75 | 4 | 0.5 | 4 | 1 | 10 | 0.9 | 6 | 0.5 | 24 | 0.75 |
| 8_2754387( | 8 | 0.5 | 9 | 0.6667 | 6 | 0.5 | 6 | 0.3333 | 29 | 0.5172 | 4 | 0.5 | 5 | 0.8 | 8 | 0.625 | 5 | 0.6 | 22 | 0.6364 |
| 8_2754390 | 6 | 0.1667 | 5 | 0 | 5 | 0.2 | 5 | 0 | 21 | 0.0952 | 4 | 0 | 5 | 0 | 8 | 0 | 5 | 0.4 | 22 | 0.0909 |
| 8_2754393 | 9 | 0.3333 | 9 | 0.5556 | 7 | 0.5714 | 5 | 0.6 | 30 | 0.5 | 4 | 1 | 6 | 0.5 | 9 | 0.6667 | 6 | 0.6667 | 25 | 0.68 |
| 8_2754402( | 10 | 0.4 | 11 | 0.7273 | 6 | 0.1667 | 7 | 0.5714 | 34 | 0.5 | 4 | 0.75 | 6 | 0.6667 | 8 | 0.625 | 6 | 0.6667 | 24 | 0.6667 |
| 8_2754411: | 9 | 0.3333 | 12 | 0.1667 | 5 | 0 | 6 | 0 | 32 | 0.1562 | 4 | 0.5 | 5 | 0.2 | 8 | 0.125 | 7 | 0.1429 | 24 | 0.2083 |
| 8_2754415 | 11 | 0.6364 | 11 | 0.5455 | 6 | 0.3333 | 7 | 0.4286 | 35 | 0.5143 | 4 | 0.75 | 6 | 0.5 | 10 | 0.3 | 7 | 0.5714 | 27 | 0.4815 |
| 8_2754427 | 11 | 0.6364 | 13 | 0.3846 | 8 | 0.375 | 7 | 0.5714 | 39 | 0.4872 | 4 | 0.75 | 7 | 0.7143 | 11 | 0.5455 | 6 | 0.1667 | 28 | 0.5357 |
| 8_2754428 | 11 | 0.5455 | 15 | 0.6 | 8 | 0.375 | 7 | 0.7143 | 41 | 0.561 | 3 | 1 | 7 | 1 | 12 | 0.4167 | 6 | 0.1667 | 28 | 0.5714 |
| 8_2754446 | 12 | 0.1667 | 13 | 0.2308 | 8 | 0.125 | 8 | 0 | 41 | 0.1463 | 4 | 0.25 | 7 | 0.2857 | 12 | 0.0833 | 7 | 0 | 30 | 0.1333 |
| 8_2761104 | 1 | 1 | 2 | 0.5 | NA | NA | NA | NA | 3 | 0.6667 | NA | NA | NA | NA | NA | NA | 1 | 1 | 1 | 1 |
| 8_2761138 | 1 | 1 | 2 | 1 | NA | NA | NA | NA | 3 | 1 | NA | NA | NA | NA | NA | NA | 1 | 1 | 1 | 1 |
| 8_2761144 | 1 | 1 | 2 | 0.5 | NA | NA | NA | NA | 3 | 0.6667 | NA | NA | NA | NA | NA | NA | 1 | 1 | 1 | 1 |
| 8_2761203 | 1 | 0 | 2 | 0.5 | NA | NA | NA | NA | 3 | 0.3333 | NA | NA | NA | NA | NA | NA | 1 | 1 | 1 | 1 |
| 8_2761225( | 1 | 0 | 1 | 1 | NA | NA | NA | NA | 2 | 0.5 | NA | NA | 1 | 1 | NA | NA | 1 | 0 | 2 | 0.5 |
| 8_2761348 | 1 | 1 | NA | NA | 1 | 1 | NA | NA | 2 | 1 | 1 | 1 | 1 | 1 | 1 | 1 | NA | NA | 3 | 1 |
| 8_2761376 | NA | NA | 1 | 1 | NA | NA | NA | NA | 1 | 1 | 1 | 1 | NA | NA | 1 | 0 | NA | NA | 2 | 0.5 |
| 8_2761379 | 1 | 0 | 1 | 1 | NA | NA | NA | NA | 2 | 0.5 | 1 | 1 | 1 | 0 | 1 | 0 | NA | NA | 3 | 0.3333 |
| 8_2761389 | 4 | 0 | 3 | 0.3333 | 2 | 0.5 | 1 | 0 | 10 | 0.2 | 1 | 0 | 3 | 0.3333 | 2 | 0.5 | 1 | 1 | 7 | 0.4286 |
| 8_2761398( | 4 | 0.5 | 2 | 0.5 | 2 | 1 | 1 | 1 | 9 | 0.6667 | 1 | 1 | 3 | 0.3333 | 3 | 0.6667 | 1 | 1 | 8 | 0.625 |
| 8_2761404 | 4 | 0.75 | 2 | 1 | 2 | 1 | 1 | 0 | 9 | 0.7778 | 1 | 1 | 3 | 1 | 3 | 0.3333 | 1 | 0 | 8 | 0.625 |
| 8_2761407 | 4 | 0.75 | 3 | 0.3333 | 2 | 0 | 1 | 1 | 10 | 0.5 | 1 | 1 | 3 | 0.3333 | 3 | 0.3333 | 1 | 0 | 8 | 0.375 |
| 8_2761418: | 5 | 0.8 | 1 | 1 | NA | NA | 1 | 1 | 7 | 0.8571 | 1 | 1 | 1 | 1 | 3 | 1 | 1 | 1 | 6 | 1 |
| 8_2761418 | 5 | 0.6 | 3 | 0.3333 | 1 | 0 | 1 | 0 | 10 | 0.4 | 1 | 1 | 3 | 0.6667 | 3 | 0.6667 | 1 | 0 | 8 | 0.625 |
| 8_2761419 | 5 | 0.4 | 3 | 0 | 1 | 0 | 1 | 0 | 10 | 0.2 | 1 | 1 | 2 | 0.5 | 3 | 0.3333 | 1 | 0 | 7 | 0.4286 |
| 8_2761432 | 4 | 0.5 | 3 | 0.6667 | NA | NA | 1 | 1 | 8 | 0.625 | 1 | 1 | NA | NA | 3 | 0.6667 | NA | NA | 4 | 0.75 |
| 8_2761441 | 4 | 0.75 | 2 | 0.5 | 1 | 1 | 1 | 1 | 8 | 0.75 | 1 | 1 | 3 | 1 | 2 | 1 | 1 | 1 | 7 | 1 |
| 8_2761453 | 5 | 1 | 2 | 0.5 | NA | NA | 1 | 1 | 8 | 0.875 | NA | NA | 3 | 0.3333 | 3 | 1 | 1 | 1 | 7 | 0.7143 |
| 8_2761594 | 5 | 0.8 | 2 | 1 | 1 | 1 | 1 | 1 | 9 | 0.8889 | NA | NA | 3 | 1 | 3 | 1 | NA | NA | 6 | 1 |
| 8_2776513 | 1 | 0 | NA | NA | NA | NA | 1 | 1 | 2 | 0.5 | NA | NA | 1 | 1 | 1 | 0 | 1 | 0 | 3 | 0.3333 |
| 8_2776536( | 1 | 0 | NA | NA | NA | NA | 1 | 1 | 2 | 0.5 | NA | NA | 1 | 1 | 1 | 0 | 1 | 1 | 3 | 0.6667 |
| 8_2776540 | 1 | 1 | NA | NA | NA | NA | 1 | 1 | 2 | 1 | NA | NA | 1 | 1 | 1 | 1 | 1 | 1 | 3 | 1 |
| 8_2913699 | 2 | 1 | 3 | 0.3333 | 2 | 1 | 3 | 0.6667 | 10 | 0.7 | 4 | 0.5 | 1 | 1 | 1 | 1 | 1 | 1 | 7 | 0.7143 |
| 8_2913712 | 2 | 0.5 | 2 | 0 | 1 | 0 | 4 | 0.25 | 9 | 0.2222 | 3 | 0.3333 | 1 | 0 | NA | NA | 1 | 0 | 5 | 0.2 |
| 8_2913848 | 1 | 1 | 2 | 0.5 | NA | NA | 2 | 1 | 5 | 0.8 | 2 | 1 | NA | NA | NA | NA | NA | NA | 2 | 1 |
| 8_2913876 | NA | NA | 2 | 1 | NA | NA | 4 | 0.25 | 6 | 0.5 | 2 | 0 | NA | NA | NA | NA | 1 | 1 | 3 | 0.3333 |
| 8_2913886 | NA | NA | 3 | 1 | NA | NA | 3 | 1 | 6 | 1 | 2 | 0.5 | NA | NA | NA | NA | NA | NA | 2 | 0.5 |
| 8_2913889 | NA | NA | 2 | 1 | NA | NA | 3 | 1 | 5 | 1 | 2 | 0.5 | NA | NA | NA | NA | 1 | 0 | 3 | 0.3333 |
| 8_2914074 | NA | NA | 3 | 0 | NA | NA | 2 | 0.5 | 5 | 0.2 | 2 | 0.5 | NA | NA | NA | NA | 1 | 0 | 3 | 0.3333 |
| 8_2914132 | NA | NA | 2 | 0.5 | NA | NA | 2 | 1 | 4 | 0.75 | 1 | 1 | NA | NA | NA | NA | NA | NA | 1 | 1 |
| 8_3158179 | NA | NA | NA | NA | 1 | 1 | NA | NA | 1 | 1 | NA | NA | 3 | 0.6667 | 1 | 0 | NA | NA | 4 | 0.5 |
| 8_3158194 | 10 | 0.1 | 10 | 0.1 | 10 | 0.2 | 7 | 0 | 37 | 0.1081 | 9 | 0.1111 | 6 | 0.1667 | 8 | 0.25 | 8 | 0 | 31 | 0.129 |
| 8_3158641 | 10 | 0.9 | 12 | 0.75 | 10 | 0.8 | 8 | 0.75 | 40 | 0.8 | 8 | 0.75 | 9 | 0.8889 | 7 | 1 | 9 | 0.8889 | 33 | 0.8788 |
| 8_3158695( | 11 | 0.3636 | 13 | 0.2308 | 13 | 0.3846 | 11 | 0.2727 | 48 | 0.3125 | 10 | 0.5 | 13 | 0.4615 | 15 | 0.4 | 10 | 0.4 | 48 | 0.4375 |
| 8_3161871 | NA | NA | NA | NA | 1 | 0 | 2 | 0.5 | 3 | 0.3333 | 1 | 1 | 1 | 1 | 2 | 0 | 2 | 0 | 6 | 0.3333 |
| 8_3161873 | NA | NA | 1 | 0 | 1 | 1 | 2 | 0.5 | 4 | 0.5 | 1 | 1 | 1 | 1 | 2 | 1 | 2 | 0.5 | 6 | 0.8333 |
| 8_3161875 | NA | NA | 1 | 0 | 1 | 1 | 2 | 0.5 | 4 | 0.5 | NA | NA | 1 | 1 | 1 | 1 | 2 | 0.5 | 4 | 0.75 |
| 8_3161879 | NA | NA | 1 | 0 | 1 | 1 | 1 | 1 | 3 | 0.6667 | NA | NA | 1 | 1 | 1 | 1 | 2 | 1 | 4 | 1 |
| 8_3161886 | NA | NA | 1 | 0 | 1 | 0 | 2 | 0.5 | 4 | 0.25 | 1 | 0 | 1 | 0 | 1 | 0 | 2 | 0.5 | 5 | 0.2 |
| 8_3161893( | NA | NA | 1 | 1 | 1 | 1 | 2 | 0.5 | 4 | 0.75 | 1 | 1 | 1 | 1 | 2 | 0.5 | 1 | 1 | 5 | 0.8 |
| 8_3161895 | NA | NA | NA | NA | 1 | 1 | 1 | 1 | 2 | 1 | 1 | 1 | 1 | 1 | 2 | 1 | 2 | 1 | 6 | 1 |
| 8_3161901( | NA | NA | 1 | 1 | 1 | 1 | 2 | 0.5 | 4 | 0.75 | 1 | 0 | 1 | 1 | NA | NA | 1 | 0 | 3 | 0.3333 |
| 8_3161904 | NA | NA | 1 | 1 | 1 | 1 | 2 | 0 | 4 | 0.5 | 1 | 0 | 1 | 1 | 2 | 0.5 | 1 | 1 | 5 | 0.6 |
| 8_3161927( | NA | NA | 1 | 1 | 1 | 0 | 2 | 0.5 | 4 | 0.5 | 1 | 0 | 1 | 0 | 2 | 0 | 2 | 0.5 | 6 | 0.1667 |
| 8_3164056 | 2 | 0.5 | NA | NA | 2 | 0 | NA | NA | 4 | 0.25 | 2 | 0 | 1 | 0 | 1 | 0 | 4 | 0.25 | 8 | 0.125 |
| 8_3164059 | 2 | 0 | NA | NA | 2 | 0.5 | NA | NA | 4 | 0.25 | 2 | 0 | 1 | 0 | 1 | 1 | 3 | 0.6667 | 7 | 0.4286 |
| 8_3164088 | 1 | 0 | 1 | 0 | 2 | 0.5 | NA | NA | 4 | 0.25 | 2 | 0.5 | 1 | 0 | 1 | 0 | 5 | 0.2 | 9 | 0.2222 |
| 8_3164100( | 2 | 0 | 1 | 1 | 2 | 0 | NA | NA | 5 | 0.2 | 2 | 0 | 2 | 0 | 2 | 0.5 | 6 | 0.1667 | 12 | 0.1667 |

| | | | | | | | | | | | | | | | | | | | | |
|---|---|---|---|---|---|---|---|---|---|---|---|---|---|---|---|---|---|---|---|---|
| 8_31641324 | 2 | 0.5 | 1 | 0 | 2 | 0 | NA | NA | 5 | 0.2 | 2 | 0 | 2 | 0 | 3 | 0 | 6 | 0.3333 | 13 | 0.1538 |
| 8_31641370 | 2 | 0.5 | NA | NA | 2 | 1 | NA | NA | 4 | 0.75 | 2 | 1 | 2 | 0.5 | 2 | 1 | 5 | 0.8 | 11 | 0.8182 |
| 8_31649639 | 2 | 0.5 | 5 | 0.4 | 4 | 0.25 | 3 | 0.3333 | 14 | 0.3571 | 5 | 0.6 | 4 | 0.5 | 3 | 0.6667 | 3 | 0 | 15 | 0.4667 |
| 8_31649693 | 4 | 0 | 7 | 0.8571 | 5 | 0.2 | 4 | 0.25 | 20 | 0.4 | 6 | 0.1667 | 5 | 0.6 | 3 | 0 | 5 | 0.2 | 19 | 0.2632 |
| 8_31649728 | 4 | 0 | 7 | 0.1429 | 4 | 0 | 4 | 0 | 19 | 0.0526 | 6 | 0 | 5 | 0 | 3 | 0 | 5 | 0.2 | 19 | 0.0526 |
| 8_31649767 | 4 | 0.25 | 7 | 0.7143 | 3 | 1 | 3 | 0.6667 | 17 | 0.6471 | 5 | 0.2 | 5 | 0.8 | 3 | 0.6667 | 5 | 0.4 | 18 | 0.5 |
| 8_31649784 | 4 | 0 | 4 | 0 | 4 | 0.25 | 4 | 0 | 16 | 0.0625 | 6 | 0 | 4 | 0 | 1 | 0 | 3 | 0.3333 | 14 | 0.0714 |
| 8_31649819 | 4 | 0.75 | 6 | 1 | 4 | 1 | 4 | 0.75 | 18 | 0.8889 | 6 | 0.8333 | 5 | 1 | 3 | 1 | 5 | 0.8 | 19 | 0.8947 |
| 8_31649871 | 3 | 0.3333 | 7 | 0 | 4 | 0 | 2 | 0 | 16 | 0.0625 | 6 | 0 | 3 | 0 | 3 | 0 | 5 | 0.2 | 17 | 0.0588 |
| 8_31650137 | 4 | 0.75 | 7 | 0.1429 | 4 | 0.5 | 2 | 0.5 | 17 | 0.4118 | 5 | 0.6 | 4 | 0.75 | 4 | 0.75 | 4 | 0 | 17 | 0.5294 |
| 8_31650498 | 3 | 0.3333 | 5 | 0.8 | 4 | 0.5 | 2 | 0.5 | 14 | 0.5714 | 3 | 0.6667 | 4 | 0.75 | 4 | 1 | 4 | 0.5 | 15 | 0.7333 |
| 8_31651175 | NA | NA | NA | NA | 1 | 1 | NA | NA | 1 | 1 | NA | NA | NA | NA | 1 | 1 | 1 | 1 | 2 | 1 |
| 8_31651965 | NA | NA | NA | NA | 2 | 0.5 | 2 | 0 | 4 | 0.25 | 1 | 0 | NA | NA | 2 | 0.5 | NA | NA | 3 | 0.3333 |
| 8_31652057 | NA | NA | NA | NA | 2 | 0.5 | 2 | 1 | 4 | 0.75 | 1 | 1 | 1 | 1 | 2 | 0.5 | NA | NA | 4 | 0.75 |
| 8_31652303 | NA | NA | NA | NA | 2 | 0.5 | 2 | 0.5 | 4 | 0.5 | 1 | 1 | 1 | 1 | 2 | 0 | NA | NA | 4 | 0.5 |
| 8_31685187 | 2 | 0.5 | NA | NA | NA | NA | 1 | 0 | 3 | 0.3333 | NA | NA | NA | NA | 1 | 1 | 1 | 0 | 2 | 0.5 |
| 8_31691430 | 2 | 1 | NA | NA | NA | NA | 2 | 0.5 | 4 | 0.75 | NA | NA | NA | NA | 1 | 0 | 1 | 1 | 2 | 0.5 |
| 8_31725860 | 2 | 0 | NA | NA | NA | NA | 2 | 0.5 | 4 | 0.25 | NA | NA | NA | NA | 1 | 1 | NA | NA | 1 | 1 |
| 8_31725870 | 2 | 1 | NA | NA | NA | NA | 1 | 1 | 3 | 1 | NA | NA | NA | NA | 1 | 1 | NA | NA | 1 | 1 |
| 8_31725905 | 2 | 1 | NA | NA | NA | NA | 2 | 1 | 4 | 1 | NA | NA | NA | NA | 1 | 1 | NA | NA | 1 | 1 |
| 8_31726263 | 1 | 1 | NA | NA | NA | NA | 1 | 1 | 2 | 1 | NA | NA | NA | NA | 1 | 1 | NA | NA | 1 | 1 |
| 8_31726376 | 2 | 1 | NA | NA | NA | NA | 2 | 1 | 4 | 1 | NA | NA | NA | NA | 1 | 1 | NA | NA | 1 | 1 |
| 8_31726388 | 2 | 0.5 | NA | NA | NA | NA | 2 | 0.5 | 4 | 0.5 | NA | NA | NA | NA | 1 | 1 | NA | NA | 1 | 1 |
| 8_31726992 | 2 | 0.5 | NA | NA | NA | NA | 1 | 0 | 3 | 0.3333 | NA | NA | NA | NA | 1 | 1 | NA | NA | 1 | 1 |
| 8_31750157 | NA | NA | NA | NA | NA | NA | 1 | 1 | 1 | 1 | NA | NA | NA | NA | NA | NA | 1 | 1 | 1 | 1 |
| 8_31750379 | 1 | 1 | NA | NA | NA | NA | 1 | 1 | 2 | 1 | NA | NA | NA | NA | NA | NA | 1 | 1 | 1 | 1 |
| 8_31750547 | 1 | 1 | NA | NA | NA | NA | NA | NA | 1 | 1 | NA | NA | NA | NA | NA | NA | 1 | 1 | 1 | 1 |
| 8_31750573 | 1 | 1 | NA | NA | NA | NA | 1 | 1 | 2 | 1 | NA | NA | NA | NA | NA | NA | 1 | 1 | 1 | 1 |
| 8_31751320 | 1 | 1 | 1 | 1 | NA | NA | 1 | 1 | 3 | 1 | NA | NA | NA | NA | NA | NA | 1 | 1 | 1 | 1 |
| 8_31757052 | 1 | 1 | NA | NA | NA | NA | NA | NA | 1 | 1 | NA | NA | 1 | 1 | NA | NA | NA | NA | 1 | 1 |
| 8_33725908 | NA | NA | 1 | 1 | NA | NA | 1 | 0 | 2 | 0.5 | 1 | 1 | 1 | 1 | 1 | 1 | NA | NA | 3 | 1 |
| 8_33726028 | NA | NA | 1 | 1 | NA | NA | 1 | 0 | 2 | 0.5 | 1 | 1 | 1 | 1 | 1 | 1 | NA | NA | 3 | 1 |
| 8_33726171 | NA | NA | 1 | 1 | NA | NA | NA | NA | 1 | 1 | 1 | 1 | NA | NA | 1 | 1 | NA | NA | 2 | 1 |
| 8_33876459 | NA | NA | 2 | 0.5 | NA | NA | NA | NA | 2 | 0.5 | NA | NA | 1 | 0 | NA | NA | 1 | 1 | 2 | 0.5 |
| 8_33876569 | NA | NA | 2 | 1 | NA | NA | NA | NA | 2 | 1 | NA | NA | 1 | 1 | NA | NA | 1 | 1 | 2 | 1 |
| 8_33876665 | NA | NA | 2 | 0.5 | NA | NA | NA | NA | 2 | 0.5 | NA | NA | 1 | 1 | NA | NA | 1 | 1 | 2 | 1 |
| 8_33876722 | NA | NA | 2 | 1 | NA | NA | NA | NA | 2 | 1 | 1 | 1 | 1 | 1 | NA | NA | 1 | 1 | 3 | 1 |
| 8_33876893 | NA | NA | 1 | 1 | NA | NA | NA | NA | 1 | 1 | NA | NA | NA | NA | NA | NA | 1 | 1 | 1 | 1 |
| 8_33877165 | NA | NA | 2 | 1 | NA | NA | NA | NA | 2 | 1 | 1 | 0 | 1 | 0 | NA | NA | 1 | 1 | 3 | 0.3333 |
| 8_33878934 | NA | NA | 1 | 1 | NA | NA | NA | NA | 1 | 1 | NA | NA | NA | NA | 1 | 1 | NA | NA | 1 | 1 |
| 8_33906675 | 3 | 0 | 4 | 0.25 | 2 | 0 | 6 | 0 | 15 | 0.0667 | 2 | 0 | 3 | 0.3333 | NA | NA | 5 | 0.2 | 10 | 0.2 |
| 8_33906692 | 2 | 1 | 6 | 0.3333 | 2 | 0.5 | 6 | 0.1667 | 16 | 0.375 | 4 | 0.25 | 3 | 0.3333 | 2 | 1 | 6 | 0.1667 | 15 | 0.3333 |
| 8_33906745 | 3 | 0.3333 | 5 | 0.4 | 2 | 0.5 | 6 | 0.3333 | 16 | 0.375 | 4 | 0.25 | 2 | 0.5 | 1 | 0 | 5 | 0.2 | 12 | 0.25 |
| 8_33906874 | 2 | 0 | 5 | 0 | 3 | 0.3333 | 5 | 0.2 | 15 | 0.1333 | 3 | 0 | 2 | 0.5 | 1 | 0 | 5 | 0 | 11 | 0.0909 |
| 8_33906882 | 3 | 0.3333 | 4 | 0.5 | 2 | 1 | 5 | 0.4 | 14 | 0.5 | 4 | 0.25 | 3 | 0.6667 | 1 | 1 | 2 | 1 | 10 | 0.6 |
| 8_33906992 | 3 | 0 | 6 | 0.1667 | 4 | 0.5 | 7 | 0.1429 | 20 | 0.2 | 4 | 0.25 | 4 | 0 | 2 | 0.5 | 7 | 0.1429 | 17 | 0.1765 |
| 8_33907238 | 4 | 0.5 | 5 | 0.2 | 4 | 0 | 7 | 0.1429 | 20 | 0.2 | 5 | 0 | 4 | 0.25 | 3 | 0 | 7 | 0.1429 | 19 | 0.1053 |
| 8_33907391 | 3 | 0 | 6 | 0 | 4 | 0.5 | 8 | 0.125 | 21 | 0.1429 | 3 | 0 | 3 | 0.3333 | 2 | 0 | 6 | 0 | 14 | 0.0714 |
| 8_33907480 | 2 | 0 | 4 | 0 | 3 | 0 | 4 | 0.25 | 13 | 0.0769 | 3 | 0.3333 | 3 | 0 | NA | NA | 5 | 0 | 11 | 0.0909 |
| 8_34106059 | 1 | 1 | NA | NA | NA | NA | NA | NA | 1 | 1 | NA | NA | 1 | 1 | NA | NA | NA | NA | 1 | 1 |
| 8_34106204 | 1 | 1 | NA | NA | NA | NA | NA | NA | 1 | 1 | NA | NA | 1 | 1 | NA | NA | NA | NA | 1 | 1 |
| 8_34109366 | 16 | 0.0625 | 19 | 0.0526 | 21 | 0.0952 | 31 | 0.1613 | 87 | 0.1034 | 18 | 0.1667 | 23 | 0.0435 | 18 | 0.1667 | 23 | 0.1739 | 82 | 0.1341 |
| 8_34109410 | 15 | 0.0667 | 21 | 0.0952 | 22 | 0.0455 | 32 | 0.0938 | 90 | 0.0778 | 20 | 0.15 | 25 | 0.04 | 15 | 0.2 | 28 | 0.0714 | 88 | 0.1023 |
| 8_34111090 | 3 | 0.3333 | 1 | 0 | 2 | 0 | 3 | 0 | 9 | 0.1111 | 2 | 0 | 2 | 0 | 2 | 0 | 7 | 0.2857 | 13 | 0.1538 |
| 8_34111150 | 2 | 0 | 2 | 0.5 | 3 | 0.3333 | 3 | 0 | 10 | 0.2 | 2 | 0.5 | 2 | 0 | 3 | 0 | 6 | 0.5 | 13 | 0.3077 |
| 8_34111187 | 3 | 1 | 2 | 1 | 3 | 1 | 4 | 0.25 | 12 | 0.75 | 3 | 0.3333 | 2 | 0 | 3 | 0.6667 | 7 | 0.8571 | 15 | 0.6 |
| 8_34111260 | 3 | 0.6667 | NA | NA | 3 | 0.6667 | 4 | 1 | 10 | 0.8 | 2 | 1 | 1 | 0 | 2 | 0.5 | 6 | 0.6667 | 11 | 0.6364 |
| 8_34111335 | 3 | 0.3333 | 1 | 0 | 2 | 0.5 | 3 | 0 | 9 | 0.2222 | 2 | 0 | 1 | 0 | 3 | 0 | 7 | 0.2857 | 13 | 0.1538 |
| 8_34111435 | 3 | 0.3333 | 2 | 0.5 | 3 | 0.3333 | 4 | 0 | 12 | 0.25 | 3 | 0 | 1 | 0 | 4 | 0 | 6 | 0.3333 | 14 | 0.1429 |
| 8_34111601 | 3 | 0.6667 | 2 | 0.5 | 2 | 1 | 3 | 0.6667 | 10 | 0.7 | 2 | 1 | 1 | 0 | 1 | 0 | 4 | 0.75 | 8 | 0.625 |
| 8_34111619 | 3 | 0.3333 | 1 | 0 | 3 | 0.3333 | 4 | 0 | 11 | 0.1818 | 1 | 0 | 1 | 0 | 2 | 0.5 | 5 | 0 | 9 | 0.1111 |
| 8_34111678 | 3 | 0.6667 | 2 | 0.5 | 3 | 0.3333 | 4 | 0 | 12 | 0.3333 | 3 | 1 | 1 | 0 | 2 | 0.5 | 5 | 0.8 | 11 | 0.7273 |
| 8_34187471 | 4 | 0 | 11 | 0.1818 | 4 | 0 | 8 | 0.125 | 27 | 0.1111 | 5 | 0.2 | 6 | 0 | 5 | 0.2 | 3 | 0 | 19 | 0.1053 |
| 8_34232236 | 1 | 1 | 1 | 0 | NA | NA | 4 | 0.25 | 6 | 0.3333 | 3 | 0.3333 | 3 | 0.3333 | 3 | 0.3333 | NA | NA | 9 | 0.3333 |
| 8_34232245 | 1 | 1 | NA | NA | NA | NA | 3 | 1 | 4 | 1 | 3 | 1 | 2 | 1 | 3 | 0.6667 | NA | NA | 8 | 0.875 |

| | | | | | | | | | | | | | | | | | | | | |
|---|---|---|---|---|---|---|---|---|---|---|---|---|---|---|---|---|---|---|---|---|
| 8_3423225 | 1 | 0 | 1 | 0 | NA | NA | 3 | 0.6667 | 5 | 0.4 | 3 | 0.3333 | 3 | 0 | 2 | 0.5 | NA | NA | 8 | 0.25 |
| 8_3423237 | 1 | 0 | 1 | 0 | NA | NA | 3 | 0.3333 | 5 | 0.2 | 4 | 0.5 | 2 | 0 | 1 | 0 | NA | NA | 7 | 0.2857 |
| 8_3423240 | 1 | 1 | 1 | 1 | NA | NA | 3 | 1 | 5 | 1 | 4 | 0.75 | 2 | 1 | 3 | 1 | NA | NA | 9 | 0.8889 |
| 8_3423249 | 1 | 1 | NA | NA | NA | NA | 3 | 1 | 4 | 1 | 3 | 1 | 3 | 1 | 3 | 1 | NA | NA | 9 | 1 |
| 8_3423252 | 1 | 0 | 1 | 1 | NA | NA | 3 | 0 | 5 | 0.2 | 3 | 0 | 2 | 0 | 3 | 0.3333 | NA | NA | 8 | 0.125 |
| 8_3426654 | 6 | 0 | 5 | 0 | 2 | 0.5 | 4 | 0 | 17 | 0.0588 | 1 | 0 | 4 | 0.25 | 2 | 0.5 | 6 | 0.1667 | 13 | 0.2308 |
| 8_3426657 | 8 | 0.625 | 7 | 0.4286 | 4 | 0 | 4 | 0.75 | 23 | 0.4783 | 3 | 0.3333 | 4 | 0.5 | 2 | 0 | 7 | 0.7143 | 16 | 0.5 |
| 8_3426660 | 7 | 0.1429 | 6 | 0.1667 | 3 | 0 | 5 | 0.4 | 21 | 0.1905 | 1 | 0 | 4 | 0 | 2 | 0.5 | 6 | 0.3333 | 13 | 0.2308 |
| 8_3426662 | 8 | 0.125 | 6 | 0 | 4 | 0.25 | 5 | 0.2 | 23 | 0.1304 | 3 | 0.3333 | 6 | 0 | 2 | 0 | 7 | 0 | 18 | 0.0556 |
| 8_3426666 | 8 | 0.625 | 7 | 0.2857 | 4 | 0.25 | 4 | 0.75 | 23 | 0.4783 | 3 | 0 | 5 | 0.2 | 2 | 0.5 | 9 | 0.2222 | 19 | 0.2105 |
| 8_3426674 | 11 | 0.0909 | 7 | 0.2857 | 6 | 0.1667 | 5 | 0 | 29 | 0.1379 | 2 | 0 | 7 | 0 | 2 | 0.5 | 9 | 0.2222 | 20 | 0.15 |
| 8_3426683 | 5 | 0 | 4 | 0.5 | 3 | 0 | 4 | 0.25 | 16 | 0.1875 | 3 | 0 | 7 | 0.2857 | 1 | 0 | 7 | 0.1429 | 18 | 0.1667 |
| 8_3455776 | 22 | 0.3636 | 17 | 0.3529 | 19 | 0.5263 | 16 | 0.625 | 74 | 0.4595 | 18 | 0.5 | 12 | 0.6667 | 16 | 0.375 | 4 | 0.25 | 50 | 0.48 |
| 8_3455782 | 23 | 0.087 | 19 | 0.1053 | 17 | 0.1765 | 16 | 0.0625 | 75 | 0.1067 | 19 | 0.1053 | 14 | 0.2857 | 16 | 0.125 | 5 | 0.2 | 54 | 0.1667 |
| 8_3457827 | 5 | 0 | 5 | 0.2 | 4 | 0 | 4 | 0.25 | 18 | 0.1111 | 3 | 0 | 2 | 0 | 3 | 0.3333 | 4 | 0 | 12 | 0.0833 |
| 8_3460294 | 11 | 0.7273 | 9 | 0.5556 | 8 | 0.5 | 13 | 0.6923 | 41 | 0.6341 | 4 | 0.75 | 13 | 0.4615 | 10 | 0.8 | 7 | 0.4286 | 34 | 0.5882 |
| 8_3460306 | 10 | 0.6 | 10 | 0.2 | 7 | 0.4286 | 14 | 0.3571 | 41 | 0.3902 | 6 | 0.1667 | 10 | 0.3 | 11 | 0.7273 | 8 | 0.375 | 35 | 0.4286 |
| 8_3460399 | 11 | 0 | 7 | 0.1429 | 6 | 0 | 12 | 0.1667 | 36 | 0.0833 | 5 | 0 | 11 | 0 | 10 | 0.1 | 5 | 0.2 | 31 | 0.0645 |
| 8_3460452 | 9 | 0.3333 | 9 | 0.3333 | 7 | 0 | 14 | 0 | 39 | 0.1538 | 5 | 0.4 | 14 | 0.0714 | 10 | 0.1 | 6 | 0 | 35 | 0.1143 |
| 8_3460466 | 6 | 1 | 6 | 1 | 6 | 0.8333 | 7 | 1 | 25 | 0.96 | 3 | 0.6667 | 12 | 0.9167 | 8 | 1 | 3 | 1 | 26 | 0.9231 |
| 8_3460472 | 8 | 1 | 8 | 0.875 | 8 | 0.875 | 14 | 0.8571 | 38 | 0.8947 | 5 | 0.8 | 15 | 1 | 8 | 1 | 6 | 0.8333 | 34 | 0.9412 |
| 8_3460484 | 8 | 0.875 | 8 | 0.875 | 6 | 0.8333 | 12 | 0.9167 | 34 | 0.8824 | 6 | 0.8333 | 11 | 1 | 8 | 1 | 5 | 0.6 | 30 | 0.9 |
| 8_3460486 | 9 | 0.8889 | 7 | 0.7143 | 7 | 0.7143 | 13 | 0.6154 | 36 | 0.7222 | 6 | 0.6667 | 12 | 0.5833 | 9 | 0.8889 | 7 | 0.8571 | 34 | 0.7353 |
| 8_3463234 | 25 | 0.24 | 37 | 0.1081 | 28 | 0.1071 | 39 | 0.1026 | 129 | 0.1318 | 20 | 0.2 | 25 | 0.12 | 34 | 0.0588 | 38 | 0.1053 | 117 | 0.1111 |
| 8_3463244 | 24 | 0.0833 | 39 | 0.1282 | 31 | 0.0645 | 37 | 0.0811 | 131 | 0.0916 | 22 | 0.1818 | 28 | 0.1429 | 32 | 0.0938 | 44 | 0.2273 | 126 | 0.1667 |
| 8_3463750 | 31 | 0.0968 | 39 | 0.0769 | 33 | 0.0303 | 42 | 0.0476 | 145 | 0.0621 | 27 | 0 | 28 | 0.0714 | 39 | 0.0769 | 39 | 0.1026 | 133 | 0.0677 |
| 8_3463756 | 31 | 0.4194 | 51 | 0.2549 | 38 | 0.2895 | 47 | 0.3191 | 167 | 0.3114 | 29 | 0.2759 | 35 | 0.4857 | 47 | 0.3191 | 50 | 0.42 | 161 | 0.3789 |
| 8_3463760 | 30 | 0.2667 | 51 | 0.2353 | 38 | 0.2368 | 47 | 0.1915 | 166 | 0.2289 | 28 | 0.1786 | 35 | 0.2857 | 45 | 0.1333 | 49 | 0.2449 | 157 | 0.2102 |
| 8_3463775 | 37 | 0.1892 | 57 | 0.193 | 37 | 0.1081 | 50 | 0.14 | 181 | 0.1602 | 32 | 0.1562 | 37 | 0.1892 | 51 | 0.2157 | 49 | 0.2653 | 169 | 0.213 |
| 8_3479554 | NA | NA | NA | NA | 1 | 1 | NA | NA | 1 | 1 | 1 | 1 | 1 | 1 | NA | NA | NA | NA | 2 | 1 |
| 8_3528593 | NA | NA | 1 | 1 | NA | NA | 1 | 1 | 2 | 1 | NA | NA | 1 | 1 | NA | NA | NA | NA | 1 | 1 |
| 8_3528601 | NA | NA | 1 | 1 | NA | NA | NA | NA | 1 | 1 | NA | NA | 1 | 1 | NA | NA | NA | NA | 1 | 1 |
| 8_3528689 | 1 | 0 | 1 | 0 | 1 | 1 | 2 | 0.5 | 5 | 0.4 | NA | NA | 1 | 1 | NA | NA | NA | NA | 1 | 1 |
| 8_3528851 | 2 | 0 | 3 | 0.3333 | 1 | 0 | 2 | 0 | 8 | 0.125 | NA | NA | 1 | 1 | NA | NA | NA | NA | 1 | 1 |
| 8_3528891 | 2 | 0.5 | 3 | 0 | 1 | 0 | 2 | 0.5 | 8 | 0.25 | 1 | 1 | 1 | 0 | NA | NA | NA | NA | 2 | 0.5 |
| 8_3529374 | 7 | 0.2857 | 12 | 0 | 5 | 0 | 10 | 0.1 | 34 | 0.0882 | 7 | 0 | 6 | 0.1667 | 11 | 0 | 10 | 0.1 | 34 | 0.0588 |
| 8_3529530 | 2 | 0 | 8 | 0 | 4 | 0.25 | 6 | 0.3333 | 20 | 0.15 | 3 | 0 | 6 | 0 | 6 | 0 | 5 | 0.2 | 20 | 0.05 |
| 8_3529687 | 6 | 0 | 6 | 0 | 3 | 0 | 4 | 0.25 | 19 | 0.0526 | 3 | 0 | 5 | 0.2 | 3 | 0 | 5 | 0 | 16 | 0.0625 |
| 8_3529827 | 4 | 0 | 6 | 0.1667 | 2 | 0 | 4 | 0.25 | 16 | 0.125 | 2 | 0 | 5 | 0.2 | 2 | 0.5 | 3 | 0 | 12 | 0.1667 |
| 8_3529842 | 3 | 0.6667 | 6 | 0.1667 | 2 | 0 | 4 | 0.25 | 15 | 0.2667 | 2 | 0 | 5 | 0.2 | 3 | 0 | 2 | 0 | 12 | 0.0833 |
| 8_3529857 | 3 | 0 | 5 | 0.4 | 1 | 0 | 4 | 0.25 | 13 | 0.2308 | 2 | 0.5 | 4 | 0.25 | 2 | 0 | 3 | 0 | 11 | 0.1818 |
| 8_3529865 | 3 | 0.3333 | 3 | 0 | 2 | 0 | 3 | 0.6667 | 11 | 0.2727 | 2 | 0 | 3 | 0.3333 | 2 | 0.5 | 3 | 0 | 10 | 0.2 |
| 8_3593606 | 4 | 0 | 1 | 0 | 4 | 0.25 | 1 | 1 | 10 | 0.2 | 1 | 0 | NA | NA | 4 | 0.25 | 2 | 0 | 7 | 0.1429 |
| 8_3593628 | 4 | 0.75 | 2 | 0 | 4 | 0.5 | 1 | 0 | 11 | 0.4545 | 1 | 0 | NA | NA | 4 | 0.5 | 2 | 0.5 | 7 | 0.4286 |
| 8_3593638 | 2 | 0.5 | 1 | 0 | 2 | 0.5 | 1 | 0 | 6 | 0.3333 | 1 | 1 | NA | NA | 3 | 0.6667 | 2 | 0 | 6 | 0.5 |
| 8_3593643 | 3 | 0.6667 | NA | NA | 2 | 0.5 | 1 | 0 | 6 | 0.5 | 1 | 1 | NA | NA | 3 | 0.3333 | 1 | 0 | 5 | 0.4 |
| 8_3605538 | NA | NA | NA | NA | 1 | 1 | NA | NA | 1 | 1 | NA | NA | NA | NA | NA | NA | 1 | 1 | 1 | 1 |
| 8_3605570 | NA | NA | NA | NA | 1 | 1 | NA | NA | 1 | 1 | NA | NA | NA | NA | NA | NA | 2 | 1 | 2 | 1 |
| 8_3605663 | NA | NA | 1 | 1 | 1 | 0 | NA | NA | 2 | 0.5 | NA | NA | NA | NA | NA | NA | 2 | 1 | 2 | 1 |
| 8_3605703 | NA | NA | 1 | 1 | 1 | 1 | NA | NA | 2 | 1 | NA | NA | NA | NA | NA | NA | 3 | 1 | 3 | 1 |
| 8_3605713 | NA | NA | 1 | 1 | 1 | 1 | NA | NA | 2 | 1 | NA | NA | NA | NA | NA | NA | 3 | 1 | 3 | 1 |
| 8_3605717 | NA | NA | 1 | 1 | 1 | 1 | 1 | 1 | 3 | 1 | NA | NA | NA | NA | NA | NA | 2 | 1 | 2 | 1 |
| 8_3605726 | NA | NA | 1 | 0 | 1 | 1 | 1 | 1 | 3 | 0.6667 | NA | NA | NA | NA | NA | NA | 2 | 1 | 2 | 1 |
| 8_3605731 | NA | NA | NA | NA | 1 | 1 | NA | NA | 1 | 1 | NA | NA | NA | NA | NA | NA | 2 | 0.5 | 2 | 0.5 |
| 8_3605751 | 1 | 1 | 1 | 1 | 1 | 0 | 1 | 0 | 4 | 0.5 | NA | NA | 1 | 1 | NA | NA | 3 | 1 | 4 | 1 |
| 8_3605756 | 1 | 1 | 1 | 1 | 1 | 1 | 1 | 1 | 4 | 1 | NA | NA | 1 | 1 | NA | NA | 2 | 1 | 3 | 1 |
| 8_3605760 | 1 | 1 | 1 | 1 | 1 | 1 | 1 | 0 | 4 | 0.75 | NA | NA | 1 | 0 | NA | NA | 3 | 0.3333 | 4 | 0.25 |
| 8_3605760 | 1 | 0 | 1 | 0 | 1 | 0 | 1 | 1 | 4 | 0.25 | NA | NA | 1 | 0 | NA | NA | 3 | 0.3333 | 4 | 0.25 |
| 8_3605776 | 1 | 0 | 2 | 1 | 1 | 1 | 1 | 1 | 5 | 0.8 | NA | NA | NA | NA | NA | NA | 1 | 1 | 1 | 1 |
| 8_3605778 | NA | NA | 2 | 1 | NA | NA | 1 | 1 | 3 | 1 | NA | NA | 1 | 1 | NA | NA | 2 | 1 | 3 | 1 |
| 8_3605796 | 2 | 1 | 2 | 1 | 1 | 1 | 1 | 1 | 6 | 1 | NA | NA | NA | NA | 1 | 0 | 2 | 0.5 | 3 | 0.3333 |
| 8_3605799 | 1 | 1 | 2 | 1 | 2 | 1 | 1 | 1 | 6 | 1 | NA | NA | 1 | 1 | 1 | 1 | 2 | 1 | 4 | 1 |
| 8_3605814 | 2 | 0 | 2 | 0.5 | 2 | 1 | 1 | 0 | 7 | 0.4286 | NA | NA | 1 | 1 | 1 | 1 | 3 | 0.6667 | 5 | 0.8 |
| 8_3605830 | 2 | 0.5 | 2 | 0 | 1 | 0 | 2 | 0 | 7 | 0.1429 | NA | NA | 1 | 0 | 1 | 0 | 3 | 0.3333 | 5 | 0.2 |
| 8_3605837 | 2 | 0.5 | 2 | 1 | 1 | 1 | 2 | 1 | 7 | 0.8571 | NA | NA | 1 | 1 | 1 | 1 | 2 | 0 | 4 | 0.5 |

| | | | | | | | | | | | | | | | | | | | | |
|---|---|---|---|---|---|---|---|---|---|---|---|---|---|---|---|---|---|---|---|---|
| 8_36058409 | 2 | 1 | 2 | 1 | 1 | 1 | 2 | 0.5 | 7 | 0.8571 | NA | NA | 1 | 1 | 1 | 0 | 4 | 0.75 | 6 | 0.6667 |
| 8_36058439 | 2 | 1 | 2 | 0.5 | 1 | 1 | 1 | 1 | 6 | 0.8333 | NA | NA | 1 | 1 | NA | NA | 3 | 1 | 4 | 1 |
| 8_36143963 | 1 | 1 | 4 | 0 | 1 | 1 | 2 | 0 | 8 | 0.25 | 1 | 0 | 1 | 0 | 2 | 1 | 4 | 0.25 | 8 | 0.375 |
| 8_36143973 | 2 | 0 | 4 | 0.25 | 2 | 0.5 | 2 | 0.5 | 10 | 0.3 | 1 | 0 | NA | NA | 2 | 0 | 4 | 0.25 | 7 | 0.1429 |
| 8_36144069 | 2 | 1 | 4 | 1 | 2 | 1 | 2 | 1 | 10 | 1 | 1 | 1 | 2 | 1 | 2 | 1 | 4 | 1 | 9 | 1 |
| 8_36144145 | 2 | 1 | 5 | 1 | 1 | 1 | 1 | 0 | 9 | 0.8889 | 1 | 1 | 2 | 1 | 1 | 1 | 3 | 1 | 7 | 1 |
| 8_36144595 | 2 | 0 | 5 | 0 | 1 | 1 | 3 | 0 | 11 | 0.0909 | 3 | 0.3333 | 3 | 0 | 2 | 0 | 3 | 0 | 11 | 0.0909 |
| 8_36145179 | 3 | 0.6667 | 5 | 0.4 | 2 | 1 | 3 | 0.3333 | 13 | 0.5385 | 3 | 1 | 3 | 0.3333 | 2 | 0 | 4 | 0.75 | 12 | 0.5833 |
| 8_36146055 | 2 | 0 | 4 | 0 | 2 | 0.5 | 5 | 0.2 | 13 | 0.1538 | 2 | 0 | 4 | 0.25 | 2 | 0 | 4 | 0.25 | 12 | 0.1667 |
| 8_36146377 | 2 | 0 | 6 | 0.1667 | 4 | 0 | 3 | 0 | 15 | 0.0667 | 2 | 0.5 | 3 | 0 | 2 | 0 | 3 | 0 | 10 | 0.1 |
| 8_36146418 | 3 | 0.6667 | 8 | 0.625 | 4 | 0.5 | 5 | 0.8 | 20 | 0.65 | 2 | 1 | 3 | 0.3333 | 2 | 0.5 | 4 | 1 | 11 | 0.7273 |
| 8_36570691 | NA | NA | NA | NA | 1 | 1 | NA | NA | 1 | 1 | NA | NA | NA | NA | 2 | 1 | NA | NA | 2 | 1 |
| 8_36570958 | NA | NA | NA | NA | 1 | 1 | NA | NA | 1 | 1 | NA | NA | NA | NA | 2 | 1 | NA | NA | 2 | 1 |
| 8_36606782 | NA | NA | NA | NA | 2 | 0.5 | NA | NA | 2 | 0.5 | NA | NA | NA | NA | 2 | 0.5 | NA | NA | 2 | 0.5 |
| 8_36606794 | NA | NA | NA | NA | 2 | 1 | NA | NA | 2 | 1 | NA | NA | NA | NA | 1 | 1 | NA | NA | 1 | 1 |
| 8_36607137 | 1 | 1 | NA | NA | 2 | 1 | NA | NA | 3 | 1 | NA | NA | NA | NA | 3 | 0.6667 | NA | NA | 3 | 0.6667 |
| 8_36613532 | 1 | 1 | NA | NA | 1 | 1 | NA | NA | 2 | 1 | NA | NA | NA | NA | 3 | 1 | NA | NA | 3 | 1 |
| 8_36613598 | 2 | 0.5 | NA | NA | 2 | 0.5 | NA | NA | 4 | 0.5 | NA | NA | NA | NA | 3 | 0.3333 | NA | NA | 3 | 0.3333 |
| 8_36613643 | 3 | 0.6667 | NA | NA | 2 | 1 | NA | NA | 5 | 0.8 | NA | NA | NA | NA | 3 | 0.3333 | 1 | 1 | 4 | 0.5 |
| 8_36613681 | 2 | 1 | NA | NA | 2 | 1 | NA | NA | 4 | 1 | NA | NA | NA | NA | 3 | 1 | 1 | 1 | 4 | 1 |
| 8_36613703 | 2 | 1 | 1 | 1 | 2 | 1 | NA | NA | 5 | 1 | NA | NA | NA | NA | 3 | 1 | 1 | 0 | 4 | 0.75 |
| 8_36613820 | 3 | 0.3333 | 1 | 0 | 2 | 0.5 | NA | NA | 6 | 0.3333 | NA | NA | NA | NA | 3 | 0.6667 | 1 | 1 | 4 | 0.75 |
| 8_36614207 | 2 | 1 | 1 | 1 | 2 | 1 | NA | NA | 5 | 1 | 1 | 1 | NA | NA | 3 | 0.6667 | 1 | 1 | 5 | 0.8 |
| 8_36614465 | 1 | 1 | 1 | 1 | 2 | 1 | NA | NA | 4 | 1 | NA | NA | NA | NA | 4 | 0.5 | NA | NA | 4 | 0.5 |
| 8_36614484 | 2 | 1 | 1 | 1 | 2 | 0.5 | NA | NA | 5 | 0.8 | NA | NA | NA | NA | 2 | 1 | 1 | 1 | 3 | 1 |
| 8_36614492 | NA | NA | NA | NA | 1 | 1 | NA | NA | 1 | 1 | 1 | 1 | NA | NA | 4 | 0.75 | 1 | 1 | 6 | 0.8333 |
| 8_36614657 | 1 | 1 | 1 | 1 | 2 | 1 | NA | NA | 4 | 1 | NA | NA | NA | NA | 3 | 1 | 1 | 1 | 4 | 1 |
| 8_36614709 | 2 | 1 | 1 | 1 | 2 | 1 | 1 | 1 | 6 | 1 | 1 | 1 | NA | NA | 2 | 1 | 1 | 1 | 4 | 1 |
| 8_36614848 | 3 | 0.6667 | 1 | 1 | 4 | 1 | 1 | 1 | 9 | 0.8889 | 1 | 1 | NA | NA | 4 | 1 | 2 | 1 | 7 | 1 |
| 8_36614896 | 3 | 0.3333 | NA | NA | 4 | 0 | 1 | 0 | 8 | 0.125 | NA | NA | NA | NA | 4 | 0.5 | 2 | 0.5 | 6 | 0.5 |
| 8_36683316 | 7 | 0 | 3 | 0.3333 | 4 | 0 | 3 | 0 | 17 | 0.0588 | 4 | 0 | 4 | 0 | 3 | 0 | 4 | 0.25 | 15 | 0.0667 |
| 8_36683464 | 7 | 0 | 4 | 0.25 | 4 | 0 | 1 | 1 | 16 | 0.125 | 4 | 0.25 | 5 | 0.2 | 3 | 0 | 3 | 0.3333 | 15 | 0.2 |
| 8_36683672 | 5 | 0 | 3 | 0 | 5 | 0.2 | 2 | 0 | 15 | 0.0667 | 1 | 0 | 2 | 0 | 2 | 0 | 4 | 0.5 | 9 | 0.2222 |
| 8_36683812 | 6 | 0 | 3 | 0 | 5 | 0.4 | 3 | 0 | 17 | 0.1176 | 3 | 0 | 3 | 0 | 2 | 0 | 4 | 0.25 | 12 | 0.0833 |
| 8_36683828 | 5 | 0.8 | 2 | 1 | 5 | 0.6 | 2 | 0.5 | 14 | 0.7143 | 2 | 1 | 3 | 0 | 2 | 1 | 3 | 0.6667 | 10 | 0.6 |
| 8_36683878 | 6 | 0.1667 | 3 | 0 | 5 | 0 | 3 | 0 | 17 | 0.0588 | 3 | 0 | 3 | 0 | 2 | 0 | 4 | 0.25 | 12 | 0.0833 |
| 8_36683991 | 5 | 0.2 | 3 | 0.3333 | 3 | 0.3333 | 3 | 0.3333 | 14 | 0.2857 | 3 | 0.6667 | 2 | 0 | 2 | 0.5 | 4 | 0.5 | 11 | 0.4545 |
| 8_36684021 | 6 | 0.1667 | 3 | 0 | 5 | 0 | 3 | 0 | 17 | 0.0588 | 3 | 0 | 3 | 0 | 2 | 0 | 4 | 0.25 | 12 | 0.0833 |
| 8_36684280 | 6 | 0.1667 | NA | NA | 5 | 0.2 | 3 | 0 | 14 | 0.1429 | 3 | 0.3333 | NA | NA | NA | NA | 2 | 0 | 5 | 0.2 |
| 8_36684362 | 4 | 1 | 2 | 1 | 4 | 1 | 3 | 1 | 13 | 1 | 2 | 1 | 1 | 1 | 2 | 1 | 2 | 1 | 7 | 1 |
| 8_36684423 | 6 | 0.1667 | 2 | 0 | 3 | 0 | 2 | 0 | 13 | 0.0769 | 3 | 0 | NA | NA | 2 | 0 | 2 | 0.5 | 7 | 0.1429 |
| 8_36684440 | 5 | 0.8 | 1 | 0 | 4 | 0.25 | 2 | 1 | 12 | 0.5833 | 3 | 1 | 1 | 0 | 2 | 0.5 | 2 | 1 | 8 | 0.75 |
| 8_36684531 | 4 | 0.25 | 2 | 0.5 | 3 | 0.6667 | 3 | 0.6667 | 12 | 0.5 | 3 | 0.6667 | 1 | 1 | 1 | 0 | 1 | 1 | 6 | 0.6667 |
| 8_36684593 | 5 | 1 | 1 | 1 | 3 | 0.6667 | 2 | 1 | 11 | 0.9091 | 1 | 1 | NA | NA | 1 | 1 | 2 | 1 | 4 | 1 |
| 8_36684610 | 6 | 0.6667 | 2 | 1 | 4 | 0.75 | 2 | 1 | 14 | 0.7857 | 3 | 1 | 1 | 1 | 2 | 1 | 2 | 1 | 8 | 1 |
| 8_36716140 | 2 | 0.5 | 1 | 1 | 3 | 0.6667 | NA | NA | 6 | 0.6667 | 1 | 1 | NA | NA | NA | NA | 1 | 1 | 2 | 1 |
| 8_36978857 | 1 | 1 | 1 | 0 | 1 | 0 | 2 | 0 | 5 | 0.2 | 2 | 0 | 2 | 0 | NA | NA | 1 | 1 | 5 | 0.2 |
| 8_36979039 | NA | NA | 1 | 1 | 1 | 1 | 2 | 1 | 4 | 1 | 3 | 1 | 1 | 1 | NA | NA | 1 | 0 | 5 | 0.8 |
| 8_36979270 | 1 | 0 | 1 | 0 | 1 | 0 | 2 | 0.5 | 5 | 0.2 | 3 | 0 | 1 | 0 | NA | NA | 1 | 1 | 5 | 0.2 |
| 8_36979304 | 1 | 1 | 1 | 1 | NA | NA | 2 | 1 | 4 | 1 | NA | NA | 3 | 1 | NA | NA | 1 | 1 | 4 | 1 |
| 8_36979346 | 2 | 1 | 1 | 1 | NA | NA | 2 | 1 | 5 | 1 | 3 | 1 | 2 | 1 | NA | NA | 1 | 1 | 6 | 1 |
| 8_36979394 | 2 | 1 | 1 | 1 | 1 | 1 | 2 | 1 | 6 | 1 | 2 | 1 | 1 | 1 | NA | NA | NA | NA | 3 | 1 |
| 8_36979441 | 2 | 0.5 | 1 | 1 | 1 | 0 | 1 | 1 | 5 | 0.6 | 2 | 0.5 | 2 | 1 | NA | NA | 1 | 0 | 5 | 0.6 |
| 8_36979459 | 1 | 1 | 1 | 1 | 1 | 1 | 2 | 1 | 5 | 1 | 3 | 1 | 2 | 1 | NA | NA | NA | NA | 5 | 1 |
| 8_36979693 | 1 | 0 | 1 | 0 | 1 | 1 | 2 | 0 | 5 | 0.2 | 3 | 0 | 3 | 0.3333 | NA | NA | NA | NA | 6 | 0.1667 |
| 8_36979702 | 1 | 1 | NA | NA | 1 | 1 | 1 | 1 | 3 | 1 | 1 | 0 | 2 | 0.5 | NA | NA | NA | NA | 3 | 0.3333 |
| 8_36979874 | 2 | 0.5 | 1 | 0 | NA | NA | 2 | 0 | 5 | 0.2 | 4 | 0.25 | 3 | 0 | NA | NA | 1 | 1 | 8 | 0.25 |
| 8_36979982 | 2 | 1 | 1 | 1 | 1 | 1 | 2 | 1 | 6 | 1 | 3 | 1 | 4 | 1 | NA | NA | 1 | 1 | 8 | 1 |
| 8_36979987 | 2 | 1 | 1 | 1 | NA | NA | 2 | 1 | 5 | 1 | 2 | 1 | 4 | 0.75 | NA | NA | 2 | 1 | 8 | 0.875 |
| 8_37035073 | 1 | 1 | 2 | 1 | 3 | 1 | 1 | 1 | 7 | 1 | 1 | 1 | NA | NA | NA | NA | 3 | 1 | 4 | 1 |
| 8_37035084 | 1 | 1 | 2 | 0.5 | 2 | 0.5 | 2 | 1 | 7 | 0.7143 | 1 | 1 | NA | NA | 1 | 0 | 3 | 0.3333 | 5 | 0.4 |
| 8_37035142 | 1 | 0 | 3 | 1 | 3 | 0.3333 | 1 | 1 | 8 | 0.625 | 1 | 1 | NA | NA | 2 | 1 | 3 | 0.3333 | 6 | 0.6667 |
| 8_37035459 | 1 | 0 | 2 | 0 | 2 | 0.5 | 2 | 0 | 7 | 0.1429 | 1 | 0 | 1 | 0 | 2 | 0 | 3 | 0.6667 | 7 | 0.2857 |
| 8_37035963 | NA | NA | 3 | 0.3333 | 2 | 0 | 3 | 0 | 8 | 0.125 | 1 | 0 | 3 | 0.3333 | NA | NA | 3 | 0.3333 | 7 | 0.2857 |
| 8_37037064 | NA | NA | 2 | 0 | 1 | 1 | 3 | 0 | 6 | 0.1667 | 2 | 0 | 2 | 0 | NA | NA | 3 | 0.3333 | 7 | 0.1429 |

| | | | | | | | | | | | | | | | | | | | | |
|---|---|---|---|---|---|---|---|---|---|---|---|---|---|---|---|---|---|---|---|---|
| 8_37037196 | NA | NA | 3 | 0.6667 | 1 | 0 | 2 | 0.5 | 6 | 0.5 | 1 | 0 | 1 | 1 | NA | NA | 3 | 0.6667 | 5 | 0.6 |
| 8_37037273 | NA | NA | 1 | 1 | 1 | 0 | 1 | 1 | 3 | 0.6667 | 2 | 1 | 1 | 1 | NA | NA | 3 | 0.3333 | 6 | 0.6667 |
| 8_37037340 | NA | NA | 2 | 1 | 1 | 1 | 1 | 0 | 4 | 0.75 | 2 | 1 | 1 | 1 | NA | NA | 2 | 1 | 5 | 1 |
| 8_37037453 | NA | NA | 2 | 0 | 1 | 1 | NA | NA | 3 | 0.3333 | 1 | 1 | 1 | 0 | NA | NA | 2 | 0.5 | 4 | 0.5 |
| 8_37037521 | NA | NA | 2 | 0.5 | 1 | 0 | 1 | 1 | 4 | 0.5 | 2 | 1 | 1 | 1 | NA | NA | NA | NA | 3 | 1 |
| 8_37041203 | NA | NA | 1 | 1 | 1 | 1 | 1 | 0 | 3 | 0.6667 | 2 | 0 | 1 | 0 | NA | NA | 2 | 0.5 | 5 | 0.2 |
| 8_37050872 | NA | NA | NA | NA | 1 | 0 | 1 | 1 | 2 | 0.5 | 1 | 1 | NA | NA | NA | NA | 1 | 1 | 2 | 1 |
| 8_37050884 | NA | NA | 1 | 0 | 1 | 1 | 1 | 0 | 3 | 0.3333 | 1 | 1 | NA | NA | NA | NA | 1 | 0 | 2 | 0.5 |
| 8_37050983 | NA | NA | 1 | 1 | 1 | 1 | 1 | 0 | 3 | 0.6667 | 1 | 1 | NA | NA | NA | NA | 1 | 0 | 2 | 0.5 |
| 8_37051175 | NA | NA | NA | NA | 1 | 1 | 1 | 0 | 2 | 0.5 | NA | NA | NA | NA | NA | NA | 1 | 1 | 1 | 1 |
| 8_37051547 | NA | NA | 1 | 1 | NA | NA | 1 | 0 | 2 | 0.5 | 1 | 1 | NA | NA | NA | NA | NA | NA | 1 | 1 |
| 8_37051801 | NA | NA | NA | NA | NA | NA | 1 | 1 | 1 | 1 | 1 | 1 | NA | NA | NA | NA | NA | NA | 1 | 1 |
| 8_39138288 | NA | NA | 1 | 1 | NA | NA | NA | NA | 1 | 1 | 1 | 1 | NA | NA | 1 | 0 | NA | NA | 2 | 0.5 |
| 8_39618085 | 46 | 0.1739 | 39 | 0.2051 | 67 | 0.194 | 56 | 0.2143 | 208 | 0.1971 | 35 | 0.1714 | 56 | 0.0714 | 63 | 0.1429 | 51 | 0.3137 | 205 | 0.1707 |
| 8_40760587 | 3 | 1 | 1 | 0 | NA | NA | 2 | 1 | 6 | 0.8333 | NA | NA | 2 | 1 | 4 | 1 | NA | NA | 6 | 1 |
| 8_40760609 | 3 | 0.6667 | 1 | 1 | NA | NA | 2 | 1 | 6 | 0.8333 | NA | NA | 2 | 1 | 3 | 1 | NA | NA | 5 | 1 |
| 8_40760809 | 3 | 0.6667 | 1 | 1 | NA | NA | 1 | 0 | 5 | 0.6 | NA | NA | 1 | 0 | 3 | 0.3333 | NA | NA | 4 | 0.25 |
| 8_40837821 | 10 | 0.1 | 12 | 0.1667 | 10 | 0.2 | 15 | 0.3333 | 47 | 0.2128 | 4 | 0.25 | 11 | 0.1818 | 4 | 0.5 | 11 | 0.1818 | 30 | 0.2333 |
| 8_40837879 | 10 | 0.2 | 13 | 0.0769 | 8 | 0.25 | 19 | 0.2105 | 50 | 0.18 | 6 | 0.3333 | 12 | 0.25 | 9 | 0 | 12 | 0.1667 | 39 | 0.1795 |
| 8_40837935 | 10 | 0.3 | 15 | 0.3333 | 9 | 0.5556 | 17 | 0.3529 | 51 | 0.3725 | 4 | 0.5 | 12 | 0.0833 | 10 | 0.7 | 11 | 0.2727 | 37 | 0.3514 |
| 8_40837970 | 10 | 0.8 | 12 | 0.9167 | 10 | 0.6 | 16 | 0.8125 | 48 | 0.7917 | 6 | 0.8333 | 9 | 0.8889 | 8 | 0.625 | 10 | 0.6 | 33 | 0.7273 |
| 8_40936915 | 5 | 0 | 1 | 0 | 5 | 0.2 | 5 | 0 | 16 | 0.0625 | 4 | 0.25 | 4 | 0 | 9 | 0 | 4 | 0.25 | 21 | 0.0952 |
| 8_40936938 | 4 | 0 | 2 | 0.5 | 4 | 0 | 6 | 0 | 16 | 0.0625 | 4 | 0 | 5 | 0.4 | 7 | 0 | 3 | 0 | 19 | 0.1053 |
| 8_40937247 | 5 | 0.2 | 3 | 0 | 6 | 0.1667 | 6 | 0.1667 | 20 | 0.15 | 5 | 0.4 | 4 | 0.5 | 7 | 0.1429 | 4 | 0 | 20 | 0.25 |
| 8_40937390 | 6 | 0.3333 | 2 | 0.5 | 6 | 0.3333 | 6 | 0.1667 | 20 | 0.3 | 6 | 0.1667 | 5 | 0 | 10 | 0.2 | 5 | 0.2 | 26 | 0.1538 |
| 8_40937426 | 5 | 0.2 | 3 | 0.3333 | 6 | 0.1667 | 6 | 0.3333 | 20 | 0.25 | 6 | 0 | 5 | 0 | 11 | 0.1818 | 6 | 0 | 28 | 0.0714 |
| 8_40937585 | 6 | 0.1667 | 3 | 0.6667 | 7 | 0.1429 | 7 | 0.2857 | 23 | 0.2609 | 6 | 0.3333 | 5 | 0.2 | 11 | 0.0909 | 7 | 0 | 29 | 0.1379 |
| 8_40938231 | 4 | 0 | 4 | 0 | 8 | 0.25 | 7 | 0.2857 | 23 | 0.1739 | 10 | 0.3 | 7 | 0.2857 | 12 | 0.0833 | 7 | 0.2857 | 36 | 0.2222 |
| 8_40945797 | 26 | 0.0769 | 19 | 0.3684 | 14 | 0.2857 | 26 | 0.2692 | 85 | 0.2353 | 14 | 0.1429 | 21 | 0.2857 | 20 | 0.1 | 13 | 0.2308 | 68 | 0.1912 |
| 8_40945890 | 24 | 0.1667 | 18 | 0.2778 | 12 | 0.4167 | 27 | 0.2593 | 81 | 0.2593 | 13 | 0.1538 | 20 | 0.1 | 20 | 0.15 | 12 | 0.3333 | 65 | 0.1692 |
| 8_40954263 | 2 | 0 | 1 | 0 | NA | NA | 2 | 0.5 | 5 | 0.2 | 2 | 0 | 3 | 0 | 2 | 1 | 4 | 0.25 | 11 | 0.2727 |
| 8_40955092 | 1 | 0 | 1 | 0 | NA | NA | 2 | 0.5 | 4 | 0.25 | 1 | 0 | 2 | 0.5 | 2 | 0 | 3 | 0 | 8 | 0.125 |
| 8_40960573 | 12 | 0.4167 | 15 | 0.2 | 8 | 0.25 | 18 | 0.5 | 53 | 0.3585 | 7 | 0.2857 | 14 | 0.5 | 12 | 0.75 | 6 | 0.1667 | 39 | 0.4872 |
| 8_40960598 | 13 | 0.0769 | 14 | 0.2857 | 7 | 0.1429 | 20 | 0.2 | 54 | 0.1852 | 8 | 0.125 | 15 | 0.2667 | 13 | 0.0769 | 7 | 0.1429 | 43 | 0.1628 |
| 8_40999355 | NA | NA | 5 | 0 | NA | NA | 5 | 0.6 | 10 | 0.3 | 3 | 0 | 4 | 0 | 3 | 0.3333 | 6 | 0.1667 | 16 | 0.125 |
| 8_40999388 | NA | NA | 5 | 0.4 | NA | NA | 3 | 0.6667 | 8 | 0.5 | 3 | 0 | 4 | 0.25 | 3 | 0 | 4 | 0.5 | 14 | 0.2143 |
| 8_40999455 | NA | NA | 4 | 0.5 | NA | NA | 4 | 0.25 | 8 | 0.375 | 3 | 0 | 4 | 0.25 | 3 | 0.3333 | 5 | 0.2 | 15 | 0.2 |
| 8_40999479 | NA | NA | 5 | 0.2 | NA | NA | 5 | 0.4 | 10 | 0.3 | 3 | 0.3333 | 4 | 0 | 3 | 0.3333 | 6 | 0.1667 | 16 | 0.1875 |
| 8_41003379 | 1 | 0 | 4 | 0.75 | 2 | 1 | 1 | 0 | 8 | 0.625 | 2 | 0.5 | 6 | 0.5 | 8 | 0.875 | 3 | 0.3333 | 19 | 0.6316 |
| 8_41004209 | 8 | 0.625 | 14 | 0.4286 | 11 | 0.4545 | 10 | 0.5 | 43 | 0.4884 | 8 | 0.375 | 11 | 0.5455 | 11 | 0.5455 | 10 | 0.2 | 40 | 0.425 |
| 8_41004247 | 7 | 1 | 14 | 0.7857 | 10 | 0.9 | 9 | 0.8889 | 40 | 0.875 | 8 | 1 | 9 | 1 | 13 | 0.9231 | 10 | 1 | 40 | 0.975 |
| 8_41004319 | 8 | 0 | 13 | 0.2308 | 11 | 0.2727 | 10 | 0.1 | 42 | 0.1667 | 7 | 0.2857 | 9 | 0.2222 | 12 | 0 | 10 | 0.2 | 38 | 0.1579 |
| 8_41004376 | 8 | 0.25 | 13 | 0.2308 | 9 | 0.4444 | 10 | 0.4 | 40 | 0.325 | 7 | 0.4286 | 11 | 0.4545 | 11 | 0.5455 | 10 | 0.5 | 39 | 0.4872 |
| 8_41004406 | 7 | 0.4286 | 12 | 0.75 | 8 | 0.5 | 8 | 0.625 | 35 | 0.6 | 4 | 0.5 | 10 | 0.7 | 9 | 0.4444 | 8 | 0.75 | 31 | 0.6129 |
| 8_41004486 | 8 | 0 | 10 | 0.1 | 8 | 0.125 | 9 | 0.1111 | 35 | 0.0857 | 6 | 0.3333 | 7 | 0 | 9 | 0 | 8 | 0.25 | 30 | 0.1333 |
| 8_41004555 | 6 | 0.1667 | 10 | 0.5 | 7 | 0.5714 | 10 | 0.1 | 33 | 0.3333 | 6 | 0.5 | 8 | 0.625 | 13 | 0.6154 | 8 | 0.25 | 35 | 0.5143 |
| 8_41004563 | 7 | 0.5714 | 10 | 0.6 | 8 | 0.75 | 7 | 0.8571 | 32 | 0.6875 | 4 | 1 | 9 | 1 | 12 | 0.8333 | 3 | 1 | 28 | 0.9286 |
| 8_41004623 | 6 | 0.8333 | 11 | 0.6364 | 9 | 0.6667 | 9 | 0.7778 | 35 | 0.7143 | 6 | 0.8333 | 10 | 0.9 | 9 | 0.7778 | 8 | 0.75 | 33 | 0.8182 |
| 8_41004629 | 6 | 0.1667 | 11 | 0 | 9 | 0.2222 | 10 | 0 | 36 | 0.0833 | 7 | 0 | 8 | 0.375 | 9 | 0.4444 | 8 | 0.125 | 32 | 0.25 |
| 8_41004752 | 7 | 0.5714 | 9 | 0.6667 | 9 | 0.4444 | 10 | 0.3 | 35 | 0.4857 | 5 | 0.4 | 8 | 0.875 | 9 | 0.6667 | 6 | 0.8333 | 28 | 0.7143 |
| 8_41305904 | NA | NA | NA | NA | NA | NA | 1 | 1 | 1 | 1 | NA | NA | NA | NA | 3 | 0.3333 | NA | NA | 3 | 0.3333 |
| 8_41306452 | NA | NA | 1 | 1 | NA | NA | 1 | 1 | 2 | 1 | NA | NA | NA | NA | 3 | 1 | NA | NA | 3 | 1 |
| 8_41306832 | NA | NA | 1 | 1 | NA | NA | NA | NA | 1 | 1 | NA | NA | NA | NA | 3 | 1 | NA | NA | 3 | 1 |
| 8_41307067 | NA | NA | 1 | 1 | NA | NA | NA | NA | 1 | 1 | NA | NA | NA | NA | 3 | 0.3333 | NA | NA | 3 | 0.3333 |
| 8_41307497 | NA | NA | 1 | 1 | NA | NA | 1 | 1 | 2 | 1 | NA | NA | NA | NA | 3 | 0.3333 | NA | NA | 3 | 0.3333 |
| 8_41307985 | NA | NA | 1 | 1 | NA | NA | NA | NA | 1 | 1 | NA | NA | NA | NA | 3 | 0.6667 | NA | NA | 3 | 0.6667 |
| 8_41308031 | NA | NA | 1 | 1 | NA | NA | NA | NA | 1 | 1 | NA | NA | NA | NA | 2 | 1 | NA | NA | 2 | 1 |
| 8_41369665 | NA | NA | 1 | 1 | 1 | 1 | NA | NA | 2 | 1 | 1 | 1 | 1 | 1 | 1 | 1 | 1 | 1 | 4 | 1 |
| 8_41374363 | 2 | 0.5 | 3 | 0.3333 | 1 | 1 | 4 | 0 | 10 | 0.3 | 3 | 0.6667 | 3 | 0.6667 | 2 | 0.5 | 4 | 0.75 | 12 | 0.6667 |
| 8_41374581 | 2 | 0 | 4 | 0.25 | 1 | 0 | 4 | 0 | 11 | 0.0909 | 6 | 0.1667 | 6 | 0 | 4 | 0 | 4 | 0.25 | 20 | 0.1 |
| 8_41374877 | 2 | 0.5 | 4 | 0.25 | 1 | 0 | 4 | 0 | 11 | 0.1818 | 4 | 0 | 3 | 0.3333 | 4 | 0 | 4 | 0 | 15 | 0.0667 |
| 8_41374995 | 1 | 0 | 4 | 0.25 | NA | NA | 3 | 0.3333 | 8 | 0.25 | 3 | 0 | 2 | 0 | 2 | 0 | 3 | 0.3333 | 10 | 0.1 |
| 8_41445793 | 7 | 0.1429 | 6 | 0.1667 | 4 | 0.25 | 10 | 0.1 | 27 | 0.1481 | 2 | 0 | 5 | 0 | 7 | 0.1429 | 7 | 0.2857 | 21 | 0.1429 |
| 8_41446058 | 5 | 0.2 | 4 | 0.75 | 4 | 0 | 10 | 0.8 | 23 | 0.5217 | 1 | 0 | 2 | 1 | 6 | 0.6667 | 9 | 0.6667 | 18 | 0.6667 |
| 8_41446088 | 6 | 0.1667 | 5 | 0.4 | 5 | 0.2 | 13 | 0.2308 | 29 | 0.2414 | 2 | 0 | 3 | 0.3333 | 6 | 0 | 10 | 0.2 | 21 | 0.1429 |

| | | | | | | | | | | | | | | | | | | | | |
|---|---|---|---|---|---|---|---|---|---|---|---|---|---|---|---|---|---|---|---|---|
| 8_4144614 | 5 | 0.2 | 4 | 0 | 4 | 0 | 11 | 0.0909 | 24 | 0.0833 | 1 | 0 | 3 | 0.3333 | 6 | 0.1667 | 8 | 0.25 | 18 | 0.2222 |
| 8_4144618 | 5 | 0.2 | 6 | 0.1667 | 4 | 0 | 14 | 0 | 29 | 0.069 | 2 | 0 | 3 | 0.3333 | 6 | 0.1667 | 9 | 0.1111 | 20 | 0.15 |
| 8_4144622 | 6 | 0.6667 | 6 | 1 | 6 | 0.8333 | 13 | 0.9231 | 31 | 0.871 | 2 | 0.5 | 3 | 1 | 6 | 0.6667 | 8 | 0.875 | 19 | 0.7895 |
| 8_4144623 | 6 | 1 | 6 | 0.8333 | 4 | 1 | 13 | 1 | 29 | 0.9655 | 2 | 1 | 3 | 1 | 5 | 1 | 7 | 1 | 17 | 1 |
| 8_4144624 | 4 | 0.25 | 5 | 0 | 5 | 0 | 9 | 0.1111 | 23 | 0.087 | 2 | 0 | 3 | 0 | 6 | 0.3333 | 3 | 0.3333 | 14 | 0.2143 |
| 8_4144627 | 5 | 0.8 | 5 | 0.6 | 4 | 0.25 | 15 | 0.7333 | 29 | 0.6552 | 2 | 1 | 3 | 0.3333 | 7 | 0.4286 | 9 | 0.7778 | 21 | 0.619 |
| 8_4144637 | 5 | 0.2 | 5 | 0.2 | 4 | 0.75 | 13 | 0.2308 | 27 | 0.2963 | 1 | 0 | 2 | 0 | 5 | 0.4 | 7 | 0 | 15 | 0.1333 |
| 8_4144640 | 5 | 1 | 5 | 1 | 4 | 1 | 14 | 1 | 28 | 1 | 2 | 1 | 3 | 1 | 7 | 0.8571 | 6 | 1 | 18 | 0.9444 |
| 8_4144643 | 6 | 0.3333 | 6 | 1 | 5 | 0.4 | 13 | 0.6154 | 30 | 0.6 | 2 | 1 | 3 | 1 | 5 | 0.8 | 6 | 0.5 | 16 | 0.75 |
| 8_4144658 | 5 | 0.2 | 5 | 0 | 6 | 0 | 13 | 0.3077 | 29 | 0.1724 | 2 | 0.5 | 3 | 0.3333 | 6 | 0 | 7 | 0.1429 | 18 | 0.1667 |
| 8_4144660 | 6 | 0.6667 | 4 | 0.5 | 6 | 0.3333 | 12 | 0.4167 | 28 | 0.4643 | 2 | 0.5 | NA | NA | 4 | 0.5 | 7 | 0.2857 | 13 | 0.3846 |
| 8_4145542 | 4 | 0 | 4 | 0.25 | 5 | 0 | 12 | 0.0833 | 25 | 0.08 | 1 | 0 | 1 | 0 | 3 | 0.3333 | 6 | 0 | 11 | 0.0909 |
| 8_4149457 | 2 | 0 | 3 | 0.3333 | 1 | 1 | 9 | 0.2222 | 15 | 0.2667 | 1 | 0 | NA | NA | 3 | 0.3333 | 5 | 0.2 | 9 | 0.2222 |
| 8_4153599 | 1 | 1 | NA | NA | NA | NA | 1 | 1 | 2 | 1 | NA | NA | NA | NA | NA | NA | 1 | 1 | 1 | 1 |
| 8_4153616 | 1 | 1 | NA | NA | 1 | 0 | 1 | 1 | 3 | 0.6667 | NA | NA | NA | NA | NA | NA | 1 | 1 | 1 | 1 |
| 8_4153671 | 1 | 0 | NA | NA | 1 | 0 | 1 | 1 | 3 | 0.3333 | NA | NA | NA | NA | NA | NA | 1 | 1 | 1 | 1 |
| 8_4153704 | NA | NA | NA | NA | 1 | 1 | NA | NA | 1 | 1 | NA | NA | NA | NA | NA | NA | 1 | 1 | 1 | 1 |
| 8_4174671 | 2 | 0.5 | 1 | 1 | 4 | 0.25 | 4 | 0.5 | 11 | 0.4545 | 1 | 0 | 3 | 0 | 5 | 0.2 | 5 | 0.2 | 14 | 0.1429 |
| 8_4174673 | 2 | 0.5 | NA | NA | 5 | 0.2 | 3 | 0 | 10 | 0.2 | 1 | 0 | 2 | 0.5 | 5 | 0 | 5 | 0.2 | 13 | 0.1538 |
| 8_4174673 | 2 | 0 | 1 | 0 | 5 | 0.2 | 3 | 0 | 11 | 0.0909 | 1 | 0 | 3 | 0.3333 | 5 | 0 | 3 | 0 | 12 | 0.0833 |
| 8_4174678 | 2 | 0 | 1 | 0 | 6 | 0.3333 | 4 | 0 | 13 | 0.1538 | 1 | 0 | 3 | 0.3333 | 5 | 0 | 5 | 0 | 14 | 0.0714 |
| 8_4177492 | 2 | 0 | 2 | 0 | 7 | 0.1429 | 4 | 0 | 15 | 0.0667 | NA | NA | 4 | 0 | 5 | 0 | 7 | 0.1429 | 16 | 0.0625 |
| 8_4178189 | 3 | 0.3333 | 7 | 0.4286 | 9 | 0.4444 | 5 | 0.6 | 24 | 0.4583 | 1 | 0 | 5 | 0.2 | 7 | 0.1429 | 8 | 0.5 | 21 | 0.2857 |
| 8_4178510 | 6 | 0 | 8 | 0 | 10 | 0.3 | 5 | 0 | 29 | 0.1034 | 1 | 0 | 6 | 0 | 6 | 0.1667 | 9 | 0.1111 | 22 | 0.0909 |
| 8_4178645 | 6 | 0.1667 | 11 | 0 | 9 | 0.4444 | 6 | 0.1667 | 32 | 0.1875 | 1 | 0 | 4 | 0.25 | 5 | 0.4 | 4 | 0.25 | 14 | 0.2857 |
| 8_4178655 | 6 | 0.1667 | 13 | 0.0769 | 10 | 0 | 7 | 0 | 36 | 0.0556 | 1 | 0 | 4 | 0.25 | 6 | 0.1667 | 5 | 0 | 16 | 0.125 |
| 8_4179442 | 5 | 0 | 9 | 0.2222 | 6 | 0 | 5 | 0 | 25 | 0.08 | 7 | 0.1429 | 6 | 0 | 7 | 0.1429 | 9 | 0.2222 | 29 | 0.1379 |
| 8_4179451 | 6 | 0.5 | 9 | 0.7778 | 5 | 0.4 | 7 | 0.5714 | 27 | 0.5926 | 8 | 0.875 | 5 | 1 | 7 | 0.7143 | 8 | 1 | 28 | 0.8929 |
| 8_4180002 | 5 | 0.2 | 6 | 0.3333 | 4 | 0 | 5 | 0 | 20 | 0.15 | 6 | 0 | 3 | 0 | 5 | 0 | 8 | 0.25 | 22 | 0.0909 |
| 8_4548444 | 1 | 0 | 1 | 1 | 1 | 0 | NA | NA | 3 | 0.3333 | NA | NA | 2 | 0 | NA | NA | 1 | 1 | 3 | 0.3333 |
| 8_4548998 | 2 | 0.5 | 1 | 1 | 1 | 1 | NA | NA | 4 | 0.75 | NA | NA | 2 | 0.5 | 1 | 0 | 2 | 1 | 5 | 0.6 |
| 8_4549377 | 4 | 0.5 | 1 | 0 | 2 | 0 | 1 | 1 | 8 | 0.375 | NA | NA | 2 | 1 | 1 | 0 | 1 | 0 | 4 | 0.5 |
| 8_4549381 | 2 | 0 | 2 | 0 | 1 | 1 | 1 | 1 | 6 | 0.3333 | NA | NA | 1 | 1 | 1 | 1 | 2 | 0 | 4 | 0.5 |
| 8_4549712 | 6 | 0.8333 | 1 | 1 | 2 | 1 | 1 | 1 | 10 | 0.9 | 2 | 1 | 2 | 1 | 1 | 1 | 4 | 0.75 | 9 | 0.8889 |
| 8_4549728 | 6 | 0.5 | 3 | 1 | 3 | 1 | 2 | 1 | 14 | 0.7857 | 2 | 1 | 2 | 1 | 2 | 1 | 4 | 1 | 10 | 1 |
| 8_4549730 | 4 | 0 | 2 | 0 | 3 | 0.3333 | 1 | 0 | 10 | 0.1 | 2 | 0 | 1 | 1 | 2 | 0 | 4 | 0.25 | 9 | 0.2222 |
| 8_4549731 | 4 | 0.5 | 2 | 0.5 | 3 | 1 | 1 | 1 | 10 | 0.7 | 2 | 0.5 | NA | NA | 1 | 1 | 3 | 0.6667 | 6 | 0.6667 |
| 8_4550383 | 6 | 0.1667 | 3 | 0 | 3 | 0 | 3 | 0 | 15 | 0.0667 | 3 | 0 | 2 | 0 | 7 | 0 | 5 | 0.2 | 17 | 0.0588 |
| 8_4550390 | 5 | 0.4 | 2 | 0.5 | 1 | 0 | 4 | 0 | 12 | 0.25 | 2 | 0 | 1 | 0 | 7 | 0.4286 | 3 | 0 | 13 | 0.2308 |
| 8_4550394 | 6 | 0.8333 | 2 | 0.5 | NA | NA | 4 | 0.75 | 12 | 0.75 | 1 | 0 | 2 | 0.5 | 4 | 0.75 | 5 | 0.8 | 12 | 0.6667 |
| 8_4550396 | 7 | 0.7143 | 2 | 1 | 3 | 0.6667 | 4 | 0.5 | 16 | 0.6875 | 3 | 0.3333 | 2 | 0.5 | 6 | 0.5 | 4 | 0.5 | 15 | 0.4667 |
| 8_4550398 | 7 | 0.7143 | 3 | 1 | 1 | 1 | 3 | 0.6667 | 14 | 0.7857 | 1 | 1 | 2 | 1 | 6 | 0.8333 | 4 | 1 | 13 | 0.9231 |
| 8_4550400 | 7 | 0.4286 | 2 | 0.5 | 3 | 0 | 4 | 0.25 | 16 | 0.3125 | 2 | 0.5 | 2 | 1 | 6 | 0.5 | 4 | 0.25 | 14 | 0.5 |
| 8_4550402 | 7 | 0.7143 | 2 | 1 | 3 | 0.3333 | 4 | 0.75 | 16 | 0.6875 | 2 | 1 | 1 | 0 | 5 | 0.6 | 5 | 0.8 | 13 | 0.6923 |
| 8_4550407 | 7 | 0.7143 | 2 | 0.5 | 3 | 0.3333 | 3 | 0.6667 | 15 | 0.6 | 3 | 0.6667 | 2 | 0.5 | 6 | 0.6667 | 5 | 0.8 | 16 | 0.6875 |
| 8_4550417 | 8 | 0.25 | 3 | 0 | 2 | 0 | 4 | 0 | 17 | 0.1176 | 3 | 0 | 2 | 0 | 8 | 0.125 | 4 | 0.25 | 17 | 0.1176 |
| 8_4550422 | 3 | 0.3333 | 3 | 0 | 2 | 0.5 | 1 | 0 | 9 | 0.2222 | 3 | 0.6667 | 2 | 0 | 4 | 0.5 | 4 | 0.5 | 13 | 0.4615 |
| 8_4550425 | 9 | 0.3333 | 2 | 1 | 3 | 0.6667 | 4 | 0 | 18 | 0.3889 | 3 | 0 | 2 | 0.5 | 8 | 0.25 | 5 | 0 | 18 | 0.1667 |
| 8_4550445 | 7 | 0 | 3 | 0 | 4 | 0 | 3 | 0.3333 | 17 | 0.0588 | 2 | 0 | 3 | 0 | 7 | 0 | 4 | 0.25 | 16 | 0.0625 |
| 8_4550447 | 9 | 1 | 3 | 1 | 4 | 1 | 4 | 1 | 20 | 1 | 3 | 1 | 3 | 1 | 7 | 1 | 6 | 0.8333 | 19 | 0.9474 |
| 8_4550448 | 9 | 0.5556 | 3 | 0.3333 | 3 | 0.3333 | 2 | 0.5 | 17 | 0.4706 | 3 | 1 | 3 | 0.3333 | 7 | 0.5714 | 3 | 0.6667 | 16 | 0.625 |
| 8_4550459 | 7 | 0.2857 | 2 | 0 | 4 | 0.25 | 1 | 0 | 14 | 0.2143 | 2 | 0 | 3 | 0 | 7 | 0.1429 | 4 | 0 | 16 | 0.0625 |
| 8_4550462 | 7 | 0.8571 | 3 | 1 | 3 | 1 | 4 | 1 | 17 | 0.9412 | 1 | 0 | 3 | 1 | 6 | 1 | 6 | 0.8333 | 16 | 0.875 |
| 8_4550506 | 8 | 0.25 | 6 | 0.1667 | 3 | 0 | 5 | 0.2 | 22 | 0.1818 | 3 | 0.3333 | 3 | 0.6667 | 9 | 0.1111 | 5 | 0.2 | 20 | 0.25 |
| 8_4550510 | 7 | 0 | 5 | 0 | 3 | 0 | 5 | 0.2 | 20 | 0.05 | 3 | 0 | 4 | 0 | 7 | 0.1429 | 6 | 0 | 20 | 0.05 |
| 8_4579176 | NA | NA | 3 | 0.3333 | 1 | 0 | NA | NA | 4 | 0.25 | NA | NA | NA | NA | NA | NA | 2 | 0.5 | 2 | 0.5 |
| 8_4579231 | 2 | 1 | 2 | 0.5 | 2 | 0.5 | NA | NA | 6 | 0.6667 | NA | NA | 2 | 0.5 | 2 | 0.5 | 4 | 0.75 | 8 | 0.625 |
| 8_4585003 | NA | NA | 2 | 1 | 1 | 1 | 2 | 1 | 5 | 1 | 1 | 1 | NA | NA | 1 | 0 | 1 | 1 | 3 | 0.6667 |
| 8_4585042 | 1 | 0 | 2 | 0 | 1 | 0 | 4 | 0.25 | 8 | 0.125 | 1 | 0 | NA | NA | 1 | 1 | 1 | 0 | 3 | 0.3333 |
| 8_4585051 | 1 | 1 | 2 | 0 | 1 | 1 | 3 | 0.3333 | 7 | 0.4286 | 1 | 0 | NA | NA | 1 | 0 | 1 | 1 | 3 | 0.3333 |
| 8_4585057 | NA | NA | 1 | 1 | NA | NA | 3 | 0.6667 | 4 | 0.75 | 1 | 1 | NA | NA | NA | NA | 1 | 1 | 2 | 1 |
| 8_4585059 | NA | NA | 2 | 1 | 1 | 1 | 4 | 1 | 7 | 1 | 1 | 1 | NA | NA | 1 | 0 | 1 | 0 | 3 | 0.3333 |
| 8_4585065 | 1 | 1 | 1 | 0 | 1 | 1 | 4 | 0.25 | 7 | 0.4286 | 1 | 1 | NA | NA | 1 | 1 | 1 | 0 | 3 | 0.6667 |
| 8_4585078 | NA | NA | 1 | 0 | NA | NA | 2 | 0.5 | 3 | 0.3333 | 1 | 1 | NA | NA | NA | NA | 1 | 0 | 2 | 0.5 |
| 8_4585098 | NA | NA | 1 | 0 | 1 | 1 | 2 | 0 | 4 | 0.25 | 1 | 1 | NA | NA | 1 | 0 | 1 | 0 | 3 | 0.3333 |

| | | | | | | | | | | | | | | | | | | | | |
|---|---|---|---|---|---|---|---|---|---|---|---|---|---|---|---|---|---|---|---|---|
| 8_4585108 | NA | NA | 2 | 0 | 1 | 0 | 3 | 0.3333 | 6 | 0.1667 | 1 | 1 | NA | NA | 1 | 0 | 1 | 0 | 3 | 0.3333 |
| 8_4585109 | 1 | 0 | 2 | 0 | 1 | 0 | 4 | 0.25 | 8 | 0.125 | 1 | 1 | NA | NA | 1 | 0 | 1 | 1 | 3 | 0.6667 |
| 8_4585111 | NA | NA | 2 | 1 | 1 | 1 | 4 | 0.75 | 7 | 0.8571 | 1 | 1 | NA | NA | 1 | 1 | 1 | 1 | 3 | 1 |
| 8_4585114 | 1 | 1 | 1 | 0 | 1 | 0 | 4 | 0.5 | 7 | 0.4286 | 1 | 1 | NA | NA | 1 | 0 | 1 | 0 | 3 | 0.3333 |
| 8_4585132 | NA | NA | 2 | 0 | 1 | 1 | 2 | 0.5 | 5 | 0.4 | 1 | 1 | NA | NA | NA | NA | NA | NA | 1 | 1 |
| 8_4585135 | NA | NA | 1 | 1 | 1 | 1 | 3 | 0 | 5 | 0.4 | 1 | 1 | NA | NA | 1 | 1 | 1 | 1 | 3 | 1 |
| 8_4585144 | NA | NA | 1 | 1 | 1 | 1 | 3 | 0.6667 | 5 | 0.8 | 1 | 1 | NA | NA | 1 | 1 | NA | NA | 2 | 1 |
| 8_4585169 | 1 | 1 | 2 | 0 | 1 | 0 | 3 | 0.3333 | 7 | 0.2857 | 1 | 1 | NA | NA | 1 | 0 | 1 | 0 | 3 | 0.3333 |
| 8_4585171 | NA | NA | 2 | 0 | 1 | 1 | 3 | 0.6667 | 6 | 0.5 | NA | NA | NA | NA | NA | NA | 1 | 1 | 1 | 1 |
| 8_4585172 | NA | NA | 2 | 0 | 1 | 0 | 3 | 0.3333 | 6 | 0.1667 | 1 | 1 | NA | NA | 1 | 0 | 1 | 0 | 3 | 0.3333 |
| 8_4585193 | NA | NA | 1 | 1 | 1 | 1 | 3 | 1 | 5 | 1 | 1 | 1 | NA | NA | NA | NA | 1 | 1 | 2 | 1 |
| 8_4585216 | NA | NA | 1 | 0 | 1 | 1 | 3 | 0.3333 | 5 | 0.4 | 1 | 0 | NA | NA | 1 | 1 | 1 | 0 | 3 | 0.3333 |
| 8_4585589 | 1 | 1 | 1 | 1 | 1 | 1 | 2 | 1 | 5 | 1 | 1 | 0 | NA | NA | 1 | 1 | 1 | 1 | 3 | 0.6667 |
| 8_4624813 | 1 | 1 | 3 | 0.3333 | 2 | 0.5 | 4 | 0.25 | 10 | 0.4 | NA | NA | 2 | 0 | 4 | 1 | 2 | 0.5 | 8 | 0.625 |
| 8_4624848 | 1 | 0 | 3 | 1 | 2 | 0.5 | 4 | 0.75 | 10 | 0.7 | NA | NA | 2 | 0.5 | 3 | 0 | NA | NA | 5 | 0.2 |
| 8_4624896 | 1 | 1 | 3 | 0 | 2 | 0 | 4 | 0.25 | 10 | 0.2 | 1 | 1 | 3 | 0.3333 | 4 | 0.5 | 3 | 0.3333 | 11 | 0.4545 |
| 8_4624913 | 1 | 1 | 4 | 0 | 3 | 0 | 4 | 0 | 12 | 0.0833 | 1 | 0 | 1 | 0 | 5 | 0 | 2 | 1 | 9 | 0.2222 |
| 8_4624924 | 1 | 1 | 5 | 0.6 | 3 | 0.3333 | 3 | 0.6667 | 12 | 0.5833 | 1 | 1 | 3 | 0.6667 | 2 | 0.5 | 4 | 0.75 | 10 | 0.7 |
| 8_4624928 | 1 | 1 | 5 | 0.6 | 3 | 0.3333 | 6 | 0.6667 | 15 | 0.6 | 1 | 1 | 3 | 0.6667 | 7 | 0.7143 | 3 | 0.6667 | 14 | 0.7143 |
| 8_4624929 | 1 | 0 | 5 | 0.6 | 3 | 0.3333 | 5 | 0.2 | 14 | 0.3571 | 1 | 0 | 1 | 1 | 7 | 0.1429 | 4 | 0.5 | 13 | 0.3077 |
| 8_4624937 | 1 | 0 | 3 | 0 | 3 | 0.6667 | 4 | 0.75 | 11 | 0.4545 | 1 | 0 | 1 | 1 | 3 | 0.3333 | 2 | 1 | 7 | 0.5714 |
| 8_4624954 | 2 | 0 | 6 | 0.1667 | 2 | 0.5 | 6 | 0.3333 | 16 | 0.25 | 1 | 0 | 2 | 0 | 7 | 0.5714 | 4 | 0 | 14 | 0.2857 |
| 8_4624956 | 2 | 1 | 6 | 1 | 2 | 1 | 6 | 0.8333 | 16 | 0.9375 | 1 | 1 | 2 | 1 | 6 | 1 | 4 | 1 | 13 | 1 |
| 8_4627884 | 7 | 0.2857 | 12 | 0.4167 | 7 | 0.4286 | 10 | 0 | 36 | 0.2778 | 3 | 0 | 9 | 0.1111 | 9 | 0.1111 | 10 | 0.1 | 31 | 0.0968 |
| 8_4627909 | 5 | 1 | 11 | 1 | 6 | 0.6667 | 7 | 1 | 29 | 0.931 | 3 | 1 | 6 | 0.8333 | 6 | 1 | 7 | 0.8571 | 22 | 0.9091 |
| 8_4627912 | 7 | 0.8571 | 14 | 0.8571 | 7 | 0.5714 | 10 | 1 | 38 | 0.8421 | 3 | 1 | 7 | 0.7143 | 11 | 0.9091 | 10 | 0.9 | 31 | 0.871 |
| 8_4627919 | 9 | 0.6667 | 15 | 1 | 7 | 0.7143 | 11 | 0.7273 | 42 | 0.8095 | 4 | 1 | 8 | 0.875 | 11 | 1 | 11 | 0.9091 | 34 | 0.9412 |
| 8_4627920 | 10 | 0.8 | 15 | 0.8667 | 8 | 0.75 | 12 | 0.6667 | 45 | 0.7778 | 4 | 0.5 | 9 | 0.7778 | 11 | 0.8182 | 11 | 0.9091 | 35 | 0.8 |
| 8_4627922 | 10 | 0.4 | 14 | 0.0714 | 7 | 0.4286 | 10 | 0.4 | 41 | 0.2927 | 4 | 0 | 7 | 0.1429 | 12 | 0.1667 | 11 | 0.2727 | 34 | 0.1765 |
| 8_4627958 | 12 | 0.75 | 14 | 0.7143 | 10 | 0.6 | 12 | 0.8333 | 48 | 0.7292 | 4 | 0.75 | 10 | 0.5 | 12 | 0.5833 | 13 | 0.6154 | 39 | 0.5897 |
| 8_4627960 | 8 | 0.125 | 12 | 0.0833 | 10 | 0.1 | 12 | 0.3333 | 42 | 0.1667 | 4 | 0 | 10 | 0 | 12 | 0.25 | 12 | 0.25 | 38 | 0.1579 |
| 8_4627963 | 10 | 0.2 | 13 | 0.0769 | 10 | 0 | 11 | 0.0909 | 44 | 0.0909 | 5 | 0 | 10 | 0 | 11 | 0 | 14 | 0.2857 | 40 | 0.1 |
| 8_4637228 | 1 | 1 | NA | NA | 1 | 0 | 3 | 0 | 5 | 0.2 | 1 | 1 | NA | NA | 1 | 0 | NA | NA | 2 | 0.5 |
| 8_4648659 | 13 | 0 | 7 | 0.4286 | 13 | 0 | 7 | 0.1429 | 40 | 0.1 | 5 | 0.2 | 10 | 0 | 9 | 0.1111 | 7 | 0 | 31 | 0.0645 |
| 8_4648662 | 9 | 0.2222 | 5 | 0.4 | 11 | 0.1818 | 6 | 0.1667 | 31 | 0.2258 | 5 | 0.4 | 8 | 0 | 8 | 0.25 | 6 | 0.1667 | 27 | 0.1852 |
| 8_4648673 | 7 | 0.2857 | 8 | 0.375 | 11 | 0.3636 | 5 | 0.6 | 31 | 0.3871 | 3 | 0.6667 | 8 | 0.125 | 9 | 0.3333 | 6 | 0.3333 | 26 | 0.3077 |
| 8_4660491 | 4 | 0.25 | 2 | 0 | 4 | 0 | 2 | 0 | 12 | 0.0833 | 1 | 1 | 4 | 0 | 3 | 0 | 3 | 0.3333 | 11 | 0.1818 |
| 8_4662275 | 2 | 1 | 1 | 1 | 2 | 1 | NA | NA | 5 | 1 | 2 | 1 | 4 | 1 | 1 | 1 | 3 | 1 | 10 | 1 |
| 8_4662276 | 2 | 0 | 1 | 1 | 2 | 0 | NA | NA | 5 | 0.2 | 2 | 0 | 3 | 0.3333 | 2 | 0 | 3 | 0 | 10 | 0.1 |
| 8_4662283 | 2 | 0 | 1 | 1 | 1 | 0 | NA | NA | 4 | 0.25 | 3 | 0.3333 | 4 | 0.75 | 2 | 1 | 2 | 1 | 11 | 0.7273 |
| 8_4662290 | 2 | 0 | 1 | 0 | 2 | 0.5 | NA | NA | 5 | 0.2 | 3 | 0.3333 | 3 | 0.3333 | 2 | 0 | 2 | 0.5 | 10 | 0.3 |
| 8_4662291 | 2 | 0 | 1 | 0 | 2 | 0.5 | NA | NA | 5 | 0.2 | 1 | 0 | 3 | 0 | 1 | 0 | 2 | 0.5 | 7 | 0.1429 |
| 8_4662300 | 2 | 1 | 1 | 1 | 2 | 0.5 | NA | NA | 5 | 0.8 | 3 | 1 | 3 | 1 | 2 | 1 | 3 | 1 | 11 | 1 |
| 8_4662321 | 2 | 1 | 2 | 0 | 2 | 1 | NA | NA | 6 | 0.6667 | 1 | 0 | 4 | 0.5 | 2 | 0 | 3 | 0.6667 | 10 | 0.4 |
| 8_4662336 | 2 | 1 | 2 | 1 | 2 | 1 | NA | NA | 6 | 1 | 4 | 1 | 3 | 1 | 2 | 1 | 3 | 1 | 12 | 1 |
| 8_4662337 | 3 | 0.3333 | 2 | 0 | 2 | 0 | NA | NA | 7 | 0.1429 | 4 | 0.25 | 4 | 0.25 | 2 | 0 | 3 | 0 | 13 | 0.1538 |
| 8_4662358 | 3 | 1 | 2 | 1 | 2 | 1 | NA | NA | 7 | 1 | 4 | 1 | 3 | 1 | 2 | 1 | 2 | 1 | 11 | 1 |
| 8_4662366 | 3 | 0.3333 | 2 | 0.5 | 2 | 0.5 | NA | NA | 7 | 0.4286 | 3 | 0.3333 | 4 | 0.5 | 2 | 0 | 2 | 1 | 11 | 0.4545 |
| 8_4663489 | 3 | 0 | 2 | 0.5 | 1 | 0 | NA | NA | 6 | 0.1667 | 3 | 1 | 4 | 0.75 | 2 | 0.5 | 2 | 0 | 11 | 0.6364 |
| 8_4664566 | 3 | 0 | 2 | 0 | 2 | 0.5 | NA | NA | 7 | 0.1429 | 4 | 0 | 4 | 0 | 2 | 0.5 | 2 | 0 | 12 | 0.0833 |
| 8_4664568 | 3 | 0.3333 | 2 | 0.5 | 1 | 0 | NA | NA | 6 | 0.3333 | 4 | 0.25 | 6 | 0 | 2 | 0 | 2 | 0 | 14 | 0.0714 |
| 8_4664608 | 3 | 0.3333 | 2 | 0 | 2 | 0 | NA | NA | 7 | 0.1429 | 3 | 0 | 5 | 0.2 | 2 | 0 | 3 | 0 | 13 | 0.0769 |
| 8_4666042 | 129 | 0.062 | 140 | 0.0357 | 121 | 0.1322 | 139 | 0.0647 | 529 | 0.0718 | 88 | 0.125 | 142 | 0.0704 | 108 | 0.0648 | 118 | 0.0763 | 456 | 0.0811 |
| 8_4666045 | 149 | 0.1007 | 150 | 0.04 | 139 | 0.1079 | 160 | 0.0688 | 598 | 0.0786 | 102 | 0.1176 | 163 | 0.1166 | 123 | 0.0976 | 140 | 0.1 | 528 | 0.108 |
| 8_4666225 | 117 | 0.812 | 104 | 0.8077 | 92 | 0.837 | 119 | 0.8319 | 432 | 0.8218 | 78 | 0.8333 | 115 | 0.8435 | 100 | 0.86 | 100 | 0.85 | 393 | 0.8473 |
| 8_4698746 | 2 | 0 | 6 | 0 | 1 | 0 | 5 | 0.2 | 14 | 0.0714 | 2 | 0 | 6 | 0 | 4 | 0 | 4 | 0.25 | 16 | 0.0625 |
| 8_4698749 | NA | NA | 9 | 0.7778 | 2 | 0.5 | 5 | 0.8 | 16 | 0.75 | 2 | 0.5 | 7 | 0.7143 | 4 | 0.75 | 2 | 1 | 15 | 0.7333 |
| 8_4698763 | 1 | 1 | 8 | 1 | 1 | 1 | 6 | 0.8333 | 16 | 0.9375 | 2 | 1 | 7 | 0.8571 | 5 | 0.6 | 4 | 1 | 18 | 0.8333 |
| 8_4698768 | 2 | 1 | 8 | 0.875 | 2 | 0.5 | 5 | 0.8 | 17 | 0.8235 | 2 | 0.5 | 5 | 0.8 | 5 | 0.6 | 4 | 0.75 | 16 | 0.6875 |
| 8_4698772 | 2 | 1 | 8 | 0.5 | 2 | 0 | 6 | 0.6667 | 18 | 0.5556 | 2 | 0.5 | 6 | 0.5 | 4 | 0.5 | 4 | 0.75 | 16 | 0.5625 |
| 8_4698775 | 2 | 0 | 9 | 0.2222 | 2 | 0 | 5 | 0 | 18 | 0.1111 | 2 | 0 | 6 | 0.1667 | 5 | 0.2 | 4 | 0.25 | 17 | 0.1765 |
| 8_4698779 | 2 | 0.5 | 9 | 0.6667 | 2 | 1 | 5 | 0.8 | 18 | 0.7222 | 2 | 0.5 | 7 | 0.4286 | 4 | 1 | 4 | 0.75 | 17 | 0.6471 |
| 8_4698784 | 1 | 0 | 5 | 0.4 | 2 | 1 | 4 | 0 | 12 | 0.3333 | 1 | 0 | 4 | 0.5 | 3 | 0 | 4 | 0.5 | 12 | 0.3333 |
| 8_4698786 | 2 | 1 | 9 | 0.8889 | 2 | 1 | 5 | 1 | 18 | 0.9444 | 1 | 1 | 7 | 1 | 5 | 1 | 6 | 1 | 19 | 1 |
| 8_4698796 | 1 | 0 | 8 | 0.375 | 2 | 0 | 4 | 0.25 | 15 | 0.2667 | 1 | 0 | 6 | 0.1667 | 5 | 0.4 | 4 | 0.5 | 16 | 0.3125 |

| | | | | | | | | | | | | | | | | | | | | |
|---|---|---|---|---|---|---|---|---|---|---|---|---|---|---|---|---|---|---|---|---|
| 8_4698799 | 2 | 0 | 10 | 0.4 | 2 | 0.5 | 5 | 0.4 | 19 | 0.3684 | NA | NA | 6 | 0.3333 | 4 | 0.25 | 5 | 1 | 15 | 0.5333 |
| 8_4698815 | 3 | 0 | 9 | 0.1111 | 1 | 0 | 5 | 0 | 18 | 0.0556 | 2 | 0 | 5 | 0.4 | 5 | 0 | 6 | 0.1667 | 18 | 0.1667 |
| 8_4698816 | 3 | 0 | 9 | 0.4444 | 2 | 0.5 | 5 | 0.4 | 19 | 0.3684 | 2 | 0.5 | 7 | 0.4286 | 5 | 0.6 | 4 | 0.5 | 18 | 0.5 |
| 8_4698822 | 2 | 0.5 | 8 | 0.5 | 3 | 1 | 4 | 0.5 | 17 | 0.5882 | 2 | 0.5 | 7 | 0.4286 | 5 | 0.2 | 5 | 1 | 19 | 0.5263 |
| 8_4698829 | 2 | 0.5 | 10 | 0.5 | 3 | 0.6667 | 6 | 0.3333 | 21 | 0.4762 | 2 | 0.5 | 7 | 0.4286 | 5 | 0.4 | 6 | 0.5 | 20 | 0.45 |
| 8_4698855 | 4 | 0 | 8 | 0 | 3 | 0 | 5 | 0.2 | 20 | 0.05 | 2 | 0 | 7 | 0.1429 | 3 | 0 | 4 | 0.25 | 16 | 0.125 |
| 8_4698859 | 4 | 1 | 7 | 0.8571 | 3 | 0.6667 | 4 | 0.75 | 18 | 0.8333 | 2 | 1 | 7 | 1 | 4 | 0.75 | 6 | 1 | 19 | 0.9474 |
| 8_4698884 | 2 | 0 | 6 | 0.3333 | 3 | 0 | 6 | 0 | 17 | 0.1176 | 3 | 0 | 2 | 0 | 4 | 0 | 6 | 0.3333 | 15 | 0.1333 |
| 8_4702894 | NA | NA | NA | NA | 2 | 0.5 | NA | NA | 2 | 0.5 | NA | NA | NA | NA | 1 | 1 | NA | NA | 1 | 1 |
| 8_4702917 | NA | NA | NA | NA | 2 | 1 | NA | NA | 2 | 1 | NA | NA | NA | NA | 1 | 1 | NA | NA | 1 | 1 |
| 8_4702933 | NA | NA | NA | NA | 2 | 1 | NA | NA | 2 | 1 | NA | NA | NA | NA | 1 | 1 | NA | NA | 1 | 1 |
| 8_4702942 | NA | NA | NA | NA | 2 | 1 | NA | NA | 2 | 1 | NA | NA | NA | NA | 1 | 1 | NA | NA | 1 | 1 |
| 8_4702965 | NA | NA | NA | NA | 2 | 1 | NA | NA | 2 | 1 | NA | NA | NA | NA | 1 | 1 | NA | NA | 1 | 1 |
| 8_4709238 | NA | NA | 3 | 0.3333 | NA | NA | 1 | 0 | 4 | 0.25 | 2 | 0 | 1 | 1 | NA | NA | 1 | 0 | 4 | 0.25 |
| 8_4719282 | NA | NA | 1 | 1 | 1 | 0 | NA | NA | 2 | 0.5 | NA | NA | 1 | 0 | 1 | 1 | NA | NA | 2 | 0.5 |
| 8_4729913 | 2 | 0 | 4 | 0.75 | 1 | 0 | NA | NA | 7 | 0.4286 | NA | NA | 5 | 0.6 | 6 | 0.3333 | 3 | 1 | 14 | 0.5714 |
| 8_4729926 | 2 | 0 | 4 | 0.5 | 1 | 1 | 1 | 0 | 8 | 0.375 | 1 | 0 | 4 | 0 | 6 | 0.1667 | 3 | 0 | 14 | 0.0714 |
| 8_4729943 | 2 | 0.5 | 5 | 0.4 | 1 | 1 | 1 | 1 | 9 | 0.5556 | 1 | 0 | 5 | 0.6 | 7 | 0.5714 | 3 | 1 | 16 | 0.625 |
| 8_4729952 | 2 | 0 | 6 | 0.3333 | 1 | 1 | 1 | 0 | 10 | 0.3 | 1 | 0 | 4 | 0.75 | 6 | 0.3333 | 3 | 0.3333 | 14 | 0.4286 |
| 8_4729961 | 2 | 0 | 6 | 0.5 | 1 | 0 | 1 | 1 | 10 | 0.4 | 1 | 1 | 5 | 0 | 8 | 0.125 | 3 | 0.6667 | 17 | 0.2353 |
| 8_4730008 | 3 | 0.6667 | 7 | 0.8571 | 1 | 1 | 1 | 1 | 12 | 0.8333 | NA | NA | 6 | 0.8333 | 6 | 0.6667 | 3 | 0.6667 | 15 | 0.7333 |
| 8_4730015 | 3 | 0.3333 | 6 | 0.1667 | NA | NA | 1 | 0 | 10 | 0.2 | 1 | 0 | 4 | 0 | 7 | 0.1429 | 3 | 1 | 15 | 0.2667 |
| 8_4754629 | NA | NA | NA | NA | 1 | 1 | NA | NA | 1 | 1 | NA | NA | NA | NA | 1 | 1 | NA | NA | 1 | 1 |
| 8_4763643 | NA | NA | 5 | 0.4 | 4 | 0.5 | 7 | 0 | 16 | 0.25 | 2 | 0 | 6 | 0.3333 | 11 | 0.2727 | 9 | 0.3333 | 28 | 0.2857 |
| 8_4763682 | 2 | 0 | 3 | 0 | 4 | 0 | 6 | 0.1667 | 15 | 0.0667 | 1 | 0 | 7 | 0 | 10 | 0.1 | 5 | 0.2 | 23 | 0.087 |
| 8_4763814 | 1 | 0 | 3 | 0 | 5 | 0.2 | 5 | 0.2 | 14 | 0.1429 | NA | NA | 5 | 0.2 | 7 | 0 | 3 | 0 | 15 | 0.0667 |
| 8_4763901 | 1 | 0 | 2 | 0 | 2 | 0 | 3 | 0.6667 | 8 | 0.25 | NA | NA | 4 | 0.25 | 6 | 0 | 4 | 0 | 14 | 0.0714 |
| 8_4763911 | 1 | 0 | 2 | 0 | 2 | 0.5 | 3 | 0 | 8 | 0.125 | NA | NA | 4 | 0.25 | 8 | 0.25 | 4 | 0.25 | 16 | 0.25 |
| 8_4763932 | 1 | 0 | 2 | 0.5 | 2 | 0 | 1 | 0 | 6 | 0.1667 | NA | NA | 4 | 0.25 | 6 | 0.1667 | 4 | 0.25 | 14 | 0.2143 |
| 8_4764489 | 1 | 0 | 1 | 0 | 2 | 0.5 | 2 | 1 | 6 | 0.5 | NA | NA | 3 | 0.6667 | 4 | 0.5 | 2 | 1 | 9 | 0.6667 |
| 8_4764492 | 1 | 1 | 1 | 0 | 1 | 0 | 2 | 0.5 | 5 | 0.4 | NA | NA | 2 | 0.5 | 3 | 0 | 3 | 0 | 8 | 0.125 |
| 8_4764497 | 1 | 0 | 1 | 1 | 2 | 1 | 2 | 1 | 6 | 0.8333 | NA | NA | 3 | 0.6667 | 6 | 0.8333 | 2 | 1 | 11 | 0.8182 |
| 8_4764502 | 1 | 1 | 1 | 1 | 2 | 1 | 2 | 0.5 | 6 | 0.8333 | NA | NA | 3 | 0.6667 | 6 | 0.6667 | 2 | 0 | 11 | 0.5455 |
| 8_4764514 | 1 | 0 | 1 | 1 | NA | NA | 2 | 0.5 | 4 | 0.5 | NA | NA | 4 | 0.25 | 6 | 0.5 | 3 | 0.3333 | 13 | 0.3846 |
| 8_4764523 | 1 | 0 | 1 | 0 | 1 | 1 | 2 | 0.5 | 5 | 0.4 | NA | NA | 2 | 0.5 | 4 | 0.5 | 2 | 0 | 8 | 0.375 |
| 8_4764535 | 1 | 0 | 1 | 0 | 1 | 1 | 2 | 0 | 5 | 0.2 | NA | NA | 4 | 0.5 | 4 | 0.5 | 2 | 0.5 | 10 | 0.5 |
| 8_4764538 | 1 | 0 | 1 | 0 | 1 | 1 | 2 | 0 | 5 | 0.2 | NA | NA | 4 | 0.25 | 4 | 0 | 3 | 0 | 11 | 0.0909 |
| 8_4764557 | NA | NA | 1 | 0 | 1 | 0 | 2 | 1 | 4 | 0.5 | NA | NA | 2 | 0 | 4 | 0.25 | 3 | 0.3333 | 9 | 0.2222 |
| 8_4764559 | 1 | 1 | 1 | 0 | 1 | 1 | 2 | 0 | 5 | 0.4 | NA | NA | 2 | 0 | 4 | 0.5 | 3 | 0.3333 | 9 | 0.3333 |
| 8_4764569 | 1 | 1 | 1 | 0 | 1 | 0 | 1 | 0 | 4 | 0.25 | NA | NA | 2 | 0.5 | 3 | 0.3333 | 1 | 0 | 6 | 0.3333 |
| 8_4764593 | 1 | 1 | 1 | 0 | 1 | 0 | 1 | 0 | 4 | 0.25 | NA | NA | 2 | 0.5 | 2 | 0 | 2 | 0 | 6 | 0.1667 |
| 8_4764678 | NA | NA | NA | NA | 1 | 0 | 1 | 1 | 2 | 0.5 | NA | NA | 1 | 0 | 2 | 0.5 | 1 | 0 | 4 | 0.25 |
| 8_4764679 | NA | NA | NA | NA | 1 | 0 | 1 | 1 | 2 | 0.5 | NA | NA | 1 | 1 | 2 | 0 | 2 | 0 | 5 | 0.2 |
| 8_4764689 | 1 | 1 | NA | NA | 1 | 1 | 1 | 0 | 3 | 0.6667 | NA | NA | 1 | 1 | 2 | 1 | 3 | 1 | 6 | 1 |
| 8_4764700 | 1 | 0 | 1 | 1 | 1 | 0 | 1 | 1 | 4 | 0.5 | NA | NA | 1 | 1 | 1 | 1 | 2 | 0 | 4 | 0.5 |
| 8_4764703 | 1 | 1 | NA | NA | 1 | 0 | 1 | 1 | 3 | 0.6667 | NA | NA | 1 | 1 | 2 | 0.5 | 2 | 1 | 5 | 0.8 |
| 8_4774098 | 1 | 1 | NA | NA | 1 | 0 | 1 | 1 | 3 | 0.6667 | NA | NA | NA | NA | 1 | 1 | 2 | 0 | 3 | 0.3333 |
| 8_4774150 | NA | NA | 1 | 0 | 1 | 0 | 1 | 1 | 3 | 0.3333 | 1 | 1 | 1 | 0 | NA | NA | 1 | 0 | 3 | 0.3333 |
| 8_4774199 | NA | NA | 1 | 1 | 3 | 1 | 3 | 1 | 7 | 1 | 1 | 1 | 1 | 0 | NA | NA | 1 | 1 | 3 | 0.6667 |
| 8_4774246 | NA | NA | 1 | 1 | 2 | 0.5 | 1 | 1 | 4 | 0.75 | 1 | 1 | 1 | 0 | NA | NA | 1 | 0 | 3 | 0.3333 |
| 8_4786740 | NA | NA | NA | NA | NA | NA | 1 | 1 | 1 | 1 | NA | NA | 1 | 1 | NA | NA | NA | NA | 1 | 1 |
| 8_4786870 | 1 | 1 | NA | NA | NA | NA | NA | NA | 1 | 1 | NA | NA | NA | NA | NA | NA | 1 | 1 | 1 | 1 |
| 8_4791196 | NA | NA | NA | NA | NA | NA | 1 | 1 | 1 | 1 | NA | NA | NA | NA | NA | NA | 1 | 1 | 1 | 1 |
| 8_4791197 | NA | NA | NA | NA | NA | NA | 1 | 1 | 1 | 1 | NA | NA | NA | NA | NA | NA | 1 | 1 | 1 | 1 |
| 8_4794371 | 5 | 0.6 | 3 | 0.3333 | 5 | 0.4 | 8 | 0.625 | 21 | 0.5238 | 4 | 0.25 | 9 | 0.4444 | 2 | 0 | 4 | 0.25 | 19 | 0.3158 |
| 8_4794376 | 5 | 0.2 | 5 | 0.2 | 6 | 0.3333 | 9 | 0.2222 | 25 | 0.24 | 4 | 0.25 | 8 | 0.125 | 8 | 0.25 | 4 | 0.25 | 24 | 0.2083 |
| 8_4794379 | 5 | 1 | 6 | 0.8333 | 6 | 0.8333 | 10 | 0.6 | 27 | 0.7778 | 4 | 0.75 | 9 | 0.8889 | 7 | 0.8571 | 4 | 0.75 | 24 | 0.8333 |
| 8_4800386 | 4 | 0.75 | NA | NA | 4 | 0 | 5 | 0.4 | 13 | 0.3846 | 1 | 0 | 5 | 0.6 | 5 | 0.2 | 2 | 0.5 | 13 | 0.3846 |
| 8_4800394 | 3 | 0.3333 | NA | NA | 3 | 0.3333 | 3 | 0.6667 | 9 | 0.4444 | 1 | 0 | 2 | 0 | 4 | 0.5 | 1 | 0 | 8 | 0.25 |
| 8_4800410 | 1 | 1 | NA | NA | 2 | 1 | 3 | 0.3333 | 6 | 0.6667 | NA | NA | 1 | 1 | 3 | 0.6667 | 2 | 1 | 6 | 0.8333 |
| 8_4800433 | 3 | 0.3333 | NA | NA | 2 | 0 | 3 | 0 | 8 | 0.125 | NA | NA | 2 | 0.5 | 3 | 0 | 2 | 0.5 | 7 | 0.2857 |
| 8_4812052 | 2 | 1 | 3 | 0.6667 | NA | NA | 3 | 1 | 8 | 0.875 | 2 | 1 | 2 | 0.5 | 1 | 1 | 2 | 1 | 7 | 0.8571 |
| 8_4812103 | 4 | 0.75 | 2 | 0 | NA | NA | 2 | 0 | 8 | 0.375 | NA | NA | 2 | 0 | 2 | 0 | 2 | 0.5 | 6 | 0.1667 |
| 8_4812116 | 3 | 0.3333 | 1 | 0 | NA | NA | 1 | 1 | 5 | 0.4 | NA | NA | 2 | 1 | 2 | 1 | 2 | 0.5 | 6 | 0.8333 |
| 8_4812145 | 1 | 1 | 3 | 0.6667 | NA | NA | 1 | 1 | 5 | 0.8 | 1 | 1 | NA | NA | 2 | 1 | 1 | 1 | 4 | 1 |

| | | | | | | | | | | | | | | | | | | | | |
|---|---|---|---|---|---|---|---|---|---|---|---|---|---|---|---|---|---|---|---|---|
| 8_4812149 | 2 | 0 | 4 | 0.25 | NA | NA | 2 | 0 | 8 | 0.125 | 1 | 1 | 2 | 0 | 2 | 0.5 | 2 | 0 | 7 | 0.2857 |
| 8_4812161 | 3 | 0.6667 | 5 | 0.4 | 3 | 0.3333 | 2 | 0.5 | 13 | 0.4615 | 1 | 0 | 1 | 1 | 3 | 0.6667 | 1 | 1 | 6 | 0.6667 |
| 8_4812172 | 5 | 0.4 | 7 | 0.7143 | 4 | 0.5 | 3 | 1 | 19 | 0.6316 | 1 | 0 | 2 | 0.5 | 5 | 0.2 | 2 | 1 | 10 | 0.4 |
| 8_4812174 | 5 | 1 | 7 | 1 | 4 | 1 | 2 | 1 | 18 | 1 | 1 | 1 | 2 | 1 | 5 | 1 | 2 | 1 | 10 | 1 |
| 8_4812176 | 6 | 0.1667 | 7 | 0.1429 | 2 | 0.5 | 3 | 0.6667 | 18 | 0.2778 | 1 | 0 | 3 | 0 | 5 | 0.2 | 2 | 0.5 | 11 | 0.1818 |
| 8_4812180 | 5 | 0.8 | 6 | 0.6667 | 2 | 0.5 | 3 | 0.3333 | 16 | 0.625 | 1 | 1 | 1 | 1 | 4 | 0.25 | 1 | 1 | 7 | 0.5714 |
| 8_4812185 | 3 | 1 | 6 | 0.8333 | 2 | 1 | 2 | 1 | 13 | 0.9231 | NA | NA | 1 | 1 | 3 | 1 | NA | NA | 4 | 1 |
| 8_4812185 | 6 | 0.6667 | 7 | 0.4286 | 3 | 1 | 3 | 0.6667 | 19 | 0.6316 | 1 | 1 | 3 | 1 | 5 | 0.6 | 2 | 0.5 | 11 | 0.7273 |
| 8_4812205 | 6 | 0.3333 | 3 | 0.3333 | 3 | 0.3333 | 2 | 0 | 14 | 0.2857 | 1 | 0 | 2 | 0 | 4 | 0 | 2 | 0.5 | 9 | 0.1111 |
| 8_4812207 | 6 | 0.8333 | 5 | 0.8 | 3 | 0.6667 | 1 | 0 | 15 | 0.7333 | 1 | 0 | 2 | 1 | 4 | 0 | 2 | 0.5 | 9 | 0.3333 |
| 8_4812762 | 4 | 0 | 3 | 0.3333 | 1 | 0 | 3 | 0 | 11 | 0.0909 | NA | NA | 2 | 0 | 3 | 0.3333 | 1 | 0 | 6 | 0.1667 |
| 8_4816330 | 2 | 0 | 3 | 0 | 3 | 0 | 5 | 0.2 | 13 | 0.0769 | 2 | 0 | 4 | 0 | 3 | 0 | 4 | 0.25 | 13 | 0.0769 |
| 8_4816395 | NA | NA | 3 | 0.3333 | 2 | 1 | 1 | 1 | 6 | 0.6667 | 3 | 0.3333 | 4 | 0.75 | 3 | 0.3333 | 3 | 1 | 13 | 0.6154 |
| 8_4816397 | NA | NA | 2 | 0.5 | 1 | 0 | 2 | 0 | 5 | 0.2 | 2 | 1 | 4 | 0.25 | 3 | 1 | 2 | 0.5 | 11 | 0.6364 |
| 8_4816430 | NA | NA | 1 | 1 | NA | NA | NA | NA | 1 | 1 | 1 | 1 | 2 | 1 | NA | NA | NA | NA | 3 | 1 |
| 8_4816443 | NA | NA | 1 | 1 | 1 | 1 | 2 | 1 | 4 | 1 | 3 | 1 | 1 | 1 | 1 | 1 | 3 | 1 | 8 | 1 |
| 8_4816465 | NA | NA | 3 | 0.3333 | 2 | 0 | 3 | 0 | 8 | 0.125 | 3 | 0 | 2 | 0 | 3 | 0.3333 | 3 | 0 | 11 | 0.0909 |
| 8_4816479 | NA | NA | 3 | 0.6667 | 1 | 0 | 3 | 0.6667 | 7 | 0.5714 | 3 | 0.3333 | 1 | 0 | 2 | 0 | 2 | 0.5 | 8 | 0.25 |
| 8_4816494 | NA | NA | 3 | 0.3333 | 2 | 0 | 3 | 0.6667 | 8 | 0.375 | 3 | 0 | 2 | 0 | 2 | 0.5 | NA | NA | 7 | 0.1429 |
| 8_4827976 | 2 | 0 | 3 | 0 | 4 | 0.25 | 3 | 0.3333 | 12 | 0.1667 | 6 | 0 | 4 | 0 | 2 | 0.5 | 2 | 0 | 14 | 0.0714 |
| 8_4828029 | 1 | 0 | 3 | 0 | 5 | 0.2 | 4 | 0.25 | 13 | 0.1538 | 8 | 0.125 | 6 | 0.1667 | 4 | 0.25 | 2 | 0 | 20 | 0.15 |
| 8_4828076 | NA | NA | 2 | 0 | 4 | 0 | 4 | 0.5 | 10 | 0.2 | 7 | 0.1429 | 6 | 0.1667 | 1 | 0 | 2 | 1 | 16 | 0.25 |
| 8_4828110 | 1 | 1 | 2 | 0.5 | 4 | 0.5 | 3 | 0.3333 | 10 | 0.5 | 4 | 0.5 | 3 | 0.3333 | 1 | 0 | 2 | 0.5 | 10 | 0.4 |
| 8_4828135 | 2 | 0 | 3 | 0 | 4 | 0 | 4 | 0.5 | 13 | 0.1538 | 7 | 0.1429 | 6 | 0 | 1 | 0 | 1 | 0 | 15 | 0.0667 |
| 8_4828142 | 2 | 0 | 3 | 0 | 4 | 0.25 | 2 | 1 | 11 | 0.2727 | 4 | 0.5 | 4 | 1 | 1 | 0 | 1 | 1 | 10 | 0.7 |
| 8_4828169 | 1 | 0 | 3 | 0.6667 | 3 | 1 | 2 | 1 | 9 | 0.7778 | 5 | 0.8 | 5 | 1 | 1 | 1 | NA | NA | 11 | 0.9091 |
| 8_4832161 | 1 | 0 | 1 | 1 | 1 | 1 | 1 | 0 | 4 | 0.5 | NA | NA | 1 | 1 | NA | NA | NA | NA | 1 | 1 |
| 8_4832165 | 1 | 1 | 1 | 1 | NA | NA | 2 | 1 | 4 | 1 | 1 | 1 | 1 | 1 | NA | NA | NA | NA | 2 | 1 |
| 8_4832167 | 1 | 0 | 1 | 1 | 1 | 0 | 2 | 0.5 | 5 | 0.4 | 1 | 1 | NA | NA | NA | NA | NA | NA | 1 | 1 |
| 8_4859358 | 1 | 1 | NA | NA | 1 | 1 | 2 | 0.5 | 4 | 0.75 | NA | NA | NA | NA | 2 | 1 | 3 | 1 | 5 | 1 |
| 8_4859370 | 1 | 0 | NA | NA | 1 | 1 | 2 | 0 | 4 | 0.25 | NA | NA | NA | NA | 2 | 0.5 | 3 | 1 | 5 | 0.8 |
| 8_4859393 | 1 | 1 | NA | NA | NA | NA | 2 | 0 | 3 | 0.3333 | NA | NA | NA | NA | 2 | 0 | 3 | 0.3333 | 5 | 0.2 |
| 8_4867902 | 13 | 0.1538 | 15 | 0.2667 | 10 | 0.2 | 4 | 0.25 | 42 | 0.2143 | 8 | 0.125 | 1 | 1 | 8 | 0.125 | 13 | 0.2308 | 30 | 0.2 |
| 8_4867918 | 11 | 0.8182 | 13 | 0.8462 | 7 | 0.4286 | 4 | 0.75 | 35 | 0.7429 | 8 | 0.625 | 1 | 1 | 5 | 1 | 12 | 0.75 | 26 | 0.7692 |
| 8_4867932 | 10 | 0.2 | 12 | 0.0833 | 8 | 0 | 5 | 0.4 | 35 | 0.1429 | 5 | 0 | 1 | 0 | 8 | 0.125 | 14 | 0.2143 | 28 | 0.1429 |
| 8_4867933 | 10 | 0.2 | 12 | 0.1667 | 6 | 0.1667 | 4 | 0 | 32 | 0.1562 | 6 | 0 | 1 | 1 | 6 | 0.1667 | 13 | 0 | 26 | 0.0769 |
| 8_4867939 | 6 | 0.3333 | 8 | 0.5 | 5 | 0.6 | 4 | 0.75 | 23 | 0.5217 | 4 | 0.5 | 1 | 1 | 7 | 0.7143 | 13 | 0.4615 | 25 | 0.56 |
| 8_4868021 | 6 | 0.3333 | 6 | 0.1667 | 4 | 0 | 1 | 0 | 17 | 0.1765 | 1 | 0 | 1 | 0 | 4 | 0 | 10 | 0.1 | 16 | 0.0625 |
| 8_4868030 | 6 | 0.3333 | 5 | 0.2 | 3 | 0 | 2 | 0 | 16 | 0.1875 | 2 | 0 | 1 | 0 | 4 | 0 | 9 | 0.1111 | 16 | 0.0625 |
| 8_4868090 | 6 | 0.3333 | 7 | 0.2857 | 4 | 0.5 | 2 | 0.5 | 19 | 0.3684 | 5 | 0.2 | NA | NA | 8 | 0.5 | 9 | 0.4444 | 22 | 0.4091 |
| 8_4868098 | 7 | 1 | 5 | 0.8 | 3 | 1 | 4 | 1 | 19 | 0.9474 | 5 | 1 | NA | NA | 7 | 1 | 11 | 0.8182 | 23 | 0.913 |
| 8_4868126 | 4 | 1 | 6 | 1 | 2 | 1 | 2 | 1 | 14 | 1 | 5 | 1 | 1 | 1 | 7 | 1 | 9 | 1 | 22 | 1 |
| 8_4868132 | 4 | 0 | 7 | 0.1429 | 1 | 1 | 3 | 0.3333 | 15 | 0.2 | 5 | 0.2 | 1 | 0 | 5 | 0.4 | 7 | 0 | 18 | 0.1667 |
| 8_4868134 | 6 | 0.5 | 7 | 0.5714 | 4 | 0.75 | 2 | 1 | 19 | 0.6316 | 4 | 0.5 | 1 | 1 | 6 | 0.3333 | 8 | 0.875 | 19 | 0.6316 |
| 8_4868138 | 6 | 0.1667 | 8 | 0.375 | 4 | 0.25 | 3 | 0.3333 | 21 | 0.2857 | 5 | 0.4 | 1 | 0 | 6 | 0.3333 | 7 | 0.2857 | 19 | 0.3158 |
| 8_4868140 | 6 | 0.3333 | 8 | 0.5 | 4 | 0.5 | 3 | 0 | 21 | 0.381 | 5 | 0.6 | 1 | 0 | 5 | 0.2 | 6 | 0.6667 | 17 | 0.4706 |
| 8_4868143 | 5 | 1 | 8 | 1 | 4 | 1 | 2 | 0.5 | 19 | 0.9474 | 5 | 1 | 1 | 1 | 6 | 1 | 7 | 1 | 19 | 1 |
| 8_4868181 | 6 | 1 | 8 | 1 | 3 | 1 | 2 | 1 | 19 | 1 | 5 | 1 | 1 | 1 | 6 | 1 | 7 | 1 | 19 | 1 |
| 8_4868194 | 6 | 1 | 7 | 0.5714 | 3 | 0.6667 | 2 | 1 | 18 | 0.7778 | 5 | 1 | 1 | 1 | 5 | 0.6 | 5 | 0.8 | 16 | 0.8125 |
| 8_4868202 | 5 | 0.6 | 7 | 0.5714 | 2 | 1 | 3 | 1 | 17 | 0.7059 | 4 | 1 | 1 | 1 | 4 | 1 | 4 | 1 | 13 | 1 |
| 8_4868840 | 6 | 0 | 6 | 0.3333 | 3 | 0.3333 | 2 | 0 | 17 | 0.1765 | 5 | 0.4 | 1 | 0 | 6 | 0.3333 | 5 | 0.6 | 17 | 0.4118 |
| 8_4868843 | 6 | 0.8333 | 5 | 0.6 | 2 | 1 | 2 | 1 | 15 | 0.8 | 4 | 0.75 | 1 | 1 | 6 | 0.8333 | 4 | 1 | 15 | 0.8667 |
| 8_4868844 | 5 | 0.8 | 5 | 0.2 | 3 | 0.3333 | 1 | 0 | 14 | 0.4286 | 4 | 0.25 | 1 | 0 | 4 | 0.75 | 4 | 0.25 | 13 | 0.3846 |
| 8_4868846 | 6 | 0 | 6 | 0.3333 | 3 | 0 | 2 | 0 | 17 | 0.1176 | 4 | 0.25 | 1 | 0 | 4 | 0 | 3 | 0 | 12 | 0.0833 |
| 8_4868855 | 6 | 0.1667 | 5 | 0.2 | 3 | 0 | 3 | 0.3333 | 17 | 0.1765 | 4 | 0.25 | 1 | 0 | 3 | 0 | 4 | 0.25 | 12 | 0.1667 |
| 8_4868863 | 4 | 0.75 | 4 | 0.5 | 3 | 0 | 1 | 0 | 12 | 0.4167 | 4 | 0.25 | 1 | 0 | 4 | 0.5 | 2 | 1 | 11 | 0.4545 |
| 8_4868870 | 2 | 0.5 | 5 | 0.6 | 2 | 0.5 | 3 | 0 | 12 | 0.4167 | 4 | 0.75 | 1 | 1 | 5 | 0.4 | 4 | 0.75 | 14 | 0.6429 |
| 8_4868880 | 5 | 0.8 | 2 | 0.5 | 2 | 0 | 2 | 1 | 11 | 0.6364 | 3 | 0 | 1 | 1 | 4 | 0.5 | 3 | 1 | 11 | 0.5455 |
| 8_4868886 | 4 | 0.75 | 4 | 0.5 | 2 | 0.5 | NA | NA | 10 | 0.6 | 4 | 0.5 | 1 | 0 | 3 | 0.3333 | 2 | 1 | 10 | 0.5 |
| 8_4868913 | 5 | 0.8 | 3 | 0.6667 | 2 | 0.5 | 1 | 0 | 11 | 0.6364 | 3 | 1 | 1 | 1 | 5 | 0.4 | 1 | 1 | 10 | 0.7 |
| 8_4868931 | 3 | 0.6667 | 2 | 0 | 2 | 1 | 2 | 1 | 9 | 0.6667 | 2 | 0 | 1 | 1 | 3 | 0.6667 | 3 | 0.6667 | 9 | 0.5556 |
| 8_4868948 | 2 | 0.5 | 3 | 0.3333 | 2 | 0 | 2 | 0.5 | 9 | 0.3333 | 3 | 0 | 1 | 0 | 3 | 0.6667 | 2 | 0.5 | 9 | 0.3333 |
| 8_4868953 | 3 | 1 | 3 | 0.6667 | 2 | 0.5 | 2 | 0.5 | 10 | 0.7 | 2 | 0 | 1 | 0 | 4 | 0.25 | 1 | 0 | 8 | 0.125 |
| 8_4869335 | 2 | 0 | 2 | 1 | NA | NA | 1 | 1 | 5 | 0.6 | 1 | 0 | 1 | 0 | 1 | 1 | 1 | 0 | 4 | 0.25 |
| 8_4870761 | 3 | 0.3333 | 2 | 0.5 | NA | NA | 1 | 0 | 6 | 0.3333 | 1 | 0 | NA | NA | 3 | 0.3333 | 1 | 1 | 5 | 0.4 |

| | | | | | | | | | | | | | | | | | | | | |
|---|---|---|---|---|---|---|---|---|---|---|---|---|---|---|---|---|---|---|---|---|
| 8_4872914 | 1 | 1 | 3 | 1 | NA | NA | NA | NA | 4 | 1 | 2 | 1 | NA | NA | 1 | 1 | 1 | 1 | 4 | 1 |
| 8_4872928 | 1 | 0 | 2 | 0 | NA | NA | 1 | 1 | 4 | 0.25 | 2 | 0.5 | NA | NA | 3 | 0 | 1 | 1 | 6 | 0.3333 |
| 8_4872930 | 1 | 0 | 1 | 0 | NA | NA | 1 | 1 | 3 | 0.3333 | 2 | 0.5 | NA | NA | 3 | 0.3333 | 1 | 0 | 6 | 0.3333 |
| 8_4872933 | 1 | 0 | 1 | 1 | NA | NA | 1 | 0 | 3 | 0.3333 | 2 | 0 | NA | NA | 3 | 0.6667 | NA | NA | 5 | 0.4 |
| 8_4872943 | 1 | 1 | NA | NA | NA | NA | 2 | 0.5 | 3 | 0.6667 | 2 | 1 | NA | NA | 1 | 1 | NA | NA | 3 | 1 |
| 8_4872944 | 1 | 1 | NA | NA | NA | NA | 1 | 1 | 2 | 1 | 2 | 0.5 | NA | NA | 1 | 1 | NA | NA | 3 | 0.6667 |
| 8_4872958 | NA | NA | 1 | 1 | NA | NA | 1 | 1 | 2 | 1 | 2 | 1 | NA | NA | NA | NA | 1 | 0 | 3 | 0.6667 |
| 8_5136939 | NA | NA | NA | NA | 1 | 1 | NA | NA | 1 | 1 | NA | NA | NA | NA | 1 | 1 | NA | NA | 1 | 1 |
| 8_5463918 | NA | NA | NA | NA | NA | NA | 1 | 1 | 1 | 1 | NA | NA | NA | NA | NA | NA | 1 | 1 | 1 | 1 |
| 8_5463919 | NA | NA | NA | NA | NA | NA | 1 | 1 | 1 | 1 | NA | NA | NA | NA | NA | NA | 1 | 1 | 1 | 1 |
| 8_5463928 | NA | NA | NA | NA | NA | NA | 1 | 1 | 1 | 1 | NA | NA | NA | NA | NA | NA | 1 | 1 | 1 | 1 |
| 8_5496605 | NA | NA | NA | NA | NA | NA | 1 | 1 | 1 | 1 | 1 | 1 | NA | NA | NA | NA | 1 | 1 | 2 | 1 |
| 8_5496607 | NA | NA | NA | NA | NA | NA | 1 | 1 | 1 | 1 | 1 | 1 | NA | NA | NA | NA | 1 | 1 | 2 | 1 |
| 8_5496617 | NA | NA | NA | NA | NA | NA | 1 | 1 | 1 | 1 | 1 | 1 | NA | NA | NA | NA | 1 | 1 | 2 | 1 |
| 8_5496636 | NA | NA | NA | NA | NA | NA | 1 | 1 | 1 | 1 | 1 | 1 | NA | NA | NA | NA | 1 | 1 | 2 | 1 |
| 8_5496678 | NA | NA | NA | NA | NA | NA | 1 | 1 | 1 | 1 | 1 | 1 | NA | NA | NA | NA | 1 | 1 | 2 | 1 |
| 8_5497384 | 11 | 0.0909 | 14 | 0.3571 | 8 | 0.25 | 15 | 0.2 | 48 | 0.2292 | 12 | 0.0833 | 20 | 0.4 | 7 | 0.4286 | 9 | 0.1111 | 48 | 0.2708 |
| 8_5497611 | 5 | 0.2 | 9 | 0.1111 | 5 | 0.4 | 11 | 0.0909 | 30 | 0.1667 | 8 | 0.25 | 16 | 0.0625 | 4 | 0 | 6 | 0.1667 | 34 | 0.1176 |
| 8_5497634 | 5 | 0.6 | 6 | 0.3333 | 4 | 0.25 | 8 | 0 | 23 | 0.2609 | 3 | 0 | 14 | 0.1429 | 4 | 0.5 | 8 | 0.625 | 29 | 0.3103 |
| 8_5508282 | 5 | 0.2 | 7 | 0.1429 | 3 | 0.3333 | 7 | 0 | 22 | 0.1364 | 2 | 0 | 6 | 0.3333 | 7 | 0.1429 | 2 | 0 | 17 | 0.1765 |
| 8_5508285 | 4 | 0 | 7 | 0.4286 | 3 | 0 | 7 | 0.2857 | 21 | 0.2381 | 3 | 0.6667 | 6 | 0.6667 | 7 | 0.5714 | 3 | 0.6667 | 19 | 0.6316 |
| 8_5508545 | 4 | 0 | 7 | 0.1429 | 4 | 0.25 | 7 | 0.1429 | 22 | 0.1364 | 3 | 0 | 6 | 0.1667 | 7 | 0 | 3 | 0 | 19 | 0.0526 |
| 8_5540819 | 65 | 0.7538 | 60 | 0.7333 | 60 | 0.6333 | 63 | 0.6825 | 248 | 0.7016 | 46 | 0.8261 | 59 | 0.7458 | 57 | 0.7368 | 69 | 0.7826 | 231 | 0.7706 |
| 8_5551210 | 107 | 0.1869 | 123 | 0.1382 | 100 | 0.13 | 107 | 0.1402 | 437 | 0.1487 | 83 | 0.1205 | 106 | 0.2075 | 101 | 0.1188 | 109 | 0.211 | 399 | 0.1679 |
| 8_5551240 | 154 | 0.2338 | 164 | 0.2561 | 136 | 0.2059 | 146 | 0.2192 | 600 | 0.23 | 109 | 0.2752 | 144 | 0.2639 | 139 | 0.1799 | 140 | 0.2643 | 532 | 0.2444 |
| 8_5551242 | 159 | 0.1824 | 167 | 0.1317 | 139 | 0.0935 | 150 | 0.1 | 615 | 0.1285 | 114 | 0.1842 | 151 | 0.1854 | 139 | 0.1655 | 147 | 0.1905 | 551 | 0.1815 |
| 8_5551276 | 150 | 0.3933 | 158 | 0.4304 | 139 | 0.3957 | 141 | 0.4184 | 588 | 0.4099 | 105 | 0.5143 | 153 | 0.4837 | 128 | 0.3984 | 137 | 0.4672 | 523 | 0.4646 |
| 8_5551297 | 176 | 0.0852 | 201 | 0.0597 | 162 | 0.1173 | 166 | 0.1145 | 705 | 0.0922 | 134 | 0.0672 | 179 | 0.1173 | 165 | 0.0667 | 157 | 0.1401 | 635 | 0.0992 |
| 8_5551321 | 192 | 0.0573 | 216 | 0.088 | 187 | 0.0588 | 176 | 0.0739 | 771 | 0.07 | 156 | 0.0897 | 174 | 0.0517 | 178 | 0.0674 | 166 | 0.0723 | 674 | 0.0697 |
| 8_5552666 | NA | NA | 2 | 0 | NA | NA | 3 | 0.3333 | 5 | 0.2 | NA | NA | 2 | 0 | NA | NA | 2 | 0.5 | 4 | 0.25 |
| 8_5552994 | 2 | 0 | 6 | 0.1667 | 6 | 0 | 3 | 0 | 17 | 0.0588 | 1 | 0 | 4 | 0 | NA | NA | 7 | 0.2857 | 12 | 0.1667 |
| 8_5658267 | 2 | 0 | NA | NA | 1 | 1 | NA | NA | 3 | 0.3333 | 1 | 1 | NA | NA | 1 | 1 | 1 | 0 | 3 | 0.6667 |
| 8_5704459 | 2 | 0 | 9 | 0.4444 | 5 | 0.4 | 8 | 0.25 | 24 | 0.3333 | 4 | 0.25 | 1 | 0 | 2 | 0 | 1 | 0 | 8 | 0.125 |
| 8_5704461 | 2 | 0.5 | 7 | 0.2857 | 4 | 0 | 8 | 0.125 | 21 | 0.1905 | 4 | 0.25 | 1 | 1 | 2 | 0 | 1 | 0 | 8 | 0.25 |
| 8_5704467 | 3 | 0 | 8 | 0.375 | 5 | 0.6 | 6 | 0.3333 | 22 | 0.3636 | 2 | 0 | 2 | 0 | 1 | 0 | 1 | 1 | 6 | 0.1667 |
| 8_5704471 | 3 | 1 | 8 | 0.375 | 5 | 0.4 | 6 | 0.5 | 22 | 0.5 | 4 | 0.25 | 2 | 1 | 1 | 1 | 1 | 1 | 8 | 0.625 |
| 8_5704471 | 3 | 0 | 7 | 0.2857 | 4 | 0 | 7 | 0.2857 | 21 | 0.1905 | 3 | 0.3333 | 2 | 0 | 2 | 0 | 1 | 0 | 8 | 0.125 |
| 8_5704478 | 3 | 0.6667 | 7 | 0.1429 | 6 | 0.5 | 7 | 0.1429 | 23 | 0.3043 | 3 | 0 | 2 | 0 | 2 | 0.5 | 1 | 0 | 8 | 0.125 |
| 8_5704485 | 3 | 0.6667 | 9 | 0.2222 | 6 | 0.5 | 8 | 0.875 | 26 | 0.5385 | 5 | 0.4 | 2 | 0.5 | 1 | 1 | 1 | 1 | 9 | 0.5556 |
| 8_5793049 | 6 | 0 | 6 | 0 | 17 | 0.1176 | 8 | 0.125 | 37 | 0.0811 | 9 | 0 | 11 | 0.1818 | 9 | 0 | 8 | 0 | 37 | 0.0541 |
| 8_5793063 | 1 | 1 | 1 | 0 | 5 | 0.2 | 2 | 0 | 9 | 0.2222 | 2 | 0.5 | NA | NA | 1 | 0 | NA | NA | 3 | 0.3333 |
| 8_5793230 | NA | NA | 1 | 1 | 6 | 0.5 | 1 | 0 | 8 | 0.5 | 2 | 0.5 | 2 | 0 | 2 | 0 | NA | NA | 6 | 0.1667 |
| 8_5797895 | NA | NA | NA | NA | 1 | 1 | NA | NA | 1 | 1 | NA | NA | 1 | 1 | NA | NA | NA | NA | 1 | 1 |
| 8_5797896 | NA | NA | NA | NA | 1 | 1 | NA | NA | 1 | 1 | NA | NA | 1 | 1 | NA | NA | NA | NA | 1 | 1 |
| 8_5797899 | NA | NA | NA | NA | 1 | 1 | NA | NA | 1 | 1 | NA | NA | 1 | 1 | NA | NA | NA | NA | 1 | 1 |
| 8_5823025 | 1 | 1 | NA | NA | NA | NA | NA | NA | 1 | 1 | NA | NA | 1 | 1 | NA | NA | NA | NA | 1 | 1 |
| 8_5823028 | 1 | 1 | NA | NA | 1 | 0 | NA | NA | 2 | 0.5 | NA | NA | 1 | 1 | NA | NA | NA | NA | 1 | 1 |
| 8_5936589 | 1 | 0 | NA | NA | 1 | 1 | 2 | 0 | 4 | 0.25 | NA | NA | NA | NA | NA | NA | 1 | 1 | 1 | 1 |
| 8_5936605 | 1 | 1 | NA | NA | NA | NA | 1 | 1 | 2 | 1 | NA | NA | NA | NA | 1 | 0 | 1 | 1 | 2 | 0.5 |
| 8_5936615 | 1 | 1 | NA | NA | 1 | 0 | 2 | 0 | 4 | 0.25 | NA | NA | NA | NA | NA | NA | 1 | 1 | 1 | 1 |
| 8_5936625 | 1 | 1 | NA | NA | 3 | 0.6667 | 3 | 1 | 7 | 0.8571 | NA | NA | NA | NA | 1 | 1 | 2 | 1 | 3 | 1 |
| 8_5936636 | NA | NA | NA | NA | 3 | 0.6667 | 2 | 1 | 5 | 0.8 | NA | NA | NA | NA | NA | NA | 1 | 1 | 1 | 1 |
| 8_5936639 | 1 | 1 | NA | NA | 3 | 0.6667 | 3 | 1 | 7 | 0.8571 | NA | NA | NA | NA | 1 | 0 | 2 | 1 | 3 | 0.6667 |
| 8_5936645 | 1 | 1 | NA | NA | 2 | 0.5 | 3 | 0 | 6 | 0.3333 | NA | NA | 1 | 1 | 1 | 0 | 1 | 0 | 3 | 0.3333 |
| 8_5936650 | 1 | 1 | NA | NA | 3 | 0.6667 | 2 | 0.5 | 6 | 0.6667 | NA | NA | 1 | 1 | NA | NA | 1 | 1 | 2 | 1 |
| 8_5936663 | 1 | 0 | NA | NA | 2 | 0.5 | 4 | 0.25 | 7 | 0.2857 | NA | NA | NA | NA | 1 | 0 | 2 | 0.5 | 3 | 0.3333 |
| 8_5936676 | 1 | 1 | NA | NA | 3 | 0 | 4 | 0.5 | 8 | 0.375 | NA | NA | 1 | 1 | 1 | 0 | 2 | 0.5 | 4 | 0.5 |
| 8_5936681 | 1 | 0 | NA | NA | 3 | 0.3333 | 3 | 0.3333 | 7 | 0.2857 | NA | NA | NA | NA | NA | NA | 1 | 1 | 1 | 1 |
| 8_5936684 | 1 | 1 | NA | NA | 2 | 0.5 | 3 | 1 | 6 | 0.8333 | NA | NA | NA | NA | 1 | 1 | 2 | 1 | 3 | 1 |
| 8_6112430 | 3 | 0 | 4 | 0.5 | 1 | 0 | 2 | 0 | 10 | 0.2 | 1 | 0 | 1 | 0 | 3 | 0.3333 | 1 | 1 | 6 | 0.3333 |
| 8_6112433 | 3 | 1 | 4 | 0.25 | 2 | 0.5 | 2 | 0.5 | 11 | 0.5455 | 1 | 1 | NA | NA | 3 | 0.6667 | 1 | 1 | 5 | 0.8 |
| 8_6112440 | 3 | 0.6667 | 3 | 0.3333 | 2 | 1 | 2 | 0.5 | 10 | 0.6 | 1 | 0 | 1 | 1 | 3 | 1 | NA | NA | 5 | 0.8 |
| 8_6112441 | 4 | 1 | 4 | 1 | 2 | 0.5 | 2 | 1 | 12 | 0.9167 | 1 | 1 | 1 | 1 | 3 | 0.6667 | 1 | 1 | 6 | 0.8333 |
| 8_6112442 | 3 | 1 | 3 | 1 | 2 | 1 | 1 | 0 | 9 | 0.8889 | 1 | 1 | 1 | 1 | 2 | 1 | 1 | 1 | 5 | 1 |
| 8_6112450 | 2 | 1 | 3 | 1 | 1 | 1 | NA | NA | 6 | 1 | 1 | 1 | 1 | 1 | NA | NA | 1 | 1 | 3 | 1 |

| | | | | | | | | | | | | | | | | | | | | |
|---|---|---|---|---|---|---|---|---|---|---|---|---|---|---|---|---|---|---|---|---|
| 8_61124664 | 3 | 0.3333 | 4 | 0 | 2 | 0 | 3 | 0 | 12 | 0.0833 | 1 | 0 | 1 | 1 | 4 | 0.25 | 1 | 0 | 7 | 0.2857 |
| 8_61124767 | 3 | 1 | 4 | 0.5 | 2 | 1 | 4 | 0.75 | 13 | 0.7692 | 1 | 1 | 1 | 1 | 3 | 0.6667 | 1 | 1 | 6 | 0.8333 |
| 8_61124812 | 4 | 0.5 | 4 | 0.25 | 2 | 0.5 | 4 | 0 | 14 | 0.2857 | 1 | 0 | 1 | 0 | 3 | 0.3333 | 1 | 0 | 6 | 0.1667 |
| 8_61124825 | 4 | 1 | 5 | 1 | 2 | 0.5 | 4 | 0.5 | 15 | 0.8 | 1 | 1 | 1 | 1 | 4 | 1 | 1 | 1 | 7 | 1 |
| 8_61124844 | 4 | 0.25 | 5 | 0.2 | 2 | 0 | 4 | 0 | 15 | 0.1333 | 1 | 1 | 1 | 0 | 4 | 0 | 1 | 1 | 7 | 0.2857 |
| 8_61124905 | 3 | 1 | 5 | 1 | 2 | 1 | 4 | 1 | 14 | 1 | 1 | 1 | 1 | 1 | 4 | 1 | 1 | 1 | 7 | 1 |
| 8_61124945 | 3 | 0 | 5 | 0.4 | 2 | 0.5 | 4 | 0 | 14 | 0.2143 | 1 | 1 | 1 | 0 | 4 | 0 | 1 | 1 | 7 | 0.2857 |
| 8_61124952 | 4 | 0.25 | 5 | 0.4 | 2 | 0 | 4 | 0 | 15 | 0.2 | 1 | 0 | 1 | 0 | 4 | 0.25 | 1 | 0 | 7 | 0.1429 |
| 8_61125149 | 3 | 0.6667 | 5 | 0.4 | 2 | 0 | 4 | 0.75 | 14 | 0.5 | 1 | 1 | 1 | 1 | 3 | 0.6667 | NA | NA | 5 | 0.8 |
| 8_61125197 | 4 | 1 | 5 | 1 | 1 | 1 | 5 | 1 | 15 | 1 | 1 | 1 | 1 | 1 | 4 | 1 | NA | NA | 6 | 1 |
| 8_61125236 | 4 | 1 | 5 | 1 | 2 | 1 | 4 | 1 | 15 | 1 | NA | NA | 1 | 1 | 3 | 1 | 1 | 1 | 5 | 1 |
| 8_61125245 | 4 | 0.5 | 5 | 0.6 | 2 | 0.5 | 5 | 0.2 | 16 | 0.4375 | 1 | 1 | 1 | 1 | 4 | 0.75 | 1 | 1 | 7 | 0.8571 |
| 8_61126764 | 4 | 0 | 5 | 0.2 | 6 | 0.1667 | 6 | 0.1667 | 21 | 0.1429 | 1 | 0 | 1 | 0 | 3 | 0.3333 | 1 | 0 | 6 | 0.1667 |
| 8_61127333 | 5 | 0.2 | 4 | 0.25 | 6 | 0 | 6 | 0 | 21 | 0.0952 | 2 | 0.5 | 1 | 0 | 5 | 0 | 1 | 0 | 9 | 0.1111 |
| 8_61128438 | 6 | 0 | 7 | 0.2857 | 9 | 0.3333 | 8 | 0 | 30 | 0.1667 | 4 | 0.25 | 3 | 0.3333 | 5 | 0.2 | 3 | 0.6667 | 15 | 0.3333 |
| 8_61129511 | 5 | 0.2 | 8 | 0 | 9 | 0 | 10 | 0.1 | 32 | 0.0625 | 6 | 0.1667 | 5 | 0 | 4 | 0 | 3 | 0 | 18 | 0.0556 |
| 8_61355787 | NA | NA | NA | NA | 1 | 1 | NA | NA | 1 | 1 | NA | NA | 1 | 1 | 1 | 0 | NA | NA | 2 | 0.5 |
| 8_61360539 | 10 | 0.1 | 15 | 0.0667 | 17 | 0 | 10 | 0.1 | 52 | 0.0577 | 3 | 0 | 14 | 0.1429 | 13 | 0 | 8 | 0 | 38 | 0.0526 |
| 8_61363716 | 9 | 0.1111 | 19 | 0.1053 | 16 | 0.0625 | 23 | 0.1739 | 67 | 0.1194 | 11 | 0.1818 | 8 | 0.125 | 15 | 0.0667 | 13 | 0.0769 | 47 | 0.1064 |
| 8_61363938 | 13 | 0.2308 | 19 | 0.2105 | 14 | 0.2857 | 25 | 0.2 | 71 | 0.2254 | 9 | 0.5556 | 11 | 0.3636 | 17 | 0.3529 | 12 | 0.1667 | 49 | 0.3469 |
| 8_61365050 | 6 | 0.1667 | 12 | 0.4167 | 8 | 0.125 | 17 | 0.3529 | 43 | 0.3023 | 7 | 0.5714 | 10 | 0.2 | 10 | 0.4 | 10 | 0.3 | 37 | 0.3514 |
| 8_61365180 | 6 | 0.5 | 10 | 0.1 | 11 | 0.3636 | 17 | 0.1765 | 44 | 0.25 | 7 | 0.4286 | 9 | 0 | 7 | 0.1429 | 10 | 0.6 | 33 | 0.303 |
| 8_61365288 | 5 | 0.2 | 12 | 0.25 | 13 | 0.3077 | 14 | 0 | 44 | 0.1818 | 8 | 0.625 | 9 | 0.2222 | 10 | 0.3 | 11 | 0.2727 | 38 | 0.3421 |
| 8_61365318 | 6 | 0.1667 | 11 | 0.1818 | 11 | 0.2727 | 13 | 0.3077 | 41 | 0.2439 | 7 | 0.1429 | 9 | 0.3333 | 10 | 0.2 | 9 | 0.1111 | 35 | 0.2 |
| 8_61365323 | 6 | 0.3333 | 12 | 0.1667 | 12 | 0.0833 | 15 | 0 | 45 | 0.1111 | 7 | 0.1429 | 9 | 0 | 9 | 0.3333 | 11 | 0.1818 | 36 | 0.1667 |
| 8_61365661 | 5 | 1 | 5 | 1 | 11 | 1 | 11 | 1 | 32 | 1 | 7 | 1 | 8 | 0.875 | 7 | 1 | 7 | 1 | 29 | 0.9655 |
| 8_61365690 | 3 | 1 | 5 | 1 | 12 | 0.6667 | 13 | 0.6923 | 33 | 0.7576 | 5 | 1 | 8 | 0.75 | 9 | 0.7778 | 6 | 1 | 28 | 0.8571 |
| 8_61365718 | 5 | 0.4 | 9 | 0.7778 | 11 | 0.4545 | 14 | 0.5714 | 39 | 0.5641 | 7 | 0.4286 | 8 | 0.875 | 8 | 0.625 | 11 | 0.6364 | 34 | 0.6471 |
| 8_61365746 | NA | NA | 1 | 1 | NA | NA | 1 | 1 | 2 | 1 | 1 | 0 | NA | NA | NA | NA | 1 | 1 | 2 | 0.5 |
| 8_61365891 | 3 | 0.6667 | 9 | 0.8889 | 8 | 1 | 11 | 0.9091 | 31 | 0.9032 | 5 | 1 | 8 | 0.75 | 6 | 0.6667 | 9 | 0.8889 | 28 | 0.8214 |
| 8_61366042 | 2 | 1 | 7 | 1 | 7 | 1 | 12 | 0.9167 | 28 | 0.9643 | 7 | 1 | 9 | 0.8889 | 6 | 1 | 10 | 0.9 | 32 | 0.9375 |
| 8_61366103 | 4 | 0.5 | 9 | 0.4444 | 6 | 0.3333 | 11 | 0.3636 | 30 | 0.4 | 2 | 0.5 | 7 | 0 | 6 | 0.3333 | 9 | 0.3333 | 24 | 0.25 |
| 8_61375947 | 4 | 0.25 | 8 | 0.125 | 4 | 0.25 | 9 | 0 | 25 | 0.12 | 4 | 0 | 7 | 0 | 6 | 0.3333 | 9 | 0 | 26 | 0.0769 |
| 8_61375955 | 5 | 0.4 | 7 | 0.5714 | 5 | 0.2 | 7 | 0.8571 | 24 | 0.5417 | 4 | 0.5 | 7 | 0.4286 | 6 | 0.5 | 9 | 0.4444 | 26 | 0.4615 |
| 8_61382306 | 1 | 1 | 7 | 1 | 3 | 0.6667 | 4 | 1 | 15 | 0.9333 | 2 | 0.5 | 4 | 1 | 4 | 0.5 | 6 | 0.8333 | 16 | 0.75 |
| 8_61382501 | 3 | 0.6667 | 5 | 1 | 2 | 0.5 | 3 | 1 | 13 | 0.8462 | 2 | 1 | 3 | 0.6667 | 4 | 1 | 5 | 1 | 14 | 0.9286 |
| 8_61386270 | 3 | 0.3333 | 4 | 0.5 | 3 | 0 | 3 | 0 | 13 | 0.2308 | 3 | 0.3333 | 3 | 0.3333 | 2 | 0 | 5 | 0 | 13 | 0.1538 |
| 8_61559922 | 1 | 0 | 1 | 1 | 3 | 0 | 3 | 0 | 8 | 0.125 | 1 | 0 | NA | NA | 1 | 1 | 1 | 0 | 3 | 0.3333 |
| 8_61583236 | 1 | 0 | 2 | 0 | 4 | 0.5 | 4 | 0 | 11 | 0.1818 | 1 | 0 | 1 | 0 | 1 | 0 | 2 | 0.5 | 5 | 0.2 |
| 8_61679119 | 1 | 1 | NA | NA | 1 | 1 | NA | NA | 2 | 1 | NA | NA | 1 | 1 | NA | NA | 1 | 0 | 2 | 0.5 |
| 8_61846745 | 2 | 0.5 | 1 | 0 | 2 | 0.5 | 1 | 1 | 6 | 0.5 | NA | NA | 1 | 1 | 2 | 0 | NA | NA | 3 | 0.3333 |
| 8_61846751 | 1 | 1 | 1 | 1 | 1 | 1 | NA | NA | 3 | 1 | NA | NA | 1 | 1 | 2 | 0 | 1 | 0 | 4 | 0.25 |
| 8_61846881 | NA | NA | 1 | 0 | 2 | 0.5 | 1 | 0 | 4 | 0.25 | NA | NA | 1 | 1 | 2 | 0.5 | 1 | 1 | 4 | 0.75 |
| 8_61846914 | 2 | 0 | 1 | 0 | 2 | 0 | 1 | 1 | 6 | 0.1667 | 1 | 1 | 1 | 0 | 2 | 0 | 1 | 0 | 5 | 0.2 |
| 8_61846958 | 1 | 1 | 1 | 1 | 1 | 1 | 1 | 1 | 4 | 1 | NA | NA | 1 | 0 | 2 | 1 | NA | NA | 3 | 0.6667 |
| 8_61847045 | 2 | 0.5 | 1 | 0 | 2 | 0.5 | 1 | 1 | 6 | 0.5 | NA | NA | 2 | 0 | 2 | 1 | 1 | 1 | 5 | 0.6 |
| 8_61847112 | 1 | 1 | NA | NA | 2 | 1 | NA | NA | 3 | 1 | 1 | 1 | 2 | 1 | 2 | 1 | 1 | 1 | 6 | 1 |
| 8_61847161 | 2 | 1 | 1 | 1 | 2 | 1 | 1 | 1 | 6 | 1 | 1 | 1 | 2 | 0.5 | NA | NA | 1 | 0 | 4 | 0.5 |
| 8_61847210 | 1 | 0 | 1 | 0 | 1 | 0 | 1 | 1 | 4 | 0.25 | 3 | 0.3333 | 2 | 0 | 1 | 0 | 1 | 0 | 7 | 0.1429 |
| 8_61847335 | 2 | 0 | 1 | 0 | 3 | 0.6667 | 1 | 1 | 7 | 0.4286 | 2 | 0.5 | 2 | 1 | 2 | 0 | 1 | 1 | 7 | 0.5714 |
| 8_61847447 | 1 | 0 | 1 | 0 | 2 | 0 | 1 | 1 | 5 | 0.2 | 3 | 0.3333 | 1 | 1 | 1 | 0 | 1 | 1 | 6 | 0.5 |
| 8_61847619 | 2 | 0 | 1 | 0 | 2 | 0.5 | NA | NA | 5 | 0.2 | 2 | 0.5 | 2 | 0.5 | 2 | 0 | NA | NA | 6 | 0.3333 |
| 8_61847902 | 2 | 0 | 1 | 0 | 2 | 1 | 1 | 1 | 6 | 0.5 | 2 | 0 | 3 | 0.3333 | 2 | 0 | 1 | 0 | 8 | 0.125 |
| 8_61847944 | 2 | 0 | 1 | 0 | 2 | 0 | 1 | 1 | 6 | 0.1667 | 3 | 0 | 3 | 0.3333 | 2 | 0 | NA | NA | 8 | 0.125 |
| 8_61848171 | 2 | 0.5 | 1 | 1 | 3 | 0.3333 | 1 | 1 | 7 | 0.5714 | 3 | 0 | 3 | 0.6667 | 2 | 0.5 | 1 | 1 | 9 | 0.4444 |
| 8_61848475 | 4 | 0.5 | 1 | 1 | 5 | 0 | 1 | 0 | 11 | 0.2727 | 2 | 0.5 | 3 | 0.3333 | 2 | 1 | 1 | 1 | 8 | 0.625 |
| 8_61848679 | 5 | 0 | 3 | 0 | 5 | 0.4 | 3 | 0 | 16 | 0.125 | 3 | 0 | 3 | 0 | 2 | 0.5 | 2 | 0 | 10 | 0.1 |
| 8_61849063 | 3 | 0.6667 | 3 | 0.6667 | 5 | 0.8 | 3 | 0.6667 | 14 | 0.7143 | 4 | 0.25 | 3 | 0 | 1 | 1 | 2 | 0 | 10 | 0.2 |
| 8_61956147 | 5 | 0.8 | 3 | 0 | 2 | 0 | 3 | 0.6667 | 13 | 0.4615 | 1 | 1 | 8 | 0.75 | 4 | 0.75 | 2 | 0 | 15 | 0.6667 |
| 8_61956306 | 3 | 0.6667 | 3 | 0.6667 | 3 | 0.6667 | 2 | 0.5 | 11 | 0.6364 | 1 | 1 | 7 | 0.2857 | 4 | 0.5 | 2 | 0 | 14 | 0.3571 |
| 8_61956327 | 5 | 0.6 | 3 | 0 | 3 | 0.3333 | 4 | 0.5 | 15 | 0.4 | 2 | 0.5 | 6 | 0.5 | 4 | 0.5 | 2 | 0.5 | 14 | 0.5 |
| 8_61956388 | 5 | 0.6 | 2 | 0.5 | 3 | 0.3333 | 2 | 1 | 12 | 0.5833 | 2 | 1 | 7 | 0.8571 | 4 | 0.75 | 1 | 0 | 14 | 0.7857 |
| 8_61956412 | 5 | 0.2 | 3 | 0 | 3 | 0 | 3 | 0 | 14 | 0.0714 | 2 | 0 | 7 | 0 | 4 | 0.25 | 2 | 0 | 15 | 0.0667 |
| 8_61956455 | 4 | 0.75 | 3 | 0.6667 | 4 | 0.5 | 4 | 1 | 15 | 0.7333 | 2 | 1 | 8 | 0.875 | 4 | 1 | 2 | 1 | 16 | 0.9375 |
| 8_63798757 | 14 | 0.2143 | 13 | 0.3077 | 14 | 0.1429 | 23 | 0.1304 | 64 | 0.1875 | 3 | 0 | 30 | 0.2667 | 28 | 0.2143 | 17 | 0.1176 | 78 | 0.2051 |

| | | | | | | | | | | | | | | | | | | | | |
|---|---|---|---|---|---|---|---|---|---|---|---|---|---|---|---|---|---|---|---|---|
| 8_6379887( | 14 | 0.0714 | 13 | 0.0769 | 14 | 0.1429 | 21 | 0.1905 | 62 | 0.129 | 2 | 0 | 27 | 0.0741 | 28 | 0.1786 | 17 | 0.0588 | 74 | 0.1081 |
| 8_6380825; | 16 | 0.25 | 15 | 0.2 | 15 | 0.1333 | 22 | 0.2727 | 68 | 0.2206 | 3 | 0 | 28 | 0.1786 | 28 | 0.1786 | 16 | 0.0625 | 75 | 0.1467 |
| 8_6380829( | 16 | 0.5 | 12 | 0.3333 | 17 | 0.4118 | 22 | 0.3636 | 67 | 0.403 | 3 | 0.6667 | 32 | 0.375 | 30 | 0.2667 | 18 | 0.3889 | 83 | 0.3494 |
| 8_6380949( | 26 | 0.2692 | 19 | 0.0526 | 23 | 0.0435 | 30 | 0.1333 | 98 | 0.1327 | 9 | 0 | 39 | 0.0256 | 36 | 0.0833 | 21 | 0.1429 | 105 | 0.0667 |
| 8_6446883( | 1 | 1 | NA | NA | NA | NA | 1 | 1 | 2 | 1 | NA | NA | NA | NA | 1 | 1 | NA | NA | 1 | 1 |
| 8_6446893( | 1 | 1 | NA | NA | NA | NA | 1 | 1 | 2 | 1 | NA | NA | NA | NA | 1 | 1 | NA | NA | 1 | 1 |
| 8_6446901 | 1 | 1 | NA | NA | NA | NA | 1 | 1 | 2 | 1 | NA | NA | NA | NA | 1 | 1 | NA | NA | 1 | 1 |
| 8_6446908 | 1 | 1 | NA | NA | NA | NA | 1 | 0 | 2 | 0.5 | NA | NA | NA | NA | 1 | 1 | NA | NA | 1 | 1 |
| 8_6504611 | 360 | 0.1417 | 333 | 0.1321 | 313 | 0.1821 | 281 | 0.1495 | 1287 | 0.1507 | 302 | 0.096 | 275 | 0.1164 | 292 | 0.1267 | 280 | 0.1393 | 1149 | 0.1192 |
| 8_6517152 | 13 | 0.6923 | 18 | 0.6111 | 16 | 0.5625 | 14 | 0.2857 | 61 | 0.541 | 16 | 0.5 | 17 | 0.4706 | 10 | 0.6 | 10 | 0.5 | 53 | 0.5094 |
| 8_6517156 | 11 | 0.4545 | 13 | 0.6154 | 14 | 0.2857 | 16 | 0.3125 | 54 | 0.4074 | 15 | 0.2667 | 16 | 0.4375 | 9 | 0.3333 | 8 | 0.25 | 48 | 0.3333 |
| 8_6517163 | 13 | 0 | 16 | 0.0625 | 15 | 0.0667 | 16 | 0.0625 | 60 | 0.05 | 16 | 0.1875 | 19 | 0.1579 | 11 | 0 | 9 | 0.2222 | 55 | 0.1455 |
| 8_6517176 | 11 | 0.6364 | 16 | 0.6875 | 14 | 0.5714 | 16 | 0.5 | 57 | 0.5965 | 16 | 0.5 | 18 | 0.4444 | 12 | 0.3333 | 8 | 0.5 | 54 | 0.4444 |
| 8_6517186 | 10 | 0.2 | 12 | 0.3333 | 13 | 0.0769 | 12 | 0 | 47 | 0.1489 | 11 | 0 | 18 | 0.0556 | 7 | 0.2857 | 8 | 0.25 | 44 | 0.1136 |
| 8_6517196 | 10 | 0.9 | 13 | 0.8462 | 16 | 0.875 | 16 | 0.75 | 55 | 0.8364 | 15 | 0.9333 | 17 | 0.9412 | 12 | 0.9167 | 8 | 0.625 | 52 | 0.8846 |
| 8_6517208 | 8 | 0.625 | 15 | 0.8 | 17 | 0.5882 | 15 | 0.7333 | 55 | 0.6909 | 13 | 0.6154 | 19 | 0.6316 | 10 | 0.9 | 7 | 1 | 49 | 0.7347 |
| 8_6517209 | 8 | 0.375 | 13 | 0.6923 | 12 | 0.5833 | 13 | 0.6154 | 46 | 0.587 | 14 | 0.8571 | 13 | 0.6923 | 9 | 0.4444 | 7 | 0.1429 | 43 | 0.6047 |
| 8_6517213 | 7 | 0.2857 | 7 | 0.5714 | 12 | 0.4167 | 12 | 0.3333 | 38 | 0.3947 | 9 | 0.6667 | 8 | 0.75 | 6 | 0.5 | 8 | 0.5 | 31 | 0.6129 |
| 8_6518078 | 7 | 0 | 8 | 0 | 12 | 0.25 | 8 | 0.125 | 35 | 0.1143 | 9 | 0.2222 | 8 | 0.25 | 5 | 0.4 | 6 | 0 | 28 | 0.2143 |
| 8_6518085 | 8 | 0.375 | 13 | 0.0769 | 13 | 0.1538 | 11 | 0.2727 | 45 | 0.2 | 10 | 0.4 | 15 | 0.4 | 9 | 0.3333 | 8 | 0.375 | 42 | 0.381 |
| 8_6519367 | 40 | 0.125 | 26 | 0.3462 | 34 | 0.2647 | 28 | 0.3929 | 128 | 0.2656 | 17 | 0.2941 | 36 | 0.1944 | 41 | 0.2683 | 35 | 0.1143 | 129 | 0.2093 |
| 8_6519370 | 29 | 0.0345 | 22 | 0.0455 | 24 | 0.125 | 23 | 0.087 | 98 | 0.0714 | 10 | 0.3 | 29 | 0.1034 | 34 | 0.0882 | 28 | 0.2857 | 101 | 0.1683 |
| 8_6519370 | 5 | 0.2 | 4 | 0.25 | 6 | 0.3333 | 2 | 0 | 17 | 0.2353 | 1 | 0 | 4 | 0 | 2 | 0 | 4 | 0.25 | 11 | 0.0909 |
| 8_6519375 | 41 | 0.1463 | 28 | 0.1429 | 33 | 0.2727 | 28 | 0.2857 | 130 | 0.2077 | 18 | 0.2222 | 40 | 0.2 | 41 | 0.2439 | 37 | 0.2703 | 136 | 0.2353 |
| 8_6519376 | 35 | 0.5714 | 21 | 0.619 | 29 | 0.6207 | 29 | 0.6207 | 114 | 0.6053 | 14 | 0.7857 | 32 | 0.75 | 36 | 0.6389 | 30 | 0.6333 | 112 | 0.6875 |
| 8_6519377 | 40 | 0.2 | 29 | 0.1724 | 29 | 0.3103 | 29 | 0.3448 | 127 | 0.252 | 15 | 0.3333 | 38 | 0.2105 | 41 | 0.2439 | 32 | 0.2188 | 126 | 0.2381 |
| 8_6519391 | 33 | 0.4848 | 22 | 0.5909 | 31 | 0.4194 | 27 | 0.5556 | 113 | 0.5044 | 12 | 0.6667 | 27 | 0.5185 | 36 | 0.6667 | 33 | 0.5152 | 108 | 0.5833 |
| 8_6519397 | 37 | 0.4595 | 27 | 0.4444 | 30 | 0.3 | 26 | 0.4615 | 120 | 0.4167 | 16 | 0.5625 | 35 | 0.4857 | 40 | 0.425 | 38 | 0.4211 | 129 | 0.4574 |
| 8_6542148 | 2 | 1 | 2 | 1 | 3 | 1 | 1 | 0 | 8 | 0.875 | 9 | 0.6667 | 5 | 0.8 | 5 | 1 | 6 | 1 | 25 | 0.84 |
| 8_6542152 | 2 | 1 | 3 | 0.6667 | 3 | 0.3333 | 1 | 1 | 9 | 0.6667 | 9 | 0.6667 | 5 | 0.2 | 5 | 0.2 | 6 | 0.5 | 25 | 0.44 |
| 8_6542153 | NA | NA | 3 | 0.3333 | 2 | 0 | 1 | 0 | 6 | 0.1667 | 8 | 0 | 4 | 0.25 | 4 | 0.25 | 7 | 0.1429 | 23 | 0.1304 |
| 8_6692105 | NA | NA | 1 | 1 | NA | NA | NA | NA | 1 | 1 | NA | NA | 2 | 1 | NA | NA | 1 | 0 | 3 | 0.6667 |
| 8_6692106 | NA | NA | 1 | 1 | NA | NA | NA | NA | 1 | 1 | NA | NA | 1 | 1 | NA | NA | 1 | 1 | 2 | 1 |
| 8_6692112 | NA | NA | 1 | 1 | NA | NA | NA | NA | 1 | 1 | NA | NA | 2 | 0.5 | NA | NA | 1 | 1 | 3 | 0.6667 |
| 8_6692119 | NA | NA | 1 | 1 | NA | NA | NA | NA | 1 | 1 | NA | NA | 2 | 1 | NA | NA | 1 | 0 | 3 | 0.6667 |
| 8_6692145 | NA | NA | 1 | 1 | NA | NA | NA | NA | 1 | 1 | NA | NA | 2 | 1 | NA | NA | 1 | 1 | 3 | 1 |
| 8_6692149 | NA | NA | 1 | 1 | NA | NA | NA | NA | 1 | 1 | NA | NA | 2 | 1 | NA | NA | 1 | 1 | 3 | 1 |
| 8_6692166 | NA | NA | 1 | 1 | NA | NA | NA | NA | 1 | 1 | NA | NA | 1 | 1 | NA | NA | 1 | 1 | 2 | 1 |
| 8_6692293 | 2 | 0.5 | 1 | 0 | 1 | 0 | NA | NA | 4 | 0.25 | 1 | 0 | 1 | 0 | NA | NA | 2 | 1 | 4 | 0.5 |
| 8_6692322 | NA | NA | 1 | 1 | 1 | 0 | NA | NA | 2 | 0.5 | NA | NA | 1 | 1 | NA | NA | 2 | 0 | 3 | 0.3333 |
| 8_6692327 | 1 | 0 | 1 | 1 | 1 | 0 | NA | NA | 3 | 0.3333 | NA | NA | 1 | 1 | NA | NA | 2 | 0 | 3 | 0.3333 |
| 8_6692636 | NA | NA | NA | NA | NA | NA | 2 | 1 | 2 | 1 | NA | NA | 1 | 0 | NA | NA | 1 | 1 | 2 | 0.5 |
| 8_6692675 | NA | NA | NA | NA | NA | NA | 2 | 0.5 | 2 | 0.5 | NA | NA | 1 | 0 | NA | NA | 1 | 1 | 2 | 0.5 |
| 8_6692723 | NA | NA | NA | NA | NA | NA | 1 | 1 | 1 | 1 | NA | NA | 1 | 1 | NA | NA | NA | NA | 1 | 1 |
| 8_6692738 | NA | NA | NA | NA | NA | NA | 1 | 1 | 1 | 1 | NA | NA | 2 | 0.5 | NA | NA | NA | NA | 2 | 0.5 |
| 8_6692771 | NA | NA | NA | NA | NA | NA | 1 | 1 | 1 | 1 | NA | NA | 1 | 1 | NA | NA | NA | NA | 1 | 1 |
| 8_6692907 | 1 | 0 | 1 | 1 | 1 | 0 | NA | NA | 3 | 0.3333 | NA | NA | 1 | 1 | NA | NA | 1 | 0 | 2 | 0.5 |
| 8_6692943 | 1 | 1 | 1 | 1 | 1 | 0 | NA | NA | 3 | 0.6667 | 1 | 1 | 1 | 1 | 1 | 0 | 1 | 0 | 4 | 0.5 |
| 8_6713288 | 1 | 0 | NA | NA | 1 | 1 | 5 | 0.2 | 7 | 0.2857 | NA | NA | 4 | 0 | 2 | 0.5 | NA | NA | 6 | 0.1667 |
| 8_6713295 | 1 | 0 | NA | NA | 2 | 0.5 | 4 | 0.75 | 7 | 0.5714 | NA | NA | 3 | 0.3333 | 1 | 0 | NA | NA | 4 | 0.25 |
| 8_6713335 | 1 | 1 | NA | NA | NA | NA | 2 | 1 | 3 | 1 | NA | NA | 3 | 0 | 2 | 0.5 | NA | NA | 5 | 0.2 |
| 8_6713342 | 1 | 1 | NA | NA | NA | NA | 2 | 1 | 3 | 1 | NA | NA | 3 | 0.6667 | 2 | 1 | NA | NA | 5 | 0.8 |
| 8_6713362 | 1 | 1 | NA | NA | NA | NA | 1 | 1 | 2 | 1 | NA | NA | 2 | 1 | 1 | 0 | NA | NA | 3 | 0.6667 |
| 8_6713392 | 1 | 1 | NA | NA | NA | NA | 1 | 0 | 2 | 0.5 | NA | NA | 3 | 0.3333 | 2 | 0 | NA | NA | 5 | 0.2 |
| 8_6713418 | NA | NA | NA | NA | NA | NA | 1 | 1 | 1 | 1 | NA | NA | 2 | 1 | 2 | 0.5 | NA | NA | 4 | 0.75 |
| 8_6713423 | NA | NA | NA | NA | NA | NA | 1 | 1 | 1 | 1 | NA | NA | 3 | 1 | 2 | 1 | NA | NA | 5 | 1 |
| 8_6714955 | NA | NA | 1 | 1 | NA | NA | NA | NA | 1 | 1 | 1 | 1 | NA | NA | NA | NA | NA | NA | 1 | 1 |
| 8_6714973 | NA | NA | 1 | 1 | NA | NA | NA | NA | 1 | 1 | 1 | 1 | NA | NA | NA | NA | NA | NA | 1 | 1 |
| 8_6714975 | NA | NA | 1 | 1 | NA | NA | NA | NA | 1 | 1 | 1 | 1 | NA | NA | NA | NA | NA | NA | 1 | 1 |
| 8_6715653 | 2 | 1 | 5 | 0.4 | 2 | 0.5 | 1 | 0 | 10 | 0.5 | 3 | 0.3333 | 1 | 1 | 4 | 0.75 | 3 | 1 | 11 | 0.7273 |
| 8_6715654 | 3 | 0.6667 | 2 | 0 | 2 | 0 | 1 | 0 | 8 | 0.25 | 3 | 0.6667 | 1 | 0 | 2 | 0.5 | 3 | 0.3333 | 9 | 0.4444 |
| 8_6715658 | 2 | 1 | 4 | 1 | 1 | 1 | 1 | 1 | 8 | 1 | 2 | 1 | 1 | 1 | 4 | 1 | 3 | 1 | 10 | 1 |
| 8_6715667 | 2 | 1 | 4 | 1 | 2 | 1 | 1 | 1 | 9 | 1 | 3 | 1 | 1 | 1 | 4 | 0.75 | 3 | 0.6667 | 11 | 0.8182 |
| 8_6715683 | 4 | 0.5 | 5 | 0.4 | 2 | 1 | 1 | 1 | 12 | 0.5833 | 3 | 0.6667 | 2 | 0.5 | 4 | 0.75 | 3 | 1 | 12 | 0.75 |
| 8_6715688 | 3 | 1 | 5 | 1 | 2 | 1 | 1 | 0 | 11 | 0.9091 | 2 | 1 | 1 | 1 | 3 | 1 | 2 | 1 | 8 | 1 |

| | | | | | | | | | | | | | | | | | | | | |
|---|---|---|---|---|---|---|---|---|---|---|---|---|---|---|---|---|---|---|---|---|
| 8_6715692 | 3 | 0.3333 | 2 | 0.5 | 2 | 0.5 | 1 | 1 | 8 | 0.5 | 3 | 0.6667 | 1 | 0 | 4 | 0.75 | 2 | 0.5 | 10 | 0.6 |
| 8_6715702 | 4 | 0 | 5 | 0.2 | 1 | 0 | 1 | 0 | 11 | 0.0909 | 3 | 0 | 2 | 0 | 5 | 0 | 4 | 0.25 | 14 | 0.0714 |
| 8_6715703 | 4 | 1 | 5 | 1 | 2 | 1 | 1 | 1 | 12 | 1 | 3 | 0.6667 | 1 | 1 | 5 | 0.8 | 4 | 1 | 13 | 0.8462 |
| 8_6715715 | 4 | 1 | 5 | 0.2 | 3 | 0.6667 | 1 | 1 | 13 | 0.6154 | 3 | 1 | 2 | 0 | 5 | 0.2 | 4 | 0.75 | 14 | 0.5 |
| 8_6715719 | 4 | 0.5 | 4 | 0.25 | 2 | 0 | 1 | 0 | 11 | 0.2727 | 3 | 0 | 2 | 0 | 5 | 0.2 | 3 | 0 | 13 | 0.0769 |
| 8_6715742 | 4 | 0.75 | 4 | 0.75 | 3 | 0.3333 | 1 | 0 | 12 | 0.5833 | 3 | 0 | 2 | 0.5 | 6 | 0.3333 | 3 | 0.3333 | 14 | 0.2857 |
| 8_6715743 | 4 | 0.75 | 5 | 0.8 | 2 | 0 | NA | NA | 11 | 0.6364 | 2 | 0.5 | 2 | 0.5 | 2 | 0.5 | 4 | 0.75 | 10 | 0.6 |
| 8_6715765 | 5 | 0.2 | 5 | 0.2 | 2 | 0.5 | 2 | 0 | 14 | 0.2143 | 2 | 0 | 1 | 0 | 5 | 0.4 | 3 | 0.3333 | 11 | 0.2727 |
| 8_6715767 | 5 | 1 | 5 | 1 | 3 | 1 | 2 | 1 | 15 | 1 | 3 | 1 | 2 | 1 | 5 | 1 | 4 | 1 | 14 | 1 |
| 8_6715770 | 4 | 1 | 5 | 0.8 | 3 | 0.6667 | 2 | 1 | 14 | 0.8571 | 2 | 1 | 1 | 1 | 6 | 0.8333 | 3 | 1 | 12 | 0.9167 |
| 8_6715782 | 4 | 0.75 | 4 | 1 | 3 | 0.6667 | 2 | 0.5 | 13 | 0.7692 | 3 | 1 | 1 | 1 | 3 | 0.6667 | 4 | 0.75 | 11 | 0.8182 |
| 8_6715795 | 5 | 0.6 | 6 | 0.8333 | 3 | 0.6667 | 2 | 1 | 16 | 0.75 | 4 | 0.25 | 2 | 0 | 6 | 0.6667 | 5 | 1 | 17 | 0.5882 |
| 8_6715797 | 5 | 0.4 | 5 | 0.2 | 3 | 0.3333 | 2 | 0 | 15 | 0.2667 | 4 | 0.25 | 2 | 0 | 5 | 0.2 | 4 | 0.5 | 15 | 0.2667 |
| 8_6715837 | 5 | 0 | 6 | 0.1667 | 3 | 0.3333 | 2 | 0 | 16 | 0.125 | 5 | 0.2 | 2 | 0 | 6 | 0.1667 | 5 | 0.4 | 18 | 0.2222 |
| 8_6715852 | 5 | 0.4 | 5 | 0.2 | 3 | 0.3333 | 2 | 0.5 | 15 | 0.3333 | 4 | 0.25 | 2 | 0 | 7 | 0.2857 | 5 | 0.4 | 18 | 0.2778 |
| 8_6715891 | 7 | 0.5714 | 6 | 0.8333 | 3 | 0 | 2 | 0 | 18 | 0.5 | 4 | 0.5 | 2 | 0.5 | 8 | 0.375 | 6 | 0.3333 | 20 | 0.4 |
| 8_6715911 | 7 | 0.5714 | 7 | 0.2857 | 2 | 0 | 2 | 0 | 18 | 0.3333 | 5 | 0.2 | 2 | 1 | 8 | 0.125 | 7 | 0.2857 | 22 | 0.2727 |
| 8_6731332 | 1 | 0 | 2 | 0.5 | 1 | 0 | NA | NA | 4 | 0.25 | 1 | 0 | 1 | 0 | 4 | 0.25 | 3 | 0.3333 | 9 | 0.2222 |
| 8_6731358 | 2 | 0.5 | 1 | 0 | 1 | 0 | NA | NA | 4 | 0.25 | 1 | 0 | 1 | 1 | 4 | 0.25 | 2 | 0 | 8 | 0.25 |
| 8_6731367 | 2 | 0 | 2 | 0.5 | 3 | 0 | NA | NA | 7 | 0.1429 | 1 | 0 | 1 | 0 | 4 | 0 | 3 | 0.3333 | 9 | 0.1111 |
| 8_6731368 | 2 | 0 | 1 | 0 | 3 | 0.3333 | 1 | 0 | 7 | 0.1429 | 1 | 1 | 1 | 0 | 3 | 0.3333 | 3 | 0 | 8 | 0.25 |
| 8_6794368 | NA | NA | 2 | 0.5 | NA | NA | NA | NA | 2 | 0.5 | 1 | 1 | NA | NA | 1 | 1 | NA | NA | 2 | 1 |
| 8_6794378 | NA | NA | 2 | 1 | NA | NA | NA | NA | 2 | 1 | 1 | 1 | NA | NA | 1 | 1 | NA | NA | 2 | 1 |
| 8_6794411 | NA | NA | 2 | 0.5 | NA | NA | NA | NA | 2 | 0.5 | 1 | 1 | NA | NA | 1 | 1 | NA | NA | 2 | 1 |
| 8_6794448 | NA | NA | 2 | 0.5 | NA | NA | NA | NA | 2 | 0.5 | 1 | 1 | NA | NA | 1 | 1 | NA | NA | 2 | 1 |
| 8_6794479 | NA | NA | 1 | 1 | NA | NA | NA | NA | 1 | 1 | NA | NA | NA | NA | 1 | 1 | NA | NA | 1 | 1 |
| 8_6796671 | 1 | 1 | 1 | 1 | NA | NA | NA | NA | 2 | 1 | 1 | 1 | NA | NA | 1 | 1 | 1 | 1 | 3 | 1 |
| 8_6814256 | 22 | 0.1818 | 26 | 0.1923 | 24 | 0.125 | 25 | 0.16 | 97 | 0.1649 | 11 | 0.0909 | 29 | 0.069 | 20 | 0.05 | 25 | 0.12 | 85 | 0.0824 |
| 8_6814888 | 3 | 0 | 4 | 0.25 | 2 | 0.5 | 2 | 0 | 11 | 0.1818 | NA | NA | 5 | 0.2 | 1 | 0 | NA | NA | 6 | 0.1667 |
| 8_6814905 | 3 | 0 | 4 | 0.25 | 1 | 1 | NA | NA | 8 | 0.25 | NA | NA | 5 | 0.2 | 1 | 0 | 1 | 0 | 7 | 0.1429 |
| 8_6814943 | 3 | 0.3333 | 4 | 0.75 | NA | NA | 1 | 0 | 8 | 0.5 | NA | NA | 4 | 0.25 | NA | NA | NA | NA | 4 | 0.25 |
| 8_6814975 | 2 | 0 | 4 | 0.5 | NA | NA | NA | NA | 6 | 0.3333 | NA | NA | 5 | 0.2 | 1 | 0 | NA | NA | 6 | 0.1667 |
| 8_6846075 | NA | NA | 1 | 1 | NA | NA | NA | NA | 1 | 1 | NA | NA | NA | NA | NA | NA | 1 | 1 | 1 | 1 |
| 8_6846097 | NA | NA | 1 | 1 | NA | NA | NA | NA | 1 | 1 | NA | NA | NA | NA | NA | NA | 1 | 1 | 1 | 1 |
| 8_6846102 | NA | NA | 1 | 1 | NA | NA | NA | NA | 1 | 1 | NA | NA | NA | NA | NA | NA | 1 | 1 | 1 | 1 |
| 8_6846126 | NA | NA | 1 | 1 | NA | NA | NA | NA | 1 | 1 | NA | NA | NA | NA | NA | NA | 1 | 1 | 1 | 1 |
| 8_6880961 | 1 | 0 | 3 | 0.3333 | 4 | 0.25 | 4 | 0.5 | 12 | 0.3333 | 5 | 0.8 | 2 | 0 | 6 | 0.1667 | 3 | 0.3333 | 16 | 0.375 |
| 8_6880989 | 3 | 0.3333 | 4 | 0.5 | 5 | 0 | 4 | 0.25 | 16 | 0.25 | 4 | 0 | 4 | 1 | 6 | 0.1667 | 5 | 0.2 | 19 | 0.3158 |
| 8_6880999 | 3 | 0.6667 | 3 | 0.3333 | 2 | 0 | 1 | 0 | 9 | 0.3333 | 4 | 0.5 | 3 | 0 | 6 | 0.3333 | 4 | 0 | 17 | 0.2353 |
| 8_6881002 | 3 | 0.6667 | 4 | 0.25 | 3 | 0.3333 | 3 | 0.3333 | 13 | 0.3846 | 4 | 0.5 | 4 | 0.25 | 6 | 0.3333 | 4 | 0.75 | 18 | 0.4444 |
| 8_6881005 | 2 | 0.5 | 2 | 0 | 2 | 0 | 3 | 0.3333 | 9 | 0.2222 | 3 | 0 | 1 | 0 | 2 | 0.5 | 3 | 0.3333 | 9 | 0.2222 |
| 8_6881009 | 3 | 0.6667 | 4 | 1 | 3 | 0.6667 | 2 | 1 | 12 | 0.8333 | 4 | 1 | 2 | 0.5 | 6 | 0.6667 | 4 | 0.75 | 16 | 0.75 |
| 8_6881011 | 3 | 0 | 4 | 0.5 | 3 | 0.6667 | 3 | 0 | 13 | 0.3077 | 4 | 0.25 | 3 | 0 | 6 | 0.3333 | 3 | 0.3333 | 16 | 0.25 |
| 8_6881015 | 2 | 1 | 4 | 0.75 | 2 | 0.5 | 2 | 1 | 10 | 0.8 | 3 | 1 | 1 | 1 | 5 | 0.6 | 4 | 0.75 | 13 | 0.7692 |
| 8_6881016 | 2 | 0 | 4 | 0.5 | 3 | 0 | 3 | 0.3333 | 12 | 0.25 | 4 | 0.25 | 2 | 0 | 6 | 0 | 4 | 0 | 16 | 0.0625 |
| 8_6881022 | 2 | 1 | 4 | 1 | 3 | 1 | 3 | 1 | 12 | 1 | 3 | 1 | 2 | 1 | 6 | 0.6667 | 3 | 1 | 14 | 0.8571 |
| 8_6881059 | 2 | 0.5 | 4 | 0 | 3 | 0 | 2 | 0.5 | 11 | 0.1818 | 1 | 0 | 3 | 0 | 4 | 0 | 4 | 0.25 | 12 | 0.0833 |
| 8_6881065 | 3 | 0.3333 | 4 | 0.25 | 3 | 0.3333 | 3 | 0.6667 | 13 | 0.3846 | 1 | 0 | 2 | 1 | 5 | 0.2 | 4 | 0.75 | 12 | 0.5 |
| 8_6881071 | 3 | 0 | 3 | 0.3333 | 1 | 0 | 3 | 0 | 10 | 0.1 | 1 | 0 | 3 | 0 | 4 | 0.25 | 4 | 0 | 12 | 0.0833 |
| 8_6881077 | 3 | 0.6667 | 4 | 0.25 | 3 | 0.6667 | 2 | 0 | 12 | 0.4167 | 2 | 1 | 3 | 0 | 4 | 0.25 | 4 | 0 | 13 | 0.2308 |
| 8_6881091 | 3 | 0 | 3 | 1 | 3 | 0.3333 | 3 | 0.3333 | 12 | 0.4167 | 2 | 1 | 3 | 0.6667 | 5 | 0.4 | 4 | 0.25 | 14 | 0.5 |
| 8_6881098 | 2 | 0.5 | 3 | 0 | 3 | 0.3333 | 3 | 1 | 11 | 0.4545 | 2 | 0.5 | 2 | 1 | 4 | 1 | 4 | 0.75 | 12 | 0.8333 |
| 8_6881105 | 3 | 0.6667 | 1 | 1 | 3 | 0.6667 | 3 | 0.6667 | 10 | 0.7 | 1 | 0 | 3 | 0.6667 | 4 | 1 | 3 | 1 | 11 | 0.8182 |
| 8_6881108 | 3 | 1 | 2 | 1 | 3 | 0.6667 | 3 | 0.6667 | 11 | 0.8182 | 2 | 0.5 | 3 | 0.3333 | 5 | 0.4 | 4 | 0.5 | 14 | 0.4286 |
| 8_6881122 | 1 | 1 | 3 | 1 | 2 | 0 | 2 | 1 | 8 | 0.75 | 2 | 1 | NA | NA | 4 | 0.5 | 4 | 1 | 10 | 0.8 |
| 8_6881126 | 3 | 1 | 3 | 0.6667 | 3 | 1 | 3 | 1 | 12 | 0.9167 | 2 | 0.5 | 2 | 0.5 | 4 | 0.75 | 4 | 0.75 | 12 | 0.6667 |
| 8_6891368 | 2 | 0 | 3 | 0 | 2 | 0 | 3 | 0.3333 | 10 | 0.1 | 1 | 0 | 3 | 0.3333 | 3 | 0 | 2 | 0 | 9 | 0.1111 |
| 8_6891372 | 2 | 0 | 3 | 0.3333 | 1 | 0 | 3 | 0.3333 | 9 | 0.2222 | 1 | 0 | 3 | 0.3333 | 3 | 0 | 2 | 0 | 9 | 0.1111 |
| 8_6891382 | 2 | 0.5 | 3 | 0 | 2 | 1 | 2 | 0.5 | 9 | 0.4444 | 1 | 0 | 3 | 0.6667 | 3 | 0.6667 | 2 | 0 | 9 | 0.4444 |
| 8_6891383 | 1 | 1 | 3 | 0.3333 | 1 | 1 | 2 | 0 | 7 | 0.4286 | 1 | 1 | 2 | 1 | 3 | 0 | 2 | 0.5 | 8 | 0.5 |
| 8_6891387 | 1 | 0 | 3 | 0.3333 | 2 | 0 | 3 | 0 | 9 | 0.1111 | NA | NA | 3 | 0.3333 | 3 | 0 | 2 | 0 | 8 | 0.125 |
| 8_6891391 | 2 | 0.5 | 3 | 0.3333 | 2 | 0 | 3 | 0.3333 | 10 | 0.3 | NA | NA | 3 | 0 | 3 | 0.6667 | 2 | 0.5 | 8 | 0.375 |
| 8_6891395 | 2 | 0.5 | 3 | 0.6667 | 2 | 1 | 1 | 0 | 8 | 0.625 | NA | NA | 3 | 0.6667 | 3 | 0.6667 | 2 | 0.5 | 8 | 0.625 |
| 8_6891406 | 2 | 0.5 | 3 | 0.3333 | 2 | 0 | 3 | 1 | 10 | 0.5 | NA | NA | 3 | 0 | 3 | 0 | 2 | 0.5 | 8 | 0.125 |
| 8_6891407 | 2 | 1 | 2 | 1 | 2 | 1 | 3 | 1 | 9 | 1 | NA | NA | 2 | 0.5 | 2 | 1 | 1 | 1 | 5 | 0.8 |

| | | | | | | | | | | | | | | | | | | | | |
|---|---|---|---|---|---|---|---|---|---|---|---|---|---|---|---|---|---|---|---|---|
| 8_6891423 | 1 | 0 | 2 | 1 | 2 | 1 | 3 | 0.6667 | 8 | 0.75 | NA | NA | 3 | 0.6667 | 2 | 1 | 1 | 1 | 6 | 0.8333 |
| 8_6891431 | 1 | 1 | 3 | 0.3333 | 2 | 0.5 | 3 | 0.6667 | 9 | 0.5556 | NA | NA | 3 | 0.3333 | 3 | 0.3333 | 2 | 0.5 | 8 | 0.375 |
| 8_6891435 | 1 | 1 | 3 | 0 | 2 | 0.5 | 3 | 0 | 9 | 0.2222 | NA | NA | 3 | 0 | 3 | 0.3333 | 2 | 0 | 8 | 0.125 |
| 8_6891436 | 1 | 0 | 3 | 0 | 2 | 0 | 3 | 0.6667 | 9 | 0.2222 | NA | NA | 3 | 0.3333 | 3 | 0 | 1 | 0 | 7 | 0.1429 |
| 8_6891442 | 1 | 1 | 3 | 0.6667 | 2 | 0.5 | 2 | 0.5 | 8 | 0.625 | NA | NA | 3 | 0.6667 | 3 | 0.6667 | 2 | 0 | 8 | 0.5 |
| 8_6927997 | 7 | 0.5714 | 8 | 0.25 | 5 | 0.2 | 4 | 0 | 24 | 0.2917 | 4 | 0 | 8 | 0 | 8 | 0.25 | 9 | 0.3333 | 29 | 0.1724 |
| 8_6928007 | 7 | 0.1429 | 7 | 0.4286 | 9 | 0.3333 | 5 | 0.2 | 28 | 0.2857 | 5 | 0.2 | 10 | 0.3 | 9 | 0.2222 | 10 | 0.3 | 34 | 0.2647 |
| 8_6928008 | 7 | 0 | 9 | 0.1111 | 9 | 0.1111 | 5 | 0 | 30 | 0.0667 | 6 | 0.1667 | 11 | 0.0909 | 8 | 0 | 10 | 0 | 35 | 0.0571 |
| 8_6928022 | 6 | 0 | 7 | 0.1429 | 8 | 0.125 | 4 | 0 | 25 | 0.08 | 6 | 0.1667 | 11 | 0.0909 | 6 | 0 | 10 | 0.2 | 33 | 0.1212 |
| 8_6928031 | 3 | 0 | 2 | 0 | 2 | 0 | 1 | 1 | 8 | 0.125 | NA | NA | 5 | 0 | 1 | 1 | 4 | 0 | 10 | 0.1 |
| 8_6928050 | 2 | 1 | 1 | 0 | 1 | 0 | NA | NA | 4 | 0.5 | NA | NA | 2 | 0.5 | 3 | 0.3333 | 5 | 0.6 | 10 | 0.5 |
| 8_6928053 | 3 | 0.3333 | 1 | 0 | 2 | 0 | NA | NA | 6 | 0.1667 | 1 | 0 | 3 | 0.6667 | 3 | 0.3333 | 4 | 0 | 11 | 0.2727 |
| 8_6928068 | 4 | 1 | 1 | 1 | 1 | 0 | NA | NA | 6 | 0.8333 | 1 | 1 | 2 | 1 | 3 | 1 | 5 | 0.8 | 11 | 0.9091 |
| 8_6928147 | 2 | 0.5 | NA | NA | 3 | 0 | NA | NA | 5 | 0.2 | 1 | 0 | 1 | 1 | 3 | 0 | 1 | 0 | 6 | 0.1667 |
| 8_6928182 | 2 | 0.5 | NA | NA | 3 | 0 | NA | NA | 5 | 0.2 | 1 | 0 | NA | NA | 3 | 0.3333 | 1 | 0 | 5 | 0.2 |
| 8_6928186 | 2 | 0.5 | NA | NA | 3 | 0.3333 | NA | NA | 5 | 0.4 | 1 | 1 | 1 | 1 | 3 | 0 | 1 | 1 | 6 | 0.5 |
| 8_6928195 | 2 | 0.5 | NA | NA | 3 | 0 | NA | NA | 5 | 0.2 | 1 | 0 | 1 | 0 | 3 | 0 | 1 | 1 | 6 | 0.1667 |
| 8_6935899 | 4 | 0.25 | 2 | 0 | NA | NA | 8 | 0.125 | 14 | 0.1429 | 2 | 0.5 | 6 | 0.3333 | 2 | 0 | 3 | 0 | 13 | 0.2308 |
| 8_6956509 | 52 | 0.0577 | 82 | 0.0488 | 43 | 0.0698 | 58 | 0.0862 | 235 | 0.0638 | 35 | 0.0857 | 68 | 0.1471 | 57 | 0.0526 | 47 | 0.0638 | 207 | 0.0918 |
| 8_6956511 | 32 | 0.2188 | 63 | 0.2698 | 29 | 0.2414 | 46 | 0.1522 | 170 | 0.2235 | 23 | 0.087 | 60 | 0.2333 | 39 | 0.1538 | 37 | 0.2703 | 159 | 0.2013 |
| 8_6956520 | 55 | 0.2182 | 78 | 0.1026 | 40 | 0.125 | 50 | 0.2 | 223 | 0.157 | 35 | 0.2 | 60 | 0.2167 | 55 | 0.2545 | 44 | 0.2273 | 194 | 0.2268 |
| 8_6956591 | 74 | 0.1757 | 101 | 0.1782 | 60 | 0.1833 | 72 | 0.1528 | 307 | 0.1726 | 50 | 0.24 | 81 | 0.2099 | 75 | 0.16 | 55 | 0.2364 | 261 | 0.2069 |
| 8_6958549 | 22 | 0.9545 | 18 | 1 | 12 | 1 | 16 | 1 | 68 | 0.9853 | 17 | 0.9412 | 19 | 1 | 24 | 0.9583 | 19 | 1 | 79 | 0.9747 |
| 8_6958554 | 26 | 0.9231 | 17 | 1 | 14 | 1 | 15 | 0.9333 | 72 | 0.9583 | 17 | 0.9412 | 19 | 0.8947 | 27 | 1 | 22 | 0.9545 | 85 | 0.9529 |
| 8_6958559 | 26 | 0.3846 | 17 | 0.2353 | 15 | 0.2 | 17 | 0.4118 | 75 | 0.32 | 21 | 0.2857 | 23 | 0.6522 | 26 | 0.2692 | 22 | 0.3636 | 92 | 0.3913 |
| 8_6958561 | 26 | 0.1923 | 17 | 0.2353 | 14 | 0.0714 | 16 | 0.25 | 73 | 0.1918 | 18 | 0.0556 | 24 | 0.0833 | 26 | 0.4231 | 21 | 0.1905 | 89 | 0.2022 |
| 8_6958569 | 24 | 0.2917 | 17 | 0.2353 | 15 | 0.3333 | 14 | 0.2143 | 70 | 0.2714 | 18 | 0.2778 | 21 | 0.2381 | 22 | 0.3182 | 21 | 0.381 | 82 | 0.3049 |
| 8_6958573 | 22 | 0.0909 | 15 | 0.0667 | 14 | 0 | 14 | 0.1429 | 65 | 0.0769 | 18 | 0.0556 | 23 | 0.1304 | 22 | 0.0455 | 19 | 0.1053 | 82 | 0.0854 |
| 8_6958587 | 21 | 0.0476 | 13 | 0.1538 | 12 | 0.0833 | 12 | 0.1667 | 58 | 0.1034 | 15 | 0.0667 | 21 | 0.0476 | 18 | 0.1111 | 19 | 0.2105 | 73 | 0.1096 |
| 8_6958593 | 21 | 0.2381 | 14 | 0.2143 | 13 | 0.1538 | 13 | 0.1538 | 61 | 0.1967 | 17 | 0.3529 | 22 | 0.1364 | 21 | 0.2381 | 20 | 0.45 | 80 | 0.2875 |
| 8_6958593 | 14 | 0.0714 | 10 | 0.2 | 9 | 0.4444 | 10 | 0.1 | 43 | 0.186 | 11 | 0.0909 | 11 | 0.1818 | 15 | 0 | 18 | 0.1667 | 55 | 0.1091 |
| 8_6958636 | 22 | 0.0455 | 14 | 0.0714 | 10 | 0 | 11 | 0.1818 | 57 | 0.0702 | 15 | 0 | 22 | 0.1364 | 20 | 0.1 | 15 | 0 | 72 | 0.0694 |
| 8_6958761 | 8 | 0.25 | 4 | 0.25 | 2 | 0 | 5 | 0.2 | 19 | 0.2105 | 5 | 0.2 | 11 | 0.3636 | 7 | 0.5714 | 8 | 0.625 | 31 | 0.4516 |
| 8_6958771 | 8 | 0 | 6 | 0.3333 | 3 | 0.3333 | 5 | 0.4 | 22 | 0.2273 | 6 | 0 | 13 | 0.1538 | 11 | 0 | 7 | 0 | 37 | 0.0541 |
| 8_6958781 | 7 | 0.2857 | 4 | 0.25 | 4 | 0.25 | 7 | 0.4286 | 22 | 0.3182 | 5 | 0 | 13 | 0.0769 | 11 | 0.3636 | 7 | 0.1429 | 36 | 0.1667 |
| 8_6958781 | 7 | 0.8571 | 5 | 0.8 | 3 | 1 | 6 | 1 | 21 | 0.9048 | 6 | 1 | 11 | 1 | 10 | 1 | 6 | 0.6667 | 33 | 0.9394 |
| 8_6958787 | 5 | 0.2 | 5 | 0 | 5 | 0 | 7 | 0.1429 | 22 | 0.0909 | 7 | 0.1429 | 9 | 0.1111 | 8 | 0 | 6 | 0.3333 | 30 | 0.1333 |
| 8_6958787 | 1 | 1 | NA | NA | NA | NA | NA | NA | 1 | 1 | NA | NA | 2 | 0.5 | 2 | 0.5 | NA | NA | 4 | 0.5 |
| 8_6958808 | 5 | 0.8 | 5 | 1 | 2 | 1 | 6 | 1 | 18 | 0.9444 | 7 | 1 | 12 | 1 | 9 | 1 | 6 | 1 | 34 | 1 |
| 8_6958812 | 2 | 0.5 | 5 | 0.6 | 3 | 0.6667 | 5 | 0.2 | 15 | 0.4667 | 6 | 0.6667 | 11 | 0.5455 | 9 | 0.8889 | 5 | 0.6 | 31 | 0.6774 |
| 8_6958812 | 5 | 0.6 | 5 | 0.2 | 2 | 0 | 6 | 0.5 | 18 | 0.3889 | 6 | 0.5 | 9 | 0.6667 | 6 | 0 | 4 | 0.5 | 25 | 0.44 |
| 8_6958817 | 5 | 1 | 3 | 1 | 3 | 1 | 5 | 1 | 16 | 1 | 7 | 0.8571 | 8 | 1 | 7 | 1 | 5 | 0.8 | 27 | 0.9259 |
| 8_6958820 | 6 | 0.5 | 2 | 0.5 | 2 | 0 | 6 | 0.1667 | 16 | 0.3125 | 7 | 0.4286 | 9 | 0.4444 | 8 | 0.5 | 5 | 0.4 | 29 | 0.4483 |
| 8_6958822 | 6 | 0 | 3 | 0.3333 | 3 | 0 | 7 | 0.4286 | 19 | 0.2105 | 6 | 0.1667 | 7 | 0.1429 | 7 | 0.1429 | 5 | 0.2 | 25 | 0.16 |
| 8_6958845 | 5 | 1 | 2 | 1 | 3 | 1 | 4 | 1 | 14 | 1 | 6 | 1 | 7 | 0.8571 | 5 | 0.6 | 5 | 0.8 | 23 | 0.8261 |
| 8_6958855 | 4 | 0 | 2 | 0 | 3 | 0 | 5 | 0.2 | 14 | 0.0714 | 6 | 0.1667 | 7 | 0.2857 | 7 | 0.1429 | 4 | 0.25 | 24 | 0.2083 |
| 8_6958857 | 5 | 0.4 | 1 | 0 | 2 | 0.5 | 4 | 0.75 | 12 | 0.5 | 5 | 0.8 | 5 | 0.6 | 6 | 0.5 | 3 | 1 | 19 | 0.6842 |
| 8_6958860 | 5 | 0.4 | 2 | 1 | 3 | 0.3333 | 4 | 1 | 14 | 0.6429 | 5 | 0.6 | 7 | 0.7143 | 5 | 0.6 | 4 | 0.25 | 21 | 0.5714 |
| 8_6958865 | 5 | 0.8 | 2 | 1 | 3 | 1 | 6 | 0.5 | 16 | 0.75 | 5 | 0.6 | 6 | 0.6667 | 7 | 0.8571 | 4 | 0.5 | 22 | 0.6818 |
| 8_6959123 | 2 | 0.5 | 2 | 1 | 3 | 0.6667 | 4 | 0.5 | 11 | 0.6364 | 4 | 0.5 | 2 | 0.5 | 5 | 0.8 | 2 | 1 | 13 | 0.6923 |
| 8_6959124 | 4 | 0.75 | 3 | 0.6667 | 3 | 0.3333 | 5 | 0.8 | 15 | 0.6667 | 5 | 0.6 | 4 | 0.75 | 6 | 0.3333 | 3 | 0 | 18 | 0.4444 |
| 8_6959132 | 4 | 0.5 | 3 | 0 | 3 | 0 | 4 | 0.25 | 14 | 0.2143 | 4 | 0.25 | 5 | 0.4 | 5 | 0 | 3 | 0 | 17 | 0.1765 |
| 8_6959148 | 4 | 1 | 2 | 1 | 2 | 1 | 4 | 1 | 12 | 1 | 2 | 1 | 5 | 0.8 | 3 | 1 | 2 | 1 | 12 | 0.9167 |
| 8_6959161 | 3 | 0.3333 | 2 | 0.5 | 3 | 0 | 5 | 0.6 | 13 | 0.3846 | 1 | 1 | 5 | 0.4 | 5 | 0.2 | 2 | 0 | 13 | 0.3077 |
| 8_6959162 | 3 | 0 | 2 | 0 | 2 | 0 | 4 | 0.25 | 11 | 0.0909 | 3 | 0.3333 | 5 | 0 | 6 | 0.3333 | 2 | 0 | 16 | 0.1875 |
| 8_6959341 | 3 | 0 | 2 | 0 | 2 | 0.5 | 4 | 0 | 11 | 0.0909 | 3 | 0 | 3 | 0 | 5 | 0.2 | 2 | 0 | 13 | 0.0769 |
| 8_6959346 | 3 | 0.3333 | 1 | 0 | 3 | 0.3333 | 4 | 1 | 11 | 0.5455 | 2 | 1 | 3 | 0.3333 | 5 | 0.6 | 1 | 0 | 11 | 0.5455 |
| 8_6968067 | 1 | 1 | NA | NA | 1 | 0 | 2 | 0.5 | 4 | 0.5 | NA | NA | NA | NA | 2 | 0.5 | NA | NA | 2 | 0.5 |
| 8_6976666 | NA | NA | 2 | 1 | NA | NA | 1 | 1 | 3 | 1 | NA | NA | 1 | 1 | NA | NA | NA | NA | 1 | 1 |
| 8_6976685 | NA | NA | 2 | 0.5 | NA | NA | 2 | 1 | 4 | 0.75 | NA | NA | 1 | 1 | NA | NA | 1 | 1 | 2 | 1 |
| 8_6976691 | NA | NA | 2 | 1 | NA | NA | 2 | 0.5 | 4 | 0.75 | NA | NA | NA | NA | NA | NA | 1 | 1 | 1 | 1 |
| 8_6976846 | 1 | 1 | 2 | 0.5 | 1 | 1 | NA | NA | 4 | 0.75 | 1 | 1 | 1 | 1 | 1 | 1 | 2 | 1 | 5 | 1 |
| 8_6976851 | 1 | 1 | 2 | 1 | 1 | 1 | 3 | 0.6667 | 7 | 0.8571 | 1 | 1 | 1 | 1 | 1 | 1 | 2 | 0 | 5 | 0.6 |
| 8_6976863 | NA | NA | 2 | 0.5 | 1 | 1 | 1 | 0 | 4 | 0.5 | 1 | 0 | 1 | 1 | 1 | 0 | 3 | 0 | 6 | 0.1667 |
| 8_6976868 | 1 | 1 | 2 | 1 | 1 | 0 | 2 | 0 | 6 | 0.5 | 1 | 0 | 1 | 0 | 2 | 0.5 | NA | NA | 4 | 0.25 |

| | | | | | | | | | | | | | | | | | | | | |
|---|---|---|---|---|---|---|---|---|---|---|---|---|---|---|---|---|---|---|---|---|
| 8_6976875 | 1 | 0 | 2 | 0 | 1 | 0 | 3 | 0.3333 | 7 | 0.1429 | 1 | 0 | NA | NA | 2 | 0 | 3 | 0.3333 | 6 | 0.1667 |
| 8_69769018 | NA | NA | 2 | 1 | NA | NA | 2 | 0.5 | 4 | 0.75 | 1 | 1 | 1 | 1 | 2 | 0.5 | 2 | 0.5 | 6 | 0.6667 |
| 8_6976905 | 1 | 1 | 2 | 0.5 | 1 | 0 | 3 | 0 | 7 | 0.2857 | NA | NA | NA | NA | 2 | 0.5 | 2 | 0.5 | 4 | 0.5 |
| 8_6976906 | 1 | 1 | 3 | 0.3333 | 1 | 0 | 3 | 0.3333 | 8 | 0.375 | 1 | 1 | 1 | 0 | 2 | 0 | 3 | 0 | 7 | 0.1429 |
| 8_6976938 | NA | NA | 3 | 0.6667 | 1 | 0 | 4 | 0.5 | 8 | 0.5 | 1 | 0 | 1 | 0 | 2 | 0.5 | 3 | 0.3333 | 7 | 0.2857 |
| 8_6977056 | NA | NA | 1 | 1 | NA | NA | NA | NA | 1 | 1 | NA | NA | NA | NA | NA | NA | 1 | 1 | 1 | 1 |
| 8_6977149 | NA | NA | 2 | 0.5 | NA | NA | NA | NA | 2 | 0.5 | NA | NA | NA | NA | NA | NA | 2 | 0.5 | 2 | 0.5 |
| 8_6977178 | NA | NA | 2 | 0.5 | NA | NA | NA | NA | 2 | 0.5 | NA | NA | NA | NA | NA | NA | 2 | 1 | 2 | 1 |
| 8_6977227 | NA | NA | 2 | 1 | NA | NA | NA | NA | 2 | 1 | NA | NA | NA | NA | NA | NA | 2 | 1 | 2 | 1 |
| 8_6977228 | NA | NA | 2 | 0.5 | NA | NA | NA | NA | 2 | 0.5 | NA | NA | NA | NA | NA | NA | 2 | 1 | 2 | 1 |
| 8_6977230 | NA | NA | 2 | 1 | NA | NA | NA | NA | 2 | 1 | NA | NA | NA | NA | NA | NA | 2 | 1 | 2 | 1 |
| 8_6977232 | NA | NA | 2 | 1 | NA | NA | NA | NA | 2 | 1 | NA | NA | NA | NA | NA | NA | 2 | 1 | 2 | 1 |
| 8_6977233 | NA | NA | 2 | 1 | NA | NA | NA | NA | 2 | 1 | NA | NA | NA | NA | NA | NA | 2 | 1 | 2 | 1 |
| 8_6982310 | NA | NA | 2 | 0.5 | NA | NA | NA | NA | 2 | 0.5 | 1 | 1 | NA | NA | NA | NA | 1 | 0 | 2 | 0.5 |
| 8_6982376 | NA | NA | 1 | 1 | 1 | 0 | 2 | 0 | 4 | 0.25 | 2 | 0.5 | 1 | 0 | 1 | 0 | 2 | 0.5 | 6 | 0.3333 |
| 8_6982384 | NA | NA | 1 | 1 | 1 | 1 | 2 | 1 | 4 | 1 | 3 | 0.6667 | 1 | 0 | 1 | 0 | 2 | 1 | 7 | 0.5714 |
| 8_6982396 | NA | NA | NA | NA | 1 | 0 | 2 | 0.5 | 3 | 0.3333 | 3 | 0 | 1 | 1 | NA | NA | 2 | 0 | 6 | 0.1667 |
| 8_6982398 | NA | NA | 1 | 1 | NA | NA | 1 | 1 | 2 | 1 | 3 | 0.6667 | 1 | 0 | 1 | 0 | 2 | 0.5 | 7 | 0.4286 |
| 8_6982435 | NA | NA | 1 | 1 | 1 | 0 | 1 | 1 | 3 | 0.6667 | 2 | 0 | 1 | 0 | 1 | 0 | 2 | 0.5 | 6 | 0.1667 |
| 8_6982498 | NA | NA | 1 | 1 | 1 | 0 | NA | NA | 2 | 0.5 | 2 | 1 | 1 | 0 | 1 | 0 | 2 | 1 | 6 | 0.6667 |
| 8_6982506 | NA | NA | 1 | 1 | 1 | 0 | NA | NA | 2 | 0.5 | 2 | 0.5 | 1 | 0 | 1 | 0 | 2 | 0.5 | 6 | 0.3333 |
| 8_6982605 | NA | NA | 1 | 1 | 1 | 1 | NA | NA | 2 | 1 | 1 | 1 | 1 | 0 | 1 | 0 | 1 | 0 | 4 | 0.25 |
| 8_6982606 | NA | NA | 1 | 1 | 1 | 0 | NA | NA | 2 | 0.5 | 1 | 1 | 1 | 1 | NA | NA | 1 | 0 | 3 | 0.6667 |
| 8_6982612 | NA | NA | 1 | 0 | 1 | 1 | NA | NA | 2 | 0.5 | 1 | 1 | 1 | 0 | 1 | 0 | 1 | 0 | 4 | 0.25 |
| 8_6982614 | NA | NA | 1 | 1 | 1 | 0 | NA | NA | 2 | 0.5 | 1 | 1 | 1 | 0 | 1 | 0 | 1 | 1 | 4 | 0.5 |
| 8_6982617 | NA | NA | 1 | 1 | 1 | 1 | NA | NA | 2 | 1 | 1 | 1 | NA | NA | 1 | 1 | 1 | 1 | 3 | 1 |
| 8_6982629 | NA | NA | 1 | 1 | 1 | 1 | NA | NA | 2 | 1 | NA | NA | NA | NA | 1 | 1 | NA | NA | 1 | 1 |
| 8_6982630 | NA | NA | 1 | 1 | 1 | 0 | NA | NA | 2 | 0.5 | 1 | 1 | 1 | 1 | 1 | 1 | 1 | 1 | 4 | 1 |
| 8_6982637 | NA | NA | 1 | 1 | 1 | 1 | NA | NA | 2 | 1 | 1 | 1 | 1 | 1 | 1 | 1 | NA | NA | 3 | 1 |
| 8_7006361 | 3 | 0.6667 | 3 | 1 | 2 | 1 | 2 | 1 | 10 | 0.9 | 4 | 0.75 | 1 | 1 | 2 | 1 | NA | NA | 7 | 0.8571 |
| 8_7006364 | 3 | 0.6667 | 8 | 0.875 | 2 | 1 | 1 | 1 | 14 | 0.8571 | 2 | 0.5 | 1 | 0 | 1 | 1 | NA | NA | 4 | 0.5 |
| 8_7006374 | 4 | 0.75 | 8 | 0.875 | 4 | 0.5 | 2 | 1 | 18 | 0.7778 | 5 | 0.6 | 2 | 1 | 2 | 1 | 1 | 1 | 10 | 0.8 |
| 8_7006393 | 4 | 1 | 8 | 1 | 4 | 0.5 | 2 | 1 | 18 | 0.8889 | 5 | 1 | 2 | 1 | 2 | 1 | 1 | 1 | 10 | 1 |
| 8_7006396 | 4 | 0 | 8 | 0.125 | 4 | 0 | 2 | 0 | 18 | 0.0556 | 5 | 0 | 2 | 0.5 | 2 | 0 | 1 | 0 | 10 | 0.1 |
| 8_7006398 | 4 | 0 | 7 | 0.7143 | 3 | 0 | 2 | 0.5 | 16 | 0.375 | 4 | 0.5 | 2 | 0.5 | 2 | 0.5 | 1 | 0 | 9 | 0.4444 |
| 8_7006399 | 4 | 0.25 | 8 | 0.25 | 4 | 0 | 2 | 0.5 | 18 | 0.2222 | 5 | 0.4 | 2 | 0.5 | 2 | 0 | 1 | 0 | 10 | 0.3 |
| 8_7006401 | 4 | 0 | 6 | 0.1667 | 4 | 0.25 | 1 | 0 | 15 | 0.1333 | 5 | 0.2 | 2 | 0.5 | 1 | 0 | 1 | 0 | 9 | 0.2222 |
| 8_7006402 | 3 | 0 | 7 | 0.4286 | 4 | 0 | 1 | 0 | 15 | 0.2 | 5 | 0.2 | 1 | 0 | 2 | 0.5 | 1 | 0 | 9 | 0.2222 |
| 8_7006414 | 4 | 0.75 | 8 | 0.5 | 2 | 0 | 2 | 1 | 16 | 0.5625 | 5 | 0.4 | 2 | 0 | 2 | 0 | 1 | 0 | 10 | 0.2 |
| 8_7006419 | 3 | 0.3333 | 5 | 0.4 | 3 | 0.6667 | 2 | 0 | 13 | 0.3846 | 5 | 0.2 | 1 | 1 | 2 | 0.5 | 1 | 1 | 9 | 0.4444 |
| 8_7006439 | 1 | 0 | 5 | 1 | 2 | 0 | 2 | 0.5 | 10 | 0.6 | 4 | 0.75 | 1 | 1 | 1 | 0 | NA | NA | 6 | 0.6667 |
| 8_7006443 | 4 | 0.25 | 7 | 0.5714 | 3 | 0 | 2 | 1 | 16 | 0.4375 | 5 | 0.4 | 2 | 0.5 | 1 | 0 | 1 | 0 | 9 | 0.3333 |
| 8_7006447 | 3 | 1 | 8 | 0.875 | 1 | 0 | 2 | 1 | 14 | 0.8571 | 3 | 0.6667 | 1 | 1 | 1 | 1 | NA | NA | 5 | 0.8 |
| 8_7006454 | 3 | 1 | 6 | 1 | 2 | 0.5 | 2 | 1 | 13 | 0.9231 | 4 | 0.5 | 1 | 1 | 1 | 1 | 1 | 0 | 7 | 0.5714 |
| 8_7006456 | 4 | 0.25 | 8 | 0.25 | 3 | 0 | 2 | 0.5 | 17 | 0.2353 | 5 | 0 | 2 | 0.5 | 1 | 1 | 1 | 0 | 9 | 0.2222 |
| 8_7006462 | 4 | 0 | 7 | 0 | 2 | 0.5 | 2 | 0 | 15 | 0.0667 | 3 | 0.3333 | 2 | 0 | 1 | 0 | 1 | 0 | 7 | 0.1429 |
| 8_7006466 | 1 | 0 | 8 | 0.375 | 3 | 0.3333 | 2 | 0 | 14 | 0.2857 | 2 | 0 | 1 | 0 | 1 | 1 | 1 | 1 | 5 | 0.4 |
| 8_7006470 | 4 | 0 | 8 | 0 | 3 | 0 | 2 | 0.5 | 17 | 0.0588 | 4 | 0 | 2 | 0.5 | 1 | 0 | 1 | 0 | 8 | 0.125 |
| 8_7006481 | 3 | 0.3333 | 7 | 0 | 2 | 0 | 2 | 0 | 14 | 0.0714 | 3 | 0 | 2 | 0.5 | 1 | 0 | 1 | 0 | 7 | 0.1429 |
| 8_7006495 | 4 | 1 | 7 | 1 | 3 | 0.6667 | 2 | 1 | 16 | 0.9375 | 3 | 1 | 2 | 0.5 | 1 | 1 | 1 | 1 | 7 | 0.8571 |
| 8_7006500 | 4 | 0.25 | 7 | 0 | 3 | 0 | 2 | 0 | 16 | 0.0625 | 4 | 0 | 1 | 1 | 1 | 0 | 1 | 0 | 7 | 0.1429 |
| 8_7015536 | NA | NA | NA | NA | 1 | 1 | NA | NA | 1 | 1 | NA | NA | NA | NA | 1 | 1 | 1 | 1 | 2 | 1 |
| 8_7015581 | NA | NA | NA | NA | 1 | 1 | NA | NA | 1 | 1 | NA | NA | NA | NA | 1 | 0 | 1 | 1 | 2 | 0.5 |
| 8_7015792 | 7 | 0.4286 | 2 | 0 | 7 | 0.2857 | 2 | 0.5 | 18 | 0.3333 | 1 | 1 | 4 | 0.75 | 3 | 0 | 4 | 0 | 12 | 0.3333 |
| 8_7015794 | 6 | 0.6667 | 2 | 0.5 | 6 | 1 | 2 | 1 | 16 | 0.8125 | 1 | 0 | 4 | 0.75 | 4 | 0.75 | 5 | 0.8 | 14 | 0.7143 |
| 8_7015835 | 8 | 0.25 | 2 | 0 | 6 | 0.5 | 2 | 0.5 | 18 | 0.3333 | 2 | 0 | 5 | 0.2 | 4 | 0.25 | 5 | 0.6 | 16 | 0.3125 |
| 8_7015866 | 5 | 0 | 2 | 0 | 8 | 0.25 | 2 | 0.5 | 17 | 0.1765 | 1 | 0 | 4 | 0.25 | 3 | 0.3333 | 3 | 0.3333 | 11 | 0.2727 |
| 8_7015890 | 3 | 0.3333 | 2 | 0.5 | 7 | 0.1429 | 2 | 0 | 14 | 0.2143 | 1 | 0 | 5 | 0 | 4 | 0 | 3 | 0.6667 | 13 | 0.1538 |
| 8_7015894 | 5 | 0 | 1 | 1 | 8 | 0.125 | 1 | 0 | 15 | 0.1333 | 2 | 0 | 4 | 0.25 | 4 | 0 | 3 | 0 | 13 | 0.0769 |
| 8_7015896 | 5 | 0.2 | 2 | 0 | 8 | 0.125 | 2 | 0 | 17 | 0.1176 | 2 | 0 | 4 | 0.25 | 4 | 0.25 | 4 | 0.25 | 14 | 0.2143 |
| 8_7015897 | 3 | 0.3333 | 2 | 0.5 | 6 | 0.1667 | 2 | 1 | 13 | 0.3846 | 2 | 0.5 | 4 | 0.25 | 4 | 0.75 | 3 | 0.3333 | 13 | 0.4615 |
| 8_7015902 | 4 | 1 | 1 | 1 | 6 | 1 | 2 | 1 | 13 | 1 | 2 | 1 | 4 | 1 | 3 | 1 | 4 | 1 | 13 | 1 |
| 8_7015905 | 3 | 0.6667 | 2 | 0.5 | 7 | 0.5714 | 2 | 0.5 | 14 | 0.5714 | 2 | 0.5 | 4 | 0.75 | 4 | 0.5 | 4 | 0.75 | 14 | 0.6429 |
| 8_7015909 | 5 | 0.2 | 1 | 0 | 8 | 0.625 | 2 | 1 | 16 | 0.5 | 2 | 1 | 4 | 1 | 4 | 1 | 4 | 0.75 | 14 | 0.9286 |
| 8_7015917 | 5 | 0 | 2 | 0 | 5 | 0.2 | 2 | 0.5 | 14 | 0.1429 | 2 | 1 | 4 | 0.5 | 2 | 0.5 | 3 | 0.3333 | 11 | 0.5455 |

| | | | | | | | | | | | | | | | | | | | | |
|---|---|---|---|---|---|---|---|---|---|---|---|---|---|---|---|---|---|---|---|---|
| 8_7015919 | 5 | 0.2 | 2 | 0 | 7 | 0.1429 | 2 | 0 | 16 | 0.125 | 2 | 0 | 5 | 0.2 | 4 | 0.25 | 4 | 0 | 15 | 0.1333 |
| 8_7015928 | 3 | 0.3333 | 2 | 0 | 6 | 0.1667 | 2 | 0.5 | 13 | 0.2308 | 2 | 0 | 5 | 0.2 | 4 | 0 | 4 | 0 | 15 | 0.0667 |
| 8_7015929 | 5 | 0.2 | 2 | 0.5 | 3 | 0 | 2 | 0 | 12 | 0.1667 | 2 | 0.5 | 4 | 0.25 | 2 | 0 | 3 | 0 | 11 | 0.1818 |
| 8_7015938 | 4 | 0.75 | 1 | 1 | 6 | 1 | 1 | 0 | 12 | 0.8333 | 2 | 0.5 | 3 | 1 | 3 | 1 | 4 | 0.75 | 12 | 0.8333 |
| 8_7015945 | 5 | 0.2 | 2 | 0.5 | 6 | 0.1667 | 1 | 0 | 14 | 0.2143 | 2 | 0 | 4 | 0 | 4 | 0.5 | 4 | 0 | 14 | 0.1429 |
| 8_7015946 | 4 | 0.25 | 2 | 0 | 6 | 0.5 | NA | NA | 12 | 0.3333 | 2 | 0.5 | 4 | 0 | 2 | 0.5 | 3 | 0.3333 | 11 | 0.2727 |
| 8_7015947 | 5 | 0.6 | 2 | 0.5 | 6 | 0 | 1 | 1 | 14 | 0.3571 | 2 | 1 | 4 | 0.5 | 3 | 0.3333 | 3 | 0.6667 | 12 | 0.5833 |
| 8_7015961 | 4 | 0 | 1 | 0 | 6 | 0.1667 | 1 | 0 | 12 | 0.0833 | 2 | 0 | 4 | 0 | 3 | 0 | 3 | 0.3333 | 12 | 0.0833 |
| 8_7015965 | 5 | 0.4 | 2 | 1 | 7 | 0.5714 | 1 | 1 | 15 | 0.6 | 2 | 0 | 4 | 0.75 | 3 | 0.6667 | 3 | 1 | 12 | 0.6667 |
| 8_7015977 | 4 | 0 | 2 | 0 | 4 | 0.25 | 1 | 0 | 11 | 0.0909 | 2 | 0.5 | 4 | 0.5 | 3 | 0 | 2 | 0 | 11 | 0.2727 |
| 8_7015993 | 4 | 1 | 2 | 1 | 5 | 1 | 1 | 1 | 12 | 1 | 2 | 1 | 3 | 1 | 4 | 1 | 3 | 1 | 12 | 1 |
| 8_7016003 | 2 | 0.5 | 1 | 0 | 4 | 0.75 | NA | NA | 7 | 0.5714 | 2 | 1 | 3 | 0.6667 | 2 | 0.5 | 1 | 1 | 8 | 0.75 |
| 8_7016004 | 1 | 0 | 1 | 1 | 1 | 0 | NA | NA | 3 | 0.3333 | NA | NA | NA | NA | NA | NA | 2 | 0.5 | 2 | 0.5 |
| 8_7016016 | 5 | 0.4 | 2 | 0.5 | 6 | 0.5 | 1 | 1 | 14 | 0.5 | 2 | 1 | 4 | 0.75 | 3 | 1 | 3 | 1 | 12 | 0.9167 |
| 8_7016018 | 5 | 1 | 2 | 0.5 | 6 | 1 | 1 | 1 | 14 | 0.9286 | 2 | 1 | 3 | 1 | 2 | 1 | 3 | 1 | 10 | 1 |
| 8_7016019 | 4 | 0.25 | 2 | 0.5 | 5 | 0.2 | 1 | 1 | 12 | 0.3333 | 2 | 1 | 4 | 0.75 | 3 | 0.6667 | 3 | 1 | 12 | 0.8333 |
| 8_7016022 | 4 | 0.25 | 1 | 0 | 6 | 0.5 | 1 | 0 | 12 | 0.3333 | 1 | 0 | 4 | 0 | 3 | 0.3333 | 2 | 0 | 10 | 0.1 |
| 8_7019504 | 2 | 1 | NA | NA | NA | NA | NA | NA | 2 | 1 | NA | NA | NA | NA | 1 | 1 | NA | NA | 1 | 1 |
| 8_7019507 | 2 | 1 | NA | NA | NA | NA | NA | NA | 2 | 1 | NA | NA | NA | NA | 1 | 1 | NA | NA | 1 | 1 |
| 8_7019508 | 2 | 0.5 | NA | NA | NA | NA | NA | NA | 2 | 0.5 | NA | NA | NA | NA | 1 | 0 | 1 | 1 | 2 | 0.5 |
| 8_7019533 | 2 | 1 | NA | NA | NA | NA | NA | NA | 2 | 1 | NA | NA | NA | NA | 1 | 1 | 1 | 0 | 2 | 0.5 |
| 8_7019575 | 1 | 1 | NA | NA | NA | NA | NA | NA | 1 | 1 | NA | NA | NA | NA | 1 | 0 | 1 | 1 | 2 | 0.5 |
| 8_7019576 | 2 | 0.5 | NA | NA | NA | NA | NA | NA | 2 | 0.5 | NA | NA | NA | NA | 1 | 0 | 1 | 1 | 2 | 0.5 |
| 8_7019585 | 2 | 1 | NA | NA | NA | NA | NA | NA | 2 | 1 | NA | NA | NA | NA | 1 | 1 | 1 | 1 | 2 | 1 |
| 8_7019602 | 2 | 0.5 | NA | NA | NA | NA | NA | NA | 2 | 0.5 | NA | NA | NA | NA | 1 | 1 | 1 | 1 | 2 | 1 |
| 8_7019604 | 2 | 1 | NA | NA | NA | NA | NA | NA | 2 | 1 | NA | NA | NA | NA | 1 | 1 | 1 | 1 | 2 | 1 |
| 8_7019615 | 2 | 0.5 | NA | NA | NA | NA | NA | NA | 2 | 0.5 | NA | NA | NA | NA | 1 | 0 | 1 | 1 | 2 | 0.5 |
| 8_7021749 | NA | NA | NA | NA | 1 | 1 | NA | NA | 1 | 1 | NA | NA | NA | NA | 3 | 0.6667 | 1 | 0 | 4 | 0.5 |
| 8_7021769 | NA | NA | NA | NA | 1 | 1 | NA | NA | 1 | 1 | NA | NA | 1 | 0 | 3 | 0.3333 | 1 | 1 | 5 | 0.4 |
| 8_7021803 | NA | NA | NA | NA | 1 | 1 | NA | NA | 1 | 1 | NA | NA | 1 | 1 | 2 | 1 | 1 | 1 | 4 | 1 |
| 8_7021815 | NA | NA | 1 | 1 | NA | NA | NA | NA | 1 | 1 | NA | NA | NA | NA | 3 | 1 | 1 | 1 | 4 | 1 |
| 8_7021856 | NA | NA | NA | NA | 1 | 1 | NA | NA | 1 | 1 | NA | NA | NA | NA | 3 | 1 | 1 | 1 | 4 | 1 |
| 8_7021865 | NA | NA | 1 | 1 | 1 | 1 | NA | NA | 2 | 1 | NA | NA | NA | NA | 2 | 1 | 1 | 1 | 3 | 1 |
| 8_7021874 | NA | NA | 1 | 0 | 1 | 1 | NA | NA | 2 | 0.5 | NA | NA | NA | NA | 3 | 1 | NA | NA | 3 | 1 |
| 8_7021882 | NA | NA | 1 | 1 | 1 | 1 | NA | NA | 2 | 1 | NA | NA | NA | NA | 2 | 1 | NA | NA | 2 | 1 |
| 8_7021905 | NA | NA | 1 | 1 | 1 | 0 | NA | NA | 2 | 0.5 | NA | NA | NA | NA | 3 | 0.3333 | 1 | 1 | 4 | 0.5 |
| 8_7021915 | NA | NA | 1 | 1 | 1 | 1 | NA | NA | 2 | 1 | NA | NA | NA | NA | 3 | 1 | 1 | 1 | 4 | 1 |
| 8_7021922 | NA | NA | 1 | 1 | 1 | 0 | NA | NA | 2 | 0.5 | NA | NA | NA | NA | 2 | 0.5 | 1 | 0 | 3 | 0.3333 |
| 8_7021930 | NA | NA | 1 | 1 | 1 | 1 | NA | NA | 2 | 1 | NA | NA | NA | NA | 3 | 1 | 1 | 1 | 4 | 1 |
| 8_7021933 | NA | NA | 1 | 1 | 1 | 1 | NA | NA | 2 | 1 | NA | NA | NA | NA | 3 | 0.6667 | 1 | 1 | 4 | 0.75 |
| 8_7025997 | 2 | 0 | 9 | 0 | 2 | 0 | 3 | 0.3333 | 16 | 0.0625 | 5 | 0.2 | 7 | 0 | 2 | 0 | 6 | 0.1667 | 20 | 0.1 |
| 8_7026010 | 2 | 0 | 9 | 0.1111 | 2 | 0 | 3 | 0 | 16 | 0.0625 | 5 | 0.4 | 6 | 0.3333 | 2 | 0 | 6 | 0.5 | 19 | 0.3684 |
| 8_7026015 | 3 | 0 | 9 | 0 | 2 | 0 | 4 | 0.25 | 18 | 0.0556 | 6 | 0 | 6 | 0 | 3 | 0.3333 | 5 | 0 | 20 | 0.05 |
| 8_7026030 | 3 | 0.6667 | 8 | 0.375 | 4 | 0.25 | 4 | 0.5 | 19 | 0.4211 | 5 | 0.4 | 7 | 0.7143 | 3 | 0 | 9 | 0.3333 | 24 | 0.4167 |
| 8_7026031 | 2 | 0.5 | 9 | 0.3333 | 4 | 0.5 | 4 | 0.25 | 19 | 0.3684 | 7 | 0.4286 | 7 | 0.1429 | 3 | 0 | 8 | 0.125 | 25 | 0.2 |
| 8_7027326 | NA | NA | NA | NA | 1 | 0 | 1 | 1 | 2 | 0.5 | NA | NA | NA | NA | NA | NA | 1 | 1 | 1 | 1 |
| 8_7027357 | NA | NA | NA | NA | 1 | 0 | 1 | 1 | 2 | 0.5 | 1 | 0 | NA | NA | NA | NA | 1 | 1 | 2 | 0.5 |
| 8_7027365 | NA | NA | NA | NA | 2 | 0.5 | 1 | 0 | 3 | 0.3333 | 1 | 1 | NA | NA | NA | NA | NA | NA | 1 | 1 |
| 8_7028534 | 3 | 0 | 3 | 0 | 1 | 1 | 1 | 0 | 8 | 0.125 | 2 | 0.5 | 4 | 0.25 | 1 | 0 | 2 | 0 | 9 | 0.2222 |
| 8_7032476 | 3 | 0 | 6 | 0.1667 | 1 | 1 | 5 | 0 | 15 | 0.1333 | 3 | 0.3333 | 7 | 0.1429 | 4 | 0 | 4 | 0.25 | 18 | 0.1667 |
| 8_7032488 | 2 | 0 | 6 | 0.1667 | 1 | 0 | 5 | 0.2 | 14 | 0.1429 | 3 | 0 | 7 | 0 | 4 | 0 | 4 | 0.25 | 18 | 0.0556 |
| 8_7032489 | 2 | 1 | 6 | 0.6667 | 1 | 0 | 4 | 0.5 | 13 | 0.6154 | 3 | 0.3333 | 5 | 0.8 | 4 | 0.5 | 4 | 0.75 | 16 | 0.625 |
| 8_7036126 | NA | NA | 1 | 1 | 3 | 0 | 1 | 0 | 5 | 0.2 | 1 | 1 | NA | NA | 1 | 0 | 1 | 1 | 3 | 0.6667 |
| 8_7036139 | 6 | 0.1667 | 3 | 0 | 5 | 0.4 | 1 | 0 | 15 | 0.2 | 3 | 0.3333 | 3 | 0 | 3 | 0 | 3 | 0 | 12 | 0.0833 |
| 8_7036169 | 7 | 0.4286 | 4 | 0.5 | 4 | 0.5 | 1 | 0 | 16 | 0.4375 | 3 | 0.3333 | 3 | 0.6667 | 3 | 0.3333 | 1 | 0 | 10 | 0.4 |
| 8_7036221 | 6 | 0.6667 | 3 | 1 | 1 | 1 | 1 | 1 | 11 | 0.8182 | 1 | 1 | 2 | 0.5 | 2 | 1 | 1 | 0 | 6 | 0.6667 |
| 8_7036226 | 3 | 0.6667 | 3 | 1 | 1 | 1 | 1 | 1 | 8 | 0.875 | 2 | 1 | NA | NA | 3 | 1 | 1 | 1 | 6 | 1 |
| 8_7036232 | 6 | 0.1667 | 3 | 0 | 2 | 1 | 1 | 0 | 12 | 0.25 | 1 | 0 | 1 | 1 | 3 | 0.3333 | 1 | 0 | 6 | 0.3333 |
| 8_7036242 | 4 | 1 | 2 | 1 | 2 | 1 | 1 | 0 | 9 | 0.8889 | 2 | 1 | 2 | 0.5 | 2 | 1 | 1 | 1 | 7 | 0.8571 |
| 8_7036243 | 4 | 0 | 2 | 0 | 1 | 1 | 1 | 1 | 8 | 0.25 | 2 | 1 | 2 | 0 | 2 | 0 | 1 | 0 | 7 | 0.2857 |
| 8_7046725 | 1 | 0 | 2 | 0 | 2 | 0 | 1 | 1 | 6 | 0.1667 | 1 | 1 | 1 | 0 | 3 | 0 | 1 | 0 | 6 | 0.1667 |
| 8_7046739 | 1 | 1 | 1 | 0 | 2 | 0 | 1 | 1 | 5 | 0.4 | 1 | 0 | 1 | 1 | 3 | 0.6667 | 1 | 0 | 6 | 0.5 |
| 8_7047123 | 5 | 0.2 | 3 | 0.3333 | 1 | 0 | 6 | 0 | 15 | 0.1333 | 2 | 0 | 4 | 0 | 4 | 0 | 6 | 0.1667 | 16 | 0.0625 |
| 8_7047142 | 7 | 1 | 5 | 0.4 | 3 | 1 | 6 | 0.8333 | 21 | 0.8095 | 2 | 1 | 5 | 1 | 4 | 0.75 | 5 | 1 | 16 | 0.9375 |
| 8_7047145 | 8 | 0.25 | 4 | 0.25 | 3 | 0.3333 | 7 | 0.1429 | 22 | 0.2273 | 3 | 0.3333 | 6 | 0 | 7 | 0.1429 | 7 | 0 | 23 | 0.087 |

| | | | | | | | | | | | | | | | | | | | | |
|---|---|---|---|---|---|---|---|---|---|---|---|---|---|---|---|---|---|---|---|---|
| 8_7047156 | 8 | 0.125 | 2 | 0.5 | 3 | 0 | 6 | 0.5 | 19 | 0.2632 | 2 | 0 | 4 | 0 | 6 | 0.1667 | 4 | 0.25 | 16 | 0.125 |
| 8_7047160 | 8 | 0.25 | 3 | 0 | 2 | 0.5 | 7 | 0.2857 | 20 | 0.25 | 3 | 0 | 6 | 0 | 7 | 0.1429 | 5 | 0.2 | 21 | 0.0952 |
| 8_7047161 | 5 | 0.6 | 3 | 0.6667 | 3 | 1 | 6 | 0.8333 | 17 | 0.7647 | 3 | 1 | 5 | 0.8 | 7 | 0.7143 | 4 | 0.75 | 19 | 0.7895 |
| 8_7047166 | 8 | 0.625 | 4 | 0.25 | 3 | 0.6667 | 6 | 0.8333 | 21 | 0.619 | 2 | 0 | 6 | 0.5 | 7 | 0.5714 | 6 | 0.5 | 21 | 0.4762 |
| 8_7047174 | 8 | 0.25 | 4 | 0 | 3 | 0 | 7 | 0.1429 | 22 | 0.1364 | 2 | 0.5 | 6 | 0.1667 | 7 | 0.2857 | 6 | 0.1667 | 21 | 0.2381 |
| 8_7047183 | 8 | 0.5 | 2 | 1 | 2 | 1 | 7 | 0.7143 | 19 | 0.6842 | 3 | 0.6667 | 5 | 1 | 6 | 0.8333 | 6 | 0.8333 | 20 | 0.85 |
| 8_7047184 | 8 | 0 | 2 | 0.5 | 3 | 0.6667 | 7 | 0.2857 | 20 | 0.25 | 3 | 0 | 6 | 0.1667 | 7 | 0.4286 | 7 | 0.2857 | 23 | 0.2609 |
| 8_7047185 | 5 | 0.4 | 2 | 0.5 | 2 | 0.5 | 6 | 1 | 15 | 0.6667 | 3 | 0.3333 | 4 | 0.5 | 5 | 0.2 | 6 | 0.6667 | 18 | 0.4444 |
| 8_7047196 | 4 | 0 | 2 | 0.5 | 2 | 0 | 6 | 0.3333 | 14 | 0.2143 | 3 | 0.3333 | 4 | 0 | 6 | 0.3333 | 7 | 0.5714 | 20 | 0.35 |
| 8_7047201 | 6 | 0.1667 | 2 | 0 | 3 | 0 | 6 | 0 | 17 | 0.0588 | NA | NA | 5 | 0 | 5 | 0.2 | 7 | 0 | 17 | 0.0588 |
| 8_7047205 | 8 | 0.375 | 2 | 1 | 3 | 0.3333 | 7 | 0.4286 | 20 | 0.45 | 2 | 0 | 6 | 0.5 | 6 | 0.6667 | 6 | 0.5 | 20 | 0.5 |
| 8_7047207 | 8 | 0.875 | 2 | 1 | 2 | 1 | 7 | 0.8571 | 19 | 0.8947 | 3 | 1 | 6 | 1 | 6 | 1 | 7 | 0.8571 | 22 | 0.9545 |
| 8_7047221 | 7 | 0.2857 | 2 | 0 | 3 | 0 | 4 | 0 | 16 | 0.125 | 2 | 0 | 3 | 0.3333 | 6 | 0 | 7 | 0.1429 | 18 | 0.1111 |
| 8_7047226 | 8 | 0.125 | 2 | 0.5 | 3 | 0 | 5 | 0.4 | 18 | 0.2222 | 2 | 0 | 6 | 0.1667 | 7 | 0.1429 | 7 | 0 | 22 | 0.0909 |
| 8_7051780 | 3 | 0.3333 | 7 | 0.1429 | 5 | 0 | 4 | 0 | 19 | 0.1053 | 9 | 0.1111 | 7 | 0.1429 | 4 | 0 | 5 | 0 | 25 | 0.08 |
| 8_7051781 | 4 | 0.25 | 7 | 0 | 6 | 0.1667 | 5 | 0 | 22 | 0.0909 | 9 | 0 | 6 | 0.3333 | 3 | 0.3333 | 5 | 0 | 23 | 0.1304 |
| 8_7052452 | 4 | 0.75 | 10 | 0.6 | 5 | 0.6 | 5 | 0.6 | 24 | 0.625 | 12 | 0.6667 | 9 | 0.5556 | 5 | 0.6 | 9 | 0.7778 | 35 | 0.6571 |
| 8_7052471 | NA | NA | 3 | 1 | 2 | 0.5 | NA | NA | 5 | 0.8 | 3 | 0.3333 | NA | NA | NA | NA | 2 | 0.5 | 5 | 0.4 |
| 8_7052478 | NA | NA | 3 | 0.3333 | 2 | 0.5 | 1 | 1 | 6 | 0.5 | 4 | 0.5 | NA | NA | NA | NA | 2 | 0.5 | 6 | 0.5 |
| 8_7053255 | 1 | 1 | 1 | 0 | NA | NA | NA | NA | 2 | 0.5 | 2 | 1 | NA | NA | NA | NA | NA | NA | 2 | 1 |
| 8_7053263 | 1 | 1 | 1 | 0 | NA | NA | NA | NA | 2 | 0.5 | 2 | 0.5 | NA | NA | NA | NA | NA | NA | 2 | 0.5 |
| 8_7053959 | 1 | 0 | 2 | 0 | 1 | 0 | 2 | 0.5 | 6 | 0.1667 | 1 | 0 | 1 | 1 | 1 | 0 | 1 | 1 | 4 | 0.5 |
| 8_7054097 | 2 | 0.5 | 3 | 0 | 2 | 0.5 | 3 | 0.6667 | 10 | 0.4 | NA | NA | NA | NA | 1 | 1 | 1 | 0 | 2 | 0.5 |
| 8_7054115 | 2 | 0 | 4 | 0 | 2 | 0.5 | 3 | 0 | 11 | 0.0909 | NA | NA | 1 | 0 | 1 | 1 | 2 | 0 | 4 | 0.25 |
| 8_7054127 | 2 | 1 | 4 | 1 | 2 | 1 | 3 | 1 | 11 | 1 | NA | NA | 1 | 1 | 2 | 1 | 2 | 1 | 5 | 1 |
| 8_7054130 | 3 | 1 | 4 | 1 | 2 | 1 | 2 | 1 | 11 | 1 | 1 | 1 | 1 | 1 | 1 | 1 | 2 | 1 | 5 | 1 |
| 8_7054133 | 3 | 1 | 4 | 1 | 2 | 1 | 2 | 1 | 11 | 1 | NA | NA | 1 | 1 | 1 | 1 | 2 | 1 | 4 | 1 |
| 8_7054142 | 3 | 0.3333 | 2 | 0.5 | 2 | 0 | 3 | 0.6667 | 10 | 0.4 | NA | NA | 1 | 1 | 1 | 1 | 2 | 1 | 4 | 1 |
| 8_7054147 | 2 | 1 | 3 | 0.6667 | 1 | 1 | 2 | 1 | 8 | 0.875 | NA | NA | NA | NA | NA | NA | 2 | 1 | 2 | 1 |
| 8_7054151 | 3 | 0 | 4 | 0.5 | 2 | 0 | 3 | 0.3333 | 12 | 0.25 | 1 | 0 | 1 | 0 | 2 | 0 | 2 | 0.5 | 6 | 0.1667 |
| 8_7054153 | 3 | 1 | 5 | 1 | 2 | 1 | 4 | 1 | 14 | 1 | 1 | 1 | 1 | 1 | 2 | 0.5 | 3 | 1 | 7 | 0.8571 |
| 8_7054155 | 3 | 0.3333 | 5 | 0.6 | 2 | 0.5 | 4 | 0.5 | 14 | 0.5 | 1 | 1 | 1 | 0 | 1 | 1 | 2 | 0 | 5 | 0.4 |
| 8_7054158 | 3 | 0.3333 | 4 | 0.5 | 2 | 0 | 3 | 0.6667 | 12 | 0.4167 | 1 | 1 | 1 | 0 | 2 | 0.5 | 3 | 1 | 7 | 0.7143 |
| 8_7054167 | 3 | 0.3333 | 5 | 0 | 2 | 0.5 | 4 | 0 | 14 | 0.1429 | 1 | 0 | 1 | 1 | 1 | 1 | 3 | 0 | 6 | 0.3333 |
| 8_7054170 | 3 | 0.3333 | 5 | 0.2 | 2 | 0 | 4 | 0.25 | 14 | 0.2143 | 1 | 0 | 1 | 1 | 2 | 0 | 3 | 0 | 7 | 0.1429 |
| 8_7054184 | 3 | 0 | 4 | 0.5 | 3 | 0.6667 | 2 | 0 | 12 | 0.3333 | 1 | 0 | 1 | 0 | 2 | 0.5 | 1 | 0 | 5 | 0.2 |
| 8_7054194 | 1 | 0 | 7 | 0 | 3 | 0.3333 | 4 | 0.5 | 15 | 0.2 | 1 | 1 | 1 | 0 | NA | NA | 3 | 0.6667 | 5 | 0.6 |
| 8_7054203 | 4 | 0.25 | 7 | 0.1429 | 4 | 0 | 5 | 0.4 | 20 | 0.2 | NA | NA | 1 | 1 | 3 | 0 | 3 | 0 | 7 | 0.1429 |
| 8_7054222 | 4 | 1 | 10 | 0.4 | 5 | 1 | 4 | 1 | 23 | 0.7391 | 2 | 1 | 1 | 0 | 3 | 1 | 3 | 1 | 9 | 0.8889 |
| 8_7054253 | 3 | 0.6667 | 7 | 0.4286 | 4 | 0.5 | 4 | 0.5 | 18 | 0.5 | NA | NA | 2 | 0 | 3 | 0.6667 | 3 | 0.3333 | 8 | 0.375 |
| 8_7054255 | 3 | 0.6667 | 8 | 0.125 | 5 | 0 | 5 | 0.2 | 21 | 0.1905 | 2 | 0 | 2 | 0.5 | 3 | 0 | 2 | 0 | 9 | 0.1111 |
| 8_7054262 | 5 | 1 | 10 | 0.9 | 5 | 0.6 | 5 | 1 | 25 | 0.88 | 2 | 1 | 2 | 1 | 2 | 1 | 3 | 1 | 9 | 1 |
| 8_7054266 | 5 | 0.4 | 10 | 0.1 | 5 | 0 | 5 | 0 | 25 | 0.12 | 1 | 0 | 2 | 0 | 1 | 0 | 3 | 0.3333 | 7 | 0.1429 |
| 8_7054285 | 5 | 0.2 | 12 | 0.3333 | 5 | 0.2 | 5 | 0 | 27 | 0.2222 | 1 | 0 | 2 | 0 | 2 | 1 | 3 | 0 | 8 | 0.25 |
| 8_7054305 | 5 | 0.4 | 11 | 0.4545 | 3 | 0 | 5 | 0 | 24 | 0.2917 | 2 | 0 | 1 | 0 | 5 | 0.4 | 3 | 0 | 11 | 0.1818 |
| 8_7054577 | 15 | 0.2 | 23 | 0.087 | 26 | 0.1538 | 13 | 0.0769 | 77 | 0.1299 | 19 | 0.1053 | 20 | 0.1 | 20 | 0 | 18 | 0.0556 | 77 | 0.0649 |
| 8_7054595 | 8 | 0.25 | 12 | 0.25 | 10 | 0.3 | 6 | 0 | 36 | 0.2222 | 7 | 0.7143 | 12 | 0 | 15 | 0.3333 | 10 | 0.2 | 44 | 0.2727 |
| 8_7054766 | 7 | 0.2857 | 7 | 0.2857 | 11 | 0.5455 | 10 | 0.4 | 35 | 0.4 | 8 | 0.25 | 12 | 0.5 | 11 | 0.1818 | 7 | 0.4286 | 38 | 0.3421 |
| 8_7054804 | 4 | 1 | 8 | 0.375 | 6 | 0.5 | 8 | 0.375 | 26 | 0.5 | 5 | 0.4 | 9 | 0.6667 | 9 | 0.4444 | 7 | 0.8571 | 30 | 0.6 |
| 8_7054807 | 4 | 0 | 8 | 0 | 8 | 0.125 | 7 | 0.1429 | 27 | 0.0741 | 10 | 0.1 | 13 | 0.0769 | 10 | 0 | 10 | 0.3 | 43 | 0.1163 |
| 8_7054828 | 5 | 0.4 | 7 | 0 | 8 | 0.375 | 7 | 0.2857 | 27 | 0.2593 | 8 | 0.5 | 12 | 0.5833 | 9 | 0.2222 | 10 | 0.7 | 39 | 0.5128 |
| 8_7054845 | 4 | 0.25 | 10 | 0.1 | 5 | 0.2 | 5 | 0.4 | 24 | 0.2083 | 6 | 0.3333 | 12 | 0.25 | 7 | 0.1429 | 7 | 0.7143 | 32 | 0.3438 |
| 8_7054860 | 6 | 0.3333 | 9 | 0 | 6 | 0 | 6 | 0 | 27 | 0.0741 | 7 | 0.1429 | 13 | 0.0769 | 8 | 0 | 9 | 0.2222 | 37 | 0.1081 |
| 8_7054865 | 5 | 0.4 | 7 | 0.1429 | 4 | 0 | 5 | 0.2 | 21 | 0.1905 | 6 | 0.5 | 11 | 0 | 7 | 0.1429 | 8 | 0.125 | 32 | 0.1562 |
| 8_7054887 | 5 | 0.2 | 7 | 0.2857 | 3 | 0.6667 | 5 | 0.6 | 20 | 0.4 | 3 | 0.6667 | 13 | 0.4615 | 6 | 0.5 | 7 | 0.7143 | 29 | 0.5517 |
| 8_7054891 | 6 | 0.3333 | 9 | 0.1111 | 6 | 0.3333 | 6 | 0.1667 | 27 | 0.2222 | 4 | 0.5 | 11 | 0.1818 | 7 | 0.2857 | 9 | 0.2222 | 31 | 0.2581 |
| 8_7056020 | 1 | 1 | 4 | 0 | 4 | 0.25 | 1 | 0 | 10 | 0.2 | 1 | 0 | 6 | 0 | 4 | 0.5 | 7 | 0.1429 | 18 | 0.1667 |
| 8_7056021 | 2 | 0.5 | 4 | 0 | 4 | 0.25 | 2 | 0.5 | 12 | 0.25 | 1 | 0 | 6 | 0.3333 | 3 | 0.6667 | 7 | 0.1429 | 17 | 0.2941 |
| 8_7056037 | 2 | 0.5 | 3 | 0 | 2 | 0 | 1 | 0 | 8 | 0.125 | 1 | 0 | 6 | 0.1667 | 4 | 0 | 7 | 0 | 18 | 0.0556 |
| 8_7056040 | 1 | 0 | 4 | 0 | 3 | 0.3333 | 2 | 0 | 10 | 0.1 | 1 | 0 | 5 | 0.2 | 2 | 0 | 7 | 0.1429 | 15 | 0.1333 |
| 8_7056059 | 2 | 1 | 4 | 1 | 3 | 1 | 2 | 1 | 11 | 1 | 1 | 0 | 4 | 0.75 | 3 | 1 | 6 | 0.8333 | 14 | 0.7857 |
| 8_7056065 | 2 | 0 | 3 | 0.3333 | 3 | 0 | 1 | 0 | 9 | 0.1111 | NA | NA | 5 | 0.6 | 3 | 0 | 4 | 0.25 | 12 | 0.3333 |
| 8_7056067 | 2 | 1 | 3 | 1 | 3 | 1 | 2 | 1 | 10 | 1 | 1 | 0 | 4 | 1 | 2 | 0 | 6 | 0.6667 | 13 | 0.6154 |
| 8_7056069 | 2 | 1 | 4 | 1 | 3 | 1 | 1 | 1 | 10 | 1 | 1 | 0 | 5 | 0.8 | 3 | 0.6667 | 5 | 0.8 | 14 | 0.7143 |
| 8_7056075 | 2 | 1 | 4 | 1 | 3 | 1 | 1 | 1 | 10 | 1 | 1 | 1 | 5 | 1 | 3 | 0.6667 | 4 | 1 | 13 | 0.9231 |

| | | | | | | | | | | | | | | | | | | | | |
|---|---|---|---|---|---|---|---|---|---|---|---|---|---|---|---|---|---|---|---|---|
| 8_7056080 | NA | NA | 4 | 0.25 | 3 | 0.3333 | 2 | 0 | 9 | 0.2222 | 1 | 0 | 4 | 0.5 | 3 | 0 | 4 | 0 | 12 | 0.1667 |
| 8_7056105 | NA | NA | 2 | 0 | 3 | 0 | 2 | 0.5 | 7 | 0.1429 | NA | NA | 5 | 0 | 2 | 0 | 4 | 0.25 | 11 | 0.0909 |
| 8_7056299 | 1 | 0 | 1 | 1 | 2 | 1 | 1 | 1 | 5 | 0.8 | NA | NA | 4 | 0.5 | NA | NA | 3 | 0.3333 | 7 | 0.4286 |
| 8_7056300 | 1 | 1 | 2 | 0 | 2 | 0 | 1 | 0 | 6 | 0.1667 | NA | NA | 4 | 0 | 1 | 1 | 3 | 0.3333 | 8 | 0.25 |
| 8_7056300 | 1 | 0 | 2 | 0 | 2 | 0.5 | 1 | 0 | 6 | 0.1667 | NA | NA | 5 | 0.2 | 1 | 0 | 3 | 0 | 9 | 0.1111 |
| 8_7056787 | NA | NA | 2 | 0 | 1 | 1 | 1 | 1 | 4 | 0.5 | NA | NA | 3 | 0.3333 | 1 | 1 | 2 | 0 | 6 | 0.3333 |
| 8_7058364 | NA | NA | 1 | 1 | NA | NA | 1 | 1 | 2 | 1 | NA | NA | 2 | 1 | 1 | 1 | 2 | 1 | 5 | 1 |
| 8_7058388 | 1 | 0 | NA | NA | NA | NA | 1 | 1 | 2 | 0.5 | 1 | 1 | 3 | 0.3333 | 1 | 0 | 1 | 1 | 6 | 0.5 |
| 8_7058398 | NA | NA | 1 | 1 | NA | NA | 1 | 1 | 2 | 1 | 1 | 0 | 3 | 0.3333 | 1 | 1 | 1 | 1 | 6 | 0.5 |
| 8_7058402 | 1 | 0 | 1 | 1 | NA | NA | NA | NA | 2 | 0.5 | 1 | 0 | 2 | 0.5 | 1 | 0 | NA | NA | 4 | 0.25 |
| 8_7058407 | 1 | 0 | 1 | 1 | NA | NA | 1 | 1 | 3 | 0.6667 | 1 | 1 | 3 | 1 | 1 | 1 | 2 | 0.5 | 7 | 0.8571 |
| 8_7058422 | NA | NA | NA | NA | NA | NA | 1 | 1 | 1 | 1 | 1 | 0 | 3 | 0.6667 | NA | NA | NA | NA | 4 | 0.5 |
| 8_7058423 | 1 | 1 | NA | NA | NA | NA | 1 | 1 | 2 | 1 | NA | NA | 3 | 0 | 1 | 0 | 2 | 0.5 | 6 | 0.1667 |
| 8_7058438 | 1 | 1 | NA | NA | NA | NA | 1 | 1 | 2 | 1 | NA | NA | 1 | 1 | 1 | 1 | 2 | 1 | 4 | 1 |
| 8_7058447 | 1 | 1 | NA | NA | NA | NA | NA | NA | 1 | 1 | 1 | 0 | 3 | 0.6667 | 1 | 0 | NA | NA | 5 | 0.4 |
| 8_7058452 | 1 | 1 | NA | NA | NA | NA | 1 | 1 | 2 | 1 | NA | NA | 2 | 1 | 1 | 1 | 1 | 1 | 4 | 1 |
| 8_7058455 | 1 | 1 | NA | NA | NA | NA | NA | NA | 1 | 1 | 1 | 1 | 3 | 0 | NA | NA | 2 | 0 | 6 | 0.1667 |
| 8_7058457 | 1 | 1 | NA | NA | NA | NA | 1 | 1 | 2 | 1 | 1 | 1 | 2 | 1 | 1 | 1 | 1 | 0 | 5 | 0.8 |
| 8_7058553 | 1 | 1 | 1 | 0 | NA | NA | 1 | 0 | 3 | 0.3333 | 1 | 1 | 1 | 0 | NA | NA | NA | NA | 2 | 0.5 |
| 8_7058567 | 1 | 1 | 1 | 0 | NA | NA | 1 | 0 | 3 | 0.3333 | 1 | 1 | 1 | 1 | NA | NA | NA | NA | 2 | 1 |
| 8_7058592 | 2 | 0.5 | 1 | 0 | NA | NA | 1 | 0 | 4 | 0.25 | NA | NA | 2 | 1 | 1 | 0 | NA | NA | 3 | 0.6667 |
| 8_7058600 | 4 | 0.5 | 3 | 0.6667 | 4 | 0 | 4 | 0.75 | 15 | 0.4667 | 3 | 0 | NA | NA | 6 | 0.3333 | 6 | 0.5 | 15 | 0.3333 |
| 8_7058601 | 1 | 0 | 2 | 0.5 | NA | NA | 1 | 1 | 4 | 0.5 | 1 | 0 | 2 | 1 | 1 | 1 | NA | NA | 4 | 0.75 |
| 8_7058607 | 3 | 0.3333 | 3 | 0.3333 | 3 | 0 | 3 | 0 | 12 | 0.1667 | 2 | 1 | NA | NA | 4 | 0.25 | 2 | 0 | 8 | 0.375 |
| 8_7058616 | 4 | 0.75 | 5 | 0.4 | 5 | 0.2 | 4 | 0.25 | 18 | 0.3889 | 3 | 0.6667 | NA | NA | 6 | 0.5 | 6 | 0.5 | 15 | 0.5333 |
| 8_7059769 | 4 | 0 | 5 | 0 | 6 | 0.1667 | 3 | 0 | 18 | 0.0556 | 5 | 0 | 4 | 0 | 4 | 0.25 | 3 | 0 | 16 | 0.0625 |
| 8_7059788 | 3 | 0 | 2 | 0 | 3 | 0.3333 | 4 | 0 | 12 | 0.0833 | 4 | 0.25 | 4 | 0 | 4 | 0 | 2 | 0 | 14 | 0.0714 |
| 8_7059791 | 2 | 0.5 | 2 | 0.5 | 5 | 0.4 | 2 | 0.5 | 11 | 0.4545 | 3 | 0.3333 | 6 | 0.3333 | 3 | 0.3333 | 3 | 0.3333 | 15 | 0.3333 |
| 8_7059791 | 2 | 0 | 2 | 0 | 6 | 0 | 3 | 0.6667 | 13 | 0.1538 | 4 | 1 | 6 | 0.1667 | 5 | 0.4 | 3 | 0.6667 | 18 | 0.5 |
| 8_7059792 | 4 | 0 | 2 | 0 | 5 | 0.2 | 3 | 0.6667 | 14 | 0.2143 | 4 | 0 | 6 | 0.3333 | 4 | 0 | 3 | 0.6667 | 17 | 0.2353 |
| 8_7059800 | 4 | 0.25 | 1 | 0 | 6 | 0 | 4 | 0 | 15 | 0.0667 | 5 | 0 | 6 | 0 | 5 | 0.2 | 4 | 0 | 20 | 0.05 |
| 8_7059810 | 4 | 0.25 | 2 | 0 | 3 | 0.6667 | 3 | 1 | 12 | 0.5 | 3 | 0.6667 | 6 | 0.6667 | 4 | 0.5 | 4 | 0.75 | 17 | 0.6471 |
| 8_7059816 | 3 | 0.3333 | 2 | 0 | 4 | 0 | 4 | 0 | 13 | 0.0769 | 5 | 0.2 | 6 | 0.1667 | 4 | 0 | 2 | 0 | 17 | 0.1176 |
| 8_7059817 | 3 | 0 | 1 | 0 | 5 | 0.2 | 3 | 0 | 12 | 0.0833 | 5 | 0.2 | 5 | 0 | 5 | 0 | 1 | 0 | 16 | 0.0625 |
| 8_7059990 | NA | NA | 4 | 0.25 | 1 | 0 | 1 | 0 | 6 | 0.1667 | 1 | 0 | 1 | 0 | 1 | 0 | 1 | 1 | 4 | 0.25 |
| 8_7059999 | NA | NA | 2 | 0.5 | NA | NA | 1 | 0 | 3 | 0.3333 | 1 | 0 | NA | NA | 1 | 0 | 1 | 1 | 3 | 0.3333 |
| 8_7063483 | 2 | 1 | 1 | 1 | 4 | 0.75 | 3 | 1 | 10 | 0.9 | 2 | 0.5 | 5 | 0.8 | 5 | 0.6 | 4 | 0.75 | 16 | 0.6875 |
| 8_7063920 | NA | NA | 1 | 1 | 1 | 1 | 1 | 1 | 3 | 1 | 1 | 1 | 3 | 1 | NA | NA | 3 | 1 | 7 | 1 |
| 8_7067310 | 2 | 0 | 7 | 0.2857 | 5 | 0 | 3 | 0 | 17 | 0.1176 | 2 | 0.5 | 3 | 0 | 6 | 0.1667 | 2 | 0 | 13 | 0.1538 |
| 8_7067329 | 2 | 0 | 7 | 0 | 6 | 0 | 4 | 0.25 | 19 | 0.0526 | 2 | 0 | 3 | 0.3333 | 5 | 0.2 | 3 | 0 | 13 | 0.1538 |
| 8_7067330 | 3 | 0.6667 | 5 | 0.2 | 6 | 0.6667 | 4 | 0.5 | 18 | 0.5 | 2 | 0 | 2 | 0.5 | 4 | 1 | 4 | 0.75 | 12 | 0.6667 |
| 8_7067335 | 3 | 0.3333 | 6 | 0.6667 | 5 | 0.2 | 4 | 0.75 | 18 | 0.5 | 1 | 0 | 3 | 0 | 4 | 0.5 | 3 | 0.6667 | 11 | 0.3636 |
| 8_7067341 | 2 | 0 | 5 | 0.2 | 4 | 0.25 | 4 | 0.25 | 15 | 0.2 | 2 | 0 | 3 | 0 | 5 | 0.2 | 3 | 0 | 13 | 0.0769 |
| 8_7067775 | 2 | 0.5 | 6 | 0.5 | 5 | 0.6 | 3 | 0.6667 | 16 | 0.5625 | 2 | 0.5 | 3 | 1 | 5 | 0.6 | 3 | 0.3333 | 13 | 0.6154 |
| 8_7067782 | 3 | 0.6667 | 5 | 0.6 | 3 | 0.3333 | 3 | 0.6667 | 14 | 0.5714 | 1 | 1 | 3 | 0.3333 | 3 | 0.3333 | 2 | 0.5 | 9 | 0.4444 |
| 8_7067790 | 3 | 0.3333 | 5 | 0 | 6 | 0.1667 | 3 | 0.3333 | 17 | 0.1765 | 1 | 1 | 2 | 0.5 | 4 | 0 | 3 | 0 | 10 | 0.2 |
| 8_7069518 | 1 | 0 | 2 | 0 | 1 | 1 | 6 | 0 | 10 | 0.1 | 1 | 0 | 2 | 0 | 3 | 0.3333 | 1 | 0 | 7 | 0.1429 |
| 8_7069536 | 1 | 1 | 1 | 0 | 1 | 1 | 5 | 0.2 | 8 | 0.375 | 1 | 1 | 2 | 0 | 3 | 0.3333 | 1 | 0 | 7 | 0.2857 |
| 8_7069540 | 1 | 1 | 2 | 0.5 | 1 | 1 | 6 | 0.6667 | 10 | 0.7 | 1 | 1 | 2 | 1 | 3 | 1 | 1 | 1 | 7 | 1 |
| 8_7069542 | NA | NA | 2 | 1 | 1 | 1 | 5 | 0.8 | 8 | 0.875 | 1 | 1 | 1 | 1 | 2 | 1 | 1 | 1 | 5 | 1 |
| 8_7069558 | 1 | 0 | 2 | 0.5 | 1 | 0 | 5 | 0.2 | 9 | 0.2222 | 1 | 0 | 1 | 0 | NA | NA | 1 | 1 | 3 | 0.3333 |
| 8_7069560 | 1 | 1 | 2 | 1 | 2 | 1 | 5 | 0.6 | 10 | 0.8 | 1 | 1 | 2 | 1 | 3 | 1 | 1 | 1 | 7 | 1 |
| 8_7069584 | 2 | 0.5 | 2 | 0 | 1 | 1 | 7 | 0.1429 | 12 | 0.25 | 1 | 0 | 2 | 0.5 | 3 | 0 | 1 | 1 | 7 | 0.2857 |
| 8_7069590 | 2 | 0.5 | 2 | 0 | 2 | 0.5 | 7 | 0.1429 | 13 | 0.2308 | 1 | 0 | 1 | 1 | 2 | 0.5 | 1 | 0 | 5 | 0.4 |
| 8_7069591 | 2 | 1 | 2 | 0.5 | 2 | 0 | 6 | 0.3333 | 12 | 0.4167 | 1 | 1 | 2 | 0 | 3 | 0 | 1 | 0 | 7 | 0.1429 |
| 8_7069605 | 2 | 0 | 2 | 0 | 1 | 0 | 4 | 0.25 | 9 | 0.1111 | NA | NA | 2 | 0.5 | 3 | 0 | 1 | 0 | 6 | 0.1667 |
| 8_7069606 | 1 | 1 | 2 | 1 | NA | NA | 6 | 0.3333 | 9 | 0.5556 | 1 | 1 | 2 | 0.5 | 3 | 0.6667 | NA | NA | 6 | 0.6667 |
| 8_7069607 | 3 | 0.3333 | 1 | 0 | 2 | 0.5 | 5 | 0 | 11 | 0.1818 | 1 | 1 | 2 | 0.5 | 3 | 0 | 1 | 0 | 7 | 0.2857 |
| 8_7069624 | 3 | 0.3333 | 3 | 0.3333 | 1 | 0 | 6 | 0 | 13 | 0.1538 | 1 | 1 | NA | NA | 3 | 0.3333 | 1 | 0 | 5 | 0.4 |
| 8_7069629 | 3 | 0.3333 | 3 | 0 | 2 | 0.5 | 8 | 0.125 | 16 | 0.1875 | 1 | 0 | 2 | 0.5 | 3 | 0.3333 | 1 | 0 | 7 | 0.2857 |
| 8_7069635 | 3 | 1 | 2 | 1 | 2 | 1 | 6 | 1 | 13 | 1 | 1 | 1 | 2 | 1 | 2 | 1 | 1 | 1 | 6 | 1 |
| 8_7069638 | 3 | 0 | 2 | 0 | 2 | 1 | 5 | 0.2 | 12 | 0.25 | 1 | 0 | 2 | 0 | 3 | 0 | 1 | 1 | 7 | 0.1429 |
| 8_7069651 | 3 | 0.3333 | 2 | 0 | 2 | 0 | 7 | 0 | 14 | 0.0714 | 2 | 0 | 1 | 1 | 3 | 0 | 1 | 0 | 7 | 0.1429 |
| 8_7069658 | 3 | 0 | 1 | 0 | 1 | 0 | 5 | 0.2 | 10 | 0.1 | 1 | 0 | 2 | 0.5 | 3 | 0 | 1 | 1 | 7 | 0.2857 |
| 8_7070451 | 4 | 0 | 5 | 0 | 2 | 0 | 7 | 0.1429 | 18 | 0.0556 | 2 | 0.5 | 3 | 0 | 4 | 0 | 3 | 0 | 12 | 0.0833 |

| | | | | | | | | | | | | | | | | | | | | |
|---|---|---|---|---|---|---|---|---|---|---|---|---|---|---|---|---|---|---|---|---|
| 8_70704620 | 5 | 0.2 | 6 | 0.5 | 3 | 0 | 8 | 0.125 | 22 | 0.2273 | 3 | 0 | 5 | 0.2 | 4 | 0.75 | 4 | 0.75 | 16 | 0.4375 |
| 8_70709837 | 4 | 0 | 5 | 0.2 | 3 | 0 | 5 | 0.2 | 17 | 0.1176 | 2 | 0 | 6 | 0.1667 | 4 | 0.5 | 3 | 0 | 15 | 0.2 |
| 8_70709881 | 3 | 0 | 6 | 0.1667 | 3 | 0.3333 | 6 | 0 | 18 | 0.1111 | 2 | 0.5 | 5 | 0.4 | 4 | 0.25 | 4 | 0.25 | 15 | 0.3333 |
| 8_70713444 | NA | NA | 2 | 0.5 | 1 | 0 | 2 | 1 | 5 | 0.6 | NA | NA | 2 | 0.5 | 1 | 0 | NA | NA | 3 | 0.3333 |
| 8_70713527 | NA | NA | 4 | 0 | 1 | 1 | 2 | 0 | 7 | 0.1429 | NA | NA | 2 | 0 | 1 | 1 | NA | NA | 3 | 0.3333 |
| 8_70713698 | NA | NA | 4 | 0 | 1 | 1 | 4 | 0 | 9 | 0.1111 | NA | NA | 3 | 0.3333 | NA | NA | NA | NA | 3 | 0.3333 |
| 8_70715543 | 3 | 0 | 10 | 0.1 | 4 | 0.25 | 10 | 0 | 27 | 0.0741 | 5 | 0 | 6 | 0.1667 | 5 | 0.2 | 7 | 0.1429 | 23 | 0.1304 |
| 8_70717858 | 2 | 0.5 | NA | NA | 3 | 0 | NA | NA | 5 | 0.2 | 2 | 0.5 | 1 | 0 | 1 | 0 | 3 | 0 | 7 | 0.1429 |
| 8_70745877 | 2 | 0.5 | NA | NA | 1 | 1 | 1 | 0 | 4 | 0.5 | 1 | 0 | NA | NA | 3 | 0.3333 | 1 | 0 | 5 | 0.2 |
| 8_70745994 | NA | NA | 1 | 1 | NA | NA | 2 | 1 | 3 | 1 | NA | NA | NA | NA | 2 | 0.5 | NA | NA | 2 | 0.5 |
| 8_70746099 | NA | NA | 1 | 1 | NA | NA | NA | NA | 1 | 1 | NA | NA | 1 | 1 | NA | NA | NA | NA | 1 | 1 |
| 8_70746162 | 5 | 0 | 7 | 0 | 6 | 0.1667 | 4 | 0.25 | 22 | 0.0909 | 6 | 0.1667 | 4 | 0.25 | 2 | 0.5 | 4 | 0.75 | 16 | 0.375 |
| 8_70746222 | NA | NA | NA | NA | 1 | 0 | 2 | 1 | 3 | 0.6667 | 1 | 1 | 1 | 0 | 2 | 0 | NA | NA | 4 | 0.25 |
| 8_70746381 | NA | NA | 1 | 1 | 2 | 1 | 2 | 1 | 5 | 1 | NA | NA | 1 | 1 | 2 | 1 | NA | NA | 3 | 1 |
| 8_70746746 | NA | NA | 1 | 0 | 2 | 0 | 2 | 1 | 5 | 0.4 | 1 | 0 | 1 | 0 | 1 | 1 | NA | NA | 3 | 0.3333 |
| 8_70746828 | NA | NA | 1 | 1 | 2 | 1 | 2 | 1 | 5 | 1 | 1 | 1 | 1 | 1 | NA | NA | NA | NA | 2 | 1 |
| 8_70759054 | NA | NA | 1 | 0 | NA | NA | 1 | 1 | 2 | 0.5 | NA | NA | 1 | 0 | NA | NA | 1 | 1 | 2 | 0.5 |
| 8_70768625 | 8 | 0 | 6 | 0.1667 | 7 | 0.2857 | 8 | 0.125 | 29 | 0.1379 | 2 | 0 | 6 | 0.1667 | 3 | 0 | 2 | 0 | 13 | 0.0769 |
| 8_70774253 | 5 | 0 | 6 | 0.1667 | 9 | 0 | 13 | 0.1538 | 33 | 0.0909 | 5 | 0 | 7 | 0.1429 | 3 | 0 | 5 | 0.2 | 20 | 0.1 |
| 8_70782261 | 12 | 0.9167 | 11 | 0.4545 | 10 | 0.7 | 23 | 0.6087 | 56 | 0.6607 | 11 | 0.5455 | 15 | 0.2667 | 10 | 0.5 | 8 | 0.5 | 44 | 0.4318 |
| 8_70782304 | 11 | 0.0909 | 12 | 0.0833 | 7 | 0.1429 | 25 | 0.04 | 55 | 0.0727 | 8 | 0.125 | 14 | 0.1429 | 10 | 0.1 | 8 | 0.25 | 40 | 0.15 |
| 8_70782383 | 13 | 0.7692 | 12 | 0.75 | 10 | 0.9 | 23 | 0.6522 | 58 | 0.7414 | 11 | 0.8182 | 15 | 0.6667 | 10 | 0.6 | 12 | 0.9167 | 48 | 0.75 |
| 8_70782428 | 9 | 0.8889 | 11 | 0.7273 | 5 | 0.6 | 11 | 1 | 36 | 0.8333 | 8 | 0.875 | 9 | 0.8889 | 8 | 0.875 | 10 | 1 | 35 | 0.9143 |
| 8_70782484 | 11 | 0.7273 | 11 | 0.7273 | 10 | 0.6 | 19 | 0.7368 | 51 | 0.7059 | 8 | 0.875 | 14 | 0.7143 | 8 | 1 | 9 | 0.6667 | 39 | 0.7949 |
| 8_70782540 | 14 | 0.6429 | 11 | 0.5455 | 9 | 0.6667 | 22 | 0.6818 | 56 | 0.6429 | 10 | 0.5 | 17 | 0.3529 | 11 | 0.3636 | 11 | 0.2727 | 49 | 0.3673 |
| 8_70782604 | 12 | 0.5 | 12 | 0.5833 | 6 | 0.5 | 24 | 0.625 | 54 | 0.5741 | 10 | 0.6 | 12 | 0.6667 | 11 | 0.4545 | 10 | 0.6 | 43 | 0.5814 |
| 8_70782614 | 9 | 0.7778 | 8 | 0.875 | 7 | 0.7143 | 18 | 0.7778 | 42 | 0.7857 | 4 | 0.75 | 7 | 0.7143 | 4 | 0.75 | 6 | 1 | 21 | 0.8095 |
| 8_70782616 | 1 | 1 | NA | NA | NA | NA | NA | NA | 1 | 1 | NA | NA | NA | NA | 1 | 1 | NA | NA | 1 | 1 |
| 8_70783405 | 11 | 0.3636 | 9 | 0.3333 | 11 | 0.8182 | 21 | 0.5714 | 52 | 0.5385 | 11 | 0.2727 | 10 | 0.5 | 12 | 0.5833 | 11 | 0.4545 | 44 | 0.4545 |
| 8_70783609 | 15 | 0.6667 | 12 | 0.8333 | 11 | 0.7273 | 24 | 0.7083 | 62 | 0.7258 | 12 | 0.6667 | 11 | 0.5455 | 12 | 0.75 | 13 | 0.6923 | 48 | 0.6667 |
| 8_70783730 | 16 | 0.25 | 16 | 0.3125 | 15 | 0.3333 | 22 | 0.1364 | 69 | 0.2464 | 16 | 0.0625 | 17 | 0.2353 | 14 | 0.2857 | 16 | 0.125 | 63 | 0.1746 |
| 8_70783929 | 16 | 0.1875 | 16 | 0 | 14 | 0.0714 | 23 | 0.087 | 69 | 0.087 | 18 | 0.1111 | 12 | 0.0833 | 14 | 0.0714 | 16 | 0 | 60 | 0.0667 |
| 8_70784137 | 18 | 0.2222 | 17 | 0.1176 | 16 | 0.125 | 23 | 0.1739 | 74 | 0.1622 | 13 | 0.3846 | 16 | 0.125 | 13 | 0.2308 | 16 | 0.0625 | 58 | 0.1897 |
| 8_70784151 | 19 | 0.2632 | 18 | 0.2778 | 14 | 0 | 25 | 0.32 | 76 | 0.2368 | 16 | 0.125 | 17 | 0.2353 | 14 | 0.1429 | 15 | 0.0667 | 62 | 0.1452 |
| 8_70784173 | 18 | 0.8333 | 14 | 0.7857 | 14 | 0.7857 | 22 | 0.7727 | 68 | 0.7941 | 13 | 1 | 17 | 0.5882 | 15 | 0.8 | 15 | 0.6667 | 60 | 0.75 |
| 8_70784221 | 11 | 0 | 13 | 0.1538 | 11 | 0.0909 | 15 | 0 | 50 | 0.06 | 11 | 0.1818 | 11 | 0 | 10 | 0.1 | 12 | 0 | 44 | 0.0682 |
| 8_70784234 | 6 | 0.3333 | 11 | 0.6364 | 7 | 0.4286 | 15 | 0.6 | 39 | 0.5385 | 8 | 0.5 | 15 | 0.5333 | 9 | 0.2222 | 7 | 0.4286 | 39 | 0.4359 |
| 8_70784368 | 7 | 0.8571 | 9 | 0.3333 | 8 | 0.25 | 14 | 0.6429 | 38 | 0.5263 | 9 | 0.5556 | 12 | 0.3333 | 9 | 0.3333 | 6 | 0.3333 | 36 | 0.3889 |
| 8_70784505 | 8 | 0.5 | 9 | 0.3333 | 9 | 0.4444 | 14 | 0.1429 | 40 | 0.325 | 9 | 0.4444 | 11 | 0.6364 | 7 | 0.5714 | 6 | 0.5 | 33 | 0.5455 |
| 8_70784510 | 9 | 0.5556 | 11 | 0.3636 | 11 | 0.3636 | 15 | 0.4 | 46 | 0.413 | 8 | 0.375 | 15 | 0.5333 | 8 | 0.375 | 6 | 0.5 | 37 | 0.4595 |
| 8_70784770 | 8 | 0.75 | 8 | 0.625 | 8 | 0.75 | 10 | 0.6 | 34 | 0.6765 | 4 | 0.5 | 13 | 0.8462 | 6 | 0.6667 | 5 | 1 | 28 | 0.7857 |
| 8_70805594 | 2 | 0.5 | 2 | 0.5 | 1 | 0 | 2 | 1 | 7 | 0.5714 | 1 | 0 | 2 | 1 | 2 | 0 | 1 | 1 | 6 | 0.5 |
| 8_70837209 | 10 | 0.2 | 7 | 0.2857 | 11 | 0.1818 | 7 | 0 | 35 | 0.1714 | 3 | 0 | 17 | 0.1765 | 7 | 0 | 8 | 0 | 35 | 0.0857 |
| 8_70837229 | 10 | 0.6 | 6 | 0.1667 | 9 | 0.5556 | 8 | 0.5 | 33 | 0.4848 | 4 | 0.5 | 15 | 0.6 | 6 | 0.5 | 6 | 0.1667 | 31 | 0.4839 |
| 8_70838062 | 6 | 0 | 3 | 0.3333 | 5 | 0.2 | 6 | 0.3333 | 20 | 0.2 | NA | NA | 9 | 0.2222 | 4 | 0 | 5 | 0.4 | 18 | 0.2222 |
| 8_70838127 | 7 | 0.7143 | 2 | 0.5 | 7 | 1 | 6 | 0.3333 | 22 | 0.6818 | 2 | 1 | 11 | 0.4545 | 5 | 0.2 | 4 | 0.75 | 22 | 0.5 |
| 8_70838207 | 10 | 0.1 | 3 | 0.3333 | 8 | 0.125 | 5 | 0.2 | 26 | 0.1538 | 6 | 0 | 14 | 0.0714 | 6 | 0 | 4 | 0.25 | 30 | 0.0667 |
| 8_70838217 | 10 | 0.1 | 4 | 0.5 | 10 | 0.1 | 4 | 0.25 | 28 | 0.1786 | 7 | 0.1429 | 16 | 0.1875 | 7 | 0 | 5 | 0 | 35 | 0.1143 |
| 8_70838784 | 7 | 0.4286 | 4 | 0 | 5 | 1 | 2 | 1 | 18 | 0.5556 | 4 | 0 | 10 | 0.6 | 5 | 1 | 5 | 0.6 | 24 | 0.5833 |
| 8_70838799 | 8 | 0.375 | 4 | 0 | 6 | 0.5 | 4 | 0.25 | 22 | 0.3182 | 6 | 0.1667 | 14 | 0.2857 | 6 | 0.1667 | 5 | 0 | 31 | 0.1935 |
| 8_70838852 | 9 | 0.7778 | 4 | 0.75 | 8 | 0.875 | 5 | 0.6 | 26 | 0.7692 | 5 | 0.4 | 13 | 0.9231 | 7 | 0.8571 | 5 | 0.6 | 30 | 0.7667 |
| 8_70838951 | 8 | 0.25 | 1 | 0 | 7 | 0.7143 | 4 | 0.5 | 20 | 0.45 | 2 | 0 | 9 | 0.3333 | 4 | 0 | 3 | 1 | 18 | 0.3333 |
| 8_70902414 | NA | NA | 2 | 0.5 | NA | NA | 1 | 0 | 3 | 0.3333 | 1 | 0 | 3 | 0.3333 | NA | NA | NA | NA | 4 | 0.25 |
| 8_70903140 | 5 | 0.8 | 7 | 0.8571 | 7 | 0.7143 | 8 | 0.5 | 27 | 0.7037 | 7 | 0.5714 | 5 | 0.2 | 3 | 1 | 4 | 0.5 | 19 | 0.5263 |
| 8_70903488 | 3 | 0 | 10 | 0.1 | 6 | 0 | 9 | 0.1111 | 28 | 0.0714 | 7 | 0.2857 | 7 | 0.4286 | 3 | 0 | 4 | 0.25 | 21 | 0.2857 |
| 8_70903571 | 3 | 0.3333 | 10 | 0.2 | 6 | 0.3333 | 7 | 0.1429 | 26 | 0.2308 | 6 | 0.3333 | 8 | 0.25 | 2 | 0 | 2 | 0 | 18 | 0.2222 |
| 8_70903670 | 3 | 0.3333 | 8 | 0.625 | 6 | 0.3333 | 8 | 0.75 | 25 | 0.56 | 6 | 0.6667 | 6 | 0.3333 | 4 | 0.5 | 4 | 0.5 | 20 | 0.5 |
| 8_70903730 | 3 | 0 | 6 | 0.1667 | 4 | 0 | 6 | 0.1667 | 19 | 0.1053 | 6 | 0 | 7 | 0.1429 | 1 | 0 | 4 | 0.25 | 18 | 0.1111 |
| 8_70903751 | 1 | 0 | 6 | 0.3333 | 5 | 0 | 7 | 0.2857 | 19 | 0.2105 | 6 | 0.3333 | 7 | 0.4286 | 3 | 0.3333 | 3 | 0.3333 | 19 | 0.3684 |
| 8_70903772 | 2 | 1 | 11 | 0.9091 | 6 | 0.8333 | 9 | 1 | 28 | 0.9286 | 6 | 1 | 8 | 1 | 3 | 0.6667 | 4 | 0.75 | 21 | 0.9048 |
| 8_70903830 | 2 | 0 | 8 | 0.125 | 5 | 0.6 | 7 | 0.2857 | 22 | 0.2727 | 6 | 0.1667 | 6 | 0.1667 | 3 | 0.3333 | 3 | 0.3333 | 18 | 0.2222 |
| 8_70903835 | 2 | 0.5 | 11 | 0.1818 | 6 | 0 | 9 | 0 | 28 | 0.1071 | 7 | 0 | 7 | 0.1429 | 4 | 0.5 | 4 | 0 | 22 | 0.1364 |
| 8_70903881 | 1 | 1 | 7 | 0.8571 | 4 | 1 | 5 | 0.6 | 17 | 0.8235 | 5 | 0.8 | 8 | 0.875 | 3 | 0.6667 | 2 | 0.5 | 18 | 0.7778 |
| 8_70904164 | 2 | 0 | 9 | 0.5556 | 6 | 0.3333 | 8 | 0.75 | 25 | 0.52 | 5 | 0.4 | 7 | 0.7143 | 3 | 0.3333 | 3 | 0.3333 | 18 | 0.5 |
| 8_70904221 | 2 | 0 | 4 | 0 | 4 | 0 | 5 | 0.4 | 15 | 0.1333 | 3 | 0 | 5 | 0.6 | 1 | 0 | 2 | 0.5 | 11 | 0.3636 |

| | | | | | | | | | | | | | | | | | | | | |
|---|---|---|---|---|---|---|---|---|---|---|---|---|---|---|---|---|---|---|---|---|
| 8_70904318 | 2 | 0.5 | 9 | 0.6667 | 6 | 0.5 | 7 | 0.5714 | 24 | 0.5833 | 6 | 0.1667 | 4 | 0 | 3 | 0.3333 | 3 | 0.3333 | 16 | 0.1875 |
| 8_70904374 | 2 | 0.5 | 6 | 0.8333 | 6 | 0.6667 | 5 | 0.8 | 19 | 0.7368 | 5 | 0.8 | 3 | 0.6667 | 3 | 0.6667 | 3 | 0.3333 | 14 | 0.6429 |
| 8_70904383 | 2 | 1 | 8 | 0.5 | 4 | 0.75 | 6 | 1 | 20 | 0.75 | 4 | 1 | 6 | 1 | 2 | 1 | 1 | 1 | 13 | 1 |
| 8_70904515 | 1 | 1 | 7 | 0.4286 | 3 | 0.6667 | 5 | 0.8 | 16 | 0.625 | 2 | 1 | 4 | 0.25 | 1 | 0 | 2 | 1 | 9 | 0.5556 |
| 8_70904623 | 2 | 0.5 | 6 | 0 | 4 | 0.75 | 5 | 0 | 17 | 0.2353 | 6 | 0.3333 | 4 | 0 | 1 | 0 | 3 | 0.3333 | 14 | 0.2143 |
| 8_70904854 | 2 | 0 | 8 | 0.125 | 2 | 0.5 | 1 | 1 | 13 | 0.2308 | 2 | 0.5 | 4 | 0.25 | 2 | 0.5 | 2 | 0.5 | 10 | 0.4 |
| 8_70904892 | 2 | 1 | 1 | 1 | 3 | 1 | 3 | 0.6667 | 9 | 0.8889 | 2 | 1 | 2 | 1 | 1 | 1 | 2 | 1 | 7 | 1 |
| 8_70905137 | 2 | 0 | 5 | 0.2 | 4 | 0.5 | 4 | 0 | 15 | 0.2 | 5 | 0 | 5 | 0.6 | 2 | 0 | 1 | 0 | 13 | 0.2308 |
| 8_70905148 | 2 | 0.5 | 5 | 0.4 | 4 | 0.5 | 5 | 0.4 | 16 | 0.4375 | 3 | 1 | 4 | 0.75 | 2 | 1 | 1 | 0 | 10 | 0.8 |
| 8_70905150 | NA | NA | 3 | 0.6667 | 2 | 0.5 | 4 | 0.75 | 9 | 0.6667 | 3 | 1 | 1 | 1 | 2 | 1 | 1 | 0 | 7 | 0.8571 |
| 8_70905262 | 2 | 0.5 | 3 | 0.3333 | 3 | 0 | 5 | 0.2 | 13 | 0.2308 | 4 | 0.25 | 5 | 0.2 | 2 | 0 | 1 | 0 | 12 | 0.1667 |
| 8_70905337 | 2 | 1 | 3 | 0.6667 | 4 | 0.75 | 4 | 0.75 | 13 | 0.7692 | 4 | 1 | 5 | 1 | 2 | 1 | 1 | 1 | 12 | 1 |
| 8_70905475 | 2 | 0.5 | 3 | 0.6667 | 4 | 0.75 | 5 | 0.4 | 14 | 0.5714 | 2 | 0.5 | 5 | 0.8 | 2 | 0.5 | 1 | 1 | 10 | 0.7 |
| 8_70905544 | 1 | 1 | 3 | 1 | 3 | 0.6667 | 4 | 0.25 | 11 | 0.6364 | 2 | 0 | 3 | 1 | 1 | 1 | 1 | 0 | 7 | 0.5714 |
| 8_70905550 | 2 | 0.5 | 3 | 0.3333 | 3 | 0 | 4 | 0.5 | 12 | 0.3333 | 3 | 0 | 4 | 0.25 | 2 | 0.5 | 1 | 0 | 10 | 0.2 |
| 8_70936587 | 61 | 0.082 | 50 | 0.08 | 38 | 0.0526 | 48 | 0.125 | 197 | 0.0863 | 42 | 0.0714 | 32 | 0.0625 | 35 | 0.0571 | 52 | 0.1346 | 161 | 0.087 |
| 8_70936729 | 53 | 0.0566 | 50 | 0.12 | 31 | 0.0645 | 42 | 0.119 | 176 | 0.0909 | 39 | 0.0769 | 28 | 0.0714 | 26 | 0.0769 | 45 | 0.0667 | 138 | 0.0725 |
| 8_70936802 | 58 | 0.0517 | 48 | 0.1458 | 31 | 0.1935 | 45 | 0.0889 | 182 | 0.1099 | 41 | 0.1463 | 32 | 0.0938 | 34 | 0.0588 | 47 | 0.1915 | 154 | 0.1299 |
| 8_70960809 | NA | NA | 3 | 0.3333 | NA | NA | 1 | 0 | 4 | 0.25 | NA | NA | 2 | 1 | NA | NA | 1 | 0 | 3 | 0.6667 |
| 8_70960897 | NA | NA | 2 | 0.5 | NA | NA | 1 | 0 | 3 | 0.3333 | NA | NA | 2 | 0.5 | NA | NA | 1 | 0 | 3 | 0.3333 |
| 8_70966063 | 4 | 0.25 | 13 | 0.3077 | 6 | 0.6667 | 2 | 0.5 | 25 | 0.4 | 7 | 0.2857 | 5 | 0.2 | 4 | 0.5 | 4 | 0.25 | 20 | 0.3 |
| 8_70966180 | 5 | 0.2 | 13 | 0.2308 | 8 | 0.25 | 3 | 0 | 29 | 0.2069 | 6 | 0 | 5 | 0.2 | 4 | 0.25 | 7 | 0.2857 | 22 | 0.1818 |
| 8_70966379 | 5 | 0.2 | 10 | 0.4 | 3 | 0.3333 | 3 | 0 | 21 | 0.2857 | 3 | 0 | 2 | 0 | 3 | 0.3333 | 4 | 0.25 | 12 | 0.1667 |
| 8_70966456 | 5 | 0 | 7 | 0.1429 | 6 | 0.3333 | 2 | 0 | 20 | 0.15 | 7 | 0.2857 | 5 | 0 | 3 | 0.3333 | 6 | 0.3333 | 21 | 0.2381 |
| 8_70987745 | 43 | 0.2558 | 52 | 0.4423 | 47 | 0.3404 | 46 | 0.3696 | 188 | 0.3564 | 41 | 0.3902 | 57 | 0.386 | 44 | 0.2045 | 32 | 0.375 | 174 | 0.3391 |
| 8_71038321 | NA | NA | NA | NA | 2 | 0.5 | 4 | 0 | 6 | 0.1667 | 2 | 0 | 3 | 0 | 2 | 0.5 | 1 | 0 | 8 | 0.125 |
| 8_71038341 | NA | NA | NA | NA | 3 | 0 | 4 | 0.25 | 7 | 0.1429 | 2 | 0 | 3 | 0.3333 | 2 | 0 | 1 | 0 | 8 | 0.125 |
| 8_71038454 | NA | NA | 1 | 0 | 3 | 0 | 4 | 0.25 | 8 | 0.125 | 2 | 0.5 | 3 | 0 | 2 | 0 | NA | NA | 7 | 0.1429 |
| 8_71038517 | NA | NA | 1 | 0 | 2 | 0.5 | 4 | 0.25 | 7 | 0.2857 | 1 | 1 | 3 | 0.3333 | 3 | 0.6667 | 1 | 1 | 8 | 0.625 |
| 8_71039247 | NA | NA | NA | NA | NA | NA | 3 | 0.3333 | 3 | 0.3333 | NA | NA | NA | NA | 1 | 1 | 2 | 0.5 | 3 | 0.6667 |
| 8_71039374 | NA | NA | NA | NA | NA | NA | 3 | 0.3333 | 3 | 0.3333 | NA | NA | NA | NA | 1 | 1 | 2 | 0.5 | 3 | 0.6667 |
| 8_71039459 | NA | NA | NA | NA | NA | NA | 3 | 0.3333 | 3 | 0.3333 | NA | NA | NA | NA | 1 | 0 | 2 | 0.5 | 3 | 0.3333 |
| 8_71039507 | NA | NA | NA | NA | NA | NA | 1 | 1 | 1 | 1 | NA | NA | NA | NA | 1 | 1 | 2 | 0.5 | 3 | 0.6667 |
| 8_71039530 | NA | NA | NA | NA | NA | NA | 1 | 1 | 1 | 1 | NA | NA | NA | NA | 1 | 1 | 2 | 1 | 3 | 1 |
| 8_71039540 | NA | NA | NA | NA | NA | NA | 2 | 1 | 2 | 1 | NA | NA | NA | NA | 1 | 0 | 2 | 1 | 3 | 0.6667 |
| 8_71039610 | NA | NA | NA | NA | NA | NA | 2 | 0.5 | 2 | 0.5 | NA | NA | NA | NA | 1 | 0 | 2 | 0.5 | 3 | 0.3333 |
| 8_71039617 | NA | NA | NA | NA | NA | NA | 1 | 1 | 1 | 1 | NA | NA | NA | NA | 1 | 0 | 2 | 1 | 3 | 0.6667 |
| 8_71039746 | NA | NA | NA | NA | NA | NA | 1 | 1 | 1 | 1 | NA | NA | NA | NA | 1 | 1 | 2 | 1 | 3 | 1 |
| 8_71039753 | NA | NA | NA | NA | NA | NA | 1 | 1 | 1 | 1 | NA | NA | NA | NA | 1 | 1 | 2 | 1 | 3 | 1 |
| 8_71040220 | NA | NA | NA | NA | NA | NA | 1 | 1 | 1 | 1 | NA | NA | NA | NA | NA | NA | 2 | 1 | 2 | 1 |
| 8_71044316 | 3 | 1 | 1 | 1 | 2 | 0.5 | 5 | 1 | 11 | 0.9091 | 1 | 1 | 3 | 0.6667 | 5 | 0.8 | 2 | 0.5 | 11 | 0.7273 |
| 8_71044341 | 3 | 0.6667 | 2 | 0.5 | 4 | 0 | 4 | 0 | 13 | 0.2308 | 1 | 0 | 5 | 0 | 5 | 0.2 | 2 | 0 | 13 | 0.0769 |
| 8_71045177 | 3 | 0 | 2 | 0 | 2 | 0 | 4 | 0.25 | 11 | 0.0909 | 2 | 0 | 6 | 0.1667 | 5 | 0.2 | 2 | 0 | 15 | 0.1333 |
| 8_71045323 | 2 | 0 | 3 | 0 | 3 | 0.3333 | 4 | 0 | 12 | 0.0833 | 2 | 0 | 6 | 0 | 5 | 0.6 | 2 | 0 | 15 | 0.2 |
| 8_71048655 | 4 | 0 | 11 | 0 | 4 | 0 | 6 | 0.3333 | 25 | 0.08 | 2 | 0 | 7 | 0.2857 | 8 | 0.125 | 9 | 0.3333 | 26 | 0.2308 |
| 8_71048790 | 4 | 0.25 | 9 | 0.1111 | 6 | 0.1667 | 6 | 0.6667 | 25 | 0.28 | 2 | 0 | 7 | 0.1429 | 8 | 0.875 | 7 | 0.4286 | 24 | 0.4583 |
| 8_71049423 | 4 | 0 | 12 | 0.0833 | 8 | 0 | 6 | 0.1667 | 30 | 0.0667 | 4 | 0 | 10 | 0.1 | 9 | 0.1111 | 14 | 0.0714 | 37 | 0.0811 |
| 8_71049429 | 4 | 0.25 | 10 | 0 | 7 | 0.2857 | 5 | 0 | 26 | 0.1154 | 4 | 0.25 | 10 | 0 | 8 | 0.125 | 14 | 0 | 36 | 0.0556 |
| 8_71050206 | 27 | 0.1852 | 17 | 0.2941 | 29 | 0.1379 | 25 | 0.08 | 98 | 0.1633 | 20 | 0.05 | 24 | 0.0417 | 38 | 0.0789 | 18 | 0.0556 | 100 | 0.06 |
| 8_71050231 | 23 | 0.3478 | 17 | 0.4118 | 27 | 0.1852 | 18 | 0.4444 | 85 | 0.3294 | 17 | 0.3529 | 22 | 0.4091 | 32 | 0.4375 | 14 | 0.1429 | 85 | 0.3647 |
| 8_71050279 | 24 | 0.7083 | 20 | 0.75 | 29 | 0.7241 | 22 | 0.7273 | 95 | 0.7263 | 17 | 0.6471 | 24 | 0.6667 | 37 | 0.7297 | 17 | 0.7647 | 95 | 0.7053 |
| 8_71050321 | 21 | 0.2857 | 16 | 0.25 | 24 | 0.1667 | 20 | 0.15 | 81 | 0.2099 | 12 | 0.3333 | 20 | 0.1 | 30 | 0.2333 | 13 | 0.2308 | 75 | 0.2133 |
| 8_71101945 | 1 | 0 | NA | NA | 2 | 0.5 | NA | NA | 3 | 0.3333 | NA | NA | 1 | 0 | 1 | 0 | 1 | 1 | 3 | 0.3333 |
| 8_71101980 | 1 | 1 | NA | NA | 1 | 1 | NA | NA | 2 | 1 | NA | NA | 1 | 1 | 1 | 1 | 1 | 1 | 3 | 1 |
| 8_71102024 | 1 | 1 | NA | NA | 2 | 0.5 | NA | NA | 3 | 0.6667 | NA | NA | 1 | 0 | 1 | 1 | 1 | 1 | 3 | 0.6667 |
| 8_71102108 | 1 | 1 | NA | NA | 2 | 1 | NA | NA | 3 | 1 | NA | NA | 1 | 1 | 1 | 1 | 1 | 1 | 3 | 1 |
| 8_71102165 | 1 | 1 | NA | NA | 2 | 1 | NA | NA | 3 | 1 | NA | NA | NA | NA | 1 | 1 | 1 | 1 | 2 | 1 |
| 8_71102199 | 1 | 1 | NA | NA | 2 | 1 | NA | NA | 3 | 1 | NA | NA | 1 | 1 | 1 | 1 | 1 | 1 | 3 | 1 |
| 8_71102294 | 1 | 1 | NA | NA | 2 | 0 | NA | NA | 3 | 0.3333 | NA | NA | 1 | 1 | 1 | 1 | NA | NA | 2 | 1 |
| 8_71102454 | 1 | 0 | NA | NA | 2 | 1 | NA | NA | 3 | 0.6667 | NA | NA | NA | NA | 1 | 1 | 1 | 1 | 2 | 1 |
| 8_71102557 | 3 | 0 | 1 | 0 | 2 | 0.5 | NA | NA | 6 | 0.1667 | 2 | 0.5 | 1 | 0 | NA | NA | 2 | 0.5 | 5 | 0.4 |
| 8_71103034 | 2 | 0.5 | 1 | 1 | 2 | 1 | NA | NA | 5 | 0.8 | 2 | 0.5 | 1 | 1 | 1 | 1 | 2 | 1 | 6 | 0.8333 |
| 8_71103127 | 1 | 1 | 1 | 1 | 2 | 1 | NA | NA | 4 | 1 | 2 | 1 | 1 | 1 | 1 | 1 | 2 | 1 | 6 | 1 |
| 8_71103175 | NA | NA | NA | NA | 1 | 1 | NA | NA | 1 | 1 | 1 | 1 | NA | NA | NA | NA | NA | NA | 1 | 1 |
| 8_71103984 | 2 | 0.5 | 1 | 0 | 2 | 0 | NA | NA | 5 | 0.2 | 1 | 0 | NA | NA | 1 | 0 | 2 | 0.5 | 4 | 0.25 |

| | | | | | | | | | | | | | | | | | | | | |
|---|---|---|---|---|---|---|---|---|---|---|---|---|---|---|---|---|---|---|---|---|
| 8_7114743( | 1 | 1 | 1 | 1 | NA | NA | NA | NA | 2 | 1 | 1 | 0 | NA | NA | 1 | 1 | NA | NA | 2 | 0.5 |
| 8_7114847! | 1 | 0 | 2 | 0.5 | 1 | 0 | NA | NA | 4 | 0.25 | 2 | 0.5 | 2 | 0.5 | 1 | 0 | 1 | 0 | 6 | 0.3333 |
| 8_7114850( | 1 | 1 | 2 | 0.5 | 1 | 1 | NA | NA | 4 | 0.75 | 1 | 0 | 2 | 0.5 | 1 | 0 | 1 | 1 | 5 | 0.4 |
| 8_7114853' | NA | NA | 2 | 0.5 | NA | NA | NA | NA | 2 | 0.5 | 2 | 0.5 | 2 | 0.5 | 1 | 0 | 1 | 0 | 6 | 0.3333 |
| 8_7114884( | 1 | 1 | 2 | 0.5 | 1 | 0 | NA | NA | 4 | 0.5 | 1 | 0 | 1 | 0 | 1 | 1 | 1 | 0 | 4 | 0.25 |
| 8_7114983! | 1 | 1 | 2 | 1 | NA | NA | NA | NA | 3 | 1 | 2 | 1 | 1 | 1 | NA | NA | 1 | 1 | 4 | 1 |
| 8_7115078( | 1 | 1 | NA | NA | NA | NA | 1 | 1 | 2 | 1 | 3 | 0.6667 | 1 | 0 | 4 | 1 | 1 | 0 | 9 | 0.6667 |
| 8_7115203 | 1 | 1 | NA | NA | NA | NA | 1 | 0 | 2 | 0.5 | 3 | 0.6667 | 1 | 0 | 4 | 0 | NA | NA | 8 | 0.25 |
| 8_7115207 | 1 | 1 | NA | NA | NA | NA | 1 | 1 | 2 | 1 | 1 | 0 | 1 | 1 | 2 | 0.5 | NA | NA | 4 | 0.5 |
| 8_7115214 | 5 | 0.4 | 27 | 0.1852 | 29 | 0.2069 | 6 | 0 | 67 | 0.194 | 22 | 0.4091 | 26 | 0.2308 | 31 | 0.2581 | 21 | 0.1905 | 100 | 0.27 |
| 8_7115247! | 8 | 0.875 | 29 | 0.931 | 32 | 0.875 | 6 | 0.8333 | 75 | 0.8933 | 17 | 0.7059 | 20 | 0.8 | 27 | 0.7778 | 19 | 0.8421 | 83 | 0.7831 |
| 8_7115260 | 6 | 0.1667 | 23 | 0.0435 | 17 | 0.1765 | 4 | 0 | 50 | 0.1 | 17 | 0.1176 | 14 | 0.1429 | 24 | 0.0417 | 15 | 0 | 70 | 0.0714 |
| 8_7115267' | 1 | 1 | NA | NA | NA | NA | NA | NA | 1 | 1 | 1 | 1 | NA | NA | 1 | 0 | NA | NA | 2 | 0.5 |
| 8_7115275 | 7 | 1 | 31 | 0.8387 | 35 | 0.9429 | 6 | 1 | 79 | 0.9114 | 24 | 0.9167 | 29 | 0.9655 | 32 | 0.9688 | 21 | 0.8571 | 106 | 0.934 |
| 8_7115278 | 7 | 1 | 32 | 0.9688 | 35 | 0.9714 | 7 | 1 | 81 | 0.9753 | 25 | 0.92 | 28 | 1 | 32 | 1 | 21 | 1 | 106 | 0.9811 |
| 8_7115279( | 6 | 0 | 30 | 0.2333 | 28 | 0.25 | 5 | 0.2 | 69 | 0.2174 | 21 | 0.0952 | 22 | 0.1364 | 31 | 0.1613 | 15 | 0.2667 | 89 | 0.1573 |
| 8_7115281' | 5 | 1 | 34 | 0.9706 | 39 | 0.8974 | 7 | 0.8571 | 85 | 0.9294 | 26 | 0.8846 | 30 | 0.9333 | 34 | 0.9118 | 20 | 0.95 | 110 | 0.9182 |
| 8_7115282 | 7 | 0 | 30 | 0.1667 | 30 | 0.1 | 4 | 0.25 | 71 | 0.1268 | 20 | 0.1 | 27 | 0.1481 | 30 | 0.2 | 19 | 0.1579 | 96 | 0.1562 |
| 8_7115283 | 5 | 0.2 | 24 | 0.0833 | 17 | 0.0588 | 4 | 0.25 | 50 | 0.1 | 16 | 0.1875 | 14 | 0 | 16 | 0.125 | 12 | 0.0833 | 58 | 0.1034 |
| 8_7115304( | 6 | 0.1667 | 27 | 0.1111 | 33 | 0.0303 | 5 | 0.2 | 71 | 0.0845 | 21 | 0.0952 | 26 | 0.0385 | 32 | 0.0625 | 19 | 0.0526 | 98 | 0.0612 |
| 8_7121105 | 120 | 0.3417 | 148 | 0.223 | 108 | 0.25 | 96 | 0.3646 | 472 | 0.2881 | 97 | 0.2887 | 94 | 0.4043 | 108 | 0.3056 | 112 | 0.3393 | 411 | 0.3333 |
| 8_7121105 | 2 | 0.5 | 3 | 0.3333 | 3 | 0.3333 | 3 | 0.6667 | 11 | 0.4545 | NA | NA | 1 | 1 | 1 | 0 | 1 | 1 | 3 | 0.6667 |
| 8_7121124 | 115 | 0.1826 | 188 | 0.1649 | 130 | 0.2 | 114 | 0.1842 | 547 | 0.181 | 136 | 0.1618 | 108 | 0.2222 | 120 | 0.1667 | 129 | 0.1395 | 493 | 0.1704 |
| 8_7121149 | 7 | 0.8571 | 7 | 0.5714 | 8 | 0.5 | 6 | 0.1667 | 28 | 0.5357 | 7 | 1 | 8 | 0.5 | 6 | 0.6667 | 4 | 0.5 | 25 | 0.68 |
| 8_7121298 | 1 | 1 | NA | NA | NA | NA | 1 | 0 | 2 | 0.5 | NA | NA | NA | NA | 1 | 1 | NA | NA | 1 | 1 |
| 8_7121299 | 1 | 1 | NA | NA | NA | NA | 1 | 0 | 2 | 0.5 | NA | NA | NA | NA | 1 | 1 | NA | NA | 1 | 1 |
| 8_7121320 | 1 | 1 | 1 | 1 | NA | NA | 1 | 0 | 3 | 0.6667 | NA | NA | 1 | 1 | NA | NA | 1 | 1 | 2 | 1 |
| 8_7121326 | 1 | 1 | NA | NA | NA | NA | 1 | 0 | 2 | 0.5 | 1 | 1 | 1 | 0 | 1 | 0 | NA | NA | 3 | 0.3333 |
| 8_7121339 | 7 | 0.1429 | 4 | 0.25 | 6 | 0 | 3 | 0 | 20 | 0.1 | 5 | 0 | 6 | 0.1667 | 6 | 0 | 3 | 0 | 20 | 0.05 |
| 8_7122096 | 14 | 0.5714 | 11 | 0.1818 | 8 | 0.375 | 9 | 0.2222 | 42 | 0.3571 | 20 | 0.3 | 10 | 0.2 | 25 | 0.28 | 12 | 0.3333 | 67 | 0.2836 |
| 8_7122099 | 8 | 0.375 | 9 | 0.3333 | 6 | 0 | 7 | 0.2857 | 30 | 0.2667 | 15 | 0.2667 | 6 | 0.5 | 18 | 0.2222 | 9 | 0.2222 | 48 | 0.2708 |
| 8_7122112 | 14 | 0.6429 | 10 | 0.5 | 3 | 0.6667 | 5 | 0.8 | 32 | 0.625 | 9 | 0.4444 | 5 | 0.6 | 21 | 0.619 | 9 | 0.3333 | 44 | 0.5227 |
| 8_7122772 | 11 | 0.2727 | 2 | 1 | 1 | 0 | 1 | 1 | 15 | 0.4 | 11 | 0.4545 | 4 | 0.5 | 7 | 0.4286 | 8 | 0.25 | 30 | 0.4 |
| 8_7122779 | 10 | 0.5 | 2 | 0.5 | 1 | 0 | 1 | 0 | 14 | 0.4286 | 10 | 0.5 | 5 | 0.4 | 6 | 0 | 6 | 0.1667 | 27 | 0.2963 |
| 8_7122781( | 9 | 0.2222 | 1 | 0 | NA | NA | 1 | 0 | 11 | 0.1818 | 11 | 0.0909 | 4 | 0 | 5 | 0.2 | 6 | 0.3333 | 26 | 0.1538 |
| 8_7122961 | 2 | 0.5 | NA | NA | 1 | 1 | 1 | 0 | 4 | 0.5 | 1 | 0 | 2 | 0.5 | 1 | 0 | NA | NA | 4 | 0.25 |
| 8_7122962 | 2 | 0.5 | NA | NA | NA | NA | 1 | 0 | 3 | 0.3333 | 1 | 1 | 2 | 1 | 1 | 0 | 1 | 1 | 5 | 0.8 |
| 8_7122999 | 1 | 1 | NA | NA | 1 | 1 | NA | NA | 2 | 1 | 1 | 1 | NA | NA | 2 | 1 | 1 | 1 | 4 | 1 |
| 8_7122999 | 1 | 1 | NA | NA | 1 | 1 | NA | NA | 2 | 1 | 1 | 0 | 1 | 1 | 2 | 0.5 | 1 | 1 | 5 | 0.6 |
| 8_7123102 | 31 | 0.1613 | 23 | 0.087 | 30 | 0.0667 | 25 | 0.04 | 109 | 0.0917 | 23 | 0.1304 | 27 | 0.1481 | 30 | 0.0667 | 27 | 0.037 | 107 | 0.0935 |
| 8_7123114 | 34 | 0.2059 | 19 | 0.1579 | 31 | 0.1613 | 24 | 0.25 | 108 | 0.1944 | 22 | 0.1364 | 25 | 0.16 | 29 | 0.1034 | 23 | 0.3478 | 99 | 0.1818 |
| 8_7123115 | 28 | 0.5714 | 20 | 0.45 | 26 | 0.6154 | 25 | 0.52 | 99 | 0.5455 | 18 | 0.6667 | 24 | 0.5417 | 29 | 0.5862 | 24 | 0.3333 | 95 | 0.5263 |
| 8_7123123( | 31 | 0.0645 | 22 | 0.1364 | 29 | 0.0345 | 27 | 0.1111 | 109 | 0.0826 | 21 | 0.0476 | 25 | 0.08 | 31 | 0.129 | 27 | 0.0741 | 104 | 0.0865 |
| 8_7123130 | 27 | 0.1111 | 18 | 0.1111 | 28 | 0.25 | 24 | 0.125 | 97 | 0.1546 | 20 | 0.05 | 23 | 0.1739 | 31 | 0.1935 | 27 | 0.1481 | 101 | 0.1485 |
| 8_7123137 | 32 | 0.1562 | 22 | 0 | 27 | 0.1481 | 23 | 0.087 | 104 | 0.1058 | 22 | 0.0909 | 22 | 0.1364 | 29 | 0.069 | 24 | 0.25 | 97 | 0.134 |
| 8_7123144 | 31 | 0.7419 | 18 | 0.8333 | 29 | 0.8621 | 25 | 0.72 | 103 | 0.7864 | 20 | 0.6 | 22 | 0.8636 | 26 | 0.7308 | 25 | 0.92 | 93 | 0.7849 |
| 8_7123151 | 31 | 0.4839 | 17 | 0.4706 | 25 | 0.44 | 25 | 0.48 | 98 | 0.4694 | 20 | 0.5 | 21 | 0.619 | 27 | 0.5556 | 24 | 0.375 | 92 | 0.5109 |
| 8_7123154( | 31 | 0.129 | 16 | 0.125 | 25 | 0.12 | 25 | 0.2 | 97 | 0.1443 | 21 | 0.1905 | 23 | 0.1739 | 23 | 0.1304 | 23 | 0.087 | 90 | 0.1444 |
| 8_7123185 | 29 | 0.8621 | 15 | 0.8667 | 20 | 0.95 | 25 | 0.76 | 89 | 0.8539 | 16 | 0.875 | 21 | 0.9524 | 20 | 0.85 | 21 | 0.9048 | 78 | 0.8974 |
| 8_7123192 | 26 | 0.6923 | 12 | 0.8333 | 21 | 0.8571 | 23 | 0.7391 | 82 | 0.7683 | 19 | 0.7895 | 22 | 0.7727 | 22 | 0.7727 | 23 | 0.7391 | 86 | 0.7674 |
| 8_7123197 | 23 | 1 | 9 | 1 | 20 | 1 | 20 | 0.95 | 72 | 0.9861 | 14 | 1 | 20 | 1 | 20 | 0.95 | 19 | 1 | 73 | 0.9863 |
| 8_7123206 | 23 | 0.6522 | 9 | 0.7778 | 17 | 0.8235 | 22 | 0.8182 | 71 | 0.7606 | 17 | 0.9412 | 23 | 0.9565 | 19 | 0.8947 | 21 | 0.7619 | 80 | 0.8875 |
| 8_7123207 | 24 | 0.5 | 10 | 0.5 | 19 | 0.4737 | 22 | 0.5455 | 75 | 0.5067 | 16 | 0.5 | 22 | 0.5 | 21 | 0.4762 | 20 | 0.6 | 79 | 0.519 |
| 8_7123247 | 13 | 0 | 6 | 0.3333 | 16 | 0.0625 | 12 | 0.0833 | 47 | 0.0851 | 10 | 0 | 13 | 0.1538 | 16 | 0.1875 | 13 | 0 | 52 | 0.0962 |
| 8_7128784 | 1 | 0 | 7 | 0.2857 | 5 | 0 | 3 | 0 | 16 | 0.125 | 4 | 0 | 6 | 0.1667 | 6 | 0.1667 | 3 | 0.3333 | 19 | 0.1579 |
| 8_7128803 | 2 | 0 | 6 | 0 | 3 | 0 | 2 | 0.5 | 13 | 0.0769 | 2 | 0 | 4 | 0.25 | 3 | 0.6667 | 2 | 0 | 11 | 0.2727 |
| 8_7128827 | 2 | 0 | 5 | 0.2 | 5 | 0 | 3 | 0 | 15 | 0.0667 | 4 | 0 | 4 | 0 | 4 | 0 | 3 | 0.3333 | 15 | 0.0667 |
| 8_7128833 | 2 | 0.5 | 6 | 0.8333 | 5 | 0.2 | 3 | 0.6667 | 16 | 0.5625 | 4 | 1 | 3 | 0.6667 | 3 | 1 | 2 | 0.5 | 12 | 0.8333 |
| 8_7128836 | 2 | 0.5 | 5 | 0.6 | 5 | 0.2 | 3 | 0.3333 | 15 | 0.4 | 3 | 0.6667 | 5 | 0.4 | 5 | 0.4 | 2 | 0 | 15 | 0.4 |
| 8_7128845 | 2 | 1 | 6 | 0.3333 | 4 | 0.5 | 3 | 0.6667 | 15 | 0.5333 | 3 | 0.3333 | 4 | 0.25 | 5 | 0.2 | 2 | 0.5 | 14 | 0.2857 |
| 8_7128851 | 1 | 0 | 5 | 0.2 | 4 | 0 | 3 | 0 | 13 | 0.0769 | 4 | 0 | 4 | 0 | 5 | 0.2 | 2 | 0 | 15 | 0.0667 |
| 8_7129601 | NA | NA | 3 | 0.3333 | 2 | 0.5 | 1 | 0 | 6 | 0.3333 | 2 | 0.5 | 1 | 0 | NA | NA | 2 | 0 | 5 | 0.2 |
| 8_7129604 | NA | NA | 2 | 0 | 2 | 0.5 | 2 | 0 | 6 | 0.1667 | 2 | 0 | 2 | 0 | 2 | 0 | 2 | 0.5 | 8 | 0.125 |
| 8_7129632 | NA | NA | 2 | 0.5 | 2 | 0.5 | 1 | 1 | 5 | 0.6 | 2 | 1 | 2 | 1 | 2 | 1 | 2 | 0.5 | 8 | 0.875 |
| 8_7129640 | NA | NA | 2 | 0.5 | 2 | 0 | 2 | 1 | 6 | 0.5 | 1 | 0 | 2 | 0.5 | 2 | 0 | 1 | 0 | 6 | 0.1667 |

| | | | | | | | | | | | | | | | | | | | | |
|---|---|---|---|---|---|---|---|---|---|---|---|---|---|---|---|---|---|---|---|---|
| 8_7129658 | 1 | 0 | 2 | 1 | 1 | 1 | 1 | 1 | 5 | 0.8 | 1 | 1 | 2 | 1 | 2 | 1 | 1 | 0 | 6 | 0.8333 |
| 8_7129663 | 1 | 0 | 2 | 0.5 | 2 | 0.5 | 1 | 0 | 6 | 0.3333 | 2 | 1 | 2 | 1 | 2 | 0.5 | 1 | 0 | 7 | 0.7143 |
| 8_7129684 | 1 | 0 | 2 | 1 | 2 | 1 | 2 | 0.5 | 7 | 0.7143 | 1 | 1 | 2 | 1 | 1 | 1 | NA | NA | 4 | 1 |
| 8_7129685 | 1 | 1 | 2 | 0.5 | NA | NA | 2 | 0.5 | 5 | 0.6 | 1 | 0 | 2 | 1 | 1 | 0 | NA | NA | 4 | 0.5 |
| 8_7129688 | 1 | 0 | 2 | 1 | 2 | 0.5 | 2 | 0.5 | 7 | 0.5714 | 2 | 0.5 | 2 | 1 | 1 | 1 | NA | NA | 5 | 0.8 |
| 8_7129689 | 1 | 0 | 1 | 0 | 2 | 0 | 1 | 1 | 5 | 0.2 | 2 | 0 | 2 | 0.5 | NA | NA | NA | NA | 4 | 0.25 |
| 8_7129694 | 1 | 0 | 2 | 1 | 2 | 0.5 | 1 | 0 | 6 | 0.5 | 1 | 1 | 2 | 1 | 1 | 0 | NA | NA | 4 | 0.75 |
| 8_7129697 | NA | NA | 2 | 0.5 | 2 | 0.5 | 2 | 1 | 6 | 0.6667 | 2 | 0.5 | 2 | 1 | 1 | 1 | NA | NA | 5 | 0.8 |
| 8_7130531 | 1 | 1 | 1 | 0 | 1 | 1 | 2 | 0 | 5 | 0.4 | 2 | 0.5 | 2 | 0.5 | 1 | 1 | NA | NA | 5 | 0.6 |
| 8_7133591 | 3 | 0.3333 | 4 | 0.25 | 4 | 0 | 3 | 0.3333 | 14 | 0.2143 | 3 | 0 | 1 | 1 | 7 | 0.1429 | NA | NA | 11 | 0.1818 |
| 8_7133592 | 3 | 0 | 2 | 0 | 5 | 0.6 | 3 | 0.6667 | 13 | 0.3846 | 4 | 0.5 | 1 | 0 | 7 | 0.2857 | NA | NA | 12 | 0.3333 |
| 8_7133593 | 2 | 0.5 | 3 | 1 | 4 | 1 | 3 | 0.6667 | 12 | 0.8333 | 2 | 1 | 1 | 1 | 6 | 0.6667 | 1 | 1 | 10 | 0.8 |
| 8_7133599 | 2 | 0.5 | 4 | 0.25 | 4 | 0.5 | 3 | 0.6667 | 13 | 0.4615 | 3 | 0.6667 | 1 | 1 | 6 | 0.3333 | 2 | 0.5 | 12 | 0.5 |
| 8_7133628 | 3 | 0.3333 | 4 | 0.25 | 5 | 0 | 3 | 0.6667 | 15 | 0.2667 | 3 | 0.6667 | 1 | 0 | 3 | 0 | 1 | 0 | 8 | 0.25 |
| 8_7133631 | 3 | 0 | 4 | 0.5 | 5 | 0 | 3 | 0.6667 | 15 | 0.2667 | 3 | 0.3333 | 1 | 0 | 5 | 0 | NA | NA | 9 | 0.1111 |
| 8_7133644 | 3 | 0 | 4 | 0 | 5 | 0.4 | 2 | 0.5 | 14 | 0.2143 | 2 | 0.5 | 1 | 1 | 5 | 0.4 | 1 | 0 | 9 | 0.4444 |
| 8_7134802 | 72 | 0.0694 | 55 | 0.0727 | 65 | 0.1231 | 89 | 0.0787 | 281 | 0.0854 | 92 | 0.0652 | 99 | 0.0505 | 94 | 0.0638 | 90 | 0.0556 | 375 | 0.0587 |
| 8_7136561 | NA | NA | 5 | 0.2 | 2 | 0 | 1 | 0 | 8 | 0.125 | 6 | 0.1667 | 6 | 0 | 2 | 0.5 | 3 | 0.3333 | 17 | 0.1765 |
| 8_7136602 | 1 | 0 | 6 | 0 | 2 | 0 | 3 | 0.3333 | 12 | 0.0833 | 6 | 0.1667 | 8 | 0.125 | 3 | 0 | 2 | 0.5 | 19 | 0.1579 |
| 8_7136603 | 1 | 1 | 4 | 1 | 2 | 0.5 | 3 | 0.6667 | 10 | 0.8 | 7 | 0.5714 | 7 | 0.5714 | 3 | 1 | 2 | 0 | 19 | 0.5789 |
| 8_7136628 | 1 | 1 | 5 | 0.2 | 2 | 0.5 | 5 | 0.6 | 13 | 0.4615 | 9 | 0.3333 | 10 | 0.5 | 3 | 0.3333 | 3 | 0.6667 | 25 | 0.44 |
| 8_7136648 | 2 | 0 | 7 | 0.1429 | 2 | 0 | 5 | 0.2 | 16 | 0.125 | 8 | 0.375 | 8 | 0.125 | 2 | 0 | 2 | 0 | 20 | 0.2 |
| 8_7136650 | 1 | 1 | 6 | 0.3333 | 2 | 0.5 | 3 | 0.6667 | 12 | 0.5 | 9 | 0.8889 | 8 | 0.75 | 4 | 0.75 | 2 | 0.5 | 23 | 0.7826 |
| 8_7136660 | 1 | 0 | 6 | 0.6667 | 2 | 1 | 2 | 1 | 11 | 0.7273 | 9 | 0.5556 | 9 | 0.4444 | 4 | 0.75 | 4 | 0.5 | 26 | 0.5385 |
| 8_7136676 | 1 | 0 | 6 | 0.1667 | 2 | 0 | 5 | 0.2 | 14 | 0.1429 | 9 | 0.2222 | 8 | 0 | 3 | 0.3333 | 4 | 0.25 | 24 | 0.1667 |
| 8_7136709 | 3 | 0.6667 | 10 | 0.4 | 3 | 0.3333 | 4 | 0.25 | 20 | 0.4 | 9 | 0.2222 | 11 | 0.3636 | 2 | 0.5 | 4 | 0.5 | 26 | 0.3462 |
| 8_7136716 | 4 | 0.5 | 8 | 0.5 | 2 | 0 | 5 | 0.8 | 19 | 0.5263 | 6 | 0.8333 | 11 | 0.7273 | 4 | 0.5 | 3 | 0.3333 | 24 | 0.6667 |
| 8_7136719 | 4 | 0.5 | 9 | 0.4444 | 1 | 0 | 3 | 0.3333 | 17 | 0.4118 | 7 | 0.4286 | 7 | 0.4286 | 1 | 1 | 4 | 0.5 | 19 | 0.4737 |
| 8_7136724 | 3 | 0.6667 | 8 | 0.875 | 2 | 1 | 4 | 0.75 | 17 | 0.8235 | 7 | 0.4286 | 8 | 0.625 | 2 | 0.5 | 4 | 1 | 21 | 0.619 |
| 8_7136730 | 3 | 0 | 9 | 0 | 2 | 0.5 | 4 | 0 | 18 | 0.0556 | 8 | 0 | 10 | 0.1 | 2 | 0.5 | 4 | 0 | 24 | 0.0833 |
| 8_7136736 | 3 | 0.3333 | 9 | 0.2222 | 2 | 0.5 | 3 | 0.3333 | 17 | 0.2941 | 3 | 0.3333 | 9 | 0.4444 | 4 | 0 | 3 | 0 | 19 | 0.2632 |
| 8_7136749 | 4 | 0.25 | 9 | 0 | 3 | 0 | 6 | 0.1667 | 22 | 0.0909 | 7 | 0 | 9 | 0.1111 | 4 | 0.25 | 3 | 0 | 23 | 0.087 |
| 8_7136751 | 4 | 0.25 | 7 | 0.1429 | 3 | 0 | 6 | 0.1667 | 20 | 0.15 | 6 | 0.3333 | 8 | 0 | 3 | 0 | 4 | 0 | 21 | 0.0952 |
| 8_7136999 | 7 | 0 | 9 | 0.2222 | 6 | 0.1667 | 7 | 0 | 29 | 0.1034 | 12 | 0.3333 | 10 | 0.3 | 7 | 0.4286 | 6 | 0 | 35 | 0.2857 |
| 8_7180141 | NA | NA | 1 | 1 | 4 | 0.25 | NA | NA | 5 | 0.4 | 1 | 1 | 2 | 0 | 3 | 0 | NA | NA | 6 | 0.1667 |
| 8_7180142 | 1 | 1 | 1 | 1 | 4 | 0.75 | NA | NA | 6 | 0.8333 | 1 | 0 | 2 | 0.5 | 4 | 0.25 | 1 | 0 | 8 | 0.25 |
| 8_7180143 | 1 | 1 | 1 | 1 | 4 | 1 | NA | NA | 6 | 1 | 1 | 1 | 1 | 1 | 3 | 1 | 1 | 0 | 6 | 0.8333 |
| 8_7181245 | 3 | 0.3333 | 2 | 0.5 | 4 | 0.25 | 2 | 0 | 11 | 0.2727 | 1 | 0 | 1 | 1 | 6 | 0 | 4 | 0.5 | 12 | 0.25 |
| 8_7181252 | 3 | 1 | 4 | 1 | 4 | 0.5 | 2 | 1 | 13 | 0.8462 | 1 | 1 | 2 | 0.5 | 6 | 0.8333 | 3 | 0.3333 | 12 | 0.6667 |
| 8_7181254 | 2 | 0.5 | 3 | 0.3333 | 4 | 0.5 | 2 | 0 | 11 | 0.3636 | 1 | 1 | 2 | 1 | 6 | 0.5 | 4 | 0.25 | 13 | 0.5385 |
| 8_7181256 | 3 | 1 | 3 | 0.6667 | 4 | 0.5 | 2 | 0.5 | 12 | 0.6667 | 1 | 1 | 2 | 0 | 5 | 0.6 | 3 | 0.6667 | 11 | 0.5455 |
| 8_7181262 | 3 | 0.6667 | 4 | 0.25 | 4 | 0.5 | 2 | 0 | 13 | 0.3846 | 1 | 1 | 1 | 1 | 5 | 0 | 3 | 0.3333 | 10 | 0.3 |
| 8_7181262 | 3 | 0.3333 | 2 | 1 | 4 | 0.25 | 2 | 0.5 | 11 | 0.4545 | 1 | 0 | 2 | 0 | 5 | 0.4 | 3 | 0 | 11 | 0.1818 |
| 8_7181265 | 3 | 0.6667 | 2 | 1 | 3 | 1 | 1 | 1 | 9 | 0.8889 | 1 | 1 | NA | NA | 5 | 0.8 | 4 | 0.5 | 10 | 0.7 |
| 8_7181275 | 3 | 0.3333 | 5 | 0.6 | 4 | 0.75 | 2 | 0.5 | 14 | 0.5714 | 1 | 1 | 2 | 0.5 | 7 | 0.4286 | 3 | 0.3333 | 13 | 0.4615 |
| 8_7181282 | 3 | 0.6667 | 2 | 0.5 | 3 | 1 | 1 | 1 | 9 | 0.7778 | 1 | 1 | 1 | 1 | 5 | 0.6 | 2 | 1 | 9 | 0.7778 |
| 8_7181288 | 3 | 0.3333 | 5 | 0 | 4 | 0.25 | 2 | 0 | 14 | 0.1429 | 2 | 1 | 2 | 1 | 6 | 0.1667 | 3 | 0.6667 | 13 | 0.5385 |
| 8_7181291 | 3 | 0.6667 | 4 | 0.75 | 3 | 1 | 3 | 0.6667 | 13 | 0.7692 | 2 | 1 | 2 | 0.5 | 5 | 0.8 | 3 | 0 | 12 | 0.5833 |
| 8_7181292 | 2 | 0 | 3 | 0 | 3 | 0.6667 | 3 | 0 | 11 | 0.1818 | 2 | 0 | NA | NA | 6 | 0.1667 | 4 | 0 | 12 | 0.0833 |
| 8_7181294 | NA | NA | 2 | 0.5 | 1 | 0 | 2 | 0.5 | 5 | 0.4 | 1 | 0 | 1 | 1 | 3 | 0 | NA | NA | 5 | 0.2 |
| 8_7181297 | 3 | 0.6667 | 5 | 0.6 | 4 | 1 | 2 | 1 | 14 | 0.7857 | 2 | 1 | NA | NA | 5 | 1 | 4 | 0.5 | 11 | 0.8182 |
| 8_7181299 | 2 | 0 | 5 | 0.2 | 4 | 0.75 | 2 | 0 | 13 | 0.3077 | 2 | 0.5 | 2 | 0.5 | 7 | 0.2857 | 4 | 0.25 | 15 | 0.3333 |
| 8_7181302 | 2 | 0.5 | 3 | 1 | 1 | 1 | 2 | 1 | 8 | 0.875 | 1 | 1 | 1 | 0 | 4 | 0.5 | 3 | 0.6667 | 9 | 0.5556 |
| 8_7181308 | 3 | 0 | 5 | 0 | 4 | 0 | 3 | 0.3333 | 15 | 0.0667 | 2 | 0 | 2 | 0 | 7 | 0.1429 | 3 | 0 | 14 | 0.0714 |
| 8_7181312 | 3 | 0 | 4 | 0.5 | 4 | 0.25 | 3 | 0 | 14 | 0.2143 | 2 | 0.5 | 2 | 0 | 7 | 0.1429 | 2 | 0 | 13 | 0.1538 |
| 8_7181321 | 3 | 0.3333 | 5 | 0.8 | 4 | 0.5 | 3 | 1 | 15 | 0.6667 | 2 | 1 | 2 | 0.5 | 7 | 0.5714 | 4 | 0.25 | 15 | 0.5333 |
| 8_7181322 | 3 | 0.3333 | 3 | 0 | 4 | 0.25 | 3 | 0.6667 | 13 | 0.3077 | 2 | 0 | 1 | 0 | 5 | 0 | 3 | 0.3333 | 11 | 0.0909 |
| 8_7181968 | 4 | 0 | 1 | 0 | NA | NA | 1 | 1 | 6 | 0.1667 | 1 | 1 | 1 | 0 | 2 | 1 | 1 | 0 | 5 | 0.6 |
| 8_7181969 | 5 | 0.2 | 2 | 0 | NA | NA | 1 | 1 | 8 | 0.25 | 1 | 1 | 1 | 1 | 2 | 0.5 | 1 | 0 | 5 | 0.6 |
| 8_7181978 | 1 | 1 | NA | NA | NA | NA | 1 | 0 | 2 | 0.5 | 1 | 1 | 1 | 1 | 1 | 1 | NA | NA | 3 | 1 |
| 8_7182167 | 1 | 0 | 1 | 1 | 3 | 0.3333 | 1 | 0 | 6 | 0.3333 | 1 | 0 | 3 | 0.6667 | NA | NA | NA | NA | 4 | 0.5 |
| 8_7182702 | 4 | 1 | 6 | 0.6667 | 5 | 0.8 | 5 | 1 | 20 | 0.85 | 10 | 0.8 | 6 | 0.8333 | 4 | 1 | 6 | 0.6667 | 26 | 0.8077 |
| 8_7182704 | 2 | 0.5 | 5 | 0.6 | 5 | 0.8 | 7 | 0.4286 | 19 | 0.5789 | 8 | 0.625 | 7 | 0.5714 | 5 | 0 | 5 | 0.4 | 25 | 0.44 |
| 8_7182709 | 1 | 0 | 5 | 0.4 | 3 | 0 | 5 | 0.2 | 14 | 0.2143 | 9 | 0.1111 | 4 | 0 | 3 | 0.3333 | 5 | 0.4 | 21 | 0.1905 |
| 8_7182710 | 3 | 0.6667 | 6 | 0.1667 | 5 | 0.4 | 8 | 0.5 | 22 | 0.4091 | 9 | 0.5556 | 6 | 0.5 | 5 | 0.6 | 6 | 0.1667 | 26 | 0.4615 |

| | | | | | | | | | | | | | | | | | | | | |
|---|---|---|---|---|---|---|---|---|---|---|---|---|---|---|---|---|---|---|---|---|
| 8_7182717( | NA | NA | 2 | 0.5 | 2 | 0 | 3 | 0.3333 | 7 | 0.2857 | 5 | 0.4 | 4 | 0.25 | 2 | 0 | 5 | 0.2 | 16 | 0.25 |
| 8_7182777 | NA | NA | 2 | 0.5 | 1 | 0 | 1 | 1 | 4 | 0.5 | 1 | 0 | NA | NA | 1 | 1 | 1 | 1 | 3 | 0.6667 |
| 8_7182817 | NA | NA | 2 | 0.5 | 1 | 0 | 1 | 1 | 4 | 0.5 | 1 | 0 | NA | NA | 1 | 0 | 1 | 1 | 3 | 0.3333 |
| 8_7182849 | 4 | 0.5 | 6 | 0.5 | 5 | 0 | 6 | 0.3333 | 21 | 0.3333 | 9 | 0.3333 | 7 | 0.1429 | 2 | 0.5 | 6 | 0.5 | 24 | 0.3333 |
| 8_7182881 | 1 | 1 | NA | NA | NA | NA | NA | NA | 1 | 1 | 1 | 1 | 1 | 1 | NA | NA | NA | NA | 2 | 1 |
| 8_7182909 | 1 | 1 | NA | NA | NA | NA | NA | NA | 1 | 1 | 1 | 1 | 1 | 1 | NA | NA | NA | NA | 2 | 1 |
| 8_7182934 | 1 | 1 | NA | NA | NA | NA | NA | NA | 1 | 1 | 1 | 1 | NA | NA | NA | NA | NA | NA | 1 | 1 |
| 8_7182957 | 1 | 1 | NA | NA | NA | NA | NA | NA | 1 | 1 | 1 | 1 | 1 | 0 | NA | NA | NA | NA | 2 | 0.5 |
| 8_7182991 | 1 | 1 | NA | NA | NA | NA | NA | NA | 1 | 1 | 1 | 1 | NA | NA | NA | NA | NA | NA | 1 | 1 |
| 8_7183006 | 1 | 1 | NA | NA | 1 | 1 | NA | NA | 2 | 1 | 1 | 1 | 1 | 1 | NA | NA | NA | NA | 2 | 1 |
| 8_7183009 | 1 | 1 | NA | NA | NA | NA | NA | NA | 1 | 1 | NA | NA | 1 | 1 | NA | NA | NA | NA | 1 | 1 |
| 8_7183022 | 1 | 1 | NA | NA | 1 | 1 | NA | NA | 2 | 1 | 1 | 1 | 1 | 1 | NA | NA | NA | NA | 2 | 1 |
| 8_7183027 | 1 | 1 | NA | NA | 1 | 0 | NA | NA | 2 | 0.5 | 1 | 1 | 1 | 0 | NA | NA | NA | NA | 2 | 0.5 |
| 8_7183065 | 1 | 1 | NA | NA | 1 | 0 | NA | NA | 2 | 0.5 | 1 | 0 | 1 | 1 | NA | NA | 1 | 0 | 3 | 0.3333 |
| 8_7183123 | 1 | 1 | NA | NA | NA | NA | NA | NA | 1 | 1 | 1 | 1 | NA | NA | NA | NA | NA | NA | 1 | 1 |
| 8_7183133 | 2 | 0.5 | NA | NA | 1 | 0 | NA | NA | 3 | 0.3333 | 1 | 1 | 1 | 0 | NA | NA | 1 | 0 | 3 | 0.3333 |
| 8_7183134 | NA | NA | NA | NA | 1 | 1 | NA | NA | 1 | 1 | NA | NA | NA | NA | NA | NA | 1 | 1 | 1 | 1 |
| 8_7183147 | 2 | 1 | NA | NA | NA | NA | NA | NA | 2 | 1 | 1 | 1 | NA | NA | NA | NA | 1 | 1 | 2 | 1 |
| 8_7183153 | 1 | 1 | NA | NA | 1 | 1 | NA | NA | 2 | 1 | 1 | 1 | 1 | 1 | NA | NA | 1 | 1 | 3 | 1 |
| 8_7183172 | 2 | 1 | NA | NA | 1 | 1 | NA | NA | 3 | 1 | 1 | 1 | 1 | 1 | NA | NA | 1 | 0 | 3 | 0.6667 |
| 8_7183178 | 2 | 1 | NA | NA | NA | NA | NA | NA | 2 | 1 | NA | NA | NA | NA | NA | NA | 1 | 1 | 1 | 1 |
| 8_7183181 | 2 | 0.5 | NA | NA | 1 | 1 | NA | NA | 3 | 0.6667 | 1 | 0 | 1 | 1 | NA | NA | 1 | 1 | 3 | 0.6667 |
| 8_7183182 | 2 | 0.5 | NA | NA | NA | NA | NA | NA | 2 | 0.5 | 1 | 0 | 1 | 1 | NA | NA | 1 | 0 | 3 | 0.3333 |
| 8_7183373 | 2 | 0 | 5 | 0.2 | 4 | 0 | 6 | 0.1667 | 17 | 0.1176 | 7 | 0.2857 | 6 | 0.1667 | 2 | 0 | 2 | 0 | 17 | 0.1765 |
| 8_7183374 | 3 | 0.6667 | 3 | 1 | 3 | 0.6667 | 4 | 0.5 | 13 | 0.6923 | 6 | 1 | 7 | 0.5714 | 2 | 0.5 | 2 | 0 | 17 | 0.6471 |
| 8_7183395 | 3 | 1 | 5 | 0.8 | 2 | 0.5 | 4 | 0.25 | 14 | 0.6429 | 5 | 0.4 | 6 | 0.6667 | 2 | 0 | 2 | 1 | 15 | 0.5333 |
| 8_7183406 | 1 | 1 | 3 | 0.3333 | 2 | 0.5 | 1 | 1 | 7 | 0.5714 | 6 | 0.6667 | 4 | 0.25 | 1 | 0 | NA | NA | 11 | 0.4545 |
| 8_7183408 | 3 | 0 | 3 | 0 | 2 | 1 | 3 | 0 | 11 | 0.1818 | 7 | 0.1429 | 6 | 0 | 2 | 0 | 2 | 0 | 17 | 0.0588 |
| 8_7183417 | 3 | 0.6667 | 3 | 0.3333 | 4 | 0.25 | 2 | 0 | 12 | 0.3333 | 6 | 0.1667 | 4 | 0 | 2 | 0 | 1 | 0 | 13 | 0.0769 |
| 8_7183420 | NA | NA | 3 | 0.3333 | 2 | 0 | 3 | 0.6667 | 8 | 0.375 | 5 | 0.2 | 2 | 0.5 | NA | NA | 1 | 0 | 8 | 0.25 |
| 8_7183420 | 3 | 0.3333 | 3 | 0.3333 | 3 | 0.3333 | 3 | 0.3333 | 12 | 0.3333 | 5 | 0.6 | 5 | 0.6 | 1 | 1 | 1 | 1 | 12 | 0.6667 |
| 8_7183438 | 2 | 0 | 3 | 0.6667 | 2 | 0 | 3 | 0 | 10 | 0.2 | 6 | 0.6667 | 5 | 0 | 1 | 0 | 2 | 0 | 14 | 0.2857 |
| 8_7185583 | NA | NA | 1 | 1 | NA | NA | NA | NA | 1 | 1 | 2 | 1 | NA | NA | 1 | 1 | 2 | 1 | 5 | 1 |
| 8_7185684 | 15 | 0.0667 | 12 | 0.25 | 12 | 0.0833 | 9 | 0.2222 | 48 | 0.1458 | 10 | 0.1 | 9 | 0.1111 | 18 | 0 | 5 | 0.2 | 42 | 0.0714 |
| 8_7185686 | 10 | 0.1 | 10 | 0.1 | 11 | 0 | 8 | 0 | 39 | 0.0513 | 6 | 0.1667 | 9 | 0.1111 | 14 | 0.0714 | 4 | 0 | 33 | 0.0909 |
| 8_7185704 | 11 | 0.2727 | 11 | 0.4545 | 10 | 0.1 | 6 | 0.3333 | 38 | 0.2895 | 14 | 0.2857 | 7 | 0.5714 | 17 | 0.4706 | 2 | 0.5 | 40 | 0.425 |
| 8_7185715 | 13 | 1 | 13 | 0.9231 | 15 | 0.6667 | 10 | 0.8 | 51 | 0.8431 | 15 | 0.7333 | 8 | 0.875 | 17 | 0.7647 | 6 | 0.6667 | 46 | 0.7609 |
| 8_7190945 | 45 | 0.1556 | 43 | 0.0698 | 41 | 0.0976 | 35 | 0.0286 | 164 | 0.0915 | 32 | 0.125 | 40 | 0.075 | 20 | 0.05 | 29 | 0.1724 | 121 | 0.1074 |
| 8_7190969 | 43 | 0.0698 | 38 | 0.1579 | 42 | 0.1429 | 32 | 0.0938 | 155 | 0.1161 | 28 | 0.0714 | 43 | 0.1163 | 27 | 0.0741 | 34 | 0.1765 | 132 | 0.1136 |
| 8_7190978 | 37 | 0.1892 | 37 | 0.1892 | 38 | 0.1579 | 31 | 0.3226 | 143 | 0.2098 | 31 | 0.0968 | 40 | 0.15 | 27 | 0.2222 | 28 | 0.25 | 126 | 0.1746 |
| 8_7190981 | 45 | 0.2444 | 38 | 0.0789 | 42 | 0.1667 | 36 | 0.1944 | 161 | 0.1739 | 33 | 0.303 | 43 | 0.2326 | 25 | 0.2 | 34 | 0.1471 | 135 | 0.2222 |
| 8_7191406 | 36 | 0.0833 | 31 | 0.2581 | 32 | 0.25 | 34 | 0.2059 | 133 | 0.1955 | 25 | 0.24 | 32 | 0.0938 | 19 | 0.0526 | 28 | 0.1786 | 104 | 0.1442 |
| 8_7191418 | 33 | 0.0909 | 29 | 0.1034 | 32 | 0.125 | 34 | 0.0882 | 128 | 0.1016 | 25 | 0.16 | 30 | 0.1 | 19 | 0.0526 | 23 | 0.1304 | 97 | 0.1134 |
| 8_7191428 | 37 | 0.2162 | 26 | 0.0769 | 33 | 0.0909 | 30 | 0.1333 | 126 | 0.1349 | 24 | 0.125 | 31 | 0.0968 | 20 | 0.1 | 26 | 0.0769 | 101 | 0.099 |
| 8_7191456 | 18 | 0.1111 | 13 | 0.0769 | 15 | 0.1333 | 24 | 0.0833 | 70 | 0.1 | 16 | 0.3125 | 22 | 0.2273 | 13 | 0.2308 | 22 | 0.2727 | 73 | 0.2603 |
| 8_7191813 | 4 | 0 | 10 | 0.3 | 4 | 0.5 | 11 | 0.3636 | 29 | 0.3103 | 14 | 0.5714 | 4 | 0.25 | 8 | 0.625 | 4 | 0.75 | 30 | 0.5667 |
| 8_7191832 | 3 | 0 | 11 | 0 | 6 | 0.3333 | 14 | 0.0714 | 34 | 0.0882 | 15 | 0.1333 | 4 | 0.25 | 11 | 0 | 7 | 0.1429 | 37 | 0.1081 |
| 8_7192741 | 12 | 0.25 | 20 | 0.45 | 13 | 0.0769 | 14 | 0.3571 | 59 | 0.3051 | 10 | 0.4 | 5 | 0 | 14 | 0.2143 | 8 | 0.5 | 37 | 0.2973 |
| 8_7192768 | 12 | 0.5833 | 19 | 0.6842 | 11 | 0.4545 | 14 | 0.8571 | 56 | 0.6607 | 10 | 0.5 | 6 | 0.8333 | 15 | 0.8 | 8 | 0.75 | 39 | 0.7179 |
| 8_7192772 | 10 | 0.5 | 19 | 0.4737 | 13 | 0.3846 | 11 | 0.3636 | 53 | 0.434 | 9 | 0.1111 | 9 | 0.1111 | 13 | 0.5385 | 10 | 0.5 | 41 | 0.3415 |
| 8_7192777 | 5 | 0.2 | 10 | 0.5 | 5 | 0.2 | 10 | 0.2 | 30 | 0.3 | 3 | 0.3333 | 5 | 0.4 | 10 | 0.4 | 6 | 0.1667 | 24 | 0.3333 |
| 8_7192783 | 6 | 0.1667 | 10 | 0.2 | 4 | 0 | 10 | 0.1 | 30 | 0.1333 | 1 | 0 | 4 | 0 | 9 | 0.1111 | 7 | 0.2857 | 21 | 0.1429 |
| 8_7192794 | 4 | 0.25 | 7 | 0.2857 | 5 | 0.2 | 9 | 0.3333 | 25 | 0.28 | 2 | 0.5 | 2 | 0 | 7 | 0.2857 | 4 | 0.25 | 15 | 0.2667 |
| 8_7192799 | 5 | 1 | 7 | 0.8571 | 5 | 0.8 | 10 | 0.8 | 27 | 0.8519 | 2 | 0.5 | 4 | 1 | 8 | 0.625 | 7 | 1 | 21 | 0.8095 |
| 8_7192818 | 5 | 0 | 8 | 0 | 7 | 0.1429 | 11 | 0.0909 | 31 | 0.0645 | 2 | 0 | 4 | 0.5 | 7 | 0.1429 | 5 | 0 | 18 | 0.1667 |
| 8_7192924 | 4 | 0 | 5 | 0.2 | 2 | 0 | 2 | 0 | 13 | 0.0769 | 1 | 0 | 7 | 0.1429 | 7 | 0.2857 | 3 | 0 | 18 | 0.1667 |
| 8_7192929 | 4 | 0 | 5 | 0.4 | 2 | 0 | 4 | 0.25 | 15 | 0.2 | 3 | 0 | 6 | 0.1667 | 7 | 0.2857 | 3 | 0 | 19 | 0.1579 |
| 8_7192933 | 3 | 0 | 5 | 0.4 | 2 | 0 | 4 | 0 | 14 | 0.1429 | 4 | 0.5 | 6 | 0.1667 | 7 | 0.2857 | 3 | 0 | 20 | 0.25 |
| 8_7193320 | 2 | 0 | 4 | 0.25 | 2 | 0 | 3 | 0 | 11 | 0.0909 | 4 | 0 | 5 | 0.2 | 6 | 0.5 | 2 | 0.5 | 17 | 0.2941 |
| 8_7193327 | 2 | 1 | 3 | 0 | 2 | 1 | 4 | 0.25 | 11 | 0.4545 | 4 | 0.5 | 3 | 0.6667 | 3 | 0.3333 | 2 | 0.5 | 12 | 0.5 |
| 8_7193347 | 2 | 0 | 3 | 0 | 1 | 0 | 2 | 1 | 8 | 0.25 | 4 | 0.5 | 4 | 0.25 | 5 | 0.4 | 1 | 1 | 14 | 0.4286 |
| 8_7194538 | 3 | 0.6667 | 2 | 1 | NA | NA | NA | NA | 5 | 0.8 | 2 | 1 | 3 | 1 | 1 | 1 | 2 | 1 | 8 | 1 |
| 8_7194542 | 3 | 0.6667 | 2 | 0.5 | NA | NA | NA | NA | 5 | 0.6 | 3 | 1 | 3 | 0.6667 | 1 | 0 | 2 | 0 | 9 | 0.5556 |
| 8_7194552 | 3 | 0.6667 | 2 | 1 | NA | NA | NA | NA | 5 | 0.8 | 3 | 0.3333 | 5 | 0.8 | 1 | 1 | 2 | 0.5 | 11 | 0.6364 |
| 8_7194569 | 4 | 0.5 | 2 | 0.5 | NA | NA | NA | NA | 6 | 0.5 | 3 | 1 | 2 | 0.5 | 1 | 0 | 2 | 0 | 8 | 0.5 |

| | | | | | | | | | | | | | | | | | | | | |
|---|---|---|---|---|---|---|---|---|---|---|---|---|---|---|---|---|---|---|---|---|
| 8_7194950 | NA | NA | NA | NA | NA | NA | 1 | 1 | 1 | 1 | 1 | 1 | NA | NA | NA | NA | NA | NA | 1 | 1 |
| 8_7199819 | 3 | 0 | 1 | 0 | NA | NA | 4 | 0.25 | 8 | 0.125 | NA | NA | 2 | 0 | 5 | 0.4 | 3 | 0 | 10 | 0.2 |
| 8_7202342 | 6 | 0.1667 | NA | NA | 6 | 0 | 3 | 0 | 15 | 0.0667 | 5 | 0.2 | 6 | 0 | 4 | 0 | 1 | 0 | 16 | 0.0625 |
| 8_7203219 | 7 | 0.2857 | 2 | 0.5 | 6 | 0.1667 | 4 | 0 | 19 | 0.2105 | 4 | 0.5 | 7 | 0.2857 | 4 | 0 | NA | NA | 15 | 0.2667 |
| 8_7203220 | 6 | 0.1667 | 3 | 0 | 6 | 0.1667 | 6 | 0 | 21 | 0.0952 | 5 | 0 | 7 | 0.1429 | 5 | 0.2 | 1 | 0 | 18 | 0.1111 |
| 8_7203230 | 6 | 0.6667 | 3 | 0.6667 | 8 | 0.5 | 5 | 0.4 | 22 | 0.5455 | 6 | 0.3333 | 4 | 0 | 5 | 0.4 | 1 | 0 | 16 | 0.25 |
| 8_7203872 | 10 | 0.4 | 11 | 0.5455 | 13 | 0.6154 | 13 | 0.4615 | 47 | 0.5106 | 7 | 0.7143 | 11 | 0.5455 | 9 | 0.7778 | 9 | 0.5556 | 36 | 0.6389 |
| 8_7203878 | 10 | 0.1 | 11 | 0.4545 | 11 | 0.5455 | 8 | 0.75 | 40 | 0.45 | 10 | 0.3 | 9 | 0.5556 | 3 | 0.3333 | 10 | 0.4 | 32 | 0.4062 |
| 8_7203882 | 12 | 0.25 | 10 | 0.1 | 13 | 0.2308 | 12 | 0.25 | 47 | 0.2128 | 8 | 0.125 | 10 | 0.5 | 10 | 0 | 10 | 0.5 | 38 | 0.2895 |
| 8_7203954 | 8 | 0.125 | 3 | 0 | 6 | 0.1667 | 3 | 0.3333 | 20 | 0.15 | 1 | 0 | 1 | 0 | 3 | 0 | 7 | 0.1429 | 12 | 0.0833 |
| 8_7203964 | 8 | 0.125 | 4 | 0 | 6 | 0.6667 | 4 | 0.5 | 22 | 0.3182 | 2 | 0 | 1 | 0 | 3 | 0 | 7 | 0.2857 | 13 | 0.1538 |
| 8_7204137 | NA | NA | NA | NA | NA | NA | 1 | 1 | 1 | 1 | NA | NA | 1 | 0 | 1 | 0 | 1 | 1 | 3 | 0.3333 |
| 8_7204145 | NA | NA | NA | NA | NA | NA | 1 | 1 | 1 | 1 | NA | NA | NA | NA | 2 | 0.5 | 1 | 1 | 3 | 0.6667 |
| 8_7207468 | 1 | 0 | 2 | 0 | 1 | 1 | NA | NA | 4 | 0.25 | NA | NA | 1 | 1 | 1 | 0 | NA | NA | 2 | 0.5 |
| 8_7207625 | 4 | 0.25 | 3 | 0.3333 | 3 | 0.3333 | 3 | 0.3333 | 13 | 0.3077 | 5 | 0 | 6 | 0.3333 | 5 | 0 | 6 | 0 | 22 | 0.0909 |
| 8_7207628 | 2 | 1 | 5 | 0.6 | 2 | 1 | 2 | 0.5 | 11 | 0.7273 | 6 | 0.6667 | 4 | 0.5 | 4 | 0.5 | 3 | 0.6667 | 17 | 0.5882 |
| 8_7207635 | 3 | 0.6667 | 5 | 0.8 | 3 | 1 | 2 | 1 | 13 | 0.8462 | 6 | 0.3333 | 4 | 0.5 | 3 | 0.6667 | 5 | 1 | 18 | 0.6111 |
| 8_7207637 | 3 | 0.3333 | 3 | 0.3333 | 2 | 0 | NA | NA | 8 | 0.25 | 4 | 0.25 | 4 | 0.25 | 5 | 0 | 5 | 0 | 18 | 0.1111 |
| 8_7207646 | 3 | 0.3333 | 3 | 0.6667 | 1 | 1 | 3 | 0.3333 | 10 | 0.5 | 5 | 0.8 | 3 | 1 | 3 | 1 | 5 | 0.8 | 16 | 0.875 |
| 8_7207670 | 4 | 0.5 | 4 | 0.5 | 2 | 0.5 | 3 | 0 | 13 | 0.3846 | 6 | 0.5 | 4 | 0.5 | 5 | 0 | 4 | 0.25 | 19 | 0.3158 |
| 8_7207700 | 1 | 1 | 3 | 1 | NA | NA | NA | NA | 4 | 1 | 1 | 0 | 1 | 1 | NA | NA | NA | NA | 2 | 0.5 |
| 8_7207720 | 6 | 0.1667 | 5 | 0 | 2 | 0 | 6 | 0 | 19 | 0.0526 | 6 | 0.1667 | 4 | 0 | 4 | 0 | 3 | 0 | 17 | 0.0588 |
| 8_7207744 | 5 | 0 | 6 | 0 | 5 | 0.2 | 7 | 0.2857 | 23 | 0.1304 | 4 | 0 | 3 | 0 | 3 | 0.3333 | 8 | 0.25 | 18 | 0.1667 |
| 8_7207746 | 5 | 0.4 | 4 | 0.25 | 5 | 0.2 | 5 | 0.2 | 19 | 0.2632 | 3 | 0 | 6 | 0 | 2 | 1 | 6 | 0.1667 | 17 | 0.1765 |
| 8_7208155 | 3 | 0.6667 | 5 | 0.4 | 6 | 0.8333 | 5 | 1 | 19 | 0.7368 | 4 | 0.75 | 3 | 1 | 6 | 0.5 | 2 | 1 | 15 | 0.7333 |
| 8_7208211 | 3 | 0.3333 | 8 | 0.25 | 6 | 0.5 | 4 | 0.5 | 21 | 0.381 | 3 | 0 | 5 | 0.4 | 9 | 0.2222 | 1 | 1 | 18 | 0.2778 |
| 8_7208231 | 3 | 0 | 7 | 0 | 4 | 0.25 | 4 | 0.5 | 18 | 0.1667 | 3 | 0.3333 | 5 | 0.2 | 7 | 0.2857 | NA | NA | 15 | 0.2667 |
| 8_7214491 | NA | NA | 1 | 0 | NA | NA | 1 | 1 | 2 | 0.5 | NA | NA | 1 | 1 | NA | NA | NA | NA | 1 | 1 |
| 8_7214558 | NA | NA | NA | NA | NA | NA | 1 | 1 | 1 | 1 | NA | NA | 1 | 1 | NA | NA | NA | NA | 1 | 1 |
| 8_7215967 | NA | NA | 1 | 0 | 2 | 0.5 | 3 | 0.6667 | 6 | 0.5 | 3 | 0.3333 | 1 | 1 | 2 | 0 | NA | NA | 6 | 0.3333 |
| 8_7216015 | 1 | 1 | 1 | 1 | 2 | 0.5 | 3 | 1 | 7 | 0.8571 | NA | NA | 1 | 0 | 2 | 1 | NA | NA | 3 | 0.6667 |
| 8_7216039 | 1 | 1 | 1 | 1 | 1 | 1 | 3 | 1 | 6 | 1 | 2 | 0.5 | NA | NA | 1 | 0 | NA | NA | 3 | 0.3333 |
| 8_7216039 | 1 | 1 | 1 | 0 | 2 | 0 | 3 | 0 | 7 | 0.1429 | 1 | 0 | NA | NA | 2 | 0.5 | NA | NA | 3 | 0.3333 |
| 8_7216044 | 1 | 0 | 1 | 1 | 1 | 1 | 2 | 0 | 5 | 0.4 | 1 | 1 | 1 | 0 | 2 | 0 | NA | NA | 4 | 0.25 |
| 8_7216073 | 1 | 1 | 1 | 0 | 1 | 1 | 3 | 0 | 6 | 0.3333 | 2 | 0.5 | 1 | 1 | 2 | 0 | NA | NA | 5 | 0.4 |
| 8_7264278 | 1 | 1 | NA | NA | NA | NA | 2 | 0.5 | 3 | 0.6667 | NA | NA | NA | NA | 1 | 0 | 1 | 1 | 2 | 0.5 |
| 8_7264295 | 1 | 0 | NA | NA | 1 | 1 | 3 | 0 | 5 | 0.2 | NA | NA | NA | NA | 1 | 1 | 1 | 0 | 2 | 0.5 |
| 8_7264305 | NA | NA | NA | NA | 1 | 1 | 3 | 0.6667 | 4 | 0.75 | NA | NA | NA | NA | 2 | 1 | 1 | 1 | 3 | 1 |
| 8_7264329 | 1 | 1 | NA | NA | 1 | 0 | 2 | 0.5 | 4 | 0.5 | NA | NA | NA | NA | 2 | 0.5 | 1 | 0 | 3 | 0.3333 |
| 8_7264339 | 1 | 1 | NA | NA | 1 | 0 | 3 | 1 | 5 | 0.8 | NA | NA | NA | NA | 2 | 1 | 1 | 1 | 3 | 1 |
| 8_7264354 | NA | NA | NA | NA | 1 | 1 | 3 | 0.6667 | 4 | 0.75 | NA | NA | NA | NA | 2 | 1 | 1 | 1 | 3 | 1 |
| 8_7264356 | 1 | 1 | NA | NA | 1 | 1 | 2 | 1 | 4 | 1 | NA | NA | NA | NA | 2 | 1 | 2 | 1 | 4 | 1 |
| 8_7264389 | 1 | 1 | NA | NA | 1 | 1 | 3 | 1 | 5 | 1 | 1 | 1 | NA | NA | 2 | 1 | 2 | 1 | 5 | 1 |
| 8_7264395 | 1 | 0 | NA | NA | 1 | 1 | 2 | 0 | 4 | 0.25 | 1 | 0 | NA | NA | NA | NA | 2 | 1 | 3 | 0.6667 |
| 8_7264412 | 1 | 0 | NA | NA | 2 | 0.5 | 2 | 0.5 | 5 | 0.4 | 1 | 1 | NA | NA | 1 | 1 | 1 | 1 | 3 | 1 |
| 8_7264414 | NA | NA | NA | NA | NA | NA | 1 | 1 | 1 | 1 | 1 | 1 | NA | NA | 1 | 1 | 2 | 1 | 4 | 1 |
| 8_7264416 | 1 | 1 | NA | NA | 1 | 0 | 3 | 1 | 5 | 0.8 | 1 | 1 | NA | NA | 2 | 1 | 2 | 1 | 5 | 1 |
| 8_7264420 | 1 | 1 | NA | NA | 2 | 1 | 3 | 1 | 6 | 1 | 1 | 1 | NA | NA | 2 | 1 | 2 | 1 | 5 | 1 |
| 8_7264430 | 1 | 1 | NA | NA | 2 | 1 | 1 | 0 | 4 | 0.75 | 1 | 1 | NA | NA | NA | NA | 2 | 0.5 | 3 | 0.6667 |
| 8_7264431 | 1 | 1 | NA | NA | 1 | 1 | 1 | 0 | 3 | 0.6667 | 1 | 0 | NA | NA | 1 | 0 | 1 | 1 | 3 | 0.3333 |
| 8_7264443 | 1 | 1 | NA | NA | 2 | 1 | 2 | 1 | 5 | 1 | 1 | 1 | NA | NA | 2 | 1 | 2 | 1 | 5 | 1 |
| 8_7264473 | 1 | 1 | NA | NA | 2 | 0.5 | 3 | 0 | 6 | 0.3333 | 1 | 0 | NA | NA | 2 | 0.5 | 2 | 0 | 5 | 0.2 |
| 8_7264857 | NA | NA | NA | NA | 1 | 0 | 1 | 1 | 2 | 0.5 | NA | NA | 1 | 0 | 2 | 0 | 1 | 1 | 4 | 0.25 |
| 8_7266849 | NA | NA | 1 | 1 | NA | NA | NA | NA | 1 | 1 | NA | NA | NA | NA | 1 | 1 | 1 | 1 | 2 | 1 |
| 8_7268604 | 2 | 1 | 3 | 0.3333 | 4 | 0.25 | 3 | 0.3333 | 12 | 0.4167 | 1 | 0 | 2 | 0.5 | 1 | 0 | 2 | 0 | 6 | 0.1667 |
| 8_7268607 | 2 | 0 | 2 | 0 | 4 | 0.25 | 2 | 0.5 | 10 | 0.2 | 1 | 0 | 3 | 0.3333 | 1 | 0 | 2 | 0 | 7 | 0.1429 |
| 8_7268609 | 2 | 1 | 2 | 1 | 4 | 1 | 3 | 0.6667 | 11 | 0.9091 | 1 | 1 | 2 | 1 | 1 | 0 | 2 | 1 | 6 | 0.8333 |
| 8_7268614 | 2 | 0.5 | 2 | 0 | 3 | 0 | 2 | 0 | 9 | 0.1111 | 1 | 0 | 1 | 0 | 1 | 0 | 2 | 0.5 | 5 | 0.2 |
| 8_7268628 | 1 | 1 | 3 | 0.6667 | 4 | 0.75 | 1 | 1 | 9 | 0.7778 | 2 | 1 | 3 | 1 | 1 | 0 | 2 | 1 | 8 | 0.875 |
| 8_7268673 | 2 | 0.5 | 3 | 0.3333 | 3 | 0.6667 | 3 | 0.3333 | 11 | 0.4545 | 1 | 0 | 3 | 1 | NA | NA | 2 | 1 | 6 | 0.8333 |
| 8_7268683 | 3 | 0 | 4 | 0.25 | 4 | 0.25 | 4 | 0.25 | 15 | 0.2 | 3 | 0 | 3 | 0.3333 | 1 | 0 | 2 | 0 | 9 | 0.1111 |
| 8_7268698 | 2 | 0.5 | 3 | 0 | 4 | 0.25 | 4 | 0 | 13 | 0.1538 | 3 | 0.3333 | 3 | 0.3333 | 1 | 1 | 2 | 0.5 | 9 | 0.4444 |
| 8_7268709 | 3 | 0 | 4 | 0.25 | 4 | 0 | 4 | 0 | 15 | 0.0667 | 3 | 0.3333 | 3 | 0 | 1 | 0 | 2 | 0 | 9 | 0.1111 |
| 8_7268715 | 3 | 0.6667 | 3 | 0.3333 | 4 | 0.25 | 4 | 0 | 14 | 0.2857 | 3 | 0.6667 | 3 | 0.6667 | 1 | 0 | 2 | 0.5 | 9 | 0.5556 |
| 8_7268732 | 3 | 0.3333 | 4 | 0.5 | 3 | 0.6667 | 4 | 0.5 | 14 | 0.5 | 4 | 0.75 | 3 | 0.6667 | 1 | 0 | 3 | 0.6667 | 11 | 0.6364 |

| | | | | | | | | | | | | | | | | | | | | |
|---|---|---|---|---|---|---|---|---|---|---|---|---|---|---|---|---|---|---|---|---|
| 8_7268734 | 3 | 0.6667 | 4 | 0.5 | 4 | 0.5 | 4 | 0.75 | 15 | 0.6 | 4 | 1 | 3 | 0.6667 | 1 | 0 | 3 | 0.6667 | 11 | 0.7273 |
| 8_7268746 | 3 | 0 | 4 | 0.25 | 4 | 0 | 4 | 0 | 15 | 0.0667 | 4 | 0 | 3 | 0.3333 | 1 | 0 | 3 | 0 | 11 | 0.0909 |
| 8_7268750 | 3 | 0.3333 | 5 | 0 | 4 | 0 | 4 | 0 | 16 | 0.0625 | 4 | 0 | 3 | 0.3333 | 1 | 0 | 3 | 0 | 11 | 0.0909 |
| 8_7268756 | 3 | 0 | 4 | 0 | 5 | 0.2 | 3 | 0 | 15 | 0.0667 | 4 | 0.25 | 2 | 0 | 1 | 0 | 3 | 0 | 10 | 0.1 |
| 8_7268782 | 3 | 0 | 5 | 0.2 | 5 | 0.6 | 6 | 0.1667 | 19 | 0.2632 | 4 | 0.25 | 3 | 0.6667 | 1 | 0 | 2 | 0 | 10 | 0.3 |
| 8_7268839 | 4 | 0 | 5 | 0.6 | 5 | 0.4 | 8 | 0.25 | 22 | 0.3182 | 4 | 0.75 | 4 | 0.75 | 1 | 0 | 2 | 0.5 | 11 | 0.6364 |
| 8_7272189 | NA | NA | NA | NA | 1 | 0 | 3 | 0.3333 | 4 | 0.25 | 2 | 0.5 | 2 | 1 | NA | NA | 1 | 1 | 5 | 0.8 |
| 8_7272219 | NA | NA | NA | NA | 1 | 1 | 3 | 1 | 4 | 1 | 2 | 1 | 2 | 0.5 | NA | NA | 1 | 1 | 5 | 0.8 |
| 8_7272224 | NA | NA | NA | NA | 1 | 0 | 4 | 0.75 | 5 | 0.6 | 2 | 0.5 | 2 | 0.5 | NA | NA | 1 | 1 | 5 | 0.6 |
| 8_7272231 | NA | NA | NA | NA | 1 | 1 | 4 | 1 | 5 | 1 | 2 | 1 | 2 | 1 | NA | NA | 1 | 1 | 5 | 1 |
| 8_7272244 | NA | NA | NA | NA | 1 | 1 | 3 | 1 | 4 | 1 | 1 | 1 | 1 | 1 | NA | NA | 1 | 1 | 3 | 1 |
| 8_7272247 | NA | NA | NA | NA | 1 | 1 | 4 | 0.75 | 5 | 0.8 | 2 | 1 | 2 | 1 | NA | NA | 1 | 1 | 5 | 1 |
| 8_7272258 | NA | NA | NA | NA | 1 | 0 | 4 | 0.5 | 5 | 0.4 | 2 | 0 | 2 | 0.5 | NA | NA | 1 | 0 | 5 | 0.2 |
| 8_7272300 | 1 | 1 | NA | NA | 1 | 0 | 4 | 0.75 | 6 | 0.6667 | 2 | 0 | 2 | 0.5 | NA | NA | NA | NA | 4 | 0.25 |
| 8_7272305 | 1 | 1 | NA | NA | 1 | 1 | 4 | 0.75 | 6 | 0.8333 | 1 | 1 | 2 | 0.5 | NA | NA | 1 | 1 | 4 | 0.75 |
| 8_7272322 | 1 | 0 | NA | NA | 1 | 0 | 4 | 0.5 | 6 | 0.3333 | 1 | 0 | 2 | 0.5 | NA | NA | 1 | 0 | 4 | 0.25 |
| 8_7272330 | 2 | 0 | NA | NA | 1 | 0 | 3 | 1 | 6 | 0.5 | 2 | 1 | 2 | 0 | NA | NA | 1 | 0 | 5 | 0.4 |
| 8_7272342 | 2 | 1 | NA | NA | 1 | 1 | 3 | 1 | 6 | 1 | 2 | 1 | 2 | 1 | NA | NA | 1 | 1 | 5 | 1 |
| 8_7290565 | 1 | 1 | 6 | 0 | 8 | 0.25 | 7 | 0.1429 | 22 | 0.1818 | 4 | 0 | 3 | 0.3333 | 4 | 0.25 | 5 | 0.4 | 16 | 0.25 |
| 8_7290634 | 4 | 0.25 | 5 | 0 | 8 | 0.125 | 6 | 0 | 23 | 0.087 | 4 | 0.25 | 3 | 0 | 5 | 0.2 | 5 | 0 | 17 | 0.1176 |
| 8_7290686 | 4 | 0.25 | 5 | 0 | 8 | 0 | 8 | 0.125 | 25 | 0.08 | 7 | 0.2857 | 4 | 0 | 6 | 0 | 7 | 0 | 24 | 0.0833 |
| 8_7293398 | 3 | 0 | 3 | 0.3333 | 2 | 0 | 10 | 0.3 | 18 | 0.2222 | 4 | 0 | 2 | 0.5 | 3 | 0 | 4 | 0.5 | 13 | 0.2308 |
| 8_7293402 | 3 | 0.6667 | 3 | 1 | 3 | 0.6667 | 9 | 0.2222 | 18 | 0.5 | 3 | 0.3333 | 2 | 1 | 3 | 0.6667 | 3 | 0.6667 | 11 | 0.6364 |
| 8_7293406 | 3 | 0 | 2 | 1 | 4 | 0.5 | 8 | 0.5 | 17 | 0.4706 | 3 | 0.6667 | 3 | 0 | 3 | 0.6667 | 4 | 0.5 | 13 | 0.4615 |
| 8_7293408 | 3 | 1 | 3 | 1 | 4 | 1 | 10 | 0.9 | 20 | 0.95 | 3 | 1 | 3 | 0.3333 | 3 | 0.6667 | 4 | 1 | 13 | 0.7692 |
| 8_7293435 | 2 | 0 | 4 | 0.5 | 3 | 0.3333 | 8 | 0.25 | 17 | 0.2941 | 5 | 0.6 | 2 | 1 | 4 | 0 | 4 | 0.5 | 15 | 0.4667 |
| 8_7293439 | 3 | 0.3333 | 4 | 0.5 | 4 | 0.5 | 8 | 0.25 | 19 | 0.3684 | 5 | 0.2 | 3 | 0.6667 | 4 | 0 | 3 | 0.3333 | 15 | 0.2667 |
| 8_7293440 | 2 | 0 | 4 | 0 | 2 | 0.5 | 9 | 0 | 17 | 0.0588 | 6 | 0 | 4 | 0 | 4 | 0.25 | 3 | 0 | 17 | 0.0588 |
| 8_7293444 | 3 | 0 | 3 | 0.3333 | 4 | 0 | 10 | 0.1 | 20 | 0.1 | 5 | 0.2 | 3 | 0.3333 | 4 | 0.25 | 3 | 0.3333 | 15 | 0.2667 |
| 8_7293448 | 3 | 0 | 4 | 0.5 | 4 | 0.25 | 8 | 0.125 | 19 | 0.2105 | 6 | 0.1667 | 4 | 0.25 | 3 | 0.3333 | 3 | 0.3333 | 16 | 0.25 |
| 8_7293453 | 3 | 0.6667 | 1 | 1 | 4 | 0.75 | 7 | 0.7143 | 15 | 0.7333 | 4 | 0.5 | 2 | 0.5 | 3 | 0.6667 | 3 | 0.3333 | 12 | 0.5 |
| 8_7293460 | 4 | 0 | 4 | 0 | 3 | 0 | 9 | 0.1111 | 20 | 0.05 | 5 | 0.2 | 4 | 0 | 3 | 0 | 4 | 0 | 16 | 0.0625 |
| 8_7293475 | 4 | 0.75 | 4 | 0 | 5 | 0.2 | 4 | 0 | 17 | 0.2353 | 3 | 0.3333 | 2 | 0 | 3 | 0 | 3 | 0.3333 | 11 | 0.1818 |
| 8_7293486 | 2 | 0 | 4 | 0.5 | 5 | 0 | 5 | 0 | 16 | 0.125 | 5 | 0 | 3 | 0 | 4 | 0 | 4 | 0.25 | 16 | 0.0625 |
| 8_7293487 | 4 | 0.5 | 4 | 0.25 | 5 | 0.6 | 6 | 0.3333 | 19 | 0.4211 | 5 | 0.6 | 1 | 1 | 4 | 0.5 | 2 | 0.5 | 12 | 0.5833 |
| 8_7293496 | 4 | 0 | 3 | 0 | 7 | 0 | 6 | 0.1667 | 20 | 0.05 | 4 | 0.25 | 2 | 0 | 5 | 0 | 4 | 0 | 15 | 0.0667 |
| 8_7297584 | 1 | 0 | NA | NA | NA | NA | 1 | 1 | 2 | 0.5 | NA | NA | 1 | 0 | 1 | 1 | NA | NA | 2 | 0.5 |
| 8_7297589 | 1 | 1 | NA | NA | 1 | 1 | 1 | 1 | 3 | 1 | 1 | 1 | 1 | 0 | 1 | 1 | NA | NA | 3 | 0.6667 |
| 8_7297594 | 1 | 0 | NA | NA | 1 | 1 | 1 | 0 | 3 | 0.3333 | 1 | 1 | 1 | 0 | 1 | 0 | NA | NA | 3 | 0.3333 |
| 8_7297602 | 10 | 0.4 | 9 | 0.1111 | 9 | 0.2222 | 12 | 0.3333 | 40 | 0.275 | 6 | 0.3333 | 7 | 0.2857 | 10 | 0.2 | 3 | 0 | 26 | 0.2308 |
| 8_7297608 | 6 | 0.3333 | 5 | 0.4 | 10 | 0.4 | 8 | 0.25 | 29 | 0.3448 | 4 | 0.25 | 7 | 0.1429 | 8 | 0.125 | 1 | 0 | 20 | 0.15 |
| 8_7297621 | 13 | 0.9231 | 9 | 0.6667 | 12 | 0.8333 | 13 | 0.6923 | 47 | 0.7872 | 7 | 0.8571 | 8 | 0.875 | 11 | 0.8182 | 3 | 0.6667 | 29 | 0.8276 |
| 8_7297628 | 12 | 0.8333 | 9 | 0.6667 | 9 | 0.7778 | 12 | 0.75 | 42 | 0.7619 | 6 | 0.6667 | 9 | 0.7778 | 11 | 0.9091 | 1 | 1 | 27 | 0.8148 |
| 8_7297639 | 13 | 0.0769 | 8 | 0.375 | 10 | 0.2 | 10 | 0.4 | 41 | 0.2439 | 6 | 0.3333 | 7 | 0.5714 | 10 | 0.4 | 3 | 0.6667 | 26 | 0.4615 |
| 8_7297641 | 5 | 0.2 | 6 | 0.1667 | 5 | 0 | 7 | 0 | 23 | 0.087 | 6 | 0.1667 | 7 | 0.1429 | 6 | 0.3333 | 2 | 0 | 21 | 0.1905 |
| 8_7297645 | 11 | 0.5455 | 8 | 0.5 | 9 | 0.5556 | 10 | 0.5 | 38 | 0.5263 | 6 | 0.3333 | 8 | 0.5 | 8 | 0.25 | 2 | 0 | 24 | 0.3333 |
| 8_7297650 | 13 | 0.1538 | 9 | 0 | 12 | 0.1667 | 11 | 0.3636 | 45 | 0.1778 | 7 | 0 | 8 | 0.5 | 11 | 0.1818 | 3 | 0 | 29 | 0.2069 |
| 8_7297650 | 13 | 0.8462 | 11 | 0.8182 | 12 | 0.6667 | 11 | 0.6364 | 47 | 0.7447 | 8 | 0.75 | 8 | 0.75 | 11 | 0.9091 | 3 | 1 | 30 | 0.8333 |
| 8_7297655 | 9 | 0.7778 | 9 | 0.5556 | 12 | 0.75 | 9 | 0.7778 | 39 | 0.7179 | 7 | 0.8571 | 7 | 0.7143 | 9 | 0.6667 | 2 | 1 | 25 | 0.76 |
| 8_7297669 | 13 | 0.0769 | 10 | 0.3 | 12 | 0.3333 | 10 | 0.1 | 45 | 0.2 | 8 | 0.25 | 8 | 0.25 | 9 | 0.3333 | 3 | 0.3333 | 28 | 0.2857 |
| 8_7297674 | 12 | 1 | 12 | 0.9167 | 13 | 0.8462 | 10 | 1 | 47 | 0.9362 | 8 | 0.875 | 9 | 0.8889 | 10 | 0.9 | 3 | 1 | 30 | 0.9 |
| 8_7297675 | 12 | 0.1667 | 12 | 0.1667 | 10 | 0.3 | 12 | 0.4167 | 46 | 0.2609 | 7 | 0.4286 | 9 | 0.1111 | 11 | 0.2727 | 3 | 0 | 30 | 0.2333 |
| 8_7297690 | 8 | 0.5 | 8 | 0.375 | 6 | 0.3333 | 6 | 0.5 | 28 | 0.4286 | 4 | 0.5 | 3 | 0.3333 | 8 | 0.5 | 4 | 0.5 | 19 | 0.4737 |
| 8_7297696 | 11 | 0.7273 | 9 | 0.5556 | 8 | 0.625 | 7 | 1 | 35 | 0.7143 | 6 | 0.6667 | 3 | 0.3333 | 8 | 0.5 | 6 | 0.5 | 23 | 0.5217 |
| 8_7297701 | 10 | 0.8 | 7 | 0.7143 | 7 | 0.7143 | 5 | 1 | 29 | 0.7931 | 6 | 1 | 3 | 0.6667 | 8 | 0.625 | 6 | 0.3333 | 23 | 0.6522 |
| 8_7297715 | 12 | 0.3333 | 9 | 0.2222 | 7 | 0.1429 | 8 | 0.5 | 36 | 0.3056 | 8 | 0.5 | 3 | 0 | 9 | 0.3333 | 6 | 0.5 | 26 | 0.3846 |
| 8_7297727 | 1 | 0 | 4 | 0.5 | 2 | 0.5 | 5 | 0.6 | 12 | 0.5 | 3 | 0.6667 | NA | NA | 1 | 1 | 3 | 0.3333 | 7 | 0.5714 |
| 8_7297755 | 3 | 0.6667 | 5 | 0.2 | 2 | 0 | 3 | 0 | 13 | 0.2308 | 2 | 0.5 | 1 | 0 | 2 | 0 | 4 | 0.25 | 9 | 0.2222 |
| 8_7297757 | 3 | 0 | 5 | 0.2 | 2 | 0.5 | 2 | 0 | 12 | 0.1667 | 2 | 0 | 1 | 1 | 2 | 0 | 2 | 0.5 | 7 | 0.2857 |
| 8_7297823 | 5 | 0.4 | 5 | 0 | 1 | 1 | 3 | 0 | 14 | 0.2143 | 2 | 0.5 | 3 | 0.3333 | 5 | 0.2 | 2 | 0.5 | 12 | 0.3333 |
| 8_7297828 | 5 | 0.2 | 1 | 0 | 2 | 0 | 2 | 0 | 10 | 0.1 | 2 | 0 | 2 | 0 | 4 | 0.25 | 3 | 0 | 11 | 0.0909 |
| 8_7297854 | 5 | 0.2 | 5 | 0 | 2 | 0 | 3 | 0 | 15 | 0.0667 | 2 | 0 | 2 | 0.5 | 6 | 0 | 3 | 0 | 13 | 0.0769 |
| 8_7299072 | NA | NA | NA | NA | 3 | 0.3333 | 1 | 0 | 4 | 0.25 | NA | NA | 1 | 0 | 3 | 0.3333 | NA | NA | 4 | 0.25 |
| 8_7299077 | NA | NA | NA | NA | 3 | 0.3333 | 1 | 1 | 4 | 0.5 | NA | NA | 1 | 0 | 3 | 0.3333 | NA | NA | 4 | 0.25 |
| 8_7299157 | NA | NA | 1 | 1 | 3 | 0.3333 | 1 | 0 | 5 | 0.4 | NA | NA | 1 | 1 | 3 | 0.3333 | 1 | 0 | 5 | 0.4 |

| | | | | | | | | | | | | | | | | | | | | |
|---|---|---|---|---|---|---|---|---|---|---|---|---|---|---|---|---|---|---|---|---|
| 8_7299214 | NA | NA | 1 | 0 | 3 | 0 | 1 | 1 | 5 | 0.2 | NA | NA | 2 | 0 | 2 | 0 | 1 | 1 | 5 | 0.2 |
| 8_7301068 | 6 | 0 | 6 | 0.1667 | 2 | 0 | 5 | 0 | 19 | 0.0526 | 1 | 0 | 6 | 0.1667 | 6 | 0 | 2 | 0 | 15 | 0.0667 |
| 8_7301072 | 12 | 0.5 | 10 | 0.3 | 5 | 0.4 | 8 | 0.375 | 35 | 0.4 | 4 | 0.75 | 10 | 0.6 | 5 | 0.8 | 6 | 0.6667 | 25 | 0.68 |
| 8_7301079 | 14 | 0.3571 | 9 | 0.3333 | 4 | 0 | 6 | 0 | 33 | 0.2424 | 4 | 0.25 | 10 | 0.2 | 5 | 0.2 | 8 | 0 | 27 | 0.1481 |
| 8_7301368 | NA | NA | NA | NA | NA | NA | 1 | 1 | 1 | 1 | NA | NA | 1 | 1 | NA | NA | NA | NA | 1 | 1 |
| 8_7307333 | 4 | 0.25 | 2 | 0.5 | 2 | 0 | 5 | 0.2 | 13 | 0.2308 | 4 | 0.75 | 3 | 0.6667 | 4 | 0.25 | NA | NA | 11 | 0.5455 |
| 8_7307362 | 8 | 0.25 | 2 | 0 | 3 | 0 | 7 | 0.1429 | 20 | 0.15 | 6 | 0 | 2 | 0 | 5 | 0.4 | 1 | 0 | 14 | 0.1429 |
| 8_7307370 | 6 | 0 | 1 | 1 | 4 | 0 | 7 | 0 | 18 | 0.0556 | 5 | 0.2 | 5 | 0 | 4 | 0 | NA | NA | 14 | 0.0714 |
| 8_7307468 | 9 | 0.2222 | 4 | 0 | 4 | 0.5 | 9 | 0.3333 | 26 | 0.2692 | 7 | 0.2857 | 5 | 0.4 | 5 | 0.2 | 2 | 0 | 19 | 0.2632 |
| 8_7307483 | 9 | 1 | 2 | 1 | 3 | 1 | 8 | 1 | 22 | 1 | 7 | 0.8571 | 4 | 1 | 4 | 1 | 1 | 0 | 16 | 0.875 |
| 8_7307485 | 9 | 0.8889 | 3 | 1 | 5 | 0.6 | 5 | 1 | 22 | 0.8636 | 7 | 0.8571 | 3 | 1 | 4 | 1 | 1 | 1 | 15 | 0.9333 |
| 8_7307486 | 7 | 0.5714 | 2 | 0 | 5 | 0.6 | 8 | 0.5 | 22 | 0.5 | 8 | 0.5 | 5 | 0.4 | 6 | 0.8333 | 1 | 0 | 20 | 0.55 |
| 8_7307487 | 9 | 0.2222 | 2 | 0.5 | 4 | 0 | 8 | 0.125 | 23 | 0.1739 | 8 | 0 | 2 | 0 | 4 | 0 | 1 | 1 | 15 | 0.0667 |
| 8_7307489 | 9 | 0.7778 | 3 | 0 | 5 | 0.6 | 8 | 0.375 | 25 | 0.52 | 7 | 0.4286 | 5 | 0.4 | 6 | 0.3333 | 1 | 1 | 19 | 0.4211 |
| 8_7307492 | 8 | 0.875 | 3 | 1 | 5 | 0.8 | 9 | 0.6667 | 25 | 0.8 | 6 | 0.5 | 5 | 0.6 | 5 | 0.6 | 1 | 1 | 17 | 0.5882 |
| 8_7307498 | 5 | 1 | 4 | 1 | 5 | 0.8 | 7 | 0.7143 | 21 | 0.8571 | 4 | 0.5 | 3 | 1 | 3 | 0.6667 | 2 | 1 | 12 | 0.75 |
| 8_7307509 | 10 | 0.3 | 4 | 0.25 | 5 | 0 | 9 | 0.4444 | 28 | 0.2857 | 9 | 0.2222 | 5 | 0 | 4 | 0 | 1 | 0 | 19 | 0.1053 |
| 8_7309512 | 2 | 0.5 | 9 | 0.4444 | 8 | 0.375 | 8 | 0.5 | 27 | 0.4444 | 7 | 0.5714 | 5 | 0.4 | 5 | 0.8 | 5 | 0.2 | 22 | 0.5 |
| 8_7309515 | 2 | 1 | 9 | 0.6667 | 7 | 0.8571 | 7 | 0.4286 | 25 | 0.68 | 6 | 0.6667 | 5 | 0.4 | 4 | 0.5 | 4 | 0.5 | 19 | 0.5263 |
| 8_7319914 | 1 | 0 | NA | NA | 1 | 0 | 1 | 1 | 3 | 0.3333 | NA | NA | 1 | 1 | NA | NA | NA | NA | 1 | 1 |
| 8_7320925 | NA | NA | NA | NA | 2 | 1 | NA | NA | 2 | 1 | NA | NA | 2 | 0.5 | NA | NA | NA | NA | 2 | 0.5 |
| 8_7320925 | NA | NA | NA | NA | 1 | 1 | NA | NA | 1 | 1 | NA | NA | 2 | 0.5 | NA | NA | NA | NA | 2 | 0.5 |
| 8_7320932 | NA | NA | NA | NA | 2 | 0.5 | NA | NA | 2 | 0.5 | NA | NA | 2 | 1 | NA | NA | NA | NA | 2 | 1 |
| 8_7321405 | NA | NA | NA | NA | 2 | 1 | 1 | 1 | 3 | 1 | NA | NA | 2 | 1 | NA | NA | NA | NA | 2 | 1 |
| 8_7321412 | NA | NA | NA | NA | 3 | 1 | NA | NA | 3 | 1 | NA | NA | 2 | 1 | NA | NA | NA | NA | 2 | 1 |
| 8_7321416 | NA | NA | NA | NA | 3 | 1 | 1 | 1 | 4 | 1 | NA | NA | 2 | 1 | NA | NA | NA | NA | 2 | 1 |
| 8_7321817 | 2 | 0 | 5 | 0.2 | 2 | 0 | 6 | 0 | 15 | 0.0667 | NA | NA | 5 | 0.2 | 3 | 0.3333 | 3 | 0 | 11 | 0.1818 |
| 8_7321818 | 2 | 0.5 | 7 | 0.2857 | 3 | 0 | 7 | 0.2857 | 19 | 0.2632 | NA | NA | 8 | 0.25 | 5 | 0.4 | 6 | 0 | 19 | 0.2105 |
| 8_7323506 | 7 | 0.2857 | 5 | 0.2 | 5 | 0 | 3 | 0.3333 | 20 | 0.2 | 3 | 0 | 3 | 0 | 4 | 0.5 | 5 | 0 | 15 | 0.1333 |
| 8_7323556 | 5 | 0.4 | 5 | 0.6 | 4 | 0.25 | 7 | 0.5714 | 21 | 0.4762 | 3 | 0.3333 | 4 | 0.75 | 5 | 1 | 6 | 0.6667 | 18 | 0.7222 |
| 8_7323603 | 5 | 0 | 4 | 0.25 | 3 | 0 | 6 | 0 | 18 | 0.0556 | 2 | 0 | 4 | 0 | 3 | 0.3333 | 5 | 0 | 14 | 0.0714 |
| 8_7323702 | 7 | 1 | 1 | 1 | 2 | 1 | 5 | 0.8 | 15 | 0.9333 | 1 | 0 | 3 | 1 | 3 | 1 | 1 | 1 | 8 | 0.875 |
| 8_7323710 | 6 | 0.1667 | 1 | 0 | 2 | 0.5 | 5 | 0.2 | 14 | 0.2143 | 2 | 0 | 3 | 0.6667 | 3 | 0 | 2 | 0 | 10 | 0.2 |
| 8_7323723 | 6 | 0.6667 | 1 | 1 | 1 | 0 | 5 | 1 | 13 | 0.7692 | 2 | 0 | 1 | 0 | 3 | 0.6667 | 2 | 1 | 8 | 0.5 |
| 8_7323730 | 4 | 0.5 | 1 | 1 | 1 | 1 | 4 | 0.75 | 10 | 0.7 | 1 | 1 | 1 | 1 | 3 | 0.6667 | 2 | 1 | 7 | 0.8571 |
| 8_7323731 | 6 | 0.6667 | NA | NA | 2 | 1 | 5 | 0.6 | 13 | 0.6923 | 2 | 0.5 | 2 | 0.5 | 4 | 0.75 | 2 | 0 | 10 | 0.5 |
| 8_7323733 | 5 | 1 | 1 | 1 | NA | NA | 2 | 1 | 8 | 1 | 1 | 1 | 2 | 1 | 2 | 1 | 2 | 0.5 | 7 | 0.8571 |
| 8_7323741 | 4 | 0.5 | 1 | 1 | 1 | 1 | 2 | 0.5 | 8 | 0.625 | 1 | 0 | 2 | 1 | 3 | 0.6667 | 1 | 0 | 7 | 0.5714 |
| 8_7323744 | 3 | 0.3333 | 1 | 1 | 2 | 1 | 3 | 0.6667 | 9 | 0.6667 | 1 | 0 | 1 | 0 | 2 | 0.5 | 2 | 0.5 | 6 | 0.3333 |
| 8_7323745 | 6 | 0.8333 | 1 | 1 | 2 | 1 | 5 | 1 | 14 | 0.9286 | NA | NA | 2 | 1 | 3 | 0.6667 | 2 | 1 | 7 | 0.8571 |
| 8_7323753 | 4 | 0.5 | NA | NA | 2 | 0.5 | 4 | 0.75 | 10 | 0.6 | 1 | 0 | 1 | 1 | 1 | 1 | 2 | 0.5 | 5 | 0.6 |
| 8_7323754 | 3 | 0.6667 | 1 | 1 | 1 | 1 | 4 | 0.5 | 9 | 0.6667 | 1 | 1 | 2 | 0 | 3 | 1 | 2 | 0.5 | 8 | 0.625 |
| 8_7323764 | 5 | 0.6 | 1 | 1 | 2 | 0 | 3 | 0.3333 | 11 | 0.4545 | 1 | 0 | 2 | 0 | 2 | 0.5 | 2 | 0 | 7 | 0.1429 |
| 8_7323770 | 5 | 0.2 | 1 | 1 | 2 | 0.5 | 5 | 0 | 13 | 0.2308 | 1 | 0 | 2 | 0 | 2 | 0.5 | NA | NA | 5 | 0.2 |
| 8_7324933 | 1 | 1 | NA | NA | 2 | 1 | 1 | 1 | 4 | 1 | 1 | 1 | NA | NA | 1 | 1 | 3 | 1 | 5 | 1 |
| 8_7324935 | NA | NA | NA | NA | 2 | 1 | 1 | 0 | 3 | 0.6667 | 1 | 1 | NA | NA | 1 | 1 | 3 | 0 | 5 | 0.4 |
| 8_7324940 | 1 | 1 | NA | NA | 1 | 1 | NA | NA | 2 | 1 | 1 | 1 | NA | NA | 1 | 0 | 3 | 1 | 5 | 0.8 |
| 8_7324947 | 1 | 1 | NA | NA | 1 | 1 | 1 | 1 | 3 | 1 | 1 | 1 | NA | NA | NA | NA | 3 | 1 | 4 | 1 |
| 8_7324951 | 1 | 0 | NA | NA | 1 | 1 | 1 | 0 | 3 | 0.3333 | 1 | 0 | NA | NA | 1 | 0 | 2 | 0.5 | 4 | 0.25 |
| 8_7324970 | NA | NA | NA | NA | 2 | 1 | 1 | 0 | 3 | 0.6667 | 1 | 1 | NA | NA | NA | NA | 2 | 1 | 3 | 1 |
| 8_7324976 | 1 | 1 | NA | NA | 2 | 0.5 | 1 | 1 | 4 | 0.75 | 1 | 1 | NA | NA | 1 | 1 | 2 | 0.5 | 4 | 0.75 |
| 8_7324983 | NA | NA | NA | NA | 2 | 1 | 1 | 0 | 3 | 0.6667 | 1 | 1 | NA | NA | 1 | 0 | 1 | 0 | 3 | 0.3333 |
| 8_7324992 | NA | NA | NA | NA | 1 | 1 | 1 | 1 | 2 | 1 | NA | NA | NA | NA | 1 | 1 | 1 | 1 | 2 | 1 |
| 8_7324999 | NA | NA | NA | NA | 2 | 1 | 1 | 1 | 3 | 1 | NA | NA | NA | NA | 1 | 1 | 1 | 1 | 2 | 1 |
| 8_7325026 | NA | NA | NA | NA | 2 | 0.5 | NA | NA | 2 | 0.5 | 1 | 0 | NA | NA | NA | NA | 1 | 1 | 2 | 0.5 |
| 8_7330284 | NA | NA | NA | NA | 2 | 1 | NA | NA | 2 | 1 | NA | NA | NA | NA | NA | NA | 1 | 1 | 1 | 1 |
| 8_7330306 | 2 | 0.5 | NA | NA | NA | NA | NA | NA | 2 | 0.5 | NA | NA | 1 | 1 | NA | NA | NA | NA | 1 | 1 |
| 8_7330382 | 2 | 0.5 | NA | NA | 2 | 0.5 | NA | NA | 4 | 0.5 | NA | NA | 1 | 1 | 2 | 0 | 1 | 1 | 4 | 0.5 |
| 8_7330404 | 2 | 0.5 | 1 | 0 | 2 | 0.5 | NA | NA | 5 | 0.4 | NA | NA | 1 | 0 | 2 | 0 | 1 | 1 | 4 | 0.25 |
| 8_7331913 | 3 | 0.3333 | 3 | 0.3333 | 1 | 0 | NA | NA | 7 | 0.2857 | 1 | 1 | 2 | 0 | 3 | 1 | 3 | 0 | 9 | 0.4444 |
| 8_7331976 | 5 | 0.2 | 3 | 0 | 4 | 0 | 4 | 0 | 16 | 0.0625 | 5 | 0 | 6 | 0 | 5 | 0.2 | 4 | 0 | 20 | 0.05 |
| 8_7331988 | 15 | 0.1333 | 8 | 0 | 8 | 0 | 10 | 0.1 | 41 | 0.0732 | 9 | 0 | 11 | 0 | 8 | 0.125 | 9 | 0.1111 | 37 | 0.0541 |
| 8_7332061 | 11 | 0.0909 | 4 | 0.25 | 8 | 0.125 | 8 | 0.125 | 31 | 0.129 | 5 | 0 | 8 | 0.125 | 4 | 0 | 8 | 0.25 | 25 | 0.12 |
| 8_7332080 | 19 | 0.0526 | 10 | 0.2 | 12 | 0 | 15 | 0 | 56 | 0.0536 | 9 | 0.3333 | 15 | 0.0667 | 9 | 0.1111 | 8 | 0.25 | 41 | 0.1707 |
| 8_7332125 | 14 | 0.1429 | 8 | 0 | 10 | 0 | 8 | 0 | 40 | 0.05 | 7 | 0.1429 | 10 | 0 | 6 | 0.1667 | 9 | 0.2222 | 32 | 0.125 |

| | | | | | | | | | | | | | | | | | | | | |
|---|---|---|---|---|---|---|---|---|---|---|---|---|---|---|---|---|---|---|---|---|
| 8_7332143 | 21 | 0.1429 | 12 | 0 | 15 | 0.1333 | 14 | 0.1429 | 62 | 0.1129 | 9 | 0.1111 | 16 | 0 | 9 | 0.3333 | 10 | 0 | 44 | 0.0909 |
| 8_7332145 | 15 | 0.8 | 11 | 0.4545 | 14 | 0.7143 | 12 | 0.6667 | 52 | 0.6731 | 8 | 0.5 | 15 | 0.5333 | 7 | 0.2857 | 9 | 0.6667 | 39 | 0.5128 |
| 8_7332154 | 15 | 0.1333 | 8 | 0.125 | 12 | 0.0833 | 13 | 0.0769 | 48 | 0.1042 | 9 | 0.3333 | 14 | 0.0714 | 6 | 0.1667 | 8 | 0 | 37 | 0.1351 |
| 8_7333984 | 13 | 0.3846 | 7 | 0.4286 | 16 | 0.4375 | 19 | 0.4211 | 55 | 0.4182 | 4 | 0.75 | 10 | 0.4 | 10 | 0.5 | 11 | 0.4545 | 35 | 0.4857 |
| 8_7333997 | 14 | 1 | 13 | 1 | 17 | 0.9412 | 20 | 1 | 64 | 0.9844 | 8 | 1 | 12 | 1 | 11 | 1 | 13 | 0.9231 | 44 | 0.9773 |
| 8_7333998 | 11 | 0.2727 | 13 | 0.3077 | 16 | 0.4375 | 19 | 0.3684 | 59 | 0.3559 | 8 | 0.125 | 8 | 0.25 | 12 | 0.25 | 9 | 0.6667 | 37 | 0.3243 |
| 8_7333999 | 11 | 0.4545 | 12 | 0.8333 | 16 | 0.4375 | 21 | 0.6667 | 60 | 0.6 | 7 | 0.5714 | 10 | 0.6 | 14 | 0.6429 | 12 | 0.9167 | 43 | 0.6977 |
| 8_7334029 | 14 | 0.0714 | 12 | 0 | 13 | 0.2308 | 17 | 0.0588 | 56 | 0.0893 | 6 | 0.1667 | 8 | 0 | 12 | 0 | 13 | 0.1538 | 39 | 0.0769 |
| 8_7334038 | 14 | 0 | 10 | 0.3 | 13 | 0.3077 | 17 | 0.0588 | 54 | 0.1481 | 5 | 0 | 8 | 0.25 | 11 | 0 | 8 | 0.125 | 32 | 0.0938 |
| 8_7334054 | 9 | 0.1111 | 12 | 0.0833 | 12 | 0.0833 | 16 | 0.125 | 49 | 0.102 | 3 | 0 | 7 | 0 | 12 | 0.25 | 11 | 0.0909 | 33 | 0.1212 |
| 8_7343427 | 2 | 1 | 1 | 1 | 1 | 0 | 2 | 0.5 | 6 | 0.6667 | NA | NA | 1 | 0 | 2 | 1 | 2 | 0.5 | 5 | 0.6 |
| 8_7343854 | 2 | 0.5 | 2 | 0.5 | 2 | 0 | 3 | 1 | 9 | 0.5556 | NA | NA | 2 | 1 | 1 | 1 | NA | NA | 3 | 1 |
| 8_7343858 | 1 | 1 | 2 | 1 | 1 | 0 | 3 | 0.3333 | 7 | 0.5714 | NA | NA | 2 | 1 | 1 | 1 | 1 | 0 | 4 | 0.75 |
| 8_7343871 | 2 | 0.5 | 1 | 0 | NA | NA | 3 | 0.3333 | 6 | 0.3333 | NA | NA | 2 | 0.5 | 1 | 1 | 1 | 1 | 4 | 0.75 |
| 8_7343886 | 1 | 0 | 1 | 0 | 1 | 1 | 2 | 0 | 5 | 0.2 | NA | NA | 2 | 0 | 1 | 1 | 2 | 0 | 5 | 0.2 |
| 8_7344183 | 4 | 0 | 4 | 0.5 | 2 | 0 | 5 | 0 | 15 | 0.1333 | 4 | 0.25 | 2 | 0 | 3 | 0.6667 | 3 | 0.3333 | 12 | 0.3333 |
| 8_7344185 | 3 | 0 | 2 | 0.5 | 2 | 0 | 3 | 0.6667 | 10 | 0.3 | 1 | 1 | 1 | 1 | 3 | 0.3333 | 2 | 0 | 7 | 0.4286 |
| 8_7344208 | 4 | 0 | 2 | 0 | 3 | 0 | 2 | 0.5 | 11 | 0.0909 | 4 | 0 | 2 | 0.5 | 3 | 0.3333 | 3 | 0.3333 | 12 | 0.25 |
| 8_7344340 | 7 | 0.4286 | 6 | 0.3333 | 3 | 0.6667 | 6 | 0.3333 | 22 | 0.4091 | 4 | 0.5 | 2 | 0.5 | 5 | 0.2 | 6 | 0.5 | 17 | 0.4118 |
| 8_7344402 | NA | NA | 1 | 1 | NA | NA | 1 | 0 | 2 | 0.5 | NA | NA | NA | NA | NA | NA | 1 | 1 | 1 | 1 |
| 8_7347127 | 2 | 0 | 1 | 0 | NA | NA | 2 | 0.5 | 5 | 0.2 | 2 | 0.5 | 3 | 0 | 2 | 0 | 3 | 0 | 10 | 0.1 |
| 8_7347657 | 2 | 0 | 1 | 1 | NA | NA | 3 | 0 | 6 | 0.1667 | 2 | 0 | 3 | 0.3333 | 4 | 0.25 | 6 | 0.1667 | 15 | 0.2 |
| 8_7347710 | 2 | 0 | 3 | 0.3333 | NA | NA | 3 | 0 | 8 | 0.125 | 2 | 0 | 4 | 0 | 2 | 0 | 7 | 0.1429 | 15 | 0.0667 |
| 8_7348387 | 3 | 0.3333 | 5 | 0.2 | 2 | 0 | 5 | 0.2 | 15 | 0.2 | 2 | 0 | 6 | 0.3333 | 7 | 0.4286 | 7 | 0.1429 | 22 | 0.2727 |
| 8_7348390 | 1 | 1 | 6 | 0 | 3 | 0.3333 | 5 | 0.2 | 15 | 0.2 | 2 | 0 | 4 | 0.25 | 7 | 0.2857 | 8 | 0.375 | 21 | 0.2857 |
| 8_7348396 | 4 | 0 | 5 | 0.4 | 3 | 0 | 6 | 0.1667 | 18 | 0.1667 | 3 | 0 | 6 | 0.3333 | 7 | 0.1429 | 8 | 0.125 | 24 | 0.1667 |
| 8_7348403 | 3 | 0.3333 | 2 | 0 | 3 | 0 | 1 | 0 | 9 | 0.1111 | 2 | 0 | 3 | 0 | 8 | 0 | 6 | 0.1667 | 19 | 0.0526 |
| 8_7348416 | 4 | 0 | 6 | 0 | 3 | 0 | 6 | 0.3333 | 19 | 0.1053 | 4 | 0 | 7 | 0 | 8 | 0.125 | 9 | 0.1111 | 28 | 0.0714 |
| 8_7348419 | 5 | 0.2 | 4 | 0 | 3 | 0.3333 | 6 | 0.3333 | 18 | 0.2222 | 4 | 0.5 | 6 | 0.3333 | 7 | 0.1429 | 7 | 0.2857 | 24 | 0.2917 |
| 8_7348428 | 6 | 0.5 | 6 | 0.3333 | 3 | 0 | 6 | 0 | 21 | 0.2381 | 4 | 0.25 | 7 | 0.4286 | 7 | 0.1429 | 9 | 0 | 27 | 0.1852 |
| 8_7354123 | 5 | 0.8 | 14 | 0.7143 | 6 | 0.6667 | 9 | 0.5556 | 34 | 0.6765 | 9 | 0.7778 | 12 | 0.5833 | 12 | 0.6667 | 18 | 0.6111 | 51 | 0.6471 |
| 8_7354132 | 8 | 0 | 21 | 0.1905 | 8 | 0 | 17 | 0.1176 | 54 | 0.1111 | 13 | 0 | 14 | 0.1429 | 10 | 0.2 | 16 | 0.125 | 53 | 0.1132 |
| 8_7354137 | 10 | 0.6 | 16 | 0.8125 | 9 | 0.6667 | 16 | 0.5 | 51 | 0.6471 | 15 | 0.8 | 16 | 0.6875 | 12 | 0.75 | 19 | 0.5263 | 62 | 0.6774 |
| 8_7354137 | 11 | 0.0909 | 17 | 0.1176 | 10 | 0 | 15 | 0.0667 | 53 | 0.0755 | 15 | 0.1333 | 17 | 0.4118 | 13 | 0.0769 | 19 | 0.1579 | 64 | 0.2031 |
| 8_7354140 | 10 | 0.3 | 16 | 0.3125 | 10 | 0.1 | 15 | 0.3333 | 51 | 0.2745 | 16 | 0.3125 | 15 | 0.4 | 11 | 0.2727 | 20 | 0.2 | 62 | 0.2903 |
| 8_7354142 | 12 | 0.8333 | 12 | 0.75 | 10 | 0.6 | 14 | 0.8571 | 48 | 0.7708 | 13 | 0.8462 | 14 | 0.8571 | 12 | 0.6667 | 13 | 0.7692 | 52 | 0.7885 |
| 8_7354144 | 10 | 1 | 14 | 1 | 9 | 1 | 14 | 1 | 47 | 1 | 15 | 1 | 15 | 0.9333 | 10 | 1 | 18 | 1 | 58 | 0.9828 |
| 8_7354152 | 10 | 0.2 | 15 | 0.0667 | 9 | 0 | 15 | 0.2 | 49 | 0.1224 | 15 | 0.1333 | 16 | 0.25 | 9 | 0 | 16 | 0.0625 | 56 | 0.125 |
| 8_7354153 | 8 | 0.75 | 11 | 1 | 7 | 0.4286 | 16 | 0.875 | 42 | 0.8095 | 12 | 0.8333 | 13 | 0.9231 | 9 | 0.8889 | 13 | 0.4615 | 47 | 0.766 |
| 8_7354156 | 5 | 0.4 | 9 | 0.6667 | 7 | 0.4286 | 11 | 0.7273 | 32 | 0.5938 | 10 | 0.8 | 10 | 0.4 | 10 | 0.3 | 9 | 0.5556 | 39 | 0.5128 |
| 8_7354156 | 8 | 0.875 | 14 | 0.3571 | 10 | 0.5 | 13 | 0.9231 | 45 | 0.6444 | 12 | 0.6667 | 14 | 0.6429 | 11 | 0.8182 | 16 | 0.6875 | 53 | 0.6981 |
| 8_7354161 | 10 | 0 | 15 | 0.6 | 8 | 0.375 | 14 | 0.4286 | 47 | 0.383 | 12 | 0.4167 | 17 | 0.4706 | 10 | 0.3 | 14 | 0.3571 | 53 | 0.3962 |
| 8_7354165 | 9 | 1 | 13 | 0.8462 | 5 | 0.8 | 13 | 1 | 40 | 0.925 | 13 | 0.9231 | 12 | 0.75 | 12 | 1 | 13 | 0.8462 | 50 | 0.88 |
| 8_7354170 | 8 | 0.875 | 15 | 0.9333 | 6 | 0.8333 | 8 | 0.875 | 37 | 0.8919 | 12 | 0.9167 | 13 | 0.8462 | 11 | 0.9091 | 18 | 0.8889 | 54 | 0.8889 |
| 8_7354176 | 6 | 0.5 | 12 | 0.5833 | 8 | 0.375 | 10 | 0.6 | 36 | 0.5278 | 13 | 0.3077 | 12 | 0.6667 | 9 | 0.5556 | 12 | 0.4167 | 46 | 0.4783 |
| 8_7354200 | 6 | 1 | 13 | 1 | 8 | 0.875 | 10 | 1 | 37 | 0.973 | 13 | 0.9231 | 13 | 1 | 9 | 1 | 15 | 1 | 50 | 0.98 |
| 8_7354201 | 7 | 1 | 14 | 1 | 7 | 1 | 10 | 1 | 38 | 1 | 13 | 1 | 14 | 1 | 8 | 0.875 | 15 | 1 | 50 | 0.98 |
| 8_7354203 | 7 | 0.7143 | 15 | 0.5333 | 8 | 0.625 | 12 | 0.6667 | 42 | 0.619 | 15 | 0.6667 | 18 | 0.6111 | 8 | 0.375 | 15 | 0.3333 | 56 | 0.5179 |
| 8_7354204 | 6 | 0 | 12 | 0.0833 | 7 | 0.1429 | 11 | 0 | 36 | 0.0556 | 14 | 0 | 14 | 0.1429 | 9 | 0.1111 | 11 | 0 | 48 | 0.0625 |
| 8_7354205 | 6 | 0.5 | 15 | 0.7333 | 8 | 0.5 | 11 | 0.9091 | 40 | 0.7 | 14 | 0.8571 | 16 | 0.5 | 10 | 0.6 | 17 | 0.7647 | 57 | 0.6842 |
| 8_7354206 | 4 | 1 | 13 | 1 | 7 | 1 | 11 | 1 | 35 | 1 | 11 | 1 | 11 | 1 | 6 | 1 | 12 | 0.9167 | 40 | 0.975 |
| 8_7354219 | 7 | 0.2857 | 14 | 0.1429 | 7 | 0 | 12 | 0.5 | 40 | 0.25 | 13 | 0.1538 | 15 | 0.2667 | 9 | 0.1111 | 13 | 0 | 50 | 0.14 |
| 8_7354225 | 7 | 0.8571 | 13 | 0.8462 | 6 | 0.5 | 12 | 0.75 | 38 | 0.7632 | 12 | 0.5 | 13 | 0.6154 | 9 | 0.4444 | 16 | 0.6875 | 50 | 0.58 |
| 8_7354228 | 8 | 0.25 | 12 | 0.1667 | 7 | 0.2857 | 11 | 0.2727 | 38 | 0.2368 | 13 | 0.1538 | 16 | 0.4375 | 10 | 0.2 | 17 | 0.0588 | 56 | 0.2143 |
| 8_7354228 | 6 | 1 | 12 | 0.8333 | 7 | 0.7143 | 7 | 0.8571 | 32 | 0.8438 | 9 | 0.5556 | 12 | 0.75 | 8 | 0.875 | 8 | 0.625 | 37 | 0.7027 |
| 8_7407928 | 2 | 0.5 | 3 | 1 | 1 | 0 | 5 | 0.8 | 11 | 0.7273 | 2 | 1 | 7 | 0.4286 | 2 | 0 | 7 | 0.5714 | 18 | 0.5 |
| 8_7572735 | 2 | 0.5 | NA | NA | NA | NA | NA | NA | 2 | 0.5 | 1 | 0 | 1 | 1 | NA | NA | NA | NA | 2 | 0.5 |
| 8_7572755 | 2 | 1 | NA | NA | NA | NA | NA | NA | 2 | 1 | 1 | 1 | 1 | 1 | NA | NA | NA | NA | 2 | 1 |
| 8_7572795 | 1 | 1 | NA | NA | NA | NA | NA | NA | 1 | 1 | 1 | 0 | 1 | 1 | NA | NA | NA | NA | 2 | 0.5 |
| 8_7579629 | 5 | 0 | 3 | 0.3333 | 4 | 0 | 3 | 0 | 15 | 0.0667 | 5 | 0.2 | 1 | 0 | 1 | 0 | 5 | 0 | 12 | 0.0833 |
| 8_7582667 | 1 | 1 | NA | NA | NA | NA | 1 | 0 | 2 | 0.5 | 1 | 1 | 1 | 0 | NA | NA | 1 | 1 | 3 | 0.6667 |
| 8_7585413 | NA | NA | NA | NA | 1 | 1 | 2 | 0.5 | 3 | 0.6667 | NA | NA | 1 | 1 | 1 | 1 | NA | NA | 2 | 1 |
| 8_7594522 | 28 | 0.3929 | 16 | 0.125 | 23 | 0.1739 | 25 | 0.32 | 92 | 0.2717 | 11 | 0.3636 | 33 | 0.4242 | 26 | 0.2692 | 14 | 0.2143 | 84 | 0.3333 |
| 8_7594533 | 34 | 0.3529 | 18 | 0.4444 | 23 | 0.4348 | 32 | 0.25 | 107 | 0.3551 | 12 | 0.5 | 36 | 0.4444 | 27 | 0.3704 | 9 | 0.2222 | 84 | 0.4048 |
| 8_7594849 | 36 | 0.0556 | 18 | 0.1111 | 23 | 0.087 | 33 | 0.0909 | 110 | 0.0818 | 9 | 0 | 39 | 0.1795 | 36 | 0.0833 | 16 | 0.1875 | 100 | 0.13 |

| | | | | | | | | | | | | | | | | | | | | |
|---|---|---|---|---|---|---|---|---|---|---|---|---|---|---|---|---|---|---|---|---|
| 8_7594856 | 38 | 0.1842 | 19 | 0.4211 | 23 | 0.2609 | 33 | 0.1818 | 113 | 0.2389 | 12 | 0.25 | 42 | 0.2143 | 37 | 0.2162 | 18 | 0.5 | 109 | 0.2661 |
| 8_7594860 | 40 | 0.6 | 17 | 0.4706 | 27 | 0.7778 | 36 | 0.5278 | 120 | 0.6 | 13 | 0.6923 | 50 | 0.62 | 36 | 0.5833 | 17 | 0.6471 | 116 | 0.6207 |
| 8_7594885 | 43 | 0.3953 | 20 | 0.45 | 31 | 0.1935 | 40 | 0.4 | 134 | 0.3582 | 16 | 0.375 | 47 | 0.4894 | 43 | 0.2791 | 19 | 0.4211 | 125 | 0.392 |
| 8_7594888 | 40 | 0.375 | 20 | 0.2 | 27 | 0.2963 | 41 | 0.4634 | 128 | 0.3594 | 15 | 0.3333 | 50 | 0.36 | 40 | 0.5 | 20 | 0.55 | 125 | 0.432 |
| 8_7594900 | 36 | 0.3056 | 17 | 0.2353 | 29 | 0.4828 | 37 | 0.2162 | 119 | 0.3109 | 15 | 0.1333 | 50 | 0.28 | 38 | 0.2632 | 19 | 0.1579 | 122 | 0.2377 |
| 8_7594906 | 47 | 0.1064 | 21 | 0.2381 | 37 | 0.1622 | 48 | 0.2292 | 153 | 0.1765 | 16 | 0.125 | 60 | 0.3 | 49 | 0.1633 | 22 | 0.2273 | 147 | 0.2245 |
| 8_7595016 | 60 | 0.1667 | 28 | 0.0714 | 49 | 0.0816 | 63 | 0.127 | 200 | 0.12 | 20 | 0 | 67 | 0.1045 | 66 | 0.1061 | 35 | 0.0857 | 188 | 0.0904 |
| 8_7595061 | 80 | 0.0625 | 33 | 0.0303 | 51 | 0.0588 | 65 | 0.0923 | 229 | 0.0655 | 23 | 0.2174 | 84 | 0.131 | 74 | 0.0541 | 42 | 0.1429 | 223 | 0.1166 |
| 8_7597146 | 1 | 1 | NA | NA | NA | NA | NA | NA | 1 | 1 | NA | NA | 1 | 1 | NA | NA | NA | NA | 1 | 1 |
| 8_7613211 | 6 | 0.3333 | 5 | 0 | 9 | 0 | 3 | 0 | 23 | 0.087 | 4 | 0 | 1 | 0 | 6 | 0.1667 | 2 | 0 | 13 | 0.0769 |
| 8_7756857 | 1 | 1 | NA | NA | 1 | 1 | 1 | 1 | 3 | 1 | NA | NA | NA | NA | 1 | 1 | NA | NA | 1 | 1 |
| 8_7756877 | NA | NA | NA | NA | 1 | 1 | 1 | 1 | 2 | 1 | NA | NA | NA | NA | 1 | 1 | NA | NA | 1 | 1 |
| 8_7763516 | 2 | 0 | 1 | 1 | NA | NA | NA | NA | 3 | 0.3333 | NA | NA | 1 | 0 | 1 | 1 | NA | NA | 2 | 0.5 |
| 8_7763518 | 2 | 1 | 1 | 1 | 1 | 1 | NA | NA | 4 | 1 | NA | NA | 1 | 1 | 1 | 1 | NA | NA | 2 | 1 |
| 8_7763540 | 1 | 1 | NA | NA | 1 | 0 | NA | NA | 2 | 0.5 | NA | NA | 1 | 1 | 1 | 1 | NA | NA | 2 | 1 |
| 8_7763540 | 2 | 0.5 | NA | NA | 1 | 1 | NA | NA | 3 | 0.6667 | NA | NA | 1 | 1 | 1 | 0 | NA | NA | 2 | 0.5 |
| 8_7763606 | 2 | 0.5 | 1 | 0 | 1 | 0 | NA | NA | 4 | 0.25 | NA | NA | 2 | 0 | 2 | 0.5 | NA | NA | 4 | 0.25 |
| 8_7763629 | 2 | 1 | 1 | 1 | 1 | 0 | NA | NA | 4 | 0.75 | NA | NA | 2 | 1 | 3 | 0.3333 | NA | NA | 5 | 0.6 |
| 8_7763632 | 2 | 0.5 | 1 | 0 | 1 | 1 | NA | NA | 4 | 0.5 | NA | NA | 1 | 0 | 3 | 0.3333 | NA | NA | 4 | 0.25 |
| 8_7763646 | 2 | 1 | NA | NA | 1 | 1 | NA | NA | 3 | 1 | NA | NA | 1 | 1 | 2 | 0.5 | NA | NA | 3 | 0.6667 |
| 8_7763652 | 2 | 0 | 1 | 1 | NA | NA | NA | NA | 3 | 0.3333 | NA | NA | 2 | 0.5 | 2 | 1 | NA | NA | 4 | 0.75 |
| 8_7791253 | 1 | 1 | 1 | 1 | 1 | 0 | NA | NA | 3 | 0.6667 | NA | NA | 2 | 0.5 | 1 | 0 | NA | NA | 3 | 0.3333 |
| 8_7796918 | 2 | 1 | 1 | 1 | 3 | 1 | NA | NA | 6 | 1 | NA | NA | 4 | 0.75 | 4 | 0.75 | 2 | 0.5 | 10 | 0.7 |
| 8_7796922 | 2 | 0.5 | NA | NA | 3 | 0.6667 | NA | NA | 5 | 0.6 | NA | NA | 3 | 1 | 3 | 1 | 2 | 0.5 | 8 | 0.875 |
| 8_7796926 | 2 | 0 | 2 | 1 | 3 | 0 | NA | NA | 7 | 0.2857 | NA | NA | 3 | 0.3333 | 4 | 0.25 | 2 | 0.5 | 9 | 0.3333 |
| 8_7796935 | 2 | 0.5 | 1 | 0 | 2 | 0.5 | NA | NA | 5 | 0.4 | NA | NA | 4 | 0.25 | 4 | 0.5 | 2 | 0.5 | 10 | 0.4 |
| 8_7796936 | 2 | 1 | 2 | 0.5 | 1 | 1 | NA | NA | 5 | 0.8 | NA | NA | 4 | 1 | 4 | 1 | 2 | 1 | 10 | 1 |
| 8_7796943 | 1 | 0 | 2 | 0.5 | 3 | 0.3333 | NA | NA | 6 | 0.3333 | NA | NA | 2 | 1 | 4 | 0.25 | 1 | 1 | 7 | 0.5714 |
| 8_7796952 | 2 | 0.5 | 2 | 1 | 3 | 1 | NA | NA | 7 | 0.8571 | NA | NA | 4 | 1 | 4 | 0.75 | 2 | 1 | 10 | 0.9 |
| 8_7796953 | 1 | 0 | 2 | 1 | 3 | 1 | NA | NA | 6 | 0.8333 | NA | NA | 4 | 0.75 | 4 | 0.25 | 2 | 1 | 10 | 0.6 |
| 8_7796968 | 1 | 1 | 1 | 1 | 2 | 0.5 | NA | NA | 4 | 0.75 | NA | NA | 2 | 0 | 2 | 0 | 2 | 1 | 6 | 0.3333 |
| 8_7797724 | 5 | 0.4 | 5 | 0 | 3 | 0.6667 | 4 | 0 | 17 | 0.2353 | 2 | 0.5 | 2 | 1 | 2 | 0 | 6 | 0.5 | 12 | 0.5 |
| 8_7797733 | 5 | 0.8 | 4 | 0.75 | 3 | 0.3333 | 4 | 0.75 | 16 | 0.6875 | 1 | 1 | 2 | 1 | 3 | 0.6667 | 5 | 0.8 | 11 | 0.8182 |
| 8_7797740 | 5 | 0.6 | 5 | 0.8 | 3 | 1 | 3 | 0 | 16 | 0.625 | 2 | 1 | 2 | 1 | 3 | 0.3333 | 6 | 0.5 | 13 | 0.6154 |
| 8_7800383 | 3 | 0.3333 | 1 | 0 | NA | NA | 3 | 0 | 7 | 0.1429 | 1 | 0 | 2 | 0 | 2 | 0 | 5 | 0.2 | 10 | 0.1 |
| 8_7829877 | 2 | 0.5 | 2 | 1 | 2 | 1 | 1 | 1 | 7 | 0.8571 | 5 | 1 | 2 | 1 | 2 | 0.5 | 2 | 1 | 11 | 0.9091 |
| 8_7829880 | 2 | 0 | 3 | 0.6667 | 2 | 0.5 | NA | NA | 7 | 0.4286 | 3 | 0.6667 | 2 | 1 | 2 | 0.5 | 2 | 0.5 | 9 | 0.6667 |
| 8_7829882 | 2 | 0.5 | 3 | 0.3333 | 2 | 0 | 1 | 0 | 8 | 0.25 | 6 | 0 | 2 | 0 | 2 | 0 | 2 | 0.5 | 12 | 0.0833 |
| 8_7829885 | 2 | 0.5 | 3 | 0.6667 | 2 | 0 | 1 | 0 | 8 | 0.375 | 6 | 0.3333 | 2 | 1 | 2 | 1 | 2 | 0 | 12 | 0.5 |
| 8_7829900 | 2 | 1 | 2 | 1 | 2 | 0.5 | NA | NA | 6 | 0.8333 | 5 | 1 | 2 | 1 | 1 | 0 | 2 | 1 | 10 | 0.9 |
| 8_7829902 | 2 | 0 | 2 | 1 | 1 | 1 | 2 | 0.5 | 7 | 0.5714 | 5 | 0.4 | 1 | 1 | 2 | 1 | 2 | 1 | 10 | 0.7 |
| 8_7829904 | 2 | 0 | 3 | 0 | 2 | 0.5 | 2 | 0 | 9 | 0.1111 | 6 | 0.1667 | 2 | 0 | 2 | 0 | 1 | 0 | 11 | 0.0909 |
| 8_7829910 | 1 | 1 | 1 | 1 | 1 | 1 | 2 | 1 | 5 | 1 | 5 | 1 | 2 | 1 | 2 | 1 | 1 | 0 | 10 | 0.9 |
| 8_7829916 | 2 | 0.5 | 3 | 0.3333 | 2 | 0.5 | 2 | 1 | 9 | 0.5556 | 6 | 0.5 | 2 | 0.5 | 2 | 0.5 | 2 | 0 | 12 | 0.4167 |
| 8_7829920 | 2 | 0.5 | 3 | 0.3333 | 2 | 0 | 2 | 0 | 9 | 0.2222 | 5 | 0 | 2 | 0.5 | 2 | 0 | 2 | 0 | 11 | 0.0909 |
| 8_7830807 | 1 | 1 | NA | NA | NA | NA | NA | NA | 1 | 1 | NA | NA | NA | NA | 2 | 0.5 | 1 | 0 | 3 | 0.3333 |
| 8_7832269 | 6 | 1 | 5 | 0.8 | 12 | 0.9167 | 12 | 0.8333 | 35 | 0.8857 | 3 | 0.6667 | 5 | 1 | 12 | 1 | 6 | 0.6667 | 26 | 0.8846 |
| 8_7832283 | 8 | 0 | 4 | 0.25 | 9 | 0.2222 | 12 | 0.25 | 33 | 0.1818 | 3 | 0.6667 | 7 | 0.2857 | 11 | 0.0909 | 6 | 0.3333 | 27 | 0.2593 |
| 8_7832299 | 10 | 0.8 | 6 | 0.8333 | 9 | 1 | 12 | 0.6667 | 37 | 0.8108 | 3 | 0.6667 | 6 | 1 | 9 | 0.8889 | 6 | 1 | 24 | 0.9167 |
| 8_7832314 | 8 | 0.375 | 6 | 0.1667 | 9 | 0.1111 | 12 | 0.1667 | 35 | 0.2 | 3 | 0 | 6 | 0.1667 | 10 | 0.1 | 6 | 0.5 | 25 | 0.2 |
| 8_7832318 | 8 | 0.375 | 4 | 0.5 | 9 | 0.5556 | 11 | 0.2727 | 32 | 0.4062 | 2 | 0.5 | 4 | 0.25 | 9 | 0.5556 | 6 | 0.5 | 21 | 0.4762 |
| 8_7832322 | 8 | 0.25 | 6 | 0.5 | 10 | 0.1 | 12 | 0.1667 | 36 | 0.2222 | 3 | 0 | 5 | 0.2 | 9 | 0 | 5 | 0.4 | 22 | 0.1364 |
| 8_7832322 | 4 | 0.25 | 3 | 0.3333 | 10 | 0.7 | 10 | 0.8 | 27 | 0.6296 | 2 | 0.5 | 5 | 0.6 | 9 | 0.3333 | 3 | 1 | 19 | 0.5263 |
| 8_7832328 | 7 | 0.7143 | 6 | 0.3333 | 10 | 0.7 | 11 | 0.5455 | 34 | 0.5882 | 2 | 0 | 3 | 0.6667 | 9 | 0.6667 | 5 | 0.4 | 19 | 0.5263 |
| 8_7832336 | 10 | 0.1 | 3 | 0.6667 | 9 | 0.1111 | 13 | 0.1538 | 35 | 0.1714 | 2 | 0 | 5 | 0.4 | 10 | 0.1 | 5 | 0.2 | 22 | 0.1818 |
| 8_7832336 | 9 | 0.4444 | 2 | 0 | 8 | 0.5 | 12 | 0.1667 | 31 | 0.3226 | 3 | 0 | 6 | 0.3333 | 9 | 0.4444 | 4 | 0.25 | 22 | 0.3182 |
| 8_7832339 | 4 | 0 | 3 | 0.3333 | 6 | 0.1667 | 10 | 0 | 23 | 0.087 | 2 | 0 | 5 | 0.2 | 6 | 0 | 4 | 0 | 17 | 0.0588 |
| 8_7832340 | 8 | 0.125 | 4 | 0 | 8 | 0.125 | 8 | 0.375 | 28 | 0.1786 | 2 | 0 | 4 | 0 | 8 | 0.25 | 5 | 0.6 | 19 | 0.2632 |
| 8_7832346 | 9 | 0 | 2 | 0 | 9 | 0.1111 | 13 | 0.0769 | 33 | 0.0606 | 2 | 0 | 6 | 0 | 10 | 0.1 | 5 | 0.2 | 23 | 0.087 |
| 8_7832348 | 10 | 0.3 | 4 | 0.25 | 8 | 0.125 | 13 | 0.1538 | 35 | 0.2 | 3 | 0 | 4 | 0.5 | 8 | 0 | 4 | 0.5 | 19 | 0.2105 |
| 8_7832352 | 8 | 0.25 | 4 | 0 | 7 | 0.4286 | 11 | 0.0909 | 30 | 0.2 | 3 | 0.3333 | 5 | 0 | 9 | 0.2222 | 5 | 0 | 22 | 0.1364 |
| 8_7832365 | 8 | 0.75 | 4 | 1 | 9 | 0.6667 | 9 | 0.6667 | 30 | 0.7333 | 3 | 0.6667 | 5 | 0.8 | 7 | 0.8571 | 4 | 1 | 19 | 0.8421 |
| 8_7833712 | 5 | 0.2 | 3 | 0.3333 | 7 | 0 | 6 | 0 | 21 | 0.0952 | 2 | 0 | 4 | 0.25 | 5 | 0 | 3 | 0.3333 | 14 | 0.1429 |
| 8_7923195 | 13 | 0.0769 | 12 | 0.1667 | 10 | 0 | 11 | 0 | 46 | 0.0652 | 13 | 0.1538 | 7 | 0 | 10 | 0.1 | 16 | 0.125 | 46 | 0.1087 |
| 8_7923209 | 14 | 1 | 12 | 1 | 10 | 1 | 10 | 0.9 | 46 | 0.9783 | 13 | 0.9231 | 8 | 1 | 11 | 1 | 16 | 0.9375 | 48 | 0.9583 |

| | | | | | | | | | | | | | | | | | | | | |
|---|---|---|---|---|---|---|---|---|---|---|---|---|---|---|---|---|---|---|---|---|
| 8_79232163 | 14 | 0.9286 | 11 | 1 | 8 | 0.875 | 10 | 1 | 43 | 0.9535 | 12 | 1 | 8 | 1 | 10 | 0.9 | 14 | 1 | 44 | 0.9773 |
| 8_79232377 | 12 | 0.0833 | 12 | 0.0833 | 9 | 0 | 10 | 0.2 | 43 | 0.093 | 13 | 0.1538 | 7 | 0.1429 | 10 | 0.1 | 14 | 0.0714 | 44 | 0.1136 |
| 8_79232482 | 14 | 0.0714 | 11 | 0 | 7 | 0.1429 | 9 | 0.1111 | 41 | 0.0732 | 13 | 0.0769 | 6 | 0.1667 | 8 | 0 | 16 | 0.1875 | 43 | 0.1163 |
| 8_79232539 | 14 | 0.2857 | 9 | 0.4444 | 9 | 0.2222 | 8 | 0.25 | 40 | 0.3 | 13 | 0.2308 | 5 | 0.4 | 9 | 0.2222 | 12 | 0.3333 | 39 | 0.2821 |
| 8_79232916 | 10 | 0.4 | 10 | 0.1 | 7 | 0.1429 | 6 | 0.1667 | 33 | 0.2121 | 10 | 0.4 | 3 | 0.6667 | 8 | 0.375 | 11 | 0.2727 | 32 | 0.375 |
| 8_79232941 | 6 | 1 | 6 | 0.8333 | 6 | 1 | 5 | 1 | 23 | 0.9565 | 5 | 1 | 3 | 1 | 4 | 1 | 8 | 1 | 20 | 1 |
| 8_79233217 | 12 | 0.25 | 8 | 0 | 7 | 0.1429 | 9 | 0.1111 | 36 | 0.1389 | 9 | 0.2222 | 4 | 0.25 | 9 | 0.1111 | 12 | 0 | 34 | 0.1176 |
| 8_79234896 | 1 | 1 | NA | NA | NA | NA | NA | NA | 1 | 1 | NA | NA | NA | NA | NA | NA | 1 | 1 | 1 | 1 |
| 8_79235081 | 1 | 1 | NA | NA | NA | NA | NA | NA | 1 | 1 | NA | NA | NA | NA | NA | NA | 1 | 1 | 1 | 1 |
| 8_79459415 | NA | NA | 1 | 1 | NA | NA | NA | NA | 1 | 1 | NA | NA | 2 | 0.5 | NA | NA | NA | NA | 2 | 0.5 |
| 8_79459491 | NA | NA | NA | NA | 1 | 1 | NA | NA | 1 | 1 | NA | NA | 2 | 0.5 | NA | NA | NA | NA | 2 | 0.5 |
| 8_79459759 | NA | NA | 1 | 0 | 1 | 1 | NA | NA | 2 | 0.5 | NA | NA | 2 | 0.5 | NA | NA | NA | NA | 2 | 0.5 |
| 8_79533330 | 2 | 1 | 2 | 0.5 | 5 | 0.6 | 3 | 0.6667 | 12 | 0.6667 | 2 | 0.5 | 1 | 1 | 3 | 0.6667 | 1 | 1 | 7 | 0.7143 |
| 8_79534401 | 1 | 0 | 2 | 0 | 2 | 0.5 | 3 | 0 | 8 | 0.125 | 1 | 0 | 2 | 0 | 3 | 0 | 2 | 0.5 | 8 | 0.125 |
| 8_79916580 | 3 | 1 | NA | NA | NA | NA | NA | NA | 3 | 1 | 2 | 1 | 2 | 1 | 1 | 1 | 1 | 1 | 6 | 1 |
| 8_79916721 | 2 | 1 | NA | NA | NA | NA | NA | NA | 2 | 1 | 2 | 1 | 1 | 1 | 1 | 1 | 1 | 1 | 5 | 1 |
| 8_79916937 | 3 | 0.3333 | NA | NA | NA | NA | NA | NA | 3 | 0.3333 | 2 | 0 | 1 | 0 | 1 | 1 | 3 | 0 | 7 | 0.1429 |
| 8_79916949 | 3 | 1 | NA | NA | NA | NA | NA | NA | 3 | 1 | 2 | 1 | 2 | 1 | 1 | 1 | 2 | 1 | 7 | 1 |
| 8_79917035 | 3 | 1 | NA | NA | NA | NA | NA | NA | 3 | 1 | 2 | 1 | 2 | 1 | 1 | 1 | 2 | 1 | 7 | 1 |
| 8_79917088 | 3 | 1 | NA | NA | NA | NA | NA | NA | 3 | 1 | 2 | 1 | 1 | 1 | 1 | 1 | 2 | 1 | 6 | 1 |
| 8_79917107 | 3 | 1 | NA | NA | NA | NA | NA | NA | 3 | 1 | 1 | 1 | 2 | 0.5 | 1 | 1 | 3 | 1 | 7 | 0.8571 |
| 8_79917118 | 3 | 0.3333 | NA | NA | NA | NA | NA | NA | 3 | 0.3333 | 2 | 0.5 | 2 | 0.5 | 1 | 1 | 2 | 0.5 | 7 | 0.5714 |
| 8_79917148 | 3 | 0.6667 | NA | NA | NA | NA | NA | NA | 3 | 0.6667 | 2 | 1 | 2 | 1 | 1 | 1 | 3 | 0.3333 | 8 | 0.75 |
| 8_79917154 | 3 | 0.3333 | NA | NA | NA | NA | NA | NA | 3 | 0.3333 | 2 | 0.5 | 2 | 1 | 1 | 0 | 4 | 1 | 9 | 0.7778 |
| 8_79917179 | 3 | 0.6667 | NA | NA | NA | NA | NA | NA | 3 | 0.6667 | 2 | 0.5 | 2 | 1 | 1 | 0 | 4 | 0.25 | 9 | 0.4444 |
| 8_79917241 | 3 | 0.3333 | NA | NA | NA | NA | NA | NA | 3 | 0.3333 | 2 | 0 | 2 | 0.5 | 1 | 1 | 4 | 0.25 | 9 | 0.3333 |
| 8_79918855 | 4 | 0.5 | NA | NA | NA | NA | 1 | 0 | 5 | 0.4 | 2 | 0 | 4 | 0.25 | 3 | 0.6667 | 8 | 0.25 | 17 | 0.2941 |
| 8_79919019 | 5 | 0.6 | NA | NA | NA | NA | 1 | 1 | 6 | 0.6667 | 2 | 1 | 4 | 0.75 | 3 | 0.6667 | 9 | 0.7778 | 18 | 0.7778 |
| 8_79919237 | 7 | 0.1429 | NA | NA | NA | NA | 1 | 0 | 8 | 0.125 | 2 | 0 | 4 | 0.25 | 3 | 0 | 10 | 0.4 | 19 | 0.2632 |
| 8_79919300 | 6 | 0.1667 | NA | NA | 1 | 0 | 1 | 0 | 8 | 0.125 | 2 | 0.5 | 3 | 0 | 2 | 0 | 10 | 0 | 17 | 0.0588 |
| 8_79919309 | 5 | 0.2 | NA | NA | 1 | 0 | 1 | 0 | 7 | 0.1429 | 2 | 0.5 | 3 | 0 | 3 | 0 | 10 | 0.2 | 18 | 0.1667 |
| 8_79919319 | 4 | 0.75 | NA | NA | NA | NA | 1 | 1 | 5 | 0.8 | 2 | 0.5 | 4 | 0.25 | 2 | 0.5 | 3 | 0.6667 | 11 | 0.4545 |
| 8_79994563 | 2 | 0.5 | 3 | 0.3333 | 5 | 0.2 | 4 | 0 | 14 | 0.2143 | 5 | 0 | 7 | 0.1429 | 3 | 0 | 4 | 0 | 19 | 0.0526 |
| 8_79994591 | 4 | 0.25 | 4 | 0 | 5 | 0 | 4 | 0 | 17 | 0.0588 | 5 | 0 | 7 | 0.1429 | 3 | 0 | 4 | 0 | 19 | 0.0526 |
| 8_79994737 | 4 | 0 | 4 | 0 | 5 | 0.2 | 4 | 0.25 | 17 | 0.1176 | 4 | 0.25 | 7 | 0.2857 | 3 | 0 | 4 | 0 | 18 | 0.1667 |
| 8_79994795 | 4 | 0 | 4 | 0.5 | 4 | 0 | 4 | 0 | 16 | 0.125 | 5 | 0 | 7 | 0.1429 | 3 | 0.3333 | 4 | 0 | 19 | 0.1053 |
| 8_80007957 | 3 | 0.3333 | NA | NA | NA | NA | 2 | 0.5 | 5 | 0.4 | 3 | 0.6667 | 7 | 0.4286 | 3 | 0.6667 | 4 | 0.5 | 17 | 0.5294 |
| 8_80007981 | 2 | 1 | NA | NA | 2 | 0.5 | 3 | 1 | 7 | 0.8571 | 4 | 0.5 | 7 | 1 | 4 | 1 | 5 | 0.2 | 20 | 0.7 |
| 8_80008014 | 2 | 1 | NA | NA | 1 | 1 | 3 | 1 | 6 | 1 | 4 | 0.5 | 5 | 0.8 | 3 | 1 | 5 | 0.8 | 17 | 0.7647 |
| 8_80008027 | 3 | 0.3333 | NA | NA | 2 | 0.5 | 3 | 0.3333 | 8 | 0.375 | 4 | 0.25 | 7 | 0.1429 | 1 | 1 | 5 | 0 | 17 | 0.1765 |
| 8_80008055 | 2 | 0.5 | NA | NA | 1 | 0 | 3 | 0.6667 | 6 | 0.5 | 4 | 0.75 | 7 | 0.1429 | 3 | 0.3333 | 3 | 0 | 17 | 0.2941 |
| 8_80008101 | 2 | 0.5 | NA | NA | 2 | 0.5 | 3 | 1 | 7 | 0.7143 | 4 | 0 | 5 | 0.4 | 2 | 0.5 | 2 | 0.5 | 13 | 0.3077 |
| 8_80008111 | 3 | 0.3333 | NA | NA | 1 | 1 | 3 | 0.3333 | 7 | 0.4286 | 4 | 0.25 | 7 | 0.2857 | 3 | 0.3333 | 4 | 0.75 | 18 | 0.3889 |
| 8_80011010 | 1 | 1 | NA | NA | NA | NA | 2 | 0.5 | 3 | 0.6667 | 3 | 0 | 2 | 0 | 3 | 0.6667 | 2 | 1 | 10 | 0.4 |
| 8_80066004 | 2 | 0 | 5 | 0 | 4 | 0.75 | 3 | 0.3333 | 14 | 0.2857 | 3 | 0.3333 | 2 | 1 | 1 | 0 | 2 | 0 | 8 | 0.375 |
| 8_80066048 | 2 | 1 | 5 | 0.6 | 4 | 1 | 3 | 1 | 14 | 0.8571 | 3 | 1 | 1 | 1 | 1 | 1 | 2 | 1 | 7 | 1 |
| 8_80066060 | 2 | 0 | 5 | 0.2 | 4 | 0 | 3 | 0 | 14 | 0.0714 | 3 | 0.3333 | 2 | 0 | 1 | 0 | 1 | 0 | 7 | 0.1429 |
| 8_80066092 | 2 | 0.5 | 5 | 0.4 | 4 | 0 | 3 | 1 | 14 | 0.4286 | 1 | 0 | 2 | 0.5 | 1 | 0 | 1 | 1 | 5 | 0.4 |
| 8_80066113 | 2 | 0 | 5 | 0.2 | 4 | 0.25 | 3 | 0.3333 | 14 | 0.2143 | 3 | 0.6667 | 1 | 1 | 1 | 1 | 2 | 0.5 | 7 | 0.7143 |
| 8_80066140 | 2 | 0.5 | 5 | 1 | 4 | 0.5 | 3 | 0.6667 | 14 | 0.7143 | 3 | 0.3333 | 1 | 1 | NA | NA | 1 | 1 | 5 | 0.6 |
| 8_80066169 | 2 | 0 | 5 | 0 | 2 | 0.5 | 3 | 0 | 12 | 0.0833 | 2 | 0.5 | 2 | 0 | 1 | 0 | NA | NA | 5 | 0.2 |
| 8_80066218 | 2 | 0 | 4 | 0.75 | 2 | 1 | 3 | 1 | 11 | 0.7273 | 2 | 1 | 2 | 1 | 1 | 1 | 2 | 1 | 7 | 1 |
| 8_80066250 | 2 | 0 | 5 | 0.2 | 3 | 0.6667 | 3 | 0 | 13 | 0.2308 | 3 | 0.3333 | 2 | 0.5 | 1 | 0 | NA | NA | 6 | 0.3333 |
| 8_80066274 | 2 | 0.5 | 5 | 0.6 | 4 | 0.5 | 3 | 1 | 14 | 0.6429 | 3 | 0.3333 | 2 | 0.5 | 1 | 0 | 2 | 0.5 | 8 | 0.375 |
| 8_80098507 | NA | NA | 1 | 1 | 1 | 0 | 2 | 1 | 4 | 0.75 | 1 | 0 | NA | NA | NA | NA | 1 | 1 | 2 | 0.5 |
| 8_80098636 | NA | NA | 1 | 0 | 3 | 0.3333 | 1 | 1 | 5 | 0.4 | 1 | 1 | NA | NA | NA | NA | 1 | 0 | 2 | 0.5 |
| 8_80098765 | NA | NA | 1 | 1 | 3 | 0.3333 | 3 | 0.6667 | 7 | 0.5714 | 2 | 1 | NA | NA | NA | NA | NA | NA | 2 | 1 |
| 8_80098827 | NA | NA | 1 | 0 | 3 | 0.3333 | 1 | 1 | 5 | 0.4 | 1 | 1 | NA | NA | NA | NA | NA | NA | 1 | 1 |
| 8_80365894 | NA | NA | NA | NA | NA | NA | 1 | 1 | 1 | 1 | NA | NA | 2 | 1 | 1 | 1 | NA | NA | 3 | 1 |
| 8_80399551 | 2 | 1 | 5 | 1 | 1 | 1 | 2 | 0.5 | 10 | 0.9 | 1 | 1 | 1 | 1 | 3 | 1 | 2 | 1 | 7 | 1 |
| 8_80399630 | 2 | 1 | 5 | 0.2 | NA | NA | 2 | 1 | 9 | 0.5556 | 1 | 1 | 1 | 0 | 3 | 0.3333 | 2 | 0.5 | 7 | 0.4286 |
| 8_80399690 | 2 | 1 | 5 | 1 | 2 | 1 | 2 | 1 | 11 | 1 | 1 | 1 | 1 | 1 | 3 | 1 | 2 | 0.5 | 7 | 0.8571 |
| 8_80399711 | 2 | 0.5 | 6 | 1 | NA | NA | NA | NA | 8 | 0.875 | 1 | 1 | 1 | 1 | 3 | 0.3333 | 1 | 0 | 6 | 0.5 |
| 8_80399776 | 2 | 0.5 | 6 | 0.6667 | 2 | 0.5 | 2 | 0.5 | 12 | 0.5833 | 1 | 0 | 1 | 0 | 2 | 0.5 | 2 | 1 | 6 | 0.5 |
| 8_80400115 | 2 | 1 | 6 | 0.8333 | 2 | 1 | 3 | 0.6667 | 13 | 0.8462 | 1 | 0 | 1 | 1 | 3 | 1 | 1 | 1 | 6 | 0.8333 |

| | | | | | | | | | | | | | | | | | | | | |
|---|---|---|---|---|---|---|---|---|---|---|---|---|---|---|---|---|---|---|---|---|
| 8_8040021 | 2 | 1 | 6 | 1 | 3 | 0.6667 | 3 | 1 | 14 | 0.9286 | 2 | 0.5 | 1 | 1 | 3 | 1 | 2 | 0.5 | 8 | 0.75 |
| 8_8040026 | 2 | 1 | 4 | 1 | 2 | 0.5 | 2 | 0.5 | 10 | 0.8 | 1 | 1 | 1 | 1 | 3 | 1 | 2 | 1 | 7 | 1 |
| 8_8040032 | 2 | 0.5 | 6 | 0.3333 | 3 | 0.3333 | 3 | 0 | 14 | 0.2857 | 2 | 1 | 1 | 0 | 3 | 0.3333 | 2 | 0 | 8 | 0.375 |
| 8_8040040 | 2 | 0.5 | 5 | 0.4 | 2 | 0.5 | 2 | 0 | 11 | 0.3636 | 2 | 0 | 1 | 0 | 3 | 0.3333 | 2 | 0 | 8 | 0.125 |
| 8_8040059 | 2 | 0 | 5 | 0.6 | 3 | 0.3333 | 3 | 0.3333 | 13 | 0.3846 | 2 | 0.5 | 1 | 0 | 3 | 0 | 2 | 0 | 8 | 0.125 |
| 8_8040070 | 1 | 1 | 5 | 0.4 | 3 | 1 | 2 | 1 | 11 | 0.7273 | 2 | 1 | 1 | 1 | 3 | 0.6667 | 2 | 1 | 8 | 0.875 |
| 8_8040075 | 2 | 0.5 | 5 | 0.6 | 1 | 1 | 1 | 1 | 9 | 0.6667 | 2 | 0.5 | 1 | 1 | 3 | 0.6667 | 1 | 1 | 7 | 0.7143 |
| 8_8040135 | 3 | 0 | 5 | 0.4 | 4 | 0 | 4 | 0 | 16 | 0.125 | 2 | 0.5 | 2 | 0 | 1 | 0 | 2 | 0 | 7 | 0.1429 |
| 8_8040181 | 3 | 0 | 6 | 0.5 | 4 | 0 | 4 | 0 | 17 | 0.1765 | 1 | 0 | 1 | 0 | 1 | 1 | 1 | 0 | 4 | 0.25 |
| 8_8041054 | 8 | 0.375 | 12 | 0.25 | 8 | 0.375 | 10 | 0.2 | 38 | 0.2895 | 5 | 0 | 14 | 0.1429 | 9 | 0.1111 | 16 | 0.3125 | 44 | 0.1818 |
| 8_8041184 | 7 | 0 | 10 | 0.1 | 4 | 0.25 | 9 | 0.1111 | 30 | 0.1 | 4 | 0.5 | 10 | 0.1 | 5 | 0 | 14 | 0.0714 | 33 | 0.1212 |
| 8_8050179 | 4 | 0.5 | 4 | 0.75 | 1 | 0 | NA | NA | 9 | 0.5556 | 1 | 0 | 1 | 0 | 1 | 1 | 3 | 0.3333 | 6 | 0.3333 |
| 8_8050180 | 3 | 0.6667 | 4 | 1 | NA | NA | NA | NA | 7 | 0.8571 | NA | NA | 2 | 1 | NA | NA | 3 | 0.6667 | 5 | 0.8 |
| 8_8146572 | NA | NA | 1 | 0 | 1 | 0 | 5 | 0.2 | 7 | 0.1429 | NA | NA | 1 | 0 | 6 | 0.1667 | NA | NA | 7 | 0.1429 |
| 8_8149140 | 2 | 0 | 3 | 0.3333 | NA | NA | 2 | 0 | 7 | 0.1429 | 4 | 0.25 | 2 | 0 | 2 | 0 | 4 | 0.5 | 12 | 0.25 |
| 8_8149221 | 1 | 0 | 3 | 1 | NA | NA | 2 | 1 | 6 | 0.8333 | 4 | 0.75 | 2 | 0.5 | 2 | 0.5 | 3 | 0 | 11 | 0.4545 |
| 8_8149226 | 1 | 1 | 1 | 0 | NA | NA | 2 | 0 | 4 | 0.25 | 4 | 0.25 | 2 | 0.5 | 2 | 0.5 | 3 | 0.3333 | 11 | 0.3636 |
| 8_8149235 | NA | NA | 3 | 0 | NA | NA | 2 | 0.5 | 5 | 0.2 | 4 | 0 | 1 | 0 | 2 | 0 | 4 | 0.25 | 11 | 0.0909 |
| 8_8149238 | 1 | 0 | 2 | 0.5 | NA | NA | 2 | 0 | 5 | 0.2 | 4 | 0.25 | 1 | 0 | 1 | 0 | 4 | 0 | 10 | 0.1 |
| 8_8149245 | 1 | 0 | 2 | 0.5 | NA | NA | NA | NA | 3 | 0.3333 | 3 | 0.3333 | 1 | 0 | 1 | 0 | 4 | 0 | 9 | 0.1111 |
| 8_8151139 | NA | NA | 2 | 1 | NA | NA | 1 | 1 | 3 | 1 | 2 | 1 | NA | NA | NA | NA | 4 | 0.5 | 6 | 0.6667 |
| 8_8151140 | NA | NA | 2 | 0 | NA | NA | 1 | 1 | 3 | 0.3333 | 3 | 0.6667 | NA | NA | NA | NA | 4 | 0 | 7 | 0.2857 |
| 8_8151521 | NA | NA | 2 | 0.5 | NA | NA | 1 | 0 | 3 | 0.3333 | 2 | 0.5 | NA | NA | NA | NA | 4 | 0.5 | 6 | 0.5 |
| 8_8151533 | NA | NA | 1 | 1 | NA | NA | NA | NA | 1 | 1 | 2 | 1 | NA | NA | NA | NA | 3 | 0.3333 | 5 | 0.6 |
| 8_8151544 | NA | NA | 1 | 1 | NA | NA | 1 | 1 | 2 | 1 | 3 | 1 | NA | NA | NA | NA | 3 | 1 | 6 | 1 |
| 8_8170779 | 5 | 0.4 | 3 | 0.3333 | 2 | 0 | 7 | 0.1429 | 17 | 0.2353 | 3 | 0.3333 | 3 | 0.3333 | 2 | 0.5 | 2 | 0.5 | 10 | 0.4 |
| 8_8170782 | 4 | 0.25 | 3 | 0 | 2 | 0.5 | 6 | 0 | 15 | 0.1333 | 2 | 0 | 3 | 0.3333 | 2 | 0 | 1 | 0 | 8 | 0.125 |
| 8_8170789 | 5 | 0 | 3 | 0 | 2 | 0 | 6 | 0.1667 | 16 | 0.0625 | 2 | 0 | 3 | 0.3333 | 2 | 0 | 2 | 0 | 9 | 0.1111 |
| 8_8170843 | 4 | 0.5 | 2 | 0.5 | 4 | 0.5 | 6 | 0.6667 | 16 | 0.5625 | 3 | 0.3333 | 3 | 0.6667 | 1 | 0 | 2 | 1 | 9 | 0.5556 |
| 8_8170846 | 4 | 0.25 | 3 | 0.3333 | 6 | 0.1667 | 6 | 0 | 19 | 0.1579 | 3 | 0.3333 | 4 | 0.25 | 1 | 0 | 3 | 0 | 11 | 0.1818 |
| 8_8171116 | 4 | 0 | 3 | 0 | 6 | 0 | 6 | 0.5 | 19 | 0.1579 | 3 | 0 | 4 | 0 | 1 | 0 | 3 | 0.3333 | 11 | 0.0909 |
| 8_8171116 | 3 | 1 | 2 | 0 | 4 | 0.25 | 6 | 0.1667 | 15 | 0.3333 | 3 | 0.3333 | 3 | 0 | 1 | 0 | 3 | 0 | 10 | 0.1 |
| 8_8171121 | 4 | 0.75 | 3 | 0 | 5 | 0.2 | 6 | 0.1667 | 18 | 0.2778 | 3 | 0.3333 | 4 | 0.25 | 1 | 0 | 3 | 0.3333 | 11 | 0.2727 |
| 8_8171127 | 4 | 0.25 | 3 | 0.3333 | 6 | 0.1667 | 6 | 0.1667 | 19 | 0.2105 | 3 | 0.3333 | 3 | 0.3333 | 1 | 1 | 2 | 0 | 9 | 0.3333 |
| 8_8171128 | 4 | 0.25 | 2 | 0 | 5 | 0 | 6 | 0 | 17 | 0.0588 | 3 | 0.3333 | 4 | 0.25 | 1 | 0 | 3 | 0 | 11 | 0.1818 |
| 8_8171132 | 4 | 1 | 3 | 1 | 5 | 0.8 | 5 | 1 | 17 | 0.9412 | 3 | 1 | 2 | 1 | 1 | 1 | 3 | 1 | 9 | 1 |
| 8_8171133 | 3 | 0 | 2 | 0 | 4 | 0 | 5 | 0.4 | 14 | 0.1429 | 1 | 1 | 4 | 0.25 | 1 | 0 | 2 | 0.5 | 8 | 0.375 |
| 8_8171154 | 4 | 1 | 3 | 1 | 5 | 1 | 6 | 1 | 18 | 1 | 3 | 1 | 4 | 1 | 1 | 1 | 3 | 1 | 11 | 1 |
| 8_8171157 | 4 | 0.5 | 2 | 0 | 5 | 0.2 | 6 | 0.6667 | 17 | 0.4118 | 2 | 0.5 | 4 | 0.25 | 1 | 0 | 3 | 0.3333 | 10 | 0.3 |
| 8_8171159 | 4 | 0 | 3 | 0.3333 | 5 | 0.4 | 6 | 0.1667 | 18 | 0.2222 | 2 | 0.5 | 3 | 0.3333 | 1 | 0 | 3 | 1 | 9 | 0.5556 |
| 8_8171164 | 4 | 1 | 2 | 1 | 3 | 1 | 5 | 1 | 14 | 1 | 3 | 1 | 3 | 1 | 1 | 1 | 2 | 1 | 9 | 1 |
| 8_8171168 | 4 | 0.25 | 2 | 0 | 3 | 0 | 6 | 0 | 15 | 0.0667 | 3 | 0 | 3 | 0.6667 | 1 | 0 | 3 | 0.3333 | 10 | 0.3 |
| 8_8171178 | 3 | 0.3333 | 3 | 0 | 4 | 0 | 5 | 0.4 | 15 | 0.2 | 3 | 0 | 3 | 0.3333 | 1 | 0 | 3 | 0.3333 | 10 | 0.2 |
| 8_8171183 | 3 | 0.3333 | 3 | 0 | 5 | 0 | 5 | 0 | 16 | 0.0625 | 3 | 0.3333 | 3 | 0 | 1 | 0 | 2 | 0 | 9 | 0.1111 |
| 8_8171186 | 3 | 0 | 3 | 0 | 4 | 0 | 4 | 0.25 | 14 | 0.0714 | 3 | 0 | 3 | 0.3333 | 1 | 0 | 2 | 0 | 9 | 0.1111 |
| 8_8171187 | 4 | 0.5 | 3 | 0 | 4 | 0 | 4 | 0 | 15 | 0.1333 | 3 | 0.3333 | 2 | 0 | 1 | 0 | 2 | 0 | 8 | 0.125 |
| 8_8171187 | 4 | 1 | 2 | 1 | 4 | 1 | 5 | 1 | 15 | 1 | 3 | 1 | 3 | 1 | 1 | 1 | 3 | 1 | 10 | 1 |
| 8_8171224 | 3 | 1 | 3 | 1 | 5 | 1 | 5 | 1 | 16 | 1 | 2 | 1 | 1 | 1 | 1 | 1 | 3 | 1 | 7 | 1 |
| 8_8171228 | 3 | 0.3333 | 2 | 0 | 5 | 0.2 | 5 | 0 | 15 | 0.1333 | 3 | 0 | 2 | 0.5 | 1 | 0 | 3 | 0.6667 | 9 | 0.3333 |
| 8_8171236 | 3 | 0.6667 | 2 | 1 | 5 | 0.8 | 4 | 1 | 14 | 0.8571 | 3 | 0.6667 | 2 | 1 | 1 | 0 | 3 | 0.3333 | 9 | 0.5556 |
| 8_8171989 | 2 | 0 | 2 | 0 | 3 | 0 | 3 | 0.3333 | 10 | 0.1 | 1 | 0 | 2 | 0.5 | 1 | 0 | 3 | 0 | 7 | 0.1429 |
| 8_8174050 | 1 | 0 | NA | NA | 1 | 0 | 3 | 0.3333 | 5 | 0.2 | NA | NA | 1 | 0 | NA | NA | 1 | 1 | 2 | 0.5 |
| 8_8174058 | 1 | 0 | 1 | 1 | 2 | 0.5 | 4 | 0 | 8 | 0.25 | 2 | 0.5 | 1 | 0 | 1 | 0 | 3 | 0 | 7 | 0.1429 |
| 8_8174075 | NA | NA | NA | NA | NA | NA | 2 | 0.5 | 2 | 0.5 | 2 | 0.5 | 2 | 0.5 | NA | NA | 2 | 1 | 6 | 0.6667 |
| 8_8174077 | 1 | 0 | NA | NA | 2 | 0.5 | 4 | 0 | 7 | 0.1429 | 2 | 0 | 2 | 0 | 1 | 1 | 3 | 0 | 8 | 0.125 |
| 8_8174081 | 1 | 0 | NA | NA | 2 | 0.5 | 4 | 0 | 7 | 0.1429 | 2 | 0.5 | 2 | 0 | 1 | 0 | 2 | 0 | 7 | 0.1429 |
| 8_8174140 | 1 | 0 | NA | NA | NA | NA | 2 | 0.5 | 3 | 0.3333 | 1 | 0 | 2 | 0 | 1 | 0 | 3 | 0.3333 | 7 | 0.1429 |
| 8_8284806 | 2 | 0.5 | NA | NA | 1 | 1 | 2 | 1 | 5 | 0.8 | NA | NA | 2 | 1 | NA | NA | 1 | 0 | 3 | 0.6667 |
| 8_8284914 | 2 | 0 | 1 | 0 | 2 | 0.5 | 2 | 0 | 7 | 0.1429 | NA | NA | 1 | 1 | NA | NA | 1 | 0 | 2 | 0.5 |
| 8_8349090 | 1 | 0 | 6 | 0.1667 | 5 | 0.6 | 6 | 0.5 | 18 | 0.3889 | 4 | 0.5 | 2 | 0.5 | 1 | 0 | 4 | 0.75 | 11 | 0.5455 |
| 8_8349091 | NA | NA | 6 | 0 | 4 | 0.25 | 4 | 0 | 14 | 0.0714 | 4 | 0.25 | 2 | 0 | 1 | 0 | 3 | 0 | 10 | 0.1 |
| 8_8349095 | 1 | 1 | 6 | 0.1667 | 5 | 0.4 | 6 | 0.1667 | 18 | 0.2778 | 4 | 0 | 2 | 0.5 | 1 | 1 | 4 | 0.25 | 11 | 0.2727 |
| 8_8381009 | NA | NA | 4 | 0.5 | NA | NA | 1 | 0 | 5 | 0.4 | 2 | 0.5 | NA | NA | 1 | 0 | NA | NA | 3 | 0.3333 |
| 8_8381014 | NA | NA | 4 | 0.75 | NA | NA | 1 | 0 | 5 | 0.6 | 2 | 0.5 | NA | NA | NA | NA | NA | NA | 2 | 0.5 |
| 8_8381016 | NA | NA | 3 | 0 | NA | NA | 1 | 1 | 4 | 0.25 | 2 | 1 | NA | NA | NA | NA | NA | NA | 2 | 1 |

| | | | | | | | | | | | | | | | | | | | | |
|---|---|---|---|---|---|---|---|---|---|---|---|---|---|---|---|---|---|---|---|---|
| 8_83810198 | NA | NA | 3 | 1 | NA | NA | 1 | 1 | 4 | 1 | 2 | 1 | NA | NA | NA | NA | NA | NA | 2 | 1 |
| 8_83810249 | NA | NA | 4 | 1 | NA | NA | 1 | 1 | 5 | 1 | 2 | 1 | NA | NA | NA | NA | NA | NA | 2 | 1 |
| 8_83810439 | NA | NA | 2 | 0.5 | NA | NA | 1 | 1 | 3 | 0.6667 | 1 | 1 | NA | NA | 1 | 0 | NA | NA | 2 | 0.5 |
| 8_83810451 | NA | NA | 3 | 0.3333 | NA | NA | NA | NA | 3 | 0.3333 | 1 | 1 | NA | NA | 1 | 0 | NA | NA | 2 | 0.5 |
| 8_83810472 | NA | NA | 4 | 0.25 | NA | NA | 1 | 0 | 5 | 0.2 | 2 | 0.5 | NA | NA | NA | NA | NA | NA | 2 | 0.5 |
| 8_83810708 | NA | NA | 4 | 0.5 | NA | NA | 2 | 1 | 6 | 0.6667 | 2 | 0.5 | NA | NA | NA | NA | NA | NA | 2 | 0.5 |
| 8_83810919 | NA | NA | 4 | 0.5 | NA | NA | 4 | 0.75 | 8 | 0.625 | 2 | 1 | NA | NA | NA | NA | NA | NA | 2 | 1 |
| 8_83810987 | NA | NA | 4 | 0.75 | 1 | 1 | 4 | 0.75 | 9 | 0.7778 | 2 | 0.5 | NA | NA | NA | NA | NA | NA | 2 | 0.5 |
| 8_83815732 | 4 | 0.5 | 9 | 0.5556 | 7 | 0.1429 | 4 | 0.75 | 24 | 0.4583 | 2 | 0.5 | 3 | 0.6667 | 6 | 0.5 | 3 | 0 | 14 | 0.4286 |
| 8_83815752 | 4 | 0 | 9 | 0.4444 | 7 | 0 | 3 | 1 | 23 | 0.3043 | 2 | 0.5 | 2 | 0.5 | 4 | 0.25 | 3 | 0 | 11 | 0.2727 |
| 8_83815888 | 4 | 0.25 | 8 | 0.625 | 6 | 0.3333 | 2 | 0.5 | 20 | 0.45 | 2 | 0.5 | 3 | 0.3333 | 6 | 0.6667 | 2 | 0 | 13 | 0.4615 |
| 8_83816504 | 7 | 0.2857 | 9 | 0.3333 | 6 | 0 | 4 | 0.5 | 26 | 0.2692 | 5 | 0.4 | 7 | 0.4286 | 7 | 0.4286 | 5 | 0 | 24 | 0.3333 |
| 8_83817294 | 8 | 0.75 | 11 | 0.7273 | 10 | 0.7 | 7 | 1 | 36 | 0.7778 | 6 | 0.6667 | 7 | 0.5714 | 10 | 0.5 | 8 | 0.75 | 31 | 0.6129 |
| 8_83817601 | 8 | 0.5 | 12 | 0.5833 | 10 | 0.3 | 7 | 0.7143 | 37 | 0.5135 | 6 | 0.3333 | 8 | 0.75 | 12 | 0.3333 | 8 | 0.5 | 34 | 0.4706 |
| 8_83897446 | 4 | 0 | 8 | 0.375 | 6 | 0.1667 | 3 | 0.3333 | 21 | 0.2381 | 7 | 0.2857 | 3 | 0 | 5 | 0 | 1 | 1 | 16 | 0.1875 |
| 8_83981421 | 3 | 0.6667 | 2 | 0 | NA | NA | NA | NA | 5 | 0.4 | 1 | 1 | 2 | 0.5 | 3 | 0.3333 | NA | NA | 6 | 0.5 |
| 8_83997797 | 3 | 1 | 6 | 0.8333 | 1 | 1 | NA | NA | 10 | 0.9 | NA | NA | 3 | 0.6667 | 4 | 1 | NA | NA | 7 | 0.8571 |
| 8_83997833 | 3 | 0.6667 | 6 | 1 | 1 | 1 | 1 | 1 | 11 | 0.9091 | 3 | 1 | 2 | 1 | 5 | 1 | 1 | 1 | 11 | 1 |
| 8_83997840 | 3 | 0.3333 | 4 | 0.25 | 1 | 0 | 2 | 0 | 10 | 0.2 | 4 | 0 | 3 | 0.3333 | 5 | 0 | NA | NA | 12 | 0.0833 |
| 8_83997848 | 2 | 1 | 6 | 0.8333 | 1 | 1 | NA | NA | 9 | 0.8889 | 3 | 0.3333 | 2 | 1 | 4 | 0.75 | NA | NA | 9 | 0.6667 |
| 8_83997863 | 3 | 0.6667 | 7 | 0.7143 | 1 | 1 | 2 | 1 | 13 | 0.7692 | 3 | 0.6667 | 3 | 1 | 5 | 0.4 | 1 | 0 | 12 | 0.5833 |
| 8_83997965 | 3 | 0.6667 | 6 | 0.1667 | NA | NA | 1 | 1 | 10 | 0.4 | 3 | 0.3333 | 2 | 0 | 4 | 0.25 | 1 | 1 | 10 | 0.3 |
| 8_83997969 | 2 | 0 | 5 | 0.2 | 1 | 0 | 2 | 0 | 10 | 0.1 | 3 | 0.3333 | 2 | 0 | 5 | 0.2 | 1 | 1 | 11 | 0.2727 |
| 8_83997983 | 1 | 1 | 6 | 1 | 1 | 1 | 2 | 1 | 10 | 1 | 3 | 1 | 3 | 1 | 4 | 1 | NA | NA | 10 | 1 |
| 8_83997995 | 1 | 1 | 7 | 0.2857 | 1 | 1 | 2 | 0.5 | 11 | 0.4545 | 2 | 1 | 3 | 0.6667 | 5 | 0.4 | 1 | 1 | 11 | 0.6364 |
| 8_83998061 | 3 | 0.3333 | 7 | 0.8571 | 1 | 1 | 1 | 0 | 12 | 0.6667 | 3 | 1 | 3 | 0.6667 | 5 | 0.6 | 1 | 0 | 12 | 0.6667 |
| 8_83998211 | 3 | 1 | 3 | 0.3333 | NA | NA | 2 | 0.5 | 8 | 0.625 | 2 | 1 | 4 | 0.75 | 4 | 0.5 | 1 | 1 | 11 | 0.7273 |
| 8_83998840 | 5 | 0.4 | 8 | 0.125 | 3 | 0.3333 | 5 | 0 | 21 | 0.1905 | 4 | 1 | 4 | 0 | 8 | 0 | 1 | 1 | 17 | 0.2941 |
| 8_84040534 | 1 | 0 | 1 | 1 | NA | NA | 1 | 1 | 3 | 0.6667 | NA | NA | 2 | 0 | NA | NA | 2 | 0.5 | 4 | 0.25 |
| 8_84161581 | 16 | 0.0625 | 25 | 0.04 | 34 | 0.0588 | 28 | 0.0714 | 103 | 0.0583 | 22 | 0.1364 | 23 | 0.0435 | 22 | 0.1364 | 29 | 0 | 96 | 0.0729 |
| 8_84161658 | 17 | 0.1176 | 32 | 0.0625 | 36 | 0.0278 | 29 | 0.1034 | 114 | 0.0702 | 23 | 0.1739 | 26 | 0.1538 | 24 | 0.125 | 28 | 0.0357 | 101 | 0.1188 |
| 8_84162318 | 13 | 0.0769 | 25 | 0.24 | 34 | 0.1765 | 24 | 0.0417 | 96 | 0.1458 | 22 | 0.1364 | 23 | 0.1304 | 21 | 0.0952 | 26 | 0.1538 | 92 | 0.1304 |
| 8_84162599 | 11 | 0.4545 | 24 | 0.2083 | 30 | 0.2333 | 22 | 0.0909 | 87 | 0.2184 | 18 | 0.3333 | 22 | 0.2727 | 17 | 0.2941 | 21 | 0.3333 | 78 | 0.3077 |
| 8_84298216 | 39 | 0.0513 | 28 | 0.0357 | 36 | 0.0833 | 32 | 0.0625 | 135 | 0.0593 | 38 | 0.0263 | 26 | 0.0385 | 34 | 0.1176 | 27 | 0.0741 | 125 | 0.064 |
| 8_84334971 | 4 | 0.5 | 4 | 0.25 | 5 | 0.4 | 1 | 0 | 14 | 0.3571 | 8 | 0.25 | 4 | 0 | 3 | 0.3333 | 3 | 0.6667 | 18 | 0.2778 |
| 8_84336608 | 6 | 1 | 6 | 1 | 7 | 1 | 1 | 1 | 20 | 1 | 13 | 1 | 6 | 1 | 5 | 1 | 5 | 1 | 29 | 1 |
| 8_84336665 | 5 | 1 | 6 | 1 | 6 | 0.8333 | 1 | 1 | 18 | 0.9444 | 13 | 0.9231 | 4 | 1 | 3 | 1 | 3 | 1 | 23 | 0.9565 |
| 8_84336877 | 8 | 0.125 | 7 | 0.4286 | 8 | 0.125 | 1 | 1 | 24 | 0.25 | 11 | 0.4545 | 6 | 0.3333 | 6 | 0.5 | 4 | 0.5 | 27 | 0.4444 |
| 8_84336953 | 6 | 0.1667 | 6 | 0 | 5 | 0.6 | 1 | 1 | 18 | 0.2778 | 12 | 0.25 | 6 | 0.5 | 5 | 0 | 5 | 0.2 | 28 | 0.25 |
| 8_84337468 | 11 | 1 | 7 | 1 | 9 | 1 | 2 | 1 | 29 | 1 | 11 | 1 | 6 | 1 | 8 | 1 | 4 | 1 | 29 | 1 |
| 8_84337504 | 11 | 0 | 9 | 0.3333 | 9 | 0 | 2 | 0 | 31 | 0.0968 | 13 | 0 | 7 | 0.1429 | 5 | 0.2 | 5 | 0 | 30 | 0.0667 |
| 8_84337607 | 8 | 0.25 | 8 | 0.625 | 8 | 0 | 1 | 0 | 25 | 0.28 | 11 | 0.1818 | 4 | 0.5 | 7 | 0.5714 | 4 | 0.25 | 26 | 0.3462 |
| 8_84337666 | 11 | 0.5455 | 8 | 0.875 | 6 | 0.8333 | 4 | 1 | 29 | 0.7586 | 10 | 0.5 | 6 | 0.8333 | 7 | 0.7143 | 5 | 1 | 28 | 0.7143 |
| 8_84337722 | 11 | 1 | 8 | 1 | 8 | 1 | 3 | 1 | 30 | 1 | 11 | 0.9091 | 6 | 1 | 7 | 1 | 3 | 1 | 27 | 0.963 |
| 8_84337761 | 11 | 0.2727 | 9 | 0.1111 | 8 | 0.25 | 4 | 0.25 | 32 | 0.2188 | 10 | 0.3 | 5 | 0.2 | 6 | 0 | 4 | 0 | 25 | 0.16 |
| 8_84337921 | 10 | 0.8 | 10 | 0.9 | 7 | 0.7143 | 3 | 0.6667 | 30 | 0.8 | 10 | 0.7 | 5 | 0.2 | 7 | 0.7143 | 4 | 0.5 | 26 | 0.5769 |
| 8_84337960 | 12 | 0.25 | 9 | 0 | 7 | 0 | 4 | 0 | 32 | 0.0938 | 10 | 0.1 | 6 | 0.1667 | 7 | 0 | 5 | 0 | 28 | 0.0714 |
| 8_84338034 | 12 | 0.1667 | 10 | 0 | 8 | 0.25 | 3 | 0 | 33 | 0.1212 | 11 | 0.1818 | 5 | 0 | 6 | 0.1667 | 1 | 0 | 23 | 0.1304 |
| 8_84338181 | 14 | 0.7857 | 7 | 0.7143 | 7 | 0.5714 | 4 | 0.25 | 32 | 0.6562 | 10 | 0.3 | 4 | 0.25 | 6 | 0.6667 | 5 | 0.8 | 25 | 0.48 |
| 8_84338188 | 14 | 0 | 9 | 0.1111 | 8 | 0.125 | 4 | 0 | 35 | 0.0571 | 9 | 0.3333 | 6 | 0 | 6 | 0.1667 | 3 | 0 | 24 | 0.1667 |
| 8_84338516 | 13 | 0.3846 | 7 | 0.5714 | 7 | 0.5714 | 5 | 0.6 | 32 | 0.5 | 12 | 0.25 | 3 | 0.6667 | 7 | 0.8571 | 6 | 0.3333 | 28 | 0.4643 |
| 8_84388624 | 9 | 0 | 9 | 0.3333 | 6 | 0.1667 | 12 | 0 | 36 | 0.1111 | 7 | 0.1429 | 2 | 0 | 7 | 0.1429 | 2 | 0 | 18 | 0.1111 |
| 8_84390900 | 8 | 0.5 | 12 | 0.5 | 6 | 0.6667 | 9 | 0.3333 | 35 | 0.4857 | 8 | 0.25 | 3 | 0.3333 | 7 | 0.4286 | 3 | 0.3333 | 21 | 0.3333 |
| 8_84390906 | 9 | 0.3333 | 11 | 0.4545 | 8 | 0.5 | 12 | 0.5833 | 40 | 0.475 | 8 | 0.625 | 2 | 0 | 8 | 0.5 | 3 | 0.3333 | 21 | 0.4762 |
| 8_84390917 | 9 | 0.4444 | 11 | 0.2727 | 7 | 0.5714 | 14 | 0.5 | 41 | 0.439 | 7 | 0.8571 | 2 | 0.5 | 6 | 0.1667 | 2 | 0.5 | 17 | 0.5294 |
| 8_84393285 | 2 | 1 | NA | NA | 2 | 1 | 1 | 1 | 5 | 1 | 2 | 1 | 2 | 0.5 | 3 | 1 | 1 | 0 | 8 | 0.75 |
| 8_84393972 | 1 | 0 | NA | NA | 2 | 1 | 1 | 0 | 4 | 0.5 | 2 | 0.5 | 1 | 1 | 3 | 0.6667 | NA | NA | 6 | 0.6667 |
| 8_84394675 | 2 | 1 | NA | NA | 2 | 1 | NA | NA | 4 | 1 | 3 | 0.6667 | 4 | 0.75 | 3 | 0.6667 | NA | NA | 10 | 0.7 |
| 8_84394756 | 2 | 1 | NA | NA | 2 | 0 | 1 | 0 | 5 | 0.4 | 2 | 0.5 | 1 | 1 | 2 | 1 | NA | NA | 5 | 0.8 |
| 8_84395071 | 2 | 0 | NA | NA | 1 | 1 | 1 | 0 | 4 | 0.25 | 3 | 0.3333 | 3 | 0 | 2 | 0.5 | NA | NA | 8 | 0.25 |
| 8_84395621 | 1 | 1 | NA | NA | 2 | 0.5 | NA | NA | 3 | 0.6667 | 3 | 0 | 1 | 0 | 4 | 0.5 | NA | NA | 8 | 0.25 |
| 8_84396120 | 1 | 1 | NA | NA | 1 | 0 | NA | NA | 2 | 0.5 | 2 | 0.5 | 1 | 1 | 2 | 1 | NA | NA | 5 | 0.8 |
| 8_84396153 | 1 | 1 | NA | NA | 1 | 1 | NA | NA | 2 | 1 | 2 | 1 | NA | NA | 2 | 0.5 | NA | NA | 4 | 0.75 |
| 8_84396181 | 2 | 0.5 | NA | NA | 1 | 1 | 1 | 0 | 4 | 0.5 | 2 | 0 | 1 | 1 | 2 | 0 | NA | NA | 5 | 0.2 |
| 8_84396216 | 1 | 1 | NA | NA | 1 | 1 | 1 | 1 | 3 | 1 | 2 | 1 | 1 | 1 | 2 | 1 | NA | NA | 5 | 1 |

| | | | | | | | | | | | | | | | | | | | | |
|---|---|---|---|---|---|---|---|---|---|---|---|---|---|---|---|---|---|---|---|---|
| 8_8439631 | 1 | 1 | NA | NA | 1 | 1 | 1 | 1 | 3 | 1 | 2 | 1 | 1 | 1 | 2 | 1 | NA | NA | 5 | 1 |
| 8_8439642 | 2 | 0.5 | NA | NA | NA | NA | NA | NA | 2 | 0.5 | 2 | 0.5 | 1 | 0 | 1 | 0 | NA | NA | 4 | 0.25 |
| 8_8439664 | 1 | 1 | NA | NA | 1 | 1 | 1 | 1 | 3 | 1 | 2 | 1 | NA | NA | 1 | 1 | NA | NA | 3 | 1 |
| 8_8439665 | 1 | 1 | NA | NA | 1 | 1 | NA | NA | 2 | 1 | 2 | 1 | NA | NA | 1 | 1 | NA | NA | 3 | 1 |
| 8_8439669 | 1 | 1 | NA | NA | 1 | 1 | 1 | 1 | 3 | 1 | 2 | 0.5 | NA | NA | 1 | 1 | NA | NA | 3 | 0.6667 |
| 8_8445017 | 2 | 1 | 3 | 0.6667 | NA | NA | 1 | 1 | 6 | 0.8333 | 1 | 1 | 2 | 0.5 | NA | NA | 3 | 1 | 6 | 0.8333 |
| 8_8461416 | NA | NA | 1 | 1 | NA | NA | NA | NA | 1 | 1 | NA | NA | 1 | 1 | NA | NA | NA | NA | 1 | 1 |
| 8_8461442 | NA | NA | 1 | 1 | NA | NA | NA | NA | 1 | 1 | NA | NA | 1 | 1 | NA | NA | NA | NA | 1 | 1 |
| 8_8465623 | 4 | 0 | 8 | 0.125 | 4 | 0 | 5 | 0.2 | 21 | 0.0952 | 4 | 0.25 | NA | NA | 3 | 0 | 4 | 0.25 | 11 | 0.1818 |
| 8_8465624 | 3 | 0 | 6 | 0.3333 | 4 | 0.25 | 3 | 0.3333 | 16 | 0.25 | 4 | 0.25 | NA | NA | 5 | 0.4 | 2 | 0 | 11 | 0.2727 |
| 8_8465626 | 3 | 0 | 7 | 0.4286 | 4 | 0.75 | 5 | 0.8 | 19 | 0.5263 | 5 | 0.6 | NA | NA | 4 | 0 | 4 | 0.75 | 13 | 0.4615 |
| 8_8465630 | 3 | 0.6667 | 5 | 0.4 | 3 | 0.3333 | 5 | 0.4 | 16 | 0.4375 | 4 | 0.5 | NA | NA | 3 | 0.6667 | 4 | 0.75 | 11 | 0.6364 |
| 8_8465642 | 3 | 0 | 8 | 0.125 | 6 | 0.6667 | 7 | 0.1429 | 24 | 0.25 | 5 | 0.2 | NA | NA | 5 | 0 | 4 | 0.75 | 14 | 0.2857 |
| 8_8465651 | 4 | 0.5 | 7 | 0.8571 | 6 | 1 | 7 | 0.8571 | 24 | 0.8333 | 6 | 0.5 | NA | NA | 5 | 0.6 | 4 | 1 | 15 | 0.6667 |
| 8_8465671 | 4 | 1 | 6 | 1 | 4 | 1 | 5 | 0.6 | 19 | 0.8947 | 5 | 1 | NA | NA | 5 | 0.6 | 5 | 0.8 | 15 | 0.8 |
| 8_8465681 | 3 | 0 | 7 | 0 | 6 | 0.3333 | 5 | 0 | 21 | 0.0952 | 4 | 0 | 1 | 0 | 6 | 0.1667 | 6 | 0.1667 | 17 | 0.1176 |
| 8_8466501 | 6 | 0 | 15 | 0.0667 | 13 | 0.1538 | 14 | 0.1429 | 48 | 0.1042 | 10 | 0.1 | 10 | 0.2 | 16 | 0 | 12 | 0.25 | 48 | 0.125 |
| 8_8466526 | 6 | 0.1667 | 16 | 0.3125 | 13 | 0.1538 | 14 | 0.2143 | 49 | 0.2245 | 8 | 0.375 | 11 | 0.2727 | 20 | 0.2 | 13 | 0 | 52 | 0.1923 |
| 8_8466529 | 5 | 0.8 | 12 | 0.75 | 5 | 0.8 | 12 | 0.5 | 34 | 0.6765 | 4 | 1 | 5 | 0.4 | 10 | 0.7 | 7 | 0.5714 | 26 | 0.6538 |
| 8_8466543 | 7 | 0.8571 | 18 | 0.8333 | 15 | 0.8667 | 12 | 0.75 | 52 | 0.8269 | 11 | 0.7273 | 12 | 0.6667 | 19 | 0.7895 | 14 | 0.7857 | 56 | 0.75 |
| 8_8466546 | 9 | 0.8889 | 17 | 0.9412 | 11 | 0.9091 | 10 | 0.9 | 47 | 0.9149 | 8 | 1 | 11 | 1 | 18 | 0.9444 | 10 | 0.9 | 47 | 0.9574 |
| 8_8466548 | 9 | 0.7778 | 18 | 0.8333 | 14 | 0.8571 | 13 | 0.7692 | 54 | 0.8148 | 8 | 0.75 | 11 | 0.9091 | 13 | 0.7692 | 13 | 0.3846 | 45 | 0.6889 |
| 8_8466572 | 11 | 0.4545 | 14 | 0.2857 | 19 | 0.6316 | 17 | 0.4706 | 61 | 0.4754 | 12 | 0.6667 | 15 | 0.4667 | 20 | 0.55 | 15 | 0.6667 | 62 | 0.5806 |
| 8_8466578 | 11 | 0.7273 | 22 | 0.8636 | 17 | 1 | 15 | 0.8 | 65 | 0.8615 | 12 | 0.5833 | 13 | 0.9231 | 23 | 0.8261 | 18 | 0.8333 | 66 | 0.803 |
| 8_8466585 | 11 | 0.2727 | 19 | 0.3684 | 14 | 0.2143 | 12 | 0.6667 | 56 | 0.375 | 12 | 0.4167 | 12 | 0.25 | 19 | 0.4737 | 9 | 0.3333 | 52 | 0.3846 |
| 8_8466603 | 11 | 0 | 21 | 0 | 18 | 0.3333 | 15 | 0.0667 | 65 | 0.1077 | 10 | 0.3 | 13 | 0 | 21 | 0.1429 | 19 | 0.1053 | 63 | 0.127 |
| 8_8466613 | 9 | 0.6667 | 18 | 0.3333 | 15 | 0.5333 | 12 | 0.5 | 54 | 0.4815 | 7 | 0.5714 | 13 | 0.5385 | 15 | 0.4667 | 11 | 0.3636 | 46 | 0.4783 |
| 8_8466615 | 10 | 0.4 | 19 | 0.1579 | 12 | 0.1667 | 16 | 0.1875 | 57 | 0.2105 | 9 | 0.2222 | 9 | 0.1111 | 18 | 0.1667 | 14 | 0.3571 | 50 | 0.22 |
| 8_8466628 | 14 | 0.1429 | 22 | 0.2273 | 17 | 0.4118 | 16 | 0.1875 | 69 | 0.2464 | 16 | 0.4375 | 16 | 0.1875 | 25 | 0.32 | 19 | 0.1579 | 76 | 0.2763 |
| 8_8466662 | 13 | 0.2308 | 22 | 0.1818 | 19 | 0.3684 | 16 | 0.375 | 70 | 0.2857 | 16 | 0.375 | 21 | 0.0476 | 25 | 0.28 | 19 | 0.2105 | 81 | 0.2222 |
| 8_8466668 | 13 | 0.5385 | 19 | 0.5789 | 17 | 0.4118 | 18 | 0.7222 | 67 | 0.5672 | 17 | 0.3529 | 18 | 0.4444 | 25 | 0.56 | 21 | 0.3333 | 81 | 0.4321 |
| 8_8466670 | 13 | 0.1538 | 15 | 0 | 12 | 0 | 15 | 0.2 | 55 | 0.0909 | 15 | 0.0667 | 14 | 0.1429 | 22 | 0.1364 | 18 | 0.1111 | 69 | 0.1159 |
| 8_8466673 | 14 | 0.7857 | 20 | 0.8 | 18 | 0.7778 | 20 | 0.95 | 72 | 0.8333 | 18 | 0.9444 | 20 | 0.85 | 26 | 0.6538 | 21 | 0.6667 | 85 | 0.7647 |
| 8_8466679 | 17 | 0.2353 | 22 | 0.4091 | 18 | 0.3889 | 19 | 0.6842 | 76 | 0.4342 | 18 | 0.5556 | 20 | 0.6 | 25 | 0.36 | 21 | 0.5714 | 84 | 0.5119 |
| 8_8466683 | 11 | 0.0909 | 20 | 0.1 | 16 | 0.0625 | 18 | 0.1667 | 65 | 0.1077 | 15 | 0.1333 | 17 | 0.0588 | 21 | 0.1905 | 17 | 0.1765 | 70 | 0.1429 |
| 8_8466704 | 16 | 0.5 | 23 | 0.5217 | 15 | 0.4 | 18 | 0.7222 | 72 | 0.5417 | 16 | 0.5 | 21 | 0.3333 | 19 | 0.4211 | 22 | 0.4545 | 78 | 0.4231 |
| 8_8466779 | 22 | 0.5 | 28 | 0.5714 | 21 | 0.4286 | 20 | 0.75 | 91 | 0.5604 | 20 | 0.65 | 31 | 0.5806 | 19 | 0.6842 | 28 | 0.5357 | 98 | 0.602 |
| 8_8466784 | 30 | 0.3667 | 43 | 0.3953 | 30 | 0.3667 | 30 | 0.3667 | 133 | 0.3759 | 25 | 0.48 | 41 | 0.2927 | 31 | 0.3871 | 37 | 0.5946 | 134 | 0.4328 |
| 8_8466791 | 19 | 0.0526 | 19 | 0.0526 | 20 | 0.05 | 13 | 0.0769 | 71 | 0.0563 | 16 | 0.0625 | 23 | 0.0435 | 24 | 0.0833 | 24 | 0.0417 | 87 | 0.0575 |
| 8_8471751 | 11 | 0.3636 | 11 | 0.4545 | 16 | 0.5 | 10 | 0.2 | 48 | 0.3958 | 7 | 0.7143 | 15 | 0.3333 | 24 | 0.2917 | 8 | 0.5 | 54 | 0.3889 |
| 8_8472452 | 1 | 0 | NA | NA | 1 | 1 | 3 | 0.3333 | 5 | 0.4 | 2 | 0.5 | NA | NA | 2 | 0 | 1 | 1 | 5 | 0.4 |
| 8_8472564 | 6 | 0 | NA | NA | 5 | 0.2 | 5 | 0.2 | 16 | 0.125 | 3 | 0 | 1 | 0 | 6 | 0 | 2 | 0.5 | 12 | 0.0833 |
| 8_8472653 | 7 | 0 | NA | NA | 6 | 0.1667 | 4 | 0 | 17 | 0.0588 | 2 | 0.5 | 3 | 0 | 6 | 0 | 4 | 0 | 15 | 0.0667 |
| 8_8472660 | 8 | 0.5 | NA | NA | 3 | 0 | 2 | 0 | 13 | 0.3077 | 1 | 0 | 1 | 0 | 4 | 0.25 | 2 | 0.5 | 8 | 0.25 |
| 8_8472670 | 9 | 0.4444 | NA | NA | 7 | 0.1429 | 6 | 0.5 | 22 | 0.3636 | 5 | 0.2 | 3 | 0 | 9 | 0.3333 | 5 | 0.2 | 22 | 0.2273 |
| 8_8472674 | 7 | 0.4286 | NA | NA | 7 | 0.4286 | 4 | 0.5 | 18 | 0.4444 | 4 | 0.5 | 4 | 0.25 | 8 | 0.75 | 6 | 0.5 | 22 | 0.5455 |
| 8_8479031 | NA | NA | NA | NA | NA | NA | 1 | 1 | 1 | 1 | 1 | 1 | NA | NA | NA | NA | NA | NA | 1 | 1 |
| 8_8479035 | 1 | 1 | NA | NA | NA | NA | 1 | 1 | 2 | 1 | 1 | 1 | NA | NA | 1 | 1 | NA | NA | 2 | 1 |
| 8_8479078 | 1 | 1 | NA | NA | NA | NA | 1 | 1 | 2 | 1 | NA | NA | NA | NA | 1 | 1 | NA | NA | 1 | 1 |
| 8_8479200 | 1 | 0 | NA | NA | NA | NA | 1 | 1 | 2 | 0.5 | NA | NA | NA | NA | 1 | 1 | NA | NA | 1 | 1 |
| 8_8482255 | 1 | 0 | NA | NA | 3 | 0 | 2 | 0.5 | 6 | 0.1667 | 2 | 0.5 | 2 | 0.5 | NA | NA | 1 | 0 | 5 | 0.4 |
| 8_8482293 | 3 | 0.6667 | NA | NA | 3 | 0.3333 | 2 | 0.5 | 8 | 0.5 | 4 | 0.75 | 3 | 1 | 1 | 1 | 1 | 0 | 9 | 0.7778 |
| 8_8482396 | 6 | 0.6667 | 7 | 0.8571 | 2 | 0.5 | 5 | 0.4 | 20 | 0.65 | 1 | 1 | NA | NA | 2 | 1 | 3 | 0.6667 | 6 | 0.8333 |
| 8_8482457 | 3 | 0 | 5 | 0 | 2 | 0 | 4 | 0.25 | 14 | 0.0714 | NA | NA | 2 | 0 | 1 | 0 | 3 | 0.3333 | 6 | 0.1667 |
| 8_8482701 | NA | NA | 1 | 0 | NA | NA | 1 | 1 | 2 | 0.5 | NA | NA | NA | NA | NA | NA | 2 | 0.5 | 2 | 0.5 |
| 8_8482709 | NA | NA | 1 | 1 | NA | NA | 1 | 1 | 2 | 1 | NA | NA | NA | NA | NA | NA | 2 | 1 | 2 | 1 |
| 8_8485948 | 3 | 0 | 7 | 0.2857 | 2 | 0 | 4 | 0 | 16 | 0.125 | 3 | 0.3333 | 4 | 0.5 | 3 | 0.3333 | 2 | 0 | 12 | 0.3333 |
| 8_8485952 | 5 | 0.4 | 7 | 0.7143 | 4 | 0.5 | 6 | 0.6667 | 22 | 0.5909 | 3 | 0.6667 | 5 | 0.8 | 3 | 1 | 3 | 0.6667 | 14 | 0.7857 |
| 8_8485972 | 4 | 0.5 | 8 | 0.375 | 5 | 0.4 | 4 | 0.25 | 21 | 0.381 | 2 | 1 | 6 | 0.5 | 4 | 0.75 | 2 | 0 | 14 | 0.5714 |
| 8_8485976 | 4 | 0.25 | 7 | 0.2857 | 5 | 0.4 | 5 | 0.4 | 21 | 0.3333 | 1 | 0 | 4 | 0.25 | 4 | 0.25 | 2 | 1 | 11 | 0.3636 |
| 8_8485978 | 4 | 0 | 10 | 0.3 | 4 | 0.5 | 5 | 0.6 | 23 | 0.3478 | 2 | 0.5 | 4 | 1 | NA | NA | 1 | 0 | 7 | 0.7143 |
| 8_8489367 | 1 | 1 | NA | NA | NA | NA | NA | NA | 1 | 1 | NA | NA | NA | NA | 1 | 1 | 1 | 1 | 2 | 1 |
| 8_8489397 | NA | NA | 1 | 1 | NA | NA | NA | NA | 1 | 1 | NA | NA | NA | NA | 1 | 1 | 1 | 1 | 2 | 1 |
| 8_8489408 | 1 | 1 | 1 | 1 | NA | NA | NA | NA | 2 | 1 | NA | NA | NA | NA | 1 | 1 | 1 | 0 | 2 | 0.5 |
| 8_8489417 | 1 | 0 | 1 | 1 | NA | NA | NA | NA | 2 | 0.5 | NA | NA | NA | NA | 1 | 1 | 1 | 0 | 2 | 0.5 |

| | | | | | | | | | | | | | | | | | | | | |
|---|---|---|---|---|---|---|---|---|---|---|---|---|---|---|---|---|---|---|---|---|
| 8_8493830( | 1 | 1 | 2 | 1 | 1 | 0 | 3 | 1 | 7 | 0.8571 | 1 | 1 | 2 | 1 | 1 | 1 | 1 | 1 | 5 | 1 |
| 8_8493842( | NA | NA | 3 | 1 | 1 | 1 | 3 | 1 | 7 | 1 | 1 | 1 | 3 | 1 | 1 | 1 | 1 | 1 | 6 | 1 |
| 8_8493843! | NA | NA | 4 | 0 | 1 | 0 | 3 | 0.3333 | 8 | 0.125 | 1 | 0 | 3 | 0 | 1 | 1 | 1 | 0 | 6 | 0.1667 |
| 8_8493857' | NA | NA | 2 | 1 | NA | NA | 1 | 1 | 3 | 1 | 1 | 1 | 2 | 1 | NA | NA | 1 | 0 | 4 | 0.75 |
| 8_8493869' | NA | NA | 3 | 1 | 1 | 0 | 2 | 1 | 6 | 0.8333 | 1 | 0 | 3 | 0.3333 | 1 | 1 | 1 | 1 | 6 | 0.5 |
| 8_8493872: | NA | NA | 3 | 0.6667 | 1 | 1 | 1 | 1 | 5 | 0.8 | 1 | 1 | 3 | 0.6667 | 1 | 1 | 1 | 0 | 6 | 0.6667 |
| 8_8493882( | NA | NA | 2 | 1 | 1 | 1 | 1 | 1 | 4 | 1 | 1 | 1 | 2 | 1 | 1 | 1 | 1 | 1 | 5 | 1 |
| 8_8493888( | NA | NA | 1 | 1 | 1 | 1 | 1 | 0 | 3 | 0.6667 | 1 | 1 | 3 | 1 | NA | NA | 1 | 1 | 5 | 1 |
| 8_8497310( | 11 | 0.6364 | 9 | 0.3333 | 13 | 0.4615 | 16 | 0.5 | 49 | 0.4898 | 6 | 0.6667 | 11 | 0.4545 | 11 | 0.2727 | 20 | 0.2 | 48 | 0.3333 |
| 8_8497314' | 13 | 0.6154 | 14 | 0.4286 | 17 | 0.5882 | 19 | 0.5789 | 63 | 0.5556 | 8 | 0.125 | 17 | 0.7059 | 17 | 0.4706 | 24 | 0.4167 | 66 | 0.4697 |
| 8_8497317' | 18 | 0.6111 | 15 | 0.6 | 18 | 0.7778 | 21 | 0.8571 | 72 | 0.7222 | 10 | 0.8 | 18 | 0.7222 | 17 | 0.6471 | 26 | 0.7308 | 71 | 0.7183 |
| 8_8498574' | 1 | 0 | 2 | 0.5 | NA | NA | NA | NA | 3 | 0.3333 | 1 | 0 | 1 | 1 | 3 | 0.3333 | 1 | 0 | 6 | 0.3333 |
| 8_8532811' | 2 | 0 | 2 | 0 | 3 | 0.3333 | 7 | 0.1429 | 14 | 0.1429 | 1 | 0 | 3 | 0 | 3 | 0.3333 | 1 | 0 | 8 | 0.125 |
| 8_8533973' | NA | NA | NA | NA | NA | NA | 2 | 0.5 | 2 | 0.5 | NA | NA | 2 | 0.5 | NA | NA | NA | NA | 2 | 0.5 |
| 8_8536619' | 6 | 0.3333 | 10 | 0.3 | 11 | 0.2727 | 11 | 0.3636 | 38 | 0.3158 | 7 | 0.2857 | 8 | 0.375 | 12 | 0.0833 | 6 | 0.5 | 33 | 0.2727 |
| 8_8536639' | 5 | 0.2 | 8 | 0.125 | 11 | 0 | 12 | 0.25 | 36 | 0.1389 | 3 | 0 | 9 | 0.6667 | 10 | 0.5 | 5 | 0.4 | 27 | 0.4815 |
| 8_8536654! | 6 | 0.1667 | 15 | 0 | 12 | 0.0833 | 12 | 0.0833 | 45 | 0.0667 | 7 | 0 | 11 | 0 | 11 | 0.0909 | 7 | 0.1429 | 36 | 0.0556 |
| 8_8536663 | 7 | 0.7143 | 14 | 0.7143 | 12 | 0.75 | 12 | 0.75 | 45 | 0.7333 | 6 | 0.8333 | 9 | 0.5556 | 12 | 0.8333 | 8 | 0.875 | 35 | 0.7714 |
| 8_8536681' | 8 | 0.375 | 12 | 0.5 | 17 | 0.3529 | 15 | 0.3333 | 52 | 0.3846 | 8 | 0.25 | 9 | 0.4444 | 13 | 0.4615 | 8 | 0 | 38 | 0.3158 |
| 8_8538818 | 2 | 1 | 1 | 0 | 2 | 0.5 | 1 | 1 | 6 | 0.6667 | 1 | 1 | NA | NA | 1 | 1 | 2 | 0.5 | 4 | 0.75 |
| 8_8538823! | 2 | 1 | 1 | 1 | 1 | 0 | 1 | 1 | 5 | 0.8 | NA | NA | NA | NA | NA | NA | 2 | 0.5 | 2 | 0.5 |
| 8_8538828: | 2 | 0 | 1 | 0 | 2 | 1 | 1 | 0 | 6 | 0.3333 | NA | NA | NA | NA | 1 | 0 | 2 | 0.5 | 3 | 0.3333 |
| 8_8538829: | 2 | 1 | 1 | 1 | 1 | 1 | 1 | 1 | 5 | 1 | NA | NA | NA | NA | 1 | 1 | 2 | 1 | 3 | 1 |
| 8_8538837( | 1 | 1 | NA | NA | 2 | 0.5 | 1 | 0 | 4 | 0.5 | NA | NA | NA | NA | 1 | 1 | 2 | 0.5 | 3 | 0.6667 |
| 8_8538853: | 2 | 1 | NA | NA | 1 | 1 | 1 | 1 | 4 | 1 | NA | NA | NA | NA | NA | NA | 2 | 1 | 2 | 1 |
| 8_8538859' | NA | NA | NA | NA | 2 | 1 | 1 | 1 | 3 | 1 | NA | NA | NA | NA | 1 | 1 | 2 | 1 | 3 | 1 |
| 8_8538882( | 2 | 0 | NA | NA | 2 | 0.5 | 1 | 0 | 5 | 0.2 | NA | NA | NA | NA | 1 | 0 | 2 | 0.5 | 3 | 0.3333 |
| 8_8538897! | 2 | 0.5 | NA | NA | 1 | 1 | 1 | 1 | 4 | 0.75 | 1 | 1 | NA | NA | 1 | 0 | 2 | 0.5 | 4 | 0.5 |
| 8_8539890' | 5 | 0.4 | 8 | 0.125 | 9 | 0 | 3 | 0 | 25 | 0.12 | 10 | 0 | 5 | 0 | 8 | 0.25 | 6 | 0 | 29 | 0.069 |
| 8_8539892( | 4 | 0 | 10 | 0 | 10 | 0.2 | 3 | 0.3333 | 27 | 0.1111 | 11 | 0.2727 | 4 | 0 | 10 | 0 | 7 | 0 | 32 | 0.0938 |
| 8_8539905( | 5 | 0 | 9 | 0 | 6 | 0.1667 | 2 | 0.5 | 22 | 0.0909 | 7 | 0.1429 | 5 | 0.6 | 9 | 0.1111 | 7 | 0 | 28 | 0.1786 |
| 8_8539944! | 4 | 0.25 | 9 | 0.3333 | 6 | 0.3333 | 3 | 0.6667 | 22 | 0.3636 | 9 | 0.3333 | 4 | 0 | 7 | 0.2857 | 5 | 0.6 | 25 | 0.32 |
| 8_8539957( | 2 | 0 | 9 | 0.3333 | 7 | 0.1429 | 2 | 1 | 20 | 0.3 | 5 | 0.8 | 4 | 0.25 | 8 | 0.75 | 4 | 1 | 21 | 0.7143 |
| 8_8539975( | 2 | 0 | 7 | 0.4286 | 5 | 0.4 | 1 | 1 | 15 | 0.4 | 4 | 0 | 1 | 0 | 6 | 0 | 3 | 0.3333 | 14 | 0.0714 |
| 8_8540331 | 2 | 0.5 | 5 | 0.4 | 2 | 0 | 1 | 0 | 10 | 0.3 | 5 | 0.2 | 2 | 0 | 2 | 0 | 4 | 0 | 13 | 0.0769 |
| 8_8540335( | 3 | 1 | 5 | 1 | 3 | 1 | NA | NA | 11 | 1 | 7 | 1 | 2 | 1 | 4 | 1 | 4 | 0.75 | 17 | 0.9412 |
| 8_8540337: | 3 | 0 | 5 | 0.2 | 3 | 0 | 1 | 0 | 12 | 0.0833 | 7 | 0 | 2 | 0 | 4 | 0 | 4 | 0.25 | 17 | 0.0588 |
| 8_8540342( | 1 | 1 | 5 | 0 | 2 | 0 | 1 | 1 | 9 | 0.2222 | 7 | 0.1429 | 1 | 0 | 1 | 0 | 2 | 0.5 | 11 | 0.1818 |
| 8_8540358 | 2 | 0.5 | 4 | 0 | 3 | 0 | 1 | 0 | 10 | 0.1 | 6 | 0 | 2 | 0 | 3 | 0.3333 | 3 | 0 | 14 | 0.0714 |
| 8_8540370' | 1 | 0 | 4 | 0.5 | 2 | 0.5 | 1 | 1 | 8 | 0.5 | 5 | 1 | NA | NA | 3 | 1 | 3 | 1 | 11 | 1 |
| 8_8540379: | 1 | 0 | 3 | 1 | 2 | 1 | 1 | 1 | 7 | 0.8571 | 6 | 0.8333 | 2 | 0.5 | 3 | 1 | 3 | 0.6667 | 14 | 0.7857 |
| 8_8541748: | 4 | 0.25 | 4 | 0 | 1 | 0 | 3 | 0.3333 | 12 | 0.1667 | 6 | 0 | 1 | 0 | 3 | 0 | 2 | 0.5 | 12 | 0.0833 |
| 8_8541755( | 3 | 0.3333 | 4 | 0 | 3 | 0 | 3 | 0 | 13 | 0.0769 | 7 | 0.1429 | NA | NA | 3 | 0 | 3 | 0.3333 | 13 | 0.1538 |
| 8_8542616( | NA | NA | NA | NA | NA | NA | 1 | 1 | 1 | 1 | 1 | 1 | NA | NA | 2 | 0 | NA | NA | 3 | 0.3333 |
| 8_8542634! | 6 | 0 | 5 | 0 | 4 | 0 | 4 | 0.25 | 19 | 0.0526 | 10 | 0 | NA | NA | 4 | 0 | 4 | 0.25 | 18 | 0.0556 |
| 8_8543086( | NA | NA | 1 | 1 | 1 | 1 | NA | NA | 2 | 1 | NA | NA | NA | NA | NA | NA | 1 | 1 | 1 | 1 |
| 8_8543105! | NA | NA | 1 | 0 | 1 | 1 | NA | NA | 2 | 0.5 | NA | NA | NA | NA | NA | NA | 1 | 1 | 1 | 1 |
| 8_8543193' | 9 | 0.1111 | 11 | 0 | 8 | 0.125 | 6 | 0.1667 | 34 | 0.0882 | 11 | 0.0909 | 8 | 0.25 | 8 | 0.125 | 9 | 0 | 36 | 0.1111 |
| 8_8543408' | 5 | 0 | 4 | 0.25 | 4 | 0 | 4 | 0 | 17 | 0.0588 | 6 | 0.1667 | 2 | 0 | 4 | 0 | 3 | 0 | 15 | 0.0667 |
| 8_8543426! | 5 | 0.6 | 3 | 0.6667 | 3 | 0.6667 | 4 | 1 | 15 | 0.7333 | 5 | 0.2 | 2 | 1 | 5 | 0.8 | 3 | 0 | 15 | 0.4667 |
| 8_8543454! | 5 | 0.6 | 3 | 0.3333 | 3 | 0.6667 | 3 | 1 | 14 | 0.6429 | 2 | 0.5 | 2 | 0 | 2 | 0.5 | 2 | 0 | 8 | 0.25 |
| 8_8543471' | 4 | 0.25 | 3 | 0.3333 | 4 | 0 | 3 | 0 | 14 | 0.1429 | 5 | 0.4 | 2 | 0 | 2 | 0.5 | 3 | 0 | 12 | 0.25 |
| 8_8543480! | 5 | 0.8 | 3 | 1 | 4 | 0.75 | 4 | 1 | 16 | 0.875 | 4 | 0.75 | 2 | 0.5 | 4 | 1 | 3 | 1 | 13 | 0.8462 |
| 8_8543482! | 5 | 0 | 3 | 0 | 3 | 0.3333 | 4 | 0.25 | 15 | 0.1333 | 4 | 0.25 | 2 | 0 | 3 | 0 | 3 | 0 | 12 | 0.0833 |
| 8_8543487 | 4 | 1 | 3 | 1 | 4 | 1 | 4 | 0.5 | 15 | 0.8667 | 5 | 0.8 | 2 | 0.5 | 4 | 0.75 | 2 | 1 | 13 | 0.7692 |
| 8_8543497! | 4 | 1 | 3 | 1 | 4 | 1 | 2 | 1 | 13 | 1 | 5 | 1 | 1 | 1 | 3 | 1 | 3 | 1 | 12 | 1 |
| 8_8543500: | 5 | 0.2 | 2 | 0.5 | 4 | 0.5 | 4 | 0.25 | 15 | 0.3333 | 4 | 0 | 2 | 0.5 | 4 | 0.5 | 3 | 0.3333 | 13 | 0.3077 |
| 8_8543501! | 5 | 1 | 3 | 1 | 4 | 0.75 | 3 | 1 | 15 | 0.9333 | 5 | 0.6 | 2 | 0.5 | 4 | 1 | 3 | 1 | 14 | 0.7857 |
| 8_8543516( | 5 | 0.2 | 3 | 0 | 3 | 0 | 4 | 0 | 15 | 0.0667 | 5 | 0 | 2 | 0.5 | 4 | 0 | 3 | 0 | 14 | 0.0714 |
| 8_8549845( | 3 | 0.3333 | 2 | 0 | 2 | 1 | 3 | 0.3333 | 10 | 0.4 | 3 | 0 | 1 | 0 | 2 | 0.5 | 2 | 0.5 | 8 | 0.25 |
| 8_8549858: | 3 | 0.3333 | 2 | 0.5 | 3 | 0.3333 | 4 | 0.5 | 12 | 0.4167 | 3 | 0.6667 | 1 | 0 | 2 | 0 | 2 | 0 | 8 | 0.25 |
| 8_8549870: | 3 | 0.6667 | 2 | 1 | 3 | 1 | 4 | 0.5 | 12 | 0.75 | 2 | 1 | 1 | 1 | 2 | 1 | 2 | 1 | 7 | 1 |
| 8_8554222: | 3 | 0.3333 | 5 | 0.4 | 1 | 0 | NA | NA | 9 | 0.3333 | 1 | 0 | 2 | 0 | 1 | 1 | 3 | 1 | 7 | 0.5714 |
| 8_8554226' | 3 | 0 | 4 | 0 | 2 | 0.5 | 1 | 0 | 10 | 0.1 | 1 | 0 | 2 | 0 | 1 | 0 | 3 | 0.3333 | 7 | 0.1429 |
| 8_8554229' | 2 | 1 | 3 | 1 | NA | NA | NA | NA | 5 | 1 | 1 | 1 | 2 | 1 | 1 | 1 | 2 | 1 | 6 | 1 |

| | | | | | | | | | | | | | | | | | | | | |
|---|---|---|---|---|---|---|---|---|---|---|---|---|---|---|---|---|---|---|---|---|
| 8_85542802 | 2 | 0.5 | 2 | 0 | 2 | 0 | 1 | 0 | 7 | 0.1429 | 2 | 0.5 | 2 | 1 | 1 | 0 | 3 | 0.3333 | 8 | 0.5 |
| 8_85542879 | 3 | 0.3333 | 3 | 0 | 2 | 1 | 1 | 0 | 9 | 0.3333 | 3 | 0.6667 | 2 | 0 | 1 | 0 | 4 | 0.75 | 10 | 0.5 |
| 8_85542889 | 3 | 0.3333 | 4 | 0.5 | 1 | 0 | 1 | 1 | 9 | 0.4444 | 3 | 0.6667 | 2 | 0 | 1 | 0 | 4 | 0.5 | 10 | 0.4 |
| 8_85559413 | 1 | 0 | NA | NA | NA | NA | 1 | 1 | 2 | 0.5 | NA | NA | 2 | 1 | NA | NA | NA | NA | 2 | 1 |
| 8_85560892 | 5 | 0.4 | 8 | 0.125 | 5 | 0 | 2 | 0 | 20 | 0.15 | 5 | 0.2 | 3 | 0 | 6 | 0.1667 | 4 | 0 | 18 | 0.1111 |
| 8_85560917 | 6 | 0.3333 | 6 | 0.5 | 2 | 0.5 | 4 | 0 | 18 | 0.3333 | 6 | 0.1667 | 2 | 0 | 6 | 0.1667 | 4 | 0.25 | 18 | 0.1667 |
| 8_85560928 | 7 | 0.1429 | 8 | 0.375 | 5 | 0.4 | 4 | 0.5 | 24 | 0.3333 | 6 | 0.5 | 2 | 0.5 | 7 | 0.2857 | 5 | 0.2 | 20 | 0.35 |
| 8_85561024 | 7 | 0.4286 | 9 | 0.5556 | 5 | 1 | 4 | 0.75 | 25 | 0.64 | 7 | 1 | 3 | 0.6667 | 8 | 0.5 | 7 | 0.2857 | 25 | 0.6 |
| 8_85561122 | 2 | 1 | 1 | 0 | 1 | 1 | 1 | 1 | 5 | 0.8 | 2 | 0.5 | 5 | 0.4 | 1 | 1 | NA | NA | 8 | 0.5 |
| 8_85561197 | 3 | 0 | 1 | 0 | 1 | 1 | NA | NA | 5 | 0.2 | 2 | 0.5 | 3 | 0.6667 | 1 | 0 | NA | NA | 6 | 0.5 |
| 8_85561216 | 2 | 1 | 1 | 1 | NA | NA | NA | NA | 3 | 1 | 2 | 1 | 5 | 1 | 1 | 1 | NA | NA | 8 | 1 |
| 8_85561355 | 2 | 0 | 1 | 0 | 3 | 0.6667 | 1 | 0 | 7 | 0.2857 | 2 | 0.5 | 3 | 0 | 1 | 0 | 1 | 0 | 7 | 0.1429 |
| 8_85561549 | 3 | 0.3333 | 2 | 0 | 3 | 0 | 1 | 0 | 9 | 0.1111 | 2 | 0 | 4 | 0.25 | 1 | 0 | 1 | 0 | 8 | 0.125 |
| 8_85561989 | 4 | 1 | 2 | 0.5 | 5 | 1 | 1 | 1 | 12 | 0.9167 | 2 | 0.5 | 7 | 1 | 1 | 1 | 2 | 1 | 12 | 0.9167 |
| 8_85564761 | 8 | 0 | 7 | 0 | 10 | 0 | 7 | 0.2857 | 32 | 0.0625 | 5 | 0.2 | 8 | 0.125 | 7 | 0 | 10 | 0.1 | 30 | 0.1 |
| 8_85568816 | 63 | 0.2698 | 74 | 0.2568 | 54 | 0.1296 | 55 | 0.1818 | 246 | 0.2154 | 67 | 0.1791 | 65 | 0.2154 | 66 | 0.1212 | 66 | 0.1667 | 264 | 0.1705 |
| 8_85569182 | 54 | 0.9074 | 70 | 0.8571 | 44 | 0.9091 | 61 | 0.9016 | 229 | 0.8908 | 60 | 0.8833 | 57 | 0.9474 | 61 | 0.9016 | 58 | 0.931 | 236 | 0.9153 |
| 8_85569190 | 55 | 0.0182 | 75 | 0.08 | 51 | 0.098 | 62 | 0.0645 | 243 | 0.0658 | 64 | 0.1562 | 59 | 0.0508 | 64 | 0.0625 | 57 | 0.1579 | 244 | 0.1066 |
| 8_85569247 | 48 | 0.5833 | 71 | 0.6056 | 45 | 0.6 | 61 | 0.5902 | 225 | 0.5956 | 58 | 0.5172 | 61 | 0.5902 | 57 | 0.5263 | 62 | 0.6452 | 238 | 0.5714 |
| 8_85613021 | 1 | 0 | 2 | 0.5 | 1 | 0 | 2 | 0.5 | 6 | 0.3333 | NA | NA | 2 | 0.5 | NA | NA | NA | NA | 2 | 0.5 |
| 8_85613306 | 7 | 0.4286 | 6 | 0.3333 | 8 | 0.875 | 7 | 0.4286 | 28 | 0.5357 | 4 | 0 | 8 | 0.5 | 2 | 1 | 2 | 0.5 | 16 | 0.4375 |
| 8_85613377 | 7 | 0.4286 | 7 | 0.4286 | 8 | 0.75 | 7 | 0.5714 | 29 | 0.5517 | 4 | 0.25 | 8 | 0.375 | 2 | 0.5 | 2 | 0 | 16 | 0.3125 |
| 8_85613489 | 8 | 0 | 6 | 0.1667 | 7 | 0.1429 | 7 | 0 | 28 | 0.0714 | 3 | 0.3333 | 6 | 0.3333 | 1 | 0 | 3 | 0 | 13 | 0.2308 |
| 8_85613506 | 6 | 0.6667 | 6 | 0.8333 | 9 | 0.6667 | 6 | 0.8333 | 27 | 0.7407 | 4 | 0.25 | 7 | 0.5714 | 1 | 1 | 3 | 0.6667 | 15 | 0.5333 |
| 8_85613552 | 7 | 0.4286 | 5 | 0.4 | 7 | 0.8571 | 6 | 0.6667 | 25 | 0.6 | 4 | 0 | 7 | 0.7143 | 1 | 1 | 3 | 1 | 15 | 0.6 |
| 8_85617743 | 3 | 0 | 4 | 0 | 5 | 0.4 | 5 | 0 | 17 | 0.1176 | 4 | 0 | 4 | 0 | NA | NA | 2 | 0.5 | 10 | 0.1 |
| 8_85626577 | 1 | 0 | 1 | 1 | 1 | 1 | 3 | 0.3333 | 6 | 0.5 | 2 | 0 | 1 | 1 | NA | NA | 2 | 0.5 | 5 | 0.4 |
| 8_85626662 | 1 | 1 | 1 | 1 | 1 | 0 | 3 | 0.6667 | 6 | 0.6667 | 2 | 0.5 | 1 | 1 | NA | NA | 2 | 1 | 5 | 0.8 |
| 8_85638671 | 4 | 0 | 11 | 0.0909 | 7 | 0.2857 | 3 | 0.3333 | 25 | 0.16 | 3 | 0 | 5 | 0 | 6 | 0.1667 | 4 | 0 | 18 | 0.0556 |
| 8_85638697 | 4 | 0 | 9 | 0.3333 | 6 | 0.5 | 2 | 0 | 21 | 0.2857 | 3 | 0.3333 | 4 | 0.25 | 4 | 0.5 | 4 | 0.5 | 15 | 0.4 |
| 8_85638819 | 3 | 0 | 9 | 0.1111 | 7 | 0.1429 | 2 | 0 | 21 | 0.0952 | 2 | 0 | 2 | 0.5 | 3 | 0.3333 | 4 | 0.25 | 11 | 0.2727 |
| 8_85638846 | 4 | 0.5 | 9 | 0.7778 | 5 | 0.4 | 3 | 0.6667 | 21 | 0.619 | 3 | 0.3333 | 4 | 0.75 | 5 | 0.4 | 3 | 0.6667 | 15 | 0.5333 |
| 8_85638939 | 4 | 0 | 11 | 0.2727 | 7 | 0.1429 | 3 | 0 | 25 | 0.16 | 3 | 0 | 4 | 0.25 | 5 | 0.2 | 3 | 0 | 15 | 0.1333 |
| 8_85638958 | 4 | 0.25 | 11 | 0.5455 | 7 | 0.1429 | 3 | 0.3333 | 25 | 0.36 | 3 | 0 | 3 | 0.3333 | 4 | 0.25 | 3 | 0.3333 | 13 | 0.2308 |
| 8_85639067 | 2 | 0 | 7 | 0.1429 | 7 | 0.1429 | 2 | 0 | 18 | 0.1111 | 2 | 0 | 3 | 0 | 4 | 0.25 | 3 | 0.3333 | 12 | 0.1667 |
| 8_85639074 | 4 | 0.5 | 5 | 0.4 | 6 | 0.3333 | 3 | 0 | 18 | 0.3333 | 2 | 0.5 | 2 | 0 | 3 | 0.3333 | 1 | 1 | 8 | 0.375 |
| 8_85639139 | 3 | 0.6667 | 7 | 0.8571 | 6 | 0.6667 | 2 | 0.5 | 18 | 0.7222 | 1 | 0 | 4 | 1 | 5 | 0.6 | 3 | 0.3333 | 13 | 0.6154 |
| 8_85639184 | 4 | 0.25 | 10 | 0.5 | 6 | 0.3333 | 3 | 0 | 23 | 0.3478 | 1 | 0 | 4 | 0 | 5 | 0.2 | 3 | 0 | 13 | 0.0769 |
| 8_85639266 | 3 | 0.6667 | 11 | 0.4545 | 7 | 0.2857 | 3 | 0.6667 | 24 | 0.4583 | 2 | 0.5 | 4 | 0.75 | 5 | 0.4 | 3 | 0.3333 | 14 | 0.5 |
| 8_85639294 | 3 | 0.3333 | 11 | 0.5455 | 7 | 0.4286 | 4 | 0 | 25 | 0.4 | 1 | 1 | 4 | 0.5 | 5 | 0 | 3 | 0.6667 | 13 | 0.3846 |
| 8_85639432 | 2 | 1 | 10 | 1 | 5 | 1 | 2 | 1 | 19 | 1 | 2 | 1 | 4 | 1 | 4 | 1 | 3 | 1 | 13 | 1 |
| 8_85639452 | 2 | 1 | 10 | 1 | 5 | 0.6 | 3 | 0.6667 | 20 | 0.85 | 2 | 1 | 4 | 0.75 | 3 | 0.6667 | 3 | 0.3333 | 12 | 0.6667 |
| 8_85639515 | 1 | 0 | 7 | 0.1429 | 4 | 0.25 | 3 | 0.3333 | 15 | 0.2 | 1 | 0 | 3 | 0 | 3 | 0.3333 | 1 | 0 | 8 | 0.125 |
| 8_85649207 | NA | NA | NA | NA | NA | NA | 1 | 1 | 1 | 1 | NA | NA | 1 | 1 | NA | NA | NA | NA | 1 | 1 |
| 8_85683652 | 1 | 0 | 1 | 1 | NA | NA | NA | NA | 2 | 0.5 | 1 | 0 | 2 | 0.5 | 1 | 0 | NA | NA | 4 | 0.25 |
| 8_85684291 | NA | NA | 1 | 1 | NA | NA | NA | NA | 1 | 1 | NA | NA | 1 | 0 | 1 | 1 | NA | NA | 2 | 0.5 |
| 8_85684488 | 3 | 0.3333 | 4 | 0.25 | 2 | 0 | 5 | 0.2 | 14 | 0.2143 | 3 | 0 | 4 | 0.25 | 1 | 1 | 4 | 0.5 | 12 | 0.3333 |
| 8_85686434 | 1 | 0 | 3 | 0.3333 | 3 | 0.3333 | 5 | 0 | 12 | 0.1667 | NA | NA | 4 | 0 | 2 | 0 | 3 | 0.3333 | 9 | 0.1111 |
| 8_85701005 | 1 | 0 | 3 | 1 | 4 | 0.75 | NA | NA | 8 | 0.75 | NA | NA | NA | NA | NA | NA | 1 | 1 | 1 | 1 |
| 8_85701078 | 1 | 0 | 1 | 1 | 4 | 0.25 | 1 | 0 | 7 | 0.2857 | NA | NA | NA | NA | 1 | 1 | 1 | 0 | 2 | 0.5 |
| 8_85703805 | 7 | 1 | 13 | 1 | 6 | 1 | 7 | 1 | 33 | 1 | 6 | 1 | 10 | 1 | 16 | 0.9375 | 4 | 1 | 36 | 0.9722 |
| 8_85703900 | 6 | 1 | 9 | 1 | 5 | 0.8 | 8 | 0.875 | 28 | 0.9286 | 4 | 1 | 10 | 1 | 15 | 0.9333 | 6 | 1 | 35 | 0.9714 |
| 8_85704001 | 3 | 1 | 6 | 1 | 4 | 1 | 6 | 1 | 19 | 1 | 4 | 1 | 7 | 1 | 9 | 0.8889 | 5 | 1 | 25 | 0.96 |
| 8_85704036 | 7 | 1 | 12 | 0.8333 | 6 | 0.8333 | 9 | 0.6667 | 34 | 0.8235 | 5 | 0.6 | 10 | 0.7 | 16 | 0.5625 | 7 | 0.8571 | 38 | 0.6579 |
| 8_85704175 | 8 | 0 | 10 | 0.1 | 5 | 0.2 | 8 | 0.125 | 31 | 0.0968 | 4 | 0 | 9 | 0.4444 | 17 | 0.1176 | 7 | 0.1429 | 37 | 0.1892 |
| 8_85704458 | 7 | 0.4286 | 8 | 0.25 | 4 | 0.25 | 7 | 0.4286 | 26 | 0.3462 | 5 | 0 | 6 | 0 | 14 | 0.2143 | 7 | 0 | 32 | 0.0938 |
| 8_85704738 | 7 | 1 | 10 | 0.8 | 5 | 1 | 3 | 1 | 25 | 0.92 | 5 | 0.8 | 8 | 0.75 | 11 | 0.9091 | 5 | 0.8 | 29 | 0.8276 |
| 8_85704960 | 5 | 0.2 | 8 | 0.25 | 2 | 0.5 | 4 | 0 | 19 | 0.2105 | 5 | 0.2 | 6 | 0.1667 | 8 | 0.125 | 7 | 0.4286 | 26 | 0.2308 |
| 8_85705041 | 6 | 0 | 7 | 0 | NA | NA | 5 | 0.4 | 18 | 0.1111 | 4 | 0 | 7 | 0.1429 | 6 | 0 | 7 | 0.1429 | 24 | 0.0833 |
| 8_85705087 | 5 | 0.2 | 9 | 0.1111 | 3 | 0 | 6 | 0 | 23 | 0.087 | 5 | 0 | 8 | 0.125 | 3 | 0 | 7 | 0.1429 | 23 | 0.087 |
| 8_85729782 | NA | NA | NA | NA | NA | NA | 1 | 1 | 1 | 1 | NA | NA | 1 | 1 | 3 | 0.3333 | 1 | 0 | 5 | 0.4 |
| 8_85753685 | 14 | 0 | 26 | 0.0385 | 16 | 0.125 | 16 | 0.125 | 72 | 0.0694 | 20 | 0 | 16 | 0.125 | 14 | 0.0714 | 9 | 0.1111 | 59 | 0.0678 |
| 8_85756611 | 11 | 0.0909 | 26 | 0.0769 | 17 | 0.0588 | 17 | 0.0588 | 71 | 0.0704 | 24 | 0.0833 | 16 | 0 | 17 | 0.1176 | 14 | 0.0714 | 71 | 0.0704 |
| 8_85756639 | 15 | 0.6667 | 26 | 0.6538 | 14 | 0.7143 | 17 | 0.3529 | 72 | 0.5972 | 19 | 0.3684 | 19 | 0.5789 | 15 | 0.4 | 12 | 0.1667 | 65 | 0.4 |
| 8_85756696 | 12 | 0.8333 | 21 | 0.7619 | 14 | 0.8571 | 17 | 0.7647 | 64 | 0.7969 | 21 | 0.7143 | 15 | 0.6667 | 16 | 0.875 | 14 | 0.7143 | 66 | 0.7424 |

| | | | | | | | | | | | | | | | | | | | | |
|---|---|---|---|---|---|---|---|---|---|---|---|---|---|---|---|---|---|---|---|---|
| 8_8575675 | 14 | 0.7143 | 26 | 0.7692 | 19 | 0.8421 | 16 | 0.625 | 75 | 0.7467 | 22 | 0.8182 | 18 | 0.5556 | 17 | 0.6471 | 15 | 0.8667 | 72 | 0.7222 |
| 8_8575678 | 16 | 0.25 | 25 | 0.24 | 17 | 0.4118 | 18 | 0.3889 | 76 | 0.3158 | 22 | 0.3182 | 18 | 0.1667 | 17 | 0.3529 | 16 | 0.125 | 73 | 0.2466 |
| 8_8578058 | 1 | 1 | NA | NA | 1 | 0 | NA | NA | 2 | 0.5 | 1 | 1 | 1 | 0 | 1 | 0 | NA | NA | 3 | 0.3333 |
| 8_8578062 | 1 | 1 | NA | NA | 1 | 0 | NA | NA | 2 | 0.5 | 1 | 1 | 1 | 1 | 2 | 0.5 | NA | NA | 4 | 0.75 |
| 8_8578067 | 1 | 1 | NA | NA | NA | NA | NA | NA | 1 | 1 | 1 | 1 | 1 | 0 | 2 | 0.5 | NA | NA | 4 | 0.5 |
| 8_8578070 | 1 | 1 | NA | NA | 1 | 0 | NA | NA | 2 | 0.5 | 1 | 0 | 1 | 1 | 2 | 0.5 | NA | NA | 4 | 0.5 |
| 8_8578085 | 1 | 1 | NA | NA | 1 | 1 | NA | NA | 2 | 1 | 1 | 0 | 1 | 1 | 2 | 0 | NA | NA | 4 | 0.25 |
| 8_8578326 | 9 | 0.2222 | 3 | 0.6667 | 6 | 0.1667 | 5 | 0.4 | 23 | 0.3043 | 3 | 0.6667 | 7 | 0.4286 | 2 | 0 | 4 | 0 | 16 | 0.3125 |
| 8_8578329 | 8 | 0.25 | 1 | 0 | 4 | 0.5 | 2 | 0 | 15 | 0.2667 | 2 | 0 | 5 | 0 | NA | NA | 8 | 0.125 | 15 | 0.0667 |
| 8_8578354 | 9 | 0 | 8 | 0.125 | 10 | 0.1 | 8 | 0 | 35 | 0.0571 | 8 | 0.125 | 10 | 0.1 | 2 | 0 | 11 | 0.0909 | 31 | 0.0968 |
| 8_8578403 | 8 | 0.375 | 13 | 0.4615 | 8 | 0.25 | 11 | 0.2727 | 40 | 0.35 | 12 | 0.4167 | 13 | 0.2308 | 8 | 0.375 | 14 | 0.3571 | 47 | 0.3404 |
| 8_8579333 | NA | NA | 1 | 1 | 1 | 0 | 4 | 0.25 | 6 | 0.3333 | NA | NA | 5 | 0.4 | 1 | 1 | 1 | 0 | 7 | 0.4286 |
| 8_8579821 | NA | NA | NA | NA | 1 | 1 | NA | NA | 1 | 1 | NA | NA | NA | NA | NA | NA | 1 | 1 | 1 | 1 |
| 8_8579939 | NA | NA | 1 | 0 | 3 | 0.3333 | NA | NA | 4 | 0.25 | 1 | 1 | 2 | 0.5 | 1 | 0 | 1 | 0 | 5 | 0.4 |
| 8_8580167 | 4 | 0.25 | 4 | 0.25 | 12 | 0.0833 | 7 | 0 | 27 | 0.1111 | 3 | 0 | 6 | 0.3333 | 6 | 0 | 8 | 0.125 | 23 | 0.1304 |
| 8_8580178 | 1 | 1 | 3 | 0 | 5 | 0 | 4 | 0 | 13 | 0.0769 | 6 | 0 | 5 | 0 | 2 | 0 | 4 | 0.25 | 17 | 0.0588 |
| 8_8580181 | 1 | 0 | 4 | 0.25 | 5 | 0 | 4 | 0 | 14 | 0.0714 | 6 | 0.1667 | 5 | 0.2 | 3 | 0 | 3 | 0.3333 | 17 | 0.1765 |
| 8_8580726 | NA | NA | 2 | 0.5 | 3 | 0 | 2 | 0 | 7 | 0.1429 | 4 | 0 | 5 | 0.2 | 2 | 0.5 | NA | NA | 11 | 0.1818 |
| 8_8580776 | 4 | 0.25 | 2 | 0 | 3 | 0 | 3 | 0.6667 | 12 | 0.25 | 1 | 0 | 3 | 0.3333 | 4 | 0 | NA | NA | 8 | 0.125 |
| 8_8580866 | 16 | 0.1875 | 13 | 0.1538 | 9 | 0.2222 | 12 | 0.25 | 50 | 0.2 | 6 | 0.1667 | 14 | 0.3571 | 18 | 0.2222 | 6 | 0.3333 | 44 | 0.2727 |
| 8_8581210 | 1 | 1 | 2 | 0.5 | 2 | 0 | 3 | 0 | 8 | 0.25 | 1 | 0 | 2 | 0.5 | 2 | 1 | 1 | 0 | 6 | 0.5 |
| 8_8583607 | NA | NA | 1 | 0 | 1 | 1 | NA | NA | 2 | 0.5 | 1 | 1 | NA | NA | NA | NA | NA | NA | 1 | 1 |
| 8_8583621 | NA | NA | 1 | 0 | 2 | 0.5 | NA | NA | 3 | 0.3333 | 1 | 1 | NA | NA | NA | NA | NA | NA | 1 | 1 |
| 8_8583624 | NA | NA | 1 | 1 | 2 | 1 | NA | NA | 3 | 1 | 1 | 1 | NA | NA | NA | NA | NA | NA | 1 | 1 |
| 8_8583628 | NA | NA | 1 | 1 | 2 | 1 | NA | NA | 3 | 1 | 1 | 1 | NA | NA | NA | NA | NA | NA | 1 | 1 |
| 8_8583629 | NA | NA | 1 | 1 | 2 | 1 | NA | NA | 3 | 1 | 1 | 1 | NA | NA | NA | NA | NA | NA | 1 | 1 |
| 8_8583650 | NA | NA | 1 | 1 | 1 | 1 | NA | NA | 2 | 1 | 1 | 1 | NA | NA | NA | NA | NA | NA | 1 | 1 |
| 8_8583675 | NA | NA | 1 | 1 | 1 | 1 | NA | NA | 2 | 1 | 1 | 1 | NA | NA | NA | NA | NA | NA | 1 | 1 |
| 8_8583687 | NA | NA | 1 | 1 | 1 | 1 | NA | NA | 2 | 1 | 1 | 1 | NA | NA | NA | NA | NA | NA | 1 | 1 |
| 8_8583747 | NA | NA | 1 | 1 | NA | NA | NA | NA | 1 | 1 | 1 | 1 | NA | NA | NA | NA | NA | NA | 1 | 1 |
| 8_8583765 | NA | NA | 1 | 1 | 1 | 1 | NA | NA | 2 | 1 | 1 | 1 | NA | NA | NA | NA | NA | NA | 1 | 1 |
| 8_8598718 | 24 | 0.1667 | 30 | 0.4 | 17 | 0.1765 | 28 | 0.2857 | 99 | 0.2727 | 16 | 0.1875 | 17 | 0.2353 | 31 | 0.3871 | 20 | 0.55 | 84 | 0.3571 |
| 8_8598720 | 22 | 0.6818 | 32 | 0.7188 | 16 | 0.625 | 27 | 0.7407 | 97 | 0.701 | 15 | 0.7333 | 16 | 0.625 | 26 | 0.7308 | 22 | 0.6818 | 79 | 0.6962 |
| 8_8598755 | 20 | 0 | 31 | 0.0645 | 11 | 0.1818 | 26 | 0.1538 | 88 | 0.0909 | 15 | 0.0667 | 13 | 0.1538 | 24 | 0.0417 | 16 | 0.125 | 68 | 0.0882 |
| 8_8598760 | 20 | 0.2 | 30 | 0.2 | 13 | 0.1538 | 28 | 0.3214 | 91 | 0.2308 | 13 | 0.0769 | 14 | 0.2143 | 28 | 0.2857 | 18 | 0.3333 | 73 | 0.2466 |
| 8_8599005 | 13 | 0 | 25 | 0.16 | 11 | 0.0909 | 19 | 0 | 68 | 0.0735 | 12 | 0.0833 | 10 | 0 | 24 | 0 | 13 | 0.1538 | 59 | 0.0508 |
| 8_8603447 | 3 | 1 | 5 | 0.8 | 1 | 1 | 6 | 0.8333 | 15 | 0.8667 | 4 | 1 | 5 | 1 | 3 | 0.6667 | 2 | 1 | 14 | 0.9286 |
| 8_8603450 | 4 | 0.25 | 6 | 0.3333 | 1 | 0 | 6 | 0.1667 | 17 | 0.2353 | 4 | 0.75 | 6 | 0 | 4 | 0 | 2 | 0 | 16 | 0.1875 |
| 8_8603453 | 4 | 0.25 | 5 | 0 | 1 | 0 | 7 | 0.1429 | 17 | 0.1176 | 3 | 0 | 6 | 0.1667 | 3 | 0 | 2 | 0.5 | 14 | 0.1429 |
| 8_8603460 | 3 | 0 | 6 | 0 | 1 | 0 | 7 | 0.2857 | 17 | 0.1176 | 3 | 0.3333 | 6 | 0 | 4 | 0 | 2 | 0 | 15 | 0.0667 |
| 8_8613609 | 3 | 0.3333 | 1 | 0 | 3 | 0 | 6 | 0 | 13 | 0.0769 | 3 | 0 | 3 | 0 | 4 | 0 | 1 | 1 | 11 | 0.0909 |
| 8_8613615 | 4 | 0.5 | 1 | 0 | 3 | 0 | 6 | 0.1667 | 14 | 0.2143 | 3 | 0.6667 | 4 | 0.25 | 4 | 0 | 3 | 0 | 14 | 0.2143 |
| 8_8613623 | 4 | 0.5 | 1 | 0 | 2 | 0 | 5 | 0.6 | 12 | 0.4167 | 3 | 0.3333 | 4 | 0.5 | 4 | 0 | 3 | 0.3333 | 14 | 0.2857 |
| 8_8613631 | 3 | 0.6667 | 1 | 1 | 3 | 0.6667 | 4 | 0.25 | 11 | 0.5455 | 3 | 0.3333 | 3 | 1 | 4 | 0.5 | 3 | 0 | 13 | 0.4615 |
| 8_8613632 | 4 | 1 | NA | NA | 3 | 1 | 5 | 0.6 | 12 | 0.8333 | 2 | 0.5 | 4 | 0.5 | 3 | 0 | 2 | 1 | 11 | 0.4545 |
| 8_8613638 | 3 | 0.3333 | 1 | 0 | 1 | 0 | 5 | 0.2 | 10 | 0.2 | 2 | 0 | 4 | 0.25 | 3 | 0.3333 | 2 | 0 | 11 | 0.1818 |
| 8_8613641 | 3 | 1 | 1 | 1 | 1 | 1 | 5 | 1 | 10 | 1 | 3 | 1 | 4 | 1 | 4 | 0.75 | 2 | 1 | 13 | 0.9231 |
| 8_8621577 | 1 | 1 | 1 | 1 | NA | NA | NA | NA | 2 | 1 | 1 | 1 | NA | NA | 1 | 1 | 1 | 1 | 3 | 1 |
| 8_8621736 | 3 | 0 | 2 | 0.5 | 2 | 0.5 | NA | NA | 7 | 0.2857 | 2 | 0.5 | NA | NA | NA | NA | 1 | 0 | 3 | 0.3333 |
| 8_8621738 | 2 | 0.5 | 2 | 0.5 | 2 | 0.5 | NA | NA | 6 | 0.5 | 1 | 1 | NA | NA | NA | NA | 1 | 1 | 2 | 1 |
| 8_8621739 | 2 | 0.5 | 1 | 1 | 2 | 0 | NA | NA | 5 | 0.4 | 2 | 1 | NA | NA | NA | NA | 1 | 0 | 3 | 0.6667 |
| 8_8621741 | 2 | 0 | 2 | 0.5 | 1 | 0 | NA | NA | 5 | 0.2 | 2 | 0.5 | NA | NA | NA | NA | 1 | 1 | 3 | 0.6667 |
| 8_8621742 | 2 | 0 | 2 | 0 | 2 | 0.5 | NA | NA | 6 | 0.1667 | 2 | 0.5 | NA | NA | NA | NA | 1 | 0 | 3 | 0.3333 |
| 8_8621804 | 2 | 0 | 2 | 0 | 2 | 0.5 | NA | NA | 6 | 0.1667 | 3 | 0.6667 | 1 | 0 | NA | NA | NA | NA | 4 | 0.5 |
| 8_8621853 | 1 | 0 | 2 | 0 | 1 | 1 | 1 | 0 | 5 | 0.2 | 3 | 0.6667 | 1 | 0 | NA | NA | 1 | 0 | 5 | 0.4 |
| 8_8621906 | 1 | 0 | NA | NA | 1 | 1 | NA | NA | 2 | 0.5 | 1 | 0 | 1 | 1 | NA | NA | 1 | 1 | 3 | 0.6667 |
| 8_8621915 | 1 | 1 | NA | NA | 1 | 1 | NA | NA | 2 | 1 | 1 | 1 | 1 | 1 | NA | NA | 1 | 0 | 3 | 0.6667 |
| 8_8621940 | 1 | 1 | NA | NA | 1 | 1 | NA | NA | 2 | 1 | 1 | 1 | 1 | 1 | NA | NA | 1 | 1 | 3 | 1 |
| 8_8621948 | 1 | 0 | NA | NA | 1 | 1 | NA | NA | 2 | 0.5 | 1 | 1 | 1 | 0 | NA | NA | 1 | 0 | 3 | 0.3333 |
| 8_8621955 | 1 | 0 | NA | NA | 1 | 1 | NA | NA | 2 | 0.5 | 1 | 0 | 1 | 0 | NA | NA | 1 | 1 | 3 | 0.3333 |
| 8_8621970 | 1 | 1 | NA | NA | 1 | 1 | NA | NA | 2 | 1 | 1 | 0 | NA | NA | NA | NA | 1 | 1 | 2 | 0.5 |
| 8_8625231 | 4 | 0.75 | 5 | 0.8 | 6 | 0.8333 | 7 | 1 | 22 | 0.8636 | 14 | 0.9286 | 2 | 0.5 | 9 | 0.4444 | 6 | 1 | 31 | 0.7742 |
| 8_8625237 | 4 | 0 | 5 | 0.2 | 6 | 0 | 6 | 0.1667 | 21 | 0.0952 | 13 | 0.2308 | 2 | 0.5 | 8 | 0.125 | 5 | 0.6 | 28 | 0.2857 |
| 8_8625264 | 6 | 0.1667 | 3 | 0.6667 | 5 | 0 | 7 | 0.1429 | 21 | 0.1905 | 16 | 0.25 | 3 | 0 | 9 | 0.3333 | 7 | 0.1429 | 35 | 0.2286 |
| 8_8625273 | 3 | 0 | 5 | 0.4 | 3 | 0.3333 | 6 | 0.3333 | 17 | 0.2941 | 16 | 0.1875 | 2 | 0 | 8 | 0.25 | 4 | 0.25 | 30 | 0.2 |

| | | | | | | | | | | | | | | | | | | | | |
|---|---|---|---|---|---|---|---|---|---|---|---|---|---|---|---|---|---|---|---|---|
| 8_8625314 | 7 | 0.1429 | 4 | 0 | 6 | 0.1667 | 7 | 0 | 24 | 0.0833 | 16 | 0.125 | 3 | 0.3333 | 10 | 0.1 | 7 | 0 | 36 | 0.1111 |
| 8_8625404 | 6 | 0.3333 | 7 | 0.4286 | 4 | 0.25 | 7 | 0.2857 | 24 | 0.3333 | 14 | 0.2857 | 4 | 0.25 | 9 | 0.4444 | 6 | 0.3333 | 33 | 0.3333 |
| 8_8626484 | 7 | 0.1429 | 7 | 0.1429 | 5 | 0.4 | 4 | 0 | 23 | 0.1739 | 4 | 0.5 | 4 | 0.25 | 5 | 0.6 | 4 | 0.75 | 17 | 0.5294 |
| 8_8626521 | 12 | 0.1667 | 12 | 0.0833 | 6 | 0 | 9 | 0 | 39 | 0.0769 | 5 | 0 | 11 | 0 | 9 | 0 | 8 | 0.25 | 33 | 0.0606 |
| 8_8626562 | 29 | 0.069 | 33 | 0.2424 | 20 | 0.1 | 32 | 0.1562 | 114 | 0.1491 | 18 | 0.1111 | 26 | 0.1154 | 25 | 0.08 | 30 | 0.2 | 99 | 0.1313 |
| 8_8626575 | 33 | 0.7879 | 26 | 0.8077 | 19 | 0.7895 | 34 | 0.7353 | 112 | 0.7768 | 19 | 0.8421 | 23 | 0.6522 | 28 | 0.6786 | 31 | 0.8065 | 101 | 0.7426 |
| 8_8626576 | 34 | 0.7059 | 27 | 0.6667 | 19 | 0.4211 | 31 | 0.6452 | 111 | 0.6306 | 17 | 0.4118 | 23 | 0.3478 | 27 | 0.7407 | 30 | 0.5 | 97 | 0.5155 |
| 8_8626587 | 31 | 0.6774 | 24 | 0.75 | 13 | 0.6154 | 27 | 0.8148 | 95 | 0.7263 | 17 | 0.6471 | 20 | 0.65 | 24 | 0.7083 | 29 | 0.7931 | 90 | 0.7111 |
| 8_8636442 | 3 | 0 | 11 | 0.1818 | 3 | 0 | 11 | 0.0909 | 28 | 0.1071 | 4 | 0.25 | 11 | 0.1818 | 12 | 0 | 5 | 0 | 32 | 0.0938 |
| 8_8636591 | 2 | 0 | 10 | 0.6 | 4 | 0.25 | 7 | 0.2857 | 23 | 0.3913 | 4 | 0.5 | 5 | 0 | 8 | 0.25 | 2 | 0.5 | 19 | 0.2632 |
| 8_8636620 | 2 | 0 | 10 | 0.2 | 4 | 0.25 | 6 | 0.3333 | 22 | 0.2273 | 4 | 0 | 6 | 0.1667 | 10 | 0.2 | 2 | 0 | 22 | 0.1364 |
| 8_8636624 | 1 | 0 | 9 | 0.1111 | 2 | 0.5 | 3 | 0 | 15 | 0.1333 | 2 | 0 | 5 | 0 | 7 | 0.1429 | 1 | 0 | 15 | 0.0667 |
| 8_8636699 | 2 | 0 | 11 | 0.1818 | 3 | 0 | 6 | 0.5 | 22 | 0.2273 | 5 | 0 | 6 | 0.1667 | 7 | 0.2857 | 2 | 0.5 | 20 | 0.2 |
| 8_8636715 | 2 | 0 | 10 | 0.5 | 4 | 0.5 | 7 | 0.4286 | 23 | 0.4348 | 4 | 0.5 | 5 | 0.4 | 6 | 0 | 2 | 0 | 17 | 0.2353 |
| 8_8636725 | 2 | 1 | 9 | 1 | 2 | 1 | 7 | 0.8571 | 20 | 0.95 | 4 | 1 | 6 | 1 | 6 | 1 | 2 | 1 | 18 | 1 |
| 8_8636732 | 2 | 1 | 9 | 0.5556 | 2 | 1 | 5 | 0.2 | 18 | 0.5556 | 3 | 0.6667 | 3 | 0.6667 | 7 | 0.8571 | 2 | 1 | 15 | 0.8 |
| 8_8636736 | 1 | 0 | 8 | 0.625 | 2 | 1 | 6 | 0.6667 | 17 | 0.6471 | 3 | 0.3333 | 5 | 0.6 | 6 | 0.8333 | 2 | 1 | 16 | 0.6875 |
| 8_8636738 | 2 | 1 | 8 | 0.25 | 2 | 0 | 6 | 0.3333 | 18 | 0.3333 | 5 | 1 | 5 | 0.6 | 5 | 0.4 | 2 | 1 | 17 | 0.7059 |
| 8_8636748 | 1 | 0 | 4 | 0.25 | 1 | 1 | 6 | 1 | 12 | 0.6667 | 5 | 0.4 | 6 | 0.8333 | 6 | 0.5 | 2 | 1 | 19 | 0.6316 |
| 8_8636754 | 1 | 1 | 8 | 1 | 1 | 1 | 5 | 1 | 15 | 1 | 4 | 1 | 4 | 1 | 6 | 0.6667 | 2 | 1 | 16 | 0.875 |
| 8_8636758 | 2 | 0.5 | 6 | 0.8333 | NA | NA | 3 | 1 | 11 | 0.8182 | 4 | 0.75 | 2 | 1 | 3 | 1 | 1 | 1 | 10 | 0.9 |
| 8_8636759 | 2 | 0.5 | 9 | 0.6667 | 2 | 0.5 | 6 | 0.8333 | 19 | 0.6842 | 5 | 1 | 6 | 0.8333 | 4 | 0.75 | 2 | 0.5 | 17 | 0.8235 |
| 8_8636769 | 2 | 1 | 6 | 1 | 2 | 1 | 6 | 0.6667 | 16 | 0.875 | 5 | 0.8 | 5 | 0.8 | 6 | 1 | 2 | 1 | 18 | 0.8889 |
| 8_8636770 | 1 | 1 | 6 | 0.5 | 1 | 0 | 4 | 0.75 | 12 | 0.5833 | 3 | 0.3333 | 6 | 0.5 | 4 | 1 | 2 | 1 | 15 | 0.6667 |
| 8_8636772 | 2 | 1 | 5 | 0.8 | 2 | 1 | 5 | 1 | 14 | 0.9286 | 5 | 0.8 | 5 | 1 | 5 | 1 | 2 | 1 | 17 | 0.9412 |
| 8_8636781 | NA | NA | 1 | 1 | NA | NA | 1 | 1 | 2 | 1 | 1 | 1 | 1 | 0 | NA | NA | NA | NA | 2 | 0.5 |
| 8_8642796 | NA | NA | NA | NA | NA | NA | 1 | 1 | 1 | 1 | NA | NA | 1 | 1 | NA | NA | 1 | 1 | 2 | 1 |
| 8_8642798 | NA | NA | 1 | 1 | NA | NA | 1 | 0 | 2 | 0.5 | NA | NA | 1 | 1 | 1 | 0 | 1 | 0 | 3 | 0.3333 |
| 8_8642800 | NA | NA | NA | NA | NA | NA | 1 | 1 | 1 | 1 | NA | NA | 1 | 1 | NA | NA | NA | NA | 1 | 1 |
| 8_8642804 | NA | NA | 1 | 1 | NA | NA | NA | NA | 1 | 1 | NA | NA | 1 | 1 | 1 | 1 | 1 | 1 | 3 | 1 |
| 8_8642818 | NA | NA | 1 | 1 | NA | NA | 1 | 0 | 2 | 0.5 | NA | NA | NA | NA | 1 | 1 | 1 | 1 | 2 | 1 |
| 8_8642823 | NA | NA | 1 | 1 | NA | NA | NA | NA | 1 | 1 | NA | NA | NA | NA | 1 | 1 | 1 | 1 | 2 | 1 |
| 8_8642893 | NA | NA | 1 | 1 | NA | NA | NA | NA | 1 | 1 | NA | NA | NA | NA | NA | NA | 1 | 1 | 1 | 1 |
| 8_8643280 | NA | NA | 1 | 1 | NA | NA | NA | NA | 1 | 1 | NA | NA | NA | NA | NA | NA | 1 | 1 | 1 | 1 |
| 8_8643387 | NA | NA | 1 | 1 | NA | NA | 1 | 1 | 2 | 1 | NA | NA | NA | NA | NA | NA | 1 | 1 | 1 | 1 |
| 8_8644471 | 40 | 0.225 | 20 | 0.4 | 24 | 0.4167 | 35 | 0.3429 | 119 | 0.3277 | 22 | 0.2727 | 38 | 0.1842 | 25 | 0.48 | 26 | 0.3846 | 111 | 0.3153 |
| 8_8644499 | 37 | 0.0541 | 17 | 0 | 23 | 0.087 | 34 | 0.0588 | 111 | 0.0541 | 15 | 0.0667 | 35 | 0.0857 | 22 | 0 | 24 | 0.0417 | 96 | 0.0521 |
| 8_8678605 | 2 | 0.5 | NA | NA | 4 | 0.75 | 5 | 0.4 | 11 | 0.5455 | 3 | 1 | 2 | 0 | 4 | 0.75 | 5 | 0.2 | 14 | 0.5 |
| 8_8678620 | 1 | 1 | NA | NA | 4 | 0.25 | 4 | 0.25 | 9 | 0.3333 | 2 | 0 | 3 | 0.3333 | 4 | 0.25 | 4 | 0.25 | 13 | 0.2308 |
| 8_8678645 | 1 | 1 | NA | NA | 3 | 1 | 3 | 0.6667 | 7 | 0.8571 | 2 | 1 | 1 | 1 | 3 | 0.3333 | 3 | 0.6667 | 9 | 0.6667 |
| 8_8678659 | 2 | 0 | NA | NA | 6 | 0.1667 | 4 | 0 | 12 | 0.0833 | 3 | 0.3333 | 3 | 0 | 4 | 0 | 5 | 0 | 15 | 0.0667 |
| 8_8678666 | 2 | 0 | NA | NA | 5 | 0 | 4 | 0.25 | 11 | 0.0909 | 3 | 0.3333 | 3 | 0 | 2 | 0 | 5 | 0 | 13 | 0.0769 |
| 8_8723181 | NA | NA | NA | NA | 1 | 1 | 1 | 1 | 2 | 1 | 1 | 0 | NA | NA | NA | NA | 1 | 1 | 2 | 0.5 |
| 8_8723185 | NA | NA | NA | NA | 1 | 1 | 1 | 0 | 2 | 0.5 | NA | NA | NA | NA | NA | NA | 1 | 1 | 1 | 1 |
| 8_8735091 | 2 | 0 | 5 | 0.2 | 6 | 0 | 2 | 0 | 15 | 0.0667 | 7 | 0.1429 | 5 | 0 | 5 | 0 | 3 | 0 | 20 | 0.05 |
| 8_8736835 | 3 | 0.6667 | 6 | 0.1667 | 8 | 0.25 | 3 | 0 | 20 | 0.25 | 6 | 0.5 | 8 | 0.125 | 9 | 0.1111 | 6 | 0.1667 | 29 | 0.2069 |
| 8_8743998 | 6 | 0.1667 | 9 | 0 | 8 | 0 | 4 | 0.25 | 27 | 0.0741 | 10 | 0 | 10 | 0.1 | 9 | 0 | 6 | 0.1667 | 35 | 0.0571 |
| 8_8744309 | 8 | 0 | 10 | 0.2 | 11 | 0.0909 | 7 | 0.2857 | 36 | 0.1389 | 12 | 0.0833 | 12 | 0.0833 | 11 | 0 | 10 | 0.1 | 45 | 0.0667 |
| 8_8744310 | 5 | 0 | 9 | 0 | 4 | 0 | 6 | 0.3333 | 24 | 0.0833 | 8 | 0 | 8 | 0 | 13 | 0.0769 | 8 | 0.125 | 37 | 0.0541 |
| 8_8745154 | 3 | 0 | NA | NA | 5 | 0.2 | 5 | 0 | 13 | 0.0769 | 3 | 0.3333 | 2 | 0 | 2 | 0.5 | 2 | 0 | 9 | 0.2222 |
| 8_8745158 | 5 | 0.2 | NA | NA | 6 | 0.3333 | 6 | 0 | 17 | 0.1765 | 4 | 0 | 3 | 0.3333 | 1 | 0 | 4 | 0.5 | 12 | 0.25 |
| 8_8745165 | 4 | 0.25 | NA | NA | 6 | 0.1667 | 4 | 0.25 | 14 | 0.2143 | 3 | 0.3333 | 2 | 0.5 | NA | NA | 3 | 0.3333 | 8 | 0.375 |
| 8_8745171 | 5 | 0.6 | NA | NA | 4 | 1 | 4 | 1 | 13 | 0.8462 | 2 | 1 | 3 | 1 | NA | NA | 2 | 1 | 7 | 1 |
| 8_8745178 | 3 | 0.6667 | NA | NA | 5 | 0 | 4 | 0.25 | 12 | 0.25 | 3 | 0.6667 | 2 | 0.5 | 1 | 0 | 3 | 0.3333 | 9 | 0.4444 |
| 8_8745184 | 4 | 0 | NA | NA | 4 | 0.25 | 4 | 0.25 | 12 | 0.1667 | 3 | 0.3333 | 2 | 0 | 1 | 0 | 4 | 0 | 10 | 0.1 |
| 8_8745195 | 4 | 1 | NA | NA | 5 | 0.4 | 3 | 0.3333 | 12 | 0.5833 | 2 | 1 | 2 | 1 | 1 | 1 | 3 | 0.6667 | 8 | 0.875 |
| 8_8745197 | 3 | 1 | NA | NA | 5 | 0.8 | 3 | 1 | 11 | 0.9091 | 3 | 1 | 3 | 1 | 1 | 1 | 4 | 1 | 11 | 1 |
| 8_8745216 | 4 | 0.75 | NA | NA | 5 | 0.6 | 2 | 0.5 | 11 | 0.6364 | 3 | 1 | 1 | 0 | NA | NA | 3 | 0.6667 | 7 | 0.7143 |
| 8_8745227 | 2 | 0 | NA | NA | 3 | 0 | 3 | 0.3333 | 8 | 0.125 | 1 | 1 | 2 | 0.5 | NA | NA | 3 | 0.6667 | 6 | 0.6667 |
| 8_8745234 | 4 | 0.5 | NA | NA | 5 | 0.6 | 4 | 0.25 | 13 | 0.4615 | 3 | 0.3333 | 3 | 0.3333 | NA | NA | 4 | 0.75 | 10 | 0.5 |
| 8_8745238 | 4 | 1 | NA | NA | 5 | 0.4 | 4 | 0.25 | 13 | 0.5385 | 3 | 0 | 1 | 1 | NA | NA | 4 | 0.5 | 8 | 0.375 |
| 8_8757071 | 1 | 1 | 1 | 0 | 2 | 0 | NA | NA | 4 | 0.25 | 1 | 0 | 3 | 0 | 1 | 0 | 2 | 0.5 | 7 | 0.1429 |
| 8_8757108 | 2 | 0 | 5 | 0.4 | 3 | 0.3333 | 2 | 0.5 | 12 | 0.3333 | NA | NA | 4 | 0.25 | 3 | 0 | 3 | 0 | 10 | 0.1 |
| 8_8757112 | 3 | 0.3333 | 5 | 0.2 | 3 | 0.6667 | 1 | 1 | 12 | 0.4167 | NA | NA | 5 | 0 | 3 | 0.3333 | 3 | 0.3333 | 11 | 0.1818 |
| 8_8757126 | 1 | 1 | 4 | 0.75 | 3 | 0.3333 | 2 | 0 | 10 | 0.5 | NA | NA | 5 | 1 | 2 | 0 | 3 | 0 | 10 | 0.5 |

| | | | | | | | | | | | | | | | | | | | | |
|---|---|---|---|---|---|---|---|---|---|---|---|---|---|---|---|---|---|---|---|---|
| 8_8757128 | 3 | 0.3333 | 4 | 0.75 | 2 | 0.5 | 1 | 0 | 10 | 0.5 | 1 | 1 | 3 | 0 | 2 | 1 | 2 | 0.5 | 8 | 0.5 |
| 8_8757131 | 2 | 1 | 4 | 1 | 2 | 1 | 1 | 1 | 9 | 1 | 1 | 1 | 3 | 1 | 2 | 1 | 3 | 1 | 9 | 1 |
| 8_8757134 | 2 | 1 | 5 | 0.2 | 2 | 0 | 2 | 1 | 11 | 0.4545 | NA | NA | 5 | 0.6 | 2 | 0 | 3 | 0.6667 | 10 | 0.5 |
| 8_8757140 | 3 | 0.6667 | 4 | 0.75 | 3 | 0.3333 | 2 | 1 | 12 | 0.6667 | 1 | 1 | 3 | 0.3333 | 2 | 0.5 | 3 | 0.6667 | 9 | 0.5556 |
| 8_8758753 | 2 | 0 | 3 | 0.3333 | 2 | 0.5 | 1 | 1 | 8 | 0.375 | 1 | 0 | 2 | 0 | 1 | 0 | 2 | 0.5 | 6 | 0.1667 |
| 8_8758755 | 1 | 0 | 1 | 1 | 2 | 1 | 1 | 0 | 5 | 0.6 | NA | NA | 2 | 0.5 | 1 | 1 | NA | NA | 3 | 0.6667 |
| 8_8758759 | 1 | 1 | 3 | 1 | 2 | 1 | 1 | 1 | 7 | 1 | NA | NA | 2 | 1 | 1 | 1 | 1 | 1 | 4 | 1 |
| 8_8758775 | 1 | 0 | 1 | 1 | 2 | 0.5 | 1 | 1 | 5 | 0.6 | 1 | 1 | NA | NA | 1 | 0 | 1 | 1 | 3 | 0.6667 |
| 8_8758818 | 1 | 1 | 2 | 0.5 | NA | NA | 1 | 1 | 4 | 0.75 | 1 | 1 | 2 | 0.5 | 1 | 0 | 1 | 0 | 5 | 0.4 |
| 8_8758827 | 2 | 0 | 3 | 0 | NA | NA | 1 | 1 | 6 | 0.1667 | 1 | 0 | 2 | 0 | 1 | 0 | 1 | 1 | 5 | 0.2 |
| 8_8758841 | 2 | 1 | 2 | 1 | NA | NA | 1 | 1 | 5 | 1 | 1 | 0 | 2 | 0.5 | 1 | 1 | 1 | 1 | 5 | 0.6 |
| 8_8761144 | NA | NA | 1 | 1 | NA | NA | NA | NA | 1 | 1 | NA | NA | NA | NA | 1 | 1 | NA | NA | 1 | 1 |
| 8_8819662 | 1 | 1 | NA | NA | NA | NA | 1 | 1 | 2 | 1 | NA | NA | 1 | 1 | NA | NA | NA | NA | 1 | 1 |
| 8_8825212 | NA | NA | 1 | 0 | NA | NA | 1 | 1 | 2 | 0.5 | 1 | 1 | NA | NA | 1 | 0 | 1 | 0 | 3 | 0.3333 |
| 8_8850708 | 1 | 1 | 3 | 0.6667 | 5 | 0.8 | 2 | 1 | 11 | 0.8182 | 2 | 1 | 1 | 1 | 3 | 0.6667 | 2 | 1 | 8 | 0.875 |
| 8_8850714 | 1 | 1 | 3 | 0.6667 | 5 | 0.6 | 2 | 1 | 11 | 0.7273 | 2 | 1 | 1 | 1 | 3 | 0.3333 | 1 | 0 | 7 | 0.5714 |
| 8_8850721 | 1 | 0 | 2 | 0 | 3 | 0.3333 | 2 | 0 | 8 | 0.125 | 2 | 0 | 1 | 0 | 3 | 0.3333 | 2 | 0 | 8 | 0.125 |
| 8_8850723 | 1 | 1 | 3 | 0.3333 | 5 | 0.2 | 1 | 0 | 10 | 0.3 | 1 | 0 | 1 | 0 | 3 | 0 | 1 | 1 | 6 | 0.1667 |
| 8_8850750 | 1 | 1 | 1 | 1 | 4 | 1 | NA | NA | 6 | 1 | 3 | 1 | 1 | 0 | 2 | 1 | 2 | 1 | 8 | 0.875 |
| 8_8850818 | 1 | 0 | 1 | 1 | 4 | 0.75 | 1 | 0 | 7 | 0.5714 | 1 | 1 | 1 | 1 | 1 | 1 | 2 | 0 | 5 | 0.6 |
| 8_8850831 | 1 | 1 | 2 | 0 | 3 | 0.3333 | 1 | 1 | 7 | 0.4286 | 3 | 1 | 1 | 1 | 2 | 0.5 | 2 | 0.5 | 8 | 0.75 |
| 8_8850906 | 1 | 1 | NA | NA | 4 | 0.75 | 1 | 1 | 6 | 0.8333 | 3 | 1 | NA | NA | 1 | 1 | 1 | 1 | 5 | 1 |
| 8_8886054 | 3 | 0.3333 | 6 | 0.3333 | 6 | 0.1667 | 3 | 0.3333 | 18 | 0.2778 | 2 | 0.5 | 4 | 1 | 6 | 0.6667 | 4 | 0.75 | 16 | 0.75 |
| 8_8886069 | 4 | 0.75 | 7 | 0.8571 | 5 | 0.6 | 4 | 0.5 | 20 | 0.7 | 2 | 1 | 4 | 0.75 | 5 | 0.8 | 4 | 0.75 | 15 | 0.8 |
| 8_8886073 | 5 | 0 | 8 | 0.125 | 4 | 0 | 4 | 0.5 | 21 | 0.1429 | 2 | 0.5 | 5 | 0.2 | 4 | 0.25 | 4 | 0 | 15 | 0.2 |
| 8_8886083 | 4 | 1 | 8 | 1 | 6 | 0.8333 | 2 | 1 | 20 | 0.95 | 2 | 1 | 5 | 1 | 4 | 1 | 4 | 1 | 15 | 1 |
| 8_8886089 | 4 | 0 | 7 | 0.2857 | 6 | 0 | 5 | 0.4 | 22 | 0.1818 | 1 | 1 | 4 | 0.25 | 2 | 0 | 3 | 0.3333 | 10 | 0.3 |
| 8_8886097 | 5 | 0.4 | 6 | 0.1667 | 6 | 0.3333 | 6 | 0.1667 | 23 | 0.2609 | 2 | 0 | 4 | 0.75 | 4 | 0.75 | 6 | 0.1667 | 16 | 0.4375 |
| 8_8886153 | 7 | 0.1429 | 7 | 0.1429 | 8 | 0 | 6 | 0 | 28 | 0.0714 | 3 | 0 | 5 | 0 | 5 | 0 | 7 | 0.1429 | 20 | 0.05 |
| 8_8889758 | 5 | 0.2 | 8 | 0 | 3 | 0 | 3 | 0 | 19 | 0.0526 | NA | NA | 3 | 0.3333 | 1 | 0 | 2 | 0 | 6 | 0.1667 |
| 8_8889759 | 5 | 0.8 | 7 | 0.8571 | 5 | 0.6 | 2 | 0.5 | 19 | 0.7368 | NA | NA | 2 | 1 | 1 | 1 | 2 | 1 | 5 | 1 |
| 8_8889760 | 5 | 0.4 | 8 | 0 | 5 | 0.2 | 3 | 0 | 21 | 0.1429 | NA | NA | 3 | 0.3333 | NA | NA | 3 | 0 | 6 | 0.1667 |
| 8_8889762 | 4 | 0 | 6 | 0.1667 | 4 | 0 | 3 | 0 | 17 | 0.0588 | NA | NA | 3 | 0.3333 | NA | NA | 3 | 0 | 6 | 0.1667 |
| 8_8892679 | 2 | 1 | NA | NA | NA | NA | NA | NA | 2 | 1 | 2 | 0.5 | 1 | 0 | 2 | 0.5 | 4 | 0.75 | 9 | 0.5556 |
| 8_8892700 | 2 | 0.5 | NA | NA | NA | NA | NA | NA | 2 | 0.5 | 2 | 1 | 1 | 0 | 2 | 1 | 5 | 0.6 | 10 | 0.7 |
| 8_8898386 | 2 | 0 | 4 | 0.5 | 1 | 0 | 2 | 0.5 | 9 | 0.3333 | 1 | 0 | 4 | 0 | 4 | 0.25 | 5 | 0.2 | 14 | 0.1429 |
| 8_8898392 | 3 | 0.6667 | 3 | 0.3333 | 1 | 1 | 2 | 0.5 | 9 | 0.5556 | 1 | 1 | 4 | 1 | 4 | 0.75 | 4 | 1 | 13 | 0.9231 |
| 8_8898395 | 3 | 1 | 3 | 0.6667 | 1 | 1 | 1 | 1 | 8 | 0.875 | 1 | 0 | 3 | 0.6667 | 3 | 0.6667 | 5 | 0.6 | 12 | 0.5833 |
| 8_8898396 | 2 | 0.5 | 3 | 1 | 1 | 1 | 2 | 0 | 8 | 0.625 | 2 | 1 | 4 | 1 | 3 | 1 | 5 | 0.8 | 14 | 0.9286 |
| 8_8898405 | 4 | 0.25 | 4 | 0.25 | 1 | 0 | 2 | 0 | 11 | 0.1818 | 2 | 0 | 4 | 0.5 | 4 | 0.75 | 5 | 0.6 | 15 | 0.5333 |
| 8_8898555 | 5 | 1 | 3 | 0.6667 | 3 | 0 | 3 | 0.6667 | 14 | 0.6429 | 2 | 0.5 | 5 | 0.4 | 6 | 0.3333 | 7 | 0.7143 | 20 | 0.5 |
| 8_8905550 | NA | NA | NA | NA | 1 | 1 | NA | NA | 1 | 1 | NA | NA | 1 | 1 | 2 | 1 | 1 | 1 | 4 | 1 |
| 8_8905551 | NA | NA | NA | NA | 1 | 1 | NA | NA | 1 | 1 | NA | NA | 1 | 0 | 2 | 1 | 1 | 1 | 4 | 0.75 |
| 8_8905621 | NA | NA | NA | NA | 1 | 1 | 1 | 1 | 2 | 1 | NA | NA | 1 | 1 | 1 | 1 | 1 | 1 | 3 | 1 |
| 8_8905638 | NA | NA | 1 | 1 | 1 | 0 | 1 | 0 | 3 | 0.3333 | NA | NA | NA | NA | 3 | 0 | 1 | 1 | 4 | 0.25 |
| 8_8905894 | 4 | 0 | 2 | 0 | 6 | 0.1667 | 9 | 0.1111 | 21 | 0.0952 | 6 | 0 | 6 | 0 | 5 | 0.2 | 2 | 0 | 19 | 0.0526 |
| 8_8905900 | 4 | 0.25 | 3 | 0.3333 | 6 | 0 | 4 | 0.25 | 17 | 0.1765 | 5 | 0.2 | 6 | 0 | 2 | 0 | 3 | 0.3333 | 16 | 0.125 |
| 8_8905906 | 9 | 0.1111 | 3 | 0.3333 | 8 | 0 | 11 | 0.0909 | 31 | 0.0968 | 7 | 0.4286 | 7 | 0 | 5 | 0.2 | 2 | 0 | 21 | 0.1905 |
| 8_8905962 | 7 | 0.1429 | 3 | 0 | 8 | 0.125 | 11 | 0 | 29 | 0.069 | 6 | 0.1667 | 7 | 0 | 7 | 0.1429 | 4 | 0 | 24 | 0.0833 |
| 8_8908796 | NA | NA | NA | NA | NA | NA | 1 | 1 | 1 | 1 | NA | NA | 1 | 1 | NA | NA | NA | NA | 1 | 1 |
| 8_8908797 | NA | NA | NA | NA | NA | NA | 1 | 1 | 1 | 1 | NA | NA | 1 | 1 | NA | NA | NA | NA | 1 | 1 |
| 8_8931922 | NA | NA | NA | NA | NA | NA | 1 | 1 | 1 | 1 | NA | NA | NA | NA | NA | NA | 1 | 1 | 1 | 1 |
| 8_8935354 | NA | NA | 1 | 1 | NA | NA | NA | NA | 1 | 1 | NA | NA | 1 | 1 | NA | NA | NA | NA | 1 | 1 |
| 8_8935367 | NA | NA | 1 | 1 | NA | NA | NA | NA | 1 | 1 | NA | NA | 1 | 1 | NA | NA | NA | NA | 1 | 1 |
| 8_8939129 | NA | NA | NA | NA | NA | NA | 1 | 1 | 1 | 1 | NA | NA | NA | NA | 1 | 1 | NA | NA | 1 | 1 |
| 8_8941335 | NA | NA | NA | NA | NA | NA | 1 | 1 | 1 | 1 | NA | NA | NA | NA | 1 | 1 | NA | NA | 1 | 1 |
| 8_8941379 | 1 | 1 | NA | NA | NA | NA | 2 | 1 | 3 | 1 | NA | NA | NA | NA | 1 | 1 | NA | NA | 1 | 1 |
| 8_8941445 | 1 | 1 | NA | NA | NA | NA | 1 | 1 | 2 | 1 | NA | NA | NA | NA | 1 | 1 | NA | NA | 1 | 1 |
| 8_8941451 | 1 | 1 | NA | NA | NA | NA | 2 | 0.5 | 3 | 0.6667 | NA | NA | NA | NA | 1 | 1 | NA | NA | 1 | 1 |
| 8_8941478 | 1 | 1 | NA | NA | NA | NA | 1 | 1 | 2 | 1 | NA | NA | NA | NA | 1 | 1 | NA | NA | 1 | 1 |
| 8_8941482 | 1 | 0 | NA | NA | NA | NA | 2 | 0.5 | 3 | 0.3333 | NA | NA | NA | NA | 1 | 1 | NA | NA | 1 | 1 |
| 8_8941485 | 1 | 1 | NA | NA | NA | NA | 2 | 0.5 | 3 | 0.6667 | NA | NA | NA | NA | 1 | 1 | NA | NA | 1 | 1 |
| 8_8943191 | NA | NA | 3 | 0.6667 | 6 | 1 | 4 | 1 | 13 | 0.9231 | NA | NA | 4 | 0.75 | 7 | 0.8571 | 2 | 1 | 13 | 0.8462 |
| 8_8943209 | 2 | 0.5 | 2 | 0 | 6 | 0.1667 | 4 | 0 | 14 | 0.1429 | NA | NA | 4 | 0.25 | 9 | 0.2222 | 4 | 0.25 | 17 | 0.2353 |
| 8_8943210 | 3 | 0 | 3 | 0.3333 | 6 | 0 | 6 | 0.5 | 18 | 0.2222 | NA | NA | 5 | 0 | 9 | 0.3333 | 4 | 0.25 | 18 | 0.2222 |

| | | | | | | | | | | | | | | | | | | | | |
|---|---|---|---|---|---|---|---|---|---|---|---|---|---|---|---|---|---|---|---|---|
| 8_8943214 | 3 | 1 | 2 | 1 | 5 | 0.6 | 4 | 0.75 | 14 | 0.7857 | NA | NA | 5 | 0.6 | 9 | 0.7778 | 2 | 0.5 | 16 | 0.6875 |
| 8_8943235 | 2 | 0 | 3 | 0.6667 | 7 | 0.7143 | 3 | 1 | 15 | 0.6667 | NA | NA | 4 | 0.5 | 9 | 0.4444 | 4 | 0.75 | 17 | 0.5294 |
| 8_8943378 | 1 | 1 | 2 | 1 | 6 | 0.8333 | 6 | 1 | 15 | 0.9333 | NA | NA | 5 | 0.6 | 11 | 0.9091 | 4 | 1 | 20 | 0.85 |
| 8_8943381 | 2 | 0.5 | 3 | 0 | 5 | 0.2 | 5 | 0.2 | 15 | 0.2 | NA | NA | 4 | 0.25 | 11 | 0.3636 | 3 | 0.3333 | 18 | 0.3333 |
| 8_8943382 | 3 | 0.3333 | 2 | 0.5 | 5 | 0.2 | 5 | 0.4 | 15 | 0.3333 | NA | NA | 3 | 0.3333 | 10 | 0.1 | 3 | 0.6667 | 16 | 0.25 |
| 8_8943395 | 3 | 0.6667 | 3 | 0 | 6 | 0 | 5 | 0 | 17 | 0.1176 | NA | NA | 3 | 0.3333 | 11 | 0 | 4 | 0 | 18 | 0.0556 |
| 8_8945617 | 3 | 0 | 4 | 0.5 | 6 | 0.1667 | 7 | 0 | 20 | 0.15 | NA | NA | 4 | 0 | 12 | 0.25 | 4 | 0.25 | 20 | 0.2 |
| 8_8945645 | 3 | 0 | 3 | 0 | 6 | 0 | 8 | 0.125 | 20 | 0.05 | NA | NA | 4 | 0 | 11 | 0.2727 | 6 | 0.1667 | 21 | 0.1905 |
| 8_8945739 | 3 | 0 | 2 | 0 | 6 | 0.1667 | 8 | 0 | 19 | 0.0526 | NA | NA | 4 | 0 | 8 | 0.25 | 5 | 0 | 17 | 0.1176 |
| 8_8947189 | 6 | 0 | 8 | 0.25 | 9 | 0.1111 | 14 | 0 | 37 | 0.0811 | 1 | 0 | 8 | 0.125 | 18 | 0.0556 | 7 | 0.1429 | 34 | 0.0882 |
| 8_8947191 | 4 | 0 | 7 | 0.1429 | 9 | 0.1111 | 13 | 0 | 33 | 0.0606 | 1 | 0 | 8 | 0 | 18 | 0.1111 | 7 | 0 | 34 | 0.0588 |
| 8_8947355 | 6 | 0.5 | 6 | 0.6667 | 10 | 0.1 | 13 | 0.2308 | 35 | 0.3143 | 1 | 0 | 8 | 0.5 | 19 | 0.4211 | 8 | 0.5 | 36 | 0.4444 |
| 8_8947358 | 7 | 0 | 8 | 0.25 | 12 | 0.3333 | 11 | 0.0909 | 38 | 0.1842 | 1 | 0 | 5 | 0.4 | 19 | 0.2105 | 7 | 0.2857 | 32 | 0.25 |
| 8_8975400 | 1 | 1 | 3 | 0.6667 | 2 | 0.5 | 3 | 0.3333 | 9 | 0.5556 | 3 | 0.6667 | 1 | 1 | 2 | 0.5 | 2 | 0.5 | 8 | 0.625 |
| 8_8975451 | 1 | 0 | 3 | 0 | 1 | 0 | 1 | 1 | 6 | 0.1667 | 3 | 0.3333 | 2 | 0.5 | 1 | 1 | 1 | 0 | 7 | 0.4286 |
| 8_8975452 | 1 | 0 | 3 | 0 | 2 | 0.5 | 1 | 1 | 7 | 0.2857 | 2 | 0.5 | 2 | 1 | 2 | 0.5 | 2 | 0 | 8 | 0.5 |
| 8_8975477 | 1 | 1 | 3 | 0.3333 | 2 | 0 | 4 | 0.75 | 10 | 0.5 | 4 | 0.5 | 2 | 0.5 | 2 | 0 | 2 | 0 | 10 | 0.3 |
| 8_8975478 | 1 | 1 | NA | NA | NA | NA | 3 | 1 | 4 | 1 | 4 | 1 | 2 | 1 | 2 | 0 | 2 | 1 | 10 | 0.8 |
| 8_8975490 | 1 | 0 | 4 | 0 | 2 | 0 | 4 | 0.25 | 11 | 0.0909 | 4 | 0.25 | 2 | 0 | 2 | 0.5 | 2 | 0 | 10 | 0.2 |
| 8_8975491 | 1 | 1 | 3 | 0.6667 | 2 | 0.5 | 3 | 0.3333 | 9 | 0.5556 | 3 | 0 | 2 | 0.5 | 2 | 0 | 2 | 1 | 9 | 0.3333 |
| 8_8975492 | 1 | 1 | 4 | 1 | 2 | 1 | 4 | 0.75 | 11 | 0.9091 | 4 | 1 | 2 | 1 | 2 | 0.5 | 2 | 1 | 10 | 0.9 |
| 8_8975503 | 1 | 0 | 4 | 0.75 | 2 | 0.5 | 3 | 0.3333 | 10 | 0.5 | 3 | 0.6667 | 2 | 0.5 | 2 | 0.5 | 2 | 0.5 | 9 | 0.5556 |
| 8_8975512 | 1 | 1 | 1 | 0 | 2 | 0.5 | 3 | 0 | 7 | 0.2857 | 2 | 0.5 | 1 | 1 | 2 | 0.5 | 2 | 0 | 7 | 0.4286 |
| 8_8975664 | 1 | 0 | 3 | 0.3333 | 2 | 0.5 | 3 | 0.3333 | 9 | 0.3333 | 3 | 0.3333 | 2 | 1 | 1 | 1 | 2 | 0 | 8 | 0.5 |
| 8_8975671 | NA | NA | 3 | 0 | 1 | 0 | 3 | 0.3333 | 7 | 0.1429 | 4 | 0.25 | 2 | 1 | NA | NA | 2 | 0 | 8 | 0.375 |
| 8_8975681 | 1 | 1 | 3 | 1 | 1 | 1 | 4 | 0.75 | 9 | 0.8889 | 4 | 0.75 | 1 | 1 | 2 | 1 | 2 | 1 | 9 | 0.8889 |
| 8_8975689 | 1 | 0 | 3 | 0.3333 | 2 | 0.5 | 2 | 0.5 | 8 | 0.375 | 3 | 0.3333 | 1 | 1 | 2 | 0.5 | 2 | 0.5 | 8 | 0.5 |
| 8_8975702 | 1 | 0 | 3 | 0.3333 | 2 | 0 | 3 | 0.3333 | 9 | 0.2222 | 4 | 0.5 | 1 | 0 | 2 | 1 | 2 | 0.5 | 9 | 0.5556 |
| 8_8975705 | 1 | 1 | 3 | 1 | 2 | 1 | 3 | 1 | 9 | 1 | 2 | 1 | 1 | 1 | 2 | 1 | 2 | 1 | 7 | 1 |
| 8_8975710 | 1 | 0 | 2 | 0 | 2 | 0 | 3 | 0.3333 | 8 | 0.125 | 1 | 0 | 1 | 1 | 2 | 0 | 2 | 0 | 6 | 0.1667 |
| 8_8975712 | 1 | 1 | 3 | 0.3333 | 2 | 0.5 | 3 | 0.3333 | 9 | 0.4444 | 4 | 0.5 | 1 | 0 | 2 | 0.5 | 2 | 0 | 9 | 0.3333 |
| 8_8975719 | 1 | 1 | 3 | 1 | 2 | 1 | 2 | 1 | 8 | 1 | 4 | 1 | 1 | 1 | 2 | 1 | 1 | 1 | 8 | 1 |
| 8_8975792 | 1 | 0 | 1 | 1 | 2 | 0 | NA | NA | 4 | 0.25 | 1 | 1 | NA | NA | NA | NA | 2 | 0 | 3 | 0.3333 |
| 8_8975807 | 1 | 1 | 2 | 1 | 2 | 1 | 3 | 0.6667 | 8 | 0.875 | 1 | 1 | NA | NA | NA | NA | 2 | 1 | 3 | 1 |
| 8_8975817 | 1 | 1 | 3 | 0.6667 | NA | NA | 2 | 1 | 6 | 0.8333 | NA | NA | NA | NA | NA | NA | 2 | 0.5 | 2 | 0.5 |
| 8_8975831 | 1 | 1 | 3 | 0.3333 | 1 | 1 | 2 | 0 | 7 | 0.4286 | 1 | 1 | NA | NA | 1 | 1 | 2 | 1 | 4 | 1 |
| 8_8975854 | 1 | 1 | 1 | 0 | 1 | 0 | 2 | 1 | 5 | 0.6 | NA | NA | NA | NA | 1 | 1 | 1 | 1 | 2 | 1 |
| 8_9097370 | 2 | 0.5 | 3 | 0.3333 | 1 | 1 | NA | NA | 6 | 0.5 | 3 | 0.3333 | 1 | 0 | 1 | 0 | 1 | 1 | 6 | 0.3333 |
| 8_9097436 | 4 | 0.75 | 3 | 1 | 1 | 1 | 1 | 1 | 9 | 0.8889 | 5 | 1 | NA | NA | 1 | 1 | 2 | 1 | 8 | 1 |
| 8_9097451 | 4 | 1 | 3 | 1 | 1 | 1 | NA | NA | 8 | 1 | 5 | 0.8 | NA | NA | 1 | 1 | 2 | 1 | 8 | 0.875 |
| 8_9097460 | 3 | 1 | 3 | 1 | 1 | 1 | 1 | 1 | 8 | 1 | 5 | 1 | NA | NA | 1 | 1 | 1 | 1 | 7 | 1 |
| 8_9097461 | 4 | 1 | 3 | 1 | 1 | 1 | 1 | 1 | 9 | 1 | 5 | 0.8 | NA | NA | 1 | 1 | 2 | 1 | 8 | 0.875 |
| 8_9097476 | 3 | 1 | 2 | 1 | 1 | 1 | 1 | 1 | 7 | 1 | 4 | 1 | NA | NA | 1 | 1 | 1 | 1 | 6 | 1 |
| 8_9097483 | 4 | 0.75 | 3 | 1 | 1 | 1 | 1 | 1 | 9 | 0.8889 | 4 | 0.5 | NA | NA | 1 | 1 | 2 | 1 | 7 | 0.7143 |
| 8_9097484 | 3 | 0.3333 | 1 | 0 | 1 | 1 | 1 | 0 | 6 | 0.3333 | 5 | 0.2 | NA | NA | 1 | 0 | 1 | 1 | 7 | 0.2857 |
| 8_9097485 | 3 | 1 | 3 | 1 | NA | NA | 1 | 1 | 7 | 1 | 5 | 1 | NA | NA | 1 | 1 | 2 | 1 | 8 | 1 |
| 8_9097493 | 4 | 0.75 | 3 | 0.6667 | 1 | 1 | 1 | 1 | 9 | 0.7778 | 5 | 0.4 | NA | NA | 1 | 1 | 2 | 0.5 | 8 | 0.5 |
| 8_9097494 | 4 | 0.75 | 2 | 1 | 1 | 1 | 1 | 1 | 8 | 0.875 | 3 | 0.6667 | NA | NA | 1 | 0 | 2 | 1 | 6 | 0.6667 |
| 8_9097540 | 3 | 0.3333 | 3 | 1 | 1 | 0 | NA | NA | 7 | 0.5714 | 3 | 0 | NA | NA | 1 | 1 | 1 | 0 | 5 | 0.2 |
| 8_9097546 | 4 | 0.75 | 3 | 0 | 1 | 1 | 1 | 1 | 9 | 0.5556 | 3 | 0.3333 | NA | NA | 1 | 0 | NA | NA | 4 | 0.25 |
| 8_9097551 | 4 | 1 | 3 | 0.6667 | 1 | 0 | NA | NA | 8 | 0.75 | 4 | 1 | NA | NA | 1 | 1 | 1 | 1 | 6 | 1 |
| 8_9097553 | 3 | 0 | 3 | 0.3333 | 1 | 0 | 1 | 0 | 8 | 0.125 | 4 | 0.75 | NA | NA | NA | NA | 1 | 0 | 5 | 0.6 |
| 8_9107452 | 2 | 0 | 1 | 1 | 1 | 0 | 1 | 0 | 5 | 0.2 | 2 | 0.5 | NA | NA | NA | NA | NA | NA | 2 | 0.5 |
| 8_9161537 | 2 | 0.5 | 5 | 0 | 3 | 0.3333 | 1 | 0 | 11 | 0.1818 | 1 | 0 | 2 | 0 | 3 | 0.3333 | 2 | 0 | 8 | 0.125 |
| 8_9161570 | 2 | 0 | 7 | 0 | 4 | 0 | 2 | 0.5 | 15 | 0.0667 | 2 | 0 | 3 | 0.3333 | 4 | 0.25 | 3 | 0.3333 | 12 | 0.25 |
| 8_9161573 | 2 | 0.5 | 7 | 0.5714 | 7 | 0.7143 | 3 | 0.6667 | 19 | 0.6316 | 2 | 0.5 | 4 | 0.5 | 4 | 1 | 3 | 0 | 13 | 0.5385 |
| 8_9161579 | 3 | 0.3333 | 5 | 0.2 | 6 | 0 | 2 | 0 | 16 | 0.125 | 1 | 0 | 4 | 0 | 4 | 0.25 | 3 | 0.3333 | 12 | 0.1667 |
| 8_9161580 | 1 | 0 | 5 | 0.2 | 3 | 0 | 1 | 0 | 10 | 0.1 | 2 | 0 | 3 | 0.3333 | 2 | 0 | 2 | 0 | 9 | 0.1111 |
| 8_9161594 | 3 | 1 | 7 | 0.7143 | 7 | 0.8571 | 3 | 1 | 20 | 0.85 | 2 | 1 | 3 | 0.6667 | 5 | 0.8 | 3 | 1 | 13 | 0.8462 |
| 8_9161599 | 2 | 0 | 7 | 0.4286 | 6 | 0.5 | 2 | 1 | 17 | 0.4706 | 2 | 0 | 5 | 0.4 | 4 | 0.75 | 3 | 0.3333 | 14 | 0.4286 |
| 8_9161610 | 3 | 0 | 6 | 0 | 6 | 0.1667 | 4 | 0 | 19 | 0.0526 | 1 | 0 | 6 | 0.1667 | 4 | 0 | 2 | 0 | 13 | 0.0769 |
| 8_9161613 | 3 | 0.3333 | 7 | 0.1429 | 7 | 0.1429 | 3 | 0.3333 | 20 | 0.2 | 3 | 0.3333 | 6 | 0 | 5 | 0 | 3 | 0 | 17 | 0.0588 |
| 8_9161639 | 2 | 0.5 | 6 | 0.1667 | 6 | 0.3333 | 2 | 0 | 16 | 0.25 | 4 | 0.5 | 5 | 0.4 | 6 | 0.1667 | 3 | 0 | 18 | 0.2778 |
| 8_9161662 | 2 | 0 | 8 | 0.5 | 8 | 0 | 4 | 0.25 | 22 | 0.2273 | 7 | 0.1429 | 6 | 0.5 | 5 | 0.2 | 4 | 0.5 | 22 | 0.3182 |
| 8_9161663 | 2 | 0.5 | 8 | 0.25 | 6 | 0 | NA | NA | 16 | 0.1875 | 6 | 0.3333 | 7 | 0.2857 | 4 | 0.25 | 4 | 0.25 | 21 | 0.2857 |

| | | | | | | | | | | | | | | | | | | | | |
|---|---|---|---|---|---|---|---|---|---|---|---|---|---|---|---|---|---|---|---|---|
| 8_9161677 | 3 | 1 | 9 | 0.8889 | 5 | 1 | 3 | 1 | 20 | 0.95 | 5 | 1 | 8 | 0.875 | 6 | 1 | 4 | 1 | 23 | 0.9565 |
| 8_9161678 | 5 | 0.4 | 8 | 0.25 | 7 | 0.1429 | 4 | 0 | 24 | 0.2083 | 8 | 0.25 | 8 | 0.25 | 5 | 0 | 5 | 0.2 | 26 | 0.1923 |
| 8_9161679 | 5 | 0.2 | 8 | 0.125 | 7 | 0.1429 | 4 | 0.25 | 24 | 0.1667 | 8 | 0 | 8 | 0.25 | 6 | 0.1667 | 6 | 0.1667 | 28 | 0.1429 |
| 8_9161681 | 5 | 0 | 9 | 0.1111 | 6 | 0.1667 | 2 | 0.5 | 22 | 0.1364 | 6 | 0.3333 | 7 | 0.1429 | 6 | 0.3333 | 5 | 0.2 | 24 | 0.25 |
| 8_9161698 | 4 | 0.25 | 9 | 0.2222 | 8 | 0.25 | 4 | 0.25 | 25 | 0.24 | 9 | 0.5556 | 7 | 0.2857 | 8 | 0.125 | 6 | 0.3333 | 30 | 0.3333 |
| 8_9161706 | 6 | 0 | 9 | 0 | 9 | 0.1111 | 5 | 0.2 | 29 | 0.069 | 9 | 0.2222 | 8 | 0 | 8 | 0 | 9 | 0 | 34 | 0.0588 |
| 8_9161715 | 3 | 0.6667 | 10 | 0.8 | 9 | 0.7778 | 6 | 0.8333 | 28 | 0.7857 | 9 | 0.6667 | 7 | 0.8571 | 6 | 0.8333 | 8 | 0.625 | 30 | 0.7333 |
| 8_9161744 | 7 | 0 | 9 | 0.3333 | 8 | 0.375 | 6 | 0.3333 | 30 | 0.2667 | 8 | 0 | 8 | 0 | 6 | 0.3333 | 9 | 0.4444 | 31 | 0.1935 |
| 8_9161750 | 7 | 0.4286 | 8 | 0.5 | 8 | 0.5 | 4 | 0.5 | 27 | 0.4815 | 8 | 0.25 | 5 | 0.4 | 7 | 0.8571 | 9 | 0.7778 | 29 | 0.5862 |
| 8_9161783 | 8 | 0.125 | 13 | 0.3846 | 8 | 0.625 | 6 | 0.3333 | 35 | 0.3714 | 11 | 0.1818 | 10 | 0.3 | 9 | 0.2222 | 12 | 0.4167 | 42 | 0.2857 |
| 8_9161784 | 9 | 0.5556 | 13 | 0.5385 | 8 | 0.625 | 9 | 0.2222 | 39 | 0.4872 | 11 | 0.4545 | 10 | 0.7 | 9 | 0.4444 | 12 | 0.5833 | 42 | 0.5476 |
| 8_9161793 | 10 | 0.2 | 13 | 0 | 7 | 0.2857 | 7 | 0 | 37 | 0.1081 | 11 | 0.1818 | 8 | 0.625 | 9 | 0.3333 | 13 | 0.3077 | 41 | 0.3415 |
| 8_9161797 | 11 | 0.3636 | 11 | 0.1818 | 8 | 0.375 | 8 | 0.125 | 38 | 0.2632 | 12 | 0.0833 | 9 | 0.1111 | 9 | 0.1111 | 13 | 0.1538 | 43 | 0.1163 |
| 8_9161824 | 13 | 0.0769 | 16 | 0.125 | 12 | 0.0833 | 7 | 0.1429 | 48 | 0.1042 | 13 | 0.0769 | 8 | 0.125 | 11 | 0.1818 | 16 | 0.125 | 48 | 0.125 |
| 8_9176002 | 2 | 0.5 | NA | NA | NA | NA | NA | NA | 2 | 0.5 | NA | NA | NA | NA | NA | NA | 3 | 0.3333 | 3 | 0.3333 |
| 8_9176019 | 2 | 0.5 | NA | NA | NA | NA | 1 | 1 | 3 | 0.6667 | NA | NA | NA | NA | 1 | 1 | 2 | 0 | 3 | 0.3333 |
| 8_9176022 | 2 | 0 | NA | NA | NA | NA | 1 | 1 | 3 | 0.3333 | NA | NA | NA | NA | 1 | 1 | 3 | 0.6667 | 4 | 0.75 |
| 8_9176028 | 2 | 1 | NA | NA | NA | NA | 1 | 1 | 3 | 1 | NA | NA | NA | NA | 1 | 1 | 3 | 0.6667 | 4 | 0.75 |
| 8_9176046 | 2 | 0 | NA | NA | NA | NA | 1 | 1 | 3 | 0.3333 | NA | NA | NA | NA | 1 | 0 | 2 | 0.5 | 3 | 0.3333 |
| 8_9177797 | 3 | 1 | 2 | 1 | NA | NA | NA | NA | 5 | 1 | NA | NA | NA | NA | 1 | 1 | 2 | 1 | 3 | 1 |
| 8_9177801 | 3 | 0.6667 | 2 | 0 | NA | NA | 1 | 1 | 6 | 0.5 | NA | NA | NA | NA | 2 | 0.5 | 2 | 1 | 4 | 0.75 |
| 8_9177802 | 4 | 0.5 | 1 | 0 | NA | NA | 1 | 0 | 6 | 0.3333 | NA | NA | NA | NA | 2 | 0 | 3 | 0.6667 | 5 | 0.4 |
| 8_9177809 | 4 | 1 | 2 | 1 | NA | NA | 1 | 1 | 7 | 1 | NA | NA | NA | NA | 2 | 1 | 5 | 1 | 7 | 1 |
| 8_9177810 | 1 | 1 | 2 | 0.5 | NA | NA | 1 | 1 | 4 | 0.75 | NA | NA | NA | NA | 1 | 1 | 4 | 0.75 | 5 | 0.8 |
| 8_9177814 | 3 | 0.6667 | 1 | 0 | NA | NA | NA | NA | 4 | 0.5 | NA | NA | NA | NA | 2 | 1 | 5 | 0.2 | 7 | 0.4286 |
| 8_9177817 | 2 | 1 | 2 | 1 | NA | NA | 1 | 1 | 5 | 1 | NA | NA | NA | NA | 1 | 1 | 6 | 0.5 | 7 | 0.5714 |
| 8_9177841 | 4 | 0.25 | 2 | 0.5 | NA | NA | 1 | 0 | 7 | 0.2857 | NA | NA | 1 | 0 | 2 | 0 | 6 | 0.1667 | 9 | 0.1111 |
| 8_9177848 | 4 | 1 | 1 | 0 | NA | NA | 1 | 1 | 6 | 0.8333 | NA | NA | 1 | 1 | 1 | 1 | 5 | 1 | 7 | 1 |
| 8_9177850 | 3 | 0.3333 | NA | NA | NA | NA | 1 | 0 | 4 | 0.25 | NA | NA | NA | NA | 2 | 0 | 4 | 0.25 | 6 | 0.1667 |
| 8_9177857 | 4 | 1 | 2 | 1 | NA | NA | 1 | 1 | 7 | 1 | NA | NA | 1 | 0 | 3 | 1 | 4 | 0.75 | 8 | 0.75 |
| 8_9177859 | 4 | 0.5 | NA | NA | NA | NA | 1 | 0 | 5 | 0.4 | NA | NA | 1 | 0 | 3 | 0.6667 | 6 | 0.5 | 10 | 0.5 |
| 8_9177862 | 4 | 0.75 | 1 | 0 | NA | NA | NA | NA | 5 | 0.6 | NA | NA | 1 | 1 | 3 | 1 | 4 | 0.75 | 8 | 0.875 |
| 8_9177865 | 4 | 0.75 | 1 | 1 | NA | NA | NA | NA | 5 | 0.8 | NA | NA | 1 | 1 | 3 | 0.3333 | 5 | 0.8 | 9 | 0.6667 |
| 8_9177878 | 5 | 0.2 | 2 | 0 | NA | NA | 1 | 1 | 8 | 0.25 | 1 | 0 | 1 | 0 | 4 | 0.5 | 6 | 0.5 | 12 | 0.4167 |
| 8_9177948 | 5 | 0.2 | 4 | 0 | NA | NA | 1 | 1 | 10 | 0.2 | NA | NA | 3 | 0.3333 | 4 | 0.25 | 4 | 0 | 11 | 0.1818 |
| 8_9178004 | 4 | 0.25 | 4 | 0 | NA | NA | 2 | 0 | 10 | 0.1 | 2 | 0 | 3 | 0 | 6 | 0 | 7 | 0.1429 | 18 | 0.0556 |
| 8_9178008 | 4 | 0 | 4 | 0.5 | NA | NA | 2 | 0.5 | 10 | 0.3 | 1 | 0 | 3 | 0.6667 | 4 | 0.75 | 5 | 0.6 | 13 | 0.6154 |
| 8_9183299 | 1 | 1 | NA | NA | 1 | 1 | 1 | 1 | 3 | 1 | NA | NA | NA | NA | NA | NA | 1 | 1 | 1 | 1 |
| 8_9183314 | 2 | 0.5 | 2 | 0.5 | 5 | 0.2 | 3 | 0.3333 | 12 | 0.3333 | 3 | 0 | 3 | 0 | NA | NA | 3 | 0.3333 | 9 | 0.1111 |
| 8_9183316 | 1 | 1 | NA | NA | 1 | 0 | 1 | 1 | 3 | 0.6667 | NA | NA | NA | NA | NA | NA | 1 | 1 | 1 | 1 |
| 8_9183322 | 1 | 1 | NA | NA | NA | NA | 1 | 1 | 2 | 1 | NA | NA | NA | NA | NA | NA | 1 | 1 | 1 | 1 |
| 8_9183337 | 1 | 1 | NA | NA | 1 | 1 | 1 | 1 | 3 | 1 | NA | NA | NA | NA | NA | NA | 1 | 1 | 1 | 1 |
| 8_9183348 | 1 | 1 | NA | NA | 1 | 1 | 1 | 1 | 3 | 1 | NA | NA | NA | NA | NA | NA | 1 | 1 | 1 | 1 |
| 8_9183374 | 1 | 0 | 3 | 0.3333 | 4 | 0.25 | 2 | 0 | 10 | 0.2 | 3 | 0 | 4 | 0.25 | NA | NA | 3 | 0 | 10 | 0.1 |
| 8_9183376 | 2 | 0.5 | 3 | 0 | 4 | 0 | 3 | 0 | 12 | 0.0833 | 2 | 0 | 5 | 0 | NA | NA | 4 | 0.25 | 11 | 0.0909 |
| 8_9183441 | 1 | 1 | NA | NA | 1 | 1 | 1 | 1 | 3 | 1 | NA | NA | NA | NA | NA | NA | 2 | 1 | 2 | 1 |
| 8_9183444 | 1 | 1 | NA | NA | 1 | 0 | 1 | 1 | 3 | 0.6667 | NA | NA | NA | NA | NA | NA | 2 | 0.5 | 2 | 0.5 |
| 8_9183466 | 1 | 0 | NA | NA | 1 | 1 | 1 | 1 | 3 | 0.6667 | NA | NA | NA | NA | NA | NA | 2 | 1 | 2 | 1 |
| 8_9183470 | 1 | 1 | NA | NA | 1 | 0 | 1 | 1 | 3 | 0.6667 | NA | NA | NA | NA | NA | NA | 2 | 0.5 | 2 | 0.5 |
| 8_9183471 | NA | NA | NA | NA | 1 | 1 | 1 | 1 | 2 | 1 | NA | NA | NA | NA | NA | NA | 2 | 1 | 2 | 1 |
| 8_9183476 | 1 | 1 | NA | NA | 1 | 1 | 1 | 1 | 3 | 1 | NA | NA | NA | NA | NA | NA | 2 | 0.5 | 2 | 0.5 |
| 8_9183883 | 1 | 1 | NA | NA | NA | NA | 2 | 0.5 | 3 | 0.6667 | NA | NA | 1 | 0 | 2 | 1 | 1 | 0 | 4 | 0.5 |
| 8_9183901 | 1 | 1 | NA | NA | NA | NA | 1 | 1 | 2 | 1 | NA | NA | 1 | 1 | 1 | 1 | 1 | 0 | 3 | 0.6667 |
| 8_9183906 | 1 | 1 | NA | NA | NA | NA | 1 | 1 | 2 | 1 | NA | NA | NA | NA | 2 | 1 | 1 | 1 | 3 | 1 |
| 8_9184890 | 2 | 1 | 3 | 0.3333 | 5 | 0.2 | 3 | 0 | 13 | 0.3077 | 4 | 0 | 6 | 0 | 1 | 1 | 5 | 0 | 16 | 0.0625 |
| 8_9184909 | 2 | 0.5 | 3 | 0 | 5 | 0.2 | 3 | 0.3333 | 13 | 0.2308 | 5 | 0.2 | 7 | 0 | 1 | 0 | 5 | 0.2 | 18 | 0.1111 |
| 8_9184910 | 2 | 1 | 1 | 1 | 4 | 1 | 2 | 1 | 9 | 1 | 5 | 1 | 6 | 0.6667 | 1 | 1 | 4 | 0.5 | 16 | 0.75 |
| 8_9184912 | 2 | 1 | 2 | 0.5 | 5 | 0.6 | 4 | 0.25 | 13 | 0.5385 | 5 | 0.4 | 6 | 0.6667 | 1 | 1 | 5 | 0 | 17 | 0.4118 |
| 8_9184913 | 2 | 0.5 | 3 | 0.3333 | 5 | 1 | 4 | 1 | 14 | 0.7857 | 5 | 0.8 | 6 | 1 | 1 | 1 | 5 | 0.6 | 17 | 0.8235 |
| 8_9184925 | 1 | 0 | 2 | 0.5 | 4 | 0 | 4 | 0 | 11 | 0.0909 | 5 | 0 | 4 | 0 | 2 | 0 | 5 | 0.2 | 16 | 0.0625 |
| 8_9184931 | 2 | 0.5 | 2 | 0 | 4 | 0.5 | 4 | 0 | 12 | 0.25 | 3 | 0.6667 | 4 | 0.25 | 1 | 1 | 3 | 0.3333 | 11 | 0.4545 |
| 8_9184937 | 3 | 0 | 3 | 0 | 5 | 0.2 | 5 | 0 | 16 | 0.0625 | 5 | 0 | 7 | 0 | 2 | 0 | 5 | 0.2 | 19 | 0.0526 |
| 8_9184939 | 3 | 0 | 3 | 0 | 4 | 0.25 | 4 | 0 | 14 | 0.0714 | 5 | 0 | 6 | 0 | 1 | 0 | 5 | 0.4 | 17 | 0.1176 |
| 8_9184947 | 3 | 0.3333 | 3 | 0 | 3 | 0.3333 | 5 | 0 | 14 | 0.1429 | 5 | 0.2 | 7 | 0.1429 | 2 | 0 | 6 | 0.1667 | 20 | 0.15 |
| 8_9184955 | 2 | 0 | 3 | 0 | 5 | 0.4 | 4 | 0 | 14 | 0.1429 | 5 | 0.2 | 7 | 0 | 1 | 0 | 5 | 0 | 18 | 0.0556 |

| | | | | | | | | | | | | | | | | | | | | |
|---|---|---|---|---|---|---|---|---|---|---|---|---|---|---|---|---|---|---|---|---|
| 8_9184962 | 3 | 0 | 2 | 0 | 5 | 0.4 | 4 | 0 | 14 | 0.1429 | 5 | 0.6 | 7 | 0.1429 | 1 | 1 | 5 | 0.2 | 18 | 0.3333 |
| 8_9184992 | 3 | 0 | 3 | 0 | 5 | 0.4 | 5 | 0.2 | 16 | 0.1875 | 5 | 0 | 7 | 0 | 2 | 0.5 | 6 | 0 | 20 | 0.05 |
| 8_9185047 | 9 | 0.2222 | 4 | 0 | 7 | 0 | 11 | 0 | 31 | 0.0645 | 12 | 0.0833 | 8 | 0.125 | 8 | 0 | 4 | 0 | 32 | 0.0625 |
| 8_9185089 | 19 | 0.3158 | 11 | 0 | 20 | 0.2 | 15 | 0.1333 | 65 | 0.1846 | 25 | 0.12 | 19 | 0.1579 | 15 | 0.0667 | 9 | 0 | 68 | 0.1029 |
| 8_9185109 | 17 | 0.1176 | 10 | 0.2 | 18 | 0 | 16 | 0 | 61 | 0.0656 | 21 | 0.0952 | 18 | 0.1111 | 13 | 0.0769 | 10 | 0 | 62 | 0.0806 |
| 8_9185112 | 17 | 0.4118 | 12 | 0.3333 | 18 | 0.2778 | 16 | 0.625 | 63 | 0.4127 | 25 | 0.48 | 17 | 0.7059 | 13 | 0.3077 | 10 | 0.4 | 65 | 0.4923 |
| 8_9185115 | 16 | 0.6875 | 12 | 1 | 15 | 0.8667 | 16 | 0.875 | 59 | 0.8475 | 24 | 0.875 | 19 | 0.6842 | 15 | 0.9333 | 9 | 1 | 67 | 0.8507 |
| 8_9194514 | 3 | 0.3333 | 2 | 0.5 | NA | NA | 2 | 0 | 7 | 0.2857 | 1 | 0 | 1 | 1 | 2 | 0 | 2 | 0.5 | 6 | 0.3333 |
| 8_9194522 | 2 | 0.5 | 2 | 0 | NA | NA | 2 | 1 | 6 | 0.5 | NA | NA | 2 | 0.5 | 2 | 1 | 2 | 0.5 | 6 | 0.6667 |
| 8_9194681 | 1 | 1 | 1 | 1 | NA | NA | 2 | 0.5 | 4 | 0.75 | NA | NA | 1 | 1 | 1 | 0 | NA | NA | 2 | 0.5 |
| 8_9194687 | 1 | 1 | 1 | 0 | NA | NA | 1 | 1 | 3 | 0.6667 | NA | NA | 1 | 1 | 1 | 0 | NA | NA | 2 | 0.5 |
| 8_9201440 | NA | NA | NA | NA | 1 | 1 | 1 | 1 | 2 | 1 | 1 | 1 | NA | NA | 1 | 1 | 1 | 0 | 3 | 0.6667 |
| 8_9201444 | NA | NA | NA | NA | 1 | 1 | 1 | 1 | 2 | 1 | 1 | 1 | NA | NA | 1 | 1 | NA | NA | 2 | 1 |
| 8_9201474 | NA | NA | NA | NA | 1 | 0 | 1 | 1 | 2 | 0.5 | 1 | 1 | NA | NA | 1 | 0 | NA | NA | 2 | 0.5 |
| 8_9213606 | 12 | 0.5833 | 9 | 0.6667 | 2 | 0 | NA | NA | 23 | 0.5652 | 3 | 0 | 9 | 0.5556 | 6 | 0.3333 | 3 | 0 | 21 | 0.3333 |
| 8_9213615 | 11 | 0 | 11 | 0 | 5 | 0.4 | 2 | 0.5 | 29 | 0.1034 | 6 | 0.1667 | 10 | 0 | 6 | 0.3333 | 4 | 0.25 | 26 | 0.1538 |
| 8_9213619 | 15 | 0.4 | 13 | 0.4615 | 5 | 0.4 | 1 | 1 | 34 | 0.4412 | 6 | 0.5 | 11 | 0.2727 | 8 | 0.375 | 4 | 0.25 | 29 | 0.3448 |
| 8_9213629 | 15 | 0.3333 | 10 | 0.7 | 5 | 0.4 | 2 | 0.5 | 32 | 0.4688 | 7 | 0.4286 | 12 | 0.75 | 10 | 0.8 | 4 | 0 | 33 | 0.6061 |
| 8_9213632 | 14 | 0.3571 | 13 | 0.1538 | 5 | 0.2 | 3 | 0.3333 | 35 | 0.2571 | 7 | 0.4286 | 10 | 0.2 | 9 | 0.2222 | 6 | 0 | 32 | 0.2188 |
| 8_9239312 | 19 | 0.4737 | 13 | 0.1538 | 7 | 0.2857 | 6 | 0.5 | 45 | 0.3556 | 7 | 0.2857 | 15 | 0.3333 | 10 | 0.2 | 6 | 0.5 | 38 | 0.3158 |
| 8_9239314 | 23 | 0.1739 | 11 | 0 | 6 | 0 | 7 | 0.1429 | 47 | 0.1064 | 10 | 0.1 | 18 | 0.2222 | 11 | 0.0909 | 7 | 0 | 46 | 0.1304 |
| 8_9239324 | 22 | 0.0909 | 16 | 0 | 10 | 0 | 9 | 0.1111 | 57 | 0.0526 | 9 | 0.1111 | 20 | 0.05 | 15 | 0 | 8 | 0.125 | 52 | 0.0577 |
| 8_9239344 | 27 | 0.1111 | 17 | 0.0588 | 9 | 0 | 11 | 0.0909 | 64 | 0.0781 | 9 | 0.1111 | 21 | 0.0476 | 15 | 0.0667 | 8 | 0 | 53 | 0.0566 |
| 8_9239345 | 25 | 0.2 | 16 | 0 | 8 | 0.125 | 10 | 0.1 | 59 | 0.1186 | 10 | 0 | 21 | 0.0952 | 14 | 0 | 8 | 0.375 | 53 | 0.0943 |
| 8_9239398 | 28 | 0.4643 | 17 | 0.4118 | 10 | 0.2 | 8 | 0.625 | 63 | 0.4286 | 11 | 0.6364 | 19 | 0.2632 | 17 | 0.4706 | 10 | 0.3 | 57 | 0.4035 |
| 8_9340649 | NA | NA | NA | NA | NA | NA | 1 | 1 | 1 | 1 | 1 | 1 | NA | NA | NA | NA | NA | NA | 1 | 1 |
| 8_9340664 | NA | NA | NA | NA | NA | NA | 1 | 1 | 1 | 1 | 1 | 1 | NA | NA | NA | NA | NA | NA | 1 | 1 |
| 8_9357962 | 2 | 1 | 1 | 1 | 2 | 1 | 1 | 1 | 6 | 1 | 1 | 1 | NA | NA | NA | NA | NA | NA | 1 | 1 |
| 8_9357983 | 1 | 1 | NA | NA | 2 | 0 | 1 | 1 | 4 | 0.5 | 1 | 1 | NA | NA | 1 | 0 | NA | NA | 2 | 0.5 |
| 8_9453755 | 20 | 0.15 | 21 | 0.0952 | 21 | 0.1905 | 21 | 0 | 83 | 0.1084 | 18 | 0.2778 | 16 | 0.1875 | 19 | 0.2105 | 27 | 0.1852 | 80 | 0.2125 |
| 8_9468279 | 2 | 0 | 5 | 0.4 | 6 | 0.1667 | 4 | 0.75 | 17 | 0.3529 | 6 | 0.1667 | 1 | 0 | 6 | 0.5 | 5 | 0.6 | 18 | 0.3889 |
| 8_9468287 | 2 | 1 | 5 | 0.8 | 7 | 1 | 4 | 1 | 18 | 0.9444 | 7 | 1 | 1 | 1 | 6 | 1 | 3 | 0.6667 | 17 | 0.9412 |
| 8_9468292 | 2 | 1 | 4 | 0.75 | 5 | 0.6 | 3 | 0.3333 | 14 | 0.6429 | 7 | 0.8571 | 1 | 0 | 6 | 0.5 | 4 | 0.75 | 18 | 0.6667 |
| 8_9468295 | 1 | 0 | 3 | 0.3333 | 6 | 0.5 | 4 | 0.25 | 14 | 0.3571 | 5 | 0.6 | 1 | 0 | 3 | 0.6667 | 3 | 0 | 12 | 0.4167 |
| 8_9468304 | 2 | 1 | 4 | 0.75 | 8 | 0.75 | 4 | 0.5 | 18 | 0.7222 | 6 | 0.8333 | 1 | 1 | 5 | 0.6 | 4 | 0.5 | 16 | 0.6875 |
| 8_9468305 | 1 | 0 | 2 | 0.5 | 6 | 0 | 4 | 0 | 13 | 0.0769 | 5 | 0 | NA | NA | 3 | 0.3333 | 4 | 0 | 12 | 0.0833 |
| 8_9468309 | 2 | 0 | 5 | 0 | 8 | 0.125 | 4 | 0.25 | 19 | 0.1053 | 7 | 0.1429 | 1 | 0 | 6 | 0.1667 | 4 | 0 | 18 | 0.1111 |
| 8_9468339 | 2 | 0.5 | 4 | 0.75 | 6 | 0.8333 | 3 | 0.6667 | 15 | 0.7333 | 5 | 1 | 1 | 1 | 7 | 0.7143 | 5 | 0.6 | 18 | 0.7778 |
| 8_9468352 | 2 | 0 | 2 | 0.5 | 7 | 0.2857 | 3 | 0.3333 | 14 | 0.2857 | 6 | 0.6667 | 1 | 0 | 6 | 0.1667 | 5 | 0.6 | 18 | 0.4444 |
| 8_9468353 | 1 | 0 | 4 | 0 | 8 | 0.125 | 4 | 0 | 17 | 0.0588 | 7 | 0.1429 | 1 | 0 | 7 | 0.1429 | 6 | 0.1667 | 21 | 0.1429 |
| 8_9468403 | 1 | 0 | 1 | 0 | 3 | 0 | 4 | 0.5 | 9 | 0.2222 | 3 | 0 | 1 | 0 | 5 | 0.2 | 6 | 0 | 15 | 0.0667 |
| 8_9468431 | 3 | 0 | 4 | 0.5 | 6 | 0.1667 | 4 | 0 | 17 | 0.1765 | 5 | 0.2 | 2 | 0.5 | 8 | 0.375 | 5 | 0 | 20 | 0.25 |
| 8_9468536 | 3 | 0.3333 | 6 | 0.1667 | 8 | 0 | 2 | 0 | 19 | 0.1053 | 7 | 0.1429 | 2 | 0 | 10 | 0.2 | 6 | 0 | 25 | 0.12 |
| 8_9468544 | 4 | 0 | 3 | 0 | 3 | 0.3333 | 2 | 0 | 12 | 0.0833 | 2 | 0 | 1 | 0 | 3 | 0.3333 | 4 | 0.5 | 10 | 0.3 |
| 8_9468554 | 2 | 0 | 5 | 0.2 | 4 | 0.25 | 4 | 0 | 15 | 0.1333 | 7 | 0 | 1 | 0 | 6 | 0.1667 | 5 | 0.6 | 19 | 0.2105 |
| 8_9468560 | 8 | 0 | 6 | 0 | 8 | 0.125 | 5 | 0.2 | 27 | 0.0741 | 5 | 0.4 | 3 | 0.3333 | 8 | 0.5 | 8 | 0.375 | 24 | 0.4167 |
| 8_9468566 | 4 | 0 | 6 | 0.3333 | 9 | 0 | 6 | 0.1667 | 25 | 0.12 | 6 | 0.1667 | 3 | 0.3333 | 11 | 0.1818 | 6 | 0.1667 | 26 | 0.1923 |
| 8_9468570 | 7 | 0.1429 | 5 | 0.4 | 6 | 0 | 4 | 0 | 22 | 0.1364 | 6 | 0 | 3 | 0 | 6 | 0.3333 | 7 | 0 | 22 | 0.0909 |
| 8_9468582 | 4 | 0.25 | 6 | 0.1667 | 8 | 0.125 | 5 | 0 | 23 | 0.1304 | 9 | 0.1111 | 3 | 0.3333 | 10 | 0.3 | 10 | 0.3 | 32 | 0.25 |
| 8_9468589 | 6 | 0 | 8 | 0.125 | 10 | 0.1 | 6 | 0.1667 | 30 | 0.1 | 9 | 0.1111 | 3 | 0 | 11 | 0.0909 | 10 | 0 | 33 | 0.0606 |
| 8_9468592 | 7 | 0.2857 | 7 | 0.5714 | 8 | 0.25 | 6 | 0.3333 | 28 | 0.3571 | 7 | 0.1429 | 1 | 0 | 10 | 0.3 | 7 | 0.4286 | 25 | 0.28 |
| 8_9468595 | 8 | 0.25 | 5 | 0.2 | 8 | 0.125 | 6 | 0.1667 | 27 | 0.1852 | 8 | 0.125 | 2 | 0 | 10 | 0.3 | 8 | 0.375 | 28 | 0.25 |
| 8_9468701 | 7 | 0 | 10 | 0.1 | 13 | 0.1538 | 7 | 0 | 37 | 0.0811 | 9 | 0 | 3 | 0.3333 | 13 | 0.2308 | 11 | 0.0909 | 36 | 0.1389 |
| 8_9468804 | NA | NA | 2 | 0.5 | 3 | 0.3333 | 4 | 0.5 | 9 | 0.4444 | NA | NA | 1 | 1 | NA | NA | NA | NA | 1 | 1 |
| 8_9468833 | NA | NA | 2 | 0 | 3 | 0 | 4 | 0.25 | 9 | 0.1111 | NA | NA | 1 | 1 | NA | NA | NA | NA | 1 | 1 |
| 8_9469391 | 23 | 0 | 22 | 0.0909 | 29 | 0.1034 | 46 | 0.0652 | 120 | 0.0667 | 20 | 0.15 | 25 | 0.12 | 25 | 0.08 | 25 | 0 | 95 | 0.0842 |
| 8_9469467 | 50 | 0.2 | 59 | 0.322 | 46 | 0.2391 | 71 | 0.2676 | 226 | 0.2611 | 36 | 0.3333 | 56 | 0.2679 | 55 | 0.3455 | 51 | 0.3137 | 198 | 0.3131 |
| 8_9469477 | 67 | 0.0896 | 71 | 0.0563 | 57 | 0.0526 | 85 | 0.0941 | 280 | 0.075 | 44 | 0.1136 | 64 | 0.0156 | 63 | 0.0317 | 63 | 0.0794 | 234 | 0.0556 |
| 8_9469896 | 25 | 0.2 | 27 | 0.1481 | 18 | 0.0556 | 25 | 0.16 | 95 | 0.1474 | 30 | 0.1333 | 20 | 0.1 | 31 | 0.129 | 22 | 0.4545 | 103 | 0.1942 |
| 8_9469925 | 21 | 0.0952 | 23 | 0.2609 | 14 | 0.0714 | 23 | 0.087 | 81 | 0.1358 | 21 | 0.0476 | 17 | 0.1765 | 22 | 0.1818 | 21 | 0.0476 | 81 | 0.1111 |
| 8_9469955 | 22 | 0.8636 | 21 | 0.9524 | 13 | 0.7692 | 17 | 0.9412 | 73 | 0.8904 | 21 | 0.8571 | 15 | 1 | 21 | 0.7619 | 22 | 0.8636 | 79 | 0.8608 |
| 8_9469956 | 21 | 0.1429 | 21 | 0.0952 | 12 | 0.0833 | 19 | 0.2105 | 73 | 0.137 | 19 | 0.3158 | 12 | 0.0833 | 20 | 0.15 | 22 | 0.2727 | 73 | 0.2192 |
| 8_9469962 | 20 | 0.2 | 20 | 0.25 | 13 | 0.3077 | 23 | 0.3478 | 76 | 0.2763 | 23 | 0.2174 | 12 | 0.5 | 20 | 0.2 | 18 | 0.2222 | 73 | 0.2603 |
| 8_9469966 | 23 | 0.6522 | 22 | 0.7727 | 15 | 0.9333 | 21 | 0.7619 | 81 | 0.7654 | 23 | 0.6957 | 14 | 1 | 23 | 0.6957 | 21 | 0.7143 | 81 | 0.7531 |
| 8_9469980 | 22 | 0.2273 | 15 | 0.0667 | 14 | 0.1429 | 19 | 0.1579 | 70 | 0.1571 | 24 | 0.0833 | 15 | 0.1333 | 23 | 0.0435 | 23 | 0.1304 | 85 | 0.0941 |

| | | | | | | | | | | | | | | | | | | | | |
|---|---|---|---|---|---|---|---|---|---|---|---|---|---|---|---|---|---|---|---|---|
| 8_94699897 | 19 | 0.1579 | 20 | 0.15 | 12 | 0 | 18 | 0.1667 | 69 | 0.1304 | 21 | 0.0952 | 13 | 0.1538 | 20 | 0.15 | 20 | 0.3 | 74 | 0.1757 |
| 8_94765698 | 1 | 0 | 2 | 0 | 2 | 0 | 4 | 0.25 | 9 | 0.1111 | 1 | 0 | 6 | 0.1667 | 4 | 0 | 3 | 0.3333 | 14 | 0.1429 |
| 8_94784823 | 1 | 0 | 2 | 0.5 | 3 | 0 | 3 | 0.3333 | 9 | 0.2222 | 1 | 0 | 8 | 0.25 | 4 | 0.25 | 3 | 0.3333 | 16 | 0.25 |
| 8_94784880 | 2 | 1 | 2 | 0.5 | 4 | 0.75 | 3 | 0.3333 | 11 | 0.6364 | NA | NA | 9 | 0.7778 | 4 | 0.5 | 4 | 1 | 17 | 0.7647 |
| 8_94790943 | 3 | 0.6667 | 3 | 0.3333 | 4 | 0.25 | 4 | 0.5 | 14 | 0.4286 | 1 | 1 | 11 | 0.4545 | 3 | 0.3333 | 4 | 0.25 | 19 | 0.4211 |
| 8_94791015 | 3 | 0 | 3 | 0.3333 | 5 | 0.2 | 5 | 0.2 | 16 | 0.1875 | NA | NA | 12 | 0.3333 | 4 | 0.25 | 3 | 0 | 19 | 0.2632 |
| 8_94791685 | 4 | 0.5 | 5 | 0.2 | 5 | 0.4 | 5 | 0.4 | 19 | 0.3684 | 1 | 0 | 10 | 0.4 | 5 | 0.2 | 4 | 0.5 | 20 | 0.35 |
| 8_94792041 | 5 | 0.4 | 7 | 0 | 5 | 0 | 5 | 0 | 22 | 0.0909 | 2 | 0 | 13 | 0.0769 | 7 | 0 | 5 | 0.2 | 27 | 0.0741 |
| 8_94792078 | 5 | 0.2 | 7 | 0 | 4 | 0 | 4 | 0 | 20 | 0.05 | 2 | 0 | 9 | 0.2222 | 6 | 0 | 3 | 0 | 20 | 0.1 |
| 8_94792984 | 5 | 0 | 6 | 0.1667 | 5 | 0 | 4 | 0 | 20 | 0.05 | 1 | 0 | 10 | 0.2 | 9 | 0 | 5 | 0.2 | 25 | 0.12 |
| 8_94793035 | 5 | 0 | 8 | 0.125 | 5 | 0.2 | 7 | 0.1429 | 25 | 0.12 | 2 | 0 | 13 | 0.0769 | 12 | 0.0833 | 7 | 0.1429 | 34 | 0.0882 |
| 8_94794431 | 4 | 0.25 | 9 | 0 | 7 | 0.2857 | 9 | 0.3333 | 29 | 0.2069 | 9 | 0 | 10 | 0.4 | 11 | 0.1818 | 6 | 0.3333 | 36 | 0.2222 |
| 8_94794780 | 4 | 0.25 | 1 | 0 | 2 | 0 | 6 | 0 | 13 | 0.0769 | 2 | 0 | 6 | 0.1667 | 1 | 0 | 2 | 0 | 11 | 0.0909 |
| 8_94803310 | NA | NA | 1 | 1 | NA | NA | NA | NA | 1 | 1 | 1 | 1 | NA | NA | NA | NA | NA | NA | 1 | 1 |
| 8_94803485 | NA | NA | 1 | 1 | NA | NA | NA | NA | 1 | 1 | 1 | 1 | NA | NA | NA | NA | NA | NA | 1 | 1 |
| 8_94880418 | 3 | 0 | NA | NA | 6 | 0.1667 | 4 | 0.25 | 13 | 0.1538 | 2 | 0 | 3 | 0.3333 | 1 | 0 | 3 | 0 | 9 | 0.1111 |
| 8_94880479 | 3 | 0 | NA | NA | 7 | 0.1429 | 4 | 0.25 | 14 | 0.1429 | 2 | 0 | 3 | 0.3333 | 2 | 0 | 3 | 0 | 10 | 0.1 |
| 8_94880574 | 262 | 0.1107 | 265 | 0.1283 | 247 | 0.081 | 222 | 0.1351 | 996 | 0.1135 | 232 | 0.1379 | 238 | 0.1513 | 220 | 0.1182 | 253 | 0.1304 | 943 | 0.1347 |
| 8_95041348 | 3 | 0.3333 | 3 | 0 | 6 | 0.1667 | 4 | 0 | 16 | 0.125 | 2 | 0 | 2 | 1 | 6 | 0.1667 | 2 | 1 | 12 | 0.4167 |
| 8_95051485 | 3 | 0 | 2 | 0.5 | 1 | 0 | 5 | 0 | 11 | 0.0909 | NA | NA | 2 | 0.5 | 1 | 0 | 3 | 0 | 6 | 0.1667 |
| 8_95052898 | 4 | 0.25 | 3 | 0 | 3 | 0.3333 | 4 | 0 | 14 | 0.1429 | NA | NA | 2 | 0 | 4 | 0.25 | 2 | 0 | 8 | 0.125 |
| 8_95118975 | 11 | 0.0909 | 10 | 0 | 12 | 0.25 | 7 | 0 | 40 | 0.1 | 12 | 0.1667 | 15 | 0.0667 | 8 | 0.125 | 9 | 0 | 44 | 0.0909 |
| 8_95119035 | 12 | 0.1667 | 9 | 0 | 12 | 0.25 | 9 | 0.2222 | 42 | 0.1667 | 12 | 0.0833 | 15 | 0.1333 | 8 | 0.125 | 8 | 0 | 43 | 0.093 |
| 8_95119049 | 10 | 0.1 | 9 | 0.3333 | 11 | 0.1818 | 8 | 0.125 | 38 | 0.1842 | 11 | 0.4545 | 15 | 0.1333 | 8 | 0 | 8 | 0.25 | 42 | 0.2143 |
| 8_95121333 | 10 | 0.6 | 9 | 0.5556 | 13 | 0.6154 | 8 | 0.375 | 40 | 0.55 | 11 | 0.3636 | 13 | 0.3846 | 8 | 0.625 | 10 | 0.3 | 42 | 0.4048 |
| 8_95121424 | 12 | 0.6667 | 11 | 0.7273 | 11 | 0.5455 | 9 | 0.6667 | 43 | 0.6512 | 11 | 0.8182 | 14 | 0.5714 | 10 | 0.5 | 9 | 0.4444 | 44 | 0.5909 |
| 8_95121467 | 9 | 0.6667 | 9 | 0.3333 | 11 | 0.7273 | 9 | 0.5556 | 38 | 0.5789 | 11 | 0.6364 | 12 | 0.5833 | 11 | 0.8182 | 9 | 0.6667 | 43 | 0.6744 |
| 8_95121530 | 12 | 0.0833 | 8 | 0.125 | 9 | 0 | 7 | 0 | 36 | 0.0556 | 10 | 0 | 11 | 0.0909 | 9 | 0 | 8 | 0.125 | 38 | 0.0526 |
| 8_95121536 | 11 | 0.1818 | 11 | 0.0909 | 11 | 0.1818 | 8 | 0.25 | 41 | 0.1707 | 13 | 0.1538 | 13 | 0.3077 | 11 | 0.0909 | 11 | 0.1818 | 48 | 0.1875 |
| 8_95121555 | 13 | 0.2308 | 9 | 0.7778 | 11 | 0.5455 | 7 | 0.7143 | 40 | 0.525 | 10 | 0.5 | 12 | 0.4167 | 10 | 0.4 | 9 | 0.3333 | 41 | 0.4146 |
| 8_95121852 | 15 | 0.1333 | 12 | 0.0833 | 13 | 0 | 10 | 0 | 50 | 0.06 | 14 | 0.1429 | 17 | 0.1765 | 12 | 0 | 12 | 0.25 | 55 | 0.1455 |
| 8_95296839 | 3 | 0.3333 | 3 | 0 | 2 | 0 | 5 | 0.2 | 13 | 0.1538 | 3 | 0.3333 | 4 | 0 | 5 | 0.2 | 2 | 0 | 14 | 0.1429 |
| 8_95301716 | 4 | 0.75 | 3 | 0.6667 | 5 | 0.6 | 5 | 0.6 | 17 | 0.6471 | 7 | 0.7143 | 8 | 0.625 | 7 | 0.5714 | 6 | 0.3333 | 28 | 0.5714 |
| 8_95301780 | 5 | 0 | 4 | 0 | 5 | 0 | 4 | 0.25 | 18 | 0.0556 | 9 | 0.2222 | 7 | 0 | 8 | 0.125 | 6 | 0.3333 | 30 | 0.1667 |
| 8_95301805 | 1 | 0 | 3 | 0.3333 | 1 | 0 | 5 | 0 | 10 | 0.1 | 7 | 0 | 5 | 0.2 | 6 | 0 | 4 | 0.25 | 22 | 0.0909 |
| 8_95302003 | 5 | 0 | 4 | 0 | 5 | 0.2 | 2 | 0.5 | 16 | 0.125 | 7 | 0.1429 | 7 | 0 | 6 | 0.3333 | 5 | 0.2 | 25 | 0.16 |
| 8_95302014 | 3 | 1 | 4 | 0.5 | 5 | 0.8 | 3 | 0.6667 | 15 | 0.7333 | 6 | 0.6667 | 5 | 0.8 | 5 | 0.8 | 5 | 1 | 21 | 0.8095 |
| 8_95302160 | 4 | 1 | 3 | 1 | 6 | 0.5 | 3 | 0.6667 | 16 | 0.75 | 7 | 0.4286 | 8 | 0.625 | 5 | 0.6 | 8 | 0.75 | 28 | 0.6071 |
| 8_95302190 | 4 | 0.25 | 2 | 0 | 3 | 0 | 2 | 0 | 11 | 0.0909 | 5 | 0.4 | 3 | 0.6667 | 4 | 0.5 | 4 | 0.25 | 16 | 0.4375 |
| 8_95328857 | 2 | 0.5 | NA | NA | 1 | 0 | 1 | 0 | 4 | 0.25 | NA | NA | 2 | 1 | NA | NA | NA | NA | 2 | 1 |
| 8_95328862 | NA | NA | NA | NA | 1 | 1 | 1 | 1 | 2 | 1 | NA | NA | 2 | 0.5 | NA | NA | NA | NA | 2 | 0.5 |
| 8_95349605 | 1 | 1 | NA | NA | 1 | 0 | 1 | 1 | 3 | 0.6667 | NA | NA | 4 | 0.75 | 1 | 0 | NA | NA | 5 | 0.6 |
| 8_95349677 | 1 | 1 | NA | NA | 2 | 1 | 1 | 1 | 4 | 1 | NA | NA | 4 | 0.25 | 1 | 0 | NA | NA | 5 | 0.2 |
| 8_95349710 | 1 | 1 | NA | NA | 3 | 1 | NA | NA | 4 | 1 | NA | NA | 4 | 1 | 1 | 1 | NA | NA | 5 | 1 |
| 8_95349849 | 2 | 1 | NA | NA | 3 | 0.6667 | 1 | 0 | 6 | 0.6667 | NA | NA | 4 | 0.75 | 1 | 1 | NA | NA | 5 | 0.8 |
| 8_95349880 | 2 | 0.5 | NA | NA | 3 | 0 | 1 | 0 | 6 | 0.1667 | NA | NA | 3 | 0.3333 | 1 | 0 | NA | NA | 4 | 0.25 |
| 8_95349930 | 1 | 1 | NA | NA | 2 | 0 | 1 | 1 | 4 | 0.5 | NA | NA | 4 | 0.5 | 1 | 1 | 1 | 0 | 6 | 0.5 |
| 8_95349950 | 2 | 1 | NA | NA | 2 | 0.5 | 1 | 1 | 5 | 0.8 | NA | NA | 4 | 1 | 1 | 1 | NA | NA | 5 | 1 |
| 8_95385451 | 3 | 0.6667 | NA | NA | 5 | 0.8 | 1 | 1 | 9 | 0.7778 | 2 | 0.5 | 4 | 0.75 | 1 | 0 | 1 | 1 | 8 | 0.625 |
| 8_95385534 | 3 | 0 | NA | NA | 5 | 0.2 | NA | NA | 8 | 0.125 | 1 | 0 | 4 | 0.25 | 1 | 0 | 1 | 0 | 7 | 0.1429 |
| 8_95385899 | 4 | 0 | NA | NA | 4 | 0 | 1 | 1 | 9 | 0.1111 | 2 | 0 | 3 | 0.3333 | NA | NA | 1 | 0 | 6 | 0.1667 |
| 8_95399825 | 21 | 0.0476 | 14 | 0.0714 | 24 | 0.0417 | 15 | 0.1333 | 74 | 0.0676 | 10 | 0.1 | 9 | 0 | 10 | 0.1 | 10 | 0 | 39 | 0.0513 |
| 8_95399844 | 21 | 0.5238 | 15 | 0.4 | 26 | 0.3462 | 15 | 0.2667 | 77 | 0.3896 | 12 | 0.5 | 11 | 0.3636 | 12 | 0.4167 | 10 | 0.4 | 45 | 0.4222 |
| 8_95399906 | 20 | 0.1 | 14 | 0 | 25 | 0.16 | 16 | 0.125 | 75 | 0.1067 | 11 | 0.0909 | 10 | 0 | 11 | 0.1818 | 9 | 0 | 41 | 0.0732 |
| 8_95399929 | 19 | 0.1053 | 15 | 0.1333 | 23 | 0.087 | 15 | 0.0667 | 72 | 0.0972 | 10 | 0.1 | 7 | 0 | 12 | 0.3333 | 10 | 0.1 | 39 | 0.1538 |
| 8_95400921 | 24 | 0.125 | 17 | 0.0588 | 23 | 0.0435 | 21 | 0.0476 | 85 | 0.0706 | 14 | 0.1429 | 13 | 0.0769 | 21 | 0.0476 | 12 | 0 | 60 | 0.0667 |
| 8_95402355 | 5 | 0.2 | 6 | 0 | 12 | 0.1667 | 7 | 0.1429 | 30 | 0.1333 | 4 | 0.25 | 8 | 0.25 | 7 | 0 | 9 | 0 | 28 | 0.1071 |
| 8_95402385 | 6 | 0 | 6 | 0 | 14 | 0.1429 | 5 | 0.2 | 31 | 0.0968 | 4 | 0 | 8 | 0 | 6 | 0.3333 | 8 | 0 | 26 | 0.0769 |
| 8_95402697 | 3 | 0.3333 | 7 | 0.4286 | 12 | 0.25 | 3 | 0.3333 | 25 | 0.32 | 3 | 0 | 4 | 0 | 5 | 0.2 | 7 | 0.1429 | 19 | 0.1053 |
| 8_95402724 | 5 | 0.6 | 6 | 0.5 | 13 | 0.6154 | 7 | 1 | 31 | 0.6774 | 4 | 0.75 | 6 | 0.8333 | 6 | 0.6667 | 9 | 0.5556 | 25 | 0.68 |
| 8_95406741 | NA | NA | 2 | 0.5 | NA | NA | NA | NA | 2 | 0.5 | NA | NA | NA | NA | 1 | 1 | NA | NA | 1 | 1 |
| 8_95506319 | 37 | 0.8108 | 42 | 0.7619 | 25 | 0.76 | 25 | 0.68 | 129 | 0.7597 | 32 | 0.7812 | 38 | 0.7632 | 57 | 0.7018 | 35 | 0.9143 | 162 | 0.7778 |
| 8_95506356 | 33 | 0.9394 | 44 | 0.8864 | 27 | 0.9259 | 26 | 0.9615 | 130 | 0.9231 | 33 | 0.9091 | 39 | 0.9231 | 59 | 0.9322 | 31 | 0.871 | 162 | 0.9136 |
| 8_95506416 | 32 | 1 | 34 | 0.9706 | 20 | 1 | 19 | 1 | 105 | 0.9905 | 32 | 0.9375 | 35 | 0.9429 | 51 | 0.9608 | 25 | 0.88 | 143 | 0.9371 |
| 8_95506420 | 29 | 0.1724 | 32 | 0.1875 | 20 | 0.2 | 23 | 0.3913 | 104 | 0.2308 | 23 | 0.1304 | 28 | 0.2143 | 38 | 0.2895 | 21 | 0.0476 | 110 | 0.1909 |

| | | | | | | | | | | | | | | | | | | | | |
|---|---|---|---|---|---|---|---|---|---|---|---|---|---|---|---|---|---|---|---|---|
| 8_95506423 | 1 | 1 | 4 | 0 | 2 | 0.5 | 1 | 0 | 8 | 0.25 | NA | NA | 3 | 0 | 3 | 0 | 3 | 0.6667 | 9 | 0.2222 |
| 8_95506447 | 37 | 0.7838 | 45 | 0.6667 | 29 | 0.5862 | 28 | 0.7143 | 139 | 0.6906 | 32 | 0.5312 | 46 | 0.8478 | 57 | 0.7193 | 31 | 0.5484 | 166 | 0.6867 |
| 8_95506456 | 29 | 0.7586 | 47 | 0.8511 | 30 | 0.8 | 27 | 0.7778 | 133 | 0.8045 | 35 | 0.8571 | 45 | 0.9111 | 54 | 0.9259 | 34 | 0.7647 | 168 | 0.875 |
| 8_95506749 | 38 | 0.6053 | 51 | 0.6471 | 31 | 0.7097 | 25 | 0.8 | 145 | 0.6759 | 35 | 0.6286 | 46 | 0.6957 | 62 | 0.6129 | 34 | 0.5588 | 177 | 0.6271 |
| 8_95506754 | 44 | 0.0909 | 55 | 0.2364 | 31 | 0.0645 | 33 | 0.0909 | 163 | 0.135 | 37 | 0.1081 | 49 | 0.0612 | 62 | 0.0806 | 38 | 0.0789 | 186 | 0.0806 |
| 8_95506959 | 43 | 0.186 | 58 | 0.2759 | 34 | 0.1765 | 38 | 0.2105 | 173 | 0.2197 | 40 | 0.125 | 51 | 0.1569 | 71 | 0.1972 | 41 | 0.122 | 203 | 0.1576 |
| 8_95507224 | 37 | 0.2703 | 50 | 0.26 | 35 | 0.2857 | 35 | 0.3429 | 157 | 0.2866 | 38 | 0.3421 | 50 | 0.38 | 62 | 0.1774 | 34 | 0.2059 | 184 | 0.2717 |
| 8_95507302 | 37 | 0.1351 | 58 | 0.1724 | 38 | 0.1579 | 42 | 0.1905 | 175 | 0.1657 | 38 | 0.1316 | 58 | 0.2069 | 68 | 0.1324 | 37 | 0.1892 | 201 | 0.1642 |
| 8_95507392 | 45 | 0.4889 | 66 | 0.5303 | 42 | 0.4762 | 44 | 0.4091 | 197 | 0.4822 | 43 | 0.3256 | 62 | 0.4194 | 83 | 0.4458 | 48 | 0.4583 | 236 | 0.4195 |
| 8_95507484 | 36 | 0.3333 | 54 | 0.4815 | 38 | 0.4737 | 33 | 0.3636 | 161 | 0.4224 | 42 | 0.619 | 48 | 0.5625 | 66 | 0.5303 | 35 | 0.5429 | 191 | 0.5602 |
| 8_95507820 | 37 | 0.1081 | 80 | 0.1 | 47 | 0.0638 | 46 | 0.1957 | 210 | 0.1143 | 50 | 0.18 | 64 | 0.2031 | 89 | 0.1124 | 45 | 0.2 | 248 | 0.1653 |
| 8_95508043 | 31 | 0.9032 | 70 | 0.9143 | 43 | 0.9767 | 43 | 0.9302 | 187 | 0.9305 | 35 | 0.9429 | 50 | 0.92 | 73 | 0.9452 | 42 | 0.881 | 200 | 0.925 |
| 8_95508047 | 29 | 0.1379 | 78 | 0.1026 | 42 | 0.0714 | 44 | 0.0909 | 193 | 0.0984 | 44 | 0.0227 | 57 | 0.0877 | 79 | 0.1392 | 40 | 0.125 | 220 | 0.1 |
| 8_95508147 | 42 | 0.1667 | 77 | 0.2208 | 50 | 0.16 | 47 | 0.2766 | 216 | 0.2083 | 55 | 0.1091 | 66 | 0.1515 | 90 | 0.0889 | 45 | 0.1556 | 256 | 0.1211 |
| 8_95508430 | 32 | 0.4688 | 77 | 0.4416 | 45 | 0.4667 | 37 | 0.4054 | 191 | 0.445 | 49 | 0.3673 | 54 | 0.4074 | 85 | 0.3765 | 36 | 0.3889 | 224 | 0.3839 |
| 8_95508494 | 40 | 0.1 | 90 | 0.0556 | 61 | 0.0328 | 45 | 0.0222 | 236 | 0.0508 | 64 | 0.0156 | 71 | 0.0704 | 101 | 0.0891 | 48 | 0.125 | 284 | 0.0739 |
| 8_95508510 | 33 | 0.4545 | 84 | 0.4048 | 58 | 0.5 | 34 | 0.3824 | 209 | 0.4354 | 58 | 0.4138 | 57 | 0.4737 | 87 | 0.4368 | 42 | 0.5476 | 244 | 0.459 |
| 8_95508532 | 41 | 0.2439 | 91 | 0.2308 | 68 | 0.3088 | 49 | 0.1837 | 249 | 0.245 | 59 | 0.1864 | 61 | 0.3279 | 107 | 0.2617 | 48 | 0.2708 | 275 | 0.2618 |
| 8_95508690 | 41 | 0.3902 | 88 | 0.4659 | 61 | 0.623 | 49 | 0.5918 | 239 | 0.5188 | 64 | 0.4688 | 74 | 0.5811 | 110 | 0.4818 | 52 | 0.4615 | 300 | 0.5 |
| 8_95508881 | 42 | 0.0952 | 93 | 0.0968 | 68 | 0.0882 | 54 | 0.1111 | 257 | 0.0973 | 82 | 0.061 | 81 | 0.0864 | 110 | 0.1 | 59 | 0.1017 | 332 | 0.0873 |
| 8_95508895 | 44 | 0.9318 | 100 | 0.89 | 70 | 0.9429 | 53 | 0.9623 | 267 | 0.9251 | 79 | 0.9241 | 82 | 0.9146 | 116 | 0.8879 | 64 | 0.9375 | 341 | 0.912 |
| 8_95508951 | 45 | 0.6 | 102 | 0.6373 | 75 | 0.5867 | 57 | 0.6842 | 279 | 0.6272 | 87 | 0.6092 | 85 | 0.7294 | 119 | 0.6639 | 65 | 0.6923 | 356 | 0.6713 |
| 8_95544652 | NA | NA | 4 | 0.75 | 2 | 0.5 | 1 | 1 | 7 | 0.7143 | 3 | 1 | NA | NA | 1 | 1 | NA | NA | 4 | 1 |
| 8_95546531 | 2 | 1 | 3 | 0.6667 | 3 | 0.6667 | NA | NA | 8 | 0.75 | NA | NA | 4 | 0.75 | 1 | 1 | 3 | 1 | 8 | 0.875 |
| 8_95548286 | 3 | 0 | 2 | 0 | 1 | 1 | 1 | 0 | 7 | 0.1429 | 2 | 0.5 | 3 | 0 | NA | NA | 1 | 0 | 6 | 0.1667 |
| 8_95549109 | 5 | 0.4 | 2 | 0.5 | 4 | 0.5 | 2 | 0 | 13 | 0.3846 | 2 | 0 | 1 | 0 | 2 | 0 | 3 | 0.3333 | 8 | 0.125 |
| 8_95549386 | 8 | 0.875 | 6 | 0.5 | 6 | 0.6667 | 7 | 0.4286 | 27 | 0.6296 | 4 | 1 | 4 | 1 | 3 | 0.6667 | 4 | 1 | 15 | 0.9333 |
| 8_95549483 | 6 | 0.8333 | 5 | 0.8 | 6 | 0.5 | 7 | 0.4286 | 24 | 0.625 | 4 | 0.5 | 4 | 0.5 | 3 | 0.6667 | 4 | 0.5 | 15 | 0.5333 |
| 8_95549528 | 8 | 0.25 | 4 | 0 | 7 | 0.2857 | 7 | 0.2857 | 26 | 0.2308 | 6 | 0 | 4 | 0 | 2 | 0 | 4 | 0.25 | 16 | 0.0625 |
| 8_95549544 | 8 | 0.25 | 4 | 0 | 7 | 0.2857 | 6 | 0 | 25 | 0.16 | 4 | 0 | 4 | 0 | 3 | 0.3333 | 5 | 0 | 16 | 0.0625 |
| 8_95549628 | 8 | 0.125 | 5 | 0 | 6 | 0.1667 | 6 | 0.1667 | 25 | 0.12 | 5 | 0 | 3 | 0.6667 | 3 | 0 | 4 | 0.25 | 15 | 0.2 |
| 8_95580880 | 5 | 0 | 12 | 0.0833 | 7 | 0.1429 | 9 | 0.1111 | 33 | 0.0909 | 7 | 0.2857 | 7 | 0 | 11 | 0.1818 | 9 | 0.1111 | 34 | 0.1471 |
| 8_95580946 | 8 | 0.625 | 11 | 0.7273 | 6 | 0.6667 | 8 | 0.875 | 33 | 0.7273 | 6 | 1 | 6 | 1 | 11 | 0.7273 | 9 | 0.7778 | 32 | 0.8438 |
| 8_95581022 | 6 | 0.3333 | 13 | 0.3077 | 9 | 0.1111 | 8 | 0.125 | 36 | 0.2222 | 7 | 0.4286 | 6 | 0.5 | 13 | 0.3846 | 7 | 0.4286 | 33 | 0.4242 |
| 8_95581028 | 7 | 1 | 12 | 0.75 | 8 | 0.625 | 7 | 0.8571 | 34 | 0.7941 | 7 | 1 | 6 | 0.6667 | 11 | 0.5455 | 7 | 0.5714 | 31 | 0.6774 |
| 8_95581067 | 9 | 0.3333 | 12 | 0.0833 | 9 | 0.1111 | 8 | 0.75 | 38 | 0.2895 | 7 | 0.1429 | 7 | 0.1429 | 11 | 0.0909 | 10 | 0.2 | 35 | 0.1429 |
| 8_95581171 | 9 | 0.3333 | 12 | 0.1667 | 9 | 0.3333 | 8 | 0.375 | 38 | 0.2895 | 8 | 0.375 | 7 | 0 | 14 | 0.0714 | 11 | 0.2727 | 40 | 0.175 |
| 8_95581358 | 6 | 0.3333 | 10 | 0.1 | 7 | 0.1429 | 8 | 0.125 | 31 | 0.1613 | 7 | 0 | 6 | 0 | 15 | 0.2 | 11 | 0.1818 | 39 | 0.1282 |
| 8_95581427 | 6 | 0.5 | 8 | 0.5 | 6 | 0.6667 | 6 | 0.6667 | 26 | 0.5769 | 9 | 0.4444 | 5 | 0.8 | 7 | 0.7143 | 9 | 0.6667 | 30 | 0.6333 |
| 8_95590177 | 12 | 0.0833 | 13 | 0.0769 | 13 | 0.2308 | 10 | 0.3 | 48 | 0.1667 | 10 | 0 | 13 | 0.2308 | 10 | 0.4 | 14 | 0.1429 | 47 | 0.1915 |
| 8_95590268 | 9 | 0.2222 | 11 | 0.1818 | 8 | 0.125 | 8 | 0.5 | 36 | 0.25 | 12 | 0.4167 | 9 | 0.3333 | 9 | 0.4444 | 13 | 0.5385 | 43 | 0.4419 |
| 8_95590931 | 1 | 0 | 6 | 0.1667 | 4 | 0.25 | 3 | 0 | 14 | 0.1429 | 3 | 0.3333 | 1 | 1 | 2 | 0 | 1 | 0 | 7 | 0.2857 |
| 8_95591080 | 1 | 0 | 6 | 0.1667 | 4 | 0 | 3 | 0 | 14 | 0.0714 | 3 | 0.3333 | 1 | 0 | 2 | 0 | 1 | 0 | 7 | 0.1429 |
| 8_95591327 | 1 | 1 | 7 | 0.1429 | 4 | 0 | 3 | 0.3333 | 15 | 0.2 | 3 | 0.3333 | 1 | 0 | 2 | 0.5 | NA | NA | 6 | 0.3333 |
| 8_95591554 | 1 | 1 | 6 | 0.8333 | 3 | 0 | 3 | 1 | 13 | 0.6923 | 3 | 0.6667 | 1 | 1 | 1 | 0 | 1 | 1 | 6 | 0.6667 |
| 8_95591687 | 1 | 0 | 5 | 0.6 | 4 | 0.25 | 3 | 0.6667 | 13 | 0.4615 | 3 | 0.6667 | 1 | 1 | 1 | 1 | 1 | 1 | 6 | 0.8333 |
| 8_95591739 | 1 | 1 | 5 | 0.2 | 3 | 0.6667 | 4 | 0.25 | 13 | 0.3846 | 2 | 0.5 | 1 | 1 | 1 | 1 | 1 | 0 | 5 | 0.6 |
| 8_95740350 | 10 | 0.1 | 15 | 0.2 | 12 | 0.0833 | 13 | 0 | 50 | 0.1 | 6 | 0.1667 | 7 | 0.1429 | 14 | 0.0714 | 9 | 0 | 36 | 0.0833 |
| 8_95740638 | 10 | 0.6 | 14 | 0.7857 | 10 | 0.6 | 12 | 0.6667 | 46 | 0.6739 | 7 | 0.7143 | 8 | 0.5 | 12 | 0.3333 | 10 | 0.3 | 37 | 0.4324 |
| 8_95825410 | 5 | 1 | 8 | 0.75 | 3 | 0.6667 | 6 | 1 | 22 | 0.8636 | 6 | 1 | 5 | 0.8 | 4 | 0.5 | 1 | 1 | 16 | 0.8125 |
| 8_95825565 | 6 | 1 | 6 | 1 | 2 | 0.5 | 8 | 0.875 | 22 | 0.9091 | 6 | 0.6667 | 4 | 1 | 5 | 1 | 1 | 0 | 16 | 0.8125 |
| 8_95825677 | 4 | 0.5 | 3 | 0.3333 | 1 | 1 | 6 | 0.3333 | 14 | 0.4286 | 5 | 0.2 | 4 | 0.75 | 3 | 1 | 1 | 1 | 13 | 0.6154 |
| 8_95825762 | 1 | 1 | NA | NA | NA | NA | 2 | 1 | 3 | 1 | 1 | 1 | 1 | 1 | 1 | 1 | NA | NA | 3 | 1 |
| 8_95825857 | 1 | 0 | NA | NA | NA | NA | 2 | 1 | 3 | 0.6667 | 1 | 0 | 1 | 1 | 1 | 0 | NA | NA | 3 | 0.3333 |
| 8_95825973 | 1 | 0 | NA | NA | NA | NA | 1 | 1 | 2 | 0.5 | 1 | 0 | 1 | 0 | 1 | 1 | NA | NA | 3 | 0.3333 |
| 8_95826152 | NA | NA | NA | NA | NA | NA | 1 | 1 | 1 | 1 | 1 | 1 | 1 | 0 | 1 | 0 | NA | NA | 3 | 0.3333 |
| 8_95826789 | 2 | 0 | 6 | 0.1667 | 3 | 0 | 2 | 0 | 13 | 0.0769 | 6 | 0.1667 | 4 | 0.5 | 2 | 1 | 4 | 0.5 | 16 | 0.4375 |
| 8_95830529 | 1 | 1 | NA | NA | NA | NA | 1 | 1 | 2 | 1 | NA | NA | NA | NA | NA | NA | 1 | 1 | 1 | 1 |
| 8_96052603 | NA | NA | 1 | 1 | NA | NA | NA | NA | 1 | 1 | 1 | 0 | 1 | 1 | 3 | 0 | NA | NA | 5 | 0.2 |
| 8_96053006 | 1 | 0 | 6 | 0.3333 | 1 | 1 | 1 | 0 | 9 | 0.3333 | 1 | 0 | 3 | 0 | 7 | 0.1429 | 1 | 0 | 12 | 0.0833 |
| 8_96053012 | 1 | 0 | 4 | 0.25 | 1 | 0 | 1 | 0 | 7 | 0.1429 | 1 | 0 | 3 | 0.3333 | 6 | 0 | 1 | 0 | 11 | 0.0909 |
| 8_96053037 | NA | NA | 5 | 0.4 | NA | NA | 1 | 0 | 6 | 0.3333 | 2 | 0 | 4 | 0.25 | 6 | 0.1667 | 1 | 0 | 13 | 0.1538 |
| 8_96053064 | 1 | 0 | 5 | 1 | 1 | 1 | 1 | 1 | 8 | 0.875 | 2 | 1 | 3 | 1 | 7 | 0.7143 | 1 | 0 | 13 | 0.7692 |
| 8_96053266 | 1 | 0 | 3 | 0.3333 | 1 | 0 | 1 | 0 | 6 | 0.1667 | 2 | 1 | 3 | 0.3333 | 7 | 0.2857 | 1 | 0 | 13 | 0.3846 |
| 8_96054008 | 1 | 0 | 2 | 0.5 | 1 | 0 | NA | NA | 4 | 0.25 | NA | NA | 3 | 0.6667 | 5 | 0.2 | 1 | 0 | 9 | 0.3333 |

| | | | | | | | | | | | | | | | | | | | | |
|---|---|---|---|---|---|---|---|---|---|---|---|---|---|---|---|---|---|---|---|---|
| 8_96054189 | 1 | 1 | 1 | 1 | NA | NA | NA | NA | 2 | 1 | NA | NA | 2 | 0.5 | 5 | 0.8 | 1 | 1 | 8 | 0.75 |
| 8_96054231 | 1 | 1 | 1 | 1 | NA | NA | NA | NA | 2 | 1 | NA | NA | 2 | 1 | 5 | 1 | 1 | 1 | 8 | 1 |
| 8_96054358 | 1 | 1 | 1 | 1 | NA | NA | NA | NA | 2 | 1 | NA | NA | 2 | 1 | 5 | 1 | 1 | 1 | 8 | 1 |
| 8_96054736 | 1 | 0 | 2 | 0.5 | NA | NA | NA | NA | 3 | 0.3333 | NA | NA | 1 | 1 | 4 | 1 | 1 | 1 | 6 | 1 |
| 8_96055016 | 1 | 1 | 1 | 1 | NA | NA | NA | NA | 2 | 1 | NA | NA | 1 | 1 | 4 | 1 | NA | NA | 5 | 1 |
| 8_96056074 | 1 | 0 | 2 | 1 | NA | NA | NA | NA | 3 | 0.6667 | NA | NA | 1 | 1 | NA | NA | NA | NA | 1 | 1 |
| 8_96056123 | 1 | 1 | 2 | 1 | NA | NA | NA | NA | 3 | 1 | NA | NA | 1 | 1 | NA | NA | NA | NA | 1 | 1 |
| 8_96056171 | NA | NA | 2 | 0.5 | NA | NA | NA | NA | 2 | 0.5 | NA | NA | 1 | 1 | NA | NA | NA | NA | 1 | 1 |
| 8_96098826 | NA | NA | 2 | 1 | 3 | 1 | 1 | 1 | 6 | 1 | NA | NA | 1 | 1 | 1 | 1 | 1 | 1 | 3 | 1 |
| 8_96099005 | NA | NA | 1 | 1 | 4 | 1 | 2 | 1 | 7 | 1 | NA | NA | NA | NA | 1 | 0 | 2 | 1 | 3 | 0.6667 |
| 8_96099190 | NA | NA | 3 | 0.6667 | 4 | 0 | 2 | 1 | 9 | 0.4444 | NA | NA | 1 | 1 | 1 | 0 | 2 | 1 | 4 | 0.75 |
| 8_96099511 | NA | NA | 2 | 1 | 4 | 0.75 | 3 | 1 | 9 | 0.8889 | NA | NA | 1 | 1 | 1 | 1 | 2 | 1 | 4 | 1 |
| 8_96099617 | NA | NA | 3 | 0 | 5 | 0 | 2 | 0.5 | 10 | 0.1 | NA | NA | 1 | 0 | 2 | 0 | 3 | 0.3333 | 6 | 0.1667 |
| 8_96100588 | NA | NA | 4 | 0.25 | 2 | 0.5 | 3 | 0.3333 | 9 | 0.3333 | 1 | 0 | 1 | 1 | 2 | 0 | 4 | 0.25 | 8 | 0.25 |
| 8_96100826 | 1 | 1 | 3 | 0.6667 | 2 | 0.5 | 3 | 1 | 9 | 0.7778 | NA | NA | 1 | 1 | 2 | 0.5 | 2 | 1 | 5 | 0.8 |
| 8_96100974 | NA | NA | NA | NA | NA | NA | 2 | 0.5 | 2 | 0.5 | 1 | 1 | NA | NA | 1 | 0 | 1 | 0 | 3 | 0.3333 |
| 8_96101133 | NA | NA | NA | NA | NA | NA | 2 | 0.5 | 2 | 0.5 | 1 | 0 | NA | NA | 1 | 0 | 1 | 1 | 3 | 0.3333 |
| 8_96147214 | 2 | 0.5 | 9 | 0.3333 | 4 | 0 | 8 | 0.5 | 23 | 0.3478 | 4 | 0.5 | 5 | 0 | 5 | 0.6 | 3 | 0 | 17 | 0.2941 |
| 8_96147215 | 2 | 0 | 2 | 0.5 | 2 | 1 | NA | NA | 6 | 0.5 | 1 | 0 | NA | NA | 1 | 0 | 2 | 1 | 4 | 0.5 |
| 8_96147602 | NA | NA | 1 | 0 | 3 | 0.3333 | 1 | 1 | 5 | 0.4 | 2 | 0.5 | NA | NA | NA | NA | NA | NA | 2 | 0.5 |
| 8_96148640 | NA | NA | 4 | 0.25 | 3 | 0 | 2 | 0 | 9 | 0.1111 | 1 | 1 | NA | NA | NA | NA | NA | NA | 1 | 1 |
| 8_96174710 | 8 | 0.625 | 6 | 0.3333 | 3 | 0.3333 | 4 | 0.25 | 21 | 0.4286 | 6 | 0.5 | 3 | 1 | 6 | 0.5 | 4 | 0.5 | 19 | 0.5789 |
| 8_96179119 | 2 | 0.5 | 2 | 0 | 1 | 0 | 1 | 0 | 6 | 0.1667 | 1 | 1 | 2 | 0.5 | 2 | 0 | 1 | 0 | 6 | 0.3333 |
| 8_96179521 | 2 | 0 | 1 | 1 | 1 | 1 | 1 | 0 | 5 | 0.4 | NA | NA | 2 | 0.5 | 2 | 0.5 | 1 | 1 | 5 | 0.6 |
| 8_96179764 | 1 | 1 | 1 | 1 | NA | NA | 1 | 0 | 3 | 0.6667 | NA | NA | 2 | 1 | 1 | 1 | 1 | 1 | 4 | 1 |
| 8_96179859 | 1 | 1 | 1 | 0 | NA | NA | 2 | 0 | 4 | 0.25 | NA | NA | 3 | 0.3333 | 1 | 1 | 1 | 0 | 5 | 0.4 |
| 8_96179929 | 1 | 1 | 1 | 1 | NA | NA | 2 | 0 | 4 | 0.5 | NA | NA | 2 | 0.5 | NA | NA | NA | NA | 2 | 0.5 |
| 8_96214999 | 3 | 0 | 5 | 0.6 | 1 | 1 | 2 | 0.5 | 11 | 0.4545 | 3 | 0.3333 | 3 | 0.3333 | 3 | 0 | 1 | 0 | 10 | 0.2 |
| 8_96272620 | 2 | 0 | 3 | 0 | NA | NA | 1 | 1 | 6 | 0.1667 | NA | NA | 1 | 0 | NA | NA | 1 | 1 | 2 | 0.5 |
| 8_96355554 | NA | NA | NA | NA | NA | NA | 1 | 1 | 1 | 1 | NA | NA | 1 | 1 | NA | NA | 1 | 1 | 2 | 1 |
| 8_96370142 | 1 | 0 | 1 | 1 | 1 | 1 | NA | NA | 3 | 0.6667 | 1 | 1 | NA | NA | NA | NA | NA | NA | 1 | 1 |
| 8_96370261 | 1 | 1 | 1 | 1 | 1 | 1 | NA | NA | 3 | 1 | 1 | 1 | NA | NA | NA | NA | NA | NA | 1 | 1 |
| 8_96370373 | 1 | 1 | 1 | 1 | 1 | 1 | NA | NA | 3 | 1 | 1 | 1 | NA | NA | NA | NA | NA | NA | 1 | 1 |
| 8_96370406 | 1 | 1 | 1 | 1 | 1 | 1 | NA | NA | 3 | 1 | 1 | 1 | NA | NA | NA | NA | NA | NA | 1 | 1 |
| 8_96370946 | 1 | 1 | 1 | 0 | 1 | 1 | NA | NA | 3 | 0.6667 | 1 | 1 | NA | NA | NA | NA | NA | NA | 1 | 1 |
| 8_96371051 | 1 | 1 | 1 | 1 | 1 | 1 | NA | NA | 3 | 1 | 1 | 1 | NA | NA | NA | NA | NA | NA | 1 | 1 |
| 8_96371305 | 1 | 1 | 1 | 1 | 1 | 1 | NA | NA | 3 | 1 | NA | NA | NA | NA | NA | NA | 1 | 1 | 1 | 1 |
| 8_96399863 | NA | NA | 1 | 1 | NA | NA | NA | NA | 1 | 1 | 1 | 1 | NA | NA | NA | NA | NA | NA | 1 | 1 |
| 8_96401600 | NA | NA | NA | NA | NA | NA | 1 | 1 | 1 | 1 | 1 | 1 | NA | NA | 1 | 1 | NA | NA | 2 | 1 |
| 8_96401743 | NA | NA | NA | NA | NA | NA | 1 | 1 | 1 | 1 | 1 | 1 | NA | NA | NA | NA | NA | NA | 1 | 1 |
| 8_96440123 | 249 | 0.0643 | 212 | 0.0566 | 213 | 0.0704 | 267 | 0.0861 | 941 | 0.0701 | 165 | 0.0788 | 270 | 0.0741 | 270 | 0.063 | 205 | 0.1122 | 910 | 0.0802 |
| 8_96440355 | 247 | 0.0931 | 225 | 0.08 | 220 | 0.0818 | 273 | 0.0476 | 965 | 0.0746 | 164 | 0.0366 | 264 | 0.0871 | 288 | 0.0833 | 229 | 0.0917 | 945 | 0.0783 |
| 8_96440675 | 317 | 0.0915 | 253 | 0.0593 | 284 | 0.0704 | 331 | 0.0363 | 1185 | 0.0641 | 191 | 0.0733 | 337 | 0.0593 | 335 | 0.0657 | 251 | 0.0717 | 1114 | 0.0664 |
| 8_96441227 | 352 | 0.0824 | 274 | 0.1131 | 297 | 0.101 | 366 | 0.0847 | 1289 | 0.0939 | 226 | 0.0841 | 350 | 0.0971 | 377 | 0.1247 | 268 | 0.1269 | 1221 | 0.1097 |
| 8_96443391 | 3 | 0.3333 | NA | NA | NA | NA | 2 | 0.5 | 5 | 0.4 | 3 | 0 | 1 | 0 | NA | NA | 1 | 1 | 5 | 0.2 |
| 8_96445276 | 3 | 0.6667 | NA | NA | NA | NA | 3 | 0.6667 | 6 | 0.6667 | 3 | 1 | 1 | 1 | 2 | 0 | 2 | 0 | 8 | 0.5 |
| 8_96445294 | 3 | 0.6667 | NA | NA | 1 | 1 | 3 | 0.6667 | 7 | 0.7143 | 2 | 1 | 1 | 1 | 2 | 0.5 | 3 | 1 | 8 | 0.875 |
| 8_96445397 | 2 | 1 | NA | NA | 1 | 0 | 3 | 0.6667 | 6 | 0.6667 | 3 | 0 | 1 | 1 | 2 | 0.5 | 2 | 0.5 | 8 | 0.375 |
| 8_96445472 | 3 | 0.3333 | NA | NA | 1 | 1 | 3 | 1 | 7 | 0.7143 | 4 | 0.75 | NA | NA | 2 | 0 | 2 | 0.5 | 8 | 0.5 |
| 8_96446853 | 8 | 0 | 13 | 0.1538 | 15 | 0 | 16 | 0.0625 | 52 | 0.0577 | 8 | 0 | 11 | 0.0909 | 13 | 0.1538 | 5 | 0.2 | 37 | 0.1081 |
| 8_96446957 | 8 | 0.125 | 12 | 0.0833 | 13 | 0.0769 | 14 | 0 | 47 | 0.0638 | 10 | 0 | 10 | 0 | 13 | 0 | 6 | 0.3333 | 39 | 0.0513 |
| 8_96446974 | 9 | 0 | 8 | 0.375 | 12 | 0.0833 | 15 | 0 | 44 | 0.0909 | 10 | 0.3 | 13 | 0.2308 | 14 | 0.1429 | 5 | 0 | 42 | 0.1905 |
| 8_96447030 | 7 | 0.4286 | 13 | 0.3846 | 15 | 0.6667 | 15 | 0.4 | 50 | 0.48 | 9 | 0.2222 | 12 | 0.25 | 14 | 0.4286 | 6 | 0.8333 | 41 | 0.3902 |
| 8_96468503 | 3 | 0 | 5 | 0 | 4 | 0.25 | 7 | 0 | 19 | 0.0526 | 6 | 0 | 4 | 0 | 4 | 0.25 | 3 | 0.3333 | 17 | 0.1176 |
| 8_96468563 | 1 | 0 | 5 | 0.2 | 4 | 0 | 7 | 0.1429 | 17 | 0.1176 | 6 | 0.1667 | 4 | 0.25 | 7 | 0.4286 | 3 | 0.3333 | 20 | 0.3 |
| 8_96479249 | 1 | 0 | 2 | 0 | 2 | 0 | 6 | 0.3333 | 11 | 0.1818 | 5 | 0.8 | 2 | 1 | 3 | 1 | 2 | 0 | 12 | 0.75 |
| 8_96552520 | 11 | 0.0909 | 13 | 0.1538 | 5 | 0 | 12 | 0.25 | 41 | 0.1463 | 7 | 0.2857 | 3 | 0.3333 | 14 | 0.0714 | 3 | 0.3333 | 27 | 0.1852 |
| 8_96552559 | 10 | 0 | 14 | 0 | 5 | 0.2 | 11 | 0.0909 | 40 | 0.05 | 7 | 0.1429 | 5 | 0.6 | 14 | 0 | 2 | 0 | 28 | 0.1429 |
| 8_96552585 | 12 | 0.6667 | 13 | 0.5385 | 2 | 0.5 | 9 | 0.5556 | 36 | 0.5833 | 4 | 0.5 | 4 | 0.5 | 8 | 0.375 | 3 | 0.6667 | 19 | 0.4737 |
| 8_96562562 | 6 | 0 | 5 | 0 | 6 | 0.1667 | 3 | 0.3333 | 20 | 0.1 | 4 | 0.5 | 4 | 0 | NA | NA | 1 | 0 | 9 | 0.2222 |
| 8_96562697 | 6 | 0.1667 | 6 | 0 | 6 | 0.1667 | 3 | 0.3333 | 21 | 0.1429 | 4 | 0.25 | 4 | 0 | NA | NA | 1 | 1 | 9 | 0.2222 |
| 8_96563581 | 6 | 0.5 | 5 | 0.4 | 5 | 0.4 | 2 | 0.5 | 18 | 0.4444 | 3 | 0 | 3 | 0.3333 | NA | NA | 1 | 0 | 7 | 0.1429 |
| 8_96563634 | 4 | 0.75 | 4 | 0.5 | 5 | 0.8 | 2 | 1 | 15 | 0.7333 | 3 | 0.6667 | 3 | 0.6667 | NA | NA | 1 | 1 | 7 | 0.7143 |
| 8_96563743 | 5 | 0.6 | 5 | 0.8 | 4 | 0.75 | 2 | 0 | 16 | 0.625 | 3 | 0.3333 | 3 | 0.3333 | NA | NA | 1 | 1 | 7 | 0.4286 |
| 8_96563813 | 6 | 1 | 4 | 1 | 5 | 1 | 2 | 1 | 17 | 1 | 4 | 1 | 2 | 1 | NA | NA | 1 | 1 | 7 | 1 |

| | | | | | | | | | | | | | | | | | | | | |
|---|---|---|---|---|---|---|---|---|---|---|---|---|---|---|---|---|---|---|---|---|
| 8_9656387 | 6 | 0.3333 | 4 | 0.5 | 4 | 0.75 | 2 | 0.5 | 16 | 0.5 | 3 | 0.3333 | 3 | 0.6667 | NA | NA | NA | NA | 6 | 0.5 |
| 8_9656388 | 6 | 0.3333 | 5 | 0.4 | 5 | 0 | 2 | 0 | 18 | 0.2222 | 3 | 0.3333 | 3 | 0 | NA | NA | 1 | 0 | 7 | 0.1429 |
| 8_9656393 | 5 | 0.8 | 4 | 1 | 3 | 0.6667 | NA | NA | 12 | 0.8333 | 3 | 1 | 2 | 1 | NA | NA | NA | NA | 5 | 1 |
| 8_9656407 | 4 | 0 | 3 | 0.6667 | 4 | 0 | 2 | 0 | 13 | 0.1538 | 3 | 0.3333 | 2 | 0 | NA | NA | 1 | 1 | 6 | 0.3333 |
| 8_9656408 | 4 | 0.75 | 3 | 1 | 4 | 1 | 1 | 1 | 12 | 0.9167 | 2 | 0.5 | 3 | 1 | NA | NA | 1 | 1 | 6 | 0.8333 |
| 8_9656409 | 6 | 0.8333 | 2 | 1 | 4 | 1 | 2 | 1 | 14 | 0.9286 | 3 | 1 | 3 | 1 | NA | NA | 1 | 1 | 7 | 1 |
| 8_9656417 | 4 | 0 | 4 | 0.25 | 5 | 0 | 2 | 0 | 15 | 0.0667 | 3 | 0.3333 | 3 | 0 | NA | NA | 1 | 0 | 7 | 0.1429 |
| 8_9656419 | 5 | 0.4 | 3 | 0 | 5 | 0.2 | 2 | 0 | 15 | 0.2 | 3 | 1 | 2 | 0.5 | NA | NA | NA | NA | 5 | 0.8 |
| 8_9657525 | 2 | 0 | 4 | 0.25 | 2 | 0 | 1 | 0 | 9 | 0.1111 | 1 | 0 | 2 | 0.5 | NA | NA | 1 | 0 | 4 | 0.25 |
| 8_9659085 | 32 | 0.0625 | 57 | 0.0526 | 39 | 0.0769 | 29 | 0.1379 | 157 | 0.0764 | 30 | 0.1333 | 28 | 0.2143 | 41 | 0.0488 | 33 | 0.1212 | 132 | 0.1212 |
| 8_9659096 | 25 | 0.16 | 53 | 0.2642 | 31 | 0.2581 | 23 | 0.4348 | 132 | 0.2727 | 28 | 0.3929 | 22 | 0.4545 | 35 | 0.3714 | 25 | 0.44 | 110 | 0.4091 |
| 8_9659123 | 30 | 0.2667 | 48 | 0.3125 | 38 | 0.3684 | 26 | 0.3462 | 142 | 0.3239 | 28 | 0.4286 | 30 | 0.4333 | 36 | 0.3333 | 26 | 0.3462 | 120 | 0.3833 |
| 8_9659130 | 30 | 0.0333 | 58 | 0.1379 | 36 | 0.1389 | 27 | 0.1111 | 151 | 0.1126 | 32 | 0.125 | 32 | 0.125 | 39 | 0.1282 | 33 | 0.0909 | 136 | 0.1176 |
| 8_9659135 | 27 | 0 | 51 | 0.0784 | 35 | 0.0286 | 25 | 0.08 | 138 | 0.0507 | 28 | 0.0714 | 29 | 0.1379 | 32 | 0.0625 | 28 | 0.0714 | 117 | 0.0855 |
| 8_9659156 | 29 | 0.2414 | 54 | 0.1481 | 36 | 0.25 | 26 | 0.0385 | 145 | 0.1724 | 27 | 0.2222 | 29 | 0.069 | 36 | 0.25 | 28 | 0.3214 | 120 | 0.2167 |
| 8_9659162 | 22 | 0.0455 | 36 | 0.1111 | 28 | 0.1429 | 18 | 0.2222 | 104 | 0.125 | 19 | 0.2105 | 21 | 0.1429 | 24 | 0.25 | 16 | 0.0625 | 80 | 0.175 |
| 8_9659173 | 26 | 0.1923 | 48 | 0.2083 | 30 | 0.2667 | 26 | 0.2692 | 130 | 0.2308 | 20 | 0.2 | 28 | 0.2143 | 28 | 0.1071 | 22 | 0.4545 | 98 | 0.2347 |
| 8_9659173 | 26 | 0 | 43 | 0.186 | 29 | 0.069 | 20 | 0.05 | 118 | 0.0932 | 22 | 0 | 24 | 0.1667 | 22 | 0.0909 | 26 | 0.1923 | 94 | 0.117 |
| 8_9975554 | 1 | 0 | 1 | 0 | NA | NA | 1 | 1 | 3 | 0.3333 | 2 | 0.5 | 2 | 0 | 1 | 0 | NA | NA | 5 | 0.2 |
| 8_9975606 | 2 | 0.5 | 1 | 1 | NA | NA | 1 | 0 | 4 | 0.5 | 1 | 0 | 5 | 0.4 | 3 | 1 | NA | NA | 9 | 0.5556 |
| 8_9975639 | 2 | 0 | 1 | 1 | NA | NA | 3 | 0 | 6 | 0.1667 | 1 | 1 | 6 | 0.5 | 3 | 0.3333 | NA | NA | 10 | 0.5 |
| 8_9975643 | 2 | 0 | 1 | 0 | NA | NA | 2 | 0.5 | 5 | 0.2 | 1 | 0 | 6 | 0.1667 | 3 | 0.3333 | NA | NA | 10 | 0.2 |
| 8_9975660 | 1 | 0 | NA | NA | NA | NA | 2 | 0.5 | 3 | 0.3333 | 1 | 1 | 2 | 1 | 2 | 1 | NA | NA | 5 | 1 |
| 8_9975669 | 1 | 0 | NA | NA | NA | NA | 2 | 0.5 | 3 | 0.3333 | 1 | 0 | 2 | 0.5 | 2 | 0.5 | NA | NA | 5 | 0.4 |
| 8_9975681 | 1 | 1 | NA | NA | NA | NA | 2 | 1 | 3 | 1 | 1 | 0 | 2 | 1 | 3 | 0.6667 | NA | NA | 6 | 0.6667 |
| 8_9975682 | 1 | 1 | NA | NA | NA | NA | 2 | 1 | 3 | 1 | 1 | 1 | 1 | 1 | 3 | 1 | NA | NA | 5 | 1 |
| 8_9975720 | 1 | 1 | NA | NA | NA | NA | 1 | 0 | 2 | 0.5 | 1 | 0 | 1 | 0 | 3 | 0.3333 | NA | NA | 5 | 0.2 |
| 8_9975721 | 1 | 0 | NA | NA | NA | NA | 2 | 0.5 | 3 | 0.3333 | 1 | 1 | NA | NA | 3 | 0 | NA | NA | 4 | 0.25 |
| 8_9975724 | 1 | 1 | NA | NA | NA | NA | 2 | 1 | 3 | 1 | 1 | 1 | 1 | 1 | 3 | 0.6667 | NA | NA | 5 | 0.8 |
| 8_9975727 | 1 | 1 | NA | NA | NA | NA | 2 | 0 | 3 | 0.3333 | 1 | 0 | 1 | 1 | 3 | 0 | NA | NA | 5 | 0.2 |
| 8_9975729 | 1 | 1 | NA | NA | NA | NA | 1 | 1 | 2 | 1 | NA | NA | 1 | 1 | 3 | 0.6667 | NA | NA | 4 | 0.75 |
| 8_9975737 | NA | NA | NA | NA | NA | NA | 2 | 1 | 2 | 1 | NA | NA | 1 | 1 | 3 | 0.6667 | NA | NA | 4 | 0.75 |
| 8_9975755 | 2 | 0 | 2 | 0 | NA | NA | 3 | 0.3333 | 7 | 0.1429 | NA | NA | 4 | 0.25 | 3 | 0 | NA | NA | 7 | 0.1429 |
| 8_9975989 | 2 | 0.5 | 2 | 0 | NA | NA | 2 | 0.5 | 6 | 0.3333 | NA | NA | 3 | 0.6667 | 2 | 0.5 | NA | NA | 5 | 0.6 |
| 8_9982553 | 2 | 0.5 | 2 | 0 | NA | NA | 2 | 0.5 | 6 | 0.3333 | NA | NA | 4 | 0.5 | 1 | 0 | NA | NA | 5 | 0.4 |
| 8_1000061 | 1 | 1 | 2 | 1 | NA | NA | 1 | 1 | 4 | 1 | NA | NA | 3 | 1 | 1 | 0 | NA | NA | 4 | 0.75 |
| 8_1000062 | 1 | 0 | 2 | 0.5 | NA | NA | 1 | 0 | 4 | 0.25 | NA | NA | 2 | 0.5 | 2 | 0.5 | NA | NA | 4 | 0.5 |
| 8_1001274 | 1 | 0 | 2 | 0.5 | NA | NA | 1 | 1 | 4 | 0.5 | NA | NA | 2 | 0.5 | 1 | 0 | NA | NA | 3 | 0.3333 |
| 8_1001275 | 1 | 1 | 1 | 1 | NA | NA | 1 | 1 | 3 | 1 | NA | NA | 2 | 1 | NA | NA | NA | NA | 2 | 1 |
| 8_1001275 | 1 | 1 | 2 | 1 | NA | NA | 1 | 1 | 4 | 1 | NA | NA | 1 | 1 | 1 | 1 | NA | NA | 2 | 1 |
| 8_1001276 | 1 | 1 | 1 | 1 | NA | NA | 1 | 0 | 3 | 0.6667 | NA | NA | 2 | 0.5 | 1 | 1 | NA | NA | 3 | 0.6667 |
| 8_1033596 | 4 | 0.25 | 1 | 1 | NA | NA | 1 | 0 | 6 | 0.3333 | 1 | 0 | 2 | 0.5 | 2 | 0 | 1 | 0 | 6 | 0.1667 |
| 8_1033597 | 4 | 0.25 | 1 | 0 | NA | NA | 1 | 0 | 6 | 0.1667 | 1 | 1 | 2 | 0 | 3 | 0.3333 | NA | NA | 6 | 0.3333 |
| 8_1033598 | 5 | 0.2 | 1 | 0 | NA | NA | 1 | 0 | 7 | 0.1429 | 1 | 0 | 1 | 0 | 2 | 0.5 | NA | NA | 4 | 0.25 |
| 8_1033604 | 8 | 0 | 5 | 0 | 7 | 0.2857 | 5 | 0 | 25 | 0.08 | 3 | 0 | 9 | 0.1111 | 5 | 0.4 | 2 | 0 | 19 | 0.1579 |
| 8_1033605 | 10 | 0.8 | 5 | 1 | 7 | 1 | 5 | 1 | 27 | 0.9259 | 4 | 1 | 8 | 1 | 5 | 1 | 3 | 1 | 20 | 1 |
| 8_1033605 | 11 | 0.3636 | 5 | 0.6 | 5 | 0.4 | 5 | 0.4 | 26 | 0.4231 | 2 | 0 | 7 | 0.2857 | 5 | 0.2 | 3 | 0.6667 | 17 | 0.2941 |
| 8_1033606 | 12 | 0.1667 | 5 | 0.4 | 5 | 0 | 5 | 0.2 | 27 | 0.1852 | 3 | 0.3333 | 7 | 0.4286 | 4 | 0 | 3 | 0 | 17 | 0.2353 |
| 8_1033606 | 10 | 0.6 | 3 | 0.6667 | 5 | 0.2 | 4 | 0.75 | 22 | 0.5455 | 3 | 0.3333 | 6 | 1 | 5 | 0.6 | 2 | 1 | 16 | 0.75 |
| 8_1033607 | 9 | 1 | 3 | 1 | 3 | 1 | 5 | 1 | 20 | 1 | 2 | 0.5 | 6 | 1 | 3 | 1 | 3 | 1 | 14 | 0.9286 |
| 8_1033608 | 10 | 0.6 | 3 | 0.3333 | 3 | 0.6667 | 5 | 0.6 | 21 | 0.5714 | 3 | 0 | 8 | 0.125 | 3 | 0.3333 | 2 | 0.5 | 16 | 0.1875 |
| 8_1033608 | 10 | 0 | 5 | 0.2 | 3 | 0.3333 | 4 | 0 | 22 | 0.0909 | 3 | 0 | 8 | 0.375 | 4 | 0 | 2 | 0.5 | 17 | 0.2353 |
| 8_1033608 | 9 | 0 | 4 | 0 | 2 | 0 | 2 | 0.5 | 17 | 0.0588 | 1 | 0 | 6 | 0 | 3 | 0 | 3 | 0.3333 | 13 | 0.0769 |
| 8_1033609 | 9 | 1 | 4 | 1 | 4 | 1 | 5 | 1 | 22 | 1 | 3 | 1 | 7 | 1 | 4 | 1 | 2 | 1 | 16 | 1 |
| 8_1033609 | 8 | 1 | 4 | 1 | 3 | 0.6667 | 5 | 1 | 20 | 0.95 | 2 | 1 | 8 | 1 | 2 | 1 | 2 | 1 | 14 | 1 |
| 8_1033609 | 11 | 0.6364 | 4 | 0.75 | 5 | 1 | 5 | 1 | 25 | 0.8 | 3 | 0.6667 | 8 | 0.875 | 4 | 1 | 3 | 1 | 18 | 0.8889 |
| 8_1033609 | 11 | 0.4545 | 3 | 0.6667 | 5 | 0.2 | 5 | 0.2 | 24 | 0.375 | 3 | 0 | 8 | 0.5 | 4 | 0.5 | 3 | 0.3333 | 18 | 0.3889 |
| 8_1033609 | 11 | 0.5455 | 3 | 0.6667 | 4 | 0.5 | 4 | 1 | 22 | 0.6364 | 3 | 0.6667 | 5 | 0.8 | 3 | 0 | 2 | 0.5 | 13 | 0.5385 |
| 8_1033610 | 11 | 0.4545 | 4 | 0.5 | 4 | 0.25 | 5 | 0.4 | 24 | 0.4167 | 3 | 0 | 8 | 0.25 | 4 | 0.25 | 3 | 0 | 18 | 0.1667 |
| 8_1033610 | 9 | 0.5556 | 3 | 1 | 4 | 0.25 | 5 | 0.4 | 21 | 0.5238 | 3 | 0.6667 | 7 | 0.2857 | 4 | 0.75 | 3 | 0.3333 | 17 | 0.4706 |
| 8_1033610 | 10 | 0.8 | 3 | 1 | 3 | 1 | 5 | 1 | 21 | 0.9048 | 3 | 1 | 7 | 0.8571 | 4 | 1 | 2 | 1 | 16 | 0.9375 |
| 8_1033610 | 10 | 0.2 | 4 | 0.75 | 5 | 0.6 | 5 | 0.6 | 24 | 0.4583 | 3 | 0 | 8 | 0 | 4 | 0 | 3 | 0.3333 | 18 | 0.0556 |
| 8_1033611 | 10 | 0.7 | 2 | 1 | 5 | 0.8 | 4 | 0.75 | 21 | 0.7619 | 3 | 0.6667 | 4 | 0.75 | 3 | 1 | 3 | 0.6667 | 13 | 0.7692 |
| 8_1033613 | 10 | 0 | 3 | 0.3333 | 3 | 0 | 4 | 0 | 20 | 0.05 | 1 | 0 | 7 | 0.1429 | 4 | 0 | 3 | 0 | 15 | 0.0667 |
| 8_1033849 | 8 | 0 | 4 | 0 | 3 | 0 | 3 | 0.3333 | 18 | 0.0556 | 2 | 0 | 4 | 0.25 | 2 | 0 | 1 | 0 | 9 | 0.1111 |

| | | | | | | | | | | | | | | | | | | | | |
|---|---|---|---|---|---|---|---|---|---|---|---|---|---|---|---|---|---|---|---|---|
| 8_1033912 | 8 | 0.125 | 3 | 0.3333 | 3 | 0 | 3 | 0 | 17 | 0.1176 | 1 | 0 | 4 | 0.25 | 2 | 0 | 2 | 0 | 9 | 0.1111 |
| 8_1034063 | 6 | 0.8333 | 2 | 1 | 3 | 0.6667 | 1 | 1 | 12 | 0.8333 | 2 | 0 | 3 | 0.3333 | NA | NA | 1 | 0 | 6 | 0.1667 |
| 8_1048693 | 1 | 1 | 1 | 1 | NA | NA | 1 | 1 | 3 | 1 | NA | NA | NA | NA | NA | NA | 1 | 1 | 1 | 1 |
| 8_1048696 | 2 | 1 | 1 | 1 | NA | NA | 1 | 0 | 4 | 0.75 | NA | NA | NA | NA | NA | NA | 1 | 1 | 1 | 1 |
| 8_1048696 | 2 | 1 | 2 | 0.5 | NA | NA | 1 | 0 | 5 | 0.6 | NA | NA | NA | NA | NA | NA | 1 | 1 | 1 | 1 |
| 8_1048697 | 1 | 1 | 2 | 1 | NA | NA | NA | NA | 3 | 1 | NA | NA | NA | NA | NA | NA | 1 | 1 | 1 | 1 |
| 8_1048699 | 2 | 1 | 2 | 0.5 | NA | NA | 1 | 1 | 5 | 0.8 | NA | NA | NA | NA | NA | NA | 1 | 1 | 1 | 1 |
| 8_1048702 | 2 | 0 | 2 | 0.5 | NA | NA | 1 | 1 | 5 | 0.4 | NA | NA | NA | NA | NA | NA | 2 | 1 | 2 | 1 |
| 8_1048708 | 2 | 0.5 | 3 | 0.6667 | 1 | 0 | 1 | 0 | 7 | 0.4286 | NA | NA | 1 | 0 | NA | NA | 2 | 0.5 | 3 | 0.3333 |
| 8_1049534 | 3 | 0 | 3 | 0.3333 | 6 | 0.3333 | 2 | 1 | 14 | 0.3571 | 1 | 0 | 2 | 0.5 | 1 | 1 | 3 | 0.3333 | 7 | 0.4286 |
| 8_1049534 | 5 | 0.4 | 4 | 0.75 | 7 | 0.2857 | 3 | 0.6667 | 19 | 0.4737 | 3 | 0 | 4 | 0 | 2 | 0.5 | 2 | 0 | 11 | 0.0909 |
| 8_1049537 | 4 | 0.25 | 3 | 0.3333 | 2 | 0 | 3 | 0.6667 | 12 | 0.3333 | 2 | 0.5 | 2 | 0.5 | 1 | 0 | 3 | 0 | 8 | 0.25 |
| 8_1049537 | 5 | 1 | 4 | 0.5 | 7 | 0.7143 | 3 | 0.3333 | 19 | 0.6842 | 2 | 0.5 | 2 | 1 | 2 | 0.5 | 4 | 0.75 | 10 | 0.7 |
| 8_1049537 | 4 | 0 | 4 | 0.25 | 6 | 0.1667 | 2 | 0 | 16 | 0.125 | 2 | 0 | 2 | 0 | 2 | 0.5 | 3 | 0 | 9 | 0.1111 |
| 8_1049537 | 5 | 1 | 2 | 1 | 7 | 1 | 3 | 0.6667 | 17 | 0.9412 | 2 | 1 | 3 | 1 | 2 | 0.5 | 3 | 1 | 10 | 0.9 |
| 8_1049538 | 5 | 0.2 | 5 | 0.2 | 8 | 0.125 | 3 | 0 | 21 | 0.1429 | 2 | 0.5 | 3 | 0.6667 | 2 | 0 | 3 | 0 | 10 | 0.3 |
| 8_1049538 | 5 | 0 | 5 | 0.4 | 8 | 0.25 | 2 | 0 | 20 | 0.2 | 2 | 0 | 3 | 0.3333 | 2 | 0 | 3 | 0 | 10 | 0.1 |
| 8_1049539 | 3 | 0 | 1 | 0 | 7 | 0.1429 | 3 | 0 | 14 | 0.0714 | 2 | 0 | 3 | 0.3333 | 2 | 0 | 3 | 0 | 10 | 0.1 |
| 8_1049539 | 5 | 0.2 | 4 | 0.5 | 5 | 0 | 3 | 0 | 17 | 0.1765 | 2 | 0.5 | 2 | 0.5 | 2 | 0 | 3 | 0.3333 | 9 | 0.3333 |
| 8_1049541 | 3 | 0.3333 | 2 | 0.5 | 8 | 0.625 | 2 | 0 | 15 | 0.4667 | 2 | 0.5 | 3 | 0 | 2 | 1 | 3 | 1 | 10 | 0.6 |
| 8_1049541 | 5 | 0 | 3 | 0.6667 | 8 | 0.25 | 3 | 0 | 19 | 0.2105 | 2 | 0 | 3 | 0 | 2 | 0.5 | 3 | 0 | 10 | 0.1 |
| 8_1049541 | 4 | 0 | 3 | 0 | 8 | 0 | 3 | 0.3333 | 18 | 0.0556 | 2 | 0 | 3 | 0 | 2 | 0 | 3 | 0.3333 | 10 | 0.1 |
| 8_1049542 | 4 | 0 | 4 | 0.5 | 7 | 0.1429 | 2 | 0 | 17 | 0.1765 | 2 | 0.5 | 2 | 0 | 1 | 1 | 3 | 0.3333 | 8 | 0.375 |
| 8_1049542 | 3 | 0 | 3 | 0.3333 | 6 | 0 | 3 | 0 | 15 | 0.0667 | 2 | 0 | 3 | 0 | 1 | 0 | 3 | 0.3333 | 9 | 0.1111 |
| 8_1049544 | 4 | 0.75 | 3 | 0.6667 | 7 | 0.7143 | 3 | 0 | 17 | 0.5882 | 2 | 1 | 2 | 1 | 1 | 0 | 2 | 1 | 7 | 0.8571 |
| 8_1049545 | 4 | 0 | 3 | 0 | 7 | 0.2857 | 3 | 0.6667 | 17 | 0.2353 | 2 | 0 | 3 | 0.3333 | 2 | 0 | 2 | 0.5 | 9 | 0.2222 |
| 8_1049545 | 3 | 0 | 3 | 0.3333 | 5 | 0.4 | 3 | 0.6667 | 14 | 0.3571 | 2 | 0 | 3 | 0.3333 | 1 | 0 | 2 | 0.5 | 8 | 0.25 |
| 8_1049546 | 1 | 1 | NA | NA | 5 | 1 | 3 | 0.3333 | 9 | 0.7778 | 1 | 1 | 3 | 0.6667 | 1 | 0 | 1 | 1 | 6 | 0.6667 |
| 8_1049548 | 4 | 0 | 4 | 0.5 | 7 | 0.2857 | 3 | 0.3333 | 18 | 0.2778 | 2 | 0.5 | 2 | 0.5 | 2 | 0 | 2 | 0 | 8 | 0.25 |
| 8_1049840 | NA | NA | 1 | 1 | 1 | 1 | NA | NA | 2 | 1 | NA | NA | NA | NA | 1 | 1 | NA | NA | 1 | 1 |
| 8_1050740 | NA | NA | 1 | 0 | 1 | 0 | 3 | 0.3333 | 5 | 0.2 | 3 | 0.3333 | NA | NA | 1 | 0 | 3 | 0.3333 | 7 | 0.2857 |
| 8_1050751 | 12 | 0.1667 | 16 | 0 | 14 | 0.1429 | 10 | 0.2 | 52 | 0.1154 | 4 | 0 | 10 | 0 | 11 | 0.2727 | 7 | 0.1429 | 32 | 0.125 |
| 8_1050751 | 10 | 0 | 18 | 0.0556 | 14 | 0.1429 | 7 | 0 | 49 | 0.0612 | 4 | 0 | 12 | 0.0833 | 11 | 0 | 6 | 0.1667 | 33 | 0.0606 |
| 8_1051438 | 2 | 0 | 2 | 0.5 | 1 | 0 | 2 | 0 | 7 | 0.1429 | NA | NA | 1 | 1 | NA | NA | 1 | 0 | 2 | 0.5 |
| 8_1051438 | NA | NA | 2 | 0 | 1 | 0 | 3 | 0.3333 | 6 | 0.1667 | NA | NA | 3 | 0.3333 | NA | NA | 1 | 0 | 4 | 0.25 |
| 8_1051438 | 2 | 0.5 | 1 | 0 | 1 | 0 | 2 | 0.5 | 6 | 0.3333 | NA | NA | 3 | 1 | NA | NA | NA | NA | 3 | 1 |
| 8_1051441 | 2 | 0.5 | 2 | 0 | NA | NA | 2 | 0.5 | 6 | 0.3333 | NA | NA | 3 | 0.3333 | NA | NA | NA | NA | 3 | 0.3333 |
| 8_1051444 | 27 | 0.037 | 29 | 0.069 | 19 | 0.1053 | 22 | 0.0455 | 97 | 0.0619 | 20 | 0 | 18 | 0.0556 | 14 | 0.0714 | 19 | 0.2105 | 71 | 0.0845 |
| 8_1051445 | 23 | 0.5217 | 29 | 0.4828 | 19 | 0.3158 | 24 | 0.375 | 95 | 0.4316 | 17 | 0.2353 | 23 | 0.2174 | 15 | 0.1333 | 22 | 0.2727 | 77 | 0.2208 |
| 8_1051453 | 14 | 0.2143 | 22 | 0.2727 | 10 | 0.1 | 16 | 0.375 | 62 | 0.2581 | 11 | 0.3636 | 17 | 0.3529 | 9 | 0.1111 | 13 | 0.3846 | 50 | 0.32 |
| 8_1051458 | 11 | 0.2727 | 19 | 0 | 8 | 0.125 | 13 | 0 | 51 | 0.0784 | 13 | 0.1538 | 16 | 0 | 8 | 0.125 | 11 | 0 | 48 | 0.0625 |
| 8_1051460 | 12 | 0.25 | 19 | 0.2632 | 5 | 0.2 | 12 | 0.25 | 48 | 0.25 | 9 | 0.4444 | 12 | 0.3333 | 6 | 0.1667 | 8 | 0 | 35 | 0.2571 |
| 8_1051470 | 10 | 0.6 | 10 | 0.8 | 4 | 0.5 | 5 | 0.2 | 29 | 0.5862 | 6 | 0.5 | 7 | 0.4286 | 4 | 0.5 | 6 | 0.8333 | 23 | 0.5652 |
| 8_1051470 | 9 | 0.3333 | 14 | 0.5714 | 7 | 0.4286 | 7 | 0.4286 | 37 | 0.4595 | 5 | 0.6 | 5 | 0.6 | 3 | 0.3333 | 7 | 0.4286 | 20 | 0.5 |
| 8_1051472 | 11 | 0.0909 | 15 | 0 | 6 | 0.1667 | 8 | 0 | 40 | 0.05 | 3 | 0.3333 | 6 | 0 | 2 | 0 | 8 | 0 | 19 | 0.0526 |
| 8_1051473 | 11 | 0.3636 | 12 | 0.3333 | 5 | 0 | 10 | 0.3 | 38 | 0.2895 | 4 | 0.5 | 6 | 0.1667 | 2 | 0.5 | 7 | 0.2857 | 19 | 0.3158 |
| 8_1052765 | NA | NA | 4 | 0 | 4 | 0.25 | 2 | 0 | 10 | 0.1 | 3 | 0 | 5 | 0.4 | 1 | 0 | 3 | 0.3333 | 12 | 0.25 |
| 8_1052767 | NA | NA | 3 | 0.3333 | 4 | 0.75 | 2 | 0.5 | 9 | 0.5556 | 1 | 0 | 5 | 0.6 | NA | NA | 3 | 0.6667 | 9 | 0.5556 |
| 8_1053208 | 1 | 1 | NA | NA | 1 | 1 | NA | NA | 2 | 1 | 2 | 1 | 1 | 1 | NA | NA | NA | NA | 3 | 1 |
| 8_1053211 | NA | NA | NA | NA | 1 | 1 | NA | NA | 1 | 1 | 2 | 1 | 1 | 1 | NA | NA | NA | NA | 3 | 1 |
| 8_1053213 | NA | NA | NA | NA | 1 | 1 | NA | NA | 1 | 1 | 1 | 1 | 1 | 1 | NA | NA | NA | NA | 2 | 1 |
| 8_1053214 | NA | NA | NA | NA | 1 | 1 | NA | NA | 1 | 1 | 2 | 1 | 1 | 1 | NA | NA | NA | NA | 3 | 1 |
| 8_1053214 | NA | NA | NA | NA | 1 | 1 | NA | NA | 1 | 1 | 2 | 1 | 1 | 0 | NA | NA | NA | NA | 3 | 0.6667 |
| 8_1053215 | NA | NA | NA | NA | 1 | 1 | NA | NA | 1 | 1 | 1 | 1 | 1 | 1 | NA | NA | NA | NA | 2 | 1 |
| 8_1053217 | NA | NA | NA | NA | 1 | 1 | NA | NA | 1 | 1 | 2 | 0.5 | 1 | 1 | NA | NA | NA | NA | 3 | 0.6667 |
| 8_1053220 | 1 | 0 | NA | NA | 1 | 1 | NA | NA | 2 | 0.5 | 2 | 0.5 | 1 | 0 | NA | NA | NA | NA | 3 | 0.3333 |
| 8_1053231 | NA | NA | 2 | 0 | NA | NA | 3 | 0.3333 | 5 | 0.2 | 3 | 0.3333 | 1 | 0 | NA | NA | NA | NA | 4 | 0.25 |
| 8_1053247 | 1 | 0 | 2 | 0 | 1 | 0 | 3 | 0.3333 | 7 | 0.1429 | 3 | 0.3333 | 2 | 0 | NA | NA | 1 | 0 | 6 | 0.1667 |
| 8_1053252 | 1 | 0 | 2 | 0 | 1 | 1 | 3 | 0.6667 | 7 | 0.4286 | 4 | 0.5 | 2 | 0.5 | NA | NA | 1 | 1 | 7 | 0.5714 |
| 8_1059426 | 3 | 0.3333 | 3 | 0 | 2 | 0 | 1 | 0 | 9 | 0.1111 | 2 | 0.5 | 1 | 0 | NA | NA | 3 | 0.3333 | 6 | 0.3333 |
| 8_1059427 | 3 | 0 | 5 | 0.8 | 2 | 0 | 1 | 0 | 11 | 0.3636 | 2 | 0 | NA | NA | NA | NA | 3 | 0.6667 | 5 | 0.4 |
| 8_1059783 | 2 | 0 | 4 | 0.5 | 1 | 0 | 7 | 0.1429 | 14 | 0.2143 | 1 | 0 | 1 | 0 | 3 | 0 | 6 | 0.1667 | 11 | 0.0909 |
| 8_1059783 | 4 | 0.5 | 4 | 0.5 | 1 | 0 | 7 | 0.4286 | 16 | 0.4375 | 2 | 0.5 | 1 | 0 | 3 | 0 | 6 | 0.5 | 12 | 0.3333 |
| 8_1059784 | 4 | 0.5 | 4 | 0.5 | 1 | 0 | 7 | 0.2857 | 16 | 0.375 | 2 | 0 | NA | NA | 3 | 0 | 6 | 0.3333 | 11 | 0.1818 |
| 8_1059784 | 4 | 0.75 | 4 | 0.25 | 1 | 1 | 7 | 0.7143 | 16 | 0.625 | 2 | 0.5 | 1 | 1 | 3 | 0.6667 | 7 | 0.5714 | 13 | 0.6154 |

| | | | | | | | | | | | | | | | | | | | | |
|---|---|---|---|---|---|---|---|---|---|---|---|---|---|---|---|---|---|---|---|---|
| 8_1059785 | 3 | 0 | 6 | 0 | 1 | 0 | 7 | 0.4286 | 17 | 0.1765 | 2 | 0.5 | NA | NA | 2 | 1 | 4 | 0.25 | 8 | 0.5 |
| 8_1059786 | 4 | 0.25 | 5 | 0.2 | 1 | 0 | 7 | 0.2857 | 17 | 0.2353 | 2 | 0 | 1 | 0 | 3 | 0.3333 | 8 | 0.375 | 14 | 0.2857 |
| 8_1059791 | 5 | 0.2 | 4 | 0 | 2 | 0 | 6 | 0 | 17 | 0.0588 | 2 | 0 | 1 | 0 | 5 | 0.2 | 6 | 0 | 14 | 0.0714 |
| 8_1059794 | 5 | 0 | 6 | 0.1667 | 3 | 0.3333 | 7 | 0.1429 | 21 | 0.1429 | 2 | 0 | 3 | 0 | 3 | 0.3333 | 7 | 0.1429 | 15 | 0.1333 |
| 8_1059795 | 6 | 0.3333 | 4 | 0.25 | 2 | 0 | 7 | 0.5714 | 19 | 0.3684 | 2 | 0.5 | 2 | 0 | 3 | 0.3333 | 5 | 0.6 | 12 | 0.4167 |
| 8_1059795 | 6 | 0 | 5 | 0.8 | 2 | 0 | 7 | 0.8571 | 20 | 0.5 | 3 | 1 | 2 | 0 | 4 | 1 | 6 | 0.5 | 15 | 0.6667 |
| 8_1059853 | NA | NA | NA | NA | 1 | 1 | 1 | 1 | 2 | 1 | 2 | 1 | 1 | 1 | NA | NA | 1 | 1 | 4 | 1 |
| 8_1059853 | 1 | 0 | NA | NA | 1 | 1 | 2 | 1 | 4 | 0.75 | 2 | 1 | 1 | 0 | 1 | 1 | 1 | 1 | 5 | 0.8 |
| 8_1059853 | NA | NA | NA | NA | 1 | 1 | 2 | 1 | 3 | 1 | 2 | 0.5 | 1 | 1 | 1 | 1 | 1 | 1 | 5 | 0.8 |
| 8_1059854 | 1 | 0 | NA | NA | NA | NA | 2 | 0.5 | 3 | 0.3333 | 2 | 1 | 1 | 1 | NA | NA | NA | NA | 3 | 1 |
| 8_1059856 | 1 | 1 | NA | NA | NA | NA | 2 | 1 | 3 | 1 | 2 | 1 | 1 | 1 | 1 | 1 | 1 | 1 | 5 | 1 |
| 8_1059861 | NA | NA | NA | NA | 1 | 0 | 2 | 0.5 | 3 | 0.3333 | 2 | 1 | 1 | 0 | 1 | 0 | 1 | 1 | 5 | 0.6 |
| 8_1059862 | NA | NA | NA | NA | NA | NA | 1 | 1 | 1 | 1 | 1 | 1 | NA | NA | NA | NA | NA | NA | 1 | 1 |
| 8_1059863 | NA | NA | NA | NA | 1 | 0 | 2 | 0.5 | 3 | 0.3333 | 2 | 1 | NA | NA | NA | NA | NA | NA | 2 | 1 |
| 8_1060024 | 1 | 0 | 3 | 0 | 6 | 0.3333 | 6 | 0 | 16 | 0.125 | 2 | 0 | 2 | 0.5 | 6 | 0 | 1 | 0 | 11 | 0.0909 |
| 8_1060066 | 6 | 0 | 4 | 0 | 5 | 0.2 | 2 | 0 | 17 | 0.0588 | 7 | 0.1429 | 3 | 0 | 2 | 0 | 1 | 0 | 13 | 0.0769 |
| 8_1060066 | NA | NA | 1 | 1 | 1 | 1 | NA | NA | 2 | 1 | 1 | 1 | NA | NA | 1 | 1 | NA | NA | 2 | 1 |
| 8_1060080 | 1 | 0 | 1 | 1 | NA | NA | NA | NA | 2 | 0.5 | 1 | 0 | 2 | 0.5 | NA | NA | 1 | 0 | 4 | 0.25 |
| 8_1060084 | 2 | 0.5 | NA | NA | NA | NA | 3 | 0.6667 | 5 | 0.6 | 3 | 0.3333 | 3 | 0.3333 | 1 | 1 | NA | NA | 7 | 0.4286 |
| 8_1060085 | 2 | 0 | 1 | 1 | NA | NA | NA | NA | 3 | 0.3333 | 1 | 0 | 2 | 0.5 | NA | NA | 1 | 0 | 4 | 0.25 |
| 8_1060085 | 2 | 0.5 | 2 | 0 | NA | NA | 3 | 0 | 7 | 0.1429 | 2 | 0 | NA | NA | 1 | 1 | NA | NA | 3 | 0.3333 |
| 8_1060086 | 2 | 0.5 | 1 | 0 | NA | NA | 3 | 0.3333 | 6 | 0.3333 | 2 | 0.5 | 2 | 1 | 1 | 0 | NA | NA | 5 | 0.6 |
| 8_1060087 | 2 | 1 | 2 | 0.5 | NA | NA | 2 | 0.5 | 6 | 0.6667 | 2 | 1 | 1 | 1 | NA | NA | 1 | 0 | 4 | 0.75 |
| 8_1060122 | 2 | 0.5 | 1 | 0 | NA | NA | 1 | 0 | 4 | 0.25 | 2 | 0.5 | 1 | 1 | NA | NA | 1 | 1 | 4 | 0.75 |
| 8_1060124 | 2 | 0.5 | 1 | 1 | NA | NA | NA | NA | 3 | 0.6667 | 2 | 1 | 1 | 1 | NA | NA | NA | NA | 3 | 1 |
| 8_1060125 | 1 | 1 | NA | NA | NA | NA | 1 | 0 | 2 | 0.5 | 1 | 1 | NA | NA | NA | NA | NA | NA | 1 | 1 |
| 8_1060133 | 1 | 1 | NA | NA | NA | NA | 1 | 0 | 2 | 0.5 | 1 | 1 | NA | NA | NA | NA | NA | NA | 1 | 1 |
| 8_1060615 | NA | NA | 1 | 1 | 3 | 0.3333 | NA | NA | 4 | 0.5 | 1 | 1 | NA | NA | 1 | 1 | NA | NA | 2 | 1 |
| 8_1060757 | 2 | 0.5 | 1 | 1 | NA | NA | NA | NA | 3 | 0.6667 | NA | NA | NA | NA | 1 | 1 | 1 | 1 | 2 | 1 |
| 8_1060833 | 6 | 0.6667 | 2 | 0 | 2 | 0.5 | 4 | 1 | 14 | 0.6429 | NA | NA | 1 | 1 | 2 | 0 | 1 | 0 | 4 | 0.25 |
| 8_1060947 | 7 | 0.1429 | 2 | 0 | 2 | 0 | 2 | 0.5 | 13 | 0.1538 | 1 | 0 | 2 | 0.5 | 3 | 0 | 3 | 0 | 9 | 0.1111 |
| 8_1060950 | 8 | 0.625 | 3 | 1 | 2 | 0 | 6 | 0.6667 | 19 | 0.6316 | 1 | 0 | 6 | 0.8333 | 6 | 0.6667 | 5 | 0.6 | 18 | 0.6667 |
| 8_1060951 | 8 | 0.75 | 2 | 0.5 | 2 | 0.5 | 5 | 0.4 | 17 | 0.5882 | 1 | 1 | 6 | 1 | 6 | 0.5 | 4 | 0.5 | 17 | 0.7059 |
| 8_1060952 | 9 | 0.5556 | 2 | 1 | 2 | 0 | 6 | 0.6667 | 19 | 0.5789 | 1 | 0 | 6 | 1 | 4 | 0.25 | 5 | 0.8 | 16 | 0.6875 |
| 8_1060953 | 12 | 0.3333 | 3 | 0.6667 | 3 | 0 | 4 | 0 | 22 | 0.2727 | 1 | 0 | 6 | 0.5 | 6 | 0.5 | 6 | 0.5 | 19 | 0.4737 |
| 8_1060954 | 11 | 0.5455 | 3 | 0.6667 | 3 | 0 | 6 | 0.5 | 23 | 0.4783 | 1 | 0 | 6 | 0.3333 | 7 | 0.4286 | 5 | 0.6 | 19 | 0.4211 |
| 8_1060957 | 12 | 0.25 | 2 | 0 | 2 | 0 | 3 | 0 | 19 | 0.1579 | 1 | 1 | 5 | 0.4 | 8 | 0.25 | 8 | 0.25 | 22 | 0.3182 |
| 8_1061942 | 9 | 0.2222 | 6 | 0 | 11 | 0 | 12 | 0.0833 | 38 | 0.0789 | 8 | 0.125 | 10 | 0.1 | 8 | 0 | 5 | 0 | 31 | 0.0645 |
| 8_1061943 | 7 | 0.5714 | 6 | 0.6667 | 12 | 0.5 | 14 | 0.5 | 39 | 0.5385 | 8 | 0.25 | 10 | 0.6 | 8 | 0.25 | 5 | 0.2 | 31 | 0.3548 |
| 8_1061990 | 3 | 0 | 2 | 0 | 3 | 0 | 6 | 0.1667 | 14 | 0.0714 | 1 | 0 | 4 | 0.25 | 5 | 0 | 2 | 0 | 12 | 0.0833 |
| 8_1061991 | 3 | 0 | 3 | 0 | 6 | 0.1667 | 9 | 0.2222 | 21 | 0.1429 | 1 | 1 | 3 | 0 | 5 | 0.2 | 3 | 0.3333 | 12 | 0.25 |
| 8_1061991 | 3 | 0.3333 | 2 | 0 | 5 | 0.4 | 9 | 0.2222 | 19 | 0.2632 | 1 | 0 | 7 | 0 | 3 | 0 | 2 | 0.5 | 13 | 0.0769 |
| 8_1061991 | 3 | 0 | 3 | 0 | 6 | 0.3333 | 10 | 0.1 | 22 | 0.1364 | 1 | 0 | 8 | 0 | 5 | 0.2 | 4 | 0.5 | 18 | 0.1667 |
| 8_1061992 | 3 | 0.3333 | 3 | 0.3333 | 4 | 0.5 | 9 | 0.4444 | 19 | 0.4211 | 1 | 0 | 7 | 0.4286 | 5 | 1 | 2 | 0.5 | 15 | 0.6 |
| 8_1061996 | 1 | 0 | 2 | 0 | 3 | 0.6667 | 7 | 0 | 13 | 0.1538 | 2 | 0 | 6 | 0.1667 | 2 | 0 | 3 | 0.3333 | 13 | 0.1538 |
| 8_1062513 | 23 | 0.0435 | 47 | 0.0213 | 35 | 0.1429 | 35 | 0.0571 | 140 | 0.0643 | 39 | 0.0769 | 24 | 0 | 25 | 0.08 | 32 | 0.0312 | 120 | 0.05 |
| 8_1063442 | 2 | 1 | 2 | 0.5 | 1 | 1 | 5 | 1 | 10 | 0.9 | NA | NA | 1 | 1 | 2 | 0.5 | 3 | 0.3333 | 6 | 0.5 |
| 8_1063442 | 2 | 1 | 4 | 0.75 | 1 | 1 | 4 | 1 | 11 | 0.9091 | NA | NA | 1 | 1 | 2 | 1 | 2 | 1 | 5 | 1 |
| 8_1063446 | 3 | 0.3333 | 4 | 0 | 1 | 0 | 3 | 0 | 11 | 0.0909 | 1 | 0 | 1 | 0 | 3 | 0 | 2 | 0.5 | 7 | 0.1429 |
| 8_1063485 | 4 | 1 | 7 | 0 | 10 | 0.3 | 8 | 0.375 | 29 | 0.3448 | 6 | 0.8333 | 9 | 0.5556 | 9 | 0.3333 | 10 | 0.2 | 34 | 0.4412 |
| 8_1063904 | 3 | 0 | 4 | 0.25 | 2 | 0 | 3 | 0.3333 | 12 | 0.1667 | NA | NA | 3 | 0 | NA | NA | 3 | 0.3333 | 6 | 0.1667 |
| 8_1063907 | 4 | 1 | 3 | 1 | 2 | 0.5 | 5 | 0.8 | 14 | 0.8571 | NA | NA | 2 | 1 | NA | NA | 3 | 0.6667 | 5 | 0.8 |
| 8_1063909 | 4 | 0.25 | 4 | 0.25 | 2 | 0 | 4 | 0.25 | 14 | 0.2143 | NA | NA | 1 | 1 | NA | NA | 2 | 0 | 3 | 0.3333 |
| 8_1063909 | 5 | 0.2 | 3 | 0.6667 | 2 | 0 | 5 | 0 | 15 | 0.2 | NA | NA | 2 | 0 | NA | NA | 4 | 0.25 | 6 | 0.1667 |
| 8_1063909 | 5 | 0 | 4 | 0.25 | 2 | 0.5 | 5 | 0.2 | 16 | 0.1875 | NA | NA | 2 | 0.5 | NA | NA | 4 | 0.25 | 6 | 0.3333 |
| 8_1064081 | 7 | 0.1429 | 5 | 0 | 4 | 0.75 | 6 | 0.1667 | 22 | 0.2273 | 3 | 0 | 4 | 0.25 | 4 | 0.25 | 3 | 0.6667 | 14 | 0.2857 |
| 8_1064082 | 8 | 0.25 | 5 | 0.2 | 6 | 0.1667 | 5 | 0.2 | 24 | 0.2083 | 4 | 0.25 | 3 | 0.3333 | 4 | 0.25 | 3 | 0.3333 | 14 | 0.2857 |
| 8_1064082 | 5 | 0.8 | 3 | 1 | 3 | 0.3333 | 2 | 0.5 | 13 | 0.6923 | 2 | 1 | 2 | 0.5 | 1 | 1 | 1 | 0 | 6 | 0.6667 |
| 8_1064085 | 13 | 0.0769 | 5 | 0 | 6 | 0 | 6 | 0.3333 | 30 | 0.1 | 4 | 0 | 6 | 0.3333 | 4 | 0 | 2 | 0 | 16 | 0.125 |
| 8_1064086 | 12 | 0.25 | 6 | 0 | 6 | 0.1667 | 7 | 0.5714 | 31 | 0.2581 | 5 | 0.2 | 6 | 0 | 4 | 0.25 | 3 | 0 | 18 | 0.1111 |
| 8_1064089 | 12 | 0.25 | 6 | 0.1667 | 5 | 0 | 7 | 0.2857 | 30 | 0.2 | 5 | 0.2 | 5 | 0.2 | 4 | 0.25 | 3 | 0 | 17 | 0.1765 |
| 8_1064254 | 2 | 0 | 5 | 0 | 5 | 0.2 | 3 | 0 | 15 | 0.0667 | 2 | 0 | 7 | 0.1429 | 3 | 0 | 2 | 0 | 14 | 0.0714 |
| 8_1064293 | 7 | 0.1429 | 8 | 0.5 | 3 | 0.3333 | 6 | 0.1667 | 24 | 0.2917 | 1 | 0 | 2 | 0.5 | 4 | 0.5 | 4 | 0.25 | 11 | 0.3636 |
| 8_1064296 | 6 | 0.3333 | 7 | 0.1429 | 6 | 0.3333 | 5 | 0.6 | 24 | 0.3333 | 2 | 0 | 4 | 0.5 | 4 | 0.25 | 5 | 0.2 | 15 | 0.2667 |
| 8_1064296 | 9 | 0.8889 | 10 | 0.9 | 5 | 0.8 | 6 | 0.8333 | 30 | 0.8667 | 1 | 1 | 4 | 0.5 | 4 | 1 | 5 | 1 | 14 | 0.8571 |

| | | | | | | | | | | | | | | | | | | | | |
|---|---|---|---|---|---|---|---|---|---|---|---|---|---|---|---|---|---|---|---|---|
| 8_10642978 | 8 | 0.25 | 9 | 0 | 5 | 0.4 | 5 | 0.4 | 27 | 0.2222 | 3 | 0 | 3 | 0.3333 | 6 | 0 | 5 | 0 | 17 | 0.0588 |
| 8_10642980 | 9 | 0.1111 | 11 | 0.1818 | 6 | 0.1667 | 6 | 0 | 32 | 0.125 | 4 | 0.25 | 4 | 0 | 6 | 0 | 5 | 0.2 | 19 | 0.1053 |
| 8_10642987 | 8 | 0.375 | 9 | 0.3333 | 4 | 0 | 7 | 0.2857 | 28 | 0.2857 | 2 | 0 | 4 | 0.5 | 5 | 0 | 5 | 0.2 | 16 | 0.1875 |
| 8_10643003 | 9 | 0.1111 | 10 | 0.1 | 6 | 0.1667 | 6 | 0.5 | 31 | 0.1935 | 3 | 0 | 4 | 0.25 | 6 | 0 | 5 | 0.2 | 18 | 0.1111 |
| 8_10643004 | 6 | 0.8333 | 6 | 1 | 5 | 1 | 6 | 0.8333 | 23 | 0.913 | 3 | 1 | 4 | 1 | 4 | 0.75 | 3 | 0.6667 | 14 | 0.8571 |
| 8_10643010 | 7 | 0 | 9 | 0 | 4 | 0 | 5 | 0.4 | 25 | 0.08 | 3 | 0.3333 | 2 | 0 | 5 | 0.2 | 4 | 0 | 14 | 0.1429 |
| 8_10643039 | 1 | 0 | NA | NA | NA | NA | 2 | 0.5 | 3 | 0.3333 | 1 | 0 | 1 | 0 | NA | NA | 2 | 0.5 | 4 | 0.25 |
| 8_10643122 | NA | NA | 1 | 1 | NA | NA | NA | NA | 1 | 1 | NA | NA | NA | NA | NA | NA | 1 | 1 | 1 | 1 |
| 8_10643130 | NA | NA | 1 | 1 | NA | NA | NA | NA | 1 | 1 | NA | NA | NA | NA | NA | NA | 1 | 1 | 1 | 1 |
| 8_10644148 | NA | NA | NA | NA | NA | NA | 1 | 1 | 1 | 1 | NA | NA | NA | NA | 1 | 1 | NA | NA | 1 | 1 |
| 8_10644295 | 2 | 0.5 | 2 | 0.5 | 1 | 1 | 5 | 0.4 | 10 | 0.5 | NA | NA | 2 | 0.5 | 2 | 0 | 6 | 0.3333 | 10 | 0.3 |
| 8_10644332 | 2 | 0 | 1 | 0 | 1 | 0 | 5 | 0.8 | 9 | 0.4444 | NA | NA | 2 | 0.5 | 2 | 0 | 5 | 0.2 | 9 | 0.2222 |
| 8_10644345 | 2 | 0.5 | 1 | 0 | 1 | 0 | 4 | 0.75 | 8 | 0.5 | NA | NA | 2 | 1 | 2 | 0 | 5 | 0.2 | 9 | 0.3333 |
| 8_10644351 | 2 | 1 | 1 | 1 | 1 | 1 | 4 | 1 | 8 | 1 | NA | NA | 2 | 1 | 2 | 1 | 5 | 1 | 9 | 1 |
| 8_10644369 | 2 | 1 | 1 | 1 | 1 | 1 | 3 | 1 | 7 | 1 | NA | NA | 3 | 1 | 2 | 1 | 5 | 1 | 10 | 1 |
| 8_10644372 | 2 | 1 | 1 | 0 | 1 | 0 | 3 | 0.3333 | 7 | 0.4286 | NA | NA | 3 | 1 | 2 | 1 | 4 | 0.25 | 9 | 0.6667 |
| 8_10644383 | 2 | 0.5 | 1 | 0 | 1 | 0 | 2 | 0 | 6 | 0.1667 | NA | NA | 3 | 0.6667 | 1 | 0 | 5 | 0.2 | 9 | 0.3333 |
| 8_10644388 | 1 | 1 | 1 | 1 | 1 | 1 | 3 | 0.3333 | 6 | 0.6667 | NA | NA | 2 | 1 | 2 | 1 | 4 | 0.5 | 8 | 0.75 |
| 8_10644393 | 2 | 0 | 1 | 1 | 1 | 0 | 1 | 0 | 5 | 0.2 | NA | NA | 2 | 0.5 | 2 | 0.5 | 5 | 0.4 | 9 | 0.4444 |
| 8_10644397 | 2 | 0.5 | NA | NA | 1 | 1 | 3 | 1 | 6 | 0.8333 | NA | NA | 2 | 0.5 | NA | NA | 5 | 0.4 | 7 | 0.4286 |
| 8_10644399 | 2 | 1 | NA | NA | 1 | 1 | 3 | 1 | 6 | 1 | NA | NA | 2 | 1 | 1 | 1 | 5 | 0.8 | 8 | 0.875 |
| 8_10645484 | 2 | 0 | 1 | 0 | 1 | 0 | 2 | 0.5 | 6 | 0.1667 | NA | NA | 2 | 0.5 | 1 | 0 | 3 | 0 | 6 | 0.1667 |
| 8_10649506 | 6 | 0 | 3 | 0.6667 | 4 | 0 | 5 | 0.8 | 18 | 0.3333 | 3 | 0.6667 | 5 | 0.4 | 4 | 0 | 1 | 1 | 13 | 0.3846 |
| 8_10649521 | 7 | 0 | 2 | 0.5 | 3 | 0 | 5 | 0 | 17 | 0.0588 | 3 | 0 | 4 | 0 | 3 | 0 | 1 | 1 | 11 | 0.0909 |
| 8_10649525 | 6 | 0.3333 | 2 | 0 | 4 | 0.25 | 5 | 0.2 | 17 | 0.2353 | 3 | 0.3333 | 4 | 0.25 | 4 | 0.75 | 1 | 0 | 12 | 0.4167 |
| 8_10649532 | 8 | 0.5 | 2 | 0.5 | 4 | 1 | 4 | 1 | 18 | 0.7222 | 2 | 1 | 3 | 1 | 4 | 1 | 1 | 1 | 10 | 1 |
| 8_10649537 | 7 | 1 | 2 | 0 | 4 | 1 | 5 | 0.8 | 18 | 0.8333 | 3 | 1 | 4 | 0.75 | 4 | 1 | 1 | 1 | 12 | 0.9167 |
| 8_10649539 | 7 | 0.2857 | 2 | 0 | 4 | 0 | 5 | 0 | 18 | 0.1111 | 3 | 0.3333 | 4 | 0 | 2 | 0 | 1 | 1 | 10 | 0.2 |
| 8_10649541 | 7 | 0 | 3 | 0 | 4 | 0.25 | 5 | 0 | 19 | 0.0526 | 3 | 0 | 4 | 0.25 | 4 | 0 | 1 | 0 | 12 | 0.0833 |
| 8_10649558 | 7 | 0.1429 | 1 | 0 | 4 | 0.25 | 5 | 0 | 17 | 0.1176 | 3 | 0 | 4 | 0.25 | 4 | 0.25 | 1 | 1 | 12 | 0.25 |
| 8_10649578 | 6 | 0.1667 | 1 | 0 | 3 | 0.3333 | 5 | 0.4 | 15 | 0.2667 | 3 | 0 | 4 | 0.25 | 4 | 0.25 | 1 | 0 | 12 | 0.1667 |
| 8_10649595 | 6 | 0 | NA | NA | 3 | 0.3333 | 5 | 0.2 | 14 | 0.1429 | 2 | 1 | 3 | 0.3333 | 4 | 0.25 | NA | NA | 9 | 0.4444 |
| 8_10649614 | 6 | 0.1667 | NA | NA | 3 | 0.3333 | 4 | 0.25 | 13 | 0.2308 | 3 | 0.6667 | 3 | 0.6667 | 4 | 0.5 | NA | NA | 10 | 0.6 |
| 8_10649675 | 5 | 0 | NA | NA | 2 | 0.5 | 3 | 0 | 10 | 0.1 | 1 | 1 | 3 | 0.3333 | 4 | 0.25 | NA | NA | 8 | 0.375 |
| 8_10649679 | 2 | 1 | NA | NA | 3 | 1 | 3 | 1 | 8 | 1 | NA | NA | 4 | 0.5 | 4 | 1 | NA | NA | 8 | 0.75 |
| 8_10649682 | 5 | 0.4 | NA | NA | 2 | 0 | 3 | 0.6667 | 10 | 0.4 | NA | NA | 4 | 0.25 | 3 | 0.6667 | NA | NA | 7 | 0.4286 |
| 8_10649693 | 5 | 0.8 | NA | NA | 2 | 1 | 3 | 0.6667 | 10 | 0.8 | NA | NA | 3 | 1 | 3 | 0.3333 | NA | NA | 6 | 0.6667 |
| 8_10649701 | 4 | 0 | NA | NA | 3 | 0 | 3 | 0.6667 | 10 | 0.2 | NA | NA | 2 | 0 | 3 | 0.3333 | NA | NA | 5 | 0.2 |
| 8_10649701 | 5 | 0.2 | NA | NA | 3 | 0.3333 | 3 | 0.6667 | 11 | 0.3636 | NA | NA | 3 | 0 | 3 | 0.3333 | NA | NA | 6 | 0.1667 |
| 8_10649705 | 5 | 1 | NA | NA | 3 | 1 | 3 | 1 | 11 | 1 | NA | NA | 3 | 1 | 1 | 1 | NA | NA | 4 | 1 |
| 8_10649714 | 4 | 0.25 | NA | NA | 3 | 0 | 1 | 1 | 8 | 0.25 | NA | NA | 3 | 0.3333 | 2 | 0 | NA | NA | 5 | 0.2 |
| 8_10649813 | 2 | 0.5 | NA | NA | 1 | 0 | 1 | 1 | 4 | 0.5 | NA | NA | 3 | 0.3333 | 2 | 0.5 | NA | NA | 5 | 0.4 |
| 8_10657131 | 2 | 0 | 4 | 0.25 | 2 | 0.5 | 3 | 0.3333 | 11 | 0.2727 | NA | NA | 3 | 0.3333 | 3 | 0.3333 | NA | NA | 6 | 0.3333 |
| 8_10658294 | 9 | 0.1111 | 6 | 0 | 4 | 0.25 | 9 | 0.1111 | 28 | 0.1071 | 4 | 0.25 | 5 | 0.6 | 6 | 0.1667 | 8 | 0 | 23 | 0.2174 |
| 8_10658300 | 8 | 0.125 | 6 | 0.5 | 4 | 0 | 9 | 0.3333 | 27 | 0.2593 | 4 | 0 | 5 | 0.2 | 5 | 0.2 | 6 | 0.1667 | 20 | 0.15 |
| 8_10658301 | 10 | 0.6 | 6 | 0.8333 | 4 | 1 | 10 | 0.5 | 30 | 0.6667 | 3 | 0.6667 | 5 | 0.8 | 5 | 0.4 | 7 | 0.5714 | 20 | 0.6 |
| 8_10658302 | 10 | 0.7 | 6 | 1 | 4 | 0.75 | 9 | 1 | 29 | 0.8621 | 4 | 0.75 | 6 | 1 | 4 | 1 | 8 | 0.875 | 22 | 0.9091 |
| 8_10658308 | 9 | 0.8889 | 6 | 0.6667 | 4 | 0.75 | 8 | 1 | 27 | 0.8519 | 2 | 1 | 6 | 1 | 5 | 1 | 9 | 0.8889 | 22 | 0.9545 |
| 8_10658311 | 10 | 1 | 7 | 1 | 4 | 0.75 | 6 | 1 | 27 | 0.963 | 3 | 1 | 5 | 0.8 | 5 | 1 | 6 | 1 | 19 | 0.9474 |
| 8_10658318 | 4 | 0.5 | 4 | 0.75 | 2 | 0.5 | 5 | 0.6 | 15 | 0.6 | 2 | 1 | 3 | 0.6667 | 2 | 0 | 3 | 1 | 10 | 0.7 |
| 8_10658318 | 2 | 0.5 | 2 | 0.5 | 1 | 1 | 7 | 0.5714 | 12 | 0.5833 | 2 | 0 | 1 | 0 | 2 | 0.5 | 3 | 0.3333 | 8 | 0.25 |
| 8_10658321 | 7 | 1 | 5 | 1 | 3 | 0.6667 | 8 | 0.875 | 23 | 0.913 | 2 | 1 | 4 | 1 | 5 | 1 | 8 | 0.875 | 19 | 0.9474 |
| 8_10658322 | 9 | 1 | 5 | 1 | 2 | 1 | 9 | 1 | 25 | 1 | 3 | 1 | 7 | 1 | 6 | 1 | 8 | 1 | 24 | 1 |
| 8_10658327 | 9 | 0.3333 | 5 | 0.4 | 3 | 0.3333 | 10 | 0.6 | 27 | 0.4444 | 4 | 1 | 6 | 0.3333 | 6 | 0.6667 | 8 | 0.625 | 24 | 0.625 |
| 8_10658336 | 9 | 0.8889 | 5 | 1 | 4 | 0.5 | 9 | 0.8889 | 27 | 0.8519 | 4 | 0.5 | 6 | 1 | 5 | 0.6 | 10 | 0.6 | 25 | 0.68 |
| 8_10658339 | 10 | 0 | 6 | 0.3333 | 4 | 0.25 | 9 | 0.1111 | 29 | 0.1379 | 3 | 0 | 7 | 0 | 4 | 0.25 | 11 | 0.1818 | 25 | 0.12 |
| 8_10658344 | 7 | 0.1429 | 7 | 0 | 5 | 0.2 | 8 | 0.125 | 27 | 0.1111 | 5 | 0.2 | 7 | 0 | 6 | 0 | 9 | 0.2222 | 27 | 0.1111 |
| 8_10658347 | 10 | 0.8 | 6 | 0.5 | 5 | 0.6 | 9 | 0.3333 | 30 | 0.5667 | 5 | 0.4 | 7 | 0.2857 | 6 | 0.3333 | 11 | 0.4545 | 29 | 0.3793 |
| 8_10660576 | 9 | 0 | 14 | 0.2857 | 12 | 0.1667 | 7 | 0.2857 | 42 | 0.1905 | 9 | 0.2222 | 6 | 0 | 11 | 0.3636 | 11 | 0 | 37 | 0.1622 |
| 8_10661586 | 3 | 0.3333 | 3 | 0.6667 | 2 | 0 | 4 | 0 | 12 | 0.25 | 1 | 0 | 3 | 0 | 4 | 0.25 | 2 | 0 | 10 | 0.1 |
| 8_10666620 | 6 | 0.1667 | 3 | 0 | 6 | 0 | 2 | 0 | 17 | 0.0588 | 6 | 0 | 2 | 0.5 | 2 | 0 | 6 | 0 | 16 | 0.0625 |
| 8_10666629 | 8 | 0.25 | 3 | 0 | 4 | 0.5 | 2 | 0.5 | 17 | 0.2941 | 6 | 0.5 | 2 | 0 | 2 | 0 | 6 | 0.5 | 16 | 0.375 |
| 8_10666661 | 6 | 0.3333 | 3 | 0.3333 | 6 | 0.1667 | 2 | 0 | 17 | 0.2353 | 5 | 0.2 | 2 | 0.5 | 1 | 0 | 5 | 0.2 | 13 | 0.2308 |
| 8_10667025 | 2 | 0 | 3 | 0.3333 | 2 | 0 | 3 | 0 | 10 | 0.1 | 2 | 0.5 | 2 | 0 | 2 | 0 | 1 | 0 | 7 | 0.1429 |
| 8_10667047 | 3 | 0.3333 | 3 | 0 | 2 | 0.5 | 2 | 0.5 | 10 | 0.3 | 3 | 0.3333 | 2 | 0 | 2 | 0 | 1 | 0 | 8 | 0.125 |

| | | | | | | | | | | | | | | | | | | | | |
|---|---|---|---|---|---|---|---|---|---|---|---|---|---|---|---|---|---|---|---|---|
| 8_10667052 | 3 | 0.6667 | 2 | 1 | 2 | 0 | 2 | 0.5 | 9 | 0.5556 | 2 | 0.5 | 2 | 0.5 | 1 | 1 | 1 | 1 | 6 | 0.6667 |
| 8_10667054 | 3 | 0.3333 | 3 | 0.3333 | 2 | 1 | 2 | 0.5 | 10 | 0.5 | 2 | 0.5 | 2 | 0.5 | 2 | 0 | 1 | 0 | 7 | 0.2857 |
| 8_10673630 | 1 | 0 | NA | NA | 2 | 0.5 | 7 | 0.5714 | 10 | 0.5 | 1 | 1 | 3 | 0.6667 | 3 | 0.6667 | 2 | 0 | 9 | 0.5556 |
| 8_10673639 | 1 | 1 | NA | NA | 3 | 0.6667 | 5 | 0.8 | 9 | 0.7778 | NA | NA | 2 | 1 | 3 | 0.6667 | 1 | 0 | 6 | 0.6667 |
| 8_10673645 | 2 | 0.5 | NA | NA | 2 | 0.5 | 8 | 0.75 | 12 | 0.6667 | 1 | 1 | 3 | 0.6667 | 4 | 0.75 | 1 | 0 | 9 | 0.6667 |
| 8_10673647 | 2 | 0.5 | NA | NA | 3 | 0.6667 | 7 | 0.7143 | 12 | 0.6667 | 1 | 0 | 3 | 0.3333 | 4 | 0.25 | 2 | 1 | 10 | 0.4 |
| 8_10673650 | 2 | 0.5 | NA | NA | 2 | 0.5 | 4 | 0.75 | 8 | 0.625 | 1 | 1 | 3 | 0.3333 | 4 | 0.25 | 1 | 0 | 9 | 0.3333 |
| 8_10673667 | 2 | 0.5 | NA | NA | 3 | 0 | 3 | 0 | 8 | 0.125 | 1 | 0 | 3 | 0.3333 | 4 | 0.5 | 1 | 0 | 9 | 0.3333 |
| 8_10673669 | 2 | 0.5 | NA | NA | 1 | 0 | 3 | 0.3333 | 6 | 0.3333 | 1 | 0 | 3 | 0 | 2 | 0.5 | 2 | 0.5 | 8 | 0.25 |
| 8_10673682 | 1 | 0 | NA | NA | 1 | 0 | 3 | 0.3333 | 5 | 0.2 | 1 | 0 | 2 | 0.5 | 2 | 0 | NA | NA | 5 | 0.2 |
| 8_10673686 | 2 | 1 | NA | NA | 2 | 0.5 | 4 | 0.5 | 8 | 0.625 | 1 | 1 | 3 | 0.6667 | 4 | 0.5 | 2 | 1 | 10 | 0.7 |
| 8_10673689 | 2 | 0.5 | NA | NA | 2 | 0 | 4 | 0 | 8 | 0.125 | 1 | 0 | 3 | 0 | 4 | 0 | 1 | 1 | 9 | 0.1111 |
| 8_10673691 | 2 | 1 | NA | NA | 1 | 0 | 5 | 0.4 | 8 | 0.5 | 1 | 0 | NA | NA | 3 | 0.6667 | 2 | 0.5 | 6 | 0.5 |
| 8_10673694 | 2 | 0.5 | NA | NA | 2 | 1 | 5 | 0.8 | 9 | 0.7778 | 1 | 1 | 2 | 1 | 4 | 0.75 | 1 | 1 | 8 | 0.875 |
| 8_10673701 | 1 | 0 | NA | NA | 1 | 0 | 5 | 0.2 | 7 | 0.1429 | NA | NA | 2 | 0 | 4 | 0 | 2 | 0.5 | 8 | 0.125 |
| 8_10673715 | 1 | 1 | NA | NA | 3 | 0.3333 | 5 | 0.6 | 9 | 0.5556 | NA | NA | 1 | 0 | 3 | 0.6667 | 2 | 0.5 | 6 | 0.5 |
| 8_10673716 | 2 | 0.5 | NA | NA | 2 | 0 | 4 | 0.25 | 8 | 0.25 | NA | NA | NA | NA | 4 | 0.25 | 2 | 0.5 | 6 | 0.3333 |
| 8_10673734 | 1 | 0 | NA | NA | 1 | 0 | 3 | 0.3333 | 5 | 0.2 | NA | NA | 1 | 0 | 3 | 0 | 2 | 1 | 6 | 0.3333 |
| 8_10673746 | 2 | 1 | NA | NA | 1 | 1 | 5 | 0.8 | 8 | 0.875 | NA | NA | 2 | 1 | 4 | 0.5 | 1 | 1 | 7 | 0.7143 |
| 8_10673766 | 2 | 0.5 | NA | NA | 2 | 0.5 | 4 | 0.5 | 8 | 0.5 | NA | NA | 2 | 1 | 3 | 0.3333 | 2 | 0 | 7 | 0.4286 |
| 8_10678012 | NA | NA | 4 | 0.75 | 3 | 0 | NA | NA | 7 | 0.4286 | 1 | 1 | 1 | 1 | 1 | 0 | 3 | 0.3333 | 6 | 0.5 |
| 8_10678018 | NA | NA | 5 | 1 | 2 | 1 | 1 | 1 | 8 | 1 | 2 | 1 | 1 | 1 | 1 | 0 | 2 | 1 | 6 | 0.8333 |
| 8_10678021 | NA | NA | 5 | 0.4 | 3 | 0 | 1 | 0 | 9 | 0.2222 | 3 | 0.3333 | 1 | 0 | 2 | 0 | 3 | 0 | 9 | 0.1111 |
| 8_10678023 | NA | NA | 5 | 1 | 3 | 0.6667 | 1 | 1 | 9 | 0.8889 | 3 | 1 | 1 | 1 | 1 | 0 | 1 | 0 | 6 | 0.6667 |
| 8_10678044 | NA | NA | 4 | 1 | 3 | 0.6667 | 1 | 0 | 8 | 0.75 | 3 | 1 | NA | NA | 2 | 0 | 2 | 0.5 | 7 | 0.5714 |
| 8_10680571 | 1 | 1 | NA | NA | NA | NA | NA | NA | 1 | 1 | NA | NA | NA | NA | NA | NA | 1 | 1 | 1 | 1 |
| 8_10680583 | 1 | 1 | NA | NA | NA | NA | NA | NA | 1 | 1 | NA | NA | NA | NA | NA | NA | 1 | 1 | 1 | 1 |
| 8_10680613 | 1 | 1 | NA | NA | NA | NA | NA | NA | 1 | 1 | NA | NA | NA | NA | NA | NA | 1 | 1 | 1 | 1 |
| 8_10683485 | 1 | 1 | NA | NA | NA | NA | NA | NA | 1 | 1 | NA | NA | NA | NA | NA | NA | 1 | 1 | 1 | 1 |
| 8_10683553 | 1 | 1 | NA | NA | 1 | 0 | NA | NA | 2 | 0.5 | NA | NA | NA | NA | NA | NA | 1 | 1 | 1 | 1 |
| 8_10685469 | 1 | 1 | NA | NA | 1 | 1 | NA | NA | 2 | 1 | NA | NA | NA | NA | NA | NA | 2 | 1 | 2 | 1 |
| 8_10685476 | 1 | 1 | NA | NA | 1 | 1 | NA | NA | 2 | 1 | NA | NA | NA | NA | NA | NA | 2 | 1 | 2 | 1 |
| 8_10685584 | 2 | 1 | 1 | 0 | 3 | 1 | NA | NA | 6 | 0.8333 | NA | NA | NA | NA | 1 | 1 | 3 | 0.6667 | 4 | 0.75 |
| 8_10688961 | NA | NA | 3 | 1 | NA | NA | 1 | 1 | 4 | 1 | 2 | 1 | 2 | 1 | 3 | 1 | 1 | 1 | 8 | 1 |
| 8_10688966 | NA | NA | 3 | 0.6667 | 2 | 0.5 | 1 | 0 | 6 | 0.5 | 1 | 1 | 2 | 0 | 5 | 0.2 | 1 | 1 | 9 | 0.3333 |
| 8_10688974 | 1 | 1 | 4 | 0.25 | 2 | 0.5 | 1 | 1 | 8 | 0.5 | 2 | 0.5 | 1 | 1 | 4 | 0.75 | 1 | 0 | 8 | 0.625 |
| 8_10688978 | NA | NA | 3 | 0.6667 | 2 | 0 | 1 | 0 | 6 | 0.3333 | 2 | 0.5 | 2 | 0 | 5 | 0.6 | 1 | 1 | 10 | 0.5 |
| 8_10688996 | 1 | 0 | 3 | 0.3333 | 2 | 0 | 1 | 1 | 7 | 0.2857 | NA | NA | 2 | 0 | 4 | 0.25 | 1 | 0 | 7 | 0.1429 |
| 8_10689000 | 1 | 0 | 3 | 0.3333 | 2 | 0 | 1 | 1 | 7 | 0.2857 | 2 | 1 | 3 | 0 | 3 | 0.6667 | 1 | 1 | 9 | 0.5556 |
| 8_10689002 | NA | NA | 1 | 1 | 1 | 1 | NA | NA | 2 | 1 | 1 | 1 | 1 | 1 | 3 | 1 | NA | NA | 5 | 1 |
| 8_10689016 | 1 | 1 | 4 | 1 | 2 | 1 | 1 | 1 | 8 | 1 | 2 | 1 | 2 | 0.5 | 5 | 1 | 1 | 1 | 10 | 0.9 |
| 8_10689017 | 1 | 1 | 3 | 0.3333 | 1 | 1 | 2 | 0.5 | 7 | 0.5714 | 2 | 1 | 3 | 1 | 5 | 0.4 | 1 | 0 | 11 | 0.6364 |
| 8_10689020 | 1 | 1 | 4 | 0.25 | 2 | 0.5 | 2 | 0 | 9 | 0.3333 | 2 | 0 | 2 | 0 | 4 | 0.25 | 1 | 1 | 9 | 0.2222 |
| 8_10689025 | NA | NA | 4 | 0 | 2 | 0 | 2 | 0.5 | 8 | 0.125 | 2 | 0 | 3 | 0 | 5 | 0.2 | 1 | 0 | 11 | 0.0909 |
| 8_10689026 | 1 | 1 | 3 | 1 | 2 | 1 | 2 | 1 | 8 | 1 | 1 | 1 | 3 | 1 | 4 | 1 | 1 | 1 | 9 | 1 |
| 8_10689047 | 1 | 1 | 5 | 0.8 | 1 | 1 | 2 | 1 | 9 | 0.8889 | 2 | 1 | 4 | 0.75 | 2 | 1 | 1 | 1 | 9 | 0.8889 |
| 8_10689062 | NA | NA | 4 | 0.75 | 2 | 1 | 2 | 1 | 8 | 0.875 | 3 | 0.6667 | 2 | 1 | 4 | 0.75 | 1 | 1 | 10 | 0.8 |
| 8_10689062 | 1 | 1 | 3 | 0.3333 | 2 | 0 | 2 | 0.5 | 8 | 0.375 | 1 | 0 | 3 | 0 | 3 | 0.3333 | NA | NA | 7 | 0.1429 |
| 8_10689063 | 1 | 0 | 5 | 0.4 | 1 | 0 | 2 | 0.5 | 9 | 0.3333 | 3 | 0.3333 | 4 | 0 | 4 | 0 | 1 | 0 | 12 | 0.0833 |
| 8_10689084 | 1 | 0 | 6 | 0.1667 | 1 | 0 | 2 | 0.5 | 10 | 0.2 | 1 | 1 | 3 | 0 | 5 | 0 | 1 | 0 | 10 | 0.1 |
| 8_10689111 | 1 | 0 | 4 | 0.5 | 2 | 0 | 1 | 0 | 8 | 0.25 | 3 | 0.3333 | 1 | 0 | 2 | 1 | 1 | 0 | 7 | 0.4286 |
| 8_10689122 | 1 | 0 | 7 | 0.5714 | 3 | 0 | 3 | 0 | 14 | 0.2857 | NA | NA | 4 | 0.25 | 5 | 0.4 | 1 | 0 | 10 | 0.3 |
| 8_10692332 | 1 | 0 | NA | NA | 2 | 0.5 | 4 | 0.25 | 7 | 0.2857 | NA | NA | 1 | 0 | 2 | 0.5 | NA | NA | 3 | 0.3333 |
| 8_10692353 | 1 | 1 | NA | NA | 3 | 0.3333 | 3 | 0.6667 | 7 | 0.5714 | 1 | 1 | 1 | 0 | 2 | 0 | NA | NA | 4 | 0.25 |
| 8_10692376 | 1 | 0 | NA | NA | 3 | 0.6667 | 1 | 0 | 5 | 0.4 | 1 | 0 | 1 | 0 | 1 | 1 | NA | NA | 3 | 0.3333 |
| 8_10692383 | 2 | 0.5 | NA | NA | 3 | 0 | 3 | 0 | 8 | 0.125 | 1 | 1 | 1 | 0 | 2 | 0.5 | NA | NA | 4 | 0.5 |
| 8_10692387 | 2 | 1 | NA | NA | 4 | 0.5 | 4 | 0.5 | 10 | 0.6 | 1 | 0 | NA | NA | 2 | 0.5 | NA | NA | 3 | 0.3333 |
| 8_10692399 | 1 | 1 | NA | NA | 4 | 0.75 | 3 | 0.3333 | 8 | 0.625 | 1 | 0 | NA | NA | 2 | 0.5 | NA | NA | 3 | 0.3333 |
| 8_10692452 | 3 | 0 | NA | NA | 3 | 0 | 4 | 0.25 | 10 | 0.1 | 1 | 0 | 1 | 0 | 3 | 0.3333 | NA | NA | 5 | 0.2 |
| 8_10692457 | 1 | 1 | NA | NA | 3 | 0.6667 | 4 | 0.25 | 8 | 0.5 | 1 | 1 | 1 | 0 | 4 | 0 | NA | NA | 6 | 0.1667 |
| 8_10692459 | 3 | 0 | NA | NA | 3 | 0 | 4 | 0.25 | 10 | 0.1 | 1 | 0 | 1 | 0 | 4 | 0.25 | NA | NA | 6 | 0.1667 |
| 8_10692464 | NA | NA | NA | NA | 1 | 0 | 1 | 1 | 2 | 0.5 | 1 | 1 | NA | NA | 2 | 1 | NA | NA | 3 | 1 |
| 8_10692472 | NA | NA | NA | NA | 1 | 1 | 1 | 1 | 2 | 1 | 1 | 1 | NA | NA | 2 | 1 | NA | NA | 3 | 1 |
| 8_10692479 | NA | NA | NA | NA | 1 | 1 | 1 | 1 | 2 | 1 | 1 | 1 | NA | NA | 2 | 0.5 | NA | NA | 3 | 0.6667 |
| 8_10692487 | NA | NA | NA | NA | 1 | 1 | 1 | 1 | 2 | 1 | 1 | 0 | NA | NA | 2 | 1 | NA | NA | 3 | 0.6667 |

| | | | | | | | | | | | | | | | | | | | | |
|---|---|---|---|---|---|---|---|---|---|---|---|---|---|---|---|---|---|---|---|---|
| 8_1069251 | NA | NA | NA | NA | 1 | 1 | 1 | 1 | 2 | 1 | 1 | 1 | NA | NA | 2 | 0.5 | NA | NA | 3 | 0.6667 |
| 8_1069252 | NA | NA | NA | NA | 2 | 0.5 | 1 | 1 | 3 | 0.6667 | 1 | 1 | NA | NA | 2 | 1 | NA | NA | 3 | 1 |
| 8_1069271 | 4 | 0 | 3 | 0.3333 | 3 | 0 | 5 | 0.2 | 15 | 0.1333 | 1 | 0 | 5 | 0.2 | 2 | 0 | 3 | 0 | 11 | 0.0909 |
| 8_1069271 | 4 | 0 | 3 | 0 | 4 | 0.25 | 3 | 0 | 14 | 0.0714 | 1 | 0 | 5 | 0.2 | 2 | 0 | 3 | 0 | 11 | 0.0909 |
| 8_1069272 | 3 | 0 | 3 | 0 | 4 | 0 | 4 | 0.25 | 14 | 0.0714 | 1 | 0 | 4 | 0.25 | 2 | 0 | 4 | 0.25 | 11 | 0.1818 |
| 8_1069275 | 5 | 0.6 | 6 | 1 | 3 | 0.3333 | 5 | 0.4 | 19 | 0.6316 | 4 | 1 | 4 | 1 | 3 | 1 | 2 | 1 | 13 | 1 |
| 8_1069275 | 5 | 0 | 6 | 0.1667 | 4 | 0 | 2 | 0 | 17 | 0.0588 | 4 | 0 | 4 | 0.25 | 3 | 0 | 3 | 0.6667 | 14 | 0.2143 |
| 8_1069373 | 2 | 1 | 1 | 1 | 2 | 0.5 | 2 | 1 | 7 | 0.8571 | 1 | 1 | 2 | 1 | NA | NA | 1 | 1 | 4 | 1 |
| 8_1069783 | 7 | 0.4286 | 6 | 0 | 4 | 0 | 4 | 0.25 | 21 | 0.1905 | 6 | 0.1667 | 4 | 0.25 | 4 | 0 | 4 | 0.5 | 18 | 0.2222 |
| 8_1069783 | NA | NA | 1 | 1 | NA | NA | NA | NA | 1 | 1 | NA | NA | NA | NA | NA | NA | 2 | 1 | 2 | 1 |
| 8_1069785 | NA | NA | 1 | 1 | NA | NA | NA | NA | 1 | 1 | NA | NA | NA | NA | NA | NA | 2 | 0.5 | 2 | 0.5 |
| 8_1069786 | NA | NA | 1 | 1 | NA | NA | NA | NA | 1 | 1 | NA | NA | NA | NA | NA | NA | 2 | 1 | 2 | 1 |
| 8_1069795 | 13 | 0 | 24 | 0.0833 | 6 | 0.3333 | 14 | 0.0714 | 57 | 0.0877 | 10 | 0.1 | 18 | 0.0556 | 18 | 0.1111 | 11 | 0.1818 | 57 | 0.1053 |
| 8_1069796 | 13 | 0.0769 | 19 | 0.1579 | 7 | 0.4286 | 16 | 0 | 55 | 0.1273 | 10 | 0 | 17 | 0.1765 | 18 | 0 | 12 | 0.1667 | 57 | 0.0877 |
| 8_1069804 | 11 | 0.6364 | 14 | 0.5 | 4 | 0.25 | 11 | 0.5455 | 40 | 0.525 | 5 | 0.6 | 12 | 0.4167 | 14 | 0.2857 | 12 | 0.4167 | 43 | 0.3953 |
| 8_1069809 | 10 | 0 | 13 | 0.0769 | 6 | 0.1667 | 8 | 0 | 37 | 0.0541 | 5 | 0 | 14 | 0.1429 | 11 | 0.0909 | 11 | 0 | 41 | 0.0732 |
| 8_1069816 | 5 | 1 | 9 | 0.7778 | 3 | 1 | 8 | 0.625 | 25 | 0.8 | 2 | 1 | 8 | 0.75 | 8 | 0.75 | 7 | 1 | 25 | 0.84 |
| 8_1069816 | 7 | 0.4286 | 9 | 0.7778 | 3 | 0.3333 | 6 | 0.1667 | 25 | 0.48 | NA | NA | 8 | 0.125 | 6 | 0.1667 | 6 | 0.3333 | 20 | 0.2 |
| 8_1069817 | 6 | 0.3333 | 7 | 0.4286 | 4 | 0.5 | 6 | 0.1667 | 23 | 0.3478 | 3 | 0 | 8 | 0.375 | 9 | 0.1111 | 8 | 0.125 | 28 | 0.1786 |
| 8_1069914 | 4 | 0.25 | 7 | 0.1429 | 1 | 0 | 3 | 0 | 15 | 0.1333 | 1 | 0 | 5 | 0 | 6 | 0 | 4 | 0.5 | 16 | 0.125 |
| 8_1069914 | 4 | 1 | 4 | 1 | 3 | 1 | 5 | 1 | 16 | 1 | 1 | 1 | 6 | 1 | 6 | 0.8333 | 5 | 1 | 18 | 0.9444 |
| 8_1069916 | 1 | 1 | 5 | 0.4 | 1 | 1 | 2 | 0.5 | 9 | 0.5556 | 1 | 0 | 7 | 0.2857 | 6 | 0.1667 | 5 | 0.2 | 19 | 0.2105 |
| 8_1069918 | 3 | 0.3333 | 6 | 0.3333 | 3 | 1 | 3 | 0.6667 | 15 | 0.5333 | 1 | 1 | 7 | 0.5714 | 5 | 1 | 4 | 1 | 17 | 0.8235 |
| 8_1069918 | 4 | 0.5 | 4 | 0 | 1 | 0 | 1 | 0 | 10 | 0.2 | 1 | 0 | 8 | 0.125 | 7 | 0 | 3 | 0.3333 | 19 | 0.1053 |
| 8_1069919 | 1 | 1 | 6 | 1 | 2 | 1 | 2 | 0.5 | 11 | 0.9091 | 1 | 0 | 4 | 0.5 | 6 | 0.6667 | 3 | 1 | 14 | 0.6429 |
| 8_1069922 | 4 | 0 | 5 | 0.2 | 2 | 0 | 3 | 0 | 14 | 0.0714 | 1 | 0 | 7 | 0.1429 | 4 | 0.5 | 4 | 0.5 | 16 | 0.3125 |
| 8_1069925 | 3 | 1 | 4 | 0.75 | 2 | 1 | 3 | 0.6667 | 12 | 0.8333 | 1 | 1 | 5 | 0.8 | 4 | 0.75 | 2 | 1 | 12 | 0.8333 |
| 8_1069926 | 2 | 0.5 | 2 | 0.5 | 1 | 0 | 3 | 0.3333 | 8 | 0.375 | 1 | 1 | 5 | 0.4 | 2 | 0.5 | 3 | 1 | 11 | 0.6364 |
| 8_1069926 | 3 | 0 | 3 | 0 | 1 | 1 | 2 | 0 | 9 | 0.1111 | 1 | 0 | 4 | 0.25 | 3 | 0.3333 | 3 | 0 | 11 | 0.1818 |
| 8_1071505 | 1 | 1 | NA | NA | NA | NA | NA | NA | 1 | 1 | NA | NA | 1 | 0 | NA | NA | 1 | 1 | 2 | 0.5 |
| 8_1071507 | 1 | 1 | NA | NA | NA | NA | 1 | 0 | 2 | 0.5 | NA | NA | 2 | 0 | NA | NA | 1 | 1 | 3 | 0.3333 |
| 8_1071508 | 1 | 1 | NA | NA | NA | NA | 1 | 1 | 2 | 1 | 1 | 0 | 2 | 1 | NA | NA | 1 | 1 | 4 | 0.75 |
| 8_1071508 | 1 | 1 | NA | NA | NA | NA | 1 | 1 | 2 | 1 | 1 | 1 | 2 | 1 | NA | NA | 1 | 1 | 4 | 1 |
| 8_1071508 | 1 | 0 | NA | NA | NA | NA | 1 | 1 | 2 | 0.5 | 1 | 0 | 2 | 0.5 | NA | NA | 1 | 0 | 4 | 0.25 |
| 8_1071508 | 1 | 1 | NA | NA | NA | NA | 1 | 0 | 2 | 0.5 | 1 | 1 | 2 | 0 | NA | NA | 1 | 0 | 4 | 0.25 |
| 8_1071508 | 1 | 1 | NA | NA | NA | NA | 1 | 1 | 2 | 1 | 1 | 1 | 2 | 0.5 | NA | NA | 1 | 0 | 4 | 0.5 |
| 8_1071510 | NA | NA | NA | NA | NA | NA | 1 | 1 | 1 | 1 | NA | NA | 2 | 1 | NA | NA | 1 | 1 | 3 | 1 |
| 8_1071514 | 1 | 1 | NA | NA | NA | NA | 2 | 0.5 | 3 | 0.6667 | NA | NA | 2 | 1 | NA | NA | 1 | 0 | 3 | 0.6667 |
| 8_1071516 | NA | NA | NA | NA | NA | NA | 1 | 1 | 1 | 1 | NA | NA | 1 | 1 | NA | NA | 1 | 1 | 2 | 1 |
| 8_1071516 | 1 | 1 | NA | NA | NA | NA | 2 | 0.5 | 3 | 0.6667 | NA | NA | 2 | 0.5 | NA | NA | 1 | 1 | 3 | 0.6667 |
| 8_1071519 | 1 | 0 | 1 | 1 | NA | NA | 2 | 0 | 4 | 0.25 | 1 | 0 | 3 | 1 | NA | NA | 1 | 1 | 5 | 0.8 |
| 8_1071520 | 1 | 1 | 1 | 1 | NA | NA | 2 | 0 | 4 | 0.5 | 1 | 0 | 2 | 0.5 | NA | NA | 1 | 0 | 4 | 0.25 |
| 8_1071521 | NA | NA | 1 | 1 | NA | NA | 1 | 0 | 2 | 0.5 | NA | NA | 2 | 0.5 | NA | NA | 1 | 1 | 3 | 0.6667 |
| 8_1071522 | 1 | 0 | 1 | 1 | NA | NA | 1 | 0 | 3 | 0.3333 | 1 | 1 | 2 | 0.5 | NA | NA | 1 | 0 | 4 | 0.5 |
| 8_1071522 | 2 | 1 | 2 | 1 | NA | NA | 2 | 1 | 6 | 1 | 1 | 1 | 2 | 1 | 1 | 1 | 3 | 1 | 7 | 1 |
| 8_1071523 | 2 | 0.5 | 2 | 1 | NA | NA | 2 | 0 | 6 | 0.5 | 2 | 0.5 | 3 | 0.3333 | 1 | 0 | 3 | 0 | 9 | 0.2222 |
| 8_1073951 | 1 | 0 | NA | NA | NA | NA | 1 | 1 | 2 | 0.5 | 1 | 1 | NA | NA | NA | NA | NA | NA | 1 | 1 |
| 8_1074058 | 1 | 1 | 1 | 1 | NA | NA | NA | NA | 2 | 1 | NA | NA | 1 | 1 | NA | NA | NA | NA | 1 | 1 |
| 8_1074060 | 1 | 1 | 1 | 0 | NA | NA | NA | NA | 2 | 0.5 | NA | NA | 1 | 1 | NA | NA | NA | NA | 1 | 1 |
| 8_1074070 | 1 | 1 | 2 | 0.5 | NA | NA | NA | NA | 3 | 0.6667 | NA | NA | 1 | 1 | NA | NA | NA | NA | 1 | 1 |
| 8_1074157 | 2 | 0.5 | NA | NA | NA | NA | NA | NA | 2 | 0.5 | NA | NA | NA | NA | 1 | 1 | NA | NA | 1 | 1 |
| 8_1075769 | 2 | 0 | 2 | 0.5 | 1 | 0 | 2 | 1 | 7 | 0.4286 | NA | NA | 1 | 0 | 2 | 0.5 | NA | NA | 3 | 0.3333 |
| 8_1075772 | 2 | 0.5 | 1 | 0 | NA | NA | 2 | 0.5 | 5 | 0.4 | NA | NA | NA | NA | 1 | 1 | NA | NA | 1 | 1 |
| 8_1075773 | 2 | 0 | 2 | 0.5 | 1 | 0 | 1 | 0 | 6 | 0.1667 | NA | NA | 1 | 0 | 2 | 0.5 | NA | NA | 3 | 0.3333 |
| 8_1076070 | NA | NA | 1 | 1 | NA | NA | NA | NA | 1 | 1 | NA | NA | NA | NA | NA | NA | 1 | 1 | 1 | 1 |
| 8_1076072 | NA | NA | 1 | 1 | 1 | 1 | NA | NA | 2 | 1 | NA | NA | 1 | 1 | NA | NA | NA | NA | 1 | 1 |
| 8_1076074 | NA | NA | 1 | 1 | NA | NA | NA | NA | 1 | 1 | NA | NA | 1 | 1 | NA | NA | 1 | 1 | 2 | 1 |
| 8_1076074 | NA | NA | 1 | 0 | 1 | 1 | NA | NA | 2 | 0.5 | NA | NA | 1 | 1 | NA | NA | 1 | 1 | 2 | 1 |
| 8_1076075 | NA | NA | 1 | 1 | 1 | 1 | NA | NA | 2 | 1 | NA | NA | 1 | 1 | NA | NA | 1 | 1 | 2 | 1 |
| 8_1076075 | NA | NA | 1 | 1 | 1 | 1 | NA | NA | 2 | 1 | NA | NA | 1 | 1 | NA | NA | 1 | 1 | 2 | 1 |
| 8_1076106 | 11 | 0.1818 | 8 | 0 | 6 | 0.1667 | 10 | 0 | 35 | 0.0857 | 10 | 0 | 10 | 0.1 | 6 | 0.1667 | 15 | 0.2667 | 41 | 0.1463 |
| 8_1076106 | 16 | 0 | 9 | 0.2222 | 4 | 0.25 | 8 | 0.25 | 37 | 0.1351 | 10 | 0.2 | 7 | 0.2857 | 8 | 0 | 15 | 0.1333 | 40 | 0.15 |
| 8_1076107 | 12 | 0.75 | 8 | 0.375 | 5 | 0.4 | 11 | 0.3636 | 36 | 0.5 | 10 | 0.5 | 9 | 0.6667 | 7 | 0.4286 | 14 | 0.6429 | 40 | 0.575 |
| 8_1076107 | 14 | 0.8571 | 7 | 0.7143 | 6 | 1 | 12 | 0.9167 | 39 | 0.8718 | 12 | 0.6667 | 10 | 0.9 | 8 | 0.875 | 16 | 0.9375 | 46 | 0.8478 |
| 8_1076108 | 9 | 0.1111 | 3 | 0.3333 | 3 | 0 | 7 | 0 | 22 | 0.0909 | 9 | 0.3333 | 6 | 0.1667 | 5 | 0 | 10 | 0 | 30 | 0.1333 |

| | | | | | | | | | | | | | | | | | | | | |
|---|---|---|---|---|---|---|---|---|---|---|---|---|---|---|---|---|---|---|---|---|
| 8_10761088 | 14 | 0.5 | 9 | 0.2222 | 6 | 0.3333 | 11 | 0.3636 | 40 | 0.375 | 11 | 0.1818 | 8 | 0.25 | 9 | 0.3333 | 16 | 0.4375 | 44 | 0.3182 |
| 8_10761101 | 9 | 1 | 7 | 0.7143 | 5 | 1 | 7 | 0.7143 | 28 | 0.8571 | 7 | 0.8571 | 7 | 1 | 8 | 0.875 | 11 | 0.8182 | 33 | 0.8788 |
| 8_10761103 | 14 | 0.7857 | 6 | 0.1667 | 5 | 0.2 | 10 | 0.4 | 35 | 0.4857 | 11 | 0.4545 | 8 | 0.875 | 8 | 0.625 | 12 | 0.8333 | 39 | 0.6923 |
| 8_10761114 | 16 | 0.75 | 6 | 0.3333 | 6 | 1 | 7 | 0.7143 | 35 | 0.7143 | 9 | 0.4444 | 7 | 0.7143 | 8 | 0.75 | 12 | 0.5833 | 36 | 0.6111 |
| 8_10761144 | 14 | 0.0714 | 7 | 0 | 6 | 0 | 10 | 0.2 | 37 | 0.0811 | 9 | 0 | 9 | 0 | 7 | 0.1429 | 15 | 0.0667 | 40 | 0.05 |
| 8_10761206 | 11 | 0.1818 | 5 | 0 | 5 | 0 | 6 | 0 | 27 | 0.0741 | 7 | 0 | 8 | 0 | 4 | 0 | 10 | 0.2 | 29 | 0.069 |
| 8_10761247 | 12 | 0.75 | 5 | 0.8 | 5 | 0.6 | 6 | 0.6667 | 28 | 0.7143 | 8 | 0.5 | 6 | 0.5 | 3 | 0.3333 | 9 | 0.6667 | 26 | 0.5385 |
| 8_10761267 | 11 | 0.2727 | 5 | 0.2 | 3 | 0.3333 | 5 | 0 | 24 | 0.2083 | 8 | 0.125 | 7 | 0.2857 | 4 | 0.5 | 9 | 0.2222 | 28 | 0.25 |
| 8_10761281 | 10 | 1 | 5 | 0.8 | 2 | 1 | 4 | 1 | 21 | 0.9524 | 6 | 1 | 5 | 1 | 4 | 1 | 8 | 0.875 | 23 | 0.9565 |
| 8_10761289 | 8 | 1 | 4 | 1 | 1 | 1 | 4 | 1 | 17 | 1 | 4 | 1 | 5 | 1 | 2 | 1 | 5 | 1 | 16 | 1 |
| 8_10761301 | 10 | 0.2 | 5 | 0 | 3 | 0.3333 | 4 | 0 | 22 | 0.1364 | 5 | 0.2 | 5 | 0 | 4 | 0.25 | 7 | 0 | 21 | 0.0952 |
| 8_10761303 | 10 | 0.6 | 5 | 0.8 | 3 | 0.6667 | 4 | 0.5 | 22 | 0.6364 | 5 | 0.8 | 4 | 0 | 4 | 0.5 | 5 | 0.2 | 18 | 0.3889 |
| 8_10761305 | 10 | 1 | 5 | 1 | 3 | 1 | 5 | 0.6 | 23 | 0.913 | 5 | 0.8 | 5 | 1 | 3 | 0.6667 | 6 | 0.5 | 19 | 0.7368 |
| 8_10763097 | 4 | 0 | 5 | 0.2 | 2 | 0 | 1 | 0 | 12 | 0.0833 | 2 | 0.5 | 3 | 0 | 1 | 0 | 1 | 0 | 7 | 0.1429 |
| 8_10764964 | 6 | 0.5 | 6 | 0.5 | 2 | 0.5 | 4 | 0.5 | 18 | 0.5 | 1 | 0 | 5 | 0.4 | 1 | 1 | NA | NA | 7 | 0.4286 |
| 8_10777175 | 17 | 0.7647 | 16 | 0.5 | 14 | 0.7143 | 15 | 0.5333 | 62 | 0.629 | 7 | 0.8571 | 14 | 0.5714 | 7 | 0.7143 | 11 | 0.6364 | 39 | 0.6667 |
| 8_10777176 | 13 | 0.0769 | 15 | 0.0667 | 12 | 0.1667 | 15 | 0.0667 | 55 | 0.0909 | 4 | 0.25 | 12 | 0.25 | 6 | 0.5 | 9 | 0.3333 | 31 | 0.3226 |
| 8_10777180 | 16 | 0 | 17 | 0 | 15 | 0.2667 | 15 | 0 | 63 | 0.0635 | 8 | 0.125 | 16 | 0.0625 | 8 | 0 | 12 | 0.0833 | 44 | 0.0682 |
| 8_10777182 | 15 | 0.6667 | 14 | 0.5 | 16 | 0.75 | 13 | 0.5385 | 58 | 0.6207 | 6 | 0.8333 | 14 | 0.5714 | 8 | 0.875 | 9 | 0.6667 | 37 | 0.7027 |
| 8_10777195 | 15 | 0.6 | 13 | 0.6923 | 15 | 0.6667 | 16 | 0.6875 | 59 | 0.661 | 10 | 0.5 | 14 | 0.5714 | 9 | 0.5556 | 10 | 0.7 | 43 | 0.5814 |
| 8_10777196 | 17 | 0.5294 | 16 | 0.6875 | 16 | 0.6875 | 18 | 0.5556 | 67 | 0.6119 | 10 | 0.8 | 15 | 0.4 | 9 | 0.3333 | 12 | 0.25 | 46 | 0.4348 |
| 8_10777216 | 20 | 0.3 | 18 | 0.3333 | 14 | 0.4286 | 17 | 0.1765 | 69 | 0.3043 | 8 | 0.25 | 15 | 0.4667 | 8 | 0.5 | 9 | 0.4444 | 40 | 0.425 |
| 8_10777232 | 17 | 0.3529 | 11 | 0.2727 | 11 | 0.4545 | 12 | 0.25 | 51 | 0.3333 | 6 | 0.1667 | 9 | 0.2222 | 5 | 0.8 | 6 | 0.1667 | 26 | 0.3077 |
| 8_10777327 | 4 | 0.5 | 6 | 0.8333 | 5 | 0.6 | 2 | 1 | 17 | 0.7059 | 2 | 0.5 | 5 | 0.8 | 9 | 0.4444 | 8 | 0.375 | 24 | 0.5 |
| 8_10777330 | 4 | 1 | 5 | 0.6 | 6 | 0.6667 | 2 | 1 | 17 | 0.7647 | 2 | 0.5 | 5 | 0.8 | 8 | 0.625 | 8 | 0.875 | 23 | 0.7391 |
| 8_10777337 | 4 | 0.25 | 4 | 0.5 | 5 | 0.2 | 2 | 0 | 15 | 0.2667 | 2 | 0 | 5 | 0.4 | 9 | 0.3333 | 6 | 0.5 | 22 | 0.3636 |
| 8_10777348 | 4 | 0.75 | 6 | 0.1667 | 6 | 0 | 1 | 0 | 17 | 0.2353 | 2 | 0 | 6 | 0.3333 | 7 | 0.1429 | 7 | 0.4286 | 22 | 0.2727 |
| 8_10777353 | 3 | 0 | 4 | 0 | 5 | 0.2 | 3 | 0 | 15 | 0.0667 | 2 | 0 | 6 | 0.1667 | 7 | 0.5714 | 6 | 0 | 21 | 0.2381 |
| 8_10777357 | 5 | 0.6 | 4 | 0.75 | 5 | 0.8 | 4 | 0.75 | 18 | 0.7222 | 2 | 1 | 4 | 1 | 10 | 1 | 7 | 1 | 23 | 1 |
| 8_10777359 | 6 | 0 | 5 | 0.2 | 5 | 0 | 3 | 0 | 19 | 0.0526 | 2 | 0 | 6 | 0.1667 | 11 | 0.1818 | 8 | 0.25 | 27 | 0.1852 |
| 8_10777365 | 6 | 0.1667 | 5 | 0.2 | 5 | 0.4 | 4 | 0.75 | 20 | 0.35 | 1 | 1 | 5 | 0.2 | 8 | 0.25 | 8 | 0.25 | 22 | 0.2727 |
| 8_10777370 | 6 | 0.3333 | 5 | 0.6 | 6 | 0.6667 | 4 | 0.25 | 21 | 0.4762 | 2 | 0 | 4 | 0.25 | 10 | 0.1 | 6 | 0 | 22 | 0.0909 |
| 8_10777374 | 5 | 0 | 3 | 0 | 5 | 0.6 | 3 | 0 | 16 | 0.1875 | 2 | 0 | 4 | 0.25 | 9 | 0 | 4 | 0 | 19 | 0.0526 |
| 8_10777375 | 4 | 0.75 | 3 | 0.6667 | 5 | 0.4 | 3 | 0 | 15 | 0.4667 | 2 | 1 | 4 | 0.5 | 10 | 0.5 | 8 | 0.875 | 24 | 0.6667 |
| 8_10777378 | 6 | 1 | 2 | 1 | 5 | 0.6 | 4 | 1 | 17 | 0.8824 | 2 | 1 | 3 | 1 | 10 | 1 | 5 | 1 | 20 | 1 |
| 8_10777379 | 4 | 1 | 4 | 1 | 5 | 0.8 | 3 | 1 | 16 | 0.9375 | 1 | 1 | 4 | 1 | 7 | 1 | 7 | 1 | 19 | 1 |
| 8_10777385 | 4 | 0.25 | 2 | 0 | 3 | 0 | 3 | 0.3333 | 12 | 0.1667 | 2 | 0 | 5 | 0.2 | 6 | 0.1667 | 6 | 0.1667 | 19 | 0.1579 |
| 8_10777562 | 3 | 0 | NA | NA | 2 | 0 | 3 | 0.3333 | 8 | 0.125 | 2 | 0.5 | 2 | 0.5 | 1 | 0 | 1 | 0 | 6 | 0.3333 |
| 8_10777566 | 1 | 0 | 1 | 1 | 1 | 1 | 1 | 1 | 4 | 0.75 | 1 | 1 | 1 | 0 | 1 | 0 | 1 | 0 | 4 | 0.25 |
| 8_10777896 | NA | NA | 1 | 1 | 2 | 0 | 3 | 0.3333 | 6 | 0.3333 | 2 | 1 | 1 | 0 | 3 | 0 | 2 | 0.5 | 8 | 0.375 |
| 8_10777907 | NA | NA | 1 | 0 | 2 | 0.5 | 3 | 1 | 6 | 0.6667 | 2 | 1 | 1 | 0 | 3 | 0.6667 | 2 | 0.5 | 8 | 0.625 |
| 8_10777923 | NA | NA | 1 | 1 | 2 | 0.5 | 3 | 1 | 6 | 0.8333 | 2 | 1 | 1 | 1 | 2 | 1 | 2 | 1 | 7 | 1 |
| 8_10777926 | NA | NA | 1 | 1 | 2 | 1 | 2 | 1 | 5 | 1 | 1 | 1 | 1 | 1 | 2 | 1 | 2 | 1 | 6 | 1 |
| 8_10777936 | NA | NA | 1 | 0 | 2 | 0.5 | 3 | 0 | 6 | 0.1667 | 2 | 0 | 1 | 0 | 2 | 0.5 | 2 | 0 | 7 | 0.1429 |
| 8_10777960 | NA | NA | 1 | 1 | 2 | 0.5 | NA | NA | 3 | 0.6667 | 2 | 0.5 | 1 | 0 | 2 | 0.5 | 1 | 1 | 6 | 0.5 |
| 8_10778511 | 4 | 0 | 4 | 0 | 4 | 0 | 4 | 0.5 | 16 | 0.125 | 2 | 0 | 3 | 0.3333 | 6 | 0 | 6 | 0 | 17 | 0.0588 |
| 8_10778514 | 4 | 0.25 | 4 | 0 | 5 | 0.2 | 4 | 0.5 | 17 | 0.2353 | 3 | 0 | 3 | 0.6667 | 4 | 0.25 | 7 | 0.1429 | 17 | 0.2353 |
| 8_10778516 | 4 | 1 | 4 | 0.75 | 5 | 0.4 | 4 | 1 | 17 | 0.7647 | 2 | 0.5 | 3 | 1 | 5 | 1 | 4 | 0.75 | 14 | 0.8571 |
| 8_10778520 | 2 | 0.5 | 4 | 0.5 | 4 | 0 | 4 | 0 | 14 | 0.2143 | 2 | 0.5 | 3 | 0.3333 | 6 | 0.1667 | 4 | 0.5 | 15 | 0.3333 |
| 8_10778538 | 3 | 0 | 4 | 0 | 5 | 0 | 3 | 0.3333 | 15 | 0.0667 | 3 | 0 | 2 | 0 | 6 | 0 | 7 | 0.1429 | 18 | 0.0556 |
| 8_10778557 | 4 | 0.25 | 4 | 0.25 | 4 | 0.5 | 5 | 0.2 | 17 | 0.2941 | 3 | 0.3333 | 3 | 0.3333 | 7 | 0.4286 | 6 | 0.5 | 19 | 0.4211 |
| 8_10778711 | 5 | 0 | 3 | 0.3333 | 1 | 0 | 1 | 0 | 10 | 0.1 | 3 | 0 | 4 | 0.25 | 3 | 0 | 3 | 0 | 13 | 0.0769 |
| 8_10778742 | 7 | 0.2857 | 4 | 0 | 2 | 0 | 1 | 0 | 14 | 0.1429 | 3 | 0.3333 | 2 | 0.5 | 2 | 0 | 4 | 0 | 11 | 0.1818 |
| 8_10779616 | 6 | 0 | 6 | 0.1667 | 7 | 0.1429 | 5 | 0.2 | 24 | 0.125 | 4 | 0 | 4 | 0.25 | 5 | 0.4 | 4 | 0.25 | 17 | 0.2353 |
| 8_10779633 | 5 | 0.6 | 6 | 0.3333 | 7 | 1 | 5 | 0.8 | 23 | 0.6957 | 2 | 0.5 | 4 | 0.5 | 4 | 0.5 | 3 | 0.6667 | 13 | 0.5385 |
| 8_10779642 | 5 | 0 | 3 | 0 | 5 | 0.2 | 4 | 0 | 17 | 0.0588 | 2 | 0 | 4 | 0 | 4 | 0 | 3 | 0.3333 | 13 | 0.0769 |
| 8_10779651 | 4 | 0.5 | 5 | 0 | 5 | 0.2 | 5 | 0.2 | 19 | 0.2105 | 3 | 0.3333 | 5 | 0 | 5 | 0.2 | 4 | 0.25 | 17 | 0.1765 |
| 8_10779653 | 4 | 0.5 | 1 | 1 | 2 | 1 | 1 | 0 | 8 | 0.625 | 2 | 0.5 | 3 | 0.6667 | 3 | 1 | 2 | 1 | 10 | 0.8 |
| 8_10779653 | NA | NA | 1 | 1 | 1 | 0 | 1 | 0 | 3 | 0.3333 | NA | NA | 2 | 1 | NA | NA | 1 | 1 | 3 | 1 |
| 8_10779680 | 5 | 0.6 | 4 | 0.5 | 7 | 0.5714 | 4 | 0.5 | 20 | 0.55 | 2 | 0.5 | 4 | 0.75 | 4 | 1 | 4 | 0.25 | 14 | 0.6429 |
| 8_10779685 | 5 | 0.6 | 5 | 0.8 | 6 | 0.6667 | 5 | 1 | 21 | 0.7619 | 3 | 0.6667 | 5 | 1 | 5 | 1 | 4 | 1 | 17 | 0.9412 |
| 8_10779695 | 4 | 0.75 | 5 | 0.6 | 7 | 0.4286 | 5 | 0.6 | 21 | 0.5714 | 3 | 1 | 5 | 0.8 | 5 | 0.6 | 2 | 0.5 | 15 | 0.7333 |
| 8_10779700 | 3 | 0 | 5 | 0.4 | 7 | 0 | 4 | 0 | 19 | 0.1053 | 3 | 0 | 5 | 0 | 3 | 0.3333 | 2 | 0.5 | 13 | 0.1538 |
| 8_10779701 | 4 | 0.5 | 6 | 0.5 | 7 | 0.5714 | 3 | 0.3333 | 20 | 0.5 | 2 | 0.5 | 5 | 0.6 | 4 | 1 | 2 | 1 | 13 | 0.7692 |
| 8_10779704 | 4 | 0.75 | 5 | 1 | 4 | 0.75 | 4 | 1 | 17 | 0.8824 | 3 | 1 | 2 | 1 | 3 | 1 | 2 | 0.5 | 10 | 0.9 |

| | | | | | | | | | | | | | | | | | | | | |
|---|---|---|---|---|---|---|---|---|---|---|---|---|---|---|---|---|---|---|---|---|
| 8_10779705 | 5 | 0 | 6 | 0.3333 | 6 | 0.3333 | 2 | 0 | 19 | 0.2105 | 3 | 0.3333 | 3 | 0.3333 | 3 | 0 | 1 | 0 | 10 | 0.2 |
| 8_10779706 | 5 | 0.2 | 6 | 0 | 7 | 0 | 3 | 0.3333 | 21 | 0.0952 | 3 | 0 | 5 | 0.4 | 4 | 0.25 | 2 | 0 | 14 | 0.2143 |
| 8_10780335 | 2 | 0 | 5 | 0.2 | 4 | 0 | 3 | 0.3333 | 14 | 0.1429 | 1 | 0 | 1 | 0 | 1 | 1 | NA | NA | 3 | 0.3333 |
| 8_10780337 | 2 | 0 | 4 | 0 | 4 | 0 | 3 | 0.3333 | 13 | 0.0769 | 1 | 0 | 3 | 0 | 2 | 0.5 | 1 | 0 | 7 | 0.1429 |
| 8_10780346 | 2 | 0 | 4 | 0 | 4 | 0.25 | 4 | 0 | 14 | 0.0714 | 2 | 0 | 4 | 0.5 | 4 | 0.25 | 1 | 0 | 11 | 0.2727 |
| 8_10780355 | 2 | 0.5 | 3 | 0 | 3 | 0.6667 | 3 | 0.3333 | 11 | 0.3636 | 1 | 0 | 6 | 0 | 4 | 0.25 | 1 | 1 | 12 | 0.1667 |
| 8_10780359 | NA | NA | 4 | 0.25 | 4 | 0.75 | 4 | 0.25 | 12 | 0.4167 | 1 | 0 | 6 | 0.1667 | 4 | 0 | 1 | 0 | 12 | 0.0833 |
| 8_10791035 | 6 | 0.3333 | 5 | 0.8 | 7 | 0.7143 | 2 | 1 | 20 | 0.65 | 3 | 0.3333 | 8 | 0.5 | 5 | 0.6 | 6 | 1 | 22 | 0.6364 |
| 8_10791057 | 5 | 0 | 4 | 0 | 7 | 0.1429 | 5 | 0.2 | 21 | 0.0952 | 4 | 0 | 8 | 0.125 | 4 | 0 | 9 | 0.2222 | 25 | 0.12 |
| 8_10791061 | 8 | 0.25 | 6 | 0.1667 | 6 | 0.1667 | 5 | 0.6 | 25 | 0.28 | 3 | 0.3333 | 7 | 0.4286 | 6 | 0.3333 | 9 | 0.4444 | 25 | 0.4 |
| 8_10791117 | 9 | 0.5556 | 7 | 0.5714 | 9 | 0.5556 | 8 | 0.625 | 33 | 0.5758 | 4 | 0.5 | 6 | 0.6667 | 9 | 0.4444 | 7 | 0.8571 | 26 | 0.6154 |
| 8_10791124 | 10 | 0 | 7 | 0.2857 | 8 | 0.125 | 6 | 0.6667 | 31 | 0.2258 | 4 | 0.25 | 9 | 0.2222 | 7 | 0.1429 | 6 | 0 | 26 | 0.1538 |
| 8_10791144 | 10 | 0 | 9 | 0.2222 | 10 | 0 | 8 | 0.125 | 37 | 0.0811 | 4 | 0 | 9 | 0.3333 | 9 | 0 | 9 | 0.1111 | 31 | 0.129 |
| 8_10791153 | 11 | 0.3636 | 9 | 0.2222 | 10 | 0.3 | 8 | 0.375 | 38 | 0.3158 | 4 | 0.25 | 10 | 0.6 | 11 | 0.4545 | 9 | 0.4444 | 34 | 0.4706 |
| 8_10791198 | 11 | 0 | 10 | 0 | 12 | 0.1667 | 8 | 0.125 | 41 | 0.0732 | 3 | 0 | 12 | 0.1667 | 9 | 0.1111 | 8 | 0.125 | 32 | 0.125 |
| 8_10791240 | 11 | 0.1818 | 13 | 0.0769 | 15 | 0.2 | 7 | 0 | 46 | 0.1304 | 5 | 0.2 | 15 | 0.0667 | 12 | 0.1667 | 9 | 0.1111 | 41 | 0.122 |
| 8_10791256 | 11 | 0 | 16 | 0.125 | 12 | 0.0833 | 10 | 0.1 | 49 | 0.0816 | 4 | 0 | 14 | 0.1429 | 12 | 0 | 9 | 0.1111 | 39 | 0.0769 |
| 8_10791309 | 9 | 0.1111 | 12 | 0.1667 | 12 | 0.1667 | 11 | 0.0909 | 44 | 0.1364 | 4 | 0.5 | 6 | 0.1667 | 7 | 0.4286 | 9 | 0.5556 | 26 | 0.4231 |
| 8_10791337 | 9 | 0.8889 | 16 | 1 | 16 | 0.9375 | 9 | 0.8889 | 50 | 0.94 | 2 | 1 | 11 | 0.8182 | 10 | 0.9 | 13 | 0.8462 | 36 | 0.8611 |
| 8_10791338 | 10 | 0 | 18 | 0.1111 | 19 | 0.1053 | 11 | 0.5455 | 58 | 0.1724 | 4 | 0 | 13 | 0.0769 | 11 | 0.1818 | 13 | 0.3077 | 41 | 0.1707 |
| 8_10791358 | 12 | 0 | 20 | 0.25 | 20 | 0.05 | 10 | 0.1 | 62 | 0.1129 | 4 | 0 | 12 | 0 | 11 | 0.0909 | 13 | 0.2308 | 40 | 0.1 |
| 8_10791363 | 11 | 0.9091 | 19 | 1 | 19 | 1 | 11 | 1 | 60 | 0.9833 | 4 | 1 | 13 | 1 | 12 | 1 | 14 | 1 | 43 | 1 |
| 8_10809385 | NA | NA | 2 | 0.5 | NA | NA | NA | NA | 2 | 0.5 | NA | NA | NA | NA | NA | NA | 2 | 0.5 | 2 | 0.5 |
| 8_10809441 | NA | NA | 4 | 0.5 | NA | NA | NA | NA | 4 | 0.5 | NA | NA | NA | NA | NA | NA | 2 | 0.5 | 2 | 0.5 |
| 8_10809783 | 1 | 1 | 5 | 1 | 1 | 0 | NA | NA | 7 | 0.8571 | NA | NA | 1 | 0 | NA | NA | 3 | 0.6667 | 4 | 0.5 |
| 8_10809786 | 1 | 0 | 5 | 0.8 | 1 | 1 | NA | NA | 7 | 0.7143 | NA | NA | 1 | 1 | NA | NA | 3 | 0.3333 | 4 | 0.5 |
| 8_10809787 | 1 | 1 | 5 | 0.2 | 1 | 0 | NA | NA | 7 | 0.2857 | NA | NA | 1 | 1 | NA | NA | 3 | 0 | 4 | 0.25 |
| 8_10809830 | 1 | 1 | 4 | 0 | 1 | 0 | NA | NA | 6 | 0.1667 | NA | NA | 1 | 0 | NA | NA | 3 | 0.3333 | 4 | 0.25 |
| 8_10810061 | NA | NA | 4 | 0.5 | 1 | 0 | 1 | 0 | 6 | 0.3333 | NA | NA | 1 | 1 | 1 | 0 | 5 | 0 | 7 | 0.1429 |
| 8_10810071 | NA | NA | 4 | 0.25 | 1 | 0 | 1 | 0 | 6 | 0.1667 | NA | NA | 1 | 1 | 2 | 0.5 | 6 | 0 | 9 | 0.2222 |
| 8_10811501 | 1 | 1 | 3 | 0.6667 | NA | NA | NA | NA | 4 | 0.75 | 1 | 0 | NA | NA | NA | NA | 1 | 1 | 2 | 0.5 |
| 8_10811508 | 1 | 1 | 3 | 0.6667 | NA | NA | NA | NA | 4 | 0.75 | 1 | 1 | NA | NA | NA | NA | 1 | 1 | 2 | 1 |
| 8_10811539 | 1 | 0 | 3 | 0.6667 | NA | NA | NA | NA | 4 | 0.5 | 1 | 1 | NA | NA | NA | NA | NA | NA | 1 | 1 |
| 8_10813919 | 7 | 0.8571 | 6 | 1 | 4 | 1 | 8 | 1 | 25 | 0.96 | 2 | 1 | 1 | 1 | 7 | 0.8571 | 3 | 1 | 13 | 0.9231 |
| 8_10813920 | 3 | 0.6667 | 4 | 1 | 5 | 0.4 | 7 | 0.5714 | 19 | 0.6316 | 2 | 0.5 | 1 | 0 | 5 | 0.8 | 2 | 1 | 10 | 0.7 |
| 8_10813925 | 5 | 0.4 | 5 | 0.6 | 5 | 0.8 | 5 | 0.8 | 20 | 0.65 | 3 | 0.3333 | 2 | 1 | 5 | 0.6 | 2 | 1 | 12 | 0.6667 |
| 8_10813935 | 8 | 0.5 | 7 | 0.7143 | 4 | 0.75 | 4 | 0.75 | 23 | 0.6522 | 3 | 0 | 1 | 1 | 4 | 1 | 3 | 1 | 11 | 0.7273 |
| 8_10813939 | 6 | 1 | 6 | 0.6667 | 6 | 0.6667 | 5 | 1 | 23 | 0.8261 | 3 | 0.6667 | 1 | 1 | 7 | 0.8571 | 2 | 1 | 13 | 0.8462 |
| 8_10813945 | 4 | 0 | 6 | 0 | 6 | 0 | 5 | 0.4 | 21 | 0.0952 | 2 | 0.5 | 2 | 0 | 3 | 0 | 1 | 0 | 8 | 0.125 |
| 8_10813954 | 5 | 0.6 | 5 | 0.2 | 5 | 0.4 | 6 | 0.5 | 21 | 0.4286 | 3 | 0.3333 | 2 | 0 | 4 | 0.5 | 3 | 0.3333 | 12 | 0.3333 |
| 8_10814334 | 1 | 0 | 2 | 0 | 1 | 1 | 1 | 0 | 5 | 0.2 | NA | NA | NA | NA | 2 | 0 | 1 | 1 | 3 | 0.3333 |
| 8_10828373 | 1 | 1 | 6 | 0.5 | 3 | 1 | 1 | 1 | 11 | 0.7273 | 3 | 0.6667 | 1 | 1 | 1 | 1 | 3 | 0.3333 | 8 | 0.625 |
| 8_10828375 | NA | NA | 4 | 0.25 | 2 | 0 | NA | NA | 6 | 0.1667 | 3 | 0.3333 | 1 | 1 | 1 | 0 | 4 | 0 | 9 | 0.2222 |
| 8_10828378 | 1 | 1 | 7 | 0.1429 | 3 | 0 | 1 | 1 | 12 | 0.25 | 3 | 0.3333 | 1 | 1 | NA | NA | 4 | 0 | 8 | 0.25 |
| 8_10828384 | 1 | 1 | 4 | 1 | 1 | 0 | 1 | 1 | 7 | 0.8571 | 2 | 0.5 | 1 | 1 | 1 | 1 | 4 | 0.25 | 8 | 0.5 |
| 8_10828390 | 1 | 1 | 2 | 1 | 1 | 0 | 1 | 1 | 5 | 0.8 | 1 | 0 | 1 | 1 | NA | NA | 1 | 0 | 3 | 0.3333 |
| 8_10828405 | 1 | 1 | 2 | 0 | 1 | 0 | NA | NA | 4 | 0.25 | 1 | 1 | 1 | 0 | 1 | 1 | 3 | 0.6667 | 6 | 0.6667 |
| 8_10830709 | 29 | 0.3793 | 23 | 0.4348 | 28 | 0.3929 | 27 | 0.4074 | 107 | 0.4019 | 14 | 0.3571 | 10 | 0.2 | 20 | 0.35 | 19 | 0.4211 | 63 | 0.3492 |
| 8_10830712 | 22 | 0.7273 | 21 | 0.6667 | 19 | 0.7368 | 23 | 0.6957 | 85 | 0.7059 | 15 | 0.5333 | 10 | 0.8 | 18 | 0.6667 | 16 | 0.875 | 59 | 0.7119 |
| 8_10830713 | 27 | 0.4444 | 22 | 0.2727 | 26 | 0.1538 | 26 | 0.3846 | 101 | 0.3168 | 14 | 0.2857 | 14 | 0.3571 | 19 | 0.2632 | 19 | 0.2105 | 66 | 0.2727 |
| 8_10830728 | 22 | 0.0455 | 20 | 0.05 | 19 | 0.1053 | 20 | 0.1 | 81 | 0.0741 | 10 | 0 | 9 | 0.2222 | 14 | 0.0714 | 16 | 0 | 49 | 0.0612 |
| 8_10830737 | 28 | 0.5357 | 18 | 0.7778 | 23 | 0.6522 | 23 | 0.5652 | 92 | 0.6196 | 11 | 0.6364 | 11 | 0.7273 | 18 | 0.5 | 18 | 0.7778 | 58 | 0.6552 |
| 8_10830742 | 25 | 0.12 | 19 | 0.4211 | 22 | 0.1818 | 20 | 0.05 | 86 | 0.186 | 7 | 0.1429 | 10 | 0.5 | 12 | 0.1667 | 15 | 0.1333 | 44 | 0.2273 |
| 8_10830747 | 19 | 0.0526 | 16 | 0.0625 | 15 | 0.0667 | 20 | 0.1 | 70 | 0.0714 | 6 | 0 | 8 | 0.125 | 14 | 0 | 12 | 0.0833 | 40 | 0.05 |
| 8_10830761 | 23 | 0.2609 | 17 | 0.2353 | 22 | 0.0909 | 21 | 0 | 83 | 0.1446 | 9 | 0.2222 | 11 | 0.0909 | 15 | 0.2667 | 16 | 0.25 | 51 | 0.2157 |
| 8_10830788 | 19 | 0.6842 | 13 | 1 | 22 | 0.9091 | 21 | 0.9524 | 75 | 0.88 | 6 | 0.8333 | 12 | 0.75 | 11 | 0.8182 | 17 | 0.8235 | 46 | 0.8043 |
| 8_10830792 | 19 | 0.0526 | 10 | 0.1 | 20 | 0.1 | 20 | 0.2 | 69 | 0.1159 | 6 | 0.1667 | 11 | 0.0909 | 15 | 0.3333 | 15 | 0.2 | 47 | 0.2128 |
| 8_10925431 | NA | NA | 1 | 1 | NA | NA | NA | NA | 1 | 1 | 1 | 1 | NA | NA | 2 | 0 | NA | NA | 3 | 0.3333 |
| 8_10925437 | NA | NA | 1 | 1 | NA | NA | NA | NA | 1 | 1 | 1 | 1 | NA | NA | 2 | 0.5 | NA | NA | 3 | 0.6667 |
| 8_10925448 | NA | NA | 1 | 1 | NA | NA | NA | NA | 1 | 1 | 1 | 1 | NA | NA | 2 | 0.5 | NA | NA | 3 | 0.6667 |
| 8_10925485 | NA | NA | 1 | 1 | NA | NA | NA | NA | 1 | 1 | 1 | 1 | NA | NA | 1 | 1 | NA | NA | 2 | 1 |
| 8_10925493 | NA | NA | 1 | 1 | NA | NA | NA | NA | 1 | 1 | 1 | 1 | NA | NA | 2 | 1 | NA | NA | 3 | 1 |
| 8_10968807 | 2 | 0.5 | NA | NA | 2 | 0.5 | 3 | 1 | 7 | 0.7143 | NA | NA | 2 | 0.5 | 1 | 0 | NA | NA | 3 | 0.3333 |
| 8_10968811 | 2 | 1 | NA | NA | 2 | 1 | 3 | 1 | 7 | 1 | NA | NA | 1 | 1 | 1 | 1 | NA | NA | 2 | 1 |
| 8_10987208 | 1 | 1 | NA | NA | 1 | 0 | NA | NA | 2 | 0.5 | 1 | 1 | NA | NA | NA | NA | NA | NA | 1 | 1 |

| | | | | | | | | | | | | | | | | | | | | |
|---|---|---|---|---|---|---|---|---|---|---|---|---|---|---|---|---|---|---|---|---|
| 8_1098721 | 1 | 0 | NA | NA | 1 | 1 | NA | NA | 2 | 0.5 | 1 | 1 | NA | NA | NA | NA | NA | NA | 1 | 1 |
| 8_1098721 | 1 | 1 | NA | NA | 1 | 0 | NA | NA | 2 | 0.5 | 1 | 1 | NA | NA | NA | NA | NA | NA | 1 | 1 |
| 8_1098722 | NA | NA | NA | NA | 1 | 1 | NA | NA | 1 | 1 | 1 | 1 | NA | NA | NA | NA | NA | NA | 1 | 1 |
| 8_1098722 | 1 | 1 | NA | NA | 1 | 0 | NA | NA | 2 | 0.5 | 1 | 1 | NA | NA | NA | NA | NA | NA | 1 | 1 |
| 8_1102747 | 3 | 0.3333 | 1 | 1 | 5 | 0.6 | 1 | 1 | 10 | 0.6 | 2 | 0.5 | 6 | 0.5 | 3 | 1 | 5 | 0.4 | 16 | 0.5625 |
| 8_1102749 | 3 | 0.6667 | 1 | 0 | 5 | 0.8 | 1 | 1 | 10 | 0.7 | 2 | 0.5 | 5 | 0.8 | 2 | 1 | 5 | 0.4 | 14 | 0.6429 |
| 8_1102750 | 1 | 1 | 1 | 0 | 4 | 0.75 | 1 | 1 | 7 | 0.7143 | 2 | 0 | 3 | 0.6667 | 2 | 0.5 | 4 | 0 | 11 | 0.2727 |
| 8_1102892 | NA | NA | NA | NA | NA | NA | 1 | 1 | 1 | 1 | 1 | 1 | NA | NA | NA | NA | NA | NA | 1 | 1 |
| 8_1102955 | 2 | 0.5 | 2 | 0.5 | 1 | 1 | 1 | 0 | 6 | 0.5 | 1 | 1 | 1 | 0 | 1 | 1 | 3 | 0.3333 | 6 | 0.5 |
| 8_1102995 | 2 | 1 | 4 | 1 | 3 | 0.6667 | 2 | 0.5 | 11 | 0.8182 | 1 | 1 | 3 | 1 | 1 | 0 | 3 | 1 | 8 | 0.875 |
| 8_1102997 | 1 | 0 | 2 | 1 | 3 | 0.6667 | 2 | 0.5 | 8 | 0.625 | 1 | 0 | 3 | 0.6667 | 1 | 1 | 1 | 1 | 6 | 0.6667 |
| 8_1102997 | 1 | 1 | 4 | 1 | 3 | 0.6667 | 2 | 1 | 10 | 0.9 | NA | NA | 3 | 0.6667 | 1 | 1 | 1 | 1 | 5 | 0.8 |
| 8_1102997 | 2 | 0 | 4 | 0.75 | 3 | 0.3333 | 2 | 1 | 11 | 0.5455 | 1 | 0 | 1 | 0 | 1 | 0 | 2 | 0.5 | 5 | 0.2 |
| 8_1102998 | 2 | 0.5 | 3 | 0.3333 | 2 | 0.5 | 2 | 0 | 9 | 0.3333 | 1 | 1 | 2 | 1 | 1 | 0 | 3 | 0 | 7 | 0.4286 |
| 8_1103000 | 2 | 0 | 2 | 0.5 | 3 | 1 | 2 | 0 | 9 | 0.4444 | 1 | 0 | 1 | 0 | 1 | 0 | 3 | 0.3333 | 6 | 0.1667 |
| 8_1103000 | 2 | 1 | 4 | 1 | 2 | 1 | 2 | 1 | 10 | 1 | 1 | 1 | 3 | 0.6667 | 1 | 1 | 3 | 1 | 8 | 0.875 |
| 8_1103001 | 2 | 0 | 4 | 0 | 3 | 0.6667 | 2 | 0 | 11 | 0.1818 | 1 | 0 | 3 | 0 | 1 | 0 | 3 | 0.3333 | 8 | 0.125 |
| 8_1103005 | 2 | 0 | 3 | 0.3333 | 4 | 0.5 | 2 | 0.5 | 11 | 0.3636 | 1 | 1 | 3 | 0.3333 | 1 | 0 | 4 | 0.5 | 9 | 0.4444 |
| 8_1103202 | NA | NA | 3 | 1 | 6 | 0.6667 | 4 | 0.25 | 13 | 0.6154 | 5 | 0.8 | 3 | 0.6667 | 6 | 0.6667 | 1 | 1 | 15 | 0.7333 |
| 8_1103204 | 1 | 1 | 1 | 1 | 6 | 0.6667 | 5 | 0.4 | 13 | 0.6154 | 5 | 0 | 4 | 0.5 | 3 | 0.3333 | 2 | 1 | 14 | 0.3571 |
| 8_1103205 | 1 | 0 | 3 | 0.3333 | 5 | 0.4 | 4 | 0.25 | 13 | 0.3077 | 3 | 0 | 5 | 0.4 | 5 | 0 | 3 | 0.3333 | 16 | 0.1875 |
| 8_1103206 | 1 | 0 | 2 | 0 | 5 | 0.6 | 4 | 0 | 12 | 0.25 | 3 | 0.3333 | 5 | 0.4 | 4 | 0.25 | 2 | 0 | 14 | 0.2857 |
| 8_1103921 | NA | NA | NA | NA | 1 | 1 | NA | NA | 1 | 1 | NA | NA | 1 | 1 | NA | NA | NA | NA | 1 | 1 |
| 8_1103982 | 16 | 0.3125 | 16 | 0.375 | 10 | 0.7 | 13 | 0.4615 | 55 | 0.4364 | 5 | 0 | 9 | 0.6667 | 6 | 0.3333 | 11 | 0.3636 | 31 | 0.3871 |
| 8_1103985 | 10 | 0.2 | 15 | 0.2667 | 5 | 0.4 | 8 | 0.375 | 38 | 0.2895 | 5 | 0 | 13 | 0.4615 | 6 | 0 | 12 | 0.0833 | 36 | 0.1944 |
| 8_1103986 | 14 | 0.7143 | 19 | 0.8421 | 8 | 0.75 | 13 | 0.7692 | 54 | 0.7778 | 5 | 1 | 12 | 0.75 | 7 | 0.5714 | 13 | 0.7692 | 37 | 0.7568 |
| 8_1103987 | 13 | 0.6923 | 17 | 0.8235 | 8 | 0.875 | 14 | 0.8571 | 52 | 0.8077 | 4 | 0.75 | 13 | 0.6154 | 7 | 0.8571 | 13 | 0.6154 | 37 | 0.6757 |
| 8_1103988 | 14 | 0.9286 | 19 | 0.9474 | 6 | 1 | 12 | 0.9167 | 51 | 0.9412 | 7 | 1 | 11 | 1 | 6 | 0.6667 | 11 | 0.9091 | 35 | 0.9143 |
| 8_1103988 | 10 | 1 | 19 | 0.9474 | 7 | 1 | 12 | 0.75 | 48 | 0.9167 | 6 | 1 | 12 | 0.9167 | 7 | 1 | 11 | 0.8182 | 36 | 0.9167 |
| 8_1103990 | 13 | 0.7692 | 17 | 0.8235 | 5 | 1 | 13 | 0.7692 | 48 | 0.8125 | 6 | 1 | 12 | 1 | 7 | 0.8571 | 12 | 0.75 | 37 | 0.8919 |
| 8_1103990 | 9 | 0.3333 | 10 | 0.6 | 3 | 1 | 5 | 0.8 | 27 | 0.5926 | 3 | 1 | 6 | 0.1667 | 1 | 0 | 8 | 0.625 | 18 | 0.5 |
| 8_1103990 | 14 | 0.5714 | 18 | 0.8333 | 5 | 0.8 | 11 | 0.5455 | 48 | 0.6875 | 4 | 0.5 | 13 | 0.7692 | 7 | 0.5714 | 12 | 0.75 | 36 | 0.6944 |
| 8_1103991 | 12 | 0.9167 | 19 | 0.9474 | 5 | 1 | 10 | 0.7 | 46 | 0.8913 | 5 | 0.6 | 12 | 0.8333 | 7 | 0.7143 | 12 | 0.8333 | 36 | 0.7778 |
| 8_1104506 | 6 | 0.6667 | 7 | 0.5714 | 3 | 0.6667 | 5 | 1 | 21 | 0.7143 | 2 | 0.5 | 5 | 1 | 3 | 0 | 3 | 1 | 13 | 0.6923 |
| 8_1104509 | 5 | 0.4 | 6 | 0.1667 | 3 | 0 | 6 | 0.1667 | 20 | 0.2 | 4 | 0 | 5 | 0 | 5 | 0.4 | 2 | 0 | 16 | 0.125 |
| 8_1104510 | 5 | 0.2 | 7 | 0.1429 | 4 | 0 | 7 | 0.1429 | 23 | 0.1304 | 5 | 0.2 | 4 | 0 | 5 | 0.2 | 3 | 0 | 17 | 0.1176 |
| 8_1104510 | 6 | 1 | 7 | 0.8571 | 3 | 0.6667 | 6 | 1 | 22 | 0.9091 | 5 | 0.8 | 4 | 1 | 5 | 0.8 | 3 | 1 | 17 | 0.8824 |
| 8_1104511 | 6 | 0.3333 | 9 | 0.4444 | 4 | 0.25 | 7 | 0.1429 | 26 | 0.3077 | 5 | 0.2 | 5 | 0.2 | 6 | 0.1667 | 3 | 0.6667 | 19 | 0.2632 |
| 8_1104511 | 3 | 1 | 7 | 1 | 4 | 0.75 | 5 | 0.8 | 19 | 0.8947 | 2 | 1 | 5 | 1 | 4 | 1 | 2 | 1 | 13 | 1 |
| 8_1104511 | 3 | 0.3333 | 7 | 0.7143 | 4 | 0.5 | 5 | 0.6 | 19 | 0.5789 | 3 | 0 | 2 | 0.5 | 5 | 0.8 | 2 | 1 | 12 | 0.5833 |
| 8_1104514 | 6 | 0.1667 | 7 | 0.1429 | 4 | 0.25 | 6 | 0 | 23 | 0.1304 | 5 | 0.2 | 5 | 0 | 5 | 0 | 4 | 0 | 19 | 0.0526 |
| 8_1104514 | 4 | 0.25 | 8 | 0.625 | 5 | 0.8 | 6 | 0.5 | 23 | 0.5652 | 4 | 0.25 | 4 | 0.75 | 4 | 0.5 | 3 | 0.3333 | 15 | 0.4667 |
| 8_1104514 | 5 | 1 | 5 | 1 | 3 | 1 | 6 | 0.8333 | 19 | 0.9474 | 5 | 0.8 | 1 | 0 | 4 | 1 | 3 | 1 | 13 | 0.8462 |
| 8_1104514 | 6 | 0.5 | 7 | 0.8571 | 5 | 1 | 6 | 0.8333 | 24 | 0.7917 | 5 | 0.8 | 5 | 0.6 | 4 | 1 | 4 | 1 | 18 | 0.8333 |
| 8_1104515 | 2 | 1 | 1 | 0 | NA | NA | 2 | 1 | 5 | 0.8 | 1 | 0 | 4 | 0.25 | NA | NA | 1 | 0 | 6 | 0.1667 |
| 8_1104531 | 5 | 0 | 1 | 0 | 3 | 0.3333 | 2 | 0 | 11 | 0.0909 | 5 | 0.2 | 2 | 0 | 7 | 0.1429 | 4 | 0.5 | 18 | 0.2222 |
| 8_1104581 | 2 | 0.5 | NA | NA | 1 | 1 | 1 | 1 | 4 | 0.75 | 1 | 1 | NA | NA | NA | NA | NA | NA | 1 | 1 |
| 8_1104584 | 3 | 0.3333 | NA | NA | 1 | 1 | 1 | 1 | 5 | 0.6 | 1 | 1 | NA | NA | NA | NA | NA | NA | 1 | 1 |
| 8_1104584 | 3 | 1 | NA | NA | NA | NA | 1 | 1 | 4 | 1 | 1 | 1 | NA | NA | NA | NA | NA | NA | 1 | 1 |
| 8_1104585 | 3 | 0.3333 | NA | NA | NA | NA | 1 | 1 | 4 | 0.5 | 1 | 1 | NA | NA | NA | NA | NA | NA | 1 | 1 |
| 8_1104585 | 3 | 1 | NA | NA | 1 | 1 | NA | NA | 4 | 1 | 1 | 1 | NA | NA | NA | NA | NA | NA | 1 | 1 |
| 8_1105867 | 6 | 0.5 | 5 | 0 | 4 | 0.25 | 5 | 0.6 | 20 | 0.35 | 3 | 0.3333 | 5 | 0.2 | NA | NA | 7 | 0 | 15 | 0.1333 |
| 8_1105869 | 6 | 1 | 3 | 1 | 3 | 1 | 7 | 1 | 19 | 1 | 3 | 1 | 5 | 0.8 | 2 | 1 | 8 | 0.875 | 18 | 0.8889 |
| 8_1105869 | 6 | 0.5 | 5 | 0 | 4 | 0.25 | 7 | 0.1429 | 22 | 0.2273 | 5 | 0 | 7 | 0.1429 | 2 | 0.5 | 9 | 0.2222 | 23 | 0.1739 |
| 8_1105869 | 3 | 1 | 4 | 1 | 4 | 1 | 7 | 1 | 18 | 1 | 5 | 1 | 8 | 0.875 | 2 | 1 | 8 | 0.875 | 23 | 0.913 |
| 8_1105871 | 4 | 1 | 6 | 0.8333 | 4 | 1 | 6 | 0.8333 | 20 | 0.9 | 4 | 1 | 7 | 1 | 2 | 1 | 9 | 0.7778 | 22 | 0.9091 |
| 8_1105871 | 5 | 0.2 | 6 | 0.1667 | 4 | 0 | 7 | 0.1429 | 22 | 0.1364 | 5 | 0.2 | 8 | 0.375 | 2 | 0 | 8 | 0.25 | 23 | 0.2609 |
| 8_1105872 | 5 | 1 | 4 | 1 | 4 | 1 | 5 | 0.8 | 18 | 0.9444 | 4 | 0.75 | 7 | 1 | 2 | 1 | 7 | 1 | 20 | 0.95 |
| 8_1105873 | 4 | 0.75 | 5 | 1 | 1 | 0 | 6 | 0.8333 | 16 | 0.8125 | 4 | 1 | 5 | 1 | NA | NA | 6 | 0.8333 | 15 | 0.9333 |
| 8_1105874 | 4 | 0 | 4 | 0 | 3 | 0 | 5 | 0.4 | 16 | 0.125 | 3 | 0 | 6 | 0.5 | 3 | 0.3333 | 7 | 0.2857 | 19 | 0.3158 |
| 8_1105876 | 8 | 0 | 6 | 0.1667 | 4 | 0.25 | 7 | 0.1429 | 25 | 0.12 | 5 | 0.2 | 7 | 0 | 4 | 0.25 | 9 | 0.2222 | 25 | 0.16 |
| 8_1105881 | 1 | 0 | 3 | 0 | 2 | 0.5 | 2 | 0 | 8 | 0.125 | 2 | 0.5 | 5 | 0 | 4 | 0 | 3 | 0 | 14 | 0.0714 |
| 8_1105886 | 1 | 1 | 4 | 0.25 | 1 | 1 | 4 | 0 | 10 | 0.3 | 2 | 0 | 5 | 0.6 | 3 | 0.3333 | 2 | 0 | 12 | 0.3333 |
| 8_1105889 | 1 | 1 | 4 | 0.5 | 1 | 1 | 3 | 0.6667 | 9 | 0.6667 | 2 | 1 | 5 | 0.4 | 3 | 0.3333 | 3 | 0.3333 | 13 | 0.4615 |
| 8_1105889 | 1 | 1 | 4 | 0.75 | 1 | 1 | 3 | 0.6667 | 9 | 0.7778 | 2 | 0.5 | 5 | 0.8 | 3 | 0.6667 | 3 | 0.3333 | 13 | 0.6154 |

| | | | | | | | | | | | | | | | | | | | | |
|---|---|---|---|---|---|---|---|---|---|---|---|---|---|---|---|---|---|---|---|---|
| 8_1105891 | 1 | 1 | 3 | 1 | NA | NA | 2 | 1 | 6 | 1 | 2 | 1 | 6 | 1 | 3 | 1 | 3 | 0.3333 | 14 | 0.8571 |
| 8_1105891 | 2 | 0.5 | 4 | 0.75 | 1 | 1 | 3 | 1 | 10 | 0.8 | 1 | 1 | 5 | 0.2 | 1 | 1 | 3 | 0.3333 | 10 | 0.4 |
| 8_1105893 | 1 | 0 | 3 | 0.3333 | 2 | 0 | 2 | 0 | 8 | 0.125 | 1 | 0 | 7 | 0.2857 | 4 | 0 | 2 | 0 | 14 | 0.1429 |
| 8_1105894 | 1 | 1 | 3 | 1 | 2 | 1 | 3 | 1 | 9 | 1 | 2 | 1 | 7 | 0.8571 | 3 | 1 | 2 | 0.5 | 14 | 0.8571 |
| 8_1105894 | 2 | 0.5 | 4 | 0.5 | 1 | 1 | 3 | 1 | 10 | 0.7 | 2 | 0.5 | 6 | 0.5 | 2 | 0.5 | 2 | 0.5 | 12 | 0.5 |
| 8_1105898 | 2 | 0 | 4 | 0.25 | 2 | 0 | 2 | 0.5 | 10 | 0.2 | 3 | 0.6667 | 9 | 0.3333 | 4 | 0.75 | 4 | 0.5 | 20 | 0.5 |
| 8_1105900 | 4 | 0.25 | 5 | 0 | 1 | 0 | 3 | 0.3333 | 13 | 0.1538 | 4 | 0 | 5 | 0 | 2 | 0 | 4 | 0.25 | 15 | 0.0667 |
| 8_1105901 | 4 | 0.75 | 6 | 0.3333 | 1 | 0 | 2 | 0.5 | 13 | 0.4615 | 3 | 0.3333 | 8 | 0.25 | 4 | 0.25 | 4 | 0 | 19 | 0.2105 |
| 8_1105902 | 5 | 0.6 | 6 | 0.6667 | 2 | 0 | 4 | 0.25 | 17 | 0.4706 | 5 | 0.2 | 7 | 0.1429 | 3 | 0.3333 | 6 | 0.3333 | 21 | 0.2381 |
| 8_1105902 | 3 | 0 | 7 | 0.2857 | 1 | 0 | 4 | 0.25 | 15 | 0.2 | 2 | 0 | 8 | 0.625 | 1 | 1 | 3 | 0.3333 | 14 | 0.5 |
| 8_1105903 | 5 | 0 | 9 | 0.2222 | 2 | 0 | 5 | 0.2 | 21 | 0.1429 | 5 | 0 | 9 | 0.1111 | 4 | 0 | 7 | 0.2857 | 25 | 0.12 |
| 8_1105904 | 6 | 0.3333 | 10 | 0.2 | 2 | 0 | 6 | 0.1667 | 24 | 0.2083 | 5 | 0.2 | 9 | 0.3333 | 6 | 0 | 7 | 0.4286 | 27 | 0.2593 |
| 8_1106687 | NA | NA | NA | NA | 1 | 0 | 4 | 0.25 | 5 | 0.2 | NA | NA | 1 | 0 | 4 | 0.25 | 1 | 0 | 6 | 0.1667 |
| 8_1106697 | 2 | 0.5 | NA | NA | NA | NA | 5 | 0.4 | 7 | 0.4286 | 1 | 0 | 2 | 0 | 8 | 0.25 | 2 | 0 | 13 | 0.1538 |
| 8_1106703 | 4 | 0.25 | 1 | 0 | 1 | 0 | 5 | 0 | 11 | 0.0909 | 2 | 0 | 1 | 1 | 10 | 0.1 | 3 | 0 | 16 | 0.125 |
| 8_1106704 | 5 | 0.2 | 1 | 0 | 2 | 0 | 5 | 0 | 13 | 0.0769 | 2 | 0 | 2 | 0 | 9 | 0 | 3 | 0.3333 | 16 | 0.0625 |
| 8_1106705 | 3 | 0.3333 | NA | NA | 2 | 0.5 | 4 | 0 | 9 | 0.2222 | 1 | 1 | 1 | 0 | 8 | 0.25 | 2 | 0 | 12 | 0.25 |
| 8_1106705 | 3 | 0.3333 | 1 | 0 | 2 | 0 | 6 | 0 | 12 | 0.0833 | 1 | 0 | 2 | 0.5 | 8 | 0 | 1 | 0 | 12 | 0.0833 |
| 8_1106706 | 4 | 0.25 | 1 | 0 | 2 | 0 | 6 | 0 | 13 | 0.0769 | 2 | 0 | 2 | 0 | 9 | 0.1111 | 2 | 0 | 15 | 0.0667 |
| 8_1106708 | 3 | 0.3333 | 1 | 1 | 2 | 0.5 | 7 | 0 | 13 | 0.2308 | NA | NA | 1 | 1 | 10 | 0.1 | 2 | 0.5 | 13 | 0.2308 |
| 8_1106709 | 5 | 0.2 | 3 | 0 | 2 | 0 | 7 | 0 | 17 | 0.0588 | 2 | 0 | 2 | 0 | 10 | 0.1 | 3 | 0 | 17 | 0.0588 |
| 8_1107561 | 1 | 1 | NA | NA | NA | NA | NA | NA | 1 | 1 | NA | NA | 1 | 1 | NA | NA | NA | NA | 1 | 1 |
| 8_1107562 | 1 | 1 | NA | NA | NA | NA | NA | NA | 1 | 1 | NA | NA | 1 | 1 | NA | NA | NA | NA | 1 | 1 |
| 8_1107562 | 1 | 1 | NA | NA | NA | NA | NA | NA | 1 | 1 | NA | NA | 1 | 1 | NA | NA | NA | NA | 1 | 1 |
| 8_1107563 | 1 | 1 | NA | NA | NA | NA | NA | NA | 1 | 1 | NA | NA | 1 | 1 | NA | NA | NA | NA | 1 | 1 |
| 8_1107567 | 1 | 1 | NA | NA | NA | NA | NA | NA | 1 | 1 | NA | NA | 1 | 1 | NA | NA | NA | NA | 1 | 1 |
| 8_1107570 | 1 | 1 | NA | NA | NA | NA | NA | NA | 1 | 1 | NA | NA | 1 | 1 | NA | NA | NA | NA | 1 | 1 |
| 8_1108151 | NA | NA | NA | NA | 1 | 0 | 1 | 1 | 2 | 0.5 | NA | NA | 2 | 0.5 | NA | NA | NA | NA | 2 | 0.5 |
| 8_1108151 | NA | NA | NA | NA | 1 | 1 | NA | NA | 1 | 1 | NA | NA | 2 | 1 | NA | NA | NA | NA | 2 | 1 |
| 8_1108152 | NA | NA | NA | NA | 1 | 1 | 1 | 0 | 2 | 0.5 | NA | NA | 2 | 0.5 | NA | NA | NA | NA | 2 | 0.5 |
| 8_1108155 | NA | NA | NA | NA | 1 | 1 | 1 | 0 | 2 | 0.5 | NA | NA | 2 | 0.5 | NA | NA | NA | NA | 2 | 0.5 |
| 8_1108157 | 1 | 1 | NA | NA | 1 | 0 | 1 | 1 | 3 | 0.6667 | NA | NA | 2 | 1 | NA | NA | NA | NA | 2 | 1 |
| 8_1108158 | 1 | 0 | NA | NA | 1 | 1 | 1 | 1 | 3 | 0.6667 | NA | NA | 2 | 1 | NA | NA | 1 | 1 | 3 | 1 |
| 8_1108160 | 2 | 0.5 | NA | NA | 1 | 0 | 1 | 1 | 4 | 0.5 | NA | NA | 2 | 0.5 | NA | NA | 1 | 1 | 3 | 0.6667 |
| 8_1108166 | NA | NA | NA | NA | 1 | 1 | 2 | 1 | 3 | 1 | NA | NA | 2 | 0.5 | NA | NA | 2 | 1 | 4 | 0.75 |
| 8_1108167 | 1 | 1 | NA | NA | NA | NA | 1 | 1 | 2 | 1 | NA | NA | 1 | 1 | NA | NA | 1 | 1 | 2 | 1 |
| 8_1108168 | 1 | 1 | NA | NA | NA | NA | 2 | 0 | 3 | 0.3333 | NA | NA | 3 | 0.3333 | NA | NA | 1 | 0 | 4 | 0.25 |
| 8_1108175 | 2 | 0.5 | NA | NA | 1 | 0 | 2 | 0 | 5 | 0.2 | NA | NA | 4 | 0.25 | NA | NA | 3 | 0 | 7 | 0.1429 |
| 8_1108177 | 2 | 1 | NA | NA | 1 | 0 | 2 | 0.5 | 5 | 0.6 | NA | NA | 3 | 0 | NA | NA | 3 | 0.3333 | 6 | 0.1667 |
| 8_1108177 | 2 | 0.5 | 2 | 0 | 1 | 0 | 2 | 0 | 7 | 0.1429 | NA | NA | 2 | 0 | NA | NA | 3 | 0.3333 | 5 | 0.2 |
| 8_1108177 | 2 | 0 | 2 | 0 | 1 | 1 | 2 | 0 | 7 | 0.1429 | NA | NA | 3 | 0.3333 | NA | NA | 3 | 0 | 6 | 0.1667 |
| 8_1108180 | 2 | 0 | 2 | 0 | 1 | 1 | 2 | 0.5 | 7 | 0.2857 | NA | NA | 3 | 0.3333 | NA | NA | 3 | 0.3333 | 6 | 0.3333 |
| 8_1109483 | NA | NA | NA | NA | NA | NA | 2 | 1 | 2 | 1 | NA | NA | 1 | 0 | 1 | 1 | NA | NA | 2 | 0.5 |
| 8_1109485 | NA | NA | NA | NA | NA | NA | 2 | 0.5 | 2 | 0.5 | NA | NA | 1 | 1 | 1 | 1 | NA | NA | 2 | 1 |
| 8_1109486 | NA | NA | NA | NA | NA | NA | 2 | 0.5 | 2 | 0.5 | NA | NA | 1 | 0 | 1 | 1 | NA | NA | 2 | 0.5 |
| 8_1109486 | NA | NA | NA | NA | NA | NA | 1 | 1 | 1 | 1 | NA | NA | 1 | 0 | 1 | 1 | NA | NA | 2 | 0.5 |
| 8_1109488 | NA | NA | NA | NA | NA | NA | 2 | 1 | 2 | 1 | NA | NA | 1 | 1 | 1 | 1 | NA | NA | 2 | 1 |
| 8_1109488 | NA | NA | NA | NA | NA | NA | 2 | 1 | 2 | 1 | NA | NA | NA | NA | 1 | 1 | NA | NA | 1 | 1 |
| 8_1109489 | NA | NA | NA | NA | NA | NA | 1 | 1 | 1 | 1 | NA | NA | 1 | 1 | 1 | 1 | NA | NA | 2 | 1 |
| 8_1109491 | NA | NA | NA | NA | NA | NA | 2 | 0.5 | 2 | 0.5 | NA | NA | 1 | 1 | 1 | 1 | NA | NA | 2 | 1 |
| 8_1109492 | NA | NA | NA | NA | NA | NA | 2 | 1 | 2 | 1 | NA | NA | 1 | 1 | 1 | 1 | NA | NA | 2 | 1 |
| 8_1109495 | NA | NA | NA | NA | NA | NA | 1 | 1 | 1 | 1 | NA | NA | 1 | 1 | 1 | 1 | NA | NA | 2 | 1 |
| 8_1109497 | NA | NA | NA | NA | NA | NA | 2 | 0.5 | 2 | 0.5 | NA | NA | 1 | 1 | 1 | 1 | NA | NA | 2 | 1 |
| 8_1109498 | NA | NA | NA | NA | NA | NA | 2 | 1 | 2 | 1 | NA | NA | 1 | 1 | 1 | 1 | NA | NA | 2 | 1 |
| 8_1109500 | NA | NA | NA | NA | NA | NA | 2 | 1 | 2 | 1 | NA | NA | 1 | 1 | 1 | 1 | NA | NA | 2 | 1 |
| 8_1109500 | NA | NA | NA | NA | NA | NA | 2 | 1 | 2 | 1 | NA | NA | 1 | 0 | 1 | 1 | NA | NA | 2 | 0.5 |
| 8_1109501 | 1 | 1 | NA | NA | NA | NA | 2 | 1 | 3 | 1 | NA | NA | 1 | 1 | 1 | 1 | NA | NA | 2 | 1 |
| 8_1109503 | 1 | 1 | NA | NA | NA | NA | 1 | 0 | 2 | 0.5 | NA | NA | NA | NA | 1 | 1 | NA | NA | 1 | 1 |
| 8_1109504 | 1 | 1 | NA | NA | NA | NA | 2 | 0.5 | 3 | 0.6667 | NA | NA | 1 | 1 | 1 | 1 | NA | NA | 2 | 1 |
| 8_1109505 | 1 | 1 | NA | NA | NA | NA | 2 | 0 | 3 | 0.3333 | NA | NA | 1 | 0 | 1 | 1 | NA | NA | 2 | 0.5 |
| 8_1114464 | 4 | 0 | 2 | 0.5 | 4 | 0.25 | 2 | 0.5 | 12 | 0.25 | 4 | 0.25 | 3 | 0.3333 | 1 | 1 | 2 | 1 | 10 | 0.5 |
| 8_1114465 | 4 | 0 | 2 | 0 | 4 | 0.5 | 3 | 0 | 13 | 0.1538 | 5 | 0.2 | 4 | 0.5 | 1 | 1 | 2 | 0.5 | 12 | 0.4167 |
| 8_1114466 | 2 | 0 | 2 | 0 | 4 | 0 | 3 | 0.3333 | 11 | 0.0909 | 5 | 0.4 | 4 | 0.75 | NA | NA | 2 | 0.5 | 11 | 0.5455 |
| 8_1114467 | 3 | 0.6667 | 2 | 0.5 | 4 | 0.25 | 3 | 0.6667 | 12 | 0.5 | 4 | 0 | 4 | 0.25 | 1 | 0 | 1 | 1 | 10 | 0.2 |
| 8_1114467 | 4 | 0.5 | 1 | 1 | 4 | 0.25 | 3 | 0 | 12 | 0.3333 | 4 | 0 | 5 | 0.2 | 1 | 0 | 3 | 0 | 13 | 0.0769 |

| | | | | | | | | | | | | | | | | | | | | |
|---|---|---|---|---|---|---|---|---|---|---|---|---|---|---|---|---|---|---|---|---|
| 8_1114468 | 4 | 0.5 | 1 | 0 | 4 | 0 | 4 | 0.25 | 13 | 0.2308 | 4 | 0.25 | 4 | 0 | 1 | 0 | 2 | 0.5 | 11 | 0.1818 |
| 8_1114468 | 4 | 0.75 | 2 | 0 | 4 | 0.25 | 4 | 0.25 | 14 | 0.3571 | 4 | 0.25 | 4 | 0.75 | 2 | 0 | 2 | 0 | 12 | 0.3333 |
| 8_1114469 | 3 | 0.3333 | 2 | 0.5 | 4 | 0 | 3 | 0.6667 | 12 | 0.3333 | 5 | 0.2 | 5 | 0.4 | 2 | 0 | 3 | 0.3333 | 15 | 0.2667 |
| 8_1114650 | 2 | 0 | 9 | 0 | 2 | 0 | 5 | 0.2 | 18 | 0.0556 | 7 | 0 | 9 | 0.1111 | 3 | 0 | 6 | 0.1667 | 25 | 0.08 |
| 8_1114653 | 2 | 0.5 | 13 | 0.3077 | 7 | 0.2857 | 9 | 0.1111 | 31 | 0.2581 | 8 | 0.25 | 10 | 0.2 | 7 | 0 | 7 | 0.2857 | 32 | 0.1875 |
| 8_1114656 | 2 | 0 | 12 | 0.0833 | 7 | 0.1429 | 10 | 0 | 31 | 0.0645 | 8 | 0 | 9 | 0.2222 | 9 | 0 | 8 | 0 | 34 | 0.0588 |
| 8_1114656 | 2 | 1 | 14 | 0.5714 | 7 | 0.2857 | 10 | 0.5 | 33 | 0.5152 | 9 | 0.4444 | 10 | 0.5 | 7 | 0.2857 | 8 | 0.5 | 34 | 0.4412 |
| 8_1114657 | 2 | 0.5 | 14 | 0 | 5 | 0 | 9 | 0.1111 | 30 | 0.0667 | 8 | 0 | 8 | 0 | 9 | 0.1111 | 8 | 0.125 | 33 | 0.0606 |
| 8_1114658 | 2 | 0 | 12 | 0.1667 | 7 | 0.2857 | 12 | 0.0833 | 33 | 0.1515 | 8 | 0.125 | 10 | 0.1 | 9 | 0 | 7 | 0.1429 | 34 | 0.0882 |
| 8_1114658 | 3 | 0.6667 | 12 | 0.4167 | 6 | 0.1667 | 9 | 0.1111 | 30 | 0.3 | 8 | 0.375 | 4 | 0 | 7 | 0.4286 | 7 | 0.1429 | 26 | 0.2692 |
| 8_1114659 | 3 | 0.6667 | 16 | 0.5625 | 6 | 0.8333 | 9 | 0.5556 | 34 | 0.6176 | 8 | 1 | 10 | 0.5 | 10 | 0.7 | 7 | 0.4286 | 35 | 0.6571 |
| 8_1114661 | 2 | 0.5 | 17 | 0.5882 | 8 | 0.625 | 8 | 0.375 | 35 | 0.5429 | 11 | 0.2727 | 11 | 0.1818 | 10 | 0.2 | 9 | 0.2222 | 41 | 0.2195 |
| 8_1114664 | 4 | 0.25 | 13 | 0.3077 | 7 | 0.5714 | 11 | 0.1818 | 35 | 0.3143 | 8 | 0.5 | 14 | 0.3571 | 8 | 0.125 | 9 | 0.3333 | 39 | 0.3333 |
| 8_1114665 | 3 | 0 | 17 | 0.1176 | 8 | 0.125 | 10 | 0.3 | 38 | 0.1579 | 10 | 0.2 | 10 | 0 | 8 | 0 | 5 | 0.2 | 33 | 0.0909 |
| 8_1114676 | 7 | 0.5714 | 22 | 0.1818 | 10 | 0.5 | 12 | 0.25 | 51 | 0.3137 | 13 | 0.1538 | 13 | 0.1538 | 8 | 0.375 | 9 | 0.2222 | 43 | 0.2093 |
| 8_1114679 | 1 | 1 | 1 | 0 | 2 | 0 | 1 | 0 | 5 | 0.2 | 4 | 0 | 7 | 0.1429 | NA | NA | 5 | 0 | 16 | 0.0625 |
| 8_1114685 | 9 | 0.4444 | 5 | 0.4 | 9 | 0.3333 | 8 | 0.25 | 31 | 0.3548 | 5 | 0.4 | 13 | 0.5385 | 9 | 0.6667 | 10 | 0.3 | 37 | 0.4865 |
| 8_1114685 | 11 | 0.0909 | 8 | 0.125 | 10 | 0.3 | 10 | 0.4 | 39 | 0.2308 | 9 | 0.5556 | 15 | 0.2667 | 13 | 0.2308 | 13 | 0.5385 | 50 | 0.38 |
| 8_1114688 | 10 | 0 | 7 | 0.1429 | 9 | 0.1111 | 8 | 0 | 34 | 0.0588 | 10 | 0.2 | 14 | 0 | 13 | 0.1538 | 11 | 0 | 48 | 0.0833 |
| 8_1114688 | 12 | 0.25 | 8 | 0.125 | 7 | 0.1429 | 8 | 0.25 | 35 | 0.2 | 8 | 0.375 | 14 | 0.1429 | 13 | 0.2308 | 9 | 0.1111 | 44 | 0.2045 |
| 8_1114820 | 9 | 0.7778 | 7 | 0.4286 | 4 | 0.25 | 5 | 0.6 | 25 | 0.56 | 8 | 0.375 | 10 | 0.5 | 6 | 0.3333 | 8 | 0.25 | 32 | 0.375 |
| 8_1114821 | 8 | 0.25 | 5 | 0 | 1 | 0 | 5 | 0 | 19 | 0.1053 | 7 | 0.2857 | 8 | 0.25 | 6 | 0 | 5 | 0.2 | 26 | 0.1923 |
| 8_1115374 | 5 | 0.2 | 11 | 0.6364 | 6 | 0 | 6 | 0.3333 | 28 | 0.3571 | 9 | 0.7778 | 8 | 0.25 | 6 | 0.5 | 13 | 0.1538 | 36 | 0.3889 |
| 8_1115374 | 5 | 0.8 | 12 | 0.75 | 6 | 0.6667 | 8 | 0.5 | 31 | 0.6774 | 12 | 0.75 | 9 | 0.4444 | 6 | 0.8333 | 16 | 0.875 | 43 | 0.7442 |
| 8_1115375 | 6 | 0.1667 | 11 | 0.1818 | 6 | 0 | 6 | 0.1667 | 29 | 0.1379 | 13 | 0.0769 | 11 | 0.1818 | 6 | 0.1667 | 16 | 0 | 46 | 0.087 |
| 8_1116084 | 3 | 0 | 8 | 0 | NA | NA | 5 | 0.4 | 16 | 0.125 | 9 | 0.3333 | 7 | 0 | 6 | 0 | 3 | 0.3333 | 25 | 0.16 |
| 8_1116085 | 3 | 0.3333 | 9 | 0.5556 | 2 | 0.5 | 5 | 0.6 | 19 | 0.5263 | 10 | 0.4 | 7 | 0.7143 | 6 | 0.5 | 3 | 0.6667 | 26 | 0.5385 |
| 8_1116085 | 3 | 0.6667 | 9 | 0.6667 | 2 | 0 | 5 | 0.8 | 19 | 0.6316 | 6 | 0.5 | 2 | 0.5 | 3 | 0.3333 | 1 | 1 | 12 | 0.5 |
| 8_1116087 | 2 | 1 | 8 | 1 | 2 | 1 | 5 | 0.6 | 17 | 0.8824 | 10 | 0.8 | 6 | 0.6667 | 5 | 0.8 | 3 | 0.6667 | 24 | 0.75 |
| 8_1116088 | 3 | 0.3333 | 10 | 0.2 | 1 | 1 | 4 | 0.25 | 18 | 0.2778 | 10 | 0.5 | 9 | 0.2222 | 6 | 0.3333 | 3 | 0.6667 | 28 | 0.3929 |
| 8_1116088 | 2 | 0 | 8 | 0.125 | 2 | 0 | 5 | 0.2 | 17 | 0.1176 | 11 | 0.0909 | 8 | 0.25 | 7 | 0 | 2 | 0 | 28 | 0.1071 |
| 8_1116093 | 2 | 1 | 1 | 1 | 2 | 0 | 1 | 1 | 6 | 0.6667 | 1 | 1 | 2 | 0.5 | NA | NA | 1 | 0 | 4 | 0.5 |
| 8_1116837 | 3 | 0 | 3 | 0.3333 | 1 | 0 | NA | NA | 7 | 0.1429 | 1 | 0 | 2 | 0 | 2 | 0 | 3 | 0.3333 | 8 | 0.125 |
| 8_1116838 | 5 | 1 | 3 | 1 | 2 | 1 | 1 | 1 | 11 | 1 | 1 | 1 | 2 | 1 | 1 | 1 | 3 | 0.6667 | 7 | 0.8571 |
| 8_1116839 | 5 | 0.4 | 4 | 0.25 | 2 | 0.5 | 1 | 0 | 12 | 0.3333 | 1 | 0 | 2 | 0 | 2 | 0 | 3 | 0.3333 | 8 | 0.125 |
| 8_1116839 | 5 | 0.4 | 4 | 0.25 | 1 | 0 | NA | NA | 10 | 0.3 | 1 | 0 | 2 | 0 | 1 | 0 | 3 | 0.6667 | 7 | 0.2857 |
| 8_1116839 | 4 | 0.25 | 3 | 0 | 1 | 0 | 1 | 0 | 9 | 0.1111 | NA | NA | 1 | 1 | 2 | 0 | 3 | 0.3333 | 6 | 0.3333 |
| 8_1116839 | 4 | 0.5 | 3 | 0.6667 | 2 | 1 | 1 | 1 | 10 | 0.7 | 1 | 1 | 1 | 1 | 2 | 1 | 3 | 0.6667 | 7 | 0.8571 |
| 8_1116840 | 5 | 1 | 2 | 1 | 2 | 1 | 1 | 1 | 10 | 1 | 1 | 0 | 2 | 1 | 2 | 0.5 | 3 | 0.3333 | 8 | 0.5 |
| 8_1116840 | 5 | 1 | 3 | 1 | 2 | 1 | 1 | 1 | 11 | 1 | NA | NA | 1 | 1 | 2 | 1 | 4 | 1 | 7 | 1 |
| 8_1116841 | 5 | 0.8 | 3 | 1 | 2 | 1 | 1 | 1 | 11 | 0.9091 | NA | NA | 2 | 1 | 1 | 1 | 3 | 0.3333 | 6 | 0.6667 |
| 8_1116841 | 5 | 0.4 | 4 | 0.5 | 1 | 0 | 1 | 0 | 11 | 0.3636 | 1 | 0 | 2 | 0.5 | 2 | 0.5 | 4 | 0.75 | 9 | 0.5556 |
| 8_1116844 | 5 | 1 | 5 | 1 | 3 | 1 | 2 | 1 | 15 | 1 | 1 | 1 | 4 | 1 | 2 | 1 | 6 | 1 | 13 | 1 |
| 8_1116845 | 7 | 1 | 6 | 0.5 | 3 | 0.6667 | 2 | 1 | 18 | 0.7778 | 1 | 1 | 4 | 0.75 | 2 | 0.5 | 6 | 0.5 | 13 | 0.6154 |
| 8_1116969 | 2 | 1 | 6 | 0.6667 | 4 | 1 | 9 | 0.8889 | 21 | 0.8571 | 3 | 0.6667 | 4 | 0.75 | 4 | 0.75 | 5 | 0.6 | 16 | 0.6875 |
| 8_1116969 | 8 | 1 | 10 | 1 | 5 | 1 | 8 | 0.875 | 31 | 0.9677 | 4 | 1 | 4 | 1 | 3 | 1 | 6 | 1 | 17 | 1 |
| 8_1116971 | 11 | 0.0909 | 10 | 0.1 | 6 | 0 | 10 | 0 | 37 | 0.0541 | 3 | 0 | 2 | 0.5 | 3 | 0 | 5 | 0 | 13 | 0.0769 |
| 8_1116971 | 7 | 0.8571 | 9 | 0.6667 | 5 | 0.6 | 10 | 0.8 | 31 | 0.7419 | 5 | 1 | 2 | 1 | 6 | 1 | 6 | 1 | 19 | 1 |
| 8_1116972 | 10 | 0.2 | 12 | 0.0833 | 7 | 0.1429 | 11 | 0.2727 | 40 | 0.175 | 5 | 0.6 | 3 | 0.3333 | 7 | 0.1429 | 6 | 0.1667 | 21 | 0.2857 |
| 8_1116975 | 6 | 0.5 | 10 | 0.9 | 8 | 0.625 | 11 | 0.6364 | 35 | 0.6857 | 6 | 0.8333 | 8 | 1 | 2 | 1 | 4 | 0.25 | 20 | 0.8 |
| 8_1116976 | 10 | 1 | 12 | 1 | 10 | 0.9 | 11 | 1 | 43 | 0.9767 | 5 | 1 | 6 | 1 | 6 | 1 | 6 | 1 | 23 | 1 |
| 8_1116977 | 9 | 0.1111 | 11 | 0.0909 | 11 | 0 | 11 | 0.0909 | 42 | 0.0714 | 5 | 0.4 | 5 | 0.2 | 7 | 0.1429 | 4 | 0 | 21 | 0.1905 |
| 8_1116980 | 6 | 0 | 11 | 0.1818 | 11 | 0 | 12 | 0 | 40 | 0.05 | 4 | 0 | 4 | 0 | 6 | 0.1667 | 4 | 0 | 18 | 0.0556 |
| 8_1116981 | 10 | 0.5 | 13 | 0.5385 | 12 | 0.4167 | 15 | 0.2667 | 50 | 0.42 | 6 | 0.3333 | 10 | 0.7 | 7 | 0.5714 | 6 | 0.1667 | 29 | 0.4828 |
| 8_1116985 | 3 | 0.3333 | 5 | 0 | 2 | 0 | 1 | 0 | 11 | 0.0909 | 1 | 1 | 2 | 1 | 2 | 0 | 1 | 0 | 6 | 0.5 |
| 8_1117018 | 4 | 0 | 5 | 0.2 | 5 | 0 | 3 | 0 | 17 | 0.0588 | 2 | 0 | 3 | 0 | 10 | 0.1 | 5 | 0 | 20 | 0.05 |
| 8_1117023 | 4 | 0 | 5 | 0.2 | 4 | 0.25 | 4 | 0.5 | 17 | 0.2353 | 3 | 0.6667 | 5 | 0.2 | 8 | 0.125 | 4 | 0 | 20 | 0.2 |
| 8_1117025 | 4 | 0.75 | 4 | 0.75 | 4 | 0.5 | 4 | 0.75 | 16 | 0.6875 | 3 | 1 | 3 | 0.6667 | 8 | 0.625 | 5 | 0.6 | 19 | 0.6842 |
| 8_1117025 | 4 | 1 | 6 | 0.3333 | 4 | 0.25 | 3 | 0 | 17 | 0.4118 | 4 | 0.75 | 5 | 0.2 | 8 | 0.375 | 3 | 0.3333 | 20 | 0.4 |
| 8_1117025 | 4 | 0 | 6 | 0.1667 | 4 | 0 | 3 | 0.3333 | 17 | 0.1176 | 4 | 0.25 | 4 | 0.25 | 8 | 0.5 | 4 | 0.25 | 20 | 0.35 |
| 8_1117308 | 2 | 0.5 | 2 | 0 | 1 | 0 | 2 | 0 | 7 | 0.1429 | NA | NA | NA | NA | 3 | 0.3333 | 3 | 0 | 6 | 0.1667 |
| 8_1117343 | NA | NA | 1 | 1 | NA | NA | NA | NA | 1 | 1 | NA | NA | NA | NA | 1 | 0 | 2 | 0.5 | 3 | 0.3333 |
| 8_1117343 | NA | NA | 1 | 1 | NA | NA | 1 | 0 | 2 | 0.5 | NA | NA | NA | NA | 1 | 0 | 2 | 0.5 | 3 | 0.3333 |
| 8_1117346 | NA | NA | 1 | 1 | NA | NA | 1 | 1 | 2 | 1 | NA | NA | NA | NA | NA | NA | 2 | 0.5 | 2 | 0.5 |
| 8_1117347 | NA | NA | 1 | 1 | NA | NA | 1 | 1 | 2 | 1 | 2 | 1 | NA | NA | 1 | 1 | 2 | 1 | 5 | 1 |

| | | | | | | | | | | | | | | | | | | | | |
|---|---|---|---|---|---|---|---|---|---|---|---|---|---|---|---|---|---|---|---|---|
| 8_11173476 | NA | NA | 1 | 1 | NA | NA | 2 | 0.5 | 3 | 0.6667 | 1 | 1 | NA | NA | 1 | 1 | 2 | 0.5 | 4 | 0.75 |
| 8_11173478 | NA | NA | 1 | 0 | NA | NA | 2 | 1 | 3 | 0.6667 | 2 | 1 | NA | NA | 1 | 1 | NA | NA | 3 | 1 |
| 8_11178173 | 13 | 0.1538 | 12 | 0.0833 | 13 | 0.2308 | 11 | 0.1818 | 49 | 0.1633 | 13 | 0.1538 | 12 | 0.1667 | 15 | 0.0667 | 14 | 0.0714 | 54 | 0.1111 |
| 8_11178183 | 11 | 0.2727 | 11 | 0.7273 | 12 | 0.4167 | 7 | 0.5714 | 41 | 0.4878 | 11 | 0.6364 | 8 | 0.125 | 16 | 0.5 | 11 | 0.8182 | 46 | 0.5435 |
| 8_11178194 | 15 | 0.2 | 13 | 0.2308 | 12 | 0.1667 | 11 | 0.1818 | 51 | 0.1961 | 13 | 0.3077 | 12 | 0.25 | 18 | 0.1667 | 15 | 0.0667 | 58 | 0.1897 |
| 8_11178218 | 12 | 0.5 | 14 | 0.3571 | 9 | 0.5556 | 14 | 0.3571 | 49 | 0.4286 | 11 | 0.4545 | 13 | 0.3846 | 16 | 0.5 | 13 | 0.4615 | 53 | 0.4528 |
| 8_11178248 | NA | NA | NA | NA | NA | NA | 1 | 1 | 1 | 1 | 1 | 1 | NA | NA | NA | NA | 1 | 0 | 2 | 0.5 |
| 8_11178271 | NA | NA | NA | NA | NA | NA | 1 | 1 | 1 | 1 | 1 | 1 | NA | NA | NA | NA | 1 | 1 | 2 | 1 |
| 8_11178281 | NA | NA | NA | NA | NA | NA | 1 | 1 | 1 | 1 | 1 | 1 | NA | NA | NA | NA | 1 | 1 | 2 | 1 |
| 8_11178337 | 3 | 0 | 3 | 0.3333 | NA | NA | 2 | 1 | 8 | 0.375 | NA | NA | 2 | 0.5 | 1 | 0 | 1 | 1 | 4 | 0.5 |
| 8_11178370 | 3 | 0 | 2 | 0.5 | 1 | 0 | 2 | 0.5 | 8 | 0.25 | 3 | 1 | NA | NA | 2 | 1 | 2 | 0 | 7 | 0.7143 |
| 8_11178371 | 2 | 1 | 3 | 0.6667 | NA | NA | 2 | 1 | 7 | 0.8571 | 3 | 1 | 2 | 1 | 3 | 0.6667 | 2 | 1 | 10 | 0.9 |
| 8_11178385 | 2 | 1 | 4 | 0.5 | 1 | 1 | 2 | 1 | 9 | 0.7778 | 3 | 1 | 2 | 1 | 2 | 1 | 3 | 1 | 10 | 1 |
| 8_11178388 | 2 | 0.5 | 3 | 0.3333 | NA | NA | 2 | 0.5 | 7 | 0.4286 | 2 | 0 | 1 | 1 | 3 | 0.3333 | 3 | 0.6667 | 9 | 0.4444 |
| 8_11178396 | 3 | 0.3333 | 2 | 1 | 1 | 1 | 1 | 1 | 7 | 0.7143 | 2 | 1 | 2 | 1 | 3 | 0.6667 | 3 | 0.6667 | 10 | 0.8 |
| 8_11178405 | 3 | 1 | 3 | 1 | 1 | 1 | 2 | 1 | 9 | 1 | 3 | 0.6667 | 2 | 0.5 | 3 | 1 | 3 | 1 | 11 | 0.8182 |
| 8_11178411 | 3 | 1 | 3 | 1 | 2 | 1 | 2 | 0.5 | 10 | 0.9 | 3 | 1 | 2 | 0.5 | 3 | 1 | 3 | 1 | 11 | 0.9091 |
| 8_11178415 | 4 | 1 | 3 | 0.6667 | 1 | 0 | 1 | 1 | 9 | 0.7778 | 4 | 1 | 2 | 1 | 2 | 1 | 3 | 0.6667 | 11 | 0.9091 |
| 8_11178417 | 4 | 0.75 | 3 | 1 | 2 | 1 | 2 | 0.5 | 11 | 0.8182 | 3 | 0.6667 | 2 | 1 | 3 | 1 | 3 | 1 | 11 | 0.9091 |
| 8_11178418 | 3 | 1 | 3 | 0.6667 | 2 | 1 | 3 | 1 | 11 | 0.9091 | 4 | 0.75 | NA | NA | 2 | 1 | 2 | 1 | 8 | 0.875 |
| 8_11178423 | 4 | 0.5 | 3 | 0 | 1 | 0 | 3 | 0 | 11 | 0.1818 | 3 | 0 | 1 | 0 | 2 | 0.5 | 1 | 0 | 7 | 0.1429 |
| 8_11186124 | 2 | 0 | 4 | 0 | 3 | 0.3333 | 1 | 0 | 10 | 0.1 | 1 | 0 | NA | NA | NA | NA | 2 | 0.5 | 3 | 0.3333 |
| 8_11186133 | 1 | 1 | 4 | 0.5 | 3 | 0 | NA | NA | 8 | 0.375 | 1 | 0 | NA | NA | NA | NA | 2 | 0.5 | 3 | 0.3333 |
| 8_11186136 | 2 | 0 | 4 | 0.5 | 5 | 0 | NA | NA | 11 | 0.1818 | 1 | 0 | NA | NA | NA | NA | 2 | 0.5 | 3 | 0.3333 |
| 8_11186147 | 1 | 1 | 3 | 1 | 4 | 0.75 | NA | NA | 8 | 0.875 | NA | NA | NA | NA | NA | NA | 1 | 1 | 1 | 1 |
| 8_11186153 | 1 | 1 | 4 | 1 | 5 | 0.6 | NA | NA | 10 | 0.8 | 1 | 1 | NA | NA | NA | NA | 1 | 1 | 2 | 1 |
| 8_11186154 | 2 | 1 | 4 | 1 | 5 | 0.8 | 1 | 1 | 12 | 0.9167 | NA | NA | NA | NA | NA | NA | 1 | 1 | 1 | 1 |
| 8_11186495 | 3 | 0 | 4 | 0 | 6 | 0.1667 | 4 | 0.25 | 17 | 0.1176 | 2 | 1 | NA | NA | 1 | 0 | 1 | 0 | 4 | 0.5 |
| 8_11186622 | 6 | 0.3333 | 5 | 0.2 | 5 | 0.2 | 5 | 0.2 | 21 | 0.2381 | 3 | 0.3333 | 2 | 0.5 | NA | NA | 5 | 0.4 | 10 | 0.4 |
| 8_11186628 | 5 | 0.4 | 6 | 0.5 | 5 | 0.4 | 3 | 1 | 19 | 0.5263 | 2 | 0.5 | 2 | 0 | 1 | 1 | 6 | 0.3333 | 11 | 0.3636 |
| 8_11188109 | 12 | 0.1667 | 11 | 0.1818 | 13 | 0.0769 | 18 | 0.1111 | 54 | 0.1296 | 4 | 0 | 15 | 0.1333 | 16 | 0.1875 | 10 | 0 | 45 | 0.1111 |
| 8_11188378 | 5 | 0.4 | 2 | 0.5 | 5 | 0.8 | 12 | 0.6667 | 24 | 0.625 | 4 | 0.25 | 9 | 0.3333 | 4 | 0.5 | 4 | 0.25 | 21 | 0.3333 |
| 8_11188394 | 6 | 0.5 | 2 | 0.5 | 6 | 0.6667 | 14 | 0.4286 | 28 | 0.5 | 5 | 0.2 | 8 | 0.625 | 4 | 0.25 | 4 | 0.25 | 21 | 0.381 |
| 8_11188396 | 5 | 0.4 | 1 | 0 | 6 | 0.6667 | 14 | 0.4286 | 26 | 0.4615 | 3 | 0.6667 | 8 | 0.625 | 5 | 0.6 | 3 | 0.3333 | 19 | 0.5789 |
| 8_11188401 | 7 | 0 | 2 | 0 | 7 | 0.1429 | 12 | 0.1667 | 28 | 0.1071 | 3 | 0 | 10 | 0.3 | 4 | 0.25 | 4 | 0 | 21 | 0.1905 |
| 8_11188405 | 6 | 0.5 | 1 | 1 | 8 | 0.625 | 14 | 0.7143 | 29 | 0.6552 | 3 | 0.3333 | 10 | 0.9 | 5 | 0.8 | 5 | 0.2 | 23 | 0.6522 |
| 8_11188410 | 2 | 0 | 1 | 0 | 3 | 0 | 10 | 0.1 | 16 | 0.0625 | 2 | 0.5 | 7 | 0 | 4 | 0 | 4 | 0 | 17 | 0.0588 |
| 8_11188410 | 7 | 0 | 2 | 1 | 7 | 0.4286 | 14 | 0.2857 | 30 | 0.3 | 3 | 0 | 10 | 0.3 | 5 | 0.6 | 3 | 0 | 21 | 0.2857 |
| 8_11188412 | 5 | 0.2 | 2 | 0.5 | 6 | 0.5 | 13 | 0.5385 | 26 | 0.4615 | 3 | 0.3333 | 8 | 0.375 | 5 | 0.4 | 3 | 0.3333 | 19 | 0.3684 |
| 8_11188416 | 4 | 0.25 | 2 | 0.5 | 7 | 0.7143 | 14 | 0.6429 | 27 | 0.5926 | 2 | 0.5 | 9 | 0.7778 | 4 | 0.75 | 3 | 0.6667 | 18 | 0.7222 |
| 8_11188424 | 4 | 1 | 1 | 0 | 3 | 1 | 7 | 0.7143 | 15 | 0.8 | NA | NA | 6 | 1 | 3 | 0.6667 | 3 | 1 | 12 | 0.9167 |
| 8_11188426 | 6 | 0.3333 | 2 | 0.5 | 6 | 0.5 | 12 | 0.3333 | 26 | 0.3846 | 2 | 0 | 9 | 0.4444 | 5 | 0.6 | 4 | 0.5 | 20 | 0.45 |
| 8_11188437 | 6 | 0.6667 | 2 | 0.5 | 8 | 0.375 | 10 | 0.4 | 26 | 0.4615 | 2 | 0.5 | 9 | 0.3333 | 4 | 0.5 | 3 | 0.6667 | 18 | 0.4444 |
| 8_11188453 | 6 | 0.6667 | 1 | 0 | 7 | 0.4286 | 10 | 0.8 | 24 | 0.625 | 1 | 0 | 8 | 0.75 | 4 | 0.5 | 3 | 1 | 16 | 0.6875 |
| 8_11199796 | 4 | 0 | 7 | 0.1429 | 3 | 0.3333 | 3 | 0 | 17 | 0.1176 | 2 | 0.5 | 1 | 0 | 2 | 0 | 2 | 0 | 7 | 0.1429 |
| 8_11199955 | 2 | 1 | 4 | 1 | 1 | 1 | 1 | 1 | 8 | 1 | 2 | 1 | NA | NA | 1 | 1 | 2 | 1 | 5 | 1 |
| 8_11199963 | 2 | 0 | 4 | 0 | NA | NA | 1 | 1 | 7 | 0.1429 | 2 | 0 | NA | NA | 1 | 1 | 2 | 0 | 5 | 0.2 |
| 8_11199967 | 2 | 0.5 | 3 | 0.6667 | 1 | 1 | 1 | 1 | 7 | 0.7143 | 2 | 0.5 | 1 | 0 | 1 | 0 | 2 | 0.5 | 6 | 0.3333 |
| 8_11200295 | 2 | 0 | 3 | 0.3333 | 1 | 0 | 1 | 0 | 7 | 0.1429 | 2 | 0.5 | 1 | 0 | 1 | 0 | 2 | 0 | 6 | 0.1667 |
| 8_11202408 | 2 | 1 | 2 | 0.5 | 1 | 1 | NA | NA | 5 | 0.8 | NA | NA | NA | NA | 1 | 1 | 2 | 1 | 3 | 1 |
| 8_11202415 | 2 | 0.5 | 2 | 0.5 | 1 | 1 | 1 | 0 | 6 | 0.5 | NA | NA | NA | NA | 1 | 0 | 1 | 1 | 2 | 0.5 |
| 8_11207192 | 2 | 0 | 6 | 0.1667 | 6 | 0 | 5 | 0 | 19 | 0.0526 | 1 | 0 | 3 | 0 | 3 | 0.3333 | 3 | 0 | 10 | 0.1 |
| 8_11207219 | 2 | 0 | 4 | 0.25 | 4 | 0.25 | 5 | 0.2 | 15 | 0.2 | 3 | 0 | 2 | 0 | 3 | 0.3333 | 2 | 0.5 | 10 | 0.2 |
| 8_11217548 | 2 | 0 | 1 | 0 | 2 | 0 | 1 | 1 | 6 | 0.1667 | 1 | 0 | 4 | 0.25 | 3 | 0 | 1 | 0 | 9 | 0.1111 |
| 8_11217660 | 2 | 0 | 2 | 0 | 2 | 0 | 1 | 1 | 7 | 0.1429 | NA | NA | 4 | 0.25 | NA | NA | NA | NA | 4 | 0.25 |
| 8_11217684 | 3 | 0.3333 | 3 | 0 | 2 | 0 | 1 | 1 | 9 | 0.2222 | 1 | 0 | 5 | 0.4 | 1 | 0 | NA | NA | 7 | 0.2857 |
| 8_11217691 | 3 | 0.3333 | 3 | 0.6667 | 1 | 0 | 1 | 1 | 8 | 0.5 | 1 | 1 | 5 | 0.6 | 1 | 0 | NA | NA | 7 | 0.5714 |
| 8_11217695 | 3 | 1 | 3 | 1 | 2 | 1 | 1 | 1 | 9 | 1 | 1 | 1 | 6 | 1 | 1 | 1 | NA | NA | 8 | 1 |
| 8_11217701 | 3 | 0 | 4 | 0.75 | 3 | 0.6667 | 2 | 1 | 12 | 0.5833 | 1 | 0 | 3 | 0.6667 | 1 | 1 | 1 | 1 | 6 | 0.6667 |
| 8_11217707 | 2 | 1 | 4 | 1 | 2 | 1 | 2 | 1 | 10 | 1 | 1 | 1 | 5 | 0.8 | 1 | 1 | 1 | 1 | 8 | 0.875 |
| 8_11217712 | 1 | 1 | 3 | 1 | 4 | 1 | 2 | 1 | 10 | 1 | 1 | 1 | 5 | 0.8 | 1 | 1 | 1 | 1 | 8 | 0.875 |
| 8_11217733 | 3 | 0.6667 | 3 | 1 | 3 | 1 | 1 | 1 | 10 | 0.9 | NA | NA | 5 | 0.8 | 1 | 1 | 1 | 1 | 7 | 0.8571 |
| 8_11217743 | 3 | 1 | 3 | 1 | 4 | 1 | 1 | 1 | 11 | 1 | 1 | 0 | 4 | 1 | 1 | 1 | 1 | 1 | 7 | 0.8571 |
| 8_11217746 | 3 | 1 | 3 | 0.6667 | 4 | 1 | 1 | 1 | 11 | 0.9091 | 1 | 0 | 5 | 0.8 | 1 | 1 | 1 | 1 | 8 | 0.75 |
| 8_11217747 | 2 | 0.5 | 1 | 1 | 2 | 0.5 | 1 | 1 | 6 | 0.6667 | 1 | 1 | 5 | 0.8 | 1 | 0 | NA | NA | 7 | 0.7143 |

| | | | | | | | | | | | | | | | | | | | | |
|---|---|---|---|---|---|---|---|---|---|---|---|---|---|---|---|---|---|---|---|---|
| 8_1121775 | 3 | 0.6667 | 3 | 0.6667 | 4 | 0.5 | 1 | 1 | 11 | 0.6364 | 1 | 0 | 5 | 0.8 | 1 | 1 | 1 | 1 | 8 | 0.75 |
| 8_1122635 | 2 | 1 | 11 | 0.7273 | 4 | 0.5 | 5 | 0.2 | 22 | 0.5909 | 2 | 0.5 | 7 | 0.1429 | 8 | 0.75 | 1 | 0 | 18 | 0.4444 |
| 8_1122635 | 3 | 0.6667 | 12 | 0.75 | 4 | 0.5 | 5 | 0.6 | 24 | 0.6667 | 2 | 1 | 8 | 0.875 | 6 | 0.6667 | 2 | 0 | 18 | 0.7222 |
| 8_1122636 | 5 | 0 | 9 | 0.3333 | 3 | 0.3333 | 4 | 0.5 | 21 | 0.2857 | 3 | 0 | 9 | 0 | 9 | 0.1111 | 2 | 0.5 | 23 | 0.087 |
| 8_1122640 | 5 | 0.6 | 12 | 0.0833 | 8 | 0.125 | 8 | 0.125 | 33 | 0.1818 | 4 | 0.25 | 13 | 0.2308 | 13 | 0.2308 | 5 | 0 | 35 | 0.2 |
| 8_1123483 | 2 | 0 | 3 | 0 | 3 | 0.3333 | 5 | 0.2 | 13 | 0.1538 | 1 | 0 | 7 | 0 | 6 | 0.1667 | 3 | 0 | 17 | 0.0588 |
| 8_1123486 | 2 | 0.5 | 3 | 0 | 5 | 0 | 4 | 0 | 14 | 0.0714 | 2 | 0.5 | 6 | 0 | 8 | 0.125 | 4 | 0 | 20 | 0.1 |
| 8_1123491 | 10 | 0.9 | 9 | 0.6667 | 13 | 0.9231 | 5 | 0.8 | 37 | 0.8378 | 7 | 0.8571 | 14 | 0.5 | 17 | 0.7647 | 9 | 1 | 47 | 0.7447 |
| 8_1123492 | 11 | 0.4545 | 14 | 0.9286 | 14 | 0.7143 | 9 | 0.7778 | 48 | 0.7292 | 7 | 0.4286 | 16 | 0.625 | 16 | 0.8125 | 7 | 0.4286 | 46 | 0.6304 |
| 8_1123493 | 12 | 1 | 14 | 0.9286 | 15 | 1 | 6 | 1 | 47 | 0.9787 | 9 | 1 | 16 | 0.9375 | 21 | 0.8571 | 7 | 1 | 53 | 0.9245 |
| 8_1123493 | 11 | 0.3636 | 11 | 0.1818 | 16 | 0.3125 | 10 | 0.4 | 48 | 0.3125 | 8 | 0.375 | 18 | 0.5556 | 18 | 0.2778 | 7 | 0.1429 | 51 | 0.3725 |
| 8_1123494 | 11 | 0.9091 | 12 | 0.9167 | 14 | 0.9286 | 7 | 0.8571 | 44 | 0.9091 | 7 | 1 | 17 | 0.8824 | 18 | 1 | 8 | 0.875 | 50 | 0.94 |
| 8_1123494 | 12 | 0.1667 | 13 | 0.0769 | 16 | 0.3125 | 8 | 0 | 49 | 0.1633 | 5 | 0.2 | 16 | 0.375 | 16 | 0.25 | 7 | 0.2857 | 44 | 0.2955 |
| 8_1123494 | 14 | 0.7857 | 16 | 0.625 | 17 | 0.7059 | 10 | 0.9 | 57 | 0.7368 | 9 | 0.6667 | 18 | 0.6667 | 23 | 0.6957 | 10 | 0.4 | 60 | 0.6333 |
| 8_1123494 | 9 | 0.3333 | 11 | 0.3636 | 9 | 0.3333 | 8 | 0.625 | 37 | 0.4054 | 1 | 1 | 9 | 0.5556 | 16 | 0.625 | 7 | 0.4286 | 33 | 0.5758 |
| 8_1123494 | 13 | 0.5385 | 15 | 0.4 | 17 | 0.2941 | 10 | 0.2 | 55 | 0.3636 | 6 | 0.3333 | 19 | 0.4737 | 22 | 0.3636 | 7 | 0.2857 | 54 | 0.3889 |
| 8_1123495 | 12 | 0.0833 | 14 | 0.0714 | 13 | 0 | 11 | 0.2727 | 50 | 0.1 | 6 | 0.3333 | 13 | 0 | 20 | 0.1 | 8 | 0.25 | 47 | 0.1277 |
| 8_1123495 | 8 | 0.375 | 13 | 0.4615 | 16 | 0.1875 | 10 | 0.4 | 47 | 0.3404 | 10 | 0.5 | 20 | 0.55 | 20 | 0.3 | 7 | 0.2857 | 57 | 0.4211 |
| 8_1123495 | 13 | 0.3077 | 16 | 0.0625 | 14 | 0.0714 | 8 | 0.25 | 51 | 0.1569 | 9 | 0.1111 | 16 | 0.125 | 19 | 0.0526 | 8 | 0.125 | 52 | 0.0962 |
| 8_1123500 | 8 | 0.375 | 13 | 0.2308 | 9 | 0.4444 | 8 | 0.625 | 38 | 0.3947 | 8 | 0.25 | 6 | 0.3333 | 14 | 0.5 | 4 | 0.5 | 32 | 0.4062 |
| 8_1123501 | 10 | 0.1 | 12 | 0.0833 | 13 | 0.1538 | 10 | 0 | 45 | 0.0889 | 9 | 0.2222 | 9 | 0.3333 | 14 | 0.1429 | 5 | 0.2 | 37 | 0.2162 |
| 8_1123502 | 9 | 0.3333 | 15 | 0.4 | 15 | 0.4 | 11 | 0.3636 | 50 | 0.38 | 11 | 0.2727 | 10 | 0.6 | 17 | 0.2353 | 5 | 0.2 | 43 | 0.3256 |
| 8_1123511 | 3 | 0.6667 | 8 | 0.75 | 8 | 0.5 | 10 | 0.6 | 29 | 0.6207 | 9 | 0.5556 | 10 | 0.3 | 10 | 0.6 | 7 | 0.4286 | 36 | 0.4722 |
| 8_1123511 | 4 | 0 | 8 | 0.125 | 12 | 0.3333 | 12 | 0.3333 | 36 | 0.25 | 12 | 0.1667 | 11 | 0.0909 | 14 | 0.1429 | 6 | 0.3333 | 43 | 0.1628 |
| 8_1123512 | 5 | 0.2 | 8 | 0 | 11 | 0.0909 | 7 | 0 | 31 | 0.0645 | 10 | 0.1 | 10 | 0.1 | 10 | 0.2 | 7 | 0.5714 | 37 | 0.2162 |
| 8_1123534 | 2 | 0.5 | 1 | 1 | NA | NA | NA | NA | 3 | 0.6667 | NA | NA | 1 | 1 | 1 | 0 | NA | NA | 2 | 0.5 |
| 8_1123962 | 1 | 1 | 5 | 0.8 | 3 | 0 | 5 | 0.6 | 14 | 0.5714 | 3 | 0.6667 | 3 | 0.6667 | 8 | 0.5 | 1 | 0 | 15 | 0.5333 |
| 8_1123962 | 1 | 1 | 5 | 1 | 3 | 1 | 4 | 1 | 13 | 1 | 3 | 1 | 3 | 1 | 8 | 0.875 | NA | NA | 14 | 0.9286 |
| 8_1123962 | 1 | 0 | 4 | 0.25 | 3 | 0.6667 | 5 | 0.2 | 13 | 0.3077 | 3 | 0.3333 | 3 | 0 | 8 | 0.125 | 1 | 0 | 15 | 0.1333 |
| 8_1123963 | 1 | 1 | 5 | 0.8 | 2 | 1 | 4 | 1 | 12 | 0.9167 | 3 | 1 | 2 | 1 | 6 | 1 | 2 | 0.5 | 13 | 0.9231 |
| 8_1123963 | 1 | 1 | 5 | 0.6 | 3 | 0.6667 | 5 | 0.8 | 14 | 0.7143 | 2 | 1 | 2 | 1 | 7 | 0.8571 | 2 | 0.5 | 13 | 0.8462 |
| 8_1123963 | 1 | 1 | 3 | 0.6667 | 2 | 1 | 4 | 1 | 10 | 0.9 | 3 | 0.6667 | 2 | 0.5 | 7 | 0.8571 | 1 | 1 | 13 | 0.7692 |
| 8_1123964 | 1 | 0 | 5 | 1 | 2 | 1 | 4 | 1 | 12 | 0.9167 | 3 | 0.6667 | 2 | 1 | 6 | 1 | 2 | 0.5 | 13 | 0.8462 |
| 8_1123970 | 1 | 1 | 3 | 0.6667 | 3 | 1 | 2 | 0.5 | 9 | 0.7778 | 1 | 1 | 3 | 1 | 7 | 0.7143 | 2 | 1 | 13 | 0.8462 |
| 8_1123972 | NA | NA | 5 | 1 | 2 | 1 | 5 | 1 | 12 | 1 | 3 | 1 | 4 | 1 | 10 | 0.9 | NA | NA | 17 | 0.9412 |
| 8_1123974 | 1 | 1 | 5 | 1 | 2 | 1 | 5 | 1 | 13 | 1 | 3 | 1 | 7 | 1 | 10 | 0.9 | 2 | 1 | 22 | 0.9545 |
| 8_1124371 | 9 | 1 | 6 | 0.5 | 3 | 0.6667 | 13 | 0.5385 | 31 | 0.6774 | 8 | 0.625 | 4 | 0.5 | 9 | 0.8889 | 3 | 0.6667 | 24 | 0.7083 |
| 8_1124374 | 10 | 0.2 | 6 | 0.5 | 3 | 0 | 11 | 0.0909 | 30 | 0.2 | 5 | 0.4 | 4 | 0 | 8 | 0.125 | 4 | 0.5 | 21 | 0.2381 |
| 8_1124375 | 11 | 0.9091 | 6 | 0.3333 | 6 | 0.6667 | 14 | 0.7857 | 37 | 0.7297 | 4 | 0.75 | 4 | 0.5 | 8 | 0.625 | 3 | 1 | 19 | 0.6842 |
| 8_1124375 | 9 | 0.5556 | 6 | 0.1667 | 5 | 1 | 12 | 0.5833 | 32 | 0.5625 | 3 | 1 | 2 | 0 | 7 | 0.7143 | 3 | 0.3333 | 15 | 0.6 |
| 8_1124375 | 10 | 0.6 | 3 | 0 | 5 | 0.6 | 12 | 0.5833 | 30 | 0.5333 | 4 | 1 | 4 | 1 | 8 | 0.625 | 3 | 1 | 19 | 0.8421 |
| 8_1124376 | 11 | 0.1818 | 5 | 0 | 5 | 0 | 12 | 0.0833 | 33 | 0.0909 | 3 | 0 | 4 | 0 | 9 | 0.1111 | 4 | 0 | 20 | 0.05 |
| 8_1124376 | 10 | 0.6 | 6 | 0.1667 | 4 | 0 | 14 | 0.1429 | 34 | 0.2647 | 3 | 0.3333 | 3 | 0.3333 | 9 | 0.1111 | 3 | 0 | 18 | 0.1667 |
| 8_1124377 | 9 | 0.8889 | 4 | 0.75 | 5 | 0.8 | 11 | 1 | 29 | 0.8966 | 4 | 1 | 2 | 1 | 5 | 0.8 | 3 | 0.6667 | 14 | 0.8571 |
| 8_1124378 | 7 | 0.2857 | 3 | 0 | 5 | 0.2 | 13 | 0.0769 | 28 | 0.1429 | 3 | 0.3333 | 3 | 0.6667 | 3 | 0 | 2 | 0 | 11 | 0.2727 |
| 8_1124379 | 8 | 0.125 | 5 | 0.2 | 5 | 0 | 12 | 0.25 | 30 | 0.1667 | 2 | 0.5 | 3 | 0.6667 | 5 | 0 | 2 | 0 | 12 | 0.25 |
| 8_1124399 | 9 | 0.2222 | 5 | 0.2 | 5 | 0.4 | 12 | 0.25 | 31 | 0.2581 | 2 | 0.5 | 4 | 0.25 | 4 | 0 | 2 | 0 | 12 | 0.1667 |
| 8_1124400 | 6 | 0.5 | 4 | 0.5 | 4 | 0.25 | 13 | 0.1538 | 27 | 0.2963 | 3 | 0.3333 | 3 | 0.3333 | 6 | 0.6667 | 3 | 0.3333 | 15 | 0.4667 |
| 8_1124401 | 6 | 1 | 4 | 0.75 | 4 | 0.75 | 7 | 1 | 21 | 0.9048 | 2 | 0 | 3 | 1 | 6 | 1 | 2 | 1 | 13 | 0.8462 |
| 8_1124402 | 7 | 0.1429 | 2 | 0.5 | 4 | 0.25 | 9 | 0.2222 | 22 | 0.2273 | 3 | 0 | 3 | 0.3333 | 5 | 0 | 1 | 0 | 12 | 0.0833 |
| 8_1124403 | 6 | 0 | 4 | 0.25 | 5 | 0.4 | 10 | 0 | 25 | 0.12 | 2 | 0 | 4 | 0.25 | 6 | 0.1667 | 3 | 0.3333 | 15 | 0.2 |
| 8_1124404 | 7 | 0.8571 | 3 | 0.6667 | 4 | 0.5 | 10 | 0.7 | 24 | 0.7083 | 3 | 1 | 2 | 0.5 | 4 | 0.75 | 3 | 1 | 12 | 0.8333 |
| 8_1124405 | 5 | 0.8 | 3 | 0.3333 | 5 | 0.4 | 5 | 0.6 | 18 | 0.5556 | 3 | 0.6667 | NA | NA | 4 | 0.75 | 2 | 0 | 9 | 0.5556 |
| 8_1124406 | 8 | 0.625 | 3 | 1 | 3 | 0.6667 | 6 | 1 | 20 | 0.8 | 2 | 1 | 2 | 0.5 | 5 | 1 | 1 | 1 | 10 | 0.9 |
| 8_1124408 | 6 | 0.8333 | 1 | 0 | 3 | 1 | 5 | 0.6 | 15 | 0.7333 | 1 | 1 | 1 | 0 | 4 | 0.5 | 1 | 1 | 7 | 0.5714 |
| 8_1124474 | 3 | 0 | 2 | 0.5 | 1 | 0 | 6 | 0.3333 | 12 | 0.25 | 2 | 0.5 | 1 | 0 | 2 | 0 | 1 | 1 | 6 | 0.3333 |
| 8_1124475 | 2 | 0 | 2 | 0 | 2 | 0 | 6 | 0.3333 | 12 | 0.1667 | 2 | 0.5 | 1 | 0 | 3 | 0 | 1 | 0 | 7 | 0.1429 |
| 8_1124477 | 2 | 0 | 2 | 0 | 2 | 0 | 6 | 0.1667 | 12 | 0.0833 | 2 | 0 | 1 | 0 | 2 | 0.5 | 1 | 0 | 6 | 0.1667 |
| 8_1124952 | 20 | 0.25 | 20 | 0.05 | 10 | 0 | 9 | 0.1111 | 59 | 0.1186 | 13 | 0 | 15 | 0.2 | 22 | 0.0909 | 12 | 0.1667 | 62 | 0.1129 |
| 8_1125899 | NA | NA | 2 | 0.5 | NA | NA | 2 | 0 | 4 | 0.25 | 1 | 1 | NA | NA | NA | NA | 1 | 0 | 2 | 0.5 |
| 8_1125925 | NA | NA | 1 | 1 | NA | NA | 1 | 1 | 2 | 1 | 1 | 1 | NA | NA | NA | NA | NA | NA | 1 | 1 |
| 8_1125925 | NA | NA | 1 | 1 | NA | NA | 1 | 1 | 2 | 1 | 1 | 1 | NA | NA | NA | NA | NA | NA | 1 | 1 |
| 8_1125929 | NA | NA | NA | NA | NA | NA | 1 | 1 | 1 | 1 | 1 | 1 | NA | NA | NA | NA | NA | NA | 1 | 1 |
| 8_1125929 | NA | NA | 1 | 1 | NA | NA | 1 | 1 | 2 | 1 | 1 | 1 | NA | NA | NA | NA | NA | NA | 1 | 1 |
| 8_1126392 | 3 | 0.3333 | 4 | 0 | 2 | 0 | 5 | 0 | 14 | 0.0714 | NA | NA | 2 | 0 | 1 | 0 | 3 | 0.3333 | 6 | 0.1667 |

| | | | | | | | | | | | | | | | | | | | | |
|---|---|---|---|---|---|---|---|---|---|---|---|---|---|---|---|---|---|---|---|---|
| 8_11263999 | 1 | 1 | 4 | 0.75 | 2 | 1 | 5 | 1 | 12 | 0.9167 | 1 | 1 | 3 | 0.6667 | NA | NA | 2 | 0 | 6 | 0.5 |
| 8_11264004 | 1 | 1 | 1 | 0 | 1 | 1 | 5 | 0.6 | 8 | 0.625 | 1 | 0 | 1 | 0 | NA | NA | 2 | 0.5 | 4 | 0.25 |
| 8_11264005 | 1 | 1 | 3 | 1 | 1 | 1 | 3 | 1 | 8 | 1 | 1 | 1 | 2 | 1 | NA | NA | 2 | 1 | 5 | 1 |
| 8_11264041 | NA | NA | 2 | 1 | 1 | 1 | 4 | 1 | 7 | 1 | NA | NA | 1 | 1 | NA | NA | 2 | 1 | 3 | 1 |
| 8_11264043 | 1 | 0 | 3 | 0.6667 | 1 | 0 | 5 | 0.4 | 10 | 0.4 | 1 | 0 | 2 | 0.5 | NA | NA | 2 | 0.5 | 5 | 0.4 |
| 8_11264046 | 1 | 1 | 3 | 1 | 2 | 1 | 5 | 0.8 | 11 | 0.9091 | 1 | 1 | 2 | 1 | NA | NA | 2 | 0.5 | 5 | 0.8 |
| 8_11264062 | 1 | 0 | 3 | 0 | 1 | 0 | 3 | 0.3333 | 8 | 0.125 | 1 | 0 | 2 | 0 | NA | NA | 2 | 0.5 | 5 | 0.2 |
| 8_11267947 | 57 | 0.0702 | 53 | 0.1698 | 45 | 0.1111 | 44 | 0.0909 | 199 | 0.1106 | 36 | 0.0833 | 42 | 0.119 | 44 | 0.1136 | 44 | 0.2045 | 166 | 0.1325 |
| 8_11268190 | 9 | 0.1111 | 4 | 0.25 | 5 | 0 | 10 | 0 | 28 | 0.0714 | 7 | 0.1429 | 9 | 0.1111 | 6 | 0 | 6 | 0 | 28 | 0.0714 |
| 8_11269486 | 1 | 1 | 1 | 1 | NA | NA | 1 | 1 | 3 | 1 | NA | NA | 1 | 1 | NA | NA | 1 | 0 | 2 | 0.5 |
| 8_11269489 | NA | NA | 1 | 1 | NA | NA | 1 | 1 | 2 | 1 | NA | NA | 1 | 1 | NA | NA | 1 | 1 | 2 | 1 |
| 8_11269501 | 1 | 0 | 1 | 1 | NA | NA | 1 | 0 | 3 | 0.3333 | NA | NA | 1 | 0 | NA | NA | 1 | 1 | 2 | 0.5 |
| 8_11269503 | NA | NA | 1 | 1 | NA | NA | 1 | 1 | 2 | 1 | NA | NA | 1 | 1 | NA | NA | 1 | 1 | 2 | 1 |
| 8_11270902 | 1 | 1 | NA | NA | NA | NA | NA | NA | 1 | 1 | NA | NA | 1 | 0 | 1 | 1 | 1 | 1 | 3 | 0.6667 |
| 8_11270903 | 1 | 1 | NA | NA | NA | NA | NA | NA | 1 | 1 | NA | NA | 1 | 1 | NA | NA | 1 | 1 | 2 | 1 |
| 8_11270930 | 1 | 1 | NA | NA | NA | NA | NA | NA | 1 | 1 | NA | NA | 1 | 0 | 1 | 1 | 1 | 0 | 3 | 0.3333 |
| 8_11273864 | 9 | 0.2222 | 3 | 0 | 3 | 0.3333 | 7 | 0.1429 | 22 | 0.1818 | 3 | 0 | 5 | 0.2 | 5 | 0.2 | 6 | 0.1667 | 19 | 0.1579 |
| 8_11274454 | 9 | 0.1111 | 5 | 0 | 3 | 0.3333 | 9 | 0.1111 | 26 | 0.1154 | 4 | 0 | 6 | 0 | 7 | 0.1429 | 8 | 0.125 | 25 | 0.08 |
| 8_11274460 | 10 | 0.3 | 5 | 0.6 | 2 | 0.5 | 8 | 0.25 | 25 | 0.36 | 3 | 0.3333 | 6 | 0.5 | 7 | 0.5714 | 6 | 0.1667 | 22 | 0.4091 |
| 8_11274468 | 8 | 1 | 4 | 1 | 3 | 0.6667 | 8 | 0.75 | 23 | 0.8696 | 3 | 0.6667 | 4 | 0.75 | 6 | 0.8333 | 7 | 0.7143 | 20 | 0.75 |
| 8_11274475 | 11 | 0 | 5 | 0 | 3 | 0 | 10 | 0.2 | 29 | 0.069 | 5 | 0 | 7 | 0.2857 | 7 | 0.2857 | 8 | 0 | 27 | 0.1481 |
| 8_11274476 | 11 | 0.5455 | 4 | 0.75 | 2 | 0.5 | 9 | 0.5556 | 26 | 0.5769 | 4 | 0.5 | 5 | 0.6 | 8 | 0.5 | 7 | 0.5714 | 24 | 0.5417 |
| 8_11274487 | 11 | 0.0909 | 4 | 0.25 | 4 | 0.25 | 9 | 0.2222 | 28 | 0.1786 | 4 | 0 | 7 | 0.1429 | 8 | 0.125 | 9 | 0.2222 | 28 | 0.1429 |
| 8_11274491 | 10 | 0.1 | 3 | 0.3333 | 4 | 0 | 10 | 0.2 | 27 | 0.1481 | 4 | 0.25 | 7 | 0.1429 | 8 | 0.125 | 8 | 0 | 27 | 0.1111 |
| 8_11274501 | 10 | 0.5 | 3 | 1 | 3 | 1 | 9 | 0.7778 | 25 | 0.72 | 7 | 0.8571 | 5 | 1 | 6 | 1 | 9 | 0.8889 | 27 | 0.9259 |
| 8_11274511 | 10 | 1 | 3 | 1 | 4 | 0.75 | 9 | 1 | 26 | 0.9615 | 5 | 0.8 | 7 | 1 | 8 | 1 | 8 | 1 | 28 | 0.9643 |
| 8_11274518 | 9 | 0.4444 | 5 | 0.2 | 4 | 0 | 10 | 0.2 | 28 | 0.25 | 7 | 0.2857 | 7 | 0.1429 | 8 | 0.25 | 9 | 0 | 31 | 0.1613 |
| 8_11274531 | 10 | 0 | 5 | 0.2 | 4 | 0.5 | 10 | 0.1 | 29 | 0.1379 | 7 | 0.2857 | 8 | 0.125 | 8 | 0.125 | 8 | 0.25 | 31 | 0.1935 |
| 8_11274533 | 11 | 0.0909 | 5 | 0.6 | 4 | 0 | 11 | 0.1818 | 31 | 0.1935 | 7 | 0 | 8 | 0.25 | 8 | 0.375 | 8 | 0 | 31 | 0.1613 |
| 8_11274541 | 9 | 0.1111 | 7 | 0 | 4 | 0.25 | 12 | 0.25 | 32 | 0.1562 | 5 | 0 | 8 | 0.375 | 7 | 0 | 6 | 0 | 26 | 0.1154 |
| 8_11274566 | 12 | 0.0833 | 6 | 0.1667 | 3 | 0.3333 | 10 | 0.1 | 31 | 0.129 | 7 | 0.2857 | 8 | 0 | 7 | 0 | 9 | 0 | 31 | 0.0645 |
| 8_11274609 | 1 | 0 | NA | NA | 2 | 0.5 | 2 | 0.5 | 5 | 0.4 | NA | NA | 3 | 0 | 1 | 0 | 4 | 0.25 | 8 | 0.125 |
| 8_11274643 | 1 | 0 | NA | NA | 1 | 1 | 3 | 0 | 5 | 0.2 | 1 | 0 | 3 | 0 | 1 | 1 | 4 | 0.25 | 9 | 0.2222 |
| 8_11274686 | 1 | 0 | 1 | 1 | 2 | 0 | 3 | 0 | 7 | 0.1429 | 1 | 0 | 4 | 0.25 | 1 | 0 | 3 | 0 | 9 | 0.1111 |
| 8_11274768 | NA | NA | 1 | 1 | NA | NA | NA | NA | 1 | 1 | NA | NA | 1 | 1 | NA | NA | NA | NA | 1 | 1 |
| 8_11356919 | 2 | 0.5 | 2 | 0 | 1 | 0 | NA | NA | 5 | 0.2 | NA | NA | 1 | 0 | 1 | 0 | 3 | 0.3333 | 5 | 0.2 |
| 8_11360843 | 4 | 0.75 | 2 | 0.5 | 2 | 0.5 | NA | NA | 8 | 0.625 | 1 | 0 | 1 | 1 | 2 | 0.5 | 4 | 0.75 | 8 | 0.625 |
| 8_11360853 | 3 | 0.3333 | 2 | 0 | 2 | 0 | NA | NA | 7 | 0.1429 | 1 | 0 | 2 | 0 | 2 | 0.5 | 4 | 0.25 | 9 | 0.2222 |
| 8_11360858 | 4 | 1 | 4 | 0.75 | 1 | 1 | NA | NA | 9 | 0.8889 | 1 | 1 | NA | NA | 3 | 1 | 5 | 1 | 9 | 1 |
| 8_11360880 | 4 | 0.25 | 7 | 0.1429 | 1 | 1 | NA | NA | 12 | 0.25 | 2 | 0 | 2 | 0.5 | 4 | 0.75 | 4 | 0.5 | 12 | 0.5 |
| 8_11360885 | 3 | 0.6667 | 5 | 0.6 | 2 | 1 | NA | NA | 10 | 0.7 | 2 | 0.5 | 2 | 1 | 4 | 1 | 5 | 0.8 | 13 | 0.8462 |
| 8_11360886 | 3 | 1 | 5 | 0.8 | 2 | 1 | NA | NA | 10 | 0.9 | 2 | 1 | 2 | 1 | 3 | 1 | 5 | 1 | 12 | 1 |
| 8_11360889 | 4 | 0 | 7 | 0 | 2 | 0.5 | NA | NA | 13 | 0.0769 | 2 | 0 | 2 | 0 | 4 | 0.25 | 4 | 0 | 12 | 0.0833 |
| 8_11436280 | 1 | 1 | 3 | 0 | 1 | 1 | NA | NA | 5 | 0.4 | 2 | 1 | 2 | 0.5 | NA | NA | NA | NA | 4 | 0.75 |
| 8_11436455 | 1 | 0 | 3 | 0 | 1 | 1 | NA | NA | 5 | 0.2 | 2 | 0.5 | 1 | 0 | 1 | 0 | NA | NA | 4 | 0.25 |
| 8_11436522 | 1 | 0 | 3 | 0 | 1 | 1 | NA | NA | 5 | 0.2 | 1 | 1 | 2 | 0 | NA | NA | NA | NA | 3 | 0.3333 |
| 8_11436533 | 1 | 0 | 3 | 0.3333 | NA | NA | NA | NA | 4 | 0.25 | 1 | 0 | 2 | 0 | 1 | 1 | NA | NA | 4 | 0.25 |
| 8_11436538 | 1 | 0 | 3 | 0.3333 | NA | NA | NA | NA | 4 | 0.25 | 1 | 0 | 2 | 0.5 | 1 | 1 | NA | NA | 4 | 0.5 |
| 8_11436539 | 1 | 1 | 4 | 1 | 1 | 1 | NA | NA | 6 | 1 | 1 | 1 | 2 | 1 | 1 | 1 | NA | NA | 4 | 1 |
| 8_11436559 | 1 | 0 | 4 | 0.75 | 1 | 1 | NA | NA | 6 | 0.6667 | 2 | 1 | 2 | 0.5 | 1 | 1 | NA | NA | 5 | 0.8 |
| 8_11436561 | 1 | 1 | 4 | 0.25 | 1 | 0 | NA | NA | 6 | 0.3333 | 2 | 0.5 | 1 | 1 | 1 | 0 | NA | NA | 4 | 0.5 |
| 8_11436875 | 2 | 1 | 4 | 1 | 1 | 1 | NA | NA | 7 | 1 | 1 | 1 | 3 | 1 | 2 | 0.5 | 2 | 1 | 8 | 0.875 |
| 8_11436877 | 2 | 0.5 | 4 | 0 | 1 | 0 | NA | NA | 7 | 0.1429 | 1 | 0 | 3 | 0 | 2 | 0.5 | 1 | 0 | 7 | 0.1429 |
| 8_11436897 | 2 | 0.5 | 3 | 0.6667 | NA | NA | NA | NA | 5 | 0.6 | 1 | 0 | 3 | 0.3333 | 2 | 0.5 | 3 | 1 | 9 | 0.5556 |
| 8_11436927 | 1 | 0 | 4 | 0.25 | NA | NA | NA | NA | 5 | 0.2 | 1 | 0 | 3 | 0 | 2 | 0 | 3 | 0.6667 | 9 | 0.2222 |
| 8_11436935 | 2 | 0.5 | 4 | 1 | NA | NA | NA | NA | 6 | 0.8333 | 1 | 0 | 3 | 1 | 2 | 1 | 2 | 1 | 8 | 0.875 |
| 8_11436952 | 2 | 0.5 | 3 | 0.6667 | NA | NA | NA | NA | 5 | 0.6 | 2 | 0.5 | 3 | 0.6667 | 2 | 1 | 3 | 0.6667 | 10 | 0.7 |
| 8_11436956 | 2 | 0.5 | 4 | 0 | NA | NA | NA | NA | 6 | 0.1667 | 2 | 0 | 1 | 1 | 1 | 0 | 3 | 0.3333 | 7 | 0.2857 |
| 8_11436957 | 1 | 1 | 4 | 1 | NA | NA | NA | NA | 5 | 1 | 2 | 1 | 2 | 1 | 2 | 1 | 2 | 1 | 8 | 1 |
| 8_11436967 | 2 | 1 | 4 | 1 | NA | NA | NA | NA | 6 | 1 | 1 | 1 | 1 | 1 | 1 | 1 | 3 | 1 | 6 | 1 |
| 8_11487849 | 2 | 0.5 | 2 | 0.5 | NA | NA | NA | NA | 4 | 0.5 | 1 | 1 | 2 | 1 | 1 | 0 | 3 | 0.6667 | 7 | 0.7143 |
| 8_11487858 | 2 | 1 | 2 | 0.5 | NA | NA | NA | NA | 4 | 0.75 | 1 | 1 | 2 | 0.5 | 2 | 0.5 | 3 | 1 | 8 | 0.75 |
| 8_11487859 | 2 | 0 | 2 | 1 | NA | NA | NA | NA | 4 | 0.5 | 1 | 0 | 2 | 0 | 2 | 1 | 1 | 1 | 6 | 0.5 |
| 8_11487872 | 2 | 0.5 | 2 | 0.5 | NA | NA | NA | NA | 4 | 0.5 | 1 | 1 | 2 | 1 | 2 | 0 | 3 | 0.6667 | 8 | 0.625 |
| 8_11493260 | 6 | 0 | 1 | 1 | 1 | 1 | 4 | 0.25 | 12 | 0.25 | NA | NA | 3 | 0.6667 | 2 | 0.5 | 2 | 1 | 7 | 0.7143 |

| | | | | | | | | | | | | | | | | | | | | |
|---|---|---|---|---|---|---|---|---|---|---|---|---|---|---|---|---|---|---|---|---|
| 8_1150985 | 5 | 0 | 2 | 0.5 | 3 | 0 | 5 | 0 | 15 | 0.0667 | 4 | 0.25 | 6 | 0 | 4 | 0.25 | 2 | 0 | 16 | 0.125 |
| 8_1160694 | 1 | 1 | 2 | 0 | 3 | 0.3333 | NA | NA | 6 | 0.3333 | NA | NA | 2 | 0.5 | NA | NA | NA | NA | 2 | 0.5 |
| 8_1160785 | 1 | 1 | 3 | 0 | 1 | 0 | 1 | 0 | 6 | 0.1667 | NA | NA | 3 | 0 | 3 | 0.3333 | NA | NA | 6 | 0.1667 |
| 8_1160787 | 1 | 0 | 3 | 0 | 1 | 0 | 2 | 0.5 | 7 | 0.1429 | NA | NA | 3 | 0.3333 | 2 | 0 | NA | NA | 5 | 0.2 |
| 8_1160787 | 1 | 0 | 4 | 0.25 | NA | NA | 2 | 0.5 | 7 | 0.2857 | NA | NA | 3 | 0 | 3 | 0 | 1 | 1 | 7 | 0.1429 |
| 8_1164109 | NA | NA | NA | NA | NA | NA | 2 | 0.5 | 2 | 0.5 | NA | NA | NA | NA | 2 | 0.5 | 2 | 0 | 4 | 0.25 |
| 8_1164111 | NA | NA | NA | NA | NA | NA | 1 | 1 | 1 | 1 | NA | NA | NA | NA | 2 | 0.5 | 2 | 1 | 4 | 0.75 |
| 8_1164111 | NA | NA | NA | NA | NA | NA | 1 | 1 | 1 | 1 | NA | NA | NA | NA | 1 | 1 | 2 | 0.5 | 3 | 0.6667 |
| 8_1164112 | NA | NA | NA | NA | NA | NA | 2 | 1 | 2 | 1 | NA | NA | NA | NA | 2 | 0.5 | 1 | 1 | 3 | 0.6667 |
| 8_1164304 | 3 | 0.6667 | 3 | 0.6667 | 1 | 0 | NA | NA | 7 | 0.5714 | NA | NA | NA | NA | NA | NA | 2 | 1 | 2 | 1 |
| 8_1164306 | 2 | 1 | 4 | 1 | NA | NA | NA | NA | 6 | 1 | NA | NA | NA | NA | NA | NA | 2 | 1 | 2 | 1 |
| 8_1164311 | 1 | 0 | 3 | 0.3333 | NA | NA | NA | NA | 4 | 0.25 | NA | NA | NA | NA | NA | NA | 2 | 0.5 | 2 | 0.5 |
| 8_1164312 | 2 | 0 | 3 | 0.3333 | NA | NA | NA | NA | 5 | 0.2 | NA | NA | NA | NA | NA | NA | 2 | 0.5 | 2 | 0.5 |
| 8_1164313 | 2 | 1 | 3 | 1 | NA | NA | NA | NA | 5 | 1 | NA | NA | NA | NA | NA | NA | 2 | 1 | 2 | 1 |
| 8_1164314 | 1 | 1 | 3 | 1 | NA | NA | NA | NA | 4 | 1 | NA | NA | NA | NA | NA | NA | 2 | 0.5 | 2 | 0.5 |
| 8_1164314 | 2 | 0 | 3 | 0.3333 | NA | NA | NA | NA | 5 | 0.2 | NA | NA | NA | NA | NA | NA | 1 | 1 | 1 | 1 |
| 8_1164314 | 2 | 0.5 | 2 | 1 | NA | NA | NA | NA | 4 | 0.75 | NA | NA | NA | NA | NA | NA | 2 | 1 | 2 | 1 |
| 8_1164316 | 2 | 0 | 2 | 0.5 | NA | NA | NA | NA | 4 | 0.25 | NA | NA | NA | NA | NA | NA | 1 | 1 | 1 | 1 |
| 8_1164321 | 1 | 1 | 1 | 1 | NA | NA | NA | NA | 2 | 1 | NA | NA | NA | NA | NA | NA | 1 | 1 | 1 | 1 |
| 8_1164335 | 2 | 1 | 1 | 0 | NA | NA | 1 | 0 | 4 | 0.5 | NA | NA | NA | NA | 2 | 0.5 | 1 | 0 | 3 | 0.3333 |
| 8_1164335 | 2 | 1 | 1 | 1 | NA | NA | 1 | 1 | 4 | 1 | NA | NA | NA | NA | 2 | 0.5 | 1 | 0 | 3 | 0.3333 |
| 8_1173013 | NA | NA | NA | NA | NA | NA | 1 | 1 | 1 | 1 | 2 | 0.5 | NA | NA | NA | NA | NA | NA | 2 | 0.5 |
| 8_1173015 | NA | NA | NA | NA | NA | NA | 1 | 1 | 1 | 1 | 2 | 0.5 | NA | NA | NA | NA | NA | NA | 2 | 0.5 |
| 8_1173019 | NA | NA | NA | NA | NA | NA | 1 | 1 | 1 | 1 | 2 | 0.5 | NA | NA | NA | NA | NA | NA | 2 | 0.5 |
| 8_1176162 | 1 | 1 | 2 | 0.5 | 2 | 0.5 | 1 | 1 | 6 | 0.6667 | 5 | 0.6 | 5 | 0.8 | 1 | 1 | 2 | 0.5 | 13 | 0.6923 |
| 8_1176162 | 1 | 0 | 2 | 0 | 1 | 0 | 3 | 0.3333 | 7 | 0.1429 | 5 | 0.2 | 6 | 0 | 1 | 0 | 3 | 0 | 15 | 0.0667 |
| 8_1176163 | 1 | 0 | 2 | 1 | 1 | 0 | 3 | 1 | 7 | 0.7143 | 5 | 0.6 | 5 | 0.8 | 1 | 0 | 2 | 0 | 13 | 0.5385 |
| 8_1176164 | 2 | 0.5 | NA | NA | 2 | 0 | 3 | 0 | 7 | 0.1429 | 5 | 0.2 | 5 | 0 | 1 | 0 | 2 | 0 | 13 | 0.0769 |
| 8_1176164 | 1 | 1 | 1 | 1 | 2 | 1 | 2 | 1 | 6 | 1 | 5 | 1 | 5 | 1 | 1 | 1 | 3 | 1 | 14 | 1 |
| 8_1176650 | NA | NA | NA | NA | 1 | 1 | NA | NA | 1 | 1 | NA | NA | 1 | 0 | NA | NA | 1 | 1 | 2 | 0.5 |
| 8_1176831 | 1 | 0 | 3 | 0 | 3 | 0.3333 | 5 | 0.2 | 12 | 0.1667 | NA | NA | 5 | 0.4 | 3 | 0 | 2 | 0.5 | 10 | 0.3 |
| 8_1176832 | 1 | 0 | 3 | 0 | 3 | 0 | 5 | 0.2 | 12 | 0.0833 | NA | NA | 7 | 0 | 4 | 0.25 | 2 | 0 | 13 | 0.0769 |
| 8_1176833 | 1 | 1 | 3 | 1 | 3 | 1 | 5 | 1 | 12 | 1 | NA | NA | 7 | 1 | 3 | 1 | 1 | 1 | 11 | 1 |
| 8_1176834 | NA | NA | 2 | 1 | 3 | 1 | 5 | 1 | 10 | 1 | NA | NA | 4 | 1 | 3 | 1 | 3 | 0.6667 | 10 | 0.9 |
| 8_1176835 | 1 | 0 | 3 | 0 | 4 | 0.25 | 6 | 0.3333 | 14 | 0.2143 | NA | NA | 7 | 0.1429 | 4 | 0.25 | 3 | 0.3333 | 14 | 0.2143 |
| 8_1176835 | 1 | 0 | 4 | 0 | 5 | 0.2 | 6 | 0 | 16 | 0.0625 | NA | NA | 7 | 0.1429 | 4 | 0 | 3 | 0 | 14 | 0.0714 |
| 8_1176836 | 1 | 1 | 3 | 1 | 5 | 1 | 6 | 0.6667 | 15 | 0.8667 | NA | NA | 5 | 1 | 3 | 0.6667 | 3 | 0.6667 | 11 | 0.8182 |
| 8_1176836 | 1 | 0 | 3 | 0 | 5 | 0.2 | 6 | 0 | 15 | 0.0667 | NA | NA | 6 | 0.1667 | 4 | 0.25 | 2 | 0 | 12 | 0.1667 |
| 8_1176837 | 1 | 0 | 4 | 0.25 | 5 | 0 | 7 | 0 | 17 | 0.0588 | NA | NA | 7 | 0.1429 | 4 | 0.25 | 3 | 0 | 14 | 0.1429 |
| 8_1176838 | 1 | 0 | 4 | 0 | 4 | 0 | 7 | 0.1429 | 16 | 0.0625 | 1 | 1 | 7 | 0 | 3 | 0.3333 | 2 | 0 | 13 | 0.1538 |
| 8_1176838 | 1 | 1 | 4 | 0.75 | 3 | 1 | 7 | 1 | 15 | 0.9333 | 1 | 1 | 4 | 1 | 4 | 1 | 2 | 1 | 11 | 1 |
| 8_1176839 | 1 | 0 | 4 | 0.25 | 6 | 0 | 7 | 0.2857 | 18 | 0.1667 | 1 | 0 | 7 | 0.2857 | 6 | 0.1667 | 3 | 0 | 17 | 0.1765 |
| 8_1176839 | 1 | 1 | 4 | 0.75 | 5 | 0.8 | 8 | 0.875 | 18 | 0.8333 | 1 | 1 | 6 | 0.6667 | 6 | 1 | 2 | 1 | 15 | 0.8667 |
| 8_1176840 | 1 | 0 | 4 | 0 | 5 | 0 | 7 | 0.1429 | 17 | 0.0588 | 1 | 0 | 6 | 0 | 6 | 0.1667 | 4 | 0 | 17 | 0.0588 |
| 8_1176840 | 1 | 0 | 4 | 0 | 6 | 0 | 7 | 0.1429 | 18 | 0.0556 | 1 | 0 | 7 | 0 | 6 | 0 | 4 | 0.25 | 18 | 0.0556 |
| 8_1176841 | 1 | 0 | 4 | 0.5 | 4 | 1 | 7 | 0 | 16 | 0.375 | 1 | 0 | 6 | 0 | 5 | 0.2 | 3 | 0.3333 | 15 | 0.1333 |
| 8_1176842 | 1 | 1 | 4 | 0.75 | 2 | 0 | 8 | 0.375 | 15 | 0.4667 | 1 | 1 | 6 | 0.3333 | 5 | 0.6 | 4 | 0 | 16 | 0.375 |
| 8_1176842 | 1 | 1 | 4 | 0.75 | 5 | 0.6 | 5 | 0.8 | 15 | 0.7333 | NA | NA | 7 | 0.7143 | 5 | 0.6 | 3 | 0.6667 | 15 | 0.6667 |
| 8_1176843 | 1 | 0 | 5 | 0.4 | 4 | 0 | 7 | 0.2857 | 17 | 0.2353 | 1 | 0 | 7 | 0.4286 | 6 | 0.1667 | 4 | 0.25 | 18 | 0.2778 |
| 8_1176844 | 1 | 0 | 5 | 0.2 | 6 | 0.3333 | 10 | 0 | 22 | 0.1364 | 1 | 0 | 5 | 0.2 | 5 | 0 | 4 | 0.25 | 15 | 0.1333 |
| 8_1176844 | 1 | 1 | 5 | 0.8 | 6 | 1 | 8 | 1 | 20 | 0.95 | 1 | 1 | 6 | 1 | 4 | 1 | 4 | 1 | 15 | 1 |
| 8_1176844 | 2 | 0.5 | 5 | 0.2 | 5 | 0 | 10 | 0.3 | 22 | 0.2273 | 1 | 0 | 7 | 0.2857 | 5 | 0.6 | 3 | 0 | 16 | 0.3125 |
| 8_1176845 | 2 | 0 | 5 | 0.2 | 5 | 0 | 10 | 0.1 | 22 | 0.0909 | 1 | 0 | 7 | 0.1429 | 7 | 0 | 4 | 0 | 19 | 0.0526 |
| 8_1176845 | 3 | 0.6667 | 5 | 0.6 | 4 | 0.75 | 8 | 0.875 | 20 | 0.75 | 1 | 0 | 7 | 0.1429 | 7 | 0.7143 | 4 | 0.25 | 19 | 0.3684 |
| 8_1176845 | NA | NA | 5 | 0 | 4 | 0 | 8 | 0.125 | 17 | 0.0588 | NA | NA | 7 | 0.2857 | 7 | 0.2857 | 4 | 0 | 18 | 0.2222 |
| 8_1176846 | 1 | 0 | 5 | 0 | 5 | 0 | 8 | 0.375 | 19 | 0.1579 | 1 | 0 | 5 | 0.4 | 5 | 0.2 | 4 | 0 | 15 | 0.2 |
| 8_1176846 | 1 | 0 | 4 | 0.5 | 6 | 0.3333 | 7 | 0.4286 | 18 | 0.3889 | 1 | 0 | 7 | 0.2857 | 6 | 0 | 3 | 0 | 17 | 0.1176 |
| 8_1176846 | 1 | 0 | 4 | 0.25 | 6 | 0 | 10 | 0.5 | 21 | 0.2857 | 1 | 0 | 6 | 0.6667 | 7 | 0.1429 | 4 | 0.25 | 18 | 0.3333 |
| 8_1176846 | 2 | 0 | 4 | 0.75 | 6 | 0.6667 | 9 | 0.4444 | 21 | 0.5238 | 1 | 0 | 7 | 0.7143 | 7 | 0.2857 | 3 | 0.6667 | 18 | 0.5 |
| 8_1176846 | 2 | 0 | 4 | 0.25 | 4 | 0.25 | 11 | 0.2727 | 21 | 0.2381 | 1 | 1 | 7 | 0 | 7 | 0.1429 | 3 | 0.3333 | 18 | 0.1667 |
| 8_1176846 | 1 | 0 | 4 | 0.5 | 5 | 0.6 | 10 | 0.3 | 20 | 0.4 | 1 | 0 | 5 | 0.6 | 6 | 0.3333 | 3 | 0 | 15 | 0.3333 |
| 8_1176847 | 2 | 0 | 5 | 0.2 | 4 | 0 | 9 | 0.2222 | 20 | 0.15 | 1 | 0 | 6 | 0.1667 | 5 | 0.2 | 4 | 0.5 | 16 | 0.25 |
| 8_1176848 | 2 | 0 | 3 | 0.3333 | 5 | 0.2 | 10 | 0.2 | 20 | 0.2 | 1 | 0 | 6 | 0 | 6 | 0.6667 | 5 | 0.4 | 18 | 0.3333 |
| 8_1176848 | 2 | 1 | 5 | 0.8 | 5 | 1 | 11 | 0.9091 | 23 | 0.913 | NA | NA | 7 | 1 | 5 | 0.8 | 4 | 1 | 16 | 0.9375 |
| 8_1176850 | 1 | 0 | 4 | 0.5 | 4 | 0.5 | 9 | 0.5556 | 18 | 0.5 | 1 | 1 | 4 | 0.25 | 4 | 0.5 | 3 | 0.6667 | 12 | 0.5 |

| | | | | | | | | | | | | | | | | | | | | |
|---|---|---|---|---|---|---|---|---|---|---|---|---|---|---|---|---|---|---|---|---|
| 8_1176853 | 3 | 0 | 5 | 0.4 | 4 | 0 | 9 | 0.2222 | 21 | 0.1905 | 1 | 1 | 5 | 0 | 4 | 0 | 3 | 0 | 13 | 0.0769 |
| 8_1176860 | 5 | 0.4 | 3 | 0.6667 | 5 | 0.4 | 6 | 0.1667 | 19 | 0.3684 | 2 | 0.5 | 1 | 0 | 1 | 0 | 1 | 0 | 5 | 0.2 |
| 8_1176861 | 6 | 0.5 | 8 | 0 | 6 | 0.3333 | 11 | 0.0909 | 31 | 0.1935 | 3 | 0.3333 | 4 | 0.25 | 3 | 0.3333 | 4 | 0.5 | 14 | 0.3571 |
| 8_1176866 | 7 | 0 | 6 | 0.1667 | 6 | 0.1667 | 9 | 0 | 28 | 0.0714 | 2 | 0 | 4 | 0.25 | 3 | 0.3333 | 5 | 0.2 | 14 | 0.2143 |
| 8_1176867 | 9 | 0.1111 | 9 | 0 | 7 | 0.1429 | 11 | 0.0909 | 36 | 0.0833 | 4 | 0.25 | 4 | 0.25 | 3 | 0 | 6 | 0 | 17 | 0.1176 |
| 8_1176868 | 7 | 0 | 10 | 0.1 | 7 | 0.1429 | 14 | 0 | 38 | 0.0526 | 4 | 0 | 6 | 0.1667 | 4 | 0.25 | 6 | 0.1667 | 20 | 0.15 |
| 8_1176868 | 7 | 0.2857 | 7 | 0.2857 | 5 | 0.8 | 12 | 0.6667 | 31 | 0.5161 | 3 | 0.6667 | 5 | 0.6 | 4 | 0.5 | 3 | 0.6667 | 15 | 0.6 |
| 8_1176870 | 9 | 0 | 8 | 0 | 7 | 0.4286 | 12 | 0.3333 | 36 | 0.1944 | 5 | 0.2 | 6 | 0.1667 | 3 | 0 | 7 | 0.2857 | 21 | 0.1905 |
| 8_1176897 | 1 | 0 | NA | NA | 1 | 1 | 1 | 1 | 3 | 0.6667 | 1 | 1 | 1 | 0 | 2 | 0.5 | 1 | 0 | 5 | 0.4 |
| 8_1176897 | 1 | 0 | NA | NA | 1 | 0 | 1 | 1 | 3 | 0.3333 | 1 | 0 | 1 | 0 | 2 | 0.5 | 1 | 1 | 5 | 0.4 |
| 8_1176899 | 1 | 0 | NA | NA | 1 | 0 | 1 | 1 | 3 | 0.3333 | 1 | 0 | 1 | 0 | 2 | 0.5 | 1 | 0 | 5 | 0.2 |
| 8_1176969 | NA | NA | 1 | 1 | NA | NA | NA | NA | 1 | 1 | NA | NA | 2 | 1 | 1 | 0 | 2 | 1 | 5 | 0.8 |
| 8_1176970 | NA | NA | 1 | 1 | NA | NA | NA | NA | 1 | 1 | 1 | 1 | 2 | 1 | 1 | 1 | 2 | 0.5 | 6 | 0.8333 |
| 8_1176972 | NA | NA | 1 | 1 | NA | NA | NA | NA | 1 | 1 | 1 | 1 | 2 | 1 | 1 | 1 | 2 | 1 | 6 | 1 |
| 8_1176973 | NA | NA | 1 | 1 | NA | NA | NA | NA | 1 | 1 | NA | NA | 2 | 0.5 | NA | NA | 1 | 0 | 3 | 0.3333 |
| 8_1176975 | NA | NA | 1 | 1 | NA | NA | NA | NA | 1 | 1 | 1 | 1 | 1 | 1 | NA | NA | 1 | 1 | 3 | 1 |
| 8_1176975 | NA | NA | 1 | 1 | NA | NA | NA | NA | 1 | 1 | 1 | 1 | 2 | 1 | NA | NA | 3 | 1 | 6 | 1 |
| 8_1176978 | NA | NA | 1 | 1 | NA | NA | NA | NA | 1 | 1 | 1 | 1 | NA | NA | NA | NA | 3 | 1 | 4 | 1 |
| 8_1176980 | NA | NA | 1 | 1 | NA | NA | NA | NA | 1 | 1 | 2 | 1 | 1 | 1 | NA | NA | 3 | 1 | 6 | 1 |
| 8_1176982 | NA | NA | 1 | 1 | NA | NA | NA | NA | 1 | 1 | 2 | 1 | 1 | 1 | NA | NA | 3 | 1 | 6 | 1 |
| 8_1176983 | NA | NA | 1 | 1 | NA | NA | NA | NA | 1 | 1 | 2 | 1 | 1 | 1 | NA | NA | 3 | 1 | 6 | 1 |
| 8_1176987 | NA | NA | 1 | 1 | NA | NA | NA | NA | 1 | 1 | 2 | 0.5 | 1 | 0 | NA | NA | 2 | 0.5 | 5 | 0.4 |
| 8_1177089 | 10 | 0 | 9 | 0 | 14 | 0.0714 | 13 | 0.2308 | 46 | 0.087 | 17 | 0 | 15 | 0.2 | 10 | 0 | 12 | 0.1667 | 54 | 0.0926 |
| 8_1177092 | 8 | 0 | 9 | 0.1111 | 10 | 0.1 | 13 | 0 | 40 | 0.05 | 15 | 0.2667 | 14 | 0.2143 | 11 | 0.1818 | 10 | 0.1 | 50 | 0.2 |
| 8_1177092 | 9 | 0.3333 | 8 | 0.5 | 10 | 0.8 | 12 | 0.25 | 39 | 0.4615 | 17 | 0.5294 | 15 | 0.4 | 12 | 0.4167 | 13 | 0.3846 | 57 | 0.4386 |
| 8_1179421 | NA | NA | 2 | 0 | 3 | 0.3333 | NA | NA | 5 | 0.2 | 1 | 1 | NA | NA | 1 | 0 | 1 | 0 | 3 | 0.3333 |
| 8_1181635 | 2 | 0 | 6 | 0 | 3 | 0.3333 | 5 | 0.2 | 16 | 0.125 | 4 | 0 | 5 | 0 | 4 | 0.25 | 11 | 0.3636 | 24 | 0.2083 |
| 8_1181863 | 7 | 0 | 14 | 0.1429 | 7 | 0.1429 | 10 | 0 | 38 | 0.0789 | 7 | 0.1429 | 12 | 0.0833 | 9 | 0 | 15 | 0.0667 | 43 | 0.0698 |
| 8_1181864 | 1 | 0 | 13 | 0.6923 | 5 | 0.4 | 7 | 0.4286 | 26 | 0.5385 | 6 | 0.6667 | 8 | 0.25 | 8 | 0.625 | 10 | 0.5 | 32 | 0.5 |
| 8_1183889 | NA | NA | NA | NA | NA | NA | 1 | 1 | 1 | 1 | NA | NA | 1 | 1 | 1 | 0 | NA | NA | 2 | 0.5 |
| 8_1183891 | NA | NA | NA | NA | NA | NA | 1 | 1 | 1 | 1 | NA | NA | 1 | 1 | NA | NA | NA | NA | 1 | 1 |
| 8_1183895 | NA | NA | NA | NA | NA | NA | 1 | 1 | 1 | 1 | NA | NA | 1 | 0 | 1 | 1 | NA | NA | 2 | 0.5 |
| 8_1183896 | NA | NA | NA | NA | NA | NA | 1 | 1 | 1 | 1 | NA | NA | 1 | 1 | 1 | 0 | NA | NA | 2 | 0.5 |
| 8_1183903 | NA | NA | NA | NA | NA | NA | 1 | 1 | 1 | 1 | NA | NA | 1 | 0 | 1 | 1 | NA | NA | 2 | 0.5 |
| 8_1185185 | 7 | 0.2857 | 3 | 0.3333 | 6 | 0.1667 | 1 | 0 | 17 | 0.2353 | 5 | 0.2 | 1 | 0 | 2 | 0 | 1 | 0 | 9 | 0.1111 |
| 8_1185187 | 6 | 0.3333 | 3 | 0 | 5 | 0.2 | 1 | 0 | 15 | 0.2 | 4 | 0 | 2 | 0 | 2 | 0 | 1 | 1 | 9 | 0.1111 |
| 8_1185187 | 6 | 0.3333 | 3 | 0 | 5 | 0 | 1 | 0 | 15 | 0.1333 | 4 | 0 | 2 | 0.5 | 2 | 0 | 1 | 0 | 9 | 0.1111 |
| 8_1185188 | 7 | 0 | 3 | 0.3333 | 6 | 0 | 1 | 0 | 17 | 0.0588 | 4 | 0.25 | 1 | 0 | 2 | 0 | 1 | 0 | 8 | 0.125 |
| 8_1185189 | 7 | 0.1429 | 2 | 0 | 4 | 0.25 | 1 | 1 | 14 | 0.2143 | 4 | 0.5 | 1 | 1 | 2 | 0 | 1 | 0 | 8 | 0.375 |
| 8_1190113 | NA | NA | NA | NA | 1 | 1 | NA | NA | 1 | 1 | NA | NA | 1 | 1 | NA | NA | NA | NA | 1 | 1 |
| 8_1200392 | 5 | 0 | 7 | 0 | 3 | 0.3333 | 2 | 0 | 17 | 0.0588 | 5 | 0 | 4 | 0.5 | 7 | 0.1429 | 5 | 0.2 | 21 | 0.1905 |
| 8_1200408 | 2 | 0 | 3 | 0 | 1 | 1 | 1 | 0 | 7 | 0.1429 | 3 | 0 | 1 | 0 | 3 | 0 | 4 | 0.5 | 11 | 0.1818 |
| 8_1200502 | 4 | 0 | 3 | 0 | 5 | 0.2 | 2 | 0 | 14 | 0.0714 | 5 | 0.4 | 5 | 0 | 6 | 0.1667 | 7 | 0 | 23 | 0.1304 |
| 8_1200503 | 4 | 0.5 | 8 | 0.375 | 4 | 0 | 4 | 0 | 20 | 0.25 | 5 | 0 | 5 | 0.2 | 8 | 0.375 | 8 | 0.25 | 26 | 0.2308 |
| 8_1200503 | 5 | 0.2 | 9 | 0.1111 | 5 | 0.2 | 4 | 0.25 | 23 | 0.1739 | 4 | 0 | 7 | 0.4286 | 9 | 0 | 5 | 0.2 | 25 | 0.16 |
| 8_1200504 | 2 | 1 | 8 | 0.875 | 4 | 1 | 3 | 0.3333 | 17 | 0.8235 | 6 | 1 | 7 | 0.8571 | 10 | 0.9 | 9 | 0.7778 | 32 | 0.875 |
| 8_1200504 | 5 | 0.6 | 9 | 0.6667 | 5 | 0.8 | 4 | 1 | 23 | 0.7391 | 6 | 0.1667 | 6 | 0.3333 | 11 | 0.3636 | 9 | 0.5556 | 32 | 0.375 |
| 8_1200504 | 3 | 0.6667 | 7 | 0.4286 | 5 | 0.4 | 3 | 0.3333 | 18 | 0.4444 | 6 | 0.5 | 5 | 0.4 | 9 | 0.3333 | 8 | 0.375 | 28 | 0.3929 |
| 8_1200507 | 3 | 0 | 9 | 0.1111 | 3 | 0 | 6 | 0.3333 | 21 | 0.1429 | 7 | 0.2857 | 8 | 0.25 | 8 | 0.375 | 9 | 0 | 32 | 0.2188 |
| 8_1200508 | 4 | 0.25 | 9 | 0.2222 | 6 | 0 | 4 | 0.5 | 23 | 0.2174 | 7 | 0.1429 | 9 | 0.2222 | 7 | 0.1429 | 8 | 0.25 | 31 | 0.1935 |
| 8_1200516 | 4 | 0.25 | 2 | 0 | 3 | 0.3333 | 4 | 0.25 | 13 | 0.2308 | 2 | 0 | 4 | 0.25 | 7 | 0.1429 | 3 | 0.3333 | 16 | 0.1875 |
| 8_1200748 | 52 | 0.1154 | 58 | 0.0172 | 25 | 0 | 40 | 0.125 | 175 | 0.0686 | 41 | 0.0732 | 40 | 0.075 | 47 | 0.1277 | 40 | 0.125 | 168 | 0.1012 |
| 8_1200751 | 31 | 0.1613 | 44 | 0.2045 | 23 | 0.1304 | 35 | 0.0857 | 133 | 0.1504 | 31 | 0.0645 | 29 | 0.1724 | 32 | 0.1875 | 31 | 0.2258 | 123 | 0.1626 |
| 8_1200752 | 33 | 0.1212 | 47 | 0.234 | 25 | 0.32 | 36 | 0.1111 | 141 | 0.1915 | 31 | 0.1613 | 32 | 0.125 | 32 | 0.125 | 31 | 0.2258 | 126 | 0.1587 |
| 8_1201368 | 5 | 0.6 | 3 | 0.6667 | NA | NA | 5 | 1 | 13 | 0.7692 | 3 | 0.3333 | 2 | 0 | 3 | 0.6667 | 4 | 0.25 | 12 | 0.3333 |
| 8_1201368 | 4 | 0.5 | 3 | 0.3333 | NA | NA | 3 | 0.3333 | 10 | 0.4 | 4 | 0.25 | 2 | 0 | 2 | 0 | 4 | 0.25 | 12 | 0.1667 |
| 8_1201371 | 6 | 0 | 5 | 0.4 | NA | NA | 6 | 0.1667 | 17 | 0.1765 | 4 | 0.25 | 2 | 0.5 | 5 | 0.2 | 6 | 0 | 17 | 0.1765 |
| 8_1201372 | 7 | 0.2857 | 4 | 0 | NA | NA | 5 | 0.4 | 16 | 0.25 | 2 | 0 | 3 | 0 | 4 | 0.25 | 5 | 0.4 | 14 | 0.2143 |
| 8_1201372 | 4 | 0.75 | 3 | 0.6667 | NA | NA | 5 | 1 | 12 | 0.8333 | 4 | 0.75 | 3 | 0.6667 | 4 | 0.75 | 8 | 0.875 | 19 | 0.7895 |
| 8_1201374 | 7 | 1 | 3 | 1 | NA | NA | 4 | 1 | 14 | 1 | 3 | 1 | 3 | 1 | 5 | 1 | 5 | 1 | 16 | 1 |
| 8_1201375 | 7 | 0.5714 | 2 | 0 | NA | NA | 6 | 0.8333 | 15 | 0.6 | 5 | 0.2 | 5 | 0.4 | 5 | 0.8 | 9 | 0.4444 | 24 | 0.4583 |
| 8_1201375 | 8 | 0.125 | 5 | 0 | NA | NA | 8 | 0.125 | 21 | 0.0952 | 6 | 0.1667 | 6 | 0 | 4 | 0.25 | 9 | 0.2222 | 25 | 0.16 |
| 8_1201376 | 6 | 0.3333 | 5 | 0.4 | NA | NA | 7 | 0.7143 | 18 | 0.5 | 5 | 0.6 | 5 | 1 | 4 | 0.5 | 8 | 0.375 | 22 | 0.5909 |
| 8_1201376 | 7 | 0 | 5 | 0.2 | NA | NA | 6 | 0.3333 | 18 | 0.1667 | 5 | 0.4 | 6 | 0.1667 | 5 | 0 | 9 | 0.1111 | 25 | 0.16 |
| 8_1201377 | 8 | 0 | 2 | 0 | NA | NA | 7 | 0.1429 | 17 | 0.0588 | 6 | 0.3333 | 5 | 0.2 | 5 | 0 | 8 | 0 | 24 | 0.125 |

| | | | | | | | | | | | | | | | | | | | | |
|---|---|---|---|---|---|---|---|---|---|---|---|---|---|---|---|---|---|---|---|---|
| 8_1201989 | 29 | 0.2414 | 18 | 0.5556 | 26 | 0.2692 | 42 | 0.1905 | 115 | 0.2783 | 13 | 0.4615 | 31 | 0.3226 | 47 | 0.2979 | 23 | 0.2609 | 114 | 0.3158 |
| 8_1201990 | 29 | 0.1379 | 16 | 0.375 | 26 | 0.1538 | 41 | 0.2683 | 112 | 0.2232 | 15 | 0.2 | 32 | 0.25 | 44 | 0.2273 | 22 | 0.1364 | 113 | 0.2124 |
| 8_1202350 | 13 | 0.7692 | 10 | 0.9 | 11 | 0.7273 | 12 | 0.6667 | 46 | 0.7609 | 4 | 0.75 | 9 | 0.6667 | 8 | 0.75 | 2 | 1 | 23 | 0.7391 |
| 8_1202350 | 9 | 0.8889 | 5 | 0.8 | 4 | 1 | 8 | 1 | 26 | 0.9231 | 3 | 0.6667 | 8 | 0.625 | 5 | 1 | 1 | 1 | 17 | 0.7647 |
| 8_1202350 | 14 | 0 | 9 | 0.1111 | 11 | 0.2727 | 12 | 0.0833 | 46 | 0.1087 | 4 | 0.25 | 9 | 0.1111 | 9 | 0.2222 | 2 | 0 | 24 | 0.1667 |
| 8_1202351 | 12 | 1 | 9 | 1 | 11 | 1 | 13 | 0.8462 | 45 | 0.9556 | 4 | 1 | 8 | 0.625 | 8 | 0.875 | 1 | 1 | 21 | 0.8095 |
| 8_1202352 | 13 | 0.3846 | 9 | 0.2222 | 9 | 0.2222 | 15 | 0.0667 | 46 | 0.2174 | 4 | 0 | 10 | 0.3 | 8 | 0.5 | 1 | 0 | 23 | 0.3043 |
| 8_1202352 | 13 | 0.9231 | 9 | 0.8889 | 9 | 1 | 14 | 0.9286 | 45 | 0.9333 | 4 | 0.75 | 10 | 0.7 | 8 | 0.75 | 1 | 1 | 23 | 0.7391 |
| 8_1202352 | 11 | 0.6364 | 8 | 0.125 | 9 | 0.2222 | 12 | 0.1667 | 40 | 0.3 | 4 | 0.75 | 8 | 0.125 | 8 | 0.25 | 1 | 1 | 21 | 0.3333 |
| 8_1202352 | 11 | 0.6364 | 9 | 0.6667 | 8 | 0.5 | 12 | 0.5833 | 40 | 0.6 | 3 | 0 | 10 | 0.5 | 7 | 0.7143 | 1 | 1 | 21 | 0.5238 |
| 8_1202353 | 12 | 0.1667 | 9 | 0 | 8 | 0.625 | 14 | 0.2143 | 43 | 0.2326 | 3 | 0 | 9 | 0.1111 | 8 | 0 | 1 | 1 | 21 | 0.0952 |
| 8_1202354 | 10 | 0.3 | 9 | 0.1111 | 10 | 0.3 | 16 | 0.0625 | 45 | 0.1778 | 4 | 0.5 | 11 | 0.0909 | 6 | 0.1667 | 1 | 0 | 22 | 0.1818 |
| 8_1202355 | 9 | 0.2222 | 9 | 0.2222 | 10 | 0.2 | 14 | 0.2143 | 42 | 0.2143 | 4 | 0.5 | 8 | 0.125 | 6 | 0.1667 | 1 | 1 | 19 | 0.2632 |
| 8_1202743 | 3 | 0.6667 | 1 | 0 | 1 | 0 | 4 | 0.25 | 9 | 0.3333 | 1 | 1 | NA | NA | 3 | 0.3333 | NA | NA | 4 | 0.5 |
| 8_1202744 | 4 | 0.5 | 1 | 0 | 1 | 0 | 4 | 0.5 | 10 | 0.4 | 1 | 1 | NA | NA | 2 | 0.5 | NA | NA | 3 | 0.6667 |
| 8_1202744 | 5 | 1 | 2 | 1 | 1 | 1 | 4 | 1 | 12 | 1 | NA | NA | NA | NA | 2 | 0.5 | NA | NA | 2 | 0.5 |
| 8_1202744 | 5 | 0.4 | 2 | 0.5 | 1 | 0 | 5 | 0.4 | 13 | 0.3846 | 1 | 1 | NA | NA | 2 | 0.5 | NA | NA | 3 | 0.6667 |
| 8_1202745 | 5 | 0.2 | 2 | 1 | 1 | 1 | 5 | 0.6 | 13 | 0.5385 | 1 | 0 | NA | NA | 2 | 0.5 | NA | NA | 3 | 0.3333 |
| 8_1202887 | 9 | 0 | 7 | 0.1429 | 11 | 0.2727 | 7 | 0.1429 | 34 | 0.1471 | 8 | 0 | 5 | 0 | 10 | 0.2 | 3 | 0 | 26 | 0.0769 |
| 8_1202888 | 8 | 0.375 | 7 | 0.8571 | 9 | 0.3333 | 7 | 0.4286 | 31 | 0.4839 | 7 | 0.2857 | 5 | 0.2 | 9 | 0.3333 | 2 | 0.5 | 23 | 0.3043 |
| 8_1202889 | 8 | 0.125 | 5 | 0 | 11 | 0.1818 | 8 | 0 | 32 | 0.0938 | 8 | 0.125 | 5 | 0.2 | 11 | 0.0909 | 3 | 0 | 27 | 0.1111 |
| 8_1202889 | 8 | 0.25 | 4 | 0.25 | 9 | 0.2222 | 8 | 0 | 29 | 0.1724 | 8 | 0.625 | 6 | 0.5 | 9 | 0.5556 | 3 | 0 | 26 | 0.5 |
| 8_1202890 | 7 | 0.4286 | 5 | 0.4 | 8 | 0.125 | 8 | 0.25 | 28 | 0.2857 | 7 | 0 | 5 | 0.6 | 8 | 0.25 | 2 | 0 | 22 | 0.2273 |
| 8_1202890 | 7 | 0.2857 | 4 | 0.25 | 11 | 0 | 6 | 0.5 | 28 | 0.2143 | 6 | 0 | 5 | 0.4 | 8 | 0.125 | 3 | 0 | 22 | 0.1364 |
| 8_1202891 | 8 | 0.375 | 6 | 0.1667 | 11 | 0.1818 | 7 | 0.1429 | 32 | 0.2188 | 6 | 0.1667 | 6 | 0.3333 | 9 | 0.1111 | 3 | 0.3333 | 24 | 0.2083 |
| 8_1202894 | 5 | 0.2 | 5 | 0 | 10 | 0.1 | 7 | 0.2857 | 27 | 0.1481 | 5 | 0.4 | 3 | 0.6667 | 9 | 0.3333 | 2 | 0.5 | 19 | 0.4211 |
| 8_1202900 | 5 | 0.4 | 5 | 0.2 | 7 | 0.1429 | 6 | 0 | 23 | 0.1739 | 5 | 0.2 | 4 | 0.5 | 6 | 0.1667 | 2 | 0 | 17 | 0.2353 |
| 8_1202902 | 3 | 0 | 4 | 0.25 | 6 | 0 | 5 | 0.2 | 18 | 0.1111 | 3 | 0.6667 | 3 | 0 | 5 | 0.2 | 2 | 0.5 | 13 | 0.3077 |
| 8_1202903 | 4 | 0 | 4 | 0.75 | 6 | 0.5 | 4 | 0.25 | 18 | 0.3889 | 4 | 0.5 | 3 | 0 | 5 | 0.2 | 3 | 0.6667 | 15 | 0.3333 |
| 8_1202905 | 5 | 0.6 | 5 | 0.4 | 5 | 0.8 | 3 | 1 | 18 | 0.6667 | 3 | 1 | 5 | 0.4 | 2 | 1 | 3 | 1 | 13 | 0.7692 |
| 8_1202906 | 1 | 1 | 5 | 0.8 | 3 | 0.6667 | 4 | 0.5 | 13 | 0.6923 | 3 | 1 | 2 | 1 | 3 | 1 | 2 | 0 | 10 | 0.8 |
| 8_1202907 | 3 | 0.3333 | 5 | 0.2 | 5 | 0 | 4 | 0.25 | 17 | 0.1765 | 4 | 0 | 4 | 0.25 | 3 | 0 | 3 | 0 | 14 | 0.0714 |
| 8_1202930 | 2 | 0.5 | 2 | 0.5 | 3 | 1 | 3 | 0 | 10 | 0.5 | 3 | 0.3333 | 3 | 0.3333 | 1 | 0 | 3 | 0.3333 | 10 | 0.3 |
| 8_1202930 | 2 | 0.5 | 2 | 0 | 3 | 0 | 4 | 0 | 11 | 0.0909 | 2 | 0 | 4 | 0.25 | 1 | 1 | 3 | 0 | 10 | 0.2 |
| 8_1203247 | 2 | 1 | 2 | 1 | 2 | 1 | 1 | 1 | 7 | 1 | NA | NA | NA | NA | 2 | 1 | 1 | 1 | 3 | 1 |
| 8_1203247 | 3 | 0.6667 | 3 | 1 | 2 | 1 | 1 | 1 | 9 | 0.8889 | NA | NA | NA | NA | 2 | 0.5 | 1 | 1 | 3 | 0.6667 |
| 8_1203252 | 2 | 1 | 3 | 0.3333 | 1 | 1 | NA | NA | 6 | 0.6667 | NA | NA | 1 | 0 | 1 | 0 | 1 | 1 | 3 | 0.3333 |
| 8_1203253 | 2 | 0.5 | 3 | 0.6667 | 1 | 0 | 1 | 1 | 7 | 0.5714 | NA | NA | NA | NA | 1 | 0 | 1 | 1 | 2 | 0.5 |
| 8_1203254 | 1 | 0 | 3 | 0.3333 | 1 | 0 | 1 | 0 | 6 | 0.1667 | NA | NA | NA | NA | 1 | 0 | 1 | 1 | 2 | 0.5 |
| 8_1203254 | 2 | 0.5 | 3 | 0.6667 | 1 | 1 | 1 | 0 | 7 | 0.5714 | NA | NA | NA | NA | 1 | 0 | 1 | 1 | 2 | 0.5 |
| 8_1203254 | 2 | 1 | 3 | 1 | 1 | 1 | 1 | 1 | 7 | 1 | NA | NA | NA | NA | 1 | 1 | 1 | 1 | 2 | 1 |
| 8_1203254 | 2 | 0 | 3 | 1 | 1 | 1 | 1 | 1 | 7 | 0.7143 | NA | NA | 1 | 1 | 1 | 0 | 1 | 1 | 3 | 0.6667 |
| 8_1203256 | 2 | 1 | 2 | 1 | NA | NA | 1 | 1 | 5 | 1 | NA | NA | 1 | 1 | NA | NA | 1 | 1 | 2 | 1 |
| 8_1203257 | 2 | 1 | 3 | 1 | 1 | 1 | 1 | 1 | 7 | 1 | NA | NA | 1 | 1 | NA | NA | 1 | 1 | 2 | 1 |
| 8_1203258 | 1 | 1 | 3 | 0 | 1 | 0 | 1 | 0 | 6 | 0.1667 | NA | NA | NA | NA | NA | NA | 1 | 1 | 1 | 1 |
| 8_1203259 | 2 | 0 | 3 | 0 | 1 | 1 | 1 | 0 | 7 | 0.1429 | NA | NA | 1 | 1 | NA | NA | 1 | 0 | 2 | 0.5 |
| 8_1204146 | 3 | 0.3333 | 4 | 0.5 | 3 | 0 | 4 | 0.25 | 14 | 0.2857 | 4 | 0.25 | 2 | 0.5 | 2 | 0.5 | 2 | 0 | 10 | 0.3 |
| 8_1204869 | 2 | 0 | 3 | 0.3333 | NA | NA | 1 | 0 | 6 | 0.1667 | 1 | 0 | 3 | 0.3333 | 3 | 0.3333 | NA | NA | 7 | 0.2857 |
| 8_1204882 | 1 | 0 | 1 | 0 | NA | NA | 1 | 1 | 3 | 0.3333 | NA | NA | 3 | 0.6667 | 2 | 1 | NA | NA | 5 | 0.8 |
| 8_1204884 | 1 | 1 | 2 | 0.5 | NA | NA | 1 | 1 | 4 | 0.75 | NA | NA | 2 | 0 | 2 | 1 | NA | NA | 4 | 0.5 |
| 8_1205364 | 8 | 0 | 17 | 0.1176 | 2 | 0 | 9 | 0.3333 | 36 | 0.1389 | 7 | 0.2857 | 9 | 0 | 10 | 0.1 | 13 | 0.2308 | 39 | 0.1538 |
| 8_1205367 | 7 | 0.1429 | 17 | 0.3529 | 2 | 0.5 | 6 | 0.1667 | 32 | 0.2812 | 7 | 0.4286 | 8 | 0.125 | 9 | 0.2222 | 10 | 0.2 | 34 | 0.2353 |
| 8_1205970 | 1 | 1 | NA | NA | NA | NA | 1 | 0 | 2 | 0.5 | 1 | 1 | 1 | 0 | NA | NA | 1 | 1 | 3 | 0.6667 |
| 8_1205973 | 1 | 1 | NA | NA | NA | NA | NA | NA | 1 | 1 | 2 | 0.5 | 1 | 1 | NA | NA | 1 | 0 | 4 | 0.5 |
| 8_1206030 | 1 | 0 | NA | NA | NA | NA | 1 | 1 | 2 | 0.5 | 2 | 1 | 1 | 0 | NA | NA | 1 | 0 | 4 | 0.5 |
| 8_1206033 | NA | NA | NA | NA | NA | NA | 1 | 1 | 1 | 1 | 2 | 1 | 1 | 1 | NA | NA | 1 | 1 | 4 | 1 |
| 8_1206055 | NA | NA | 2 | 0.5 | NA | NA | NA | NA | 2 | 0.5 | 2 | 1 | NA | NA | NA | NA | NA | NA | 2 | 1 |
| 8_1206077 | NA | NA | 2 | 0.5 | NA | NA | 2 | 0 | 4 | 0.25 | 2 | 0 | 1 | 0 | NA | NA | 2 | 0.5 | 5 | 0.2 |
| 8_1206682 | 3 | 1 | 2 | 0.5 | 1 | 1 | 1 | 0 | 7 | 0.7143 | 1 | 1 | 2 | 0.5 | NA | NA | 4 | 1 | 7 | 0.8571 |
| 8_1206683 | 4 | 1 | 2 | 1 | 1 | 1 | 1 | 0 | 8 | 0.875 | 1 | 1 | 2 | 1 | 1 | 1 | 5 | 1 | 9 | 1 |
| 8_1206683 | 4 | 0.75 | 2 | 1 | 1 | 1 | 1 | 1 | 8 | 0.875 | 1 | 1 | 2 | 1 | 1 | 1 | 4 | 1 | 8 | 1 |
| 8_1206683 | 2 | 0.5 | 3 | 1 | NA | NA | NA | NA | 5 | 0.8 | NA | NA | 3 | 1 | 1 | 1 | 4 | 0.5 | 8 | 0.75 |
| 8_1206684 | 4 | 0.5 | 4 | 0.25 | 1 | 0 | 1 | 1 | 10 | 0.4 | NA | NA | 2 | 0.5 | 1 | 0 | 4 | 0.5 | 7 | 0.4286 |
| 8_1206684 | 4 | 0.5 | 4 | 0.75 | NA | NA | 1 | 1 | 9 | 0.6667 | 1 | 0 | 3 | 0.6667 | 1 | 1 | 6 | 0.5 | 11 | 0.5455 |
| 8_1206685 | 4 | 0.5 | 4 | 0.75 | 1 | 0 | NA | NA | 9 | 0.5556 | 1 | 1 | 3 | 0.6667 | 1 | 1 | 4 | 0.25 | 9 | 0.5556 |

| | | | | | | | | | | | | | | | | | | | | |
|---|---|---|---|---|---|---|---|---|---|---|---|---|---|---|---|---|---|---|---|---|
| 8_1206685( | 3 | 1 | 3 | 0.6667 | 1 | 0 | 1 | 1 | 8 | 0.75 | 1 | 1 | 3 | 0.3333 | NA | NA | 4 | 0.75 | 8 | 0.625 |
| 8_1206685! | 3 | 1 | 1 | 0 | NA | NA | 1 | 0 | 5 | 0.6 | 1 | 0 | 3 | 0.6667 | 1 | 0 | 6 | 0.1667 | 11 | 0.2727 |
| 8_1206686: | 4 | 0.75 | 3 | 0.6667 | 1 | 0 | 1 | 1 | 9 | 0.6667 | 1 | 1 | 3 | 0.6667 | 1 | 1 | 6 | 0.8333 | 11 | 0.8182 |
| 8_1206687( | 4 | 0.25 | 4 | 0 | 1 | 0 | 1 | 0 | 10 | 0.1 | 1 | 0 | 3 | 0 | 1 | 0 | 5 | 0.2 | 10 | 0.1 |
| 8_1206835! | 7 | 0.5714 | 9 | 0.4444 | 3 | 0.6667 | 4 | 0.25 | 23 | 0.4783 | 4 | 0.5 | 3 | 0.3333 | 7 | 0.7143 | 7 | 0.5714 | 21 | 0.5714 |
| 8_1206836( | 7 | 0.2857 | 4 | 0.25 | 2 | 0 | 3 | 0 | 16 | 0.1875 | 4 | 0 | 3 | 0 | 8 | 0.25 | 6 | 0.3333 | 21 | 0.1905 |
| 8_1206836( | 7 | 0.1429 | 6 | 0.3333 | 3 | 0 | 2 | 0.5 | 18 | 0.2222 | 4 | 0.25 | 4 | 0.25 | 7 | 0 | 8 | 0.125 | 23 | 0.1304 |
| 8_1206836! | 8 | 1 | 7 | 1 | 2 | 0.5 | 1 | 1 | 18 | 0.9444 | 4 | 1 | 4 | 1 | 6 | 1 | 6 | 0.8333 | 20 | 0.95 |
| 8_1206841: | 7 | 0.1429 | 8 | 0.125 | 6 | 0 | 2 | 0 | 23 | 0.087 | 3 | 0 | 4 | 0 | 7 | 0.1429 | 6 | 0.3333 | 20 | 0.15 |
| 8_1206841( | 6 | 1 | 8 | 0.875 | 5 | 0.8 | 3 | 1 | 22 | 0.9091 | 6 | 0.6667 | 5 | 0.8 | 7 | 0.8571 | 7 | 1 | 25 | 0.84 |
| 8_1208080( | 7 | 0.1429 | 7 | 0.1429 | 6 | 0.1667 | 4 | 0 | 24 | 0.125 | 4 | 0 | 4 | 0.25 | 4 | 0 | 4 | 0 | 16 | 0.0625 |
| 8_1208080( | 9 | 0.3333 | 8 | 0.125 | 6 | 0 | 4 | 0 | 27 | 0.1481 | 4 | 0.25 | 4 | 0 | 3 | 0.6667 | 5 | 0 | 16 | 0.1875 |
| 8_1208081: | 7 | 0.5714 | 5 | 0.4 | 5 | 0.8 | 3 | 0.3333 | 20 | 0.55 | 3 | 0.3333 | 4 | 0.5 | 4 | 0.5 | 3 | 0.6667 | 14 | 0.5 |
| 8_1208081: | 9 | 0 | 5 | 0.4 | 4 | 0 | 2 | 0 | 20 | 0.1 | 2 | 0 | 4 | 0.5 | 4 | 0 | 5 | 0 | 15 | 0.1333 |
| 8_1208082' | 9 | 0.5556 | 8 | 0.5 | 6 | 0.6667 | 4 | 0.75 | 27 | 0.5926 | 4 | 0.5 | 4 | 1 | 5 | 1 | 5 | 0.8 | 18 | 0.8333 |
| 8_1208083: | 8 | 0.625 | 7 | 0.8571 | 5 | 0.4 | 3 | 0 | 23 | 0.5652 | 4 | 0.75 | 4 | 0.75 | 3 | 0.6667 | 5 | 0.6 | 16 | 0.6875 |
| 8_1208086: | 7 | 0.1429 | 7 | 0.1429 | 4 | 0 | 1 | 0 | 19 | 0.1053 | 4 | 0.25 | 4 | 0 | 4 | 0 | 5 | 0 | 17 | 0.0588 |
| 8_1208087: | 7 | 0 | 8 | 0.125 | 3 | 0.3333 | 3 | 0 | 21 | 0.0952 | 2 | 0 | 1 | 0 | 4 | 0.25 | 4 | 0 | 11 | 0.0909 |
| 8_1208088! | 5 | 0.6 | 2 | 0.5 | 2 | 0.5 | 2 | 0.5 | 11 | 0.5455 | 1 | 0 | 2 | 1 | 3 | 0 | 3 | 0.3333 | 9 | 0.3333 |
| 8_1208089' | 5 | 0.6 | 3 | 0.6667 | 1 | 0 | 1 | 1 | 10 | 0.6 | NA | NA | 3 | 1 | 4 | 0.5 | 3 | 0.3333 | 10 | 0.6 |
| 8_1208089: | 6 | 1 | 7 | 1 | 3 | 1 | 4 | 1 | 20 | 1 | 2 | 1 | 3 | 0.6667 | 5 | 1 | 5 | 1 | 15 | 0.9333 |
| 8_1208091( | 5 | 1 | 5 | 1 | NA | NA | 2 | 1 | 12 | 1 | 2 | 0.5 | 3 | 1 | 5 | 1 | 4 | 1 | 14 | 0.9286 |
| 8_1208092! | 6 | 0.5 | 5 | 0 | 2 | 0 | 2 | 0.5 | 15 | 0.2667 | 2 | 0 | 3 | 0 | 3 | 0 | 4 | 0.75 | 12 | 0.25 |
| 8_1208093' | 6 | 0.5 | 5 | 0 | 2 | 0 | 3 | 0 | 16 | 0.1875 | 2 | 0.5 | NA | NA | 3 | 0 | 4 | 0.5 | 9 | 0.3333 |
| 8_1208145! | 4 | 0.5 | 4 | 0.5 | 1 | 0 | 1 | 0 | 10 | 0.4 | 1 | 0 | 1 | 0 | 2 | 0 | 4 | 0.25 | 8 | 0.125 |
| 8_1208719: | 4 | 0.25 | 4 | 0 | 3 | 0 | 2 | 0.5 | 13 | 0.1538 | 2 | 0 | NA | NA | 4 | 0.25 | 1 | 0 | 7 | 0.1429 |
| 8_1208721' | 5 | 0.6 | 3 | 0.6667 | 4 | 1 | 2 | 0.5 | 14 | 0.7143 | 3 | 0.3333 | NA | NA | 5 | 0.6 | 2 | 1 | 10 | 0.6 |
| 8_1208721' | 6 | 0.5 | 4 | 0.5 | 4 | 1 | 2 | 0 | 16 | 0.5625 | 3 | 0.3333 | NA | NA | 5 | 0.8 | 2 | 0.5 | 10 | 0.6 |
| 8_1208880! | 11 | 0.5455 | 5 | 0 | 9 | 0.2222 | 4 | 0 | 29 | 0.2759 | 5 | 0.4 | 4 | 0.5 | 6 | 0.3333 | 6 | 0.1667 | 21 | 0.3333 |
| 8_1208881' | 11 | 0.8182 | 8 | 1 | 9 | 0.6667 | 3 | 0.6667 | 31 | 0.8065 | 4 | 0.75 | 4 | 1 | 6 | 0.5 | 5 | 0.8 | 19 | 0.7368 |
| 8_1208881! | 12 | 0.3333 | 9 | 0.4444 | 8 | 0.25 | 4 | 0 | 33 | 0.303 | 4 | 0 | 4 | 0.25 | 6 | 0.6667 | 6 | 0 | 20 | 0.25 |
| 8_1208882: | 11 | 0.3636 | 5 | 0.2 | 7 | 0 | 6 | 0 | 29 | 0.1724 | 1 | 0 | 6 | 0.5 | 4 | 0.25 | 3 | 0.3333 | 14 | 0.3571 |
| 8_1208883( | 13 | 0.3846 | 9 | 0.5556 | 9 | 0.5556 | 6 | 0.6667 | 37 | 0.5135 | 5 | 0.8 | 5 | 0.4 | 5 | 0.4 | 6 | 0.5 | 21 | 0.5238 |
| 8_1208883! | 10 | 0.7 | 5 | 1 | 7 | 0.7143 | 5 | 0.8 | 27 | 0.7778 | 4 | 1 | 4 | 1 | 4 | 0.5 | 7 | 0.8571 | 19 | 0.8421 |
| 8_1208884( | 13 | 0.3077 | 10 | 0.3 | 8 | 0.25 | 5 | 0.4 | 36 | 0.3056 | 6 | 0.1667 | 5 | 0.6 | 6 | 0.6667 | 6 | 0.5 | 23 | 0.4783 |
| 8_1208884: | 12 | 0.5833 | 10 | 0.5 | 9 | 0.3333 | 7 | 0.7143 | 38 | 0.5263 | 6 | 0.6667 | 6 | 0.5 | 8 | 0.75 | 7 | 0.4286 | 27 | 0.5926 |
| 8_1208884! | 12 | 0.5 | 6 | 1 | 9 | 0.8889 | 5 | 0.8 | 32 | 0.75 | 6 | 0.8333 | 5 | 1 | 7 | 0.8571 | 7 | 0.8571 | 25 | 0.88 |
| 8_1208884! | 14 | 0.8571 | 9 | 0.8889 | 6 | 1 | 4 | 0.75 | 33 | 0.8788 | 7 | 1 | 5 | 1 | 8 | 0.75 | 6 | 1 | 26 | 0.9231 |
| 8_1208885( | 13 | 0.2308 | 10 | 0.1 | 7 | 0.1429 | 8 | 0.5 | 38 | 0.2368 | 7 | 0.4286 | 5 | 0.2 | 8 | 0.5 | 6 | 0.5 | 26 | 0.4231 |
| 8_1208885! | 12 | 0.75 | 9 | 0.7778 | 7 | 0.8571 | 7 | 0.4286 | 35 | 0.7143 | 6 | 0.6667 | 6 | 0.5 | 7 | 0.7143 | 6 | 1 | 25 | 0.72 |
| 8_1208887' | 13 | 0 | 9 | 0 | 8 | 0.375 | 8 | 0.5 | 38 | 0.1842 | 6 | 0.1667 | 5 | 0.2 | 9 | 0.1111 | 4 | 0.25 | 24 | 0.1667 |
| 8_1208887: | 11 | 1 | 9 | 0.6667 | 8 | 0.625 | 8 | 0.5 | 36 | 0.7222 | 7 | 0.4286 | 4 | 0.75 | 5 | 1 | 5 | 0.6 | 21 | 0.6667 |
| 8_1208887! | 14 | 0.0714 | 11 | 0.1818 | 10 | 0 | 9 | 0 | 44 | 0.0682 | 7 | 0 | 4 | 0 | 9 | 0 | 5 | 0.4 | 25 | 0.08 |
| 8_1208888' | 17 | 0.3529 | 11 | 0.1818 | 9 | 0.2222 | 8 | 0.125 | 45 | 0.2444 | 7 | 0.2857 | 5 | 0.2 | 9 | 0.2222 | 6 | 0 | 27 | 0.1852 |
| 8_1208889: | 15 | 0.2667 | 10 | 0.5 | 12 | 0.3333 | 7 | 0.1429 | 44 | 0.3182 | 12 | 0.1667 | 5 | 0.2 | 8 | 0.375 | 5 | 0.2 | 30 | 0.2333 |
| 8_1208889: | 10 | 0.9 | 3 | 0.3333 | 7 | 1 | 1 | 1 | 21 | 0.8571 | 2 | 0.5 | 6 | 0.8333 | 4 | 1 | 1 | 0 | 13 | 0.7692 |
| 8_1208893( | 16 | 0.1875 | 11 | 0.1818 | 10 | 0.3 | 8 | 0.375 | 45 | 0.2444 | 11 | 0 | 7 | 0.1429 | 10 | 0.1 | 4 | 0.5 | 32 | 0.125 |
| 8_1208894: | 13 | 0.6154 | 8 | 0.5 | 6 | 1 | 7 | 0.7143 | 34 | 0.6765 | 11 | 0.7273 | 7 | 0.5714 | 10 | 0.6 | 4 | 0.5 | 32 | 0.625 |
| 8_1208895! | 14 | 0.8571 | 7 | 0.8571 | 9 | 0.8889 | 7 | 1 | 37 | 0.8919 | 11 | 0.7273 | 6 | 0.8333 | 8 | 0.625 | 5 | 0.8 | 30 | 0.7333 |
| 8_1208895( | 14 | 0.5 | 11 | 0.7273 | 11 | 0.8182 | 9 | 0.8889 | 45 | 0.7111 | 12 | 0.5 | 8 | 0.625 | 10 | 0.7 | 5 | 0.8 | 35 | 0.6286 |
| 8_1208895: | 14 | 0.7857 | 10 | 1 | 10 | 1 | 8 | 0.875 | 42 | 0.9048 | 11 | 0.7273 | 6 | 0.6667 | 7 | 0.8571 | 2 | 0.5 | 26 | 0.7308 |
| 8_1208902! | 15 | 0.1333 | 10 | 0 | 14 | 0 | 10 | 0.1 | 49 | 0.0612 | 12 | 0 | 10 | 0 | 11 | 0.0909 | 5 | 0.2 | 38 | 0.0526 |
| 8_1208903! | 16 | 0.3125 | 11 | 0 | 14 | 0.2143 | 14 | 0.0714 | 55 | 0.1636 | 16 | 0.1875 | 10 | 0 | 12 | 0.0833 | 5 | 0.4 | 43 | 0.1395 |
| 8_1208904' | 19 | 0.5263 | 12 | 0.5 | 12 | 0.75 | 12 | 0.5833 | 55 | 0.5818 | 15 | 0.7333 | 10 | 0.6 | 12 | 0.6667 | 5 | 0.8 | 42 | 0.6905 |
| 8_1212132' | NA | NA | NA | NA | NA | NA | 1 | 1 | 1 | 1 | 1 | 1 | NA | NA | NA | NA | NA | NA | 1 | 1 |
| 8_1212157: | NA | NA | 1 | 1 | NA | NA | NA | NA | 1 | 1 | 1 | 1 | 1 | 0 | NA | NA | NA | NA | 2 | 0.5 |
| 8_1212975! | 2 | 1 | 1 | 1 | NA | NA | NA | NA | 3 | 1 | 2 | 1 | 2 | 1 | 3 | 1 | 3 | 1 | 10 | 1 |
| 8_1212978: | 2 | 0.5 | 1 | 0 | NA | NA | NA | NA | 3 | 0.3333 | 3 | 0.6667 | 3 | 0.3333 | 3 | 0 | 3 | 0.6667 | 12 | 0.4167 |
| 8_1212978: | 2 | 1 | 1 | 1 | NA | NA | NA | NA | 3 | 1 | 1 | 1 | 3 | 0.6667 | 3 | 1 | 3 | 1 | 10 | 0.9 |
| 8_1213000! | NA | NA | 1 | 1 | NA | NA | NA | NA | 1 | 1 | NA | NA | 2 | 1 | NA | NA | NA | NA | 2 | 1 |
| 8_1213013( | NA | NA | 1 | 1 | NA | NA | NA | NA | 1 | 1 | NA | NA | 2 | 1 | NA | NA | NA | NA | 2 | 1 |
| 8_1213015( | NA | NA | 1 | 1 | NA | NA | NA | NA | 1 | 1 | NA | NA | 2 | 1 | NA | NA | 1 | 1 | 3 | 1 |
| 8_1213019' | 2 | 0.5 | 2 | 1 | 2 | 0 | 1 | 0 | 7 | 0.4286 | 2 | 0.5 | 2 | 0.5 | 3 | 0.3333 | 4 | 0 | 11 | 0.2727 |
| 8_1213031: | 2 | 0.5 | 4 | 0.75 | 1 | 1 | 2 | 0 | 9 | 0.5556 | 3 | 0 | 4 | 0.5 | 1 | 0 | 4 | 0.5 | 12 | 0.3333 |
| 8_1213031! | 2 | 0 | 5 | 0 | 2 | 0.5 | 2 | 0.5 | 11 | 0.1818 | 2 | 0 | 4 | 0 | 2 | 0.5 | 4 | 0.25 | 12 | 0.1667 |

| | | | | | | | | | | | | | | | | | | | | |
|---|---|---|---|---|---|---|---|---|---|---|---|---|---|---|---|---|---|---|---|---|
| 8_12130536 | 2 | 0 | 3 | 0.3333 | 2 | 0 | 1 | 0 | 8 | 0.125 | 2 | 1 | 3 | 0.6667 | 3 | 0.3333 | 5 | 0.6 | 13 | 0.6154 |
| 8_12130553 | 3 | 1 | 4 | 1 | 3 | 1 | 3 | 1 | 13 | 1 | 2 | 1 | 4 | 1 | 4 | 1 | 6 | 1 | 16 | 1 |
| 8_12130573 | 3 | 1 | 4 | 0.5 | 2 | 1 | 3 | 1 | 12 | 0.8333 | 3 | 1 | 4 | 1 | 4 | 1 | 7 | 1 | 18 | 1 |
| 8_12130586 | 1 | 0 | 3 | 0.6667 | 1 | 0 | 2 | 0 | 7 | 0.2857 | 2 | 1 | 3 | 0.3333 | 2 | 0 | 6 | 0 | 13 | 0.2308 |
| 8_12130636 | 1 | 0 | 6 | 0.3333 | 5 | 0.2 | 3 | 0.6667 | 15 | 0.3333 | 2 | 0 | 5 | 0.4 | 4 | 0 | 8 | 0.125 | 19 | 0.1579 |
| 8_12130651 | 3 | 0.3333 | 6 | 0 | 4 | 0 | 2 | 0 | 15 | 0.0667 | 2 | 0.5 | 4 | 0 | 3 | 0 | 5 | 0 | 14 | 0.0714 |
| 8_12130753 | 6 | 0 | 11 | 0.0909 | 7 | 0.1429 | 2 | 0 | 26 | 0.0769 | 4 | 0 | 7 | 0 | 5 | 0.2 | 7 | 0.1429 | 23 | 0.087 |
| 8_12133538 | 4 | 0 | 3 | 0.3333 | 2 | 0 | 4 | 0 | 13 | 0.0769 | 2 | 0.5 | 5 | 0 | 2 | 0 | 2 | 0 | 11 | 0.0909 |
| 8_12133543 | 5 | 1 | 3 | 0.6667 | 2 | 1 | 4 | 1 | 14 | 0.9286 | 2 | 1 | 6 | 0.5 | 2 | 0.5 | 2 | 1 | 12 | 0.6667 |
| 8_12133546 | 4 | 0.5 | 2 | 0.5 | 2 | 0.5 | 2 | 1 | 10 | 0.6 | 2 | 0.5 | 3 | 0.3333 | 1 | 1 | 2 | 1 | 8 | 0.625 |
| 8_12138431 | 2 | 0.5 | 2 | 1 | 1 | 0 | NA | NA | 5 | 0.6 | NA | NA | 1 | 1 | 1 | 1 | 1 | 1 | 3 | 1 |
| 8_12138460 | 5 | 0 | 6 | 0.3333 | 1 | 0 | 3 | 0 | 15 | 0.1333 | 4 | 0 | 2 | 0 | 2 | 0.5 | 2 | 0 | 10 | 0.1 |
| 8_12138469 | 6 | 1 | 8 | 0.875 | 4 | 1 | 5 | 1 | 23 | 0.9565 | 7 | 0.8571 | 2 | 1 | 4 | 1 | 2 | 1 | 15 | 0.9333 |
| 8_12138482 | 5 | 0.4 | 8 | 0.25 | 6 | 0.3333 | 5 | 0.4 | 24 | 0.3333 | 6 | 0.1667 | 2 | 0 | 4 | 0 | 2 | 0.5 | 14 | 0.1429 |
| 8_12138486 | 6 | 0 | 4 | 0 | 5 | 0 | 4 | 0.25 | 19 | 0.0526 | 7 | 0 | 1 | 0 | 2 | 0.5 | 2 | 0 | 12 | 0.0833 |
| 8_12138533 | 6 | 0.1667 | 6 | 0 | 4 | 0 | 4 | 0 | 20 | 0.05 | 4 | 0 | 1 | 0 | 4 | 0 | 2 | 0.5 | 11 | 0.0909 |
| 8_12140081 | 206 | 0.034 | 200 | 0.08 | 205 | 0.0683 | 188 | 0.0372 | 799 | 0.0551 | 190 | 0.0632 | 201 | 0.0647 | 171 | 0.076 | 181 | 0.0608 | 743 | 0.0659 |
| 8_12181963 | 1 | 1 | NA | NA | NA | NA | NA | NA | 1 | 1 | NA | NA | NA | NA | 1 | 1 | NA | NA | 1 | 1 |
| 8_12181973 | 1 | 1 | NA | NA | NA | NA | NA | NA | 1 | 1 | NA | NA | NA | NA | 1 | 1 | NA | NA | 1 | 1 |
| 8_12181989 | 1 | 1 | NA | NA | NA | NA | NA | NA | 1 | 1 | NA | NA | NA | NA | 1 | 1 | NA | NA | 1 | 1 |
| 8_12181997 | 1 | 1 | NA | NA | NA | NA | NA | NA | 1 | 1 | NA | NA | NA | NA | 1 | 1 | NA | NA | 1 | 1 |
| 8_12181998 | 1 | 1 | NA | NA | NA | NA | NA | NA | 1 | 1 | NA | NA | NA | NA | 1 | 1 | NA | NA | 1 | 1 |
| 8_12182052 | 1 | 1 | NA | NA | NA | NA | NA | NA | 1 | 1 | NA | NA | NA | NA | 1 | 1 | NA | NA | 1 | 1 |
| 8_12182060 | 1 | 1 | NA | NA | NA | NA | NA | NA | 1 | 1 | NA | NA | NA | NA | 1 | 1 | NA | NA | 1 | 1 |
| 8_12182554 | 1 | 1 | NA | NA | NA | NA | 1 | 0 | 2 | 0.5 | NA | NA | 1 | 1 | NA | NA | 3 | 1 | 4 | 1 |
| 8_12227635 | 4 | 0.25 | 10 | 0.5 | 6 | 0.6667 | 10 | 0.5 | 30 | 0.5 | 4 | 0.75 | 8 | 0.625 | 9 | 0.3333 | 8 | 0.375 | 29 | 0.4828 |
| 8_12227652 | 6 | 0.3333 | 12 | 0.1667 | 9 | 0.2222 | 11 | 0 | 38 | 0.1579 | 9 | 0.1111 | 11 | 0.0909 | 9 | 0.2222 | 8 | 0.125 | 37 | 0.1351 |
| 8_12227661 | 5 | 0.6 | 11 | 0.1818 | 8 | 0.375 | 15 | 0.2667 | 39 | 0.3077 | 8 | 0.125 | 13 | 0.0769 | 9 | 0.2222 | 9 | 0.2222 | 39 | 0.1538 |
| 8_12227664 | 6 | 0.6667 | 10 | 0.5 | 7 | 0.4286 | 12 | 0.6667 | 35 | 0.5714 | 7 | 0.8571 | 12 | 0.6667 | 9 | 0.5556 | 7 | 0.4286 | 35 | 0.6286 |
| 8_12227666 | 5 | 0 | 11 | 0.1818 | 7 | 0.1429 | 11 | 0 | 34 | 0.0882 | 8 | 0.125 | 13 | 0.1538 | 9 | 0.1111 | 8 | 0.25 | 38 | 0.1579 |
| 8_12227668 | 6 | 0.8333 | 10 | 0.8 | 9 | 0.7778 | 15 | 0.7333 | 40 | 0.775 | 9 | 0.8889 | 14 | 0.9286 | 9 | 0.7778 | 8 | 1 | 40 | 0.9 |
| 8_12227673 | 4 | 0.75 | 5 | 0.8 | 8 | 0.5 | 12 | 0.8333 | 29 | 0.7241 | 7 | 0.8571 | 11 | 0.1818 | 8 | 0.5 | 7 | 0.4286 | 33 | 0.4545 |
| 8_12227677 | 5 | 0.2 | 3 | 0.6667 | 4 | 0 | 9 | 0.3333 | 21 | 0.2857 | 6 | 0.1667 | 12 | 0.4167 | 6 | 0.1667 | 4 | 0 | 28 | 0.25 |
| 8_12227704 | 5 | 0.4 | 9 | 0.3333 | 6 | 0 | 13 | 0.3846 | 33 | 0.303 | 8 | 0.125 | 9 | 0.2222 | 7 | 0 | 6 | 0.3333 | 30 | 0.1667 |
| 8_12227718 | 4 | 0.25 | 9 | 0 | 8 | 0.125 | 13 | 0.0769 | 34 | 0.0882 | 6 | 0.3333 | 11 | 0.2727 | 8 | 0 | 7 | 0.1429 | 32 | 0.1875 |
| 8_12227725 | 3 | 0.3333 | 4 | 0.5 | 6 | 0.5 | 10 | 0.5 | 23 | 0.4783 | 5 | 0.6 | 7 | 0.2857 | 7 | 0.2857 | 6 | 0.3333 | 25 | 0.36 |
| 8_12227881 | 8 | 0.125 | 13 | 0 | 8 | 0 | 15 | 0.1333 | 44 | 0.0682 | 14 | 0.1429 | 13 | 0.2308 | 11 | 0 | 12 | 0.0833 | 50 | 0.12 |
| 8_12227898 | 4 | 0 | 11 | 0 | 7 | 0.2857 | 10 | 0 | 32 | 0.0625 | 9 | 0.2222 | 11 | 0.0909 | 10 | 0.2 | 9 | 0.2222 | 39 | 0.1795 |
| 8_12227910 | 6 | 0 | 16 | 0.0625 | 7 | 0.1429 | 11 | 0.0909 | 40 | 0.075 | 14 | 0 | 10 | 0.2 | 8 | 0.125 | 11 | 0 | 43 | 0.0698 |
| 8_12228682 | 3 | 0 | 7 | 0.1429 | 5 | 0.6 | 10 | 0.2 | 25 | 0.24 | 11 | 0.0909 | 11 | 0.4545 | 6 | 0 | 7 | 0 | 35 | 0.1714 |
| 8_12232349 | 27 | 0.2593 | 35 | 0.2571 | 36 | 0.3611 | 32 | 0.3438 | 130 | 0.3077 | 28 | 0.3214 | 21 | 0.4762 | 27 | 0.2222 | 29 | 0.3793 | 105 | 0.3429 |
| 8_12232351 | 36 | 0.4167 | 37 | 0.4054 | 44 | 0.3409 | 39 | 0.4615 | 156 | 0.4038 | 32 | 0.375 | 21 | 0.2381 | 27 | 0.5185 | 29 | 0.3448 | 109 | 0.3761 |
| 8_12232363 | 40 | 0.575 | 44 | 0.5682 | 43 | 0.5581 | 38 | 0.6053 | 165 | 0.5758 | 33 | 0.7273 | 22 | 0.5 | 29 | 0.5862 | 27 | 0.5185 | 111 | 0.5946 |
| 8_12232364 | 38 | 0.2105 | 43 | 0.1163 | 46 | 0.0652 | 34 | 0.1176 | 161 | 0.1242 | 32 | 0.125 | 21 | 0.0476 | 28 | 0.0357 | 29 | 0.069 | 110 | 0.0727 |
| 8_12232450 | 35 | 0.4286 | 42 | 0.3333 | 37 | 0.3784 | 34 | 0.5588 | 148 | 0.4189 | 31 | 0.5484 | 23 | 0.3478 | 28 | 0.3929 | 30 | 0.5333 | 112 | 0.4643 |
| 8_12232570 | 12 | 0.75 | 11 | 0.9091 | 11 | 0.7273 | 14 | 0.7857 | 48 | 0.7917 | 5 | 1 | 13 | 1 | 16 | 0.75 | 11 | 0.6364 | 45 | 0.8222 |
| 8_12232575 | 11 | 0.3636 | 9 | 0.5556 | 10 | 0.5 | 13 | 0.4615 | 43 | 0.4651 | 3 | 0.6667 | 12 | 0.4167 | 14 | 0.4286 | 9 | 0.2222 | 38 | 0.3947 |
| 8_12232601 | 12 | 0.1667 | 11 | 0.1818 | 10 | 0.2 | 12 | 0.0833 | 45 | 0.1556 | 4 | 0.25 | 12 | 0.4167 | 14 | 0.1429 | 11 | 0.0909 | 41 | 0.2195 |
| 8_12232603 | 10 | 0 | 10 | 0.4 | 10 | 0.4 | 12 | 0.25 | 42 | 0.2619 | 4 | 0.25 | 12 | 0.3333 | 15 | 0.2667 | 11 | 0.0909 | 42 | 0.2381 |
| 8_12232605 | 9 | 0 | 9 | 0 | 11 | 0.0909 | 13 | 0.1538 | 42 | 0.0714 | 3 | 0 | 12 | 0 | 11 | 0.0909 | 10 | 0.1 | 36 | 0.0556 |
| 8_12232607 | 11 | 0.3636 | 12 | 0.5833 | 12 | 0.5 | 14 | 0.6429 | 49 | 0.5306 | 4 | 0.5 | 12 | 0.6667 | 14 | 0.5 | 11 | 0.0909 | 41 | 0.439 |
| 8_12232621 | 9 | 0 | 9 | 0.1111 | 5 | 0 | 8 | 0.125 | 31 | 0.0645 | 2 | 0 | 7 | 0.1429 | 9 | 0.1111 | 7 | 0 | 25 | 0.08 |
| 8_12232628 | 5 | 0 | 6 | 0 | 7 | 0.2857 | 13 | 0.1538 | 31 | 0.129 | 4 | 0.25 | 8 | 0.25 | 9 | 0.1111 | 10 | 0.2 | 31 | 0.1935 |
| 8_12232641 | 10 | 0.2 | 11 | 0.3636 | 9 | 0.5556 | 10 | 0.4 | 40 | 0.375 | 3 | 0.3333 | 12 | 0.5 | 11 | 0.4545 | 10 | 0.5 | 36 | 0.4722 |
| 8_12232644 | 11 | 0.2727 | 11 | 0.2727 | 10 | 0.4 | 10 | 0.4 | 42 | 0.3333 | 4 | 0.25 | 10 | 0.3 | 9 | 0.3333 | 7 | 0.2857 | 30 | 0.3 |
| 8_12232654 | 9 | 0.2222 | 11 | 0.1818 | 8 | 0.125 | 14 | 0.1429 | 42 | 0.1667 | 3 | 0 | 11 | 0.0909 | 14 | 0.0714 | 11 | 0.0909 | 39 | 0.0769 |
| 8_12232664 | 7 | 1 | 11 | 0.9091 | 8 | 0.75 | 14 | 0.9286 | 40 | 0.9 | 3 | 1 | 11 | 0.9091 | 12 | 1 | 11 | 0.9091 | 37 | 0.9459 |
| 8_12232668 | 5 | 0.2 | 6 | 0.3333 | 6 | 0.5 | 8 | 0.375 | 25 | 0.36 | 1 | 0 | 8 | 0.375 | 8 | 0.5 | 5 | 0.4 | 22 | 0.4091 |
| 8_12232671 | 7 | 0 | 10 | 0.2 | 9 | 0.2222 | 11 | 0 | 37 | 0.1081 | 3 | 0.6667 | 11 | 0.2727 | 14 | 0.0714 | 8 | 0 | 36 | 0.1667 |
| 8_12232683 | 4 | 0.75 | 10 | 0.6 | 9 | 0.7778 | 13 | 0.6923 | 36 | 0.6944 | 2 | 1 | 9 | 0.6667 | 13 | 0.6154 | 10 | 0.3 | 34 | 0.5588 |
| 8_12232701 | 7 | 0.2857 | 9 | 0.1111 | 7 | 0 | 13 | 0.3077 | 36 | 0.1944 | 2 | 0.5 | 11 | 0 | 13 | 0.2308 | 9 | 0.2222 | 35 | 0.1714 |
| 8_12232711 | 7 | 0.7143 | 8 | 0.875 | 6 | 1 | 12 | 0.75 | 33 | 0.8182 | 2 | 1 | 8 | 0.875 | 12 | 0.5833 | 6 | 0.1667 | 28 | 0.6071 |
| 8_12232719 | 6 | 0 | 8 | 0.25 | 7 | 0.2857 | 12 | 0.3333 | 33 | 0.2424 | 2 | 0 | 12 | 0 | 10 | 0.2 | 10 | 0.3 | 34 | 0.1471 |
| 8_12232727 | 7 | 0.2857 | 7 | 0.4286 | 8 | 0.25 | 12 | 0.25 | 34 | 0.2941 | 2 | 0 | 11 | 0.0909 | 9 | 0.2222 | 6 | 0.6667 | 28 | 0.25 |
| 8_12232730 | 5 | 0.4 | 6 | 0.6667 | 8 | 0.625 | 10 | 0.5 | 29 | 0.5517 | 2 | 0 | 8 | 0.75 | 11 | 0.6364 | 5 | 0.6 | 26 | 0.6154 |

| | | | | | | | | | | | | | | | | | | | | |
|---|---|---|---|---|---|---|---|---|---|---|---|---|---|---|---|---|---|---|---|---|
| 8_12232733 | 4 | 1 | 7 | 1 | 5 | 1 | 7 | 1 | 23 | 1 | 1 | 1 | 8 | 1 | 10 | 1 | 6 | 1 | 25 | 1 |
| 8_12232735 | 6 | 0.1667 | 7 | 0.1429 | 9 | 0.3333 | 9 | 0 | 31 | 0.1613 | 1 | 0 | 12 | 0.1667 | 10 | 0.3 | 7 | 0.2857 | 30 | 0.2333 |
| 8_12232755 | 4 | 0 | 6 | 0.3333 | 7 | 0 | 10 | 0.3 | 27 | 0.1852 | 2 | 0.5 | 11 | 0.0909 | 7 | 0.2857 | 7 | 0 | 27 | 0.1481 |
| 8_12232760 | 5 | 1 | 5 | 1 | 9 | 1 | 10 | 1 | 29 | 1 | 2 | 1 | 9 | 1 | 8 | 1 | 7 | 1 | 26 | 1 |
| 8_12232772 | 5 | 0.2 | 3 | 0 | 6 | 0.3333 | 10 | 0.1 | 24 | 0.1667 | 2 | 0 | 8 | 0.25 | 7 | 0.2857 | 3 | 0 | 20 | 0.2 |
| 8_12232776 | 4 | 0.75 | 4 | 0.75 | 7 | 1 | 10 | 1 | 25 | 0.92 | 2 | 1 | 11 | 1 | 9 | 1 | 3 | 1 | 25 | 1 |
| 8_12232789 | 5 | 0 | 3 | 0.3333 | 4 | 0 | 10 | 0.1 | 22 | 0.0909 | 1 | 0 | 8 | 0.25 | 9 | 0 | 2 | 0 | 20 | 0.1 |
| 8_12232796 | 5 | 0.2 | 8 | 0.5 | 9 | 0.4444 | 9 | 0.1111 | 31 | 0.3226 | 2 | 0 | 9 | 0.6667 | 9 | 0.4444 | 4 | 0.25 | 24 | 0.4583 |
| 8_12232796 | 5 | 1 | 7 | 1 | 6 | 1 | 11 | 1 | 29 | 1 | 2 | 1 | 9 | 1 | 7 | 1 | 5 | 1 | 23 | 1 |
| 8_12232797 | 3 | 0.3333 | 5 | 0 | 9 | 0.2222 | 11 | 0.1818 | 28 | 0.1786 | 1 | 0 | 8 | 0.125 | 7 | 0.4286 | 4 | 0.25 | 20 | 0.25 |
| 8_12232799 | 4 | 0 | 5 | 0.2 | 8 | 0.125 | 11 | 0.0909 | 28 | 0.1071 | 2 | 0 | 11 | 0.0909 | 8 | 0 | 5 | 0.2 | 26 | 0.0769 |
| 8_12232801 | 5 | 0 | 7 | 0.1429 | 7 | 0 | 10 | 0.3 | 29 | 0.1379 | 2 | 0.5 | 11 | 0.0909 | 8 | 0.125 | 3 | 0.3333 | 24 | 0.1667 |
| 8_12232802 | 5 | 1 | 7 | 0.8571 | 8 | 0.5 | 11 | 0.6364 | 31 | 0.7097 | 2 | 0.5 | 10 | 0.5 | 9 | 0.4444 | 4 | 0.25 | 25 | 0.44 |
| 8_12232805 | 5 | 0 | 7 | 0.2857 | 8 | 0 | 10 | 0 | 30 | 0.0667 | 2 | 0 | 11 | 0.1818 | 7 | 0.2857 | 3 | 0.3333 | 23 | 0.2174 |
| 8_12232811 | 4 | 0.5 | 5 | 0.4 | 7 | 0.5714 | 9 | 0.6667 | 25 | 0.56 | 2 | 1 | 9 | 0.6667 | 7 | 0.8571 | 4 | 0.75 | 22 | 0.7727 |
| 8_12232815 | 4 | 0.5 | 6 | 0.3333 | 6 | 0.3333 | 8 | 0.375 | 24 | 0.375 | 2 | 0.5 | 9 | 0.4444 | 7 | 0.2857 | 4 | 0.75 | 22 | 0.4545 |
| 8_12232835 | 4 | 0.5 | 6 | 0.6667 | 7 | 0.2857 | 9 | 0.7778 | 26 | 0.5769 | NA | NA | 10 | 0.4 | 8 | 0.75 | 3 | 0.3333 | 21 | 0.5238 |
| 8_12232843 | 4 | 0.75 | 6 | 1 | 6 | 0.8333 | 7 | 1 | 23 | 0.913 | NA | NA | 7 | 1 | 6 | 1 | 4 | 1 | 17 | 1 |
| 8_12232845 | 4 | 0.75 | 5 | 0.2 | 8 | 0.375 | 9 | 0.3333 | 26 | 0.3846 | NA | NA | 11 | 0.7273 | 8 | 0.25 | 4 | 0.25 | 23 | 0.4783 |
| 8_12232846 | 4 | 0.75 | 4 | 0.25 | 5 | 0.6 | 8 | 0.75 | 21 | 0.619 | NA | NA | 7 | 1 | 7 | 0.7143 | 3 | 1 | 17 | 0.8824 |
| 8_12232858 | 5 | 0.6 | 6 | 0.3333 | 5 | 0.4 | 7 | 0.7143 | 23 | 0.5217 | NA | NA | 11 | 0.3636 | 6 | 0.5 | 4 | 0.5 | 21 | 0.4286 |
| 8_12232875 | 5 | 0.6 | 5 | 0.8 | 3 | 0.6667 | 7 | 0.5714 | 20 | 0.65 | NA | NA | 11 | 0.7273 | 8 | 0.5 | 4 | 0.75 | 23 | 0.6522 |
| 8_12232880 | 4 | 0.25 | 6 | 0.1667 | 4 | 0.25 | 8 | 0 | 22 | 0.1364 | NA | NA | 9 | 0.2222 | 7 | 0 | 3 | 0.6667 | 19 | 0.2105 |
| 8_12232884 | 3 | 0 | 5 | 0 | 5 | 0 | 6 | 0.1667 | 19 | 0.0526 | NA | NA | 10 | 0.2 | 6 | 0 | 2 | 0 | 18 | 0.1111 |
| 8_12232895 | NA | NA | 4 | 1 | 3 | 0.6667 | 7 | 0.4286 | 14 | 0.6429 | NA | NA | 9 | 0.7778 | 6 | 0.5 | 1 | 1 | 16 | 0.6875 |
| 8_12232898 | 2 | 0 | 4 | 0.5 | 2 | 0 | 7 | 0.1429 | 15 | 0.2 | NA | NA | 7 | 0 | 6 | 0.3333 | 2 | 0.5 | 15 | 0.2 |
| 8_12232901 | 3 | 0.6667 | 5 | 0.8 | 4 | 0.75 | 8 | 1 | 20 | 0.85 | NA | NA | 10 | 0.5 | 7 | 0.8571 | 3 | 0.6667 | 20 | 0.65 |
| 8_12232902 | 2 | 0 | 5 | 0.2 | 4 | 0 | 8 | 0.125 | 19 | 0.1053 | NA | NA | 11 | 0.1818 | 6 | 0 | 2 | 0 | 19 | 0.1053 |
| 8_12233056 | 2 | 1 | 6 | 0.3333 | 5 | 0.2 | 7 | 0.4286 | 20 | 0.4 | NA | NA | 10 | 0.4 | 7 | 0.4286 | 2 | 0 | 19 | 0.3684 |
| 8_12233058 | 3 | 0.3333 | 6 | 0.3333 | 4 | 0.5 | 6 | 0.1667 | 19 | 0.3158 | NA | NA | 10 | 0.1 | 7 | 0 | 3 | 0 | 20 | 0.05 |
| 8_12233071 | 2 | 1 | 4 | 0.5 | 5 | 0.6 | 5 | 0.6 | 16 | 0.625 | NA | NA | 8 | 0.625 | 7 | 0.8571 | 3 | 0.3333 | 18 | 0.6667 |
| 8_12233076 | 3 | 0 | 6 | 0.1667 | 5 | 0.2 | 7 | 0.1429 | 21 | 0.1429 | NA | NA | 11 | 0.2727 | 8 | 0.375 | 3 | 0 | 22 | 0.2727 |
| 8_12233107 | 2 | 0 | 5 | 0 | 5 | 0.2 | 3 | 0 | 15 | 0.0667 | NA | NA | 6 | 0.1667 | 4 | 0.25 | 2 | 0 | 12 | 0.1667 |
| 8_12233132 | 3 | 0.6667 | 5 | 1 | 4 | 1 | 4 | 0.75 | 16 | 0.875 | NA | NA | 6 | 0.8333 | 3 | 1 | 2 | 1 | 11 | 0.9091 |
| 8_12233132 | 3 | 0 | 4 | 0.5 | 4 | 0 | 4 | 0.25 | 15 | 0.2 | NA | NA | 4 | 0.25 | 4 | 0.25 | 2 | 0 | 10 | 0.2 |
| 8_12233137 | 4 | 0 | 5 | 0.2 | 5 | 0 | 4 | 0.25 | 18 | 0.1111 | NA | NA | 6 | 0.5 | 4 | 0 | 2 | 1 | 12 | 0.4167 |
| 8_12233146 | 3 | 0.6667 | 4 | 0.25 | 4 | 0.75 | 4 | 0.5 | 15 | 0.5333 | NA | NA | 6 | 0.6667 | 3 | 1 | 2 | 0 | 11 | 0.6364 |
| 8_12233149 | 4 | 0.75 | 4 | 1 | 3 | 0.6667 | 3 | 1 | 14 | 0.8571 | NA | NA | 5 | 0.8 | 3 | 0.6667 | 2 | 1 | 10 | 0.8 |
| 8_12233153 | 1 | 0 | 2 | 1 | 2 | 0.5 | 2 | 0.5 | 7 | 0.5714 | NA | NA | 1 | 1 | 1 | 1 | 1 | 1 | 3 | 1 |
| 8_12233159 | 2 | 1 | 5 | 0.2 | 4 | 0 | 3 | 0.3333 | 14 | 0.2857 | NA | NA | 5 | 0.2 | 4 | 0.25 | 2 | 0.5 | 11 | 0.2727 |
| 8_12233165 | 2 | 1 | 5 | 0.6 | 3 | 0.6667 | 2 | 0.5 | 12 | 0.6667 | NA | NA | 5 | 0.6 | 4 | 0.75 | 2 | 0.5 | 11 | 0.6364 |
| 8_12233168 | 3 | 0 | 4 | 0 | 2 | 0 | 2 | 0.5 | 11 | 0.0909 | NA | NA | 4 | 0 | 2 | 1 | 1 | 0 | 7 | 0.2857 |
| 8_12237888 | 1 | 1 | 2 | 1 | 1 | 1 | 2 | 0 | 6 | 0.6667 | NA | NA | 2 | 1 | 1 | 1 | NA | NA | 3 | 1 |
| 8_12245714 | 3 | 0.6667 | 3 | 0.3333 | 8 | 0.25 | 5 | 0.4 | 19 | 0.3684 | 4 | 0 | 3 | 0.3333 | 7 | 0.2857 | 2 | 0.5 | 16 | 0.25 |
| 8_12245770 | 11 | 0.0909 | 11 | 0.0909 | 10 | 0 | 12 | 0.0833 | 44 | 0.0682 | 7 | 0.1429 | 7 | 0 | 17 | 0.1176 | 9 | 0.1111 | 40 | 0.1 |
| 8_12247980 | 13 | 0.1538 | 14 | 0.2143 | 12 | 0.25 | 14 | 0 | 53 | 0.1509 | 7 | 0 | 9 | 0.1111 | 18 | 0.2222 | 11 | 0.0909 | 45 | 0.1333 |
| 8_12247993 | 13 | 0.1538 | 16 | 0.125 | 10 | 0.1 | 14 | 0 | 53 | 0.0943 | 8 | 0.25 | 8 | 0.625 | 20 | 0.15 | 10 | 0.3 | 46 | 0.2826 |
| 8_12247996 | 14 | 0.8571 | 15 | 1 | 12 | 1 | 10 | 0.7 | 51 | 0.902 | 8 | 1 | 7 | 1 | 16 | 0.9375 | 11 | 0.9091 | 42 | 0.9524 |
| 8_12248030 | 13 | 0.1538 | 15 | 0.2 | 13 | 0.3846 | 14 | 0.2857 | 55 | 0.2545 | 7 | 0.5714 | 9 | 0.2222 | 17 | 0.1176 | 8 | 0.125 | 41 | 0.2195 |
| 8_12251136 | 16 | 0.25 | 17 | 0.0588 | 17 | 0.0588 | 19 | 0.0526 | 69 | 0.1014 | 8 | 0 | 12 | 0 | 21 | 0.1905 | 15 | 0.2 | 56 | 0.125 |
| 8_12251149 | 18 | 0.1667 | 21 | 0.0952 | 20 | 0.1 | 19 | 0.1053 | 78 | 0.1154 | 8 | 0.5 | 13 | 0 | 25 | 0.08 | 17 | 0.1176 | 63 | 0.127 |
| 8_12251155 | 17 | 0.1765 | 19 | 0.2105 | 16 | 0.1875 | 15 | 0 | 67 | 0.1493 | 8 | 0.125 | 10 | 0.3 | 22 | 0.1818 | 15 | 0.1333 | 55 | 0.1818 |
| 8_12251158 | 17 | 0.9412 | 19 | 0.7895 | 17 | 0.9412 | 18 | 0.8889 | 71 | 0.8873 | 9 | 1 | 12 | 1 | 19 | 0.8947 | 13 | 0.9231 | 53 | 0.9434 |
| 8_12251167 | 15 | 0.9333 | 20 | 1 | 19 | 0.9474 | 17 | 1 | 71 | 0.9718 | 9 | 1 | 12 | 1 | 23 | 0.913 | 20 | 1 | 64 | 0.9688 |
| 8_12251171 | 18 | 0.0556 | 23 | 0.1304 | 17 | 0.0588 | 18 | 0 | 76 | 0.0658 | 11 | 0.0909 | 13 | 0.0769 | 25 | 0.04 | 15 | 0.2 | 64 | 0.0938 |
| 8_12251176 | 16 | 0.9375 | 17 | 1 | 14 | 0.9286 | 11 | 1 | 58 | 0.9655 | 8 | 1 | 11 | 1 | 21 | 1 | 13 | 1 | 53 | 1 |
| 8_12251193 | 15 | 0 | 16 | 0.0625 | 11 | 0.0909 | 16 | 0.1875 | 58 | 0.0862 | 7 | 0.4286 | 13 | 0.2308 | 15 | 0.1333 | 9 | 0.3333 | 44 | 0.25 |
| 8_12251196 | 16 | 0.875 | 24 | 0.9167 | 18 | 0.9444 | 18 | 0.9444 | 76 | 0.9211 | 10 | 1 | 21 | 0.9048 | 25 | 0.96 | 16 | 1 | 72 | 0.9583 |
| 8_12251198 | 20 | 0.1 | 24 | 0.0833 | 19 | 0.1053 | 21 | 0.2381 | 84 | 0.131 | 12 | 0.1667 | 20 | 0.1 | 26 | 0.1154 | 19 | 0.0526 | 77 | 0.1039 |
| 8_12251202 | 23 | 0 | 26 | 0.1923 | 20 | 0 | 24 | 0.1667 | 93 | 0.0968 | 12 | 0 | 20 | 0.05 | 29 | 0.069 | 20 | 0.1 | 81 | 0.0617 |
| 8_12251208 | 23 | 0.1304 | 26 | 0 | 20 | 0.1 | 25 | 0.16 | 94 | 0.0957 | 12 | 0.25 | 21 | 0.1905 | 29 | 0.1379 | 21 | 0.2381 | 83 | 0.1928 |
| 8_12251620 | 20 | 0.45 | 25 | 0.32 | 17 | 0.4118 | 21 | 0.4286 | 83 | 0.3976 | 12 | 0.4167 | 19 | 0.4211 | 28 | 0.5 | 19 | 0.3684 | 78 | 0.4359 |
| 8_12251626 | NA | NA | NA | NA | 1 | 1 | 1 | 1 | 2 | 1 | NA | NA | NA | NA | 1 | 1 | NA | NA | 1 | 1 |
| 8_12251630 | 17 | 0.4118 | 21 | 0.2857 | 21 | 0.2857 | 15 | 0.5333 | 74 | 0.3649 | 9 | 0.4444 | 16 | 0.5 | 27 | 0.2593 | 15 | 0.4667 | 67 | 0.3881 |
| 8_12251651 | 23 | 0.4783 | 23 | 0.4783 | 23 | 0.3043 | 21 | 0.3333 | 90 | 0.4 | 14 | 0.6429 | 20 | 0.7 | 34 | 0.5882 | 21 | 0.4286 | 89 | 0.5843 |

| | | | | | | | | | | | | | | | | | | | | |
|---|---|---|---|---|---|---|---|---|---|---|---|---|---|---|---|---|---|---|---|---|
| 8_1225167 | 31 | 0.0645 | 28 | 0.1429 | 27 | 0.1481 | 24 | 0.125 | 110 | 0.1182 | 15 | 0.1333 | 24 | 0.1667 | 34 | 0.1765 | 26 | 0.1923 | 99 | 0.1717 |
| 8_1225169 | 31 | 0.2581 | 23 | 0.2174 | 22 | 0.4545 | 21 | 0.2857 | 97 | 0.299 | 14 | 0.3571 | 23 | 0.4783 | 27 | 0.2963 | 23 | 0.2174 | 87 | 0.3333 |
| 8_1225172 | 31 | 0.2581 | 33 | 0.1212 | 30 | 0.0667 | 28 | 0.0714 | 122 | 0.1311 | 16 | 0.1875 | 26 | 0.2692 | 36 | 0.2778 | 26 | 0.1923 | 104 | 0.2404 |
| 8_1225174 | 38 | 0.4737 | 35 | 0.6286 | 33 | 0.5758 | 26 | 0.4615 | 132 | 0.5379 | 20 | 0.7 | 30 | 0.4333 | 34 | 0.6176 | 31 | 0.4839 | 115 | 0.5478 |
| 8_1225203 | 2 | 0 | 3 | 0 | 2 | 0.5 | 1 | 0 | 8 | 0.125 | 5 | 0.2 | 1 | 0 | 4 | 0.25 | 2 | 0 | 12 | 0.1667 |
| 8_1225206 | 2 | 0.5 | 4 | 0.5 | 2 | 0.5 | 1 | 1 | 9 | 0.5556 | 5 | 0.2 | 2 | 0 | 4 | 0 | 1 | 1 | 12 | 0.1667 |
| 8_1225206 | 3 | 0.3333 | 3 | 0.3333 | 4 | 0.5 | 2 | 0.5 | 12 | 0.4167 | 5 | 0.4 | 2 | 0.5 | 3 | 0.3333 | 2 | 0 | 12 | 0.3333 |
| 8_1225207 | 2 | 0 | 2 | 0.5 | 3 | 0.6667 | 1 | 0 | 8 | 0.375 | 4 | 0.25 | 1 | 1 | 4 | 0.75 | 2 | 1 | 11 | 0.6364 |
| 8_1225214 | NA | NA | NA | NA | 1 | 1 | NA | NA | 1 | 1 | 1 | 0 | NA | NA | 1 | 1 | 1 | 0 | 3 | 0.3333 |
| 8_1225215 | NA | NA | NA | NA | 1 | 1 | NA | NA | 1 | 1 | 1 | 1 | NA | NA | 1 | 0 | 1 | 0 | 3 | 0.3333 |
| 8_1225217 | NA | NA | NA | NA | 2 | 0.5 | NA | NA | 2 | 0.5 | NA | NA | NA | NA | 1 | 0 | 1 | 1 | 2 | 0.5 |
| 8_1225232 | 4 | 1 | 1 | 1 | 5 | 0.6 | 1 | 0 | 11 | 0.7273 | NA | NA | NA | NA | 2 | 0 | 4 | 0.75 | 6 | 0.5 |
| 8_1225233 | 4 | 0 | 1 | 0 | 6 | 0.1667 | 1 | 0 | 12 | 0.0833 | 1 | 1 | 1 | 0 | 1 | 0 | 5 | 0.4 | 8 | 0.375 |
| 8_1225234 | 4 | 0 | 2 | 0 | 7 | 0.4286 | 2 | 0.5 | 15 | 0.2667 | 2 | 0 | 1 | 0 | 2 | 0.5 | 4 | 0.5 | 9 | 0.3333 |
| 8_1225241 | 2 | 0.5 | NA | NA | 1 | 1 | NA | NA | 3 | 0.6667 | NA | NA | NA | NA | NA | NA | 2 | 0.5 | 2 | 0.5 |
| 8_1225246 | 3 | 0.3333 | NA | NA | 5 | 0.8 | 1 | 1 | 9 | 0.6667 | 1 | 1 | NA | NA | 2 | 0.5 | 3 | 0.6667 | 6 | 0.6667 |
| 8_1225247 | 5 | 0.8 | 1 | 1 | 5 | 0.4 | 2 | 0 | 13 | 0.5385 | NA | NA | NA | NA | 2 | 0.5 | 2 | 0 | 4 | 0.25 |
| 8_1225248 | 4 | 0.25 | 1 | 1 | 5 | 0.4 | 1 | 1 | 11 | 0.4545 | NA | NA | 1 | 1 | 1 | 0 | 4 | 0.75 | 6 | 0.6667 |
| 8_1225655 | 1 | 1 | 2 | 1 | NA | NA | NA | NA | 3 | 1 | 1 | 1 | NA | NA | 1 | 0 | NA | NA | 2 | 0.5 |
| 8_1225657 | 1 | 1 | 2 | 1 | NA | NA | NA | NA | 3 | 1 | 1 | 1 | NA | NA | NA | NA | NA | NA | 1 | 1 |
| 8_1225657 | 1 | 1 | 2 | 1 | NA | NA | NA | NA | 3 | 1 | 1 | 1 | NA | NA | NA | NA | NA | NA | 1 | 1 |
| 8_1225658 | 1 | 1 | 2 | 0 | NA | NA | NA | NA | 3 | 0.3333 | 1 | 1 | NA | NA | NA | NA | NA | NA | 1 | 1 |
| 8_1225658 | 1 | 1 | 2 | 0.5 | NA | NA | NA | NA | 3 | 0.6667 | 1 | 1 | NA | NA | NA | NA | NA | NA | 1 | 1 |
| 8_1225665 | 1 | 1 | 2 | 0.5 | NA | NA | NA | NA | 3 | 0.6667 | 1 | 1 | NA | NA | 1 | 0 | NA | NA | 2 | 0.5 |
| 8_1225667 | 1 | 1 | 2 | 1 | NA | NA | NA | NA | 3 | 1 | 1 | 0 | NA | NA | 2 | 0.5 | NA | NA | 3 | 0.3333 |
| 8_1226085 | 6 | 0.3333 | 9 | 0 | 5 | 0.6 | 7 | 0 | 27 | 0.1852 | 2 | 0.5 | 7 | 0.1429 | 2 | 0 | 8 | 0.25 | 19 | 0.2105 |
| 8_1226088 | 7 | 0 | 8 | 0.375 | 5 | 0.2 | 7 | 0 | 27 | 0.1481 | 2 | 0 | 7 | 0.1429 | 2 | 0.5 | 10 | 0.2 | 21 | 0.1905 |
| 8_1226091 | 5 | 0 | 8 | 0.5 | 4 | 0 | 7 | 0.4286 | 24 | 0.2917 | 2 | 0 | 8 | 0.125 | 2 | 0.5 | 8 | 0.125 | 20 | 0.15 |
| 8_1226091 | 6 | 0.8333 | 8 | 1 | 4 | 1 | 7 | 0.7143 | 25 | 0.88 | 1 | 1 | 7 | 1 | 2 | 1 | 9 | 0.8889 | 19 | 0.9474 |
| 8_1226092 | 5 | 0.8 | 7 | 0.8571 | 4 | 0.75 | 6 | 0.5 | 22 | 0.7273 | NA | NA | 7 | 0.8571 | 2 | 1 | 8 | 0.875 | 17 | 0.8824 |
| 8_1226092 | 4 | 0.5 | 7 | 0.7143 | 5 | 0.8 | 7 | 0.4286 | 23 | 0.6087 | 1 | 0 | 6 | 0.5 | 2 | 0 | 9 | 0.4444 | 18 | 0.3889 |
| 8_1226092 | 5 | 0 | 8 | 0.125 | 5 | 0 | 7 | 0.1429 | 25 | 0.08 | 1 | 0 | 8 | 0 | 2 | 0 | 9 | 0.2222 | 20 | 0.1 |
| 8_1226093 | 5 | 0.4 | 7 | 0.1429 | 3 | 0 | 5 | 0 | 20 | 0.15 | 1 | 0 | 7 | 0.1429 | 2 | 0.5 | 9 | 0.5556 | 19 | 0.3684 |
| 8_1226095 | 4 | 0.75 | 5 | 0.8 | 3 | 0.6667 | 7 | 0.2857 | 19 | 0.5789 | 1 | 1 | 6 | 0.5 | 2 | 0.5 | 9 | 0.5556 | 18 | 0.5556 |
| 8_1226095 | 3 | 0.6667 | 4 | 0.5 | 3 | 0.6667 | 5 | 0 | 15 | 0.4 | 1 | 0 | 4 | 0.25 | 2 | 0.5 | 7 | 0.2857 | 14 | 0.2857 |
| 8_1226095 | 5 | 1 | 7 | 1 | 3 | 1 | 6 | 1 | 21 | 1 | 1 | 1 | 6 | 0.8333 | 2 | 1 | 8 | 0.875 | 17 | 0.8824 |
| 8_1226096 | 5 | 0.4 | 5 | 0.6 | 2 | 1 | 5 | 0.4 | 17 | 0.5294 | NA | NA | 6 | 0.6667 | 2 | 0.5 | 9 | 0.5556 | 17 | 0.5882 |
| 8_1226340 | 3 | 1 | 1 | 1 | 3 | 1 | 5 | 0.8 | 12 | 0.9167 | NA | NA | 1 | 1 | NA | NA | 4 | 1 | 5 | 1 |
| 8_1227526 | 2 | 0.5 | 3 | 0.3333 | 1 | 1 | 2 | 1 | 8 | 0.625 | 1 | 0 | 2 | 0.5 | 1 | 0 | 3 | 1 | 7 | 0.5714 |
| 8_1227528 | 3 | 1 | 3 | 0.6667 | 1 | 1 | 1 | 0 | 8 | 0.75 | 1 | 1 | 1 | 1 | NA | NA | 3 | 1 | 5 | 1 |
| 8_1230622 | 1 | 0 | 1 | 1 | 2 | 0.5 | 3 | 0 | 7 | 0.2857 | 2 | 0 | 2 | 0 | 1 | 1 | 2 | 0 | 7 | 0.1429 |
| 8_1230622 | NA | NA | 1 | 0 | 1 | 0 | 4 | 0.25 | 6 | 0.1667 | 2 | 0 | 2 | 0.5 | 1 | 0 | 1 | 0 | 6 | 0.1667 |
| 8_1230623 | 2 | 0.5 | NA | NA | 1 | 1 | 4 | 0 | 7 | 0.2857 | 4 | 0 | 2 | 0 | 1 | 1 | 3 | 0.6667 | 10 | 0.3 |
| 8_1230624 | NA | NA | 1 | 1 | 1 | 1 | 4 | 0.25 | 6 | 0.5 | 3 | 0.3333 | 1 | 1 | 1 | 0 | 2 | 0 | 7 | 0.2857 |
| 8_1230624 | 1 | 1 | 1 | 1 | 2 | 1 | 3 | 0.6667 | 7 | 0.8571 | 4 | 0.75 | 1 | 1 | NA | NA | 2 | 1 | 7 | 0.8571 |
| 8_1230625 | 2 | 1 | 1 | 1 | 2 | 1 | 4 | 1 | 9 | 1 | 4 | 1 | 2 | 1 | 1 | 1 | 3 | 1 | 10 | 1 |
| 8_1230627 | 1 | 1 | 1 | 1 | NA | NA | 3 | 0.6667 | 5 | 0.8 | 2 | 1 | 2 | 0.5 | 1 | 1 | 3 | 1 | 8 | 0.875 |
| 8_1230628 | 2 | 0 | 1 | 0 | 2 | 0.5 | 4 | 0 | 9 | 0.1111 | 4 | 0.5 | 2 | 0.5 | 1 | 0 | 3 | 0.6667 | 10 | 0.5 |
| 8_1230631 | 2 | 0.5 | 2 | 0 | 3 | 0 | 5 | 0 | 12 | 0.0833 | 3 | 0.3333 | 2 | 0 | 2 | 0 | 4 | 0.25 | 11 | 0.1818 |
| 8_1230633 | 1 | 1 | 2 | 0.5 | 4 | 0.5 | 3 | 0 | 10 | 0.4 | 3 | 1 | 2 | 0 | 1 | 1 | 3 | 0.6667 | 9 | 0.6667 |
| 8_1230637 | 1 | 1 | 3 | 0.6667 | 4 | 0.5 | 3 | 0.3333 | 11 | 0.5455 | 4 | 0.75 | 3 | 1 | 1 | 1 | 4 | 1 | 12 | 0.9167 |
| 8_1230639 | 1 | 1 | 2 | 1 | 3 | 0.6667 | 4 | 1 | 10 | 0.9 | 4 | 0.75 | 2 | 0.5 | 2 | 1 | 3 | 1 | 11 | 0.8182 |
| 8_1230639 | 1 | 1 | 2 | 0 | 4 | 1 | 4 | 0.25 | 11 | 0.5455 | 3 | 0.6667 | 2 | 0 | 2 | 1 | 2 | 0.5 | 9 | 0.5556 |
| 8_1230639 | 1 | 0 | NA | NA | 3 | 1 | 4 | 1 | 8 | 0.875 | 4 | 1 | 2 | 0.5 | NA | NA | 2 | 1 | 8 | 0.875 |
| 8_1230641 | 1 | 1 | NA | NA | NA | NA | NA | NA | 1 | 1 | 2 | 1 | NA | NA | 1 | 1 | 1 | 0 | 4 | 0.75 |
| 8_1230642 | 1 | 0 | 1 | 1 | NA | NA | NA | NA | 2 | 0.5 | 2 | 0.5 | NA | NA | NA | NA | 1 | 1 | 3 | 0.6667 |
| 8_1230645 | 1 | 0 | 1 | 1 | NA | NA | 1 | 0 | 3 | 0.3333 | 2 | 1 | NA | NA | NA | NA | 1 | 0 | 3 | 0.6667 |
| 8_1230657 | NA | NA | 1 | 1 | NA | NA | NA | NA | 1 | 1 | 1 | 1 | NA | NA | NA | NA | NA | NA | 1 | 1 |
| 8_1230803 | NA | NA | NA | NA | NA | NA | 1 | 1 | 1 | 1 | 1 | 1 | NA | NA | NA | NA | NA | NA | 1 | 1 |
| 8_1230803 | NA | NA | NA | NA | NA | NA | 1 | 1 | 1 | 1 | 1 | 1 | NA | NA | NA | NA | NA | NA | 1 | 1 |
| 8_1230804 | NA | NA | NA | NA | NA | NA | 1 | 1 | 1 | 1 | 1 | 1 | NA | NA | NA | NA | NA | NA | 1 | 1 |
| 8_1230822 | 2 | 0 | 2 | 0 | 1 | 0 | 1 | 1 | 6 | 0.1667 | 2 | 0.5 | 1 | 0 | 2 | 0 | 2 | 0 | 7 | 0.1429 |
| 8_1230825 | 1 | 0 | 2 | 0.5 | 2 | 0.5 | 2 | 0 | 7 | 0.2857 | 2 | 0 | 1 | 1 | 2 | 0 | 2 | 0.5 | 7 | 0.2857 |
| 8_1230827 | 2 | 0 | 2 | 1 | 2 | 0.5 | 2 | 0.5 | 8 | 0.5 | 2 | 0 | NA | NA | NA | NA | 2 | 0.5 | 4 | 0.25 |
| 8_1230828 | 2 | 0.5 | 3 | 0.3333 | 2 | 0 | 1 | 0 | 8 | 0.25 | 1 | 0 | 1 | 0 | 1 | 0 | 3 | 0.3333 | 6 | 0.1667 |

| | | | | | | | | | | | | | | | | | | | | |
|---|---|---|---|---|---|---|---|---|---|---|---|---|---|---|---|---|---|---|---|---|
| 8_12308288 | 2 | 0.5 | 2 | 0.5 | 3 | 0 | 2 | 0.5 | 9 | 0.3333 | 1 | 0 | 1 | 1 | 1 | 0 | 2 | 1 | 5 | 0.6 |
| 8_12308297 | 2 | 1 | 3 | 1 | 3 | 1 | 2 | 1 | 10 | 1 | 1 | 1 | 1 | 1 | 2 | 1 | 3 | 0.6667 | 7 | 0.8571 |
| 8_12308297 | 2 | 0 | 3 | 0.6667 | 3 | 0 | 2 | 1 | 10 | 0.4 | 1 | 0 | 1 | 0 | 1 | 0 | 3 | 1 | 6 | 0.5 |
| 8_12308299 | 2 | 1 | 1 | 1 | 2 | 1 | 2 | 1 | 7 | 1 | NA | NA | 1 | 1 | 2 | 1 | 3 | 1 | 6 | 1 |
| 8_12308302 | 2 | 1 | 3 | 1 | 3 | 1 | 2 | 1 | 10 | 1 | NA | NA | 1 | 1 | 2 | 1 | 3 | 1 | 6 | 1 |
| 8_12308304 | 2 | 0 | 3 | 0.6667 | 3 | 0 | 2 | 1 | 10 | 0.4 | NA | NA | 1 | 1 | 2 | 0 | 3 | 0.3333 | 6 | 0.3333 |
| 8_12308312 | 2 | 0.5 | 3 | 1 | 3 | 1 | 2 | 1 | 10 | 0.9 | NA | NA | 1 | 1 | 2 | 1 | 3 | 1 | 6 | 1 |
| 8_12308314 | 1 | 1 | 2 | 0.5 | 2 | 0 | 1 | 0 | 6 | 0.3333 | NA | NA | 1 | 1 | 2 | 0.5 | 3 | 0.6667 | 6 | 0.6667 |
| 8_12308318 | 1 | 1 | 3 | 0.6667 | 3 | 0.6667 | 2 | 1 | 9 | 0.7778 | NA | NA | 1 | 1 | 2 | 0 | 3 | 0.3333 | 6 | 0.3333 |
| 8_12308334 | 2 | 1 | 3 | 0.6667 | 3 | 0.6667 | 1 | 1 | 9 | 0.7778 | NA | NA | 1 | 1 | 2 | 1 | 1 | 1 | 4 | 1 |
| 8_12308345 | 1 | 0 | 1 | 0 | 3 | 0.6667 | NA | NA | 5 | 0.4 | NA | NA | NA | NA | 1 | 0 | 2 | 0.5 | 3 | 0.3333 |
| 8_12308347 | 2 | 1 | 3 | 1 | 3 | 1 | 1 | 1 | 9 | 1 | NA | NA | 1 | 1 | 2 | 1 | 2 | 1 | 5 | 1 |
| 8_12308369 | 1 | 1 | 3 | 1 | 2 | 1 | NA | NA | 6 | 1 | NA | NA | NA | NA | 2 | 1 | 1 | 1 | 3 | 1 |
| 8_12308372 | 1 | 1 | 2 | 1 | 3 | 1 | NA | NA | 6 | 1 | NA | NA | 1 | 1 | 2 | 1 | 2 | 1 | 5 | 1 |
| 8_12308382 | 1 | 0 | 1 | 0 | 3 | 0.6667 | NA | NA | 5 | 0.4 | NA | NA | 1 | 0 | 2 | 1 | 2 | 0 | 5 | 0.4 |
| 8_12308384 | 1 | 1 | 2 | 1 | 3 | 1 | NA | NA | 6 | 1 | NA | NA | 1 | 1 | 2 | 1 | 2 | 1 | 5 | 1 |
| 8_12308394 | 1 | 1 | 2 | 1 | 2 | 1 | NA | NA | 5 | 1 | NA | NA | 1 | 1 | 2 | 1 | 2 | 0.5 | 5 | 0.8 |
| 8_12308403 | 1 | 1 | 2 | 1 | 3 | 1 | NA | NA | 6 | 1 | NA | NA | 1 | 1 | 1 | 1 | 1 | 1 | 3 | 1 |
| 8_12308541 | 1 | 1 | 1 | 1 | 3 | 1 | NA | NA | 5 | 1 | NA | NA | NA | NA | 2 | 0.5 | 1 | 1 | 3 | 0.6667 |
| 8_12308716 | 1 | 1 | 1 | 1 | 2 | 1 | NA | NA | 4 | 1 | NA | NA | NA | NA | 2 | 0.5 | 1 | 1 | 3 | 0.6667 |
| 8_12308773 | NA | NA | 1 | 1 | 2 | 0.5 | NA | NA | 3 | 0.6667 | NA | NA | NA | NA | 1 | 1 | 1 | 1 | 2 | 1 |
| 8_12308798 | NA | NA | 1 | 1 | 2 | 1 | NA | NA | 3 | 1 | NA | NA | NA | NA | 1 | 1 | 1 | 1 | 2 | 1 |
| 8_12311033 | 1 | 1 | NA | NA | NA | NA | 1 | 1 | 2 | 1 | 1 | 0 | NA | NA | 2 | 0.5 | NA | NA | 3 | 0.3333 |
| 8_12311050 | 1 | 0 | NA | NA | NA | NA | 1 | 1 | 2 | 0.5 | 1 | 1 | NA | NA | 2 | 0.5 | NA | NA | 3 | 0.6667 |
| 8_12311052 | 1 | 1 | NA | NA | NA | NA | 1 | 1 | 2 | 1 | 1 | 0 | NA | NA | 2 | 1 | NA | NA | 3 | 0.6667 |
| 8_12314343 | 4 | 0.5 | 2 | 0.5 | 3 | 1 | 5 | 0.8 | 14 | 0.7143 | 4 | 0.25 | 2 | 0.5 | 5 | 0.4 | 4 | 1 | 15 | 0.5333 |
| 8_12314361 | 4 | 0.5 | 2 | 0 | 3 | 0.6667 | 5 | 0.2 | 14 | 0.3571 | 4 | 0 | 3 | 0 | 5 | 0.6 | 4 | 0.25 | 16 | 0.25 |
| 8_12314365 | 2 | 0.5 | 2 | 0 | 2 | 0 | 2 | 0.5 | 8 | 0.25 | 4 | 0.5 | 3 | 1 | 5 | 0.6 | 3 | 0.3333 | 15 | 0.6 |
| 8_12314365 | 4 | 0.5 | 2 | 0.5 | 3 | 0 | 5 | 0 | 14 | 0.2143 | 4 | 0.25 | 3 | 0 | 5 | 0.4 | 4 | 0.75 | 16 | 0.375 |
| 8_12314368 | 4 | 0 | 2 | 0.5 | 3 | 0.3333 | 5 | 0.2 | 14 | 0.2143 | 4 | 0.25 | 3 | 0 | 6 | 0.1667 | 4 | 0.25 | 17 | 0.1765 |
| 8_12314380 | 5 | 0.4 | 2 | 0 | 2 | 0 | 5 | 0.2 | 14 | 0.2143 | 4 | 0.25 | 3 | 0.3333 | 7 | 0.2857 | 3 | 0.6667 | 17 | 0.3529 |
| 8_12314381 | 4 | 0.25 | 2 | 0.5 | 2 | 0.5 | 3 | 0.3333 | 11 | 0.3636 | 4 | 0 | 2 | 0 | 7 | 0.2857 | 4 | 0.25 | 17 | 0.1765 |
| 8_12314382 | 5 | 0.8 | 2 | 1 | 2 | 0.5 | 3 | 1 | 12 | 0.8333 | 4 | 0.5 | 2 | 0.5 | 5 | 1 | 2 | 0.5 | 13 | 0.6923 |
| 8_12314392 | 4 | 0.25 | 3 | 0 | 3 | 0.3333 | 6 | 0 | 16 | 0.125 | 4 | 0 | 2 | 0 | 6 | 0.1667 | 4 | 0.25 | 16 | 0.125 |
| 8_12316061 | 3 | 0.3333 | 2 | 0.5 | 4 | 0.75 | 2 | 0.5 | 11 | 0.5455 | 2 | 0.5 | 1 | 1 | 1 | 0 | NA | NA | 4 | 0.5 |
| 8_12316078 | 2 | 0 | 1 | 1 | 3 | 0.6667 | NA | NA | 6 | 0.5 | 1 | 0 | NA | NA | 1 | 1 | NA | NA | 2 | 0.5 |
| 8_12316327 | 3 | 0 | NA | NA | 4 | 0.25 | 2 | 0 | 9 | 0.1111 | 2 | 0.5 | 1 | 0 | 3 | 0 | NA | NA | 6 | 0.1667 |
| 8_12319361 | 10 | 0 | 5 | 0.2 | 6 | 0.1667 | 2 | 0 | 23 | 0.087 | 6 | 0.1667 | 9 | 0.1111 | 3 | 0 | 6 | 0.3333 | 24 | 0.1667 |
| 8_12319391 | 8 | 0.25 | 4 | 0 | 4 | 0.25 | 1 | 1 | 17 | 0.2353 | 6 | 0 | 6 | 0.1667 | 2 | 0.5 | 5 | 0.4 | 19 | 0.2105 |
| 8_12320873 | 1 | 1 | 1 | 0 | NA | NA | 1 | 1 | 3 | 0.6667 | 1 | 0 | 1 | 1 | NA | NA | NA | NA | 2 | 0.5 |
| 8_12329144 | NA | NA | NA | NA | NA | NA | 1 | 1 | 1 | 1 | NA | NA | NA | NA | 1 | 1 | NA | NA | 1 | 1 |
| 8_12329154 | NA | NA | NA | NA | NA | NA | 1 | 1 | 1 | 1 | NA | NA | NA | NA | 1 | 1 | NA | NA | 1 | 1 |
| 8_12329161 | NA | NA | NA | NA | NA | NA | 1 | 1 | 1 | 1 | NA | NA | NA | NA | 1 | 1 | NA | NA | 1 | 1 |
| 8_12329211 | NA | NA | NA | NA | NA | NA | 1 | 1 | 1 | 1 | NA | NA | NA | NA | 1 | 1 | NA | NA | 1 | 1 |
| 8_12329364 | NA | NA | NA | NA | NA | NA | 1 | 1 | 1 | 1 | NA | NA | 1 | 0 | 1 | 1 | 2 | 0 | 4 | 0.25 |
| 8_12329371 | NA | NA | NA | NA | NA | NA | 1 | 1 | 1 | 1 | NA | NA | 1 | 0 | 1 | 1 | 2 | 0 | 4 | 0.25 |
| 8_12329385 | NA | NA | NA | NA | NA | NA | 1 | 1 | 1 | 1 | NA | NA | 1 | 0 | 1 | 0 | 2 | 1 | 4 | 0.5 |
| 8_12329397 | NA | NA | NA | NA | NA | NA | 1 | 1 | 1 | 1 | NA | NA | NA | NA | 1 | 1 | 2 | 0.5 | 3 | 0.6667 |
| 8_12329417 | NA | NA | NA | NA | NA | NA | 1 | 1 | 1 | 1 | NA | NA | NA | NA | 1 | 1 | NA | NA | 1 | 1 |
| 8_12330157 | 2 | 0.5 | 5 | 0.2 | 6 | 0.5 | 5 | 0.2 | 18 | 0.3333 | 6 | 0 | 9 | 0.1111 | 4 | 0.5 | 4 | 0.5 | 23 | 0.2174 |
| 8_12330166 | 2 | 0.5 | 6 | 0.1667 | 6 | 0.1667 | 7 | 0 | 21 | 0.1429 | 6 | 0.3333 | 8 | 0.25 | 4 | 0.25 | 4 | 0.5 | 22 | 0.3182 |
| 8_12330518 | NA | NA | 2 | 1 | 2 | 1 | 1 | 1 | 5 | 1 | 1 | 1 | NA | NA | NA | NA | NA | NA | 1 | 1 |
| 8_12330525 | NA | NA | 2 | 1 | 1 | 1 | 1 | 0 | 4 | 0.75 | 1 | 1 | NA | NA | 1 | 1 | NA | NA | 2 | 1 |
| 8_12333996 | 12 | 0.1667 | 14 | 0 | 12 | 0.0833 | 15 | 0.2 | 53 | 0.1132 | 6 | 0.1667 | 12 | 0.1667 | 16 | 0.25 | 6 | 0.1667 | 40 | 0.2 |
| 8_12335237 | NA | NA | NA | NA | 1 | 1 | 1 | 1 | 2 | 1 | 1 | 0 | NA | NA | NA | NA | 1 | 1 | 2 | 0.5 |
| 8_12335265 | NA | NA | NA | NA | 1 | 0 | 1 | 1 | 2 | 0.5 | NA | NA | 1 | 1 | NA | NA | 1 | 0 | 2 | 0.5 |
| 8_12335569 | 1 | 0 | 4 | 0.5 | 5 | 0.2 | 6 | 0.1667 | 16 | 0.25 | 4 | 0.5 | 2 | 0.5 | 2 | 0 | 3 | 0 | 11 | 0.2727 |
| 8_12335570 | 1 | 0 | 2 | 0.5 | 3 | 0.6667 | 5 | 0.4 | 11 | 0.4545 | 4 | 0.75 | 1 | 1 | 2 | 0.5 | 1 | 1 | 8 | 0.75 |
| 8_12335596 | 1 | 0 | 4 | 0.25 | 3 | 0 | 4 | 0.25 | 12 | 0.1667 | 4 | 0.25 | 1 | 1 | 2 | 0 | 2 | 0.5 | 9 | 0.3333 |
| 8_12335609 | 1 | 0 | 4 | 0.25 | 5 | 0 | 4 | 0.5 | 14 | 0.2143 | 4 | 0.25 | 1 | 0 | 2 | 0 | 3 | 0.3333 | 10 | 0.2 |
| 8_12335611 | NA | NA | 4 | 0 | 2 | 0 | 3 | 0.3333 | 9 | 0.1111 | 2 | 0 | 1 | 0 | 1 | 1 | 3 | 0.3333 | 7 | 0.2857 |
| 8_12335613 | 1 | 0 | 2 | 1 | 5 | 0.4 | 4 | 0 | 12 | 0.3333 | 3 | 0.3333 | 1 | 0 | 2 | 0.5 | 1 | 0 | 7 | 0.2857 |
| 8_12335614 | 1 | 1 | 4 | 1 | 2 | 0.5 | 2 | 0.5 | 9 | 0.7778 | 4 | 1 | NA | NA | 2 | 1 | 3 | 0.3333 | 9 | 0.7778 |
| 8_12335622 | 1 | 1 | 4 | 1 | 5 | 1 | 4 | 1 | 14 | 1 | 4 | 1 | NA | NA | 2 | 1 | 3 | 1 | 9 | 1 |
| 8_12335658 | NA | NA | 4 | 0 | 5 | 0.2 | 4 | 0.25 | 13 | 0.1538 | 3 | 0.3333 | NA | NA | 2 | 0 | 2 | 0 | 7 | 0.1429 |

| | | | | | | | | | | | | | | | | | | | | |
|---|---|---|---|---|---|---|---|---|---|---|---|---|---|---|---|---|---|---|---|---|
| 8_12335674 | NA | NA | 4 | 0.75 | 4 | 0.5 | 4 | 0.5 | 12 | 0.5833 | 1 | 0 | NA | NA | 4 | 0.75 | 1 | 1 | 6 | 0.6667 |
| 8_12335688 | NA | NA | 5 | 1 | 4 | 1 | 2 | 1 | 11 | 1 | 3 | 1 | NA | NA | 3 | 1 | NA | NA | 6 | 1 |
| 8_12335692 | NA | NA | 3 | 1 | 4 | 0.75 | 2 | 1 | 9 | 0.8889 | 3 | 1 | NA | NA | 3 | 1 | 2 | 1 | 8 | 1 |
| 8_12335693 | NA | NA | 2 | 1 | 3 | 1 | 3 | 1 | 8 | 1 | 4 | 1 | NA | NA | NA | NA | 2 | 1 | 6 | 1 |
| 8_12335699 | NA | NA | 4 | 0.75 | 3 | 0.3333 | 1 | 0 | 8 | 0.5 | 1 | 1 | NA | NA | 2 | 1 | 2 | 1 | 5 | 1 |
| 8_12335707 | NA | NA | 5 | 0 | 3 | 0 | 4 | 0.25 | 12 | 0.0833 | 3 | 0 | NA | NA | 2 | 0.5 | 1 | 0 | 6 | 0.1667 |
| 8_12335731 | NA | NA | 5 | 0.8 | 3 | 0.6667 | 2 | 0 | 10 | 0.6 | 3 | 1 | NA | NA | 3 | 0.6667 | 1 | 1 | 7 | 0.8571 |
| 8_12335755 | NA | NA | 5 | 0.8 | 3 | 0.3333 | 2 | 1 | 10 | 0.7 | 2 | 0.5 | NA | NA | 2 | 1 | NA | NA | 4 | 0.75 |
| 8_12335762 | NA | NA | 4 | 0.5 | 3 | 1 | 4 | 0.5 | 11 | 0.6364 | 1 | 1 | NA | NA | 3 | 0.6667 | 1 | 1 | 5 | 0.8 |
| 8_12345186 | 1 | 0 | 4 | 0.5 | NA | NA | 2 | 0.5 | 7 | 0.4286 | 2 | 1 | 4 | 0.75 | NA | NA | 1 | 0 | 7 | 0.7143 |
| 8_12345226 | NA | NA | 2 | 1 | NA | NA | 2 | 0.5 | 4 | 0.75 | NA | NA | 3 | 0.3333 | NA | NA | NA | NA | 3 | 0.3333 |
| 8_12345279 | 1 | 1 | 2 | 0.5 | NA | NA | 1 | 1 | 4 | 0.75 | 2 | 1 | 4 | 0.5 | NA | NA | 1 | 1 | 7 | 0.7143 |
| 8_12345948 | 1 | 1 | NA | NA | NA | NA | NA | NA | 1 | 1 | NA | NA | 1 | 1 | NA | NA | NA | NA | 1 | 1 |
| 8_12350687 | 1 | 0 | 1 | 0 | 2 | 0.5 | 2 | 0.5 | 6 | 0.3333 | 3 | 0.6667 | NA | NA | 1 | 1 | NA | NA | 4 | 0.75 |
| 8_12350689 | 1 | 0 | 1 | 0 | 2 | 0.5 | 2 | 0.5 | 6 | 0.3333 | 3 | 0.3333 | NA | NA | 1 | 0 | NA | NA | 4 | 0.25 |
| 8_12350691 | 1 | 1 | 1 | 1 | 2 | 1 | 1 | 1 | 5 | 1 | 3 | 1 | NA | NA | 1 | 1 | NA | NA | 4 | 1 |
| 8_12350712 | 1 | 1 | 1 | 1 | NA | NA | 1 | 0 | 3 | 0.6667 | 2 | 1 | NA | NA | NA | NA | NA | NA | 2 | 1 |
| 8_12354449 | 1 | 1 | 6 | 0.3333 | 3 | 0 | 3 | 0.3333 | 13 | 0.3077 | 4 | 0.5 | 1 | 0 | 1 | 0 | 2 | 0.5 | 8 | 0.375 |
| 8_12359369 | NA | NA | NA | NA | 2 | 0.5 | 1 | 1 | 3 | 0.6667 | NA | NA | 1 | 0 | 4 | 0.25 | 1 | 0 | 6 | 0.1667 |
| 8_12359371 | NA | NA | NA | NA | 2 | 0.5 | 1 | 1 | 3 | 0.6667 | NA | NA | 1 | 1 | 3 | 0.3333 | 1 | 1 | 5 | 0.6 |
| 8_12359374 | NA | NA | NA | NA | 2 | 0 | 1 | 1 | 3 | 0.3333 | NA | NA | 1 | 0 | 3 | 0.3333 | 1 | 0 | 5 | 0.2 |
| 8_12359375 | NA | NA | NA | NA | 2 | 1 | 1 | 1 | 3 | 1 | NA | NA | 1 | 1 | 4 | 1 | 1 | 1 | 6 | 1 |
| 8_12359389 | NA | NA | NA | NA | 2 | 0 | 1 | 1 | 3 | 0.3333 | NA | NA | 1 | 0 | 3 | 0.3333 | 1 | 0 | 5 | 0.2 |
| 8_12359393 | NA | NA | NA | NA | 1 | 1 | 1 | 1 | 2 | 1 | NA | NA | 1 | 1 | 4 | 0.25 | 1 | 1 | 6 | 0.5 |
| 8_12359397 | NA | NA | NA | NA | 2 | 0.5 | 1 | 1 | 3 | 0.6667 | NA | NA | 1 | 1 | 3 | 0.3333 | 1 | 1 | 5 | 0.6 |
| 8_12361036 | 9 | 0.8889 | 14 | 0.9286 | 6 | 1 | 11 | 0.9091 | 40 | 0.925 | 5 | 1 | 8 | 0.875 | 10 | 1 | 8 | 0.875 | 31 | 0.9355 |
| 8_12361038 | 9 | 0.1111 | 14 | 0.1429 | 7 | 0 | 13 | 0.1538 | 43 | 0.1163 | 4 | 0.25 | 8 | 0.125 | 11 | 0.0909 | 9 | 0.2222 | 32 | 0.1562 |
| 8_12361042 | 9 | 0.8889 | 11 | 1 | 5 | 1 | 12 | 0.9167 | 37 | 0.9459 | 5 | 1 | 7 | 0.8571 | 10 | 1 | 8 | 1 | 30 | 0.9667 |
| 8_12361050 | 10 | 0.6 | 12 | 1 | 7 | 1 | 10 | 0.9 | 39 | 0.8718 | 5 | 1 | 8 | 0.875 | 11 | 0.9091 | 7 | 1 | 31 | 0.9355 |
| 8_12361063 | 9 | 0.1111 | 10 | 0 | 3 | 0 | 12 | 0.25 | 34 | 0.1176 | 3 | 0 | 5 | 0 | 10 | 0.1 | 7 | 0.1429 | 25 | 0.08 |
| 8_12361064 | 10 | 0.2 | 14 | 0 | 3 | 0 | 13 | 0.1538 | 40 | 0.1 | 2 | 0 | 6 | 0 | 10 | 0.4 | 9 | 0.1111 | 27 | 0.1852 |
| 8_12361065 | 8 | 0.5 | 10 | 0.7 | 2 | 0.5 | 12 | 0.6667 | 32 | 0.625 | 2 | 0.5 | 5 | 0.6 | 10 | 0.7 | 7 | 0.5714 | 24 | 0.625 |
| 8_12361068 | 9 | 0.4444 | 13 | 0.0769 | 3 | 0.3333 | 12 | 0.0833 | 37 | 0.1892 | 2 | 0.5 | 7 | 0.4286 | 10 | 0.4 | 8 | 0.25 | 27 | 0.3704 |
| 8_12361077 | 10 | 0.9 | 14 | 0.9286 | 2 | 0.5 | 12 | 0.9167 | 38 | 0.8947 | 4 | 1 | 8 | 0.875 | 11 | 0.9091 | 9 | 1 | 32 | 0.9375 |
| 8_12361086 | 10 | 0.8 | 13 | 0.8462 | 3 | 1 | 12 | 0.75 | 38 | 0.8158 | 3 | 0.3333 | 5 | 0.6 | 11 | 0.7273 | 7 | 0.7143 | 26 | 0.6538 |
| 8_12361655 | 6 | 0.1667 | 12 | 0.25 | 3 | 0 | 13 | 0.3077 | 34 | 0.2353 | 2 | 0 | 5 | 0 | 7 | 0.1429 | 6 | 0 | 20 | 0.05 |
| 8_12361676 | 7 | 0.2857 | 8 | 0.25 | 2 | 0 | 11 | 0.1818 | 28 | 0.2143 | 2 | 1 | 5 | 0.2 | 8 | 0.125 | 7 | 0.4286 | 22 | 0.3182 |
| 8_12361700 | 6 | 0.8333 | 7 | 0.1429 | 2 | 0 | 10 | 0.7 | 25 | 0.52 | 2 | 0.5 | 4 | 0.75 | 8 | 0.5 | 5 | 0.6 | 19 | 0.5789 |
| 8_12361704 | 6 | 0 | 8 | 0 | 2 | 0 | 9 | 0.2222 | 25 | 0.08 | 2 | 0 | 4 | 0.25 | 8 | 0 | 4 | 0 | 18 | 0.0556 |
| 8_12361753 | 5 | 0 | 8 | 0.25 | 1 | 0 | 10 | 0.3 | 24 | 0.2083 | 1 | 1 | 2 | 0 | 7 | 0 | 3 | 0 | 13 | 0.0769 |
| 8_12361757 | 5 | 0 | 7 | 0.1429 | 1 | 0 | 7 | 0.2857 | 20 | 0.15 | 2 | 0 | 2 | 0 | 6 | 0 | 4 | 0.25 | 14 | 0.0714 |
| 8_12361758 | 2 | 0 | 6 | 0.5 | 1 | 1 | 9 | 0.1111 | 18 | 0.2778 | 2 | 1 | 2 | 1 | 5 | 0.2 | 3 | 0.6667 | 12 | 0.5833 |
| 8_12361770 | 5 | 0 | 2 | 0 | 1 | 0 | 8 | 0.125 | 16 | 0.0625 | 1 | 1 | 2 | 0 | 5 | 0 | 4 | 0 | 12 | 0.0833 |
| 8_12361779 | 5 | 0.4 | 8 | 0.125 | 1 | 1 | 9 | 0.2222 | 23 | 0.2609 | 1 | 0 | 3 | 0.6667 | 6 | 0 | 4 | 0 | 14 | 0.1429 |
| 8_12361797 | 4 | 0.75 | 5 | 0.8 | 1 | 1 | 6 | 1 | 16 | 0.875 | 1 | 1 | 2 | 0 | 5 | 0.4 | 2 | 0.5 | 10 | 0.4 |
| 8_12361811 | 4 | 0.5 | 6 | 0.5 | 1 | 1 | 9 | 0.4444 | 20 | 0.5 | NA | NA | 3 | 0.6667 | 6 | 0.3333 | 2 | 0 | 11 | 0.3636 |
| 8_12361814 | 3 | 0.6667 | 5 | 0.4 | 1 | 1 | 7 | 1 | 16 | 0.75 | NA | NA | 2 | 0.5 | 4 | 0.75 | 1 | 1 | 7 | 0.7143 |
| 8_12361824 | 5 | 0.4 | 6 | 0.3333 | 1 | 1 | 8 | 0.625 | 20 | 0.5 | NA | NA | 2 | 0 | 6 | 0.3333 | 2 | 0 | 10 | 0.2 |
| 8_12361827 | 3 | 0.3333 | 5 | 0.2 | 1 | 0 | 8 | 0.625 | 17 | 0.4118 | NA | NA | 3 | 0 | 6 | 0.5 | 1 | 1 | 10 | 0.4 |
| 8_12361836 | 3 | 0.3333 | 4 | 0.25 | 1 | 0 | 9 | 0.2222 | 17 | 0.2353 | NA | NA | 3 | 0.3333 | 4 | 0 | 2 | 0 | 9 | 0.1111 |
| 8_12361841 | 5 | 0.4 | 4 | 0.25 | 1 | 0 | 7 | 0 | 17 | 0.1765 | NA | NA | 3 | 0.3333 | 4 | 0.25 | 2 | 0.5 | 9 | 0.3333 |
| 8_12361844 | 2 | 1 | 5 | 0.4 | 1 | 1 | 7 | 0.4286 | 15 | 0.5333 | NA | NA | 2 | 1 | 4 | 0.75 | 2 | 1 | 8 | 0.875 |
| 8_12361880 | 1 | 0 | 1 | 0 | NA | NA | 7 | 0.5714 | 9 | 0.4444 | NA | NA | 2 | 0 | 2 | 0.5 | 2 | 0.5 | 6 | 0.3333 |
| 8_12361891 | 4 | 0.5 | 1 | 0 | NA | NA | 7 | 0.4286 | 12 | 0.4167 | NA | NA | 1 | 1 | 4 | 0.5 | 1 | 1 | 6 | 0.6667 |
| 8_12361913 | 3 | 0.3333 | 1 | 0 | NA | NA | 9 | 0.3333 | 13 | 0.3077 | NA | NA | 2 | 0.5 | 5 | 0 | 2 | 0 | 9 | 0.1111 |
| 8_12361936 | 3 | 0.3333 | NA | NA | NA | NA | 7 | 0.1429 | 10 | 0.2 | NA | NA | 1 | 1 | 5 | 0.2 | 2 | 0 | 8 | 0.25 |
| 8_12361959 | 1 | 1 | NA | NA | NA | NA | 6 | 1 | 7 | 1 | NA | NA | 2 | 1 | 4 | 0.75 | NA | NA | 6 | 0.8333 |
| 8_12361977 | 1 | 0 | NA | NA | NA | NA | 5 | 0.2 | 6 | 0.1667 | NA | NA | 1 | 0 | 5 | 0.2 | 1 | 0 | 7 | 0.1429 |
| 8_12361979 | 2 | 0.5 | NA | NA | NA | NA | 4 | 0.25 | 6 | 0.3333 | NA | NA | NA | NA | 3 | 0 | 1 | 1 | 4 | 0.25 |
| 8_12362015 | 3 | 0.3333 | NA | NA | NA | NA | 6 | 0.1667 | 9 | 0.2222 | NA | NA | 1 | 0 | 4 | 0.25 | 1 | 0 | 6 | 0.1667 |
| 8_12362104 | 1 | 1 | NA | NA | NA | NA | 4 | 0.75 | 5 | 0.8 | NA | NA | 2 | 0.5 | 4 | 1 | NA | NA | 6 | 0.8333 |
| 8_12362118 | 1 | 0 | NA | NA | NA | NA | 3 | 0.3333 | 4 | 0.25 | NA | NA | 2 | 0 | 3 | 0.6667 | NA | NA | 5 | 0.4 |
| 8_12362120 | NA | NA | NA | NA | NA | NA | 3 | 1 | 3 | 1 | NA | NA | 2 | 1 | 2 | 1 | NA | NA | 4 | 1 |
| 8_12362165 | NA | NA | NA | NA | NA | NA | 4 | 0.5 | 4 | 0.5 | NA | NA | 1 | 1 | 2 | 1 | NA | NA | 3 | 1 |
| 8_12362207 | NA | NA | NA | NA | NA | NA | 4 | 0.75 | 4 | 0.75 | NA | NA | 1 | 1 | 1 | 0 | NA | NA | 2 | 0.5 |

| | | | | | | | | | | | | | | | | | | | | |
|---|---|---|---|---|---|---|---|---|---|---|---|---|---|---|---|---|---|---|---|---|
| 8_12362225 | NA | NA | NA | NA | NA | NA | 4 | 0.25 | 4 | 0.25 | NA | NA | 2 | 0.5 | 3 | 0.3333 | NA | NA | 5 | 0.4 |
| 8_12362282 | NA | NA | NA | NA | NA | NA | 4 | 0.75 | 4 | 0.75 | NA | NA | 1 | 1 | 2 | 0.5 | NA | NA | 3 | 0.6667 |
| 8_12362397 | 2 | 0 | 4 | 0.25 | 5 | 0.4 | 6 | 0.1667 | 17 | 0.2353 | 1 | 0 | 2 | 0 | 4 | 0.5 | 5 | 0.2 | 12 | 0.25 |
| 8_12362404 | 1 | 0 | 5 | 0 | 6 | 0.3333 | 3 | 0.3333 | 15 | 0.2 | NA | NA | 1 | 0 | 3 | 0 | 3 | 0.3333 | 7 | 0.1429 |
| 8_12362502 | NA | NA | 1 | 0 | 2 | 1 | 5 | 0.2 | 8 | 0.375 | NA | NA | NA | NA | 3 | 0 | 3 | 0.3333 | 6 | 0.1667 |
| 8_12362520 | NA | NA | 1 | 0 | 5 | 0.2 | 4 | 0 | 10 | 0.1 | NA | NA | NA | NA | 3 | 0 | 3 | 0.6667 | 6 | 0.3333 |
| 8_12362526 | NA | NA | 1 | 1 | 3 | 0.3333 | 4 | 0.5 | 8 | 0.5 | NA | NA | 1 | 1 | 2 | 1 | 3 | 0.6667 | 6 | 0.8333 |
| 8_12362537 | NA | NA | 1 | 1 | 4 | 0 | 5 | 0 | 10 | 0.1 | NA | NA | 2 | 0 | 5 | 0.2 | 2 | 0 | 9 | 0.1111 |
| 8_12376957 | 2 | 0.5 | 2 | 1 | 2 | 1 | 1 | 1 | 7 | 0.8571 | 3 | 1 | 2 | 1 | 3 | 1 | 1 | 1 | 9 | 1 |
| 8_12376963 | 2 | 0.5 | 3 | 0.3333 | 1 | 0 | 1 | 0 | 7 | 0.2857 | 2 | 1 | 1 | 1 | 2 | 0.5 | 1 | 1 | 6 | 0.8333 |
| 8_12376969 | 2 | 0.5 | 3 | 0.6667 | 2 | 0 | 1 | 1 | 8 | 0.5 | 2 | 0.5 | 2 | 1 | 4 | 0.25 | 1 | 0 | 9 | 0.4444 |
| 8_12376975 | 3 | 0.6667 | 3 | 1 | 2 | 0 | 1 | 0 | 9 | 0.5556 | 2 | 0.5 | 3 | 0.6667 | 4 | 1 | 1 | 1 | 10 | 0.8 |
| 8_12376989 | 2 | 0.5 | 3 | 0.3333 | 2 | 0 | 1 | 0 | 8 | 0.25 | 2 | 0.5 | 3 | 0.3333 | 3 | 0 | 1 | 1 | 9 | 0.3333 |
| 8_12376993 | 2 | 1 | 3 | 1 | 1 | 1 | NA | NA | 6 | 1 | 2 | 1 | 2 | 0.5 | 3 | 1 | 1 | 1 | 8 | 0.875 |
| 8_12377023 | 2 | 0.5 | 3 | 0.6667 | 3 | 1 | 1 | 0 | 9 | 0.6667 | 3 | 0.6667 | 3 | 1 | 4 | 0.25 | 3 | 1 | 13 | 0.6923 |
| 8_12377025 | 2 | 0 | 2 | 0.5 | 2 | 0.5 | 1 | 0 | 7 | 0.2857 | 1 | 0 | 2 | 0 | 3 | 0.3333 | 3 | 0 | 9 | 0.1111 |
| 8_12377032 | 2 | 0.5 | 3 | 0.6667 | 3 | 0.6667 | 1 | 1 | 9 | 0.6667 | 3 | 1 | 3 | 1 | 5 | 0.4 | 3 | 0.3333 | 14 | 0.6429 |
| 8_12377081 | 3 | 1 | 3 | 0.6667 | 3 | 0 | 1 | 0 | 10 | 0.5 | 3 | 0.3333 | 1 | 0 | 5 | 0.6 | 3 | 1 | 12 | 0.5833 |
| 8_12377082 | 2 | 0 | 3 | 0.3333 | 2 | 0 | 1 | 0 | 8 | 0.125 | 3 | 0 | 2 | 0 | 5 | 0.2 | 2 | 0 | 12 | 0.0833 |
| 8_12377086 | 3 | 0.6667 | 3 | 0.6667 | 3 | 1 | 1 | 0 | 10 | 0.7 | 3 | 0.3333 | 2 | 0.5 | 5 | 0.8 | 3 | 1 | 13 | 0.6923 |
| 8_12377087 | 3 | 0.3333 | 3 | 0.6667 | 1 | 0 | 1 | 0 | 8 | 0.375 | 2 | 0.5 | 2 | 0.5 | 4 | 0.5 | 2 | 0 | 10 | 0.4 |
| 8_12377093 | 2 | 1 | 3 | 1 | 2 | 1 | 1 | 1 | 8 | 1 | 2 | 1 | 1 | 1 | 6 | 1 | 2 | 1 | 11 | 1 |
| 8_12377110 | 3 | 0.6667 | 3 | 1 | 3 | 0.3333 | 1 | 1 | 10 | 0.7 | 3 | 0.3333 | 3 | 0.6667 | 6 | 0.5 | 3 | 1 | 15 | 0.6 |
| 8_12377113 | 2 | 1 | NA | NA | 2 | 1 | 1 | 1 | 5 | 1 | 2 | 1 | 3 | 1 | 5 | 0.6 | 3 | 1 | 13 | 0.8462 |
| 8_12377116 | 3 | 1 | 3 | 1 | 2 | 1 | 1 | 1 | 9 | 1 | 3 | 1 | 3 | 1 | 6 | 1 | 2 | 1 | 14 | 1 |
| 8_12377129 | 2 | 0.5 | 3 | 0.6667 | 2 | 1 | 1 | 0 | 8 | 0.625 | 3 | 0.6667 | 3 | 0 | 7 | 0 | 3 | 0.3333 | 16 | 0.1875 |
| 8_12382417 | 4 | 0 | 7 | 0.2857 | 4 | 0.75 | 4 | 0.75 | 19 | 0.4211 | 4 | 0.25 | 3 | 0 | 8 | 0.375 | 1 | 0 | 16 | 0.25 |
| 8_12382426 | 7 | 0 | 5 | 0 | 2 | 0 | 5 | 0.2 | 19 | 0.0526 | 3 | 0 | 3 | 0 | 10 | 0.1 | 2 | 0 | 18 | 0.0556 |
| 8_12382435 | 7 | 0 | 6 | 0.1667 | 6 | 0 | 6 | 0.6667 | 25 | 0.2 | 5 | 0.4 | 4 | 0.75 | 7 | 0.1429 | 3 | 0.3333 | 19 | 0.3684 |
| 8_12382440 | 5 | 0.4 | 8 | 0.375 | 4 | 0.5 | 4 | 0 | 21 | 0.3333 | 4 | 0.5 | 3 | 0.6667 | 9 | 0.2222 | 2 | 0 | 18 | 0.3333 |
| 8_12382444 | 7 | 0 | 9 | 0.4444 | 5 | 0.6 | 7 | 0.2857 | 28 | 0.3214 | 6 | 0.3333 | 5 | 0.2 | 10 | 0.3 | 2 | 0 | 23 | 0.2609 |
| 8_12382461 | NA | NA | NA | NA | NA | NA | 1 | 1 | 1 | 1 | 1 | 1 | NA | NA | NA | NA | NA | NA | 1 | 1 |
| 8_12385338 | 1 | 0 | 6 | 0 | 3 | 0.3333 | NA | NA | 10 | 0.1 | 2 | 0 | 2 | 0 | 1 | 0 | 4 | 0.25 | 9 | 0.1111 |
| 8_12386138 | 4 | 0.5 | 11 | 0.2727 | 3 | 0.3333 | 5 | 0.2 | 23 | 0.3043 | 5 | 0.2 | 4 | 0.25 | 2 | 0.5 | 5 | 0.2 | 16 | 0.25 |
| 8_12386173 | 4 | 0.25 | 9 | 0.2222 | 4 | 0.25 | 3 | 0.3333 | 20 | 0.25 | 8 | 0.25 | 3 | 0.3333 | 3 | 0.3333 | 8 | 0.5 | 22 | 0.3636 |
| 8_12386179 | 5 | 0 | 10 | 0 | 4 | 0.25 | 5 | 0.2 | 24 | 0.0833 | 9 | 0.2222 | 4 | 0 | 2 | 0 | 9 | 0.1111 | 24 | 0.125 |
| 8_12393143 | 15 | 0.5333 | 19 | 0.8421 | 7 | 0.7143 | 16 | 0.8125 | 57 | 0.7368 | 3 | 1 | 17 | 0.7647 | 6 | 0.8333 | 8 | 0.75 | 34 | 0.7941 |
| 8_12393144 | 15 | 0.0667 | 18 | 0 | 8 | 0 | 16 | 0.125 | 57 | 0.0526 | 4 | 0.25 | 18 | 0.2778 | 8 | 0.125 | 8 | 0.25 | 38 | 0.2368 |
| 8_12393150 | 15 | 0.1333 | 21 | 0.1429 | 9 | 0.2222 | 16 | 0.25 | 61 | 0.1803 | 5 | 0 | 18 | 0.1667 | 10 | 0.3 | 10 | 0.3 | 43 | 0.2093 |
| 8_12393164 | 14 | 0.1429 | 19 | 0.1053 | 9 | 0 | 15 | 0 | 57 | 0.0702 | 2 | 0.5 | 19 | 0.1579 | 9 | 0 | 10 | 0 | 40 | 0.1 |
| 8_12393166 | 11 | 0.5455 | 17 | 0.3529 | 9 | 0.5556 | 14 | 0.2857 | 51 | 0.4118 | 4 | 0.5 | 19 | 0.5789 | 7 | 0.5714 | 8 | 0.125 | 38 | 0.4737 |
| 8_12393171 | 12 | 0.0833 | 20 | 0.1 | 9 | 0.1111 | 14 | 0 | 55 | 0.0727 | 4 | 0 | 19 | 0.1579 | 9 | 0.2222 | 9 | 0.1111 | 41 | 0.1463 |
| 8_12393176 | 10 | 0.9 | 17 | 1 | 8 | 0.875 | 10 | 0.9 | 45 | 0.9333 | 2 | 1 | 12 | 1 | 9 | 1 | 10 | 0.9 | 33 | 0.9697 |
| 8_12393183 | 13 | 0.5385 | 19 | 0.3158 | 9 | 0.3333 | 13 | 0.4615 | 54 | 0.4074 | 4 | 0.25 | 17 | 0.5294 | 10 | 0.5 | 9 | 0.3333 | 40 | 0.45 |
| 8_12393196 | 12 | 0.0833 | 17 | 0.1176 | 9 | 0.1111 | 12 | 0 | 50 | 0.08 | 2 | 0 | 13 | 0.2308 | 8 | 0 | 6 | 0 | 29 | 0.1034 |
| 8_12393200 | 13 | 1 | 18 | 0.9444 | 9 | 1 | 11 | 0.9091 | 51 | 0.9608 | 4 | 1 | 15 | 0.8667 | 7 | 0.8571 | 5 | 1 | 31 | 0.9032 |
| 8_12393202 | 8 | 0 | 15 | 0.2 | 6 | 0 | 9 | 0 | 38 | 0.0789 | 3 | 0 | 13 | 0 | 7 | 0.1429 | 6 | 0.1667 | 29 | 0.069 |
| 8_12393207 | 9 | 0.1111 | 16 | 0.1875 | 7 | 0.1429 | 9 | 0.1111 | 41 | 0.1463 | 4 | 0 | 13 | 0.1538 | 5 | 0 | 4 | 0 | 26 | 0.0769 |
| 8_12393214 | 12 | 0.1667 | 20 | 0.2 | 9 | 0.2222 | 12 | 0.25 | 53 | 0.2075 | 4 | 0.25 | 14 | 0.2143 | 8 | 0.625 | 6 | 0.5 | 32 | 0.375 |
| 8_12393217 | 12 | 1 | 19 | 0.8947 | 8 | 0.875 | 12 | 0.8333 | 51 | 0.902 | 3 | 1 | 15 | 0.7333 | 6 | 0.6667 | 4 | 0.75 | 28 | 0.75 |
| 8_12393233 | 11 | 0.9091 | 17 | 0.8235 | 8 | 0.625 | 10 | 0.7 | 46 | 0.7826 | 3 | 1 | 11 | 0.7273 | 6 | 0.8333 | 3 | 1 | 23 | 0.8261 |
| 8_12395163 | 4 | 0.25 | 4 | 0.25 | 3 | 0 | 2 | 0 | 13 | 0.1538 | 4 | 0.25 | 2 | 0 | 3 | 0 | 1 | 0 | 10 | 0.1 |
| 8_12395526 | 1 | 1 | NA | NA | 1 | 1 | NA | NA | 2 | 1 | NA | NA | NA | NA | NA | NA | 1 | 1 | 1 | 1 |
| 8_12395539 | 1 | 1 | NA | NA | NA | NA | NA | NA | 1 | 1 | NA | NA | NA | NA | NA | NA | 1 | 1 | 1 | 1 |
| 8_12395571 | 1 | 1 | NA | NA | 1 | 1 | NA | NA | 2 | 1 | NA | NA | NA | NA | NA | NA | 1 | 1 | 1 | 1 |
| 8_12395657 | NA | NA | NA | NA | 1 | 1 | NA | NA | 1 | 1 | NA | NA | NA | NA | NA | NA | 1 | 1 | 1 | 1 |
| 8_12395861 | 6 | 0.5 | 7 | 0.2857 | 7 | 0 | 5 | 0.2 | 25 | 0.24 | 7 | 0.2857 | 2 | 0 | 5 | 0.4 | 2 | 0.5 | 16 | 0.3125 |
| 8_12395875 | 6 | 0.3333 | 6 | 0.1667 | 8 | 0.5 | 5 | 0.6 | 25 | 0.4 | 5 | 0 | 1 | 1 | 6 | 0.6667 | 2 | 0.5 | 14 | 0.4286 |
| 8_12395883 | 6 | 0 | 7 | 0.1429 | 9 | 0.2222 | 5 | 0 | 27 | 0.1111 | 6 | 0.1667 | 2 | 0 | 8 | 0.125 | 2 | 0 | 18 | 0.1111 |
| 8_12395890 | 7 | 0 | 7 | 0.1429 | 7 | 0.4286 | 6 | 0 | 27 | 0.1481 | 6 | 0 | 2 | 0 | 9 | 0.4444 | 2 | 0 | 19 | 0.2105 |
| 8_12395896 | 5 | 0.6 | 5 | 0.6 | 9 | 0.6667 | 6 | 0.3333 | 25 | 0.56 | 5 | 0.6 | 2 | 0 | 9 | 0.5556 | 2 | 0 | 18 | 0.4444 |
| 8_12395905 | 6 | 0.6667 | 7 | 0.8571 | 5 | 0 | 6 | 0.1667 | 24 | 0.4583 | 3 | 1 | 5 | 0.2 | 9 | 0.7778 | 3 | 1 | 20 | 0.7 |
| 8_12395910 | 8 | 0.25 | 9 | 0.1111 | 6 | 0.1667 | 7 | 0.1429 | 30 | 0.1667 | 3 | 0 | 7 | 0 | 10 | 0.1 | 8 | 0.125 | 28 | 0.0714 |
| 8_12395965 | 6 | 0.1667 | 6 | 0.1667 | 5 | 0.2 | 4 | 0.25 | 21 | 0.1905 | 1 | 0 | 5 | 0.4 | 9 | 0.4444 | 5 | 0.4 | 20 | 0.4 |
| 8_12395970 | 5 | 1 | 4 | 0.5 | 4 | 0.75 | 4 | 0.75 | 17 | 0.7647 | 1 | 1 | 2 | 1 | 6 | 0.6667 | 4 | 1 | 13 | 0.8462 |

| | | | | | | | | | | | | | | | | | | | | |
|---|---|---|---|---|---|---|---|---|---|---|---|---|---|---|---|---|---|---|---|---|
| 8_1239598 | 7 | 0.7143 | 7 | 0.5714 | 6 | 0.6667 | 4 | 0.5 | 24 | 0.625 | 1 | 0 | 5 | 0.8 | 7 | 0.7143 | 2 | 1 | 15 | 0.7333 |
| 8_1239599 | 7 | 1 | 6 | 1 | 6 | 1 | 5 | 0.8 | 24 | 0.9583 | 1 | 1 | 3 | 1 | 9 | 1 | 4 | 1 | 17 | 1 |
| 8_1239599 | 6 | 1 | 5 | 1 | 5 | 1 | 5 | 1 | 21 | 1 | 1 | 1 | 4 | 0.75 | 9 | 0.6667 | 3 | 1 | 17 | 0.7647 |
| 8_1239599 | 7 | 0.7143 | 6 | 0.6667 | 5 | 0.6 | 5 | 0.4 | 23 | 0.6087 | 1 | 0 | 3 | 0.3333 | 8 | 0.5 | 3 | 0.6667 | 15 | 0.4667 |
| 8_1239600 | 3 | 0.3333 | 5 | 0.2 | 2 | 0 | 2 | 0 | 12 | 0.1667 | NA | NA | 2 | 1 | 4 | 0.5 | 3 | 0.3333 | 9 | 0.5556 |
| 8_1239603 | 7 | 0 | 7 | 0 | 5 | 0.2 | 6 | 0.3333 | 25 | 0.12 | 1 | 0 | 4 | 0 | 8 | 0.375 | 4 | 0 | 17 | 0.1765 |
| 8_1239603 | 7 | 0 | 6 | 0 | 5 | 0.4 | 6 | 0 | 24 | 0.0833 | 1 | 0 | 2 | 0 | 7 | 0.1429 | 4 | 0.25 | 14 | 0.1429 |
| 8_1239604 | 4 | 0.75 | 4 | 1 | 4 | 1 | 3 | 1 | 15 | 0.9333 | NA | NA | 3 | 0.6667 | 5 | 0.8 | 4 | 1 | 12 | 0.8333 |
| 8_1239605 | 6 | 1 | 4 | 1 | 5 | 0.8 | 4 | 1 | 19 | 0.9474 | NA | NA | 2 | 0.5 | 5 | 0.8 | 2 | 1 | 9 | 0.7778 |
| 8_1239605 | 7 | 0.4286 | 6 | 0.3333 | 4 | 0.75 | 5 | 0.2 | 22 | 0.4091 | NA | NA | 2 | 0.5 | 6 | 0.5 | 3 | 1 | 11 | 0.6364 |
| 8_1239606 | 7 | 0.7143 | 6 | 0.5 | 4 | 0.75 | 6 | 0.5 | 23 | 0.6087 | NA | NA | 2 | 0 | 7 | 0.5714 | 4 | 0.75 | 13 | 0.5385 |
| 8_1239607 | 7 | 0.1429 | 5 | 0.2 | 3 | 0.3333 | 3 | 0 | 18 | 0.1667 | NA | NA | 2 | 0 | 6 | 0.3333 | 3 | 0 | 11 | 0.1818 |
| 8_1239608 | 7 | 0.1429 | 4 | 0.25 | 3 | 0 | 4 | 0 | 18 | 0.1111 | NA | NA | 2 | 0 | 5 | 0.2 | 4 | 0.25 | 11 | 0.1818 |
| 8_1239625 | 5 | 0.4 | 5 | 0 | 3 | 0.6667 | 3 | 0.6667 | 16 | 0.375 | NA | NA | 1 | 0 | 6 | 0.3333 | 3 | 0.3333 | 10 | 0.3 |
| 8_1239692 | 4 | 0 | 2 | 0 | 4 | 0 | 3 | 0.3333 | 13 | 0.0769 | 7 | 0 | 11 | 0.0909 | 8 | 0.25 | 3 | 0.3333 | 29 | 0.1379 |
| 8_1239695 | 2 | 1 | 2 | 0.5 | 3 | 1 | 1 | 1 | 8 | 0.875 | 6 | 1 | 7 | 0.8571 | 5 | 0.6 | 2 | 1 | 20 | 0.85 |
| 8_1239697 | 3 | 0 | 2 | 0.5 | 2 | 0.5 | 2 | 0.5 | 9 | 0.3333 | 4 | 0.25 | 7 | 0.4286 | 4 | 0.75 | 1 | 1 | 16 | 0.5 |
| 8_1239734 | 3 | 0.3333 | 2 | 0 | 2 | 0 | 2 | 0.5 | 9 | 0.2222 | 1 | 0 | 11 | 0.2727 | 5 | 0.4 | NA | NA | 17 | 0.2941 |
| 8_1239735 | 2 | 0.5 | 2 | 0 | 2 | 0.5 | 2 | 0.5 | 8 | 0.375 | 1 | 0 | 10 | 0.3 | 3 | 0 | 1 | 0 | 15 | 0.2 |
| 8_1239735 | 3 | 0.3333 | 2 | 0 | 2 | 0 | 2 | 0 | 9 | 0.1111 | 1 | 0 | 11 | 0.2727 | 5 | 0.4 | 1 | 0 | 18 | 0.2778 |
| 8_1239736 | 3 | 0 | 1 | 0 | 1 | 1 | 2 | 0.5 | 7 | 0.2857 | 1 | 1 | 9 | 0.1111 | 5 | 0.2 | 1 | 1 | 16 | 0.25 |
| 8_1239737 | 2 | 0 | 2 | 0.5 | 2 | 0 | 2 | 0.5 | 8 | 0.25 | 1 | 1 | 11 | 0.5455 | 4 | 0.75 | 1 | 0 | 17 | 0.5882 |
| 8_1239833 | NA | NA | NA | NA | 1 | 0 | 2 | 0.5 | 3 | 0.3333 | NA | NA | 1 | 1 | NA | NA | NA | NA | 1 | 1 |
| 8_1239945 | NA | NA | 2 | 0.5 | 1 | 1 | 2 | 0.5 | 5 | 0.6 | NA | NA | 2 | 1 | 1 | 1 | NA | NA | 3 | 1 |
| 8_1239945 | NA | NA | 3 | 0.3333 | 1 | 1 | 2 | 0.5 | 6 | 0.5 | NA | NA | 3 | 0.3333 | 1 | 1 | NA | NA | 4 | 0.5 |
| 8_1239946 | NA | NA | 3 | 0.6667 | NA | NA | 2 | 0.5 | 5 | 0.6 | NA | NA | 3 | 0.3333 | 1 | 0 | NA | NA | 4 | 0.25 |
| 8_1239947 | NA | NA | 3 | 1 | 1 | 1 | 2 | 0.5 | 6 | 0.8333 | NA | NA | 3 | 1 | NA | NA | NA | NA | 3 | 1 |
| 8_1239948 | NA | NA | 3 | 0.6667 | 1 | 0 | 2 | 0.5 | 6 | 0.5 | NA | NA | 2 | 0.5 | 2 | 1 | NA | NA | 4 | 0.75 |
| 8_1239948 | NA | NA | 2 | 0 | 1 | 1 | 2 | 0.5 | 5 | 0.4 | NA | NA | 3 | 0.6667 | 2 | 1 | NA | NA | 5 | 0.8 |
| 8_1239953 | NA | NA | 1 | 1 | NA | NA | 2 | 0.5 | 3 | 0.6667 | NA | NA | 3 | 0.3333 | 1 | 0 | NA | NA | 4 | 0.25 |
| 8_1239954 | NA | NA | 2 | 1 | NA | NA | 2 | 0.5 | 4 | 0.75 | NA | NA | 3 | 0.3333 | 1 | 1 | NA | NA | 4 | 0.5 |
| 8_1239958 | NA | NA | 3 | 1 | NA | NA | 2 | 0.5 | 5 | 0.8 | NA | NA | 2 | 0.5 | 2 | 0.5 | NA | NA | 4 | 0.5 |
| 8_1239959 | NA | NA | 2 | 0.5 | NA | NA | 2 | 0 | 4 | 0.25 | NA | NA | 3 | 0.6667 | 2 | 1 | NA | NA | 5 | 0.8 |
| 8_1239960 | NA | NA | 3 | 0.6667 | NA | NA | NA | NA | 3 | 0.6667 | NA | NA | 3 | 0.6667 | 1 | 1 | NA | NA | 4 | 0.75 |
| 8_1239961 | NA | NA | 3 | 0.3333 | NA | NA | 2 | 0 | 5 | 0.2 | NA | NA | 3 | 0.6667 | 1 | 0 | NA | NA | 4 | 0.5 |
| 8_1239962 | NA | NA | 3 | 1 | NA | NA | 2 | 1 | 5 | 1 | NA | NA | 4 | 1 | 2 | 1 | NA | NA | 6 | 1 |
| 8_1239963 | NA | NA | 3 | 1 | NA | NA | 2 | 0.5 | 5 | 0.8 | NA | NA | 4 | 0.75 | 2 | 1 | NA | NA | 6 | 0.8333 |
| 8_1239963 | NA | NA | 2 | 1 | NA | NA | 2 | 1 | 4 | 1 | NA | NA | 4 | 1 | 2 | 1 | NA | NA | 6 | 1 |
| 8_1240961 | 1 | 0 | 3 | 0.3333 | 2 | 0.5 | 1 | 0 | 7 | 0.2857 | 2 | 0 | 3 | 0.6667 | 2 | 1 | 1 | 1 | 8 | 0.625 |
| 8_1240961 | 1 | 1 | 3 | 0.6667 | 2 | 0.5 | 1 | 1 | 7 | 0.7143 | 2 | 0 | 2 | 0.5 | 2 | 0 | 1 | 1 | 7 | 0.2857 |
| 8_1241279 | 3 | 0.3333 | 7 | 0.4286 | 2 | 0 | 4 | 0 | 16 | 0.25 | 4 | 0.25 | 2 | 0.5 | 4 | 0.5 | NA | NA | 10 | 0.4 |
| 8_1241279 | 3 | 0 | 6 | 0.6667 | 3 | 0.3333 | 4 | 0.25 | 16 | 0.375 | 2 | 0.5 | 2 | 0 | 4 | 0.75 | 1 | 0 | 9 | 0.4444 |
| 8_1241279 | 3 | 0 | 6 | 0.1667 | 2 | 0.5 | 2 | 0.5 | 13 | 0.2308 | 2 | 1 | 1 | 0 | 2 | 0.5 | 1 | 0 | 6 | 0.5 |
| 8_1241283 | 2 | 0 | 3 | 0.3333 | 3 | 0 | NA | NA | 8 | 0.125 | 1 | 0 | 1 | 1 | 1 | 0 | NA | NA | 3 | 0.3333 |
| 8_1241290 | 4 | 0.5 | 6 | 0 | 3 | 0 | 2 | 0.5 | 15 | 0.2 | 3 | 0.3333 | 2 | 0 | 1 | 0 | 1 | 0 | 7 | 0.1429 |
| 8_1241294 | 4 | 0 | 5 | 0 | 6 | 0.1667 | 2 | 0 | 17 | 0.0588 | 4 | 0.25 | 3 | 0.3333 | 2 | 0.5 | 1 | 0 | 10 | 0.3 |
| 8_1241298 | 6 | 0.1667 | 5 | 0.2 | 4 | 0.5 | 1 | 1 | 16 | 0.3125 | 3 | 0 | 3 | 0.3333 | 3 | 0.3333 | 1 | 1 | 10 | 0.3 |
| 8_1241299 | 47 | 0.1489 | 62 | 0.1129 | 60 | 0.05 | 33 | 0.1515 | 202 | 0.1089 | 55 | 0.0909 | 42 | 0.1429 | 43 | 0.0233 | 44 | 0.1136 | 184 | 0.0924 |
| 8_1241301 | 24 | 0.7083 | 26 | 0.5 | 29 | 0.6207 | 15 | 0.6667 | 94 | 0.617 | 27 | 0.7407 | 17 | 0.7647 | 26 | 0.7308 | 20 | 0.8 | 90 | 0.7556 |
| 8_1241301 | 54 | 0.0741 | 67 | 0.0597 | 64 | 0.0469 | 35 | 0.1714 | 220 | 0.0773 | 67 | 0.0448 | 45 | 0.0889 | 52 | 0.0962 | 48 | 0.0833 | 212 | 0.0755 |
| 8_1241305 | 49 | 0.3878 | 57 | 0.386 | 57 | 0.3158 | 33 | 0.3333 | 196 | 0.3571 | 63 | 0.2857 | 46 | 0.3913 | 48 | 0.2917 | 42 | 0.5476 | 199 | 0.3668 |
| 8_1241306 | 8 | 0.625 | 4 | 0.5 | NA | NA | 1 | 0 | 13 | 0.5385 | 1 | 0 | 4 | 0.5 | 1 | 0 | 3 | 0 | 9 | 0.2222 |
| 8_1241306 | 6 | 0.1667 | 4 | 0.25 | 4 | 0.75 | 3 | 0 | 17 | 0.2941 | 1 | 0 | 5 | 0.2 | 3 | 0.3333 | 4 | 0.25 | 13 | 0.2308 |
| 8_1241308 | 6 | 0.1667 | 3 | 0 | 3 | 0 | 3 | 0 | 15 | 0.0667 | 1 | 0 | 7 | 0.1429 | 3 | 0 | 3 | 0 | 14 | 0.0714 |
| 8_1241474 | 37 | 0.4054 | 58 | 0.5862 | 36 | 0.3333 | 35 | 0.4571 | 166 | 0.4639 | 42 | 0.4286 | 25 | 0.44 | 34 | 0.4412 | 38 | 0.3947 | 139 | 0.4245 |
| 8_1241474 | 36 | 0.4722 | 51 | 0.5294 | 36 | 0.4444 | 37 | 0.4324 | 160 | 0.475 | 41 | 0.5366 | 26 | 0.5385 | 30 | 0.5 | 38 | 0.4737 | 135 | 0.5111 |
| 8_1241475 | 24 | 0.125 | 36 | 0.1389 | 22 | 0.2273 | 23 | 0.2174 | 105 | 0.1714 | 22 | 0.0909 | 21 | 0.1429 | 24 | 0.0833 | 28 | 0.1786 | 95 | 0.1263 |
| 8_1241475 | 40 | 0.375 | 50 | 0.54 | 34 | 0.5 | 36 | 0.6111 | 160 | 0.5063 | 40 | 0.65 | 21 | 0.5238 | 32 | 0.5938 | 40 | 0.6 | 133 | 0.6015 |
| 8_1241477 | 50 | 0.48 | 57 | 0.4211 | 45 | 0.4 | 38 | 0.4474 | 190 | 0.4368 | 42 | 0.3333 | 32 | 0.5938 | 34 | 0.4412 | 47 | 0.5957 | 155 | 0.4903 |
| 8_1241478 | 37 | 0.1351 | 55 | 0.1455 | 40 | 0.175 | 33 | 0.1515 | 165 | 0.1515 | 42 | 0.2619 | 32 | 0.0938 | 33 | 0.1818 | 43 | 0.2558 | 150 | 0.2067 |
| 8_1241479 | 44 | 0.1136 | 61 | 0.1639 | 42 | 0.1667 | 40 | 0.125 | 187 | 0.1444 | 48 | 0.1042 | 34 | 0.1471 | 40 | 0.175 | 45 | 0.2222 | 167 | 0.1617 |
| 8_1241480 | 54 | 0.0741 | 78 | 0.1282 | 48 | 0.0417 | 45 | 0.0667 | 225 | 0.0844 | 52 | 0.0769 | 40 | 0.075 | 48 | 0.0417 | 50 | 0.04 | 190 | 0.0579 |
| 8_1241485 | 64 | 0.3594 | 92 | 0.4457 | 65 | 0.2923 | 66 | 0.3636 | 287 | 0.3728 | 73 | 0.4247 | 48 | 0.3333 | 47 | 0.3617 | 58 | 0.3621 | 226 | 0.3761 |
| 8_1241863 | 3 | 0 | 5 | 0 | 4 | 0 | 5 | 0.2 | 17 | 0.0588 | 3 | 0.3333 | 3 | 0 | 7 | 0 | 2 | 0 | 15 | 0.0667 |
| 8_1241882 | 3 | 0 | 7 | 0.1429 | 5 | 0.2 | 7 | 0.4286 | 22 | 0.2273 | 3 | 0 | 2 | 0.5 | 8 | 0.5 | 2 | 0.5 | 15 | 0.4 |

| | | | | | | | | | | | | | | | | | | | | |
|---|---|---|---|---|---|---|---|---|---|---|---|---|---|---|---|---|---|---|---|---|
| 8_1241883( | 3 | 0.3333 | 5 | 0 | 3 | 0.3333 | 6 | 0.3333 | 17 | 0.2353 | 4 | 0.5 | NA | NA | 6 | 0 | 1 | 0 | 11 | 0.1818 |
| 8_1241884! | 4 | 0.5 | 6 | 0.5 | 4 | 0.25 | 4 | 0 | 18 | 0.3333 | 4 | 0.75 | 3 | 0.3333 | 8 | 0.25 | 3 | 0.3333 | 18 | 0.3889 |
| 8_1241884( | 3 | 1 | 6 | 1 | 4 | 1 | 4 | 0.75 | 17 | 0.9412 | 4 | 0.75 | 3 | 1 | 8 | 0.875 | 2 | 1 | 17 | 0.8824 |
| 8_1241884! | 4 | 0.5 | 7 | 0.4286 | 6 | 0.1667 | 4 | 0.75 | 21 | 0.4286 | 4 | 0.5 | 2 | 0.5 | 8 | 0.5 | 3 | 0.6667 | 17 | 0.5294 |
| 8_1241885 | 3 | 1 | 7 | 0.5714 | 4 | 0.75 | 4 | 0.25 | 18 | 0.6111 | 3 | 0.6667 | 2 | 0 | 5 | 0.8 | 3 | 1 | 13 | 0.6923 |
| 8_1241886( | 4 | 0.25 | 6 | 0.3333 | 5 | 0.2 | 3 | 0.3333 | 18 | 0.2778 | 1 | 0 | 3 | 0.3333 | 7 | 0.1429 | 3 | 0.3333 | 14 | 0.2143 |
| 8_1241887 | 4 | 0.25 | 7 | 0.1429 | 6 | 0.5 | 5 | 0.4 | 22 | 0.3182 | 4 | 0.5 | 4 | 0.5 | 8 | 0 | 3 | 0 | 19 | 0.2105 |
| 8_1241887( | 4 | 0.25 | 7 | 0.1429 | 6 | 0.3333 | 4 | 0 | 21 | 0.1905 | 4 | 0.25 | 4 | 0.5 | 8 | 0.125 | 3 | 0 | 19 | 0.2105 |
| 8_1241888( | 3 | 0 | 7 | 0.1429 | 6 | 0 | 4 | 0 | 20 | 0.05 | 3 | 0.3333 | 4 | 0 | 7 | 0 | 3 | 0.3333 | 17 | 0.1176 |
| 8_1241896( | 4 | 0.25 | 6 | 0.3333 | 6 | 0.1667 | 3 | 0 | 19 | 0.2105 | 4 | 0 | 4 | 0 | 7 | 0.2857 | 5 | 0.2 | 20 | 0.15 |
| 8_1241896( | 5 | 0 | 5 | 0.2 | 7 | 0.1429 | 5 | 0 | 22 | 0.0909 | 3 | 0.3333 | 2 | 0 | 7 | 0 | 4 | 0 | 16 | 0.0625 |
| 8_1241909( | 2 | 0 | 2 | 0.5 | 2 | 0 | 1 | 0 | 7 | 0.1429 | 2 | 0.5 | 1 | 1 | 1 | 1 | NA | NA | 4 | 0.75 |
| 8_1241910 | 2 | 0.5 | 2 | 1 | 2 | 0.5 | 2 | 0.5 | 8 | 0.625 | 2 | 1 | 1 | 1 | NA | NA | NA | NA | 3 | 1 |
| 8_1241937 | 2 | 0 | 5 | 0 | 4 | 0.25 | 3 | 0 | 14 | 0.0714 | 3 | 0.3333 | 1 | 0 | 1 | 0 | 1 | 0 | 6 | 0.1667 |
| 8_1241937! | 3 | 0.6667 | 6 | 0.8333 | 4 | 0.75 | 3 | 0.6667 | 16 | 0.75 | 2 | 1 | 1 | 1 | 1 | 1 | 1 | 1 | 5 | 1 |
| 8_1241937! | 4 | 0 | 5 | 0.2 | 4 | 0.25 | 7 | 0.2857 | 20 | 0.2 | 3 | 0.3333 | NA | NA | 1 | 1 | 2 | 0 | 6 | 0.3333 |
| 8_1241938 | 3 | 0.3333 | 6 | 0.5 | 4 | 1 | 7 | 0.5714 | 20 | 0.6 | 3 | 0.6667 | 1 | 1 | NA | NA | 2 | 1 | 6 | 0.8333 |
| 8_1241938 | 3 | 0 | 6 | 0 | 2 | 0.5 | 7 | 0.2857 | 18 | 0.1667 | 3 | 0.3333 | 1 | 0 | 1 | 0 | 1 | 0 | 6 | 0.1667 |
| 8_1241939! | 3 | 0.6667 | 4 | 1 | 7 | 1 | 4 | 1 | 18 | 0.9444 | 2 | 0.5 | NA | NA | 1 | 1 | 2 | 1 | 5 | 0.8 |
| 8_1241947 | 1 | 1 | 2 | 0.5 | NA | NA | NA | NA | 3 | 0.6667 | 1 | 0 | 1 | 1 | NA | NA | NA | NA | 2 | 0.5 |
| 8_1242287 | NA | NA | 3 | 1 | 4 | 0.5 | 2 | 0 | 9 | 0.5556 | 1 | 1 | 2 | 0 | 1 | 1 | NA | NA | 4 | 0.5 |
| 8_1242288! | 2 | 1 | 4 | 0.25 | 5 | 0.4 | 4 | 0 | 15 | 0.3333 | NA | NA | 1 | 0 | 2 | 0.5 | NA | NA | 3 | 0.3333 |
| 8_1242298 | 1 | 0 | 2 | 0.5 | 3 | 0.3333 | 1 | 0 | 7 | 0.2857 | 2 | 1 | 1 | 0 | 2 | 0 | NA | NA | 5 | 0.4 |
| 8_1242304 | 3 | 0 | 2 | 0.5 | 3 | 0.3333 | 1 | 0 | 9 | 0.2222 | 2 | 0.5 | 1 | 0 | 1 | 0 | NA | NA | 4 | 0.25 |
| 8_1242324 | NA | NA | 3 | 0.3333 | 1 | 1 | 1 | 1 | 5 | 0.6 | 4 | 1 | 3 | 0.6667 | 3 | 1 | 1 | 1 | 11 | 0.9091 |
| 8_1242327 | NA | NA | 3 | 0 | 2 | 0.5 | 1 | 1 | 6 | 0.3333 | 4 | 0.5 | 3 | 0.3333 | 4 | 0.5 | 1 | 0 | 12 | 0.4167 |
| 8_1242327! | NA | NA | 1 | 0 | 2 | 0.5 | 2 | 0 | 5 | 0.2 | 4 | 0 | 2 | 0 | 4 | 0.25 | 1 | 0 | 11 | 0.0909 |
| 8_1242329 | NA | NA | 3 | 0 | NA | NA | 1 | 1 | 4 | 0.25 | 4 | 0.25 | 3 | 0.3333 | 2 | 0.5 | 1 | 0 | 10 | 0.3 |
| 8_1242331! | 1 | 0 | 3 | 0.3333 | 1 | 0 | 2 | 0 | 7 | 0.1429 | 4 | 0.25 | 2 | 0.5 | 2 | 0 | 1 | 0 | 9 | 0.2222 |
| 8_1242333 | 1 | 1 | 1 | 0 | 1 | 1 | 2 | 0.5 | 5 | 0.6 | 3 | 0.6667 | 3 | 0.3333 | 3 | 0.6667 | 1 | 1 | 10 | 0.6 |
| 8_1242333! | 1 | 1 | 3 | 1 | 2 | 1 | 1 | 0 | 7 | 0.8571 | 4 | 0.75 | 2 | 0.5 | 3 | 1 | 1 | 1 | 10 | 0.8 |
| 8_1242333 | 1 | 1 | 3 | 0.6667 | 2 | 1 | 1 | 1 | 7 | 0.8571 | 4 | 1 | 3 | 0.6667 | 3 | 0.6667 | 1 | 0 | 11 | 0.7273 |
| 8_1242334! | 1 | 0 | 3 | 0.6667 | 1 | 0 | 1 | 0 | 6 | 0.3333 | 3 | 0.6667 | 3 | 0.3333 | 2 | 0.5 | 1 | 1 | 9 | 0.5556 |
| 8_1242334( | 1 | 1 | 3 | 0.3333 | 2 | 1 | 1 | 0 | 7 | 0.5714 | 4 | 0.5 | 3 | 0.3333 | 3 | 0.3333 | 1 | 0 | 11 | 0.3636 |
| 8_1242361( | 1 | 1 | NA | NA | NA | NA | NA | NA | 1 | 1 | 1 | 1 | NA | NA | 1 | 0 | NA | NA | 2 | 0.5 |
| 8_1242362 | 1 | 1 | NA | NA | NA | NA | NA | NA | 1 | 1 | 1 | 1 | NA | NA | 2 | 0 | NA | NA | 3 | 0.3333 |
| 8_1242365! | 1 | 1 | NA | NA | NA | NA | NA | NA | 1 | 1 | 1 | 1 | NA | NA | 2 | 0.5 | NA | NA | 3 | 0.6667 |
| 8_1242379! | NA | NA | 1 | 1 | NA | NA | NA | NA | 1 | 1 | 1 | 0 | NA | NA | 2 | 0.5 | NA | NA | 3 | 0.3333 |
| 8_1242632 | 2 | 1 | NA | NA | 1 | 0 | 2 | 0 | 5 | 0.4 | 2 | 0 | NA | NA | 2 | 0.5 | 1 | 0 | 5 | 0.2 |
| 8_1243877! | 2 | 0 | 4 | 0.25 | 2 | 0.5 | 4 | 0 | 12 | 0.1667 | 2 | 0 | 1 | 0 | 8 | 0.25 | 2 | 0 | 13 | 0.1538 |
| 8_1243877( | 1 | 0 | 2 | 0 | 1 | 0 | 4 | 0.25 | 8 | 0.125 | 1 | 0 | 2 | 0.5 | 7 | 0.1429 | 3 | 0.3333 | 13 | 0.2308 |
| 8_1243878( | 1 | 1 | 2 | 1 | 2 | 0 | 3 | 0.6667 | 8 | 0.625 | 1 | 0 | 2 | 1 | 3 | 0.3333 | 2 | 1 | 8 | 0.625 |
| 8_1243878( | 2 | 0 | 3 | 0 | 2 | 0.5 | 3 | 0.6667 | 10 | 0.3 | NA | NA | 1 | 1 | 6 | 0 | 3 | 0 | 10 | 0.1 |
| 8_1243878! | 2 | 0.5 | 4 | 0 | 2 | 0 | 3 | 1 | 11 | 0.3636 | 1 | 1 | 2 | 1 | 7 | 0.7143 | 3 | 0.3333 | 13 | 0.6923 |
| 8_1243879! | 1 | 1 | 3 | 1 | 2 | 0.5 | 4 | 0.75 | 10 | 0.8 | NA | NA | 2 | 0.5 | 8 | 0.625 | 3 | 0.6667 | 13 | 0.6154 |
| 8_1243879! | 2 | 0 | 3 | 0 | 1 | 0 | 4 | 0.25 | 10 | 0.1 | 1 | 0 | 1 | 0 | 8 | 0.125 | 3 | 0.3333 | 13 | 0.1538 |
| 8_1243880 | 2 | 0.5 | 4 | 0.5 | 2 | 0.5 | 3 | 0.6667 | 11 | 0.5455 | 2 | 0.5 | 2 | 0.5 | 8 | 0.5 | 3 | 0.3333 | 15 | 0.4667 |
| 8_1243880! | 2 | 0.5 | 3 | 0.6667 | 2 | 1 | 3 | 0.6667 | 10 | 0.7 | 2 | 1 | 2 | 1 | 7 | 0.4286 | 2 | 0.5 | 13 | 0.6154 |
| 8_1243883 | 2 | 0.5 | 4 | 0.25 | 2 | 0.5 | 4 | 0.5 | 12 | 0.4167 | 7 | 0.2857 | 2 | 0 | 8 | 0.375 | 4 | 0.5 | 21 | 0.3333 |
| 8_1243883! | 2 | 0.5 | 4 | 0.5 | 2 | 1 | 1 | 0 | 9 | 0.5556 | 6 | 0.6667 | 1 | 1 | 7 | 0.5714 | 4 | 0.75 | 18 | 0.6667 |
| 8_1243887 | 2 | 0 | 5 | 0 | 2 | 0.5 | 3 | 0 | 12 | 0.0833 | 7 | 0.1429 | 1 | 1 | 8 | 0 | 4 | 0 | 20 | 0.1 |
| 8_1243887 | 1 | 0 | 4 | 0 | 2 | 0.5 | 4 | 0.25 | 11 | 0.1818 | 6 | 0.3333 | 1 | 1 | 6 | 0 | 3 | 0.3333 | 16 | 0.25 |
| 8_1243889 | 3 | 0.3333 | 3 | 0 | 1 | 0 | 3 | 0 | 10 | 0.1 | 6 | 0 | 1 | 0 | 6 | 0.1667 | 2 | 0 | 15 | 0.0667 |
| 8_1243896( | 8 | 0.625 | 6 | 0.5 | 2 | 1 | 6 | 0.6667 | 22 | 0.6364 | 7 | 0.2857 | 2 | 1 | 7 | 0.5714 | 7 | 0.4286 | 23 | 0.4783 |
| 8_1243896 | 8 | 0.25 | 6 | 0.1667 | 2 | 0.5 | 5 | 0.2 | 21 | 0.2381 | 6 | 0.3333 | 2 | 0 | 8 | 0.125 | 7 | 0 | 23 | 0.1304 |
| 8_1243896( | 8 | 0.875 | 6 | 0.6667 | 2 | 1 | 4 | 1 | 20 | 0.85 | 7 | 0.8571 | 1 | 1 | 8 | 0.75 | 7 | 1 | 23 | 0.8696 |
| 8_1243897( | 8 | 0.125 | 7 | 0.2857 | 2 | 0 | 6 | 0.1667 | 23 | 0.1739 | 8 | 0.25 | 2 | 0 | 8 | 0 | 7 | 0.2857 | 25 | 0.16 |
| 8_1243899 | 8 | 0.125 | 8 | 0.125 | 5 | 0 | 5 | 0 | 26 | 0.0769 | 10 | 0.2 | 3 | 0.3333 | 7 | 0 | 6 | 0 | 26 | 0.1154 |
| 8_1243899( | 8 | 0.25 | 7 | 0.1429 | 6 | 0.1667 | 6 | 0.5 | 27 | 0.2593 | 10 | 0.2 | 3 | 0.3333 | 9 | 0.3333 | 6 | 0.6667 | 28 | 0.3571 |
| 8_1243900 | 7 | 1 | 7 | 1 | 5 | 1 | 4 | 1 | 23 | 1 | 8 | 1 | 2 | 1 | 8 | 0.875 | 4 | 1 | 22 | 0.9545 |
| 8_1243900 | NA | NA | NA | NA | NA | NA | 1 | 1 | 1 | 1 | 1 | 1 | NA | NA | NA | NA | NA | NA | 1 | 1 |
| 8_1243900! | 6 | 0.8333 | 7 | 1 | 4 | 1 | 4 | 1 | 21 | 0.9524 | 9 | 0.7778 | 2 | 1 | 7 | 1 | 5 | 1 | 23 | 0.913 |
| 8_1243901 | 7 | 0.5714 | 4 | 0.75 | 5 | 0.4 | 3 | 0.3333 | 19 | 0.5263 | 7 | 0.5714 | 3 | 0 | 7 | 0.4286 | 5 | 0.4 | 22 | 0.4091 |
| 8_1243904 | 7 | 0.2857 | 6 | 0.5 | 5 | 0.2 | 6 | 0.3333 | 24 | 0.3333 | 11 | 0.0909 | 2 | 1 | 6 | 0.3333 | 4 | 0.75 | 23 | 0.3478 |
| 8_1243905! | 6 | 0.6667 | 7 | 0.4286 | 6 | 0.3333 | 5 | 0.4 | 24 | 0.4583 | 10 | 0.4 | 2 | 0.5 | 7 | 0.4286 | 7 | 0.4286 | 26 | 0.4231 |

| | | | | | | | | | | | | | | | | | | | | |
|---|---|---|---|---|---|---|---|---|---|---|---|---|---|---|---|---|---|---|---|---|
| 8_1243906 | 4 | 0.5 | 3 | 0 | 2 | 0.5 | 2 | 0 | 11 | 0.2727 | 7 | 0.1429 | 2 | 0 | 6 | 0.1667 | 4 | 0.5 | 19 | 0.2105 |
| 8_1245615 | 8 | 0 | 10 | 0.2 | 12 | 0 | 14 | 0.0714 | 44 | 0.0682 | 9 | 0.1111 | 8 | 0 | 8 | 0.125 | 15 | 0 | 40 | 0.05 |
| 8_1245615 | 7 | 0.4286 | 11 | 0.4545 | 9 | 0.3333 | 11 | 0.3636 | 38 | 0.3947 | 6 | 0.3333 | 6 | 0 | 8 | 0.75 | 12 | 0.5833 | 32 | 0.4688 |
| 8_1245618 | 8 | 0 | 10 | 0.2 | 11 | 0.0909 | 14 | 0.5714 | 43 | 0.2558 | 7 | 0.1429 | 8 | 0.375 | 10 | 0.1 | 15 | 0.2 | 40 | 0.2 |
| 8_1245619 | 7 | 0.2857 | 11 | 0.3636 | 12 | 0.3333 | 14 | 0.2857 | 44 | 0.3182 | 8 | 0.375 | 6 | 0.3333 | 9 | 0.5556 | 15 | 0.2 | 38 | 0.3421 |
| 8_1245663 | NA | NA | NA | NA | 1 | 1 | NA | NA | 1 | 1 | NA | NA | 1 | 0 | NA | NA | 1 | 1 | 2 | 0.5 |
| 8_1245670 | 5 | 0.2 | NA | NA | 4 | 0.75 | 3 | 0.6667 | 12 | 0.5 | 3 | 0 | 10 | 0.4 | 3 | 0.6667 | 1 | 1 | 17 | 0.4118 |
| 8_1245671 | 4 | 0.25 | NA | NA | 4 | 0 | 3 | 0.3333 | 11 | 0.0909 | 2 | 0 | 6 | 0.1667 | 3 | 0.3333 | 1 | 0 | 12 | 0.1667 |
| 8_1245672 | 4 | 0 | 1 | 0 | 4 | 0.25 | 3 | 0 | 12 | 0.0833 | 3 | 0.3333 | 9 | 0 | 4 | 0 | 1 | 0 | 17 | 0.0588 |
| 8_1245677 | 7 | 0.5714 | 3 | 0 | 7 | 0.5714 | 7 | 0.1429 | 24 | 0.375 | 10 | 0.3 | 14 | 0.4286 | 14 | 0.4286 | 8 | 0.875 | 46 | 0.4783 |
| 8_1245677 | 7 | 0.5714 | 7 | 0.7143 | 8 | 0.625 | 8 | 0.5 | 30 | 0.6 | 9 | 0.5556 | 16 | 0.625 | 16 | 0.5 | 8 | 1 | 49 | 0.6327 |
| 8_1245677 | 7 | 0.1429 | 4 | 0 | 4 | 0.25 | 8 | 0.125 | 23 | 0.1304 | 9 | 0 | 16 | 0.1875 | 15 | 0.2 | 8 | 0.125 | 48 | 0.1458 |
| 8_1245677 | 7 | 0.5714 | 5 | 0.6 | 7 | 0.2857 | 8 | 0.375 | 27 | 0.4444 | 9 | 0.3333 | 15 | 0.4667 | 14 | 0.2857 | 8 | 0.625 | 46 | 0.413 |
| 8_1245679 | 6 | 0.3333 | 7 | 0.1429 | 7 | 0.5714 | 8 | 0.375 | 28 | 0.3571 | 9 | 0.4444 | 15 | 0.1333 | 15 | 0.2667 | 8 | 0.125 | 47 | 0.234 |
| 8_1245679 | 6 | 0 | 5 | 0 | 6 | 0.3333 | 8 | 0 | 25 | 0.08 | 8 | 0.25 | 15 | 0.2 | 14 | 0.2143 | 8 | 0.125 | 45 | 0.2 |
| 8_1245681 | 4 | 1 | 7 | 1 | 7 | 1 | 7 | 0.7143 | 25 | 0.92 | 5 | 1 | 13 | 0.6923 | 11 | 0.7273 | 6 | 1 | 35 | 0.8 |
| 8_1245682 | 6 | 0.5 | 5 | 0.2 | 7 | 0.2857 | 8 | 0 | 26 | 0.2308 | 6 | 0.5 | 10 | 0.1 | 13 | 0.2308 | 7 | 0.2857 | 36 | 0.25 |
| 8_1245682 | 4 | 1 | 4 | 0.75 | 5 | 1 | 4 | 1 | 17 | 0.9412 | 5 | 0.8 | 11 | 1 | 12 | 1 | 7 | 1 | 35 | 0.9714 |
| 8_1245682 | 6 | 0.8333 | 6 | 0.5 | 7 | 0.5714 | 7 | 0.4286 | 26 | 0.5769 | 8 | 0.5 | 12 | 0.5 | 13 | 0.6154 | 8 | 0.25 | 41 | 0.4878 |
| 8_1245684 | 4 | 1 | 6 | 0.6667 | 7 | 0.7143 | 5 | 0.8 | 22 | 0.7727 | 7 | 0.1429 | 15 | 0.2667 | 10 | 0.4 | 7 | 0.8571 | 39 | 0.3846 |
| 8_1245685 | 4 | 0.25 | 6 | 0 | 8 | 0.625 | 6 | 0 | 24 | 0.25 | 5 | 0 | 16 | 0.375 | 11 | 0.3636 | 9 | 0.3333 | 41 | 0.3171 |
| 8_1245685 | 4 | 0.75 | 4 | 0.5 | 6 | 0.3333 | 6 | 0.6667 | 20 | 0.55 | 5 | 0 | 11 | 0.2727 | 9 | 0.2222 | 3 | 0.3333 | 28 | 0.2143 |
| 8_1245685 | 5 | 0.8 | 5 | 0.6 | 6 | 0.5 | 6 | 0.3333 | 22 | 0.5455 | 7 | 0.2857 | 15 | 0.7333 | 10 | 0.8 | 9 | 0.5556 | 41 | 0.6341 |
| 8_1245686 | 4 | 1 | 4 | 1 | 6 | 1 | 5 | 1 | 19 | 1 | 6 | 1 | 15 | 1 | 8 | 1 | 5 | 1 | 34 | 1 |
| 8_1245688 | 5 | 0 | 4 | 0 | 5 | 0.2 | 6 | 0.3333 | 20 | 0.15 | 6 | 0 | 16 | 0.125 | 11 | 0.2727 | 9 | 0.2222 | 42 | 0.1667 |
| 8_1245689 | 5 | 0 | 4 | 0 | 5 | 0.4 | 5 | 0.2 | 19 | 0.1579 | 5 | 0.2 | 14 | 0.0714 | 9 | 0.1111 | 8 | 0.125 | 36 | 0.1111 |
| 8_1245689 | 4 | 0.5 | 4 | 0.75 | 3 | 0.6667 | 4 | 0.25 | 15 | 0.5333 | 4 | 0.75 | 12 | 0.75 | 5 | 0.8 | 6 | 1 | 27 | 0.8148 |
| 8_1245690 | 3 | 1 | 2 | 1 | 3 | 1 | 4 | 1 | 12 | 1 | 3 | 1 | 12 | 0.9167 | 9 | 1 | 8 | 0.875 | 32 | 0.9375 |
| 8_1245691 | 5 | 0 | 5 | 0 | 4 | 0.25 | 5 | 0 | 19 | 0.0526 | 5 | 0.2 | 16 | 0.125 | 11 | 0.1818 | 9 | 0 | 41 | 0.122 |
| 8_1245691 | 4 | 0 | 5 | 0.4 | 4 | 0 | 5 | 0 | 18 | 0.1111 | 5 | 0 | 14 | 0.1429 | 11 | 0.0909 | 9 | 0 | 39 | 0.0769 |
| 8_1245719 | 5 | 0 | 4 | 0 | 4 | 0 | 5 | 0.4 | 18 | 0.1111 | 4 | 0.5 | 13 | 0.3077 | 7 | 0.1429 | 8 | 0.5 | 32 | 0.3438 |
| 8_1245719 | 4 | 0 | 4 | 0 | 3 | 0.3333 | 4 | 0 | 15 | 0.0667 | 3 | 0.3333 | 10 | 0.2 | 8 | 0 | 6 | 0 | 27 | 0.1111 |
| 8_1245721 | 5 | 0.6 | 4 | 0.75 | 4 | 0.25 | 1 | 0 | 14 | 0.5 | 3 | 1 | 9 | 0.3333 | 9 | 0.5556 | 6 | 0.5 | 27 | 0.5185 |
| 8_1245859 | 4 | 0.5 | 3 | 0.3333 | 2 | 0 | 4 | 0.5 | 13 | 0.3846 | 2 | 0 | 9 | 0.1111 | 8 | 0.5 | 5 | 0 | 24 | 0.2083 |
| 8_1246186 | 4 | 0.25 | 3 | 0 | 1 | 0 | 2 | 0.5 | 10 | 0.2 | 2 | 0.5 | 3 | 0.3333 | NA | NA | 2 | 0 | 7 | 0.2857 |
| 8_1246194 | 2 | 0 | 3 | 0.3333 | NA | NA | NA | NA | 5 | 0.2 | 1 | 1 | NA | NA | NA | NA | 1 | 0 | 2 | 0.5 |
| 8_1246795 | 4 | 0.5 | 1 | 0 | 6 | 0.5 | 3 | 0 | 14 | 0.3571 | 2 | 0.5 | 8 | 1 | 2 | 1 | 2 | 1 | 14 | 0.9286 |
| 8_1246797 | 4 | 0.25 | 2 | 0.5 | 6 | 0.5 | 3 | 0.6667 | 15 | 0.4667 | 2 | 0.5 | 8 | 0.75 | 4 | 0.5 | 2 | 0.5 | 16 | 0.625 |
| 8_1246800 | 3 | 0.3333 | 1 | 0 | 5 | 0.2 | 2 | 0 | 11 | 0.1818 | 2 | 0 | 7 | 0.1429 | 4 | 0.25 | 2 | 0 | 15 | 0.1333 |
| 8_1246807 | 3 | 0 | 1 | 0 | 4 | 0.25 | 2 | 0 | 10 | 0.1 | 1 | 0 | 6 | 0.5 | 3 | 1 | 2 | 0 | 12 | 0.5 |
| 8_1246809 | 3 | 0.6667 | 2 | 0.5 | 3 | 0.6667 | 2 | 1 | 10 | 0.7 | 1 | 1 | 7 | 0.5714 | 3 | 0.6667 | 2 | 0.5 | 13 | 0.6154 |
| 8_1246809 | 3 | 0.3333 | 2 | 0.5 | 3 | 0 | 2 | 0 | 10 | 0.2 | 1 | 1 | 5 | 0.2 | 2 | 0 | 2 | 0 | 10 | 0.2 |
| 8_1246810 | 2 | 1 | 1 | 1 | 2 | 1 | 2 | 1 | 7 | 1 | 1 | 1 | 7 | 1 | 2 | 1 | 2 | 1 | 12 | 1 |
| 8_1246810 | 1 | 1 | NA | NA | 3 | 0.3333 | 1 | 1 | 5 | 0.6 | 1 | 0 | 3 | 0.3333 | 2 | 1 | 1 | 1 | 7 | 0.5714 |
| 8_1246811 | 2 | 0.5 | 1 | 0 | 2 | 0.5 | 1 | 0 | 6 | 0.3333 | NA | NA | 2 | 1 | 2 | 0 | 2 | 0 | 6 | 0.3333 |
| 8_1246811 | 2 | 0.5 | 1 | 1 | 1 | 1 | 1 | 0 | 5 | 0.6 | 1 | 1 | 3 | 1 | 2 | 0.5 | 2 | 1 | 8 | 0.875 |
| 8_1246811 | 3 | 0 | 1 | 0 | 2 | 0.5 | 2 | 0.5 | 8 | 0.25 | 1 | 1 | 7 | 0.5714 | 3 | 0.6667 | 2 | 0.5 | 13 | 0.6154 |
| 8_1247232 | 1 | 1 | NA | NA | 3 | 0.6667 | 2 | 1 | 6 | 0.8333 | 3 | 0.6667 | 5 | 0.2 | 3 | 0.6667 | 7 | 0.4286 | 18 | 0.4444 |
| 8_1247233 | NA | NA | NA | NA | 3 | 0.6667 | 3 | 0.6667 | 6 | 0.6667 | 3 | 0.6667 | 5 | 0.6 | 3 | 1 | 7 | 0.2857 | 18 | 0.5556 |
| 8_1247235 | 1 | 0 | NA | NA | 3 | 0.3333 | 3 | 0 | 7 | 0.1429 | 3 | 0.3333 | 6 | 0.5 | 3 | 0.3333 | 6 | 0.3333 | 18 | 0.3889 |
| 8_1247239 | 1 | 1 | NA | NA | 3 | 0.6667 | 1 | 1 | 5 | 0.8 | 3 | 1 | 5 | 1 | 1 | 1 | 5 | 0.8 | 14 | 0.9286 |
| 8_1247240 | 1 | 0 | NA | NA | 3 | 0 | 1 | 1 | 5 | 0.2 | 3 | 0 | 6 | 0.3333 | 2 | 0 | 6 | 0 | 17 | 0.1176 |
| 8_1247240 | 1 | 0 | NA | NA | 3 | 0.3333 | NA | NA | 4 | 0.25 | 3 | 1 | 5 | 0.4 | 2 | 0.5 | 6 | 0.5 | 16 | 0.5625 |
| 8_1247240 | 1 | 1 | NA | NA | 3 | 0.3333 | 1 | 0 | 5 | 0.4 | 3 | 0 | 5 | 0.4 | 2 | 0.5 | 5 | 0.4 | 15 | 0.3333 |
| 8_1247242 | 1 | 1 | NA | NA | 2 | 0.5 | 1 | 1 | 4 | 0.75 | 2 | 0 | 6 | 0.3333 | 2 | 0.5 | 5 | 0.4 | 15 | 0.3333 |
| 8_1247242 | NA | NA | NA | NA | 3 | 0.6667 | 1 | 1 | 4 | 0.75 | 3 | 0.6667 | 4 | 0.75 | 2 | 0.5 | 5 | 0.8 | 14 | 0.7143 |
| 8_1247242 | 1 | 0 | NA | NA | 3 | 0.3333 | 1 | 0 | 5 | 0.2 | 3 | 0 | 6 | 0.3333 | 1 | 0 | 6 | 0.1667 | 16 | 0.1875 |
| 8_1247242 | 1 | 1 | NA | NA | 2 | 1 | 1 | 1 | 4 | 1 | 1 | 1 | 5 | 1 | 1 | 1 | 3 | 1 | 10 | 1 |
| 8_1247243 | NA | NA | NA | NA | 2 | 1 | 1 | 0 | 3 | 0.6667 | 3 | 1 | 2 | 0.5 | 2 | 0.5 | 4 | 0.75 | 11 | 0.7273 |
| 8_1247243 | 1 | 1 | NA | NA | 3 | 1 | 1 | 1 | 5 | 1 | 2 | 1 | 4 | 1 | 2 | 1 | 5 | 0.8 | 13 | 0.9231 |
| 8_1247243 | 1 | 1 | NA | NA | 3 | 1 | NA | NA | 4 | 1 | 3 | 1 | 6 | 1 | 2 | 1 | 5 | 1 | 16 | 1 |
| 8_1247244 | 1 | 1 | NA | NA | 2 | 0 | 1 | 0 | 4 | 0.25 | 1 | 0 | 6 | 0.1667 | 2 | 0 | 5 | 0 | 14 | 0.0714 |
| 8_1247244 | NA | NA | NA | NA | 2 | 1 | NA | NA | 2 | 1 | NA | NA | 1 | 1 | 2 | 1 | 4 | 1 | 7 | 1 |
| 8_1247244 | 1 | 0 | NA | NA | 3 | 0.3333 | 1 | 0 | 5 | 0.2 | 1 | 0 | 5 | 0.4 | 2 | 0 | 5 | 0.2 | 13 | 0.2308 |
| 8_1247245 | 1 | 1 | NA | NA | 3 | 1 | 1 | 1 | 5 | 1 | 3 | 1 | 6 | 1 | 2 | 1 | 5 | 1 | 16 | 1 |

| | | | | | | | | | | | | | | | | | | | | |
|---|---|---|---|---|---|---|---|---|---|---|---|---|---|---|---|---|---|---|---|---|
| 8_12472454 | 1 | 1 | NA | NA | 3 | 0.3333 | 1 | 0 | 5 | 0.4 | 2 | 1 | 4 | 0.75 | NA | NA | 5 | 0.4 | 11 | 0.6364 |
| 8_12472464 | 1 | 0 | NA | NA | 3 | 0.6667 | 1 | 1 | 5 | 0.6 | 3 | 1 | 6 | 0.8333 | 2 | 1 | 5 | 0.8 | 16 | 0.875 |
| 8_12472467 | NA | NA | NA | NA | 3 | 1 | 1 | 1 | 4 | 1 | 3 | 1 | 4 | 1 | 2 | 1 | 5 | 0.8 | 14 | 0.9286 |
| 8_12472470 | NA | NA | NA | NA | 3 | 1 | 1 | 1 | 4 | 1 | 3 | 0.6667 | 5 | 1 | 2 | 1 | 5 | 1 | 15 | 0.9333 |
| 8_12472984 | NA | NA | NA | NA | 3 | 1 | 1 | 0 | 4 | 0.75 | 2 | 0 | 4 | 0.25 | NA | NA | 4 | 0 | 10 | 0.1 |
| 8_12472988 | NA | NA | NA | NA | 3 | 0.6667 | 1 | 0 | 4 | 0.5 | 1 | 0 | 5 | 0.4 | 2 | 0 | 2 | 0 | 10 | 0.2 |
| 8_12472993 | 1 | 0 | NA | NA | 3 | 0 | 1 | 1 | 5 | 0.2 | 2 | 0.5 | 4 | 0.5 | 1 | 0 | 4 | 0 | 11 | 0.2727 |
| 8_12473007 | NA | NA | NA | NA | 1 | 1 | 1 | 0 | 2 | 0.5 | 2 | 0 | 6 | 0.8333 | 2 | 0.5 | 4 | 1 | 14 | 0.7143 |
| 8_12473024 | NA | NA | NA | NA | 3 | 1 | 1 | 1 | 4 | 1 | 2 | 1 | 5 | 1 | 2 | 0.5 | 4 | 0.5 | 13 | 0.7692 |
| 8_12475590 | NA | NA | 1 | 1 | NA | NA | NA | NA | 1 | 1 | 1 | 1 | NA | NA | 2 | 1 | NA | NA | 3 | 1 |
| 8_12475634 | NA | NA | 1 | 1 | NA | NA | NA | NA | 1 | 1 | 1 | 1 | 1 | 1 | 2 | 1 | 1 | 1 | 5 | 1 |
| 8_12475643 | NA | NA | 1 | 1 | NA | NA | NA | NA | 1 | 1 | NA | NA | 1 | 1 | 2 | 1 | 1 | 1 | 4 | 1 |
| 8_12475760 | NA | NA | 1 | 1 | 1 | 1 | NA | NA | 2 | 1 | 3 | 0.6667 | 1 | 1 | 3 | 1 | 1 | 1 | 8 | 0.875 |
| 8_12475774 | NA | NA | 1 | 1 | 1 | 1 | NA | NA | 2 | 1 | 3 | 0.6667 | NA | NA | 1 | 1 | 1 | 1 | 5 | 0.8 |
| 8_12477400 | NA | NA | 1 | 1 | 1 | 1 | NA | NA | 2 | 1 | 4 | 0.75 | 1 | 1 | 3 | 0.6667 | 2 | 1 | 10 | 0.8 |
| 8_12477408 | NA | NA | 1 | 0 | 1 | 1 | NA | NA | 2 | 0.5 | 4 | 0.5 | 1 | 0 | 3 | 0 | 1 | 0 | 9 | 0.2222 |
| 8_12477468 | NA | NA | 1 | 1 | 1 | 0 | NA | NA | 2 | 0.5 | 4 | 0.25 | NA | NA | 4 | 0.75 | 1 | 1 | 9 | 0.5556 |
| 8_12507026 | 4 | 1 | 7 | 0.8571 | 4 | 0.75 | 6 | 0.8333 | 21 | 0.8571 | 2 | 1 | 4 | 1 | 4 | 1 | 3 | 0.6667 | 13 | 0.9231 |
| 8_12507028 | 3 | 1 | 6 | 0.6667 | 3 | 0.6667 | 8 | 0.375 | 20 | 0.6 | 2 | 0.5 | 3 | 1 | 3 | 0.6667 | 3 | 0.3333 | 11 | 0.6364 |
| 8_12507031 | 2 | 0 | 6 | 0.1667 | 3 | 0.3333 | 6 | 0.6667 | 17 | 0.3529 | 2 | 0.5 | 4 | 0.5 | 4 | 0.5 | 3 | 0.6667 | 13 | 0.5385 |
| 8_12507039 | 3 | 1 | 9 | 0.8889 | 3 | 0.3333 | 9 | 1 | 24 | 0.875 | 2 | 1 | 5 | 1 | 5 | 0.8 | 3 | 0.6667 | 15 | 0.8667 |
| 8_12507047 | 2 | 0 | 7 | 0 | 3 | 0 | 6 | 0.5 | 18 | 0.1667 | 1 | 1 | 4 | 0.25 | 4 | 0 | 2 | 0.5 | 11 | 0.2727 |
| 8_12507050 | 4 | 0.25 | 8 | 0.125 | 2 | 0.5 | 5 | 0 | 19 | 0.1579 | 2 | 0 | 4 | 0.25 | 5 | 0 | 3 | 0 | 14 | 0.0714 |
| 8_12507059 | 3 | 0 | 9 | 0.1111 | 4 | 0 | 7 | 0.1429 | 23 | 0.087 | 2 | 0 | 4 | 0.25 | 5 | 0 | 2 | 0 | 13 | 0.0769 |
| 8_12507076 | 4 | 0.25 | 9 | 0.3333 | 3 | 0.3333 | 8 | 0.25 | 24 | 0.2917 | 2 | 0.5 | 4 | 0.5 | 5 | 0.2 | 2 | 1 | 13 | 0.4615 |
| 8_12507079 | 4 | 0.75 | 6 | 1 | 4 | 0.75 | 8 | 0.875 | 22 | 0.8636 | 1 | 1 | 3 | 1 | 5 | 0.8 | 4 | 0.75 | 13 | 0.8462 |
| 8_12507080 | 3 | 1 | 11 | 0.9091 | 5 | 1 | 10 | 1 | 29 | 0.9655 | 2 | 1 | 6 | 1 | 4 | 1 | 4 | 0.75 | 16 | 0.9375 |
| 8_12507117 | 5 | 0.2 | 10 | 0.3 | 5 | 0.4 | 11 | 0.3636 | 31 | 0.3226 | 2 | 0 | 5 | 0.2 | 6 | 0.3333 | 4 | 0.25 | 17 | 0.2353 |
| 8_12507119 | 5 | 0.2 | 11 | 0.3636 | 5 | 0.4 | 9 | 0.5556 | 30 | 0.4 | 3 | 0.3333 | 4 | 1 | 7 | 0.2857 | 3 | 0 | 17 | 0.4118 |
| 8_12507120 | 3 | 0 | 7 | 0 | 4 | 0 | 8 | 0.25 | 22 | 0.0909 | 2 | 0.5 | 5 | 0.4 | 5 | 0.2 | 4 | 0.25 | 16 | 0.3125 |
| 8_12507167 | 5 | 0 | 9 | 0.4444 | 5 | 0 | 11 | 0.1818 | 30 | 0.2 | 3 | 0 | 4 | 0.25 | 7 | 0.1429 | 5 | 0.2 | 19 | 0.1579 |
| 8_12507174 | 6 | 0.1667 | 7 | 0 | 5 | 0.2 | 8 | 0.125 | 26 | 0.1154 | 4 | 0 | 7 | 0.1429 | 5 | 0.4 | 5 | 0 | 21 | 0.1429 |
| 8_12507185 | 5 | 0 | 9 | 0.2222 | 5 | 0 | 8 | 0 | 27 | 0.0741 | 3 | 0.3333 | 7 | 0 | 7 | 0.1429 | 4 | 0 | 21 | 0.0952 |
| 8_12509644 | 12 | 0.1667 | 21 | 0.3333 | 23 | 0.1739 | 11 | 0 | 67 | 0.194 | 20 | 0.05 | 12 | 0.0833 | 23 | 0.1739 | 15 | 0.2 | 70 | 0.1286 |
| 8_12509679 | 11 | 0 | 19 | 0.1053 | 16 | 0.375 | 10 | 0.1 | 56 | 0.1607 | 10 | 0 | 10 | 0.2 | 14 | 0 | 12 | 0.0833 | 46 | 0.0652 |
| 8_12509683 | 14 | 0.2857 | 16 | 0.375 | 18 | 0.1111 | 11 | 0.0909 | 59 | 0.2203 | 20 | 0.15 | 13 | 0.3077 | 19 | 0.3158 | 15 | 0.2667 | 67 | 0.2537 |
| 8_12509687 | 10 | 0.4 | 13 | 0.1538 | 12 | 0.4167 | 6 | 0.1667 | 41 | 0.2927 | 17 | 0.4118 | 11 | 0.5455 | 15 | 0.4 | 9 | 0.3333 | 52 | 0.4231 |
| 8_12509698 | 13 | 0.4615 | 14 | 0.5714 | 17 | 0.1765 | 6 | 0.6667 | 50 | 0.42 | 14 | 0.5714 | 9 | 1 | 19 | 0.4737 | 12 | 0.5833 | 54 | 0.6111 |
| 8_12509705 | 13 | 0.2308 | 18 | 0.2222 | 15 | 0.4 | 9 | 0.4444 | 55 | 0.3091 | 17 | 0.2941 | 10 | 0.4 | 19 | 0.3158 | 15 | 0.3333 | 61 | 0.3279 |
| 8_12509706 | 12 | 0.0833 | 19 | 0.1053 | 16 | 0.0625 | 10 | 0.1 | 57 | 0.0877 | 17 | 0.2353 | 12 | 0.1667 | 21 | 0.1429 | 16 | 0 | 66 | 0.1364 |
| 8_12509720 | 13 | 0.7692 | 17 | 0.6471 | 13 | 0.6923 | 10 | 0.8 | 53 | 0.717 | 17 | 0.3529 | 13 | 0.4615 | 17 | 0.4706 | 15 | 0.6 | 62 | 0.4677 |
| 8_12509722 | 13 | 0.8462 | 18 | 0.7222 | 15 | 0.6 | 8 | 0.875 | 54 | 0.7407 | 18 | 0.8333 | 12 | 1 | 15 | 0.7333 | 14 | 0.8571 | 59 | 0.8475 |
| 8_12509724 | 14 | 0.9286 | 19 | 0.7895 | 16 | 0.6875 | 8 | 0.75 | 57 | 0.7895 | 15 | 0.8667 | 12 | 0.8333 | 18 | 0.7778 | 15 | 0.8 | 60 | 0.8167 |
| 8_12509727 | 10 | 0.8 | 18 | 0.8333 | 13 | 0.9231 | 9 | 1 | 50 | 0.88 | 18 | 0.9444 | 13 | 0.9231 | 12 | 0.75 | 14 | 0.9286 | 57 | 0.8947 |
| 8_12509728 | 2 | 1 | 2 | 1 | 2 | 1 | NA | NA | 6 | 1 | NA | NA | NA | NA | 2 | 1 | 1 | 0 | 3 | 0.6667 |
| 8_12516075 | 2 | 0 | 10 | 0.3 | 4 | 0 | 1 | 0 | 17 | 0.1765 | 4 | 0 | 8 | 0 | 5 | 0.2 | 4 | 0.25 | 21 | 0.0952 |
| 8_12527838 | 5 | 0.6 | 3 | 1 | 2 | 0.5 | NA | NA | 10 | 0.7 | 2 | 0 | NA | NA | 2 | 0 | 3 | 1 | 7 | 0.4286 |
| 8_12527855 | 5 | 0.4 | 3 | 0.3333 | 2 | 0.5 | 2 | 1 | 12 | 0.5 | 3 | 0 | 2 | 0.5 | 2 | 0 | 6 | 0.3333 | 13 | 0.2308 |
| 8_12527969 | NA | NA | 1 | 1 | NA | NA | NA | NA | 1 | 1 | NA | NA | 1 | 0 | NA | NA | 1 | 1 | 2 | 0.5 |
| 8_12527987 | NA | NA | 1 | 1 | NA | NA | NA | NA | 1 | 1 | NA | NA | NA | NA | NA | NA | 1 | 1 | 1 | 1 |
| 8_12528038 | NA | NA | 1 | 1 | NA | NA | NA | NA | 1 | 1 | 1 | 1 | 1 | 0 | NA | NA | 1 | 0 | 3 | 0.3333 |
| 8_12528039 | NA | NA | 1 | 1 | NA | NA | NA | NA | 1 | 1 | 1 | 0 | 1 | 1 | NA | NA | 1 | 1 | 3 | 0.6667 |
| 8_12528039 | NA | NA | 1 | 1 | NA | NA | NA | NA | 1 | 1 | 1 | 1 | 1 | 1 | NA | NA | 1 | 1 | 3 | 1 |
| 8_12528052 | NA | NA | 1 | 1 | NA | NA | NA | NA | 1 | 1 | NA | NA | 1 | 1 | NA | NA | 2 | 1 | 3 | 1 |
| 8_12528063 | NA | NA | 1 | 1 | NA | NA | NA | NA | 1 | 1 | 1 | 1 | 1 | 1 | NA | NA | 2 | 0.5 | 4 | 0.75 |
| 8_12528064 | NA | NA | 1 | 1 | NA | NA | NA | NA | 1 | 1 | 1 | 1 | 1 | 1 | NA | NA | 2 | 1 | 4 | 1 |
| 8_12528079 | NA | NA | 1 | 1 | NA | NA | NA | NA | 1 | 1 | 1 | 1 | NA | NA | NA | NA | 2 | 0.5 | 3 | 0.6667 |
| 8_12528103 | NA | NA | 1 | 1 | 1 | 1 | NA | NA | 2 | 1 | 1 | 0 | 1 | 1 | NA | NA | 2 | 0.5 | 4 | 0.5 |
| 8_12528113 | NA | NA | 1 | 1 | 1 | 1 | NA | NA | 2 | 1 | 1 | 0 | 1 | 0 | NA | NA | 2 | 0.5 | 4 | 0.25 |
| 8_12528116 | NA | NA | 1 | 1 | NA | NA | NA | NA | 1 | 1 | 1 | 0 | 1 | 1 | NA | NA | 2 | 0.5 | 4 | 0.5 |
| 8_12528122 | NA | NA | 1 | 1 | 1 | 1 | NA | NA | 2 | 1 | 1 | 1 | 1 | 1 | NA | NA | 2 | 0 | 4 | 0.5 |
| 8_12528124 | NA | NA | 1 | 1 | 1 | 0 | NA | NA | 2 | 0.5 | 1 | 1 | 1 | 1 | NA | NA | 2 | 0 | 4 | 0.5 |
| 8_12528372 | 1 | 1 | NA | NA | 2 | 0 | 2 | 0.5 | 5 | 0.4 | NA | NA | 2 | 0.5 | 2 | 0 | 1 | 0 | 5 | 0.2 |
| 8_12529052 | 2 | 0.5 | NA | NA | 2 | 1 | 2 | 1 | 6 | 0.8333 | NA | NA | 1 | 1 | 1 | 1 | NA | NA | 2 | 1 |
| 8_12529079 | 2 | 1 | NA | NA | 2 | 1 | 2 | 1 | 6 | 1 | NA | NA | 1 | 1 | 2 | 1 | NA | NA | 3 | 1 |

| | | | | | | | | | | | | | | | | | | | | |
|---|---|---|---|---|---|---|---|---|---|---|---|---|---|---|---|---|---|---|---|---|
| 8_1252908 | 2 | 0.5 | NA | NA | 2 | 0 | 2 | 0.5 | 6 | 0.3333 | NA | NA | 1 | 0 | 2 | 0.5 | NA | NA | 3 | 0.3333 |
| 8_1253629 | 6 | 0.8333 | 8 | 1 | 4 | 0.5 | 10 | 0.6 | 28 | 0.75 | 8 | 0.875 | 3 | 1 | 5 | 0.6 | 14 | 1 | 30 | 0.9 |
| 8_1253631 | 5 | 0 | 7 | 0.1429 | 4 | 0 | 9 | 0.2222 | 25 | 0.12 | 8 | 0.125 | 3 | 0.3333 | 4 | 0 | 14 | 0.1429 | 29 | 0.1379 |
| 8_1253636 | 6 | 0.1667 | 6 | 0.3333 | 4 | 0.25 | 9 | 0.1111 | 25 | 0.2 | 8 | 0.25 | 3 | 0.3333 | 5 | 0.2 | 16 | 0.25 | 32 | 0.25 |
| 8_1253641 | 5 | 0 | 4 | 0 | 4 | 0.25 | 5 | 0 | 18 | 0.0556 | 6 | 0 | 3 | 0.3333 | 4 | 0 | 13 | 0.0769 | 26 | 0.0769 |
| 8_1253642 | 5 | 0.2 | 5 | 0.2 | 4 | 0 | 7 | 0.1429 | 21 | 0.1429 | 7 | 0 | 3 | 0.3333 | 5 | 0 | 14 | 0.0714 | 29 | 0.069 |
| 8_1253643 | 5 | 0.2 | 5 | 0 | 3 | 0 | 7 | 0 | 20 | 0.05 | 6 | 0 | 2 | 0.5 | 5 | 0.2 | 14 | 0 | 27 | 0.0741 |
| 8_1253646 | 5 | 0.2 | 4 | 0.25 | 2 | 0 | 8 | 0.125 | 19 | 0.1579 | 6 | 0.1667 | 3 | 0.6667 | 2 | 0 | 12 | 0.1667 | 23 | 0.2174 |
| 8_1253660 | 4 | 0.25 | 5 | 0 | 3 | 0 | 6 | 0 | 18 | 0.0556 | 6 | 0 | 2 | 0 | 4 | 0 | 11 | 0.1818 | 23 | 0.087 |
| 8_1253983 | 3 | 0 | 5 | 0.2 | 7 | 0.1429 | 8 | 0.125 | 23 | 0.1304 | 4 | 0.25 | 2 | 0 | 2 | 0 | 4 | 0 | 12 | 0.0833 |
| 8_1254587 | NA | NA | 1 | 1 | NA | NA | NA | NA | 1 | 1 | NA | NA | 3 | 0.3333 | NA | NA | 1 | 1 | 4 | 0.5 |
| 8_1254587 | NA | NA | 1 | 1 | NA | NA | NA | NA | 1 | 1 | NA | NA | 3 | 1 | NA | NA | NA | NA | 3 | 1 |
| 8_1254589 | NA | NA | 1 | 0 | 1 | 1 | NA | NA | 2 | 0.5 | NA | NA | 3 | 0.6667 | NA | NA | 1 | 1 | 4 | 0.75 |
| 8_1254593 | 1 | 1 | 2 | 1 | 1 | 1 | NA | NA | 4 | 1 | NA | NA | 3 | 0 | NA | NA | 1 | 1 | 4 | 0.25 |
| 8_1254598 | NA | NA | 2 | 0.5 | 1 | 0 | NA | NA | 3 | 0.3333 | NA | NA | NA | NA | NA | NA | 1 | 1 | 1 | 1 |
| 8_1254601 | 1 | 1 | 1 | 0 | 1 | 1 | NA | NA | 3 | 0.6667 | NA | NA | 3 | 0 | NA | NA | 1 | 1 | 4 | 0.25 |
| 8_1254606 | 1 | 0 | 2 | 0.5 | 1 | 0 | NA | NA | 4 | 0.25 | 1 | 0 | 2 | 0.5 | NA | NA | 2 | 0.5 | 5 | 0.4 |
| 8_1255184 | 1 | 1 | 1 | 1 | NA | NA | NA | NA | 2 | 1 | NA | NA | 1 | 1 | NA | NA | NA | NA | 1 | 1 |
| 8_1255185 | 1 | 1 | 1 | 0 | NA | NA | NA | NA | 2 | 0.5 | NA | NA | 1 | 1 | NA | NA | NA | NA | 1 | 1 |
| 8_1255188 | 1 | 1 | 1 | 1 | NA | NA | NA | NA | 2 | 1 | NA | NA | 1 | 1 | NA | NA | NA | NA | 1 | 1 |
| 8_1255192 | 1 | 0 | 1 | 1 | NA | NA | NA | NA | 2 | 0.5 | NA | NA | 1 | 1 | NA | NA | NA | NA | 1 | 1 |
| 8_1255194 | 1 | 1 | 1 | 1 | NA | NA | NA | NA | 2 | 1 | NA | NA | 1 | 1 | NA | NA | NA | NA | 1 | 1 |
| 8_1255196 | 2 | 1 | 1 | 0 | NA | NA | NA | NA | 3 | 0.6667 | NA | NA | 1 | 1 | NA | NA | NA | NA | 1 | 1 |
| 8_1255202 | 2 | 1 | 1 | 1 | NA | NA | NA | NA | 3 | 1 | NA | NA | 1 | 1 | NA | NA | NA | NA | 1 | 1 |
| 8_1255207 | 2 | 0.5 | 1 | 1 | NA | NA | NA | NA | 3 | 0.6667 | NA | NA | 1 | 1 | NA | NA | NA | NA | 1 | 1 |
| 8_1255209 | 2 | 1 | 1 | 1 | NA | NA | NA | NA | 3 | 1 | NA | NA | 1 | 1 | NA | NA | NA | NA | 1 | 1 |
| 8_1255209 | 2 | 1 | 1 | 1 | NA | NA | NA | NA | 3 | 1 | NA | NA | 1 | 1 | NA | NA | NA | NA | 1 | 1 |
| 8_1255662 | 2 | 0.5 | 1 | 0 | 1 | 0 | 3 | 0.3333 | 7 | 0.2857 | 1 | 0 | 2 | 0.5 | NA | NA | 1 | 1 | 4 | 0.5 |
| 8_1255665 | 2 | 0.5 | 1 | 0 | 2 | 0.5 | 5 | 0.6 | 10 | 0.5 | 1 | 0 | 2 | 0 | 1 | 1 | 1 | 0 | 5 | 0.2 |
| 8_1255665 | 2 | 0.5 | 1 | 1 | 2 | 0 | 5 | 0.2 | 10 | 0.3 | NA | NA | 2 | 0.5 | 1 | 1 | 1 | 1 | 4 | 0.75 |
| 8_1255665 | 2 | 0.5 | NA | NA | 1 | 0 | 3 | 0 | 6 | 0.1667 | 1 | 0 | NA | NA | NA | NA | 1 | 1 | 2 | 0.5 |
| 8_1255878 | 2 | 0.5 | NA | NA | 2 | 0 | 3 | 0.3333 | 7 | 0.2857 | 1 | 0 | 1 | 1 | 1 | 0 | 1 | 0 | 4 | 0.25 |
| 8_1255879 | 2 | 0 | NA | NA | 2 | 1 | 2 | 0.5 | 6 | 0.5 | 1 | 1 | 1 | 0 | 1 | 0 | 1 | 0 | 4 | 0.25 |
| 8_1256070 | 2 | 0 | 7 | 0 | 10 | 0.1 | 4 | 0.25 | 23 | 0.087 | 1 | 0 | 7 | 0.2857 | 5 | 0 | 5 | 0 | 18 | 0.1111 |
| 8_1256101 | 3 | 0.3333 | 6 | 0.3333 | 9 | 0.3333 | 4 | 0.25 | 22 | 0.3182 | 2 | 1 | 6 | 0.6667 | 6 | 0.3333 | 5 | 0.8 | 19 | 0.6316 |
| 8_1256160 | 3 | 0 | 6 | 0.3333 | 8 | 0 | 5 | 0 | 22 | 0.0909 | 5 | 0.2 | 8 | 0.125 | 9 | 0 | 5 | 0 | 27 | 0.0741 |
| 8_1256161 | 4 | 0.75 | 7 | 0.7143 | 9 | 0.7778 | 6 | 0.3333 | 26 | 0.6538 | 5 | 0.4 | 7 | 0.7143 | 9 | 0.5556 | 8 | 0.75 | 29 | 0.6207 |
| 8_1256164 | 5 | 0.2 | 5 | 0 | 7 | 0.1429 | 5 | 0 | 22 | 0.0909 | 3 | 0.3333 | 8 | 0.375 | 6 | 0.3333 | 6 | 0.3333 | 23 | 0.3478 |
| 8_1256164 | 5 | 0.2 | 5 | 0.2 | 6 | 0.3333 | 6 | 0.3333 | 22 | 0.2727 | 2 | 0.5 | 7 | 0.2857 | 7 | 0.1429 | 5 | 0.2 | 21 | 0.2381 |
| 8_1256165 | 5 | 0.2 | 8 | 0.25 | 8 | 0.375 | 7 | 0.2857 | 28 | 0.2857 | 4 | 0 | 9 | 0.1111 | 9 | 0.2222 | 8 | 0.125 | 30 | 0.1333 |
| 8_1256166 | 3 | 0.3333 | 7 | 0.1429 | 5 | 0 | 6 | 0.6667 | 21 | 0.2857 | 2 | 0.5 | 9 | 0.3333 | 8 | 0.125 | 7 | 0.1429 | 26 | 0.2308 |
| 8_1256167 | NA | NA | 4 | 0.25 | 2 | 0.5 | 4 | 0.25 | 10 | 0.3 | 3 | 0.3333 | 7 | 0.1429 | 6 | 0.3333 | 6 | 0.6667 | 22 | 0.3636 |
| 8_1256170 | NA | NA | 2 | 0.5 | 3 | 0 | 2 | 0 | 7 | 0.1429 | 1 | 1 | 1 | 0 | 1 | 1 | NA | NA | 3 | 0.6667 |
| 8_1256171 | 1 | 0 | 2 | 0 | 2 | 0.5 | 2 | 0.5 | 7 | 0.2857 | 1 | 1 | 1 | 0 | 3 | 0 | NA | NA | 5 | 0.2 |
| 8_1256175 | NA | NA | 1 | 1 | 3 | 0.3333 | 1 | 0 | 5 | 0.4 | NA | NA | 1 | 0 | 3 | 0.3333 | NA | NA | 4 | 0.25 |
| 8_1256199 | 5 | 0 | 5 | 0.6 | 7 | 0.2857 | 11 | 0.0909 | 28 | 0.2143 | 5 | 0.2 | 2 | 0.5 | 9 | 0.3333 | 3 | 0.3333 | 19 | 0.3158 |
| 8_1256213 | 3 | 0 | 4 | 0 | 4 | 0 | 4 | 0.25 | 15 | 0.0667 | 2 | 0 | 2 | 0 | 7 | 0.1429 | 1 | 0 | 12 | 0.0833 |
| 8_1256215 | 3 | 0 | 2 | 0.5 | 3 | 0.3333 | 2 | 0.5 | 10 | 0.3 | 3 | 0 | 1 | 0 | 6 | 0.8333 | 1 | 0 | 11 | 0.4545 |
| 8_1256215 | 3 | 0 | 2 | 1 | 3 | 0.3333 | 4 | 0.75 | 12 | 0.5 | 2 | 1 | 2 | 1 | 7 | 0.5714 | 1 | 0 | 12 | 0.6667 |
| 8_1256216 | 4 | 0 | 3 | 0.3333 | 3 | 0 | 3 | 0 | 13 | 0.0769 | 2 | 0.5 | 4 | 0 | 4 | 0 | 1 | 0 | 11 | 0.0909 |
| 8_1256219 | 4 | 0.5 | 2 | 0.5 | NA | NA | 3 | 0.6667 | 9 | 0.5556 | 3 | 0.3333 | 3 | 0 | 6 | 0.3333 | NA | NA | 12 | 0.25 |
| 8_1256220 | 2 | 1 | 3 | 1 | 1 | 1 | 3 | 1 | 9 | 1 | 2 | 1 | 3 | 1 | 5 | 1 | NA | NA | 10 | 1 |
| 8_1256221 | 3 | 0 | 3 | 0 | 2 | 0 | 3 | 0.6667 | 11 | 0.1818 | 3 | 0 | 5 | 0.2 | 6 | 0 | 1 | 0 | 15 | 0.0667 |
| 8_1256221 | 3 | 0.3333 | 3 | 0.3333 | 2 | 0 | 3 | 0.3333 | 11 | 0.2727 | 3 | 0.6667 | 5 | 0.8 | 7 | 0.7143 | NA | NA | 15 | 0.7333 |
| 8_1256222 | 3 | 0 | 3 | 0.3333 | 2 | 0 | 2 | 0 | 10 | 0.1 | 2 | 0.5 | 5 | 0.2 | 7 | 0.2857 | NA | NA | 14 | 0.2857 |
| 8_1256223 | 2 | 0.5 | 3 | 0 | 1 | 1 | 1 | 1 | 7 | 0.4286 | 2 | 1 | 2 | 1 | 6 | 1 | NA | NA | 10 | 1 |
| 8_1256223 | 3 | 0 | 3 | 0 | 1 | 0 | 3 | 0.3333 | 10 | 0.1 | 3 | 0 | 5 | 0 | 6 | 0.1667 | NA | NA | 14 | 0.0714 |
| 8_1256224 | 3 | 1 | 1 | 1 | 1 | 1 | 3 | 1 | 8 | 1 | 1 | 0 | 2 | 0.5 | 2 | 0.5 | NA | NA | 5 | 0.4 |
| 8_1256224 | 2 | 1 | 2 | 1 | 2 | 1 | 3 | 1 | 9 | 1 | 3 | 1 | 4 | 1 | 4 | 1 | NA | NA | 11 | 1 |
| 8_1256225 | 3 | 1 | 2 | 1 | 1 | 1 | 2 | 1 | 8 | 1 | 2 | 0.5 | 4 | 0.75 | 3 | 1 | NA | NA | 9 | 0.7778 |
| 8_1256225 | 3 | 0.3333 | 2 | 0.5 | 1 | 0 | 2 | 0.5 | 8 | 0.375 | 3 | 0.3333 | 5 | 0.6 | 3 | 0.6667 | NA | NA | 11 | 0.5455 |
| 8_1256225 | 1 | 0 | 2 | 1 | 1 | 0 | 3 | 0 | 7 | 0.2857 | 2 | 0.5 | 4 | 0.25 | 5 | 0.2 | 1 | 1 | 12 | 0.3333 |
| 8_1256225 | 3 | 0.3333 | 1 | 0 | 1 | 1 | 3 | 0.3333 | 8 | 0.375 | 2 | 0.5 | 2 | 0.5 | 4 | 1 | NA | NA | 8 | 0.75 |
| 8_1256227 | 2 | 0 | 1 | 0 | 1 | 1 | 3 | 0.3333 | 7 | 0.2857 | 2 | 0 | 5 | 0.2 | 5 | 0.4 | NA | NA | 12 | 0.25 |
| 8_1256229 | 3 | 1 | 2 | 1 | 1 | 1 | 3 | 1 | 9 | 1 | 3 | 1 | 4 | 1 | 4 | 1 | NA | NA | 11 | 1 |

| | | | | | | | | | | | | | | | | | | | | |
|---|---|---|---|---|---|---|---|---|---|---|---|---|---|---|---|---|---|---|---|---|
| 8_1256233 | 3 | 0.3333 | 1 | 0 | NA | NA | 3 | 0.3333 | 7 | 0.2857 | 2 | 0.5 | 4 | 0.5 | 3 | 1 | 1 | 1 | 10 | 0.7 |
| 8_1256233 | 3 | 0.6667 | 2 | 1 | NA | NA | 3 | 1 | 8 | 0.875 | 2 | 0.5 | 3 | 0.6667 | 3 | 1 | NA | NA | 8 | 0.75 |
| 8_1256233 | 3 | 1 | 2 | 1 | NA | NA | 3 | 1 | 8 | 1 | 1 | 1 | 2 | 0.5 | 3 | 1 | NA | NA | 6 | 0.8333 |
| 8_1256234 | 3 | 0.6667 | NA | NA | NA | NA | 3 | 0 | 6 | 0.3333 | 1 | 1 | 3 | 0 | 2 | 0 | 1 | 0 | 7 | 0.1429 |
| 8_1256236 | 2 | 0.5 | 1 | 0 | NA | NA | 2 | 1 | 5 | 0.6 | 2 | 0.5 | 2 | 0 | 2 | 0 | 1 | 0 | 7 | 0.1429 |
| 8_1256296 | NA | NA | NA | NA | 1 | 1 | 1 | 1 | 2 | 1 | 1 | 1 | 1 | 1 | NA | NA | NA | NA | 2 | 1 |
| 8_1256297 | NA | NA | NA | NA | 1 | 1 | NA | NA | 1 | 1 | 1 | 0 | 1 | 1 | NA | NA | NA | NA | 2 | 0.5 |
| 8_1256299 | NA | NA | NA | NA | 1 | 1 | NA | NA | 1 | 1 | 1 | 0 | 1 | 0 | NA | NA | 1 | 1 | 3 | 0.3333 |
| 8_1256299 | NA | NA | NA | NA | 1 | 1 | 1 | 1 | 2 | 1 | NA | NA | 1 | 1 | NA | NA | 1 | 0 | 2 | 0.5 |
| 8_1256299 | NA | NA | NA | NA | 1 | 0 | 1 | 1 | 2 | 0.5 | 1 | 1 | 1 | 1 | NA | NA | 1 | 1 | 3 | 1 |
| 8_1256301 | NA | NA | NA | NA | 1 | 1 | 1 | 1 | 2 | 1 | 1 | 0 | NA | NA | NA | NA | 1 | 1 | 2 | 0.5 |
| 8_1256302 | NA | NA | NA | NA | 1 | 0 | 1 | 1 | 2 | 0.5 | 1 | 0 | 1 | 1 | NA | NA | 1 | 1 | 3 | 0.6667 |
| 8_1256302 | NA | NA | NA | NA | 1 | 1 | 1 | 0 | 2 | 0.5 | 1 | 1 | 1 | 0 | NA | NA | NA | NA | 2 | 0.5 |
| 8_1256313 | 1 | 0 | NA | NA | 1 | 0 | 2 | 0.5 | 4 | 0.25 | 2 | 0 | 1 | 1 | NA | NA | 2 | 0 | 5 | 0.2 |
| 8_1256315 | 1 | 0 | NA | NA | NA | NA | 2 | 0.5 | 3 | 0.3333 | 2 | 0 | 1 | 1 | NA | NA | 2 | 0 | 5 | 0.2 |
| 8_1256354 | 17 | 0.2941 | 16 | 0.4375 | 18 | 0.2778 | 17 | 0.1765 | 68 | 0.2941 | 14 | 0.1429 | 21 | 0.0952 | 22 | 0.3182 | 12 | 0.1667 | 69 | 0.1884 |
| 8_1256355 | 14 | 0.2857 | 16 | 0.0625 | 15 | 0.2 | 17 | 0.2353 | 62 | 0.1935 | 11 | 0.2727 | 17 | 0.2941 | 19 | 0.5263 | 11 | 0.2727 | 58 | 0.3621 |
| 8_1256358 | 13 | 0.5385 | 16 | 0.3125 | 11 | 0.3636 | 12 | 0.4167 | 52 | 0.4038 | 9 | 0.3333 | 19 | 0.4211 | 19 | 0.3158 | 11 | 0.8182 | 58 | 0.4483 |
| 8_1256364 | 12 | 0 | 10 | 0.1 | 8 | 0 | 6 | 0.1667 | 36 | 0.0556 | 4 | 0.25 | 12 | 0.25 | 10 | 0.1 | 8 | 0.125 | 34 | 0.1765 |
| 8_1256364 | 15 | 0.0667 | 12 | 0.1667 | 9 | 0 | 12 | 0.0833 | 48 | 0.0833 | 9 | 0 | 17 | 0 | 18 | 0.1111 | 12 | 0.0833 | 56 | 0.0536 |
| 8_1256365 | 12 | 0.25 | 11 | 0.1818 | 8 | 0.125 | 9 | 0.5556 | 40 | 0.275 | 8 | 0.125 | 15 | 0.5333 | 15 | 0.3333 | 8 | 0.5 | 46 | 0.3913 |
| 8_1256366 | 16 | 0.0625 | 14 | 0.1429 | 8 | 0 | 10 | 0.2 | 48 | 0.1042 | 9 | 0 | 15 | 0.1333 | 16 | 0 | 11 | 0.0909 | 51 | 0.0588 |
| 8_1256366 | 11 | 0.9091 | 7 | 0.7143 | 5 | 0.6 | 9 | 0.7778 | 32 | 0.7812 | 6 | 0.6667 | 11 | 0.9091 | 9 | 0.7778 | 6 | 0.8333 | 32 | 0.8125 |
| 8_1256367 | 11 | 0.1818 | 10 | 0.2 | 5 | 0 | 10 | 0.6 | 36 | 0.2778 | 5 | 0.2 | 10 | 0.4 | 15 | 0.1333 | 9 | 0 | 39 | 0.1795 |
| 8_1256370 | 13 | 0.6154 | 14 | 0.7857 | 8 | 0.5 | 9 | 0.7778 | 44 | 0.6818 | 8 | 0.625 | 12 | 0.5 | 17 | 0.7059 | 6 | 0.5 | 43 | 0.6047 |
| 8_1256371 | 15 | 0.1333 | 13 | 0.0769 | 6 | 0.5 | 8 | 0.125 | 42 | 0.1667 | 7 | 0.2857 | 16 | 0.1875 | 14 | 0.2143 | 6 | 0.3333 | 43 | 0.2326 |
| 8_1256371 | 15 | 0.0667 | 15 | 0.2667 | 7 | 0 | 10 | 0 | 47 | 0.1064 | 8 | 0.125 | 17 | 0.0588 | 18 | 0 | 6 | 0.1667 | 49 | 0.0612 |
| 8_1256371 | 14 | 0.7143 | 11 | 0.6364 | 7 | 0.4286 | 9 | 0.6667 | 41 | 0.6341 | 7 | 0.5714 | 15 | 0.4667 | 17 | 0.5294 | 4 | 1 | 43 | 0.5581 |
| 8_1256372 | 13 | 0.5385 | 12 | 0.75 | 4 | 0.75 | 10 | 0.7 | 39 | 0.6667 | 7 | 0.7143 | 14 | 0.5 | 15 | 0.4667 | 5 | 1 | 41 | 0.5854 |
| 8_1256372 | 14 | 0.9286 | 13 | 1 | 6 | 1 | 10 | 0.9 | 43 | 0.9535 | 5 | 1 | 14 | 1 | 17 | 0.9412 | 6 | 1 | 42 | 0.9762 |
| 8_1256373 | 13 | 0.4615 | 11 | 0.2727 | 6 | 0.6667 | 8 | 0.75 | 38 | 0.5 | 8 | 0.375 | 12 | 0.5833 | 11 | 0.0909 | 7 | 0.2857 | 38 | 0.3421 |
| 8_1256373 | 10 | 0.1 | 11 | 0.1818 | 5 | 0 | 7 | 0 | 33 | 0.0909 | 7 | 0.1429 | 14 | 0.2857 | 13 | 0.1538 | 5 | 0.2 | 39 | 0.2051 |
| 8_1256373 | 14 | 0.7143 | 12 | 0.6667 | 5 | 0.4 | 6 | 0.3333 | 37 | 0.5946 | 8 | 0.875 | 14 | 0.2857 | 13 | 0.3846 | 7 | 0.8571 | 42 | 0.5238 |
| 8_1256751 | 9 | 0.7778 | 9 | 0.2222 | 6 | 0.8333 | 8 | 0.375 | 32 | 0.5312 | 6 | 0.8333 | 9 | 0.5556 | 13 | 0.6923 | 6 | 0.6667 | 34 | 0.6765 |
| 8_1256751 | 10 | 0.1 | 10 | 0 | 5 | 0.2 | 9 | 0.2222 | 34 | 0.1176 | 7 | 0.1429 | 9 | 0.1111 | 14 | 0.2857 | 6 | 0 | 36 | 0.1667 |
| 8_1256756 | 8 | 0.625 | 4 | 0.5 | 5 | 0.6 | 4 | 0.25 | 21 | 0.5238 | 7 | 0.4286 | 3 | 0.3333 | 10 | 0.6 | 1 | 1 | 21 | 0.5238 |
| 8_1257424 | 14 | 0.0714 | 16 | 0.0625 | 11 | 0.0909 | 16 | 0.0625 | 57 | 0.0702 | 11 | 0 | 21 | 0.1905 | 20 | 0.05 | 14 | 0.2143 | 66 | 0.1212 |
| 8_1257593 | 15 | 0 | 17 | 0.0588 | 13 | 0.1538 | 18 | 0.1111 | 63 | 0.0794 | 8 | 0.125 | 24 | 0.0833 | 21 | 0.0476 | 13 | 0.0769 | 66 | 0.0758 |
| 8_1257593 | 24 | 0.0833 | 24 | 0.0417 | 22 | 0.1818 | 20 | 0.05 | 90 | 0.0889 | 15 | 0.0667 | 25 | 0.08 | 25 | 0.12 | 16 | 0.125 | 81 | 0.0988 |
| 8_1257595 | 24 | 0.1667 | 30 | 0.0667 | 23 | 0.1304 | 24 | 0.0417 | 101 | 0.099 | 15 | 0.0667 | 29 | 0.1034 | 29 | 0.1724 | 21 | 0.1429 | 94 | 0.1277 |
| 8_1257596 | 25 | 0.2 | 30 | 0.1667 | 24 | 0.2083 | 23 | 0.1739 | 102 | 0.1863 | 13 | 0.3846 | 31 | 0.129 | 29 | 0.3448 | 21 | 0.1905 | 94 | 0.2447 |
| 8_1257606 | 30 | 0.0667 | 35 | 0.2286 | 30 | 0.0667 | 26 | 0.0385 | 121 | 0.1074 | 19 | 0.2105 | 35 | 0.0571 | 36 | 0.0278 | 26 | 0.1538 | 116 | 0.0948 |
| 8_1257607 | 32 | 0.1875 | 32 | 0.0625 | 30 | 0.1333 | 24 | 0 | 118 | 0.1017 | 21 | 0.1905 | 35 | 0.1714 | 33 | 0.1818 | 28 | 0.1429 | 117 | 0.1709 |
| 8_1261453 | 1 | 0 | 7 | 0 | 2 | 0 | 3 | 0.3333 | 13 | 0.0769 | 8 | 0.25 | 4 | 0 | 2 | 0 | 2 | 0 | 16 | 0.125 |
| 8_1261455 | 2 | 0 | 6 | 0.3333 | 2 | 1 | 2 | 0 | 12 | 0.3333 | 8 | 0.375 | 4 | 0.25 | 2 | 1 | 2 | 1 | 16 | 0.5 |
| 8_1261456 | 1 | 0 | 4 | 0.5 | 2 | 0 | 2 | 0 | 9 | 0.2222 | 4 | 0.5 | 4 | 0.75 | 1 | 0 | 1 | 0 | 10 | 0.5 |
| 8_1261910 | NA | NA | 2 | 0.5 | NA | NA | NA | NA | 2 | 0.5 | 2 | 0 | NA | NA | 1 | 0 | 2 | 0.5 | 5 | 0.2 |
| 8_1263972 | NA | NA | NA | NA | 1 | 1 | 2 | 0.5 | 3 | 0.6667 | NA | NA | NA | NA | 1 | 1 | NA | NA | 1 | 1 |
| 8_1263974 | NA | NA | NA | NA | 1 | 1 | 2 | 0.5 | 3 | 0.6667 | NA | NA | NA | NA | 1 | 1 | NA | NA | 1 | 1 |
| 8_1263975 | NA | NA | NA | NA | 1 | 0 | 2 | 0.5 | 3 | 0.3333 | NA | NA | NA | NA | 1 | 1 | NA | NA | 1 | 1 |
| 8_1263980 | NA | NA | NA | NA | 1 | 1 | 2 | 1 | 3 | 1 | NA | NA | NA | NA | 1 | 1 | NA | NA | 1 | 1 |
| 8_1263981 | NA | NA | NA | NA | 1 | 1 | 2 | 1 | 3 | 1 | NA | NA | NA | NA | 1 | 1 | NA | NA | 1 | 1 |
| 8_1263985 | NA | NA | NA | NA | 1 | 1 | 2 | 1 | 3 | 1 | NA | NA | NA | NA | 1 | 1 | NA | NA | 1 | 1 |
| 8_1263989 | NA | NA | NA | NA | 2 | 0.5 | 2 | 1 | 4 | 0.75 | NA | NA | 1 | 1 | 1 | 1 | NA | NA | 2 | 1 |
| 8_1263991 | NA | NA | NA | NA | 3 | 0 | 2 | 1 | 5 | 0.4 | NA | NA | 1 | 0 | 1 | 1 | NA | NA | 2 | 0.5 |
| 8_1263991 | NA | NA | NA | NA | 3 | 0.6667 | 2 | 1 | 5 | 0.8 | NA | NA | 1 | 1 | 1 | 1 | NA | NA | 2 | 1 |
| 8_1263992 | NA | NA | NA | NA | 2 | 0.5 | 2 | 1 | 4 | 0.75 | NA | NA | 1 | 1 | 1 | 0 | NA | NA | 2 | 0.5 |
| 8_1263997 | NA | NA | NA | NA | 2 | 0 | 2 | 0.5 | 4 | 0.25 | NA | NA | NA | NA | 2 | 0.5 | NA | NA | 2 | 0.5 |
| 8_1264746 | 8 | 0.125 | 5 | 0 | 4 | 0 | 3 | 0 | 20 | 0.05 | 1 | 0 | 3 | 0 | 1 | 0 | 6 | 0.1667 | 11 | 0.0909 |
| 8_1264786 | 2 | 1 | 2 | 1 | 6 | 0.8333 | 5 | 0.4 | 15 | 0.7333 | 3 | 1 | 1 | 1 | 7 | 0.8571 | 3 | 0.6667 | 14 | 0.8571 |
| 8_1264789 | 2 | 0 | 3 | 0.3333 | 5 | 0.2 | 5 | 0 | 15 | 0.1333 | 4 | 0.25 | 1 | 0 | 7 | 0 | 2 | 0.5 | 14 | 0.1429 |
| 8_1264802 | 3 | 1 | 2 | 1 | 2 | 0.5 | 2 | 0.5 | 9 | 0.7778 | 3 | 0.3333 | 1 | 1 | 4 | 1 | 3 | 0.3333 | 11 | 0.6364 |
| 8_1266138 | 1 | 1 | 1 | 1 | NA | NA | NA | NA | 2 | 1 | 1 | 0 | NA | NA | 2 | 0 | 1 | 1 | 4 | 0.25 |
| 8_1266139 | 1 | 1 | NA | NA | NA | NA | NA | NA | 1 | 1 | 1 | 1 | NA | NA | 2 | 1 | 1 | 0 | 4 | 0.75 |
| 8_1267206 | NA | NA | 1 | 1 | 1 | 1 | NA | NA | 2 | 1 | 1 | 1 | 1 | 0 | NA | NA | NA | NA | 2 | 0.5 |

| | | | | | | | | | | | | | | | | | | | | |
|---|---|---|---|---|---|---|---|---|---|---|---|---|---|---|---|---|---|---|---|---|
| 8_1267207( | NA | NA | NA | NA | 1 | 1 | NA | NA | 1 | 1 | 1 | 1 | 1 | 1 | NA | NA | NA | NA | 2 | 1 |
| 8_1267207: | NA | NA | 1 | 1 | NA | NA | NA | NA | 1 | 1 | 1 | 1 | 1 | 0 | NA | NA | NA | NA | 2 | 0.5 |
| 8_1267207: | NA | NA | 1 | 1 | NA | NA | NA | NA | 1 | 1 | 1 | 1 | 1 | 1 | NA | NA | NA | NA | 2 | 1 |
| 8_1267208 | NA | NA | 1 | 1 | NA | NA | NA | NA | 1 | 1 | 1 | 1 | 1 | 1 | NA | NA | NA | NA | 2 | 1 |
| 8_1267222( | NA | NA | NA | NA | 1 | 1 | NA | NA | 1 | 1 | 1 | 1 | 1 | 1 | NA | NA | NA | NA | 2 | 1 |
| 8_1267225( | 17 | 0.2941 | 20 | 0.7 | 21 | 0.4286 | 16 | 0.4375 | 74 | 0.473 | 12 | 0.3333 | 3 | 0.3333 | 12 | 0.1667 | 18 | 0.2222 | 45 | 0.2444 |
| 8_1267520: | 22 | 0.0909 | 26 | 0.0769 | 22 | 0.4091 | 19 | 0.2105 | 89 | 0.191 | 16 | 0 | 6 | 0 | 13 | 0.1538 | 24 | 0.125 | 59 | 0.0847 |
| 8_1267520: | 21 | 0.2381 | 23 | 0.2609 | 23 | 0.4783 | 19 | 0.3158 | 86 | 0.3256 | 17 | 0.4706 | 4 | 0 | 13 | 0.3846 | 24 | 0.375 | 58 | 0.3793 |
| 8_1267520( | 23 | 0.3913 | 25 | 0.36 | 24 | 0.375 | 17 | 0.4706 | 89 | 0.3933 | 15 | 0.5333 | 5 | 0.6 | 15 | 0.2 | 16 | 0.375 | 51 | 0.3922 |
| 8_1267565( | 23 | 0.8261 | 23 | 0.6957 | 23 | 0.6087 | 16 | 0.8125 | 85 | 0.7294 | 17 | 0.8235 | 6 | 0.5 | 14 | 0.8571 | 20 | 0.75 | 57 | 0.7719 |
| 8_1267565 | 21 | 0.6667 | 23 | 0.5652 | 26 | 0.7308 | 18 | 0.6111 | 88 | 0.6477 | 18 | 0.5 | 7 | 0.5714 | 14 | 0.5714 | 23 | 0.6087 | 62 | 0.5645 |
| 8_1267566( | 19 | 0.9474 | 18 | 0.8889 | 18 | 1 | 14 | 0.9286 | 69 | 0.942 | 12 | 1 | 5 | 1 | 14 | 0.9286 | 18 | 0.9444 | 49 | 0.9592 |
| 8_1267567( | 25 | 0.72 | 24 | 0.5 | 25 | 0.52 | 19 | 0.6842 | 93 | 0.6022 | 18 | 0.6667 | 6 | 0.6667 | 16 | 0.5 | 21 | 0.6667 | 61 | 0.623 |
| 8_1267572 | 29 | 0.7241 | 34 | 0.7059 | 29 | 0.7931 | 21 | 0.5714 | 113 | 0.708 | 21 | 0.8095 | 8 | 1 | 19 | 0.6842 | 27 | 0.5556 | 75 | 0.7067 |
| 8_1268622: | NA | NA | NA | NA | 1 | 1 | NA | NA | 1 | 1 | NA | NA | 1 | 1 | NA | NA | NA | NA | 1 | 1 |
| 8_1268642( | NA | NA | NA | NA | 1 | 1 | NA | NA | 1 | 1 | 1 | 1 | NA | NA | 2 | 0.5 | NA | NA | 3 | 0.6667 |
| 8_1268643 | NA | NA | NA | NA | 1 | 1 | NA | NA | 1 | 1 | 1 | 1 | NA | NA | 2 | 0 | 1 | 1 | 4 | 0.5 |
| 8_1268649: | NA | NA | NA | NA | 1 | 1 | NA | NA | 1 | 1 | 1 | 0 | NA | NA | 2 | 0.5 | 1 | 1 | 4 | 0.5 |
| 8_1268649! | NA | NA | NA | NA | 1 | 1 | NA | NA | 1 | 1 | 1 | 0 | NA | NA | 2 | 1 | 1 | 1 | 4 | 0.75 |
| 8_1268652 | NA | NA | NA | NA | 1 | 1 | NA | NA | 1 | 1 | 1 | 1 | NA | NA | 1 | 1 | 1 | 1 | 3 | 1 |
| 8_1268661! | NA | NA | NA | NA | 1 | 1 | NA | NA | 1 | 1 | 1 | 1 | NA | NA | NA | NA | NA | NA | 1 | 1 |
| 8_1268664( | 4 | 0.5 | 2 | 0 | 3 | 0 | 1 | 0 | 10 | 0.2 | NA | NA | 1 | 0 | 1 | 0 | 1 | 1 | 3 | 0.3333 |
| 8_1271089: | 6 | 0.3333 | NA | NA | 4 | 0.5 | 1 | 0 | 11 | 0.3636 | 1 | 0 | 1 | 0 | 1 | 1 | NA | NA | 3 | 0.3333 |
| 8_1271213 | 9 | 0.1111 | 1 | 0 | 4 | 0.25 | 1 | 0 | 15 | 0.1333 | 1 | 0 | 3 | 0 | 2 | 0 | 1 | 1 | 7 | 0.1429 |
| 8_1271214 | 8 | 0 | NA | NA | 5 | 0.2 | 1 | 0 | 14 | 0.0714 | 1 | 0 | 2 | 0 | 2 | 0 | 1 | 1 | 6 | 0.1667 |
| 8_1271214: | 2 | 0.5 | NA | NA | 4 | 0.75 | NA | NA | 6 | 0.6667 | 1 | 1 | 2 | 1 | NA | NA | 1 | 1 | 4 | 1 |
| 8_1271215 | 8 | 0 | 2 | 0.5 | 3 | 0.6667 | NA | NA | 13 | 0.2308 | 1 | 0 | 3 | 0.3333 | 1 | 0 | 1 | 0 | 6 | 0.1667 |
| 8_1271215: | 10 | 0 | 2 | 0 | 6 | 0.1667 | 1 | 0 | 19 | 0.0526 | NA | NA | 3 | 0 | 2 | 0 | 1 | 1 | 6 | 0.1667 |
| 8_1271216( | 9 | 0 | 2 | 0.5 | 7 | 0.2857 | 1 | 0 | 19 | 0.1579 | 1 | 0 | 2 | 0 | 1 | 0 | 1 | 1 | 5 | 0.2 |
| 8_1271217 | 9 | 0.1111 | 2 | 0 | 7 | 0 | 1 | 0 | 19 | 0.0526 | NA | NA | 3 | 0.3333 | 2 | 0 | 1 | 0 | 6 | 0.1667 |
| 8_1271218( | 6 | 0.3333 | 1 | 0 | 5 | 0 | 1 | 0 | 13 | 0.1538 | 1 | 0 | 2 | 0.5 | 1 | 0 | 1 | 0 | 5 | 0.2 |
| 8_1271223( | 12 | 0.3333 | 5 | 0.2 | 8 | 0.125 | 2 | 0.5 | 27 | 0.2593 | 1 | 0 | 3 | 0.3333 | 2 | 0 | 1 | 0 | 7 | 0.1429 |
| 8_1271407( | 1 | 0 | NA | NA | 1 | 1 | 2 | 0.5 | 4 | 0.5 | NA | NA | 1 | 1 | NA | NA | NA | NA | 1 | 1 |
| 8_1271408 | 1 | 0 | NA | NA | 1 | 1 | 2 | 0.5 | 4 | 0.5 | NA | NA | 2 | 0.5 | NA | NA | NA | NA | 2 | 0.5 |
| 8_1271518 | 10 | 1 | 5 | 0.8 | 7 | 0.8571 | 8 | 0.875 | 30 | 0.9 | 4 | 1 | 6 | 0.8333 | 6 | 0.8333 | 5 | 0.8 | 21 | 0.8571 |
| 8_1271518( | 8 | 0.75 | 5 | 0.8 | 6 | 0.8333 | 8 | 0.875 | 27 | 0.8148 | 4 | 0.75 | 7 | 0.7143 | 7 | 0.8571 | 5 | 0.8 | 23 | 0.7826 |
| 8_1271519( | 10 | 0 | 6 | 0 | 8 | 0.125 | 8 | 0.125 | 32 | 0.0625 | 5 | 0.2 | 7 | 0 | 8 | 0.25 | 5 | 0.4 | 25 | 0.2 |
| 8_1271520( | 11 | 0.8182 | 5 | 0.8 | 7 | 0.8571 | 7 | 1 | 30 | 0.8667 | 3 | 1 | 7 | 1 | 6 | 0.8333 | 6 | 1 | 22 | 0.9545 |
| 8_1271536! | 1 | 1 | NA | NA | NA | NA | NA | NA | 1 | 1 | NA | NA | NA | NA | 1 | 1 | NA | NA | 1 | 1 |
| 8_1273159! | 4 | 0 | 3 | 0 | 3 | 0 | 7 | 0.1429 | 17 | 0.0588 | 5 | 0.2 | 2 | 0 | 5 | 0 | 8 | 0 | 20 | 0.05 |
| 8_1273177: | 4 | 0.25 | 6 | 0.6667 | NA | NA | 11 | 0.1818 | 21 | 0.3333 | 5 | 0.4 | 1 | 0 | 6 | 0.1667 | 3 | 0 | 15 | 0.2 |
| 8_1273178( | 4 | 1 | 5 | 1 | NA | NA | 10 | 1 | 19 | 1 | 3 | 1 | 1 | 1 | 6 | 1 | 3 | 1 | 13 | 1 |
| 8_1273178 | 4 | 0.25 | 6 | 0.3333 | NA | NA | 12 | 0.1667 | 22 | 0.2273 | 3 | 0.3333 | 1 | 0 | 5 | 0.2 | 2 | 0 | 11 | 0.1818 |
| 8_1273178! | 3 | 0.3333 | 4 | 0.25 | 1 | 0 | 11 | 0.6364 | 19 | 0.4737 | 3 | 0.6667 | 1 | 1 | 6 | 0.3333 | 3 | 0 | 13 | 0.3846 |
| 8_1273178! | 4 | 0.75 | 6 | 1 | 1 | 1 | 11 | 0.6364 | 22 | 0.7727 | 3 | 1 | 1 | 1 | 4 | 1 | 3 | 1 | 11 | 1 |
| 8_1273181: | 3 | 0.6667 | 5 | 1 | NA | NA | 9 | 0.8889 | 17 | 0.8824 | 4 | 1 | 1 | 1 | 2 | 0.5 | 2 | 0.5 | 9 | 0.7778 |
| 8_1273181( | 4 | 1 | 5 | 1 | NA | NA | 9 | 1 | 18 | 1 | 4 | 1 | 1 | 1 | 5 | 1 | 2 | 1 | 12 | 1 |
| 8_1273182: | 2 | 1 | 5 | 0.8 | NA | NA | 6 | 1 | 13 | 0.9231 | 3 | 1 | 1 | 1 | 3 | 1 | 2 | 0 | 9 | 0.7778 |
| 8_1273182! | 4 | 0.25 | 4 | 0.25 | NA | NA | 7 | 0.1429 | 15 | 0.2 | 4 | 0.25 | 1 | 0 | 5 | 0 | 1 | 0 | 11 | 0.0909 |
| 8_1273183 | 1 | 1 | 4 | 0.25 | NA | NA | 8 | 0.5 | 13 | 0.4615 | 2 | 1 | 1 | 1 | 5 | 0.4 | 2 | 0.5 | 10 | 0.6 |
| 8_1273183( | 3 | 0.3333 | 3 | 0.6667 | NA | NA | 8 | 1 | 14 | 0.7857 | NA | NA | 1 | 1 | 2 | 1 | 2 | 1 | 5 | 1 |
| 8_1273185! | 1 | 0 | 4 | 0 | NA | NA | 10 | 0.2 | 15 | 0.1333 | 2 | 0.5 | 1 | 0 | 4 | 0.25 | 1 | 0 | 8 | 0.25 |
| 8_1273193 | NA | NA | 3 | 0 | NA | NA | 9 | 0.1111 | 12 | 0.0833 | 2 | 1 | 1 | 0 | 4 | 0.25 | 1 | 0 | 8 | 0.375 |
| 8_1273193 | 1 | 1 | 4 | 1 | NA | NA | 8 | 0.875 | 13 | 0.9231 | 2 | 1 | 1 | 1 | 3 | 0.6667 | 1 | 1 | 7 | 0.8571 |
| 8_1273196( | NA | NA | 1 | 0 | NA | NA | 6 | 0.3333 | 7 | 0.2857 | NA | NA | 1 | 0 | 3 | 0.3333 | NA | NA | 4 | 0.25 |
| 8_1273199 | 1 | 1 | 3 | 0.3333 | NA | NA | 8 | 0.625 | 12 | 0.5833 | 2 | 1 | 1 | 1 | 2 | 0.5 | 2 | 0.5 | 7 | 0.7143 |
| 8_1273200( | NA | NA | 2 | 1 | NA | NA | 8 | 0.625 | 10 | 0.7 | 2 | 0.5 | NA | NA | 2 | 0.5 | 2 | 0.5 | 6 | 0.5 |
| 8_1276616: | 34 | 0.3235 | 39 | 0.0769 | 29 | 0.1379 | 54 | 0.1111 | 156 | 0.1538 | 24 | 0.1667 | 53 | 0.1509 | 37 | 0.0811 | 48 | 0.2292 | 162 | 0.1605 |
| 8_1276616( | 33 | 0.4545 | 37 | 0.3784 | 28 | 0.5 | 47 | 0.2553 | 145 | 0.3793 | 27 | 0.5185 | 45 | 0.4889 | 30 | 0.5667 | 42 | 0.6429 | 144 | 0.5556 |
| 8_1276740( | 3 | 0.6667 | 2 | 0.5 | NA | NA | 4 | 1 | 9 | 0.7778 | 1 | 1 | NA | NA | 2 | 1 | NA | NA | 3 | 1 |
| 8_1276759( | 3 | 0.3333 | 3 | 0.3333 | 1 | 0 | 2 | 0.5 | 9 | 0.3333 | 3 | 0 | 3 | 0.3333 | 2 | 0 | 1 | 0 | 9 | 0.1111 |
| 8_1276760: | 3 | 1 | 2 | 0 | 1 | 1 | 3 | 0.3333 | 9 | 0.5556 | 3 | 0.6667 | 3 | 0.3333 | 2 | 0.5 | 1 | 1 | 9 | 0.5556 |
| 8_1276761 | 2 | 1 | 2 | 0.5 | 1 | 1 | 2 | 1 | 7 | 0.8571 | 2 | 1 | 3 | 1 | 1 | 1 | 1 | 1 | 7 | 1 |
| 8_1276762( | 3 | 0 | 3 | 0 | 1 | 0 | 2 | 0.5 | 9 | 0.1111 | 2 | 1 | 3 | 0 | 3 | 0 | NA | NA | 8 | 0.25 |
| 8_1281272! | NA | NA | NA | NA | 1 | 0 | 1 | 1 | 2 | 0.5 | NA | NA | 2 | 0.5 | NA | NA | 1 | 0 | 3 | 0.3333 |

| | | | | | | | | | | | | | | | | | | | | |
|---|---|---|---|---|---|---|---|---|---|---|---|---|---|---|---|---|---|---|---|---|
| 8_1283372 | NA | NA | NA | NA | 1 | 1 | 1 | 1 | 2 | 1 | 1 | 1 | 2 | 1 | 2 | 1 | 2 | 1 | 7 | 1 |
| 8_1283372 | NA | NA | 1 | 1 | 1 | 1 | 1 | 1 | 3 | 1 | 1 | 1 | 6 | 1 | 2 | 0.5 | 2 | 1 | 11 | 0.9091 |
| 8_1283373 | NA | NA | 1 | 0 | 1 | 1 | 1 | 1 | 3 | 0.6667 | 1 | 1 | 5 | 0.6 | 3 | 1 | 4 | 1 | 13 | 0.8462 |
| 8_1283373 | NA | NA | 1 | 1 | NA | NA | NA | NA | 1 | 1 | 1 | 1 | 4 | 0.75 | 3 | 0 | 1 | 1 | 9 | 0.5556 |
| 8_1283374 | NA | NA | 1 | 0 | 1 | 1 | 1 | 1 | 3 | 0.6667 | 1 | 1 | 5 | 0.4 | 3 | 0.3333 | 4 | 0.75 | 13 | 0.5385 |
| 8_1283375 | NA | NA | 1 | 1 | 1 | 1 | 1 | 1 | 3 | 1 | 1 | 1 | 6 | 1 | 3 | 1 | 3 | 1 | 13 | 1 |
| 8_1283376 | NA | NA | 1 | 1 | 1 | 1 | 2 | 1 | 4 | 1 | 1 | 0 | 7 | 0.7143 | 1 | 1 | 3 | 1 | 12 | 0.75 |
| 8_1283376 | NA | NA | NA | NA | 1 | 1 | 2 | 1 | 3 | 1 | 1 | 1 | 4 | 0.75 | 1 | 1 | 2 | 1 | 8 | 0.875 |
| 8_1283377 | NA | NA | 1 | 0 | 1 | 1 | 2 | 1 | 4 | 0.75 | 1 | 1 | 7 | 0.1429 | 1 | 0 | 2 | 1 | 11 | 0.3636 |
| 8_1283379 | NA | NA | 1 | 0 | 1 | 0 | 2 | 0.5 | 4 | 0.25 | 1 | 1 | 7 | 0.2857 | 3 | 0 | 3 | 0.6667 | 14 | 0.3571 |
| 8_1283384 | NA | NA | 2 | 0.5 | 1 | 0 | 2 | 0 | 5 | 0.2 | 1 | 0 | 6 | 0 | 2 | 0 | 3 | 0.3333 | 12 | 0.0833 |
| 8_1283387 | NA | NA | 2 | 0 | 2 | 0.5 | 2 | 0 | 6 | 0.1667 | 1 | 0 | 7 | 0 | 5 | 0.2 | 3 | 0 | 16 | 0.0625 |
| 8_1292291 | NA | NA | 1 | 0 | 1 | 1 | NA | NA | 2 | 0.5 | 2 | 0 | 2 | 1 | 3 | 0 | 1 | 0 | 8 | 0.25 |
| 8_1292292 | NA | NA | 1 | 1 | 1 | 0 | NA | NA | 2 | 0.5 | 2 | 1 | 2 | 1 | 2 | 0.5 | 1 | 1 | 7 | 0.8571 |
| 8_1292292 | NA | NA | 1 | 1 | 1 | 1 | NA | NA | 2 | 1 | 2 | 1 | 1 | 1 | 3 | 0.6667 | 1 | 1 | 7 | 0.8571 |
| 8_1292292 | NA | NA | NA | NA | 1 | 1 | NA | NA | 1 | 1 | 2 | 1 | NA | NA | 1 | 1 | 1 | 1 | 4 | 1 |
| 8_1292293 | NA | NA | 1 | 1 | NA | NA | NA | NA | 1 | 1 | 2 | 1 | 2 | 1 | 3 | 1 | 1 | 1 | 8 | 1 |
| 8_1292293 | NA | NA | 1 | 1 | NA | NA | NA | NA | 1 | 1 | 2 | 1 | 2 | 1 | 3 | 1 | 1 | 1 | 8 | 1 |
| 8_1292295 | NA | NA | 1 | 1 | 1 | 0 | NA | NA | 2 | 0.5 | 2 | 1 | 1 | 1 | 2 | 0.5 | 1 | 1 | 6 | 0.8333 |
| 8_1292295 | NA | NA | NA | NA | 1 | 1 | NA | NA | 1 | 1 | 2 | 1 | 1 | 0 | 4 | 0.75 | 1 | 1 | 8 | 0.75 |
| 8_1292296 | NA | NA | 1 | 0 | 1 | 1 | NA | NA | 2 | 0.5 | 2 | 0.5 | 2 | 0 | 4 | 0.25 | 1 | 1 | 9 | 0.3333 |
| 8_1292297 | NA | NA | 1 | 1 | 1 | 1 | NA | NA | 2 | 1 | 1 | 1 | 1 | 1 | 3 | 1 | 1 | 1 | 6 | 1 |
| 8_1292297 | NA | NA | 1 | 0 | 1 | 1 | NA | NA | 2 | 0.5 | 2 | 0 | 1 | 1 | 3 | 0.3333 | 1 | 0 | 7 | 0.2857 |
| 8_1292299 | NA | NA | 1 | 1 | 1 | 0 | NA | NA | 2 | 0.5 | 2 | 1 | 2 | 1 | 4 | 0.75 | 1 | 1 | 9 | 0.8889 |
| 8_1292300 | NA | NA | 1 | 1 | NA | NA | NA | NA | 1 | 1 | 2 | 0 | 3 | 0 | 4 | 0.5 | 1 | 0 | 10 | 0.2 |
| 8_1292304 | 1 | 1 | 1 | 0 | 1 | 0 | NA | NA | 3 | 0.3333 | 3 | 0.3333 | 3 | 0.3333 | 3 | 0.3333 | 1 | 0 | 10 | 0.3 |
| 8_1292305 | NA | NA | 1 | 0 | 1 | 1 | NA | NA | 2 | 0.5 | 3 | 0 | 3 | 0.3333 | 4 | 0 | 1 | 0 | 11 | 0.0909 |
| 8_1292310 | 1 | 1 | 1 | 0 | 1 | 0 | NA | NA | 3 | 0.3333 | 3 | 0 | 3 | 0.3333 | 4 | 0 | 1 | 0 | 11 | 0.0909 |
| 8_1292316 | 2 | 0.5 | 1 | 0 | 1 | 0 | 1 | 0 | 5 | 0.2 | 3 | 0.6667 | 2 | 0 | 4 | 0.25 | 2 | 0 | 11 | 0.2727 |
| 8_1294493 | 5 | 0.2 | 6 | 0.3333 | 2 | 0 | 2 | 0.5 | 15 | 0.2667 | 1 | 0 | 4 | 0.25 | 5 | 0 | 4 | 0.25 | 14 | 0.1429 |
| 8_1294508 | 5 | 0.2 | 5 | 0 | 2 | 0 | 3 | 0 | 15 | 0.0667 | 1 | 0 | 2 | 0 | 6 | 0.1667 | 3 | 0 | 12 | 0.0833 |
| 8_1294587 | 8 | 0.25 | 8 | 0.125 | 2 | 0.5 | 2 | 0 | 20 | 0.2 | 3 | 0.6667 | 5 | 0.4 | 6 | 0.6667 | 5 | 0.8 | 19 | 0.6316 |
| 8_1294588 | 8 | 0.125 | 8 | 0.125 | 2 | 0.5 | 2 | 0 | 20 | 0.15 | 2 | 0.5 | 5 | 0 | 7 | 0.5714 | 2 | 0.5 | 16 | 0.375 |
| 8_1294589 | 7 | 0.1429 | 9 | 0 | 1 | 0 | 2 | 0 | 19 | 0.0526 | 2 | 0 | 5 | 0.2 | 7 | 0 | 4 | 0.25 | 18 | 0.1111 |
| 8_1299435 | 3 | 0.3333 | 3 | 0.3333 | 6 | 0 | 3 | 0.3333 | 15 | 0.2 | NA | NA | 4 | 0 | 2 | 0 | 1 | 1 | 7 | 0.1429 |
| 8_1299437 | 2 | 0 | 3 | 0 | 6 | 0.3333 | 3 | 0.3333 | 14 | 0.2143 | NA | NA | 5 | 0.4 | 2 | 0 | 3 | 0 | 10 | 0.2 |
| 8_1299438 | 1 | 1 | 3 | 1 | 6 | 1 | 3 | 1 | 13 | 1 | NA | NA | 6 | 1 | 2 | 1 | 3 | 1 | 11 | 1 |
| 8_1299438 | 1 | 1 | 2 | 1 | 4 | 0.25 | 3 | 0.3333 | 10 | 0.5 | NA | NA | 4 | 0.5 | NA | NA | 2 | 0.5 | 6 | 0.5 |
| 8_1299438 | 1 | 1 | 2 | 0.5 | 6 | 0.6667 | 3 | 0.3333 | 12 | 0.5833 | NA | NA | 5 | 0.4 | 2 | 0.5 | 3 | 0.6667 | 10 | 0.5 |
| 8_1299439 | 1 | 0 | 3 | 0 | 5 | 0.2 | 3 | 0 | 12 | 0.0833 | NA | NA | 5 | 0 | 2 | 0 | 3 | 0.3333 | 10 | 0.1 |
| 8_1299441 | 1 | 0 | 3 | 0.6667 | 5 | 0.2 | 3 | 0.3333 | 12 | 0.3333 | NA | NA | 5 | 0.4 | 2 | 0.5 | 3 | 0.3333 | 10 | 0.4 |
| 8_1299441 | 1 | 1 | 2 | 1 | 5 | 1 | 3 | 1 | 11 | 1 | NA | NA | 6 | 0.6667 | 2 | 0.5 | 3 | 1 | 11 | 0.7273 |
| 8_1299441 | 1 | 1 | 4 | 0.25 | 6 | 0.8333 | 2 | 0.5 | 13 | 0.6154 | NA | NA | 6 | 0.5 | 2 | 0.5 | 3 | 0.6667 | 11 | 0.5455 |
| 8_1299441 | 1 | 1 | 3 | 0.6667 | 5 | 0.8 | 2 | 1 | 11 | 0.8182 | NA | NA | 5 | 1 | 2 | 1 | 3 | 0.6667 | 10 | 0.9 |
| 8_1299441 | 1 | 1 | 4 | 0.75 | 4 | 0.5 | 2 | 1 | 11 | 0.7273 | NA | NA | 5 | 1 | 2 | 1 | 3 | 0.3333 | 10 | 0.8 |
| 8_1299444 | NA | NA | 2 | 0.5 | 3 | 0 | 1 | 1 | 6 | 0.3333 | NA | NA | 4 | 0.5 | 1 | 1 | 2 | 1 | 7 | 0.7143 |
| 8_1299445 | NA | NA | 2 | 0.5 | 3 | 1 | 1 | 1 | 6 | 0.8333 | NA | NA | 3 | 1 | 2 | 1 | 2 | 0.5 | 7 | 0.8571 |
| 8_1299445 | NA | NA | 2 | 1 | 3 | 0.3333 | 1 | 1 | 6 | 0.6667 | NA | NA | 3 | 1 | 1 | 1 | 2 | 0.5 | 6 | 0.8333 |
| 8_1299448 | NA | NA | 1 | 0 | 3 | 0.3333 | 1 | 0 | 5 | 0.2 | NA | NA | 4 | 0.25 | 1 | 0 | 2 | 0 | 7 | 0.1429 |
| 8_1299449 | NA | NA | 2 | 0 | 3 | 0.3333 | 1 | 0 | 6 | 0.1667 | NA | NA | 3 | 0.3333 | 1 | 0 | 2 | 0 | 6 | 0.1667 |
| 8_1299451 | NA | NA | 2 | 0.5 | 3 | 0.3333 | 1 | 1 | 6 | 0.5 | NA | NA | 3 | 0.6667 | NA | NA | 2 | 0.5 | 5 | 0.6 |
| 8_1299455 | NA | NA | 1 | 1 | 3 | 0.6667 | 1 | 1 | 5 | 0.8 | NA | NA | 3 | 0.3333 | 1 | 1 | 2 | 0 | 6 | 0.3333 |
| 8_1299455 | NA | NA | 1 | 1 | 4 | 0.5 | 1 | 1 | 6 | 0.6667 | NA | NA | 4 | 1 | 1 | 1 | 2 | 1 | 7 | 1 |
| 8_1299459 | NA | NA | 1 | 1 | 3 | 0.6667 | 1 | 1 | 5 | 0.8 | NA | NA | 4 | 0.75 | NA | NA | 2 | 0.5 | 6 | 0.6667 |
| 8_1299980 | NA | NA | NA | NA | NA | NA | 1 | 1 | 1 | 1 | NA | NA | 1 | 1 | NA | NA | NA | NA | 1 | 1 |
| 8_1300114 | 1 | 1 | NA | NA | NA | NA | NA | NA | 1 | 1 | NA | NA | NA | NA | NA | NA | 1 | 1 | 1 | 1 |
| 8_1300115 | 2 | 1 | NA | NA | NA | NA | NA | NA | 2 | 1 | 1 | 0 | NA | NA | NA | NA | 2 | 1 | 3 | 0.6667 |
| 9_3389763 | 1 | 0 | 3 | 0.3333 | NA | NA | 6 | 0 | 10 | 0.1 | NA | NA | 1 | 0 | 2 | 0 | 5 | 0.2 | 8 | 0.125 |
| 9_3389858 | NA | NA | NA | NA | 1 | 1 | 3 | 0.6667 | 4 | 0.75 | NA | NA | NA | NA | 2 | 0 | 3 | 0.6667 | 5 | 0.4 |
| 9_3389885 | NA | NA | 3 | 0 | NA | NA | 6 | 0.5 | 9 | 0.3333 | NA | NA | 1 | 0 | 4 | 0.5 | 2 | 0.5 | 7 | 0.4286 |
| 9_3390711 | NA | NA | 2 | 0.5 | NA | NA | 5 | 0 | 7 | 0.1429 | NA | NA | 1 | 0 | 2 | 1 | 2 | 0.5 | 5 | 0.6 |
| 9_3391102 | NA | NA | 3 | 0 | NA | NA | 5 | 0.2 | 8 | 0.125 | NA | NA | 2 | 0.5 | 2 | 0.5 | 2 | 1 | 6 | 0.6667 |
| 9_3430499 | 1 | 0 | NA | NA | 2 | 0.5 | NA | NA | 3 | 0.3333 | 1 | 0 | 2 | 0 | 2 | 0 | 2 | 0.5 | 7 | 0.1429 |
| 9_3430681 | 1 | 0 | NA | NA | 2 | 0.5 | NA | NA | 3 | 0.3333 | 1 | 1 | 1 | 1 | 2 | 1 | 2 | 1 | 6 | 1 |
| 9_3430859 | 1 | 1 | NA | NA | 1 | 1 | NA | NA | 2 | 1 | 1 | 1 | 1 | 0 | 1 | 1 | 3 | 1 | 6 | 0.8333 |

| | | | | | | | | | | | | | | | | | | | | |
|---|---|---|---|---|---|---|---|---|---|---|---|---|---|---|---|---|---|---|---|---|
| 9_3430880 | 1 | 1 | NA | NA | 1 | 0 | NA | NA | 2 | 0.5 | 1 | 1 | 2 | 0 | 1 | 0 | 3 | 0.3333 | 7 | 0.2857 |
| 9_3478240 | 3 | 0 | 1 | 1 | 2 | 0.5 | 1 | 1 | 7 | 0.4286 | 1 | 1 | 3 | 0.3333 | 1 | 0 | 5 | 0.4 | 10 | 0.4 |
| 9_3634544 | NA | NA | 1 | 1 | NA | NA | NA | NA | 1 | 1 | NA | NA | NA | NA | 2 | 1 | NA | NA | 2 | 1 |
| 9_3634585 | NA | NA | 1 | 1 | NA | NA | NA | NA | 1 | 1 | NA | NA | NA | NA | 2 | 0.5 | NA | NA | 2 | 0.5 |
| 9_3634776 | NA | NA | 1 | 1 | NA | NA | NA | NA | 1 | 1 | NA | NA | NA | NA | 2 | 1 | NA | NA | 2 | 1 |
| 9_3759485 | NA | NA | 1 | 1 | NA | NA | NA | NA | 1 | 1 | NA | NA | NA | NA | 2 | 1 | 1 | 1 | 3 | 1 |
| 9_3759500 | NA | NA | 1 | 1 | NA | NA | NA | NA | 1 | 1 | NA | NA | NA | NA | 2 | 0.5 | 1 | 1 | 3 | 0.6667 |
| 9_3759624 | NA | NA | 1 | 1 | NA | NA | NA | NA | 1 | 1 | NA | NA | NA | NA | 1 | 1 | 1 | 1 | 2 | 1 |
| 9_3759766 | NA | NA | 1 | 1 | NA | NA | NA | NA | 1 | 1 | NA | NA | NA | NA | 2 | 0.5 | 1 | 0 | 3 | 0.3333 |
| 9_3759833 | NA | NA | 1 | 1 | NA | NA | NA | NA | 1 | 1 | NA | NA | NA | NA | 2 | 1 | 1 | 1 | 3 | 1 |
| 9_3895991 | NA | NA | 1 | 0 | 1 | 1 | NA | NA | 2 | 0.5 | NA | NA | NA | NA | 1 | 0 | 2 | 0.5 | 3 | 0.3333 |
| 9_3897853 | 3 | 0 | 1 | 0 | 1 | 1 | 2 | 0 | 7 | 0.1429 | NA | NA | NA | NA | 1 | 0 | 2 | 0.5 | 3 | 0.3333 |
| 9_3905167 | 9 | 0 | 12 | 0 | 6 | 0.3333 | 13 | 0.2308 | 40 | 0.125 | 2 | 0 | 5 | 0.2 | 6 | 0.1667 | 4 | 0 | 17 | 0.1176 |
| 9_4286577 | 2 | 0.5 | 1 | 1 | NA | NA | 1 | 1 | 4 | 0.75 | 1 | 1 | NA | NA | 2 | 1 | 2 | 0.5 | 5 | 0.8 |
| 9_4288129 | 1 | 0 | 1 | 0 | NA | NA | 1 | 1 | 3 | 0.3333 | 1 | 0 | 1 | 0 | 2 | 0.5 | 1 | 0 | 5 | 0.2 |
| 9_4296334 | 3 | 0.3333 | 2 | 0 | 6 | 0 | 6 | 0.3333 | 17 | 0.1765 | 4 | 0 | 5 | 0 | 2 | 0.5 | 7 | 0 | 18 | 0.0556 |
| 9_4296350 | 3 | 0.3333 | 3 | 0.3333 | 6 | 0.1667 | 5 | 0.6 | 17 | 0.3529 | 5 | 0.6 | 5 | 0.2 | 1 | 0 | 9 | 0.6667 | 20 | 0.5 |
| 9_4296379 | 2 | 0 | 3 | 0.3333 | 5 | 0.2 | 4 | 0.25 | 14 | 0.2143 | 5 | 0.2 | 5 | 0.2 | 2 | 0 | 9 | 0.3333 | 21 | 0.2381 |
| 9_4296487 | 3 | 0.3333 | 3 | 0.6667 | 4 | 0.25 | 6 | 0.5 | 16 | 0.4375 | 4 | 0.5 | 4 | 0 | 1 | 0 | 8 | 0.375 | 17 | 0.2941 |
| 9_4296568 | 3 | 0.3333 | 3 | 0 | 6 | 0 | 6 | 0.3333 | 18 | 0.1667 | 5 | 0.2 | 5 | 0 | 2 | 0 | 8 | 0.125 | 20 | 0.1 |
| 9_4296727 | 2 | 1 | 2 | 0.5 | 3 | 1 | 8 | 0.625 | 15 | 0.7333 | 5 | 1 | 5 | 1 | 3 | 0.6667 | 9 | 1 | 22 | 0.9545 |
| 9_4302540 | 1 | 0 | NA | NA | 1 | 1 | 6 | 0.1667 | 8 | 0.25 | 1 | 1 | 3 | 0 | 2 | 0 | 3 | 0.3333 | 9 | 0.2222 |
| 9_4308762 | 1 | 0 | 2 | 0 | 4 | 0.25 | 8 | 0.125 | 15 | 0.1333 | 1 | 0 | 5 | 0.4 | 3 | 0 | 6 | 0.1667 | 15 | 0.2 |
| 9_4308813 | 3 | 0.3333 | 2 | 0 | 4 | 0.25 | 8 | 0.375 | 17 | 0.2941 | 3 | 0 | 5 | 0.4 | 3 | 0.3333 | 8 | 0.5 | 19 | 0.3684 |
| 9_4309210 | 1 | 1 | 1 | 0 | 1 | 1 | 2 | 0.5 | 5 | 0.6 | NA | NA | 1 | 1 | NA | NA | NA | NA | 1 | 1 |
| 9_4316943 | 3 | 0.6667 | 1 | 1 | NA | NA | 1 | 1 | 5 | 0.8 | 1 | 1 | 3 | 0 | 2 | 0 | 1 | 1 | 7 | 0.2857 |
| 9_4330043 | 1 | 1 | 2 | 1 | 1 | 0 | 1 | 0 | 5 | 0.6 | 1 | 1 | 3 | 0.6667 | 3 | 0 | 2 | 1 | 9 | 0.5556 |
| 9_4330181 | 1 | 1 | 1 | 1 | NA | NA | NA | NA | 2 | 1 | NA | NA | NA | NA | NA | NA | 1 | 1 | 1 | 1 |
| 9_4330373 | 1 | 1 | 1 | 1 | NA | NA | NA | NA | 2 | 1 | NA | NA | NA | NA | NA | NA | 1 | 1 | 1 | 1 |
| 9_4330412 | 1 | 1 | NA | NA | NA | NA | NA | NA | 1 | 1 | NA | NA | NA | NA | NA | NA | 1 | 1 | 1 | 1 |
| 9_4330571 | NA | NA | 1 | 1 | NA | NA | NA | NA | 1 | 1 | NA | NA | NA | NA | NA | NA | 1 | 1 | 1 | 1 |
| 9_4383587 | 4 | 0 | 7 | 0.2857 | 9 | 0.3333 | 8 | 0.625 | 28 | 0.3571 | 3 | 0.3333 | 7 | 0.1429 | 2 | 0.5 | 3 | 0 | 15 | 0.2 |
| 9_4383729 | 5 | 0.6 | 7 | 0.4286 | 9 | 0.3333 | 9 | 0.6667 | 30 | 0.5 | 4 | 0.25 | 6 | 0.5 | 3 | 0.6667 | 3 | 1 | 16 | 0.5625 |
| 9_4383735 | 6 | 0 | 7 | 0.2857 | 9 | 0.1111 | 9 | 0.2222 | 31 | 0.1613 | 4 | 0 | 7 | 0.2857 | 3 | 0 | 3 | 0 | 17 | 0.1176 |
| 9_4383873 | 4 | 0.25 | 7 | 0.2857 | 9 | 0.3333 | 8 | 0.125 | 28 | 0.25 | 4 | 0.75 | 5 | 0.4 | 3 | 0 | 2 | 0 | 14 | 0.3571 |
| 9_4383904 | 5 | 0.8 | 6 | 1 | 8 | 0.75 | 6 | 0.8333 | 25 | 0.84 | 3 | 1 | 7 | 0.7143 | 3 | 1 | 1 | 1 | 14 | 0.8571 |
| 9_4383994 | 3 | 0.3333 | 7 | 0.5714 | 8 | 0.125 | 9 | 0.4444 | 27 | 0.3704 | 3 | 0.3333 | 7 | 0.2857 | 2 | 0.5 | 2 | 0.5 | 14 | 0.3571 |
| 9_4384893 | 5 | 0.4 | 5 | 0.2 | 6 | 0 | 9 | 0.4444 | 25 | 0.28 | 4 | 0.25 | 5 | 0.4 | 3 | 0.3333 | 3 | 0.3333 | 15 | 0.3333 |
| 9_4385007 | 6 | 0.1667 | 9 | 0.1111 | 9 | 0.2222 | 10 | 0.5 | 34 | 0.2647 | 4 | 0.25 | 6 | 0.5 | 3 | 0.6667 | 3 | 0.3333 | 16 | 0.4375 |
| 9_4385151 | 6 | 1 | 10 | 0.7 | 10 | 0.5 | 8 | 0.625 | 34 | 0.6765 | 4 | 0.5 | 8 | 0.75 | 2 | 0.5 | 2 | 0 | 16 | 0.5625 |
| 9_4385173 | 7 | 0 | 11 | 0.0909 | 10 | 0.1 | 9 | 0.1111 | 37 | 0.0811 | 4 | 0 | 8 | 0 | 3 | 0.3333 | 3 | 0 | 18 | 0.0556 |
| 9_4385281 | 7 | 0.8571 | 11 | 1 | 10 | 1 | 8 | 1 | 36 | 0.9722 | 4 | 0.75 | 7 | 1 | 2 | 1 | 3 | 1 | 16 | 0.9375 |
| 9_4385314 | 5 | 1 | 8 | 1 | 9 | 1 | 6 | 1 | 28 | 1 | 4 | 1 | 7 | 1 | 2 | 1 | 2 | 1 | 15 | 1 |
| 9_4385656 | 9 | 0.3333 | 12 | 0.5 | 11 | 0.6364 | 10 | 0.4 | 42 | 0.4762 | 3 | 0.6667 | 8 | 0.5 | 3 | 0.3333 | 4 | 0.75 | 18 | 0.5556 |
| 9_4385665 | 7 | 1 | 10 | 1 | 9 | 1 | 9 | 1 | 35 | 1 | 4 | 1 | 8 | 1 | 3 | 1 | 4 | 1 | 19 | 1 |
| 9_4385807 | 8 | 0.375 | 11 | 0.0909 | 12 | 0.5 | 7 | 0.2857 | 38 | 0.3158 | 3 | 0.6667 | 8 | 0.375 | 2 | 0 | 4 | 0 | 17 | 0.2941 |
| 9_4386006 | 8 | 0.25 | 10 | 0.2 | 11 | 0.7273 | 11 | 0.3636 | 40 | 0.4 | 5 | 0 | 8 | 0.25 | 3 | 0.6667 | 5 | 0.4 | 21 | 0.2857 |
| 9_4386298 | 7 | 0.2857 | 11 | 0.3636 | 12 | 0.3333 | 12 | 0.0833 | 42 | 0.2619 | 5 | 0.2 | 7 | 0.2857 | 3 | 0.3333 | 6 | 0.6667 | 21 | 0.381 |
| 9_4418188 | 5 | 0.4 | 3 | 0 | 6 | 0.6667 | 3 | 0 | 17 | 0.3529 | 2 | 1 | 2 | 0 | 3 | 0 | 3 | 0.3333 | 10 | 0.3 |
| 9_4418257 | 5 | 0.2 | 4 | 0.25 | 5 | 0.6 | 3 | 0 | 17 | 0.2941 | 2 | 0 | 2 | 0 | 3 | 0.3333 | 3 | 0.3333 | 10 | 0.2 |
| 9_4419231 | 4 | 0 | 3 | 0 | 5 | 0.4 | 1 | 0 | 13 | 0.1538 | 2 | 0 | 2 | 0 | 3 | 0.3333 | 2 | 0 | 9 | 0.1111 |
| 9_4419312 | 4 | 0.5 | 2 | 0 | 5 | 0.4 | 1 | 0 | 12 | 0.3333 | 2 | 0.5 | 2 | 0.5 | 3 | 1 | 2 | 0.5 | 9 | 0.6667 |
| 9_4419474 | 3 | 0 | 3 | 0.6667 | 5 | 0.6 | 1 | 1 | 12 | 0.5 | 2 | 0.5 | 2 | 0 | 3 | 0 | 2 | 0.5 | 9 | 0.2222 |
| 9_4419841 | 3 | 0 | 2 | 0.5 | 5 | 0.4 | 1 | 0 | 11 | 0.2727 | 2 | 0 | 2 | 0 | 2 | 0.5 | 2 | 0 | 8 | 0.125 |
| 9_4419862 | 2 | 0.5 | 2 | 1 | 5 | 1 | 1 | 1 | 10 | 0.9 | 2 | 1 | 2 | 1 | 3 | 0.3333 | 2 | 0.5 | 9 | 0.6667 |
| 9_4419978 | 2 | 0.5 | NA | NA | 4 | 0 | 1 | 0 | 7 | 0.1429 | 2 | 0 | 1 | 0 | 3 | 0.6667 | 2 | 0.5 | 8 | 0.375 |
| 9_4420014 | 2 | 0.5 | 1 | 1 | 4 | 1 | 1 | 1 | 8 | 0.875 | 2 | 1 | 1 | 0 | 3 | 0.3333 | 2 | 0 | 8 | 0.375 |
| 9_4420030 | 1 | 0 | 2 | 0 | 4 | 0.5 | 1 | 0 | 8 | 0.25 | 2 | 1 | 2 | 0 | 2 | 0.5 | 2 | 0 | 8 | 0.375 |
| 9_4420044 | 2 | 0.5 | 1 | 1 | 3 | 0.3333 | 1 | 0 | 7 | 0.4286 | 1 | 0 | 2 | 1 | 2 | 0.5 | 1 | 1 | 6 | 0.6667 |
| 9_4420082 | 2 | 0 | 2 | 0.5 | 2 | 1 | NA | NA | 6 | 0.5 | 2 | 0 | 1 | 0 | 1 | 0 | 2 | 0.5 | 6 | 0.1667 |
| 9_4420107 | 1 | 1 | 2 | 1 | 3 | 1 | 1 | 1 | 7 | 1 | 2 | 1 | 2 | 1 | 2 | 1 | 2 | 1 | 8 | 1 |
| 9_4420119 | 2 | 0.5 | 2 | 0.5 | 4 | 1 | 1 | 1 | 9 | 0.7778 | 2 | 1 | 2 | 0.5 | 2 | 0.5 | 2 | 1 | 8 | 0.75 |
| 9_4420162 | 1 | 0 | 2 | 1 | 4 | 0 | 1 | 1 | 8 | 0.375 | 2 | 0.5 | 2 | 0.5 | 2 | 0 | 2 | 1 | 8 | 0.5 |
| 9_4420205 | 2 | 0.5 | 2 | 0.5 | 4 | 1 | NA | NA | 8 | 0.75 | 2 | 0.5 | 2 | 0.5 | 2 | 1 | 1 | 1 | 7 | 0.7143 |
| 9_4432837 | NA | NA | 1 | 1 | 1 | 0 | 1 | 0 | 3 | 0.3333 | 1 | 0 | NA | NA | NA | NA | 1 | 1 | 2 | 0.5 |

| | | | | | | | | | | | | | | | | | | | | |
|---|---|---|---|---|---|---|---|---|---|---|---|---|---|---|---|---|---|---|---|---|
| 9_4477706 | 1 | 1 | 6 | 0.3333 | 2 | 0.5 | 3 | 0.6667 | 12 | 0.5 | 4 | 0.5 | 4 | 0.5 | 5 | 0.6 | 7 | 0.4286 | 20 | 0.5 |
| 9_4478594 | 2 | 0 | 5 | 0 | 2 | 0.5 | 2 | 0 | 11 | 0.0909 | 1 | 0 | 2 | 0.5 | 2 | 0.5 | 6 | 0.1667 | 11 | 0.2727 |
| 9_4478655 | 2 | 0.5 | 5 | 0.6 | 2 | 1 | 2 | 1 | 11 | 0.7273 | 3 | 0.3333 | 3 | 1 | 3 | 0.3333 | 7 | 0.7143 | 16 | 0.625 |
| 9_4478696 | 2 | 1 | 4 | 0.75 | 2 | 1 | 2 | 1 | 10 | 0.9 | 3 | 1 | 3 | 1 | 4 | 0.75 | 7 | 0.8571 | 17 | 0.8824 |
| 9_4664967 | 1 | 0 | 1 | 0 | 1 | 0 | 2 | 0.5 | 5 | 0.2 | 1 | 0 | 3 | 0 | 3 | 1 | 3 | 0.6667 | 10 | 0.5 |
| 9_4796138 | 2 | 0.5 | 1 | 0 | NA | NA | NA | NA | 3 | 0.3333 | 1 | 0 | 1 | 1 | 1 | 0 | 3 | 0 | 6 | 0.1667 |
| 9_6377056 | NA | NA | 1 | 0 | NA | NA | 2 | 1 | 3 | 0.6667 | NA | NA | NA | NA | NA | NA | 1 | 1 | 1 | 1 |
| 9_6929511 | 3 | 0 | NA | NA | NA | NA | 4 | 0.25 | 7 | 0.1429 | 1 | 0 | 2 | 0 | 4 | 0 | 1 | 1 | 8 | 0.125 |
| 9_7183479 | NA | NA | 2 | 0.5 | NA | NA | 5 | 0 | 7 | 0.1429 | 3 | 0 | NA | NA | 1 | 0 | 1 | 1 | 5 | 0.2 |
| 9_7183491 | NA | NA | 1 | 0 | NA | NA | 5 | 0.6 | 6 | 0.5 | 2 | 0.5 | NA | NA | 1 | 0 | 1 | 1 | 4 | 0.5 |
| 9_7183505 | NA | NA | 2 | 0 | NA | NA | 4 | 0.25 | 6 | 0.1667 | 3 | 0.3333 | NA | NA | NA | NA | 1 | 1 | 4 | 0.5 |
| 9_7183661 | NA | NA | 2 | 0.5 | NA | NA | 4 | 0.5 | 6 | 0.5 | 1 | 0 | NA | NA | 1 | 0 | 1 | 1 | 3 | 0.3333 |
| 9_7183806 | NA | NA | 2 | 1 | NA | NA | 4 | 0.75 | 6 | 0.8333 | 3 | 1 | NA | NA | 1 | 1 | 1 | 1 | 5 | 1 |
| 9_7183902 | NA | NA | 2 | 0.5 | NA | NA | 5 | 0.2 | 7 | 0.2857 | 3 | 0.3333 | NA | NA | 1 | 0 | 1 | 1 | 5 | 0.4 |
| 9_7183915 | NA | NA | 2 | 0.5 | NA | NA | 5 | 0.2 | 7 | 0.2857 | 3 | 0.3333 | NA | NA | 1 | 0 | 1 | 0 | 5 | 0.2 |
| 9_7183950 | NA | NA | 1 | 1 | NA | NA | 5 | 0.8 | 6 | 0.8333 | 1 | 1 | NA | NA | 1 | 1 | NA | NA | 2 | 1 |
| 9_7184071 | NA | NA | 1 | 0 | NA | NA | 5 | 0.2 | 6 | 0.1667 | 3 | 0.3333 | NA | NA | 1 | 1 | 1 | 0 | 5 | 0.4 |
| 9_7184098 | NA | NA | 2 | 1 | NA | NA | 4 | 0.75 | 6 | 0.8333 | 1 | 1 | NA | NA | 1 | 1 | 1 | 1 | 3 | 1 |
| 9_7184125 | NA | NA | 2 | 1 | NA | NA | 3 | 1 | 5 | 1 | 3 | 1 | NA | NA | 1 | 1 | 1 | 1 | 5 | 1 |
| 9_7184137 | NA | NA | 2 | 1 | NA | NA | 5 | 1 | 7 | 1 | 3 | 1 | NA | NA | 1 | 1 | 1 | 1 | 5 | 1 |
| 9_7184157 | NA | NA | 2 | 1 | NA | NA | 3 | 1 | 5 | 1 | 2 | 0.5 | NA | NA | 1 | 0 | NA | NA | 3 | 0.3333 |
| 9_7206781 | 5 | 0.2 | 7 | 0.1429 | 8 | 0 | 6 | 0.1667 | 26 | 0.1154 | 5 | 0 | 7 | 0.1429 | 5 | 0 | 2 | 0 | 19 | 0.0526 |
| 9_7206866 | 2 | 0.5 | 5 | 0.4 | 6 | 0.5 | 6 | 0.8333 | 19 | 0.5789 | 5 | 0.8 | 7 | 0.7143 | 6 | 0.8333 | 3 | 0.3333 | 21 | 0.7143 |
| 9_7206950 | 5 | 0.2 | 5 | 0.2 | 8 | 0.625 | 7 | 0.4286 | 25 | 0.4 | 8 | 0.75 | 6 | 0.1667 | 5 | 0.4 | 4 | 0.25 | 23 | 0.4348 |
| 9_7206961 | 5 | 0 | 7 | 0 | 9 | 0.1111 | 6 | 0.1667 | 27 | 0.0741 | 8 | 0.125 | 7 | 0 | 6 | 0.1667 | 4 | 0 | 25 | 0.08 |
| 9_7219555 | 1 | 1 | NA | NA | 4 | 0.25 | 2 | 0.5 | 7 | 0.4286 | 1 | 0 | 1 | 0 | 3 | 0.6667 | 2 | 0 | 7 | 0.2857 |
| 9_7752240 | 2 | 1 | NA | NA | 1 | 1 | 4 | 0.25 | 7 | 0.5714 | 2 | 1 | NA | NA | 3 | 0.6667 | 3 | 1 | 8 | 0.875 |
| 9_7793682 | NA | NA | 2 | 0.5 | 1 | 0 | NA | NA | 3 | 0.3333 | 4 | 0 | NA | NA | 2 | 0 | 2 | 0.5 | 8 | 0.125 |
| 9_7818509 | 6 | 0.1667 | 7 | 0.1429 | 3 | 0 | 5 | 0 | 21 | 0.0952 | 4 | 0 | 1 | 0 | 1 | 0 | 6 | 0.3333 | 12 | 0.1667 |
| 9_7818561 | 5 | 0.4 | 9 | 0.7778 | 3 | 0.3333 | 5 | 0.6 | 22 | 0.5909 | 4 | 0.5 | 1 | 0 | 1 | 1 | 5 | 0.6 | 11 | 0.5455 |
| 9_7818667 | 5 | 0.8 | 6 | 0.5 | 3 | 1 | 5 | 1 | 19 | 0.7895 | 3 | 1 | 1 | 1 | 1 | 1 | 5 | 1 | 10 | 1 |
| 9_7818693 | 4 | 0.5 | 5 | 0.4 | 3 | 0 | 3 | 0.3333 | 15 | 0.3333 | 3 | 0.3333 | 1 | 0 | 1 | 0 | 2 | 0 | 7 | 0.1429 |
| 9_7818722 | 5 | 0.4 | 6 | 0.6667 | 2 | 1 | 5 | 0.4 | 18 | 0.5556 | 2 | 0.5 | NA | NA | NA | NA | 4 | 0.25 | 6 | 0.3333 |
| 9_7818774 | 2 | 1 | 7 | 0.8571 | 3 | 0.6667 | 5 | 1 | 17 | 0.8824 | 2 | 1 | 1 | 1 | 1 | 1 | 4 | 0.75 | 8 | 0.875 |
| 9_7818786 | 4 | 0 | 7 | 0 | 3 | 0.3333 | 5 | 0 | 19 | 0.0526 | 2 | 0 | 1 | 0 | NA | NA | 4 | 0.25 | 7 | 0.1429 |
| 9_7818881 | 4 | 1 | 8 | 0.5 | 2 | 1 | 3 | 0.6667 | 17 | 0.7059 | 2 | 1 | 1 | 1 | 1 | 1 | 4 | 1 | 8 | 1 |
| 9_7833678 | 3 | 0.3333 | 2 | 1 | 2 | 0.5 | 2 | 0.5 | 9 | 0.5556 | 1 | 1 | 1 | 1 | 1 | 1 | 4 | 0.75 | 7 | 0.8571 |
| 9_7833775 | 3 | 0.3333 | 4 | 0.5 | 1 | 1 | 2 | 0.5 | 10 | 0.5 | 1 | 1 | 1 | 1 | 1 | 1 | 4 | 0.75 | 7 | 0.8571 |
| 9_7833823 | 3 | 0.3333 | 4 | 0.25 | 1 | 1 | 2 | 0 | 10 | 0.3 | 1 | 1 | 1 | 0 | 1 | 0 | 4 | 0.25 | 7 | 0.2857 |
| 9_7833995 | 3 | 0.3333 | 3 | 0.3333 | 1 | 0 | 2 | 0.5 | 9 | 0.3333 | 1 | 1 | 1 | 0 | 1 | 0 | 3 | 0.3333 | 6 | 0.3333 |
| 9_7834137 | 3 | 1 | 4 | 0.75 | 1 | 1 | 2 | 0.5 | 10 | 0.8 | 1 | 1 | NA | NA | 1 | 0 | 4 | 0.75 | 6 | 0.6667 |
| 9_7834168 | 3 | 0.3333 | 4 | 0.25 | 1 | 1 | 2 | 0.5 | 10 | 0.4 | 1 | 1 | NA | NA | 1 | 1 | 4 | 0 | 6 | 0.3333 |
| 9_7834190 | 3 | 0.6667 | 4 | 0.75 | NA | NA | 3 | 1 | 10 | 0.8 | 1 | 1 | NA | NA | 1 | 1 | 3 | 1 | 5 | 1 |
| 9_7834342 | 1 | 1 | 2 | 1 | NA | NA | 2 | 0.5 | 5 | 0.8 | NA | NA | NA | NA | 1 | 1 | 2 | 0.5 | 3 | 0.6667 |
| 9_7834355 | 2 | 0.5 | 3 | 0.3333 | 1 | 0 | 3 | 0 | 9 | 0.2222 | NA | NA | NA | NA | 1 | 0 | 4 | 0.25 | 5 | 0.2 |
| 9_7834456 | 3 | 0.6667 | 4 | 0.5 | 1 | 1 | 3 | 0.6667 | 11 | 0.6364 | 1 | 0 | NA | NA | 1 | 0 | 4 | 0.5 | 6 | 0.3333 |
| 9_7834475 | 1 | 1 | 2 | 1 | 1 | 1 | 2 | 0.5 | 6 | 0.8333 | NA | NA | NA | NA | 1 | 1 | 3 | 0.6667 | 4 | 0.75 |
| 9_7835091 | 3 | 0.6667 | 3 | 0.6667 | 1 | 0 | 2 | 0.5 | 9 | 0.5556 | 1 | 1 | NA | NA | 1 | 0 | 4 | 0.75 | 6 | 0.6667 |
| 9_7835309 | 2 | 0 | 3 | 0.3333 | 1 | 1 | 2 | 0.5 | 8 | 0.375 | 1 | 0 | NA | NA | 1 | 0 | 4 | 0.25 | 6 | 0.1667 |
| 9_7835317 | 2 | 1 | 2 | 0 | 1 | 1 | 2 | 1 | 7 | 0.7143 | 1 | 0 | NA | NA | 1 | 0 | 4 | 1 | 6 | 0.6667 |
| 9_7932181 | 4 | 0 | 5 | 0.2 | 2 | 0 | 2 | 0 | 13 | 0.0769 | 5 | 0.4 | 1 | 1 | 3 | 0 | 1 | 1 | 10 | 0.4 |
| 9_7932376 | 2 | 0.5 | 5 | 0.2 | 3 | 0 | 2 | 0 | 12 | 0.1667 | 6 | 0.1667 | 1 | 0 | 3 | 0.3333 | 1 | 1 | 11 | 0.2727 |
| 9_7934029 | 1 | 1 | 1 | 0 | 2 | 1 | NA | NA | 4 | 0.75 | 1 | 0 | NA | NA | 4 | 0.5 | 1 | 1 | 6 | 0.5 |
| 9_7934135 | NA | NA | 1 | 0 | 2 | 0.5 | NA | NA | 3 | 0.3333 | 2 | 1 | NA | NA | 2 | 1 | 1 | 1 | 5 | 1 |
| 9_7934150 | 1 | 1 | 1 | 1 | 1 | 1 | NA | NA | 3 | 1 | 2 | 1 | NA | NA | 4 | 1 | 1 | 1 | 7 | 1 |
| 9_7934157 | 1 | 1 | 1 | 1 | 1 | 1 | NA | NA | 3 | 1 | 2 | 0.5 | NA | NA | 3 | 1 | NA | NA | 5 | 0.8 |
| 9_7934165 | NA | NA | 1 | 1 | 2 | 0.5 | NA | NA | 3 | 0.6667 | 2 | 1 | NA | NA | 4 | 0 | NA | NA | 6 | 0.3333 |
| 9_7934271 | 1 | 0 | 1 | 0 | 1 | 1 | NA | NA | 3 | 0.3333 | 2 | 0.5 | NA | NA | 2 | 0.5 | 1 | 0 | 5 | 0.4 |
| 9_7934287 | 1 | 0 | 1 | 0 | 2 | 1 | NA | NA | 4 | 0.5 | 2 | 0.5 | NA | NA | 3 | 0.6667 | 1 | 0 | 6 | 0.5 |
| 9_7934300 | 1 | 1 | 1 | 0 | 2 | 0 | NA | NA | 4 | 0.25 | 2 | 0.5 | NA | NA | 4 | 0.5 | 1 | 1 | 7 | 0.5714 |
| 9_7934649 | 1 | 1 | 1 | 1 | 2 | 0.5 | NA | NA | 4 | 0.75 | 2 | 1 | NA | NA | 3 | 1 | 1 | 1 | 6 | 1 |
| 9_7934659 | 1 | 0 | 1 | 1 | 2 | 0 | NA | NA | 4 | 0.25 | 2 | 1 | NA | NA | 3 | 0 | 1 | 1 | 6 | 0.5 |
| 9_8100338 | 2 | 1 | NA | NA | 2 | 0 | 1 | 0 | 5 | 0.4 | 1 | 1 | NA | NA | 1 | 1 | NA | NA | 2 | 1 |
| 9_8100428 | 2 | 1 | NA | NA | 1 | 1 | NA | NA | 3 | 1 | 1 | 0 | NA | NA | 1 | 1 | NA | NA | 2 | 0.5 |
| 9_8101284 | 2 | 0 | NA | NA | 1 | 1 | NA | NA | 3 | 0.3333 | 1 | 1 | NA | NA | 1 | 1 | NA | NA | 2 | 1 |

| | | | | | | | | | | | | | | | | | | | | |
|---|---|---|---|---|---|---|---|---|---|---|---|---|---|---|---|---|---|---|---|---|
| 9_8101451 | 2 | 0.5 | NA | NA | NA | NA | NA | NA | 2 | 0.5 | 1 | 1 | NA | NA | 1 | 0 | NA | NA | 2 | 0.5 |
| 9_8101648 | 2 | 0.5 | NA | NA | 1 | 1 | 1 | 1 | 4 | 0.75 | 1 | 1 | NA | NA | 1 | 1 | NA | NA | 2 | 1 |
| 9_8899856 | NA | NA | 1 | 1 | 1 | 1 | 1 | 1 | 3 | 1 | NA | NA | NA | NA | 2 | 0 | 1 | 1 | 3 | 0.3333 |
| 9_8900086 | NA | NA | 1 | 1 | 1 | 1 | 2 | 1 | 4 | 1 | NA | NA | NA | NA | 2 | 1 | 1 | 1 | 3 | 1 |
| 9_8900103 | NA | NA | 1 | 1 | 1 | 1 | 2 | 1 | 4 | 1 | NA | NA | NA | NA | 2 | 1 | 1 | 1 | 3 | 1 |
| 9_8900131 | NA | NA | 1 | 0 | 1 | 1 | 1 | 0 | 3 | 0.3333 | NA | NA | NA | NA | 2 | 0.5 | 1 | 0 | 3 | 0.3333 |
| 9_8900169 | NA | NA | 1 | 1 | 1 | 1 | 2 | 1 | 4 | 1 | NA | NA | NA | NA | 2 | 1 | 1 | 1 | 3 | 1 |
| 9_8900544 | NA | NA | 1 | 1 | 1 | 1 | NA | NA | 2 | 1 | NA | NA | NA | NA | 1 | 0 | 1 | 1 | 2 | 0.5 |
| 9_8900573 | NA | NA | 1 | 1 | 1 | 0 | NA | NA | 2 | 0.5 | NA | NA | NA | NA | 1 | 1 | NA | NA | 1 | 1 |
| 9_8900658 | NA | NA | 1 | 0 | 1 | 1 | NA | NA | 2 | 0.5 | NA | NA | NA | NA | 1 | 0 | 1 | 1 | 2 | 0.5 |
| 9_8900812 | NA | NA | 1 | 1 | 1 | 1 | NA | NA | 2 | 1 | NA | NA | NA | NA | 1 | 1 | 1 | 1 | 2 | 1 |
| 9_8900894 | NA | NA | NA | NA | 1 | 1 | NA | NA | 1 | 1 | NA | NA | NA | NA | 1 | 1 | 1 | 1 | 2 | 1 |
| 9_8900932 | NA | NA | 1 | 1 | NA | NA | NA | NA | 1 | 1 | NA | NA | NA | NA | NA | NA | 1 | 1 | 1 | 1 |
| 9_8901274 | NA | NA | 1 | 1 | NA | NA | NA | NA | 1 | 1 | NA | NA | NA | NA | 1 | 0 | 1 | 1 | 2 | 0.5 |
| 9_8952672 | NA | NA | NA | NA | NA | NA | 2 | 0.5 | 2 | 0.5 | NA | NA | NA | NA | NA | NA | 2 | 0.5 | 2 | 0.5 |
| 9_8994855 | 3 | 1 | NA | NA | 1 | 1 | NA | NA | 4 | 1 | 1 | 1 | NA | NA | NA | NA | 2 | 1 | 3 | 1 |
| 9_8997463 | 1 | 1 | NA | NA | NA | NA | NA | NA | 1 | 1 | 2 | 0.5 | NA | NA | NA | NA | 1 | 1 | 3 | 0.6667 |
| 9_8997512 | 1 | 1 | NA | NA | NA | NA | NA | NA | 1 | 1 | 2 | 0.5 | NA | NA | NA | NA | 1 | 0 | 3 | 0.3333 |
| 9_9661172 | NA | NA | 1 | 1 | 2 | 1 | 1 | 0 | 4 | 0.75 | NA | NA | NA | NA | 1 | 0 | 1 | 1 | 2 | 0.5 |
| 9_9661257 | NA | NA | NA | NA | 1 | 1 | 2 | 0.5 | 3 | 0.6667 | NA | NA | NA | NA | 1 | 0 | 1 | 1 | 2 | 0.5 |
| 9_9661291 | NA | NA | NA | NA | 1 | 0 | 2 | 1 | 3 | 0.6667 | NA | NA | NA | NA | 1 | 1 | 1 | 1 | 2 | 1 |
| 9_1324377 | 1 | 1 | NA | NA | NA | NA | 3 | 0 | 4 | 0.25 | 1 | 1 | 2 | 0 | NA | NA | 3 | 0.3333 | 6 | 0.3333 |
| 9_1324411 | NA | NA | NA | NA | 2 | 0 | 3 | 0.3333 | 5 | 0.2 | 1 | 0 | 1 | 1 | NA | NA | 3 | 0 | 5 | 0.2 |
| 9_1324415 | 1 | 1 | NA | NA | 1 | 1 | 3 | 1 | 5 | 1 | 1 | 1 | 2 | 1 | NA | NA | 3 | 1 | 6 | 1 |
| 9_1324418 | 1 | 0 | NA | NA | 2 | 0 | 3 | 0.3333 | 6 | 0.1667 | 1 | 1 | 2 | 0 | NA | NA | 3 | 0 | 6 | 0.1667 |
| 9_1324421 | NA | NA | NA | NA | 2 | 1 | 2 | 1 | 4 | 1 | 1 | 1 | 1 | 1 | NA | NA | 3 | 1 | 5 | 1 |
| 9_1324425 | 1 | 1 | NA | NA | 2 | 1 | 1 | 1 | 4 | 1 | 1 | 1 | 2 | 1 | NA | NA | 3 | 1 | 6 | 1 |
| 9_1324435 | 1 | 1 | NA | NA | 1 | 0 | 4 | 1 | 6 | 0.8333 | 1 | 1 | 2 | 0.5 | NA | NA | 3 | 0.3333 | 6 | 0.5 |
| 9_1324439 | 1 | 0 | NA | NA | 2 | 0 | 4 | 0.75 | 7 | 0.4286 | 1 | 0 | 2 | 0.5 | NA | NA | 4 | 1 | 7 | 0.7143 |
| 9_1324440 | 1 | 1 | NA | NA | 2 | 1 | 4 | 1 | 7 | 1 | 1 | 1 | 2 | 1 | NA | NA | 3 | 1 | 6 | 1 |
| 9_1324446 | 1 | 1 | NA | NA | 2 | 1 | 3 | 1 | 6 | 1 | 1 | 1 | 1 | 1 | NA | NA | 3 | 1 | 5 | 1 |
| 9_1324453 | 1 | 1 | NA | NA | 2 | 1 | 3 | 1 | 6 | 1 | 1 | 1 | 2 | 1 | NA | NA | 3 | 1 | 6 | 1 |
| 9_1324498 | NA | NA | NA | NA | 2 | 1 | 3 | 0.3333 | 5 | 0.6 | NA | NA | 2 | 0.5 | NA | NA | NA | NA | 2 | 0.5 |
| 9_1324499 | 1 | 1 | NA | NA | 2 | 0.5 | 3 | 0.6667 | 6 | 0.6667 | 1 | 0 | 2 | 1 | NA | NA | 2 | 1 | 5 | 0.8 |
| 9_1324509 | 1 | 1 | NA | NA | 2 | 0.5 | 4 | 0.5 | 7 | 0.5714 | 1 | 0 | 2 | 0.5 | NA | NA | 3 | 0.6667 | 6 | 0.5 |
| 9_1324512 | NA | NA | NA | NA | 2 | 1 | 4 | 1 | 6 | 1 | NA | NA | NA | NA | NA | NA | 2 | 1 | 2 | 1 |
| 9_1324521 | 1 | 0 | NA | NA | 2 | 0 | 4 | 0.5 | 7 | 0.2857 | 1 | 1 | 1 | 0 | NA | NA | 2 | 0 | 4 | 0.25 |
| 9_1324536 | 1 | 1 | NA | NA | 2 | 0.5 | 4 | 0.75 | 7 | 0.7143 | 1 | 1 | 1 | 1 | NA | NA | 3 | 1 | 5 | 1 |
| 9_1324542 | 1 | 0 | NA | NA | 2 | 1 | 4 | 0.5 | 7 | 0.5714 | 1 | 0 | 1 | 1 | NA | NA | 2 | 0.5 | 4 | 0.5 |
| 9_1324562 | 1 | 0 | NA | NA | 2 | 0.5 | 3 | 0.6667 | 6 | 0.5 | 1 | 1 | 1 | 1 | NA | NA | 1 | 0 | 3 | 0.6667 |
| 9_1324588 | 1 | 1 | NA | NA | 2 | 0.5 | 4 | 1 | 7 | 0.8571 | 1 | 1 | 1 | 1 | NA | NA | 2 | 0.5 | 4 | 0.75 |
| 9_1325179 | 2 | 0.5 | 3 | 0.6667 | 1 | 0 | 1 | 1 | 7 | 0.5714 | 1 | 1 | NA | NA | 2 | 1 | 2 | 1 | 5 | 1 |
| 9_1325185 | 2 | 0.5 | 3 | 0.3333 | 1 | 0 | 1 | 0 | 7 | 0.2857 | 1 | 0 | NA | NA | 2 | 1 | 2 | 1 | 5 | 0.8 |
| 9_1325189 | 2 | 0 | 3 | 0.6667 | 1 | 0 | NA | NA | 6 | 0.3333 | 1 | 0 | NA | NA | 2 | 0.5 | 2 | 0.5 | 5 | 0.4 |
| 9_1325194 | 2 | 1 | 2 | 0 | 1 | 0 | NA | NA | 5 | 0.4 | 1 | 1 | NA | NA | 1 | 0 | NA | NA | 2 | 0.5 |
| 9_1325206 | 1 | 0 | 3 | 0.3333 | NA | NA | 1 | 0 | 5 | 0.2 | 1 | 1 | NA | NA | 2 | 0 | 2 | 0.5 | 5 | 0.4 |
| 9_1325220 | 1 | 0 | 3 | 0 | 2 | 0.5 | 1 | 0 | 7 | 0.1429 | 1 | 0 | NA | NA | 2 | 0.5 | 2 | 0.5 | 5 | 0.4 |
| 9_1325222 | 2 | 0 | 2 | 0 | 2 | 0.5 | 1 | 1 | 7 | 0.2857 | 1 | 1 | NA | NA | 2 | 0.5 | 2 | 0 | 5 | 0.4 |
| 9_1325285 | 2 | 0.5 | 3 | 0.3333 | 2 | 0.5 | 1 | 1 | 8 | 0.5 | 1 | 0 | NA | NA | 2 | 0.5 | 2 | 0.5 | 5 | 0.4 |
| 9_1325286 | 2 | 0.5 | 3 | 0.3333 | 2 | 0.5 | 1 | 1 | 8 | 0.5 | 1 | 0 | NA | NA | 2 | 1 | 1 | 1 | 4 | 0.75 |
| 9_1325389 | 1 | 0 | 1 | 0 | 1 | 0 | 1 | 1 | 4 | 0.25 | NA | NA | NA | NA | 2 | 0.5 | 1 | 1 | 3 | 0.6667 |
| 9_1329063 | 1 | 0 | 1 | 1 | 1 | 1 | 1 | 1 | 4 | 0.75 | 1 | 1 | 1 | 1 | 1 | 1 | NA | NA | 3 | 1 |
| 9_1329064 | 1 | 1 | 1 | 1 | 1 | 1 | 1 | 1 | 4 | 1 | 1 | 1 | 1 | 1 | 1 | 1 | NA | NA | 3 | 1 |
| 9_1329077 | 1 | 1 | 1 | 1 | 1 | 1 | 1 | 0 | 4 | 0.75 | 1 | 0 | 1 | 1 | NA | NA | NA | NA | 2 | 0.5 |
| 9_1329087 | 1 | 0 | 1 | 0 | NA | NA | 1 | 1 | 3 | 0.3333 | NA | NA | 1 | 0 | 1 | 1 | NA | NA | 2 | 0.5 |
| 9_1329088 | NA | NA | 1 | 0 | 1 | 1 | 1 | 1 | 3 | 0.6667 | NA | NA | 1 | 1 | 1 | 0 | NA | NA | 2 | 0.5 |
| 9_1329090 | 1 | 1 | NA | NA | 1 | 1 | 1 | 1 | 3 | 1 | 1 | 1 | 1 | 1 | NA | NA | NA | NA | 2 | 1 |
| 9_1329105 | 1 | 0 | 1 | 1 | 2 | 0.5 | 1 | 1 | 5 | 0.6 | 1 | 1 | 1 | 1 | 1 | 1 | NA | NA | 3 | 1 |
| 9_1329119 | 1 | 0 | 1 | 1 | 2 | 0.5 | 1 | 0 | 5 | 0.4 | 1 | 0 | 1 | 1 | 1 | 1 | NA | NA | 3 | 0.6667 |
| 9_1329139 | 2 | 1 | 1 | 1 | 2 | 0 | 1 | 0 | 6 | 0.5 | 1 | 0 | 1 | 0 | 1 | 1 | NA | NA | 3 | 0.3333 |
| 9_1329141 | 1 | 1 | 1 | 0 | 1 | 1 | 1 | 1 | 4 | 0.75 | 1 | 1 | 1 | 1 | 1 | 1 | NA | NA | 3 | 1 |
| 9_1329147 | 1 | 1 | 1 | 1 | 2 | 1 | 1 | 0 | 5 | 0.8 | 1 | 0 | 1 | 1 | 1 | 1 | NA | NA | 3 | 0.6667 |
| 9_1361751 | 1 | 1 | 2 | 0.5 | NA | NA | NA | NA | 3 | 0.6667 | NA | NA | NA | NA | NA | NA | 1 | 1 | 1 | 1 |
| 9_1362002 | 1 | 1 | 2 | 1 | NA | NA | 2 | 1 | 5 | 1 | 2 | 0.5 | NA | NA | NA | NA | 1 | 0 | 3 | 0.3333 |
| 9_1371680 | 4 | 0.5 | 6 | 0.3333 | 6 | 0 | 7 | 0.1429 | 23 | 0.2174 | 3 | 0 | 3 | 0.3333 | 4 | 0 | 7 | 0.4286 | 17 | 0.2353 |

| | | | | | | | | | | | | | | | | | | | | |
|---|---|---|---|---|---|---|---|---|---|---|---|---|---|---|---|---|---|---|---|---|
| 9_1371682 | 5 | 0.2 | 5 | 0 | 7 | 0.2857 | 7 | 0.2857 | 24 | 0.2083 | 4 | 0.25 | 3 | 0 | 5 | 0.2 | 7 | 0.4286 | 19 | 0.2632 |
| 9_1371683 | 3 | 1 | 3 | 1 | 6 | 1 | 6 | 1 | 18 | 1 | 4 | 1 | 2 | 1 | 4 | 1 | 6 | 1 | 16 | 1 |
| 9_1371686 | 4 | 0.5 | 5 | 0.2 | 7 | 0.2857 | 8 | 0.125 | 24 | 0.25 | 4 | 0 | 3 | 0.3333 | 4 | 0.5 | 6 | 0 | 17 | 0.1765 |
| 9_1371686 | 5 | 0 | 6 | 0 | 4 | 0 | 7 | 0.2857 | 22 | 0.0909 | 4 | 0 | 3 | 0.6667 | 5 | 0.4 | 6 | 0.3333 | 18 | 0.3333 |
| 9_1371699 | 5 | 0 | 6 | 0.3333 | 4 | 0 | 7 | 0 | 22 | 0.0909 | 4 | 0.25 | 3 | 0 | 5 | 0.4 | 8 | 0.125 | 20 | 0.2 |
| 9_1371757 | 6 | 0 | 5 | 0.4 | 7 | 0.2857 | 7 | 0.1429 | 25 | 0.2 | 4 | 1 | 4 | 0.75 | 8 | 0.625 | 6 | 0 | 22 | 0.5455 |
| 9_1371792 | 2 | 0 | 1 | 1 | 2 | 0 | 3 | 0.3333 | 8 | 0.25 | 1 | 0 | NA | NA | 2 | 0 | 3 | 0.3333 | 6 | 0.1667 |
| 9_1371945 | 1 | 0 | 4 | 0.25 | 2 | 0 | 3 | 0 | 10 | 0.1 | 2 | 0 | 1 | 0 | 5 | 0.2 | 5 | 0 | 13 | 0.0769 |
| 9_1371984 | 1 | 1 | 2 | 0.5 | 2 | 0.5 | 2 | 1 | 7 | 0.7143 | 1 | 0 | NA | NA | 4 | 0 | 5 | 0.4 | 10 | 0.2 |
| 9_1371985 | 1 | 1 | 2 | 0.5 | 1 | 1 | 3 | 1 | 7 | 0.8571 | 1 | 1 | 1 | 1 | 3 | 1 | 5 | 1 | 10 | 1 |
| 9_1371995 | NA | NA | 3 | 1 | 2 | 0 | 3 | 1 | 8 | 0.75 | 1 | 0 | NA | NA | 4 | 0.75 | 4 | 0.5 | 9 | 0.5556 |
| 9_1372003 | 4 | 0 | 3 | 0 | 3 | 0.3333 | 5 | 0.2 | 15 | 0.1333 | 1 | 0 | NA | NA | 3 | 0.3333 | 3 | 0 | 7 | 0.1429 |
| 9_1372008 | 1 | 1 | 3 | 0.3333 | 2 | 0 | 3 | 0.3333 | 9 | 0.3333 | 1 | 0 | 1 | 0 | 4 | 0 | 5 | 0.6 | 11 | 0.2727 |
| 9_1375563 | 5 | 0.2 | 2 | 0.5 | 1 | 0 | 1 | 0 | 9 | 0.2222 | 1 | 0 | 1 | 0 | 1 | 1 | 1 | 0 | 4 | 0.25 |
| 9_1423705 | 3 | 1 | 7 | 0.5714 | 3 | 0.6667 | 2 | 0 | 15 | 0.6 | 2 | 0.5 | 2 | 0.5 | 9 | 0.5556 | 6 | 0.3333 | 19 | 0.4737 |
| 9_1423734 | 2 | 0 | 7 | 0.5714 | 4 | 0.25 | 3 | 0.6667 | 16 | 0.4375 | 2 | 0 | 3 | 0.6667 | 7 | 0.2857 | 3 | 0 | 15 | 0.2667 |
| 9_1424079 | 7 | 0 | 4 | 0 | 9 | 0.2222 | 10 | 0 | 30 | 0.0667 | 8 | 0.125 | 10 | 0.2 | 9 | 0.1111 | 7 | 0.2857 | 34 | 0.1765 |
| 9_1424285 | 25 | 0.16 | 7 | 0.1429 | 13 | 0 | 20 | 0.05 | 65 | 0.0923 | 9 | 0 | 16 | 0.125 | 19 | 0.1053 | 15 | 0.1333 | 59 | 0.1017 |
| 9_1424288 | 26 | 0.3077 | 7 | 0.4286 | 15 | 0.4 | 23 | 0.1739 | 71 | 0.2958 | 11 | 0.4545 | 16 | 0.6875 | 24 | 0.4167 | 16 | 0.1875 | 67 | 0.4328 |
| 9_1424295 | 26 | 0.7308 | 8 | 0.75 | 14 | 0.8571 | 23 | 0.6957 | 71 | 0.7465 | 11 | 0.9091 | 17 | 0.8235 | 21 | 0.7619 | 15 | 0.8 | 64 | 0.8125 |
| 9_1424305 | 27 | 0.037 | 8 | 0.125 | 14 | 0.0714 | 24 | 0.0417 | 73 | 0.0548 | 11 | 0.0909 | 18 | 0.1111 | 22 | 0.1818 | 14 | 0 | 65 | 0.1077 |
| 9_1424315 | 24 | 0.875 | 6 | 1 | 12 | 0.8333 | 23 | 0.7391 | 65 | 0.8308 | 10 | 0.9 | 18 | 0.9444 | 20 | 1 | 13 | 0.9231 | 61 | 0.9508 |
| 9_1424320 | 24 | 0.4583 | 7 | 0.2857 | 13 | 0.3077 | 24 | 0.2083 | 68 | 0.3235 | 10 | 0.2 | 16 | 0.375 | 21 | 0.4762 | 12 | 0.5833 | 59 | 0.4237 |
| 9_1424324 | 25 | 0.04 | 8 | 0 | 16 | 0.0625 | 24 | 0.0833 | 73 | 0.0548 | 12 | 0.0833 | 18 | 0.0556 | 23 | 0.1739 | 11 | 0.0909 | 64 | 0.1094 |
| 9_1424332 | 25 | 0.44 | 9 | 0.5556 | 15 | 0.4 | 27 | 0.3704 | 76 | 0.4211 | 12 | 0.5833 | 19 | 0.6842 | 22 | 0.4545 | 17 | 0.4706 | 70 | 0.5429 |
| 9_1424336 | 26 | 0.5 | 7 | 0 | 12 | 0.3333 | 25 | 0.24 | 70 | 0.3286 | 11 | 0.4545 | 18 | 0.7222 | 24 | 0.5417 | 17 | 0.6471 | 70 | 0.6 |
| 9_1424343 | 29 | 0.3448 | 10 | 0.1 | 15 | 0.2 | 24 | 0.25 | 78 | 0.2564 | 11 | 0.1818 | 17 | 0.2353 | 24 | 0.2083 | 13 | 0.3077 | 65 | 0.2308 |
| 9_1424346 | 24 | 0.125 | 6 | 0 | 16 | 0.125 | 22 | 0.1364 | 68 | 0.1176 | 10 | 0.1 | 19 | 0.2105 | 26 | 0.0769 | 15 | 0.0667 | 70 | 0.1143 |
| 9_1424367 | 29 | 0.2759 | 10 | 0.3 | 16 | 0.3125 | 22 | 0.3182 | 77 | 0.2987 | 11 | 0.3636 | 15 | 0.2667 | 26 | 0.5 | 14 | 0.3571 | 66 | 0.3939 |
| 9_1424418 | 34 | 0.2059 | 13 | 0.1538 | 18 | 0.2222 | 27 | 0.3333 | 92 | 0.2391 | 15 | 0.2667 | 22 | 0.2727 | 30 | 0.3333 | 18 | 0.2778 | 85 | 0.2941 |
| 9_1426539 | 10 | 1 | 15 | 0.4667 | 10 | 0.4 | 7 | 0.8571 | 42 | 0.6429 | 7 | 0.7143 | 8 | 0.75 | 4 | 1 | 6 | 0.6667 | 25 | 0.76 |
| 9_1426544 | 15 | 0.1333 | 18 | 0.0556 | 14 | 0 | 8 | 0.125 | 55 | 0.0727 | 7 | 0 | 10 | 0.1 | 5 | 0.2 | 7 | 0.2857 | 29 | 0.1379 |
| 9_1426550 | 9 | 0.7778 | 19 | 0.5263 | 10 | 0.3 | 9 | 0.2222 | 47 | 0.4681 | 10 | 0.1 | 8 | 0.625 | 5 | 0.6 | 8 | 0.5 | 31 | 0.4194 |
| 9_1426551 | NA | NA | 1 | 1 | 1 | 1 | 2 | 0.5 | 4 | 0.75 | NA | NA | 1 | 1 | NA | NA | NA | NA | 1 | 1 |
| 9_1426568 | 11 | 0.0909 | 13 | 0.1538 | 13 | 0.0769 | 12 | 0.1667 | 49 | 0.1224 | 8 | 0.125 | 11 | 0.0909 | 6 | 0.3333 | 7 | 0.1429 | 32 | 0.1562 |
| 9_1426580 | 10 | 0 | 16 | 0.125 | 13 | 0.0769 | 11 | 0.0909 | 50 | 0.08 | 8 | 0.125 | 12 | 0.0833 | 5 | 0 | 7 | 0 | 32 | 0.0625 |
| 9_1426590 | 11 | 0.1818 | 16 | 0.1875 | 12 | 0 | 12 | 0.1667 | 51 | 0.1373 | 6 | 0.1667 | 11 | 0.1818 | 5 | 0 | 7 | 0.2857 | 29 | 0.1724 |
| 9_1426606 | 12 | 0.25 | 13 | 0.0769 | 11 | 0.1818 | 12 | 0.25 | 48 | 0.1875 | 8 | 0.125 | 10 | 0.2 | 5 | 0.4 | 6 | 0 | 29 | 0.1724 |
| 9_1426622 | 10 | 0.1 | 12 | 0.25 | 12 | 0 | 10 | 0.1 | 44 | 0.1136 | 8 | 0.125 | 7 | 0 | 5 | 0.4 | 6 | 0.3333 | 26 | 0.1923 |
| 9_1426623 | 7 | 0.1429 | 12 | 0.0833 | 12 | 0.0833 | 8 | 0.375 | 39 | 0.1538 | 8 | 0.375 | 8 | 0 | 4 | 0 | 5 | 0.2 | 25 | 0.16 |
| 9_1426636 | 8 | 0.125 | 13 | 0.1538 | 8 | 0.375 | 9 | 0.3333 | 38 | 0.2368 | 4 | 0.5 | 7 | 0.1429 | 3 | 0 | 5 | 0.2 | 19 | 0.2105 |
| 9_1426648 | 10 | 0.2 | 10 | 0.4 | 5 | 0.2 | 10 | 0.2 | 35 | 0.2571 | 5 | 0.4 | 8 | 0.25 | 4 | 0.5 | 6 | 0 | 23 | 0.2609 |
| 9_1426654 | 8 | 0.125 | 12 | 0.25 | 8 | 0.125 | 11 | 0.1818 | 39 | 0.1795 | 6 | 0.1667 | 7 | 0.1429 | 4 | 0.25 | 6 | 0.1667 | 23 | 0.1739 |
| 9_1426659 | 7 | 0 | 11 | 0.2727 | 8 | 0.375 | 12 | 0.25 | 38 | 0.2368 | 5 | 0.2 | 8 | 0 | 3 | 0 | 3 | 0.3333 | 19 | 0.1053 |
| 9_1426671 | 9 | 0.3333 | 12 | 0.25 | 9 | 0.1111 | 11 | 0.2727 | 41 | 0.2439 | 5 | 0.2 | 7 | 0.1429 | 4 | 0.25 | 6 | 0.1667 | 22 | 0.1818 |
| 9_1426773 | 7 | 1 | 11 | 0.9091 | 7 | 0.8571 | 13 | 0.9231 | 38 | 0.9211 | 6 | 0.8333 | 8 | 1 | 5 | 1 | 7 | 0.8571 | 26 | 0.9231 |
| 9_1426797 | 3 | 0.6667 | 13 | 0.7692 | 8 | 0.75 | 13 | 0.8462 | 37 | 0.7838 | 6 | 0.6667 | 7 | 0.7143 | 6 | 1 | 6 | 0.5 | 25 | 0.72 |
| 9_1426799 | 3 | 0.6667 | 12 | 0.3333 | 7 | 0.4286 | 10 | 0.4 | 32 | 0.4062 | 4 | 0.25 | 4 | 0.25 | 6 | 0.1667 | 4 | 0.75 | 18 | 0.3333 |
| 9_1426807 | 2 | 0.5 | 10 | 0.4 | 6 | 0.6667 | 9 | 0.3333 | 27 | 0.4444 | 6 | 0.6667 | 7 | 0.2857 | 6 | 0.8333 | 6 | 0.6667 | 25 | 0.6 |
| 9_1426812 | 3 | 0.6667 | 12 | 1 | 6 | 1 | 12 | 1 | 33 | 0.9697 | 6 | 0.8333 | 6 | 1 | 6 | 1 | 6 | 1 | 24 | 0.9583 |
| 9_1426830 | 1 | 1 | 12 | 0.9167 | 5 | 1 | 10 | 1 | 28 | 0.9643 | 5 | 1 | 6 | 1 | 3 | 1 | 7 | 1 | 21 | 1 |
| 9_1426837 | 1 | 1 | 9 | 1 | 3 | 1 | 8 | 0.75 | 21 | 0.9048 | 3 | 0.6667 | 4 | 0.75 | 4 | 0.5 | 1 | 1 | 12 | 0.6667 |
| 9_1426849 | 1 | 1 | 8 | 0.875 | 5 | 1 | 9 | 1 | 23 | 0.9565 | 4 | 0.75 | 4 | 0.75 | 3 | 1 | 5 | 1 | 16 | 0.875 |
| 9_1426868 | 1 | 0 | 9 | 0.2222 | 6 | 0 | 11 | 0.2727 | 27 | 0.1852 | 6 | 0 | 5 | 0 | 5 | 0.2 | 4 | 0 | 20 | 0.05 |
| 9_1426875 | 1 | 1 | 9 | 0.5556 | 5 | 0.6 | 10 | 0.6 | 25 | 0.6 | 4 | 0.75 | 4 | 0.75 | 6 | 0.6667 | 4 | 0.75 | 18 | 0.7222 |
| 9_1426884 | 1 | 1 | 6 | 0.8333 | 5 | 1 | 10 | 0.7 | 22 | 0.8182 | 4 | 1 | 3 | 0.6667 | 5 | 1 | 3 | 0.6667 | 15 | 0.8667 |
| 9_1426890 | 1 | 1 | 9 | 0.6667 | 5 | 0.6 | 10 | 0.7 | 25 | 0.68 | 4 | 0.75 | 4 | 0.5 | 6 | 0.3333 | 4 | 0.5 | 18 | 0.5 |
| 9_1426893 | 1 | 1 | 9 | 0.8889 | 4 | 1 | 11 | 0.7273 | 25 | 0.84 | 4 | 1 | 4 | 1 | 6 | 1 | 4 | 1 | 18 | 1 |
| 9_1426898 | 1 | 0 | 8 | 0.25 | 4 | 0 | 10 | 0.1 | 23 | 0.1304 | 3 | 0 | 4 | 0 | 6 | 0 | 4 | 0.25 | 17 | 0.0588 |
| 9_1426905 | 1 | 1 | 9 | 0.7778 | 3 | 1 | 8 | 1 | 21 | 0.9048 | 2 | 0.5 | 4 | 0.75 | 6 | 1 | 4 | 1 | 16 | 0.875 |
| 9_1426907 | 1 | 0 | 6 | 0.3333 | 4 | 0 | 11 | 0.1818 | 22 | 0.1818 | 3 | 0 | 3 | 0 | 6 | 0.1667 | 3 | 0 | 15 | 0.0667 |
| 9_1426907 | 1 | 0 | 8 | 0.875 | 4 | 0.75 | 9 | 0.6667 | 22 | 0.7273 | 3 | 0.6667 | 4 | 1 | 6 | 0.6667 | 4 | 1 | 17 | 0.8235 |
| 9_1426911 | NA | NA | 9 | 0.1111 | 4 | 0.75 | 10 | 0.4 | 23 | 0.3478 | 3 | 0 | 3 | 0.6667 | 6 | 0.1667 | 4 | 0.25 | 16 | 0.25 |
| 9_1436160 | NA | NA | NA | NA | 1 | 1 | NA | NA | 1 | 1 | NA | NA | 1 | 1 | NA | NA | NA | NA | 1 | 1 |
| 9_1436195 | NA | NA | NA | NA | 1 | 1 | NA | NA | 1 | 1 | NA | NA | 1 | 1 | NA | NA | NA | NA | 1 | 1 |

| | | | | | | | | | | | | | | | | | | | | |
|---|---|---|---|---|---|---|---|---|---|---|---|---|---|---|---|---|---|---|---|---|
| 9_1437404 | 1 | 1 | NA | NA | NA | NA | 1 | 0 | 2 | 0.5 | NA | NA | 1 | 1 | NA | NA | NA | NA | 1 | 1 |
| 9_1437425 | NA | NA | NA | NA | NA | NA | 1 | 1 | 1 | 1 | NA | NA | 1 | 1 | NA | NA | NA | NA | 1 | 1 |
| 9_1437439 | NA | NA | NA | NA | NA | NA | 1 | 1 | 1 | 1 | NA | NA | 1 | 1 | NA | NA | NA | NA | 1 | 1 |
| 9_1437481 | NA | NA | NA | NA | NA | NA | 1 | 1 | 1 | 1 | NA | NA | 1 | 1 | NA | NA | NA | NA | 1 | 1 |
| 9_1437482 | NA | NA | NA | NA | NA | NA | 1 | 1 | 1 | 1 | NA | NA | 1 | 1 | NA | NA | NA | NA | 1 | 1 |
| 9_1437517 | NA | NA | NA | NA | NA | NA | 1 | 1 | 1 | 1 | NA | NA | 1 | 1 | NA | NA | NA | NA | 1 | 1 |
| 9_1437527 | NA | NA | NA | NA | NA | NA | 1 | 1 | 1 | 1 | NA | NA | 1 | 1 | NA | NA | NA | NA | 1 | 1 |
| 9_1445343 | 1 | 1 | 4 | 0.25 | 1 | 1 | 8 | 0.625 | 14 | 0.5714 | 2 | 0.5 | 4 | 0.75 | 3 | 0.6667 | 6 | 0.5 | 15 | 0.6 |
| 9_1445347 | 2 | 0 | 4 | 0 | 1 | 1 | 7 | 0.1429 | 14 | 0.1429 | 2 | 0 | 3 | 0 | 5 | 0.4 | 6 | 0 | 16 | 0.125 |
| 9_1445897 | NA | NA | NA | NA | 1 | 1 | 1 | 1 | 2 | 1 | 1 | 1 | 1 | 1 | NA | NA | NA | NA | 2 | 1 |
| 9_1445898 | NA | NA | NA | NA | 1 | 1 | 1 | 0 | 2 | 0.5 | 1 | 0 | 1 | 1 | 1 | 0 | NA | NA | 3 | 0.3333 |
| 9_1450425 | NA | NA | NA | NA | 1 | 1 | 1 | 0 | 2 | 0.5 | 1 | 1 | NA | NA | NA | NA | NA | NA | 1 | 1 |
| 9_1469131 | NA | NA | 1 | 1 | NA | NA | 2 | 1 | 3 | 1 | 1 | 0 | 3 | 1 | 6 | 1 | NA | NA | 10 | 0.9 |
| 9_1469184 | NA | NA | 1 | 1 | NA | NA | 1 | 0 | 2 | 0.5 | 1 | 1 | 3 | 0 | 4 | 0.25 | NA | NA | 8 | 0.25 |
| 9_1469211 | NA | NA | 1 | 1 | NA | NA | 2 | 0 | 3 | 0.3333 | 1 | 0 | 3 | 0 | 6 | 0.1667 | NA | NA | 10 | 0.1 |
| 9_1469244 | NA | NA | 1 | 1 | NA | NA | 2 | 1 | 3 | 1 | 1 | 1 | 3 | 1 | 5 | 1 | NA | NA | 9 | 1 |
| 9_1469259 | NA | NA | 1 | 0 | NA | NA | 2 | 0.5 | 3 | 0.3333 | 1 | 1 | 3 | 0.6667 | 5 | 0.6 | NA | NA | 9 | 0.6667 |
| 9_1469260 | NA | NA | 1 | 0 | NA | NA | 2 | 1 | 3 | 0.6667 | 1 | 1 | 3 | 0.6667 | 3 | 1 | NA | NA | 7 | 0.8571 |
| 9_1469260 | NA | NA | 1 | 0 | NA | NA | 2 | 1 | 3 | 0.6667 | 1 | 0 | 3 | 0.6667 | 4 | 0.75 | NA | NA | 8 | 0.625 |
| 9_1469273 | NA | NA | 1 | 0 | NA | NA | 2 | 1 | 3 | 0.6667 | 1 | 1 | 3 | 0.6667 | 4 | 0.25 | NA | NA | 8 | 0.5 |
| 9_1469275 | NA | NA | 1 | 1 | NA | NA | 2 | 1 | 3 | 1 | 1 | 1 | 3 | 1 | 4 | 1 | NA | NA | 8 | 1 |
| 9_1469406 | NA | NA | 1 | 1 | NA | NA | 2 | 1 | 3 | 1 | 1 | 1 | 2 | 0 | 5 | 0.8 | NA | NA | 8 | 0.625 |
| 9_1469411 | NA | NA | 1 | 0 | NA | NA | 2 | 1 | 3 | 0.6667 | NA | NA | 2 | 1 | 5 | 1 | NA | NA | 7 | 1 |
| 9_1474497 | NA | NA | 1 | 1 | NA | NA | NA | NA | 1 | 1 | 2 | 0.5 | NA | NA | NA | NA | 2 | 0 | 4 | 0.25 |
| 9_1474502 | NA | NA | 1 | 1 | NA | NA | NA | NA | 1 | 1 | 2 | 0 | NA | NA | 1 | 0 | 2 | 0.5 | 5 | 0.2 |
| 9_1474519 | NA | NA | 1 | 1 | NA | NA | NA | NA | 1 | 1 | 1 | 1 | NA | NA | 1 | 0 | 2 | 0.5 | 4 | 0.5 |
| 9_1474522 | NA | NA | 1 | 1 | NA | NA | NA | NA | 1 | 1 | 1 | 1 | NA | NA | 1 | 0 | 2 | 0.5 | 4 | 0.5 |
| 9_1474723 | NA | NA | 1 | 1 | NA | NA | NA | NA | 1 | 1 | NA | NA | NA | NA | 1 | 0 | 1 | 1 | 2 | 0.5 |
| 9_1474786 | NA | NA | 2 | 0.5 | NA | NA | NA | NA | 2 | 0.5 | NA | NA | NA | NA | 1 | 0 | 1 | 1 | 2 | 0.5 |
| 9_1477173 | 4 | 0.5 | 5 | 1 | 5 | 1 | 5 | 1 | 19 | 0.8947 | NA | NA | 7 | 0.8571 | 9 | 0.7778 | 7 | 0.8571 | 23 | 0.8261 |
| 9_1477182 | 4 | 0 | 5 | 0 | 6 | 0.3333 | 4 | 0 | 19 | 0.1053 | NA | NA | 8 | 0.5 | 7 | 0 | 8 | 0 | 23 | 0.1739 |
| 9_1477198 | 3 | 0 | 4 | 0 | 4 | 0 | 7 | 0.1429 | 18 | 0.0556 | NA | NA | 6 | 0.1667 | 6 | 0.1667 | 5 | 0.2 | 17 | 0.1765 |
| 9_1477199 | 2 | 0.5 | 5 | 0.2 | 4 | 0.5 | 4 | 0.25 | 15 | 0.3333 | NA | NA | 5 | 0.2 | 8 | 0.125 | 5 | 0.4 | 18 | 0.2222 |
| 9_1477947 | 5 | 0.4 | 4 | 0.25 | 6 | 0 | 13 | 0.4615 | 28 | 0.3214 | 1 | 0 | 9 | 0.5556 | 12 | 0.25 | 8 | 0.125 | 30 | 0.3 |
| 9_1477950 | 4 | 0.25 | 6 | 0 | 6 | 0.1667 | 16 | 0.1875 | 32 | 0.1562 | 1 | 0 | 11 | 0.1818 | 15 | 0.1333 | 8 | 0.125 | 35 | 0.1429 |
| 9_1477954 | 5 | 0 | 5 | 0 | 7 | 0.2857 | 17 | 0.1176 | 34 | 0.1176 | 1 | 0 | 12 | 0.3333 | 16 | 0.125 | 8 | 0.25 | 37 | 0.2162 |
| 9_1477987 | 4 | 1 | 6 | 0.8333 | 5 | 1 | 16 | 1 | 31 | 0.9677 | 1 | 1 | 14 | 1 | 12 | 0.8333 | 9 | 1 | 36 | 0.9444 |
| 9_1477997 | 4 | 1 | 7 | 0.7143 | 6 | 0.8333 | 18 | 0.8889 | 35 | 0.8571 | 1 | 1 | 18 | 0.9444 | 15 | 0.9333 | 10 | 1 | 44 | 0.9545 |
| 9_1478014 | 5 | 0 | 9 | 0.1111 | 5 | 0 | 20 | 0.15 | 39 | 0.1026 | 1 | 0 | 15 | 0.0667 | 18 | 0.2222 | 11 | 0.1818 | 45 | 0.1556 |
| 9_1478023 | 5 | 0 | 7 | 0.1429 | 7 | 0.1429 | 17 | 0.1176 | 36 | 0.1111 | 1 | 0 | 13 | 0.1538 | 18 | 0.1667 | 8 | 0 | 40 | 0.125 |
| 9_1478038 | 6 | 0.5 | 9 | 0.4444 | 8 | 0.5 | 21 | 0.619 | 44 | 0.5455 | 1 | 1 | 17 | 0.7059 | 18 | 0.5 | 10 | 0.5 | 46 | 0.587 |
| 9_1478043 | 6 | 0 | 5 | 0.4 | 8 | 0.375 | 16 | 0.0625 | 35 | 0.1714 | 1 | 0 | 12 | 0.25 | 11 | 0.2727 | 8 | 0.375 | 32 | 0.2812 |
| 9_1478051 | 6 | 0.1667 | 9 | 0 | 10 | 0.3 | 21 | 0.2857 | 46 | 0.2174 | 2 | 0 | 19 | 0.2105 | 19 | 0.2105 | 12 | 0.0833 | 52 | 0.1731 |
| 9_1478101 | 7 | 0.4286 | 9 | 0.5556 | 11 | 0.1818 | 21 | 0.4762 | 48 | 0.4167 | 2 | 0.5 | 18 | 0.4444 | 16 | 0.5625 | 13 | 0.7692 | 49 | 0.5714 |
| 9_1478183 | 11 | 0.0909 | 10 | 0.1 | 15 | 0.0667 | 22 | 0.1364 | 58 | 0.1034 | 2 | 0 | 28 | 0.2143 | 18 | 0.2222 | 15 | 0.2667 | 63 | 0.2222 |
| 9_1478199 | 13 | 0.0769 | 11 | 0.1818 | 16 | 0 | 25 | 0.04 | 65 | 0.0615 | 4 | 0.25 | 26 | 0.0385 | 23 | 0.0435 | 18 | 0.0556 | 71 | 0.0563 |
| 9_1489763 | 1 | 1 | NA | NA | NA | NA | NA | NA | 1 | 1 | NA | NA | 1 | 1 | NA | NA | 2 | 1 | 3 | 1 |
| 9_1489778 | 1 | 1 | NA | NA | NA | NA | NA | NA | 1 | 1 | NA | NA | 1 | 1 | NA | NA | 2 | 1 | 3 | 1 |
| 9_1489798 | 1 | 1 | NA | NA | NA | NA | NA | NA | 1 | 1 | NA | NA | 1 | 1 | NA | NA | 2 | 0.5 | 3 | 0.6667 |
| 9_1489800 | 1 | 1 | NA | NA | NA | NA | NA | NA | 1 | 1 | NA | NA | 1 | 1 | NA | NA | 2 | 1 | 3 | 1 |
| 9_1489808 | 1 | 1 | NA | NA | NA | NA | NA | NA | 1 | 1 | NA | NA | 1 | 1 | NA | NA | 1 | 1 | 2 | 1 |
| 9_1489814 | 1 | 1 | NA | NA | NA | NA | NA | NA | 1 | 1 | NA | NA | NA | NA | NA | NA | 1 | 1 | 1 | 1 |
| 9_1491715 | 4 | 0 | 4 | 0.25 | 7 | 0.4286 | 6 | 0.1667 | 21 | 0.2381 | 1 | 0 | 2 | 0 | 3 | 0.6667 | 3 | 0.3333 | 9 | 0.3333 |
| 9_1491717 | 4 | 1 | 3 | 0.6667 | 5 | 1 | 4 | 0.75 | 16 | 0.875 | 1 | 1 | 1 | 1 | 3 | 1 | 3 | 0.3333 | 8 | 0.75 |
| 9_1491740 | 4 | 0.25 | 3 | 0 | 5 | 0.2 | 5 | 0.2 | 17 | 0.1765 | 1 | 0 | 2 | 0.5 | 2 | 0 | 2 | 0 | 7 | 0.1429 |
| 9_1491873 | 4 | 0.5 | 4 | 0.25 | 6 | 0.6667 | 5 | 0.6 | 19 | 0.5263 | 1 | 1 | 2 | 0.5 | 3 | 0.3333 | 2 | 0.5 | 8 | 0.5 |
| 9_1491880 | 2 | 0.5 | 3 | 1 | 6 | 0.6667 | 4 | 0.25 | 15 | 0.6 | 1 | 1 | 1 | 1 | 2 | 1 | 3 | 1 | 7 | 1 |
| 9_1491884 | 3 | 0 | 3 | 0.3333 | 3 | 0.6667 | 3 | 0.3333 | 12 | 0.3333 | 1 | 0 | 2 | 0 | 3 | 0 | 3 | 0.3333 | 9 | 0.1111 |
| 9_1491888 | 4 | 0 | 3 | 0 | 5 | 0 | 3 | 0.3333 | 15 | 0.0667 | 1 | 0 | 1 | 0 | 3 | 0.3333 | 3 | 0.3333 | 8 | 0.25 |
| 9_1491895 | 4 | 0.5 | 3 | 1 | 4 | 1 | 3 | 1 | 14 | 0.8571 | 1 | 1 | 2 | 1 | 3 | 1 | 3 | 1 | 9 | 1 |
| 9_1491920 | 4 | 0.25 | 3 | 0.6667 | 5 | 0.6 | 4 | 0.25 | 16 | 0.4375 | 1 | 0 | 2 | 0.5 | 2 | 0 | 3 | 0.3333 | 8 | 0.25 |
| 9_1491924 | 4 | 1 | 2 | 1 | 5 | 1 | 4 | 0.75 | 15 | 0.9333 | 1 | 1 | 1 | 1 | NA | NA | 3 | 1 | 5 | 1 |
| 9_1495629 | 3 | 0.3333 | 1 | 0 | 4 | 0 | 3 | 0 | 11 | 0.0909 | 1 | 1 | 1 | 0 | 1 | 1 | 2 | 0.5 | 5 | 0.6 |
| 9_1516955 | NA | NA | NA | NA | 1 | 1 | 6 | 0.5 | 7 | 0.5714 | 3 | 0.6667 | 3 | 0.3333 | 2 | 0.5 | 3 | 0.3333 | 11 | 0.4545 |
| 9_1516968 | NA | NA | NA | NA | 1 | 0 | 5 | 0.2 | 6 | 0.1667 | 2 | 1 | 3 | 0 | 3 | 0.3333 | 3 | 0 | 11 | 0.2727 |

| | | | | | | | | | | | | | | | | | | | | |
|---|---|---|---|---|---|---|---|---|---|---|---|---|---|---|---|---|---|---|---|---|
| 9_1516976 | NA | NA | NA | NA | 1 | 1 | 6 | 0.8333 | 7 | 0.8571 | 3 | 1 | 5 | 0.6 | 2 | 1 | 2 | 1 | 12 | 0.8333 |
| 9_1516977 | NA | NA | NA | NA | 1 | 1 | 4 | 0.75 | 5 | 0.8 | 3 | 0.6667 | 3 | 1 | 2 | 0.5 | 3 | 0 | 11 | 0.5455 |
| 9_1517003 | 2 | 1 | NA | NA | 1 | 0 | 6 | 0.6667 | 9 | 0.6667 | 3 | 0.6667 | 5 | 1 | 3 | 1 | 3 | 1 | 14 | 0.9286 |
| 9_1517050 | 2 | 0 | NA | NA | NA | NA | 4 | 0.25 | 6 | 0.1667 | 1 | 1 | 1 | 0 | 1 | 0 | 3 | 0.3333 | 6 | 0.3333 |
| 9_1517055 | 1 | 0 | NA | NA | 1 | 0 | 7 | 0.2857 | 9 | 0.2222 | 3 | 0.3333 | 4 | 0.25 | 3 | 0.3333 | 3 | 0.3333 | 13 | 0.3077 |
| 9_1517342 | 5 | 0.6 | 8 | 0.625 | 2 | 0.5 | 1 | 0 | 16 | 0.5625 | 2 | 0.5 | 2 | 0 | 4 | 0.75 | 5 | 0.2 | 13 | 0.3846 |
| 9_1517349 | 4 | 1 | 8 | 0.875 | 2 | 1 | 1 | 1 | 15 | 0.9333 | 3 | 1 | 2 | 1 | 3 | 1 | 5 | 1 | 13 | 1 |
| 9_1517361 | 5 | 0 | 9 | 0.1111 | NA | NA | 1 | 0 | 15 | 0.0667 | 4 | 0.25 | 2 | 0 | 4 | 0.25 | 4 | 0 | 14 | 0.1429 |
| 9_1518168 | 4 | 0.5 | 7 | 0 | 1 | 0 | 1 | 0 | 13 | 0.1538 | 5 | 0.2 | 3 | 0 | 3 | 0.3333 | 2 | 0 | 13 | 0.1538 |
| 9_1519498 | 3 | 0 | 5 | 0.4 | 3 | 0 | 3 | 0 | 14 | 0.1429 | 2 | 0 | 6 | 0 | 1 | 0 | 3 | 0.3333 | 12 | 0.0833 |
| 9_1519526 | 2 | 0 | 4 | 0.25 | 3 | 0.6667 | 3 | 0.6667 | 12 | 0.4167 | 2 | 0 | 5 | 0.4 | 1 | 0 | 3 | 0 | 11 | 0.1818 |
| 9_1520202 | NA | NA | 2 | 0.5 | NA | NA | 2 | 0 | 4 | 0.25 | 1 | 0 | 1 | 1 | 1 | 0 | NA | NA | 3 | 0.3333 |
| 9_1521994 | NA | NA | NA | NA | NA | NA | 5 | 0.2 | 5 | 0.2 | 1 | 1 | 1 | 0 | NA | NA | 3 | 0 | 5 | 0.2 |
| 9_1522336 | NA | NA | NA | NA | NA | NA | 1 | 1 | 1 | 1 | NA | NA | 1 | 1 | 1 | 1 | 2 | 0.5 | 4 | 0.75 |
| 9_1523390 | 1 | 1 | NA | NA | 1 | 1 | 1 | 0 | 3 | 0.6667 | NA | NA | NA | NA | 1 | 1 | NA | NA | 1 | 1 |
| 9_1523402 | 1 | 1 | NA | NA | 2 | 1 | 1 | 1 | 4 | 1 | NA | NA | 2 | 1 | 1 | 1 | 1 | 1 | 4 | 1 |
| 9_1523409 | 1 | 1 | NA | NA | 2 | 0.5 | 1 | 1 | 4 | 0.75 | NA | NA | 2 | 1 | 2 | 0.5 | 1 | 1 | 5 | 0.8 |
| 9_1523425 | 1 | 1 | NA | NA | 2 | 1 | 1 | 1 | 4 | 1 | NA | NA | 2 | 0 | 2 | 1 | 1 | 1 | 5 | 0.6 |
| 9_1523698 | 1 | 0 | NA | NA | 2 | 0.5 | 1 | 0 | 4 | 0.25 | NA | NA | 2 | 0 | 1 | 1 | 1 | 0 | 4 | 0.25 |
| 9_1524363 | 1 | 1 | NA | NA | NA | NA | 1 | 1 | 2 | 1 | NA | NA | 2 | 0.5 | 1 | 1 | NA | NA | 3 | 0.6667 |
| 9_1524365 | 1 | 1 | NA | NA | NA | NA | 1 | 1 | 2 | 1 | NA | NA | 1 | 1 | 2 | 1 | NA | NA | 3 | 1 |
| 9_1524414 | 1 | 1 | NA | NA | NA | NA | 1 | 1 | 2 | 1 | NA | NA | 2 | 0.5 | 2 | 0.5 | NA | NA | 4 | 0.5 |
| 9_1524421 | 1 | 1 | NA | NA | NA | NA | 1 | 0 | 2 | 0.5 | NA | NA | 2 | 0.5 | 2 | 0.5 | NA | NA | 4 | 0.5 |
| 9_1524562 | 1 | 1 | NA | NA | NA | NA | 1 | 0 | 2 | 0.5 | NA | NA | NA | NA | 1 | 1 | NA | NA | 1 | 1 |
| 9_1524588 | 1 | 0 | NA | NA | NA | NA | 1 | 1 | 2 | 0.5 | NA | NA | NA | NA | 1 | 1 | NA | NA | 1 | 1 |
| 9_1524617 | 1 | 1 | NA | NA | NA | NA | 1 | 1 | 2 | 1 | NA | NA | NA | NA | 1 | 1 | NA | NA | 1 | 1 |
| 9_1564949 | 3 | 1 | 7 | 0.7143 | 2 | 1 | 1 | 1 | 13 | 0.8462 | 2 | 1 | NA | NA | 2 | 1 | 1 | 1 | 5 | 1 |
| 9_1564952 | 2 | 1 | 4 | 0.5 | 2 | 0.5 | 2 | 1 | 10 | 0.7 | 1 | 0 | NA | NA | 2 | 0.5 | 1 | 1 | 4 | 0.5 |
| 9_1564964 | 3 | 0.6667 | 6 | 1 | 1 | 1 | 1 | 1 | 11 | 0.9091 | 2 | 1 | NA | NA | 2 | 1 | 1 | 1 | 5 | 1 |
| 9_1564969 | 3 | 0 | 7 | 0.1429 | 2 | 0 | 2 | 0.5 | 14 | 0.1429 | 2 | 0 | 2 | 1 | 2 | 0 | 1 | 1 | 7 | 0.4286 |
| 9_1564976 | 2 | 1 | 4 | 0.75 | 2 | 1 | 1 | 1 | 9 | 0.8889 | 1 | 0 | 2 | 1 | 2 | 0.5 | 1 | 1 | 6 | 0.6667 |
| 9_1564980 | 1 | 1 | 3 | 1 | NA | NA | 2 | 1 | 6 | 1 | 2 | 1 | 2 | 1 | 2 | 1 | 1 | 1 | 7 | 1 |
| 9_1564981 | 2 | 1 | 5 | 0.8 | 2 | 1 | 1 | 1 | 10 | 0.9 | 2 | 0.5 | 1 | 1 | 1 | 1 | 1 | 1 | 5 | 0.8 |
| 9_1564981 | 2 | 0.5 | 6 | 0.8333 | 2 | 0.5 | 2 | 1 | 12 | 0.75 | 2 | 1 | 2 | 1 | 2 | 0.5 | NA | NA | 6 | 0.8333 |
| 9_1564984 | 3 | 0.3333 | 7 | 0.1429 | 1 | 1 | 2 | 0.5 | 13 | 0.3077 | 2 | 0 | 1 | 1 | 2 | 1 | 1 | 1 | 6 | 0.6667 |
| 9_1564987 | 2 | 0 | 6 | 0.1667 | 2 | 0.5 | 2 | 0 | 12 | 0.1667 | 2 | 0.5 | 2 | 0.5 | 2 | 0.5 | 1 | 0 | 7 | 0.4286 |
| 9_1565117 | 3 | 0.3333 | 5 | 0 | 2 | 0.5 | 2 | 0 | 12 | 0.1667 | 3 | 0.3333 | 4 | 0.5 | 2 | 0 | 2 | 0 | 11 | 0.2727 |
| 9_1565124 | 5 | 0 | 5 | 0.2 | 2 | 0 | 3 | 0.3333 | 15 | 0.1333 | 3 | 0 | 5 | 0.4 | 2 | 0 | 2 | 0.5 | 12 | 0.25 |
| 9_1565128 | 4 | 0.75 | 5 | 0.2 | 1 | 1 | 2 | 1 | 12 | 0.5833 | 4 | 0.75 | 4 | 1 | 2 | 0.5 | 2 | 0.5 | 12 | 0.75 |
| 9_1565139 | 5 | 0 | 6 | 0.1667 | 2 | 0 | 3 | 0 | 16 | 0.0625 | 2 | 0 | 5 | 0.2 | 2 | 0 | 2 | 0 | 11 | 0.0909 |
| 9_1565141 | 4 | 0.75 | 5 | 0.8 | 2 | 0.5 | 3 | 1 | 14 | 0.7857 | 4 | 0.75 | 3 | 1 | 2 | 0 | 2 | 1 | 11 | 0.7273 |
| 9_1565143 | 4 | 0.25 | 6 | 0.3333 | 2 | 0.5 | 3 | 0 | 15 | 0.2667 | 2 | 0.5 | 5 | 0.8 | 2 | 0.5 | 2 | 0 | 11 | 0.5455 |
| 9_1565162 | 6 | 0 | 6 | 0 | 3 | 0 | 4 | 0.25 | 19 | 0.0526 | 5 | 0 | 5 | 0 | 2 | 0 | 2 | 0.5 | 14 | 0.0714 |
| 9_1565254 | 7 | 0.4286 | 9 | 0.6667 | 3 | 0.3333 | 4 | 0.25 | 23 | 0.4783 | 5 | 0.2 | 4 | 0.75 | 5 | 0.2 | 2 | 0 | 16 | 0.3125 |
| 9_1565341 | 6 | 0.1667 | 9 | 0.1111 | 3 | 0 | 7 | 0.1429 | 25 | 0.12 | 4 | 0 | 5 | 0.2 | 5 | 0 | 4 | 0.25 | 18 | 0.1111 |
| 9_1565410 | NA | NA | 1 | 1 | NA | NA | NA | NA | 1 | 1 | NA | NA | NA | NA | 1 | 1 | 1 | 0 | 2 | 0.5 |
| 9_1565419 | NA | NA | 1 | 1 | NA | NA | 1 | 1 | 2 | 1 | NA | NA | NA | NA | 1 | 1 | 1 | 1 | 2 | 1 |
| 9_1565435 | NA | NA | 1 | 1 | NA | NA | 1 | 1 | 2 | 1 | NA | NA | NA | NA | 2 | 1 | 1 | 1 | 3 | 1 |
| 9_1565442 | NA | NA | 1 | 1 | NA | NA | 1 | 1 | 2 | 1 | NA | NA | NA | NA | 2 | 1 | NA | NA | 2 | 1 |
| 9_1565444 | NA | NA | 1 | 0 | NA | NA | 1 | 1 | 2 | 0.5 | NA | NA | NA | NA | 2 | 0.5 | NA | NA | 2 | 0.5 |
| 9_1565445 | NA | NA | 1 | 1 | NA | NA | 1 | 0 | 2 | 0.5 | NA | NA | NA | NA | 2 | 0.5 | 1 | 1 | 3 | 0.6667 |
| 9_1565460 | NA | NA | 1 | 1 | NA | NA | 1 | 1 | 2 | 1 | NA | NA | NA | NA | 2 | 0.5 | 1 | 0 | 3 | 0.3333 |
| 9_1582204 | 2 | 1 | 5 | 0.8 | 2 | 0 | 4 | 0.25 | 13 | 0.5385 | 4 | 1 | 2 | 1 | 3 | 0.6667 | 4 | 0.75 | 13 | 0.8462 |
| 9_1582207 | 1 | 0 | 4 | 0.5 | 2 | 0.5 | 5 | 0.2 | 12 | 0.3333 | 4 | 0.75 | 2 | 0 | 2 | 0.5 | 4 | 0 | 12 | 0.3333 |
| 9_1582242 | 1 | 0 | 5 | 0 | 2 | 0 | 4 | 0.25 | 12 | 0.0833 | 3 | 0.3333 | 3 | 0 | 4 | 0 | 3 | 0 | 13 | 0.0769 |
| 9_1582563 | 1 | 0 | 2 | 1 | NA | NA | 3 | 0.3333 | 6 | 0.5 | 2 | 0 | NA | NA | 4 | 0.25 | NA | NA | 6 | 0.1667 |
| 9_1582607 | 2 | 1 | 1 | 1 | NA | NA | 3 | 1 | 6 | 1 | 2 | 1 | NA | NA | 4 | 1 | NA | NA | 6 | 1 |
| 9_1582609 | 2 | 1 | NA | NA | NA | NA | 3 | 1 | 5 | 1 | 2 | 1 | NA | NA | 4 | 1 | NA | NA | 6 | 1 |
| 9_1582613 | 2 | 0 | 1 | 0 | NA | NA | 3 | 0.3333 | 6 | 0.1667 | 2 | 0 | NA | NA | 3 | 0.3333 | NA | NA | 5 | 0.2 |
| 9_1582615 | 2 | 1 | 1 | 0 | NA | NA | 3 | 0.3333 | 6 | 0.5 | 2 | 1 | NA | NA | 2 | 0.5 | NA | NA | 4 | 0.75 |
| 9_1582624 | 2 | 1 | 1 | 1 | NA | NA | 3 | 0.3333 | 6 | 0.6667 | 2 | 1 | NA | NA | 2 | 1 | NA | NA | 4 | 1 |
| 9_1582626 | 2 | 0.5 | 1 | 0 | NA | NA | 3 | 0.6667 | 6 | 0.5 | 2 | 0.5 | NA | NA | 2 | 1 | NA | NA | 4 | 0.75 |
| 9_1582637 | 2 | 1 | 1 | 1 | NA | NA | 3 | 1 | 6 | 1 | 2 | 1 | NA | NA | 3 | 1 | NA | NA | 5 | 1 |
| 9_1582647 | 1 | 1 | 1 | 1 | NA | NA | 2 | 0.5 | 4 | 0.75 | 2 | 1 | NA | NA | 2 | 1 | NA | NA | 4 | 1 |
| 9_1582650 | NA | NA | 1 | 1 | NA | NA | 2 | 0.5 | 3 | 0.6667 | 1 | 0 | NA | NA | 2 | 0.5 | NA | NA | 3 | 0.3333 |

| | | | | | | | | | | | | | | | | | | | | |
|---|---|---|---|---|---|---|---|---|---|---|---|---|---|---|---|---|---|---|---|---|
| 9_1582662 | 1 | 1 | 1 | 0 | NA | NA | 2 | 1 | 4 | 0.75 | 2 | 0.5 | NA | NA | 2 | 0.5 | NA | NA | 4 | 0.5 |
| 9_1582669 | 1 | 1 | 1 | 1 | NA | NA | 3 | 1 | 5 | 1 | 2 | 0.5 | NA | NA | 3 | 1 | NA | NA | 5 | 0.8 |
| 9_1582679 | 1 | 0 | 1 | 0 | NA | NA | 3 | 0.6667 | 5 | 0.4 | 2 | 0.5 | NA | NA | 3 | 0.6667 | NA | NA | 5 | 0.6 |
| 9_1583405 | 1 | 1 | NA | NA | NA | NA | 1 | 0 | 2 | 0.5 | 1 | 0 | NA | NA | 3 | 0.6667 | NA | NA | 4 | 0.5 |
| 9_1584945 | 1 | 0 | NA | NA | NA | NA | 1 | 1 | 2 | 0.5 | 2 | 0.5 | NA | NA | NA | NA | NA | NA | 2 | 0.5 |
| 9_1587157 | NA | NA | NA | NA | NA | NA | 1 | 1 | 1 | 1 | 1 | 1 | NA | NA | NA | NA | NA | NA | 1 | 1 |
| 9_1587166 | 1 | 1 | NA | NA | NA | NA | 1 | 0 | 2 | 0.5 | 2 | 0.5 | NA | NA | NA | NA | NA | NA | 2 | 0.5 |
| 9_1588114 | 1 | 1 | NA | NA | NA | NA | NA | NA | 1 | 1 | 2 | 0.5 | NA | NA | NA | NA | NA | NA | 2 | 0.5 |
| 9_1590736 | 1 | 1 | NA | NA | NA | NA | NA | NA | 1 | 1 | 1 | 1 | NA | NA | NA | NA | NA | NA | 1 | 1 |
| 9_1590737 | 1 | 1 | NA | NA | NA | NA | NA | NA | 1 | 1 | 1 | 1 | NA | NA | NA | NA | NA | NA | 1 | 1 |
| 9_1590749 | 1 | 1 | NA | NA | NA | NA | NA | NA | 1 | 1 | 1 | 1 | NA | NA | NA | NA | NA | NA | 1 | 1 |
| 9_1822460 | 8 | 0.125 | 14 | 0.0714 | 8 | 0.125 | 8 | 0.125 | 38 | 0.1053 | 7 | 0.2857 | 8 | 0.125 | 13 | 0.2308 | 8 | 0.25 | 36 | 0.2222 |
| 9_1822490 | 7 | 0.1429 | 15 | 0.0667 | 10 | 0 | 6 | 0 | 38 | 0.0526 | 4 | 0.25 | 8 | 0.125 | 10 | 0 | 6 | 0.1667 | 28 | 0.1071 |
| 9_1822680 | 2 | 0 | 2 | 0.5 | 3 | 0 | 1 | 0 | 8 | 0.125 | 1 | 0 | 3 | 0 | 6 | 0.1667 | 1 | 0 | 11 | 0.0909 |
| 9_1822836 | 2 | 0.5 | NA | NA | 1 | 0 | 1 | 0 | 4 | 0.25 | 2 | 0 | 2 | 0 | 3 | 0.3333 | NA | NA | 7 | 0.1429 |
| 9_1955833 | NA | NA | 4 | 0.5 | 3 | 0.6667 | 3 | 0.3333 | 10 | 0.5 | 1 | 1 | 1 | 1 | 1 | 1 | 2 | 0 | 5 | 0.6 |
| 9_1955835 | NA | NA | 4 | 0.5 | 4 | 0.75 | 2 | 1 | 10 | 0.7 | 2 | 0.5 | 1 | 0 | 2 | 1 | 2 | 1 | 7 | 0.7143 |
| 9_1955853 | NA | NA | 4 | 1 | 4 | 0.75 | 3 | 1 | 11 | 0.9091 | 2 | 1 | 2 | 1 | 2 | 1 | 2 | 0.5 | 8 | 0.875 |
| 9_1955857 | NA | NA | 3 | 0.3333 | 3 | 0 | 3 | 0 | 9 | 0.1111 | 2 | 0 | 1 | 0 | 2 | 0 | 2 | 0.5 | 7 | 0.1429 |
| 9_1955883 | NA | NA | 1 | 1 | 2 | 1 | 1 | 1 | 4 | 1 | 2 | 1 | 1 | 1 | NA | NA | 2 | 1 | 5 | 1 |
| 9_1955893 | NA | NA | 4 | 0.75 | 4 | 0.75 | 3 | 1 | 11 | 0.8182 | 3 | 0.3333 | 2 | 1 | 3 | 1 | 2 | 0.5 | 10 | 0.7 |
| 9_1955938 | NA | NA | 5 | 0.6 | 4 | 0.5 | 4 | 0.25 | 13 | 0.4615 | 3 | 0.6667 | 2 | 0 | 3 | 0.3333 | 1 | 1 | 9 | 0.4444 |
| 9_1955942 | NA | NA | 6 | 0 | 5 | 0.4 | 4 | 0 | 15 | 0.1333 | 3 | 0.3333 | 2 | 0 | 2 | 0.5 | 2 | 0.5 | 9 | 0.3333 |
| 9_1955946 | NA | NA | 5 | 0.2 | 5 | 0 | 4 | 0.5 | 14 | 0.2143 | 2 | 0.5 | 2 | 0.5 | 3 | 0.3333 | 2 | 0 | 9 | 0.3333 |
| 9_1955947 | NA | NA | 6 | 1 | 4 | 1 | 4 | 1 | 14 | 1 | 3 | 1 | 2 | 1 | 3 | 1 | 2 | 0 | 10 | 0.8 |
| 9_1955950 | NA | NA | 6 | 0.6667 | 5 | 1 | 3 | 1 | 14 | 0.8571 | 3 | 0.6667 | 2 | 1 | 3 | 1 | 1 | 1 | 9 | 0.8889 |
| 9_1955953 | NA | NA | 4 | 1 | 5 | 1 | 4 | 1 | 13 | 1 | 3 | 1 | 2 | 1 | 3 | 1 | 2 | 1 | 10 | 1 |
| 9_1955963 | NA | NA | 4 | 0.25 | 2 | 0.5 | 4 | 0.5 | 10 | 0.4 | 3 | 0 | 2 | 0.5 | 3 | 0.3333 | 2 | 0.5 | 10 | 0.3 |
| 9_1955972 | NA | NA | 4 | 0.25 | 6 | 0 | 3 | 0 | 13 | 0.0769 | 2 | 0.5 | 2 | 0 | 3 | 0.3333 | 2 | 0 | 9 | 0.2222 |
| 9_1955975 | NA | NA | 3 | 0.6667 | 6 | 0.3333 | 2 | 1 | 11 | 0.5455 | 2 | 1 | 1 | 1 | 1 | 0 | 2 | 0.5 | 6 | 0.6667 |
| 9_1956005 | NA | NA | 6 | 0.3333 | 6 | 0 | 4 | 0.25 | 16 | 0.1875 | 3 | 0.3333 | 1 | 1 | 3 | 0.6667 | 3 | 0.3333 | 10 | 0.5 |
| 9_1956063 | 1 | 0 | 6 | 0.1667 | 6 | 0 | 5 | 0 | 18 | 0.0556 | 4 | 0 | 2 | 0.5 | 3 | 0 | 5 | 0.2 | 14 | 0.1429 |
| 9_1956078 | 1 | 0 | 7 | 0 | 6 | 0.1667 | 4 | 0.25 | 18 | 0.1111 | 4 | 0 | 2 | 0 | 3 | 0.3333 | 5 | 0 | 14 | 0.0714 |
| 9_1956096 | 2 | 0.5 | 5 | 0.2 | 5 | 0 | 5 | 0.2 | 17 | 0.1765 | 3 | 0 | 2 | 0 | 2 | 0 | 4 | 0.25 | 11 | 0.0909 |
| 9_1956098 | 1 | 0 | 7 | 0.4286 | 6 | 0.1667 | 5 | 0.4 | 19 | 0.3158 | 3 | 0.3333 | 3 | 0 | 2 | 0.5 | 2 | 1 | 10 | 0.4 |
| 9_2025795 | NA | NA | NA | NA | NA | NA | 1 | 1 | 1 | 1 | NA | NA | 1 | 1 | NA | NA | 1 | 0 | 2 | 0.5 |
| 9_2026394 | 3 | 0.3333 | 3 | 1 | 1 | 1 | 2 | 0.5 | 9 | 0.6667 | 1 | 1 | 2 | 1 | 6 | 0.8333 | 1 | 1 | 10 | 0.9 |
| 9_2026398 | 4 | 0 | 3 | 0.3333 | 2 | 0 | 2 | 0 | 11 | 0.0909 | 1 | 0 | 2 | 0 | 5 | 0.2 | NA | NA | 8 | 0.125 |
| 9_2026404 | 4 | 0 | 2 | 0.5 | 1 | 0 | 2 | 0.5 | 9 | 0.2222 | 2 | 0.5 | 2 | 0 | 6 | 0 | 1 | 0 | 11 | 0.0909 |
| 9_2034442 | 1 | 0 | 1 | 1 | NA | NA | 3 | 0.3333 | 5 | 0.4 | NA | NA | NA | NA | NA | NA | 2 | 0.5 | 2 | 0.5 |
| 9_2034492 | 3 | 0.3333 | 2 | 0.5 | 1 | 0 | 4 | 0.25 | 10 | 0.3 | NA | NA | NA | NA | NA | NA | 1 | 1 | 1 | 1 |
| 9_2038065 | 4 | 0 | 3 | 0 | 3 | 0 | 4 | 0.25 | 14 | 0.0714 | NA | NA | NA | NA | 5 | 0.4 | 2 | 0 | 7 | 0.2857 |
| 9_2038164 | 3 | 1 | 3 | 1 | 2 | 0.5 | 2 | 1 | 10 | 0.9 | NA | NA | NA | NA | 5 | 1 | 2 | 1 | 7 | 1 |
| 9_2038180 | 3 | 0 | 3 | 0.3333 | 3 | 0.3333 | 3 | 0.3333 | 12 | 0.25 | NA | NA | NA | NA | 5 | 0.8 | 2 | 0.5 | 7 | 0.7143 |
| 9_2038197 | 1 | 1 | 3 | 0.6667 | 2 | 0.5 | 3 | 0.3333 | 9 | 0.5556 | NA | NA | NA | NA | 3 | 0.6667 | 2 | 1 | 5 | 0.8 |
| 9_2038214 | 2 | 1 | 3 | 0.6667 | 3 | 0.6667 | 2 | 0.5 | 10 | 0.7 | NA | NA | NA | NA | 5 | 0.8 | 1 | 1 | 6 | 0.8333 |
| 9_2038216 | 2 | 0.5 | 3 | 0 | 3 | 0.6667 | 3 | 0.6667 | 11 | 0.4545 | NA | NA | NA | NA | 4 | 0.25 | 2 | 0 | 6 | 0.1667 |
| 9_2038217 | 2 | 1 | 3 | 0 | 3 | 0 | 3 | 0 | 11 | 0.1818 | NA | NA | NA | NA | 5 | 0.2 | 2 | 0.5 | 7 | 0.2857 |
| 9_2038220 | 2 | 0.5 | 3 | 0 | 3 | 0 | 3 | 0.3333 | 11 | 0.1818 | NA | NA | NA | NA | 5 | 0.4 | 1 | 0 | 6 | 0.3333 |
| 9_2038222 | 2 | 0.5 | 3 | 0.3333 | 2 | 1 | 2 | 0 | 9 | 0.4444 | NA | NA | NA | NA | 5 | 0.4 | 2 | 0 | 7 | 0.2857 |
| 9_2038236 | 1 | 0 | 2 | 0.5 | 2 | 0 | 2 | 1 | 7 | 0.4286 | NA | NA | NA | NA | 5 | 0.2 | 2 | 0 | 7 | 0.1429 |
| 9_2038238 | 2 | 0 | 2 | 0.5 | 2 | 0 | 1 | 0 | 7 | 0.1429 | NA | NA | NA | NA | 3 | 0 | 2 | 0.5 | 5 | 0.2 |
| 9_2038240 | 2 | 0.5 | 3 | 0.6667 | 2 | 1 | 2 | 1 | 9 | 0.7778 | NA | NA | NA | NA | 5 | 0.8 | 2 | 1 | 7 | 0.8571 |
| 9_2038246 | 2 | 0 | 3 | 0.3333 | 2 | 0.5 | 2 | 0 | 9 | 0.2222 | NA | NA | NA | NA | 4 | 0.25 | 2 | 0.5 | 6 | 0.3333 |
| 9_2038247 | 1 | 0 | 3 | 0 | 2 | 0.5 | 2 | 0 | 8 | 0.125 | NA | NA | NA | NA | 4 | 0 | 2 | 0.5 | 6 | 0.1667 |
| 9_2040397 | 1 | 1 | 1 | 0 | NA | NA | 3 | 0 | 5 | 0.2 | NA | NA | NA | NA | 3 | 0 | 2 | 0.5 | 5 | 0.2 |
| 9_2041129 | 1 | 1 | NA | NA | 3 | 1 | 1 | 1 | 5 | 1 | 1 | 1 | 1 | 1 | 2 | 1 | 1 | 1 | 5 | 1 |
| 9_2041139 | 1 | 0 | NA | NA | 3 | 0.6667 | 1 | 1 | 5 | 0.6 | 1 | 0 | 1 | 0 | 2 | 0 | 1 | 1 | 5 | 0.2 |
| 9_2041147 | 1 | 0 | NA | NA | 3 | 1 | NA | NA | 4 | 0.75 | 1 | 1 | 1 | 1 | 2 | 1 | 1 | 1 | 5 | 1 |
| 9_2041153 | NA | NA | NA | NA | 3 | 0.3333 | 1 | 1 | 4 | 0.5 | 1 | 1 | 1 | 0 | 2 | 0.5 | 1 | 1 | 5 | 0.6 |
| 9_2041157 | 1 | 0 | NA | NA | 3 | 1 | 1 | 1 | 5 | 0.8 | 1 | 1 | NA | NA | 2 | 1 | 1 | 1 | 4 | 1 |
| 9_2041161 | NA | NA | NA | NA | 3 | 1 | 1 | 1 | 4 | 1 | 1 | 1 | 1 | 0 | 2 | 1 | NA | NA | 4 | 0.75 |
| 9_2041162 | 1 | 0 | NA | NA | 1 | 0 | 1 | 1 | 3 | 0.3333 | 1 | 1 | NA | NA | 1 | 1 | 1 | 1 | 3 | 1 |
| 9_2050456 | NA | NA | 2 | 0.5 | 1 | 0 | 1 | 0 | 4 | 0.25 | NA | NA | 1 | 1 | 3 | 0.3333 | 3 | 0.3333 | 7 | 0.4286 |
| 9_2050457 | NA | NA | 1 | 1 | 1 | 0 | 1 | 0 | 3 | 0.3333 | NA | NA | 1 | 1 | 4 | 0.5 | 2 | 0.5 | 7 | 0.5714 |

| | | | | | | | | | | | | | | | | | | | | |
|---|---|---|---|---|---|---|---|---|---|---|---|---|---|---|---|---|---|---|---|---|
| 9_2050476; | NA | NA | 2 | 0.5 | 1 | 1 | NA | NA | 3 | 0.6667 | NA | NA | 1 | 1 | 3 | 0.3333 | 2 | 1 | 6 | 0.6667 |
| 9_2050477; | NA | NA | 1 | 1 | 1 | 1 | 1 | 1 | 3 | 1 | NA | NA | 1 | 0 | 3 | 1 | 2 | 1 | 6 | 0.8333 |
| 9_2050484‹ | NA | NA | 2 | 0.5 | NA | NA | 1 | 0 | 3 | 0.3333 | NA | NA | 1 | 0 | 2 | 0 | 3 | 0.3333 | 6 | 0.1667 |
| 9_2050495 | NA | NA | 2 | 0 | 2 | 0 | 1 | 1 | 5 | 0.2 | NA | NA | 1 | 0 | 4 | 0.25 | 3 | 0 | 8 | 0.125 |
| 9_2050504‹ | NA | NA | 2 | 0 | 2 | 0.5 | 1 | 1 | 5 | 0.4 | NA | NA | 1 | 1 | 4 | 0 | 3 | 0.6667 | 8 | 0.375 |
| 9_2050511' | NA | NA | 2 | 0.5 | 1 | 1 | NA | NA | 3 | 0.6667 | NA | NA | 1 | 1 | 3 | 0.6667 | 3 | 1 | 7 | 0.8571 |
| 9_2050512( | NA | NA | 1 | 1 | 1 | 0 | 1 | 0 | 3 | 0.3333 | NA | NA | 1 | 0 | 3 | 0 | 3 | 0.3333 | 7 | 0.1429 |
| 9_2050528; | NA | NA | 1 | 1 | 2 | 0.5 | 1 | 1 | 4 | 0.75 | NA | NA | 1 | 0 | 4 | 0.5 | 2 | 0.5 | 7 | 0.4286 |
| 9_2050545( | NA | NA | 1 | 0 | 1 | 1 | 2 | 1 | 4 | 0.75 | NA | NA | 1 | 0 | 4 | 1 | 2 | 1 | 7 | 0.8571 |
| 9_2050560: | NA | NA | 1 | 1 | 1 | 1 | 2 | 1 | 4 | 1 | NA | NA | 1 | 1 | 4 | 0.75 | 2 | 0.5 | 7 | 0.7143 |
| 9_2050577; | NA | NA | 2 | 0 | 2 | 0 | 1 | 1 | 5 | 0.2 | NA | NA | 1 | 0 | 3 | 0.3333 | 3 | 0 | 7 | 0.1429 |
| 9_2050585! | NA | NA | 2 | 0.5 | 1 | 0 | NA | NA | 3 | 0.3333 | NA | NA | 1 | 1 | 4 | 0.25 | 3 | 0 | 8 | 0.25 |
| 9_2050586' | NA | NA | 2 | 1 | 1 | 1 | 1 | 1 | 4 | 1 | NA | NA | 1 | 1 | 3 | 1 | 3 | 1 | 7 | 1 |
| 9_2050593( | NA | NA | 2 | 1 | 1 | 1 | 2 | 1 | 5 | 1 | NA | NA | NA | NA | 3 | 1 | 3 | 1 | 6 | 1 |
| 9_2050596( | NA | NA | 1 | 1 | 1 | 1 | 2 | 0.5 | 4 | 0.75 | NA | NA | 1 | 1 | 4 | 1 | 2 | 1 | 7 | 1 |
| 9_2050600! | NA | NA | 2 | 0 | 1 | 0 | 2 | 0.5 | 5 | 0.2 | NA | NA | 1 | 1 | 4 | 0.5 | 3 | 0 | 8 | 0.375 |
| 9_2050602( | NA | NA | 1 | 1 | 1 | 0 | 1 | 1 | 3 | 0.6667 | NA | NA | 1 | 0 | 3 | 0.6667 | 3 | 0.6667 | 7 | 0.5714 |
| 9_2050616; | NA | NA | 2 | 0.5 | 1 | 0 | 2 | 0.5 | 5 | 0.4 | NA | NA | 1 | 1 | 2 | 0 | 3 | 0.6667 | 6 | 0.5 |
| 9_2050628( | NA | NA | 1 | 0 | NA | NA | 2 | 0.5 | 3 | 0.3333 | NA | NA | NA | NA | 4 | 0.25 | 3 | 0.6667 | 7 | 0.4286 |
| 9_2050640' | NA | NA | 2 | 0.5 | 1 | 1 | 2 | 0 | 5 | 0.4 | NA | NA | 1 | 0 | 3 | 0.3333 | 3 | 0.6667 | 7 | 0.4286 |
| 9_2052600( | NA | NA | 2 | 0.5 | 1 | 1 | NA | NA | 3 | 0.6667 | NA | NA | 2 | 1 | NA | NA | 1 | 0 | 3 | 0.6667 |
| 9_2052607! | NA | NA | 2 | 0.5 | 1 | 0 | NA | NA | 3 | 0.3333 | NA | NA | 3 | 0.6667 | NA | NA | 1 | 1 | 4 | 0.75 |
| 9_2052612: | NA | NA | 2 | 1 | 1 | 1 | NA | NA | 3 | 1 | NA | NA | 3 | 1 | NA | NA | 1 | 1 | 4 | 1 |
| 9_2052621; | NA | NA | 2 | 0.5 | 2 | 0 | NA | NA | 4 | 0.25 | NA | NA | 1 | 1 | NA | NA | 1 | 1 | 2 | 1 |
| 9_2052648( | NA | NA | 1 | 1 | NA | NA | NA | NA | 1 | 1 | NA | NA | 3 | 1 | NA | NA | 1 | 1 | 4 | 1 |
| 9_2052674! | NA | NA | 1 | 1 | NA | NA | NA | NA | 1 | 1 | NA | NA | 3 | 1 | NA | NA | 1 | 1 | 4 | 1 |
| 9_2052682: | NA | NA | 1 | 1 | NA | NA | NA | NA | 1 | 1 | NA | NA | 2 | 1 | NA | NA | 1 | 1 | 3 | 1 |
| 9_2052683: | NA | NA | 1 | 1 | NA | NA | NA | NA | 1 | 1 | NA | NA | 2 | 1 | NA | NA | 1 | 0 | 3 | 0.6667 |
| 9_2052696. | NA | NA | 1 | 1 | NA | NA | NA | NA | 1 | 1 | NA | NA | 3 | 1 | NA | NA | 1 | 1 | 4 | 1 |
| 9_2052708. | NA | NA | 1 | 1 | NA | NA | NA | NA | 1 | 1 | NA | NA | 2 | 0.5 | NA | NA | 1 | 1 | 3 | 0.6667 |
| 9_2052717' | NA | NA | 1 | 1 | NA | NA | NA | NA | 1 | 1 | NA | NA | 2 | 1 | NA | NA | 1 | 1 | 3 | 1 |
| 9_2054938' | 1 | 0 | 2 | 0 | NA | NA | 3 | 0.3333 | 6 | 0.1667 | 1 | 0 | 1 | 0 | 2 | 0.5 | NA | NA | 4 | 0.25 |
| 9_2054955' | 1 | 0 | 4 | 0 | NA | NA | 4 | 0.25 | 9 | 0.1111 | 1 | 1 | 1 | 0 | 3 | 0 | 1 | 0 | 6 | 0.1667 |
| 9_2055017( | 1 | 0 | 3 | 0.6667 | NA | NA | 2 | 0.5 | 6 | 0.5 | 1 | 1 | 1 | 1 | 2 | 0.5 | 1 | 1 | 5 | 0.8 |
| 9_2055027; | 1 | 0 | 3 | 0 | NA | NA | 2 | 0.5 | 6 | 0.1667 | 1 | 1 | 1 | 0 | 2 | 1 | 1 | 1 | 5 | 0.8 |
| 9_2055356( | 1 | 0 | 2 | 0.5 | NA | NA | NA | NA | 3 | 0.3333 | NA | NA | NA | NA | 1 | 1 | NA | NA | 1 | 1 |
| 9_2055575. | NA | NA | 2 | 0.5 | NA | NA | 1 | 1 | 3 | 0.6667 | 1 | 1 | NA | NA | 1 | 1 | NA | NA | 2 | 1 |
| 9_2057454( | 21 | 0.1905 | 18 | 0 | 17 | 0 | 9 | 0.2222 | 65 | 0.0923 | 19 | 0.1579 | 19 | 0.1053 | 16 | 0.0625 | 15 | 0.1333 | 69 | 0.1159 |
| 9_2057467: | 25 | 0.24 | 14 | 0.0714 | 18 | 0.2222 | 9 | 0.1111 | 66 | 0.1818 | 16 | 0.125 | 18 | 0.1667 | 17 | 0.0588 | 12 | 0.1667 | 63 | 0.127 |
| 9_2057606: | 2 | 0 | NA | NA | NA | NA | 1 | 1 | 3 | 0.3333 | 2 | 0.5 | NA | NA | 3 | 0 | NA | NA | 5 | 0.2 |
| 9_2057893! | NA | NA | 1 | 1 | NA | NA | NA | NA | 1 | 1 | 1 | 0 | NA | NA | 1 | 1 | NA | NA | 2 | 0.5 |
| 9_2057941( | 8 | 0.25 | 15 | 0 | 10 | 0.1 | 10 | 0.1 | 43 | 0.093 | 8 | 0.125 | 8 | 0 | 8 | 0.25 | 10 | 0 | 34 | 0.0882 |
| 9_2057968: | 8 | 0.625 | 13 | 0.6923 | 10 | 0.5 | 10 | 0.2 | 41 | 0.5122 | 3 | 0.3333 | 11 | 0.3636 | 6 | 0.1667 | 9 | 0.4444 | 29 | 0.3448 |
| 9_2057971( | 9 | 0.2222 | 11 | 0.0909 | 9 | 0 | 10 | 0 | 39 | 0.0769 | 4 | 0 | 8 | 0.125 | 6 | 0.1667 | 8 | 0 | 26 | 0.0769 |
| 9_2057979. | 7 | 1 | 9 | 1 | 6 | 1 | 9 | 0.7778 | 31 | 0.9355 | 5 | 1 | 8 | 1 | 5 | 0.8 | 5 | 1 | 23 | 0.9565 |
| 9_2057980: | 8 | 0.875 | 10 | 0.9 | 8 | 0.75 | 9 | 0.4444 | 35 | 0.7429 | 4 | 0.75 | 7 | 0.8571 | 5 | 0.4 | 7 | 0.7143 | 23 | 0.6957 |
| 9_2058000' | 7 | 0.1429 | 13 | 0.1538 | 7 | 0.1429 | 8 | 0.375 | 35 | 0.2 | 3 | 0 | 6 | 0.3333 | 3 | 0.3333 | 7 | 0 | 19 | 0.1579 |
| 9_2063813( | 2 | 0.5 | 3 | 0.6667 | 3 | 0.6667 | 4 | 0.5 | 12 | 0.5833 | 1 | 1 | 3 | 0.6667 | NA | NA | 3 | 1 | 7 | 0.8571 |
| 9_2078133! | NA | NA | 2 | 0.5 | NA | NA | NA | NA | 2 | 0.5 | 3 | 0.6667 | 1 | 0 | 4 | 0.5 | NA | NA | 8 | 0.5 |
| 9_2078137! | NA | NA | 2 | 0.5 | NA | NA | NA | NA | 2 | 0.5 | 2 | 0.5 | 1 | 0 | 4 | 0 | NA | NA | 7 | 0.1429 |
| 9_2078169; | NA | NA | 2 | 0.5 | 1 | 0 | NA | NA | 3 | 0.3333 | 3 | 0.3333 | 1 | 0 | 5 | 0 | NA | NA | 9 | 0.1111 |
| 9_2078218: | 1 | 0 | 3 | 0.3333 | 1 | 0 | NA | NA | 5 | 0.2 | 3 | 0.6667 | 1 | 1 | 4 | 0.25 | NA | NA | 8 | 0.5 |
| 9_2078498 | 1 | 0 | 3 | 0.3333 | 5 | 0.2 | 1 | 0 | 10 | 0.2 | 4 | 0.5 | 8 | 0.125 | 5 | 0.2 | 1 | 0 | 18 | 0.2222 |
| 9_2078500( | 3 | 0.3333 | 10 | 0.4 | 5 | 0.2 | 2 | 0.5 | 20 | 0.35 | 7 | 0.4286 | 9 | 0.5556 | 10 | 0.1 | 1 | 0 | 27 | 0.3333 |
| 9_2078506. | 3 | 0.3333 | 10 | 0.1 | 6 | 0.1667 | 2 | 0 | 21 | 0.1429 | 7 | 0.1429 | 7 | 0.1429 | 11 | 0.2727 | 2 | 0.5 | 27 | 0.2222 |
| 9_2078509; | 2 | 0.5 | 8 | 0.375 | 6 | 0 | 2 | 0.5 | 18 | 0.2778 | 5 | 0 | 9 | 0 | 11 | 0.0909 | 2 | 0.5 | 27 | 0.0741 |
| 9_2078509( | 2 | 0 | 9 | 0.2222 | 6 | 0 | 2 | 0 | 19 | 0.1053 | 8 | 0.125 | 10 | 0 | 11 | 0.0909 | 2 | 0 | 31 | 0.0645 |
| 9_2078515' | 4 | 0 | 7 | 0.1429 | 5 | 0 | 1 | 0 | 17 | 0.0588 | 7 | 0 | 7 | 0 | 10 | 0.3 | 2 | 0.5 | 26 | 0.1538 |
| 9_2078517‹ | 4 | 1 | 9 | 0.8889 | 6 | 0.8333 | 2 | 1 | 21 | 0.9048 | 6 | 0.8333 | 10 | 0.9 | 7 | 1 | 2 | 1 | 25 | 0.92 |
| 9_2078517( | 4 | 0.25 | NA | NA | 1 | 1 | 2 | 0 | 7 | 0.2857 | 2 | 0.5 | 1 | 0 | 1 | 0 | 2 | 0.5 | 6 | 0.3333 |
| 9_2078524( | 4 | 0.75 | 8 | 0.25 | 5 | 0.4 | 1 | 1 | 18 | 0.4444 | 8 | 0.5 | 10 | 0.5 | 11 | 0.2727 | 2 | 1 | 31 | 0.4516 |
| 9_2080331' | 4 | 0 | NA | NA | 3 | 0.3333 | 3 | 0.3333 | 10 | 0.2 | 3 | 0 | 3 | 0 | 2 | 0.5 | 1 | 0 | 9 | 0.1111 |
| 9_2080336; | 4 | 0.75 | NA | NA | 3 | 0.3333 | 4 | 0.75 | 11 | 0.6364 | 2 | 1 | 4 | 0.5 | 2 | 0.5 | 1 | 1 | 9 | 0.6667 |
| 9_2080343! | 4 | 0 | NA | NA | 3 | 0.3333 | 4 | 0 | 11 | 0.0909 | 2 | 0.5 | 2 | 0 | 1 | 0 | 1 | 0 | 6 | 0.1667 |
| 9_2080346. | 2 | 0.5 | NA | NA | 1 | 0 | 2 | 0 | 5 | 0.2 | 2 | 0 | 4 | 0 | 2 | 0 | 1 | 1 | 9 | 0.1111 |

| | | | | | | | | | | | | | | | | | | | | |
|---|---|---|---|---|---|---|---|---|---|---|---|---|---|---|---|---|---|---|---|---|
| 9_2081882 | 5 | 0.2 | 7 | 0 | 8 | 0 | 4 | 0.25 | 24 | 0.0833 | 7 | 0.1429 | 3 | 0 | 3 | 0.3333 | 1 | 0 | 14 | 0.1429 |
| 9_2087765 | NA | NA | NA | NA | NA | NA | 1 | 1 | 1 | 1 | NA | NA | 1 | 1 | NA | NA | NA | NA | 1 | 1 |
| 9_2091988 | 11 | 0.2727 | 16 | 0.0625 | 11 | 0 | 13 | 0 | 51 | 0.0784 | 7 | 0 | 12 | 0.0833 | 17 | 0.1765 | 11 | 0 | 47 | 0.0851 |
| 9_2091992 | 7 | 0.1429 | 15 | 0.1333 | 11 | 0.0909 | 13 | 0.0769 | 46 | 0.1087 | 6 | 0 | 15 | 0 | 19 | 0.2105 | 10 | 0 | 50 | 0.08 |
| 9_2091995 | 9 | 0.2222 | 14 | 0.1429 | 6 | 0 | 10 | 0.1 | 39 | 0.1282 | 6 | 0.1667 | 8 | 0 | 16 | 0.1875 | 8 | 0.125 | 38 | 0.1316 |
| 9_2092003 | 11 | 0.1818 | 11 | 0.0909 | 9 | 0 | 13 | 0.2308 | 44 | 0.1364 | 6 | 0.1667 | 14 | 0.1429 | 18 | 0.2222 | 9 | 0.2222 | 47 | 0.1915 |
| 9_2092012 | 1 | 1 | NA | NA | 1 | 1 | 4 | 0.75 | 6 | 0.8333 | 3 | 0.6667 | 3 | 0.3333 | 3 | 1 | 2 | 0 | 11 | 0.5455 |
| 9_2093940 | 1 | 1 | NA | NA | NA | NA | NA | NA | 1 | 1 | NA | NA | NA | NA | 1 | 1 | NA | NA | 1 | 1 |
| 9_2093993 | 2 | 0.5 | NA | NA | NA | NA | NA | NA | 2 | 0.5 | NA | NA | NA | NA | 1 | 1 | NA | NA | 1 | 1 |
| 9_2094383 | 4 | 0 | 3 | 0 | 3 | 0.3333 | 4 | 0.25 | 14 | 0.1429 | 1 | 1 | NA | NA | NA | NA | 1 | 0 | 2 | 0.5 |
| 9_2094397 | 4 | 0.25 | 3 | 0.6667 | 3 | 0.3333 | 3 | 0 | 13 | 0.3077 | 1 | 1 | NA | NA | NA | NA | 1 | 0 | 2 | 0.5 |
| 9_2094408 | 6 | 0.1667 | 3 | 0 | 3 | 0.3333 | 5 | 0.4 | 17 | 0.2353 | 1 | 1 | 1 | 0 | NA | NA | 1 | 0 | 3 | 0.3333 |
| 9_2094506 | 20 | 0.25 | 11 | 0.1818 | 15 | 0.1333 | 17 | 0.1765 | 63 | 0.1905 | 17 | 0.2353 | 18 | 0.2778 | 15 | 0.1333 | 15 | 0.0667 | 65 | 0.1846 |
| 9_2094589 | 16 | 0.125 | 10 | 0.2 | 17 | 0.0588 | 17 | 0.1176 | 60 | 0.1167 | 16 | 0.125 | 19 | 0.1579 | 20 | 0.1 | 13 | 0.2308 | 68 | 0.1471 |
| 9_2094592 | 20 | 0.75 | 12 | 0.5 | 19 | 0.4737 | 17 | 0.6471 | 68 | 0.6029 | 15 | 0.6 | 16 | 0.625 | 19 | 0.6316 | 13 | 0.4615 | 63 | 0.5873 |
| 9_2094612 | 21 | 0.1429 | 13 | 0 | 22 | 0.0909 | 18 | 0 | 74 | 0.0676 | 17 | 0 | 19 | 0.1579 | 19 | 0.0526 | 17 | 0.0588 | 72 | 0.0694 |
| 9_2094797 | 30 | 0.3333 | 18 | 0.1111 | 25 | 0.12 | 26 | 0.3846 | 99 | 0.2525 | 25 | 0.28 | 27 | 0.2963 | 27 | 0.1852 | 23 | 0.2609 | 102 | 0.2549 |
| 9_2094799 | 29 | 0.5172 | 20 | 0.45 | 23 | 0.2609 | 24 | 0.375 | 96 | 0.4062 | 22 | 0.4545 | 23 | 0.5217 | 29 | 0.4828 | 19 | 0.5789 | 93 | 0.5054 |
| 9_2097884 | 15 | 0.8667 | 11 | 0.7273 | 8 | 1 | 6 | 1 | 40 | 0.875 | 8 | 0.875 | 13 | 0.7692 | 10 | 0.9 | 14 | 0.8571 | 45 | 0.8444 |
| 9_2097886 | 16 | 0.75 | 10 | 0.6 | 7 | 0.8571 | 6 | 0.6667 | 39 | 0.7179 | 6 | 0.6667 | 13 | 0.6923 | 10 | 0.5 | 13 | 0.6154 | 42 | 0.619 |
| 9_2097897 | 14 | 0.3571 | 12 | 0.1667 | 7 | 0.1429 | 6 | 0.5 | 39 | 0.2821 | 7 | 0 | 12 | 0.1667 | 11 | 0.0909 | 11 | 0.1818 | 41 | 0.122 |
| 9_2097899 | 14 | 0.2857 | 9 | 0.4444 | 7 | 0.5714 | 5 | 0.4 | 35 | 0.4 | 7 | 0.5714 | 9 | 0.5556 | 11 | 0.5455 | 12 | 0.4167 | 39 | 0.5128 |
| 9_2097903 | 12 | 0.5 | 11 | 0.1818 | 4 | 0.5 | 5 | 0.6 | 32 | 0.4062 | 6 | 0.3333 | 13 | 0.2308 | 12 | 0.0833 | 12 | 0.25 | 43 | 0.2093 |
| 9_2097905 | 11 | 0.5455 | 10 | 0.2 | 6 | 0 | 4 | 0 | 31 | 0.2581 | 6 | 0.1667 | 7 | 0.2857 | 8 | 0.625 | 10 | 0.3 | 31 | 0.3548 |
| 9_2098584 | 1 | 0 | 5 | 0.4 | 4 | 0.5 | 4 | 0.25 | 14 | 0.3571 | 3 | 0 | 10 | 0.5 | 6 | 0.1667 | 4 | 0 | 23 | 0.2609 |
| 9_2098625 | 2 | 0.5 | 6 | 0.3333 | 6 | 0.5 | 4 | 0 | 18 | 0.3333 | 3 | 0.3333 | 8 | 0.25 | 6 | 0 | 6 | 0.1667 | 23 | 0.1739 |
| 9_2098650 | 2 | 1 | 6 | 0.8333 | 5 | 1 | 3 | 0.6667 | 16 | 0.875 | 2 | 1 | 7 | 0.7143 | 5 | 0.8 | 6 | 1 | 20 | 0.85 |
| 9_2099078 | 1 | 0 | 3 | 0.3333 | 4 | 0 | 2 | 0 | 10 | 0.1 | 1 | 1 | 5 | 0 | 5 | 0 | 4 | 0 | 15 | 0.0667 |
| 9_2099767 | NA | NA | 3 | 0.6667 | 2 | 1 | 1 | 1 | 6 | 0.8333 | 2 | 1 | 2 | 0 | 2 | 0.5 | 3 | 0.6667 | 9 | 0.5556 |
| 9_2099772 | 1 | 0 | 3 | 0 | 1 | 1 | 1 | 0 | 6 | 0.1667 | 2 | 0 | 2 | 0 | 4 | 0.5 | 3 | 0 | 11 | 0.1818 |
| 9_2099774 | 1 | 1 | 2 | 1 | 1 | 1 | 1 | 1 | 5 | 1 | 2 | 1 | 1 | 0 | 4 | 0.25 | 3 | 0.6667 | 10 | 0.5 |
| 9_2101621 | 1 | 1 | 2 | 0 | 2 | 0.5 | 1 | 0 | 6 | 0.3333 | 2 | 0.5 | 1 | 1 | 1 | 1 | 1 | 0 | 5 | 0.6 |
| 9_2105097 | 30 | 0.3 | 31 | 0.3226 | 19 | 0.3158 | 18 | 0.2222 | 98 | 0.2959 | 19 | 0.2105 | 19 | 0.4737 | 21 | 0.4286 | 19 | 0.5263 | 78 | 0.4103 |
| 9_2107717 | NA | NA | 2 | 0.5 | 1 | 0 | NA | NA | 3 | 0.3333 | NA | NA | 1 | 1 | 2 | 0 | NA | NA | 3 | 0.3333 |
| 9_2107737 | NA | NA | 1 | 0 | 1 | 1 | NA | NA | 2 | 0.5 | 2 | 0 | NA | NA | 3 | 0.3333 | NA | NA | 5 | 0.2 |
| 9_2111751 | 6 | 0.1667 | 10 | 0.3 | 12 | 0.0833 | 3 | 0 | 31 | 0.1613 | 6 | 0.1667 | 4 | 0.25 | 9 | 0 | 3 | 0.3333 | 22 | 0.1364 |
| 9_2112226 | 6 | 0.1667 | 17 | 0.3529 | 9 | 0.1111 | 3 | 0.3333 | 35 | 0.2571 | 8 | 0.75 | 5 | 0.2 | 6 | 0.1667 | 5 | 0 | 24 | 0.3333 |
| 9_2112230 | 2 | 0.5 | 4 | 0.75 | 4 | 0.25 | 2 | 1 | 12 | 0.5833 | 2 | 0.5 | 3 | 1 | 2 | 1 | 2 | 0.5 | 9 | 0.7778 |
| 9_2112243 | 6 | 0 | 17 | 0.0588 | 13 | 0.0769 | 5 | 0.2 | 41 | 0.0732 | 7 | 0 | 5 | 0 | 9 | 0.3333 | 5 | 0 | 26 | 0.1154 |
| 9_2112244 | 7 | 0.5714 | 16 | 0.5 | 11 | 0.5455 | 4 | 1 | 38 | 0.5789 | 7 | 0.7143 | 5 | 0.6 | 6 | 0.5 | 3 | 1 | 21 | 0.6667 |
| 9_2112245 | 4 | 1 | 18 | 0.8889 | 12 | 0.9167 | 5 | 1 | 39 | 0.9231 | 6 | 1 | 4 | 1 | 9 | 1 | 4 | 0.5 | 23 | 0.913 |
| 9_2112253 | 6 | 1 | 18 | 0.8889 | 14 | 0.8571 | 6 | 0.6667 | 44 | 0.8636 | 9 | 0.7778 | 6 | 0.8333 | 10 | 0.9 | 4 | 0.5 | 29 | 0.7931 |
| 9_2112257 | 5 | 0.8 | 18 | 0.7222 | 13 | 0.6923 | 5 | 1 | 41 | 0.7561 | 7 | 0.7143 | 6 | 1 | 11 | 0.8182 | 4 | 1 | 28 | 0.8571 |
| 9_2112258 | 5 | 0.2 | 21 | 0.5238 | 14 | 0.2857 | 5 | 0.2 | 45 | 0.3778 | 10 | 0.3 | 4 | 0 | 11 | 0.3636 | 5 | 0 | 30 | 0.2333 |
| 9_2112283 | 7 | 0.5714 | 19 | 0.2105 | 13 | 0 | 6 | 0.3333 | 45 | 0.2222 | 6 | 0.6667 | 6 | 0.5 | 11 | 0.4545 | 5 | 0.6 | 28 | 0.5357 |
| 9_2112304 | 7 | 0.8571 | 22 | 0.7727 | 11 | 0.3636 | 5 | 0.4 | 45 | 0.6444 | 11 | 0.6364 | 8 | 0.75 | 12 | 0.75 | 10 | 0.5 | 41 | 0.6585 |
| 9_2112330 | 7 | 1 | 22 | 0.9545 | 14 | 1 | 6 | 0.8333 | 49 | 0.9592 | 9 | 0.8889 | 9 | 0.7778 | 11 | 0.9091 | 9 | 0.7778 | 38 | 0.8421 |
| 9_2112413 | 6 | 0.1667 | 16 | 0.0625 | 10 | 0.1 | 3 | 0 | 35 | 0.0857 | 6 | 0.1667 | 7 | 0.1429 | 7 | 0.1429 | 8 | 0 | 28 | 0.1071 |
| 9_2112435 | 7 | 0.5714 | 23 | 0.5652 | 9 | 0.3333 | 5 | 0.8 | 44 | 0.5455 | 11 | 0.5455 | 7 | 0.4286 | 11 | 0.4545 | 12 | 0.6667 | 41 | 0.5366 |
| 9_2114103 | 5 | 0 | 8 | 0.75 | 6 | 0.5 | 5 | 0.6 | 24 | 0.5 | 8 | 0.375 | 2 | 0.5 | 4 | 0.25 | 5 | 0.4 | 19 | 0.3684 |
| 9_2114155 | 10 | 0.4 | 12 | 0.5833 | 13 | 0.2308 | 7 | 0.1429 | 42 | 0.3571 | 6 | 0.5 | 6 | 0.5 | 6 | 0.8333 | 11 | 0.3636 | 29 | 0.5172 |
| 9_2114156 | 7 | 0.2857 | 11 | 0.3636 | 11 | 0.6364 | 5 | 0.2 | 34 | 0.4118 | 9 | 0.5556 | 6 | 0.3333 | 4 | 0 | 9 | 0.5556 | 28 | 0.4286 |
| 9_2114160 | 8 | 0.125 | 9 | 0.2222 | 8 | 0.125 | 5 | 0.6 | 30 | 0.2333 | 5 | 0.2 | 6 | 0.5 | 5 | 0.8 | 8 | 0.25 | 24 | 0.4167 |
| 9_2114161 | 8 | 0.5 | 8 | 0.25 | 10 | 0.2 | 6 | 0.1667 | 32 | 0.2812 | 6 | 0.6667 | 6 | 0.3333 | 5 | 0.4 | 10 | 0.3 | 27 | 0.4074 |
| 9_2114168 | 4 | 0 | 8 | 0 | 7 | 0 | 6 | 0.3333 | 25 | 0.08 | 4 | 0 | 6 | 0.3333 | 4 | 0 | 8 | 0.125 | 22 | 0.1364 |
| 9_2114170 | 9 | 0 | 8 | 0.25 | 9 | 0 | 5 | 0.2 | 31 | 0.0968 | 5 | 0.2 | 4 | 0 | 3 | 0.3333 | 7 | 0.2857 | 19 | 0.2105 |
| 9_2114180 | 8 | 0.75 | 7 | 0.8571 | 7 | 1 | 5 | 0.6 | 27 | 0.8148 | 3 | 1 | 6 | 0.8333 | 3 | 0.6667 | 6 | 0.3333 | 18 | 0.6667 |
| 9_2114181 | 9 | 1 | 7 | 1 | 8 | 0.875 | 7 | 1 | 31 | 0.9677 | 5 | 1 | 6 | 0.8333 | 4 | 1 | 9 | 0.8889 | 24 | 0.9167 |
| 9_2114195 | 7 | 0.1429 | 6 | 0 | 4 | 0.25 | 6 | 0 | 23 | 0.087 | 3 | 0.3333 | 5 | 0.4 | 4 | 0 | 5 | 0.2 | 17 | 0.2353 |
| 9_2114196 | 9 | 0.7778 | 6 | 0.5 | 8 | 0.75 | 6 | 0.6667 | 29 | 0.6897 | 4 | 1 | 5 | 0.6 | 3 | 0.6667 | 7 | 0.4286 | 19 | 0.6316 |
| 9_2114201 | 7 | 0.7143 | 7 | 0.2857 | 7 | 0.7143 | 5 | 0.2 | 26 | 0.5 | 5 | 0.6 | 3 | 0.6667 | 2 | 0.5 | 6 | 0 | 16 | 0.375 |
| 9_2114211 | 9 | 0.3333 | 6 | 0.3333 | 6 | 0.1667 | 6 | 0.5 | 27 | 0.3333 | 4 | 0 | 5 | 0.2 | 2 | 1 | 4 | 0 | 15 | 0.2 |
| 9_2114213 | 7 | 0.1429 | 5 | 0 | 8 | 0.25 | 5 | 0.2 | 25 | 0.16 | 4 | 0 | 4 | 0 | 2 | 0 | 5 | 0.2 | 15 | 0.0667 |
| 9_2114482 | 6 | 1 | 7 | 0.7143 | 5 | 1 | 3 | 0.6667 | 21 | 0.8571 | 3 | 1 | 2 | 1 | 1 | 1 | 6 | 1 | 12 | 1 |
| 9_2114484 | 6 | 0.3333 | 6 | 0.1667 | 4 | 0 | 3 | 0 | 19 | 0.1579 | 4 | 0 | 2 | 0.5 | 1 | 0 | 5 | 0 | 12 | 0.0833 |

| | | | | | | | | | | | | | | | | | | | | |
|---|---|---|---|---|---|---|---|---|---|---|---|---|---|---|---|---|---|---|---|---|
| 9_21145113 | 4 | 1 | 6 | 1 | 4 | 1 | 3 | 0.6667 | 17 | 0.9412 | 3 | 1 | 2 | 1 | 2 | 1 | 6 | 1 | 13 | 1 |
| 9_21148746 | 4 | 1 | 6 | 1 | 1 | 1 | 3 | 1 | 14 | 1 | 1 | 1 | 1 | 1 | 1 | 1 | 3 | 1 | 6 | 1 |
| 9_21154587 | NA | NA | 4 | 0.5 | 2 | 1 | 2 | 0.5 | 8 | 0.625 | NA | NA | 2 | 0 | 2 | 0.5 | NA | NA | 4 | 0.25 |
| 9_21154624 | NA | NA | 4 | 0.75 | 3 | 0.6667 | 3 | 1 | 10 | 0.8 | NA | NA | 2 | 0.5 | 2 | 0.5 | NA | NA | 4 | 0.5 |
| 9_21154662 | NA | NA | 3 | 0.3333 | 2 | 0 | 2 | 0 | 7 | 0.1429 | NA | NA | 2 | 0.5 | 2 | 0 | NA | NA | 4 | 0.25 |
| 9_21154811 | NA | NA | 4 | 0 | 3 | 0.3333 | 3 | 0 | 10 | 0.1 | NA | NA | 2 | 0.5 | 2 | 0 | NA | NA | 4 | 0.25 |
| 9_21154847 | NA | NA | 3 | 1 | 3 | 1 | 2 | 1 | 8 | 1 | NA | NA | 2 | 1 | 2 | 0.5 | NA | NA | 4 | 0.75 |
| 9_21155063 | NA | NA | 2 | 1 | 2 | 1 | 3 | 1 | 7 | 1 | NA | NA | 1 | 1 | 2 | 1 | NA | NA | 3 | 1 |
| 9_21155091 | NA | NA | 3 | 0.3333 | 3 | 0 | 3 | 0 | 9 | 0.1111 | 1 | 0 | 2 | 0.5 | 1 | 0 | NA | NA | 4 | 0.25 |
| 9_21155140 | NA | NA | 1 | 0 | 3 | 0.3333 | 2 | 0 | 6 | 0.1667 | NA | NA | 2 | 0.5 | 1 | 0 | NA | NA | 3 | 0.3333 |
| 9_21155195 | NA | NA | 1 | 0 | 3 | 1 | 2 | 1 | 6 | 0.8333 | NA | NA | 2 | 0.5 | 1 | 1 | NA | NA | 3 | 0.6667 |
| 9_21155226 | NA | NA | 1 | 1 | 3 | 1 | 2 | 0.5 | 6 | 0.8333 | NA | NA | 2 | 0.5 | 1 | 1 | NA | NA | 3 | 0.6667 |
| 9_21155295 | NA | NA | 2 | 0.5 | 3 | 0 | 3 | 0 | 8 | 0.125 | NA | NA | 2 | 0 | 2 | 0.5 | NA | NA | 4 | 0.25 |
| 9_21155430 | NA | NA | 2 | 0.5 | 2 | 1 | 3 | 0.3333 | 7 | 0.5714 | NA | NA | 1 | 0 | 2 | 1 | NA | NA | 3 | 0.6667 |
| 9_21155627 | NA | NA | 2 | 0 | 3 | 1 | 2 | 0 | 7 | 0.4286 | NA | NA | 1 | 1 | 2 | 0.5 | NA | NA | 3 | 0.6667 |
| 9_21155904 | NA | NA | 2 | 1 | 3 | 1 | 3 | 0 | 8 | 0.625 | NA | NA | 1 | 0 | 2 | 1 | NA | NA | 3 | 0.6667 |
| 9_21156002 | NA | NA | 2 | 1 | 3 | 1 | 1 | 0 | 6 | 0.8333 | NA | NA | NA | NA | 1 | 1 | NA | NA | 1 | 1 |
| 9_21156180 | NA | NA | 2 | 1 | 2 | 1 | NA | NA | 4 | 1 | NA | NA | 1 | 1 | 1 | 1 | NA | NA | 2 | 1 |
| 9_21156264 | NA | NA | 1 | 1 | 2 | 1 | NA | NA | 3 | 1 | NA | NA | NA | NA | 1 | 1 | NA | NA | 1 | 1 |
| 9_21184295 | NA | NA | 2 | 0 | 3 | 0.6667 | NA | NA | 5 | 0.4 | NA | NA | 1 | 1 | 1 | 0 | NA | NA | 2 | 0.5 |
| 9_21184716 | 6 | 0.8333 | 7 | 1 | 8 | 1 | 7 | 1 | 28 | 0.9643 | 1 | 1 | 5 | 1 | 3 | 1 | 4 | 1 | 13 | 1 |
| 9_21184830 | 3 | 0 | 3 | 0.3333 | 2 | 0 | 1 | 0 | 9 | 0.1111 | 3 | 0 | 4 | 0.25 | NA | NA | 4 | 0.25 | 11 | 0.1818 |
| 9_21184838 | 4 | 0 | 3 | 0 | 2 | 0 | 1 | 1 | 10 | 0.1 | 2 | 0 | 3 | 0.3333 | 1 | 0 | 2 | 0 | 8 | 0.125 |
| 9_21184953 | 3 | 1 | 3 | 0.6667 | NA | NA | 1 | 1 | 7 | 0.8571 | 3 | 1 | 1 | 0 | NA | NA | 3 | 0.3333 | 12 | 0.8333 |
| 9_21184953 | 3 | 1 | 3 | 0.6667 | NA | NA | 1 | 1 | 7 | 0.8571 | 3 | 1 | 6 | 1 | NA | NA | 3 | 0.3333 | 12 | 0.8333 |
| 9_21184979 | 4 | 0.25 | 3 | 0 | 3 | 0.3333 | 1 | 0 | 11 | 0.1818 | 2 | 0 | 4 | 0.5 | 1 | 0 | 4 | 0 | 11 | 0.1818 |
| 9_21184987 | 4 | 1 | 3 | 1 | 3 | 1 | 1 | 1 | 11 | 1 | 2 | 0.5 | 6 | 1 | 1 | 1 | 6 | 1 | 15 | 0.9333 |
| 9_21184996 | 3 | 1 | 2 | 0.5 | 1 | 1 | 1 | 1 | 7 | 0.8571 | 2 | 1 | 5 | 1 | 1 | 1 | 3 | 0.6667 | 11 | 0.9091 |
| 9_21185001 | 3 | 0.3333 | 3 | 0.6667 | 2 | 0 | 1 | 1 | 9 | 0.4444 | 2 | 0 | 5 | 0 | NA | NA | 5 | 0.4 | 12 | 0.1667 |
| 9_21185011 | 5 | 0.2 | 6 | 0 | 7 | 0.2857 | 7 | 0 | 25 | 0.12 | 2 | 0 | 3 | 0 | 3 | 0 | 6 | 0.1667 | 14 | 0.0714 |
| 9_21185015 | 3 | 0.3333 | 7 | 0.2857 | 7 | 0.2857 | 7 | 0.1429 | 24 | 0.25 | 2 | 0.5 | 3 | 0.6667 | 3 | 0.3333 | 6 | 0 | 14 | 0.2857 |
| 9_21185074 | 3 | 0.3333 | 3 | 0.3333 | 2 | 0 | 1 | 0 | 9 | 0.2222 | 3 | 0 | 5 | 0.2 | 1 | 0 | 4 | 0.25 | 13 | 0.1538 |
| 9_21200076 | 6 | 0.3333 | 6 | 0.3333 | 6 | 0.1667 | 7 | 0.1429 | 25 | 0.24 | 13 | 0.4615 | 11 | 0.4545 | 9 | 0.2222 | 12 | 0.1667 | 45 | 0.3333 |
| 9_21200212 | 8 | 0.75 | 10 | 0.5 | 8 | 0.25 | 6 | 0.6667 | 32 | 0.5312 | 15 | 0.4667 | 11 | 0.4545 | 11 | 0.3636 | 10 | 0.4 | 47 | 0.4255 |
| 9_21200303 | 7 | 1 | 11 | 1 | 5 | 1 | 7 | 1 | 30 | 1 | 17 | 1 | 10 | 1 | 12 | 0.9167 | 13 | 1 | 52 | 0.9808 |
| 9_21200459 | 3 | 0.3333 | 6 | 0.8333 | 8 | 0.125 | 4 | 0.25 | 21 | 0.381 | 8 | 0.5 | 5 | 0.4 | 7 | 0.4286 | 8 | 0.5 | 28 | 0.4643 |
| 9_21265045 | 7 | 0.1429 | 4 | 0.5 | 3 | 0.3333 | 5 | 0 | 19 | 0.2105 | 4 | 0.25 | 3 | 0 | 3 | 0 | 2 | 0 | 12 | 0.0833 |
| 9_21265165 | 7 | 0.4286 | 4 | 0.75 | 2 | 0.5 | 6 | 1 | 19 | 0.6842 | 4 | 0.75 | 3 | 0.3333 | 3 | 0.3333 | 1 | 1 | 11 | 0.5455 |
| 9_21265230 | 4 | 1 | 3 | 1 | 2 | 1 | 6 | 0.8333 | 15 | 0.9333 | 3 | 1 | 3 | 1 | 3 | 0.6667 | 2 | 1 | 11 | 0.9091 |
| 9_21265296 | 5 | 1 | 4 | 0.25 | 2 | 1 | 3 | 0.6667 | 14 | 0.7143 | 4 | 0.5 | 3 | 0.3333 | 2 | 0 | 2 | 1 | 11 | 0.4545 |
| 9_21265439 | 6 | 1 | 2 | 0.5 | 2 | 0.5 | 5 | 1 | 15 | 0.8667 | 4 | 0.75 | 3 | 1 | 3 | 0.6667 | 3 | 0.6667 | 13 | 0.7692 |
| 9_21265471 | 6 | 0.5 | 4 | 0 | 2 | 0.5 | 5 | 0.4 | 17 | 0.3529 | 4 | 0.25 | NA | NA | 2 | 0.5 | 2 | 0 | 8 | 0.25 |
| 9_21265498 | 7 | 0.1429 | 4 | 0 | 2 | 0.5 | 6 | 0 | 19 | 0.1053 | 4 | 0.25 | 3 | 0 | 4 | 0.25 | 2 | 0 | 13 | 0.1538 |
| 9_21265579 | 7 | 0.1429 | 4 | 0 | 3 | 0.3333 | 5 | 0 | 19 | 0.1053 | 3 | 0.3333 | 3 | 0 | 3 | 0 | 2 | 0 | 11 | 0.0909 |
| 9_21265621 | 8 | 0.75 | 4 | 0.5 | 1 | 1 | 5 | 0.6 | 18 | 0.6667 | 3 | 0.3333 | 3 | 0.6667 | 3 | 1 | 1 | 0 | 10 | 0.6 |
| 9_21265710 | 6 | 0.5 | 4 | 0.25 | 1 | 1 | 6 | 0.5 | 17 | 0.4706 | 4 | 0.25 | 3 | 0.3333 | 3 | 0.3333 | 3 | 0.6667 | 13 | 0.3846 |
| 9_21265736 | 8 | 0 | 4 | 0.5 | 2 | 0 | 6 | 0.1667 | 20 | 0.15 | 4 | 0 | 3 | 0.6667 | 4 | 0 | 3 | 0 | 14 | 0.1429 |
| 9_21265847 | 8 | 0.25 | 4 | 0.75 | 2 | 0.5 | 5 | 1 | 19 | 0.5789 | 3 | 0.6667 | 3 | 0.6667 | 2 | 0.5 | 3 | 0.3333 | 11 | 0.5455 |
| 9_21265854 | 7 | 0.1429 | 4 | 0 | 3 | 0 | 7 | 0.4286 | 21 | 0.1905 | 4 | 0.25 | 3 | 0.3333 | 4 | 0.25 | 3 | 0.3333 | 14 | 0.2857 |
| 9_21265869 | 7 | 0.4286 | 3 | 0 | 3 | 0.3333 | 7 | 0.1429 | 20 | 0.25 | 3 | 0.3333 | 3 | 0 | 4 | 0 | 3 | 0 | 13 | 0.0769 |
| 9_21265962 | 8 | 0.125 | 4 | 0.5 | 2 | 0 | 6 | 0.1667 | 20 | 0.2 | 5 | 0.4 | 2 | 0 | 5 | 0 | 4 | 0 | 16 | 0.125 |
| 9_21265993 | 8 | 0.625 | 4 | 1 | 2 | 0.5 | 7 | 0.5714 | 21 | 0.6667 | 4 | 1 | 2 | 1 | 5 | 0.6 | 2 | 0.5 | 13 | 0.7692 |
| 9_21311933 | 1 | 1 | NA | NA | 1 | 1 | 2 | 0 | 4 | 0.5 | 2 | 0 | 3 | 0 | 2 | 0.5 | NA | NA | 7 | 0.1429 |
| 9_21312158 | 1 | 0 | 1 | 1 | NA | NA | 2 | 0 | 4 | 0.25 | 1 | 0 | 3 | 0.3333 | 2 | 0 | NA | NA | 6 | 0.1667 |
| 9_21312354 | 1 | 1 | 1 | 0 | NA | NA | 2 | 0.5 | 4 | 0.5 | 1 | 1 | 3 | 0.6667 | 3 | 0.6667 | NA | NA | 7 | 0.7143 |
| 9_21312361 | NA | NA | 1 | 1 | NA | NA | NA | NA | 1 | 1 | 2 | 1 | 3 | 0.6667 | 3 | 0.6667 | NA | NA | 8 | 0.75 |
| 9_21316112 | 6 | 0 | 2 | 1 | 3 | 0 | 12 | 0.25 | 23 | 0.2174 | 1 | 1 | 7 | 0.1429 | 4 | 0 | 6 | 0.6667 | 18 | 0.3333 |
| 9_21316289 | 8 | 0.375 | 4 | 0.75 | 8 | 0.5 | 16 | 0.25 | 36 | 0.3889 | 3 | 0 | 9 | 0.2222 | 6 | 0.1667 | 8 | 0.5 | 26 | 0.2692 |
| 9_21316324 | 5 | 1 | 3 | 1 | 7 | 0.8571 | 13 | 0.7692 | 28 | 0.8571 | 1 | 1 | 10 | 0.9 | 8 | 1 | 8 | 0.875 | 27 | 0.9259 |
| 9_21331502 | 3 | 0.3333 | 1 | 1 | 2 | 1 | 2 | 1 | 8 | 0.75 | 2 | 0.5 | 1 | 0 | NA | NA | NA | NA | 3 | 0.3333 |
| 9_21417749 | 7 | 0 | 2 | 0 | 5 | 0.2 | 4 | 0.5 | 18 | 0.1667 | 5 | 0.2 | 5 | 0.6 | 3 | 0 | 2 | 0 | 15 | 0.2667 |
| 9_21417844 | 4 | 0.25 | 3 | 0.3333 | 3 | 0.3333 | 5 | 0.4 | 15 | 0.3333 | 3 | 0.3333 | 7 | 0.2857 | 2 | 0 | 1 | 0 | 13 | 0.2308 |
| 9_21417935 | 7 | 0 | 3 | 0.6667 | 6 | 0.1667 | 4 | 0.25 | 20 | 0.2 | 5 | 0.2 | 6 | 0.1667 | 3 | 0 | 2 | 0 | 16 | 0.125 |
| 9_21417986 | 7 | 0.2857 | 3 | 0.3333 | 6 | 0.1667 | 7 | 0.2857 | 23 | 0.2609 | 4 | 0 | 8 | 0.25 | 1 | 0 | 1 | 0 | 14 | 0.1429 |
| 9_21418034 | 7 | 0.1429 | 3 | 0.3333 | 6 | 0.1667 | 6 | 0.1667 | 22 | 0.1818 | 6 | 0.3333 | 9 | 0.1111 | 3 | 0 | 2 | 0 | 20 | 0.15 |

| | | | | | | | | | | | | | | | | | | | | |
|---|---|---|---|---|---|---|---|---|---|---|---|---|---|---|---|---|---|---|---|---|
| 9_2141805 | 5 | 0.6 | 4 | 0.5 | 5 | 0.6 | 6 | 0.3333 | 20 | 0.5 | 6 | 0 | 7 | 0.4286 | 3 | 0 | 1 | 1 | 17 | 0.2353 |
| 9_2141826 | 6 | 0.3333 | 3 | 0.3333 | 6 | 0.8333 | 8 | 0.375 | 23 | 0.4783 | 3 | 0 | 7 | 0.5714 | 3 | 0 | 2 | 0.5 | 15 | 0.3333 |
| 9_2141904 | 1 | 0 | NA | NA | 4 | 0.5 | 1 | 0 | 6 | 0.3333 | 2 | 0.5 | 1 | 1 | 3 | 0.3333 | 2 | 0.5 | 8 | 0.5 |
| 9_2141940 | 1 | 0 | NA | NA | 6 | 0.3333 | 1 | 0 | 8 | 0.25 | 1 | 0 | 3 | 0.3333 | 3 | 0.3333 | 2 | 0 | 9 | 0.2222 |
| 9_2141941 | 1 | 0 | NA | NA | 6 | 0.5 | 1 | 0 | 8 | 0.375 | 1 | 0 | 3 | 0.3333 | 3 | 0.6667 | 2 | 0.5 | 9 | 0.4444 |
| 9_2141942 | 1 | 0 | NA | NA | 4 | 0.25 | 1 | 0 | 6 | 0.1667 | 1 | 0 | 3 | 0.6667 | 3 | 1 | 2 | 0 | 9 | 0.5556 |
| 9_2149962 | 8 | 0.5 | 4 | 0.5 | 3 | 1 | 4 | 0.75 | 19 | 0.6316 | 6 | 0.8333 | 6 | 1 | 7 | 0.7143 | 3 | 0.3333 | 22 | 0.7727 |
| 9_2149962 | 9 | 0.8889 | 6 | 1 | 4 | 1 | 7 | 1 | 26 | 0.9615 | 7 | 1 | 7 | 1 | 11 | 1 | 2 | 1 | 27 | 1 |
| 9_2149993 | 11 | 0.4545 | 6 | 0.3333 | 3 | 0.3333 | 5 | 0.2 | 25 | 0.36 | 7 | 0.1429 | 8 | 0.25 | 8 | 0.25 | 4 | 0.5 | 27 | 0.2593 |
| 9_2150019 | 2 | 0.5 | 4 | 0 | 1 | 0 | 1 | 0 | 8 | 0.125 | 1 | 1 | NA | NA | NA | NA | NA | NA | 1 | 1 |
| 9_2150026 | 4 | 0 | 1 | 0 | 5 | 0.2 | 5 | 0 | 15 | 0.0667 | 7 | 0 | 6 | 0.1667 | 2 | 0 | NA | NA | 15 | 0.0667 |
| 9_2150066 | 2 | 0.5 | 2 | 0.5 | 2 | 1 | NA | NA | 6 | 0.6667 | NA | NA | NA | NA | 1 | 1 | 2 | 0.5 | 3 | 0.6667 |
| 9_2150079 | 3 | 0.3333 | 1 | 1 | 1 | 0 | NA | NA | 5 | 0.4 | NA | NA | NA | NA | 1 | 1 | 3 | 0 | 4 | 0.25 |
| 9_2150394 | NA | NA | NA | NA | 1 | 1 | NA | NA | 1 | 1 | NA | NA | 1 | 1 | NA | NA | NA | NA | 1 | 1 |
| 9_2150447 | 2 | 0 | 4 | 0.25 | 2 | 0.5 | 4 | 0.5 | 12 | 0.3333 | NA | NA | 7 | 0.2857 | 3 | 0 | 4 | 0 | 14 | 0.1429 |
| 9_2150472 | 4 | 0 | 6 | 0.1667 | 1 | 0 | 8 | 0.25 | 19 | 0.1579 | NA | NA | 10 | 0.2 | 4 | 0 | 5 | 0 | 19 | 0.1053 |
| 9_2150482 | 5 | 0.6 | 6 | 0.8333 | 2 | 1 | 7 | 1 | 20 | 0.85 | 1 | 1 | 8 | 0.75 | 4 | 0.75 | 5 | 1 | 18 | 0.8333 |
| 9_2150517 | 7 | 0.7143 | 6 | 0.6667 | 3 | 0.6667 | 9 | 0.3333 | 25 | 0.56 | 2 | 0.5 | 14 | 0.5714 | 3 | 0 | 5 | 0.6 | 24 | 0.5 |
| 9_2150527 | 9 | 0.5556 | 7 | 0.4286 | 3 | 0.3333 | 7 | 0.4286 | 26 | 0.4615 | 2 | 0.5 | 14 | 0.5714 | 4 | 0.25 | 5 | 0.6 | 25 | 0.52 |
| 9_2150531 | 9 | 1 | 6 | 0.8333 | 3 | 1 | 9 | 1 | 27 | 0.963 | 2 | 1 | 11 | 0.9091 | 5 | 0.8 | 6 | 0.8333 | 24 | 0.875 |
| 9_2150565 | 11 | 0.2727 | 7 | 0 | 5 | 0.4 | 9 | 0 | 32 | 0.1562 | 2 | 0.5 | 14 | 0 | 5 | 0 | 8 | 0.25 | 29 | 0.1034 |
| 9_2150590 | 9 | 0 | 8 | 0 | 5 | 0.2 | 9 | 0.1111 | 31 | 0.0645 | 2 | 0 | 15 | 0.0667 | 8 | 0 | 7 | 0.1429 | 32 | 0.0625 |
| 9_2150596 | 11 | 0.8182 | 8 | 0.625 | 3 | 1 | 6 | 0.5 | 28 | 0.7143 | 2 | 0.5 | 12 | 0.5833 | 7 | 0.5714 | 5 | 1 | 26 | 0.6538 |
| 9_2150608 | 12 | 0.1667 | 9 | 0.3333 | 4 | 0 | 9 | 0 | 34 | 0.1471 | 2 | 0.5 | 12 | 0.3333 | 8 | 0.25 | 6 | 0.3333 | 28 | 0.3214 |
| 9_2150641 | 12 | 0.25 | 9 | 0.1111 | 5 | 0.2 | 10 | 0.1 | 36 | 0.1667 | 2 | 0 | 14 | 0 | 7 | 0.2857 | 8 | 0 | 31 | 0.0645 |
| 9_2159709 | 1 | 0 | NA | NA | 2 | 0.5 | NA | NA | 3 | 0.3333 | 1 | 0 | 1 | 0 | 1 | 1 | 1 | 0 | 4 | 0.25 |
| 9_2161436 | 18 | 0.3889 | 14 | 0.2857 | 13 | 0.1538 | 17 | 0.0588 | 62 | 0.2258 | 15 | 0.1333 | 9 | 0.1111 | 18 | 0.3333 | 13 | 0.3077 | 55 | 0.2364 |
| 9_2165968 | 1 | 0 | NA | NA | 2 | 0.5 | 1 | 1 | 4 | 0.5 | NA | NA | NA | NA | 1 | 1 | NA | NA | 1 | 1 |
| 9_2165985 | 1 | 1 | NA | NA | 1 | 1 | NA | NA | 2 | 1 | NA | NA | NA | NA | 2 | 1 | NA | NA | 2 | 1 |
| 9_2165992 | 1 | 1 | NA | NA | 2 | 1 | 1 | 1 | 4 | 1 | NA | NA | NA | NA | 2 | 1 | 1 | 1 | 3 | 1 |
| 9_2165999 | NA | NA | NA | NA | 1 | 1 | 1 | 1 | 2 | 1 | NA | NA | NA | NA | 2 | 1 | 1 | 1 | 3 | 1 |
| 9_2166009 | 1 | 1 | NA | NA | 3 | 0 | NA | NA | 4 | 0.25 | NA | NA | NA | NA | 2 | 0.5 | 1 | 0 | 3 | 0.3333 |
| 9_2166035 | NA | NA | NA | NA | 3 | 1 | 1 | 1 | 4 | 1 | NA | NA | NA | NA | 2 | 0.5 | 1 | 1 | 3 | 0.6667 |
| 9_2166075 | 3 | 0.3333 | NA | NA | 5 | 0.8 | 1 | 0 | 9 | 0.5556 | NA | NA | NA | NA | 2 | 0.5 | 1 | 0 | 3 | 0.3333 |
| 9_2166081 | 2 | 0 | NA | NA | 5 | 0.2 | 1 | 0 | 8 | 0.125 | NA | NA | NA | NA | 2 | 0.5 | 1 | 0 | 3 | 0.3333 |
| 9_2166089 | 3 | 1 | NA | NA | 4 | 0.5 | NA | NA | 7 | 0.7143 | NA | NA | NA | NA | 2 | 0.5 | 1 | 1 | 3 | 0.6667 |
| 9_2166745 | NA | NA | NA | NA | 2 | 1 | NA | NA | 2 | 1 | NA | NA | NA | NA | 1 | 1 | NA | NA | 1 | 1 |
| 9_2166755 | NA | NA | NA | NA | 2 | 0.5 | NA | NA | 2 | 0.5 | NA | NA | NA | NA | 1 | 1 | 1 | 1 | 2 | 1 |
| 9_2183235 | 6 | 0 | 10 | 0.2 | 8 | 0.375 | 4 | 0 | 28 | 0.1786 | 6 | 0 | 7 | 0 | 11 | 0.1818 | 4 | 0.5 | 28 | 0.1429 |
| 9_2184062 | NA | NA | 1 | 1 | NA | NA | NA | NA | 1 | 1 | 1 | 1 | 1 | 0 | 1 | 0 | NA | NA | 3 | 0.3333 |
| 9_2184710 | NA | NA | 3 | 0.3333 | NA | NA | NA | NA | 3 | 0.3333 | 3 | 0.3333 | 1 | 0 | 1 | 1 | NA | NA | 5 | 0.4 |
| 9_2185908 | 9 | 0.2222 | 13 | 0.1538 | 11 | 0 | 7 | 0.1429 | 40 | 0.125 | 7 | 0.2857 | 11 | 0.0909 | 13 | 0.0769 | 13 | 0 | 44 | 0.0909 |
| 9_2185915 | 14 | 0.5714 | 16 | 0.3125 | 13 | 0.3846 | 11 | 0.0909 | 54 | 0.3519 | 12 | 0.1667 | 13 | 0.3077 | 21 | 0.381 | 14 | 0.3571 | 60 | 0.3167 |
| 9_2185917 | 12 | 0.6667 | 16 | 0.875 | 12 | 0.6667 | 10 | 0.5 | 50 | 0.7 | 10 | 0.8 | 13 | 0.7692 | 17 | 0.7059 | 13 | 0.6154 | 53 | 0.717 |
| 9_2186000 | 20 | 0 | 27 | 0.1111 | 18 | 0.1667 | 15 | 0.0667 | 80 | 0.0875 | 19 | 0 | 13 | 0.0769 | 25 | 0.08 | 15 | 0.2 | 72 | 0.0833 |
| 9_2186831 | 6 | 0.3333 | 1 | 0 | 3 | 1 | NA | NA | 10 | 0.5 | NA | NA | 2 | 0.5 | 2 | 0.5 | NA | NA | 4 | 0.5 |
| 9_2186836 | 3 | 1 | 2 | 1 | 3 | 1 | NA | NA | 8 | 1 | NA | NA | 2 | 1 | 2 | 1 | 1 | 1 | 5 | 1 |
| 9_2186848 | 5 | 0.4 | 2 | 0 | 2 | 1 | NA | NA | 9 | 0.4444 | NA | NA | 2 | 0.5 | 1 | 1 | NA | NA | 3 | 0.6667 |
| 9_2186854 | 6 | 0.1667 | 2 | 0.5 | 3 | 0.3333 | NA | NA | 11 | 0.2727 | NA | NA | 3 | 0 | 3 | 0.6667 | 1 | 1 | 7 | 0.4286 |
| 9_2186856 | 6 | 0.5 | 1 | 0 | 3 | 0.3333 | NA | NA | 10 | 0.4 | NA | NA | 3 | 0.3333 | 3 | 0.6667 | 1 | 0 | 7 | 0.4286 |
| 9_2186864 | 5 | 0.2 | 2 | 0 | 2 | 0.5 | NA | NA | 9 | 0.2222 | NA | NA | 3 | 0 | 2 | 0.5 | 1 | 0 | 6 | 0.1667 |
| 9_2186866 | 5 | 0.4 | 2 | 0 | 3 | 0.3333 | NA | NA | 10 | 0.3 | NA | NA | 3 | 0.3333 | 3 | 0.3333 | 1 | 0 | 7 | 0.2857 |
| 9_2186868 | 5 | 0.8 | 2 | 1 | 3 | 1 | NA | NA | 10 | 0.9 | NA | NA | 3 | 0.6667 | 3 | 0.3333 | 1 | 0 | 7 | 0.4286 |
| 9_2186924 | 6 | 0.1667 | 3 | 0 | 2 | 0.5 | NA | NA | 11 | 0.1818 | NA | NA | 3 | 0 | 4 | 0.25 | 1 | 0 | 8 | 0.125 |
| 9_2186940 | 6 | 0 | 3 | 0 | 3 | 0.3333 | NA | NA | 12 | 0.0833 | NA | NA | 3 | 0 | 4 | 0 | 1 | 1 | 8 | 0.125 |
| 9_2186941 | 5 | 0.2 | 2 | 0 | 2 | 0 | 1 | 0 | 10 | 0.1 | NA | NA | 3 | 0.3333 | 4 | 0 | 1 | 0 | 8 | 0.125 |
| 9_2187035 | 2 | 0 | 1 | 1 | 1 | 1 | NA | NA | 4 | 0.5 | 1 | 1 | 2 | 1 | NA | NA | 1 | 0 | 4 | 0.75 |
| 9_2187074 | 2 | 0 | 1 | 1 | NA | NA | 1 | 0 | 4 | 0.25 | 1 | 1 | 2 | 0 | NA | NA | 1 | 0 | 4 | 0.25 |
| 9_2187084 | 2 | 0 | 1 | 0 | 1 | 1 | 1 | 0 | 5 | 0.2 | 1 | 0 | 1 | 0 | NA | NA | 1 | 1 | 3 | 0.3333 |
| 9_2192648 | 6 | 0 | 17 | 0.2353 | 11 | 0.0909 | 11 | 0.3636 | 45 | 0.2 | 10 | 0.3 | 20 | 0.25 | 28 | 0.3929 | 4 | 0.25 | 62 | 0.3226 |
| 9_2192731 | 6 | 0 | 16 | 0.0625 | 10 | 0.1 | 12 | 0.25 | 44 | 0.1136 | 5 | 0.2 | 22 | 0.0909 | 28 | 0.0714 | 2 | 0 | 57 | 0.0877 |
| 9_2192741 | 5 | 0.8 | 12 | 0.6667 | 7 | 0.5714 | 10 | 0.3 | 34 | 0.5588 | 4 | 0.5 | 18 | 0.6667 | 20 | 0.55 | 3 | 0.6667 | 45 | 0.6 |
| 9_2192751 | 3 | 0 | 9 | 0.1111 | 7 | 0 | 8 | 0.25 | 27 | 0.1111 | 4 | 0.25 | 17 | 0.1176 | 19 | 0.0526 | 2 | 0 | 42 | 0.0952 |
| 9_2192755 | 6 | 0.5 | 12 | 0.5 | 7 | 0.5714 | 8 | 0.375 | 33 | 0.4848 | 4 | 0.5 | 20 | 0.45 | 18 | 0.2778 | 3 | 0.3333 | 45 | 0.3778 |
| 9_2192785 | 6 | 0.1667 | 11 | 0.2727 | 6 | 0.1667 | 8 | 0 | 31 | 0.1613 | 4 | 0 | 18 | 0.1111 | 19 | 0.1053 | 2 | 0 | 43 | 0.093 |

| | | | | | | | | | | | | | | | | | | | | |
|---|---|---|---|---|---|---|---|---|---|---|---|---|---|---|---|---|---|---|---|---|
| 9_2192888 | 4 | 0 | 8 | 0.125 | 4 | 0.25 | 5 | 0.2 | 21 | 0.1429 | 3 | 0 | 13 | 0.0769 | 18 | 0.0556 | 3 | 0 | 37 | 0.0541 |
| 9_2192998 | 1 | 1 | 2 | 1 | 2 | 0 | 3 | 1 | 8 | 0.75 | 1 | 0 | 4 | 1 | 11 | 0.6364 | 3 | 0.6667 | 19 | 0.6842 |
| 9_2193762 | NA | NA | 2 | 0 | 3 | 0.6667 | 4 | 0 | 9 | 0.2222 | 1 | 0 | 2 | 0 | 11 | 0.1818 | 4 | 0.25 | 18 | 0.1667 |
| 9_2196680 | 2 | 0.5 | 6 | 0.3333 | 2 | 1 | 1 | 0 | 11 | 0.4545 | 4 | 0.5 | NA | NA | 2 | 0.5 | 1 | 0 | 7 | 0.4286 |
| 9_2196696 | 2 | 0.5 | 6 | 0.5 | 1 | 1 | 1 | 1 | 10 | 0.6 | 4 | 0.75 | NA | NA | 1 | 1 | 2 | 0.5 | 7 | 0.7143 |
| 9_2196699 | 2 | 0 | 5 | 0.4 | 1 | 0 | 1 | 1 | 9 | 0.3333 | 4 | 0.75 | NA | NA | 1 | 1 | 2 | 0.5 | 7 | 0.7143 |
| 9_2196706 | 2 | 1 | 3 | 0.6667 | 1 | 0 | 1 | 1 | 7 | 0.7143 | 3 | 0.3333 | NA | NA | 2 | 0.5 | 1 | 0 | 6 | 0.3333 |
| 9_2196884 | 2 | 0 | 3 | 0.6667 | NA | NA | NA | NA | 5 | 0.4 | 2 | 0.5 | NA | NA | 2 | 0 | 1 | 1 | 5 | 0.4 |
| 9_2196896 | 2 | 0 | 4 | 0.5 | 1 | 0 | NA | NA | 7 | 0.2857 | 2 | 0.5 | NA | NA | 2 | 0 | 1 | 1 | 5 | 0.4 |
| 9_2198359 | 14 | 0.1429 | 9 | 0.1111 | 6 | 0 | 12 | 0.0833 | 41 | 0.0976 | 10 | 0.1 | 5 | 0.2 | 11 | 0.0909 | 7 | 0.1429 | 33 | 0.1212 |
| 9_2198766 | 10 | 0.5 | 6 | 0.6667 | 4 | 0.5 | 7 | 1 | 27 | 0.6667 | 4 | 0.5 | 4 | 0.75 | 6 | 0.3333 | 2 | 1 | 16 | 0.5625 |
| 9_2198771 | 8 | 0.75 | 5 | 0.4 | 4 | 1 | 6 | 1 | 23 | 0.7826 | 4 | 0.5 | 2 | 0.5 | 6 | 0.3333 | 3 | 0.6667 | 15 | 0.4667 |
| 9_2198778 | 7 | 0 | 5 | 0 | 4 | 0.25 | 6 | 0.3333 | 22 | 0.1364 | 4 | 0 | 3 | 0 | 5 | 0.2 | 4 | 0 | 16 | 0.0625 |
| 9_2199547 | 7 | 0.1429 | 5 | 0 | 3 | 0 | 4 | 0 | 19 | 0.0526 | 2 | 0.5 | 4 | 0 | 6 | 0.3333 | 3 | 0 | 15 | 0.2 |
| 9_2202446 | 13 | 0.2308 | 10 | 0.2 | 11 | 0.1818 | 12 | 0.0833 | 46 | 0.1739 | 12 | 0.1667 | 12 | 0.5833 | 10 | 0.3 | 8 | 0.375 | 42 | 0.3571 |
| 9_2202447 | 16 | 0.1875 | 11 | 0.0909 | 11 | 0.2727 | 12 | 0.25 | 50 | 0.2 | 13 | 0.1538 | 12 | 0.1667 | 11 | 0.2727 | 8 | 0.25 | 44 | 0.2045 |
| 9_2202463 | 14 | 0.1429 | 9 | 0 | 11 | 0.0909 | 11 | 0.2727 | 45 | 0.1333 | 12 | 0.1667 | 10 | 0.2 | 12 | 0.1667 | 8 | 0.125 | 42 | 0.1667 |
| 9_2202471 | 12 | 0.5833 | 9 | 0.6667 | 9 | 0.7778 | 9 | 0.7778 | 39 | 0.6923 | 11 | 0.7273 | 9 | 0.5556 | 12 | 0.6667 | 7 | 0.5714 | 39 | 0.641 |
| 9_2202478 | 8 | 1 | 7 | 1 | 10 | 0.9 | 9 | 1 | 34 | 0.9706 | 8 | 0.75 | 8 | 0.875 | 10 | 0.9 | 4 | 0.75 | 30 | 0.8333 |
| 9_2202482 | 12 | 0.1667 | 8 | 0.125 | 11 | 0.1818 | 9 | 0.4444 | 40 | 0.225 | 9 | 0.3333 | 9 | 0.4444 | 12 | 0.5 | 7 | 0.1429 | 37 | 0.3784 |
| 9_2206914 | 6 | 0.5 | 9 | 0.4444 | 5 | 0.4 | 4 | 0.5 | 24 | 0.4583 | 10 | 0.6 | 7 | 0.4286 | 3 | 0.3333 | 3 | 0.6667 | 23 | 0.5217 |
| 9_2206919 | 5 | 0.2 | 9 | 0.1111 | 7 | 0 | 5 | 0 | 26 | 0.0769 | 10 | 0.1 | 7 | 0 | 6 | 0 | 4 | 0.25 | 27 | 0.0741 |
| 9_2206923 | 7 | 0.7143 | 6 | 0.8333 | 3 | 1 | 3 | 1 | 19 | 0.8421 | 6 | 0.6667 | 5 | 1 | 4 | 0.75 | 2 | 1 | 17 | 0.8235 |
| 9_2206926 | 7 | 0.5714 | 8 | 0.5 | 5 | 1 | 4 | 0.75 | 24 | 0.6667 | 10 | 0.7 | 7 | 0.2857 | 5 | 0.4 | 4 | 0.5 | 26 | 0.5 |
| 9_2206947 | 6 | 0.6667 | 9 | 0.5556 | 5 | 0.4 | 3 | 1 | 23 | 0.6087 | 11 | 0.9091 | 7 | 0.2857 | 4 | 0.5 | 3 | 0.6667 | 25 | 0.64 |
| 9_2206949 | 4 | 0.25 | 9 | 0.1111 | 6 | 0.3333 | 5 | 0 | 24 | 0.1667 | 8 | 0.375 | 8 | 0.375 | 6 | 0 | 3 | 0 | 25 | 0.24 |
| 9_2206950 | 4 | 0 | 7 | 0.2857 | 5 | 0 | 4 | 0 | 20 | 0.1 | 8 | 0 | 6 | 0.1667 | 3 | 0 | 1 | 0 | 18 | 0.0556 |
| 9_2209685 | 36 | 0.1389 | 28 | 0.1071 | 30 | 0.1333 | 35 | 0.0857 | 129 | 0.1163 | 36 | 0.25 | 39 | 0.1538 | 37 | 0.1081 | 34 | 0.1765 | 146 | 0.1712 |
| 9_2214998 | NA | NA | NA | NA | 1 | 1 | 6 | 1 | 7 | 1 | NA | NA | 1 | 1 | 2 | 1 | 1 | 1 | 4 | 1 |
| 9_2215003 | NA | NA | 1 | 0 | 1 | 1 | 4 | 0.5 | 6 | 0.5 | NA | NA | 1 | 1 | 2 | 0.5 | 1 | 1 | 4 | 0.75 |
| 9_2215012 | NA | NA | 1 | 0 | 1 | 0 | 5 | 0.2 | 7 | 0.1429 | NA | NA | 1 | 0 | 2 | 0.5 | 1 | 0 | 4 | 0.25 |
| 9_2215020 | NA | NA | 1 | 0 | NA | NA | 5 | 0.4 | 6 | 0.3333 | NA | NA | 1 | 1 | 1 | 1 | NA | NA | 2 | 1 |
| 9_2215035 | NA | NA | 1 | 0 | 1 | 0 | 6 | 0.1667 | 8 | 0.125 | NA | NA | 1 | 0 | 2 | 0 | 1 | 1 | 4 | 0.25 |
| 9_2215054 | NA | NA | 1 | 1 | 1 | 0 | 3 | 0 | 5 | 0.2 | NA | NA | 1 | 0 | 1 | 1 | 1 | 0 | 3 | 0.3333 |
| 9_2215073 | NA | NA | NA | NA | 1 | 0 | 5 | 0.2 | 6 | 0.1667 | NA | NA | 1 | 0 | 2 | 0.5 | 1 | 0 | 4 | 0.25 |
| 9_2215074 | NA | NA | NA | NA | 1 | 1 | 5 | 1 | 6 | 1 | NA | NA | 1 | 1 | 2 | 1 | 1 | 1 | 4 | 1 |
| 9_2215081 | NA | NA | NA | NA | NA | NA | 5 | 1 | 5 | 1 | NA | NA | 1 | 1 | 2 | 1 | 1 | 1 | 4 | 1 |
| 9_2215083 | NA | NA | NA | NA | NA | NA | 4 | 0.75 | 4 | 0.75 | NA | NA | 1 | 1 | 2 | 0.5 | 1 | 0 | 4 | 0.5 |
| 9_2215086 | NA | NA | NA | NA | NA | NA | 4 | 1 | 4 | 1 | NA | NA | 1 | 0 | 2 | 0.5 | 1 | 0 | 4 | 0.25 |
| 9_2215091 | 1 | 1 | NA | NA | NA | NA | 4 | 0.25 | 5 | 0.4 | NA | NA | 1 | 0 | 2 | 1 | 1 | 0 | 4 | 0.5 |
| 9_2215104 | 1 | 1 | NA | NA | NA | NA | 4 | 1 | 5 | 1 | NA | NA | 1 | 0 | 2 | 0.5 | 1 | 0 | 4 | 0.25 |
| 9_2215363 | 1 | 0 | NA | NA | 3 | 0.3333 | 6 | 0.1667 | 10 | 0.2 | NA | NA | 5 | 0.2 | 2 | 0 | 3 | 0 | 10 | 0.1 |
| 9_2215438 | 1 | 0 | NA | NA | 2 | 0.5 | 7 | 1 | 10 | 0.8 | 1 | 1 | 4 | 0.5 | 2 | 1 | 4 | 0.5 | 11 | 0.6364 |
| 9_2215442 | 1 | 0 | 2 | 1 | 3 | 1 | 9 | 1 | 15 | 0.9333 | 1 | 1 | 6 | 1 | 2 | 1 | 4 | 1 | 13 | 1 |
| 9_2215444 | NA | NA | 2 | 1 | 3 | 1 | 9 | 0.8889 | 14 | 0.9286 | 1 | 1 | 6 | 0.8333 | 3 | 0.6667 | 3 | 1 | 13 | 0.8462 |
| 9_2215450 | NA | NA | 2 | 1 | 4 | 0.5 | 8 | 0.625 | 14 | 0.6429 | 1 | 1 | 6 | 0.5 | 3 | 1 | 4 | 0.25 | 14 | 0.5714 |
| 9_2215459 | NA | NA | 1 | 1 | 3 | 0.3333 | 9 | 0.3333 | 13 | 0.3846 | 1 | 0 | 5 | 0.6 | 2 | 0.5 | 2 | 0 | 10 | 0.4 |
| 9_2215917 | 1 | 1 | NA | NA | NA | NA | NA | NA | 1 | 1 | NA | NA | NA | NA | 2 | 0.5 | 1 | 1 | 3 | 0.6667 |
| 9_2215919 | 1 | 1 | NA | NA | NA | NA | NA | NA | 1 | 1 | NA | NA | NA | NA | 1 | 1 | 2 | 1 | 3 | 1 |
| 9_2215920 | 1 | 0 | NA | NA | NA | NA | 1 | 1 | 2 | 0.5 | NA | NA | NA | NA | 1 | 1 | 2 | 0.5 | 3 | 0.6667 |
| 9_2215922 | 1 | 0 | NA | NA | NA | NA | 1 | 1 | 2 | 0.5 | NA | NA | NA | NA | 2 | 1 | 2 | 1 | 4 | 1 |
| 9_2215923 | 1 | 1 | NA | NA | NA | NA | 1 | 0 | 2 | 0.5 | NA | NA | NA | NA | 2 | 0.5 | 1 | 1 | 3 | 0.6667 |
| 9_2215923 | 1 | 1 | NA | NA | NA | NA | 1 | 0 | 2 | 0.5 | NA | NA | NA | NA | 2 | 1 | 1 | 1 | 3 | 1 |
| 9_2215999 | NA | NA | NA | NA | NA | NA | 1 | 1 | 1 | 1 | NA | NA | NA | NA | 1 | 1 | 1 | 1 | 2 | 1 |
| 9_2216019 | NA | NA | NA | NA | NA | NA | 1 | 1 | 1 | 1 | NA | NA | NA | NA | 1 | 1 | 1 | 1 | 2 | 1 |
| 9_2218831 | 2 | 0 | 4 | 0.25 | NA | NA | 1 | 0 | 7 | 0.1429 | 1 | 0 | 2 | 0 | 3 | 0 | 4 | 0.25 | 10 | 0.1 |
| 9_2218838 | 1 | 0 | 4 | 0.25 | NA | NA | 1 | 1 | 6 | 0.3333 | 1 | 0 | 1 | 0 | 3 | 0.3333 | 4 | 0.25 | 9 | 0.2222 |
| 9_2218905 | 1 | 1 | 4 | 1 | NA | NA | 1 | 1 | 6 | 1 | 1 | 1 | 2 | 1 | 3 | 0.6667 | 3 | 1 | 9 | 0.8889 |
| 9_2218934 | NA | NA | 2 | 1 | NA | NA | 1 | 1 | 3 | 1 | 1 | 0 | 2 | 0.5 | 3 | 0.3333 | 2 | 0.5 | 8 | 0.375 |
| 9_2218954 | 1 | 0 | 3 | 0.6667 | NA | NA | 1 | 0 | 5 | 0.4 | 1 | 0 | 2 | 0.5 | 3 | 0.3333 | 2 | 0.5 | 8 | 0.375 |
| 9_2218964 | NA | NA | 2 | 1 | NA | NA | NA | NA | 2 | 1 | 1 | 0 | 1 | 1 | 1 | 1 | 2 | 0.5 | 5 | 0.6 |
| 9_2218967 | 1 | 1 | 3 | 0.6667 | NA | NA | 1 | 0 | 5 | 0.6 | NA | NA | 2 | 0.5 | 3 | 0.3333 | 2 | 0.5 | 7 | 0.4286 |
| 9_2218996 | 1 | 0 | 3 | 1 | NA | NA | 1 | 1 | 5 | 0.8 | 1 | 1 | 2 | 1 | 2 | 1 | 1 | 1 | 6 | 1 |
| 9_2219004 | NA | NA | 2 | 1 | NA | NA | 1 | 1 | 3 | 1 | 1 | 0 | 2 | 0.5 | 2 | 0.5 | 2 | 1 | 7 | 0.5714 |
| 9_2219013 | NA | NA | 2 | 0.5 | NA | NA | 1 | 1 | 3 | 0.6667 | 1 | 0 | 2 | 1 | 3 | 1 | 2 | 1 | 8 | 0.875 |

| | | | | | | | | | | | | | | | | | | | | |
|---|---|---|---|---|---|---|---|---|---|---|---|---|---|---|---|---|---|---|---|---|
| 9_2219018 | NA | NA | 3 | 1 | NA | NA | 1 | 1 | 4 | 1 | NA | NA | 2 | 1 | 3 | 1 | 2 | 0.5 | 7 | 0.8571 |
| 9_2219027 | NA | NA | 1 | 1 | NA | NA | 1 | 1 | 2 | 1 | 1 | 0 | 2 | 1 | 3 | 1 | 2 | 0.5 | 8 | 0.75 |
| 9_2219044 | NA | NA | 2 | 1 | NA | NA | 1 | 1 | 3 | 1 | NA | NA | 2 | 0.5 | 2 | 1 | 2 | 1 | 6 | 0.8333 |
| 9_2224369 | 5 | 0 | 5 | 0 | 2 | 0 | 6 | 0.1667 | 18 | 0.0556 | 4 | 0 | 1 | 0 | 3 | 0 | 6 | 0.1667 | 14 | 0.0714 |
| 9_2224431 | 3 | 0.6667 | 4 | 0.5 | 3 | 1 | 4 | 0.75 | 14 | 0.7143 | 4 | 0.5 | NA | NA | 3 | 0.6667 | 4 | 1 | 11 | 0.7273 |
| 9_2233552 | 2 | 0.5 | 1 | 1 | NA | NA | 1 | 1 | 4 | 0.75 | 1 | 1 | NA | NA | 1 | 1 | NA | NA | 2 | 1 |
| 9_2233570 | 1 | 1 | NA | NA | NA | NA | 1 | 1 | 2 | 1 | 1 | 0 | NA | NA | 2 | 0.5 | NA | NA | 3 | 0.3333 |
| 9_2233592 | 2 | 0 | 2 | 0.5 | NA | NA | 1 | 0 | 5 | 0.2 | 1 | 1 | NA | NA | 2 | 0.5 | NA | NA | 3 | 0.6667 |
| 9_2233595 | 1 | 0 | 1 | 1 | NA | NA | 1 | 1 | 3 | 0.6667 | 1 | 1 | NA | NA | 2 | 0.5 | NA | NA | 3 | 0.6667 |
| 9_2233600 | 2 | 0.5 | 2 | 1 | NA | NA | 1 | 1 | 5 | 0.8 | 1 | 1 | NA | NA | 2 | 1 | NA | NA | 3 | 1 |
| 9_2235966 | 7 | 0.7143 | 6 | 0.8333 | 7 | 0.7143 | 7 | 0.8571 | 27 | 0.7778 | 9 | 0.6667 | 3 | 1 | 9 | 0.6667 | 4 | 0.25 | 25 | 0.64 |
| 9_2235968 | 9 | 0.1111 | 8 | 0.125 | 10 | 0.1 | 10 | 0 | 37 | 0.0811 | 13 | 0 | 4 | 0 | 12 | 0.0833 | 6 | 0.1667 | 35 | 0.0571 |
| 9_2235973 | 9 | 0 | 7 | 0 | 10 | 0.1 | 11 | 0.1818 | 37 | 0.0811 | 11 | 0.0909 | 4 | 0.25 | 14 | 0.1429 | 7 | 0.1429 | 36 | 0.1389 |
| 9_2235978 | 8 | 0.125 | 7 | 0.4286 | 11 | 0.0909 | 11 | 0.4545 | 37 | 0.2703 | 10 | 0.1 | 5 | 0.4 | 11 | 0.2727 | 7 | 0.2857 | 33 | 0.2424 |
| 9_2235989 | 7 | 0 | 9 | 0 | 7 | 0.4286 | 9 | 0.2222 | 32 | 0.1562 | 11 | 0.1818 | 6 | 0.1667 | 12 | 0.25 | 5 | 0.2 | 34 | 0.2059 |
| 9_2235990 | 8 | 0.625 | 9 | 0.4444 | 9 | 0.7778 | 11 | 0.6364 | 37 | 0.6216 | 12 | 0.5 | 6 | 0 | 11 | 0.7273 | 6 | 0.5 | 35 | 0.4857 |
| 9_2235992 | 5 | 0 | 8 | 0.125 | 8 | 0.125 | 10 | 0.1 | 31 | 0.0968 | 11 | 0.0909 | 6 | 0.3333 | 13 | 0.1538 | 7 | 0.4286 | 37 | 0.2162 |
| 9_2279924 | 1 | 1 | 1 | 1 | NA | NA | 1 | 0 | 3 | 0.6667 | NA | NA | 2 | 0.5 | 2 | 0 | 1 | 0 | 5 | 0.2 |
| 9_2331070 | NA | NA | NA | NA | 2 | 0 | 3 | 0.3333 | 5 | 0.2 | NA | NA | 1 | 1 | NA | NA | NA | NA | 1 | 1 |
| 9_2331074 | NA | NA | NA | NA | 2 | 0.5 | 2 | 1 | 4 | 0.75 | NA | NA | 1 | 1 | NA | NA | NA | NA | 1 | 1 |
| 9_2339534 | NA | NA | 1 | 1 | 2 | 1 | 3 | 0.3333 | 6 | 0.6667 | NA | NA | 1 | 1 | NA | NA | NA | NA | 1 | 1 |
| 9_2339539 | NA | NA | NA | NA | 3 | 0.6667 | 3 | 0.6667 | 6 | 0.6667 | NA | NA | 1 | 1 | NA | NA | NA | NA | 1 | 1 |
| 9_2432395 | 6 | 0.5 | 11 | 0.4545 | 2 | 0.5 | 9 | 0.2222 | 28 | 0.3929 | 7 | 0.4286 | 14 | 0.4286 | 6 | 0.1667 | 9 | 0.6667 | 36 | 0.4444 |
| 9_2432401 | 7 | 0.5714 | 12 | 0.25 | 3 | 0.3333 | 10 | 0.1 | 32 | 0.2812 | 8 | 0.625 | 14 | 0.3571 | 6 | 0.1667 | 6 | 0.3333 | 34 | 0.3824 |
| 9_2432403 | 6 | 0.3333 | 11 | 0.1818 | 3 | 0.3333 | 8 | 0.375 | 28 | 0.2857 | 8 | 0.25 | 13 | 0.5385 | 6 | 0.6667 | 9 | 0.7778 | 36 | 0.5556 |
| 9_2432411 | 7 | 0.2857 | 12 | 0.0833 | 3 | 0.3333 | 7 | 0.1429 | 29 | 0.1724 | 8 | 0.125 | 8 | 0.125 | 4 | 0.25 | 8 | 0.125 | 28 | 0.1429 |
| 9_2432413 | 5 | 0.4 | 11 | 0.2727 | 3 | 0 | 10 | 0.2 | 29 | 0.2414 | 7 | 0.2857 | 13 | 0.3846 | 5 | 1 | 7 | 0.4286 | 32 | 0.4688 |
| 9_2432415 | 5 | 0 | 14 | 0.0714 | 3 | 0.6667 | 9 | 0.2222 | 31 | 0.1613 | 8 | 0.125 | 13 | 0.2308 | 6 | 0.1667 | 7 | 0.1429 | 34 | 0.1765 |
| 9_2432524 | 1 | 1 | 3 | 0 | NA | NA | 1 | 0 | 5 | 0.2 | NA | NA | 2 | 0.5 | NA | NA | 1 | 0 | 3 | 0.3333 |
| 9_2432542 | 1 | 1 | 4 | 0.25 | 1 | 0 | 1 | 0 | 7 | 0.2857 | 1 | 0 | 5 | 0.2 | 1 | 0 | 2 | 0 | 9 | 0.1111 |
| 9_2432547 | 1 | 1 | 4 | 0.75 | NA | NA | 1 | 1 | 6 | 0.8333 | NA | NA | 4 | 0.75 | 1 | 1 | 1 | 1 | 6 | 0.8333 |
| 9_2432556 | 4 | 0 | 11 | 0.1818 | 3 | 0 | 4 | 0.25 | 22 | 0.1364 | 8 | 0 | 14 | 0.1429 | 6 | 0.3333 | 7 | 0 | 35 | 0.1143 |
| 9_2501250 | 4 | 0.25 | 6 | 0.3333 | 2 | 0 | 4 | 0.5 | 16 | 0.3125 | 12 | 0.5 | 10 | 0.4 | 13 | 0.4615 | 6 | 0.5 | 41 | 0.4634 |
| 9_2501262 | 3 | 0 | 5 | 0.6 | 1 | 0 | 1 | 0 | 10 | 0.3 | 7 | 0.4286 | 8 | 0.375 | 12 | 0.4167 | 3 | 0 | 30 | 0.3667 |
| 9_2501997 | 3 | 0 | 11 | 0.2727 | 8 | 0.625 | 4 | 0.5 | 26 | 0.3846 | 3 | 0 | 9 | 0.3333 | 5 | 0.2 | 6 | 0.5 | 23 | 0.3043 |
| 9_2502013 | 3 | 1 | 6 | 0.3333 | 6 | 0.6667 | 4 | 0 | 19 | 0.4737 | 1 | 1 | 8 | 0.5 | 4 | 0.75 | 4 | 1 | 17 | 0.7059 |
| 9_2502028 | 5 | 0 | 11 | 0.4545 | 5 | 0.6 | 5 | 0.2 | 26 | 0.3462 | 2 | 0.5 | 7 | 0.2857 | 4 | 0.5 | 3 | 0 | 16 | 0.3125 |
| 9_2502181 | 4 | 0.25 | 10 | 0.2 | 7 | 0.1429 | 4 | 0.25 | 25 | 0.2 | 1 | 0 | 6 | 0.5 | 4 | 0.5 | 2 | 0.5 | 13 | 0.4615 |
| 9_2506216 | 3 | 0.3333 | 4 | 0.5 | 6 | 1 | 3 | 0.6667 | 16 | 0.6875 | 1 | 1 | 6 | 0.8333 | 3 | 1 | 4 | 1 | 14 | 0.9286 |
| 9_2506217 | 3 | 1 | 3 | 0.6667 | 3 | 0.6667 | 3 | 1 | 12 | 0.8333 | 1 | 1 | 4 | 0.75 | 3 | 0.3333 | 1 | 1 | 9 | 0.6667 |
| 9_2506219 | 3 | 0 | 3 | 0.6667 | 5 | 0.4 | 4 | 0.25 | 15 | 0.3333 | 1 | 0 | 6 | 0.1667 | 4 | 0.25 | 4 | 0 | 15 | 0.1333 |
| 9_2506223 | 3 | 0 | 3 | 0.6667 | 5 | 0.4 | 3 | 0 | 14 | 0.2857 | 1 | 1 | 8 | 0.25 | 3 | 0.6667 | 4 | 0 | 16 | 0.3125 |
| 9_2506225 | 2 | 1 | 3 | 0.6667 | 4 | 0.5 | 3 | 0.3333 | 12 | 0.5833 | NA | NA | 6 | 0.8333 | 4 | 1 | 2 | 0.5 | 12 | 0.8333 |
| 9_2506228 | 3 | 0 | 4 | 0.25 | 1 | 0 | 3 | 0 | 11 | 0.0909 | NA | NA | 6 | 0.1667 | 3 | 0 | 3 | 0 | 12 | 0.0833 |
| 9_2506230 | 1 | 1 | 2 | 1 | 5 | 0.4 | 2 | 1 | 10 | 0.7 | 1 | 0 | 8 | 0.75 | 4 | 0.5 | 3 | 0.6667 | 16 | 0.625 |
| 9_2506234 | 3 | 0.6667 | 4 | 0.5 | 5 | 0.6 | 4 | 0.5 | 16 | 0.5625 | 1 | 1 | 6 | 0.3333 | 2 | 1 | 4 | 0.25 | 13 | 0.4615 |
| 9_2506237 | 3 | 0.6667 | 3 | 0.3333 | 5 | 0.8 | 4 | 0.5 | 15 | 0.6 | 1 | 0 | 7 | 0.8571 | 4 | 0.75 | 4 | 0.5 | 16 | 0.6875 |
| 9_2506247 | 2 | 0.5 | 2 | 0 | 5 | 0.2 | 4 | 0 | 13 | 0.1538 | 1 | 0 | 6 | 0 | 3 | 0 | 4 | 0.25 | 14 | 0.0714 |
| 9_2516272 | 1 | 1 | 1 | 1 | 1 | 1 | 1 | 1 | 4 | 1 | NA | NA | NA | NA | NA | NA | 1 | 1 | 1 | 1 |
| 9_2516289 | 1 | 1 | 1 | 1 | 1 | 1 | 1 | 0 | 4 | 0.75 | NA | NA | NA | NA | 1 | 1 | 1 | 1 | 2 | 1 |
| 9_2516304 | 1 | 1 | 1 | 0 | 1 | 1 | 1 | 1 | 4 | 0.75 | NA | NA | NA | NA | 1 | 1 | 1 | 1 | 2 | 1 |
| 9_2516316 | 1 | 1 | 1 | 0 | 1 | 1 | 1 | 1 | 4 | 0.75 | NA | NA | NA | NA | 1 | 1 | 1 | 1 | 2 | 1 |
| 9_2516321 | 1 | 1 | NA | NA | 1 | 1 | 1 | 1 | 3 | 1 | NA | NA | NA | NA | 1 | 1 | NA | NA | 1 | 1 |
| 9_2516326 | 1 | 1 | NA | NA | 1 | 1 | 1 | 0 | 3 | 0.6667 | NA | NA | NA | NA | 1 | 1 | 1 | 0 | 2 | 0.5 |
| 9_2516345 | 1 | 1 | NA | NA | 1 | 1 | 1 | 1 | 3 | 1 | NA | NA | NA | NA | 1 | 1 | 1 | 0 | 2 | 0.5 |
| 9_2516350 | 1 | 1 | NA | NA | NA | NA | 1 | 0 | 2 | 0.5 | NA | NA | NA | NA | 1 | 1 | 1 | 0 | 2 | 0.5 |
| 9_2518679 | NA | NA | 1 | 1 | NA | NA | NA | NA | 1 | 1 | NA | NA | NA | NA | 1 | 1 | NA | NA | 1 | 1 |
| 9_2518683 | NA | NA | 2 | 1 | NA | NA | NA | NA | 2 | 1 | NA | NA | 1 | 1 | 2 | 1 | NA | NA | 3 | 1 |
| 9_2518686 | NA | NA | 2 | 1 | NA | NA | NA | NA | 2 | 1 | NA | NA | 1 | 1 | 2 | 1 | NA | NA | 3 | 1 |
| 9_2518762 | NA | NA | 1 | 1 | NA | NA | NA | NA | 1 | 1 | NA | NA | 1 | 0 | 2 | 0.5 | NA | NA | 3 | 0.3333 |
| 9_2521889 | 17 | 0.2353 | 37 | 0.1351 | 44 | 0.1591 | 21 | 0.4286 | 119 | 0.2101 | 23 | 0.1304 | 29 | 0.2414 | 34 | 0.2647 | 28 | 0.2857 | 114 | 0.2368 |
| 9_2521919 | 26 | 0.3462 | 43 | 0.5814 | 42 | 0.4048 | 29 | 0.3793 | 140 | 0.4429 | 25 | 0.56 | 34 | 0.3235 | 40 | 0.575 | 37 | 0.6216 | 136 | 0.5221 |
| 9_2539388 | 2 | 0 | 3 | 0.3333 | NA | NA | 1 | 1 | 6 | 0.3333 | NA | NA | 1 | 1 | 1 | 1 | 1 | 1 | 3 | 1 |
| 9_2539389 | 2 | 0.5 | 3 | 0.6667 | NA | NA | NA | NA | 5 | 0.6 | 1 | 0 | 1 | 1 | 1 | 0 | NA | NA | 3 | 0.3333 |
| 9_2539400 | 3 | 0.3333 | 3 | 0.3333 | NA | NA | 1 | 0 | 7 | 0.2857 | 1 | 1 | 1 | 0 | 1 | 0 | 1 | 1 | 4 | 0.5 |

| | | | | | | | | | | | | | | | | | | | | |
|---|---|---|---|---|---|---|---|---|---|---|---|---|---|---|---|---|---|---|---|---|
| 9_2539405 | 3 | 0.3333 | 3 | 0.3333 | NA | NA | 1 | 1 | 7 | 0.4286 | 1 | 0 | 1 | 0 | 1 | 1 | 1 | 0 | 4 | 0.25 |
| 9_2539422 | 3 | 1 | 4 | 0.5 | NA | NA | 2 | 1 | 9 | 0.7778 | NA | NA | 1 | 1 | 1 | 1 | NA | NA | 2 | 1 |
| 9_2539426 | 4 | 1 | 4 | 1 | NA | NA | 1 | 1 | 9 | 1 | 1 | 1 | 1 | 1 | 1 | 1 | 1 | 1 | 4 | 1 |
| 9_2539430 | 4 | 0.5 | 3 | 0.6667 | NA | NA | 2 | 1 | 9 | 0.6667 | 1 | 1 | 1 | 1 | 1 | 1 | NA | NA | 3 | 1 |
| 9_2539431 | 3 | 0 | 3 | 0.3333 | NA | NA | 1 | 0 | 7 | 0.1429 | 1 | 0 | 1 | 1 | 1 | 0 | 1 | 0 | 4 | 0.25 |
| 9_2539442 | 4 | 0.75 | 4 | 0.75 | NA | NA | 2 | 1 | 10 | 0.8 | 1 | 0 | 1 | 1 | 1 | 1 | 1 | 1 | 4 | 0.75 |
| 9_2539443 | 4 | 0 | 3 | 0.6667 | NA | NA | 1 | 1 | 8 | 0.375 | 1 | 1 | 1 | 1 | 1 | 1 | NA | NA | 3 | 1 |
| 9_2539453 | 3 | 0.6667 | 4 | 0.25 | NA | NA | 2 | 0.5 | 9 | 0.4444 | NA | NA | 1 | 0 | NA | NA | 1 | 1 | 2 | 0.5 |
| 9_2551482 | 4 | 0.25 | 4 | 0.75 | 1 | 0 | 4 | 1 | 13 | 0.6154 | 1 | 1 | 1 | 0 | 1 | 0 | 3 | 0 | 6 | 0.1667 |
| 9_2551500 | 4 | 1 | 3 | 0.3333 | 1 | 1 | 3 | 1 | 11 | 0.8182 | NA | NA | 2 | 0.5 | 1 | 0 | 2 | 0.5 | 5 | 0.4 |
| 9_2551502 | 4 | 0 | 3 | 0 | 1 | 1 | 4 | 0.5 | 12 | 0.25 | 1 | 0 | 2 | 0 | NA | NA | 3 | 0.3333 | 6 | 0.1667 |
| 9_2551504 | 3 | 0.6667 | 1 | 0 | NA | NA | 1 | 1 | 5 | 0.6 | NA | NA | 2 | 0.5 | 1 | 0 | 2 | 1 | 5 | 0.6 |
| 9_2551511 | 4 | 0.5 | 4 | 0.5 | 1 | 0 | 4 | 0.25 | 13 | 0.3846 | 1 | 1 | 2 | 0 | 1 | 0 | 3 | 0.6667 | 7 | 0.4286 |
| 9_2551519 | 3 | 0 | 3 | 0.6667 | 1 | 0 | 3 | 0.6667 | 10 | 0.4 | NA | NA | 2 | 0.5 | NA | NA | 2 | 0 | 4 | 0.25 |
| 9_2666287 | 11 | 0.2727 | 10 | 0.6 | 2 | 0.5 | 5 | 0.6 | 28 | 0.4643 | 10 | 0.2 | 3 | 0 | 5 | 0.6 | 13 | 0.1538 | 31 | 0.2258 |
| 9_2666312 | 9 | 0.2222 | 8 | 0.125 | 2 | 0.5 | 5 | 0 | 24 | 0.1667 | 7 | 0.2857 | 2 | 0 | 6 | 0.1667 | 10 | 0.2 | 25 | 0.2 |
| 9_2666656 | 1 | 0 | 2 | 0.5 | 1 | 0 | 1 | 0 | 5 | 0.2 | 3 | 0.3333 | 1 | 0 | 1 | 0 | 2 | 0 | 7 | 0.1429 |
| 9_2666678 | 1 | 1 | 1 | 0 | 1 | 0 | 1 | 0 | 4 | 0.25 | 3 | 0 | 1 | 1 | 1 | 0 | 4 | 0.25 | 9 | 0.2222 |
| 9_2666721 | 11 | 0.0909 | 11 | 0 | 4 | 0 | 5 | 0.4 | 31 | 0.0968 | 8 | 0.25 | 3 | 0 | 6 | 0.1667 | 11 | 0.0909 | 28 | 0.1429 |
| 9_2666732 | 9 | 0.2222 | 9 | 0.1111 | 3 | 0.6667 | 2 | 0 | 23 | 0.2174 | 8 | 0.125 | 4 | 0.75 | 6 | 0.3333 | 8 | 0.125 | 26 | 0.2692 |
| 9_2666732 | 1 | 1 | NA | NA | NA | NA | NA | NA | 1 | 1 | NA | NA | NA | NA | NA | NA | 1 | 1 | 1 | 1 |
| 9_2667058 | 16 | 0.375 | 12 | 0.3333 | 6 | 0.8333 | 6 | 0.5 | 40 | 0.45 | 13 | 0.5385 | 6 | 0.3333 | 9 | 0.4444 | 12 | 0.5833 | 40 | 0.5 |
| 9_2667065 | 15 | 0.0667 | 15 | 0 | 9 | 0.4444 | 6 | 0 | 45 | 0.1111 | 12 | 0.0833 | 5 | 0.2 | 9 | 0 | 14 | 0.2143 | 40 | 0.125 |
| 9_2667087 | 17 | 0.7647 | 16 | 0.5625 | 10 | 0.7 | 6 | 0.5 | 49 | 0.6531 | 15 | 0.6667 | 6 | 0.6667 | 8 | 1 | 17 | 0.8824 | 46 | 0.8043 |
| 9_2667100 | 16 | 0.125 | 16 | 0.125 | 10 | 0.1 | 7 | 0.1429 | 49 | 0.1224 | 16 | 0.25 | 6 | 0.1667 | 8 | 0 | 15 | 0.3333 | 45 | 0.2222 |
| 9_2667126 | 19 | 0.1579 | 13 | 0.2308 | 11 | 0.2727 | 8 | 0.125 | 51 | 0.1961 | 16 | 0.25 | 6 | 0 | 7 | 0.2857 | 14 | 0.0714 | 43 | 0.1628 |
| 9_2667215 | 27 | 0.1111 | 21 | 0.0476 | 19 | 0.1053 | 10 | 0.1 | 77 | 0.0909 | 16 | 0.125 | 10 | 0.2 | 14 | 0.2143 | 22 | 0.2273 | 62 | 0.1935 |
| 9_2667236 | 23 | 0.2609 | 23 | 0.3913 | 17 | 0.3529 | 9 | 0.2222 | 72 | 0.3194 | 19 | 0.3684 | 10 | 0.7 | 14 | 0.2143 | 21 | 0.4286 | 64 | 0.4062 |
| 9_2667241 | 22 | 0.2273 | 17 | 0.1176 | 15 | 0.0667 | 9 | 0 | 63 | 0.127 | 15 | 0.2667 | 10 | 0.3 | 11 | 0.1818 | 18 | 0.1667 | 54 | 0.2222 |
| 9_2688632 | 2 | 0 | 2 | 0 | 2 | 0.5 | 3 | 0 | 9 | 0.1111 | 3 | 0 | 2 | 0 | 1 | 1 | 1 | 0 | 7 | 0.1429 |
| 9_2688633 | 2 | 0 | 2 | 0.5 | 2 | 0 | 3 | 0 | 9 | 0.1111 | 4 | 0.25 | 2 | 0 | 1 | 0 | 1 | 0 | 8 | 0.125 |
| 9_2688657 | 5 | 0 | 7 | 0.1429 | 4 | 0.5 | 7 | 0 | 23 | 0.1304 | 5 | 0 | 5 | 0 | 3 | 0.6667 | 4 | 0 | 17 | 0.1176 |
| 9_2688662 | 4 | 0.75 | 6 | 0.1667 | 3 | 0 | 3 | 0.6667 | 16 | 0.375 | 4 | 0.25 | 4 | 0.25 | 2 | 0 | 4 | 0 | 14 | 0.1429 |
| 9_2688675 | 4 | 0.75 | 8 | 0.375 | 4 | 0.25 | 5 | 0 | 21 | 0.3333 | 4 | 0 | 5 | 0.2 | 2 | 0 | 4 | 0 | 15 | 0.0667 |
| 9_2688684 | 4 | 0.5 | 6 | 0 | 3 | 0.3333 | 6 | 0.5 | 19 | 0.3158 | 3 | 0.6667 | 4 | 0.5 | 2 | 0.5 | 4 | 0 | 13 | 0.3846 |
| 9_2688688 | 3 | 1 | 5 | 1 | 4 | 0.75 | 6 | 0.6667 | 18 | 0.8333 | 4 | 0.5 | 4 | 0.75 | 2 | 0.5 | 3 | 0.6667 | 13 | 0.6154 |
| 9_2690261 | 1 | 0 | 4 | 0 | 3 | 0 | 3 | 0.3333 | 11 | 0.0909 | 1 | 0 | 2 | 0.5 | 2 | 0 | 4 | 0.25 | 9 | 0.2222 |
| 9_2691056 | 1 | 1 | NA | NA | NA | NA | NA | NA | 1 | 1 | NA | NA | 1 | 1 | NA | NA | NA | NA | 1 | 1 |
| 9_2691693 | 3 | 0 | 5 | 0 | 2 | 0 | 7 | 0.2857 | 17 | 0.1176 | 7 | 0.1429 | 6 | 0 | 9 | 0.1111 | 2 | 0.5 | 24 | 0.125 |
| 9_2691822 | 2 | 0 | 7 | 0 | 3 | 0.3333 | 9 | 0.2222 | 21 | 0.1429 | 5 | 0.4 | 7 | 0.1429 | 8 | 0.25 | 2 | 0 | 22 | 0.2273 |
| 9_2691970 | 10 | 0.1 | 14 | 0.2143 | 9 | 0.2222 | 7 | 0.2857 | 40 | 0.2 | 6 | 0.1667 | 8 | 0.375 | 9 | 0.1111 | 10 | 0.5 | 33 | 0.303 |
| 9_2691975 | 9 | 0.2222 | 13 | 0.3846 | 10 | 0.4 | 6 | 0.3333 | 38 | 0.3421 | 7 | 0.2857 | 9 | 0.4444 | 11 | 0.1818 | 9 | 0.1111 | 36 | 0.25 |
| 9_2692058 | 10 | 0.5 | 15 | 0.4 | 9 | 0.2222 | 7 | 0.4286 | 41 | 0.3902 | 4 | 0 | 9 | 0.3333 | 14 | 0.2143 | 7 | 0.1429 | 34 | 0.2059 |
| 9_2692076 | 8 | 0.875 | 14 | 0.6429 | 10 | 0.2 | 6 | 0.6667 | 38 | 0.5789 | 3 | 0.6667 | 9 | 0.3333 | 12 | 0.5 | 6 | 0.3333 | 30 | 0.4333 |
| 9_2692102 | 4 | 0 | 11 | 0.3636 | 7 | 0.2857 | 5 | 0.4 | 27 | 0.2963 | 3 | 0 | 8 | 0.375 | 8 | 0.375 | 6 | 0.1667 | 25 | 0.28 |
| 9_2694083 | 3 | 0 | 8 | 0.125 | 3 | 0 | 4 | 0 | 18 | 0.0556 | 2 | 0 | 5 | 0 | 9 | 0.1111 | 4 | 0 | 20 | 0.05 |
| 9_2700621 | NA | NA | 2 | 0 | 1 | 0 | 5 | 0.2 | 8 | 0.125 | 1 | 0 | 3 | 0.6667 | 1 | 0 | 1 | 0 | 6 | 0.3333 |
| 9_2700908 | 2 | 0.5 | 3 | 0 | 7 | 0.1429 | 4 | 0 | 16 | 0.125 | 3 | 0 | 4 | 0 | 2 | 0.5 | 5 | 0 | 14 | 0.0714 |
| 9_2700922 | 2 | 0.5 | 4 | 0.5 | 7 | 0.4286 | 4 | 0.5 | 17 | 0.4706 | 3 | 0.3333 | 4 | 0.5 | 2 | 0 | 2 | 0.5 | 11 | 0.3636 |
| 9_2700927 | 2 | 0.5 | 4 | 0 | 7 | 0 | 4 | 0 | 17 | 0.0588 | 1 | 0 | 4 | 0.5 | 2 | 0.5 | 5 | 0.4 | 12 | 0.4167 |
| 9_2700934 | 2 | 0 | 5 | 0 | 7 | 0 | 4 | 0.5 | 18 | 0.1111 | 3 | 0 | 4 | 0 | 2 | 0.5 | 5 | 0 | 14 | 0.0714 |
| 9_2700937 | 2 | 1 | 5 | 0.8 | 7 | 0.7143 | 3 | 1 | 17 | 0.8235 | 3 | 0.6667 | 3 | 1 | 2 | 1 | 5 | 1 | 13 | 0.9231 |
| 9_2700945 | 1 | 1 | 4 | 0.75 | 5 | 1 | 3 | 0.6667 | 13 | 0.8462 | 3 | 1 | 3 | 0.6667 | 1 | 0 | 5 | 0.8 | 12 | 0.75 |
| 9_2700946 | 1 | 0 | 4 | 0 | 5 | 0.2 | 4 | 0 | 14 | 0.0714 | 3 | 0 | 4 | 0.25 | 1 | 0 | 5 | 0 | 13 | 0.0769 |
| 9_2700953 | 3 | 0.3333 | 5 | 0 | 7 | 0 | 2 | 0 | 17 | 0.0588 | 2 | 0 | 3 | 0 | 2 | 0 | 5 | 0.2 | 12 | 0.0833 |
| 9_2883145 | 6 | 0.5 | 2 | 0 | 4 | 0 | 1 | 0 | 13 | 0.2308 | NA | NA | 4 | 0 | 2 | 0.5 | 4 | 0.5 | 10 | 0.3 |
| 9_2883153 | 4 | 0.5 | 2 | 1 | 3 | 1 | 1 | 1 | 10 | 0.8 | 1 | 1 | 3 | 0.3333 | 2 | 1 | 5 | 0.8 | 11 | 0.7273 |
| 9_2883269 | 6 | 0.6667 | 4 | 0.5 | 7 | 0 | 2 | 0 | 19 | 0.3158 | 2 | 0.5 | 7 | 0.2857 | 4 | 0.5 | 9 | 0.4444 | 22 | 0.4091 |
| 9_2883288 | 7 | 0.5714 | 5 | 0.8 | 5 | 0.4 | 2 | 0.5 | 19 | 0.5789 | 1 | 0 | 7 | 0.4286 | 5 | 0.4 | 10 | 0.6 | 23 | 0.4783 |
| 9_2883299 | 10 | 0.4 | 5 | 0.2 | 6 | 0.5 | 2 | 0 | 23 | 0.3478 | 2 | 0.5 | 5 | 0.2 | 5 | 0.4 | 7 | 0.7143 | 19 | 0.4737 |
| 9_2883318 | 11 | 0.4545 | 5 | 0.2 | 7 | 0.4286 | 2 | 0.5 | 25 | 0.4 | 2 | 1 | 6 | 0.3333 | 5 | 0.4 | 9 | 0.5556 | 22 | 0.5 |
| 9_2883335 | 11 | 0.7273 | 3 | 0 | 7 | 0.7143 | 1 | 0 | 22 | 0.5909 | NA | NA | 6 | 0.6667 | 3 | 0.3333 | 8 | 0.625 | 17 | 0.5882 |
| 9_2883343 | 9 | 0.6667 | 5 | 0.6 | 6 | 0.6667 | 2 | 1 | 22 | 0.6818 | 2 | 0.5 | 8 | 0.75 | 4 | 0.5 | 8 | 0.75 | 22 | 0.6818 |
| 9_2883348 | 11 | 0.1818 | 7 | 0.1429 | 7 | 0.5714 | 1 | 1 | 26 | 0.3077 | 3 | 0.3333 | 7 | 0.2857 | 5 | 0.4 | 10 | 0.5 | 25 | 0.4 |
| 9_2883353 | 8 | 0.375 | 7 | 0 | 5 | 0.2 | 2 | 0 | 22 | 0.1818 | 2 | 0 | 6 | 0 | 4 | 0.25 | 9 | 0.2222 | 21 | 0.1429 |

| | | | | | | | | | | | | | | | | | | | | |
|---|---|---|---|---|---|---|---|---|---|---|---|---|---|---|---|---|---|---|---|---|
| 9_2883354 | 10 | 1 | 7 | 0.5714 | 6 | 0.6667 | 2 | 0.5 | 25 | 0.76 | 3 | 0.6667 | 8 | 1 | 3 | 1 | 9 | 0.8889 | 23 | 0.913 |
| 9_2883365 | 11 | 0.0909 | 4 | 0 | 6 | 0.1667 | 3 | 0 | 24 | 0.0833 | 3 | 0.6667 | 6 | 0.1667 | 5 | 0.2 | 9 | 0.5556 | 23 | 0.3913 |
| 9_2883379 | 12 | 0.1667 | 6 | 0 | 7 | 0 | 2 | 0 | 27 | 0.0741 | 4 | 0 | 8 | 0.125 | 5 | 0 | 11 | 0.1818 | 28 | 0.1071 |
| 9_2883382 | 14 | 0.5714 | 6 | 0.3333 | 6 | 0.5 | 3 | 1 | 29 | 0.5517 | 3 | 1 | 7 | 0.8571 | 4 | 0.25 | 10 | 0.5 | 24 | 0.625 |
| 9_2883385 | 10 | 0 | 3 | 0 | 6 | 0 | 1 | 1 | 20 | 0.05 | 3 | 0.6667 | 5 | 0.2 | 3 | 0.6667 | 10 | 0.1 | 21 | 0.2857 |
| 9_2883398 | 13 | 0.2308 | 6 | 0.1667 | 7 | 0.1429 | 3 | 0.3333 | 29 | 0.2069 | 3 | 0.3333 | 7 | 0.4286 | 6 | 0.3333 | 10 | 0.3 | 26 | 0.3462 |
| 9_2883406 | 13 | 0.1538 | 7 | 0 | 7 | 0.1429 | 3 | 0.3333 | 30 | 0.1333 | 3 | 0 | 7 | 0.1429 | 5 | 0 | 10 | 0.2 | 25 | 0.12 |
| 9_2883410 | 11 | 0.8182 | 5 | 0.4 | 6 | 0.8333 | 3 | 0.6667 | 25 | 0.72 | 4 | 0.75 | 5 | 1 | 5 | 0.4 | 9 | 0.8889 | 23 | 0.7826 |
| 9_2883416 | 11 | 0.9091 | 7 | 1 | 6 | 1 | 3 | 1 | 27 | 0.963 | 4 | 0.75 | 8 | 0.875 | 6 | 1 | 12 | 0.8333 | 30 | 0.8667 |
| 9_2883432 | 13 | 0.2308 | 7 | 0.1429 | 7 | 0.4286 | 3 | 0 | 30 | 0.2333 | 4 | 0.5 | 8 | 0.5 | 5 | 0.2 | 11 | 0.3636 | 28 | 0.3929 |
| 9_2883435 | 12 | 0.25 | 6 | 0.3333 | 7 | 0.2857 | 4 | 0.5 | 29 | 0.3103 | 4 | 0.5 | 7 | 0.1429 | 6 | 0.1667 | 10 | 0.3 | 27 | 0.2593 |
| 9_2883439 | 12 | 0.5833 | 5 | 0.6 | 7 | 0.5714 | 3 | 0.6667 | 27 | 0.5926 | 4 | 0.75 | 5 | 0.8 | 6 | 0.5 | 10 | 0.8 | 25 | 0.72 |
| 9_2883443 | 12 | 0.0833 | 6 | 0 | 6 | 0.1667 | 3 | 0 | 27 | 0.0741 | 4 | 0 | 5 | 0 | 5 | 0 | 11 | 0.1818 | 25 | 0.08 |
| 9_2883597 | 22 | 0.3636 | 10 | 0.4 | 12 | 0.0833 | 6 | 0.3333 | 50 | 0.3 | 10 | 0 | 9 | 0.3333 | 8 | 0.375 | 19 | 0.2105 | 46 | 0.2174 |
| 9_2883604 | 19 | 0.1579 | 10 | 0.1 | 13 | 0.0769 | 8 | 0.125 | 50 | 0.12 | 8 | 0 | 8 | 0.125 | 8 | 0.125 | 19 | 0.1053 | 43 | 0.093 |
| 9_2883639 | 20 | 0.2 | 10 | 0.2 | 13 | 0.2308 | 9 | 0.1111 | 52 | 0.1923 | 11 | 0.2727 | 6 | 0 | 9 | 0 | 17 | 0.1176 | 43 | 0.1163 |
| 9_2890615 | 18 | 0.1111 | 26 | 0.3462 | 22 | 0.1818 | 7 | 0 | 73 | 0.2055 | 27 | 0.1852 | 23 | 0.087 | 15 | 0.0667 | 17 | 0.3529 | 82 | 0.1707 |
| 9_2890627 | 16 | 0.1875 | 26 | 0.1923 | 19 | 0.3158 | 6 | 0.1667 | 67 | 0.2239 | 23 | 0.1739 | 19 | 0.2105 | 14 | 0.2857 | 16 | 0.1875 | 72 | 0.2083 |
| 9_2890628 | 20 | 0.25 | 27 | 0.1852 | 23 | 0.1739 | 8 | 0.125 | 78 | 0.1923 | 28 | 0.1786 | 21 | 0.1905 | 19 | 0.2105 | 18 | 0.1111 | 86 | 0.1744 |
| 9_2890636 | 18 | 0.8889 | 29 | 0.8276 | 19 | 0.8421 | 8 | 0.625 | 74 | 0.8243 | 26 | 1 | 20 | 0.9 | 18 | 0.8889 | 16 | 0.9375 | 80 | 0.9375 |
| 9_2890647 | 18 | 0.3333 | 31 | 0.2258 | 20 | 0.35 | 8 | 0.25 | 77 | 0.2857 | 26 | 0.1538 | 23 | 0.4783 | 18 | 0.3889 | 15 | 0.3333 | 82 | 0.3293 |
| 9_2890649 | 17 | 0.8235 | 27 | 0.6667 | 20 | 0.65 | 8 | 0.25 | 72 | 0.6528 | 24 | 0.625 | 21 | 0.619 | 19 | 0.4737 | 13 | 0.9231 | 77 | 0.6364 |
| 9_2890662 | 18 | 0.0556 | 26 | 0.0769 | 18 | 0.0556 | 8 | 0 | 70 | 0.0571 | 24 | 0.0833 | 21 | 0 | 16 | 0.0625 | 15 | 0.0667 | 76 | 0.0526 |
| 9_2890679 | 17 | 0.0588 | 23 | 0.1304 | 16 | 0.0625 | 8 | 0 | 64 | 0.0781 | 23 | 0.0435 | 18 | 0.1111 | 13 | 0 | 13 | 0.1538 | 67 | 0.0746 |
| 9_2890700 | 13 | 0 | 21 | 0.1429 | 15 | 0.0667 | 7 | 0 | 56 | 0.0714 | 20 | 0.2 | 19 | 0.1053 | 14 | 0 | 12 | 0 | 65 | 0.0923 |
| 9_2890700 | 9 | 0.1111 | 16 | 0.125 | 12 | 0.1667 | 7 | 0 | 44 | 0.1136 | 19 | 0 | 14 | 0.1429 | 12 | 0.1667 | 10 | 0 | 55 | 0.0727 |
| 9_2890700 | 2 | 0 | 5 | 0.2 | 2 | 0 | NA | NA | 9 | 0.1111 | 1 | 0 | 2 | 0 | 1 | 1 | 1 | 0 | 5 | 0.2 |
| 9_2890710 | 12 | 0.9167 | 20 | 0.9 | 14 | 1 | 5 | 0.6 | 51 | 0.902 | 18 | 0.9444 | 13 | 1 | 14 | 1 | 12 | 0.9167 | 57 | 0.9649 |
| 9_2890720 | 14 | 0.0714 | 15 | 0 | 12 | 0.0833 | 6 | 0.1667 | 47 | 0.0638 | 19 | 0.1579 | 11 | 0.0909 | 13 | 0 | 10 | 0.1 | 53 | 0.0943 |
| 9_2890736 | 14 | 0.0714 | 11 | 0.0909 | 12 | 0.1667 | 4 | 0.25 | 41 | 0.122 | 14 | 0.2143 | 12 | 0.3333 | 13 | 0 | 5 | 0.4 | 44 | 0.2045 |
| 9_2890742 | 13 | 0.3846 | 19 | 0.1579 | 14 | 0.1429 | 6 | 0.3333 | 52 | 0.2308 | 17 | 0.1765 | 15 | 0.2667 | 15 | 0.5333 | 9 | 0.1111 | 56 | 0.2857 |
| 9_2890747 | 9 | 0.3333 | 14 | 0.4286 | 10 | 0 | 5 | 0.4 | 38 | 0.2895 | 12 | 0.5833 | 13 | 0.6154 | 10 | 0.3 | 7 | 0.2857 | 42 | 0.4762 |
| 9_2890753 | 13 | 0.1538 | 17 | 0.2941 | 15 | 0.0667 | 6 | 0.1667 | 51 | 0.1765 | 16 | 0.125 | 10 | 0.3 | 14 | 0.1429 | 10 | 0 | 50 | 0.14 |
| 9_2890756 | 12 | 0.5833 | 20 | 0.6 | 15 | 0.7333 | 6 | 0.5 | 53 | 0.6226 | 18 | 0.6111 | 14 | 0.7143 | 14 | 0.7143 | 9 | 0.7778 | 55 | 0.6909 |
| 9_2890769 | 8 | 0.375 | 16 | 0.4375 | 13 | 0.3077 | 5 | 0.4 | 42 | 0.381 | 16 | 0.1875 | 12 | 0.25 | 12 | 0.25 | 9 | 0.3333 | 49 | 0.2449 |
| 9_2890771 | 7 | 0.1429 | 17 | 0.5294 | 15 | 0.3333 | 7 | 0.4286 | 46 | 0.3913 | 16 | 0.375 | 12 | 0.3333 | 12 | 0.3333 | 9 | 0.3333 | 49 | 0.3469 |
| 9_2890789 | 7 | 0.2857 | 16 | 0.1875 | 11 | 0.2727 | 4 | 0 | 38 | 0.2105 | 16 | 0.5 | 9 | 0.3333 | 10 | 0.4 | 7 | 0.5714 | 42 | 0.4524 |
| 9_2892542 | 7 | 0 | 17 | 0.2353 | 6 | 0 | 6 | 0 | 36 | 0.1111 | 14 | 0.1429 | 8 | 0.125 | 11 | 0.0909 | 8 | 0.25 | 41 | 0.1463 |
| 9_3033876 | NA | NA | 1 | 0 | NA | NA | 1 | 1 | 2 | 0.5 | 1 | 0 | NA | NA | NA | NA | 2 | 1 | 3 | 0.6667 |
| 9_3033919 | NA | NA | 2 | 0 | NA | NA | 2 | 0.5 | 4 | 0.25 | NA | NA | NA | NA | NA | NA | 2 | 0.5 | 2 | 0.5 |
| 9_3033931 | NA | NA | 2 | 0.5 | NA | NA | NA | NA | 2 | 0.5 | 1 | 1 | NA | NA | NA | NA | 2 | 0.5 | 3 | 0.6667 |
| 9_3033951 | NA | NA | 2 | 1 | NA | NA | 2 | 0.5 | 4 | 0.75 | 1 | 1 | NA | NA | NA | NA | 3 | 0.6667 | 4 | 0.75 |
| 9_3033956 | 1 | 0 | 2 | 1 | NA | NA | 2 | 0 | 5 | 0.4 | 1 | 1 | NA | NA | NA | NA | 3 | 1 | 4 | 1 |
| 9_3033970 | NA | NA | 1 | 1 | NA | NA | 2 | 0 | 3 | 0.3333 | 1 | 1 | NA | NA | NA | NA | 1 | 0 | 2 | 0.5 |
| 9_3033998 | 3 | 0.3333 | 2 | 0.5 | 2 | 0.5 | 2 | 0 | 9 | 0.3333 | 1 | 1 | NA | NA | NA | NA | 3 | 0 | 4 | 0.25 |
| 9_3033999 | 4 | 0.25 | 2 | 0 | 2 | 0 | 3 | 0 | 11 | 0.0909 | 2 | 0 | NA | NA | NA | NA | 3 | 0.3333 | 5 | 0.2 |
| 9_3034001 | 3 | 0 | 2 | 1 | 2 | 1 | 3 | 0.3333 | 10 | 0.5 | 2 | 0.5 | NA | NA | NA | NA | 3 | 1 | 5 | 0.8 |
| 9_3034017 | 3 | 1 | 2 | 1 | 2 | 1 | 3 | 0.6667 | 10 | 0.9 | 2 | 0.5 | NA | NA | NA | NA | 2 | 1 | 4 | 0.75 |
| 9_3034018 | 4 | 0.5 | 2 | 0.5 | 2 | 1 | 3 | 0.3333 | 11 | 0.5455 | 2 | 1 | NA | NA | NA | NA | 3 | 0.3333 | 5 | 0.6 |
| 9_3034032 | 2 | 0 | 2 | 0 | 2 | 0.5 | 3 | 0 | 9 | 0.1111 | 2 | 0.5 | NA | NA | NA | NA | 3 | 0 | 5 | 0.2 |
| 9_3034051 | 5 | 0.4 | 2 | 0.5 | 2 | 0.5 | 3 | 0 | 12 | 0.3333 | 2 | 0.5 | NA | NA | 1 | 1 | 3 | 0.6667 | 6 | 0.6667 |
| 9_3037595 | 6 | 0.6667 | 1 | 1 | 4 | 0.5 | 2 | 0 | 13 | 0.5385 | 1 | 0 | 3 | 0.6667 | 1 | 0 | 4 | 0 | 9 | 0.2222 |
| 9_3037604 | 4 | 0.25 | 3 | 0 | 5 | 0.2 | 4 | 0 | 16 | 0.125 | 3 | 0 | 3 | 0 | 2 | 0 | 6 | 0.3333 | 14 | 0.1429 |
| 9_3037608 | 6 | 0.5 | 1 | 0 | 5 | 0.4 | 2 | 0 | 14 | 0.3571 | 3 | 0 | 4 | 0.25 | 1 | 0 | 5 | 0 | 13 | 0.0769 |
| 9_3037623 | 7 | 0.8571 | 3 | 1 | 6 | 0.8333 | 3 | 1 | 19 | 0.8947 | 4 | 1 | 4 | 1 | 2 | 1 | 6 | 0.6667 | 16 | 0.875 |
| 9_3037626 | 7 | 1 | 3 | 0.6667 | 6 | 1 | 2 | 1 | 18 | 0.9444 | 4 | 0.75 | 4 | 1 | 2 | 1 | 6 | 0.8333 | 16 | 0.875 |
| 9_3037634 | 6 | 0.3333 | 2 | 0 | 4 | 0 | 3 | 0.3333 | 15 | 0.2 | 3 | 0 | 4 | 0.25 | 2 | 0 | 5 | 0.2 | 14 | 0.1429 |
| 9_3037698 | 4 | 0 | 3 | 0 | 6 | 0.1667 | 5 | 0.4 | 18 | 0.1667 | 4 | 0.25 | 5 | 0.2 | NA | NA | 6 | 0 | 15 | 0.1333 |
| 9_3037782 | 2 | 1 | 1 | 0 | 3 | 0 | 2 | 0 | 8 | 0.25 | 3 | 0.3333 | 2 | 0 | 1 | 0 | NA | NA | 6 | 0.1667 |
| 9_3081077 | NA | NA | 3 | 1 | NA | NA | NA | NA | 3 | 1 | 2 | 0.5 | NA | NA | 1 | 1 | NA | NA | 3 | 0.6667 |
| 9_3081092 | NA | NA | 3 | 0.3333 | NA | NA | NA | NA | 3 | 0.3333 | 2 | 0.5 | NA | NA | 1 | 0 | NA | NA | 3 | 0.3333 |
| 9_3096467 | 1 | 1 | NA | NA | 1 | 0 | NA | NA | 2 | 0.5 | 2 | 1 | 2 | 0.5 | NA | NA | NA | NA | 4 | 0.75 |
| 9_3096470 | 1 | 1 | NA | NA | NA | NA | NA | NA | 1 | 1 | 2 | 0 | 2 | 0.5 | NA | NA | NA | NA | 4 | 0.25 |
| 9_3096482 | 1 | 1 | NA | NA | 1 | 1 | NA | NA | 2 | 1 | 2 | 1 | 2 | 1 | NA | NA | NA | NA | 4 | 1 |
| 9_3096528 | 1 | 1 | NA | NA | 1 | 1 | NA | NA | 2 | 1 | 2 | 1 | 2 | 1 | NA | NA | NA | NA | 4 | 1 |

| | | | | | | | | | | | | | | | | | | | | |
|---|---|---|---|---|---|---|---|---|---|---|---|---|---|---|---|---|---|---|---|---|
| 9_3096534 | 1 | 1 | NA | NA | 1 | 1 | NA | NA | 2 | 1 | NA | NA | 1 | 1 | NA | NA | NA | NA | 1 | 1 |
| 9_3098023 | 1 | 1 | NA | NA | 1 | 1 | NA | NA | 2 | 1 | 2 | 1 | 2 | 0.5 | NA | NA | NA | NA | 4 | 0.75 |
| 9_3098434 | 3 | 0.3333 | 1 | 0 | 3 | 0.3333 | 2 | 0 | 9 | 0.2222 | 2 | 0.5 | NA | NA | 4 | 0 | NA | NA | 6 | 0.1667 |
| 9_3098635 | 1 | 0 | 1 | 0 | 2 | 0 | 2 | 0.5 | 6 | 0.1667 | 2 | 0 | 1 | 1 | 7 | 0.1429 | 3 | 0 | 13 | 0.1538 |
| 9_3098651 | 2 | 0 | NA | NA | 4 | 0.25 | 2 | 0 | 8 | 0.125 | 3 | 0 | 1 | 0 | 8 | 0.125 | 3 | 0 | 15 | 0.0667 |
| 9_3098658 | 2 | 0 | 1 | 0 | 4 | 0 | 2 | 0.5 | 9 | 0.1111 | 3 | 0 | 1 | 0 | 7 | 0.2857 | 3 | 0 | 14 | 0.1429 |
| 9_3098676 | 2 | 0 | 1 | 0 | 5 | 0.2 | 3 | 0 | 11 | 0.0909 | 3 | 0 | 1 | 0 | 8 | 0.125 | 4 | 0 | 16 | 0.0625 |
| 9_3106122 | 50 | 0.18 | 44 | 0.2045 | 45 | 0.1333 | 50 | 0.16 | 189 | 0.1693 | 43 | 0.2326 | 39 | 0.2308 | 34 | 0.3235 | 48 | 0.25 | 164 | 0.2561 |
| 9_3106203 | 47 | 0.3617 | 36 | 0.2222 | 46 | 0.3478 | 47 | 0.2128 | 176 | 0.2898 | 40 | 0.25 | 37 | 0.2703 | 40 | 0.3 | 42 | 0.3571 | 159 | 0.2956 |
| 9_3106209 | 42 | 0.0714 | 33 | 0.0303 | 38 | 0.0789 | 46 | 0.1522 | 159 | 0.0881 | 37 | 0.027 | 30 | 0.0667 | 39 | 0.0769 | 41 | 0.1463 | 147 | 0.0816 |
| 9_3106216 | 46 | 0.1957 | 32 | 0.2188 | 44 | 0.2273 | 48 | 0.125 | 170 | 0.1882 | 35 | 0.2571 | 35 | 0.2286 | 39 | 0.2821 | 45 | 0.2222 | 154 | 0.2468 |
| 9_3128974 | 1 | 1 | NA | NA | 2 | 0.5 | 2 | 0 | 5 | 0.4 | NA | NA | 1 | 1 | NA | NA | 1 | 1 | 2 | 1 |
| 9_3128976 | NA | NA | NA | NA | 2 | 0.5 | 2 | 0.5 | 4 | 0.5 | NA | NA | 1 | 0 | NA | NA | 1 | 1 | 2 | 0.5 |
| 9_3128995 | 1 | 0 | NA | NA | 2 | 0 | 2 | 0.5 | 5 | 0.2 | NA | NA | 1 | 1 | NA | NA | 1 | 1 | 2 | 1 |
| 9_3129001 | 1 | 1 | NA | NA | 2 | 1 | 2 | 1 | 5 | 1 | NA | NA | 1 | 1 | NA | NA | 1 | 1 | 2 | 1 |
| 9_3129008 | 1 | 0 | NA | NA | 2 | 1 | 2 | 0.5 | 5 | 0.6 | NA | NA | 1 | 1 | NA | NA | 1 | 1 | 2 | 1 |
| 9_3129023 | 1 | 1 | NA | NA | 2 | 1 | 2 | 0.5 | 5 | 0.8 | NA | NA | 1 | 1 | NA | NA | 1 | 1 | 2 | 1 |
| 9_3129159 | 3 | 0.3333 | 1 | 0 | 3 | 0 | 2 | 0 | 9 | 0.1111 | NA | NA | 2 | 0 | 2 | 0.5 | 2 | 0.5 | 6 | 0.3333 |
| 9_3129182 | 3 | 0.3333 | 1 | 0 | 3 | 0 | 2 | 0 | 9 | 0.1111 | NA | NA | 2 | 0.5 | 2 | 0 | 2 | 0 | 6 | 0.1667 |
| 9_3129186 | 3 | 1 | 1 | 1 | 3 | 1 | 2 | 0.5 | 9 | 0.8889 | NA | NA | 2 | 1 | 2 | 1 | 3 | 0.6667 | 7 | 0.8571 |
| 9_3129193 | 3 | 0 | 1 | 1 | 2 | 0 | 3 | 0 | 9 | 0.1111 | NA | NA | 2 | 0.5 | 2 | 0 | 3 | 0 | 7 | 0.1429 |
| 9_3129196 | 3 | 0 | 1 | 0 | 3 | 0.3333 | 3 | 0 | 10 | 0.1 | NA | NA | 2 | 0 | 2 | 0 | 3 | 0.3333 | 7 | 0.1429 |
| 9_3129278 | 3 | 0 | 1 | 0 | 3 | 0.3333 | 3 | 0 | 10 | 0.1 | 1 | 0 | NA | NA | 4 | 0 | 4 | 0.25 | 9 | 0.1111 |
| 9_3132547 | NA | NA | 4 | 0.25 | 2 | 0 | NA | NA | 6 | 0.1667 | NA | NA | 2 | 0.5 | 2 | 0 | NA | NA | 4 | 0.25 |
| 9_3132552 | NA | NA | 4 | 0.5 | 1 | 1 | NA | NA | 5 | 0.6 | NA | NA | 2 | 1 | 2 | 1 | NA | NA | 4 | 1 |
| 9_3132570 | NA | NA | 4 | 1 | 2 | 1 | NA | NA | 6 | 1 | NA | NA | 2 | 1 | 2 | 0.5 | NA | NA | 4 | 0.75 |
| 9_3132591 | NA | NA | 4 | 1 | 1 | 1 | NA | NA | 5 | 1 | NA | NA | 1 | 1 | 2 | 1 | 1 | 1 | 4 | 1 |
| 9_3132597 | NA | NA | 4 | 0.5 | 2 | 1 | NA | NA | 6 | 0.6667 | NA | NA | 2 | 0.5 | 1 | 0 | NA | NA | 3 | 0.3333 |
| 9_3132601 | NA | NA | 4 | 0.25 | 2 | 0 | NA | NA | 6 | 0.1667 | NA | NA | 2 | 0.5 | 2 | 0.5 | NA | NA | 4 | 0.5 |
| 9_3133251 | NA | NA | 2 | 0 | 3 | 0.6667 | 2 | 0.5 | 7 | 0.4286 | NA | NA | 2 | 0.5 | 2 | 1 | 1 | 0 | 5 | 0.6 |
| 9_3216669 | NA | NA | NA | NA | NA | NA | 2 | 1 | 2 | 1 | 2 | 1 | 2 | 1 | 3 | 1 | 1 | 1 | 8 | 1 |
| 9_3216685 | NA | NA | NA | NA | NA | NA | 2 | 0.5 | 2 | 0.5 | 2 | 0 | 1 | 0 | 3 | 0.6667 | 1 | 1 | 7 | 0.4286 |
| 9_3216705 | NA | NA | NA | NA | NA | NA | 2 | 1 | 2 | 1 | 2 | 0.5 | 1 | 1 | 2 | 1 | 1 | 1 | 6 | 0.8333 |
| 9_3216729 | NA | NA | NA | NA | NA | NA | 2 | 0.5 | 2 | 0.5 | 1 | 0 | 1 | 0 | 2 | 0.5 | 1 | 0 | 5 | 0.2 |
| 9_3216816 | NA | NA | 1 | 0 | NA | NA | 2 | 0.5 | 3 | 0.3333 | 2 | 0 | 2 | 0.5 | 3 | 0 | 1 | 0 | 8 | 0.125 |
| 9_3216831 | NA | NA | 1 | 1 | NA | NA | 2 | 1 | 3 | 1 | 1 | 1 | 2 | 1 | 2 | 1 | 1 | 1 | 6 | 1 |
| 9_3216832 | NA | NA | 1 | 0 | NA | NA | 2 | 0.5 | 3 | 0.3333 | 2 | 0 | 2 | 0 | 2 | 0 | 1 | 1 | 7 | 0.1429 |
| 9_3216838 | NA | NA | 1 | 1 | NA | NA | 2 | 0 | 3 | 0.3333 | 1 | 0 | 2 | 0.5 | 2 | 0.5 | 1 | 0 | 6 | 0.3333 |
| 9_3216841 | NA | NA | 1 | 0 | NA | NA | 2 | 0.5 | 3 | 0.3333 | 2 | 0.5 | 2 | 0 | 3 | 0 | 1 | 0 | 8 | 0.125 |
| 9_3216858 | NA | NA | 2 | 0 | NA | NA | 4 | 0.5 | 6 | 0.3333 | 2 | 0.5 | 2 | 0 | 1 | 0 | 2 | 0 | 7 | 0.1429 |
| 9_3216866 | NA | NA | 1 | 1 | NA | NA | 4 | 1 | 5 | 1 | 2 | 0.5 | 2 | 1 | 2 | 1 | 2 | 1 | 8 | 0.875 |
| 9_3216867 | NA | NA | 2 | 0 | NA | NA | 3 | 0.3333 | 5 | 0.2 | 2 | 0 | 2 | 1 | 2 | 0 | 2 | 0 | 8 | 0.25 |
| 9_3216870 | NA | NA | 2 | 0 | NA | NA | 4 | 0.25 | 6 | 0.1667 | 2 | 0.5 | 2 | 0.5 | 2 | 0.5 | 2 | 0.5 | 8 | 0.5 |
| 9_3217032 | 1 | 1 | 2 | 0.5 | NA | NA | 3 | 1 | 6 | 0.8333 | 2 | 1 | 2 | 1 | 1 | 1 | 2 | 1 | 7 | 1 |
| 9_3217043 | 3 | 1 | 2 | 1 | NA | NA | 5 | 1 | 10 | 1 | 2 | 1 | 2 | 1 | 4 | 1 | 2 | 1 | 10 | 1 |
| 9_3217065 | 3 | 1 | 2 | 1 | NA | NA | 5 | 0.8 | 10 | 0.9 | 2 | 1 | 2 | 1 | 3 | 0.6667 | 2 | 1 | 9 | 0.8889 |
| 9_3217067 | 3 | 0 | 2 | 0 | NA | NA | 5 | 0.6 | 10 | 0.3 | 2 | 0 | 2 | 0.5 | 4 | 0.75 | 2 | 0.5 | 10 | 0.5 |
| 9_3217067 | 2 | 1 | 2 | 0 | NA | NA | 4 | 0 | 8 | 0.25 | 2 | 0.5 | 1 | 0 | 3 | 0.3333 | 2 | 0 | 8 | 0.25 |
| 9_3217082 | 2 | 0.5 | 2 | 0.5 | NA | NA | 4 | 1 | 8 | 0.75 | 1 | 0 | NA | NA | 3 | 0.3333 | 2 | 0.5 | 6 | 0.3333 |
| 9_3217090 | 3 | 0.6667 | 2 | 0.5 | NA | NA | 5 | 0.4 | 10 | 0.5 | 2 | 0.5 | 2 | 0 | 5 | 0.4 | 2 | 0 | 11 | 0.2727 |
| 9_3217098 | 4 | 0.75 | 2 | 1 | NA | NA | 5 | 0.6 | 11 | 0.7273 | 2 | 0.5 | 3 | 0.3333 | 5 | 0.8 | 2 | 1 | 12 | 0.6667 |
| 9_3217107 | 3 | 0 | 2 | 0.5 | NA | NA | 5 | 0.6 | 10 | 0.4 | 2 | 0 | 3 | 0.3333 | 5 | 0.4 | 2 | 0.5 | 12 | 0.3333 |
| 9_3217108 | 3 | 0 | 1 | 1 | NA | NA | 3 | 0 | 7 | 0.1429 | 2 | 0.5 | 2 | 0 | 4 | 0.5 | 2 | 0 | 10 | 0.3 |
| 9_3217124 | 4 | 0.75 | 2 | 1 | NA | NA | 5 | 1 | 11 | 0.9091 | 2 | 1 | 3 | 0.6667 | 4 | 1 | 3 | 0.6667 | 12 | 0.8333 |
| 9_3217135 | 5 | 0.2 | 2 | 0.5 | 1 | 0 | 6 | 0.3333 | 14 | 0.2857 | 2 | 0 | 4 | 0.25 | 5 | 0.2 | 3 | 0.3333 | 14 | 0.2143 |
| 9_3217141 | 3 | 0.6667 | 3 | 0.3333 | 1 | 0 | 5 | 0.6 | 12 | 0.5 | 2 | 0.5 | 4 | 0.5 | 5 | 0.2 | 2 | 0.5 | 13 | 0.3846 |
| 9_3217142 | 5 | 0.2 | 3 | 0 | 1 | 0 | 6 | 0.6667 | 15 | 0.3333 | 2 | 0 | 3 | 0 | 5 | 0.4 | 3 | 0.6667 | 13 | 0.3077 |
| 9_3217153 | 4 | 0.5 | 3 | 0.3333 | 1 | 0 | 6 | 0.5 | 14 | 0.4286 | 2 | 0 | 3 | 0.3333 | 5 | 0 | 4 | 0.25 | 14 | 0.1429 |
| 9_3217158 | 4 | 0.75 | 2 | 1 | 1 | 1 | 6 | 1 | 13 | 0.9231 | 1 | 1 | 3 | 1 | 3 | 1 | 3 | 0.6667 | 10 | 0.9 |
| 9_3217177 | 4 | 0.5 | 3 | 0 | 2 | 0 | 6 | 0.3333 | 15 | 0.2667 | 2 | 0 | 5 | 0 | 5 | 0.2 | 3 | 0 | 15 | 0.0667 |
| 9_3217186 | 3 | 1 | 1 | 1 | 1 | 1 | 6 | 1 | 11 | 1 | 2 | 0 | 4 | 1 | 4 | 0.75 | 1 | 1 | 11 | 0.7273 |
| 9_3217186 | 1 | 1 | NA | NA | NA | NA | 1 | 1 | 2 | 1 | NA | NA | NA | NA | NA | NA | 1 | 1 | 1 | 1 |
| 9_3217192 | 5 | 0 | 3 | 0 | 3 | 0 | 7 | 0.2857 | 18 | 0.1111 | 3 | 0 | 5 | 0.2 | 7 | 0 | 4 | 0.25 | 19 | 0.1053 |
| 9_3217194 | 2 | 1 | 2 | 0.5 | 1 | 1 | 4 | 0.25 | 9 | 0.5556 | 1 | 1 | 2 | 0.5 | 3 | 0.6667 | NA | NA | 6 | 0.6667 |
| 9_3217196 | 6 | 0.3333 | 3 | 0.3333 | 1 | 0 | 4 | 0 | 14 | 0.2143 | 3 | 0 | 4 | 0.25 | 7 | 0 | 3 | 0.3333 | 17 | 0.1176 |

| | | | | | | | | | | | | | | | | | | | | |
|---|---|---|---|---|---|---|---|---|---|---|---|---|---|---|---|---|---|---|---|---|
| 9_3217202( | 4 | 1 | 2 | 1 | 4 | 1 | 5 | 0.8 | 15 | 0.9333 | 1 | 0 | 1 | 1 | 5 | 0.8 | 3 | 1 | 10 | 0.8 |
| 9_3217206; | 5 | 0 | 3 | 0 | 4 | 0.25 | 7 | 0 | 19 | 0.0526 | 3 | 0 | 6 | 0.3333 | 7 | 0 | 4 | 0 | 20 | 0.1 |
| 9_3217207- | 6 | 1 | 4 | 1 | 4 | 1 | 7 | 0.8571 | 21 | 0.9524 | 3 | 1 | 6 | 1 | 6 | 0.8333 | 3 | 1 | 18 | 0.9444 |
| 9_3217209; | 6 | 0.5 | 4 | 0.25 | 3 | 0.3333 | 6 | 0.3333 | 19 | 0.3684 | 2 | 0 | 5 | 0 | 8 | 0.125 | 3 | 0 | 18 | 0.0556 |
| 9_3217215( | 6 | 0 | 4 | 0.25 | 4 | 0 | 6 | 0 | 20 | 0.05 | 1 | 0 | 7 | 0.1429 | 5 | 0 | 2 | 0 | 15 | 0.0667 |
| 9_3217216; | 3 | 0 | 4 | 0 | 3 | 0 | 7 | 0.2857 | 17 | 0.1176 | 3 | 0 | 7 | 0.2857 | 4 | 0.5 | 4 | 0 | 18 | 0.2222 |
| 9_3217217! | 6 | 0.5 | 4 | 0.75 | 4 | 0.75 | 6 | 0.5 | 20 | 0.6 | 3 | 0.6667 | 7 | 0.4286 | 7 | 0.5714 | 4 | 0.25 | 21 | 0.4762 |
| 9_3217220! | 6 | 0 | 3 | 0 | 4 | 0 | 6 | 0.1667 | 19 | 0.0526 | 3 | 0.3333 | 7 | 0.1429 | 7 | 0 | 4 | 0 | 21 | 0.0952 |
| 9_3217223; | 4 | 0.5 | 4 | 0.75 | 4 | 0.75 | 5 | 0.6 | 17 | 0.6471 | 3 | 1 | 6 | 1 | 4 | 0.75 | 4 | 0.75 | 17 | 0.8824 |
| 9_3217235! | 6 | 1 | 4 | 0.75 | 3 | 1 | 5 | 0.8 | 18 | 0.8889 | 3 | 1 | 7 | 1 | 7 | 1 | 4 | 1 | 21 | 1 |
| 9_3217248; | 3 | 0.6667 | 3 | 1 | 5 | 0.6 | 8 | 0.875 | 19 | 0.7895 | 4 | 1 | 9 | 0.8889 | 5 | 1 | 3 | 1 | 21 | 0.9524 |
| 9_3217248! | 6 | 1 | 4 | 1 | 5 | 0.8 | 7 | 1 | 22 | 0.9545 | 4 | 1 | 8 | 1 | 6 | 1 | 4 | 1 | 22 | 1 |
| 9_3217274! | 6 | 0.1667 | 4 | 0.5 | 5 | 0.2 | 5 | 0 | 20 | 0.2 | 4 | 0 | 9 | 0 | 5 | 0.2 | 4 | 0.25 | 22 | 0.0909 |
| 9_3217275- | 6 | 1 | 5 | 1 | 5 | 1 | 6 | 0.8333 | 22 | 0.9545 | 4 | 1 | 11 | 1 | 5 | 1 | 4 | 1 | 24 | 1 |
| 9_3217277! | 4 | 0 | 4 | 0 | 5 | 0.4 | 6 | 0 | 19 | 0.1053 | 2 | 0 | 10 | 0.1 | 5 | 0 | 4 | 0.25 | 21 | 0.0952 |
| 9_3217290( | 7 | 0.5714 | 5 | 0.4 | 5 | 0.4 | 5 | 0.6 | 22 | 0.5 | 5 | 0.6 | 7 | 0.5714 | 3 | 1 | 4 | 1 | 19 | 0.7368 |
| 9_3217318! | 6 | 0.1667 | 4 | 0 | 5 | 0.4 | 6 | 0 | 21 | 0.1429 | 5 | 0.2 | 11 | 0.0909 | 3 | 0.3333 | 5 | 0 | 24 | 0.125 |
| 9_3217361- | 9 | 0.5556 | 6 | 0.3333 | 6 | 0.5 | 7 | 0.5714 | 28 | 0.5 | 5 | 0.2 | 12 | 0.4167 | 7 | 0.4286 | 7 | 0.4286 | 31 | 0.3871 |
| 9_3217484! | 11 | 0.2727 | 8 | 0.125 | 6 | 0.1667 | 9 | 0.1111 | 34 | 0.1765 | 7 | 0.2857 | 11 | 0.0909 | 8 | 0.125 | 12 | 0.0833 | 38 | 0.1316 |
| 9_3217493! | 11 | 0 | 8 | 0.125 | 5 | 0 | 8 | 0.125 | 32 | 0.0625 | 9 | 0.1111 | 10 | 0.1 | 9 | 0.1111 | 10 | 0.1 | 38 | 0.1053 |
| 9_3217589- | 17 | 0.0588 | 18 | 0.1111 | 10 | 0.3 | 11 | 0.0909 | 56 | 0.125 | 15 | 0.0667 | 11 | 0.2727 | 15 | 0.0667 | 12 | 0.0833 | 53 | 0.1132 |
| 9_3217606; | 1 | 0 | NA | NA | 3 | 0.3333 | NA | NA | 4 | 0.25 | 3 | 0.3333 | 1 | 0 | NA | NA | 2 | 0 | 6 | 0.1667 |
| 9_3217648- | 1 | 1 | NA | NA | 3 | 0.6667 | NA | NA | 4 | 0.75 | 2 | 0 | 1 | 0 | NA | NA | 2 | 0.5 | 5 | 0.2 |
| 9_3217662; | 1 | 1 | NA | NA | 3 | 0.6667 | NA | NA | 4 | 0.75 | 3 | 1 | 1 | 0 | NA | NA | 2 | 0.5 | 6 | 0.6667 |
| 9_3217698 | 1 | 0 | NA | NA | 3 | 0.3333 | NA | NA | 4 | 0.25 | 3 | 0.3333 | 2 | 0 | NA | NA | 2 | 0 | 7 | 0.1429 |
| 9_3442163 | NA | NA | NA | NA | NA | NA | 1 | 1 | 1 | 1 | NA | NA | 1 | 1 | NA | NA | NA | NA | 1 | 1 |
| 9_3494534! | NA | NA | 1 | 0 | NA | NA | 1 | 1 | 2 | 0.5 | NA | NA | 1 | 0 | 1 | 1 | 2 | 0.5 | 4 | 0.5 |
| 9_3494631! | 3 | 1 | 3 | 0.6667 | 3 | 1 | 1 | 1 | 10 | 0.9 | 3 | 1 | 3 | 0.6667 | 4 | 1 | 4 | 1 | 14 | 0.9286 |
| 9_3494641( | 5 | 0.4 | 3 | 0 | 2 | 0.5 | 4 | 0.25 | 14 | 0.2857 | 3 | 0.6667 | 4 | 0 | 4 | 0.25 | 8 | 0.25 | 19 | 0.2632 |
| 9_3494652; | 6 | 0.8333 | 1 | 0 | 1 | 1 | 3 | 1 | 11 | 0.8182 | 1 | 0 | 2 | 0.5 | 4 | 0.75 | 4 | 1 | 11 | 0.7273 |
| 9_3494656! | 6 | 0.5 | 4 | 0.25 | 3 | 0.3333 | 4 | 0.25 | 17 | 0.3529 | 3 | 0.6667 | 5 | 0.2 | 5 | 0.2 | 5 | 0.2 | 18 | 0.2778 |
| 9_3494657! | 6 | 0.6667 | 4 | 0.25 | 2 | 0.5 | 2 | 0.5 | 14 | 0.5 | 2 | 0.5 | 3 | 0.6667 | 5 | 0.2 | 6 | 0.3333 | 16 | 0.375 |
| 9_3494661( | 6 | 1 | 2 | 1 | 3 | 1 | 4 | 1 | 15 | 1 | 4 | 0.5 | 5 | 0.6 | 4 | 1 | 7 | 1 | 20 | 0.8 |
| 9_3494666; | 5 | 0.8 | 4 | 0.5 | 3 | 1 | 4 | 0.75 | 16 | 0.75 | 4 | 1 | 5 | 1 | 5 | 1 | 7 | 0.7143 | 21 | 0.9048 |
| 9_3494667 | 4 | 0.75 | 3 | 1 | 1 | 1 | 5 | 1 | 13 | 0.9231 | 2 | 1 | 4 | 0.75 | 3 | 0.6667 | 4 | 1 | 13 | 0.8462 |
| 9_3494671( | 6 | 0.1667 | 2 | 0 | 3 | 0.3333 | 5 | 0 | 16 | 0.125 | 3 | 0 | 4 | 0.25 | 6 | 0 | 7 | 0 | 20 | 0.05 |
| 9_3494687( | 7 | 1 | 4 | 1 | 3 | 1 | 5 | 1 | 19 | 1 | 4 | 1 | 5 | 1 | 6 | 1 | 5 | 1 | 20 | 1 |
| 9_3494688( | 7 | 0.8571 | 3 | 1 | 2 | 1 | 6 | 1 | 18 | 0.9444 | 3 | 1 | 3 | 1 | 5 | 1 | 7 | 0.8571 | 18 | 0.9444 |
| 9_3494697( | 7 | 1 | 2 | 1 | 3 | 1 | 6 | 1 | 18 | 1 | 4 | 1 | 5 | 1 | 6 | 1 | 7 | 1 | 22 | 1 |
| 9_3494714; | 7 | 1 | 5 | 1 | 2 | 1 | 7 | 1 | 21 | 1 | 5 | 0.8 | 4 | 1 | 5 | 1 | 8 | 0.875 | 22 | 0.9091 |
| 9_3494717( | 7 | 1 | 5 | 1 | 3 | 0.6667 | 6 | 1 | 21 | 0.9524 | 4 | 1 | 4 | 1 | 6 | 1 | 7 | 0.7143 | 21 | 0.9048 |
| 9_3494722! | 5 | 0.6 | 5 | 0.6 | 3 | 0.3333 | 4 | 0.75 | 17 | 0.5882 | 5 | 0.8 | 3 | 0.6667 | 4 | 1 | 6 | 0.5 | 18 | 0.7222 |
| 9_3494739( | 8 | 0.625 | 5 | 0.2 | 2 | 0.5 | 6 | 0.5 | 21 | 0.4762 | 6 | 0.5 | 5 | 0.4 | 4 | 0.75 | 7 | 0.5714 | 22 | 0.5455 |
| 9_3494746! | 7 | 0.2857 | 5 | 0.8 | 4 | 0.25 | 6 | 0.3333 | 22 | 0.4091 | 6 | 0.3333 | 5 | 0.8 | 6 | 0.8333 | 7 | 0.4286 | 24 | 0.5833 |
| 9_3494749 | 7 | 0.2857 | 4 | 0.25 | 4 | 0.25 | 4 | 0.25 | 19 | 0.2632 | 6 | 0.1667 | 5 | 0.4 | 7 | 0.5714 | 6 | 0 | 24 | 0.2917 |
| 9_3494751! | 6 | 0.5 | 5 | 0.4 | 2 | 0.5 | 6 | 0.5 | 19 | 0.4737 | 5 | 0.8 | 6 | 0.8333 | 6 | 0.8333 | 8 | 0.875 | 25 | 0.84 |
| 9_3494753( | 6 | 0.5 | 6 | 0.3333 | 4 | 0.5 | 7 | 0.4286 | 23 | 0.4348 | 6 | 0.5 | 6 | 0.6667 | 5 | 0.8 | 7 | 0.5714 | 24 | 0.625 |
| 9_3495827( | 3 | 0.3333 | 2 | 0 | 3 | 0 | 5 | 0.2 | 13 | 0.1538 | 9 | 0 | 4 | 0.5 | 4 | 0.25 | 1 | 0 | 18 | 0.1667 |
| 9_3495829! | 1 | 1 | 3 | 0 | 2 | 0 | 4 | 0.25 | 10 | 0.2 | 6 | 0 | 4 | 0 | 3 | 0.3333 | 1 | 1 | 14 | 0.1429 |
| 9_3495863( | 2 | 0.5 | 2 | 0 | 2 | 0.5 | 3 | 0.3333 | 9 | 0.3333 | 6 | 0.3333 | 1 | 1 | 1 | 0 | 1 | 0 | 9 | 0.3333 |
| 9_3511579- | 5 | 0 | 6 | 0.3333 | 6 | 0.3333 | 1 | 0 | 18 | 0.2222 | 1 | 0 | 5 | 0.4 | 3 | 0.3333 | 5 | 0.2 | 14 | 0.2857 |
| 9_3511582( | 4 | 0.25 | 6 | 0 | 5 | 0.2 | 2 | 0 | 17 | 0.1176 | 1 | 0 | 5 | 0.4 | 3 | 0 | 4 | 0.25 | 13 | 0.2308 |
| 9_3511588; | 3 | 0 | 5 | 0.6 | 4 | 0.5 | 2 | 0 | 14 | 0.3571 | 1 | 0 | 5 | 0.2 | 3 | 0.3333 | 5 | 0.2 | 14 | 0.2143 |
| 9_3511595( | 5 | 0.2 | 4 | 0.75 | 4 | 0.5 | 1 | 0 | 14 | 0.4286 | 1 | 0 | 5 | 0.2 | 3 | 0.3333 | 6 | 0.1667 | 15 | 0.2 |
| 9_3511601; | 4 | 0 | 5 | 0.4 | 4 | 0.5 | 1 | 0 | 14 | 0.2857 | NA | NA | 4 | 0.25 | 2 | 0 | 5 | 0.2 | 11 | 0.1818 |
| 9_3511660! | 4 | 0 | 5 | 0 | 6 | 0 | 1 | 1 | 16 | 0.0625 | 1 | 0 | 5 | 0.2 | 2 | 0 | 5 | 0.2 | 13 | 0.1538 |
| 9_3512259; | 2 | 0 | 2 | 0.5 | 4 | 0 | 5 | 0 | 13 | 0.0769 | 6 | 0 | 5 | 0.2 | 3 | 0 | NA | NA | 14 | 0.0714 |
| 9_3512789; | 6 | 0 | 5 | 0 | 8 | 0.125 | 11 | 0.0909 | 30 | 0.0667 | 7 | 0.2857 | 9 | 0.2222 | 8 | 0.125 | 5 | 0 | 29 | 0.1724 |
| 9_3512871- | 3 | 0.6667 | 2 | 0 | 4 | 0.25 | 5 | 0 | 14 | 0.2143 | 3 | 1 | 1 | 1 | 5 | 0.6 | 4 | 0.75 | 13 | 0.7692 |
| 9_3512877( | 3 | 0.3333 | 1 | 0 | 5 | 0.4 | 4 | 0 | 13 | 0.2308 | 3 | 0.3333 | 1 | 0 | 5 | 0.2 | 4 | 0.25 | 13 | 0.2308 |
| 9_3512878 | 3 | 0 | 1 | 1 | 5 | 0 | 6 | 0.1667 | 15 | 0.1333 | 4 | 0.5 | 1 | 0 | 4 | 0.25 | 4 | 0.25 | 13 | 0.3077 |
| 9_3512880! | 3 | 0.3333 | 2 | 0 | 5 | 0.2 | 6 | 0 | 16 | 0.125 | 4 | 0.5 | 1 | 1 | 5 | 0.2 | 3 | 0.3333 | 13 | 0.3846 |
| 9_3512895- | 3 | 0 | 2 | 0.5 | 5 | 0 | 6 | 0.1667 | 16 | 0.125 | 3 | 0 | NA | NA | 5 | 0.2 | 3 | 0 | 11 | 0.0909 |
| 9_3513583 | NA | NA | 1 | 1 | NA | NA | 1 | 0 | 2 | 0.5 | NA | NA | 1 | 0 | NA | NA | 1 | 1 | 2 | 0.5 |
| 9_3513623! | NA | NA | 2 | 1 | NA | NA | 1 | 1 | 3 | 1 | NA | NA | 1 | 1 | NA | NA | NA | NA | 1 | 1 |

| | | | | | | | | | | | | | | | | | | | | |
|---|---|---|---|---|---|---|---|---|---|---|---|---|---|---|---|---|---|---|---|---|
| 9_3513649 | NA | NA | 2 | 0.5 | NA | NA | 1 | 1 | 3 | 0.6667 | NA | NA | 1 | 1 | 1 | 1 | NA | NA | 2 | 1 |
| 9_3513671 | NA | NA | 2 | 1 | NA | NA | 1 | 0 | 3 | 0.6667 | NA | NA | 1 | 1 | NA | NA | NA | NA | 1 | 1 |
| 9_3513685 | NA | NA | 2 | 1 | NA | NA | 1 | 1 | 3 | 1 | NA | NA | 1 | 1 | NA | NA | NA | NA | 1 | 1 |
| 9_3513708 | NA | NA | 2 | 1 | NA | NA | 1 | 1 | 3 | 1 | NA | NA | 1 | 1 | 1 | 1 | NA | NA | 2 | 1 |
| 9_3513719 | NA | NA | 2 | 1 | NA | NA | 1 | 1 | 3 | 1 | NA | NA | 1 | 1 | 1 | 1 | NA | NA | 2 | 1 |
| 9_3513727 | NA | NA | 2 | 0.5 | NA | NA | 1 | 0 | 3 | 0.3333 | NA | NA | 1 | 1 | NA | NA | NA | NA | 1 | 1 |
| 9_3513865 | 2 | 1 | 2 | 0 | 3 | 0.6667 | 3 | 0.3333 | 10 | 0.5 | 1 | 0 | 1 | 0 | 1 | 0 | 1 | 1 | 4 | 0.25 |
| 9_3518667 | 2 | 0 | NA | NA | 2 | 0 | 3 | 0.3333 | 7 | 0.1429 | 2 | 0 | 1 | 0 | 2 | 0.5 | 1 | 0 | 6 | 0.1667 |
| 9_3518672 | 2 | 1 | NA | NA | 2 | 0.5 | 3 | 0.6667 | 7 | 0.7143 | 2 | 0.5 | 1 | 1 | 2 | 1 | 1 | 1 | 6 | 0.8333 |
| 9_3518679 | 2 | 0 | NA | NA | 2 | 0.5 | 2 | 0 | 6 | 0.1667 | 1 | 0 | 1 | 0 | 2 | 0 | 1 | 1 | 5 | 0.2 |
| 9_3518684 | 2 | 1 | NA | NA | 2 | 0.5 | 2 | 1 | 6 | 0.8333 | 2 | 1 | NA | NA | 2 | 1 | 1 | 1 | 5 | 1 |
| 9_3518688 | 2 | 1 | NA | NA | 2 | 1 | 2 | 1 | 6 | 1 | 2 | 0.5 | 1 | 1 | 2 | 1 | 1 | 1 | 6 | 0.8333 |
| 9_3518712 | 2 | 0 | NA | NA | 2 | 0 | 3 | 0.3333 | 7 | 0.1429 | 2 | 0.5 | 1 | 0 | 1 | 1 | 1 | 0 | 5 | 0.4 |
| 9_3541543 | 1 | 1 | NA | NA | NA | NA | 3 | 1 | 4 | 1 | 1 | 1 | NA | NA | NA | NA | NA | NA | 1 | 1 |
| 9_3541560 | 1 | 1 | NA | NA | NA | NA | 3 | 1 | 4 | 1 | 1 | 1 | NA | NA | NA | NA | NA | NA | 1 | 1 |
| 9_3541681 | 1 | 1 | NA | NA | NA | NA | 3 | 0.3333 | 4 | 0.5 | 1 | 1 | NA | NA | NA | NA | NA | NA | 1 | 1 |
| 9_3541777 | 2 | 0.5 | NA | NA | 1 | 1 | 2 | 0 | 5 | 0.4 | 1 | 1 | NA | NA | NA | NA | NA | NA | 1 | 1 |
| 9_3544814 | NA | NA | 1 | 0 | 1 | 1 | 1 | 0 | 3 | 0.3333 | 1 | 0 | NA | NA | 3 | 0.3333 | 1 | 1 | 5 | 0.4 |
| 9_3544825 | NA | NA | 1 | 0 | 1 | 1 | 1 | 0 | 3 | 0.3333 | 1 | 0 | NA | NA | 3 | 0.3333 | 2 | 0 | 6 | 0.1667 |
| 9_3545315 | 4 | 0 | 6 | 0 | 4 | 0.25 | 5 | 0 | 19 | 0.0526 | 8 | 0.125 | 6 | 0.1667 | 8 | 0.25 | 4 | 0.25 | 26 | 0.1923 |
| 9_3545318 | 10 | 0 | 6 | 0.1667 | 6 | 0 | 10 | 0.1 | 32 | 0.0625 | 8 | 0.125 | 7 | 0.1429 | 8 | 0.375 | 4 | 0 | 27 | 0.1852 |
| 9_3545320 | 7 | 0.1429 | 5 | 0.2 | 4 | 0.25 | 9 | 0.2222 | 25 | 0.2 | 5 | 0.2 | 4 | 0.5 | 6 | 0.3333 | 4 | 0.25 | 19 | 0.3158 |
| 9_3545826 | 6 | 0.1667 | 6 | 0.1667 | 5 | 0.2 | 9 | 0.1111 | 26 | 0.1538 | 7 | 0 | 8 | 0.25 | 5 | 0 | 4 | 0 | 24 | 0.0833 |
| 9_3547236 | 2 | 1 | 1 | 1 | 1 | 1 | NA | NA | 4 | 1 | NA | NA | NA | NA | 1 | 1 | NA | NA | 1 | 1 |
| 9_3547270 | 2 | 1 | NA | NA | 1 | 1 | NA | NA | 3 | 1 | NA | NA | NA | NA | 1 | 1 | NA | NA | 1 | 1 |
| 9_3547622 | 3 | 0.3333 | 1 | 1 | NA | NA | 4 | 0 | 8 | 0.25 | NA | NA | NA | NA | 1 | 0 | 1 | 1 | 2 | 0.5 |
| 9_3547678 | 3 | 1 | 1 | 1 | NA | NA | 3 | 0.6667 | 7 | 0.8571 | NA | NA | 1 | 1 | 1 | 1 | 2 | 1 | 4 | 1 |
| 9_3547687 | 3 | 0 | 1 | 1 | NA | NA | 3 | 0.3333 | 7 | 0.2857 | NA | NA | 1 | 0 | NA | NA | 2 | 0.5 | 3 | 0.3333 |
| 9_3547692 | 3 | 1 | 1 | 1 | NA | NA | 3 | 1 | 7 | 1 | NA | NA | 1 | 0 | 1 | 1 | 2 | 1 | 4 | 0.75 |
| 9_3547784 | 3 | 0 | 1 | 1 | 1 | 0 | 3 | 0 | 8 | 0.125 | 1 | 0 | 1 | 0 | 1 | 0 | 3 | 0.6667 | 6 | 0.3333 |
| 9_3547788 | 4 | 1 | 1 | 1 | NA | NA | 1 | 1 | 6 | 1 | 1 | 0 | NA | NA | 1 | 1 | 2 | 1 | 4 | 0.75 |
| 9_3547791 | 4 | 0.5 | 1 | 1 | 1 | 0 | 2 | 0 | 8 | 0.375 | 1 | 0 | 1 | 0 | 1 | 0 | 3 | 0.3333 | 6 | 0.1667 |
| 9_3547803 | 4 | 0.75 | 1 | 1 | 1 | 1 | 3 | 0.6667 | 9 | 0.7778 | 1 | 1 | NA | NA | 1 | 1 | 4 | 0.75 | 6 | 0.8333 |
| 9_3547812 | 3 | 1 | 1 | 1 | 1 | 1 | 3 | 0.3333 | 8 | 0.75 | 1 | 1 | 1 | 0 | 1 | 1 | 4 | 0.75 | 7 | 0.7143 |
| 9_3664250 | 2 | 0 | NA | NA | NA | NA | 2 | 0.5 | 4 | 0.25 | 1 | 0 | 3 | 0 | 1 | 1 | 3 | 0.3333 | 8 | 0.25 |
| 9_3664384 | 4 | 0.5 | 4 | 0.25 | 2 | 0 | 1 | 0 | 11 | 0.2727 | 2 | 0.5 | 3 | 0.6667 | 3 | 0.3333 | 5 | 0.4 | 13 | 0.4615 |
| 9_3664408 | 3 | 0.6667 | 3 | 0.3333 | 2 | 0 | 1 | 0 | 9 | 0.3333 | 2 | 0.5 | 2 | 1 | 2 | 0 | 3 | 0.6667 | 9 | 0.5556 |
| 9_3664413 | 4 | 0.25 | 2 | 0 | 1 | 0 | NA | NA | 7 | 0.1429 | 1 | 0 | 2 | 0.5 | 1 | 1 | 4 | 0 | 8 | 0.25 |
| 9_3664422 | 6 | 0.1667 | 4 | 0 | 2 | 0 | 1 | 1 | 13 | 0.1538 | 2 | 0 | 4 | 0.25 | 1 | 0 | 4 | 0 | 11 | 0.0909 |
| 9_3669060 | 19 | 0 | 23 | 0.087 | 16 | 0.125 | 12 | 0.25 | 70 | 0.1 | 13 | 0.0769 | 14 | 0.2857 | 7 | 0.1429 | 9 | 0 | 43 | 0.1395 |
| 9_3669116 | 17 | 0.4118 | 17 | 0.2941 | 11 | 0.4545 | 11 | 0.2727 | 56 | 0.3571 | 12 | 0.6667 | 8 | 0.5 | 7 | 0.5714 | 7 | 0.8571 | 34 | 0.6471 |
| 9_3675512 | 20 | 0.3 | 24 | 0.4167 | 22 | 0.3636 | 24 | 0.4583 | 90 | 0.3889 | 22 | 0.5455 | 25 | 0.32 | 23 | 0.6522 | 24 | 0.4167 | 94 | 0.4787 |
| 9_3675518 | 25 | 0.2 | 25 | 0.16 | 22 | 0.1364 | 24 | 0.125 | 96 | 0.1562 | 22 | 0.3636 | 29 | 0.2759 | 28 | 0.3214 | 26 | 0.1538 | 105 | 0.2762 |
| 9_3675529 | 23 | 0.2609 | 24 | 0.125 | 22 | 0.1364 | 23 | 0.1304 | 92 | 0.163 | 23 | 0.1304 | 24 | 0.125 | 25 | 0.12 | 22 | 0.0455 | 94 | 0.1064 |
| 9_3675551 | NA | NA | NA | NA | 1 | 1 | NA | NA | 1 | 1 | NA | NA | NA | NA | NA | NA | 1 | 1 | 1 | 1 |
| 9_3675901 | 1 | 0 | NA | NA | 1 | 1 | NA | NA | 2 | 0.5 | NA | NA | 1 | 1 | NA | NA | 2 | 1 | 3 | 1 |
| 9_3675920 | 1 | 0 | NA | NA | 1 | 1 | NA | NA | 2 | 0.5 | NA | NA | 1 | 0 | 1 | 0 | 1 | 1 | 3 | 0.3333 |
| 9_3680243 | 2 | 0 | 9 | 0.1111 | 7 | 0.1429 | 2 | 0.5 | 20 | 0.15 | 5 | 0 | 3 | 0 | 8 | 0.125 | 4 | 0.25 | 20 | 0.1 |
| 9_3680255 | 2 | 0.5 | 7 | 0 | 7 | 0.1429 | 2 | 0.5 | 18 | 0.1667 | 5 | 0.2 | 3 | 0 | 9 | 0.5556 | 4 | 0 | 21 | 0.2857 |
| 9_3680292 | 1 | 1 | 7 | 0.7143 | 3 | 1 | 3 | 0.6667 | 14 | 0.7857 | 5 | 0.6 | 2 | 0.5 | 7 | 0.7143 | 4 | 0.75 | 18 | 0.6667 |
| 9_3680302 | 1 | 1 | 5 | 0.6 | 5 | 1 | 3 | 0.3333 | 14 | 0.7143 | 4 | 0.75 | 3 | 0.6667 | 6 | 1 | 3 | 0.6667 | 16 | 0.8125 |
| 9_3680322 | 2 | 0.5 | 6 | 0.1667 | 6 | 0 | 3 | 0 | 17 | 0.1176 | 6 | 0 | 3 | 0 | 8 | 0.125 | 4 | 0.25 | 21 | 0.0952 |
| 9_3680327 | 2 | 0 | 5 | 0.2 | 6 | 0.3333 | 2 | 0 | 15 | 0.2 | 6 | 0.5 | 3 | 0 | 8 | 0.375 | 3 | 0 | 20 | 0.3 |
| 9_3680333 | 2 | 1 | 5 | 0.6 | 3 | 0.6667 | 3 | 1 | 13 | 0.7692 | 5 | 0.8 | 2 | 1 | 6 | 0.6667 | 3 | 1 | 16 | 0.8125 |
| 9_3680346 | 1 | 1 | 3 | 0 | 5 | 0 | 3 | 0.3333 | 12 | 0.1667 | 4 | 1 | 3 | 0.3333 | 7 | 0.4286 | 4 | 0.25 | 18 | 0.5 |
| 9_3680349 | 2 | 1 | 3 | 1 | 4 | 1 | 2 | 1 | 11 | 1 | 2 | 1 | 2 | 1 | 5 | 1 | 4 | 1 | 13 | 1 |
| 9_3680352 | 1 | 0 | 4 | 0.25 | 4 | 0 | 3 | 0 | 12 | 0.0833 | 5 | 0.2 | 3 | 0.3333 | 5 | 0.4 | 4 | 0.5 | 17 | 0.3529 |
| 9_3680355 | 2 | 1 | 5 | 0.6 | 5 | 1 | 3 | 1 | 15 | 0.8667 | 5 | 0.6 | 3 | 0.6667 | 5 | 1 | 4 | 1 | 17 | 0.8235 |
| 9_3680364 | 1 | 1 | 4 | 0.75 | 5 | 0.8 | 3 | 1 | 13 | 0.8462 | 4 | 1 | 3 | 0.6667 | 7 | 1 | 2 | 1 | 16 | 0.9375 |
| 9_3680365 | 1 | 1 | 5 | 0 | 4 | 1 | 3 | 0.6667 | 13 | 0.5385 | 4 | 0.25 | 2 | 1 | 5 | 0.6 | 3 | 1 | 14 | 0.6429 |
| 9_3680372 | 1 | 0 | 4 | 0.5 | 3 | 0 | 3 | 0.3333 | 11 | 0.2727 | 3 | 0.3333 | 2 | 0.5 | 5 | 0.4 | 3 | 0.3333 | 13 | 0.3846 |
| 9_3680374 | 1 | 0 | 5 | 0.2 | 3 | 0 | 3 | 0.6667 | 12 | 0.25 | 2 | 0 | 3 | 0.3333 | 6 | 0.1667 | 1 | 1 | 12 | 0.25 |
| 9_3680376 | 2 | 0.5 | 5 | 0.4 | 4 | 0.75 | 2 | 0.5 | 13 | 0.5385 | 2 | 0 | 3 | 0.3333 | 5 | 0.8 | 3 | 1 | 13 | 0.6154 |
| 9_3680380 | 2 | 1 | 5 | 1 | 3 | 1 | 3 | 1 | 13 | 1 | 3 | 1 | 2 | 1 | 5 | 0.8 | 3 | 1 | 13 | 0.9231 |
| 9_3680384 | 2 | 1 | 5 | 0.2 | 4 | 0.25 | 2 | 0.5 | 13 | 0.3846 | 3 | 1 | 3 | 0 | 5 | 0.4 | 3 | 0.6667 | 14 | 0.5 |

| | | | | | | | | | | | | | | | | | | | | |
|---|---|---|---|---|---|---|---|---|---|---|---|---|---|---|---|---|---|---|---|---|
| 9_36803915 | 2 | 0 | 3 | 0.3333 | 4 | 0 | 3 | 0.3333 | 12 | 0.1667 | 3 | 0.3333 | 3 | 0.6667 | 5 | 0 | 3 | 0 | 14 | 0.2143 |
| 9_36804134 | 2 | 0 | 5 | 0.2 | 5 | 0.4 | 3 | 0.3333 | 15 | 0.2667 | 2 | 0.5 | 3 | 0.3333 | 6 | 0.1667 | 3 | 0.3333 | 14 | 0.2857 |
| 9_36804142 | 1 | 0 | 3 | 0 | 4 | 0.5 | 3 | 0.6667 | 11 | 0.3636 | 3 | 0 | 3 | 0.3333 | 3 | 0 | 3 | 0.3333 | 12 | 0.1667 |
| 9_36834998 | 2 | 0 | 4 | 0 | 1 | 0 | 3 | 0.3333 | 10 | 0.1 | 1 | 0 | 1 | 0 | 2 | 0.5 | 1 | 0 | 5 | 0.2 |
| 9_37057328 | NA | NA | 1 | 1 | NA | NA | NA | NA | 1 | 1 | 1 | 1 | NA | NA | NA | NA | NA | NA | 1 | 1 |
| 9_37057976 | NA | NA | 1 | 1 | NA | NA | NA | NA | 1 | 1 | 1 | 1 | NA | NA | NA | NA | NA | NA | 1 | 1 |
| 9_37058430 | NA | NA | 1 | 1 | NA | NA | NA | NA | 1 | 1 | 1 | 1 | NA | NA | NA | NA | NA | NA | 1 | 1 |
| 9_37058623 | NA | NA | 1 | 1 | NA | NA | NA | NA | 1 | 1 | 1 | 1 | NA | NA | NA | NA | NA | NA | 1 | 1 |
| 9_37060039 | NA | NA | 1 | 1 | NA | NA | NA | NA | 1 | 1 | 1 | 1 | NA | NA | NA | NA | NA | NA | 1 | 1 |
| 9_37060090 | NA | NA | 1 | 1 | NA | NA | NA | NA | 1 | 1 | 1 | 1 | NA | NA | NA | NA | NA | NA | 1 | 1 |
| 9_37133931 | 10 | 0.6 | 2 | 0.5 | 5 | 0.4 | 1 | 1 | 18 | 0.5556 | 4 | 0.75 | 3 | 0.6667 | 1 | 1 | 4 | 1 | 12 | 0.8333 |
| 9_37133994 | 10 | 0 | 2 | 0 | 5 | 0.4 | 1 | 0 | 18 | 0.1111 | 4 | 0.25 | 3 | 0 | 2 | 0 | 4 | 0 | 13 | 0.0769 |
| 9_37134051 | 10 | 0.2 | 2 | 1 | 5 | 0.4 | 1 | 0 | 18 | 0.3333 | 4 | 0.5 | 3 | 0.6667 | 2 | 0.5 | 4 | 0.5 | 13 | 0.5385 |
| 9_37134121 | 10 | 0 | 2 | 0 | 5 | 0.2 | 1 | 0 | 18 | 0.0556 | 4 | 0 | 3 | 0.3333 | 2 | 0 | 3 | 0.3333 | 12 | 0.1667 |
| 9_37134155 | 8 | 0.25 | NA | NA | 4 | 0.25 | 1 | 1 | 13 | 0.3077 | NA | NA | 2 | 0.5 | 1 | 1 | 2 | 0 | 5 | 0.4 |
| 9_37296045 | 8 | 0.5 | 18 | 0.4444 | 12 | 0.5833 | 9 | 0.4444 | 47 | 0.4894 | 9 | 0.2222 | 7 | 0.8571 | 6 | 0.6667 | 11 | 0.5455 | 33 | 0.5455 |
| 9_37296357 | 8 | 0.5 | 19 | 0.2632 | 10 | 0.3 | 9 | 0.4444 | 46 | 0.3478 | 12 | 0.3333 | 9 | 0.2222 | 9 | 0.3333 | 11 | 0.3636 | 41 | 0.3171 |
| 9_37296583 | 4 | 0.25 | 13 | 0 | 9 | 0 | 5 | 0.2 | 31 | 0.0645 | 8 | 0.125 | 8 | 0 | 10 | 0 | 10 | 0.1 | 36 | 0.0556 |
| 9_37296648 | 5 | 0.6 | 15 | 0.6667 | 9 | 0.2222 | 5 | 0 | 34 | 0.4412 | 9 | 0.5556 | 10 | 0.3 | 10 | 0.4 | 9 | 0.4444 | 38 | 0.4211 |
| 9_37296844 | 5 | 0.4 | 16 | 0.625 | 10 | 0.7 | 6 | 1 | 37 | 0.6757 | 8 | 0.25 | 10 | 0.5 | 11 | 0.4545 | 10 | 0.5 | 39 | 0.4359 |
| 9_37296892 | 4 | 0.25 | 16 | 0.1875 | 8 | 0.375 | 4 | 0.5 | 32 | 0.2812 | 8 | 0.5 | 10 | 0.2 | 11 | 0.2727 | 10 | 0.3 | 39 | 0.3077 |
| 9_37296925 | 4 | 0.5 | 15 | 0.1333 | 10 | 0.2 | 6 | 0.3333 | 35 | 0.2286 | 9 | 0.3333 | 10 | 0.5 | 11 | 0.1818 | 9 | 0.4444 | 39 | 0.359 |
| 9_37297132 | 4 | 0.5 | 15 | 0.2 | 9 | 0.1111 | 7 | 0.5714 | 35 | 0.2857 | 10 | 0.1 | 9 | 0.1111 | 9 | 0.2222 | 11 | 0.4545 | 39 | 0.2308 |
| 9_37297139 | 4 | 0.25 | 15 | 0.2667 | 8 | 0.25 | 8 | 0.125 | 35 | 0.2286 | 7 | 0.1429 | 12 | 0.25 | 10 | 0.2 | 10 | 0.2 | 39 | 0.2051 |
| 9_37325017 | NA | NA | 1 | 1 | NA | NA | NA | NA | 1 | 1 | 2 | 0.5 | 2 | 0.5 | 1 | 1 | 3 | 0.6667 | 8 | 0.625 |
| 9_37325052 | 1 | 1 | 2 | 1 | NA | NA | NA | NA | 3 | 1 | 2 | 0 | 2 | 1 | 1 | 0 | 3 | 0.3333 | 8 | 0.375 |
| 9_37325084 | 1 | 1 | 1 | 1 | NA | NA | NA | NA | 2 | 1 | 2 | 1 | 3 | 1 | 1 | 1 | 2 | 1 | 8 | 1 |
| 9_37325218 | 1 | 0 | 1 | 1 | NA | NA | NA | NA | 2 | 0.5 | 2 | 0 | 2 | 0 | 1 | 1 | 2 | 0 | 7 | 0.1429 |
| 9_37325292 | 1 | 1 | 2 | 0.5 | NA | NA | NA | NA | 3 | 0.6667 | 2 | 1 | 3 | 0.6667 | 1 | 0 | 2 | 0.5 | 8 | 0.625 |
| 9_37325307 | NA | NA | 1 | 1 | NA | NA | NA | NA | 1 | 1 | 1 | 0 | 2 | 0.5 | 1 | 0 | 2 | 0.5 | 6 | 0.3333 |
| 9_37434046 | 3 | 1 | 1 | 1 | 3 | 1 | 2 | 1 | 9 | 1 | 1 | 1 | 1 | 1 | 5 | 0.6 | 2 | 1 | 9 | 0.7778 |
| 9_37434066 | 3 | 0 | NA | NA | 3 | 0.3333 | 2 | 0.5 | 8 | 0.25 | 1 | 0 | 1 | 0 | 4 | 0.25 | 1 | 0 | 7 | 0.1429 |
| 9_37434258 | 3 | 1 | 1 | 1 | 2 | 1 | 2 | 1 | 8 | 1 | 1 | 1 | 1 | 1 | 5 | 1 | 2 | 0.5 | 9 | 0.8889 |
| 9_37434336 | 2 | 1 | 1 | 1 | 3 | 1 | 2 | 1 | 8 | 1 | 1 | 1 | NA | NA | 5 | 0.8 | 2 | 1 | 8 | 0.875 |
| 9_37434492 | 3 | 1 | 1 | 1 | 2 | 1 | 1 | 1 | 7 | 1 | 1 | 1 | NA | NA | 4 | 1 | 2 | 0.5 | 7 | 0.8571 |
| 9_37434608 | 2 | 1 | 1 | 1 | 3 | 1 | 1 | 1 | 7 | 1 | 1 | 1 | 1 | 1 | 4 | 1 | 1 | 1 | 7 | 1 |
| 9_37434699 | 2 | 0.5 | 1 | 1 | 4 | 0.75 | 1 | 1 | 8 | 0.75 | 1 | 1 | 1 | 1 | 5 | 0.8 | 2 | 1 | 9 | 0.8889 |
| 9_37434928 | 3 | 0.6667 | 1 | 1 | 3 | 0.6667 | 2 | 1 | 9 | 0.7778 | 1 | 1 | 1 | 1 | 4 | 0.75 | 1 | 1 | 7 | 0.8571 |
| 9_37434985 | 2 | 1 | 1 | 1 | 3 | 1 | 1 | 1 | 7 | 1 | 1 | 1 | NA | NA | 4 | 1 | 2 | 1 | 7 | 1 |
| 9_37435141 | 3 | 0.6667 | 1 | 0 | 4 | 0.75 | 1 | 1 | 9 | 0.6667 | 1 | 0 | NA | NA | 5 | 1 | 2 | 1 | 8 | 0.875 |
| 9_37435719 | NA | NA | NA | NA | 1 | 1 | NA | NA | 1 | 1 | NA | NA | NA | NA | NA | NA | 1 | 1 | 1 | 1 |
| 9_37435738 | NA | NA | NA | NA | 1 | 1 | NA | NA | 1 | 1 | NA | NA | NA | NA | NA | NA | 1 | 1 | 1 | 1 |
| 9_37435964 | NA | NA | NA | NA | 1 | 1 | NA | NA | 1 | 1 | NA | NA | NA | NA | NA | NA | 1 | 1 | 1 | 1 |
| 9_37448943 | 1 | 1 | 5 | 0.2 | 3 | 0 | 1 | 1 | 10 | 0.3 | NA | NA | 2 | 0.5 | 4 | 0.25 | 2 | 0 | 8 | 0.25 |
| 9_37449148 | 3 | 0.6667 | 5 | 1 | 2 | 0.5 | 1 | 1 | 11 | 0.8182 | NA | NA | 2 | 1 | 5 | 0.8 | 1 | 1 | 8 | 0.875 |
| 9_37449236 | 3 | 0.6667 | 5 | 0.2 | 3 | 0 | 1 | 1 | 12 | 0.3333 | NA | NA | 3 | 0 | 4 | 0.25 | 2 | 0 | 9 | 0.1111 |
| 9_37449256 | 3 | 0.6667 | 4 | 0.5 | 3 | 0.6667 | 1 | 1 | 11 | 0.6364 | NA | NA | 3 | 0.6667 | 5 | 0.4 | 1 | 0 | 9 | 0.4444 |
| 9_37449273 | 2 | 1 | 4 | 0.75 | 2 | 0.5 | 1 | 1 | 9 | 0.7778 | NA | NA | 1 | 1 | 3 | 1 | 1 | 1 | 5 | 1 |
| 9_37456497 | 75 | 0.12 | 142 | 0.1127 | 106 | 0.1887 | 68 | 0.2794 | 391 | 0.1637 | 83 | 0.241 | 70 | 0.1429 | 68 | 0.1618 | 90 | 0.1444 | 311 | 0.1736 |
| 9_37557659 | 1 | 0 | 1 | 0 | NA | NA | 4 | 0.25 | 6 | 0.1667 | 1 | 0 | 2 | 0.5 | NA | NA | NA | NA | 3 | 0.3333 |
| 9_37557823 | 2 | 1 | NA | NA | NA | NA | 2 | 1 | 4 | 1 | 2 | 0.5 | 3 | 1 | NA | NA | NA | NA | 5 | 0.8 |
| 9_37557884 | 2 | 0.5 | NA | NA | NA | NA | 2 | 1 | 4 | 0.75 | 2 | 0.5 | 3 | 0.3333 | NA | NA | NA | NA | 5 | 0.4 |
| 9_37558152 | 2 | 0.5 | NA | NA | NA | NA | 2 | 1 | 4 | 0.75 | 2 | 0 | 3 | 0.6667 | NA | NA | NA | NA | 5 | 0.4 |
| 9_37558171 | 2 | 1 | NA | NA | NA | NA | 3 | 1 | 5 | 1 | 1 | 0 | 3 | 0.6667 | NA | NA | NA | NA | 4 | 0.5 |
| 9_37559919 | 2 | 0.5 | NA | NA | NA | NA | NA | NA | 2 | 0.5 | NA | NA | 1 | 1 | NA | NA | 1 | 0 | 2 | 0.5 |
| 9_37560296 | 2 | 0.5 | NA | NA | NA | NA | NA | NA | 2 | 0.5 | NA | NA | 1 | 1 | NA | NA | 1 | 1 | 2 | 1 |
| 9_38653853 | 2 | 0 | 2 | 0.5 | 1 | 0 | 2 | 1 | 7 | 0.4286 | 2 | 1 | NA | NA | NA | NA | 1 | 1 | 3 | 1 |
| 9_38653918 | 4 | 0.25 | 1 | 0 | 1 | 1 | 1 | 0 | 7 | 0.2857 | 2 | 0.5 | NA | NA | NA | NA | NA | NA | 2 | 0.5 |
| 9_40122316 | NA | NA | NA | NA | 1 | 1 | NA | NA | 1 | 1 | NA | NA | NA | NA | 1 | 1 | 1 | 1 | 2 | 1 |
| 9_40122473 | NA | NA | NA | NA | 1 | 1 | NA | NA | 1 | 1 | NA | NA | NA | NA | 1 | 1 | NA | NA | 1 | 1 |
| 9_40122837 | NA | NA | NA | NA | 1 | 1 | NA | NA | 1 | 1 | NA | NA | NA | NA | 1 | 1 | NA | NA | 1 | 1 |
| 9_40123240 | NA | NA | NA | NA | 1 | 1 | NA | NA | 1 | 1 | NA | NA | NA | NA | 1 | 1 | 1 | 1 | 2 | 1 |
| 9_40123401 | NA | NA | NA | NA | 1 | 1 | NA | NA | 1 | 1 | NA | NA | NA | NA | 1 | 1 | NA | NA | 1 | 1 |
| 9_40123548 | NA | NA | NA | NA | 1 | 1 | NA | NA | 1 | 1 | NA | NA | NA | NA | 1 | 1 | 1 | 1 | 2 | 1 |
| 9_40124085 | NA | NA | NA | NA | 1 | 1 | NA | NA | 1 | 1 | NA | NA | NA | NA | 1 | 0 | 1 | 1 | 2 | 0.5 |

| | | | | | | | | | | | | | | | | | | | | |
|---|---|---|---|---|---|---|---|---|---|---|---|---|---|---|---|---|---|---|---|---|
| 9_4018076( | 5 | 0 | 8 | 0 | 4 | 0.25 | 1 | 1 | 18 | 0.1111 | 1 | 0 | 2 | 0 | 9 | 0.3333 | 4 | 0 | 16 | 0.1875 |
| 9_4019981( | NA | NA | 5 | 0.4 | NA | NA | 1 | 0 | 6 | 0.3333 | NA | NA | NA | NA | 1 | 1 | 2 | 0.5 | 3 | 0.6667 |
| 9_4020003' | 5 | 0.8 | 9 | 1 | 3 | 1 | 3 | 0.6667 | 20 | 0.9 | 1 | 1 | 1 | 1 | 8 | 1 | 9 | 1 | 19 | 1 |
| 9_4020020: | 6 | 0.8333 | 9 | 0.8889 | 3 | 1 | 3 | 0.6667 | 21 | 0.8571 | 1 | 1 | 1 | 0 | 11 | 0.9091 | 9 | 0.8889 | 22 | 0.8636 |
| 9_4020021. | 7 | 0.5714 | 8 | 0.375 | 3 | 0.3333 | 3 | 0.3333 | 21 | 0.4286 | 1 | 0 | 1 | 0 | 10 | 0.3 | 8 | 0.25 | 20 | 0.25 |
| 9_4020024' | 6 | 0.5 | 10 | 0 | 4 | 0 | 3 | 0 | 23 | 0.1304 | 1 | 1 | 1 | 0 | 8 | 0.5 | 8 | 0.125 | 18 | 0.3333 |
| 9_4020026: | 7 | 0.7143 | 11 | 0.7273 | 3 | 1 | 3 | 0.6667 | 24 | 0.75 | 1 | 1 | 1 | 1 | 9 | 1 | 9 | 1 | 20 | 1 |
| 9_4020034: | 8 | 0.625 | 9 | 0.2222 | 3 | 1 | 3 | 0.6667 | 23 | 0.5217 | 2 | 0.5 | 1 | 1 | 11 | 0.7273 | 9 | 0.7778 | 23 | 0.7391 |
| 9_4020055! | 10 | 0.1 | 13 | 0.2308 | 5 | 0.6 | 2 | 0.5 | 30 | 0.2667 | 4 | 0.25 | 3 | 0.3333 | 10 | 0.7 | 10 | 0.6 | 27 | 0.5556 |
| 9_4020064! | 7 | 0 | 12 | 0.1667 | 5 | 0.4 | 3 | 0 | 27 | 0.1481 | 2 | 0 | 2 | 0 | 7 | 0.4286 | 10 | 0.1 | 21 | 0.1905 |
| 9_4020082. | 9 | 0.5556 | 12 | 0.1667 | 3 | 0.6667 | 2 | 0.5 | 26 | 0.3846 | 4 | 0.25 | 4 | 0.75 | 10 | 0.8 | 10 | 0.6 | 28 | 0.6429 |
| 9_4020084( | 7 | 0.2857 | 7 | 0.1429 | 3 | 0 | 2 | 0 | 19 | 0.1579 | 3 | 0 | 3 | 0 | 9 | 0.1111 | 7 | 0.1429 | 22 | 0.0909 |
| 9_4020086' | 8 | 0.25 | 11 | 0.0909 | 4 | 0.25 | 2 | 0 | 25 | 0.16 | 4 | 0 | 4 | 0 | 11 | 0.1818 | 11 | 0.0909 | 30 | 0.1 |
| 9_4020092: | 8 | 1 | 13 | 0.8462 | 4 | 1 | 3 | 1 | 28 | 0.9286 | 4 | 1 | 5 | 0.8 | 9 | 1 | 9 | 0.8889 | 27 | 0.9259 |
| 9_4020099! | 9 | 0.5556 | 15 | 0.3333 | 3 | 0.6667 | 3 | 1 | 30 | 0.5 | 4 | 0.5 | 4 | 0.25 | 13 | 0.6154 | 11 | 0.7273 | 32 | 0.5938 |
| 9_4020104! | 9 | 0.1111 | 15 | 0.3333 | 4 | 0.25 | 3 | 0.6667 | 31 | 0.2903 | 3 | 0 | 5 | 0.4 | 11 | 0.3636 | 12 | 0.25 | 31 | 0.2903 |
| 9_4020110' | 9 | 0.6667 | 11 | 0.4545 | 4 | 0.5 | 3 | 0.6667 | 27 | 0.5556 | 2 | 1 | 5 | 0.8 | 13 | 0.6923 | 11 | 0.9091 | 31 | 0.8065 |
| 9_4020179: | 9 | 0 | 17 | 0.1176 | 5 | 0.4 | 3 | 0.3333 | 34 | 0.1471 | 3 | 0.3333 | 5 | 0 | 14 | 0.2143 | 13 | 0 | 35 | 0.1143 |
| 9_4020492: | 9 | 0.5556 | 29 | 0.4828 | 20 | 0.45 | 7 | 0.5714 | 65 | 0.4923 | 15 | 0.4 | 12 | 0.6667 | 13 | 0.6154 | 13 | 0.4615 | 53 | 0.5283 |
| 9_4020501! | 10 | 0.3 | 30 | 0.5 | 21 | 0.5238 | 9 | 0.4444 | 70 | 0.4714 | 15 | 0.2667 | 10 | 0.4 | 17 | 0.2353 | 15 | 0.4 | 57 | 0.3158 |
| 9_4020506: | 6 | 0.8333 | 24 | 0.4583 | 16 | 0.625 | 6 | 0.6667 | 52 | 0.5769 | 9 | 0.8889 | 9 | 0.5556 | 11 | 0.6364 | 14 | 0.5 | 43 | 0.6279 |
| 9_4020506! | 9 | 0 | 26 | 0.1154 | 18 | 0.3333 | 9 | 0.2222 | 62 | 0.1774 | 9 | 0.2222 | 10 | 0.4 | 12 | 0.5 | 11 | 0.0909 | 42 | 0.3095 |
| 9_4020516( | 10 | 0.9 | 25 | 0.96 | 20 | 0.9 | 9 | 0.8889 | 64 | 0.9219 | 15 | 0.8 | 11 | 0.9091 | 18 | 1 | 13 | 1 | 57 | 0.9298 |
| 9_4020528. | 8 | 0.25 | 23 | 0.3043 | 18 | 0.7778 | 8 | 0.125 | 57 | 0.4211 | 15 | 0.4667 | 11 | 0.6364 | 17 | 0.4118 | 12 | 0.5 | 55 | 0.4909 |
| 9_4020566( | 7 | 0.1429 | 17 | 0.2941 | 13 | 0.5385 | 6 | 0.3333 | 43 | 0.3488 | 9 | 0.1111 | 7 | 0.1429 | 12 | 0.25 | 9 | 0.1111 | 37 | 0.1622 |
| 9_4020568: | 6 | 0.3333 | 17 | 0.2941 | 14 | 0.3571 | 8 | 0.125 | 45 | 0.2889 | 6 | 0.3333 | 7 | 0.1429 | 11 | 0 | 10 | 0.3 | 34 | 0.1765 |
| 9_4020585' | 2 | 0 | 8 | 0 | 3 | 0.3333 | 2 | 0 | 15 | 0.0667 | 6 | 0 | 3 | 0.6667 | 11 | 0.3636 | 2 | 0 | 22 | 0.2727 |
| 9_4020869( | 4 | 0.25 | 3 | 0 | 2 | 0 | 1 | 0 | 10 | 0.1 | 4 | 0.25 | 5 | 0 | 6 | 0.3333 | 2 | 0.5 | 17 | 0.2353 |
| 9_4020923: | 2 | 0.5 | 7 | 0.1429 | 3 | 0 | 2 | 0.5 | 14 | 0.2143 | 6 | 0.5 | 4 | 0.25 | 7 | 0 | 2 | 0 | 19 | 0.2105 |
| 9_4020927( | 4 | 0.25 | 9 | 0.3333 | 6 | 0.3333 | 1 | 0 | 20 | 0.3 | 6 | 0.3333 | 5 | 0.2 | 6 | 0.3333 | 4 | 0.25 | 21 | 0.2857 |
| 9_4069379! | 2 | 0 | 1 | 1 | 2 | 0 | NA | NA | 5 | 0.2 | NA | NA | 3 | 0.6667 | 2 | 0 | 5 | 0.4 | 10 | 0.4 |
| 9_4069390( | 1 | 0 | 1 | 0 | 2 | 0.5 | NA | NA | 4 | 0.25 | NA | NA | 3 | 0 | 2 | 0 | 4 | 0.25 | 9 | 0.1111 |
| 9_4069391: | 1 | 1 | 1 | 1 | 2 | 1 | NA | NA | 4 | 1 | NA | NA | 3 | 1 | 1 | 1 | 5 | 1 | 9 | 1 |
| 9_4069403: | 2 | 1 | 1 | 0 | 2 | 0.5 | NA | NA | 5 | 0.6 | NA | NA | 3 | 0.6667 | 2 | 0 | 6 | 0.6667 | 11 | 0.5455 |
| 9_4069437! | 3 | 0.6667 | 1 | 1 | 1 | 0 | 1 | 1 | 6 | 0.6667 | 1 | 0 | 2 | 1 | 1 | 0 | 6 | 1 | 10 | 0.8 |
| 9_4069444( | 3 | 0.3333 | 1 | 0 | 1 | 0 | 1 | 0 | 6 | 0.1667 | NA | NA | 1 | 1 | 1 | 0 | 6 | 0.3333 | 8 | 0.375 |
| 9_4069451! | 3 | 1 | 1 | 1 | 1 | 1 | 1 | 1 | 6 | 1 | 1 | 1 | 1 | 1 | 1 | 0 | 6 | 1 | 9 | 0.8889 |
| 9_4069456! | 3 | 0.3333 | 1 | 0 | 1 | 0 | 1 | 0 | 6 | 0.1667 | 1 | 0 | 2 | 0.5 | NA | NA | 5 | 0 | 8 | 0.125 |
| 9_4069457! | 1 | 0 | 1 | 1 | 1 | 0 | NA | NA | 3 | 0.3333 | NA | NA | NA | NA | NA | NA | 4 | 0.5 | 4 | 0.5 |
| 9_4069460: | 2 | 0.5 | 1 | 1 | 1 | 0 | 1 | 1 | 5 | 0.6 | 1 | 1 | 1 | 1 | 1 | 1 | 6 | 0.6667 | 9 | 0.7778 |
| 9_4069470! | 1 | 0 | NA | NA | NA | NA | 1 | 1 | 2 | 0.5 | 1 | 1 | NA | NA | NA | NA | 5 | 0.8 | 6 | 0.8333 |
| 9_4069472. | 2 | 0.5 | 1 | 1 | 1 | 1 | 1 | 1 | 5 | 0.8 | 1 | 1 | 1 | 1 | NA | NA | 6 | 1 | 8 | 1 |
| 9_4069479: | 2 | 0.5 | 1 | 1 | 1 | 1 | 1 | 1 | 5 | 0.8 | 1 | 1 | NA | NA | NA | NA | 3 | 0.6667 | 4 | 0.75 |
| 9_4069489: | 2 | 0.5 | 1 | 1 | 2 | 0.5 | 1 | 1 | 6 | 0.6667 | 1 | 1 | 1 | 1 | 1 | 1 | 4 | 0.5 | 7 | 0.7143 |
| 9_4069494: | 3 | 0.6667 | NA | NA | 2 | 1 | NA | NA | 5 | 0.8 | 1 | 1 | 1 | 1 | 2 | 1 | 6 | 1 | 10 | 1 |
| 9_4069500: | 2 | 0.5 | 1 | 1 | 2 | 1 | 1 | 1 | 6 | 0.8333 | 1 | 1 | 2 | 1 | 1 | 1 | 5 | 1 | 9 | 1 |
| 9_4069514! | 3 | 0.6667 | 1 | 1 | 2 | 0.5 | 1 | 1 | 7 | 0.7143 | 1 | 1 | 1 | 1 | 2 | 1 | 6 | 1 | 10 | 1 |
| 9_4069537! | 2 | 0.5 | 1 | 0 | 3 | 0 | 1 | 1 | 7 | 0.2857 | 1 | 1 | 1 | 0 | 2 | 0.5 | 5 | 0.6 | 9 | 0.5556 |
| 9_4069546' | 1 | 0 | 3 | 0.3333 | 2 | 0 | 1 | 0 | 7 | 0.1429 | 1 | 0 | 2 | 0.5 | 2 | 0.5 | 5 | 0 | 10 | 0.2 |
| 9_4069558: | 3 | 0.6667 | 2 | 1 | 1 | 1 | 2 | 1 | 8 | 0.875 | 1 | 1 | 1 | 1 | 2 | 1 | 7 | 0.7143 | 11 | 0.8182 |
| 9_4069582. | 2 | 1 | 5 | 1 | 3 | 0.3333 | 1 | 0 | 11 | 0.7273 | 2 | 0.5 | 1 | 0 | 3 | 0.3333 | 8 | 0.375 | 14 | 0.3571 |
| 9_4069589: | 2 | 0 | 3 | 0.3333 | 3 | 0 | 2 | 0 | 10 | 0.1 | 2 | 0.5 | NA | NA | 2 | 0 | 7 | 0 | 11 | 0.0909 |
| 9_4069622! | 2 | 0 | 5 | 0.2 | 3 | 0 | 2 | 0 | 12 | 0.0833 | 3 | 0.3333 | NA | NA | 2 | 0 | 6 | 0 | 11 | 0.0909 |
| 9_4069624: | 3 | 0 | 4 | 0.5 | 1 | 0 | 2 | 0 | 10 | 0.2 | 3 | 0 | 1 | 0 | 2 | 0.5 | 7 | 0 | 13 | 0.0769 |
| 9_4069641( | 4 | 0.25 | 6 | 0.3333 | 3 | 0.6667 | 2 | 0 | 15 | 0.3333 | 4 | 0.5 | 1 | 0 | 1 | 1 | 8 | 0.125 | 14 | 0.2857 |
| 9_4069649: | 2 | 0 | 4 | 0.75 | 5 | 0.4 | 2 | 0.5 | 13 | 0.4615 | 4 | 0 | 1 | 0 | 2 | 0.5 | 7 | 0.1429 | 14 | 0.1429 |
| 9_4071458( | 247 | 0.166 | 330 | 0.1242 | 242 | 0.1074 | 234 | 0.1368 | 1053 | 0.133 | 217 | 0.1336 | 255 | 0.1804 | 235 | 0.1447 | 270 | 0.163 | 977 | 0.1566 |
| 9_4071464! | 250 | 0.176 | 342 | 0.1433 | 249 | 0.1245 | 251 | 0.1554 | 1092 | 0.1493 | 226 | 0.1372 | 250 | 0.184 | 239 | 0.1506 | 273 | 0.1758 | 988 | 0.163 |
| 9_4071468! | 243 | 0.07 | 300 | 0.06 | 229 | 0.0699 | 229 | 0.0786 | 1001 | 0.0689 | 205 | 0.0683 | 247 | 0.085 | 225 | 0.0622 | 252 | 0.0992 | 929 | 0.0797 |
| 9_4092513( | 2 | 0.5 | 3 | 0.3333 | 2 | 0.5 | 2 | 0.5 | 9 | 0.4444 | 3 | 0 | 1 | 1 | 3 | 0.3333 | NA | NA | 7 | 0.2857 |
| 9_4092535: | 2 | 0 | 3 | 0.3333 | 2 | 0.5 | 1 | 0 | 8 | 0.25 | 3 | 0.3333 | 1 | 1 | 3 | 0.6667 | NA | NA | 7 | 0.5714 |
| 9_4092537! | NA | NA | 2 | 1 | 2 | 0.5 | 2 | 0.5 | 6 | 0.6667 | 3 | 0.6667 | 1 | 1 | 3 | 0.6667 | NA | NA | 7 | 0.7143 |
| 9_4092538( | 2 | 0.5 | 3 | 0.3333 | 1 | 0 | 2 | 0 | 8 | 0.25 | 3 | 0.3333 | 1 | 0 | 3 | 0 | NA | NA | 7 | 0.1429 |
| 9_4092543( | 2 | 0.5 | 2 | 0.5 | 2 | 1 | 2 | 0.5 | 8 | 0.625 | 2 | 1 | 1 | 1 | 3 | 1 | NA | NA | 6 | 1 |
| 9_4092582! | 2 | 1 | 1 | 1 | 2 | 1 | 1 | 0 | 6 | 0.8333 | 1 | 1 | 1 | 1 | 3 | 1 | NA | NA | 5 | 1 |

| | | | | | | | | | | | | | | | | | | | | |
|---|---|---|---|---|---|---|---|---|---|---|---|---|---|---|---|---|---|---|---|---|
| 9_4092598 | 3 | 0.6667 | 2 | 1 | 1 | 1 | 1 | 1 | 7 | 0.8571 | 1 | 1 | NA | NA | 3 | 1 | NA | NA | 4 | 1 |
| 9_4092610 | 2 | 0.5 | 2 | 0.5 | 2 | 0 | 1 | 1 | 7 | 0.4286 | 1 | 0 | 1 | 0 | 2 | 1 | NA | NA | 4 | 0.5 |
| 9_4092617 | 1 | 1 | 2 | 0.5 | 1 | 1 | 1 | 0 | 5 | 0.6 | 1 | 0 | 1 | 1 | 2 | 0.5 | NA | NA | 4 | 0.5 |
| 9_4150131 | 3 | 0.3333 | 4 | 0.25 | 2 | 0 | 2 | 0 | 11 | 0.1818 | 1 | 0 | NA | NA | 3 | 0.3333 | NA | NA | 4 | 0.25 |
| 9_4150156 | 2 | 0 | 3 | 0.3333 | 2 | 1 | 3 | 0.3333 | 10 | 0.4 | NA | NA | 1 | 1 | 2 | 0.5 | 1 | 0 | 4 | 0.5 |
| 9_4150172 | 1 | 1 | 4 | 0.5 | 2 | 0 | 1 | 0 | 8 | 0.375 | NA | NA | NA | NA | 2 | 0.5 | 1 | 1 | 3 | 0.6667 |
| 9_4150329 | 4 | 0 | 6 | 0.1667 | 2 | 0 | 5 | 0.2 | 17 | 0.1176 | 1 | 0 | 3 | 0 | 3 | 0 | 3 | 0.6667 | 10 | 0.2 |
| 9_4150359 | 3 | 0 | 4 | 0 | 2 | 0.5 | 5 | 0.2 | 14 | 0.1429 | 1 | 0 | 3 | 0.3333 | 3 | 0 | 5 | 0 | 12 | 0.0833 |
| 9_4152620 | NA | NA | 1 | 1 | 1 | 0 | NA | NA | 2 | 0.5 | NA | NA | NA | NA | NA | NA | 2 | 0.5 | 2 | 0.5 |
| 9_4152628 | NA | NA | 2 | 0.5 | 1 | 0 | 1 | 0 | 4 | 0.25 | NA | NA | NA | NA | NA | NA | 2 | 0.5 | 2 | 0.5 |
| 9_4156642 | NA | NA | 2 | 1 | 1 | 1 | NA | NA | 3 | 1 | 1 | 1 | NA | NA | NA | NA | NA | NA | 1 | 1 |
| 9_4156681 | NA | NA | 2 | 0.5 | 1 | 1 | NA | NA | 3 | 0.6667 | 1 | 1 | NA | NA | NA | NA | NA | NA | 1 | 1 |
| 9_4156797 | NA | NA | 1 | 1 | 2 | 0.5 | 1 | 1 | 4 | 0.75 | 1 | 1 | NA | NA | NA | NA | 1 | 0 | 2 | 0.5 |
| 9_4156803 | NA | NA | 1 | 1 | 2 | 0.5 | 1 | 0 | 4 | 0.5 | 1 | 1 | NA | NA | NA | NA | 2 | 1 | 3 | 1 |
| 9_4156804 | NA | NA | 2 | 0.5 | 1 | 0 | 1 | 0 | 4 | 0.25 | 2 | 1 | NA | NA | NA | NA | 2 | 1 | 4 | 1 |
| 9_4156805 | NA | NA | 2 | 0 | 2 | 0.5 | NA | NA | 4 | 0.25 | 2 | 0 | NA | NA | NA | NA | 2 | 0.5 | 4 | 0.25 |
| 9_4156806 | NA | NA | 2 | 0 | 2 | 0.5 | 1 | 1 | 5 | 0.4 | 2 | 0.5 | NA | NA | NA | NA | 2 | 0 | 4 | 0.25 |
| 9_4156812 | NA | NA | 2 | 0 | 2 | 0.5 | NA | NA | 4 | 0.25 | 2 | 1 | NA | NA | NA | NA | 2 | 1 | 4 | 1 |
| 9_4156813 | NA | NA | 2 | 0 | 2 | 0.5 | 1 | 0 | 5 | 0.2 | 2 | 0 | NA | NA | NA | NA | 2 | 0.5 | 4 | 0.25 |
| 9_4156860 | NA | NA | 2 | 0 | 3 | 0.3333 | NA | NA | 5 | 0.2 | 1 | 1 | NA | NA | 1 | 1 | 1 | 1 | 3 | 1 |
| 9_4156873 | NA | NA | 2 | 0.5 | 3 | 0.6667 | 1 | 1 | 6 | 0.6667 | 2 | 1 | NA | NA | 1 | 0 | 2 | 0.5 | 5 | 0.6 |
| 9_4156884 | NA | NA | 2 | 0.5 | 3 | 0.6667 | NA | NA | 5 | 0.6 | 2 | 1 | NA | NA | 1 | 1 | 2 | 0.5 | 5 | 0.8 |
| 9_4156897 | NA | NA | 2 | 0.5 | 2 | 0.5 | 1 | 1 | 5 | 0.6 | 2 | 1 | NA | NA | 1 | 0 | 2 | 1 | 5 | 0.8 |
| 9_4187639 | 17 | 0.7059 | 22 | 0.5455 | 17 | 0.4706 | 9 | 0.4444 | 65 | 0.5538 | 13 | 0.7692 | 17 | 0.4706 | 17 | 0.4706 | 12 | 0.75 | 59 | 0.5932 |
| 9_4187640 | 17 | 0.4706 | 26 | 0.5 | 16 | 0.5 | 11 | 0.2727 | 70 | 0.4571 | 13 | 0.4615 | 18 | 0.5 | 18 | 0.3889 | 14 | 0.3571 | 63 | 0.4286 |
| 9_4187646 | 15 | 0.9333 | 21 | 0.9524 | 16 | 1 | 9 | 1 | 61 | 0.9672 | 12 | 0.9167 | 16 | 1 | 18 | 1 | 13 | 1 | 59 | 0.9831 |
| 9_4187647 | 15 | 0.5333 | 22 | 0.7727 | 14 | 0.7143 | 10 | 0.7 | 61 | 0.6885 | 11 | 0.7273 | 14 | 0.6429 | 18 | 0.6667 | 14 | 0.6429 | 57 | 0.6667 |
| 9_4187658 | 19 | 0.4211 | 25 | 0.16 | 15 | 0.4 | 11 | 0.3636 | 70 | 0.3143 | 10 | 0.3 | 18 | 0.1111 | 19 | 0.1579 | 15 | 0.3333 | 62 | 0.2097 |
| 9_4187667 | 16 | 0.75 | 25 | 0.8 | 13 | 0.8462 | 10 | 0.7 | 64 | 0.7812 | 7 | 0.7143 | 17 | 0.8824 | 17 | 0.7647 | 12 | 0.5833 | 53 | 0.7547 |
| 9_4187670 | 17 | 0.2941 | 24 | 0.4167 | 14 | 0.2143 | 12 | 0.25 | 67 | 0.3134 | 11 | 0.1818 | 17 | 0.3529 | 18 | 0.2778 | 13 | 0.4615 | 59 | 0.322 |
| 9_4187688 | 17 | 0.3529 | 22 | 0.2273 | 11 | 0.1818 | 10 | 0 | 60 | 0.2167 | 10 | 0.1 | 17 | 0 | 16 | 0.1875 | 14 | 0.1429 | 57 | 0.1053 |
| 9_4187748 | 15 | 0.6 | 16 | 0.3125 | 14 | 0.3571 | 10 | 0.1 | 55 | 0.3636 | 11 | 0.2727 | 16 | 0.3125 | 15 | 0.2667 | 13 | 0.2308 | 55 | 0.2727 |
| 9_4187766 | 12 | 0 | 16 | 0.1875 | 12 | 0.1667 | 7 | 0.1429 | 47 | 0.1277 | 11 | 0 | 15 | 0.2 | 12 | 0.0833 | 10 | 0.1 | 48 | 0.1042 |
| 9_4187850 | 9 | 0.2222 | 7 | 0 | 12 | 0.0833 | 6 | 0 | 34 | 0.0882 | 5 | 0.2 | 10 | 0.1 | 7 | 0 | 11 | 0.0909 | 33 | 0.0909 |
| 9_4187866 | 10 | 0.7 | 10 | 0.7 | 11 | 0.7273 | 7 | 0.4286 | 38 | 0.6579 | 6 | 0.5 | 9 | 0.6667 | 9 | 0.8889 | 10 | 0.9 | 34 | 0.7647 |
| 9_4187927 | 6 | 0.1667 | 11 | 0 | 10 | 0.3 | 4 | 0 | 31 | 0.129 | 5 | 0.6 | 6 | 0 | 5 | 0.4 | 11 | 0.3636 | 27 | 0.3333 |
| 9_4187941 | 5 | 0 | 9 | 0.1111 | 10 | 0.1 | 4 | 0 | 28 | 0.0714 | 5 | 0 | 4 | 0 | 4 | 0 | 10 | 0.2 | 23 | 0.087 |
| 9_4187972 | 4 | 0.5 | 17 | 0.4118 | 9 | 0.4444 | 3 | 0.3333 | 33 | 0.4242 | 7 | 0.4286 | 5 | 0.6 | 10 | 0.2 | 9 | 0.4444 | 31 | 0.3871 |
| 9_4187974 | 8 | 0.5 | 20 | 0.15 | 13 | 0.0769 | 6 | 0 | 47 | 0.1702 | 8 | 0 | 9 | 0.2222 | 9 | 0.2222 | 7 | 0.4286 | 33 | 0.2121 |
| 9_4188536 | 7 | 0.1429 | 15 | 0.1333 | 12 | 0.0833 | 3 | 0.3333 | 37 | 0.1351 | 7 | 0 | 8 | 0.125 | 8 | 0 | 8 | 0.125 | 31 | 0.0645 |
| 9_4194305 | NA | NA | 1 | 0 | 2 | 0.5 | NA | NA | 3 | 0.3333 | 2 | 0 | NA | NA | NA | NA | 3 | 0.3333 | 5 | 0.2 |
| 9_4216512 | 1 | 0 | 2 | 0.5 | 3 | 1 | 3 | 0 | 9 | 0.4444 | 2 | 0 | NA | NA | 1 | 1 | 3 | 0.6667 | 6 | 0.5 |
| 9_4216574 | 4 | 0 | 3 | 0 | 8 | 0.25 | 7 | 0.1429 | 22 | 0.1364 | 4 | 0.25 | 2 | 0.5 | 3 | 0 | 3 | 0.3333 | 12 | 0.25 |
| 9_4216620 | 8 | 0.375 | 3 | 1 | 10 | 0.8 | 10 | 0.4 | 31 | 0.5806 | 7 | 0.7143 | 4 | 1 | 4 | 0.5 | 2 | 0.5 | 17 | 0.7059 |
| 9_4216628 | 7 | 0.4286 | 2 | 1 | 7 | 0 | 6 | 0.3333 | 22 | 0.3182 | 6 | 0.5 | 4 | 0.75 | 4 | 0.25 | 2 | 0 | 16 | 0.4375 |
| 9_4216644 | 7 | 0.1429 | 3 | 0 | 8 | 0.125 | 8 | 0 | 26 | 0.0769 | 7 | 0 | 4 | 0.25 | 4 | 0 | 2 | 0 | 17 | 0.0588 |
| 9_4216661 | 5 | 0 | 3 | 0.6667 | 6 | 0.1667 | 10 | 0.1 | 24 | 0.1667 | 5 | 0.2 | 4 | 0.75 | 4 | 0.5 | 2 | 0.5 | 15 | 0.4667 |
| 9_4216672 | 5 | 0 | 1 | 0 | 7 | 0.1429 | 10 | 0.1 | 23 | 0.087 | 7 | 0.1429 | 4 | 0 | 4 | 0.25 | 2 | 0.5 | 17 | 0.1765 |
| 9_4216681 | 5 | 1 | 2 | 1 | 5 | 0.8 | 6 | 0.8333 | 18 | 0.8889 | 7 | 0.8571 | 3 | 1 | 3 | 1 | 1 | 1 | 14 | 0.9286 |
| 9_4216682 | 5 | 0.2 | 3 | 0.3333 | 5 | 0.2 | 8 | 0.125 | 21 | 0.1905 | 7 | 0 | 3 | 1 | 4 | 0 | 1 | 0 | 15 | 0.2 |
| 9_4216692 | 4 | 0.75 | 3 | 0.6667 | 5 | 1 | 8 | 0.625 | 20 | 0.75 | 7 | 0.8571 | 3 | 1 | 4 | 0.5 | 1 | 1 | 15 | 0.8 |
| 9_4232377 | 7 | 0.1429 | 6 | 0.1667 | 2 | 0.5 | 6 | 0.1667 | 21 | 0.1905 | 4 | 0.5 | 5 | 0.4 | 4 | 0.25 | 4 | 0.25 | 17 | 0.3529 |
| 9_4232378 | 7 | 0 | 6 | 0.1667 | 2 | 0 | 5 | 0 | 20 | 0.05 | 4 | 0.5 | 5 | 0 | 5 | 0.4 | 4 | 0 | 18 | 0.2222 |
| 9_4232384 | 6 | 0.1667 | 4 | 0 | NA | NA | 4 | 0 | 14 | 0.0714 | 4 | 0.25 | 4 | 0.25 | 6 | 0.1667 | 4 | 0 | 18 | 0.1667 |
| 9_4232410 | 5 | 0.2 | 4 | 0.25 | 2 | 0 | 4 | 0 | 15 | 0.1333 | 4 | 0.25 | 4 | 0 | 6 | 0.5 | 4 | 0.25 | 18 | 0.2778 |
| 9_4232425 | 6 | 0.3333 | 5 | 0 | 2 | 0 | 6 | 0 | 19 | 0.1053 | 4 | 0.5 | 4 | 0.25 | 6 | 0.1667 | 4 | 0.25 | 18 | 0.2778 |
| 9_4232465 | 6 | 0.1667 | 3 | 0 | 2 | 0 | 5 | 0 | 16 | 0.0625 | 3 | 0.3333 | 3 | 0 | 2 | 0.5 | 4 | 0 | 12 | 0.1667 |
| 9_4232471 | 7 | 0.1429 | 5 | 0 | 2 | 0 | 6 | 0.1667 | 20 | 0.1 | 4 | 0 | 3 | 0.3333 | 4 | 0.25 | 3 | 0 | 14 | 0.1429 |
| 9_4232512 | 7 | 0.2857 | 5 | 0.2 | 1 | 0 | 4 | 0 | 17 | 0.1765 | 2 | 0 | 2 | 0 | 4 | 0 | 3 | 0.3333 | 11 | 0.0909 |
| 9_4232532 | 6 | 0.1667 | 4 | 0.75 | 1 | 1 | 4 | 0.25 | 15 | 0.4 | 4 | 0.25 | 3 | 0.3333 | 1 | 0 | 4 | 0 | 12 | 0.1667 |
| 9_4232587 | 6 | 0 | 5 | 0.2 | 1 | 0 | 4 | 0 | 16 | 0.0625 | 4 | 0 | 2 | 0 | 2 | 0 | 3 | 0.3333 | 11 | 0.0909 |
| 9_4232596 | 6 | 0.6667 | 4 | 0.5 | 1 | 1 | 3 | 1 | 14 | 0.7143 | 4 | 0.25 | 3 | 0.6667 | 2 | 1 | 4 | 0 | 13 | 0.3846 |
| 9_4232608 | 6 | 0.5 | 4 | 0.25 | 1 | 0 | 2 | 0 | 13 | 0.3077 | 2 | 0 | 2 | 0.5 | 3 | 0.6667 | 1 | 0 | 8 | 0.375 |
| 9_4232639 | 4 | 0.5 | 3 | 0.3333 | 1 | 0 | 4 | 0 | 12 | 0.25 | 4 | 0.25 | 3 | 0 | NA | NA | 3 | 0 | 10 | 0.1 |
| 9_4232644 | 4 | 0.5 | 4 | 0 | 1 | 0 | 4 | 0 | 13 | 0.1538 | 4 | 0.25 | 3 | 0 | 1 | 0 | 3 | 0 | 11 | 0.0909 |

| | | | | | | | | | | | | | | | | | | | | |
|---|---|---|---|---|---|---|---|---|---|---|---|---|---|---|---|---|---|---|---|---|
| 9_4232675 | 3 | 0 | 2 | 0 | 1 | 0 | 3 | 0.3333 | 9 | 0.1111 | 1 | 0 | 2 | 1 | 1 | 0 | 3 | 0 | 7 | 0.2857 |
| 9_4232704 | 3 | 1 | 4 | 1 | 1 | 1 | 3 | 0.6667 | 11 | 0.9091 | 1 | 1 | 2 | 0.5 | 1 | 1 | 3 | 1 | 7 | 0.8571 |
| 9_4232709 | 3 | 1 | 2 | 1 | 1 | 0 | 2 | 1 | 8 | 0.875 | 1 | 1 | 2 | 1 | 1 | 1 | 2 | 1 | 6 | 1 |
| 9_4232710 | 3 | 1 | 3 | 1 | NA | NA | 3 | 0.3333 | 9 | 0.7778 | 1 | 1 | 2 | 1 | NA | NA | 3 | 0.6667 | 6 | 0.8333 |
| 9_4232728 | 3 | 0 | 3 | 0.6667 | 1 | 0 | 3 | 0 | 10 | 0.2 | NA | NA | 2 | 0 | 1 | 1 | 3 | 0 | 6 | 0.1667 |
| 9_4232730 | 3 | 0.3333 | 2 | 0 | 1 | 0 | 2 | 0 | 8 | 0.125 | 1 | 0 | 2 | 1 | 1 | 0 | 3 | 0 | 7 | 0.2857 |
| 9_4232739 | 3 | 0.6667 | 3 | 0 | 1 | 1 | 3 | 0 | 10 | 0.3 | 1 | 0 | 2 | 0.5 | 1 | 0 | 3 | 0 | 7 | 0.1429 |
| 9_4235425 | 1 | 0 | 2 | 0.5 | 1 | 0 | 1 | 0 | 5 | 0.2 | NA | NA | 2 | 0 | NA | NA | 2 | 1 | 4 | 0.5 |
| 9_4243190 | 3 | 1 | 1 | 1 | NA | NA | 1 | 1 | 5 | 1 | 1 | 1 | 1 | 0 | 1 | 1 | NA | NA | 3 | 0.6667 |
| 9_4243203 | 3 | 0.6667 | 1 | 1 | NA | NA | NA | NA | 4 | 0.75 | 1 | 1 | 1 | 1 | NA | NA | NA | NA | 2 | 1 |
| 9_4243210 | 3 | 1 | 1 | 1 | NA | NA | 1 | 1 | 5 | 1 | 1 | 1 | 1 | 1 | NA | NA | NA | NA | 2 | 1 |
| 9_4243228 | 3 | 1 | 1 | 1 | NA | NA | NA | NA | 4 | 1 | 1 | 1 | NA | NA | NA | NA | NA | NA | 1 | 1 |
| 9_4287531 | 23 | 0.2174 | 21 | 0.4762 | 15 | 0.1333 | 23 | 0.3478 | 82 | 0.3049 | 23 | 0.087 | 19 | 0.3684 | 23 | 0.3478 | 18 | 0.3889 | 83 | 0.2892 |
| 9_4287579 | 17 | 0.0588 | 21 | 0.2381 | 11 | 0.3636 | 13 | 0.0769 | 62 | 0.1774 | 16 | 0.1875 | 15 | 0.1333 | 17 | 0.1176 | 13 | 0.1538 | 61 | 0.1475 |
| 9_4287596 | 21 | 0.4286 | 18 | 0.5 | 11 | 0.4545 | 16 | 0.625 | 66 | 0.5 | 18 | 0.5 | 14 | 0.4286 | 16 | 0.5625 | 16 | 0.5625 | 64 | 0.5156 |
| 9_4287661 | 16 | 0.5625 | 16 | 0.6875 | 9 | 0.4444 | 15 | 0.8667 | 56 | 0.6607 | 16 | 0.75 | 14 | 0.4286 | 18 | 0.6111 | 11 | 0.6364 | 59 | 0.6102 |
| 9_4287668 | 13 | 0.1538 | 15 | 0.2 | 7 | 0 | 16 | 0.1875 | 51 | 0.1569 | 14 | 0.2857 | 14 | 0.0714 | 15 | 0.1333 | 13 | 0.1538 | 56 | 0.1607 |
| 9_4287698 | 13 | 0.8462 | 13 | 0.6154 | 8 | 0.75 | 13 | 0.7692 | 47 | 0.7447 | 13 | 0.6154 | 10 | 0.5 | 16 | 0.5625 | 11 | 0.6364 | 50 | 0.58 |
| 9_4287708 | 11 | 0.4545 | 13 | 0.5385 | 6 | 0.1667 | 12 | 0.25 | 42 | 0.381 | 13 | 0.3846 | 11 | 0.1818 | 15 | 0.2667 | 10 | 0.6 | 49 | 0.3469 |
| 9_4287731 | 11 | 0.5455 | 10 | 0.6 | 5 | 0.2 | 14 | 0.7143 | 40 | 0.575 | 9 | 0.6667 | 9 | 0 | 15 | 0.5333 | 10 | 0.4 | 43 | 0.4186 |
| 9_4287733 | 10 | 0.3 | 10 | 0.2 | 8 | 0 | 12 | 0.1667 | 40 | 0.175 | 12 | 0.0833 | 8 | 0 | 13 | 0.2308 | 7 | 0.1429 | 40 | 0.125 |
| 9_4287744 | 8 | 0.5 | 13 | 0.3846 | 9 | 0.3333 | 13 | 0.3077 | 43 | 0.3721 | 10 | 0.3 | 11 | 0.0909 | 9 | 0.2222 | 5 | 0.2 | 35 | 0.2 |
| 9_4287784 | 9 | 0.3333 | 12 | 0.25 | 5 | 0.6 | 13 | 0.5385 | 39 | 0.4103 | 11 | 0.4545 | 9 | 0.4444 | 14 | 0.5 | 6 | 0.3333 | 40 | 0.45 |
| 9_4287894 | 2 | 0 | 3 | 0.3333 | 2 | 0 | 10 | 0 | 17 | 0.0588 | 5 | 0 | 5 | 0 | 7 | 0.1429 | 4 | 0.25 | 21 | 0.0952 |
| 9_4287921 | 6 | 0.1667 | 6 | 0.1667 | 5 | 0.2 | 13 | 0.3077 | 30 | 0.2333 | 5 | 0.4 | 9 | 0.1111 | 10 | 0.2 | 5 | 0.2 | 29 | 0.2069 |
| 9_4287959 | 3 | 0 | 2 | 0.5 | 1 | 0 | 5 | 0 | 11 | 0.0909 | 3 | 0.3333 | 3 | 0.3333 | 5 | 0.2 | NA | NA | 11 | 0.2727 |
| 9_4287995 | 2 | 0 | 2 | 0 | NA | NA | 4 | 0.5 | 8 | 0.25 | 4 | 0.5 | 3 | 0.3333 | 4 | 0 | NA | NA | 11 | 0.2727 |
| 9_4288032 | 3 | 1 | 2 | 1 | NA | NA | 3 | 1 | 8 | 1 | 3 | 0.3333 | 2 | 0.5 | 3 | 1 | NA | NA | 8 | 0.625 |
| 9_4288039 | 1 | 0 | 2 | 0 | NA | NA | 2 | 0.5 | 5 | 0.2 | 2 | 0 | 1 | 0 | 4 | 0.25 | NA | NA | 7 | 0.1429 |
| 9_4288249 | 1 | 0 | 2 | 0.5 | NA | NA | 1 | 0 | 4 | 0.25 | 1 | 0 | 2 | 0 | 3 | 0.3333 | NA | NA | 6 | 0.1667 |
| 9_4288345 | 1 | 0 | 2 | 0.5 | NA | NA | 2 | 0.5 | 5 | 0.4 | 2 | 0 | 2 | 0.5 | 2 | 1 | NA | NA | 6 | 0.5 |
| 9_4288354 | 2 | 1 | 2 | 1 | NA | NA | 2 | 1 | 6 | 1 | NA | NA | 2 | 0.5 | 3 | 0.6667 | NA | NA | 5 | 0.6 |
| 9_4288357 | 2 | 0.5 | 2 | 1 | NA | NA | 1 | 1 | 5 | 0.8 | NA | NA | 1 | 1 | 3 | 1 | NA | NA | 4 | 1 |
| 9_4288359 | 2 | 1 | 2 | 1 | NA | NA | 1 | 0 | 5 | 0.8 | 2 | 0 | 2 | 0 | 3 | 0.6667 | NA | NA | 7 | 0.2857 |
| 9_4288363 | 3 | 0 | 2 | 1 | NA | NA | 2 | 0.5 | 7 | 0.4286 | 2 | 1 | 2 | 0.5 | 3 | 0.6667 | NA | NA | 7 | 0.7143 |
| 9_4302054 | 1 | 0 | 2 | 0 | 4 | 0.25 | 1 | 0 | 8 | 0.125 | 2 | 0 | 1 | 1 | 3 | 0.3333 | NA | NA | 6 | 0.3333 |
| 9_4302196 | 1 | 1 | NA | NA | 3 | 0.3333 | 1 | 0 | 5 | 0.4 | 2 | 0.5 | 1 | 0 | 2 | 0 | NA | NA | 5 | 0.2 |
| 9_4302203 | NA | NA | 1 | 1 | 2 | 0.5 | 1 | 0 | 4 | 0.5 | 1 | 0 | 1 | 0 | 2 | 1 | NA | NA | 4 | 0.5 |
| 9_4302206 | NA | NA | 1 | 1 | 1 | 0 | 1 | 0 | 3 | 0.3333 | 1 | 1 | NA | NA | 2 | 0 | NA | NA | 3 | 0.3333 |
| 9_4302208 | 1 | 0 | 1 | 0 | 3 | 0.3333 | 1 | 0 | 6 | 0.1667 | 1 | 0 | 1 | 1 | 1 | 0 | NA | NA | 3 | 0.3333 |
| 9_4302218 | NA | NA | 1 | 0 | 2 | 0.5 | 1 | 1 | 4 | 0.5 | NA | NA | NA | NA | 2 | 1 | NA | NA | 2 | 1 |
| 9_4302230 | 1 | 1 | 1 | 0 | 3 | 0 | 2 | 0 | 7 | 0.1429 | 1 | 1 | NA | NA | 2 | 0.5 | NA | NA | 3 | 0.6667 |
| 9_4302287 | 2 | 1 | 1 | 1 | 3 | 1 | 1 | 1 | 7 | 1 | 1 | 1 | NA | NA | 2 | 1 | NA | NA | 3 | 1 |
| 9_4312697 | NA | NA | NA | NA | 1 | 1 | NA | NA | 1 | 1 | NA | NA | NA | NA | NA | NA | 1 | 1 | 1 | 1 |
| 9_4312728 | NA | NA | NA | NA | 1 | 1 | NA | NA | 1 | 1 | NA | NA | NA | NA | NA | NA | 1 | 1 | 1 | 1 |
| 9_4313333 | 5 | 0.8 | 3 | 1 | 6 | 0.8333 | NA | NA | 14 | 0.8571 | 2 | 1 | 3 | 1 | 2 | 1 | 2 | 1 | 9 | 1 |
| 9_4313348 | 4 | 0.5 | 4 | 0 | 6 | 0.1667 | NA | NA | 14 | 0.2143 | 2 | 0 | 5 | 0.2 | 3 | 0.6667 | 4 | 0.25 | 14 | 0.2857 |
| 9_4313349 | 4 | 0.25 | NA | NA | 4 | 0 | NA | NA | 8 | 0.125 | 3 | 0.3333 | 3 | 0 | 1 | 1 | 2 | 0 | 9 | 0.2222 |
| 9_4313355 | 3 | 0.6667 | 4 | 0.5 | 5 | 0.8 | NA | NA | 12 | 0.6667 | 2 | 0 | 5 | 0.2 | 2 | 1 | 3 | 0.3333 | 12 | 0.3333 |
| 9_4313373 | 2 | 1 | 4 | 1 | 6 | 0.8333 | NA | NA | 12 | 0.9167 | 2 | 1 | 2 | 1 | NA | NA | 3 | 1 | 7 | 1 |
| 9_4313385 | 5 | 0 | 4 | 0 | 6 | 0.1667 | NA | NA | 15 | 0.0667 | 3 | 0 | 6 | 0.1667 | 1 | 0 | 5 | 0.2 | 15 | 0.1333 |
| 9_4313386 | 5 | 0 | 4 | 0 | 6 | 0.1667 | NA | NA | 15 | 0.0667 | 3 | 0.3333 | 6 | 0 | 1 | 0 | 4 | 0 | 14 | 0.0714 |
| 9_4313387 | 5 | 0.8 | 4 | 0.5 | 5 | 0.4 | NA | NA | 14 | 0.5714 | 3 | 0.6667 | 4 | 0.25 | 1 | 1 | 4 | 0.25 | 12 | 0.4167 |
| 9_4313392 | 4 | 0.75 | 2 | 0.5 | 5 | 0.6 | NA | NA | 11 | 0.6364 | 2 | 0.5 | 5 | 0.2 | 1 | 0 | 3 | 0.3333 | 11 | 0.2727 |
| 9_4313413 | 3 | 0 | 4 | 0.5 | 4 | 0 | NA | NA | 11 | 0.1818 | 3 | 0 | 4 | 0 | NA | NA | 3 | 0.3333 | 10 | 0.1 |
| 9_4342529 | NA | NA | NA | NA | NA | NA | 2 | 0.5 | 2 | 0.5 | 1 | 1 | 1 | 0 | NA | NA | 1 | 0 | 3 | 0.3333 |
| 9_4371520 | NA | NA | 3 | 0.3333 | NA | NA | NA | NA | 3 | 0.3333 | 4 | 0.5 | NA | NA | NA | NA | 3 | 0.6667 | 7 | 0.5714 |
| 9_4371521 | NA | NA | 4 | 1 | NA | NA | NA | NA | 4 | 1 | 6 | 1 | NA | NA | NA | NA | 2 | 1 | 8 | 1 |
| 9_4371524 | NA | NA | 7 | 1 | NA | NA | NA | NA | 7 | 1 | 7 | 1 | NA | NA | NA | NA | 3 | 1 | 10 | 1 |
| 9_4371657 | NA | NA | 8 | 0 | 2 | 0.5 | NA | NA | 10 | 0.1 | 7 | 0.1429 | NA | NA | 3 | 0 | 7 | 0 | 17 | 0.0588 |
| 9_4371659 | NA | NA | 8 | 0.125 | 2 | 0 | NA | NA | 10 | 0.1 | 5 | 0 | NA | NA | 3 | 0 | 5 | 0.2 | 13 | 0.0769 |
| 9_4371772 | 1 | 0 | 10 | 0.1 | 2 | 0 | NA | NA | 13 | 0.0769 | 9 | 0.1111 | NA | NA | 2 | 0 | 7 | 0 | 18 | 0.0556 |
| 9_4371798 | 1 | 0 | 17 | 0.1176 | 3 | 0 | 2 | 0.5 | 23 | 0.1304 | 13 | 0.1538 | 1 | 0 | 5 | 0 | 9 | 0.1111 | 28 | 0.1071 |
| 9_4371823 | 1 | 1 | 16 | 0.25 | 4 | 0 | 2 | 0 | 23 | 0.2174 | 15 | 0.4667 | 1 | 0 | 5 | 0.4 | 9 | 0.2222 | 30 | 0.3667 |
| 9_4371833 | 3 | 0 | 19 | 0.2105 | 4 | 0 | 2 | 0 | 28 | 0.1429 | 14 | 0.0714 | 1 | 0 | 5 | 0.4 | 10 | 0.2 | 30 | 0.1667 |

| | | | | | | | | | | | | | | | | | | | | |
|---|---|---|---|---|---|---|---|---|---|---|---|---|---|---|---|---|---|---|---|---|
| 9_4395873 | 17 | 0.2941 | 85 | 0.3059 | 62 | 0.4194 | 24 | 0.375 | 188 | 0.3511 | 52 | 0.3077 | 55 | 0.2909 | 52 | 0.3462 | 53 | 0.3208 | 212 | 0.316 |
| 9_4395874 | 19 | 0.2105 | 84 | 0.2381 | 63 | 0.2381 | 27 | 0.2593 | 193 | 0.2383 | 45 | 0.3778 | 60 | 0.2833 | 51 | 0.2549 | 54 | 0.2593 | 210 | 0.2905 |
| 9_4395885 | 25 | 0.04 | 86 | 0.0814 | 63 | 0.1429 | 28 | 0.0714 | 202 | 0.0941 | 49 | 0.1633 | 53 | 0.0755 | 55 | 0.0545 | 50 | 0.14 | 207 | 0.1063 |
| 9_4395915 | 26 | 0.1923 | 95 | 0.1684 | 80 | 0.1625 | 24 | 0.125 | 225 | 0.1644 | 52 | 0.1538 | 61 | 0.1803 | 59 | 0.1525 | 53 | 0.1887 | 225 | 0.1689 |
| 9_4398702 | NA | NA | 1 | 0 | 2 | 1 | NA | NA | 3 | 0.6667 | NA | NA | 3 | 1 | 2 | 0 | 3 | 1 | 8 | 0.75 |
| 9_4398709 | NA | NA | 2 | 0.5 | 2 | 0.5 | NA | NA | 4 | 0.5 | NA | NA | 4 | 0.25 | 2 | 0 | 3 | 0.3333 | 9 | 0.2222 |
| 9_4398716 | NA | NA | 2 | 0.5 | 1 | 1 | NA | NA | 3 | 0.6667 | NA | NA | 4 | 0.5 | 2 | 0.5 | 2 | 0.5 | 8 | 0.5 |
| 9_4398720 | NA | NA | 1 | 0 | 2 | 0.5 | NA | NA | 3 | 0.3333 | NA | NA | 3 | 0.6667 | 1 | 1 | 3 | 0.3333 | 7 | 0.5714 |
| 9_4398733 | NA | NA | 2 | 1 | 1 | 1 | NA | NA | 3 | 1 | NA | NA | 3 | 0.6667 | 2 | 1 | 3 | 0.6667 | 8 | 0.75 |
| 9_4398734 | NA | NA | 2 | 1 | 1 | 1 | NA | NA | 3 | 1 | NA | NA | 4 | 1 | 1 | 1 | 2 | 1 | 7 | 1 |
| 9_4400513 | 8 | 0.625 | 7 | 0.5714 | 7 | 0.5714 | 10 | 0.5 | 32 | 0.5625 | 4 | 1 | 10 | 0.6 | 6 | 0.5 | 5 | 0.4 | 25 | 0.6 |
| 9_4400524 | 8 | 0.125 | 8 | 0 | 10 | 0 | 10 | 0.1 | 36 | 0.0556 | 4 | 0 | 7 | 0.2857 | 4 | 0 | 7 | 0.1429 | 22 | 0.1364 |
| 9_4400526 | 7 | 0.1429 | 7 | 0 | 6 | 0 | 7 | 0.1429 | 27 | 0.0741 | 3 | 0.3333 | 5 | 0 | 6 | 0 | 5 | 0 | 19 | 0.0526 |
| 9_4400542 | 9 | 1 | 7 | 0.8571 | 10 | 1 | 8 | 1 | 34 | 0.9706 | 4 | 1 | 9 | 1 | 3 | 1 | 5 | 1 | 21 | 1 |
| 9_4400543 | 7 | 0.1429 | 9 | 0 | 9 | 0.3333 | 7 | 0.1429 | 32 | 0.1562 | 3 | 0 | 8 | 0.125 | 4 | 0 | 4 | 0 | 19 | 0.0526 |
| 9_4400554 | 8 | 0.125 | 8 | 0 | 11 | 0.3636 | 8 | 0.25 | 35 | 0.2 | 5 | 0 | 10 | 0.3 | 5 | 0.2 | 7 | 0.2857 | 27 | 0.2222 |
| 9_4400581 | 5 | 0.4 | 7 | 0.2857 | 12 | 0.4167 | 7 | 0.4286 | 31 | 0.3871 | 3 | 0 | 9 | 0.3333 | 3 | 0 | 4 | 0.5 | 19 | 0.2632 |
| 9_4400587 | 10 | 0.7 | 8 | 0.625 | 10 | 0.7 | 7 | 0.8571 | 35 | 0.7143 | 7 | 0.8571 | 13 | 0.7692 | 4 | 0.75 | 9 | 0.5556 | 33 | 0.7273 |
| 9_4400648 | 9 | 0.2222 | 8 | 0.25 | 10 | 0.7 | 5 | 1 | 32 | 0.5 | 7 | 0.7143 | 15 | 0.4 | 7 | 0.4286 | 12 | 0.3333 | 41 | 0.439 |
| 9_4400674 | 10 | 0.1 | 13 | 0.1538 | 13 | 0 | 9 | 0.1111 | 45 | 0.0889 | 12 | 0 | 19 | 0.1053 | 9 | 0.1111 | 16 | 0 | 56 | 0.0536 |
| 9_4401997 | 5 | 0 | 4 | 0 | 4 | 0.5 | 2 | 0 | 15 | 0.1333 | 4 | 0.25 | 1 | 0 | 3 | 0 | 2 | 0 | 10 | 0.1 |
| 9_4401998 | 5 | 0.8 | 5 | 0.2 | 3 | 0.3333 | 2 | 0.5 | 15 | 0.4667 | 4 | 0 | 1 | 1 | 3 | 0.3333 | 2 | 0 | 10 | 0.2 |
| 9_4402004 | 5 | 0.2 | 6 | 0.1667 | 4 | 0.25 | 2 | 0 | 17 | 0.1765 | 5 | 0 | 1 | 0 | 4 | 0.25 | 2 | 0 | 12 | 0.0833 |
| 9_4402021 | 2 | 0.5 | 5 | 0.2 | 2 | 0 | 2 | 0.5 | 11 | 0.2727 | 3 | 0 | 1 | 0 | 3 | 0.3333 | 2 | 0 | 9 | 0.1111 |
| 9_4402031 | 4 | 0.75 | 7 | 0.2857 | 4 | 0.75 | 3 | 0.3333 | 18 | 0.5 | 5 | 0.4 | 1 | 1 | 3 | 0.3333 | 2 | 0 | 11 | 0.3636 |
| 9_4402036 | 4 | 0.25 | 7 | 0.1429 | 4 | 0 | 2 | 0.5 | 17 | 0.1765 | 4 | 0.25 | 1 | 1 | 4 | 0.25 | 2 | 0 | 11 | 0.2727 |
| 9_4402211 | 3 | 0.6667 | 5 | 0.6 | 3 | 0.3333 | 4 | 0.5 | 15 | 0.5333 | 2 | 0.5 | 5 | 0.2 | 1 | 0 | 2 | 0 | 10 | 0.2 |
| 9_4402231 | 3 | 0.6667 | 5 | 0.2 | 2 | 0 | 3 | 0.3333 | 13 | 0.3077 | 3 | 0 | 3 | 0.3333 | 1 | 0 | 4 | 0.5 | 11 | 0.2727 |
| 9_4402237 | 5 | 0.4 | 5 | 0.4 | 2 | 0 | 4 | 0 | 16 | 0.25 | 3 | 0.6667 | 4 | 0 | 1 | 0 | 3 | 0 | 11 | 0.1818 |
| 9_4402239 | 4 | 0.5 | 5 | 0.2 | 3 | 0.6667 | 4 | 0.5 | 16 | 0.4375 | 2 | 0 | 5 | 0.2 | 1 | 0 | 4 | 0.25 | 12 | 0.1667 |
| 9_4402243 | 5 | 0.4 | 5 | 0 | 3 | 0 | 3 | 0 | 16 | 0.125 | 3 | 0 | 4 | 0 | 1 | 0 | 4 | 0.5 | 12 | 0.1667 |
| 9_4402248 | 5 | 0.2 | 2 | 0 | 1 | 0 | 4 | 0 | 12 | 0.0833 | 3 | 0 | 5 | 0.2 | 1 | 0 | 3 | 0 | 12 | 0.0833 |
| 9_4402254 | 5 | 0.2 | 4 | 0.5 | 2 | 0 | 4 | 0.25 | 15 | 0.2667 | 3 | 0.3333 | 4 | 0.5 | 1 | 0 | 4 | 0.25 | 12 | 0.3333 |
| 9_4402270 | 4 | 0.25 | 4 | 0.25 | 3 | 0.6667 | 2 | 1 | 13 | 0.4615 | 3 | 0.6667 | 4 | 1 | 1 | 1 | 2 | 0.5 | 10 | 0.8 |
| 9_4402279 | 5 | 0.6 | 3 | 0.3333 | 1 | 1 | 3 | 1 | 12 | 0.6667 | 2 | 1 | 4 | 1 | 1 | 1 | 3 | 0.6667 | 10 | 0.9 |
| 9_4402280 | 5 | 0.8 | 4 | 0.5 | 2 | 0 | 3 | 1 | 14 | 0.6429 | 3 | 1 | 4 | 0.75 | 1 | 1 | 4 | 1 | 12 | 0.9167 |
| 9_4402292 | 5 | 1 | 3 | 1 | 2 | 1 | 2 | 0.5 | 12 | 0.9167 | 3 | 0.6667 | 5 | 0.6 | NA | NA | 3 | 1 | 11 | 0.7273 |
| 9_4402296 | 4 | 0.75 | 2 | 1 | 2 | 0.5 | 3 | 1 | 11 | 0.8182 | 2 | 1 | 4 | 0.75 | 1 | 1 | 2 | 0.5 | 9 | 0.7778 |
| 9_4402312 | 4 | 0.25 | 2 | 0 | 2 | 0 | 3 | 0 | 11 | 0.0909 | 3 | 0.3333 | 4 | 0.25 | 1 | 1 | 2 | 0 | 10 | 0.3 |
| 9_4402323 | 4 | 0.5 | 3 | 0 | 2 | 0.5 | 3 | 0 | 12 | 0.25 | 3 | 0 | 4 | 0.25 | 1 | 1 | 2 | 0.5 | 10 | 0.3 |
| 9_4402327 | 5 | 0 | 3 | 0.6667 | 1 | 0 | 3 | 0 | 12 | 0.1667 | 3 | 0.6667 | 4 | 0 | 1 | 0 | 3 | 0.3333 | 11 | 0.2727 |
| 9_4402350 | 2 | 0.5 | 2 | 0 | NA | NA | 2 | 0.5 | 6 | 0.3333 | 1 | 0 | 3 | 0.3333 | 1 | 0 | 2 | 0.5 | 7 | 0.2857 |
| 9_4402363 | 4 | 0.75 | 2 | 1 | 1 | 1 | 2 | 0.5 | 9 | 0.7778 | 2 | 1 | 3 | 0.3333 | 1 | 1 | 3 | 0.6667 | 9 | 0.6667 |
| 9_4402376 | 5 | 1 | 2 | 1 | NA | NA | 2 | 0.5 | 9 | 0.8889 | 1 | 1 | 3 | 1 | 1 | 1 | 3 | 1 | 8 | 1 |
| 9_4402401 | 4 | 1 | NA | NA | 1 | 1 | 1 | 1 | 6 | 1 | 2 | 1 | 2 | 0.5 | 1 | 1 | 3 | 1 | 8 | 0.875 |
| 9_4402420 | 5 | 1 | 1 | 1 | 1 | 0 | 2 | 1 | 9 | 0.8889 | 1 | 1 | 2 | 1 | 1 | 1 | 1 | 1 | 5 | 1 |
| 9_4402426 | 5 | 0.6 | 1 | 1 | 1 | 1 | 2 | 0 | 9 | 0.5556 | 2 | 0.5 | 2 | 1 | 1 | 1 | 2 | 0.5 | 7 | 0.7143 |
| 9_4405457 | 7 | 0.5714 | 8 | 0.625 | 7 | 0.8571 | 5 | 0.8 | 27 | 0.7037 | 6 | 0.5 | 13 | 0.6154 | 7 | 0.8571 | 6 | 0.5 | 32 | 0.625 |
| 9_4405530 | 7 | 0.2857 | 6 | 0.5 | 7 | 0.2857 | 2 | 0 | 22 | 0.3182 | 4 | 0 | 12 | 0.1667 | 7 | 0.1429 | 7 | 0.4286 | 30 | 0.2 |
| 9_4405587 | 5 | 0.6 | 4 | 0 | 6 | 0.5 | 2 | 1 | 17 | 0.4706 | 4 | 0.25 | 11 | 0.4545 | 9 | 0.3333 | 4 | 0.5 | 28 | 0.3929 |
| 9_4405627 | 4 | 0 | 5 | 0 | 6 | 0.3333 | 2 | 0.5 | 17 | 0.1765 | 4 | 0.25 | 10 | 0.1 | 6 | 0.3333 | 4 | 0 | 24 | 0.1667 |
| 9_4405628 | 4 | 0.5 | 2 | 0 | 4 | 0 | 1 | 0 | 11 | 0.1818 | 3 | 0.3333 | 7 | 0 | 6 | 0.1667 | 4 | 0.25 | 20 | 0.15 |
| 9_4405633 | 5 | 0 | 5 | 0.2 | 6 | 0 | 2 | 0 | 18 | 0.0556 | 4 | 0 | 10 | 0.1 | 8 | 0.125 | 7 | 0 | 29 | 0.069 |
| 9_4405753 | 5 | 0.8 | 4 | 1 | 4 | 1 | 1 | 1 | 14 | 0.9286 | 2 | 0.5 | 6 | 1 | 4 | 1 | 4 | 0.25 | 16 | 0.75 |
| 9_4405765 | 3 | 0.6667 | 2 | 0.5 | 3 | 1 | 2 | 1 | 10 | 0.8 | 2 | 0.5 | 7 | 0.8571 | 3 | 1 | 5 | 0.4 | 17 | 0.7059 |
| 9_4406268 | 1 | 1 | NA | NA | NA | NA | NA | NA | 1 | 1 | NA | NA | NA | NA | 2 | 1 | 2 | 1 | 4 | 1 |
| 9_4406271 | NA | NA | NA | NA | NA | NA | 1 | 1 | 1 | 1 | NA | NA | NA | NA | 2 | 1 | 2 | 1 | 4 | 1 |
| 9_4406284 | 1 | 1 | NA | NA | NA | NA | 1 | 1 | 2 | 1 | NA | NA | NA | NA | 2 | 1 | 2 | 1 | 4 | 1 |
| 9_4416416 | NA | NA | 2 | 1 | NA | NA | NA | NA | 2 | 1 | 1 | 1 | NA | NA | 1 | 1 | NA | NA | 2 | 1 |
| 9_4416428 | NA | NA | 1 | 1 | NA | NA | NA | NA | 1 | 1 | 1 | 1 | NA | NA | 1 | 1 | NA | NA | 2 | 1 |
| 9_4416431 | NA | NA | 2 | 0.5 | NA | NA | NA | NA | 2 | 0.5 | 1 | 1 | NA | NA | 1 | 1 | NA | NA | 2 | 1 |
| 9_4416432 | NA | NA | 1 | 1 | NA | NA | NA | NA | 1 | 1 | 1 | 1 | NA | NA | 1 | 1 | NA | NA | 2 | 1 |
| 9_4416439 | NA | NA | 1 | 1 | NA | NA | NA | NA | 1 | 1 | 1 | 0 | NA | NA | 1 | 1 | NA | NA | 2 | 0.5 |
| 9_4416450 | NA | NA | 2 | 0.5 | NA | NA | NA | NA | 2 | 0.5 | 1 | 1 | NA | NA | 1 | 1 | NA | NA | 2 | 1 |
| 9_4416452 | NA | NA | 1 | 1 | NA | NA | NA | NA | 1 | 1 | 1 | 1 | NA | NA | 1 | 1 | NA | NA | 2 | 1 |

| | | | | | | | | | | | | | | | | | | | | |
|---|---|---|---|---|---|---|---|---|---|---|---|---|---|---|---|---|---|---|---|---|
| 9_4417421 | NA | NA | 1 | 1 | NA | NA | NA | NA | 1 | 1 | NA | NA | NA | NA | 1 | 1 | NA | NA | 1 | 1 |
| 9_4418460 | 6 | 0.3333 | 4 | 0.5 | 6 | 0.5 | 11 | 0.5455 | 27 | 0.4815 | 9 | 0.3333 | 6 | 0.5 | 4 | 0.5 | 7 | 0.4286 | 26 | 0.4231 |
| 9_4418470 | 6 | 0.1667 | 5 | 0 | 5 | 0 | 8 | 0.125 | 24 | 0.0833 | 6 | 0.1667 | 6 | 0.1667 | 4 | 0 | 7 | 0.1429 | 23 | 0.1304 |
| 9_4418497 | 6 | 0.1667 | 5 | 0.2 | 6 | 0.5 | 10 | 0.6 | 27 | 0.4074 | 9 | 0.1111 | 6 | 0.3333 | 5 | 0 | 7 | 0.1429 | 27 | 0.1481 |
| 9_4418522 | 5 | 0.4 | 4 | 0.5 | 5 | 0.2 | 8 | 0.25 | 22 | 0.3182 | 8 | 0.125 | 5 | 0 | 4 | 0 | 6 | 0.1667 | 23 | 0.087 |
| 9_4418537 | 4 | 0.75 | 2 | 0 | 5 | 0.2 | 5 | 0.4 | 16 | 0.375 | 8 | 0.375 | 3 | 0 | 2 | 0 | 2 | 0 | 15 | 0.2 |
| 9_4418539 | 5 | 1 | 1 | 1 | 3 | 1 | 6 | 0.8333 | 15 | 0.9333 | 8 | 1 | 4 | 1 | 4 | 1 | 4 | 0.75 | 20 | 0.95 |
| 9_4418545 | 3 | 1 | 4 | 0.25 | 5 | 0.6 | 5 | 0.4 | 17 | 0.5294 | 7 | 0.2857 | 5 | 0.4 | 4 | 0.5 | 5 | 0.4 | 21 | 0.381 |
| 9_4418547 | 3 | 1 | 3 | 1 | 5 | 1 | 9 | 1 | 20 | 1 | 6 | 1 | 4 | 0.5 | 4 | 1 | 5 | 0.4 | 19 | 0.7368 |
| 9_4418556 | 2 | 0 | 2 | 0.5 | 4 | 0.25 | 5 | 0 | 13 | 0.1538 | 3 | 0 | 2 | 0 | 4 | 0.5 | 4 | 0 | 13 | 0.1538 |
| 9_4418558 | 5 | 0.4 | 3 | 0 | 5 | 0.4 | 8 | 0.25 | 21 | 0.2857 | 5 | 0.2 | 4 | 0.5 | 3 | 0.3333 | 5 | 0.2 | 17 | 0.2941 |
| 9_4418566 | 3 | 0.3333 | 1 | 0 | 3 | 1 | 2 | 0.5 | 9 | 0.5556 | 3 | 0.3333 | 2 | 0 | 2 | 0.5 | 5 | 0.6 | 12 | 0.4167 |
| 9_4418586 | 4 | 0.75 | 1 | 1 | 4 | 0.25 | 6 | 0.6667 | 15 | 0.6 | 5 | 0.4 | 1 | 1 | 3 | 1 | 2 | 0.5 | 11 | 0.6364 |
| 9_4418601 | 5 | 1 | 4 | 1 | 3 | 1 | 4 | 1 | 16 | 1 | 4 | 1 | 4 | 0.5 | 3 | 0.6667 | 3 | 1 | 14 | 0.7857 |
| 9_4420225 | 1 | 1 | NA | NA | NA | NA | NA | NA | 1 | 1 | NA | NA | 1 | 0 | NA | NA | 1 | 1 | 2 | 0.5 |
| 9_4420714 | 3 | 0 | NA | NA | 3 | 0.6667 | NA | NA | 6 | 0.3333 | NA | NA | NA | NA | 1 | 1 | NA | NA | 1 | 1 |
| 9_4420731 | 1 | 1 | NA | NA | 2 | 1 | NA | NA | 3 | 1 | NA | NA | NA | NA | 1 | 1 | NA | NA | 1 | 1 |
| 9_4420769 | 3 | 0 | NA | NA | 2 | 0.5 | NA | NA | 5 | 0.2 | NA | NA | NA | NA | 1 | 1 | NA | NA | 1 | 1 |
| 9_4422162 | 25 | 0.04 | 29 | 0.0345 | 29 | 0.0345 | 38 | 0.1316 | 121 | 0.0661 | 11 | 0.0909 | 28 | 0.1071 | 30 | 0.1 | 27 | 0 | 96 | 0.0729 |
| 9_4422209 | 17 | 0.5882 | 23 | 0.5217 | 26 | 0.4615 | 34 | 0.5 | 100 | 0.51 | 8 | 0.25 | 25 | 0.48 | 28 | 0.5 | 27 | 0.4074 | 88 | 0.4432 |
| 9_4422461 | 1 | 1 | 1 | 0 | NA | NA | 1 | 0 | 3 | 0.3333 | 1 | 1 | NA | NA | 1 | 1 | NA | NA | 2 | 1 |
| 9_4422478 | 1 | 1 | NA | NA | 1 | 1 | 1 | 1 | 3 | 1 | NA | NA | 1 | 1 | 1 | 1 | NA | NA | 2 | 1 |
| 9_4422750 | 7 | 0 | 14 | 0 | 22 | 0.0909 | 18 | 0.2222 | 61 | 0.0984 | 5 | 0.2 | 16 | 0 | 19 | 0.1053 | 16 | 0.125 | 56 | 0.0893 |
| 9_4423088 | 5 | 0 | 4 | 0.25 | 15 | 0.0667 | 8 | 0.25 | 32 | 0.125 | 2 | 0 | 12 | 0.0833 | 7 | 0.1429 | 8 | 0 | 29 | 0.069 |
| 9_4423117 | 7 | 0.2857 | 2 | 1 | 13 | 0.8462 | 9 | 0.4444 | 31 | 0.6129 | 2 | 1 | 12 | 0.75 | 7 | 0.7143 | 8 | 0.75 | 29 | 0.7586 |
| 9_4424457 | 3 | 0 | 2 | 1 | 1 | 0 | 2 | 0.5 | 8 | 0.375 | 2 | 0.5 | 3 | 0.6667 | 1 | 0 | 1 | 0 | 7 | 0.4286 |
| 9_4424458 | 3 | 1 | 3 | 1 | 1 | 1 | 2 | 1 | 9 | 1 | 3 | 1 | 3 | 0.6667 | 1 | 1 | 2 | 0.5 | 9 | 0.7778 |
| 9_4424460 | 3 | 1 | 4 | 0.75 | 1 | 1 | 2 | 1 | 10 | 0.9 | 3 | 1 | 2 | 1 | 1 | 1 | 1 | 1 | 7 | 1 |
| 9_4424488 | 2 | 0 | 2 | 0 | 1 | 1 | 2 | 0.5 | 7 | 0.2857 | 4 | 0.5 | 3 | 0 | 1 | 1 | 2 | 0.5 | 10 | 0.4 |
| 9_4424691 | 5 | 1 | 4 | 0.75 | 2 | 1 | 7 | 0.8571 | 18 | 0.8889 | 7 | 0.7143 | 9 | 0.6667 | 14 | 0.8571 | 5 | 1 | 35 | 0.8 |
| 9_4424692 | 6 | 0.8333 | 5 | 0.4 | 3 | 0.6667 | 5 | 1 | 19 | 0.7368 | 2 | 1 | 7 | 0.8571 | 13 | 0.9231 | 3 | 0.6667 | 25 | 0.88 |
| 9_4424700 | 6 | 0 | 5 | 0.2 | 3 | 0 | 6 | 0 | 20 | 0.05 | 7 | 0 | 8 | 0.125 | 14 | 0.0714 | 5 | 0.2 | 34 | 0.0882 |
| 9_4424706 | 6 | 0.1667 | 6 | 0 | 2 | 0 | 7 | 0.1429 | 21 | 0.0952 | 7 | 0 | 9 | 0.4444 | 16 | 0.3125 | 5 | 0 | 37 | 0.2432 |
| 9_4424711 | 1 | 1 | 1 | 1 | NA | NA | NA | NA | 2 | 1 | 1 | 1 | NA | NA | 1 | 1 | NA | NA | 2 | 1 |
| 9_4424712 | 6 | 0.5 | 5 | 0.4 | 3 | 0.6667 | 5 | 0.8 | 19 | 0.5789 | 8 | 0.625 | 8 | 0.625 | 17 | 0.6471 | 5 | 0.4 | 38 | 0.6053 |
| 9_4424725 | 5 | 1 | 7 | 1 | 3 | 1 | 6 | 1 | 21 | 1 | 10 | 1 | 8 | 1 | 18 | 0.9444 | 4 | 1 | 40 | 0.975 |
| 9_4424727 | 3 | 0.3333 | 7 | 0.8571 | 1 | 0 | 5 | 0.6 | 16 | 0.625 | 6 | 0.5 | 9 | 0.4444 | 8 | 0.75 | 5 | 0.6 | 28 | 0.5714 |
| 9_4424728 | 5 | 0.8 | 5 | 0.8 | 3 | 0.6667 | 7 | 1 | 20 | 0.85 | 8 | 1 | 8 | 0.75 | 16 | 0.875 | 5 | 0.8 | 37 | 0.8649 |
| 9_4424730 | 5 | 0 | 6 | 0.1667 | 3 | 0 | 6 | 0 | 20 | 0.05 | 9 | 0 | 10 | 0 | 15 | 0.2 | 6 | 0.1667 | 40 | 0.1 |
| 9_4424784 | 4 | 0.25 | 2 | 0.5 | 6 | 0.5 | 2 | 0.5 | 14 | 0.4286 | 5 | 0.4 | 6 | 0.3333 | 1 | 0 | 1 | 1 | 13 | 0.3846 |
| 9_4424786 | 3 | 1 | 3 | 0.6667 | 7 | 0.7143 | 2 | 0.5 | 15 | 0.7333 | 5 | 0.8 | 3 | 0.3333 | 2 | 0 | 1 | 1 | 11 | 0.5455 |
| 9_4425937 | 1 | 0 | 4 | 1 | 1 | 0 | 6 | 0.5 | 12 | 0.5833 | 2 | 0 | 1 | 1 | 4 | 0.75 | 1 | 1 | 8 | 0.625 |
| 9_4425938 | 1 | 1 | 6 | 1 | 2 | 1 | 6 | 1 | 15 | 1 | 3 | 1 | 2 | 1 | 6 | 1 | 2 | 1 | 13 | 1 |
| 9_4425939 | 3 | 0 | 10 | 0.8 | 8 | 0.625 | 13 | 0.6923 | 34 | 0.6471 | 7 | 0.7143 | 5 | 0.8 | 4 | 0.75 | 2 | 0.5 | 18 | 0.7222 |
| 9_4425949 | 7 | 1 | 12 | 0.6667 | 9 | 0.8889 | 14 | 0.9286 | 42 | 0.8571 | 7 | 0.8571 | 5 | 0.8 | 9 | 0.7778 | 2 | 1 | 23 | 0.8261 |
| 9_4425960 | 7 | 1 | 10 | 0.7 | 8 | 0.125 | 12 | 0.75 | 37 | 0.6486 | 7 | 1 | 5 | 0.6 | 8 | 1 | 4 | 1 | 24 | 0.9167 |
| 9_4425961 | 8 | 0.375 | 13 | 0.3846 | 7 | 0.2857 | 13 | 0.3077 | 41 | 0.3415 | 9 | 0.3333 | 4 | 0.25 | 8 | 0.375 | 5 | 0.4 | 26 | 0.3462 |
| 9_4425972 | 6 | 0 | 9 | 0.1111 | 6 | 0 | 11 | 0.2727 | 32 | 0.125 | 9 | 0 | 4 | 0.25 | 4 | 0 | 4 | 0.5 | 21 | 0.1429 |
| 9_4425974 | 8 | 0 | 11 | 0.0909 | 6 | 0.1667 | 13 | 0 | 38 | 0.0526 | 8 | 0.125 | 5 | 0 | 8 | 0.125 | 4 | 0.5 | 25 | 0.16 |
| 9_4425975 | 7 | 0.4286 | 10 | 0.1 | 6 | 0 | 9 | 0.3333 | 32 | 0.2188 | 6 | 0.5 | 5 | 0.2 | 3 | 0 | 3 | 0.3333 | 17 | 0.2941 |
| 9_4425984 | 5 | 0.2 | 9 | 0.2222 | 5 | 0.2 | 9 | 0 | 28 | 0.1429 | 6 | 0.1667 | 3 | 0 | 5 | 0 | 4 | 0.25 | 18 | 0.1111 |
| 9_4430060 | 6 | 0.1667 | 7 | 0 | 3 | 0 | 5 | 0.2 | 21 | 0.0952 | 6 | 0.1667 | 2 | 0 | 6 | 0.1667 | 6 | 0.1667 | 20 | 0.15 |
| 9_4430234 | 10 | 0.3 | 10 | 0.2 | 6 | 0.1667 | 7 | 0.2857 | 33 | 0.2424 | 9 | 0.2222 | 7 | 0.2857 | 7 | 0.1429 | 7 | 0.2857 | 30 | 0.2333 |
| 9_4430290 | 9 | 0.2222 | 11 | 0.3636 | 6 | 0.3333 | 10 | 0.7 | 36 | 0.4167 | 12 | 0.4167 | 7 | 0.4286 | 10 | 0.2 | 8 | 0.375 | 37 | 0.3514 |
| 9_4430301 | 8 | 0 | 11 | 0 | 6 | 0.3333 | 7 | 0.1429 | 32 | 0.0938 | 14 | 0.2143 | 7 | 0 | 12 | 0.25 | 9 | 0 | 42 | 0.1429 |
| 9_4431381 | NA | NA | 3 | 0.6667 | 2 | 0 | 1 | 0 | 6 | 0.3333 | NA | NA | 3 | 0.3333 | 1 | 1 | 2 | 1 | 6 | 0.6667 |
| 9_4431397 | 1 | 0 | 4 | 0.25 | 2 | 0 | 1 | 0 | 8 | 0.125 | NA | NA | 3 | 0.3333 | 2 | 0.5 | 2 | 0 | 7 | 0.2857 |
| 9_4438830 | NA | NA | 1 | 0 | NA | NA | 1 | 1 | 2 | 0.5 | 1 | 0 | NA | NA | NA | NA | 1 | 1 | 2 | 0.5 |
| 9_4438948 | 2 | 1 | 1 | 1 | 2 | 0.5 | NA | NA | 5 | 0.8 | 1 | 1 | 1 | 1 | NA | NA | 2 | 1 | 4 | 1 |
| 9_4438953 | 1 | 1 | 3 | 0.6667 | 2 | 1 | NA | NA | 6 | 0.8333 | 1 | 0 | 1 | 1 | NA | NA | 2 | 1 | 4 | 0.75 |
| 9_4438953 | 2 | 0.5 | 2 | 0 | 1 | 1 | NA | NA | 5 | 0.4 | NA | NA | NA | NA | NA | NA | 1 | 1 | 1 | 1 |
| 9_4438955 | 2 | 0.5 | 1 | 1 | 2 | 1 | NA | NA | 5 | 0.8 | 1 | 1 | NA | NA | NA | NA | 1 | 1 | 2 | 1 |
| 9_4438958 | 1 | 1 | 2 | 0 | 2 | 0 | NA | NA | 5 | 0.2 | 1 | 1 | 1 | 1 | NA | NA | 2 | 1 | 4 | 1 |
| 9_4438965 | NA | NA | 2 | 0.5 | 1 | 1 | NA | NA | 3 | 0.6667 | NA | NA | 1 | 1 | NA | NA | 1 | 0 | 2 | 0.5 |
| 9_4438999 | 1 | 1 | 1 | 1 | 3 | 1 | NA | NA | 5 | 1 | NA | NA | 1 | 1 | NA | NA | 2 | 1 | 3 | 1 |

| | | | | | | | | | | | | | | | | | | | | |
|---|---|---|---|---|---|---|---|---|---|---|---|---|---|---|---|---|---|---|---|---|
| 9_4439024 | NA | NA | 1 | 1 | 2 | 0 | NA | NA | 3 | 0.3333 | NA | NA | NA | NA | NA | NA | 1 | 1 | 1 | 1 |
| 9_4439029 | NA | NA | NA | NA | 1 | 1 | NA | NA | 1 | 1 | NA | NA | NA | NA | NA | NA | 1 | 1 | 1 | 1 |
| 9_4439038 | NA | NA | 1 | 0 | 1 | 1 | NA | NA | 2 | 0.5 | NA | NA | NA | NA | NA | NA | 1 | 1 | 1 | 1 |
| 9_4439039 | NA | NA | 1 | 1 | 1 | 1 | NA | NA | 2 | 1 | NA | NA | NA | NA | NA | NA | 1 | 1 | 1 | 1 |
| 9_4439048 | NA | NA | 1 | 1 | NA | NA | NA | NA | 1 | 1 | NA | NA | NA | NA | NA | NA | 1 | 1 | 1 | 1 |
| 9_4439050 | 4 | 0.25 | 1 | 0 | NA | NA | NA | NA | 5 | 0.2 | NA | NA | 1 | 0 | NA | NA | 2 | 0.5 | 3 | 0.3333 |
| 9_4439060 | NA | NA | 1 | 1 | 1 | 0 | NA | NA | 2 | 0.5 | NA | NA | NA | NA | NA | NA | 1 | 1 | 1 | 1 |
| 9_4439061 | 4 | 0.25 | 1 | 0 | NA | NA | NA | NA | 5 | 0.2 | NA | NA | 1 | 1 | NA | NA | 2 | 0 | 3 | 0.3333 |
| 9_4439078 | NA | NA | NA | NA | 1 | 1 | NA | NA | 1 | 1 | NA | NA | NA | NA | NA | NA | 1 | 1 | 1 | 1 |
| 9_4439079 | NA | NA | NA | NA | 1 | 1 | NA | NA | 1 | 1 | NA | NA | NA | NA | NA | NA | 1 | 1 | 1 | 1 |
| 9_4439309 | 4 | 0.5 | 1 | 1 | NA | NA | NA | NA | 5 | 0.6 | NA | NA | 2 | 1 | NA | NA | 1 | 1 | 3 | 1 |
| 9_4441214 | 1 | 1 | NA | NA | NA | NA | NA | NA | 1 | 1 | NA | NA | 1 | 0 | NA | NA | 3 | 0.3333 | 4 | 0.25 |
| 9_4441708 | 1 | 1 | NA | NA | 1 | 0 | NA | NA | 2 | 0.5 | NA | NA | 1 | 1 | 2 | 1 | 2 | 0.5 | 5 | 0.8 |
| 9_4441721 | NA | NA | NA | NA | 1 | 1 | NA | NA | 1 | 1 | NA | NA | 1 | 1 | 2 | 0.5 | 2 | 0.5 | 5 | 0.6 |
| 9_4441753 | 1 | 0 | NA | NA | 1 | 1 | NA | NA | 2 | 0.5 | NA | NA | 1 | 0 | 3 | 1 | 3 | 1 | 7 | 0.8571 |
| 9_4441768 | 1 | 1 | NA | NA | 1 | 1 | NA | NA | 2 | 1 | NA | NA | 1 | 1 | 3 | 1 | 3 | 1 | 7 | 1 |
| 9_4441793 | 2 | 0.5 | NA | NA | 1 | 1 | NA | NA | 3 | 0.6667 | NA | NA | 1 | 0 | 4 | 0.75 | 3 | 0.6667 | 8 | 0.625 |
| 9_4441796 | 2 | 0.5 | NA | NA | 1 | 0 | NA | NA | 3 | 0.3333 | NA | NA | NA | NA | 4 | 0 | 1 | 1 | 5 | 0.2 |
| 9_4441815 | 2 | 0.5 | NA | NA | NA | NA | NA | NA | 2 | 0.5 | NA | NA | 1 | 0 | 3 | 0.3333 | 2 | 1 | 6 | 0.5 |
| 9_4441819 | 2 | 0 | NA | NA | 1 | 1 | NA | NA | 3 | 0.3333 | NA | NA | 1 | 1 | 4 | 0.5 | 3 | 0 | 8 | 0.375 |
| 9_4441830 | 1 | 1 | NA | NA | NA | NA | NA | NA | 1 | 1 | NA | NA | NA | NA | 2 | 1 | 3 | 1 | 5 | 1 |
| 9_4441838 | 1 | 1 | NA | NA | 1 | 1 | NA | NA | 2 | 1 | NA | NA | 1 | 0 | 3 | 0 | 3 | 0.3333 | 7 | 0.1429 |
| 9_4441839 | 1 | 0 | NA | NA | 1 | 1 | NA | NA | 2 | 0.5 | NA | NA | 2 | 0 | 4 | 0.25 | 3 | 0.6667 | 9 | 0.3333 |
| 9_4441851 | NA | NA | NA | NA | 1 | 1 | NA | NA | 1 | 1 | NA | NA | 2 | 0 | 4 | 0.25 | 2 | 0.5 | 8 | 0.25 |
| 9_4441860 | 1 | 1 | NA | NA | 1 | 1 | NA | NA | 2 | 1 | NA | NA | 2 | 0.5 | 4 | 1 | 3 | 1 | 9 | 0.8889 |
| 9_4441862 | 2 | 1 | NA | NA | 1 | 1 | NA | NA | 3 | 1 | NA | NA | 2 | 1 | 4 | 1 | 3 | 1 | 9 | 1 |
| 9_4441883 | 2 | 0 | NA | NA | 1 | 1 | NA | NA | 3 | 0.3333 | NA | NA | 2 | 0.5 | 4 | 0 | 3 | 0 | 9 | 0.1111 |
| 9_4442006 | 2 | 1 | NA | NA | 1 | 1 | 1 | 1 | 4 | 1 | NA | NA | NA | NA | 5 | 0.4 | 3 | 0.3333 | 8 | 0.375 |
| 9_4442016 | 1 | 1 | 1 | 0 | 1 | 1 | 1 | 1 | 4 | 0.75 | NA | NA | 3 | 0.6667 | 7 | 0.5714 | 3 | 0.6667 | 13 | 0.6154 |
| 9_4442018 | 2 | 0.5 | NA | NA | 1 | 0 | NA | NA | 3 | 0.3333 | NA | NA | 2 | 1 | 4 | 0 | 3 | 0.6667 | 9 | 0.4444 |
| 9_4442037 | 2 | 0.5 | 1 | 0 | 1 | 1 | 3 | 0 | 7 | 0.2857 | NA | NA | 3 | 0.3333 | 6 | 0.3333 | 3 | 0 | 12 | 0.25 |
| 9_4442043 | 1 | 0 | 1 | 0 | 1 | 1 | 2 | 0.5 | 5 | 0.4 | NA | NA | NA | NA | 6 | 0.5 | 3 | 0.3333 | 9 | 0.4444 |
| 9_4442066 | 2 | 0.5 | NA | NA | 1 | 1 | 3 | 0 | 6 | 0.3333 | 1 | 0 | 3 | 1 | 6 | 0.3333 | 3 | 0.3333 | 13 | 0.4615 |
| 9_4442076 | 2 | 0.5 | NA | NA | NA | NA | NA | NA | 2 | 0.5 | 1 | 1 | 1 | 0 | 6 | 0.5 | 2 | 0.5 | 10 | 0.5 |
| 9_4442078 | 2 | 0.5 | 1 | 1 | 1 | 1 | 2 | 0 | 6 | 0.5 | 1 | 1 | 4 | 0.25 | 6 | 0.8333 | 3 | 0.6667 | 14 | 0.6429 |
| 9_4442102 | 2 | 0.5 | 2 | 1 | 1 | 1 | 1 | 0 | 6 | 0.6667 | 1 | 1 | 3 | 1 | 7 | 0.5714 | 2 | 1 | 13 | 0.7692 |
| 9_4442110 | 2 | 0 | 2 | 0 | 2 | 0.5 | 3 | 0.3333 | 9 | 0.2222 | 2 | 0.5 | 5 | 0 | 6 | 0.1667 | 3 | 0.3333 | 16 | 0.1875 |
| 9_4442131 | 2 | 0 | 1 | 0 | 1 | 1 | 4 | 0 | 8 | 0.125 | 3 | 0 | 4 | 0.25 | 7 | 0 | 2 | 0 | 16 | 0.0625 |
| 9_4442145 | 2 | 0.5 | 2 | 0.5 | 1 | 1 | 2 | 0 | 7 | 0.4286 | 1 | 1 | 5 | 0.8 | 7 | 0.2857 | 2 | 0 | 15 | 0.4667 |
| 9_4442185 | 1 | 0 | 2 | 0.5 | 2 | 0.5 | 5 | 0.2 | 10 | 0.3 | 3 | 0.6667 | 1 | 0 | 6 | 0.3333 | 5 | 0.2 | 15 | 0.3333 |
| 9_4442215 | 2 | 0 | 3 | 0 | 2 | 1 | 5 | 0.4 | 12 | 0.3333 | 3 | 0.3333 | 1 | 0 | 6 | 0.6667 | 4 | 0.25 | 14 | 0.4286 |
| 9_4455025 | NA | NA | NA | NA | 1 | 1 | 1 | 0 | 2 | 0.5 | NA | NA | 1 | 1 | NA | NA | NA | NA | 1 | 1 |
| 9_4455134 | 14 | 0.1429 | 16 | 0.25 | 16 | 0.1875 | 17 | 0.1176 | 63 | 0.1746 | 9 | 0.1111 | 20 | 0.05 | 18 | 0.1111 | 9 | 0.1111 | 56 | 0.0893 |
| 9_4455151 | 15 | 0.2667 | 9 | 0.3333 | 15 | 0.0667 | 19 | 0.1579 | 58 | 0.1897 | 9 | 0.1111 | 16 | 0.0625 | 17 | 0.1176 | 11 | 0.4545 | 53 | 0.1698 |
| 9_4455191 | 29 | 0.1034 | 17 | 0 | 26 | 0.0385 | 32 | 0.0625 | 104 | 0.0577 | 19 | 0.2632 | 20 | 0.2 | 26 | 0.0769 | 21 | 0.0476 | 86 | 0.1395 |
| 9_4459790 | 3 | 0.3333 | 10 | 0.2 | 2 | 0 | 5 | 0.4 | 20 | 0.25 | 5 | 0 | 1 | 0 | 2 | 0 | 8 | 0.375 | 16 | 0.1875 |
| 9_4459848 | 4 | 0 | 7 | 0.4286 | 3 | 0 | 3 | 0.3333 | 17 | 0.2353 | 7 | 0.2857 | 2 | 0.5 | 2 | 0.5 | 8 | 0.5 | 19 | 0.4211 |
| 9_4459881 | 4 | 0.25 | 5 | 0.4 | 3 | 0.6667 | 3 | 1 | 15 | 0.5333 | 7 | 0.4286 | 1 | 0 | 1 | 0 | 6 | 0.6667 | 15 | 0.4667 |
| 9_4459885 | 3 | 1 | 7 | 0.2857 | 3 | 0.3333 | 4 | 1 | 17 | 0.5882 | 3 | 0.3333 | 2 | 0.5 | 2 | 0.5 | 2 | 0.5 | 9 | 0.4444 |
| 9_4462674 | 1 | 1 | 4 | 0.75 | 2 | 1 | 3 | 0.3333 | 10 | 0.7 | 2 | 1 | 1 | 0 | NA | NA | 1 | 0 | 4 | 0.5 |
| 9_4462689 | 1 | 1 | 4 | 0.75 | 1 | 1 | 3 | 0.6667 | 9 | 0.7778 | 1 | 0 | 1 | 1 | NA | NA | 1 | 1 | 3 | 0.6667 |
| 9_4462692 | 1 | 1 | 3 | 0.6667 | 2 | 1 | 3 | 1 | 9 | 0.8889 | 2 | 1 | 1 | 1 | NA | NA | 1 | 0 | 4 | 0.75 |
| 9_4462694 | 1 | 0 | 3 | 1 | 2 | 0.5 | 3 | 1 | 9 | 0.7778 | 2 | 1 | 1 | 1 | NA | NA | 1 | 1 | 4 | 1 |
| 9_4462701 | 1 | 1 | 4 | 0.5 | 2 | 1 | 1 | 1 | 8 | 0.75 | 2 | 1 | 1 | 1 | NA | NA | 1 | 0 | 4 | 0.75 |
| 9_4462707 | 1 | 0 | 3 | 0 | 1 | 0 | 3 | 0.3333 | 8 | 0.125 | 1 | 0 | NA | NA | NA | NA | 1 | 1 | 2 | 0.5 |
| 9_4462708 | 1 | 1 | 3 | 0.6667 | 1 | 1 | 2 | 1 | 7 | 0.8571 | 1 | 1 | NA | NA | NA | NA | 1 | 0 | 2 | 0.5 |
| 9_4462715 | NA | NA | 3 | 0.3333 | NA | NA | 3 | 0.3333 | 6 | 0.3333 | 2 | 1 | NA | NA | NA | NA | 1 | 0 | 3 | 0.6667 |
| 9_4462724 | NA | NA | 4 | 0.25 | 2 | 0 | 2 | 0 | 8 | 0.125 | 2 | 0.5 | 1 | 0 | NA | NA | 1 | 0 | 4 | 0.25 |
| 9_4462744 | NA | NA | 2 | 0 | 1 | 1 | 1 | 1 | 4 | 0.5 | 1 | 1 | 1 | 1 | NA | NA | 1 | 0 | 3 | 0.6667 |
| 9_4462761 | 1 | 0 | 2 | 0 | 1 | 0 | 2 | 0.5 | 6 | 0.1667 | 2 | 0.5 | 1 | 0 | NA | NA | 1 | 0 | 4 | 0.25 |
| 9_4462769 | NA | NA | 2 | 0 | 2 | 0.5 | NA | NA | 4 | 0.25 | 2 | 0.5 | 1 | 1 | NA | NA | 1 | 0 | 4 | 0.5 |
| 9_4462785 | 1 | 1 | 2 | 0 | 2 | 0 | 2 | 0.5 | 7 | 0.2857 | 2 | 0.5 | NA | NA | NA | NA | 1 | 0 | 3 | 0.3333 |
| 9_4464597 | NA | NA | NA | NA | 1 | 0 | 1 | 1 | 2 | 0.5 | 1 | 1 | NA | NA | NA | NA | NA | NA | 1 | 1 |
| 9_4464608 | NA | NA | NA | NA | 1 | 0 | 1 | 1 | 2 | 0.5 | 1 | 1 | NA | NA | NA | NA | NA | NA | 1 | 1 |
| 9_4464635 | NA | NA | NA | NA | 1 | 0 | 1 | 1 | 2 | 0.5 | 1 | 1 | NA | NA | NA | NA | NA | NA | 1 | 1 |

| | | | | | | | | | | | | | | | | | | | | |
|---|---|---|---|---|---|---|---|---|---|---|---|---|---|---|---|---|---|---|---|---|
| 9_4464649 | NA | NA | NA | NA | 2 | 0.5 | 1 | 1 | 3 | 0.6667 | 1 | 1 | NA | NA | NA | NA | NA | NA | 1 | 1 |
| 9_4464714 | NA | NA | NA | NA | 1 | 0 | 1 | 1 | 2 | 0.5 | 1 | 1 | NA | NA | NA | NA | NA | NA | 1 | 1 |
| 9_4465342 | 4 | 0 | 5 | 0.4 | 2 | 0 | 4 | 0.5 | 15 | 0.2667 | 3 | 0.6667 | 3 | 0 | 3 | 0.3333 | 4 | 0.25 | 13 | 0.3077 |
| 9_4465502 | 3 | 0 | 8 | 0.125 | 10 | 0.1 | 9 | 0 | 30 | 0.0667 | 6 | 0 | 8 | 0 | 10 | 0.2 | 18 | 0.1111 | 42 | 0.0952 |
| 9_4465504 | 4 | 1 | 9 | 1 | 10 | 1 | 9 | 0.8889 | 32 | 0.9688 | 7 | 1 | 9 | 1 | 10 | 0.8 | 18 | 0.9444 | 44 | 0.9318 |
| 9_4465507 | 4 | 0.5 | 8 | 0.25 | 9 | 0.4444 | 11 | 0.4545 | 32 | 0.4062 | 6 | 0.1667 | 10 | 0.1 | 9 | 0.4444 | 20 | 0.6 | 45 | 0.4 |
| 9_4465524 | 2 | 0.5 | 9 | 0.3333 | 9 | 0.1111 | 8 | 0 | 28 | 0.1786 | 5 | 0 | 8 | 0.125 | 9 | 0.1111 | 18 | 0.2222 | 40 | 0.15 |
| 9_4467133 | 2 | 0.5 | 6 | 0 | 8 | 0.125 | 7 | 0 | 23 | 0.087 | 3 | 0.3333 | 8 | 0.25 | 6 | 0 | 14 | 0 | 31 | 0.0968 |
| 9_4467140 | 1 | 0 | 6 | 0 | 7 | 0 | 4 | 0.5 | 18 | 0.1111 | 3 | 0 | 7 | 0.1429 | 5 | 0.2 | 14 | 0.0714 | 29 | 0.1034 |
| 9_4470189 | 3 | 1 | 6 | 0.3333 | 5 | 0.4 | 4 | 0 | 18 | 0.3889 | 6 | 0.6667 | 6 | 0.5 | 5 | 0.4 | 5 | 0.4 | 22 | 0.5 |
| 9_4470209 | 3 | 0 | 7 | 0.2857 | 6 | 0 | 5 | 0.2 | 21 | 0.1429 | 4 | 0 | 5 | 0.6 | 2 | 0.5 | 5 | 0.4 | 16 | 0.375 |
| 9_4470211 | 3 | 0 | 5 | 0.2 | 6 | 0.3333 | 3 | 0 | 17 | 0.1765 | 5 | 0.2 | 6 | 0 | 4 | 0 | 5 | 0 | 20 | 0.05 |
| 9_4470212 | 3 | 0.3333 | 7 | 0 | 6 | 0 | 4 | 0.5 | 20 | 0.15 | 5 | 0.2 | 6 | 0 | 3 | 0 | 3 | 0.3333 | 17 | 0.1176 |
| 9_4470213 | 2 | 0 | 6 | 0.1667 | 6 | 0.1667 | 3 | 0.3333 | 17 | 0.1765 | 5 | 0.6 | 6 | 0.3333 | 3 | 0 | 4 | 0.25 | 18 | 0.3333 |
| 9_4470214 | 3 | 0.6667 | 7 | 0.4286 | 8 | 0.125 | 4 | 0.25 | 22 | 0.3182 | 3 | 0.3333 | 6 | 0.1667 | 4 | 1 | 4 | 0.25 | 17 | 0.4118 |
| 9_4470525 | 6 | 0.3333 | 5 | 0.4 | 5 | 0.6 | 1 | 0 | 17 | 0.4118 | 4 | 0.25 | 3 | 0.3333 | 1 | 1 | 1 | 0 | 9 | 0.3333 |
| 9_4470573 | 6 | 0.1667 | 7 | 0 | 4 | 0 | 3 | 0 | 20 | 0.05 | 3 | 0 | 2 | 0 | 2 | 0 | 5 | 0.4 | 12 | 0.1667 |
| 9_4470604 | 5 | 0 | 7 | 0 | 2 | 0.5 | 3 | 0.3333 | 17 | 0.1176 | 2 | 0 | 3 | 0 | 2 | 0.5 | 4 | 0 | 11 | 0.0909 |
| 9_4470623 | 5 | 0.8 | 7 | 0.5714 | 1 | 1 | 3 | 0.3333 | 16 | 0.625 | 2 | 0 | 1 | 1 | 2 | 1 | 4 | 0.75 | 9 | 0.6667 |
| 9_4470777 | 4 | 0.25 | 5 | 0 | 2 | 0 | 3 | 0 | 14 | 0.0714 | 2 | 0 | 2 | 0.5 | 2 | 0 | 3 | 0.3333 | 9 | 0.2222 |
| 9_4471478 | 9 | 0 | 11 | 0.1818 | 6 | 0.1667 | 14 | 0.2857 | 40 | 0.175 | 5 | 0.2 | 5 | 0.2 | 13 | 0.2308 | 7 | 0.1429 | 30 | 0.2 |
| 9_4471584 | 4 | 0.75 | 5 | 0.8 | 3 | 1 | 11 | 0.7273 | 23 | 0.7826 | 2 | 1 | 5 | 1 | 9 | 0.8889 | 5 | 1 | 21 | 0.9524 |
| 9_4471592 | NA | NA | 1 | 0 | NA | NA | 2 | 0.5 | 3 | 0.3333 | NA | NA | NA | NA | 1 | 1 | 2 | 0 | 3 | 0.3333 |
| 9_4471592 | 1 | 0 | NA | NA | 2 | 0.5 | 5 | 0.2 | 8 | 0.25 | 1 | 1 | NA | NA | 2 | 1 | 1 | 1 | 4 | 1 |
| 9_4471592 | 2 | 0 | 3 | 0 | 1 | 0 | 4 | 0.5 | 10 | 0.2 | NA | NA | 1 | 1 | 4 | 0.5 | 2 | 1 | 7 | 0.7143 |
| 9_4471601 | 4 | 0.25 | 5 | 0.2 | 4 | 0.25 | 14 | 0.0714 | 27 | 0.1481 | 2 | 0 | 4 | 0.25 | 8 | 0.25 | 6 | 0.3333 | 20 | 0.25 |
| 9_4471614 | 4 | 0.5 | 3 | 0 | 2 | 1 | 8 | 0.375 | 17 | 0.4118 | 2 | 0.5 | 3 | 0.3333 | 9 | 0.5556 | 4 | 0.75 | 18 | 0.5556 |
| 9_4471625 | 4 | 0.5 | 2 | 0 | 3 | 0 | 11 | 0.2727 | 20 | 0.25 | 1 | 0 | 3 | 0.6667 | 9 | 0.2222 | 6 | 0.5 | 19 | 0.3684 |
| 9_4471632 | 4 | 1 | 4 | 1 | 3 | 1 | 12 | 1 | 23 | 1 | 2 | 1 | 3 | 0.6667 | 8 | 1 | 6 | 1 | 19 | 0.9474 |
| 9_4471694 | 3 | 0.3333 | 3 | 0.3333 | 4 | 0.25 | 13 | 0.4615 | 23 | 0.3913 | 1 | 0 | 4 | 1 | 7 | 0.7143 | 6 | 0.5 | 18 | 0.6667 |
| 9_4471703 | 3 | 0.3333 | 2 | 0 | 3 | 0.3333 | 13 | 0.3077 | 21 | 0.2857 | 1 | 0 | 4 | 0.5 | 6 | 0.3333 | 6 | 0.3333 | 17 | 0.3529 |
| 9_4471705 | 3 | 1 | 3 | 1 | 2 | 0.5 | 11 | 0.9091 | 19 | 0.8947 | 1 | 1 | 2 | 1 | 5 | 1 | 3 | 1 | 11 | 1 |
| 9_4471706 | 2 | 0 | 2 | 0 | 3 | 0.3333 | 11 | 0.0909 | 18 | 0.1111 | 1 | 0 | 3 | 0.6667 | 7 | 0.2857 | 6 | 0.1667 | 17 | 0.2941 |
| 9_4471711 | 3 | 1 | 3 | 0.6667 | 3 | 0.6667 | 12 | 0.8333 | 21 | 0.8095 | 1 | 1 | 3 | 1 | 8 | 0.875 | 6 | 0.8333 | 18 | 0.8889 |
| 9_4471712 | 2 | 0.5 | 3 | 0.6667 | 2 | 0.5 | 8 | 0.5 | 15 | 0.5333 | 1 | 0 | 4 | 1 | 6 | 0.6667 | 4 | 0.5 | 15 | 0.6667 |
| 9_4471726 | 2 | 0 | 2 | 0 | 3 | 0 | 11 | 0.1818 | 18 | 0.1111 | 1 | 0 | 3 | 0 | 7 | 0.1429 | 5 | 0 | 16 | 0.0625 |
| 9_4471752 | 2 | 0.5 | 1 | 1 | 3 | 0.3333 | 11 | 0.6364 | 17 | 0.5882 | 1 | 0 | 4 | 0.25 | 7 | 0.8571 | 3 | 1 | 15 | 0.6667 |
| 9_4471805 | 2 | 0.5 | 2 | 0 | 2 | 0 | 9 | 0 | 15 | 0.0667 | 1 | 0 | 2 | 0 | 6 | 0 | 3 | 0.3333 | 12 | 0.0833 |
| 9_4471895 | 2 | 0 | 2 | 0.5 | 2 | 0.5 | 3 | 0.3333 | 9 | 0.3333 | 1 | 0 | 1 | 0 | 4 | 1 | 3 | 0.6667 | 9 | 0.6667 |
| 9_4471913 | 2 | 1 | 2 | 1 | 1 | 1 | 3 | 1 | 8 | 1 | NA | NA | 2 | 1 | 6 | 1 | 3 | 0.6667 | 11 | 0.9091 |
| 9_4471916 | 2 | 1 | 2 | 0 | 2 | 0.5 | 5 | 0.8 | 11 | 0.6364 | 1 | 0 | 2 | 0 | 5 | 0.2 | 3 | 0 | 11 | 0.0909 |
| 9_4471920 | 2 | 0.5 | 1 | 0 | 2 | 0 | 5 | 0.2 | 10 | 0.2 | 1 | 1 | 2 | 0.5 | 4 | 1 | 3 | 0.3333 | 10 | 0.7 |
| 9_4471928 | 2 | 1 | 1 | 1 | 2 | 0 | 5 | 0.4 | 10 | 0.5 | 1 | 0 | 1 | 0 | 6 | 0.3333 | 3 | 0 | 11 | 0.1818 |
| 9_4472964 | 2 | 0 | 2 | 0.5 | 1 | 0 | 3 | 0 | 8 | 0.125 | NA | NA | 1 | 0 | 4 | 0.5 | 2 | 0.5 | 7 | 0.4286 |
| 9_4472982 | 1 | 1 | 1 | 1 | NA | NA | 2 | 1 | 4 | 1 | NA | NA | 1 | 0 | 4 | 1 | 2 | 0 | 7 | 0.5714 |
| 9_4472998 | 2 | 0 | 2 | 0.5 | 1 | 0 | 2 | 0 | 7 | 0.1429 | NA | NA | 1 | 1 | 3 | 0.3333 | 2 | 0.5 | 6 | 0.5 |
| 9_4473002 | 1 | 1 | 2 | 0 | 1 | 0 | 2 | 0 | 6 | 0.1667 | NA | NA | 1 | 1 | 4 | 0.25 | 2 | 0 | 7 | 0.2857 |
| 9_4473008 | 2 | 1 | 1 | 1 | NA | NA | 2 | 0.5 | 5 | 0.8 | NA | NA | 1 | 1 | 4 | 1 | 1 | 1 | 6 | 1 |
| 9_4473014 | 2 | 0.5 | 2 | 0 | 1 | 0 | 3 | 0 | 8 | 0.125 | NA | NA | 1 | 0 | 3 | 0.3333 | 1 | 0 | 5 | 0.2 |
| 9_4473021 | 2 | 0.5 | 2 | 0.5 | 1 | 1 | 2 | 1 | 7 | 0.7143 | NA | NA | 1 | 0 | 3 | 0.3333 | 2 | 0 | 6 | 0.1667 |
| 9_4473048 | 1 | 1 | 1 | 1 | NA | NA | 1 | 1 | 3 | 1 | NA | NA | NA | NA | 2 | 0.5 | 2 | 0.5 | 4 | 0.5 |
| 9_4473062 | 2 | 1 | 2 | 0 | NA | NA | 1 | 0 | 5 | 0.4 | NA | NA | 1 | 1 | 2 | 0.5 | 1 | 1 | 4 | 0.75 |
| 9_4473067 | 2 | 1 | 2 | 1 | NA | NA | 1 | 1 | 5 | 1 | NA | NA | 1 | 1 | 2 | 1 | 2 | 0.5 | 5 | 0.8 |
| 9_4473092 | 2 | 1 | 1 | 1 | NA | NA | 1 | 1 | 4 | 1 | NA | NA | 1 | 1 | 3 | 1 | 1 | 1 | 5 | 1 |
| 9_4473105 | 2 | 0.5 | 1 | 0 | NA | NA | 1 | 1 | 4 | 0.5 | NA | NA | 1 | 0 | 3 | 0.3333 | 1 | 1 | 5 | 0.4 |
| 9_4473160 | 2 | 1 | NA | NA | NA | NA | NA | NA | 2 | 1 | NA | NA | 1 | 1 | 2 | 0.5 | NA | NA | 3 | 0.6667 |
| 9_4473168 | 1 | 1 | NA | NA | NA | NA | NA | NA | 1 | 1 | NA | NA | 1 | 0 | 3 | 1 | 1 | 1 | 5 | 0.8 |
| 9_4473177 | 2 | 1 | NA | NA | NA | NA | NA | NA | 2 | 1 | NA | NA | 1 | 1 | 3 | 1 | 1 | 1 | 5 | 1 |
| 9_4473215 | 1 | 1 | NA | NA | NA | NA | NA | NA | 1 | 1 | NA | NA | 1 | 1 | 2 | 0.5 | 1 | 1 | 4 | 0.75 |
| 9_4473226 | 2 | 1 | NA | NA | NA | NA | NA | NA | 2 | 1 | NA | NA | 1 | 1 | 1 | 1 | 1 | 1 | 3 | 1 |
| 9_4473329 | 2 | 0.5 | NA | NA | NA | NA | NA | NA | 2 | 0.5 | NA | NA | NA | NA | 1 | 0 | 1 | 1 | 2 | 0.5 |
| 9_4479304 | 2 | 0.5 | 3 | 0.3333 | NA | NA | NA | NA | 5 | 0.4 | 1 | 1 | NA | NA | NA | NA | NA | NA | 1 | 1 |
| 9_4482460 | 31 | 0.1613 | 37 | 0.1081 | 46 | 0.087 | 41 | 0.122 | 155 | 0.1161 | 37 | 0.1622 | 39 | 0.1538 | 45 | 0.1333 | 35 | 0.0286 | 156 | 0.1218 |
| 9_4483459 | 1 | 0 | 4 | 0.25 | 5 | 0 | 9 | 0.1111 | 19 | 0.1053 | 4 | 0 | 5 | 0 | 1 | 0 | 8 | 0.125 | 18 | 0.0556 |
| 9_4483476 | 3 | 0.3333 | 5 | 0.6 | 4 | 0.5 | 7 | 0.1429 | 19 | 0.3684 | 3 | 0 | 4 | 0.5 | 1 | 1 | 8 | 0.25 | 16 | 0.3125 |

| | | | | | | | | | | | | | | | | | | | | |
|---|---|---|---|---|---|---|---|---|---|---|---|---|---|---|---|---|---|---|---|---|
| 9_4483484 | 3 | 0 | 5 | 0 | 5 | 0 | 8 | 0.25 | 21 | 0.0952 | 3 | 0 | 4 | 0 | 1 | 0 | 8 | 0.125 | 16 | 0.0625 |
| 9_4483488 | 4 | 0 | 5 | 0 | 5 | 0.2 | 9 | 0.2222 | 23 | 0.1304 | 2 | 0 | 4 | 0.25 | 1 | 0 | 7 | 0 | 14 | 0.0714 |
| 9_4483494 | 2 | 0 | 5 | 0.2 | 3 | 0 | 9 | 0.8889 | 19 | 0.4737 | 3 | 0 | 4 | 0.75 | 1 | 1 | 7 | 0.4286 | 15 | 0.4667 |
| 9_4483497 | 3 | 0.6667 | 4 | 0.75 | 5 | 0.6 | 8 | 0.625 | 20 | 0.65 | 3 | 1 | 3 | 1 | 1 | 1 | 5 | 0.4 | 12 | 0.75 |
| 9_4483503 | 2 | 1 | 2 | 1 | 3 | 0.6667 | 9 | 0.8889 | 16 | 0.875 | 2 | 0.5 | 4 | 1 | 1 | 1 | 6 | 1 | 13 | 0.9231 |
| 9_4483504 | 3 | 0 | 5 | 1 | 3 | 0.3333 | 8 | 0.5 | 19 | 0.5263 | 3 | 0.3333 | 4 | 0.25 | 1 | 0 | 5 | 0.6 | 13 | 0.3846 |
| 9_4483507 | 2 | 1 | 5 | 0.8 | 3 | 0 | 9 | 0.6667 | 19 | 0.6316 | 1 | 1 | 3 | 1 | 1 | 1 | 6 | 0.8333 | 11 | 0.9091 |
| 9_4483516 | 2 | 0.5 | 3 | 0.6667 | 3 | 0.6667 | 7 | 0.5714 | 15 | 0.6 | 2 | 1 | 2 | 0.5 | 1 | 1 | 6 | 0.6667 | 11 | 0.7273 |
| 9_4483526 | 2 | 0 | 5 | 0.4 | 3 | 0.3333 | 7 | 0.5714 | 17 | 0.4118 | 3 | 0.3333 | 3 | 0.6667 | NA | NA | 5 | 0.6 | 11 | 0.5455 |
| 9_4483548 | 2 | 0.5 | 5 | 0.4 | 3 | 0.6667 | 9 | 0.3333 | 19 | 0.4211 | 3 | 0.6667 | 4 | 0.75 | 1 | 1 | 6 | 0.1667 | 14 | 0.5 |
| 9_4483559 | 3 | 0 | 4 | 0 | 2 | 0 | 9 | 0.1111 | 18 | 0.0556 | 3 | 0 | 4 | 0.25 | 1 | 0 | 6 | 0 | 14 | 0.0714 |
| 9_4483713 | 2 | 0 | 2 | 0 | 4 | 0.25 | 3 | 0.3333 | 11 | 0.1818 | 1 | 0 | 4 | 0.25 | 1 | 0 | 3 | 0 | 9 | 0.1111 |
| 9_4485771 | NA | NA | NA | NA | 3 | 0 | 2 | 1 | 5 | 0.4 | 1 | 1 | 3 | 0.3333 | NA | NA | 3 | 0.3333 | 7 | 0.4286 |
| 9_4485941 | NA | NA | NA | NA | 3 | 0.3333 | 2 | 0.5 | 5 | 0.4 | NA | NA | 4 | 0.5 | NA | NA | 2 | 0 | 6 | 0.3333 |
| 9_4498820 | NA | NA | NA | NA | NA | NA | 1 | 1 | 1 | 1 | NA | NA | 2 | 0.5 | 1 | 1 | NA | NA | 3 | 0.6667 |
| 9_4498849 | NA | NA | NA | NA | NA | NA | 1 | 1 | 1 | 1 | NA | NA | 2 | 1 | 1 | 1 | 1 | 0 | 4 | 0.75 |
| 9_4498860 | NA | NA | NA | NA | NA | NA | 1 | 1 | 1 | 1 | NA | NA | 2 | 0.5 | 1 | 1 | 1 | 1 | 4 | 0.75 |
| 9_4498865 | NA | NA | NA | NA | NA | NA | 1 | 1 | 1 | 1 | NA | NA | 2 | 0 | 1 | 1 | 1 | 0 | 4 | 0.25 |
| 9_4503767 | 12 | 0.0833 | 9 | 0.1111 | 6 | 0.1667 | 8 | 0 | 35 | 0.0857 | 4 | 0.25 | 2 | 0 | 8 | 0 | 6 | 0 | 20 | 0.05 |
| 9_4503771 | 13 | 0.0769 | 8 | 0 | 5 | 0 | 8 | 0.25 | 34 | 0.0882 | 3 | 0.6667 | 4 | 0 | 9 | 0 | 6 | 0.1667 | 22 | 0.1364 |
| 9_4503779 | 9 | 0.1111 | 5 | 0.2 | 4 | 0.5 | 8 | 0 | 26 | 0.1538 | 4 | 0.25 | 3 | 0.3333 | 6 | 0 | 4 | 0.25 | 17 | 0.1765 |
| 9_4503786 | 12 | 0.5833 | 10 | 0.6 | 6 | 0.3333 | 10 | 0.3 | 38 | 0.4737 | 5 | 0.6 | 5 | 0.4 | 9 | 0.1111 | 7 | 0.7143 | 26 | 0.4231 |
| 9_4503796 | 12 | 0.0833 | 9 | 0.2222 | 6 | 0.3333 | 10 | 0.2 | 37 | 0.1892 | 4 | 0.25 | 3 | 0 | 8 | 0.25 | 8 | 0.25 | 23 | 0.2174 |
| 9_4503803 | 11 | 0.0909 | 8 | 0 | 6 | 0 | 8 | 0.125 | 33 | 0.0606 | 5 | 0.2 | 7 | 0.2857 | 8 | 0 | 6 | 0.1667 | 26 | 0.1538 |
| 9_4503819 | 14 | 0 | 11 | 0 | 6 | 0.1667 | 9 | 0.3333 | 40 | 0.1 | 7 | 0 | 8 | 0.125 | 8 | 0 | 9 | 0.1111 | 32 | 0.0625 |
| 9_4503832 | 16 | 0.0625 | 11 | 0.0909 | 7 | 0.2857 | 11 | 0.0909 | 45 | 0.1111 | 7 | 0 | 9 | 0 | 11 | 0.0909 | 9 | 0.3333 | 36 | 0.1111 |
| 9_4503833 | 10 | 0 | 9 | 0.1111 | 6 | 0.1667 | 11 | 0.1818 | 36 | 0.1111 | 7 | 0.2857 | 8 | 0 | 11 | 0.1818 | 9 | 0 | 35 | 0.1143 |
| 9_4503842 | 11 | 0 | 6 | 0 | 6 | 0.3333 | 7 | 0.1429 | 30 | 0.1 | 6 | 0 | 8 | 0 | 10 | 0 | 8 | 0.375 | 32 | 0.0938 |
| 9_4503845 | 14 | 0.2143 | 9 | 0.1111 | 8 | 0.375 | 10 | 0.1 | 41 | 0.1951 | 6 | 0.1667 | 9 | 0.1111 | 11 | 0.2727 | 7 | 0.1429 | 33 | 0.1818 |
| 9_4503872 | 18 | 0.0556 | 12 | 0.0833 | 9 | 0.1111 | 12 | 0.1667 | 51 | 0.098 | 9 | 0.2222 | 11 | 0 | 13 | 0.2308 | 10 | 0.1 | 43 | 0.1395 |
| 9_4503982 | 25 | 0.2 | 18 | 0.0556 | 18 | 0.1667 | 24 | 0.0833 | 85 | 0.1294 | 11 | 0.2727 | 24 | 0 | 23 | 0.087 | 15 | 0.0667 | 73 | 0.0822 |
| 9_4504023 | 33 | 0.1212 | 24 | 0.125 | 23 | 0.087 | 27 | 0 | 107 | 0.0841 | 15 | 0.0667 | 31 | 0.2258 | 35 | 0.1714 | 25 | 0.16 | 106 | 0.1698 |
| 9_4504064 | 36 | 0.8889 | 30 | 0.8333 | 31 | 0.8387 | 31 | 0.871 | 128 | 0.8594 | 21 | 0.6667 | 36 | 0.8333 | 38 | 0.8421 | 26 | 0.8077 | 121 | 0.8017 |
| 9_4504067 | 34 | 0.5 | 30 | 0.3333 | 27 | 0.6296 | 30 | 0.5 | 121 | 0.4876 | 16 | 0.5 | 36 | 0.3611 | 40 | 0.475 | 30 | 0.5 | 122 | 0.4508 |
| 9_4504092 | 35 | 0.2857 | 27 | 0.2593 | 37 | 0.3514 | 34 | 0.3824 | 133 | 0.3233 | 20 | 0.25 | 38 | 0.3421 | 42 | 0.4048 | 35 | 0.2571 | 135 | 0.3259 |
| 9_4505033 | 4 | 0.25 | 2 | 0 | 3 | 0 | 3 | 0 | 12 | 0.0833 | 3 | 0 | 3 | 0.6667 | 3 | 0 | NA | NA | 9 | 0.2222 |
| 9_4506062 | 6 | 0.3333 | 2 | 0 | 7 | 0.1429 | 3 | 0 | 18 | 0.1667 | 3 | 0.6667 | 3 | 0 | 4 | 0.25 | NA | NA | 10 | 0.3 |
| 9_4506189 | 7 | 0.1429 | 2 | 0 | 7 | 0.1429 | 4 | 0.5 | 20 | 0.2 | 2 | 0 | 3 | 0 | 6 | 0.1667 | NA | NA | 11 | 0.0909 |
| 9_4506191 | 8 | 0.5 | 2 | 0.5 | 6 | 0.1667 | 4 | 0.25 | 20 | 0.35 | 3 | 0.6667 | 5 | 0.8 | 7 | 0.2857 | NA | NA | 15 | 0.5333 |
| 9_4506196 | 7 | 0.2857 | 2 | 0.5 | 7 | 0.4286 | 4 | 0 | 20 | 0.3 | 3 | 0.6667 | 5 | 0.4 | 6 | 0.1667 | NA | NA | 14 | 0.3571 |
| 9_4506259 | 7 | 0 | 3 | 0 | 10 | 0.1 | 7 | 0.1429 | 27 | 0.0741 | 6 | 0.1667 | 7 | 0 | 7 | 0.1429 | 3 | 0 | 23 | 0.087 |
| 9_4506272 | 10 | 0.1 | 3 | 0 | 9 | 0.3333 | 8 | 0 | 30 | 0.1333 | 6 | 0.3333 | 7 | 0.1429 | 8 | 0.125 | 4 | 0.5 | 25 | 0.24 |
| 9_4506307 | 10 | 0.1 | 3 | 0 | 14 | 0 | 7 | 0.1429 | 34 | 0.0588 | 6 | 0 | 9 | 0.2222 | 7 | 0 | 7 | 0 | 29 | 0.069 |
| 9_4506337 | 13 | 0.0769 | 3 | 0 | 13 | 0.1538 | 10 | 0 | 39 | 0.0769 | 8 | 0.25 | 13 | 0.0769 | 10 | 0 | 6 | 0 | 37 | 0.0811 |
| 9_4506454 | 19 | 0.2105 | 8 | 0.125 | 14 | 0.2143 | 14 | 0.0714 | 55 | 0.1636 | 10 | 0.4 | 23 | 0.1739 | 19 | 0.1579 | 10 | 0.2 | 62 | 0.2097 |
| 9_4506500 | 31 | 0.0968 | 17 | 0.0588 | 32 | 0 | 49 | 0.0816 | 129 | 0.062 | 19 | 0.1579 | 48 | 0.1667 | 48 | 0.0833 | 26 | 0.0769 | 141 | 0.1206 |
| 9_4506771 | 2 | 0 | 2 | 1 | NA | NA | NA | NA | 4 | 0.5 | NA | NA | 1 | 0 | 1 | 0 | 2 | 0.5 | 4 | 0.25 |
| 9_4530676 | 10 | 0.5 | 11 | 0 | 8 | 0.625 | 5 | 0.6 | 34 | 0.3824 | 8 | 0.375 | 18 | 0.3889 | 10 | 0.4 | 14 | 0.4286 | 50 | 0.4 |
| 9_4530705 | 9 | 0 | 15 | 0.0667 | 7 | 0.1429 | 6 | 0 | 37 | 0.0541 | 12 | 0.0833 | 18 | 0.1667 | 7 | 0.1429 | 16 | 0.125 | 53 | 0.1321 |
| 9_4530738 | 9 | 0.2222 | 13 | 0.0769 | 7 | 0.4286 | 6 | 0 | 35 | 0.1714 | 13 | 0.1538 | 19 | 0.2105 | 6 | 0 | 16 | 0.125 | 54 | 0.1481 |
| 9_4558146 | 3 | 0 | 2 | 0 | 2 | 0 | 1 | 1 | 8 | 0.125 | 1 | 0 | NA | NA | 1 | 1 | 2 | 0 | 4 | 0.25 |
| 9_4558591 | 2 | 1 | 2 | 0.5 | 2 | 1 | NA | NA | 6 | 0.8333 | NA | NA | NA | NA | 1 | 1 | 1 | 0 | 2 | 0.5 |
| 9_4566346 | 2 | 0 | 3 | 0.3333 | 4 | 0 | 4 | 0 | 13 | 0.0769 | NA | NA | 3 | 0 | 1 | 1 | 6 | 0 | 10 | 0.1 |
| 9_4566410 | 4 | 0.25 | 3 | 0 | 4 | 0.25 | 4 | 0 | 15 | 0.1333 | NA | NA | 3 | 0 | 3 | 0.3333 | 8 | 0.125 | 14 | 0.1429 |
| 9_4566414 | 4 | 0.25 | 3 | 0.3333 | 4 | 0 | 4 | 0.5 | 15 | 0.2667 | NA | NA | 2 | 0.5 | 3 | 0.3333 | 7 | 0.5714 | 12 | 0.5 |
| 9_4566421 | 3 | 1 | 2 | 0.5 | 2 | 1 | 4 | 1 | 11 | 0.9091 | NA | NA | 3 | 1 | 3 | 1 | 7 | 0.8571 | 13 | 0.9231 |
| 9_4566432 | 3 | 0 | 2 | 1 | 1 | 0 | NA | NA | 6 | 0.3333 | NA | NA | 2 | 0.5 | 2 | 0.5 | 6 | 0 | 10 | 0.2 |
| 9_4566437 | 3 | 1 | 3 | 1 | 4 | 0.75 | 4 | 1 | 14 | 0.9286 | NA | NA | 3 | 1 | 2 | 1 | 8 | 0.75 | 13 | 0.8462 |
| 9_4566449 | 4 | 0.5 | 3 | 0.3333 | 3 | 0.3333 | 3 | 0.6667 | 13 | 0.4615 | NA | NA | 3 | 1 | 2 | 1 | 7 | 0.2857 | 12 | 0.5833 |
| 9_4566455 | 4 | 0.5 | 3 | 0.3333 | 4 | 0.25 | 3 | 0.3333 | 14 | 0.3571 | NA | NA | 3 | 0 | 3 | 0 | 8 | 0.25 | 14 | 0.1429 |
| 9_4566460 | 2 | 0.5 | 3 | 0 | 2 | 0.5 | 1 | 1 | 8 | 0.375 | NA | NA | 2 | 0.5 | 2 | 1 | 6 | 0.5 | 10 | 0.6 |
| 9_4566462 | 4 | 1 | 3 | 0.6667 | 4 | 0.75 | 4 | 1 | 15 | 0.8667 | NA | NA | 3 | 0.6667 | 1 | 1 | 8 | 0.625 | 12 | 0.6667 |
| 9_4566474 | 4 | 0 | 4 | 0.25 | 4 | 0 | 4 | 0.5 | 16 | 0.1875 | NA | NA | 3 | 0 | 2 | 0.5 | 7 | 0.1429 | 12 | 0.1667 |
| 9_4566480 | 4 | 0 | 4 | 0 | 4 | 0.25 | 4 | 0.25 | 16 | 0.125 | NA | NA | 3 | 0 | 2 | 0 | 5 | 0.2 | 10 | 0.1 |
| 9_4566488 | 3 | 0.3333 | 5 | 0.4 | 4 | 0.25 | 3 | 0.6667 | 15 | 0.4 | 1 | 0 | 3 | 0.6667 | 3 | 0.6667 | 6 | 0 | 13 | 0.3077 |

| | | | | | | | | | | | | | | | | | | | | |
|---|---|---|---|---|---|---|---|---|---|---|---|---|---|---|---|---|---|---|---|---|
| 9_4567928 | 3 | 1 | 1 | 1 | NA | NA | 2 | 0.5 | 6 | 0.8333 | NA | NA | NA | NA | 1 | 1 | NA | NA | 1 | 1 |
| 9_4567934 | 2 | 0.5 | 1 | 1 | NA | NA | 2 | 1 | 5 | 0.8 | NA | NA | NA | NA | 1 | 1 | NA | NA | 1 | 1 |
| 9_4567937 | 2 | 1 | 1 | 1 | NA | NA | 2 | 1 | 5 | 1 | NA | NA | NA | NA | 1 | 1 | NA | NA | 1 | 1 |
| 9_4567944 | 3 | 1 | 1 | 1 | NA | NA | 2 | 1 | 6 | 1 | NA | NA | NA | NA | 1 | 1 | NA | NA | 1 | 1 |
| 9_4567957 | 2 | 1 | 1 | 0 | NA | NA | NA | NA | 3 | 0.6667 | NA | NA | NA | NA | 1 | 1 | NA | NA | 1 | 1 |
| 9_4575019 | 2 | 0 | 1 | 0 | 4 | 0.5 | 2 | 0 | 9 | 0.2222 | 3 | 0.3333 | 1 | 0 | 4 | 0.5 | NA | NA | 8 | 0.375 |
| 9_4575029 | 2 | 0.5 | 3 | 0.3333 | 4 | 0.75 | 1 | 1 | 10 | 0.6 | 2 | 0.5 | 3 | 0.6667 | 2 | 1 | NA | NA | 7 | 0.7143 |
| 9_4575051 | 2 | 0 | 4 | 0.25 | 4 | 0 | 2 | 0 | 12 | 0.0833 | 4 | 0 | 5 | 0.2 | 4 | 0 | 2 | 0 | 15 | 0.0667 |
| 9_4576845 | 5 | 0.2 | 2 | 0 | 2 | 0 | 4 | 0.25 | 13 | 0.1538 | NA | NA | 1 | 0 | 5 | 0 | 4 | 0.25 | 10 | 0.1 |
| 9_4576977 | 4 | 0.75 | 1 | 1 | 2 | 1 | 3 | 1 | 10 | 0.9 | NA | NA | NA | NA | 3 | 0.6667 | 3 | 0.6667 | 6 | 0.6667 |
| 9_4576981 | 4 | 0.75 | 1 | 1 | 1 | 0 | 1 | 1 | 7 | 0.7143 | NA | NA | NA | NA | 2 | 1 | 3 | 0.6667 | 5 | 0.8 |
| 9_4576999 | 3 | 1 | 1 | 0 | 2 | 0.5 | 2 | 1 | 8 | 0.75 | NA | NA | NA | NA | 2 | 1 | 3 | 1 | 5 | 1 |
| 9_4577013 | 3 | 0 | 1 | 0 | 1 | 0 | 2 | 0.5 | 7 | 0.1429 | NA | NA | NA | NA | 2 | 0 | 3 | 0.3333 | 5 | 0.2 |
| 9_4577021 | 2 | 0.5 | 1 | 0 | 1 | 0 | 1 | 1 | 5 | 0.4 | NA | NA | NA | NA | 1 | 1 | 3 | 0 | 4 | 0.25 |
| 9_4577029 | 3 | 1 | 1 | 0 | 1 | 1 | 1 | 1 | 6 | 0.8333 | NA | NA | NA | NA | 2 | 1 | 1 | 1 | 3 | 1 |
| 9_4577033 | 3 | 1 | 1 | 1 | 1 | 1 | 2 | 1 | 7 | 1 | NA | NA | NA | NA | 2 | 1 | 3 | 1 | 5 | 1 |
| 9_4577063 | 2 | 1 | 1 | 0 | NA | NA | 2 | 1 | 5 | 0.8 | NA | NA | NA | NA | 1 | 1 | 3 | 0.6667 | 4 | 0.75 |
| 9_4577063 | 2 | 0.5 | 1 | 0 | NA | NA | 2 | 0 | 5 | 0.2 | NA | NA | NA | NA | 1 | 1 | 2 | 0 | 3 | 0.3333 |
| 9_4577077 | 1 | 0 | 1 | 1 | NA | NA | 2 | 0 | 4 | 0.25 | NA | NA | NA | NA | 1 | 0 | 1 | 1 | 2 | 0.5 |
| 9_4577115 | NA | NA | 1 | 0 | NA | NA | 2 | 0.5 | 3 | 0.3333 | NA | NA | NA | NA | 1 | 1 | 2 | 0.5 | 3 | 0.6667 |
| 9_4577115 | NA | NA | 1 | 0 | NA | NA | 1 | 1 | 2 | 0.5 | NA | NA | NA | NA | 1 | 0 | 2 | 0.5 | 3 | 0.3333 |
| 9_4577141 | 3 | 1 | 6 | 0.3333 | 5 | 0.2 | 5 | 0 | 19 | 0.3158 | 7 | 0.1429 | 7 | 0.4286 | 6 | 0.5 | 5 | 0.4 | 25 | 0.36 |
| 9_4577156 | 3 | 1 | 4 | 0.5 | 5 | 0.6 | 4 | 0.75 | 16 | 0.6875 | 6 | 0.5 | 8 | 0.75 | 5 | 0.8 | 5 | 0.8 | 24 | 0.7083 |
| 9_4577162 | 3 | 0.3333 | 5 | 0.2 | 3 | 0 | 2 | 0.5 | 13 | 0.2308 | 5 | 0.2 | 9 | 0.1111 | 6 | 0.3333 | 5 | 0.2 | 25 | 0.2 |
| 9_4577170 | 4 | 0.75 | 6 | 1 | 3 | 1 | 6 | 1 | 19 | 0.9474 | 7 | 0.7143 | 8 | 0.875 | 7 | 1 | 4 | 0.25 | 26 | 0.7692 |
| 9_4577173 | 4 | 0 | 6 | 0 | 4 | 0.25 | 4 | 0 | 18 | 0.0556 | 6 | 0.1667 | 7 | 0 | 7 | 0.2857 | 5 | 0 | 25 | 0.12 |
| 9_4577174 | 3 | 0.6667 | 5 | 0.6 | 2 | 0 | 5 | 0.6 | 15 | 0.5333 | 3 | 0 | 6 | 0.5 | 5 | 0.6 | 3 | 0 | 17 | 0.3529 |
| 9_4577178 | 5 | 1 | 5 | 1 | 5 | 1 | 6 | 0.8333 | 21 | 0.9524 | 8 | 1 | 6 | 1 | 7 | 1 | 5 | 1 | 26 | 1 |
| 9_4577183 | 4 | 0 | 6 | 0.1667 | 4 | 0 | 3 | 0 | 17 | 0.0588 | 7 | 0.1429 | 6 | 0 | 5 | 0 | 4 | 0.5 | 22 | 0.1364 |
| 9_4577184 | 4 | 0.75 | 6 | 0.3333 | 5 | 0.8 | 4 | 0.25 | 19 | 0.5263 | 8 | 0.75 | 8 | 0.625 | 7 | 0.5714 | 5 | 0.4 | 28 | 0.6071 |
| 9_4577190 | NA | NA | NA | NA | NA | NA | 2 | 0.5 | 2 | 0.5 | NA | NA | NA | NA | 1 | 0 | 2 | 0.5 | 3 | 0.3333 |
| 9_4577207 | 4 | 0.75 | 6 | 0.5 | 5 | 0.4 | 5 | 0.4 | 20 | 0.5 | 6 | 0.5 | 8 | 0.25 | 6 | 0.1667 | 5 | 0.4 | 25 | 0.32 |
| 9_4577208 | 4 | 0 | 5 | 0.4 | 5 | 0 | 7 | 0.1429 | 21 | 0.1429 | 7 | 0.5714 | 8 | 0.125 | 8 | 0.125 | 5 | 0 | 28 | 0.2143 |
| 9_4577210 | 4 | 1 | 6 | 1 | 5 | 1 | 5 | 0.8 | 20 | 0.95 | 7 | 0.7143 | 7 | 0.7143 | 8 | 0.75 | 3 | 0.6667 | 25 | 0.72 |
| 9_4577213 | 4 | 0.25 | 6 | 0.1667 | 5 | 0 | 5 | 0 | 20 | 0.1 | 6 | 0 | 8 | 0.125 | 7 | 0.1429 | 4 | 0 | 25 | 0.08 |
| 9_4577219 | 4 | 0 | 6 | 0 | 4 | 0.25 | 5 | 0 | 19 | 0.0526 | 8 | 0.125 | 8 | 0.125 | 5 | 0 | 5 | 0 | 26 | 0.0769 |
| 9_4577229 | 4 | 1 | 5 | 0.8 | 4 | 0.25 | 5 | 0.6 | 18 | 0.6667 | 7 | 1 | 6 | 0.6667 | 8 | 0.625 | 3 | 0.3333 | 24 | 0.7083 |
| 9_4577233 | 4 | 0.5 | 6 | 0.3333 | 2 | 0 | 7 | 0.2857 | 19 | 0.3158 | 8 | 0.375 | 8 | 0.375 | 7 | 0.4286 | 5 | 0.8 | 28 | 0.4643 |
| 9_4577242 | 4 | 0.5 | 6 | 0.3333 | 2 | 0 | 6 | 0.1667 | 18 | 0.2778 | 7 | 0.5714 | 7 | 0 | 7 | 0 | 5 | 0.2 | 26 | 0.1923 |
| 9_4577253 | 4 | 0.5 | 7 | 0.1429 | 5 | 0 | 7 | 0 | 23 | 0.1304 | 8 | 0.125 | 7 | 0 | 9 | 0.1111 | 5 | 0 | 29 | 0.069 |
| 9_4577319 | 3 | 0.3333 | 7 | 0.7143 | 5 | 0.4 | 7 | 0.5714 | 22 | 0.5455 | 7 | 0.5714 | 10 | 0.6 | 10 | 0.9 | 3 | 0.6667 | 30 | 0.7 |
| 9_4577321 | 3 | 1 | 4 | 0.75 | 4 | 1 | 4 | 1 | 15 | 0.9333 | 3 | 0.3333 | 6 | 0.5 | 10 | 0.8 | 5 | 0.6 | 24 | 0.625 |
| 9_4577336 | 2 | 0.5 | 3 | 0.3333 | 3 | 0.3333 | 1 | 1 | 9 | 0.4444 | 4 | 0.25 | 3 | 1 | 3 | 1 | 5 | 0.2 | 15 | 0.5333 |
| 9_4577338 | 3 | 0.3333 | 5 | 0.4 | 5 | 0.2 | 7 | 0.2857 | 20 | 0.3 | 4 | 0.25 | 6 | 0.3333 | 9 | 0.2222 | 6 | 0.5 | 25 | 0.32 |
| 9_4577340 | 2 | 0.5 | 5 | 0 | 5 | 0 | 6 | 0.3333 | 18 | 0.1667 | 4 | 0 | 5 | 0 | 7 | 0.4286 | 5 | 0.4 | 21 | 0.2381 |
| 9_4577348 | 3 | 1 | 5 | 0.4 | 5 | 0.6 | 4 | 0.75 | 17 | 0.6471 | 4 | 0.5 | 4 | 0.75 | 7 | 0.4286 | 4 | 0.75 | 19 | 0.5789 |
| 9_4577355 | 3 | 1 | 4 | 0.25 | 4 | 0.25 | 6 | 0.3333 | 17 | 0.4118 | 4 | 0.75 | 6 | 0.5 | 8 | 0.5 | 6 | 0.3333 | 24 | 0.5 |
| 9_4577366 | 4 | 0.5 | 4 | 0.25 | 4 | 0.5 | 5 | 0.6 | 17 | 0.4706 | 5 | 0.4 | 5 | 0.4 | 8 | 0.625 | 6 | 0.5 | 24 | 0.5 |
| 9_4577376 | 3 | 0.3333 | 1 | 0 | 1 | 1 | 3 | 0.3333 | 8 | 0.375 | 2 | 0.5 | 1 | 0 | 1 | 0 | 1 | 0 | 5 | 0.2 |
| 9_4577385 | 3 | 0 | 1 | 0 | 2 | 0 | 4 | 0.25 | 10 | 0.1 | 2 | 0.5 | 2 | 0 | 3 | 0.3333 | NA | NA | 7 | 0.2857 |
| 9_4577527 | 5 | 0.2 | 4 | 0 | 4 | 0 | 3 | 0.6667 | 16 | 0.1875 | 2 | 0 | 3 | 0.3333 | 1 | 1 | 3 | 0.3333 | 9 | 0.3333 |
| 9_4577560 | 2 | 0 | 2 | 0.5 | NA | NA | 3 | 0 | 7 | 0.1429 | 1 | 1 | 4 | 0.25 | 3 | 0.3333 | NA | NA | 8 | 0.375 |
| 9_4577616 | 8 | 0 | 4 | 0 | 4 | 0.25 | 2 | 0 | 18 | 0.0556 | 2 | 0 | 2 | 0 | 1 | 0 | 7 | 0.1429 | 12 | 0.0833 |
| 9_4581105 | 1 | 0 | 3 | 0.3333 | 2 | 0 | NA | NA | 6 | 0.1667 | NA | NA | NA | NA | 1 | 0 | 3 | 0.3333 | 4 | 0.25 |
| 9_4581115 | 1 | 1 | 2 | 0.5 | 2 | 0.5 | NA | NA | 5 | 0.6 | NA | NA | 1 | 1 | 1 | 1 | 2 | 1 | 4 | 1 |
| 9_4581130 | 2 | 0 | 3 | 0 | 1 | 1 | NA | NA | 6 | 0.1667 | NA | NA | NA | NA | 1 | 1 | 3 | 0 | 4 | 0.25 |
| 9_4581133 | 2 | 0.5 | 2 | 0.5 | 1 | 1 | NA | NA | 5 | 0.6 | NA | NA | NA | NA | 1 | 1 | 3 | 0.6667 | 4 | 0.75 |
| 9_4581797 | NA | NA | NA | NA | 1 | 1 | NA | NA | 1 | 1 | NA | NA | NA | NA | NA | NA | 4 | 0.5 | 4 | 0.5 |
| 9_4583984 | 1 | 0 | 1 | 1 | 1 | 0 | NA | NA | 3 | 0.3333 | 1 | 0 | 2 | 0 | 2 | 1 | 2 | 1 | 7 | 0.5714 |
| 9_4583996 | 1 | 1 | 3 | 0.3333 | 2 | 0.5 | 2 | 0 | 8 | 0.375 | 1 | 0 | 2 | 0 | 3 | 0.6667 | 3 | 0.6667 | 9 | 0.4444 |
| 9_4583999 | 1 | 1 | 3 | 0.3333 | 2 | 0 | 2 | 0.5 | 8 | 0.375 | NA | NA | 2 | 0.5 | 4 | 0.25 | 4 | 0 | 10 | 0.2 |
| 9_4584002 | 1 | 1 | 3 | 1 | 2 | 1 | 2 | 1 | 8 | 1 | 1 | 1 | 1 | 1 | 3 | 0.6667 | 4 | 1 | 9 | 0.8889 |
| 9_4584017 | 1 | 0 | 3 | 0.3333 | 1 | 0 | NA | NA | 5 | 0.2 | 1 | 0 | 2 | 0 | 3 | 0 | 2 | 0.5 | 8 | 0.125 |
| 9_4584022 | 2 | 0 | 3 | 0 | 2 | 1 | 2 | 0 | 9 | 0.2222 | 1 | 0 | 2 | 0 | 3 | 0.3333 | 4 | 0.25 | 10 | 0.2 |
| 9_4584023 | 2 | 0 | 3 | 0.3333 | 2 | 0.5 | 2 | 1 | 9 | 0.4444 | 1 | 0 | 2 | 0 | 4 | 0.5 | 4 | 0.5 | 11 | 0.3636 |

| | | | | | | | | | | | | | | | | | | | | |
|---|---|---|---|---|---|---|---|---|---|---|---|---|---|---|---|---|---|---|---|---|
| 9_4584030 | 2 | 1 | 2 | 1 | NA | NA | 2 | 1 | 6 | 1 | 1 | 1 | 2 | 1 | 3 | 1 | 4 | 1 | 10 | 1 |
| 9_4584033 | 2 | 0 | 2 | 0.5 | NA | NA | 2 | 1 | 6 | 0.5 | 1 | 0 | 2 | 0.5 | 2 | 0 | 4 | 0.75 | 9 | 0.4444 |
| 9_4584042 | 2 | 0.5 | 2 | 0.5 | 1 | 0 | 2 | 1 | 7 | 0.5714 | 1 | 0 | 2 | 0.5 | 4 | 0.5 | 3 | 0.6667 | 10 | 0.5 |
| 9_4584051 | 1 | 1 | 1 | 0 | 2 | 0 | 1 | 0 | 5 | 0.2 | 1 | 0 | 1 | 0 | 5 | 0.4 | 4 | 0 | 11 | 0.1818 |
| 9_4584053 | 2 | 1 | 3 | 0.3333 | 2 | 1 | 1 | 1 | 8 | 0.75 | 1 | 1 | 2 | 0.5 | 6 | 1 | 4 | 1 | 13 | 0.9231 |
| 9_4584067 | 2 | 0.5 | 3 | 0.3333 | 2 | 0.5 | 2 | 0 | 9 | 0.3333 | NA | NA | 2 | 0 | 4 | 0.5 | 2 | 0.5 | 8 | 0.375 |
| 9_4584073 | 2 | 0.5 | 3 | 0.3333 | 2 | 0.5 | 2 | 1 | 9 | 0.5556 | 1 | 0 | 2 | 1 | 6 | 0.8333 | 4 | 0.75 | 13 | 0.7692 |
| 9_4584074 | 3 | 0 | 3 | 0 | 1 | 0 | 2 | 0.5 | 9 | 0.1111 | 1 | 0 | 2 | 0 | 6 | 0.1667 | 5 | 0 | 14 | 0.0714 |
| 9_4584085 | 2 | 0.5 | 3 | 0.3333 | 1 | 0 | 2 | 0 | 8 | 0.25 | NA | NA | 2 | 1 | 6 | 0.3333 | 5 | 0.4 | 13 | 0.4615 |
| 9_4584955 | 9 | 0.4444 | 7 | 0.4286 | 7 | 0.2857 | 3 | 0.3333 | 26 | 0.3846 | 6 | 0 | 7 | 0.1429 | 5 | 0.2 | 6 | 0 | 24 | 0.0833 |
| 9_4584970 | 9 | 0.1111 | 7 | 0.1429 | 6 | 0 | 3 | 0 | 25 | 0.08 | 7 | 0 | 8 | 0 | 6 | 0 | 7 | 0.2857 | 28 | 0.0714 |
| 9_4584984 | 8 | 0.25 | 7 | 0.8571 | 7 | 0.4286 | 2 | 0 | 24 | 0.4583 | 5 | 0.4 | 7 | 0.4286 | 6 | 0.5 | 5 | 0.8 | 23 | 0.5217 |
| 9_4584998 | 8 | 0.5 | 6 | 0.3333 | 7 | 0.4286 | 1 | 0 | 22 | 0.4091 | 7 | 0.5714 | 7 | 0.5714 | 4 | 0.25 | 7 | 0.7143 | 25 | 0.56 |
| 9_4585006 | 7 | 0 | 4 | 0 | 6 | 0.1667 | 1 | 0 | 18 | 0.0556 | 7 | 0 | 6 | 0 | 5 | 0.4 | 6 | 0 | 24 | 0.0833 |
| 9_4585014 | 8 | 0.375 | 7 | 0 | 6 | 0.1667 | 2 | 0 | 23 | 0.1739 | 7 | 0.1429 | 7 | 0 | 4 | 0 | 8 | 0.5 | 26 | 0.1923 |
| 9_4585032 | 8 | 0.75 | 7 | 0.4286 | 4 | 0.25 | 2 | 0 | 21 | 0.4762 | 7 | 0.1429 | 7 | 0.1429 | 4 | 0 | 8 | 0.25 | 26 | 0.1538 |
| 9_4587706 | 19 | 0.2105 | 28 | 0.3214 | 21 | 0.1429 | 25 | 0.16 | 93 | 0.2151 | 13 | 0.0769 | 16 | 0.125 | 9 | 0 | 17 | 0 | 55 | 0.0545 |
| 9_4587977 | 15 | 0.6667 | 15 | 0.4667 | 17 | 0.4706 | 11 | 0.7273 | 58 | 0.569 | 11 | 0.4545 | 12 | 0.5833 | 9 | 0.7778 | 17 | 0.5882 | 49 | 0.5918 |
| 9_4587998 | 16 | 0.25 | 13 | 0.6154 | 13 | 0.2308 | 10 | 0.8 | 52 | 0.4423 | 12 | 0.4167 | 10 | 0.2 | 9 | 0.5556 | 12 | 0.5 | 43 | 0.4186 |
| 9_4588004 | 9 | 1 | 12 | 1 | 14 | 0.9286 | 11 | 1 | 46 | 0.9783 | 9 | 1 | 10 | 1 | 7 | 1 | 11 | 0.9091 | 37 | 0.973 |
| 9_4611353 | 9 | 0.4444 | 10 | 0.3 | 5 | 0.4 | 15 | 0.4 | 39 | 0.3846 | 6 | 0.5 | 4 | 0 | 17 | 0.3529 | 5 | 0.6 | 32 | 0.375 |
| 9_4611358 | 9 | 0.6667 | 9 | 0.5556 | 3 | 1 | 14 | 0.7857 | 35 | 0.7143 | 7 | 0.4286 | 3 | 0.3333 | 16 | 0.625 | 5 | 1 | 31 | 0.6129 |
| 9_4611370 | 5 | 1 | 8 | 1 | 4 | 1 | 14 | 0.9286 | 31 | 0.9677 | 6 | 1 | 4 | 1 | 14 | 0.9286 | 5 | 1 | 29 | 0.9655 |
| 9_4611373 | 9 | 0.2222 | 11 | 0.1818 | 4 | 0.25 | 14 | 0.0714 | 38 | 0.1579 | 5 | 0.2 | 6 | 0 | 15 | 0.0667 | 6 | 0 | 32 | 0.0625 |
| 9_4611376 | 11 | 0.7273 | 11 | 0.7273 | 5 | 0.8 | 15 | 0.8667 | 42 | 0.7857 | 7 | 0.8571 | 5 | 0.8 | 16 | 0.5625 | 5 | 1 | 33 | 0.7273 |
| 9_4611377 | 9 | 0.8889 | 9 | 0.8889 | 5 | 1 | 13 | 0.8462 | 36 | 0.8889 | 7 | 1 | 6 | 0.8333 | 17 | 0.9412 | 6 | 1 | 36 | 0.9444 |
| 9_4612184 | 4 | 0.25 | 1 | 0 | 1 | 1 | 9 | 0.1111 | 15 | 0.2 | 3 | 0 | 5 | 0 | 3 | 0.3333 | 8 | 0.5 | 19 | 0.2632 |
| 9_4612303 | 4 | 0.25 | 3 | 0 | 6 | 0 | 10 | 0.3 | 23 | 0.1739 | 2 | 0 | 9 | 0.3333 | 3 | 0 | 9 | 0 | 23 | 0.1304 |
| 9_4612321 | 3 | 0.3333 | 5 | 0 | 6 | 0 | 9 | 0.1111 | 23 | 0.087 | 3 | 0.6667 | 7 | 0.1429 | 2 | 0 | 11 | 0 | 23 | 0.1304 |
| 9_4612325 | 5 | 1 | 5 | 1 | 6 | 0.8333 | 11 | 0.9091 | 27 | 0.9259 | 3 | 0.6667 | 11 | 0.8182 | 4 | 1 | 13 | 0.9231 | 31 | 0.871 |
| 9_4612329 | 4 | 0.75 | 4 | 1 | 6 | 0.6667 | 9 | 1 | 23 | 0.8696 | 3 | 0.6667 | 8 | 0.625 | 3 | 0.6667 | 9 | 0.7778 | 23 | 0.6957 |
| 9_4612341 | 3 | 0.6667 | 3 | 0.3333 | 6 | 0.6667 | 10 | 0.5 | 22 | 0.5455 | 3 | 0.6667 | 10 | 0.5 | 1 | 1 | 11 | 0.4545 | 25 | 0.52 |
| 9_4612345 | 5 | 0.8 | 5 | 0.6 | 8 | 0.75 | 11 | 0.3636 | 29 | 0.5862 | 5 | 0.4 | 11 | 0.6364 | 5 | 0.6 | 12 | 0.75 | 33 | 0.6364 |
| 9_4612349 | 5 | 0.6 | 4 | 0.25 | 11 | 0.5455 | 7 | 0.2857 | 27 | 0.4444 | 5 | 0.2 | 10 | 0.4 | 5 | 0.4 | 13 | 0.3846 | 33 | 0.3636 |
| 9_4612364 | 5 | 0.8 | 3 | 0.3333 | 11 | 0.6364 | 8 | 0.5 | 27 | 0.5926 | 5 | 0.6 | 13 | 0.6154 | 4 | 0.25 | 15 | 0.6 | 37 | 0.5676 |
| 9_4612369 | 6 | 0 | 4 | 0 | 9 | 0 | 12 | 0.1667 | 31 | 0.0645 | 4 | 0 | 11 | 0 | 4 | 0 | 11 | 0.1818 | 30 | 0.0667 |
| 9_4612375 | 7 | 0.1429 | 5 | 0.4 | 10 | 0.4 | 12 | 0.1667 | 34 | 0.2647 | 5 | 0.4 | 12 | 0.4167 | 6 | 0 | 14 | 0.3571 | 37 | 0.3243 |
| 9_4612377 | 7 | 0.5714 | 4 | 0.75 | 10 | 0.6 | 10 | 0.7 | 31 | 0.6452 | 3 | 0.6667 | 8 | 0.75 | 5 | 0.4 | 11 | 0.8182 | 27 | 0.7037 |
| 9_4612381 | 6 | 0 | 5 | 0 | 11 | 0.2727 | 12 | 0.25 | 34 | 0.1765 | 5 | 0.2 | 11 | 0.1818 | 6 | 0 | 14 | 0.0714 | 36 | 0.1111 |
| 9_4613352 | 11 | 0.8182 | 6 | 0.6667 | 12 | 0.5 | 15 | 0.5333 | 44 | 0.6136 | 4 | 0.5 | 17 | 0.4118 | 6 | 0.1667 | 19 | 0.7368 | 46 | 0.5217 |
| 9_4613395 | 4 | 0.25 | 3 | 0 | 5 | 0.2 | 8 | 0.25 | 20 | 0.2 | 2 | 0 | 7 | 0 | 2 | 0.5 | 5 | 0.4 | 16 | 0.1875 |
| 9_4613449 | 12 | 0.0833 | 11 | 0 | 14 | 0.0714 | 16 | 0.0625 | 53 | 0.0566 | 6 | 0.3333 | 17 | 0.0588 | 8 | 0.125 | 19 | 0 | 50 | 0.08 |
| 9_4613481 | 11 | 0.6364 | 12 | 0.5833 | 15 | 0.4 | 18 | 0.4444 | 56 | 0.5 | 6 | 0.3333 | 16 | 0.5625 | 5 | 0.2 | 17 | 0.7059 | 44 | 0.5455 |
| 9_4613501 | 14 | 0.0714 | 14 | 0 | 15 | 0 | 17 | 0.1176 | 60 | 0.05 | 8 | 0.125 | 20 | 0.15 | 7 | 0 | 21 | 0.0476 | 56 | 0.0893 |
| 9_4613521 | 14 | 0.2857 | 17 | 0.2353 | 15 | 0.2667 | 19 | 0.3158 | 65 | 0.2769 | 6 | 0.8333 | 17 | 0.3529 | 8 | 0.25 | 21 | 0.2857 | 52 | 0.3654 |
| 9_4613544 | 12 | 0.25 | 11 | 0 | 15 | 0 | 11 | 0 | 49 | 0.0612 | 4 | 0 | 18 | 0.0556 | 6 | 0.1667 | 14 | 0.0714 | 42 | 0.0714 |
| 9_4618465 | 4 | 0.25 | 3 | 0 | 4 | 0.75 | 2 | 0 | 13 | 0.3077 | 4 | 0.25 | 5 | 0.4 | 2 | 0 | 4 | 0.25 | 15 | 0.2667 |
| 9_4619253 | 5 | 0.4 | 3 | 0.3333 | 5 | 0.6 | 3 | 0 | 16 | 0.375 | 3 | 0.3333 | 5 | 0.2 | 3 | 0.6667 | 4 | 0.25 | 15 | 0.3333 |
| 9_4619276 | 6 | 0.6667 | 3 | 0.6667 | 6 | 0.6667 | 4 | 0.5 | 19 | 0.6316 | 5 | 0.8 | 7 | 0.2857 | 5 | 0.6 | 6 | 0.5 | 23 | 0.5217 |
| 9_4619282 | 7 | 0.4286 | 4 | 0.25 | 7 | 0.4286 | 6 | 0.3333 | 24 | 0.375 | 6 | 0.3333 | 7 | 0.4286 | 5 | 0.8 | 6 | 0.1667 | 24 | 0.4167 |
| 9_4619294 | NA | NA | NA | NA | 1 | 1 | NA | NA | 1 | 1 | NA | NA | 1 | 1 | NA | NA | NA | NA | 1 | 1 |
| 9_4619948 | NA | NA | 1 | 1 | 1 | 1 | 4 | 1 | 6 | 1 | NA | NA | 1 | 1 | 1 | 1 | 1 | 1 | 3 | 1 |
| 9_4619954 | NA | NA | 1 | 0 | 1 | 1 | 4 | 1 | 6 | 0.8333 | NA | NA | 1 | 1 | 1 | 1 | 2 | 0 | 4 | 0.5 |
| 9_4620994 | NA | NA | 1 | 0 | 1 | 0 | 5 | 0.2 | 7 | 0.1429 | NA | NA | 1 | 0 | 3 | 0.3333 | 1 | 0 | 5 | 0.2 |
| 9_4621000 | NA | NA | 2 | 0.5 | NA | NA | 5 | 0 | 7 | 0.1429 | NA | NA | 3 | 0.3333 | 3 | 0.3333 | 2 | 0.5 | 8 | 0.375 |
| 9_4621024 | NA | NA | NA | NA | NA | NA | 1 | 1 | 1 | 1 | NA | NA | NA | NA | 2 | 1 | NA | NA | 2 | 1 |
| 9_4621032 | NA | NA | NA | NA | NA | NA | 1 | 1 | 1 | 1 | NA | NA | NA | NA | 2 | 1 | NA | NA | 2 | 1 |
| 9_4621039 | NA | NA | NA | NA | NA | NA | 1 | 1 | 1 | 1 | NA | NA | NA | NA | 1 | 1 | NA | NA | 1 | 1 |
| 9_4621049 | NA | NA | NA | NA | NA | NA | 1 | 1 | 1 | 1 | NA | NA | NA | NA | 2 | 0.5 | NA | NA | 2 | 0.5 |
| 9_4621071 | NA | NA | NA | NA | NA | NA | 1 | 1 | 1 | 1 | NA | NA | NA | NA | 2 | 0.5 | NA | NA | 2 | 0.5 |
| 9_4621082 | NA | NA | NA | NA | NA | NA | 1 | 1 | 1 | 1 | NA | NA | NA | NA | 2 | 1 | NA | NA | 2 | 1 |
| 9_4683197 | 3 | 0.3333 | 7 | 0 | 2 | 0 | 10 | 0.2 | 22 | 0.1364 | 1 | 0 | 5 | 0 | 1 | 0 | 3 | 0.6667 | 10 | 0.2 |
| 9_4683203 | 4 | 0.5 | 7 | 0.4286 | 3 | 0 | 10 | 0.2 | 24 | 0.2917 | 2 | 0 | 6 | 0.3333 | 1 | 1 | 5 | 0.6 | 14 | 0.4286 |
| 9_4683261 | 5 | 0.4 | 7 | 0.2857 | 4 | 0 | 10 | 0 | 26 | 0.1538 | 2 | 0.5 | 6 | 0.3333 | 4 | 0.75 | 4 | 0.25 | 16 | 0.4375 |
| 9_4683263 | 8 | 0.25 | 8 | 0 | 6 | 0.3333 | 11 | 0.2727 | 33 | 0.2121 | 3 | 0.3333 | 7 | 0 | 5 | 0.4 | 5 | 0.2 | 20 | 0.2 |

| | | | | | | | | | | | | | | | | | | | | |
|---|---|---|---|---|---|---|---|---|---|---|---|---|---|---|---|---|---|---|---|---|
| 9_4683265 | 8 | 0.125 | 9 | 0.1111 | 6 | 0 | 11 | 0.0909 | 34 | 0.0882 | 3 | 0 | 7 | 0 | 5 | 0.2 | 6 | 0.1667 | 21 | 0.0952 |
| 9_4683269 | 7 | 1 | 8 | 0.875 | 6 | 1 | 12 | 0.9167 | 33 | 0.9394 | 1 | 1 | 5 | 1 | 4 | 0.75 | 5 | 0.8 | 15 | 0.8667 |
| 9_4683282 | 7 | 0.1429 | 7 | 0 | 5 | 0.4 | 13 | 0 | 32 | 0.0938 | 2 | 0.5 | 7 | 0 | 5 | 0 | 6 | 0.1667 | 20 | 0.1 |
| 9_4683284 | 5 | 1 | 7 | 0.7143 | 4 | 0.75 | 11 | 0.9091 | 27 | 0.8519 | 2 | 0.5 | 6 | 1 | 5 | 1 | 4 | 0.75 | 17 | 0.8824 |
| 9_4683290 | 9 | 0.3333 | 8 | 0.25 | 6 | 0 | 12 | 0.4167 | 35 | 0.2857 | 1 | 0 | 6 | 0.3333 | 6 | 0.3333 | 6 | 0.3333 | 19 | 0.3158 |
| 9_4683296 | 7 | 1 | 7 | 0.7143 | 6 | 1 | 13 | 0.9231 | 33 | 0.9091 | 3 | 1 | 6 | 1 | 6 | 1 | 6 | 0.8333 | 21 | 0.9524 |
| 9_4683299 | 9 | 0.4444 | 9 | 0 | 6 | 0.3333 | 13 | 0.2308 | 37 | 0.2432 | 3 | 0 | 7 | 0 | 5 | 0 | 6 | 0.5 | 21 | 0.1429 |
| 9_4683302 | 9 | 0.7778 | 7 | 0.7143 | 6 | 0.8333 | 9 | 0.7778 | 31 | 0.7742 | 3 | 1 | 7 | 0.8571 | 5 | 1 | 5 | 1 | 20 | 0.95 |
| 9_4683327 | 6 | 0.3333 | 7 | 0 | 5 | 0.8 | 10 | 0.6 | 28 | 0.4286 | 3 | 0.6667 | 6 | 0.1667 | 4 | 0.25 | 5 | 0.6 | 18 | 0.3889 |
| 9_4683638 | NA | NA | NA | NA | NA | NA | 1 | 1 | 1 | 1 | 1 | 1 | 1 | 0 | NA | NA | 1 | 0 | 3 | 0.3333 |
| 9_4683782 | 1 | 0 | 2 | 1 | NA | NA | 2 | 0 | 5 | 0.4 | 1 | 1 | 4 | 0.25 | 2 | 0 | 2 | 0.5 | 9 | 0.3333 |
| 9_4683783 | 1 | 0 | 2 | 1 | NA | NA | NA | NA | 3 | 0.6667 | 1 | 0 | 3 | 0.6667 | 1 | 0 | 1 | 1 | 6 | 0.5 |
| 9_4683786 | 1 | 1 | 2 | 1 | NA | NA | 2 | 0.5 | 5 | 0.8 | 1 | 1 | 4 | 1 | 2 | 1 | 2 | 0.5 | 9 | 0.8889 |
| 9_4683816 | 1 | 0 | 2 | 0 | NA | NA | 2 | 1 | 5 | 0.4 | 1 | 1 | 3 | 0 | 2 | 0 | 2 | 0 | 8 | 0.125 |
| 9_4683831 | 1 | 1 | 2 | 1 | NA | NA | 2 | 0.5 | 5 | 0.8 | 1 | 1 | 3 | 1 | 1 | 1 | 2 | 0.5 | 7 | 0.8571 |
| 9_4683835 | 1 | 1 | 1 | 1 | NA | NA | 2 | 0.5 | 4 | 0.75 | 1 | 1 | 4 | 0.5 | 1 | 1 | 2 | 0.5 | 8 | 0.625 |
| 9_4772522 | 2 | 0.5 | 10 | 0.1 | 9 | 0 | 10 | 0.1 | 31 | 0.0968 | 9 | 0.3333 | 8 | 0.125 | 11 | 0.1818 | 20 | 0.25 | 48 | 0.2292 |
| 9_4775727 | NA | NA | NA | NA | 1 | 1 | 1 | 1 | 2 | 1 | NA | NA | 1 | 1 | NA | NA | NA | NA | 1 | 1 |
| 9_4775740 | NA | NA | NA | NA | 1 | 1 | 1 | 1 | 2 | 1 | NA | NA | 1 | 1 | NA | NA | NA | NA | 1 | 1 |
| 9_4775762 | NA | NA | NA | NA | NA | NA | 1 | 1 | 1 | 1 | NA | NA | 1 | 1 | NA | NA | NA | NA | 1 | 1 |
| 9_4776174 | 5 | 0.4 | 9 | 0.1111 | 10 | 0.7 | 13 | 0.6923 | 37 | 0.5135 | 11 | 0.5455 | 9 | 0.5556 | 11 | 0.4545 | 19 | 0.5263 | 50 | 0.52 |
| 9_4776175 | 6 | 0.5 | 9 | 0.3333 | 10 | 0.3 | 12 | 0.3333 | 37 | 0.3514 | 9 | 0.4444 | 7 | 0.1429 | 9 | 0.5556 | 16 | 0.1875 | 41 | 0.3171 |
| 9_4776243 | 5 | 0.2 | 10 | 0.1 | 8 | 0 | 14 | 0 | 37 | 0.0541 | 13 | 0.0769 | 7 | 0 | 6 | 0.1667 | 10 | 0 | 36 | 0.0556 |
| 9_4776864 | NA | NA | NA | NA | NA | NA | 1 | 1 | 1 | 1 | NA | NA | 1 | 0 | 1 | 1 | 1 | 0 | 3 | 0.3333 |
| 9_4776883 | NA | NA | NA | NA | NA | NA | 1 | 1 | 1 | 1 | NA | NA | 1 | 0 | 1 | 1 | 1 | 0 | 3 | 0.3333 |
| 9_4776915 | NA | NA | NA | NA | NA | NA | 1 | 1 | 1 | 1 | NA | NA | 1 | 0 | 1 | 1 | 1 | 0 | 3 | 0.3333 |
| 9_4776946 | NA | NA | NA | NA | NA | NA | 1 | 1 | 1 | 1 | NA | NA | 1 | 0 | 1 | 1 | 1 | 0 | 3 | 0.3333 |
| 9_4777042 | NA | NA | NA | NA | NA | NA | 1 | 1 | 1 | 1 | NA | NA | NA | NA | 2 | 1 | 1 | 0 | 3 | 0.6667 |
| 9_4777323 | NA | NA | 2 | 1 | NA | NA | 1 | 1 | 3 | 1 | NA | NA | 2 | 0.5 | 1 | 0 | 1 | 0 | 4 | 0.25 |
| 9_4830791 | NA | NA | 1 | 1 | NA | NA | 1 | 0 | 2 | 0.5 | 1 | 1 | NA | NA | NA | NA | NA | NA | 1 | 1 |
| 9_4830989 | NA | NA | 2 | 1 | NA | NA | 1 | 1 | 3 | 1 | 1 | 1 | NA | NA | NA | NA | NA | NA | 1 | 1 |
| 9_4830997 | NA | NA | 2 | 1 | NA | NA | 1 | 0 | 3 | 0.6667 | 1 | 1 | NA | NA | NA | NA | NA | NA | 1 | 1 |
| 9_4831027 | NA | NA | 2 | 1 | NA | NA | 1 | 1 | 3 | 1 | 1 | 1 | NA | NA | NA | NA | NA | NA | 1 | 1 |
| 9_4831095 | NA | NA | 2 | 0 | NA | NA | 1 | 1 | 3 | 0.3333 | 1 | 1 | NA | NA | NA | NA | NA | NA | 1 | 1 |
| 9_4838000 | 5 | 0.2 | 9 | 0.1111 | 3 | 0 | 4 | 0.25 | 21 | 0.1429 | 4 | 0 | 7 | 0 | 10 | 0.2 | 9 | 0 | 30 | 0.0667 |
| 9_4838013 | 5 | 0 | 9 | 0.2222 | 5 | 0 | 6 | 0 | 25 | 0.08 | 3 | 0.3333 | 5 | 0 | 6 | 0.5 | 7 | 0.1429 | 21 | 0.2381 |
| 9_4838531 | 1 | 0 | 1 | 0 | 1 | 0 | 2 | 0.5 | 5 | 0.2 | NA | NA | 6 | 0 | 2 | 0 | 1 | 1 | 9 | 0.1111 |
| 9_4838552 | 1 | 0 | 1 | 0 | 1 | 0 | 2 | 1 | 5 | 0.4 | NA | NA | 5 | 0.2 | 2 | 1 | 1 | 1 | 8 | 0.5 |
| 9_4838604 | NA | NA | 1 | 0 | 1 | 0 | 1 | 1 | 3 | 0.3333 | 1 | 1 | 4 | 0.5 | 1 | 0 | 1 | 1 | 7 | 0.5714 |
| 9_4838606 | NA | NA | 1 | 1 | 1 | 1 | 1 | 1 | 3 | 1 | 1 | 1 | 4 | 1 | 2 | 1 | 1 | 1 | 8 | 1 |
| 9_4838611 | NA | NA | 2 | 1 | NA | NA | 1 | 0 | 3 | 0.6667 | 1 | 1 | 3 | 0.6667 | NA | NA | 1 | 0 | 5 | 0.6 |
| 9_4838625 | NA | NA | 2 | 0.5 | 1 | 1 | NA | NA | 3 | 0.6667 | NA | NA | 3 | 1 | 2 | 1 | NA | NA | 5 | 1 |
| 9_4840076 | 10 | 0.3 | 6 | 0.5 | 5 | 0.2 | 2 | 0.5 | 23 | 0.3478 | 3 | 0.3333 | 7 | 0.1429 | 3 | 0.3333 | 10 | 0.1 | 23 | 0.1739 |
| 9_4840091 | 10 | 0.6 | 4 | 0.75 | 2 | 1 | 4 | 1 | 20 | 0.75 | 3 | 1 | 7 | 0.7143 | 7 | 0.7143 | 9 | 0.7778 | 26 | 0.7692 |
| 9_4840096 | 7 | 0.5714 | 6 | 0.5 | 5 | 0.8 | 1 | 1 | 19 | 0.6316 | 2 | 1 | 7 | 0.5714 | 5 | 0.6 | 10 | 0.9 | 24 | 0.75 |
| 9_4840105 | 10 | 0.2 | 2 | 0.5 | 6 | 0 | 3 | 0.3333 | 21 | 0.1905 | 3 | 0 | 7 | 0.1429 | 7 | 0 | 12 | 0.0833 | 29 | 0.069 |
| 9_4840109 | 9 | 0.5556 | 4 | 0.25 | 5 | 0.6 | 3 | 0.3333 | 21 | 0.4762 | 3 | 0.3333 | 6 | 0.3333 | 7 | 0.7143 | 13 | 0.6923 | 29 | 0.5862 |
| 9_4840112 | 10 | 0.3 | 5 | 0.4 | 6 | 0 | 3 | 0.3333 | 24 | 0.25 | 3 | 0 | 7 | 0.1429 | 7 | 0.4286 | 13 | 0.3077 | 30 | 0.2667 |
| 9_4840719 | 2 | 0.5 | NA | NA | 1 | 1 | 1 | 0 | 4 | 0.5 | 2 | 1 | 1 | 1 | 1 | 1 | 1 | 0 | 5 | 0.8 |
| 9_4856334 | 6 | 0.1667 | 11 | 0 | 3 | 0 | 6 | 0.1667 | 26 | 0.0769 | 6 | 0.1667 | 11 | 0.0909 | 5 | 0 | 6 | 0.1667 | 28 | 0.1071 |
| 9_4856835 | 1 | 1 | 3 | 0.3333 | NA | NA | 1 | 0 | 5 | 0.4 | NA | NA | 1 | 1 | 2 | 1 | 1 | 0 | 4 | 0.75 |
| 9_4874321 | NA | NA | 2 | 0.5 | NA | NA | 1 | 0 | 3 | 0.3333 | NA | NA | 1 | 1 | 1 | 1 | 1 | 0 | 3 | 0.6667 |
| 9_4874336 | NA | NA | 3 | 0 | NA | NA | 1 | 1 | 4 | 0.25 | NA | NA | 1 | 0 | 1 | 0 | 1 | 1 | 3 | 0.3333 |
| 9_4874366 | NA | NA | NA | NA | NA | NA | 2 | 1 | 2 | 1 | NA | NA | 1 | 1 | NA | NA | 1 | 1 | 2 | 1 |
| 9_4874368 | NA | NA | 1 | 1 | NA | NA | 2 | 1 | 3 | 1 | NA | NA | 1 | 1 | 1 | 1 | NA | NA | 2 | 1 |
| 9_4874370 | NA | NA | 2 | 0.5 | NA | NA | 2 | 1 | 4 | 0.75 | NA | NA | 1 | 1 | 1 | 0 | 1 | 1 | 3 | 0.6667 |
| 9_4874376 | NA | NA | 2 | 0.5 | NA | NA | 2 | 0 | 4 | 0.25 | NA | NA | 1 | 0 | 1 | 0 | 1 | 1 | 3 | 0.3333 |
| 9_4874421 | NA | NA | 2 | 1 | NA | NA | 2 | 0.5 | 4 | 0.75 | NA | NA | NA | NA | 1 | 1 | 1 | 1 | 2 | 1 |
| 9_4881110 | NA | NA | NA | NA | 1 | 1 | 2 | 0.5 | 3 | 0.6667 | NA | NA | NA | NA | 1 | 0 | 1 | 1 | 2 | 0.5 |
| 9_4895286 | 2 | 0.5 | 2 | 1 | 1 | 1 | 2 | 1 | 7 | 0.8571 | NA | NA | 3 | 0.6667 | NA | NA | 1 | 0 | 4 | 0.5 |
| 9_4895291 | 3 | 0.3333 | 1 | 0 | 3 | 0 | 3 | 0.3333 | 10 | 0.2 | NA | NA | 3 | 0 | 1 | 0 | 1 | 1 | 5 | 0.2 |
| 9_4898964 | 1 | 1 | 5 | 0.8 | 2 | 1 | 4 | 1 | 12 | 0.9167 | 2 | 1 | 2 | 1 | 3 | 0.3333 | 4 | 0.75 | 11 | 0.7273 |
| 9_4898975 | 2 | 0 | 4 | 0.75 | 3 | 0.6667 | 3 | 0.6667 | 12 | 0.5833 | 3 | 0.6667 | 1 | 1 | 3 | 0.3333 | 1 | 0 | 8 | 0.5 |
| 9_4898978 | 2 | 0.5 | 5 | 0.8 | 2 | 0.5 | 1 | 1 | 10 | 0.7 | 3 | 0.6667 | 2 | 0.5 | 3 | 0.3333 | 4 | 0.5 | 12 | 0.5 |
| 9_4898978 | 2 | 1 | 4 | 0.75 | 3 | 0.6667 | 2 | 0.5 | 11 | 0.7273 | 3 | 1 | 1 | 1 | 3 | 1 | 4 | 0.75 | 11 | 0.9091 |

| | | | | | | | | | | | | | | | | | | | | |
|---|---|---|---|---|---|---|---|---|---|---|---|---|---|---|---|---|---|---|---|---|
| 9_4898982 | 1 | 0 | 5 | 0.8 | 3 | 0.6667 | 4 | 0.25 | 13 | 0.5385 | 3 | 0.6667 | 2 | 1 | 1 | 1 | 4 | 1 | 10 | 0.9 |
| 9_4930696 | 5 | 0 | 2 | 0 | 3 | 0.3333 | 2 | 0.5 | 12 | 0.1667 | 4 | 0 | 10 | 0 | 9 | 0.2222 | 3 | 0 | 26 | 0.0769 |
| 9_4931630 | 11 | 0.0909 | 7 | 0.2857 | 10 | 0 | 10 | 0.3 | 38 | 0.1579 | 6 | 0.1667 | 16 | 0.1875 | 13 | 0.0769 | 11 | 0.1818 | 46 | 0.1522 |
| 9_4931826 | 11 | 0.4545 | 8 | 0.25 | 8 | 0.25 | 10 | 0.4 | 37 | 0.3514 | 5 | 0.6 | 15 | 0.4 | 19 | 0.2632 | 10 | 0.2 | 49 | 0.3265 |
| 9_4931845 | 11 | 0.1818 | 8 | 0.25 | 10 | 0.2 | 13 | 0.1538 | 42 | 0.1905 | 6 | 0.1667 | 17 | 0.0588 | 19 | 0.2632 | 12 | 0.0833 | 54 | 0.1481 |
| 9_4931849 | 13 | 1 | 8 | 0.875 | 8 | 1 | 14 | 0.9286 | 43 | 0.9535 | 4 | 0.75 | 19 | 1 | 21 | 1 | 11 | 0.9091 | 55 | 0.9636 |
| 9_4931859 | 13 | 0.0769 | 9 | 0.1111 | 9 | 0.2222 | 11 | 0.0909 | 42 | 0.119 | 6 | 0 | 22 | 0.0909 | 22 | 0.0909 | 13 | 0.2308 | 63 | 0.1111 |
| 9_4931881 | 14 | 0.7857 | 9 | 0.7778 | 10 | 0.9 | 13 | 0.9231 | 46 | 0.8478 | 6 | 0.8333 | 19 | 0.7895 | 21 | 0.8095 | 12 | 0.8333 | 58 | 0.8103 |
| 9_4931891 | 13 | 0.2308 | 9 | 0.2222 | 12 | 0.3333 | 15 | 0.2 | 49 | 0.2449 | 6 | 0 | 21 | 0.381 | 23 | 0.3478 | 12 | 0.3333 | 62 | 0.3226 |
| 9_4931892 | 14 | 0.2143 | 9 | 0.1111 | 11 | 0.4545 | 14 | 0.3571 | 48 | 0.2917 | 5 | 0.4 | 19 | 0.3158 | 24 | 0.3333 | 14 | 0.2857 | 62 | 0.3226 |
| 9_4931908 | 16 | 0.9375 | 8 | 1 | 13 | 1 | 14 | 1 | 51 | 0.9804 | 6 | 1 | 20 | 0.9 | 21 | 0.8571 | 13 | 0.8462 | 60 | 0.8833 |
| 9_4931915 | 15 | 0.8 | 4 | 1 | 11 | 0.8182 | 15 | 0.5333 | 45 | 0.7333 | 4 | 0.5 | 18 | 0.8889 | 21 | 0.9048 | 11 | 0.7273 | 54 | 0.8333 |
| 9_4931918 | 9 | 0.4444 | 8 | 0.625 | 10 | 0.8 | 10 | 0.6 | 37 | 0.6216 | 3 | 0.3333 | 16 | 0.625 | 19 | 0.5263 | 13 | 0.6923 | 51 | 0.5882 |
| 9_4931921 | 16 | 0.25 | 8 | 0.125 | 12 | 0.5833 | 14 | 0.2857 | 50 | 0.32 | 4 | 0 | 17 | 0.1176 | 21 | 0.3333 | 12 | 0.5 | 54 | 0.2778 |
| 9_4931924 | 12 | 0.4167 | 7 | 0.2857 | 8 | 0.5 | 11 | 0.3636 | 38 | 0.3947 | 5 | 0.6 | 11 | 0.1818 | 17 | 0.5882 | 8 | 0.75 | 41 | 0.5122 |
| 9_4934857 | 1 | 1 | NA | NA | NA | NA | 1 | 1 | 2 | 1 | NA | NA | NA | NA | 1 | 1 | 1 | 1 | 2 | 1 |
| 9_4934869 | 1 | 1 | NA | NA | NA | NA | 1 | 1 | 2 | 1 | NA | NA | NA | NA | 1 | 0 | 1 | 1 | 2 | 0.5 |
| 9_4934892 | 1 | 1 | NA | NA | NA | NA | 1 | 1 | 2 | 1 | NA | NA | NA | NA | 1 | 1 | 1 | 1 | 2 | 1 |
| 9_4941386 | 12 | 0 | 18 | 0.1111 | 12 | 0 | 17 | 0.1765 | 59 | 0.0847 | 12 | 0.25 | 11 | 0.0909 | 25 | 0.04 | 23 | 0.087 | 71 | 0.0986 |
| 9_4941415 | 8 | 0 | 15 | 0.2 | 9 | 0.1111 | 10 | 0 | 42 | 0.0952 | 8 | 0 | 11 | 0 | 19 | 0.1579 | 15 | 0.2 | 53 | 0.1132 |
| 9_4941420 | 12 | 0.0833 | 16 | 0.125 | 10 | 0.1 | 18 | 0.1111 | 56 | 0.1071 | 11 | 0.2727 | 11 | 0.3636 | 17 | 0.2941 | 18 | 0.2778 | 57 | 0.2982 |
| 9_4941461 | 10 | 0.2 | 16 | 0.0625 | 8 | 0 | 15 | 0.0667 | 49 | 0.0816 | 6 | 0 | 11 | 0.0909 | 20 | 0.15 | 15 | 0 | 52 | 0.0769 |
| 9_4941549 | 5 | 0.2 | 9 | 0.1111 | 2 | 0 | 6 | 0.1667 | 22 | 0.1364 | 6 | 0.5 | 9 | 0.1111 | 15 | 0.2 | 13 | 0.2308 | 43 | 0.2326 |
| 9_4941621 | 2 | 0 | 7 | 0.2857 | 1 | 0 | 7 | 0 | 17 | 0.1176 | 4 | 0.25 | 8 | 0 | 14 | 0.0714 | 8 | 0 | 34 | 0.0588 |
| 9_4941644 | 1 | 0 | 7 | 0.2857 | NA | NA | 6 | 0 | 14 | 0.1429 | 5 | 0 | 8 | 0.375 | 12 | 0.0833 | 6 | 0 | 31 | 0.129 |
| 9_4941830 | NA | NA | 2 | 0.5 | NA | NA | 2 | 0 | 4 | 0.25 | NA | NA | 3 | 0 | 3 | 0 | 4 | 0.25 | 10 | 0.1 |
| 9_4941832 | NA | NA | 2 | 1 | NA | NA | NA | NA | 2 | 1 | NA | NA | 3 | 0.6667 | 2 | 0.5 | 3 | 1 | 8 | 0.75 |
| 9_4942687 | 6 | 0.3333 | 6 | 0 | 7 | 0 | 6 | 0 | 25 | 0.08 | 3 | 0 | 4 | 0.25 | 9 | 0 | 5 | 0.2 | 21 | 0.0952 |
| 9_4942714 | 5 | 0 | 5 | 0 | 5 | 0.2 | 4 | 0 | 19 | 0.0526 | 2 | 0.5 | 3 | 0 | 6 | 0 | 4 | 0 | 15 | 0.0667 |
| 9_4942748 | 3 | 1 | 5 | 0.2 | 6 | 0 | 4 | 0.25 | 18 | 0.2778 | 3 | 0 | 3 | 0.3333 | 6 | 0.1667 | 5 | 0.2 | 17 | 0.1765 |
| 9_4942879 | 1 | 0 | 1 | 1 | 5 | 0 | 1 | 0 | 8 | 0.125 | 1 | 0 | 2 | 0 | 10 | 0 | 6 | 0.1667 | 19 | 0.0526 |
| 9_4942882 | 8 | 0 | 7 | 0.1429 | 8 | 0 | 9 | 0.1111 | 32 | 0.0625 | 6 | 0.1667 | 5 | 0 | 18 | 0.0556 | 10 | 0.2 | 39 | 0.1026 |
| 9_5043314 | 5 | 0.4 | 5 | 0.2 | 5 | 0.4 | 2 | 0 | 17 | 0.2941 | 4 | 0 | 4 | 0.25 | 5 | 0.2 | 5 | 0 | 18 | 0.1111 |
| 9_5043325 | 6 | 0.1667 | 7 | 0.1429 | 5 | 0.2 | 4 | 0 | 22 | 0.1364 | 5 | 0.2 | 6 | 0.1667 | 6 | 0.1667 | 8 | 0.125 | 25 | 0.16 |
| 9_5049290 | 4 | 0.75 | 4 | 0.25 | 6 | 0 | 5 | 0.2 | 19 | 0.2632 | 5 | 0.2 | 3 | 0.3333 | 6 | 0.5 | 2 | 0 | 16 | 0.3125 |
| 9_5050802 | 26 | 0.1154 | 21 | 0.1905 | 18 | 0.1667 | 14 | 0.0714 | 79 | 0.1392 | 11 | 0 | 14 | 0.0714 | 19 | 0.1053 | 19 | 0.2632 | 63 | 0.127 |
| 9_5050814 | 19 | 0.0526 | 20 | 0.1 | 13 | 0.0769 | 11 | 0.0909 | 63 | 0.0794 | 7 | 0.1429 | 12 | 0.0833 | 15 | 0.0667 | 15 | 0.0667 | 49 | 0.0816 |
| 9_5050814 | 22 | 0.1818 | 22 | 0.1364 | 16 | 0.1875 | 12 | 0.1667 | 72 | 0.1667 | 11 | 0.0909 | 14 | 0.2143 | 18 | 0 | 21 | 0.1905 | 64 | 0.125 |
| 9_5051643 | 44 | 0.7045 | 42 | 0.6429 | 32 | 0.6875 | 29 | 0.5172 | 147 | 0.6463 | 28 | 0.75 | 28 | 0.5 | 28 | 0.7143 | 42 | 0.6429 | 126 | 0.6508 |
| 9_5051751 | NA | NA | 1 | 0 | 2 | 0 | 2 | 0.5 | 5 | 0.2 | 1 | 0 | 1 | 1 | 1 | 0 | NA | NA | 3 | 0.3333 |
| 9_5051839 | NA | NA | 1 | 1 | 2 | 0 | 2 | 0 | 5 | 0.2 | 1 | 1 | 1 | 1 | 1 | 0 | NA | NA | 3 | 0.6667 |
| 9_5051847 | NA | NA | 1 | 0 | 2 | 0.5 | 2 | 1 | 5 | 0.6 | NA | NA | 1 | 0 | 2 | 1 | NA | NA | 3 | 0.6667 |
| 9_5051870 | NA | NA | 1 | 0 | 1 | 1 | 1 | 0 | 3 | 0.3333 | 1 | 1 | 1 | 1 | 1 | 0 | NA | NA | 3 | 0.6667 |
| 9_5051878 | NA | NA | NA | NA | 2 | 0.5 | 1 | 1 | 3 | 0.6667 | 1 | 0 | NA | NA | 2 | 0.5 | NA | NA | 3 | 0.3333 |
| 9_5051889 | NA | NA | 1 | 1 | 2 | 1 | 1 | 0 | 4 | 0.75 | 1 | 0 | 1 | 1 | 1 | 1 | NA | NA | 3 | 0.6667 |
| 9_5051907 | NA | NA | 1 | 1 | 2 | 1 | 2 | 1 | 5 | 1 | 1 | 1 | 1 | 0 | 2 | 0.5 | NA | NA | 4 | 0.5 |
| 9_5054602 | 4 | 0.5 | 7 | 0.4286 | 5 | 0.2 | 2 | 0.5 | 18 | 0.3889 | 2 | 0 | 5 | 0.4 | 7 | 0.7143 | 2 | 0 | 16 | 0.4375 |
| 9_5054605 | 4 | 0.25 | 7 | 0 | 6 | 0 | 2 | 0 | 19 | 0.0526 | 2 | 0 | 5 | 0 | 6 | 0.1667 | 2 | 0 | 15 | 0.0667 |
| 9_5054621 | 6 | 0.1667 | 8 | 0.125 | 8 | 0 | 2 | 0 | 24 | 0.0833 | 3 | 0 | 6 | 0 | 7 | 0.1429 | 4 | 0.5 | 20 | 0.15 |
| 9_5054642 | 6 | 0.3333 | 9 | 0.1111 | 9 | 0 | 2 | 0 | 26 | 0.1154 | 3 | 0 | 6 | 0.1667 | 7 | 0.2857 | 4 | 0 | 20 | 0.15 |
| 9_5054660 | 6 | 0 | 7 | 0.1429 | 10 | 0.2 | 4 | 0 | 27 | 0.1111 | 3 | 0.3333 | 6 | 0 | 7 | 0.1429 | 4 | 0 | 20 | 0.1 |
| 9_5054699 | 5 | 0.4 | 7 | 0.4286 | 8 | 0.375 | 3 | 0.6667 | 23 | 0.4348 | 2 | 0 | 5 | 0.2 | 6 | 0.5 | 3 | 0.3333 | 16 | 0.3125 |
| 9_5054703 | 5 | 0 | 7 | 0.1429 | 9 | 0.2222 | 3 | 0.3333 | 24 | 0.1667 | 2 | 0.5 | 7 | 0.2857 | 8 | 0.375 | 4 | 0.5 | 21 | 0.381 |
| 9_5054752 | 9 | 0.1111 | 9 | 0.1111 | 12 | 0.0833 | 8 | 0.125 | 38 | 0.1053 | 8 | 0.125 | 8 | 0 | 13 | 0.1538 | 11 | 0 | 40 | 0.075 |
| 9_5057446 | 8 | 0.25 | 5 | 0.2 | 4 | 0.5 | 6 | 0.3333 | 23 | 0.3043 | 3 | 0 | 8 | 0 | 9 | 0.4444 | 6 | 0.1667 | 26 | 0.1923 |
| 9_5057532 | NA | NA | NA | NA | 1 | 1 | NA | NA | 1 | 1 | 1 | 1 | NA | NA | NA | NA | NA | NA | 1 | 1 |
| 9_5057553 | 4 | 0 | 6 | 0.1667 | 4 | 0 | 5 | 0 | 19 | 0.0526 | 3 | 0 | 6 | 0.1667 | 6 | 0 | 4 | 0 | 19 | 0.0526 |
| 9_5057569 | 5 | 0.2 | 6 | 0.1667 | 4 | 0.25 | 9 | 0.1111 | 24 | 0.1667 | 5 | 0.4 | 5 | 0.6 | 8 | 0.375 | 6 | 0.5 | 24 | 0.4583 |
| 9_5057613 | 3 | 0 | 7 | 0.4286 | 3 | 0.3333 | 9 | 0.1111 | 22 | 0.2273 | 6 | 0.5 | 2 | 0.5 | 5 | 0 | 5 | 0.6 | 18 | 0.3889 |
| 9_5057622 | 3 | 0 | 6 | 0.3333 | 3 | 0.3333 | 10 | 0.1 | 22 | 0.1818 | 6 | 0.1667 | 4 | 0 | 5 | 0.2 | 6 | 0 | 21 | 0.0952 |
| 9_5057627 | 2 | 0.5 | 7 | 0.1429 | 4 | 0 | 6 | 0 | 19 | 0.1053 | 5 | 0 | 3 | 0 | 4 | 0.25 | 5 | 0 | 17 | 0.0588 |
| 9_5057630 | 4 | 0 | 8 | 0.25 | 4 | 0.25 | 10 | 0.2 | 26 | 0.1923 | 6 | 0.6667 | 4 | 0.5 | 6 | 0.3333 | 6 | 0.1667 | 22 | 0.4091 |
| 9_5057632 | 4 | 0.75 | 7 | 0.4286 | 3 | 1 | 8 | 0.375 | 22 | 0.5455 | 5 | 0.2 | 3 | 0.3333 | 5 | 0.4 | 4 | 0.5 | 17 | 0.3529 |
| 9_5057656 | 4 | 1 | 6 | 0.5 | 3 | 0 | 6 | 0.5 | 19 | 0.5263 | 5 | 0.6 | 3 | 1 | 6 | 1 | 3 | 1 | 17 | 0.8824 |
| 9_5057668 | 4 | 0.25 | 8 | 0 | 2 | 0 | 6 | 0 | 20 | 0.05 | 5 | 0.2 | 4 | 0 | 5 | 0.2 | 5 | 0.2 | 19 | 0.1579 |

| | | | | | | | | | | | | | | | | | | | | |
|---|---|---|---|---|---|---|---|---|---|---|---|---|---|---|---|---|---|---|---|---|
| 9_5057672 | 3 | 0.3333 | 7 | 0.1429 | 2 | 0 | 5 | 0.2 | 17 | 0.1765 | 5 | 0.2 | 4 | 0 | 6 | 0.1667 | 5 | 0.4 | 20 | 0.2 |
| 9_5057675 | 4 | 0.5 | 7 | 0.2857 | 2 | 0.5 | 7 | 0.1429 | 20 | 0.3 | 5 | 0.6 | 3 | 0 | 5 | 0.4 | 5 | 0.8 | 18 | 0.5 |
| 9_5057679 | 3 | 1 | 6 | 0.5 | 1 | 1 | 6 | 0.5 | 16 | 0.625 | 3 | 0.3333 | 3 | 0.3333 | 6 | 0.6667 | 4 | 0.5 | 16 | 0.5 |
| 9_5057683 | 2 | 1 | 8 | 1 | 2 | 1 | 5 | 0.8 | 17 | 0.9412 | 3 | 1 | 3 | 1 | 6 | 1 | 4 | 1 | 16 | 1 |
| 9_5057685 | 2 | 0.5 | 6 | 0.6667 | 1 | 0 | 5 | 0.6 | 14 | 0.5714 | 3 | 1 | 3 | 0.3333 | 5 | 1 | 4 | 0.75 | 15 | 0.8 |
| 9_5057697 | 2 | 1 | 6 | 0.3333 | 2 | 0.5 | 4 | 0.25 | 14 | 0.4286 | 1 | 0 | 3 | 0.6667 | 4 | 0.5 | 5 | 0.6 | 13 | 0.5385 |
| 9_5057698 | 4 | 1 | 7 | 0.5714 | 1 | 0 | 6 | 0.8333 | 18 | 0.7222 | 5 | 0.4 | 4 | 0.25 | 6 | 0.5 | 4 | 1 | 19 | 0.5263 |
| 9_5057707 | 4 | 0.75 | 6 | 0.3333 | 3 | 0.6667 | 6 | 0.5 | 19 | 0.5263 | 5 | 0.8 | 4 | 0.5 | 5 | 0.2 | 5 | 0.4 | 19 | 0.4737 |
| 9_5057718 | 3 | 0 | 5 | 0.2 | 2 | 0.5 | 7 | 0 | 17 | 0.1176 | 3 | 0 | 4 | 0.25 | 5 | 0 | 5 | 0 | 17 | 0.0588 |
| 9_5057725 | 3 | 1 | 3 | 1 | 2 | 1 | 5 | 1 | 13 | 1 | 5 | 0.8 | 3 | 0.3333 | 3 | 1 | 5 | 1 | 16 | 0.8125 |
| 9_5057728 | 4 | 0 | 3 | 0.3333 | 2 | 0.5 | 6 | 0.5 | 15 | 0.3333 | 5 | 0.4 | 3 | 0.3333 | 3 | 0.6667 | 3 | 0 | 14 | 0.3571 |
| 9_5057745 | 4 | 0.75 | 4 | 1 | 3 | 1 | 5 | 1 | 16 | 0.9375 | 3 | 1 | 4 | 1 | 2 | 1 | 4 | 1 | 13 | 1 |
| 9_5061011 | 1 | 0 | 1 | 0 | NA | NA | 1 | 1 | 3 | 0.3333 | NA | NA | 2 | 0.5 | NA | NA | 2 | 1 | 4 | 0.75 |
| 9_5066443 | 5 | 0 | 9 | 0.1111 | 9 | 0.3333 | 12 | 0.1667 | 35 | 0.1714 | 13 | 0.1538 | 7 | 0 | 5 | 0 | 6 | 0.1667 | 31 | 0.0968 |
| 9_5066458 | 25 | 0.12 | 27 | 0.1111 | 23 | 0.087 | 18 | 0 | 93 | 0.086 | 24 | 0.0833 | 22 | 0.0909 | 8 | 0.25 | 10 | 0.2 | 64 | 0.125 |
| 9_5066770 | 31 | 0.0645 | 32 | 0.0625 | 23 | 0.087 | 21 | 0 | 107 | 0.0561 | 26 | 0.1154 | 27 | 0.0741 | 11 | 0.0909 | 15 | 0.0667 | 79 | 0.0886 |
| 9_5067510 | 7 | 0.2857 | 8 | 0 | 1 | 0 | 5 | 0 | 21 | 0.0952 | 7 | 0.1429 | 6 | 0 | 10 | 0.1 | 3 | 0 | 26 | 0.0769 |
| 9_5067926 | 3 | 0.6667 | 3 | 0.6667 | NA | NA | 2 | 0 | 8 | 0.5 | 6 | 0.1667 | 3 | 0 | 8 | 0 | 3 | 0 | 20 | 0.05 |
| 9_5068278 | 1 | 0 | 1 | 1 | NA | NA | NA | NA | 2 | 0.5 | NA | NA | 3 | 0.3333 | NA | NA | 1 | 0 | 4 | 0.25 |
| 9_5068280 | 1 | 1 | 1 | 1 | NA | NA | NA | NA | 2 | 1 | NA | NA | 2 | 0.5 | NA | NA | 1 | 0 | 3 | 0.3333 |
| 9_5068312 | 1 | 0 | 1 | 1 | NA | NA | NA | NA | 2 | 0.5 | NA | NA | 3 | 0.6667 | NA | NA | NA | NA | 3 | 0.6667 |
| 9_5068315 | 1 | 1 | 1 | 1 | NA | NA | NA | NA | 2 | 1 | NA | NA | 3 | 0.6667 | NA | NA | 1 | 1 | 4 | 0.75 |
| 9_5068321 | 1 | 1 | 1 | 1 | NA | NA | NA | NA | 2 | 1 | NA | NA | 3 | 1 | NA | NA | 1 | 1 | 4 | 1 |
| 9_5068331 | 1 | 1 | 1 | 1 | NA | NA | NA | NA | 2 | 1 | NA | NA | 3 | 0.3333 | NA | NA | 1 | 1 | 4 | 0.5 |
| 9_5068350 | 1 | 1 | 1 | 1 | NA | NA | NA | NA | 2 | 1 | NA | NA | 3 | 0.6667 | NA | NA | 1 | 1 | 4 | 0.75 |
| 9_5068350 | 1 | 1 | 1 | 1 | NA | NA | NA | NA | 2 | 1 | NA | NA | 3 | 1 | NA | NA | 1 | 1 | 4 | 1 |
| 9_5068355 | 1 | 1 | 1 | 1 | NA | NA | NA | NA | 2 | 1 | NA | NA | 3 | 0.6667 | NA | NA | 1 | 1 | 4 | 0.75 |
| 9_5068368 | NA | NA | 1 | 1 | NA | NA | NA | NA | 1 | 1 | NA | NA | 2 | 1 | NA | NA | 1 | 1 | 3 | 1 |
| 9_5068377 | 1 | 1 | 1 | 1 | NA | NA | NA | NA | 2 | 1 | NA | NA | 3 | 1 | NA | NA | 1 | 1 | 4 | 1 |
| 9_5068379 | NA | NA | 1 | 1 | NA | NA | NA | NA | 1 | 1 | NA | NA | 1 | 1 | NA | NA | 1 | 1 | 2 | 1 |
| 9_5068381 | 1 | 1 | 1 | 1 | NA | NA | NA | NA | 2 | 1 | NA | NA | 3 | 0.6667 | NA | NA | 1 | 1 | 4 | 0.75 |
| 9_5068432 | 1 | 0 | NA | NA | NA | NA | 1 | 1 | 2 | 0.5 | NA | NA | 3 | 0.3333 | NA | NA | NA | NA | 3 | 0.3333 |
| 9_5068439 | 1 | 0 | NA | NA | NA | NA | 1 | 1 | 2 | 0.5 | NA | NA | 3 | 0.6667 | NA | NA | NA | NA | 3 | 0.6667 |
| 9_5068649 | 2 | 0.5 | 1 | 0 | NA | NA | 1 | 0 | 4 | 0.25 | 1 | 1 | NA | NA | NA | NA | NA | NA | 1 | 1 |
| 9_5075417 | 6 | 0.3333 | NA | NA | 1 | 0 | NA | NA | 7 | 0.2857 | 4 | 0.25 | 2 | 0 | 2 | 0 | NA | NA | 8 | 0.125 |
| 9_5075447 | 7 | 0.4286 | NA | NA | 1 | 1 | NA | NA | 8 | 0.5 | 3 | 0.6667 | 2 | 1 | 2 | 0.5 | NA | NA | 7 | 0.7143 |
| 9_5075448 | 5 | 0.8 | NA | NA | 1 | 0 | NA | NA | 6 | 0.6667 | 3 | 0.6667 | 2 | 0 | 2 | 1 | NA | NA | 7 | 0.5714 |
| 9_5075455 | 7 | 0.7143 | NA | NA | NA | NA | NA | NA | 7 | 0.7143 | 3 | 0.6667 | 1 | 0 | 2 | 0.5 | NA | NA | 6 | 0.5 |
| 9_5075469 | 6 | 1 | NA | NA | 1 | 0 | NA | NA | 7 | 0.8571 | 4 | 0.75 | NA | NA | 3 | 0.6667 | NA | NA | 7 | 0.7143 |
| 9_5075474 | 6 | 1 | NA | NA | 1 | 1 | NA | NA | 7 | 1 | 2 | 1 | 1 | 1 | 3 | 1 | NA | NA | 6 | 1 |
| 9_5079130 | 1 | 1 | 2 | 1 | NA | NA | 2 | 0.5 | 5 | 0.8 | 6 | 0.3333 | 1 | 1 | 1 | 0 | 1 | 1 | 9 | 0.4444 |
| 9_5079132 | 1 | 1 | 1 | 1 | NA | NA | 1 | 1 | 3 | 1 | 5 | 0.6 | NA | NA | 1 | 1 | 1 | 1 | 7 | 0.7143 |
| 9_5079149 | 1 | 1 | 2 | 0 | NA | NA | 1 | 0 | 4 | 0.25 | 6 | 0.1667 | NA | NA | 1 | 0 | 1 | 0 | 8 | 0.125 |
| 9_5079151 | 1 | 1 | 2 | 0.5 | NA | NA | 2 | 1 | 5 | 0.8 | 6 | 0.5 | 1 | 0 | 1 | 1 | 2 | 1 | 10 | 0.6 |
| 9_5079161 | 1 | 0 | 2 | 0.5 | NA | NA | 1 | 0 | 4 | 0.25 | 6 | 0.5 | 1 | 0 | 1 | 1 | 2 | 0.5 | 10 | 0.5 |
| 9_5079164 | NA | NA | 2 | 1 | NA | NA | 1 | 0 | 3 | 0.6667 | 6 | 1 | 1 | 1 | 1 | 1 | 1 | 1 | 9 | 1 |
| 9_5079176 | 1 | 1 | 2 | 1 | NA | NA | 2 | 1 | 5 | 1 | 3 | 0.6667 | 1 | 1 | 1 | 1 | 2 | 1 | 7 | 0.8571 |
| 9_5079185 | NA | NA | 2 | 0.5 | NA | NA | 2 | 0.5 | 4 | 0.5 | 4 | 1 | 1 | 0 | 1 | 1 | 1 | 1 | 7 | 0.8571 |
| 9_5079207 | 1 | 1 | 1 | 0 | NA | NA | 2 | 0 | 4 | 0.25 | 6 | 0 | NA | NA | 1 | 0 | 2 | 0.5 | 9 | 0.1111 |
| 9_5079235 | 1 | 1 | 2 | 0 | NA | NA | 2 | 0 | 5 | 0.2 | 4 | 0.5 | 1 | 0 | NA | NA | 2 | 0.5 | 7 | 0.4286 |
| 9_5080414 | 3 | 0 | 3 | 0.3333 | 2 | 0 | 3 | 0.6667 | 11 | 0.2727 | 1 | 0 | 3 | 0.6667 | 4 | 0.25 | 2 | 1 | 10 | 0.5 |
| 9_5080433 | 3 | 0.6667 | 3 | 0.3333 | 2 | 0 | 1 | 1 | 9 | 0.4444 | NA | NA | 3 | 1 | 5 | 1 | 1 | 1 | 9 | 1 |
| 9_5080452 | 3 | 0.3333 | 3 | 0.6667 | 2 | 0.5 | 2 | 0 | 10 | 0.4 | 1 | 1 | 2 | 1 | 4 | 0.75 | 2 | 1 | 9 | 0.8889 |
| 9_5080464 | 3 | 0.6667 | 3 | 0.3333 | 1 | 1 | 2 | 0.5 | 9 | 0.5556 | NA | NA | 3 | 0.6667 | 5 | 0.4 | 2 | 1 | 10 | 0.6 |
| 9_5080474 | 2 | 0.5 | 3 | 0.3333 | 1 | 1 | 1 | 0 | 7 | 0.4286 | 1 | 1 | 3 | 0.6667 | 3 | 0.6667 | 1 | 1 | 8 | 0.75 |
| 9_5080502 | 2 | 0.5 | 2 | 0.5 | NA | NA | 2 | 0.5 | 6 | 0.5 | 1 | 0 | 3 | 0.3333 | 3 | 0.3333 | NA | NA | 7 | 0.2857 |
| 9_5080512 | 2 | 0.5 | 2 | 1 | NA | NA | 2 | 1 | 6 | 0.8333 | 1 | 1 | 3 | 1 | 3 | 0.6667 | NA | NA | 7 | 0.8571 |
| 9_5080515 | 2 | 0 | 2 | 0.5 | NA | NA | 2 | 0 | 6 | 0.1667 | 1 | 0 | 2 | 0.5 | 3 | 0 | NA | NA | 6 | 0.1667 |
| 9_5080669 | 2 | 1 | 1 | 1 | NA | NA | 3 | 0.3333 | 6 | 0.6667 | NA | NA | 1 | 0 | 3 | 0.6667 | 1 | 0 | 5 | 0.4 |
| 9_5080673 | 2 | 0 | 1 | 0 | NA | NA | 4 | 0.25 | 7 | 0.1429 | NA | NA | 2 | 0 | 3 | 0.3333 | 1 | 0 | 6 | 0.1667 |
| 9_5080674 | 2 | 0.5 | 1 | 1 | NA | NA | 4 | 0 | 7 | 0.2857 | NA | NA | 2 | 1 | 3 | 0.6667 | 1 | 0 | 6 | 0.6667 |
| 9_5080676 | 2 | 0.5 | 1 | 0 | NA | NA | 3 | 0.6667 | 6 | 0.5 | NA | NA | 2 | 1 | 3 | 0.6667 | 1 | 0 | 6 | 0.6667 |
| 9_5080686 | 2 | 0 | 1 | 0 | NA | NA | 4 | 0.5 | 7 | 0.2857 | NA | NA | 2 | 1 | 3 | 0.6667 | 1 | 0 | 6 | 0.6667 |
| 9_5118192 | 1 | 1 | NA | NA | NA | NA | NA | NA | 1 | 1 | 1 | 1 | NA | NA | NA | NA | NA | NA | 1 | 1 |
| 9_5118193 | 1 | 1 | NA | NA | NA | NA | NA | NA | 1 | 1 | 1 | 1 | NA | NA | NA | NA | NA | NA | 1 | 1 |

| | | | | | | | | | | | | | | | | | | | | |
|---|---|---|---|---|---|---|---|---|---|---|---|---|---|---|---|---|---|---|---|---|
| 9_51759991 | 1 | 1 | 7 | 1 | 1 | 1 | 8 | 1 | 17 | 1 | 1 | 1 | 2 | 1 | NA | NA | 2 | 0.5 | 5 | 0.8 |
| 9_51760066 | 1 | 1 | 7 | 0.1429 | 1 | 1 | 8 | 0.25 | 17 | 0.2941 | 1 | 1 | 2 | 0 | NA | NA | 1 | 0 | 4 | 0.25 |
| 9_51760276 | 1 | 0 | 6 | 0.5 | 1 | 1 | 7 | 0.5714 | 15 | 0.5333 | 1 | 1 | 2 | 1 | 1 | 1 | 2 | 0.5 | 6 | 0.8333 |
| 9_51760512 | 1 | 0 | 7 | 0.5714 | 1 | 1 | 7 | 0.4286 | 16 | 0.5 | 1 | 0 | 2 | 0.5 | 1 | 0 | 2 | 0.5 | 6 | 0.3333 |
| 9_51760521 | 1 | 1 | 7 | 0.5714 | 1 | 0 | 7 | 0.7143 | 16 | 0.625 | 1 | 1 | 2 | 1 | 1 | 1 | 2 | 0.5 | 6 | 0.8333 |
| 9_51760931 | 2 | 1 | 7 | 0.8571 | 1 | 1 | 8 | 1 | 18 | 0.9444 | 1 | 1 | 3 | 0.6667 | 1 | 1 | 2 | 0 | 7 | 0.5714 |
| 9_51761014 | 2 | 0.5 | 8 | 0.25 | 1 | 1 | 8 | 0.5 | 19 | 0.4211 | 1 | 1 | 3 | 0.3333 | 1 | 0 | 2 | 0.5 | 7 | 0.4286 |
| 9_51761039 | 3 | 0 | 7 | 0.5714 | 1 | 0 | 7 | 0 | 18 | 0.2222 | 1 | 0 | 3 | 0 | 1 | 0 | 2 | 0.5 | 7 | 0.1429 |
| 9_51761125 | 3 | 0.6667 | 7 | 0.2857 | 1 | 1 | 8 | 0.25 | 19 | 0.3684 | 1 | 0 | 2 | 1 | 1 | 1 | 2 | 1 | 6 | 0.8333 |
| 9_51761293 | 2 | 0.5 | 8 | 1 | 1 | 1 | 3 | 1 | 14 | 0.9286 | 1 | 1 | 3 | 1 | 1 | 1 | 2 | 1 | 7 | 1 |
| 9_51761428 | 3 | 0.6667 | 10 | 0.5 | 2 | 0 | 8 | 0.625 | 23 | 0.5217 | 1 | 1 | 3 | 0.6667 | 1 | 1 | 2 | 0.5 | 7 | 0.7143 |
| 9_51761447 | 3 | 1 | 9 | 1 | 2 | 1 | 8 | 1 | 22 | 1 | NA | NA | 3 | 1 | 1 | 1 | 2 | 1 | 6 | 1 |
| 9_51761491 | 3 | 1 | 8 | 0.875 | 3 | 1 | 8 | 1 | 22 | 0.9545 | 1 | 1 | 2 | 1 | 1 | 1 | 1 | 1 | 5 | 1 |
| 9_51761537 | 2 | 1 | 9 | 0.7778 | 3 | 1 | 7 | 0.5714 | 21 | 0.7619 | 1 | 1 | 3 | 0.6667 | 1 | 1 | 2 | 0.5 | 7 | 0.7143 |
| 9_51761688 | 3 | 0 | 8 | 0.875 | 2 | 1 | 7 | 1 | 20 | 0.8 | 2 | 1 | 3 | 1 | 1 | 1 | 2 | 1 | 8 | 1 |
| 9_51761706 | 3 | 0.3333 | 7 | 0.5714 | 3 | 0.6667 | 8 | 0.75 | 21 | 0.619 | 2 | 0 | 4 | 0.75 | 1 | 1 | 2 | 0.5 | 9 | 0.5556 |
| 9_51761764 | 3 | 0.6667 | 7 | 0.7143 | 3 | 0.6667 | 8 | 0.75 | 21 | 0.7143 | 2 | 1 | 5 | 0.6 | 1 | 1 | 2 | 0.5 | 10 | 0.7 |
| 9_51761779 | 3 | 0.3333 | 9 | 0 | 3 | 0 | 5 | 0.2 | 20 | 0.1 | 2 | 0 | 4 | 0 | 1 | 1 | 2 | 0 | 9 | 0.1111 |
| 9_51761785 | 1 | 0 | 8 | 0.625 | 3 | 0.6667 | 6 | 0.1667 | 18 | 0.4444 | 1 | 1 | 4 | 0.75 | 1 | 0 | 2 | 0.5 | 8 | 0.625 |
| 9_51761838 | 4 | 0.75 | 8 | 0.875 | 4 | 1 | 8 | 1 | 24 | 0.9167 | 2 | 1 | 4 | 1 | 1 | 1 | 2 | 1 | 9 | 1 |
| 9_51763599 | 5 | 0.4 | 10 | 0.4 | 7 | 0 | 12 | 0.3333 | 34 | 0.2941 | 4 | 0.5 | 10 | 0.2 | 7 | 0.4286 | 5 | 0.4 | 26 | 0.3462 |
| 9_51763840 | 5 | 0 | 10 | 0.1 | 7 | 0 | 12 | 0.25 | 34 | 0.1176 | 5 | 0.2 | 10 | 0.2 | 7 | 0 | 5 | 0 | 27 | 0.1111 |
| 9_51764028 | 6 | 0.5 | 12 | 0.25 | 7 | 0 | 13 | 0 | 38 | 0.1579 | 4 | 0.75 | 10 | 0.1 | 7 | 0.1429 | 5 | 0.2 | 26 | 0.2308 |
| 9_51764299 | 8 | 0.125 | 10 | 0.1 | 6 | 0.1667 | 15 | 0.1333 | 39 | 0.1282 | 5 | 0.2 | 12 | 0.1667 | 8 | 0.125 | 5 | 0 | 30 | 0.1333 |
| 9_51764368 | 7 | 0.2857 | 4 | 0.25 | 5 | 0.4 | 6 | 0.1667 | 22 | 0.2727 | 3 | 0.3333 | 4 | 0 | 4 | 0.25 | 4 | 0.5 | 15 | 0.2667 |
| 9_51765001 | 7 | 0.8571 | 9 | 0.2222 | 7 | 0.4286 | 8 | 0.25 | 31 | 0.4194 | 3 | 0 | 6 | 0 | 5 | 0.2 | 3 | 0.6667 | 17 | 0.1765 |
| 9_51846710 | NA | NA | NA | NA | 1 | 1 | NA | NA | 1 | 1 | NA | NA | NA | NA | 1 | 1 | NA | NA | 1 | 1 |
| 9_51854908 | 3 | 0.3333 | 8 | 0 | 2 | 0 | 4 | 0.25 | 17 | 0.1176 | 1 | 0 | 2 | 0 | 5 | 0.2 | 4 | 0.75 | 12 | 0.3333 |
| 9_51911421 | NA | NA | NA | NA | 1 | 1 | NA | NA | 1 | 1 | 1 | 1 | NA | NA | NA | NA | NA | NA | 1 | 1 |
| 9_51911763 | NA | NA | NA | NA | 1 | 1 | NA | NA | 1 | 1 | 1 | 1 | NA | NA | NA | NA | NA | NA | 1 | 1 |
| 9_51913544 | NA | NA | NA | NA | 1 | 1 | NA | NA | 1 | 1 | 1 | 1 | 1 | 1 | NA | NA | NA | NA | 2 | 1 |
| 9_51913563 | NA | NA | NA | NA | 1 | 1 | NA | NA | 1 | 1 | NA | NA | 1 | 1 | NA | NA | NA | NA | 1 | 1 |
| 9_51913605 | NA | NA | 1 | 1 | 1 | 1 | NA | NA | 2 | 1 | 1 | 1 | 1 | 1 | NA | NA | NA | NA | 2 | 1 |
| 9_51913638 | NA | NA | 1 | 0 | 1 | 1 | NA | NA | 2 | 0.5 | 1 | 1 | 1 | 0 | NA | NA | NA | NA | 2 | 0.5 |
| 9_51914023 | NA | NA | 1 | 1 | 1 | 1 | NA | NA | 2 | 1 | 1 | 1 | 1 | 1 | NA | NA | NA | NA | 2 | 1 |
| 9_51914466 | NA | NA | NA | NA | 1 | 1 | NA | NA | 1 | 1 | NA | NA | 1 | 1 | NA | NA | NA | NA | 1 | 1 |
| 9_51977052 | 10 | 0.1 | 11 | 0.0909 | 5 | 0.4 | 6 | 0 | 32 | 0.125 | 2 | 0.5 | 5 | 0.2 | 5 | 0 | 5 | 0 | 17 | 0.1176 |
| 9_51997916 | 5 | 0.4 | 7 | 0.2857 | 4 | 0.5 | 2 | 0 | 18 | 0.3333 | 4 | 0.5 | 2 | 0.5 | 4 | 0.25 | 3 | 0.3333 | 13 | 0.3846 |
| 9_51997997 | 8 | 0.125 | 12 | 0.25 | 4 | 0 | 5 | 0.4 | 29 | 0.2069 | 5 | 0 | 5 | 0 | 4 | 0 | 3 | 0.3333 | 17 | 0.0588 |
| 9_51998138 | 8 | 0.375 | 11 | 0.3636 | 4 | 0.75 | 5 | 0.4 | 28 | 0.4286 | 5 | 0.4 | 6 | 0.5 | 4 | 0.5 | 3 | 0.3333 | 18 | 0.4444 |
| 9_51998781 | 10 | 0 | 12 | 0.0833 | 4 | 0 | 6 | 0.1667 | 32 | 0.0625 | 5 | 0 | 6 | 0.1667 | 5 | 0 | 3 | 0 | 19 | 0.0526 |
| 9_51998894 | 8 | 0.375 | 9 | 0.5556 | 3 | 0.3333 | 6 | 0.5 | 26 | 0.4615 | 4 | 0 | 5 | 0.4 | 5 | 0.4 | 3 | 0.6667 | 17 | 0.3529 |
| 9_51999432 | 12 | 0.3333 | 10 | 0.7 | 5 | 0.6 | 8 | 0.75 | 35 | 0.5714 | 5 | 0.8 | 4 | 0.75 | 4 | 0.5 | 4 | 0.75 | 17 | 0.7059 |
| 9_51999452 | 6 | 0.3333 | 9 | 0.4444 | 3 | 0.3333 | 9 | 0.5556 | 27 | 0.4444 | 5 | 0.2 | 2 | 0.5 | 4 | 0.5 | 5 | 0.2 | 16 | 0.3125 |
| 9_51999471 | 12 | 0.1667 | 13 | 0.2308 | 6 | 0.1667 | 9 | 0.3333 | 40 | 0.225 | 5 | 0.4 | 6 | 0.1667 | 7 | 0.4286 | 5 | 0 | 23 | 0.2609 |
| 9_51999503 | 10 | 0 | 12 | 0.1667 | 3 | 0 | 6 | 0 | 31 | 0.0645 | 4 | 0 | 4 | 0 | 7 | 0.1429 | 4 | 0 | 19 | 0.0526 |
| 9_51999624 | 11 | 0.1818 | 13 | 0.0769 | 6 | 0.1667 | 9 | 0.1111 | 39 | 0.1282 | 6 | 0.1667 | 5 | 0 | 6 | 0 | 5 | 0.2 | 22 | 0.0909 |
| 9_52014179 | 2 | 0.5 | 7 | 0.1429 | NA | NA | 6 | 0.3333 | 15 | 0.2667 | NA | NA | 2 | 0 | 4 | 0.5 | 4 | 0.25 | 10 | 0.3 |
| 9_52014347 | 1 | 1 | 6 | 0 | NA | NA | 3 | 0.3333 | 10 | 0.2 | NA | NA | 2 | 0.5 | 5 | 0 | 3 | 0 | 10 | 0.1 |
| 9_52014353 | 2 | 0 | 8 | 0.375 | NA | NA | 5 | 0 | 15 | 0.2 | 1 | 0 | 2 | 0 | 5 | 0.4 | 3 | 0.3333 | 11 | 0.2727 |
| 9_52014427 | NA | NA | 4 | 0 | NA | NA | 1 | 1 | 5 | 0.2 | 1 | 0 | 1 | 1 | 3 | 0.3333 | 2 | 0 | 7 | 0.2857 |
| 9_52014517 | 2 | 0.5 | 8 | 1 | 1 | 1 | 4 | 1 | 15 | 0.9333 | NA | NA | 2 | 1 | 4 | 1 | 4 | 1 | 10 | 1 |
| 9_52014540 | 2 | 0 | 9 | 0 | 2 | 0 | 4 | 0.25 | 17 | 0.0588 | NA | NA | 2 | 0.5 | 5 | 0.4 | 4 | 0 | 11 | 0.2727 |
| 9_52014589 | 3 | 0 | 8 | 0.125 | 2 | 0.5 | 6 | 0 | 19 | 0.1053 | 1 | 0 | 2 | 0 | 5 | 0.2 | 4 | 0.25 | 12 | 0.1667 |
| 9_52022956 | 2 | 0.5 | 2 | 0.5 | 1 | 0 | 1 | 0 | 6 | 0.3333 | 1 | 1 | NA | NA | 1 | 0 | 1 | 0 | 3 | 0.3333 |
| 9_52026146 | 2 | 0.5 | 2 | 0 | NA | NA | NA | NA | 4 | 0.25 | 1 | 1 | NA | NA | NA | NA | NA | NA | 1 | 1 |
| 9_52026818 | 2 | 0.5 | 1 | 0 | NA | NA | NA | NA | 3 | 0.3333 | 1 | 0 | 1 | 1 | NA | NA | NA | NA | 2 | 0.5 |
| 9_52026851 | 2 | 1 | 1 | 1 | NA | NA | NA | NA | 3 | 1 | 1 | 1 | NA | NA | NA | NA | NA | NA | 1 | 1 |
| 9_52027630 | 1 | 1 | 1 | 1 | NA | NA | NA | NA | 2 | 1 | NA | NA | 1 | 1 | NA | NA | NA | NA | 1 | 1 |
| 9_52584565 | 13 | 0.3846 | 11 | 0.2727 | 10 | 0.3 | 16 | 0.1875 | 50 | 0.28 | 4 | 0 | 12 | 0.1667 | 12 | 0.25 | 12 | 0.4167 | 40 | 0.25 |
| 9_52584588 | 13 | 0.1538 | 11 | 0.0909 | 11 | 0.2727 | 15 | 0.0667 | 50 | 0.14 | 4 | 0.5 | 13 | 0.1538 | 10 | 0.2 | 12 | 0.1667 | 39 | 0.2051 |
| 9_52584846 | 11 | 0.3636 | 10 | 0.5 | 11 | 0.4545 | 15 | 0.5333 | 47 | 0.4681 | 2 | 1 | 11 | 0.7273 | 11 | 0.7273 | 11 | 0.5455 | 35 | 0.6857 |
| 9_52585090 | 9 | 0.7778 | 9 | 0.7778 | 8 | 0.5 | 14 | 0.7143 | 40 | 0.7 | 2 | 1 | 11 | 0.4545 | 11 | 0.6364 | 9 | 0.2222 | 33 | 0.4848 |
| 9_52585340 | 11 | 0 | 9 | 0.2222 | 9 | 0.1111 | 13 | 0.2308 | 42 | 0.1429 | 1 | 0 | 8 | 0 | 11 | 0.1818 | 11 | 0.0909 | 31 | 0.0968 |
| 9_52585559 | 8 | 0.75 | 9 | 0.8889 | 7 | 0.4286 | 13 | 0.4615 | 37 | 0.6216 | 2 | 1 | 9 | 0.3333 | 11 | 0.7273 | 9 | 0.5556 | 31 | 0.5806 |

| | | | | | | | | | | | | | | | | | | | | |
|---|---|---|---|---|---|---|---|---|---|---|---|---|---|---|---|---|---|---|---|---|
| 9_5258566 | 8 | 0.5 | 8 | 0.875 | 8 | 0.875 | 14 | 0.8571 | 38 | 0.7895 | 2 | 0.5 | 7 | 0.7143 | 10 | 0.7 | 10 | 0.6 | 29 | 0.6552 |
| 9_5258859 | 10 | 0.1 | 5 | 0.4 | 6 | 0.3333 | 11 | 0.0909 | 32 | 0.1875 | 1 | 0 | 6 | 0.3333 | 7 | 0.4286 | 13 | 0.0769 | 27 | 0.2222 |
| 9_5258901 | 9 | 1 | 2 | 1 | 5 | 1 | 12 | 1 | 28 | 1 | 3 | 1 | 5 | 1 | 6 | 1 | 13 | 1 | 27 | 1 |
| 9_5258916 | 8 | 1 | 3 | 1 | 4 | 1 | 16 | 1 | 31 | 1 | 4 | 1 | 6 | 1 | 6 | 0.8333 | 13 | 1 | 29 | 0.9655 |
| 9_5258942 | 8 | 0.625 | 3 | 0.3333 | 3 | 0.6667 | 16 | 0.8125 | 30 | 0.7 | 2 | 1 | 5 | 0.8 | 7 | 0.7143 | 9 | 0.7778 | 23 | 0.7826 |
| 9_5258951 | 9 | 1 | 3 | 1 | 3 | 0.6667 | 12 | 1 | 27 | 0.963 | 2 | 1 | 5 | 1 | 6 | 1 | 9 | 1 | 22 | 1 |
| 9_5258952 | 9 | 0.1111 | 3 | 0 | 3 | 0 | 10 | 0.1 | 25 | 0.08 | 3 | 0 | 4 | 0.25 | 6 | 0.1667 | 4 | 0 | 17 | 0.1176 |
| 9_5258963 | 9 | 0.1111 | 3 | 0.3333 | 5 | 0 | 13 | 0 | 30 | 0.0667 | 1 | 0 | 6 | 0.1667 | 7 | 0 | 8 | 0.375 | 22 | 0.1818 |
| 9_5258977 | 9 | 0.7778 | 1 | 0 | 3 | 0.6667 | 8 | 1 | 21 | 0.8095 | 1 | 1 | 4 | 0.75 | 5 | 0.6 | 6 | 0.5 | 16 | 0.625 |
| 9_5301038 | 2 | 0.5 | 2 | 0.5 | 5 | 0.4 | 2 | 0.5 | 11 | 0.4545 | NA | NA | NA | NA | NA | NA | 2 | 0.5 | 2 | 0.5 |
| 9_5301038 | 2 | 0 | 2 | 0.5 | 5 | 0 | 2 | 0 | 11 | 0.0909 | NA | NA | NA | NA | 1 | 0 | 2 | 0.5 | 3 | 0.3333 |
| 9_5311543 | 2 | 0.5 | 2 | 0 | 4 | 0.25 | 2 | 0.5 | 10 | 0.3 | NA | NA | NA | NA | NA | NA | 2 | 0.5 | 2 | 0.5 |
| 9_5311561 | 2 | 0 | 1 | 0 | 4 | 0.5 | 3 | 0.3333 | 10 | 0.3 | NA | NA | NA | NA | 1 | 0 | 2 | 1 | 3 | 0.6667 |
| 9_5331133 | NA | NA | 1 | 1 | 2 | 0 | 2 | 0.5 | 5 | 0.4 | NA | NA | 1 | 1 | NA | NA | NA | NA | 1 | 1 |
| 9_5331190 | NA | NA | 1 | 0 | 2 | 0 | 2 | 1 | 5 | 0.4 | 1 | 1 | NA | NA | NA | NA | NA | NA | 1 | 1 |
| 9_5331211 | NA | NA | 1 | 0 | 2 | 0 | 2 | 0.5 | 5 | 0.2 | 2 | 0 | 1 | 1 | NA | NA | NA | NA | 3 | 0.3333 |
| 9_5334910 | 1 | 0 | 2 | 0.5 | 1 | 0 | 2 | 0 | 6 | 0.1667 | 1 | 0 | 2 | 0.5 | 1 | 0 | NA | NA | 4 | 0.25 |
| 9_5335102 | NA | NA | 1 | 0 | 1 | 1 | NA | NA | 2 | 0.5 | 1 | 1 | NA | NA | NA | NA | NA | NA | 1 | 1 |
| 9_5335112 | NA | NA | 2 | 0.5 | 2 | 0.5 | NA | NA | 4 | 0.5 | 1 | 0 | 1 | 1 | NA | NA | NA | NA | 2 | 0.5 |
| 9_5347360 | 1 | 1 | NA | NA | NA | NA | NA | NA | 1 | 1 | NA | NA | 1 | 1 | 2 | 0.5 | NA | NA | 3 | 0.6667 |
| 9_5347373 | 1 | 1 | NA | NA | 1 | 1 | NA | NA | 2 | 1 | NA | NA | 1 | 1 | 2 | 1 | NA | NA | 3 | 1 |
| 9_5347395 | 1 | 1 | NA | NA | NA | NA | NA | NA | 1 | 1 | NA | NA | 1 | 0 | 1 | 1 | NA | NA | 2 | 0.5 |
| 9_5347405 | 1 | 1 | 1 | 0 | 1 | 1 | NA | NA | 3 | 0.6667 | NA | NA | 1 | 1 | 2 | 0.5 | NA | NA | 3 | 0.6667 |
| 9_5347480 | 1 | 1 | 1 | 1 | 1 | 0 | NA | NA | 3 | 0.6667 | NA | NA | 1 | 0 | 2 | 0.5 | 1 | 0 | 4 | 0.25 |
| 9_5347482 | NA | NA | 1 | 0 | 1 | 1 | NA | NA | 2 | 0.5 | NA | NA | 1 | 0 | 2 | 1 | 1 | 0 | 4 | 0.5 |
| 9_5348138 | 3 | 0.6667 | 1 | 0 | 1 | 0 | NA | NA | 5 | 0.4 | NA | NA | 1 | 0 | 3 | 0.3333 | 2 | 0 | 6 | 0.1667 |
| 9_5348167 | 3 | 0.3333 | 1 | 1 | 1 | 1 | NA | NA | 5 | 0.6 | NA | NA | NA | NA | 3 | 0.6667 | 2 | 0 | 5 | 0.4 |
| 9_5348195 | 4 | 0.5 | 1 | 1 | 2 | 1 | NA | NA | 7 | 0.7143 | NA | NA | 1 | 1 | 3 | 1 | 2 | 1 | 6 | 1 |
| 9_5348206 | 4 | 0.25 | 1 | 1 | 2 | 0.5 | NA | NA | 7 | 0.4286 | NA | NA | 1 | 0 | 3 | 0.3333 | 1 | 1 | 5 | 0.4 |
| 9_5348212 | 4 | 0.25 | NA | NA | 1 | 0 | NA | NA | 5 | 0.2 | NA | NA | 1 | 0 | 3 | 0.3333 | 2 | 0 | 6 | 0.1667 |
| 9_5348221 | 4 | 0.25 | 1 | 0 | 2 | 0.5 | NA | NA | 7 | 0.2857 | NA | NA | 1 | 1 | 2 | 0.5 | 2 | 0.5 | 5 | 0.6 |
| 9_5348231 | 4 | 0.75 | 1 | 0 | 2 | 0 | NA | NA | 7 | 0.4286 | NA | NA | 1 | 1 | 3 | 0.6667 | 2 | 0.5 | 6 | 0.6667 |
| 9_5348391 | 4 | 0 | 1 | 1 | 1 | 0 | NA | NA | 6 | 0.1667 | NA | NA | 1 | 0 | 2 | 0.5 | 1 | 0 | 4 | 0.25 |
| 9_5348396 | 4 | 0.75 | 1 | 1 | 2 | 0.5 | NA | NA | 7 | 0.7143 | NA | NA | 1 | 1 | 4 | 0.25 | 2 | 0 | 7 | 0.2857 |
| 9_5348403 | 4 | 0.25 | 1 | 1 | 2 | 0 | NA | NA | 7 | 0.2857 | NA | NA | 1 | 1 | 4 | 0.75 | 2 | 0 | 7 | 0.5714 |
| 9_5348405 | 3 | 0.6667 | 1 | 1 | 1 | 0 | NA | NA | 5 | 0.6 | NA | NA | 1 | 0 | 3 | 0.6667 | 2 | 0.5 | 6 | 0.5 |
| 9_5348412 | 4 | 1 | 1 | 1 | 3 | 1 | NA | NA | 8 | 1 | NA | NA | 1 | 0 | 3 | 0.6667 | 2 | 1 | 6 | 0.6667 |
| 9_5348415 | 3 | 0.3333 | 1 | 0 | 3 | 0.3333 | NA | NA | 7 | 0.2857 | NA | NA | 1 | 0 | 4 | 0.25 | 2 | 0 | 7 | 0.1429 |
| 9_5348475 | 4 | 0.25 | 1 | 0 | 4 | 0 | NA | NA | 9 | 0.1111 | NA | NA | 1 | 0 | 4 | 0.25 | 2 | 0 | 7 | 0.1429 |
| 9_5348503 | 3 | 1 | 1 | 1 | 2 | 0.5 | 1 | 1 | 7 | 0.8571 | NA | NA | 1 | 1 | 3 | 0.6667 | 2 | 0.5 | 6 | 0.6667 |
| 9_5349303 | 1 | 1 | 2 | 0 | 2 | 1 | 2 | 0 | 7 | 0.4286 | 5 | 0.6 | 1 | 0 | 4 | 0 | 1 | 0 | 11 | 0.2727 |
| 9_5349304 | 4 | 0 | 4 | 0.25 | 5 | 0 | 7 | 0 | 20 | 0.05 | 7 | 0.2857 | 3 | 0.3333 | 9 | 0.1111 | 4 | 0 | 23 | 0.1739 |
| 9_5349310 | 5 | 1 | 4 | 0.75 | 6 | 0.6667 | 7 | 0.7143 | 22 | 0.7727 | 5 | 0.4 | 4 | 1 | 6 | 0.6667 | 3 | 0.6667 | 18 | 0.6667 |
| 9_5349334 | 3 | 0 | 4 | 0 | 4 | 0.25 | 8 | 0.125 | 19 | 0.1053 | 7 | 0.2857 | 4 | 0 | 7 | 0 | 3 | 0.3333 | 21 | 0.1429 |
| 9_5352641 | 9 | 0 | 7 | 0.4286 | 6 | 0 | 7 | 0.1429 | 29 | 0.1379 | 2 | 0.5 | 4 | 0.25 | 8 | 0.25 | 6 | 0.1667 | 20 | 0.25 |
| 9_5352734 | 3 | 0 | 6 | 0.1667 | 2 | 0 | 5 | 0 | 16 | 0.0625 | 2 | 0 | 3 | 0 | 5 | 0 | 7 | 0.2857 | 17 | 0.1176 |
| 9_5352784 | 3 | 0.3333 | 6 | 0.1667 | 2 | 0 | 3 | 0.3333 | 14 | 0.2143 | 2 | 0.5 | 3 | 0 | 5 | 0.4 | 5 | 0.2 | 15 | 0.2667 |
| 9_5352851 | 2 | 0 | 5 | 0.8 | 5 | 0.2 | 5 | 0.4 | 17 | 0.4118 | 4 | 0.25 | 4 | 0.5 | 7 | 0.4286 | 7 | 0.4286 | 22 | 0.4091 |
| 9_5352887 | NA | NA | 7 | 0.8571 | 3 | 1 | 3 | 1 | 13 | 0.9231 | 2 | 1 | 3 | 0.6667 | 5 | 0.8 | 5 | 1 | 15 | 0.8667 |
| 9_5352901 | 3 | 0.3333 | 6 | 0.5 | 3 | 0.3333 | 4 | 0.5 | 16 | 0.4375 | 4 | 0.75 | 3 | 0.6667 | 4 | 0.25 | 4 | 0.25 | 15 | 0.4667 |
| 9_5352903 | 3 | 1 | 7 | 0.8571 | 3 | 0.6667 | 5 | 1 | 18 | 0.8889 | 4 | 1 | 3 | 1 | 5 | 1 | 5 | 1 | 17 | 1 |
| 9_5352925 | 2 | 0.5 | 5 | 0.4 | 2 | 0 | 5 | 1 | 14 | 0.5714 | 3 | 0.3333 | 2 | 0.5 | 5 | 0.4 | 5 | 0.4 | 15 | 0.4 |
| 9_5359564 | NA | NA | NA | NA | 1 | 1 | NA | NA | 1 | 1 | NA | NA | NA | NA | NA | NA | 1 | 1 | 1 | 1 |
| 9_5372420 | 1 | 1 | NA | NA | 1 | 1 | NA | NA | 2 | 1 | 1 | 1 | NA | NA | NA | NA | NA | NA | 1 | 1 |
| 9_5381885 | 63 | 0.2063 | 62 | 0.3226 | 33 | 0.303 | 63 | 0.1587 | 221 | 0.2398 | 26 | 0.2308 | 64 | 0.2031 | 80 | 0.25 | 52 | 0.3654 | 222 | 0.2613 |
| 9_5381920 | 60 | 0.0833 | 60 | 0 | 35 | 0.0857 | 58 | 0.0517 | 213 | 0.0516 | 23 | 0 | 63 | 0.0317 | 72 | 0.0972 | 51 | 0.0392 | 209 | 0.0526 |
| 9_5381937 | 57 | 0.1228 | 56 | 0.1607 | 38 | 0.0789 | 56 | 0.0357 | 207 | 0.1014 | 26 | 0.1923 | 66 | 0.1212 | 68 | 0.1618 | 52 | 0.1154 | 212 | 0.1415 |
| 9_5381950 | 60 | 0.0833 | 57 | 0.2281 | 39 | 0.2564 | 55 | 0.2 | 211 | 0.1848 | 26 | 0.0385 | 72 | 0.1389 | 65 | 0.1846 | 47 | 0.1702 | 210 | 0.1476 |
| 9_5381954 | 60 | 0.0667 | 57 | 0.0702 | 39 | 0.1538 | 55 | 0.0545 | 211 | 0.0806 | 26 | 0.0769 | 64 | 0.0469 | 64 | 0.0938 | 46 | 0.0652 | 200 | 0.07 |
| 9_5381990 | 49 | 0.102 | 53 | 0.0566 | 40 | 0.1 | 50 | 0.1 | 192 | 0.0885 | 20 | 0.05 | 61 | 0.0656 | 54 | 0.1296 | 46 | 0.1304 | 181 | 0.0994 |
| 9_5381997 | 33 | 0.0606 | 38 | 0.0263 | 33 | 0.2424 | 44 | 0.1364 | 148 | 0.1149 | 17 | 0.2353 | 47 | 0.1702 | 44 | 0.0909 | 39 | 0.1026 | 147 | 0.1361 |
| 9_5382012 | 42 | 0.3095 | 41 | 0.3659 | 37 | 0.3243 | 42 | 0.3571 | 162 | 0.3395 | 16 | 0.125 | 56 | 0.3036 | 48 | 0.25 | 39 | 0.2564 | 159 | 0.2579 |
| 9_5384145 | 42 | 0.119 | 39 | 0.0769 | 32 | 0.0625 | 35 | 0.0571 | 148 | 0.0811 | 17 | 0.2353 | 51 | 0.0784 | 47 | 0.0638 | 39 | 0.1282 | 154 | 0.1039 |
| 9_5393095 | 4 | 0.5 | 5 | 0.4 | 1 | 0 | 1 | 1 | 11 | 0.4545 | 1 | 0 | 3 | 0.3333 | NA | NA | 3 | 0.6667 | 7 | 0.4286 |
| 9_5393155 | 3 | 0.3333 | 5 | 0.4 | 1 | 0 | NA | NA | 9 | 0.3333 | 1 | 0 | 3 | 0.3333 | NA | NA | 4 | 0.5 | 8 | 0.375 |

| | | | | | | | | | | | | | | | | | | | | |
|---|---|---|---|---|---|---|---|---|---|---|---|---|---|---|---|---|---|---|---|---|
| 9_5393168 | 3 | 0.3333 | 4 | 0.25 | 2 | 1 | 1 | 1 | 10 | 0.5 | 1 | 1 | 3 | 0.3333 | NA | NA | 4 | 0.25 | 8 | 0.375 |
| 9_5393181 | 4 | 0 | 4 | 0.25 | 2 | 0 | 1 | 0 | 11 | 0.0909 | 1 | 0 | 3 | 0.3333 | NA | NA | 4 | 0 | 8 | 0.125 |
| 9_5393201 | 4 | 0.5 | 4 | 0 | 2 | 0 | NA | NA | 10 | 0.2 | NA | NA | 2 | 0.5 | NA | NA | 4 | 0.25 | 6 | 0.3333 |
| 9_5393202 | 4 | 0.75 | 4 | 0.5 | 2 | 1 | NA | NA | 10 | 0.7 | 1 | 1 | 2 | 1 | NA | NA | 4 | 0.75 | 7 | 0.8571 |
| 9_5393204 | 4 | 0.25 | 5 | 0.2 | 1 | 1 | NA | NA | 10 | 0.3 | 1 | 0 | 2 | 1 | NA | NA | 4 | 0.5 | 7 | 0.5714 |
| 9_5393205 | 2 | 0.5 | 5 | 0 | NA | NA | NA | NA | 7 | 0.1429 | 1 | 0 | 2 | 0.5 | NA | NA | 2 | 0 | 5 | 0.2 |
| 9_5393209 | 4 | 0 | 4 | 0.25 | 2 | 0 | NA | NA | 10 | 0.1 | 1 | 1 | 2 | 0 | NA | NA | 4 | 0 | 7 | 0.1429 |
| 9_5393343 | 5 | 0 | 3 | 0.3333 | 2 | 0 | 2 | 0 | 12 | 0.0833 | 1 | 0 | 4 | 0 | NA | NA | 4 | 0.5 | 9 | 0.2222 |
| 9_5393348 | 5 | 1 | 3 | 0.6667 | 2 | 1 | 2 | 1 | 12 | 0.9167 | 1 | 1 | 4 | 0.75 | NA | NA | 2 | 1 | 7 | 0.8571 |
| 9_5393360 | 4 | 1 | 2 | 0 | 1 | 1 | 2 | 0.5 | 9 | 0.6667 | 1 | 1 | 4 | 0.75 | NA | NA | 4 | 1 | 9 | 0.8889 |
| 9_5393362 | 4 | 0.75 | 3 | 1 | 2 | 1 | 2 | 0.5 | 11 | 0.8182 | 1 | 1 | 3 | 0.3333 | NA | NA | 4 | 0.75 | 8 | 0.625 |
| 9_5393391 | 2 | 1 | 1 | 1 | 1 | 1 | 1 | 1 | 5 | 1 | 1 | 1 | 3 | 0.6667 | NA | NA | 2 | 1 | 6 | 0.8333 |
| 9_5393392 | 4 | 0.5 | 3 | 0.6667 | 1 | 0 | 2 | 0.5 | 10 | 0.5 | 1 | 1 | 4 | 0.75 | NA | NA | 4 | 0.75 | 9 | 0.7778 |
| 9_5393395 | 4 | 1 | 2 | 1 | 1 | 1 | 2 | 1 | 9 | 1 | 1 | 0 | 4 | 0.75 | NA | NA | 3 | 1 | 8 | 0.75 |
| 9_5393396 | 4 | 0 | 2 | 0 | 1 | 0 | 2 | 0.5 | 9 | 0.1111 | 1 | 1 | 2 | 0 | NA | NA | 3 | 0.3333 | 6 | 0.3333 |
| 9_5393396 | 4 | 1 | 3 | 1 | NA | NA | 1 | 0 | 8 | 0.875 | 1 | 1 | 4 | 1 | NA | NA | 4 | 1 | 9 | 1 |
| 9_5393400 | 4 | 1 | 3 | 1 | 1 | 1 | 2 | 1 | 10 | 1 | 1 | 1 | 4 | 0.75 | NA | NA | 4 | 1 | 9 | 0.8889 |
| 9_5393406 | 3 | 0.3333 | 3 | 0.3333 | NA | NA | 2 | 0.5 | 8 | 0.375 | 1 | 0 | 3 | 0 | NA | NA | 3 | 0.3333 | 7 | 0.1429 |
| 9_5393417 | 4 | 1 | 2 | 1 | 1 | 1 | 2 | 1 | 9 | 1 | 1 | 1 | 2 | 1 | NA | NA | 3 | 1 | 6 | 1 |
| 9_5393422 | 2 | 0 | 3 | 0.6667 | 1 | 0 | 2 | 0 | 8 | 0.25 | 1 | 0 | 1 | 0 | NA | NA | 4 | 0.5 | 6 | 0.3333 |
| 9_5393422 | 3 | 0 | 2 | 0 | 1 | 1 | 1 | 0 | 7 | 0.1429 | 1 | 1 | 2 | 0.5 | NA | NA | 3 | 0 | 6 | 0.3333 |
| 9_5393424 | 1 | 0 | 3 | 0.6667 | 1 | 1 | 1 | 1 | 6 | 0.6667 | 1 | 0 | 1 | 1 | NA | NA | 3 | 0.6667 | 5 | 0.6 |
| 9_5393433 | 2 | 1 | 2 | 1 | 1 | 1 | 2 | 1 | 7 | 1 | 1 | 1 | 2 | 1 | NA | NA | 4 | 1 | 7 | 1 |
| 9_5393435 | 2 | 0.5 | 3 | 0 | 1 | 0 | 1 | 0 | 7 | 0.1429 | 1 | 1 | 2 | 0 | NA | NA | 4 | 0.5 | 7 | 0.4286 |
| 9_5393450 | 3 | 0.3333 | NA | NA | 1 | 0 | 2 | 0 | 6 | 0.1667 | 1 | 1 | 2 | 0 | NA | NA | 4 | 0 | 7 | 0.1429 |
| 9_5393453 | 2 | 1 | NA | NA | NA | NA | 2 | 1 | 4 | 1 | 1 | 1 | 1 | 1 | NA | NA | 3 | 1 | 5 | 1 |
| 9_5393454 | 3 | 0.6667 | NA | NA | 1 | 1 | 2 | 1 | 6 | 0.8333 | 1 | 1 | 2 | 0.5 | NA | NA | 4 | 0.75 | 7 | 0.7143 |
| 9_5393463 | 3 | 0.3333 | NA | NA | 1 | 0 | 1 | 0 | 5 | 0.2 | NA | NA | 2 | 0 | NA | NA | 4 | 0.25 | 6 | 0.1667 |
| 9_5427308 | 14 | 0.1429 | 13 | 0.1538 | 11 | 0.1818 | 3 | 0.3333 | 41 | 0.1707 | 8 | 0.125 | 9 | 0.3333 | 12 | 0.3333 | 8 | 0 | 37 | 0.2162 |
| 9_5427357 | 15 | 0.1333 | 15 | 0.0667 | 12 | 0.0833 | 7 | 0 | 49 | 0.0816 | 8 | 0.375 | 8 | 0.125 | 12 | 0.1667 | 8 | 0.125 | 36 | 0.1944 |
| 9_5427371 | 12 | 0.1667 | 12 | 0.25 | 12 | 0.1667 | 5 | 0 | 41 | 0.1707 | 8 | 0 | 7 | 0 | 11 | 0.0909 | 9 | 0.2222 | 35 | 0.0857 |
| 9_5427379 | 11 | 0.2727 | 12 | 0.0833 | 12 | 0.0833 | 6 | 0.5 | 41 | 0.1951 | 7 | 0.1429 | 7 | 0 | 12 | 0 | 9 | 0.3333 | 35 | 0.1143 |
| 9_5430823 | 2 | 0 | 4 | 0 | 4 | 0 | 3 | 0.6667 | 13 | 0.1538 | 1 | 1 | 1 | 0 | 3 | 0.3333 | 4 | 0.5 | 9 | 0.4444 |
| 9_5430841 | 2 | 0.5 | 4 | 0 | 6 | 0 | 2 | 0 | 14 | 0.0714 | 3 | 0 | 1 | 0 | 4 | 0 | 3 | 0.3333 | 11 | 0.0909 |
| 9_5430844 | 2 | 0 | 4 | 0.75 | 5 | 0.6 | 2 | 0 | 13 | 0.4615 | 3 | 0.6667 | NA | NA | 3 | 0.3333 | 3 | 0.3333 | 9 | 0.4444 |
| 9_5430854 | 1 | 0 | 5 | 0.2 | 6 | 0 | 2 | 0 | 14 | 0.0714 | 2 | 0.5 | 1 | 0 | 2 | 0 | 2 | 0 | 7 | 0.1429 |
| 9_5430859 | 1 | 0 | 4 | 1 | 6 | 1 | 2 | 1 | 13 | 0.9231 | 3 | 1 | 1 | 1 | 4 | 1 | 2 | 1 | 10 | 1 |
| 9_5430860 | 1 | 0 | 5 | 0.2 | 6 | 0.5 | 1 | 1 | 13 | 0.3846 | 3 | 0.3333 | 1 | 0 | 4 | 0.25 | 1 | 0 | 9 | 0.2222 |
| 9_5430897 | 1 | 0 | 3 | 0.3333 | 6 | 0.3333 | 3 | 0.3333 | 13 | 0.3077 | 2 | 1 | 1 | 0 | 3 | 0 | 1 | 0 | 7 | 0.2857 |
| 9_5430900 | 1 | 1 | 3 | 0 | 4 | 0.25 | 3 | 0.3333 | 11 | 0.2727 | 3 | 0 | 1 | 1 | 4 | 0 | 1 | 0 | 9 | 0.1111 |
| 9_5430913 | 1 | 0 | 3 | 0 | 5 | 0.4 | 2 | 0 | 11 | 0.1818 | 2 | 0.5 | 1 | 0 | 4 | 0.25 | 1 | 0 | 8 | 0.25 |
| 9_5430914 | 1 | 1 | 1 | 0 | 4 | 0.25 | 3 | 0.3333 | 9 | 0.3333 | 2 | 0 | 1 | 0 | 3 | 0.3333 | 1 | 0 | 7 | 0.1429 |
| 9_5430917 | 1 | 1 | 3 | 1 | 3 | 0.3333 | 3 | 0.6667 | 10 | 0.7 | 2 | 1 | 1 | 1 | 3 | 1 | 1 | 0 | 7 | 0.8571 |
| 9_5432285 | 1 | 0 | 2 | 0 | 4 | 0 | 3 | 0.3333 | 10 | 0.1 | 2 | 0 | NA | NA | 2 | 0.5 | NA | NA | 4 | 0.25 |
| 9_5432288 | 1 | 1 | 2 | 1 | 4 | 0.75 | 2 | 1 | 9 | 0.8889 | 2 | 1 | NA | NA | 2 | 1 | 1 | 1 | 5 | 1 |
| 9_5432301 | 1 | 0 | 2 | 0.5 | 5 | 0.6 | 3 | 0.3333 | 11 | 0.4545 | 2 | 0 | NA | NA | 2 | 0.5 | 1 | 1 | 5 | 0.4 |
| 9_5432304 | NA | NA | 1 | 1 | 5 | 0.6 | 3 | 0.6667 | 9 | 0.6667 | 2 | 0.5 | NA | NA | 1 | 1 | NA | NA | 3 | 0.6667 |
| 9_5432306 | 1 | 0 | 2 | 0.5 | 4 | 0.5 | 3 | 0 | 10 | 0.3 | 2 | 0 | NA | NA | 2 | 0 | 1 | 1 | 5 | 0.2 |
| 9_5432311 | 1 | 1 | 2 | 1 | 5 | 1 | 3 | 1 | 11 | 1 | 2 | 1 | NA | NA | 2 | 0.5 | NA | NA | 4 | 0.75 |
| 9_5432319 | 1 | 1 | 2 | 0.5 | 5 | 0.4 | 3 | 0.3333 | 11 | 0.4545 | 2 | 0 | NA | NA | 2 | 0.5 | 1 | 1 | 5 | 0.4 |
| 9_5432331 | 1 | 1 | 2 | 0.5 | 4 | 0.75 | 3 | 0.6667 | 10 | 0.7 | 2 | 1 | NA | NA | 2 | 1 | 1 | 1 | 5 | 1 |
| 9_5432350 | 1 | 1 | 2 | 0.5 | 4 | 0.5 | 3 | 0.6667 | 10 | 0.6 | 1 | 0 | NA | NA | NA | NA | 1 | 1 | 2 | 0.5 |
| 9_5432367 | 1 | 1 | 2 | 1 | 3 | 0.6667 | 3 | 1 | 9 | 0.8889 | 1 | 1 | NA | NA | 1 | 1 | 1 | 1 | 3 | 1 |
| 9_5432379 | 1 | 0 | 2 | 0 | 4 | 0.25 | 3 | 0 | 10 | 0.1 | 1 | 1 | NA | NA | 1 | 1 | 1 | 0 | 3 | 0.6667 |
| 9_5432386 | NA | NA | 2 | 1 | 4 | 0.75 | 3 | 0.6667 | 9 | 0.7778 | 1 | 1 | NA | NA | 1 | 1 | 1 | 1 | 3 | 1 |
| 9_5432424 | 1 | 0 | 1 | 0 | 3 | 0.3333 | 3 | 0 | 8 | 0.125 | 1 | 0 | NA | NA | 1 | 1 | 1 | 0 | 3 | 0.3333 |
| 9_5435376 | 1 | 1 | 1 | 0 | 2 | 0.5 | 1 | 0 | 5 | 0.4 | NA | NA | 1 | 1 | NA | NA | 1 | 0 | 2 | 0.5 |
| 9_5435391 | 1 | 1 | 1 | 1 | 2 | 1 | 3 | 1 | 7 | 1 | NA | NA | NA | NA | 1 | 1 | 1 | 1 | 2 | 1 |
| 9_5435393 | 1 | 1 | 1 | 1 | 2 | 0.5 | 3 | 1 | 7 | 0.8571 | NA | NA | NA | NA | 1 | 1 | 1 | 0 | 2 | 0.5 |
| 9_5435405 | NA | NA | 1 | 1 | 2 | 0 | 2 | 0 | 5 | 0.2 | NA | NA | NA | NA | 1 | 0 | 1 | 1 | 2 | 0.5 |
| 9_5445262 | 3 | 0.6667 | 4 | 1 | 2 | 1 | 3 | 0.6667 | 12 | 0.8333 | 2 | 0.5 | 2 | 1 | 2 | 1 | 7 | 0.5714 | 13 | 0.6923 |
| 9_5451063 | 33 | 0.3333 | 40 | 0.25 | 35 | 0.2286 | 37 | 0.2432 | 145 | 0.2621 | 25 | 0.2 | 26 | 0.1538 | 30 | 0.2333 | 25 | 0.2 | 106 | 0.1981 |
| 9_5451090 | 32 | 0.0625 | 43 | 0.1395 | 33 | 0.1212 | 41 | 0.0488 | 149 | 0.094 | 27 | 0.1852 | 27 | 0.0741 | 27 | 0.1852 | 26 | 0.1923 | 107 | 0.1589 |
| 9_5451154 | 13 | 0.0769 | 16 | 0.1875 | 22 | 0.2273 | 17 | 0.0588 | 68 | 0.1471 | 17 | 0.0588 | 17 | 0.1176 | 14 | 0.1429 | 14 | 0.1429 | 62 | 0.1129 |
| 9_5451183 | 23 | 0.087 | 24 | 0.0833 | 24 | 0 | 19 | 0.1579 | 90 | 0.0778 | 19 | 0.0526 | 21 | 0.0476 | 17 | 0.1176 | 18 | 0.0556 | 75 | 0.0667 |

| | | | | | | | | | | | | | | | | | | | | |
|---|---|---|---|---|---|---|---|---|---|---|---|---|---|---|---|---|---|---|---|---|
| 9_5462176 | 5 | 0 | 7 | 0 | 7 | 0.2857 | 8 | 0.125 | 27 | 0.1111 | 3 | 0 | 7 | 0.1429 | 5 | 0 | 5 | 0 | 20 | 0.05 |
| 9_5462198 | 7 | 1 | 11 | 1 | 8 | 1 | 12 | 1 | 38 | 1 | 6 | 1 | 8 | 1 | 12 | 1 | 8 | 1 | 34 | 1 |
| 9_5462222 | 10 | 0.1 | 12 | 0.0833 | 6 | 0 | 13 | 0.1538 | 41 | 0.0976 | 5 | 0 | 8 | 0 | 13 | 0.0769 | 8 | 0.125 | 34 | 0.0588 |
| 9_5462245 | 9 | 0.1111 | 11 | 0 | 6 | 0 | 13 | 0.0769 | 39 | 0.0513 | 6 | 0 | 8 | 0.125 | 15 | 0.0667 | 7 | 0 | 36 | 0.0556 |
| 9_5462252 | 10 | 0.1 | 9 | 0.1111 | 6 | 0.1667 | 11 | 0 | 36 | 0.0833 | 4 | 0.25 | 3 | 0 | 11 | 0 | 6 | 0.1667 | 24 | 0.0833 |
| 9_5462338 | 7 | 0.4286 | 10 | 0.1 | 6 | 0 | 9 | 0 | 32 | 0.125 | 7 | 0 | 7 | 0.1429 | 14 | 0.2143 | 7 | 0 | 35 | 0.1143 |
| 9_5462345 | 6 | 0.1667 | 10 | 0 | 5 | 0.2 | 11 | 0 | 32 | 0.0625 | 7 | 0.2857 | 7 | 0 | 14 | 0 | 7 | 0 | 35 | 0.0571 |
| 9_5463164 | 7 | 0.4286 | 2 | 0.5 | 4 | 0.5 | 7 | 0.2857 | 20 | 0.4 | 9 | 0.1111 | 6 | 0.3333 | 6 | 0.3333 | 6 | 0.1667 | 27 | 0.2222 |
| 9_5481700 | 3 | 0.6667 | 2 | 0.5 | 3 | 1 | 4 | 0.5 | 12 | 0.6667 | NA | NA | 3 | 0.6667 | 5 | 0.4 | 3 | 0.3333 | 11 | 0.4545 |
| 9_5481705 | 3 | 0 | 2 | 0 | 3 | 0.3333 | 6 | 0.3333 | 14 | 0.2143 | NA | NA | 3 | 0.6667 | 5 | 0.2 | 3 | 0.3333 | 11 | 0.3636 |
| 9_5481708 | 3 | 0 | 2 | 0 | 3 | 0.3333 | 5 | 0.4 | 13 | 0.2308 | NA | NA | 2 | 0 | 3 | 0.6667 | 3 | 0 | 8 | 0.25 |
| 9_5481710 | 2 | 0.5 | 2 | 1 | 3 | 1 | 6 | 0.6667 | 13 | 0.7692 | NA | NA | 3 | 0.6667 | 5 | 0.4 | 2 | 0.5 | 10 | 0.5 |
| 9_5481716 | 3 | 0.6667 | 2 | 0.5 | 3 | 1 | 6 | 0.8333 | 14 | 0.7857 | NA | NA | 3 | 0.6667 | 4 | 0.5 | 3 | 0.6667 | 10 | 0.6 |
| 9_5481721 | 3 | 0.3333 | 2 | 0 | 4 | 0 | 6 | 0 | 15 | 0.0667 | NA | NA | 3 | 0 | 4 | 0.25 | 3 | 0 | 10 | 0.1 |
| 9_5481728 | 1 | 0 | NA | NA | 3 | 0 | 4 | 0.25 | 8 | 0.125 | NA | NA | 2 | 0 | 5 | 0.4 | 2 | 0 | 9 | 0.2222 |
| 9_5481920 | 4 | 0 | 5 | 0.2 | 6 | 0 | 11 | 0.0909 | 26 | 0.0769 | 2 | 0 | 4 | 0.25 | 8 | 0.125 | 3 | 0 | 17 | 0.1176 |
| 9_5481953 | 4 | 0 | 6 | 0.8333 | 6 | 0.5 | 12 | 0.5 | 28 | 0.5 | 3 | 0 | 4 | 0.5 | 8 | 0.5 | 4 | 0.25 | 19 | 0.3684 |
| 9_5485381 | NA | NA | NA | NA | 1 | 1 | NA | NA | 1 | 1 | NA | NA | NA | NA | 1 | 1 | NA | NA | 1 | 1 |
| 9_5486507 | 22 | 0 | 15 | 0.0667 | 15 | 0.2 | 21 | 0.1429 | 73 | 0.0959 | 17 | 0.1765 | 20 | 0.05 | 19 | 0.2105 | 11 | 0.1818 | 67 | 0.1493 |
| 9_5486519 | 19 | 0.8421 | 17 | 0.8824 | 17 | 0.7059 | 21 | 0.7619 | 74 | 0.7973 | 16 | 0.6875 | 19 | 0.5263 | 19 | 0.5789 | 11 | 0.8182 | 65 | 0.6308 |
| 9_5486525 | 11 | 0.7273 | 14 | 0.5714 | 14 | 0.6429 | 17 | 0.4706 | 56 | 0.5893 | 16 | 0.6875 | 22 | 0.5455 | 18 | 0.7222 | 10 | 0.6 | 66 | 0.6364 |
| 9_5505595 | 1 | 1 | NA | NA | NA | NA | NA | NA | 1 | 1 | 2 | 1 | 2 | 0.5 | 1 | 1 | NA | NA | 5 | 0.8 |
| 9_5505599 | 1 | 1 | NA | NA | NA | NA | NA | NA | 1 | 1 | 2 | 1 | 2 | 1 | 1 | 1 | NA | NA | 5 | 1 |
| 9_5505600 | 1 | 1 | NA | NA | NA | NA | NA | NA | 1 | 1 | 2 | 1 | 2 | 1 | 1 | 1 | NA | NA | 5 | 1 |
| 9_5511329 | 6 | 0 | 7 | 0.1429 | 8 | 0 | 12 | 0.0833 | 33 | 0.0606 | 4 | 0 | 8 | 0.125 | 9 | 0 | 10 | 0.1 | 31 | 0.0645 |
| 9_5511346 | 6 | 0.6667 | 7 | 0.8571 | 7 | 0.8571 | 11 | 0.8182 | 31 | 0.8065 | 2 | 0 | 8 | 0.75 | 8 | 0.75 | 9 | 1 | 27 | 0.7778 |
| 9_5511408 | 7 | 0 | 7 | 0 | 9 | 0 | 13 | 0.1538 | 36 | 0.0556 | 5 | 0.2 | 9 | 0.1111 | 8 | 0.25 | 10 | 0.1 | 32 | 0.1562 |
| 9_5511427 | 9 | 0 | 7 | 0.1429 | 10 | 0 | 13 | 0.0769 | 39 | 0.0513 | 6 | 0 | 10 | 0.2 | 9 | 0 | 12 | 0.1667 | 37 | 0.1081 |
| 9_5513056 | 6 | 1 | 6 | 0.8333 | 8 | 1 | 2 | 1 | 22 | 0.9545 | 2 | 0.5 | 4 | 0.75 | 2 | 1 | 5 | 0.8 | 13 | 0.7692 |
| 9_5513067 | 5 | 0.2 | 6 | 0 | 9 | 0.2222 | 1 | 1 | 21 | 0.1905 | 2 | 0 | 4 | 0.25 | 3 | 0 | 3 | 0 | 12 | 0.0833 |
| 9_5513084 | 7 | 1 | 7 | 0.4286 | 9 | 1 | 1 | 1 | 24 | 0.8333 | 3 | 1 | 3 | 1 | 3 | 1 | 6 | 0.8333 | 15 | 0.9333 |
| 9_5513088 | 7 | 0.5714 | 8 | 0.375 | 9 | 0.3333 | 2 | 1 | 26 | 0.4615 | 3 | 0.6667 | 4 | 0 | 3 | 0.3333 | 6 | 0.3333 | 16 | 0.3125 |
| 9_5513093 | 5 | 0.2 | 5 | 0 | 9 | 0.1111 | 2 | 0 | 21 | 0.0952 | 2 | 0 | 4 | 0.25 | 2 | 0 | 5 | 0 | 13 | 0.0769 |
| 9_5513097 | 5 | 1 | 4 | 1 | 7 | 1 | 1 | 1 | 17 | 1 | 2 | 1 | 4 | 1 | 1 | 1 | 5 | 1 | 12 | 1 |
| 9_5513111 | 7 | 0.2857 | 8 | 0.125 | 11 | 0.3636 | 2 | 0 | 28 | 0.25 | 3 | 0.3333 | 6 | 0.1667 | 4 | 0.25 | 4 | 0.5 | 17 | 0.2941 |
| 9_5513127 | 6 | 0.5 | 8 | 0.25 | 12 | 0.4167 | 3 | 0.3333 | 29 | 0.3793 | 2 | 0.5 | 7 | 0.4286 | 4 | 0.25 | 7 | 0.2857 | 20 | 0.35 |
| 9_5513149 | 6 | 0.5 | 8 | 0 | 12 | 0.4167 | 3 | 0.3333 | 29 | 0.3103 | 4 | 0.25 | 7 | 0.1429 | 3 | 0.3333 | 6 | 0.3333 | 20 | 0.25 |
| 9_5513154 | 8 | 0 | 7 | 0.1429 | 13 | 0.0769 | 3 | 0 | 31 | 0.0645 | 4 | 0.25 | 6 | 0.3333 | 4 | 0 | 8 | 0.125 | 22 | 0.1818 |
| 9_5513163 | 8 | 0.125 | 9 | 0 | 14 | 0.0714 | 3 | 0 | 34 | 0.0588 | 4 | 0 | 7 | 0.2857 | 4 | 0.25 | 8 | 0.25 | 23 | 0.2174 |
| 9_5534461 | 1 | 1 | 1 | 1 | NA | NA | 3 | 1 | 5 | 1 | 1 | 1 | NA | NA | NA | NA | 1 | 1 | 2 | 1 |
| 9_5534471 | 1 | 0 | 1 | 1 | NA | NA | 3 | 0.3333 | 5 | 0.4 | NA | NA | NA | NA | NA | NA | 1 | 1 | 1 | 1 |
| 9_5534474 | 1 | 1 | 1 | 0 | NA | NA | 3 | 0.6667 | 5 | 0.6 | 1 | 1 | NA | NA | NA | NA | 1 | 1 | 2 | 1 |
| 9_5534551 | 2 | 1 | 2 | 1 | NA | NA | 3 | 0.6667 | 7 | 0.8571 | 2 | 1 | NA | NA | NA | NA | 2 | 1 | 4 | 1 |
| 9_5545792 | 2 | 0 | 5 | 0.2 | 4 | 0.5 | 5 | 0 | 16 | 0.1875 | 2 | 0 | 3 | 0.3333 | 3 | 0.3333 | 4 | 0.25 | 12 | 0.25 |
| 9_5549359 | 2 | 0 | 3 | 0 | 3 | 0.3333 | 2 | 0.5 | 10 | 0.2 | 3 | 0 | 4 | 0.25 | 3 | 0.3333 | 2 | 0 | 12 | 0.1667 |
| 9_5594936 | 8 | 0.375 | 18 | 0.3333 | 14 | 0.5 | 21 | 0.5238 | 61 | 0.4426 | 11 | 0.0909 | 13 | 0.3846 | 14 | 0.2857 | 21 | 0.2857 | 59 | 0.2712 |
| 9_5594994 | 7 | 0.2857 | 19 | 0.3684 | 11 | 0.4545 | 21 | 0.3333 | 58 | 0.3621 | 13 | 0.2308 | 14 | 0.2857 | 15 | 0.2 | 18 | 0.2778 | 60 | 0.25 |
| 9_5596541 | 12 | 0.0833 | 20 | 0 | 16 | 0.0625 | 26 | 0.1538 | 74 | 0.0811 | 20 | 0.2 | 17 | 0.0588 | 20 | 0.15 | 25 | 0.08 | 82 | 0.122 |
| 9_5596554 | 13 | 0 | 20 | 0 | 17 | 0.2353 | 24 | 0.0833 | 74 | 0.0811 | 21 | 0.0476 | 18 | 0.0556 | 20 | 0.05 | 25 | 0.16 | 84 | 0.0833 |
| 9_5596566 | 11 | 0.0909 | 21 | 0.0952 | 14 | 0.1429 | 25 | 0.28 | 71 | 0.169 | 20 | 0 | 18 | 0.1667 | 18 | 0.2778 | 25 | 0.04 | 81 | 0.1111 |
| 9_5596595 | 15 | 0.0667 | 25 | 0.32 | 19 | 0.1053 | 26 | 0.3077 | 85 | 0.2235 | 23 | 0.1304 | 20 | 0.25 | 23 | 0.2609 | 29 | 0.2759 | 95 | 0.2316 |
| 9_5604341 | 87 | 0.1264 | 61 | 0.082 | 52 | 0.1923 | 73 | 0.1507 | 273 | 0.1355 | 48 | 0.1458 | 60 | 0.0833 | 67 | 0.1493 | 52 | 0.1731 | 227 | 0.1366 |
| 9_5604358 | 58 | 0.0862 | 46 | 0.0435 | 31 | 0.0645 | 53 | 0.1321 | 188 | 0.0851 | 32 | 0.0938 | 49 | 0.0612 | 48 | 0.0833 | 32 | 0.1875 | 161 | 0.0994 |
| 9_5604363 | 77 | 0.5325 | 59 | 0.5593 | 49 | 0.5102 | 75 | 0.56 | 260 | 0.5423 | 43 | 0.5349 | 58 | 0.5172 | 60 | 0.4833 | 48 | 0.5625 | 209 | 0.5215 |
| 9_5610865 | 3 | 0 | 4 | 0 | 1 | 0 | 6 | 0.1667 | 14 | 0.0714 | 1 | 0 | 5 | 0.2 | 5 | 0 | 2 | 0 | 13 | 0.0769 |
| 9_5610874 | 3 | 0.3333 | 5 | 0.4 | 2 | 0 | 5 | 0.2 | 15 | 0.2667 | NA | NA | 4 | 0.25 | 5 | 0.2 | 1 | 0 | 10 | 0.2 |
| 9_5610935 | 2 | 0.5 | 6 | 0 | 1 | 1 | 4 | 0 | 13 | 0.1538 | 1 | 0 | 4 | 0.25 | 5 | 0.2 | 1 | 0 | 11 | 0.1818 |
| 9_5611052 | 1 | 0 | 5 | 0.4 | 1 | 1 | 2 | 0 | 9 | 0.3333 | 1 | 0 | 3 | 0.3333 | 4 | 0.25 | NA | NA | 8 | 0.25 |
| 9_5611137 | 1 | 0 | 3 | 0 | 1 | 0 | 2 | 0.5 | 7 | 0.1429 | NA | NA | 2 | 0.5 | 1 | 1 | NA | NA | 3 | 0.6667 |
| 9_5611170 | 1 | 0 | 5 | 0.2 | NA | NA | 2 | 0 | 8 | 0.125 | NA | NA | 2 | 0.5 | 2 | 0 | NA | NA | 4 | 0.25 |
| 9_5611240 | 1 | 0 | 2 | 0 | 1 | 1 | 2 | 0.5 | 6 | 0.3333 | NA | NA | 1 | 0 | 2 | 0.5 | NA | NA | 3 | 0.3333 |
| 9_5611251 | NA | NA | 3 | 1 | 1 | 1 | 1 | 1 | 5 | 1 | NA | NA | 3 | 0.3333 | 2 | 1 | NA | NA | 5 | 0.6 |
| 9_5611257 | NA | NA | 3 | 0.3333 | 2 | 0.5 | 2 | 1 | 7 | 0.5714 | NA | NA | 2 | 1 | 2 | 0.5 | NA | NA | 4 | 0.75 |
| 9_5611260 | NA | NA | 1 | 1 | 2 | 1 | 2 | 1 | 5 | 1 | NA | NA | 2 | 1 | 2 | 1 | NA | NA | 4 | 1 |
| 9_5611267 | NA | NA | 2 | 0.5 | 2 | 0.5 | 2 | 0.5 | 6 | 0.5 | NA | NA | 3 | 0 | 1 | 1 | NA | NA | 4 | 0.25 |

| | | | | | | | | | | | | | | | | | | | | |
|---|---|---|---|---|---|---|---|---|---|---|---|---|---|---|---|---|---|---|---|---|
| 9_5611278( | NA | NA | 1 | 1 | 2 | 0 | 2 | 0.5 | 5 | 0.4 | NA | NA | 2 | 0.5 | 2 | 0 | NA | NA | 4 | 0.25 |
| 9_5611289' | NA | NA | 1 | 0 | 2 | 0 | 1 | 1 | 4 | 0.25 | NA | NA | 2 | 0 | 2 | 0.5 | NA | NA | 4 | 0.25 |
| 9_5611292: | NA | NA | 1 | 0 | 1 | 1 | 1 | 1 | 3 | 0.6667 | NA | NA | NA | NA | 2 | 1 | NA | NA | 2 | 1 |
| 9_5611318( | NA | NA | 1 | 0 | 2 | 1 | 2 | 1 | 5 | 0.8 | NA | NA | 2 | 1 | 1 | 1 | NA | NA | 3 | 1 |
| 9_5611321( | NA | NA | 1 | 1 | 2 | 1 | 2 | 1 | 5 | 1 | NA | NA | 2 | 1 | 1 | 1 | NA | NA | 3 | 1 |
| 9_5611325 | NA | NA | 1 | 1 | 2 | 0.5 | 2 | 0.5 | 5 | 0.6 | NA | NA | 2 | 1 | 1 | 1 | NA | NA | 3 | 1 |
| 9_5611337! | NA | NA | 1 | 0 | 1 | 1 | 1 | 0 | 3 | 0.3333 | NA | NA | 2 | 0.5 | 1 | 0 | NA | NA | 3 | 0.3333 |
| 9_5611377: | NA | NA | NA | NA | 1 | 0 | 1 | 1 | 2 | 0.5 | NA | NA | 2 | 0.5 | 1 | 1 | NA | NA | 3 | 0.6667 |
| 9_5611381' | NA | NA | NA | NA | 1 | 1 | 2 | 1 | 3 | 1 | NA | NA | 2 | 1 | 1 | 1 | NA | NA | 3 | 1 |
| 9_5611383l | NA | NA | NA | NA | 1 | 1 | 2 | 1 | 3 | 1 | NA | NA | 2 | 0.5 | 1 | 1 | NA | NA | 3 | 0.6667 |
| 9_5611389! | NA | NA | NA | NA | 1 | 1 | 1 | 0 | 2 | 0.5 | NA | NA | 2 | 0.5 | NA | NA | NA | NA | 2 | 0.5 |
| 9_5611441' | NA | NA | NA | NA | NA | NA | 1 | 1 | 1 | 1 | NA | NA | 2 | 0.5 | 1 | 1 | NA | NA | 3 | 0.6667 |
| 9_5611468l | NA | NA | NA | NA | NA | NA | 1 | 1 | 1 | 1 | NA | NA | 1 | 1 | 1 | 1 | NA | NA | 2 | 1 |
| 9_5613456: | NA | NA | NA | NA | NA | NA | 1 | 1 | 1 | 1 | NA | NA | 2 | 0.5 | 1 | 1 | NA | NA | 3 | 0.6667 |
| 9_5613483( | NA | NA | NA | NA | NA | NA | 1 | 1 | 1 | 1 | NA | NA | 1 | 1 | NA | NA | NA | NA | 1 | 1 |
| 9_5613484! | NA | NA | NA | NA | NA | NA | 1 | 1 | 1 | 1 | NA | NA | 1 | 1 | NA | NA | NA | NA | 1 | 1 |
| 9_5616505l | NA | NA | NA | NA | NA | NA | 1 | 1 | 1 | 1 | NA | NA | 1 | 1 | 1 | 1 | NA | NA | 2 | 1 |
| 9_5640187! | NA | NA | 2 | 0 | 5 | 0.6 | 5 | 0.4 | 12 | 0.4167 | NA | NA | 7 | 0.4286 | 1 | 0 | 1 | 0 | 9 | 0.3333 |
| 9_5640188: | NA | NA | 2 | 0 | 2 | 0.5 | 4 | 0.5 | 8 | 0.375 | 1 | 0 | 6 | 0.6667 | 1 | 1 | 1 | 0 | 9 | 0.5556 |
| 9_5640190' | NA | NA | 2 | 1 | 3 | 0 | 3 | 0 | 8 | 0.25 | 1 | 0 | 11 | 0.0909 | 1 | 1 | 3 | 0 | 16 | 0.125 |
| 9_5640199' | 2 | 1 | NA | NA | 4 | 1 | 4 | 1 | 10 | 1 | 1 | 1 | 7 | 1 | 1 | 1 | 3 | 1 | 12 | 1 |
| 9_5640212l | 2 | 0.5 | 2 | 0.5 | 4 | 0 | 5 | 0.6 | 13 | 0.3846 | 1 | 1 | 11 | 0.7273 | 2 | 0.5 | 3 | 0.3333 | 17 | 0.6471 |
| 9_5640223: | 1 | 1 | 1 | 1 | 6 | 1 | 6 | 1 | 14 | 1 | 1 | 1 | 12 | 1 | 1 | 1 | 3 | 1 | 17 | 1 |
| 9_5640227l | 2 | 0 | 2 | 0.5 | 5 | 0 | 6 | 0.5 | 15 | 0.2667 | 1 | 0 | 9 | 0.3333 | 2 | 0 | 2 | 0 | 14 | 0.2143 |
| 9_5640230· | 1 | 1 | 2 | 1 | 6 | 1 | 5 | 1 | 14 | 1 | 1 | 1 | 13 | 1 | 2 | 1 | 3 | 1 | 19 | 1 |
| 9_5640236: | 2 | 1 | 2 | 1 | 6 | 0.8333 | 5 | 1 | 15 | 0.9333 | 1 | 1 | 13 | 1 | 2 | 1 | 2 | 1 | 18 | 1 |
| 9_5640237: | 1 | 1 | NA | NA | 4 | 0.5 | 6 | 0.3333 | 11 | 0.4545 | 1 | 1 | 11 | 0.4545 | 2 | 1 | 2 | 0.5 | 16 | 0.5625 |
| 9_5640248· | 2 | 0.5 | 2 | 1 | 6 | 0.8333 | 6 | 1 | 16 | 0.875 | 1 | 1 | 10 | 0.9 | 3 | 1 | 4 | 1 | 18 | 0.9444 |
| 9_5640269' | 2 | 0.5 | 3 | 0.3333 | 6 | 0.3333 | 6 | 0.3333 | 17 | 0.3529 | 1 | 0 | 11 | 0.5455 | 3 | 0 | 3 | 0.3333 | 18 | 0.3889 |
| 9_5640270: | 2 | 0 | 2 | 0.5 | 5 | 0.6 | 5 | 0.2 | 14 | 0.3571 | 1 | 0 | 9 | 0.4444 | 3 | 0.3333 | 3 | 0 | 16 | 0.3125 |
| 9_5640275: | 3 | 0.6667 | 4 | 1 | 6 | 0.6667 | 5 | 1 | 18 | 0.8333 | 1 | 1 | 10 | 0.8 | 2 | 1 | 4 | 0.5 | 17 | 0.7647 |
| 9_5640279l | 3 | 1 | 4 | 1 | 6 | 1 | 6 | 1 | 19 | 1 | 1 | 1 | 13 | 1 | 3 | 1 | 4 | 1 | 21 | 1 |
| 9_5640286· | 4 | 0.75 | 4 | 0.5 | 6 | 0.8333 | 6 | 0.8333 | 20 | 0.75 | 2 | 1 | 12 | 0.6667 | 3 | 1 | 4 | 0.5 | 21 | 0.7143 |
| 9_5640331( | 5 | 0.2 | 6 | 0 | 6 | 0.3333 | 11 | 0.0909 | 28 | 0.1429 | 5 | 0 | 11 | 0.0909 | 5 | 0 | 5 | 0.2 | 26 | 0.0769 |
| 9_5640345 | 6 | 0 | 4 | 0 | 2 | 0 | 8 | 0.125 | 20 | 0.05 | 5 | 0 | 9 | 0.1111 | 4 | 0 | 3 | 0.3333 | 21 | 0.0952 |
| 9_5652581· | 6 | 0 | 19 | 0.3158 | 8 | 0 | 2 | 0 | 35 | 0.1714 | 8 | 0 | 8 | 0.125 | 8 | 0.375 | 9 | 0 | 33 | 0.1212 |
| 9_5652611! | 6 | 0.1667 | 14 | 0.4286 | 7 | 0.4286 | 3 | 0 | 30 | 0.3333 | 4 | 0.5 | 7 | 0.2857 | 2 | 1 | 7 | 0.8571 | 20 | 0.6 |
| 9_5652636! | 7 | 0.1429 | 19 | 0.3684 | 9 | 0.4444 | 3 | 0 | 38 | 0.3158 | 6 | 0.3333 | 6 | 0.3333 | 8 | 0.25 | 9 | 0.3333 | 29 | 0.3103 |
| 9_5652639l | 3 | 0.6667 | 14 | 0.7143 | 8 | 0.625 | 3 | 0.3333 | 28 | 0.6429 | 7 | 0.4286 | 5 | 0.4 | 6 | 0.6667 | 5 | 0.4 | 23 | 0.4783 |
| 9_5652640( | 1 | 0 | 1 | 1 | 1 | 1 | NA | NA | 3 | 0.6667 | 2 | 1 | NA | NA | 1 | 0 | 1 | 1 | 4 | 0.75 |
| 9_5652642: | 7 | 0.5714 | 24 | 0.3333 | 14 | 0.5 | 3 | 0.3333 | 48 | 0.4167 | 13 | 0.2308 | 7 | 0.5714 | 10 | 0.5 | 13 | 0.2308 | 43 | 0.3488 |
| 9_5652645 | 9 | 0.2222 | 25 | 0.12 | 16 | 0.1875 | 5 | 0.2 | 55 | 0.1636 | 14 | 0.0714 | 8 | 0.125 | 14 | 0.0714 | 18 | 0.1667 | 54 | 0.1111 |
| 9_5652646! | 7 | 0.7143 | 20 | 0.25 | 17 | 0.4706 | 3 | 0.6667 | 47 | 0.4255 | 11 | 0.1818 | 8 | 0.5 | 11 | 0.2727 | 14 | 0.2857 | 44 | 0.2955 |
| 9_5652651( | 8 | 0 | 22 | 0.1818 | 18 | 0.0556 | 5 | 0.2 | 53 | 0.1132 | 17 | 0 | 7 | 0.1429 | 15 | 0.2 | 15 | 0.1333 | 54 | 0.1111 |
| 9_5652655( | 9 | 0.3333 | 27 | 0.4444 | 18 | 0.3333 | 5 | 0.2 | 59 | 0.3729 | 16 | 0.3125 | 7 | 0.1429 | 14 | 0.2857 | 18 | 0.3889 | 55 | 0.3091 |
| 9_5652669( | 2 | 0.5 | 9 | 0.4444 | 5 | 0.4 | NA | NA | 16 | 0.4375 | 7 | 0.1429 | 1 | 0 | 4 | 0.25 | 2 | 0 | 14 | 0.1429 |
| 9_5652671( | 6 | 0.3333 | 21 | 0.8095 | 17 | 0.5294 | 4 | 0.75 | 48 | 0.6458 | 11 | 0.7273 | 6 | 0.6667 | 11 | 0.9091 | 16 | 0.625 | 44 | 0.7273 |
| 9_5652674( | 1 | 1 | 3 | 0.6667 | NA | NA | NA | NA | 4 | 0.75 | 2 | 0.5 | NA | NA | 2 | 0.5 | NA | NA | 4 | 0.5 |
| 9_5652702· | 5 | 0 | 23 | 0.1739 | 18 | 0.1111 | 4 | 0.25 | 50 | 0.14 | 11 | 0.0909 | 6 | 0.3333 | 12 | 0 | 14 | 0.1429 | 43 | 0.1163 |
| 9_5652704 | 5 | 0 | 19 | 0.1579 | 17 | 0.0588 | 3 | 0 | 44 | 0.0909 | 11 | 0.2727 | 3 | 0 | 14 | 0.0714 | 9 | 0.1111 | 37 | 0.1351 |
| 9_5652713· | 6 | 0.8333 | 20 | 0.75 | 15 | 0.8 | 5 | 0.8 | 46 | 0.7826 | 7 | 0.5714 | 3 | 0.6667 | 12 | 1 | 13 | 0.6154 | 35 | 0.7429 |
| 9_5652732: | 6 | 0.3333 | 24 | 0.2917 | 16 | 0.25 | 4 | 0.5 | 50 | 0.3 | 7 | 0.1429 | 4 | 0 | 13 | 0.2308 | 10 | 0.1 | 34 | 0.1471 |
| 9_5652738( | 5 | 1 | 23 | 1 | 18 | 1 | 3 | 1 | 49 | 1 | 12 | 1 | 4 | 1 | 10 | 1 | 12 | 1 | 38 | 1 |
| 9_5652748! | 5 | 0.4 | 17 | 0.4706 | 13 | 0.0769 | 4 | 0.25 | 39 | 0.3077 | 8 | 0.5 | 5 | 0.6 | 8 | 0.25 | 12 | 0.3333 | 33 | 0.3939 |
| 9_5652750( | 5 | 1 | 16 | 0.9375 | 14 | 0.9286 | 5 | 0.8 | 40 | 0.925 | 7 | 0.5714 | 4 | 1 | 9 | 1 | 11 | 0.9091 | 31 | 0.871 |
| 9_5652778( | 4 | 0.25 | 18 | 0.3333 | 13 | 0.3846 | 4 | 0.5 | 39 | 0.359 | 9 | 0.4444 | 5 | 0.6 | 11 | 0.4545 | 11 | 0.2727 | 36 | 0.4167 |
| 9_5652784l | 3 | 1 | 19 | 0.7368 | 13 | 0.6923 | 3 | 1 | 38 | 0.7632 | 10 | 0.5 | 4 | 1 | 10 | 0.7 | 11 | 0.4545 | 35 | 0.6 |
| 9_5652791' | 4 | 0.75 | 19 | 0.9474 | 13 | 0.9231 | 4 | 0.5 | 40 | 0.875 | 10 | 1 | 5 | 0.8 | 11 | 0.7273 | 10 | 0.9 | 36 | 0.8611 |
| 9_5652816( | 1 | 0 | 15 | 0.3333 | 10 | 0.2 | 4 | 0 | 30 | 0.2333 | 7 | 0.1429 | 4 | 0.25 | 11 | 0.1818 | 4 | 0.25 | 26 | 0.1923 |
| 9_5652823! | 2 | 1 | 14 | 0.8571 | 12 | 0.9167 | 3 | 1 | 31 | 0.9032 | 6 | 1 | 4 | 1 | 7 | 0.7143 | 7 | 1 | 24 | 0.9167 |
| 9_5652830: | 2 | 0.5 | 17 | 0.4118 | 11 | 0.0909 | 4 | 0 | 34 | 0.2647 | 7 | 0.2857 | 4 | 0.25 | 9 | 0.4444 | 6 | 0.3333 | 26 | 0.3462 |
| 9_5652835: | 3 | 0 | 13 | 0 | 8 | 0.25 | 1 | 0 | 25 | 0.08 | 5 | 0 | 3 | 0 | 8 | 0.125 | 4 | 0 | 20 | 0.05 |
| 9_5652837( | 3 | 0.3333 | 16 | 0.4375 | 12 | 0.1667 | 3 | 0.6667 | 34 | 0.3529 | 7 | 0.4286 | 4 | 0.5 | 11 | 0.4545 | 4 | 0.25 | 26 | 0.4231 |
| 9_5652838! | 3 | 0 | 15 | 0.3333 | 11 | 0.1818 | 2 | 1 | 31 | 0.2903 | 6 | 0.3333 | 3 | 0.3333 | 10 | 0.2 | 5 | 0.4 | 24 | 0.2917 |
| 9_5652845' | 3 | 0.3333 | 12 | 0.1667 | 12 | 0.1667 | 3 | 0.6667 | 30 | 0.2333 | 7 | 0.2857 | 3 | 0 | 12 | 0.5 | 5 | 0.2 | 27 | 0.3333 |

| | | | | | | | | | | | | | | | | | | | | |
|---|---|---|---|---|---|---|---|---|---|---|---|---|---|---|---|---|---|---|---|---|
| 9_5654251( | 2 | 0.5 | 5 | 0.4 | 6 | 0.5 | 1 | 0 | 14 | 0.4286 | 5 | 0.4 | NA | NA | 4 | 0.5 | 4 | 0.5 | 13 | 0.4615 |
| 9_5654252 | 1 | 0 | 5 | 0.2 | 5 | 0.2 | 3 | 0 | 14 | 0.1429 | 6 | 0.1667 | NA | NA | 7 | 0 | 4 | 0 | 17 | 0.0588 |
| 9_5654266( | 2 | 0.5 | 5 | 0 | 7 | 0.2857 | 3 | 0 | 17 | 0.1765 | 6 | 0 | 1 | 0 | 5 | 0.2 | 4 | 0.5 | 16 | 0.1875 |
| 9_5679585 | NA | NA | NA | NA | 1 | 1 | NA | NA | 1 | 1 | NA | NA | 1 | 1 | NA | NA | NA | NA | 1 | 1 |
| 9_5680455 | NA | NA | 1 | 1 | 1 | 1 | NA | NA | 2 | 1 | 1 | 1 | NA | NA | 1 | 1 | NA | NA | 2 | 1 |
| 9_5680462 | 1 | 1 | 1 | 0 | 1 | 1 | NA | NA | 3 | 0.6667 | 1 | 0 | 1 | 0 | 1 | 1 | NA | NA | 3 | 0.3333 |
| 9_5680463 | 1 | 1 | 1 | 1 | 1 | 1 | NA | NA | 3 | 1 | NA | NA | 1 | 1 | 1 | 1 | NA | NA | 2 | 1 |
| 9_5680464 | 1 | 0 | 1 | 1 | 1 | 1 | NA | NA | 3 | 0.6667 | 1 | 1 | 1 | 1 | 1 | 0 | NA | NA | 3 | 0.6667 |
| 9_5680479( | 1 | 0 | 1 | 0 | 2 | 0.5 | NA | NA | 4 | 0.25 | 1 | 0 | 1 | 1 | 2 | 0.5 | NA | NA | 4 | 0.5 |
| 9_5680487 | 1 | 1 | 1 | 1 | 2 | 1 | NA | NA | 4 | 1 | 1 | 0 | 1 | 1 | 1 | 1 | NA | NA | 3 | 0.6667 |
| 9_5680487 | 1 | 0 | 1 | 1 | 2 | 1 | NA | NA | 4 | 0.75 | 1 | 1 | 1 | 0 | 1 | 0 | NA | NA | 3 | 0.3333 |
| 9_5680546 | 1 | 0 | 1 | 0 | 2 | 0.5 | 1 | 1 | 5 | 0.4 | 1 | 1 | 1 | 0 | 1 | 1 | NA | NA | 3 | 0.6667 |
| 9_5680553 | 1 | 1 | NA | NA | 1 | 0 | 1 | 1 | 3 | 0.6667 | 1 | 1 | 1 | 1 | NA | NA | NA | NA | 2 | 1 |
| 9_5680555 | 1 | 1 | 1 | 0 | 2 | 0.5 | 1 | 0 | 5 | 0.4 | 1 | 0 | 1 | 1 | 2 | 0.5 | NA | NA | 4 | 0.5 |
| 9_5680556 | 1 | 1 | 1 | 1 | 1 | 1 | 1 | 1 | 4 | 1 | 1 | 0 | 1 | 1 | 1 | 1 | NA | NA | 3 | 0.6667 |
| 9_5680562 | 1 | 1 | 1 | 1 | 2 | 0.5 | 1 | 1 | 5 | 0.8 | 1 | 1 | 1 | 1 | 2 | 1 | NA | NA | 4 | 1 |
| 9_5680568 | 1 | 1 | 1 | 1 | 2 | 1 | 1 | 1 | 5 | 1 | 1 | 1 | 1 | 1 | 1 | 1 | NA | NA | 3 | 1 |
| 9_5680602 | 1 | 1 | 1 | 1 | 2 | 1 | 2 | 1 | 6 | 1 | 1 | 1 | 1 | 1 | 2 | 1 | NA | NA | 4 | 1 |
| 9_5680611 | 1 | 1 | 1 | 1 | NA | NA | 1 | 1 | 3 | 1 | NA | NA | NA | NA | 1 | 1 | NA | NA | 1 | 1 |
| 9_5680613 | NA | NA | 1 | 0 | 2 | 1 | 1 | 1 | 4 | 0.75 | 1 | 1 | 1 | 1 | 1 | 1 | NA | NA | 3 | 1 |
| 9_5680627 | 1 | 1 | 1 | 0 | 1 | 1 | 2 | 1 | 5 | 0.8 | 1 | 1 | 1 | 1 | 1 | 1 | NA | NA | 3 | 1 |
| 9_5680645 | 1 | 1 | 1 | 0 | 2 | 0.5 | 2 | 0.5 | 6 | 0.5 | 1 | 0 | 1 | 0 | 1 | 1 | 1 | 1 | 4 | 0.5 |
| 9_5680648 | 1 | 0 | 1 | 1 | 2 | 0.5 | 1 | 0 | 5 | 0.4 | 1 | 0 | 1 | 1 | 1 | 1 | NA | NA | 3 | 0.6667 |
| 9_5680677 | 1 | 1 | 1 | 1 | 1 | 1 | 2 | 1 | 5 | 1 | 1 | 1 | 1 | 1 | 1 | 1 | 1 | 1 | 4 | 1 |
| 9_5680683 | 1 | 0 | 1 | 0 | 2 | 0.5 | 2 | 0 | 6 | 0.1667 | 1 | 0 | 1 | 0 | 1 | 0 | 1 | 1 | 4 | 0.25 |
| 9_5680698 | 1 | 1 | 1 | 1 | 2 | 0 | 2 | 0 | 6 | 0.3333 | 1 | 0 | NA | NA | 1 | 1 | 1 | 0 | 3 | 0.3333 |
| 9_5680699 | NA | NA | 1 | 0 | 1 | 1 | 2 | 0.5 | 4 | 0.5 | NA | NA | 1 | 1 | NA | NA | 1 | 0 | 2 | 0.5 |
| 9_5682501 | 5 | 0.6 | 1 | 0 | NA | NA | NA | NA | 6 | 0.5 | 1 | 1 | 2 | 0 | 2 | 0 | NA | NA | 5 | 0.2 |
| 9_5682517 | 4 | 1 | NA | NA | NA | NA | NA | NA | 4 | 1 | 1 | 1 | 2 | 1 | 1 | 1 | NA | NA | 4 | 1 |
| 9_5682519 | 3 | 0.6667 | NA | NA | NA | NA | NA | NA | 3 | 0.6667 | 1 | 0 | 2 | 0.5 | 1 | 0 | NA | NA | 4 | 0.25 |
| 9_5682520 | 3 | 0.3333 | NA | NA | NA | NA | NA | NA | 3 | 0.3333 | 1 | 1 | 2 | 1 | 1 | 1 | NA | NA | 4 | 1 |
| 9_5682528 | 4 | 0.25 | NA | NA | NA | NA | NA | NA | 4 | 0.25 | 1 | 1 | 1 | 0 | 1 | 0 | NA | NA | 3 | 0.3333 |
| 9_5682534 | 3 | 1 | NA | NA | NA | NA | NA | NA | 3 | 1 | 1 | 1 | 2 | 1 | 1 | 1 | NA | NA | 4 | 1 |
| 9_5682538 | 3 | 0.3333 | NA | NA | NA | NA | NA | NA | 3 | 0.3333 | 1 | 1 | 2 | 0 | 1 | 0 | NA | NA | 4 | 0.25 |
| 9_5682542 | 3 | 0.3333 | NA | NA | NA | NA | NA | NA | 3 | 0.3333 | 1 | 0 | 2 | 1 | 1 | 1 | NA | NA | 4 | 0.75 |
| 9_5682543 | 2 | 1 | NA | NA | NA | NA | NA | NA | 2 | 1 | 1 | 1 | 2 | 1 | NA | NA | NA | NA | 3 | 1 |
| 9_5682556 | 3 | 0.3333 | NA | NA | NA | NA | NA | NA | 3 | 0.3333 | 1 | 0 | 2 | 0 | 1 | 1 | NA | NA | 4 | 0.25 |
| 9_5682561 | 3 | 1 | NA | NA | NA | NA | NA | NA | 3 | 1 | 1 | 0 | 2 | 1 | 1 | 0 | NA | NA | 4 | 0.5 |
| 9_5682573 | 3 | 1 | NA | NA | NA | NA | NA | NA | 3 | 1 | 1 | 1 | 2 | 1 | 1 | 1 | NA | NA | 4 | 1 |
| 9_5683272 | 2 | 1 | NA | NA | NA | NA | NA | NA | 2 | 1 | 1 | 0 | 1 | 1 | NA | NA | NA | NA | 2 | 0.5 |
| 9_5683277 | 2 | 1 | NA | NA | NA | NA | NA | NA | 2 | 1 | 1 | 1 | 2 | 1 | NA | NA | NA | NA | 3 | 1 |
| 9_5683336 | 1 | 1 | NA | NA | NA | NA | NA | NA | 1 | 1 | NA | NA | 2 | 0.5 | NA | NA | NA | NA | 2 | 0.5 |
| 9_5683343 | 1 | 1 | NA | NA | NA | NA | NA | NA | 1 | 1 | NA | NA | 2 | 1 | NA | NA | NA | NA | 2 | 1 |
| 9_5683386 | 1 | 1 | NA | NA | NA | NA | NA | NA | 1 | 1 | NA | NA | 1 | 1 | NA | NA | NA | NA | 1 | 1 |
| 9_5683425 | 1 | 1 | NA | NA | NA | NA | NA | NA | 1 | 1 | NA | NA | 2 | 1 | NA | NA | NA | NA | 2 | 1 |
| 9_5683449 | 1 | 1 | NA | NA | NA | NA | NA | NA | 1 | 1 | NA | NA | 2 | 1 | NA | NA | NA | NA | 2 | 1 |
| 9_5683497( | 1 | 1 | NA | NA | NA | NA | NA | NA | 1 | 1 | NA | NA | 1 | 1 | NA | NA | NA | NA | 1 | 1 |
| 9_5683506 | 1 | 1 | NA | NA | NA | NA | NA | NA | 1 | 1 | NA | NA | 1 | 1 | NA | NA | NA | NA | 1 | 1 |
| 9_5684503 | 8 | 0.75 | 12 | 0.25 | 14 | 0.3571 | 8 | 0.5 | 42 | 0.4286 | 5 | 0.6 | 18 | 0.6111 | 7 | 0.7143 | 5 | 0.2 | 35 | 0.5714 |
| 9_5684507 | 8 | 0.625 | 13 | 0.5385 | 14 | 0.7143 | 9 | 0.2222 | 44 | 0.5455 | 6 | 0.3333 | 20 | 0.55 | 8 | 0.625 | 5 | 0.2 | 39 | 0.4872 |
| 9_5684525 | 7 | 0.7143 | 14 | 0.7143 | 12 | 0.6667 | 8 | 0.875 | 41 | 0.7317 | 7 | 0.8571 | 16 | 0.6875 | 9 | 0.7778 | 7 | 0.8571 | 39 | 0.7692 |
| 9_5684527 | 8 | 0.375 | 14 | 0.5 | 14 | 0.5714 | 6 | 0.5 | 42 | 0.5 | 6 | 0.3333 | 21 | 0.4762 | 5 | 0.6 | 6 | 0.5 | 38 | 0.4737 |
| 9_5684532 | 10 | 0.5 | 15 | 0.2 | 15 | 0.2667 | 8 | 0.125 | 48 | 0.2708 | 6 | 0.3333 | 20 | 0.35 | 7 | 0.4286 | 8 | 0.125 | 41 | 0.3171 |
| 9_5684538 | 8 | 0 | 10 | 0.1 | 14 | 0.2857 | 8 | 0 | 40 | 0.125 | 5 | 0 | 14 | 0.1429 | 7 | 0 | 6 | 0.3333 | 32 | 0.125 |
| 9_5684544 | 9 | 1 | 14 | 1 | 15 | 0.9333 | 8 | 0.875 | 46 | 0.9565 | 6 | 0.8333 | 17 | 1 | 8 | 1 | 7 | 1 | 38 | 0.9737 |
| 9_5684545 | 8 | 0.875 | 14 | 1 | 9 | 1 | 7 | 1 | 38 | 0.9737 | 7 | 1 | 14 | 1 | 7 | 0.8571 | 6 | 0.8333 | 34 | 0.9412 |
| 9_5684547 | 11 | 0.7273 | 15 | 0.8667 | 13 | 0.6154 | 8 | 0.75 | 47 | 0.7447 | 6 | 0.8333 | 15 | 0.6 | 9 | 0.5556 | 6 | 0.6667 | 36 | 0.6389 |
| 9_5689025 | 2 | 1 | 2 | 0.5 | 2 | 0.5 | 1 | 1 | 7 | 0.7143 | 1 | 1 | 2 | 1 | 2 | 1 | 3 | 1 | 8 | 1 |
| 9_5689027 | 2 | 0 | 2 | 0.5 | 2 | 0.5 | 1 | 0 | 7 | 0.2857 | 1 | 0 | 2 | 0.5 | 1 | 0 | 3 | 0.6667 | 7 | 0.4286 |
| 9_5689029 | 2 | 1 | 2 | 0.5 | 2 | 1 | 1 | 0 | 7 | 0.7143 | 1 | 1 | 2 | 1 | 2 | 0.5 | 3 | 0.6667 | 8 | 0.75 |
| 9_5689031 | 2 | 0.5 | 2 | 0 | 2 | 0 | 1 | 1 | 7 | 0.2857 | 1 | 0 | 2 | 0.5 | 2 | 0 | 3 | 0 | 8 | 0.125 |
| 9_5689037 | 2 | 0.5 | 2 | 0 | 2 | 0.5 | 1 | 0 | 7 | 0.2857 | 1 | 1 | 2 | 1 | 2 | 0.5 | 2 | 1 | 7 | 0.8571 |
| 9_5689961 | NA | NA | NA | NA | 1 | 1 | NA | NA | 1 | 1 | NA | NA | NA | NA | NA | NA | 1 | 1 | 1 | 1 |
| 9_5689971 | NA | NA | NA | NA | 1 | 1 | NA | NA | 1 | 1 | NA | NA | NA | NA | NA | NA | 1 | 1 | 1 | 1 |
| 9_5689990 | NA | NA | NA | NA | 2 | 1 | NA | NA | 2 | 1 | NA | NA | NA | NA | NA | NA | 1 | 1 | 1 | 1 |

| | | | | | | | | | | | | | | | | | | | | |
|---|---|---|---|---|---|---|---|---|---|---|---|---|---|---|---|---|---|---|---|---|
| 9_5690005 | NA | NA | NA | NA | 2 | 1 | NA | NA | 2 | 1 | NA | NA | NA | NA | NA | NA | 1 | 1 | 1 | 1 |
| 9_5690256 | NA | NA | 1 | 1 | 2 | 0.5 | 4 | 0.75 | 7 | 0.7143 | NA | NA | 2 | 1 | 1 | 1 | NA | NA | 3 | 1 |
| 9_5694395 | 1 | 1 | 2 | 0 | 2 | 0.5 | 4 | 0.5 | 9 | 0.4444 | 1 | 1 | 3 | 0.3333 | 1 | 1 | NA | NA | 5 | 0.6 |
| 9_5696831 | 3 | 0.3333 | 2 | 0 | 2 | 0.5 | 5 | 0 | 12 | 0.1667 | 2 | 0 | 5 | 0.2 | 4 | 0 | 2 | 0.5 | 13 | 0.1538 |
| 9_5696844 | 3 | 0.6667 | 2 | 0 | 2 | 0.5 | 5 | 0.4 | 12 | 0.4167 | 3 | 0.3333 | 5 | 0.6 | 2 | 0 | 2 | 0 | 12 | 0.3333 |
| 9_5696848 | 3 | 0 | 2 | 0 | 2 | 0.5 | 4 | 0 | 11 | 0.0909 | 3 | 0 | 5 | 0.2 | 3 | 0 | 2 | 0 | 13 | 0.0769 |
| 9_5696849 | 2 | 1 | 2 | 0.5 | 2 | 1 | 5 | 1 | 11 | 0.9091 | 2 | 1 | 4 | 1 | 3 | 1 | 2 | 0.5 | 11 | 0.9091 |
| 9_5696861 | 3 | 0.3333 | 2 | 0.5 | 2 | 0 | 4 | 0.5 | 11 | 0.3636 | 2 | 0.5 | 4 | 0.25 | 3 | 0 | 1 | 0 | 10 | 0.2 |
| 9_5696867 | 4 | 0.75 | 2 | 1 | 2 | 1 | 3 | 1 | 11 | 0.9091 | 3 | 1 | 5 | 1 | 3 | 1 | 2 | 1 | 13 | 1 |
| 9_5696878 | 2 | 0.5 | 2 | 0.5 | 1 | 0 | 3 | 0.6667 | 8 | 0.5 | 2 | 0 | 1 | 1 | 2 | 0 | 1 | 0 | 6 | 0.1667 |
| 9_5696913 | 2 | 0.5 | 2 | 0 | 3 | 0.6667 | 6 | 0.3333 | 13 | 0.3846 | 2 | 0 | 3 | 0.3333 | 2 | 1 | 1 | 1 | 8 | 0.5 |
| 9_5696925 | 3 | 0.3333 | 2 | 0.5 | 2 | 0 | 5 | 0.6 | 12 | 0.4167 | 3 | 0 | 5 | 0.4 | 3 | 0 | 2 | 1 | 13 | 0.3077 |
| 9_5696999 | 4 | 0.5 | 3 | 0 | 3 | 0 | 5 | 0.6 | 15 | 0.3333 | 7 | 0 | 5 | 0.4 | 7 | 0.2857 | 3 | 0.3333 | 22 | 0.2273 |
| 9_5701800 | 3 | 0.3333 | NA | NA | NA | NA | 2 | 0.5 | 5 | 0.4 | 2 | 0.5 | 4 | 0.75 | 2 | 1 | 2 | 1 | 10 | 0.8 |
| 9_5701825 | 4 | 0.25 | 1 | 0 | NA | NA | 2 | 0.5 | 7 | 0.2857 | 2 | 0 | 4 | 0 | 2 | 0.5 | 2 | 0 | 10 | 0.1 |
| 9_5703449 | 5 | 0.4 | 4 | 0.5 | NA | NA | 2 | 0.5 | 11 | 0.4545 | 1 | 0 | 5 | 0 | 2 | 0.5 | 3 | 0.3333 | 11 | 0.1818 |
| 9_5703459 | 6 | 1 | 4 | 1 | 1 | 1 | 3 | 1 | 14 | 1 | 2 | 0.5 | 5 | 0.8 | 3 | 0.6667 | 4 | 1 | 14 | 0.7857 |
| 9_5703486 | 3 | 0.6667 | 3 | 0.6667 | 1 | 1 | 2 | 1 | 9 | 0.7778 | 1 | 1 | 5 | 0.6 | 2 | 1 | 4 | 1 | 12 | 0.8333 |
| 9_5703492 | 6 | 0.6667 | 5 | 1 | 1 | 1 | 3 | 1 | 15 | 0.8667 | 2 | 0.5 | 5 | 0.6 | 4 | 0.5 | 5 | 0.6 | 16 | 0.5625 |
| 9_5703510 | 6 | 0 | 5 | 0.2 | 1 | 0 | 2 | 0 | 14 | 0.0714 | 1 | 0 | 5 | 0 | 4 | 0.25 | 5 | 0 | 15 | 0.0667 |
| 9_5703516 | 7 | 0 | 5 | 0.2 | 1 | 0 | 3 | 0.3333 | 16 | 0.125 | 2 | 0 | 4 | 0 | 3 | 0 | 5 | 0.2 | 14 | 0.0714 |
| 9_5703527 | 6 | 0.1667 | 5 | 0.4 | 1 | 0 | 3 | 0.6667 | 15 | 0.3333 | 2 | 0.5 | 4 | 0.25 | 3 | 0.3333 | 6 | 0.5 | 15 | 0.4 |
| 9_5705021 | 1 | 0 | NA | NA | 2 | 0.5 | 1 | 0 | 4 | 0.25 | 2 | 0 | NA | NA | NA | NA | 1 | 1 | 3 | 0.3333 |
| 9_5705044 | 2 | 0 | NA | NA | 2 | 0.5 | 1 | 0 | 5 | 0.2 | 2 | 0 | NA | NA | NA | NA | 1 | 1 | 3 | 0.3333 |
| 9_5706505 | NA | NA | NA | NA | 1 | 1 | NA | NA | 1 | 1 | 1 | 0 | NA | NA | 2 | 0.5 | NA | NA | 3 | 0.3333 |
| 9_5706528 | NA | NA | NA | NA | 1 | 1 | NA | NA | 1 | 1 | 1 | 1 | NA | NA | 1 | 1 | NA | NA | 2 | 1 |
| 9_5716052 | NA | NA | NA | NA | 1 | 1 | NA | NA | 1 | 1 | 1 | 1 | NA | NA | 1 | 0 | NA | NA | 2 | 0.5 |
| 9_5716059 | NA | NA | NA | NA | 1 | 1 | NA | NA | 1 | 1 | 2 | 0.5 | 1 | 0 | 1 | 0 | NA | NA | 4 | 0.25 |
| 9_5716069 | NA | NA | NA | NA | 1 | 1 | NA | NA | 1 | 1 | 2 | 1 | NA | NA | 1 | 1 | NA | NA | 3 | 1 |
| 9_5716076 | NA | NA | NA | NA | 1 | 1 | NA | NA | 1 | 1 | 2 | 0 | 1 | 1 | 1 | 0 | NA | NA | 4 | 0.25 |
| 9_5716080 | NA | NA | NA | NA | 1 | 1 | NA | NA | 1 | 1 | 2 | 0.5 | NA | NA | 1 | 0 | NA | NA | 3 | 0.3333 |
| 9_5716083 | NA | NA | NA | NA | 1 | 1 | NA | NA | 1 | 1 | 2 | 1 | NA | NA | 1 | 1 | NA | NA | 3 | 1 |
| 9_5716104 | NA | NA | NA | NA | 1 | 1 | NA | NA | 1 | 1 | NA | NA | NA | NA | 1 | 1 | NA | NA | 1 | 1 |
| 9_5716111 | NA | NA | NA | NA | 1 | 1 | NA | NA | 1 | 1 | 3 | 1 | NA | NA | 1 | 1 | 1 | 1 | 5 | 1 |
| 9_5716112 | NA | NA | NA | NA | 1 | 1 | NA | NA | 1 | 1 | 3 | 1 | NA | NA | 1 | 1 | NA | NA | 4 | 1 |
| 9_5716120 | NA | NA | NA | NA | 1 | 1 | NA | NA | 1 | 1 | 3 | 1 | 1 | 1 | 1 | 1 | 1 | 1 | 6 | 1 |
| 9_5716402 | NA | NA | NA | NA | 1 | 1 | NA | NA | 1 | 1 | 1 | 1 | 1 | 1 | NA | NA | 1 | 1 | 3 | 1 |
| 9_5716416 | NA | NA | NA | NA | 1 | 1 | NA | NA | 1 | 1 | 1 | 1 | 1 | 0 | NA | NA | 1 | 1 | 3 | 0.6667 |
| 9_5716434 | NA | NA | NA | NA | 1 | 1 | NA | NA | 1 | 1 | 1 | 1 | NA | NA | NA | NA | 1 | 1 | 2 | 1 |
| 9_5729257 | 1 | 0 | 1 | 1 | NA | NA | NA | NA | 2 | 0.5 | NA | NA | NA | NA | 2 | 0.5 | NA | NA | 2 | 0.5 |
| 9_5732005 | NA | NA | NA | NA | 1 | 1 | NA | NA | 1 | 1 | 1 | 1 | NA | NA | NA | NA | NA | NA | 1 | 1 |
| 9_5732016 | NA | NA | NA | NA | 1 | 1 | NA | NA | 1 | 1 | 1 | 1 | 2 | 1 | NA | NA | NA | NA | 3 | 1 |
| 9_5732018 | NA | NA | NA | NA | 1 | 1 | NA | NA | 1 | 1 | 1 | 1 | 2 | 1 | NA | NA | NA | NA | 3 | 1 |
| 9_5732140 | 1 | 0 | 1 | 0 | 2 | 0.5 | NA | NA | 4 | 0.25 | 2 | 0 | 1 | 0 | 2 | 0.5 | 1 | 0 | 6 | 0.1667 |
| 9_5732162 | 1 | 1 | 1 | 0 | 2 | 0.5 | NA | NA | 4 | 0.5 | 1 | 0 | 1 | 0 | 1 | 1 | 1 | 1 | 4 | 0.5 |
| 9_5732163 | 1 | 0 | 1 | 1 | 1 | 1 | NA | NA | 3 | 0.6667 | 1 | 1 | 1 | 1 | NA | NA | NA | NA | 2 | 1 |
| 9_5732167 | 1 | 1 | NA | NA | 2 | 1 | NA | NA | 3 | 1 | 1 | 0 | 1 | 1 | NA | NA | 1 | 0 | 3 | 0.3333 |
| 9_5735091 | 24 | 0.125 | 22 | 0.1364 | 20 | 0.15 | 18 | 0.1667 | 84 | 0.1429 | 20 | 0.3 | 15 | 0.0667 | 25 | 0.2 | 13 | 0 | 73 | 0.1644 |
| 9_5735100 | 25 | 0.12 | 21 | 0.0476 | 22 | 0.0909 | 19 | 0 | 87 | 0.069 | 25 | 0.16 | 18 | 0 | 28 | 0.2857 | 18 | 0.0556 | 89 | 0.1461 |
| 9_5738556 | 1 | 1 | NA | NA | NA | NA | NA | NA | 1 | 1 | NA | NA | NA | NA | 1 | 1 | NA | NA | 1 | 1 |
| 9_5738567 | 1 | 1 | NA | NA | NA | NA | NA | NA | 1 | 1 | NA | NA | NA | NA | 1 | 1 | NA | NA | 1 | 1 |
| 9_5741137 | 4 | 0 | 2 | 0.5 | 3 | 0 | 2 | 0.5 | 11 | 0.1818 | 2 | 0 | 2 | 0 | 1 | 0 | 4 | 0.25 | 9 | 0.1111 |
| 9_5741159 | 4 | 0 | 2 | 1 | 3 | 0.6667 | 1 | 1 | 10 | 0.5 | 2 | 0 | 3 | 0 | 1 | 1 | 4 | 0.5 | 10 | 0.3 |
| 9_5741210 | 3 | 0 | 2 | 0.5 | 1 | 0 | 2 | 0.5 | 8 | 0.25 | 5 | 0.4 | 2 | 1 | 2 | 0 | 3 | 1 | 12 | 0.5833 |
| 9_5741211 | 3 | 0.6667 | 4 | 1 | 3 | 0.6667 | 3 | 0.6667 | 13 | 0.7692 | 5 | 0.4 | 3 | 0.3333 | 2 | 0.5 | 5 | 0.2 | 15 | 0.3333 |
| 9_5741223 | 3 | 0 | 4 | 0.25 | 3 | 0 | 3 | 0 | 13 | 0.0769 | 3 | 0.3333 | 3 | 0 | 3 | 0 | 4 | 0.25 | 13 | 0.1538 |
| 9_5741232 | 4 | 0.25 | 4 | 0 | 3 | 0.3333 | 2 | 0 | 13 | 0.1538 | 5 | 0.2 | 2 | 0 | 3 | 0.3333 | 4 | 0.25 | 14 | 0.2143 |
| 9_5741233 | 3 | 1 | 4 | 1 | 3 | 0.6667 | 2 | 1 | 12 | 0.9167 | 4 | 1 | 3 | 0.6667 | 3 | 1 | 4 | 0.75 | 14 | 0.8571 |
| 9_5741241 | 4 | 0.75 | 4 | 1 | 3 | 0.6667 | 3 | 0.6667 | 14 | 0.7857 | 5 | 1 | 3 | 1 | 2 | 1 | 5 | 1 | 15 | 1 |
| 9_5741243 | 3 | 0 | 4 | 0.25 | 3 | 0.6667 | 4 | 0 | 14 | 0.2143 | 4 | 0.5 | 3 | 0.3333 | 3 | 0 | 4 | 0.5 | 14 | 0.3571 |
| 9_5741247 | 4 | 0.25 | 4 | 0.25 | 3 | 0 | 4 | 0 | 15 | 0.1333 | 4 | 0 | 3 | 0 | 3 | 0 | 4 | 0.25 | 14 | 0.0714 |
| 9_5741256 | 4 | 0.5 | 4 | 0.5 | 3 | 0.6667 | 4 | 0 | 15 | 0.4 | 5 | 0.2 | 3 | 0.3333 | 3 | 0.6667 | 5 | 0.4 | 16 | 0.375 |
| 9_5741265 | 5 | 0.4 | 3 | 0.6667 | 2 | 0 | 4 | 0.5 | 14 | 0.4286 | 3 | 0.3333 | 3 | 0 | 2 | 0.5 | 5 | 0.2 | 13 | 0.2308 |
| 9_5741266 | 5 | 0.2 | 4 | 0 | 2 | 0 | 5 | 0 | 16 | 0.0625 | 4 | 0.25 | 3 | 0.3333 | 3 | 0.3333 | 4 | 0.25 | 14 | 0.2857 |
| 9_5744839 | NA | NA | 1 | 1 | NA | NA | NA | NA | 1 | 1 | 1 | 1 | 1 | 0 | 1 | 1 | NA | NA | 3 | 0.6667 |

| | | | | | | | | | | | | | | | | | | | | |
|---|---|---|---|---|---|---|---|---|---|---|---|---|---|---|---|---|---|---|---|---|
| 9_57451610 | 7 | 0.1429 | 5 | 0 | 1 | 0 | 4 | 0.25 | 17 | 0.1176 | 4 | 0 | 4 | 0.5 | 10 | 0 | 2 | 0.5 | 20 | 0.15 |
| 9_57451844 | 5 | 0 | 3 | 0 | 2 | 0.5 | 2 | 0.5 | 12 | 0.1667 | 4 | 0 | 3 | 0.3333 | 11 | 0.3636 | 2 | 0 | 20 | 0.25 |
| 9_57452000 | 7 | 0.1429 | 4 | 0 | 5 | 0.2 | 5 | 0.2 | 21 | 0.1429 | 5 | 0 | 4 | 0 | 15 | 0 | 4 | 0.5 | 28 | 0.0714 |
| 9_57452149 | 6 | 0.5 | 4 | 0 | 3 | 0.3333 | 4 | 0.25 | 17 | 0.2941 | 2 | 0.5 | 3 | 0 | 13 | 0.3846 | 4 | 0.5 | 22 | 0.3636 |
| 9_57453826 | 3 | 0 | 4 | 0.25 | 3 | 0 | 2 | 0 | 12 | 0.0833 | 1 | 0 | 2 | 0.5 | 8 | 0 | 3 | 0.3333 | 14 | 0.1429 |
| 9_57494924 | 4 | 0.25 | 3 | 0 | 4 | 0 | 1 | 0 | 12 | 0.0833 | 5 | 0.2 | 2 | 0 | 2 | 0 | 2 | 0 | 11 | 0.0909 |
| 9_57495019 | 4 | 0.5 | 3 | 0 | 4 | 0 | 1 | 0 | 12 | 0.1667 | 6 | 0.1667 | 2 | 0 | 2 | 0.5 | 2 | 0.5 | 12 | 0.25 |
| 9_57495025 | 3 | 1 | 3 | 0.6667 | 4 | 1 | 1 | 1 | 11 | 0.9091 | 6 | 0.6667 | 2 | 0.5 | 2 | 1 | 2 | 1 | 12 | 0.75 |
| 9_57495125 | 4 | 0.75 | 3 | 0.3333 | 3 | 1 | 1 | 0 | 11 | 0.6364 | 4 | 0.25 | 1 | 0 | 2 | 1 | 2 | 0.5 | 9 | 0.4444 |
| 9_57495149 | 4 | 0.75 | 3 | 1 | 4 | 0.75 | 1 | 1 | 12 | 0.8333 | 2 | 1 | NA | NA | 2 | 1 | 2 | 1 | 6 | 1 |
| 9_57495507 | 5 | 0.4 | 1 | 1 | 4 | 0 | 1 | 0 | 11 | 0.2727 | 3 | 0 | 2 | 0 | 2 | 0.5 | 1 | 0 | 8 | 0.125 |
| 9_57502323 | 3 | 0.6667 | 4 | 0.25 | 2 | 0.5 | NA | NA | 9 | 0.4444 | 1 | 1 | 4 | 0.5 | 3 | 0.6667 | 3 | 0 | 11 | 0.4545 |
| 9_57502439 | 2 | 0.5 | 4 | 0.25 | 3 | 0.6667 | NA | NA | 9 | 0.4444 | NA | NA | 5 | 0.2 | 2 | 0.5 | 3 | 0 | 10 | 0.2 |
| 9_57502455 | 3 | 0.3333 | 3 | 0.6667 | 3 | 0.6667 | NA | NA | 9 | 0.5556 | NA | NA | 3 | 0.3333 | 3 | 1 | 3 | 0.3333 | 9 | 0.5556 |
| 9_57502485 | 2 | 1 | 4 | 0.75 | 3 | 0.6667 | NA | NA | 9 | 0.7778 | 1 | 0 | 5 | 0.8 | 3 | 0.3333 | 3 | 0.3333 | 12 | 0.5 |
| 9_57502540 | 3 | 1 | 4 | 1 | 3 | 1 | NA | NA | 10 | 1 | 1 | 1 | 5 | 1 | 3 | 1 | 3 | 0.3333 | 12 | 0.8333 |
| 9_57502630 | 2 | 1 | 3 | 1 | 3 | 1 | NA | NA | 8 | 1 | NA | NA | 5 | 1 | 3 | 1 | 3 | 1 | 11 | 1 |
| 9_57502650 | 3 | 1 | 4 | 1 | 3 | 1 | NA | NA | 10 | 1 | NA | NA | 4 | 1 | 2 | 1 | 2 | 1 | 8 | 1 |
| 9_57502684 | 3 | 0 | 4 | 0 | 3 | 0.3333 | NA | NA | 10 | 0.1 | NA | NA | 4 | 0.25 | 3 | 0 | 3 | 0 | 10 | 0.1 |
| 9_57502738 | 3 | 0.3333 | 3 | 0.6667 | NA | NA | NA | NA | 6 | 0.5 | NA | NA | 5 | 0.4 | 3 | 0.6667 | 3 | 0 | 11 | 0.3636 |
| 9_57502889 | 2 | 0.5 | 4 | 0.25 | 3 | 0 | NA | NA | 9 | 0.2222 | NA | NA | 4 | 0 | 1 | 1 | 3 | 0.3333 | 8 | 0.25 |
| 9_57502986 | 2 | 0.5 | 4 | 0 | 2 | 0.5 | NA | NA | 8 | 0.25 | 1 | 1 | 5 | 0.2 | 2 | 0.5 | 3 | 0 | 11 | 0.2727 |
| 9_57508968 | 1 | 1 | 3 | 0.6667 | 3 | 0.6667 | 1 | 1 | 8 | 0.75 | NA | NA | 1 | 1 | NA | NA | 1 | 1 | 2 | 1 |
| 9_57509416 | 1 | 1 | 2 | 0.5 | 2 | 0.5 | 1 | 1 | 6 | 0.6667 | NA | NA | NA | NA | NA | NA | 2 | 0.5 | 2 | 0.5 |
| 9_57509581 | 1 | 1 | 4 | 0 | 3 | 0 | 1 | 0 | 9 | 0.1111 | NA | NA | 1 | 0 | NA | NA | 2 | 0.5 | 3 | 0.3333 |
| 9_57533937 | 2 | 0.5 | 3 | 0 | 2 | 0 | NA | NA | 7 | 0.1429 | 3 | 0.6667 | 1 | 0 | 3 | 0 | 3 | 0 | 10 | 0.2 |
| 9_57534415 | 2 | 1 | 7 | 1 | 5 | 1 | 3 | 1 | 17 | 1 | 5 | 1 | 5 | 1 | 6 | 0.6667 | 2 | 0.5 | 18 | 0.8333 |
| 9_57534447 | 4 | 0.25 | 7 | 0.1429 | 4 | 0 | 4 | 0 | 19 | 0.1053 | 5 | 0 | 5 | 0 | 9 | 0.2222 | 4 | 0 | 23 | 0.087 |
| 9_57534512 | 1 | 1 | 5 | 1 | 3 | 1 | 3 | 0.6667 | 12 | 0.9167 | 3 | 1 | 5 | 1 | 7 | 0.8571 | 1 | 1 | 16 | 0.9375 |
| 9_57534518 | 2 | 0 | 7 | 0.4286 | 4 | 0.25 | 3 | 0.6667 | 16 | 0.375 | 5 | 0.6 | 4 | 0.75 | 6 | 0.6667 | 3 | 0.6667 | 18 | 0.6667 |
| 9_57534569 | 3 | 0 | 5 | 0.4 | 1 | 0 | 4 | 0 | 13 | 0.1538 | 3 | 0 | 4 | 0 | 6 | 0.3333 | 2 | 0 | 15 | 0.1333 |
| 9_57634592 | 2 | 0.5 | 3 | 1 | 1 | 1 | 1 | 1 | 7 | 0.8571 | NA | NA | NA | NA | 2 | 0.5 | NA | NA | 2 | 0.5 |
| 9_57655601 | 2 | 0.5 | 3 | 0 | 1 | 0 | 1 | 0 | 7 | 0.1429 | 1 | 0 | NA | NA | 3 | 0.3333 | NA | NA | 4 | 0.25 |
| 9_57655684 | 3 | 0.6667 | 6 | 0.1667 | 1 | 0 | 2 | 0 | 12 | 0.25 | 1 | 1 | 1 | 0 | 4 | 0.5 | NA | NA | 6 | 0.5 |
| 9_57655798 | 3 | 0.3333 | 4 | 0.75 | 2 | 0 | 2 | 0.5 | 11 | 0.4545 | 1 | 1 | 1 | 0 | 4 | 0.75 | NA | NA | 6 | 0.6667 |
| 9_57655867 | 5 | 0.6 | 4 | 0.75 | 2 | 1 | 2 | 0.5 | 13 | 0.6923 | 1 | 1 | 2 | 1 | 4 | 1 | NA | NA | 7 | 1 |
| 9_57655876 | 5 | 0.6 | 4 | 0.5 | 2 | 0 | 2 | 0.5 | 13 | 0.4615 | NA | NA | 2 | 0 | 3 | 0.6667 | NA | NA | 5 | 0.4 |
| 9_57656069 | 4 | 0.25 | 4 | 0.5 | 2 | 0.5 | 2 | 0.5 | 12 | 0.4167 | 1 | 1 | 3 | 0 | 3 | 0.3333 | NA | NA | 7 | 0.2857 |
| 9_57657103 | 4 | 0 | 6 | 0.6667 | 3 | 0.3333 | 3 | 0 | 16 | 0.3125 | 2 | 0.5 | 3 | 0.3333 | 4 | 0 | NA | NA | 9 | 0.2222 |
| 9_57657129 | 4 | 0.25 | 6 | 0.1667 | 3 | 0 | 3 | 0.3333 | 16 | 0.1875 | 3 | 0.3333 | 3 | 0.6667 | 3 | 0 | NA | NA | 9 | 0.3333 |
| 9_57657148 | 4 | 0.25 | 4 | 0 | 3 | 0 | 3 | 0 | 14 | 0.0714 | 3 | 0 | 3 | 0.3333 | 4 | 0 | NA | NA | 10 | 0.1 |
| 9_57657228 | 3 | 0.3333 | 5 | 0.2 | 3 | 0 | 3 | 0 | 14 | 0.1429 | 4 | 0.25 | 3 | 0 | 3 | 0.3333 | NA | NA | 10 | 0.2 |
| 9_57657282 | 3 | 0 | 5 | 0 | 3 | 0 | 3 | 0.6667 | 14 | 0.1429 | 3 | 0.3333 | 3 | 0 | 4 | 0 | NA | NA | 10 | 0.1 |
| 9_57658126 | 7 | 0.8571 | 2 | 0.5 | 2 | 1 | 3 | 0.6667 | 14 | 0.7857 | 3 | 1 | 4 | 0.25 | 4 | 0.5 | 1 | 0 | 12 | 0.5 |
| 9_57658190 | 7 | 0.7143 | 4 | 0.75 | 5 | 0.8 | 2 | 0.5 | 18 | 0.7222 | 2 | 1 | 5 | 0.4 | 3 | 0.3333 | NA | NA | 10 | 0.5 |
| 9_57741001 | 1 | 1 | NA | NA | NA | NA | 1 | 0 | 2 | 0.5 | NA | NA | 1 | 1 | 1 | 1 | NA | NA | 2 | 1 |
| 9_57837052 | 21 | 0 | 18 | 0.1111 | 14 | 0.2143 | 13 | 0 | 66 | 0.0758 | 17 | 0.1176 | 14 | 0.0714 | 12 | 0.0833 | 12 | 0.0833 | 55 | 0.0909 |
| 9_57837134 | 14 | 0.2143 | 17 | 0.1176 | 13 | 0.2308 | 11 | 0.1818 | 55 | 0.1818 | 18 | 0.1111 | 15 | 0.3333 | 11 | 0.2727 | 11 | 0.1818 | 55 | 0.2182 |
| 9_57869085 | NA | NA | 1 | 1 | NA | NA | 1 | 1 | 2 | 1 | 1 | 0 | NA | NA | 3 | 0.6667 | NA | NA | 4 | 0.5 |
| 9_57869263 | NA | NA | 2 | 0.5 | NA | NA | 1 | 1 | 3 | 0.6667 | 1 | 0 | NA | NA | 2 | 0.5 | NA | NA | 3 | 0.3333 |
| 9_57869803 | 1 | 0 | 9 | 0.1111 | 3 | 0.3333 | 1 | 0 | 14 | 0.1429 | 5 | 0.2 | 4 | 0.5 | 8 | 0.25 | 1 | 0 | 18 | 0.2778 |
| 9_57869917 | NA | NA | 11 | 0.0909 | 4 | 0 | 1 | 0 | 16 | 0.0625 | 5 | 0 | 3 | 0 | 8 | 0.125 | 2 | 0.5 | 18 | 0.1111 |
| 9_57913076 | NA | NA | NA | NA | 1 | 1 | NA | NA | 1 | 1 | NA | NA | 1 | 1 | NA | NA | NA | NA | 1 | 1 |
| 9_57913307 | NA | NA | 1 | 1 | 1 | 0 | NA | NA | 2 | 0.5 | NA | NA | 1 | 1 | NA | NA | NA | NA | 1 | 1 |
| 9_57913370 | NA | NA | 1 | 0 | 1 | 1 | NA | NA | 2 | 0.5 | NA | NA | 1 | 1 | NA | NA | NA | NA | 1 | 1 |
| 9_57913410 | NA | NA | NA | NA | 1 | 1 | NA | NA | 1 | 1 | NA | NA | 1 | 1 | NA | NA | NA | NA | 1 | 1 |
| 9_57913593 | NA | NA | 1 | 1 | 1 | 1 | NA | NA | 2 | 1 | NA | NA | 1 | 1 | NA | NA | NA | NA | 1 | 1 |
| 9_57913967 | NA | NA | 2 | 1 | NA | NA | NA | NA | 2 | 1 | NA | NA | 1 | 1 | NA | NA | NA | NA | 1 | 1 |
| 9_58060744 | 1 | 0 | 5 | 0 | 2 | 0 | 6 | 0.5 | 14 | 0.2143 | 1 | 1 | NA | NA | 1 | 0 | 4 | 0 | 6 | 0.1667 |
| 9_58063998 | 9 | 0.1111 | 7 | 0 | 4 | 0.25 | 3 | 0 | 23 | 0.087 | 4 | 0 | 1 | 0 | 1 | 0 | 3 | 0.3333 | 9 | 0.1111 |
| 9_58064090 | 9 | 0.4444 | 7 | 0.5714 | 3 | 0.3333 | 3 | 0.3333 | 22 | 0.4545 | 3 | 0 | 1 | 0 | 1 | 0 | 2 | 0.5 | 7 | 0.1429 |
| 9_58064559 | 8 | 1 | 5 | 1 | 3 | 0.6667 | NA | NA | 16 | 0.9375 | 2 | 1 | 1 | 1 | NA | NA | 2 | 1 | 5 | 1 |
| 9_58064723 | 7 | 0.8571 | 5 | 1 | 2 | 1 | 1 | 1 | 15 | 0.9333 | 2 | 1 | 1 | 0 | 1 | 1 | 2 | 1 | 6 | 0.8333 |
| 9_58064889 | 4 | 1 | 4 | 0.25 | 2 | 0 | NA | NA | 10 | 0.5 | NA | NA | NA | NA | 1 | 0 | 2 | 1 | 3 | 0.6667 |
| 9_58064929 | 6 | 0 | 4 | 0 | 1 | 1 | NA | NA | 11 | 0.0909 | NA | NA | NA | NA | 1 | 0 | 2 | 0.5 | 3 | 0.3333 |

| | | | | | | | | | | | | | | | | | | | | |
|---|---|---|---|---|---|---|---|---|---|---|---|---|---|---|---|---|---|---|---|---|
| 9_5806511 | 5 | 0.6 | 2 | 0.5 | 2 | 0 | NA | NA | 9 | 0.4444 | NA | NA | NA | NA | 1 | 1 | 2 | 0.5 | 3 | 0.6667 |
| 9_5806540 | 5 | 0.8 | 1 | 1 | 2 | 0.5 | NA | NA | 8 | 0.75 | NA | NA | NA | NA | NA | NA | 2 | 1 | 2 | 1 |
| 9_5806543 | 5 | 0.6 | 2 | 1 | 1 | 1 | NA | NA | 8 | 0.75 | NA | NA | NA | NA | 1 | 0 | 2 | 1 | 3 | 0.6667 |
| 9_5810412 | 8 | 0.875 | 6 | 0.8333 | 6 | 1 | 5 | 0.8 | 25 | 0.88 | 5 | 0.6 | 6 | 0.8333 | 3 | 0.6667 | 6 | 1 | 20 | 0.8 |
| 9_5810421 | 8 | 0.125 | 5 | 0 | 5 | 0 | 5 | 0.2 | 23 | 0.087 | 5 | 0 | 6 | 0.3333 | 3 | 0 | 5 | 0.2 | 19 | 0.1579 |
| 9_5810456 | 6 | 0.5 | 3 | 0.6667 | 4 | 1 | 4 | 0.25 | 17 | 0.5882 | 5 | 0.8 | 5 | 1 | 1 | 1 | 5 | 0.8 | 16 | 0.875 |
| 9_5810459 | 6 | 0.6667 | 5 | 1 | 5 | 1 | 4 | 0.75 | 20 | 0.85 | 4 | 0.75 | 4 | 1 | 2 | 0.5 | 5 | 0.8 | 15 | 0.8 |
| 9_5810467 | 7 | 0.1429 | 4 | 0.5 | 3 | 0.3333 | 5 | 0.2 | 19 | 0.2632 | 5 | 0.2 | 6 | 0 | 2 | 0 | 5 | 0 | 18 | 0.0556 |
| 9_5810468 | 4 | 1 | 4 | 1 | 1 | 1 | 5 | 0.6 | 14 | 0.8571 | 3 | 0.6667 | 4 | 1 | NA | NA | 4 | 1 | 11 | 0.9091 |
| 9_5810493 | 6 | 1 | 5 | 1 | 2 | 1 | 6 | 0.8333 | 19 | 0.9474 | 4 | 1 | 6 | 1 | 3 | 1 | 3 | 1 | 16 | 1 |
| 9_5810495 | 7 | 1 | 5 | 1 | 2 | 1 | 6 | 1 | 20 | 1 | 3 | 1 | 4 | 1 | 2 | 1 | 4 | 1 | 13 | 1 |
| 9_5810519 | 3 | 0 | 5 | 0.4 | 2 | 0.5 | 3 | 0 | 13 | 0.2308 | 4 | 0.25 | 5 | 0.2 | 1 | 0 | 1 | 0 | 11 | 0.1818 |
| 9_5810520 | 5 | 0 | 5 | 0 | 2 | 0 | 5 | 0.2 | 17 | 0.0588 | 2 | 0 | 6 | 0.1667 | 2 | 0 | 3 | 0 | 13 | 0.0769 |
| 9_5810535 | 4 | 1 | 5 | 0.8 | 2 | 1 | 6 | 0.6667 | 17 | 0.8235 | 4 | 0.75 | 4 | 0.5 | 2 | 0.5 | 3 | 1 | 13 | 0.6923 |
| 9_5810550 | 5 | 1 | 3 | 1 | 2 | 1 | 4 | 1 | 14 | 1 | 3 | 1 | 2 | 1 | 2 | 1 | 3 | 1 | 10 | 1 |
| 9_5810587 | 3 | 0.3333 | 4 | 0.5 | 1 | 1 | 3 | 0.3333 | 11 | 0.4545 | 2 | 0.5 | 1 | 0 | 2 | 0 | 3 | 1 | 8 | 0.5 |
| 9_5810609 | 2 | 0.5 | 3 | 0 | 1 | 0 | 3 | 0 | 9 | 0.1111 | 2 | 0 | 2 | 0 | 1 | 1 | 2 | 0 | 7 | 0.1429 |
| 9_5810632 | 2 | 0 | 1 | 0 | 1 | 1 | 4 | 0.25 | 8 | 0.25 | 2 | 0 | 2 | 0 | 1 | 0 | 2 | 0.5 | 7 | 0.1429 |
| 9_5810633 | 3 | 0.3333 | 1 | 0 | 1 | 0 | 4 | 0.25 | 9 | 0.2222 | 2 | 1 | 2 | 0.5 | NA | NA | 2 | 0 | 6 | 0.5 |
| 9_5810711 | NA | NA | 1 | 1 | NA | NA | 4 | 0.75 | 5 | 0.8 | 2 | 0 | 1 | 1 | NA | NA | 1 | 0 | 4 | 0.25 |
| 9_5810715 | NA | NA | 1 | 1 | NA | NA | 4 | 0.75 | 5 | 0.8 | 1 | 0 | 1 | 1 | NA | NA | 2 | 0.5 | 4 | 0.5 |
| 9_5810871 | NA | NA | NA | NA | NA | NA | 1 | 1 | 1 | 1 | 1 | 0 | 1 | 1 | NA | NA | NA | NA | 2 | 0.5 |
| 9_5812579 | NA | NA | NA | NA | 1 | 1 | 1 | 1 | 2 | 1 | NA | NA | NA | NA | 3 | 1 | NA | NA | 3 | 1 |
| 9_5812606 | 3 | 0.6667 | 1 | 0 | 2 | 0.5 | 1 | 0 | 7 | 0.4286 | NA | NA | NA | NA | 5 | 0.4 | 1 | 1 | 6 | 0.5 |
| 9_5812606 | 2 | 0 | 1 | 0 | 2 | 0.5 | 1 | 0 | 6 | 0.1667 | NA | NA | NA | NA | 6 | 0.1667 | 1 | 1 | 7 | 0.2857 |
| 9_5812715 | NA | NA | 1 | 1 | 1 | 1 | NA | NA | 2 | 1 | NA | NA | NA | NA | 6 | 0.8333 | 1 | 1 | 7 | 0.8571 |
| 9_5812743 | 1 | 1 | 1 | 1 | 2 | 1 | 1 | 1 | 5 | 1 | NA | NA | NA | NA | 5 | 0.8 | 1 | 1 | 6 | 0.8333 |
| 9_5812746 | 1 | 1 | 1 | 0 | 2 | 1 | 1 | 1 | 5 | 0.8 | NA | NA | NA | NA | 5 | 1 | 1 | 1 | 6 | 1 |
| 9_5812754 | NA | NA | 1 | 1 | 2 | 0 | 1 | 0 | 4 | 0.25 | NA | NA | NA | NA | 5 | 0.4 | 1 | 0 | 6 | 0.3333 |
| 9_5812761 | NA | NA | 1 | 1 | 2 | 0.5 | 1 | 1 | 4 | 0.75 | NA | NA | NA | NA | 5 | 0.4 | 1 | 1 | 6 | 0.5 |
| 9_5812766 | NA | NA | NA | NA | 2 | 0.5 | 1 | 1 | 3 | 0.6667 | NA | NA | NA | NA | 5 | 0.2 | 1 | 1 | 6 | 0.3333 |
| 9_5814195 | 1 | 0 | NA | NA | 2 | 0.5 | NA | NA | 3 | 0.3333 | NA | NA | NA | NA | 3 | 0.6667 | NA | NA | 3 | 0.6667 |
| 9_5814202 | 1 | 0 | NA | NA | 2 | 0.5 | NA | NA | 3 | 0.3333 | NA | NA | NA | NA | 3 | 0.6667 | NA | NA | 3 | 0.6667 |
| 9_5814208 | 1 | 1 | NA | NA | NA | NA | NA | NA | 1 | 1 | NA | NA | NA | NA | 3 | 0.6667 | NA | NA | 3 | 0.6667 |
| 9_5815465 | 1 | 0 | NA | NA | 2 | 0.5 | NA | NA | 3 | 0.3333 | NA | NA | NA | NA | 3 | 0.3333 | NA | NA | 3 | 0.3333 |
| 9_5816899 | 5 | 0 | 1 | 0 | 7 | 0.1429 | 4 | 0 | 17 | 0.0588 | 8 | 0.25 | 8 | 0 | 7 | 0.4286 | 1 | 0 | 24 | 0.2083 |
| 9_5816902 | 2 | 1 | NA | NA | 5 | 0.6 | 1 | 1 | 8 | 0.75 | 3 | 0.6667 | 4 | 0.5 | 3 | 0.6667 | NA | NA | 10 | 0.6 |
| 9_5816914 | 6 | 0.1667 | 1 | 0 | 7 | 0.2857 | 3 | 0 | 17 | 0.1765 | 7 | 0.5714 | 8 | 0.375 | 8 | 0.625 | 1 | 0 | 24 | 0.5 |
| 9_5816915 | 6 | 0.3333 | 1 | 0 | 7 | 0.1429 | 3 | 0.6667 | 17 | 0.2941 | 8 | 0.25 | 8 | 0.125 | 8 | 0.375 | 1 | 1 | 25 | 0.28 |
| 9_5816921 | 4 | 0.25 | 1 | 0 | 5 | 0.2 | 4 | 0 | 14 | 0.1429 | 6 | 0 | 6 | 0.3333 | 8 | 0.25 | 1 | 0 | 21 | 0.1905 |
| 9_5816926 | 5 | 0 | NA | NA | 6 | 0.3333 | 4 | 0.25 | 15 | 0.2 | 8 | 0.25 | 7 | 0.1429 | 7 | 0.1429 | 1 | 0 | 23 | 0.1739 |
| 9_5816933 | 6 | 1 | 1 | 1 | 7 | 0.7143 | 4 | 1 | 18 | 0.8889 | 9 | 1 | 8 | 1 | 8 | 0.875 | 1 | 1 | 26 | 0.9615 |
| 9_5816938 | 4 | 0.25 | NA | NA | 7 | 0.1429 | 2 | 0 | 13 | 0.1538 | 8 | 0.25 | 9 | 0.2222 | 7 | 0 | 1 | 0 | 25 | 0.16 |
| 9_5816939 | 6 | 0 | 1 | 0 | 6 | 0.3333 | 4 | 0.25 | 17 | 0.1765 | 7 | 0.2857 | 8 | 0.125 | 6 | 0 | 1 | 1 | 22 | 0.1818 |
| 9_5816951 | 6 | 0.3333 | 1 | 0 | 8 | 0.375 | 2 | 0.5 | 17 | 0.3529 | 9 | 0.2222 | 8 | 0.25 | 5 | 0.4 | 1 | 1 | 23 | 0.3043 |
| 9_5816960 | 5 | 0 | 1 | 0 | 8 | 0.25 | 4 | 0.25 | 18 | 0.1667 | 9 | 0.4444 | 7 | 0 | 8 | 0.25 | 2 | 0 | 26 | 0.2308 |
| 9_5816963 | 4 | 0.75 | 1 | 1 | 4 | 1 | 4 | 0.75 | 13 | 0.8462 | 8 | 0.625 | 7 | 1 | 7 | 0.4286 | 2 | 1 | 24 | 0.7083 |
| 9_5816966 | 7 | 0.2857 | 1 | 0 | 8 | 0.125 | 4 | 0 | 20 | 0.15 | 9 | 0.1111 | 8 | 0.25 | 8 | 0 | 2 | 0 | 27 | 0.1111 |
| 9_5816978 | 2 | 0.5 | NA | NA | 6 | 0.5 | 3 | 0.3333 | 11 | 0.4545 | 8 | 0.5 | 6 | 0.5 | 6 | 0.6667 | 1 | 1 | 21 | 0.5714 |
| 9_5819604 | 2 | 1 | NA | NA | NA | NA | 1 | 1 | 3 | 1 | NA | NA | NA | NA | 2 | 1 | NA | NA | 2 | 1 |
| 9_5821946 | 1 | 1 | NA | NA | NA | NA | 2 | 1 | 3 | 1 | NA | NA | NA | NA | 1 | 1 | NA | NA | 1 | 1 |
| 9_5821957 | 2 | 0.5 | NA | NA | NA | NA | 2 | 0.5 | 4 | 0.5 | NA | NA | NA | NA | 1 | 1 | NA | NA | 1 | 1 |
| 9_5821976 | 2 | 1 | NA | NA | NA | NA | NA | NA | 2 | 1 | NA | NA | NA | NA | 1 | 1 | NA | NA | 1 | 1 |
| 9_5821984 | 2 | 0.5 | NA | NA | NA | NA | NA | NA | 2 | 0.5 | NA | NA | NA | NA | 1 | 1 | NA | NA | 1 | 1 |
| 9_5822028 | 2 | 1 | NA | NA | NA | NA | NA | NA | 2 | 1 | NA | NA | NA | NA | 1 | 1 | NA | NA | 1 | 1 |
| 9_5839588 | 15 | 0.3333 | 5 | 0.2 | 16 | 0.3125 | 6 | 0.3333 | 42 | 0.3095 | 12 | 0.25 | 11 | 0.3636 | 10 | 0.3 | 6 | 0.3333 | 39 | 0.3077 |
| 9_5839592 | 12 | 0.3333 | 6 | 0.3333 | 11 | 0.4545 | 6 | 0.1667 | 35 | 0.3429 | 8 | 0.125 | 12 | 0.3333 | 6 | 0.1667 | 4 | 0.25 | 30 | 0.2333 |
| 9_5839597 | 15 | 0.6667 | 6 | 0.6667 | 15 | 0.5333 | 6 | 0.6667 | 42 | 0.619 | 12 | 0.8333 | 14 | 0.5 | 11 | 0.9091 | 5 | 1 | 42 | 0.7619 |
| 9_5839600 | 16 | 0.3125 | 6 | 0.1667 | 13 | 0.1538 | 5 | 0.4 | 40 | 0.25 | 11 | 0.2727 | 13 | 0.5385 | 12 | 0.1667 | 6 | 0 | 42 | 0.2857 |
| 9_5839605 | 7 | 0.1429 | 4 | 0 | 11 | 0.1818 | 4 | 0.25 | 26 | 0.1538 | 6 | 0.1667 | 9 | 0.2222 | 5 | 0.2 | 4 | 0 | 24 | 0.1667 |
| 9_5839609 | 14 | 0.8571 | 4 | 1 | 15 | 0.8667 | 5 | 0.8 | 38 | 0.8684 | 13 | 0.8462 | 11 | 0.7273 | 10 | 0.9 | 6 | 1 | 40 | 0.85 |
| 9_5839611 | 16 | 0.125 | 3 | 0.3333 | 14 | 0.0714 | 7 | 0.1429 | 40 | 0.125 | 10 | 0.1 | 13 | 0.0769 | 13 | 0.0769 | 4 | 0 | 40 | 0.075 |
| 9_5839706 | 22 | 0.6818 | 8 | 0.5 | 21 | 0.5238 | 9 | 0.4444 | 60 | 0.5667 | 19 | 0.4737 | 19 | 0.4737 | 13 | 0.4615 | 10 | 0.5 | 61 | 0.4754 |
| 9_5839708 | 16 | 0.125 | 8 | 0.125 | 16 | 0.0625 | 9 | 0.1111 | 49 | 0.102 | 17 | 0.1176 | 17 | 0.1176 | 13 | 0 | 4 | 0.25 | 51 | 0.098 |
| 9_5839730 | 20 | 0.85 | 9 | 0.7778 | 20 | 0.85 | 10 | 0.9 | 59 | 0.8475 | 20 | 0.85 | 17 | 0.7647 | 15 | 0.7333 | 10 | 0.8 | 62 | 0.7903 |

| | | | | | | | | | | | | | | | | | | | | |
|---|---|---|---|---|---|---|---|---|---|---|---|---|---|---|---|---|---|---|---|---|
| 9_58397524 | 13 | 0.3846 | 5 | 0.4 | 15 | 0.4667 | 8 | 0.5 | 41 | 0.439 | 12 | 0.6667 | 12 | 0.4167 | 8 | 0.625 | 10 | 0.5 | 42 | 0.5476 |
| 9_58397542 | 26 | 0.1154 | 11 | 0.1818 | 22 | 0.1364 | 11 | 0.2727 | 70 | 0.1571 | 23 | 0.1739 | 19 | 0.2632 | 15 | 0.2 | 15 | 0.4 | 72 | 0.25 |
| 9_58397645 | 23 | 0.0435 | 11 | 0.0909 | 19 | 0.1053 | 9 | 0.2222 | 62 | 0.0968 | 21 | 0 | 11 | 0 | 14 | 0.0714 | 13 | 0.1538 | 59 | 0.0508 |
| 9_58406279 | 23 | 0.0435 | 8 | 0 | 21 | 0.0952 | 11 | 0.0909 | 63 | 0.0635 | 18 | 0.0556 | 17 | 0.1765 | 14 | 0.0714 | 16 | 0.125 | 65 | 0.1077 |
| 9_58406292 | 16 | 0.125 | 7 | 0.5714 | 15 | 0.4 | 9 | 0.1111 | 47 | 0.2766 | 16 | 0.25 | 14 | 0.2857 | 11 | 0.0909 | 11 | 0.2727 | 52 | 0.2308 |
| 9_58406387 | 23 | 1 | 11 | 1 | 20 | 0.85 | 12 | 1 | 66 | 0.9545 | 20 | 0.95 | 19 | 0.9474 | 15 | 0.9333 | 16 | 0.9375 | 70 | 0.9429 |
| 9_58406414 | 20 | 0.15 | 11 | 0.1818 | 23 | 0.087 | 11 | 0 | 65 | 0.1077 | 22 | 0.0455 | 18 | 0.0556 | 13 | 0.1538 | 15 | 0.2667 | 68 | 0.1176 |
| 9_58406459 | 18 | 0.6667 | 12 | 0.8333 | 18 | 0.8333 | 9 | 0.8889 | 57 | 0.7895 | 19 | 0.3684 | 16 | 0.75 | 9 | 0.8889 | 12 | 0.5833 | 56 | 0.6071 |
| 9_58406564 | 22 | 0.4545 | 12 | 0.4167 | 23 | 0.4348 | 11 | 0.3636 | 68 | 0.4265 | 19 | 0.2105 | 19 | 0.3158 | 14 | 0.2143 | 15 | 0.4667 | 67 | 0.2985 |
| 9_58406623 | 21 | 1 | 14 | 1 | 23 | 1 | 12 | 1 | 70 | 1 | 23 | 1 | 17 | 1 | 16 | 1 | 14 | 1 | 70 | 1 |
| 9_58406657 | 23 | 0.0435 | 13 | 0.2308 | 25 | 0.16 | 11 | 0 | 72 | 0.1111 | 22 | 0.2273 | 21 | 0.1429 | 16 | 0.1875 | 17 | 0.2353 | 76 | 0.1974 |
| 9_58406882 | 16 | 0.125 | 12 | 0.25 | 24 | 0.2083 | 10 | 0.3 | 62 | 0.2097 | 17 | 0.1765 | 14 | 0.5 | 11 | 0.0909 | 13 | 0.5385 | 55 | 0.3273 |
| 9_58406896 | 19 | 0.3158 | 13 | 0.0769 | 22 | 0.4091 | 12 | 0.3333 | 66 | 0.303 | 21 | 0.4286 | 17 | 0.4118 | 14 | 0.3571 | 18 | 0.3333 | 70 | 0.3857 |
| 9_58406906 | 22 | 0.0909 | 16 | 0.0625 | 29 | 0 | 12 | 0.0833 | 79 | 0.0506 | 23 | 0.087 | 21 | 0.0476 | 13 | 0 | 18 | 0.2778 | 75 | 0.1067 |
| 9_58406920 | 21 | 0 | 13 | 0 | 22 | 0.1364 | 7 | 0.1429 | 63 | 0.0635 | 18 | 0.1111 | 18 | 0.1667 | 13 | 0 | 16 | 0 | 65 | 0.0769 |
| 9_58406966 | 18 | 0.3889 | 8 | 0.75 | 21 | 0.7619 | 7 | 0.7143 | 54 | 0.6296 | 15 | 0.6 | 11 | 0.7273 | 10 | 0.6 | 15 | 0.8 | 51 | 0.6863 |
| 9_58407316 | 7 | 0 | 3 | 0.3333 | 2 | 0 | NA | NA | 12 | 0.0833 | 3 | 0.6667 | 1 | 1 | 3 | 0.3333 | 2 | 0.5 | 9 | 0.5556 |
| 9_58408450 | 8 | 0.25 | 4 | 0 | 3 | 0.3333 | 1 | 0 | 16 | 0.1875 | 4 | 0.25 | NA | NA | 2 | 0 | 2 | 0 | 8 | 0.125 |
| 9_58408764 | 9 | 0.1111 | 5 | 0.4 | 3 | 0 | NA | NA | 17 | 0.1765 | 7 | 0.1429 | 1 | 0 | 4 | 0.25 | 2 | 0 | 14 | 0.1429 |
| 9_58408798 | 8 | 0.125 | 5 | 0.2 | 2 | 1 | 1 | 0 | 16 | 0.25 | 7 | 0.1429 | 2 | 0 | 4 | 0.25 | 2 | 0.5 | 15 | 0.2 |
| 9_58408983 | 9 | 0.1111 | 6 | 0.3333 | 2 | 0 | 1 | 0 | 18 | 0.1667 | 7 | 0.4286 | 2 | 0 | 4 | 0.25 | 2 | 0 | 15 | 0.2667 |
| 9_58409140 | 9 | 0.2222 | 5 | 0 | 3 | 0.6667 | 1 | 0 | 18 | 0.2222 | 7 | 0 | 2 | 0.5 | 4 | 0.75 | 2 | 0 | 15 | 0.2667 |
| 9_58411091 | 2 | 0.5 | 2 | 0 | NA | NA | NA | NA | 4 | 0.25 | 1 | 0 | 1 | 1 | NA | NA | NA | NA | 2 | 0.5 |
| 9_58431631 | 3 | 0 | 1 | 0 | 1 | 0 | 3 | 0.3333 | 8 | 0.125 | 1 | 1 | 2 | 1 | 1 | 1 | 2 | 0 | 6 | 0.6667 |
| 9_58431652 | 3 | 0.6667 | 1 | 0 | 1 | 0 | 3 | 0.6667 | 8 | 0.5 | 1 | 0 | 3 | 0.6667 | 1 | 1 | 3 | 0 | 8 | 0.375 |
| 9_58431686 | 3 | 0.3333 | 1 | 0 | 1 | 0 | 4 | 0.5 | 9 | 0.3333 | 1 | 0 | 3 | 0.3333 | 2 | 0.5 | 3 | 0.3333 | 9 | 0.3333 |
| 9_58431739 | 3 | 0 | 1 | 0 | 1 | 0 | 3 | 0.3333 | 8 | 0.125 | 1 | 0 | 3 | 0.3333 | 2 | 0.5 | 3 | 0 | 9 | 0.2222 |
| 9_58431762 | 3 | 1 | 1 | 1 | NA | NA | 3 | 1 | 7 | 1 | NA | NA | 1 | 1 | 1 | 1 | 3 | 1 | 5 | 1 |
| 9_58431834 | 2 | 0.5 | NA | NA | NA | NA | 2 | 0.5 | 4 | 0.5 | 1 | 1 | 3 | 0 | 2 | 0.5 | 2 | 0 | 8 | 0.25 |
| 9_58431855 | 3 | 0.3333 | NA | NA | 1 | 0 | 3 | 0.6667 | 7 | 0.4286 | 1 | 0 | 2 | 0.5 | 2 | 0 | 2 | 0.5 | 7 | 0.2857 |
| 9_58431956 | 3 | 0.3333 | NA | NA | 1 | 0 | 3 | 0.6667 | 7 | 0.4286 | NA | NA | 1 | 0 | 1 | 1 | 3 | 0.3333 | 5 | 0.4 |
| 9_58432021 | 3 | 1 | NA | NA | 1 | 1 | 2 | 1 | 6 | 1 | 1 | 1 | 3 | 1 | 1 | 1 | 3 | 1 | 8 | 1 |
| 9_58432054 | 3 | 1 | NA | NA | 1 | 1 | 2 | 0 | 6 | 0.6667 | 1 | 1 | 3 | 1 | 1 | 1 | 3 | 1 | 8 | 1 |
| 9_58432091 | 3 | 0 | NA | NA | 1 | 0 | 3 | 0.3333 | 7 | 0.1429 | 1 | 0 | 3 | 0 | 2 | 1 | 3 | 0 | 9 | 0.2222 |
| 9_58432183 | 2 | 1 | NA | NA | 1 | 1 | 2 | 0.5 | 5 | 0.8 | 1 | 0 | 3 | 0 | 1 | 0 | 3 | 0.3333 | 8 | 0.125 |
| 9_58432201 | 2 | 1 | NA | NA | 1 | 0 | 1 | 0 | 4 | 0.5 | 1 | 1 | 3 | 0.6667 | 1 | 0 | 3 | 0 | 8 | 0.375 |
| 9_58432277 | 2 | 0.5 | NA | NA | 1 | 0 | 3 | 0.6667 | 6 | 0.5 | 1 | 0 | 3 | 0.6667 | 1 | 0 | 3 | 0.6667 | 8 | 0.5 |
| 9_58432660 | 1 | 1 | NA | NA | 1 | 0 | 3 | 0 | 5 | 0.2 | 2 | 0.5 | 2 | 0 | 1 | 0 | 3 | 0 | 8 | 0.125 |
| 9_58432720 | 1 | 1 | NA | NA | 1 | 1 | 2 | 1 | 4 | 1 | 1 | 0 | 2 | 0.5 | 1 | 1 | 3 | 1 | 7 | 0.7143 |
| 9_58433230 | 1 | 0 | NA | NA | 1 | 1 | 1 | 0 | 3 | 0.3333 | 1 | 1 | 2 | 0 | 1 | 0 | 3 | 0 | 7 | 0.1429 |
| 9_58433238 | NA | NA | NA | NA | 1 | 0 | 2 | 1 | 3 | 0.6667 | 1 | 1 | 2 | 1 | 1 | 1 | 3 | 1 | 7 | 1 |
| 9_58433285 | NA | NA | NA | NA | 1 | 1 | 2 | 1 | 3 | 1 | 1 | 1 | 2 | 1 | 1 | 0 | 3 | 1 | 7 | 0.8571 |
| 9_58433315 | NA | NA | NA | NA | 1 | 0 | 2 | 0.5 | 3 | 0.3333 | NA | NA | 2 | 0 | 1 | 0 | 3 | 0.3333 | 6 | 0.1667 |
| 9_58433384 | NA | NA | NA | NA | 1 | 0 | 2 | 1 | 3 | 0.6667 | 1 | 1 | 2 | 0 | 1 | 1 | 3 | 0.3333 | 7 | 0.4286 |
| 9_58433463 | NA | NA | NA | NA | 1 | 1 | 1 | 1 | 2 | 1 | 1 | 1 | 2 | 1 | 1 | 1 | 2 | 0.5 | 6 | 0.8333 |
| 9_58444587 | NA | NA | NA | NA | 1 | 1 | 1 | 1 | 2 | 1 | NA | NA | 2 | 0 | 1 | 0 | 1 | 1 | 4 | 0.25 |
| 9_58526183 | 25 | 0.44 | 42 | 0.4762 | 34 | 0.5 | 45 | 0.4889 | 146 | 0.4795 | 23 | 0.4348 | 47 | 0.617 | 38 | 0.6053 | 46 | 0.4783 | 154 | 0.5455 |
| 9_58526191 | 28 | 0.3929 | 42 | 0.5238 | 28 | 0.4286 | 37 | 0.5405 | 135 | 0.4815 | 22 | 0.3636 | 43 | 0.3256 | 36 | 0.4167 | 39 | 0.3077 | 140 | 0.35 |
| 9_58559676 | 75 | 0.0533 | 103 | 0.0485 | 89 | 0.0674 | 94 | 0.0638 | 361 | 0.0582 | 54 | 0 | 85 | 0.1176 | 92 | 0.0326 | 75 | 0.0533 | 306 | 0.0556 |
| 9_58559694 | 81 | 0.0741 | 108 | 0.0463 | 91 | 0.033 | 96 | 0.0833 | 376 | 0.0585 | 58 | 0.0345 | 90 | 0.0333 | 101 | 0.0792 | 79 | 0.0759 | 328 | 0.0579 |
| 9_58782625 | 1 | 0 | 2 | 0.5 | 8 | 0.25 | 1 | 0 | 12 | 0.25 | 6 | 0.3333 | 3 | 0.3333 | 4 | 0.5 | 5 | 0 | 18 | 0.2778 |
| 9_58782684 | 2 | 0 | 2 | 0.5 | 6 | 0 | 2 | 0 | 12 | 0.0833 | 6 | 0 | 4 | 0 | 5 | 0.2 | 4 | 0 | 19 | 0.0526 |
| 9_58783821 | 2 | 0.5 | 1 | 0 | 6 | 0.1667 | 2 | 0 | 11 | 0.1818 | 3 | 0 | 4 | 0 | 6 | 0.6667 | 4 | 0.5 | 17 | 0.3529 |
| 9_58783992 | 6 | 0.3333 | 4 | 0.25 | 8 | 0.125 | 2 | 0 | 20 | 0.2 | 1 | 1 | 4 | 0.5 | 4 | 0.25 | 8 | 0.375 | 17 | 0.4118 |
| 9_58784097 | 5 | 0.4 | 5 | 0.8 | 10 | 0.9 | 4 | 1 | 24 | 0.7917 | 2 | 1 | 6 | 0.5 | 4 | 0.75 | 7 | 0.4286 | 19 | 0.5789 |
| 9_58784104 | 6 | 0.1667 | 6 | 0 | 9 | 0 | 4 | 0.25 | 25 | 0.08 | 3 | 0.3333 | 5 | 0 | 4 | 0 | 7 | 0.1429 | 19 | 0.1053 |
| 9_58784151 | 5 | 0.2 | 3 | 0 | 7 | 0 | 2 | 0 | 17 | 0.0588 | 3 | 0 | 4 | 0 | 4 | 0.25 | 6 | 0 | 17 | 0.0588 |
| 9_58784200 | 6 | 0.3333 | 6 | 0.3333 | 8 | 0.5 | 3 | 0.6667 | 23 | 0.4348 | 3 | 0.3333 | 6 | 0.1667 | 4 | 1 | 7 | 0.2857 | 20 | 0.4 |
| 9_58784323 | 6 | 1 | 6 | 1 | 7 | 0.8571 | 3 | 1 | 22 | 0.9545 | 3 | 1 | 4 | 1 | 4 | 1 | 7 | 1 | 18 | 1 |
| 9_58784467 | 5 | 0.2 | 4 | 0.5 | 7 | 0.5714 | 3 | 1 | 19 | 0.5263 | 2 | 1 | 5 | 0.4 | 3 | 0.6667 | 5 | 0.6 | 15 | 0.6 |
| 9_58784508 | 5 | 0.6 | 4 | 0.5 | 9 | 0.4444 | 3 | 0.6667 | 21 | 0.5238 | 3 | 0.6667 | 4 | 1 | 4 | 0.5 | 5 | 0.4 | 16 | 0.625 |
| 9_58784542 | 4 | 0.25 | 5 | 0 | 8 | 0.125 | 3 | 0 | 20 | 0.1 | 2 | 0 | 4 | 0 | 4 | 0.25 | 5 | 0.2 | 15 | 0.1333 |
| 9_58784582 | 5 | 0 | 5 | 0 | 7 | 0.7143 | 3 | 0.3333 | 20 | 0.3 | 2 | 0.5 | 5 | 0.4 | 4 | 0.25 | 7 | 0.2857 | 18 | 0.3333 |
| 9_58784593 | 5 | 0.4 | 5 | 0.2 | 8 | 0.75 | 3 | 0.3333 | 21 | 0.4762 | 2 | 0 | 3 | 1 | 4 | 0.5 | 4 | 0.25 | 13 | 0.4615 |
| 9_58796407 | 4 | 0 | 4 | 0 | 7 | 0.1429 | 1 | 0 | 16 | 0.0625 | 2 | 0 | 4 | 0.25 | 2 | 0 | 6 | 0 | 14 | 0.0714 |

| | | | | | | | | | | | | | | | | | | | | |
|---|---|---|---|---|---|---|---|---|---|---|---|---|---|---|---|---|---|---|---|---|
| 9_58892957 | 2 | 0 | 1 | 1 | 1 | 0 | NA | NA | 4 | 0.25 | 1 | 0 | 2 | 0 | 1 | 1 | 1 | 0 | 5 | 0.2 |
| 9_58897651 | 3 | 0.6667 | 1 | 0 | NA | NA | NA | NA | 4 | 0.5 | 1 | 0 | 1 | 1 | 1 | 0 | 2 | 0.5 | 5 | 0.4 |
| 9_59222256 | 4 | 0.75 | 5 | 0.8 | 1 | 1 | 1 | 1 | 11 | 0.8182 | 3 | 1 | 2 | 0.5 | 2 | 1 | 2 | 1 | 9 | 0.8889 |
| 9_59222308 | 3 | 1 | 5 | 1 | 1 | 1 | 1 | 1 | 10 | 1 | 3 | 0.6667 | 2 | 0.5 | 2 | 1 | 4 | 1 | 11 | 0.8182 |
| 9_59222336 | 3 | 0 | 4 | 0.25 | 1 | 0 | 1 | 1 | 9 | 0.2222 | 2 | 0.5 | 2 | 0 | 1 | 0 | 2 | 0 | 7 | 0.1429 |
| 9_59222414 | 4 | 0 | 5 | 0.2 | NA | NA | 1 | 1 | 10 | 0.2 | 2 | 0 | 2 | 0.5 | 2 | 0 | 4 | 0.5 | 10 | 0.3 |
| 9_59222500 | 5 | 1 | 1 | 1 | 1 | 1 | 1 | 1 | 8 | 1 | 1 | 1 | 1 | 1 | 2 | 1 | 1 | 1 | 5 | 1 |
| 9_59222629 | 5 | 0.2 | 5 | 0.2 | 1 | 0 | 2 | 0.5 | 13 | 0.2308 | 3 | 0.3333 | 2 | 0 | 2 | 0.5 | 4 | 0.25 | 11 | 0.2727 |
| 9_59222650 | 3 | 0.6667 | 4 | 0.5 | 1 | 0 | 1 | 1 | 9 | 0.5556 | 3 | 0.6667 | 2 | 0.5 | 2 | 0.5 | 3 | 0.3333 | 10 | 0.5 |
| 9_59222739 | 5 | 0.4 | 5 | 0.4 | 1 | 0 | 2 | 0 | 13 | 0.3077 | 2 | 0 | 2 | 0.5 | 2 | 0 | 1 | 0 | 7 | 0.1429 |
| 9_59222780 | 5 | 1 | 6 | 1 | 1 | 0 | 2 | 1 | 14 | 0.9286 | 3 | 1 | 2 | 0.5 | 2 | 1 | 4 | 1 | 11 | 0.9091 |
| 9_59222853 | 5 | 0.2 | 5 | 0.4 | 1 | 0 | 2 | 0.5 | 13 | 0.3077 | 3 | 0.3333 | 2 | 0.5 | 2 | 0 | 4 | 0 | 11 | 0.1818 |
| 9_59222990 | 5 | 0.8 | 6 | 0.3333 | 1 | 0 | 2 | 1 | 14 | 0.5714 | 3 | 0.6667 | 2 | 0 | 3 | 0 | 3 | 0.6667 | 11 | 0.3636 |
| 9_59223087 | 4 | 1 | 6 | 0.8333 | NA | NA | 2 | 0.5 | 12 | 0.8333 | 3 | 0.6667 | 2 | 0.5 | 4 | 0.25 | 4 | 0.75 | 13 | 0.5385 |
| 9_59223173 | 3 | 1 | 4 | 1 | 1 | 1 | 2 | 1 | 10 | 1 | 2 | 1 | 2 | 1 | 4 | 1 | 4 | 0.5 | 12 | 0.8333 |
| 9_59223254 | 5 | 1 | 6 | 0.8333 | 1 | 0 | 2 | 0.5 | 14 | 0.7857 | 2 | 1 | 2 | 0.5 | 4 | 0.75 | 4 | 0.75 | 12 | 0.75 |
| 9_59229628 | 5 | 0 | 3 | 0 | 5 | 0.4 | 8 | 0.125 | 21 | 0.1429 | 6 | 0.5 | 9 | 0.3333 | 4 | 0.25 | 4 | 0 | 23 | 0.3043 |
| 9_59229778 | 4 | 0.25 | 3 | 0.3333 | 1 | 0 | 8 | 0 | 16 | 0.125 | 5 | 0.4 | 10 | 0.5 | 4 | 0.5 | 4 | 0.75 | 23 | 0.5217 |
| 9_59229848 | 6 | 0.1667 | 3 | 0.6667 | 2 | 0.5 | 9 | 0.3333 | 20 | 0.35 | 5 | 0.4 | 11 | 0.7273 | 4 | 0.5 | 5 | 0.2 | 25 | 0.52 |
| 9_59300555 | 9 | 0.8889 | 5 | 1 | 2 | 1 | 6 | 0.8333 | 22 | 0.9091 | 3 | 0.6667 | 9 | 0.7778 | 3 | 1 | 5 | 0.8 | 20 | 0.8 |
| 9_59300575 | 11 | 0.0909 | 6 | 0 | 3 | 0 | 7 | 0.1429 | 27 | 0.0741 | 3 | 0 | 9 | 0 | 4 | 0.5 | 5 | 0.2 | 21 | 0.1429 |
| 9_59300581 | 10 | 0.5 | 4 | 0.75 | 3 | 0.3333 | 7 | 0.5714 | 24 | 0.5417 | 3 | 0.6667 | 9 | 0.8889 | 4 | 0.5 | 3 | 0.3333 | 19 | 0.6842 |
| 9_59393361 | 5 | 0 | 1 | 0 | 1 | 0 | 4 | 0.25 | 11 | 0.0909 | 1 | 1 | 3 | 0 | 2 | 0 | 1 | 0 | 7 | 0.1429 |
| 9_59393626 | 5 | 0.2 | 1 | 1 | 1 | 0 | 4 | 0.5 | 11 | 0.3636 | 1 | 1 | 2 | 0 | 1 | 1 | 2 | 0 | 6 | 0.3333 |
| 9_59426343 | 1 | 1 | NA | NA | NA | NA | NA | NA | 1 | 1 | 1 | 1 | NA | NA | NA | NA | NA | NA | 1 | 1 |
| 9_59513353 | 3 | 0.3333 | NA | NA | 1 | 0 | 4 | 0.25 | 8 | 0.25 | 1 | 1 | 4 | 0 | 4 | 0 | 3 | 0 | 12 | 0.0833 |
| 9_59513445 | 3 | 0.3333 | 1 | 0 | 2 | 0 | 4 | 0 | 10 | 0.1 | 1 | 0 | 4 | 0.25 | 6 | 0.3333 | 3 | 0.3333 | 14 | 0.2857 |
| 9_59513565 | 1 | 1 | 1 | 1 | 1 | 1 | 4 | 0.75 | 7 | 0.8571 | 1 | 1 | 6 | 0.8333 | 6 | 1 | 3 | 1 | 16 | 0.9375 |
| 9_59517700 | NA | NA | 2 | 0.5 | NA | NA | NA | NA | 2 | 0.5 | NA | NA | NA | NA | NA | NA | 1 | 1 | 1 | 1 |
| 9_59517716 | NA | NA | 2 | 0.5 | NA | NA | NA | NA | 2 | 0.5 | NA | NA | NA | NA | 1 | 1 | 1 | 1 | 2 | 1 |
| 9_59517785 | NA | NA | 1 | 1 | NA | NA | NA | NA | 1 | 1 | NA | NA | NA | NA | 1 | 1 | 1 | 1 | 2 | 1 |
| 9_59517961 | NA | NA | 2 | 1 | NA | NA | NA | NA | 2 | 1 | NA | NA | NA | NA | 1 | 0 | 1 | 1 | 2 | 0.5 |
| 9_59518062 | NA | NA | 1 | 1 | NA | NA | NA | NA | 1 | 1 | NA | NA | NA | NA | 1 | 0 | 1 | 1 | 2 | 0.5 |
| 9_59519404 | NA | NA | 2 | 0.5 | NA | NA | NA | NA | 2 | 0.5 | NA | NA | NA | NA | 1 | 0 | 1 | 1 | 2 | 0.5 |
| 9_59519741 | NA | NA | 2 | 1 | NA | NA | NA | NA | 2 | 1 | NA | NA | NA | NA | 1 | 1 | 1 | 1 | 2 | 1 |
| 9_59525888 | 13 | 0.0769 | 9 | 0.1111 | 8 | 0.25 | 9 | 0 | 39 | 0.1026 | 4 | 0 | 14 | 0.1429 | 14 | 0.0714 | 8 | 0.125 | 40 | 0.1 |
| 9_59557289 | 21 | 0.1905 | 24 | 0.125 | 15 | 0.0667 | 13 | 0.0769 | 73 | 0.1233 | 13 | 0.1538 | 19 | 0 | 19 | 0.1579 | 18 | 0.2222 | 69 | 0.1304 |
| 9_59557305 | 21 | 0.1429 | 19 | 0.1579 | 12 | 0 | 13 | 0.2308 | 65 | 0.1385 | 8 | 0 | 15 | 0.2 | 15 | 0.0667 | 12 | 0.25 | 50 | 0.14 |
| 9_59557369 | 18 | 0.2778 | 21 | 0.381 | 15 | 0.2 | 13 | 0.4615 | 67 | 0.3284 | 12 | 0.3333 | 19 | 0.2105 | 19 | 0.1579 | 15 | 0.5333 | 65 | 0.2923 |
| 9_59557563 | 6 | 0 | 4 | 0.25 | 6 | 0.6667 | 5 | 0 | 21 | 0.2381 | 3 | 0.3333 | 8 | 0.25 | 6 | 0.1667 | 7 | 0.1429 | 24 | 0.2083 |
| 9_59623590 | 1 | 1 | NA | NA | 1 | 0 | NA | NA | 2 | 0.5 | NA | NA | NA | NA | 2 | 0.5 | NA | NA | 2 | 0.5 |
| 9_59623704 | 1 | 1 | NA | NA | 1 | 1 | NA | NA | 2 | 1 | NA | NA | NA | NA | 2 | 0.5 | NA | NA | 2 | 0.5 |
| 9_59623722 | 1 | 0 | NA | NA | 1 | 1 | NA | NA | 2 | 0.5 | NA | NA | NA | NA | 2 | 0.5 | NA | NA | 2 | 0.5 |
| 9_59643793 | NA | NA | NA | NA | 1 | 1 | 1 | 1 | 2 | 1 | NA | NA | 1 | 1 | 1 | 1 | 3 | 1 | 5 | 1 |
| 9_59643799 | NA | NA | NA | NA | 1 | 1 | NA | NA | 1 | 1 | NA | NA | 1 | 0 | 1 | 1 | 2 | 1 | 4 | 0.75 |
| 9_59643848 | NA | NA | NA | NA | 1 | 1 | 1 | 1 | 2 | 1 | NA | NA | 1 | 1 | NA | NA | 2 | 1 | 3 | 1 |
| 9_59643880 | NA | NA | NA | NA | 1 | 1 | 1 | 1 | 2 | 1 | NA | NA | 1 | 1 | 1 | 0 | 2 | 0.5 | 4 | 0.5 |
| 9_59643921 | NA | NA | NA | NA | 1 | 0 | 1 | 1 | 2 | 0.5 | NA | NA | 1 | 0 | 1 | 1 | 1 | 0 | 3 | 0.3333 |
| 9_59644230 | NA | NA | NA | NA | 1 | 0 | 1 | 1 | 2 | 0.5 | NA | NA | NA | NA | 1 | 1 | NA | NA | 1 | 1 |
| 9_59644243 | NA | NA | NA | NA | 1 | 1 | 1 | 1 | 2 | 1 | NA | NA | 1 | 1 | 1 | 1 | NA | NA | 2 | 1 |
| 9_59644257 | NA | NA | NA | NA | 1 | 1 | 1 | 1 | 2 | 1 | NA | NA | 1 | 0 | 1 | 1 | NA | NA | 2 | 0.5 |
| 9_59777770 | 1 | 1 | NA | NA | 1 | 0 | 2 | 0 | 4 | 0.25 | 1 | 1 | NA | NA | 1 | 0 | 2 | 0.5 | 4 | 0.5 |
| 9_59777922 | 1 | 1 | NA | NA | 1 | 1 | 2 | 0.5 | 4 | 0.75 | 1 | 1 | NA | NA | 1 | 0 | 2 | 0 | 4 | 0.25 |
| 9_59777982 | 1 | 0 | NA | NA | 1 | 1 | 2 | 0 | 4 | 0.25 | 1 | 0 | NA | NA | 1 | 0 | 2 | 0.5 | 4 | 0.25 |
| 9_59778244 | NA | NA | 1 | 0 | NA | NA | 1 | 1 | 2 | 0.5 | 1 | 1 | NA | NA | NA | NA | 1 | 1 | 2 | 1 |
| 9_59778440 | 1 | 0 | 1 | 0 | 1 | 1 | 2 | 1 | 5 | 0.6 | 1 | 1 | 1 | 0 | 1 | 1 | 2 | 0.5 | 5 | 0.6 |
| 9_59778501 | 1 | 1 | 1 | 0 | 1 | 1 | 1 | 1 | 4 | 0.75 | 1 | 1 | 1 | 1 | 1 | 1 | 2 | 1 | 5 | 1 |
| 9_59778878 | 2 | 0.5 | NA | NA | 1 | 1 | 1 | 0 | 4 | 0.5 | NA | NA | NA | NA | 2 | 1 | 1 | 1 | 3 | 1 |
| 9_59779121 | 2 | 0.5 | 1 | 0 | NA | NA | 3 | 0 | 6 | 0.1667 | 1 | 0 | 1 | 1 | 2 | 0.5 | 2 | 1 | 6 | 0.6667 |
| 9_59779180 | 1 | 1 | 1 | 0 | 1 | 1 | 4 | 1 | 7 | 0.8571 | 1 | 1 | 1 | 1 | 1 | 1 | 2 | 0.5 | 5 | 0.8 |
| 9_59831941 | 6 | 0.1667 | 3 | 0 | 3 | 0.3333 | 8 | 0.625 | 20 | 0.35 | 2 | 0 | 2 | 0 | 5 | 0.2 | 4 | 0 | 13 | 0.0769 |
| 9_59831950 | 5 | 0 | 3 | 0.3333 | 4 | 0 | 7 | 0.1429 | 19 | 0.1053 | 2 | 0 | 2 | 0.5 | 5 | 0 | 4 | 0 | 13 | 0.0769 |
| 9_59831986 | 6 | 0.1667 | 3 | 0 | 4 | 0.25 | 6 | 0.3333 | 19 | 0.2105 | 2 | 0 | 2 | 0.5 | 5 | 0 | 4 | 0.25 | 13 | 0.1538 |
| 9_59832086 | 6 | 0.6667 | 4 | 0.75 | 4 | 1 | 7 | 0.8571 | 21 | 0.8095 | 1 | 1 | 1 | 0 | 5 | 1 | 3 | 1 | 10 | 0.9 |
| 9_59832141 | 5 | 0.4 | 3 | 0.6667 | 4 | 0.25 | 5 | 0.8 | 17 | 0.5294 | 1 | 1 | 2 | 0.5 | 5 | 0 | 3 | 0.3333 | 11 | 0.2727 |

| | | | | | | | | | | | | | | | | | | | | |
|---|---|---|---|---|---|---|---|---|---|---|---|---|---|---|---|---|---|---|---|---|
| 9_5983217 | 5 | 0.8 | 2 | 0.5 | 2 | 1 | 6 | 1 | 15 | 0.8667 | 2 | 1 | 1 | 1 | 5 | 0.8 | 3 | 1 | 11 | 0.9091 |
| 9_5983218 | 6 | 0.8333 | 3 | 0.3333 | 3 | 0.3333 | 7 | 0.8571 | 19 | 0.6842 | 2 | 1 | 2 | 1 | 4 | 0.5 | 4 | 0.75 | 12 | 0.75 |
| 9_5983221 | 6 | 0 | 4 | 0.5 | 4 | 0 | 7 | 0.2857 | 21 | 0.1905 | 1 | 0 | 2 | 0 | 5 | 0.4 | 4 | 0 | 12 | 0.1667 |
| 9_5983222 | 6 | 0.8333 | 3 | 0 | 4 | 0.75 | 6 | 0.6667 | 19 | 0.6316 | 1 | 1 | 1 | 0 | 1 | 1 | 3 | 0.3333 | 6 | 0.5 |
| 9_5983224 | 6 | 0.5 | 2 | 0.5 | 3 | 0.3333 | 8 | 0.375 | 19 | 0.4211 | 2 | 0.5 | 2 | 1 | 6 | 0.5 | 3 | 0.3333 | 13 | 0.5385 |
| 9_5983225 | 5 | 0.8 | 4 | 1 | 3 | 1 | 5 | 0.8 | 17 | 0.8824 | 2 | 1 | 2 | 1 | 4 | 1 | 3 | 0.6667 | 11 | 0.9091 |
| 9_5983225 | 3 | 0.3333 | 3 | 0 | 4 | 0.5 | 7 | 0.4286 | 17 | 0.3529 | 2 | 0 | 2 | 0.5 | 6 | 0.3333 | 3 | 0.3333 | 13 | 0.3077 |
| 9_5983231 | 5 | 1 | 4 | 1 | 4 | 1 | 7 | 1 | 20 | 1 | 2 | 1 | 2 | 1 | 5 | 1 | 4 | 1 | 13 | 1 |
| 9_5983241 | 6 | 0 | 4 | 0.5 | 3 | 0 | 9 | 0.2222 | 22 | 0.1818 | 2 | 0.5 | 1 | 1 | 6 | 0.1667 | 4 | 0 | 13 | 0.2308 |
| 9_5983252 | 6 | 0 | 3 | 0 | 5 | 0.2 | 4 | 0 | 18 | 0.0556 | 1 | 0 | 1 | 1 | 2 | 0 | 2 | 0 | 6 | 0.1667 |
| 9_5983253 | 7 | 0.8571 | 4 | 0.75 | 5 | 1 | 9 | 0.7778 | 25 | 0.84 | 2 | 1 | 2 | 1 | 6 | 0.8333 | 4 | 1 | 14 | 0.9286 |
| 9_5983272 | 7 | 0 | 4 | 0 | 5 | 0.2 | 9 | 0.1111 | 25 | 0.08 | 2 | 0.5 | 2 | 0.5 | 6 | 0.1667 | 4 | 0.25 | 14 | 0.2857 |
| 9_5983311 | 7 | 0.1429 | 4 | 0 | 8 | 0.125 | 9 | 0.1111 | 28 | 0.1071 | 2 | 1 | 2 | 0 | 5 | 0.6 | 4 | 0.25 | 13 | 0.4615 |
| 9_5983332 | 7 | 0.2857 | 4 | 0 | 8 | 0 | 10 | 0.1 | 29 | 0.1034 | 3 | 0.6667 | 3 | 0 | 7 | 0.5714 | 5 | 0.2 | 18 | 0.3889 |
| 9_5983389 | 9 | 0.1111 | 4 | 0.25 | 7 | 0 | 13 | 0.3077 | 33 | 0.1818 | 3 | 0.3333 | 3 | 0.3333 | 7 | 0.2857 | 3 | 0 | 16 | 0.25 |
| 9_5983407 | 10 | 0.1 | 4 | 0 | 6 | 0 | 10 | 0.1 | 30 | 0.0667 | 3 | 0 | 2 | 0 | 8 | 0 | 3 | 0.3333 | 16 | 0.0625 |
| 9_5983520 | 9 | 0.2222 | 4 | 0 | 7 | 0 | 11 | 0 | 31 | 0.0645 | 1 | 0 | 4 | 0.5 | 6 | 0 | 5 | 0 | 16 | 0.125 |
| 9_5983560 | 9 | 0.1111 | 4 | 0.25 | 6 | 0 | 7 | 0.1429 | 26 | 0.1154 | 1 | 0 | 3 | 0.6667 | 5 | 0 | 3 | 0 | 12 | 0.1667 |
| 9_5983583 | 11 | 0.2727 | 5 | 0 | 8 | 0.5 | 11 | 0.3636 | 35 | 0.3143 | 2 | 1 | 2 | 0 | 8 | 0.25 | 6 | 0.5 | 18 | 0.3889 |
| 9_5983590 | 11 | 0 | 5 | 0 | 8 | 0.125 | 10 | 0.1 | 34 | 0.0588 | 3 | 0 | 4 | 0.25 | 7 | 0.1429 | 6 | 0.5 | 20 | 0.25 |
| 9_5983596 | 12 | 0.25 | 6 | 0 | 6 | 0.1667 | 8 | 0.375 | 32 | 0.2188 | 2 | 0.5 | 2 | 0 | 7 | 0.4286 | 3 | 0.6667 | 14 | 0.4286 |
| 9_5987495 | NA | NA | NA | NA | 1 | 0 | 2 | 0.5 | 3 | 0.3333 | 1 | 1 | NA | NA | NA | NA | 1 | 0 | 2 | 0.5 |
| 9_5987526 | NA | NA | NA | NA | NA | NA | 2 | 1 | 2 | 1 | 1 | 1 | 1 | 0 | NA | NA | 1 | 1 | 3 | 0.6667 |
| 9_6049461 | 4 | 0.5 | 8 | 0.5 | 10 | 0.7 | 3 | 0.3333 | 25 | 0.56 | 9 | 0.5556 | 14 | 0.4286 | 10 | 0.5 | 2 | 0.5 | 35 | 0.4857 |
| 9_6049479 | 2 | 1 | 8 | 0.625 | 10 | 0.7 | 3 | 1 | 23 | 0.7391 | 8 | 0.625 | 13 | 0.7692 | 10 | 0.7 | 3 | 1 | 34 | 0.7353 |
| 9_6049489 | 3 | 0.3333 | 7 | 0 | 9 | 0.2222 | 3 | 0 | 22 | 0.1364 | 7 | 0.2857 | 14 | 0.0714 | 8 | 0 | 2 | 0.5 | 31 | 0.129 |
| 9_6049495 | 3 | 1 | 7 | 0.4286 | 7 | 0.2857 | 3 | 1 | 20 | 0.55 | 7 | 0.5714 | 15 | 0.4667 | 8 | 0.75 | 2 | 1 | 32 | 0.5938 |
| 9_6049498 | 3 | 0 | 7 | 0 | 6 | 0.1667 | 3 | 0.3333 | 19 | 0.1053 | 5 | 0.2 | 15 | 0 | 8 | 0.125 | 2 | 0 | 30 | 0.0667 |
| 9_6049500 | 3 | 0.3333 | 6 | 0.3333 | 7 | 0.1429 | 2 | 0 | 18 | 0.2222 | 7 | 0.1429 | 15 | 0.2 | 7 | 0.2857 | 2 | 0 | 31 | 0.1935 |
| 9_6059571 | NA | NA | NA | NA | NA | NA | 1 | 1 | 1 | 1 | 1 | 1 | NA | NA | NA | NA | NA | NA | 1 | 1 |
| 9_6064434 | 4 | 0.25 | 4 | 0 | 2 | 0 | 3 | 0.3333 | 13 | 0.1538 | 1 | 0 | 3 | 0 | 3 | 0 | 2 | 0.5 | 9 | 0.1111 |
| 9_6064436 | 6 | 0.1667 | 4 | 0.25 | 1 | 0 | 5 | 0 | 16 | 0.125 | 2 | 0 | 5 | 0.2 | 2 | 0 | 3 | 0.3333 | 12 | 0.1667 |
| 9_6064440 | 4 | 0.25 | 5 | 0.2 | NA | NA | 5 | 0.2 | 14 | 0.2143 | 1 | 1 | 2 | 0 | NA | NA | 3 | 0.6667 | 6 | 0.5 |
| 9_6064452 | 6 | 0.8333 | 6 | 1 | 2 | 1 | 6 | 1 | 20 | 0.95 | 2 | 1 | 5 | 1 | 2 | 1 | 4 | 0.75 | 13 | 0.9231 |
| 9_6064477 | 6 | 0 | 7 | 0.2857 | 2 | 0 | 5 | 0 | 20 | 0.1 | 2 | 0 | 5 | 0.4 | 4 | 0 | 5 | 0.2 | 16 | 0.1875 |
| 9_6064482 | 4 | 0 | 4 | 0 | 1 | 0 | 2 | 0.5 | 11 | 0.0909 | NA | NA | 6 | 0 | 3 | 0.3333 | 4 | 0 | 13 | 0.0769 |
| 9_6064483 | 7 | 0.7143 | 7 | 0.4286 | 2 | 0.5 | 5 | 0.8 | 21 | 0.619 | 1 | 0 | 6 | 0.6667 | 2 | 1 | 4 | 1 | 13 | 0.7692 |
| 9_6064507 | 9 | 0.7778 | 8 | 0.625 | 2 | 0.5 | 4 | 0.25 | 23 | 0.6087 | 2 | 1 | 5 | 0.6 | 4 | 0.75 | 4 | 0.5 | 15 | 0.6667 |
| 9_6064507 | 10 | 0.2 | 8 | 0 | 3 | 0.6667 | 5 | 0.6 | 26 | 0.2692 | 2 | 0 | 7 | 0.2857 | 4 | 0.25 | 4 | 0 | 17 | 0.1765 |
| 9_6064518 | 7 | 1 | 7 | 1 | 2 | 1 | 7 | 1 | 23 | 1 | 2 | 1 | 7 | 1 | 5 | 0.8 | 5 | 1 | 19 | 0.9474 |
| 9_6064523 | 9 | 0.1111 | 5 | 0.4 | 3 | 0.3333 | 7 | 0.1429 | 24 | 0.2083 | 2 | 1 | 7 | 0.2857 | 4 | 0.5 | 5 | 0 | 18 | 0.3333 |
| 9_6064531 | 9 | 1 | 5 | 1 | 2 | 1 | 5 | 1 | 21 | 1 | 2 | 1 | 6 | 1 | 3 | 1 | 4 | 1 | 15 | 1 |
| 9_6064549 | 7 | 1 | 6 | 0.6667 | 2 | 0.5 | 8 | 1 | 23 | 0.8696 | 3 | 0.6667 | 6 | 0.8333 | 2 | 0.5 | 5 | 1 | 16 | 0.8125 |
| 9_6064561 | 6 | 0.5 | 4 | 0.25 | 3 | 0.3333 | 4 | 0.5 | 17 | 0.4118 | 2 | 0.5 | 7 | 0.2857 | 5 | 0.6 | 4 | 0.25 | 18 | 0.3889 |
| 9_6064564 | 9 | 0.5556 | 8 | 0.5 | 2 | 0.5 | 8 | 0.625 | 27 | 0.5556 | 4 | 1 | 7 | 0.4286 | 5 | 0.4 | 4 | 0.75 | 20 | 0.6 |
| 9_6064566 | 7 | 0.4286 | 6 | 0.3333 | 2 | 0 | 7 | 0.7143 | 22 | 0.4545 | 3 | 0 | 5 | 0.6 | 4 | 0.25 | 5 | 1 | 17 | 0.5294 |
| 9_6064577 | 8 | 0.625 | 7 | 0.7143 | 4 | 0.75 | 8 | 0.625 | 27 | 0.6667 | 4 | 0.75 | 7 | 0.5714 | 7 | 0.7143 | 4 | 1 | 22 | 0.7273 |
| 9_6077698 | NA | NA | 1 | 1 | 1 | 0 | NA | NA | 2 | 0.5 | NA | NA | NA | NA | 1 | 1 | NA | NA | 1 | 1 |
| 9_6077703 | NA | NA | 1 | 0 | 1 | 1 | NA | NA | 2 | 0.5 | NA | NA | NA | NA | 1 | 1 | NA | NA | 1 | 1 |
| 9_6077746 | NA | NA | 1 | 1 | NA | NA | NA | NA | 1 | 1 | 1 | 1 | NA | NA | 2 | 0 | NA | NA | 3 | 0.3333 |
| 9_6077809 | NA | NA | NA | NA | 2 | 1 | NA | NA | 2 | 1 | 1 | 1 | NA | NA | 2 | 0 | NA | NA | 3 | 0.3333 |
| 9_6077818 | NA | NA | 2 | 0 | 2 | 1 | NA | NA | 4 | 0.5 | 1 | 1 | NA | NA | 2 | 0.5 | NA | NA | 3 | 0.6667 |
| 9_6077863 | NA | NA | 2 | 0.5 | 1 | 1 | NA | NA | 3 | 0.6667 | 1 | 0 | NA | NA | 2 | 1 | NA | NA | 3 | 0.6667 |
| 9_6077936 | NA | NA | 2 | 0 | 1 | 1 | NA | NA | 3 | 0.3333 | 1 | 1 | NA | NA | 2 | 1 | NA | NA | 3 | 1 |
| 9_6128010 | 1 | 1 | NA | NA | NA | NA | 1 | 0 | 2 | 0.5 | NA | NA | 2 | 0.5 | NA | NA | NA | NA | 2 | 0.5 |
| 9_6132400 | 5 | 1 | 7 | 1 | 2 | 1 | 3 | 0.6667 | 17 | 0.9412 | 1 | 1 | 2 | 0.5 | 2 | 1 | NA | NA | 5 | 0.8 |
| 9_6132416 | 5 | 0.4 | 7 | 0.2857 | 2 | 1 | 4 | 0.5 | 18 | 0.4444 | 2 | 0 | 2 | 0.5 | 2 | 0 | 1 | 1 | 7 | 0.2857 |
| 9_6132417 | 4 | 0.75 | 7 | 1 | 1 | 1 | 3 | 1 | 15 | 0.9333 | 2 | 1 | 2 | 1 | 1 | 1 | 1 | 1 | 6 | 1 |
| 9_6132426 | 4 | 0 | 7 | 0 | 2 | 0.5 | 4 | 0.25 | 17 | 0.1176 | 1 | 0 | 2 | 0 | 1 | 1 | 1 | 0 | 5 | 0.2 |
| 9_6132440 | 3 | 0 | 3 | 0.3333 | 2 | 0.5 | 5 | 0.2 | 13 | 0.2308 | 1 | 0 | 4 | 0.25 | 1 | 0 | 1 | 0 | 7 | 0.1429 |
| 9_6132442 | 4 | 0.25 | 6 | 0.1667 | 2 | 1 | 4 | 0.75 | 16 | 0.4375 | 2 | 0 | 4 | 0.5 | 1 | 0 | 1 | 1 | 8 | 0.375 |
| 9_6132466 | 4 | 0 | 8 | 0.625 | 3 | 0.6667 | 4 | 0.75 | 19 | 0.5263 | 2 | 0.5 | 6 | 0.3333 | NA | NA | 2 | 1 | 10 | 0.5 |
| 9_6132474 | 2 | 1 | 7 | 0.2857 | NA | NA | 2 | 0 | 11 | 0.3636 | 2 | 0 | 3 | 0.3333 | NA | NA | 2 | 0 | 7 | 0.1429 |
| 9_6132544 | 1 | 0 | 3 | 0 | 3 | 0 | 1 | 1 | 8 | 0.125 | 2 | 0.5 | 3 | 0.3333 | NA | NA | 3 | 0 | 8 | 0.25 |
| 9_6132554 | 1 | 0 | 5 | 0.4 | 4 | 0 | 3 | 0 | 13 | 0.1538 | 3 | 0.6667 | 3 | 0.3333 | 1 | 0 | 2 | 0.5 | 9 | 0.4444 |

| | | | | | | | | | | | | | | | | | | | | |
|---|---|---|---|---|---|---|---|---|---|---|---|---|---|---|---|---|---|---|---|---|
| 9_6196529; | 2 | 0.5 | NA | NA | 1 | 0 | 2 | 0 | 5 | 0.2 | NA | NA | 8 | 0.5 | 2 | 0 | 2 | 0.5 | 12 | 0.4167 |
| 9_6196558( | 1 | 1 | 1 | 0 | 3 | 0.3333 | 2 | 0 | 7 | 0.2857 | NA | NA | 6 | 0.3333 | 3 | 0 | 3 | 0.6667 | 12 | 0.3333 |
| 9_6196621( | 1 | 0 | NA | NA | 3 | 0 | 2 | 0.5 | 6 | 0.1667 | NA | NA | 3 | 0.3333 | 1 | 0 | 3 | 0 | 7 | 0.1429 |
| 9_6196646' | 2 | 0.5 | NA | NA | 2 | 0 | 2 | 0 | 6 | 0.1667 | NA | NA | 3 | 0.3333 | 1 | 0 | 3 | 0 | 7 | 0.1429 |
| 9_6196674! | 1 | 1 | NA | NA | 2 | 0.5 | 2 | 0.5 | 5 | 0.6 | NA | NA | 1 | 0 | 1 | 0 | 3 | 0.3333 | 5 | 0.2 |
| 9_6196685! | 1 | 1 | NA | NA | 2 | 1 | 2 | 0.5 | 5 | 0.8 | NA | NA | NA | NA | 1 | 1 | 3 | 1 | 4 | 1 |
| 9_6196714! | 1 | 1 | NA | NA | 3 | 0 | 1 | 0 | 5 | 0.2 | NA | NA | 2 | 0.5 | NA | NA | 3 | 0 | 5 | 0.2 |
| 9_6196717' | 1 | 1 | NA | NA | 3 | 0.6667 | 2 | 1 | 6 | 0.8333 | NA | NA | 1 | 0 | NA | NA | 3 | 0.3333 | 4 | 0.25 |
| 9_6196741 | 1 | 1 | NA | NA | 2 | 1 | 2 | 1 | 5 | 1 | NA | NA | 2 | 1 | NA | NA | 3 | 1 | 5 | 1 |
| 9_6196756; | 1 | 0 | NA | NA | 2 | 0.5 | 2 | 0.5 | 5 | 0.4 | NA | NA | 2 | 1 | NA | NA | 3 | 0.3333 | 5 | 0.6 |
| 9_6196757' | 1 | 0 | NA | NA | 2 | 1 | 2 | 0.5 | 5 | 0.6 | NA | NA | NA | NA | NA | NA | 2 | 0.5 | 2 | 0.5 |
| 9_6196766: | NA | NA | NA | NA | 2 | 0.5 | 2 | 1 | 4 | 0.75 | NA | NA | 2 | 1 | NA | NA | 3 | 0.6667 | 5 | 0.8 |
| 9_6196775! | 1 | 1 | NA | NA | 1 | 1 | 2 | 1 | 4 | 1 | NA | NA | 1 | 1 | NA | NA | 1 | 1 | 2 | 1 |
| 9_6196791! | 1 | 1 | NA | NA | 1 | 1 | 2 | 1 | 4 | 1 | NA | NA | NA | NA | NA | NA | 1 | 1 | 1 | 1 |
| 9_6196808! | 1 | 0 | NA | NA | 2 | 0.5 | 2 | 0.5 | 5 | 0.4 | NA | NA | 2 | 0 | NA | NA | 2 | 0.5 | 4 | 0.25 |
| 9_6196811' | 1 | 0 | NA | NA | 2 | 0.5 | 2 | 0.5 | 5 | 0.4 | NA | NA | 2 | 0.5 | NA | NA | 1 | 0 | 3 | 0.3333 |
| 9_6196814( | 1 | 1 | NA | NA | 2 | 0.5 | 2 | 0.5 | 5 | 0.6 | NA | NA | 2 | 1 | NA | NA | 2 | 1 | 4 | 1 |
| 9_6196814( | NA | NA | NA | NA | 2 | 1 | 2 | 0.5 | 4 | 0.75 | NA | NA | 2 | 0.5 | NA | NA | 2 | 1 | 4 | 0.75 |
| 9_6196827! | 1 | 1 | NA | NA | 1 | 1 | 2 | 0.5 | 4 | 0.75 | NA | NA | 2 | 0.5 | NA | NA | 2 | 1 | 4 | 0.75 |
| 9_6197756( | 1 | 1 | NA | NA | 1 | 1 | 1 | 1 | 3 | 1 | NA | NA | 1 | 1 | NA | NA | 2 | 1 | 3 | 1 |
| 9_6228586( | 23 | 0.1739 | 12 | 0.1667 | 8 | 0.375 | 14 | 0.1429 | 57 | 0.193 | 15 | 0.2 | 12 | 0.25 | 10 | 0.2 | 9 | 0 | 46 | 0.1739 |
| 9_6232690: | 11 | 0.3636 | 19 | 0.2105 | 25 | 0.12 | 19 | 0.1579 | 74 | 0.1892 | 16 | 0.25 | 37 | 0.2703 | 23 | 0.1739 | 29 | 0.069 | 105 | 0.1905 |
| 9_6232706! | 12 | 0.5 | 15 | 0.6667 | 21 | 0.6667 | 19 | 0.6316 | 67 | 0.6269 | 13 | 0.4615 | 36 | 0.3889 | 22 | 0.5455 | 30 | 0.4333 | 101 | 0.4455 |
| 9_6232713! | 8 | 0 | 17 | 0.1176 | 19 | 0.1579 | 18 | 0.1111 | 62 | 0.1129 | 15 | 0.0667 | 26 | 0.0385 | 15 | 0.0667 | 24 | 0.125 | 80 | 0.075 |
| 9_6232798' | 4 | 0 | 10 | 0.4 | 9 | 0.1111 | 11 | 0.2727 | 34 | 0.2353 | 6 | 0.3333 | 17 | 0.4706 | 9 | 0.2222 | 21 | 0.3333 | 53 | 0.3585 |
| 9_6232816' | 11 | 0.9091 | 14 | 0.6429 | 12 | 0.5 | 17 | 0.7059 | 54 | 0.6852 | 9 | 0.7778 | 26 | 0.8077 | 16 | 0.75 | 21 | 0.619 | 72 | 0.7361 |
| 9_6232823( | 8 | 0.75 | 12 | 0.4167 | 12 | 0.5833 | 17 | 0.6471 | 49 | 0.5918 | 8 | 0.5 | 23 | 0.5652 | 10 | 0.6 | 19 | 0.5789 | 60 | 0.5667 |
| 9_6232830! | 8 | 0.5 | 14 | 0.2143 | 11 | 0.0909 | 16 | 0.125 | 49 | 0.2041 | 9 | 0 | 22 | 0.2727 | 16 | 0.125 | 20 | 0 | 67 | 0.1194 |
| 9_6232835! | 11 | 0 | 12 | 0 | 12 | 0.1667 | 17 | 0.1765 | 52 | 0.0962 | 8 | 0.25 | 25 | 0.16 | 14 | 0.2143 | 22 | 0.0455 | 69 | 0.1449 |
| 9_6232838' | 6 | 0.6667 | 12 | 0.5833 | 10 | 0.9 | 11 | 0.8182 | 39 | 0.7436 | 6 | 0.8333 | 20 | 0.85 | 8 | 1 | 19 | 0.6842 | 53 | 0.8113 |
| 9_6232842' | 7 | 1 | 12 | 0.9167 | 8 | 0.875 | 14 | 0.8571 | 41 | 0.9024 | 9 | 0.8889 | 22 | 1 | 10 | 1 | 20 | 0.95 | 61 | 0.9672 |
| 9_6232843! | 10 | 0.2 | 12 | 0.0833 | 11 | 0.0909 | 16 | 0.125 | 49 | 0.1224 | 9 | 0.3333 | 25 | 0.12 | 12 | 0.0833 | 22 | 0.0909 | 68 | 0.1324 |
| 9_6232846! | 7 | 0.1429 | 10 | 0.1 | 10 | 0 | 14 | 0.1429 | 41 | 0.0976 | 6 | 0.3333 | 20 | 0.25 | 13 | 0.3077 | 20 | 0.25 | 59 | 0.2712 |
| 9_6232857' | 9 | 0.8889 | 10 | 1 | 8 | 0.875 | 17 | 0.8235 | 44 | 0.8864 | 6 | 0.6667 | 21 | 0.8095 | 11 | 0.9091 | 20 | 0.85 | 58 | 0.8276 |
| 9_6232860( | 8 | 0 | 12 | 0.0833 | 9 | 0.3333 | 17 | 0.4706 | 46 | 0.2609 | 6 | 0 | 22 | 0.3182 | 12 | 0.5 | 19 | 0.2632 | 59 | 0.3051 |
| 9_6269008' | NA | NA | NA | NA | 1 | 0 | 1 | 1 | 2 | 0.5 | NA | NA | 1 | 1 | NA | NA | NA | NA | 1 | 1 |
| 9_6269016( | NA | NA | NA | NA | 1 | 1 | 1 | 1 | 2 | 1 | NA | NA | 1 | 1 | NA | NA | NA | NA | 1 | 1 |
| 9_6269020' | 1 | 1 | NA | NA | 1 | 1 | NA | NA | 2 | 1 | NA | NA | 1 | 1 | NA | NA | NA | NA | 1 | 1 |
| 9_6269037' | 1 | 1 | NA | NA | 1 | 1 | NA | NA | 2 | 1 | NA | NA | 1 | 1 | NA | NA | NA | NA | 1 | 1 |
| 9_6269055( | 1 | 1 | NA | NA | 1 | 0 | NA | NA | 2 | 0.5 | NA | NA | 1 | 1 | NA | NA | NA | NA | 1 | 1 |
| 9_6270009( | 13 | 0.0769 | 10 | 0 | 5 | 0.2 | 9 | 0.2222 | 37 | 0.1081 | 9 | 0 | 10 | 0 | 14 | 0.0714 | 18 | 0.1111 | 51 | 0.0588 |
| 9_6270026! | 10 | 0.1 | 9 | 0.2222 | 3 | 0 | 9 | 0 | 31 | 0.0968 | 7 | 0.1429 | 12 | 0.0833 | 12 | 0 | 11 | 0.1818 | 42 | 0.0952 |
| 9_6270029' | 11 | 0 | 4 | 0 | 3 | 0.3333 | 5 | 0.4 | 23 | 0.1304 | 6 | 0.1667 | 11 | 0.1818 | 11 | 0.1818 | 10 | 0.4 | 38 | 0.2368 |
| 9_6270247' | 9 | 0.1111 | 4 | 0 | 1 | 0 | 4 | 0 | 18 | 0.0556 | 2 | 0 | 6 | 0.1667 | 6 | 0 | 5 | 0 | 19 | 0.0526 |
| 9_6270274: | 7 | 0.4286 | 4 | 0.75 | 1 | 1 | 5 | 0.8 | 17 | 0.6471 | 1 | 1 | 5 | 0.8 | 6 | 0.6667 | 4 | 1 | 16 | 0.8125 |
| 9_6270287: | 6 | 1 | 4 | 1 | 1 | 1 | 4 | 1 | 15 | 1 | 1 | 1 | 5 | 1 | 4 | 1 | 4 | 1 | 14 | 1 |
| 9_6270296( | 7 | 0.8571 | 1 | 0 | 1 | 1 | 4 | 1 | 13 | 0.8462 | NA | NA | 2 | 1 | 2 | 1 | 2 | 1 | 6 | 1 |
| 9_6270297' | 7 | 0.2857 | 5 | 0.8 | 1 | 0 | 5 | 0.8 | 18 | 0.5556 | NA | NA | 3 | 1 | 3 | 1 | 4 | 1 | 10 | 1 |
| 9_6270316: | 8 | 1 | 4 | 1 | 1 | 1 | 4 | 1 | 17 | 1 | 1 | 1 | 5 | 1 | 3 | 1 | 3 | 1 | 12 | 1 |
| 9_6270317! | 7 | 0 | 1 | 0 | NA | NA | 5 | 0.2 | 13 | 0.0769 | 1 | 0 | 5 | 0 | 1 | 0 | 3 | 0.3333 | 10 | 0.1 |
| 9_6270331' | 7 | 0.5714 | 4 | 0.75 | 1 | 1 | 5 | 0.8 | 17 | 0.7059 | 1 | 0 | 5 | 0.4 | 2 | 1 | 3 | 1 | 11 | 0.6364 |
| 9_6270332' | 7 | 1 | 3 | 1 | 1 | 1 | 7 | 1 | 18 | 1 | 1 | 1 | 5 | 1 | 3 | 1 | 3 | 1 | 12 | 1 |
| 9_6270337' | 7 | 1 | 2 | 1 | 1 | 1 | 7 | 1 | 17 | 1 | 1 | 0 | 4 | 1 | 2 | 1 | 3 | 1 | 10 | 0.9 |
| 9_6270340' | 7 | 0.8571 | 3 | 1 | 1 | 1 | 7 | 1 | 18 | 0.9444 | 1 | 1 | 4 | 1 | 3 | 1 | 3 | 1 | 11 | 1 |
| 9_6270343: | 5 | 0.8 | 4 | 1 | NA | NA | 7 | 0.8571 | 16 | 0.875 | 1 | 1 | 1 | 1 | 2 | 1 | 2 | 1 | 6 | 1 |
| 9_6270348! | 4 | 0 | 3 | 0.3333 | NA | NA | 5 | 0.6 | 12 | 0.3333 | NA | NA | 2 | 0 | 1 | 0 | 2 | 1 | 5 | 0.4 |
| 9_6270352( | 8 | 0 | 3 | 0.3333 | 1 | 0 | 7 | 0.1429 | 19 | 0.1053 | 1 | 0 | 2 | 0 | 4 | 0.25 | 3 | 0 | 10 | 0.1 |
| 9_6270353( | 8 | 0.375 | 3 | 0.3333 | 1 | 0 | 3 | 0.3333 | 15 | 0.3333 | 1 | 0 | 3 | 0 | 4 | 0.5 | 3 | 0 | 11 | 0.1818 |
| 9_6270360! | 8 | 0 | 4 | 0 | 1 | 1 | 7 | 0.1429 | 20 | 0.1 | NA | NA | 3 | 0 | 4 | 0.25 | 1 | 0 | 8 | 0.125 |
| 9_6270367: | 5 | 0.6 | 2 | 1 | NA | NA | 3 | 1 | 10 | 0.8 | 1 | 1 | NA | NA | 2 | 1 | 1 | 1 | 4 | 1 |
| 9_6270370( | 8 | 1 | 2 | 1 | 1 | 1 | 4 | 1 | 15 | 1 | NA | NA | 3 | 1 | 2 | 1 | 2 | 1 | 7 | 1 |
| 9_6270382( | 8 | 0 | 1 | 1 | 1 | 0 | 6 | 0 | 16 | 0.0625 | NA | NA | 3 | 0 | 4 | 0.5 | 2 | 0.5 | 9 | 0.3333 |
| 9_6270399' | 6 | 0.5 | 1 | 1 | 1 | 1 | 3 | 1 | 11 | 0.7273 | 1 | 0 | 2 | 1 | 2 | 0 | 3 | 1 | 8 | 0.625 |
| 9_6270434: | 7 | 0.8571 | 1 | 1 | 1 | 1 | 4 | 0.75 | 13 | 0.8462 | NA | NA | 2 | 1 | 2 | 1 | 3 | 0.6667 | 7 | 0.8571 |
| 9_6270434! | 5 | 0.2 | 3 | 0 | 1 | 0 | 4 | 0.25 | 13 | 0.1538 | NA | NA | 1 | 1 | NA | NA | 2 | 0.5 | 3 | 0.6667 |

| | | | | | | | | | | | | | | | | | | | | |
|---|---|---|---|---|---|---|---|---|---|---|---|---|---|---|---|---|---|---|---|---|
| 9_6270446 | 5 | 0.4 | 1 | 1 | NA | NA | 3 | 0.3333 | 9 | 0.4444 | 1 | 0 | 2 | 0.5 | 3 | 0.3333 | 3 | 0.6667 | 9 | 0.4444 |
| 9_6270455 | 6 | 1 | 3 | 1 | 1 | 1 | 2 | 1 | 12 | 1 | 1 | 1 | 1 | 1 | 2 | 1 | 2 | 0.5 | 6 | 0.8333 |
| 9_6270464 | 6 | 0.8333 | 3 | 1 | 1 | 1 | 3 | 1 | 13 | 0.9231 | NA | NA | 3 | 1 | 2 | 1 | 3 | 1 | 8 | 1 |
| 9_6270465 | 6 | 0.3333 | 1 | 0 | 1 | 1 | 4 | 0.25 | 12 | 0.3333 | 1 | 0 | 2 | 1 | 2 | 0.5 | 3 | 0.6667 | 8 | 0.625 |
| 9_6270485 | 6 | 1 | 2 | 1 | 1 | 1 | 3 | 1 | 12 | 1 | 1 | 1 | 3 | 1 | 2 | 1 | 2 | 1 | 8 | 1 |
| 9_6270486 | 5 | 0.8 | 1 | 1 | NA | NA | 4 | 0.75 | 10 | 0.8 | 1 | 0 | 1 | 0 | 3 | 0.6667 | 3 | 0.6667 | 8 | 0.5 |
| 9_6270496 | 6 | 0.5 | 3 | 0.6667 | 1 | 1 | 4 | 0.25 | 14 | 0.5 | 1 | 1 | 3 | 0.6667 | 2 | 0.5 | 3 | 0.6667 | 9 | 0.6667 |
| 9_6275823 | 2 | 1 | 2 | 0 | 1 | 1 | 4 | 0.25 | 9 | 0.4444 | NA | NA | NA | NA | 3 | 0.6667 | 2 | 0.5 | 5 | 0.6 |
| 9_6275834 | 1 | 0 | 1 | 0 | 1 | 1 | 4 | 0 | 7 | 0.1429 | NA | NA | NA | NA | 3 | 0.6667 | 2 | 0 | 5 | 0.4 |
| 9_6275845 | 3 | 0.3333 | 2 | 0 | 2 | 0 | 4 | 0 | 11 | 0.0909 | NA | NA | NA | NA | 3 | 0.3333 | 2 | 0 | 5 | 0.2 |
| 9_6278587 | 6 | 0 | 5 | 0.2 | 5 | 0 | 4 | 0 | 20 | 0.05 | 3 | 0.3333 | 3 | 0 | 5 | 0 | 4 | 0.25 | 15 | 0.1333 |
| 9_6278608 | 4 | 0.25 | 4 | 0.25 | 4 | 0.5 | 4 | 0.5 | 16 | 0.375 | 2 | 0 | 3 | 0 | 8 | 0.375 | 4 | 0 | 17 | 0.1765 |
| 9_6278676 | 6 | 0 | 4 | 0 | 2 | 0 | 6 | 0.1667 | 18 | 0.0556 | 1 | 0 | 3 | 0.3333 | 7 | 0 | 3 | 0 | 14 | 0.0714 |
| 9_6278812 | 4 | 0 | 8 | 0.375 | 3 | 0 | 8 | 0.125 | 23 | 0.1739 | 3 | 0 | 5 | 0.2 | 3 | 0 | 3 | 0.3333 | 14 | 0.1429 |
| 9_6278894 | 2 | 0 | 7 | 0 | 2 | 1 | 4 | 0 | 15 | 0.1333 | 4 | 0.5 | 3 | 0 | 2 | 0 | 3 | 0 | 12 | 0.1667 |
| 9_6278902 | 3 | 0 | 7 | 0 | 2 | 0.5 | 5 | 0 | 17 | 0.0588 | 3 | 0.3333 | 3 | 0 | 3 | 0 | 3 | 0.3333 | 12 | 0.1667 |
| 9_6278904 | 3 | 1 | 7 | 0.8571 | 2 | 1 | 4 | 0.5 | 16 | 0.8125 | 3 | 0.6667 | 3 | 0.6667 | 3 | 0.6667 | 3 | 0.6667 | 12 | 0.6667 |
| 9_6278908 | 2 | 0.5 | 6 | 0 | 2 | 0 | 3 | 0 | 13 | 0.0769 | 4 | 0 | 4 | 0.25 | 3 | 0 | 2 | 0 | 13 | 0.0769 |
| 9_6278919 | 2 | 1 | 7 | 0.8571 | 2 | 1 | 5 | 0.8 | 16 | 0.875 | 3 | 1 | 3 | 0.6667 | 5 | 0.8 | 5 | 1 | 16 | 0.875 |
| 9_6278920 | 2 | 0 | 7 | 0.1429 | 1 | 0 | 6 | 0.1667 | 16 | 0.125 | 4 | 0.5 | 4 | 0.75 | 6 | 0.5 | 4 | 0.5 | 18 | 0.5556 |
| 9_6278925 | 2 | 1 | 7 | 1 | 2 | 1 | 6 | 0.8333 | 17 | 0.9412 | 3 | 1 | 3 | 1 | 4 | 0.75 | 4 | 1 | 14 | 0.9286 |
| 9_6278926 | 2 | 1 | 6 | 0.6667 | 2 | 0 | 6 | 0.3333 | 16 | 0.5 | 4 | 0.5 | 4 | 0.5 | 6 | 0.5 | 3 | 0.3333 | 17 | 0.4706 |
| 9_6285915 | NA | NA | 3 | 0.6667 | NA | NA | 3 | 0.6667 | 6 | 0.6667 | 1 | 0 | 4 | 0.25 | 6 | 0.6667 | 2 | 0.5 | 13 | 0.4615 |
| 9_6285916 | NA | NA | 2 | 0 | 1 | 0 | 5 | 0.4 | 8 | 0.25 | 3 | 0 | 4 | 0.25 | 5 | 0 | 2 | 0 | 14 | 0.0714 |
| 9_6285928 | NA | NA | 3 | 1 | 1 | 1 | 5 | 1 | 9 | 1 | 3 | 0.6667 | 4 | 0.5 | 5 | 0.8 | 2 | 1 | 14 | 0.7143 |
| 9_6285933 | NA | NA | 3 | 1 | NA | NA | 2 | 1 | 5 | 1 | 3 | 0.6667 | 3 | 0.6667 | 2 | 1 | 2 | 1 | 10 | 0.8 |
| 9_6285939 | NA | NA | 2 | 1 | NA | NA | 5 | 1 | 7 | 1 | 2 | 1 | 4 | 1 | 5 | 0.6 | 2 | 0.5 | 13 | 0.7692 |
| 9_6307135 | 2 | 0 | 8 | 0 | 4 | 0 | 5 | 0.2 | 19 | 0.0526 | 4 | 0 | 5 | 0.2 | 5 | 0 | 2 | 0 | 16 | 0.0625 |
| 9_6328473 | 2 | 0.5 | 5 | 0.4 | 2 | 1 | 3 | 0.6667 | 12 | 0.5833 | 2 | 0.5 | 4 | 0.5 | 2 | 0.5 | 2 | 1 | 10 | 0.6 |
| 9_6328479 | 2 | 0 | 4 | 0.25 | 2 | 0 | 3 | 0.3333 | 11 | 0.1818 | 3 | 0 | 4 | 0.5 | 2 | 0 | 1 | 0 | 10 | 0.2 |
| 9_6328484 | 2 | 0.5 | 5 | 0.6 | 2 | 1 | 2 | 0.5 | 11 | 0.6364 | 2 | 0 | 4 | 1 | 1 | 0 | 2 | 1 | 9 | 0.6667 |
| 9_6330188 | 7 | 0.1429 | 5 | 0.4 | 2 | 0 | 6 | 0.3333 | 20 | 0.25 | 8 | 0.5 | 5 | 0.6 | 6 | 0.5 | 5 | 0.6 | 24 | 0.5417 |
| 9_6330190 | 8 | 1 | 5 | 1 | 3 | 1 | 6 | 0.8333 | 22 | 0.9545 | 8 | 0.875 | 6 | 0.8333 | 6 | 1 | 5 | 0.8 | 25 | 0.88 |
| 9_6330193 | 5 | 0.4 | 4 | 0.75 | 3 | 0.6667 | 2 | 0.5 | 14 | 0.5714 | 4 | 0.5 | 5 | 0.6 | 4 | 1 | 2 | 0.5 | 15 | 0.6667 |
| 9_6330454 | NA | NA | 1 | 1 | NA | NA | NA | NA | 1 | 1 | NA | NA | NA | NA | 1 | 1 | NA | NA | 1 | 1 |
| 9_6330588 | NA | NA | 1 | 1 | NA | NA | NA | NA | 1 | 1 | NA | NA | 2 | 1 | 1 | 1 | NA | NA | 3 | 1 |
| 9_6330635 | 4 | 0.25 | 3 | 0.3333 | 3 | 0.3333 | 5 | 0.6 | 15 | 0.4 | 4 | 0.75 | 5 | 0.2 | 4 | 0.5 | 2 | 1 | 15 | 0.5333 |
| 9_6354758 | 4 | 0 | 3 | 0 | 3 | 0.3333 | 1 | 1 | 11 | 0.1818 | 4 | 0.25 | 7 | 0.1429 | 3 | 0 | 2 | 0 | 16 | 0.125 |
| 9_6354759 | 6 | 0.3333 | 5 | 0 | 5 | 0.2 | 1 | 0 | 17 | 0.1765 | 4 | 0.25 | 7 | 0.2857 | 3 | 0.3333 | 3 | 0.3333 | 17 | 0.2941 |
| 9_6354768 | 5 | 0.6 | 4 | 0.75 | 5 | 0.6 | 2 | 0.5 | 16 | 0.625 | 4 | 0 | 6 | 0.6667 | 3 | 0.6667 | 3 | 1 | 16 | 0.5625 |
| 9_6354775 | 4 | 0.75 | 4 | 0.75 | 4 | 1 | 2 | 0.5 | 14 | 0.7857 | 4 | 0.5 | 7 | 0.8571 | 3 | 0.3333 | 3 | 1 | 17 | 0.7059 |
| 9_6354777 | 7 | 0 | 5 | 0 | 5 | 0.2 | 2 | 0 | 19 | 0.0526 | 5 | 0.4 | 7 | 0.2857 | 4 | 0 | 3 | 0.3333 | 19 | 0.2632 |
| 9_6354781 | 6 | 0.1667 | 4 | 0 | 5 | 0.4 | 2 | 0 | 17 | 0.1765 | 5 | 0.4 | 6 | 0.1667 | 4 | 0.5 | 3 | 0 | 18 | 0.2778 |
| 9_6354785 | 7 | 0 | 5 | 0.2 | 5 | 0 | 2 | 0 | 19 | 0.0526 | 5 | 0 | 6 | 0 | 3 | 0 | 3 | 0.3333 | 17 | 0.0588 |
| 9_6354877 | 9 | 0.2222 | 9 | 0 | 6 | 0 | 2 | 0 | 26 | 0.0769 | 6 | 0.3333 | 7 | 0.1429 | 5 | 0 | 4 | 0 | 22 | 0.1364 |
| 9_6354912 | 7 | 0.2857 | 8 | 0 | 6 | 0.1667 | 2 | 0 | 23 | 0.1304 | 5 | 0 | 8 | 0.375 | 6 | 0.1667 | 5 | 0 | 24 | 0.1667 |
| 9_6355426 | 8 | 0.875 | 3 | 1 | 3 | 1 | 10 | 1 | 24 | 0.9583 | 6 | 1 | 7 | 1 | 17 | 0.9412 | 7 | 1 | 37 | 0.973 |
| 9_6355445 | 10 | 0.1 | 3 | 0 | 2 | 0 | 6 | 0.1667 | 21 | 0.0952 | 6 | 0.3333 | 7 | 0 | 17 | 0.1176 | 6 | 0.1667 | 36 | 0.1389 |
| 9_6355448 | 8 | 0.625 | 2 | 1 | 2 | 1 | 9 | 0.8889 | 21 | 0.8095 | 6 | 0.3333 | 7 | 0.7143 | 18 | 0.6111 | 8 | 0.875 | 39 | 0.641 |
| 9_6355454 | 10 | 0.4 | 3 | 0 | 2 | 0 | 9 | 0.5556 | 24 | 0.375 | 4 | 0.25 | 7 | 0.2857 | 15 | 0.4 | 7 | 0.2857 | 33 | 0.3333 |
| 9_6355457 | 9 | 0.3333 | 3 | 0.3333 | 2 | 0 | 7 | 0.2857 | 21 | 0.2857 | 4 | 0.5 | 6 | 0.1667 | 14 | 0.2143 | 5 | 0 | 29 | 0.2069 |
| 9_6355466 | 9 | 0.5556 | NA | NA | 2 | 0 | 9 | 0.4444 | 20 | 0.45 | 4 | 0.75 | 7 | 0.5714 | 12 | 0.5833 | 5 | 0.6 | 28 | 0.6071 |
| 9_6355487 | 9 | 0.1111 | 3 | 0.3333 | 3 | 0 | 10 | 0.1 | 25 | 0.12 | 4 | 0 | 7 | 0.2857 | 17 | 0 | 7 | 0 | 35 | 0.0571 |
| 9_6355499 | 8 | 0.25 | 3 | 0 | 3 | 0 | 6 | 0.1667 | 20 | 0.15 | 4 | 0.25 | 5 | 0 | 11 | 0.0909 | 8 | 0 | 28 | 0.0714 |
| 9_6355520 | 9 | 0.2222 | 3 | 0 | 3 | 0 | 10 | 0 | 25 | 0.08 | 4 | 0.25 | 6 | 0 | 11 | 0 | 6 | 0.1667 | 27 | 0.0741 |
| 9_6355521 | 7 | 0 | 2 | 0 | 3 | 0.3333 | 9 | 0.2222 | 21 | 0.1429 | 3 | 0.3333 | 7 | 0.2857 | 11 | 0.1818 | 5 | 0 | 26 | 0.1923 |
| 9_6355524 | 8 | 0.625 | 3 | 1 | 2 | 1 | 9 | 0.6667 | 22 | 0.7273 | 3 | 0.6667 | 7 | 0.7143 | 9 | 0.6667 | 5 | 1 | 24 | 0.75 |
| 9_6355527 | 8 | 0.125 | 1 | 0 | 3 | 0.3333 | 7 | 0.1429 | 19 | 0.1579 | 4 | 0.25 | 6 | 0.3333 | 10 | 0.2 | 7 | 0.2857 | 27 | 0.2593 |
| 9_6355557 | 8 | 0.25 | 1 | 0 | 2 | 0.5 | 7 | 0.1429 | 18 | 0.2222 | 1 | 1 | 6 | 0.3333 | 9 | 0.5556 | 2 | 0.5 | 18 | 0.5 |
| 9_6355585 | 7 | 0.5714 | 3 | 0.6667 | 2 | 1 | 8 | 0.75 | 20 | 0.7 | 2 | 0.5 | 7 | 0.5714 | 10 | 0.6 | 6 | 0.5 | 25 | 0.56 |
| 9_6355599 | 8 | 0.375 | 2 | 0 | 3 | 0.3333 | 6 | 0.3333 | 19 | 0.3158 | 2 | 0.5 | 7 | 0 | 7 | 0 | 4 | 0.25 | 20 | 0.1 |
| 9_6355602 | 6 | 0.6667 | 3 | 0.3333 | 3 | 0.6667 | 7 | 0.4286 | 19 | 0.5263 | 2 | 1 | 6 | 0.6667 | 10 | 0.6 | 6 | 0.5 | 24 | 0.625 |
| 9_6355605 | 7 | 1 | 3 | 1 | 3 | 1 | 6 | 1 | 19 | 1 | 2 | 1 | 7 | 1 | 7 | 0.8571 | 6 | 1 | 22 | 0.9545 |
| 9_6355608 | 4 | 0.5 | 2 | 0.5 | 1 | 1 | 5 | 0.4 | 12 | 0.5 | 2 | 1 | 5 | 0.6 | 9 | 1 | 6 | 0.8333 | 22 | 0.8636 |
| 9_6355610 | 7 | 0.5714 | 3 | 0.3333 | 3 | 0 | 6 | 0.6667 | 19 | 0.4737 | 2 | 0.5 | 6 | 0.3333 | 10 | 0.5 | 6 | 0.8333 | 24 | 0.5417 |

| | | | | | | | | | | | | | | | | | | | | |
|---|---|---|---|---|---|---|---|---|---|---|---|---|---|---|---|---|---|---|---|---|
| 9_6355617 | 7 | 0.1429 | 3 | 0.3333 | 3 | 0 | 6 | 0.1667 | 19 | 0.1579 | 2 | 0.5 | 5 | 0 | 10 | 0.3 | 7 | 0.1429 | 24 | 0.2083 |
| 9_6355624 | 5 | 1 | 2 | 1 | 2 | 0.5 | 5 | 0.8 | 14 | 0.8571 | 1 | 1 | 4 | 0.75 | 9 | 0.7778 | 4 | 1 | 18 | 0.8333 |
| 9_6355641 | 4 | 0.25 | 2 | 0.5 | 3 | 0 | 6 | 0.1667 | 15 | 0.2 | 1 | 0 | 3 | 0 | 4 | 0.25 | 2 | 0 | 10 | 0.1 |
| 9_6355651 | 3 | 0.3333 | 3 | 0.3333 | 3 | 0.6667 | 5 | 0.2 | 14 | 0.3571 | 1 | 1 | 1 | 0 | 10 | 0.6 | 5 | 0.2 | 17 | 0.4706 |
| 9_6355652 | 2 | 1 | 1 | 1 | 2 | 1 | 3 | 0.6667 | 8 | 0.875 | 2 | 0.5 | 2 | 0 | 6 | 0.3333 | 4 | 0.5 | 14 | 0.3571 |
| 9_6355666 | 4 | 0.25 | 3 | 0.3333 | 3 | 0.3333 | 6 | 0.1667 | 16 | 0.25 | 2 | 0 | 1 | 0 | 10 | 0.1 | 6 | 0 | 19 | 0.0526 |
| 9_6355673 | 3 | 1 | 2 | 1 | 2 | 1 | 4 | 1 | 11 | 1 | 1 | 1 | 2 | 1 | 9 | 1 | 4 | 1 | 16 | 1 |
| 9_6355677 | 1 | 0 | 1 | 0 | 2 | 0.5 | 4 | 0.25 | 8 | 0.25 | 1 | 0 | 2 | 0.5 | 7 | 0.2857 | 3 | 0 | 13 | 0.2308 |
| 9_6355713 | 3 | 0.3333 | 1 | 0 | 2 | 0 | 4 | 0.25 | 10 | 0.2 | 1 | 1 | 1 | 0 | 9 | 0.5556 | 4 | 0.25 | 15 | 0.4667 |
| 9_6355726 | 3 | 0 | 2 | 0 | 2 | 0 | 4 | 0.75 | 11 | 0.2727 | 2 | 0 | 1 | 1 | 7 | 0.5714 | 4 | 0.25 | 14 | 0.4286 |
| 9_6355727 | 4 | 0.5 | 2 | 0 | 2 | 0 | 4 | 0 | 12 | 0.1667 | 2 | 0.5 | 1 | 0 | 8 | 0.125 | 5 | 0 | 16 | 0.125 |
| 9_6355730 | 2 | 1 | 1 | 1 | 1 | 1 | NA | NA | 4 | 1 | NA | NA | 1 | 1 | 2 | 0.5 | NA | NA | 3 | 0.6667 |
| 9_6355737 | 4 | 0.25 | 2 | 1 | 2 | 1 | 4 | 1 | 12 | 0.75 | 1 | 1 | 1 | 0 | 8 | 1 | 4 | 0.5 | 14 | 0.7857 |
| 9_6355745 | 3 | 0 | 2 | 0.5 | 2 | 0.5 | 3 | 0.6667 | 10 | 0.4 | 1 | 1 | NA | NA | 7 | 0.4286 | 6 | 0.3333 | 14 | 0.4286 |
| 9_6355747 | 3 | 0.6667 | 2 | 0 | 1 | 0 | 3 | 0.3333 | 9 | 0.3333 | NA | NA | 1 | 0 | 6 | 0.8333 | 5 | 0.8 | 12 | 0.75 |
| 9_6355749 | 3 | 0 | 1 | 0 | 1 | 1 | 4 | 0 | 9 | 0.1111 | 1 | 0 | 1 | 0 | 7 | 0.1429 | 5 | 0 | 14 | 0.0714 |
| 9_6355754 | 3 | 0.6667 | NA | NA | 1 | 1 | 4 | 0.5 | 8 | 0.625 | 1 | 0 | 1 | 1 | 7 | 0.5714 | 6 | 0.3333 | 15 | 0.4667 |
| 9_6355761 | 3 | 0.3333 | 1 | 0 | 1 | 0 | 4 | 0.25 | 9 | 0.2222 | 1 | 0 | NA | NA | 7 | 0.2857 | 6 | 0.6667 | 14 | 0.4286 |
| 9_6366505 | NA | NA | NA | NA | 1 | 1 | NA | NA | 1 | 1 | 1 | 1 | 1 | 0 | NA | NA | NA | NA | 2 | 0.5 |
| 9_6366510 | NA | NA | NA | NA | 1 | 1 | NA | NA | 1 | 1 | 1 | 1 | 1 | 1 | NA | NA | NA | NA | 2 | 1 |
| 9_6366564 | NA | NA | NA | NA | 1 | 1 | NA | NA | 1 | 1 | 1 | 1 | 1 | 1 | NA | NA | NA | NA | 2 | 1 |
| 9_6366565 | NA | NA | NA | NA | 1 | 1 | NA | NA | 1 | 1 | NA | NA | 1 | 1 | NA | NA | NA | NA | 1 | 1 |
| 9_6366568 | NA | NA | NA | NA | 1 | 1 | NA | NA | 1 | 1 | 1 | 1 | 1 | 1 | NA | NA | NA | NA | 2 | 1 |
| 9_6366597 | NA | NA | NA | NA | 1 | 1 | NA | NA | 1 | 1 | 1 | 1 | 1 | 0 | NA | NA | NA | NA | 2 | 0.5 |
| 9_6366599 | NA | NA | NA | NA | 1 | 1 | NA | NA | 1 | 1 | 1 | 1 | 1 | 1 | NA | NA | NA | NA | 2 | 1 |
| 9_6366602 | NA | NA | NA | NA | 1 | 1 | NA | NA | 1 | 1 | 1 | 1 | 1 | 0 | NA | NA | NA | NA | 2 | 0.5 |
| 9_6386073 | 1 | 1 | 3 | 0.6667 | 1 | 0 | NA | NA | 5 | 0.6 | NA | NA | 1 | 1 | 1 | 0 | 2 | 1 | 4 | 0.75 |
| 9_6386074 | 1 | 1 | 2 | 1 | 1 | 1 | NA | NA | 4 | 1 | NA | NA | 1 | 0 | 1 | 0 | 2 | 1 | 4 | 0.5 |
| 9_6386087 | 1 | 1 | 3 | 0.6667 | 1 | 1 | NA | NA | 5 | 0.8 | NA | NA | 1 | 0 | 1 | 1 | 2 | 0.5 | 4 | 0.5 |
| 9_6392773 | NA | NA | 1 | 1 | NA | NA | NA | NA | 1 | 1 | NA | NA | 1 | 0 | 1 | 1 | 1 | 0 | 3 | 0.3333 |
| 9_6392795 | 1 | 1 | 2 | 1 | 1 | 1 | NA | NA | 4 | 1 | NA | NA | 1 | 1 | 1 | 1 | 2 | 1 | 4 | 1 |
| 9_6392836 | 1 | 0 | 3 | 0.6667 | 1 | 1 | NA | NA | 5 | 0.6 | NA | NA | 1 | 0 | NA | NA | 2 | 0.5 | 3 | 0.3333 |
| 9_6392855 | 1 | 1 | 3 | 0 | NA | NA | NA | NA | 4 | 0.25 | NA | NA | 1 | 1 | NA | NA | 2 | 0.5 | 3 | 0.6667 |
| 9_6392887 | 1 | 1 | 2 | 0.5 | NA | NA | NA | NA | 3 | 0.6667 | NA | NA | NA | NA | NA | NA | 2 | 1 | 2 | 1 |
| 9_6392896 | 1 | 1 | 2 | 0 | NA | NA | NA | NA | 3 | 0.3333 | NA | NA | 1 | 1 | NA | NA | 2 | 1 | 3 | 1 |
| 9_6408576 | 106 | 0.0755 | 121 | 0.0579 | 97 | 0.0412 | 92 | 0.0435 | 416 | 0.0553 | 89 | 0.0674 | 120 | 0.0583 | 138 | 0.029 | 103 | 0.0583 | 450 | 0.0511 |
| 9_6408581 | 4 | 0.75 | 3 | 0.6667 | 1 | 0 | 2 | 0 | 10 | 0.5 | 1 | 1 | 1 | 0 | 1 | 0 | 2 | 0 | 5 | 0.2 |
| 9_6408958 | 5 | 0 | 2 | 0 | 4 | 0 | 2 | 0.5 | 13 | 0.0769 | 3 | 0 | 4 | 0.25 | 5 | 0.4 | 4 | 0 | 16 | 0.1875 |
| 9_6408964 | 5 | 0.2 | 2 | 0 | 4 | 0.25 | 4 | 0 | 15 | 0.1333 | 4 | 0 | 6 | 0.1667 | 8 | 0.25 | 4 | 0.25 | 22 | 0.1818 |
| 9_6408984 | 6 | 0.1667 | 5 | 0.2 | 4 | 0 | 4 | 0.25 | 19 | 0.1579 | 3 | 0 | 6 | 0 | 8 | 0.25 | 4 | 0 | 21 | 0.0952 |
| 9_6409345 | 57 | 0.0702 | 41 | 0.1707 | 40 | 0.075 | 63 | 0.0635 | 201 | 0.0896 | 34 | 0.0588 | 68 | 0.1324 | 59 | 0.0339 | 45 | 0.1111 | 206 | 0.0874 |
| 9_6409361 | 54 | 0.5185 | 39 | 0.4615 | 38 | 0.6579 | 60 | 0.5 | 191 | 0.5288 | 36 | 0.6667 | 62 | 0.5 | 55 | 0.5455 | 40 | 0.6 | 193 | 0.5648 |
| 9_6409365 | 49 | 0.0408 | 42 | 0.0238 | 39 | 0.1538 | 53 | 0.0566 | 183 | 0.0656 | 32 | 0.0625 | 62 | 0.0806 | 53 | 0.0755 | 48 | 0.0208 | 195 | 0.0615 |
| 9_6409371 | 47 | 0.2128 | 40 | 0.075 | 40 | 0.1 | 57 | 0.0702 | 184 | 0.1141 | 29 | 0.1379 | 59 | 0.1356 | 54 | 0.1111 | 47 | 0 | 189 | 0.0952 |
| 9_6421173 | 3 | 0 | 6 | 0.3333 | 1 | 0 | 1 | 0 | 11 | 0.1818 | 1 | 0 | 1 | 0 | 1 | 1 | NA | NA | 3 | 0.3333 |
| 9_6421174 | 5 | 0.2 | 6 | 0 | 1 | 0 | 1 | 0 | 13 | 0.0769 | 1 | 1 | 1 | 0 | 1 | 0 | NA | NA | 3 | 0.3333 |
| 9_6421174 | 3 | 0.6667 | 5 | 0 | 1 | 0 | 1 | 0 | 10 | 0.2 | 1 | 0 | 1 | 0 | 1 | 0 | 1 | 1 | 4 | 0.25 |
| 9_6421188 | 5 | 0 | 6 | 0.1667 | 1 | 0 | 1 | 0 | 13 | 0.0769 | 1 | 0 | 1 | 0 | 1 | 1 | 1 | 0 | 4 | 0.25 |
| 9_6421436 | 5 | 0.4 | 4 | 0.25 | 1 | 1 | 1 | 0 | 11 | 0.3636 | 2 | 0.5 | 3 | 0.3333 | 3 | 0 | 4 | 0.25 | 12 | 0.25 |
| 9_6421453 | 4 | 0.25 | 4 | 0.25 | 2 | 0.5 | 1 | 0 | 11 | 0.2727 | NA | NA | 2 | 0.5 | 3 | 0 | 4 | 0.25 | 9 | 0.2222 |
| 9_6421873 | 1 | 0 | 3 | 0.3333 | 2 | 0 | 1 | 1 | 7 | 0.2857 | 1 | 0 | 1 | 1 | 3 | 0.3333 | 3 | 0.3333 | 8 | 0.375 |
| 9_6421902 | 1 | 0 | 2 | 0.5 | 2 | 0 | 1 | 0 | 6 | 0.1667 | NA | NA | 1 | 0 | 2 | 0.5 | 4 | 0.25 | 7 | 0.2857 |
| 9_6421910 | 1 | 1 | 3 | 0.3333 | 1 | 1 | 1 | 0 | 6 | 0.5 | NA | NA | 1 | 1 | 2 | 0.5 | 3 | 0.3333 | 6 | 0.5 |
| 9_6422507 | 1 | 0 | 2 | 0.5 | 1 | 0 | 2 | 0 | 6 | 0.1667 | NA | NA | 1 | 1 | 1 | 0 | 3 | 0 | 5 | 0.2 |
| 9_6460774 | 1 | 1 | NA | NA | NA | NA | NA | NA | 1 | 1 | 1 | 1 | NA | NA | NA | NA | NA | NA | 1 | 1 |
| 9_6461343 | 1 | 1 | NA | NA | NA | NA | NA | NA | 1 | 1 | 1 | 0 | 1 | 0 | NA | NA | 1 | 1 | 3 | 0.3333 |
| 9_6461378 | 1 | 1 | NA | NA | NA | NA | NA | NA | 1 | 1 | 1 | 1 | 1 | 1 | NA | NA | NA | NA | 2 | 1 |
| 9_6461411 | 1 | 1 | NA | NA | NA | NA | 1 | 0 | 2 | 0.5 | 1 | 1 | 1 | 1 | 1 | 1 | 2 | 1 | 5 | 1 |
| 9_6461420 | 1 | 1 | NA | NA | NA | NA | 1 | 0 | 2 | 0.5 | NA | NA | 1 | 1 | 1 | 0 | 1 | 0 | 3 | 0.3333 |
| 9_6462352 | 9 | 0.2222 | 10 | 0.2 | 5 | 0.2 | 15 | 0.2 | 39 | 0.2051 | 7 | 0 | 10 | 0.3 | 6 | 0.1667 | 7 | 0 | 30 | 0.1333 |
| 9_6474974 | NA | NA | 1 | 1 | NA | NA | 1 | 0 | 2 | 0.5 | 1 | 0 | NA | NA | 1 | 1 | 1 | 0 | 3 | 0.3333 |
| 9_6477914 | 1 | 0 | 1 | 0 | NA | NA | 1 | 1 | 3 | 0.3333 | 1 | 0 | 1 | 1 | 1 | 0 | 1 | 0 | 4 | 0.25 |
| 9_6480444 | 1 | 0 | 2 | 1 | 1 | 1 | 2 | 1 | 6 | 0.8333 | 1 | 1 | 3 | 0.6667 | 3 | 0.6667 | 1 | 1 | 8 | 0.75 |
| 9_6480449 | 1 | 0 | 2 | 0.5 | 1 | 1 | 2 | 0.5 | 6 | 0.5 | 1 | 1 | 3 | 0 | 2 | 0 | 1 | 1 | 7 | 0.2857 |
| 9_6480449 | NA | NA | 2 | 0.5 | 1 | 0 | 2 | 0 | 5 | 0.2 | 1 | 0 | 3 | 0 | 3 | 0 | 1 | 1 | 8 | 0.125 |

| | | | | | | | | | | | | | | | | | | | | |
|---|---|---|---|---|---|---|---|---|---|---|---|---|---|---|---|---|---|---|---|---|
| 9_6481742 | 1 | 0 | 4 | 0 | 2 | 0 | 5 | 0.2 | 12 | 0.0833 | 1 | 1 | 5 | 0 | 3 | 0 | 2 | 0 | 11 | 0.0909 |
| 9_6482524 | 1 | 0 | 4 | 0.25 | 1 | 1 | 7 | 0.5714 | 13 | 0.4615 | 1 | 0 | 3 | 0.6667 | 4 | 0.5 | 1 | 0 | 9 | 0.4444 |
| 9_6482668 | 1 | 0 | 3 | 0.3333 | 2 | 0 | 2 | 0 | 8 | 0.125 | 2 | 0 | 4 | 0 | 5 | 0.2 | 1 | 0 | 12 | 0.0833 |
| 9_6486922 | 2 | 0 | 1 | 0 | 1 | 1 | 1 | 1 | 5 | 0.4 | NA | NA | 1 | 1 | 1 | 0 | 2 | 1 | 4 | 0.75 |
| 9_6486925 | 2 | 0.5 | 1 | 0 | 1 | 0 | 1 | 0 | 5 | 0.2 | NA | NA | 1 | 0 | 3 | 0.3333 | 2 | 0.5 | 6 | 0.3333 |
| 9_6489335 | 2 | 0 | 1 | 1 | 1 | 1 | 2 | 1 | 6 | 0.6667 | 2 | 1 | 2 | 0 | 4 | 0.25 | 3 | 0 | 11 | 0.2727 |
| 9_6489359 | 3 | 1 | 2 | 0 | 1 | 1 | 3 | 0.6667 | 9 | 0.6667 | 1 | 1 | 2 | 1 | 4 | 1 | 4 | 1 | 11 | 1 |
| 9_6489368 | 2 | 1 | 2 | 0.5 | 1 | 1 | 2 | 1 | 7 | 0.8571 | 2 | 1 | 3 | 1 | 4 | 1 | 4 | 1 | 13 | 1 |
| 9_6489373 | 3 | 1 | 3 | 0.3333 | 1 | 1 | 3 | 0.6667 | 10 | 0.7 | 2 | 0.5 | 3 | 0.3333 | 5 | 0.4 | 3 | 0.6667 | 13 | 0.4615 |
| 9_6489378 | 2 | 0.5 | 2 | 0 | 1 | 0 | 3 | 0.3333 | 8 | 0.25 | 2 | 0.5 | 3 | 0 | 5 | 0.4 | 3 | 0 | 13 | 0.2308 |
| 9_6489384 | 4 | 1 | 2 | 0 | 1 | 1 | 4 | 0.5 | 11 | 0.6364 | 2 | 0.5 | 4 | 0.25 | 6 | 0 | 4 | 0 | 16 | 0.125 |
| 9_6489441 | 26 | 0.3846 | 25 | 0.32 | 22 | 0.4091 | 14 | 0.1429 | 87 | 0.3333 | 12 | 0.25 | 19 | 0.2632 | 18 | 0.3889 | 16 | 0.375 | 65 | 0.3231 |
| 9_6489452 | 31 | 0.0323 | 30 | 0.0667 | 27 | 0.1111 | 18 | 0.1667 | 106 | 0.0849 | 18 | 0.0556 | 21 | 0.0952 | 19 | 0.1053 | 19 | 0.0526 | 77 | 0.0779 |
| 9_6489456 | 30 | 0.2 | 28 | 0.0714 | 28 | 0.0714 | 19 | 0.0526 | 105 | 0.1048 | 17 | 0.1176 | 23 | 0.1739 | 17 | 0.1176 | 19 | 0.0526 | 76 | 0.1184 |
| 9_6489473 | 27 | 0.3333 | 25 | 0.08 | 25 | 0.24 | 14 | 0.3571 | 91 | 0.2418 | 17 | 0.2941 | 19 | 0.1579 | 18 | 0.3889 | 19 | 0.3158 | 73 | 0.2877 |
| 9_6489530 | 19 | 0.2105 | 23 | 0.1304 | 18 | 0.2222 | 13 | 0.2308 | 73 | 0.1918 | 14 | 0.2857 | 19 | 0.2632 | 14 | 0.2143 | 15 | 0.3333 | 62 | 0.2742 |
| 9_6498806 | 8 | 0.375 | 9 | 0.3333 | 3 | 0.6667 | 6 | 0.5 | 26 | 0.4231 | 4 | 0.75 | 7 | 0.2857 | 7 | 0.1429 | 8 | 0.75 | 26 | 0.4615 |
| 9_6498815 | 8 | 0.375 | 12 | 0.3333 | 4 | 0.75 | 8 | 0.75 | 32 | 0.5 | 4 | 0.75 | 7 | 0.7143 | 5 | 0.6 | 10 | 0.3 | 26 | 0.5385 |
| 9_6498821 | 9 | 0.1111 | 13 | 0 | 4 | 0.25 | 8 | 0.125 | 34 | 0.0882 | 5 | 0 | 8 | 0.125 | 7 | 0.1429 | 11 | 0.2727 | 31 | 0.1613 |
| 9_6498832 | 9 | 0.2222 | 12 | 0.25 | 4 | 0.5 | 7 | 0.1429 | 32 | 0.25 | 3 | 0.3333 | 7 | 0.1429 | 7 | 0.2857 | 11 | 0.2727 | 28 | 0.25 |
| 9_6498833 | 8 | 0.25 | 12 | 0.1667 | 4 | 0 | 9 | 0 | 33 | 0.1212 | 4 | 0.25 | 7 | 0 | 7 | 0.1429 | 13 | 0 | 31 | 0.0645 |
| 9_6498835 | 2 | 0.5 | 5 | 0 | 1 | 0 | 6 | 0.1667 | 14 | 0.1429 | 1 | 0 | 6 | 0 | 4 | 0 | 7 | 0.1429 | 18 | 0.0556 |
| 9_6498844 | 8 | 0.875 | 12 | 0.5 | 4 | 1 | 9 | 0.5556 | 33 | 0.6667 | 6 | 0.6667 | 9 | 0.4444 | 6 | 0.5 | 13 | 0.5385 | 34 | 0.5294 |
| 9_6498845 | 9 | 0.2222 | 12 | 0.3333 | 3 | 0.3333 | 7 | 0.5714 | 31 | 0.3548 | 6 | 0.6667 | 9 | 0.5556 | 6 | 0.5 | 12 | 0.4167 | 33 | 0.5152 |
| 9_6498861 | 10 | 0.1 | 13 | 0.0769 | 5 | 0 | 9 | 0 | 37 | 0.0541 | 6 | 0.1667 | 8 | 0.125 | 7 | 0.1429 | 13 | 0 | 34 | 0.0882 |
| 9_6499044 | 12 | 0.0833 | 19 | 0.0526 | 10 | 0.2 | 19 | 0 | 60 | 0.0667 | 7 | 0.1429 | 8 | 0.25 | 3 | 0.3333 | 10 | 0.1 | 28 | 0.1786 |
| 9_6504253 | NA | NA | 1 | 0 | 1 | 1 | 1 | 1 | 3 | 0.6667 | 2 | 0.5 | 1 | 0 | 1 | 0 | 2 | 0 | 6 | 0.1667 |
| 9_6504296 | 1 | 0 | 1 | 0 | 1 | 0 | 3 | 0.6667 | 6 | 0.3333 | 1 | 0 | 1 | 1 | 1 | 0 | 3 | 0 | 6 | 0.1667 |
| 9_6504391 | 2 | 0 | 1 | 1 | 3 | 0 | 2 | 0 | 8 | 0.125 | 2 | 0 | 2 | 0.5 | 2 | 0 | 2 | 0 | 8 | 0.125 |
| 9_6523737 | 6 | 0.6667 | 8 | 0.875 | 4 | 0.5 | 7 | 0.8571 | 25 | 0.76 | 3 | 0.6667 | 4 | 0.5 | 7 | 0.7143 | 5 | 0.6 | 19 | 0.6316 |
| 9_6526620 | 7 | 0.4286 | 1 | 0 | 4 | 0.75 | 4 | 0.5 | 16 | 0.5 | 2 | 1 | 7 | 0.2857 | 1 | 1 | 5 | 0.8 | 15 | 0.6 |
| 9_6526651 | 7 | 0.4286 | 2 | 0 | 5 | 0.2 | 5 | 0 | 19 | 0.2105 | 2 | 0 | 9 | 0.3333 | 1 | 0 | 5 | 0 | 17 | 0.1765 |
| 9_6526668 | 5 | 0.2 | 2 | 0.5 | 5 | 0.2 | 5 | 0 | 17 | 0.1765 | 3 | 0.3333 | 7 | 0.1429 | 2 | 0 | 6 | 0.1667 | 18 | 0.1667 |
| 9_6528707 | 1 | 1 | NA | NA | NA | NA | NA | NA | 1 | 1 | NA | NA | NA | NA | NA | NA | 2 | 0.5 | 2 | 0.5 |
| 9_6528710 | 1 | 1 | NA | NA | NA | NA | NA | NA | 1 | 1 | NA | NA | NA | NA | 1 | 0 | 2 | 0.5 | 3 | 0.3333 |
| 9_6531814 | NA | NA | 1 | 1 | NA | NA | NA | NA | 1 | 1 | 1 | 1 | NA | NA | 1 | 1 | NA | NA | 2 | 1 |
| 9_6531844 | NA | NA | 1 | 1 | NA | NA | NA | NA | 1 | 1 | 1 | 1 | NA | NA | 1 | 0 | NA | NA | 2 | 0.5 |
| 9_6531845 | NA | NA | 1 | 1 | NA | NA | NA | NA | 1 | 1 | 1 | 1 | NA | NA | 1 | 1 | NA | NA | 2 | 1 |
| 9_6531877 | NA | NA | 1 | 1 | NA | NA | NA | NA | 1 | 1 | 1 | 0 | NA | NA | 1 | 1 | NA | NA | 2 | 0.5 |
| 9_6531898 | NA | NA | 1 | 1 | NA | NA | NA | NA | 1 | 1 | 1 | 1 | NA | NA | 1 | 0 | NA | NA | 2 | 0.5 |
| 9_6531914 | NA | NA | 1 | 1 | NA | NA | NA | NA | 1 | 1 | 1 | 1 | NA | NA | 1 | 1 | NA | NA | 2 | 1 |
| 9_6531986 | NA | NA | 1 | 1 | NA | NA | NA | NA | 1 | 1 | 1 | 1 | NA | NA | 1 | 1 | NA | NA | 2 | 1 |
| 9_6535941 | 1 | 0 | 1 | 0 | 3 | 0 | 2 | 1 | 7 | 0.2857 | 2 | 1 | 2 | 0 | 2 | 1 | NA | NA | 6 | 0.6667 |
| 9_6535949 | 2 | 0 | 1 | 0 | 3 | 0.6667 | 2 | 0.5 | 8 | 0.375 | 2 | 1 | 2 | 0.5 | 2 | 0.5 | NA | NA | 6 | 0.6667 |
| 9_6535950 | 2 | 0 | 1 | 1 | 2 | 1 | 2 | 1 | 7 | 0.7143 | 2 | 1 | 2 | 0.5 | 2 | 0.5 | NA | NA | 6 | 0.6667 |
| 9_6535963 | 2 | 1 | 1 | 1 | 1 | 1 | 3 | 1 | 7 | 1 | 1 | 1 | 2 | 1 | 2 | 1 | NA | NA | 5 | 1 |
| 9_6535978 | 1 | 1 | 1 | 1 | NA | NA | 2 | 1 | 4 | 1 | NA | NA | 1 | 1 | 1 | 0 | NA | NA | 2 | 0.5 |
| 9_6535995 | 1 | 1 | 1 | 1 | NA | NA | 2 | 0.5 | 4 | 0.75 | NA | NA | 1 | 1 | NA | NA | 1 | 1 | 2 | 1 |
| 9_6536002 | NA | NA | 1 | 1 | NA | NA | 1 | 1 | 2 | 1 | NA | NA | 1 | 1 | 1 | 1 | 1 | 1 | 3 | 1 |
| 9_6536019 | 1 | 1 | 1 | 1 | NA | NA | 1 | 1 | 3 | 1 | NA | NA | 1 | 0 | 1 | 0 | 1 | 1 | 3 | 0.3333 |
| 9_6539432 | 5 | 1 | 8 | 1 | 7 | 0.8571 | 4 | 1 | 24 | 0.9583 | 8 | 0.75 | 9 | 0.8889 | 10 | 0.8 | 6 | 1 | 33 | 0.8485 |
| 9_6539437 | 4 | 0 | 7 | 0.2857 | 7 | 0 | 4 | 0.5 | 22 | 0.1818 | 8 | 0.125 | 8 | 0.125 | 9 | 0.2222 | 5 | 0.2 | 30 | 0.1667 |
| 9_6539442 | 3 | 1 | 6 | 0.6667 | 6 | 0.8333 | 4 | 1 | 19 | 0.8421 | 7 | 0.4286 | 8 | 0.875 | 10 | 0.9 | 5 | 0.8 | 30 | 0.7667 |
| 9_6539443 | 5 | 1 | 7 | 0.8571 | 7 | 0.8571 | 4 | 0.75 | 23 | 0.8696 | 6 | 0.6667 | 8 | 0.625 | 10 | 0.9 | 5 | 0.6 | 29 | 0.7241 |
| 9_6546323 | 1 | 1 | NA | NA | 1 | 1 | NA | NA | 2 | 1 | 2 | 1 | NA | NA | NA | NA | NA | NA | 2 | 1 |
| 9_6546327 | 1 | 1 | NA | NA | NA | NA | NA | NA | 1 | 1 | 2 | 1 | NA | NA | NA | NA | NA | NA | 2 | 1 |
| 9_6546343 | 1 | 1 | NA | NA | 1 | 1 | NA | NA | 2 | 1 | 2 | 1 | NA | NA | NA | NA | NA | NA | 2 | 1 |
| 9_6546356 | NA | NA | NA | NA | 1 | 1 | NA | NA | 1 | 1 | 1 | 1 | NA | NA | NA | NA | NA | NA | 1 | 1 |
| 9_6546369 | NA | NA | NA | NA | 1 | 1 | NA | NA | 1 | 1 | 2 | 1 | NA | NA | NA | NA | NA | NA | 2 | 1 |
| 9_6546374 | NA | NA | NA | NA | 1 | 1 | NA | NA | 1 | 1 | 2 | 1 | NA | NA | NA | NA | NA | NA | 2 | 1 |
| 9_6546403 | NA | NA | NA | NA | 1 | 1 | NA | NA | 1 | 1 | 2 | 1 | NA | NA | NA | NA | NA | NA | 2 | 1 |
| 9_6546408 | NA | NA | NA | NA | 1 | 1 | NA | NA | 1 | 1 | 1 | 1 | NA | NA | NA | NA | NA | NA | 1 | 1 |
| 9_6559629 | 2 | 1 | 1 | 0 | 1 | 1 | NA | NA | 4 | 0.75 | NA | NA | NA | NA | 2 | 1 | 1 | 0 | 3 | 0.6667 |
| 9_6559650 | 29 | 0.3448 | 22 | 0.3636 | 31 | 0.4839 | 21 | 0.3333 | 103 | 0.3883 | 20 | 0.45 | 18 | 0.2222 | 19 | 0.3684 | 13 | 0.3846 | 70 | 0.3571 |
| 9_6559660 | 28 | 0.2857 | 14 | 0.3571 | 26 | 0.2308 | 19 | 0.3684 | 87 | 0.2989 | 15 | 0.2 | 16 | 0.1875 | 17 | 0.1765 | 12 | 0.1667 | 60 | 0.1833 |

| | | | | | | | | | | | | | | | | | | | | |
|---|---|---|---|---|---|---|---|---|---|---|---|---|---|---|---|---|---|---|---|---|
| 9_6559674 | 26 | 0.0769 | 22 | 0.2273 | 32 | 0.25 | 22 | 0.1818 | 102 | 0.1863 | 20 | 0.25 | 20 | 0.25 | 19 | 0.3158 | 13 | 0.1538 | 72 | 0.25 |
| 9_6559743 | 31 | 0.129 | 28 | 0.2143 | 37 | 0.1892 | 25 | 0.12 | 121 | 0.1653 | 20 | 0.1 | 19 | 0.2105 | 21 | 0.1905 | 19 | 0.1053 | 79 | 0.1519 |
| 9_6560266 | NA | NA | NA | NA | 1 | 1 | NA | NA | 1 | 1 | 1 | 1 | 5 | 0.8 | NA | NA | NA | NA | 6 | 0.8333 |
| 9_6560277 | NA | NA | NA | NA | 1 | 1 | 2 | 1 | 3 | 1 | 1 | 1 | 3 | 1 | 1 | 1 | NA | NA | 5 | 1 |
| 9_6560834 | NA | NA | NA | NA | 1 | 1 | NA | NA | 1 | 1 | NA | NA | 1 | 1 | 1 | 0 | NA | NA | 2 | 0.5 |
| 9_6560970 | NA | NA | NA | NA | 1 | 0 | 1 | 1 | 2 | 0.5 | NA | NA | 1 | 1 | NA | NA | NA | NA | 1 | 1 |
| 9_6561023 | NA | NA | NA | NA | 1 | 0 | 1 | 1 | 2 | 0.5 | NA | NA | 1 | 1 | NA | NA | NA | NA | 1 | 1 |
| 9_6561030 | NA | NA | NA | NA | 1 | 1 | 1 | 1 | 2 | 1 | NA | NA | 1 | 1 | NA | NA | NA | NA | 1 | 1 |
| 9_6573760 | 1 | 1 | NA | NA | NA | NA | NA | NA | 1 | 1 | NA | NA | NA | NA | NA | NA | 2 | 0.5 | 2 | 0.5 |
| 9_6573763 | 1 | 1 | NA | NA | NA | NA | NA | NA | 1 | 1 | NA | NA | NA | NA | NA | NA | 2 | 0.5 | 2 | 0.5 |
| 9_6573923 | 4 | 0.25 | 2 | 0 | 4 | 0 | 4 | 0.5 | 14 | 0.2143 | 3 | 0.3333 | 2 | 0 | 2 | 0.5 | 1 | 0 | 8 | 0.25 |
| 9_6573937 | 4 | 0.25 | 3 | 0 | 5 | 0 | 3 | 0 | 15 | 0.0667 | 2 | 0.5 | 2 | 0 | 1 | 1 | 2 | 0 | 7 | 0.2857 |
| 9_6574064 | 4 | 0.25 | 1 | 0 | 3 | 0.3333 | 2 | 0 | 10 | 0.2 | 4 | 0.25 | 1 | 1 | 1 | 0 | 2 | 0.5 | 8 | 0.375 |
| 9_6586575 | NA | NA | 1 | 0 | 2 | 1 | NA | NA | 3 | 0.6667 | 1 | 1 | 1 | 0 | NA | NA | NA | NA | 2 | 0.5 |
| 9_6586580 | NA | NA | 1 | 1 | 2 | 0.5 | NA | NA | 3 | 0.6667 | 1 | 1 | 1 | 1 | NA | NA | NA | NA | 2 | 1 |
| 9_6586582 | NA | NA | 1 | 1 | 2 | 0 | NA | NA | 3 | 0.3333 | 1 | 1 | 1 | 0 | NA | NA | NA | NA | 2 | 0.5 |
| 9_6594694 | NA | NA | 2 | 0 | 2 | 0.5 | NA | NA | 4 | 0.25 | 1 | 0 | 3 | 0 | NA | NA | 1 | 1 | 5 | 0.2 |
| 9_6594699 | NA | NA | 2 | 0 | 1 | 1 | NA | NA | 3 | 0.3333 | NA | NA | 1 | 1 | NA | NA | 1 | 1 | 2 | 1 |
| 9_6594717 | NA | NA | 2 | 0.5 | 1 | 1 | NA | NA | 3 | 0.6667 | 1 | 1 | 3 | 1 | 1 | 1 | 1 | 1 | 6 | 1 |
| 9_6594721 | NA | NA | 2 | 0 | 2 | 0.5 | NA | NA | 4 | 0.25 | 1 | 0 | 3 | 0.6667 | NA | NA | 1 | 0 | 5 | 0.4 |
| 9_6594722 | NA | NA | 2 | 1 | 2 | 1 | NA | NA | 4 | 1 | NA | NA | 3 | 1 | 1 | 1 | 1 | 1 | 5 | 1 |
| 9_6594765 | NA | NA | 3 | 0.3333 | 1 | 1 | NA | NA | 4 | 0.5 | 1 | 0 | 3 | 0.6667 | 1 | 0 | 1 | 1 | 6 | 0.5 |
| 9_6594782 | NA | NA | 3 | 1 | 2 | 1 | NA | NA | 5 | 1 | 1 | 1 | 3 | 0.3333 | 1 | 1 | NA | NA | 5 | 0.6 |
| 9_6594805 | NA | NA | 3 | 0.6667 | 2 | 0 | NA | NA | 5 | 0.4 | 1 | 0 | 3 | 0.6667 | 1 | 1 | NA | NA | 5 | 0.6 |
| 9_6594814 | NA | NA | 3 | 1 | 2 | 1 | NA | NA | 5 | 1 | 1 | 1 | 3 | 1 | 1 | 0 | NA | NA | 5 | 0.8 |
| 9_6594816 | NA | NA | 3 | 1 | 2 | 1 | NA | NA | 5 | 1 | 1 | 1 | 2 | 1 | 1 | 1 | NA | NA | 4 | 1 |
| 9_6594817 | NA | NA | 2 | 1 | 2 | 1 | NA | NA | 4 | 1 | 1 | 1 | 3 | 0.6667 | 1 | 0 | NA | NA | 5 | 0.6 |
| 9_6594819 | NA | NA | 2 | 1 | 2 | 0.5 | NA | NA | 4 | 0.75 | 1 | 0 | 3 | 0.6667 | 1 | 1 | NA | NA | 5 | 0.6 |
| 9_6594823 | 1 | 0 | 3 | 0 | 2 | 1 | NA | NA | 6 | 0.3333 | NA | NA | 3 | 0.6667 | NA | NA | NA | NA | 3 | 0.6667 |
| 9_6594853 | 2 | 1 | 3 | 0.3333 | 2 | 0.5 | NA | NA | 7 | 0.5714 | 1 | 0 | 3 | 0.6667 | 1 | 1 | NA | NA | 5 | 0.6 |
| 9_6594891 | 1 | 0 | 2 | 0.5 | 1 | 1 | NA | NA | 4 | 0.5 | NA | NA | 2 | 0.5 | 1 | 0 | NA | NA | 3 | 0.3333 |
| 9_6595128 | 3 | 0 | 4 | 0.25 | 1 | 1 | 1 | 0 | 9 | 0.2222 | 2 | 0 | 6 | 0 | 3 | 0.3333 | 2 | 0 | 13 | 0.0769 |
| 9_6595213 | 3 | 1 | 3 | 1 | 3 | 1 | 2 | 1 | 11 | 1 | 2 | 1 | 9 | 1 | 4 | 1 | 4 | 1 | 19 | 1 |
| 9_6595223 | 3 | 0 | 4 | 0.25 | 3 | 0 | 2 | 0 | 12 | 0.0833 | 2 | 0 | 9 | 0 | 5 | 0.4 | 4 | 0 | 20 | 0.1 |
| 9_6597431 | NA | NA | NA | NA | 2 | 0.5 | NA | NA | 2 | 0.5 | NA | NA | 1 | 0 | 1 | 1 | 1 | 0 | 3 | 0.3333 |
| 9_6599570 | 16 | 0.125 | 16 | 0.3125 | 12 | 0.3333 | 15 | 0.4 | 59 | 0.2881 | 11 | 0.2727 | 9 | 0.3333 | 7 | 0.5714 | 10 | 0 | 37 | 0.2703 |
| 9_6599573 | 15 | 0.2667 | 16 | 0 | 13 | 0.0769 | 16 | 0 | 60 | 0.0833 | 8 | 0 | 11 | 0.0909 | 7 | 0 | 11 | 0.0909 | 37 | 0.0541 |
| 9_6635543 | 1 | 0 | 1 | 1 | NA | NA | 2 | 1 | 4 | 0.75 | NA | NA | 1 | 1 | NA | NA | NA | NA | 1 | 1 |
| 9_6635546 | 1 | 0 | 1 | 1 | NA | NA | 2 | 1 | 4 | 0.75 | NA | NA | 1 | 1 | NA | NA | NA | NA | 1 | 1 |
| 9_6635562 | 1 | 0 | 1 | 1 | NA | NA | 2 | 0 | 4 | 0.25 | 1 | 0 | 1 | 1 | NA | NA | NA | NA | 2 | 0.5 |
| 9_6635563 | 1 | 1 | 1 | 1 | NA | NA | 2 | 0.5 | 4 | 0.75 | 1 | 1 | 1 | 1 | NA | NA | NA | NA | 2 | 1 |
| 9_6635819 | 2 | 1 | 2 | 1 | NA | NA | 2 | 1 | 6 | 1 | 1 | 1 | 2 | 0.5 | NA | NA | NA | NA | 3 | 0.6667 |
| 9_6635838 | 2 | 1 | 2 | 0 | NA | NA | 1 | 1 | 5 | 0.6 | 1 | 1 | 2 | 0 | NA | NA | NA | NA | 3 | 0.3333 |
| 9_6635855 | 2 | 0.5 | 2 | 0.5 | NA | NA | 2 | 1 | 6 | 0.6667 | NA | NA | 2 | 1 | NA | NA | NA | NA | 2 | 1 |
| 9_6641554 | 7 | 0.4286 | 12 | 0.25 | 7 | 0 | 10 | 0.5 | 36 | 0.3056 | 2 | 0.5 | 12 | 0.25 | 4 | 0.5 | 8 | 0.25 | 26 | 0.3077 |
| 9_6641557 | 5 | 0.4 | 7 | 0.2857 | 12 | 0.25 | 9 | 0.4444 | 33 | 0.3333 | 5 | 0.4 | 13 | 0.3077 | 4 | 0.25 | 10 | 0.7 | 32 | 0.4375 |
| 9_6641559 | 7 | 0.1429 | 11 | 0 | 11 | 0 | 9 | 0.2222 | 38 | 0.0789 | 4 | 0 | 13 | 0 | 2 | 0 | 11 | 0.1818 | 30 | 0.0667 |
| 9_6641560 | 7 | 0.4286 | 12 | 0.5833 | 9 | 0.4444 | 9 | 0.2222 | 37 | 0.4324 | 4 | 0.5 | 10 | 0.6 | 3 | 0.3333 | 12 | 0.5 | 29 | 0.5172 |
| 9_6642175 | NA | NA | NA | NA | 1 | 1 | NA | NA | 1 | 1 | NA | NA | 1 | 1 | NA | NA | NA | NA | 1 | 1 |
| 9_6642205 | NA | NA | NA | NA | 1 | 1 | NA | NA | 1 | 1 | NA | NA | 1 | 1 | NA | NA | NA | NA | 1 | 1 |
| 9_6642388 | 4 | 0.25 | 5 | 0.2 | 4 | 0.25 | 4 | 0.25 | 17 | 0.2353 | 4 | 0.25 | 6 | 0 | 5 | 0 | 2 | 0 | 17 | 0.0588 |
| 9_6642514 | 6 | 0.3333 | 5 | 0.2 | 2 | 0 | 2 | 0 | 15 | 0.2 | 2 | 0 | 5 | 0 | 5 | 0.4 | 3 | 0 | 15 | 0.1333 |
| 9_6642526 | 7 | 0.2857 | 6 | 0.3333 | 2 | 0 | 4 | 0.25 | 19 | 0.2632 | 3 | 0.3333 | 5 | 0.2 | 5 | 0.2 | 2 | 1 | 15 | 0.3333 |
| 9_6642566 | 4 | 1 | 6 | 0.1667 | 2 | 0 | 4 | 0.25 | 16 | 0.375 | 3 | 0 | 6 | 0.3333 | 4 | 0 | 3 | 0.3333 | 16 | 0.1875 |
| 9_6642570 | 5 | 0.6 | 5 | 0.6 | 2 | 1 | 4 | 0.75 | 16 | 0.6875 | 2 | 0 | 6 | 1 | 6 | 0.8333 | 3 | 1 | 17 | 0.8235 |
| 9_6642576 | 5 | 0.2 | 5 | 0 | 2 | 0 | 3 | 0 | 15 | 0.0667 | 3 | 0 | 6 | 0 | 5 | 0.2 | 3 | 0 | 17 | 0.0588 |
| 9_6642578 | 5 | 0.4 | 5 | 0.6 | 2 | 0 | 4 | 0.5 | 16 | 0.4375 | 3 | 0 | 4 | 1 | 6 | 0.1667 | 3 | 0.6667 | 16 | 0.4375 |
| 9_6649965 | 1 | 1 | NA | NA | NA | NA | NA | NA | 1 | 1 | NA | NA | NA | NA | 2 | 0.5 | NA | NA | 2 | 0.5 |
| 9_6650020 | NA | NA | NA | NA | NA | NA | 1 | 1 | 1 | 1 | NA | NA | 1 | 1 | 3 | 0.6667 | 1 | 1 | 5 | 0.8 |
| 9_6650036 | NA | NA | NA | NA | NA | NA | 1 | 1 | 1 | 1 | NA | NA | 1 | 1 | 3 | 0.6667 | 1 | 1 | 5 | 0.8 |
| 9_6650039 | NA | NA | NA | NA | NA | NA | 1 | 1 | 1 | 1 | NA | NA | 1 | 1 | 2 | 1 | 1 | 1 | 4 | 1 |
| 9_6650054 | NA | NA | NA | NA | NA | NA | 1 | 1 | 1 | 1 | NA | NA | 1 | 1 | 3 | 1 | 1 | 1 | 5 | 1 |
| 9_6650063 | NA | NA | NA | NA | NA | NA | 1 | 1 | 1 | 1 | NA | NA | 1 | 0 | 2 | 0.5 | 1 | 1 | 4 | 0.5 |
| 9_6650076 | NA | NA | NA | NA | NA | NA | 1 | 1 | 1 | 1 | NA | NA | 1 | 1 | 3 | 1 | 1 | 1 | 5 | 1 |
| 9_6650081 | NA | NA | NA | NA | NA | NA | 1 | 1 | 1 | 1 | NA | NA | NA | NA | 1 | 1 | NA | NA | 1 | 1 |

| | | | | | | | | | | | | | | | | | | | | |
|---|---|---|---|---|---|---|---|---|---|---|---|---|---|---|---|---|---|---|---|---|
| 9_66500860 | NA | NA | NA | NA | NA | NA | 1 | 1 | 1 | 1 | NA | NA | NA | NA | 2 | 0.5 | 1 | 0 | 3 | 0.3333 |
| 9_66671672 | 2 | 0.5 | 2 | 0 | 3 | 0 | 1 | 0 | 8 | 0.125 | 1 | 0 | NA | NA | 3 | 0.3333 | 5 | 0.2 | 9 | 0.2222 |
| 9_66671684 | 2 | 0 | 1 | 0 | 3 | 1 | 1 | 0 | 7 | 0.4286 | 1 | 0 | NA | NA | 3 | 0.6667 | 5 | 0.4 | 9 | 0.4444 |
| 9_66671708 | 2 | 1 | 2 | 0.5 | 3 | 0.6667 | 1 | 1 | 8 | 0.75 | 1 | 0 | NA | NA | 3 | 0.6667 | 5 | 1 | 9 | 0.7778 |
| 9_66671764 | 2 | 0.5 | 1 | 1 | 1 | 0 | 1 | 0 | 5 | 0.4 | 1 | 0 | NA | NA | 3 | 0 | 4 | 0.25 | 8 | 0.125 |
| 9_66671840 | 2 | 1 | 1 | 1 | 2 | 1 | 1 | 1 | 6 | 1 | 2 | 1 | NA | NA | 2 | 0.5 | 2 | 0.5 | 6 | 0.6667 |
| 9_66671845 | 1 | 0 | 1 | 0 | 2 | 0.5 | 1 | 0 | 5 | 0.2 | 1 | 0 | NA | NA | 3 | 0 | 4 | 0.5 | 8 | 0.25 |
| 9_66672641 | 2 | 0 | 1 | 0 | 2 | 0.5 | 2 | 0 | 7 | 0.1429 | 2 | 0.5 | 1 | 1 | 7 | 0.1429 | 6 | 0.3333 | 16 | 0.3125 |
| 9_66673193 | 2 | 0.5 | 2 | 0 | 3 | 0 | 2 | 0 | 9 | 0.1111 | 2 | 0.5 | 1 | 0 | 7 | 0 | 5 | 0.4 | 15 | 0.2 |
| 9_66673359 | 2 | 1 | 1 | 1 | 2 | 1 | 2 | 0.5 | 7 | 0.8571 | 2 | 0.5 | 1 | 1 | 7 | 1 | 4 | 0.5 | 14 | 0.7857 |
| 9_66673394 | 3 | 0 | 2 | 0.5 | 4 | 0.5 | 2 | 0 | 11 | 0.2727 | 2 | 0 | 1 | 0 | 6 | 0.1667 | 6 | 0 | 15 | 0.0667 |
| 9_66673805 | 4 | 0 | 1 | 0 | 4 | 0.25 | 4 | 0 | 13 | 0.0769 | 3 | 0 | 2 | 0.5 | 9 | 0 | 5 | 0 | 19 | 0.0526 |
| 9_66673875 | 4 | 0 | 2 | 0.5 | 3 | 0.3333 | 4 | 0 | 13 | 0.1538 | 3 | 0 | 2 | 0 | 8 | 0.375 | 6 | 0 | 19 | 0.1579 |
| 9_66685638 | NA | NA | 2 | 0.5 | 1 | 0 | 3 | 0.3333 | 6 | 0.3333 | 2 | 0.5 | 3 | 0.6667 | 3 | 0.3333 | 2 | 0.5 | 10 | 0.5 |
| 9_66686234 | 2 | 1 | 1 | 1 | 2 | 0.5 | 3 | 0.6667 | 8 | 0.75 | 2 | 1 | 3 | 0.6667 | 4 | 0.5 | 2 | 1 | 11 | 0.7273 |
| 9_66686258 | 2 | 0.5 | 1 | 1 | 2 | 1 | 3 | 0.6667 | 8 | 0.75 | 4 | 0.5 | 4 | 0 | 3 | 0.3333 | 2 | 1 | 13 | 0.3846 |
| 9_66686373 | 1 | 0 | 1 | 1 | 1 | 1 | 4 | 1 | 7 | 0.8571 | 4 | 1 | 1 | 1 | 2 | 1 | 1 | 1 | 8 | 1 |
| 9_66686388 | 2 | 1 | 1 | 1 | 2 | 0.5 | 4 | 1 | 9 | 0.8889 | 4 | 1 | 3 | 1 | 2 | 1 | NA | NA | 9 | 1 |
| 9_66686404 | 1 | 1 | 1 | 1 | 2 | 0 | 4 | 0.75 | 8 | 0.625 | 3 | 0.6667 | 3 | 1 | 4 | 0.75 | 1 | 1 | 11 | 0.8182 |
| 9_66686420 | 2 | 0 | 2 | 0 | 2 | 0.5 | 3 | 0 | 9 | 0.1111 | 3 | 0 | 3 | 0.3333 | 4 | 0 | 2 | 0.5 | 12 | 0.1667 |
| 9_66686552 | 2 | 0 | 2 | 0 | 2 | 0 | 2 | 0.5 | 8 | 0.125 | 3 | 0 | 4 | 0 | 3 | 0 | 2 | 0.5 | 12 | 0.0833 |
| 9_66726315 | NA | NA | 3 | 1 | NA | NA | 3 | 0.3333 | 6 | 0.6667 | 1 | 1 | NA | NA | NA | NA | NA | NA | 1 | 1 |
| 9_66726339 | NA | NA | 3 | 1 | 1 | 0 | 3 | 1 | 7 | 0.8571 | 1 | 1 | NA | NA | 1 | 0 | NA | NA | 2 | 0.5 |
| 9_66726363 | NA | NA | 3 | 1 | 1 | 1 | 3 | 1 | 7 | 1 | 1 | 1 | NA | NA | NA | NA | NA | NA | 1 | 1 |
| 9_66726469 | NA | NA | 2 | 1 | 1 | 1 | 1 | 1 | 4 | 1 | 1 | 1 | NA | NA | 1 | 1 | NA | NA | 2 | 1 |
| 9_66726485 | NA | NA | 3 | 0.6667 | 1 | 1 | 3 | 1 | 7 | 0.8571 | 1 | 1 | NA | NA | NA | NA | NA | NA | 1 | 1 |
| 9_66751093 | 5 | 0.2 | 5 | 0 | 5 | 0.2 | 2 | 0 | 17 | 0.1176 | 4 | 0.5 | 7 | 0 | 5 | 0.2 | NA | NA | 16 | 0.1875 |
| 9_66751167 | 5 | 0.2 | 4 | 0 | 5 | 0.2 | 2 | 0 | 16 | 0.125 | 3 | 0.3333 | 8 | 0.25 | 4 | 0.5 | NA | NA | 15 | 0.3333 |
| 9_66751187 | 5 | 0.2 | 5 | 0 | 3 | 0 | 2 | 0 | 15 | 0.0667 | 4 | 0 | 8 | 0.125 | 4 | 0 | NA | NA | 16 | 0.0625 |
| 9_66751220 | 5 | 0.4 | 4 | 0.25 | 4 | 0.75 | 1 | 1 | 14 | 0.5 | 3 | 0.3333 | 8 | 0.625 | 4 | 0.75 | NA | NA | 15 | 0.6 |
| 9_66752121 | 5 | 0.2 | 4 | 0 | 2 | 0 | NA | NA | 11 | 0.0909 | 3 | 0 | 6 | 0 | 3 | 0.3333 | NA | NA | 12 | 0.0833 |
| 9_66754354 | 2 | 1 | 3 | 0.6667 | 1 | 0 | NA | NA | 6 | 0.6667 | 2 | 0 | 2 | 0.5 | 2 | 0.5 | NA | NA | 6 | 0.3333 |
| 9_66754680 | 2 | 0 | 3 | 0.3333 | 1 | 0 | NA | NA | 6 | 0.1667 | 1 | 0 | 2 | 0.5 | 2 | 0 | NA | NA | 5 | 0.2 |
| 9_66754778 | 1 | 1 | 3 | 0.3333 | 1 | 1 | NA | NA | 5 | 0.6 | 2 | 1 | 2 | 0.5 | 1 | 0 | NA | NA | 5 | 0.6 |
| 9_66754826 | 2 | 1 | 2 | 0.5 | NA | NA | NA | NA | 4 | 0.75 | 1 | 1 | 2 | 1 | 2 | 1 | NA | NA | 5 | 1 |
| 9_66754979 | 2 | 1 | 3 | 1 | 1 | 1 | NA | NA | 6 | 1 | 2 | 1 | 2 | 1 | 2 | 1 | NA | NA | 6 | 1 |
| 9_66754997 | 1 | 1 | 2 | 0.5 | 1 | 0 | NA | NA | 4 | 0.5 | 2 | 1 | 2 | 0.5 | 2 | 1 | NA | NA | 6 | 0.8333 |
| 9_66862759 | 1 | 1 | 8 | 0.125 | 4 | 0 | 1 | 0 | 14 | 0.1429 | 2 | 0 | 4 | 0.25 | 1 | 1 | 3 | 0 | 10 | 0.2 |
| 9_66862961 | 4 | 0.5 | 8 | 0.125 | 4 | 0.25 | 1 | 1 | 17 | 0.2941 | 2 | 0 | 4 | 0.25 | 1 | 0 | 4 | 0.5 | 11 | 0.2727 |
| 9_66862970 | 3 | 0.3333 | 8 | 0.125 | 3 | 0.3333 | 1 | 0 | 15 | 0.2 | 2 | 0.5 | 3 | 0.3333 | 1 | 0 | 4 | 0.75 | 10 | 0.5 |
| 9_66863090 | 4 | 1 | 6 | 0.6667 | 4 | 1 | 1 | 1 | 15 | 0.8667 | 1 | 1 | 3 | 0.6667 | 1 | 1 | 2 | 1 | 7 | 0.8571 |
| 9_66863095 | 3 | 0 | 5 | 0.6 | 4 | 0.25 | 1 | 0 | 13 | 0.3077 | 2 | 1 | 3 | 0.6667 | 1 | 1 | 2 | 1 | 8 | 0.875 |
| 9_66863268 | 4 | 0.5 | 6 | 0.3333 | 4 | 0.5 | 1 | 1 | 15 | 0.4667 | 2 | 0.5 | 4 | 0.25 | 1 | 1 | 4 | 0.5 | 11 | 0.4545 |
| 9_66863286 | 5 | 0 | 7 | 0.1429 | 4 | 0.25 | 1 | 0 | 17 | 0.1176 | 2 | 0.5 | 4 | 0 | 1 | 0 | 2 | 0 | 9 | 0.1111 |
| 9_66863327 | 4 | 0.75 | 8 | 0.875 | 4 | 1 | 1 | 1 | 17 | 0.8824 | 1 | 1 | 4 | 1 | 1 | 1 | 4 | 1 | 10 | 1 |
| 9_66878072 | 2 | 0 | 5 | 0.8 | 3 | 0.3333 | 1 | 0 | 11 | 0.4545 | 1 | 1 | 2 | 0.5 | 1 | 0 | 2 | 0 | 6 | 0.3333 |
| 9_66878157 | 3 | 0.3333 | 7 | 0.2857 | 2 | 0 | 1 | 0 | 13 | 0.2308 | 1 | 0 | 4 | 0.5 | NA | NA | 2 | 0 | 7 | 0.2857 |
| 9_66930667 | 12 | 0.0833 | 6 | 0 | 9 | 0.1111 | 6 | 0 | 33 | 0.0606 | 10 | 0 | 16 | 0.0625 | 15 | 0.0667 | 18 | 0.0556 | 59 | 0.0508 |
| 9_66931502 | 23 | 0.1304 | 16 | 0.125 | 13 | 0.2308 | 20 | 0 | 72 | 0.1111 | 14 | 0.1429 | 30 | 0.1333 | 21 | 0.3333 | 22 | 0.0455 | 87 | 0.1609 |
| 9_66931519 | 24 | 0.3333 | 15 | 0.3333 | 14 | 0.2143 | 19 | 0.4211 | 72 | 0.3333 | 13 | 0.3846 | 33 | 0.3939 | 18 | 0.3333 | 18 | 0.2778 | 82 | 0.3537 |
| 9_66931724 | 17 | 0.4118 | 13 | 0.2308 | 15 | 0.1333 | 19 | 0.4211 | 64 | 0.3125 | 11 | 0.0909 | 30 | 0.3 | 17 | 0.3529 | 22 | 0.5 | 80 | 0.3375 |
| 9_66935312 | 12 | 0.0833 | 11 | 0.0909 | 14 | 0.1429 | 10 | 0.2 | 47 | 0.1277 | 7 | 0 | 11 | 0.0909 | 9 | 0.2222 | 7 | 0.1429 | 34 | 0.1176 |
| 9_67124873 | 3 | 0 | 4 | 0 | 6 | 0.1667 | 9 | 0.2222 | 22 | 0.1364 | 2 | 1 | 4 | 0 | 8 | 0.25 | 4 | 0.25 | 18 | 0.2778 |
| 9_67126804 | 3 | 0 | 1 | 1 | 5 | 0.2 | 3 | 0 | 12 | 0.1667 | 1 | 0 | 1 | 1 | 2 | 0 | 2 | 0 | 6 | 0.1667 |
| 9_67126899 | 3 | 0 | 1 | 0 | 5 | 0.2 | 4 | 0.25 | 13 | 0.1538 | 1 | 0 | 2 | 0.5 | 5 | 0 | 1 | 0 | 9 | 0.1111 |
| 9_67128191 | NA | NA | 4 | 0.75 | 3 | 0.3333 | 4 | 0.5 | 11 | 0.5455 | NA | NA | 4 | 0.75 | 2 | 0 | NA | NA | 6 | 0.5 |
| 9_67128205 | 4 | 0 | 8 | 0.125 | 6 | 0.1667 | 9 | 0.1111 | 27 | 0.1111 | 3 | 0 | 7 | 0 | 4 | 0.25 | 6 | 0 | 20 | 0.05 |
| 9_67128226 | 7 | 0.8571 | 9 | 0.8889 | 6 | 0.8333 | 9 | 0.8889 | 31 | 0.871 | 3 | 0.6667 | 7 | 1 | 5 | 1 | 5 | 1 | 20 | 0.95 |
| 9_67128268 | 5 | 0.8 | 7 | 1 | 7 | 1 | 10 | 1 | 29 | 0.9655 | 2 | 1 | 6 | 1 | 4 | 0.75 | 6 | 1 | 18 | 0.9444 |
| 9_67128309 | 7 | 0.5714 | 5 | 0.4 | 7 | 0.1429 | 12 | 0.6667 | 31 | 0.4839 | 2 | 0 | 7 | 0.2857 | 5 | 0.2 | 4 | 0.25 | 18 | 0.2222 |
| 9_67128350 | 4 | 0.75 | 8 | 0.75 | 6 | 0.6667 | 11 | 0.8182 | 29 | 0.7586 | 3 | 0.6667 | 5 | 0.8 | 4 | 0.75 | 4 | 1 | 16 | 0.8125 |
| 9_67128417 | 5 | 0.8 | 5 | 1 | 6 | 1 | 11 | 0.9091 | 27 | 0.9259 | 1 | 1 | 6 | 0.8333 | 4 | 0.75 | 4 | 0.75 | 15 | 0.8 |
| 9_67128437 | 5 | 0.4 | 7 | 0.1429 | 7 | 0.1429 | 9 | 0.6667 | 28 | 0.3571 | 2 | 0 | 8 | 0.375 | 4 | 0 | 5 | 0.6 | 19 | 0.3158 |
| 9_67128574 | 6 | 0 | 8 | 0.125 | 4 | 0 | 12 | 0.1667 | 30 | 0.1 | 2 | 0 | 6 | 0 | 4 | 0.25 | 5 | 0 | 17 | 0.0588 |
| 9_67128585 | 5 | 0.2 | 8 | 0.125 | 4 | 0.25 | 12 | 0 | 29 | 0.1034 | 1 | 0 | 8 | 0.25 | 4 | 0 | 6 | 0 | 19 | 0.1053 |

| | | | | | | | | | | | | | | | | | | | | |
|---|---|---|---|---|---|---|---|---|---|---|---|---|---|---|---|---|---|---|---|---|
| 9_6712861 | 4 | 0.5 | 6 | 0.8333 | 2 | 1 | 12 | 0.5 | 24 | 0.625 | 2 | 1 | 6 | 0.5 | 3 | 1 | 2 | 1 | 13 | 0.7692 |
| 9_6712861 | 2 | 1 | 4 | 0.75 | 1 | 0 | 8 | 0.75 | 15 | 0.7333 | 1 | 1 | 6 | 1 | 3 | 0.3333 | 3 | 0.6667 | 13 | 0.7692 |
| 9_6766524 | NA | NA | 1 | 1 | NA | NA | NA | NA | 1 | 1 | NA | NA | 1 | 1 | NA | NA | NA | NA | 1 | 1 |
| 9_6789070 | 2 | 0.5 | 1 | 1 | 2 | 1 | 4 | 0.25 | 9 | 0.5556 | 1 | 1 | NA | NA | 2 | 0 | 2 | 0.5 | 5 | 0.4 |
| 9_6789108 | 2 | 0 | 1 | 1 | 2 | 1 | 3 | 0.6667 | 8 | 0.625 | 1 | 0 | NA | NA | 2 | 0 | 2 | 0.5 | 5 | 0.2 |
| 9_6789230 | 2 | 1 | 3 | 0.3333 | 1 | 0 | 6 | 0.1667 | 12 | 0.3333 | 2 | 0 | 1 | 0 | 2 | 0.5 | 5 | 0.6 | 10 | 0.4 |
| 9_6789297 | 3 | 0 | 4 | 0 | 1 | 0 | 6 | 0.1667 | 14 | 0.0714 | 1 | 0 | 1 | 0 | 2 | 0.5 | 5 | 0.2 | 9 | 0.2222 |
| 9_6789361 | 4 | 0.75 | 6 | 0.5 | 1 | 1 | 6 | 0.8333 | 17 | 0.7059 | 3 | 0.3333 | 2 | 1 | 3 | 0.3333 | 6 | 0.5 | 14 | 0.5 |
| 9_6789362 | 4 | 0 | 3 | 0.6667 | 1 | 1 | 4 | 0.75 | 12 | 0.5 | 2 | 0.5 | 1 | 0 | 2 | 0.5 | 5 | 0.2 | 10 | 0.3 |
| 9_6789375 | 4 | 0.5 | 5 | 0.6 | 1 | 0 | 4 | 0.25 | 14 | 0.4286 | 3 | 0.3333 | 2 | 1 | 3 | 0 | 6 | 0.5 | 14 | 0.4286 |
| 9_6789377 | 5 | 0 | 4 | 0.25 | 1 | 0 | 4 | 0 | 14 | 0.0714 | 2 | 0 | 1 | 0 | 3 | 0 | 4 | 0.25 | 10 | 0.1 |
| 9_6789379 | 4 | 0.25 | 5 | 0.4 | 1 | 1 | 4 | 1 | 14 | 0.5714 | 3 | 0.3333 | 1 | 1 | 3 | 0.6667 | 4 | 0.5 | 11 | 0.5455 |
| 9_6789393 | 5 | 0 | 5 | 0.4 | 2 | 0.5 | 6 | 0.3333 | 18 | 0.2778 | 2 | 0.5 | 2 | 0.5 | 2 | 0.5 | 6 | 0.5 | 12 | 0.5 |
| 9_6789404 | 4 | 0.25 | 4 | 0.25 | NA | NA | 6 | 0.5 | 14 | 0.3571 | 3 | 0 | 2 | 0 | 3 | 0.6667 | 6 | 0.1667 | 14 | 0.2143 |
| 9_6789415 | 4 | 0.5 | 6 | 0.3333 | 2 | 0.5 | 5 | 0.2 | 17 | 0.3529 | 3 | 0 | 1 | 0 | 3 | 0.6667 | 6 | 0.1667 | 13 | 0.2308 |
| 9_6928646 | NA | NA | NA | NA | NA | NA | 1 | 1 | 1 | 1 | NA | NA | NA | NA | NA | NA | 1 | 1 | 1 | 1 |
| 9_6928651 | NA | NA | NA | NA | NA | NA | 1 | 1 | 1 | 1 | NA | NA | NA | NA | NA | NA | 1 | 1 | 1 | 1 |
| 9_6928661 | NA | NA | NA | NA | NA | NA | 1 | 1 | 1 | 1 | NA | NA | NA | NA | NA | NA | 1 | 1 | 1 | 1 |
| 9_6928689 | NA | NA | 1 | 1 | NA | NA | 1 | 0 | 2 | 0.5 | NA | NA | NA | NA | NA | NA | 1 | 1 | 1 | 1 |
| 9_6929002 | 2 | 0 | 2 | 0 | 1 | 1 | 4 | 0.25 | 9 | 0.2222 | NA | NA | 1 | 1 | 2 | 0.5 | 1 | 0 | 4 | 0.5 |
| 9_6929402 | 2 | 0.5 | 5 | 0.2 | 2 | 0 | 5 | 0 | 14 | 0.1429 | 2 | 0.5 | NA | NA | 1 | 0 | NA | NA | 3 | 0.3333 |
| 9_6929450 | 2 | 1 | 7 | 0.5714 | 2 | 0 | 4 | 0.5 | 15 | 0.5333 | 2 | 1 | NA | NA | 1 | 0 | NA | NA | 3 | 0.6667 |
| 9_6929515 | 3 | 0.3333 | 7 | 0.7143 | 3 | 0.6667 | 5 | 0.4 | 18 | 0.5556 | 1 | 1 | 2 | 1 | 1 | 0 | 1 | 0 | 5 | 0.6 |
| 9_6929521 | 3 | 0 | 7 | 0.1429 | 3 | 0 | 3 | 0 | 16 | 0.0625 | 1 | 0 | 1 | 1 | 1 | 0 | 1 | 0 | 4 | 0.25 |
| 9_6932267 | NA | NA | 1 | 1 | 1 | 1 | NA | NA | 2 | 1 | NA | NA | 1 | 1 | NA | NA | NA | NA | 1 | 1 |
| 9_6932304 | NA | NA | 1 | 1 | 1 | 1 | NA | NA | 2 | 1 | NA | NA | 1 | 1 | NA | NA | NA | NA | 1 | 1 |
| 9_6932312 | NA | NA | 1 | 0 | 1 | 1 | NA | NA | 2 | 0.5 | NA | NA | 1 | 1 | NA | NA | NA | NA | 1 | 1 |
| 9_6933951 | NA | NA | 1 | 0 | 3 | 1 | NA | NA | 4 | 0.75 | 1 | 1 | NA | NA | 1 | 1 | NA | NA | 2 | 1 |
| 9_6933977 | 1 | 0 | 1 | 0 | 3 | 0.3333 | NA | NA | 5 | 0.2 | 1 | 1 | 1 | 0 | 1 | 0 | NA | NA | 3 | 0.3333 |
| 9_6991452 | 5 | 0.6 | 11 | 0.7273 | 6 | 1 | 2 | 1 | 24 | 0.7917 | 4 | 0.75 | 4 | 0.75 | 4 | 1 | 6 | 1 | 18 | 0.8889 |
| 9_6991453 | 7 | 0.2857 | 12 | 0.5 | 6 | 0.8333 | 2 | 0.5 | 27 | 0.5185 | 4 | 0.5 | 3 | 0.3333 | 4 | 0.5 | 7 | 0.2857 | 18 | 0.3889 |
| 9_6991511 | 8 | 0 | 10 | 0.3 | 8 | 0.125 | 3 | 0.3333 | 29 | 0.1724 | 4 | 0 | 3 | 0 | 2 | 0 | 6 | 0.1667 | 15 | 0.0667 |
| 9_6991514 | 7 | 0 | 12 | 0.1667 | 6 | 0.6667 | 3 | 0.3333 | 28 | 0.25 | 3 | 0.3333 | 4 | 0 | 3 | 0.3333 | 6 | 0.6667 | 16 | 0.375 |
| 9_6991532 | 8 | 0.125 | 13 | 0.0769 | 8 | 0 | 3 | 0 | 32 | 0.0625 | 4 | 0.25 | 3 | 0 | 3 | 0.3333 | 5 | 0.6 | 15 | 0.3333 |
| 9_6991722 | NA | NA | NA | NA | NA | NA | 1 | 1 | 1 | 1 | NA | NA | 1 | 1 | NA | NA | NA | NA | 1 | 1 |
| 9_6992981 | 14 | 0.3571 | 5 | 0.6 | 10 | 0.5 | 4 | 1 | 33 | 0.5152 | 10 | 0.2 | 9 | 0.7778 | 7 | 0.5714 | 6 | 0.6667 | 32 | 0.5312 |
| 9_6999622 | 20 | 0.15 | 34 | 0.2353 | 15 | 0.2667 | 8 | 0.125 | 77 | 0.2078 | 22 | 0.2273 | 11 | 0.3636 | 11 | 0.2727 | 23 | 0.087 | 67 | 0.209 |
| 9_6999625 | 16 | 0.25 | 25 | 0.32 | 12 | 0.25 | 7 | 0.4286 | 60 | 0.3 | 19 | 0.2105 | 10 | 0.1 | 13 | 0.2308 | 19 | 0.3158 | 61 | 0.2295 |
| 9_6999652 | 18 | 0.1667 | 36 | 0.1667 | 16 | 0.1875 | 7 | 0.2857 | 77 | 0.1818 | 21 | 0.1905 | 12 | 0.1667 | 12 | 0.0833 | 22 | 0.1818 | 67 | 0.1642 |
| 9_6999664 | 19 | 0.0526 | 31 | 0.0645 | 16 | 0.1875 | 10 | 0 | 76 | 0.0789 | 20 | 0.3 | 12 | 0 | 11 | 0.1818 | 22 | 0.0909 | 65 | 0.1538 |
| 9_6999691 | 15 | 0.0667 | 30 | 0.1333 | 13 | 0 | 6 | 0 | 64 | 0.0781 | 20 | 0.05 | 10 | 0.2 | 11 | 0.0909 | 20 | 0.1 | 61 | 0.0984 |
| 9_6999775 | 10 | 0 | 23 | 0.087 | 14 | 0.1429 | 8 | 0 | 55 | 0.0727 | 20 | 0.15 | 8 | 0 | 5 | 0 | 17 | 0.1765 | 50 | 0.12 |
| 9_6999801 | 7 | 0.2857 | 22 | 0.4545 | 11 | 0.3636 | 5 | 0.8 | 45 | 0.4444 | 16 | 0.375 | 8 | 0.25 | 6 | 0.6667 | 18 | 0.2778 | 48 | 0.3542 |
| 9_6999803 | 8 | 0.375 | 22 | 0.2273 | 13 | 0.3077 | 5 | 0.6 | 48 | 0.3125 | 17 | 0.3529 | 7 | 0.1429 | 6 | 0.5 | 18 | 0.3333 | 48 | 0.3333 |
| 9_6999810 | 9 | 0.2222 | 20 | 0.1 | 14 | 0 | 7 | 0.1429 | 50 | 0.1 | 16 | 0.0625 | 7 | 0.1429 | 6 | 0.1667 | 20 | 0.15 | 49 | 0.1224 |
| 9_7030612 | 1 | 0 | 3 | 0.3333 | NA | NA | NA | NA | 4 | 0.25 | NA | NA | 4 | 0.5 | NA | NA | 2 | 0 | 6 | 0.3333 |
| 9_7030613 | 1 | 1 | 3 | 0.3333 | NA | NA | NA | NA | 4 | 0.5 | NA | NA | 4 | 0.25 | NA | NA | 3 | 0.6667 | 7 | 0.4286 |
| 9_7030614 | 1 | 1 | 3 | 0 | NA | NA | NA | NA | 4 | 0.25 | NA | NA | 4 | 0.25 | NA | NA | 3 | 0 | 7 | 0.1429 |
| 9_7030625 | 2 | 0 | 3 | 0.6667 | NA | NA | NA | NA | 5 | 0.4 | NA | NA | 3 | 0.6667 | NA | NA | 3 | 0.3333 | 6 | 0.5 |
| 9_7030636 | 2 | 0 | 3 | 0.6667 | NA | NA | 1 | 0 | 6 | 0.3333 | NA | NA | 3 | 0.3333 | NA | NA | 2 | 0.5 | 5 | 0.4 |
| 9_7030654 | 2 | 1 | 3 | 1 | NA | NA | 1 | 0 | 6 | 0.8333 | NA | NA | 3 | 0.6667 | 1 | 0 | 2 | 0 | 6 | 0.3333 |
| 9_7030663 | 2 | 0.5 | NA | NA | NA | NA | 1 | 0 | 3 | 0.3333 | NA | NA | 4 | 0.25 | NA | NA | 2 | 1 | 6 | 0.5 |
| 9_7033343 | 6 | 0.1667 | 3 | 0 | 3 | 0 | 5 | 0 | 17 | 0.0588 | 2 | 0 | 1 | 1 | 7 | 0.1429 | 7 | 0.1429 | 17 | 0.1765 |
| 9_7033348 | 9 | 0.1111 | 4 | 0.25 | 4 | 0 | 5 | 0 | 22 | 0.0909 | 2 | 0 | 2 | 0 | 6 | 0.3333 | 7 | 0 | 17 | 0.1176 |
| 9_7033412 | 7 | 0 | 4 | 0.25 | 4 | 0 | 4 | 0 | 19 | 0.0526 | 2 | 0 | 2 | 0 | 4 | 0.25 | 7 | 0.1429 | 15 | 0.1333 |
| 9_7033415 | 7 | 0 | 3 | 0 | 4 | 0.25 | 4 | 0 | 18 | 0.0556 | 2 | 0 | 1 | 0 | 3 | 0 | 6 | 0.1667 | 12 | 0.0833 |
| 9_7033416 | 7 | 0.1429 | 4 | 0.25 | 3 | 0 | 4 | 0 | 18 | 0.1111 | 2 | 0 | 1 | 0 | 4 | 0.25 | 6 | 0.1667 | 13 | 0.1538 |
| 9_7033425 | 6 | 0.8333 | 3 | 1 | 4 | 1 | 5 | 1 | 18 | 0.9444 | 2 | 1 | 3 | 1 | 4 | 0.5 | 6 | 0.8333 | 15 | 0.8 |
| 9_7033427 | 8 | 0 | 4 | 0.5 | 4 | 0.5 | 5 | 0.2 | 21 | 0.2381 | 2 | 0.5 | 3 | 0 | 4 | 0.25 | 7 | 0.2857 | 16 | 0.25 |
| 9_7033430 | 5 | 0.2 | 3 | 0 | 3 | 0 | 5 | 0 | 16 | 0.0625 | 2 | 0.5 | 2 | 0 | 4 | 0 | 6 | 0.1667 | 14 | 0.1429 |
| 9_7035074 | 5 | 0 | 3 | 0.3333 | 3 | 0 | 5 | 0.2 | 16 | 0.125 | 1 | 0 | 2 | 0 | 5 | 0.2 | 4 | 0 | 12 | 0.0833 |
| 9_7035095 | 5 | 0.2 | 3 | 0 | 3 | 0 | 4 | 0 | 15 | 0.0667 | 1 | 0 | 3 | 0 | 5 | 0 | 5 | 0.2 | 14 | 0.0714 |
| 9_7038333 | 1 | 0 | 1 | 0 | 2 | 1 | 3 | 0.3333 | 7 | 0.4286 | 1 | 0 | NA | NA | 3 | 0.3333 | 1 | 1 | 5 | 0.4 |
| 9_7038347 | 1 | 1 | 1 | 1 | 2 | 1 | 4 | 0.75 | 8 | 0.875 | 1 | 0 | NA | NA | 2 | 0.5 | 1 | 1 | 4 | 0.5 |
| 9_7038354 | 2 | 0 | 1 | 0 | 2 | 0.5 | 2 | 0.5 | 7 | 0.2857 | 1 | 1 | NA | NA | 1 | 1 | 1 | 0 | 3 | 0.6667 |

| | | | | | | | | | | | | | | | | | | | | |
|---|---|---|---|---|---|---|---|---|---|---|---|---|---|---|---|---|---|---|---|---|
| 9_7038361 | 1 | 1 | 1 | 1 | 1 | 1 | 4 | 1 | 7 | 1 | 1 | 1 | NA | NA | 2 | 1 | 1 | 1 | 4 | 1 |
| 9_7038370 | 1 | 1 | NA | NA | 2 | 0.5 | 3 | 0.6667 | 6 | 0.6667 | 1 | 0 | NA | NA | 1 | 0 | 1 | 1 | 3 | 0.3333 |
| 9_7048104 | 2 | 0 | 3 | 0 | 2 | 0 | 3 | 0.3333 | 10 | 0.1 | 1 | 1 | 3 | 0 | 6 | 0.1667 | 5 | 0 | 15 | 0.1333 |
| 9_7049415 | 8 | 0.125 | 5 | 0.4 | 3 | 0.3333 | 4 | 0.5 | 20 | 0.3 | 4 | 0.5 | 4 | 0.25 | 11 | 0 | 8 | 0.375 | 27 | 0.2222 |
| 9_7049903 | 8 | 0.25 | 7 | 0 | 3 | 0.3333 | 4 | 0.25 | 22 | 0.1818 | 2 | 0.5 | 2 | 0 | 9 | 0.4444 | 9 | 0.2222 | 22 | 0.3182 |
| 9_7049905 | 7 | 0 | 6 | 0.1667 | 1 | 0 | 2 | 0 | 16 | 0.0625 | 4 | 0.5 | 3 | 0.3333 | 10 | 0.4 | 9 | 0.2222 | 26 | 0.3462 |
| 9_7049907 | 7 | 0.5714 | 6 | 0.3333 | 3 | 0 | 5 | 0.2 | 21 | 0.3333 | 4 | 0.75 | 5 | 0.6 | 11 | 0.2727 | 8 | 0.25 | 28 | 0.3929 |
| 9_7049915 | 9 | 0.2222 | 7 | 0 | 4 | 0 | 6 | 0 | 26 | 0.0769 | 5 | 0.2 | 6 | 0.1667 | 10 | 0.1 | 9 | 0 | 30 | 0.1 |
| 9_7050652 | 4 | 0 | 13 | 0.3077 | 6 | 0.1667 | 3 | 0 | 26 | 0.1923 | 6 | 0.1667 | 4 | 0.25 | 4 | 0.5 | 4 | 0.75 | 18 | 0.3889 |
| 9_7050660 | 3 | 0.3333 | 12 | 0.1667 | 5 | 0 | 4 | 0.25 | 24 | 0.1667 | 6 | 0.1667 | 5 | 0.4 | 5 | 0.6 | 4 | 0.5 | 20 | 0.4 |
| 9_7050685 | 3 | 0.3333 | 13 | 0.0769 | 5 | 0 | 4 | 0 | 25 | 0.08 | 6 | 0 | 5 | 0.2 | 5 | 0 | 3 | 0.3333 | 19 | 0.1053 |
| 9_7050697 | 4 | 0 | 12 | 0.1667 | 5 | 0 | 4 | 0.25 | 25 | 0.12 | 5 | 0 | 4 | 0.25 | 4 | 0 | 3 | 0.3333 | 16 | 0.125 |
| 9_7050712 | 4 | 0 | 11 | 0 | 5 | 0 | 4 | 0.5 | 24 | 0.0833 | 5 | 0.2 | 4 | 0.25 | 5 | 0.4 | 4 | 0.5 | 18 | 0.3333 |
| 9_7050727 | 3 | 0.3333 | 12 | 0.25 | 5 | 0.2 | 4 | 0 | 24 | 0.2083 | 4 | 0.25 | 4 | 0.75 | 5 | 0.4 | 4 | 0.25 | 17 | 0.4118 |
| 9_7051018 | NA | NA | 7 | 0 | 3 | 0 | 1 | 1 | 11 | 0.0909 | 3 | 0 | 3 | 0 | 2 | 0.5 | NA | NA | 8 | 0.125 |
| 9_7051191 | NA | NA | 3 | 0.3333 | 4 | 0.25 | 2 | 0 | 9 | 0.2222 | 3 | 0.3333 | 2 | 0 | 2 | 0 | NA | NA | 7 | 0.1429 |
| 9_7051203 | NA | NA | 4 | 0.75 | 4 | 0.5 | 2 | 0 | 10 | 0.5 | 3 | 0.3333 | 3 | 0.6667 | 2 | 0.5 | NA | NA | 8 | 0.5 |
| 9_7051219 | NA | NA | 3 | 0.3333 | 4 | 0 | 2 | 0.5 | 9 | 0.2222 | 1 | 0 | 2 | 0.5 | 2 | 0.5 | NA | NA | 5 | 0.4 |
| 9_7051225 | NA | NA | 3 | 0.6667 | 3 | 0.6667 | 2 | 1 | 8 | 0.75 | 1 | 1 | 2 | 0.5 | 2 | 1 | NA | NA | 5 | 0.8 |
| 9_7051248 | NA | NA | 3 | 0.3333 | 4 | 0.25 | 3 | 0.3333 | 10 | 0.3 | 2 | 0.5 | 2 | 0 | 2 | 0 | NA | NA | 6 | 0.1667 |
| 9_7056367 | NA | NA | 2 | 1 | 3 | 0.3333 | 1 | 0 | 6 | 0.5 | 1 | 0 | NA | NA | 2 | 0.5 | NA | NA | 3 | 0.3333 |
| 9_7056387 | NA | NA | 1 | 1 | 1 | 1 | 1 | 1 | 3 | 1 | 1 | 1 | NA | NA | 1 | 1 | NA | NA | 2 | 1 |
| 9_7067613 | 3 | 0 | 3 | 0 | 5 | 0 | 3 | 0.3333 | 14 | 0.0714 | 1 | 1 | 3 | 0 | 4 | 0 | 2 | 0 | 10 | 0.1 |
| 9_7068536 | 4 | 0.25 | 5 | 0.4 | 7 | 0.4286 | 3 | 0 | 19 | 0.3158 | 1 | 1 | 4 | 1 | 9 | 0.3333 | 2 | 0.5 | 16 | 0.5625 |
| 9_7068542 | 4 | 0 | 6 | 0 | 7 | 0.1429 | 3 | 0 | 20 | 0.05 | 1 | 1 | 4 | 0 | 8 | 0 | 2 | 0 | 15 | 0.0667 |
| 9_7068560 | 5 | 0.4 | 4 | 0.75 | 8 | 0.625 | 1 | 0 | 18 | 0.5556 | NA | NA | 4 | 0.5 | 4 | 0.25 | 2 | 0.5 | 10 | 0.4 |
| 9_7068570 | 6 | 0.6667 | 5 | 0.6 | 9 | 0.3333 | 5 | 1 | 25 | 0.6 | 3 | 1 | 4 | 1 | 7 | 0.8571 | 3 | 0.3333 | 17 | 0.8235 |
| 9_7068571 | 3 | 1 | 4 | 1 | 7 | 0.4286 | 5 | 0.8 | 19 | 0.7368 | 1 | 0 | 3 | 1 | 5 | 0.8 | 2 | 1 | 11 | 0.8182 |
| 9_7068694 | 10 | 0.1 | 10 | 0.1 | 13 | 0.0769 | 7 | 0.2857 | 40 | 0.125 | 5 | 0 | 4 | 0.25 | 12 | 0 | 4 | 0.5 | 25 | 0.12 |
| 9_7068708 | 9 | 0.4444 | 10 | 0.7 | 12 | 0.3333 | 3 | 0 | 34 | 0.4412 | 3 | 0 | 3 | 0.3333 | 14 | 0.4286 | 5 | 0.4 | 25 | 0.36 |
| 9_7068728 | 13 | 0.2308 | 8 | 0.5 | 13 | 0.3077 | 8 | 0.5 | 42 | 0.3571 | 3 | 0.3333 | 3 | 0.6667 | 12 | 0.3333 | 4 | 0.5 | 22 | 0.4091 |
| 9_7076131 | 1 | 1 | NA | NA | NA | NA | NA | NA | 1 | 1 | NA | NA | 1 | 1 | NA | NA | 1 | 1 | 2 | 1 |
| 9_7076142 | 1 | 0 | 1 | 0 | NA | NA | 1 | 1 | 3 | 0.3333 | NA | NA | 1 | 1 | NA | NA | 1 | 1 | 2 | 1 |
| 9_7138603 | 34 | 0.6176 | 28 | 0.7143 | 17 | 0.7059 | 29 | 0.7931 | 108 | 0.7037 | 19 | 0.6316 | 24 | 0.7917 | 17 | 0.3529 | 20 | 0.75 | 80 | 0.65 |
| 9_7138611 | 36 | 0.0556 | 30 | 0.1667 | 19 | 0.2632 | 35 | 0.0571 | 120 | 0.1167 | 23 | 0.087 | 24 | 0.2917 | 18 | 0.0556 | 18 | 0.1667 | 83 | 0.1566 |
| 9_7138614 | 38 | 0.2105 | 32 | 0.125 | 21 | 0.1905 | 37 | 0.2703 | 128 | 0.2031 | 24 | 0.1667 | 26 | 0.0769 | 21 | 0.1905 | 21 | 0.1905 | 92 | 0.1522 |
| 9_7138619 | 39 | 0.0769 | 35 | 0.0857 | 20 | 0.15 | 33 | 0.0303 | 127 | 0.0787 | 23 | 0.087 | 22 | 0 | 22 | 0.1364 | 19 | 0.1579 | 86 | 0.093 |
| 9_7138623 | 37 | 0.7297 | 33 | 0.7879 | 21 | 0.9524 | 34 | 0.7941 | 125 | 0.8 | 22 | 0.8182 | 25 | 0.88 | 22 | 0.7727 | 22 | 0.9091 | 91 | 0.8462 |
| 9_7138648 | 33 | 0.2424 | 28 | 0.2857 | 18 | 0.0556 | 34 | 0.1471 | 113 | 0.1947 | 17 | 0.2353 | 17 | 0.2353 | 16 | 0.125 | 12 | 0.3333 | 62 | 0.2258 |
| 9_7138653 | 33 | 0.1212 | 31 | 0.1935 | 20 | 0.15 | 32 | 0.1562 | 116 | 0.1552 | 23 | 0.1304 | 20 | 0.2 | 18 | 0 | 14 | 0.2143 | 75 | 0.1333 |
| 9_7138657 | 34 | 0.5588 | 28 | 0.6786 | 14 | 0.6429 | 30 | 0.7667 | 106 | 0.6604 | 22 | 0.8182 | 23 | 0.7391 | 19 | 0.6316 | 16 | 0.375 | 80 | 0.6625 |
| 9_7138659 | 32 | 0.0625 | 32 | 0.0938 | 17 | 0.2941 | 34 | 0.1176 | 115 | 0.1217 | 22 | 0.1364 | 22 | 0.1818 | 20 | 0.05 | 17 | 0.1176 | 81 | 0.1235 |
| 9_7138662 | 23 | 0.4348 | 29 | 0.6207 | 19 | 0.3684 | 30 | 0.4667 | 101 | 0.4851 | 22 | 0.2273 | 19 | 0.4211 | 17 | 0.7059 | 14 | 0.4286 | 72 | 0.4306 |
| 9_7138666 | 29 | 0.2069 | 30 | 0.3333 | 17 | 0.1176 | 33 | 0.2424 | 109 | 0.2385 | 22 | 0.3182 | 23 | 0.1304 | 19 | 0.2632 | 14 | 0.1429 | 78 | 0.2179 |
| 9_7138678 | 26 | 0.5385 | 26 | 0.7692 | 18 | 0.7222 | 35 | 0.5429 | 105 | 0.6286 | 23 | 0.5217 | 24 | 0.5833 | 20 | 0.45 | 17 | 0.7059 | 84 | 0.5595 |
| 9_7139054 | 20 | 0.25 | 17 | 0.1765 | 12 | 0.0833 | 21 | 0.1429 | 70 | 0.1714 | 13 | 0.0769 | 20 | 0.05 | 15 | 0.3333 | 10 | 0 | 58 | 0.1207 |
| 9_7139055 | 21 | 0.1429 | 20 | 0.05 | 16 | 0.1875 | 30 | 0.1667 | 87 | 0.1379 | 20 | 0.2 | 24 | 0.2917 | 18 | 0.1667 | 12 | 0.0833 | 74 | 0.2027 |
| 9_7139077 | 24 | 0.5 | 19 | 0.6316 | 15 | 0.5333 | 30 | 0.6 | 88 | 0.5682 | 18 | 0.4444 | 18 | 0.4444 | 16 | 0.4375 | 10 | 0.7 | 62 | 0.4839 |
| 9_7139080 | 26 | 0.1923 | 19 | 0.1053 | 14 | 0.2143 | 33 | 0.2424 | 92 | 0.1957 | 19 | 0.2105 | 22 | 0.2727 | 16 | 0.3125 | 11 | 0.3636 | 68 | 0.2794 |
| 9_7139084 | 24 | 0.4583 | 20 | 0.4 | 13 | 0.7692 | 33 | 0.6364 | 90 | 0.5556 | 20 | 0.3 | 24 | 0.5 | 17 | 0.4706 | 11 | 0.4545 | 72 | 0.4306 |
| 9_7139086 | 24 | 0.3333 | 20 | 0.5 | 15 | 0.4 | 30 | 0.3333 | 89 | 0.382 | 20 | 0.45 | 21 | 0.3333 | 14 | 0.3571 | 12 | 0.4167 | 67 | 0.3881 |
| 9_7139095 | 20 | 0.4 | 18 | 0.5 | 12 | 0.3333 | 30 | 0.4 | 80 | 0.4125 | 20 | 0.6 | 22 | 0.4545 | 16 | 0.5 | 12 | 0.25 | 70 | 0.4714 |
| 9_7139098 | 20 | 0.8 | 18 | 0.6667 | 14 | 0.7143 | 32 | 0.7188 | 84 | 0.7262 | 11 | 0.7273 | 18 | 0.8889 | 11 | 0.9091 | 11 | 1 | 51 | 0.8824 |
| 9_7175209 | 8 | 0 | 4 | 0 | 8 | 0.25 | 4 | 0 | 24 | 0.0833 | 3 | 0.3333 | 3 | 0 | 5 | 0.2 | 5 | 0 | 16 | 0.125 |
| 9_7175222 | 7 | 0.2857 | 3 | 0 | 9 | 0.2222 | 6 | 0.1667 | 25 | 0.2 | 4 | 0 | 5 | 0 | 5 | 0 | 5 | 0.2 | 19 | 0.0526 |
| 9_7175226 | 7 | 0.7143 | 5 | 0.8 | 8 | 0.625 | 5 | 1 | 25 | 0.76 | 3 | 0.6667 | 4 | 0.5 | 5 | 0.6 | 6 | 0.5 | 18 | 0.5556 |
| 9_7175226 | 9 | 0.2222 | 5 | 0 | 9 | 0 | 6 | 0 | 29 | 0.069 | 4 | 0.25 | 5 | 0 | 5 | 0 | 6 | 0.1667 | 20 | 0.1 |
| 9_7175229 | 7 | 0.2857 | 4 | 0.75 | 9 | 0.6667 | 5 | 0.4 | 25 | 0.52 | 4 | 1 | 5 | 0.4 | 3 | 0 | 6 | 0.8333 | 18 | 0.6111 |
| 9_7175347 | 7 | 1 | 1 | 0 | 7 | 0.7143 | 7 | 0.8571 | 22 | 0.8182 | 4 | 1 | 4 | 0.75 | 2 | 1 | 4 | 1 | 14 | 0.9286 |
| 9_7175351 | 8 | 0.125 | 4 | 0 | 7 | 0.2857 | 6 | 0.1667 | 25 | 0.16 | 4 | 0.25 | 4 | 0 | 4 | 0.25 | 4 | 0 | 16 | 0.125 |
| 9_7175352 | 8 | 0.125 | 4 | 0.75 | 7 | 0.2857 | 7 | 0.5714 | 26 | 0.3846 | 4 | 0 | 4 | 0.25 | 4 | 0.25 | 4 | 0.5 | 16 | 0.25 |
| 9_7175357 | 6 | 1 | 2 | 1 | 7 | 0.8571 | 3 | 1 | 18 | 0.9444 | 3 | 1 | 4 | 1 | 4 | 0.75 | 4 | 0.75 | 15 | 0.8667 |
| 9_7175364 | 7 | 0 | 4 | 0.25 | 7 | 0 | 6 | 0.3333 | 24 | 0.125 | 4 | 0 | 5 | 0.2 | 3 | 0 | 4 | 0.25 | 16 | 0.125 |
| 9_7175371 | 6 | 0.6667 | 3 | 1 | 6 | 0.8333 | 5 | 1 | 20 | 0.85 | 3 | 1 | 4 | 0.5 | 3 | 0 | 4 | 1 | 14 | 0.6429 |
| 9_7175373 | 6 | 0 | 2 | 0 | 5 | 0.2 | 5 | 0.2 | 18 | 0.1111 | 4 | 0 | 4 | 0 | 3 | 0 | 4 | 0.25 | 15 | 0.0667 |

| | | | | | | | | | | | | | | | | | | | | |
|---|---|---|---|---|---|---|---|---|---|---|---|---|---|---|---|---|---|---|---|---|
| 9_7175378 | 4 | 0.5 | 3 | 0.3333 | 6 | 0 | 4 | 0 | 17 | 0.1765 | 4 | 0.75 | 3 | 0.3333 | 2 | 0 | 4 | 0 | 13 | 0.3077 |
| 9_7226934 | 3 | 0 | 5 | 0 | 10 | 0.2 | 1 | 0 | 19 | 0.1053 | 3 | 0 | 3 | 0.3333 | 2 | 1 | 2 | 0 | 10 | 0.3 |
| 9_7226937 | 2 | 0 | 5 | 0.2 | 10 | 0.3 | 2 | 0.5 | 19 | 0.2632 | 3 | 0 | 2 | 0 | 2 | 0 | 2 | 0.5 | 9 | 0.1111 |
| 9_7226944 | 3 | 0 | 6 | 0.3333 | 11 | 0.2727 | 2 | 0.5 | 22 | 0.2727 | 2 | 1 | 3 | 0.3333 | 2 | 0.5 | 2 | 0.5 | 9 | 0.5556 |
| 9_7226947 | 3 | 1 | 5 | 1 | 7 | 1 | 2 | 1 | 17 | 1 | 3 | 1 | 3 | 0.6667 | 2 | 1 | 2 | 1 | 10 | 0.9 |
| 9_7226949 | 3 | 1 | 6 | 0.8333 | 11 | 0.8182 | 2 | 1 | 22 | 0.8636 | 3 | 1 | 3 | 1 | 2 | 0.5 | 2 | 1 | 10 | 0.9 |
| 9_7226955 | 1 | 1 | 6 | 1 | 7 | 0.7143 | 2 | 0.5 | 16 | 0.8125 | 2 | 0.5 | 2 | 1 | 2 | 1 | 2 | 0.5 | 8 | 0.75 |
| 9_7226958 | 3 | 0.6667 | 6 | 0.1667 | 9 | 0.1111 | 2 | 0 | 20 | 0.2 | 3 | 0 | 4 | 0.25 | 2 | 0 | 2 | 0.5 | 11 | 0.1818 |
| 9_7226960 | 3 | 0.3333 | 5 | 0.6 | 8 | 0.125 | 2 | 0 | 18 | 0.2778 | 2 | 0 | 3 | 0.3333 | 2 | 0.5 | 2 | 0 | 9 | 0.2222 |
| 9_7226969 | 4 | 0.25 | 6 | 0.1667 | 10 | 0.1 | 2 | 0 | 22 | 0.1364 | 3 | 0.3333 | 4 | 0 | 2 | 0 | 2 | 0.5 | 11 | 0.1818 |
| 9_7226975 | 4 | 0.5 | 7 | 0.7143 | 11 | 0.3636 | 2 | 0.5 | 24 | 0.5 | 3 | 0.3333 | 4 | 0.25 | 2 | 1 | 2 | 0 | 11 | 0.3636 |
| 9_7226983 | 2 | 0.5 | 3 | 0.3333 | 7 | 0 | 1 | 0 | 13 | 0.1538 | 2 | 0 | 2 | 0 | 2 | 0.5 | 2 | 0 | 8 | 0.125 |
| 9_7226999 | 4 | 0.25 | 7 | 0.2857 | 11 | 0.0909 | 2 | 0 | 24 | 0.1667 | 3 | 0 | 4 | 0 | 3 | 0.3333 | 2 | 0 | 12 | 0.0833 |
| 9_7227010 | 4 | 0 | 7 | 0.1429 | 11 | 0.0909 | 2 | 0.5 | 24 | 0.125 | 3 | 0 | 4 | 0 | 3 | 0 | 2 | 0.5 | 12 | 0.0833 |
| 9_7227072 | 4 | 0.5 | 9 | 0.2222 | 11 | 0 | 4 | 0 | 28 | 0.1429 | 3 | 0 | 4 | 0.25 | 5 | 0.2 | 3 | 0 | 15 | 0.1333 |
| 9_7236724 | 1 | 0 | NA | NA | NA | NA | 1 | 1 | 2 | 0.5 | 2 | 0 | NA | NA | 2 | 0.5 | 3 | 0 | 7 | 0.1429 |
| 9_7236827 | 1 | 0 | NA | NA | NA | NA | 1 | 1 | 2 | 0.5 | 1 | 0 | NA | NA | 2 | 0.5 | 2 | 0 | 5 | 0.2 |
| 9_7251919 | 1 | 1 | 1 | 0 | NA | NA | NA | NA | 2 | 0.5 | NA | NA | NA | NA | 1 | 1 | NA | NA | 1 | 1 |
| 9_7252404 | NA | NA | 1 | 1 | 2 | 1 | NA | NA | 3 | 1 | NA | NA | NA | NA | 1 | 1 | 1 | 1 | 2 | 1 |
| 9_7252447 | 1 | 0 | 1 | 0 | 2 | 0.5 | NA | NA | 4 | 0.25 | NA | NA | NA | NA | 1 | 1 | 1 | 0 | 2 | 0.5 |
| 9_7252470 | 1 | 0 | 2 | 0.5 | 1 | 1 | NA | NA | 4 | 0.5 | NA | NA | NA | NA | 1 | 1 | 1 | 1 | 2 | 1 |
| 9_7252475 | 1 | 1 | 2 | 0.5 | 1 | 1 | NA | NA | 4 | 0.75 | NA | NA | NA | NA | 1 | 0 | 1 | 1 | 2 | 0.5 |
| 9_7252490 | 1 | 0 | 2 | 1 | 2 | 0 | NA | NA | 5 | 0.4 | NA | NA | NA | NA | NA | NA | 1 | 1 | 1 | 1 |
| 9_7252511 | 1 | 1 | 3 | 0.3333 | 2 | 1 | NA | NA | 6 | 0.6667 | NA | NA | 1 | 1 | 1 | 0 | NA | NA | 2 | 0.5 |
| 9_7252529 | 1 | 0 | 3 | 1 | 2 | 0.5 | NA | NA | 6 | 0.6667 | NA | NA | 1 | 1 | 1 | 0 | 1 | 1 | 3 | 0.6667 |
| 9_7252539 | 1 | 1 | 3 | 0.3333 | 1 | 1 | NA | NA | 5 | 0.6 | NA | NA | NA | NA | 1 | 1 | 1 | 1 | 2 | 1 |
| 9_7252545 | 1 | 1 | 1 | 1 | NA | NA | NA | NA | 2 | 1 | NA | NA | 1 | 1 | 1 | 1 | NA | NA | 2 | 1 |
| 9_7252596 | 1 | 1 | 3 | 0.3333 | 2 | 0 | 1 | 0 | 7 | 0.2857 | NA | NA | 1 | 0 | 1 | 1 | 1 | 1 | 3 | 0.6667 |
| 9_7252612 | 1 | 1 | 1 | 1 | 1 | 1 | 1 | 1 | 4 | 1 | NA | NA | 1 | 0 | 1 | 0 | 1 | 1 | 3 | 0.3333 |
| 9_7252615 | 1 | 0 | 3 | 0 | 2 | 0.5 | 1 | 1 | 7 | 0.2857 | NA | NA | 1 | 1 | 1 | 0 | 1 | 0 | 3 | 0.3333 |
| 9_7252626 | NA | NA | 1 | 0 | 2 | 0.5 | 1 | 1 | 4 | 0.5 | NA | NA | 1 | 1 | 1 | 1 | 1 | 1 | 3 | 1 |
| 9_7252649 | 1 | 1 | 3 | 0.6667 | 2 | 1 | NA | NA | 6 | 0.8333 | NA | NA | NA | NA | 2 | 1 | 1 | 1 | 3 | 1 |
| 9_7252654 | NA | NA | 2 | 1 | 2 | 0 | NA | NA | 4 | 0.5 | NA | NA | 2 | 1 | 1 | 1 | 1 | 1 | 4 | 1 |
| 9_7252657 | 1 | 1 | 3 | 0.6667 | 2 | 0 | 1 | 0 | 7 | 0.4286 | NA | NA | 2 | 1 | NA | NA | 2 | 1 | 4 | 1 |
| 9_7252658 | 1 | 0 | 2 | 0 | 2 | 0.5 | 1 | 1 | 6 | 0.3333 | NA | NA | 2 | 0.5 | 2 | 0 | 2 | 0.5 | 6 | 0.3333 |
| 9_7252667 | NA | NA | 3 | 0.3333 | 2 | 0 | 1 | 1 | 6 | 0.3333 | NA | NA | 3 | 0.3333 | 2 | 0 | 2 | 0.5 | 7 | 0.2857 |
| 9_7252683 | 1 | 1 | 2 | 0.5 | 1 | 1 | NA | NA | 4 | 0.75 | NA | NA | 3 | 0.6667 | 3 | 0.3333 | 1 | 0 | 7 | 0.4286 |
| 9_7252686 | 1 | 1 | 2 | 1 | 1 | 1 | 1 | 1 | 5 | 1 | NA | NA | 2 | 1 | 3 | 1 | 1 | 1 | 6 | 1 |
| 9_7252700 | 1 | 0 | 2 | 1 | 2 | 1 | 1 | 1 | 6 | 0.8333 | NA | NA | 3 | 0.3333 | 3 | 0.6667 | 2 | 1 | 8 | 0.625 |
| 9_7252701 | NA | NA | 2 | 0 | 2 | 0 | 1 | 1 | 5 | 0.2 | NA | NA | 3 | 0 | 3 | 0 | 2 | 1 | 8 | 0.25 |
| 9_7252704 | 1 | 1 | 3 | 0.6667 | 2 | 1 | 1 | 1 | 7 | 0.8571 | NA | NA | 2 | 1 | 3 | 1 | 1 | 0 | 6 | 0.8333 |
| 9_7252712 | 1 | 1 | 2 | 1 | 2 | 1 | 1 | 1 | 6 | 1 | NA | NA | 2 | 1 | 2 | 1 | 2 | 1 | 6 | 1 |
| 9_7252712 | NA | NA | 2 | 0.5 | 2 | 0.5 | 1 | 0 | 5 | 0.4 | NA | NA | 3 | 0.3333 | 3 | 0.3333 | 2 | 0.5 | 8 | 0.375 |
| 9_7252716 | 1 | 1 | 2 | 1 | 2 | 0.5 | 1 | 1 | 6 | 0.8333 | NA | NA | 3 | 0.6667 | 3 | 0.3333 | 2 | 1 | 8 | 0.625 |
| 9_7252729 | 1 | 0 | 2 | 0 | 2 | 0 | 1 | 1 | 6 | 0.1667 | NA | NA | 3 | 1 | 3 | 0.3333 | 2 | 0 | 8 | 0.5 |
| 9_7252736 | NA | NA | 3 | 0.3333 | 1 | 0 | 1 | 0 | 5 | 0.2 | NA | NA | 3 | 0.3333 | 2 | 0.5 | 2 | 0 | 7 | 0.2857 |
| 9_7252771 | 1 | 0 | 2 | 0.5 | 1 | 0 | NA | NA | 4 | 0.25 | NA | NA | 2 | 0.5 | 2 | 0 | 2 | 0 | 6 | 0.1667 |
| 9_7252917 | 2 | 0.5 | 4 | 0.75 | 4 | 0 | 1 | 0 | 11 | 0.3636 | NA | NA | 3 | 0 | 4 | 0.25 | 4 | 0.25 | 11 | 0.1818 |
| 9_7252977 | 1 | 1 | 7 | 0.2857 | 3 | 0 | 2 | 0 | 13 | 0.2308 | NA | NA | 3 | 0.3333 | 4 | 0.25 | 3 | 0 | 10 | 0.2 |
| 9_7252982 | 2 | 0 | 5 | 0.4 | 4 | 0.5 | 2 | 0.5 | 13 | 0.3846 | NA | NA | 4 | 0.75 | 3 | 0 | 4 | 0.5 | 11 | 0.4545 |
| 9_7252982 | 1 | 1 | 5 | 1 | 2 | 0.5 | 1 | 1 | 9 | 0.8889 | NA | NA | 2 | 1 | 3 | 1 | 4 | 1 | 9 | 1 |
| 9_7265445 | 17 | 0.3529 | 30 | 0.1 | 16 | 0.1875 | 33 | 0.3333 | 96 | 0.2396 | 13 | 0.3077 | 28 | 0.1429 | 23 | 0.2174 | 12 | 0.25 | 76 | 0.2105 |
| 9_7265464 | 22 | 0.4091 | 33 | 0.3333 | 18 | 0.2778 | 35 | 0.5143 | 108 | 0.3981 | 18 | 0.5 | 32 | 0.4062 | 27 | 0.2593 | 18 | 0.2222 | 95 | 0.3474 |
| 9_7285206 | 2 | 0 | 2 | 0 | 2 | 0.5 | 1 | 0 | 7 | 0.1429 | NA | NA | NA | NA | 1 | 0 | 1 | 1 | 2 | 0.5 |
| 9_7285225 | 2 | 1 | 2 | 1 | 2 | 1 | 1 | 1 | 7 | 1 | NA | NA | NA | NA | 1 | 0 | 1 | 1 | 2 | 0.5 |
| 9_7285231 | 1 | 0 | 2 | 0.5 | 2 | 1 | 1 | 0 | 6 | 0.5 | NA | NA | NA | NA | 1 | 1 | 1 | 1 | 2 | 1 |
| 9_7285231 | 2 | 1 | 2 | 1 | 2 | 1 | 1 | 1 | 7 | 1 | NA | NA | NA | NA | 1 | 1 | 1 | 1 | 2 | 1 |
| 9_7285253 | NA | NA | 2 | 0.5 | 2 | 1 | 1 | 1 | 5 | 0.8 | NA | NA | NA | NA | NA | NA | 1 | 1 | 1 | 1 |
| 9_7285271 | 2 | 1 | 2 | 1 | 2 | 0.5 | 1 | 1 | 7 | 0.8571 | NA | NA | 1 | 1 | 1 | 1 | 1 | 1 | 3 | 1 |
| 9_7285293 | 2 | 0.5 | 2 | 0 | 2 | 0 | 1 | 0 | 7 | 0.1429 | NA | NA | 1 | 1 | 1 | 1 | 1 | 1 | 3 | 1 |
| 9_7285298 | 4 | 0.25 | 2 | 0.5 | 1 | 0 | 1 | 0 | 8 | 0.25 | NA | NA | 1 | 0 | 1 | 1 | 1 | 1 | 3 | 0.6667 |
| 9_7285312 | 3 | 0 | 1 | 1 | 2 | 0 | 1 | 0 | 7 | 0.1429 | NA | NA | 1 | 0 | 1 | 0 | 1 | 1 | 3 | 0.3333 |
| 9_7285326 | 4 | 0 | 2 | 0.5 | 1 | 1 | 1 | 0 | 8 | 0.25 | NA | NA | 2 | 1 | 1 | 0 | 1 | 0 | 4 | 0.5 |
| 9_7285584 | 1 | 0 | 5 | 0.2 | 1 | 1 | NA | NA | 7 | 0.2857 | NA | NA | 2 | 1 | NA | NA | 1 | 1 | 3 | 1 |
| 9_7285830 | 1 | 0 | 4 | 0.5 | 1 | 0 | 1 | 0 | 7 | 0.2857 | 1 | 1 | 2 | 1 | 2 | 0 | 3 | 0 | 8 | 0.375 |

| | | | | | | | | | | | | | | | | | | | | |
|---|---|---|---|---|---|---|---|---|---|---|---|---|---|---|---|---|---|---|---|---|
| 9_72858386 | NA | NA | 7 | 0.1429 | 1 | 0 | 2 | 0 | 10 | 0.1 | 1 | 0 | 3 | 0.3333 | 2 | 0 | 5 | 0 | 11 | 0.0909 |
| 9_72858442 | 1 | 0 | 6 | 0.5 | 1 | 0 | 1 | 1 | 9 | 0.4444 | NA | NA | 4 | 1 | 2 | 0 | 6 | 0.6667 | 12 | 0.6667 |
| 9_72858487 | 1 | 0 | 7 | 0.5714 | 1 | 1 | 2 | 0.5 | 11 | 0.5455 | 1 | 0 | 3 | 0.6667 | 2 | 0 | 5 | 0.8 | 11 | 0.5455 |
| 9_72858565 | 1 | 0 | 6 | 0.1667 | 2 | 0 | 2 | 0 | 11 | 0.0909 | 1 | 0 | 3 | 0.3333 | 2 | 0 | 6 | 0 | 12 | 0.0833 |
| 9_72858689 | NA | NA | 6 | 0.3333 | 1 | 1 | 2 | 0 | 9 | 0.3333 | 1 | 0 | 1 | 1 | 2 | 1 | 4 | 0.5 | 8 | 0.625 |
| 9_72858703 | 1 | 0 | 7 | 1 | 1 | 1 | 1 | 0 | 10 | 0.8 | NA | NA | 3 | 1 | 1 | 1 | 6 | 0.8333 | 10 | 0.9 |
| 9_72858712 | 1 | 1 | 7 | 1 | 2 | 1 | 2 | 1 | 12 | 1 | 1 | 1 | 4 | 1 | 2 | 1 | 6 | 1 | 13 | 1 |
| 9_72858759 | 1 | 0 | 5 | 0.4 | 2 | 0 | 2 | 0 | 10 | 0.2 | 1 | 0 | 5 | 0.4 | 2 | 0.5 | 7 | 0.5714 | 15 | 0.4667 |
| 9_72858768 | 1 | 0 | 6 | 0.6667 | 2 | 1 | 2 | 0 | 11 | 0.5455 | 1 | 0 | 3 | 1 | NA | NA | 6 | 0.8333 | 10 | 0.8 |
| 9_72858961 | 2 | 1 | 7 | 1 | 2 | 1 | 2 | 1 | 13 | 1 | 2 | 1 | 5 | 0.8 | 1 | 1 | 6 | 1 | 14 | 0.9286 |
| 9_72858985 | 2 | 0 | 5 | 0.4 | 2 | 0 | 2 | 0.5 | 11 | 0.2727 | NA | NA | 5 | 0.2 | 2 | 0 | 6 | 0.5 | 13 | 0.3077 |
| 9_72858996 | NA | NA | 3 | 1 | NA | NA | 1 | 1 | 4 | 1 | NA | NA | 2 | 1 | 1 | 0 | 1 | 0 | 4 | 0.5 |
| 9_72859100 | 1 | 1 | 7 | 0 | 2 | 0.5 | 1 | 0 | 11 | 0.1818 | NA | NA | 4 | 0.25 | 1 | 0 | 6 | 0.5 | 11 | 0.3636 |
| 9_72859750 | 1 | 1 | NA | NA | 1 | 0 | NA | NA | 2 | 0.5 | NA | NA | NA | NA | NA | NA | 1 | 1 | 1 | 1 |
| 9_72919537 | 4 | 0.75 | 5 | 0.6 | 6 | 0.6667 | 3 | 0.6667 | 18 | 0.6667 | 4 | 0.75 | 1 | 0 | 5 | 0.6 | 4 | 0.75 | 14 | 0.6429 |
| 9_72919580 | 4 | 0.25 | 6 | 0.3333 | 6 | 0.5 | 2 | 0 | 18 | 0.3333 | 5 | 0.4 | 2 | 0 | 6 | 0 | 4 | 0.25 | 17 | 0.1765 |
| 9_72919638 | 3 | 0.6667 | 5 | 0 | 5 | 0.2 | 3 | 0 | 16 | 0.1875 | 5 | 0.2 | 2 | 0 | 4 | 0 | 6 | 0 | 17 | 0.0588 |
| 9_72919733 | 3 | 0.6667 | 8 | 0.75 | 6 | 0.8333 | 3 | 0.6667 | 20 | 0.75 | 5 | 1 | 2 | 1 | 6 | 0.8333 | 7 | 0.5714 | 20 | 0.8 |
| 9_72919845 | 4 | 0.75 | 7 | 0.7143 | 4 | 1 | 4 | 0.5 | 19 | 0.7368 | 4 | 0.75 | 2 | 0.5 | 5 | 0.6 | 5 | 0.6 | 16 | 0.625 |
| 9_72920070 | 8 | 0 | 9 | 0.1111 | 6 | 0.1667 | 4 | 0.25 | 27 | 0.1111 | 6 | 0 | 1 | 0 | 7 | 0 | 7 | 0.2857 | 21 | 0.0952 |
| 9_72920236 | 7 | 0.1429 | 9 | 0.1111 | 6 | 0.1667 | 4 | 0 | 26 | 0.1154 | 6 | 0 | 2 | 0 | 8 | 0.125 | 8 | 0.375 | 24 | 0.1667 |
| 9_72920294 | 5 | 0.6 | 2 | 1 | 5 | 0.2 | 3 | 0.3333 | 15 | 0.4667 | 4 | 0.25 | 1 | 1 | 5 | 0.6 | 6 | 0.5 | 16 | 0.5 |
| 9_72920344 | 7 | 0.4286 | 5 | 0.4 | 7 | 0.7143 | 3 | 0.6667 | 22 | 0.5455 | 3 | 0.3333 | 1 | 1 | 8 | 0.375 | 6 | 0.8333 | 18 | 0.5556 |
| 9_72920381 | 3 | 1 | 5 | 0.4 | 3 | 0.6667 | 1 | 1 | 12 | 0.6667 | 3 | 1 | NA | NA | 4 | 0.75 | 6 | 0.6667 | 13 | 0.7692 |
| 9_72920388 | 7 | 0.8571 | 7 | 1 | 7 | 0.7143 | 3 | 1 | 24 | 0.875 | 4 | 0.75 | 1 | 0 | 9 | 0.7778 | 7 | 0.8571 | 21 | 0.7619 |
| 9_72920408 | 7 | 0.5714 | 5 | 0.4 | 6 | 0.8333 | 2 | 1 | 20 | 0.65 | 4 | 0.5 | 1 | 1 | 8 | 0.625 | 5 | 0.8 | 18 | 0.6667 |
| 9_72920453 | 9 | 0.1111 | 5 | 0.4 | 5 | 0 | 3 | 0 | 22 | 0.1364 | 5 | 0.2 | 1 | 0 | 7 | 0 | 7 | 0.1429 | 20 | 0.1 |
| 9_72920570 | 5 | 0.8 | 5 | 0 | 2 | 0.5 | 2 | 1 | 14 | 0.5 | 3 | 0 | NA | NA | 4 | 1 | 5 | 0.8 | 12 | 0.6667 |
| 9_72920572 | 1 | 1 | NA | NA | 1 | 1 | NA | NA | 2 | 1 | NA | NA | NA | NA | 1 | 1 | 1 | 0 | 2 | 0.5 |
| 9_72920603 | 8 | 0.75 | 7 | 0.4286 | 6 | 0.5 | 3 | 0 | 24 | 0.5 | 6 | 0.3333 | 1 | 0 | 7 | 0.7143 | 7 | 0.5714 | 21 | 0.5238 |
| 9_72920738 | 2 | 1 | NA | NA | 1 | 1 | NA | NA | 3 | 1 | 1 | 0 | NA | NA | NA | NA | 1 | 1 | 2 | 0.5 |
| 9_72920756 | 8 | 0 | 8 | 0.125 | 6 | 0.1667 | 3 | 0 | 25 | 0.08 | 4 | 0 | 1 | 0 | 7 | 0.1429 | 5 | 0 | 17 | 0.0588 |
| 9_72920969 | 8 | 0.25 | 7 | 0 | 6 | 0.1667 | 3 | 0 | 24 | 0.125 | 5 | 0.2 | 1 | 0 | 7 | 0.2857 | 5 | 0.2 | 18 | 0.2222 |
| 9_72921119 | 3 | 0.3333 | NA | NA | 1 | 0 | NA | NA | 4 | 0.25 | 1 | 0 | NA | NA | NA | NA | 1 | 1 | 2 | 0.5 |
| 9_72921126 | 3 | 0 | NA | NA | 1 | 1 | NA | NA | 4 | 0.25 | 1 | 1 | NA | NA | NA | NA | 1 | 0 | 2 | 0.5 |
| 9_72921158 | 1 | 0 | NA | NA | 1 | 1 | NA | NA | 2 | 0.5 | 1 | 1 | NA | NA | NA | NA | 1 | 1 | 2 | 1 |
| 9_72921176 | 2 | 0.5 | NA | NA | 1 | 1 | NA | NA | 3 | 0.6667 | 1 | 1 | NA | NA | NA | NA | 1 | 1 | 2 | 1 |
| 9_72921181 | 3 | 0.6667 | NA | NA | 1 | 1 | NA | NA | 4 | 0.75 | 1 | 1 | NA | NA | NA | NA | 1 | 0 | 2 | 0.5 |
| 9_72921239 | 3 | 1 | NA | NA | NA | NA | NA | NA | 3 | 1 | 1 | 1 | NA | NA | NA | NA | 1 | 0 | 2 | 0.5 |
| 9_72921496 | 3 | 0.3333 | NA | NA | 1 | 0 | NA | NA | 4 | 0.25 | 1 | 0 | NA | NA | NA | NA | 1 | 1 | 2 | 0.5 |
| 9_72921647 | 3 | 0 | NA | NA | 1 | 1 | NA | NA | 4 | 0.25 | 1 | 1 | NA | NA | NA | NA | NA | NA | 1 | 1 |
| 9_72922770 | 5 | 0.2 | 3 | 0 | 5 | 0.2 | 2 | 0.5 | 15 | 0.2 | 3 | 0 | 1 | 0 | 1 | 1 | 3 | 0 | 8 | 0.125 |
| 9_72922868 | 5 | 0.4 | 3 | 0 | 3 | 0 | 3 | 0.3333 | 14 | 0.2143 | 5 | 0.2 | 1 | 0 | 1 | 0 | 3 | 0 | 10 | 0.1 |
| 9_72923021 | 2 | 1 | NA | NA | 1 | 1 | NA | NA | 3 | 1 | 1 | 1 | NA | NA | 1 | 0 | NA | NA | 2 | 0.5 |
| 9_72923112 | 3 | 1 | 5 | 0.8 | 5 | 0.4 | 3 | 0.3333 | 16 | 0.625 | 7 | 0.2857 | 2 | 0.5 | 1 | 0 | 5 | 0.8 | 15 | 0.4667 |
| 9_72923221 | 5 | 0 | 10 | 0.6 | 5 | 0.6 | 1 | 1 | 21 | 0.4762 | 8 | 0.25 | 4 | 0.5 | 1 | 0 | 4 | 0.25 | 17 | 0.2941 |
| 9_72923260 | 3 | 0 | 9 | 0.3333 | 5 | 0.2 | 2 | 0.5 | 19 | 0.2632 | 7 | 0 | 4 | 0 | 1 | 1 | 4 | 0.25 | 16 | 0.125 |
| 9_72923285 | 3 | 0.3333 | NA | NA | 1 | 1 | NA | NA | 4 | 0.5 | 1 | 1 | NA | NA | 1 | 0 | NA | NA | 2 | 0.5 |
| 9_72923334 | NA | NA | 2 | 1 | NA | NA | 1 | 1 | 3 | 1 | NA | NA | 2 | 0.5 | 1 | 1 | 1 | 1 | 4 | 0.75 |
| 9_72923391 | 3 | 0.3333 | NA | NA | 1 | 1 | NA | NA | 4 | 0.5 | 1 | 1 | NA | NA | 1 | 0 | NA | NA | 2 | 0.5 |
| 9_72923424 | 3 | 1 | NA | NA | 1 | 1 | NA | NA | 4 | 1 | 1 | 0 | NA | NA | 1 | 1 | NA | NA | 2 | 0.5 |
| 9_72923433 | 3 | 0.6667 | NA | NA | 1 | 1 | NA | NA | 4 | 0.75 | 1 | 1 | NA | NA | 1 | 0 | NA | NA | 2 | 0.5 |
| 9_72923491 | 1 | 0 | 3 | 0.3333 | NA | NA | 1 | 0 | 5 | 0.2 | NA | NA | 1 | 1 | 1 | 0 | 1 | 0 | 3 | 0.3333 |
| 9_72923594 | 1 | 0 | 3 | 0.3333 | NA | NA | 1 | 1 | 5 | 0.4 | NA | NA | 2 | 0 | 1 | 0 | 1 | 1 | 4 | 0.25 |
| 9_72949874 | 4 | 0.75 | 3 | 0.3333 | NA | NA | 1 | 1 | 8 | 0.625 | 1 | 1 | 3 | 0.3333 | 1 | 1 | 2 | 1 | 7 | 0.7143 |
| 9_72949959 | 6 | 0.6667 | 3 | 0.6667 | NA | NA | 3 | 1 | 12 | 0.75 | 2 | 0.5 | 6 | 0.5 | 1 | 1 | 2 | 1 | 11 | 0.6364 |
| 9_72950021 | 7 | 0.4286 | 3 | 0 | 1 | 0 | 3 | 0 | 14 | 0.2143 | 2 | 0.5 | 6 | 0.3333 | 2 | 0 | 2 | 0 | 12 | 0.25 |
| 9_73015321 | NA | NA | 1 | 1 | NA | NA | NA | NA | 1 | 1 | 1 | 0 | NA | NA | NA | NA | 1 | 1 | 2 | 0.5 |
| 9_73016620 | NA | NA | 1 | 1 | NA | NA | NA | NA | 1 | 1 | 1 | 0 | 1 | 0 | NA | NA | 1 | 1 | 3 | 0.3333 |
| 9_73029666 | 10 | 0 | 10 | 0.1 | 9 | 0 | 6 | 0.3333 | 35 | 0.0857 | 11 | 0.1818 | 8 | 0.25 | 8 | 0.125 | 5 | 0 | 32 | 0.1562 |
| 9_73029856 | 7 | 0.8571 | 9 | 0.8889 | 7 | 0.5714 | 6 | 0.8333 | 29 | 0.7931 | 11 | 0.8182 | 5 | 1 | 3 | 0.6667 | 6 | 1 | 25 | 0.88 |
| 9_73029977 | 10 | 0.2 | 12 | 0 | 8 | 0 | 6 | 0 | 36 | 0.0556 | 10 | 0 | 8 | 0.125 | 8 | 0.125 | 6 | 0.1667 | 32 | 0.0938 |
| 9_73387754 | 7 | 0 | 1 | 1 | 1 | 0 | 8 | 0 | 17 | 0.0588 | 1 | 0 | 10 | 0.1 | 11 | 0.0909 | 6 | 0 | 28 | 0.0714 |
| 9_73388001 | 5 | 0 | NA | NA | 1 | 0 | 3 | 0.6667 | 9 | 0.2222 | 1 | 0 | 5 | 0.2 | 6 | 0.1667 | 4 | 0 | 16 | 0.125 |
| 9_73388113 | 5 | 0.4 | 1 | 1 | 1 | 0 | 9 | 0.6667 | 16 | 0.5625 | 1 | 1 | 10 | 0.5 | 8 | 0.875 | 7 | 0.5714 | 26 | 0.6538 |

| | | | | | | | | | | | | | | | | | | | | |
|---|---|---|---|---|---|---|---|---|---|---|---|---|---|---|---|---|---|---|---|---|
| 9_7338814 | 5 | 0.6 | 1 | 1 | 1 | 0 | 10 | 0.7 | 17 | 0.6471 | 1 | 1 | 10 | 0.5 | 9 | 1 | 7 | 0.8571 | 27 | 0.7778 |
| 9_7338819 | 4 | 0.5 | 1 | 0 | 1 | 1 | 9 | 0.6667 | 15 | 0.6 | 1 | 1 | 9 | 0.6667 | 9 | 0.7778 | 6 | 0.5 | 25 | 0.68 |
| 9_7338822 | 4 | 0.25 | 1 | 0 | 2 | 0 | 8 | 0.125 | 15 | 0.1333 | 1 | 0 | 9 | 0.2222 | 8 | 0.125 | 7 | 0.2857 | 25 | 0.2 |
| 9_7338825 | 3 | 0 | 1 | 1 | 2 | 0 | 9 | 0.6667 | 15 | 0.4667 | 1 | 0 | 8 | 0.375 | 6 | 0.5 | 6 | 0.3333 | 21 | 0.381 |
| 9_7338834 | 3 | 0 | NA | NA | 1 | 0 | 10 | 0.1 | 14 | 0.0714 | 1 | 0 | 6 | 0 | 7 | 0 | 6 | 0.3333 | 20 | 0.1 |
| 9_7338838 | 5 | 0.4 | 1 | 0 | 1 | 1 | 10 | 0.3 | 17 | 0.3529 | 1 | 1 | 9 | 0.2222 | 7 | 0.4286 | 6 | 0.8333 | 23 | 0.4783 |
| 9_7338840 | 5 | 0 | 1 | 0 | 2 | 0 | 9 | 0.1111 | 17 | 0.0588 | 1 | 0 | 6 | 0.1667 | 5 | 0.2 | 7 | 0.1429 | 19 | 0.1579 |
| 9_7383806 | NA | NA | 1 | 1 | 2 | 0.5 | 3 | 0 | 6 | 0.3333 | NA | NA | 4 | 0.75 | 3 | 0.3333 | 2 | 0.5 | 9 | 0.5556 |
| 9_7383815 | NA | NA | NA | NA | 2 | 0.5 | 3 | 0 | 5 | 0.2 | NA | NA | 4 | 0.75 | 3 | 1 | 2 | 0.5 | 9 | 0.7778 |
| 9_7383818 | NA | NA | NA | NA | 2 | 0.5 | 3 | 0.3333 | 5 | 0.4 | NA | NA | 4 | 1 | 3 | 1 | 2 | 0 | 9 | 0.7778 |
| 9_7383826 | NA | NA | NA | NA | 2 | 0 | 3 | 0.3333 | 5 | 0.2 | NA | NA | 4 | 0.25 | 3 | 0.3333 | 2 | 0 | 9 | 0.2222 |
| 9_7383835 | NA | NA | NA | NA | 1 | 1 | 3 | 0.6667 | 4 | 0.75 | NA | NA | 4 | 0.75 | 3 | 0.3333 | 2 | 0.5 | 9 | 0.5556 |
| 9_7383849 | NA | NA | NA | NA | 1 | 1 | 2 | 1 | 3 | 1 | NA | NA | 4 | 0.75 | 2 | 0.5 | 2 | 1 | 8 | 0.75 |
| 9_7383856 | NA | NA | NA | NA | 1 | 0 | 3 | 0.6667 | 4 | 0.5 | NA | NA | 3 | 0.3333 | 3 | 0.3333 | 2 | 0.5 | 8 | 0.375 |
| 9_7383867 | NA | NA | NA | NA | 1 | 0 | 3 | 0.3333 | 4 | 0.25 | NA | NA | 3 | 0 | 1 | 0 | 1 | 1 | 5 | 0.2 |
| 9_7383873 | NA | NA | NA | NA | 1 | 0 | 3 | 0.6667 | 4 | 0.5 | NA | NA | 3 | 0.3333 | 3 | 0 | 1 | 0 | 7 | 0.1429 |
| 9_7383887 | NA | NA | NA | NA | 1 | 1 | 2 | 1 | 3 | 1 | NA | NA | 3 | 1 | 3 | 0.6667 | 2 | 1 | 8 | 0.875 |
| 9_7383911 | NA | NA | NA | NA | 1 | 1 | NA | NA | 1 | 1 | NA | NA | 3 | 0.6667 | 2 | 0 | 2 | 0 | 7 | 0.2857 |
| 9_7383915 | NA | NA | NA | NA | NA | NA | 1 | 1 | 1 | 1 | NA | NA | NA | NA | 3 | 0 | 2 | 1 | 5 | 0.4 |
| 9_7383992 | NA | NA | NA | NA | 1 | 1 | NA | NA | 1 | 1 | NA | NA | 1 | 0 | 3 | 0.3333 | NA | NA | 4 | 0.25 |
| 9_7384093 | NA | NA | NA | NA | 1 | 1 | NA | NA | 1 | 1 | NA | NA | 1 | 0 | 1 | 1 | NA | NA | 2 | 0.5 |
| 9_7486126 | NA | NA | NA | NA | 1 | 1 | NA | NA | 1 | 1 | 1 | 0 | 1 | 1 | NA | NA | NA | NA | 2 | 0.5 |
| 9_7491622 | 1 | 1 | NA | NA | 4 | 0.75 | 1 | 0 | 6 | 0.6667 | 2 | 1 | NA | NA | 1 | 1 | NA | NA | 3 | 1 |
| 9_7491649 | 3 | 0.6667 | NA | NA | 5 | 0.8 | 1 | 1 | 9 | 0.7778 | 2 | 1 | 1 | 1 | 2 | 1 | NA | NA | 5 | 1 |
| 9_7491658 | 1 | 0 | NA | NA | 4 | 0 | 2 | 0.5 | 7 | 0.1429 | 2 | 0 | 1 | 0 | 2 | 0.5 | NA | NA | 5 | 0.2 |
| 9_7491722 | 3 | 0 | NA | NA | 3 | 0.6667 | 1 | 0 | 7 | 0.2857 | 2 | 0.5 | 1 | 1 | 4 | 0 | NA | NA | 7 | 0.2857 |
| 9_7491733 | 5 | 0.4 | NA | NA | 4 | 0 | 4 | 0.25 | 13 | 0.2308 | 3 | 0 | 2 | 1 | 4 | 0.25 | 2 | 0.5 | 11 | 0.3636 |
| 9_7491734 | 4 | 0.25 | NA | NA | 2 | 0 | 3 | 0 | 9 | 0.1111 | 2 | 0.5 | 1 | 1 | 3 | 0.3333 | NA | NA | 6 | 0.5 |
| 9_7491746 | 3 | 0.3333 | NA | NA | 4 | 0.75 | 4 | 0.75 | 11 | 0.6364 | 3 | 0.6667 | 2 | 0.5 | 3 | 1 | 2 | 0.5 | 10 | 0.7 |
| 9_7491748 | 5 | 0.4 | NA | NA | 5 | 0.6 | 3 | 0.6667 | 13 | 0.5385 | 3 | 1 | 2 | 0.5 | 4 | 0.5 | 2 | 0 | 11 | 0.5455 |
| 9_7491754 | 4 | 0.75 | NA | NA | 5 | 0.4 | 3 | 1 | 12 | 0.6667 | 3 | 0.6667 | 2 | 0.5 | 4 | 0.25 | 1 | 0 | 10 | 0.4 |
| 9_7491767 | 5 | 0.6 | NA | NA | 5 | 1 | 2 | 1 | 12 | 0.8333 | 3 | 0.6667 | 2 | 1 | 4 | 0.75 | 1 | 1 | 10 | 0.8 |
| 9_7492950 | 2 | 0.5 | NA | NA | NA | NA | 1 | 0 | 3 | 0.3333 | 1 | 0 | 1 | 0 | 2 | 0.5 | NA | NA | 4 | 0.25 |
| 9_7493889 | 7 | 0.7143 | 3 | 0.3333 | 6 | 0.5 | 8 | 0.375 | 24 | 0.5 | 2 | 1 | 7 | 0.2857 | 9 | 0.3333 | 3 | 1 | 21 | 0.4762 |
| 9_7493895 | 6 | 0.5 | 4 | 0.5 | 7 | 0.2857 | 11 | 0.4545 | 28 | 0.4286 | 4 | 0.25 | 7 | 0.2857 | 10 | 0.3 | 3 | 0.3333 | 24 | 0.2917 |
| 9_7493906 | 7 | 0.1429 | 3 | 0.3333 | 6 | 0.1667 | 10 | 0.4 | 26 | 0.2692 | 4 | 0.25 | 7 | 0.1429 | 11 | 0.2727 | 3 | 0 | 25 | 0.2 |
| 9_7493908 | 6 | 0.5 | 4 | 0.75 | 6 | 1 | 9 | 0.3333 | 25 | 0.6 | 4 | 0.75 | 7 | 0.7143 | 10 | 0.5 | 3 | 0.6667 | 24 | 0.625 |
| 9_7493912 | 7 | 1 | 4 | 1 | 7 | 0.4286 | 11 | 0.8182 | 29 | 0.7931 | 4 | 0.75 | 6 | 0.8333 | 11 | 0.7273 | 3 | 1 | 24 | 0.7917 |
| 9_7493917 | 6 | 1 | 4 | 1 | 5 | 1 | 10 | 0.8 | 25 | 0.92 | 2 | 1 | 7 | 1 | 8 | 1 | 3 | 1 | 20 | 1 |
| 9_7495935 | NA | NA | 3 | 0 | 1 | 0 | 4 | 0.25 | 8 | 0.125 | NA | NA | 2 | 0 | 2 | 0 | 4 | 0.25 | 8 | 0.125 |
| 9_7495944 | 49 | 0.0612 | 27 | 0.1111 | 40 | 0.075 | 49 | 0.0816 | 165 | 0.0788 | 18 | 0.1111 | 65 | 0.0462 | 54 | 0.0556 | 54 | 0.0556 | 191 | 0.0576 |
| 9_7496393 | 68 | 0.1029 | 54 | 0.0741 | 50 | 0.02 | 68 | 0.0882 | 240 | 0.075 | 37 | 0.0811 | 86 | 0.0233 | 69 | 0.087 | 73 | 0.0685 | 265 | 0.0604 |
| 9_7496403 | 64 | 0.0938 | 46 | 0.087 | 49 | 0.0408 | 60 | 0.1 | 219 | 0.0822 | 32 | 0.0938 | 75 | 0.12 | 58 | 0.1379 | 69 | 0.087 | 234 | 0.1111 |
| 9_7496410 | 56 | 0.6786 | 43 | 0.6047 | 52 | 0.5385 | 61 | 0.6393 | 212 | 0.6179 | 36 | 0.6389 | 72 | 0.6528 | 58 | 0.6897 | 63 | 0.6825 | 229 | 0.6681 |
| 9_7496413 | 58 | 0.2241 | 39 | 0.1026 | 37 | 0.1622 | 52 | 0.2115 | 186 | 0.1828 | 26 | 0.1923 | 70 | 0.1571 | 55 | 0.1455 | 57 | 0.1404 | 208 | 0.1538 |
| 9_7496418 | 62 | 0.1935 | 47 | 0.2553 | 43 | 0.186 | 52 | 0.2115 | 204 | 0.2108 | 24 | 0.2083 | 70 | 0.2 | 60 | 0.2667 | 61 | 0.2787 | 215 | 0.2419 |
| 9_7496428 | 55 | 0.5636 | 42 | 0.5952 | 38 | 0.6316 | 52 | 0.6731 | 187 | 0.615 | 26 | 0.5385 | 70 | 0.6714 | 55 | 0.6727 | 59 | 0.678 | 210 | 0.6571 |
| 9_7496431 | 50 | 0.04 | 41 | 0.0732 | 34 | 0.0588 | 49 | 0.1837 | 174 | 0.092 | 24 | 0 | 67 | 0.0448 | 52 | 0.0769 | 53 | 0.0755 | 196 | 0.0561 |
| 9_7496444 | 61 | 0.1475 | 49 | 0.1224 | 46 | 0.1522 | 63 | 0.1587 | 219 | 0.1461 | 26 | 0.0385 | 76 | 0.1447 | 73 | 0.137 | 62 | 0.2581 | 237 | 0.1603 |
| 9_7496457 | 52 | 0.4038 | 40 | 0.3 | 31 | 0.2903 | 54 | 0.3704 | 177 | 0.3503 | 21 | 0.3333 | 65 | 0.3846 | 58 | 0.3448 | 63 | 0.4603 | 207 | 0.3913 |
| 9_7496458 | 62 | 0.9194 | 44 | 0.8864 | 37 | 0.8649 | 59 | 0.8983 | 202 | 0.896 | 27 | 0.963 | 73 | 0.9315 | 67 | 0.8955 | 61 | 0.9344 | 228 | 0.9254 |
| 9_7496466 | 64 | 0.5469 | 48 | 0.5208 | 50 | 0.46 | 63 | 0.4444 | 225 | 0.4933 | 32 | 0.375 | 80 | 0.4375 | 77 | 0.5455 | 72 | 0.5278 | 261 | 0.4866 |
| 9_7496467 | 65 | 0.3846 | 47 | 0.2128 | 49 | 0.2449 | 64 | 0.3438 | 225 | 0.3067 | 33 | 0.303 | 80 | 0.4125 | 71 | 0.3239 | 71 | 0.3239 | 255 | 0.349 |
| 9_7496469 | 63 | 0.0476 | 50 | 0.08 | 47 | 0.1489 | 60 | 0.05 | 220 | 0.0773 | 31 | 0.0645 | 78 | 0.0128 | 74 | 0.0811 | 73 | 0.1233 | 256 | 0.0703 |
| 9_7496716 | 8 | 0.125 | 6 | 0.3333 | 11 | 0.2727 | 10 | 0.1 | 35 | 0.2 | 11 | 0.0909 | 14 | 0.2143 | 22 | 0.3182 | 13 | 0.3846 | 60 | 0.2667 |
| 9_7496724 | 10 | 0.4 | 6 | 0.3333 | 11 | 0.3636 | 10 | 0.2 | 37 | 0.3243 | 8 | 0.125 | 10 | 0.2 | 14 | 0.2857 | 11 | 0.3636 | 43 | 0.2558 |
| 9_7496743 | 13 | 0.1538 | 8 | 0.25 | 10 | 0.2 | 13 | 0.1538 | 44 | 0.1818 | 12 | 0 | 15 | 0.2 | 23 | 0.0435 | 13 | 0.1538 | 63 | 0.0952 |
| 9_7505424 | 6 | 0 | 2 | 0 | NA | NA | 6 | 0.1667 | 14 | 0.0714 | 3 | 0 | 6 | 0 | 1 | 1 | 5 | 0 | 15 | 0.0667 |
| 9_7512532 | 14 | 0.7143 | 7 | 0.5714 | 13 | 0.7692 | 21 | 0.7619 | 55 | 0.7273 | 8 | 0.625 | 11 | 0.6364 | 13 | 0.7692 | 15 | 0.8 | 47 | 0.7234 |
| 9_7512545 | 22 | 0.0455 | 13 | 0.3846 | 23 | 0.3913 | 23 | 0.1739 | 81 | 0.2346 | 10 | 0.3 | 20 | 0.2 | 23 | 0.087 | 20 | 0.35 | 73 | 0.2192 |
| 9_7512557 | 23 | 0.3043 | 12 | 0.75 | 21 | 0.2857 | 24 | 0.625 | 80 | 0.4625 | 11 | 0.5455 | 22 | 0.5 | 22 | 0.3636 | 21 | 0.4762 | 76 | 0.4605 |
| 9_7512564 | 16 | 0.9375 | 9 | 0.8889 | 19 | 0.8421 | 22 | 0.9545 | 66 | 0.9091 | 11 | 1 | 18 | 0.8889 | 18 | 0.8889 | 21 | 0.9524 | 68 | 0.9265 |
| 9_7512565 | 15 | 0 | 8 | 0.375 | 19 | 0 | 21 | 0.0476 | 63 | 0.0635 | 11 | 0.0909 | 16 | 0.125 | 17 | 0 | 18 | 0.0556 | 62 | 0.0645 |
| 9_7512566 | 21 | 0.5714 | 12 | 0.8333 | 22 | 0.6818 | 27 | 0.6296 | 82 | 0.6585 | 11 | 0.7273 | 21 | 0.5714 | 19 | 0.5263 | 22 | 0.6818 | 73 | 0.6164 |
| 9_7512573 | 22 | 0.0909 | 12 | 0.0833 | 22 | 0.1818 | 29 | 0.2414 | 85 | 0.1647 | 10 | 0 | 19 | 0.1053 | 19 | 0.1053 | 25 | 0.16 | 73 | 0.1096 |

| | | | | | | | | | | | | | | | | | | | | |
|---|---|---|---|---|---|---|---|---|---|---|---|---|---|---|---|---|---|---|---|---|
| 9_7512575 | 20 | 0.3 | 12 | 0.75 | 19 | 0.5789 | 26 | 0.5769 | 77 | 0.5325 | 10 | 0.5 | 20 | 0.65 | 20 | 0.6 | 19 | 0.6842 | 69 | 0.6232 |
| 9_7512576 | 12 | 0.25 | 11 | 0.3636 | 18 | 0.1667 | 17 | 0.2941 | 58 | 0.2586 | 8 | 0.125 | 16 | 0.25 | 11 | 0.5455 | 19 | 0.3684 | 54 | 0.3333 |
| 9_7512578 | 10 | 0.2 | 10 | 0.2 | 15 | 0.1333 | 19 | 0.0526 | 54 | 0.1296 | 7 | 0 | 15 | 0.2 | 13 | 0.2308 | 19 | 0.0526 | 54 | 0.1296 |
| 9_7512592 | 10 | 0.8 | 12 | 1 | 22 | 0.8182 | 19 | 0.7895 | 63 | 0.8413 | 9 | 0.7778 | 16 | 0.8125 | 12 | 0.9167 | 21 | 0.9524 | 58 | 0.8793 |
| 9_7512593 | 11 | 0.0909 | 11 | 0.2727 | 19 | 0.2632 | 18 | 0.1667 | 59 | 0.2034 | 8 | 0.125 | 15 | 0 | 7 | 0 | 20 | 0.1 | 50 | 0.06 |
| 9_7512625 | 12 | 0.0833 | 14 | 0.3571 | 21 | 0.2857 | 23 | 0.3478 | 70 | 0.2857 | 10 | 0.5 | 19 | 0.2105 | 14 | 0.4286 | 19 | 0.3158 | 62 | 0.3387 |
| 9_7513071 | 17 | 0.3529 | 27 | 0.4444 | 41 | 0.3171 | 23 | 0.3478 | 108 | 0.3611 | 13 | 0.3846 | 22 | 0.5 | 22 | 0.5 | 18 | 0.5556 | 75 | 0.4933 |
| 9_7513085 | 21 | 0.2381 | 39 | 0.0769 | 51 | 0.1176 | 30 | 0.1333 | 141 | 0.1277 | 20 | 0.1 | 31 | 0.0968 | 31 | 0.0323 | 30 | 0 | 112 | 0.0536 |
| 9_7521803 | 2 | 1 | 2 | 0 | 2 | 0.5 | 1 | 1 | 7 | 0.5714 | 2 | 0 | 1 | 1 | NA | NA | NA | NA | 3 | 0.3333 |
| 9_7525243 | 33 | 0.1818 | 22 | 0 | 22 | 0 | 35 | 0.1429 | 112 | 0.0982 | 23 | 0.087 | 38 | 0.0789 | 26 | 0 | 26 | 0.1538 | 113 | 0.0796 |
| 9_7525246 | 27 | 0.7407 | 19 | 0.6842 | 14 | 0.7857 | 33 | 0.5455 | 93 | 0.6667 | 18 | 0.5556 | 35 | 0.7714 | 28 | 0.75 | 22 | 0.8182 | 103 | 0.7379 |
| 9_7525258 | 37 | 0.2432 | 20 | 0.35 | 22 | 0.5 | 38 | 0.3684 | 117 | 0.3504 | 23 | 0.3478 | 35 | 0.4571 | 28 | 0.2857 | 29 | 0.3103 | 115 | 0.3565 |
| 9_7525260 | 34 | 0.8824 | 19 | 0.8421 | 17 | 0.8235 | 34 | 0.8235 | 104 | 0.8462 | 22 | 0.8636 | 34 | 0.8529 | 27 | 0.9259 | 27 | 0.9259 | 110 | 0.8909 |
| 9_7525264 | 36 | 0.8333 | 21 | 0.8095 | 21 | 0.8571 | 37 | 0.7297 | 115 | 0.8 | 19 | 0.6842 | 35 | 0.7714 | 26 | 0.6538 | 26 | 0.9615 | 106 | 0.7736 |
| 9_7525272 | 35 | 0.1429 | 21 | 0.1429 | 25 | 0.12 | 41 | 0.122 | 122 | 0.1311 | 22 | 0.2273 | 37 | 0.0811 | 28 | 0.0714 | 31 | 0.0968 | 118 | 0.1102 |
| 9_7525284 | 27 | 0.2593 | 12 | 0.25 | 20 | 0.2 | 21 | 0.2381 | 80 | 0.2375 | 20 | 0.2 | 26 | 0.1923 | 24 | 0.1667 | 26 | 0.1538 | 96 | 0.1771 |
| 9_7525286 | 25 | 0.24 | 13 | 0.0769 | 18 | 0.1667 | 20 | 0.05 | 76 | 0.1447 | 18 | 0.0556 | 20 | 0.2 | 19 | 0.1053 | 26 | 0.0769 | 83 | 0.1084 |
| 9_7529532 | 8 | 0 | 9 | 0 | 6 | 0 | 4 | 0.5 | 27 | 0.0741 | 3 | 0 | 8 | 0.375 | 7 | 0.1429 | 5 | 0 | 23 | 0.1739 |
| 9_7529533 | 6 | 0 | 9 | 0 | 5 | 0.4 | 4 | 0.25 | 24 | 0.125 | 2 | 0.5 | 7 | 0.4286 | 4 | 0.5 | 6 | 0 | 19 | 0.3158 |
| 9_7529557 | 4 | 0 | 5 | 0 | 4 | 0.25 | 4 | 0.25 | 17 | 0.1176 | 2 | 0 | 7 | 0.5714 | 4 | 0.25 | 5 | 0.4 | 18 | 0.3889 |
| 9_7529561 | 4 | 0.5 | 6 | 0.3333 | 4 | 0.25 | 4 | 0 | 18 | 0.2778 | 2 | 0 | 5 | 0.2 | 6 | 0.1667 | 3 | 0.3333 | 16 | 0.1875 |
| 9_7529572 | 3 | 0 | 6 | 0 | 5 | 0.2 | 3 | 0 | 17 | 0.0588 | 1 | 0 | 7 | 0 | 4 | 0.25 | 2 | 0.5 | 14 | 0.1429 |
| 9_7529590 | 6 | 0 | 5 | 0 | 4 | 0 | 4 | 0.25 | 19 | 0.0526 | 2 | 0 | 6 | 0.5 | 6 | 0 | 4 | 0.25 | 18 | 0.2222 |
| 9_7529592 | 6 | 0.1667 | 5 | 0 | 5 | 0 | 4 | 0 | 20 | 0.05 | 2 | 0 | 6 | 0 | 6 | 0.1667 | 4 | 0 | 18 | 0.0556 |
| 9_7529619 | 6 | 0 | 6 | 0.3333 | 4 | 0 | 3 | 0 | 19 | 0.1053 | 1 | 0 | 5 | 0 | 5 | 0.2 | 3 | 0.3333 | 14 | 0.1429 |
| 9_7530507 | 5 | 0.8 | 5 | 0.2 | 3 | 0.3333 | 3 | 0.6667 | 16 | 0.5 | 2 | 1 | 4 | 0.5 | 6 | 0.5 | 1 | 1 | 13 | 0.6154 |
| 9_7530520 | 4 | 0.25 | 4 | 0 | 3 | 0 | 3 | 0 | 14 | 0.0714 | 2 | 0 | 3 | 0 | 6 | 0.3333 | 2 | 0 | 13 | 0.1538 |
| 9_7530529 | 4 | 1 | 6 | 1 | 3 | 0.3333 | 3 | 1 | 16 | 0.875 | 1 | 1 | 4 | 0.75 | 5 | 0.8 | 2 | 1 | 12 | 0.8333 |
| 9_7530530 | 5 | 0 | 6 | 0 | 3 | 0.3333 | 3 | 0 | 17 | 0.0588 | 2 | 0 | 4 | 0.25 | 4 | 0 | 2 | 0 | 12 | 0.0833 |
| 9_7530535 | 4 | 1 | 6 | 1 | 2 | 1 | 3 | 1 | 15 | 1 | 2 | 0.5 | 2 | 1 | 6 | 1 | 2 | 1 | 12 | 0.9167 |
| 9_7530538 | 3 | 1 | 4 | 1 | 2 | 1 | 3 | 1 | 12 | 1 | 1 | 1 | 2 | 0.5 | 2 | 1 | 2 | 0.5 | 7 | 0.7143 |
| 9_7530558 | 4 | 0 | 5 | 0.2 | 2 | 0.5 | 1 | 0 | 12 | 0.1667 | 2 | 0.5 | 3 | 0 | 5 | 0.2 | 2 | 0 | 12 | 0.1667 |
| 9_7530565 | 4 | 0.75 | 5 | 1 | 3 | 0.6667 | 2 | 0.5 | 14 | 0.7857 | 1 | 1 | 3 | 0.6667 | 5 | 1 | 1 | 1 | 10 | 0.9 |
| 9_7535263 | 3 | 0.6667 | 1 | 1 | 3 | 0.3333 | NA | NA | 7 | 0.5714 | NA | NA | 3 | 0.6667 | 1 | 1 | 4 | 0.5 | 8 | 0.625 |
| 9_7535269 | 3 | 0 | 1 | 0 | 3 | 0.3333 | NA | NA | 7 | 0.1429 | 1 | 0 | 3 | 0.6667 | 1 | 1 | 3 | 0.6667 | 8 | 0.625 |
| 9_7535277 | 3 | 0.3333 | NA | NA | 4 | 0.25 | NA | NA | 7 | 0.2857 | 1 | 1 | 3 | 0.3333 | 1 | 0 | 4 | 0 | 9 | 0.2222 |
| 9_7537358 | 3 | 0 | 2 | 0 | 6 | 0.1667 | 2 | 0 | 13 | 0.0769 | 1 | 0 | 4 | 0 | 3 | 0 | 5 | 0.2 | 13 | 0.0769 |
| 9_7540592 | 8 | 0 | 1 | 1 | 4 | 0 | 3 | 0 | 16 | 0.0625 | NA | NA | 1 | 0 | 5 | 0.2 | 4 | 0 | 10 | 0.1 |
| 9_7540706 | 4 | 0.25 | 1 | 0 | 4 | 0.25 | 3 | 0 | 12 | 0.1667 | NA | NA | NA | NA | 4 | 0.25 | 4 | 0.25 | 8 | 0.25 |
| 9_7540731 | 6 | 0.3333 | 2 | 0 | 4 | 0 | 3 | 0.6667 | 15 | 0.2667 | NA | NA | NA | NA | 5 | 0.2 | 2 | 0.5 | 7 | 0.2857 |
| 9_7542826 | 4 | 1 | 3 | 1 | NA | NA | 5 | 0.8 | 12 | 0.9167 | 4 | 1 | 2 | 0.5 | 6 | 1 | 3 | 0.6667 | 15 | 0.8667 |
| 9_7542862 | 5 | 0.2 | 1 | 0 | NA | NA | 6 | 0.1667 | 12 | 0.1667 | 4 | 0 | 2 | 0 | 5 | 0.2 | 3 | 0 | 14 | 0.0714 |
| 9_7547303 | 35 | 0.0571 | 19 | 0.0526 | 32 | 0.0312 | 34 | 0.0588 | 120 | 0.05 | 16 | 0.1875 | 21 | 0.1905 | 30 | 0.1 | 27 | 0.1111 | 94 | 0.1383 |
| 9_7547322 | 33 | 0.0606 | 22 | 0 | 32 | 0.0312 | 33 | 0.1212 | 120 | 0.0583 | 18 | 0.1111 | 22 | 0.0909 | 25 | 0.08 | 29 | 0.0345 | 94 | 0.0745 |
| 9_7547365 | 30 | 0.1333 | 15 | 0.0667 | 30 | 0.0333 | 29 | 0.1379 | 104 | 0.0962 | 15 | 0.1333 | 17 | 0.0588 | 24 | 0.125 | 22 | 0.0909 | 78 | 0.1026 |
| 9_7547647 | 3 | 0 | 7 | 0.1429 | 6 | 0 | 8 | 0.25 | 24 | 0.125 | 3 | 0 | 2 | 0.5 | 8 | 0 | 5 | 0 | 18 | 0.0556 |
| 9_7547664 | 4 | 0 | 6 | 0 | 8 | 0.25 | 11 | 0 | 29 | 0.069 | 5 | 0 | 5 | 0.2 | 10 | 0.1 | 7 | 0 | 27 | 0.0741 |
| 9_7547779 | 2 | 0 | 4 | 0 | 5 | 0 | 6 | 0.1667 | 17 | 0.0588 | 2 | 0 | 2 | 0 | 7 | 0.2857 | 6 | 0 | 17 | 0.1176 |
| 9_7547907 | 3 | 0 | 2 | 0.5 | 6 | 0 | 9 | 0 | 20 | 0.05 | 1 | 0 | 2 | 0 | 5 | 0 | 5 | 0.2 | 13 | 0.0769 |
| 9_7547917 | 2 | 0 | 2 | 0 | 6 | 0 | 7 | 0.2857 | 17 | 0.1176 | NA | NA | NA | NA | 6 | 0 | 4 | 0.25 | 10 | 0.1 |
| 9_7547988 | NA | NA | NA | NA | 4 | 0 | 7 | 0.1429 | 11 | 0.0909 | 2 | 0 | 2 | 0 | 5 | 0.4 | 5 | 0 | 14 | 0.1429 |
| 9_7547990 | 1 | 0 | 1 | 0 | 5 | 0.2 | 8 | 0 | 15 | 0.0667 | 2 | 0 | 2 | 0 | 3 | 0.6667 | 6 | 0.5 | 13 | 0.3846 |
| 9_7548035 | 2 | 0 | 4 | 0 | 6 | 0.1667 | 8 | 0 | 20 | 0.05 | 4 | 0 | 4 | 0 | 7 | 0.1429 | 7 | 0.1429 | 22 | 0.0909 |
| 9_7548117 | 2 | 0.5 | 6 | 0 | 8 | 0 | 6 | 0.1667 | 22 | 0.0909 | 6 | 0 | 4 | 0.5 | 10 | 0.3 | 2 | 0 | 22 | 0.2273 |
| 9_7548124 | 2 | 0 | 5 | 0.4 | 7 | 0 | 5 | 0 | 19 | 0.1053 | 6 | 0.1667 | 4 | 0.25 | 8 | 0 | 5 | 0.2 | 23 | 0.1304 |
| 9_7548129 | 1 | 0 | 5 | 0 | 10 | 0 | 5 | 0.4 | 21 | 0.0952 | 6 | 0.3333 | 5 | 0.2 | 9 | 0.1111 | 4 | 0.25 | 24 | 0.2083 |
| 9_7548895 | 4 | 0.25 | 2 | 0.5 | 1 | 0 | 2 | 1 | 9 | 0.4444 | 2 | 0 | 3 | 0.6667 | 2 | 0 | NA | NA | 7 | 0.2857 |
| 9_7548910 | 3 | 0 | 3 | 0.3333 | 1 | 1 | 2 | 0 | 9 | 0.2222 | 1 | 1 | 3 | 0 | 3 | 0.3333 | 1 | 0 | 8 | 0.25 |
| 9_7550223 | 4 | 0.5 | 6 | 0 | 9 | 0.1111 | 7 | 0 | 26 | 0.1154 | 8 | 0 | 7 | 0.2857 | 11 | 0 | 7 | 0 | 33 | 0.0606 |
| 9_7553386 | NA | NA | 1 | 1 | NA | NA | NA | NA | 1 | 1 | 1 | 0 | NA | NA | 1 | 1 | 1 | 1 | 3 | 0.6667 |
| 9_7554269 | 8 | 0.125 | 14 | 0.2857 | 7 | 0.4286 | 8 | 0.375 | 37 | 0.2973 | 7 | 0.1429 | 10 | 0.3 | 6 | 0.1667 | 6 | 0.5 | 29 | 0.2759 |
| 9_7554270 | 7 | 0.4286 | 11 | 0.7273 | 7 | 0.2857 | 8 | 0.5 | 33 | 0.5152 | 7 | 0.1429 | 9 | 0.4444 | 6 | 0.6667 | 4 | 0.75 | 26 | 0.4615 |
| 9_7554277 | 8 | 1 | 14 | 1 | 7 | 1 | 8 | 1 | 37 | 1 | 8 | 1 | 13 | 0.9231 | 7 | 1 | 5 | 1 | 33 | 0.9697 |
| 9_7554280 | 7 | 0.1429 | 14 | 0.0714 | 7 | 0.1429 | 8 | 0.25 | 36 | 0.1389 | 5 | 0.2 | 13 | 0.1538 | 7 | 0.2857 | 5 | 0.4 | 30 | 0.2333 |
| 9_7554285 | 8 | 0.375 | 10 | 0.2 | 7 | 0.2857 | 6 | 0 | 31 | 0.2258 | 6 | 0.5 | 12 | 0.4167 | 5 | 0.2 | 5 | 0.2 | 28 | 0.3571 |

| | | | | | | | | | | | | | | | | | | | | |
|---|---|---|---|---|---|---|---|---|---|---|---|---|---|---|---|---|---|---|---|---|
| 9_7554294( | 8 | 0.375 | 13 | 0.3846 | 8 | 0.5 | 8 | 0.375 | 37 | 0.4054 | 8 | 0.25 | 13 | 0.2308 | 6 | 0.3333 | 6 | 0.3333 | 33 | 0.2727 |
| 9_7554294! | 8 | 0.5 | 14 | 0.5714 | 6 | 0.5 | 6 | 1 | 34 | 0.6176 | 5 | 0.4 | 12 | 0.3333 | 6 | 0.8333 | 5 | 0.2 | 28 | 0.4286 |
| 9_7554453! | 7 | 0.1429 | 13 | 0.0769 | 8 | 0 | 8 | 0 | 36 | 0.0556 | 5 | 0.2 | 12 | 0 | 6 | 0 | 5 | 0.4 | 28 | 0.1071 |
| 9_7554455 | 8 | 0.25 | 14 | 0.3571 | 6 | 0.6667 | 7 | 0 | 35 | 0.3143 | 7 | 0.1429 | 14 | 0.1429 | 7 | 0.4286 | 7 | 0.2857 | 35 | 0.2286 |
| 9_7554457 | 9 | 0 | 14 | 0.2857 | 8 | 0 | 8 | 0.125 | 39 | 0.1282 | 5 | 0 | 14 | 0.0714 | 8 | 0.125 | 9 | 0 | 36 | 0.0556 |
| 9_7554473! | 10 | 0.1 | 13 | 0 | 8 | 0.125 | 8 | 0 | 39 | 0.0513 | 7 | 0.1429 | 13 | 0.0769 | 8 | 0 | 8 | 0 | 36 | 0.0556 |
| 9_7554481 | 10 | 0.7 | 15 | 0.6667 | 8 | 0.75 | 9 | 0.7778 | 42 | 0.7143 | 8 | 0.625 | 15 | 0.8667 | 8 | 0.5 | 8 | 0.75 | 39 | 0.7179 |
| 9_7555083! | 30 | 0.6 | 26 | 0.5769 | 35 | 0.5429 | 25 | 0.48 | 116 | 0.5517 | 32 | 0.5 | 23 | 0.6087 | 29 | 0.5862 | 24 | 0.75 | 108 | 0.6019 |
| 9_7555120 | 31 | 0.0968 | 25 | 0.04 | 31 | 0.0645 | 24 | 0.0833 | 111 | 0.0721 | 25 | 0.04 | 22 | 0.0455 | 24 | 0.125 | 26 | 0 | 97 | 0.0515 |
| 9_7604109 | NA | NA | 2 | 0 | 2 | 0.5 | 2 | 0 | 6 | 0.1667 | 5 | 0.2 | NA | NA | 3 | 0.3333 | 3 | 0.3333 | 11 | 0.2727 |
| 9_7604127( | NA | NA | 1 | 0 | 2 | 0 | 2 | 1 | 5 | 0.4 | 5 | 0.4 | 1 | 0 | 3 | 0.3333 | 3 | 0.3333 | 12 | 0.3333 |
| 9_7604135 | NA | NA | 2 | 0 | 1 | 0 | 2 | 1 | 5 | 0.4 | 5 | 0.6 | 1 | 1 | 4 | 0.75 | 4 | 0.75 | 14 | 0.7143 |
| 9_7604140! | NA | NA | 2 | 0.5 | 2 | 1 | 1 | 1 | 5 | 0.8 | 6 | 1 | 1 | 1 | 4 | 1 | 3 | 0.6667 | 14 | 0.9286 |
| 9_7604142( | NA | NA | 2 | 1 | 1 | 1 | 2 | 1 | 5 | 1 | 5 | 0.8 | 1 | 1 | 4 | 1 | 4 | 0.75 | 14 | 0.8571 |
| 9_7604160 | 1 | 1 | 2 | 0.5 | 2 | 0 | 2 | 0.5 | 7 | 0.4286 | 5 | 0.4 | 1 | 1 | 4 | 0.25 | 4 | 0.5 | 14 | 0.4286 |
| 9_7604165 | NA | NA | 2 | 0.5 | 2 | 1 | 1 | 1 | 5 | 0.8 | 4 | 1 | 1 | 1 | 3 | 1 | 3 | 0.6667 | 11 | 0.9091 |
| 9_7604167! | 1 | 0 | 2 | 0 | 2 | 0 | 2 | 0.5 | 7 | 0.1429 | 6 | 0.1667 | 1 | 0 | 4 | 0 | 3 | 0 | 14 | 0.0714 |
| 9_7604169 | NA | NA | 2 | 0.5 | 2 | 1 | 2 | 1 | 6 | 0.8333 | 7 | 0.7143 | 1 | 1 | 3 | 1 | 4 | 1 | 15 | 0.8667 |
| 9_7604185 | NA | NA | 2 | 0 | 2 | 0.5 | 2 | 0 | 6 | 0.1667 | 6 | 0 | 1 | 0 | 4 | 0.25 | 4 | 0 | 15 | 0.0667 |
| 9_7604189( | NA | NA | 1 | 0 | 1 | 1 | 2 | 0.5 | 4 | 0.5 | 5 | 0.2 | 1 | 1 | 3 | 0.6667 | 3 | 0.3333 | 12 | 0.4167 |
| 9_7604190( | NA | NA | 1 | 0 | 2 | 0 | 1 | 1 | 4 | 0.25 | 6 | 0.1667 | 1 | 0 | 4 | 0 | 3 | 0 | 14 | 0.0714 |
| 9_7604220 | NA | NA | 2 | 0 | 2 | 0 | 1 | 1 | 5 | 0.2 | 3 | 0.3333 | 1 | 0 | 3 | 0 | 2 | 0 | 9 | 0.1111 |
| 9_7604248! | NA | NA | NA | NA | 2 | 0.5 | NA | NA | 2 | 0.5 | 3 | 0 | NA | NA | 3 | 0.3333 | NA | NA | 6 | 0.1667 |
| 9_7613494( | 1 | 1 | 1 | 1 | NA | NA | NA | NA | 2 | 1 | 1 | 1 | NA | NA | NA | NA | 1 | 1 | 2 | 1 |
| 9_7613494 | NA | NA | 1 | 1 | NA | NA | NA | NA | 1 | 1 | 1 | 1 | NA | NA | NA | NA | 1 | 1 | 2 | 1 |
| 9_7613496 | 1 | 1 | NA | NA | NA | NA | NA | NA | 1 | 1 | 1 | 1 | NA | NA | NA | NA | 1 | 1 | 2 | 1 |
| 9_7613508 | 1 | 0 | 1 | 1 | NA | NA | NA | NA | 2 | 0.5 | 1 | 1 | NA | NA | NA | NA | 1 | 1 | 2 | 1 |
| 9_7613519 | 1 | 0 | 1 | 1 | NA | NA | NA | NA | 2 | 0.5 | 1 | 1 | NA | NA | NA | NA | 1 | 0 | 2 | 0.5 |
| 9_7613528 | 1 | 1 | 1 | 1 | NA | NA | NA | NA | 2 | 1 | 1 | 1 | NA | NA | NA | NA | 1 | 1 | 2 | 1 |
| 9_7613532 | NA | NA | 1 | 1 | NA | NA | NA | NA | 1 | 1 | 1 | 1 | NA | NA | NA | NA | 1 | 1 | 2 | 1 |
| 9_7623068! | NA | NA | 1 | 1 | NA | NA | NA | NA | 1 | 1 | NA | NA | NA | NA | 1 | 1 | 2 | 0.5 | 3 | 0.6667 |
| 9_7713691( | NA | NA | NA | NA | 3 | 1 | 1 | 0 | 4 | 0.75 | NA | NA | NA | NA | NA | NA | 1 | 1 | 1 | 1 |
| 9_7713697 | NA | NA | NA | NA | 3 | 0.6667 | 1 | 1 | 4 | 0.75 | 2 | 0.5 | 1 | 1 | 1 | 1 | 1 | 0 | 5 | 0.6 |
| 9_7713712 | NA | NA | NA | NA | 3 | 0.6667 | 1 | 1 | 4 | 0.75 | 1 | 1 | NA | NA | 1 | 0 | 1 | 1 | 3 | 0.6667 |
| 9_7713724 | NA | NA | NA | NA | 3 | 0.3333 | 1 | 0 | 4 | 0.25 | 2 | 0.5 | 1 | 0 | 1 | 0 | 1 | 0 | 5 | 0.2 |
| 9_7713725! | NA | NA | NA | NA | 2 | 1 | 1 | 1 | 3 | 1 | 2 | 1 | 1 | 1 | 1 | 0 | 1 | 0 | 5 | 0.6 |
| 9_7713734 | NA | NA | NA | NA | NA | NA | 1 | 1 | 1 | 1 | 1 | 1 | 1 | 0 | 1 | 0 | 1 | 1 | 4 | 0.5 |
| 9_7733811( | NA | NA | 3 | 0.3333 | 3 | 0.3333 | NA | NA | 6 | 0.3333 | NA | NA | 1 | 0 | 1 | 1 | 1 | 1 | 3 | 0.6667 |
| 9_7733839 | 1 | 0 | 5 | 0.2 | 5 | 0.2 | 2 | 0 | 13 | 0.1538 | NA | NA | 1 | 1 | 2 | 1 | 3 | 0.3333 | 6 | 0.6667 |
| 9_7733843 | 1 | 1 | 4 | 0.5 | 5 | 0.8 | 2 | 1 | 12 | 0.75 | NA | NA | 1 | 1 | 2 | 1 | 1 | 1 | 4 | 1 |
| 9_7733863 | 1 | 0 | 2 | 0 | 4 | 0 | 2 | 0.5 | 9 | 0.1111 | NA | NA | 2 | 0.5 | 2 | 0 | 1 | 0 | 5 | 0.2 |
| 9_7733906( | NA | NA | 3 | 0.6667 | 4 | 1 | 2 | 1 | 9 | 0.8889 | NA | NA | 2 | 1 | 2 | 1 | 1 | 1 | 5 | 1 |
| 9_7733908 | NA | NA | 3 | 0 | 4 | 0 | 2 | 1 | 9 | 0.2222 | NA | NA | 2 | 0.5 | 2 | 0.5 | 1 | 1 | 5 | 0.6 |
| 9_7733909 | NA | NA | 4 | 1 | 4 | 1 | 1 | 1 | 9 | 1 | NA | NA | 2 | 1 | 2 | 1 | 1 | 1 | 5 | 1 |
| 9_7733946 | NA | NA | 3 | 1 | 5 | 0.8 | 1 | 1 | 9 | 0.8889 | NA | NA | 2 | 1 | 2 | 1 | 2 | 0.5 | 6 | 0.8333 |
| 9_7770033 | 5 | 0 | 10 | 0.1 | 3 | 0 | 9 | 0.2222 | 27 | 0.1111 | 1 | 0 | 5 | 0 | 6 | 0.3333 | 4 | 0.25 | 16 | 0.1875 |
| 9_7770039 | 5 | 0.6 | 7 | 0.2857 | 5 | 0.4 | 6 | 0.3333 | 23 | 0.3913 | 1 | 0 | 5 | 0.6 | 5 | 0.6 | 3 | 0 | 14 | 0.4286 |
| 9_7770077 | 7 | 0 | 10 | 0 | 5 | 0.2 | 8 | 0.125 | 30 | 0.0667 | 1 | 0 | 5 | 0 | 6 | 0.3333 | 5 | 0 | 17 | 0.1176 |
| 9_7770100 | 5 | 0.2 | 7 | 0.4286 | 7 | 0.2857 | 6 | 0.3333 | 25 | 0.32 | NA | NA | 3 | 0.3333 | 7 | 0.2857 | 4 | 0.25 | 14 | 0.2857 |
| 9_7770113 | 9 | 0.1111 | 10 | 0.1 | 7 | 0 | 11 | 0.0909 | 37 | 0.0811 | 1 | 1 | 6 | 0 | 6 | 0 | 6 | 0.1667 | 19 | 0.1053 |
| 9_7786830 | 4 | 0 | 4 | 0 | 7 | 0.1429 | NA | NA | 15 | 0.0667 | 4 | 0 | 7 | 0 | 3 | 0 | 6 | 0.1667 | 20 | 0.05 |
| 9_7789020! | 17 | 0.2941 | 10 | 0.1 | 9 | 0.1111 | 10 | 0.4 | 46 | 0.2391 | 7 | 0.1429 | 14 | 0.3571 | 9 | 0.1111 | 13 | 0.1538 | 43 | 0.2093 |
| 9_7789021 | 18 | 0.2778 | 10 | 0.2 | 11 | 0.1818 | 9 | 0.2222 | 48 | 0.2292 | 7 | 0 | 13 | 0.3077 | 10 | 0.2 | 12 | 0.4167 | 42 | 0.2619 |
| 9_7789031 | 13 | 0.0769 | 9 | 0.1111 | 13 | 0.3077 | 10 | 0.1 | 45 | 0.1556 | 6 | 0.5 | 11 | 0.4545 | 9 | 0.1111 | 13 | 0.3846 | 39 | 0.359 |
| 9_7789128 | 13 | 0.1538 | 15 | 0.4 | 14 | 0.3571 | 12 | 0.1667 | 54 | 0.2778 | 8 | 0.25 | 17 | 0.4706 | 11 | 0.1818 | 12 | 0.0833 | 48 | 0.2708 |
| 9_7789134 | 18 | 0.1667 | 13 | 0.0769 | 12 | 0.1667 | 14 | 0.2143 | 57 | 0.1579 | 9 | 0.1111 | 14 | 0.1429 | 12 | 0.3333 | 13 | 0.1538 | 48 | 0.1875 |
| 9_7807503 | 2 | 0.5 | 3 | 0.6667 | 1 | 1 | NA | NA | 6 | 0.6667 | 1 | 0 | NA | NA | 1 | 1 | NA | NA | 2 | 0.5 |
| 9_7807692( | NA | NA | 2 | 0.5 | 1 | 0 | NA | NA | 3 | 0.3333 | 1 | 0 | 1 | 1 | NA | NA | NA | NA | 2 | 0.5 |
| 9_7807833 | NA | NA | 2 | 0 | 1 | 1 | NA | NA | 3 | 0.3333 | 1 | 0 | NA | NA | NA | NA | 1 | 1 | 2 | 0.5 |
| 9_7808549 | 2 | 0 | 1 | 1 | NA | NA | NA | NA | 3 | 0.3333 | NA | NA | 1 | 1 | NA | NA | NA | NA | 1 | 1 |
| 9_7808591( | 1 | 1 | 1 | 0 | NA | NA | NA | NA | 2 | 0.5 | NA | NA | 1 | 1 | NA | NA | 1 | 1 | 2 | 1 |
| 9_7808598 | 3 | 0.6667 | 1 | 0 | NA | NA | NA | NA | 4 | 0.5 | NA | NA | 1 | 1 | NA | NA | NA | NA | 1 | 1 |
| 9_7808606 | 3 | 0.6667 | 1 | 0 | NA | NA | NA | NA | 4 | 0.5 | NA | NA | 1 | 1 | NA | NA | 1 | 0 | 2 | 0.5 |
| 9_7811644 | 14 | 0.2857 | 7 | 0.1429 | 6 | 0.6667 | 5 | 0.4 | 32 | 0.3438 | 7 | 0.8571 | 7 | 0.2857 | 5 | 0.8 | 4 | 0.75 | 23 | 0.6522 |
| 9_7836009 | 2 | 0 | 2 | 1 | 4 | 0 | 2 | 0 | 10 | 0.2 | 1 | 0 | 3 | 0.6667 | 2 | 0 | NA | NA | 6 | 0.3333 |

| | | | | | | | | | | | | | | | | | | | | |
|---|---|---|---|---|---|---|---|---|---|---|---|---|---|---|---|---|---|---|---|---|
| 9_7836021 | 2 | 0.5 | 2 | 0.5 | 5 | 0.8 | 2 | 0.5 | 11 | 0.6364 | 1 | 0 | 3 | 0 | 3 | 0.3333 | 1 | 0 | 8 | 0.125 |
| 9_7838119 | 2 | 0.5 | 3 | 0 | 5 | 0.4 | 3 | 0 | 13 | 0.2308 | 3 | 0 | 1 | 0 | 6 | 0.3333 | 2 | 0 | 12 | 0.1667 |
| 9_7838121 | 2 | 0 | 3 | 0.3333 | 3 | 0.6667 | 3 | 0.6667 | 11 | 0.4545 | 3 | 0 | 2 | 0 | 4 | 0.25 | 1 | 0 | 10 | 0.1 |
| 9_7838586 | 510 | 0.1392 | 438 | 0.1438 | 433 | 0.1132 | 497 | 0.1268 | 1878 | 0.131 | 357 | 0.1289 | 459 | 0.098 | 521 | 0.1344 | 412 | 0.1262 | 1749 | 0.1218 |
| 9_7838591 | 553 | 0.6799 | 462 | 0.7035 | 450 | 0.7222 | 550 | 0.7055 | 2015 | 0.7017 | 408 | 0.6642 | 503 | 0.6461 | 559 | 0.6923 | 452 | 0.708 | 1922 | 0.6779 |
| 9_7838595 | 502 | 0.6076 | 416 | 0.6202 | 418 | 0.5885 | 504 | 0.5952 | 1840 | 0.6027 | 374 | 0.5267 | 454 | 0.5242 | 494 | 0.5607 | 390 | 0.6026 | 1712 | 0.5532 |
| 9_7847480 | 1 | 0 | NA | NA | 1 | 1 | NA | NA | 2 | 0.5 | 2 | 1 | 1 | 0 | NA | NA | NA | NA | 3 | 0.6667 |
| 9_7847483 | 1 | 1 | NA | NA | 1 | 1 | NA | NA | 2 | 1 | 1 | 1 | 1 | 1 | NA | NA | NA | NA | 2 | 1 |
| 9_7847491 | 1 | 1 | NA | NA | 1 | 1 | NA | NA | 2 | 1 | 2 | 1 | 1 | 0 | NA | NA | NA | NA | 3 | 0.6667 |
| 9_7847534 | 2 | 1 | NA | NA | 1 | 1 | NA | NA | 3 | 1 | 1 | 1 | NA | NA | NA | NA | NA | NA | 1 | 1 |
| 9_7847543 | 2 | 0.5 | NA | NA | 1 | 1 | NA | NA | 3 | 0.6667 | 1 | 1 | NA | NA | 1 | 1 | NA | NA | 2 | 1 |
| 9_7847559 | 2 | 1 | NA | NA | 1 | 1 | NA | NA | 3 | 1 | 1 | 1 | NA | NA | NA | NA | NA | NA | 1 | 1 |
| 9_7847639 | 2 | 1 | NA | NA | 1 | 1 | NA | NA | 3 | 1 | 1 | 1 | NA | NA | NA | NA | NA | NA | 1 | 1 |
| 9_7847662 | 2 | 1 | NA | NA | 1 | 1 | NA | NA | 3 | 1 | 1 | 1 | NA | NA | NA | NA | NA | NA | 1 | 1 |
| 9_7967594 | 6 | 0.5 | 5 | 0.2 | 8 | 0.5 | 7 | 0.2857 | 26 | 0.3846 | 3 | 0.3333 | 7 | 0.2857 | 6 | 0.6667 | 8 | 0.5 | 24 | 0.4583 |
| 9_7967600 | 6 | 0 | 6 | 0.1667 | 8 | 0.125 | 6 | 0 | 26 | 0.0769 | 2 | 0 | 8 | 0 | 6 | 0.1667 | 7 | 0.1429 | 23 | 0.087 |
| 9_7967637 | 43 | 0.093 | 50 | 0.06 | 35 | 0.0571 | 47 | 0.1277 | 175 | 0.0857 | 22 | 0.1818 | 36 | 0.0833 | 36 | 0.0556 | 45 | 0.1333 | 139 | 0.1079 |
| 9_7967653 | 43 | 0.8837 | 49 | 0.9388 | 37 | 0.973 | 46 | 0.913 | 175 | 0.9257 | 21 | 0.9524 | 35 | 0.9429 | 40 | 0.9 | 46 | 1 | 142 | 0.9507 |
| 9_7967671 | 35 | 0.2857 | 40 | 0.15 | 25 | 0.32 | 38 | 0.2632 | 138 | 0.2464 | 15 | 0.1333 | 28 | 0.1429 | 31 | 0.0968 | 38 | 0.2895 | 112 | 0.1786 |
| 9_7967674 | 36 | 0.9722 | 45 | 0.9333 | 30 | 0.9667 | 40 | 0.9 | 151 | 0.9404 | 14 | 0.9286 | 29 | 0.931 | 36 | 0.8333 | 37 | 0.973 | 116 | 0.9138 |
| 9_7967689 | 37 | 0.2162 | 43 | 0.1395 | 28 | 0.2143 | 35 | 0.3143 | 143 | 0.2168 | 15 | 0.2667 | 27 | 0.2222 | 30 | 0.0333 | 35 | 0.1143 | 107 | 0.1402 |
| 9_7967698 | 39 | 0.0513 | 44 | 0.1136 | 31 | 0.0968 | 40 | 0.075 | 154 | 0.0844 | 14 | 0 | 31 | 0.0968 | 37 | 0.0811 | 41 | 0.0488 | 123 | 0.065 |
| 9_7967810 | 26 | 0.0769 | 38 | 0.0789 | 25 | 0.16 | 33 | 0.0909 | 122 | 0.0984 | 14 | 0.1429 | 28 | 0.1786 | 28 | 0.1071 | 33 | 0.0909 | 103 | 0.1262 |
| 9_7967859 | 14 | 0.0714 | 30 | 0.1 | 17 | 0.1176 | 21 | 0.1905 | 82 | 0.122 | 9 | 0 | 18 | 0.1667 | 21 | 0.1429 | 26 | 0.0769 | 74 | 0.1081 |
| 9_7970048 | 17 | 0.1176 | 25 | 0.24 | 17 | 0.1765 | 21 | 0.2857 | 80 | 0.2125 | 7 | 0.1429 | 18 | 0.1111 | 21 | 0.2381 | 27 | 0.1852 | 73 | 0.1781 |
| 9_7972235 | 1 | 0 | 1 | 0 | 2 | 0 | 6 | 0.1667 | 10 | 0.1 | 2 | 0 | 3 | 0 | 4 | 0.25 | 2 | 0 | 11 | 0.0909 |
| 9_7972255 | 1 | 1 | 1 | 0 | 2 | 1 | 6 | 0.1667 | 10 | 0.4 | 3 | 0 | 3 | 0.3333 | 3 | 0 | 3 | 0.3333 | 12 | 0.1667 |
| 9_7972259 | 1 | 1 | 1 | 0 | 3 | 0.3333 | 5 | 0.4 | 10 | 0.4 | 2 | 0 | 5 | 0.4 | 2 | 0.5 | 1 | 0 | 10 | 0.3 |
| 9_7972304 | NA | NA | NA | NA | 1 | 1 | 4 | 0 | 5 | 0.2 | 2 | 0 | 3 | 0.3333 | 2 | 1 | 3 | 1 | 10 | 0.6 |
| 9_7972548 | NA | NA | 1 | 1 | NA | NA | 4 | 1 | 5 | 1 | 1 | 0 | 2 | 1 | 2 | 1 | 1 | 1 | 6 | 0.8333 |
| 9_7972551 | NA | NA | 1 | 0 | 1 | 1 | 6 | 1 | 8 | 0.875 | 1 | 1 | 3 | 1 | 2 | 1 | 2 | 1 | 8 | 1 |
| 9_7972598 | NA | NA | NA | NA | 1 | 0 | 5 | 0.4 | 6 | 0.3333 | 1 | 1 | 3 | 0.3333 | 2 | 1 | 2 | 0.5 | 8 | 0.625 |
| 9_7972603 | NA | NA | 1 | 0 | NA | NA | 5 | 0.6 | 6 | 0.5 | 2 | 0.5 | 2 | 0 | 1 | 0 | 1 | 0 | 6 | 0.1667 |
| 9_7972666 | NA | NA | NA | NA | NA | NA | 1 | 1 | 1 | 1 | 2 | 0 | 1 | 0 | 2 | 0.5 | 1 | 1 | 6 | 0.3333 |
| 9_7972670 | NA | NA | NA | NA | NA | NA | 2 | 1 | 2 | 1 | 1 | 1 | 2 | 0.5 | 2 | 0.5 | 1 | 0 | 6 | 0.5 |
| 9_7997425 | NA | NA | NA | NA | NA | NA | 1 | 1 | 1 | 1 | 1 | 0 | 1 | 1 | NA | NA | NA | NA | 2 | 0.5 |
| 9_8002116 | 1 | 0 | 6 | 0.1667 | 1 | 0 | 8 | 0 | 16 | 0.0625 | 1 | 0 | 4 | 0 | 3 | 0.3333 | 3 | 0 | 11 | 0.0909 |
| 9_8002117 | 1 | 0 | 5 | 0 | 1 | 0 | 9 | 0.3333 | 16 | 0.1875 | 2 | 0 | 4 | 0 | 4 | 0.25 | 5 | 0.2 | 15 | 0.1333 |
| 9_8002118 | 1 | 1 | 4 | 0 | 1 | 0 | 6 | 0.3333 | 12 | 0.25 | 2 | 0.5 | 5 | 0.4 | 3 | 0 | 4 | 0.25 | 14 | 0.2857 |
| 9_8002120 | 1 | 0 | 7 | 0.8571 | 1 | 1 | 9 | 0.5556 | 18 | 0.6667 | 1 | 1 | 5 | 0.6 | 3 | 0.6667 | 4 | 0.25 | 13 | 0.5385 |
| 9_8002123 | 1 | 0 | 6 | 0.3333 | 1 | 0 | 7 | 0.2857 | 15 | 0.2667 | 1 | 0 | 5 | 0.4 | 3 | 0 | 4 | 0 | 13 | 0.1538 |
| 9_8004379 | 1 | 1 | 9 | 0.4444 | 2 | 0 | 6 | 0.1667 | 18 | 0.3333 | 1 | 0 | 5 | 0 | 6 | 0 | 5 | 0.2 | 17 | 0.0588 |
| 9_8004383 | 1 | 0 | 7 | 0 | 5 | 0 | 7 | 0.1429 | 20 | 0.05 | 2 | 0 | 5 | 0 | 5 | 0 | 3 | 0.3333 | 15 | 0.0667 |
| 9_8004385 | 1 | 1 | 10 | 0.3 | 5 | 0 | 9 | 0.3333 | 25 | 0.28 | 1 | 0 | 6 | 0.5 | 6 | 0.1667 | 5 | 0.4 | 18 | 0.3333 |
| 9_8004389 | 1 | 0 | 11 | 0.2727 | 4 | 0 | 9 | 0.4444 | 25 | 0.28 | 2 | 0 | 6 | 0.5 | 6 | 0.1667 | 5 | 0.4 | 19 | 0.3158 |
| 9_8004395 | 1 | 0 | 10 | 0.3 | 5 | 0.4 | 8 | 0.375 | 24 | 0.3333 | 1 | 1 | 5 | 0.2 | 6 | 0.6667 | 2 | 0 | 14 | 0.4286 |
| 9_8005097 | 3 | 0.6667 | 8 | 0.375 | 7 | 0.7143 | 11 | 0.4545 | 29 | 0.5172 | 2 | 0.5 | 5 | 0.6 | 8 | 0.625 | 4 | 0.75 | 19 | 0.6316 |
| 9_8005107 | 4 | 0.75 | 12 | 0.75 | 8 | 0.75 | 10 | 0.6 | 34 | 0.7059 | 2 | 1 | 6 | 0.5 | 8 | 0.5 | 5 | 0.4 | 21 | 0.5238 |
| 9_8005118 | 3 | 0.3333 | 14 | 0.1429 | 7 | 0.1429 | 11 | 0.2727 | 35 | 0.2 | 1 | 1 | 6 | 0 | 9 | 0.3333 | 5 | 0.6 | 21 | 0.3333 |
| 9_8005118 | 3 | 0.3333 | 12 | 0.1667 | 8 | 0 | 11 | 0.0909 | 34 | 0.1176 | 2 | 0 | 3 | 0 | 7 | 0.2857 | 5 | 0 | 17 | 0.1176 |
| 9_8005121 | 3 | 0 | 14 | 0.2143 | 6 | 0.1667 | 12 | 0.4167 | 35 | 0.2571 | 2 | 0.5 | 5 | 0.8 | 9 | 0.3333 | 4 | 0 | 20 | 0.4 |
| 9_8005124 | 5 | 1 | 13 | 1 | 9 | 0.6667 | 12 | 0.75 | 39 | 0.8462 | 2 | 1 | 5 | 0.8 | 8 | 1 | 5 | 0.8 | 20 | 0.9 |
| 9_8021515 | 4 | 0 | 7 | 0 | 7 | 0.1429 | 6 | 0.1667 | 24 | 0.0833 | 4 | 0 | 6 | 0 | 4 | 0 | 4 | 0.25 | 18 | 0.0556 |
| 9_8021525 | 2 | 0 | 6 | 0.1667 | 6 | 0 | 6 | 0.1667 | 20 | 0.1 | 6 | 0.3333 | 6 | 0 | 4 | 0.5 | 6 | 0.5 | 22 | 0.3182 |
| 9_8021532 | 4 | 0 | 6 | 0 | 9 | 0.3333 | 7 | 0.4286 | 26 | 0.2308 | 5 | 0 | 7 | 0.1429 | 6 | 0.1667 | 6 | 0.1667 | 24 | 0.125 |
| 9_8021546 | 3 | 0.3333 | 7 | 0.1429 | 7 | 0 | 6 | 0 | 23 | 0.087 | 5 | 0.4 | 6 | 0 | 4 | 0 | 6 | 0 | 21 | 0.0952 |
| 9_8021594 | 7 | 0.7143 | 7 | 0 | 9 | 0.3333 | 6 | 0.5 | 29 | 0.3793 | 6 | 0.6667 | 5 | 0.4 | 5 | 0.8 | 5 | 0.2 | 21 | 0.5238 |
| 9_8025256 | 1 | 1 | NA | NA | 1 | 1 | NA | NA | 2 | 1 | 1 | 1 | 1 | 1 | NA | NA | 1 | 1 | 3 | 1 |
| 9_8025287 | 1 | 1 | NA | NA | NA | NA | NA | NA | 1 | 1 | 1 | 1 | 1 | 1 | NA | NA | 2 | 0.5 | 4 | 0.75 |
| 9_8151050 | 6 | 0.1667 | 4 | 0.5 | 2 | 0.5 | 11 | 0.1818 | 23 | 0.2609 | 2 | 0 | 6 | 0.5 | 2 | 0.5 | 3 | 0.6667 | 13 | 0.4615 |
| 9_8151061 | 6 | 0.1667 | 4 | 0 | 2 | 0.5 | 12 | 0.1667 | 24 | 0.1667 | 1 | 0 | 5 | 0.2 | 2 | 0 | 3 | 0 | 11 | 0.0909 |
| 9_8151083 | 6 | 0 | 3 | 0 | 2 | 0 | 9 | 0.2222 | 20 | 0.1 | 1 | 0 | 4 | 0.25 | 1 | 0 | 3 | 0 | 9 | 0.1111 |
| 9_8151089 | 6 | 0.3333 | 3 | 0.6667 | 2 | 0 | 11 | 0.3636 | 22 | 0.3636 | 1 | 0 | 5 | 0 | 2 | 0.5 | 3 | 0 | 11 | 0.0909 |
| 9_8151102 | 5 | 0.4 | 2 | 0 | 1 | 0 | 10 | 0.1 | 18 | 0.1667 | 1 | 0 | 5 | 0 | 2 | 0 | 3 | 0.3333 | 11 | 0.0909 |
| 9_8151150 | 3 | 0.6667 | 3 | 0.3333 | 2 | 0 | 13 | 0.1538 | 21 | 0.2381 | 2 | 0 | 6 | 0.3333 | 6 | 0.3333 | 5 | 0 | 19 | 0.2105 |

| | | | | | | | | | | | | | | | | | | | | |
|---|---|---|---|---|---|---|---|---|---|---|---|---|---|---|---|---|---|---|---|---|
| 9_8151236( | 3 | 0.3333 | 3 | 0 | 2 | 0.5 | 8 | 0.25 | 16 | 0.25 | 1 | 1 | 4 | 0 | 5 | 0 | 4 | 0 | 14 | 0.0714 |
| 9_8151243( | 3 | 0.3333 | 3 | 0 | 2 | 0 | 10 | 0.1 | 18 | 0.1111 | 2 | 0.5 | 3 | 0 | 4 | 0 | 4 | 0 | 13 | 0.0769 |
| 9_8151264( | 2 | 0 | 2 | 0 | 2 | 0 | 10 | 0.1 | 16 | 0.0625 | 2 | 0 | 3 | 0 | 1 | 0 | 3 | 0.3333 | 9 | 0.1111 |
| 9_8151295 | 2 | 0 | 3 | 0 | 2 | 0 | 9 | 0.2222 | 16 | 0.125 | 1 | 0 | 2 | 0 | 3 | 0 | 2 | 0.5 | 8 | 0.125 |
| 9_8151298( | 2 | 1 | 3 | 1 | 1 | 1 | 9 | 0.8889 | 15 | 0.9333 | 1 | 1 | 2 | 1 | 3 | 1 | 2 | 1 | 8 | 1 |
| 9_8151300 | 3 | 0 | 3 | 0.3333 | 1 | 0 | 6 | 0.8333 | 13 | 0.4615 | 1 | 1 | 2 | 0.5 | 2 | 0 | 2 | 0 | 7 | 0.2857 |
| 9_8151301 | 3 | 1 | 1 | 1 | 2 | 1 | 7 | 1 | 13 | 1 | 1 | 1 | 1 | 1 | 2 | 1 | 2 | 1 | 6 | 1 |
| 9_8151314 | 3 | 0.3333 | NA | NA | 1 | 1 | 9 | 0.1111 | 13 | 0.2308 | 1 | 0 | 2 | 0.5 | 1 | 0 | 1 | 0 | 5 | 0.2 |
| 9_8151325 | 2 | 0.5 | 1 | 0 | 2 | 0.5 | 7 | 0.2857 | 12 | 0.3333 | NA | NA | 3 | 0.6667 | 2 | 0 | 1 | 1 | 6 | 0.5 |
| 9_8151340 | 3 | 1 | 1 | 1 | 2 | 1 | 7 | 1 | 13 | 1 | NA | NA | 1 | 1 | 2 | 1 | 1 | 1 | 4 | 1 |
| 9_8151344 | 2 | 1 | NA | NA | 2 | 1 | 5 | 1 | 9 | 1 | NA | NA | 2 | 0.5 | 3 | 0.6667 | 1 | 1 | 6 | 0.6667 |
| 9_8151347 | 2 | 1 | 1 | 1 | 2 | 1 | 6 | 1 | 11 | 1 | NA | NA | 1 | 1 | 3 | 1 | 1 | 1 | 5 | 1 |
| 9_8151349 | 3 | 1 | 1 | 1 | 2 | 1 | 5 | 1 | 11 | 1 | NA | NA | 3 | 1 | 3 | 1 | 1 | 1 | 7 | 1 |
| 9_8151356 | 3 | 0.3333 | 1 | 1 | 1 | 1 | 5 | 0.6 | 10 | 0.6 | NA | NA | 2 | 0.5 | 1 | 1 | NA | NA | 3 | 0.6667 |
| 9_8151370 | 3 | 0.3333 | 1 | 0 | 2 | 0.5 | 6 | 0.1667 | 12 | 0.25 | NA | NA | 2 | 0 | 2 | 0.5 | NA | NA | 4 | 0.25 |
| 9_8151384 | 1 | 1 | 1 | 1 | 2 | 1 | 6 | 1 | 10 | 1 | NA | NA | 1 | 1 | 3 | 1 | NA | NA | 4 | 1 |
| 9_8151389 | 3 | 0 | 1 | 0 | 2 | 0.5 | 6 | 0.3333 | 12 | 0.25 | NA | NA | 1 | 0 | 3 | 0.3333 | NA | NA | 4 | 0.25 |
| 9_8151402 | 3 | 1 | NA | NA | 2 | 1 | 5 | 1 | 10 | 1 | NA | NA | 2 | 0.5 | 3 | 1 | NA | NA | 5 | 0.8 |
| 9_8151413 | 3 | 1 | 1 | 0 | 2 | 1 | 4 | 1 | 10 | 0.9 | NA | NA | NA | NA | 3 | 1 | NA | NA | 3 | 1 |
| 9_8151418 | 3 | 0 | 1 | 0 | 1 | 0 | 4 | 0.25 | 9 | 0.1111 | NA | NA | 2 | 0.5 | 3 | 0.3333 | NA | NA | 5 | 0.4 |
| 9_8151427 | 2 | 0.5 | NA | NA | 2 | 0.5 | 1 | 1 | 5 | 0.6 | NA | NA | 1 | 1 | 1 | 1 | NA | NA | 2 | 1 |
| 9_8151438 | 3 | 0.3333 | 1 | 0 | 2 | 0 | 5 | 0 | 11 | 0.0909 | NA | NA | 3 | 0.3333 | 1 | 0 | NA | NA | 4 | 0.25 |
| 9_8151449 | 3 | 0.3333 | NA | NA | 2 | 0.5 | 4 | 0.5 | 9 | 0.4444 | NA | NA | 2 | 1 | 1 | 1 | NA | NA | 3 | 1 |
| 9_8151472( | 2 | 1 | NA | NA | 2 | 1 | 5 | 1 | 9 | 1 | NA | NA | 1 | 1 | NA | NA | NA | NA | 1 | 1 |
| 9_8151490 | 2 | 0.5 | NA | NA | 1 | 0 | 3 | 1 | 6 | 0.6667 | NA | NA | 2 | 0.5 | NA | NA | NA | NA | 2 | 0.5 |
| 9_8152616 | 1 | 1 | NA | NA | 1 | 1 | 3 | 0.6667 | 5 | 0.8 | 1 | 1 | 1 | 1 | NA | NA | NA | NA | 2 | 1 |
| 9_8152625 | 1 | 0 | NA | NA | 1 | 0 | 3 | 0.6667 | 5 | 0.4 | 2 | 1 | 1 | 1 | 1 | 0 | NA | NA | 4 | 0.75 |
| 9_8180945 | NA | NA | NA | NA | NA | NA | 1 | 1 | 1 | 1 | NA | NA | NA | NA | 1 | 1 | NA | NA | 1 | 1 |
| 9_8190781 | NA | NA | NA | NA | NA | NA | 1 | 1 | 1 | 1 | NA | NA | 1 | 1 | 1 | 0 | NA | NA | 2 | 0.5 |
| 9_8272869 | 3 | 0.6667 | 3 | 0 | 6 | 0.3333 | 3 | 0.6667 | 15 | 0.4 | 4 | 0.25 | 6 | 0.6667 | 3 | 0.6667 | 7 | 0.2857 | 20 | 0.45 |
| 9_8272873 | 3 | 0 | 4 | 0 | 7 | 0.2857 | 3 | 0 | 17 | 0.1176 | 3 | 0.3333 | 5 | 0.4 | 3 | 0.3333 | 6 | 0.3333 | 17 | 0.3529 |
| 9_8272874 | 2 | 0 | 1 | 0 | 3 | 0.3333 | 3 | 0 | 9 | 0.1111 | 1 | 1 | 3 | 0.3333 | 2 | 0 | 5 | 0.8 | 11 | 0.5455 |
| 9_8272892 | 3 | 0.6667 | 5 | 0.8 | 6 | 0.6667 | 4 | 1 | 18 | 0.7778 | 4 | 0.75 | 5 | 1 | 3 | 0.6667 | 6 | 0.8333 | 18 | 0.8333 |
| 9_8274861 | 3 | 0.3333 | 7 | 0.5714 | 6 | 0.3333 | 12 | 0.4167 | 28 | 0.4286 | 4 | 0.25 | 11 | 0.3636 | 8 | 0.5 | 8 | 0.375 | 31 | 0.3871 |
| 9_8274863 | 1 | 0 | 8 | 0.125 | 5 | 0.4 | 15 | 0.3333 | 29 | 0.2759 | 3 | 0.3333 | 11 | 0.1818 | 6 | 0.1667 | 9 | 0.1111 | 29 | 0.1724 |
| 9_8275127( | NA | NA | 4 | 0.25 | 1 | 0 | 5 | 0 | 10 | 0.1 | 2 | 0 | 4 | 0 | 2 | 0.5 | 6 | 0.3333 | 14 | 0.2143 |
| 9_8275165 | NA | NA | 3 | 0 | 1 | 0 | 6 | 0.1667 | 10 | 0.1 | 2 | 0.5 | 3 | 0 | 2 | 0.5 | 5 | 0 | 12 | 0.1667 |
| 9_8275256 | NA | NA | 2 | 0 | 2 | 0 | 7 | 0.2857 | 11 | 0.1818 | 1 | 0 | 3 | 0 | 2 | 0.5 | 5 | 0.2 | 11 | 0.1818 |
| 9_8275263( | NA | NA | 1 | 1 | NA | NA | 6 | 0 | 7 | 0.1429 | 1 | 0 | 3 | 0 | 1 | 1 | 4 | 0.25 | 9 | 0.2222 |
| 9_8275274 | NA | NA | 1 | 0 | 2 | 0.5 | 5 | 0.4 | 8 | 0.375 | 1 | 1 | 3 | 0.3333 | 2 | 0 | 5 | 0 | 11 | 0.1818 |
| 9_8275300 | NA | NA | 1 | 0 | 1 | 1 | 5 | 0 | 7 | 0.1429 | 1 | 0 | 3 | 0 | 2 | 0.5 | 4 | 0.25 | 10 | 0.2 |
| 9_8275314 | NA | NA | 1 | 1 | 1 | 1 | 2 | 1 | 4 | 1 | 1 | 1 | 3 | 0.6667 | 2 | 0.5 | 4 | 1 | 10 | 0.8 |
| 9_8275327 | NA | NA | 1 | 0 | 1 | 0 | 4 | 0.25 | 6 | 0.1667 | 1 | 0 | 2 | 0 | 2 | 0 | 4 | 0.25 | 9 | 0.1111 |
| 9_8275331 | NA | NA | 1 | 1 | 1 | 1 | 1 | 1 | 3 | 1 | 1 | 1 | 2 | 1 | 1 | 1 | 3 | 0.6667 | 7 | 0.8571 |
| 9_8275334 | NA | NA | 1 | 1 | NA | NA | 2 | 1 | 3 | 1 | 1 | 0 | 2 | 1 | 1 | 1 | 4 | 0.75 | 8 | 0.75 |
| 9_8275335 | NA | NA | 1 | 0 | 1 | 0 | 3 | 0.3333 | 5 | 0.2 | 1 | 0 | 2 | 0 | 2 | 0 | 3 | 0.6667 | 8 | 0.25 |
| 9_8275336 | NA | NA | 1 | 1 | 1 | 1 | 4 | 0.25 | 6 | 0.5 | 1 | 1 | 1 | 0 | 2 | 0 | 3 | 0 | 7 | 0.1429 |
| 9_8275337 | NA | NA | 1 | 0 | NA | NA | 4 | 0.5 | 5 | 0.4 | 1 | 1 | 1 | 1 | 2 | 0.5 | 3 | 0.6667 | 7 | 0.7143 |
| 9_8275342( | NA | NA | 1 | 1 | 1 | 1 | 4 | 0.75 | 6 | 0.8333 | NA | NA | 2 | 1 | 2 | 1 | 2 | 1 | 6 | 1 |
| 9_8275344 | NA | NA | 1 | 1 | NA | NA | 2 | 1 | 3 | 1 | 1 | 1 | 2 | 1 | 1 | 1 | 1 | 1 | 5 | 1 |
| 9_8275346 | NA | NA | 1 | 1 | 1 | 1 | 3 | 0.6667 | 5 | 0.8 | 1 | 1 | 2 | 1 | 2 | 1 | 3 | 1 | 8 | 1 |
| 9_8275355 | NA | NA | 1 | 1 | 1 | 1 | 3 | 1 | 5 | 1 | 1 | 1 | 2 | 1 | 2 | 1 | 2 | 1 | 7 | 1 |
| 9_8275357 | NA | NA | 1 | 1 | NA | NA | 3 | 0 | 4 | 0.25 | 1 | 0 | 2 | 0.5 | 2 | 0.5 | 2 | 0 | 7 | 0.2857 |
| 9_8275365 | NA | NA | NA | NA | 1 | 1 | 4 | 0.5 | 5 | 0.6 | 1 | 1 | 2 | 1 | 2 | 0 | 2 | 1 | 7 | 0.7143 |
| 9_8275372 | NA | NA | NA | NA | 1 | 0 | 2 | 1 | 3 | 0.6667 | 1 | 1 | 2 | 0.5 | 2 | 0.5 | 2 | 0.5 | 7 | 0.5714 |
| 9_8275382 | NA | NA | NA | NA | 1 | 0 | 3 | 0.3333 | 4 | 0.25 | NA | NA | 2 | 0.5 | 2 | 0 | 3 | 0 | 7 | 0.1429 |
| 9_8299216 | 7 | 0.4286 | 8 | 0.5 | 7 | 0.4286 | 5 | 0.8 | 27 | 0.5185 | 6 | 0.5 | 9 | 0.4444 | 12 | 0.4167 | 8 | 0.5 | 35 | 0.4571 |
| 9_8299230 | 8 | 0.375 | 9 | 0.8889 | 8 | 0.75 | 5 | 1 | 30 | 0.7333 | 6 | 1 | 11 | 0.9091 | 12 | 0.6667 | 7 | 0.7143 | 36 | 0.8056 |
| 9_8299234 | 7 | 0.8571 | 9 | 0.8889 | 8 | 0.75 | 5 | 1 | 29 | 0.8621 | 6 | 1 | 11 | 1 | 12 | 0.9167 | 8 | 0.5 | 37 | 0.8649 |
| 9_8299240 | 8 | 0 | 8 | 0.125 | 8 | 0.375 | 5 | 0 | 29 | 0.1379 | 6 | 0.3333 | 10 | 0.2 | 13 | 0.3077 | 7 | 0 | 36 | 0.2222 |
| 9_8327497( | NA | NA | NA | NA | NA | NA | 1 | 1 | 1 | 1 | NA | NA | NA | NA | NA | NA | 1 | 1 | 1 | 1 |
| 9_8327529 | NA | NA | NA | NA | NA | NA | 1 | 1 | 1 | 1 | NA | NA | NA | NA | NA | NA | 1 | 1 | 1 | 1 |
| 9_8327565 | 2 | 0.5 | NA | NA | NA | NA | 1 | 1 | 3 | 0.6667 | NA | NA | 1 | 1 | 1 | 1 | 1 | 1 | 3 | 1 |
| 9_8327593 | 2 | 0.5 | NA | NA | NA | NA | 1 | 0 | 3 | 0.3333 | NA | NA | 1 | 1 | 1 | 0 | 1 | 0 | 3 | 0.3333 |
| 9_8327667 | 1 | 1 | NA | NA | NA | NA | 1 | 0 | 2 | 0.5 | NA | NA | 1 | 1 | NA | NA | NA | NA | 1 | 1 |

| | | | | | | | | | | | | | | | | | | | | |
|---|---|---|---|---|---|---|---|---|---|---|---|---|---|---|---|---|---|---|---|---|
| 9_8327722 | 1 | 0 | NA | NA | NA | NA | 1 | 1 | 2 | 0.5 | NA | NA | 1 | 1 | 1 | 1 | NA | NA | 2 | 1 |
| 9_8327725 | 1 | 1 | NA | NA | NA | NA | 1 | 1 | 2 | 1 | NA | NA | NA | NA | 1 | 1 | 1 | 1 | 2 | 1 |
| 9_8327731 | 1 | 1 | NA | NA | NA | NA | 1 | 0 | 2 | 0.5 | NA | NA | NA | NA | 1 | 0 | 1 | 1 | 2 | 0.5 |
| 9_8327742 | 1 | 1 | NA | NA | NA | NA | 1 | 1 | 2 | 1 | NA | NA | NA | NA | 1 | 1 | 1 | 1 | 2 | 1 |
| 9_8327770 | 1 | 1 | NA | NA | NA | NA | 1 | 0 | 2 | 0.5 | NA | NA | NA | NA | 1 | 1 | 1 | 1 | 2 | 1 |
| 9_8327776 | 1 | 1 | NA | NA | NA | NA | 1 | 1 | 2 | 1 | NA | NA | NA | NA | 1 | 1 | 1 | 1 | 2 | 1 |
| 9_8327795 | 1 | 1 | NA | NA | NA | NA | 1 | 1 | 2 | 1 | NA | NA | NA | NA | 1 | 0 | 1 | 1 | 2 | 0.5 |
| 9_8327796 | 1 | 1 | NA | NA | NA | NA | 1 | 0 | 2 | 0.5 | NA | NA | NA | NA | 1 | 0 | 1 | 1 | 2 | 0.5 |
| 9_8330521 | 1 | 1 | NA | NA | NA | NA | 1 | 1 | 2 | 1 | NA | NA | NA | NA | 1 | 1 | NA | NA | 1 | 1 |
| 9_8347943 | 2 | 1 | 7 | 1 | 1 | 1 | 1 | 1 | 11 | 1 | NA | NA | 1 | 1 | 2 | 1 | 6 | 1 | 9 | 1 |
| 9_8347945 | 2 | 1 | 8 | 1 | 1 | 1 | 2 | 1 | 13 | 1 | NA | NA | 1 | 1 | 3 | 1 | 6 | 0.8333 | 10 | 0.9 |
| 9_8347957 | 2 | 1 | 5 | 1 | 1 | 1 | 1 | 1 | 9 | 1 | NA | NA | 1 | 1 | 2 | 1 | 7 | 1 | 10 | 1 |
| 9_8347961 | 1 | 0 | 7 | 0.2857 | NA | NA | 2 | 0 | 10 | 0.2 | NA | NA | 1 | 0 | 2 | 0 | 5 | 0.4 | 8 | 0.25 |
| 9_8347963 | 2 | 1 | 5 | 1 | 1 | 1 | 2 | 1 | 10 | 1 | NA | NA | 1 | 1 | 3 | 0.6667 | 6 | 1 | 10 | 0.9 |
| 9_8347965 | 2 | 0 | 7 | 0.1429 | 1 | 0 | 2 | 0 | 12 | 0.0833 | NA | NA | 1 | 1 | 3 | 0 | 7 | 0.2857 | 11 | 0.2727 |
| 9_8347975 | 1 | 1 | 7 | 0.2857 | 1 | 0 | 2 | 1 | 11 | 0.4545 | NA | NA | 1 | 0 | 2 | 0.5 | 6 | 0.3333 | 9 | 0.3333 |
| 9_8347975 | 2 | 1 | 5 | 1 | 1 | 1 | 2 | 1 | 10 | 1 | NA | NA | NA | NA | 2 | 1 | 6 | 1 | 8 | 1 |
| 9_8348002 | 3 | 0 | 7 | 0 | 2 | 0.5 | 2 | 0 | 14 | 0.0714 | NA | NA | 1 | 0 | 3 | 0 | 6 | 0.1667 | 10 | 0.1 |
| 9_8348027 | 4 | 0.25 | 6 | 0.5 | 2 | 0.5 | 2 | 0 | 14 | 0.3571 | NA | NA | 1 | 0 | 3 | 0 | 7 | 0.1429 | 11 | 0.0909 |
| 9_8348086 | 4 | 1 | 7 | 0.4286 | 2 | 0.5 | 3 | 0.3333 | 16 | 0.5625 | NA | NA | 1 | 0 | 2 | 0 | 6 | 0.3333 | 9 | 0.2222 |
| 9_8348256 | 1 | 0 | 6 | 0.5 | 3 | 0.6667 | 1 | 1 | 11 | 0.5455 | NA | NA | 2 | 0.5 | 4 | 0.25 | 6 | 0.3333 | 12 | 0.3333 |
| 9_8348275 | 3 | 0 | 4 | 0 | 2 | 0.5 | 3 | 0 | 12 | 0.0833 | NA | NA | 2 | 0.5 | 4 | 0 | 5 | 0 | 11 | 0.0909 |
| 9_8348304 | 4 | 0.25 | 7 | 0 | 5 | 0 | 3 | 0 | 19 | 0.0526 | NA | NA | 2 | 0 | 5 | 0 | 5 | 0.2 | 12 | 0.0833 |
| 9_8348315 | 5 | 0.2 | 8 | 0.5 | 6 | 0.8333 | 3 | 0 | 22 | 0.4545 | NA | NA | 2 | 0.5 | 8 | 0.375 | 7 | 0.4286 | 17 | 0.4118 |
| 9_8366201 | 9 | 0.1111 | 13 | 0.0769 | 9 | 0.1111 | 4 | 0 | 35 | 0.0857 | 5 | 0 | 7 | 0.1429 | 7 | 0 | 10 | 0.1 | 29 | 0.069 |
| 9_8366258 | 9 | 0.6667 | 13 | 0.3077 | 8 | 0.25 | 3 | 0 | 33 | 0.3636 | 2 | 0.5 | 5 | 0.4 | 6 | 0.3333 | 8 | 0.5 | 21 | 0.4286 |
| 9_8366272 | 7 | 0.4286 | 12 | 0.0833 | 7 | 0 | 4 | 0.25 | 30 | 0.1667 | 2 | 0 | 3 | 0 | 5 | 0.4 | 9 | 0.3333 | 19 | 0.2632 |
| 9_8400612 | 10 | 0.1 | 7 | 0.1429 | 3 | 0.3333 | 4 | 0.5 | 24 | 0.2083 | 4 | 0.25 | 3 | 0 | 4 | 0.5 | 5 | 0.4 | 16 | 0.3125 |
| 9_8400623 | 10 | 0.1 | 8 | 0 | 4 | 0 | 5 | 0.2 | 27 | 0.0741 | 4 | 0 | 3 | 0 | 6 | 0.3333 | 6 | 0 | 19 | 0.1053 |
| 9_8520261 | NA | NA | NA | NA | 1 | 1 | NA | NA | 1 | 1 | 1 | 1 | NA | NA | 1 | 0 | 2 | 1 | 4 | 0.75 |
| 9_8556991 | 4 | 0 | 3 | 0.3333 | 2 | 0 | 1 | 0 | 10 | 0.1 | 1 | 1 | 1 | 1 | 1 | 0 | 3 | 0.3333 | 6 | 0.5 |
| 9_8557010 | 4 | 0.25 | 3 | 0.6667 | 2 | 0.5 | 2 | 0 | 11 | 0.3636 | NA | NA | 1 | 0 | 1 | 0 | 3 | 0.3333 | 5 | 0.2 |
| 9_8560307 | 1 | 1 | NA | NA | 1 | 0 | 2 | 0 | 4 | 0.25 | NA | NA | 1 | 1 | NA | NA | 2 | 0.5 | 3 | 0.6667 |
| 9_8560328 | 1 | 1 | NA | NA | 1 | 1 | 1 | 1 | 3 | 1 | NA | NA | 1 | 0 | NA | NA | 2 | 1 | 3 | 0.6667 |
| 9_8560331 | 1 | 0 | NA | NA | 1 | 1 | 1 | 1 | 3 | 0.6667 | NA | NA | 1 | 1 | NA | NA | 2 | 0.5 | 3 | 0.6667 |
| 9_8562554 | NA | NA | NA | NA | NA | NA | 1 | 1 | 1 | 1 | NA | NA | 1 | 1 | NA | NA | 2 | 1 | 3 | 1 |
| 9_8562578 | NA | NA | NA | NA | NA | NA | 1 | 1 | 1 | 1 | NA | NA | 1 | 0 | NA | NA | 1 | 1 | 2 | 0.5 |
| 9_8562598 | NA | NA | NA | NA | NA | NA | 1 | 1 | 1 | 1 | NA | NA | 1 | 1 | NA | NA | 1 | 1 | 2 | 1 |
| 9_8563233 | 1 | 1 | NA | NA | NA | NA | NA | NA | 1 | 1 | 1 | 1 | 1 | 1 | 1 | 1 | 2 | 1 | 5 | 1 |
| 9_8572513 | NA | NA | 1 | 1 | NA | NA | 1 | 1 | 2 | 1 | 1 | 1 | NA | NA | NA | NA | 1 | 1 | 2 | 1 |
| 9_8572517 | NA | NA | 1 | 0 | NA | NA | 1 | 1 | 2 | 0.5 | 1 | 1 | NA | NA | NA | NA | 1 | 1 | 2 | 1 |
| 9_8572531 | NA | NA | NA | NA | NA | NA | 1 | 1 | 1 | 1 | 1 | 1 | NA | NA | NA | NA | 1 | 1 | 2 | 1 |
| 9_8572673 | 5 | 1 | 7 | 1 | 6 | 1 | 4 | 1 | 22 | 1 | 9 | 1 | 9 | 1 | 2 | 1 | 5 | 1 | 25 | 1 |
| 9_8572676 | 4 | 0.5 | 6 | 0.3333 | 6 | 0.6667 | 5 | 0.8 | 21 | 0.5714 | 9 | 0.6667 | 10 | 0.6 | 2 | 0.5 | 6 | 0.3333 | 27 | 0.5556 |
| 9_8572677 | 3 | 0.3333 | 6 | 0 | 5 | 0 | 2 | 0.5 | 16 | 0.125 | 6 | 0.3333 | 8 | 0.125 | 1 | 0 | 6 | 0.5 | 21 | 0.2857 |
| 9_8572682 | 3 | 0.3333 | 5 | 0 | 5 | 0 | 5 | 0 | 18 | 0.0556 | 4 | 0 | 7 | 0.1429 | 2 | 0 | 6 | 0 | 19 | 0.0526 |
| 9_8572685 | 5 | 0.6 | 6 | 0.8333 | 7 | 0.5714 | 4 | 1 | 22 | 0.7273 | 9 | 0.5556 | 8 | 0.375 | 1 | 1 | 6 | 0.1667 | 24 | 0.4167 |
| 9_8572692 | 6 | 0.6667 | 5 | 0.6 | 8 | 0.875 | 5 | 0.8 | 24 | 0.75 | 9 | 1 | 10 | 0.8 | 2 | 1 | 6 | 0.8333 | 27 | 0.8889 |
| 9_8572698 | 6 | 1 | 7 | 1 | 6 | 1 | 5 | 1 | 24 | 1 | 9 | 1 | 7 | 0.8571 | 1 | 1 | 3 | 0.6667 | 20 | 0.9 |
| 9_8572707 | 6 | 1 | 7 | 1 | 8 | 1 | 6 | 1 | 27 | 1 | 9 | 1 | 10 | 1 | 3 | 1 | 7 | 1 | 29 | 1 |
| 9_8572711 | 6 | 0.5 | 7 | 0.5714 | 8 | 0.375 | 6 | 0.1667 | 27 | 0.4074 | 9 | 0.4444 | 7 | 0.4286 | 3 | 1 | 7 | 0.5714 | 26 | 0.5385 |
| 9_8572718 | 6 | 0.3333 | 7 | 0.4286 | 8 | 0.125 | 6 | 0.1667 | 27 | 0.2593 | 7 | 0.2857 | 8 | 0.25 | 3 | 0 | 7 | 0.2857 | 25 | 0.24 |
| 9_8572727 | 6 | 0.8333 | 7 | 0.7143 | 8 | 0.625 | 6 | 0.8333 | 27 | 0.7407 | 9 | 0.8889 | 9 | 0.7778 | 3 | 1 | 7 | 0.5714 | 28 | 0.7857 |
| 9_8572732 | 6 | 0 | 7 | 0 | 9 | 0.4444 | 6 | 0.1667 | 28 | 0.1786 | 8 | 0.25 | 10 | 0.2 | 3 | 0.3333 | 6 | 0.1667 | 27 | 0.2222 |
| 9_8572733 | 6 | 1 | 6 | 0.8333 | 7 | 1 | 6 | 0.8333 | 25 | 0.92 | 8 | 1 | 9 | 1 | 4 | 1 | 7 | 1 | 28 | 1 |
| 9_8572735 | 5 | 0.4 | 7 | 0.7143 | 9 | 0.4444 | 6 | 0.3333 | 27 | 0.4815 | 8 | 0.625 | 9 | 0.5556 | 3 | 0.6667 | 8 | 0.5 | 28 | 0.5714 |
| 9_8572742 | 6 | 1 | 7 | 0.7143 | 9 | 0.5556 | 6 | 0.6667 | 28 | 0.7143 | 8 | 0.875 | 10 | 0.9 | 4 | 0.75 | 8 | 0.875 | 30 | 0.8667 |
| 9_8572747 | 7 | 0.8571 | 8 | 0.625 | 8 | 0.75 | 6 | 0.8333 | 29 | 0.7586 | 9 | 0.5556 | 8 | 0.625 | 4 | 0.75 | 8 | 0.5 | 29 | 0.5862 |
| 9_8572757 | 7 | 1 | 10 | 0.6 | 9 | 0.5556 | 7 | 0.8571 | 33 | 0.7273 | 9 | 0.6667 | 10 | 0.7 | 4 | 0.75 | 9 | 0.6667 | 32 | 0.6875 |
| 9_8572776 | 6 | 1 | 7 | 1 | 7 | 0.8571 | 2 | 1 | 22 | 0.9545 | 6 | 0.8333 | 8 | 0.75 | 4 | 1 | 4 | 1 | 22 | 0.8636 |
| 9_8572786 | 8 | 0.125 | 13 | 0.1538 | 11 | 0 | 8 | 0.125 | 40 | 0.1 | 9 | 0.1111 | 10 | 0 | 5 | 0.2 | 10 | 0.3 | 34 | 0.1471 |
| 9_8618978 | NA | NA | NA | NA | NA | NA | 2 | 0.5 | 2 | 0.5 | 1 | 1 | NA | NA | NA | NA | 1 | 1 | 2 | 1 |
| 9_8639476 | 1 | 1 | NA | NA | NA | NA | NA | NA | 1 | 1 | NA | NA | NA | NA | NA | NA | 1 | 1 | 1 | 1 |
| 9_8639498 | 2 | 0.5 | NA | NA | NA | NA | NA | NA | 2 | 0.5 | NA | NA | 1 | 0 | NA | NA | 2 | 0.5 | 3 | 0.3333 |
| 9_8639505 | 2 | 1 | NA | NA | NA | NA | NA | NA | 2 | 1 | NA | NA | 1 | 0 | 1 | 1 | 2 | 0 | 4 | 0.25 |

| | | | | | | | | | | | | | | | | | | | | |
|---|---|---|---|---|---|---|---|---|---|---|---|---|---|---|---|---|---|---|---|---|
| 9_8639598 | 2 | 1 | NA | NA | 1 | 0 | NA | NA | 3 | 0.6667 | NA | NA | 1 | 1 | 1 | 0 | 2 | 0 | 4 | 0.25 |
| 9_8639599 | 2 | 0.5 | 1 | 1 | 1 | 0 | NA | NA | 4 | 0.5 | NA | NA | 1 | 0 | 1 | 1 | 2 | 0.5 | 4 | 0.5 |
| 9_8640202 | 2 | 1 | NA | NA | 1 | 1 | NA | NA | 3 | 1 | NA | NA | 2 | 1 | 1 | 1 | 2 | 1 | 5 | 1 |
| 9_8640207 | 2 | 0.5 | 1 | 0 | 1 | 0 | NA | NA | 4 | 0.25 | NA | NA | 2 | 0 | 1 | 0 | 2 | 0.5 | 5 | 0.2 |
| 9_8640253 | 2 | 1 | 1 | 1 | 1 | 0 | NA | NA | 4 | 0.75 | NA | NA | 2 | 1 | 1 | 0 | 1 | 1 | 4 | 0.75 |
| 9_8640278 | 2 | 1 | 1 | 0 | 1 | 1 | NA | NA | 4 | 0.75 | NA | NA | 2 | 0.5 | 1 | 1 | 2 | 0.5 | 5 | 0.6 |
| 9_8640279 | 2 | 0.5 | 1 | 1 | NA | NA | NA | NA | 3 | 0.6667 | NA | NA | 1 | 1 | 1 | 0 | 2 | 0 | 4 | 0.25 |
| 9_8640293 | 3 | 0.6667 | 1 | 0 | 2 | 1 | NA | NA | 6 | 0.6667 | NA | NA | 2 | 1 | 1 | 1 | NA | NA | 3 | 1 |
| 9_8640309 | 3 | 0.3333 | 1 | 1 | 2 | 0.5 | NA | NA | 6 | 0.5 | NA | NA | 2 | 1 | 2 | 1 | 2 | 0.5 | 6 | 0.8333 |
| 9_8640322 | 3 | 0.6667 | 1 | 1 | 2 | 0 | NA | NA | 6 | 0.5 | NA | NA | 2 | 0 | 2 | 0 | 3 | 0.3333 | 7 | 0.1429 |
| 9_8640327 | 3 | 0.3333 | NA | NA | 2 | 1 | NA | NA | 5 | 0.6 | NA | NA | 1 | 1 | 1 | 0 | 2 | 1 | 4 | 0.75 |
| 9_8640350 | 3 | 0.3333 | 1 | 1 | 2 | 0.5 | NA | NA | 6 | 0.5 | NA | NA | 1 | 0 | 2 | 0.5 | 3 | 0.3333 | 6 | 0.3333 |
| 9_8643609 | 7 | 0.1429 | 1 | 0 | 2 | 0 | 5 | 0 | 15 | 0.0667 | 2 | 0.5 | 6 | 0 | 3 | 0 | 4 | 0 | 15 | 0.0667 |
| 9_8643622 | 5 | 0.2 | NA | NA | 2 | 0 | 5 | 0 | 12 | 0.0833 | 2 | 0 | 6 | 0 | 2 | 0 | 5 | 0.2 | 15 | 0.0667 |
| 9_8643627 | 7 | 0.4286 | NA | NA | 2 | 0.5 | 5 | 0.2 | 14 | 0.3571 | 2 | 1 | 5 | 0.6 | 3 | 0.3333 | 4 | 0.25 | 14 | 0.5 |
| 9_8643634 | 5 | 0.8 | 1 | 1 | 3 | 0.6667 | 5 | 1 | 14 | 0.8571 | 1 | 1 | 6 | 1 | 3 | 0.3333 | 5 | 0.8 | 15 | 0.8 |
| 9_8643636 | 7 | 0.2857 | 1 | 1 | 3 | 0.3333 | 4 | 0.75 | 15 | 0.4667 | 2 | 0.5 | 6 | 0.6667 | 4 | 0.5 | 4 | 1 | 16 | 0.6875 |
| 9_8643718 | 4 | 0 | 3 | 0.3333 | 3 | 0 | NA | NA | 10 | 0.1 | NA | NA | 2 | 0 | 1 | 1 | 1 | 0 | 4 | 0.25 |
| 9_8645615 | 1 | 0 | 3 | 1 | 1 | 1 | 5 | 0.8 | 10 | 0.8 | NA | NA | 2 | 0.5 | 1 | 1 | 2 | 0.5 | 5 | 0.6 |
| 9_8645628 | 2 | 0.5 | 3 | 0 | 1 | 0 | 4 | 0.25 | 10 | 0.2 | NA | NA | 4 | 0 | 5 | 0.2 | 2 | 0 | 11 | 0.0909 |
| 9_8645633 | 2 | 0.5 | 3 | 0.6667 | 1 | 0 | 5 | 0.4 | 11 | 0.4545 | NA | NA | 4 | 0.75 | 5 | 0.8 | 2 | 0 | 11 | 0.6364 |
| 9_8645637 | 2 | 0.5 | 3 | 1 | 1 | 1 | 5 | 1 | 11 | 0.9091 | NA | NA | 4 | 1 | 4 | 0.75 | 1 | 1 | 9 | 0.8889 |
| 9_8645651 | 2 | 0.5 | 1 | 1 | 1 | 0 | 5 | 0.2 | 9 | 0.3333 | NA | NA | 4 | 0.25 | 3 | 0.6667 | 1 | 1 | 8 | 0.5 |
| 9_8645658 | 2 | 1 | 2 | 0.5 | NA | NA | 6 | 0.6667 | 10 | 0.7 | 1 | 1 | 3 | 0 | 5 | 0.6 | 1 | 0 | 10 | 0.4 |
| 9_8645661 | 2 | 0 | 4 | 0.25 | 1 | 0 | 6 | 0.1667 | 13 | 0.1538 | 1 | 0 | 3 | 0 | 5 | 0.2 | 1 | 0 | 10 | 0.1 |
| 9_8645678 | 2 | 0 | 4 | 0 | 2 | 0 | 7 | 0.1429 | 15 | 0.0667 | 1 | 1 | 4 | 0 | 6 | 0 | 2 | 0 | 13 | 0.0769 |
| 9_8663877 | NA | NA | 1 | 1 | NA | NA | NA | NA | 1 | 1 | NA | NA | NA | NA | NA | NA | 1 | 1 | 1 | 1 |
| 9_8663879 | NA | NA | 1 | 1 | NA | NA | NA | NA | 1 | 1 | NA | NA | NA | NA | NA | NA | 1 | 1 | 1 | 1 |
| 9_8664850 | 37 | 0.0811 | 42 | 0.0476 | 29 | 0.069 | 49 | 0.0204 | 157 | 0.051 | 17 | 0.2353 | 26 | 0.0769 | 27 | 0.0741 | 28 | 0.1786 | 98 | 0.1327 |
| 9_8664898 | 27 | 0.5556 | 27 | 0.7778 | 24 | 0.5 | 42 | 0.7143 | 120 | 0.65 | 13 | 0.5385 | 16 | 0.625 | 23 | 0.5652 | 24 | 0.5833 | 76 | 0.5789 |
| 9_8664900 | 27 | 0.0741 | 28 | 0.0714 | 24 | 0.1667 | 37 | 0.0811 | 116 | 0.0948 | 13 | 0.0769 | 22 | 0.0455 | 24 | 0.125 | 23 | 0 | 82 | 0.061 |
| 9_8664923 | 22 | 0.1818 | 24 | 0.2083 | 21 | 0.1905 | 42 | 0.2619 | 109 | 0.2202 | 13 | 0.2308 | 19 | 0.1579 | 20 | 0.2 | 24 | 0.2083 | 76 | 0.1974 |
| 9_8664937 | 22 | 0.0455 | 26 | 0.0769 | 21 | 0 | 42 | 0.0714 | 111 | 0.0541 | 13 | 0.0769 | 20 | 0.05 | 21 | 0.0952 | 25 | 0.04 | 79 | 0.0633 |
| 9_8676147 | 12 | 0.5 | 7 | 0.2857 | 5 | 0.4 | 11 | 0.2727 | 35 | 0.3714 | 6 | 0 | 8 | 0.125 | 3 | 0.3333 | 9 | 0.2222 | 26 | 0.1538 |
| 9_8690906 | 3 | 0 | 6 | 0 | 4 | 0.25 | 4 | 0 | 17 | 0.0588 | 2 | 0 | 6 | 0 | 2 | 0 | 5 | 0.2 | 15 | 0.0667 |
| 9_8695909 | 7 | 0.1429 | 9 | 0.1111 | 4 | 0.25 | 4 | 0 | 24 | 0.125 | 4 | 0 | 7 | 0.1429 | 3 | 0 | 5 | 0 | 19 | 0.0526 |
| 9_8695916 | 8 | 0.625 | 10 | 0.7 | 6 | 0.5 | 4 | 0.25 | 28 | 0.5714 | 4 | 0.5 | 8 | 0.375 | 3 | 0 | 5 | 0.4 | 20 | 0.35 |
| 9_8706463 | 2 | 1 | 3 | 0.6667 | NA | NA | 2 | 0.5 | 7 | 0.7143 | NA | NA | 2 | 0.5 | 2 | 0 | NA | NA | 4 | 0.25 |
| 9_8706464 | 2 | 0.5 | 4 | 1 | NA | NA | 3 | 1 | 9 | 0.8889 | NA | NA | 1 | 1 | 2 | 0.5 | NA | NA | 3 | 0.6667 |
| 9_8706477 | 2 | 0 | 3 | 0 | NA | NA | 3 | 0.3333 | 8 | 0.125 | NA | NA | 2 | 0 | 1 | 1 | NA | NA | 3 | 0.3333 |
| 9_8706487 | 2 | 1 | 4 | 0.25 | NA | NA | 2 | 0 | 8 | 0.375 | NA | NA | 2 | 0 | 2 | 0.5 | NA | NA | 4 | 0.25 |
| 9_8706488 | NA | NA | 3 | 1 | NA | NA | 3 | 1 | 6 | 1 | NA | NA | 2 | 1 | 1 | 1 | NA | NA | 3 | 1 |
| 9_8706488 | 2 | 1 | 4 | 0.75 | NA | NA | 3 | 1 | 9 | 0.8889 | NA | NA | 2 | 1 | 2 | 1 | NA | NA | 4 | 1 |
| 9_8706556 | 1 | 0 | 5 | 0.4 | 2 | 0 | 2 | 0 | 10 | 0.2 | NA | NA | 2 | 0 | 2 | 0.5 | 1 | 0 | 5 | 0.2 |
| 9_8706560 | 2 | 0 | 5 | 0.4 | 1 | 0 | NA | NA | 8 | 0.25 | NA | NA | 1 | 0 | 1 | 1 | 1 | 0 | 3 | 0.3333 |
| 9_8706566 | 2 | 0 | 6 | 0.1667 | 2 | 0.5 | 3 | 0.3333 | 13 | 0.2308 | NA | NA | 2 | 0 | 2 | 0.5 | 1 | 0 | 5 | 0.2 |
| 9_8706577 | 2 | 0.5 | 6 | 0.1667 | 2 | 0.5 | 3 | 0.3333 | 13 | 0.3077 | NA | NA | 2 | 0 | 2 | 0.5 | 1 | 0 | 5 | 0.2 |
| 9_8706597 | 2 | 0.5 | 5 | 0.4 | 2 | 0 | 2 | 0 | 11 | 0.2727 | NA | NA | 2 | 0.5 | 2 | 0.5 | 1 | 1 | 5 | 0.6 |
| 9_8707177 | 1 | 0 | 2 | 0.5 | NA | NA | 2 | 0 | 5 | 0.2 | NA | NA | NA | NA | 2 | 0.5 | NA | NA | 2 | 0.5 |
| 9_8707191 | 1 | 0 | 2 | 0 | NA | NA | 2 | 0.5 | 5 | 0.2 | NA | NA | NA | NA | 2 | 0.5 | NA | NA | 2 | 0.5 |
| 9_8707455 | 1 | 1 | 1 | 1 | NA | NA | 1 | 1 | 3 | 1 | NA | NA | 1 | 1 | 2 | 1 | 1 | 0 | 4 | 0.75 |
| 9_8707465 | 1 | 0 | 2 | 0.5 | NA | NA | 1 | 1 | 4 | 0.5 | NA | NA | 1 | 0 | 2 | 0.5 | 1 | 0 | 4 | 0.25 |
| 9_8707470 | NA | NA | 2 | 0.5 | NA | NA | 1 | 0 | 3 | 0.3333 | NA | NA | 1 | 1 | 2 | 0.5 | 1 | 0 | 4 | 0.5 |
| 9_8707473 | 1 | 1 | 2 | 0.5 | NA | NA | 1 | 0 | 4 | 0.5 | NA | NA | 1 | 0 | 2 | 0.5 | 1 | 0 | 4 | 0.25 |
| 9_8707476 | 1 | 0 | 1 | 1 | NA | NA | NA | NA | 2 | 0.5 | NA | NA | NA | NA | 2 | 1 | 1 | 0 | 3 | 0.6667 |
| 9_8707486 | 1 | 1 | 1 | 1 | NA | NA | 1 | 1 | 3 | 1 | NA | NA | 1 | 1 | 1 | 1 | 1 | 1 | 3 | 1 |
| 9_8708582 | NA | NA | 1 | 1 | NA | NA | NA | NA | 1 | 1 | NA | NA | NA | NA | 2 | 0.5 | NA | NA | 2 | 0.5 |
| 9_8708585 | NA | NA | 1 | 1 | NA | NA | NA | NA | 1 | 1 | NA | NA | 1 | 0 | 2 | 0.5 | NA | NA | 3 | 0.3333 |
| 9_8708621 | NA | NA | 1 | 1 | NA | NA | NA | NA | 1 | 1 | NA | NA | 1 | 0 | 2 | 0.5 | NA | NA | 3 | 0.3333 |
| 9_8708622 | NA | NA | 1 | 1 | NA | NA | NA | NA | 1 | 1 | NA | NA | 1 | 0 | 2 | 0.5 | NA | NA | 3 | 0.3333 |
| 9_8758681 | NA | NA | 1 | 1 | NA | NA | NA | NA | 1 | 1 | NA | NA | NA | NA | NA | NA | 1 | 1 | 1 | 1 |
| 9_8758696 | NA | NA | 1 | 1 | NA | NA | NA | NA | 1 | 1 | NA | NA | NA | NA | NA | NA | 1 | 1 | 1 | 1 |
| 9_8758701 | NA | NA | 1 | 1 | NA | NA | NA | NA | 1 | 1 | NA | NA | NA | NA | NA | NA | 1 | 1 | 1 | 1 |
| 9_8758703 | NA | NA | 1 | 1 | NA | NA | NA | NA | 1 | 1 | NA | NA | NA | NA | NA | NA | 1 | 1 | 1 | 1 |
| 9_8758740 | NA | NA | 1 | 1 | NA | NA | NA | NA | 1 | 1 | NA | NA | NA | NA | NA | NA | 1 | 1 | 1 | 1 |

| | | | | | | | | | | | | | | | | | | | | |
|---|---|---|---|---|---|---|---|---|---|---|---|---|---|---|---|---|---|---|---|---|
| 9_8758773( | NA | NA | 1 | 1 | NA | NA | NA | NA | 1 | 1 | NA | NA | NA | NA | NA | NA | 1 | 1 | 1 | 1 |
| 9_88234354 | NA | NA | 1 | 0 | 3 | 0.6667 | NA | NA | 4 | 0.5 | 1 | 0 | 2 | 0 | 3 | 0.3333 | NA | NA | 6 | 0.1667 |
| 9_8823444 | 1 | 0 | 1 | 0 | 3 | 0.3333 | 1 | 0 | 6 | 0.1667 | 1 | 0 | 2 | 0 | 3 | 0.3333 | NA | NA | 6 | 0.1667 |
| 9_8825250( | 3 | 1 | 2 | 0 | 2 | 0 | 2 | 1 | 9 | 0.5556 | 2 | 1 | 3 | 0.6667 | 4 | 0.75 | 1 | 1 | 10 | 0.8 |
| 9_8825341( | 2 | 0 | 2 | 0 | 2 | 0 | 3 | 0.3333 | 9 | 0.1111 | 2 | 0.5 | 4 | 0.5 | 3 | 0.3333 | 1 | 0 | 10 | 0.4 |
| 9_8825381 | 2 | 0 | 3 | 0.3333 | 3 | 0.6667 | 4 | 0.25 | 12 | 0.3333 | 2 | 1 | 4 | 0.5 | 5 | 0.8 | 1 | 0 | 12 | 0.6667 |
| 9_8825393 | 3 | 0 | 3 | 0 | 3 | 0.3333 | 5 | 0 | 14 | 0.0714 | 2 | 0.5 | 5 | 0 | 5 | 0 | 2 | 0 | 14 | 0.0714 |
| 9_8833330( | 6 | 0 | 1 | 0 | 2 | 0.5 | 2 | 0 | 11 | 0.0909 | 5 | 0 | 3 | 0.3333 | 3 | 0 | 4 | 0 | 15 | 0.0667 |
| 9_8833614( | 6 | 0.8333 | 7 | 1 | 7 | 1 | 5 | 1 | 25 | 0.96 | 4 | 1 | 5 | 1 | 4 | 1 | 7 | 1 | 20 | 1 |
| 9_8833615 | 6 | 0.1667 | 8 | 0.125 | 7 | 0 | 4 | 0 | 25 | 0.08 | 3 | 0 | 5 | 0 | 4 | 0.25 | 6 | 0 | 18 | 0.0556 |
| 9_8833615 | 6 | 0.8333 | 8 | 0.875 | 7 | 0.8571 | 4 | 0.5 | 25 | 0.8 | 3 | 1 | 5 | 0.8 | 5 | 0.8 | 6 | 0.5 | 19 | 0.7368 |
| 9_8833630 | 5 | 0.2 | 6 | 0.1667 | 5 | 0.6 | 5 | 0.2 | 21 | 0.2857 | 5 | 0 | 4 | 0 | 4 | 0.25 | 6 | 0.1667 | 19 | 0.1053 |
| 9_8833816 | 1 | 1 | 2 | 0.5 | 3 | 0.6667 | 1 | 1 | 7 | 0.7143 | NA | NA | 2 | 0.5 | 2 | 0 | NA | NA | 4 | 0.25 |
| 9_8833831 | 1 | 0 | 1 | 1 | 2 | 1 | 2 | 0.5 | 6 | 0.6667 | 1 | 0 | 2 | 0.5 | 2 | 0.5 | NA | NA | 5 | 0.4 |
| 9_8840482 | 1 | 0 | 1 | 0 | 2 | 0.5 | 2 | 0 | 6 | 0.1667 | 1 | 0 | 1 | 1 | NA | NA | 1 | 0 | 3 | 0.3333 |
| 9_8843037 | 1 | 1 | 1 | 1 | NA | NA | NA | NA | 2 | 1 | NA | NA | 2 | 0.5 | NA | NA | 3 | 0.3333 | 5 | 0.4 |
| 9_8845083 | 2 | 0 | 2 | 1 | NA | NA | 1 | 0 | 5 | 0.4 | NA | NA | 3 | 1 | 2 | 0 | 3 | 0.3333 | 8 | 0.5 |
| 9_8845093 | 1 | 1 | 1 | 0 | NA | NA | 1 | 0 | 3 | 0.3333 | NA | NA | 1 | 1 | NA | NA | NA | NA | 1 | 1 |
| 9_8845108 | NA | NA | 1 | 1 | NA | NA | NA | NA | 1 | 1 | NA | NA | 2 | 0.5 | NA | NA | 2 | 0.5 | 4 | 0.5 |
| 9_8845109 | NA | NA | 2 | 0 | NA | NA | 1 | 1 | 3 | 0.3333 | NA | NA | 1 | 1 | NA | NA | 2 | 0.5 | 3 | 0.6667 |
| 9_8845228 | 2 | 0 | 1 | 1 | 1 | 0 | NA | NA | 4 | 0.25 | NA | NA | 1 | 1 | NA | NA | 1 | 0 | 2 | 0.5 |
| 9_8847022 | 1 | 1 | NA | NA | NA | NA | 1 | 1 | 2 | 1 | NA | NA | NA | NA | NA | NA | 1 | 1 | 1 | 1 |
| 9_8847037 | 1 | 0 | NA | NA | NA | NA | 1 | 1 | 2 | 0.5 | NA | NA | NA | NA | NA | NA | 1 | 1 | 1 | 1 |
| 9_8857075 | 3 | 0.3333 | 4 | 0.25 | 2 | 0.5 | 5 | 0.4 | 14 | 0.3571 | 1 | 0 | 1 | 0 | 2 | 0.5 | 3 | 0 | 7 | 0.1429 |
| 9_8857103 | 2 | 0.5 | 3 | 0 | 2 | 0 | 7 | 0 | 14 | 0.0714 | 1 | 0 | 1 | 0 | 2 | 0.5 | 5 | 0 | 9 | 0.1111 |
| 9_8859764 | NA | NA | 3 | 0.3333 | NA | NA | 1 | 0 | 4 | 0.25 | NA | NA | NA | NA | 1 | 1 | 4 | 0.75 | 5 | 0.8 |
| 9_8859771 | NA | NA | 2 | 0.5 | NA | NA | 2 | 0 | 4 | 0.25 | NA | NA | NA | NA | 1 | 1 | 3 | 0.6667 | 4 | 0.75 |
| 9_8958128 | 2 | 0.5 | 5 | 0.2 | 2 | 0 | 5 | 0 | 14 | 0.1429 | 5 | 0 | 10 | 0.3 | 2 | 0 | 2 | 0 | 19 | 0.1579 |
| 9_8958143 | 1 | 1 | 5 | 0 | 3 | 0 | 5 | 0.2 | 14 | 0.1429 | 4 | 0.25 | 8 | 0 | 2 | 0 | 2 | 0 | 16 | 0.0625 |
| 9_8958147 | 2 | 0 | 3 | 0.3333 | 3 | 0 | 5 | 0.2 | 13 | 0.1538 | 3 | 0.3333 | 6 | 0 | NA | NA | 2 | 0 | 11 | 0.0909 |
| 9_8958153 | 2 | 0.5 | 4 | 0 | 1 | 1 | 5 | 0.2 | 12 | 0.25 | 6 | 0 | 9 | 0.1111 | 2 | 0 | 1 | 0 | 18 | 0.0556 |
| 9_8958189 | 2 | 0 | 4 | 0 | 3 | 0.3333 | 5 | 0.6 | 14 | 0.2857 | 4 | 0 | 9 | 0.2222 | 2 | 0.5 | 1 | 0 | 16 | 0.1875 |
| 9_8960747 | 1 | 0 | 1 | 1 | NA | NA | NA | NA | 2 | 0.5 | 1 | 1 | NA | NA | NA | NA | NA | NA | 1 | 1 |
| 9_8979209 | 9 | 0.2222 | 10 | 0.2 | 10 | 0.2 | 15 | 0.1333 | 44 | 0.1818 | 13 | 0.3077 | 4 | 0 | 7 | 0.2857 | 8 | 0.125 | 32 | 0.2188 |
| 9_8979215 | 9 | 0.2222 | 10 | 0 | 7 | 0.1429 | 15 | 0.0667 | 41 | 0.0976 | 10 | 0.1 | 4 | 0.25 | 7 | 0.4286 | 9 | 0 | 30 | 0.1667 |
| 9_8979359 | 1 | 1 | 3 | 0.3333 | 1 | 1 | 1 | 1 | 6 | 0.6667 | NA | NA | NA | NA | NA | NA | 1 | 1 | 1 | 1 |
| 9_8979646( | NA | NA | 1 | 0 | NA | NA | 1 | 1 | 2 | 0.5 | NA | NA | 2 | 0 | NA | NA | 3 | 0.3333 | 5 | 0.2 |
| 9_8979688 | 2 | 1 | 1 | 1 | NA | NA | 1 | 1 | 4 | 1 | NA | NA | 3 | 1 | 1 | 1 | 5 | 1 | 9 | 1 |
| 9_8979693 | 2 | 0 | 1 | 1 | NA | NA | 2 | 0.5 | 5 | 0.4 | NA | NA | 3 | 0.6667 | 1 | 0 | 4 | 0.75 | 8 | 0.625 |
| 9_8979702 | 1 | 0 | 1 | 0 | NA | NA | 1 | 1 | 3 | 0.3333 | 1 | 0 | 2 | 0.5 | 1 | 0 | 3 | 0.6667 | 7 | 0.4286 |
| 9_8979723( | 1 | 0 | 1 | 0 | NA | NA | 2 | 1 | 4 | 0.5 | 1 | 0 | 4 | 0.25 | 1 | 0 | 5 | 0.4 | 11 | 0.2727 |
| 9_8979738 | 1 | 1 | 1 | 1 | NA | NA | 2 | 1 | 4 | 1 | NA | NA | 3 | 0.3333 | NA | NA | 3 | 0.6667 | 6 | 0.5 |
| 9_8979756 | 3 | 0.3333 | 3 | 0 | 3 | 0 | 3 | 0 | 12 | 0.0833 | NA | NA | 3 | 0 | 1 | 0 | 5 | 0.2 | 9 | 0.1111 |
| 9_8979774 | 3 | 0.3333 | 2 | 0 | 2 | 0.5 | 4 | 0.25 | 11 | 0.2727 | NA | NA | 3 | 0.6667 | 1 | 1 | 3 | 0.3333 | 7 | 0.5714 |
| 9_8987691( | 22 | 0.0909 | 19 | 0.2105 | 21 | 0.1429 | 27 | 0.1852 | 89 | 0.1573 | 26 | 0.0385 | 16 | 0.3125 | 22 | 0.0455 | 21 | 0.1429 | 85 | 0.1176 |
| 9_8987699( | 19 | 0.1053 | 14 | 0.1429 | 15 | 0.1333 | 24 | 0.0833 | 72 | 0.1111 | 17 | 0.1176 | 13 | 0.0769 | 18 | 0.0556 | 22 | 0 | 70 | 0.0571 |
| 9_8995794 | 2 | 0 | 2 | 0 | 5 | 0.2 | 1 | 0 | 10 | 0.1 | 3 | 0 | 1 | 1 | 1 | 0 | NA | NA | 5 | 0.2 |
| 9_8995795 | 2 | 0 | 2 | 0.5 | 5 | 0.2 | 1 | 0 | 10 | 0.2 | 3 | 0.3333 | 1 | 0 | 1 | 0 | NA | NA | 5 | 0.2 |
| 9_8995804 | 2 | 1 | 2 | 0 | 5 | 0.8 | 1 | 0 | 10 | 0.6 | 2 | 1 | 1 | 0 | 1 | 0 | NA | NA | 4 | 0.5 |
| 9_8997414 | 74 | 0.0405 | 74 | 0.0541 | 53 | 0.0755 | 60 | 0.05 | 261 | 0.0536 | 27 | 0.1111 | 63 | 0.0635 | 57 | 0.0175 | 55 | 0.0545 | 202 | 0.0545 |
| 9_8997415 | 75 | 0.0933 | 75 | 0.0667 | 54 | 0.0185 | 61 | 0.0984 | 265 | 0.0717 | 28 | 0.0714 | 62 | 0.0323 | 59 | 0.0678 | 59 | 0.0678 | 208 | 0.0577 |
| 9_9008420 | 1 | 1 | NA | NA | 1 | 0 | 1 | 0 | 3 | 0.3333 | 2 | 0 | 1 | 0 | 2 | 0 | 1 | 1 | 6 | 0.1667 |
| 9_9008595( | NA | NA | NA | NA | 2 | 1 | 1 | 1 | 3 | 1 | 1 | 0 | NA | NA | NA | NA | 1 | 1 | 2 | 0.5 |
| 9_9008600 | 1 | 1 | NA | NA | 2 | 0 | 1 | 0 | 4 | 0.25 | 1 | 0 | 1 | 1 | NA | NA | 1 | 1 | 3 | 0.6667 |
| 9_9008609 | 1 | 1 | NA | NA | 2 | 0.5 | NA | NA | 3 | 0.6667 | 1 | 1 | 1 | 1 | NA | NA | 1 | 1 | 3 | 1 |
| 9_9008610 | 1 | 1 | NA | NA | 1 | 1 | 1 | 1 | 3 | 1 | 1 | 1 | 1 | 1 | NA | NA | 1 | 1 | 3 | 1 |
| 9_9008629 | 1 | 1 | NA | NA | 2 | 0 | 1 | 0 | 4 | 0.25 | 1 | 0 | NA | NA | NA | NA | 1 | 1 | 2 | 0.5 |
| 9_9008634( | 1 | 1 | NA | NA | 2 | 0 | 1 | 1 | 4 | 0.5 | 1 | 1 | 1 | 1 | NA | NA | 1 | 1 | 3 | 1 |
| 9_9008652 | 1 | 1 | NA | NA | 2 | 0 | 1 | 1 | 4 | 0.5 | 1 | 1 | 1 | 0 | NA | NA | 1 | 0 | 3 | 0.3333 |
| 9_9008656( | 1 | 1 | NA | NA | 2 | 1 | 1 | 1 | 4 | 1 | 1 | 1 | 1 | 1 | NA | NA | 1 | 1 | 3 | 1 |
| 9_9008665 | 1 | 1 | NA | NA | 2 | 0 | 1 | 0 | 4 | 0.25 | 1 | 1 | 1 | 0 | NA | NA | 1 | 0 | 3 | 0.3333 |
| 9_9008677 | 1 | 1 | NA | NA | 2 | 0.5 | NA | NA | 3 | 0.6667 | 1 | 1 | 1 | 1 | NA | NA | NA | NA | 2 | 1 |
| 9_9008683 | 1 | 1 | NA | NA | 2 | 1 | 1 | 1 | 4 | 1 | 1 | 1 | 1 | 1 | NA | NA | 1 | 1 | 3 | 1 |
| 9_9124053 | 3 | 0 | 3 | 0 | 4 | 0.25 | 4 | 0 | 14 | 0.0714 | NA | NA | 2 | 1 | 4 | 0 | 1 | 0 | 7 | 0.2857 |
| 9_9124259 | 3 | 0.3333 | 2 | 0.5 | 7 | 0 | 3 | 0 | 15 | 0.1333 | 1 | 0 | 2 | 0 | 4 | 0.25 | 3 | 0 | 10 | 0.1 |

| | | | | | | | | | | | | | | | | | | | | |
|---|---|---|---|---|---|---|---|---|---|---|---|---|---|---|---|---|---|---|---|---|
| 9_9124343 | 4 | 0.75 | NA | NA | 6 | 0.5 | 2 | 1 | 12 | 0.6667 | NA | NA | 2 | 1 | 1 | 1 | 2 | 1 | 5 | 1 |
| 9_9124358 | 3 | 1 | NA | NA | 6 | 0.8333 | 1 | 1 | 10 | 0.9 | NA | NA | 1 | 1 | 1 | 1 | 2 | 1 | 4 | 1 |
| 9_9124620 | 4 | 0.5 | NA | NA | 2 | 0 | 1 | 1 | 7 | 0.4286 | NA | NA | NA | NA | NA | NA | 2 | 1 | 2 | 1 |
| 9_9124646 | 4 | 0.25 | NA | NA | 2 | 0.5 | 1 | 1 | 7 | 0.4286 | NA | NA | NA | NA | 1 | 0 | 2 | 0.5 | 3 | 0.3333 |
| 9_9124652 | 4 | 0 | NA | NA | 2 | 0.5 | 1 | 0 | 7 | 0.1429 | NA | NA | NA | NA | NA | NA | 2 | 0.5 | 2 | 0.5 |
| 9_9124656 | 4 | 1 | NA | NA | 1 | 1 | 1 | 1 | 6 | 1 | NA | NA | NA | NA | NA | NA | 2 | 1 | 2 | 1 |
| 9_9124677 | 3 | 1 | NA | NA | 1 | 1 | NA | NA | 4 | 1 | NA | NA | NA | NA | NA | NA | 1 | 1 | 1 | 1 |
| 9_9124736 | 2 | 0.5 | NA | NA | 1 | 1 | NA | NA | 3 | 0.6667 | NA | NA | NA | NA | NA | NA | 1 | 1 | 1 | 1 |
| 9_9237355 | 1 | 0 | 1 | 1 | NA | NA | 1 | 1 | 3 | 0.6667 | NA | NA | 1 | 1 | NA | NA | 1 | 0 | 2 | 0.5 |
| 9_9440037 | 4 | 0 | 4 | 0 | 3 | 0.3333 | 9 | 0.1111 | 20 | 0.1 | 2 | 0 | 6 | 0.1667 | 3 | 0 | 4 | 0 | 15 | 0.0667 |
| 9_9440042 | 3 | 0 | 4 | 0 | 2 | 0.5 | 8 | 0.25 | 17 | 0.1765 | 2 | 0 | 3 | 0 | 3 | 0 | 4 | 0.25 | 12 | 0.0833 |
| 9_9440214 | 1 | 1 | 4 | 1 | 3 | 1 | 6 | 1 | 14 | 1 | 1 | 1 | 2 | 1 | 1 | 1 | 3 | 1 | 7 | 1 |
| 9_9440221 | 1 | 1 | 4 | 1 | 4 | 0.75 | 6 | 1 | 15 | 0.9333 | 2 | 1 | 2 | 1 | 1 | 1 | 3 | 1 | 8 | 1 |
| 9_9440225 | 1 | 1 | 4 | 1 | 4 | 1 | 6 | 1 | 15 | 1 | 1 | 1 | 2 | 1 | 1 | 1 | 3 | 1 | 7 | 1 |
| 9_9440230 | 1 | 1 | 1 | 1 | 2 | 1 | 2 | 1 | 6 | 1 | 1 | 1 | NA | NA | NA | NA | NA | NA | 1 | 1 |
| 9_9440232 | 1 | 0 | 2 | 0 | 2 | 0.5 | 3 | 1 | 8 | 0.5 | 2 | 0.5 | 2 | 0.5 | 1 | 0 | 2 | 1 | 7 | 0.5714 |
| 9_9440232 | 1 | 1 | 4 | 0.5 | 4 | 0.25 | 6 | 0.1667 | 15 | 0.3333 | NA | NA | 1 | 1 | 1 | 0 | 3 | 0.3333 | 5 | 0.4 |
| 9_9440241 | 1 | 1 | 3 | 1 | 3 | 1 | 4 | 1 | 11 | 1 | 2 | 1 | 2 | 1 | 1 | 1 | 2 | 1 | 7 | 1 |
| 9_9440252 | 1 | 1 | 4 | 1 | 2 | 1 | 6 | 1 | 13 | 1 | 2 | 1 | 2 | 1 | 1 | 1 | 2 | 1 | 7 | 1 |
| 9_9440256 | NA | NA | 4 | 0.5 | 3 | 0.6667 | 6 | 0 | 13 | 0.3077 | 2 | 0.5 | 2 | 0.5 | 1 | 0 | 2 | 0 | 7 | 0.2857 |
| 9_9440259 | NA | NA | 4 | 0.75 | 2 | 1 | 6 | 0.6667 | 12 | 0.75 | 2 | 0 | 2 | 0.5 | 1 | 0 | 2 | 0.5 | 7 | 0.2857 |
| 9_9440657 | NA | NA | 3 | 1 | 2 | 1 | 4 | 1 | 9 | 1 | 2 | 1 | 1 | 1 | 1 | 1 | 2 | 0.5 | 6 | 0.8333 |
| 9_9441933 | NA | NA | 4 | 0.25 | 3 | 0 | 6 | 0.1667 | 13 | 0.1538 | 2 | 1 | 1 | 0 | 1 | 0 | 1 | 0 | 5 | 0.4 |
| 9_9441936 | NA | NA | 4 | 0.25 | 3 | 0 | 3 | 0.3333 | 10 | 0.2 | 2 | 1 | NA | NA | 1 | 0 | 1 | 1 | 4 | 0.75 |
| 9_9441943 | NA | NA | 4 | 0.25 | 3 | 0 | 5 | 0.2 | 12 | 0.1667 | 2 | 0.5 | 1 | 0 | 1 | 0 | 1 | 0 | 5 | 0.2 |
| 9_9441967 | NA | NA | 3 | 0.3333 | 2 | 0.5 | 3 | 0.6667 | 8 | 0.5 | NA | NA | NA | NA | 1 | 0 | 1 | 1 | 2 | 0.5 |
| 9_9511111 | NA | NA | 3 | 1 | 2 | 1 | 1 | 1 | 6 | 1 | 2 | 1 | NA | NA | 1 | 1 | 1 | 1 | 4 | 1 |
| 9_9511113 | NA | NA | 3 | 0.3333 | 2 | 0 | 1 | 1 | 6 | 0.3333 | 2 | 0.5 | 1 | 1 | 1 | 0 | 1 | 0 | 5 | 0.4 |
| 9_9511122 | NA | NA | 2 | 0.5 | 2 | 0 | 1 | 0 | 5 | 0.2 | 2 | 0 | 1 | 1 | 1 | 0 | 1 | 0 | 5 | 0.2 |
| 9_9511123 | NA | NA | 3 | 1 | 1 | 1 | 1 | 1 | 5 | 1 | 2 | 1 | 1 | 0 | 1 | 1 | 1 | 1 | 5 | 0.8 |
| 9_9511148 | NA | NA | 2 | 0.5 | 3 | 0.3333 | 1 | 0 | 6 | 0.3333 | 1 | 0 | 1 | 0 | 1 | 1 | NA | NA | 3 | 0.3333 |
| 9_9511157 | NA | NA | 3 | 0.6667 | 4 | 0 | 1 | 1 | 8 | 0.375 | 2 | 0.5 | 1 | 0 | 1 | 0 | 1 | 1 | 5 | 0.4 |
| 9_9528144 | NA | NA | 1 | 1 | 2 | 0 | 1 | 1 | 4 | 0.5 | 1 | 1 | 1 | 1 | 2 | 0.5 | 1 | 0 | 5 | 0.6 |
| 9_9528157 | NA | NA | NA | NA | 1 | 1 | NA | NA | 1 | 1 | 1 | 0 | 1 | 0 | NA | NA | 1 | 1 | 3 | 0.3333 |
| 9_9528322 | 6 | 0.3333 | 5 | 0.4 | 4 | 0 | 8 | 0 | 23 | 0.1739 | 4 | 0.25 | 3 | 1 | 7 | 0.2857 | 1 | 0 | 15 | 0.4 |
| 9_9528501 | 3 | 0.3333 | 3 | 0 | 2 | 0 | 4 | 0.25 | 12 | 0.1667 | 2 | 0 | 1 | 0 | 3 | 0.3333 | 3 | 0.6667 | 9 | 0.3333 |
| 9_9528573 | 4 | 1 | 4 | 0.75 | 2 | 0 | 4 | 0.75 | 14 | 0.7143 | 2 | 0 | 2 | 0 | 3 | 0.3333 | 2 | 0.5 | 9 | 0.2222 |
| 9_9528580 | 4 | 1 | 4 | 0.5 | 2 | 0.5 | 4 | 0.75 | 14 | 0.7143 | 2 | 0.5 | 2 | 1 | 2 | 0.5 | 2 | 1 | 8 | 0.75 |
| 9_9528620 | 27 | 0.1852 | 24 | 0 | 19 | 0 | 36 | 0.0833 | 106 | 0.0755 | 24 | 0.0417 | 17 | 0.1176 | 25 | 0.08 | 27 | 0.1481 | 93 | 0.0968 |
| 9_9528647 | 24 | 0.2083 | 24 | 0.0417 | 22 | 0.1818 | 30 | 0.1333 | 100 | 0.14 | 26 | 0.1923 | 14 | 0.0714 | 27 | 0.0741 | 28 | 0.2857 | 95 | 0.1684 |
| 9_9528657 | 13 | 0.2308 | 13 | 0.0769 | 18 | 0.2222 | 20 | 0.05 | 64 | 0.1406 | 17 | 0.0588 | 11 | 0 | 14 | 0.1429 | 15 | 0.0667 | 57 | 0.0702 |
| 9_9528673 | 20 | 0.25 | 19 | 0.1053 | 16 | 0.0625 | 26 | 0.1154 | 81 | 0.1358 | 23 | 0.1304 | 14 | 0.2143 | 22 | 0.0455 | 23 | 0.1739 | 82 | 0.1341 |
| 9_9528677 | 21 | 0.381 | 20 | 0.05 | 17 | 0.4706 | 30 | 0.3333 | 88 | 0.3068 | 22 | 0.3182 | 15 | 0.4 | 23 | 0.3913 | 24 | 0.375 | 84 | 0.369 |
| 9_9528678 | 22 | 0.1364 | 20 | 0.2 | 20 | 0 | 32 | 0.1562 | 94 | 0.1277 | 24 | 0 | 14 | 0.0714 | 26 | 0.1154 | 28 | 0.1071 | 92 | 0.0761 |
| 9_9528689 | 20 | 0.05 | 18 | 0.1667 | 19 | 0.1579 | 30 | 0.1 | 87 | 0.1149 | 21 | 0.0476 | 14 | 0.1429 | 23 | 0.1739 | 24 | 0.2083 | 82 | 0.1463 |
| 9_9528697 | 18 | 0.0556 | 13 | 0 | 16 | 0.1875 | 24 | 0 | 71 | 0.0563 | 18 | 0.0556 | 12 | 0 | 19 | 0.0526 | 21 | 0.1429 | 70 | 0.0714 |
| 9_9528697 | 14 | 0.1429 | 13 | 0.0769 | 15 | 0.2667 | 23 | 0.087 | 65 | 0.1385 | 19 | 0.1053 | 9 | 0 | 15 | 0.1333 | 20 | 0.05 | 63 | 0.0794 |
| 9_9528702 | 13 | 1 | 8 | 0.75 | 9 | 1 | 15 | 0.8667 | 45 | 0.9111 | 10 | 1 | 6 | 1 | 8 | 1 | 9 | 1 | 33 | 1 |
| 9_9528704 | 21 | 0.0952 | 16 | 0.1875 | 20 | 0.2 | 29 | 0.1379 | 86 | 0.1512 | 23 | 0.1304 | 14 | 0.1429 | 25 | 0.12 | 24 | 0.2083 | 86 | 0.1512 |
| 9_9528709 | 21 | 0.5714 | 17 | 0.4706 | 19 | 0.6842 | 30 | 0.6667 | 87 | 0.6092 | 25 | 0.56 | 13 | 0.8462 | 25 | 0.52 | 22 | 0.6364 | 85 | 0.6118 |
| 9_9528710 | 22 | 0.1364 | 18 | 0.1667 | 18 | 0.1111 | 25 | 0.2 | 83 | 0.1566 | 23 | 0.1739 | 13 | 0.2308 | 25 | 0.04 | 20 | 0.15 | 81 | 0.1358 |
| 9_9528718 | 17 | 0.5294 | 12 | 0.3333 | 13 | 0.6923 | 16 | 0.375 | 58 | 0.4828 | 17 | 0.5882 | 11 | 0.4545 | 19 | 0.5263 | 20 | 0.55 | 67 | 0.5373 |
| 9_9528737 | 17 | 0.4706 | 12 | 0.5833 | 15 | 0.5333 | 27 | 0.4815 | 71 | 0.507 | 15 | 0.5333 | 10 | 0.6 | 22 | 0.2727 | 20 | 0.4 | 67 | 0.4179 |
| 9_9528752 | 18 | 0.9444 | 15 | 1 | 13 | 0.9231 | 27 | 1 | 73 | 0.9726 | 20 | 0.9 | 9 | 1 | 24 | 0.9167 | 21 | 0.9048 | 74 | 0.9189 |
| 9_9528769 | 19 | 0.9474 | 13 | 0.8462 | 12 | 0.9167 | 20 | 0.95 | 64 | 0.9219 | 19 | 0.9474 | 8 | 1 | 21 | 1 | 20 | 0.95 | 68 | 0.9706 |
| 9_9528773 | 12 | 1 | 6 | 1 | 9 | 1 | 15 | 1 | 42 | 1 | 11 | 1 | 7 | 1 | 11 | 1 | 11 | 1 | 40 | 1 |
| 9_9528778 | 16 | 0.625 | 14 | 0.2857 | 14 | 0.2857 | 23 | 0.3478 | 67 | 0.3881 | 19 | 0.6842 | 9 | 0.3333 | 20 | 0.25 | 17 | 0.1176 | 65 | 0.3538 |
| 9_9528786 | 16 | 0.5 | 11 | 0.3636 | 9 | 0.3333 | 23 | 0.3913 | 59 | 0.4068 | 14 | 0.4286 | 9 | 0.4444 | 18 | 0.5556 | 16 | 0.5625 | 57 | 0.5088 |
| 9_9528798 | 18 | 1 | 10 | 1 | 13 | 1 | 23 | 0.9565 | 64 | 0.9844 | 15 | 1 | 7 | 0.8571 | 15 | 1 | 18 | 0.9444 | 55 | 0.9636 |
| 9_9528812 | 14 | 1 | 10 | 1 | 13 | 1 | 21 | 1 | 58 | 1 | 13 | 1 | 8 | 1 | 14 | 0.9286 | 16 | 0.9375 | 51 | 0.9608 |
| 9_9528856 | 1 | 1 | NA | NA | NA | NA | 4 | 0.5 | 5 | 0.6 | NA | NA | 2 | 0.5 | NA | NA | 2 | 0.5 | 4 | 0.5 |
| 9_9528860 | 1 | 1 | NA | NA | NA | NA | 3 | 0.6667 | 4 | 0.75 | NA | NA | 2 | 0.5 | NA | NA | 2 | 0.5 | 4 | 0.5 |
| 9_9528868 | 9 | 0.1111 | 3 | 0 | 8 | 0 | 9 | 0.2222 | 29 | 0.1034 | 3 | 0.6667 | 3 | 1 | 12 | 0.0833 | 10 | 0.1 | 28 | 0.25 |
| 9_9528878 | 11 | 0 | 6 | 0 | 9 | 0.2222 | 6 | 0.1667 | 32 | 0.0938 | 6 | 0.3333 | 6 | 0 | 7 | 0 | 11 | 0.0909 | 30 | 0.1 |
| 9_9528890 | 15 | 0 | 6 | 0.1667 | 11 | 0 | 17 | 0.1176 | 49 | 0.0612 | 11 | 0.0909 | 6 | 0.3333 | 14 | 0.1429 | 14 | 0.0714 | 45 | 0.1333 |

| | | | | | | | | | | | | | | | | | | | | |
|---|---|---|---|---|---|---|---|---|---|---|---|---|---|---|---|---|---|---|---|---|
| 9_95319429 | NA | NA | NA | NA | NA | NA | 2 | 1 | 2 | 1 | NA | NA | NA | NA | 1 | 1 | NA | NA | 1 | 1 |
| 9_95442475 | 3 | 1 | 3 | 0.6667 | 1 | 0 | NA | NA | 7 | 0.7143 | 2 | 0 | 4 | 0.75 | NA | NA | 2 | 1 | 8 | 0.625 |
| 9_95442607 | 7 | 1 | 3 | 1 | 1 | 1 | NA | NA | 11 | 1 | 2 | 1 | 5 | 1 | 2 | 1 | 2 | 1 | 11 | 1 |
| 9_95443088 | 6 | 1 | 4 | 1 | NA | NA | 2 | 1 | 12 | 1 | 1 | 1 | 4 | 1 | 2 | 1 | 1 | 1 | 8 | 1 |
| 9_95443111 | 4 | 0.5 | 3 | 0.6667 | 1 | 0 | 1 | 0 | 9 | 0.4444 | 1 | 0 | 3 | 0.3333 | 1 | 0 | 2 | 0.5 | 7 | 0.2857 |
| 9_95443381 | 7 | 0.8571 | 2 | 1 | 1 | 0 | 2 | 1 | 12 | 0.8333 | 3 | 0.6667 | 4 | 0.75 | 2 | 1 | 2 | 1 | 11 | 0.8182 |
| 9_95443413 | 6 | 0.5 | 4 | 0.25 | 1 | 0 | 2 | 0.5 | 13 | 0.3846 | 2 | 0 | 6 | 0.1667 | 2 | 1 | 2 | 0 | 12 | 0.25 |
| 9_95443601 | 5 | 0.6 | 2 | 0 | NA | NA | 2 | 0.5 | 9 | 0.4444 | 4 | 0.25 | 4 | 0.5 | 1 | 1 | 2 | 0.5 | 11 | 0.4545 |
| 9_95443607 | 6 | 0.1667 | 5 | 0 | NA | NA | 2 | 0 | 13 | 0.0769 | 3 | 0.3333 | 4 | 0 | 2 | 0.5 | 2 | 0 | 11 | 0.1818 |
| 9_95443628 | 6 | 0.3333 | 4 | 0.25 | 1 | 0 | 2 | 0 | 13 | 0.2308 | 3 | 0.6667 | 6 | 0.1667 | 2 | 0 | 3 | 0 | 14 | 0.2143 |
| 9_95443646 | 8 | 0.125 | 4 | 0.25 | 1 | 0 | 3 | 0 | 16 | 0.125 | 4 | 0 | 6 | 0 | 2 | 0 | 3 | 0.6667 | 15 | 0.1333 |
| 9_95443801 | 3 | 0.6667 | 2 | 0.5 | 1 | 1 | 3 | 1 | 9 | 0.7778 | 3 | 1 | 5 | 0.8 | NA | NA | 3 | 0.6667 | 11 | 0.8182 |
| 9_95443806 | 6 | 0.5 | 4 | 0.5 | 1 | 0 | 4 | 0.5 | 15 | 0.4667 | 3 | 0.3333 | 6 | 0.5 | 1 | 1 | 3 | 1 | 13 | 0.6154 |
| 9_95443888 | 8 | 0.375 | 4 | 0.5 | 2 | 0 | 4 | 0.25 | 18 | 0.3333 | 4 | 0.25 | 6 | 0.5 | 2 | 0 | 3 | 0.3333 | 15 | 0.3333 |
| 9_95443898 | 9 | 0.2222 | 4 | 0 | 2 | 0 | 4 | 0 | 19 | 0.1053 | 4 | 0 | 6 | 0.1667 | 2 | 0 | 2 | 0 | 14 | 0.0714 |
| 9_95443909 | 8 | 0.375 | 3 | 0.3333 | 2 | 0.5 | 4 | 1 | 17 | 0.5294 | 3 | 0 | 7 | 0.7143 | 2 | 0.5 | 3 | 0.3333 | 15 | 0.4667 |
| 9_95443920 | 10 | 0.2 | 5 | 0 | 2 | 0 | 3 | 0.3333 | 20 | 0.15 | 4 | 0.25 | 7 | 0.2857 | 1 | 0 | 3 | 0.6667 | 15 | 0.3333 |
| 9_95444944 | 2 | 0 | 1 | 0 | NA | NA | 2 | 0.5 | 5 | 0.2 | 1 | 1 | 2 | 0 | NA | NA | 1 | 0 | 4 | 0.25 |
| 9_95447499 | 1 | 0 | 2 | 0.5 | 1 | 0 | 6 | 0 | 10 | 0.1 | 1 | 0 | 5 | 0.2 | 2 | 0 | 2 | 0 | 10 | 0.1 |
| 9_95448749 | 2 | 0.5 | 3 | 0 | 3 | 0 | 5 | 0 | 13 | 0.0769 | 1 | 0 | 5 | 0 | 2 | 0.5 | 3 | 0 | 11 | 0.0909 |
| 9_95448897 | 2 | 0 | 3 | 0 | 3 | 0 | 6 | 0.3333 | 14 | 0.1429 | 1 | 0 | 4 | 0.25 | 2 | 0 | 2 | 0 | 9 | 0.1111 |
| 9_95449196 | 2 | 0.5 | 3 | 0 | 4 | 0.25 | 6 | 0.1667 | 15 | 0.2 | 2 | 0 | 5 | 0.6 | 2 | 0.5 | 3 | 0.3333 | 12 | 0.4167 |
| 9_95449504 | 2 | 1 | 3 | 1 | 5 | 0.8 | 6 | 1 | 16 | 0.9375 | 2 | 1 | 5 | 1 | 2 | 1 | 3 | 1 | 12 | 1 |
| 9_95449553 | 1 | 0 | 2 | 0 | 5 | 0.2 | 5 | 0.2 | 13 | 0.1538 | 3 | 0 | 2 | 0 | 2 | 0 | 3 | 0.6667 | 10 | 0.2 |
| 9_95449693 | 2 | 0 | 3 | 0 | 5 | 0.2 | 6 | 0.1667 | 16 | 0.125 | 3 | 0.3333 | 3 | 0 | 3 | 0.6667 | 4 | 0.25 | 13 | 0.3077 |
| 9_95449714 | NA | NA | 1 | 1 | 2 | 0.5 | 3 | 0.6667 | 6 | 0.6667 | 1 | 1 | 2 | 0.5 | NA | NA | 1 | 1 | 4 | 0.75 |
| 9_95449718 | 2 | 0.5 | NA | NA | 3 | 0.3333 | 3 | 0.3333 | 8 | 0.375 | 1 | 0 | 1 | 1 | 2 | 1 | NA | NA | 4 | 0.75 |
| 9_95449768 | 3 | 1 | 3 | 1 | 5 | 0.8 | 6 | 1 | 17 | 0.9412 | 3 | 1 | 5 | 1 | 2 | 1 | 3 | 1 | 13 | 1 |
| 9_95449902 | 2 | 0.5 | 1 | 1 | 5 | 0.4 | 5 | 0.8 | 13 | 0.6154 | 2 | 0 | 2 | 1 | 3 | 0.6667 | 3 | 1 | 10 | 0.7 |
| 9_95449984 | 2 | 0 | 4 | 0.25 | 5 | 0.4 | 7 | 0 | 18 | 0.1667 | 2 | 0 | 5 | 0.2 | 2 | 0 | 4 | 0 | 13 | 0.0769 |
| 9_95577174 | 1 | 1 | NA | NA | 2 | 1 | NA | NA | 3 | 1 | 1 | 1 | NA | NA | NA | NA | 2 | 0.5 | 3 | 0.6667 |
| 9_95587534 | 4 | 0.25 | 5 | 0.2 | 5 | 0 | 12 | 0.0833 | 26 | 0.1154 | 1 | 0 | 2 | 0.5 | 5 | 0 | 5 | 0 | 13 | 0.0769 |
| 9_95588744 | 3 | 0 | 5 | 0.6 | 4 | 0.5 | 8 | 0.625 | 20 | 0.5 | 1 | 1 | 2 | 0.5 | 7 | 1 | 7 | 1 | 17 | 0.9412 |
| 9_95588828 | 3 | 0.6667 | 6 | 1 | 4 | 0.75 | 8 | 0.875 | 21 | 0.8571 | 1 | 1 | 3 | 0 | 7 | 0.7143 | 7 | 0.8571 | 18 | 0.6667 |
| 9_95588886 | 2 | 0.5 | 4 | 0.75 | 3 | 0.3333 | 7 | 0.1429 | 16 | 0.375 | 1 | 1 | 3 | 0 | 5 | 0.4 | 4 | 0.5 | 13 | 0.3846 |
| 9_95592264 | 3 | 0 | 2 | 0.5 | 4 | 0 | 6 | 0 | 15 | 0.0667 | 1 | 0 | 2 | 0 | 3 | 0 | 6 | 0.1667 | 12 | 0.0833 |
| 9_95603426 | 3 | 1 | 3 | 0.6667 | 3 | 0 | 5 | 0.6 | 14 | 0.5714 | 1 | 1 | 1 | 0 | 5 | 0.2 | 4 | 0.5 | 11 | 0.3636 |
| 9_95603472 | 2 | 0.5 | 2 | 0 | 2 | 0 | 5 | 0 | 11 | 0.0909 | 1 | 0 | 1 | 0 | 4 | 0.25 | 2 | 0 | 8 | 0.125 |
| 9_95631864 | 2 | 0 | 3 | 0 | 2 | 0.5 | 4 | 0.25 | 11 | 0.1818 | 1 | 0 | 2 | 0 | 2 | 0 | 2 | 1 | 7 | 0.2857 |
| 9_95632151 | 2 | 0 | 3 | 0 | 2 | 0 | 3 | 0.3333 | 10 | 0.1 | NA | NA | 2 | 0 | 1 | 0 | 3 | 0.3333 | 6 | 0.1667 |
| 9_95636364 | 2 | 0.5 | 2 | 0.5 | 1 | 0 | 1 | 1 | 6 | 0.5 | 2 | 0.5 | 1 | 1 | 1 | 1 | 1 | 0 | 5 | 0.6 |
| 9_95936911 | 2 | 0 | 2 | 0 | 2 | 0.5 | 3 | 0 | 9 | 0.1111 | NA | NA | 1 | 0 | 2 | 0.5 | NA | NA | 3 | 0.3333 |
| 9_95936932 | 1 | 1 | 1 | 1 | 2 | 1 | 3 | 0.6667 | 7 | 0.8571 | NA | NA | 1 | 1 | 2 | 1 | NA | NA | 3 | 1 |
| 9_95937090 | 2 | 1 | 2 | 0.5 | 2 | 1 | 3 | 0.3333 | 9 | 0.6667 | NA | NA | 1 | 1 | 2 | 0.5 | NA | NA | 3 | 0.6667 |
| 9_95937275 | 2 | 0.5 | 3 | 0.3333 | 1 | 1 | 3 | 0.3333 | 9 | 0.4444 | NA | NA | 2 | 0.5 | 1 | 0 | NA | NA | 3 | 0.3333 |
| 9_95939687 | 3 | 0 | 4 | 0.5 | 3 | 0 | 2 | 1 | 12 | 0.3333 | 1 | 1 | 2 | 0.5 | 2 | 0 | 2 | 0.5 | 7 | 0.4286 |
| 9_96199575 | NA | NA | NA | NA | 1 | 1 | NA | NA | 1 | 1 | NA | NA | NA | NA | 2 | 0.5 | NA | NA | 2 | 0.5 |
| 9_96200037 | NA | NA | NA | NA | 2 | 1 | NA | NA | 2 | 1 | NA | NA | 1 | 1 | 2 | 1 | NA | NA | 3 | 1 |
| 9_96215000 | 16 | 0.125 | 21 | 0.1429 | 13 | 0 | 11 | 0 | 61 | 0.082 | 7 | 0.2857 | 12 | 0 | 9 | 0 | 11 | 0.0909 | 39 | 0.0769 |
| 9_96215048 | 17 | 0.0588 | 20 | 0.1 | 12 | 0.1667 | 8 | 0 | 57 | 0.0877 | 7 | 0.2857 | 13 | 0.0769 | 8 | 0 | 12 | 0.1667 | 40 | 0.125 |
| 9_96215095 | 18 | 0 | 20 | 0.15 | 13 | 0 | 11 | 0.0909 | 62 | 0.0645 | 7 | 0.1429 | 13 | 0 | 10 | 0.3 | 12 | 0.0833 | 42 | 0.119 |
| 9_96215171 | 24 | 0.25 | 26 | 0.1923 | 17 | 0.1765 | 13 | 0.2308 | 80 | 0.2125 | 11 | 0.1818 | 15 | 0.1333 | 15 | 0.4667 | 17 | 0.0588 | 58 | 0.2069 |
| 9_96215233 | 14 | 0.2143 | 20 | 0.15 | 14 | 0.0714 | 14 | 0.0714 | 62 | 0.129 | 7 | 0.2857 | 12 | 0.0833 | 11 | 0.1818 | 11 | 0 | 41 | 0.122 |
| 9_96215356 | 22 | 0.3182 | 26 | 0.2692 | 15 | 0.2 | 14 | 0.4286 | 77 | 0.2987 | 10 | 0.4 | 13 | 0.2308 | 14 | 0.2857 | 16 | 0.25 | 53 | 0.283 |
| 9_96215381 | 21 | 0.0952 | 25 | 0.12 | 16 | 0.1875 | 11 | 0.0909 | 73 | 0.1233 | 10 | 0.2 | 13 | 0.3077 | 13 | 0 | 15 | 0.0667 | 51 | 0.1373 |
| 9_96215408 | 16 | 0.3125 | 22 | 0.3636 | 12 | 0.1667 | 10 | 0.5 | 60 | 0.3333 | 6 | 0.3333 | 9 | 0.2222 | 8 | 0.25 | 10 | 0.2 | 33 | 0.2424 |
| 9_96215456 | 23 | 0.1304 | 24 | 0.0417 | 15 | 0.0667 | 12 | 0.0833 | 74 | 0.0811 | 9 | 0.2222 | 16 | 0.0625 | 12 | 0 | 14 | 0 | 51 | 0.0588 |
| 9_96215520 | 21 | 0.4286 | 24 | 0.625 | 14 | 0.5 | 11 | 0.6364 | 70 | 0.5429 | 9 | 0.6667 | 14 | 0.7857 | 13 | 0.6923 | 13 | 0.6154 | 49 | 0.6939 |
| 9_96215647 | 21 | 0.1429 | 25 | 0 | 14 | 0 | 12 | 0.1667 | 72 | 0.0694 | 9 | 0 | 15 | 0.1333 | 14 | 0.1429 | 14 | 0.1429 | 52 | 0.1154 |
| 9_96236975 | NA | NA | 1 | 1 | 1 | 1 | NA | NA | 2 | 1 | 1 | 0 | 2 | 1 | NA | NA | NA | NA | 3 | 0.6667 |
| 9_96237042 | NA | NA | NA | NA | 1 | 1 | NA | NA | 1 | 1 | 1 | 0 | 2 | 0.5 | NA | NA | NA | NA | 3 | 0.3333 |
| 9_96245500 | 2 | 1 | 4 | 1 | 1 | 1 | NA | NA | 7 | 1 | 2 | 1 | 2 | 1 | 3 | 1 | 1 | 1 | 8 | 1 |
| 9_96245786 | 2 | 1 | 4 | 1 | 1 | 1 | NA | NA | 7 | 1 | 2 | 1 | 2 | 1 | 2 | 1 | 1 | 1 | 7 | 1 |
| 9_96245817 | 1 | 1 | 4 | 1 | NA | NA | NA | NA | 5 | 1 | 1 | 1 | 3 | 1 | 3 | 0.6667 | 1 | 1 | 8 | 0.875 |
| 9_96245871 | 2 | 1 | 4 | 0.75 | 1 | 1 | NA | NA | 7 | 0.8571 | 2 | 0.5 | 2 | 1 | 3 | 0.6667 | 1 | 1 | 8 | 0.75 |

| | | | | | | | | | | | | | | | | | | | | |
|---|---|---|---|---|---|---|---|---|---|---|---|---|---|---|---|---|---|---|---|---|
| 9_9624588 | 2 | 1 | 4 | 1 | 1 | 1 | NA | NA | 7 | 1 | 2 | 1 | 3 | 1 | 3 | 1 | 1 | 1 | 9 | 1 |
| 9_9624619 | 15 | 0 | 20 | 0.05 | 10 | 0.2 | 7 | 0 | 52 | 0.0577 | 9 | 0.1111 | 13 | 0.0769 | 7 | 0.1429 | 12 | 0.0833 | 41 | 0.0976 |
| 9_9635318 | 18 | 0.2222 | 28 | 0.3929 | 11 | 0.2727 | 18 | 0.1111 | 75 | 0.2667 | 8 | 0.125 | 22 | 0.3636 | 12 | 0.3333 | 14 | 0.2857 | 56 | 0.3036 |
| 9_9635329 | 13 | 0.0769 | 27 | 0.037 | 9 | 0.1111 | 18 | 0.1111 | 67 | 0.0746 | 9 | 0 | 18 | 0 | 11 | 0.1818 | 12 | 0.1667 | 50 | 0.08 |
| 9_9635368 | 16 | 0.0625 | 27 | 0.1852 | 10 | 0 | 16 | 0.0625 | 69 | 0.1014 | 13 | 0.0769 | 25 | 0 | 14 | 0.2857 | 7 | 0.1429 | 59 | 0.1017 |
| 9_9635374 | 18 | 0.2222 | 26 | 0.2308 | 9 | 0.4444 | 17 | 0.2353 | 70 | 0.2571 | 13 | 0.2308 | 26 | 0.2308 | 13 | 0.1538 | 10 | 0.1 | 62 | 0.1935 |
| 9_9635381 | 15 | 0.2 | 29 | 0.1034 | 11 | 0.2727 | 18 | 0.1667 | 73 | 0.1644 | 12 | 0 | 29 | 0.1034 | 14 | 0.2143 | 11 | 0.0909 | 66 | 0.1061 |
| 9_9642166 | 1 | 0 | 1 | 1 | 2 | 0 | 2 | 1 | 6 | 0.5 | 1 | 0 | 2 | 0 | 1 | 0 | 1 | 1 | 5 | 0.2 |
| 9_9642434 | 1 | 0 | 1 | 1 | 1 | 0 | NA | NA | 3 | 0.3333 | NA | NA | 2 | 0.5 | NA | NA | NA | NA | 2 | 0.5 |
| 9_9642441 | NA | NA | 1 | 1 | 1 | 1 | 1 | 1 | 3 | 1 | NA | NA | 3 | 0.6667 | NA | NA | NA | NA | 3 | 0.6667 |
| 9_9642447 | 1 | 0 | 1 | 1 | 1 | 0 | 1 | 0 | 4 | 0.25 | NA | NA | 3 | 0.3333 | NA | NA | NA | NA | 3 | 0.3333 |
| 9_9656498 | 8 | 0.25 | 10 | 0.2 | 11 | 0.2727 | 7 | 0.4286 | 36 | 0.2778 | 7 | 0 | 4 | 0.25 | 9 | 0.3333 | 11 | 0.3636 | 31 | 0.2581 |
| 9_9656500 | 5 | 0 | 8 | 0.125 | 8 | 0.25 | 6 | 0.1667 | 27 | 0.1481 | 5 | 0 | 4 | 0.25 | 7 | 0.5714 | 9 | 0.2222 | 25 | 0.28 |
| 9_9656530 | 6 | 1 | 7 | 0.4286 | 10 | 0.7 | 6 | 0.5 | 29 | 0.6552 | 6 | 0.8333 | 3 | 0.6667 | 8 | 0.625 | 11 | 0.9091 | 28 | 0.7857 |
| 9_9656556 | 6 | 0.1667 | 7 | 0 | 8 | 0.25 | 6 | 0.3333 | 27 | 0.1852 | 3 | 0 | 4 | 0.25 | 7 | 0.1429 | 8 | 0 | 22 | 0.0909 |
| 9_9656569 | 4 | 1 | 7 | 0.4286 | 8 | 0.5 | 6 | 0.5 | 25 | 0.56 | 4 | 0.5 | 3 | 0 | 6 | 0.5 | 8 | 0.875 | 21 | 0.5714 |
| 9_9656582 | 3 | 0.6667 | 4 | 0.25 | 6 | 0.8333 | 5 | 0.4 | 18 | 0.5556 | 4 | 0.5 | 3 | 0.6667 | 6 | 0.6667 | 6 | 0.6667 | 19 | 0.6316 |
| 9_9656589 | 1 | 0 | 2 | 0 | 5 | 0.2 | 3 | 0 | 11 | 0.0909 | 4 | 0.5 | 1 | 0 | 4 | 0.25 | 4 | 0 | 13 | 0.2308 |
| 9_9656589 | 2 | 0 | 5 | 0.2 | 7 | 0 | 5 | 0 | 19 | 0.0526 | 4 | 0 | 2 | 0 | 5 | 0 | 6 | 0.1667 | 17 | 0.0588 |
| 9_9656598 | 2 | 0 | 5 | 0.2 | 8 | 0 | 5 | 0.4 | 20 | 0.15 | 4 | 0.25 | 2 | 0 | 5 | 0.4 | 5 | 0.4 | 16 | 0.3125 |
| 9_9656606 | 2 | 0.5 | 2 | 0.5 | 6 | 0.1667 | 4 | 0.25 | 14 | 0.2857 | 2 | 1 | 2 | 0 | 5 | 0.2 | 6 | 0.1667 | 15 | 0.2667 |
| 9_9656615 | 1 | 0 | 4 | 0.25 | 6 | 0.1667 | 4 | 0 | 15 | 0.1333 | 4 | 0.5 | 2 | 0 | 5 | 0.2 | 5 | 0 | 16 | 0.1875 |
| 9_9656666 | NA | NA | 4 | 0.25 | 2 | 0 | 3 | 0 | 9 | 0.1111 | 2 | 0.5 | 2 | 0 | 3 | 0 | 4 | 0 | 11 | 0.0909 |
| 9_9656674 | NA | NA | 4 | 0 | 5 | 0 | 2 | 0.5 | 11 | 0.0909 | 3 | 0.3333 | 2 | 0 | 4 | 0 | 5 | 0.4 | 14 | 0.2143 |
| 9_9656677 | NA | NA | 4 | 0.75 | 6 | 0 | 2 | 0 | 12 | 0.25 | 3 | 0.3333 | 2 | 0 | 4 | 0.5 | 5 | 0.4 | 14 | 0.3571 |
| 9_9656710 | 1 | 1 | 4 | 1 | 5 | 1 | 1 | 1 | 11 | 1 | 2 | 1 | 1 | 1 | 4 | 1 | 4 | 1 | 11 | 1 |
| 9_9656717 | 1 | 1 | 3 | 0 | 4 | 0.25 | 1 | 0 | 9 | 0.2222 | 2 | 0.5 | NA | NA | 4 | 0.25 | 4 | 0.5 | 10 | 0.4 |
| 9_9656748 | NA | NA | 3 | 0 | 4 | 0.25 | NA | NA | 7 | 0.1429 | 2 | 0.5 | 1 | 1 | 2 | 0 | 4 | 0 | 9 | 0.2222 |
| 9_9656753 | NA | NA | 2 | 0.5 | 3 | 0 | NA | NA | 5 | 0.2 | 2 | 0 | 1 | 0 | 2 | 0 | 3 | 0.3333 | 8 | 0.125 |
| 9_9656753 | NA | NA | 2 | 0.5 | 3 | 0.3333 | NA | NA | 5 | 0.4 | 2 | 0.5 | 1 | 0 | 1 | 0 | 4 | 0.25 | 8 | 0.25 |
| 9_9656762 | NA | NA | 2 | 1 | 1 | 0 | NA | NA | 3 | 0.6667 | 2 | 1 | 1 | 0 | 2 | 0 | 3 | 0.6667 | 8 | 0.5 |
| 9_9656763 | NA | NA | 2 | 1 | 1 | 0 | NA | NA | 3 | 0.6667 | 2 | 1 | 1 | 1 | 1 | 1 | 4 | 1 | 8 | 1 |
| 9_9656774 | NA | NA | 2 | 0.5 | 1 | 1 | NA | NA | 3 | 0.6667 | 2 | 0.5 | 1 | 0 | 1 | 0 | 3 | 0 | 7 | 0.1429 |
| 9_9656847 | NA | NA | 1 | 1 | 1 | 1 | NA | NA | 2 | 1 | 1 | 1 | 1 | 1 | 1 | 1 | 2 | 1 | 5 | 1 |
| 9_9656864 | NA | NA | 1 | 0 | 1 | 1 | NA | NA | 2 | 0.5 | 2 | 0 | 1 | 1 | 1 | 0 | 2 | 0 | 6 | 0.1667 |
| 9_9656939 | NA | NA | NA | NA | 1 | 1 | NA | NA | 1 | 1 | NA | NA | 1 | 1 | NA | NA | 1 | 1 | 2 | 1 |
| 9_9656945 | NA | NA | 1 | 1 | 2 | 0.5 | NA | NA | 3 | 0.6667 | 2 | 1 | 1 | 1 | NA | NA | 1 | 1 | 4 | 1 |
| 9_9656967 | NA | NA | 1 | 1 | 1 | 1 | NA | NA | 2 | 1 | 2 | 0.5 | 1 | 1 | NA | NA | 1 | 0 | 4 | 0.5 |
| 9_9670648 | 2 | 0.5 | NA | NA | 1 | 0 | 3 | 0.6667 | 6 | 0.5 | NA | NA | 1 | 1 | 1 | 0 | 2 | 0.5 | 4 | 0.5 |
| 9_9670677 | 2 | 0.5 | NA | NA | 1 | 0 | 2 | 0 | 5 | 0.2 | NA | NA | NA | NA | NA | NA | 2 | 0.5 | 2 | 0.5 |
| 9_9670682 | 2 | 1 | NA | NA | 1 | 1 | 2 | 1 | 5 | 1 | NA | NA | 1 | 1 | NA | NA | 2 | 1 | 3 | 1 |
| 9_9670687 | 2 | 0.5 | NA | NA | 1 | 0 | 3 | 0.6667 | 6 | 0.5 | NA | NA | 1 | 0 | NA | NA | 2 | 0.5 | 3 | 0.3333 |
| 9_9670693 | 2 | 1 | NA | NA | NA | NA | 2 | 1 | 4 | 1 | NA | NA | 1 | 1 | NA | NA | 2 | 1 | 3 | 1 |
| 9_9670733 | 2 | 1 | NA | NA | NA | NA | 2 | 0.5 | 4 | 0.75 | NA | NA | 1 | 1 | NA | NA | 2 | 1 | 3 | 1 |
| 9_9670736 | 2 | 0.5 | NA | NA | NA | NA | 2 | 1 | 4 | 0.75 | NA | NA | 1 | 1 | NA | NA | 2 | 1 | 3 | 1 |
| 9_9670750 | 2 | 0.5 | NA | NA | NA | NA | 1 | 1 | 3 | 0.6667 | NA | NA | 1 | 1 | NA | NA | 2 | 1 | 3 | 1 |
| 9_9677819 | 124 | 0.1452 | 117 | 0.1624 | 83 | 0.1205 | 97 | 0.1546 | 421 | 0.1473 | 106 | 0.1792 | 105 | 0.1429 | 105 | 0.0762 | 90 | 0.2333 | 406 | 0.1552 |
| 9_9677821 | 140 | 0.05 | 133 | 0.0827 | 90 | 0.0111 | 113 | 0.0796 | 476 | 0.0588 | 118 | 0.0678 | 123 | 0.065 | 102 | 0.098 | 102 | 0.0784 | 445 | 0.0764 |
| 9_9677840 | 2 | 0.5 | 1 | 0 | 4 | 0 | 1 | 0 | 8 | 0.125 | 5 | 0.2 | 4 | 0 | 3 | 0 | NA | NA | 12 | 0.0833 |
| 9_9695787 | 2 | 0 | 2 | 0.5 | 5 | 0.2 | 3 | 0.6667 | 12 | 0.3333 | 1 | 1 | 9 | 0.2222 | 7 | 0 | 3 | 0.3333 | 20 | 0.2 |
| 9_9695817 | 2 | 0 | 1 | 0 | 7 | 0.1429 | 3 | 0 | 13 | 0.0769 | 1 | 0 | 8 | 0.125 | 7 | 0 | 3 | 0.6667 | 19 | 0.1579 |
| 9_9695841 | 2 | 0.5 | 2 | 0 | 6 | 0.5 | 2 | 0.5 | 12 | 0.4167 | 1 | 0 | 8 | 0.375 | 5 | 0.2 | 2 | 0.5 | 16 | 0.3125 |
| 9_9695850 | 2 | 0.5 | 2 | 1 | 6 | 0.6667 | 2 | 1 | 12 | 0.75 | NA | NA | 7 | 0.5714 | 5 | 0.4 | 3 | 0.3333 | 15 | 0.4667 |
| 9_9696045 | 1 | 0 | 2 | 0.5 | 3 | 0 | NA | NA | 6 | 0.1667 | 1 | 0 | 8 | 0 | 5 | 0.2 | 2 | 0 | 16 | 0.0625 |
| 9_9696100 | 1 | 0 | 1 | 1 | 2 | 0 | 1 | 0 | 5 | 0.2 | 1 | 0 | 7 | 0 | 6 | 0.1667 | 2 | 0 | 16 | 0.0625 |
| 9_9696101 | 1 | 1 | 1 | 1 | 4 | 0.75 | NA | NA | 6 | 0.8333 | 1 | 1 | 5 | 0.6 | 2 | 0.5 | 2 | 1 | 10 | 0.7 |
| 9_9696104 | 1 | 0 | 2 | 0.5 | 2 | 1 | 1 | 0 | 6 | 0.5 | 1 | 0 | 7 | 1 | 3 | 0.6667 | 2 | 1 | 13 | 0.8462 |
| 9_9696112 | NA | NA | NA | NA | 2 | 1 | NA | NA | 2 | 1 | NA | NA | 4 | 1 | 2 | 1 | 1 | 1 | 7 | 1 |
| 9_9696126 | 1 | 1 | 2 | 1 | 4 | 0.75 | 1 | 1 | 8 | 0.875 | 1 | 1 | 5 | 1 | 4 | 1 | 1 | 1 | 11 | 1 |
| 9_9696236 | 1 | 1 | 2 | 1 | 3 | 0.3333 | 1 | 0 | 7 | 0.5714 | 1 | 1 | 4 | 0.5 | 6 | 0.6667 | 1 | 1 | 12 | 0.6667 |
| 9_9696242 | 1 | 1 | 1 | 1 | 4 | 0.5 | 1 | 0 | 7 | 0.5714 | 1 | 1 | 6 | 1 | 6 | 0.8333 | 1 | 0 | 14 | 0.8571 |
| 9_9696253 | 1 | 1 | 1 | 0 | 4 | 0 | 1 | 0 | 7 | 0.1429 | 1 | 1 | 6 | 0.1667 | 5 | 0 | 1 | 0 | 13 | 0.1538 |
| 9_9732704 | 4 | 0 | 7 | 0.2857 | 9 | 0.1111 | 2 | 0 | 22 | 0.1364 | 2 | 0 | 3 | 0 | 5 | 0 | 2 | 0.5 | 12 | 0.0833 |
| 9_9732723 | 3 | 0 | 7 | 0.2857 | 10 | 0.4 | 2 | 0.5 | 22 | 0.3182 | 2 | 0.5 | 3 | 0 | 5 | 0.2 | 2 | 1 | 12 | 0.3333 |
| 9_9732736 | 1 | 0 | 5 | 0.2 | 7 | 0.5714 | 2 | 0 | 15 | 0.3333 | 1 | 1 | 2 | 0 | 3 | 0.3333 | 2 | 1 | 8 | 0.5 |

| | | | | | | | | | | | | | | | | | | | | |
|---|---|---|---|---|---|---|---|---|---|---|---|---|---|---|---|---|---|---|---|---|
| 9_97327648 | 2 | 0 | 4 | 0 | 11 | 0.0909 | 2 | 0 | 19 | 0.0526 | 2 | 0 | 2 | 0 | 3 | 0.3333 | 1 | 0 | 8 | 0.125 |
| 9_98292286 | 1 | 1 | NA | NA | 1 | 1 | NA | NA | 2 | 1 | 1 | 1 | NA | NA | NA | NA | 2 | 0 | 3 | 0.3333 |
| 9_98292486 | 1 | 1 | NA | NA | 1 | 1 | NA | NA | 2 | 1 | NA | NA | NA | NA | 1 | 1 | 3 | 0.6667 | 4 | 0.75 |
| 9_98307623 | 11 | 0 | 4 | 0.25 | 10 | 0.2 | 10 | 0.1 | 35 | 0.1143 | 7 | 0.2857 | 7 | 0.1429 | 10 | 0.1 | 10 | 0 | 34 | 0.1176 |
| 9_98470260 | 14 | 0.1429 | 5 | 0.2 | 11 | 0.7273 | 9 | 0.3333 | 39 | 0.359 | 13 | 0.4615 | 10 | 0.3 | 6 | 0.1667 | 13 | 0.2308 | 42 | 0.3095 |
| 9_98883629 | NA | NA | 4 | 0.5 | 1 | 0 | 1 | 0 | 6 | 0.3333 | 4 | 0.5 | 1 | 0 | NA | NA | 4 | 0 | 9 | 0.2222 |
| 9_98883791 | 1 | 0 | 5 | 0 | 3 | 0 | 1 | 1 | 10 | 0.1 | 5 | 0.2 | 3 | 0 | 2 | 0 | 4 | 0 | 14 | 0.0714 |
| 9_98883928 | 1 | 1 | 5 | 0.4 | 5 | 0 | 1 | 0 | 12 | 0.25 | 3 | 0.3333 | 3 | 0.3333 | 2 | 0 | 5 | 0.6 | 13 | 0.3846 |
| 9_98918855 | NA | NA | 1 | 1 | NA | NA | NA | NA | 1 | 1 | NA | NA | 1 | 1 | NA | NA | NA | NA | 1 | 1 |
| 9_98920611 | 4 | 0.25 | 5 | 0 | 2 | 0 | 1 | 0 | 12 | 0.0833 | 2 | 0 | 3 | 1 | 3 | 0 | NA | NA | 8 | 0.375 |
| 9_98920675 | 3 | 0 | 7 | 0.1429 | 2 | 0 | 1 | 0 | 13 | 0.0769 | 3 | 0 | 4 | 0.25 | 3 | 0.3333 | 1 | 0 | 11 | 0.1818 |
| 9_98921307 | 2 | 0 | 4 | 1 | 3 | 1 | 1 | 1 | 10 | 0.8 | 3 | 0.3333 | 3 | 0.6667 | 1 | 0 | 1 | 0 | 8 | 0.375 |
| 9_98921360 | 2 | 0 | 6 | 0.3333 | 2 | 0.5 | NA | NA | 10 | 0.3 | 3 | 0 | 2 | 0.5 | 2 | 0 | 1 | 0 | 8 | 0.125 |
| 9_98921506 | 2 | 1 | 3 | 1 | 2 | 1 | 1 | 1 | 8 | 1 | 2 | 1 | 3 | 1 | 2 | 1 | 1 | 1 | 8 | 1 |
| 9_98921573 | 2 | 0.5 | 5 | 0.8 | 1 | 1 | NA | NA | 8 | 0.75 | 2 | 0 | 2 | 0.5 | 3 | 1 | 1 | 1 | 8 | 0.625 |
| 9_98921696 | 2 | 0.5 | 4 | 0 | 2 | 0 | 1 | 0 | 9 | 0.1111 | 2 | 0 | 2 | 0 | 2 | 0.5 | 1 | 0 | 7 | 0.1429 |
| 9_98921701 | 2 | 0 | 3 | 1 | 2 | 0.5 | 1 | 1 | 8 | 0.625 | 2 | 0.5 | 1 | 1 | 2 | 0 | NA | NA | 5 | 0.4 |
| 9_98921779 | 2 | 0 | 5 | 0.4 | 2 | 0.5 | 1 | 0 | 10 | 0.3 | 2 | 0 | 3 | 0.6667 | 3 | 0.3333 | 1 | 0 | 9 | 0.3333 |
| 9_98921806 | 1 | 0 | 4 | 0.25 | 2 | 1 | 1 | 1 | 8 | 0.5 | 2 | 1 | 3 | 0.6667 | 2 | 1 | 1 | 0 | 8 | 0.75 |
| 9_98921819 | 1 | 1 | 4 | 1 | 2 | 1 | 1 | 1 | 8 | 1 | 2 | 1 | 3 | 1 | 1 | 0 | 1 | 1 | 7 | 0.8571 |
| 9_98955756 | 1 | 0 | 4 | 0.25 | 2 | 0 | 1 | 0 | 8 | 0.125 | 2 | 0.5 | 2 | 0 | NA | NA | 1 | 0 | 5 | 0.2 |
| 9_99145756 | 2 | 1 | 2 | 0.5 | 2 | 0.5 | 1 | 1 | 7 | 0.7143 | 2 | 0.5 | 1 | 0 | 2 | 0 | 3 | 0.3333 | 8 | 0.25 |
| 9_99181884 | 1 | 1 | 3 | 0 | 3 | 0 | NA | NA | 7 | 0.1429 | NA | NA | 2 | 0 | 1 | 1 | NA | NA | 3 | 0.3333 |
| 9_99182122 | 1 | 0 | 3 | 0.3333 | 3 | 0 | NA | NA | 7 | 0.1429 | 1 | 1 | 2 | 0.5 | 2 | 0.5 | 1 | 0 | 6 | 0.5 |
| 9_99183397 | NA | NA | NA | NA | 1 | 1 | NA | NA | 1 | 1 | 1 | 1 | NA | NA | NA | NA | NA | NA | 1 | 1 |
| 9_99269486 | 6 | 0 | 12 | 0 | 5 | 0.4 | 7 | 0.4286 | 30 | 0.1667 | 8 | 0.125 | 9 | 0.3333 | 11 | 0.0909 | 7 | 0 | 35 | 0.1429 |
| 9_99269589 | 3 | 0 | 8 | 0 | 2 | 0 | 7 | 0.4286 | 20 | 0.15 | 7 | 0 | 5 | 0 | 6 | 0.1667 | 4 | 0.25 | 22 | 0.0909 |
| 9_99269636 | 5 | 0.6 | 10 | 0.4 | 5 | 0.2 | 7 | 0.2857 | 27 | 0.3704 | 7 | 0.5714 | 6 | 0.6667 | 10 | 0.5 | 6 | 0.3333 | 29 | 0.5172 |
| 9_99270243 | 10 | 0.2 | 9 | 0 | 8 | 0 | 9 | 0 | 36 | 0.0556 | 6 | 0.3333 | 12 | 0.3333 | 11 | 0.0909 | 8 | 0 | 37 | 0.1892 |
| 9_99270352 | 5 | 1 | 7 | 0.8571 | 5 | 0.8 | 9 | 0.8889 | 26 | 0.8846 | 6 | 0.8333 | 12 | 0.4167 | 6 | 0.8333 | 6 | 0.8333 | 30 | 0.6667 |
| 9_99270479 | 10 | 0.2 | 8 | 0 | 7 | 0 | 8 | 0.125 | 33 | 0.0909 | 6 | 0.1667 | 9 | 0.1111 | 9 | 0 | 6 | 0 | 30 | 0.0667 |
| 9_99360848 | 2 | 0 | 6 | 0 | 3 | 0.3333 | 3 | 0.6667 | 14 | 0.2143 | 2 | 0 | 8 | 0.125 | 5 | 0.2 | 2 | 0 | 17 | 0.1176 |
| 9_99360925 | 3 | 0.3333 | 6 | 0.1667 | 2 | 0 | 4 | 0 | 15 | 0.1333 | 1 | 0 | 8 | 0 | 6 | 0.1667 | 2 | 0.5 | 17 | 0.1176 |
| 9_99361082 | 4 | 0 | 5 | 0 | 2 | 0.5 | 4 | 0 | 15 | 0.0667 | 2 | 0 | 7 | 0.1429 | 5 | 0 | 1 | 0 | 15 | 0.0667 |
| 9_99361105 | 4 | 0 | 5 | 0.2 | 2 | 0 | 4 | 0 | 15 | 0.0667 | 2 | 0 | 7 | 0.1429 | 5 | 0.2 | 2 | 0 | 16 | 0.125 |
| 9_99361114 | 1 | 0 | 6 | 0 | 2 | 0.5 | 2 | 0 | 11 | 0.0909 | 2 | 0 | 7 | 0 | 4 | 0.25 | 1 | 0 | 14 | 0.0714 |
| 9_99362528 | 2 | 0 | 4 | 0.25 | 1 | 0 | 5 | 0.4 | 12 | 0.25 | 3 | 0 | 5 | 0 | 5 | 0 | 3 | 0.3333 | 16 | 0.0625 |
| 9_99362562 | 2 | 1 | 4 | 0.75 | 2 | 1 | 4 | 1 | 12 | 0.9167 | 3 | 1 | 6 | 0.8333 | 5 | 1 | 5 | 1 | 19 | 0.9474 |
| 9_99362603 | 2 | 0.5 | 3 | 0.3333 | 2 | 1 | 5 | 0.8 | 12 | 0.6667 | 3 | 1 | 5 | 0.8 | 5 | 0.6 | 5 | 0.2 | 18 | 0.6111 |
| 9_99362635 | 1 | 0 | 3 | 0.3333 | 1 | 0 | 3 | 0 | 8 | 0.125 | 2 | 0.5 | 3 | 0.3333 | 4 | 0.25 | 4 | 0.25 | 13 | 0.3077 |
| 9_99362696 | 1 | 1 | NA | NA | NA | NA | NA | NA | 1 | 1 | 2 | 1 | 3 | 0.6667 | 3 | 1 | 3 | 1 | 11 | 0.9091 |
| 9_99401104 | 2 | 0.5 | 3 | 0.3333 | 2 | 0 | 2 | 1 | 9 | 0.4444 | 1 | 0 | 6 | 0.5 | 2 | 0 | 1 | 0 | 10 | 0.3 |
| 9_99401387 | 3 | 0.3333 | 4 | 0.5 | 2 | 0 | 1 | 1 | 10 | 0.4 | 1 | 0 | 5 | 0.8 | 2 | 0.5 | 1 | 0 | 9 | 0.5556 |
| 9_99402390 | 7 | 0.2857 | 7 | 0 | 1 | 0 | 5 | 0.4 | 20 | 0.2 | 2 | 1 | 6 | 1 | 5 | 0.4 | 4 | 0.5 | 17 | 0.7059 |
| 9_99402446 | 7 | 1 | 6 | 1 | 1 | 0 | 4 | 1 | 18 | 0.9444 | 2 | 1 | 6 | 1 | 3 | 1 | 2 | 0.5 | 13 | 0.9231 |
| 9_99465636 | 2 | 1 | 2 | 0.5 | 2 | 0 | 3 | 0.6667 | 9 | 0.5556 | 1 | 1 | 2 | 1 | 2 | 0.5 | NA | NA | 5 | 0.8 |
| 9_99465676 | 2 | 0 | 2 | 0 | 2 | 0.5 | 3 | 0.3333 | 9 | 0.2222 | 1 | 1 | NA | NA | 2 | 0 | NA | NA | 3 | 0.3333 |
| 9_99465775 | 2 | 0 | 2 | 0 | 1 | 0 | 4 | 0.5 | 9 | 0.2222 | 1 | 0 | 2 | 1 | 2 | 1 | NA | NA | 5 | 0.8 |
| 9_99465781 | 2 | 0.5 | 2 | 0.5 | 2 | 0.5 | 3 | 0.3333 | 9 | 0.4444 | 1 | 1 | 1 | 0 | 1 | 1 | NA | NA | 3 | 0.6667 |
| 9_99465850 | NA | NA | 3 | 0.6667 | 2 | 0.5 | 4 | 0.25 | 9 | 0.4444 | 1 | 1 | 2 | 0.5 | 2 | 0 | NA | NA | 5 | 0.4 |
| 9_99465952 | 2 | 1 | 3 | 1 | 2 | 1 | 3 | 1 | 10 | 1 | 1 | 1 | NA | NA | 2 | 1 | NA | NA | 3 | 1 |
| 9_99466078 | 2 | 0.5 | 3 | 0.3333 | 2 | 0 | 3 | 0.3333 | 10 | 0.3 | 1 | 1 | 1 | 1 | 1 | 1 | NA | NA | 3 | 1 |
| 9_99466089 | 2 | 1 | 1 | 1 | 1 | 1 | 4 | 1 | 8 | 1 | 1 | 1 | 2 | 1 | 2 | 1 | NA | NA | 5 | 1 |
| 9_99466158 | 2 | 1 | 2 | 1 | 2 | 1 | 3 | 0.6667 | 9 | 0.8889 | 2 | 1 | 2 | 1 | 2 | 1 | NA | NA | 6 | 1 |
| 9_99466349 | 2 | 0 | 3 | 0 | 3 | 0 | 3 | 1 | 11 | 0.2727 | 1 | 0 | 2 | 1 | 2 | 0.5 | 1 | 1 | 6 | 0.6667 |
| 9_99519779 | NA | NA | 1 | 1 | NA | NA | NA | NA | 1 | 1 | 1 | 1 | 2 | 1 | 1 | 0 | NA | NA | 4 | 0.75 |
| 9_99550081 | 2 | 0.5 | 2 | 0.5 | 3 | 0 | 1 | 1 | 8 | 0.375 | 2 | 0 | 3 | 0.3333 | 3 | 0.6667 | 2 | 0.5 | 10 | 0.4 |
| 9_99550216 | 1 | 1 | 2 | 1 | 2 | 0 | 1 | 1 | 6 | 0.6667 | 2 | 0.5 | 2 | 0 | 3 | 0 | 2 | 0.5 | 9 | 0.2222 |
| 9_99550795 | 2 | 1 | 2 | 0.5 | 6 | 0.5 | 1 | 1 | 11 | 0.6364 | 2 | 0.5 | 4 | 0.25 | 4 | 0.75 | 1 | 0 | 11 | 0.4545 |
| 9_99551013 | 2 | 0.5 | 1 | 0 | 2 | 0 | 1 | 1 | 6 | 0.3333 | 2 | 0 | 4 | 0.25 | 2 | 0.5 | 2 | 0.5 | 10 | 0.3 |
| 9_10037712 | 4 | 0 | 5 | 0.2 | 6 | 0 | 1 | 0 | 16 | 0.0625 | 4 | 0.25 | 3 | 0.6667 | 2 | 0 | 2 | 0.5 | 11 | 0.3636 |
| 9_10037741 | 2 | 0 | 2 | 0.5 | 6 | 0.1667 | 1 | 0 | 11 | 0.1818 | 4 | 0.25 | 3 | 0.3333 | 2 | 0.5 | 2 | 1 | 11 | 0.4545 |
| 9_10037747 | 2 | 1 | 4 | 0.5 | 5 | 1 | 1 | 1 | 12 | 0.8333 | 4 | 0.75 | 2 | 0.5 | 1 | 1 | 1 | 1 | 8 | 0.75 |
| 9_10037751 | 2 | 1 | 3 | 1 | 5 | 1 | NA | NA | 10 | 1 | 4 | 1 | 1 | 1 | 2 | 1 | 2 | 1 | 9 | 1 |
| 9_10037760 | 2 | 1 | 4 | 0 | 3 | 0 | 1 | 0 | 10 | 0.2 | 2 | 1 | 1 | 0 | 2 | 0.5 | 1 | 0 | 6 | 0.5 |

| | | | | | | | | | | | | | | | | | | | | |
|---|---|---|---|---|---|---|---|---|---|---|---|---|---|---|---|---|---|---|---|---|
| 9_10037761 | 2 | 0.5 | 3 | 0 | 6 | 0.5 | 1 | 0 | 12 | 0.3333 | 4 | 0.5 | 3 | 0 | 2 | 0 | 2 | 0 | 11 | 0.1818 |
| 9_10037765 | 2 | 1 | 4 | 1 | 6 | 1 | 1 | 1 | 13 | 1 | 4 | 1 | 3 | 1 | 2 | 1 | 2 | 1 | 11 | 1 |
| 9_10037947 | 1 | 0 | 4 | 0 | 5 | 0.2 | 1 | 0 | 11 | 0.0909 | 4 | 0.25 | 3 | 0.3333 | 1 | 0 | 2 | 0 | 10 | 0.2 |
| 9_10037949 | 1 | 0 | 3 | 0.6667 | 3 | 0 | NA | NA | 7 | 0.2857 | 3 | 0 | 3 | 0.3333 | 1 | 0 | 2 | 0.5 | 9 | 0.2222 |
| 9_10037951 | 1 | 1 | 4 | 0.25 | 5 | 0.4 | 1 | 1 | 11 | 0.4545 | 3 | 0.3333 | 3 | 0.3333 | 1 | 1 | 1 | 0 | 8 | 0.375 |
| 9_10037962 | 1 | 1 | 3 | 0.6667 | 5 | 0.8 | NA | NA | 9 | 0.7778 | 4 | 0.75 | 3 | 1 | 2 | 1 | 1 | 1 | 10 | 0.9 |
| 9_10037975 | 1 | 0 | 4 | 0.25 | 2 | 0 | 1 | 0 | 8 | 0.125 | 3 | 0 | 2 | 0.5 | 2 | 0.5 | 1 | 0 | 8 | 0.25 |
| 9_10037976 | 1 | 1 | 4 | 0 | 5 | 0.2 | 1 | 0 | 11 | 0.1818 | 3 | 0.6667 | 1 | 0 | NA | NA | NA | NA | 4 | 0.5 |
| 9_10037980 | 1 | 1 | 3 | 0.6667 | 4 | 0.75 | 1 | 0 | 9 | 0.6667 | 3 | 1 | 3 | 0.3333 | 2 | 0.5 | 2 | 1 | 10 | 0.7 |
| 9_10037985 | 1 | 0 | 3 | 0.3333 | 5 | 0.8 | 1 | 0 | 10 | 0.5 | 4 | 0.5 | 3 | 0.6667 | 2 | 0.5 | 2 | 0.5 | 11 | 0.5455 |
| 9_10043444 | 2 | 1 | 5 | 0.8 | 5 | 1 | 3 | 1 | 15 | 0.9333 | 4 | 0.75 | 3 | 1 | 1 | 1 | 3 | 1 | 11 | 0.9091 |
| 9_10043465 | 3 | 1 | 5 | 1 | 6 | 1 | 1 | 1 | 15 | 1 | 3 | 0.6667 | 1 | 1 | 1 | 1 | 1 | 1 | 6 | 0.8333 |
| 9_10043478 | 2 | 0 | 4 | 0 | 4 | 0.25 | 2 | 0.5 | 12 | 0.1667 | 3 | 0.3333 | 2 | 0.5 | NA | NA | 3 | 0 | 8 | 0.25 |
| 9_10043479 | 2 | 1 | 1 | 1 | 5 | 1 | 2 | 1 | 10 | 1 | 2 | 1 | 2 | 1 | 1 | 1 | 2 | 1 | 7 | 1 |
| 9_10043481 | 3 | 0.6667 | 4 | 0.5 | 6 | 0.1667 | 1 | 0 | 14 | 0.3571 | 3 | 0.3333 | 2 | 0.5 | 1 | 0 | 3 | 0.6667 | 9 | 0.4444 |
| 9_10043488 | 3 | 1 | 3 | 1 | 5 | 1 | 1 | 1 | 12 | 1 | 3 | 1 | 2 | 1 | 1 | 1 | 3 | 1 | 9 | 1 |
| 9_10043493 | 3 | 1 | 4 | 0.75 | 5 | 1 | 2 | 1 | 14 | 0.9286 | 3 | 1 | 2 | 1 | NA | NA | 3 | 1 | 8 | 1 |
| 9_10043500 | 3 | 0.3333 | 4 | 0 | 6 | 0.5 | 2 | 1 | 15 | 0.4 | 3 | 0 | 2 | 0 | 1 | 0 | 3 | 0.3333 | 9 | 0.1111 |
| 9_10043508 | 3 | 1 | 3 | 0.3333 | 5 | 0.8 | 1 | 1 | 12 | 0.75 | 3 | 1 | 2 | 1 | 1 | 1 | 2 | 0.5 | 8 | 0.875 |
| 9_10043526 | 1 | 1 | 3 | 1 | 3 | 1 | 1 | 1 | 8 | 1 | 2 | 1 | 2 | 1 | 1 | 1 | 2 | 1 | 7 | 1 |
| 9_10043538 | 2 | 0 | 2 | 0 | 2 | 0.5 | NA | NA | 6 | 0.1667 | 1 | 1 | NA | NA | 1 | 1 | 2 | 0.5 | 4 | 0.75 |
| 9_10043540 | 2 | 1 | 3 | 1 | 5 | 0.8 | 2 | 1 | 12 | 0.9167 | 3 | 0.6667 | 2 | 1 | 1 | 1 | 3 | 1 | 9 | 0.8889 |
| 9_10043545 | 2 | 0.5 | 4 | 0.75 | 4 | 1 | 2 | 1 | 12 | 0.8333 | 3 | 0.6667 | 2 | 1 | 1 | 1 | 3 | 0.6667 | 9 | 0.7778 |
| 9_10043547 | 2 | 0.5 | 4 | 0.5 | 4 | 0.25 | 2 | 0.5 | 12 | 0.4167 | 3 | 0 | 2 | 0 | 1 | 0 | 3 | 0.3333 | 9 | 0.1111 |
| 9_10083881 | 1 | 0 | 4 | 0.5 | 2 | 0 | 2 | 0 | 9 | 0.2222 | 2 | 0 | 2 | 0 | 4 | 0.25 | 1 | 0 | 9 | 0.1111 |
| 9_10083885 | 1 | 1 | 4 | 0.25 | 2 | 0 | 2 | 0.5 | 9 | 0.3333 | 1 | 0 | 1 | 0 | 3 | 0.3333 | NA | NA | 5 | 0.2 |
| 9_10084025 | 1 | 1 | 4 | 1 | 2 | 0.5 | 2 | 0.5 | 9 | 0.7778 | 2 | 0.5 | 1 | 1 | 2 | 0 | 1 | 0 | 6 | 0.3333 |
| 9_10086463 | 4 | 0.75 | 10 | 0.6 | 8 | 0.625 | 1 | 1 | 23 | 0.6522 | 7 | 0.5714 | 6 | 0.3333 | 9 | 0.6667 | 8 | 0.5 | 30 | 0.5333 |
| 9_10086472 | 4 | 0.5 | 11 | 0.3636 | 9 | 0.2222 | 2 | 1 | 26 | 0.3846 | 5 | 0.4 | 6 | 0.3333 | 9 | 0 | 7 | 0 | 27 | 0.1481 |
| 9_10095691 | 1 | 0 | 1 | 0 | 1 | 1 | 1 | 0 | 4 | 0.25 | 1 | 1 | NA | NA | 1 | 1 | NA | NA | 2 | 1 |
| 9_10095698 | 1 | 0 | 1 | 1 | 1 | 0 | 1 | 1 | 4 | 0.5 | 2 | 0.5 | NA | NA | 1 | 0 | NA | NA | 3 | 0.3333 |
| 9_10095707 | 2 | 1 | 1 | 0 | 1 | 1 | NA | NA | 4 | 0.75 | 2 | 1 | NA | NA | 1 | 1 | 1 | 1 | 4 | 1 |
| 9_10095712 | 3 | 0.6667 | 1 | 1 | 1 | 1 | 1 | 1 | 6 | 0.8333 | 2 | 1 | NA | NA | 1 | 1 | 3 | 1 | 6 | 1 |
| 9_10095727 | 1 | 0 | 1 | 1 | 1 | 1 | 1 | 0 | 4 | 0.5 | 2 | 1 | NA | NA | 2 | 1 | 4 | 0.5 | 8 | 0.75 |
| 9_10097788 | 2 | 1 | 1 | 1 | 1 | 1 | 1 | 1 | 5 | 1 | 2 | 1 | NA | NA | 2 | 1 | 4 | 1 | 8 | 1 |
| 9_10097794 | 1 | 1 | 1 | 1 | 1 | 1 | 1 | 1 | 4 | 1 | 3 | 1 | NA | NA | 2 | 1 | 4 | 0.75 | 9 | 0.8889 |
| 9_10097799 | NA | NA | 1 | 1 | 1 | 1 | 1 | 1 | 3 | 1 | 1 | 1 | NA | NA | 2 | 0.5 | 4 | 1 | 7 | 0.8571 |
| 9_10097830 | 1 | 1 | NA | NA | 1 | 1 | 1 | 1 | 3 | 1 | 2 | 1 | NA | NA | 2 | 0.5 | 3 | 0.6667 | 7 | 0.7143 |
| 9_10097836 | 1 | 1 | 1 | 1 | 1 | 1 | NA | NA | 3 | 1 | 3 | 0.6667 | NA | NA | 2 | 0.5 | 5 | 0.6 | 10 | 0.6 |
| 9_10097841 | 1 | 1 | 1 | 1 | 1 | 1 | 1 | 1 | 4 | 1 | 2 | 1 | NA | NA | 2 | 0.5 | 3 | 0.6667 | 7 | 0.7143 |
| 9_10097846 | 1 | 1 | 1 | 1 | 1 | 1 | 1 | 1 | 4 | 1 | 3 | 1 | NA | NA | 2 | 1 | 4 | 1 | 9 | 1 |
| 9_10097853 | 1 | 1 | 1 | 1 | 1 | 1 | 1 | 1 | 4 | 1 | 2 | 1 | NA | NA | 1 | 1 | 4 | 1 | 7 | 1 |
| 9_10097867 | 3 | 0.3333 | NA | NA | 1 | 0 | 2 | 0.5 | 6 | 0.3333 | 2 | 0.5 | NA | NA | 2 | 0.5 | 5 | 0.2 | 9 | 0.3333 |
| 9_10097874 | 2 | 0.5 | 1 | 0 | 1 | 1 | 2 | 1 | 6 | 0.6667 | 3 | 1 | NA | NA | 2 | 1 | 4 | 0.75 | 9 | 0.8889 |
| 9_10097882 | 3 | 0.6667 | 1 | 1 | 1 | 1 | 2 | 1 | 7 | 0.8571 | 3 | 1 | NA | NA | 1 | 1 | 5 | 0.8 | 9 | 0.8889 |
| 9_10097886 | 3 | 0.3333 | 1 | 0 | 1 | 0 | 2 | 0.5 | 7 | 0.2857 | 2 | 0 | NA | NA | 3 | 0.3333 | 5 | 0.4 | 10 | 0.3 |
| 9_10097891 | 3 | 0.6667 | 1 | 1 | 1 | 1 | 2 | 1 | 7 | 0.8571 | 2 | 1 | NA | NA | 3 | 0.6667 | 5 | 0.8 | 10 | 0.8 |
| 9_10097897 | 3 | 1 | 1 | 1 | 1 | 1 | 2 | 0.5 | 7 | 0.8571 | 2 | 1 | NA | NA | 3 | 1 | 5 | 1 | 10 | 1 |
| 9_10097912 | 4 | 0.5 | 1 | 0 | 1 | 0 | 2 | 1 | 8 | 0.5 | 3 | 0.6667 | NA | NA | 3 | 0 | 5 | 0.4 | 11 | 0.3636 |
| 9_10097922 | 4 | 1 | NA | NA | 1 | 1 | 1 | 1 | 6 | 1 | 3 | 1 | NA | NA | 3 | 1 | 5 | 1 | 11 | 1 |
| 9_10097962 | 3 | 0.3333 | 1 | 1 | 2 | 0 | 3 | 0.6667 | 9 | 0.4444 | 3 | 1 | NA | NA | 3 | 0.3333 | 2 | 0.5 | 8 | 0.625 |
| 9_10098061 | 8 | 0.125 | 3 | 0.6667 | 3 | 0 | 4 | 0.5 | 18 | 0.2778 | 6 | 0.1667 | 3 | 0 | 2 | 0 | 7 | 0.5714 | 18 | 0.2778 |
| 9_10098081 | 6 | 0 | 3 | 0.3333 | 4 | 0 | 4 | 0 | 17 | 0.0588 | 6 | 0 | 1 | 1 | 2 | 0.5 | 6 | 0 | 15 | 0.1333 |
| 9_10098143 | 5 | 0.6 | NA | NA | 2 | 0 | 4 | 0.75 | 11 | 0.5455 | 4 | 0.75 | 2 | 0.5 | NA | NA | 3 | 0.6667 | 9 | 0.6667 |
| 9_10098177 | NA | NA | 1 | 0 | 1 | 1 | 2 | 0.5 | 4 | 0.5 | 2 | 0.5 | NA | NA | NA | NA | 1 | 0 | 3 | 0.3333 |
| 9_10098277 | 3 | 0 | 3 | 0 | 5 | 0.6 | 6 | 0 | 17 | 0.1765 | 5 | 0.2 | 1 | 0 | 3 | 0.3333 | 6 | 0.5 | 15 | 0.3333 |
| 9_10098282 | 2 | 0 | 2 | 0 | 3 | 0 | 7 | 0.2857 | 14 | 0.1429 | 1 | 1 | 1 | 0 | 4 | 0 | 5 | 0 | 11 | 0.0909 |
| 9_10098298 | 1 | 1 | 4 | 0 | 5 | 0.4 | 7 | 0.1429 | 17 | 0.2353 | 3 | 0.3333 | 1 | 0 | 3 | 0 | 6 | 0 | 13 | 0.0769 |
| 9_10098344 | NA | NA | 4 | 0 | 4 | 0.25 | 5 | 0.4 | 13 | 0.2308 | 4 | 0.25 | NA | NA | 3 | 0 | 6 | 0.3333 | 13 | 0.2308 |
| 9_10098391 | 2 | 0 | 4 | 0 | 4 | 0 | 6 | 0.1667 | 16 | 0.0625 | 3 | 0 | 1 | 0 | 3 | 0.3333 | 7 | 0.1429 | 14 | 0.1429 |
| 9_10098393 | 2 | 0.5 | 3 | 0 | 4 | 0.5 | 6 | 0.5 | 15 | 0.4 | 3 | 0.3333 | 1 | 0 | 3 | 0 | 7 | 0.4286 | 14 | 0.2857 |
| 9_10098420 | 1 | 1 | 4 | 0 | 3 | 0.3333 | 6 | 0.3333 | 14 | 0.2857 | 3 | 0.3333 | 1 | 0 | 4 | 0 | 6 | 0.1667 | 14 | 0.1429 |
| 9_10098426 | 2 | 0 | 4 | 0.25 | 3 | 0 | 6 | 0.5 | 15 | 0.2667 | 2 | 0.5 | 1 | 1 | 3 | 0 | 7 | 0.1429 | 13 | 0.2308 |
| 9_10098440 | 1 | 0 | 3 | 0 | 2 | 0 | 5 | 0.2 | 11 | 0.0909 | 3 | 0.3333 | 2 | 0.5 | 2 | 0 | 6 | 0 | 13 | 0.1538 |
| 9_10098442 | 1 | 1 | 4 | 0.75 | 3 | 1 | 6 | 0.6667 | 14 | 0.7857 | 4 | 0.75 | 2 | 0.5 | 3 | 0.3333 | 7 | 0.7143 | 16 | 0.625 |

| | | | | | | | | | | | | | | | | | | | | |
|---|---|---|---|---|---|---|---|---|---|---|---|---|---|---|---|---|---|---|---|---|
| 9_10098443 | 1 | 1 | 3 | 0.3333 | 3 | 0.6667 | 6 | 0.5 | 13 | 0.5385 | 3 | 0.6667 | 2 | 0.5 | 1 | 0 | 4 | 0.25 | 10 | 0.4 |
| 9_10098451 | 1 | 1 | 4 | 0 | 3 | 0 | 6 | 0.1667 | 14 | 0.1429 | 4 | 0.25 | 2 | 0 | 2 | 0 | 5 | 0 | 13 | 0.0769 |
| 9_10098455 | 1 | 1 | 2 | 0.5 | 3 | 0.6667 | 4 | 0.75 | 10 | 0.7 | 3 | 0.6667 | 1 | 0 | 3 | 0 | 5 | 0.8 | 12 | 0.5 |
| 9_10098456 | 1 | 1 | 3 | 0.6667 | 3 | 0.6667 | 6 | 0.3333 | 13 | 0.5385 | 2 | 0.5 | 2 | 1 | 3 | 0.3333 | 5 | 0.4 | 12 | 0.5 |
| 9_10098459 | 1 | 0 | 4 | 0 | 2 | 0.5 | 5 | 0.2 | 12 | 0.1667 | 4 | 0 | 1 | 1 | 2 | 0 | 5 | 0 | 12 | 0.0833 |
| 9_10098463 | 1 | 1 | 2 | 0 | 3 | 1 | 4 | 0.25 | 10 | 0.5 | 3 | 1 | 1 | 1 | 3 | 0 | 6 | 0.5 | 13 | 0.5385 |
| 9_10098465 | 2 | 0.5 | 2 | 0.5 | 2 | 0.5 | 4 | 0.25 | 10 | 0.4 | 3 | 0.6667 | 1 | 1 | 3 | 0.3333 | 6 | 0.5 | 13 | 0.5385 |
| 9_10098477 | 1 | 1 | 3 | 0 | 3 | 0 | 6 | 0 | 13 | 0.0769 | 3 | 0.3333 | 1 | 1 | 3 | 0 | 6 | 0 | 13 | 0.1538 |
| 9_10098479 | 1 | 0 | 1 | 0 | 3 | 0.6667 | 6 | 0 | 11 | 0.1818 | 4 | 0 | 1 | 0 | 3 | 0 | 6 | 0.1667 | 14 | 0.0714 |
| 9_10098494 | 1 | 0 | 2 | 0.5 | 2 | 0.5 | 6 | 0.5 | 11 | 0.4545 | 3 | 0 | 1 | 0 | 3 | 0 | 6 | 0.3333 | 13 | 0.1538 |
| 9_10098512 | 1 | 0 | 3 | 0 | 3 | 0 | 6 | 0.1667 | 13 | 0.0769 | 3 | 0.3333 | 1 | 0 | 3 | 0 | 5 | 0 | 12 | 0.0833 |
| 9_10107146 | 1 | 0 | 2 | 0 | 1 | 1 | 4 | 0.25 | 8 | 0.25 | NA | NA | 1 | 0 | 1 | 1 | 3 | 0 | 5 | 0.2 |
| 9_10108858 | 1 | 0 | 2 | 1 | 2 | 0 | 3 | 1 | 8 | 0.625 | NA | NA | NA | NA | 1 | 1 | 1 | 1 | 2 | 1 |
| 9_10108868 | NA | NA | 2 | 0 | 2 | 0.5 | 3 | 0.3333 | 7 | 0.2857 | NA | NA | 1 | 1 | 1 | 1 | 2 | 0 | 4 | 0.5 |
| 9_10108901 | 1 | 1 | 2 | 1 | 2 | 0.5 | 3 | 0.3333 | 8 | 0.625 | NA | NA | 1 | 1 | 1 | 0 | 2 | 0.5 | 4 | 0.5 |
| 9_10108916 | 1 | 0 | 1 | 0 | 2 | 0.5 | 2 | 1 | 6 | 0.5 | NA | NA | 1 | 1 | 1 | 1 | 1 | 0 | 3 | 0.6667 |
| 9_10108923 | 1 | 1 | 1 | 1 | 2 | 0.5 | 3 | 0.3333 | 7 | 0.5714 | NA | NA | 1 | 1 | 1 | 1 | NA | NA | 2 | 1 |
| 9_10108928 | 1 | 0 | 1 | 1 | 2 | 0.5 | 3 | 0.3333 | 7 | 0.4286 | NA | NA | 1 | 0 | 1 | 1 | 2 | 0.5 | 4 | 0.5 |
| 9_10108934 | NA | NA | 1 | 1 | 2 | 0.5 | 3 | 0.3333 | 6 | 0.5 | NA | NA | 1 | 1 | 1 | 1 | 1 | 0 | 3 | 0.6667 |
| 9_10108939 | 1 | 0 | 1 | 0 | 1 | 0 | 3 | 0.3333 | 6 | 0.1667 | NA | NA | 1 | 0 | 1 | 0 | 2 | 0.5 | 4 | 0.25 |
| 9_10108962 | NA | NA | 1 | 0 | 2 | 0.5 | 3 | 0.3333 | 6 | 0.3333 | NA | NA | 1 | 1 | NA | NA | 2 | 0.5 | 3 | 0.6667 |
| 9_10108964 | NA | NA | NA | NA | 2 | 0.5 | 3 | 0.6667 | 5 | 0.6 | NA | NA | 1 | 0 | 1 | 1 | 2 | 0.5 | 4 | 0.5 |
| 9_10108993 | NA | NA | NA | NA | 2 | 0.5 | 3 | 0.3333 | 5 | 0.4 | NA | NA | NA | NA | 1 | 1 | 2 | 0 | 3 | 0.3333 |
| 9_10108999 | NA | NA | NA | NA | 2 | 1 | 3 | 0.6667 | 5 | 0.8 | NA | NA | NA | NA | NA | NA | 2 | 0.5 | 2 | 0.5 |
| 9_10109019 | NA | NA | NA | NA | 2 | 0.5 | 2 | 0 | 4 | 0.25 | NA | NA | NA | NA | NA | NA | 2 | 1 | 2 | 1 |
| 9_10109030 | NA | NA | NA | NA | 1 | 1 | 2 | 1 | 3 | 1 | NA | NA | NA | NA | 1 | 0 | 2 | 0.5 | 3 | 0.3333 |
| 9_10109069 | NA | NA | NA | NA | 2 | 0.5 | 2 | 0 | 4 | 0.25 | NA | NA | NA | NA | 1 | 1 | 2 | 0 | 3 | 0.3333 |
| 9_10109632 | NA | NA | NA | NA | NA | NA | 1 | 1 | 1 | 1 | NA | NA | NA | NA | NA | NA | 1 | 1 | 1 | 1 |
| 9_10179954 | 4 | 1 | 6 | 1 | 2 | 1 | 1 | 1 | 13 | 1 | 2 | 1 | 2 | 1 | 4 | 1 | 3 | 1 | 11 | 1 |
| 9_10179956 | 4 | 0.75 | 6 | 0.8333 | 3 | 1 | 1 | 0 | 14 | 0.7857 | 2 | 0.5 | 1 | 1 | 5 | 0.8 | 2 | 0.5 | 10 | 0.7 |
| 9_10179967 | 4 | 0 | 6 | 0.1667 | 2 | 0 | 1 | 0 | 13 | 0.0769 | 2 | 0.5 | 2 | 0 | 6 | 0.1667 | 2 | 0 | 12 | 0.1667 |
| 9_10179992 | 4 | 1 | 4 | 1 | 2 | 1 | 1 | 1 | 11 | 1 | 2 | 1 | 2 | 1 | 5 | 1 | 3 | 1 | 12 | 1 |
| 9_10180013 | 4 | 1 | 6 | 1 | 3 | 1 | 1 | 1 | 14 | 1 | 2 | 1 | 2 | 1 | 4 | 1 | 1 | 1 | 9 | 1 |
| 9_10180020 | 4 | 0.25 | 6 | 0.1667 | 4 | 0 | 1 | 0 | 15 | 0.1333 | 2 | 0.5 | 2 | 0.5 | 5 | 0 | 2 | 0.5 | 11 | 0.2727 |
| 9_10180028 | 4 | 0.5 | 6 | 0.6667 | 4 | 0.75 | NA | NA | 14 | 0.6429 | 1 | 1 | 3 | 0.6667 | 4 | 0.25 | 3 | 0.6667 | 11 | 0.5455 |
| 9_10180032 | 4 | 0.25 | 6 | 0.1667 | 4 | 0.25 | 1 | 0 | 15 | 0.2 | 1 | 0 | 3 | 0.3333 | 5 | 0 | 4 | 0 | 13 | 0.0769 |
| 9_10180037 | 3 | 0.6667 | 5 | 0.6 | 3 | 0.6667 | 2 | 0 | 13 | 0.5385 | 1 | 1 | 3 | 0.6667 | 5 | 0.6 | 4 | 0.75 | 13 | 0.6923 |
| 9_10180040 | 4 | 0 | 5 | 0.2 | 4 | 0.25 | 2 | 0.5 | 15 | 0.2 | 1 | 1 | 3 | 0.3333 | 5 | 0.2 | 3 | 0 | 12 | 0.25 |
| 9_10180046 | 4 | 0.5 | 6 | 0.8333 | 3 | 0.6667 | 1 | 0 | 14 | 0.6429 | 1 | 1 | 2 | 0.5 | 4 | 1 | 4 | 0.75 | 11 | 0.8182 |
| 9_10180054 | 4 | 1 | 6 | 0.8333 | 3 | 1 | 1 | 1 | 14 | 0.9286 | NA | NA | 3 | 1 | 5 | 0.8 | 2 | 1 | 10 | 0.9 |
| 9_10180054 | 2 | 1 | 4 | 1 | 2 | 1 | 1 | 0 | 9 | 0.8889 | 1 | 1 | 1 | 1 | 2 | 1 | 2 | 0 | 6 | 0.6667 |
| 9_10180064 | 4 | 0.5 | 6 | 0.1667 | 4 | 0.25 | 1 | 1 | 15 | 0.3333 | 1 | 1 | 2 | 1 | 3 | 0.3333 | 4 | 0.5 | 10 | 0.6 |
| 9_10180068 | 4 | 0 | 6 | 1 | 4 | 0.25 | 2 | 0.5 | 16 | 0.5 | 1 | 0 | 3 | 0.3333 | 4 | 0.25 | 2 | 0.5 | 10 | 0.3 |
| 9_10180661 | 4 | 0.25 | 6 | 0.3333 | 4 | 0.25 | 1 | 0 | 15 | 0.2667 | 1 | 1 | 2 | 1 | 4 | 0 | 3 | 0 | 10 | 0.3 |
| 9_10191834 | 2 | 0.5 | 3 | 0 | NA | NA | 1 | 0 | 6 | 0.1667 | 1 | 1 | 1 | 0 | 1 | 0 | 2 | 0.5 | 5 | 0.4 |
| 9_10207206 | 2 | 1 | 3 | 0.3333 | 1 | 0 | NA | NA | 6 | 0.5 | 1 | 0 | NA | NA | 1 | 1 | 2 | 0 | 4 | 0.25 |
| 9_10207215 | 2 | 0.5 | 3 | 0 | 1 | 0 | 2 | 0 | 8 | 0.125 | 1 | 0 | NA | NA | 2 | 0 | 2 | 0.5 | 5 | 0.2 |
| 9_10207236 | 2 | 0.5 | 3 | 0.6667 | 1 | 0 | 1 | 0 | 7 | 0.4286 | 1 | 1 | NA | NA | 2 | 0 | 2 | 0.5 | 5 | 0.4 |
| 9_10207242 | 2 | 0.5 | 2 | 0.5 | NA | NA | 1 | 0 | 5 | 0.4 | 1 | 1 | NA | NA | 2 | 0.5 | 2 | 0 | 5 | 0.4 |
| 9_10207242 | 2 | 0 | 3 | 1 | NA | NA | 1 | 0 | 6 | 0.5 | 1 | 0 | NA | NA | 2 | 1 | 2 | 1 | 5 | 0.8 |
| 9_10207249 | 1 | 1 | 3 | 0.6667 | 1 | 0 | 1 | 0 | 6 | 0.5 | 1 | 1 | NA | NA | 2 | 1 | 2 | 0.5 | 5 | 0.8 |
| 9_10242309 | 1 | 1 | NA | NA | NA | NA | NA | NA | 1 | 1 | NA | NA | NA | NA | 1 | 0 | 1 | 1 | 2 | 0.5 |
| 9_10246389 | 1 | 0 | 2 | 0 | 1 | 0 | 2 | 1 | 6 | 0.3333 | 2 | 0.5 | 4 | 0 | 9 | 0.2222 | 7 | 0.2857 | 22 | 0.2273 |
| 9_10246391 | 1 | 0 | 5 | 0.4 | 2 | 0 | 2 | 0.5 | 10 | 0.3 | 2 | 0.5 | 6 | 0.3333 | 8 | 0.25 | 6 | 0.5 | 22 | 0.3636 |
| 9_10246393 | 1 | 0 | 5 | 0.8 | 2 | 1 | 2 | 1 | 10 | 0.8 | 1 | 1 | 6 | 0.8333 | 8 | 0.875 | 8 | 0.875 | 23 | 0.8696 |
| 9_10246396 | 1 | 0 | 5 | 0.4 | 2 | 0.5 | 2 | 0.5 | 10 | 0.4 | NA | NA | 3 | 0 | 9 | 0.3333 | 7 | 0.5714 | 19 | 0.3684 |
| 9_10246410 | 1 | 0 | 5 | 0.2 | 2 | 0 | 1 | 1 | 9 | 0.2222 | 2 | 0 | 5 | 0.2 | 9 | 0 | 8 | 0.25 | 24 | 0.125 |
| 9_10251128 | 5 | 0 | 7 | 0.1429 | 4 | 0.25 | 3 | 0.3333 | 19 | 0.1579 | 6 | 0 | 2 | 0 | 6 | 0.1667 | 2 | 0.5 | 16 | 0.125 |
| 9_10259742 | 2 | 1 | NA | NA | NA | NA | NA | NA | 2 | 1 | 1 | 1 | NA | NA | NA | NA | 1 | 0 | 2 | 0.5 |
| 9_10259752 | 2 | 0.5 | NA | NA | NA | NA | NA | NA | 2 | 0.5 | 1 | 0 | NA | NA | NA | NA | 1 | 1 | 2 | 0.5 |
| 9_10260907 | 3 | 1 | 4 | 1 | 4 | 1 | 1 | 1 | 12 | 1 | 1 | 1 | 3 | 1 | 2 | 1 | 1 | 1 | 7 | 1 |
| 9_10260933 | 5 | 0.4 | 5 | 0.4 | 3 | 0.6667 | 1 | 0 | 14 | 0.4286 | 1 | 0 | 3 | 0.6667 | 1 | 1 | 1 | 0 | 6 | 0.5 |
| 9_10260935 | 2 | 1 | 3 | 1 | 4 | 0.75 | 2 | 1 | 11 | 0.9091 | 1 | 1 | 3 | 1 | 2 | 0.5 | 1 | 1 | 7 | 0.8571 |
| 9_10260949 | 5 | 0.4 | 5 | 0 | 4 | 0 | 2 | 0 | 16 | 0.125 | NA | NA | 2 | 0 | 2 | 0.5 | 1 | 0 | 5 | 0.2 |
| 9_10260972 | 4 | 1 | 4 | 0.75 | 4 | 0.5 | 1 | 1 | 13 | 0.7692 | 1 | 0 | 2 | 0.5 | 2 | 0.5 | 1 | 1 | 6 | 0.5 |

| | | | | | | | | | | | | | | | | | | | | |
|---|---|---|---|---|---|---|---|---|---|---|---|---|---|---|---|---|---|---|---|---|
| 9_1026097 | 3 | 0.6667 | 4 | 0.5 | 4 | 0.75 | 2 | 0.5 | 13 | 0.6154 | NA | NA | 2 | 0.5 | 2 | 1 | 1 | 0 | 5 | 0.6 |
| 9_1026097 | 3 | 1 | 4 | 0.75 | 5 | 0.8 | 1 | 0 | 13 | 0.7692 | NA | NA | 3 | 1 | 3 | 0.3333 | 1 | 1 | 7 | 0.7143 |
| 9_1026098 | 5 | 0.2 | 4 | 0.25 | 5 | 0 | 2 | 0 | 16 | 0.125 | 1 | 0 | 3 | 0.3333 | 3 | 0 | 1 | 0 | 8 | 0.125 |
| 9_1026098 | 5 | 1 | 6 | 0.8333 | 4 | 0.75 | 2 | 0.5 | 17 | 0.8235 | 1 | 0 | 3 | 1 | 2 | 0.5 | 1 | 0 | 7 | 0.5714 |
| 9_1026099 | 5 | 1 | 5 | 1 | 4 | 1 | 2 | 0.5 | 16 | 0.9375 | 1 | 1 | 2 | 1 | 2 | 0.5 | 1 | 1 | 6 | 0.8333 |
| 9_1026100 | 5 | 1 | 5 | 1 | 4 | 1 | 1 | 1 | 15 | 1 | 1 | 1 | 2 | 1 | 1 | 0 | 2 | 1 | 6 | 0.8333 |
| 9_1026100 | 5 | 0 | 6 | 0.1667 | 5 | 0.2 | 2 | 0.5 | 18 | 0.1667 | 1 | 0 | 3 | 0.3333 | 3 | 0.3333 | 1 | 0 | 8 | 0.25 |
| 9_1027759 | 5 | 0 | 4 | 0 | 8 | 0 | 7 | 0.5714 | 24 | 0.1667 | 1 | 0 | 5 | 0.2 | 3 | 0 | 5 | 0 | 14 | 0.0714 |
| 9_1027759 | 6 | 0.1667 | 3 | 0.3333 | 6 | 0 | 6 | 0.3333 | 21 | 0.1905 | 1 | 0 | 1 | 0 | 3 | 0 | 5 | 0.2 | 10 | 0.1 |
| 9_1027833 | 6 | 0.1667 | 3 | 0.6667 | 7 | 0 | 6 | 0 | 22 | 0.1364 | 1 | 0 | 3 | 0.3333 | 6 | 0 | 4 | 0.75 | 14 | 0.2857 |
| 9_1027844 | 6 | 0.1667 | 3 | 0.3333 | 8 | 0.125 | 7 | 0.4286 | 24 | 0.25 | NA | NA | 5 | 0.4 | 5 | 0.6 | 7 | 0.5714 | 17 | 0.5294 |
| 9_1027844 | 4 | 0.75 | 4 | 0.75 | 8 | 0.5 | 7 | 0.4286 | 23 | 0.5652 | 1 | 1 | 4 | 0 | 4 | 0.5 | 8 | 0.125 | 17 | 0.2353 |
| 9_1027851 | 4 | 0.25 | 3 | 0.3333 | 4 | 0.25 | 3 | 0.3333 | 14 | 0.2857 | 3 | 0.3333 | 4 | 0.25 | 3 | 0.3333 | 3 | 0.6667 | 13 | 0.3846 |
| 9_1030588 | 30 | 0.0333 | 35 | 0.0857 | 34 | 0.0588 | 40 | 0.025 | 139 | 0.0504 | 23 | 0.087 | 23 | 0.087 | 31 | 0.0645 | 30 | 0.0333 | 107 | 0.0654 |
| 9_1030591 | 30 | 0.2333 | 37 | 0.1081 | 33 | 0.2424 | 38 | 0.2368 | 138 | 0.2029 | 24 | 0.2917 | 26 | 0.1154 | 34 | 0.2353 | 28 | 0.1786 | 112 | 0.2054 |
| 9_1030593 | 35 | 0.1429 | 44 | 0.1364 | 42 | 0.0952 | 53 | 0.1887 | 174 | 0.1437 | 28 | 0.0714 | 36 | 0.1667 | 37 | 0.1892 | 33 | 0.2121 | 134 | 0.1642 |
| 9_1030594 | 33 | 0.2424 | 44 | 0.0682 | 39 | 0.1026 | 52 | 0.0769 | 168 | 0.1131 | 27 | 0.1852 | 37 | 0.1081 | 40 | 0.25 | 38 | 0.1316 | 142 | 0.169 |
| 9_1030598 | 43 | 0.0465 | 50 | 0.06 | 38 | 0.1316 | 60 | 0.0333 | 191 | 0.0628 | 29 | 0.0345 | 39 | 0.0513 | 46 | 0.0652 | 39 | 0.1538 | 153 | 0.0784 |
| 9_1030600 | 42 | 0.2143 | 54 | 0.1852 | 42 | 0.1905 | 62 | 0.2097 | 200 | 0.2 | 32 | 0.1875 | 42 | 0.1429 | 46 | 0.1522 | 41 | 0.122 | 161 | 0.1491 |
| 9_1030600 | 50 | 0.1 | 55 | 0.1273 | 49 | 0.0612 | 67 | 0.0448 | 221 | 0.0814 | 34 | 0.1765 | 46 | 0.0217 | 54 | 0.0556 | 42 | 0.0714 | 176 | 0.0739 |
| 9_1030601 | 49 | 0.1633 | 56 | 0.1071 | 47 | 0.1489 | 66 | 0.1364 | 218 | 0.1376 | 37 | 0.0541 | 45 | 0.0667 | 53 | 0.0189 | 42 | 0.0714 | 177 | 0.0508 |
| 9_1030606 | 45 | 0.1333 | 53 | 0.0943 | 40 | 0.125 | 54 | 0.1852 | 192 | 0.1354 | 41 | 0.1463 | 45 | 0.2 | 50 | 0.14 | 39 | 0.1026 | 175 | 0.1486 |
| 9_1030621 | 67 | 0.3433 | 77 | 0.3896 | 61 | 0.3443 | 70 | 0.4286 | 275 | 0.3782 | 49 | 0.3878 | 56 | 0.375 | 66 | 0.3333 | 58 | 0.2931 | 229 | 0.345 |
| 9_1030623 | 74 | 0.0811 | 91 | 0.1319 | 81 | 0.037 | 89 | 0.0787 | 335 | 0.0836 | 64 | 0.0625 | 64 | 0.0469 | 87 | 0.069 | 68 | 0.0735 | 283 | 0.0636 |
| 9_1030657 | NA | NA | 2 | 0 | 1 | 1 | NA | NA | 3 | 0.3333 | NA | NA | NA | NA | 1 | 1 | 1 | 0 | 2 | 0.5 |
| 9_1030663 | 1 | 0 | 6 | 0.1667 | 1 | 0 | 2 | 0 | 10 | 0.1 | 3 | 0 | 2 | 0 | 3 | 0.3333 | 4 | 0 | 12 | 0.0833 |
| 9_1030673 | 6 | 1 | 4 | 1 | 4 | 1 | 8 | 1 | 22 | 1 | 4 | 1 | 4 | 1 | 6 | 1 | 3 | 1 | 17 | 1 |
| 9_1030673 | 7 | 0.2857 | 3 | 0.6667 | 3 | 1 | 7 | 0.2857 | 20 | 0.45 | 2 | 0 | 4 | 0.25 | 5 | 0 | 4 | 0 | 15 | 0.0667 |
| 9_1030674 | 7 | 0.7143 | 4 | 0.25 | 5 | 0.4 | 8 | 0.625 | 24 | 0.5417 | 3 | 0.3333 | 4 | 0.75 | 6 | 0.6667 | 4 | 0.25 | 17 | 0.5294 |
| 9_1030674 | 6 | 0.3333 | 5 | 0.4 | 5 | 0.2 | 8 | 0.25 | 24 | 0.2917 | 3 | 0.3333 | 4 | 1 | 4 | 0.5 | 4 | 0.5 | 15 | 0.6 |
| 9_1030675 | 4 | 0 | 5 | 0 | 4 | 0 | 6 | 0.1667 | 19 | 0.0526 | 2 | 0 | 4 | 0 | 4 | 0 | 4 | 0.25 | 14 | 0.0714 |
| 9_1032272 | 1 | 1 | NA | NA | 4 | 0 | 2 | 0 | 7 | 0.1429 | NA | NA | 2 | 0 | 1 | 1 | 1 | 0 | 4 | 0.25 |
| 9_1032305 | 10 | 0 | 11 | 0.0909 | 7 | 0.1429 | 7 | 0.1429 | 35 | 0.0857 | 9 | 0 | 6 | 0 | 8 | 0 | 11 | 0.1818 | 34 | 0.0588 |
| 9_1032306 | 10 | 0.1 | 11 | 0.1818 | 8 | 0.125 | 6 | 0 | 35 | 0.1143 | 9 | 0 | 7 | 0.1429 | 6 | 0 | 11 | 0.1818 | 33 | 0.0909 |
| 9_1032307 | 10 | 0 | 11 | 0.0909 | 6 | 0.3333 | 7 | 0.1429 | 34 | 0.1176 | 8 | 0.25 | 6 | 0.5 | 5 | 0.2 | 12 | 0.0833 | 31 | 0.2258 |
| 9_1032313 | 5 | 0.4 | 7 | 0 | 4 | 0 | 4 | 0.25 | 20 | 0.15 | 7 | 0.1429 | 6 | 0 | 4 | 0.25 | 10 | 0 | 27 | 0.0741 |
| 9_1032313 | 5 | 0.2 | 5 | 0.2 | 4 | 0.5 | 3 | 1 | 17 | 0.4118 | 4 | 0 | 4 | 0.25 | 2 | 0 | 8 | 0.125 | 18 | 0.1111 |
| 9_1032313 | 6 | 0.3333 | 8 | 0.5 | 3 | 0.6667 | 5 | 0.2 | 22 | 0.4091 | 8 | 0.25 | 6 | 0.3333 | 4 | 0.75 | 11 | 0.3636 | 29 | 0.3793 |
| 9_1032314 | 6 | 0.1667 | 7 | 0.1429 | 5 | 0 | 4 | 0.25 | 22 | 0.1364 | 8 | 0.125 | 5 | 0.2 | 4 | 0.25 | 9 | 0.1111 | 26 | 0.1538 |
| 9_1032318 | 4 | 0.25 | 5 | 0.2 | 2 | 0 | 2 | 0 | 13 | 0.1538 | 5 | 0.4 | 1 | 0 | 3 | 0.3333 | 8 | 0.25 | 17 | 0.2941 |
| 9_1032318 | 4 | 0.5 | 5 | 0 | 2 | 0 | 2 | 0 | 13 | 0.1538 | 5 | 0.4 | 2 | 0 | 3 | 0.6667 | 8 | 0.375 | 18 | 0.3889 |
| 9_1032320 | 1 | 0 | 5 | 0.2 | 1 | 0 | 1 | 0 | 8 | 0.125 | 5 | 0 | NA | NA | 3 | 0.3333 | 6 | 0.1667 | 14 | 0.1429 |
| 9_1032322 | 1 | 1 | 1 | 1 | 2 | 1 | NA | NA | 4 | 1 | 5 | 1 | NA | NA | 2 | 1 | 4 | 1 | 11 | 1 |
| 9_1032322 | 2 | 0 | 3 | 0 | 2 | 0.5 | 1 | 0 | 8 | 0.125 | 3 | 0.3333 | NA | NA | 2 | 0.5 | 5 | 0.2 | 10 | 0.3 |
| 9_1032322 | 2 | 1 | 3 | 0.3333 | 2 | 0 | 1 | 0 | 8 | 0.375 | 3 | 0.3333 | 1 | 1 | 1 | 0 | 5 | 0.2 | 10 | 0.3 |
| 9_1032327 | 2 | 0.5 | 2 | 0 | 2 | 0 | NA | NA | 6 | 0.1667 | 2 | 0.5 | NA | NA | NA | NA | 3 | 0.3333 | 5 | 0.4 |
| 9_1032328 | 1 | 0 | NA | NA | 2 | 0.5 | NA | NA | 3 | 0.3333 | 2 | 0.5 | NA | NA | 1 | 0 | 3 | 0.3333 | 6 | 0.3333 |
| 9_1032328 | 2 | 0.5 | NA | NA | 1 | 1 | NA | NA | 3 | 0.6667 | 2 | 1 | NA | NA | NA | NA | 3 | 0.6667 | 5 | 0.8 |
| 9_1032329 | 1 | 0 | NA | NA | 2 | 0.5 | NA | NA | 3 | 0.3333 | 2 | 1 | NA | NA | 1 | 0 | 3 | 0.6667 | 6 | 0.6667 |
| 9_1032329 | 2 | 0.5 | NA | NA | 2 | 0.5 | NA | NA | 4 | 0.5 | 2 | 0.5 | NA | NA | NA | NA | 2 | 1 | 4 | 0.75 |
| 9_1032330 | 2 | 1 | NA | NA | 2 | 1 | NA | NA | 4 | 1 | 1 | 1 | NA | NA | 1 | 1 | 2 | 1 | 4 | 1 |
| 9_1032331 | 2 | 0.5 | NA | NA | 2 | 0.5 | NA | NA | 4 | 0.5 | 2 | 1 | NA | NA | 1 | 0 | 2 | 1 | 5 | 0.8 |
| 9_1032900 | NA | NA | NA | NA | NA | NA | 1 | 1 | 1 | 1 | NA | NA | 1 | 1 | NA | NA | NA | NA | 1 | 1 |
| 9_1033570 | NA | NA | NA | NA | 1 | 1 | NA | NA | 1 | 1 | NA | NA | 1 | 1 | 1 | 0 | 1 | 1 | 3 | 0.6667 |
| 9_1033761 | NA | NA | 3 | 0.6667 | 1 | 1 | NA | NA | 4 | 0.75 | 1 | 1 | NA | NA | NA | NA | 1 | 1 | 2 | 1 |
| 9_1033763 | NA | NA | 3 | 0.6667 | 1 | 0 | NA | NA | 4 | 0.5 | 1 | 0 | NA | NA | NA | NA | 1 | 1 | 2 | 0.5 |
| 9_1033764 | NA | NA | 3 | 1 | 1 | 1 | NA | NA | 4 | 1 | NA | NA | NA | NA | NA | NA | 1 | 1 | 1 | 1 |
| 9_1033766 | NA | NA | 3 | 1 | 1 | 0 | NA | NA | 4 | 0.75 | NA | NA | NA | NA | NA | NA | 1 | 1 | 1 | 1 |
| 9_1039191 | NA | NA | 3 | 0.3333 | 1 | 0 | 1 | 0 | 5 | 0.2 | NA | NA | NA | NA | NA | NA | 2 | 0.5 | 2 | 0.5 |
| 9_1039202 | NA | NA | 2 | 0.5 | 1 | 1 | NA | NA | 3 | 0.6667 | NA | NA | NA | NA | NA | NA | 1 | 1 | 1 | 1 |
| 9_1039239 | 2 | 0.5 | 11 | 0.6364 | 3 | 0 | 5 | 0.4 | 21 | 0.4762 | 4 | 0.75 | 5 | 0.6 | 4 | 0.5 | 8 | 0.5 | 21 | 0.5714 |
| 9_1039240 | 3 | 0.3333 | 9 | 0.2222 | 3 | 0 | 4 | 0.25 | 19 | 0.2105 | 3 | 0 | 4 | 0 | 4 | 0 | 6 | 0.1667 | 17 | 0.0588 |
| 9_1039244 | 5 | 0.6 | 10 | 0.9 | 3 | 0.6667 | 7 | 0.7143 | 25 | 0.76 | 4 | 0.75 | 8 | 0.25 | 7 | 0.7143 | 10 | 0.8 | 29 | 0.6207 |
| 9_1039244 | 2 | 0 | 9 | 0.6667 | 4 | 0.75 | 7 | 0.8571 | 22 | 0.6818 | 4 | 0.75 | 7 | 0.7143 | 5 | 0.4 | 7 | 0.8571 | 23 | 0.6957 |
| 9_1039245 | 4 | 0.25 | 8 | 0.375 | 2 | 0.5 | 5 | 0.2 | 19 | 0.3158 | 3 | 0.3333 | 7 | 0.2857 | 6 | 0.6667 | 7 | 0.2857 | 23 | 0.3913 |

| | | | | | | | | | | | | | | | | | | | | |
|---|---|---|---|---|---|---|---|---|---|---|---|---|---|---|---|---|---|---|---|---|
| 9_10392454 | 3 | 1 | 8 | 0.75 | 4 | 1 | 8 | 0.75 | 23 | 0.8261 | 2 | 0.5 | 5 | 0.8 | 6 | 1 | 6 | 0.8333 | 19 | 0.8421 |
| 9_10392467 | 5 | 0.2 | 9 | 0.4444 | 4 | 0.5 | 7 | 0.2857 | 25 | 0.36 | 3 | 0.6667 | 7 | 0.2857 | 7 | 0.2857 | 8 | 0.25 | 25 | 0.32 |
| 9_10392472 | 4 | 0 | 5 | 0.2 | 2 | 0 | 4 | 0.75 | 15 | 0.2667 | 2 | 0.5 | 6 | 0 | 7 | 0.1429 | 9 | 0.2222 | 24 | 0.1667 |
| 9_10392473 | 6 | 0.1667 | 11 | 0.9091 | 3 | 1 | 5 | 0.8 | 25 | 0.72 | 3 | 0.6667 | 7 | 0.5714 | 6 | 1 | 9 | 0.5556 | 25 | 0.68 |
| 9_10392474 | 5 | 0 | 10 | 0.5 | 3 | 1 | 6 | 0.5 | 24 | 0.4583 | 2 | 0.5 | 6 | 0.3333 | 6 | 0.5 | 9 | 0.2222 | 23 | 0.3478 |
| 9_10394009 | 1 | 1 | 7 | 0 | 1 | 0 | 4 | 0.5 | 13 | 0.2308 | 3 | 0 | 5 | 0.2 | 4 | 0 | 4 | 0 | 16 | 0.0625 |
| 9_10394048 | NA | NA | NA | NA | NA | NA | 1 | 1 | 1 | 1 | NA | NA | NA | NA | NA | NA | 1 | 1 | 1 | 1 |
| 9_10402915 | 5 | 0 | 9 | 0.2222 | 5 | 0.2 | 6 | 0 | 25 | 0.12 | 5 | 0 | 5 | 0 | 3 | 0.3333 | 3 | 0.3333 | 16 | 0.125 |
| 9_10402918 | 5 | 0.6 | 8 | 0.125 | 4 | 0.25 | 4 | 0.75 | 21 | 0.381 | 5 | 0 | 6 | 0.3333 | 3 | 0 | 2 | 0 | 16 | 0.125 |
| 9_10402922 | 5 | 0.2 | 6 | 0.1667 | 5 | 0.2 | 7 | 0 | 23 | 0.1304 | 5 | 0.2 | 3 | 0 | 2 | 0 | 3 | 0 | 13 | 0.0769 |
| 9_10402924 | 5 | 0.6 | 8 | 0.5 | 5 | 0.6 | 6 | 0.6667 | 24 | 0.5833 | 5 | 0.6 | 5 | 0.4 | 3 | 0 | 5 | 0.8 | 18 | 0.5 |
| 9_10402938 | 3 | 1 | 5 | 0.8 | 3 | 1 | 3 | 0.6667 | 14 | 0.8571 | 4 | 0.5 | 4 | 1 | 3 | 0.6667 | 2 | 1 | 13 | 0.7692 |
| 9_10402944 | 4 | 0 | 8 | 0.25 | 5 | 0.6 | 6 | 0.1667 | 23 | 0.2609 | 5 | 0 | 5 | 0.6 | 3 | 0 | 4 | 0.25 | 17 | 0.2353 |
| 9_10403312 | NA | NA | 2 | 0.5 | 1 | 0 | NA | NA | 3 | 0.3333 | NA | NA | NA | NA | 2 | 0.5 | NA | NA | 2 | 0.5 |
| 9_10474994 | 2 | 0 | 1 | 0 | 1 | 0 | 3 | 0.3333 | 7 | 0.1429 | NA | NA | 2 | 0.5 | 2 | 0 | NA | NA | 4 | 0.25 |
| 9_10491000 | 7 | 0.2857 | 4 | 0.25 | 3 | 0 | 2 | 0 | 16 | 0.1875 | 2 | 0 | 1 | 0 | 2 | 0.5 | 1 | 0 | 6 | 0.1667 |
| 9_10491006 | 5 | 0 | 6 | 0.3333 | 4 | 0 | 4 | 0 | 19 | 0.1053 | 2 | 0 | 2 | 0.5 | 3 | 0 | 1 | 0 | 8 | 0.125 |
| 9_10491028 | 6 | 0.3333 | 6 | 0.1667 | 3 | 0 | 6 | 0.3333 | 21 | 0.2381 | 3 | 0 | 2 | 1 | 3 | 0.3333 | 1 | 1 | 9 | 0.4444 |
| 9_10491048 | 7 | 0.2857 | 6 | 0.1667 | 4 | 0.25 | 6 | 0.1667 | 23 | 0.2174 | 3 | 0 | 3 | 0 | 2 | 0 | 1 | 1 | 9 | 0.1111 |
| 9_10491071 | 4 | 0 | 4 | 0.25 | 3 | 0 | 3 | 0 | 14 | 0.0714 | 3 | 0 | 2 | 0.5 | 4 | 0.25 | 2 | 0 | 11 | 0.1818 |
| 9_10491081 | 4 | 0 | 3 | 0 | 4 | 0.25 | 5 | 0 | 16 | 0.0625 | 3 | 0.3333 | 3 | 0 | 3 | 0 | 1 | 0 | 10 | 0.1 |
| 9_10491141 | 5 | 0 | 5 | 0 | 4 | 0.25 | 6 | 0 | 20 | 0.05 | 2 | 0 | 5 | 0.2 | 4 | 0 | 3 | 0 | 14 | 0.0714 |
| 9_10493053 | 7 | 0.1429 | 7 | 0 | 6 | 0 | 6 | 0.1667 | 26 | 0.0769 | 3 | 0 | 8 | 0.125 | 3 | 0 | 6 | 0 | 20 | 0.05 |
| 9_10495278 | NA | NA | 1 | 0 | 1 | 0 | 2 | 0.5 | 4 | 0.25 | 2 | 0 | 1 | 1 | 1 | 0 | 3 | 0.3333 | 7 | 0.2857 |
| 9_10495286 | 1 | 0 | 1 | 0 | 1 | 1 | 2 | 0.5 | 5 | 0.4 | 2 | 0 | 1 | 1 | 2 | 0 | 2 | 0 | 7 | 0.1429 |
| 9_10495288 | NA | NA | 1 | 0 | NA | NA | 2 | 0.5 | 3 | 0.3333 | 2 | 0.5 | 1 | 1 | 2 | 0 | 3 | 0.6667 | 8 | 0.5 |
| 9_10495328 | 1 | 0 | 1 | 0 | 1 | 1 | 2 | 0.5 | 5 | 0.4 | 2 | 0 | 1 | 0 | 3 | 0 | 2 | 0.5 | 8 | 0.125 |
| 9_10495419 | 10 | 0.3 | 11 | 0.0909 | 6 | 0.1667 | 10 | 0.5 | 37 | 0.2703 | 2 | 0 | 7 | 0.5714 | 8 | 0.375 | 8 | 0.375 | 25 | 0.4 |
| 9_10495436 | 11 | 0.9091 | 10 | 0.9 | 8 | 0.875 | 12 | 1 | 41 | 0.9268 | 2 | 1 | 9 | 0.8889 | 8 | 0.875 | 7 | 0.8571 | 26 | 0.8846 |
| 9_10495438 | 10 | 0 | 10 | 0.3 | 7 | 0.1429 | 10 | 0.3 | 37 | 0.1892 | 2 | 0 | 9 | 0.6667 | 8 | 0.25 | 8 | 0.25 | 27 | 0.3704 |
| 9_10495445 | 10 | 0.6 | 10 | 0.7 | 7 | 0.4286 | 13 | 0.5385 | 40 | 0.575 | 2 | 0 | 6 | 0.8333 | 9 | 1 | 7 | 0.5714 | 24 | 0.75 |
| 9_10495453 | 12 | 0.0833 | 10 | 0.2 | 9 | 0.1111 | 12 | 0.0833 | 43 | 0.1163 | 4 | 0.25 | 7 | 0 | 8 | 0.25 | 7 | 0 | 26 | 0.1154 |
| 9_10495638 | 1 | 0 | 1 | 0 | NA | NA | 1 | 1 | 3 | 0.3333 | NA | NA | NA | NA | 1 | 0 | 3 | 0.3333 | 4 | 0.25 |
| 9_10500747 | NA | NA | 3 | 0.6667 | NA | NA | 4 | 0.5 | 7 | 0.5714 | 1 | 0 | 2 | 0.5 | 1 | 0 | 1 | 1 | 5 | 0.4 |
| 9_10500751 | NA | NA | 3 | 1 | NA | NA | 3 | 0.6667 | 6 | 0.8333 | 1 | 0 | 1 | 1 | 2 | 1 | 1 | 1 | 5 | 0.8 |
| 9_10500752 | NA | NA | 3 | 0 | NA | NA | 4 | 0.5 | 7 | 0.2857 | NA | NA | 2 | 0.5 | 2 | 0.5 | 1 | 1 | 5 | 0.6 |
| 9_10500755 | NA | NA | 3 | 0 | NA | NA | 4 | 0.25 | 7 | 0.1429 | 1 | 0 | 2 | 0.5 | 1 | 0 | 1 | 0 | 5 | 0.2 |
| 9_10500758 | NA | NA | 3 | 1 | NA | NA | 4 | 0.5 | 7 | 0.7143 | 1 | 1 | 2 | 0.5 | 1 | 1 | 1 | 1 | 5 | 0.8 |
| 9_10500850 | NA | NA | 3 | 0 | NA | NA | 2 | 1 | 5 | 0.4 | 1 | 0 | 1 | 1 | 2 | 0.5 | NA | NA | 4 | 0.5 |
| 9_10527419 | 1 | 1 | 1 | 1 | NA | NA | 1 | 1 | 3 | 1 | NA | NA | NA | NA | 1 | 1 | NA | NA | 1 | 1 |
| 9_10527424 | 1 | 1 | NA | NA | NA | NA | 1 | 1 | 2 | 1 | NA | NA | NA | NA | 1 | 1 | NA | NA | 1 | 1 |
| 9_10527446 | 1 | 1 | 1 | 1 | NA | NA | 1 | 1 | 3 | 1 | NA | NA | NA | NA | 1 | 1 | NA | NA | 1 | 1 |
| 9_10527452 | 1 | 1 | 1 | 1 | NA | NA | NA | NA | 2 | 1 | NA | NA | NA | NA | 1 | 1 | NA | NA | 1 | 1 |
| 9_10527480 | 1 | 1 | NA | NA | NA | NA | 1 | 1 | 2 | 1 | NA | NA | NA | NA | 1 | 1 | NA | NA | 1 | 1 |
| 9_10527488 | 1 | 1 | 1 | 1 | NA | NA | 1 | 1 | 3 | 1 | NA | NA | NA | NA | 1 | 1 | NA | NA | 1 | 1 |
| 9_10528230 | 1 | 0 | 1 | 1 | NA | NA | 1 | 0 | 3 | 0.3333 | NA | NA | 1 | 1 | 2 | 1 | NA | NA | 3 | 1 |
| 9_10528245 | 1 | 0 | 1 | 1 | 1 | 0 | 2 | 0 | 5 | 0.2 | NA | NA | 1 | 1 | 2 | 0.5 | NA | NA | 3 | 0.6667 |
| 9_10528249 | 1 | 0 | 1 | 1 | 1 | 0 | 2 | 0.5 | 5 | 0.4 | NA | NA | 1 | 1 | 2 | 0.5 | NA | NA | 3 | 0.6667 |
| 9_10529002 | 22 | 0.2273 | 16 | 0.25 | 17 | 0.1176 | 10 | 0.1 | 65 | 0.1846 | 7 | 0.1429 | 17 | 0.2353 | 11 | 0.1818 | 11 | 0.0909 | 46 | 0.1739 |
| 9_10529010 | 19 | 0.6316 | 13 | 0.6154 | 13 | 0.4615 | 14 | 0.5 | 59 | 0.5593 | 6 | 0.8333 | 16 | 0.8125 | 11 | 0.6364 | 13 | 0.7692 | 46 | 0.7609 |
| 9_10529011 | 23 | 0.6522 | 19 | 0.5789 | 16 | 0.4375 | 11 | 0.6364 | 69 | 0.5797 | 8 | 0.75 | 17 | 0.6471 | 13 | 0.5385 | 13 | 0.6154 | 51 | 0.6275 |
| 9_10529014 | 22 | 0.2273 | 19 | 0.1579 | 16 | 0.0625 | 11 | 0.1818 | 68 | 0.1618 | 7 | 0.5714 | 17 | 0.2353 | 15 | 0.3333 | 13 | 0.3077 | 52 | 0.3269 |
| 9_10529016 | 23 | 0.7391 | 18 | 0.8889 | 15 | 0.6 | 13 | 0.7692 | 69 | 0.7536 | 8 | 0.75 | 18 | 0.8333 | 14 | 0.6429 | 11 | 0.7273 | 51 | 0.7451 |
| 9_10539841 | NA | NA | 1 | 1 | NA | NA | NA | NA | 1 | 1 | 1 | 0 | 3 | 0.3333 | NA | NA | NA | NA | 4 | 0.25 |
| 9_10552150 | 1 | 1 | 2 | 0 | NA | NA | 3 | 1 | 6 | 0.6667 | NA | NA | NA | NA | 3 | 0.6667 | 1 | 1 | 4 | 0.75 |
| 9_10552165 | 1 | 1 | 2 | 0.5 | NA | NA | 3 | 1 | 6 | 0.8333 | NA | NA | NA | NA | 2 | 0.5 | 1 | 1 | 3 | 0.6667 |
| 9_10552168 | 1 | 1 | 2 | 1 | NA | NA | 3 | 0.6667 | 6 | 0.8333 | NA | NA | NA | NA | 3 | 1 | 1 | 1 | 4 | 1 |
| 9_10552191 | NA | NA | 2 | 0.5 | NA | NA | 2 | 1 | 4 | 0.75 | NA | NA | NA | NA | 2 | 0 | 1 | 1 | 3 | 0.3333 |
| 9_10552201 | 1 | 0 | 2 | 0 | NA | NA | 3 | 0.3333 | 6 | 0.1667 | NA | NA | NA | NA | 2 | 0 | 1 | 1 | 3 | 0.3333 |
| 9_10552215 | 2 | 0.5 | 2 | 0 | NA | NA | 2 | 0.5 | 6 | 0.3333 | NA | NA | NA | NA | 3 | 0.3333 | 1 | 0 | 4 | 0.25 |
| 9_10552772 | 2 | 1 | 2 | 1 | NA | NA | 2 | 1 | 6 | 1 | NA | NA | 1 | 1 | 3 | 0.6667 | 2 | 0.5 | 6 | 0.6667 |
| 9_10552793 | 2 | 1 | 2 | 1 | NA | NA | 3 | 0.6667 | 7 | 0.8571 | NA | NA | 2 | 0.5 | 2 | 1 | NA | NA | 4 | 0.75 |
| 9_10552796 | 1 | 1 | 1 | 1 | NA | NA | 2 | 0.5 | 4 | 0.75 | NA | NA | 2 | 0 | 3 | 1 | 2 | 0 | 7 | 0.4286 |
| 9_10555544 | 3 | 0 | 2 | 0.5 | 1 | 0 | 3 | 0.3333 | 9 | 0.2222 | NA | NA | 5 | 0 | 6 | 0.1667 | 4 | 0 | 15 | 0.0667 |
| 9_10556440 | 6 | 0 | 5 | 0.2 | 2 | 0 | 5 | 0 | 18 | 0.0556 | 1 | 0 | 9 | 0.2222 | 7 | 0.1429 | 3 | 0 | 20 | 0.15 |

| | | | | | | | | | | | | | | | | | | | | |
|---|---|---|---|---|---|---|---|---|---|---|---|---|---|---|---|---|---|---|---|---|
| 9_1055644 | 5 | 0.2 | 6 | 0.3333 | 3 | 0.6667 | 5 | 0 | 19 | 0.2632 | 1 | 0 | 9 | 0.4444 | 7 | 0.1429 | 4 | 0.25 | 21 | 0.2857 |
| 9_1055645 | 6 | 0.8333 | 6 | 0.8333 | 3 | 0.3333 | 7 | 0.5714 | 22 | 0.6818 | 1 | 0 | 10 | 0.8 | 8 | 0.625 | 7 | 0.7143 | 26 | 0.6923 |
| 9_1055645 | 6 | 0 | 5 | 0 | 2 | 0 | 7 | 0.1429 | 20 | 0.05 | 1 | 0 | 8 | 0 | 7 | 0.2857 | 6 | 0 | 22 | 0.0909 |
| 9_1055645 | 6 | 0.5 | 5 | 0.8 | 3 | 0.6667 | 6 | 1 | 20 | 0.75 | 1 | 1 | 11 | 0.8182 | 8 | 0.875 | 7 | 0.8571 | 27 | 0.8519 |
| 9_1055646 | 7 | 0.2857 | 5 | 0.4 | 3 | 0.3333 | 7 | 0.4286 | 22 | 0.3636 | 1 | 0 | 11 | 0.3636 | 8 | 0.5 | 6 | 0.1667 | 26 | 0.3462 |
| 9_1055646 | 7 | 0.1429 | 6 | 0.1667 | 2 | 0 | 6 | 0.3333 | 21 | 0.1905 | 1 | 1 | 10 | 0.2 | 9 | 0.1111 | 7 | 0.2857 | 27 | 0.2222 |
| 9_1060485 | 3 | 0 | 6 | 0 | 2 | 0.5 | 9 | 0 | 20 | 0.05 | 3 | 0.3333 | 9 | 0 | 5 | 0.2 | 6 | 0.1667 | 23 | 0.1304 |
| 9_1060658 | 10 | 0 | 11 | 0.0909 | 5 | 0 | 12 | 0.1667 | 38 | 0.0789 | 7 | 0.1429 | 10 | 0 | 7 | 0.1429 | 12 | 0.0833 | 36 | 0.0833 |
| 9_1060658 | 12 | 0.5833 | 12 | 0.25 | 6 | 0.3333 | 13 | 0.1538 | 43 | 0.3256 | 7 | 0.2857 | 11 | 0.3636 | 7 | 0.2857 | 13 | 0.5385 | 38 | 0.3947 |
| 9_1060659 | 12 | 0.25 | 13 | 0.1538 | 4 | 0 | 12 | 0.1667 | 41 | 0.1707 | 8 | 0.125 | 9 | 0.2222 | 8 | 0.5 | 12 | 0.1667 | 37 | 0.2432 |
| 9_1060660 | 12 | 0.8333 | 12 | 0.8333 | 4 | 1 | 14 | 0.8571 | 42 | 0.8571 | 6 | 1 | 10 | 0.9 | 11 | 0.6364 | 13 | 1 | 40 | 0.875 |
| 9_1060660 | 11 | 0.1818 | 10 | 0.3 | 6 | 0.5 | 11 | 0.4545 | 38 | 0.3421 | 8 | 0.875 | 11 | 0.3636 | 10 | 0.3 | 11 | 0.5455 | 40 | 0.5 |
| 9_1060660 | 14 | 0.3571 | 13 | 0.3077 | 3 | 0.3333 | 11 | 0.1818 | 41 | 0.2927 | 8 | 0.125 | 8 | 0.25 | 11 | 0.0909 | 9 | 0.3333 | 36 | 0.1944 |
| 9_1060660 | 9 | 0.8889 | 9 | 0.6667 | 3 | 0.6667 | 11 | 0.8182 | 32 | 0.7812 | 4 | 1 | 8 | 0.625 | 8 | 0.375 | 13 | 0.3846 | 33 | 0.5152 |
| 9_1060676 | 21 | 0.0952 | 6 | 0 | 9 | 0.1111 | 16 | 0.125 | 52 | 0.0962 | 10 | 0.3 | 11 | 0.1818 | 8 | 0 | 14 | 0 | 43 | 0.1163 |
| 9_1060676 | 23 | 0.3478 | 11 | 0.3636 | 12 | 0.25 | 15 | 0.2667 | 61 | 0.3115 | 12 | 0.25 | 18 | 0.3333 | 11 | 0.1818 | 16 | 0.25 | 57 | 0.2632 |
| 9_1060676 | 20 | 0.05 | 12 | 0.0833 | 12 | 0.0833 | 17 | 0.0588 | 61 | 0.0656 | 12 | 0.0833 | 15 | 0.1333 | 8 | 0 | 12 | 0.1667 | 47 | 0.1064 |
| 9_1060676 | 18 | 0.3333 | 14 | 0.2857 | 11 | 0.1818 | 16 | 0.375 | 59 | 0.3051 | 16 | 0.4375 | 17 | 0.3529 | 9 | 0.1111 | 17 | 0.2941 | 59 | 0.322 |
| 9_1060677 | 21 | 0.4762 | 13 | 0.3077 | 11 | 0.3636 | 17 | 0.7059 | 62 | 0.4839 | 16 | 0.5625 | 17 | 0.6471 | 8 | 0.5 | 18 | 0.6667 | 59 | 0.6102 |
| 9_1060677 | 24 | 0.25 | 14 | 0.2857 | 12 | 0.5 | 20 | 0.3 | 70 | 0.3143 | 15 | 0.3333 | 17 | 0.3529 | 8 | 0.25 | 13 | 0.2308 | 53 | 0.3019 |
| 9_1060722 | 2 | 0.5 | 1 | 1 | NA | NA | 3 | 0.6667 | 6 | 0.6667 | NA | NA | NA | NA | 1 | 0 | 1 | 1 | 2 | 0.5 |
| 9_1060725 | 1 | 1 | 1 | 1 | 1 | 1 | 3 | 0.6667 | 6 | 0.8333 | NA | NA | NA | NA | NA | NA | 2 | 1 | 2 | 1 |
| 9_1060922 | 4 | 1 | 6 | 0.8333 | 2 | 0.5 | 6 | 0.8333 | 18 | 0.8333 | 3 | 0.6667 | 2 | 0.5 | 1 | 0 | 4 | 0.75 | 10 | 0.6 |
| 9_1060922 | 5 | 0.4 | 6 | 0.8333 | 1 | 0 | 5 | 0.2 | 17 | 0.4706 | 5 | 0.6 | 4 | 0.5 | 1 | 0 | 4 | 0.25 | 14 | 0.4286 |
| 9_1060922 | 5 | 0.8 | 5 | 0.8 | 1 | 0 | 5 | 0.8 | 16 | 0.75 | 4 | 0.75 | 2 | 0 | 2 | 0.5 | 5 | 0.8 | 13 | 0.6154 |
| 9_1060923 | 5 | 0.2 | 5 | 0 | 1 | 0 | 5 | 0 | 16 | 0.0625 | 7 | 0 | 5 | 0.4 | 1 | 0 | 5 | 0.2 | 18 | 0.1667 |
| 9_1060923 | 5 | 0.2 | 7 | 0.1429 | 1 | 0 | 5 | 0.4 | 18 | 0.2222 | 7 | 0 | 5 | 0.2 | 1 | 1 | 5 | 0.2 | 18 | 0.1667 |
| 9_1061152 | 9 | 0 | 7 | 0.1429 | 4 | 0 | 6 | 0.3333 | 26 | 0.1154 | 4 | 0.25 | 4 | 0.25 | 5 | 0 | 6 | 0.1667 | 19 | 0.1579 |
| 9_1062458 | 4 | 0 | 2 | 0 | 2 | 0.5 | 1 | 1 | 9 | 0.2222 | 2 | 0.5 | 3 | 0.3333 | 3 | 0 | NA | NA | 8 | 0.25 |
| 9_1062479 | 4 | 0.25 | 5 | 0.2 | 4 | 0 | 2 | 0 | 15 | 0.1333 | 5 | 0 | 5 | 0 | 5 | 0 | 4 | 0.25 | 19 | 0.0526 |
| 9_1062479 | 4 | 0 | 4 | 0 | 3 | 0.6667 | 2 | 0.5 | 13 | 0.2308 | 4 | 0 | 6 | 0 | 5 | 0.2 | 5 | 0.2 | 20 | 0.1 |
| 9_1062481 | 7 | 0.8571 | 4 | 1 | 4 | 1 | 2 | 1 | 17 | 0.9412 | 6 | 1 | 6 | 0.8333 | 5 | 0.6 | 4 | 1 | 21 | 0.8571 |
| 9_1062509 | 9 | 0.4444 | 6 | 0.5 | 6 | 0.1667 | 2 | 0.5 | 23 | 0.3913 | 7 | 0.4286 | 6 | 0.5 | 6 | 0.1667 | 4 | 0.75 | 23 | 0.4348 |
| 9_1062510 | 7 | 1 | 9 | 1 | 5 | 0.8 | 2 | 0.5 | 23 | 0.913 | 5 | 1 | 5 | 1 | 5 | 1 | 2 | 1 | 17 | 1 |
| 9_1062512 | 11 | 0.4545 | 9 | 0 | 6 | 0 | 4 | 0 | 30 | 0.1667 | 7 | 0.1429 | 6 | 0 | 5 | 0 | 6 | 0.3333 | 24 | 0.125 |
| 9_1062513 | 11 | 0.3636 | 11 | 0.3636 | 6 | 0.5 | 4 | 0.5 | 32 | 0.4062 | 8 | 0.375 | 6 | 0.5 | 6 | 0.5 | 6 | 0.1667 | 26 | 0.3846 |
| 9_1062513 | 8 | 0.875 | 9 | 0.8889 | 6 | 0.6667 | 1 | 1 | 24 | 0.8333 | 8 | 0.75 | 5 | 1 | 4 | 1 | 4 | 0.75 | 21 | 0.8571 |
| 9_1062514 | 11 | 0.9091 | 10 | 0.8 | 8 | 0.875 | 4 | 1 | 33 | 0.8788 | 5 | 1 | 8 | 0.875 | 7 | 1 | 5 | 1 | 25 | 0.96 |
| 9_1062514 | 9 | 0.6667 | 11 | 0.7273 | 6 | 1 | 3 | 0.3333 | 29 | 0.7241 | 5 | 0.8 | 7 | 0.7143 | 7 | 0.7143 | 5 | 0.4 | 24 | 0.6667 |
| 9_1062514 | 11 | 0.1818 | 10 | 0.3 | 8 | 0.25 | 4 | 0.25 | 33 | 0.2424 | 7 | 0.2857 | 8 | 0.5 | 7 | 0.1429 | 6 | 0.1667 | 28 | 0.2857 |
| 9_1063086 | 93 | 0.6129 | 66 | 0.6515 | 90 | 0.6333 | 105 | 0.6571 | 354 | 0.6384 | 78 | 0.641 | 92 | 0.663 | 103 | 0.6893 | 84 | 0.7738 | 357 | 0.6919 |
| 9_1063173 | 10 | 0.1 | 3 | 0 | 6 | 0 | 11 | 0.0909 | 30 | 0.0667 | 5 | 0 | 9 | 0 | 6 | 0.1667 | 7 | 0.1429 | 27 | 0.0741 |
| 9_1063173 | 7 | 0 | 3 | 0 | 6 | 0 | 8 | 0.25 | 24 | 0.0833 | 5 | 0.6 | 8 | 0 | 2 | 0 | 6 | 0.3333 | 21 | 0.2381 |
| 9_1063174 | 5 | 0.8 | 3 | 0.3333 | 4 | 0.25 | 10 | 0.5 | 22 | 0.5 | 6 | 0.8333 | 7 | 0.7143 | 5 | 1 | 5 | 0.6 | 23 | 0.7826 |
| 9_1063229 | 2 | 0 | NA | NA | NA | NA | 3 | 0.3333 | 5 | 0.2 | 1 | 0 | 2 | 0.5 | NA | NA | 1 | 0 | 4 | 0.25 |
| 9_1063293 | 1 | 0 | 3 | 0.3333 | 1 | 0 | 1 | 1 | 6 | 0.3333 | NA | NA | 1 | 1 | 2 | 0.5 | NA | NA | 3 | 0.6667 |
| 9_1063293 | 1 | 0 | 3 | 0.3333 | 1 | 0 | 1 | 0 | 6 | 0.1667 | NA | NA | 1 | 1 | 2 | 1 | NA | NA | 3 | 1 |
| 9_1063407 | 8 | 0.125 | 19 | 0.3158 | 11 | 0.4545 | 5 | 0.6 | 43 | 0.3488 | 11 | 0.5455 | 6 | 0 | 13 | 0.3846 | 14 | 0.5714 | 44 | 0.4318 |
| 9_1063408 | 9 | 0 | 19 | 0.1053 | 17 | 0.2941 | 9 | 0 | 54 | 0.1296 | 15 | 0.1333 | 6 | 0 | 16 | 0.0625 | 14 | 0 | 51 | 0.0588 |
| 9_1063410 | 10 | 0.2 | 21 | 0.1429 | 14 | 0.1429 | 7 | 0.2857 | 52 | 0.1731 | 15 | 0 | 7 | 0 | 16 | 0.125 | 15 | 0.0667 | 53 | 0.0566 |
| 9_1063410 | 9 | 0.4444 | 21 | 0.4286 | 14 | 0.2857 | 8 | 0.375 | 52 | 0.3846 | 16 | 0.5625 | 5 | 0.6 | 12 | 0.3333 | 12 | 0.5 | 45 | 0.4889 |
| 9_1063412 | 1 | 0 | 3 | 0.3333 | 4 | 0.25 | NA | NA | 8 | 0.25 | 1 | 0 | NA | NA | 1 | 1 | 2 | 0.5 | 4 | 0.5 |
| 9_1063413 | 1 | 1 | 3 | 0.6667 | 5 | 1 | 1 | 1 | 10 | 0.9 | 1 | 1 | NA | NA | 1 | 1 | 1 | 1 | 3 | 1 |
| 9_1063413 | NA | NA | 3 | 0.6667 | 4 | 1 | NA | NA | 7 | 0.8571 | NA | NA | NA | NA | 1 | 1 | 2 | 1 | 3 | 1 |
| 9_1063413 | 1 | 0 | 3 | 0.3333 | 3 | 1 | 1 | 1 | 8 | 0.625 | 1 | 1 | NA | NA | 1 | 1 | 2 | 0.5 | 4 | 0.75 |
| 9_1063413 | 1 | 1 | 3 | 0 | 4 | 0 | 1 | 1 | 9 | 0.2222 | NA | NA | NA | NA | 1 | 0 | 2 | 1 | 3 | 0.6667 |
| 9_1063415 | 1 | 0 | 4 | 0.25 | 2 | 1 | 1 | 0 | 8 | 0.375 | 1 | 1 | NA | NA | 1 | 0 | 2 | 1 | 4 | 0.75 |
| 9_1063416 | 1 | 0 | 3 | 0 | 2 | 0.5 | 1 | 1 | 7 | 0.2857 | 1 | 1 | NA | NA | 1 | 1 | 2 | 0 | 4 | 0.5 |
| 9_1063418 | NA | NA | 3 | 1 | 2 | 0.5 | 1 | 0 | 6 | 0.6667 | 1 | 1 | NA | NA | NA | NA | 1 | 1 | 2 | 1 |
| 9_1063419 | NA | NA | 3 | 0.3333 | 2 | 1 | NA | NA | 5 | 0.6 | 1 | 0 | NA | NA | NA | NA | 1 | 1 | 2 | 0.5 |
| 9_1063419 | 1 | 1 | 2 | 1 | 2 | 1 | NA | NA | 5 | 1 | 1 | 1 | NA | NA | NA | NA | 1 | 0 | 2 | 0.5 |
| 9_1063422 | 1 | 0 | 2 | 0.5 | NA | NA | NA | NA | 3 | 0.3333 | 1 | 1 | NA | NA | NA | NA | NA | NA | 1 | 1 |
| 9_1063426 | 1 | 0 | 2 | 1 | NA | NA | NA | NA | 3 | 0.6667 | 1 | 1 | NA | NA | NA | NA | NA | NA | 1 | 1 |
| 9_1063428 | 1 | 0 | 2 | 0.5 | NA | NA | NA | NA | 3 | 0.3333 | 1 | 1 | NA | NA | NA | NA | NA | NA | 1 | 1 |
| 9_1063430 | 1 | 1 | 2 | 1 | NA | NA | NA | NA | 3 | 1 | 1 | 1 | NA | NA | NA | NA | NA | NA | 1 | 1 |

| | | | | | | | | | | | | | | | | | | | | |
|---|---|---|---|---|---|---|---|---|---|---|---|---|---|---|---|---|---|---|---|---|
| 9_10634329 | 1 | 0 | 1 | 1 | NA | NA | NA | NA | 2 | 0.5 | 1 | 0 | 2 | 1 | 2 | 0.5 | NA | NA | 5 | 0.6 |
| 9_10634339 | 1 | 0 | 1 | 1 | NA | NA | NA | NA | 2 | 0.5 | 1 | 0 | 1 | 1 | 2 | 0.5 | NA | NA | 4 | 0.5 |
| 9_10634361 | NA | NA | 1 | 1 | NA | NA | NA | NA | 1 | 1 | 1 | 1 | 2 | 1 | 2 | 1 | NA | NA | 5 | 1 |
| 9_10634377 | 1 | 1 | NA | NA | NA | NA | NA | NA | 1 | 1 | 1 | 0 | 1 | 1 | 1 | 1 | NA | NA | 3 | 0.6667 |
| 9_10634825 | NA | NA | NA | NA | NA | NA | 1 | 1 | 1 | 1 | NA | NA | NA | NA | 2 | 1 | 3 | 1 | 5 | 1 |
| 9_10652479 | NA | NA | 2 | 0.5 | 1 | 0 | 1 | 0 | 4 | 0.25 | 1 | 0 | 1 | 0 | NA | NA | 2 | 0.5 | 4 | 0.25 |
| 9_10652500 | NA | NA | 1 | 1 | 1 | 1 | NA | NA | 2 | 1 | 1 | 0 | 1 | 1 | NA | NA | 2 | 1 | 4 | 0.75 |
| 9_10652506 | NA | NA | 1 | 1 | 2 | 1 | 1 | 1 | 4 | 1 | 1 | 1 | 1 | 1 | NA | NA | 2 | 1 | 4 | 1 |
| 9_10652509 | NA | NA | 2 | 1 | 1 | 1 | 1 | 0 | 4 | 0.75 | 1 | 1 | 1 | 1 | NA | NA | 2 | 1 | 4 | 1 |
| 9_10652527 | NA | NA | 1 | 1 | NA | NA | NA | NA | 1 | 1 | 1 | 1 | 1 | 0 | NA | NA | 1 | 1 | 3 | 0.6667 |
| 9_10652762 | NA | NA | 2 | 1 | 2 | 0.5 | NA | NA | 4 | 0.75 | 1 | 0 | 1 | 1 | NA | NA | 2 | 1 | 4 | 0.75 |
| 9_10652776 | NA | NA | 2 | 0.5 | 2 | 0 | NA | NA | 4 | 0.25 | 1 | 0 | 1 | 1 | NA | NA | 1 | 1 | 3 | 0.6667 |
| 9_10652788 | NA | NA | 2 | 1 | 1 | 0 | NA | NA | 3 | 0.6667 | NA | NA | 1 | 1 | NA | NA | 1 | 1 | 2 | 1 |
| 9_10652790 | NA | NA | 2 | 1 | 2 | 1 | NA | NA | 4 | 1 | 1 | 1 | 1 | 1 | NA | NA | 2 | 1 | 4 | 1 |
| 9_10652822 | NA | NA | 2 | 0 | 1 | 1 | NA | NA | 3 | 0.3333 | 1 | 1 | 1 | 0 | NA | NA | 2 | 1 | 4 | 0.75 |
| 9_10652823 | NA | NA | 2 | 0.5 | 2 | 1 | NA | NA | 4 | 0.75 | 1 | 1 | NA | NA | NA | NA | 2 | 0.5 | 3 | 0.6667 |
| 9_10653117 | NA | NA | 2 | 0.5 | 2 | 1 | NA | NA | 4 | 0.75 | 1 | 0 | 1 | 1 | NA | NA | 2 | 0.5 | 4 | 0.5 |
| 9_10653133 | NA | NA | 2 | 0.5 | 2 | 1 | NA | NA | 4 | 0.75 | 1 | 0 | 1 | 1 | NA | NA | 2 | 0 | 4 | 0.25 |
| 9_10653600 | 12 | 0.0833 | 13 | 0.1538 | 9 | 0.1111 | 11 | 0.0909 | 45 | 0.1111 | 10 | 0.1 | 17 | 0.0588 | 8 | 0.125 | 10 | 0.2 | 45 | 0.1111 |
| 9_10653604 | 11 | 0.8182 | 14 | 0.8571 | 10 | 1 | 12 | 0.5833 | 47 | 0.8085 | 10 | 0.9 | 16 | 0.75 | 9 | 0.8889 | 10 | 0.9 | 45 | 0.8444 |
| 9_10653618 | 9 | 0.6667 | 12 | 0.5833 | 10 | 0.9 | 9 | 0.8889 | 40 | 0.75 | 10 | 0.7 | 15 | 0.7333 | 9 | 0.6667 | 10 | 0.8 | 44 | 0.7273 |
| 9_10653631 | 11 | 0.1818 | 13 | 0.0769 | 10 | 0.2 | 11 | 0.2727 | 45 | 0.1778 | 7 | 0.1429 | 15 | 0.4667 | 9 | 0.2222 | 9 | 0.1111 | 40 | 0.275 |
| 9_10653632 | 9 | 0.2222 | 12 | 0.1667 | 8 | 0.125 | 7 | 0.1429 | 36 | 0.1667 | 8 | 0.125 | 11 | 0 | 5 | 0.2 | 8 | 0.25 | 32 | 0.125 |
| 9_10655923 | 7 | 0.8571 | 6 | 0.5 | 5 | 0.4 | 4 | 0.5 | 22 | 0.5909 | 3 | 1 | 8 | 0.625 | 3 | 0.3333 | 5 | 0.6 | 19 | 0.6316 |
| 9_10656292 | 1 | 1 | 3 | 0 | 1 | 0 | 1 | 0 | 6 | 0.1667 | 2 | 0.5 | 3 | 0 | 1 | 0 | 1 | 0 | 7 | 0.1429 |
| 9_10656543 | 2 | 0 | 1 | 0 | NA | NA | 2 | 0.5 | 5 | 0.2 | 3 | 0.3333 | NA | NA | 1 | 0 | 2 | 0 | 6 | 0.1667 |
| 9_10671553 | 1 | 1 | NA | NA | NA | NA | NA | NA | 1 | 1 | 1 | 1 | 1 | 1 | 1 | 1 | NA | NA | 3 | 1 |
| 9_10673564 | 2 | 0.5 | NA | NA | 2 | 0.5 | 2 | 0 | 6 | 0.3333 | 4 | 0.5 | 1 | 0 | 2 | 0 | NA | NA | 7 | 0.2857 |
| 9_10673568 | 2 | 0 | NA | NA | 1 | 0 | 2 | 0.5 | 5 | 0.2 | 4 | 0.25 | 1 | 0 | 2 | 0.5 | NA | NA | 7 | 0.2857 |
| 9_10673572 | 2 | 1 | NA | NA | 2 | 0.5 | 3 | 0.6667 | 7 | 0.7143 | 3 | 1 | 1 | 0 | 1 | 1 | NA | NA | 5 | 0.8 |
| 9_10673575 | 2 | 0 | NA | NA | 2 | 0.5 | 2 | 0.5 | 6 | 0.3333 | 5 | 0.2 | 1 | 0 | 2 | 0 | NA | NA | 8 | 0.125 |
| 9_10673598 | 2 | 0.5 | 1 | 0 | 2 | 0 | 3 | 0.3333 | 8 | 0.25 | 5 | 0.8 | 2 | 1 | 2 | 1 | 1 | 1 | 10 | 0.9 |
| 9_10673602 | 1 | 0 | 1 | 0 | 1 | 1 | 3 | 0 | 6 | 0.1667 | 3 | 0.3333 | 1 | 0 | 1 | 0 | NA | NA | 5 | 0.2 |
| 9_10673603 | 2 | 0 | 1 | 1 | 2 | 1 | 3 | 0 | 8 | 0.375 | 5 | 0.4 | 2 | 0 | 2 | 0 | 1 | 1 | 10 | 0.3 |
| 9_10673606 | 2 | 1 | 1 | 1 | 2 | 0.5 | 2 | 1 | 7 | 0.8571 | 3 | 0.6667 | 1 | 1 | 2 | 1 | 1 | 0 | 7 | 0.7143 |
| 9_10673611 | 2 | 0 | 1 | 1 | 2 | 0 | 3 | 0 | 8 | 0.125 | 4 | 0 | 2 | 0 | 2 | 0.5 | 1 | 0 | 9 | 0.1111 |
| 9_10673619 | 2 | 1 | 1 | 1 | 1 | 0 | 4 | 0.75 | 8 | 0.75 | 5 | 0.4 | 2 | 0.5 | 2 | 0 | 1 | 1 | 10 | 0.4 |
| 9_10675132 | 2 | 0 | 2 | 0 | 1 | 0 | 5 | 0.4 | 10 | 0.2 | 6 | 0 | 1 | 1 | 1 | 0 | 1 | 0 | 9 | 0.1111 |
| 9_10675723 | 2 | 0 | 3 | 0.6667 | 2 | 0 | 8 | 0.5 | 15 | 0.4 | 8 | 0.375 | 4 | 0 | 3 | 0 | 2 | 0 | 17 | 0.1765 |
| 9_10675729 | 2 | 0 | 3 | 0.6667 | 3 | 0.3333 | 4 | 0.25 | 12 | 0.3333 | 8 | 0.375 | 4 | 0.5 | 3 | 0 | 2 | 0.5 | 17 | 0.3529 |
| 9_10675731 | 2 | 0.5 | 3 | 0.6667 | 3 | 1 | 7 | 0.7143 | 15 | 0.7333 | 6 | 0.6667 | 3 | 1 | 1 | 1 | 2 | 0.5 | 12 | 0.75 |
| 9_10675851 | NA | NA | 2 | 0 | 3 | 0.3333 | 1 | 0 | 6 | 0.1667 | 2 | 0.5 | 2 | 0.5 | 6 | 0.1667 | 3 | 0 | 13 | 0.2308 |
| 9_10675899 | NA | NA | 2 | 0.5 | 3 | 0.6667 | 1 | 0 | 6 | 0.5 | 1 | 1 | 2 | 0.5 | 5 | 0.8 | 2 | 0.5 | 10 | 0.7 |
| 9_10676151 | 5 | 0.8 | 3 | 1 | 2 | 0.5 | 7 | 0.8571 | 17 | 0.8235 | 10 | 0.8 | 4 | 0.75 | 8 | 0.875 | 3 | 0.6667 | 25 | 0.8 |
| 9_10676153 | 6 | 0.3333 | 3 | 0.6667 | 2 | 0.5 | 9 | 0.3333 | 20 | 0.4 | 10 | 0.3 | 4 | 0.5 | 10 | 0.5 | 3 | 0 | 27 | 0.3704 |
| 9_10676156 | 7 | 1 | 3 | 1 | 2 | 1 | 8 | 1 | 20 | 1 | 11 | 1 | 4 | 1 | 9 | 1 | 3 | 1 | 27 | 1 |
| 9_10676163 | 7 | 1 | 3 | 1 | 2 | 0.5 | 10 | 1 | 22 | 0.9545 | 10 | 0.9 | 2 | 1 | 8 | 1 | 2 | 0.5 | 22 | 0.9091 |
| 9_10676168 | 5 | 0.2 | 2 | 0 | 1 | 1 | 7 | 0.1429 | 15 | 0.2 | 7 | 0.1429 | 1 | 0 | 7 | 0 | 2 | 0 | 17 | 0.0588 |
| 9_10676170 | 6 | 0.6667 | 2 | 0.5 | 2 | 0.5 | 10 | 0.9 | 20 | 0.75 | 11 | 0.5455 | 4 | 0.75 | 8 | 0.375 | 3 | 1 | 26 | 0.5769 |
| 9_10676191 | 5 | 0 | 3 | 0 | 2 | 0 | 8 | 0.125 | 18 | 0.0556 | 7 | 0.1429 | 3 | 0.3333 | 6 | 0 | 3 | 0 | 19 | 0.1053 |
| 9_10676204 | 7 | 1 | 3 | 1 | 1 | 1 | 8 | 0.875 | 19 | 0.9474 | 10 | 1 | 4 | 0.75 | 5 | 1 | 3 | 1 | 22 | 0.9545 |
| 9_10676206 | 6 | 0 | 3 | 0.3333 | 2 | 0 | 7 | 0 | 18 | 0.0556 | 8 | 0.125 | 3 | 0 | 5 | 0 | 3 | 0 | 19 | 0.0526 |
| 9_10676208 | 6 | 0.1667 | 3 | 0.6667 | 2 | 0.5 | 6 | 0.3333 | 17 | 0.3529 | 10 | 0.3 | 4 | 0.5 | 6 | 0.5 | 3 | 0.3333 | 23 | 0.3913 |
| 9_10676213 | 6 | 0.6667 | 3 | 0 | 2 | 0.5 | 8 | 0.25 | 19 | 0.3684 | 11 | 0.5455 | 4 | 0.25 | 6 | 0.1667 | 3 | 0.3333 | 24 | 0.375 |
| 9_10676215 | 5 | 0.4 | 3 | 0 | 2 | 0 | 8 | 0.25 | 18 | 0.2222 | 11 | 0.2727 | 4 | 0.5 | 6 | 0 | 3 | 0.3333 | 24 | 0.25 |
| 9_10676224 | 6 | 0 | 3 | 0.6667 | 2 | 0.5 | 6 | 0.1667 | 17 | 0.2353 | 7 | 0 | 3 | 0.6667 | 3 | 0.6667 | 3 | 0.3333 | 16 | 0.3125 |
| 9_10676232 | 3 | 0.6667 | 3 | 0.3333 | 2 | 0 | 5 | 0.4 | 13 | 0.3846 | 4 | 0.5 | 2 | 0 | 4 | 0 | 3 | 0.3333 | 13 | 0.2308 |
| 9_10676242 | 6 | 0.1667 | 2 | 0 | 2 | 0 | 8 | 0 | 18 | 0.0556 | 8 | 0.125 | 3 | 0 | 5 | 0 | 3 | 0 | 19 | 0.0526 |
| 9_10676252 | 4 | 0 | 1 | 0 | NA | NA | 4 | 0.75 | 9 | 0.3333 | 8 | 0.125 | 4 | 0.5 | 4 | 0.5 | 2 | 0 | 18 | 0.2778 |
| 9_10676253 | 5 | 1 | 2 | 1 | 1 | 1 | 6 | 0.8333 | 14 | 0.9286 | 8 | 0.75 | 4 | 0.5 | 4 | 1 | 2 | 1 | 18 | 0.7778 |
| 9_10676254 | 5 | 0.4 | 2 | 1 | 1 | 1 | 5 | 0.4 | 13 | 0.5385 | 4 | 0.75 | 3 | 1 | 5 | 0.6 | 2 | 0 | 14 | 0.6429 |
| 9_10676258 | 5 | 0 | 2 | 0 | 1 | 1 | 8 | 0.125 | 16 | 0.125 | 8 | 0 | 4 | 0 | 5 | 0 | 3 | 0.3333 | 20 | 0.05 |
| 9_10676260 | 5 | 0.4 | 2 | 0 | 1 | 0 | 6 | 0.1667 | 14 | 0.2143 | 8 | 0.125 | 2 | 0 | 3 | 0 | 3 | 0 | 16 | 0.0625 |
| 9_10676261 | 4 | 0.25 | 2 | 0.5 | 1 | 0 | 7 | 0 | 14 | 0.1429 | 5 | 0 | 2 | 0 | 4 | 0.25 | 3 | 0.3333 | 14 | 0.1429 |
| 9_10676269 | 5 | 0.4 | 2 | 0.5 | 1 | 0 | 8 | 0.25 | 16 | 0.3125 | 8 | 0.25 | 4 | 0.25 | 5 | 0.6 | 3 | 0.6667 | 20 | 0.4 |

| | | | | | | | | | | | | | | | | | | | | |
|---|---|---|---|---|---|---|---|---|---|---|---|---|---|---|---|---|---|---|---|---|
| 9_1067629( | 3 | 0.3333 | 1 | 0 | 1 | 0 | 7 | 0.7143 | 12 | 0.5 | 4 | 1 | 3 | 0.3333 | 4 | 0.75 | 2 | 1 | 13 | 0.7692 |
| 9_1067630' | 5 | 0.4 | 1 | 0 | 1 | 1 | 6 | 0.8333 | 13 | 0.6154 | 6 | 0.3333 | 2 | 0.5 | 5 | 1 | 2 | 1 | 15 | 0.6667 |
| 9_1067632( | 4 | 0.25 | 1 | 0 | 1 | 1 | 6 | 0 | 12 | 0.1667 | 4 | 0 | 3 | 0 | 4 | 0.25 | 3 | 0.6667 | 14 | 0.2143 |
| 9_1067632( | 3 | 0.6667 | 1 | 1 | 1 | 0 | 6 | 0.8333 | 11 | 0.7273 | 4 | 1 | 2 | 1 | 3 | 1 | 1 | 1 | 10 | 1 |
| 9_1067632( | 5 | 0.8 | 1 | 1 | 1 | 1 | 6 | 0.8333 | 13 | 0.8462 | 3 | 0.6667 | 3 | 1 | 2 | 1 | 2 | 1 | 10 | 0.9 |
| 9_1067636' | 4 | 0.25 | 1 | 0 | 1 | 0 | 4 | 0.25 | 10 | 0.2 | 3 | 0 | 1 | 1 | 1 | 0 | 2 | 0.5 | 7 | 0.2857 |
| 9_1067648 | 1 | 1 | NA | NA | 1 | 1 | 1 | 1 | 3 | 1 | NA | NA | NA | NA | 1 | 1 | NA | NA | 1 | 1 |
| 9_1067648 | 1 | 1 | NA | NA | 1 | 1 | 1 | 1 | 3 | 1 | NA | NA | NA | NA | 1 | 1 | NA | NA | 1 | 1 |
| 9_1067649 | 1 | 1 | NA | NA | 1 | 1 | 1 | 1 | 3 | 1 | NA | NA | NA | NA | 1 | 1 | NA | NA | 1 | 1 |
| 9_1067660( | 18 | 0.0556 | 21 | 0.1429 | 15 | 0.2667 | 11 | 0.1818 | 65 | 0.1538 | 22 | 0.0909 | 14 | 0.2143 | 18 | 0.0556 | 18 | 0.2222 | 72 | 0.1389 |
| 9_1067662! | 17 | 0.7059 | 20 | 0.65 | 12 | 0.6667 | 11 | 0.8182 | 60 | 0.7 | 15 | 0.5333 | 16 | 0.8125 | 17 | 0.6471 | 13 | 0.6923 | 61 | 0.6721 |
| 9_1067662! | 14 | 0.8571 | 14 | 0.6429 | 11 | 0.5455 | 10 | 0.7 | 49 | 0.6939 | 18 | 0.7222 | 15 | 0.6 | 16 | 0.6875 | 14 | 0.6429 | 63 | 0.6667 |
| 9_1067705 | 37 | 0.2432 | 31 | 0.3226 | 34 | 0.2353 | 29 | 0.1724 | 131 | 0.2443 | 16 | 0.4375 | 18 | 0.1667 | 29 | 0.2759 | 24 | 0.375 | 87 | 0.3103 |
| 9_1067718 | 27 | 0.8889 | 16 | 1 | 21 | 0.9524 | 18 | 0.8333 | 82 | 0.9146 | 8 | 1 | 14 | 0.9286 | 17 | 0.9412 | 14 | 1 | 53 | 0.9623 |
| 9_1067719 | 23 | 0.6087 | 14 | 0.7857 | 21 | 0.6667 | 18 | 0.6111 | 76 | 0.6579 | 6 | 0.8333 | 10 | 0.7 | 18 | 0.8333 | 11 | 0.8182 | 45 | 0.8 |
| 9_1067721' | 21 | 0.7143 | 13 | 0.8462 | 17 | 0.7647 | 17 | 0.7059 | 68 | 0.75 | 8 | 0.625 | 11 | 0.8182 | 17 | 0.7647 | 13 | 0.8462 | 49 | 0.7755 |
| 9_1067721 | 23 | 1 | 13 | 1 | 22 | 1 | 18 | 0.8889 | 76 | 0.9737 | 7 | 1 | 12 | 0.9167 | 18 | 0.9444 | 13 | 1 | 50 | 0.96 |
| 9_1067721 | 20 | 0.55 | 13 | 0.6154 | 22 | 0.5 | 16 | 0.5625 | 71 | 0.5493 | 7 | 0.4286 | 12 | 0.5833 | 17 | 0.5294 | 12 | 0.5 | 48 | 0.5208 |
| 9_1067721 | 21 | 0.8571 | 13 | 0.9231 | 17 | 0.9412 | 17 | 0.8824 | 68 | 0.8971 | 8 | 0.875 | 12 | 0.9167 | 13 | 0.9231 | 14 | 1 | 47 | 0.9362 |
| 9_1067722 | 21 | 0.381 | 11 | 0.5455 | 18 | 0.6111 | 19 | 0.3684 | 69 | 0.4638 | 9 | 0.6667 | 7 | 0.5714 | 13 | 0.5385 | 13 | 0.6154 | 42 | 0.5952 |
| 9_1067724 | 19 | 1 | 7 | 1 | 10 | 0.9 | 10 | 0.9 | 46 | 0.9565 | 5 | 1 | 10 | 1 | 11 | 1 | 10 | 1 | 36 | 1 |
| 9_1067725 | 12 | 0.25 | 10 | 0.6 | 13 | 0.3846 | 9 | 0.4444 | 44 | 0.4091 | 4 | 0.25 | 9 | 0.4444 | 8 | 0.5 | 6 | 0.6667 | 27 | 0.4815 |
| 9_1067727 | 20 | 0.8 | 10 | 1 | 14 | 0.7857 | 14 | 0.8571 | 58 | 0.8448 | 6 | 1 | 13 | 0.9231 | 13 | 0.9231 | 11 | 0.9091 | 43 | 0.9302 |
| 9_1067729 | 19 | 0.9474 | 10 | 1 | 12 | 0.8333 | 12 | 0.8333 | 53 | 0.9057 | 4 | 1 | 10 | 0.9 | 10 | 1 | 9 | 0.6667 | 33 | 0.8788 |
| 9_1067729 | 20 | 0.1 | 9 | 0.1111 | 13 | 0.0769 | 9 | 0 | 51 | 0.0784 | 4 | 0 | 10 | 0 | 12 | 0.1667 | 9 | 0 | 35 | 0.0571 |
| 9_1067729 | 17 | 0.1765 | 10 | 0.4 | 11 | 0.4545 | 14 | 0.4286 | 52 | 0.3462 | 3 | 0.6667 | 9 | 0.5556 | 11 | 0.3636 | 9 | 0.7778 | 32 | 0.5625 |
| 9_1071332( | NA | NA | NA | NA | 1 | 1 | NA | NA | 1 | 1 | 1 | 1 | NA | NA | 1 | 1 | NA | NA | 2 | 1 |
| 9_1071333 | NA | NA | NA | NA | 1 | 1 | NA | NA | 1 | 1 | 1 | 1 | NA | NA | NA | NA | NA | NA | 1 | 1 |
| 9_1071777 | NA | NA | 2 | 1 | 1 | 1 | 1 | 1 | 4 | 1 | NA | NA | NA | NA | 1 | 1 | 1 | 1 | 2 | 1 |
| 9_1071779( | NA | NA | 2 | 0.5 | 1 | 0 | 2 | 0.5 | 5 | 0.4 | NA | NA | NA | NA | NA | NA | 1 | 1 | 1 | 1 |
| 9_1071780 | NA | NA | 4 | 1 | 2 | 1 | NA | NA | 6 | 1 | NA | NA | NA | NA | 1 | 1 | 1 | 1 | 2 | 1 |
| 9_1071780! | NA | NA | 2 | 0.5 | 2 | 0.5 | 2 | 1 | 6 | 0.6667 | NA | NA | NA | NA | 1 | 0 | 1 | 1 | 2 | 0.5 |
| 9_1071781( | NA | NA | 4 | 1 | 1 | 1 | 2 | 1 | 7 | 1 | NA | NA | NA | NA | 1 | 1 | 1 | 1 | 2 | 1 |
| 9_1071782 | NA | NA | 4 | 0.25 | 2 | 0 | 2 | 0.5 | 8 | 0.25 | NA | NA | NA | NA | 1 | 0 | 1 | 1 | 2 | 0.5 |
| 9_1071783 | NA | NA | 4 | 0 | 2 | 0.5 | 1 | 0 | 7 | 0.1429 | NA | NA | NA | NA | 1 | 0 | 1 | 1 | 2 | 0.5 |
| 9_1071784' | NA | NA | 3 | 0.6667 | 1 | 1 | 2 | 1 | 6 | 0.8333 | NA | NA | NA | NA | 1 | 1 | 1 | 1 | 2 | 1 |
| 9_1071785! | NA | NA | 4 | 0.25 | 2 | 0 | 2 | 0.5 | 8 | 0.25 | NA | NA | NA | NA | 1 | 1 | 1 | 1 | 2 | 1 |
| 9_1072050 | 2 | 0.5 | 2 | 0 | 3 | 0 | 1 | 0 | 8 | 0.125 | NA | NA | 2 | 0.5 | 2 | 0 | 3 | 0.6667 | 7 | 0.4286 |
| 9_1072144( | NA | NA | NA | NA | 1 | 1 | NA | NA | 1 | 1 | NA | NA | NA | NA | NA | NA | 2 | 0.5 | 2 | 0.5 |
| 9_1072253! | 2 | 1 | NA | NA | NA | NA | NA | NA | 2 | 1 | NA | NA | 1 | 1 | NA | NA | NA | NA | 1 | 1 |
| 9_1072254 | 2 | 1 | NA | NA | NA | NA | NA | NA | 2 | 1 | 1 | 1 | 1 | 1 | NA | NA | NA | NA | 2 | 1 |
| 9_1074021 | NA | NA | 1 | 1 | 3 | 0.6667 | 2 | 0 | 6 | 0.5 | 4 | 0 | 2 | 0 | 5 | 0.6 | 4 | 0 | 15 | 0.2 |
| 9_1074047 | NA | NA | 1 | 0 | 4 | 0.75 | 7 | 0.1429 | 12 | 0.3333 | 6 | 0.1667 | 4 | 0.25 | 9 | 0.1111 | 5 | 0.4 | 24 | 0.2083 |
| 9_1074048( | NA | NA | 2 | 0.5 | 4 | 0.75 | 6 | 0.1667 | 12 | 0.4167 | 4 | 0.5 | 2 | 0 | 11 | 0.0909 | 3 | 0.3333 | 20 | 0.2 |
| 9_1074114! | 2 | 0 | 3 | 0 | 4 | 0.25 | 4 | 0 | 13 | 0.0769 | 1 | 0 | 1 | 0 | 5 | 0.2 | 4 | 0.25 | 11 | 0.1818 |
| 9_1074117! | 2 | 0 | 5 | 0.4 | 1 | 1 | 3 | 0.3333 | 11 | 0.3636 | 1 | 1 | 1 | 1 | 4 | 0.25 | 3 | 0.6667 | 9 | 0.5556 |
| 9_1074118! | 4 | 1 | 5 | 0.8 | 2 | 0.5 | 4 | 0.75 | 15 | 0.8 | 2 | 1 | 3 | 1 | 4 | 0.75 | 3 | 1 | 12 | 0.9167 |
| 9_1074118 | 4 | 0 | 4 | 0 | 3 | 0 | 3 | 0.3333 | 14 | 0.0714 | 2 | 0 | 3 | 0.6667 | 3 | 0.6667 | 3 | 0 | 11 | 0.3636 |
| 9_1074121 | 4 | 0.5 | 7 | 0.8571 | 6 | 0.3333 | 4 | 0.5 | 21 | 0.5714 | 1 | 1 | 5 | 0.4 | 5 | 0.2 | 4 | 1 | 15 | 0.5333 |
| 9_1074121 | 6 | 0.6667 | 8 | 0.625 | 4 | 0.5 | 3 | 1 | 21 | 0.6667 | 1 | 0 | 5 | 1 | 5 | 0.4 | 4 | 0.75 | 15 | 0.6667 |
| 9_1074123 | 6 | 0.1667 | 7 | 0.1429 | 6 | 0.5 | 5 | 0.6 | 24 | 0.3333 | 3 | 0.3333 | 4 | 0.75 | 4 | 0.25 | 6 | 0.1667 | 17 | 0.3529 |
| 9_1074123! | 5 | 0.2 | 6 | 0.6667 | 5 | 0.4 | 5 | 0.6 | 21 | 0.4762 | 3 | 0.6667 | 3 | 0 | 3 | 0.3333 | 6 | 0.6667 | 15 | 0.4667 |
| 9_1074151( | 3 | 1 | 4 | 1 | 3 | 1 | 5 | 1 | 15 | 1 | 3 | 1 | 3 | 1 | 2 | 1 | 6 | 1 | 14 | 1 |
| 9_1074152 | 7 | 0 | 8 | 0.25 | 6 | 0.1667 | 6 | 0 | 27 | 0.1111 | 3 | 0.3333 | 5 | 0.2 | 5 | 0 | 5 | 0 | 18 | 0.1111 |
| 9_1074153 | 7 | 0.2857 | 7 | 0 | 6 | 0.1667 | 5 | 0 | 25 | 0.12 | 1 | 0 | 4 | 0 | 5 | 0 | 7 | 0.1429 | 17 | 0.0588 |
| 9_1074153! | 6 | 0.6667 | 7 | 0.8571 | 6 | 1 | 5 | 1 | 24 | 0.875 | 3 | 0.6667 | 5 | 0.8 | 6 | 1 | 6 | 0.8333 | 20 | 0.85 |
| 9_1074154' | 6 | 0.6667 | 8 | 0.875 | 6 | 0.6667 | 6 | 0.6667 | 26 | 0.7308 | 3 | 0.6667 | 6 | 0.8333 | 6 | 0.3333 | 6 | 0.8333 | 21 | 0.6667 |
| 9_1074154 | 7 | 0.4286 | 6 | 0.5 | 5 | 0.6 | 6 | 0.3333 | 24 | 0.4583 | 3 | 0.3333 | 6 | 0.6667 | 5 | 0.4 | 6 | 0.8333 | 20 | 0.6 |
| 9_1074155! | 7 | 1 | 8 | 1 | 6 | 1 | 5 | 1 | 26 | 1 | 2 | 1 | 6 | 1 | 6 | 1 | 6 | 1 | 20 | 1 |
| 9_1074155! | 6 | 0.5 | 7 | 0.8571 | 4 | 0.75 | 4 | 1 | 21 | 0.7619 | 3 | 1 | 7 | 1 | 6 | 0.8333 | 5 | 0.8 | 21 | 0.9048 |
| 9_1074156! | 6 | 0.1667 | 6 | 0 | 7 | 0.1429 | 5 | 0 | 24 | 0.0833 | 3 | 1 | 5 | 0.2 | 6 | 0.1667 | 6 | 0 | 20 | 0.25 |
| 9_1074156 | 5 | 0.4 | 6 | 0 | 2 | 0 | 6 | 0 | 19 | 0.1053 | 2 | 0.5 | 5 | 0 | 4 | 0 | 7 | 0.1429 | 18 | 0.1111 |
| 9_1074156! | 6 | 0.5 | 7 | 0.7143 | 6 | 0.8333 | 7 | 0.4286 | 26 | 0.6154 | 2 | 0 | 7 | 0.7143 | 5 | 0.4 | 5 | 0.8 | 19 | 0.5789 |
| 9_1074157! | 6 | 0.3333 | 7 | 0.8571 | 5 | 0.6 | 4 | 0 | 22 | 0.5 | 2 | 0.5 | 6 | 0.5 | 6 | 0.3333 | 7 | 0.1429 | 21 | 0.3333 |
| 9_1074169 | 1 | 1 | 5 | 0.6 | NA | NA | 3 | 0.6667 | 9 | 0.6667 | 3 | 0.3333 | 2 | 0.5 | 2 | 0.5 | 3 | 0.6667 | 10 | 0.5 |

| | | | | | | | | | | | | | | | | | | | | |
|---|---|---|---|---|---|---|---|---|---|---|---|---|---|---|---|---|---|---|---|---|
| 9_1074169 | 2 | 1 | 5 | 0.6 | NA | NA | 1 | 1 | 8 | 0.75 | 3 | 1 | 2 | 1 | 2 | 1 | 2 | 1 | 9 | 1 |
| 9_1074170 | 2 | 1 | 5 | 0.8 | NA | NA | 3 | 1 | 10 | 0.9 | 3 | 1 | 2 | 1 | 1 | 1 | 3 | 1 | 9 | 1 |
| 9_1074172 | 1 | 0 | 2 | 0.5 | NA | NA | 1 | 0 | 4 | 0.25 | 2 | 0 | 2 | 1 | 1 | 0 | 2 | 0 | 7 | 0.2857 |
| 9_1074248 | 1 | 1 | NA | NA | NA | NA | NA | NA | 1 | 1 | NA | NA | NA | NA | 1 | 1 | NA | NA | 1 | 1 |
| 9_1074282 | 3 | 0.6667 | NA | NA | NA | NA | 2 | 0 | 5 | 0.4 | 1 | 1 | NA | NA | 1 | 0 | NA | NA | 2 | 0.5 |
| 9_1074390 | NA | NA | 2 | 0.5 | 3 | 0 | 2 | 0.5 | 7 | 0.2857 | 2 | 0 | 2 | 0.5 | 5 | 0 | 4 | 0.25 | 13 | 0.1538 |
| 9_1074421 | 12 | 0.0833 | 14 | 0.0714 | 13 | 0.0769 | 7 | 0.1429 | 46 | 0.087 | 5 | 0 | 6 | 0.3333 | 10 | 0 | 15 | 0 | 36 | 0.0556 |
| 9_1074422 | 9 | 0.5556 | 14 | 0.5714 | 13 | 0.5385 | 8 | 0.625 | 44 | 0.5682 | 6 | 0.6667 | 6 | 1 | 10 | 0.5 | 13 | 0.3077 | 35 | 0.5429 |
| 9_1074422 | 8 | 0.625 | 15 | 0.4 | 9 | 0.1111 | 5 | 0 | 37 | 0.3243 | 7 | 0.4286 | 4 | 0.5 | 7 | 0.2857 | 10 | 0.4 | 28 | 0.3929 |
| 9_1074423 | 5 | 0.6 | 8 | 0.5 | 12 | 0.25 | 8 | 0.5 | 33 | 0.4242 | 5 | 0.2 | 5 | 0.2 | 6 | 0.3333 | 8 | 0.375 | 24 | 0.2917 |
| 9_1074423 | 10 | 1 | 16 | 0.875 | 14 | 0.9286 | 10 | 0.9 | 50 | 0.92 | 6 | 0.8333 | 7 | 0.8571 | 7 | 0.8571 | 13 | 0.9231 | 33 | 0.8788 |
| 9_1074425 | 8 | 0.125 | 16 | 0.25 | 12 | 0.4167 | 5 | 0 | 41 | 0.2439 | 7 | 0.2857 | 5 | 0.2 | 6 | 0.1667 | 11 | 0.3636 | 29 | 0.2759 |
| 9_1074426 | 9 | 0.3333 | 13 | 0.4615 | 14 | 0.2857 | 9 | 0.6667 | 45 | 0.4222 | 7 | 0.4286 | 4 | 0.25 | 6 | 0.5 | 12 | 0.25 | 29 | 0.3448 |
| 9_1074426 | 10 | 0.3 | 16 | 0.125 | 16 | 0.1875 | 10 | 0.1 | 52 | 0.1731 | 8 | 0.25 | 6 | 0.3333 | 8 | 0.625 | 13 | 0.2308 | 35 | 0.3429 |
| 9_1074426 | 10 | 0.6 | 18 | 0.6111 | 15 | 0.4667 | 10 | 0.4 | 53 | 0.5283 | 9 | 0.6667 | 6 | 0.5 | 9 | 0.6667 | 11 | 0.6364 | 35 | 0.6286 |
| 9_1074427 | 10 | 0.3 | 19 | 0.2105 | 14 | 0.3571 | 9 | 0.2222 | 52 | 0.2692 | 10 | 0.5 | 7 | 0.1429 | 7 | 0 | 13 | 0.4615 | 37 | 0.3243 |
| 9_1074427 | 11 | 0.6364 | 19 | 0.8421 | 17 | 0.8235 | 9 | 0.8889 | 56 | 0.8036 | 10 | 0.8 | 6 | 1 | 7 | 1 | 14 | 0.6429 | 37 | 0.8108 |
| 9_1074428 | 10 | 0.2 | 18 | 0.0556 | 16 | 0.25 | 9 | 0.1111 | 53 | 0.1509 | 10 | 0.1 | 6 | 0.3333 | 10 | 0.1 | 13 | 0.0769 | 39 | 0.1282 |
| 9_1074473 | 2 | 1 | 2 | 0 | 1 | 0 | 1 | 1 | 6 | 0.5 | 2 | 1 | 2 | 1 | 1 | 0 | 2 | 1 | 7 | 0.8571 |
| 9_1074475 | 2 | 1 | 1 | 1 | 1 | 1 | NA | NA | 4 | 1 | NA | NA | 2 | 0.5 | NA | NA | 2 | 0 | 4 | 0.25 |
| 9_1074476 | 2 | 1 | 2 | 0.5 | 1 | 0 | 1 | 0 | 6 | 0.5 | 1 | 0 | 1 | 1 | 1 | 0 | 2 | 0.5 | 5 | 0.4 |
| 9_1074477 | 1 | 1 | 2 | 1 | 1 | 1 | NA | NA | 4 | 1 | 2 | 0.5 | 2 | 1 | 1 | 1 | 2 | 0.5 | 7 | 0.7143 |
| 9_1074477 | 2 | 0 | 2 | 1 | 1 | 0 | 1 | 1 | 6 | 0.5 | 2 | 0 | 2 | 0.5 | 1 | 0 | 2 | 0 | 7 | 0.1429 |
| 9_1074479 | 2 | 0 | 2 | 0 | 2 | 0 | 1 | 1 | 7 | 0.1429 | 1 | 0 | 3 | 0.3333 | 1 | 0 | 2 | 0.5 | 7 | 0.2857 |
| 9_1074480 | 2 | 1 | 2 | 1 | 2 | 1 | 1 | 1 | 7 | 1 | NA | NA | 3 | 1 | NA | NA | 2 | 1 | 5 | 1 |
| 9_1074495 | NA | NA | 2 | 1 | 2 | 0.5 | NA | NA | 4 | 0.75 | 1 | 1 | 3 | 1 | 2 | 1 | 2 | 1 | 8 | 1 |
| 9_1074496 | 2 | 0.5 | 2 | 0.5 | 2 | 1 | 1 | 1 | 7 | 0.7143 | 2 | 1 | 4 | 1 | 2 | 1 | 2 | 0.5 | 10 | 0.9 |
| 9_1074599 | NA | NA | 1 | 1 | NA | NA | NA | NA | 1 | 1 | NA | NA | 2 | 0.5 | NA | NA | NA | NA | 2 | 0.5 |
| 9_1074600 | NA | NA | 1 | 0 | 1 | 1 | 1 | 0 | 3 | 0.3333 | 3 | 0 | 4 | 0.25 | NA | NA | NA | NA | 7 | 0.1429 |
| 9_1074606 | 1 | 1 | NA | NA | 1 | 0 | 1 | 1 | 3 | 0.6667 | 2 | 0.5 | 4 | 0 | NA | NA | NA | NA | 6 | 0.1667 |
| 9_1074609 | 1 | 0 | 1 | 0 | 2 | 0 | 2 | 1 | 6 | 0.3333 | 2 | 0.5 | 2 | 0.5 | 1 | 1 | NA | NA | 5 | 0.6 |
| 9_1074609 | 1 | 0 | 1 | 1 | 2 | 0.5 | 3 | 1 | 7 | 0.7143 | 3 | 0.6667 | 4 | 0.25 | 2 | 0.5 | NA | NA | 9 | 0.4444 |
| 9_1074610 | 1 | 0 | NA | NA | 2 | 0.5 | 3 | 0.6667 | 6 | 0.5 | 2 | 0.5 | 3 | 1 | 2 | 0.5 | NA | NA | 7 | 0.7143 |
| 9_1074610 | 1 | 0 | 1 | 0 | 2 | 0 | 3 | 0.6667 | 7 | 0.2857 | 4 | 0 | 3 | 0.3333 | 2 | 0 | NA | NA | 9 | 0.1111 |
| 9_1074611 | 1 | 1 | 1 | 0 | 2 | 1 | 2 | 0.5 | 6 | 0.6667 | 4 | 0.5 | 4 | 0.5 | 2 | 0.5 | NA | NA | 10 | 0.5 |
| 9_1074673 | NA | NA | NA | NA | 1 | 1 | 1 | 1 | 2 | 1 | NA | NA | 1 | 1 | NA | NA | 3 | 0 | 4 | 0.25 |
| 9_1074917 | 27 | 0.1111 | 26 | 0 | 23 | 0.087 | 23 | 0 | 99 | 0.0505 | 21 | 0.0952 | 30 | 0.0667 | 20 | 0 | 28 | 0.1429 | 99 | 0.0808 |
| 9_1074917 | 27 | 0.1111 | 26 | 0.0385 | 27 | 0 | 28 | 0.1429 | 108 | 0.0741 | 30 | 0 | 34 | 0.1176 | 21 | 0.0476 | 38 | 0.1316 | 123 | 0.0813 |
| 9_1075285 | 6 | 0 | 7 | 0.1429 | 7 | 0 | 7 | 0.1429 | 27 | 0.0741 | 8 | 0.125 | 9 | 0.1111 | 5 | 0 | 7 | 0 | 29 | 0.069 |
| 9_1075289 | 5 | 0.2 | 6 | 0 | 5 | 0 | 6 | 0.1667 | 22 | 0.0909 | 5 | 0.2 | 4 | 0 | 4 | 0.5 | 3 | 0.3333 | 16 | 0.25 |
| 9_1075290 | 4 | 0.25 | 6 | 0.5 | 7 | 0.5714 | 6 | 0.3333 | 23 | 0.4348 | 6 | 0.5 | 9 | 0.4444 | 6 | 0.6667 | 7 | 0.1429 | 28 | 0.4286 |
| 9_1075592 | 3 | 0.6667 | NA | NA | 1 | 0 | 3 | 0.6667 | 7 | 0.5714 | 1 | 1 | 1 | 1 | 1 | 1 | NA | NA | 3 | 1 |
| 9_1075592 | 2 | 0.5 | NA | NA | 3 | 0.6667 | 3 | 1 | 8 | 0.75 | 1 | 0 | 2 | 0.5 | 2 | 1 | NA | NA | 5 | 0.6 |
| 9_1075594 | 2 | 0.5 | NA | NA | 1 | 1 | 3 | 1 | 6 | 0.8333 | 1 | 0 | 2 | 0.5 | 1 | 1 | NA | NA | 4 | 0.5 |
| 9_1075594 | 3 | 0.3333 | NA | NA | 2 | 0.5 | 1 | 1 | 6 | 0.5 | 1 | 1 | 1 | 1 | 1 | 1 | NA | NA | 3 | 1 |
| 9_1075595 | 3 | 0.6667 | NA | NA | 3 | 0 | 2 | 1 | 8 | 0.5 | 1 | 0 | 1 | 0 | 2 | 1 | NA | NA | 4 | 0.5 |
| 9_1075596 | 3 | 0 | NA | NA | 3 | 0.3333 | 3 | 0.3333 | 9 | 0.2222 | 1 | 0 | 2 | 0 | 2 | 0.5 | NA | NA | 5 | 0.2 |
| 9_1075596 | 3 | 0.3333 | NA | NA | 1 | 0 | 1 | 1 | 5 | 0.4 | 1 | 1 | 2 | 0.5 | 2 | 0 | NA | NA | 5 | 0.4 |
| 9_1076179 | 11 | 0.2727 | 11 | 0.0909 | 20 | 0.05 | 18 | 0.1111 | 60 | 0.1167 | 6 | 0 | 16 | 0.0625 | 18 | 0.1111 | 14 | 0 | 54 | 0.0556 |
| 9_1076179 | 8 | 0 | 11 | 0.0909 | 13 | 0.1538 | 9 | 0.2222 | 41 | 0.122 | 4 | 0.25 | 11 | 0.0909 | 15 | 0.2 | 12 | 0 | 42 | 0.119 |
| 9_1076179 | 10 | 0.2 | 8 | 0.125 | 17 | 0.1176 | 15 | 0.1333 | 50 | 0.14 | 5 | 0.2 | 16 | 0.125 | 16 | 0.1875 | 13 | 0.2308 | 50 | 0.18 |
| 9_1076349 | 2 | 0 | 5 | 0.2 | 2 | 0 | 1 | 0 | 10 | 0.1 | 1 | 0 | 1 | 1 | 3 | 0 | 6 | 0 | 11 | 0.0909 |
| 9_1077294 | 2 | 1 | 2 | 1 | 1 | 1 | 2 | 1 | 7 | 1 | NA | NA | NA | NA | 3 | 1 | 1 | 1 | 4 | 1 |
| 9_1077295 | 2 | 0.5 | 2 | 0 | 1 | 0 | 2 | 0 | 7 | 0.1429 | NA | NA | NA | NA | 3 | 0.6667 | 1 | 1 | 4 | 0.75 |
| 9_1077296 | 2 | 1 | 2 | 1 | 1 | 1 | 2 | 1 | 7 | 1 | NA | NA | NA | NA | 3 | 1 | 1 | 1 | 4 | 1 |
| 9_1077298 | 2 | 0.5 | 1 | 1 | NA | NA | 2 | 0.5 | 5 | 0.6 | NA | NA | NA | NA | 3 | 1 | 1 | 0 | 4 | 0.75 |
| 9_1077299 | 2 | 0 | 1 | 1 | NA | NA | 2 | 0.5 | 5 | 0.4 | NA | NA | NA | NA | 2 | 0.5 | NA | NA | 2 | 0.5 |
| 9_1077299 | 2 | 1 | 1 | 1 | 1 | 0 | 2 | 1 | 6 | 0.8333 | NA | NA | NA | NA | 3 | 0.6667 | NA | NA | 3 | 0.6667 |
| 9_1077302 | 2 | 1 | NA | NA | 1 | 1 | 2 | 1 | 5 | 1 | NA | NA | NA | NA | 3 | 1 | NA | NA | 3 | 1 |
| 9_1077303 | 2 | 1 | 1 | 1 | NA | NA | 2 | 1 | 5 | 1 | NA | NA | NA | NA | 3 | 0.6667 | NA | NA | 3 | 0.6667 |
| 9_1077303 | 2 | 1 | 1 | 0 | 1 | 0 | 1 | 0 | 5 | 0.4 | NA | NA | NA | NA | 2 | 0.5 | NA | NA | 2 | 0.5 |
| 9_1077304 | 2 | 1 | 1 | 1 | 1 | 0 | 2 | 1 | 6 | 0.8333 | NA | NA | NA | NA | 2 | 1 | NA | NA | 2 | 1 |
| 9_1077306 | NA | NA | 1 | 1 | NA | NA | NA | NA | 1 | 1 | NA | NA | NA | NA | 2 | 1 | NA | NA | 2 | 1 |
| 9_1077307 | NA | NA | 1 | 1 | NA | NA | 1 | 0 | 2 | 0.5 | NA | NA | NA | NA | 2 | 0.5 | NA | NA | 2 | 0.5 |
| 9_1077773 | 1 | 1 | 4 | 0.5 | 1 | 0 | 4 | 0.25 | 10 | 0.4 | 4 | 0.5 | 1 | 0 | NA | NA | 2 | 0.5 | 7 | 0.4286 |

| | | | | | | | | | | | | | | | | | | | | |
|---|---|---|---|---|---|---|---|---|---|---|---|---|---|---|---|---|---|---|---|---|
| 9_1077773 | 1 | 1 | 4 | 0 | 1 | 1 | 6 | 0.1667 | 12 | 0.25 | 4 | 0.25 | 1 | 0 | NA | NA | 2 | 0 | 7 | 0.1429 |
| 9_1077784 | NA | NA | 1 | 1 | NA | NA | 1 | 1 | 2 | 1 | NA | NA | 1 | 1 | NA | NA | 1 | 0 | 2 | 0.5 |
| 9_1077786 | 1 | 0 | 5 | 0.2 | 1 | 0 | 8 | 0 | 15 | 0.0667 | 5 | 0 | 3 | 0.3333 | 1 | 0 | 3 | 0 | 12 | 0.0833 |
| 9_1077786 | 1 | 1 | 6 | 0.8333 | 1 | 1 | 6 | 0.5 | 14 | 0.7143 | 4 | 0.75 | 3 | 0.6667 | 1 | 1 | 3 | 0.6667 | 11 | 0.7273 |
| 9_1077787 | 1 | 1 | 7 | 0.7143 | 1 | 1 | 8 | 1 | 17 | 0.8824 | 5 | 0.8 | 3 | 0.6667 | 2 | 0.5 | 3 | 0.6667 | 13 | 0.6923 |
| 9_1077788 | 1 | 1 | 7 | 0.8571 | 1 | 1 | 7 | 0.5714 | 16 | 0.75 | 5 | 1 | 2 | 1 | 2 | 1 | 3 | 0.3333 | 12 | 0.8333 |
| 9_1077789 | 1 | 0 | 8 | 0.25 | 1 | 0 | 9 | 0.2222 | 19 | 0.2105 | 3 | 0 | 4 | 0.25 | 2 | 1 | 3 | 0.3333 | 12 | 0.3333 |
| 9_1077789 | NA | NA | 6 | 0.6667 | 1 | 0 | 8 | 1 | 15 | 0.8 | 4 | 0.75 | 4 | 0.75 | 2 | 1 | 3 | 0.6667 | 13 | 0.7692 |
| 9_1077791 | 1 | 1 | 8 | 0.75 | 1 | 1 | 9 | 0.7778 | 19 | 0.7895 | 5 | 0.6 | 5 | 0.6 | 2 | 0.5 | 3 | 0.6667 | 15 | 0.6 |
| 9_1077792 | NA | NA | 1 | 0 | NA | NA | 1 | 1 | 2 | 0.5 | 1 | 1 | 1 | 1 | 2 | 0 | 1 | 0 | 5 | 0.4 |
| 9_1077799 | NA | NA | 1 | 1 | NA | NA | 1 | 1 | 2 | 1 | 1 | 0 | 1 | 1 | 1 | 1 | NA | NA | 3 | 0.6667 |
| 9_1077800 | 2 | 0 | 6 | 0.3333 | 1 | 0 | 7 | 0.4286 | 16 | 0.3125 | 4 | 0.5 | 2 | 1 | 4 | 0.25 | 3 | 0.3333 | 13 | 0.4615 |
| 9_1077802 | 2 | 0 | 5 | 0.6 | NA | NA | 9 | 0.2222 | 16 | 0.3125 | 3 | 0.3333 | 5 | 0.4 | 4 | 0.5 | 3 | 1 | 15 | 0.5333 |
| 9_1078254 | 29 | 0.8621 | 29 | 0.7241 | 25 | 0.76 | 48 | 0.8125 | 131 | 0.7939 | 29 | 0.8276 | 36 | 0.7222 | 40 | 0.85 | 27 | 0.8148 | 132 | 0.803 |
| 9_1078262 | 60 | 0.0333 | 45 | 0.1333 | 41 | 0.0976 | 73 | 0.0274 | 219 | 0.0639 | 52 | 0.0385 | 71 | 0.1127 | 81 | 0.037 | 48 | 0.0208 | 252 | 0.0556 |
| 9_1078263 | 52 | 0.0769 | 43 | 0.0465 | 37 | 0.0811 | 70 | 0.1 | 202 | 0.0792 | 42 | 0.0476 | 64 | 0.1406 | 70 | 0.0714 | 49 | 0.0612 | 225 | 0.0844 |
| 9_1078264 | 49 | 0.1633 | 44 | 0.2727 | 34 | 0.2059 | 58 | 0.2414 | 185 | 0.2216 | 42 | 0.5 | 61 | 0.2787 | 68 | 0.2794 | 46 | 0.1957 | 217 | 0.3041 |
| 9_1078270 | 4 | 0.5 | 2 | 1 | 4 | 0.75 | 1 | 1 | 11 | 0.7273 | 2 | 1 | 5 | 0.2 | 5 | 0.2 | 3 | 0.6667 | 15 | 0.4 |
| 9_1078271 | 4 | 0.75 | 3 | 1 | 3 | 0.3333 | 2 | 1 | 12 | 0.75 | 2 | 0.5 | 5 | 0.8 | 5 | 0.8 | 1 | 1 | 13 | 0.7692 |
| 9_1078482 | 2 | 0.5 | 1 | 1 | 2 | 0.5 | 2 | 0.5 | 7 | 0.5714 | 1 | 0 | 5 | 0.8 | 4 | 0.75 | 4 | 0.5 | 14 | 0.6429 |
| 9_1078485 | 2 | 1 | 1 | 1 | 3 | 1 | 1 | 1 | 7 | 1 | 1 | 0 | 5 | 1 | 4 | 1 | 4 | 1 | 14 | 0.9286 |
| 9_1078485 | 2 | 0.5 | NA | NA | 3 | 0 | 2 | 0 | 7 | 0.1429 | 1 | 0 | 5 | 0.2 | 2 | 0.5 | 5 | 0.6 | 13 | 0.3846 |
| 9_1078486 | 3 | 0.3333 | 1 | 0 | 3 | 0 | 2 | 0 | 9 | 0.1111 | 1 | 0 | 5 | 0 | 4 | 0 | 4 | 0.25 | 14 | 0.0714 |
| 9_1078614 | NA | NA | NA | NA | NA | NA | 1 | 1 | 1 | 1 | NA | NA | 1 | 1 | NA | NA | NA | NA | 1 | 1 |
| 9_1078616 | NA | NA | NA | NA | NA | NA | 1 | 1 | 1 | 1 | NA | NA | 1 | 1 | NA | NA | NA | NA | 1 | 1 |
| 9_1078619 | NA | NA | NA | NA | NA | NA | 1 | 1 | 1 | 1 | NA | NA | 1 | 1 | NA | NA | NA | NA | 1 | 1 |
| 9_1078620 | NA | NA | NA | NA | NA | NA | 1 | 1 | 1 | 1 | NA | NA | 1 | 1 | NA | NA | NA | NA | 1 | 1 |
| 9_1079227 | 6 | 0.3333 | 7 | 0.8571 | 7 | 0.7143 | 9 | 0.6667 | 29 | 0.6552 | 8 | 0.5 | 9 | 0.6667 | 8 | 0.625 | 6 | 0.6667 | 31 | 0.6129 |
| 9_1079227 | 8 | 0.5 | 12 | 0.75 | 13 | 0.6923 | 9 | 0.6667 | 42 | 0.6667 | 8 | 0.5 | 9 | 0.7778 | 8 | 0.875 | 7 | 0.8571 | 32 | 0.75 |
| 9_1079228 | 9 | 0.5556 | 13 | 0.7692 | 9 | 0.6667 | 10 | 0.4 | 41 | 0.6098 | 8 | 0.375 | 8 | 0.625 | 9 | 0.5556 | 4 | 0.75 | 29 | 0.5517 |
| 9_1079229 | 7 | 0.7143 | 7 | 1 | 8 | 0.875 | 6 | 1 | 28 | 0.8929 | 8 | 0.75 | 3 | 0.6667 | 7 | 1 | 5 | 0.8 | 23 | 0.8261 |
| 9_1079230 | 11 | 1 | 12 | 0.9167 | 10 | 1 | 8 | 1 | 41 | 0.9756 | 9 | 0.8889 | 8 | 1 | 9 | 0.8889 | 6 | 1 | 32 | 0.9375 |
| 9_1079231 | 11 | 0.0909 | 10 | 0.2 | 11 | 0.1818 | 9 | 0.1111 | 41 | 0.1463 | 10 | 0.1 | 5 | 0.2 | 7 | 0 | 8 | 0 | 30 | 0.0667 |
| 9_1079231 | 13 | 0.6154 | 11 | 0.6364 | 13 | 0.3077 | 12 | 0.5 | 49 | 0.5102 | 9 | 0.4444 | 9 | 0.3333 | 11 | 0.1818 | 9 | 0.3333 | 38 | 0.3158 |
| 9_1079232 | 12 | 0.4167 | 11 | 0.4545 | 13 | 0.2308 | 10 | 0.6 | 46 | 0.413 | 9 | 0.4444 | 9 | 0.5556 | 11 | 0.6364 | 10 | 0.2 | 39 | 0.4615 |
| 9_1079232 | 11 | 0.6364 | 10 | 0.8 | 11 | 0.9091 | 11 | 0.8182 | 43 | 0.7907 | 7 | 0.7143 | 8 | 0.75 | 8 | 1 | 8 | 0.875 | 31 | 0.8387 |
| 9_1079232 | 10 | 0.5 | 12 | 0.4167 | 14 | 0.6429 | 11 | 0.4545 | 47 | 0.5106 | 10 | 0.3 | 7 | 0.5714 | 10 | 0.6 | 9 | 0.5556 | 36 | 0.5 |
| 9_1079232 | 13 | 0.7692 | 13 | 0.8462 | 14 | 0.5714 | 9 | 0.8889 | 49 | 0.7551 | 9 | 0.6667 | 9 | 0.7778 | 7 | 1 | 9 | 0.7778 | 34 | 0.7941 |
| 9_1079233 | 10 | 0 | 10 | 0.1 | 12 | 0.0833 | 11 | 0.1818 | 43 | 0.093 | 9 | 0.1111 | 7 | 0 | 8 | 0.125 | 8 | 0.125 | 32 | 0.0938 |
| 9_1079233 | 11 | 0.1818 | 12 | 0.25 | 14 | 0.2857 | 9 | 0.3333 | 46 | 0.2609 | 9 | 0.2222 | 10 | 0.2 | 12 | 0.0833 | 9 | 0.5556 | 40 | 0.25 |
| 9_1079234 | 13 | 0.8462 | 12 | 0.9167 | 14 | 0.7143 | 8 | 0.875 | 47 | 0.8298 | 10 | 0.7 | 8 | 0.625 | 12 | 1 | 8 | 0.75 | 38 | 0.7895 |
| 9_1079235 | 13 | 0.0769 | 9 | 0.1111 | 12 | 0 | 6 | 0.1667 | 40 | 0.075 | 10 | 0.2 | 10 | 0.1 | 12 | 0 | 8 | 0.125 | 40 | 0.1 |
| 9_1079236 | 13 | 0.0769 | 10 | 0.3 | 14 | 0.2857 | 10 | 0.1 | 47 | 0.1915 | 9 | 0.2222 | 7 | 0.4286 | 9 | 0.3333 | 10 | 0.4 | 35 | 0.3429 |
| 9_1079241 | 15 | 0 | 17 | 0.4118 | 19 | 0.2105 | 16 | 0.0625 | 67 | 0.1791 | 11 | 0.0909 | 10 | 0 | 16 | 0.1875 | 10 | 0 | 47 | 0.0851 |
| 9_1079247 | 20 | 0.1 | 19 | 0.2632 | 17 | 0 | 18 | 0.1111 | 74 | 0.1216 | 18 | 0.0556 | 17 | 0.0588 | 21 | 0.0952 | 15 | 0.1333 | 71 | 0.0845 |
| 9_1079255 | 31 | 0.129 | 24 | 0.4167 | 27 | 0.1852 | 21 | 0.1429 | 103 | 0.2136 | 19 | 0.0526 | 19 | 0.1053 | 24 | 0.2083 | 18 | 0.2778 | 80 | 0.1625 |
| 9_1079256 | 33 | 0.303 | 30 | 0.4667 | 35 | 0.4286 | 32 | 0.4375 | 130 | 0.4077 | 19 | 0.3684 | 24 | 0.3333 | 27 | 0.4074 | 24 | 0.2917 | 94 | 0.3511 |
| 9_1079257 | 30 | 0.1667 | 24 | 0.0417 | 30 | 0.2 | 23 | 0.0435 | 107 | 0.1215 | 16 | 0 | 20 | 0.15 | 27 | 0.1852 | 15 | 0.1333 | 78 | 0.1282 |
| 9_1079257 | 34 | 0.3235 | 34 | 0.3824 | 35 | 0.4 | 32 | 0.3125 | 135 | 0.3556 | 23 | 0.3043 | 23 | 0.4783 | 31 | 0.5484 | 24 | 0.375 | 101 | 0.4356 |
| 9_1079285 | 1 | 1 | 1 | 1 | NA | NA | 1 | 1 | 3 | 1 | 2 | 1 | 1 | 1 | 2 | 1 | 1 | 1 | 6 | 1 |
| 9_1079287 | 2 | 0.5 | 4 | 1 | NA | NA | 1 | 1 | 7 | 0.8571 | 2 | 1 | 1 | 1 | 3 | 1 | 2 | 1 | 8 | 1 |
| 9_1079287 | 2 | 1 | 4 | 0.5 | NA | NA | 1 | 1 | 7 | 0.7143 | 2 | 1 | 1 | 1 | 3 | 0.6667 | 2 | 1 | 8 | 0.875 |
| 9_1079287 | 2 | 1 | 3 | 1 | NA | NA | 1 | 1 | 6 | 1 | 3 | 1 | 1 | 0 | 2 | 0.5 | 2 | 0.5 | 8 | 0.625 |
| 9_1079287 | 1 | 0 | 3 | 0.3333 | NA | NA | 1 | 0 | 5 | 0.2 | 3 | 0.6667 | 1 | 1 | 2 | 0 | 2 | 0 | 8 | 0.375 |
| 9_1079288 | 1 | 1 | 3 | 0.6667 | NA | NA | NA | NA | 4 | 0.75 | 1 | 1 | 1 | 1 | 3 | 1 | NA | NA | 5 | 1 |
| 9_1079288 | 2 | 0 | 2 | 0.5 | NA | NA | 1 | 1 | 5 | 0.4 | 3 | 0.3333 | 1 | 0 | 3 | 0 | 2 | 0 | 9 | 0.1111 |
| 9_1079289 | 1 | 1 | 2 | 0.5 | NA | NA | 1 | 1 | 4 | 0.75 | 2 | 1 | 1 | 0 | 1 | 1 | NA | NA | 4 | 0.75 |
| 9_1079290 | NA | NA | 3 | 0.3333 | NA | NA | 1 | 0 | 4 | 0.25 | 2 | 0 | 1 | 0 | 2 | 0.5 | NA | NA | 5 | 0.2 |
| 9_1079290 | 4 | 0.25 | 5 | 0.2 | 5 | 0.4 | 8 | 0.375 | 22 | 0.3182 | 3 | 0 | 6 | 0.3333 | 1 | 0 | 11 | 0.2727 | 21 | 0.2381 |
| 9_1079292 | NA | NA | 2 | 0.5 | NA | NA | 2 | 0 | 4 | 0.25 | 1 | 1 | NA | NA | NA | NA | 1 | 1 | 2 | 1 |
| 9_1079297 | 5 | 0.2 | 5 | 0.2 | 1 | 0 | 6 | 0.1667 | 17 | 0.1765 | 3 | 0.3333 | 5 | 0 | 1 | 0 | 7 | 0 | 16 | 0.0625 |
| 9_1079318 | NA | NA | NA | NA | NA | NA | 2 | 1 | 2 | 1 | NA | NA | NA | NA | NA | NA | 1 | 1 | 1 | 1 |
| 9_1079318 | NA | NA | NA | NA | NA | NA | 2 | 1 | 2 | 1 | NA | NA | NA | NA | NA | NA | 2 | 1 | 2 | 1 |
| 9_1079626 | NA | NA | 2 | 0.5 | 2 | 0.5 | 3 | 0 | 7 | 0.2857 | 1 | 1 | 7 | 0.5714 | 5 | 0.6 | 6 | 0.6667 | 19 | 0.6316 |
| 9_1079626 | NA | NA | 2 | 0 | 2 | 0 | 3 | 0.6667 | 7 | 0.2857 | 1 | 0 | 6 | 0.1667 | 5 | 0.2 | 6 | 0.3333 | 18 | 0.2222 |

| | | | | | | | | | | | | | | | | | | | | |
|---|---|---|---|---|---|---|---|---|---|---|---|---|---|---|---|---|---|---|---|---|
| 9_10796267 | NA | NA | 1 | 1 | 2 | 0 | 3 | 0.3333 | 6 | 0.3333 | 1 | 0 | 7 | 0.2857 | 5 | 0.6 | 4 | 0.5 | 17 | 0.4118 |
| 9_10796273 | NA | NA | 1 | 0 | 2 | 0.5 | 2 | 0.5 | 5 | 0.4 | 1 | 0 | 6 | 0.5 | 5 | 0.6 | 4 | 0.25 | 16 | 0.4375 |
| 9_10796280 | NA | NA | NA | NA | 2 | 0.5 | NA | NA | 2 | 0.5 | 1 | 0 | 4 | 0 | 4 | 0.5 | 6 | 0 | 15 | 0.1333 |
| 9_10797324 | 20 | 0.2 | 31 | 0.0645 | 23 | 0.0435 | 19 | 0.0526 | 93 | 0.086 | 24 | 0.125 | 24 | 0.25 | 15 | 0.1333 | 22 | 0.0455 | 85 | 0.1412 |
| 9_10797381 | 14 | 0 | 21 | 0.0952 | 15 | 0 | 15 | 0.1333 | 65 | 0.0615 | 18 | 0.1111 | 16 | 0.125 | 9 | 0 | 13 | 0 | 56 | 0.0714 |
| 9_10797453 | 12 | 0.1667 | 16 | 0 | 10 | 0.4 | 9 | 0.3333 | 47 | 0.1915 | 15 | 0.2 | 9 | 0.2222 | 4 | 0.25 | 12 | 0.0833 | 40 | 0.175 |
| 9_10797478 | 11 | 0.0909 | 18 | 0.0556 | 9 | 0.1111 | 12 | 0.0833 | 50 | 0.08 | 13 | 0.2308 | 16 | 0.25 | 7 | 0.1429 | 14 | 0.1429 | 50 | 0.2 |
| 9_10797488 | 11 | 0.0909 | 17 | 0 | 12 | 0 | 13 | 0.3077 | 53 | 0.0943 | 13 | 0.3077 | 14 | 0.2143 | 6 | 0.3333 | 14 | 0.2143 | 47 | 0.2553 |
| 9_10797520 | 10 | 0 | 15 | 0.1333 | 7 | 0.1429 | 14 | 0.0714 | 46 | 0.087 | 14 | 0.1429 | 12 | 0 | 6 | 0 | 13 | 0.0769 | 45 | 0.0667 |
| 9_10797592 | 6 | 0.1667 | 8 | 0.125 | 7 | 0.1429 | 4 | 0 | 25 | 0.12 | 12 | 0.0833 | 9 | 0.1111 | 3 | 0 | 9 | 0.1111 | 33 | 0.0909 |
| 9_10797596 | 7 | 0 | 11 | 0 | 10 | 0 | 10 | 0.2 | 38 | 0.0526 | 12 | 0.0833 | 9 | 0.1111 | 5 | 0.2 | 11 | 0.0909 | 37 | 0.1081 |
| 9_10797617 | 7 | 0.1429 | 11 | 0.0909 | 10 | 0.2 | 9 | 0.1111 | 37 | 0.1351 | 8 | 0.125 | 7 | 0.1429 | 7 | 0.1429 | 10 | 0.1 | 32 | 0.125 |
| 9_10797627 | 5 | 0 | 10 | 0.3 | 6 | 0 | 5 | 0.4 | 26 | 0.1923 | 5 | 0.2 | 6 | 0.3333 | 2 | 0.5 | 7 | 0.2857 | 20 | 0.3 |
| 9_10797645 | 3 | 0 | 7 | 0.1429 | 6 | 0.1667 | 4 | 0 | 20 | 0.1 | 6 | 0 | 7 | 0 | 6 | 0.1667 | 5 | 0.2 | 24 | 0.0833 |
| 9_10797655 | 4 | 1 | 9 | 0.8889 | 8 | 0.875 | 8 | 0.875 | 29 | 0.8966 | 10 | 0.6 | 5 | 0.8 | 7 | 0.8571 | 8 | 0.875 | 30 | 0.7667 |
| 9_10797656 | 3 | 0.3333 | 7 | 0.4286 | 5 | 0.4 | 6 | 0.6667 | 21 | 0.4762 | 5 | 0.8 | 6 | 0.3333 | 4 | 0.25 | 7 | 0.4286 | 22 | 0.4545 |
| 9_10797657 | 5 | 1 | 8 | 0.875 | 7 | 1 | 7 | 1 | 27 | 0.963 | 8 | 0.875 | 6 | 1 | 6 | 1 | 7 | 1 | 27 | 0.963 |
| 9_10797671 | 3 | 1 | 7 | 0.8571 | 2 | 1 | 2 | 1 | 14 | 0.9286 | 7 | 1 | 5 | 0.6 | 5 | 1 | 6 | 0.8333 | 23 | 0.8696 |
| 9_10797692 | 3 | 0.6667 | 10 | 0.2 | 4 | 0.25 | 8 | 0.375 | 25 | 0.32 | 8 | 0.5 | 6 | 0.3333 | 5 | 0.6 | 7 | 0.1429 | 26 | 0.3846 |
| 9_10797694 | 3 | 0 | 9 | 0.5556 | 3 | 1 | 8 | 0.625 | 23 | 0.5652 | 7 | 0.8571 | 6 | 0.6667 | 5 | 0.8 | 6 | 0.6667 | 24 | 0.75 |
| 9_10797696 | 3 | 0.6667 | 8 | 0.5 | 2 | 0 | 6 | 0.5 | 19 | 0.4737 | 4 | 0.75 | 5 | 0.2 | 5 | 0.8 | 3 | 0.6667 | 17 | 0.5882 |
| 9_10797697 | 3 | 0 | 9 | 0.5556 | 4 | 0 | 7 | 0.4286 | 23 | 0.3478 | 7 | 0.5714 | 5 | 0.2 | 4 | 0.25 | 6 | 0.5 | 22 | 0.4091 |
| 9_10798270 | 1 | 0 | 5 | 0.4 | 2 | 0.5 | 4 | 0.5 | 12 | 0.4167 | 6 | 0.1667 | 6 | 0 | 2 | 0 | 2 | 1 | 16 | 0.1875 |
| 9_10798277 | 1 | 1 | 5 | 0 | 3 | 0 | 4 | 0 | 13 | 0.0769 | 5 | 0 | 5 | 0 | 1 | 0 | 3 | 0.3333 | 14 | 0.0714 |
| 9_10798298 | 1 | 0 | 7 | 0.2857 | 3 | 0.3333 | 3 | 0.3333 | 14 | 0.2857 | 6 | 0.1667 | 5 | 0 | 1 | 0 | 1 | 0 | 13 | 0.0769 |
| 9_10798338 | 1 | 1 | 4 | 0.25 | 2 | 0.5 | 2 | 0 | 9 | 0.3333 | 6 | 0.3333 | 5 | 0.2 | 3 | 0.3333 | 2 | 0.5 | 16 | 0.3125 |
| 9_10798344 | 1 | 0 | 6 | 0.1667 | 3 | 0 | 3 | 0 | 13 | 0.0769 | 6 | 0.1667 | 5 | 0.6 | 3 | 0.3333 | 2 | 0 | 16 | 0.3125 |
| 9_10798361 | 1 | 0 | 6 | 0.1667 | 3 | 1 | 3 | 0.6667 | 13 | 0.4615 | 6 | 0.5 | 4 | 0.75 | 3 | 0 | 1 | 1 | 14 | 0.5 |
| 9_10798403 | 1 | 1 | 6 | 0 | 3 | 0 | 1 | 0 | 11 | 0.0909 | 7 | 0.1429 | 2 | 0 | 3 | 0 | 2 | 0 | 14 | 0.0714 |
| 9_10798404 | 1 | 0 | 4 | 0 | 3 | 0.3333 | 1 | 0 | 9 | 0.1111 | 7 | 0.1429 | 3 | 0 | 3 | 0 | 2 | 0.5 | 15 | 0.1333 |
| 9_10798405 | 1 | 0 | 6 | 0.8333 | 2 | 0.5 | 2 | 0.5 | 11 | 0.6364 | 5 | 0.4 | 3 | 1 | 3 | 1 | 2 | 1 | 13 | 0.7692 |
| 9_10798406 | 1 | 1 | 6 | 0.1667 | 2 | 0 | 2 | 0 | 11 | 0.1818 | 6 | 0 | 3 | 0 | 3 | 0 | 2 | 0.5 | 14 | 0.0714 |
| 9_10798414 | 1 | 0 | 6 | 0.8333 | 3 | 0.3333 | 2 | 0.5 | 12 | 0.5833 | 6 | 0.5 | 3 | 1 | 3 | 0.6667 | 1 | 1 | 13 | 0.6923 |
| 9_10798419 | 1 | 1 | 6 | 0.1667 | 2 | 0 | NA | NA | 9 | 0.2222 | 7 | 0.1429 | 3 | 0.3333 | 3 | 0 | 1 | 0 | 14 | 0.1429 |
| 9_10798429 | 1 | 0 | 5 | 0 | 3 | 0.3333 | 2 | 0 | 11 | 0.0909 | 4 | 0.5 | 3 | 0 | 2 | 0 | 1 | 0 | 10 | 0.2 |
| 9_10798473 | NA | NA | 6 | 1 | 2 | 1 | NA | NA | 8 | 1 | 2 | 1 | 1 | 1 | 2 | 1 | 1 | 1 | 6 | 1 |
| 9_10798481 | NA | NA | 5 | 0.2 | 2 | 0.5 | NA | NA | 7 | 0.2857 | 3 | 0.3333 | 1 | 0 | 1 | 1 | 1 | 0 | 6 | 0.3333 |
| 9_10798492 | NA | NA | 6 | 1 | 2 | 1 | NA | NA | 8 | 1 | 3 | 1 | 1 | 1 | 2 | 1 | 1 | 1 | 7 | 1 |
| 9_10798534 | NA | NA | 6 | 0.8333 | 1 | 0 | NA | NA | 7 | 0.7143 | 1 | 0 | 1 | 0 | 2 | 0.5 | 1 | 0 | 5 | 0.2 |
| 9_10798560 | NA | NA | NA | NA | 1 | 1 | NA | NA | 1 | 1 | 1 | 1 | NA | NA | 1 | 1 | NA | NA | 2 | 1 |
| 9_10798721 | NA | NA | 6 | 0.5 | NA | NA | NA | NA | 6 | 0.5 | NA | NA | NA | NA | 2 | 0 | 1 | 1 | 3 | 0.3333 |
| 9_10798751 | NA | NA | 6 | 0.1667 | NA | NA | NA | NA | 6 | 0.1667 | NA | NA | NA | NA | 1 | 0 | 1 | 1 | 2 | 0.5 |
| 9_10798755 | NA | NA | 4 | 0.5 | NA | NA | NA | NA | 4 | 0.5 | NA | NA | NA | NA | 1 | 0 | 1 | 1 | 2 | 0.5 |
| 9_10798759 | NA | NA | 4 | 0.5 | NA | NA | NA | NA | 4 | 0.5 | NA | NA | NA | NA | 1 | 1 | 1 | 0 | 2 | 0.5 |
| 9_10798781 | NA | NA | 6 | 0.1667 | NA | NA | NA | NA | 6 | 0.1667 | NA | NA | NA | NA | 1 | 0 | 1 | 1 | 2 | 0.5 |
| 9_10798787 | NA | NA | 5 | 0.4 | 1 | 1 | NA | NA | 6 | 0.5 | NA | NA | NA | NA | 1 | 1 | 1 | 0 | 2 | 0.5 |
| 9_10798818 | NA | NA | 4 | 1 | 1 | 1 | NA | NA | 5 | 1 | NA | NA | NA | NA | 1 | 0 | 1 | 1 | 2 | 0.5 |
| 9_10798890 | NA | NA | 2 | 1 | NA | NA | NA | NA | 2 | 1 | NA | NA | NA | NA | NA | NA | 1 | 1 | 1 | 1 |
| 9_10799029 | NA | NA | 1 | 1 | NA | NA | NA | NA | 1 | 1 | NA | NA | NA | NA | NA | NA | 1 | 1 | 1 | 1 |
| 9_10799145 | NA | NA | 1 | 1 | NA | NA | NA | NA | 1 | 1 | NA | NA | NA | NA | NA | NA | 1 | 1 | 1 | 1 |
| 9_10799182 | NA | NA | 1 | 1 | NA | NA | NA | NA | 1 | 1 | NA | NA | NA | NA | NA | NA | 1 | 1 | 1 | 1 |
| 9_10808221 | 16 | 0.375 | 16 | 0.5 | 13 | 0.3077 | 13 | 0.3077 | 58 | 0.3793 | 9 | 0.3333 | 18 | 0.3889 | 14 | 0.0714 | 10 | 0.3 | 51 | 0.2745 |
| 9_10808227 | 9 | 0 | 12 | 0.0833 | 10 | 0.3 | 10 | 0 | 41 | 0.0976 | 6 | 0.3333 | 16 | 0.25 | 9 | 0 | 6 | 0.3333 | 37 | 0.2162 |
| 9_10808308 | 10 | 0.4 | 11 | 0.4545 | 9 | 0.3333 | 11 | 0.1818 | 41 | 0.3415 | 6 | 0.1667 | 8 | 0.375 | 5 | 0.4 | 8 | 0.125 | 27 | 0.2593 |
| 9_10808339 | 8 | 0 | 10 | 0.2 | 9 | 0.3333 | 11 | 0.0909 | 38 | 0.1579 | 4 | 0.25 | 11 | 0 | 5 | 0.2 | 8 | 0.125 | 28 | 0.1071 |
| 9_10808351 | 7 | 0 | 11 | 0.4545 | 8 | 0.125 | 11 | 0.3636 | 37 | 0.2703 | 4 | 0 | 12 | 0.3333 | 6 | 0.5 | 6 | 0.1667 | 28 | 0.2857 |
| 9_10808360 | 9 | 0.2222 | 8 | 0.625 | 9 | 0 | 8 | 0 | 34 | 0.2059 | 5 | 0 | 10 | 0.6 | 6 | 0.5 | 6 | 0.3333 | 27 | 0.4074 |
| 9_10808367 | 7 | 0.2857 | 11 | 0.2727 | 7 | 0.1429 | 7 | 0.1429 | 32 | 0.2188 | 4 | 0 | 9 | 0.6667 | 6 | 0.3333 | 5 | 0.6 | 24 | 0.4583 |
| 9_10808385 | 8 | 0 | 10 | 0.2 | 10 | 0.3 | 7 | 0.1429 | 35 | 0.1714 | 5 | 0 | 8 | 0.375 | 3 | 0 | 4 | 0.5 | 20 | 0.25 |
| 9_10808399 | 8 | 0.75 | 9 | 0.7778 | 7 | 0.8571 | 5 | 0.8 | 29 | 0.7931 | 2 | 0.5 | 6 | 1 | 3 | 1 | 4 | 0.75 | 15 | 0.8667 |
| 9_10808407 | 5 | 0 | 10 | 0.5 | 8 | 0.125 | 7 | 0.1429 | 30 | 0.2333 | 4 | 0 | 6 | 0.3333 | 2 | 0.5 | 4 | 0.5 | 16 | 0.3125 |
| 9_10808412 | 4 | 0.25 | 7 | 0.1429 | 8 | 0.75 | 6 | 0.5 | 25 | 0.44 | 4 | 0.5 | 6 | 0.8333 | 3 | 0.6667 | 4 | 1 | 17 | 0.7647 |
| 9_10808413 | 6 | 0.1667 | 10 | 0.1 | 8 | 0 | 6 | 0 | 30 | 0.0667 | 4 | 0.25 | 7 | 0 | 2 | 0 | 3 | 0 | 16 | 0.0625 |
| 9_10808425 | 4 | 0.25 | 6 | 0.1667 | 7 | 0 | 5 | 0.6 | 22 | 0.2273 | 2 | 0 | 7 | 0.4286 | 3 | 0.3333 | 3 | 0.6667 | 15 | 0.4 |
| 9_10808426 | 6 | 0.8333 | 9 | 0.7778 | 8 | 0.75 | 5 | 0.6 | 28 | 0.75 | 4 | 0.75 | 6 | 1 | 3 | 1 | 3 | 1 | 16 | 0.9375 |

| | | | | | | | | | | | | | | | | | | | | |
|---|---|---|---|---|---|---|---|---|---|---|---|---|---|---|---|---|---|---|---|---|
| 9_1080843 | 6 | 0.1667 | 8 | 0.5 | 7 | 0.1429 | 5 | 0.2 | 26 | 0.2692 | 4 | 0.25 | 7 | 0.4286 | 2 | 0.5 | 2 | 1 | 15 | 0.4667 |
| 9_1080843 | 4 | 0 | 7 | 0.1429 | 7 | 0.2857 | 6 | 0.5 | 24 | 0.25 | 4 | 0.5 | 7 | 0.2857 | 2 | 0.5 | 3 | 0.3333 | 16 | 0.375 |
| 9_1080845 | 3 | 0.6667 | 6 | 0.1667 | 4 | 0.5 | 4 | 0.5 | 17 | 0.4118 | 3 | 0.3333 | 5 | 0.2 | 2 | 0.5 | 2 | 0.5 | 12 | 0.3333 |
| 9_1080846 | 5 | 1 | 8 | 1 | 6 | 1 | 5 | 1 | 24 | 1 | 4 | 1 | 6 | 1 | 1 | 1 | 3 | 1 | 14 | 1 |
| 9_1080847 | 4 | 0.75 | 9 | 0.8889 | 4 | 0.75 | 6 | 1 | 23 | 0.8696 | 3 | 1 | 6 | 1 | 2 | 1 | 3 | 1 | 14 | 1 |
| 9_1080847 | 4 | 0.75 | 9 | 0.6667 | 6 | 0.6667 | 4 | 0.5 | 23 | 0.6522 | 3 | 1 | 6 | 0.8333 | 1 | 1 | 2 | 1 | 12 | 0.9167 |
| 9_1080847 | 4 | 0 | 8 | 0.25 | 7 | 0 | 5 | 0 | 24 | 0.0833 | 4 | 0 | 5 | 0 | 2 | 0 | 3 | 0.3333 | 14 | 0.0714 |
| 9_1080849 | 4 | 0.25 | 7 | 0.2857 | 7 | 0 | 4 | 0.25 | 22 | 0.1818 | 4 | 0.5 | 5 | 0.6 | NA | NA | 3 | 0.6667 | 12 | 0.5833 |
| 9_1080850 | 4 | 0 | 7 | 0.2857 | 7 | 0 | 3 | 0.6667 | 21 | 0.1905 | 4 | 0 | 4 | 0.25 | 1 | 0 | 3 | 0.3333 | 12 | 0.1667 |
| 9_1080851 | 3 | 1 | 5 | 1 | 6 | 1 | 4 | 1 | 18 | 1 | 2 | 1 | 5 | 1 | NA | NA | 3 | 0.6667 | 10 | 0.9 |
| 9_1080852 | 3 | 0 | 5 | 0.2 | 7 | 0.7143 | 2 | 0 | 17 | 0.3529 | 2 | 0.5 | 4 | 0 | 1 | 0 | 3 | 0 | 10 | 0.1 |
| 9_1080852 | 4 | 0.5 | 6 | 0.3333 | 7 | 0.2857 | 4 | 0.25 | 21 | 0.3333 | 4 | 0.75 | 5 | 0.4 | NA | NA | 2 | 0 | 11 | 0.4545 |
| 9_1080852 | 3 | 0.3333 | 6 | 0.3333 | 7 | 0.5714 | 4 | 0.75 | 20 | 0.5 | 4 | 0.5 | 4 | 0.75 | 1 | 0 | 3 | 0.3333 | 12 | 0.5 |
| 9_1080853 | 2 | 1 | 4 | 0.75 | 4 | 0.25 | 3 | 0.6667 | 13 | 0.6154 | 3 | 0.3333 | 3 | 0.6667 | NA | NA | 3 | 0.6667 | 9 | 0.5556 |
| 9_1080854 | 2 | 0 | 3 | 0.3333 | 6 | 0.5 | 3 | 0.3333 | 14 | 0.3571 | 3 | 0.3333 | 4 | 0.25 | NA | NA | 2 | 0 | 9 | 0.2222 |
| 9_1080854 | 2 | 0 | 4 | 0 | 6 | 0.1667 | 3 | 0 | 15 | 0.0667 | 3 | 0 | 3 | 0.3333 | NA | NA | 1 | 0 | 7 | 0.1429 |
| 9_1080855 | 2 | 0 | 4 | 0.25 | 5 | 0 | 2 | 0 | 13 | 0.0769 | 3 | 0 | 4 | 0.25 | NA | NA | 1 | 0 | 8 | 0.125 |
| 9_1080857 | 2 | 0 | 5 | 0 | 5 | 0 | 3 | 0.3333 | 15 | 0.0667 | 3 | 0.3333 | 4 | 0 | 1 | 0 | 2 | 0 | 10 | 0.1 |
| 9_1080857 | 2 | 0 | 5 | 0.8 | 4 | 0.5 | 3 | 1 | 14 | 0.6429 | 3 | 0.6667 | 4 | 0.5 | 1 | 1 | 1 | 0 | 9 | 0.5556 |
| 9_1080859 | 2 | 0.5 | 3 | 0 | 4 | 0.25 | 1 | 0 | 10 | 0.2 | 2 | 0.5 | 3 | 0.6667 | NA | NA | 1 | 0 | 6 | 0.5 |
| 9_1080860 | 2 | 0 | 3 | 0.6667 | 4 | 0 | 2 | 0.5 | 11 | 0.2727 | 3 | 0.3333 | 4 | 0.5 | NA | NA | 2 | 0 | 9 | 0.3333 |
| 9_1080863 | 1 | 0 | 4 | 0 | 4 | 0.25 | 1 | 0 | 10 | 0.1 | 3 | 0 | 4 | 0.25 | NA | NA | 1 | 0 | 8 | 0.125 |
| 9_1080864 | 2 | 1 | 3 | 1 | 3 | 1 | 2 | 1 | 10 | 1 | 3 | 1 | 4 | 1 | NA | NA | NA | NA | 7 | 1 |
| 9_1080864 | 2 | 0.5 | 3 | 0.3333 | 3 | 0 | 2 | 0 | 10 | 0.2 | 2 | 0.5 | 3 | 0.3333 | NA | NA | 1 | 0 | 6 | 0.3333 |
| 9_1080865 | 2 | 1 | 3 | 1 | 4 | 0.75 | 1 | 1 | 10 | 0.9 | 3 | 1 | 3 | 1 | NA | NA | NA | NA | 6 | 1 |
| 9_1080953 | 1 | 0 | 2 | 0 | 3 | 0.6667 | 2 | 0 | 8 | 0.25 | 3 | 0.6667 | 3 | 0.3333 | NA | NA | 1 | 0 | 7 | 0.4286 |
| 9_1080954 | 1 | 0 | 2 | 0 | 3 | 0.3333 | 2 | 1 | 8 | 0.375 | 2 | 0 | 3 | 0.6667 | NA | NA | NA | NA | 5 | 0.4 |
| 9_1081724 | 10 | 0.2 | 9 | 0.3333 | 11 | 0.2727 | 7 | 0.5714 | 37 | 0.3243 | 7 | 0.7143 | 10 | 0.7 | 11 | 0.2727 | 11 | 0.4545 | 39 | 0.5128 |
| 9_1081729 | 6 | 0.3333 | 8 | 0.25 | 6 | 0.5 | 6 | 0.3333 | 26 | 0.3462 | 6 | 0.5 | 4 | 0.5 | 11 | 0.1818 | 9 | 0.5556 | 30 | 0.4 |
| 9_1081731 | 10 | 0.6 | 13 | 0.3846 | 10 | 0.6 | 11 | 0.8182 | 44 | 0.5909 | 7 | 0.7143 | 10 | 0.9 | 13 | 0.5385 | 13 | 0.8462 | 43 | 0.7442 |
| 9_1081732 | 10 | 0.3 | 8 | 0 | 10 | 0.1 | 9 | 0.1111 | 37 | 0.1351 | 6 | 0.1667 | 9 | 0.2222 | 13 | 0.1538 | 10 | 0 | 38 | 0.1316 |
| 9_1081734 | 11 | 0.4545 | 13 | 0.3077 | 10 | 0.5 | 11 | 0.6364 | 45 | 0.4667 | 8 | 0.5 | 11 | 0.5455 | 13 | 0.3846 | 12 | 0.5833 | 44 | 0.5 |
| 9_1081786 | 2 | 0 | 3 | 0.3333 | 3 | 0.3333 | 3 | 0 | 11 | 0.1818 | 2 | 0 | 4 | 0.25 | 1 | 0 | 1 | 0 | 8 | 0.125 |
| 9_1081792 | 3 | 0 | 3 | 0 | 3 | 0 | 3 | 0.3333 | 12 | 0.0833 | 2 | 0 | 3 | 0.3333 | 1 | 0 | NA | NA | 6 | 0.1667 |
| 9_1081793 | 2 | 0.5 | 3 | 0 | 1 | 0 | 2 | 0 | 8 | 0.125 | 1 | 0 | 3 | 0.3333 | NA | NA | 1 | 0 | 5 | 0.2 |
| 9_1081803 | 6 | 0.5 | 3 | 0.6667 | 7 | 0.4286 | 4 | 0.5 | 20 | 0.5 | 4 | 0.5 | 2 | 0 | 3 | 0.3333 | 3 | 0 | 12 | 0.25 |
| 9_1081803 | 8 | 0.5 | 4 | 0.5 | 7 | 0.5714 | 4 | 0 | 23 | 0.4348 | 5 | 0.8 | 2 | 0 | 4 | 0.5 | 3 | 0.6667 | 14 | 0.5714 |
| 9_1081809 | 8 | 0.25 | 5 | 0.2 | 6 | 0.1667 | 4 | 0.25 | 23 | 0.2174 | 5 | 0 | 2 | 0 | 4 | 0 | 3 | 0.3333 | 14 | 0.0714 |
| 9_1081811 | 7 | 0.1429 | 5 | 0.6 | 5 | 0.4 | 3 | 0.6667 | 20 | 0.4 | 5 | 0.8 | 2 | 0 | 4 | 0.5 | 3 | 0.3333 | 14 | 0.5 |
| 9_1081812 | 5 | 0.4 | 4 | 0.5 | 4 | 0 | 3 | 0 | 16 | 0.25 | 5 | 0.2 | 2 | 0.5 | 5 | 0.4 | 2 | 0 | 14 | 0.2857 |
| 9_1081813 | 6 | 0.1667 | 4 | 0.5 | 5 | 0.4 | 4 | 0.5 | 19 | 0.3684 | 4 | 0.5 | 2 | 0.5 | 4 | 0.5 | 3 | 0.3333 | 13 | 0.4615 |
| 9_1081830 | 5 | 0 | 3 | 0.3333 | 5 | 0.2 | 3 | 0 | 16 | 0.125 | 5 | 0.4 | 2 | 0 | 5 | 0 | 3 | 0 | 15 | 0.1333 |
| 9_1082148 | 24 | 0 | 16 | 0 | 25 | 0.16 | 15 | 0.0667 | 80 | 0.0625 | 15 | 0.1333 | 15 | 0.2 | 14 | 0.0714 | 14 | 0.2143 | 58 | 0.1552 |
| 9_1082149 | 17 | 0.2353 | 22 | 0.1364 | 22 | 0.1818 | 19 | 0.1579 | 80 | 0.175 | 12 | 0.0833 | 17 | 0.1765 | 14 | 0.2143 | 18 | 0.2222 | 61 | 0.1803 |
| 9_1082172 | 41 | 0.0732 | 37 | 0.1351 | 22 | 0.2273 | 20 | 0.2 | 120 | 0.1417 | 24 | 0.1667 | 27 | 0.1481 | 29 | 0.1724 | 38 | 0.1316 | 118 | 0.1525 |
| 9_1082172 | 44 | 0.1136 | 38 | 0.0526 | 22 | 0.1364 | 25 | 0.24 | 129 | 0.124 | 30 | 0.0667 | 30 | 0.2 | 29 | 0.069 | 43 | 0.093 | 132 | 0.1061 |
| 9_1082660 | NA | NA | NA | NA | 1 | 0 | 2 | 0.5 | 3 | 0.3333 | NA | NA | NA | NA | 1 | 0 | 2 | 0.5 | 3 | 0.3333 |
| 9_1082690 | 15 | 0.6667 | 11 | 0.9091 | 4 | 0.75 | 6 | 0.8333 | 36 | 0.7778 | 12 | 0.8333 | 7 | 0.7143 | 13 | 0.8462 | 12 | 0.75 | 44 | 0.7955 |
| 9_1082691 | 17 | 0.5294 | 12 | 0.75 | 5 | 0.4 | 4 | 0.5 | 38 | 0.5789 | 9 | 0.5556 | 6 | 0.8333 | 12 | 0.3333 | 12 | 0.6667 | 39 | 0.5641 |
| 9_1082691 | 16 | 0.9375 | 15 | 0.9333 | 5 | 0.8 | 6 | 1 | 42 | 0.9286 | 12 | 0.8333 | 8 | 0.875 | 13 | 0.7692 | 13 | 0.8462 | 46 | 0.8261 |
| 9_1082692 | 16 | 0.3125 | 13 | 0.4615 | 6 | 0 | 8 | 0.375 | 43 | 0.3256 | 12 | 0.4167 | 8 | 0 | 14 | 0.2857 | 14 | 0.2857 | 48 | 0.2708 |
| 9_1082693 | 16 | 0.5625 | 11 | 0.7273 | 5 | 0.8 | 5 | 0.6 | 37 | 0.6486 | 11 | 0.8182 | 6 | 1 | 13 | 0.4615 | 9 | 0.8889 | 39 | 0.7436 |
| 9_1082695 | 9 | 0.2222 | 7 | 0.2857 | 3 | 0.3333 | 3 | 0 | 22 | 0.2273 | 8 | 0.375 | 4 | 0.25 | 10 | 0.2 | 6 | 0.3333 | 28 | 0.2857 |
| 9_1082696 | 8 | 0.125 | 6 | 0.3333 | 2 | 0 | 2 | 0 | 18 | 0.1667 | 8 | 0 | 4 | 0.25 | 11 | 0.0909 | 6 | 0.5 | 29 | 0.1724 |
| 9_1082696 | 5 | 0.6 | 4 | 0.5 | 1 | 0 | 2 | 0 | 12 | 0.4167 | 7 | 0.5714 | 2 | 0.5 | 10 | 0.2 | 5 | 0.6 | 24 | 0.4167 |
| 9_1083250 | 3 | 0.3333 | 3 | 0.6667 | 1 | 1 | 1 | 0 | 8 | 0.5 | 2 | 0.5 | 2 | 0.5 | NA | NA | 1 | 1 | 5 | 0.6 |
| 9_1083283 | 2 | 0.5 | 1 | 0 | 1 | 1 | NA | NA | 4 | 0.5 | NA | NA | 1 | 1 | NA | NA | NA | NA | 1 | 1 |
| 9_1083404 | 2 | 0 | 2 | 0.5 | 3 | 0.3333 | NA | NA | 7 | 0.2857 | NA | NA | NA | NA | NA | NA | 1 | 1 | 1 | 1 |
| 9_1083405 | 3 | 0 | 3 | 0.6667 | 2 | 0.5 | 2 | 0 | 10 | 0.3 | NA | NA | NA | NA | NA | NA | 1 | 1 | 1 | 1 |
| 9_1083407 | 3 | 0.6667 | 2 | 0.5 | 2 | 0.5 | 1 | 0 | 8 | 0.5 | NA | NA | 1 | 1 | NA | NA | 1 | 0 | 2 | 0.5 |
| 9_1083408 | 3 | 0.3333 | 2 | 1 | 3 | 0.6667 | 1 | 0 | 9 | 0.5556 | NA | NA | NA | NA | NA | NA | 1 | 1 | 1 | 1 |
| 9_1083410 | 3 | 0.3333 | 3 | 0 | 2 | 1 | 1 | 1 | 9 | 0.4444 | NA | NA | 1 | 0 | NA | NA | 1 | 1 | 2 | 0.5 |
| 9_1083411 | 3 | 0.3333 | 4 | 0.25 | 3 | 0.6667 | 3 | 0.3333 | 13 | 0.3846 | NA | NA | 1 | 0 | NA | NA | 1 | 1 | 2 | 0.5 |
| 9_1083412 | 3 | 1 | 6 | 1 | 4 | 0.75 | 2 | 0.5 | 15 | 0.8667 | NA | NA | 2 | 1 | NA | NA | 1 | 1 | 3 | 1 |
| 9_1083415 | 3 | 0.3333 | 7 | 0.4286 | 5 | 0.4 | 2 | 0 | 17 | 0.3529 | NA | NA | 3 | 0.6667 | 1 | 0 | 1 | 0 | 5 | 0.4 |

| | | | | | | | | | | | | | | | | | | | | |
|---|---|---|---|---|---|---|---|---|---|---|---|---|---|---|---|---|---|---|---|---|
| 9_10834158 | 3 | 0.6667 | 6 | 0.8333 | 4 | 0.5 | 2 | 1 | 15 | 0.7333 | NA | NA | 4 | 1 | 1 | 1 | 2 | 1 | 7 | 1 |
| 9_10834160 | 3 | 0.6667 | 6 | 1 | 4 | 0.75 | 2 | 1 | 15 | 0.8667 | NA | NA | 4 | 0.75 | 1 | 1 | 2 | 1 | 7 | 0.8571 |
| 9_10834161 | 3 | 1 | 6 | 1 | 4 | 1 | 3 | 1 | 16 | 1 | NA | NA | 4 | 1 | 1 | 1 | 2 | 1 | 7 | 1 |
| 9_10834169 | 2 | 0.5 | 7 | 0 | 2 | 0 | 1 | 0 | 12 | 0.0833 | NA | NA | 4 | 0.25 | 1 | 0 | 2 | 0.5 | 7 | 0.2857 |
| 9_10834236 | 1 | 1 | 2 | 0.5 | NA | NA | NA | NA | 3 | 0.6667 | NA | NA | 2 | 1 | NA | NA | 1 | 1 | 3 | 1 |
| 9_10834725 | 1 | 0 | NA | NA | NA | NA | 1 | 1 | 2 | 0.5 | NA | NA | NA | NA | NA | NA | 1 | 1 | 1 | 1 |
| 9_10836606 | 4 | 0 | 1 | 0 | 1 | 1 | 4 | 0.25 | 10 | 0.2 | 1 | 0 | 2 | 0.5 | NA | NA | 1 | 0 | 4 | 0.25 |
| 9_10837611 | 3 | 0 | 8 | 0 | 2 | 0.5 | 6 | 0 | 19 | 0.0526 | 5 | 0 | 3 | 0 | 5 | 0.4 | 6 | 0 | 19 | 0.1053 |
| 9_10837873 | 4 | 0.5 | 1 | 0 | 2 | 0 | 6 | 0.1667 | 13 | 0.2308 | 2 | 0.5 | 2 | 0.5 | 1 | 1 | 1 | 0 | 6 | 0.5 |
| 9_10837874 | 2 | 0 | 1 | 0 | 2 | 0 | 4 | 0.25 | 9 | 0.1111 | 3 | 0.6667 | 2 | 0.5 | 2 | 1 | NA | NA | 7 | 0.7143 |
| 9_10837917 | 7 | 0.2857 | 7 | 0 | 5 | 0.2 | 5 | 0 | 24 | 0.125 | 3 | 0.6667 | 5 | 0.4 | 6 | 0.6667 | 7 | 0.5714 | 21 | 0.5714 |
| 9_10837924 | 8 | 0.5 | 9 | 0.5556 | 5 | 0.8 | 6 | 0.5 | 28 | 0.5714 | 3 | 0 | 3 | 0.3333 | 6 | 0.6667 | 7 | 0.7143 | 19 | 0.5263 |
| 9_10838612 | 1 | 0 | 2 | 0 | 4 | 0.25 | 3 | 0 | 10 | 0.1 | 1 | 1 | 1 | 0 | 1 | 0 | 4 | 0.25 | 7 | 0.2857 |
| 9_10839304 | 9 | 0.2222 | 6 | 0.1667 | 6 | 0 | 5 | 0.4 | 26 | 0.1923 | 5 | 0 | 4 | 0.25 | 1 | 0 | 7 | 0.1429 | 17 | 0.1176 |
| 9_10840579 | 5 | 0.2 | 5 | 0.2 | 6 | 0 | 6 | 0 | 22 | 0.0909 | 6 | 0 | 6 | 0.3333 | 1 | 0 | 4 | 0 | 17 | 0.1176 |
| 9_10840582 | 3 | 0 | 4 | 0.25 | 4 | 0 | 4 | 0 | 15 | 0.0667 | 4 | 0 | 4 | 0 | 1 | 1 | 3 | 0 | 12 | 0.0833 |
| 9_10840590 | 4 | 0.5 | 5 | 0.6 | 6 | 0.6667 | 3 | 0 | 18 | 0.5 | 3 | 0.6667 | 5 | 0.6 | 1 | 1 | 4 | 0.5 | 13 | 0.6154 |
| 9_10840595 | 3 | 0.6667 | 5 | 0 | 6 | 0.1667 | 5 | 0.4 | 19 | 0.2632 | 5 | 0.2 | 5 | 0.4 | 2 | 0 | 2 | 0 | 14 | 0.2143 |
| 9_10840598 | 5 | 0.4 | 5 | 0.2 | 6 | 0.3333 | 6 | 0.1667 | 22 | 0.2727 | 6 | 0.1667 | 5 | 0.2 | 2 | 0.5 | 4 | 0.25 | 17 | 0.2353 |
| 9_10840603 | 5 | 0.4 | 4 | 0.75 | 6 | 0.1667 | 6 | 0.1667 | 21 | 0.3333 | 6 | 0.3333 | 6 | 0 | 2 | 0.5 | 4 | 0.25 | 18 | 0.2222 |
| 9_10841093 | 6 | 0.3333 | 6 | 0 | 4 | 0 | 5 | 0.2 | 21 | 0.1429 | 7 | 0.1429 | 5 | 0.2 | 3 | 0.3333 | 4 | 0.25 | 19 | 0.2105 |
| 9_10841112 | 5 | 0.6 | 6 | 0.8333 | 6 | 1 | 6 | 0.8333 | 23 | 0.8261 | 8 | 0.875 | 6 | 0.8333 | 4 | 0.75 | 4 | 0.5 | 22 | 0.7727 |
| 9_10841123 | 6 | 0.8333 | 6 | 0.3333 | 6 | 0.6667 | 6 | 0.1667 | 24 | 0.5 | 10 | 0.2 | 5 | 0.4 | 4 | 0.25 | 4 | 0 | 23 | 0.2174 |
| 9_10841133 | 7 | 0.2857 | 6 | 0.5 | 6 | 0.3333 | 5 | 0.4 | 24 | 0.375 | 9 | 0.4444 | 6 | 0.3333 | 5 | 0.2 | 3 | 1 | 23 | 0.4348 |
| 9_10841134 | 7 | 0.1429 | 4 | 0.5 | 3 | 0.3333 | 4 | 0.75 | 18 | 0.3889 | 9 | 0.5556 | 4 | 0.75 | 5 | 0.4 | 3 | 0 | 21 | 0.4762 |
| 9_10841134 | 5 | 0.4 | 5 | 0.6 | 5 | 0 | 5 | 0 | 20 | 0.25 | 10 | 0.7 | 5 | 0 | 4 | 0 | 4 | 0.5 | 23 | 0.3913 |
| 9_10841137 | 5 | 0.8 | 5 | 0.2 | 6 | 0.6667 | 6 | 0.8333 | 22 | 0.6364 | 9 | 0.4444 | 5 | 0.4 | 5 | 0.6 | 6 | 0.5 | 25 | 0.48 |
| 9_10841140 | 7 | 1 | 5 | 0.4 | 4 | 0.25 | 4 | 0.5 | 20 | 0.6 | 8 | 0.5 | 2 | 1 | 2 | 0.5 | 6 | 0.8333 | 18 | 0.6667 |
| 9_10841162 | 5 | 0.2 | 7 | 0.2857 | 6 | 0.5 | 4 | 0 | 22 | 0.2727 | 8 | 0.25 | 6 | 0 | 3 | 0 | 8 | 0.25 | 25 | 0.16 |
| 9_10841171 | 7 | 0.8571 | 7 | 0.5714 | 5 | 0.6 | 6 | 0.6667 | 25 | 0.68 | 9 | 0.5556 | 4 | 0.25 | 5 | 0.2 | 8 | 0.75 | 26 | 0.5 |
| 9_10841172 | 6 | 0.3333 | 6 | 0.3333 | 6 | 0.3333 | 5 | 0 | 23 | 0.2609 | 10 | 0.1 | 4 | 0 | 4 | 0 | 8 | 0.25 | 26 | 0.1154 |
| 9_10843668 | 7 | 0.4286 | 12 | 0.5833 | 9 | 0.5556 | 9 | 1 | 37 | 0.6486 | 14 | 0.5 | 8 | 0.625 | 9 | 0.5556 | 10 | 0.7 | 41 | 0.5854 |
| 9_10843691 | 9 | 0.1111 | 13 | 0.0769 | 14 | 0.1429 | 11 | 0 | 47 | 0.0851 | 16 | 0.0625 | 13 | 0.0769 | 9 | 0.1111 | 12 | 0.1667 | 50 | 0.1 |
| 9_10843692 | 6 | 0.1667 | 13 | 0.6154 | 12 | 0.6667 | 9 | 0.7778 | 40 | 0.6 | 14 | 0.5 | 10 | 0.5 | 5 | 1 | 11 | 0.5455 | 40 | 0.575 |
| 9_10844283 | 2 | 0.5 | NA | NA | NA | NA | 2 | 0 | 4 | 0.25 | 1 | 1 | 5 | 0.4 | 2 | 0 | 2 | 0 | 10 | 0.3 |
| 9_10844319 | 8 | 0.125 | 8 | 0.125 | 17 | 0.1765 | 9 | 0.1111 | 42 | 0.1429 | 5 | 0.2 | 12 | 0.25 | 9 | 0.2222 | 5 | 0 | 31 | 0.1935 |
| 9_10844918 | 13 | 0.3077 | 8 | 0.375 | 2 | 0.5 | 6 | 0.3333 | 29 | 0.3448 | 2 | 0 | 5 | 0.4 | 8 | 0.25 | 7 | 0.4286 | 22 | 0.3182 |
| 9_10845017 | 10 | 0.3 | 11 | 0.1818 | 10 | 0.4 | 9 | 0.1111 | 40 | 0.25 | 12 | 0.3333 | 22 | 0.2273 | 20 | 0.2 | 19 | 0.2632 | 73 | 0.2466 |
| 9_10845023 | 10 | 1 | 11 | 0.9091 | 10 | 0.7 | 8 | 0.5 | 39 | 0.7949 | 8 | 0.75 | 21 | 0.6667 | 17 | 0.7647 | 15 | 0.8667 | 61 | 0.7541 |
| 9_10845029 | 10 | 0 | 8 | 0 | 9 | 0.1111 | 8 | 0.125 | 35 | 0.0571 | 10 | 0 | 17 | 0.0588 | 17 | 0.1176 | 18 | 0.0556 | 62 | 0.0645 |
| 9_10845163 | 9 | 0.2222 | 8 | 0.375 | 3 | 0.3333 | 3 | 0 | 23 | 0.2609 | 6 | 0 | 13 | 0.2308 | 9 | 0.1111 | 9 | 0.1111 | 37 | 0.1351 |
| 9_10845166 | 7 | 0.2857 | 5 | 0 | 3 | 0 | 3 | 0 | 18 | 0.1111 | 5 | 0 | 13 | 0 | 10 | 0 | 7 | 0.2857 | 35 | 0.0571 |
| 9_10845204 | 7 | 0.8571 | 4 | 0.5 | 3 | 0.6667 | 2 | 1 | 16 | 0.75 | 3 | 0.6667 | 7 | 0.7143 | 5 | 1 | 4 | 1 | 19 | 0.8421 |
| 9_10845235 | 6 | 0 | 4 | 0.25 | 4 | 0 | 2 | 0 | 16 | 0.0625 | 4 | 0.25 | 8 | 0 | 6 | 0 | 6 | 0.1667 | 24 | 0.0833 |
| 9_10845258 | 4 | 0.5 | 2 | 0 | 2 | 1 | 1 | 1 | 9 | 0.5556 | 2 | 0 | 5 | 1 | 7 | 0.4286 | 5 | 0.8 | 19 | 0.6316 |
| 9_10845266 | 5 | 1 | 3 | 0.6667 | 3 | 1 | 2 | 1 | 13 | 0.9231 | 4 | 1 | 6 | 1 | 7 | 1 | 5 | 0.8 | 22 | 0.9545 |
| 9_10845272 | 5 | 1 | 3 | 0.6667 | 4 | 0.5 | 2 | 1 | 14 | 0.7857 | 4 | 0.75 | 6 | 0.6667 | 8 | 0.75 | 2 | 1 | 20 | 0.75 |
| 9_10845278 | 4 | 1 | 2 | 0.5 | 2 | 1 | 2 | 1 | 10 | 0.9 | 4 | 1 | 5 | 0.8 | 7 | 0.8571 | 5 | 1 | 21 | 0.9048 |
| 9_10845281 | 5 | 0.6 | 3 | 0.6667 | 2 | 0.5 | 2 | 0.5 | 12 | 0.5833 | 4 | 1 | 6 | 0.5 | 7 | 0.2857 | 5 | 0.4 | 22 | 0.5 |
| 9_10845305 | 2 | 0.5 | 3 | 0.6667 | 2 | 1 | 2 | 0.5 | 9 | 0.6667 | 2 | 1 | 4 | 0.5 | 8 | 0.75 | 4 | 0.75 | 18 | 0.7222 |
| 9_10845307 | 4 | 0.5 | 1 | 0 | 1 | 0 | 2 | 0 | 8 | 0.25 | 2 | 1 | 5 | 0.4 | 7 | 0.4286 | 3 | 0.6667 | 17 | 0.5294 |
| 9_10845316 | 4 | 0.5 | 2 | 0 | NA | NA | 2 | 0 | 8 | 0.25 | 3 | 0.6667 | 5 | 0.4 | 7 | 0.5714 | 4 | 0.75 | 19 | 0.5789 |
| 9_10845329 | 2 | 0 | 2 | 0.5 | NA | NA | 2 | 0.5 | 6 | 0.3333 | 1 | 1 | 3 | 0.3333 | 6 | 0.1667 | 3 | 0 | 13 | 0.2308 |
| 9_10845336 | 3 | 0.6667 | 2 | 0.5 | NA | NA | 1 | 1 | 6 | 0.6667 | 3 | 0.6667 | 4 | 0.75 | 7 | 0.4286 | 3 | 1 | 17 | 0.6471 |
| 9_10845364 | 3 | 0.6667 | 2 | 0 | NA | NA | 2 | 0 | 7 | 0.2857 | 3 | 0.3333 | 3 | 0.6667 | 5 | 1 | 4 | 0.75 | 15 | 0.7333 |
| 9_10845374 | 3 | 0.3333 | 2 | 0.5 | NA | NA | 2 | 0.5 | 7 | 0.4286 | 2 | 0 | 4 | 0.25 | 6 | 0 | 4 | 0.25 | 16 | 0.125 |
| 9_10845621 | 9 | 0.2222 | 13 | 0.3846 | 5 | 0.4 | 15 | 0.3333 | 42 | 0.3333 | 10 | 0.5 | 14 | 0.2857 | 4 | 0.25 | 7 | 0.4286 | 35 | 0.3714 |
| 9_10845624 | 12 | 0 | 23 | 0.1304 | 10 | 0.5 | 18 | 0.0556 | 63 | 0.1429 | 13 | 0 | 13 | 0.0769 | 10 | 0.1 | 9 | 0.1111 | 45 | 0.0667 |
| 9_10845632 | 12 | 0.3333 | 18 | 0.1111 | 5 | 0.2 | 16 | 0.125 | 51 | 0.1765 | 12 | 0.1667 | 13 | 0.1538 | 7 | 0.1429 | 7 | 0 | 39 | 0.1282 |
| 9_10845634 | 9 | 1 | 19 | 0.9474 | 9 | 1 | 16 | 0.875 | 53 | 0.9434 | 11 | 0.9091 | 14 | 0.9286 | 6 | 1 | 9 | 1 | 40 | 0.95 |
| 9_10845641 | 8 | 0.5 | 18 | 0.4444 | 7 | 0.4286 | 12 | 0.3333 | 45 | 0.4222 | 10 | 0.4 | 10 | 0.3 | 7 | 0 | 6 | 0.3333 | 33 | 0.2727 |
| 9_10848205 | 1 | 1 | 2 | 0.5 | 1 | 1 | 2 | 1 | 6 | 0.8333 | 3 | 1 | 2 | 0.5 | 7 | 0.7143 | 6 | 0.6667 | 18 | 0.7222 |
| 9_10848215 | 2 | 1 | 2 | 1 | 2 | 1 | 2 | 0.5 | 8 | 0.875 | 3 | 0.6667 | 3 | 0.3333 | 5 | 0 | 6 | 0.1667 | 17 | 0.2353 |
| 9_10852137 | 2 | 0.5 | NA | NA | NA | NA | NA | NA | 2 | 0.5 | 2 | 0.5 | NA | NA | NA | NA | 1 | 0 | 3 | 0.3333 |
| 9_10856157 | 2 | 0 | 4 | 0 | 2 | 0.5 | 1 | 1 | 9 | 0.2222 | NA | NA | NA | NA | 3 | 0.3333 | NA | NA | 3 | 0.3333 |

| | | | | | | | | | | | | | | | | | | | | |
|---|---|---|---|---|---|---|---|---|---|---|---|---|---|---|---|---|---|---|---|---|
| 9_1086241 | 3 | 0 | 3 | 0.3333 | 1 | 0 | 1 | 0 | 8 | 0.125 | NA | NA | 5 | 0 | 2 | 1 | 1 | 0 | 8 | 0.25 |
| 9_1086241 | 2 | 1 | 3 | 1 | 1 | 1 | 1 | 1 | 7 | 1 | NA | NA | 5 | 0.8 | 2 | 1 | 1 | 1 | 8 | 0.875 |
| 9_1086242 | 2 | 1 | 3 | 0 | 1 | 0 | 1 | 0 | 7 | 0.2857 | NA | NA | 5 | 0.2 | 2 | 0.5 | 2 | 0 | 9 | 0.2222 |
| 9_1086248 | 3 | 0.3333 | 4 | 0.25 | 2 | 0.5 | 1 | 0 | 10 | 0.3 | 2 | 0 | 8 | 0.125 | 3 | 0 | 3 | 0 | 16 | 0.0625 |
| 9_1086251 | 3 | 0 | 3 | 0 | 2 | 0.5 | 2 | 0 | 10 | 0.1 | 2 | 0 | 8 | 0.125 | 2 | 0 | 1 | 0 | 13 | 0.0769 |
| 9_1086254 | 4 | 0 | 4 | 0 | 3 | 0.3333 | 4 | 0 | 15 | 0.0667 | 1 | 0 | 9 | 0 | 3 | 0.3333 | 3 | 0 | 16 | 0.0625 |
| 9_1086256 | 3 | 0 | 4 | 0.5 | 3 | 0 | 5 | 0 | 15 | 0.1333 | 3 | 0.3333 | 9 | 0.1111 | 4 | 0.5 | 3 | 0.6667 | 19 | 0.3158 |
| 9_1086258 | 4 | 0.25 | 4 | 0.75 | 3 | 0 | 5 | 0.2 | 16 | 0.3125 | 3 | 0.3333 | 11 | 0.1818 | 4 | 0.5 | 3 | 0.6667 | 21 | 0.3333 |
| 9_1086747 | 3 | 1 | NA | NA | 3 | 1 | 1 | 0 | 7 | 0.8571 | NA | NA | 2 | 0.5 | NA | NA | 1 | 0 | 3 | 0.3333 |
| 9_1086747 | 3 | 1 | 4 | 0.75 | 4 | 0.5 | 1 | 0 | 12 | 0.6667 | NA | NA | 3 | 0.3333 | NA | NA | NA | NA | 3 | 0.3333 |
| 9_1086748 | 3 | 0 | 3 | 0 | 3 | 0.6667 | NA | NA | 9 | 0.2222 | NA | NA | 3 | 0.3333 | NA | NA | 1 | 0 | 4 | 0.25 |
| 9_1086755 | 19 | 0.1053 | 12 | 0.0833 | 9 | 0.2222 | 11 | 0.0909 | 51 | 0.1176 | 8 | 0.25 | 14 | 0.0714 | 14 | 0.1429 | 13 | 0.0769 | 49 | 0.1224 |
| 9_1086755 | 15 | 0.3333 | 11 | 0.2727 | 9 | 0.4444 | 12 | 0.3333 | 47 | 0.3404 | 8 | 0.375 | 15 | 0.4 | 13 | 0.1538 | 14 | 0.3571 | 50 | 0.32 |
| 9_1086829 | 21 | 0.381 | 16 | 0.6875 | 10 | 0.5 | 15 | 0.4667 | 62 | 0.5 | 13 | 0.3077 | 14 | 0.7857 | 16 | 0.6875 | 13 | 0.3846 | 56 | 0.5536 |
| 9_1086830 | 26 | 0.5769 | 17 | 0.4706 | 11 | 0.3636 | 18 | 0.6111 | 72 | 0.5278 | 13 | 0.8462 | 18 | 0.6111 | 16 | 0.5 | 17 | 0.4118 | 64 | 0.5781 |
| 9_1086831 | 25 | 0.4 | 16 | 0.4375 | 9 | 0.3333 | 14 | 0.5 | 64 | 0.4219 | 12 | 0.6667 | 18 | 0.6111 | 18 | 0.3889 | 17 | 0.5294 | 65 | 0.5385 |
| 9_1086831 | 22 | 0.5 | 18 | 0.6667 | 11 | 0.4545 | 12 | 0.6667 | 63 | 0.5714 | 13 | 0.6154 | 15 | 0.8667 | 16 | 0.6875 | 15 | 0.6667 | 59 | 0.7119 |
| 9_1086832 | 22 | 0.6818 | 15 | 0.6 | 12 | 0.6667 | 14 | 0.7857 | 63 | 0.6825 | 15 | 0.5333 | 15 | 0.6667 | 13 | 0.5385 | 16 | 0.375 | 59 | 0.5254 |
| 9_1086832 | 28 | 0.1786 | 19 | 0.1579 | 12 | 0.1667 | 15 | 0.2 | 74 | 0.1757 | 16 | 0.125 | 19 | 0.2105 | 19 | 0.1053 | 15 | 0.0667 | 69 | 0.1304 |
| 9_1086832 | 28 | 0.9286 | 20 | 0.85 | 12 | 1 | 17 | 0.9412 | 77 | 0.9221 | 16 | 0.875 | 18 | 0.7778 | 20 | 0.9 | 18 | 0.9444 | 72 | 0.875 |
| 9_1086833 | 22 | 0.1818 | 11 | 0.0909 | 8 | 0.25 | 10 | 0.2 | 51 | 0.1765 | 11 | 0.1818 | 12 | 0 | 14 | 0.0714 | 10 | 0.1 | 47 | 0.0851 |
| 9_1086894 | 5 | 0 | 4 | 0.5 | 2 | 0 | 9 | 0.5556 | 20 | 0.35 | 3 | 0.3333 | 3 | 0.6667 | 3 | 0.6667 | 4 | 0.5 | 13 | 0.5385 |
| 9_1086897 | 6 | 0 | 3 | 0 | 2 | 0 | 9 | 0.1111 | 20 | 0.05 | 3 | 0 | 3 | 0 | 3 | 0.3333 | 4 | 0 | 13 | 0.0769 |
| 9_1086897 | 5 | 0.2 | 4 | 0 | 1 | 0 | 8 | 0 | 18 | 0.0556 | 3 | 0.3333 | 3 | 0 | 3 | 0 | 6 | 0 | 15 | 0.0667 |
| 9_1086948 | 9 | 0.4444 | 10 | 0.2 | 3 | 0 | 9 | 0.4444 | 31 | 0.3226 | 4 | 0.5 | 5 | 0.4 | 6 | 0.5 | 8 | 0.5 | 23 | 0.4783 |
| 9_1086949 | 10 | 0.3 | 12 | 0.3333 | 4 | 0 | 8 | 0.125 | 34 | 0.2353 | 5 | 0 | 5 | 0.4 | 5 | 0.4 | 8 | 0.125 | 23 | 0.2174 |
| 9_1086950 | 9 | 0.1111 | 9 | 0.4444 | 4 | 0.25 | 7 | 0.4286 | 29 | 0.3103 | 3 | 0.3333 | 5 | 0.4 | 5 | 0.6 | 6 | 0.1667 | 19 | 0.3684 |
| 9_1086951 | 3 | 0 | 9 | 0.1111 | NA | NA | 4 | 0 | 16 | 0.0625 | 5 | 0.2 | 4 | 0.25 | 3 | 0 | 5 | 0 | 17 | 0.1176 |
| 9_1086951 | 7 | 0.1429 | 11 | 0.1818 | 4 | 0 | 7 | 0.1429 | 29 | 0.1379 | 3 | 0.3333 | 4 | 0.25 | 4 | 0 | 7 | 0.4286 | 18 | 0.2778 |
| 9_1086954 | 10 | 0 | 11 | 0.0909 | 4 | 0.25 | 10 | 0.1 | 35 | 0.0857 | 6 | 0.1667 | 7 | 0.1429 | 5 | 0 | 8 | 0 | 26 | 0.0769 |
| 9_1086954 | 11 | 0.1818 | 11 | 0.0909 | 4 | 0.5 | 11 | 0.0909 | 37 | 0.1622 | 6 | 0.5 | 7 | 0.1429 | 6 | 0.1667 | 8 | 0.25 | 27 | 0.2593 |
| 9_1086954 | 10 | 0 | 11 | 0 | 3 | 0 | 8 | 0.25 | 32 | 0.0625 | 5 | 0 | 4 | 0.25 | 5 | 0 | 6 | 0.1667 | 20 | 0.1 |
| 9_1087259 | 2 | 0.5 | 2 | 0.5 | 2 | 0 | NA | NA | 6 | 0.3333 | 2 | 1 | 4 | 0.75 | 1 | 1 | 2 | 0.5 | 9 | 0.7778 |
| 9_1087260 | 2 | 0 | 3 | 0 | 3 | 0.3333 | NA | NA | 8 | 0.125 | 2 | 0 | 5 | 0.8 | 2 | 0 | 2 | 0 | 11 | 0.3636 |
| 9_1087261 | 2 | 1 | 3 | 1 | 2 | 1 | NA | NA | 7 | 1 | 2 | 1 | 5 | 1 | 3 | 0.6667 | 1 | 1 | 11 | 0.9091 |
| 9_1087262 | 2 | 0.5 | 4 | 1 | 3 | 1 | NA | NA | 9 | 0.8889 | 2 | 1 | 5 | 0.6 | 1 | 1 | 2 | 1 | 10 | 0.8 |
| 9_1087262 | 2 | 1 | 3 | 0.6667 | 3 | 1 | 1 | 0 | 9 | 0.7778 | 2 | 1 | 5 | 0.4 | 3 | 0.6667 | 2 | 1 | 12 | 0.6667 |
| 9_1087263 | 1 | 1 | 4 | 0.5 | 2 | 0.5 | 1 | 0 | 8 | 0.5 | 1 | 1 | 5 | 0.2 | 3 | 0.3333 | 2 | 0.5 | 11 | 0.3636 |
| 9_1087263 | 1 | 1 | 4 | 1 | 2 | 1 | 1 | 1 | 8 | 1 | 1 | 1 | 4 | 0.75 | 2 | 1 | 1 | 1 | 8 | 0.875 |
| 9_1087263 | 1 | 1 | 4 | 0.75 | 2 | 1 | NA | NA | 7 | 0.8571 | 1 | 1 | 2 | 1 | 1 | 1 | 1 | 1 | 5 | 1 |
| 9_1087263 | 2 | 0 | 3 | 0.3333 | 2 | 0 | 1 | 0 | 8 | 0.125 | 1 | 0 | 4 | 0.25 | 1 | 0 | 2 | 0 | 8 | 0.125 |
| 9_1087264 | 2 | 0.5 | 3 | 0.3333 | 1 | 1 | 1 | 0 | 7 | 0.4286 | 1 | 0 | 4 | 0.25 | 3 | 0 | 2 | 0 | 10 | 0.1 |
| 9_1087265 | 2 | 0.5 | 4 | 0.5 | 2 | 1 | 1 | 0 | 9 | 0.5556 | 2 | 1 | 4 | 0.5 | 3 | 1 | 2 | 1 | 11 | 0.8182 |
| 9_1087285 | 2 | 0 | 3 | 1 | 3 | 0.6667 | 2 | 0 | 10 | 0.5 | 1 | 1 | 7 | 0.4286 | 4 | 0.25 | 2 | 0.5 | 14 | 0.4286 |
| 9_1087287 | 2 | 0.5 | 5 | 0 | 3 | 0.3333 | 2 | 0 | 12 | 0.1667 | 2 | 0 | 7 | 0.2857 | 4 | 0 | 2 | 0 | 15 | 0.1333 |
| 9_1087288 | 2 | 0 | 3 | 0.3333 | 3 | 0 | 1 | 0 | 9 | 0.1111 | 2 | 0 | 6 | 0.1667 | 5 | 0 | 2 | 0 | 15 | 0.0667 |
| 9_1087291 | 3 | 0.3333 | 5 | 0.6 | 3 | 0 | 2 | 0 | 13 | 0.3077 | 3 | 0 | 7 | 0.1429 | 5 | 0.2 | 1 | 0 | 16 | 0.125 |
| 9_1087295 | 3 | 0 | 5 | 0 | 3 | 0.3333 | 2 | 0 | 13 | 0.0769 | 3 | 0 | 7 | 0 | 5 | 0 | 4 | 0.25 | 19 | 0.0526 |
| 9_1087394 | NA | NA | NA | NA | 1 | 1 | 1 | 1 | 2 | 1 | NA | NA | 1 | 1 | NA | NA | 3 | 0.6667 | 4 | 0.75 |
| 9_1088597 | 1 | 1 | 1 | 0 | 1 | 1 | 1 | 1 | 4 | 0.75 | 1 | 1 | NA | NA | NA | NA | NA | NA | 1 | 1 |
| 9_1088597 | 1 | 1 | NA | NA | 1 | 0 | 1 | 1 | 3 | 0.6667 | 1 | 1 | NA | NA | NA | NA | NA | NA | 1 | 1 |
| 9_1088597 | NA | NA | 1 | 1 | 1 | 1 | NA | NA | 2 | 1 | 1 | 1 | NA | NA | NA | NA | NA | NA | 1 | 1 |
| 9_1088600 | 1 | 0 | 1 | 1 | 1 | 1 | 1 | 1 | 4 | 0.75 | 1 | 1 | NA | NA | NA | NA | NA | NA | 1 | 1 |
| 9_1088601 | 1 | 1 | NA | NA | 1 | 0 | 1 | 0 | 3 | 0.3333 | 1 | 1 | NA | NA | NA | NA | NA | NA | 1 | 1 |
| 9_1088603 | 1 | 0 | 1 | 1 | 1 | 1 | 1 | 0 | 4 | 0.5 | 1 | 1 | NA | NA | NA | NA | NA | NA | 1 | 1 |
| 9_1088605 | 1 | 0 | 1 | 1 | 1 | 1 | 1 | 1 | 4 | 0.75 | 1 | 1 | NA | NA | 1 | 0 | NA | NA | 2 | 0.5 |
| 9_1088606 | 1 | 1 | 1 | 0 | 1 | 0 | 1 | 0 | 4 | 0.25 | 1 | 1 | NA | NA | 1 | 0 | 1 | 0 | 3 | 0.3333 |
| 9_1088608 | 1 | 1 | 1 | 1 | NA | NA | NA | NA | 2 | 1 | 1 | 1 | NA | NA | 1 | 1 | NA | NA | 2 | 1 |
| 9_1088857 | 6 | 0 | 5 | 0.2 | 4 | 0 | 3 | 0 | 18 | 0.0556 | 7 | 0.1429 | 6 | 0 | 5 | 0 | 8 | 0.125 | 26 | 0.0769 |
| 9_1088866 | 7 | 0.7143 | 3 | 0.3333 | 5 | 0.2 | 3 | 0 | 18 | 0.3889 | 6 | 0.3333 | 2 | 0 | 4 | 0.5 | 9 | 0 | 21 | 0.1905 |
| 9_1088870 | 5 | 0.6 | 1 | 1 | 4 | 0.75 | 2 | 1 | 12 | 0.75 | 2 | 1 | 5 | 0.8 | 3 | 0.6667 | 3 | 1 | 13 | 0.8462 |
| 9_1088870 | 5 | 1 | 1 | 1 | 4 | 1 | 2 | 1 | 12 | 1 | 2 | 1 | 5 | 1 | 3 | 1 | 3 | 1 | 13 | 1 |
| 9_1088874 | 4 | 1 | 1 | 1 | 4 | 1 | NA | NA | 9 | 1 | 2 | 1 | 4 | 0.75 | 2 | 0.5 | 3 | 1 | 11 | 0.8182 |
| 9_1088875 | 3 | 0.3333 | 1 | 1 | 4 | 0.75 | 3 | 0.6667 | 11 | 0.6364 | 2 | 0.5 | 3 | 0.3333 | 2 | 0.5 | 3 | 0.3333 | 10 | 0.4 |
| 9_1088876 | 2 | 1 | 1 | 0 | 1 | 1 | 3 | 0 | 7 | 0.4286 | 2 | 1 | 3 | 0.3333 | 2 | 0 | 3 | 0.6667 | 10 | 0.5 |

| | | | | | | | | | | | | | | | | | | | | |
|---|---|---|---|---|---|---|---|---|---|---|---|---|---|---|---|---|---|---|---|---|
| 9_1088877 | 4 | 0.25 | 1 | 0 | 4 | 0.75 | 3 | 1 | 12 | 0.5833 | 2 | 0 | 3 | 0.6667 | 2 | 0 | 2 | 0 | 9 | 0.2222 |
| 9_1088878 | 3 | 0.3333 | NA | NA | 3 | 0.6667 | 3 | 0.6667 | 9 | 0.5556 | 2 | 0.5 | 4 | 0.5 | 2 | 1 | 2 | 0.5 | 10 | 0.6 |
| 9_1088878 | 3 | 0 | 1 | 0 | 3 | 0 | 3 | 0.3333 | 10 | 0.1 | 2 | 0 | 3 | 0 | 2 | 0.5 | 2 | 0 | 9 | 0.1111 |
| 9_1088943 | 2 | 1 | 3 | 0.3333 | 1 | 0 | 2 | 0.5 | 8 | 0.5 | 3 | 1 | NA | NA | NA | NA | 2 | 0.5 | 5 | 0.8 |
| 9_1088944 | 2 | 0.5 | 3 | 0 | 2 | 0 | 2 | 0 | 9 | 0.1111 | 3 | 0.3333 | NA | NA | NA | NA | 2 | 0 | 5 | 0.2 |
| 9_1088944 | 1 | 0 | 3 | 1 | 1 | 1 | 2 | 0.5 | 7 | 0.7143 | 2 | 0.5 | NA | NA | NA | NA | 2 | 1 | 4 | 0.75 |
| 9_1088945 | 2 | 1 | 2 | 1 | 2 | 1 | 2 | 1 | 8 | 1 | 2 | 1 | NA | NA | NA | NA | 2 | 0.5 | 4 | 0.75 |
| 9_1088945 | 2 | 1 | 2 | 1 | 2 | 1 | 3 | 0.6667 | 9 | 0.8889 | 2 | 1 | NA | NA | NA | NA | 2 | 1 | 4 | 1 |
| 9_1088947 | 2 | 1 | 2 | 1 | 2 | 1 | 3 | 1 | 9 | 1 | 3 | 0.6667 | NA | NA | NA | NA | 2 | 0.5 | 5 | 0.6 |
| 9_1088948 | 2 | 0.5 | 1 | 0 | 1 | 0 | 2 | 0 | 6 | 0.1667 | 1 | 1 | NA | NA | NA | NA | 2 | 0 | 3 | 0.3333 |
| 9_1089185 | 2 | 0 | 1 | 1 | NA | NA | 1 | 1 | 4 | 0.5 | 1 | 1 | 1 | 0 | 2 | 0 | NA | NA | 4 | 0.25 |
| 9_1089208 | 2 | 1 | 1 | 1 | NA | NA | 1 | 1 | 4 | 1 | 1 | 1 | 1 | 1 | 3 | 1 | 1 | 0 | 6 | 0.8333 |
| 9_1089209 | 2 | 1 | NA | NA | 1 | 1 | 1 | 1 | 4 | 1 | 1 | 1 | 1 | 1 | 3 | 0.3333 | 1 | 1 | 6 | 0.6667 |
| 9_1089210 | 2 | 1 | 1 | 1 | 3 | 1 | 1 | 1 | 7 | 1 | NA | NA | 2 | 1 | 3 | 1 | NA | NA | 5 | 1 |
| 9_1089211 | 1 | 0 | 1 | 0 | 1 | 0 | 1 | 1 | 4 | 0.25 | NA | NA | 2 | 1 | 3 | 0 | 1 | 0 | 6 | 0.3333 |
| 9_1089212 | 2 | 1 | 1 | 1 | 2 | 1 | 1 | 1 | 6 | 1 | 1 | 1 | 1 | 1 | 3 | 1 | 1 | 1 | 6 | 1 |
| 9_1089213 | 2 | 0.5 | 2 | 0.5 | 1 | 1 | 2 | 0 | 7 | 0.4286 | 1 | 0 | 2 | 1 | 3 | 0.3333 | 1 | 1 | 7 | 0.5714 |
| 9_1089213 | 1 | 0 | 2 | 0.5 | 3 | 0.6667 | 2 | 1 | 8 | 0.625 | 1 | 0 | 1 | 0 | 3 | 0.6667 | 1 | 1 | 6 | 0.5 |
| 9_1089213 | 1 | 1 | 1 | 0 | 2 | 0 | 2 | 0.5 | 6 | 0.3333 | 1 | 0 | 1 | 0 | 1 | 0 | 1 | 1 | 4 | 0.25 |
| 9_1089214 | 2 | 0.5 | 2 | 1 | 3 | 0.6667 | 2 | 0.5 | 9 | 0.6667 | 1 | 1 | 2 | 0.5 | 3 | 0.6667 | 1 | 0 | 7 | 0.5714 |
| 9_1089295 | NA | NA | NA | NA | NA | NA | 1 | 1 | 1 | 1 | NA | NA | NA | NA | NA | NA | 1 | 1 | 1 | 1 |
| 9_1089470 | 1 | 0 | 2 | 0.5 | 2 | 0 | 4 | 0 | 9 | 0.1111 | 5 | 0 | 2 | 0 | 3 | 0.3333 | 5 | 0 | 15 | 0.0667 |
| 9_1089471 | 1 | 0 | 1 | 0 | 2 | 0.5 | 4 | 0.25 | 8 | 0.25 | 4 | 0.25 | 2 | 0.5 | 4 | 0.25 | 6 | 0.3333 | 16 | 0.3125 |
| 9_1089471 | 1 | 0 | 2 | 0.5 | 2 | 0 | 3 | 1 | 8 | 0.5 | 5 | 0.4 | 1 | 0 | 5 | 0.2 | 4 | 0.25 | 15 | 0.2667 |
| 9_1089472 | 1 | 0 | 2 | 1 | 3 | 0.3333 | 4 | 0.25 | 10 | 0.4 | 5 | 0 | 3 | 0.6667 | 4 | 0.5 | 5 | 0.2 | 17 | 0.2941 |
| 9_1089472 | 1 | 0 | 2 | 0 | 3 | 0.6667 | 4 | 0.25 | 10 | 0.3 | 5 | 0 | 1 | 1 | 5 | 0 | 6 | 0.1667 | 17 | 0.1176 |
| 9_1089472 | 1 | 1 | 2 | 0.5 | 3 | 1 | 4 | 0.25 | 10 | 0.6 | 5 | 0.2 | 3 | 0.3333 | 5 | 0.2 | 6 | 0.3333 | 19 | 0.2632 |
| 9_1089473 | 1 | 0 | 2 | 0.5 | 3 | 0.3333 | 4 | 0 | 10 | 0.2 | 5 | 0 | 3 | 0.6667 | 4 | 0 | 6 | 0.1667 | 18 | 0.1667 |
| 9_1089477 | 1 | 0 | 4 | 0.25 | 5 | 0.2 | 1 | 1 | 11 | 0.2727 | 5 | 0.4 | 4 | 0.5 | 6 | 0.8333 | 3 | 0 | 18 | 0.5 |
| 9_1093832 | 1 | 1 | 1 | 1 | NA | NA | 1 | 1 | 3 | 1 | 1 | 1 | NA | NA | NA | NA | 1 | 1 | 2 | 1 |
| 9_1093833 | 1 | 1 | 1 | 1 | NA | NA | 1 | 1 | 3 | 1 | 1 | 1 | NA | NA | NA | NA | 1 | 1 | 2 | 1 |
| 9_1093834 | 1 | 1 | 1 | 1 | NA | NA | 1 | 1 | 3 | 1 | 1 | 0 | NA | NA | NA | NA | 1 | 1 | 2 | 0.5 |
| 9_1093853 | 1 | 0 | 1 | 0 | NA | NA | 1 | 1 | 3 | 0.3333 | NA | NA | NA | NA | 2 | 0.5 | 1 | 1 | 3 | 0.6667 |
| 9_1096594 | 1 | 1 | NA | NA | 1 | 1 | 1 | 1 | 3 | 1 | NA | NA | NA | NA | NA | NA | 1 | 1 | 1 | 1 |
| 9_1096594 | 1 | 1 | NA | NA | 1 | 0 | 1 | 1 | 3 | 0.6667 | NA | NA | NA | NA | NA | NA | 1 | 1 | 1 | 1 |
| 9_1096595 | 1 | 0 | NA | NA | 1 | 1 | 1 | 0 | 3 | 0.3333 | NA | NA | NA | NA | NA | NA | 1 | 1 | 1 | 1 |
| 9_1096599 | 1 | 1 | NA | NA | 1 | 1 | 1 | 1 | 3 | 1 | NA | NA | NA | NA | NA | NA | 1 | 1 | 1 | 1 |
| 9_1096605 | 1 | 1 | NA | NA | 1 | 0 | NA | NA | 2 | 0.5 | NA | NA | NA | NA | NA | NA | 1 | 1 | 1 | 1 |
| 9_1096717 | NA | NA | 1 | 1 | NA | NA | NA | NA | 1 | 1 | 1 | 1 | 1 | 0 | 1 | 1 | NA | NA | 3 | 0.6667 |
| 9_1096719 | NA | NA | 1 | 1 | 1 | 1 | NA | NA | 2 | 1 | 1 | 1 | NA | NA | 1 | 1 | NA | NA | 2 | 1 |
| 9_1097219 | 1 | 0 | NA | NA | 1 | 1 | NA | NA | 2 | 0.5 | NA | NA | 1 | 1 | NA | NA | NA | NA | 1 | 1 |
| 9_1098318 | 12 | 0 | 21 | 0.0476 | 16 | 0.1875 | 16 | 0.0625 | 65 | 0.0769 | 10 | 0 | 16 | 0.25 | 14 | 0.0714 | 17 | 0.1176 | 57 | 0.1228 |
| 9_1098320 | 12 | 0.0833 | 21 | 0.0476 | 15 | 0.0667 | 17 | 0.0588 | 65 | 0.0615 | 12 | 0.1667 | 16 | 0.0625 | 14 | 0 | 18 | 0.1111 | 60 | 0.0833 |
| 9_1098345 | 9 | 0.2222 | 7 | 0.1429 | 4 | 0 | 3 | 0.3333 | 23 | 0.1739 | 3 | 0.3333 | 8 | 0.125 | 3 | 0 | 4 | 0.25 | 18 | 0.1667 |
| 9_1098346 | 10 | 0.1 | 7 | 0.4286 | 4 | 0.25 | 6 | 0.1667 | 27 | 0.2222 | 3 | 0.3333 | 8 | 0.25 | 3 | 0.3333 | 4 | 0.25 | 18 | 0.2778 |
| 9_1098356 | 3 | 0 | 3 | 0 | 2 | 0 | 4 | 0.25 | 12 | 0.0833 | 4 | 0 | 2 | 0.5 | 1 | 0 | 3 | 0.3333 | 10 | 0.2 |
| 9_1098636 | 6 | 0.3333 | 2 | 0 | 8 | 0.25 | 2 | 0.5 | 18 | 0.2778 | 2 | 0 | 1 | 0 | 3 | 0 | 2 | 0.5 | 8 | 0.125 |
| 9_1098638 | 4 | 1 | 2 | 1 | 7 | 0.5714 | 3 | 0.6667 | 16 | 0.75 | 4 | 0.5 | 1 | 0 | 2 | 0.5 | 2 | 0.5 | 9 | 0.4444 |
| 9_1098641 | 6 | 0.1667 | 2 | 0 | 4 | 0 | 3 | 0.6667 | 15 | 0.2 | 4 | 0.5 | 1 | 0 | 2 | 0 | 3 | 0 | 10 | 0.2 |
| 9_1098641 | 7 | 0.4286 | 3 | 0.3333 | 8 | 0.125 | 2 | 0.5 | 20 | 0.3 | 4 | 0 | 1 | 0 | 2 | 0 | 3 | 0.3333 | 10 | 0.1 |
| 9_1098641 | 6 | 0 | 2 | 0 | 6 | 0.1667 | 2 | 0 | 16 | 0.0625 | 2 | 0 | 1 | 0 | 3 | 0 | 3 | 0.3333 | 9 | 0.1111 |
| 9_1098643 | 6 | 0.1667 | 3 | 0 | 7 | 0.2857 | 3 | 0.3333 | 19 | 0.2105 | 3 | 0.3333 | 1 | 0 | 3 | 0.3333 | 2 | 0.5 | 9 | 0.3333 |
| 9_1098644 | 6 | 0.1667 | 4 | 0 | 8 | 0.25 | 3 | 0.3333 | 21 | 0.1905 | 4 | 0.25 | 1 | 0 | 3 | 0.3333 | 3 | 0 | 11 | 0.1818 |
| 9_1098814 | NA | NA | 1 | 0 | 2 | 0.5 | 2 | 0 | 5 | 0.2 | 1 | 0 | 1 | 0 | 3 | 0.3333 | 2 | 0 | 7 | 0.1429 |
| 9_1098815 | NA | NA | 2 | 0 | 2 | 0.5 | 3 | 0.3333 | 7 | 0.2857 | 1 | 0 | 1 | 0 | 2 | 0.5 | 1 | 1 | 5 | 0.4 |
| 9_1098816 | 1 | 1 | 2 | 0.5 | 1 | 1 | 3 | 0.6667 | 7 | 0.7143 | 1 | 1 | 1 | 1 | 2 | 1 | 2 | 0 | 6 | 0.6667 |
| 9_1098817 | 1 | 0 | 2 | 0 | 2 | 0.5 | 5 | 0.4 | 10 | 0.3 | 1 | 1 | 2 | 0.5 | 2 | 0.5 | 2 | 0.5 | 7 | 0.5714 |
| 9_1098818 | 1 | 0 | 3 | 0.3333 | 2 | 0 | 5 | 0 | 11 | 0.0909 | 1 | 0 | 2 | 0.5 | 3 | 0 | 2 | 0 | 8 | 0.125 |
| 9_1098821 | 1 | 1 | 3 | 0 | 2 | 0 | 5 | 0.6 | 11 | 0.3636 | 1 | 0 | 3 | 1 | 2 | 0 | 2 | 0.5 | 8 | 0.5 |
| 9_1098822 | 1 | 0 | 4 | 0 | 1 | 0 | 6 | 0.3333 | 12 | 0.1667 | 1 | 1 | 2 | 0.5 | 3 | 0.6667 | 2 | 0 | 8 | 0.5 |
| 9_1098823 | 2 | 0 | 3 | 0 | 2 | 0.5 | 7 | 0.4286 | 14 | 0.2857 | 1 | 0 | 5 | 0.6 | 3 | 0.6667 | 2 | 0 | 11 | 0.4545 |
| 9_1098933 | 3 | 0 | 3 | 0.3333 | 7 | 0.1429 | 4 | 0 | 17 | 0.1176 | 2 | 0 | 4 | 0 | 3 | 0.3333 | 3 | 0 | 12 | 0.0833 |
| 9_1099110 | 13 | 0.1538 | 14 | 0.2143 | 15 | 0.2667 | 13 | 0.1538 | 55 | 0.2 | 7 | 0.2857 | 17 | 0 | 12 | 0.1667 | 9 | 0.1111 | 45 | 0.1111 |
| 9_1099113 | 10 | 0.3 | 14 | 0.5 | 14 | 0.5 | 14 | 0.5714 | 52 | 0.4808 | 10 | 0.5 | 16 | 0.5 | 10 | 0.3 | 12 | 0.5833 | 48 | 0.4792 |
| 9_1099113 | 9 | 0.6667 | 14 | 0.6429 | 13 | 0.4615 | 14 | 0.4286 | 50 | 0.54 | 10 | 0.7 | 17 | 0.5294 | 12 | 0.5 | 12 | 0.3333 | 51 | 0.5098 |

| | | | | | | | | | | | | | | | | | | | | |
|---|---|---|---|---|---|---|---|---|---|---|---|---|---|---|---|---|---|---|---|---|
| 9_1099115 | 13 | 0.4615 | 16 | 0.625 | 14 | 0.2857 | 14 | 0.5 | 57 | 0.4737 | 12 | 0.1667 | 18 | 0.4444 | 11 | 0.4545 | 9 | 0.4444 | 50 | 0.38 |
| 9_1099119 | 8 | 0 | 16 | 0.125 | 9 | 0 | 12 | 0.0833 | 45 | 0.0667 | 11 | 0.0909 | 15 | 0.1333 | 8 | 0 | 7 | 0.1429 | 41 | 0.0976 |
| 9_1099120 | 18 | 0.1111 | 18 | 0.1111 | 19 | 0.1053 | 18 | 0.1111 | 73 | 0.1096 | 15 | 0.0667 | 22 | 0.1364 | 14 | 0.0714 | 15 | 0.0667 | 66 | 0.0909 |
| 9_1099125 | 18 | 0.0556 | 15 | 0.1333 | 18 | 0 | 18 | 0.0556 | 69 | 0.058 | 13 | 0.1538 | 23 | 0.0435 | 16 | 0.0625 | 13 | 0.1538 | 65 | 0.0923 |
| 9_1099125 | 10 | 0.2 | 13 | 0.5385 | 12 | 0.4167 | 16 | 0.4375 | 51 | 0.4118 | 11 | 0.5455 | 17 | 0.2941 | 8 | 0.625 | 9 | 0.3333 | 45 | 0.4222 |
| 9_1099126 | 14 | 0.1429 | 15 | 0.4 | 16 | 0.0625 | 16 | 0.1875 | 61 | 0.1967 | 13 | 0.1538 | 18 | 0.3333 | 13 | 0.2308 | 11 | 0.4545 | 55 | 0.2909 |
| 9_1099134 | 11 | 0.1818 | 16 | 0 | 12 | 0.0833 | 9 | 0 | 48 | 0.0625 | 9 | 0.1111 | 13 | 0 | 14 | 0.1429 | 8 | 0.125 | 44 | 0.0909 |
| 9_1099135 | 11 | 0.1818 | 16 | 0.3125 | 10 | 0.3 | 9 | 0 | 46 | 0.2174 | 11 | 0.5455 | 11 | 0.1818 | 17 | 0.0588 | 8 | 0.125 | 47 | 0.2128 |
| 9_1099136 | 9 | 0.3333 | 11 | 0.3636 | 9 | 0.2222 | 6 | 0 | 35 | 0.2571 | 9 | 0.2222 | 10 | 0.1 | 13 | 0.3077 | 8 | 0.25 | 40 | 0.225 |
| 9_1099165 | 8 | 0.125 | 9 | 0 | 6 | 0 | 5 | 0.2 | 28 | 0.0714 | 6 | 0 | 7 | 0.1429 | 8 | 0.125 | 3 | 0.3333 | 24 | 0.125 |
| 9_1099167 | 7 | 0 | 8 | 0.5 | 5 | 0.4 | 4 | 0.25 | 24 | 0.2917 | 4 | 0.5 | 5 | 0.2 | 6 | 0.1667 | 2 | 0 | 17 | 0.2353 |
| 9_1099169 | 6 | 0.8333 | 6 | 0.8333 | 4 | 1 | 1 | 1 | 17 | 0.8824 | 7 | 1 | 4 | 1 | 8 | 0.875 | 1 | 1 | 20 | 0.95 |
| 9_1099169 | 4 | 0.5 | 4 | 0.75 | NA | NA | 2 | 0.5 | 10 | 0.6 | 2 | 0.5 | 1 | 0 | 4 | 0.25 | NA | NA | 7 | 0.2857 |
| 9_1099170 | 8 | 0.25 | 4 | 0.25 | 3 | 0.3333 | 2 | 0 | 17 | 0.2353 | 4 | 0.25 | 6 | 0.3333 | 6 | 0.3333 | 3 | 0.3333 | 19 | 0.3158 |
| 9_1099171 | 7 | 0.4286 | 5 | 1 | 3 | 1 | 1 | 0 | 16 | 0.6875 | 6 | 0.6667 | 6 | 0.6667 | 6 | 0.6667 | 2 | 0.5 | 20 | 0.65 |
| 9_1099172 | 5 | 1 | 3 | 1 | 4 | 0.75 | 2 | 1 | 14 | 0.9286 | 5 | 0.4 | 3 | 0.3333 | 6 | 0.5 | 3 | 0.6667 | 17 | 0.4706 |
| 9_1100253 | 2 | 0.5 | 2 | 0 | 3 | 0 | 1 | 0 | 8 | 0.125 | 3 | 0 | 6 | 0.1667 | 5 | 0 | 2 | 0.5 | 16 | 0.125 |
| 9_1100670 | 3 | 0.6667 | 3 | 0.6667 | 4 | 0.75 | 4 | 0.75 | 14 | 0.7143 | 3 | 1 | 7 | 0.8571 | 5 | 0.2 | 3 | 0.6667 | 18 | 0.6667 |
| 9_1100671 | 3 | 0 | 3 | 0.3333 | 4 | 0 | 5 | 0.2 | 15 | 0.1333 | 3 | 0 | 7 | 0.1429 | 5 | 0.2 | 3 | 0 | 18 | 0.1111 |
| 9_1100682 | 3 | 0 | 6 | 0 | 5 | 0.2 | 5 | 0 | 19 | 0.0526 | 4 | 0 | 6 | 0.3333 | 7 | 0 | 3 | 0 | 20 | 0.1 |
| 9_1100687 | 3 | 0 | 6 | 0.1667 | 4 | 0 | 5 | 0 | 18 | 0.0556 | 6 | 0.1667 | 7 | 0.1429 | 8 | 0 | 5 | 0 | 26 | 0.0769 |
| 9_1100688 | 4 | 0 | 5 | 0.2 | 5 | 0.4 | 4 | 0.25 | 18 | 0.2222 | 5 | 0.4 | 7 | 0 | 9 | 0 | 5 | 0 | 26 | 0.0769 |
| 9_1100754 | 21 | 0.2381 | 22 | 0.5455 | 19 | 0.4737 | 8 | 0.375 | 70 | 0.4143 | 16 | 0.3125 | 21 | 0.381 | 17 | 0.4118 | 17 | 0.5882 | 71 | 0.4225 |
| 9_1100755 | 24 | 0.25 | 26 | 0.3077 | 22 | 0.3636 | 17 | 0.4118 | 89 | 0.3258 | 15 | 0.2 | 20 | 0.45 | 23 | 0.3043 | 21 | 0.4286 | 79 | 0.3544 |
| 9_1100756 | 33 | 0.7273 | 26 | 0.2692 | 20 | 0.65 | 18 | 0.3333 | 97 | 0.5155 | 21 | 0.619 | 27 | 0.4815 | 25 | 0.68 | 23 | 0.6522 | 96 | 0.6042 |
| 9_1100757 | 23 | 0.3913 | 32 | 0.3438 | 23 | 0.2174 | 20 | 0.2 | 98 | 0.2959 | 25 | 0.2 | 31 | 0.2581 | 22 | 0.3182 | 22 | 0.2273 | 100 | 0.25 |
| 9_1100757 | 19 | 0.3684 | 29 | 0.3793 | 18 | 0.1111 | 16 | 0.75 | 82 | 0.3902 | 20 | 0.6 | 27 | 0.3333 | 25 | 0.28 | 24 | 0.5 | 96 | 0.4167 |
| 9_1100758 | 35 | 0.5143 | 38 | 0.5526 | 32 | 0.5625 | 26 | 0.5769 | 131 | 0.5496 | 31 | 0.6129 | 36 | 0.4722 | 35 | 0.5714 | 32 | 0.4688 | 134 | 0.5299 |
| 9_1100759 | 37 | 0.1622 | 41 | 0.0488 | 31 | 0.1613 | 29 | 0.1034 | 138 | 0.1159 | 31 | 0.2258 | 40 | 0.175 | 34 | 0.1471 | 36 | 0.1667 | 141 | 0.1773 |
| 9_1100760 | 40 | 0.15 | 40 | 0.2 | 33 | 0.1818 | 32 | 0.3125 | 145 | 0.2069 | 31 | 0.1935 | 42 | 0.1905 | 36 | 0.1667 | 34 | 0.1176 | 143 | 0.1678 |
| 9_1100762 | 33 | 0.8485 | 32 | 0.75 | 23 | 0.6522 | 26 | 0.5769 | 114 | 0.7193 | 26 | 0.6923 | 28 | 0.6429 | 23 | 0.6957 | 27 | 0.7778 | 104 | 0.7019 |
| 9_1100762 | 23 | 0.3478 | 32 | 0.2812 | 21 | 0.1429 | 22 | 0.3182 | 98 | 0.2755 | 22 | 0.2727 | 26 | 0.2308 | 20 | 0.15 | 24 | 0.125 | 92 | 0.1957 |
| 9_1100763 | 37 | 0.5135 | 45 | 0.3778 | 34 | 0.4706 | 32 | 0.2812 | 148 | 0.4122 | 33 | 0.3939 | 45 | 0.2444 | 39 | 0.5641 | 38 | 0.4211 | 155 | 0.4 |
| 9_1100764 | 39 | 0.0513 | 45 | 0.0889 | 29 | 0.1034 | 27 | 0.037 | 140 | 0.0714 | 30 | 0.0667 | 40 | 0.075 | 37 | 0.1081 | 33 | 0.0606 | 140 | 0.0786 |
| 9_1100801 | 43 | 0.0465 | 55 | 0.0545 | 35 | 0.0286 | 32 | 0.125 | 165 | 0.0606 | 34 | 0.0882 | 41 | 0.0732 | 40 | 0.125 | 41 | 0.0488 | 156 | 0.0833 |
| 9_1100912 | 48 | 0.1042 | 66 | 0.1212 | 38 | 0.1316 | 38 | 0.0526 | 190 | 0.1053 | 34 | 0.0882 | 51 | 0.0392 | 45 | 0.1556 | 46 | 0.1087 | 176 | 0.0966 |
| 9_1100912 | 45 | 0.2889 | 61 | 0.2787 | 36 | 0.25 | 36 | 0.1944 | 178 | 0.2584 | 36 | 0.4167 | 49 | 0.2041 | 46 | 0.2609 | 46 | 0.1957 | 177 | 0.2599 |
| 9_1100912 | 44 | 0.1364 | 65 | 0.1385 | 33 | 0.1515 | 31 | 0.2581 | 173 | 0.1618 | 34 | 0.1471 | 44 | 0.2727 | 47 | 0.2128 | 41 | 0.2439 | 166 | 0.2229 |
| 9_1100913 | 47 | 0.1277 | 64 | 0.2188 | 39 | 0.2564 | 40 | 0.175 | 190 | 0.1947 | 34 | 0.2059 | 50 | 0.18 | 46 | 0.2174 | 45 | 0.1333 | 175 | 0.1829 |
| 9_1100962 | NA | NA | 1 | 1 | NA | NA | NA | NA | 1 | 1 | 2 | 0.5 | NA | NA | NA | NA | NA | NA | 2 | 0.5 |
| 9_1100963 | NA | NA | 1 | 1 | NA | NA | NA | NA | 1 | 1 | 2 | 1 | NA | NA | NA | NA | NA | NA | 2 | 1 |
| 9_1101449 | 1 | 0 | 2 | 0.5 | 1 | 0 | NA | NA | 4 | 0.25 | NA | NA | NA | NA | 2 | 0.5 | NA | NA | 2 | 0.5 |
| 9_1101500 | 2 | 1 | 3 | 0 | 1 | 0 | 2 | 0.5 | 8 | 0.375 | NA | NA | 1 | 0 | 1 | 1 | 2 | 0.5 | 4 | 0.5 |
| 9_1101500 | 2 | 0.5 | 3 | 0.3333 | 1 | 1 | 1 | 1 | 7 | 0.5714 | NA | NA | NA | NA | 1 | 1 | 1 | 0 | 2 | 0.5 |
| 9_1101500 | 2 | 0.5 | 3 | 0.6667 | NA | NA | 2 | 1 | 7 | 0.7143 | NA | NA | 1 | 1 | 2 | 0.5 | 2 | 0.5 | 5 | 0.6 |
| 9_1101500 | 1 | 0 | 3 | 0.3333 | 1 | 0 | 2 | 0 | 7 | 0.1429 | NA | NA | NA | NA | 2 | 0.5 | 2 | 0 | 4 | 0.25 |
| 9_1101501 | 2 | 1 | 3 | 0.3333 | 1 | 0 | 2 | 0.5 | 8 | 0.5 | NA | NA | 1 | 1 | 1 | 1 | 2 | 1 | 4 | 1 |
| 9_1101503 | 3 | 0 | 4 | 0.25 | 1 | 1 | 1 | 0 | 9 | 0.2222 | NA | NA | 1 | 0 | 2 | 0.5 | 1 | 0 | 4 | 0.25 |
| 9_1101505 | 1 | 1 | 1 | 0 | NA | NA | NA | NA | 2 | 0.5 | NA | NA | NA | NA | NA | NA | 1 | 1 | 1 | 1 |
| 9_1102100 | 9 | 0.1111 | 4 | 0 | 7 | 0.1429 | 11 | 0 | 31 | 0.0645 | 4 | 0 | 8 | 0 | 5 | 0.2 | 3 | 0 | 20 | 0.05 |
| 9_1102103 | 11 | 0.8182 | 4 | 0.75 | 5 | 0.8 | 11 | 0.7273 | 31 | 0.7742 | 6 | 0.8333 | 9 | 0.5556 | 4 | 0.5 | 3 | 0.6667 | 22 | 0.6364 |
| 9_1102104 | 6 | 0.1667 | 3 | 0.6667 | 5 | 0.2 | 6 | 0.1667 | 20 | 0.25 | 4 | 0.25 | 7 | 0.1429 | 6 | 0 | 3 | 0.3333 | 20 | 0.15 |
| 9_1102104 | 10 | 0.3 | 4 | 0.5 | 6 | 0 | 9 | 0.3333 | 29 | 0.2759 | 6 | 0 | 9 | 0.1111 | 8 | 0.25 | 3 | 0 | 26 | 0.1154 |
| 9_1102137 | 14 | 0 | 5 | 0.2 | 9 | 0.4444 | 12 | 0.0833 | 40 | 0.15 | 13 | 0 | 11 | 0 | 11 | 0.1818 | 4 | 0 | 39 | 0.0513 |
| 9_1102137 | 19 | 0.1579 | 3 | 1 | 9 | 0.3333 | 12 | 0.0833 | 43 | 0.2326 | 13 | 0 | 9 | 0.3333 | 12 | 0.1667 | 5 | 0.2 | 39 | 0.1538 |
| 9_1102138 | 8 | 0.75 | 4 | 0.75 | 4 | 0.5 | 9 | 0.5556 | 25 | 0.64 | 7 | 0.5714 | 8 | 0.375 | 5 | 0.8 | 3 | 0.3333 | 23 | 0.5217 |
| 9_1102141 | 4 | 0 | 5 | 0.2 | 5 | 0.4 | 2 | 0 | 16 | 0.1875 | 4 | 0 | 6 | 0.5 | 3 | 0.3333 | 6 | 0.3333 | 19 | 0.3158 |
| 9_1102142 | 4 | 0 | 7 | 0.1429 | 5 | 0 | 2 | 0.5 | 18 | 0.1111 | 4 | 0.25 | 5 | 0 | 3 | 0.3333 | 7 | 0 | 19 | 0.1053 |
| 9_1102142 | 4 | 0.5 | 7 | 0 | 5 | 0 | 2 | 0 | 18 | 0.1111 | 4 | 0 | 5 | 0.2 | 2 | 0 | 5 | 0.2 | 16 | 0.125 |
| 9_1102143 | 4 | 0 | 7 | 0 | 4 | 0.25 | NA | NA | 15 | 0.0667 | 5 | 0.2 | 5 | 0 | 3 | 0 | 7 | 0 | 20 | 0.05 |
| 9_1102143 | 4 | 0.5 | 5 | 1 | 4 | 1 | 1 | 0 | 14 | 0.7857 | 4 | 1 | 4 | 1 | 2 | 0.5 | 7 | 0.7143 | 17 | 0.8235 |
| 9_1102144 | 4 | 0.5 | 6 | 0.1667 | 1 | 1 | 2 | 0 | 13 | 0.3077 | 3 | 0.3333 | 5 | 0.2 | 2 | 1 | 7 | 0.4286 | 17 | 0.4118 |
| 9_1102144 | 2 | 1 | 5 | 0.6 | 1 | 1 | 1 | 0 | 9 | 0.6667 | 2 | 0.5 | 2 | 0.5 | 2 | 1 | 3 | 0.6667 | 9 | 0.6667 |
| 9_1102147 | 5 | 0.4 | 6 | 0.1667 | 5 | 0 | 2 | 0 | 18 | 0.1667 | 3 | 0 | 6 | 0 | 3 | 0 | 7 | 0.2857 | 19 | 0.1053 |
| 9_1102161 | NA | NA | 3 | 0 | 2 | 0.5 | 1 | 0 | 6 | 0.1667 | 1 | 1 | 1 | 0 | 1 | 0 | 5 | 0.2 | 8 | 0.25 |

| | | | | | | | | | | | | | | | | | | | | |
|---|---|---|---|---|---|---|---|---|---|---|---|---|---|---|---|---|---|---|---|---|
| 9_1102161 | 2 | 0 | 4 | 0 | 4 | 0.25 | 1 | 0 | 11 | 0.0909 | 2 | 1 | 5 | 0 | 2 | 0 | 6 | 0.1667 | 15 | 0.2 |
| 9_1102174 | 2 | 0.5 | 2 | 0 | NA | NA | NA | NA | 4 | 0.25 | 1 | 0 | 4 | 0 | 1 | 0 | 5 | 0.2 | 11 | 0.0909 |
| 9_1102551 | 4 | 0.5 | 3 | 0.6667 | 5 | 0.2 | 3 | 0 | 15 | 0.3333 | 5 | 0.4 | 1 | 0 | 3 | 0.3333 | 1 | 0 | 10 | 0.3 |
| 9_1102552 | 4 | 0.25 | 4 | 0.5 | 5 | 0.6 | 3 | 0 | 16 | 0.375 | 5 | 0.4 | 1 | 0 | 3 | 1 | 1 | 0 | 10 | 0.5 |
| 9_1102552 | 5 | 0 | 4 | 0 | 5 | 0.4 | 3 | 0 | 17 | 0.1176 | 6 | 0.3333 | 1 | 0 | 4 | 0.5 | 1 | 0 | 12 | 0.3333 |
| 9_1102556 | 5 | 0.4 | 2 | 0.5 | 5 | 0.4 | 3 | 0.6667 | 15 | 0.4667 | 3 | 0.3333 | 1 | 1 | 3 | 0.6667 | 1 | 0 | 8 | 0.5 |
| 9_1102557 | 5 | 1 | 2 | 1 | 6 | 1 | 3 | 1 | 16 | 1 | 4 | 0.5 | 1 | 1 | 2 | 1 | 1 | 0 | 8 | 0.625 |
| 9_1102558 | 5 | 0.6 | 1 | 1 | 4 | 1 | 3 | 1 | 13 | 0.8462 | 4 | 1 | 1 | 1 | 2 | 1 | NA | NA | 7 | 1 |
| 9_1102561 | 2 | 1 | 1 | 1 | 3 | 0.6667 | 3 | 1 | 9 | 0.8889 | 3 | 0.6667 | 1 | 1 | 2 | 1 | NA | NA | 6 | 0.8333 |
| 9_1102561 | 2 | 1 | 1 | 1 | 3 | 0.6667 | 3 | 1 | 9 | 0.8889 | 3 | 0.6667 | 1 | 0 | 3 | 1 | NA | NA | 7 | 0.7143 |
| 9_1102562 | 3 | 1 | 1 | 1 | 3 | 1 | 2 | 0.5 | 9 | 0.8889 | 4 | 1 | 1 | 1 | 3 | 1 | NA | NA | 8 | 1 |
| 9_1102562 | 3 | 1 | 1 | 1 | 3 | 1 | 3 | 1 | 10 | 1 | 3 | 1 | 1 | 1 | 3 | 0.6667 | NA | NA | 7 | 0.8571 |
| 9_1102564 | 2 | 1 | 1 | 1 | 1 | 1 | 2 | 0.5 | 6 | 0.8333 | 1 | 0 | 1 | 0 | 2 | 0.5 | NA | NA | 4 | 0.25 |
| 9_1102567 | NA | NA | 1 | 1 | 3 | 0 | 3 | 1 | 7 | 0.5714 | 4 | 0.75 | NA | NA | 1 | 1 | NA | NA | 5 | 0.8 |
| 9_1102568 | 1 | 0 | 1 | 1 | 2 | 0.5 | 3 | 0 | 7 | 0.2857 | 2 | 0.5 | NA | NA | 1 | 1 | NA | NA | 3 | 0.6667 |
| 9_1102568 | 2 | 0.5 | 1 | 1 | 3 | 0.3333 | 3 | 1 | 9 | 0.6667 | 3 | 1 | 1 | 1 | 1 | 1 | NA | NA | 5 | 1 |
| 9_1102569 | NA | NA | 1 | 1 | 3 | 0.6667 | 2 | 0.5 | 6 | 0.6667 | 3 | 1 | NA | NA | 2 | 0.5 | NA | NA | 5 | 0.8 |
| 9_1102571 | NA | NA | 1 | 0 | 2 | 1 | 3 | 1 | 6 | 0.8333 | 4 | 0.75 | NA | NA | 2 | 1 | NA | NA | 6 | 0.8333 |
| 9_1102572 | NA | NA | 1 | 1 | 2 | 0 | 3 | 1 | 6 | 0.6667 | 4 | 0.75 | NA | NA | 1 | 1 | NA | NA | 5 | 0.8 |
| 9_1102574 | NA | NA | 1 | 1 | 3 | 0.3333 | 2 | 0 | 6 | 0.3333 | 3 | 0 | NA | NA | 2 | 0.5 | NA | NA | 5 | 0.2 |
| 9_1102575 | NA | NA | 1 | 1 | 3 | 0.6667 | 2 | 0.5 | 6 | 0.6667 | 2 | 0.5 | NA | NA | 2 | 0.5 | NA | NA | 4 | 0.5 |
| 9_1102577 | NA | NA | 1 | 1 | 1 | 1 | 3 | 1 | 5 | 1 | 2 | 1 | NA | NA | NA | NA | NA | NA | 2 | 1 |
| 9_1102577 | NA | NA | 1 | 0 | 1 | 0 | 3 | 0.3333 | 5 | 0.2 | 2 | 0.5 | NA | NA | 1 | 0 | NA | NA | 3 | 0.3333 |
| 9_1102578 | NA | NA | NA | NA | 2 | 1 | 2 | 1 | 4 | 1 | 2 | 1 | NA | NA | 1 | 1 | NA | NA | 3 | 1 |
| 9_1103765 | NA | NA | NA | NA | NA | NA | 2 | 0.5 | 2 | 0.5 | NA | NA | 1 | 1 | 2 | 1 | 1 | 0 | 4 | 0.75 |
| 9_1103769 | NA | NA | NA | NA | NA | NA | 3 | 0.3333 | 3 | 0.3333 | NA | NA | 1 | 0 | 2 | 0.5 | 1 | 1 | 4 | 0.5 |
| 9_1103771 | NA | NA | NA | NA | NA | NA | 3 | 0.6667 | 3 | 0.6667 | NA | NA | 1 | 0 | 2 | 1 | 1 | 1 | 4 | 0.75 |
| 9_1103776 | NA | NA | NA | NA | NA | NA | 3 | 1 | 3 | 1 | NA | NA | 1 | 1 | 1 | 1 | 1 | 1 | 3 | 1 |
| 9_1103777 | NA | NA | NA | NA | NA | NA | 3 | 0.6667 | 3 | 0.6667 | NA | NA | 1 | 0 | 2 | 0.5 | 1 | 0 | 4 | 0.25 |
| 9_1103779 | NA | NA | NA | NA | NA | NA | 3 | 0.3333 | 3 | 0.3333 | NA | NA | 1 | 0 | 2 | 0.5 | NA | NA | 3 | 0.3333 |
| 9_1103780 | NA | NA | NA | NA | NA | NA | 3 | 0.6667 | 3 | 0.6667 | NA | NA | 1 | 1 | 2 | 1 | 1 | 1 | 4 | 1 |
| 9_1103782 | NA | NA | NA | NA | NA | NA | 3 | 0.6667 | 3 | 0.6667 | NA | NA | 1 | 1 | NA | NA | 1 | 1 | 2 | 1 |
| 9_1103783 | NA | NA | NA | NA | NA | NA | 3 | 0.3333 | 3 | 0.3333 | NA | NA | 1 | 0 | 2 | 0.5 | 1 | 0 | 4 | 0.25 |
| 9_1103785 | NA | NA | NA | NA | NA | NA | 3 | 0.3333 | 3 | 0.3333 | NA | NA | 1 | 1 | 4 | 0.75 | 1 | 1 | 6 | 0.8333 |
| 9_1103786 | NA | NA | NA | NA | NA | NA | 3 | 0.3333 | 3 | 0.3333 | NA | NA | 1 | 0 | 4 | 0.25 | 1 | 0 | 6 | 0.1667 |
| 9_1103791 | NA | NA | NA | NA | NA | NA | 3 | 0.3333 | 3 | 0.3333 | NA | NA | 1 | 0 | 3 | 0 | 1 | 1 | 5 | 0.2 |
| 9_1103794 | NA | NA | NA | NA | NA | NA | 3 | 0.3333 | 3 | 0.3333 | NA | NA | 1 | 0 | 3 | 0 | 2 | 0.5 | 6 | 0.1667 |
| 9_1103796 | NA | NA | NA | NA | NA | NA | 3 | 0.3333 | 3 | 0.3333 | NA | NA | 1 | 0 | 2 | 0.5 | 2 | 0 | 5 | 0.2 |
| 9_1103799 | NA | NA | NA | NA | NA | NA | 3 | 0.3333 | 3 | 0.3333 | NA | NA | 1 | 0 | 4 | 0 | 1 | 1 | 6 | 0.1667 |
| 9_1103802 | NA | NA | NA | NA | 1 | 1 | 4 | 0.75 | 5 | 0.8 | 1 | 0 | 3 | 0.6667 | 5 | 0.6 | 3 | 1 | 12 | 0.6667 |
| 9_1104026 | NA | NA | NA | NA | 1 | 0 | 6 | 0.3333 | 7 | 0.2857 | 1 | 0 | 3 | 0.3333 | 6 | 0 | 6 | 0 | 16 | 0.0625 |
| 9_1104027 | NA | NA | NA | NA | 1 | 1 | 6 | 0.6667 | 7 | 0.7143 | 1 | 0 | 3 | 0.3333 | 3 | 0.6667 | 5 | 0.6 | 12 | 0.5 |
| 9_1104029 | NA | NA | NA | NA | 1 | 1 | 4 | 1 | 5 | 1 | 1 | 1 | 3 | 1 | 5 | 1 | 6 | 1 | 15 | 1 |
| 9_1104030 | NA | NA | NA | NA | 1 | 1 | 6 | 1 | 7 | 1 | 1 | 1 | 3 | 0.6667 | 5 | 0.6 | 6 | 0.5 | 15 | 0.6 |
| 9_1104030 | NA | NA | NA | NA | NA | NA | 7 | 1 | 7 | 1 | NA | NA | 3 | 1 | 6 | 0.8333 | 5 | 1 | 14 | 0.9286 |
| 9_1104215 | NA | NA | 1 | 0 | 1 | 0 | 7 | 0.1429 | 9 | 0.1111 | 1 | 0 | 2 | 0 | 6 | 0.1667 | 4 | 0 | 13 | 0.0769 |
| 9_1104232 | NA | NA | 1 | 0 | 2 | 0.5 | 7 | 0.2857 | 10 | 0.3 | 1 | 1 | 3 | 0.3333 | 5 | 0.4 | 5 | 0.2 | 14 | 0.3571 |
| 9_1104233 | NA | NA | 1 | 0 | 2 | 0.5 | 5 | 0.6 | 8 | 0.5 | 1 | 1 | 3 | 0.6667 | 4 | 0.75 | 7 | 1 | 15 | 0.8667 |
| 9_1104234 | NA | NA | 1 | 1 | 2 | 1 | 7 | 0.8571 | 10 | 0.9 | 1 | 1 | 3 | 1 | 5 | 1 | 7 | 1 | 16 | 1 |
| 9_1104234 | NA | NA | 1 | 1 | 2 | 1 | 8 | 1 | 11 | 1 | 1 | 1 | 3 | 0.6667 | 6 | 1 | 7 | 0.7143 | 17 | 0.8235 |
| 9_1104236 | NA | NA | 1 | 0 | 2 | 0.5 | 8 | 0.375 | 11 | 0.3636 | 1 | 0 | 3 | 0.3333 | 5 | 0.6 | 6 | 0.3333 | 15 | 0.4 |
| 9_1104236 | NA | NA | 1 | 1 | 1 | 0 | 7 | 0.8571 | 9 | 0.7778 | 1 | 1 | 2 | 0.5 | 5 | 0.8 | 6 | 0.8333 | 14 | 0.7857 |
| 9_1104237 | NA | NA | 1 | 0 | 2 | 0 | 8 | 0.375 | 11 | 0.2727 | 2 | 1 | 4 | 0 | 6 | 0.6667 | 7 | 0.8571 | 19 | 0.6316 |
| 9_1104470 | NA | NA | 2 | 0 | 3 | 0.6667 | 9 | 0.1111 | 14 | 0.2143 | 3 | 0.6667 | 4 | 0.25 | 9 | 0.4444 | 11 | 0.2727 | 27 | 0.3704 |
| 9_1104474 | NA | NA | 3 | 0 | 4 | 0.5 | 10 | 0.1 | 17 | 0.1765 | 4 | 0.25 | 6 | 0.3333 | 10 | 0.2 | 11 | 0 | 31 | 0.1613 |
| 9_1104506 | NA | NA | NA | NA | 1 | 1 | NA | NA | 1 | 1 | NA | NA | NA | NA | 2 | 1 | 1 | 0 | 3 | 0.6667 |
| 9_1104526 | NA | NA | NA | NA | 1 | 1 | NA | NA | 1 | 1 | NA | NA | NA | NA | 1 | 1 | 1 | 0 | 2 | 0.5 |
| 9_1104610 | NA | NA | 2 | 1 | NA | NA | NA | NA | 2 | 1 | NA | NA | 1 | 0 | 1 | 1 | NA | NA | 2 | 0.5 |
| 9_1104676 | NA | NA | 1 | 1 | NA | NA | NA | NA | 1 | 1 | NA | NA | 1 | 1 | 1 | 1 | NA | NA | 2 | 1 |
| 9_1105403 | 11 | 0.1818 | 10 | 0.2 | 4 | 0 | 10 | 0 | 35 | 0.1143 | 5 | 0.2 | 7 | 0 | 11 | 0 | 3 | 0.3333 | 26 | 0.0769 |
| 9_1105512 | 1 | 1 | 2 | 0.5 | 1 | 0 | 1 | 1 | 5 | 0.6 | 1 | 1 | 1 | 1 | NA | NA | 2 | 0.5 | 4 | 0.75 |
| 9_1105512 | 1 | 1 | 2 | 0 | 1 | 0 | 1 | 1 | 5 | 0.4 | 1 | 1 | 1 | 1 | NA | NA | 2 | 1 | 4 | 1 |
| 9_1105512 | 1 | 1 | 2 | 1 | 1 | 0 | NA | NA | 4 | 0.75 | 1 | 1 | 1 | 1 | NA | NA | 2 | 1 | 4 | 1 |
| 9_1105513 | 1 | 1 | 2 | 1 | 1 | 1 | 1 | 1 | 5 | 1 | NA | NA | 1 | 1 | NA | NA | 2 | 1 | 3 | 1 |
| 9_1105513 | 1 | 1 | 2 | 0 | NA | NA | 1 | 0 | 4 | 0.25 | 1 | 1 | 1 | 0 | NA | NA | 1 | 0 | 3 | 0.3333 |

| | | | | | | | | | | | | | | | | | | | | |
|---|---|---|---|---|---|---|---|---|---|---|---|---|---|---|---|---|---|---|---|---|
| 9_1105515 | 1 | 1 | 2 | 1 | 1 | 1 | 1 | 1 | 5 | 1 | 1 | 1 | 1 | 0 | NA | NA | 2 | 0.5 | 4 | 0.5 |
| 9_1106948 | 2 | 0.5 | NA | NA | NA | NA | 2 | 0.5 | 4 | 0.5 | NA | NA | 1 | 1 | NA | NA | NA | NA | 1 | 1 |
| 9_1106949 | 1 | 1 | NA | NA | NA | NA | 1 | 1 | 2 | 1 | NA | NA | 1 | 1 | NA | NA | NA | NA | 1 | 1 |
| 9_1107275 | NA | NA | 1 | 1 | NA | NA | NA | NA | 1 | 1 | 1 | 0 | NA | NA | 1 | 0 | 1 | 1 | 3 | 0.3333 |
| 9_1107283 | NA | NA | 1 | 1 | NA | NA | NA | NA | 1 | 1 | 1 | 0 | 1 | 1 | 1 | 1 | 1 | 0 | 4 | 0.5 |
| 9_1107284 | NA | NA | 2 | 0.5 | NA | NA | NA | NA | 2 | 0.5 | 1 | 1 | 1 | 1 | 1 | 0 | 1 | 0 | 4 | 0.5 |
| 9_1107287 | NA | NA | 1 | 1 | NA | NA | NA | NA | 1 | 1 | 1 | 1 | NA | NA | 1 | 1 | 1 | 0 | 3 | 0.6667 |
| 9_1107290 | NA | NA | 2 | 0.5 | NA | NA | NA | NA | 2 | 0.5 | 1 | 1 | 1 | 0 | 1 | 0 | NA | NA | 3 | 0.3333 |
| 9_1110531 | 4 | 0.25 | 7 | 0.1429 | 2 | 0 | 6 | 0 | 19 | 0.1053 | 5 | 0 | 1 | 0 | 5 | 0.2 | 7 | 0 | 18 | 0.0556 |
| 9_1110531 | 5 | 0.4 | 8 | 0.125 | 3 | 0 | 5 | 0 | 21 | 0.1429 | 4 | 0.25 | 4 | 0.25 | 6 | 0.1667 | 7 | 0.1429 | 21 | 0.1905 |
| 9_1110541 | 2 | 0.5 | 2 | 0 | NA | NA | 2 | 0.5 | 6 | 0.3333 | NA | NA | 2 | 0 | 2 | 0.5 | 1 | 0 | 5 | 0.2 |
| 9_1110574 | NA | NA | 1 | 0 | 1 | 0 | 4 | 1 | 6 | 0.6667 | 1 | 1 | NA | NA | 4 | 0.75 | 3 | 0.6667 | 8 | 0.75 |
| 9_1110575 | NA | NA | 1 | 1 | 2 | 1 | 3 | 1 | 6 | 1 | 1 | 1 | NA | NA | 5 | 0.6 | 3 | 0.6667 | 9 | 0.6667 |
| 9_1110576 | NA | NA | 2 | 0.5 | 2 | 1 | 4 | 0.5 | 8 | 0.625 | 1 | 1 | NA | NA | 5 | 0.6 | 3 | 0.3333 | 9 | 0.5556 |
| 9_1110594 | NA | NA | 2 | 0 | NA | NA | 3 | 0.3333 | 5 | 0.2 | NA | NA | NA | NA | 2 | 0 | 2 | 0.5 | 4 | 0.25 |
| 9_1110594 | NA | NA | 2 | 0 | 1 | 1 | 3 | 0.6667 | 6 | 0.5 | NA | NA | NA | NA | 2 | 0.5 | 2 | 1 | 4 | 0.75 |
| 9_1110596 | NA | NA | NA | NA | NA | NA | 3 | 0.6667 | 3 | 0.6667 | NA | NA | NA | NA | 2 | 0.5 | 2 | 1 | 4 | 0.75 |
| 9_1110597 | NA | NA | NA | NA | 1 | 1 | 2 | 0.5 | 3 | 0.6667 | NA | NA | NA | NA | 2 | 1 | 1 | 0 | 3 | 0.6667 |
| 9_1111013 | 4 | 0 | 5 | 0.4 | 3 | 0 | 7 | 0.1429 | 19 | 0.1579 | 3 | 0 | 4 | 0 | 1 | 0 | 4 | 0.25 | 12 | 0.0833 |
| 9_1111014 | 5 | 0 | 5 | 0.2 | 4 | 0 | 7 | 0.1429 | 21 | 0.0952 | 4 | 0 | 5 | 0.2 | 2 | 0.5 | 5 | 0.4 | 16 | 0.25 |
| 9_1111015 | 6 | 0.8333 | 6 | 1 | 4 | 0.5 | 7 | 0.8571 | 23 | 0.8261 | 4 | 1 | 8 | 1 | 1 | 1 | 5 | 1 | 18 | 1 |
| 9_1111015 | 3 | 0.6667 | 5 | 0.6 | 2 | 0 | 7 | 0.2857 | 17 | 0.4118 | 3 | 0.6667 | 3 | 0.6667 | 1 | 1 | 4 | 0.5 | 11 | 0.6364 |
| 9_1111017 | 4 | 1 | 6 | 1 | 4 | 1 | 9 | 1 | 23 | 1 | 4 | 1 | 7 | 1 | 2 | 1 | 4 | 1 | 17 | 1 |
| 9_1111018 | 3 | 1 | 7 | 1 | 4 | 0.25 | 9 | 0.6667 | 23 | 0.7391 | 5 | 0.4 | 8 | 0.5 | 2 | 0.5 | 5 | 0.8 | 20 | 0.55 |
| 9_1111018 | 4 | 0.75 | 6 | 1 | 5 | 1 | 7 | 1 | 22 | 0.9545 | 4 | 1 | 6 | 1 | 2 | 1 | 5 | 1 | 17 | 1 |
| 9_1111019 | 5 | 0 | 7 | 0 | 5 | 0 | 6 | 0.3333 | 23 | 0.087 | 5 | 0 | 8 | 0 | 2 | 0 | 6 | 0.3333 | 21 | 0.0952 |
| 9_1111019 | 5 | 0.4 | 7 | 0.4286 | 5 | 0.4 | 9 | 0.4444 | 26 | 0.4231 | 4 | 0 | 8 | 0.25 | 2 | 0.5 | 4 | 0.75 | 18 | 0.3333 |
| 9_1111020 | 5 | 0 | 7 | 0.1429 | 6 | 0.1667 | 9 | 0.3333 | 27 | 0.1852 | 4 | 0 | 9 | 0.2222 | 2 | 0 | 6 | 0 | 21 | 0.0952 |
| 9_1111021 | 5 | 1 | 7 | 1 | 5 | 1 | 8 | 1 | 25 | 1 | 4 | 0.75 | 9 | 1 | 2 | 1 | 6 | 1 | 21 | 0.9524 |
| 9_1111022 | 5 | 0.4 | 7 | 0.1429 | 5 | 0.4 | 7 | 0 | 24 | 0.2083 | 5 | 0.8 | 9 | 0.4444 | 2 | 0.5 | 5 | 0.6 | 21 | 0.5714 |
| 9_1111023 | 4 | 0.25 | 5 | 0.8 | 4 | 0.25 | 8 | 0.125 | 21 | 0.3333 | 4 | 0.5 | 8 | 0.5 | 2 | 0.5 | 6 | 0 | 20 | 0.35 |
| 9_1111023 | 5 | 1 | 7 | 0.7143 | 6 | 1 | 9 | 0.8889 | 27 | 0.8889 | 5 | 0.8 | 7 | 0.8571 | 2 | 0.5 | 6 | 1 | 20 | 0.85 |
| 9_1111023 | 4 | 1 | 7 | 0.7143 | 5 | 0.8 | 7 | 0.7143 | 23 | 0.7826 | 5 | 0.8 | 9 | 0.7778 | 2 | 0.5 | 5 | 1 | 21 | 0.8095 |
| 9_1111024 | 6 | 1 | 7 | 1 | 6 | 1 | 8 | 1 | 27 | 1 | 3 | 1 | 10 | 1 | 2 | 1 | 6 | 1 | 21 | 1 |
| 9_1111024 | 4 | 0.75 | 6 | 0.5 | 6 | 0.5 | 8 | 0.75 | 24 | 0.625 | 5 | 0.2 | 10 | 0.5 | 2 | 0.5 | 6 | 0.6667 | 23 | 0.4783 |
| 9_1111024 | 4 | 1 | 6 | 1 | 5 | 1 | 6 | 1 | 21 | 1 | 4 | 1 | 10 | 1 | 2 | 1 | 6 | 1 | 22 | 1 |
| 9_1111024 | 4 | 0.75 | 6 | 0.8333 | 4 | 0.5 | 6 | 1 | 20 | 0.8 | 5 | 1 | 9 | 0.8889 | 2 | 1 | 6 | 0.8333 | 22 | 0.9091 |
| 9_1111026 | 4 | 1 | 6 | 0.5 | 5 | 0.6 | 8 | 0.75 | 23 | 0.6957 | 5 | 0.8 | 9 | 0.8889 | 2 | 1 | 5 | 1 | 21 | 0.9048 |
| 9_1111026 | 5 | 0.8 | 7 | 1 | 6 | 0.8333 | 9 | 1 | 27 | 0.9259 | 4 | 1 | 7 | 1 | 2 | 1 | 6 | 0.8333 | 19 | 0.9474 |
| 9_1111026 | 4 | 1 | 7 | 0.8571 | 5 | 1 | 9 | 1 | 25 | 0.96 | 5 | 1 | 9 | 1 | 2 | 1 | 4 | 1 | 20 | 1 |
| 9_1111029 | 4 | 0.75 | 4 | 1 | 5 | 1 | 9 | 1 | 22 | 0.9545 | 5 | 1 | 10 | 1 | 3 | 1 | 5 | 1 | 23 | 1 |
| 9_1111029 | 4 | 0 | 6 | 0.3333 | 7 | 0.1429 | 10 | 0 | 27 | 0.1111 | 5 | 0 | 9 | 0.2222 | 2 | 0.5 | 6 | 0 | 22 | 0.1364 |
| 9_1111030 | 5 | 1 | 7 | 1 | 4 | 1 | 8 | 1 | 24 | 1 | 4 | 1 | 9 | 0.8889 | 3 | 1 | 6 | 1 | 22 | 0.9545 |
| 9_1111030 | 3 | 0.6667 | 7 | 0.5714 | 6 | 0.6667 | 9 | 0.8889 | 25 | 0.72 | 5 | 0.2 | 4 | 0.5 | 2 | 0.5 | 5 | 0.8 | 16 | 0.5 |
| 9_1111030 | 5 | 0.6 | 7 | 0.5714 | 7 | 0.7143 | 11 | 0.6364 | 30 | 0.6333 | 5 | 0 | 7 | 0.1429 | 3 | 0.6667 | 6 | 0.6667 | 21 | 0.3333 |
| 9_1111031 | 4 | 0 | 7 | 0 | 6 | 0.3333 | 12 | 0.25 | 29 | 0.1724 | 5 | 0.4 | 10 | 0.1 | 2 | 0 | 5 | 0.4 | 22 | 0.2273 |
| 9_1111032 | 6 | 0 | 9 | 0.1111 | 6 | 0.1667 | 11 | 0.0909 | 32 | 0.0938 | 6 | 0.1667 | 10 | 0.2 | 3 | 0 | 5 | 0.4 | 24 | 0.2083 |
| 9_1111036 | 6 | 0.1667 | 9 | 0.1111 | 6 | 0.1667 | 12 | 0.25 | 33 | 0.1818 | 5 | 0.2 | 10 | 0.2 | 3 | 0 | 6 | 0 | 24 | 0.125 |
| 9_1111038 | 5 | 0 | 11 | 0.0909 | 6 | 0 | 11 | 0.1818 | 33 | 0.0909 | 5 | 0.2 | 9 | 0 | 4 | 0.25 | 6 | 0.1667 | 24 | 0.125 |
| 9_1111039 | 6 | 0.6667 | 12 | 0.5833 | 5 | 0.8 | 10 | 0.4 | 33 | 0.5758 | 5 | 0.8 | 9 | 0.3333 | 5 | 0.6 | 6 | 0.6667 | 25 | 0.56 |
| 9_1111039 | 4 | 0.25 | 12 | 0.1667 | 7 | 0.1429 | 12 | 0.25 | 35 | 0.2 | 4 | 0 | 9 | 0.2222 | 5 | 0.4 | 5 | 0 | 23 | 0.1739 |
| 9_1111039 | 6 | 0.3333 | 12 | 0.5 | 7 | 0.2857 | 10 | 0.2 | 35 | 0.3429 | 5 | 0.8 | 10 | 0.3 | 5 | 1 | 6 | 0.3333 | 26 | 0.5385 |
| 9_1111041 | 5 | 0 | 11 | 0.1818 | 5 | 0 | 10 | 0.1 | 31 | 0.0968 | 4 | 0 | 10 | 0.1 | 6 | 0.1667 | 6 | 0 | 26 | 0.0769 |
| 9_1111041 | 7 | 0.4286 | 10 | 0.5 | 6 | 0.8333 | 12 | 0.6667 | 35 | 0.6 | 5 | 0.8 | 11 | 0.6364 | 6 | 0.8333 | 4 | 0.25 | 26 | 0.6538 |
| 9_1111044 | 8 | 0.375 | 12 | 0.1667 | 5 | 0.2 | 13 | 0.1538 | 38 | 0.2105 | 5 | 0.2 | 11 | 0.2727 | 7 | 0.4286 | 7 | 0.1429 | 30 | 0.2667 |
| 9_1111050 | 10 | 0.1 | 6 | 0 | 6 | 0.1667 | 8 | 0 | 30 | 0.0667 | 4 | 0 | 8 | 0.25 | 2 | 0 | 7 | 0.1429 | 21 | 0.1429 |
| 9_1111060 | 11 | 0.0909 | 9 | 0.1111 | 7 | 0.1429 | 9 | 0.1111 | 36 | 0.1111 | 3 | 0.3333 | 9 | 0.1111 | 2 | 0 | 6 | 0.1667 | 20 | 0.15 |
| 9_1111066 | 14 | 0.1429 | 11 | 0.0909 | 9 | 0.2222 | 14 | 0.2857 | 48 | 0.1875 | 10 | 0.1 | 12 | 0.3333 | 2 | 0 | 7 | 0.2857 | 31 | 0.2258 |
| 9_1112131 | NA | NA | NA | NA | 1 | 1 | 2 | 0 | 3 | 0.3333 | NA | NA | NA | NA | NA | NA | 1 | 1 | 1 | 1 |
| 9_1112194 | NA | NA | 3 | 0.3333 | 2 | 0.5 | 3 | 1 | 8 | 0.625 | 2 | 1 | 4 | 0.75 | 3 | 1 | 4 | 0.75 | 13 | 0.8462 |
| 9_1112198 | NA | NA | 1 | 1 | 2 | 1 | 3 | 1 | 6 | 1 | 3 | 1 | 3 | 0.6667 | 4 | 0.75 | 3 | 1 | 13 | 0.8462 |
| 9_1112198 | NA | NA | 4 | 0 | 2 | 0 | 4 | 0.25 | 10 | 0.1 | 3 | 0 | 2 | 0 | 4 | 0.25 | 6 | 0.1667 | 15 | 0.1333 |
| 9_1112198 | NA | NA | 4 | 0.25 | 2 | 0 | 4 | 0.25 | 10 | 0.2 | 3 | 0.3333 | 4 | 0.5 | 4 | 0.25 | 6 | 0.5 | 17 | 0.4118 |
| 9_1112202 | 1 | 0 | 2 | 0 | 4 | 0 | 5 | 0.2 | 12 | 0.0833 | 3 | 0 | 4 | 0 | 4 | 0 | 6 | 0.1667 | 17 | 0.0588 |
| 9_1112203 | NA | NA | 4 | 0.75 | 4 | 0.75 | 4 | 0.75 | 12 | 0.75 | 2 | 0.5 | 4 | 0.5 | 4 | 0.75 | 7 | 0.8571 | 17 | 0.7059 |

| | | | | | | | | | | | | | | | | | | | | |
|---|---|---|---|---|---|---|---|---|---|---|---|---|---|---|---|---|---|---|---|---|
| 9_1112204 | 1 | 0 | 3 | 0.3333 | 2 | 0.5 | 4 | 0.25 | 10 | 0.3 | 3 | 0.6667 | 4 | 0.25 | 4 | 0.25 | 6 | 0.6667 | 17 | 0.4706 |
| 9_1112204 | 1 | 0 | 4 | 0.75 | 3 | 0.6667 | 3 | 0 | 11 | 0.4545 | 2 | 0 | 5 | 0.2 | 5 | 0.4 | 6 | 0.6667 | 18 | 0.3889 |
| 9_1112205 | 1 | 0 | 3 | 0.3333 | 4 | 0.25 | 5 | 0 | 13 | 0.1538 | 3 | 0 | 4 | 0.25 | 5 | 0.2 | 6 | 0.1667 | 18 | 0.1667 |
| 9_1112208 | 1 | 0 | 3 | 0 | 3 | 0 | 4 | 0.25 | 11 | 0.0909 | 3 | 0.3333 | 4 | 0 | 5 | 0 | 5 | 0.2 | 17 | 0.1176 |
| 9_1112209 | 2 | 1 | 3 | 0.3333 | 2 | 0.5 | 4 | 0.25 | 11 | 0.4545 | 2 | 0.5 | 4 | 0 | 3 | 0.6667 | 7 | 0.2857 | 16 | 0.3125 |
| 9_1112209 | 2 | 0 | 5 | 0.2 | 4 | 0 | 5 | 0 | 16 | 0.0625 | 3 | 0 | 4 | 0 | 4 | 0 | 6 | 0.1667 | 17 | 0.0588 |
| 9_1112209 | 2 | 0.5 | 6 | 0 | 4 | 0 | 6 | 0 | 18 | 0.0556 | 3 | 0 | 4 | 0 | 4 | 0 | 7 | 0.1429 | 18 | 0.0556 |
| 9_1112210 | 2 | 1 | 5 | 0.6 | 4 | 0.5 | 4 | 0.75 | 15 | 0.6667 | 2 | 0.5 | 5 | 0.8 | 4 | 0.75 | 8 | 0.625 | 19 | 0.6842 |
| 9_1112210 | 2 | 0.5 | 6 | 0.3333 | 5 | 0.2 | 5 | 0.6 | 18 | 0.3889 | 3 | 0.6667 | 5 | 0.6 | 5 | 0.8 | 6 | 0.6667 | 19 | 0.6842 |
| 9_1112213 | 2 | 1 | 6 | 0.6667 | 4 | 0.75 | 9 | 0.6667 | 21 | 0.7143 | 2 | 1 | 6 | 1 | 5 | 0.4 | 9 | 0.6667 | 22 | 0.7273 |
| 9_1112213 | 1 | 1 | 6 | 0.5 | 4 | 0.75 | 6 | 0.5 | 17 | 0.5882 | 3 | 0.6667 | 5 | 0.6 | 4 | 0.5 | 5 | 0.8 | 17 | 0.6471 |
| 9_1112213 | 2 | 1 | 8 | 0.75 | 5 | 0.8 | 9 | 0.5556 | 24 | 0.7083 | 3 | 1 | 6 | 0.8333 | 7 | 0.7143 | 8 | 0.75 | 24 | 0.7917 |
| 9_1112214 | 2 | 0.5 | 7 | 0.4286 | 3 | 0.3333 | 7 | 0.4286 | 19 | 0.4211 | 2 | 0.5 | 7 | 0.1429 | 6 | 1 | 7 | 0.7143 | 22 | 0.5909 |
| 9_1112214 | 2 | 1 | 7 | 0.4286 | 5 | 0.4 | 7 | 0.2857 | 21 | 0.4286 | 2 | 1 | 5 | 0.6 | 7 | 0.5714 | 7 | 0.4286 | 21 | 0.5714 |
| 9_1112216 | 2 | 0.5 | 6 | 0 | 5 | 0.2 | 8 | 0.25 | 21 | 0.1905 | 3 | 0 | 6 | 0.1667 | 7 | 0.2857 | 8 | 0.375 | 24 | 0.25 |
| 9_1112216 | 2 | 0 | 8 | 0.125 | 6 | 0.5 | 9 | 0.1111 | 25 | 0.2 | 3 | 0 | 8 | 0 | 6 | 0.1667 | 10 | 0.1 | 27 | 0.0741 |
| 9_1112217 | 2 | 1 | 4 | 0.75 | 3 | 1 | 7 | 1 | 16 | 0.9375 | 2 | 1 | 5 | 0.8 | 4 | 1 | 9 | 1 | 20 | 0.95 |
| 9_1112235 | 2 | 0 | 7 | 0.1429 | 4 | 0 | 7 | 0 | 20 | 0.05 | 1 | 0 | 6 | 0.1667 | 5 | 0 | 6 | 0.1667 | 18 | 0.1111 |
| 9_1112237 | 3 | 0 | 10 | 0.1 | 5 | 0 | 9 | 0.1111 | 27 | 0.0741 | 3 | 0 | 7 | 0.1429 | 10 | 0.1 | 13 | 0 | 33 | 0.0606 |
| 9_1112242 | 3 | 0 | 7 | 0.1429 | 4 | 0.5 | 9 | 0.3333 | 23 | 0.2609 | 5 | 0 | 8 | 0.25 | 14 | 0 | 14 | 0.2143 | 41 | 0.122 |
| 9_1112244 | 4 | 0.5 | 10 | 0.6 | 5 | 0.8 | 12 | 0.5833 | 31 | 0.6129 | 6 | 0.5 | 9 | 0.6667 | 15 | 0.5333 | 17 | 0.8235 | 47 | 0.6596 |
| 9_1112244 | 4 | 0.25 | 13 | 0.1538 | 7 | 0.4286 | 16 | 0.5625 | 40 | 0.375 | 5 | 0.2 | 12 | 0.5 | 18 | 0.3333 | 18 | 0.4444 | 53 | 0.3962 |
| 9_1112966 | 6 | 0 | 1 | 0 | 2 | 0.5 | 4 | 0 | 13 | 0.0769 | 2 | 0 | 6 | 0.1667 | 2 | 0.5 | 1 | 0 | 11 | 0.1818 |
| 9_1112971 | 5 | 0 | 2 | 0 | 2 | 0 | 4 | 0.25 | 13 | 0.0769 | 2 | 0.5 | 7 | 0 | 2 | 0 | 4 | 0 | 15 | 0.0667 |
| 9_1112974 | 8 | 0.125 | 2 | 0 | 3 | 0.3333 | 6 | 0.3333 | 19 | 0.2105 | 3 | 0.3333 | 7 | 0 | 5 | 0.2 | 5 | 0 | 20 | 0.1 |
| 9_1112975 | 9 | 0 | 2 | 0 | 3 | 0 | 6 | 0.3333 | 20 | 0.1 | 3 | 0 | 7 | 0.2857 | 7 | 0.1429 | 7 | 0.1429 | 24 | 0.1667 |
| 9_1112976 | 8 | 0.25 | 3 | 0 | 3 | 0.3333 | 6 | 0 | 20 | 0.15 | 2 | 0.5 | 6 | 0 | 4 | 0.5 | 6 | 0.1667 | 18 | 0.2222 |
| 9_1112976 | 9 | 0.2222 | 3 | 0 | 2 | 0 | 7 | 0.2857 | 21 | 0.1905 | 2 | 0.5 | 7 | 0.2857 | 6 | 0.6667 | 7 | 0.2857 | 22 | 0.4091 |
| 9_1112980 | 11 | 0.0909 | 2 | 0.5 | 5 | 0.2 | 10 | 0.1 | 28 | 0.1429 | 3 | 0 | 17 | 0.2353 | 12 | 0.1667 | 7 | 0 | 39 | 0.1538 |
| 9_1112981 | 10 | 1 | 4 | 1 | 3 | 1 | 8 | 1 | 25 | 1 | 5 | 1 | 17 | 0.8824 | 13 | 0.9231 | 7 | 1 | 42 | 0.9286 |
| 9_1112981 | 11 | 0 | 4 | 0.25 | 5 | 0.2 | 10 | 0 | 30 | 0.0667 | 4 | 0 | 19 | 0.1579 | 14 | 0.0714 | 7 | 0 | 44 | 0.0909 |
| 9_1112982 | 14 | 0.7143 | 2 | 0.5 | 4 | 0.75 | 10 | 0.6 | 30 | 0.6667 | 5 | 0.6 | 19 | 0.6842 | 15 | 0.8667 | 7 | 0.7143 | 46 | 0.7391 |
| 9_1112983 | 12 | 0.75 | 5 | 0.6 | 5 | 0.8 | 8 | 1 | 30 | 0.8 | 4 | 0.75 | 20 | 0.9 | 15 | 0.9333 | 9 | 0.5556 | 48 | 0.8333 |
| 9_1113177 | 16 | 0.125 | 2 | 0 | 2 | 0.5 | 8 | 0.125 | 28 | 0.1429 | 6 | 0 | 14 | 0.2857 | 20 | 0 | 7 | 0.4286 | 47 | 0.1489 |
| 9_1113177 | 18 | 0.1667 | 5 | 0 | 6 | 0.3333 | 14 | 0.2143 | 43 | 0.186 | 8 | 0.125 | 26 | 0.2692 | 19 | 0.0526 | 12 | 0.1667 | 65 | 0.1692 |
| 9_1113178 | 18 | 0.0556 | 6 | 0.3333 | 8 | 0.25 | 13 | 0.0769 | 45 | 0.1333 | 6 | 0.1667 | 24 | 0.0833 | 25 | 0.32 | 12 | 0.3333 | 67 | 0.2239 |
| 9_1113182 | 13 | 0.3846 | 4 | 0 | 8 | 0.375 | 10 | 0.3 | 35 | 0.3143 | 7 | 0.4286 | 21 | 0.1905 | 16 | 0.3125 | 7 | 0.5714 | 51 | 0.3137 |
| 9_1113184 | 15 | 0 | 4 | 0 | 8 | 0.125 | 11 | 0.0909 | 38 | 0.0526 | 9 | 0 | 22 | 0.0455 | 16 | 0.125 | 10 | 0.2 | 57 | 0.0877 |
| 9_1113905 | NA | NA | 1 | 0 | 2 | 0.5 | NA | NA | 3 | 0.3333 | NA | NA | 1 | 1 | 1 | 0 | NA | NA | 2 | 0.5 |
| 9_1113907 | NA | NA | 1 | 0 | 2 | 0.5 | NA | NA | 3 | 0.3333 | NA | NA | 1 | 1 | NA | NA | NA | NA | 1 | 1 |
| 9_1113909 | NA | NA | 1 | 0 | 1 | 1 | NA | NA | 2 | 0.5 | NA | NA | NA | NA | 1 | 1 | NA | NA | 1 | 1 |
| 9_1113914 | NA | NA | 1 | 1 | 2 | 0.5 | NA | NA | 3 | 0.6667 | NA | NA | 1 | 1 | NA | NA | NA | NA | 1 | 1 |
| 9_1118946 | 13 | 0.7692 | 17 | 0.7647 | 14 | 0.8571 | 13 | 0.7692 | 57 | 0.7895 | 3 | 0.6667 | 9 | 0.8889 | 13 | 0.8462 | 10 | 0.5 | 35 | 0.7429 |
| 9_1118947 | 14 | 0.7143 | 22 | 0.7727 | 16 | 0.875 | 13 | 0.8462 | 65 | 0.8 | 5 | 0.6 | 11 | 0.5455 | 11 | 0.6364 | 11 | 0.8182 | 38 | 0.6579 |
| 9_1118947 | 14 | 0.1429 | 21 | 0.0952 | 14 | 0.3571 | 13 | 0.0769 | 62 | 0.1613 | 5 | 0 | 10 | 0 | 12 | 0.1667 | 12 | 0 | 39 | 0.0513 |
| 9_1118948 | 10 | 0.9 | 22 | 0.8636 | 14 | 0.9286 | 13 | 0.4615 | 59 | 0.7966 | 4 | 1 | 7 | 1 | 12 | 0.75 | 11 | 0.7273 | 34 | 0.8235 |
| 9_1119538 | 5 | 0 | 17 | 0 | 11 | 0.2727 | 6 | 0 | 39 | 0.0769 | 3 | 0.3333 | 8 | 0 | 10 | 0.1 | 8 | 0 | 29 | 0.069 |
| 9_1120104 | 87 | 0.3678 | 64 | 0.2344 | 73 | 0.3562 | 114 | 0.2544 | 338 | 0.3018 | 35 | 0.4 | 141 | 0.3972 | 139 | 0.3597 | 78 | 0.3718 | 393 | 0.3791 |
| 9_1120105 | 68 | 0.1618 | 47 | 0.1702 | 52 | 0.0769 | 94 | 0.1809 | 261 | 0.1533 | 26 | 0.1154 | 101 | 0.1287 | 101 | 0.1089 | 66 | 0.0606 | 294 | 0.1054 |
| 9_1120111 | 90 | 0.1889 | 61 | 0.1311 | 75 | 0.1467 | 121 | 0.1488 | 347 | 0.1556 | 39 | 0.1538 | 139 | 0.2086 | 136 | 0.0735 | 72 | 0.0972 | 386 | 0.1347 |
| 9_1120116 | 79 | 0.7089 | 46 | 0.5 | 64 | 0.7188 | 100 | 0.77 | 289 | 0.699 | 27 | 0.5926 | 110 | 0.6818 | 109 | 0.7156 | 57 | 0.5439 | 303 | 0.6601 |
| 9_1120116 | 72 | 0.5833 | 44 | 0.4773 | 55 | 0.6364 | 94 | 0.5426 | 265 | 0.5623 | 28 | 0.7857 | 95 | 0.6316 | 101 | 0.5743 | 51 | 0.3922 | 275 | 0.5818 |
| 9_1120118 | 76 | 0.0658 | 46 | 0.0652 | 62 | 0.0323 | 103 | 0.0583 | 287 | 0.0557 | 32 | 0 | 117 | 0.0684 | 115 | 0.0522 | 57 | 0.0877 | 321 | 0.0592 |
| 9_1120118 | 81 | 0.1358 | 49 | 0.0408 | 67 | 0.0597 | 108 | 0.1481 | 305 | 0.1082 | 35 | 0.0286 | 124 | 0.0968 | 120 | 0.075 | 65 | 0.0769 | 344 | 0.0785 |
| 9_1120121 | 6 | 0.3333 | 4 | 0.25 | 2 | 0 | 1 | 1 | 13 | 0.3077 | NA | NA | 5 | 0.2 | 2 | 0 | 2 | 0 | 9 | 0.1111 |
| 9_1120634 | 1 | 0 | 2 | 0 | 1 | 1 | 1 | 0 | 5 | 0.2 | 1 | 1 | 3 | 0 | NA | NA | 1 | 0 | 5 | 0.2 |
| 9_1120636 | NA | NA | 2 | 0.5 | 2 | 1 | 1 | 1 | 5 | 0.8 | 1 | 0 | 3 | 0.6667 | NA | NA | 2 | 0 | 6 | 0.3333 |
| 9_1120637 | NA | NA | 2 | 0.5 | 2 | 1 | 1 | 0 | 5 | 0.6 | 1 | 0 | 3 | 0 | NA | NA | 2 | 0.5 | 6 | 0.1667 |
| 9_1120637 | NA | NA | 2 | 0.5 | 2 | 0 | 2 | 0 | 6 | 0.1667 | 1 | 0 | 3 | 0 | NA | NA | 2 | 0.5 | 6 | 0.1667 |
| 9_1120637 | NA | NA | 2 | 0 | 2 | 0.5 | 2 | 0 | 6 | 0.1667 | 1 | 0 | 1 | 1 | NA | NA | 2 | 0 | 4 | 0.25 |
| 9_1120649 | 3 | 0.3333 | 2 | 0 | 7 | 0.2857 | 2 | 0.5 | 14 | 0.2857 | NA | NA | 3 | 0 | 2 | 0 | 3 | 0.3333 | 8 | 0.125 |
| 9_1120649 | 3 | 0 | 2 | 0 | 6 | 0.3333 | 2 | 0 | 13 | 0.1538 | NA | NA | 3 | 0 | 2 | 0.5 | 4 | 0 | 9 | 0.1111 |
| 9_1120649 | 2 | 0 | 2 | 0.5 | 8 | 0.125 | 1 | 0 | 13 | 0.1538 | NA | NA | 4 | 0 | 1 | 0 | 4 | 0.25 | 9 | 0.1111 |
| 9_1134826 | 6 | 0.3333 | 8 | 0 | 2 | 0 | 3 | 0 | 19 | 0.1053 | 2 | 0.5 | 4 | 0 | 5 | 0 | 2 | 0 | 13 | 0.0769 |
| 9_1134827 | 6 | 0.1667 | 8 | 0 | 2 | 0 | 3 | 0 | 19 | 0.0526 | 2 | 0 | 5 | 0.2 | 5 | 0 | 2 | 0.5 | 14 | 0.1429 |

| | | | | | | | | | | | | | | | | | | | | |
|---|---|---|---|---|---|---|---|---|---|---|---|---|---|---|---|---|---|---|---|---|
| 9_1134829 | 5 | 0.4 | 7 | 0.5714 | 2 | 0 | 2 | 0 | 16 | 0.375 | 2 | 0 | 4 | 0.25 | 4 | 0 | 2 | 0.5 | 12 | 0.1667 |
| 9_1134829 | 6 | 0.1667 | 5 | 0.4 | 2 | 0.5 | 3 | 0.3333 | 16 | 0.3125 | 2 | 1 | 3 | 0 | 4 | 0.25 | 2 | 0.5 | 11 | 0.3636 |
| 9_1134833 | 3 | 0 | 4 | 0.75 | 3 | 0.3333 | 2 | 0.5 | 12 | 0.4167 | 1 | 0 | 2 | 0.5 | 3 | 0.3333 | 1 | 0 | 7 | 0.2857 |
| 9_1134836 | 10 | 0.5 | 9 | 0.1111 | 6 | 0 | 4 | 0.5 | 29 | 0.2759 | 5 | 0 | 8 | 0.25 | 5 | 0 | 4 | 0 | 22 | 0.0909 |
| 9_1134839 | 6 | 0.1667 | 5 | 0.2 | 4 | 0.5 | 4 | 0.25 | 19 | 0.2632 | 3 | 0 | 5 | 0.4 | 4 | 0.25 | 4 | 0.25 | 16 | 0.25 |
| 9_1134840 | 6 | 0.1667 | 10 | 0.1 | 5 | 0 | 4 | 0 | 25 | 0.08 | 4 | 0.25 | 7 | 0 | 3 | 0 | 2 | 0 | 16 | 0.0625 |
| 9_1137471 | 12 | 0.0833 | 12 | 0 | 18 | 0.2222 | 13 | 0.1538 | 55 | 0.1273 | 11 | 0.1818 | 14 | 0.0714 | 14 | 0.1429 | 13 | 0.0769 | 52 | 0.1154 |
| 9_1137472 | 16 | 0.125 | 17 | 0.3529 | 19 | 0.1053 | 18 | 0.4444 | 70 | 0.2571 | 8 | 0.125 | 18 | 0.2778 | 14 | 0.2143 | 17 | 0.1176 | 57 | 0.193 |
| 9_1137472 | 18 | 0.2778 | 16 | 0.375 | 19 | 0.3158 | 20 | 0.25 | 73 | 0.3014 | 11 | 0.3636 | 19 | 0.2632 | 16 | 0.3125 | 17 | 0.2353 | 63 | 0.2857 |
| 9_1137477 | 18 | 0 | 18 | 0.2778 | 19 | 0.0526 | 23 | 0.0435 | 78 | 0.0897 | 15 | 0.0667 | 19 | 0.0526 | 18 | 0 | 17 | 0.1176 | 69 | 0.058 |
| 9_1138044 | 6 | 0.8333 | 4 | 1 | 4 | 0.75 | 6 | 0.8333 | 20 | 0.85 | 4 | 1 | 6 | 0.8333 | 10 | 0.9 | 2 | 1 | 22 | 0.9091 |
| 9_1138044 | 7 | 0.5714 | 4 | 0.75 | 3 | 1 | 4 | 0.75 | 18 | 0.7222 | 3 | 1 | 5 | 0.6 | 6 | 0.5 | 1 | 1 | 15 | 0.6667 |
| 9_1138045 | 7 | 0 | 6 | 0.1667 | 5 | 0.2 | 4 | 0.75 | 22 | 0.2273 | 5 | 0.2 | 5 | 0.2 | 12 | 0.0833 | 3 | 0.3333 | 25 | 0.16 |
| 9_1138047 | 7 | 0.1429 | 4 | 0 | 5 | 0.2 | 5 | 0 | 21 | 0.0952 | 5 | 0.2 | 6 | 0.3333 | 13 | 0.0769 | 3 | 0 | 27 | 0.1481 |
| 9_1138047 | 4 | 0.75 | 5 | 0.6 | 4 | 1 | 6 | 0.6667 | 19 | 0.7368 | 4 | 0.75 | 5 | 1 | 10 | 0.6 | 5 | 1 | 24 | 0.7917 |
| 9_1138048 | 7 | 0.2857 | 5 | 0.4 | 5 | 0.4 | 4 | 0.25 | 21 | 0.3333 | 3 | 0.3333 | 6 | 0.3333 | 10 | 0.4 | 5 | 0.2 | 24 | 0.3333 |
| 9_1138048 | 6 | 0 | 5 | 0.2 | 4 | 0 | 2 | 0.5 | 17 | 0.1176 | 5 | 0 | 5 | 0 | 12 | 0.1667 | 5 | 0.2 | 27 | 0.1111 |
| 9_1138049 | 10 | 0.5 | 6 | 0.5 | 3 | 0.3333 | 4 | 0 | 23 | 0.3913 | 3 | 0.3333 | 4 | 0 | 8 | 0.25 | 4 | 0.25 | 19 | 0.2105 |
| 9_1138050 | 10 | 0.2 | 7 | 0 | 5 | 0 | 6 | 0 | 28 | 0.0714 | 5 | 0 | 7 | 0 | 13 | 0.0769 | 6 | 0.1667 | 31 | 0.0645 |
| 9_1138058 | 9 | 0.2222 | 8 | 0.375 | 7 | 0 | 7 | 0.1429 | 31 | 0.1935 | 5 | 0 | 8 | 0 | 8 | 0.125 | 6 | 0.1667 | 27 | 0.0741 |
| 9_1138059 | 10 | 0.1 | 12 | 0.25 | 8 | 0.125 | 6 | 0.5 | 36 | 0.2222 | 6 | 0.3333 | 10 | 0.4 | 12 | 0.0833 | 8 | 0 | 36 | 0.1944 |
| 9_1138067 | 1 | 1 | NA | NA | NA | NA | 1 | 0 | 2 | 0.5 | 1 | 1 | 1 | 1 | NA | NA | 2 | 1 | 4 | 1 |
| 9_1138070 | 1 | 1 | NA | NA | NA | NA | NA | NA | 1 | 1 | 1 | 1 | 1 | 0 | NA | NA | 2 | 0 | 4 | 0.25 |
| 9_1138074 | 6 | 0 | 7 | 0 | 3 | 0.3333 | 4 | 0 | 20 | 0.05 | 3 | 0 | 6 | 0.1667 | 2 | 0 | 3 | 0.6667 | 14 | 0.2143 |
| 9_1138075 | 6 | 0.3333 | 7 | 0.1429 | 3 | 0.3333 | 4 | 0.25 | 20 | 0.25 | 2 | 0.5 | 6 | 0.3333 | 2 | 0 | 4 | 0.25 | 14 | 0.2857 |
| 9_1138077 | 4 | 0.5 | 7 | 0.7143 | 2 | 0 | 3 | 0 | 16 | 0.4375 | 2 | 1 | 5 | 0.4 | 2 | 0.5 | 4 | 0.25 | 13 | 0.4615 |
| 9_1139271 | NA | NA | 2 | 0.5 | NA | NA | 1 | 1 | 3 | 0.6667 | NA | NA | 1 | 1 | NA | NA | NA | NA | 1 | 1 |
| 9_1139273 | NA | NA | 2 | 1 | 1 | 1 | 1 | 1 | 4 | 1 | NA | NA | 1 | 1 | NA | NA | NA | NA | 1 | 1 |
| 9_1139275 | NA | NA | 2 | 0.5 | 1 | 0 | 1 | 1 | 4 | 0.5 | NA | NA | 1 | 1 | NA | NA | NA | NA | 1 | 1 |
| 9_1142380 | 4 | 0.25 | NA | NA | NA | NA | NA | NA | 4 | 0.25 | 1 | 0 | NA | NA | 1 | 1 | 1 | 1 | 3 | 0.6667 |
| 9_1142395 | 4 | 0.25 | 1 | 0 | NA | NA | NA | NA | 5 | 0.2 | 1 | 1 | NA | NA | 1 | 0 | 2 | 0.5 | 4 | 0.5 |
| 9_1142396 | 5 | 0.4 | 1 | 0 | NA | NA | 1 | 0 | 7 | 0.2857 | 1 | 0 | NA | NA | 1 | 1 | 2 | 0 | 4 | 0.25 |
| 9_1142932 | 1 | 1 | 3 | 0 | 1 | 0 | 2 | 0 | 7 | 0.1429 | 2 | 0 | 2 | 1 | 1 | 0 | 1 | 0 | 6 | 0.3333 |
| 9_1143030 | 3 | 0.3333 | 3 | 0.3333 | 2 | 0 | 4 | 0 | 12 | 0.1667 | 3 | 0 | 2 | 0.5 | 1 | 0 | 3 | 0.3333 | 9 | 0.2222 |
| 9_1143030 | 3 | 0.3333 | 3 | 0.3333 | 2 | 0 | 3 | 0.6667 | 11 | 0.3636 | 2 | 0.5 | 2 | 0 | 1 | 1 | 3 | 0 | 8 | 0.25 |
| 9_1143031 | 3 | 0.3333 | 3 | 0.3333 | 2 | 0.5 | 4 | 0.5 | 12 | 0.4167 | 3 | 0.3333 | 2 | 0 | 1 | 0 | 4 | 0.5 | 10 | 0.3 |
| 9_1143031 | 3 | 0.6667 | 2 | 1 | 2 | 1 | 2 | 1 | 9 | 0.8889 | 1 | 1 | 2 | 1 | 1 | 1 | 4 | 1 | 8 | 1 |
| 9_1143032 | 3 | 0.3333 | 3 | 0.3333 | 3 | 0.6667 | 3 | 0.6667 | 12 | 0.5 | 2 | 1 | 1 | 1 | 1 | 1 | 4 | 1 | 8 | 1 |
| 9_1143033 | 3 | 0 | 3 | 0 | 2 | 0 | 3 | 0.3333 | 11 | 0.0909 | 2 | 0.5 | 2 | 0 | 1 | 0 | 4 | 0 | 9 | 0.1111 |
| 9_1144277 | NA | NA | 1 | 0 | 1 | 1 | 1 | 0 | 3 | 0.3333 | 2 | 0.5 | 3 | 0 | 2 | 0 | 1 | 0 | 8 | 0.125 |
| 9_1145351 | 8 | 0.25 | 5 | 0 | 8 | 0.25 | 4 | 0.5 | 25 | 0.24 | 6 | 0.3333 | 7 | 0 | 5 | 0 | 2 | 0 | 20 | 0.1 |
| 9_1145352 | 6 | 0.3333 | 6 | 0 | 5 | 0 | 4 | 0 | 21 | 0.0952 | 5 | 0.2 | 6 | 0 | 3 | 0 | 3 | 0 | 17 | 0.0588 |
| 9_1145524 | 14 | 0 | 6 | 0 | 10 | 0.2 | 9 | 0 | 39 | 0.0513 | 8 | 0.125 | 9 | 0.1111 | 8 | 0 | 8 | 0 | 33 | 0.0606 |
| 9_1145525 | 12 | 0.1667 | 7 | 0.2857 | 6 | 0.3333 | 7 | 0.1429 | 32 | 0.2188 | 4 | 0 | 9 | 0.3333 | 6 | 0.5 | 6 | 0.3333 | 25 | 0.32 |
| 9_1145526 | 15 | 0.0667 | 10 | 0.1 | 11 | 0.0909 | 10 | 0.1 | 46 | 0.087 | 6 | 0 | 9 | 0.1111 | 10 | 0.2 | 8 | 0.125 | 33 | 0.1212 |
| 9_1145527 | 14 | 0.1429 | 12 | 0.0833 | 14 | 0.1429 | 12 | 0.1667 | 52 | 0.1346 | 9 | 0.3333 | 8 | 0.125 | 10 | 0.5 | 9 | 0 | 36 | 0.25 |
| 9_1145527 | 14 | 0.1429 | 11 | 0 | 12 | 0 | 9 | 0.1111 | 46 | 0.0652 | 8 | 0.125 | 8 | 0 | 9 | 0.2222 | 9 | 0.1111 | 34 | 0.1176 |
| 9_1145528 | 15 | 0.0667 | 11 | 0.1818 | 12 | 0 | 13 | 0.1538 | 51 | 0.098 | 8 | 0.25 | 8 | 0 | 9 | 0.4444 | 9 | 0 | 34 | 0.1765 |
| 9_1145528 | 14 | 0.2857 | 10 | 0.4 | 11 | 0.3636 | 12 | 0.4167 | 47 | 0.3617 | 9 | 0.2222 | 6 | 0.3333 | 8 | 0.5 | 10 | 0.3 | 33 | 0.3333 |
| 9_1145529 | 14 | 0.0714 | 8 | 0.125 | 14 | 0.0714 | 9 | 0 | 45 | 0.0667 | 8 | 0.25 | 9 | 0.2222 | 10 | 0.1 | 11 | 0 | 38 | 0.1316 |
| 9_1145529 | 17 | 0.2941 | 13 | 0.1538 | 14 | 0.2857 | 13 | 0.4615 | 57 | 0.2982 | 10 | 0.3 | 9 | 0.1111 | 11 | 0.3636 | 12 | 0.3333 | 42 | 0.2857 |
| 9_1145530 | 20 | 0.15 | 13 | 0.1538 | 15 | 0.0667 | 14 | 0.2857 | 62 | 0.1613 | 11 | 0.1818 | 10 | 0 | 11 | 0.0909 | 12 | 0.3333 | 44 | 0.1591 |
| 9_1145532 | 17 | 0.1176 | 15 | 0.2 | 15 | 0.2 | 13 | 0.3846 | 60 | 0.2167 | 10 | 0.4 | 10 | 0.1 | 11 | 0.1818 | 11 | 0.3636 | 42 | 0.2619 |
| 9_1145757 | NA | NA | 1 | 0 | 1 | 0 | 3 | 0.3333 | 5 | 0.2 | NA | NA | 1 | 0 | 1 | 1 | 1 | 0 | 3 | 0.3333 |
| 9_1145780 | 1 | 0 | 2 | 0 | 2 | 0 | 5 | 0.4 | 10 | 0.2 | 2 | 0.5 | 1 | 0 | 2 | 0.5 | 3 | 0 | 8 | 0.25 |
| 9_1145781 | 2 | 0 | 2 | 0 | 1 | 0 | 5 | 0.2 | 10 | 0.1 | 2 | 0 | 1 | 0 | 2 | 0 | 3 | 0.3333 | 8 | 0.125 |
| 9_1145861 | 2 | 0.5 | 4 | 0 | 1 | 0 | 1 | 0 | 8 | 0.125 | NA | NA | NA | NA | 1 | 0 | 1 | 1 | 2 | 0.5 |
| 9_1145861 | 1 | 1 | 4 | 0.75 | 1 | 1 | NA | NA | 6 | 0.8333 | NA | NA | NA | NA | 1 | 1 | 1 | 1 | 2 | 1 |
| 9_1147289 | 9 | 0.2222 | 15 | 0.3333 | 8 | 0.375 | 10 | 0 | 42 | 0.2381 | 8 | 0.25 | 10 | 0.2 | 7 | 0.2857 | 9 | 0.1111 | 34 | 0.2059 |
| 9_1147290 | 8 | 0.375 | 7 | 0.1429 | 4 | 0 | 8 | 0.125 | 27 | 0.1852 | 5 | 0.2 | 8 | 0.25 | 4 | 0.5 | 5 | 0.2 | 22 | 0.2727 |
| 9_1147293 | 9 | 0.1111 | 14 | 0 | 5 | 0 | 12 | 0.1667 | 40 | 0.075 | 9 | 0.1111 | 8 | 0 | 8 | 0.125 | 8 | 0.125 | 33 | 0.0909 |
| 9_1147307 | 3 | 0.3333 | 7 | 0.4286 | 2 | 0 | 8 | 0.5 | 20 | 0.4 | 2 | 0 | 5 | 0.2 | 5 | 0.2 | 4 | 0 | 16 | 0.125 |
| 9_1147307 | 4 | 0.25 | 7 | 0.1429 | 2 | 0 | 7 | 0.4286 | 20 | 0.25 | 2 | 0 | 5 | 0 | 5 | 0.6 | 4 | 0 | 16 | 0.1875 |
| 9_1147309 | 3 | 0.3333 | 5 | 0 | 2 | 0 | 5 | 0.6 | 15 | 0.2667 | 2 | 1 | 3 | 0.3333 | 3 | 0 | 4 | 0.5 | 12 | 0.4167 |
| 9_1147311 | 3 | 0 | 6 | 0 | 1 | 0 | 5 | 0.2 | 15 | 0.0667 | 1 | 1 | 3 | 0.3333 | 5 | 0 | 2 | 0 | 11 | 0.1818 |

| | | | | | | | | | | | | | | | | | | | | |
|---|---|---|---|---|---|---|---|---|---|---|---|---|---|---|---|---|---|---|---|---|
| 9_11473139 | 3 | 0.6667 | 4 | 0.5 | 1 | 0 | 5 | 0.6 | 13 | 0.5385 | 1 | 1 | 1 | 1 | 4 | 0.5 | 1 | 0 | 7 | 0.5714 |
| 9_11504585 | 3 | 0.6667 | 1 | 1 | 4 | 1 | 5 | 0.8 | 13 | 0.8462 | 2 | 0 | 1 | 1 | 2 | 0 | NA | NA | 5 | 0.2 |
| 9_11504587 | 3 | 1 | 1 | 1 | 5 | 0.8 | 4 | 1 | 13 | 0.9231 | 2 | 1 | 1 | 1 | 2 | 0.5 | NA | NA | 5 | 0.8 |
| 9_11504592 | 3 | 0 | 1 | 1 | 5 | 0 | 5 | 0 | 14 | 0.0714 | 2 | 0 | 1 | 0 | 2 | 0.5 | 2 | 0.5 | 7 | 0.2857 |
| 9_11504602 | 3 | 1 | 1 | 1 | 5 | 1 | 5 | 0.8 | 14 | 0.9286 | 2 | 0.5 | 1 | 1 | 2 | 0.5 | 2 | 0.5 | 7 | 0.5714 |
| 9_11504604 | 3 | 1 | 1 | 0 | 4 | 0.25 | 3 | 0.6667 | 11 | 0.5455 | 2 | 1 | 1 | 1 | 1 | 1 | 2 | 0 | 6 | 0.6667 |
| 9_11506213 | 5 | 0 | 3 | 0.6667 | 4 | 0 | 6 | 0.3333 | 18 | 0.2222 | 2 | 0 | 2 | 0 | 2 | 0 | 2 | 0.5 | 8 | 0.125 |
| 9_11507182 | 10 | 0.3 | 15 | 0.1333 | 16 | 0.1875 | 18 | 0 | 59 | 0.1356 | 10 | 0.4 | 10 | 0.3 | 12 | 0.1667 | 10 | 0.3 | 42 | 0.2857 |
| 9_11507185 | 9 | 0.5556 | 13 | 0.3846 | 16 | 0.125 | 15 | 0.2667 | 53 | 0.3019 | 7 | 0.2857 | 7 | 0.5714 | 15 | 0.3333 | 11 | 0.5455 | 40 | 0.425 |
| 9_11507191 | 11 | 0.4545 | 17 | 0.2353 | 17 | 0.1765 | 19 | 0.3684 | 64 | 0.2969 | 6 | 0.3333 | 10 | 0.3 | 15 | 0.1333 | 13 | 0.4615 | 44 | 0.2955 |
| 9_11507214 | 6 | 0 | 15 | 0.2 | 11 | 0.0909 | 18 | 0.1667 | 50 | 0.14 | 6 | 0 | 9 | 0 | 15 | 0.0667 | 11 | 0.2727 | 41 | 0.0976 |
| 9_11507216 | 6 | 0.1667 | 13 | 0.2308 | 10 | 0.4 | 18 | 0.1111 | 47 | 0.2128 | 4 | 0.75 | 8 | 0.375 | 15 | 0.1333 | 6 | 0.1667 | 33 | 0.2727 |
| 9_11507219 | 7 | 0.8571 | 13 | 0.6154 | 9 | 0.2222 | 13 | 0.2308 | 42 | 0.4524 | 5 | 0.8 | 6 | 0.3333 | 14 | 0.5 | 7 | 0.5714 | 32 | 0.5312 |
| 9_11507254 | 5 | 0.8 | 16 | 0.3125 | 11 | 0.2727 | 16 | 0.4375 | 48 | 0.3958 | 5 | 0.4 | 8 | 0.625 | 12 | 0.5833 | 6 | 0.8333 | 31 | 0.6129 |
| 9_11507263 | 6 | 0 | 15 | 0.2 | 9 | 0.3333 | 14 | 0.2143 | 44 | 0.2045 | 6 | 0.1667 | 8 | 0.125 | 11 | 0.0909 | 8 | 0.25 | 33 | 0.1515 |
| 9_11507266 | 3 | 1 | 10 | 0.4 | 9 | 0.6667 | 13 | 0.4615 | 35 | 0.5429 | 5 | 0.4 | 9 | 0.5556 | 11 | 0.5455 | 8 | 0.625 | 33 | 0.5455 |
| 9_11507273 | 5 | 0.4 | 14 | 0.3571 | 10 | 0.3 | 13 | 0.2308 | 42 | 0.3095 | 4 | 0 | 9 | 0.2222 | 13 | 0.5385 | 8 | 0.375 | 34 | 0.3529 |
| 9_11507275 | 6 | 1 | 15 | 1 | 10 | 1 | 14 | 1 | 45 | 1 | 6 | 1 | 11 | 1 | 13 | 0.8462 | 7 | 0.8571 | 37 | 0.9189 |
| 9_11507283 | 2 | 0 | 15 | 0.1333 | 5 | 0.2 | 10 | 0.1 | 32 | 0.125 | 4 | 0 | 8 | 0.125 | 9 | 0.1111 | 8 | 0.125 | 29 | 0.1034 |
| 9_11507286 | 3 | 0.3333 | 13 | 0.3846 | 8 | 0.75 | 12 | 0.1667 | 36 | 0.3889 | 5 | 0.6 | 8 | 0.5 | 11 | 0.3636 | 8 | 0.375 | 32 | 0.4375 |
| 9_11507294 | 5 | 0 | 15 | 0.1333 | 9 | 0.2222 | 13 | 0.1538 | 42 | 0.1429 | 6 | 0 | 9 | 0 | 12 | 0.1667 | 10 | 0 | 37 | 0.0541 |
| 9_11507304 | 5 | 0.2 | 15 | 0.2667 | 10 | 0.3 | 12 | 0.3333 | 42 | 0.2857 | 6 | 0.5 | 9 | 0.6667 | 12 | 0.4167 | 8 | 0.625 | 35 | 0.5429 |
| 9_11510588 | 4 | 0 | 7 | 0.2857 | 6 | 0.1667 | 6 | 0.1667 | 23 | 0.1739 | 2 | 0.5 | 6 | 0 | 8 | 0.375 | 4 | 0.25 | 20 | 0.25 |
| 9_11510591 | 3 | 0.3333 | 7 | 0.4286 | 6 | 0.3333 | 4 | 0 | 20 | 0.3 | 1 | 0 | 4 | 0.5 | 8 | 0.375 | 3 | 0 | 16 | 0.3125 |
| 9_11513930 | 1 | 1 | NA | NA | NA | NA | NA | NA | 1 | 1 | 1 | 1 | NA | NA | NA | NA | NA | NA | 1 | 1 |
| 9_11513996 | 1 | 1 | NA | NA | NA | NA | NA | NA | 1 | 1 | 1 | 1 | NA | NA | NA | NA | NA | NA | 1 | 1 |
| 9_11514021 | 1 | 1 | NA | NA | NA | NA | NA | NA | 1 | 1 | 1 | 1 | NA | NA | NA | NA | NA | NA | 1 | 1 |
| 9_11514031 | 1 | 1 | NA | NA | NA | NA | 1 | 1 | 2 | 1 | 1 | 1 | NA | NA | NA | NA | NA | NA | 1 | 1 |
| 9_11514036 | 1 | 0 | NA | NA | NA | NA | 1 | 1 | 2 | 0.5 | 1 | 1 | NA | NA | NA | NA | NA | NA | 1 | 1 |
| 9_11591697 | 5 | 0 | 1 | 1 | NA | NA | 1 | 0 | 7 | 0.1429 | NA | NA | NA | NA | 3 | 0.3333 | 1 | 1 | 4 | 0.5 |
| 9_11591748 | 3 | 0 | 1 | 1 | NA | NA | 1 | 0 | 5 | 0.2 | NA | NA | NA | NA | 3 | 0.3333 | 2 | 0 | 5 | 0.2 |
| 9_11591768 | 3 | 0.3333 | 1 | 0 | NA | NA | 1 | 0 | 5 | 0.2 | NA | NA | NA | NA | 3 | 0.3333 | 1 | 0 | 4 | 0.25 |
| 9_11591783 | 3 | 0.3333 | 2 | 0 | NA | NA | 1 | 0 | 6 | 0.1667 | NA | NA | NA | NA | 2 | 0 | 2 | 0.5 | 4 | 0.25 |
| 9_11591897 | 1 | 0 | 1 | 0 | NA | NA | 2 | 0.5 | 4 | 0.25 | NA | NA | NA | NA | 3 | 0.6667 | 2 | 0.5 | 5 | 0.6 |
| 9_11591922 | 1 | 1 | NA | NA | NA | NA | 2 | 1 | 3 | 1 | NA | NA | NA | NA | 3 | 1 | 2 | 1 | 5 | 1 |
| 9_11591934 | 1 | 1 | NA | NA | NA | NA | 2 | 1 | 3 | 1 | NA | NA | NA | NA | 2 | 1 | 2 | 1 | 4 | 1 |
| 9_11591939 | 1 | 1 | NA | NA | NA | NA | 2 | 0 | 3 | 0.3333 | NA | NA | NA | NA | 3 | 0 | 2 | 1 | 5 | 0.4 |
| 9_11591945 | NA | NA | NA | NA | NA | NA | 1 | 1 | 1 | 1 | NA | NA | NA | NA | 1 | 0 | 2 | 0.5 | 3 | 0.3333 |
| 9_11593869 | 1 | 1 | 1 | 1 | NA | NA | 2 | 1 | 4 | 1 | NA | NA | NA | NA | 2 | 0.5 | 2 | 1 | 4 | 0.75 |
| 9_11640206 | 2 | 0.5 | 2 | 0 | 1 | 1 | 3 | 0.3333 | 8 | 0.375 | 1 | 0 | 1 | 1 | 4 | 0.25 | 2 | 0.5 | 8 | 0.375 |
| 9_11640238 | NA | NA | 3 | 0.3333 | 1 | 1 | 1 | 0 | 5 | 0.4 | NA | NA | NA | NA | 3 | 1 | 2 | 0 | 5 | 0.6 |
| 9_11708270 | 2 | 0.5 | 2 | 0.5 | NA | NA | NA | NA | 4 | 0.5 | 1 | 1 | 1 | 0 | NA | NA | NA | NA | 2 | 0.5 |
| 9_11708316 | 1 | 1 | NA | NA | NA | NA | 1 | 1 | 2 | 1 | NA | NA | 1 | 1 | NA | NA | NA | NA | 1 | 1 |
| 9_11708323 | 3 | 1 | 2 | 1 | NA | NA | 1 | 1 | 6 | 1 | 1 | 1 | 1 | 1 | NA | NA | NA | NA | 2 | 1 |
| 9_11708325 | 3 | 0.3333 | 1 | 0 | NA | NA | 1 | 0 | 5 | 0.2 | 1 | 1 | 1 | 0 | NA | NA | NA | NA | 2 | 0.5 |
| 9_11708329 | 2 | 1 | 1 | 1 | NA | NA | 1 | 1 | 4 | 1 | 1 | 1 | NA | NA | NA | NA | NA | NA | 1 | 1 |
| 9_11708333 | 2 | 0.5 | 2 | 0.5 | NA | NA | 1 | 1 | 5 | 0.6 | 1 | 0 | 2 | 1 | NA | NA | NA | NA | 3 | 0.6667 |
| 9_11708339 | 3 | 0 | 2 | 0.5 | NA | NA | NA | NA | 5 | 0.2 | 1 | 1 | 2 | 1 | 1 | 0 | NA | NA | 4 | 0.75 |
| 9_11708344 | 2 | 1 | 1 | 1 | NA | NA | 1 | 1 | 4 | 1 | 1 | 1 | 2 | 1 | 1 | 1 | NA | NA | 4 | 1 |
| 9_11708355 | 1 | 0 | 2 | 0.5 | NA | NA | 1 | 1 | 4 | 0.5 | 1 | 1 | NA | NA | NA | NA | NA | NA | 1 | 1 |
| 9_11708368 | 2 | 0 | 2 | 0.5 | NA | NA | 1 | 0 | 5 | 0.2 | 1 | 1 | 2 | 0.5 | 1 | 1 | NA | NA | 4 | 0.75 |
| 9_11708374 | 2 | 0 | 1 | 0 | NA | NA | 1 | 1 | 4 | 0.25 | 1 | 0 | 1 | 1 | 1 | 1 | NA | NA | 3 | 0.6667 |
| 9_11753594 | NA | NA | NA | NA | 1 | 1 | 2 | 0 | 3 | 0.3333 | NA | NA | 3 | 0 | 1 | 1 | 2 | 1 | 6 | 0.5 |
| 9_11753601 | NA | NA | NA | NA | NA | NA | 2 | 0.5 | 2 | 0.5 | NA | NA | 2 | 0 | 1 | 1 | 2 | 0.5 | 5 | 0.4 |
| 9_11789086 | 3 | 0.3333 | 3 | 1 | 5 | 0.4 | 6 | 0.6667 | 17 | 0.5882 | 2 | 0 | 5 | 0.8 | 1 | 1 | 8 | 0.5 | 16 | 0.5625 |
| 9_11789097 | 2 | 1 | 5 | 1 | 6 | 0.8333 | 6 | 0.8333 | 19 | 0.8947 | 3 | 0.6667 | 6 | 0.8333 | 3 | 1 | 12 | 0.9167 | 24 | 0.875 |
| 9_11789101 | 4 | 0.5 | 4 | 0.25 | 5 | 0.6 | 6 | 0.6667 | 19 | 0.5263 | 3 | 0.6667 | 7 | 0.8571 | 3 | 0.3333 | 9 | 0.7778 | 22 | 0.7273 |
| 9_11789109 | 4 | 0.25 | 4 | 0.25 | 5 | 0.6 | 6 | 0 | 19 | 0.2632 | 3 | 0 | 8 | 0.375 | 2 | 0 | 12 | 0.3333 | 25 | 0.28 |
| 9_11789117 | 4 | 0.75 | 4 | 1 | 6 | 0.6667 | 6 | 1 | 20 | 0.85 | 2 | 1 | 5 | 1 | 4 | 1 | 12 | 1 | 23 | 1 |
| 9_11789122 | 3 | 1 | 5 | 1 | 5 | 1 | 5 | 1 | 18 | 1 | 2 | 1 | 7 | 1 | 5 | 1 | 12 | 1 | 26 | 1 |
| 9_11789126 | 4 | 1 | 5 | 1 | 6 | 0.8333 | 5 | 1 | 20 | 0.95 | 2 | 0.5 | 6 | 1 | 4 | 1 | 12 | 1 | 24 | 0.9583 |
| 9_11789245 | 2 | 0.5 | 6 | 0.3333 | 2 | 0 | 3 | 0.3333 | 13 | 0.3077 | 3 | 0.3333 | 3 | 0.3333 | 4 | 0.5 | 14 | 0.2857 | 24 | 0.3333 |
| 9_11789251 | 3 | 0.3333 | 7 | 0.4286 | 4 | 0 | 5 | 0.4 | 19 | 0.3158 | 3 | 0.3333 | 3 | 0.3333 | 5 | 0.2 | 14 | 0.3571 | 25 | 0.32 |
| 9_11789327 | NA | NA | 1 | 1 | NA | NA | 3 | 0.6667 | 4 | 0.75 | NA | NA | 1 | 1 | NA | NA | 1 | 1 | 2 | 1 |
| 9_11789423 | 4 | 1 | 2 | 0.5 | 3 | 1 | 2 | 1 | 11 | 0.9091 | 2 | 1 | 4 | 1 | 2 | 1 | 2 | 1 | 10 | 1 |

| | | | | | | | | | | | | | | | | | | | | |
|---|---|---|---|---|---|---|---|---|---|---|---|---|---|---|---|---|---|---|---|---|
| 9_1178982 | NA | NA | 1 | 1 | 1 | 1 | 1 | 1 | 3 | 1 | 1 | 1 | 2 | 1 | NA | NA | 1 | 1 | 4 | 1 |
| 9_1178983 | NA | NA | 1 | 1 | 1 | 1 | 1 | 1 | 3 | 1 | 1 | 1 | 2 | 1 | NA | NA | 1 | 1 | 4 | 1 |
| 9_1178984 | NA | NA | 1 | 1 | 1 | 1 | 1 | 1 | 3 | 1 | 1 | 1 | 1 | 1 | NA | NA | 1 | 1 | 3 | 1 |
| 9_1183847 | NA | NA | 3 | 0.6667 | NA | NA | 1 | 0 | 4 | 0.5 | 1 | 0 | 2 | 0.5 | 1 | 0 | 1 | 0 | 5 | 0.2 |
| 9_1183848 | NA | NA | 3 | 0.6667 | NA | NA | 1 | 1 | 4 | 0.75 | NA | NA | 2 | 1 | 1 | 0 | 1 | 1 | 4 | 0.75 |
| 9_1183849 | NA | NA | 2 | 0 | NA | NA | 1 | 1 | 3 | 0.3333 | NA | NA | 1 | 0 | 1 | 1 | 1 | 0 | 3 | 0.3333 |
| 9_1183851 | NA | NA | 3 | 0.3333 | NA | NA | NA | NA | 3 | 0.3333 | 1 | 0 | 2 | 0 | NA | NA | 1 | 1 | 4 | 0.25 |
| 9_1183855 | NA | NA | 3 | 0.3333 | NA | NA | 1 | 0 | 4 | 0.25 | NA | NA | 3 | 0.3333 | 2 | 0.5 | 1 | 0 | 6 | 0.3333 |
| 9_1183865 | 1 | 0 | 2 | 0.5 | NA | NA | NA | NA | 3 | 0.3333 | NA | NA | 1 | 0 | 3 | 0.3333 | 2 | 0 | 6 | 0.1667 |
| 9_1183873 | 2 | 0.5 | 4 | 0.5 | NA | NA | 2 | 0.5 | 8 | 0.5 | 1 | 1 | 2 | 0.5 | 4 | 0.25 | 2 | 0.5 | 9 | 0.4444 |
| 9_1183874 | 2 | 0 | 5 | 0.2 | NA | NA | 2 | 0.5 | 9 | 0.2222 | 1 | 0 | 4 | 0.5 | 4 | 0.25 | 2 | 0.5 | 11 | 0.3636 |
| 9_1183874 | 2 | 0.5 | 2 | 0 | NA | NA | 2 | 0.5 | 6 | 0.3333 | 1 | 1 | 3 | 0 | 4 | 0 | 2 | 0 | 10 | 0.1 |
| 9_1183875 | 2 | 0 | 5 | 0.2 | NA | NA | 2 | 0.5 | 9 | 0.2222 | 1 | 0 | 3 | 0.6667 | 4 | 0.5 | 2 | 1 | 10 | 0.6 |
| 9_1183876 | 2 | 0 | 5 | 0.2 | NA | NA | 2 | 0 | 9 | 0.1111 | 1 | 1 | 4 | 0.5 | 4 | 0.5 | 2 | 0 | 11 | 0.4545 |
| 9_1183881 | 2 | 0.5 | 5 | 0.6 | 1 | 1 | 2 | 0.5 | 10 | 0.6 | 1 | 0 | 4 | 0.25 | 3 | 0.3333 | 2 | 1 | 10 | 0.4 |
| 9_1183882 | 2 | 0 | 4 | 0.25 | 1 | 0 | 3 | 0 | 10 | 0.1 | 2 | 0 | 2 | 0.5 | 5 | 0.2 | 2 | 0 | 11 | 0.1818 |
| 9_1183883 | 2 | 0 | 6 | 0.1667 | 2 | 0.5 | 3 | 0.3333 | 13 | 0.2308 | NA | NA | 3 | 0.3333 | 5 | 0.2 | 2 | 0.5 | 10 | 0.3 |
| 9_1183884 | 3 | 0 | 6 | 0.6667 | 2 | 0.5 | 3 | 0.3333 | 14 | 0.4286 | 1 | 1 | 2 | 0.5 | 5 | 0.2 | 2 | 0.5 | 10 | 0.4 |
| 9_1183884 | 3 | 0.6667 | 5 | 0.6 | 2 | 1 | 3 | 0.3333 | 13 | 0.6154 | NA | NA | 3 | 1 | 4 | 1 | 2 | 0.5 | 9 | 0.8889 |
| 9_1183885 | 3 | 0 | 6 | 0.5 | 3 | 0.3333 | 3 | 0.6667 | 15 | 0.4 | 2 | 0.5 | 3 | 0.6667 | 5 | 0.2 | 2 | 1 | 12 | 0.5 |
| 9_1184112 | 10 | 0.2 | 11 | 0.2727 | 6 | 0 | 3 | 0 | 30 | 0.1667 | 3 | 0.6667 | 9 | 0 | 5 | 0 | 6 | 0 | 23 | 0.087 |
| 9_1184112 | 11 | 0 | 10 | 0.2 | 6 | 0.1667 | 3 | 0 | 30 | 0.1 | 5 | 0 | 9 | 0.3333 | 5 | 0 | 6 | 0 | 25 | 0.12 |
| 9_1184113 | 9 | 0.1111 | 11 | 0.0909 | 6 | 0 | 4 | 0 | 30 | 0.0667 | 4 | 0 | 10 | 0.1 | 6 | 0.1667 | 5 | 0 | 25 | 0.08 |
| 9_1184114 | 8 | 0.25 | 11 | 0 | 5 | 0 | 4 | 0 | 28 | 0.0714 | 3 | 0 | 7 | 0.2857 | 5 | 0 | 6 | 0 | 21 | 0.0952 |
| 9_1187268 | 1 | 1 | NA | NA | NA | NA | NA | NA | 1 | 1 | NA | NA | 1 | 1 | NA | NA | NA | NA | 1 | 1 |
| 9_1187268 | 1 | 1 | NA | NA | NA | NA | NA | NA | 1 | 1 | NA | NA | 1 | 1 | NA | NA | NA | NA | 1 | 1 |
| 9_1187270 | 1 | 1 | NA | NA | NA | NA | NA | NA | 1 | 1 | NA | NA | 1 | 1 | NA | NA | NA | NA | 1 | 1 |
| 9_1187536 | 6 | 1 | 1 | 1 | 4 | 1 | 8 | 1 | 19 | 1 | 4 | 1 | 9 | 1 | 3 | 1 | 6 | 1 | 22 | 1 |
| 9_1187537 | 1 | 0 | NA | NA | 3 | 1 | 3 | 1 | 7 | 0.8571 | 4 | 1 | 2 | 1 | 1 | 1 | 3 | 1 | 10 | 1 |
| 9_1187537 | 5 | 0.8 | 1 | 1 | 4 | 0.25 | 5 | 0.2 | 15 | 0.4667 | 3 | 0 | 7 | 0.7143 | 1 | 1 | 6 | 0.6667 | 17 | 0.5882 |
| 9_1187537 | 3 | 1 | 1 | 1 | 4 | 1 | 6 | 1 | 14 | 1 | 2 | 1 | 5 | 1 | 1 | 1 | 3 | 1 | 11 | 1 |
| 9_1187538 | 2 | 1 | NA | NA | 2 | 0.5 | 3 | 0.3333 | 7 | 0.5714 | 3 | 0.3333 | 6 | 1 | 1 | 1 | 3 | 1 | 13 | 0.8462 |
| 9_1187538 | 4 | 0.25 | 1 | 0 | 2 | 0 | 6 | 0.1667 | 13 | 0.1538 | 4 | 0.5 | 8 | 0 | 3 | 0.3333 | 4 | 0.5 | 19 | 0.2632 |
| 9_1187539 | 5 | 1 | 3 | 1 | 2 | 1 | 3 | 1 | 13 | 1 | 6 | 0.8333 | 4 | 1 | 2 | 0.5 | 6 | 1 | 18 | 0.8889 |
| 9_1187541 | 5 | 0 | 3 | 0 | 2 | 0 | 4 | 0.5 | 14 | 0.1429 | 4 | 0.25 | 6 | 0 | 3 | 0 | 4 | 0 | 17 | 0.0588 |
| 9_1187542 | 4 | 0.25 | 2 | 0.5 | 2 | 0 | 2 | 0.5 | 10 | 0.3 | 5 | 0.6 | 3 | 0.6667 | 3 | 0.3333 | 3 | 0.3333 | 14 | 0.5 |
| 9_1187543 | 4 | 0 | 1 | 1 | 1 | 1 | 4 | 0 | 10 | 0.2 | 5 | 0.2 | 5 | 0 | 1 | 0 | 3 | 0.3333 | 14 | 0.1429 |
| 9_1187544 | 4 | 1 | 1 | 1 | 2 | 1 | 6 | 0.8333 | 13 | 0.9231 | 4 | 1 | 4 | 1 | 2 | 1 | 4 | 1 | 14 | 1 |
| 9_1187545 | 2 | 0.5 | 1 | 0 | NA | NA | 4 | 0.5 | 7 | 0.4286 | 1 | 1 | 3 | 0.3333 | 1 | 0 | 2 | 0.5 | 7 | 0.4286 |
| 9_1187547 | 5 | 0.2 | 2 | 0.5 | NA | NA | 4 | 1 | 11 | 0.5455 | 3 | 0.3333 | 3 | 0.6667 | 2 | 0.5 | 4 | 1 | 12 | 0.6667 |
| 9_1187548 | 3 | 0 | 2 | 0 | NA | NA | 2 | 0.5 | 7 | 0.1429 | 2 | 0.5 | 2 | 0 | 2 | 0.5 | 2 | 0.5 | 8 | 0.375 |
| 9_1187548 | 3 | 1 | 2 | 1 | NA | NA | 2 | 1 | 7 | 1 | 2 | 1 | 2 | 1 | 2 | 0 | 4 | 1 | 10 | 0.8 |
| 9_1188697 | NA | NA | NA | NA | 2 | 1 | NA | NA | 2 | 1 | NA | NA | 1 | 1 | 4 | 1 | 4 | 0.5 | 9 | 0.7778 |
| 9_1188697 | 1 | 1 | NA | NA | 2 | 0 | NA | NA | 3 | 0.3333 | NA | NA | 1 | 0 | 4 | 0 | 3 | 0.3333 | 8 | 0.125 |
| 9_1188698 | 1 | 1 | NA | NA | 1 | 0 | NA | NA | 2 | 0.5 | NA | NA | 1 | 0 | 4 | 0 | 3 | 0.3333 | 8 | 0.125 |
| 9_1188699 | NA | NA | NA | NA | NA | NA | 1 | 1 | 1 | 1 | NA | NA | 1 | 1 | 3 | 1 | 3 | 1 | 7 | 1 |
| 9_1188699 | 1 | 1 | NA | NA | 2 | 1 | 1 | 1 | 4 | 1 | NA | NA | 1 | 1 | 4 | 1 | 3 | 1 | 8 | 1 |
| 9_1188700 | 1 | 0 | NA | NA | 2 | 0 | 1 | 1 | 4 | 0.25 | NA | NA | 1 | 1 | 4 | 0.75 | 3 | 0.6667 | 8 | 0.75 |
| 9_1188702 | 1 | 1 | NA | NA | 3 | 1 | 1 | 1 | 5 | 1 | NA | NA | 1 | 1 | 3 | 1 | 4 | 0.25 | 8 | 0.625 |
| 9_1188702 | 1 | 1 | NA | NA | 2 | 1 | 1 | 0 | 4 | 0.75 | NA | NA | 1 | 0 | 4 | 0 | 3 | 0.3333 | 8 | 0.125 |
| 9_1188703 | NA | NA | NA | NA | 1 | 1 | NA | NA | 1 | 1 | NA | NA | NA | NA | 2 | 1 | 2 | 1 | 4 | 1 |
| 9_1188703 | NA | NA | NA | NA | 2 | 1 | 1 | 0 | 3 | 0.6667 | NA | NA | NA | NA | 3 | 0 | 3 | 0.6667 | 6 | 0.3333 |
| 9_1188704 | 1 | 1 | NA | NA | 2 | 0.5 | 1 | 1 | 4 | 0.75 | NA | NA | NA | NA | 4 | 1 | 3 | 0.3333 | 7 | 0.7143 |
| 9_1188705 | 1 | 0 | NA | NA | 2 | 0.5 | NA | NA | 3 | 0.3333 | NA | NA | 3 | 0 | 4 | 0.25 | 3 | 0.6667 | 10 | 0.3 |
| 9_1188707 | 1 | 1 | 1 | 1 | 3 | 0.6667 | 1 | 1 | 6 | 0.8333 | NA | NA | 3 | 1 | 4 | 0.5 | 4 | 1 | 11 | 0.8182 |
| 9_1188707 | NA | NA | NA | NA | 2 | 0.5 | 1 | 1 | 3 | 0.6667 | NA | NA | 3 | 0.6667 | 4 | 0.25 | 3 | 0.3333 | 10 | 0.4 |
| 9_1188708 | 1 | 0 | 1 | 1 | 2 | 0.5 | 1 | 0 | 5 | 0.4 | NA | NA | 1 | 1 | 3 | 0 | 4 | 0 | 8 | 0.125 |
| 9_1188709 | 1 | 1 | 1 | 1 | 3 | 1 | 1 | 1 | 6 | 1 | NA | NA | 3 | 1 | 4 | 1 | 3 | 1 | 10 | 1 |
| 9_1188709 | NA | NA | 1 | 0 | 2 | 1 | NA | NA | 3 | 0.6667 | NA | NA | NA | NA | 2 | 1 | 1 | 1 | 3 | 1 |
| 9_1188710 | 1 | 0 | 1 | 1 | 2 | 0.5 | 2 | 0 | 6 | 0.3333 | NA | NA | 3 | 0.3333 | 3 | 0.3333 | 4 | 0.25 | 10 | 0.3 |
| 9_1188710 | NA | NA | 1 | 1 | 2 | 0 | 1 | 0 | 4 | 0.25 | NA | NA | 2 | 0.5 | 4 | 0.25 | 2 | 0 | 8 | 0.25 |
| 9_1188711 | 1 | 0 | NA | NA | 3 | 0.6667 | 2 | 0 | 6 | 0.3333 | NA | NA | 3 | 0.3333 | 3 | 0 | 4 | 0 | 10 | 0.1 |
| 9_1188713 | 1 | 0 | 1 | 0 | 4 | 0.75 | 2 | 0 | 8 | 0.375 | NA | NA | 2 | 0.5 | 3 | 0.3333 | 4 | 0.25 | 9 | 0.3333 |
| 9_1188715 | NA | NA | 1 | 0 | 3 | 0.3333 | 2 | 0.5 | 6 | 0.3333 | NA | NA | 3 | 0.3333 | 4 | 0.25 | 4 | 0.25 | 11 | 0.2727 |
| 9_1188715 | 1 | 1 | 1 | 1 | 4 | 0.75 | 2 | 0.5 | 8 | 0.75 | NA | NA | 2 | 1 | 5 | 0.4 | 3 | 0.3333 | 10 | 0.5 |

| | | | | | | | | | | | | | | | | | | | | |
|---|---|---|---|---|---|---|---|---|---|---|---|---|---|---|---|---|---|---|---|---|
| 9_1188717 | 1 | 1 | 1 | 0 | 3 | 1 | 2 | 1 | 7 | 0.8571 | NA | NA | 1 | 1 | 5 | 1 | 3 | 0.6667 | 9 | 0.8889 |
| 9_1188726 | 1 | 0 | 2 | 0.5 | 4 | 0 | 2 | 0.5 | 9 | 0.2222 | 2 | 0 | 3 | 1 | 4 | 0 | 4 | 0.25 | 13 | 0.3077 |
| 9_1189008 | 1 | 0 | NA | NA | 4 | 0.25 | NA | NA | 5 | 0.2 | 1 | 0 | 1 | 1 | NA | NA | 1 | 0 | 3 | 0.3333 |
| 9_1190676 | 3 | 0.3333 | 7 | 0.2857 | 5 | 0.2 | 2 | 0 | 17 | 0.2353 | 1 | 0 | 5 | 0.2 | 3 | 0 | 5 | 0 | 14 | 0.0714 |
| 9_1190679 | 4 | 0 | 4 | 0.5 | 3 | 0 | 1 | 0 | 12 | 0.1667 | NA | NA | 4 | 0.25 | 4 | 0.5 | 4 | 0.25 | 12 | 0.3333 |
| 9_1190680 | 4 | 0.25 | 7 | 0.2857 | 5 | 0.6 | 2 | 0 | 18 | 0.3333 | NA | NA | 4 | 0.5 | 3 | 1 | 4 | 0.75 | 11 | 0.7273 |
| 9_1190685 | 3 | 0 | 6 | 0.1667 | 3 | 0.3333 | 2 | 0.5 | 14 | 0.2143 | NA | NA | 4 | 0.25 | 2 | 0 | 3 | 0.3333 | 9 | 0.2222 |
| 9_1190686 | 3 | 0 | 6 | 0.1667 | 4 | 0.25 | 2 | 0 | 15 | 0.1333 | NA | NA | 2 | 0 | 3 | 0.3333 | 2 | 0 | 7 | 0.1429 |
| 9_1190687 | 4 | 0 | 6 | 0.1667 | 4 | 0.25 | 2 | 0 | 16 | 0.125 | NA | NA | 3 | 0.3333 | 2 | 0 | 3 | 0 | 8 | 0.125 |
| 9_1190688 | 4 | 0 | 6 | 0 | 4 | 0.5 | 2 | 0 | 16 | 0.125 | NA | NA | 4 | 0 | 3 | 0.3333 | 3 | 0.3333 | 10 | 0.2 |
| 9_1190689 | 2 | 0 | 3 | 0 | 2 | 0.5 | 1 | 0 | 8 | 0.125 | NA | NA | 1 | 0 | 1 | 1 | 3 | 0 | 5 | 0.2 |
| 9_1190691 | 3 | 0 | 4 | 0 | 2 | 0.5 | 1 | 0 | 10 | 0.1 | NA | NA | 3 | 0 | 2 | 0 | 2 | 0.5 | 7 | 0.1429 |
| 9_1190691 | 3 | 1 | 2 | 1 | 4 | 0.5 | 1 | 1 | 10 | 0.8 | NA | NA | 2 | 1 | 1 | 1 | 2 | 1 | 5 | 1 |
| 9_1190692 | 3 | 1 | 5 | 1 | 5 | 1 | 1 | 1 | 14 | 1 | 1 | 1 | 2 | 1 | 2 | 1 | 3 | 1 | 8 | 1 |
| 9_1190692 | 4 | 1 | 6 | 0.8333 | 4 | 0.75 | 1 | 1 | 15 | 0.8667 | 1 | 1 | 3 | 1 | 2 | 1 | 2 | 1 | 8 | 1 |
| 9_1190693 | 3 | 0.6667 | 3 | 0.3333 | 4 | 0.75 | NA | NA | 10 | 0.6 | 1 | 1 | 2 | 1 | 2 | 1 | 2 | 1 | 7 | 1 |
| 9_1190693 | 4 | 1 | 5 | 1 | 4 | 1 | 1 | 1 | 14 | 1 | 1 | 1 | 2 | 1 | 2 | 1 | 3 | 1 | 8 | 1 |
| 9_1190694 | 3 | 0 | 4 | 0.25 | 2 | 0 | 1 | 1 | 10 | 0.2 | NA | NA | 3 | 0 | 2 | 0 | 3 | 0.3333 | 8 | 0.125 |
| 9_1190695 | 4 | 0 | 6 | 0.3333 | 4 | 0 | 1 | 0 | 15 | 0.1333 | 1 | 0 | 3 | 0.3333 | 2 | 0 | 3 | 0.6667 | 9 | 0.3333 |
| 9_1190695 | 4 | 0.25 | 5 | 0.4 | 4 | 0.75 | NA | NA | 13 | 0.4615 | 1 | 0 | 2 | 0.5 | 2 | 1 | 3 | 1 | 8 | 0.75 |
| 9_1190695 | 3 | 0 | 4 | 0.25 | 4 | 0 | 1 | 0 | 12 | 0.0833 | 1 | 0 | 2 | 0.5 | 2 | 0.5 | 2 | 0 | 7 | 0.2857 |
| 9_1190696 | 2 | 0 | 3 | 1 | 2 | 0.5 | 1 | 1 | 8 | 0.625 | 1 | 1 | 2 | 0.5 | 2 | 1 | 2 | 1 | 7 | 0.8571 |
| 9_1190698 | 3 | 1 | 6 | 0.3333 | 2 | 1 | 1 | 0 | 12 | 0.5833 | 2 | 0 | 1 | 1 | 2 | 0.5 | 3 | 0.6667 | 8 | 0.5 |
| 9_1190698 | 2 | 1 | 6 | 0.6667 | 1 | 0 | 2 | 0.5 | 11 | 0.6364 | 2 | 1 | 1 | 1 | 1 | 1 | 3 | 1 | 7 | 1 |
| 9_1190701 | 2 | 0.5 | 5 | 0.2 | 1 | 0 | 2 | 0.5 | 10 | 0.3 | 1 | 0 | 1 | 0 | 2 | 0.5 | 1 | 1 | 5 | 0.4 |
| 9_1191529 | NA | NA | NA | NA | NA | NA | 1 | 1 | 1 | 1 | NA | NA | 1 | 1 | NA | NA | NA | NA | 1 | 1 |
| 9_1191658 | 1 | 1 | NA | NA | NA | NA | 1 | 0 | 2 | 0.5 | 1 | 0 | NA | NA | 1 | 0 | 1 | 1 | 3 | 0.3333 |
| 9_1191790 | 9 | 0.1111 | 6 | 0.3333 | 8 | 0.125 | 10 | 0.1 | 33 | 0.1515 | 7 | 0.1429 | 6 | 0 | 9 | 0.2222 | 7 | 0 | 29 | 0.1034 |
| 9_1191792 | 4 | 0 | 5 | 0.2 | 4 | 0.25 | 6 | 0.3333 | 19 | 0.2105 | 6 | 0 | 5 | 0.2 | 5 | 0.2 | 5 | 0.2 | 21 | 0.1429 |
| 9_1191792 | 8 | 0.125 | 6 | 0 | 7 | 0.1429 | 9 | 0.1111 | 30 | 0.1 | 5 | 0 | 6 | 0.1667 | 8 | 0.125 | 4 | 0 | 23 | 0.087 |
| 9_1191792 | 7 | 0.2857 | 6 | 0 | 7 | 0 | 10 | 0.1 | 30 | 0.1 | 8 | 0 | 8 | 0.125 | 11 | 0 | 6 | 0.1667 | 33 | 0.0606 |
| 9_1191792 | 7 | 0.2857 | 10 | 0 | 10 | 0.1 | 11 | 0 | 38 | 0.0789 | 8 | 0.25 | 8 | 0.125 | 10 | 0.2 | 9 | 0 | 35 | 0.1429 |
| 9_1192153 | NA | NA | NA | NA | 1 | 1 | NA | NA | 1 | 1 | NA | NA | 1 | 1 | NA | NA | NA | NA | 1 | 1 |
| 9_1192153 | NA | NA | NA | NA | 1 | 1 | NA | NA | 1 | 1 | NA | NA | 1 | 1 | NA | NA | NA | NA | 1 | 1 |
| 9_1192219 | NA | NA | 1 | 0 | 1 | 1 | 1 | 0 | 3 | 0.3333 | 1 | 0 | NA | NA | NA | NA | 1 | 1 | 2 | 0.5 |
| 9_1192221 | 1 | 0 | 1 | 1 | 1 | 1 | NA | NA | 3 | 0.6667 | 1 | 0 | NA | NA | NA | NA | 1 | 1 | 2 | 0.5 |
| 9_1192222 | 1 | 1 | 1 | 0 | 1 | 0 | 1 | 0 | 4 | 0.25 | 1 | 1 | NA | NA | NA | NA | 1 | 0 | 2 | 0.5 |
| 9_1192629 | NA | NA | 1 | 1 | NA | NA | NA | NA | 1 | 1 | 1 | 1 | NA | NA | NA | NA | NA | NA | 1 | 1 |
| 9_1192629 | NA | NA | 1 | 1 | NA | NA | NA | NA | 1 | 1 | 1 | 1 | NA | NA | NA | NA | NA | NA | 1 | 1 |
| 9_1192631 | NA | NA | 1 | 1 | NA | NA | NA | NA | 1 | 1 | 1 | 1 | NA | NA | NA | NA | NA | NA | 1 | 1 |
| 9_1192632 | NA | NA | 1 | 1 | NA | NA | NA | NA | 1 | 1 | 1 | 1 | NA | NA | NA | NA | NA | NA | 1 | 1 |
| 9_1192634 | NA | NA | 1 | 1 | NA | NA | NA | NA | 1 | 1 | 1 | 1 | NA | NA | NA | NA | NA | NA | 1 | 1 |
| 9_1192699 | NA | NA | 2 | 0 | NA | NA | 3 | 0.3333 | 5 | 0.2 | 2 | 0.5 | 5 | 0.2 | 1 | 0 | 2 | 0 | 10 | 0.2 |
| 9_1192701 | NA | NA | 2 | 0 | NA | NA | 3 | 0.3333 | 5 | 0.2 | 2 | 0 | 4 | 0.5 | 1 | 0 | 2 | 0.5 | 9 | 0.3333 |
| 9_1192702 | NA | NA | 2 | 0.5 | NA | NA | 3 | 0.6667 | 5 | 0.6 | 2 | 0.5 | 5 | 0.8 | 1 | 1 | 2 | 1 | 10 | 0.8 |
| 9_1192703 | NA | NA | 2 | 0.5 | NA | NA | 1 | 0 | 3 | 0.3333 | 2 | 0 | 4 | 0.25 | 1 | 0 | 2 | 0.5 | 9 | 0.2222 |
| 9_1192916 | 4 | 0.25 | 2 | 0 | 3 | 0.6667 | 6 | 0.5 | 15 | 0.4 | 3 | 0.3333 | 3 | 0 | 3 | 0 | 3 | 1 | 12 | 0.3333 |
| 9_1192916 | 3 | 1 | 3 | 0.6667 | 2 | 1 | 3 | 0.6667 | 11 | 0.8182 | 3 | 0.6667 | 3 | 1 | 3 | 0.6667 | 3 | 0.6667 | 12 | 0.75 |
| 9_1192916 | NA | NA | NA | NA | NA | NA | 1 | 1 | 1 | 1 | 1 | 1 | NA | NA | NA | NA | NA | NA | 1 | 1 |
| 9_1192917 | 4 | 0 | 3 | 0 | 3 | 0 | 6 | 0.1667 | 16 | 0.0625 | 3 | 0.3333 | 3 | 0 | 2 | 0 | 3 | 0 | 11 | 0.0909 |
| 9_1192918 | 3 | 0.6667 | 2 | 0.5 | 2 | 1 | 5 | 0.4 | 12 | 0.5833 | 2 | 1 | 3 | 1 | 3 | 0.6667 | 3 | 1 | 11 | 0.9091 |
| 9_1192919 | 5 | 1 | 4 | 1 | 3 | 1 | 6 | 1 | 18 | 1 | 4 | 1 | 3 | 1 | 1 | 1 | 2 | 1 | 10 | 1 |
| 9_1192920 | 5 | 0.2 | 4 | 0.75 | 3 | 0.6667 | 6 | 0.1667 | 18 | 0.3889 | 4 | 0.25 | 3 | 0.3333 | 3 | 0.3333 | 3 | 0.3333 | 13 | 0.3077 |
| 9_1192920 | 5 | 0.6 | 3 | 0.3333 | 2 | 1 | 6 | 0.6667 | 16 | 0.625 | 5 | 0.8 | 3 | 0.3333 | 2 | 1 | 3 | 1 | 13 | 0.7692 |
| 9_1192921 | 5 | 1 | 4 | 1 | 3 | 1 | 6 | 1 | 18 | 1 | 5 | 1 | 3 | 1 | 3 | 1 | 3 | 1 | 14 | 1 |
| 9_1192921 | 5 | 1 | 4 | 1 | 3 | 1 | 6 | 1 | 18 | 1 | 5 | 1 | 3 | 1 | 3 | 1 | 3 | 1 | 14 | 1 |
| 9_1192922 | 5 | 0.4 | 3 | 0 | 3 | 0 | 5 | 0.4 | 16 | 0.25 | 5 | 0.4 | 3 | 0 | 2 | 0 | 3 | 0 | 13 | 0.1538 |
| 9_1192922 | 4 | 0 | 4 | 0.5 | 2 | 0 | 6 | 0.3333 | 16 | 0.25 | 3 | 0.3333 | 3 | 0 | 3 | 0.3333 | 4 | 0.25 | 13 | 0.2308 |
| 9_1192923 | 4 | 1 | 2 | 1 | 1 | 1 | 4 | 0.75 | 11 | 0.9091 | 2 | 1 | 2 | 1 | 3 | 1 | 3 | 1 | 10 | 1 |
| 9_1192924 | 6 | 1 | 4 | 1 | 3 | 1 | 5 | 1 | 18 | 1 | 1 | 1 | 2 | 1 | 3 | 1 | 3 | 1 | 9 | 1 |
| 9_1192925 | 4 | 0 | 4 | 0 | 2 | 0 | 4 | 0.25 | 14 | 0.0714 | 3 | 0.3333 | 1 | 0 | 2 | 0 | 4 | 0 | 10 | 0.1 |
| 9_1192926 | 5 | 1 | 4 | 1 | 2 | 1 | 6 | 1 | 17 | 1 | 5 | 1 | 3 | 1 | 3 | 1 | 5 | 0.6 | 16 | 0.875 |
| 9_1192927 | 4 | 1 | 4 | 1 | 3 | 1 | 6 | 1 | 17 | 1 | 4 | 1 | 3 | 1 | 3 | 1 | 4 | 1 | 14 | 1 |
| 9_1192927 | 4 | 1 | 3 | 0.6667 | 2 | 1 | 5 | 0.8 | 14 | 0.8571 | 5 | 0.4 | 3 | 0.6667 | 3 | 1 | 4 | 0.75 | 15 | 0.6667 |
| 9_1192928 | 5 | 0.6 | 2 | 0 | 3 | 0.3333 | 6 | 0.5 | 16 | 0.4375 | 4 | 0 | 3 | 0.3333 | 3 | 0.3333 | 4 | 0.5 | 14 | 0.2857 |

| | | | | | | | | | | | | | | | | | | | | |
|---|---|---|---|---|---|---|---|---|---|---|---|---|---|---|---|---|---|---|---|---|
| 9_1192928 | 6 | 1 | 2 | 1 | 4 | 1 | 4 | 1 | 16 | 1 | 2 | 1 | 1 | 1 | 4 | 0.75 | 5 | 0.8 | 12 | 0.8333 |
| 9_1192928 | 5 | 0.4 | 2 | 0.5 | 3 | 0.3333 | 5 | 0.2 | 15 | 0.3333 | 1 | 0 | 1 | 0 | 4 | 0.25 | 5 | 0.2 | 11 | 0.1818 |
| 9_1192929 | 4 | 0 | 1 | 0 | 3 | 0 | 2 | 0.5 | 10 | 0.1 | 2 | 0.5 | 1 | 0 | 3 | 0 | 4 | 0 | 10 | 0.1 |
| 9_1192932 | 5 | 0.4 | 2 | 0.5 | 4 | 0.25 | 6 | 0.1667 | 17 | 0.2941 | 2 | 0 | 1 | 0 | 4 | 0.25 | 5 | 0 | 12 | 0.0833 |
| 9_1192933 | 4 | 1 | 1 | 1 | 2 | 0 | 3 | 1 | 10 | 0.8 | NA | NA | 1 | 0 | 4 | 0.5 | 4 | 0 | 9 | 0.2222 |
| 9_1192933 | 2 | 0.5 | 2 | 0.5 | NA | NA | 1 | 1 | 5 | 0.6 | 1 | 1 | NA | NA | 1 | 0 | 2 | 0.5 | 4 | 0.5 |
| 9_1192935 | 3 | 0.3333 | 2 | 0 | NA | NA | NA | NA | 5 | 0.2 | 1 | 1 | NA | NA | 1 | 1 | 1 | 0 | 3 | 0.6667 |
| 9_1192936 | 2 | 1 | 1 | 0 | NA | NA | NA | NA | 3 | 0.6667 | NA | NA | NA | NA | 1 | 1 | 2 | 1 | 3 | 1 |
| 9_1192936 | 2 | 0.5 | 1 | 0 | NA | NA | 1 | 0 | 4 | 0.25 | NA | NA | NA | NA | 1 | 0 | 1 | 1 | 2 | 0.5 |
| 9_1192936 | 1 | 1 | NA | NA | NA | NA | NA | NA | 1 | 1 | 1 | 1 | NA | NA | 1 | 1 | NA | NA | 2 | 1 |
| 9_1192939 | 3 | 0.6667 | NA | NA | NA | NA | NA | NA | 3 | 0.6667 | 1 | 1 | NA | NA | 1 | 0 | NA | NA | 2 | 0.5 |
| 9_1192942 | 3 | 0.3333 | NA | NA | NA | NA | NA | NA | 3 | 0.3333 | 1 | 1 | NA | NA | 1 | 0 | NA | NA | 2 | 0.5 |
| 9_1192944 | 3 | 0 | NA | NA | NA | NA | 1 | 1 | 4 | 0.25 | 1 | 1 | NA | NA | 1 | 0 | NA | NA | 2 | 0.5 |
| 9_1192945 | 3 | 0.3333 | NA | NA | NA | NA | NA | NA | 3 | 0.3333 | 1 | 1 | NA | NA | 1 | 1 | NA | NA | 2 | 1 |
| 9_1193126 | 1 | 0 | 2 | 0 | 3 | 0.6667 | 3 | 0 | 9 | 0.2222 | 2 | 0 | 3 | 0.3333 | NA | NA | 1 | 0 | 6 | 0.1667 |
| 9_1193132 | NA | NA | 2 | 1 | 1 | 0 | 3 | 0.6667 | 6 | 0.6667 | 1 | 1 | 3 | 0.6667 | NA | NA | 1 | 1 | 5 | 0.8 |
| 9_1193135 | NA | NA | 2 | 0 | 2 | 0 | 3 | 0.3333 | 7 | 0.1429 | 1 | 1 | 3 | 0.3333 | NA | NA | 1 | 1 | 5 | 0.6 |
| 9_1193136 | NA | NA | 2 | 0.5 | 2 | 1 | 2 | 1 | 6 | 0.8333 | 1 | 1 | 2 | 0.5 | NA | NA | 1 | 1 | 4 | 0.75 |
| 9_1193140 | NA | NA | 2 | 0.5 | 1 | 0 | 3 | 0 | 6 | 0.1667 | 1 | 1 | 1 | 1 | NA | NA | 1 | 1 | 3 | 1 |
| 9_1193140 | NA | NA | 2 | 0.5 | NA | NA | 3 | 0 | 5 | 0.2 | 1 | 1 | 1 | 1 | NA | NA | NA | NA | 2 | 1 |
| 9_1193142 | NA | NA | 1 | 1 | 1 | 1 | 2 | 1 | 4 | 1 | 1 | 1 | 1 | 1 | NA | NA | 1 | 1 | 3 | 1 |
| 9_1193142 | NA | NA | NA | NA | 1 | 1 | 2 | 1 | 3 | 1 | 1 | 1 | NA | NA | NA | NA | 1 | 1 | 2 | 1 |
| 9_1193142 | NA | NA | NA | NA | NA | NA | 3 | 1 | 3 | 1 | 1 | 1 | 1 | 0 | NA | NA | 1 | 0 | 3 | 0.3333 |
| 9_1193143 | NA | NA | NA | NA | 1 | 1 | 3 | 1 | 4 | 1 | 1 | 1 | 1 | 1 | NA | NA | 1 | 0 | 3 | 0.6667 |
| 9_1193146 | NA | NA | NA | NA | 1 | 1 | 1 | 1 | 2 | 1 | 1 | 1 | 1 | 1 | NA | NA | NA | NA | 2 | 1 |
| 9_1193146 | NA | NA | NA | NA | NA | NA | 3 | 0.6667 | 3 | 0.6667 | 1 | 1 | 1 | 1 | NA | NA | 1 | 1 | 3 | 1 |
| 9_1193147 | NA | NA | NA | NA | 1 | 0 | 3 | 0.6667 | 4 | 0.5 | NA | NA | 1 | 1 | NA | NA | 1 | 1 | 2 | 1 |
| 9_1193148 | NA | NA | NA | NA | 1 | 0 | 3 | 0.3333 | 4 | 0.25 | 1 | 0 | 1 | 0 | NA | NA | 1 | 1 | 3 | 0.3333 |
| 9_1193148 | NA | NA | NA | NA | 1 | 1 | 3 | 1 | 4 | 1 | 1 | 1 | 1 | 1 | NA | NA | 1 | 1 | 3 | 1 |
| 9_1193151 | NA | NA | NA | NA | 1 | 1 | 3 | 1 | 4 | 1 | 1 | 1 | 1 | 1 | NA | NA | 1 | 1 | 3 | 1 |
| 9_1193151 | NA | NA | NA | NA | 1 | 1 | 3 | 0.6667 | 4 | 0.75 | 1 | 1 | NA | NA | NA | NA | 1 | 1 | 2 | 1 |
| 9_1193154 | NA | NA | NA | NA | NA | NA | 1 | 1 | 1 | 1 | 1 | 1 | 1 | 1 | NA | NA | NA | NA | 2 | 1 |
| 9_1193155 | NA | NA | NA | NA | 1 | 0 | 3 | 0.3333 | 4 | 0.25 | 1 | 0 | 1 | 1 | NA | NA | 1 | 0 | 3 | 0.3333 |
| 9_1193628 | NA | NA | NA | NA | 1 | 0 | 1 | 1 | 2 | 0.5 | 1 | 1 | NA | NA | NA | NA | NA | NA | 1 | 1 |
| 9_1197532 | NA | NA | 1 | 1 | NA | NA | 1 | 1 | 2 | 1 | 2 | 1 | NA | NA | 1 | 1 | NA | NA | 3 | 1 |
| 9_1197534 | NA | NA | 1 | 1 | NA | NA | 1 | 1 | 2 | 1 | 2 | 0.5 | NA | NA | NA | NA | NA | NA | 2 | 0.5 |
| 9_1197537 | NA | NA | 1 | 1 | NA | NA | NA | NA | 1 | 1 | 2 | 1 | NA | NA | NA | NA | NA | NA | 2 | 1 |
| 9_1197537 | 2 | 0 | 3 | 0 | 7 | 0.2857 | 2 | 1 | 14 | 0.2857 | 4 | 0 | 4 | 0.25 | 7 | 0 | 4 | 0 | 19 | 0.0526 |
| 9_1197538 | 3 | 0.3333 | 5 | 0.2 | 7 | 0.4286 | 1 | 0 | 16 | 0.3125 | 4 | 0.25 | 4 | 0 | 8 | 0.125 | 4 | 0.25 | 20 | 0.15 |
| 9_1197538 | NA | NA | 1 | 1 | NA | NA | NA | NA | 1 | 1 | 2 | 1 | NA | NA | NA | NA | NA | NA | 2 | 1 |
| 9_1197539 | 4 | 0 | 5 | 0.2 | 8 | 0.125 | 3 | 0.3333 | 20 | 0.15 | 4 | 0.25 | 4 | 0.25 | 7 | 0.2857 | 4 | 0.25 | 19 | 0.2632 |
| 9_1197540 | 5 | 0.6 | 4 | 0.5 | 7 | 0.8571 | 3 | 0.6667 | 19 | 0.6842 | 2 | 1 | 4 | 1 | 8 | 0.875 | 4 | 0.75 | 18 | 0.8889 |
| 9_1197540 | 2 | 0.5 | 5 | 0.4 | 4 | 0.75 | 1 | 0 | 12 | 0.5 | 4 | 0.5 | 3 | 0.6667 | 4 | 0.75 | 3 | 0.6667 | 14 | 0.6429 |
| 9_1197541 | 5 | 0 | 6 | 0.1667 | 8 | 0.25 | 3 | 0 | 22 | 0.1364 | 4 | 0.25 | 5 | 0.2 | 6 | 0.3333 | 3 | 0 | 18 | 0.2222 |
| 9_1197541 | 5 | 0.2 | 5 | 0 | 7 | 0.4286 | 3 | 0.3333 | 20 | 0.25 | 3 | 0.3333 | 5 | 0.2 | 5 | 0.4 | 3 | 0.3333 | 16 | 0.3125 |
| 9_1197543 | 5 | 0.2 | 6 | 0 | 6 | 0.1667 | 4 | 0 | 21 | 0.0952 | 4 | 0.25 | 5 | 0 | 7 | 0.2857 | 3 | 0 | 19 | 0.1579 |
| 9_1197543 | 7 | 0 | 6 | 0 | 7 | 0.2857 | 3 | 0 | 23 | 0.087 | 4 | 0.25 | 5 | 0 | 8 | 0 | 4 | 0.5 | 21 | 0.1429 |
| 9_1197544 | 6 | 0.8333 | 5 | 0.6 | 8 | 0.625 | 4 | 0.5 | 23 | 0.6522 | 3 | 0 | 4 | 1 | 7 | 0.7143 | 4 | 1 | 18 | 0.7222 |
| 9_1197546 | 3 | 0.3333 | 5 | 0.6 | 7 | 0.5714 | 4 | 0.5 | 19 | 0.5263 | 4 | 0.75 | 4 | 0.5 | 6 | 0.6667 | 4 | 0.75 | 18 | 0.6667 |
| 9_1197547 | NA | NA | 2 | 0.5 | NA | NA | NA | NA | 2 | 0.5 | 3 | 0.6667 | NA | NA | 1 | 0 | NA | NA | 4 | 0.5 |
| 9_1197548 | 1 | 0 | 2 | 0.5 | NA | NA | NA | NA | 3 | 0.3333 | 3 | 0.3333 | NA | NA | 1 | 0 | NA | NA | 4 | 0.25 |
| 9_1197548 | 4 | 1 | 5 | 0.6 | 7 | 0.8571 | 3 | 0.6667 | 19 | 0.7895 | 3 | 0.6667 | 4 | 0.75 | 3 | 0 | 1 | 0 | 11 | 0.4545 |
| 9_1197548 | 7 | 0.1429 | 5 | 0 | 6 | 0 | 3 | 0.3333 | 21 | 0.0952 | 4 | 0.25 | 4 | 0.5 | 4 | 0 | 3 | 0 | 15 | 0.2 |
| 9_1197548 | 5 | 0.2 | 5 | 0 | 7 | 0.1429 | 4 | 0 | 21 | 0.0952 | 3 | 0.3333 | 5 | 0 | 3 | 0 | 3 | 0 | 14 | 0.0714 |
| 9_1197549 | 1 | 0 | 2 | 0 | 1 | 1 | NA | NA | 4 | 0.25 | 4 | 0.25 | NA | NA | 1 | 1 | NA | NA | 5 | 0.4 |
| 9_1197549 | 6 | 0 | 4 | 0 | 6 | 0.3333 | 3 | 0 | 19 | 0.1053 | 3 | 0.3333 | 5 | 0.6 | 3 | 0 | 4 | 0.5 | 15 | 0.4 |
| 9_1197551 | 5 | 0 | 7 | 0.2857 | 10 | 0.1 | 5 | 0 | 27 | 0.1111 | 5 | 0.4 | 6 | 0 | 4 | 0.5 | 6 | 0 | 21 | 0.1905 |
| 9_1197553 | 5 | 0.2 | 5 | 0 | 6 | 0.3333 | 3 | 0 | 19 | 0.1579 | 4 | 0 | 2 | 0 | 3 | 0 | 6 | 0.1667 | 15 | 0.0667 |
| 9_1197560 | 2 | 0.5 | 4 | 0 | 6 | 0.1667 | 4 | 0.25 | 16 | 0.1875 | 8 | 0.25 | 4 | 0.25 | 1 | 0 | 3 | 0.3333 | 16 | 0.25 |
| 9_1197568 | 1 | 0 | 3 | 0.3333 | 3 | 0 | 4 | 0.5 | 11 | 0.2727 | 8 | 0.25 | 4 | 0.5 | 1 | 0 | 2 | 0 | 15 | 0.2667 |
| 9_1197568 | 2 | 1 | 3 | 1 | 2 | 1 | 4 | 0.5 | 11 | 0.8182 | 6 | 0.8333 | 3 | 1 | 1 | 1 | 1 | 1 | 11 | 0.9091 |
| 9_1198006 | 1 | 0 | 3 | 0.3333 | 2 | 0 | 2 | 0 | 8 | 0.125 | NA | NA | 2 | 0.5 | NA | NA | 3 | 0 | 5 | 0.2 |
| 9_1198008 | 1 | 1 | 2 | 0.5 | 1 | 1 | NA | NA | 4 | 0.75 | NA | NA | 2 | 1 | NA | NA | 3 | 0.6667 | 5 | 0.8 |
| 9_1198015 | 1 | 1 | 1 | 1 | 1 | 1 | 1 | 1 | 4 | 1 | NA | NA | NA | NA | NA | NA | 1 | 1 | 1 | 1 |
| 9_1198015 | 1 | 1 | 1 | 0 | 1 | 1 | 1 | 1 | 4 | 0.75 | NA | NA | NA | NA | NA | NA | 1 | 1 | 1 | 1 |

| | | | | | | | | | | | | | | | | | | | | |
|---|---|---|---|---|---|---|---|---|---|---|---|---|---|---|---|---|---|---|---|---|
| 9_1198015 | 1 | 1 | NA | NA | 1 | 0 | 1 | 1 | 3 | 0.6667 | NA | NA | NA | NA | NA | NA | 1 | 1 | 1 | 1 |
| 9_1198016 | NA | NA | 1 | 1 | 1 | 1 | NA | NA | 2 | 1 | NA | NA | NA | NA | NA | NA | 1 | 1 | 1 | 1 |
| 9_1198017 | NA | NA | 1 | 1 | 1 | 1 | 1 | 1 | 3 | 1 | NA | NA | NA | NA | NA | NA | 1 | 1 | 1 | 1 |
| 9_1198020 | NA | NA | 1 | 1 | 1 | 1 | 1 | 1 | 3 | 1 | NA | NA | NA | NA | NA | NA | 1 | 1 | 1 | 1 |
| 9_1198020 | NA | NA | 1 | 1 | 1 | 0 | 1 | 0 | 3 | 0.3333 | NA | NA | NA | NA | NA | NA | 1 | 1 | 1 | 1 |
| 9_1198021 | NA | NA | 1 | 1 | 1 | 1 | 1 | 1 | 3 | 1 | NA | NA | NA | NA | NA | NA | 1 | 1 | 1 | 1 |
| 9_1199528 | 6 | 0 | 3 | 0.3333 | 3 | 0 | 3 | 0 | 15 | 0.0667 | 3 | 0.3333 | 1 | 0 | 2 | 0 | 4 | 0 | 10 | 0.1 |
| 9_1199532 | 5 | 0.2 | 3 | 0.3333 | 3 | 0 | 3 | 0.3333 | 14 | 0.2143 | 3 | 0 | 1 | 0 | 3 | 0.3333 | 4 | 0 | 11 | 0.0909 |
| 9_1199532 | 4 | 0.25 | 2 | 0 | 2 | 0.5 | 2 | 0.5 | 10 | 0.3 | 3 | 0.3333 | NA | NA | 2 | 0.5 | 4 | 0.75 | 9 | 0.5556 |
| 9_1199532 | 5 | 0.4 | 3 | 0 | 3 | 0 | 3 | 0.3333 | 14 | 0.2143 | 2 | 1 | NA | NA | 3 | 0 | 3 | 0 | 8 | 0.25 |
| 9_1199541 | 1 | 0 | 1 | 1 | NA | NA | NA | NA | 2 | 0.5 | 1 | 1 | NA | NA | NA | NA | 2 | 0.5 | 3 | 0.6667 |
| 9_1199594 | 87 | 0.092 | 93 | 0.086 | 91 | 0.0659 | 95 | 0.1158 | 366 | 0.0902 | 74 | 0.0946 | 94 | 0.1277 | 96 | 0.125 | 73 | 0.1096 | 337 | 0.1157 |
| 9_1199974 | 10 | 0.2 | 15 | 0.3333 | 26 | 0.0385 | 13 | 0.1538 | 64 | 0.1562 | 15 | 0.3333 | 2 | 0 | 9 | 0 | 13 | 0.3077 | 39 | 0.2308 |
| 9_1200022 | 49 | 0.0204 | 68 | 0.0882 | 66 | 0.0758 | 48 | 0.0625 | 231 | 0.0649 | 84 | 0.0833 | 33 | 0.0303 | 44 | 0.0909 | 45 | 0.0667 | 206 | 0.0728 |
| 9_1200022 | 42 | 0.119 | 64 | 0.1094 | 63 | 0.1587 | 48 | 0.1667 | 217 | 0.1382 | 84 | 0.2262 | 31 | 0.1613 | 44 | 0.2045 | 38 | 0.1842 | 197 | 0.203 |
| 9_1200028 | 4 | 0 | 3 | 0.3333 | 3 | 0 | 5 | 0 | 15 | 0.0667 | 7 | 0 | 3 | 0 | 8 | 0.25 | 8 | 0.125 | 26 | 0.1154 |
| 9_1200031 | 2 | 0.5 | 2 | 0 | 3 | 0 | 1 | 0 | 8 | 0.125 | 6 | 0 | 2 | 0 | 5 | 0.2 | 5 | 0 | 18 | 0.0556 |
| 9_1200036 | 6 | 0.1667 | 4 | 0 | 5 | 0 | 5 | 0 | 20 | 0.05 | 11 | 0.0909 | 5 | 0.2 | 7 | 0 | 11 | 0 | 34 | 0.0588 |
| 9_1200037 | 6 | 0.6667 | 3 | 0.3333 | 6 | 0.8333 | 6 | 0.5 | 21 | 0.619 | 9 | 0.7778 | 4 | 0 | 6 | 0.5 | 8 | 0.5 | 27 | 0.5185 |
| 9_1200038 | 9 | 0.5556 | 5 | 0 | 6 | 0 | 4 | 0 | 24 | 0.2083 | 13 | 0.0769 | 5 | 0.2 | 7 | 0.2857 | 10 | 0.2 | 35 | 0.1714 |
| 9_1200045 | 13 | 0 | 6 | 0 | 8 | 0 | 7 | 0.2857 | 34 | 0.0588 | 15 | 0.0667 | 7 | 0 | 11 | 0 | 13 | 0.1538 | 46 | 0.0652 |
| 9_1200047 | 13 | 0 | 7 | 0.2857 | 10 | 0.1 | 7 | 0 | 37 | 0.0811 | 17 | 0.0588 | 9 | 0 | 12 | 0.1667 | 13 | 0 | 51 | 0.0588 |
| 9_1200087 | 48 | 0.7708 | 75 | 0.7467 | 72 | 0.8056 | 54 | 0.9259 | 249 | 0.8072 | 78 | 0.8333 | 38 | 0.6842 | 41 | 0.8537 | 46 | 0.7174 | 203 | 0.7833 |
| 9_1200088 | 37 | 0.8649 | 64 | 0.8438 | 75 | 0.7867 | 49 | 0.7755 | 225 | 0.8133 | 74 | 0.7838 | 36 | 0.8889 | 42 | 0.7143 | 42 | 0.7857 | 194 | 0.7887 |
| 9_1200090 | 44 | 0.0909 | 61 | 0.1311 | 61 | 0.1639 | 49 | 0.1224 | 215 | 0.1302 | 73 | 0.0411 | 30 | 0.0333 | 35 | 0.0571 | 38 | 0.1316 | 176 | 0.0625 |
| 9_1200095 | 47 | 0.0426 | 79 | 0.0759 | 77 | 0.1299 | 55 | 0.0727 | 258 | 0.0853 | 80 | 0.0375 | 40 | 0.15 | 46 | 0.0217 | 45 | 0.0889 | 211 | 0.0664 |
| 9_1200099 | 42 | 0.0476 | 80 | 0.05 | 76 | 0.0658 | 58 | 0.069 | 256 | 0.0586 | 71 | 0.0563 | 31 | 0.0323 | 47 | 0.0426 | 43 | 0.0698 | 192 | 0.0521 |
| 9_1200099 | 53 | 0.0377 | 86 | 0.0698 | 86 | 0.1279 | 68 | 0.0882 | 293 | 0.0853 | 84 | 0.0833 | 46 | 0.0435 | 54 | 0.0741 | 51 | 0.0588 | 235 | 0.0681 |
| 9_1200100 | 56 | 0.8393 | 83 | 0.8554 | 82 | 0.8049 | 65 | 0.8154 | 286 | 0.8287 | 83 | 0.8675 | 43 | 0.7907 | 54 | 0.8889 | 51 | 0.8431 | 231 | 0.8528 |
| 9_1200100 | 53 | 0.0755 | 90 | 0.0667 | 83 | 0.0602 | 67 | 0.0896 | 293 | 0.0717 | 82 | 0.061 | 43 | 0.0698 | 56 | 0.0179 | 54 | 0.0556 | 235 | 0.0511 |
| 9_1200101 | 57 | 0.3509 | 87 | 0.3448 | 87 | 0.3793 | 70 | 0.2143 | 301 | 0.3256 | 84 | 0.2262 | 46 | 0.2391 | 54 | 0.2778 | 57 | 0.2982 | 241 | 0.2573 |
| 9_1200105 | 4 | 0 | 8 | 0.125 | 7 | 0 | 6 | 0.1667 | 25 | 0.08 | 12 | 0 | 9 | 0 | 3 | 0 | 8 | 0.25 | 32 | 0.0625 |
| 9_1201866 | 1 | 1 | NA | NA | NA | NA | NA | NA | 1 | 1 | NA | NA | NA | NA | NA | NA | 1 | 1 | 1 | 1 |
| 9_1202615 | 3 | 0 | 2 | 0 | 2 | 0.5 | 2 | 0 | 9 | 0.1111 | 1 | 1 | 3 | 0 | NA | NA | NA | NA | 4 | 0.25 |
| 9_1202615 | 3 | 0.3333 | 2 | 0 | 1 | 0 | 2 | 0 | 8 | 0.125 | NA | NA | 6 | 0.1667 | NA | NA | 1 | 0 | 7 | 0.1429 |
| 9_1202617 | 5 | 0.4 | 1 | 1 | 2 | 0 | 2 | 0.5 | 10 | 0.4 | 1 | 0 | 6 | 0.6667 | NA | NA | 1 | 1 | 8 | 0.625 |
| 9_1202618 | 5 | 0 | 2 | 0.5 | 2 | 0 | 3 | 0.3333 | 12 | 0.1667 | 1 | 0 | 6 | 0.3333 | NA | NA | 1 | 0 | 8 | 0.25 |
| 9_1202705 | 4 | 0 | 2 | 1 | 3 | 0.6667 | 3 | 0.6667 | 12 | 0.5 | 1 | 1 | 7 | 0.7143 | NA | NA | 1 | 1 | 9 | 0.7778 |
| 9_1203022 | 5 | 0.2 | 4 | 0 | 2 | 0 | 3 | 0 | 14 | 0.0714 | 1 | 0 | 11 | 0.0909 | 2 | 0 | 1 | 0 | 15 | 0.0667 |
| 9_1203036 | 4 | 0 | 4 | 0.25 | 2 | 0 | 6 | 0 | 16 | 0.0625 | 2 | 0.5 | 9 | 0 | 3 | 0 | 2 | 0 | 16 | 0.0625 |
| 9_1203197 | NA | NA | 1 | 1 | NA | NA | NA | NA | 1 | 1 | NA | NA | NA | NA | 1 | 1 | NA | NA | 1 | 1 |
| 9_1203241 | 27 | 0.3333 | 28 | 0.3929 | 24 | 0.5 | 24 | 0.375 | 103 | 0.3981 | 24 | 0.5417 | 27 | 0.3333 | 28 | 0.3214 | 29 | 0.4483 | 108 | 0.4074 |
| 9_1203241 | 23 | 0.0435 | 21 | 0.0952 | 19 | 0.0526 | 17 | 0 | 80 | 0.05 | 18 | 0.1667 | 23 | 0.087 | 21 | 0.0476 | 26 | 0.0385 | 88 | 0.0795 |
| 9_1203955 | 2 | 0.5 | NA | NA | 3 | 0.3333 | 1 | 1 | 6 | 0.5 | 1 | 0 | 1 | 1 | 1 | 1 | 2 | 0 | 5 | 0.4 |
| 9_1203956 | 2 | 0.5 | NA | NA | 3 | 0.3333 | 1 | 0 | 6 | 0.3333 | 1 | 0 | 1 | 1 | 3 | 0.6667 | 1 | 0 | 6 | 0.5 |
| 9_1203956 | 2 | 1 | NA | NA | 3 | 1 | 2 | 1 | 7 | 1 | 1 | 1 | 1 | 1 | 3 | 0.6667 | 2 | 0.5 | 7 | 0.7143 |
| 9_1203957 | 2 | 1 | NA | NA | 3 | 1 | 1 | 1 | 6 | 1 | 1 | 1 | 1 | 1 | 3 | 1 | 2 | 0.5 | 7 | 0.8571 |
| 9_1203959 | 1 | 0 | NA | NA | 2 | 0.5 | 1 | 1 | 4 | 0.5 | 2 | 1 | 1 | 1 | 1 | 1 | 1 | 0 | 5 | 0.8 |
| 9_1203963 | 1 | 0 | NA | NA | 2 | 0.5 | 1 | 1 | 4 | 0.5 | 1 | 0 | 1 | 0 | 2 | 0.5 | 1 | 1 | 5 | 0.4 |
| 9_1203964 | 2 | 0 | NA | NA | 3 | 0.3333 | 3 | 0 | 8 | 0.125 | 2 | 0.5 | 1 | 0 | 3 | 0 | 1 | 1 | 7 | 0.2857 |
| 9_1203965 | 2 | 1 | NA | NA | 3 | 0.6667 | 3 | 0.6667 | 8 | 0.75 | 2 | 1 | NA | NA | 2 | 0.5 | 1 | 1 | 5 | 0.8 |
| 9_1203967 | 1 | 1 | NA | NA | 1 | 1 | NA | NA | 2 | 1 | 2 | 0.5 | 1 | 0 | NA | NA | 1 | 1 | 4 | 0.5 |
| 9_1204035 | NA | NA | NA | NA | NA | NA | 1 | 1 | 1 | 1 | 2 | 1 | 3 | 1 | NA | NA | NA | NA | 5 | 1 |
| 9_1204035 | NA | NA | 1 | 1 | NA | NA | 1 | 1 | 2 | 1 | 2 | 1 | 4 | 0 | NA | NA | NA | NA | 6 | 0.3333 |
| 9_1204412 | 1 | 1 | NA | NA | NA | NA | NA | NA | 1 | 1 | NA | NA | 1 | 1 | NA | NA | NA | NA | 1 | 1 |
| 9_1207846 | 31 | 0.0968 | 31 | 0.0645 | 22 | 0.2273 | 27 | 0.1111 | 111 | 0.1171 | 25 | 0.12 | 25 | 0.04 | 15 | 0.2667 | 19 | 0.1053 | 84 | 0.119 |
| 9_1207884 | 38 | 0.1579 | 40 | 0.175 | 31 | 0.1935 | 32 | 0.25 | 141 | 0.1915 | 33 | 0.2121 | 38 | 0.2105 | 26 | 0.3077 | 29 | 0.2759 | 126 | 0.246 |
| 9_1207884 | 38 | 0.1842 | 38 | 0.0526 | 30 | 0.1 | 34 | 0.1176 | 140 | 0.1143 | 31 | 0.129 | 36 | 0.0833 | 26 | 0.0769 | 29 | 0.069 | 122 | 0.0902 |
| 9_1207885 | 40 | 0.15 | 43 | 0.093 | 33 | 0.1212 | 38 | 0.3684 | 154 | 0.1818 | 39 | 0.1282 | 39 | 0.1282 | 28 | 0.25 | 29 | 0.1724 | 135 | 0.163 |
| 9_1207886 | 43 | 0.0465 | 46 | 0.087 | 35 | 0.0286 | 36 | 0.0833 | 160 | 0.0625 | 33 | 0.0303 | 36 | 0.1667 | 25 | 0.04 | 30 | 0.1667 | 124 | 0.1048 |
| 9_1207886 | 43 | 0.5814 | 47 | 0.7234 | 36 | 0.6389 | 36 | 0.6944 | 162 | 0.6605 | 32 | 0.3438 | 35 | 0.7143 | 25 | 0.8 | 33 | 0.7576 | 125 | 0.648 |
| 9_1207890 | 45 | 0.0444 | 62 | 0.1452 | 41 | 0.1463 | 44 | 0.0909 | 192 | 0.1094 | 43 | 0.0465 | 47 | 0.0851 | 37 | 0.0811 | 39 | 0.1538 | 166 | 0.0904 |
| 9_1207894 | 45 | 0.1111 | 52 | 0.1346 | 36 | 0.1111 | 29 | 0.2069 | 162 | 0.1358 | 29 | 0.069 | 38 | 0.0526 | 29 | 0.2069 | 40 | 0.2 | 136 | 0.1324 |
| 9_1212846 | 6 | 0 | 7 | 0.1429 | NA | NA | 1 | 0 | 14 | 0.0714 | 2 | 1 | 5 | 0 | 2 | 0 | 1 | 0 | 10 | 0.2 |
| 9_1212847 | 7 | 0 | 7 | 0 | NA | NA | 1 | 1 | 15 | 0.0667 | 2 | 1 | 5 | 0.4 | 2 | 0 | 3 | 0.3333 | 12 | 0.4167 |

| | | | | | | | | | | | | | | | | | | | | |
|---|---|---|---|---|---|---|---|---|---|---|---|---|---|---|---|---|---|---|---|---|
| 9_12128952 | 3 | 1 | 4 | 0.25 | 8 | 0.375 | 10 | 0.6 | 25 | 0.52 | 5 | 0 | 11 | 0.4545 | 12 | 0.3333 | 5 | 0.6 | 33 | 0.3636 |
| 9_12128966 | 2 | 0 | 1 | 0 | 3 | 0.3333 | 5 | 0.6 | 11 | 0.3636 | 1 | 0 | 4 | 0.75 | 3 | 0.3333 | 1 | 1 | 9 | 0.5556 |
| 9_12129102 | 1 | 1 | 1 | 1 | 1 | 1 | 4 | 0 | 7 | 0.4286 | NA | NA | 4 | 0.25 | 2 | 0 | NA | NA | 6 | 0.1667 |
| 9_12130138 | 4 | 0 | 8 | 0.25 | 8 | 0 | 8 | 0 | 28 | 0.0714 | 7 | 0.2857 | 9 | 0.1111 | 10 | 0.1 | 6 | 0.3333 | 32 | 0.1875 |
| 9_12130151 | 3 | 0.3333 | 4 | 0 | 8 | 0.25 | 6 | 0.1667 | 21 | 0.1905 | 7 | 0.2857 | 9 | 0.2222 | 8 | 0.25 | 2 | 0.5 | 26 | 0.2692 |
| 9_12130158 | 3 | 0 | 7 | 0.1429 | 8 | 0 | 8 | 0.125 | 26 | 0.0769 | 6 | 0.1667 | 9 | 0 | 8 | 0 | 3 | 0.3333 | 26 | 0.0769 |
| 9_12130160 | 2 | 0.5 | 7 | 0.1429 | 7 | 0.1429 | 7 | 0.4286 | 23 | 0.2609 | 6 | 0 | 8 | 0.125 | 8 | 0.25 | 6 | 0 | 28 | 0.1071 |
| 9_12130177 | 4 | 1 | 6 | 0.8333 | 5 | 1 | 4 | 1 | 19 | 0.9474 | 8 | 1 | 7 | 0.8571 | 7 | 1 | 4 | 0.75 | 26 | 0.9231 |
| 9_12130180 | 3 | 1 | 8 | 0.625 | 9 | 0.5556 | 7 | 0.7143 | 27 | 0.6667 | 8 | 0.75 | 8 | 0.5 | 11 | 0.2727 | 7 | 1 | 34 | 0.5882 |
| 9_12130183 | 4 | 0.75 | 7 | 0.4286 | 7 | 0.5714 | 5 | 0.6 | 23 | 0.5652 | 8 | 0.625 | 9 | 0.4444 | 9 | 0.7778 | 6 | 0.6667 | 32 | 0.625 |
| 9_12130184 | 5 | 0.2 | 8 | 0.125 | 8 | 0 | 5 | 0.4 | 26 | 0.1538 | 8 | 0.25 | 9 | 0.2222 | 8 | 0.125 | 6 | 0.1667 | 31 | 0.1935 |
| 9_12130185 | 5 | 0.8 | 9 | 0.8889 | 8 | 0.875 | 9 | 0.7778 | 31 | 0.8387 | 7 | 1 | 10 | 0.5 | 11 | 0.7273 | 7 | 0.7143 | 35 | 0.7143 |
| 9_12130186 | 5 | 1 | 8 | 0.75 | 6 | 0.8333 | 9 | 0.7778 | 28 | 0.8214 | 7 | 0.8571 | 8 | 0.5 | 10 | 1 | 5 | 1 | 30 | 0.8333 |
| 9_12130205 | 4 | 0.25 | 2 | 1 | 6 | 0.8333 | 7 | 1 | 19 | 0.7895 | 5 | 1 | 8 | 0.625 | 6 | 0.6667 | 6 | 1 | 25 | 0.8 |
| 9_12130219 | 5 | 0.8 | 7 | 0.2857 | 9 | 0.5556 | 9 | 0.4444 | 30 | 0.5 | 6 | 0.1667 | 11 | 0.1818 | 11 | 0.2727 | 7 | 0.5714 | 35 | 0.2857 |
| 9_12130222 | 6 | 1 | 10 | 0.9 | 10 | 1 | 9 | 1 | 35 | 0.9714 | 8 | 1 | 14 | 1 | 12 | 1 | 8 | 1 | 42 | 1 |
| 9_12130222 | 6 | 0.3333 | 9 | 0.5556 | 10 | 0.5 | 9 | 0.5556 | 34 | 0.5 | 7 | 0.7143 | 15 | 0.7333 | 12 | 0.8333 | 8 | 0.75 | 42 | 0.7619 |
| 9_12130241 | 5 | 1 | 10 | 0.6 | 7 | 1 | 8 | 1 | 30 | 0.8667 | 5 | 1 | 11 | 0.9091 | 8 | 1 | 6 | 1 | 30 | 0.9667 |
| 9_12130250 | 6 | 0.8333 | 10 | 1 | 9 | 1 | 7 | 1 | 32 | 0.9688 | 8 | 1 | 12 | 0.8333 | 11 | 1 | 7 | 1 | 38 | 0.9474 |
| 9_12130260 | 7 | 0 | 13 | 0.0769 | 10 | 0.3 | 9 | 0.2222 | 39 | 0.1538 | 9 | 0.1111 | 13 | 0 | 13 | 0.1538 | 6 | 0 | 41 | 0.0732 |
| 9_12130285 | 8 | 0.125 | 12 | 0.1667 | 11 | 0.1818 | 8 | 0.125 | 39 | 0.1538 | 8 | 0 | 16 | 0.125 | 13 | 0 | 8 | 0.5 | 45 | 0.1333 |
| 9_12130292 | 9 | 0.3333 | 14 | 0.2857 | 11 | 0.6364 | 9 | 0.2222 | 43 | 0.3721 | 8 | 0.125 | 15 | 0.1333 | 13 | 0.1538 | 8 | 0.5 | 44 | 0.2045 |
| 9_12130377 | 12 | 0.0833 | 18 | 0.5 | 15 | 0.2 | 12 | 0.25 | 57 | 0.2807 | 12 | 0.1667 | 19 | 0.2105 | 16 | 0.1875 | 9 | 0.5556 | 56 | 0.25 |
| 9_12131894 | 56 | 0.0179 | 99 | 0.0707 | 54 | 0.0741 | 14 | 0.0714 | 223 | 0.0583 | 98 | 0.051 | 21 | 0.1429 | 24 | 0.0833 | 58 | 0.0172 | 201 | 0.0547 |
| 9_12131915 | 50 | 0.08 | 90 | 0.1111 | 44 | 0.0682 | 15 | 0.1333 | 199 | 0.0955 | 84 | 0.0476 | 19 | 0.1053 | 20 | 0.15 | 52 | 0.0962 | 175 | 0.08 |
| 9_12131919 | 31 | 0 | 54 | 0.0741 | 30 | 0.1 | 9 | 0 | 124 | 0.0565 | 50 | 0.1 | 10 | 0 | 14 | 0.0714 | 36 | 0.0278 | 110 | 0.0636 |
| 9_12132360 | 2 | 0 | 4 | 0.5 | NA | NA | NA | NA | 6 | 0.3333 | 2 | 1 | NA | NA | NA | NA | 4 | 0.5 | 6 | 0.6667 |
| 9_12132382 | 4 | 0.25 | 4 | 0 | NA | NA | NA | NA | 8 | 0.125 | 2 | 0 | NA | NA | NA | NA | 3 | 0.6667 | 5 | 0.4 |
| 9_12132498 | NA | NA | 1 | 1 | NA | NA | NA | NA | 1 | 1 | 2 | 1 | NA | NA | NA | NA | NA | NA | 2 | 1 |
| 9_12149872 | NA | NA | 3 | 0.3333 | 1 | 1 | NA | NA | 4 | 0.5 | 1 | 1 | NA | NA | 1 | 1 | 2 | 1 | 4 | 1 |
| 9_12149875 | NA | NA | 3 | 1 | 1 | 0 | NA | NA | 4 | 0.75 | 1 | 1 | NA | NA | 1 | 1 | 2 | 1 | 4 | 1 |
| 9_12149882 | NA | NA | 2 | 1 | 1 | 1 | NA | NA | 3 | 1 | 1 | 1 | NA | NA | 1 | 1 | 1 | 0 | 3 | 0.6667 |
| 9_12149891 | NA | NA | 2 | 1 | 1 | 1 | NA | NA | 3 | 1 | 1 | 1 | NA | NA | 1 | 1 | 2 | 1 | 4 | 1 |
| 9_12149899 | NA | NA | 3 | 0.6667 | 1 | 1 | NA | NA | 4 | 0.75 | NA | NA | NA | NA | 1 | 0 | 1 | 1 | 2 | 0.5 |
| 9_12149918 | NA | NA | 2 | 0.5 | 1 | 0 | 1 | 0 | 4 | 0.25 | 1 | 1 | NA | NA | 1 | 0 | 2 | 0.5 | 4 | 0.5 |
| 9_12149924 | NA | NA | 3 | 0.6667 | 1 | 1 | 1 | 1 | 5 | 0.8 | 1 | 0 | NA | NA | 1 | 0 | 2 | 1 | 4 | 0.5 |
| 9_12150100 | 2 | 0.5 | 4 | 0 | 2 | 0 | 3 | 0 | 11 | 0.0909 | 2 | 0 | 2 | 0.5 | 3 | 0 | 4 | 0.25 | 11 | 0.1818 |
| 9_12150165 | 2 | 0 | 4 | 0.25 | 2 | 0 | 3 | 0 | 11 | 0.0909 | 1 | 1 | 2 | 0 | 4 | 0 | 4 | 0 | 11 | 0.0909 |
| 9_12151148 | 1 | 0 | 1 | 0 | 1 | 0 | 4 | 0.25 | 7 | 0.1429 | 1 | 0 | 2 | 0 | 3 | 0.3333 | 4 | 0 | 10 | 0.1 |
| 9_12151193 | 2 | 0 | 1 | 0 | 3 | 0.3333 | 4 | 0 | 10 | 0.1 | 1 | 1 | 3 | 0.3333 | 2 | 1 | 1 | 0 | 7 | 0.5714 |
| 9_12151208 | 2 | 0 | 1 | 0 | 3 | 0.3333 | 4 | 0 | 10 | 0.1 | 1 | 0 | 3 | 0 | 1 | 1 | 1 | 0 | 6 | 0.1667 |
| 9_12151239 | 2 | 1 | 1 | 0 | 4 | 0.25 | 4 | 0 | 11 | 0.2727 | 1 | 1 | 3 | 0.3333 | 3 | 0 | 3 | 0.3333 | 10 | 0.3 |
| 9_12151338 | 2 | 0 | 1 | 0 | 1 | 0 | 3 | 0.3333 | 7 | 0.1429 | NA | NA | 3 | 0.3333 | 1 | 0 | 1 | 0 | 5 | 0.2 |
| 9_12151339 | 2 | 0 | 1 | 0 | 1 | 0 | 2 | 0.5 | 6 | 0.1667 | NA | NA | 2 | 0 | 2 | 0.5 | 3 | 0.6667 | 7 | 0.4286 |
| 9_12151351 | 2 | 0 | 1 | 1 | 1 | 0 | 2 | 0 | 6 | 0.1667 | NA | NA | 3 | 1 | 2 | 0.5 | 2 | 0.5 | 7 | 0.7143 |
| 9_12151369 | 2 | 0.5 | NA | NA | NA | NA | 2 | 0.5 | 4 | 0.5 | NA | NA | 3 | 0.6667 | 1 | 1 | 3 | 0.3333 | 7 | 0.5714 |
| 9_12151373 | 2 | 1 | 1 | 1 | NA | NA | 3 | 0.3333 | 6 | 0.6667 | NA | NA | 3 | 1 | 1 | 1 | 2 | 0.5 | 6 | 0.8333 |
| 9_12153959 | 3 | 0.6667 | 4 | 0.75 | 6 | 0.1667 | 4 | 0.75 | 17 | 0.5294 | 1 | 0 | 5 | 0.4 | 2 | 0 | 4 | 0.5 | 12 | 0.3333 |
| 9_12154174 | 11 | 0.7273 | 13 | 0.6923 | 17 | 0.7647 | 18 | 0.6667 | 59 | 0.7119 | 10 | 0.5 | 15 | 0.8 | 9 | 0.7778 | 17 | 0.8235 | 51 | 0.7451 |
| 9_12154181 | 10 | 0.7 | 14 | 0.6429 | 17 | 0.7059 | 18 | 0.4444 | 59 | 0.6102 | 11 | 0.6364 | 15 | 0.8 | 9 | 0.7778 | 17 | 0.5882 | 52 | 0.6923 |
| 9_12154185 | 10 | 0.2 | 15 | 0.2667 | 17 | 0.1765 | 19 | 0.1579 | 61 | 0.1967 | 11 | 0.2727 | 17 | 0.3529 | 9 | 0.2222 | 17 | 0.3529 | 54 | 0.3148 |
| 9_12154190 | 8 | 0 | 13 | 0.0769 | 15 | 0.0667 | 19 | 0.0526 | 55 | 0.0545 | 9 | 0 | 13 | 0.0769 | 9 | 0.1111 | 14 | 0.0714 | 45 | 0.0667 |
| 9_12154564 | 1 | 0 | 3 | 0.3333 | 1 | 1 | 1 | 0 | 6 | 0.3333 | 1 | 1 | 3 | 0.3333 | NA | NA | NA | NA | 4 | 0.5 |
| 9_12154578 | NA | NA | 3 | 0.6667 | 1 | 0 | NA | NA | 4 | 0.5 | 1 | 0 | 2 | 1 | NA | NA | NA | NA | 3 | 0.6667 |
| 9_12154584 | 1 | 1 | 3 | 0.6667 | 1 | 1 | 1 | 1 | 6 | 0.8333 | 1 | 1 | 3 | 1 | NA | NA | NA | NA | 4 | 1 |
| 9_12157748 | 1 | 1 | 1 | 1 | 2 | 1 | 1 | 0 | 5 | 0.8 | 1 | 1 | 1 | 1 | 1 | 1 | 1 | 1 | 4 | 1 |
| 9_12157859 | 2 | 0.5 | 1 | 0 | 2 | 0 | 2 | 0 | 7 | 0.1429 | 2 | 0.5 | 1 | 0 | 2 | 1 | 2 | 0 | 7 | 0.4286 |
| 9_12157916 | 2 | 0.5 | 2 | 0.5 | 3 | 0.3333 | 2 | 0 | 9 | 0.3333 | 2 | 0 | 2 | 0.5 | 3 | 0 | 2 | 0.5 | 9 | 0.2222 |
| 9_12157932 | 2 | 0.5 | 3 | 0.3333 | 3 | 0 | 1 | 0 | 9 | 0.2222 | 1 | 0 | 3 | 1 | 5 | 1 | 2 | 1 | 11 | 0.9091 |
| 9_12157946 | 2 | 1 | 2 | 1 | 2 | 1 | 1 | 1 | 7 | 1 | 2 | 1 | 2 | 1 | 5 | 0.8 | 3 | 1 | 12 | 0.9167 |
| 9_12157949 | 2 | 0 | 3 | 0 | 3 | 0.6667 | 2 | 0 | 10 | 0.2 | 1 | 0 | 3 | 0.3333 | 3 | 0 | 3 | 0.3333 | 10 | 0.2 |
| 9_12157952 | 2 | 0.5 | 3 | 0 | 3 | 0.3333 | 2 | 0 | 10 | 0.2 | 2 | 0 | 3 | 0.6667 | 5 | 0 | 3 | 0.3333 | 13 | 0.2308 |
| 9_12158439 | 2 | 0 | 4 | 0.25 | 4 | 0 | 2 | 0 | 12 | 0.0833 | 6 | 0.3333 | 6 | 0.3333 | 9 | 0 | 4 | 0 | 25 | 0.16 |
| 9_12158526 | 4 | 0 | 6 | 0 | 8 | 0.125 | 5 | 0.2 | 23 | 0.087 | 5 | 0 | 9 | 0 | 13 | 0.1538 | 5 | 0 | 32 | 0.0625 |
| 9_12158535 | 5 | 0 | 6 | 0.1667 | 8 | 0.125 | 4 | 0 | 23 | 0.087 | 6 | 0.1667 | 10 | 0 | 14 | 0 | 5 | 0.2 | 35 | 0.0571 |

| | | | | | | | | | | | | | | | | | | | | |
|---|---|---|---|---|---|---|---|---|---|---|---|---|---|---|---|---|---|---|---|---|
| 9_1215854 | 5 | 0 | 7 | 0.5714 | 9 | 0 | 4 | 0.25 | 25 | 0.2 | 5 | 0.2 | 9 | 0.4444 | 14 | 0.3571 | 5 | 0.2 | 33 | 0.3333 |
| 9_1215875 | 1 | 1 | NA | NA | NA | NA | NA | NA | 1 | 1 | 2 | 1 | NA | NA | NA | NA | 1 | 1 | 3 | 1 |
| 9_1215878 | 1 | 0 | NA | NA | 2 | 0.5 | NA | NA | 3 | 0.3333 | 1 | 1 | NA | NA | NA | NA | 1 | 0 | 2 | 0.5 |
| 9_1215883 | 1 | 1 | NA | NA | 2 | 0.5 | NA | NA | 3 | 0.6667 | 2 | 1 | NA | NA | NA | NA | 1 | 1 | 3 | 1 |
| 9_1215883 | 2 | 0 | NA | NA | 1 | 1 | NA | NA | 3 | 0.3333 | 2 | 0.5 | NA | NA | NA | NA | NA | NA | 2 | 0.5 |
| 9_1215886 | 2 | 0.5 | NA | NA | 2 | 0 | NA | NA | 4 | 0.25 | 1 | 0 | NA | NA | NA | NA | 1 | 1 | 2 | 0.5 |
| 9_1215886 | 2 | 1 | NA | NA | 2 | 1 | NA | NA | 4 | 1 | 2 | 1 | NA | NA | NA | NA | 1 | 1 | 3 | 1 |
| 9_1215923 | 1 | 1 | NA | NA | 2 | 0 | NA | NA | 3 | 0.3333 | 3 | 0.3333 | NA | NA | NA | NA | 5 | 0.8 | 8 | 0.625 |
| 9_1215924 | 1 | 1 | NA | NA | 1 | 0 | NA | NA | 2 | 0.5 | 2 | 0.5 | NA | NA | NA | NA | 4 | 0.75 | 6 | 0.6667 |
| 9_1215925 | 1 | 0 | NA | NA | 2 | 0.5 | NA | NA | 3 | 0.3333 | 3 | 0.6667 | NA | NA | NA | NA | 4 | 0.75 | 7 | 0.7143 |
| 9_1215925 | 1 | 1 | 1 | 0 | 2 | 0.5 | NA | NA | 4 | 0.5 | 1 | 1 | NA | NA | 1 | 0 | 3 | 1 | 5 | 0.8 |
| 9_1215928 | 3 | 0.3333 | 1 | 0 | 2 | 1 | 4 | 0 | 10 | 0.3 | 1 | 0 | 2 | 0.5 | 2 | 0 | 4 | 0.5 | 9 | 0.3333 |
| 9_1215931 | 4 | 0 | 1 | 0 | 3 | 0.3333 | 5 | 0.4 | 13 | 0.2308 | 4 | 0.5 | 2 | 0 | 3 | 0.3333 | 5 | 0 | 14 | 0.2143 |
| 9_1215945 | 4 | 0 | 1 | 0 | 3 | 0 | 4 | 0.25 | 12 | 0.0833 | 3 | 0.3333 | 2 | 0 | 3 | 0.3333 | 5 | 0 | 13 | 0.1538 |
| 9_1215968 | 4 | 1 | 1 | 1 | 3 | 1 | 3 | 1 | 11 | 1 | 5 | 1 | 1 | 1 | 3 | 1 | 7 | 1 | 16 | 1 |
| 9_1215968 | 3 | 0.3333 | 1 | 0 | 4 | 0.75 | 4 | 0.25 | 12 | 0.4167 | 5 | 0.4 | NA | NA | 3 | 0.3333 | 5 | 0.6 | 13 | 0.4615 |
| 9_1215968 | 5 | 0.2 | 1 | 0 | 3 | 0.3333 | 5 | 0.2 | 14 | 0.2143 | 4 | 0.25 | 2 | 0 | 4 | 0.25 | 7 | 0.1429 | 17 | 0.1765 |
| 9_1215968 | 5 | 0.6 | 1 | 1 | 3 | 1 | 5 | 0.2 | 14 | 0.5714 | 5 | 0.4 | 2 | 1 | 3 | 0 | 5 | 0.2 | 15 | 0.3333 |
| 9_1215969 | 1 | 0 | 1 | 1 | 3 | 0.6667 | 3 | 0.3333 | 8 | 0.5 | 4 | 0.5 | 1 | 1 | 3 | 0 | 7 | 0.7143 | 15 | 0.5333 |
| 9_1215969 | 1 | 1 | 1 | 1 | 4 | 1 | 4 | 1 | 10 | 1 | 2 | 1 | 2 | 1 | 1 | 1 | 5 | 1 | 10 | 1 |
| 9_1215971 | 5 | 0.4 | 2 | 0.5 | 2 | 0.5 | 3 | 0.3333 | 12 | 0.4167 | 2 | 0.5 | 2 | 0.5 | 4 | 1 | 6 | 0.5 | 14 | 0.6429 |
| 9_1215972 | 1 | 0 | 2 | 0 | 4 | 0.5 | 5 | 0 | 12 | 0.1667 | 2 | 0.5 | 2 | 0 | 4 | 0.25 | 5 | 0.2 | 13 | 0.2308 |
| 9_1215973 | 4 | 0 | 3 | 0 | 4 | 0.5 | 7 | 0.4286 | 18 | 0.2778 | 6 | 0.1667 | 1 | 0 | 5 | 0.2 | 6 | 0.3333 | 18 | 0.2222 |
| 9_1215980 | 9 | 0.1111 | 6 | 0 | 5 | 0.2 | 6 | 0.3333 | 26 | 0.1538 | 3 | 0 | 3 | 0 | 4 | 0 | 7 | 0.1429 | 17 | 0.0588 |
| 9_1215980 | 7 | 0.2857 | 4 | 1 | 4 | 0.25 | 5 | 0 | 20 | 0.35 | 3 | 0 | 6 | 0 | 5 | 0 | 9 | 0.4444 | 23 | 0.1739 |
| 9_1216004 | NA | NA | 1 | 1 | NA | NA | 1 | 1 | 2 | 1 | NA | NA | NA | NA | 1 | 1 | NA | NA | 1 | 1 |
| 9_1216079 | 1 | 1 | NA | NA | 1 | 1 | 1 | 1 | 3 | 1 | NA | NA | 1 | 0 | NA | NA | 1 | 1 | 2 | 0.5 |
| 9_1216090 | NA | NA | NA | NA | 1 | 1 | 1 | 1 | 2 | 1 | NA | NA | 1 | 1 | NA | NA | 1 | 0 | 2 | 0.5 |
| 9_1216126 | NA | NA | NA | NA | 1 | 1 | 1 | 0 | 2 | 0.5 | NA | NA | NA | NA | NA | NA | 1 | 1 | 1 | 1 |
| 9_1216126 | 1 | 1 | NA | NA | 1 | 1 | 1 | 1 | 3 | 1 | NA | NA | 1 | 1 | NA | NA | 1 | 1 | 2 | 1 |
| 9_1216127 | 1 | 1 | NA | NA | 1 | 1 | 1 | 1 | 3 | 1 | NA | NA | NA | NA | NA | NA | 1 | 1 | 1 | 1 |
| 9_1216130 | 1 | 0 | 5 | 0 | 1 | 1 | 3 | 0.3333 | 10 | 0.2 | 3 | 0.3333 | 3 | 0 | 1 | 0 | 3 | 0.3333 | 10 | 0.2 |
| 9_1216135 | NA | NA | 1 | 1 | NA | NA | NA | NA | 1 | 1 | NA | NA | NA | NA | NA | NA | 1 | 1 | 1 | 1 |
| 9_1216146 | NA | NA | 1 | 1 | NA | NA | NA | NA | 1 | 1 | NA | NA | NA | NA | NA | NA | 1 | 1 | 1 | 1 |
| 9_1217380 | 1 | 1 | NA | NA | NA | NA | NA | NA | 1 | 1 | NA | NA | 1 | 1 | NA | NA | NA | NA | 1 | 1 |
| 9_1217382 | 1 | 1 | NA | NA | NA | NA | NA | NA | 1 | 1 | NA | NA | 1 | 1 | NA | NA | NA | NA | 1 | 1 |
| 9_1217390 | 1 | 1 | NA | NA | NA | NA | NA | NA | 1 | 1 | NA | NA | 1 | 1 | NA | NA | NA | NA | 1 | 1 |
| 9_1217393 | 1 | 1 | NA | NA | NA | NA | NA | NA | 1 | 1 | NA | NA | 1 | 1 | NA | NA | NA | NA | 1 | 1 |
| 9_1217395 | 1 | 1 | NA | NA | NA | NA | NA | NA | 1 | 1 | NA | NA | 1 | 1 | NA | NA | NA | NA | 1 | 1 |
| 9_1217396 | 1 | 1 | NA | NA | NA | NA | NA | NA | 1 | 1 | NA | NA | 1 | 1 | NA | NA | NA | NA | 1 | 1 |
| 9_1217398 | 1 | 1 | NA | NA | NA | NA | NA | NA | 1 | 1 | NA | NA | 1 | 1 | NA | NA | NA | NA | 1 | 1 |
| 9_1217400 | 1 | 1 | NA | NA | NA | NA | NA | NA | 1 | 1 | NA | NA | 1 | 1 | NA | NA | NA | NA | 1 | 1 |
| 9_1217756 | 1 | 1 | 1 | 0 | 1 | 0 | 4 | 0.25 | 7 | 0.2857 | NA | NA | NA | NA | 2 | 0.5 | 2 | 0 | 4 | 0.25 |
| 9_1217939 | 1 | 0 | NA | NA | 1 | 1 | 1 | 1 | 3 | 0.6667 | NA | NA | 1 | 1 | NA | NA | 1 | 0 | 2 | 0.5 |
| 9_1217939 | 1 | 0 | NA | NA | 1 | 0 | 1 | 1 | 3 | 0.3333 | NA | NA | 1 | 1 | NA | NA | 1 | 1 | 2 | 1 |
| 9_1217946 | 2 | 1 | NA | NA | 1 | 1 | 1 | 1 | 4 | 1 | NA | NA | 1 | 1 | NA | NA | NA | NA | 1 | 1 |
| 9_1217947 | 2 | 0 | NA | NA | 1 | 0 | 1 | 1 | 4 | 0.25 | NA | NA | 1 | 1 | NA | NA | NA | NA | 1 | 1 |
| 9_1217947 | 2 | 0.5 | NA | NA | 1 | 0 | 1 | 0 | 4 | 0.25 | NA | NA | 1 | 1 | NA | NA | NA | NA | 1 | 1 |
| 9_1217948 | 3 | 0 | NA | NA | 1 | 1 | 1 | 0 | 5 | 0.2 | NA | NA | 1 | 1 | NA | NA | NA | NA | 1 | 1 |
| 9_1217949 | 3 | 0.3333 | NA | NA | 1 | 0 | 1 | 0 | 5 | 0.2 | NA | NA | 1 | 1 | NA | NA | NA | NA | 1 | 1 |
| 9_1217949 | 3 | 1 | NA | NA | 1 | 1 | 1 | 1 | 5 | 1 | NA | NA | 1 | 1 | NA | NA | NA | NA | 1 | 1 |
| 9_1218074 | 2 | 0.5 | NA | NA | 1 | 0 | NA | NA | 3 | 0.3333 | NA | NA | 1 | 1 | NA | NA | NA | NA | 1 | 1 |
| 9_1218074 | 2 | 1 | NA | NA | 1 | 1 | 1 | 1 | 4 | 1 | NA | NA | 1 | 1 | NA | NA | 1 | 1 | 2 | 1 |
| 9_1218075 | 2 | 1 | NA | NA | 1 | 0 | NA | NA | 3 | 0.6667 | NA | NA | 1 | 0 | NA | NA | 1 | 1 | 2 | 0.5 |
| 9_1218080 | 2 | 1 | NA | NA | 1 | 0 | 1 | 1 | 4 | 0.75 | NA | NA | 1 | 1 | NA | NA | 1 | 0 | 2 | 0.5 |
| 9_1218082 | 2 | 0.5 | NA | NA | 1 | 1 | 1 | 0 | 4 | 0.5 | NA | NA | 1 | 1 | NA | NA | 1 | 0 | 2 | 0.5 |
| 9_1218082 | 2 | 1 | NA | NA | 1 | 1 | 1 | 1 | 4 | 1 | NA | NA | 1 | 1 | NA | NA | 1 | 1 | 2 | 1 |
| 9_1218083 | 2 | 1 | NA | NA | NA | NA | 1 | 0 | 3 | 0.6667 | NA | NA | 1 | 1 | NA | NA | 1 | 0 | 2 | 0.5 |
| 9_1218085 | 2 | 0.5 | NA | NA | 1 | 0 | 1 | 1 | 4 | 0.5 | NA | NA | 1 | 1 | NA | NA | NA | NA | 1 | 1 |
| 9_1218090 | 2 | 0.5 | NA | NA | 1 | 1 | 1 | 0 | 4 | 0.5 | NA | NA | 1 | 1 | NA | NA | 1 | 0 | 2 | 0.5 |
| 9_1218110 | 3 | 0.6667 | 6 | 1 | 3 | 0.6667 | 6 | 0.8333 | 18 | 0.8333 | 6 | 1 | 5 | 0.8 | 7 | 0.8571 | 4 | 0.75 | 22 | 0.8636 |
| 9_1218111 | 4 | 0.25 | 5 | 0.6 | 4 | 0.25 | 5 | 0.4 | 18 | 0.3889 | 4 | 0.5 | 5 | 0.8 | 6 | 0.3333 | 5 | 0.6 | 20 | 0.55 |
| 9_1218111 | 4 | 0.25 | 5 | 0 | 3 | 0 | 7 | 0.1429 | 19 | 0.1053 | 5 | 0 | 6 | 0 | 8 | 0.125 | 4 | 0.25 | 23 | 0.087 |
| 9_1218114 | 4 | 0.25 | 6 | 0.6667 | 4 | 0.25 | 7 | 0.4286 | 21 | 0.4286 | 6 | 0.3333 | 4 | 0.25 | 7 | 0.1429 | 5 | 0.4 | 22 | 0.2727 |
| 9_1218114 | 2 | 0.5 | 4 | 0.75 | 3 | 1 | 3 | 1 | 12 | 0.8333 | 3 | 0.6667 | 3 | 1 | 8 | 1 | 4 | 0.75 | 18 | 0.8889 |

| | | | | | | | | | | | | | | | | | | | | |
|---|---|---|---|---|---|---|---|---|---|---|---|---|---|---|---|---|---|---|---|---|
| 9_12181152 | 1 | 0 | 1 | 1 | NA | NA | 5 | 0.4 | 7 | 0.4286 | 3 | 0.6667 | NA | NA | 2 | 0.5 | 2 | 0 | 7 | 0.4286 |
| 9_12181155 | 3 | 0.6667 | 5 | 0.8 | 4 | 1 | 7 | 0.7143 | 19 | 0.7895 | 4 | 0.5 | 5 | 0.8 | 7 | 0.4286 | 4 | 1 | 20 | 0.65 |
| 9_12181158 | 4 | 0 | 5 | 0.4 | 4 | 0.5 | 5 | 0 | 18 | 0.2222 | 5 | 0.2 | 4 | 0 | 6 | 0.3333 | 5 | 0.4 | 20 | 0.25 |
| 9_12181162 | 4 | 1 | 5 | 1 | 3 | 0.6667 | 7 | 1 | 19 | 0.9474 | 6 | 1 | 4 | 1 | 7 | 0.8571 | 3 | 1 | 20 | 0.95 |
| 9_12181169 | 4 | 1 | 4 | 1 | 4 | 1 | 6 | 1 | 18 | 1 | 5 | 1 | 5 | 1 | 6 | 1 | 2 | 1 | 18 | 1 |
| 9_12181180 | 3 | 0 | 6 | 0.1667 | 2 | 0 | 7 | 0.2857 | 18 | 0.1667 | 5 | 0 | 4 | 0.25 | 4 | 0 | 3 | 0 | 16 | 0.0625 |
| 9_12181188 | 2 | 1 | 4 | 0.5 | 3 | 0.3333 | 7 | 0.4286 | 16 | 0.5 | 5 | 0.4 | 4 | 0.5 | 5 | 0.8 | 2 | 1 | 16 | 0.625 |
| 9_12181196 | 1 | 1 | 1 | 0 | 3 | 1 | 7 | 0.8571 | 12 | 0.8333 | 4 | 0.75 | 4 | 0.25 | 4 | 1 | 2 | 1 | 14 | 0.7143 |
| 9_12181211 | 4 | 0.75 | 5 | 0.6 | 2 | 0.5 | 7 | 0.2857 | 18 | 0.5 | 4 | 0.25 | 4 | 0.25 | 6 | 0.1667 | 2 | 0 | 16 | 0.1875 |
| 9_12181215 | 4 | 0.75 | 3 | 0 | 2 | 0 | 7 | 0.4286 | 16 | 0.375 | 3 | 1 | 5 | 0.6 | 4 | 0.5 | 1 | 0 | 13 | 0.6154 |
| 9_12181216 | 3 | 0 | 6 | 0.3333 | 2 | 0 | 5 | 0.4 | 16 | 0.25 | 5 | 0.2 | 5 | 0.2 | 6 | 0 | 1 | 0 | 17 | 0.1176 |
| 9_12181240 | 3 | 0.3333 | 4 | 0.25 | 2 | 0 | 5 | 0.4 | 14 | 0.2857 | 4 | 0.75 | 5 | 0.6 | 6 | 0.5 | 2 | 0 | 17 | 0.5294 |
| 9_12181242 | 4 | 0.75 | 2 | 0.5 | 2 | 0.5 | 6 | 1 | 14 | 0.7857 | 4 | 0.75 | 5 | 1 | 4 | 0.5 | 2 | 1 | 15 | 0.8 |
| 9_12181255 | 4 | 1 | 4 | 1 | 2 | 0.5 | 6 | 1 | 16 | 0.9375 | 3 | 1 | 3 | 0.6667 | 5 | 1 | 2 | 1 | 13 | 0.9231 |
| 9_12181263 | 4 | 0.25 | 4 | 0.5 | 2 | 0 | 5 | 0.2 | 15 | 0.2667 | 3 | 1 | 4 | 0.75 | 4 | 0.25 | 2 | 0 | 13 | 0.5385 |
| 9_12181266 | 3 | 0.6667 | 2 | 1 | NA | NA | 4 | 0.75 | 9 | 0.7778 | 2 | 0.5 | 2 | 0.5 | 3 | 0.6667 | 2 | 1 | 9 | 0.6667 |
| 9_12181682 | 1 | 1 | NA | NA | 1 | 1 | 2 | 0 | 4 | 0.5 | NA | NA | NA | NA | 1 | 1 | NA | NA | 1 | 1 |
| 9_12181698 | 1 | 1 | NA | NA | 1 | 0 | 2 | 0.5 | 4 | 0.5 | NA | NA | NA | NA | 1 | 1 | NA | NA | 1 | 1 |
| 9_12196613 | 2 | 0 | 2 | 1 | 2 | 0.5 | NA | NA | 6 | 0.5 | NA | NA | NA | NA | 1 | 1 | 2 | 0.5 | 3 | 0.6667 |
| 9_12196623 | 1 | 1 | 1 | 1 | 3 | 0.6667 | 1 | 0 | 6 | 0.6667 | 1 | 1 | 1 | 1 | 1 | 1 | 2 | 0.5 | 5 | 0.8 |
| 9_12196628 | 2 | 0.5 | 2 | 0 | 3 | 0 | 2 | 0 | 9 | 0.1111 | 1 | 1 | 1 | 0 | 1 | 0 | 2 | 0 | 5 | 0.2 |
| 9_12196633 | 2 | 0.5 | 1 | 1 | 3 | 0 | 2 | 1 | 8 | 0.5 | 1 | 0 | 1 | 0 | 1 | 0 | 2 | 0.5 | 5 | 0.2 |
| 9_12196635 | 2 | 1 | 2 | 0.5 | 1 | 1 | 2 | 0.5 | 7 | 0.7143 | 1 | 1 | 1 | 1 | 1 | 1 | 1 | 1 | 4 | 1 |
| 9_12196641 | 2 | 0 | 3 | 0 | 3 | 0 | 2 | 0.5 | 10 | 0.1 | 1 | 0 | 1 | 0 | 1 | 1 | 2 | 0 | 5 | 0.2 |
| 9_12196649 | 3 | 1 | 3 | 1 | 3 | 1 | 2 | 1 | 11 | 1 | 1 | 1 | 2 | 1 | 1 | 1 | 3 | 0.6667 | 7 | 0.8571 |
| 9_12196655 | 3 | 0.6667 | 3 | 0.6667 | 3 | 0.6667 | 2 | 1 | 11 | 0.7273 | 1 | 1 | 2 | 0 | 1 | 1 | 3 | 0.3333 | 7 | 0.4286 |
| 9_12199576 | 2 | 1 | 4 | 1 | 6 | 1 | 3 | 1 | 15 | 1 | 2 | 1 | 1 | 1 | 1 | 1 | 3 | 0.6667 | 7 | 0.8571 |
| 9_12199588 | 3 | 0 | 4 | 0 | 6 | 0.1667 | 2 | 0.5 | 15 | 0.1333 | 2 | 0.5 | 5 | 0 | 3 | 0 | 4 | 0.5 | 14 | 0.2143 |
| 9_12199589 | 2 | 1 | 3 | 1 | 5 | 1 | 4 | 0.75 | 14 | 0.9286 | 3 | 0.6667 | 4 | 0.75 | 3 | 1 | 2 | 0.5 | 12 | 0.75 |
| 9_12199592 | 3 | 0 | 4 | 0.25 | 6 | 0 | 3 | 0 | 16 | 0.0625 | 3 | 0 | 3 | 0 | 3 | 0.3333 | 3 | 0 | 12 | 0.0833 |
| 9_12199600 | 1 | 1 | 3 | 0.6667 | 5 | 0.4 | 2 | 0.5 | 11 | 0.5455 | 3 | 1 | 3 | 1 | 3 | 1 | 3 | 1 | 12 | 1 |
| 9_12199605 | 3 | 0.3333 | 4 | 0.5 | 5 | 0.4 | 4 | 0.25 | 16 | 0.375 | 3 | 0.6667 | 5 | 0.4 | 4 | 0.75 | 3 | 0 | 15 | 0.4667 |
| 9_12199610 | 3 | 0.3333 | 4 | 0 | 4 | 0.25 | 3 | 0 | 14 | 0.1429 | 3 | 0 | 4 | 0.5 | 5 | 0.2 | 3 | 0.3333 | 15 | 0.2667 |
| 9_12199617 | NA | NA | 4 | 0 | 5 | 0.2 | 2 | 0.5 | 11 | 0.1818 | 3 | 0 | 4 | 0.25 | 3 | 0.3333 | 4 | 0 | 14 | 0.1429 |
| 9_12199622 | 3 | 0.3333 | 5 | 0.4 | 6 | 0.5 | 4 | 0.75 | 18 | 0.5 | 3 | 0.6667 | 4 | 0.5 | 5 | 0.6 | 4 | 0.25 | 16 | 0.5 |
| 9_12199638 | 3 | 0 | 4 | 0 | 5 | 0.2 | 4 | 0.25 | 16 | 0.125 | 3 | 0.6667 | 5 | 0.6 | 4 | 0.5 | 4 | 0 | 16 | 0.4375 |
| 9_12199639 | 3 | 0 | 6 | 0 | 6 | 0.1667 | 2 | 0 | 17 | 0.0588 | 3 | 0 | 3 | 0.3333 | 4 | 0 | 4 | 0.25 | 14 | 0.1429 |
| 9_12199650 | 3 | 0 | 6 | 0.6667 | 6 | 0.3333 | 4 | 0.75 | 19 | 0.4737 | 2 | 0.5 | 4 | 0.75 | 5 | 0.2 | 4 | 0.75 | 15 | 0.5333 |
| 9_12199663 | 3 | 0.3333 | 6 | 0.3333 | 4 | 0.5 | 6 | 0.5 | 19 | 0.4211 | 2 | 1 | 2 | 1 | 5 | 0.2 | 4 | 0.25 | 13 | 0.4615 |
| 9_12199677 | 3 | 0 | 8 | 0.375 | 6 | 0.6667 | 5 | 0.6 | 22 | 0.4545 | 3 | 0 | 5 | 0.2 | 5 | 0.4 | 4 | 0.25 | 17 | 0.2353 |
| 9_12199678 | 3 | 0 | 6 | 0.3333 | 5 | 0.2 | 5 | 0.2 | 19 | 0.2105 | 1 | 0 | 3 | 0.6667 | 3 | 0 | 2 | 0 | 9 | 0.2222 |
| 9_12199686 | 3 | 0.6667 | 8 | 0.125 | 7 | 0 | 6 | 0.3333 | 24 | 0.2083 | 2 | 0 | 4 | 0 | 4 | 0.25 | 3 | 0 | 13 | 0.0769 |
| 9_12199705 | 2 | 0 | 5 | 0.2 | 9 | 0.3333 | 2 | 0.5 | 18 | 0.2778 | 2 | 0 | 5 | 0.2 | 5 | 0 | 2 | 0 | 14 | 0.0714 |
| 9_12199716 | 3 | 0.3333 | 9 | 0.3333 | 9 | 0.1111 | 4 | 0 | 25 | 0.2 | 3 | 0.3333 | 7 | 0.1429 | 5 | 0 | 4 | 0 | 19 | 0.1053 |
| 9_12199738 | 3 | 0.3333 | 9 | 0.1111 | 9 | 0.1111 | 5 | 0.2 | 26 | 0.1538 | 4 | 0 | 8 | 0.5 | 4 | 0 | 3 | 0 | 19 | 0.2105 |
| 9_12199749 | 3 | 1 | 8 | 0.875 | 10 | 0.4 | 5 | 0.6 | 26 | 0.6538 | 4 | 0.75 | 6 | 0.6667 | 6 | 0.6667 | 3 | 0.3333 | 19 | 0.6316 |
| 9_12199773 | 1 | 1 | 10 | 0.1 | 12 | 0 | 6 | 0 | 29 | 0.069 | 5 | 0.2 | 8 | 0.125 | 7 | 0 | 6 | 0.1667 | 26 | 0.1154 |
| 9_12199804 | 4 | 0.25 | 9 | 0 | 13 | 0.3077 | 6 | 0.1667 | 32 | 0.1875 | 7 | 0.2857 | 6 | 0.1667 | 7 | 0.1429 | 6 | 0.1667 | 26 | 0.1923 |
| 9_12199843 | 6 | 0 | 10 | 0.1 | 12 | 0.25 | 6 | 0 | 34 | 0.1176 | 7 | 0.2857 | 8 | 0.125 | 7 | 0 | 7 | 0 | 29 | 0.1034 |
| 9_12200527 | 2 | 0 | 2 | 0.5 | 2 | 0.5 | 2 | 0.5 | 8 | 0.375 | NA | NA | 2 | 0.5 | 1 | 0 | 6 | 0.3333 | 9 | 0.3333 |
| 9_12200539 | 2 | 0.5 | 2 | 0.5 | 2 | 0.5 | 1 | 1 | 7 | 0.5714 | NA | NA | 1 | 0 | 1 | 1 | 5 | 0.6 | 7 | 0.5714 |
| 9_12200543 | 1 | 1 | 1 | 1 | 2 | 0 | 1 | 1 | 5 | 0.6 | NA | NA | 2 | 0.5 | 1 | 0 | 6 | 0.3333 | 9 | 0.3333 |
| 9_12209029 | 1 | 1 | 2 | 1 | 2 | 1 | 1 | 0 | 6 | 0.8333 | NA | NA | 1 | 1 | NA | NA | 4 | 1 | 5 | 1 |
| 9_12209038 | 1 | 0 | 2 | 0.5 | 2 | 0 | 1 | 0 | 6 | 0.1667 | NA | NA | 1 | 0 | 1 | 0 | 6 | 0.1667 | 8 | 0.125 |
| 9_12209049 | 1 | 0 | 1 | 1 | 2 | 0.5 | 1 | 0 | 5 | 0.4 | NA | NA | NA | NA | 1 | 1 | 5 | 0.8 | 6 | 0.8333 |
| 9_12209062 | 1 | 0 | 2 | 0.5 | 1 | 1 | NA | NA | 4 | 0.5 | NA | NA | 1 | 1 | 1 | 0 | 6 | 0.8333 | 8 | 0.75 |
| 9_12219708 | NA | NA | NA | NA | 2 | 1 | 2 | 0.5 | 4 | 0.75 | NA | NA | 1 | 1 | 1 | 1 | 2 | 0.5 | 4 | 0.75 |
| 9_12219727 | NA | NA | NA | NA | 2 | 0.5 | 2 | 0.5 | 4 | 0.5 | NA | NA | 1 | 0 | 1 | 0 | 2 | 0.5 | 4 | 0.25 |
| 9_12220866 | 1 | 0 | NA | NA | 2 | 0.5 | 1 | 1 | 4 | 0.5 | NA | NA | 1 | 0 | 1 | 0 | 3 | 0.3333 | 5 | 0.2 |
| 9_12220874 | 1 | 1 | NA | NA | 2 | 0 | 3 | 0 | 6 | 0.1667 | NA | NA | 1 | 0 | 1 | 1 | 2 | 0.5 | 4 | 0.5 |
| 9_12220875 | 1 | 0 | NA | NA | 2 | 0.5 | 3 | 0.6667 | 6 | 0.5 | NA | NA | 1 | 1 | 1 | 0 | 1 | 1 | 3 | 0.6667 |
| 9_12220891 | NA | NA | NA | NA | 1 | 1 | 1 | 0 | 2 | 0.5 | NA | NA | 1 | 0 | 1 | 1 | 1 | 1 | 3 | 0.6667 |
| 9_12220973 | 1 | 1 | NA | NA | 2 | 0 | 1 | 0 | 4 | 0.25 | NA | NA | 1 | 0 | 2 | 0 | 3 | 0.3333 | 6 | 0.1667 |
| 9_12266391 | 3 | 0.3333 | 8 | 0.875 | 2 | 0.5 | 3 | 0.6667 | 16 | 0.6875 | 1 | 1 | NA | NA | 2 | 1 | 2 | 0.5 | 5 | 0.8 |
| 9_12266401 | 3 | 0 | 7 | 0.1429 | 2 | 0.5 | 3 | 0.3333 | 15 | 0.2 | 2 | 0 | NA | NA | 2 | 0.5 | 1 | 0 | 5 | 0.2 |

| | | | | | | | | | | | | | | | | | | | | |
|---|---|---|---|---|---|---|---|---|---|---|---|---|---|---|---|---|---|---|---|---|
| 9_12266407 | 4 | 0.5 | 9 | 0.7778 | 2 | 0 | 3 | 0.6667 | 18 | 0.6111 | 2 | 0 | NA | NA | 2 | 1 | 2 | 1 | 6 | 0.6667 |
| 9_12267858 | 11 | 0 | 7 | 0.4286 | 8 | 0.125 | 12 | 0.3333 | 38 | 0.2105 | 5 | 0.4 | 6 | 0.5 | 6 | 0.1667 | 8 | 0.375 | 25 | 0.36 |
| 9_12267870 | 9 | 0 | 6 | 0 | 7 | 0 | 11 | 0.1818 | 33 | 0.0606 | 5 | 0 | 5 | 0.2 | 5 | 0 | 7 | 0.1429 | 22 | 0.0909 |
| 9_12267871 | 8 | 0.5 | 6 | 0.1667 | 8 | 0.125 | 11 | 0.3636 | 33 | 0.303 | 5 | 0.2 | 6 | 0.1667 | 5 | 0.2 | 8 | 0.25 | 24 | 0.2083 |
| 9_12267882 | 7 | 0.1429 | 7 | 0 | 8 | 0 | 10 | 0.1 | 32 | 0.0625 | 4 | 0 | 6 | 0.1667 | 4 | 0 | 8 | 0.375 | 22 | 0.1818 |
| 9_12267907 | 2 | 0 | 6 | 0 | 4 | 0 | 10 | 0.2 | 22 | 0.0909 | 3 | 0.3333 | 4 | 0 | 2 | 0.5 | 6 | 0 | 15 | 0.1333 |
| 9_12267973 | 2 | 0 | 5 | 0 | 3 | 0 | 7 | 0.2857 | 17 | 0.1176 | 3 | 0 | 3 | 0.3333 | 2 | 0 | 1 | 1 | 9 | 0.2222 |
| 9_12268092 | 1 | 0 | 3 | 0 | 1 | 1 | 4 | 0 | 9 | 0.1111 | 2 | 0 | 1 | 0 | 1 | 1 | NA | NA | 4 | 0.25 |
| 9_12268101 | 1 | 0 | 3 | 0.6667 | 1 | 0 | 4 | 0.25 | 9 | 0.3333 | 2 | 0 | 1 | 1 | NA | NA | NA | NA | 3 | 0.3333 |
| 9_12268159 | NA | NA | 3 | 0.6667 | 1 | 1 | 3 | 0.6667 | 7 | 0.7143 | 1 | 0 | 1 | 1 | NA | NA | NA | NA | 2 | 0.5 |
| 9_12268193 | NA | NA | 3 | 0.3333 | 1 | 0 | 1 | 0 | 5 | 0.2 | 1 | 1 | 1 | 0 | NA | NA | NA | NA | 2 | 0.5 |
| 9_12268283 | NA | NA | 1 | 0 | 1 | 1 | 2 | 0.5 | 4 | 0.5 | 1 | 1 | NA | NA | NA | NA | NA | NA | 1 | 1 |
| 9_12271762 | 1 | 0 | 2 | 0.5 | NA | NA | NA | NA | 3 | 0.3333 | NA | NA | 1 | 1 | NA | NA | NA | NA | 1 | 1 |
| 9_12272379 | 1 | 0 | 2 | 0.5 | NA | NA | NA | NA | 3 | 0.3333 | NA | NA | 1 | 1 | NA | NA | NA | NA | 1 | 1 |
| 9_12272384 | 1 | 1 | 2 | 1 | NA | NA | NA | NA | 3 | 1 | NA | NA | 1 | 1 | NA | NA | NA | NA | 1 | 1 |
| 9_12272394 | 1 | 1 | 1 | 1 | NA | NA | NA | NA | 2 | 1 | NA | NA | 1 | 1 | NA | NA | NA | NA | 1 | 1 |
| 9_12272413 | 1 | 1 | 2 | 1 | NA | NA | NA | NA | 3 | 1 | NA | NA | 1 | 1 | NA | NA | NA | NA | 1 | 1 |
| 9_12272416 | NA | NA | 2 | 0.5 | NA | NA | NA | NA | 2 | 0.5 | NA | NA | 1 | 1 | NA | NA | NA | NA | 1 | 1 |
| 9_12272431 | 1 | 1 | 2 | 1 | NA | NA | NA | NA | 3 | 1 | NA | NA | 1 | 1 | NA | NA | NA | NA | 1 | 1 |
| 9_12272480 | NA | NA | 2 | 1 | NA | NA | NA | NA | 2 | 1 | NA | NA | 1 | 1 | NA | NA | NA | NA | 1 | 1 |
| 9_12272563 | 1 | 1 | NA | NA | NA | NA | NA | NA | 1 | 1 | NA | NA | 1 | 1 | NA | NA | NA | NA | 1 | 1 |
| 9_12272636 | 1 | 1 | NA | NA | NA | NA | NA | NA | 1 | 1 | NA | NA | 1 | 1 | NA | NA | NA | NA | 1 | 1 |
| 9_12275419 | NA | NA | 1 | 0 | 2 | 0.5 | NA | NA | 3 | 0.3333 | NA | NA | 1 | 1 | NA | NA | NA | NA | 1 | 1 |
| 9_12275914 | NA | NA | NA | NA | 1 | 1 | 1 | 1 | 2 | 1 | NA | NA | 1 | 1 | NA | NA | NA | NA | 1 | 1 |
| 9_12275950 | NA | NA | 1 | 1 | 2 | 1 | 1 | 1 | 4 | 1 | NA | NA | 1 | 1 | NA | NA | NA | NA | 1 | 1 |
| 9_12275988 | NA | NA | 1 | 1 | 1 | 0 | 1 | 1 | 3 | 0.6667 | NA | NA | 1 | 1 | NA | NA | NA | NA | 1 | 1 |
| 9_12275999 | NA | NA | 1 | 1 | 1 | 1 | 1 | 1 | 3 | 1 | NA | NA | 1 | 1 | NA | NA | NA | NA | 1 | 1 |
| 9_12277686 | 7 | 0.4286 | 6 | 0.1667 | 7 | 0.1429 | 5 | 0.4 | 25 | 0.28 | 5 | 0.2 | 4 | 0 | 5 | 0.2 | 1 | 0 | 15 | 0.1333 |
| 9_12277694 | 7 | 1 | 5 | 1 | 5 | 1 | 5 | 1 | 22 | 1 | 4 | 1 | 4 | 1 | 6 | 1 | 1 | 1 | 15 | 1 |
| 9_12277702 | 7 | 0.7143 | 5 | 0.6 | 7 | 0.7143 | 6 | 0.6667 | 25 | 0.68 | 4 | 0.75 | 4 | 0.75 | 5 | 0.6 | 1 | 0 | 14 | 0.6429 |
| 9_12285231 | 1 | 1 | 4 | 0.75 | 3 | 0.6667 | 3 | 1 | 11 | 0.8182 | NA | NA | 1 | 1 | 4 | 0.75 | 1 | 1 | 6 | 0.8333 |
| 9_12289945 | 1 | 1 | NA | NA | NA | NA | NA | NA | 1 | 1 | 1 | 1 | NA | NA | NA | NA | NA | NA | 1 | 1 |
| 9_12290346 | NA | NA | 1 | 1 | 3 | 0.6667 | 1 | 1 | 5 | 0.8 | 2 | 0.5 | 2 | 0.5 | 2 | 1 | NA | NA | 6 | 0.6667 |
| 9_12290348 | NA | NA | 2 | 0 | 3 | 0.3333 | 1 | 0 | 6 | 0.1667 | 1 | 0 | 2 | 1 | 1 | 1 | NA | NA | 4 | 0.75 |
| 9_12290354 | NA | NA | 2 | 1 | 2 | 1 | 2 | 0.5 | 6 | 0.8333 | 1 | 0 | 1 | 1 | NA | NA | NA | NA | 2 | 0.5 |
| 9_12290356 | 1 | 0 | 3 | 1 | 3 | 0.6667 | 2 | 0.5 | 9 | 0.6667 | 1 | 0 | 2 | 1 | 2 | 0.5 | NA | NA | 5 | 0.6 |
| 9_12290377 | 1 | 1 | 2 | 1 | 3 | 0.6667 | 2 | 0.5 | 8 | 0.75 | NA | NA | 2 | 1 | 1 | 1 | NA | NA | 3 | 1 |
| 9_12290386 | 1 | 1 | 2 | 0.5 | 3 | 1 | 1 | 0 | 7 | 0.7143 | 1 | 0 | 1 | 1 | 2 | 1 | NA | NA | 4 | 0.75 |
| 9_12290432 | NA | NA | 2 | 0 | 3 | 0.3333 | 1 | 0 | 6 | 0.1667 | 1 | 0 | 1 | 0 | 2 | 1 | NA | NA | 4 | 0.5 |
| 9_12290640 | 10 | 0.1 | 7 | 0.2857 | 3 | 0.3333 | 4 | 0.25 | 24 | 0.2083 | 6 | 0 | 5 | 0.2 | 8 | 0.25 | 7 | 0 | 26 | 0.1154 |
| 9_12290667 | 8 | 0.625 | 8 | 0.625 | 4 | 0 | 4 | 0.25 | 24 | 0.4583 | 7 | 0 | 1 | 0 | 8 | 0.5 | 4 | 0.5 | 20 | 0.3 |
| 9_12290671 | 7 | 1 | 8 | 0.875 | 5 | 1 | 3 | 1 | 23 | 0.9565 | 6 | 0.8333 | 2 | 1 | 8 | 1 | 7 | 1 | 23 | 0.9565 |
| 9_12290682 | 8 | 1 | 8 | 1 | 4 | 1 | 3 | 1 | 23 | 1 | 5 | 0.8 | 2 | 0.5 | 10 | 1 | 7 | 1 | 24 | 0.9167 |
| 9_12290689 | 7 | 0.8571 | 3 | 0.6667 | 1 | 1 | 3 | 0.6667 | 14 | 0.7857 | 2 | 1 | 2 | 1 | 5 | 0.8 | 4 | 1 | 13 | 0.9231 |
| 9_12290704 | 8 | 0.75 | 4 | 0.5 | 4 | 0.25 | 1 | 0 | 17 | 0.5294 | 5 | 0.8 | 2 | 0.5 | 8 | 0.25 | 7 | 0.2857 | 22 | 0.4091 |
| 9_12290714 | 4 | 0.5 | 5 | 0.4 | 2 | 0 | 3 | 1 | 14 | 0.5 | 5 | 1 | 2 | 1 | 7 | 0.8571 | 7 | 0.7143 | 21 | 0.8571 |
| 9_12290715 | 4 | 1 | 4 | 1 | 2 | 1 | 2 | 0 | 12 | 0.8333 | 5 | 0.6 | 2 | 1 | 5 | 0.8 | 6 | 0.3333 | 18 | 0.6111 |
| 9_12290722 | 3 | 1 | 6 | 0.5 | 3 | 0.3333 | 2 | 0 | 14 | 0.5 | 5 | 0.6 | 2 | 1 | 8 | 0.25 | 7 | 0.5714 | 22 | 0.5 |
| 9_12290732 | 3 | 0 | 5 | 0 | 2 | 0 | 3 | 0.6667 | 13 | 0.1538 | 3 | 0 | 2 | 0.5 | 7 | 0 | 7 | 0 | 19 | 0.0526 |
| 9_12308822 | 54 | 0.5556 | 33 | 0.3939 | 34 | 0.3824 | 64 | 0.3906 | 185 | 0.4378 | 26 | 0.5 | 67 | 0.4925 | 81 | 0.3704 | 51 | 0.4902 | 225 | 0.4489 |
| 9_12308841 | 57 | 0.1228 | 28 | 0.1429 | 29 | 0.1724 | 52 | 0.1538 | 166 | 0.1446 | 20 | 0.1 | 67 | 0.2239 | 80 | 0.1625 | 48 | 0.1875 | 215 | 0.1814 |
| 9_12308852 | 53 | 0.6226 | 27 | 0.5926 | 28 | 0.7857 | 59 | 0.661 | 167 | 0.6587 | 22 | 0.6364 | 58 | 0.6207 | 70 | 0.6429 | 51 | 0.7059 | 201 | 0.6517 |
| 9_12308863 | 51 | 0.1176 | 23 | 0.1739 | 27 | 0.1852 | 51 | 0.1176 | 152 | 0.1382 | 17 | 0.0588 | 45 | 0.2222 | 63 | 0.1587 | 40 | 0.175 | 165 | 0.1697 |
| 9_12308875 | 48 | 0.0833 | 30 | 0.1667 | 25 | 0.12 | 53 | 0.0566 | 156 | 0.0962 | 18 | 0.1111 | 57 | 0.0702 | 65 | 0.0923 | 47 | 0.1064 | 187 | 0.0909 |
| 9_12308883 | 43 | 0.7209 | 27 | 0.6296 | 25 | 0.72 | 46 | 0.587 | 141 | 0.6596 | 15 | 0.6 | 54 | 0.7037 | 62 | 0.7258 | 36 | 0.6944 | 167 | 0.7006 |
| 9_12308887 | 36 | 0.2222 | 22 | 0.2727 | 16 | 0.1875 | 34 | 0.2353 | 108 | 0.2315 | 14 | 0.0714 | 34 | 0.2059 | 54 | 0.2037 | 31 | 0.0323 | 133 | 0.1504 |
| 9_12308905 | 35 | 0.9143 | 22 | 0.8636 | 19 | 0.7895 | 46 | 0.7609 | 122 | 0.8279 | 12 | 0.6667 | 47 | 0.766 | 57 | 0.9123 | 38 | 0.8158 | 154 | 0.8247 |
| 9_12308947 | 29 | 0.6897 | 19 | 0.6316 | 12 | 0.5833 | 27 | 0.5185 | 87 | 0.6092 | 7 | 0 | 25 | 0.72 | 35 | 0.6571 | 22 | 0.6364 | 89 | 0.618 |
| 9_12308971 | 2 | 1 | NA | NA | 3 | 0.6667 | 1 | 1 | 6 | 0.8333 | NA | NA | 2 | 1 | 1 | 1 | NA | NA | 3 | 1 |
| 9_12308976 | 2 | 0.5 | NA | NA | 2 | 0.5 | 1 | 1 | 5 | 0.6 | NA | NA | 2 | 0.5 | 1 | 1 | NA | NA | 3 | 0.6667 |
| 9_12308976 | 2 | 0.5 | NA | NA | 2 | 0.5 | 1 | 0 | 5 | 0.6 | NA | NA | 2 | 0.5 | 1 | 1 | NA | NA | 3 | 0.6667 |
| 9_12308976 | 2 | 0.5 | NA | NA | 2 | 0.5 | 1 | 1 | 5 | 0.6 | NA | NA | 2 | 0.5 | 1 | 1 | NA | NA | 3 | 0.6667 |
| 9_12308976 | 2 | 0.5 | NA | NA | 2 | 0.5 | 1 | 0 | 5 | 0.6 | NA | NA | 2 | 0.5 | 1 | 1 | NA | NA | 3 | 0.6667 |
| 9_12308995 | 1 | 1 | NA | NA | 2 | 0.5 | NA | NA | 3 | 0.6667 | NA | NA | 2 | 0.5 | 1 | 1 | NA | NA | 3 | 0.6667 |
| 9_12308999 | 2 | 0.5 | NA | NA | NA | NA | 1 | 1 | 3 | 0.6667 | NA | NA | 1 | 1 | 1 | 0 | NA | NA | 2 | 0.5 |

| | | | | | | | | | | | | | | | | | | | | |
|---|---|---|---|---|---|---|---|---|---|---|---|---|---|---|---|---|---|---|---|---|
| 9_12309009 | 2 | 0 | NA | NA | 3 | 0 | 1 | 1 | 6 | 0.1667 | NA | NA | 2 | 0.5 | 1 | 0 | NA | NA | 3 | 0.3333 |
| 9_12309014 | 2 | 0.5 | NA | NA | 3 | 0.3333 | 1 | 0 | 6 | 0.3333 | NA | NA | 1 | 0 | 1 | 1 | NA | NA | 2 | 0.5 |
| 9_12309014 | 2 | 1 | NA | NA | 2 | 1 | 1 | 1 | 5 | 1 | NA | NA | 2 | 1 | 1 | 1 | NA | NA | 3 | 1 |
| 9_12319626 | NA | NA | NA | NA | 1 | 1 | NA | NA | 1 | 1 | NA | NA | 1 | 1 | NA | NA | NA | NA | 1 | 1 |
| 9_12329063 | 2 | 1 | 3 | 0.3333 | 3 | 0.3333 | 2 | 0 | 10 | 0.4 | NA | NA | 2 | 0 | 1 | 0 | 2 | 1 | 5 | 0.4 |
| 9_12329071 | 4 | 0.5 | 3 | 0.6667 | 3 | 1 | 2 | 0.5 | 12 | 0.6667 | NA | NA | 1 | 1 | 1 | 1 | 1 | 1 | 3 | 1 |
| 9_12329075 | 5 | 0.6 | 3 | 0.6667 | 2 | 0 | 2 | 1 | 12 | 0.5833 | 1 | 1 | 2 | 0 | 1 | 1 | 2 | 1 | 6 | 0.6667 |
| 9_12329082 | 7 | 0.7143 | 2 | 1 | 1 | 1 | 1 | 1 | 11 | 0.8182 | 1 | 0 | 2 | 1 | 1 | 1 | 2 | 1 | 6 | 0.8333 |
| 9_12329084 | 6 | 0.6667 | 2 | 0.5 | 3 | 0 | 2 | 0 | 13 | 0.3846 | NA | NA | 2 | 1 | 1 | 1 | 2 | 0.5 | 5 | 0.8 |
| 9_12329088 | 6 | 0.3333 | 4 | 0.5 | 1 | 0 | 2 | 0.5 | 13 | 0.3846 | 1 | 1 | 1 | 1 | 2 | 0 | 1 | 0 | 5 | 0.4 |
| 9_12329103 | 3 | 0.3333 | 1 | 0 | 2 | 0 | 1 | 0 | 7 | 0.1429 | 1 | 0 | 2 | 0 | NA | NA | 2 | 0.5 | 5 | 0.2 |
| 9_12329145 | 7 | 0.2857 | 3 | 0 | 4 | 0 | 2 | 0 | 16 | 0.125 | 1 | 0 | 3 | 0 | 1 | 0 | 2 | 0.5 | 7 | 0.1429 |
| 9_12329168 | 3 | 0.3333 | 3 | 0 | 3 | 0 | 2 | 0 | 11 | 0.0909 | 1 | 0 | 3 | 0 | 1 | 0 | 1 | 1 | 6 | 0.1667 |
| 9_12334861 | NA | NA | NA | NA | 2 | 0 | 2 | 0.5 | 4 | 0.25 | NA | NA | NA | NA | NA | NA | 1 | 1 | 1 | 1 |
| 9_12335006 | NA | NA | 1 | 1 | 2 | 0 | 2 | 0 | 5 | 0.2 | 1 | 1 | NA | NA | NA | NA | 1 | 0 | 2 | 0.5 |
| 9_12341992 | 2 | 0 | 3 | 0 | 7 | 0.4286 | 11 | 0.5455 | 23 | 0.3913 | 6 | 0.1667 | 2 | 0.5 | 10 | 0.2 | 7 | 0.1429 | 25 | 0.2 |
| 9_12341997 | 4 | 0.25 | 4 | 0 | 8 | 0.375 | 10 | 0.2 | 26 | 0.2308 | 5 | 0.4 | 7 | 0.2857 | 12 | 0.1667 | 9 | 0.1111 | 33 | 0.2121 |
| 9_12341999 | 4 | 1 | 3 | 1 | 3 | 0.6667 | 9 | 1 | 19 | 0.9474 | 2 | 1 | 6 | 0.8333 | 6 | 0.8333 | 8 | 0.875 | 22 | 0.8636 |
| 9_12342005 | 5 | 0.2 | 4 | 0.25 | 8 | 0.125 | 9 | 0.1111 | 26 | 0.1538 | 5 | 0.2 | 7 | 0 | 10 | 0.2 | 8 | 0.125 | 30 | 0.1333 |
| 9_12342154 | 8 | 0 | 4 | 0.5 | 8 | 0.125 | 13 | 0.1538 | 33 | 0.1515 | 8 | 0.25 | 14 | 0.0714 | 12 | 0.1667 | 8 | 0 | 42 | 0.119 |
| 9_12346553 | 13 | 0.3077 | 4 | 0.75 | 3 | 0.3333 | 1 | 1 | 21 | 0.4286 | 2 | 0 | 1 | 0 | 4 | 0.5 | 1 | 0 | 8 | 0.25 |
| 9_12346576 | 10 | 0.1 | 4 | 0.5 | 4 | 0.25 | 3 | 0.6667 | 21 | 0.2857 | 1 | 0 | 2 | 0 | 5 | 0.2 | 1 | 1 | 9 | 0.2222 |
| 9_12346576 | 9 | 0.1111 | 3 | 0 | 4 | 0 | 2 | 0 | 18 | 0.0556 | 1 | 0 | 2 | 0 | 5 | 0 | 1 | 1 | 9 | 0.1111 |
| 9_12346599 | 10 | 0.1 | 1 | 0 | 2 | 0 | 3 | 0.6667 | 16 | 0.1875 | 1 | 1 | 2 | 0.5 | 3 | 0.6667 | NA | NA | 6 | 0.6667 |
| 9_12346605 | 6 | 0.5 | 2 | 0.5 | 3 | 1 | 3 | 0.3333 | 14 | 0.5714 | 2 | 0.5 | 2 | 1 | 4 | 0.75 | 1 | 1 | 9 | 0.7778 |
| 9_12346611 | 10 | 0.4 | 3 | 0.3333 | 3 | 0 | 3 | 0.3333 | 19 | 0.3158 | 2 | 0.5 | 2 | 1 | 4 | 0.75 | 1 | 1 | 9 | 0.7778 |
| 9_12350740 | NA | NA | NA | NA | 1 | 1 | NA | NA | 1 | 1 | NA | NA | NA | NA | 1 | 1 | NA | NA | 1 | 1 |
| 9_12350743 | NA | NA | NA | NA | 1 | 1 | NA | NA | 1 | 1 | NA | NA | NA | NA | 1 | 1 | NA | NA | 1 | 1 |
| 9_12351811 | 3 | 0.6667 | 1 | 0 | NA | NA | 2 | 0.5 | 6 | 0.5 | NA | NA | NA | NA | NA | NA | 1 | 1 | 1 | 1 |
| 9_12351865 | 3 | 1 | 1 | 0 | NA | NA | 1 | 1 | 5 | 0.8 | NA | NA | NA | NA | NA | NA | 1 | 1 | 1 | 1 |
| 9_12352231 | 1 | 1 | NA | NA | NA | NA | NA | NA | 1 | 1 | NA | NA | NA | NA | NA | NA | 1 | 1 | 1 | 1 |
| 9_12352420 | NA | NA | 4 | 0.25 | 2 | 0 | NA | NA | 6 | 0.1667 | NA | NA | 3 | 0 | 3 | 0 | 1 | 1 | 7 | 0.1429 |
| 9_12352439 | 1 | 0 | 2 | 0 | 2 | 0.5 | 2 | 0 | 7 | 0.1429 | NA | NA | 3 | 0.3333 | 3 | 0 | NA | NA | 6 | 0.1667 |
| 9_12352447 | 1 | 0 | 2 | 0 | 2 | 0 | 2 | 0.5 | 7 | 0.1429 | NA | NA | 2 | 0.5 | 3 | 0 | NA | NA | 5 | 0.2 |
| 9_12352474 | 1 | 1 | 4 | 0.5 | 2 | 0.5 | NA | NA | 7 | 0.5714 | NA | NA | 3 | 0.3333 | 3 | 0.6667 | NA | NA | 6 | 0.5 |
| 9_12352795 | NA | NA | 1 | 0 | NA | NA | 3 | 0.6667 | 4 | 0.5 | NA | NA | NA | NA | 2 | 0.5 | NA | NA | 2 | 0.5 |
| 9_12352807 | NA | NA | 1 | 1 | NA | NA | 3 | 0.3333 | 4 | 0.5 | NA | NA | NA | NA | 2 | 0.5 | NA | NA | 2 | 0.5 |
| 9_12352914 | 2 | 0 | 8 | 0.25 | 4 | 0 | 5 | 0.2 | 19 | 0.1579 | 2 | 0 | 4 | 0 | 4 | 0.25 | 2 | 0 | 12 | 0.0833 |
| 9_12352994 | 2 | 0 | 7 | 0.1429 | 4 | 0 | 4 | 0.25 | 17 | 0.1176 | 1 | 0 | 4 | 0.25 | 4 | 0 | 2 | 0 | 11 | 0.0909 |
| 9_12361946 | 2 | 0 | 3 | 0.6667 | NA | NA | 1 | 1 | 6 | 0.5 | 2 | 0 | 6 | 0.5 | 4 | 0.75 | 2 | 1 | 14 | 0.5714 |
| 9_12361981 | 2 | 0 | 2 | 0 | NA | NA | 1 | 1 | 5 | 0.2 | 2 | 0.5 | 6 | 0 | 2 | 0 | 2 | 0 | 12 | 0.0833 |
| 9_12362013 | NA | NA | 2 | 0 | NA | NA | 1 | 1 | 3 | 0.3333 | 2 | 0 | 3 | 0.3333 | 3 | 0 | 1 | 0 | 9 | 0.1111 |
| 9_12372324 | 1 | 1 | NA | NA | NA | NA | 1 | 0 | 2 | 0.5 | 1 | 1 | NA | NA | NA | NA | 1 | 0 | 2 | 0.5 |
| 9_12392459 | 1 | 1 | NA | NA | NA | NA | NA | NA | 1 | 1 | 1 | 1 | NA | NA | 1 | 1 | NA | NA | 2 | 1 |
| 9_12392514 | 1 | 1 | NA | NA | NA | NA | NA | NA | 1 | 1 | 1 | 1 | NA | NA | 1 | 1 | NA | NA | 2 | 1 |
| 9_12392519 | 1 | 1 | 1 | 1 | NA | NA | NA | NA | 2 | 1 | 1 | 1 | NA | NA | 1 | 1 | NA | NA | 2 | 1 |
| 9_12392522 | 1 | 1 | 1 | 1 | NA | NA | NA | NA | 2 | 1 | 1 | 1 | NA | NA | 1 | 1 | NA | NA | 2 | 1 |
| 9_12392525 | 1 | 1 | 1 | 1 | NA | NA | NA | NA | 2 | 1 | 1 | 1 | NA | NA | 1 | 1 | NA | NA | 2 | 1 |
| 9_12392547 | 1 | 1 | 1 | 1 | NA | NA | NA | NA | 2 | 1 | 1 | 1 | NA | NA | 1 | 1 | NA | NA | 2 | 1 |
| 9_12392568 | 1 | 0 | 1 | 0 | 1 | 1 | NA | NA | 3 | 0.3333 | 1 | 1 | NA | NA | 1 | 1 | NA | NA | 2 | 1 |
| 9_12392609 | NA | NA | 1 | 1 | NA | NA | 1 | 0 | 2 | 0.5 | NA | NA | NA | NA | 1 | 1 | NA | NA | 1 | 1 |
| 9_12392661 | 1 | 1 | 1 | 1 | 1 | 0 | 1 | 1 | 4 | 0.75 | 1 | 1 | NA | NA | 1 | 1 | NA | NA | 2 | 1 |
| 9_12392668 | 1 | 0 | 1 | 0 | 1 | 0 | 1 | 1 | 4 | 0.25 | 1 | 1 | NA | NA | 1 | 0 | NA | NA | 2 | 0.5 |
| 9_12403165 | 2 | 0.5 | 1 | 1 | NA | NA | 1 | 0 | 4 | 0.5 | NA | NA | 1 | 1 | 1 | 0 | 1 | 1 | 3 | 0.6667 |
| 9_12403172 | 1 | 1 | 1 | 0 | 1 | 0 | 1 | 0 | 4 | 0.25 | NA | NA | 1 | 0 | 1 | 0 | 1 | 1 | 3 | 0.3333 |
| 9_12403173 | 2 | 0.5 | NA | NA | 1 | 0 | 1 | 0 | 4 | 0.25 | NA | NA | 1 | 1 | 1 | 1 | 1 | 1 | 3 | 1 |
| 9_12403180 | 2 | 1 | 1 | 1 | 1 | 1 | 1 | 0 | 5 | 0.8 | NA | NA | 1 | 1 | 1 | 1 | 2 | 1 | 4 | 1 |
| 9_12403190 | 2 | 1 | NA | NA | 1 | 1 | 1 | 0 | 4 | 0.75 | NA | NA | NA | NA | 1 | 1 | 2 | 1 | 3 | 1 |
| 9_12403201 | 2 | 1 | 1 | 1 | 1 | 1 | 1 | 1 | 5 | 1 | NA | NA | NA | NA | 1 | 1 | 2 | 1 | 3 | 1 |
| 9_12403220 | 1 | 1 | NA | NA | 1 | 1 | 1 | 1 | 3 | 1 | NA | NA | NA | NA | 1 | 1 | 2 | 1 | 3 | 1 |
| 9_12403231 | 2 | 0 | 1 | 1 | 1 | 0 | 1 | 0 | 5 | 0.2 | NA | NA | NA | NA | 1 | 0 | 1 | 1 | 2 | 0.5 |
| 9_12403247 | 2 | 1 | 1 | 0 | 1 | 0 | 2 | 0.5 | 6 | 0.5 | NA | NA | NA | NA | 1 | 1 | 2 | 0.5 | 3 | 0.6667 |
| 9_12405531 | 2 | 0.5 | NA | NA | NA | NA | 2 | 1 | 4 | 0.75 | 1 | 0 | 2 | 1 | 1 | 1 | 2 | 0.5 | 6 | 0.6667 |
| 9_12405572 | 1 | 1 | NA | NA | NA | NA | 1 | 0 | 2 | 0.5 | NA | NA | 2 | 0.5 | 1 | 1 | 2 | 0 | 5 | 0.4 |
| 9_12405605 | 1 | 1 | NA | NA | NA | NA | 1 | 0 | 2 | 0.5 | NA | NA | 2 | 1 | 1 | 1 | 2 | 0 | 5 | 0.6 |

| | | | | | | | | | | | | | | | | | | | | |
|---|---|---|---|---|---|---|---|---|---|---|---|---|---|---|---|---|---|---|---|---|
| 9_1240563 | 1 | 1 | NA | NA | NA | NA | 1 | 0 | 2 | 0.5 | NA | NA | 1 | 1 | 1 | 0 | 1 | 0 | 3 | 0.3333 |
| 9_1240565 | 1 | 1 | NA | NA | NA | NA | 1 | 1 | 2 | 1 | NA | NA | 1 | 1 | NA | NA | 2 | 1 | 3 | 1 |
| 9_1240566 | 1 | 1 | NA | NA | NA | NA | 1 | 1 | 2 | 1 | NA | NA | NA | NA | 1 | 0 | 2 | 1 | 3 | 0.6667 |
| 9_1241957 | 3 | 1 | NA | NA | NA | NA | NA | NA | 3 | 1 | 2 | 1 | 2 | 1 | 2 | 1 | NA | NA | 6 | 1 |
| 9_1242033 | NA | NA | 1 | 1 | NA | NA | NA | NA | 1 | 1 | NA | NA | NA | NA | 1 | 1 | NA | NA | 1 | 1 |
| 9_1242033 | NA | NA | 1 | 1 | NA | NA | NA | NA | 1 | 1 | NA | NA | NA | NA | 1 | 1 | NA | NA | 1 | 1 |
| MT_1006 | 5 | 0.8 | 4 | 1 | 3 | 1 | 6 | 1 | 15 | 1 | 5 | 0.6 | 4 | 0.75 | 9 | 0.7778 | 6 | 1 | 23 | 0.7826 |
| MT_2014 | 5 | 0 | 8 | 0 | 15 | 0.0667 | 11 | 0 | 11 | 0.0909 | 7 | 0 | 9 | 0.1111 | 18 | 0 | 10 | 0 | 16 | 0.0625 |
| MT_6681 | 38 | 0.0526 | 25 | 0.08 | 33 | 0.0909 | 22 | 0.1364 | 35 | 0.0571 | 22 | 0.2273 | 29 | 0.0345 | 40 | 0.05 | 38 | 0.1053 | 43 | 0.1163 |
| MT_8429 | 2 | 0 | 7 | 0.1429 | 3 | 0.3333 | 4 | 0 | 7 | 0.2857 | 3 | 0 | 4 | 0 | 2 | 0.5 | 6 | 0.3333 | 6 | 0.1667 |
| MT_9933 | 1 | 0 | 5 | 0.2 | 2 | 0 | NA | NA | 7 | 0.1429 | 1 | 1 | 3 | 0.3333 | NA | NA | 1 | 0 | 5 | 0.4 |
| MT_13602 | 3 | 0 | 1 | 0 | 1 | 1 | 3 | 0.6667 | 8 | 0.375 | NA | NA | 2 | 0.5 | 2 | 0 | 1 | 0 | 5 | 0.2 |
| X_5959718 | 3 | 0.3333 | 5 | 0.4 | 3 | 1 | 5 | 0.6 | 16 | 0.5625 | 1 | 0 | 4 | 0.75 | 2 | 0.5 | 2 | 0 | 9 | 0.4444 |
| X_5959722 | 3 | 0 | 3 | 0.3333 | 4 | 0.25 | 3 | 0.3333 | 13 | 0.2308 | 1 | 0 | 4 | 0.25 | 2 | 0 | 1 | 0 | 8 | 0.125 |
| X_5960390 | 5 | 0.2 | 6 | 0.1667 | 2 | 0 | 9 | 0.3333 | 22 | 0.2273 | 1 | 0 | 4 | 0.25 | 3 | 0 | 4 | 0 | 12 | 0.0833 |
| X_5965187 | 6 | 0 | 7 | 0 | 5 | 0.2 | 9 | 0.1111 | 27 | 0.0741 | 3 | 0 | 6 | 0 | 4 | 0.25 | 5 | 0 | 18 | 0.0556 |
| X_5965221 | 5 | 0.6 | 6 | 0.8333 | 5 | 0.4 | 11 | 0.5455 | 27 | 0.5926 | 4 | 0.75 | 7 | 0.5714 | 3 | 1 | 3 | 0.3333 | 17 | 0.6471 |
| X_5965285 | 6 | 0.8333 | 7 | 1 | 5 | 1 | 11 | 1 | 29 | 0.9655 | 4 | 1 | 8 | 1 | 5 | 1 | 6 | 1 | 23 | 1 |
| X_5965363 | 7 | 0.2857 | 7 | 0.4286 | 4 | 0.25 | 9 | 0.6667 | 27 | 0.4444 | 4 | 0.75 | 7 | 0.5714 | 7 | 0.4286 | 6 | 0.5 | 24 | 0.5417 |
| X_5965399 | 5 | 0 | 6 | 0.1667 | 5 | 0 | 10 | 0.4 | 26 | 0.1923 | 4 | 0 | 8 | 0.625 | 6 | 0 | 4 | 0 | 22 | 0.2273 |
| X_5965569 | 5 | 0.2 | 7 | 0.2857 | 5 | 0.4 | 10 | 0.1 | 27 | 0.2222 | 3 | 0.3333 | 9 | 0.1111 | 7 | 0.2857 | 6 | 0.5 | 25 | 0.28 |
| X_5966207 | 10 | 0.4 | 10 | 0.3 | 4 | 0.5 | 11 | 0.6364 | 35 | 0.4571 | 5 | 0.2 | 15 | 0.4667 | 5 | 0.6 | 6 | 0.3333 | 31 | 0.4194 |
| X_5966299 | 8 | 0 | 8 | 0.375 | 5 | 0 | 11 | 0.2727 | 32 | 0.1875 | 5 | 0 | 12 | 0 | 7 | 0 | 6 | 0.3333 | 30 | 0.0667 |
| X_5966692 | 7 | 0.1429 | 9 | 0 | 4 | 0 | 11 | 0.0909 | 31 | 0.0645 | 6 | 0.1667 | 12 | 0.0833 | 8 | 0.125 | 6 | 0.3333 | 32 | 0.1562 |
| X_6080886 | 3 | 0 | 6 | 0 | 4 | 0.5 | 1 | 0 | 14 | 0.1429 | 6 | 0.1667 | 2 | 0.5 | 4 | 0.25 | 3 | 0 | 15 | 0.2 |
| X_6081212 | 3 | 0.6667 | 5 | 0 | 5 | 0 | 1 | 0 | 14 | 0.1429 | 4 | 0.25 | 2 | 0 | 6 | 0 | 2 | 0 | 14 | 0.0714 |
| X_6081328 | 1 | 0 | 5 | 0.2 | 5 | 0.2 | 2 | 0 | 13 | 0.1538 | 5 | 0 | 2 | 0 | 6 | 0 | 3 | 0.3333 | 16 | 0.0625 |
| X_6081719 | 1 | 1 | 3 | 0.6667 | 1 | 1 | 2 | 1 | 7 | 0.8571 | 2 | 1 | 2 | 1 | 4 | 1 | 1 | 0 | 9 | 0.8889 |
| X_6081743 | 1 | 1 | 4 | 1 | 4 | 0.5 | 2 | 1 | 11 | 0.8182 | 1 | 1 | 3 | 1 | 5 | 0.8 | 3 | 1 | 12 | 0.9167 |
| X_6084432 | 1 | 1 | 3 | 0.3333 | 1 | 0 | 2 | 0 | 7 | 0.2857 | 2 | 0 | 3 | 0.6667 | 1 | 0 | 1 | 0 | 7 | 0.2857 |
| X_6084564 | 1 | 0 | 2 | 0.5 | 2 | 0 | 2 | 0 | 7 | 0.1429 | 3 | 0 | 2 | 0 | 2 | 0 | 2 | 0.5 | 9 | 0.1111 |
| X_6084613 | 1 | 1 | 3 | 0 | 2 | 0.5 | 2 | 0.5 | 8 | 0.375 | 3 | 0.3333 | 2 | 0 | 3 | 0 | 1 | 0 | 9 | 0.1111 |
| X_6084627 | 1 | 1 | 3 | 0 | 2 | 0.5 | 2 | 0 | 8 | 0.25 | 3 | 0.3333 | 2 | 0 | 3 | 0 | 2 | 0.5 | 10 | 0.2 |
| X_6085130 | NA | NA | 2 | 0.5 | 1 | 1 | 1 | 1 | 4 | 0.75 | 1 | 0 | 1 | 0 | 1 | 1 | NA | NA | 3 | 0.3333 |
| X_7025935 | NA | NA | NA | NA | 1 | 1 | 1 | 1 | 2 | 1 | NA | NA | NA | NA | 1 | 1 | NA | NA | 1 | 1 |
| X_7025993 | NA | NA | NA | NA | 1 | 0 | 1 | 1 | 2 | 0.5 | NA | NA | NA | NA | 1 | 1 | NA | NA | 1 | 1 |
| X_7238857 | 1 | 0 | 2 | 0.5 | NA | NA | 2 | 1 | 5 | 0.6 | 2 | 1 | 3 | 0 | 2 | 0 | 3 | 0.6667 | 10 | 0.4 |
| X_7239671 | NA | NA | NA | NA | NA | NA | 1 | 1 | 1 | 1 | 1 | 0 | 1 | 1 | NA | NA | NA | NA | 2 | 0.5 |
| X_7239706 | 1 | 1 | NA | NA | NA | NA | NA | NA | 1 | 1 | 1 | 1 | NA | NA | NA | NA | NA | NA | 1 | 1 |
| X_7239872 | 1 | 1 | NA | NA | NA | NA | NA | NA | 1 | 1 | 1 | 1 | NA | NA | NA | NA | NA | NA | 1 | 1 |
| X_7240054 | 1 | 1 | NA | NA | NA | NA | NA | NA | 1 | 1 | 1 | 1 | NA | NA | NA | NA | NA | NA | 1 | 1 |
| X_7240406 | 1 | 1 | NA | NA | NA | NA | NA | NA | 1 | 1 | 1 | 1 | NA | NA | NA | NA | NA | NA | 1 | 1 |
| X_7240832 | 1 | 1 | NA | NA | NA | NA | NA | NA | 1 | 1 | 1 | 1 | NA | NA | NA | NA | NA | NA | 1 | 1 |
| X_7278366 | 7 | 0.1429 | 11 | 0 | 3 | 0 | 12 | 0.0833 | 33 | 0.0606 | 14 | 0 | 14 | 0.0714 | 12 | 0 | 14 | 0.1429 | 54 | 0.0556 |
| X_7425737 | 2 | 0.5 | 5 | 0.4 | NA | NA | 3 | 0.3333 | 10 | 0.4 | 4 | 0.25 | 3 | 0.3333 | 1 | 0 | 4 | 0 | 12 | 0.1667 |
| X_7425961 | NA | NA | 4 | 0.5 | NA | NA | 3 | 0.3333 | 7 | 0.4286 | 2 | 0.5 | 4 | 0 | 1 | 1 | 3 | 0 | 10 | 0.2 |
| X_7426083 | NA | NA | 3 | 0 | NA | NA | 3 | 0.3333 | 6 | 0.1667 | 2 | 0.5 | 3 | 0.3333 | 1 | 0 | 3 | 0 | 9 | 0.2222 |
| X_7426311 | NA | NA | 4 | 0.25 | NA | NA | 2 | 0.5 | 6 | 0.3333 | 3 | 0.6667 | 3 | 0 | 1 | 1 | 4 | 0.25 | 11 | 0.3636 |
| X_7426557 | NA | NA | 5 | 1 | NA | NA | 1 | 1 | 6 | 1 | 2 | 1 | 2 | 1 | 1 | 1 | 2 | 1 | 7 | 1 |
| X_7426626 | NA | NA | 2 | 1 | NA | NA | 1 | 1 | 3 | 1 | 2 | 1 | 1 | 1 | 1 | 1 | 1 | 1 | 5 | 1 |
| X_7426671 | NA | NA | 4 | 0.5 | NA | NA | 1 | 1 | 5 | 0.6 | 1 | 0 | 2 | 0.5 | 1 | 1 | 3 | 1 | 7 | 0.7143 |
| X_7426977 | NA | NA | 4 | 1 | NA | NA | NA | NA | 4 | 1 | 2 | 1 | 2 | 1 | 1 | 1 | NA | NA | 5 | 1 |
| X_7514977 | 47 | 0.1702 | 84 | 0.2619 | 44 | 0.3182 | 57 | 0.2632 | 232 | 0.2543 | 48 | 0.4167 | 58 | 0.1897 | 49 | 0.2653 | 43 | 0.2093 | 198 | 0.2677 |
| X_7574355 | 1 | 0 | 5 | 0.2 | 1 | 0 | 4 | 0 | 11 | 0.0909 | 2 | 0 | 1 | 0 | 3 | 0.3333 | 3 | 0 | 9 | 0.1111 |
| X_7574583 | 1 | 1 | 6 | 0.3333 | 2 | 0 | 4 | 0.5 | 13 | 0.3846 | 2 | 0.5 | 1 | 1 | 3 | 0.6667 | 3 | 1 | 9 | 0.7778 |
| X_7575164 | 1 | 1 | 6 | 0.5 | 2 | 0 | 4 | 0.25 | 13 | 0.3846 | 2 | 0.5 | 2 | 0 | 4 | 0.25 | 3 | 0.3333 | 11 | 0.2727 |
| X_7575594 | 1 | 1 | 7 | 0.1429 | 2 | 0 | 5 | 0 | 15 | 0.1333 | 2 | 0.5 | 2 | 0 | 4 | 0 | 3 | 0.3333 | 11 | 0.1818 |
| X_7576369 | 1 | 0 | 5 | 0 | 2 | 0 | 5 | 0.2 | 13 | 0.0769 | 1 | 0 | 3 | 0 | 4 | 0 | 3 | 0.3333 | 11 | 0.0909 |
| X_7592203 | 3 | 0 | 7 | 0.1429 | 4 | 0 | 6 | 0 | 20 | 0.05 | 6 | 0.1667 | 3 | 0 | 6 | 0 | 2 | 0 | 17 | 0.0588 |
| X_7593610 | 4 | 0 | 7 | 0.1429 | 7 | 0.4286 | 4 | 0 | 22 | 0.1818 | 5 | 0.2 | 2 | 0 | 8 | 0.25 | 2 | 0 | 17 | 0.1765 |
| X_7593633 | 3 | 0.3333 | 5 | 0 | 4 | 0.75 | 5 | 0 | 17 | 0.2353 | 4 | 0 | 3 | 0.3333 | 7 | 0.2857 | 1 | 1 | 15 | 0.2667 |
| X_7594091 | 4 | 0 | 7 | 0.1429 | 8 | 0.375 | 7 | 0.1429 | 26 | 0.1923 | 6 | 0.3333 | 4 | 0 | 9 | 0.2222 | NA | NA | 19 | 0.2105 |
| X_7594148 | 5 | 0.2 | 7 | 0.1429 | 8 | 0.5 | 7 | 0.2857 | 27 | 0.2963 | 6 | 0.3333 | 4 | 0 | 6 | 0.1667 | 2 | 0.5 | 18 | 0.2222 |
| X_7594259 | 5 | 0.6 | 3 | 0.3333 | 4 | 1 | 6 | 0.6667 | 18 | 0.6667 | 4 | 0.5 | 4 | 0.75 | 6 | 0.3333 | 1 | 1 | 15 | 0.5333 |
| X_7594328 | 5 | 0.2 | 6 | 0.1667 | 6 | 0.5 | 6 | 0.3333 | 23 | 0.3043 | 4 | 1 | 4 | 0.25 | 7 | 0.4286 | 2 | 0.5 | 17 | 0.5294 |

| | | | | | | | | | | | | | | | | | | | | |
|---|---|---|---|---|---|---|---|---|---|---|---|---|---|---|---|---|---|---|---|---|
| X_7596760 | 18 | 0.1111 | 17 | 0.1765 | 16 | 0.1875 | 14 | 0.0714 | 65 | 0.1385 | 19 | 0.0526 | 18 | 0.2778 | 21 | 0.1905 | 28 | 0.0714 | 86 | 0.1395 |
| X_7596858 | 19 | 0.0526 | 17 | 0.1765 | 16 | 0.125 | 13 | 0.0769 | 65 | 0.1077 | 20 | 0.05 | 18 | 0.1667 | 21 | 0.1429 | 31 | 0.0645 | 90 | 0.1 |
| X_7596890 | 16 | 0.3125 | 13 | 0.2308 | 15 | 0.2667 | 11 | 0.1818 | 55 | 0.2545 | 16 | 0.1875 | 13 | 0.3077 | 17 | 0.1765 | 30 | 0.2 | 76 | 0.2105 |
| X_7596925 | 12 | 0 | 14 | 0.0714 | 11 | 0.0909 | 9 | 0.1111 | 46 | 0.0652 | 13 | 0.0769 | 15 | 0.2 | 15 | 0 | 24 | 0 | 67 | 0.0597 |
| X_7596962 | 16 | 0.0625 | 14 | 0.2143 | 14 | 0.2143 | 12 | 0.1667 | 56 | 0.1607 | 18 | 0.2778 | 14 | 0.2857 | 15 | 0.1333 | 25 | 0.24 | 72 | 0.2361 |
| X_7597005 | 19 | 0.1579 | 15 | 0.2 | 14 | 0.0714 | 14 | 0.1429 | 62 | 0.1452 | 15 | 0.3333 | 17 | 0.1765 | 20 | 0.05 | 31 | 0.2258 | 83 | 0.1928 |
| X_7597209 | 13 | 0 | 15 | 0.0667 | 14 | 0.2143 | 13 | 0.0769 | 55 | 0.0909 | 18 | 0.1111 | 15 | 0.1333 | 18 | 0 | 28 | 0 | 79 | 0.0506 |
| X_7598466 | 2 | 0.5 | NA | NA | NA | NA | 3 | 0.6667 | 5 | 0.6 | 1 | 0 | 2 | 0 | 1 | 1 | 1 | 1 | 5 | 0.4 |
| X_7598642 | 1 | 1 | NA | NA | NA | NA | 3 | 0.6667 | 4 | 0.75 | 1 | 1 | 3 | 0.6667 | 1 | 1 | 1 | 1 | 6 | 0.8333 |
| X_7599612 | 1 | 1 | NA | NA | NA | NA | 2 | 0.5 | 3 | 0.6667 | 1 | 0 | 3 | 0.6667 | 1 | 1 | 1 | 0 | 6 | 0.5 |
| X_7599957 | 1 | 1 | NA | NA | NA | NA | 2 | 0.5 | 3 | 0.6667 | 1 | 1 | 2 | 1 | NA | NA | NA | NA | 3 | 1 |
| X_7639786 | 3 | 0.6667 | 4 | 0 | 6 | 0.3333 | 2 | 0.5 | 15 | 0.3333 | 1 | 0 | 2 | 1 | 2 | 0.5 | 2 | 0.5 | 7 | 0.5714 |
| X_7639950 | 3 | 0 | 3 | 0 | 4 | 0.25 | 3 | 0 | 13 | 0.0769 | 2 | 0 | 2 | 0 | 2 | 1 | NA | NA | 6 | 0.3333 |
| X_7640013 | 4 | 1 | 4 | 0.75 | 6 | 1 | 2 | 1 | 16 | 0.9375 | 2 | 1 | 1 | 1 | 2 | 1 | 2 | 0.5 | 7 | 0.8571 |
| X_7640047 | 3 | 0.3333 | 3 | 0 | 4 | 0.25 | 3 | 0.3333 | 13 | 0.2308 | 1 | 0 | NA | NA | 1 | 1 | 2 | 0.5 | 4 | 0.5 |
| X_7640230 | 4 | 0.5 | 4 | 0.5 | 5 | 0.4 | 3 | 0.3333 | 16 | 0.4375 | 2 | 1 | 2 | 0.5 | 2 | 0.5 | 2 | 0.5 | 8 | 0.625 |
| X_7640887 | 5 | 0.6 | 4 | 0.25 | 5 | 0.2 | 3 | 0 | 17 | 0.2941 | 2 | 0.5 | 3 | 0.6667 | 6 | 0.3333 | 2 | 0.5 | 13 | 0.4615 |
| X_7673565 | NA | NA | NA | NA | 1 | 1 | NA | NA | 1 | 1 | NA | NA | NA | NA | NA | NA | 1 | 1 | 1 | 1 |
| X_7685396 | NA | NA | 3 | 0.3333 | 1 | 0 | 3 | 0 | 7 | 0.1429 | NA | NA | 2 | 0.5 | NA | NA | 2 | 0 | 4 | 0.25 |
| X_7686242 | 5 | 0 | 8 | 0.125 | 7 | 0 | 15 | 0.0667 | 35 | 0.0571 | 9 | 0 | 7 | 0 | 8 | 0.25 | 7 | 0.1429 | 31 | 0.0968 |
| X_7686315 | 5 | 0 | 9 | 0 | 6 | 0 | 17 | 0.1176 | 37 | 0.0541 | 12 | 0 | 7 | 0 | 8 | 0.125 | 7 | 0.1429 | 34 | 0.0588 |
| X_7686445 | 5 | 0.8 | 9 | 0.7778 | 7 | 0.8571 | 16 | 0.9375 | 37 | 0.8649 | 12 | 0.75 | 5 | 0.4 | 9 | 1 | 10 | 0.9 | 36 | 0.8056 |
| X_7686611 | 6 | 0 | 12 | 0 | 8 | 0 | 17 | 0.1765 | 43 | 0.0698 | 16 | 0.0625 | 6 | 0.1667 | 12 | 0.0833 | 11 | 0.0909 | 45 | 0.0889 |
| X_7686681 | 6 | 0.6667 | 12 | 0.4167 | 7 | 0 | 19 | 0.4211 | 44 | 0.3864 | 12 | 0.5 | 7 | 0.5714 | 10 | 0.6 | 10 | 0.6 | 39 | 0.5641 |
| X_7690368 | 2 | 1 | 2 | 1 | NA | NA | 1 | 1 | 5 | 1 | NA | NA | NA | NA | 1 | 1 | 1 | 1 | 2 | 1 |
| X_7690470 | 1 | 1 | 2 | 1 | NA | NA | 2 | 1 | 5 | 1 | NA | NA | NA | NA | 1 | 1 | 1 | 0 | 2 | 0.5 |
| X_7690494 | 2 | 0.5 | 2 | 0 | NA | NA | 2 | 0.5 | 6 | 0.3333 | NA | NA | NA | NA | 1 | 0 | 1 | 1 | 2 | 0.5 |
| X_7690655 | 2 | 0.5 | 2 | 1 | NA | NA | 1 | 1 | 5 | 0.8 | 1 | 1 | NA | NA | 1 | 0 | 1 | 1 | 3 | 0.6667 |
| X_7690785 | 1 | 1 | 2 | 1 | NA | NA | 2 | 1 | 5 | 1 | 1 | 1 | 1 | 1 | 1 | 1 | 1 | 1 | 4 | 1 |
| X_7702549 | 1 | 1 | NA | NA | NA | NA | 2 | 0.5 | 3 | 0.6667 | 1 | 0 | NA | NA | 2 | 0.5 | 1 | 0 | 4 | 0.25 |
| X_7702564 | 2 | 0 | 2 | 0 | NA | NA | 2 | 0.5 | 6 | 0.1667 | 1 | 1 | NA | NA | 3 | 0 | 1 | 0 | 5 | 0.2 |
| X_7702719 | 2 | 1 | 2 | 0.5 | NA | NA | 3 | 0.6667 | 7 | 0.7143 | NA | NA | 1 | 1 | 3 | 1 | 2 | 0.5 | 6 | 0.8333 |
| X_7702786 | 2 | 0.5 | 1 | 0 | NA | NA | 2 | 0 | 5 | 0.2 | 1 | 1 | NA | NA | 2 | 0 | 2 | 0 | 5 | 0.2 |
| X_7702861 | 1 | 1 | 2 | 1 | NA | NA | 3 | 0 | 6 | 0.5 | 1 | 1 | NA | NA | 4 | 0.5 | 2 | 0 | 7 | 0.4286 |
| X_7703491 | 2 | 0 | 2 | 0 | NA | NA | 3 | 0.3333 | 7 | 0.1429 | 1 | 1 | NA | NA | 3 | 0 | 2 | 0 | 6 | 0.1667 |
| X_7703609 | 1 | 0 | 2 | 1 | NA | NA | 2 | 0.5 | 5 | 0.6 | NA | NA | 1 | 0 | 4 | 1 | 1 | 1 | 6 | 0.8333 |
| X_7703700 | 2 | 0.5 | 2 | 0 | 1 | 1 | 4 | 0 | 9 | 0.2222 | 1 | 0 | NA | NA | 4 | 0.5 | 2 | 0.5 | 7 | 0.4286 |
| X_7703930 | 2 | 0.5 | NA | NA | NA | NA | 2 | 0.5 | 4 | 0.5 | 1 | 0 | NA | NA | 3 | 0.6667 | 2 | 0.5 | 6 | 0.5 |
| X_7708359 | 1 | 0 | 2 | 0.5 | NA | NA | 1 | 0 | 4 | 0.25 | 4 | 0.25 | 2 | 0 | 3 | 0.3333 | NA | NA | 9 | 0.2222 |
| X_7740597 | 4 | 0 | 15 | 0.2667 | 7 | 0.2857 | 6 | 0.3333 | 32 | 0.25 | 9 | 0 | 10 | 0.1 | 12 | 0.25 | 2 | 0 | 33 | 0.1212 |
| X_7740607 | 5 | 0.2 | 14 | 0.6429 | 7 | 0.2857 | 6 | 0.5 | 32 | 0.4688 | 8 | 0.5 | 10 | 0.5 | 13 | 0.2308 | 3 | 0.6667 | 34 | 0.4118 |
| X_7740612 | 4 | 0.25 | 12 | 0.0833 | 5 | 0 | 4 | 0 | 25 | 0.08 | 6 | 0.1667 | 9 | 0 | 10 | 0 | 2 | 0.5 | 27 | 0.0741 |
| X_7740842 | 6 | 0.6667 | 14 | 0.8571 | 8 | 0.75 | 5 | 0.6 | 33 | 0.7576 | 7 | 0.5714 | 11 | 0.8182 | 11 | 0.7273 | 2 | 0.5 | 31 | 0.7097 |
| X_7740884 | 6 | 0 | 12 | 0.0833 | 8 | 0.125 | 3 | 0.6667 | 29 | 0.1379 | 7 | 0 | 10 | 0.2 | 10 | 0.2 | 3 | 1 | 30 | 0.2333 |
| X_7740921 | 5 | 0.4 | 12 | 0.4167 | 9 | 0.5556 | 5 | 0 | 31 | 0.3871 | 8 | 0 | 10 | 0.5 | 9 | 0.2222 | 3 | 0.3333 | 30 | 0.2667 |
| X_7740972 | 5 | 0.2 | 15 | 0.1333 | 7 | 0.4286 | 6 | 0.1667 | 33 | 0.2121 | 8 | 0.125 | 10 | 0.2 | 11 | 0.4545 | 2 | 0 | 31 | 0.2581 |
| X_7741186 | 8 | 0.125 | 11 | 0.0909 | 6 | 0.1667 | 5 | 0.2 | 30 | 0.1333 | 10 | 0.1 | 5 | 0.2 | 8 | 0 | 1 | 0 | 24 | 0.0833 |
| X_7741298 | 9 | 0.3333 | 12 | 0.1667 | 6 | 0 | 6 | 0.1667 | 33 | 0.1818 | 11 | 0.0909 | 7 | 0.2857 | 10 | 0 | NA | NA | 28 | 0.1071 |
| X_7741305 | 10 | 0.2 | 11 | 0.2727 | 5 | 0 | 5 | 0.2 | 31 | 0.1935 | 10 | 0.4 | 5 | 0.4 | 5 | 0.2 | 2 | 1 | 22 | 0.4091 |
| X_7741684 | 4 | 0 | 4 | 0.25 | 2 | 0 | 2 | 0.5 | 12 | 0.1667 | 2 | 0 | 3 | 0.3333 | 5 | 0.2 | 4 | 0 | 14 | 0.1429 |
| X_7741986 | 3 | 0 | 4 | 0.25 | 2 | 0.5 | 3 | 0.3333 | 12 | 0.25 | 1 | 0 | 3 | 0.3333 | 4 | 0.5 | 3 | 0.3333 | 11 | 0.3636 |
| X_7742689 | 6 | 0.1667 | 5 | 0 | 3 | 0 | 5 | 0 | 19 | 0.0526 | 2 | 0 | 3 | 0 | 6 | 0.1667 | 5 | 0.2 | 16 | 0.125 |
| X_7742819 | 4 | 0 | 5 | 0 | 3 | 0.3333 | 6 | 0.1667 | 18 | 0.1111 | 3 | 0.3333 | 4 | 0 | 6 | 0.1667 | 4 | 0.25 | 17 | 0.1765 |
| X_7748811 | 6 | 0.1667 | 3 | 0 | 5 | 0.4 | 4 | 0.25 | 18 | 0.2222 | 5 | 0.4 | 2 | 0 | 8 | 0.5 | 2 | 0 | 17 | 0.3529 |
| X_7749424 | 10 | 0.1 | 8 | 0.125 | 12 | 0 | 6 | 0.1667 | 36 | 0.0833 | 6 | 0.1667 | 6 | 0.3333 | 8 | 0.375 | 6 | 0 | 26 | 0.2308 |
| X_7758545 | 1 | 0 | 4 | 0.75 | 3 | 0.3333 | 1 | 0 | 9 | 0.4444 | 2 | 0.5 | 2 | 0.5 | 1 | 1 | 2 | 0.5 | 7 | 0.5714 |
| X_7758830 | 3 | 0 | 4 | 0.25 | 4 | 0 | 1 | 0 | 12 | 0.0833 | 5 | 0 | 2 | 0 | 1 | 0 | 2 | 0.5 | 10 | 0.1 |
| X_7758914 | 3 | 0.3333 | 4 | 0.25 | 4 | 1 | 1 | 1 | 12 | 0.5833 | 4 | 0.5 | 3 | 0.6667 | 1 | 0 | 2 | 1 | 10 | 0.6 |
| X_7759307 | 3 | 0.6667 | 3 | 0.3333 | 6 | 0.1667 | NA | NA | 12 | 0.3333 | 3 | 0.3333 | NA | NA | NA | NA | 2 | 1 | 5 | 0.6 |
| X_7759473 | 3 | 0 | 4 | 0.5 | 6 | 0.1667 | NA | NA | 13 | 0.2308 | 3 | 0.3333 | NA | NA | NA | NA | 2 | 0 | 5 | 0.2 |
| X_7760519 | 2 | 0.5 | 3 | 0.3333 | 2 | 0 | NA | NA | 7 | 0.2857 | 1 | 0 | 3 | 0 | 1 | 0 | 1 | 1 | 6 | 0.1667 |
| X_7760954 | 7 | 0 | 9 | 0.2222 | 10 | 0.3 | 14 | 0.1429 | 40 | 0.175 | 13 | 0.3846 | 12 | 0.3333 | 18 | 0.0556 | 10 | 0.3 | 53 | 0.2453 |
| X_7762341 | 7 | 0 | 11 | 0 | 8 | 0 | 15 | 0.2 | 41 | 0.0732 | 12 | 0.0833 | 9 | 0.2222 | 12 | 0 | 8 | 0 | 41 | 0.0732 |
| X_7772893 | 1 | 0 | NA | NA | 4 | 0.25 | 5 | 0.2 | 10 | 0.2 | 2 | 0 | 4 | 0.25 | 2 | 0 | 2 | 1 | 10 | 0.3 |
| X_7796556 | 4 | 0 | 3 | 0 | NA | NA | 2 | 1 | 9 | 0.2222 | 3 | 0 | 1 | 1 | 3 | 0.3333 | 2 | 0.5 | 9 | 0.3333 |

| | | | | | | | | | | | | | | | | | | | | |
|---|---|---|---|---|---|---|---|---|---|---|---|---|---|---|---|---|---|---|---|---|
| X_7797929 | 3 | 0.6667 | 2 | 0 | NA | NA | 1 | 0 | 6 | 0.3333 | 2 | 0.5 | 1 | 0 | 3 | 0 | 4 | 0 | 10 | 0.1 |
| X_7927750 | 4 | 0.25 | 2 | 0 | 3 | 0 | 6 | 0.1667 | 15 | 0.1333 | 4 | 0.25 | 5 | 0.6 | 3 | 0 | 2 | 0 | 14 | 0.2857 |
| X_7928739 | 2 | 0.5 | NA | NA | 1 | 0 | 5 | 0.4 | 8 | 0.375 | 4 | 0.25 | 3 | 0 | 1 | 0 | 1 | 0 | 9 | 0.1111 |
| X_7928929 | 3 | 0.3333 | 1 | 0 | 2 | 0 | 5 | 0.6 | 11 | 0.3636 | 4 | 0.5 | 3 | 0 | 1 | 0 | NA | NA | 8 | 0.25 |
| X_7929118 | 1 | 0 | 1 | 0 | 2 | 0 | 3 | 0.3333 | 7 | 0.1429 | 4 | 0.75 | 2 | 0 | 1 | 0 | NA | NA | 7 | 0.4286 |
| X_7937101 | 4 | 1 | 1 | 1 | 2 | 0.5 | 3 | 0.3333 | 10 | 0.7 | 2 | 1 | 4 | 0.5 | NA | NA | NA | NA | 6 | 0.6667 |
| X_7937323 | 3 | 1 | 1 | 1 | 2 | 0 | 3 | 0.6667 | 9 | 0.6667 | 1 | 0 | 4 | 0.5 | NA | NA | NA | NA | 5 | 0.4 |
| X_7937380 | 3 | 1 | 1 | 1 | 1 | 1 | 3 | 0.6667 | 8 | 0.875 | NA | NA | 1 | 1 | NA | NA | NA | NA | 1 | 1 |
| X_7989953 | 16 | 0.3125 | 20 | 0.4 | 20 | 0.4 | 19 | 0.5789 | 75 | 0.4267 | 12 | 0.25 | 19 | 0.3158 | 15 | 0.4667 | 14 | 0.2857 | 60 | 0.3333 |
| X_7990032 | 13 | 0.2308 | 18 | 0.1111 | 16 | 0.3125 | 20 | 0.3 | 67 | 0.2388 | 12 | 0 | 17 | 0.1176 | 16 | 0.3125 | 15 | 0.2 | 60 | 0.1667 |
| X_7990235 | 16 | 0.6875 | 18 | 0.5 | 17 | 0.5882 | 18 | 0.5556 | 69 | 0.5797 | 13 | 0.3846 | 17 | 0.7647 | 16 | 0.5 | 16 | 0.75 | 62 | 0.6129 |
| X_7990331 | 17 | 0.2941 | 21 | 0.2381 | 18 | 0.3333 | 18 | 0.1667 | 74 | 0.2568 | 10 | 0.4 | 22 | 0.2727 | 14 | 0.1429 | 13 | 0.1538 | 59 | 0.2373 |
| X_7990506 | 13 | 0.3077 | 16 | 0.4375 | 13 | 0.3846 | 12 | 0.5833 | 54 | 0.4259 | 12 | 0.25 | 18 | 0.2222 | 13 | 0.3077 | 14 | 0.2143 | 57 | 0.2456 |
| X_7991636 | 9 | 0.2222 | 12 | 0.25 | 7 | 0.1429 | 10 | 0.2 | 38 | 0.2105 | 3 | 0 | 11 | 0.0909 | 8 | 0 | 11 | 0.0909 | 33 | 0.0606 |
| X_7995374 | 8 | 0.5 | 7 | 0.2857 | 4 | 0 | 4 | 0.25 | 23 | 0.3043 | 1 | 1 | 5 | 0 | 4 | 0 | 5 | 0.2 | 15 | 0.1333 |
| X_7998629 | 5 | 0.2 | 4 | 0 | 3 | 0 | 2 | 0 | 14 | 0.0714 | NA | NA | 4 | 0 | 3 | 0 | 5 | 0.2 | 12 | 0.0833 |
| X_7998797 | 3 | 0 | 3 | 0 | 3 | 0.3333 | 1 | 0 | 10 | 0.1 | NA | NA | 2 | 0 | 1 | 1 | 1 | 1 | 4 | 0.5 |
| X_8009050 | 41 | 0.1463 | 25 | 0.24 | 16 | 0.125 | 26 | 0.0385 | 108 | 0.1389 | 25 | 0.04 | 20 | 0.05 | 22 | 0.1818 | 16 | 0.0625 | 83 | 0.0843 |
| X_8020876 | 1 | 0 | NA | NA | 2 | 0.5 | 3 | 0.6667 | 6 | 0.5 | 1 | 0 | 1 | 1 | NA | NA | NA | NA | 2 | 0.5 |
| X_8020981 | 1 | 1 | NA | NA | 2 | 1 | 3 | 1 | 6 | 1 | NA | NA | 1 | 1 | 2 | 0.5 | NA | NA | 3 | 0.6667 |
| X_8021529 | NA | NA | NA | NA | 2 | 1 | 3 | 1 | 5 | 1 | 1 | 1 | 1 | 1 | NA | NA | NA | NA | 2 | 1 |
| X_8021842 | NA | NA | NA | NA | 2 | 0.5 | 2 | 1 | 4 | 0.75 | 1 | 1 | NA | NA | NA | NA | NA | NA | 1 | 1 |
| X_8021856 | NA | NA | NA | NA | 2 | 0.5 | 2 | 0.5 | 4 | 0.5 | 1 | 1 | 1 | 1 | NA | NA | NA | NA | 2 | 1 |
| X_8021974 | NA | NA | NA | NA | 2 | 1 | 2 | 1 | 4 | 1 | NA | NA | 1 | 1 | NA | NA | NA | NA | 1 | 1 |
| X_8022244 | NA | NA | NA | NA | 2 | 1 | 2 | 0 | 4 | 0.5 | NA | NA | 1 | 1 | NA | NA | NA | NA | 1 | 1 |
| X_8051740 | 5 | 0.6 | 9 | 0.6667 | 6 | 0.5 | 4 | 1 | 24 | 0.6667 | 3 | 0.6667 | 1 | 0 | 2 | 1 | 3 | 0.6667 | 9 | 0.6667 |
| X_8051748 | 4 | 0 | 6 | 0.1667 | 6 | 0 | 4 | 0 | 20 | 0.05 | 3 | 0.3333 | 1 | 0 | 2 | 0 | 2 | 0 | 8 | 0.125 |
| X_8051790 | 4 | 0 | 9 | 0.1111 | 7 | 0 | 4 | 0.25 | 24 | 0.0833 | 2 | 0 | 2 | 0 | 1 | 0 | 3 | 0.3333 | 8 | 0.125 |
| X_8051886 | 4 | 0.5 | 8 | 0.25 | 5 | 0 | 2 | 0.5 | 19 | 0.2632 | 1 | 1 | 1 | 0 | 2 | 0 | 3 | 0.6667 | 7 | 0.4286 |
| X_8106870 | 8 | 0.5 | 3 | 1 | 10 | 0.9 | 7 | 0.5714 | 28 | 0.7143 | 5 | 1 | 7 | 0.5714 | 5 | 0.6 | 2 | 1 | 19 | 0.7368 |
| X_8106967 | 6 | 0 | 3 | 1 | 9 | 0.2222 | 3 | 1 | 21 | 0.381 | 6 | 0.6667 | 6 | 0.3333 | 4 | 0 | 1 | 1 | 17 | 0.4118 |
| X_8759157 | 1 | 1 | 2 | 0.5 | 2 | 1 | 2 | 0 | 7 | 0.5714 | NA | NA | 3 | 1 | 3 | 1 | 1 | 1 | 7 | 1 |
| X_8759334 | 2 | 0.5 | 2 | 0 | 2 | 0 | 3 | 0 | 9 | 0.1111 | 4 | 0 | 5 | 0 | 8 | 0.25 | 3 | 0 | 20 | 0.1 |
| X_8759683 | 2 | 0.5 | 4 | 0.75 | 5 | 1 | 3 | 0.3333 | 14 | 0.7143 | 7 | 0.4286 | 9 | 0.6667 | 9 | 0.4444 | 7 | 0.7143 | 32 | 0.5625 |
| X_8759886 | 2 | 1 | 6 | 1 | 5 | 1 | 5 | 1 | 18 | 1 | 5 | 1 | 9 | 1 | 9 | 1 | 8 | 1 | 31 | 1 |
| X_8760265 | 3 | 0.3333 | 4 | 0.75 | 3 | 0.6667 | 3 | 1 | 13 | 0.6923 | 5 | 0.8 | 9 | 0.6667 | 7 | 0.4286 | 8 | 0.625 | 29 | 0.6207 |
| X_8760361 | 1 | 0 | 5 | 0.4 | 5 | 0.6 | 3 | 0.3333 | 14 | 0.4286 | 7 | 0.1429 | 8 | 0 | 7 | 0.2857 | 8 | 0.5 | 30 | 0.2333 |
| X_8760401 | 3 | 0.3333 | 4 | 0.25 | 4 | 0.25 | 6 | 0.3333 | 17 | 0.2941 | 8 | 0.125 | 8 | 0.125 | 9 | 0.1111 | 8 | 0.125 | 33 | 0.1212 |
| X_8760520 | 1 | 1 | 8 | 0.5 | 5 | 0.6 | 5 | 0.6 | 19 | 0.5789 | 7 | 0.5714 | 10 | 0.5 | 7 | 0.4286 | 6 | 0.5 | 30 | 0.5 |
| X_8760524 | 2 | 0.5 | 7 | 0 | 5 | 0.4 | 5 | 0 | 19 | 0.1579 | 6 | 0.3333 | 8 | 0.125 | 6 | 0 | 8 | 0.125 | 28 | 0.1429 |
| X_8760542 | 3 | 0 | 8 | 0.5 | 5 | 0.6 | 5 | 0.6 | 21 | 0.4762 | 9 | 0.5556 | 11 | 0.7273 | 10 | 0.6 | 9 | 0.6667 | 39 | 0.641 |
| X_8760549 | 3 | 0 | 8 | 0.125 | 6 | 0 | 6 | 0.1667 | 23 | 0.087 | 9 | 0.2222 | 13 | 0.0769 | 10 | 0.1 | 9 | 0 | 41 | 0.0976 |
| X_8760594 | 4 | 1 | 8 | 0.875 | 6 | 1 | 5 | 0.8 | 23 | 0.913 | 8 | 0.75 | 13 | 0.4615 | 10 | 0.7 | 9 | 0.6667 | 40 | 0.625 |
| X_8760618 | 4 | 0 | 7 | 0.1429 | 6 | 0.1667 | 2 | 0 | 19 | 0.1053 | 8 | 0.125 | 9 | 0.1111 | 8 | 0.125 | 9 | 0.2222 | 34 | 0.1471 |
| X_8760637 | 4 | 1 | 5 | 1 | 5 | 1 | 4 | 1 | 18 | 1 | 6 | 1 | 8 | 1 | 8 | 1 | 6 | 1 | 28 | 1 |
| X_8760650 | 5 | 0.8 | 8 | 1 | 5 | 1 | 6 | 1 | 24 | 0.9583 | 9 | 0.8889 | 12 | 0.9167 | 8 | 1 | 9 | 1 | 38 | 0.9474 |
| X_8760666 | 5 | 0 | 8 | 0.125 | 5 | 0 | 6 | 0.1667 | 24 | 0.0833 | 7 | 0 | 13 | 0 | 9 | 0.1111 | 9 | 0.1111 | 38 | 0.0526 |
| X_9137229 | NA | NA | 1 | 1 | 1 | 1 | NA | NA | 2 | 1 | NA | NA | 1 | 0 | 4 | 0.25 | 2 | 0 | 7 | 0.1429 |
| X_9175444 | 1 | 1 | NA | NA | 1 | 1 | 3 | 0.6667 | 5 | 0.8 | 1 | 1 | 1 | 1 | 1 | 0 | 2 | 1 | 5 | 0.8 |
| X_9175585 | 1 | 1 | NA | NA | 1 | 1 | 3 | 0.3333 | 5 | 0.6 | 1 | 1 | 1 | 1 | 1 | 1 | 2 | 1 | 5 | 1 |
| X_9175630 | 1 | 1 | NA | NA | 1 | 0 | 3 | 0.3333 | 5 | 0.4 | 1 | 0 | 1 | 1 | 1 | 0 | 2 | 0.5 | 5 | 0.4 |
| X_9175665 | NA | NA | NA | NA | 1 | 1 | 1 | 1 | 2 | 1 | 1 | 1 | 1 | 1 | NA | NA | NA | NA | 2 | 1 |
| X_9175888 | 1 | 1 | NA | NA | 1 | 1 | 1 | 1 | 3 | 1 | 1 | 1 | 1 | 1 | 1 | 1 | 3 | 0.6667 | 6 | 0.8333 |
| X_9175962 | 1 | 1 | NA | NA | 1 | 1 | 1 | 1 | 3 | 1 | 1 | 1 | NA | NA | 1 | 1 | 3 | 1 | 5 | 1 |
| X_9176050 | 1 | 1 | NA | NA | 1 | 1 | 1 | 1 | 3 | 1 | 1 | 1 | 1 | 0 | NA | NA | 3 | 0.6667 | 5 | 0.6 |
| X_9176172 | 1 | 1 | NA | NA | 1 | 0 | 1 | 0 | 3 | 0.3333 | 1 | 0 | NA | NA | NA | NA | 3 | 0.3333 | 4 | 0.25 |
| X_9179417 | 2 | 1 | 6 | 0.8333 | 2 | 0 | 2 | 0.5 | 12 | 0.6667 | 1 | 1 | 1 | 0 | 2 | 0.5 | 1 | 0 | 5 | 0.4 |
| X_9520942 | 18 | 0 | 20 | 0.25 | 26 | 0.1538 | 17 | 0 | 81 | 0.1111 | 13 | 0 | 14 | 0.2857 | 16 | 0.0625 | 16 | 0.125 | 59 | 0.1186 |
| X_9860673 | NA | NA | 1 | 1 | NA | NA | 1 | 1 | 2 | 1 | NA | NA | 2 | 1 | NA | NA | NA | NA | 2 | 1 |
| X_9860843 | NA | NA | 2 | 1 | NA | NA | NA | NA | 2 | 1 | NA | NA | 2 | 0.5 | NA | NA | NA | NA | 2 | 0.5 |
| X_9860849 | NA | NA | 2 | 1 | NA | NA | 1 | 1 | 3 | 1 | NA | NA | 2 | 1 | NA | NA | NA | NA | 2 | 1 |
| X_9870745 | NA | NA | 1 | 1 | 2 | 0.5 | 4 | 0.25 | 7 | 0.4286 | NA | NA | NA | NA | 2 | 0.5 | 3 | 0 | 5 | 0.2 |
| X_1002477 | 2 | 0 | 5 | 0.6 | 4 | 0.5 | 5 | 0.8 | 16 | 0.5625 | NA | NA | 2 | 0.5 | 3 | 0.6667 | 5 | 0.6 | 10 | 0.6 |
| X_1002478 | 3 | 0 | 4 | 0 | 4 | 0.25 | 6 | 0 | 17 | 0.0588 | 1 | 0 | 4 | 0 | 3 | 0.3333 | 4 | 0 | 12 | 0.0833 |
| X_1002484 | 3 | 0.6667 | 5 | 0.2 | 4 | 0 | 6 | 0.1667 | 18 | 0.2222 | 1 | 0 | 2 | 0 | 3 | 0 | 5 | 0.4 | 11 | 0.1818 |

| | | | | | | | | | | | | | | | | | | | | |
|---|---|---|---|---|---|---|---|---|---|---|---|---|---|---|---|---|---|---|---|---|
| X_1057986 | NA | NA | 1 | 1 | NA | NA | NA | NA | 1 | 1 | NA | NA | NA | NA | 1 | 1 | NA | NA | 1 | 1 |
| X_1058036 | NA | NA | 1 | 1 | NA | NA | NA | NA | 1 | 1 | NA | NA | NA | NA | 1 | 1 | NA | NA | 1 | 1 |
| X_1058379 | 3 | 0.3333 | 5 | 0.4 | 7 | 0.2857 | 9 | 0.2222 | 24 | 0.2917 | 1 | 0 | 5 | 0.4 | 5 | 0.4 | 3 | 0 | 14 | 0.2857 |
| X_1058454 | 7 | 0 | 12 | 0.1667 | 12 | 0 | 12 | 0.1667 | 43 | 0.093 | 5 | 0.2 | 7 | 0 | 9 | 0.1111 | 8 | 0.125 | 29 | 0.1034 |
| X_1058455 | 10 | 0.7 | 12 | 0.6667 | 11 | 0.7273 | 12 | 0.6667 | 45 | 0.6889 | 3 | 1 | 8 | 0.75 | 11 | 0.6364 | 9 | 1 | 31 | 0.8065 |
| X_1058472 | 12 | 0.9167 | 13 | 0.9231 | 15 | 1 | 13 | 0.8462 | 53 | 0.9245 | 4 | 0.75 | 7 | 1 | 12 | 1 | 10 | 0.9 | 33 | 0.9394 |
| X_1058483 | 15 | 0.8667 | 15 | 0.8 | 14 | 1 | 13 | 1 | 57 | 0.9123 | 6 | 0.6667 | 10 | 0.8 | 14 | 0.8571 | 11 | 0.8182 | 41 | 0.8049 |
| X_1058494 | 9 | 0.1111 | 13 | 0 | 13 | 0.1538 | 9 | 0 | 44 | 0.0682 | 4 | 0 | 10 | 0 | 12 | 0.0833 | 8 | 0.125 | 34 | 0.0588 |
| X_1058526 | 16 | 0.25 | 17 | 0.2353 | 12 | 0.4167 | 14 | 0.3571 | 59 | 0.3051 | 4 | 0.75 | 10 | 0.2 | 17 | 0.4706 | 13 | 0.2308 | 44 | 0.3636 |
| X_1190377 | 2 | 0.5 | 3 | 1 | 2 | 1 | 1 | 0 | 8 | 0.75 | 4 | 1 | NA | NA | 2 | 0.5 | 2 | 1 | 8 | 0.875 |
| X_1190387 | 2 | 0 | 2 | 0.5 | 1 | 0 | 2 | 0 | 7 | 0.1429 | 4 | 0 | 1 | 0 | 1 | 0 | 2 | 1 | 8 | 0.25 |
| X_1190389 | 2 | 0 | 3 | 0 | 3 | 0.3333 | 2 | 0 | 10 | 0.1 | 4 | 0.25 | 1 | 1 | 2 | 0 | 2 | 0.5 | 9 | 0.3333 |
| X_1192275 | 1 | 1 | 1 | 1 | NA | NA | 1 | 0 | 3 | 0.6667 | 2 | 0.5 | NA | NA | NA | NA | 1 | 1 | 3 | 0.6667 |
| X_1192284 | 1 | 1 | 1 | 1 | 1 | 0 | 2 | 1 | 5 | 0.8 | 2 | 0.5 | NA | NA | 2 | 1 | 1 | 1 | 5 | 0.8 |
| X_1192300 | 1 | 0 | 1 | 0 | 1 | 1 | 2 | 1 | 5 | 0.6 | NA | NA | NA | NA | 1 | 1 | NA | NA | 1 | 1 |
| X_1192309 | 1 | 0 | 1 | 1 | 1 | 0 | 2 | 0.5 | 5 | 0.4 | 2 | 0 | NA | NA | 2 | 0.5 | 1 | 0 | 5 | 0.2 |
| X_1192314 | 1 | 0 | 1 | 0 | 1 | 1 | 1 | 1 | 4 | 0.5 | 2 | 0.5 | NA | NA | 2 | 0 | 1 | 0 | 5 | 0.2 |
| X_1192356 | 1 | 1 | 1 | 1 | 1 | 1 | 2 | 1 | 5 | 1 | 1 | 0 | NA | NA | 1 | 0 | 1 | 1 | 3 | 0.3333 |
| X_1192399 | NA | NA | NA | NA | 1 | 1 | 2 | 1 | 3 | 1 | 1 | 1 | NA | NA | NA | NA | 1 | 0 | 2 | 0.5 |
| X_1199733 | NA | NA | 2 | 1 | NA | NA | 1 | 1 | 3 | 1 | 2 | 1 | 1 | 1 | NA | NA | 1 | 1 | 4 | 1 |
| X_1199736 | NA | NA | 2 | 1 | NA | NA | 1 | 0 | 3 | 0.6667 | 2 | 0.5 | 1 | 0 | NA | NA | 1 | 1 | 4 | 0.5 |
| X_1199759 | NA | NA | 2 | 0.5 | NA | NA | 1 | 0 | 3 | 0.3333 | 1 | 0 | 1 | 1 | NA | NA | 1 | 1 | 3 | 0.6667 |
| X_1199762 | NA | NA | 3 | 0 | NA | NA | 1 | 1 | 4 | 0.25 | 2 | 0.5 | 1 | 0 | NA | NA | 1 | 0 | 4 | 0.25 |
| X_1199767 | NA | NA | 1 | 1 | NA | NA | 1 | 1 | 2 | 1 | 2 | 1 | 1 | 1 | NA | NA | 1 | 1 | 4 | 1 |
| X_1199842 | NA | NA | 3 | 0.3333 | NA | NA | 1 | 0 | 4 | 0.25 | 2 | 0.5 | 1 | 1 | NA | NA | 1 | 1 | 4 | 0.75 |
| X_1200280 | 1 | 1 | NA | NA | NA | NA | NA | NA | 1 | 1 | 1 | 0 | NA | NA | 1 | 1 | 1 | 0 | 3 | 0.3333 |
| X_1200310 | 1 | 1 | 1 | 0 | NA | NA | NA | NA | 2 | 0.5 | 1 | 1 | NA | NA | NA | NA | NA | NA | 1 | 1 |
| X_1200312 | 1 | 1 | 1 | 1 | NA | NA | NA | NA | 2 | 1 | 1 | 1 | NA | NA | NA | NA | NA | NA | 1 | 1 |
| X_1200317 | 1 | 0 | 1 | 1 | NA | NA | NA | NA | 2 | 0.5 | 1 | 1 | NA | NA | NA | NA | NA | NA | 1 | 1 |
| X_1248239 | 54 | 0.0926 | 35 | 0.0571 | 43 | 0.1395 | 32 | 0.3125 | 164 | 0.1402 | 16 | 0.125 | 35 | 0.1714 | 37 | 0.1892 | 33 | 0.1818 | 121 | 0.1736 |
| X_1248251 | 57 | 0.0175 | 37 | 0.0541 | 45 | 0.0889 | 35 | 0.1143 | 174 | 0.0632 | 22 | 0.0455 | 38 | 0.0789 | 35 | 0.1429 | 38 | 0.1053 | 133 | 0.0977 |
| X_1252142 | 4 | 0.25 | NA | NA | 1 | 0 | 5 | 0.2 | 10 | 0.2 | NA | NA | 2 | 0.5 | 2 | 0 | 4 | 0.5 | 8 | 0.375 |
| X_1252164 | 4 | 0.25 | NA | NA | 2 | 0.5 | 5 | 0.2 | 11 | 0.2727 | NA | NA | 1 | 0 | 3 | 0.3333 | 3 | 0.3333 | 7 | 0.2857 |
| X_1252179 | 4 | 0.25 | NA | NA | 1 | 1 | 6 | 0.3333 | 11 | 0.3636 | NA | NA | 2 | 0.5 | 3 | 0 | 4 | 0 | 9 | 0.1111 |
| X_1252199 | 3 | 0.3333 | NA | NA | 1 | 0 | 6 | 0.3333 | 10 | 0.3 | NA | NA | 2 | 0 | 4 | 0.25 | 4 | 0.5 | 10 | 0.3 |
| X_1252255 | 2 | 0.5 | NA | NA | 1 | 1 | 4 | 0.25 | 7 | 0.4286 | NA | NA | 1 | 1 | 3 | 0.6667 | 5 | 0.4 | 9 | 0.5556 |
| X_1252271 | 1 | 0 | NA | NA | NA | NA | 4 | 0.25 | 5 | 0.2 | NA | NA | 1 | 0 | 3 | 0.3333 | 5 | 0.2 | 9 | 0.2222 |
| X_1254371 | NA | NA | NA | NA | NA | NA | 3 | 1 | 3 | 1 | NA | NA | 1 | 1 | 1 | 0 | 1 | 1 | 3 | 0.6667 |
| X_1254381 | 1 | 0 | NA | NA | NA | NA | 3 | 1 | 4 | 0.75 | NA | NA | 3 | 0.3333 | 1 | 0 | 1 | 0 | 5 | 0.2 |
| X_1254616 | 6 | 0.1667 | 7 | 0.1429 | 7 | 0.2857 | 9 | 0.2222 | 29 | 0.2069 | 5 | 0 | 5 | 0.2 | 1 | 1 | 3 | 0.3333 | 14 | 0.2143 |
| X_1254623 | 8 | 0.25 | 9 | 0.4444 | 6 | 0 | 7 | 0.4286 | 30 | 0.3 | 5 | 0 | 7 | 0.4286 | 1 | 0 | 3 | 0 | 16 | 0.1875 |
| X_1254626 | 8 | 0.375 | 11 | 0.4545 | 6 | 0.3333 | 8 | 0.375 | 33 | 0.3939 | 6 | 0.3333 | 7 | 0.4286 | 1 | 0 | 4 | 0.25 | 18 | 0.3333 |
| X_1280351 | 2 | 0.5 | NA | NA | NA | NA | 2 | 0 | 4 | 0.25 | NA | NA | NA | NA | 2 | 0.5 | NA | NA | 2 | 0.5 |
| X_1280386 | 1 | 1 | NA | NA | 1 | 1 | 2 | 1 | 4 | 1 | NA | NA | NA | NA | 1 | 1 | NA | NA | 1 | 1 |
| X_1280391 | 2 | 0.5 | NA | NA | 1 | 1 | 3 | 1 | 6 | 0.8333 | NA | NA | 1 | 1 | 2 | 1 | 1 | 1 | 4 | 1 |
| X_1280392 | 2 | 1 | NA | NA | NA | NA | 3 | 1 | 5 | 1 | NA | NA | 1 | 1 | 2 | 1 | 1 | 0 | 4 | 0.75 |
| X_1280396 | 3 | 0.3333 | NA | NA | 1 | 1 | 3 | 0.6667 | 7 | 0.5714 | NA | NA | 1 | 0 | 2 | 1 | 1 | 0 | 4 | 0.5 |
| X_1280403 | 2 | 1 | NA | NA | 1 | 1 | 3 | 0.6667 | 6 | 0.8333 | NA | NA | NA | NA | 2 | 1 | 1 | 1 | 3 | 1 |
| X_1280406 | 1 | 0 | NA | NA | 1 | 1 | 2 | 0 | 4 | 0.25 | NA | NA | 1 | 0 | 2 | 0 | 1 | 1 | 4 | 0.25 |
| X_1280415 | 3 | 1 | NA | NA | 1 | 1 | 3 | 0.6667 | 7 | 0.8571 | NA | NA | 1 | 0 | 2 | 1 | 1 | 1 | 4 | 0.75 |
| X_1280437 | 2 | 0.5 | NA | NA | 1 | 0 | 3 | 0.3333 | 6 | 0.3333 | NA | NA | 1 | 0 | 2 | 1 | 1 | 0 | 4 | 0.5 |
| X_1280448 | 3 | 0.3333 | NA | NA | 1 | 0 | 3 | 0 | 7 | 0.1429 | NA | NA | 1 | 0 | 2 | 0.5 | 1 | 0 | 4 | 0.25 |
| X_1296025 | 1 | 0 | NA | NA | 3 | 0 | 1 | 1 | 5 | 0.2 | 1 | 1 | 2 | 0 | NA | NA | 1 | 0 | 4 | 0.25 |
| X_1298957 | NA | NA | 3 | 0.6667 | 2 | 0 | 2 | 0 | 7 | 0.2857 | 1 | 0 | 4 | 0 | NA | NA | 3 | 0.3333 | 8 | 0.125 |
| X_1301146 | NA | NA | 5 | 0.2 | 4 | 0 | 6 | 0 | 15 | 0.0667 | 5 | 0 | 8 | 0 | 4 | 0 | 7 | 0.2857 | 24 | 0.0833 |
| X_1302237 | 2 | 0 | 7 | 0.1429 | 4 | 0.25 | 11 | 0 | 24 | 0.0833 | 5 | 0.4 | 9 | 0 | 6 | 0.1667 | 8 | 0 | 28 | 0.1071 |
| X_1302249 | 4 | 1 | 5 | 0.6 | 3 | 1 | 7 | 0.5714 | 19 | 0.7368 | 4 | 0.75 | 4 | 1 | 5 | 0.8 | 7 | 0.5714 | 20 | 0.75 |
| X_1302253 | 4 | 1 | 7 | 1 | 4 | 1 | 11 | 0.8182 | 26 | 0.9231 | 3 | 1 | 9 | 0.8889 | 5 | 0.6 | 8 | 0.875 | 25 | 0.84 |
| X_1302277 | 5 | 0.8 | 7 | 0.8571 | 4 | 1 | 12 | 0.6667 | 28 | 0.7857 | 5 | 1 | 8 | 0.625 | 6 | 1 | 8 | 0.75 | 27 | 0.8148 |
| X_1303565 | 7 | 0 | 14 | 0.1429 | 8 | 0.25 | 19 | 0 | 48 | 0.0833 | 8 | 0.25 | 9 | 0.1111 | 14 | 0.0714 | 9 | 0.1111 | 40 | 0.125 |
| X_1303573 | 7 | 0.2857 | 11 | 0.3636 | 7 | 0.2857 | 11 | 0.0909 | 36 | 0.25 | 6 | 0.1667 | 9 | 0.3333 | 12 | 0.5 | 9 | 0.2222 | 36 | 0.3333 |
| X_1303575 | 8 | 0.5 | 13 | 0.5385 | 8 | 0.5 | 16 | 0.75 | 45 | 0.6 | 7 | 0.5714 | 9 | 1 | 13 | 0.7692 | 12 | 0.75 | 41 | 0.7805 |
| X_1303580 | 6 | 0 | 10 | 0.1 | 8 | 0.25 | 13 | 0.0769 | 37 | 0.1081 | 6 | 0 | 8 | 0.125 | 12 | 0.1667 | 10 | 0.4 | 36 | 0.1944 |
| X_1303581 | 8 | 0.25 | 14 | 0.3571 | 8 | 0.375 | 17 | 0.4118 | 47 | 0.3617 | 8 | 0 | 10 | 0.6 | 15 | 0.3333 | 9 | 0.5556 | 42 | 0.381 |
| X_1303598 | 8 | 0 | 13 | 0.3077 | 9 | 0.1111 | 14 | 0.1429 | 44 | 0.1591 | 6 | 0.3333 | 9 | 0 | 12 | 0.1667 | 8 | 0.125 | 35 | 0.1429 |

| | | | | | | | | | | | | | | | | | | | | |
|---|---|---|---|---|---|---|---|---|---|---|---|---|---|---|---|---|---|---|---|---|
| X_1303602 | 8 | 0 | 11 | 0 | 9 | 0.1111 | 16 | 0.125 | 44 | 0.0682 | 7 | 0.1429 | 9 | 0.2222 | 12 | 0.3333 | 9 | 0.2222 | 37 | 0.2432 |
| X_1303649 | 5 | 0.2 | 11 | 0.0909 | 9 | 0.1111 | 16 | 0.1875 | 41 | 0.1463 | 6 | 0.1667 | 8 | 0.125 | 14 | 0.1429 | 9 | 0.1111 | 37 | 0.1351 |
| X_1303658 | 8 | 0 | 13 | 0.1538 | 10 | 0.1 | 16 | 0 | 47 | 0.0638 | 6 | 0 | 10 | 0.1 | 14 | 0.0714 | 8 | 0 | 38 | 0.0526 |
| X_1303672 | 8 | 0 | 15 | 0.0667 | 9 | 0 | 15 | 0.2 | 47 | 0.0851 | 6 | 0 | 11 | 0.1818 | 14 | 0.2857 | 10 | 0.1 | 41 | 0.1707 |
| X_1303724 | 4 | 0 | 5 | 0 | 4 | 0.25 | 6 | 0 | 19 | 0.0526 | 3 | 0.3333 | 4 | 0 | 6 | 0.3333 | 7 | 0.5714 | 20 | 0.35 |
| X_1303909 | 2 | 0 | 7 | 0 | 4 | 0.25 | 4 | 0 | 17 | 0.0588 | 1 | 0 | 4 | 0.25 | 5 | 0 | 4 | 0 | 14 | 0.0714 |
| X_1303910 | 2 | 0.5 | 7 | 0.2857 | 4 | 0 | 4 | 0 | 17 | 0.1765 | 2 | 0 | 4 | 0.25 | 4 | 0.25 | 5 | 0.4 | 15 | 0.2667 |
| X_1304584 | 1 | 0 | NA | NA | 1 | 1 | NA | NA | 2 | 0.5 | 1 | 0 | 1 | 1 | 2 | 1 | 1 | 1 | 5 | 0.8 |
| X_1304658 | 3 | 0 | 1 | 1 | 2 | 1 | 1 | 0 | 7 | 0.4286 | 1 | 0 | 1 | 1 | 1 | 1 | 2 | 0 | 5 | 0.4 |
| X_1304659 | 2 | 0.5 | 2 | 0 | 3 | 0 | 1 | 0 | 8 | 0.125 | 2 | 0 | 1 | 0 | 2 | 0.5 | 2 | 0 | 7 | 0.1429 |
| X_1304677 | 3 | 0.3333 | 2 | 0 | 3 | 0 | 1 | 0 | 9 | 0.1111 | 2 | 0 | 1 | 0 | 1 | 0 | 3 | 0.3333 | 7 | 0.1429 |
| X_1304682 | 3 | 0 | 2 | 0 | 3 | 0.3333 | 1 | 0 | 9 | 0.1111 | 2 | 0 | 1 | 0 | 1 | 0 | 3 | 0.3333 | 7 | 0.1429 |
| X_1304734 | 1 | 0 | 1 | 1 | 1 | 0 | 2 | 0.5 | 5 | 0.4 | 2 | 0.5 | 1 | 1 | NA | NA | 3 | 0 | 6 | 0.3333 |
| X_1305592 | 1 | 0 | 1 | 1 | NA | NA | 1 | 1 | 3 | 0.6667 | NA | NA | 1 | 1 | NA | NA | 1 | 1 | 2 | 1 |
| X_1305594 | 1 | 1 | NA | NA | NA | NA | NA | NA | 1 | 1 | NA | NA | 1 | 1 | NA | NA | 1 | 1 | 2 | 1 |
| X_1305600 | NA | NA | 2 | 0.5 | NA | NA | 1 | 0 | 3 | 0.3333 | NA | NA | 1 | 1 | NA | NA | 1 | 0 | 2 | 0.5 |
| X_1305669 | 1 | 0 | 2 | 0 | 1 | 1 | NA | NA | 4 | 0.25 | 1 | 0 | 1 | 0 | 1 | 0 | 1 | 1 | 4 | 0.25 |
| X_1315771 | 7 | 0.4286 | 12 | 0.0833 | 11 | 0.1818 | 6 | 0 | 36 | 0.1667 | 3 | 0.3333 | 8 | 0 | 3 | 0 | 8 | 0.25 | 22 | 0.1364 |
| X_1315779 | 9 | 0.3333 | 15 | 0.2667 | 14 | 0.1429 | 7 | 0.2857 | 45 | 0.2444 | 3 | 0.3333 | 8 | 0.125 | 5 | 0.2 | 15 | 0.3333 | 31 | 0.2581 |
| X_1315782 | 9 | 0.3333 | 13 | 0.3846 | 15 | 0.2667 | 7 | 0.5714 | 44 | 0.3636 | 3 | 0.6667 | 10 | 0.3 | 5 | 0.6 | 12 | 0.5 | 30 | 0.4667 |
| X_1315801 | 10 | 0 | 15 | 0 | 14 | 0 | 8 | 0.375 | 47 | 0.0638 | 3 | 0 | 11 | 0 | 4 | 0 | 14 | 0.1429 | 32 | 0.0625 |
| X_1315813 | 8 | 0 | 10 | 0.2 | 15 | 0.0667 | 8 | 0.125 | 41 | 0.0976 | 4 | 0 | 9 | 0.1111 | 4 | 0.5 | 12 | 0.1667 | 29 | 0.1724 |
| X_1338830 | 3 | 0.3333 | 1 | 1 | 3 | 0 | 7 | 0 | 14 | 0.1429 | 1 | 0 | 5 | 0.4 | 6 | 0.3333 | 3 | 0 | 15 | 0.2667 |
| X_1350554 | NA | NA | 2 | 0.5 | 1 | 0 | 3 | 0 | 6 | 0.1667 | 3 | 0 | 3 | 0.3333 | 2 | 0 | NA | NA | 8 | 0.125 |
| X_1350565 | NA | NA | 2 | 1 | NA | NA | 2 | 1 | 4 | 1 | 2 | 1 | 2 | 1 | 2 | 1 | NA | NA | 6 | 1 |
| X_1350581 | 1 | 0 | 2 | 1 | 1 | 1 | 2 | 0.5 | 6 | 0.6667 | 3 | 1 | 3 | 0.6667 | 1 | 1 | NA | NA | 7 | 0.8571 |
| X_1350584 | 1 | 0 | 2 | 0.5 | 1 | 0 | 3 | 1 | 7 | 0.5714 | 2 | 1 | 2 | 0.5 | 2 | 0 | NA | NA | 6 | 0.5 |
| X_1350586 | NA | NA | 2 | 0 | NA | NA | 2 | 0.5 | 4 | 0.25 | 2 | 1 | 3 | 0 | 2 | 0 | NA | NA | 7 | 0.2857 |
| X_1350607 | 2 | 0 | 2 | 0.5 | NA | NA | 2 | 0.5 | 6 | 0.3333 | 3 | 0.6667 | 2 | 0.5 | 2 | 0 | NA | NA | 7 | 0.4286 |
| X_1350611 | 2 | 0.5 | 2 | 0 | 1 | 0 | 3 | 0 | 8 | 0.125 | 1 | 1 | 3 | 0 | 2 | 0 | NA | NA | 6 | 0.1667 |
| X_1350653 | 2 | 1 | 3 | 0 | 1 | 1 | 3 | 0.6667 | 9 | 0.5556 | 3 | 0.3333 | 3 | 0.3333 | 2 | 0 | 1 | 0 | 9 | 0.2222 |
| X_1675212 | 1 | 0 | 2 | 0.5 | 2 | 0 | NA | NA | 5 | 0.2 | 4 | 0 | NA | NA | 1 | 0 | 2 | 0.5 | 7 | 0.1429 |
| X_1675215 | 1 | 0 | 2 | 0 | 1 | 1 | NA | NA | 4 | 0.25 | 2 | 0 | NA | NA | 1 | 0 | 2 | 0.5 | 5 | 0.2 |
| X_1675253 | 1 | 1 | 2 | 0 | 2 | 0.5 | NA | NA | 5 | 0.4 | 4 | 0.25 | NA | NA | 1 | 1 | 2 | 0 | 7 | 0.2857 |
| X_1675254 | 1 | 0 | 1 | 1 | 1 | 1 | NA | NA | 3 | 0.6667 | 4 | 1 | NA | NA | 1 | 1 | 2 | 1 | 7 | 1 |
| X_1675256 | 1 | 1 | 2 | 1 | 2 | 1 | NA | NA | 5 | 1 | 4 | 0.75 | NA | NA | 1 | 1 | 1 | 1 | 6 | 0.8333 |
| X_1675264 | NA | NA | 2 | 1 | 2 | 1 | NA | NA | 4 | 1 | 4 | 0.75 | NA | NA | 1 | 1 | 2 | 1 | 7 | 0.8571 |
| X_1675268 | 1 | 1 | 2 | 0 | 1 | 1 | NA | NA | 4 | 0.5 | 4 | 0.25 | NA | NA | 1 | 1 | 2 | 0.5 | 7 | 0.4286 |
| X_1675270 | 1 | 1 | 2 | 0.5 | 2 | 1 | NA | NA | 5 | 0.8 | 4 | 0.75 | NA | NA | 1 | 0 | 2 | 0.5 | 7 | 0.5714 |
| X_1676071 | 1 | 1 | 2 | 1 | 2 | 0.5 | NA | NA | 5 | 0.8 | 4 | 1 | NA | NA | 1 | 1 | 2 | 1 | 7 | 1 |
| X_1676073 | 1 | 0 | 1 | 1 | 1 | 0 | NA | NA | 3 | 0.3333 | 4 | 0.75 | NA | NA | NA | NA | 2 | 0.5 | 6 | 0.6667 |
| X_1676090 | 1 | 1 | 2 | 1 | 1 | 1 | NA | NA | 4 | 1 | 3 | 0.3333 | NA | NA | 1 | 1 | 1 | 0 | 5 | 0.4 |
| X_1676095 | 1 | 1 | 1 | 1 | 2 | 0.5 | NA | NA | 4 | 0.75 | 3 | 0 | NA | NA | 1 | 0 | 2 | 0.5 | 6 | 0.1667 |
| X_1725994 | NA | NA | 1 | 1 | NA | NA | NA | NA | 1 | 1 | NA | NA | 1 | 1 | NA | NA | NA | NA | 1 | 1 |
| X_1742407 | 5 | 0.4 | 6 | 0 | 6 | 0 | 8 | 0.125 | 25 | 0.12 | 4 | 0.25 | 4 | 0.25 | 4 | 0 | 4 | 0 | 16 | 0.125 |
| X_1742412 | 10 | 0.4 | 8 | 0.375 | 11 | 0.4545 | 11 | 0.2727 | 40 | 0.375 | 7 | 0.5714 | 7 | 0 | 7 | 0.7143 | 5 | 0.2 | 26 | 0.3846 |
| X_1742417 | 9 | 0.8889 | 8 | 0.875 | 10 | 0.7 | 10 | 0.7 | 37 | 0.7838 | 7 | 0.5714 | 7 | 1 | 7 | 0.5714 | 6 | 0.8333 | 27 | 0.7407 |
| X_1811712 | 2 | 1 | NA | NA | 1 | 1 | NA | NA | 3 | 1 | 1 | 0 | 1 | 0 | 3 | 1 | 2 | 1 | 7 | 0.7143 |
| X_1812036 | 2 | 1 | NA | NA | 1 | 0 | NA | NA | 3 | 0.6667 | 1 | 0 | 1 | 0 | 3 | 0.6667 | 1 | 0 | 6 | 0.3333 |
| X_1812055 | 2 | 1 | NA | NA | 1 | 1 | NA | NA | 3 | 1 | 1 | 1 | 1 | 0 | 3 | 1 | 2 | 1 | 7 | 0.8571 |
| X_1812064 | 2 | 0.5 | NA | NA | 1 | 0 | NA | NA | 3 | 0.3333 | 1 | 0 | 1 | 0 | 3 | 0.6667 | 2 | 0 | 7 | 0.2857 |
| X_1812075 | 2 | 0.5 | NA | NA | 1 | 1 | 1 | 1 | 4 | 0.75 | 1 | 0 | 1 | 0 | 3 | 0.6667 | 2 | 1 | 7 | 0.5714 |
| X_1814490 | 5 | 0.8 | 2 | 1 | 1 | 0 | 2 | 0.5 | 10 | 0.7 | 1 | 1 | 1 | 0 | 5 | 0.4 | 4 | 1 | 11 | 0.6364 |
| X_1814507 | 4 | 1 | 2 | 1 | 1 | 1 | 2 | 1 | 9 | 1 | 1 | 1 | 2 | 1 | 5 | 1 | 4 | 1 | 12 | 1 |
| X_1814509 | 5 | 0.4 | 2 | 0 | 1 | 1 | 2 | 0 | 10 | 0.3 | 1 | 0 | 2 | 1 | 5 | 0.2 | 4 | 0.25 | 12 | 0.3333 |
| X_1814515 | 5 | 0 | 3 | 0 | 1 | 0 | 1 | 1 | 10 | 0.1 | 1 | 1 | 3 | 0 | 4 | 0 | 4 | 0.5 | 12 | 0.25 |
| X_1814517 | 4 | 0.75 | 3 | 1 | 1 | 1 | 1 | 1 | 9 | 0.8889 | 1 | 1 | 3 | 0.3333 | 5 | 0.6 | 4 | 0.75 | 13 | 0.6154 |
| X_1814550 | 6 | 0.1667 | 3 | 0 | 1 | 0 | 2 | 0 | 12 | 0.0833 | 1 | 0 | 4 | 0.25 | 5 | 0.4 | 3 | 0 | 13 | 0.2308 |
| X_1901176 | 1 | 0 | NA | NA | 1 | 1 | 4 | 0.25 | 6 | 0.3333 | NA | NA | 1 | 1 | NA | NA | 1 | 0 | 2 | 0.5 |
| X_1901197 | 1 | 1 | NA | NA | 1 | 0 | 1 | 0 | 3 | 0.3333 | NA | NA | 1 | 1 | NA | NA | 1 | 0 | 2 | 0.5 |
| X_1901213 | 1 | 1 | NA | NA | NA | NA | 4 | 0.5 | 5 | 0.6 | NA | NA | 1 | 1 | NA | NA | 1 | 1 | 2 | 1 |
| X_1996540 | 5 | 0.6 | 6 | 0.8333 | 5 | 1 | 8 | 0.75 | 24 | 0.7917 | 1 | 1 | 7 | 1 | 5 | 0.8 | 3 | 0.6667 | 16 | 0.875 |
| X_1996558 | 5 | 0.2 | 6 | 0.3333 | 7 | 0.4286 | 9 | 0.3333 | 27 | 0.3333 | 1 | 1 | 9 | 0.5556 | 6 | 0 | 4 | 0.25 | 20 | 0.35 |
| X_1996570 | 7 | 0.1429 | 6 | 0.1667 | 6 | 0 | 9 | 0.1111 | 28 | 0.1071 | 2 | 0 | 7 | 0 | 3 | 0.3333 | 6 | 0.5 | 18 | 0.2222 |
| X_1996580 | 7 | 0.7143 | 7 | 0.7143 | 7 | 0.8571 | 8 | 0.875 | 29 | 0.7931 | 2 | 1 | 7 | 0.5714 | 6 | 0.6667 | 4 | 0.5 | 19 | 0.6316 |

| | | | | | | | | | | | | | | | | | | | | |
|---|---|---|---|---|---|---|---|---|---|---|---|---|---|---|---|---|---|---|---|---|
| X_1996586 | 4 | 0.25 | 5 | 0.2 | 4 | 0 | 5 | 0.2 | 18 | 0.1667 | NA | NA | 6 | 0 | 4 | 0 | 6 | 0.1667 | 16 | 0.0625 |
| X_1996588 | 5 | 0.8 | 7 | 0.4286 | 6 | 0.3333 | 9 | 0.5556 | 27 | 0.5185 | 2 | 0.5 | 8 | 0.625 | 6 | 0.1667 | 6 | 0.8333 | 22 | 0.5455 |
| X_1996593 | 5 | 0.4 | 5 | 0.8 | 6 | 0.5 | 9 | 0.3333 | 25 | 0.48 | 1 | 0 | 7 | 0.5714 | 4 | 0.25 | 6 | 0.5 | 18 | 0.4444 |
| X_1996621 | 5 | 0.2 | 6 | 0.1667 | 6 | 0.5 | 8 | 0.25 | 25 | 0.28 | 2 | 0 | 6 | 0.1667 | 5 | 0.2 | 6 | 0.3333 | 19 | 0.2105 |
| X_1996633 | 5 | 0 | 5 | 0.2 | 5 | 0.4 | 5 | 0 | 20 | 0.15 | 2 | 0 | 4 | 0 | 3 | 0.3333 | 3 | 0.3333 | 12 | 0.1667 |
| X_1996650 | 2 | 0 | 2 | 0 | 2 | 0 | 3 | 0.3333 | 9 | 0.1111 | 1 | 0 | 1 | 0 | 3 | 0 | 4 | 0.25 | 9 | 0.1111 |
| X_1996678 | 4 | 0 | 7 | 0.1429 | 7 | 0.2857 | 8 | 0 | 26 | 0.1154 | 2 | 0.5 | 5 | 0 | 4 | 0 | 6 | 0.3333 | 17 | 0.1765 |
| X_1996696 | 3 | 0 | 7 | 0.1429 | 5 | 0 | 7 | 0.2857 | 22 | 0.1364 | 2 | 0 | 5 | 0 | 4 | 0.25 | 6 | 0.1667 | 17 | 0.1176 |
| X_1996713 | 3 | 0 | 4 | 0.5 | 4 | 0 | 8 | 0.125 | 19 | 0.1579 | 1 | 0 | 4 | 0 | 3 | 0 | 5 | 0.4 | 13 | 0.1538 |
| X_1996755 | NA | NA | 3 | 0 | 5 | 0.6 | 7 | 0 | 15 | 0.2 | 2 | 0.5 | 4 | 0 | 4 | 0.25 | 3 | 0.6667 | 13 | 0.3077 |
| X_1996759 | NA | NA | 4 | 0.25 | 6 | 0.1667 | 8 | 0.375 | 18 | 0.2778 | 2 | 0 | 5 | 0.8 | 5 | 0.4 | 4 | 0.5 | 16 | 0.5 |
| X_1996775 | NA | NA | 4 | 0 | 6 | 0.3333 | 7 | 0.1429 | 17 | 0.1765 | 2 | 0 | 1 | 0 | 5 | 0.2 | 3 | 0.3333 | 11 | 0.1818 |
| X_1996792 | NA | NA | 4 | 0 | 6 | 0 | 9 | 0.1111 | 19 | 0.0526 | 2 | 0 | 5 | 0.2 | 5 | 0 | 3 | 0 | 15 | 0.0667 |
| X_1996806 | NA | NA | 4 | 0 | 4 | 0.25 | 6 | 0 | 14 | 0.0714 | 2 | 0.5 | 4 | 0 | 4 | 0 | 3 | 0 | 13 | 0.0769 |
| X_1997204 | NA | NA | 1 | 0 | 1 | 0 | 1 | 1 | 3 | 0.3333 | 1 | 0 | 2 | 0.5 | 1 | 0 | 1 | 0 | 5 | 0.2 |
| X_1997210 | NA | NA | 1 | 0 | 1 | 0 | 1 | 1 | 3 | 0.3333 | 1 | 1 | 2 | 0.5 | 2 | 0.5 | NA | NA | 5 | 0.6 |
| X_1997227 | NA | NA | 1 | 0 | NA | NA | 3 | 0.3333 | 4 | 0.25 | 1 | 0 | 1 | 0 | 2 | 0 | 1 | 1 | 5 | 0.2 |
| X_2024322 | 1 | 1 | NA | NA | 1 | 1 | 2 | 1 | 4 | 1 | NA | NA | 1 | 1 | NA | NA | NA | NA | 1 | 1 |
| X_2024328 | 1 | 1 | NA | NA | 1 | 1 | 2 | 0.5 | 4 | 0.75 | NA | NA | 1 | 1 | 1 | 0 | NA | NA | 2 | 0.5 |
| X_2024329 | 1 | 0 | NA | NA | 1 | 1 | 2 | 0 | 4 | 0.25 | NA | NA | 1 | 1 | 1 | 0 | NA | NA | 2 | 0.5 |
| X_2024337 | 1 | 1 | NA | NA | 1 | 1 | 1 | 1 | 3 | 1 | NA | NA | 1 | 1 | 1 | 1 | NA | NA | 2 | 1 |
| X_2024362 | 1 | 0 | NA | NA | 1 | 0 | 2 | 0.5 | 4 | 0.25 | 1 | 0 | 1 | 1 | NA | NA | 1 | 0 | 3 | 0.3333 |
| X_2038322 | NA | NA | NA | NA | 1 | 1 | 1 | 0 | 2 | 0.5 | NA | NA | NA | NA | 1 | 1 | 1 | 0 | 2 | 0.5 |
| X_2038334 | NA | NA | NA | NA | 1 | 1 | 1 | 0 | 2 | 0.5 | NA | NA | NA | NA | 1 | 0 | 1 | 1 | 2 | 0.5 |
| X_2038360 | NA | NA | NA | NA | 1 | 1 | 1 | 1 | 2 | 1 | NA | NA | NA | NA | 1 | 1 | 1 | 1 | 2 | 1 |
| X_2048418 | 1 | 0 | NA | NA | 4 | 0.25 | NA | NA | 5 | 0.2 | 2 | 0.5 | 1 | 0 | NA | NA | NA | NA | 3 | 0.3333 |
| X_2051698 | 10 | 0.2 | 6 | 0.1667 | 8 | 0.375 | 7 | 0.7143 | 31 | 0.3548 | 5 | 0.6 | 1 | 0 | 6 | 0.1667 | 5 | 0.2 | 17 | 0.2941 |
| X_2051706 | 9 | 0.4444 | 6 | 0 | 8 | 0.25 | 9 | 0.1111 | 32 | 0.2188 | 5 | 0 | 2 | 0 | 9 | 0.1111 | 6 | 0.1667 | 22 | 0.0909 |
| X_2056529 | 17 | 0 | 23 | 0.1304 | 10 | 0.2 | 13 | 0 | 63 | 0.0794 | 18 | 0.2778 | 11 | 0.2727 | 8 | 0.125 | 9 | 0.1111 | 46 | 0.2174 |
| X_2056568 | 26 | 0.0769 | 26 | 0.1923 | 17 | 0.1765 | 18 | 0.1111 | 87 | 0.1379 | 19 | 0.1053 | 10 | 0 | 8 | 0 | 13 | 0.1538 | 50 | 0.08 |
| X_2056569 | 20 | 0.15 | 17 | 0.1176 | 11 | 0.0909 | 14 | 0.0714 | 62 | 0.1129 | 14 | 0.1429 | 8 | 0 | 5 | 0 | 7 | 0 | 34 | 0.0588 |
| X_2056588 | 19 | 0 | 27 | 0.0741 | 17 | 0 | 17 | 0.1176 | 80 | 0.05 | 17 | 0.2941 | 13 | 0.1538 | 12 | 0 | 11 | 0.0909 | 53 | 0.1509 |
| X_2056594 | 12 | 0.25 | 24 | 0.0833 | 13 | 0 | 14 | 0.2143 | 63 | 0.127 | 17 | 0.2353 | 13 | 0.2308 | 9 | 0.1111 | 10 | 0.1 | 49 | 0.1837 |
| X_2058477 | 2 | 0 | 2 | 0 | 2 | 0.5 | 2 | 0.5 | 8 | 0.25 | NA | NA | 2 | 0 | 2 | 1 | 2 | 0.5 | 6 | 0.5 |
| X_2071832 | 16 | 0.0625 | 20 | 0.15 | 23 | 0.1304 | 27 | 0.0741 | 86 | 0.1047 | 13 | 0 | 23 | 0.0435 | 26 | 0.0769 | 27 | 0.0741 | 89 | 0.0562 |
| X_2071892 | 23 | 0.3913 | 34 | 0.5 | 36 | 0.4444 | 38 | 0.4211 | 131 | 0.4427 | 15 | 0.2667 | 26 | 0.3846 | 32 | 0.4688 | 30 | 0.4 | 103 | 0.3981 |
| X_2072642 | 1 | 0 | 3 | 0.3333 | 1 | 0 | NA | NA | 5 | 0.2 | 1 | 1 | 1 | 0 | NA | NA | 1 | 1 | 3 | 0.6667 |
| X_2072659 | 1 | 0 | 5 | 0.2 | 2 | 0 | 1 | 0 | 9 | 0.1111 | 1 | 0 | NA | NA | 1 | 1 | 1 | 0 | 3 | 0.3333 |
| X_2072672 | 1 | 0 | 4 | 0.5 | 1 | 1 | NA | NA | 6 | 0.5 | NA | NA | NA | NA | 1 | 1 | 2 | 1 | 3 | 1 |
| X_2072682 | 50 | 0.32 | 86 | 0.3488 | 72 | 0.2222 | 41 | 0.1951 | 249 | 0.2811 | 68 | 0.3235 | 64 | 0.3906 | 45 | 0.3333 | 59 | 0.3559 | 236 | 0.3517 |
| X_2072695 | 52 | 0.3077 | 85 | 0.2588 | 61 | 0.2623 | 40 | 0.275 | 238 | 0.2731 | 59 | 0.4407 | 56 | 0.3571 | 46 | 0.3261 | 56 | 0.2857 | 217 | 0.3548 |
| X_2072717 | 46 | 0.1087 | 77 | 0.1688 | 56 | 0.1607 | 35 | 0.2571 | 214 | 0.1682 | 50 | 0.1 | 46 | 0.0435 | 40 | 0.2 | 48 | 0.2083 | 184 | 0.1359 |
| X_2072872 | 5 | 0.2 | 6 | 0 | 5 | 0 | 4 | 0.25 | 20 | 0.1 | 4 | 0 | 8 | 0.25 | 3 | 0 | 7 | 0.1429 | 22 | 0.1364 |
| X_2078700 | 9 | 0.2222 | 33 | 0.0606 | 19 | 0.1579 | 12 | 0 | 73 | 0.0959 | 14 | 0.1429 | 22 | 0 | 17 | 0.0588 | 18 | 0.1111 | 71 | 0.0704 |
| X_2080026 | NA | NA | 1 | 0 | 2 | 0 | 3 | 0.3333 | 6 | 0.1667 | NA | NA | 3 | 0.3333 | 1 | 1 | 1 | 0 | 5 | 0.4 |
| X_2080027 | NA | NA | 1 | 1 | 2 | 0.5 | 2 | 1 | 5 | 0.8 | NA | NA | 3 | 0.3333 | NA | NA | 1 | 1 | 4 | 0.5 |
| X_2080030 | NA | NA | 1 | 1 | 1 | 0 | 2 | 0.5 | 4 | 0.5 | NA | NA | 3 | 0.6667 | 1 | 1 | 1 | 0 | 5 | 0.6 |
| X_2080046 | 7 | 0.2857 | 6 | 0.3333 | 10 | 0.4 | 11 | 0.4545 | 34 | 0.3824 | 3 | 0.3333 | 7 | 0 | 12 | 0.3333 | 3 | 0.3333 | 25 | 0.24 |
| X_2080058 | 7 | 0.2857 | 5 | 0 | 10 | 0.1 | 8 | 0.5 | 30 | 0.2333 | 4 | 0 | 13 | 0 | 12 | 0.1667 | 4 | 0.25 | 33 | 0.0909 |
| X_2080067 | 5 | 0.2 | 2 | 0 | 9 | 0 | 8 | 0.125 | 24 | 0.0833 | 3 | 0 | 10 | 0 | 9 | 0.2222 | 4 | 0.5 | 26 | 0.1538 |
| X_2080105 | 4 | 0 | 3 | 0 | 8 | 0.125 | 3 | 0 | 18 | 0.0556 | 4 | 0 | 7 | 0.1429 | 8 | 0 | NA | NA | 19 | 0.0526 |
| X_2080152 | 1 | 0 | 4 | 0 | 5 | 0.6 | 4 | 0 | 14 | 0.2143 | 4 | 0.25 | 8 | 0.375 | 6 | 0 | 1 | 0 | 19 | 0.2105 |
| X_2080159 | NA | NA | 3 | 0.3333 | 3 | 0.6667 | 2 | 1 | 8 | 0.625 | 2 | 0.5 | 3 | 0.3333 | 4 | 0.5 | NA | NA | 9 | 0.4444 |
| X_2080160 | 1 | 0 | 4 | 0.25 | 5 | 0.4 | 2 | 0.5 | 12 | 0.3333 | 2 | 1 | 6 | 0 | 6 | 0.3333 | 1 | 0 | 15 | 0.2667 |
| X_2080164 | 1 | 0 | 2 | 0.5 | 5 | 0.4 | 2 | 0 | 10 | 0.3 | 2 | 0 | 6 | 0.1667 | 4 | 0.25 | NA | NA | 12 | 0.1667 |
| X_2080255 | 1 | 0 | 4 | 0 | 6 | 0.1667 | 1 | 0 | 12 | 0.0833 | 3 | 0.3333 | 6 | 0.3333 | 4 | 0 | 1 | 0 | 14 | 0.2143 |
| X_2080259 | 1 | 0 | 4 | 0.25 | 6 | 0 | 1 | 0 | 12 | 0.0833 | 2 | 0 | 5 | 0 | 2 | 0 | 1 | 1 | 10 | 0.1 |
| X_2080260 | 1 | 0 | 4 | 0 | 6 | 0 | 1 | 1 | 12 | 0.0833 | 3 | 0.3333 | 5 | 0 | 3 | 0 | 1 | 0 | 12 | 0.0833 |
| X_2080364 | 1 | 0 | 1 | 0 | 5 | 0.6 | 1 | 1 | 8 | 0.5 | 3 | 0.6667 | 2 | 0 | 3 | 0.6667 | NA | NA | 8 | 0.5 |
| X_2081826 | 2 | 0 | 2 | 0.5 | 1 | 0 | 2 | 0 | 7 | 0.1429 | 1 | 0 | NA | NA | 2 | 0.5 | NA | NA | 3 | 0.3333 |
| X_2085326 | 2 | 1 | NA | NA | 2 | 0.5 | 1 | 0 | 5 | 0.6 | 1 | 1 | 1 | 1 | 1 | 0 | 1 | 0 | 4 | 0.5 |
| X_2085353 | NA | NA | NA | NA | 2 | 1 | 1 | 1 | 3 | 1 | NA | NA | 1 | 1 | NA | NA | 1 | 1 | 2 | 1 |
| X_2089535 | 1 | 1 | NA | NA | NA | NA | 1 | 0 | 2 | 0.5 | NA | NA | NA | NA | 3 | 0.3333 | 1 | 1 | 4 | 0.5 |
| X_2089551 | 1 | 1 | NA | NA | NA | NA | NA | NA | 1 | 1 | NA | NA | NA | NA | 3 | 1 | 1 | 0 | 4 | 0.75 |
| X_2089552 | 1 | 1 | NA | NA | NA | NA | 1 | 0 | 2 | 0.5 | NA | NA | NA | NA | 3 | 1 | 1 | 0 | 4 | 0.75 |

| | | | | | | | | | | | | | | | | | | | | |
|---|---|---|---|---|---|---|---|---|---|---|---|---|---|---|---|---|---|---|---|---|
| X_2089577 | 1 | 1 | NA | NA | NA | NA | NA | NA | 1 | 1 | NA | NA | 1 | 1 | 3 | 1 | 1 | 1 | 5 | 1 |
| X_2089580 | 1 | 1 | NA | NA | NA | NA | 1 | 0 | 2 | 0.5 | NA | NA | 1 | 1 | 3 | 0.6667 | NA | NA | 4 | 0.75 |
| X_2089583 | 1 | 1 | NA | NA | NA | NA | 1 | 0 | 2 | 0.5 | NA | NA | 1 | 1 | 3 | 0.6667 | 1 | 1 | 5 | 0.8 |
| X_2089585 | 1 | 1 | NA | NA | NA | NA | NA | NA | 1 | 1 | NA | NA | 1 | 0 | 3 | 0.6667 | 1 | 0 | 5 | 0.4 |
| X_2089603 | 1 | 1 | NA | NA | NA | NA | 1 | 1 | 2 | 1 | NA | NA | 1 | 1 | 3 | 1 | 1 | 1 | 5 | 1 |
| X_2089620 | 1 | 1 | NA | NA | NA | NA | 1 | 0 | 2 | 0.5 | NA | NA | 1 | 0 | 3 | 0.3333 | 1 | 0 | 5 | 0.2 |
| X_2089641 | 1 | 1 | NA | NA | NA | NA | 1 | 1 | 2 | 1 | NA | NA | 1 | 1 | 3 | 0.6667 | NA | NA | 4 | 0.75 |
| X_2089666 | 1 | 1 | NA | NA | NA | NA | NA | NA | 1 | 1 | NA | NA | 1 | 1 | 4 | 1 | NA | NA | 5 | 1 |
| X_2089683 | 1 | 1 | NA | NA | NA | NA | NA | NA | 1 | 1 | NA | NA | 1 | 1 | 4 | 1 | NA | NA | 5 | 1 |
| X_2089725 | 1 | 0 | NA | NA | NA | NA | 1 | 1 | 2 | 0.5 | NA | NA | NA | NA | 2 | 1 | NA | NA | 2 | 1 |
| X_2089732 | 1 | 1 | NA | NA | NA | NA | 1 | 1 | 2 | 1 | NA | NA | 1 | 0 | 2 | 1 | NA | NA | 3 | 0.6667 |
| X_2089742 | NA | NA | NA | NA | NA | NA | 1 | 1 | 1 | 1 | NA | NA | 1 | 1 | 3 | 0.6667 | NA | NA | 4 | 0.75 |
| X_2089777 | 1 | 1 | NA | NA | NA | NA | 1 | 1 | 2 | 1 | NA | NA | 1 | 1 | 3 | 1 | NA | NA | 4 | 1 |
| X_2122912 | NA | NA | 1 | 1 | NA | NA | NA | NA | 1 | 1 | NA | NA | NA | NA | 1 | 1 | NA | NA | 1 | 1 |
| X_2122925 | 1 | 1 | 1 | 0 | NA | NA | NA | NA | 2 | 0.5 | NA | NA | NA | NA | 1 | 1 | NA | NA | 1 | 1 |
| X_2122926 | 1 | 1 | 1 | 1 | NA | NA | NA | NA | 2 | 1 | NA | NA | NA | NA | 1 | 1 | NA | NA | 1 | 1 |
| X_2122933 | 1 | 1 | 1 | 0 | NA | NA | NA | NA | 2 | 0.5 | NA | NA | NA | NA | 1 | 1 | NA | NA | 1 | 1 |
| X_2122936 | 1 | 1 | 1 | 1 | NA | NA | NA | NA | 2 | 1 | NA | NA | NA | NA | 1 | 1 | NA | NA | 1 | 1 |
| X_2122942 | 1 | 1 | 1 | 0 | NA | NA | NA | NA | 2 | 0.5 | NA | NA | NA | NA | 1 | 1 | NA | NA | 1 | 1 |
| X_2123015 | 1 | 0 | NA | NA | 1 | 1 | NA | NA | 2 | 0.5 | NA | NA | NA | NA | 1 | 1 | NA | NA | 1 | 1 |
| X_2308661 | 9 | 0 | 8 | 0 | 17 | 0.0588 | 20 | 0.25 | 54 | 0.1111 | 12 | 0.25 | 18 | 0.2222 | 26 | 0.3846 | 17 | 0.0588 | 73 | 0.2466 |
| X_2308687 | 9 | 0.1111 | 8 | 0.125 | 15 | 0.2667 | 19 | 0.1053 | 51 | 0.1569 | 10 | 0.2 | 16 | 0.125 | 25 | 0.16 | 15 | 0.2 | 66 | 0.1667 |
| X_2308702 | 6 | 0 | 6 | 0.5 | 15 | 0.0667 | 19 | 0.2632 | 46 | 0.1957 | 10 | 0 | 16 | 0.125 | 20 | 0.15 | 15 | 0.0667 | 61 | 0.0984 |
| X_2309488 | 8 | 0.125 | 8 | 0.125 | 15 | 0 | 14 | 0.0714 | 45 | 0.0667 | 9 | 0 | 17 | 0 | 17 | 0.1176 | 14 | 0.0714 | 57 | 0.0526 |
| X_2310162 | 8 | 0.375 | 6 | 0 | 16 | 0.0625 | 13 | 0 | 43 | 0.093 | 8 | 0 | 18 | 0.1667 | 15 | 0.1333 | 15 | 0.1333 | 56 | 0.125 |
| X_2310173 | 8 | 0 | 6 | 0.3333 | 15 | 0.1333 | 13 | 0.3846 | 42 | 0.2143 | 7 | 0.1429 | 17 | 0.2941 | 16 | 0.125 | 15 | 0.3333 | 55 | 0.2364 |
| X_2310183 | 8 | 1 | 5 | 1 | 15 | 1 | 13 | 0.8462 | 41 | 0.9512 | 7 | 0.8571 | 16 | 1 | 14 | 1 | 15 | 0.9333 | 52 | 0.9615 |
| X_2310191 | 6 | 0.1667 | 3 | 0 | 13 | 0.1538 | 9 | 0.4444 | 31 | 0.2258 | 4 | 0.25 | 11 | 0.0909 | 8 | 0.125 | 9 | 0.2222 | 32 | 0.1562 |
| X_2310192 | 4 | 1 | 4 | 1 | 10 | 1 | 7 | 1 | 25 | 1 | 6 | 1 | 15 | 1 | 12 | 1 | 8 | 1 | 41 | 1 |
| X_2310206 | 7 | 0.2857 | 4 | 0.5 | 12 | 0.3333 | 10 | 0.4 | 33 | 0.3636 | 6 | 0.8333 | 14 | 0.5714 | 8 | 0.5 | 15 | 0.3333 | 43 | 0.5116 |
| X_2362275 | NA | NA | 5 | 0.8 | 1 | 1 | NA | NA | 6 | 0.8333 | NA | NA | 1 | 1 | 1 | 1 | 3 | 0.3333 | 5 | 0.6 |
| X_2362279 | NA | NA | 5 | 0.8 | 2 | 1 | 1 | 1 | 8 | 0.875 | NA | NA | 1 | 1 | 1 | 0 | 4 | 0.5 | 6 | 0.5 |
| X_2362281 | NA | NA | 5 | 0.6 | 1 | 1 | 1 | 1 | 7 | 0.7143 | NA | NA | NA | NA | 1 | 1 | 3 | 1 | 4 | 1 |
| X_2362296 | NA | NA | 5 | 0.6 | 2 | 0 | 1 | 0 | 8 | 0.375 | NA | NA | 1 | 1 | 1 | 1 | 4 | 0.25 | 6 | 0.5 |
| X_2362301 | NA | NA | 5 | 0.2 | 1 | 1 | 1 | 1 | 7 | 0.4286 | NA | NA | NA | NA | 1 | 0 | 3 | 0.3333 | 4 | 0.25 |
| X_2362308 | NA | NA | 5 | 0.6 | 2 | 0.5 | 1 | 1 | 8 | 0.625 | NA | NA | 1 | 0 | 1 | 1 | 4 | 0.75 | 6 | 0.6667 |
| X_2367197 | NA | NA | 7 | 0 | 6 | 0.1667 | 3 | 0.3333 | 16 | 0.125 | NA | NA | 2 | 0.5 | 5 | 0.2 | 7 | 0.2857 | 14 | 0.2857 |
| X_3543263 | 1 | 1 | 2 | 1 | 3 | 1 | 1 | 1 | 7 | 1 | NA | NA | 1 | 1 | 2 | 1 | 1 | 1 | 4 | 1 |
| X_3543265 | 1 | 1 | 2 | 0.5 | 3 | 1 | 1 | 1 | 7 | 0.8571 | NA | NA | NA | NA | 2 | 1 | 1 | 1 | 3 | 1 |
| X_3543270 | 1 | 1 | 3 | 1 | 3 | 1 | 1 | 1 | 8 | 1 | NA | NA | 1 | 1 | 1 | 1 | 1 | 0 | 3 | 0.6667 |
| X_3543281 | 1 | 0 | 3 | 0.3333 | 2 | 0.5 | 1 | 0 | 7 | 0.2857 | NA | NA | NA | NA | 2 | 0 | 1 | 1 | 3 | 0.3333 |
| X_3543283 | NA | NA | 3 | 1 | 3 | 1 | 1 | 1 | 7 | 1 | NA | NA | 1 | 0 | 2 | 1 | 1 | 1 | 4 | 0.75 |
| X_3543300 | NA | NA | 3 | 0 | 3 | 0.3333 | 1 | 0 | 7 | 0.1429 | NA | NA | 1 | 0 | 1 | 1 | 1 | 1 | 3 | 0.6667 |
| X_3543303 | 1 | 0 | 3 | 0.3333 | 3 | 0.6667 | 1 | 0 | 8 | 0.375 | NA | NA | 1 | 0 | 2 | 0.5 | 1 | 0 | 4 | 0.25 |
| X_3543321 | 1 | 0 | 3 | 0 | 3 | 1 | 1 | 1 | 8 | 0.5 | NA | NA | 1 | 1 | 1 | 1 | 1 | 1 | 3 | 1 |
| X_3543342 | 1 | 0 | 2 | 0.5 | 3 | 0.6667 | 1 | 1 | 7 | 0.5714 | NA | NA | 1 | 1 | 2 | 0.5 | 1 | 1 | 4 | 0.75 |
| X_3546104 | 12 | 0 | 10 | 0.1 | 12 | 0 | 22 | 0.1364 | 56 | 0.0714 | 7 | 0.1429 | 20 | 0.1 | 15 | 0 | 15 | 0 | 57 | 0.0526 |
| X_3562557 | 1 | 0 | 1 | 1 | NA | NA | 2 | 1 | 4 | 0.75 | 1 | 0 | NA | NA | 1 | 1 | NA | NA | 2 | 0.5 |
| X_3563008 | 4 | 0 | 3 | 0 | 4 | 0.25 | 4 | 0.25 | 15 | 0.1333 | 1 | 0 | 3 | 0 | 6 | 0.1667 | 3 | 0.3333 | 13 | 0.1538 |
| X_3563021 | 5 | 0 | 3 | 0 | 4 | 0.25 | 4 | 0 | 16 | 0.0625 | 1 | 0 | 3 | 0.3333 | 6 | 0 | 3 | 0 | 13 | 0.0769 |
| X_3586189 | 37 | 0.0811 | 29 | 0.1724 | 36 | 0.1667 | 47 | 0.2766 | 149 | 0.1812 | 24 | 0.1667 | 32 | 0.2188 | 29 | 0.2069 | 34 | 0.1176 | 119 | 0.1765 |
| X_3586864 | 53 | 0.0377 | 45 | 0.0667 | 44 | 0.1136 | 64 | 0.0938 | 206 | 0.0777 | 35 | 0.0571 | 44 | 0.1136 | 38 | 0.0789 | 45 | 0.0667 | 162 | 0.0802 |
| X_3586912 | 47 | 0.1915 | 46 | 0.3696 | 43 | 0.186 | 51 | 0.2353 | 187 | 0.246 | 30 | 0.2333 | 41 | 0.2195 | 38 | 0.2632 | 46 | 0.2391 | 155 | 0.2387 |
| X_3586942 | 70 | 0.1286 | 57 | 0.1053 | 58 | 0.1379 | 76 | 0.1579 | 261 | 0.1341 | 44 | 0.0682 | 58 | 0.1552 | 46 | 0.087 | 58 | 0.1207 | 206 | 0.1117 |
| X_3587292 | 3 | 0.3333 | NA | NA | NA | NA | NA | NA | 3 | 0.3333 | NA | NA | 2 | 0.5 | 4 | 0.25 | 1 | 1 | 7 | 0.4286 |
| X_3587355 | 3 | 0.6667 | 2 | 0 | NA | NA | NA | NA | 5 | 0.4 | 1 | 1 | 2 | 0 | 3 | 0 | 1 | 1 | 7 | 0.2857 |
| X_3587372 | 1 | 1 | 2 | 0.5 | NA | NA | NA | NA | 3 | 0.6667 | NA | NA | 2 | 0.5 | 3 | 0.6667 | 1 | 0 | 6 | 0.5 |
| X_3587392 | NA | NA | 2 | 1 | NA | NA | NA | NA | 2 | 1 | 1 | 1 | 2 | 1 | 1 | 1 | NA | NA | 4 | 1 |
| X_3587395 | NA | NA | 1 | 1 | NA | NA | NA | NA | 1 | 1 | 1 | 1 | 2 | 1 | 3 | 1 | NA | NA | 6 | 1 |
| X_3587416 | NA | NA | 2 | 1 | NA | NA | NA | NA | 2 | 1 | 1 | 1 | 2 | 0.5 | 2 | 0 | NA | NA | 5 | 0.4 |
| X_3587422 | NA | NA | 1 | 1 | NA | NA | NA | NA | 1 | 1 | NA | NA | NA | NA | 2 | 0.5 | NA | NA | 2 | 0.5 |
| X_3587423 | NA | NA | 2 | 1 | NA | NA | NA | NA | 2 | 1 | 1 | 0 | 2 | 1 | 3 | 0.6667 | NA | NA | 6 | 0.6667 |
| X_3587456 | NA | NA | 1 | 1 | NA | NA | NA | NA | 1 | 1 | NA | NA | 1 | 0 | 2 | 0.5 | NA | NA | 3 | 0.3333 |
| X_3587550 | NA | NA | 1 | 1 | NA | NA | NA | NA | 1 | 1 | 1 | 1 | NA | NA | 2 | 1 | NA | NA | 3 | 1 |
| X_3587560 | NA | NA | 1 | 1 | NA | NA | NA | NA | 1 | 1 | 1 | 1 | NA | NA | 2 | 1 | NA | NA | 3 | 1 |

| | | | | | | | | | | | | | | | | | | | | |
|---|---|---|---|---|---|---|---|---|---|---|---|---|---|---|---|---|---|---|---|---|
| X_3587602 | NA | NA | 1 | 1 | NA | NA | NA | NA | 1 | 1 | 1 | 0 | NA | NA | 2 | 0.5 | NA | NA | 3 | 0.3333 |
| X_3588735 | NA | NA | 1 | 1 | NA | NA | NA | NA | 1 | 1 | 1 | 1 | NA | NA | 1 | 1 | NA | NA | 2 | 1 |
| X_3588765 | NA | NA | 1 | 1 | NA | NA | NA | NA | 1 | 1 | 1 | 1 | NA | NA | 1 | 0 | NA | NA | 2 | 0.5 |
| X_3588775 | NA | NA | 1 | 1 | NA | NA | NA | NA | 1 | 1 | 1 | 1 | NA | NA | 1 | 1 | NA | NA | 2 | 1 |
| X_3606055 | 55 | 0.1091 | 60 | 0.1167 | 61 | 0.1311 | 54 | 0.0926 | 230 | 0.113 | 48 | 0.1042 | 58 | 0.1034 | 59 | 0.0847 | 49 | 0.2245 | 214 | 0.1262 |
| X_3606069 | 53 | 0.2075 | 59 | 0.1186 | 56 | 0.2321 | 52 | 0.1731 | 220 | 0.1818 | 45 | 0.2 | 51 | 0.1176 | 55 | 0.1273 | 44 | 0.0455 | 195 | 0.1231 |
| X_3606086 | 63 | 0.1429 | 65 | 0.2154 | 70 | 0.1 | 64 | 0.0938 | 262 | 0.1374 | 53 | 0.1509 | 61 | 0.0984 | 69 | 0.1449 | 59 | 0.1356 | 242 | 0.1322 |
| X_3606225 | 60 | 0.1333 | 67 | 0.0896 | 65 | 0.0308 | 63 | 0.0794 | 255 | 0.0824 | 55 | 0.0364 | 61 | 0.1148 | 67 | 0.0299 | 51 | 0.0588 | 234 | 0.0598 |
| X_3606226 | 57 | 0.1228 | 56 | 0.1429 | 56 | 0.125 | 50 | 0.26 | 219 | 0.1598 | 46 | 0.1739 | 51 | 0.1765 | 59 | 0.1525 | 46 | 0.1522 | 202 | 0.1634 |
| X_3606239 | 61 | 0.0984 | 65 | 0.1692 | 60 | 0.1 | 67 | 0.1642 | 253 | 0.1344 | 64 | 0.0469 | 61 | 0.0984 | 73 | 0.137 | 56 | 0.1429 | 254 | 0.1063 |
| X_3611286 | 4 | 0 | 5 | 0.2 | 2 | 0 | 8 | 0 | 19 | 0.0526 | 3 | 0.3333 | 5 | 0 | 6 | 0.1667 | 6 | 0 | 20 | 0.1 |
| X_3611343 | 3 | 0.3333 | 5 | 0.2 | 2 | 0 | 5 | 0.6 | 15 | 0.3333 | 2 | 0 | 5 | 0 | 3 | 0.3333 | 3 | 0.6667 | 13 | 0.2308 |
| X_3614697 | 9 | 0.1111 | 8 | 0.125 | 6 | 0 | 10 | 0 | 33 | 0.0606 | 4 | 0.25 | 4 | 0.25 | 5 | 0.2 | 9 | 0 | 22 | 0.1364 |
| X_3615138 | 1 | 0 | 3 | 1 | 4 | 0.5 | 3 | 1 | 11 | 0.7273 | 1 | 1 | 3 | 1 | 3 | 1 | 4 | 1 | 11 | 1 |
| X_3615147 | 1 | 1 | 7 | 0.5714 | 5 | 1 | 3 | 0.6667 | 16 | 0.75 | NA | NA | 5 | 1 | 2 | 0.5 | 4 | 1 | 11 | 0.9091 |
| X_3615150 | 2 | 0.5 | 4 | 1 | 4 | 1 | 3 | 1 | 13 | 0.9231 | 1 | 1 | 4 | 1 | 2 | 1 | 4 | 0.75 | 11 | 0.9091 |
| X_3615160 | 2 | 0 | 6 | 0.1667 | 5 | 0.6 | 2 | 0 | 15 | 0.2667 | NA | NA | 4 | 0.5 | 2 | 0.5 | 4 | 0.25 | 10 | 0.4 |
| X_3615224 | 1 | 1 | 3 | 1 | 5 | 1 | 2 | 0.5 | 11 | 0.9091 | NA | NA | 1 | 1 | 1 | 1 | 4 | 0.75 | 6 | 0.8333 |
| X_3615246 | 1 | 0 | 3 | 0 | 5 | 0.2 | 1 | 0 | 10 | 0.1 | NA | NA | 1 | 0 | NA | NA | 3 | 0.6667 | 4 | 0.5 |
| X_3615270 | 1 | 1 | 4 | 1 | 3 | 1 | NA | NA | 8 | 1 | NA | NA | 2 | 1 | NA | NA | 3 | 1 | 5 | 1 |
| X_3615326 | 1 | 1 | 2 | 0.5 | 1 | 1 | NA | NA | 4 | 0.75 | NA | NA | NA | NA | NA | NA | 2 | 1 | 2 | 1 |
| X_3615340 | 1 | 0 | 3 | 0.6667 | 1 | 1 | NA | NA | 5 | 0.6 | NA | NA | 1 | 1 | NA | NA | 3 | 1 | 4 | 1 |
| X_3615364 | 1 | 0 | 3 | 0.6667 | 1 | 1 | NA | NA | 5 | 0.6 | NA | NA | 1 | 1 | NA | NA | 3 | 0 | 4 | 0.25 |
| X_3615397 | 1 | 1 | 2 | 0 | NA | NA | NA | NA | 3 | 0.3333 | NA | NA | 1 | 1 | NA | NA | 3 | 0.6667 | 4 | 0.75 |
| X_3617633 | 7 | 0.7143 | 5 | 1 | 5 | 0.6 | 6 | 0.5 | 23 | 0.6957 | 4 | 0.5 | 5 | 0.2 | 1 | 1 | 3 | 0.6667 | 13 | 0.4615 |
| X_3617636 | 5 | 0.2 | 8 | 0.375 | 4 | 0 | 5 | 0.2 | 22 | 0.2273 | 5 | 0.2 | 6 | 0.1667 | 1 | 0 | 4 | 0.5 | 16 | 0.25 |
| X_3617640 | 8 | 0.75 | 8 | 1 | 7 | 0.7143 | 5 | 1 | 28 | 0.8571 | 4 | 1 | 7 | 0.5714 | 1 | 1 | 3 | 0.6667 | 15 | 0.7333 |
| X_3617648 | 7 | 0 | 7 | 0.5714 | 5 | 0.2 | 6 | 0.5 | 25 | 0.32 | 5 | 0.8 | 5 | 0.2 | 1 | 1 | 2 | 0.5 | 13 | 0.5385 |
| X_3617657 | 9 | 0.7778 | 8 | 0.875 | 7 | 0.5714 | 5 | 0.8 | 29 | 0.7586 | 5 | 1 | 5 | 0.4 | 1 | 1 | 2 | 1 | 13 | 0.7692 |
| X_3617663 | 9 | 0.3333 | 7 | 0.2857 | 7 | 0.7143 | 6 | 0.5 | 29 | 0.4483 | 3 | 1 | 5 | 0.4 | 1 | 1 | 2 | 0.5 | 11 | 0.6364 |
| X_3635598 | 4 | 0.25 | 4 | 0.5 | 3 | 0.6667 | 5 | 0.8 | 16 | 0.5625 | 3 | 1 | 6 | 0.6667 | 4 | 1 | 4 | 0.5 | 17 | 0.7647 |
| X_3635607 | 4 | 0 | 2 | 0 | 3 | 0.3333 | 3 | 0 | 12 | 0.0833 | 3 | 0 | 4 | 0.25 | 2 | 0 | 4 | 0 | 13 | 0.0769 |
| X_3635943 | 1 | 1 | 2 | 0.5 | 2 | 0.5 | 1 | 1 | 6 | 0.6667 | 3 | 0 | 5 | 0.4 | 1 | 0 | 2 | 0 | 11 | 0.1818 |
| X_3641162 | NA | NA | 1 | 1 | 2 | 1 | 3 | 1 | 6 | 1 | NA | NA | 1 | 0 | 5 | 1 | 3 | 1 | 9 | 0.8889 |
| X_3641178 | NA | NA | 1 | 1 | 1 | 0 | 3 | 0.6667 | 5 | 0.6 | 1 | 0 | 1 | 0 | 2 | 0 | 2 | 1 | 6 | 0.3333 |
| X_3645540 | NA | NA | NA | NA | 1 | 1 | 1 | 0 | 2 | 0.5 | NA | NA | 1 | 1 | 1 | 0 | 2 | 0.5 | 4 | 0.5 |
| X_3645564 | 1 | 0 | NA | NA | 1 | 1 | 1 | 1 | 3 | 0.6667 | NA | NA | 1 | 1 | 1 | 1 | 3 | 0.6667 | 5 | 0.8 |
| X_3645566 | 1 | 0 | NA | NA | 1 | 0 | 2 | 1 | 4 | 0.5 | NA | NA | 1 | 0 | 2 | 0.5 | 3 | 0.3333 | 6 | 0.3333 |
| X_3645568 | 1 | 1 | NA | NA | 1 | 0 | 2 | 0.5 | 4 | 0.5 | NA | NA | 1 | 0 | 2 | 0.5 | 3 | 0.3333 | 6 | 0.3333 |
| X_3645597 | 1 | 1 | 1 | 1 | 1 | 1 | 2 | 1 | 5 | 1 | NA | NA | 1 | 1 | 2 | 0.5 | 2 | 1 | 5 | 0.8 |
| X_3645607 | 1 | 0 | 1 | 0 | 1 | 1 | 2 | 0.5 | 5 | 0.4 | NA | NA | 1 | 1 | 1 | 1 | 3 | 0.3333 | 5 | 0.6 |
| X_3728134 | NA | NA | NA | NA | NA | NA | 1 | 1 | 1 | 1 | NA | NA | NA | NA | 1 | 1 | NA | NA | 1 | 1 |
| X_3728262 | NA | NA | NA | NA | NA | NA | 1 | 1 | 1 | 1 | 1 | 0 | 1 | 1 | 1 | 0 | 1 | 1 | 4 | 0.5 |
| X_3728282 | NA | NA | 1 | 1 | NA | NA | 1 | 1 | 2 | 1 | 1 | 1 | 1 | 1 | 1 | 1 | 1 | 1 | 4 | 1 |
| X_3728311 | NA | NA | NA | NA | NA | NA | 1 | 1 | 1 | 1 | NA | NA | 1 | 1 | 1 | 1 | 1 | 1 | 3 | 1 |
| X_3728481 | 1 | 0 | 3 | 0.3333 | 5 | 0 | 1 | 0 | 10 | 0.1 | 4 | 0.25 | 5 | 0 | 3 | 0 | 3 | 0 | 15 | 0.0667 |
| X_3728516 | 2 | 0 | 3 | 0.3333 | 5 | 0.2 | 1 | 0 | 11 | 0.1818 | 3 | 0 | 5 | 0.4 | 3 | 0 | 2 | 0.5 | 13 | 0.2308 |
| X_3728672 | 7 | 0.2857 | 2 | 0 | 1 | 0 | 9 | 0.3333 | 19 | 0.2632 | 4 | 0 | 6 | 0.1667 | 6 | 0.1667 | 6 | 0 | 22 | 0.0909 |
| X_3728718 | 6 | 0.1667 | 1 | 0 | 1 | 0 | 9 | 0 | 17 | 0.0588 | 1 | 0 | 4 | 0.25 | 6 | 0 | 6 | 0 | 17 | 0.0588 |
| X_3728749 | 5 | 0.2 | 2 | 0 | 1 | 0 | 6 | 0 | 14 | 0.0714 | 1 | 0 | 4 | 0 | 5 | 0.2 | 6 | 0.1667 | 16 | 0.125 |
| X_3728800 | 5 | 0 | 2 | 0 | 1 | 0 | 7 | 0.2857 | 15 | 0.1333 | 2 | 0 | 4 | 0 | 4 | 0.25 | 5 | 0 | 15 | 0.0667 |
| X_3728828 | 5 | 0 | NA | NA | 1 | 0 | 6 | 0.5 | 12 | 0.25 | 2 | 1 | 2 | 0 | 4 | 0.25 | 4 | 0.5 | 12 | 0.4167 |
| X_3728849 | 4 | 0.25 | NA | NA | NA | NA | 4 | 0 | 8 | 0.125 | 1 | 0 | NA | NA | 3 | 0 | 3 | 0.3333 | 7 | 0.1429 |
| X_3748879 | 5 | 0 | 4 | 0 | 5 | 0 | 4 | 0.25 | 18 | 0.0556 | 2 | 0 | 3 | 0 | 7 | 0 | 5 | 0.2 | 17 | 0.0588 |
| X_3748903 | 7 | 0.1429 | 5 | 0 | 5 | 0.2 | 4 | 0.5 | 21 | 0.1905 | 2 | 0 | 3 | 0.3333 | 7 | 0 | 4 | 0.5 | 16 | 0.1875 |
| X_3749038 | 4 | 0 | 8 | 0.125 | 6 | 0.1667 | 4 | 0 | 22 | 0.0909 | 3 | 0 | 6 | 0.3333 | 5 | 0 | 7 | 0 | 21 | 0.0952 |
| X_3749059 | 3 | 0 | 7 | 0.1429 | 5 | 0 | 4 | 0 | 19 | 0.0526 | 3 | 0 | 6 | 0 | 5 | 0 | 8 | 0.375 | 22 | 0.1364 |
| X_3749069 | 1 | 0 | 8 | 0.125 | 4 | 0.5 | 3 | 0.3333 | 16 | 0.25 | 3 | 0.6667 | 4 | 0 | 3 | 0 | 9 | 0.6667 | 19 | 0.4211 |
| X_3749081 | 2 | 0.5 | 6 | 0.8333 | 6 | 0.8333 | 3 | 0.3333 | 17 | 0.7059 | 1 | 1 | 5 | 0.2 | 4 | 0.5 | 7 | 0.7143 | 17 | 0.5294 |
| X_3749083 | 1 | 0 | 7 | 0.1429 | 5 | 0 | 4 | 0 | 17 | 0.0588 | 2 | 0.5 | 5 | 0 | 3 | 0.3333 | 5 | 0.2 | 15 | 0.2 |
| X_3749088 | 2 | 0 | 7 | 0.4286 | 6 | 0 | 4 | 0.25 | 19 | 0.2105 | 2 | 0 | 6 | 0 | 5 | 0 | 8 | 0.375 | 21 | 0.1429 |
| X_3749090 | 2 | 0.5 | 6 | 0.8333 | 3 | 1 | 3 | 0.6667 | 14 | 0.7857 | 2 | 0.5 | 4 | 0.75 | 4 | 0.75 | 6 | 0.8333 | 16 | 0.75 |
| X_3749094 | 1 | 1 | 6 | 1 | 4 | 0.75 | 4 | 0.75 | 15 | 0.8667 | 1 | 0 | 3 | 0.3333 | 3 | 1 | 4 | 0.25 | 11 | 0.4545 |
| X_3749430 | 7 | 0.1429 | 4 | 0.25 | 2 | 0 | 5 | 0.2 | 18 | 0.1667 | 2 | 0 | 2 | 0.5 | 2 | 0 | 2 | 0 | 8 | 0.125 |
| X_3750884 | 8 | 0 | 4 | 0.5 | 4 | 0 | 4 | 0 | 20 | 0.1 | 4 | 0.25 | 4 | 0 | 3 | 0 | 3 | 0.3333 | 14 | 0.1429 |

| | | | | | | | | | | | | | | | | | | | | |
|---|---|---|---|---|---|---|---|---|---|---|---|---|---|---|---|---|---|---|---|---|
| X_3751021 | 5 | 0 | 4 | 0 | 4 | 0 | 3 | 0.3333 | 16 | 0.0625 | 3 | 0 | 2 | 0.5 | 2 | 0 | 3 | 0 | 10 | 0.1 |
| X_3751059 | 5 | 0.2 | 5 | 0.4 | 3 | 0 | 4 | 0 | 17 | 0.1765 | 4 | 0.25 | 4 | 0 | 3 | 0.3333 | 2 | 0.5 | 13 | 0.2308 |
| X_3762237 | 3 | 0.3333 | 4 | 1 | 9 | 0.6667 | 6 | 0.6667 | 22 | 0.6818 | 3 | 0.6667 | 5 | 0.8 | 10 | 0.8 | 1 | 1 | 19 | 0.7895 |
| X_3762249 | 2 | 0 | 4 | 0.5 | 8 | 0.625 | 6 | 0.5 | 20 | 0.5 | 3 | 0.3333 | 6 | 0.6667 | 9 | 0.4444 | NA | NA | 18 | 0.5 |
| X_3763569 | 2 | 0.5 | 2 | 0.5 | 5 | 0.4 | 3 | 0 | 12 | 0.3333 | 2 | 0.5 | 6 | 0.1667 | 7 | 0.1429 | 1 | 1 | 16 | 0.25 |
| X_3765254 | 2 | 0 | 2 | 0 | 5 | 0 | 2 | 0.5 | 11 | 0.0909 | 1 | 0 | 5 | 0.4 | 8 | 0 | NA | NA | 14 | 0.1429 |
| X_3765267 | 2 | 0 | 2 | 0 | 4 | 0 | 3 | 0.3333 | 11 | 0.0909 | 1 | 0 | 5 | 0.2 | 7 | 0.1429 | NA | NA | 13 | 0.1538 |
| X_3765292 | 2 | 0 | 2 | 0.5 | 5 | 0 | 3 | 0 | 12 | 0.0833 | 1 | 0 | 5 | 0 | 7 | 0.1429 | NA | NA | 13 | 0.0769 |
| X_3772001 | 5 | 0 | 4 | 0 | 5 | 0 | 7 | 0.2857 | 21 | 0.0952 | 6 | 0.1667 | 7 | 0 | 3 | 0.3333 | 3 | 0 | 19 | 0.1053 |
| X_3772004 | 5 | 0.2 | 4 | 0.75 | 4 | 0.25 | 6 | 0.6667 | 19 | 0.4737 | 6 | 0.5 | 5 | 0.2 | 3 | 0.3333 | 3 | 1 | 17 | 0.4706 |
| X_3772010 | 4 | 0.5 | 5 | 1 | 5 | 0.8 | 7 | 0.8571 | 21 | 0.8095 | 3 | 1 | 5 | 1 | 3 | 1 | 3 | 0.6667 | 14 | 0.9286 |
| X_3772015 | 3 | 0 | 4 | 0 | 4 | 0.25 | 7 | 0 | 18 | 0.0556 | 4 | 0.25 | 3 | 0 | 1 | 0 | 2 | 0.5 | 10 | 0.2 |
| X_3772016 | 3 | 1 | 4 | 1 | 3 | 1 | 2 | 1 | 12 | 1 | 4 | 1 | 4 | 1 | 2 | 1 | 2 | 1 | 12 | 1 |
| X_3772022 | 5 | 0.8 | 3 | 0.3333 | 5 | 0.6 | 7 | 0.8571 | 20 | 0.7 | 5 | 0.8 | 6 | 0.6667 | 1 | 1 | 2 | 0.5 | 14 | 0.7143 |
| X_3772033 | 4 | 0 | 3 | 0.3333 | 2 | 0 | 6 | 0 | 15 | 0.0667 | 3 | 0 | 6 | 0 | 2 | 0 | 2 | 0.5 | 13 | 0.0769 |
| X_3772041 | 4 | 0.75 | 5 | 0.8 | 4 | 0.75 | 6 | 0.8333 | 19 | 0.7895 | 5 | 0.8 | 4 | 1 | 2 | 0.5 | 3 | 0.6667 | 14 | 0.7857 |
| X_3772047 | 4 | 0.25 | 4 | 0 | 5 | 0.8 | 7 | 0.2857 | 20 | 0.35 | 5 | 0 | 5 | 0 | 2 | 0 | 3 | 0.6667 | 15 | 0.1333 |
| X_3772049 | 4 | 1 | 5 | 1 | 5 | 0.8 | 7 | 1 | 21 | 0.9524 | 5 | 1 | 4 | 1 | 2 | 1 | 3 | 1 | 14 | 1 |
| X_3772059 | 5 | 0.8 | 4 | 0.25 | 5 | 0.8 | 6 | 0.6667 | 20 | 0.65 | 3 | 0.6667 | 3 | 1 | 2 | 1 | 2 | 1 | 10 | 0.9 |
| X_3772065 | 4 | 0.5 | 5 | 0.2 | 5 | 0.8 | 5 | 0.2 | 19 | 0.4211 | 5 | 0.4 | 4 | 0.25 | 2 | 0 | 3 | 0.3333 | 14 | 0.2857 |
| X_3772066 | 4 | 1 | 3 | 1 | 3 | 1 | 4 | 1 | 14 | 1 | 4 | 1 | 5 | 1 | 2 | 1 | 3 | 1 | 14 | 1 |
| X_3772073 | 4 | 0.75 | 5 | 0.4 | 5 | 0.4 | 4 | 0 | 18 | 0.3889 | 5 | 0.4 | 4 | 0.25 | 2 | 0.5 | 2 | 0.5 | 13 | 0.3846 |
| X_3772073 | 5 | 0.8 | 3 | 0.3333 | 5 | 0.8 | 5 | 0.8 | 18 | 0.7222 | 4 | 1 | 5 | 0.8 | 2 | 0 | 2 | 0.5 | 13 | 0.6923 |
| X_4076536 | 9 | 0.6667 | 15 | 0.5333 | 11 | 0.6364 | 11 | 0.6364 | 46 | 0.6087 | 8 | 0.125 | 11 | 0.6364 | 14 | 0.7143 | 17 | 0.4706 | 50 | 0.52 |
| X_4076539 | 9 | 0.3333 | 15 | 0.4667 | 12 | 0.5833 | 12 | 0.1667 | 48 | 0.3958 | 8 | 0.375 | 13 | 0.2308 | 11 | 0.4545 | 17 | 0.1765 | 49 | 0.2857 |
| X_4076539 | NA | NA | 1 | 1 | NA | NA | NA | NA | 1 | 1 | NA | NA | NA | NA | NA | NA | 1 | 1 | 1 | 1 |
| X_4076540 | 9 | 0.4444 | 16 | 0.5 | 10 | 0.5 | 13 | 0.2308 | 48 | 0.4167 | 4 | 0.75 | 16 | 0.5 | 15 | 0.4667 | 19 | 0.3158 | 54 | 0.4444 |
| X_4076548 | 9 | 1 | 17 | 1 | 15 | 0.8667 | 15 | 0.8 | 56 | 0.9107 | 8 | 1 | 15 | 0.8667 | 15 | 0.9333 | 22 | 0.9091 | 60 | 0.9167 |
| X_4076552 | 8 | 0.125 | 15 | 0.3333 | 11 | 0.1818 | 10 | 0.1 | 44 | 0.2045 | 6 | 0 | 11 | 0.2727 | 16 | 0.4375 | 17 | 0.2353 | 50 | 0.28 |
| X_4076568 | 11 | 0.2727 | 21 | 0 | 14 | 0.2143 | 15 | 0.0667 | 61 | 0.1148 | 7 | 0.1429 | 15 | 0.2 | 16 | 0.1875 | 22 | 0.0909 | 60 | 0.15 |
| X_4076575 | 10 | 0 | 19 | 0.1579 | 15 | 0 | 15 | 0.0667 | 59 | 0.0678 | 8 | 0.125 | 15 | 0.0667 | 17 | 0.1176 | 23 | 0 | 63 | 0.0635 |
| X_4076591 | 12 | 0.0833 | 20 | 0.3 | 14 | 0.2143 | 14 | 0.2857 | 60 | 0.2333 | 8 | 0.125 | 15 | 0.2667 | 18 | 0.2778 | 24 | 0.4583 | 65 | 0.3231 |
| X_4076637 | 10 | 0 | 24 | 0.25 | 17 | 0.1765 | 15 | 0.2667 | 66 | 0.197 | 10 | 0.1 | 16 | 0.1875 | 17 | 0.1765 | 22 | 0 | 65 | 0.1077 |
| X_4076643 | 11 | 0.0909 | 24 | 0.0417 | 20 | 0.15 | 16 | 0 | 71 | 0.0704 | 10 | 0.1 | 15 | 0.1333 | 17 | 0.1765 | 25 | 0.08 | 67 | 0.1194 |
| X_4076646 | 12 | 0 | 23 | 0.3478 | 20 | 0.35 | 16 | 0.25 | 71 | 0.2676 | 7 | 0 | 19 | 0.1579 | 18 | 0.2222 | 23 | 0.3043 | 67 | 0.209 |
| X_4076650 | 11 | 0.2727 | 24 | 0.0417 | 20 | 0.1 | 16 | 0.125 | 71 | 0.1127 | 10 | 0 | 19 | 0.0526 | 19 | 0.2632 | 24 | 0.0833 | 72 | 0.1111 |
| X_4076655 | 10 | 0 | 20 | 0.25 | 14 | 0.2143 | 15 | 0.2667 | 59 | 0.2034 | 8 | 0.375 | 15 | 0.2 | 15 | 0.3333 | 16 | 0.25 | 54 | 0.2778 |
| X_4076673 | 14 | 0.2857 | 22 | 0.2727 | 21 | 0.0952 | 16 | 0.125 | 73 | 0.1918 | 12 | 0.3333 | 20 | 0.2 | 21 | 0.0952 | 22 | 0.1364 | 75 | 0.1733 |
| X_4076713 | 15 | 0.0667 | 29 | 0.1034 | 24 | 0 | 17 | 0.1176 | 85 | 0.0706 | 14 | 0.0714 | 20 | 0 | 21 | 0.0476 | 23 | 0.087 | 78 | 0.0513 |
| X_4088574 | 3 | 0 | 1 | 0 | 3 | 0.6667 | 6 | 0.6667 | 13 | 0.4615 | 4 | 0.25 | 4 | 0.5 | 4 | 0.25 | 7 | 0.1429 | 19 | 0.2632 |
| X_4088635 | 2 | 0.5 | 1 | 0 | 1 | 1 | 6 | 0.8333 | 10 | 0.7 | 1 | 0 | 2 | 1 | 2 | 0.5 | 3 | 0.3333 | 8 | 0.5 |
| X_4088635 | 3 | 0 | 1 | 0 | 2 | 0.5 | 6 | 0.3333 | 12 | 0.25 | NA | NA | NA | NA | 2 | 0 | 5 | 0.2 | 7 | 0.1429 |
| X_4088639 | 2 | 0 | 1 | 0 | 3 | 0.3333 | 6 | 0.1667 | 12 | 0.1667 | 1 | 0 | 3 | 1 | 3 | 0 | 5 | 0.8 | 12 | 0.5833 |
| X_4088643 | 1 | 0 | 1 | 0 | 3 | 0.3333 | 5 | 0 | 10 | 0.1 | 2 | 0 | 3 | 0 | 2 | 0 | 5 | 0.2 | 12 | 0.0833 |
| X_4088646 | 2 | 0 | 1 | 0 | 3 | 1 | 6 | 0.8333 | 12 | 0.6667 | NA | NA | 2 | 1 | 3 | 0.6667 | 5 | 0.6 | 10 | 0.7 |
| X_4088648 | 1 | 0 | 1 | 0 | 3 | 0.3333 | 5 | 0.4 | 10 | 0.3 | 2 | 1 | 4 | 0.5 | 5 | 0.4 | 7 | 0.2857 | 18 | 0.4444 |
| X_4088658 | 2 | 0 | 1 | 0 | 3 | 0.3333 | 5 | 0.4 | 11 | 0.2727 | 2 | 0 | 3 | 0.3333 | 5 | 0 | 7 | 0 | 17 | 0.0588 |
| X_4090262 | NA | NA | NA | NA | 1 | 1 | NA | NA | 1 | 1 | NA | NA | NA | NA | NA | NA | 1 | 1 | 1 | 1 |
| X_4118330 | NA | NA | 2 | 0.5 | 2 | 0 | NA | NA | 4 | 0.25 | NA | NA | 1 | 0 | NA | NA | 1 | 1 | 2 | 0.5 |
| X_4118363 | NA | NA | 1 | 1 | 2 | 1 | NA | NA | 3 | 1 | NA | NA | 1 | 1 | NA | NA | 1 | 1 | 2 | 1 |
| X_4118367 | NA | NA | 2 | 0 | 2 | 1 | NA | NA | 4 | 0.5 | NA | NA | 1 | 1 | NA | NA | 1 | 0 | 2 | 0.5 |
| X_4118370 | NA | NA | 2 | 1 | 2 | 1 | NA | NA | 4 | 1 | NA | NA | 1 | 1 | NA | NA | 1 | 0 | 2 | 0.5 |
| X_4118379 | NA | NA | 2 | 0.5 | 2 | 0.5 | NA | NA | 4 | 0.5 | NA | NA | NA | NA | NA | NA | 1 | 1 | 1 | 1 |
| X_4118379 | NA | NA | 2 | 1 | 1 | 1 | NA | NA | 3 | 1 | NA | NA | 1 | 1 | NA | NA | 1 | 1 | 2 | 1 |
| X_4119093 | NA | NA | 3 | 0.3333 | 2 | 0 | 1 | 0 | 6 | 0.1667 | NA | NA | 2 | 1 | NA | NA | 1 | 0 | 3 | 0.6667 |
| X_4119376 | NA | NA | 4 | 0.5 | 2 | 0.5 | 1 | 0 | 7 | 0.4286 | NA | NA | 2 | 0.5 | NA | NA | 2 | 0.5 | 4 | 0.5 |
| X_4119391 | 1 | 0 | 4 | 0.5 | 3 | 0.3333 | 2 | 0 | 10 | 0.3 | 1 | 0 | 2 | 0.5 | NA | NA | 1 | 0 | 4 | 0.25 |
| X_4119455 | 1 | 0 | 4 | 0 | 6 | 0.1667 | 3 | 0 | 14 | 0.0714 | 1 | 0 | 3 | 0.3333 | 3 | 0 | 2 | 0 | 9 | 0.1111 |
| X_4136434 | 7 | 0.1429 | 4 | 0 | 7 | 0.1429 | 7 | 0.4286 | 25 | 0.2 | NA | NA | 4 | 0.25 | NA | NA | 3 | 0 | 7 | 0.1429 |
| X_4136583 | 6 | 0.5 | 8 | 0.625 | 6 | 0.1667 | 12 | 0.5 | 32 | 0.4688 | 1 | 0 | 7 | 0.4286 | 3 | 0.6667 | 4 | 0.5 | 15 | 0.4667 |
| X_4161694 | 2 | 0 | 2 | 0.5 | 2 | 0.5 | 8 | 0.375 | 14 | 0.3571 | 5 | 0.6 | 3 | 0 | 3 | 0 | 7 | 0.1429 | 18 | 0.2222 |
| X_4161695 | 2 | 0 | 1 | 0 | 3 | 0.3333 | 7 | 0.2857 | 13 | 0.2308 | 6 | 0.5 | 3 | 0.3333 | 3 | 0.3333 | 7 | 0.2857 | 19 | 0.3684 |
| X_4161714 | 1 | 0 | 2 | 1 | 1 | 1 | 8 | 0.625 | 12 | 0.6667 | 4 | 1 | 3 | 1 | 2 | 1 | 5 | 0.8 | 14 | 0.9286 |
| X_4161730 | 3 | 0 | 2 | 0.5 | NA | NA | 8 | 0.25 | 13 | 0.2308 | 5 | 0.4 | 3 | 0.3333 | 3 | 0.3333 | 6 | 0.3333 | 17 | 0.3529 |
| X_4161748 | 2 | 0.5 | 2 | 0 | NA | NA | 9 | 0 | 13 | 0.0769 | 3 | 0 | 2 | 0 | 3 | 0 | 5 | 0.4 | 13 | 0.1538 |

| | | | | | | | | | | | | | | | | | | | | |
|---|---|---|---|---|---|---|---|---|---|---|---|---|---|---|---|---|---|---|---|---|
| X_4162149 | NA | NA | 2 | 0.5 | NA | NA | NA | NA | 2 | 0.5 | NA | NA | 1 | 1 | NA | NA | 1 | 1 | 2 | 1 |
| X_4162150 | NA | NA | 2 | 1 | NA | NA | NA | NA | 2 | 1 | NA | NA | NA | NA | NA | NA | 1 | 1 | 1 | 1 |
| X_4162167 | NA | NA | 2 | 1 | NA | NA | NA | NA | 2 | 1 | NA | NA | 1 | 1 | NA | NA | NA | NA | 1 | 1 |
| X_4162180 | NA | NA | 1 | 1 | NA | NA | NA | NA | 1 | 1 | NA | NA | 1 | 1 | NA | NA | 1 | 1 | 2 | 1 |
| X_4162548 | NA | NA | 1 | 1 | NA | NA | NA | NA | 1 | 1 | NA | NA | 1 | 0 | NA | NA | 1 | 1 | 2 | 0.5 |
| X_4387599 | 9 | 0.4444 | 12 | 0.5 | 4 | 0.75 | 9 | 0.4444 | 34 | 0.5 | 3 | 0.3333 | 8 | 0.75 | 5 | 0.6 | 6 | 0.5 | 22 | 0.5909 |
| X_4387632 | 10 | 0.3 | 11 | 0.0909 | 3 | 0 | 10 | 0.2 | 34 | 0.1765 | 3 | 0 | 8 | 0.25 | 3 | 0 | 7 | 0.1429 | 21 | 0.1429 |
| X_4387684 | 10 | 0 | 9 | 0.2222 | 1 | 0 | 9 | 0.1111 | 29 | 0.1034 | 3 | 0 | 6 | 0.3333 | 3 | 0 | 5 | 0 | 17 | 0.1176 |
| X_4387687 | 9 | 0.6667 | 10 | 0.7 | 1 | 1 | 9 | 0.6667 | 29 | 0.6897 | 2 | 0.5 | 5 | 0.8 | 3 | 0.6667 | 5 | 0.6 | 15 | 0.6667 |
| X_4387706 | 9 | 1 | 11 | 0.9091 | 1 | 1 | 6 | 1 | 27 | 0.963 | 3 | 1 | 6 | 1 | 4 | 1 | 4 | 1 | 17 | 1 |
| X_4387750 | 9 | 0 | 9 | 0.2222 | 1 | 0 | 8 | 0 | 27 | 0.0741 | 3 | 0 | 6 | 0 | 3 | 0 | 4 | 0.25 | 16 | 0.0625 |
| X_4387796 | 6 | 0.8333 | 9 | 0.7778 | 1 | 1 | 9 | 1 | 25 | 0.88 | 3 | 1 | 6 | 1 | 3 | 1 | 3 | 1 | 15 | 1 |
| X_4387806 | 7 | 0.1429 | 7 | 0.1429 | 1 | 0 | 7 | 0 | 22 | 0.0909 | 2 | 0 | 6 | 0 | 3 | 0.3333 | 2 | 0 | 13 | 0.0769 |
| X_4387809 | 6 | 1 | 7 | 0.8571 | 1 | 1 | 8 | 1 | 22 | 0.9545 | 3 | 1 | 6 | 1 | 2 | 1 | 3 | 1 | 14 | 1 |
| X_4387830 | 4 | 0 | 6 | 0.1667 | 1 | 0 | 7 | 0 | 18 | 0.0556 | 2 | 0 | 3 | 0 | 2 | 0.5 | 3 | 0 | 10 | 0.1 |
| X_4387864 | 4 | 0.25 | 4 | 0.25 | 1 | 0 | 7 | 0.1429 | 16 | 0.1875 | 1 | 1 | 4 | 0.5 | 1 | 0 | 2 | 0 | 8 | 0.375 |
| X_4387886 | 2 | 0.5 | 3 | 1 | 1 | 0 | 5 | 0 | 11 | 0.3636 | 2 | 0 | 3 | 0.3333 | 2 | 0 | 1 | 0 | 8 | 0.125 |
| X_4689826 | 1 | 1 | NA | NA | 1 | 0 | 2 | 0 | 4 | 0.25 | 1 | 1 | 2 | 0 | NA | NA | NA | NA | 3 | 0.3333 |
| X_4689851 | 4 | 0.5 | NA | NA | 2 | 0.5 | 3 | 0.3333 | 9 | 0.4444 | 1 | 0 | 2 | 0 | 2 | 0.5 | 1 | 1 | 6 | 0.3333 |
| X_4689967 | 2 | 0.5 | 3 | 0.3333 | 1 | 0 | 2 | 0.5 | 8 | 0.375 | 1 | 1 | 3 | 0.6667 | 3 | 0.3333 | 2 | 0.5 | 9 | 0.5556 |
| X_4691239 | 4 | 0 | 2 | 0 | 3 | 0.3333 | 2 | 0 | 11 | 0.0909 | 2 | 0 | 7 | 0 | 4 | 0.25 | 2 | 1 | 15 | 0.2 |
| X_4703704 | 2 | 0 | 3 | 0 | 1 | 0 | 7 | 0.1429 | 13 | 0.0769 | 1 | 0 | 6 | 0 | NA | NA | 5 | 0.2 | 12 | 0.0833 |
| X_4705239 | 2 | 0 | 4 | 0 | 2 | 0.5 | 9 | 0 | 17 | 0.0588 | 1 | 0 | 5 | 0 | 2 | 0 | 8 | 0.25 | 16 | 0.125 |
| X_4705249 | 2 | 0 | 4 | 0 | 2 | 0 | 10 | 0.2 | 18 | 0.1111 | 1 | 0 | 5 | 0.2 | 2 | 0.5 | 7 | 0.2857 | 15 | 0.2667 |
| X_4705284 | 3 | 0.3333 | 5 | 0.4 | 3 | 0 | 10 | 0 | 21 | 0.1429 | 1 | 0 | 4 | 0 | 2 | 0 | 7 | 0.1429 | 14 | 0.0714 |
| X_4705296 | 3 | 1 | 5 | 0.6 | 4 | 0.5 | 9 | 0.5556 | 21 | 0.619 | 1 | 1 | 5 | 1 | 1 | 1 | 8 | 0.625 | 15 | 0.8 |
| X_4705314 | 2 | 1 | 6 | 0.8333 | 4 | 0.75 | 9 | 0.6667 | 21 | 0.7619 | 1 | 1 | 5 | 0.6 | 2 | 0.5 | 8 | 0.5 | 16 | 0.5625 |
| X_4705352 | 3 | 0.6667 | 5 | 0.4 | 3 | 0.3333 | 10 | 0.5 | 21 | 0.4762 | 1 | 1 | 6 | 0.5 | NA | NA | 9 | 0.3333 | 16 | 0.4375 |
| X_4705353 | 3 | 0.6667 | 6 | 0.6667 | 4 | 0.75 | 10 | 0.6 | 23 | 0.6522 | 1 | 0 | 6 | 0.5 | 2 | 0.5 | 6 | 0.3333 | 15 | 0.4 |
| X_4705362 | 3 | 0.3333 | 5 | 0.2 | 3 | 0 | 9 | 0.1111 | 20 | 0.15 | 1 | 1 | 3 | 0.3333 | 2 | 0.5 | 7 | 0.4286 | 13 | 0.4615 |
| X_4705365 | 3 | 0.3333 | 6 | 0.6667 | 5 | 0.4 | 10 | 0 | 24 | 0.2917 | 1 | 1 | 6 | 0.3333 | 2 | 0 | 8 | 0.125 | 17 | 0.2353 |
| X_4705369 | 2 | 0 | 4 | 0.5 | 5 | 0.2 | 10 | 0.3 | 21 | 0.2857 | 1 | 0 | 6 | 0.5 | 2 | 0 | 8 | 0.125 | 17 | 0.2353 |
| X_4705452 | 3 | 0.6667 | 8 | 0.25 | 5 | 0.2 | 13 | 0.0769 | 29 | 0.2069 | 2 | 0 | 5 | 0 | 3 | 0 | 12 | 0.25 | 22 | 0.1364 |
| X_4711417 | 1 | 1 | 2 | 1 | 2 | 0 | 1 | 0 | 6 | 0.5 | 3 | 0.6667 | 2 | 0 | 1 | 0 | 1 | 1 | 7 | 0.4286 |
| X_4711440 | 1 | 0 | 2 | 0.5 | 2 | 0.5 | 1 | 1 | 6 | 0.5 | 3 | 0 | 2 | 1 | 1 | 0 | 1 | 0 | 7 | 0.2857 |
| X_4711536 | 1 | 0 | 3 | 0 | 2 | 0.5 | 1 | 1 | 7 | 0.2857 | 2 | 0.5 | 1 | 0 | 1 | 0 | NA | NA | 4 | 0.25 |
| X_4711613 | 1 | 1 | 1 | 1 | 3 | 1 | NA | NA | 5 | 1 | 1 | 1 | 2 | 0.5 | 1 | 1 | NA | NA | 4 | 0.75 |
| X_4711617 | 1 | 0 | 2 | 1 | 1 | 1 | 1 | 1 | 5 | 0.8 | 2 | 1 | 2 | 1 | 1 | 1 | NA | NA | 5 | 1 |
| X_4711623 | NA | NA | 2 | 1 | 3 | 0.3333 | 1 | 1 | 6 | 0.6667 | 2 | 0.5 | 2 | 1 | 1 | 0 | NA | NA | 5 | 0.6 |
| X_4711623 | 1 | 1 | 2 | 1 | 3 | 0.6667 | 1 | 0 | 7 | 0.7143 | 1 | 1 | 2 | 1 | 1 | 1 | NA | NA | 4 | 1 |
| X_4711631 | 1 | 0 | 2 | 0 | 3 | 0.3333 | 1 | 0 | 7 | 0.1429 | 1 | 1 | 1 | 1 | 1 | 0 | NA | NA | 3 | 0.6667 |
| X_4711634 | 1 | 1 | 2 | 1 | 1 | 1 | NA | NA | 4 | 1 | NA | NA | 2 | 1 | NA | NA | NA | NA | 2 | 1 |
| X_4726099 | 3 | 0 | 6 | 0 | 3 | 0 | 6 | 0.1667 | 18 | 0.0556 | 2 | 0 | 5 | 0.2 | 4 | 0.25 | 6 | 0.3333 | 17 | 0.2353 |
| X_4726105 | 2 | 0 | 5 | 0.4 | 1 | 0 | 5 | 0.2 | 13 | 0.2308 | 2 | 0 | 2 | 1 | 3 | 0.3333 | 7 | 0.1429 | 14 | 0.2857 |
| X_4726203 | 3 | 0.3333 | 5 | 0 | 2 | 0 | 6 | 0 | 16 | 0.0625 | 2 | 0 | 5 | 0 | NA | NA | 5 | 0.2 | 12 | 0.0833 |
| X_4726211 | 2 | 0.5 | 4 | 0.25 | 1 | 0 | 3 | 0.3333 | 10 | 0.3 | 2 | 0.5 | 4 | 0 | NA | NA | 5 | 0 | 11 | 0.0909 |
| X_4726216 | 3 | 0 | 6 | 0.6667 | 2 | 0 | 6 | 0.5 | 17 | 0.4118 | 2 | 0.5 | 4 | 0 | NA | NA | 6 | 0.3333 | 12 | 0.25 |
| X_4726218 | 3 | 0.3333 | 5 | 0.2 | 2 | 0 | 6 | 1 | 16 | 0.5 | 2 | 0.5 | 4 | 0.5 | NA | NA | 7 | 0.4286 | 13 | 0.4615 |
| X_4726221 | 3 | 0 | 6 | 0 | 2 | 0 | 4 | 0.25 | 15 | 0.0667 | 2 | 0 | 3 | 0 | NA | NA | 4 | 0.25 | 9 | 0.1111 |
| X_4726222 | 3 | 0.3333 | 3 | 0 | 2 | 0 | 3 | 0.3333 | 11 | 0.1818 | 2 | 0 | 4 | 0.25 | NA | NA | 3 | 0 | 9 | 0.1111 |
| X_4726238 | 3 | 1 | 6 | 1 | 2 | 1 | 5 | 0.8 | 16 | 0.9375 | 2 | 0.5 | 4 | 0.75 | NA | NA | 7 | 0.7143 | 13 | 0.6923 |
| X_4726249 | 3 | 0 | 6 | 0 | 2 | 0.5 | 6 | 0 | 17 | 0.0588 | 2 | 0 | 2 | 0 | NA | NA | 6 | 0.1667 | 10 | 0.1 |
| X_4729572 | 2 | 0 | 6 | 0.3333 | 1 | 0 | 4 | 0.25 | 13 | 0.2308 | 2 | 0 | 2 | 0.5 | NA | NA | 6 | 0.5 | 10 | 0.4 |
| X_4736220 | NA | NA | NA | NA | NA | NA | 1 | 1 | 1 | 1 | 1 | 1 | NA | NA | NA | NA | NA | NA | 1 | 1 |
| X_4736229 | NA | NA | NA | NA | NA | NA | 1 | 1 | 1 | 1 | 1 | 1 | NA | NA | NA | NA | NA | NA | 1 | 1 |
| X_4745910 | 1 | 1 | NA | NA | NA | NA | NA | NA | 1 | 1 | NA | NA | NA | NA | 1 | 1 | NA | NA | 1 | 1 |
| X_4745922 | 1 | 1 | NA | NA | NA | NA | NA | NA | 1 | 1 | NA | NA | 1 | 1 | NA | NA | NA | NA | 1 | 1 |
| X_4745951 | 1 | 1 | 1 | 1 | NA | NA | NA | NA | 2 | 1 | NA | NA | 1 | 0 | 1 | 1 | NA | NA | 2 | 0.5 |
| X_4745976 | 1 | 1 | 1 | 0 | NA | NA | NA | NA | 2 | 0.5 | NA | NA | 1 | 0 | 1 | 1 | NA | NA | 2 | 0.5 |
| X_4747289 | 2 | 0.5 | 2 | 0.5 | NA | NA | NA | NA | 4 | 0.5 | 1 | 1 | 1 | 1 | 1 | 0 | NA | NA | 3 | 0.6667 |
| X_4749497 | 3 | 0.3333 | 3 | 0 | NA | NA | 1 | 1 | 7 | 0.2857 | 1 | 1 | 2 | 1 | 3 | 0 | 1 | 0 | 7 | 0.4286 |
| X_4749499 | 3 | 1 | 3 | 1 | NA | NA | 2 | 0.5 | 8 | 0.875 | 1 | 1 | 2 | 1 | 3 | 1 | 1 | 1 | 7 | 1 |
| X_4749516 | 3 | 0.3333 | 2 | 0 | NA | NA | 1 | 0 | 6 | 0.1667 | 1 | 1 | 2 | 0.5 | 3 | 0.3333 | 1 | 0 | 7 | 0.4286 |
| X_4749518 | 3 | 0.6667 | 3 | 0 | NA | NA | 1 | 0 | 7 | 0.2857 | 1 | 0 | 2 | 0.5 | 3 | 1 | NA | NA | 6 | 0.6667 |
| X_4749522 | 2 | 0.5 | 2 | 0.5 | NA | NA | NA | NA | 4 | 0.5 | NA | NA | 1 | 1 | 1 | 1 | NA | NA | 2 | 1 |

| | | | | | | | | | | | | | | | | | | | | |
|---|---|---|---|---|---|---|---|---|---|---|---|---|---|---|---|---|---|---|---|---|
| X_4749537 | 2 | 1 | 2 | 1 | NA | NA | NA | NA | 4 | 1 | 1 | 1 | 1 | 1 | 1 | 1 | NA | NA | 3 | 1 |
| X_4750348 | NA | NA | NA | NA | NA | NA | 1 | 1 | 1 | 1 | NA | NA | NA | NA | 1 | 1 | NA | NA | 1 | 1 |
| X_4756393 | 6 | 0.3333 | 3 | 0.6667 | 7 | 0.4286 | 2 | 0 | 18 | 0.3889 | 4 | 0.25 | 5 | 0.2 | 3 | 0.3333 | 4 | 0.5 | 16 | 0.3125 |
| X_4761872 | 2 | 0.5 | 1 | 0 | 6 | 0.3333 | 4 | 0.25 | 13 | 0.3077 | 4 | 0.5 | 1 | 0 | 4 | 0.5 | 6 | 0.3333 | 15 | 0.4 |
| X_4761881 | 2 | 0.5 | 1 | 0 | 4 | 0.75 | 6 | 1 | 13 | 0.7692 | 4 | 0.5 | 1 | 1 | 5 | 0.8 | 8 | 0.75 | 18 | 0.7222 |
| X_4761893 | 3 | 0.3333 | 1 | 0 | 4 | 0.5 | 6 | 0.5 | 14 | 0.4286 | 2 | 0.5 | 1 | 1 | 5 | 0.8 | 8 | 0.875 | 16 | 0.8125 |
| X_4763144 | 1 | 1 | NA | NA | 2 | 0.5 | 2 | 0.5 | 5 | 0.6 | NA | NA | 1 | 0 | 2 | 0.5 | NA | NA | 3 | 0.3333 |
| X_4775092 | 13 | 0.1538 | 15 | 0.2 | 13 | 0.1538 | 12 | 0.25 | 53 | 0.1887 | 7 | 0.4286 | 12 | 0.3333 | 19 | 0.1579 | 7 | 0 | 45 | 0.2222 |
| X_4809982 | 1 | 1 | 1 | 1 | 2 | 0.5 | 4 | 0 | 8 | 0.375 | 1 | 0 | 1 | 0 | 2 | 0 | 3 | 0.6667 | 7 | 0.2857 |
| X_4810028 | 2 | 1 | NA | NA | 1 | 1 | 4 | 0.75 | 7 | 0.8571 | 1 | 1 | 2 | 0.5 | NA | NA | 3 | 0.3333 | 6 | 0.5 |
| X_4875031 | 2 | 0.5 | NA | NA | 1 | 0 | 3 | 0.6667 | 6 | 0.5 | NA | NA | 1 | 1 | 1 | 1 | 2 | 0.5 | 4 | 0.75 |
| X_4964717 | NA | NA | NA | NA | 1 | 1 | NA | NA | 1 | 1 | NA | NA | 1 | 0 | NA | NA | 1 | 1 | 2 | 0.5 |
| X_4998349 | NA | NA | 1 | 1 | NA | NA | NA | NA | 1 | 1 | NA | NA | NA | NA | NA | NA | 1 | 1 | 1 | 1 |
| X_4998373 | NA | NA | 1 | 1 | NA | NA | NA | NA | 1 | 1 | NA | NA | NA | NA | NA | NA | 1 | 1 | 1 | 1 |
| X_4998376 | NA | NA | 1 | 1 | NA | NA | NA | NA | 1 | 1 | NA | NA | NA | NA | NA | NA | 1 | 1 | 1 | 1 |
| X_4998404 | NA | NA | 1 | 1 | NA | NA | NA | NA | 1 | 1 | NA | NA | NA | NA | NA | NA | 1 | 1 | 1 | 1 |
| X_4998409 | NA | NA | 1 | 1 | NA | NA | NA | NA | 1 | 1 | NA | NA | NA | NA | NA | NA | 1 | 1 | 1 | 1 |
| X_5009508 | 6 | 0.3333 | 3 | 0.3333 | 4 | 0.25 | 3 | 0.3333 | 16 | 0.3125 | 3 | 0 | 8 | 0.25 | 2 | 0.5 | 4 | 0.75 | 17 | 0.3529 |
| X_5009547 | 4 | 0 | 2 | 0 | 3 | 0.3333 | 4 | 0.5 | 13 | 0.2308 | 3 | 0 | 8 | 0 | 2 | 1 | 4 | 0 | 17 | 0.1176 |
| X_5009556 | 5 | 0.4 | 2 | 0 | 2 | 1 | 2 | 0.5 | 11 | 0.4545 | 3 | 0 | 7 | 0.2857 | 2 | 0.5 | 3 | 0 | 15 | 0.2 |
| X_5009558 | 4 | 0 | 3 | 0.3333 | 2 | 0 | 3 | 0.6667 | 12 | 0.25 | 1 | 0 | 6 | 0.3333 | 2 | 0.5 | 3 | 0 | 12 | 0.25 |
| X_5010394 | 3 | 1 | 2 | 0.5 | 4 | 0.75 | 3 | 1 | 12 | 0.8333 | 2 | 0.5 | 5 | 0.6 | 2 | 1 | 2 | 0 | 11 | 0.5455 |
| X_5010396 | 4 | 0.75 | 3 | 0.6667 | 4 | 1 | 3 | 1 | 14 | 0.8571 | 3 | 0.6667 | 6 | 0.5 | 2 | 1 | 3 | 1 | 14 | 0.7143 |
| X_5010407 | 3 | 0.3333 | 3 | 0.3333 | 3 | 0 | 3 | 0.3333 | 12 | 0.25 | 3 | 0 | 6 | 0.1667 | 2 | 0 | 3 | 0 | 14 | 0.0714 |
| X_5010663 | 4 | 0.25 | 3 | 0 | 3 | 0.6667 | 3 | 0.3333 | 13 | 0.3077 | 3 | 0.3333 | 6 | 0 | 1 | 0 | 3 | 0.6667 | 13 | 0.2308 |
| X_5010669 | 4 | 1 | 3 | 1 | 3 | 1 | 3 | 0.6667 | 13 | 0.9231 | 3 | 1 | 6 | 1 | 1 | 1 | 3 | 1 | 13 | 1 |
| X_5010677 | 3 | 0.3333 | 3 | 0.3333 | 3 | 1 | 3 | 0.3333 | 12 | 0.5 | 2 | 0.5 | 7 | 0.2857 | 2 | 0.5 | 3 | 0 | 14 | 0.2857 |
| X_5010693 | 2 | 0.5 | 2 | 0 | 3 | 0 | 1 | 0 | 8 | 0.125 | 2 | 0 | 4 | 0.5 | 2 | 0.5 | 3 | 0.3333 | 11 | 0.3636 |
| X_5047650 | 3 | 0 | 4 | 0 | 3 | 0.3333 | 4 | 0 | 14 | 0.0714 | 3 | 0.6667 | 8 | 0.125 | 4 | 0.25 | 4 | 0.25 | 19 | 0.2632 |
| X_5047727 | 5 | 0 | 4 | 0 | 5 | 0.2 | 3 | 0 | 17 | 0.0588 | 2 | 0 | 8 | 0.375 | 5 | 0.4 | 4 | 0 | 19 | 0.2632 |
| X_5047735 | 5 | 0.2 | 3 | 0 | 4 | 0.25 | 4 | 0 | 16 | 0.125 | 2 | 0.5 | 8 | 0.375 | 4 | 0 | 4 | 0.5 | 18 | 0.3333 |
| X_5047740 | 6 | 0.5 | 4 | 0 | 5 | 0.6 | 4 | 0.75 | 19 | 0.4737 | 2 | 0.5 | 7 | 0.2857 | 5 | 0.6 | 5 | 0 | 19 | 0.3158 |
| X_5047741 | 5 | 0.2 | 2 | 0 | 5 | 0.4 | 3 | 0.3333 | 15 | 0.2667 | 2 | 0.5 | 6 | 0.8333 | 2 | 0.5 | 3 | 0.3333 | 13 | 0.6154 |
| X_5047870 | 4 | 0 | 4 | 0.25 | 1 | 0 | 4 | 0 | 13 | 0.0769 | 1 | 0 | 7 | 0.1429 | 4 | 0 | 1 | 0 | 13 | 0.0769 |
| X_5047874 | 4 | 1 | 2 | 0 | 1 | 0 | 4 | 0.75 | 11 | 0.6364 | 1 | 1 | 5 | 0.8 | 4 | 1 | 1 | 1 | 11 | 0.9091 |
| X_5047883 | 3 | 1 | 4 | 0.75 | 2 | 1 | 4 | 1 | 13 | 0.9231 | NA | NA | 3 | 1 | 4 | 1 | 1 | 1 | 8 | 1 |
| X_5047899 | 4 | 1 | 3 | 0.6667 | 2 | 1 | 4 | 0.5 | 13 | 0.7692 | 1 | 1 | 3 | 0.6667 | 4 | 0.5 | 1 | 1 | 9 | 0.6667 |
| X_5047899 | 4 | 0.25 | 3 | 0 | 2 | 0 | 4 | 0 | 13 | 0.0769 | NA | NA | 5 | 0.4 | 2 | 0 | NA | NA | 7 | 0.2857 |
| X_5047904 | 3 | 1 | 3 | 1 | 2 | 1 | 4 | 1 | 12 | 1 | 1 | 1 | 5 | 1 | 4 | 1 | 1 | 0 | 11 | 0.9091 |
| X_5047908 | 3 | 0.6667 | 4 | 0.5 | 1 | 0 | 3 | 0 | 11 | 0.3636 | 1 | 0 | 6 | 0.5 | 3 | 0.6667 | 1 | 0 | 11 | 0.4545 |
| X_5114228 | NA | NA | NA | NA | NA | NA | 2 | 0.5 | 2 | 0.5 | 2 | 1 | NA | NA | NA | NA | NA | NA | 2 | 1 |
| X_5114231 | NA | NA | NA | NA | NA | NA | 2 | 1 | 2 | 1 | 2 | 1 | 1 | 1 | NA | NA | NA | NA | 3 | 1 |
| X_5188725 | NA | NA | 1 | 1 | NA | NA | NA | NA | 1 | 1 | NA | NA | 1 | 1 | 1 | 1 | NA | NA | 2 | 1 |
| X_5188745 | NA | NA | 1 | 1 | NA | NA | NA | NA | 1 | 1 | NA | NA | 1 | 1 | 1 | 1 | NA | NA | 2 | 1 |
| X_5188748 | NA | NA | 1 | 1 | NA | NA | NA | NA | 1 | 1 | NA | NA | 1 | 1 | NA | NA | NA | NA | 1 | 1 |
| X_5188812 | NA | NA | 1 | 1 | 1 | 0 | NA | NA | 2 | 0.5 | NA | NA | 1 | 1 | 1 | 1 | NA | NA | 2 | 1 |
| X_5204291 | NA | NA | 1 | 0 | 3 | 0.6667 | 1 | 1 | 5 | 0.6 | NA | NA | 1 | 0 | 2 | 0.5 | NA | NA | 3 | 0.3333 |
| X_5204305 | 1 | 1 | 1 | 0 | 2 | 0 | 1 | 1 | 5 | 0.4 | NA | NA | 2 | 0 | 3 | 0.3333 | NA | NA | 5 | 0.2 |
| X_5204307 | NA | NA | 2 | 0 | 3 | 0.3333 | 2 | 0 | 7 | 0.1429 | NA | NA | 2 | 0 | 3 | 0.3333 | NA | NA | 5 | 0.2 |
| X_5204308 | 1 | 0 | NA | NA | 3 | 1 | 1 | 0 | 5 | 0.6 | NA | NA | 2 | 0.5 | 1 | 0 | NA | NA | 3 | 0.3333 |
| X_5204323 | 1 | 1 | 2 | 0 | 3 | 1 | 2 | 0.5 | 8 | 0.625 | NA | NA | 2 | 1 | 3 | 0.6667 | NA | NA | 5 | 0.8 |
| X_5204334 | 1 | 1 | 2 | 1 | 3 | 1 | 2 | 1 | 8 | 1 | 1 | 1 | 3 | 0.6667 | 3 | 0.6667 | NA | NA | 7 | 0.7143 |
| X_5204339 | 1 | 0 | 2 | 0 | 3 | 0 | 1 | 1 | 7 | 0.1429 | 1 | 0 | 3 | 0 | 2 | 0.5 | NA | NA | 6 | 0.1667 |
| X_5204353 | 1 | 1 | 2 | 0.5 | 3 | 0.6667 | 2 | 1 | 8 | 0.75 | 1 | 1 | 3 | 0.6667 | 3 | 0.6667 | NA | NA | 7 | 0.7143 |
| X_5204354 | 1 | 1 | 2 | 0 | 3 | 0.3333 | 2 | 0.5 | 8 | 0.375 | NA | NA | 3 | 0 | 3 | 0.3333 | NA | NA | 6 | 0.1667 |
| X_5204358 | 1 | 1 | 2 | 0 | 2 | 0 | 2 | 0 | 7 | 0.1429 | 1 | 1 | 3 | 0.3333 | 2 | 0 | NA | NA | 6 | 0.3333 |
| X_5204368 | 1 | 0 | 2 | 0 | 2 | 0.5 | 2 | 0 | 7 | 0.1429 | 1 | 1 | 2 | 0 | 4 | 0 | NA | NA | 7 | 0.1429 |
| X_5204411 | 1 | 0 | 2 | 0 | 3 | 0 | 2 | 0.5 | 8 | 0.125 | 1 | 0 | 4 | 0.25 | 4 | 0 | NA | NA | 9 | 0.1111 |
| X_5204460 | 1 | 0 | 2 | 1 | 3 | 0.3333 | 3 | 1 | 9 | 0.6667 | 2 | 0 | 4 | 0.75 | 4 | 0.5 | 2 | 0 | 12 | 0.4167 |
| X_5243429 | 1 | 0 | 2 | 0.5 | NA | NA | 4 | 0 | 7 | 0.1429 | 1 | 1 | 1 | 0 | 1 | 0 | NA | NA | 3 | 0.3333 |
| X_5243445 | 1 | 0 | 1 | 0 | 1 | 0 | 4 | 0.25 | 7 | 0.1429 | 1 | 1 | 1 | 0 | 2 | 0 | NA | NA | 4 | 0.25 |
| X_5243466 | 1 | 1 | 1 | 0 | 1 | 1 | 3 | 0.6667 | 6 | 0.6667 | 1 | 1 | NA | NA | 2 | 0.5 | NA | NA | 3 | 0.6667 |
| X_5243478 | 1 | 1 | 1 | 1 | 1 | 1 | 4 | 1 | 7 | 1 | 1 | 0 | NA | NA | 1 | 1 | NA | NA | 2 | 0.5 |
| X_5261026 | 37 | 0.1081 | 30 | 0 | 35 | 0.0857 | 37 | 0.1081 | 139 | 0.0791 | 20 | 0.05 | 35 | 0.1429 | 42 | 0.0714 | 26 | 0.1154 | 123 | 0.0976 |
| X_5261026 | 38 | 0.1053 | 32 | 0.0625 | 36 | 0.1111 | 36 | 0.1389 | 142 | 0.1056 | 19 | 0.0526 | 38 | 0.0526 | 41 | 0.0732 | 27 | 0.037 | 125 | 0.056 |

| | | | | | | | | | | | | | | | | | | | | |
|---|---|---|---|---|---|---|---|---|---|---|---|---|---|---|---|---|---|---|---|---|
| X_5261034 | 40 | 0.7 | 28 | 0.6429 | 39 | 0.7949 | 41 | 0.561 | 148 | 0.6757 | 20 | 0.8 | 41 | 0.4634 | 48 | 0.75 | 28 | 0.6786 | 137 | 0.6569 |
| X_5261063 | 6 | 0.1667 | 4 | 0.25 | 6 | 0.6667 | 6 | 0.6667 | 22 | 0.4545 | 3 | 0.3333 | 2 | 0 | 9 | 0.2222 | 4 | 0.5 | 18 | 0.2778 |
| X_5261072 | 36 | 0.6667 | 24 | 0.625 | 38 | 0.5263 | 39 | 0.4615 | 137 | 0.562 | 14 | 0.6429 | 28 | 0.6786 | 45 | 0.6222 | 25 | 0.6 | 112 | 0.6339 |
| X_5261080 | 43 | 0.8372 | 30 | 0.9333 | 36 | 0.8611 | 39 | 0.8718 | 148 | 0.8716 | 19 | 0.6842 | 39 | 0.9231 | 49 | 0.8776 | 27 | 0.8519 | 134 | 0.8582 |
| X_5261082 | 36 | 0.5556 | 29 | 0.6897 | 37 | 0.7297 | 38 | 0.5263 | 140 | 0.6214 | 15 | 0.6667 | 31 | 0.5484 | 43 | 0.4651 | 23 | 0.6957 | 112 | 0.5625 |
| X_5261086 | 50 | 0.96 | 39 | 0.9231 | 43 | 0.9535 | 44 | 0.8864 | 176 | 0.9318 | 21 | 0.9524 | 42 | 0.9286 | 50 | 0.88 | 29 | 0.931 | 142 | 0.9155 |
| X_5261089 | 48 | 0.25 | 38 | 0.1316 | 42 | 0.119 | 44 | 0.1364 | 172 | 0.1628 | 25 | 0.12 | 42 | 0.1667 | 54 | 0.1296 | 30 | 0.2667 | 151 | 0.1656 |
| X_5261094 | 36 | 0.6389 | 30 | 0.6333 | 33 | 0.5152 | 41 | 0.4634 | 140 | 0.5571 | 18 | 0.6667 | 32 | 0.6875 | 42 | 0.5 | 23 | 0.6087 | 115 | 0.6 |
| X_5264461 | 27 | 0.7778 | 32 | 0.75 | 25 | 0.72 | 41 | 0.7805 | 125 | 0.76 | 32 | 0.75 | 28 | 0.7143 | 18 | 0.6111 | 29 | 0.8276 | 107 | 0.7383 |
| X_5264466 | 42 | 0.2619 | 42 | 0.3571 | 28 | 0.25 | 53 | 0.1509 | 165 | 0.2485 | 37 | 0.0811 | 29 | 0.2759 | 19 | 0.1053 | 39 | 0.2308 | 124 | 0.1774 |
| X_5264468 | 41 | 0.9756 | 41 | 1 | 28 | 1 | 49 | 1 | 159 | 0.9937 | 34 | 0.9412 | 31 | 0.9677 | 18 | 1 | 39 | 1 | 122 | 0.9754 |
| X_5264472 | 31 | 0.7097 | 35 | 0.9429 | 25 | 0.72 | 39 | 0.6923 | 130 | 0.7692 | 29 | 0.8966 | 25 | 0.84 | 17 | 0.7647 | 29 | 0.6207 | 100 | 0.78 |
| X_5264475 | 38 | 0.8947 | 34 | 0.7941 | 24 | 0.875 | 47 | 0.8511 | 143 | 0.8531 | 31 | 0.9032 | 29 | 0.931 | 17 | 0.9412 | 34 | 0.9412 | 111 | 0.9279 |
| X_5264482 | 36 | 0.3611 | 39 | 0.3333 | 29 | 0.2414 | 49 | 0.2653 | 153 | 0.3007 | 35 | 0.2571 | 26 | 0.3077 | 18 | 0.1667 | 38 | 0.2632 | 117 | 0.2564 |
| X_5264482 | 30 | 0.3 | 33 | 0.1818 | 22 | 0.3636 | 37 | 0.2703 | 122 | 0.2705 | 30 | 0.2 | 25 | 0.2 | 14 | 0.2143 | 34 | 0.2059 | 103 | 0.2039 |
| X_5264486 | 34 | 0.9118 | 35 | 0.7429 | 32 | 0.9062 | 51 | 0.8824 | 152 | 0.8618 | 37 | 1 | 26 | 0.9615 | 18 | 0.9444 | 37 | 0.9459 | 118 | 0.9661 |
| X_5264488 | 33 | 0.9394 | 31 | 0.871 | 25 | 0.92 | 40 | 0.9 | 129 | 0.907 | 27 | 0.7778 | 25 | 0.88 | 17 | 0.9412 | 34 | 0.9118 | 103 | 0.8738 |
| X_5264493 | 37 | 0.8108 | 37 | 0.8649 | 28 | 0.8214 | 51 | 0.7843 | 153 | 0.817 | 30 | 0.8333 | 27 | 0.7778 | 17 | 0.7647 | 36 | 0.75 | 110 | 0.7818 |
| X_5264512 | 29 | 0.2069 | 26 | 0.1154 | 22 | 0.2273 | 26 | 0.2308 | 103 | 0.1942 | 24 | 0.25 | 22 | 0.1818 | 10 | 0.1 | 22 | 0.1364 | 78 | 0.1795 |
| X_5264525 | 37 | 1 | 28 | 1 | 24 | 0.9583 | 40 | 1 | 129 | 0.9922 | 33 | 0.9697 | 25 | 0.96 | 14 | 1 | 37 | 0.973 | 109 | 0.9725 |
| X_5267093 | 2 | 1 | NA | NA | NA | NA | NA | NA | 2 | 1 | NA | NA | NA | NA | NA | NA | 1 | 1 | 1 | 1 |
| X_5267129 | 46 | 0.7391 | 37 | 0.5405 | 36 | 0.6111 | 34 | 0.7059 | 153 | 0.6536 | 30 | 0.8333 | 29 | 0.6552 | 24 | 0.5417 | 34 | 0.6471 | 117 | 0.6752 |
| X_5267134 | 55 | 0.1636 | 51 | 0.1373 | 41 | 0.1951 | 32 | 0.25 | 179 | 0.1788 | 40 | 0.2 | 41 | 0.2683 | 36 | 0.1944 | 35 | 0.2 | 152 | 0.2171 |
| X_5267136 | 54 | 0.9815 | 53 | 0.9057 | 45 | 0.9556 | 33 | 0.9697 | 185 | 0.9514 | 37 | 1 | 39 | 1 | 34 | 0.9706 | 37 | 1 | 147 | 0.9932 |
| X_5267140 | 39 | 0.9487 | 38 | 0.8684 | 28 | 0.9286 | 21 | 0.9524 | 126 | 0.9206 | 28 | 0.8214 | 29 | 1 | 29 | 0.8966 | 27 | 0.8889 | 113 | 0.9027 |
| X_5267143 | 53 | 0.9245 | 42 | 0.9048 | 39 | 0.9231 | 34 | 0.8529 | 168 | 0.9048 | 38 | 0.9211 | 35 | 0.9429 | 29 | 0.931 | 34 | 0.8824 | 136 | 0.9191 |
| X_5267150 | 57 | 0.2105 | 49 | 0.2653 | 41 | 0.2683 | 35 | 0.2 | 182 | 0.2363 | 41 | 0.2439 | 39 | 0.1795 | 34 | 0.2941 | 36 | 0.3056 | 150 | 0.2533 |
| X_5267150 | 48 | 0.2083 | 44 | 0.2727 | 36 | 0.25 | 30 | 0.1667 | 158 | 0.2278 | 35 | 0.1714 | 32 | 0.3125 | 32 | 0.1562 | 29 | 0.1724 | 128 | 0.2031 |
| X_5267154 | 53 | 0.7925 | 47 | 0.8298 | 44 | 0.8864 | 33 | 0.7273 | 177 | 0.8136 | 39 | 0.9487 | 38 | 0.8947 | 30 | 0.8 | 35 | 0.9143 | 142 | 0.8944 |
| X_5267156 | 47 | 0.8511 | 45 | 0.8667 | 39 | 0.8718 | 30 | 0.8333 | 161 | 0.8571 | 40 | 0.85 | 41 | 1 | 30 | 0.8333 | 39 | 0.9231 | 150 | 0.9067 |
| X_5267161 | 53 | 0.8491 | 48 | 0.8542 | 40 | 0.75 | 32 | 0.6562 | 173 | 0.7919 | 38 | 0.7368 | 40 | 0.75 | 32 | 0.625 | 38 | 0.8421 | 148 | 0.7432 |
| X_5267180 | 31 | 0.0968 | 29 | 0.1379 | 20 | 0.1 | 19 | 0.1579 | 99 | 0.1212 | 22 | 0.2273 | 26 | 0.2308 | 21 | 0.1905 | 23 | 0.1304 | 92 | 0.1957 |
| X_5267193 | 48 | 0.9167 | 43 | 0.9535 | 38 | 0.9474 | 29 | 0.9655 | 158 | 0.943 | 35 | 0.9714 | 39 | 1 | 27 | 0.963 | 36 | 0.9722 | 137 | 0.9781 |
| X_5267215 | 43 | 0.0698 | 37 | 0.027 | 29 | 0.069 | 26 | 0.1923 | 135 | 0.0815 | 34 | 0.0882 | 35 | 0.0857 | 25 | 0 | 34 | 0.0294 | 128 | 0.0547 |
| X_5269333 | NA | NA | 1 | 1 | 1 | 1 | 1 | 1 | 3 | 1 | NA | NA | 1 | 1 | NA | NA | NA | NA | 1 | 1 |
| X_5541992 | NA | NA | NA | NA | NA | NA | 1 | 1 | 1 | 1 | NA | NA | NA | NA | 1 | 1 | NA | NA | 1 | 1 |
| X_5542032 | 2 | 0 | 2 | 0 | 1 | 0 | 2 | 0.5 | 7 | 0.1429 | 1 | 0 | 2 | 0 | 2 | 0.5 | 1 | 0 | 6 | 0.1667 |
| X_5542160 | 3 | 1 | 2 | 1 | 1 | 1 | 2 | 1 | 8 | 1 | 1 | 1 | 1 | 1 | 2 | 1 | NA | NA | 4 | 1 |
| X_5542161 | 4 | 1 | NA | NA | 1 | 1 | 2 | 1 | 7 | 1 | 1 | 1 | 2 | 1 | 2 | 1 | 1 | 1 | 6 | 1 |
| X_5542167 | 1 | 1 | NA | NA | 1 | 1 | 1 | 1 | 3 | 1 | NA | NA | 2 | 1 | 2 | 1 | NA | NA | 4 | 1 |
| X_5542174 | 4 | 0.75 | 2 | 0.5 | NA | NA | 1 | 0 | 7 | 0.5714 | 1 | 0 | 2 | 0.5 | 2 | 0 | 1 | 0 | 6 | 0.1667 |
| X_5542179 | 4 | 1 | 2 | 1 | 1 | 1 | 2 | 1 | 9 | 1 | 1 | 1 | 1 | 1 | 1 | 1 | 1 | 1 | 4 | 1 |
| X_5542193 | 4 | 0.25 | 2 | 0 | NA | NA | 2 | 0 | 8 | 0.125 | 2 | 0.5 | 2 | 0.5 | 2 | 0 | 1 | 0 | 7 | 0.2857 |
| X_5542225 | 2 | 0.5 | 2 | 0 | 1 | 0 | 3 | 0 | 8 | 0.125 | 1 | 0 | NA | NA | 2 | 0.5 | NA | NA | 3 | 0.3333 |
| X_5542250 | 3 | 0 | 3 | 0 | 2 | 0 | 3 | 0.3333 | 11 | 0.0909 | 1 | 0 | NA | NA | 2 | 0.5 | 1 | 0 | 4 | 0.25 |
| X_5542306 | 4 | 0.25 | 5 | 0.4 | 2 | 0 | 3 | 1 | 14 | 0.4286 | 2 | 1 | 3 | 0.6667 | 1 | 0 | 1 | 1 | 7 | 0.7143 |
| X_5542470 | NA | NA | 1 | 1 | NA | NA | NA | NA | 1 | 1 | NA | NA | 1 | 1 | 1 | 1 | NA | NA | 2 | 1 |
| X_5542484 | NA | NA | 1 | 1 | NA | NA | NA | NA | 1 | 1 | NA | NA | 1 | 1 | 1 | 0 | NA | NA | 2 | 0.5 |
| X_5555243 | 5 | 0.4 | 6 | 0.1667 | 5 | 0 | 3 | 0.3333 | 19 | 0.2105 | 3 | 0.3333 | 1 | 0 | 7 | 0.1429 | 3 | 0 | 14 | 0.1429 |
| X_5555247 | 6 | 0.1667 | 8 | 0.25 | 5 | 0 | 3 | 0 | 22 | 0.1364 | 3 | 0 | 1 | 1 | 5 | 0 | 2 | 0.5 | 11 | 0.1818 |
| X_5555264 | 6 | 1 | 5 | 0.6 | 4 | 0.75 | 2 | 0 | 17 | 0.7059 | 4 | 0.75 | 1 | 1 | 9 | 0.7778 | 2 | 1 | 16 | 0.8125 |
| X_5555267 | 6 | 0.6667 | 6 | 0.8333 | 3 | 1 | 3 | 0 | 18 | 0.6667 | 3 | 0.6667 | 1 | 1 | 9 | 0.3333 | 1 | 0 | 14 | 0.4286 |
| X_5555277 | 3 | 0.6667 | 6 | 0.8333 | 4 | 0.75 | 2 | 1 | 15 | 0.8 | 3 | 1 | NA | NA | 9 | 0.5556 | 2 | 0.5 | 14 | 0.6429 |
| X_5555286 | 4 | 0 | 6 | 0.1667 | 5 | 0.2 | 2 | 0 | 17 | 0.1176 | 4 | 0.25 | 1 | 0 | 7 | 0 | 3 | 0.3333 | 15 | 0.1333 |
| X_5555293 | 6 | 0.1667 | 7 | 0.1429 | 5 | 0.2 | 3 | 0.3333 | 21 | 0.1905 | 3 | 0.3333 | 2 | 0 | 7 | 0.2857 | 2 | 0.5 | 14 | 0.2857 |
| X_5555301 | 6 | 0.8333 | 5 | 0.8 | 5 | 0.6 | 3 | 0.6667 | 19 | 0.7368 | 4 | 0.75 | 2 | 0.5 | 7 | 1 | 3 | 0.6667 | 16 | 0.8125 |
| X_5563403 | 1 | 1 | 5 | 0 | 1 | 0 | 2 | 0.5 | 9 | 0.2222 | 1 | 1 | 3 | 0 | 4 | 0 | 4 | 0.5 | 12 | 0.25 |
| X_5563447 | NA | NA | 4 | 0.25 | 2 | 0.5 | 2 | 0.5 | 8 | 0.375 | 1 | 0 | 2 | 0.5 | 2 | 0 | 3 | 1 | 8 | 0.5 |
| X_5570668 | 11 | 0.0909 | 7 | 0 | 6 | 0.1667 | 11 | 0.3636 | 35 | 0.1714 | 5 | 0.4 | 8 | 0.125 | 8 | 0.375 | 4 | 0 | 25 | 0.24 |
| X_5570671 | 12 | 0.0833 | 6 | 0.1667 | 7 | 0 | 10 | 0.3 | 35 | 0.1429 | 5 | 0 | 9 | 0 | 9 | 0.2222 | 4 | 0.25 | 27 | 0.1111 |
| X_5570683 | 11 | 0.0909 | 8 | 0.125 | 7 | 0.1429 | 12 | 0.0833 | 38 | 0.1053 | 6 | 0.1667 | 10 | 0 | 11 | 0.0909 | 4 | 0 | 31 | 0.0645 |
| X_5570697 | 11 | 0 | 7 | 0.4286 | 7 | 0 | 10 | 0.1 | 35 | 0.1143 | 8 | 0.25 | 9 | 0.1111 | 11 | 0.3636 | 4 | 0 | 32 | 0.2188 |
| X_5570727 | 8 | 0 | 6 | 0 | 6 | 0 | 10 | 0.2 | 30 | 0.0667 | 8 | 0.125 | 12 | 0.1667 | 10 | 0 | 3 | 0 | 33 | 0.0909 |
| X_5570757 | 10 | 0.2 | 9 | 0.2222 | 6 | 0.1667 | 7 | 0.1429 | 32 | 0.1875 | 7 | 0.1429 | 12 | 0.25 | 8 | 0.125 | 4 | 0 | 31 | 0.1613 |
| X_5570890 | 12 | 0 | 15 | 0 | 12 | 0.0833 | 18 | 0.2222 | 57 | 0.0877 | 8 | 0.25 | 19 | 0.1053 | 13 | 0.0769 | 7 | 0 | 47 | 0.1064 |

| | | | | | | | | | | | | | | | | | | | | |
|---|---|---|---|---|---|---|---|---|---|---|---|---|---|---|---|---|---|---|---|---|
| X_5583660 | 3 | 0 | 2 | 0.5 | 5 | 0.2 | 4 | 0.5 | 14 | 0.2857 | 1 | 0 | 4 | 0 | 2 | 0.5 | 6 | 0.1667 | 13 | 0.1538 |
| X_5583673 | 4 | 0 | 2 | 0 | 5 | 0 | 4 | 0.25 | 15 | 0.0667 | 1 | 0 | 4 | 0.25 | 2 | 0 | 6 | 0 | 13 | 0.0769 |
| X_5583724 | 8 | 0 | 7 | 0.1429 | 8 | 0.25 | 10 | 0.2 | 33 | 0.1515 | 1 | 0 | 9 | 0.1111 | 11 | 0.0909 | 8 | 0.125 | 29 | 0.1034 |
| X_5583733 | 7 | 0.2857 | 6 | 0.3333 | 14 | 0.2857 | 13 | 0.2308 | 40 | 0.275 | 2 | 0 | 12 | 0.1667 | 14 | 0.1429 | 11 | 0.3636 | 39 | 0.2051 |
| X_5583752 | 8 | 0.125 | 7 | 0.1429 | 14 | 0.2857 | 14 | 0.2143 | 43 | 0.2093 | 2 | 1 | 12 | 0.5 | 13 | 0.3077 | 10 | 0.2 | 37 | 0.3784 |
| X_5583761 | 6 | 0.1667 | 7 | 0.1429 | 13 | 0.0769 | 11 | 0.1818 | 37 | 0.1351 | 2 | 1 | 8 | 0.25 | 15 | 0.2667 | 10 | 0.5 | 35 | 0.3714 |
| X_5583761 | 8 | 0 | 8 | 0.125 | 12 | 0.0833 | 10 | 0.3 | 38 | 0.1316 | 3 | 0 | 9 | 0.1111 | 15 | 0.3333 | 11 | 0.1818 | 38 | 0.2105 |
| X_5583781 | 9 | 0.1111 | 10 | 0.1 | 14 | 0.2143 | 13 | 0.4615 | 46 | 0.2391 | 2 | 0 | 12 | 0.25 | 15 | 0.0667 | 11 | 0 | 40 | 0.1 |
| X_5583785 | 7 | 0 | 8 | 0 | 10 | 0 | 12 | 0.1667 | 37 | 0.0541 | 3 | 0 | 10 | 0.1 | 13 | 0.1538 | 9 | 0.1111 | 35 | 0.1143 |
| X_5583792 | 7 | 0.4286 | 9 | 0.2222 | 11 | 0.2727 | 10 | 0.4 | 37 | 0.3243 | 1 | 0 | 10 | 0.2 | 11 | 0.3636 | 8 | 0.375 | 30 | 0.3 |
| X_5583794 | 9 | 0.4444 | 11 | 0.1818 | 14 | 0.2857 | 17 | 0.5294 | 51 | 0.3725 | 3 | 0 | 11 | 0.4545 | 16 | 0.375 | 9 | 0.4444 | 39 | 0.3846 |
| X_5583809 | 9 | 0 | 11 | 0 | 15 | 0.0667 | 15 | 0.1333 | 50 | 0.06 | 3 | 0 | 9 | 0.1111 | 17 | 0.0588 | 11 | 0.0909 | 40 | 0.075 |
| X_5583826 | 7 | 0.4286 | 9 | 0 | 17 | 0.0588 | 14 | 0.1429 | 47 | 0.1277 | 4 | 0.25 | 10 | 0.1 | 14 | 0.2857 | 12 | 0.4167 | 40 | 0.275 |
| X_5583840 | 8 | 0.375 | 9 | 0.2222 | 18 | 0.1111 | 15 | 0.2667 | 50 | 0.22 | 6 | 0.1667 | 12 | 0.0833 | 15 | 0.3333 | 10 | 0.2 | 43 | 0.2093 |
| X_5583842 | 9 | 0.5556 | 12 | 0.1667 | 16 | 0.375 | 17 | 0.4706 | 54 | 0.3889 | 5 | 0.8 | 14 | 0.2857 | 19 | 0.4737 | 10 | 0.8 | 48 | 0.5208 |
| X_5583847 | 7 | 0.4286 | 9 | 0.2222 | 15 | 0.1333 | 16 | 0.375 | 47 | 0.2766 | 6 | 0.1667 | 11 | 0.2727 | 15 | 0.3333 | 10 | 0.6 | 42 | 0.3571 |
| X_5610247 | 1 | 1 | NA | NA | 1 | 0 | 1 | 0 | 3 | 0.3333 | 1 | 0 | 1 | 0 | 2 | 0.5 | 1 | 0 | 5 | 0.2 |
| X_5610250 | 1 | 0 | NA | NA | 1 | 0 | 1 | 1 | 3 | 0.3333 | 1 | 0 | NA | NA | 1 | 1 | NA | NA | 2 | 0.5 |
| X_5611172 | 7 | 0.7143 | 7 | 0.2857 | 7 | 0.1429 | 6 | 0.3333 | 27 | 0.3704 | 5 | 0.6 | 4 | 0.75 | 6 | 0.1667 | 9 | 0.5556 | 24 | 0.5 |
| X_5611212 | 6 | 1 | 9 | 0.8889 | 6 | 1 | 11 | 0.9091 | 32 | 0.9375 | 5 | 0.8 | 7 | 1 | 6 | 1 | 10 | 1 | 28 | 0.9643 |
| X_5611232 | 6 | 0.1667 | 10 | 0 | 8 | 0.25 | 11 | 0 | 35 | 0.0857 | 6 | 0.1667 | 7 | 0 | 5 | 0.2 | 8 | 0 | 26 | 0.0769 |
| X_5627799 | NA | NA | 2 | 0 | NA | NA | 4 | 0.25 | 6 | 0.1667 | NA | NA | 3 | 0 | NA | NA | 3 | 0.3333 | 6 | 0.1667 |
| X_5642894 | 3 | 0.3333 | 6 | 0.5 | 2 | 0 | 4 | 0 | 15 | 0.2667 | 2 | 0.5 | 5 | 0.4 | 2 | 0 | 1 | 0 | 10 | 0.3 |
| X_5643235 | 10 | 0.9 | 6 | 1 | 5 | 0.8 | 12 | 0.9167 | 33 | 0.9091 | 1 | 1 | 6 | 1 | 3 | 1 | 6 | 1 | 16 | 1 |
| X_5643236 | 8 | 0.375 | 7 | 0.1429 | 5 | 0.2 | 9 | 0.1111 | 29 | 0.2069 | 1 | 0 | 6 | 0.1667 | 4 | 0 | 3 | 0.3333 | 14 | 0.1429 |
| X_5643238 | 10 | 0.3 | 7 | 0.5714 | 6 | 0.3333 | 10 | 0.1 | 33 | 0.303 | NA | NA | 6 | 0.5 | 3 | 0 | 5 | 0.4 | 14 | 0.3571 |
| X_5643242 | 9 | 0.7778 | 6 | 0.8333 | 6 | 0.8333 | 10 | 0.8 | 31 | 0.8065 | NA | NA | 5 | 0.8 | 4 | 0.25 | 6 | 0.6667 | 15 | 0.6 |
| X_5643243 | 5 | 0.8 | 6 | 0.6667 | 4 | 0.5 | 9 | 0.3333 | 24 | 0.5417 | NA | NA | 6 | 0.8333 | 4 | 0.75 | 6 | 0.5 | 16 | 0.6875 |
| X_5643244 | 7 | 0 | 7 | 0 | 2 | 0.5 | 10 | 0.1 | 26 | 0.0769 | 1 | 0 | 5 | 0.4 | 4 | 0 | 4 | 0 | 14 | 0.1429 |
| X_5643284 | NA | NA | 2 | 0.5 | NA | NA | 1 | 1 | 3 | 0.6667 | NA | NA | 1 | 1 | 1 | 1 | NA | NA | 2 | 1 |
| X_5654291 | 1 | 1 | NA | NA | NA | NA | 3 | 0.3333 | 4 | 0.5 | NA | NA | 1 | 1 | NA | NA | 1 | 0 | 2 | 0.5 |
| X_5707619 | NA | NA | 1 | 1 | NA | NA | NA | NA | 1 | 1 | NA | NA | NA | NA | 1 | 1 | NA | NA | 1 | 1 |
| X_5707707 | NA | NA | 2 | 1 | NA | NA | NA | NA | 2 | 1 | NA | NA | NA | NA | 1 | 1 | NA | NA | 1 | 1 |
| X_5707716 | NA | NA | 2 | 1 | NA | NA | NA | NA | 2 | 1 | NA | NA | NA | NA | 1 | 1 | NA | NA | 1 | 1 |
| X_5707747 | 1 | 1 | 1 | 0 | NA | NA | 1 | 1 | 3 | 0.6667 | NA | NA | NA | NA | 1 | 1 | NA | NA | 1 | 1 |
| X_5810865 | 10 | 0 | 13 | 0.0769 | 15 | 0.0667 | 15 | 0.0667 | 53 | 0.0566 | 10 | 0.1 | 11 | 0 | 10 | 0.1 | 16 | 0.125 | 47 | 0.0851 |
| X_5817977 | 5 | 0.2 | 4 | 0 | 9 | 0.1111 | 10 | 0.4 | 28 | 0.2143 | 3 | 0 | 7 | 0.2857 | 6 | 0.3333 | 8 | 0.125 | 24 | 0.2083 |
| X_5817996 | 4 | 0.25 | 4 | 0.5 | 8 | 0.375 | 4 | 0.5 | 20 | 0.4 | 2 | 0 | 4 | 0.5 | 3 | 0.6667 | 5 | 0.6 | 14 | 0.5 |
| X_5879796 | NA | NA | NA | NA | 1 | 1 | NA | NA | 1 | 1 | NA | NA | NA | NA | 1 | 1 | NA | NA | 1 | 1 |
| X_5879808 | NA | NA | NA | NA | 1 | 1 | NA | NA | 1 | 1 | NA | NA | NA | NA | 1 | 1 | NA | NA | 1 | 1 |
| X_5927005 | 1 | 0 | NA | NA | 1 | 1 | 1 | 0 | 3 | 0.3333 | NA | NA | 2 | 0.5 | 1 | 0 | NA | NA | 3 | 0.3333 |
| X_5959186 | NA | NA | NA | NA | 1 | 1 | 2 | 1 | 3 | 1 | NA | NA | 1 | 1 | NA | NA | 1 | 1 | 2 | 1 |
| X_5959236 | NA | NA | NA | NA | NA | NA | 2 | 0.5 | 2 | 0.5 | NA | NA | NA | NA | NA | NA | 1 | 1 | 1 | 1 |
| X_5959242 | NA | NA | NA | NA | 2 | 0 | 2 | 0.5 | 4 | 0.25 | NA | NA | 1 | 1 | NA | NA | 1 | 0 | 2 | 0.5 |
| X_5959254 | NA | NA | NA | NA | 1 | 1 | 2 | 1 | 3 | 1 | NA | NA | 2 | 1 | NA | NA | NA | NA | 2 | 1 |
| X_5959263 | NA | NA | NA | NA | 1 | 1 | 2 | 0.5 | 3 | 0.6667 | NA | NA | NA | NA | NA | NA | 2 | 0.5 | 2 | 0.5 |
| X_5959264 | NA | NA | NA | NA | 1 | 0 | 2 | 0.5 | 3 | 0.3333 | NA | NA | NA | NA | NA | NA | 2 | 0.5 | 2 | 0.5 |
| X_5959267 | NA | NA | NA | NA | NA | NA | 2 | 0.5 | 2 | 0.5 | NA | NA | 2 | 0 | 1 | 0 | 1 | 1 | 4 | 0.25 |
| X_5993533 | 1 | 1 | 2 | 1 | NA | NA | NA | NA | 3 | 1 | NA | NA | 1 | 1 | NA | NA | NA | NA | 1 | 1 |
| X_5993549 | 1 | 1 | 3 | 1 | NA | NA | NA | NA | 4 | 1 | NA | NA | 1 | 1 | NA | NA | NA | NA | 1 | 1 |
| X_5993554 | 1 | 1 | 2 | 1 | NA | NA | NA | NA | 3 | 1 | NA | NA | 1 | 1 | NA | NA | NA | NA | 1 | 1 |
| X_5993559 | 1 | 0 | 2 | 1 | NA | NA | NA | NA | 3 | 0.6667 | NA | NA | 1 | 1 | NA | NA | NA | NA | 1 | 1 |
| X_6022497 | NA | NA | NA | NA | 1 | 0 | 2 | 0.5 | 3 | 0.3333 | NA | NA | NA | NA | 1 | 1 | 1 | 1 | 2 | 1 |
| X_6022989 | NA | NA | 1 | 0 | 1 | 0 | 4 | 0.75 | 6 | 0.5 | NA | NA | 1 | 1 | 1 | 0 | 1 | 0 | 3 | 0.3333 |
| X_6023017 | NA | NA | 2 | 0 | 2 | 0 | 4 | 0.25 | 8 | 0.125 | NA | NA | 1 | 1 | 3 | 0.3333 | 2 | 0 | 6 | 0.3333 |
| X_6023041 | NA | NA | 3 | 0.3333 | 2 | 0.5 | 5 | 0.2 | 10 | 0.3 | NA | NA | 1 | 0 | 3 | 0 | 3 | 0.3333 | 7 | 0.1429 |
| X_6023111 | NA | NA | 2 | 0 | 3 | 0.3333 | 4 | 0.5 | 9 | 0.3333 | NA | NA | 1 | 1 | 4 | 0.5 | 3 | 0.3333 | 8 | 0.5 |
| X_6023201 | NA | NA | 3 | 0.6667 | 2 | 0 | 7 | 0.1429 | 12 | 0.25 | 1 | 0 | 2 | 0.5 | 4 | 0 | 2 | 0 | 9 | 0.1111 |
| X_6023514 | 1 | 0 | 2 | 0 | 4 | 0 | 4 | 0.25 | 11 | 0.0909 | 1 | 0 | NA | NA | 2 | 0 | 2 | 0.5 | 5 | 0.2 |
| X_6033509 | 1 | 1 | 2 | 0.5 | NA | NA | 2 | 1 | 5 | 0.8 | NA | NA | NA | NA | NA | NA | 1 | 1 | 1 | 1 |
| X_6033532 | 1 | 1 | 2 | 0.5 | NA | NA | 2 | 0.5 | 5 | 0.6 | NA | NA | NA | NA | 1 | 1 | 1 | 1 | 2 | 1 |
| X_6033533 | 1 | 1 | 2 | 1 | NA | NA | 1 | 1 | 4 | 1 | NA | NA | NA | NA | 1 | 1 | 2 | 0.5 | 3 | 0.6667 |
| X_6075422 | NA | NA | NA | NA | 1 | 1 | NA | NA | 1 | 1 | NA | NA | NA | NA | NA | NA | 1 | 1 | 1 | 1 |
| X_6075423 | NA | NA | NA | NA | 1 | 1 | NA | NA | 1 | 1 | NA | NA | NA | NA | NA | NA | 1 | 1 | 1 | 1 |
| X_6075434 | NA | NA | NA | NA | 1 | 1 | NA | NA | 1 | 1 | 1 | 1 | NA | NA | NA | NA | 1 | 1 | 2 | 1 |

| | | | | | | | | | | | | | | | | | | | | |
|---|---|---|---|---|---|---|---|---|---|---|---|---|---|---|---|---|---|---|---|---|
| X_6075445 | NA | NA | NA | NA | 1 | 1 | NA | NA | 1 | 1 | 1 | 1 | NA | NA | NA | NA | 1 | 1 | 2 | 1 |
| X_6075447 | NA | NA | NA | NA | 1 | 1 | NA | NA | 1 | 1 | 1 | 0 | NA | NA | NA | NA | 1 | 1 | 2 | 0.5 |
| X_6330888 | 2 | 0 | 9 | 0.1111 | NA | NA | 2 | 0 | 13 | 0.0769 | 2 | 0 | 1 | 0 | 3 | 0.6667 | 3 | 0 | 9 | 0.2222 |
| X_6331108 | 1 | 0 | 6 | 0.1667 | NA | NA | 2 | 0 | 9 | 0.1111 | 2 | 0 | 1 | 0 | 1 | 1 | 5 | 0 | 9 | 0.1111 |
| X_6331309 | NA | NA | 4 | 0 | NA | NA | 1 | 1 | 5 | 0.2 | NA | NA | NA | NA | NA | NA | 4 | 0.25 | 4 | 0.25 |
| X_6331364 | NA | NA | 5 | 0.2 | 1 | 1 | 2 | 0 | 8 | 0.25 | 1 | 0 | NA | NA | 1 | 1 | 1 | 0 | 3 | 0.3333 |
| X_6331380 | NA | NA | 5 | 0.8 | 1 | 1 | 1 | 0 | 7 | 0.7143 | 1 | 1 | NA | NA | NA | NA | 3 | 0.6667 | 4 | 0.75 |
| X_6331380 | NA | NA | 1 | 1 | NA | NA | NA | NA | 1 | 1 | NA | NA | NA | NA | 1 | 1 | 1 | 0 | 2 | 0.5 |
| X_6331392 | NA | NA | 7 | 0.4286 | 1 | 0 | 2 | 0.5 | 10 | 0.4 | 1 | 0 | NA | NA | 1 | 1 | 4 | 0.5 | 6 | 0.5 |
| X_6331402 | NA | NA | 6 | 0.3333 | 1 | 1 | 1 | 0 | 8 | 0.375 | 1 | 0 | NA | NA | 1 | 1 | 3 | 0.6667 | 5 | 0.6 |
| X_6331408 | NA | NA | 6 | 0.3333 | 1 | 0 | 1 | 0 | 8 | 0.25 | 1 | 0 | NA | NA | NA | NA | 3 | 0.3333 | 4 | 0.25 |
| X_6331409 | NA | NA | 6 | 0.6667 | 1 | 1 | 2 | 1 | 9 | 0.7778 | 1 | 0 | NA | NA | 1 | 1 | 4 | 0.75 | 6 | 0.6667 |
| X_6331412 | NA | NA | 5 | 1 | 1 | 1 | 1 | 1 | 7 | 1 | 1 | 1 | NA | NA | 1 | 1 | 4 | 1 | 6 | 1 |
| X_6331413 | NA | NA | 5 | 0.4 | NA | NA | 2 | 0 | 7 | 0.2857 | 1 | 1 | NA | NA | 1 | 0 | 3 | 0.6667 | 5 | 0.6 |
| X_6331416 | NA | NA | 6 | 0.1667 | NA | NA | NA | NA | 6 | 0.1667 | 1 | 0 | NA | NA | 1 | 0 | 4 | 0.5 | 6 | 0.3333 |
| X_6331422 | NA | NA | 7 | 1 | 1 | 1 | NA | NA | 8 | 1 | 1 | 1 | NA | NA | 1 | 1 | 1 | 1 | 3 | 1 |
| X_6331425 | NA | NA | 5 | 0.2 | 1 | 0 | 1 | 0 | 7 | 0.1429 | NA | NA | NA | NA | 1 | 0 | 3 | 0.3333 | 4 | 0.25 |
| X_6331431 | NA | NA | 7 | 0 | 1 | 0 | 1 | 1 | 9 | 0.1111 | 1 | 0 | NA | NA | 1 | 1 | 4 | 0 | 6 | 0.1667 |
| X_6331442 | NA | NA | 3 | 1 | 1 | 1 | 1 | 1 | 5 | 1 | 1 | 0 | NA | NA | 1 | 1 | 2 | 1 | 4 | 0.75 |
| X_6331444 | NA | NA | 5 | 1 | NA | NA | 2 | 1 | 7 | 1 | 1 | 1 | NA | NA | NA | NA | 4 | 1 | 5 | 1 |
| X_6331448 | NA | NA | 6 | 0.5 | 1 | 0 | 1 | 1 | 8 | 0.5 | 1 | 0 | NA | NA | 1 | 1 | 2 | 0.5 | 4 | 0.5 |
| X_6331451 | NA | NA | 6 | 0.3333 | 1 | 1 | 1 | 1 | 8 | 0.5 | 1 | 0 | NA | NA | NA | NA | 4 | 0.25 | 5 | 0.2 |
| X_6331454 | NA | NA | 4 | 0.75 | 1 | 1 | 1 | 1 | 6 | 0.8333 | 1 | 1 | NA | NA | 1 | 0 | 4 | 0.75 | 6 | 0.6667 |
| X_6331462 | NA | NA | 4 | 1 | 1 | 1 | 1 | 1 | 6 | 1 | 1 | 1 | NA | NA | 1 | 1 | 2 | 1 | 4 | 1 |
| X_6331477 | NA | NA | 5 | 0.4 | 1 | 1 | 1 | 0 | 7 | 0.4286 | 1 | 0 | NA | NA | NA | NA | 3 | 0.6667 | 4 | 0.5 |
| X_6331498 | NA | NA | 5 | 0.2 | 1 | 0 | 2 | 1 | 8 | 0.375 | 1 | 0 | NA | NA | 1 | 0 | 4 | 0.5 | 6 | 0.3333 |
| X_6331499 | NA | NA | 6 | 0.6667 | 1 | 1 | 1 | 1 | 8 | 0.75 | 1 | 1 | NA | NA | NA | NA | 3 | 1 | 4 | 1 |
| X_6331506 | NA | NA | 5 | 0.8 | 1 | 1 | 1 | 1 | 7 | 0.8571 | 1 | 0 | NA | NA | 1 | 1 | 3 | 0.6667 | 5 | 0.6 |
| X_6331508 | NA | NA | 5 | 0.6 | NA | NA | 1 | 1 | 6 | 0.6667 | 1 | 1 | NA | NA | 1 | 0 | 2 | 0.5 | 4 | 0.5 |
| X_6331516 | NA | NA | 4 | 0.5 | NA | NA | NA | NA | 4 | 0.5 | 1 | 0 | NA | NA | NA | NA | 3 | 1 | 4 | 0.75 |
| X_6331522 | NA | NA | 5 | 0.4 | 1 | 1 | 2 | 1 | 8 | 0.625 | 1 | 1 | NA | NA | 1 | 0 | 4 | 0.75 | 6 | 0.6667 |
| X_6331525 | NA | NA | 5 | 0.4 | 1 | 0 | 2 | 0.5 | 8 | 0.375 | NA | NA | NA | NA | NA | NA | 3 | 0.6667 | 3 | 0.6667 |
| X_6331532 | NA | NA | 4 | 1 | 1 | 1 | 1 | 0 | 6 | 0.8333 | NA | NA | NA | NA | 1 | 1 | 4 | 1 | 5 | 1 |
| X_6331547 | NA | NA | 4 | 0.75 | 1 | 1 | 1 | 0 | 6 | 0.6667 | 1 | 1 | NA | NA | 1 | 0 | 3 | 0.3333 | 5 | 0.4 |
| X_6331548 | NA | NA | 3 | 1 | 1 | 1 | 1 | 1 | 5 | 1 | 1 | 1 | NA | NA | 1 | 1 | 4 | 1 | 6 | 1 |
| X_6331586 | NA | NA | 2 | 1 | 1 | 1 | 2 | 1 | 5 | 1 | 1 | 1 | NA | NA | 1 | 1 | 4 | 1 | 6 | 1 |
| X_6331592 | NA | NA | 3 | 0.3333 | 1 | 1 | 1 | 1 | 5 | 0.6 | 1 | 0 | NA | NA | NA | NA | 3 | 0.6667 | 4 | 0.5 |
| X_6331601 | NA | NA | 3 | 0.6667 | NA | NA | 1 | 1 | 4 | 0.75 | 1 | 0 | NA | NA | 1 | 1 | 4 | 1 | 6 | 0.8333 |
| X_6331617 | NA | NA | 2 | 0.5 | 1 | 1 | 2 | 1 | 5 | 0.8 | 1 | 0 | NA | NA | 1 | 0 | 4 | 0.5 | 6 | 0.3333 |
| X_6331619 | NA | NA | 2 | 1 | 1 | 1 | 2 | 0.5 | 5 | 0.8 | 1 | 1 | NA | NA | NA | NA | 2 | 0.5 | 3 | 0.6667 |
| X_6331627 | NA | NA | 3 | 0.3333 | 1 | 1 | 1 | 0 | 5 | 0.4 | 1 | 0 | NA | NA | 1 | 0 | 4 | 0.25 | 6 | 0.1667 |
| X_6331629 | NA | NA | 3 | 0 | 1 | 1 | 2 | 0.5 | 6 | 0.3333 | 1 | 0 | NA | NA | 1 | 0 | 4 | 0.25 | 6 | 0.1667 |
| X_6331637 | NA | NA | 2 | 0 | 1 | 1 | 1 | 0 | 4 | 0.25 | 1 | 0 | NA | NA | 1 | 0 | 2 | 1 | 4 | 0.5 |
| X_6331646 | NA | NA | 1 | 1 | 1 | 1 | NA | NA | 2 | 1 | 1 | 0 | NA | NA | 1 | 1 | 3 | 0 | 5 | 0.2 |
| X_6569759 | 1 | 0 | NA | NA | NA | NA | 1 | 1 | 2 | 0.5 | NA | NA | 1 | 0 | 1 | 0 | 1 | 1 | 3 | 0.3333 |
| X_6569764 | 1 | 1 | NA | NA | NA | NA | 2 | 1 | 3 | 1 | NA | NA | 1 | 1 | 2 | 0.5 | 1 | 0 | 4 | 0.5 |
| X_6569773 | 1 | 1 | NA | NA | NA | NA | 2 | 0.5 | 3 | 0.6667 | NA | NA | 1 | 1 | 2 | 1 | 1 | 1 | 4 | 1 |
| X_6569785 | 1 | 1 | NA | NA | NA | NA | 1 | 1 | 2 | 1 | NA | NA | NA | NA | 2 | 1 | 1 | 1 | 3 | 1 |
| X_6569789 | 1 | 1 | NA | NA | NA | NA | 2 | 1 | 3 | 1 | NA | NA | 1 | 1 | 2 | 1 | NA | NA | 3 | 1 |
| X_6569807 | 1 | 1 | NA | NA | NA | NA | 2 | 0.5 | 3 | 0.6667 | NA | NA | 1 | 1 | 2 | 0 | NA | NA | 3 | 0.3333 |
| X_6569817 | 1 | 0 | NA | NA | NA | NA | 3 | 0.3333 | 4 | 0.25 | NA | NA | NA | NA | 2 | 1 | 1 | 0 | 3 | 0.6667 |
| X_6569820 | 1 | 0 | NA | NA | NA | NA | 3 | 0.3333 | 4 | 0.25 | NA | NA | NA | NA | 2 | 1 | 1 | 0 | 3 | 0.6667 |
| X_6569845 | 1 | 1 | NA | NA | NA | NA | 3 | 1 | 4 | 1 | NA | NA | 2 | 1 | 1 | 1 | 1 | 1 | 4 | 1 |
| X_6569865 | NA | NA | NA | NA | NA | NA | 3 | 0.3333 | 3 | 0.3333 | NA | NA | 1 | 1 | 2 | 0.5 | 1 | 0 | 4 | 0.5 |
| X_6569879 | 1 | 1 | NA | NA | NA | NA | 3 | 0.3333 | 4 | 0.5 | NA | NA | 2 | 0 | 2 | 1 | 1 | 0 | 5 | 0.4 |
| X_6569884 | 1 | 1 | NA | NA | NA | NA | 3 | 1 | 4 | 1 | NA | NA | 1 | 1 | 2 | 1 | 1 | 1 | 4 | 1 |
| X_6569890 | 1 | 0 | NA | NA | NA | NA | 3 | 0.3333 | 4 | 0.25 | NA | NA | 1 | 0 | 3 | 0.3333 | 2 | 0.5 | 6 | 0.3333 |
| X_6569910 | 1 | 1 | NA | NA | 1 | 1 | 3 | 1 | 5 | 1 | NA | NA | 2 | 1 | 4 | 1 | 2 | 1 | 8 | 1 |
| X_6569913 | 1 | 1 | NA | NA | 1 | 1 | 3 | 1 | 5 | 1 | NA | NA | 2 | 1 | 3 | 1 | 2 | 1 | 7 | 1 |
| X_6569918 | 1 | 0 | NA | NA | NA | NA | 2 | 0.5 | 3 | 0.3333 | NA | NA | 2 | 0.5 | 3 | 1 | 2 | 0 | 7 | 0.5714 |
| X_6569927 | 1 | 1 | NA | NA | 1 | 1 | 3 | 1 | 5 | 1 | NA | NA | 2 | 1 | 3 | 1 | 1 | 1 | 6 | 1 |
| X_6569934 | 1 | 1 | NA | NA | 1 | 1 | 2 | 1 | 4 | 1 | NA | NA | 1 | 1 | 2 | 0 | 2 | 0 | 5 | 0.2 |
| X_6569953 | 1 | 1 | NA | NA | 1 | 0 | 3 | 0.6667 | 5 | 0.6 | NA | NA | 2 | 0.5 | 4 | 0.75 | 2 | 0.5 | 8 | 0.625 |
| X_6569966 | 1 | 1 | NA | NA | 1 | 0 | 2 | 0.5 | 4 | 0.5 | NA | NA | 1 | 1 | 3 | 1 | 2 | 1 | 6 | 1 |
| X_6569980 | 1 | 1 | NA | NA | 1 | 0 | 3 | 1 | 5 | 0.8 | NA | NA | 2 | 1 | 4 | 1 | 2 | 1 | 8 | 1 |

| | | | | | | | | | | | | | | | | | | | | |
|---|---|---|---|---|---|---|---|---|---|---|---|---|---|---|---|---|---|---|---|---|
| X_6569983 | 1 | 1 | NA | NA | NA | NA | 3 | 1 | 4 | 1 | NA | NA | 2 | 1 | 3 | 1 | 2 | 0 | 7 | 0.7143 |
| X_6569992 | 1 | 1 | NA | NA | NA | NA | 3 | 1 | 4 | 1 | NA | NA | 2 | 1 | 3 | 1 | 1 | 1 | 6 | 1 |
| X_6570003 | 1 | 1 | NA | NA | 1 | 1 | 3 | 0.3333 | 5 | 0.6 | NA | NA | 1 | 0 | 2 | 1 | 1 | 1 | 4 | 0.75 |
| X_6570009 | 1 | 1 | NA | NA | 1 | 1 | 3 | 1 | 5 | 1 | NA | NA | 2 | 1 | 4 | 1 | 2 | 1 | 8 | 1 |
| X_6570015 | NA | NA | NA | NA | 1 | 1 | 2 | 0.5 | 3 | 0.6667 | NA | NA | 2 | 1 | 4 | 0.75 | 2 | 1 | 8 | 0.875 |
| X_6570028 | 1 | 1 | NA | NA | 1 | 0 | 3 | 0.6667 | 5 | 0.6 | NA | NA | 2 | 0 | 4 | 0.25 | 2 | 0 | 8 | 0.125 |
| X_6570047 | NA | NA | NA | NA | NA | NA | 3 | 0.6667 | 3 | 0.6667 | NA | NA | 2 | 0 | 3 | 1 | NA | NA | 5 | 0.6 |
| X_6570455 | 1 | 0 | 1 | 0 | NA | NA | 2 | 0.5 | 4 | 0.25 | 3 | 0.6667 | 1 | 1 | 5 | 1 | 2 | 0.5 | 11 | 0.8182 |
| X_6775959 | 4 | 0 | 10 | 0.2 | 1 | 0 | 4 | 0 | 19 | 0.1053 | 1 | 0 | 4 | 0 | 3 | 0.3333 | 5 | 0 | 13 | 0.0769 |
| X_6891140 | NA | NA | 1 | 1 | NA | NA | NA | NA | 1 | 1 | 1 | 1 | NA | NA | NA | NA | NA | NA | 1 | 1 |
| X_6891159 | NA | NA | 1 | 1 | NA | NA | NA | NA | 1 | 1 | 1 | 1 | NA | NA | NA | NA | NA | NA | 1 | 1 |
| X_6938969 | 34 | 0.0294 | 42 | 0.1429 | 38 | 0.0789 | 40 | 0.05 | 154 | 0.0779 | 31 | 0.2258 | 38 | 0.1053 | 36 | 0.0833 | 27 | 0.1111 | 132 | 0.1288 |
| X_6938989 | 32 | 0.0625 | 38 | 0.0263 | 29 | 0.069 | 33 | 0.1212 | 132 | 0.0682 | 26 | 0.1538 | 37 | 0.1622 | 37 | 0.1622 | 27 | 0.2222 | 127 | 0.1732 |
| X_6939010 | 32 | 0.125 | 43 | 0.1395 | 36 | 0.1389 | 43 | 0.186 | 154 | 0.1494 | 31 | 0.129 | 41 | 0.0976 | 40 | 0.025 | 35 | 0.1714 | 147 | 0.102 |
| X_6939021 | 30 | 0.2667 | 43 | 0.4419 | 37 | 0.4595 | 42 | 0.4048 | 152 | 0.4013 | 30 | 0.3333 | 39 | 0.359 | 37 | 0.3514 | 30 | 0.4667 | 136 | 0.375 |
| X_6939022 | 26 | 0.2692 | 36 | 0.4167 | 31 | 0.2581 | 38 | 0.2105 | 131 | 0.2901 | 27 | 0.4074 | 37 | 0.2432 | 33 | 0.3333 | 28 | 0.1429 | 125 | 0.28 |
| X_6939025 | 28 | 0.0714 | 41 | 0.0976 | 35 | 0.2 | 40 | 0.1 | 144 | 0.1181 | 23 | 0.1739 | 36 | 0.2222 | 34 | 0.1176 | 30 | 0.2 | 123 | 0.1789 |
| X_6943165 | 1 | 1 | 2 | 0 | 4 | 0.25 | 2 | 0.5 | 9 | 0.3333 | 2 | 0 | 4 | 0.5 | 1 | 1 | 2 | 0.5 | 9 | 0.4444 |
| X_6943249 | 1 | 0 | 2 | 0.5 | 3 | 0.3333 | 2 | 0 | 8 | 0.25 | 1 | 0 | 4 | 0.25 | 2 | 0 | 2 | 0 | 9 | 0.1111 |
| X_6943269 | 1 | 1 | 2 | 0 | 3 | 0.3333 | 2 | 0.5 | 8 | 0.375 | 2 | 0 | 4 | 0.25 | 1 | 0 | 2 | 0 | 9 | 0.1111 |
| X_6943277 | 1 | 0 | 2 | 0 | 4 | 0.5 | 2 | 0.5 | 9 | 0.3333 | 2 | 0 | 3 | 0.3333 | 2 | 0 | 2 | 0 | 9 | 0.1111 |
| X_6943289 | 1 | 0 | 2 | 0 | 5 | 0.4 | 2 | 0 | 10 | 0.2 | 2 | 0 | 4 | 0.25 | 2 | 0.5 | 1 | 1 | 9 | 0.3333 |
| X_6943291 | 1 | 1 | 1 | 1 | 5 | 1 | 2 | 1 | 9 | 1 | 3 | 1 | 3 | 1 | 2 | 1 | 2 | 0.5 | 10 | 0.9 |
| X_6950377 | 11 | 0.0909 | 5 | 0 | 7 | 0.2857 | 8 | 0.125 | 31 | 0.129 | 3 | 0 | 6 | 0.1667 | 8 | 0.25 | 16 | 0.125 | 33 | 0.1515 |
| X_6950444 | 8 | 0.125 | 4 | 0.25 | 7 | 0 | 9 | 0.2222 | 28 | 0.1429 | 3 | 0.6667 | 6 | 0.1667 | 5 | 0 | 14 | 0.3571 | 28 | 0.2857 |
| X_6950446 | 10 | 0 | 4 | 0 | 7 | 0.5714 | 9 | 0 | 30 | 0.1333 | 3 | 0 | 6 | 0.1667 | 5 | 0 | 13 | 0.1538 | 27 | 0.1111 |
| X_6950449 | 10 | 0.9 | 4 | 1 | 6 | 1 | 8 | 0.875 | 28 | 0.9286 | 2 | 0.5 | 5 | 1 | 6 | 0.8333 | 13 | 0.7692 | 26 | 0.8077 |
| X_6950451 | 9 | 0.1111 | 3 | 0.3333 | 4 | 0 | 8 | 0.25 | 24 | 0.1667 | 3 | 0 | 6 | 0.3333 | 4 | 0.25 | 11 | 0.1818 | 24 | 0.2083 |
| X_6950465 | 10 | 0.3 | 3 | 0.6667 | 7 | 0.5714 | 7 | 0.2857 | 27 | 0.4074 | 3 | 0 | 5 | 0 | 4 | 0.5 | 9 | 0.5556 | 21 | 0.3333 |
| X_7016201 | NA | NA | 2 | 0.5 | NA | NA | 2 | 0 | 4 | 0.25 | 1 | 0 | 1 | 0 | NA | NA | 1 | 1 | 3 | 0.3333 |
| X_7016498 | 2 | 0 | 2 | 0 | NA | NA | 2 | 0.5 | 6 | 0.1667 | 1 | 0 | 1 | 0 | 1 | 0 | 2 | 0.5 | 5 | 0.2 |
| X_7016505 | 2 | 0.5 | 1 | 0 | NA | NA | 1 | 1 | 4 | 0.5 | 2 | 0 | NA | NA | 2 | 0.5 | 3 | 0 | 7 | 0.1429 |
| X_7019757 | 3 | 0.6667 | 3 | 0 | 1 | 0 | 2 | 0 | 9 | 0.2222 | 4 | 0 | 3 | 0.3333 | 2 | 0 | 5 | 0.6 | 14 | 0.2857 |
| X_7019831 | 4 | 0.25 | 3 | 0 | 2 | 0 | 1 | 0 | 10 | 0.1 | 3 | 0 | 3 | 0.6667 | 1 | 1 | 5 | 0.4 | 12 | 0.4167 |
| X_7019899 | 3 | 0 | 3 | 0 | 2 | 0 | 2 | 0.5 | 10 | 0.1 | 3 | 0 | 4 | 0.25 | 3 | 0 | 5 | 0 | 15 | 0.0667 |
| X_7035742 | 1 | 1 | 1 | 0 | 4 | 0.25 | 1 | 0 | 7 | 0.2857 | 1 | 0 | NA | NA | 1 | 1 | 1 | 1 | 3 | 0.6667 |
| X_7035744 | 2 | 1 | 1 | 0 | 4 | 0.25 | 1 | 1 | 8 | 0.5 | 1 | 1 | NA | NA | 1 | 1 | 1 | 1 | 3 | 1 |
| X_7035749 | 1 | 1 | 1 | 1 | 3 | 0.3333 | 1 | 0 | 6 | 0.5 | 1 | 0 | NA | NA | 1 | 1 | NA | NA | 2 | 0.5 |
| X_7035754 | 1 | 0 | NA | NA | 4 | 0.5 | 1 | 1 | 6 | 0.5 | 1 | 0 | NA | NA | 1 | 1 | 1 | 0 | 3 | 0.3333 |
| X_7046023 | 1 | 1 | 3 | 0.3333 | 1 | 0 | 1 | 0 | 6 | 0.3333 | 2 | 0.5 | 4 | 0.25 | 2 | 0 | 6 | 0.5 | 14 | 0.3571 |
| X_7046044 | 1 | 0 | 3 | 1 | 1 | 1 | 2 | 0.5 | 7 | 0.7143 | 3 | 1 | 4 | 0.75 | 3 | 0.6667 | 5 | 0.4 | 15 | 0.6667 |
| X_7046045 | 1 | 0 | 4 | 0.5 | 2 | 0.5 | 2 | 0 | 9 | 0.3333 | 3 | 1 | 3 | 0.3333 | 5 | 0.2 | 5 | 0.6 | 16 | 0.5 |
| X_7046066 | 1 | 0 | 1 | 0 | 1 | 0 | 1 | 1 | 4 | 0.25 | 1 | 1 | 3 | 0 | 2 | 0 | 4 | 0.25 | 10 | 0.2 |
| X_7046073 | 1 | 1 | 3 | 0.6667 | 2 | 0.5 | 2 | 0.5 | 8 | 0.625 | 2 | 1 | 3 | 0.3333 | 4 | 0.5 | 5 | 0.6 | 14 | 0.5714 |
| X_7046132 | 1 | 1 | 4 | 0.25 | 2 | 0 | 2 | 0 | 9 | 0.2222 | 3 | 0 | 4 | 0.5 | 3 | 0 | 6 | 0 | 16 | 0.125 |
| X_7046144 | 1 | 0 | 4 | 0.25 | 2 | 0.5 | 3 | 0 | 10 | 0.2 | 2 | 0 | 2 | 0 | 3 | 0 | 5 | 0.2 | 12 | 0.0833 |
| X_7046201 | 1 | 0 | 4 | 0.5 | 3 | 0 | 4 | 0.25 | 12 | 0.25 | 3 | 0.3333 | 5 | 0.6 | 2 | 0.5 | 6 | 0.3333 | 16 | 0.4375 |
| X_7046233 | 2 | 0 | 7 | 0 | 3 | 0.3333 | 3 | 0 | 15 | 0.0667 | 3 | 0 | 4 | 0 | 2 | 0 | 6 | 0.3333 | 15 | 0.1333 |
| X_7046240 | 2 | 0 | 7 | 0.1429 | 3 | 0.3333 | 4 | 0 | 16 | 0.125 | 4 | 0.25 | 4 | 0.25 | 1 | 0 | 6 | 0.5 | 15 | 0.3333 |
| X_7046389 | 14 | 0.1429 | 8 | 0.25 | 6 | 0.1667 | 12 | 0.0833 | 40 | 0.15 | 9 | 0.3333 | 6 | 0 | 9 | 0.3333 | 11 | 0.1818 | 35 | 0.2286 |
| X_7046403 | 13 | 0.1538 | 7 | 0.2857 | 3 | 0.3333 | 13 | 0.1538 | 36 | 0.1944 | 8 | 0.5 | 6 | 0.1667 | 9 | 0.1111 | 11 | 0.0909 | 34 | 0.2059 |
| X_7046449 | 12 | 0.3333 | 8 | 0.375 | 2 | 0 | 10 | 0.3 | 32 | 0.3125 | 6 | 0.1667 | 5 | 0.4 | 6 | 0.3333 | 10 | 0.4 | 27 | 0.3333 |
| X_7046456 | 11 | 0.0909 | 7 | 0.1429 | 2 | 0 | 9 | 0 | 29 | 0.069 | 6 | 0 | 5 | 0 | 5 | 0.2 | 8 | 0.125 | 24 | 0.0833 |
| X_7046467 | 10 | 0.2 | 7 | 0.1429 | 2 | 0.5 | 11 | 0 | 30 | 0.1333 | 6 | 0.1667 | 4 | 0.25 | 6 | 0.1667 | 11 | 0.1818 | 27 | 0.1852 |
| X_7046483 | 10 | 0.6 | 6 | 0.6667 | 2 | 0.5 | 10 | 0.4 | 28 | 0.5357 | 4 | 0.5 | 5 | 0.8 | 5 | 0.8 | 9 | 0.5556 | 23 | 0.6522 |
| X_7046488 | 10 | 0.1 | 4 | 0.25 | 2 | 0 | 11 | 0 | 27 | 0.0741 | 2 | 0 | 3 | 0 | 4 | 0 | 9 | 0.1111 | 18 | 0.0556 |
| X_7046489 | 10 | 0.1 | 6 | 0.3333 | 2 | 0 | 12 | 0.0833 | 30 | 0.1333 | 5 | 0 | 5 | 0.2 | 6 | 0 | 8 | 0.25 | 24 | 0.125 |
| X_7046498 | 5 | 0.2 | 4 | 0.75 | 1 | 1 | 6 | 0 | 16 | 0.3125 | 2 | 0.5 | 3 | 1 | 4 | 0 | 4 | 0.25 | 13 | 0.3846 |
| X_7046506 | 8 | 0.125 | 5 | 0.8 | 2 | 0.5 | 8 | 0.25 | 23 | 0.3478 | 2 | 0 | 4 | 0.25 | 4 | 0.25 | 8 | 0.625 | 18 | 0.3889 |
| X_7046544 | 4 | 0.25 | 2 | 0 | NA | NA | 6 | 0.1667 | 12 | 0.1667 | 2 | 0 | 3 | 0.3333 | 3 | 0.6667 | 2 | 0 | 10 | 0.3 |
| X_7046550 | 8 | 0.25 | 5 | 0 | 1 | 0 | 9 | 0 | 23 | 0.087 | 3 | 0.3333 | 4 | 0 | 5 | 0.2 | 6 | 0 | 18 | 0.1111 |
| X_7046607 | 4 | 0 | 3 | 0 | 1 | 0 | 7 | 0.1429 | 15 | 0.0667 | 3 | 0 | 4 | 0 | 6 | 0.1667 | 6 | 0 | 19 | 0.0526 |
| X_7046608 | 5 | 1 | 4 | 1 | 1 | 1 | 7 | 1 | 17 | 1 | 3 | 0.6667 | 4 | 0.75 | 6 | 0.8333 | 6 | 0.8333 | 19 | 0.7895 |
| X_7046611 | 5 | 0.6 | 4 | 0.5 | 1 | 1 | 7 | 0.4286 | 17 | 0.5294 | 2 | 1 | 4 | 0.75 | 3 | 0 | 5 | 0.8 | 14 | 0.6429 |
| X_7046612 | 6 | 0 | 4 | 0.25 | 1 | 0 | 6 | 0 | 17 | 0.0588 | 2 | 0.5 | 4 | 0 | 5 | 0 | 6 | 0 | 17 | 0.0588 |

| | | | | | | | | | | | | | | | | | | | | |
|---|---|---|---|---|---|---|---|---|---|---|---|---|---|---|---|---|---|---|---|---|
| X_7046617 | 5 | 0.6 | 3 | 1 | 1 | 1 | 6 | 0.3333 | 15 | 0.6 | 2 | 0.5 | 3 | 0.3333 | 5 | 0.4 | 4 | 0.5 | 14 | 0.4286 |
| X_7060359 | 10 | 0.3 | 7 | 0 | 6 | 0.1667 | 7 | 0.2857 | 30 | 0.2 | 6 | 0 | 8 | 0.125 | 7 | 0.1429 | 5 | 0 | 26 | 0.0769 |
| X_7060410 | 12 | 0.1667 | 6 | 0.1667 | 7 | 0 | 7 | 0.1429 | 32 | 0.125 | 5 | 0 | 9 | 0.1111 | 12 | 0.1667 | 4 | 0 | 30 | 0.1 |
| X_7086443 | 12 | 0 | 8 | 0.125 | 4 | 0 | 10 | 0.2 | 34 | 0.0882 | 8 | 0 | 7 | 0.1429 | 10 | 0 | 12 | 0.3333 | 37 | 0.1351 |
| X_7086494 | 9 | 0.2222 | 6 | 0 | 3 | 0 | 8 | 0 | 26 | 0.0769 | 7 | 0.1429 | 7 | 0.2857 | 8 | 0.125 | 11 | 0.1818 | 33 | 0.1818 |
| X_7101586 | 1 | 0 | 2 | 0.5 | 2 | 0 | 2 | 0 | 7 | 0.1429 | NA | NA | 1 | 0 | 1 | 1 | 1 | 0 | 3 | 0.3333 |
| X_7101591 | 1 | 1 | 2 | 1 | 1 | 1 | 2 | 1 | 6 | 1 | NA | NA | 1 | 1 | 1 | 1 | 1 | 0 | 3 | 0.6667 |
| X_7147647 | 17 | 0.0588 | 11 | 0.2727 | 7 | 0 | 13 | 0.0769 | 48 | 0.1042 | 9 | 0 | 5 | 0 | 11 | 0.1818 | 9 | 0.3333 | 34 | 0.1471 |
| X_7147679 | 13 | 0 | 4 | 0.5 | 6 | 0.1667 | 9 | 0.1111 | 32 | 0.125 | 6 | 0.5 | 4 | 0.25 | 10 | 0 | 6 | 0.1667 | 26 | 0.1923 |
| X_7147715 | 10 | 0.3 | 4 | 0 | 6 | 0.1667 | 10 | 0.1 | 30 | 0.1667 | 4 | 0.25 | 4 | 0.25 | 9 | 0.2222 | 6 | 0 | 23 | 0.1739 |
| X_7147740 | 9 | 0 | 4 | 0.25 | 4 | 0.5 | 9 | 0.1111 | 26 | 0.1538 | 3 | 0 | 4 | 0 | 10 | 0.3 | 7 | 0 | 24 | 0.125 |
| X_7147794 | 8 | 0.125 | 3 | 0.3333 | 3 | 0 | 8 | 0 | 22 | 0.0909 | 2 | 0 | 4 | 0.25 | 9 | 0 | 6 | 0.1667 | 21 | 0.0952 |
| X_7147794 | 8 | 0.25 | 3 | 0 | 3 | 0.6667 | 7 | 0.1429 | 21 | 0.2381 | 2 | 0 | 3 | 0 | 8 | 0.25 | 5 | 0 | 18 | 0.1111 |
| X_7147806 | 5 | 0 | 3 | 0.3333 | 3 | 0.6667 | 5 | 0.6 | 16 | 0.375 | 2 | 0.5 | 3 | 0 | 7 | 0.2857 | 4 | 0.5 | 16 | 0.3125 |
| X_7147807 | 8 | 0.125 | 3 | 0 | 3 | 0.3333 | 8 | 0 | 22 | 0.0909 | 2 | 0 | 4 | 0.25 | 8 | 0.25 | 5 | 0.4 | 19 | 0.2632 |
| X_7147814 | 6 | 0.3333 | 4 | 0.25 | 3 | 0 | 8 | 0 | 21 | 0.1429 | 2 | 0 | 5 | 0.4 | 9 | 0.1111 | 5 | 0.2 | 21 | 0.1905 |
| X_7147826 | 5 | 0 | 4 | 0.25 | 1 | 0 | 8 | 0 | 18 | 0.0556 | 2 | 0 | 5 | 0 | 8 | 0.125 | 3 | 0 | 18 | 0.0556 |
| X_7147829 | 7 | 0.1429 | 3 | 0 | 2 | 0 | 8 | 0.125 | 20 | 0.1 | 2 | 0.5 | 4 | 0 | 9 | 0.1111 | 3 | 0 | 18 | 0.1111 |
| X_7147836 | 6 | 0.3333 | 3 | 0 | 3 | 0 | 6 | 0 | 18 | 0.1111 | 2 | 0 | 4 | 0 | 8 | 0.125 | 3 | 0 | 17 | 0.0588 |
| X_7147841 | 1 | 0 | 2 | 0.5 | 1 | 1 | 3 | 1 | 7 | 0.7143 | 2 | 0.5 | 2 | 0.5 | 2 | 0 | 2 | 0.5 | 8 | 0.375 |
| X_7148890 | 6 | 0 | 2 | 0 | 3 | 0 | 8 | 0.25 | 19 | 0.1053 | 2 | 0 | 3 | 0 | 8 | 0.25 | 3 | 0 | 16 | 0.125 |
| X_7154479 | 5 | 0.2 | 3 | 0 | 3 | 0.6667 | 8 | 0.125 | 19 | 0.2105 | 2 | 0 | 3 | 0 | 6 | 0.1667 | 4 | 0 | 15 | 0.0667 |
| X_7195756 | 5 | 0.2 | 7 | 0.1429 | 9 | 0 | 13 | 0 | 34 | 0.0588 | 6 | 0 | 14 | 0.2143 | 5 | 0 | 13 | 0.0769 | 38 | 0.1053 |
| X_7200109 | 2 | 0 | 4 | 0.25 | 2 | 0 | 3 | 0.3333 | 11 | 0.1818 | NA | NA | 1 | 0 | 4 | 0 | 4 | 0.25 | 9 | 0.1111 |
| X_7200118 | 2 | 1 | 4 | 0.25 | 4 | 0.75 | 3 | 1 | 13 | 0.6923 | NA | NA | 2 | 0.5 | 5 | 0.6 | 5 | 0.6 | 12 | 0.5833 |
| X_7210851 | NA | NA | 1 | 1 | NA | NA | 1 | 1 | 2 | 1 | 1 | 0 | 1 | 1 | NA | NA | NA | NA | 2 | 0.5 |
| X_7210863 | NA | NA | 1 | 0 | NA | NA | 1 | 1 | 2 | 0.5 | 1 | 0 | 1 | 1 | NA | NA | NA | NA | 2 | 0.5 |
| X_7210874 | NA | NA | 1 | 1 | NA | NA | NA | NA | 1 | 1 | 1 | 0 | 1 | 1 | NA | NA | 1 | 1 | 3 | 0.6667 |
| X_7210880 | NA | NA | 1 | 1 | NA | NA | 1 | 0 | 2 | 0.5 | 1 | 0 | 1 | 1 | NA | NA | 1 | 0 | 3 | 0.3333 |
| X_7210886 | NA | NA | 1 | 1 | NA | NA | NA | NA | 1 | 1 | 1 | 0 | 1 | 0 | NA | NA | 1 | 1 | 3 | 0.3333 |
| X_7211008 | NA | NA | 2 | 1 | NA | NA | 2 | 0 | 4 | 0.5 | 1 | 0 | 1 | 1 | NA | NA | 1 | 0 | 3 | 0.3333 |
| X_7211011 | NA | NA | 1 | 1 | NA | NA | 2 | 1 | 3 | 1 | 3 | 0.6667 | 1 | 1 | NA | NA | 1 | 1 | 5 | 0.8 |
| X_7211039 | NA | NA | 2 | 0.5 | 1 | 0 | 2 | 0.5 | 5 | 0.4 | 3 | 0.6667 | 1 | 0 | NA | NA | 2 | 0 | 6 | 0.3333 |
| X_7211044 | NA | NA | 2 | 1 | 1 | 1 | 2 | 1 | 5 | 1 | 3 | 0.3333 | 1 | 1 | NA | NA | 2 | 1 | 6 | 0.6667 |
| X_7211057 | NA | NA | 2 | 0.5 | 1 | 1 | 2 | 1 | 5 | 0.8 | 3 | 0.6667 | NA | NA | NA | NA | 2 | 0.5 | 5 | 0.6 |
| X_7211081 | NA | NA | 3 | 0.3333 | 1 | 0 | 2 | 0.5 | 6 | 0.3333 | 3 | 0.3333 | NA | NA | NA | NA | 1 | 0 | 4 | 0.25 |
| X_7211134 | NA | NA | 3 | 1 | 4 | 0.75 | 2 | 0.5 | 9 | 0.7778 | 3 | 0.3333 | 1 | 1 | NA | NA | 4 | 0.5 | 8 | 0.5 |
| X_7211138 | NA | NA | 3 | 0 | 3 | 0.3333 | 2 | 0 | 8 | 0.125 | 2 | 0 | 1 | 0 | NA | NA | 4 | 0.25 | 7 | 0.1429 |
| X_7211139 | NA | NA | 3 | 0.6667 | 3 | 0.6667 | 2 | 0.5 | 8 | 0.625 | 3 | 0.3333 | NA | NA | NA | NA | 3 | 0 | 6 | 0.1667 |
| X_7211141 | NA | NA | 3 | 1 | 2 | 0.5 | 2 | 0 | 7 | 0.5714 | 2 | 0 | 1 | 0 | NA | NA | 4 | 0.25 | 7 | 0.1429 |
| X_7211145 | NA | NA | 3 | 0 | 3 | 0.3333 | 2 | 0.5 | 8 | 0.25 | 3 | 0 | 1 | 0 | NA | NA | 4 | 0.25 | 8 | 0.125 |
| X_7227198 | 2 | 0.5 | 5 | 0.8 | NA | NA | 4 | 0.25 | 11 | 0.5455 | 3 | 0.3333 | 6 | 0.3333 | 4 | 0.25 | 4 | 0.25 | 17 | 0.2941 |
| X_7227236 | 3 | 1 | 5 | 0 | NA | NA | 3 | 0 | 11 | 0.2727 | 4 | 0 | 5 | 0.4 | 4 | 0 | 4 | 0 | 17 | 0.1176 |
| X_7227263 | 4 | 0.5 | 7 | 0.2857 | NA | NA | 4 | 0 | 15 | 0.2667 | 5 | 0 | 6 | 0.3333 | 5 | 0.2 | 4 | 0.5 | 20 | 0.25 |
| X_7227268 | 3 | 0.6667 | 6 | 0.3333 | NA | NA | 2 | 0.5 | 11 | 0.4545 | 5 | 0.4 | 5 | 0.2 | 4 | 0.5 | 2 | 0.5 | 16 | 0.375 |
| X_7227294 | 1 | 1 | 6 | 1 | NA | NA | 2 | 1 | 9 | 1 | 5 | 1 | 4 | 1 | 1 | 1 | 1 | 1 | 11 | 1 |
| X_7227305 | 3 | 0.6667 | 5 | 0.6 | NA | NA | 4 | 0.5 | 12 | 0.5833 | 5 | 0.4 | 5 | 1 | 3 | 1 | 2 | 1 | 15 | 0.8 |
| X_7227310 | 4 | 0.75 | 6 | 0.8333 | NA | NA | 5 | 0.8 | 15 | 0.8 | 6 | 0.3333 | 7 | 0.8571 | 4 | 0.75 | 5 | 0.6 | 22 | 0.6364 |
| X_7227338 | 4 | 0 | 7 | 0.1429 | 1 | 0 | 5 | 0 | 17 | 0.0588 | 7 | 0.2857 | 6 | 0 | 6 | 0 | 4 | 0 | 23 | 0.087 |
| X_7227339 | 4 | 0.25 | 6 | 0 | 1 | 0 | 4 | 0.25 | 15 | 0.1333 | 6 | 0 | 7 | 0.1429 | 5 | 0.2 | 4 | 0 | 22 | 0.0909 |
| X_7239694 | 1 | 1 | 1 | 1 | NA | NA | 1 | 1 | 3 | 1 | NA | NA | 1 | 1 | 1 | 1 | 1 | 1 | 3 | 1 |
| X_7239781 | 1 | 1 | 4 | 1 | NA | NA | 2 | 1 | 7 | 1 | 1 | 1 | 1 | 1 | 3 | 1 | 1 | 1 | 6 | 1 |
| X_7239805 | NA | NA | 4 | 0.75 | NA | NA | 2 | 1 | 6 | 0.8333 | 1 | 1 | 1 | 1 | 3 | 1 | 1 | 0 | 6 | 0.8333 |
| X_7239928 | 1 | 0 | 4 | 0.5 | 2 | 0 | 2 | 0 | 9 | 0.2222 | 3 | 0.3333 | 1 | 0 | 4 | 0.25 | 1 | 0 | 9 | 0.2222 |
| X_7239942 | 1 | 0 | 4 | 0.75 | 2 | 0.5 | 2 | 0 | 9 | 0.4444 | 3 | 0 | 1 | 0 | 4 | 0.25 | 1 | 0 | 9 | 0.1111 |
| X_7239959 | 1 | 1 | 4 | 0.75 | 2 | 1 | 1 | 1 | 8 | 0.875 | 3 | 1 | 1 | 1 | 3 | 1 | 1 | 0 | 8 | 0.875 |
| X_7239961 | NA | NA | 4 | 0.5 | NA | NA | 2 | 1 | 6 | 0.6667 | 3 | 0.6667 | 2 | 1 | 4 | 0.5 | 1 | 1 | 10 | 0.7 |
| X_7239964 | 1 | 1 | 4 | 1 | 2 | 1 | 2 | 1 | 9 | 1 | 2 | 1 | 2 | 1 | 4 | 1 | 1 | 1 | 9 | 1 |
| X_7239968 | 1 | 1 | 4 | 0.25 | 2 | 0.5 | 2 | 0.5 | 9 | 0.4444 | 3 | 1 | 2 | 0.5 | 4 | 1 | 1 | 0 | 10 | 0.8 |
| X_7239979 | 1 | 0 | 4 | 0.5 | 2 | 0 | NA | NA | 7 | 0.2857 | 3 | 0 | 2 | 0 | 4 | 0.5 | 1 | 0 | 10 | 0.2 |
| X_7239986 | 1 | 0 | 3 | 0.6667 | 2 | 0.5 | 2 | 0 | 8 | 0.375 | 3 | 0.3333 | 2 | 1 | 4 | 0.25 | 1 | 0 | 10 | 0.4 |
| X_7239990 | NA | NA | 3 | 0.6667 | 2 | 0.5 | 2 | 1 | 7 | 0.7143 | 2 | 1 | 1 | 1 | 3 | 1 | NA | NA | 6 | 1 |
| X_7239992 | 1 | 1 | 4 | 1 | 2 | 1 | 2 | 0.5 | 9 | 0.8889 | 2 | 1 | 1 | 1 | 4 | 1 | NA | NA | 7 | 1 |
| X_7240014 | NA | NA | 3 | 1 | 2 | 1 | 2 | 1 | 7 | 1 | 3 | 1 | 2 | 1 | 2 | 0.5 | 1 | 1 | 8 | 0.875 |
| X_7240018 | 1 | 0 | 4 | 0.5 | 2 | 0 | 1 | 1 | 8 | 0.375 | 2 | 0.5 | 2 | 0.5 | 3 | 0.6667 | 1 | 0 | 8 | 0.5 |

| | | | | | | | | | | | | | | | | | | | | |
|---|---|---|---|---|---|---|---|---|---|---|---|---|---|---|---|---|---|---|---|---|
| X_7240156 | 3 | 0.3333 | 4 | 0.25 | 3 | 0 | 3 | 0 | 13 | 0.1538 | 4 | 0 | 4 | 0 | 4 | 0.25 | 3 | 0 | 15 | 0.0667 |
| X_7240166 | 1 | 0 | 4 | 0.25 | 2 | 1 | 1 | 0 | 8 | 0.375 | 4 | 0.25 | 3 | 0.3333 | 4 | 0.5 | 1 | 0 | 12 | 0.3333 |
| X_7240253 | 4 | 0 | 6 | 0.3333 | 4 | 0 | 2 | 0 | 16 | 0.125 | 6 | 0.1667 | 5 | 0 | 4 | 0.25 | 3 | 0 | 18 | 0.1111 |
| X_7253825 | 3 | 0 | 3 | 0.3333 | NA | NA | 1 | 0 | 7 | 0.1429 | NA | NA | 1 | 1 | 2 | 0 | 2 | 0 | 5 | 0.2 |
| X_7253922 | 2 | 0.5 | 2 | 0.5 | NA | NA | 1 | 1 | 5 | 0.6 | NA | NA | 2 | 0 | 2 | 0 | 3 | 0.3333 | 7 | 0.1429 |
| X_7259460 | 1 | 0 | NA | NA | 2 | 0 | 3 | 0.6667 | 6 | 0.3333 | 2 | 0.5 | 4 | 0.25 | 1 | 0 | 1 | 0 | 8 | 0.25 |
| X_7261394 | 1 | 0 | NA | NA | 4 | 0 | 5 | 0.2 | 10 | 0.1 | 1 | 0 | 4 | 0 | NA | NA | 1 | 1 | 6 | 0.1667 |
| X_7261416 | 1 | 0 | NA | NA | 3 | 0.3333 | 6 | 0.6667 | 10 | 0.5 | 2 | 0 | 4 | 0.5 | NA | NA | 1 | 0 | 7 | 0.2857 |
| X_7261430 | 3 | 1 | NA | NA | 4 | 1 | 6 | 1 | 13 | 1 | 2 | 1 | 2 | 0.5 | NA | NA | 1 | 1 | 5 | 0.8 |
| X_7261435 | 2 | 0 | NA | NA | 5 | 0.8 | 6 | 0.6667 | 13 | 0.6154 | 3 | 0.6667 | 4 | 0.25 | 1 | 1 | 1 | 1 | 9 | 0.5556 |
| X_7261452 | 4 | 0.75 | NA | NA | 7 | 0.1429 | 5 | 0.6 | 16 | 0.4375 | 2 | 0 | 5 | 0.2 | 1 | 1 | 1 | 0 | 9 | 0.2222 |
| X_7261472 | 4 | 0 | 2 | 0.5 | 7 | 0 | 6 | 0.3333 | 19 | 0.1579 | 3 | 0 | 7 | 0.1429 | 3 | 0 | 3 | 0 | 16 | 0.0625 |
| X_7261481 | 3 | 0.3333 | 1 | 1 | 6 | 0.3333 | 7 | 0.5714 | 17 | 0.4706 | 3 | 0.6667 | 7 | 0.7143 | 3 | 1 | 3 | 0.6667 | 16 | 0.75 |
| X_7261491 | 4 | 0 | 2 | 0 | 7 | 0.4286 | 7 | 0.5714 | 20 | 0.35 | 3 | 0.3333 | 6 | 0 | 3 | 1 | 3 | 0 | 15 | 0.2667 |
| X_7261504 | 5 | 0.4 | 3 | 0.3333 | 7 | 0.2857 | 7 | 0.2857 | 22 | 0.3182 | 3 | 0.3333 | 8 | 0.25 | 3 | 0.3333 | 3 | 0.6667 | 17 | 0.3529 |
| X_7261507 | 5 | 1 | 6 | 1 | 8 | 0.875 | 6 | 1 | 25 | 0.96 | 3 | 0.6667 | 7 | 1 | 4 | 0.75 | 4 | 1 | 18 | 0.8889 |
| X_7261512 | 5 | 0 | 5 | 0.2 | 8 | 0 | 7 | 0.4286 | 25 | 0.16 | 3 | 0.3333 | 8 | 0 | 3 | 0.3333 | 4 | 0.25 | 18 | 0.1667 |
| X_7261524 | 4 | 0 | 4 | 0.5 | 9 | 0.2222 | 6 | 0 | 23 | 0.1739 | 3 | 0 | 7 | 0 | 2 | 1 | 3 | 0 | 15 | 0.1333 |
| X_7261525 | 4 | 0.25 | 5 | 0.6 | 9 | 0.2222 | 6 | 0 | 24 | 0.25 | 3 | 0.3333 | 7 | 0.1429 | 2 | 0.5 | 4 | 0.25 | 16 | 0.25 |
| X_7261528 | 3 | 0.6667 | 7 | 0.7143 | 8 | 0.5 | 6 | 0.6667 | 24 | 0.625 | 3 | 0.3333 | 7 | 0.5714 | 3 | 0.6667 | 4 | 0.5 | 17 | 0.5294 |
| X_7261548 | 2 | 0.5 | 4 | 0 | 8 | 0.125 | 7 | 0.2857 | 21 | 0.1905 | 2 | 0 | 6 | 0.3333 | 3 | 0.3333 | 4 | 0 | 15 | 0.2 |
| X_7261551 | 4 | 0.25 | 6 | 0 | 9 | 0.3333 | 4 | 0.25 | 23 | 0.2174 | 2 | 0 | 7 | 0.2857 | 2 | 0.5 | 4 | 0.5 | 15 | 0.3333 |
| X_7261556 | 3 | 0.6667 | 8 | 0.375 | 7 | 0.5714 | 8 | 0.625 | 26 | 0.5385 | 1 | 1 | 7 | 0.5714 | 2 | 1 | 4 | 0.75 | 14 | 0.7143 |
| X_7261639 | 8 | 0 | 8 | 0.125 | 10 | 0.3 | 9 | 0.2222 | 35 | 0.1714 | 4 | 0 | 10 | 0.2 | 4 | 0.25 | 4 | 0 | 22 | 0.1364 |
| X_7261878 | NA | NA | 1 | 1 | 1 | 0 | 1 | 0 | 3 | 0.3333 | NA | NA | 1 | 1 | NA | NA | NA | NA | 1 | 1 |
| X_7269967 | 8 | 0.125 | 10 | 0.2 | 7 | 0.1429 | 14 | 0.2857 | 39 | 0.2051 | 9 | 0.1111 | 15 | 0.1333 | 10 | 0.2 | 19 | 0.3684 | 53 | 0.2264 |
| X_7269974 | 8 | 0.375 | 16 | 0.4375 | 8 | 0.375 | 18 | 0.5 | 50 | 0.44 | 13 | 0.3077 | 14 | 0.5714 | 12 | 0.5833 | 18 | 0.5 | 57 | 0.4912 |
| X_7269976 | 9 | 0 | 16 | 0.0625 | 9 | 0.1111 | 19 | 0.1579 | 53 | 0.0943 | 15 | 0.0667 | 15 | 0.0667 | 11 | 0.0909 | 17 | 0.1176 | 58 | 0.0862 |
| X_7269987 | 9 | 0.2222 | 14 | 0.1429 | 8 | 0.125 | 21 | 0.0476 | 52 | 0.1154 | 11 | 0.0909 | 16 | 0.1875 | 10 | 0.3 | 17 | 0.1176 | 54 | 0.1667 |
| X_7269996 | 6 | 0 | 16 | 0.0625 | 7 | 0.1429 | 19 | 0.0526 | 48 | 0.0625 | 9 | 0.2222 | 15 | 0 | 8 | 0 | 18 | 0.1111 | 50 | 0.08 |
| X_7270141 | 6 | 0 | 11 | 0 | 7 | 0.2857 | 15 | 0.2 | 39 | 0.1282 | 9 | 0.1111 | 13 | 0.0769 | 8 | 0 | 17 | 0.1176 | 47 | 0.0851 |
| X_7270158 | NA | NA | NA | NA | 1 | 1 | NA | NA | 1 | 1 | 1 | 1 | NA | NA | NA | NA | NA | NA | 1 | 1 |
| X_7271490 | NA | NA | NA | NA | 2 | 0.5 | NA | NA | 2 | 0.5 | NA | NA | NA | NA | NA | NA | 1 | 1 | 1 | 1 |
| X_7271525 | NA | NA | NA | NA | 1 | 1 | NA | NA | 1 | 1 | NA | NA | 1 | 1 | NA | NA | 1 | 1 | 2 | 1 |
| X_7271719 | 1 | 0 | 3 | 0.3333 | 2 | 1 | NA | NA | 6 | 0.5 | 2 | 0 | NA | NA | 2 | 0.5 | 2 | 0.5 | 6 | 0.3333 |
| X_7271755 | 1 | 1 | 6 | 0.3333 | 4 | 0 | 3 | 0 | 14 | 0.2143 | 4 | 0 | NA | NA | 4 | 0.25 | 2 | 0.5 | 10 | 0.2 |
| X_7271757 | 1 | 0 | 8 | 0.625 | 4 | 0 | 3 | 0 | 16 | 0.3125 | 3 | 0.3333 | NA | NA | 3 | 0 | 2 | 0 | 8 | 0.125 |
| X_7272582 | 1 | 0 | 10 | 0.1 | 5 | 0.6 | 2 | 0.5 | 18 | 0.2778 | 7 | 0 | 1 | 0 | 8 | 0.125 | 4 | 0.25 | 20 | 0.1 |
| X_7272599 | 2 | 1 | 10 | 0.5 | 6 | 0.3333 | 7 | 0.7143 | 25 | 0.56 | 7 | 0.8571 | 4 | 0.75 | 4 | 0.75 | 3 | 0.6667 | 18 | 0.7778 |
| X_7272998 | 11 | 0.1818 | 12 | 0.0833 | 9 | 0 | 16 | 0.125 | 48 | 0.1042 | 15 | 0.1333 | 5 | 0 | 8 | 0.25 | 12 | 0.0833 | 40 | 0.125 |
| X_7277482 | NA | NA | NA | NA | NA | NA | 1 | 1 | 1 | 1 | NA | NA | NA | NA | 1 | 1 | NA | NA | 1 | 1 |
| X_7281595 | NA | NA | NA | NA | 4 | 0.5 | 2 | 0.5 | 6 | 0.5 | 1 | 0 | 1 | 1 | 5 | 0.2 | 2 | 0 | 9 | 0.2222 |
| X_7282232 | NA | NA | NA | NA | NA | NA | 2 | 1 | 2 | 1 | 1 | 1 | NA | NA | NA | NA | NA | NA | 1 | 1 |
| X_7282264 | 25 | 0.2 | 28 | 0.3929 | 32 | 0.2188 | 28 | 0.2857 | 113 | 0.2743 | 21 | 0.2857 | 21 | 0.2857 | 16 | 0.3125 | 21 | 0.4286 | 79 | 0.3291 |
| X_7283707 | 14 | 0.1429 | 5 | 0.6 | 15 | 0.4 | 15 | 0.4 | 49 | 0.3469 | 9 | 0.2222 | 15 | 0.4667 | 14 | 0.5714 | 14 | 0.2857 | 52 | 0.4038 |
| X_7283729 | 14 | 0.1429 | 5 | 0 | 10 | 0.1 | 18 | 0.0556 | 47 | 0.0851 | 12 | 0.0833 | 20 | 0.25 | 15 | 0.0667 | 14 | 0.0714 | 61 | 0.1311 |
| X_7283767 | 7 | 0.5714 | 3 | 0 | 6 | 0.6667 | 7 | 0.7143 | 23 | 0.5652 | 3 | 0.3333 | 5 | 0.4 | 4 | 0.5 | 5 | 0.2 | 17 | 0.3529 |
| X_7283782 | 2 | 1 | 2 | 0.5 | 5 | 0.8 | 5 | 0.6 | 14 | 0.7143 | 1 | 1 | 4 | 0.75 | 4 | 0.5 | 2 | 1 | 11 | 0.7273 |
| X_7283785 | 7 | 1 | 4 | 1 | 6 | 0.6667 | 6 | 0.8333 | 23 | 0.8696 | 3 | 1 | 5 | 0.8 | 5 | 0.8 | 6 | 0.8333 | 19 | 0.8421 |
| X_7283808 | 7 | 0.5714 | 4 | 0.5 | 4 | 0.25 | 4 | 0.5 | 19 | 0.4737 | 3 | 0 | 3 | 0.6667 | 4 | 0.5 | 3 | 0.3333 | 13 | 0.3846 |
| X_7283877 | 4 | 0 | 3 | 0 | 1 | 1 | 5 | 0.6 | 13 | 0.3077 | 3 | 0 | 3 | 0.6667 | 4 | 0 | 3 | 0 | 13 | 0.1538 |
| X_7283891 | 5 | 0.2 | 4 | 0.75 | 1 | 1 | 4 | 0.5 | 14 | 0.5 | 2 | 0 | 1 | 1 | 3 | 0.6667 | 2 | 0.5 | 8 | 0.5 |
| X_7283896 | 5 | 0 | 4 | 0 | 2 | 0 | 5 | 0.2 | 16 | 0.0625 | 3 | 0.3333 | 3 | 0 | 4 | 0 | 3 | 0 | 13 | 0.0769 |
| X_7283931 | 2 | 1 | 4 | 0.75 | 2 | 1 | 4 | 1 | 12 | 0.9167 | 2 | 1 | 3 | 1 | 3 | 1 | 1 | 1 | 9 | 1 |
| X_7283944 | 2 | 1 | 4 | 0.75 | 2 | 1 | 2 | 1 | 10 | 0.9 | 3 | 1 | 1 | 1 | 2 | 0.5 | 1 | 0 | 7 | 0.7143 |
| X_7283946 | 1 | 1 | 4 | 0.25 | 2 | 0.5 | 2 | 1 | 9 | 0.5556 | 2 | 0.5 | 1 | 1 | 1 | 1 | 1 | 0 | 5 | 0.6 |
| X_7283946 | 2 | 0 | 4 | 0.25 | 1 | 0 | 3 | 0.3333 | 10 | 0.2 | 3 | 0.3333 | 1 | 0 | 1 | 1 | 1 | 0 | 6 | 0.3333 |
| X_7286840 | 1 | 1 | 1 | 1 | NA | NA | NA | NA | 2 | 1 | NA | NA | NA | NA | 1 | 1 | 1 | 0 | 2 | 0.5 |
| X_7289771 | 13 | 0.0769 | 13 | 0.0769 | 5 | 0 | 14 | 0.1429 | 45 | 0.0889 | 9 | 0 | 11 | 0.3636 | 16 | 0 | 14 | 0.1429 | 50 | 0.12 |
| X_7289815 | 9 | 0.1111 | 11 | 0.0909 | 3 | 0 | 11 | 0 | 34 | 0.0588 | 9 | 0 | 7 | 0 | 15 | 0.1333 | 12 | 0.1667 | 43 | 0.093 |
| X_7289819 | 10 | 0.4 | 11 | 0.1818 | 4 | 0.25 | 13 | 0.3077 | 38 | 0.2895 | 9 | 0.3333 | 9 | 0.2222 | 15 | 0.3333 | 11 | 0.2727 | 44 | 0.2955 |
| X_7289822 | 7 | 0.5714 | 11 | 0.2727 | 3 | 0.3333 | 12 | 0.3333 | 33 | 0.3636 | 5 | 0.2 | 5 | 0.6 | 11 | 0.5455 | 10 | 0.3 | 31 | 0.4194 |
| X_7289827 | 10 | 0.1 | 11 | 0.0909 | 4 | 0 | 13 | 0.0769 | 38 | 0.0789 | 8 | 0 | 8 | 0 | 14 | 0.1429 | 11 | 0.0909 | 41 | 0.0732 |
| X_7289846 | 8 | 0.75 | 11 | 0.7273 | 3 | 1 | 8 | 0.625 | 30 | 0.7333 | 5 | 0.6 | 7 | 1 | 10 | 0.8 | 8 | 0.75 | 30 | 0.8 |
| X_7289847 | 6 | 0.1667 | 10 | 0.1 | 4 | 0.25 | 8 | 0.375 | 28 | 0.2143 | 7 | 0.2857 | 5 | 0.4 | 12 | 0.0833 | 10 | 0.1 | 34 | 0.1765 |

| | | | | | | | | | | | | | | | | | | | | |
|---|---|---|---|---|---|---|---|---|---|---|---|---|---|---|---|---|---|---|---|---|
| X_7289848 | 10 | 0.1 | 10 | 0.2 | 2 | 0 | 7 | 0 | 29 | 0.1034 | 7 | 0.1429 | 10 | 0.2 | 12 | 0.0833 | 10 | 0 | 39 | 0.1026 |
| X_7289852 | 6 | 0.5 | 6 | 1 | 4 | 0.75 | 8 | 1 | 24 | 0.8333 | 7 | 1 | 8 | 0.875 | 7 | 1 | 6 | 0.6667 | 28 | 0.8929 |
| X_7289856 | 9 | 0.1111 | 11 | 0.0909 | 4 | 0 | 7 | 0 | 31 | 0.0645 | 8 | 0.125 | 9 | 0.1111 | 12 | 0.0833 | 11 | 0.0909 | 40 | 0.1 |
| X_7289869 | 7 | 0.5714 | 8 | 0.625 | 3 | 1 | 6 | 0.3333 | 24 | 0.5833 | 4 | 0.25 | 6 | 0.3333 | 8 | 0.5 | 8 | 0.625 | 26 | 0.4615 |
| X_7290624 | 6 | 0.1667 | 3 | 0 | 1 | 0 | 3 | 0 | 13 | 0.0769 | 1 | 0 | 1 | 0 | 2 | 0.5 | 2 | 0 | 6 | 0.1667 |
| X_7298651 | 9 | 0.1111 | 8 | 0.125 | 4 | 0.25 | 10 | 0.1 | 31 | 0.129 | 4 | 0 | 6 | 0 | 4 | 0 | 10 | 0.2 | 24 | 0.0833 |
| X_7298667 | 7 | 0.2857 | 6 | 0.5 | 4 | 0.25 | 8 | 0 | 25 | 0.24 | 2 | 0.5 | 7 | 0.2857 | 2 | 0 | 8 | 0.125 | 19 | 0.2105 |
| X_7298772 | 6 | 0.3333 | 4 | 0.25 | 3 | 0 | 7 | 0.2857 | 20 | 0.25 | 2 | 0.5 | 2 | 0 | 4 | 0 | 6 | 0.5 | 14 | 0.2857 |
| X_7298774 | 6 | 0.6667 | 7 | 1 | 3 | 0.6667 | 8 | 0.75 | 24 | 0.7917 | 2 | 1 | 3 | 1 | 4 | 0.75 | 6 | 0.8333 | 15 | 0.8667 |
| X_7298776 | 6 | 0.6667 | 5 | 0.8 | 3 | 0.3333 | 9 | 0.6667 | 23 | 0.6522 | 2 | 0.5 | 3 | 0.6667 | 4 | 0.25 | 5 | 1 | 14 | 0.6429 |
| X_7299346 | NA | NA | 2 | 0.5 | NA | NA | 5 | 0.8 | 7 | 0.7143 | NA | NA | NA | NA | 1 | 1 | NA | NA | 1 | 1 |
| X_7299363 | NA | NA | 2 | 1 | NA | NA | 4 | 0.5 | 6 | 0.6667 | NA | NA | 1 | 1 | 1 | 0 | NA | NA | 2 | 0.5 |
| X_7299383 | NA | NA | 2 | 1 | NA | NA | 2 | 0.5 | 4 | 0.75 | NA | NA | 1 | 1 | 1 | 1 | NA | NA | 2 | 1 |
| X_7299396 | NA | NA | 2 | 0.5 | NA | NA | 2 | 1 | 4 | 0.75 | NA | NA | 1 | 0 | 1 | 1 | NA | NA | 2 | 0.5 |
| X_7305811 | 2 | 0 | 6 | 0.1667 | 5 | 0 | 4 | 0 | 17 | 0.0588 | 8 | 0 | 6 | 0.1667 | 5 | 0.2 | 7 | 0 | 26 | 0.0769 |
| X_7305820 | 3 | 0 | 5 | 0.2 | 4 | 0.25 | 5 | 0.2 | 17 | 0.1765 | 8 | 0.375 | 5 | 0 | 4 | 0 | 7 | 0.1429 | 24 | 0.1667 |
| X_7305854 | 1 | 0 | 5 | 0.2 | 5 | 0.2 | 5 | 0 | 16 | 0.125 | 9 | 0 | 4 | 0.25 | 3 | 0 | 4 | 0 | 20 | 0.05 |
| X_7305857 | 2 | 0 | 5 | 0 | 5 | 0.2 | 6 | 0 | 18 | 0.0556 | 6 | 0 | 2 | 0.5 | 2 | 0 | 4 | 0 | 14 | 0.0714 |
| X_7305886 | NA | NA | 5 | 0.4 | 4 | 0.25 | 5 | 0.2 | 14 | 0.2857 | 7 | 0.2857 | 4 | 0.5 | 1 | 0 | 4 | 0.25 | 16 | 0.3125 |
| X_7305888 | 1 | 0 | 4 | 0 | 4 | 0 | 5 | 0.4 | 14 | 0.1429 | 8 | 0.125 | 3 | 0 | 2 | 0 | 5 | 0.2 | 18 | 0.1111 |
| X_7305890 | 2 | 0 | 5 | 0 | 5 | 0.2 | 7 | 0 | 19 | 0.0526 | 9 | 0 | 3 | 0 | 2 | 0 | 6 | 0.1667 | 20 | 0.05 |
| X_7305901 | 2 | 0 | 5 | 0.2 | 5 | 0.4 | 6 | 0.3333 | 18 | 0.2778 | 9 | 0.1111 | 3 | 0.6667 | 3 | 0 | 6 | 0.1667 | 21 | 0.1905 |
| X_7306103 | 2 | 0 | 3 | 0.3333 | 1 | 0 | 3 | 0 | 9 | 0.1111 | 6 | 0 | 3 | 0 | NA | NA | 4 | 0.25 | 13 | 0.0769 |
| X_7306183 | 1 | 0 | 4 | 0.5 | 4 | 0.25 | 5 | 0 | 14 | 0.2143 | 6 | 0.3333 | 3 | 0.3333 | NA | NA | 4 | 0.5 | 13 | 0.3846 |
| X_7307059 | 18 | 0.1111 | 8 | 0 | 3 | 0 | 19 | 0.0526 | 48 | 0.0625 | 8 | 0 | 15 | 0 | 12 | 0.25 | 10 | 0.1 | 45 | 0.0889 |
| X_7307099 | 12 | 0.0833 | 7 | 0.2857 | 4 | 0 | 15 | 0 | 38 | 0.0789 | 5 | 0 | 10 | 0.1 | 9 | 0.1111 | 7 | 0 | 31 | 0.0645 |
| X_7307129 | 12 | 0.0833 | 8 | 0.25 | 2 | 0 | 14 | 0.0714 | 36 | 0.1111 | 7 | 0.2857 | 11 | 0 | 12 | 0 | 6 | 0.1667 | 36 | 0.0833 |
| X_7307199 | 10 | 0 | 6 | 0 | 2 | 0.5 | 11 | 0.1818 | 29 | 0.1034 | 5 | 0.4 | 6 | 0.1667 | 9 | 0.3333 | 6 | 0 | 26 | 0.2308 |
| X_7307205 | 11 | 0.6364 | 8 | 0.5 | 2 | 1 | 12 | 0.6667 | 33 | 0.6364 | 6 | 0.5 | 7 | 0.4286 | 8 | 0.5 | 8 | 0.625 | 29 | 0.5172 |
| X_7307217 | 11 | 0 | 7 | 0 | 2 | 0 | 13 | 0.1538 | 33 | 0.0606 | 2 | 0 | 9 | 0 | 10 | 0.4 | 9 | 0 | 30 | 0.1333 |
| X_7307220 | 11 | 0.3636 | 7 | 0.5714 | 2 | 0.5 | 11 | 0.4545 | 31 | 0.4516 | 1 | 1 | 8 | 0.25 | 6 | 0.5 | 8 | 0.5 | 23 | 0.4348 |
| X_7307494 | 3 | 0.6667 | 1 | 0 | 1 | 0 | 5 | 0.4 | 10 | 0.4 | 2 | 0 | 3 | 0 | 5 | 0.6 | 4 | 0 | 14 | 0.2143 |
| X_7307851 | 1 | 0 | NA | NA | 1 | 0 | 1 | 1 | 3 | 0.3333 | NA | NA | 2 | 0.5 | 1 | 1 | 2 | 0 | 5 | 0.4 |
| X_7307892 | 2 | 0 | 1 | 0 | 1 | 0 | 2 | 0.5 | 6 | 0.1667 | 1 | 1 | 1 | 0 | 1 | 0 | 2 | 0.5 | 5 | 0.4 |
| X_7307902 | 2 | 0.5 | 1 | 0 | 3 | 0.6667 | 3 | 0.6667 | 9 | 0.5556 | 1 | 0 | 1 | 1 | 1 | 1 | 2 | 1 | 5 | 0.8 |
| X_7307905 | 2 | 0 | 1 | 1 | 3 | 0 | 3 | 0.3333 | 9 | 0.2222 | 1 | 1 | 1 | 0 | NA | NA | 2 | 0.5 | 4 | 0.5 |
| X_7307910 | 2 | 1 | 1 | 1 | 2 | 1 | 3 | 1 | 8 | 1 | 1 | 1 | 1 | 0 | 1 | 1 | 2 | 1 | 5 | 0.8 |
| X_7307917 | 2 | 0.5 | NA | NA | 2 | 1 | 2 | 0.5 | 6 | 0.6667 | NA | NA | NA | NA | 1 | 1 | 2 | 1 | 3 | 1 |
| X_7307926 | 2 | 1 | 1 | 1 | 3 | 1 | 1 | 1 | 7 | 1 | 1 | 1 | NA | NA | 1 | 1 | 2 | 1 | 4 | 1 |
| X_7307929 | 1 | 1 | 1 | 0 | 3 | 1 | 2 | 0.5 | 7 | 0.7143 | 1 | 0 | NA | NA | 1 | 1 | 2 | 0.5 | 4 | 0.5 |
| X_7307947 | 2 | 0 | NA | NA | 3 | 0.3333 | 2 | 0 | 7 | 0.1429 | 1 | 0 | NA | NA | 1 | 0 | 2 | 1 | 4 | 0.5 |
| X_7307958 | 2 | 0 | 1 | 1 | 3 | 0.6667 | 2 | 1 | 8 | 0.625 | 1 | 0 | NA | NA | 1 | 1 | 2 | 1 | 4 | 0.75 |
| X_7307982 | 2 | 0.5 | 1 | 0 | 1 | 1 | 1 | 0 | 5 | 0.4 | 1 | 0 | NA | NA | 1 | 0 | 2 | 0.5 | 4 | 0.25 |
| X_7307992 | 2 | 1 | 1 | 1 | 2 | 0.5 | 1 | 1 | 6 | 0.8333 | 1 | 1 | NA | NA | NA | NA | 2 | 1 | 3 | 1 |
| X_7308001 | 2 | 0 | 1 | 1 | 3 | 0 | NA | NA | 6 | 0.1667 | 1 | 0 | NA | NA | NA | NA | 1 | 1 | 2 | 0.5 |
| X_7308002 | 2 | 0 | 1 | 1 | 3 | 0.6667 | 1 | 1 | 7 | 0.5714 | 1 | 1 | NA | NA | 1 | 0 | 1 | 1 | 3 | 0.6667 |
| X_7326747 | 2 | 0 | 6 | 0.3333 | 3 | 0.3333 | 2 | 0 | 13 | 0.2308 | 1 | 0 | NA | NA | 4 | 0.5 | 2 | 0.5 | 7 | 0.4286 |
| X_7326850 | 2 | 0 | 4 | 0 | 3 | 0 | 2 | 0.5 | 11 | 0.0909 | 1 | 0 | 1 | 0 | 3 | 0.3333 | 1 | 0 | 6 | 0.1667 |
| X_7327948 | 1 | 1 | 1 | 1 | 1 | 0 | 1 | 1 | 4 | 0.75 | NA | NA | NA | NA | 1 | 1 | NA | NA | 1 | 1 |
| X_7332666 | 1 | 0 | NA | NA | 3 | 0.3333 | NA | NA | 4 | 0.25 | 1 | 1 | 3 | 0 | NA | NA | 4 | 0 | 8 | 0.125 |
| X_7333293 | 2 | 0 | 1 | 0 | 5 | 0.2 | 2 | 0 | 10 | 0.1 | 2 | 0.5 | 3 | 0 | 4 | 0 | 6 | 0 | 15 | 0.0667 |
| X_7333305 | 2 | 0.5 | NA | NA | 4 | 1 | 2 | 0.5 | 8 | 0.75 | 1 | 1 | 3 | 0 | 4 | 0.75 | 5 | 0.2 | 13 | 0.3846 |
| X_7333314 | 1 | 0 | 1 | 0 | 4 | 0.75 | 2 | 0 | 8 | 0.375 | 2 | 1 | 2 | 0.5 | 2 | 0 | 5 | 0.8 | 11 | 0.6364 |
| X_7333316 | 2 | 0.5 | NA | NA | 6 | 1 | 2 | 1 | 10 | 0.9 | 3 | 1 | 3 | 1 | 4 | 0.75 | 6 | 0.6667 | 16 | 0.8125 |
| X_7333330 | 2 | 0.5 | 1 | 0 | 1 | 0 | 2 | 0 | 6 | 0.1667 | 3 | 0 | 2 | 0.5 | 4 | 0.25 | 4 | 0 | 13 | 0.1538 |
| X_7333341 | 2 | 0 | 1 | 0 | 6 | 0.3333 | 2 | 0 | 11 | 0.1818 | 3 | 0 | 3 | 0 | 4 | 0 | 6 | 0.3333 | 16 | 0.125 |
| X_7334708 | 35 | 0.0857 | 50 | 0.06 | 45 | 0.0444 | 38 | 0.0526 | 168 | 0.0595 | 36 | 0.0556 | 43 | 0.093 | 39 | 0.1026 | 23 | 0.1304 | 141 | 0.0922 |
| X_7334755 | 41 | 0.0976 | 54 | 0.1111 | 47 | 0.0851 | 36 | 0.1111 | 178 | 0.1011 | 42 | 0.0714 | 45 | 0.1556 | 38 | 0.0263 | 24 | 0.0833 | 149 | 0.0872 |
| X_7334771 | 41 | 0.122 | 53 | 0.1132 | 45 | 0.0444 | 37 | 0.027 | 176 | 0.0795 | 41 | 0.0732 | 40 | 0.125 | 39 | 0.1026 | 23 | 0.0435 | 143 | 0.0909 |
| X_7334777 | 46 | 0.0435 | 52 | 0.0962 | 43 | 0.1163 | 37 | 0.027 | 178 | 0.073 | 38 | 0.0789 | 42 | 0.0714 | 38 | 0.0526 | 22 | 0.0909 | 140 | 0.0714 |
| X_7334800 | 53 | 0.2264 | 52 | 0.3269 | 54 | 0.3704 | 43 | 0.2558 | 202 | 0.297 | 44 | 0.2045 | 46 | 0.3261 | 45 | 0.2 | 23 | 0.2174 | 158 | 0.2405 |
| X_7334804 | NA | NA | NA | NA | 1 | 1 | NA | NA | 1 | 1 | 1 | 1 | NA | NA | 1 | 0 | 2 | 0.5 | 4 | 0.5 |
| X_7334805 | 50 | 0.1 | 57 | 0.0877 | 50 | 0.04 | 40 | 0.1 | 197 | 0.0812 | 46 | 0.087 | 44 | 0.0909 | 50 | 0.12 | 30 | 0.0667 | 170 | 0.0941 |
| X_7334821 | 46 | 0.2609 | 61 | 0.3279 | 54 | 0.537 | 41 | 0.4878 | 202 | 0.401 | 46 | 0.4565 | 41 | 0.3902 | 49 | 0.3878 | 30 | 0.3 | 166 | 0.3916 |
| X_7338525 | NA | NA | 1 | 0 | 1 | 1 | 2 | 0 | 4 | 0.25 | 1 | 1 | NA | NA | 2 | 0 | 1 | 0 | 4 | 0.25 |

| | | | | | | | | | | | | | | | | | | | | |
|---|---|---|---|---|---|---|---|---|---|---|---|---|---|---|---|---|---|---|---|---|
| X_7338780 | 1 | 0 | 1 | 0 | 1 | 0 | 1 | 1 | 4 | 0.25 | NA | NA | NA | NA | 2 | 0 | 1 | 1 | 3 | 0.3333 |
| X_7338790 | 1 | 0 | 1 | 1 | 1 | 1 | NA | NA | 3 | 0.6667 | 1 | 1 | NA | NA | 1 | 0 | NA | NA | 2 | 0.5 |
| X_7340953 | 12 | 0.25 | 9 | 0.3333 | 7 | 0 | 7 | 0 | 35 | 0.1714 | 6 | 0 | 8 | 0.25 | 10 | 0 | 3 | 0 | 27 | 0.0741 |
| X_7340962 | 14 | 0.1429 | 6 | 0 | 8 | 0 | 10 | 0 | 38 | 0.0526 | 6 | 0.1667 | 8 | 0 | 12 | 0.0833 | 5 | 0 | 31 | 0.0645 |
| X_7340964 | 10 | 0.3 | 5 | 0 | 7 | 0.2857 | 7 | 0 | 29 | 0.1724 | 2 | 0.5 | 6 | 0 | 10 | 0 | 5 | 0.4 | 23 | 0.1304 |
| X_7341003 | 10 | 0 | 4 | 0.25 | 4 | 0.25 | 9 | 0.1111 | 27 | 0.1111 | 5 | 0 | 6 | 0.1667 | 9 | 0 | 4 | 0.25 | 24 | 0.0833 |
| X_7341111 | 4 | 0.25 | 8 | 0.25 | 10 | 0.1 | 7 | 0.4286 | 29 | 0.2414 | 6 | 0.1667 | 7 | 0.5714 | 9 | 0.5556 | 3 | 0.3333 | 25 | 0.44 |
| X_7341316 | 1 | 0 | NA | NA | NA | NA | 1 | 1 | 2 | 0.5 | NA | NA | NA | NA | 2 | 0.5 | 1 | 0 | 3 | 0.3333 |
| X_7341385 | NA | NA | NA | NA | NA | NA | 1 | 1 | 1 | 1 | NA | NA | NA | NA | 2 | 1 | 1 | 1 | 3 | 1 |
| X_7342850 | 4 | 0.75 | 5 | 0.2 | NA | NA | 3 | 0 | 12 | 0.3333 | NA | NA | 6 | 0.5 | 1 | 0 | NA | NA | 7 | 0.4286 |
| X_7342858 | 4 | 0.25 | 5 | 0.2 | NA | NA | 3 | 0 | 12 | 0.1667 | 1 | 0 | 6 | 0.5 | 3 | 0 | 1 | 0 | 11 | 0.2727 |
| X_7343640 | NA | NA | 4 | 0.25 | NA | NA | 2 | 0 | 6 | 0.1667 | NA | NA | 6 | 0.3333 | 1 | 1 | 2 | 0 | 9 | 0.3333 |
| X_7343645 | 1 | 0 | 3 | 0.3333 | NA | NA | 2 | 0.5 | 6 | 0.3333 | NA | NA | 5 | 0.6 | 1 | 0 | 2 | 0.5 | 8 | 0.5 |
| X_7343646 | 1 | 0 | 4 | 0.25 | NA | NA | 2 | 0.5 | 7 | 0.2857 | NA | NA | 4 | 0.25 | 2 | 0 | 1 | 1 | 7 | 0.2857 |
| X_7343648 | 1 | 0 | 4 | 0.25 | NA | NA | 2 | 0 | 7 | 0.1429 | NA | NA | 5 | 0.4 | 2 | 0.5 | 2 | 0 | 9 | 0.3333 |
| X_7343653 | 1 | 0 | 4 | 0.25 | NA | NA | 2 | 0 | 7 | 0.1429 | NA | NA | 6 | 0.3333 | 2 | 0 | 2 | 0 | 10 | 0.2 |
| X_7345333 | 6 | 0.1667 | 6 | 0.1667 | NA | NA | 3 | 0 | 15 | 0.1333 | 3 | 0 | 3 | 0 | 7 | 0.1429 | 7 | 0 | 20 | 0.05 |
| X_7345340 | 5 | 0 | 6 | 0.1667 | 3 | 0.3333 | 3 | 0.3333 | 17 | 0.1765 | 4 | 0 | 6 | 0 | 8 | 0 | 7 | 0.2857 | 25 | 0.08 |
| X_7345343 | 4 | 0 | 5 | 0 | 2 | 0 | 3 | 0.3333 | 14 | 0.0714 | 2 | 0 | 4 | 0.25 | 7 | 0.1429 | 6 | 0.1667 | 19 | 0.1579 |
| X_7345345 | 2 | 0 | 4 | 0.25 | 3 | 0 | 2 | 0 | 11 | 0.0909 | 2 | 0 | 3 | 0 | 4 | 0 | 7 | 0.1429 | 16 | 0.0625 |
| X_7345370 | 4 | 0.75 | 6 | 0.5 | 1 | 0 | 3 | 0.3333 | 14 | 0.5 | 3 | 0.6667 | 6 | 0.3333 | 6 | 0.6667 | 7 | 0.8571 | 22 | 0.6364 |
| X_7345372 | 5 | 0.4 | 4 | 0.5 | 2 | 0.5 | 3 | 0.3333 | 14 | 0.4286 | 3 | 0.3333 | 6 | 0.1667 | 7 | 0.4286 | 5 | 0.2 | 21 | 0.2857 |
| X_7345408 | 3 | 0 | 6 | 0.1667 | 2 | 0 | 3 | 0.3333 | 14 | 0.1429 | 3 | 0 | 2 | 0 | 6 | 0.1667 | 6 | 0 | 17 | 0.0588 |
| X_7349192 | 1 | 1 | 1 | 0 | 1 | 0 | 2 | 1 | 5 | 0.6 | NA | NA | 1 | 1 | NA | NA | 1 | 0 | 2 | 0.5 |
| X_7349491 | 4 | 0 | 2 | 0 | 1 | 0 | 2 | 0.5 | 9 | 0.1111 | NA | NA | 3 | 0.3333 | NA | NA | 2 | 0 | 5 | 0.2 |
| X_7349504 | 4 | 0.25 | 3 | 0.3333 | 1 | 0 | 2 | 0 | 10 | 0.2 | NA | NA | 2 | 1 | NA | NA | 2 | 0 | 4 | 0.5 |
| X_7375700 | NA | NA | NA | NA | NA | NA | 1 | 1 | 1 | 1 | NA | NA | 1 | 1 | NA | NA | NA | NA | 1 | 1 |
| X_7414016 | 2 | 0.5 | NA | NA | NA | NA | NA | NA | 2 | 0.5 | NA | NA | 1 | 0 | NA | NA | 1 | 1 | 2 | 0.5 |
| X_7415435 | 8 | 0 | 7 | 0.1429 | 2 | 0.5 | 5 | 0 | 22 | 0.0909 | 3 | 0 | 3 | 0 | 4 | 0.5 | 3 | 0 | 13 | 0.1538 |
| X_7443799 | 8 | 0.375 | 9 | 0.3333 | 11 | 0.5455 | 10 | 0.1 | 38 | 0.3421 | 8 | 0.125 | 10 | 0.2 | 9 | 0.3333 | 10 | 0.1 | 37 | 0.1892 |
| X_7443806 | 8 | 0.5 | 10 | 0.2 | 9 | 0.2222 | 8 | 0.125 | 35 | 0.2571 | 9 | 0.1111 | 8 | 0.25 | 9 | 0.3333 | 10 | 0.3 | 36 | 0.25 |
| X_7444878 | 3 | 0 | 3 | 0.3333 | 3 | 0.3333 | 3 | 0.3333 | 12 | 0.25 | NA | NA | 2 | 0.5 | NA | NA | NA | NA | 2 | 0.5 |
| X_7449734 | 1 | 1 | 1 | 0 | 2 | 0 | NA | NA | 4 | 0.25 | NA | NA | NA | NA | 3 | 0.3333 | 1 | 0 | 4 | 0.25 |
| X_7457784 | 7 | 0.1429 | 9 | 0.1111 | 7 | 0.5714 | 5 | 0 | 28 | 0.2143 | 2 | 0 | 9 | 0.1111 | 4 | 0.5 | 11 | 0.3636 | 26 | 0.2692 |
| X_7457837 | 8 | 0.125 | 15 | 0.0667 | 11 | 0.0909 | 6 | 0 | 40 | 0.075 | 6 | 0 | 10 | 0.1 | 6 | 0 | 16 | 0.125 | 38 | 0.0789 |
| X_7461171 | 1 | 0 | 2 | 0 | 1 | 1 | NA | NA | 4 | 0.25 | NA | NA | NA | NA | 2 | 0.5 | 1 | 1 | 3 | 0.6667 |
| X_7461800 | 1 | 0 | 1 | 1 | 1 | 0 | 4 | 0.25 | 7 | 0.2857 | NA | NA | 2 | 0 | 4 | 0.5 | 3 | 0 | 9 | 0.2222 |
| X_7461801 | 1 | 0 | 3 | 0.3333 | 1 | 1 | 4 | 0 | 9 | 0.2222 | NA | NA | 2 | 0 | 4 | 0.25 | 3 | 0 | 9 | 0.1111 |
| X_7461810 | 2 | 0 | 3 | 0 | 1 | 0 | 3 | 0.3333 | 9 | 0.1111 | NA | NA | 2 | 0.5 | 4 | 0 | 1 | 0 | 7 | 0.1429 |
| X_7461811 | 2 | 0 | 3 | 0.3333 | 1 | 0 | 4 | 0.25 | 10 | 0.2 | NA | NA | 2 | 0 | 4 | 0.5 | 3 | 0.3333 | 9 | 0.3333 |
| X_7461814 | 2 | 0.5 | 2 | 0.5 | 1 | 0 | 4 | 0 | 9 | 0.2222 | NA | NA | 2 | 0.5 | 2 | 0.5 | 2 | 0.5 | 6 | 0.5 |
| X_7461832 | 2 | 0 | 3 | 0.6667 | 1 | 0 | 4 | 0.25 | 10 | 0.3 | NA | NA | 2 | 1 | 4 | 1 | 3 | 0.3333 | 9 | 0.7778 |
| X_7461840 | 2 | 0.5 | 3 | 0.6667 | 1 | 0 | 4 | 0.25 | 10 | 0.4 | NA | NA | 2 | 0 | 4 | 0.25 | 3 | 0.6667 | 9 | 0.3333 |
| X_7461842 | 2 | 0.5 | 3 | 0 | 1 | 1 | 4 | 0.5 | 10 | 0.4 | NA | NA | 2 | 1 | 3 | 0.6667 | 3 | 0.6667 | 8 | 0.75 |
| X_7461851 | 1 | 1 | 3 | 0.3333 | 1 | 0 | 4 | 0 | 9 | 0.2222 | NA | NA | 2 | 0 | 4 | 0.25 | 3 | 0.6667 | 9 | 0.3333 |
| X_7461856 | 1 | 0 | 3 | 0.6667 | 1 | 1 | 4 | 0.5 | 9 | 0.5556 | NA | NA | 2 | 1 | 4 | 0.5 | 3 | 0 | 9 | 0.4444 |
| X_7483026 | 4 | 1 | 7 | 0.8571 | 7 | 0.5714 | 9 | 0.6667 | 27 | 0.7407 | 8 | 0.75 | 7 | 0.5714 | 12 | 0.9167 | 10 | 0.6 | 37 | 0.7297 |
| X_7483041 | 4 | 0.25 | 8 | 0 | 5 | 0.2 | 6 | 0.1667 | 23 | 0.1304 | 7 | 0.1429 | 7 | 0.1429 | 9 | 0.1111 | 9 | 0.2222 | 32 | 0.1562 |
| X_7670343 | 3 | 0 | 2 | 0 | 3 | 0.3333 | 4 | 0 | 12 | 0.0833 | 4 | 0.25 | 6 | 0.1667 | 9 | 0.2222 | 6 | 0 | 25 | 0.16 |
| X_7670344 | 2 | 0.5 | 2 | 0.5 | 4 | 0.25 | 3 | 0.6667 | 11 | 0.4545 | 5 | 0.4 | 6 | 0.3333 | 7 | 0.2857 | 6 | 0.8333 | 24 | 0.4583 |
| X_7670366 | 3 | 1 | 2 | 0.5 | 4 | 0.25 | 3 | 0.3333 | 12 | 0.5 | 3 | 0.3333 | 8 | 0.625 | 8 | 0.375 | 6 | 1 | 25 | 0.6 |
| X_7670385 | 2 | 0 | 2 | 0.5 | 5 | 0 | 5 | 0 | 14 | 0.0714 | 4 | 0 | 7 | 0.1429 | 9 | 0.2222 | 6 | 0 | 26 | 0.1154 |
| X_7670610 | 9 | 0.1111 | 6 | 0.1667 | 7 | 0 | 8 | 0 | 30 | 0.0667 | 5 | 0 | 9 | 0 | 11 | 0.0909 | 12 | 0.0833 | 37 | 0.0541 |
| X_7670615 | 7 | 0.1429 | 6 | 0.5 | 6 | 0.1667 | 6 | 0 | 25 | 0.2 | 4 | 0.5 | 9 | 0.1111 | 10 | 0 | 12 | 0.1667 | 35 | 0.1429 |
| X_7680612 | 1 | 1 | 2 | 0 | 2 | 0 | 2 | 0 | 7 | 0.1429 | 1 | 0 | NA | NA | 1 | 0 | 1 | 1 | 3 | 0.3333 |
| X_7749490 | 1 | 1 | NA | NA | 1 | 0 | NA | NA | 2 | 0.5 | NA | NA | 2 | 1 | 1 | 1 | 2 | 0.5 | 5 | 0.8 |
| X_7749493 | 1 | 0 | NA | NA | 1 | 0 | 1 | 1 | 3 | 0.3333 | NA | NA | 2 | 1 | 1 | 1 | 3 | 0.3333 | 6 | 0.6667 |
| X_7749618 | 1 | 0 | NA | NA | 1 | 1 | NA | NA | 2 | 0.5 | NA | NA | 2 | 0.5 | NA | NA | 2 | 0.5 | 4 | 0.5 |
| X_7749660 | 1 | 1 | NA | NA | NA | NA | NA | NA | 1 | 1 | NA | NA | NA | NA | NA | NA | 1 | 1 | 1 | 1 |
| X_7749672 | 1 | 1 | NA | NA | NA | NA | NA | NA | 1 | 1 | NA | NA | 1 | 1 | NA | NA | 1 | 1 | 2 | 1 |
| X_7950152 | 3 | 0.3333 | 6 | 0.3333 | 6 | 0.6667 | 5 | 0.6 | 20 | 0.5 | 3 | 1 | 2 | 0.5 | 2 | 0 | 4 | 0.75 | 11 | 0.6364 |
| X_7950160 | 4 | 0.5 | 5 | 0.8 | 7 | 0.7143 | 5 | 0.6 | 21 | 0.6667 | 3 | 1 | 2 | 1 | 3 | 0.3333 | 4 | 1 | 12 | 0.8333 |
| X_8013821 | 8 | 0.125 | 6 | 0.3333 | 7 | 0.1429 | 13 | 0.0769 | 34 | 0.1471 | 9 | 0.1111 | 9 | 0.1111 | 13 | 0.1538 | 12 | 0.0833 | 43 | 0.1163 |
| X_8013991 | 12 | 0 | 10 | 0.4 | 8 | 0.25 | 17 | 0.1765 | 47 | 0.1915 | 10 | 0.5 | 13 | 0.0769 | 14 | 0.1429 | 17 | 0.2941 | 54 | 0.2407 |
| X_8014019 | 12 | 0.0833 | 11 | 0.0909 | 11 | 0 | 16 | 0.0625 | 50 | 0.06 | 9 | 0 | 15 | 0.0667 | 16 | 0.125 | 18 | 0.0556 | 58 | 0.069 |

| | | | | | | | | | | | | | | | | | | | | |
|---|---|---|---|---|---|---|---|---|---|---|---|---|---|---|---|---|---|---|---|---|
| X_8424773 | 6 | 0 | 4 | 0.25 | 2 | 0.5 | 2 | 0 | 14 | 0.1429 | 3 | 0 | 2 | 0 | 4 | 0.25 | 8 | 0 | 17 | 0.0588 |
| X_8424835 | 7 | 0 | 4 | 0.25 | 4 | 0 | 5 | 0 | 20 | 0.05 | 3 | 0.3333 | 3 | 0 | 4 | 0 | 7 | 0 | 17 | 0.0588 |
| X_8424846 | 7 | 0.4286 | 3 | 0 | 6 | 0 | 5 | 0.2 | 21 | 0.1905 | 4 | 0.25 | 3 | 0.3333 | 4 | 0.25 | 9 | 0.4444 | 20 | 0.35 |
| X_8465776 | NA | NA | 1 | 1 | 1 | 1 | 2 | 1 | 4 | 1 | NA | NA | NA | NA | 1 | 1 | 3 | 1 | 4 | 1 |
| X_8465778 | NA | NA | NA | NA | 1 | 1 | 2 | 1 | 3 | 1 | NA | NA | NA | NA | 1 | 1 | 3 | 1 | 4 | 1 |
| X_8465809 | NA | NA | NA | NA | NA | NA | 2 | 0.5 | 2 | 0.5 | NA | NA | NA | NA | 1 | 0 | 2 | 1 | 3 | 0.6667 |
| X_8465809 | NA | NA | NA | NA | 1 | 1 | 2 | 0.5 | 3 | 0.6667 | NA | NA | NA | NA | 1 | 1 | 3 | 0.3333 | 4 | 0.5 |
| X_8465829 | NA | NA | 1 | 1 | 1 | 1 | 2 | 1 | 4 | 1 | NA | NA | NA | NA | 1 | 1 | 2 | 0.5 | 3 | 0.6667 |
| X_8465849 | NA | NA | 1 | 1 | 1 | 1 | 2 | 1 | 4 | 1 | NA | NA | NA | NA | 1 | 0 | 2 | 0.5 | 3 | 0.3333 |
| X_8465861 | NA | NA | 2 | 0.5 | 1 | 1 | 3 | 0.6667 | 6 | 0.6667 | NA | NA | NA | NA | 1 | 1 | 3 | 0 | 4 | 0.25 |
| X_8465879 | NA | NA | 2 | 0.5 | 1 | 1 | 3 | 0.6667 | 6 | 0.6667 | NA | NA | NA | NA | 1 | 1 | 3 | 0.6667 | 4 | 0.75 |
| X_8467422 | NA | NA | 2 | 1 | 1 | 0 | 3 | 0.3333 | 6 | 0.5 | NA | NA | NA | NA | 1 | 1 | 3 | 0.3333 | 4 | 0.5 |
| X_8467660 | NA | NA | 3 | 0 | 1 | 0 | 6 | 0.1667 | 10 | 0.1 | NA | NA | NA | NA | 2 | 0 | 4 | 0.25 | 6 | 0.1667 |
| X_8474789 | NA | NA | 2 | 0 | 1 | 1 | 3 | 0.6667 | 6 | 0.5 | 2 | 1 | 1 | 0 | 2 | 0.5 | 2 | 0.5 | 7 | 0.5714 |
| X_8578755 | NA | NA | 1 | 0 | NA | NA | 2 | 0.5 | 3 | 0.3333 | NA | NA | NA | NA | 2 | 1 | 1 | 1 | 3 | 1 |
| X_8578764 | NA | NA | 1 | 1 | NA | NA | 2 | 0 | 3 | 0.3333 | NA | NA | NA | NA | 2 | 0.5 | 1 | 0 | 3 | 0.3333 |
| X_8578793 | NA | NA | 1 | 0 | NA | NA | 2 | 0.5 | 3 | 0.3333 | NA | NA | NA | NA | 2 | 0.5 | 1 | 0 | 3 | 0.3333 |
| X_8578856 | 2 | 1 | 2 | 0.5 | NA | NA | 3 | 0.6667 | 7 | 0.7143 | NA | NA | 1 | 1 | 1 | 1 | 1 | 1 | 3 | 1 |
| X_8578857 | 2 | 1 | 1 | 1 | NA | NA | 3 | 1 | 6 | 1 | NA | NA | 1 | 1 | 1 | 1 | 1 | 1 | 3 | 1 |
| X_8578865 | 2 | 0 | 2 | 0.5 | NA | NA | 3 | 0.6667 | 7 | 0.4286 | NA | NA | 1 | 0 | 1 | 1 | 1 | 0 | 3 | 0.3333 |
| X_8578867 | 2 | 0 | 2 | 0 | NA | NA | 3 | 0.3333 | 7 | 0.1429 | NA | NA | 1 | 1 | 1 | 0 | 1 | 1 | 3 | 0.6667 |
| X_8578917 | 2 | 0.5 | 1 | 1 | NA | NA | 3 | 0.3333 | 6 | 0.5 | NA | NA | 1 | 1 | 1 | 1 | 1 | 0 | 3 | 0.6667 |
| X_8579040 | 1 | 1 | NA | NA | NA | NA | 2 | 1 | 3 | 1 | NA | NA | 1 | 1 | NA | NA | 1 | 1 | 2 | 1 |
| X_8579044 | NA | NA | NA | NA | NA | NA | 2 | 0.5 | 2 | 0.5 | NA | NA | 1 | 1 | NA | NA | 1 | 0 | 2 | 0.5 |
| X_8579046 | NA | NA | NA | NA | NA | NA | 1 | 1 | 1 | 1 | NA | NA | 1 | 1 | NA | NA | 1 | 0 | 2 | 0.5 |
| X_8579058 | 1 | 1 | NA | NA | NA | NA | 2 | 1 | 3 | 1 | NA | NA | 1 | 1 | NA | NA | 1 | 1 | 2 | 1 |
| X_8579063 | 1 | 0 | NA | NA | NA | NA | 2 | 1 | 3 | 0.6667 | NA | NA | 1 | 1 | NA | NA | 1 | 1 | 2 | 1 |
| X_8579073 | 1 | 1 | NA | NA | NA | NA | 2 | 0.5 | 3 | 0.6667 | NA | NA | 1 | 1 | NA | NA | 1 | 1 | 2 | 1 |
| X_8579078 | 1 | 1 | NA | NA | NA | NA | 2 | 1 | 3 | 1 | NA | NA | 1 | 1 | NA | NA | NA | NA | 1 | 1 |
| X_8579083 | NA | NA | NA | NA | NA | NA | 2 | 1 | 2 | 1 | NA | NA | 1 | 1 | NA | NA | 1 | 1 | 2 | 1 |
| X_8579124 | 1 | 0 | NA | NA | NA | NA | 2 | 0.5 | 3 | 0.3333 | NA | NA | 1 | 0 | NA | NA | 1 | 1 | 2 | 0.5 |
| X_8579129 | 1 | 1 | NA | NA | NA | NA | 1 | 1 | 2 | 1 | NA | NA | 1 | 1 | NA | NA | 1 | 1 | 2 | 1 |
| X_8579131 | 1 | 1 | NA | NA | NA | NA | 2 | 1 | 3 | 1 | NA | NA | 1 | 1 | NA | NA | 1 | 1 | 2 | 1 |
| X_8579138 | NA | NA | NA | NA | NA | NA | 2 | 0.5 | 2 | 0.5 | NA | NA | 1 | 0 | NA | NA | 1 | 1 | 2 | 0.5 |
| X_9234160 | 4 | 0.75 | 1 | 1 | 3 | 0.6667 | 2 | 0.5 | 10 | 0.7 | 2 | 1 | 2 | 0.5 | 3 | 1 | 2 | 0.5 | 9 | 0.7778 |
| X_9234917 | 1 | 1 | 2 | 0.5 | NA | NA | NA | NA | 3 | 0.6667 | NA | NA | NA | NA | 1 | 0 | 1 | 1 | 2 | 0.5 |
| X_9278996 | NA | NA | NA | NA | 1 | 0 | 1 | 1 | 2 | 0.5 | 2 | 0 | NA | NA | NA | NA | 2 | 0.5 | 4 | 0.25 |
| X_9279023 | NA | NA | NA | NA | 2 | 0.5 | 1 | 0 | 3 | 0.3333 | 3 | 0.3333 | 1 | 0 | 1 | 0 | 2 | 0.5 | 7 | 0.2857 |
| X_9279060 | 1 | 1 | 1 | 1 | 2 | 1 | 1 | 1 | 5 | 1 | 3 | 1 | 1 | 1 | 1 | 1 | 1 | 1 | 6 | 1 |
| X_9279061 | 1 | 1 | 2 | 0 | 2 | 0 | 2 | 0 | 7 | 0.1429 | 3 | 0 | 1 | 0 | 1 | 0 | 2 | 0.5 | 7 | 0.1429 |
| X_9279067 | 1 | 1 | 1 | 1 | 2 | 0.5 | 1 | 1 | 5 | 0.8 | 3 | 0.6667 | 1 | 1 | 1 | 1 | 2 | 1 | 7 | 0.8571 |
| X_9279075 | NA | NA | 2 | 0.5 | 2 | 0 | NA | NA | 4 | 0.25 | 4 | 0.25 | 1 | 0 | NA | NA | 1 | 1 | 6 | 0.3333 |
| X_9279087 | 1 | 1 | 2 | 1 | 2 | 1 | 2 | 1 | 7 | 1 | 3 | 1 | 1 | 1 | NA | NA | 2 | 1 | 6 | 1 |
| X_9279089 | 1 | 0 | 2 | 0.5 | 3 | 0 | 3 | 0 | 9 | 0.1111 | 4 | 0.25 | 1 | 0 | NA | NA | 3 | 0.3333 | 8 | 0.25 |
| X_9279101 | NA | NA | 1 | 1 | 2 | 1 | 3 | 1 | 6 | 1 | 4 | 0.75 | 1 | 1 | 3 | 1 | 3 | 0.6667 | 11 | 0.8182 |
| X_9279109 | 1 | 1 | 2 | 0 | 3 | 0 | 3 | 0 | 9 | 0.1111 | 4 | 0.25 | 1 | 0 | 3 | 0 | 3 | 0 | 11 | 0.0909 |
| X_9279122 | NA | NA | 2 | 0 | 3 | 0.3333 | 3 | 0 | 8 | 0.125 | 4 | 0.25 | 1 | 0 | 3 | 0.3333 | 3 | 0 | 11 | 0.1818 |
| X_9279127 | 1 | 1 | 2 | 0.5 | 3 | 0.3333 | 3 | 0.3333 | 9 | 0.4444 | 5 | 0.8 | NA | NA | 3 | 0 | 3 | 0.6667 | 11 | 0.5455 |
| X_9279162 | 2 | 0.5 | 2 | 0 | 4 | 0 | 3 | 0 | 11 | 0.0909 | 4 | 0.25 | 1 | 0 | 3 | 0.6667 | 3 | 0 | 11 | 0.2727 |
| X_9279173 | 1 | 1 | 2 | 0.5 | 4 | 0.25 | 2 | 0 | 9 | 0.3333 | 4 | 0.25 | 1 | 0 | 3 | 0 | 3 | 0 | 11 | 0.0909 |
| X_9279184 | 2 | 0.5 | 2 | 0.5 | 4 | 0.25 | 4 | 0 | 12 | 0.25 | 4 | 0 | 1 | 1 | 3 | 0.3333 | 3 | 0 | 11 | 0.1818 |
| X_9279201 | 3 | 0.6667 | 2 | 0.5 | 4 | 0.25 | 4 | 0 | 13 | 0.3077 | 4 | 0.5 | 1 | 1 | 3 | 0.3333 | 4 | 0.25 | 12 | 0.4167 |
| X_9279225 | 3 | 0.6667 | 1 | 0 | 4 | 0.75 | 5 | 0 | 13 | 0.3846 | 4 | 0.25 | 1 | 1 | 2 | 0 | 4 | 0.25 | 11 | 0.2727 |
| X_9279309 | 5 | 0.2 | 4 | 0 | 3 | 0 | 4 | 0 | 16 | 0.0625 | 4 | 0.5 | 1 | 0 | 2 | 0 | 5 | 0 | 12 | 0.1667 |
| X_9279907 | 5 | 0 | 8 | 0.375 | 2 | 0 | 5 | 0.4 | 20 | 0.25 | 4 | 0.5 | 9 | 0 | 5 | 0.2 | 6 | 0.1667 | 24 | 0.1667 |
| X_9279951 | 4 | 0.75 | 6 | 0.1667 | 1 | 0 | 4 | 0 | 15 | 0.2667 | 4 | 0.75 | 6 | 0.3333 | 4 | 0.25 | 6 | 0.1667 | 20 | 0.35 |
| X_9279965 | 4 | 0 | 6 | 0 | 1 | 1 | 4 | 0.25 | 15 | 0.1333 | 4 | 0.5 | 9 | 0.2222 | 4 | 0.5 | 5 | 0.4 | 22 | 0.3636 |
| X_9279986 | 3 | 0.3333 | 5 | 0 | NA | NA | 3 | 0 | 11 | 0.0909 | 4 | 0.25 | 6 | 0 | 4 | 0 | 6 | 0 | 20 | 0.05 |
| X_9280035 | 3 | 0 | 4 | 0 | 1 | 1 | 4 | 0 | 12 | 0.0833 | 3 | 0 | 7 | 0 | 3 | 0 | 5 | 0.2 | 18 | 0.0556 |
| X_9280130 | 2 | 0 | 1 | 0 | 1 | 1 | 3 | 0 | 7 | 0.1429 | 1 | 0 | 5 | 0.2 | 2 | 0 | 2 | 0 | 10 | 0.1 |
| X_9280147 | 2 | 0 | 1 | 0 | 1 | 1 | 3 | 0 | 7 | 0.1429 | 1 | 0 | 7 | 0.1429 | 2 | 0 | 3 | 0.3333 | 13 | 0.1538 |
| X_9280837 | 6 | 0.3333 | 3 | 0.3333 | 8 | 0 | 4 | 0.5 | 21 | 0.2381 | 5 | 0.4 | 5 | 0 | 4 | 0 | 5 | 0.4 | 19 | 0.2105 |
| X_9280857 | 7 | 0.1429 | 2 | 0 | 6 | 0 | 5 | 0 | 20 | 0.05 | 5 | 0.2 | 6 | 0 | 3 | 0 | 5 | 0 | 19 | 0.0526 |
| X_9280883 | 9 | 0 | 6 | 0.1667 | 5 | 0.4 | 4 | 0.25 | 24 | 0.1667 | 7 | 0.1429 | 7 | 0.1429 | 6 | 0 | 8 | 0.375 | 28 | 0.1786 |
| X_9311842 | 1 | 1 | 1 | 1 | 5 | 0 | 2 | 0 | 9 | 0.2222 | NA | NA | NA | NA | 2 | 0.5 | 1 | 0 | 3 | 0.3333 |

| | | | | | | | | | | | | | | | | | | | | |
|---|---|---|---|---|---|---|---|---|---|---|---|---|---|---|---|---|---|---|---|---|
| X_9311865 | 2 | 0 | NA | NA | 5 | 0.2 | 2 | 0 | 9 | 0.1111 | NA | NA | NA | NA | 2 | 0.5 | 1 | 0 | 3 | 0.3333 |
| X_9311989 | 1 | 0 | 1 | 1 | 3 | 0.3333 | 3 | 0 | 8 | 0.25 | 1 | 0 | 2 | 0.5 | 1 | 0 | 2 | 0 | 6 | 0.1667 |
| X_9312021 | 1 | 0 | 1 | 0 | 3 | 0.3333 | 1 | 0 | 6 | 0.1667 | NA | NA | 1 | 1 | NA | NA | 2 | 0 | 3 | 0.3333 |
| X_9312202 | 1 | 1 | 1 | 1 | 3 | 1 | 2 | 0.5 | 7 | 0.8571 | NA | NA | NA | NA | 1 | 1 | NA | NA | 1 | 1 |
| X_9312267 | 1 | 1 | NA | NA | 2 | 1 | 2 | 1 | 5 | 1 | NA | NA | NA | NA | 1 | 1 | NA | NA | 1 | 1 |
| X_9312273 | 1 | 1 | NA | NA | 3 | 1 | 2 | 1 | 6 | 1 | NA | NA | NA | NA | 1 | 1 | NA | NA | 1 | 1 |
| X_9312274 | NA | NA | NA | NA | 3 | 1 | 2 | 0.5 | 5 | 0.8 | NA | NA | NA | NA | 1 | 1 | NA | NA | 1 | 1 |
| X_9312279 | 1 | 1 | NA | NA | 3 | 1 | 2 | 1 | 6 | 1 | NA | NA | NA | NA | 1 | 1 | NA | NA | 1 | 1 |
| X_9312292 | 1 | 0 | NA | NA | 3 | 1 | 2 | 0.5 | 6 | 0.6667 | NA | NA | NA | NA | 1 | 1 | NA | NA | 1 | 1 |
| X_9312298 | 1 | 0 | NA | NA | 3 | 1 | 2 | 1 | 6 | 0.8333 | NA | NA | NA | NA | 1 | 1 | NA | NA | 1 | 1 |
| X_9312299 | 1 | 1 | NA | NA | 3 | 0.6667 | 2 | 0.5 | 6 | 0.6667 | NA | NA | NA | NA | 1 | 1 | NA | NA | 1 | 1 |
| X_9312303 | 1 | 0 | NA | NA | 1 | 1 | 1 | 1 | 3 | 0.6667 | NA | NA | NA | NA | 1 | 1 | NA | NA | 1 | 1 |
| X_9312344 | 1 | 1 | NA | NA | 3 | 1 | 1 | 1 | 5 | 1 | NA | NA | NA | NA | 1 | 1 | NA | NA | 1 | 1 |
| X_9312356 | 1 | 0 | NA | NA | 3 | 0.3333 | 1 | 0 | 5 | 0.2 | NA | NA | NA | NA | 1 | 1 | NA | NA | 1 | 1 |
| X_9314198 | 1 | 0 | NA | NA | 2 | 0.5 | NA | NA | 3 | 0.3333 | NA | NA | NA | NA | 1 | 1 | NA | NA | 1 | 1 |
| X_9314207 | 1 | 1 | NA | NA | 2 | 1 | 1 | 0 | 4 | 0.75 | NA | NA | NA | NA | 1 | 1 | NA | NA | 1 | 1 |
| X_9323343 | 15 | 0.1333 | 19 | 0.1579 | 12 | 0.3333 | 13 | 0.2308 | 59 | 0.2034 | 6 | 0.5 | 6 | 0 | 10 | 0.1 | 13 | 0.3077 | 35 | 0.2286 |
| X_9323356 | 13 | 0 | 13 | 0 | 10 | 0.1 | 13 | 0.2308 | 49 | 0.0816 | 4 | 0 | 6 | 0.1667 | 10 | 0.1 | 11 | 0.0909 | 31 | 0.0968 |
| X_9323372 | 10 | 0 | 15 | 0.1333 | 8 | 0.125 | 10 | 0.2 | 43 | 0.1163 | 5 | 0.2 | 6 | 0.1667 | 8 | 0.25 | 11 | 0.2727 | 30 | 0.2333 |
| X_9323410 | 12 | 0.3333 | 16 | 0.125 | 10 | 0 | 14 | 0.0714 | 52 | 0.1346 | 5 | 0 | 6 | 0 | 12 | 0.0833 | 13 | 0.0769 | 36 | 0.0556 |
| X_9327882 | NA | NA | 1 | 1 | NA | NA | NA | NA | 1 | 1 | 2 | 0.5 | 2 | 1 | NA | NA | NA | NA | 4 | 0.75 |
| X_9329623 | NA | NA | 2 | 0.5 | NA | NA | NA | NA | 2 | 0.5 | 2 | 1 | 1 | 0 | NA | NA | NA | NA | 3 | 0.6667 |
| X_9329630 | 1 | 1 | 1 | 1 | NA | NA | NA | NA | 2 | 1 | 1 | 1 | 1 | 1 | 1 | 0 | NA | NA | 3 | 0.6667 |
| X_9329631 | 2 | 0.5 | 2 | 1 | NA | NA | NA | NA | 4 | 0.75 | 2 | 0 | 1 | 1 | 1 | 0 | NA | NA | 4 | 0.25 |
| X_9329632 | 2 | 1 | 2 | 1 | NA | NA | NA | NA | 4 | 1 | 2 | 1 | 2 | 1 | 1 | 1 | NA | NA | 5 | 1 |
| X_9329644 | 2 | 0.5 | 2 | 1 | NA | NA | NA | NA | 4 | 0.75 | 2 | 1 | 1 | 1 | 1 | 0 | NA | NA | 4 | 0.75 |
| X_9331606 | NA | NA | 3 | 1 | NA | NA | NA | NA | 3 | 1 | 1 | 1 | 2 | 0.5 | 1 | 0 | NA | NA | 4 | 0.5 |
| X_9331612 | 1 | 1 | 3 | 1 | NA | NA | NA | NA | 4 | 1 | 1 | 1 | 2 | 1 | 1 | 1 | NA | NA | 4 | 1 |
| X_9331618 | 1 | 1 | 3 | 1 | NA | NA | NA | NA | 4 | 1 | 1 | 1 | 2 | 1 | 1 | 1 | NA | NA | 4 | 1 |
| X_9331626 | NA | NA | 3 | 0.6667 | NA | NA | NA | NA | 3 | 0.6667 | 2 | 0.5 | 2 | 0.5 | 1 | 1 | NA | NA | 5 | 0.6 |
| X_9331632 | 1 | 1 | 3 | 0 | NA | NA | NA | NA | 4 | 0.25 | 2 | 0.5 | 2 | 0 | NA | NA | NA | NA | 4 | 0.25 |
| X_9331643 | 1 | 1 | 2 | 1 | NA | NA | NA | NA | 3 | 1 | 1 | 1 | 2 | 1 | NA | NA | NA | NA | 3 | 1 |
| X_9331651 | 1 | 1 | 2 | 1 | NA | NA | NA | NA | 3 | 1 | 2 | 1 | 2 | 1 | NA | NA | NA | NA | 4 | 1 |
| X_9331652 | 1 | 1 | 3 | 1 | NA | NA | NA | NA | 4 | 1 | 1 | 1 | 2 | 1 | 1 | 1 | NA | NA | 4 | 1 |
| X_9331661 | 1 | 1 | 3 | 1 | NA | NA | NA | NA | 4 | 1 | 2 | 1 | 2 | 1 | 2 | 1 | NA | NA | 6 | 1 |
| X_9331662 | 1 | 0 | 2 | 0.5 | NA | NA | NA | NA | 3 | 0.3333 | 2 | 0.5 | 2 | 0 | 2 | 0.5 | NA | NA | 6 | 0.3333 |
| X_9331682 | 1 | 1 | 2 | 1 | NA | NA | NA | NA | 3 | 1 | 2 | 1 | 2 | 1 | 2 | 1 | NA | NA | 6 | 1 |
| X_9331686 | 1 | 1 | 2 | 1 | NA | NA | NA | NA | 3 | 1 | 2 | 1 | 2 | 1 | 2 | 1 | NA | NA | 6 | 1 |
| X_9331703 | 1 | 1 | 3 | 1 | NA | NA | NA | NA | 4 | 1 | 2 | 1 | 2 | 1 | 2 | 1 | NA | NA | 6 | 1 |
| X_9331707 | 1 | 1 | 3 | 0.3333 | NA | NA | NA | NA | 4 | 0.5 | 2 | 0.5 | 1 | 0 | 2 | 0.5 | NA | NA | 5 | 0.4 |
| X_9331742 | 1 | 1 | 3 | 0.3333 | NA | NA | NA | NA | 4 | 0.5 | 1 | 1 | 3 | 0 | 2 | 0 | NA | NA | 6 | 0.1667 |
| X_9346064 | 14 | 0 | 7 | 0 | 5 | 0.2 | 5 | 0.2 | 31 | 0.0645 | 8 | 0 | 11 | 0 | 5 | 0 | 14 | 0.1429 | 38 | 0.0526 |
| X_9348626 | NA | NA | NA | NA | 1 | 1 | NA | NA | 1 | 1 | NA | NA | NA | NA | 2 | 1 | 1 | 1 | 3 | 1 |
| X_9358364 | 42 | 0.0476 | 61 | 0.0984 | 40 | 0.025 | 61 | 0.082 | 204 | 0.0686 | 32 | 0.0312 | 48 | 0.0625 | 61 | 0.0984 | 60 | 0.0667 | 201 | 0.0697 |
| X_9368027 | 4 | 0.5 | NA | NA | 2 | 0.5 | 1 | 1 | 7 | 0.5714 | 1 | 1 | 4 | 0.75 | 2 | 1 | 1 | 1 | 8 | 0.875 |
| X_9368053 | 4 | 0.5 | NA | NA | 2 | 1 | 4 | 0.75 | 10 | 0.7 | 2 | 1 | 4 | 0.75 | 3 | 1 | 2 | 0.5 | 11 | 0.8182 |
| X_9368057 | 5 | 0.8 | NA | NA | 3 | 0.3333 | 4 | 1 | 12 | 0.75 | 2 | 0.5 | 4 | 0.75 | 3 | 1 | 2 | 0.5 | 11 | 0.7273 |
| X_9368083 | 4 | 0.75 | NA | NA | 4 | 0.75 | 4 | 1 | 12 | 0.8333 | 2 | 1 | 4 | 0.5 | 3 | 0.3333 | 2 | 1 | 11 | 0.6364 |
| X_9368085 | 3 | 0.3333 | NA | NA | 4 | 0.75 | 4 | 0.25 | 11 | 0.4545 | 2 | 1 | 2 | 0 | 3 | 0.6667 | 2 | 0 | 9 | 0.4444 |
| X_9368098 | 4 | 0.75 | NA | NA | 4 | 0.75 | 3 | 0.6667 | 11 | 0.7273 | 2 | 1 | 4 | 0.5 | 2 | 0 | 2 | 1 | 10 | 0.6 |
| X_9368102 | 4 | 1 | NA | NA | 5 | 1 | 3 | 1 | 12 | 1 | 2 | 1 | 4 | 1 | 3 | 1 | 1 | 1 | 10 | 1 |
| X_9368105 | 4 | 1 | NA | NA | 5 | 1 | 4 | 1 | 13 | 1 | 1 | 1 | 2 | 1 | 3 | 0.6667 | 2 | 1 | 8 | 0.875 |
| X_9368105 | 4 | 0.25 | NA | NA | 5 | 0.2 | 4 | 0.5 | 13 | 0.3077 | 2 | 1 | 2 | 0.5 | 3 | 0.3333 | 2 | 0 | 9 | 0.4444 |
| X_9368131 | 5 | 0.8 | NA | NA | 3 | 1 | 4 | 0.5 | 12 | 0.75 | 2 | 0.5 | 3 | 0.6667 | 3 | 0.6667 | 2 | 1 | 10 | 0.7 |
| X_9368135 | 2 | 0.5 | NA | NA | 5 | 0.4 | 4 | 0.75 | 11 | 0.5455 | 2 | 1 | 3 | 0 | 2 | 0.5 | 1 | 1 | 8 | 0.5 |
| X_9368155 | 5 | 0.2 | NA | NA | 4 | 0.25 | 5 | 0.2 | 14 | 0.2143 | 2 | 0 | 4 | 0 | 2 | 0.5 | 2 | 0 | 10 | 0.1 |
| X_9368163 | 6 | 0.5 | NA | NA | 5 | 0.6 | 5 | 1 | 16 | 0.6875 | 2 | 0.5 | 4 | 0.75 | 3 | 0.3333 | 2 | 0 | 11 | 0.4545 |
| X_9368164 | 5 | 0.4 | NA | NA | 5 | 0.4 | 4 | 0.25 | 14 | 0.3571 | 2 | 1 | 4 | 0.5 | 2 | 0.5 | 2 | 0.5 | 10 | 0.6 |
| X_9368166 | 5 | 0.4 | NA | NA | 5 | 0 | 5 | 0.6 | 15 | 0.3333 | 2 | 0 | 4 | 0.25 | 2 | 0 | 2 | 0.5 | 10 | 0.2 |
| X_9368175 | 4 | 1 | 1 | 1 | 4 | 1 | 4 | 1 | 13 | 1 | 1 | 1 | 4 | 1 | 2 | 1 | 3 | 1 | 10 | 1 |
| X_9368180 | 5 | 0.4 | 1 | 1 | 5 | 0 | 5 | 0.2 | 16 | 0.25 | 2 | 0.5 | 4 | 0 | 3 | 0 | 2 | 0 | 11 | 0.0909 |
| X_9368183 | 5 | 0.8 | 1 | 1 | 2 | 0.5 | 4 | 1 | 12 | 0.8333 | 2 | 1 | 2 | 0.5 | 3 | 1 | 2 | 0.5 | 9 | 0.7778 |
| X_9368190 | 6 | 0.3333 | 1 | 0 | 5 | 0.4 | 5 | 0.2 | 17 | 0.2941 | 2 | 1 | 4 | 0.75 | 4 | 0.5 | 3 | 0.6667 | 13 | 0.6923 |
| X_9368196 | 4 | 1 | 1 | 1 | 5 | 0.8 | 5 | 0.8 | 15 | 0.8667 | 2 | 1 | 4 | 1 | 4 | 0.75 | 4 | 0.5 | 14 | 0.7857 |
| X_9368208 | 1 | 0 | 1 | 1 | 1 | 0 | 6 | 0 | 9 | 0.1111 | 1 | 0 | 1 | 0 | 2 | 0 | 2 | 0.5 | 6 | 0.1667 |

| | | | | | | | | | | | | | | | | | | | | |
|---|---|---|---|---|---|---|---|---|---|---|---|---|---|---|---|---|---|---|---|---|
| X_9368641 | NA | NA | NA | NA | 1 | 1 | 1 | 0 | 2 | 0.5 | NA | NA | 2 | 1 | 3 | 0 | 1 | 1 | 6 | 0.5 |
| X_9368645 | NA | NA | NA | NA | 1 | 1 | 1 | 1 | 2 | 1 | NA | NA | 2 | 0.5 | 3 | 0.3333 | 1 | 1 | 6 | 0.5 |
| X_9376893 | 1 | 1 | 2 | 1 | 1 | 1 | NA | NA | 4 | 1 | 1 | 0 | NA | NA | 1 | 0 | 1 | 1 | 3 | 0.3333 |
| X_9376942 | 2 | 0 | 3 | 0.3333 | 1 | 1 | NA | NA | 6 | 0.3333 | 1 | 0 | 3 | 0.3333 | 1 | 0 | 1 | 0 | 6 | 0.1667 |
| X_9376975 | 3 | 0.6667 | 4 | 0.5 | 3 | 0.6667 | 2 | 0.5 | 12 | 0.5833 | 1 | 1 | 3 | 1 | 1 | 0 | 1 | 1 | 6 | 0.8333 |
| X_9376984 | 4 | 0 | 3 | 0 | 3 | 0 | 2 | 0.5 | 12 | 0.0833 | 1 | 0 | 3 | 0 | 1 | 1 | NA | NA | 5 | 0.2 |
| X_9376987 | 4 | 0 | 3 | 0 | 2 | 0.5 | 2 | 0 | 11 | 0.0909 | 1 | 0 | 2 | 0.5 | 2 | 0 | 1 | 0 | 6 | 0.1667 |
| X_9377035 | 3 | 0.6667 | 3 | 0.6667 | 4 | 0.25 | 1 | 0 | 11 | 0.4545 | 2 | 0.5 | 3 | 0.6667 | 3 | 0 | 2 | 0.5 | 10 | 0.4 |
| X_9377038 | 3 | 0 | 3 | 0.6667 | 4 | 1 | 3 | 0.3333 | 13 | 0.5385 | 2 | 0.5 | 4 | 0.25 | 3 | 0.6667 | 1 | 1 | 10 | 0.5 |
| X_9377040 | 2 | 0.5 | NA | NA | 4 | 0.5 | NA | NA | 6 | 0.5 | 1 | 1 | 3 | 0.6667 | 3 | 1 | 1 | 1 | 8 | 0.875 |
| X_9377050 | 5 | 0.2 | 3 | 0.3333 | 3 | 0 | 1 | 0 | 12 | 0.1667 | 2 | 0 | 2 | 0.5 | 3 | 0 | 2 | 0 | 9 | 0.1111 |
| X_9377054 | 3 | 0 | 2 | 0 | 2 | 0.5 | 1 | 0 | 8 | 0.125 | 1 | 0 | NA | NA | 3 | 0.3333 | 2 | 0 | 6 | 0.1667 |
| X_9377058 | 3 | 1 | 4 | 1 | 3 | 0.6667 | 2 | 1 | 12 | 0.9167 | 2 | 1 | 3 | 1 | 2 | 0.5 | 2 | 0.5 | 9 | 0.7778 |
| X_9377063 | 5 | 0.8 | 4 | 0.25 | 4 | 0.75 | 3 | 0.6667 | 16 | 0.625 | 2 | 0.5 | 1 | 0 | 3 | 0.3333 | 2 | 0.5 | 8 | 0.375 |
| X_9377068 | 5 | 1 | 4 | 1 | 4 | 1 | 3 | 1 | 16 | 1 | 2 | 0 | 3 | 1 | 4 | 1 | 2 | 1 | 11 | 0.8182 |
| X_9377068 | 5 | 0.8 | 3 | 0.3333 | 3 | 0.6667 | 3 | 1 | 14 | 0.7143 | 2 | 1 | 4 | 0.75 | 4 | 0.25 | 3 | 1 | 13 | 0.6923 |
| X_9377078 | 4 | 0.5 | 4 | 0 | 3 | 0 | 3 | 0.6667 | 14 | 0.2857 | NA | NA | 3 | 0.3333 | 3 | 0.3333 | 3 | 0 | 9 | 0.2222 |
| X_9377079 | 5 | 0.6 | 4 | 0.25 | 3 | 1 | 4 | 0.75 | 16 | 0.625 | 2 | 0.5 | 3 | 0.6667 | 4 | 0.5 | 2 | 1 | 11 | 0.6364 |
| X_9377085 | 5 | 1 | 4 | 0.75 | 4 | 1 | 3 | 0.6667 | 16 | 0.875 | 2 | 1 | 3 | 0.6667 | 2 | 1 | 3 | 1 | 10 | 0.9 |
| X_9377087 | 3 | 0.6667 | 3 | 1 | 5 | 1 | 4 | 1 | 15 | 0.9333 | 1 | 1 | 3 | 1 | 4 | 1 | 1 | 1 | 9 | 1 |
| X_9377090 | 4 | 0 | 2 | 1 | 3 | 0.6667 | 3 | 0.6667 | 12 | 0.5 | 1 | 1 | 4 | 0.25 | 5 | 0.4 | 2 | 0.5 | 12 | 0.4167 |
| X_9377110 | 6 | 1 | 4 | 1 | 5 | 1 | 3 | 1 | 18 | 1 | 2 | 1 | 4 | 1 | 6 | 1 | 3 | 1 | 15 | 1 |
| X_9377111 | 5 | 1 | 4 | 1 | 5 | 1 | 3 | 0.6667 | 17 | 0.9412 | 3 | 1 | 4 | 0.75 | 4 | 1 | 2 | 1 | 13 | 0.9231 |
| X_9377117 | 4 | 1 | 3 | 1 | 4 | 1 | 4 | 1 | 15 | 1 | 2 | 1 | 4 | 0.75 | 6 | 1 | 3 | 1 | 15 | 0.9333 |
| X_9377259 | 5 | 0 | 1 | 0 | 2 | 0.5 | 3 | 0.3333 | 11 | 0.1818 | 3 | 0.3333 | 4 | 0 | 4 | 0 | 3 | 0.3333 | 14 | 0.1429 |
| X_9377268 | 6 | 0.1667 | 2 | 0.5 | 2 | 0 | 3 | 0.3333 | 13 | 0.2308 | 3 | 0 | 4 | 0 | 5 | 0.4 | 3 | 0 | 15 | 0.1333 |
| X_9383678 | 1 | 1 | 1 | 1 | 2 | 0 | NA | NA | 4 | 0.5 | NA | NA | 1 | 1 | 1 | 0 | 2 | 0 | 4 | 0.25 |
| X_9383873 | 1 | 0 | NA | NA | 2 | 0.5 | NA | NA | 3 | 0.3333 | NA | NA | 1 | 1 | NA | NA | 1 | 0 | 2 | 0.5 |
| X_9383895 | 1 | 1 | 2 | 1 | NA | NA | NA | NA | 3 | 1 | 1 | 1 | 2 | 1 | NA | NA | 3 | 1 | 6 | 1 |
| X_9383897 | 1 | 1 | 2 | 1 | NA | NA | NA | NA | 3 | 1 | 1 | 1 | 2 | 1 | NA | NA | 3 | 1 | 6 | 1 |
| X_9383919 | 1 | 1 | 2 | 1 | NA | NA | NA | NA | 3 | 1 | 1 | 1 | 2 | 1 | NA | NA | 3 | 0.6667 | 6 | 0.8333 |
| X_9383927 | 1 | 0 | 2 | 0.5 | NA | NA | NA | NA | 3 | 0.3333 | NA | NA | 2 | 1 | NA | NA | 2 | 1 | 4 | 1 |
| X_9383928 | 1 | 1 | 2 | 0.5 | NA | NA | NA | NA | 3 | 0.6667 | NA | NA | 2 | 0.5 | NA | NA | 3 | 0 | 5 | 0.2 |
| X_9383938 | 2 | 0.5 | 2 | 1 | NA | NA | NA | NA | 4 | 0.75 | 1 | 1 | 2 | 0.5 | NA | NA | 1 | 1 | 4 | 0.75 |
| X_9383939 | 2 | 0.5 | 2 | 1 | NA | NA | NA | NA | 4 | 0.75 | 1 | 1 | 2 | 1 | NA | NA | 2 | 0.5 | 5 | 0.8 |
| X_9383943 | 1 | 1 | 1 | 0 | NA | NA | NA | NA | 2 | 0.5 | 1 | 0 | 1 | 0 | NA | NA | 2 | 0.5 | 4 | 0.25 |
| X_9383949 | NA | NA | 2 | 1 | NA | NA | NA | NA | 2 | 1 | 1 | 1 | 2 | 0.5 | NA | NA | 3 | 1 | 6 | 0.8333 |
| X_9383966 | 1 | 0 | 1 | 1 | NA | NA | NA | NA | 2 | 0.5 | 1 | 1 | 1 | 1 | NA | NA | 3 | 1 | 5 | 1 |
| X_9383968 | 1 | 1 | 1 | 1 | NA | NA | NA | NA | 2 | 1 | 1 | 1 | 1 | 1 | NA | NA | 2 | 0 | 4 | 0.5 |
| X_9383971 | 1 | 0 | 2 | 1 | NA | NA | NA | NA | 3 | 0.6667 | 1 | 0 | 1 | 1 | NA | NA | 2 | 0 | 4 | 0.25 |
| X_9384064 | 1 | 1 | 1 | 0 | NA | NA | NA | NA | 2 | 0.5 | NA | NA | 1 | 1 | NA | NA | 1 | 0 | 2 | 0.5 |
| X_9384113 | 1 | 1 | 2 | 0.5 | NA | NA | NA | NA | 3 | 0.6667 | NA | NA | 1 | 1 | NA | NA | 1 | 1 | 2 | 1 |
| X_9409308 | 35 | 0.0571 | 40 | 0.1 | 25 | 0.04 | 29 | 0.0345 | 129 | 0.062 | 18 | 0.0556 | 40 | 0.175 | 26 | 0 | 27 | 0.0741 | 111 | 0.0901 |
| X_9409529 | 18 | 0.0556 | 22 | 0.2273 | 18 | 0 | 19 | 0.1053 | 77 | 0.1039 | 11 | 0.2727 | 29 | 0.0345 | 24 | 0.125 | 21 | 0.0952 | 85 | 0.1059 |
| X_9409552 | 19 | 0.3684 | 16 | 0.1875 | 15 | 0.7333 | 17 | 0.7059 | 67 | 0.4925 | 8 | 0.5 | 26 | 0.5769 | 20 | 0.35 | 18 | 0.4444 | 72 | 0.4722 |
| X_9420935 | 5 | 0.2 | 9 | 0.1111 | 7 | 0 | 6 | 0.1667 | 27 | 0.1111 | 5 | 0 | 11 | 0.1818 | 8 | 0.125 | 4 | 0 | 28 | 0.1071 |
| X_9420975 | 4 | 0.5 | 6 | 0 | 2 | 0 | 6 | 0 | 18 | 0.1111 | 5 | 0 | 8 | 0.125 | 8 | 0 | 2 | 0.5 | 23 | 0.087 |
| X_9446440 | 1 | 0 | 1 | 1 | NA | NA | 1 | 1 | 3 | 0.6667 | 1 | 1 | NA | NA | NA | NA | NA | NA | 1 | 1 |
| X_9446449 | 1 | 0 | 1 | 0 | NA | NA | 1 | 1 | 3 | 0.3333 | 1 | 1 | 1 | 1 | NA | NA | NA | NA | 2 | 1 |
| X_9446450 | 2 | 0 | 1 | 1 | NA | NA | 1 | 1 | 4 | 0.5 | 1 | 0 | 1 | 1 | NA | NA | NA | NA | 2 | 0.5 |
| X_9446465 | 2 | 0 | NA | NA | NA | NA | 1 | 1 | 3 | 0.3333 | NA | NA | 1 | 1 | NA | NA | NA | NA | 1 | 1 |
| X_9446577 | 2 | 0.5 | 1 | 1 | NA | NA | NA | NA | 3 | 0.6667 | NA | NA | 1 | 1 | NA | NA | NA | NA | 1 | 1 |
| X_9446579 | NA | NA | 1 | 1 | NA | NA | NA | NA | 1 | 1 | NA | NA | 1 | 1 | NA | NA | NA | NA | 1 | 1 |
| X_9446590 | 1 | 1 | 1 | 1 | NA | NA | NA | NA | 2 | 1 | NA | NA | 1 | 1 | NA | NA | NA | NA | 1 | 1 |
| X_9446618 | 1 | 1 | NA | NA | NA | NA | NA | NA | 1 | 1 | NA | NA | 1 | 1 | NA | NA | NA | NA | 1 | 1 |
| X_9446623 | 1 | 1 | NA | NA | NA | NA | NA | NA | 1 | 1 | NA | NA | 1 | 1 | NA | NA | NA | NA | 1 | 1 |
| X_9468297 | 1 | 0 | 1 | 0 | 3 | 0.3333 | 3 | 0 | 8 | 0.125 | 4 | 0.5 | 9 | 0.2222 | 3 | 0 | 1 | 0 | 17 | 0.2353 |
| X_9468379 | 1 | 0 | 2 | 0.5 | 2 | 0 | 1 | 0 | 6 | 0.1667 | 3 | 0 | 6 | 0.5 | 3 | 0.6667 | 1 | 1 | 13 | 0.4615 |
| X_9468382 | 1 | 0 | 1 | 0 | 2 | 0.5 | 1 | 0 | 5 | 0.2 | 2 | 0 | 6 | 0.1667 | 3 | 0.3333 | NA | NA | 11 | 0.1818 |
| X_9468387 | 1 | 1 | 2 | 0 | 2 | 0.5 | 1 | 1 | 6 | 0.5 | 2 | 0 | 2 | 0 | 3 | 0.3333 | 1 | 0 | 8 | 0.125 |
| X_9468392 | 1 | 1 | 2 | 0 | 3 | 0.6667 | 1 | 1 | 7 | 0.5714 | 2 | 1 | 5 | 0.4 | 3 | 0.3333 | NA | NA | 10 | 0.5 |
| X_9497496 | NA | NA | 1 | 0 | 1 | 1 | 2 | 0 | 4 | 0.25 | 3 | 0.6667 | 1 | 0 | 1 | 0 | NA | NA | 5 | 0.4 |
| X_9497501 | NA | NA | 2 | 0 | 3 | 0.3333 | 2 | 0 | 7 | 0.1429 | 3 | 0.6667 | 1 | 0 | 1 | 0 | NA | NA | 5 | 0.4 |
| X_9497504 | NA | NA | 2 | 0 | 3 | 0.3333 | 2 | 0 | 7 | 0.1429 | 3 | 0.3333 | 1 | 0 | 1 | 0 | NA | NA | 5 | 0.2 |
| X_9497505 | NA | NA | 2 | 0.5 | 1 | 1 | NA | NA | 3 | 0.6667 | 1 | 0 | 1 | 1 | 1 | 1 | NA | NA | 3 | 0.6667 |

| | | | | | | | | | | | | | | | | | | | | |
|---|---|---|---|---|---|---|---|---|---|---|---|---|---|---|---|---|---|---|---|---|
| X_9497519 | NA | NA | 2 | 0 | 2 | 1 | 2 | 0 | 6 | 0.3333 | 3 | 0.3333 | 1 | 0 | 2 | 0 | NA | NA | 6 | 0.1667 |
| X_9497903 | 7 | 0.1429 | 7 | 0.2857 | 3 | 0.6667 | 5 | 0.4 | 22 | 0.3182 | 4 | 0 | 4 | 0.25 | 3 | 0.3333 | 7 | 0.5714 | 18 | 0.3333 |
| X_9497904 | 6 | 0.1667 | 5 | 0 | 4 | 0 | 4 | 0 | 19 | 0.0526 | 3 | 0 | 3 | 0 | 3 | 0 | 6 | 0.1667 | 15 | 0.0667 |
| X_9497905 | 7 | 0.4286 | 4 | 0.25 | 4 | 0 | 4 | 0.75 | 19 | 0.3684 | 3 | 0.3333 | 2 | 0 | 3 | 0 | 7 | 0.1429 | 15 | 0.1333 |
| X_9497916 | 6 | 0.3333 | 6 | 0.3333 | 3 | 0.3333 | 4 | 0.25 | 19 | 0.3158 | 4 | 0 | 2 | 0.5 | 1 | 0 | 4 | 0.25 | 11 | 0.1818 |
| X_9561718 | 2 | 0 | NA | NA | 2 | 0.5 | 2 | 0 | 6 | 0.1667 | 1 | 0 | NA | NA | NA | NA | 2 | 0.5 | 3 | 0.3333 |
| X_9561748 | 1 | 0 | 1 | 0 | 2 | 0 | 2 | 0.5 | 6 | 0.1667 | 1 | 0 | NA | NA | 1 | 1 | 2 | 0.5 | 4 | 0.5 |
| X_9561792 | 2 | 0 | 1 | 1 | 2 | 1 | 2 | 1 | 7 | 0.7143 | NA | NA | NA | NA | 1 | 0 | 2 | 1 | 3 | 0.6667 |
| X_9576068 | NA | NA | 1 | 0 | NA | NA | 2 | 0.5 | 3 | 0.3333 | 1 | 0 | 2 | 0.5 | NA | NA | 1 | 0 | 4 | 0.25 |
| X_9576174 | NA | NA | 1 | 1 | NA | NA | 2 | 0 | 3 | 0.3333 | 1 | 1 | 2 | 0 | NA | NA | 1 | 0 | 4 | 0.25 |
| X_9576184 | NA | NA | 1 | 1 | NA | NA | 2 | 0.5 | 3 | 0.6667 | NA | NA | 2 | 0.5 | NA | NA | 1 | 1 | 3 | 0.6667 |
| X_9576189 | NA | NA | 1 | 1 | NA | NA | 2 | 0.5 | 3 | 0.6667 | 1 | 0 | 3 | 0.6667 | NA | NA | 1 | 0 | 5 | 0.4 |
| X_9576308 | NA | NA | 1 | 1 | NA | NA | 2 | 0 | 3 | 0.3333 | 1 | 0 | 3 | 0 | NA | NA | 1 | 1 | 5 | 0.2 |
| X_9576506 | 1 | 1 | 1 | 1 | 1 | 0 | 2 | 1 | 5 | 0.8 | 2 | 1 | 2 | 0 | NA | NA | NA | NA | 4 | 0.5 |
| X_9576507 | 1 | 1 | 1 | 0 | 1 | 0 | 2 | 0 | 5 | 0.2 | 1 | 1 | 3 | 0 | NA | NA | NA | NA | 4 | 0.25 |
| X_9576509 | 1 | 1 | NA | NA | 1 | 0 | 1 | 0 | 3 | 0.3333 | 1 | 0 | 2 | 0.5 | NA | NA | NA | NA | 3 | 0.3333 |
| X_9576510 | NA | NA | 1 | 1 | 1 | 1 | 1 | 1 | 3 | 1 | 1 | 1 | 2 | 1 | NA | NA | NA | NA | 3 | 1 |
| X_9576522 | NA | NA | 2 | 0 | 1 | 1 | 1 | 0 | 4 | 0.25 | 3 | 0.3333 | 2 | 0.5 | 1 | 1 | NA | NA | 6 | 0.5 |
| X_9736194 | 1 | 1 | 1 | 1 | NA | NA | NA | NA | 2 | 1 | NA | NA | NA | NA | 1 | 1 | NA | NA | 1 | 1 |
| X_9736212 | 1 | 1 | 1 | 1 | NA | NA | NA | NA | 2 | 1 | NA | NA | NA | NA | 1 | 1 | NA | NA | 1 | 1 |
| X_9736238 | 1 | 1 | 1 | 1 | NA | NA | NA | NA | 2 | 1 | NA | NA | NA | NA | 1 | 1 | NA | NA | 1 | 1 |
| X_9736242 | 1 | 1 | 1 | 1 | NA | NA | NA | NA | 2 | 1 | NA | NA | NA | NA | 1 | 1 | NA | NA | 1 | 1 |
| X_9736250 | 1 | 1 | 1 | 1 | NA | NA | NA | NA | 2 | 1 | NA | NA | NA | NA | 1 | 1 | NA | NA | 1 | 1 |
| X_9736362 | 1 | 1 | 1 | 0 | NA | NA | NA | NA | 2 | 0.5 | NA | NA | NA | NA | 1 | 1 | NA | NA | 1 | 1 |
| X_9736369 | 1 | 1 | 1 | 1 | NA | NA | NA | NA | 2 | 1 | NA | NA | NA | NA | 1 | 1 | NA | NA | 1 | 1 |
| X_9736376 | 1 | 1 | 1 | 1 | NA | NA | NA | NA | 2 | 1 | NA | NA | NA | NA | 1 | 1 | NA | NA | 1 | 1 |
| X_9736514 | 2 | 0 | 2 | 0.5 | 1 | 0 | 1 | 0 | 6 | 0.1667 | NA | NA | NA | NA | 1 | 0 | 3 | 0.3333 | 4 | 0.25 |
| X_9760159 | 2 | 0.5 | 1 | 1 | 2 | 0 | NA | NA | 5 | 0.4 | NA | NA | NA | NA | 1 | 1 | 3 | 0 | 4 | 0.25 |
| X_9760163 | 2 | 0 | 1 | 0 | 2 | 1 | NA | NA | 5 | 0.4 | NA | NA | 1 | 1 | 1 | 1 | 2 | 0.5 | 4 | 0.75 |
| X_9760178 | 2 | 1 | 1 | 1 | 1 | 0 | NA | NA | 4 | 0.75 | 1 | 0 | NA | NA | NA | NA | 3 | 0.6667 | 4 | 0.5 |
| X_9760187 | 2 | 0.5 | 2 | 0 | 3 | 0 | NA | NA | 7 | 0.1429 | 2 | 0.5 | 1 | 1 | NA | NA | 3 | 0 | 6 | 0.3333 |
| X_9760200 | 3 | 0.3333 | 1 | 0 | 2 | 0 | NA | NA | 6 | 0.1667 | 2 | 0.5 | 1 | 1 | NA | NA | 4 | 0.25 | 7 | 0.4286 |
| X_9798862 | 2 | 0.5 | 3 | 0.6667 | 1 | 1 | NA | NA | 6 | 0.6667 | NA | NA | 3 | 1 | 1 | 1 | 2 | 1 | 6 | 1 |
| X_9798874 | 2 | 1 | 4 | 0.75 | 3 | 1 | 2 | 0.5 | 11 | 0.8182 | 1 | 1 | 3 | 0.6667 | 1 | 1 | 2 | 0.5 | 7 | 0.7143 |
| X_9798885 | 2 | 0 | 4 | 0.25 | 3 | 0.3333 | 2 | 0.5 | 11 | 0.2727 | NA | NA | 4 | 0.25 | 1 | 0 | 1 | 0 | 6 | 0.1667 |
| X_9798951 | 2 | 0.5 | 5 | 0 | 3 | 0 | 2 | 0 | 12 | 0.0833 | 2 | 0 | 3 | 0 | 1 | 0 | 3 | 0.3333 | 9 | 0.1111 |
| X_9798984 | 1 | 0 | 5 | 0 | 3 | 0.3333 | 2 | 0 | 11 | 0.0909 | 3 | 0.3333 | 4 | 0.25 | 2 | 0 | 3 | 0.3333 | 12 | 0.25 |
| X_9799085 | 2 | 0 | 5 | 0.4 | 2 | 0 | 1 | 0 | 10 | 0.2 | 4 | 0.25 | 3 | 0.3333 | 2 | 0 | 3 | 0.6667 | 12 | 0.3333 |
| X_9799090 | 2 | 0.5 | 5 | 0 | 4 | 0 | 1 | 0 | 12 | 0.0833 | 4 | 0 | 5 | 0.4 | 2 | 0 | 3 | 0 | 14 | 0.1429 |
| X_9799096 | 2 | 0.5 | 4 | 0.25 | 1 | 0 | 1 | 0 | 8 | 0.25 | 3 | 0.6667 | 3 | 0 | 2 | 0 | 3 | 0.3333 | 11 | 0.2727 |
| X_9799106 | 2 | 0.5 | 5 | 0 | 4 | 1 | 1 | 0 | 12 | 0.4167 | 4 | 0.5 | 4 | 0.25 | 4 | 0.5 | 3 | 0 | 15 | 0.3333 |
| X_9799150 | 2 | 0 | 6 | 0.1667 | 4 | 0.25 | 1 | 0 | 13 | 0.1538 | 4 | 0.25 | 5 | 0 | 4 | 0.25 | 3 | 0 | 16 | 0.125 |
| X_9799170 | 2 | 0 | 5 | 0 | 5 | 0 | 1 | 1 | 13 | 0.0769 | 3 | 0 | 5 | 0 | 4 | 0 | 2 | 0.5 | 14 | 0.0714 |
| X_9799247 | 2 | 0 | 8 | 0.125 | 3 | 0 | 1 | 0 | 14 | 0.0714 | 1 | 0 | 6 | 0.1667 | 2 | 0 | 2 | 0 | 11 | 0.0909 |
| X_9810968 | NA | NA | 1 | 1 | NA | NA | NA | NA | 1 | 1 | NA | NA | NA | NA | 1 | 1 | NA | NA | 1 | 1 |
| X_9811676 | 1 | 1 | 2 | 0.5 | NA | NA | NA | NA | 3 | 0.6667 | 1 | 1 | NA | NA | 2 | 0.5 | NA | NA | 3 | 0.6667 |
| X_9811686 | NA | NA | 2 | 1 | NA | NA | NA | NA | 2 | 1 | NA | NA | NA | NA | 1 | 1 | NA | NA | 1 | 1 |
| X_9811688 | 1 | 1 | 2 | 0.5 | 1 | 1 | NA | NA | 4 | 0.75 | 1 | 1 | NA | NA | 2 | 0.5 | 1 | 0 | 4 | 0.5 |
| X_9811732 | 1 | 1 | 1 | 0 | 1 | 0 | NA | NA | 3 | 0.3333 | NA | NA | NA | NA | 3 | 0.3333 | 1 | 0 | 4 | 0.25 |
| X_9817981 | NA | NA | NA | NA | 1 | 1 | NA | NA | 1 | 1 | 1 | 0 | NA | NA | 1 | 1 | NA | NA | 2 | 0.5 |
| X_9818015 | NA | NA | NA | NA | 1 | 1 | NA | NA | 1 | 1 | 1 | 1 | 1 | 0 | 1 | 1 | NA | NA | 3 | 0.6667 |
| X_9819086 | NA | NA | 1 | 0 | 1 | 1 | 2 | 0 | 4 | 0.25 | 1 | 0 | 1 | 0 | 1 | 1 | NA | NA | 3 | 0.3333 |
| X_9819101 | NA | NA | 1 | 1 | 3 | 0.6667 | 2 | 1 | 6 | 0.8333 | 1 | 1 | 1 | 1 | 2 | 1 | NA | NA | 4 | 1 |
| X_9819106 | NA | NA | 1 | 0 | 3 | 1 | 2 | 1 | 6 | 0.8333 | 1 | 1 | 1 | 0 | 2 | 0 | NA | NA | 4 | 0.25 |
| X_9819139 | NA | NA | 2 | 0 | 3 | 0.3333 | 2 | 0.5 | 7 | 0.2857 | NA | NA | NA | NA | 2 | 0.5 | NA | NA | 2 | 0.5 |
| X_9819140 | NA | NA | 2 | 0 | 2 | 0.5 | 3 | 0.3333 | 7 | 0.2857 | 1 | 1 | NA | NA | 3 | 0.6667 | NA | NA | 4 | 0.75 |
| X_9819177 | NA | NA | 1 | 0 | 4 | 0.25 | 4 | 0 | 9 | 0.1111 | 1 | 0 | 1 | 0 | 4 | 0.25 | NA | NA | 6 | 0.1667 |
| X_9819230 | NA | NA | 3 | 1 | 4 | 0.5 | 4 | 0.25 | 11 | 0.5455 | 3 | 0.6667 | 1 | 0 | 3 | 0.3333 | NA | NA | 7 | 0.4286 |
| X_9850938 | 11 | 0.1818 | 15 | 0.1333 | 14 | 0.2857 | 12 | 0.25 | 52 | 0.2115 | 9 | 0.2222 | 9 | 0 | 17 | 0.2941 | 15 | 0.0667 | 50 | 0.16 |
| X_9850987 | 31 | 0.2903 | 30 | 0.2333 | 32 | 0.2812 | 25 | 0.16 | 118 | 0.2458 | 19 | 0.5789 | 19 | 0.5789 | 32 | 0.375 | 35 | 0.4857 | 105 | 0.4857 |
| X_9850992 | 27 | 0.1852 | 23 | 0.087 | 30 | 0.2667 | 22 | 0.0455 | 102 | 0.1569 | 12 | 0.25 | 20 | 0.15 | 34 | 0.1765 | 34 | 0.1471 | 100 | 0.17 |
| X_9850995 | 22 | 0.1364 | 23 | 0.1304 | 21 | 0.1429 | 20 | 0.1 | 86 | 0.1279 | 15 | 0.2 | 19 | 0 | 23 | 0.1739 | 29 | 0.2069 | 86 | 0.1512 |
| X_9851000 | 29 | 0.3793 | 26 | 0.6538 | 31 | 0.2903 | 24 | 0.2917 | 110 | 0.4 | 18 | 0.2222 | 22 | 0.3636 | 28 | 0.25 | 36 | 0.3889 | 104 | 0.3173 |
| X_9851006 | 29 | 0.2414 | 28 | 0.2143 | 33 | 0.3333 | 29 | 0.1034 | 119 | 0.2269 | 19 | 0.2105 | 21 | 0.2381 | 33 | 0.3636 | 33 | 0.4545 | 106 | 0.3396 |
| X_9851009 | 26 | 0.0385 | 29 | 0.1379 | 26 | 0.0769 | 26 | 0.1923 | 107 | 0.1121 | 17 | 0.0588 | 20 | 0.1 | 26 | 0.1154 | 31 | 0.1613 | 94 | 0.117 |

| | | | | | | | | | | | | | | | | | | | | |
|---|---|---|---|---|---|---|---|---|---|---|---|---|---|---|---|---|---|---|---|---|
| X_9851057 | 22 | 1 | 26 | 0.9231 | 25 | 0.92 | 20 | 0.95 | 93 | 0.9462 | 16 | 0.9375 | 18 | 0.9444 | 25 | 0.84 | 27 | 0.963 | 86 | 0.9186 |
| X_9851084 | 19 | 0.2632 | 25 | 0.08 | 22 | 0.1364 | 23 | 0.2174 | 89 | 0.1685 | 13 | 0.0769 | 17 | 0.1765 | 25 | 0.12 | 25 | 0.16 | 80 | 0.1375 |
| X_9851124 | 10 | 0.2 | 19 | 0.1053 | 8 | 0 | 14 | 0.2143 | 51 | 0.1373 | 10 | 0.2 | 15 | 0.1333 | 20 | 0.1 | 18 | 0 | 63 | 0.0952 |
| X_9851166 | 9 | 0.6667 | 14 | 0.5 | 5 | 0.8 | 13 | 0.5385 | 41 | 0.5854 | 4 | 0.5 | 8 | 0.375 | 12 | 0.5 | 13 | 0.5385 | 37 | 0.4865 |
| X_9851167 | 9 | 0.8889 | 14 | 0.8571 | 3 | 1 | 12 | 1 | 38 | 0.9211 | 7 | 1 | 12 | 0.9167 | 15 | 0.9333 | 12 | 1 | 46 | 0.9565 |
| X_9886472 | 1 | 0 | 2 | 1 | NA | NA | 1 | 0 | 4 | 0.5 | NA | NA | 2 | 0.5 | NA | NA | NA | NA | 2 | 0.5 |
| X_9886513 | 1 | 1 | 3 | 0.6667 | NA | NA | 1 | 1 | 5 | 0.8 | NA | NA | 2 | 0.5 | 2 | 1 | NA | NA | 4 | 0.75 |
| X_9886528 | 1 | 0 | 2 | 1 | NA | NA | 1 | 1 | 4 | 0.75 | NA | NA | 2 | 0.5 | 2 | 1 | NA | NA | 4 | 0.75 |
| X_9886530 | 1 | 0 | 3 | 0.6667 | NA | NA | 1 | 1 | 5 | 0.6 | NA | NA | 1 | 1 | 1 | 1 | NA | NA | 2 | 1 |
| X_9886532 | 1 | 1 | 2 | 1 | NA | NA | 1 | 1 | 4 | 1 | NA | NA | 2 | 1 | 2 | 1 | NA | NA | 4 | 1 |
| X_9886533 | 1 | 1 | 1 | 1 | NA | NA | 1 | 1 | 3 | 1 | NA | NA | 2 | 1 | 2 | 1 | NA | NA | 4 | 1 |
| X_9886538 | 1 | 1 | 3 | 0 | NA | NA | 1 | 0 | 5 | 0.2 | NA | NA | 2 | 0 | 1 | 1 | NA | NA | 3 | 0.3333 |
| X_9888965 | 1 | 1 | 3 | 1 | NA | NA | 1 | 0 | 5 | 0.8 | NA | NA | 2 | 0.5 | 2 | 0.5 | NA | NA | 4 | 0.5 |
| X_9888989 | 1 | 0 | 2 | 1 | NA | NA | NA | NA | 3 | 0.6667 | NA | NA | 2 | 0 | 2 | 1 | NA | NA | 4 | 0.5 |
| X_9888990 | 1 | 1 | 1 | 0 | NA | NA | 1 | 0 | 3 | 0.3333 | NA | NA | 1 | 0 | 2 | 0.5 | NA | NA | 3 | 0.3333 |
| X_9889121 | 1 | 0 | 1 | 1 | NA | NA | 3 | 0 | 5 | 0.2 | 1 | 0 | 2 | 0 | 3 | 0.3333 | 1 | 0 | 7 | 0.1429 |
| X_9889126 | 1 | 0 | 3 | 0 | NA | NA | 3 | 0.3333 | 7 | 0.1429 | 1 | 1 | 2 | 0 | 3 | 0 | 1 | 0 | 7 | 0.1429 |
| X_9889135 | 1 | 1 | 2 | 0 | NA | NA | 2 | 0.5 | 5 | 0.4 | 1 | 1 | 1 | 1 | 3 | 0.3333 | 1 | 0 | 6 | 0.5 |
| X_9889143 | 1 | 0 | 2 | 0 | NA | NA | 3 | 0.3333 | 6 | 0.1667 | 1 | 0 | 2 | 0 | 4 | 0 | 1 | 1 | 8 | 0.125 |
| X_9889148 | 1 | 1 | 2 | 0 | NA | NA | 3 | 0 | 6 | 0.1667 | NA | NA | 1 | 0 | 2 | 0.5 | NA | NA | 3 | 0.3333 |
| X_9889187 | 1 | 1 | 3 | 0.3333 | NA | NA | 3 | 0 | 7 | 0.2857 | 1 | 0 | NA | NA | 3 | 0.3333 | 2 | 0.5 | 6 | 0.3333 |
| X_9953864 | 9 | 0.1111 | 6 | 0.1667 | 3 | 0 | 4 | 0 | 22 | 0.0909 | 5 | 0 | 1 | 0 | 5 | 0 | 4 | 0.25 | 15 | 0.0667 |
| X_9953871 | 9 | 0.2222 | 6 | 0.3333 | 4 | 0 | 6 | 0.1667 | 25 | 0.2 | 5 | 0.2 | 1 | 0 | 4 | 0.25 | 5 | 0.2 | 15 | 0.2 |
| X_9953871 | 9 | 0.1111 | 7 | 0.1429 | 6 | 0 | 6 | 0 | 28 | 0.0714 | 5 | 0 | 1 | 0 | 5 | 0 | 5 | 0.2 | 16 | 0.0625 |
| X_9955933 | 8 | 0.125 | 5 | 0.2 | 7 | 0.1429 | 7 | 0.1429 | 27 | 0.1481 | 7 | 0.1429 | 2 | 1 | 5 | 0 | 7 | 0.4286 | 21 | 0.2857 |
| X_9955936 | 8 | 0.25 | 6 | 0 | 7 | 0.1429 | 7 | 0 | 28 | 0.1071 | 6 | 0.1667 | 2 | 0 | 4 | 0.25 | 6 | 0.3333 | 18 | 0.2222 |
| X_9966699 | 8 | 0.125 | 3 | 0.3333 | 4 | 0.25 | 8 | 0.25 | 23 | 0.2174 | 4 | 0.25 | 10 | 0 | 6 | 0 | 7 | 0.2857 | 27 | 0.1111 |
| X_9966701 | 7 | 0.1429 | 3 | 0 | 5 | 0.2 | 9 | 0.1111 | 24 | 0.125 | 4 | 0 | 10 | 0.1 | 6 | 0 | 6 | 0.3333 | 26 | 0.1154 |
| X_9985988 | 18 | 0.0556 | 13 | 0.0769 | 8 | 0.125 | 7 | 0 | 46 | 0.0652 | 11 | 0.0909 | 9 | 0 | 13 | 0.0769 | 11 | 0.0909 | 44 | 0.0682 |
| X_9985992 | 15 | 0.2667 | 11 | 0.3636 | 7 | 0.4286 | 7 | 0.5714 | 40 | 0.375 | 8 | 0.375 | 10 | 0.2 | 11 | 0.4545 | 9 | 0.4444 | 38 | 0.3684 |
| X_9986002 | 13 | 0.6154 | 12 | 0.6667 | 6 | 0.8333 | 6 | 0.8333 | 37 | 0.7027 | 11 | 0.4545 | 9 | 0.6667 | 11 | 0.3636 | 9 | 1 | 40 | 0.6 |
| X_9986017 | 16 | 1 | 13 | 0.9231 | 7 | 1 | 6 | 1 | 42 | 0.9762 | 11 | 1 | 9 | 1 | 10 | 0.8 | 11 | 1 | 41 | 0.9512 |
| X_9986025 | 16 | 0.8125 | 11 | 0.7273 | 9 | 0.7778 | 7 | 0.7143 | 43 | 0.7674 | 13 | 1 | 11 | 0.9091 | 13 | 0.8462 | 10 | 0.8 | 47 | 0.8936 |
| X_9986027 | 18 | 0.2222 | 13 | 0.1538 | 9 | 0.1111 | 9 | 0 | 49 | 0.1429 | 12 | 0 | 11 | 0.1818 | 14 | 0.0714 | 9 | 0.1111 | 46 | 0.087 |
| X_9986027 | 17 | 0.6471 | 13 | 0.3846 | 7 | 0.7143 | 9 | 0.4444 | 46 | 0.5435 | 10 | 0.5 | 11 | 0.5455 | 15 | 0.5333 | 11 | 0.6364 | 47 | 0.5532 |
| X_9986028 | 14 | 0.2857 | 11 | 0.2727 | 7 | 0 | 5 | 0.4 | 37 | 0.2432 | 11 | 0.1818 | 7 | 0.2857 | 13 | 0.2308 | 8 | 0.375 | 39 | 0.2564 |
| X_9986051 | 15 | 0.6667 | 12 | 0.8333 | 8 | 1 | 8 | 0.625 | 43 | 0.7674 | 9 | 0.5556 | 7 | 0.5714 | 13 | 0.8462 | 10 | 0.8 | 39 | 0.7179 |
| X_9986076 | 18 | 0.0556 | 13 | 0.1538 | 12 | 0 | 8 | 0.125 | 51 | 0.0784 | 11 | 0.1818 | 11 | 0 | 11 | 0.0909 | 12 | 0.3333 | 45 | 0.1556 |
| X_9986122 | 21 | 0.0952 | 18 | 0 | 12 | 0 | 9 | 0.1111 | 60 | 0.05 | 12 | 0.0833 | 11 | 0.2727 | 11 | 0 | 13 | 0.1538 | 47 | 0.1277 |
| X_1001231 | NA | NA | 1 | 1 | NA | NA | 1 | 1 | 2 | 1 | NA | NA | 1 | 1 | 1 | 0 | NA | NA | 2 | 0.5 |
| X_1001331 | NA | NA | NA | NA | NA | NA | 1 | 1 | 1 | 1 | NA | NA | NA | NA | 1 | 1 | NA | NA | 1 | 1 |
| X_1001331 | NA | NA | NA | NA | NA | NA | 1 | 1 | 1 | 1 | NA | NA | NA | NA | 1 | 1 | NA | NA | 1 | 1 |
| X_1001332 | NA | NA | NA | NA | NA | NA | 1 | 1 | 1 | 1 | NA | NA | NA | NA | 1 | 1 | NA | NA | 1 | 1 |
| X_1001417 | 1 | 1 | 1 | 1 | 2 | 0 | 2 | 0 | 6 | 0.3333 | 1 | 1 | 1 | 0 | 1 | 1 | 2 | 0 | 5 | 0.4 |
| X_1002573 | 2 | 0.5 | 2 | 0 | 2 | 0.5 | 1 | 1 | 7 | 0.4286 | 2 | 0 | 2 | 1 | 5 | 0.4 | 1 | 0 | 10 | 0.4 |
| X_1002983 | 1 | 1 | NA | NA | NA | NA | NA | NA | 1 | 1 | 1 | 1 | NA | NA | NA | NA | NA | NA | 1 | 1 |
| X_1003364 | NA | NA | 6 | 0.1667 | 2 | 0.5 | NA | NA | 8 | 0.25 | NA | NA | NA | NA | 2 | 0 | 3 | 0.3333 | 5 | 0.2 |
| X_1003408 | 3 | 0 | 7 | 0 | 5 | 0.2 | NA | NA | 15 | 0.0667 | NA | NA | NA | NA | 4 | 0.25 | 3 | 0 | 7 | 0.1429 |
| X_1003409 | 2 | 0 | 4 | 0.5 | 5 | 0.2 | NA | NA | 11 | 0.2727 | NA | NA | NA | NA | 2 | 0.5 | 3 | 0.6667 | 5 | 0.6 |
| X_1003457 | 3 | 0.6667 | 1 | 0 | 1 | 1 | 1 | 0 | 6 | 0.5 | 1 | 0 | 1 | 0 | 1 | 1 | 2 | 0 | 5 | 0.2 |
| X_1003615 | 2 | 0.5 | 2 | 0 | 1 | 0 | 1 | 0 | 6 | 0.1667 | 1 | 0 | 3 | 0.3333 | 3 | 0 | 4 | 0 | 11 | 0.0909 |
| X_1003616 | 2 | 0.5 | 3 | 1 | 2 | 0 | 1 | 1 | 8 | 0.625 | 2 | 0.5 | 3 | 0.3333 | 3 | 0.6667 | 4 | 0.5 | 12 | 0.5 |
| X_1003617 | 2 | 1 | 3 | 0.6667 | 2 | 1 | 1 | 1 | 8 | 0.875 | 2 | 0.5 | 2 | 1 | 4 | 0.75 | 4 | 0.75 | 12 | 0.75 |
| X_1003619 | 2 | 0.5 | 3 | 0.6667 | NA | NA | 1 | 0 | 6 | 0.5 | 2 | 0.5 | 3 | 0.6667 | 2 | 0.5 | 4 | 1 | 11 | 0.7273 |
| X_1003619 | 2 | 1 | 3 | 0.6667 | 1 | 0 | 1 | 0 | 7 | 0.5714 | 1 | 1 | 3 | 0.3333 | 2 | 0 | 2 | 0 | 8 | 0.25 |
| X_1003619 | 2 | 0.5 | 3 | 1 | 2 | 1 | 2 | 1 | 9 | 0.8889 | 2 | 1 | 4 | 1 | 3 | 1 | 4 | 0.75 | 13 | 0.9231 |
| X_1003619 | 4 | 0.5 | 4 | 0 | 2 | 0.5 | 2 | 0 | 12 | 0.25 | 2 | 0 | 4 | 0 | 2 | 0 | 4 | 0.5 | 12 | 0.1667 |
| X_1003620 | 3 | 1 | 3 | 1 | 2 | 1 | 2 | 1 | 10 | 1 | 2 | 1 | 3 | 1 | 3 | 1 | 4 | 1 | 12 | 1 |
| X_1003632 | 2 | 1 | 4 | 0.5 | 4 | 0.5 | 2 | 0 | 12 | 0.5 | 2 | 0 | 4 | 0 | 4 | 0.25 | 4 | 0.5 | 14 | 0.2143 |
| X_1003634 | 3 | 0.6667 | 4 | 0.25 | 4 | 0.25 | 3 | 0 | 14 | 0.2857 | 2 | 0 | 5 | 0.8 | 4 | 0 | 4 | 0.5 | 15 | 0.4 |
| X_1003634 | 4 | 0.75 | 5 | 0.4 | 3 | 0.6667 | 2 | 0.5 | 14 | 0.5714 | 1 | 1 | 3 | 0.3333 | 5 | 0.8 | 4 | 0.5 | 13 | 0.6154 |
| X_1003635 | 4 | 1 | 2 | 1 | 3 | 1 | 2 | 1 | 11 | 1 | 2 | 0.5 | 3 | 1 | 3 | 1 | 4 | 1 | 12 | 0.9167 |
| X_1003635 | 4 | 0.25 | 4 | 0 | 2 | 0 | 1 | 0 | 11 | 0.0909 | 2 | 0 | 3 | 0.3333 | 2 | 0.5 | 3 | 0 | 10 | 0.2 |
| X_1003635 | 4 | 0.25 | 4 | 0.75 | 2 | 0.5 | 1 | 0 | 11 | 0.4545 | NA | NA | 2 | 0.5 | 3 | 0.3333 | 5 | 0.4 | 10 | 0.4 |
| X_1003640 | 5 | 0.2 | 7 | 0.1429 | 5 | 0 | 3 | 0.3333 | 20 | 0.15 | 1 | 0 | 6 | 0 | 6 | 0.1667 | 5 | 0 | 18 | 0.0556 |

| | | | | | | | | | | | | | | | | | | | | |
|---|---|---|---|---|---|---|---|---|---|---|---|---|---|---|---|---|---|---|---|---|
| X_1003640 | 7 | 0.4286 | 7 | 0.2857 | 5 | 0.4 | 4 | 0.5 | 23 | 0.3913 | 2 | 0 | 7 | 0.1429 | 6 | 0.6667 | 6 | 0.1667 | 21 | 0.2857 |
| X_1003641 | 6 | 0.1667 | 5 | 0 | 5 | 0 | 4 | 0 | 20 | 0.05 | 1 | 1 | 6 | 0 | 6 | 0 | 5 | 0 | 18 | 0.0556 |
| X_1003641 | 7 | 0.1429 | 8 | 0.375 | 3 | 0.3333 | 2 | 1 | 20 | 0.35 | 1 | 1 | 5 | 0.4 | 6 | 0.1667 | 6 | 0.5 | 18 | 0.3889 |
| X_1003641 | 7 | 0.2857 | 7 | 0.1429 | 4 | 0.5 | 4 | 0.75 | 22 | 0.3636 | 1 | 0 | 6 | 0.6667 | 7 | 0.5714 | 5 | 0.4 | 19 | 0.5263 |
| X_1003642 | 4 | 1 | 6 | 0.3333 | 6 | 0.8333 | 4 | 0.75 | 20 | 0.7 | 2 | 0 | 8 | 0.5 | 7 | 0.4286 | 7 | 0.4286 | 24 | 0.4167 |
| X_1003642 | 5 | 0.4 | 5 | 0.8 | 4 | 0.25 | 4 | 0 | 18 | 0.3889 | 2 | 0.5 | 7 | 0.8571 | 7 | 0.4286 | 5 | 0.4 | 21 | 0.5714 |
| X_1003642 | 7 | 0.7143 | 9 | 0.2222 | 5 | 0 | 3 | 1 | 24 | 0.4167 | 2 | 1 | 8 | 0.25 | 6 | 0.1667 | 6 | 0.3333 | 22 | 0.3182 |
| X_1003643 | 5 | 0.2 | 9 | 0 | 6 | 0.1667 | 4 | 0 | 24 | 0.0833 | 2 | 0 | 8 | 0.125 | 8 | 0 | 4 | 0.5 | 22 | 0.1364 |
| X_1003644 | 7 | 1 | 11 | 1 | 6 | 1 | 4 | 1 | 28 | 1 | 2 | 1 | 6 | 0.8333 | 9 | 1 | 7 | 1 | 24 | 0.9583 |
| X_1003644 | 5 | 0.6 | 10 | 0.7 | 6 | 0.3333 | 3 | 1 | 24 | 0.625 | 2 | 1 | 6 | 0.6667 | 8 | 0.5 | 9 | 0.6667 | 25 | 0.64 |
| X_1003644 | 8 | 0.125 | 7 | 0.4286 | 3 | 0.3333 | 1 | 0 | 19 | 0.2632 | 1 | 0 | 7 | 0.5714 | 7 | 0.5714 | 9 | 0.4444 | 24 | 0.5 |
| X_1003644 | 5 | 1 | 11 | 0.9091 | 6 | 1 | 4 | 1 | 26 | 0.9615 | 1 | 1 | 8 | 0.625 | 10 | 1 | 8 | 1 | 27 | 0.8889 |
| X_1003645 | 6 | 0.5 | 10 | 0.5 | 6 | 0.8333 | 3 | 0.6667 | 25 | 0.6 | 2 | 0.5 | 6 | 1 | 11 | 0.5455 | 9 | 0.7778 | 28 | 0.7143 |
| X_1003645 | 7 | 0.7143 | 10 | 0.7 | 5 | 0.8 | 4 | 1 | 26 | 0.7692 | 3 | 0.3333 | 6 | 0.3333 | 10 | 0.5 | 9 | 0.5556 | 28 | 0.4643 |
| X_1003646 | 5 | 1 | 9 | 1 | 4 | 1 | 4 | 1 | 22 | 1 | 3 | 1 | 8 | 0.875 | 8 | 0.875 | 10 | 0.9 | 29 | 0.8966 |
| X_1003647 | 8 | 0.625 | 13 | 0.3846 | 7 | 0.2857 | 6 | 0.5 | 34 | 0.4412 | 4 | 0.25 | 9 | 0.4444 | 11 | 0.4545 | 9 | 0.3333 | 33 | 0.3939 |
| X_1003653 | 6 | 1 | 3 | 1 | 2 | 1 | 3 | 1 | 14 | 1 | 3 | 1 | 1 | 1 | NA | NA | 2 | 1 | 6 | 1 |
| X_1003653 | 7 | 0 | 6 | 0.1667 | 2 | 0.5 | 3 | 0.3333 | 18 | 0.1667 | 3 | 0.3333 | 2 | 0 | NA | NA | 4 | 0 | 9 | 0.1111 |
| X_1003654 | 5 | 0.8 | 3 | 0.6667 | NA | NA | 2 | 1 | 10 | 0.8 | 1 | 1 | 2 | 0.5 | NA | NA | 3 | 0.6667 | 6 | 0.6667 |
| X_1003654 | 5 | 1 | 5 | 1 | 3 | 1 | 3 | 1 | 16 | 1 | 3 | 1 | 1 | 1 | NA | NA | 4 | 1 | 8 | 1 |
| X_1003655 | 7 | 0.1429 | 7 | 0.1429 | 3 | 0.3333 | 3 | 0 | 20 | 0.15 | 2 | 0 | 1 | 0 | NA | NA | 6 | 0.1667 | 9 | 0.1111 |
| X_1003656 | 7 | 0.2857 | 4 | 0.5 | 2 | 0.5 | 2 | 0 | 15 | 0.3333 | 1 | 0 | 2 | 0.5 | NA | NA | 4 | 0.75 | 7 | 0.5714 |
| X_1003656 | 7 | 0.1429 | 8 | 0.375 | 3 | 0.3333 | 3 | 0 | 21 | 0.2381 | 3 | 0 | 3 | 0 | 1 | 0 | 7 | 0.1429 | 14 | 0.0714 |
| X_1003657 | 7 | 0 | 7 | 0 | 3 | 0 | 3 | 0.3333 | 20 | 0.05 | 3 | 0 | 2 | 0 | 1 | 0 | 7 | 0.2857 | 13 | 0.1538 |
| X_1003657 | 7 | 0.2857 | 6 | 0 | 3 | 0.3333 | 3 | 0 | 19 | 0.1579 | 2 | 0 | 2 | 0 | 2 | 0 | 6 | 0.5 | 12 | 0.25 |
| X_1003658 | 7 | 0.1429 | 6 | 0 | 3 | 0 | 3 | 0 | 19 | 0.0526 | 2 | 0 | 1 | 1 | 1 | 0 | 2 | 0 | 6 | 0.1667 |
| X_1003658 | 6 | 0.5 | 7 | 0.2857 | 3 | 0.3333 | 4 | 0.25 | 20 | 0.35 | 3 | 0.3333 | 4 | 0.25 | 2 | 0 | 7 | 0.4286 | 16 | 0.3125 |
| X_1003660 | 10 | 0.1 | 8 | 0.25 | 4 | 0 | 7 | 0 | 29 | 0.1034 | 3 | 0 | 3 | 0 | 2 | 0 | 8 | 0.125 | 16 | 0.0625 |
| X_1003664 | 10 | 0.3 | 11 | 0.2727 | 4 | 0 | 9 | 0.2222 | 34 | 0.2353 | 7 | 0.2857 | 3 | 0 | 3 | 0.3333 | 9 | 0.2222 | 22 | 0.2273 |
| X_1003674 | 10 | 0.2 | 14 | 0 | 8 | 0 | 10 | 0.1 | 42 | 0.0714 | 11 | 0.1818 | 6 | 0 | 5 | 0.2 | 12 | 0 | 34 | 0.0882 |
| X_1003676 | 2 | 0.5 | 4 | 0.5 | 2 | 0.5 | 3 | 0 | 11 | 0.3636 | 7 | 0.5714 | NA | NA | 3 | 0.3333 | 4 | 0.25 | 14 | 0.4286 |
| X_1003677 | 3 | 0.3333 | 4 | 0 | 3 | 0 | 2 | 0 | 12 | 0.0833 | 6 | 0.3333 | NA | NA | 3 | 0 | 4 | 0 | 13 | 0.1538 |
| X_1003682 | 4 | 0 | 5 | 0.2 | 4 | 0.25 | 3 | 0 | 16 | 0.125 | 7 | 0 | NA | NA | 5 | 0.2 | 5 | 0 | 17 | 0.0588 |
| X_1003684 | 3 | 0.6667 | 5 | 0.6 | 4 | 0.25 | 4 | 0.5 | 16 | 0.5 | 7 | 1 | NA | NA | 4 | 0.75 | 4 | 0.5 | 15 | 0.8 |
| X_1003685 | 4 | 0.5 | 5 | 0.8 | 4 | 0.5 | 4 | 0.25 | 17 | 0.5294 | 5 | 0.8 | NA | NA | 5 | 0.8 | 6 | 0.6667 | 16 | 0.75 |
| X_1003689 | 6 | 0.5 | 2 | 0 | 5 | 0.8 | 5 | 0.4 | 18 | 0.5 | 9 | 0.2222 | NA | NA | 6 | 0.3333 | 5 | 0.8 | 20 | 0.4 |
| X_1003691 | 6 | 0.1667 | 4 | 0.25 | 5 | 0.2 | 6 | 0.1667 | 21 | 0.1905 | 6 | 0.1667 | NA | NA | 5 | 0 | 4 | 0 | 15 | 0.0667 |
| X_1003694 | 5 | 0.2 | 6 | 0.5 | 7 | 0.5714 | 6 | 0.6667 | 24 | 0.5 | 8 | 0.625 | 1 | 1 | 7 | 0.5714 | 6 | 0.1667 | 22 | 0.5 |
| X_1004281 | 1 | 0 | 3 | 0.3333 | 1 | 1 | 2 | 0.5 | 7 | 0.4286 | 2 | 0 | NA | NA | 1 | 1 | 1 | 0 | 4 | 0.25 |
| X_1004281 | 1 | 1 | 2 | 1 | 1 | 1 | 2 | 0.5 | 6 | 0.8333 | 2 | 1 | NA | NA | 1 | 0 | 1 | 1 | 4 | 0.75 |
| X_1004281 | 1 | 0 | 2 | 0.5 | 1 | 1 | 2 | 1 | 6 | 0.6667 | 2 | 0 | NA | NA | 1 | 1 | 1 | 1 | 4 | 0.5 |
| X_1004283 | 1 | 1 | 2 | 1 | 1 | 1 | 2 | 1 | 6 | 1 | 2 | 1 | NA | NA | 1 | 1 | 1 | 1 | 4 | 1 |
| X_1004285 | NA | NA | 4 | 1 | 1 | 1 | 2 | 1 | 7 | 1 | 2 | 1 | NA | NA | NA | NA | 1 | 1 | 3 | 1 |
| X_1004287 | 1 | 1 | 5 | 0.2 | NA | NA | 2 | 0.5 | 8 | 0.375 | 2 | 0 | NA | NA | 1 | 1 | 1 | 1 | 4 | 0.5 |
| X_1004290 | 1 | 0 | 5 | 0.2 | 1 | 0 | 2 | 0 | 9 | 0.1111 | 1 | 1 | NA | NA | 2 | 0.5 | 1 | 0 | 4 | 0.5 |
| X_1004492 | 4 | 0 | 10 | 0.1 | 2 | 0.5 | 10 | 0.2 | 26 | 0.1538 | 5 | 0.6 | 7 | 0 | 15 | 0.0667 | 3 | 0.3333 | 30 | 0.1667 |
| X_1004493 | 4 | 0.25 | 5 | 0.2 | 3 | 0.3333 | 7 | 0.2857 | 19 | 0.2632 | 7 | 0.4286 | 7 | 0.5714 | 12 | 0.3333 | 1 | 0 | 27 | 0.4074 |
| X_1004589 | 5 | 0.2 | 3 | 0.6667 | 2 | 0 | 7 | 0.1429 | 17 | 0.2353 | 5 | 0.2 | 4 | 0.75 | 12 | 0.5 | 4 | 0.75 | 25 | 0.52 |
| X_1004600 | 2 | 0 | 2 | 0.5 | 1 | 0 | 5 | 0.2 | 10 | 0.2 | 4 | 0 | 6 | 0.1667 | 5 | 0.2 | 3 | 0 | 18 | 0.1111 |
| X_1004619 | 4 | 0.5 | 1 | 0 | NA | NA | 4 | 0 | 9 | 0.2222 | 2 | 0.5 | 5 | 0.2 | 6 | 0.1667 | 3 | 0.3333 | 16 | 0.25 |
| X_1004622 | 2 | 1 | 2 | 0.5 | NA | NA | 2 | 0.5 | 6 | 0.6667 | 1 | 1 | 3 | 0.6667 | 5 | 0.6 | 3 | 0.3333 | 12 | 0.5833 |
| X_1004623 | 4 | 0 | 1 | 1 | NA | NA | 4 | 0 | 9 | 0.1111 | 1 | 0 | 4 | 0 | 6 | 0.5 | 4 | 0 | 15 | 0.2 |
| X_1004852 | 18 | 0.0556 | 12 | 0.1667 | 9 | 0 | 26 | 0.0769 | 65 | 0.0769 | 10 | 0.2 | 7 | 0.2857 | 13 | 0.2308 | 14 | 0 | 44 | 0.1591 |
| X_1004913 | 22 | 0.3636 | 17 | 0.4706 | 11 | 0.3636 | 33 | 0.1515 | 83 | 0.3012 | 15 | 0.3333 | 12 | 0.3333 | 9 | 0.2222 | 21 | 0.4762 | 57 | 0.3684 |
| X_1004915 | 22 | 0.1818 | 20 | 0.1 | 14 | 0.2857 | 33 | 0.1212 | 89 | 0.1573 | 17 | 0.0588 | 13 | 0 | 16 | 0.3125 | 24 | 0.0833 | 70 | 0.1143 |
| X_1006430 | NA | NA | 5 | 0.2 | 2 | 0 | 4 | 0.25 | 11 | 0.1818 | NA | NA | 1 | 0 | 2 | 0.5 | 4 | 0 | 7 | 0.1429 |
| X_1006431 | NA | NA | 5 | 1 | 1 | 1 | 4 | 1 | 10 | 1 | NA | NA | 2 | 1 | 3 | 1 | 2 | 0.5 | 7 | 0.8571 |
| X_1006442 | 2 | 0 | 7 | 0 | 3 | 0 | 6 | 0.1667 | 18 | 0.0556 | NA | NA | 3 | 0 | 3 | 0 | 7 | 0.1429 | 13 | 0.0769 |
| X_1007258 | 7 | 0.2857 | 10 | 0.1 | 12 | 0.0833 | 10 | 0.1 | 39 | 0.1282 | 5 | 0 | 3 | 0.3333 | 15 | 0.1333 | 18 | 0.1111 | 41 | 0.122 |
| X_1007259 | 11 | 0.9091 | 9 | 1 | 9 | 0.7778 | 7 | 1 | 36 | 0.9167 | 3 | 1 | 3 | 1 | 14 | 0.9286 | 15 | 0.9333 | 35 | 0.9429 |
| X_1007259 | 11 | 0.4545 | 13 | 0.6923 | 12 | 0.4167 | 13 | 0.5385 | 49 | 0.5306 | 6 | 0.5 | 3 | 0.3333 | 17 | 0.5294 | 20 | 0.7 | 46 | 0.587 |
| X_1007260 | 9 | 0.7778 | 13 | 0.6923 | 12 | 0.75 | 11 | 0.6364 | 45 | 0.7111 | 4 | 0.75 | 2 | 1 | 16 | 0.8125 | 18 | 0.7222 | 40 | 0.775 |
| X_1007261 | 10 | 0.4 | 13 | 0.3846 | 15 | 0.3333 | 13 | 0.3077 | 51 | 0.3529 | 6 | 0.1667 | 3 | 0 | 18 | 0.3889 | 18 | 0.2778 | 45 | 0.2889 |
| X_1007264 | 11 | 0.6364 | 10 | 0.8 | 8 | 0.25 | 9 | 0.5556 | 38 | 0.5789 | 5 | 0.4 | 2 | 1 | 11 | 0.6364 | 13 | 0.6923 | 31 | 0.6452 |
| X_1007264 | 15 | 0.2667 | 13 | 0.3077 | 18 | 0.2222 | 15 | 0.1333 | 61 | 0.2295 | 7 | 0.1429 | 4 | 0 | 18 | 0.1667 | 21 | 0.1905 | 50 | 0.16 |

| | | | | | | | | | | | | | | | | | | | | |
|---|---|---|---|---|---|---|---|---|---|---|---|---|---|---|---|---|---|---|---|---|
| X_1007265 | 13 | 0.2308 | 14 | 0 | 16 | 0 | 13 | 0.0769 | 56 | 0.0714 | 7 | 0 | 5 | 0 | 18 | 0.0556 | 22 | 0.1364 | 52 | 0.0769 |
| X_1007265 | 16 | 0.6875 | 14 | 0.6429 | 17 | 0.7647 | 15 | 0.4667 | 62 | 0.6452 | 7 | 0.4286 | 5 | 0.4 | 17 | 0.5294 | 20 | 0.4 | 49 | 0.449 |
| X_1007266 | 15 | 0.5333 | 10 | 0.7 | 16 | 0.5 | 13 | 0.4615 | 54 | 0.537 | 8 | 0.625 | 5 | 0.2 | 18 | 0.6111 | 19 | 0.5263 | 50 | 0.54 |
| X_1007266 | 10 | 0 | 11 | 0.1818 | 10 | 0 | 7 | 0.1429 | 38 | 0.0789 | 3 | 0.3333 | 3 | 0.3333 | 12 | 0.0833 | 13 | 0.1538 | 31 | 0.1613 |
| X_1007270 | 15 | 0.4667 | 16 | 0.3125 | 18 | 0.2778 | 10 | 0.2 | 59 | 0.322 | 7 | 0.1429 | 6 | 0 | 17 | 0.4118 | 20 | 0.25 | 50 | 0.26 |
| X_1007270 | 15 | 0.6 | 17 | 0.4706 | 19 | 0.3158 | 13 | 0.6154 | 64 | 0.4844 | 6 | 0.5 | 7 | 0.4286 | 20 | 0.6 | 21 | 0.3333 | 54 | 0.463 |
| X_1007271 | 12 | 0.9167 | 15 | 0.9333 | 16 | 0.8125 | 12 | 0.75 | 55 | 0.8545 | 4 | 0.75 | 7 | 0.7143 | 13 | 0.6923 | 20 | 0.75 | 44 | 0.7273 |
| X_1007272 | 15 | 0.1333 | 13 | 0.3077 | 15 | 0.1333 | 12 | 0.0833 | 55 | 0.1636 | 7 | 0 | 7 | 0.1429 | 19 | 0 | 19 | 0.1053 | 52 | 0.0577 |
| X_1007273 | 19 | 0.2632 | 20 | 0.05 | 21 | 0.1429 | 13 | 0.0769 | 73 | 0.137 | 8 | 0.125 | 7 | 0 | 20 | 0.15 | 19 | 0.1053 | 54 | 0.1111 |
| X_1007274 | 16 | 0.75 | 15 | 0.4 | 15 | 0.6 | 8 | 0.375 | 54 | 0.5556 | 8 | 0.25 | 9 | 0.5556 | 20 | 0.5 | 19 | 0.5789 | 56 | 0.5 |
| X_1007276 | 13 | 0.1538 | 15 | 0 | 14 | 0.1429 | 12 | 0.1667 | 54 | 0.1111 | 8 | 0.125 | 10 | 0 | 17 | 0.2353 | 18 | 0.2222 | 53 | 0.1698 |
| X_1007276 | 18 | 0.4444 | 19 | 0.4211 | 19 | 0.3158 | 13 | 0.3077 | 69 | 0.3768 | 8 | 0.125 | 11 | 0.4545 | 17 | 0.3529 | 21 | 0.3333 | 57 | 0.3333 |
| X_1007277 | 19 | 0.6316 | 18 | 0.6111 | 20 | 0.55 | 12 | 0.3333 | 69 | 0.5507 | 7 | 0.7143 | 10 | 0.5 | 21 | 0.8095 | 20 | 0.45 | 58 | 0.6207 |
| X_1007278 | 20 | 0.2 | 19 | 0.2105 | 22 | 0.0455 | 12 | 0 | 73 | 0.1233 | 10 | 0.2 | 11 | 0 | 19 | 0.1579 | 22 | 0.1364 | 62 | 0.129 |
| X_1011104 | NA | NA | 1 | 1 | NA | NA | 3 | 0.6667 | 4 | 0.75 | 3 | 0.3333 | 1 | 0 | 4 | 0.75 | 2 | 1 | 10 | 0.6 |
| X_1011106 | NA | NA | NA | NA | NA | NA | 3 | 0.3333 | 3 | 0.3333 | 2 | 0.5 | 1 | 0 | 3 | 0.6667 | 2 | 0 | 8 | 0.375 |
| X_1011113 | NA | NA | NA | NA | NA | NA | 2 | 1 | 2 | 1 | 2 | 0.5 | 1 | 0 | 4 | 0.25 | NA | NA | 7 | 0.2857 |
| X_1011113 | NA | NA | 1 | 0 | NA | NA | 2 | 0.5 | 3 | 0.3333 | 2 | 1 | 1 | 1 | 3 | 1 | 1 | 1 | 7 | 1 |
| X_1011114 | NA | NA | 1 | 0 | NA | NA | 3 | 0.6667 | 4 | 0.5 | 2 | 0.5 | 1 | 0 | 3 | 0 | 1 | 1 | 7 | 0.2857 |
| X_1011115 | NA | NA | 1 | 1 | NA | NA | 2 | 1 | 3 | 1 | 2 | 0.5 | 1 | 0 | 2 | 1 | 1 | 1 | 6 | 0.6667 |
| X_1011116 | NA | NA | 1 | 1 | NA | NA | 2 | 1 | 3 | 1 | 2 | 1 | 1 | 1 | 2 | 1 | 1 | 0 | 6 | 0.8333 |
| X_1011116 | NA | NA | 1 | 1 | NA | NA | 2 | 0.5 | 3 | 0.6667 | 2 | 1 | 1 | 0 | 1 | 1 | 1 | 1 | 5 | 0.8 |
| X_1011119 | NA | NA | 1 | 1 | NA | NA | 2 | 1 | 3 | 1 | 2 | 1 | 1 | 0 | 2 | 1 | 1 | 1 | 6 | 0.8333 |
| X_1011119 | NA | NA | NA | NA | NA | NA | 1 | 1 | 1 | 1 | 1 | 1 | 1 | 1 | 2 | 1 | NA | NA | 4 | 1 |
| X_1011120 | NA | NA | 1 | 1 | NA | NA | 1 | 1 | 2 | 1 | 2 | 1 | NA | NA | 2 | 1 | 1 | 1 | 5 | 1 |
| X_1011122 | NA | NA | 1 | 1 | NA | NA | NA | NA | 1 | 1 | 1 | 1 | NA | NA | 2 | 0 | 1 | 0 | 4 | 0.25 |
| X_1011124 | NA | NA | 1 | 1 | NA | NA | 1 | 1 | 2 | 1 | 2 | 0.5 | NA | NA | 2 | 0 | 1 | 0 | 5 | 0.2 |
| X_1011135 | NA | NA | 1 | 1 | NA | NA | NA | NA | 1 | 1 | 2 | 0.5 | NA | NA | NA | NA | NA | NA | 2 | 0.5 |
| X_1011139 | NA | NA | 1 | 1 | NA | NA | 1 | 1 | 2 | 1 | 1 | 1 | NA | NA | NA | NA | NA | NA | 1 | 1 |
| X_1011141 | NA | NA | 1 | 1 | NA | NA | 1 | 1 | 2 | 1 | 1 | 1 | NA | NA | NA | NA | NA | NA | 1 | 1 |
| X_1011265 | NA | NA | 1 | 1 | NA | NA | 3 | 1 | 4 | 1 | 1 | 1 | NA | NA | NA | NA | NA | NA | 1 | 1 |
| X_1011268 | NA | NA | 1 | 0 | NA | NA | 2 | 0.5 | 3 | 0.3333 | 1 | 1 | NA | NA | NA | NA | NA | NA | 1 | 1 |
| X_1011269 | NA | NA | 1 | 1 | NA | NA | 2 | 1 | 3 | 1 | 1 | 1 | NA | NA | NA | NA | NA | NA | 1 | 1 |
| X_1011270 | NA | NA | 1 | 1 | NA | NA | 3 | 0.6667 | 4 | 0.75 | 1 | 1 | NA | NA | NA | NA | NA | NA | 1 | 1 |
| X_1011271 | NA | NA | 1 | 1 | NA | NA | 3 | 1 | 4 | 1 | 1 | 1 | NA | NA | NA | NA | NA | NA | 1 | 1 |
| X_1011278 | NA | NA | 1 | 1 | NA | NA | 3 | 1 | 4 | 1 | 2 | 1 | NA | NA | NA | NA | NA | NA | 2 | 1 |
| X_1011280 | NA | NA | 1 | 1 | 1 | 0 | 3 | 0.6667 | 5 | 0.6 | 2 | 1 | NA | NA | 1 | 0 | NA | NA | 3 | 0.6667 |
| X_1011281 | NA | NA | 1 | 1 | 1 | 1 | 4 | 0.5 | 6 | 0.6667 | 2 | 0.5 | NA | NA | 1 | 0 | NA | NA | 3 | 0.3333 |
| X_1011282 | NA | NA | NA | NA | 1 | 1 | 3 | 1 | 4 | 1 | NA | NA | NA | NA | 1 | 1 | NA | NA | 1 | 1 |
| X_1011285 | NA | NA | 1 | 1 | 2 | 1 | 3 | 1 | 6 | 1 | 1 | 0 | NA | NA | 1 | 1 | NA | NA | 2 | 0.5 |
| X_1011285 | NA | NA | 1 | 1 | 1 | 1 | 1 | 0 | 3 | 0.6667 | 1 | 1 | NA | NA | NA | NA | NA | NA | 1 | 1 |
| X_1011287 | NA | NA | 1 | 1 | 2 | 1 | 3 | 1 | 6 | 1 | 2 | 1 | NA | NA | 1 | 1 | NA | NA | 3 | 1 |
| X_1011302 | NA | NA | 2 | 0 | 2 | 0 | 4 | 0.25 | 8 | 0.125 | 2 | 0.5 | 2 | 0 | 2 | 0 | NA | NA | 6 | 0.1667 |
| X_1011306 | NA | NA | 2 | 0.5 | 2 | 0.5 | 4 | 0.75 | 8 | 0.625 | 1 | 1 | 2 | 1 | 2 | 1 | NA | NA | 5 | 1 |
| X_1011324 | 1 | 0 | 2 | 0 | 3 | 0.3333 | 7 | 0 | 13 | 0.0769 | 3 | 0 | 6 | 0.1667 | 4 | 0 | 2 | 0 | 15 | 0.0667 |
| X_1011332 | 2 | 0 | 2 | 0.5 | 3 | 0.3333 | 6 | 0 | 13 | 0.1538 | 3 | 0 | 5 | 0.2 | 4 | 0 | 3 | 0 | 15 | 0.0667 |
| X_1011351 | 4 | 0 | 3 | 0 | 4 | 0 | 10 | 0.2 | 21 | 0.0952 | 4 | 0 | 7 | 0.1429 | 5 | 0.2 | 6 | 0 | 22 | 0.0909 |
| X_1011352 | 4 | 0 | 4 | 0.25 | 4 | 0 | 8 | 0 | 20 | 0.05 | 4 | 0 | 7 | 0.1429 | 6 | 0 | 5 | 0.2 | 22 | 0.0909 |
| X_1011353 | 4 | 0.25 | 4 | 0 | 3 | 0 | 8 | 0.125 | 19 | 0.1053 | 3 | 0.3333 | 6 | 0 | 7 | 0.2857 | 5 | 0 | 21 | 0.1429 |
| X_1015584 | NA | NA | NA | NA | NA | NA | 2 | 0.5 | 2 | 0.5 | 2 | 0.5 | NA | NA | 1 | 0 | 3 | 0.3333 | 6 | 0.3333 |
| X_1015595 | NA | NA | NA | NA | 1 | 1 | 2 | 0.5 | 3 | 0.6667 | 2 | 1 | NA | NA | 2 | 0 | 1 | 0 | 5 | 0.4 |
| X_1015595 | NA | NA | NA | NA | 1 | 0 | 2 | 1 | 3 | 0.6667 | 2 | 1 | NA | NA | 1 | 0 | 1 | 0 | 4 | 0.5 |
| X_1015596 | NA | NA | NA | NA | NA | NA | 2 | 0.5 | 2 | 0.5 | 1 | 1 | NA | NA | 1 | 0 | 1 | 0 | 3 | 0.3333 |
| X_1019473 | 1 | 0 | 3 | 0 | NA | NA | 3 | 0.3333 | 7 | 0.1429 | 2 | 0.5 | 1 | 1 | 2 | 0 | NA | NA | 5 | 0.4 |
| X_1019474 | 1 | 1 | 3 | 0.3333 | NA | NA | 3 | 0.3333 | 7 | 0.4286 | 2 | 0.5 | 1 | 0 | 2 | 0 | 1 | 0 | 6 | 0.1667 |
| X_1019769 | 1 | 1 | NA | NA | NA | NA | NA | NA | 1 | 1 | NA | NA | NA | NA | 1 | 1 | NA | NA | 1 | 1 |
| X_1020574 | 3 | 0.3333 | 5 | 0 | 1 | 0 | 1 | 0 | 10 | 0.1 | NA | NA | 4 | 0 | 3 | 0 | 3 | 0.3333 | 10 | 0.1 |
| X_1020575 | 3 | 1 | 6 | 1 | 1 | 1 | 1 | 1 | 11 | 1 | NA | NA | 4 | 1 | 5 | 1 | 3 | 1 | 12 | 1 |
| X_1020578 | 2 | 0.5 | 3 | 0 | 1 | 0 | 1 | 0 | 7 | 0.1429 | NA | NA | 3 | 0 | 4 | 0.5 | 4 | 0.75 | 11 | 0.4545 |
| X_1020580 | 2 | 0 | 5 | 0.2 | 1 | 1 | 2 | 0.5 | 10 | 0.3 | NA | NA | 4 | 0.25 | 4 | 0 | 3 | 0 | 11 | 0.0909 |
| X_1020587 | 3 | 0.6667 | 5 | 0.6 | 1 | 0 | 2 | 1 | 11 | 0.6364 | NA | NA | 5 | 0.2 | 4 | 0.25 | 3 | 0.6667 | 12 | 0.3333 |
| X_1020588 | 5 | 1 | 6 | 1 | 1 | 1 | 2 | 1 | 14 | 1 | NA | NA | 6 | 1 | 4 | 1 | 4 | 1 | 14 | 1 |
| X_1020588 | 7 | 1 | 7 | 1 | 1 | 1 | 1 | 1 | 16 | 1 | 1 | 1 | 6 | 1 | 6 | 1 | 5 | 1 | 18 | 1 |
| X_1020590 | 7 | 0.8571 | 7 | 0.5714 | 2 | 0.5 | 2 | 1 | 18 | 0.7222 | NA | NA | 6 | 0.8333 | 5 | 1 | 5 | 0.4 | 16 | 0.75 |
| X_1020590 | 7 | 0.7143 | 7 | 0.8571 | 2 | 1 | 2 | 0.5 | 18 | 0.7778 | NA | NA | 3 | 0.6667 | 5 | 0.8 | 2 | 0.5 | 10 | 0.7 |

| | | | | | | | | | | | | | | | | | | | | |
|---|---|---|---|---|---|---|---|---|---|---|---|---|---|---|---|---|---|---|---|---|
| X_1020590 | 5 | 1 | 5 | 1 | NA | NA | 2 | 0.5 | 12 | 0.9167 | NA | NA | 5 | 1 | 5 | 1 | 3 | 1 | 13 | 1 |
| X_1020606 | NA | NA | 3 | 0 | 1 | 1 | 1 | 0 | 5 | 0.2 | NA | NA | 2 | 0.5 | 2 | 1 | NA | NA | 4 | 0.75 |
| X_1020609 | NA | NA | 4 | 0.25 | 1 | 1 | 1 | 0 | 6 | 0.3333 | NA | NA | 2 | 0.5 | 1 | 1 | NA | NA | 3 | 0.6667 |
| X_1021091 | NA | NA | 12 | 0.5833 | 7 | 1 | 9 | 0.7778 | 28 | 0.75 | 3 | 0.3333 | 7 | 0.7143 | 7 | 0.8571 | 5 | 0.8 | 22 | 0.7273 |
| X_1021091 | 1 | 0 | 10 | 0.3 | 5 | 0.4 | 5 | 0.8 | 21 | 0.4286 | 4 | 0.75 | 6 | 0.5 | 8 | 0.75 | 4 | 0.75 | 22 | 0.6818 |
| X_1021091 | NA | NA | 10 | 0.8 | 6 | 0.5 | 6 | 0.5 | 22 | 0.6364 | 5 | 0.8 | 6 | 0.8333 | 6 | 0.6667 | 4 | 0.75 | 21 | 0.7619 |
| X_1021092 | 1 | 1 | 12 | 0.9167 | 7 | 1 | 8 | 1 | 28 | 0.9643 | 5 | 0.8 | 8 | 1 | 8 | 1 | 5 | 1 | 26 | 0.9615 |
| X_1021092 | 1 | 1 | 13 | 1 | 7 | 1 | 8 | 1 | 29 | 1 | 5 | 1 | 7 | 1 | 8 | 1 | 6 | 1 | 26 | 1 |
| X_1021093 | 1 | 0 | 12 | 0.25 | 6 | 0.1667 | 8 | 0 | 27 | 0.1481 | 4 | 0.25 | 7 | 0.1429 | 5 | 0 | 6 | 0.1667 | 22 | 0.1364 |
| X_1021093 | NA | NA | 10 | 0.5 | 4 | 0.5 | 7 | 0.4286 | 21 | 0.4762 | 4 | 0.5 | 8 | 0.5 | 3 | 1 | 5 | 0.8 | 20 | 0.65 |
| X_1021101 | 1 | 0 | 10 | 0 | 3 | 0.3333 | 5 | 0.4 | 19 | 0.1579 | 1 | 0 | 5 | 0.2 | 6 | 0.6667 | 4 | 0 | 16 | 0.3125 |
| X_1021102 | NA | NA | 6 | 0 | 4 | 0.25 | 6 | 0.5 | 16 | 0.25 | 2 | 0 | 4 | 0 | 5 | 0.2 | 1 | 0 | 12 | 0.0833 |
| X_1021105 | 1 | 1 | 8 | 1 | 1 | 1 | 7 | 1 | 17 | 1 | 3 | 1 | 3 | 1 | 2 | 1 | 3 | 1 | 11 | 1 |
| X_1021107 | 1 | 0 | 7 | 0 | 4 | 0 | 7 | 0.1429 | 19 | 0.0526 | 3 | 0 | 6 | 0.1667 | 3 | 0.3333 | 2 | 0 | 14 | 0.1429 |
| X_1022065 | NA | NA | NA | NA | NA | NA | 1 | 1 | 1 | 1 | 1 | 0 | NA | NA | NA | NA | 2 | 0.5 | 3 | 0.3333 |
| X_1022278 | 13 | 0.3077 | 22 | 0.5 | 25 | 0.48 | 14 | 0.2857 | 74 | 0.4189 | 13 | 0.4615 | 14 | 0.4286 | 13 | 0.2308 | 20 | 0.55 | 60 | 0.4333 |
| X_1022280 | 12 | 0.0833 | 22 | 0 | 27 | 0.1111 | 20 | 0.1 | 81 | 0.0741 | 12 | 0.0833 | 14 | 0.1429 | 14 | 0.0714 | 23 | 0.1304 | 63 | 0.1111 |
| X_1022281 | 14 | 0.0714 | 20 | 0.1 | 25 | 0.08 | 20 | 0.15 | 79 | 0.1013 | 11 | 0.3636 | 11 | 0.1818 | 11 | 0 | 21 | 0.0952 | 54 | 0.1481 |
| X_1022283 | 12 | 0.1667 | 21 | 0.1429 | 23 | 0.1739 | 20 | 0.4 | 76 | 0.2237 | 10 | 0 | 13 | 0.1538 | 13 | 0.1538 | 18 | 0.2222 | 54 | 0.1481 |
| X_1022288 | 10 | 1 | 16 | 1 | 21 | 1 | 14 | 1 | 61 | 1 | 4 | 1 | 9 | 1 | 9 | 1 | 14 | 1 | 36 | 1 |
| X_1022288 | 9 | 0 | 15 | 0.0667 | 21 | 0.2381 | 16 | 0.375 | 61 | 0.1967 | 5 | 0 | 9 | 0 | 12 | 0 | 14 | 0.2143 | 40 | 0.075 |
| X_1022288 | 5 | 0.4 | 12 | 0.5 | 18 | 0.6667 | 9 | 0.5556 | 44 | 0.5682 | 3 | 0.6667 | 9 | 0.5556 | 7 | 1 | 6 | 0.5 | 25 | 0.68 |
| X_1022288 | 7 | 0.2857 | 15 | 0.4 | 20 | 0.35 | 11 | 0.2727 | 53 | 0.3396 | 5 | 0.6 | 9 | 0.4444 | 11 | 0.5455 | 16 | 0.4375 | 41 | 0.4878 |
| X_1022289 | 6 | 0.6667 | 13 | 0.1538 | 18 | 0.3333 | 14 | 0.4286 | 51 | 0.3529 | 4 | 0.75 | 8 | 0.375 | 10 | 0.2 | 9 | 0.6667 | 31 | 0.4516 |
| X_1022290 | 9 | 1 | 16 | 1 | 16 | 1 | 14 | 1 | 55 | 1 | 5 | 1 | 10 | 1 | 11 | 1 | 15 | 1 | 41 | 1 |
| X_1022291 | 9 | 0.2222 | 15 | 0.0667 | 18 | 0.1111 | 14 | 0.1429 | 56 | 0.125 | 3 | 0 | 10 | 0.2 | 9 | 0.1111 | 16 | 0 | 38 | 0.0789 |
| X_1022291 | 9 | 0.8889 | 16 | 0.9375 | 19 | 1 | 12 | 1 | 56 | 0.9643 | 4 | 1 | 10 | 1 | 11 | 1 | 15 | 0.9333 | 40 | 0.975 |
| X_1022293 | 8 | 0.875 | 14 | 1 | 19 | 1 | 16 | 1 | 57 | 0.9825 | 2 | 1 | 10 | 1 | 11 | 1 | 12 | 1 | 35 | 1 |
| X_1022293 | 10 | 0 | 16 | 0.0625 | 19 | 0.1579 | 14 | 0 | 59 | 0.0678 | 3 | 0 | 10 | 0.1 | 12 | 0.1667 | 15 | 0.0667 | 40 | 0.1 |
| X_1022293 | 7 | 0.7143 | 15 | 0.6667 | 19 | 0.7368 | 16 | 0.6875 | 57 | 0.7018 | 3 | 0.6667 | 9 | 0.8889 | 12 | 0.75 | 12 | 0.75 | 36 | 0.7778 |
| X_1022294 | 9 | 0.7778 | 16 | 0.9375 | 18 | 0.9444 | 15 | 0.8667 | 58 | 0.8966 | 3 | 0.6667 | 8 | 1 | 11 | 0.9091 | 14 | 0.7143 | 36 | 0.8333 |
| X_1022296 | 5 | 1 | 13 | 1 | 16 | 1 | 12 | 0.8333 | 46 | 0.9565 | 3 | 1 | 9 | 1 | 10 | 1 | 12 | 0.9167 | 34 | 0.9706 |
| X_1022296 | 4 | 0.5 | 14 | 0.1429 | 17 | 0.2941 | 13 | 0.3077 | 48 | 0.2708 | 3 | 0.3333 | 8 | 0 | 11 | 0.2727 | 11 | 0.2727 | 33 | 0.2121 |
| X_1022296 | 6 | 1 | 11 | 0.8182 | 15 | 0.7333 | 11 | 0.9091 | 43 | 0.8372 | 3 | 1 | 8 | 1 | 9 | 0.8889 | 12 | 0.8333 | 32 | 0.9062 |
| X_1022298 | 4 | 1 | 12 | 0.8333 | 15 | 0.7333 | 9 | 0.8889 | 40 | 0.825 | 3 | 0.6667 | 8 | 0.875 | 11 | 0.6364 | 12 | 0.9167 | 34 | 0.7941 |
| X_1024379 | 1 | 0 | 4 | 0 | 2 | 0.5 | 4 | 0 | 11 | 0.0909 | 2 | 0 | 2 | 0 | 3 | 0 | 5 | 0.2 | 12 | 0.0833 |
| X_1024380 | 1 | 0 | 4 | 0.75 | 2 | 0.5 | 4 | 0.5 | 11 | 0.5455 | 1 | 0 | 2 | 0 | 3 | 0.6667 | 4 | 0.5 | 10 | 0.4 |
| X_1024389 | 2 | 0 | 4 | 0.5 | 2 | 0.5 | 2 | 0 | 10 | 0.3 | 1 | 1 | 1 | 1 | 3 | 0 | 4 | 0 | 9 | 0.2222 |
| X_1024392 | 3 | 0 | 7 | 0.4286 | 3 | 0 | 4 | 0.5 | 17 | 0.2941 | 3 | 0.3333 | 3 | 0.3333 | 4 | 0.5 | 4 | 0.5 | 14 | 0.4286 |
| X_1024393 | 4 | 0.75 | 7 | 0.8571 | 4 | 0.75 | 4 | 1 | 19 | 0.8421 | 2 | 0.5 | 4 | 1 | 4 | 0.75 | 5 | 1 | 15 | 0.8667 |
| X_1024393 | 4 | 0.5 | 6 | 0.6667 | 3 | 0.3333 | 4 | 0.5 | 17 | 0.5294 | 3 | 0.6667 | 5 | 0.2 | 6 | 0.1667 | 4 | 0.5 | 18 | 0.3333 |
| X_1024394 | 4 | 0 | 7 | 0.1429 | 4 | 0 | 4 | 0 | 19 | 0.0526 | 2 | 0 | 3 | 0 | 5 | 0 | 4 | 0.25 | 14 | 0.0714 |
| X_1024394 | 4 | 0.75 | 5 | 0.6 | 3 | 0.6667 | 3 | 0.6667 | 15 | 0.6667 | 3 | 1 | 4 | 1 | 6 | 0.8333 | 3 | 1 | 16 | 0.9375 |
| X_1024395 | 4 | 0.75 | 6 | 0.8333 | 3 | 0.6667 | 4 | 0.75 | 17 | 0.7647 | 4 | 1 | 4 | 0.75 | 6 | 1 | 5 | 0.6 | 19 | 0.8421 |
| X_1024395 | 3 | 0.3333 | 5 | 0.4 | 5 | 0.4 | 2 | 0.5 | 15 | 0.4 | 3 | 0.3333 | 3 | 0.3333 | 2 | 0 | 4 | 0.5 | 12 | 0.3333 |
| X_1026133 | 1 | 1 | 2 | 0 | 3 | 0.3333 | 3 | 0 | 9 | 0.2222 | 1 | 1 | 3 | 0.6667 | 1 | 0 | 5 | 0.4 | 10 | 0.5 |
| X_1026136 | 1 | 1 | 3 | 0.6667 | 3 | 0.3333 | 5 | 0.2 | 12 | 0.4167 | 1 | 0 | 4 | 0.25 | 1 | 0 | 5 | 0.2 | 11 | 0.1818 |
| X_1026136 | 1 | 1 | 3 | 1 | 3 | 1 | 1 | 1 | 8 | 1 | 1 | 1 | 4 | 1 | 1 | 1 | 5 | 1 | 11 | 1 |
| X_1026136 | 1 | 1 | 3 | 1 | 3 | 0.6667 | 4 | 0.75 | 11 | 0.8182 | NA | NA | 3 | 1 | NA | NA | 6 | 1 | 9 | 1 |
| X_1026137 | 1 | 0 | 3 | 0.3333 | 3 | 0 | 5 | 0 | 12 | 0.0833 | 1 | 0 | 4 | 0 | 1 | 0 | 5 | 0.2 | 11 | 0.0909 |
| X_1026138 | 1 | 0 | 3 | 1 | 3 | 0 | 4 | 0.25 | 11 | 0.3636 | 1 | 1 | 3 | 0.3333 | 1 | 0 | 5 | 0.4 | 10 | 0.4 |
| X_1026138 | 1 | 1 | 2 | 1 | 3 | 1 | 4 | 1 | 10 | 1 | 1 | 1 | 4 | 0.75 | 1 | 1 | 5 | 1 | 11 | 0.9091 |
| X_1026139 | 1 | 0 | 3 | 0.3333 | 2 | 0 | 4 | 0.25 | 10 | 0.2 | 1 | 0 | 3 | 0 | 1 | 0 | 5 | 0.2 | 10 | 0.1 |
| X_1026139 | 1 | 1 | 3 | 1 | 1 | 1 | 3 | 1 | 8 | 1 | 1 | 1 | 3 | 1 | NA | NA | 5 | 1 | 9 | 1 |
| X_1026139 | NA | NA | 3 | 0 | 2 | 0 | 4 | 0.25 | 9 | 0.1111 | 1 | 0 | 4 | 0.25 | NA | NA | 5 | 0 | 10 | 0.1 |
| X_1026140 | 1 | 0 | 3 | 0 | 2 | 0 | 3 | 0.3333 | 9 | 0.1111 | 1 | 0 | 3 | 0.3333 | 1 | 0 | 5 | 0 | 10 | 0.1 |
| X_1026140 | 1 | 0 | 2 | 1 | 2 | 0 | 4 | 0.5 | 9 | 0.4444 | 1 | 1 | 3 | 1 | 1 | 0 | 5 | 0.4 | 10 | 0.6 |
| X_1026141 | 1 | 0 | 3 | 0.6667 | 2 | 0.5 | 4 | 0.25 | 10 | 0.4 | 1 | 1 | 3 | 0.3333 | 1 | 1 | 6 | 0.8333 | 11 | 0.7273 |
| X_1026141 | 1 | 1 | 3 | 1 | 2 | 1 | 4 | 1 | 10 | 1 | 1 | 1 | 3 | 1 | 1 | 1 | 6 | 0.6667 | 11 | 0.8182 |
| X_1026142 | 1 | 0 | 3 | 0 | 2 | 0.5 | 4 | 0.25 | 10 | 0.2 | 1 | 1 | 3 | 0.3333 | 1 | 1 | 6 | 0.3333 | 11 | 0.4545 |
| X_1026143 | 1 | 0 | 2 | 0.5 | 2 | 0 | 3 | 0 | 8 | 0.125 | 1 | 0 | 3 | 0.6667 | 1 | 1 | 5 | 0.8 | 10 | 0.7 |
| X_1027410 | 7 | 0.1429 | 5 | 0.4 | 2 | 0.5 | 5 | 0 | 19 | 0.2105 | 5 | 0.2 | 6 | 0.3333 | 6 | 0.3333 | 3 | 0 | 20 | 0.25 |
| X_1027413 | 7 | 0.2857 | 4 | 0 | 1 | 0 | 4 | 0 | 16 | 0.125 | 5 | 0 | 4 | 0 | 6 | 0.1667 | 3 | 0.3333 | 18 | 0.1111 |
| X_1027413 | 5 | 0 | 5 | 0.2 | 1 | 0 | 2 | 0 | 13 | 0.0769 | 5 | 0.2 | 4 | 0.25 | 4 | 0 | 1 | 0 | 14 | 0.1429 |
| X_1027419 | 5 | 0 | 4 | 0.25 | NA | NA | 3 | 0 | 12 | 0.0833 | 5 | 0 | 3 | 0 | 3 | 0.3333 | 5 | 0 | 16 | 0.0625 |

| | | | | | | | | | | | | | | | | | | | | |
|---|---|---|---|---|---|---|---|---|---|---|---|---|---|---|---|---|---|---|---|---|
| X_1027422 | 1 | 0 | 1 | 1 | NA | NA | NA | NA | 2 | 0.5 | 3 | 0 | 2 | 0 | 2 | 0.5 | 1 | 0 | 8 | 0.125 |
| X_1027422 | 4 | 0.25 | 3 | 1 | NA | NA | 2 | 1 | 9 | 0.6667 | 3 | 0.3333 | 2 | 1 | 4 | 1 | 4 | 1 | 13 | 0.8462 |
| X_1027423 | 3 | 0 | 3 | 0 | 1 | 0 | 2 | 0.5 | 9 | 0.1111 | 3 | 0 | 3 | 0.3333 | 4 | 0 | 5 | 0.2 | 15 | 0.1333 |
| X_1027425 | 4 | 0 | 3 | 0.3333 | NA | NA | 2 | 1 | 9 | 0.3333 | 5 | 0.2 | 3 | 0.6667 | 4 | 0.75 | 3 | 0.3333 | 15 | 0.4667 |
| X_1027428 | 4 | 0 | 3 | 0.3333 | 1 | 1 | 1 | 0 | 9 | 0.2222 | 4 | 0 | 3 | 0 | 4 | 0 | 4 | 0.5 | 15 | 0.1333 |
| X_1027429 | 3 | 0.3333 | 3 | 0 | 1 | 0 | 1 | 0 | 8 | 0.125 | 2 | 0 | 2 | 0 | 4 | 0 | 2 | 0.5 | 10 | 0.1 |
| X_1027433 | 3 | 0.3333 | 1 | 0 | 1 | 0 | NA | NA | 5 | 0.2 | 2 | 1 | 3 | 0.6667 | 3 | 0.6667 | 2 | 0.5 | 10 | 0.7 |
| X_1028651 | 2 | 0.5 | 2 | 0 | 1 | 0 | NA | NA | 5 | 0.2 | 4 | 0 | 1 | 0 | 3 | 0.3333 | 2 | 0 | 10 | 0.1 |
| X_1030011 | 10 | 0 | 7 | 0.1429 | 5 | 0 | 8 | 0.125 | 30 | 0.0667 | 4 | 0 | 6 | 0.1667 | 5 | 0 | 8 | 0.125 | 23 | 0.087 |
| X_1030011 | 10 | 0.5 | 7 | 0.5714 | 5 | 1 | 9 | 0.7778 | 31 | 0.6774 | 4 | 0.5 | 6 | 0.5 | 5 | 0.8 | 6 | 0.6667 | 21 | 0.619 |
| X_1030012 | 9 | 0.2222 | 7 | 0.1429 | 6 | 0.1667 | 7 | 0.2857 | 29 | 0.2069 | 3 | 0.3333 | 6 | 0.1667 | 5 | 0 | 7 | 0.1429 | 21 | 0.1429 |
| X_1030047 | 4 | 0 | 1 | 1 | NA | NA | NA | NA | 5 | 0.2 | 1 | 0 | 4 | 0.25 | NA | NA | 2 | 0 | 7 | 0.1429 |
| X_1030048 | 4 | 1 | 1 | 1 | NA | NA | NA | NA | 5 | 1 | 1 | 1 | 4 | 1 | NA | NA | 1 | 1 | 6 | 1 |
| X_1030050 | 3 | 1 | 1 | 0 | NA | NA | NA | NA | 4 | 0.75 | NA | NA | 3 | 0.3333 | NA | NA | 1 | 1 | 4 | 0.5 |
| X_1030050 | 4 | 0.5 | 1 | 0 | NA | NA | NA | NA | 5 | 0.4 | 1 | 1 | 4 | 1 | NA | NA | 2 | 1 | 7 | 1 |
| X_1030051 | 4 | 0.75 | 1 | 0 | NA | NA | NA | NA | 5 | 0.6 | 1 | 1 | 3 | 0.6667 | NA | NA | 2 | 0.5 | 6 | 0.6667 |
| X_1030053 | 4 | 0.25 | 1 | 1 | NA | NA | NA | NA | 5 | 0.4 | 1 | 1 | 5 | 0.4 | NA | NA | 1 | 0 | 7 | 0.4286 |
| X_1030055 | 2 | 1 | 1 | 1 | NA | NA | NA | NA | 3 | 1 | 1 | 1 | 4 | 0.5 | NA | NA | 2 | 0 | 7 | 0.4286 |
| X_1030058 | 1 | 0 | 1 | 1 | NA | NA | NA | NA | 2 | 0.5 | 1 | 1 | 4 | 1 | NA | NA | 1 | 1 | 6 | 1 |
| X_1030060 | 2 | 1 | 1 | 1 | 1 | 1 | NA | NA | 4 | 1 | 1 | 1 | 4 | 1 | NA | NA | 2 | 1 | 7 | 1 |
| X_1030061 | 2 | 0.5 | NA | NA | 1 | 1 | NA | NA | 3 | 0.6667 | 1 | 0 | 4 | 0.25 | NA | NA | 3 | 0.3333 | 8 | 0.25 |
| X_1030063 | 2 | 0.5 | NA | NA | NA | NA | NA | NA | 2 | 0.5 | 1 | 0 | 3 | 0.6667 | NA | NA | 1 | 1 | 5 | 0.6 |
| X_1030063 | 2 | 1 | 1 | 1 | NA | NA | NA | NA | 3 | 1 | 1 | 1 | 5 | 0.8 | NA | NA | 2 | 1 | 8 | 0.875 |
| X_1030066 | 2 | 0.5 | 1 | 0 | NA | NA | NA | NA | 3 | 0.3333 | NA | NA | 4 | 0.5 | NA | NA | 3 | 0.3333 | 7 | 0.4286 |
| X_1030068 | 2 | 0.5 | NA | NA | NA | NA | NA | NA | 2 | 0.5 | NA | NA | 5 | 0.4 | NA | NA | 2 | 0 | 7 | 0.2857 |
| X_1030075 | 1 | 1 | NA | NA | NA | NA | NA | NA | 1 | 1 | NA | NA | 3 | 1 | NA | NA | 2 | 0 | 5 | 0.6 |
| X_1030075 | 2 | 0.5 | NA | NA | NA | NA | NA | NA | 2 | 0.5 | NA | NA | 4 | 0.25 | NA | NA | 2 | 0 | 6 | 0.1667 |
| X_1031212 | 1 | 0 | 1 | 0 | 1 | 1 | 2 | 0 | 5 | 0.2 | 3 | 0 | 3 | 0.3333 | 1 | 1 | 2 | 0 | 9 | 0.2222 |
| X_1031217 | 1 | 0 | 2 | 0 | NA | NA | 2 | 0.5 | 5 | 0.2 | 1 | 0 | 2 | 0 | 1 | 0 | 2 | 0.5 | 6 | 0.1667 |
| X_1033263 | 1 | 0 | 5 | 0 | 6 | 0.1667 | 3 | 0.3333 | 15 | 0.1333 | 2 | 0 | 1 | 0 | 2 | 0 | 4 | 0.25 | 9 | 0.1111 |
| X_1033276 | 1 | 1 | 7 | 0.4286 | 6 | 0 | NA | NA | 14 | 0.2857 | 2 | 0.5 | 1 | 0 | 3 | 0.3333 | 3 | 0.3333 | 9 | 0.3333 |
| X_1035498 | 1 | 1 | NA | NA | NA | NA | NA | NA | 1 | 1 | NA | NA | 1 | 0 | 1 | 1 | NA | NA | 2 | 0.5 |
| X_1035846 | 1 | 0 | NA | NA | NA | NA | 2 | 0.5 | 3 | 0.3333 | NA | NA | 2 | 0.5 | NA | NA | NA | NA | 2 | 0.5 |
| X_1035846 | 1 | 1 | NA | NA | NA | NA | 2 | 1 | 3 | 1 | NA | NA | 1 | 1 | NA | NA | NA | NA | 1 | 1 |
| X_1038986 | 1 | 1 | 1 | 1 | 1 | 0 | NA | NA | 3 | 0.6667 | 1 | 0 | 4 | 0.25 | 2 | 0 | 1 | 1 | 8 | 0.25 |
| X_1038990 | 1 | 1 | 1 | 1 | NA | NA | NA | NA | 2 | 1 | NA | NA | 4 | 1 | 2 | 1 | 1 | 1 | 7 | 1 |
| X_1038990 | 1 | 0 | 2 | 0.5 | 1 | 0 | NA | NA | 4 | 0.25 | NA | NA | 4 | 0.25 | 2 | 0 | 1 | 0 | 7 | 0.1429 |
| X_1038990 | 1 | 1 | 2 | 1 | 1 | 1 | NA | NA | 4 | 1 | NA | NA | 3 | 1 | 2 | 1 | 1 | 1 | 6 | 1 |
| X_1038991 | 1 | 1 | 2 | 1 | 1 | 0 | NA | NA | 4 | 0.75 | NA | NA | 2 | 1 | 2 | 1 | 1 | 1 | 5 | 1 |
| X_1038993 | 1 | 0 | 1 | 1 | 1 | 0 | NA | NA | 3 | 0.3333 | NA | NA | 1 | 1 | 1 | 1 | 1 | 0 | 3 | 0.6667 |
| X_1038997 | 1 | 1 | 1 | 1 | 1 | 1 | NA | NA | 3 | 1 | NA | NA | 1 | 1 | NA | NA | 1 | 1 | 2 | 1 |
| X_1038999 | NA | NA | 1 | 1 | 1 | 1 | NA | NA | 2 | 1 | NA | NA | 2 | 1 | 1 | 1 | 1 | 1 | 4 | 1 |
| X_1039001 | 1 | 1 | 1 | 1 | 1 | 1 | NA | NA | 3 | 1 | NA | NA | 2 | 1 | 1 | 1 | NA | NA | 3 | 1 |
| X_1039001 | 1 | 1 | 1 | 1 | 1 | 1 | NA | NA | 3 | 1 | NA | NA | 2 | 1 | 1 | 1 | 1 | 1 | 4 | 1 |
| X_1040824 | 1 | 0 | 1 | 0 | 2 | 0 | 2 | 0.5 | 6 | 0.1667 | NA | NA | 5 | 0.2 | 2 | 0.5 | 1 | 0 | 8 | 0.25 |
| X_1040829 | 1 | 1 | 1 | 0 | 2 | 0 | 3 | 0.6667 | 7 | 0.4286 | NA | NA | 5 | 0.2 | 1 | 0 | 1 | 1 | 7 | 0.2857 |
| X_1040830 | 1 | 0 | 1 | 0 | NA | NA | 2 | 1 | 4 | 0.5 | NA | NA | 3 | 0.3333 | 1 | 1 | NA | NA | 4 | 0.5 |
| X_1040830 | 1 | 1 | 1 | 0 | 1 | 1 | 2 | 1 | 5 | 0.8 | NA | NA | 2 | 1 | 1 | 1 | 1 | 1 | 4 | 1 |
| X_1040830 | 1 | 0 | 1 | 0 | 2 | 1 | 3 | 1 | 7 | 0.7143 | NA | NA | 5 | 1 | 1 | 1 | 1 | 1 | 7 | 1 |
| X_1040830 | NA | NA | 1 | 0 | 2 | 0 | 3 | 0.6667 | 6 | 0.3333 | NA | NA | 5 | 0.2 | 1 | 0 | 1 | 0 | 7 | 0.1429 |
| X_1040832 | 1 | 1 | 1 | 0 | 1 | 0 | 3 | 0.6667 | 6 | 0.5 | NA | NA | 3 | 0.3333 | 1 | 1 | NA | NA | 4 | 0.5 |
| X_1041268 | NA | NA | 8 | 0.25 | 1 | 1 | 2 | 0.5 | 11 | 0.3636 | 1 | 0 | 1 | 1 | 2 | 0.5 | NA | NA | 4 | 0.5 |
| X_1041269 | NA | NA | 9 | 0.8889 | NA | NA | 1 | 1 | 10 | 0.9 | NA | NA | 1 | 1 | 2 | 1 | NA | NA | 3 | 1 |
| X_1041269 | NA | NA | 8 | 1 | 1 | 1 | 2 | 1 | 11 | 1 | 1 | 1 | 1 | 1 | 2 | 1 | NA | NA | 4 | 1 |
| X_1041485 | 2 | 1 | NA | NA | 1 | 0 | NA | NA | 3 | 0.6667 | NA | NA | NA | NA | NA | NA | 1 | 1 | 1 | 1 |
| X_1041597 | 1 | 1 | NA | NA | NA | NA | NA | NA | 1 | 1 | NA | NA | 1 | 0 | 1 | 0 | 1 | 1 | 3 | 0.3333 |
| X_1041644 | 3 | 0 | NA | NA | 4 | 0 | 9 | 0.2222 | 16 | 0.125 | 1 | 0 | 11 | 0 | 4 | 0 | 3 | 0.3333 | 19 | 0.0526 |
| X_1041645 | 9 | 0 | 4 | 0 | 4 | 0 | 8 | 0.25 | 25 | 0.08 | 4 | 0.5 | 14 | 0.0714 | 10 | 0 | 4 | 0.25 | 32 | 0.125 |
| X_1041648 | 14 | 0 | 7 | 0.1429 | 9 | 0.1111 | 14 | 0.3571 | 44 | 0.1591 | 5 | 0.2 | 21 | 0.1905 | 11 | 0.0909 | 9 | 0.2222 | 46 | 0.1739 |
| X_1041648 | 15 | 0.4667 | 6 | 0.6667 | 8 | 0.5 | 13 | 0.4615 | 42 | 0.5 | 5 | 0.6 | 21 | 0.3333 | 13 | 0.3077 | 8 | 0.625 | 47 | 0.4043 |
| X_1041648 | 8 | 0.75 | 3 | 0.3333 | 7 | 0.5714 | 9 | 0.7778 | 27 | 0.6667 | 4 | 1 | 19 | 0.7368 | 10 | 0.9 | 4 | 0.75 | 37 | 0.8108 |
| X_1041648 | 11 | 1 | 6 | 1 | 7 | 1 | 11 | 1 | 35 | 1 | 5 | 1 | 16 | 1 | 11 | 0.9091 | 7 | 1 | 39 | 0.9744 |
| X_1041650 | 17 | 0.7059 | 8 | 0.5 | 9 | 0.3333 | 15 | 0.6 | 49 | 0.5714 | 5 | 0.6 | 20 | 0.85 | 12 | 0.5 | 10 | 0.7 | 47 | 0.7021 |
| X_1041651 | 13 | 0.8462 | 7 | 1 | 5 | 1 | 13 | 1 | 38 | 0.9474 | 3 | 1 | 19 | 1 | 7 | 1 | 10 | 1 | 39 | 1 |
| X_1041655 | 19 | 0.8947 | 9 | 0.8889 | 9 | 0.7778 | 18 | 0.8333 | 55 | 0.8545 | 8 | 0.375 | 22 | 0.7273 | 13 | 0.8462 | 11 | 0.9091 | 54 | 0.7407 |

| | | | | | | | | | | | | | | | | | | | | |
|---|---|---|---|---|---|---|---|---|---|---|---|---|---|---|---|---|---|---|---|---|
| X_1041656 | 15 | 0.6 | 9 | 0.7778 | 6 | 0.3333 | 19 | 0.4737 | 49 | 0.551 | 7 | 0.8571 | 21 | 0.6667 | 10 | 0.6 | 11 | 0.7273 | 49 | 0.6939 |
| X_1041657 | 17 | 0.5882 | 9 | 0.5556 | 11 | 0.2727 | 16 | 0.4375 | 53 | 0.4717 | 5 | 1 | 21 | 0.2381 | 6 | 0.3333 | 10 | 0.4 | 42 | 0.381 |
| X_1041658 | 7 | 0.1429 | 8 | 0.25 | 9 | 0.1111 | 16 | 0.125 | 40 | 0.15 | 7 | 0 | 21 | 0.1429 | 13 | 0.3077 | 8 | 0.25 | 49 | 0.1837 |
| X_1041658 | 17 | 0.9412 | 11 | 1 | 12 | 0.9167 | 20 | 0.9 | 60 | 0.9333 | 7 | 0.7143 | 24 | 0.875 | 14 | 1 | 12 | 0.8333 | 57 | 0.8772 |
| X_1041659 | 16 | 0.5625 | 11 | 0.3636 | 12 | 0.4167 | 16 | 0.8125 | 55 | 0.5636 | 8 | 0.375 | 25 | 0.4 | 16 | 0.4375 | 11 | 0.5455 | 60 | 0.4333 |
| X_1041660 | 20 | 1 | 11 | 1 | 13 | 1 | 20 | 0.95 | 64 | 0.9844 | 8 | 1 | 24 | 1 | 16 | 1 | 12 | 1 | 60 | 1 |
| X_1041660 | 12 | 0.1667 | 7 | 0 | 7 | 0.4286 | 13 | 0.3077 | 39 | 0.2308 | 7 | 0 | 19 | 0.1053 | 12 | 0.1667 | 7 | 0.1429 | 45 | 0.1111 |
| X_1041660 | 16 | 0.75 | 9 | 0.5556 | 10 | 0.8 | 16 | 0.625 | 51 | 0.6863 | 9 | 0.6667 | 20 | 0.85 | 11 | 0.9091 | 9 | 0.8889 | 49 | 0.8367 |
| X_1041661 | 15 | 0.9333 | 9 | 0.8889 | 13 | 0.9231 | 19 | 0.9474 | 56 | 0.9286 | 7 | 0.7143 | 20 | 0.9 | 11 | 0.9091 | 10 | 0.9 | 48 | 0.875 |
| X_1041665 | 17 | 1 | 13 | 1 | 11 | 1 | 15 | 1 | 56 | 1 | 9 | 1 | 18 | 1 | 17 | 1 | 11 | 1 | 55 | 1 |
| X_1041667 | 22 | 0.5 | 14 | 0.8571 | 13 | 0.5385 | 29 | 0.6897 | 78 | 0.641 | 7 | 0.5714 | 24 | 0.75 | 13 | 0.6154 | 13 | 0.3846 | 57 | 0.614 |
| X_1041667 | 22 | 0.1818 | 15 | 0.2667 | 15 | 0.2 | 24 | 0.2083 | 76 | 0.2105 | 9 | 0.1111 | 25 | 0.24 | 13 | 0.1538 | 10 | 0.2 | 57 | 0.193 |
| X_1041668 | 24 | 0.125 | 13 | 0.3846 | 18 | 0.1111 | 26 | 0.1923 | 81 | 0.1852 | 10 | 0.1 | 28 | 0.3214 | 17 | 0.2353 | 13 | 0.2308 | 68 | 0.25 |
| X_1041672 | 31 | 0.1935 | 20 | 0.2 | 22 | 0.2727 | 30 | 0.2667 | 103 | 0.233 | 12 | 0.0833 | 29 | 0.1724 | 21 | 0.2381 | 13 | 0.2308 | 75 | 0.1867 |
| X_1042081 | 1 | 1 | NA | NA | 1 | 1 | NA | NA | 2 | 1 | NA | NA | NA | NA | NA | NA | 1 | 1 | 1 | 1 |
| X_1042082 | NA | NA | NA | NA | 1 | 1 | NA | NA | 1 | 1 | NA | NA | NA | NA | NA | NA | 1 | 1 | 1 | 1 |
| X_1042084 | NA | NA | NA | NA | 1 | 1 | NA | NA | 1 | 1 | NA | NA | NA | NA | NA | NA | 1 | 1 | 1 | 1 |
| X_1042090 | NA | NA | NA | NA | 1 | 1 | NA | NA | 1 | 1 | NA | NA | NA | NA | NA | NA | 1 | 1 | 1 | 1 |
| X_1043307 | NA | NA | NA | NA | NA | NA | 1 | 1 | 1 | 1 | NA | NA | 1 | 1 | NA | NA | NA | NA | 1 | 1 |
| X_1043307 | NA | NA | NA | NA | NA | NA | 2 | 0.5 | 2 | 0.5 | NA | NA | 1 | 1 | NA | NA | NA | NA | 1 | 1 |
| X_1043310 | NA | NA | NA | NA | NA | NA | 2 | 1 | 2 | 1 | NA | NA | 1 | 1 | NA | NA | NA | NA | 1 | 1 |
| X_1043311 | NA | NA | NA | NA | NA | NA | 2 | 1 | 2 | 1 | NA | NA | 1 | 1 | NA | NA | NA | NA | 1 | 1 |
| X_1048423 | 3 | 0 | 10 | 0.4 | 4 | 0.5 | 10 | 0.3 | 27 | 0.3333 | 3 | 0.3333 | 10 | 0.3 | 7 | 0.1429 | 12 | 0.4167 | 32 | 0.3125 |
| X_1048438 | 1 | 0 | 8 | 0.25 | 1 | 0 | 11 | 0.1818 | 21 | 0.1905 | 2 | 0 | 4 | 0 | 3 | 0.3333 | 9 | 0.2222 | 18 | 0.1667 |
| X_1048439 | NA | NA | 9 | 0.2222 | 2 | 0 | 10 | 0.1 | 21 | 0.1429 | 2 | 0 | 4 | 0.25 | 3 | 0 | 7 | 0.4286 | 16 | 0.25 |
| X_1048439 | 2 | 0 | 8 | 0.25 | 1 | 0 | 8 | 0.25 | 19 | 0.2105 | 2 | 0.5 | 2 | 0 | 2 | 0 | 5 | 0.2 | 11 | 0.1818 |
| X_1048617 | 1 | 0 | 8 | 0.125 | 1 | 0 | 8 | 0.25 | 18 | 0.1667 | 2 | 0 | 3 | 0.3333 | 1 | 0 | 7 | 0.1429 | 13 | 0.1538 |
| X_1049185 | NA | NA | 3 | 0.6667 | 1 | 1 | NA | NA | 4 | 0.75 | NA | NA | 1 | 1 | 1 | 1 | 4 | 0.25 | 6 | 0.5 |
| X_1049197 | NA | NA | 1 | 1 | 1 | 0 | NA | NA | 2 | 0.5 | NA | NA | NA | NA | 1 | 0 | 2 | 0.5 | 3 | 0.3333 |
| X_1049207 | NA | NA | 1 | 1 | 1 | 0 | NA | NA | 2 | 0.5 | NA | NA | NA | NA | 1 | 0 | 2 | 0.5 | 3 | 0.3333 |
| X_1049210 | NA | NA | 1 | 1 | 1 | 1 | NA | NA | 2 | 1 | NA | NA | NA | NA | 1 | 1 | 1 | 1 | 2 | 1 |
| X_1050123 | 1 | 1 | 4 | 0 | NA | NA | 6 | 0.1667 | 11 | 0.1818 | 2 | 0.5 | 2 | 0 | 3 | 0.3333 | 2 | 0 | 9 | 0.2222 |
| X_1050142 | 1 | 1 | 3 | 0 | NA | NA | 6 | 0 | 10 | 0.1 | 1 | 1 | NA | NA | NA | NA | 1 | 1 | 2 | 1 |
| X_1050145 | 1 | 1 | NA | NA | NA | NA | 5 | 0.2 | 6 | 0.3333 | 1 | 0 | NA | NA | NA | NA | 1 | 1 | 2 | 0.5 |
| X_1052017 | NA | NA | 1 | 1 | 2 | 1 | NA | NA | 3 | 1 | NA | NA | NA | NA | NA | NA | 1 | 1 | 1 | 1 |
| X_1052019 | NA | NA | 1 | 0 | 2 | 0.5 | NA | NA | 3 | 0.3333 | NA | NA | NA | NA | NA | NA | 1 | 1 | 1 | 1 |
| X_1052033 | 1 | 1 | NA | NA | NA | NA | NA | NA | 1 | 1 | NA | NA | NA | NA | NA | NA | 1 | 1 | 1 | 1 |
| X_1052033 | 1 | 0 | 1 | 1 | 1 | 0 | NA | NA | 3 | 0.3333 | NA | NA | NA | NA | NA | NA | 1 | 1 | 1 | 1 |
| X_1052034 | 1 | 0 | 1 | 1 | 2 | 1 | NA | NA | 4 | 0.75 | NA | NA | NA | NA | NA | NA | 1 | 1 | 1 | 1 |
| X_1052037 | 1 | 1 | 1 | 1 | 1 | 1 | NA | NA | 3 | 1 | NA | NA | NA | NA | NA | NA | 1 | 1 | 1 | 1 |
| X_1052038 | 1 | 1 | 1 | 1 | 2 | 1 | NA | NA | 4 | 1 | NA | NA | NA | NA | NA | NA | 1 | 1 | 1 | 1 |
| X_1052039 | 1 | 1 | 1 | 1 | 2 | 1 | NA | NA | 4 | 1 | NA | NA | NA | NA | NA | NA | 1 | 1 | 1 | 1 |
| X_1052470 | 70 | 0.1714 | 54 | 0.0741 | 54 | 0.0741 | 50 | 0.2 | 228 | 0.1316 | 57 | 0.1053 | 47 | 0.0851 | 54 | 0.1667 | 40 | 0.2 | 198 | 0.1364 |
| X_1052509 | 4 | 0 | 4 | 0 | 4 | 0.25 | 11 | 0.1818 | 23 | 0.1304 | 2 | 0 | 5 | 0.2 | 5 | 0 | 3 | 0 | 15 | 0.0667 |
| X_1052513 | 2 | 0 | 4 | 0 | 4 | 0 | 9 | 0.1111 | 19 | 0.0526 | 3 | 0.3333 | 5 | 0 | 5 | 0 | 3 | 0.3333 | 16 | 0.125 |
| X_1052513 | 3 | 0 | 3 | 0 | 4 | 0 | 10 | 0.2 | 20 | 0.1 | 3 | 0 | 5 | 0 | 3 | 0.3333 | 3 | 0.3333 | 14 | 0.1429 |
| X_1052515 | 3 | 0 | 4 | 0 | 4 | 0 | 9 | 0.1111 | 20 | 0.05 | 3 | 0 | 4 | 0 | 5 | 0.2 | 2 | 0 | 14 | 0.0714 |
| X_1052518 | 3 | 0.6667 | 4 | 0.25 | 3 | 0.6667 | 9 | 0.6667 | 19 | 0.5789 | 3 | 0.3333 | 4 | 0.75 | 2 | 0 | 2 | 1 | 11 | 0.5455 |
| X_1052518 | 2 | 0 | 2 | 0 | 4 | 0 | 6 | 0.1667 | 14 | 0.0714 | 3 | 0.3333 | 4 | 0 | 4 | 0.25 | 3 | 0.3333 | 14 | 0.2143 |
| X_1052520 | 3 | 0 | 4 | 0.5 | 4 | 0.25 | 8 | 0.375 | 19 | 0.3158 | 3 | 0.3333 | 3 | 0.3333 | 3 | 0.6667 | 2 | 0.5 | 11 | 0.4545 |
| X_1059738 | NA | NA | NA | NA | NA | NA | 1 | 1 | 1 | 1 | 1 | 1 | NA | NA | NA | NA | NA | NA | 1 | 1 |
| X_1059738 | 1 | 0 | NA | NA | NA | NA | 1 | 1 | 2 | 0.5 | 1 | 1 | NA | NA | NA | NA | NA | NA | 1 | 1 |
| X_1059741 | 1 | 1 | NA | NA | NA | NA | 1 | 1 | 2 | 1 | 1 | 1 | NA | NA | NA | NA | NA | NA | 1 | 1 |
| X_1059742 | 1 | 1 | NA | NA | NA | NA | 1 | 1 | 2 | 1 | 1 | 1 | NA | NA | NA | NA | NA | NA | 1 | 1 |
| X_1068307 | 3 | 0.3333 | 1 | 0 | 7 | 0 | 1 | 0 | 12 | 0.0833 | 3 | 0 | 3 | 0 | NA | NA | 3 | 0.3333 | 9 | 0.1111 |
| X_1077866 | 1 | 1 | NA | NA | NA | NA | NA | NA | 1 | 1 | NA | NA | NA | NA | NA | NA | 1 | 1 | 1 | 1 |
| X_1077867 | 1 | 1 | NA | NA | NA | NA | NA | NA | 1 | 1 | NA | NA | NA | NA | NA | NA | 1 | 1 | 1 | 1 |
| X_1080487 | 1 | 0 | 1 | 0 | 3 | 0 | 2 | 0.5 | 7 | 0.1429 | NA | NA | 1 | 0 | NA | NA | 4 | 0.25 | 5 | 0.2 |
| X_1080493 | 1 | 0 | NA | NA | 3 | 0.3333 | 2 | 1 | 6 | 0.5 | NA | NA | NA | NA | NA | NA | 3 | 0.3333 | 3 | 0.3333 |
| X_1082041 | 12 | 0.1667 | 9 | 0.1111 | 13 | 0 | 8 | 0.125 | 42 | 0.0952 | 4 | 0.25 | 9 | 0 | 21 | 0.2381 | 14 | 0.1429 | 48 | 0.1667 |
| X_1082049 | 10 | 0.1 | 10 | 0.1 | 18 | 0.1111 | 10 | 0 | 48 | 0.0833 | 8 | 0.125 | 11 | 0.0909 | 22 | 0.0455 | 15 | 0.1333 | 56 | 0.0893 |
| X_1082054 | 12 | 0.3333 | 6 | 0.1667 | 13 | 0.2308 | 14 | 0.0714 | 45 | 0.2 | 8 | 0 | 4 | 0.5 | 16 | 0.3125 | 13 | 0.2308 | 41 | 0.2439 |
| X_1082058 | 12 | 0 | 11 | 0 | 20 | 0.15 | 15 | 0.1333 | 58 | 0.0862 | 8 | 0.25 | 14 | 0.0714 | 21 | 0.0952 | 13 | 0.3077 | 56 | 0.1607 |
| X_1098585 | NA | NA | NA | NA | 2 | 0.5 | 1 | 0 | 3 | 0.3333 | 1 | 1 | 4 | 0.5 | 1 | 1 | 3 | 0.6667 | 9 | 0.6667 |
| X_1098588 | NA | NA | NA | NA | 1 | 1 | 1 | 0 | 2 | 0.5 | 1 | 1 | 4 | 0 | NA | NA | 4 | 0 | 9 | 0.1111 |

| | | | | | | | | | | | | | | | | | | | | |
|---|---|---|---|---|---|---|---|---|---|---|---|---|---|---|---|---|---|---|---|---|
| X_1098590 | NA | NA | NA | NA | 2 | 1 | 1 | 1 | 3 | 1 | 1 | 1 | 2 | 1 | 1 | 1 | 3 | 0.6667 | 7 | 0.8571 |
| X_1098591 | NA | NA | NA | NA | 2 | 0.5 | 1 | 1 | 3 | 0.6667 | 1 | 0 | 2 | 0 | 1 | 0 | 3 | 0.3333 | 7 | 0.1429 |
| X_1098595 | 1 | 1 | NA | NA | 2 | 1 | 1 | 1 | 4 | 1 | 1 | 1 | 5 | 0.8 | 1 | 1 | 7 | 0.8571 | 14 | 0.8571 |
| X_1098600 | 2 | 0.5 | 2 | 1 | 2 | 1 | NA | NA | 6 | 0.8333 | NA | NA | 6 | 1 | 1 | 1 | 5 | 1 | 12 | 1 |
| X_1098600 | 3 | 0 | 1 | 0 | 4 | 0 | 3 | 0.3333 | 11 | 0.0909 | 1 | 0 | 6 | 0 | 1 | 0 | 5 | 0.4 | 13 | 0.1538 |
| X_1098601 | 3 | 0.6667 | 2 | 1 | 3 | 0.3333 | 3 | 1 | 11 | 0.7273 | 1 | 1 | 8 | 0.625 | 1 | 1 | 5 | 0.6 | 15 | 0.6667 |
| X_1098603 | 3 | 1 | 2 | 1 | 4 | 1 | 5 | 1 | 14 | 1 | 1 | 1 | 7 | 1 | 1 | 1 | 6 | 1 | 15 | 1 |
| X_1098603 | 3 | 1 | 1 | 1 | 2 | 1 | 5 | 1 | 11 | 1 | 1 | 1 | 6 | 1 | 2 | 1 | 7 | 1 | 16 | 1 |
| X_1098604 | 3 | 0.6667 | 2 | 1 | 3 | 0.6667 | 4 | 0.75 | 12 | 0.75 | 1 | 1 | 7 | 0.4286 | 2 | 0 | 7 | 0.4286 | 17 | 0.4118 |
| X_1098608 | 4 | 0 | 3 | 0 | 3 | 0 | 4 | 0.25 | 14 | 0.0714 | 1 | 0 | 7 | 0.1429 | 3 | 0 | 5 | 0 | 16 | 0.0625 |
| X_1098610 | 4 | 0 | 4 | 0.5 | 5 | 0.4 | 6 | 0.6667 | 19 | 0.4211 | 1 | 0 | 9 | 0.3333 | 3 | 0.6667 | 7 | 0.1429 | 20 | 0.3 |
| X_1115422 | 1 | 0 | 2 | 0.5 | NA | NA | 1 | 0 | 4 | 0.25 | NA | NA | NA | NA | 2 | 0.5 | 1 | 0 | 3 | 0.3333 |
| X_1115423 | 1 | 0 | 2 | 0.5 | NA | NA | 1 | 1 | 4 | 0.5 | NA | NA | NA | NA | 1 | 0 | 1 | 1 | 2 | 0.5 |
| X_1115424 | 1 | 1 | 1 | 0 | NA | NA | 1 | 1 | 3 | 0.6667 | NA | NA | NA | NA | 2 | 1 | 1 | 1 | 3 | 1 |
| X_1115424 | 1 | 1 | 2 | 1 | NA | NA | 1 | 1 | 4 | 1 | NA | NA | NA | NA | 2 | 1 | 1 | 1 | 3 | 1 |
| X_1115425 | 1 | 0 | 2 | 0.5 | NA | NA | 1 | 1 | 4 | 0.5 | NA | NA | NA | NA | 1 | 1 | 1 | 1 | 2 | 1 |
| X_1115426 | 1 | 0 | 2 | 0.5 | NA | NA | 1 | 0 | 4 | 0.25 | NA | NA | NA | NA | 2 | 0 | 1 | 1 | 3 | 0.3333 |
| X_1115427 | 1 | 1 | 2 | 1 | NA | NA | NA | NA | 3 | 1 | NA | NA | NA | NA | 2 | 1 | 1 | 1 | 3 | 1 |
| X_1115428 | 1 | 1 | 2 | 1 | NA | NA | 1 | 1 | 4 | 1 | NA | NA | NA | NA | 2 | 0.5 | 1 | 1 | 3 | 0.6667 |
| X_1115428 | 1 | 0 | 2 | 0.5 | NA | NA | 1 | 0 | 4 | 0.25 | NA | NA | NA | NA | 2 | 0.5 | 1 | 1 | 3 | 0.6667 |
| X_1115429 | 1 | 1 | 2 | 1 | NA | NA | 1 | 1 | 4 | 1 | NA | NA | NA | NA | 2 | 1 | NA | NA | 2 | 1 |
| X_1115430 | 1 | 0 | 2 | 1 | NA | NA | 1 | 0 | 4 | 0.5 | NA | NA | NA | NA | 2 | 0 | 1 | 1 | 3 | 0.3333 |
| X_1115431 | 1 | 1 | NA | NA | NA | NA | 1 | 1 | 2 | 1 | NA | NA | NA | NA | 1 | 1 | 1 | 1 | 2 | 1 |
| X_1115432 | 1 | 1 | 1 | 1 | NA | NA | 1 | 1 | 3 | 1 | NA | NA | NA | NA | 1 | 1 | NA | NA | 1 | 1 |
| X_1115433 | 1 | 0 | 2 | 0.5 | NA | NA | 1 | 0 | 4 | 0.25 | NA | NA | NA | NA | 1 | 0 | 1 | 1 | 2 | 0.5 |
| X_1115437 | 1 | 0 | 2 | 0.5 | NA | NA | 1 | 0 | 4 | 0.25 | NA | NA | NA | NA | 1 | 0 | 1 | 1 | 2 | 0.5 |
| X_1115447 | NA | NA | 2 | 0.5 | 1 | 1 | 2 | 0 | 5 | 0.4 | 3 | 0.3333 | NA | NA | 2 | 0 | 1 | 0 | 6 | 0.1667 |
| X_1119505 | 1 | 0 | 6 | 0 | 6 | 0.3333 | 4 | 0 | 17 | 0.1176 | 5 | 0 | 4 | 0.25 | 6 | 0 | 9 | 0.1111 | 24 | 0.0833 |
| X_1119512 | 1 | 0 | 3 | 0 | 5 | 0.4 | 4 | 0 | 13 | 0.1538 | 2 | 0 | 2 | 0 | 5 | 0 | 8 | 0.125 | 17 | 0.0588 |
| X_1119522 | 1 | 0 | 2 | 0 | 5 | 0.2 | 4 | 0.25 | 12 | 0.1667 | 3 | 0.3333 | 1 | 1 | 3 | 0 | 7 | 0 | 14 | 0.1429 |
| X_1119526 | 1 | 0 | 2 | 0 | 4 | 0.25 | 3 | 0 | 10 | 0.1 | 3 | 0 | 2 | 0 | 4 | 0.25 | 8 | 0.25 | 17 | 0.1765 |
| X_1119528 | 1 | 1 | 3 | 1 | 3 | 1 | 4 | 1 | 11 | 1 | 4 | 0.75 | 2 | 1 | 4 | 1 | 9 | 1 | 19 | 0.9474 |
| X_1119529 | 1 | 0 | 3 | 0.6667 | 4 | 0 | 4 | 0.25 | 12 | 0.25 | 3 | 0 | 1 | 0 | 4 | 0.5 | 6 | 0.1667 | 14 | 0.2143 |
| X_1119529 | 1 | 1 | 2 | 0 | 5 | 0 | 3 | 0.3333 | 11 | 0.1818 | 4 | 0.25 | 2 | 0 | 4 | 0.75 | 8 | 0.375 | 18 | 0.3889 |
| X_1119529 | 1 | 0 | 3 | 0 | 4 | 0.25 | 2 | 0 | 10 | 0.1 | 3 | 0 | 2 | 0 | 4 | 0.25 | 8 | 0.375 | 17 | 0.2353 |
| X_1119530 | 1 | 0 | 3 | 0 | 5 | 0 | 4 | 0.5 | 13 | 0.1538 | 4 | 0 | 2 | 0.5 | 4 | 0.25 | 7 | 0 | 17 | 0.1176 |
| X_1119531 | 1 | 0 | 3 | 0.3333 | 5 | 0.4 | 4 | 0.25 | 13 | 0.3077 | 3 | 0.3333 | 1 | 0 | 4 | 0.25 | 7 | 0.4286 | 15 | 0.3333 |
| X_1119531 | 1 | 0 | 3 | 0 | 4 | 0.5 | 4 | 0 | 12 | 0.1667 | 3 | 0.3333 | 2 | 0 | 4 | 0 | 8 | 0.125 | 17 | 0.1176 |
| X_1119531 | 1 | 0 | 2 | 0.5 | 3 | 0 | 3 | 0.3333 | 9 | 0.2222 | 3 | 0 | 2 | 0 | 3 | 0 | 6 | 0.1667 | 14 | 0.0714 |
| X_1119532 | 1 | 0 | 2 | 0 | 3 | 0 | 4 | 0.25 | 10 | 0.1 | 3 | 0.3333 | 2 | 0 | 3 | 0 | 8 | 0 | 16 | 0.0625 |
| X_1120257 | 1 | 1 | 2 | 0 | 2 | 0 | 1 | 0 | 6 | 0.1667 | 3 | 0 | 2 | 0 | 2 | 0 | 6 | 0.1667 | 13 | 0.0769 |
| X_1127452 | NA | NA | NA | NA | NA | NA | 1 | 1 | 1 | 1 | NA | NA | NA | NA | NA | NA | 1 | 1 | 1 | 1 |
| X_1188386 | NA | NA | 1 | 1 | NA | NA | 1 | 1 | 2 | 1 | NA | NA | NA | NA | NA | NA | 1 | 1 | 1 | 1 |
| X_1188386 | NA | NA | 1 | 1 | NA | NA | 1 | 0 | 2 | 0.5 | NA | NA | NA | NA | NA | NA | 1 | 1 | 1 | 1 |
| X_1188386 | NA | NA | 1 | 0 | NA | NA | 1 | 1 | 2 | 0.5 | NA | NA | NA | NA | NA | NA | 1 | 1 | 1 | 1 |
| X_1188440 | NA | NA | 3 | 0.6667 | 1 | 1 | 2 | 0 | 6 | 0.5 | 1 | 0 | NA | NA | 2 | 0 | 1 | 1 | 4 | 0.25 |
| X_1188440 | NA | NA | 3 | 0.3333 | 1 | 1 | 2 | 0.5 | 6 | 0.5 | 1 | 0 | NA | NA | 2 | 0 | 1 | 1 | 4 | 0.25 |
| X_1193100 | 1 | 1 | NA | NA | NA | NA | 3 | 1 | 4 | 1 | NA | NA | NA | NA | 1 | 1 | NA | NA | 1 | 1 |
| X_1193102 | 1 | 1 | NA | NA | NA | NA | 3 | 1 | 4 | 1 | NA | NA | NA | NA | 1 | 1 | NA | NA | 1 | 1 |
| X_1193107 | 1 | 1 | NA | NA | NA | NA | 3 | 0.3333 | 4 | 0.5 | NA | NA | NA | NA | 1 | 1 | 1 | 1 | 2 | 1 |
| X_1193109 | 1 | 0 | NA | NA | NA | NA | 3 | 1 | 4 | 0.75 | NA | NA | NA | NA | 1 | 1 | 1 | 1 | 2 | 1 |
| X_1193109 | 1 | 1 | NA | NA | NA | NA | 2 | 1 | 3 | 1 | NA | NA | 1 | 1 | 1 | 0 | 1 | 0 | 3 | 0.3333 |
| X_1193114 | NA | NA | NA | NA | NA | NA | 2 | 0.5 | 2 | 0.5 | NA | NA | 1 | 0 | 1 | 1 | 1 | 1 | 3 | 0.6667 |
| X_1193114 | 1 | 0 | NA | NA | NA | NA | 2 | 0.5 | 3 | 0.3333 | NA | NA | NA | NA | 1 | 1 | 1 | 0 | 2 | 0.5 |
| X_1198201 | 4 | 0.5 | 1 | 0 | 3 | 0 | 5 | 0.6 | 13 | 0.3846 | 1 | 0 | 3 | 0.6667 | 3 | 0.3333 | 2 | 0 | 9 | 0.3333 |
| X_1213046 | 4 | 0.5 | 2 | 0 | 8 | 0.375 | 7 | 0.5714 | 21 | 0.4286 | 2 | 0.5 | 9 | 0.5556 | 8 | 0.5 | 6 | 0.5 | 25 | 0.52 |
| X_1213049 | 2 | 1 | 1 | 1 | 7 | 1 | 4 | 0.75 | 14 | 0.9286 | 1 | 1 | 7 | 0.8571 | 1 | 1 | 4 | 1 | 13 | 0.9231 |
| X_1213049 | 3 | 0 | 1 | 0 | 7 | 0.1429 | 4 | 0 | 15 | 0.0667 | 2 | 0 | 8 | 0 | 3 | 0 | 6 | 0.1667 | 19 | 0.0526 |
| X_1213049 | 3 | 0 | NA | NA | 8 | 0.125 | 4 | 0.25 | 15 | 0.1333 | 2 | 0.5 | 8 | 0.25 | 5 | 0.6 | 6 | 0.1667 | 21 | 0.3333 |
| X_1213050 | 3 | 0.3333 | 1 | 0 | 8 | 0.25 | 5 | 0.2 | 17 | 0.2353 | 1 | 0 | 9 | 0.2222 | 6 | 0.3333 | 5 | 0 | 21 | 0.1905 |
| X_1213051 | 2 | 0 | 2 | 0 | 4 | 0.25 | 6 | 0.1667 | 14 | 0.1429 | 2 | 0 | 6 | 0.3333 | 5 | 0.2 | 5 | 0.6 | 18 | 0.3333 |
| X_1213052 | 1 | 1 | NA | NA | 7 | 1 | 6 | 1 | 14 | 1 | 2 | 1 | 7 | 1 | 4 | 1 | 3 | 1 | 16 | 1 |
| X_1213053 | 2 | 1 | 2 | 1 | 7 | 1 | 5 | 1 | 16 | 1 | 2 | 1 | 7 | 1 | 6 | 1 | 4 | 1 | 19 | 1 |
| X_1213056 | 2 | 1 | 1 | 1 | 5 | 0.6 | 3 | 1 | 11 | 0.8182 | 2 | 0 | 5 | 0.4 | 6 | 1 | 6 | 0.1667 | 19 | 0.4737 |
| X_1213056 | 2 | 1 | 1 | 1 | 4 | 1 | 4 | 1 | 11 | 1 | 2 | 1 | 6 | 1 | 6 | 1 | 5 | 1 | 19 | 1 |

| | | | | | | | | | | | | | | | | | | | | |
|---|---|---|---|---|---|---|---|---|---|---|---|---|---|---|---|---|---|---|---|---|
| X_1213056 | 2 | 0 | 2 | 0 | 5 | 0.2 | 4 | 0 | 13 | 0.0769 | 2 | 0 | 7 | 0.1429 | 5 | 0.2 | 5 | 0 | 19 | 0.1053 |
| X_1213059 | 2 | 0 | 1 | 0 | 5 | 0.2 | 3 | 0 | 11 | 0.0909 | 1 | 0 | 7 | 0.1429 | 4 | 0.25 | 4 | 0 | 16 | 0.125 |
| X_1213062 | 1 | 1 | 1 | 0 | 5 | 0.6 | 2 | 0.5 | 9 | 0.5556 | 2 | 1 | 7 | 0.5714 | 4 | 0.75 | 3 | 1 | 16 | 0.75 |
| X_1213064 | 2 | 0 | 2 | 0 | 5 | 0.2 | 2 | 0 | 11 | 0.0909 | 1 | 0 | 6 | 0.3333 | 3 | 0 | 3 | 0 | 13 | 0.1538 |
| X_1213064 | 2 | 0 | 1 | 0 | 5 | 0 | 3 | 0.3333 | 11 | 0.0909 | 2 | 0 | 5 | 0 | 3 | 0 | 2 | 0.5 | 12 | 0.0833 |
| X_1213066 | 2 | 0 | 2 | 0 | 6 | 0.1667 | 3 | 0 | 13 | 0.0769 | 2 | 0.5 | 7 | 0 | 2 | 0 | 3 | 0 | 14 | 0.0714 |
| X_1293617 | NA | NA | NA | NA | 1 | 0 | 1 | 1 | 2 | 0.5 | 1 | 1 | 2 | 0 | 3 | 0 | NA | NA | 6 | 0.1667 |
| X_1293663 | 2 | 0.5 | 6 | 0 | 4 | 0 | 3 | 0.3333 | 15 | 0.1333 | 1 | 0 | 3 | 0.3333 | 7 | 0.2857 | 3 | 0 | 14 | 0.2143 |
| X_1324847 | NA | NA | 3 | 0.3333 | 3 | 0.3333 | 2 | 0 | 8 | 0.25 | 1 | 0 | 1 | 0 | 3 | 0.6667 | 5 | 0.6 | 10 | 0.5 |
| X_1324847 | 1 | 1 | 3 | 0.3333 | 3 | 0.3333 | 1 | 1 | 8 | 0.5 | 1 | 0 | 1 | 0 | 3 | 0.6667 | 6 | 0.1667 | 11 | 0.2727 |
| X_1324851 | 1 | 1 | 3 | 0.3333 | 3 | 0.3333 | 1 | 0 | 8 | 0.375 | NA | NA | NA | NA | 1 | 1 | 4 | 0 | 5 | 0.2 |
| X_1324879 | NA | NA | NA | NA | NA | NA | 1 | 1 | 1 | 1 | NA | NA | NA | NA | 1 | 1 | NA | NA | 1 | 1 |
| X_1324890 | NA | NA | NA | NA | NA | NA | 1 | 1 | 1 | 1 | NA | NA | NA | NA | 1 | 1 | NA | NA | 1 | 1 |
| X_1324891 | NA | NA | NA | NA | NA | NA | 1 | 1 | 1 | 1 | NA | NA | NA | NA | 1 | 1 | NA | NA | 1 | 1 |
| X_1325859 | NA | NA | NA | NA | NA | NA | 1 | 1 | 1 | 1 | NA | NA | NA | NA | 1 | 1 | NA | NA | 1 | 1 |
| X_1329753 | 3 | 0 | 5 | 0.4 | 2 | 0 | 1 | 0 | 11 | 0.1818 | NA | NA | 10 | 0.1 | 5 | 0.2 | 6 | 0 | 21 | 0.0952 |
| X_1329754 | 4 | 0 | 5 | 0 | 2 | 0.5 | 1 | 0 | 12 | 0.0833 | NA | NA | 11 | 0.1818 | 4 | 0 | 5 | 0.2 | 20 | 0.15 |
| X_1329754 | 4 | 0.25 | 5 | 0 | 2 | 0 | 1 | 0 | 12 | 0.0833 | NA | NA | 11 | 0.1818 | 6 | 0.1667 | 7 | 0 | 24 | 0.125 |
| X_1329851 | 4 | 0 | 6 | 0.3333 | 1 | 0 | 1 | 0 | 12 | 0.1667 | NA | NA | 8 | 0 | 6 | 0 | 6 | 0.1667 | 20 | 0.05 |
| X_1329851 | 5 | 0.2 | 5 | 0.4 | 2 | 0.5 | 1 | 0 | 13 | 0.3077 | NA | NA | 8 | 0.5 | 4 | 0.75 | 6 | 0.1667 | 18 | 0.4444 |
| X_1329853 | 3 | 0 | 3 | 0.3333 | 1 | 0 | 1 | 0 | 8 | 0.125 | NA | NA | 7 | 0.1429 | 5 | 0 | 4 | 0 | 16 | 0.0625 |
| X_1329868 | 7 | 0.4286 | 4 | 0 | 7 | 0 | 3 | 0 | 21 | 0.1429 | 2 | 0.5 | 11 | 0.1818 | 5 | 0 | 7 | 0 | 25 | 0.12 |
| X_1330499 | NA | NA | NA | NA | NA | NA | 1 | 1 | 1 | 1 | NA | NA | 1 | 1 | 1 | 1 | NA | NA | 2 | 1 |
| X_1330513 | NA | NA | NA | NA | NA | NA | 1 | 1 | 1 | 1 | NA | NA | 1 | 1 | 1 | 0 | NA | NA | 2 | 0.5 |
| X_1330516 | NA | NA | NA | NA | NA | NA | 1 | 1 | 1 | 1 | NA | NA | NA | NA | 1 | 1 | NA | NA | 1 | 1 |
| X_1331239 | 2 | 0 | 3 | 0 | 3 | 0 | 3 | 0.6667 | 11 | 0.1818 | 1 | 0 | 1 | 0 | NA | NA | 2 | 0.5 | 4 | 0.25 |
| X_1331249 | NA | NA | 2 | 0.5 | NA | NA | 2 | 0 | 4 | 0.25 | 1 | 0 | 2 | 0 | NA | NA | 2 | 1 | 5 | 0.4 |
| X_1331251 | NA | NA | 2 | 1 | NA | NA | 2 | 1 | 4 | 1 | 1 | 1 | 2 | 1 | NA | NA | 2 | 1 | 5 | 1 |
| X_1331252 | NA | NA | 1 | 1 | NA | NA | 2 | 1 | 3 | 1 | 1 | 0 | 2 | 1 | NA | NA | 2 | 0.5 | 5 | 0.6 |
| X_1331635 | NA | NA | 3 | 0.3333 | NA | NA | 2 | 0 | 5 | 0.2 | NA | NA | 1 | 0 | NA | NA | 2 | 0.5 | 3 | 0.3333 |
| X_1331635 | NA | NA | 3 | 0.3333 | NA | NA | 2 | 0 | 5 | 0.2 | NA | NA | 1 | 0 | NA | NA | 2 | 0.5 | 3 | 0.3333 |
| X_1331963 | 13 | 0.2308 | 6 | 0 | 6 | 0 | 10 | 0.3 | 35 | 0.1714 | 5 | 0.6 | 6 | 0.3333 | 15 | 0.2667 | 6 | 0.1667 | 32 | 0.3125 |
| X_1332056 | 10 | 1 | 8 | 1 | 9 | 1 | 10 | 1 | 37 | 1 | 7 | 1 | 8 | 1 | 13 | 1 | 7 | 1 | 35 | 1 |
| X_1332057 | 14 | 0.2143 | 8 | 0.25 | 9 | 0 | 10 | 0.1 | 41 | 0.1463 | 6 | 0.3333 | 6 | 0.3333 | 17 | 0.2941 | 8 | 0.25 | 37 | 0.2973 |
| X_1332058 | 14 | 0.2143 | 10 | 0.4 | 10 | 0.3 | 11 | 0.5455 | 45 | 0.3556 | 6 | 0.8333 | 9 | 0.2222 | 17 | 0.6471 | 8 | 0.375 | 40 | 0.525 |
| X_1332058 | 15 | 0.0667 | 10 | 0 | 10 | 0.2 | 11 | 0 | 46 | 0.0652 | 5 | 0 | 9 | 0.2222 | 17 | 0 | 8 | 0 | 39 | 0.0513 |
| X_1332060 | 14 | 0.3571 | 8 | 0.125 | 11 | 0.2727 | 10 | 0.2 | 43 | 0.2558 | 6 | 0.1667 | 9 | 0.3333 | 16 | 0.4375 | 9 | 0.5556 | 40 | 0.4 |
| X_1333534 | 2 | 0.5 | 1 | 0 | 1 | 0 | 4 | 0.5 | 8 | 0.375 | 1 | 0 | 6 | 0 | 1 | 0 | 3 | 0.3333 | 11 | 0.0909 |
| X_1333539 | 2 | 0 | 1 | 0 | 1 | 0 | 4 | 0.25 | 8 | 0.125 | 1 | 0 | 7 | 0.1429 | 2 | 0.5 | 4 | 0.25 | 14 | 0.2143 |
| X_1333548 | 2 | 0 | 2 | 0 | 1 | 1 | 1 | 0 | 6 | 0.1667 | 1 | 1 | 6 | 0.3333 | 3 | 0.3333 | 4 | 0.25 | 14 | 0.3571 |
| X_1333569 | 5 | 0.6 | 1 | 0 | NA | NA | 1 | 1 | 7 | 0.5714 | 1 | 0 | 5 | 0.4 | 5 | 0.2 | 6 | 0.5 | 17 | 0.3529 |
| X_1333569 | 5 | 0.2 | 1 | 0 | NA | NA | NA | NA | 6 | 0.1667 | 1 | 0 | 4 | 0 | 5 | 0 | 5 | 0.2 | 15 | 0.0667 |
| X_1334383 | 3 | 0 | 3 | 0.3333 | 6 | 0 | 3 | 0 | 15 | 0.0667 | 4 | 0 | 6 | 0 | 4 | 0.25 | 2 | 0 | 16 | 0.0625 |
| X_1334385 | 3 | 0 | 3 | 0 | 5 | 0.2 | 4 | 0 | 15 | 0.0667 | 4 | 0 | 6 | 0 | 3 | 0 | 3 | 0.3333 | 16 | 0.0625 |
| X_1334385 | 3 | 0.3333 | 2 | 0.5 | 5 | 0.6 | 3 | 0.6667 | 13 | 0.5385 | 4 | 0.75 | 6 | 0.3333 | 4 | 0.75 | 3 | 0.3333 | 17 | 0.5294 |
| X_1334385 | 3 | 0.6667 | 3 | 0.6667 | 5 | 0.2 | 4 | 1 | 15 | 0.6 | 3 | 0.3333 | 5 | 0.2 | 4 | 0.5 | 3 | 0.3333 | 15 | 0.3333 |
| X_1334387 | 2 | 0.5 | 3 | 0.3333 | 4 | 0 | 4 | 0.25 | 13 | 0.2308 | 2 | 0 | 6 | 0.5 | 4 | 0.25 | 3 | 0 | 15 | 0.2667 |
| X_1334387 | 1 | 0 | 2 | 0.5 | 2 | 0 | 3 | 0 | 8 | 0.125 | 1 | 0 | 4 | 0.25 | 3 | 0 | 3 | 0 | 11 | 0.0909 |
| X_1334387 | 2 | 1 | 3 | 1 | 4 | 0.5 | 4 | 0.75 | 13 | 0.7692 | 3 | 0.6667 | 6 | 0.3333 | 4 | 0.5 | 3 | 0.6667 | 16 | 0.5 |
| X_1334388 | 1 | 1 | 2 | 1 | 4 | 1 | 4 | 0.5 | 11 | 0.8182 | 2 | 0.5 | 6 | 0.5 | 2 | 1 | 3 | 0.6667 | 13 | 0.6154 |
| X_1334389 | 1 | 1 | 3 | 0.3333 | 3 | 0.6667 | 4 | 0.75 | 11 | 0.6364 | 3 | 0.3333 | 6 | 0.5 | 4 | 0.75 | 2 | 0.5 | 15 | 0.5333 |
| X_1334389 | 1 | 0 | 3 | 0.6667 | 3 | 0.6667 | 3 | 0 | 10 | 0.4 | 2 | 0.5 | 2 | 0.5 | 2 | 1 | 1 | 0 | 7 | 0.5714 |
| X_1334866 | 1 | 1 | 4 | 0.25 | 5 | 0.6 | 1 | 1 | 11 | 0.5455 | NA | NA | 2 | 1 | 4 | 0.25 | NA | NA | 6 | 0.5 |
| X_1334906 | NA | NA | 3 | 0.6667 | 1 | 1 | 2 | 1 | 6 | 0.8333 | NA | NA | NA | NA | NA | NA | 1 | 1 | 1 | 1 |
| X_1335058 | 5 | 0.8 | 3 | 0 | 1 | 0 | 6 | 0.5 | 15 | 0.4667 | 3 | 0.3333 | 6 | 0.3333 | 2 | 1 | 2 | 0.5 | 13 | 0.4615 |
| X_1335058 | 5 | 0.6 | 3 | 0.3333 | NA | NA | 5 | 0.2 | 13 | 0.3846 | 1 | 0 | 7 | 0.2857 | 3 | 0.3333 | 4 | 0.5 | 15 | 0.3333 |
| X_1335058 | 5 | 0.6 | 4 | 0.25 | NA | NA | 6 | 0.1667 | 15 | 0.3333 | 3 | 0.3333 | 5 | 0.4 | 3 | 0 | 4 | 0.75 | 15 | 0.4 |
| X_1335058 | 3 | 1 | 4 | 0.5 | 1 | 1 | 5 | 1 | 13 | 0.8462 | 3 | 1 | 5 | 1 | 3 | 0.6667 | 4 | 0.75 | 15 | 0.8667 |
| X_1335060 | 4 | 0.75 | 4 | 0 | NA | NA | 7 | 0.4286 | 15 | 0.4 | 3 | 0.6667 | 7 | 0.2857 | 3 | 0 | 3 | 0 | 16 | 0.25 |
| X_1335060 | 5 | 0.8 | 4 | 1 | 1 | 0 | 6 | 0.5 | 16 | 0.6875 | 2 | 1 | 6 | 0.3333 | 2 | 1 | 2 | 1 | 12 | 0.6667 |
| X_1335063 | 6 | 0 | 4 | 0.25 | NA | NA | 8 | 0 | 18 | 0.0556 | 3 | 0.3333 | 8 | 0.25 | 5 | 0 | 3 | 0 | 19 | 0.1579 |
| X_1335959 | NA | NA | 1 | 1 | NA | NA | 1 | 1 | 2 | 1 | NA | NA | 2 | 0.5 | 1 | 1 | NA | NA | 3 | 0.6667 |
| X_1335959 | 2 | 0.5 | 1 | 0 | 1 | 0 | 1 | 0 | 5 | 0.2 | NA | NA | 2 | 0 | 1 | 1 | 1 | 1 | 4 | 0.5 |
| X_1335960 | 3 | 0 | 1 | 0 | 1 | 0 | 1 | 1 | 6 | 0.1667 | NA | NA | 2 | 0 | 1 | 1 | 1 | 0 | 4 | 0.25 |
| X_1335960 | 3 | 0.3333 | 1 | 1 | 1 | 1 | 1 | 1 | 6 | 0.6667 | NA | NA | 2 | 1 | 1 | 1 | 1 | 0 | 4 | 0.75 |

| | | | | | | | | | | | | | | | | | | | | |
|---|---|---|---|---|---|---|---|---|---|---|---|---|---|---|---|---|---|---|---|---|
| X_1335964 | 3 | 0.3333 | 1 | 0 | 1 | 0 | 1 | 1 | 6 | 0.3333 | NA | NA | 2 | 0 | 1 | 1 | 3 | 0.6667 | 6 | 0.5 |
| X_1335967 | 3 | 0.6667 | 2 | 0.5 | 1 | 1 | 1 | 1 | 7 | 0.7143 | NA | NA | 2 | 0.5 | 1 | 1 | 3 | 1 | 6 | 0.8333 |
| X_1335968 | 3 | 0.6667 | 2 | 0.5 | 1 | 1 | 1 | 1 | 7 | 0.7143 | NA | NA | 3 | 0.3333 | 1 | 1 | 3 | 1 | 7 | 0.7143 |
| X_1335968 | 3 | 1 | 2 | 0.5 | 1 | 1 | 1 | 1 | 7 | 0.8571 | NA | NA | 3 | 0 | 1 | 1 | 3 | 1 | 7 | 0.5714 |
| X_1335969 | 3 | 0 | 2 | 0 | 2 | 0.5 | 1 | 1 | 8 | 0.25 | NA | NA | 3 | 0.3333 | NA | NA | 3 | 0 | 6 | 0.1667 |
| X_1335971 | 3 | 0.3333 | 3 | 0.6667 | 3 | 0.3333 | 2 | 0 | 11 | 0.3636 | 1 | 0 | 2 | 0.5 | 2 | 1 | 3 | 1 | 8 | 0.75 |
| X_1335972 | 5 | 0.2 | 2 | 0.5 | 3 | 0.3333 | 2 | 0 | 12 | 0.25 | 1 | 0 | 2 | 0 | 3 | 0.6667 | 3 | 0.6667 | 9 | 0.4444 |
| X_1335972 | 5 | 0.8 | 3 | 0.6667 | 3 | 0.3333 | 1 | 0 | 12 | 0.5833 | 1 | 0 | 2 | 0 | 1 | 1 | 2 | 0.5 | 6 | 0.3333 |
| X_1336209 | 3 | 0 | 12 | 0.4167 | 3 | 1 | 4 | 0.5 | 22 | 0.4545 | 2 | 0 | 3 | 0.3333 | 5 | 0.8 | 2 | 0.5 | 12 | 0.5 |
| X_1336210 | 6 | 0.5 | 13 | 0 | 6 | 0 | 5 | 0.4 | 30 | 0.1667 | 3 | 0 | 3 | 0.6667 | 4 | 0 | 5 | 0 | 15 | 0.1333 |
| X_1336210 | 5 | 0.2 | 14 | 0.1429 | 6 | 0 | 5 | 0.4 | 30 | 0.1667 | 3 | 0 | 3 | 0 | 4 | 0 | 5 | 0.2 | 15 | 0.0667 |
| X_1336213 | 6 | 0.1667 | 17 | 0.0588 | 8 | 0.5 | 4 | 0 | 35 | 0.1714 | 2 | 0 | 4 | 0 | 6 | 0 | 6 | 0.5 | 18 | 0.1667 |
| X_1336506 | 1 | 1 | NA | NA | 1 | 0 | NA | NA | 2 | 0.5 | NA | NA | 1 | 1 | NA | NA | 1 | 0 | 2 | 0.5 |
| X_1336506 | 1 | 0 | NA | NA | 1 | 1 | NA | NA | 2 | 0.5 | NA | NA | 1 | 1 | NA | NA | NA | NA | 1 | 1 |
| X_1336507 | 1 | 0 | NA | NA | 1 | 1 | NA | NA | 2 | 0.5 | NA | NA | 1 | 1 | NA | NA | 1 | 1 | 2 | 1 |
| X_1336507 | 1 | 1 | NA | NA | 1 | 0 | NA | NA | 2 | 0.5 | NA | NA | 1 | 0 | NA | NA | 1 | 1 | 2 | 0.5 |
| X_1336507 | 1 | 1 | NA | NA | 1 | 0 | NA | NA | 2 | 0.5 | NA | NA | 1 | 1 | NA | NA | 1 | 0 | 2 | 0.5 |
| X_1336592 | 2 | 0 | 7 | 0.4286 | 6 | 0.3333 | 5 | 0.2 | 20 | 0.3 | 6 | 0.3333 | 1 | 1 | 5 | 0.6 | 7 | 0.4286 | 19 | 0.4737 |
| X_1336592 | 1 | 1 | 5 | 0.4 | 5 | 0.6 | 3 | 0.6667 | 14 | 0.5714 | 6 | 0.5 | 1 | 0 | 5 | 0.8 | 6 | 0.3333 | 18 | 0.5 |
| X_1336593 | 2 | 0 | 6 | 0.5 | 4 | 0.25 | 4 | 0 | 16 | 0.25 | 5 | 0.2 | 1 | 1 | 5 | 0.6 | 7 | 0.1429 | 18 | 0.3333 |
| X_1336594 | 1 | 1 | 3 | 1 | 5 | 1 | 3 | 0.3333 | 12 | 0.8333 | 5 | 0.8 | 1 | 1 | 4 | 1 | 6 | 0.8333 | 16 | 0.875 |
| X_1336595 | 3 | 0.6667 | 7 | 0.5714 | 6 | 0.1667 | 4 | 0 | 20 | 0.35 | 7 | 0.2857 | 2 | 0 | 5 | 0.2 | 7 | 0.2857 | 21 | 0.2381 |
| X_1336596 | 3 | 0.3333 | 8 | 0 | 6 | 0.1667 | 5 | 0 | 22 | 0.0909 | 7 | 0.2857 | 3 | 0 | 5 | 0.4 | 9 | 0.2222 | 24 | 0.25 |
| X_1336600 | 3 | 0.3333 | 9 | 0.6667 | 8 | 0.375 | 6 | 0 | 26 | 0.3846 | 8 | 0.25 | 4 | 0.75 | 6 | 0.1667 | 8 | 0.25 | 26 | 0.3077 |
| X_1336600 | 3 | 0.3333 | 7 | 0.2857 | 7 | 0.5714 | 6 | 0.1667 | 23 | 0.3478 | 7 | 0.1429 | 4 | 0.25 | 5 | 0.2 | 7 | 0 | 23 | 0.1304 |
| X_1336601 | 2 | 1 | 7 | 0.2857 | 9 | 0.4444 | 4 | 0 | 22 | 0.3636 | 7 | 0.4286 | 4 | 0.25 | 6 | 0.1667 | 8 | 0.375 | 25 | 0.32 |
| X_1336604 | 3 | 0.3333 | 10 | 0 | 12 | 0 | 6 | 0.1667 | 31 | 0.0645 | 9 | 0.1111 | 3 | 0 | 6 | 0 | 8 | 0.125 | 26 | 0.0769 |
| X_1337055 | 5 | 0 | 12 | 0.0833 | 9 | 0.3333 | 6 | 0 | 32 | 0.125 | 15 | 0.2 | 11 | 0 | 12 | 0.1667 | 15 | 0.2 | 53 | 0.1509 |
| X_1337061 | 1 | 0 | 3 | 0 | 6 | 0.1667 | 4 | 0 | 14 | 0.0714 | 9 | 0.2222 | 6 | 0.1667 | 8 | 0.125 | 12 | 0.0833 | 35 | 0.1429 |
| X_1337074 | 1 | 0 | NA | NA | 3 | 0.3333 | 1 | 1 | 5 | 0.4 | 6 | 0.3333 | 8 | 0.125 | 3 | 0.3333 | 7 | 0.1429 | 24 | 0.2083 |
| X_1337076 | 1 | 1 | 1 | 1 | 3 | 1 | 1 | 0 | 6 | 0.8333 | 7 | 0.8571 | 5 | 0.6 | 4 | 0.5 | 5 | 0.4 | 21 | 0.619 |
| X_1337077 | 1 | 1 | 1 | 0 | 3 | 0.3333 | 1 | 0 | 6 | 0.3333 | 5 | 0.2 | 8 | 0.125 | 5 | 0.4 | 4 | 0.25 | 22 | 0.2273 |
| X_1338722 | NA | NA | 2 | 0.5 | 2 | 0 | 2 | 0 | 6 | 0.1667 | NA | NA | 3 | 0 | 6 | 0 | 3 | 0.3333 | 12 | 0.0833 |
| X_1338724 | NA | NA | 2 | 0.5 | 2 | 0.5 | 3 | 1 | 7 | 0.7143 | NA | NA | 2 | 1 | 6 | 0.8333 | 3 | 0.6667 | 11 | 0.8182 |
| X_1338727 | NA | NA | 2 | 1 | 1 | 0 | 2 | 0 | 5 | 0.4 | NA | NA | 2 | 0 | 4 | 0.25 | 1 | 1 | 7 | 0.2857 |
| X_1338728 | NA | NA | 2 | 0.5 | 1 | 1 | 2 | 0.5 | 5 | 0.6 | NA | NA | 2 | 0 | 5 | 0.4 | 3 | 0.3333 | 10 | 0.3 |
| X_1338732 | NA | NA | 1 | 1 | 2 | 1 | 1 | 1 | 4 | 1 | NA | NA | 1 | 1 | 3 | 1 | 1 | 1 | 5 | 1 |
| X_1338732 | NA | NA | 2 | 0 | 1 | 1 | 2 | 0 | 5 | 0.2 | NA | NA | 1 | 0 | 4 | 0.25 | 2 | 0.5 | 7 | 0.2857 |
| X_1338733 | NA | NA | 2 | 0.5 | NA | NA | 2 | 0.5 | 4 | 0.5 | NA | NA | 2 | 0 | 3 | 0.3333 | 2 | 0 | 7 | 0.1429 |
| X_1338738 | NA | NA | 2 | 0 | 1 | 0 | 2 | 0.5 | 5 | 0.2 | NA | NA | 1 | 0 | 3 | 0 | 2 | 0.5 | 6 | 0.1667 |
| X_1338740 | NA | NA | 2 | 0 | NA | NA | 2 | 0.5 | 4 | 0.25 | NA | NA | 2 | 0 | 3 | 0 | 2 | 0.5 | 7 | 0.1429 |
| X_1338740 | NA | NA | 1 | 1 | NA | NA | 2 | 0 | 3 | 0.3333 | NA | NA | 1 | 1 | 3 | 0.3333 | 1 | 1 | 5 | 0.6 |
| X_1338741 | NA | NA | 1 | 1 | NA | NA | 1 | 0 | 2 | 0.5 | NA | NA | 2 | 0.5 | 2 | 0.5 | 2 | 1 | 6 | 0.6667 |
| X_1338744 | NA | NA | 2 | 1 | NA | NA | 2 | 1 | 4 | 1 | NA | NA | 2 | 1 | 3 | 1 | 1 | 1 | 6 | 1 |
| X_1338745 | NA | NA | 2 | 0.5 | NA | NA | 2 | 0.5 | 4 | 0.5 | NA | NA | 2 | 0 | 3 | 0.6667 | 2 | 0 | 7 | 0.2857 |
| X_1338745 | NA | NA | 1 | 0 | NA | NA | 2 | 0.5 | 3 | 0.3333 | NA | NA | 2 | 0 | 3 | 0.3333 | 2 | 0.5 | 7 | 0.2857 |
| X_1338745 | NA | NA | 2 | 0.5 | NA | NA | 1 | 0 | 3 | 0.3333 | NA | NA | 1 | 1 | 2 | 0 | 1 | 0 | 4 | 0.25 |
| X_1341095 | 20 | 0.3 | 15 | 0.2667 | 22 | 0.1818 | 16 | 0 | 73 | 0.1918 | 6 | 0.1667 | 18 | 0.1667 | 17 | 0.1176 | 11 | 0 | 52 | 0.1154 |
| X_1341096 | 19 | 0 | 13 | 0.0769 | 23 | 0.087 | 14 | 0.0714 | 69 | 0.058 | 6 | 0 | 16 | 0.25 | 19 | 0.1579 | 11 | 0.2727 | 52 | 0.1923 |
| X_1341099 | 16 | 0.125 | 9 | 0 | 19 | 0.1053 | 13 | 0 | 57 | 0.0702 | 4 | 0 | 14 | 0.2143 | 13 | 0.1538 | 9 | 0.5556 | 40 | 0.25 |
| X_1346458 | 2 | 0.5 | NA | NA | 1 | 0 | 3 | 0 | 6 | 0.1667 | 1 | 1 | 1 | 0 | 3 | 0.6667 | 2 | 0 | 7 | 0.4286 |
| X_1346458 | 2 | 1 | NA | NA | 1 | 1 | 3 | 0.3333 | 6 | 0.6667 | 1 | 0 | 1 | 1 | 3 | 1 | 2 | 1 | 7 | 0.8571 |
| X_1346458 | 2 | 0.5 | NA | NA | 1 | 0 | 3 | 0.6667 | 6 | 0.5 | 1 | 1 | 1 | 1 | 3 | 0.3333 | 2 | 0.5 | 7 | 0.5714 |
| X_1346458 | 2 | 0.5 | NA | NA | 1 | 0 | 2 | 0 | 5 | 0.2 | 1 | 0 | 1 | 0 | 3 | 0.3333 | 2 | 0 | 7 | 0.1429 |
| X_1346461 | 1 | 1 | NA | NA | 1 | 1 | 3 | 0.3333 | 5 | 0.6 | 1 | 1 | 1 | 0 | 3 | 0.3333 | 2 | 0.5 | 7 | 0.4286 |
| X_1346461 | 2 | 0.5 | NA | NA | 1 | 0 | 3 | 0.6667 | 6 | 0.5 | NA | NA | 1 | 1 | 2 | 0.5 | 2 | 0 | 5 | 0.4 |
| X_1346462 | 2 | 0 | NA | NA | NA | NA | 1 | 1 | 3 | 0.3333 | 1 | 0 | 1 | 1 | 2 | 0.5 | NA | NA | 4 | 0.5 |
| X_1346464 | 1 | 0 | NA | NA | NA | NA | 3 | 0.3333 | 4 | 0.25 | 1 | 1 | 1 | 1 | 3 | 0.3333 | 2 | 0 | 7 | 0.4286 |
| X_1346464 | 2 | 0.5 | NA | NA | NA | NA | 2 | 0.5 | 4 | 0.5 | 1 | 1 | 1 | 0 | 3 | 0 | 1 | 0 | 6 | 0.1667 |
| X_1346465 | 1 | 1 | NA | NA | 1 | 0 | 3 | 0.6667 | 5 | 0.6 | 1 | 1 | 1 | 1 | 3 | 0.6667 | 2 | 0.5 | 7 | 0.7143 |
| X_1346468 | 2 | 0.5 | NA | NA | 1 | 1 | 3 | 0.6667 | 6 | 0.6667 | 1 | 0 | 1 | 0 | 3 | 0.3333 | 2 | 0 | 7 | 0.1429 |
| X_1346468 | 2 | 1 | NA | NA | 1 | 1 | 3 | 0.6667 | 6 | 0.8333 | 1 | 1 | 1 | 1 | 3 | 1 | 2 | 1 | 7 | 1 |
| X_1346469 | 2 | 0.5 | NA | NA | 1 | 0 | 3 | 0.6667 | 6 | 0.5 | 1 | 0 | 1 | 0 | 4 | 0.5 | 2 | 0 | 8 | 0.25 |
| X_1346469 | 2 | 0.5 | NA | NA | 2 | 0.5 | 2 | 0.5 | 6 | 0.5 | 1 | 1 | 1 | 0 | 4 | 0.75 | 2 | 0.5 | 8 | 0.625 |
| X_1346470 | 2 | 0 | NA | NA | 2 | 0.5 | 3 | 0.3333 | 7 | 0.2857 | 1 | 0 | 1 | 0 | 4 | 0.5 | 2 | 0 | 8 | 0.25 |

| | | | | | | | | | | | | | | | | | | | | |
|---|---|---|---|---|---|---|---|---|---|---|---|---|---|---|---|---|---|---|---|---|
| X_1346471 | 1 | 0 | NA | NA | 2 | 1 | 3 | 0.6667 | 6 | 0.6667 | 1 | 0 | 1 | 0 | 3 | 0.3333 | 2 | 0.5 | 7 | 0.2857 |
| X_1346472 | 2 | 0 | NA | NA | 1 | 0 | 3 | 0.3333 | 6 | 0.1667 | 1 | 0 | 1 | 0 | 3 | 0.3333 | 2 | 0 | 7 | 0.1429 |
| X_1346996 | 3 | 0 | 3 | 0.3333 | 2 | 0 | 9 | 0.3333 | 17 | 0.2353 | 6 | 0 | 6 | 0.3333 | 7 | 0.2857 | 6 | 0.5 | 25 | 0.28 |
| X_1346996 | 3 | 0 | 4 | 0.25 | 3 | 0 | 8 | 0.5 | 18 | 0.2778 | 6 | 0 | 6 | 0.3333 | 7 | 0.2857 | 4 | 0 | 23 | 0.1739 |
| X_1346998 | 4 | 0 | 6 | 0.1667 | 3 | 0 | 10 | 0.1 | 23 | 0.087 | 7 | 0.2857 | 5 | 0.2 | 8 | 0.125 | 7 | 0 | 27 | 0.1481 |
| X_1347002 | 5 | 0 | 8 | 0.25 | 3 | 0 | 13 | 0.0769 | 29 | 0.1034 | 8 | 0.125 | 10 | 0.1 | 11 | 0 | 8 | 0 | 37 | 0.0541 |
| X_1347002 | 4 | 0.5 | 8 | 0.75 | 2 | 1 | 10 | 0.5 | 24 | 0.625 | 8 | 0.75 | 10 | 0.7 | 9 | 0.8889 | 8 | 0.875 | 35 | 0.8 |
| X_1347010 | 10 | 0.1 | 12 | 0.1667 | 2 | 0 | 15 | 0 | 39 | 0.0769 | 12 | 0.0833 | 13 | 0.0769 | 17 | 0.1176 | 10 | 0.2 | 52 | 0.1154 |
| X_1347011 | 9 | 0.3333 | 11 | 0.9091 | 3 | 0.3333 | 14 | 0.7143 | 37 | 0.6486 | 11 | 0.3636 | 13 | 0.4615 | 17 | 0.4706 | 11 | 0.8182 | 52 | 0.5192 |
| X_1347015 | 16 | 0.1875 | 14 | 0 | 5 | 0 | 23 | 0.087 | 58 | 0.0862 | 18 | 0.0556 | 19 | 0.0526 | 21 | 0.0952 | 17 | 0 | 75 | 0.0533 |
| X_1347019 | 17 | 0.2353 | 18 | 0.3333 | 6 | 0.1667 | 28 | 0.2143 | 69 | 0.2464 | 16 | 0.3125 | 20 | 0.25 | 26 | 0.2308 | 21 | 0.2381 | 83 | 0.253 |
| X_1347020 | 16 | 0.1875 | 17 | 0.1765 | 6 | 0 | 22 | 0.2273 | 61 | 0.1803 | 18 | 0.1667 | 18 | 0.2778 | 23 | 0.3043 | 22 | 0.1364 | 81 | 0.2222 |
| X_1347023 | 17 | 0.9412 | 17 | 0.8235 | 7 | 0.5714 | 21 | 0.8571 | 62 | 0.8387 | 15 | 0.8667 | 21 | 0.7619 | 20 | 0.8 | 21 | 0.9048 | 77 | 0.8312 |
| X_1347023 | 19 | 0.2105 | 18 | 0.1111 | 9 | 0 | 27 | 0.1852 | 73 | 0.1507 | 17 | 0.1176 | 24 | 0.125 | 29 | 0.069 | 24 | 0.125 | 94 | 0.1064 |
| X_1347023 | 19 | 0.2105 | 19 | 0 | 8 | 0.125 | 26 | 0 | 72 | 0.0694 | 18 | 0.0556 | 25 | 0.04 | 31 | 0.129 | 20 | 0.15 | 94 | 0.0957 |
| X_1347024 | 16 | 0.8125 | 18 | 0.7778 | 9 | 0.5556 | 25 | 0.72 | 68 | 0.7353 | 17 | 0.5882 | 20 | 0.6 | 25 | 0.36 | 25 | 0.56 | 87 | 0.5172 |
| X_1347025 | 23 | 0.1304 | 22 | 0.0909 | 10 | 0 | 28 | 0 | 83 | 0.0602 | 19 | 0.0526 | 24 | 0.0833 | 31 | 0.0968 | 28 | 0.0714 | 102 | 0.0784 |
| X_1347026 | 23 | 0.0435 | 24 | 0.0417 | 9 | 0 | 31 | 0.0968 | 87 | 0.0575 | 22 | 0.0455 | 27 | 0.0741 | 31 | 0.129 | 28 | 0.0714 | 108 | 0.0833 |
| X_1347031 | 23 | 0.1304 | 23 | 0.3043 | 11 | 0.3636 | 31 | 0.1613 | 88 | 0.2159 | 23 | 0.1304 | 29 | 0.2414 | 38 | 0.2895 | 24 | 0.2917 | 114 | 0.2456 |
| X_1347036 | 35 | 0.2571 | 31 | 0.2581 | 20 | 0.15 | 43 | 0.2791 | 129 | 0.2481 | 29 | 0.2759 | 34 | 0.2647 | 45 | 0.2667 | 35 | 0.2857 | 143 | 0.2727 |
| X_1347038 | 33 | 0.1212 | 30 | 0.0667 | 20 | 0 | 39 | 0.1538 | 122 | 0.0984 | 29 | 0.1034 | 37 | 0.027 | 49 | 0.1224 | 31 | 0.0645 | 146 | 0.0822 |
| X_1347442 | 8 | 0 | 8 | 0.125 | 4 | 0 | 11 | 0.0909 | 31 | 0.0645 | 8 | 0 | 10 | 0.1 | 12 | 0.1667 | 9 | 0.1111 | 39 | 0.1026 |
| X_1347444 | 10 | 0.2 | 9 | 0.1111 | 5 | 0 | 10 | 0.1 | 34 | 0.1176 | 5 | 0.2 | 9 | 0.1111 | 13 | 0 | 8 | 0.125 | 35 | 0.0857 |
| X_1347896 | 11 | 0.7273 | 6 | 0.8333 | 7 | 0.2857 | 9 | 0.7778 | 33 | 0.6667 | 3 | 0.6667 | 15 | 0.4667 | 11 | 0.6364 | 2 | 1 | 31 | 0.5806 |
| X_1347896 | 9 | 0.1111 | 7 | 0 | 6 | 0 | 9 | 0.1111 | 31 | 0.0645 | 3 | 0 | 15 | 0.1333 | 9 | 0 | 2 | 0 | 29 | 0.069 |
| X_1347896 | 11 | 0.8182 | 6 | 0.8333 | 7 | 0.7143 | 8 | 0.625 | 32 | 0.75 | 3 | 0.3333 | 15 | 0.8 | 12 | 0.6667 | 1 | 1 | 31 | 0.7097 |
| X_1347904 | 12 | 0.1667 | 7 | 0.1429 | 11 | 0 | 15 | 0.1333 | 45 | 0.1111 | 4 | 0 | 20 | 0.15 | 14 | 0 | 4 | 0.25 | 42 | 0.0952 |
| X_1348947 | 10 | 0.2 | 6 | 0 | 8 | 0.125 | 9 | 0.2222 | 33 | 0.1515 | 5 | 0.4 | 4 | 0.25 | 13 | 0.2308 | 10 | 0 | 32 | 0.1875 |
| X_1348949 | 12 | 0.1667 | 8 | 0.5 | 8 | 0.125 | 11 | 0.0909 | 39 | 0.2051 | 6 | 0.3333 | 6 | 0.3333 | 16 | 0.125 | 10 | 0.1 | 38 | 0.1842 |
| X_1348950 | 12 | 0.25 | 9 | 0.4444 | 9 | 0.2222 | 10 | 0.4 | 40 | 0.325 | 6 | 0.1667 | 5 | 0.2 | 15 | 0.4 | 10 | 0.2 | 36 | 0.2778 |
| X_1348950 | 12 | 0.0833 | 9 | 0.1111 | 8 | 0 | 8 | 0.125 | 37 | 0.0811 | 5 | 0 | 6 | 0 | 13 | 0.0769 | 9 | 0.1111 | 33 | 0.0606 |
| X_1348951 | 11 | 1 | 9 | 1 | 8 | 1 | 8 | 1 | 36 | 1 | 3 | 1 | 6 | 1 | 12 | 1 | 9 | 0.8889 | 30 | 0.9667 |
| X_1348951 | 11 | 0.9091 | 6 | 0.8333 | 8 | 1 | 9 | 0.7778 | 34 | 0.8824 | 5 | 0.8 | 6 | 0.6667 | 12 | 0.9167 | 9 | 0.8889 | 32 | 0.8438 |
| X_1348951 | 11 | 0.8182 | 7 | 0.8571 | 9 | 0.7778 | 9 | 1 | 36 | 0.8611 | 4 | 0.75 | 5 | 1 | 14 | 0.9286 | 8 | 0.75 | 31 | 0.871 |
| X_1348952 | 12 | 0.1667 | 7 | 0.1429 | 10 | 0.2 | 13 | 0.4615 | 42 | 0.2619 | 6 | 0.3333 | 6 | 0.6667 | 14 | 0.4286 | 10 | 0.4 | 36 | 0.4444 |
| X_1348952 | 13 | 0.9231 | 9 | 1 | 9 | 1 | 12 | 0.9167 | 43 | 0.9535 | 3 | 1 | 5 | 1 | 14 | 1 | 10 | 1 | 32 | 1 |
| X_1348953 | 12 | 0.5833 | 10 | 0.4 | 8 | 0.375 | 12 | 0.25 | 42 | 0.4048 | 4 | 0.75 | 5 | 0 | 15 | 0.2667 | 11 | 0.2727 | 35 | 0.2857 |
| X_1348953 | 13 | 0.0769 | 8 | 0 | 7 | 0.1429 | 10 | 0 | 38 | 0.0526 | 2 | 0 | 3 | 0 | 13 | 0.0769 | 8 | 0.125 | 26 | 0.0769 |
| X_1348953 | 14 | 1 | 10 | 1 | 10 | 1 | 12 | 1 | 46 | 1 | 5 | 1 | 5 | 1 | 14 | 1 | 10 | 1 | 34 | 1 |
| X_1348954 | 15 | 1 | 9 | 1 | 9 | 1 | 11 | 1 | 44 | 1 | 6 | 1 | 6 | 1 | 15 | 0.8667 | 11 | 1 | 38 | 0.9474 |
| X_1348954 | 14 | 0.0714 | 9 | 0.1111 | 10 | 0 | 12 | 0.1667 | 45 | 0.0889 | 6 | 0 | 5 | 0 | 14 | 0.0714 | 12 | 0.1667 | 37 | 0.0811 |
| X_1348955 | 11 | 0.5455 | 9 | 0.6667 | 6 | 0.6667 | 12 | 0.75 | 38 | 0.6579 | 4 | 0.5 | 7 | 0.5714 | 14 | 0.3571 | 11 | 0.6364 | 36 | 0.5 |
| X_1349342 | 4 | 0 | 4 | 0.25 | 4 | 0.25 | 4 | 0.25 | 16 | 0.1875 | 2 | 0 | 6 | 0 | 8 | 0.25 | 3 | 0 | 19 | 0.1053 |
| X_1349343 | 5 | 0.2 | 4 | 0 | 4 | 0 | 4 | 0.25 | 17 | 0.1176 | 2 | 0 | 5 | 0.2 | 9 | 0.2222 | 4 | 0 | 20 | 0.15 |
| X_1349344 | 5 | 0 | 5 | 0 | 4 | 0.25 | 4 | 0 | 18 | 0.0556 | 2 | 0 | 5 | 0.2 | 9 | 0 | 4 | 0 | 20 | 0.05 |
| X_1349344 | 5 | 0.6 | 4 | 0.25 | 5 | 0.4 | 5 | 0 | 19 | 0.3158 | 2 | 0 | 6 | 0.1667 | 9 | 0.2222 | 4 | 0.75 | 21 | 0.2857 |
| X_1349345 | 6 | 0 | 5 | 0.2 | 4 | 0.25 | 5 | 0.4 | 20 | 0.2 | 2 | 1 | 5 | 0.2 | 8 | 0.125 | 4 | 0.5 | 19 | 0.3158 |
| X_1349346 | 5 | 0.2 | 3 | 0 | 2 | 0 | 3 | 0 | 13 | 0.0769 | 2 | 0 | 6 | 0.1667 | 8 | 0.125 | 4 | 0.25 | 20 | 0.15 |
| X_1349346 | 6 | 1 | 3 | 1 | 6 | 1 | 4 | 1 | 19 | 1 | 2 | 1 | 5 | 1 | 8 | 1 | 4 | 1 | 19 | 1 |
| X_1349346 | 5 | 0.6 | 2 | 1 | 3 | 0.6667 | 4 | 1 | 14 | 0.7857 | 2 | 0.5 | 5 | 0.8 | 8 | 0.875 | 4 | 1 | 19 | 0.8421 |
| X_1349347 | 5 | 1 | 4 | 1 | 5 | 1 | 4 | 1 | 18 | 1 | 1 | 1 | 6 | 1 | 7 | 0.8571 | 2 | 1 | 16 | 0.9375 |
| X_1349348 | 5 | 0.4 | 5 | 0.8 | 7 | 0.5714 | 5 | 0.2 | 22 | 0.5 | 2 | 0 | 7 | 0.5714 | 8 | 0.375 | 4 | 0.25 | 21 | 0.381 |
| X_1349348 | 6 | 1 | 5 | 1 | 7 | 1 | 5 | 0.8 | 23 | 0.9565 | 2 | 0.5 | 8 | 1 | 8 | 1 | 4 | 0.75 | 22 | 0.9091 |
| X_1349348 | 4 | 0.75 | 3 | 1 | 5 | 0.8 | 5 | 0.8 | 17 | 0.8235 | 2 | 0.5 | 6 | 0.5 | 7 | 0.4286 | 4 | 0.75 | 19 | 0.5263 |
| X_1349354 | 4 | 0.25 | 5 | 0.2 | 2 | 0 | 5 | 0.4 | 16 | 0.25 | 2 | 0 | 7 | 0.2857 | 6 | 0 | 2 | 0 | 17 | 0.1176 |
| X_1349677 | 100 | 0.11 | 79 | 0.1139 | 67 | 0.1642 | 105 | 0.1333 | 351 | 0.1282 | 49 | 0.2041 | 87 | 0.1494 | 98 | 0.1429 | 80 | 0.2125 | 314 | 0.172 |
| X_1350408 | 6 | 0.3333 | 7 | 0.2857 | 6 | 0.1667 | 9 | 0.1111 | 28 | 0.2143 | 2 | 0 | 9 | 0.1111 | 8 | 0.125 | 13 | 0.0769 | 32 | 0.0938 |
| X_1350709 | 10 | 0.2 | 15 | 0 | 14 | 0.1429 | 10 | 0.2 | 49 | 0.1224 | 10 | 0 | 11 | 0.1818 | 3 | 0 | 10 | 0.2 | 34 | 0.1176 |
| X_1350713 | 10 | 0.2 | 18 | 0.4444 | 15 | 0.4667 | 12 | 0.5 | 55 | 0.4182 | 9 | 0.4444 | 7 | 0.5714 | 8 | 0.625 | 12 | 0.6667 | 36 | 0.5833 |
| X_1350714 | 12 | 0.6667 | 18 | 0.6111 | 12 | 0.6667 | 12 | 1 | 54 | 0.7222 | 5 | 0.6 | 10 | 0.6 | 6 | 0.8333 | 12 | 0.8333 | 33 | 0.7273 |
| X_1350715 | 9 | 0.4444 | 16 | 0.0625 | 10 | 0.4 | 10 | 0.3 | 45 | 0.2667 | 4 | 0.5 | 8 | 0.25 | 6 | 0.6667 | 9 | 0.4444 | 27 | 0.4444 |
| X_1350716 | 11 | 0.0909 | 13 | 0.0769 | 10 | 0.1 | 12 | 0.0833 | 46 | 0.087 | 5 | 0.2 | 8 | 0.125 | 7 | 0 | 12 | 0.25 | 32 | 0.1562 |
| X_1351017 | 10 | 0.4 | 12 | 0.5 | 13 | 0.3846 | 12 | 0.25 | 47 | 0.383 | 10 | 0.6 | 8 | 0.375 | 12 | 0.5833 | 13 | 0.5385 | 43 | 0.5349 |
| X_1351019 | 15 | 0.4667 | 8 | 0.25 | 13 | 0.0769 | 11 | 0.1818 | 47 | 0.2553 | 10 | 0.1 | 10 | 0.3 | 15 | 0.2 | 19 | 0.1579 | 54 | 0.1852 |
| X_1351473 | 21 | 0.0952 | 17 | 0.1176 | 14 | 0.0714 | 13 | 0 | 65 | 0.0769 | 11 | 0.0909 | 22 | 0.2273 | 15 | 0.2 | 17 | 0.1176 | 65 | 0.1692 |

| | | | | | | | | | | | | | | | | | | | | |
|---|---|---|---|---|---|---|---|---|---|---|---|---|---|---|---|---|---|---|---|---|
| X_1351724 | 54 | 0.1111 | 44 | 0.1136 | 45 | 0.1333 | 42 | 0.0238 | 185 | 0.0973 | 33 | 0.1212 | 49 | 0.102 | 38 | 0.0263 | 32 | 0.0625 | 152 | 0.0789 |
| X_1351724 | 54 | 0.2222 | 43 | 0.2093 | 49 | 0.2857 | 41 | 0.1463 | 187 | 0.2193 | 32 | 0.1562 | 50 | 0.16 | 41 | 0.1951 | 34 | 0.1765 | 157 | 0.172 |
| X_1355686 | 22 | 0.2273 | 24 | 0.1667 | 17 | 0.1176 | 27 | 0.1111 | 90 | 0.1556 | 19 | 0.2632 | 16 | 0.0625 | 18 | 0.3333 | 29 | 0.2414 | 82 | 0.2317 |
| X_1355689 | 26 | 0.1538 | 34 | 0.0588 | 19 | 0.2632 | 30 | 0.1333 | 109 | 0.1376 | 21 | 0.1429 | 17 | 0 | 19 | 0.2105 | 38 | 0.0789 | 95 | 0.1053 |
| X_1355689 | 22 | 0.0909 | 32 | 0.0312 | 20 | 0.2 | 27 | 0.1481 | 101 | 0.1089 | 17 | 0.4118 | 15 | 0 | 18 | 0.1111 | 35 | 0.1714 | 85 | 0.1765 |
| X_1356098 | 5 | 1 | 4 | 1 | 1 | 1 | 6 | 1 | 16 | 1 | 3 | 1 | 6 | 1 | 4 | 1 | 7 | 1 | 20 | 1 |
| X_1356099 | 5 | 0.4 | 5 | 0.2 | 3 | 0.6667 | 7 | 0.1429 | 20 | 0.3 | 4 | 0.5 | 7 | 0.1429 | 4 | 0.5 | 7 | 0.2857 | 22 | 0.3182 |
| X_1356100 | 4 | 1 | 6 | 0.6667 | 3 | 1 | 6 | 0.8333 | 19 | 0.8421 | 2 | 1 | 4 | 0.75 | 4 | 1 | 3 | 0.6667 | 13 | 0.8462 |
| X_1356103 | 5 | 0 | 6 | 0.1667 | 4 | 0.25 | 7 | 0.1429 | 22 | 0.1364 | 5 | 0.6 | 8 | 0.25 | 3 | 0 | 7 | 0 | 23 | 0.2174 |
| X_1356348 | 31 | 0.9677 | 35 | 0.9429 | 37 | 0.8919 | 37 | 0.973 | 140 | 0.9429 | 25 | 0.96 | 39 | 0.7692 | 34 | 0.8824 | 35 | 0.8571 | 133 | 0.8571 |
| X_1356348 | 24 | 0.6667 | 33 | 0.7273 | 39 | 0.6667 | 34 | 0.7059 | 130 | 0.6923 | 22 | 0.7273 | 40 | 0.725 | 34 | 0.6471 | 35 | 0.6857 | 131 | 0.6947 |
| X_1357370 | 76 | 0.3816 | 152 | 0.5197 | 118 | 0.4661 | 119 | 0.4118 | 465 | 0.4559 | 98 | 0.5 | 69 | 0.3913 | 76 | 0.5 | 87 | 0.4368 | 330 | 0.4606 |
| X_1357373 | 75 | 0.3867 | 147 | 0.415 | 111 | 0.4234 | 117 | 0.3504 | 450 | 0.3956 | 83 | 0.3735 | 71 | 0.3803 | 68 | 0.4559 | 72 | 0.4444 | 294 | 0.4116 |
| X_1357373 | NA | NA | 3 | 0 | 1 | 0 | 2 | 0.5 | 6 | 0.1667 | 2 | 0.5 | 1 | 0 | 3 | 0.3333 | 2 | 0 | 8 | 0.25 |
| X_1357374 | 85 | 0.5765 | 177 | 0.6215 | 120 | 0.625 | 135 | 0.6889 | 517 | 0.6325 | 94 | 0.7021 | 77 | 0.7013 | 82 | 0.6098 | 98 | 0.6327 | 351 | 0.661 |
| X_1357377 | 95 | 0.2316 | 194 | 0.1804 | 145 | 0.2207 | 145 | 0.1931 | 579 | 0.2021 | 110 | 0.1909 | 89 | 0.2809 | 93 | 0.2366 | 105 | 0.1429 | 397 | 0.2091 |
| X_1357618 | 3 | 0 | 2 | 0 | NA | NA | 4 | 0.25 | 9 | 0.1111 | 1 | 0 | 1 | 0 | 2 | 0.5 | 4 | 0 | 8 | 0.125 |
| X_1357620 | 4 | 0.25 | 2 | 0 | 2 | 0 | 4 | 0.25 | 12 | 0.1667 | 2 | 0.5 | 2 | 0.5 | 3 | 0.3333 | 4 | 0.25 | 11 | 0.3636 |
| X_1357621 | 3 | 0.3333 | 2 | 0 | 2 | 0 | 3 | 0.6667 | 10 | 0.3 | 2 | 0.5 | 2 | 0 | 3 | 1 | 5 | 0.2 | 12 | 0.4167 |
| X_1357622 | 4 | 0.75 | 2 | 1 | 1 | 0 | 4 | 0.75 | 11 | 0.7273 | 1 | 1 | 1 | 0 | 1 | 1 | 5 | 1 | 8 | 0.875 |
| X_1357623 | 4 | 0.25 | 2 | 1 | 1 | 1 | 4 | 0.25 | 11 | 0.4545 | 2 | 0.5 | 1 | 0 | 3 | 0.3333 | 5 | 0.4 | 11 | 0.3636 |
| X_1358834 | 1 | 1 | NA | NA | NA | NA | 1 | 0 | 2 | 0.5 | NA | NA | NA | NA | 1 | 1 | 1 | 1 | 2 | 1 |
| X_1358835 | 1 | 1 | NA | NA | NA | NA | 1 | 0 | 2 | 0.5 | NA | NA | NA | NA | 1 | 1 | 1 | 1 | 2 | 1 |
| X_1358839 | 1 | 0 | NA | NA | NA | NA | 1 | 1 | 2 | 0.5 | NA | NA | NA | NA | 1 | 1 | 1 | 1 | 2 | 1 |
| X_1358839 | 1 | 1 | NA | NA | NA | NA | 1 | 0 | 2 | 0.5 | NA | NA | NA | NA | 1 | 1 | 1 | 1 | 2 | 1 |
| X_1358840 | 1 | 1 | NA | NA | NA | NA | 1 | 1 | 2 | 1 | NA | NA | NA | NA | 1 | 1 | 1 | 1 | 2 | 1 |
| X_1358939 | 1 | 0 | NA | NA | 2 | 0 | 1 | 1 | 4 | 0.25 | 2 | 0.5 | 4 | 0 | 4 | 0 | 1 | 0 | 11 | 0.0909 |
| X_1358953 | 25 | 0.56 | 26 | 0.5769 | 18 | 0.4444 | 22 | 0.4545 | 91 | 0.5165 | 13 | 0.6923 | 32 | 0.4375 | 21 | 0.381 | 18 | 0.6111 | 84 | 0.5 |
| X_1358955 | 32 | 0.0312 | 30 | 0 | 22 | 0.0909 | 24 | 0.125 | 108 | 0.0556 | 22 | 0.0455 | 31 | 0.0645 | 24 | 0.125 | 24 | 0.0833 | 101 | 0.0792 |
| X_1358959 | 35 | 0.0857 | 32 | 0.0938 | 24 | 0.125 | 30 | 0.1 | 121 | 0.0992 | 19 | 0.1579 | 33 | 0.0303 | 22 | 0.1818 | 25 | 0.08 | 99 | 0.101 |
| X_1358970 | 42 | 0.0714 | 41 | 0.1951 | 35 | 0.0571 | 37 | 0.1892 | 155 | 0.129 | 32 | 0.0625 | 44 | 0.1364 | 34 | 0.0294 | 35 | 0.1429 | 145 | 0.0966 |
| X_1359021 | 1 | 0 | 1 | 0 | NA | NA | 2 | 1 | 4 | 0.5 | 1 | 1 | NA | NA | 1 | 0 | NA | NA | 2 | 0.5 |
| X_1359042 | 1 | 0 | 1 | 0 | NA | NA | 1 | 1 | 3 | 0.3333 | 2 | 0.5 | 1 | 0 | 3 | 0.3333 | 2 | 0.5 | 8 | 0.375 |
| X_1359043 | 1 | 1 | 1 | 1 | NA | NA | NA | NA | 2 | 1 | 2 | 0.5 | 1 | 0 | 3 | 0 | 1 | 0 | 7 | 0.1429 |
| X_1377354 | NA | NA | NA | NA | NA | NA | 1 | 1 | 1 | 1 | NA | NA | 1 | 1 | NA | NA | NA | NA | 1 | 1 |
| X_1381353 | NA | NA | NA | NA | 1 | 1 | NA | NA | 1 | 1 | 1 | 1 | NA | NA | NA | NA | 1 | 0 | 2 | 0.5 |
| X_1381353 | NA | NA | NA | NA | 1 | 1 | NA | NA | 1 | 1 | NA | NA | NA | NA | NA | NA | 1 | 1 | 1 | 1 |
| X_1381358 | NA | NA | NA | NA | 1 | 1 | NA | NA | 1 | 1 | 1 | 1 | NA | NA | NA | NA | 1 | 1 | 2 | 1 |
| X_1381359 | NA | NA | NA | NA | 1 | 1 | NA | NA | 1 | 1 | NA | NA | NA | NA | NA | NA | 1 | 1 | 1 | 1 |
| X_1381363 | NA | NA | NA | NA | 1 | 1 | 1 | 1 | 2 | 1 | 1 | 1 | NA | NA | NA | NA | 1 | 1 | 2 | 1 |
| X_1381364 | NA | NA | NA | NA | 1 | 1 | 1 | 1 | 2 | 1 | 1 | 0 | NA | NA | NA | NA | 1 | 1 | 2 | 0.5 |
| X_1381370 | NA | NA | NA | NA | 1 | 0 | 2 | 0.5 | 3 | 0.3333 | 1 | 1 | 2 | 0.5 | NA | NA | 1 | 1 | 4 | 0.75 |
| X_1381372 | NA | NA | NA | NA | 1 | 1 | 2 | 0.5 | 3 | 0.6667 | 1 | 1 | 2 | 1 | NA | NA | 1 | 0 | 4 | 0.75 |
| X_1381374 | NA | NA | NA | NA | NA | NA | 2 | 0.5 | 2 | 0.5 | 1 | 1 | 1 | 1 | NA | NA | 1 | 1 | 3 | 1 |
| X_1381377 | NA | NA | NA | NA | 1 | 0 | 2 | 0.5 | 3 | 0.3333 | 1 | 0 | 1 | 1 | NA | NA | 1 | 0 | 3 | 0.3333 |
| X_1381377 | NA | NA | NA | NA | 1 | 0 | 2 | 0.5 | 3 | 0.3333 | 1 | 1 | 2 | 0 | NA | NA | 1 | 0 | 4 | 0.25 |
| X_1385726 | 2 | 0 | 1 | 0 | 4 | 0.5 | NA | NA | 7 | 0.2857 | 2 | 0 | 2 | 0.5 | 4 | 0.5 | 3 | 0.6667 | 11 | 0.4545 |
| X_1385726 | 3 | 0.6667 | NA | NA | 5 | 0.2 | 2 | 0.5 | 10 | 0.4 | 3 | 0.3333 | 1 | 0 | 5 | 0.4 | 3 | 0 | 12 | 0.25 |
| X_1385734 | 3 | 0 | 4 | 0 | 7 | 0.1429 | 4 | 0 | 18 | 0.0556 | 5 | 0.2 | 2 | 0 | 5 | 0 | 3 | 0.3333 | 15 | 0.1333 |
| X_1393287 | NA | NA | 1 | 1 | NA | NA | NA | NA | 1 | 1 | NA | NA | NA | NA | 1 | 1 | 1 | 1 | 2 | 1 |
| X_1393290 | NA | NA | 1 | 1 | NA | NA | NA | NA | 1 | 1 | NA | NA | NA | NA | 1 | 0 | 1 | 1 | 2 | 0.5 |
| X_1393314 | NA | NA | 1 | 1 | NA | NA | NA | NA | 1 | 1 | NA | NA | NA | NA | 2 | 0.5 | NA | NA | 2 | 0.5 |
| X_1393322 | NA | NA | 1 | 1 | NA | NA | NA | NA | 1 | 1 | NA | NA | NA | NA | 3 | 0.3333 | NA | NA | 3 | 0.3333 |
| X_1393327 | NA | NA | 1 | 1 | NA | NA | NA | NA | 1 | 1 | NA | NA | NA | NA | 2 | 0.5 | NA | NA | 2 | 0.5 |
| X_1393329 | 2 | 0.5 | 1 | 0 | 1 | 0 | NA | NA | 4 | 0.25 | NA | NA | NA | NA | 3 | 0.6667 | NA | NA | 3 | 0.6667 |
| X_1393329 | 1 | 0 | 1 | 1 | 1 | 1 | NA | NA | 3 | 0.6667 | NA | NA | NA | NA | 3 | 0.3333 | NA | NA | 3 | 0.3333 |
| X_1393336 | 2 | 0 | 1 | 0 | 1 | 1 | NA | NA | 4 | 0.25 | 1 | 0 | NA | NA | 3 | 0.3333 | NA | NA | 4 | 0.25 |
| X_1393338 | 1 | 1 | 1 | 1 | 1 | 1 | NA | NA | 3 | 1 | 1 | 1 | NA | NA | 3 | 1 | NA | NA | 4 | 1 |
| X_1393339 | 2 | 0.5 | 1 | 1 | 1 | 1 | NA | NA | 4 | 0.75 | 1 | 1 | NA | NA | 3 | 0.6667 | NA | NA | 4 | 0.75 |
| X_1393341 | 2 | 0.5 | 1 | 0 | 1 | 0 | NA | NA | 4 | 0.25 | 1 | 1 | NA | NA | 2 | 0.5 | NA | NA | 3 | 0.6667 |
| X_1393343 | 1 | 0 | 1 | 0 | 1 | 1 | NA | NA | 3 | 0.3333 | 1 | 0 | NA | NA | 3 | 0.6667 | NA | NA | 4 | 0.5 |
| X_1393345 | 2 | 0 | 1 | 0 | 1 | 1 | NA | NA | 4 | 0.25 | 1 | 0 | NA | NA | 3 | 0.3333 | NA | NA | 4 | 0.25 |
| X_1393351 | 1 | 1 | 2 | 0.5 | NA | NA | NA | NA | 3 | 0.6667 | 2 | 1 | 1 | 0 | 3 | 0.6667 | NA | NA | 6 | 0.6667 |
| X_1393762 | 13 | 0.2308 | 16 | 0.125 | 9 | 0.2222 | 12 | 0.0833 | 50 | 0.16 | 13 | 0.1538 | 19 | 0.0526 | 6 | 0 | 7 | 0 | 45 | 0.0667 |
| X_1393766 | 12 | 0 | 20 | 0.25 | 11 | 0.1818 | 13 | 0.1538 | 56 | 0.1607 | 16 | 0.25 | 19 | 0.1053 | 7 | 0.2857 | 8 | 0 | 50 | 0.16 |

| | | | | | | | | | | | | | | | | | | | | |
|---|---|---|---|---|---|---|---|---|---|---|---|---|---|---|---|---|---|---|---|---|
| X_1394412 | 28 | 0.6429 | 20 | 0.9 | 13 | 0.5385 | 18 | 0.8889 | 79 | 0.7468 | 28 | 0.9643 | 24 | 0.7917 | 24 | 0.6667 | 24 | 0.5833 | 100 | 0.76 |
| X_1394412 | 28 | 0.75 | 19 | 0.9474 | 12 | 0.75 | 24 | 0.8333 | 83 | 0.8193 | 24 | 0.7917 | 24 | 0.8333 | 23 | 0.913 | 20 | 0.7 | 91 | 0.8132 |
| X_1394414 | 30 | 0.2333 | 20 | 0.45 | 14 | 0.2143 | 25 | 0.32 | 89 | 0.3034 | 30 | 0.4333 | 27 | 0.2593 | 28 | 0.1786 | 21 | 0.1429 | 106 | 0.2642 |
| X_1394414 | 27 | 0.0741 | 19 | 0.1579 | 14 | 0.1429 | 21 | 0.0952 | 81 | 0.1111 | 29 | 0.1034 | 25 | 0 | 23 | 0.087 | 18 | 0 | 95 | 0.0526 |
| X_1394414 | 25 | 0.68 | 14 | 0.7143 | 13 | 0.5385 | 23 | 0.913 | 75 | 0.7333 | 26 | 0.7692 | 25 | 0.72 | 21 | 0.8095 | 18 | 0.7778 | 90 | 0.7667 |
| X_1394415 | 25 | 0.88 | 18 | 0.8889 | 12 | 0.75 | 19 | 0.8947 | 74 | 0.8649 | 26 | 0.8077 | 22 | 0.9545 | 26 | 0.8077 | 16 | 0.875 | 90 | 0.8556 |
| X_1394439 | 1 | 0 | 2 | 0 | 2 | 0 | 1 | 1 | 6 | 0.1667 | 1 | 0 | 3 | 0 | 1 | 1 | 3 | 0 | 8 | 0.125 |
| X_1394491 | NA | NA | NA | NA | NA | NA | 1 | 1 | 1 | 1 | NA | NA | NA | NA | 1 | 1 | NA | NA | 1 | 1 |
| X_1395789 | NA | NA | NA | NA | 1 | 1 | NA | NA | 1 | 1 | 1 | 0 | NA | NA | 3 | 0.3333 | NA | NA | 4 | 0.25 |
| X_1395791 | NA | NA | NA | NA | 1 | 1 | NA | NA | 1 | 1 | 1 | 1 | NA | NA | 3 | 1 | NA | NA | 4 | 1 |
| X_1396643 | 3 | 0.3333 | 2 | 0 | 1 | 0 | 1 | 1 | 7 | 0.2857 | NA | NA | 3 | 0 | 3 | 0.3333 | 1 | 0 | 7 | 0.1429 |
| X_1396644 | 3 | 0.3333 | 2 | 0 | 1 | 0 | NA | NA | 6 | 0.1667 | 1 | 0 | 2 | 0.5 | 4 | 0 | 1 | 0 | 8 | 0.125 |
| X_1396666 | 4 | 0.5 | 4 | 0.25 | 2 | 0 | 3 | 0.6667 | 13 | 0.3846 | 2 | 0.5 | 4 | 0.5 | 6 | 0.3333 | 3 | 0 | 15 | 0.3333 |
| X_1396671 | 4 | 0 | 4 | 0.25 | 1 | 0 | 3 | 0 | 12 | 0.0833 | 2 | 0 | 5 | 0 | 7 | 0.1429 | 3 | 0 | 17 | 0.0588 |
| X_1396675 | 5 | 0.4 | 4 | 0.25 | 1 | 0 | 3 | 1 | 13 | 0.4615 | 3 | 0.3333 | 5 | 0.4 | 7 | 0.4286 | 3 | 0 | 18 | 0.3333 |
| X_1396675 | 4 | 0 | 4 | 0.25 | 1 | 0 | 3 | 0 | 12 | 0.0833 | 3 | 0 | 5 | 0 | 7 | 0.2857 | 4 | 0 | 19 | 0.1053 |
| X_1396680 | 5 | 0.8 | 5 | 0.4 | 1 | 1 | 3 | 1 | 14 | 0.7143 | 2 | 1 | 7 | 0.4286 | 9 | 0.5556 | 5 | 0.6 | 23 | 0.5652 |
| X_1398495 | 1 | 1 | 1 | 1 | 3 | 0 | 1 | 0 | 6 | 0.3333 | 2 | 0 | 2 | 0.5 | NA | NA | NA | NA | 4 | 0.25 |
| X_1398496 | 1 | 0 | 1 | 1 | 3 | 0 | 1 | 0 | 6 | 0.1667 | 2 | 0.5 | 2 | 0.5 | NA | NA | NA | NA | 4 | 0.5 |
| X_1398497 | 3 | 0.3333 | 5 | 0 | 5 | 0.6 | 3 | 0 | 16 | 0.25 | 2 | 0 | 3 | 0 | 4 | 0.25 | 1 | 0 | 10 | 0.1 |
| X_1398499 | 4 | 0.25 | 5 | 0.4 | 4 | 0.25 | 4 | 0.5 | 17 | 0.3529 | 2 | 0.5 | 2 | 0.5 | 4 | 0 | 2 | 1 | 10 | 0.4 |
| X_1410216 | 6 | 0 | 2 | 0.5 | 5 | 0.2 | 9 | 0 | 22 | 0.0909 | 2 | 0 | 5 | 0 | 7 | 0.2857 | 6 | 0.3333 | 20 | 0.2 |
| X_1410218 | 6 | 0.1667 | 1 | 0 | 6 | 0 | 11 | 0.0909 | 24 | 0.0833 | 2 | 0 | 4 | 0 | 5 | 0 | 6 | 0.1667 | 17 | 0.0588 |
| X_1410879 | NA | NA | NA | NA | 3 | 0.3333 | NA | NA | 3 | 0.3333 | 1 | 0 | NA | NA | 1 | 1 | 1 | 1 | 3 | 0.6667 |
| X_1410880 | NA | NA | NA | NA | 1 | 1 | NA | NA | 1 | 1 | 2 | 1 | NA | NA | 1 | 1 | NA | NA | 3 | 1 |
| X_1410880 | NA | NA | NA | NA | 2 | 0.5 | NA | NA | 2 | 0.5 | 2 | 0 | NA | NA | 1 | 0 | 1 | 1 | 4 | 0.25 |
| X_1410881 | NA | NA | NA | NA | 2 | 0.5 | NA | NA | 2 | 0.5 | 2 | 0 | NA | NA | 1 | 1 | 1 | 0 | 4 | 0.25 |
| X_1410882 | NA | NA | NA | NA | 3 | 1 | NA | NA | 3 | 1 | 1 | 1 | NA | NA | 1 | 1 | 1 | 1 | 3 | 1 |
| X_1410882 | NA | NA | NA | NA | 1 | 1 | NA | NA | 1 | 1 | 2 | 1 | NA | NA | 1 | 1 | 1 | 1 | 4 | 1 |
| X_1411012 | 17 | 0.2941 | 16 | 0.25 | 10 | 0.3 | 7 | 0.2857 | 50 | 0.28 | 9 | 0.1111 | 7 | 0.2857 | 11 | 0.2727 | 12 | 0.1667 | 39 | 0.2051 |
| X_1411019 | 11 | 0.1818 | 14 | 0 | 6 | 0 | 8 | 0.125 | 39 | 0.0769 | 6 | 0.1667 | 6 | 0 | 8 | 0 | 10 | 0.1 | 30 | 0.0667 |
| X_1411024 | 12 | 0.0833 | 11 | 0.3636 | 8 | 0.25 | 9 | 0.4444 | 40 | 0.275 | 6 | 0.1667 | 5 | 0.2 | 7 | 0.4286 | 10 | 0.3 | 28 | 0.2857 |
| X_1411033 | 8 | 0.625 | 12 | 0.25 | 4 | 1 | 6 | 0.3333 | 30 | 0.4667 | 5 | 0.2 | 3 | 0.6667 | 6 | 0.1667 | 9 | 0.3333 | 23 | 0.3043 |
| X_1411034 | 8 | 0 | 11 | 0.1818 | 6 | 0.1667 | 9 | 0.3333 | 34 | 0.1765 | 5 | 0 | 2 | 0 | 5 | 0 | 9 | 0.2222 | 21 | 0.0952 |
| X_1411035 | 8 | 0 | 12 | 0.1667 | 5 | 0 | 8 | 0 | 33 | 0.0606 | 6 | 0 | 3 | 0 | 6 | 0.1667 | 8 | 0.125 | 23 | 0.087 |
| X_1411037 | 6 | 0.3333 | 9 | 0.1111 | 5 | 0.6 | 7 | 0.2857 | 27 | 0.2963 | 5 | 0 | 2 | 0.5 | 6 | 0.1667 | 8 | 0.25 | 21 | 0.1905 |
| X_1411037 | 8 | 0.125 | 12 | 0.0833 | 5 | 0 | 8 | 0 | 33 | 0.0606 | 6 | 0 | 3 | 0 | 6 | 0.3333 | 8 | 0 | 23 | 0.087 |
| X_1411340 | 3 | 0 | 7 | 0.1429 | 1 | 0 | 7 | 0.1429 | 18 | 0.1111 | 3 | 0.3333 | 2 | 0 | 3 | 0 | 6 | 0.1667 | 14 | 0.1429 |
| X_1411347 | 2 | 0.5 | 6 | 0.5 | NA | NA | 6 | 0.1667 | 14 | 0.3571 | 2 | 0 | 1 | 1 | 1 | 1 | 3 | 0.3333 | 7 | 0.4286 |
| X_1414140 | NA | NA | 1 | 1 | NA | NA | NA | NA | 1 | 1 | 1 | 1 | NA | NA | NA | NA | NA | NA | 1 | 1 |
| X_1416210 | 3 | 0 | 3 | 0.3333 | 2 | 0.5 | 1 | 0 | 9 | 0.2222 | 1 | 0 | 1 | 1 | 3 | 0 | 2 | 0 | 7 | 0.1429 |
| X_1420706 | 1 | 1 | NA | NA | NA | NA | 1 | 1 | 2 | 1 | NA | NA | 2 | 0.5 | NA | NA | NA | NA | 2 | 0.5 |
| X_1425743 | 2 | 0 | 2 | 1 | 2 | 0 | 7 | 0.4286 | 13 | 0.3846 | NA | NA | 1 | 0 | 1 | 1 | NA | NA | 2 | 0.5 |
| X_1425743 | 2 | 0.5 | 2 | 0 | 2 | 0 | 7 | 0.1429 | 13 | 0.1538 | NA | NA | 1 | 1 | 2 | 0 | NA | NA | 3 | 0.3333 |
| X_1425744 | 2 | 0 | 2 | 0.5 | 2 | 0 | 7 | 0 | 13 | 0.0769 | NA | NA | 2 | 0.5 | 2 | 0 | NA | NA | 4 | 0.25 |
| X_1425747 | 3 | 0 | 2 | 0 | 2 | 0 | 8 | 0.25 | 15 | 0.1333 | NA | NA | 2 | 0.5 | 1 | 0 | NA | NA | 3 | 0.3333 |
| X_1425750 | 4 | 0.5 | 2 | 1 | 2 | 1 | 8 | 0.75 | 16 | 0.75 | NA | NA | 2 | 0.5 | 1 | 1 | NA | NA | 3 | 0.6667 |
| X_1426390 | NA | NA | 3 | 0.3333 | 1 | 1 | 3 | 0.3333 | 7 | 0.4286 | NA | NA | NA | NA | 4 | 0.25 | 1 | 0 | 5 | 0.2 |
| X_1426391 | NA | NA | 3 | 0.3333 | 1 | 0 | 3 | 0.6667 | 7 | 0.4286 | 1 | 0 | NA | NA | 4 | 0.5 | 1 | 0 | 6 | 0.3333 |
| X_1426401 | NA | NA | 3 | 0.3333 | NA | NA | 3 | 0 | 6 | 0.1667 | 1 | 0 | NA | NA | 4 | 0.25 | NA | NA | 5 | 0.2 |
| X_1430585 | 1 | 1 | NA | NA | NA | NA | NA | NA | 1 | 1 | NA | NA | NA | NA | NA | NA | 1 | 1 | 1 | 1 |
| X_1430588 | 1 | 1 | NA | NA | NA | NA | NA | NA | 1 | 1 | NA | NA | NA | NA | NA | NA | 1 | 1 | 1 | 1 |
| X_1442295 | NA | NA | 5 | 0.2 | 2 | 0 | 1 | 0 | 8 | 0.125 | 4 | 0 | 4 | 0.25 | 2 | 0.5 | 1 | 0 | 11 | 0.1818 |
| X_1442303 | 1 | 1 | 4 | 0.25 | NA | NA | 1 | 0 | 6 | 0.3333 | 2 | 0.5 | 2 | 0.5 | 1 | 0 | 1 | 1 | 6 | 0.5 |
| X_1442304 | 1 | 0 | 5 | 0.2 | 1 | 0 | NA | NA | 7 | 0.1429 | 2 | 0.5 | 2 | 0.5 | 2 | 0 | 1 | 1 | 7 | 0.4286 |
| X_1442304 | NA | NA | 4 | 0.5 | 1 | 0 | 1 | 0 | 6 | 0.3333 | 3 | 0.6667 | 2 | 1 | 1 | 0 | NA | NA | 6 | 0.6667 |
| X_1442305 | NA | NA | 2 | 0 | 1 | 1 | 1 | 0 | 4 | 0.25 | 3 | 0.3333 | 2 | 0.5 | 1 | 0 | NA | NA | 6 | 0.3333 |
| X_1442307 | NA | NA | 4 | 0.25 | 1 | 0 | NA | NA | 5 | 0.2 | 2 | 0.5 | 2 | 0.5 | 1 | 0 | 1 | 1 | 6 | 0.5 |
| X_1457456 | 3 | 0.3333 | 4 | 1 | 1 | 1 | 1 | 0 | 9 | 0.6667 | 4 | 0.75 | 1 | 1 | 2 | 1 | NA | NA | 7 | 0.8571 |
| X_1459767 | 2 | 0 | 8 | 0.25 | 3 | 0 | 3 | 0.6667 | 16 | 0.25 | 5 | 0.4 | 3 | 0.6667 | 4 | 0.25 | 4 | 0.5 | 16 | 0.4375 |
| X_1459768 | 3 | 0.3333 | 4 | 0.5 | 3 | 0 | 3 | 0.3333 | 13 | 0.3077 | 4 | 0.5 | 3 | 0.3333 | 4 | 0 | 3 | 0 | 14 | 0.2143 |
| X_1459769 | 3 | 0.6667 | 9 | 0.6667 | 3 | 0.6667 | 3 | 0.3333 | 18 | 0.6111 | 4 | 0.25 | 3 | 0 | 2 | 0.5 | 3 | 0.6667 | 12 | 0.3333 |
| X_1459770 | NA | NA | 2 | 0.5 | 3 | 1 | 1 | 1 | 6 | 0.8333 | 5 | 1 | 5 | 0.8 | 3 | 1 | 3 | 1 | 16 | 0.9375 |
| X_1459772 | 3 | 0.3333 | 6 | 0.1667 | 2 | 0 | 5 | 0 | 16 | 0.125 | 5 | 0.2 | 3 | 0.6667 | 5 | 0 | 3 | 0.3333 | 16 | 0.25 |
| X_1459788 | 6 | 0.6667 | 14 | 0.3571 | 5 | 0.2 | 6 | 0.8333 | 31 | 0.4839 | 10 | 0.3 | 9 | 0.1111 | 12 | 0.25 | 11 | 0 | 42 | 0.1667 |

| | | | | | | | | | | | | | | | | | | | | |
|---|---|---|---|---|---|---|---|---|---|---|---|---|---|---|---|---|---|---|---|---|
| X_1459801 | 11 | 0.1818 | 22 | 0.0909 | 10 | 0.1 | 16 | 0.125 | 59 | 0.1186 | 13 | 0 | 13 | 0.0769 | 12 | 0.1667 | 13 | 0 | 51 | 0.0588 |
| X_1462539 | 5 | 0 | 2 | 0.5 | 5 | 0 | 2 | 0 | 14 | 0.0714 | 1 | 0 | 3 | 0.3333 | 5 | 0 | 9 | 0 | 18 | 0.0556 |
| X_1462550 | 5 | 0.2 | 1 | 0 | 2 | 0 | 1 | 0 | 9 | 0.1111 | NA | NA | 1 | 0 | 4 | 0.25 | 6 | 0 | 11 | 0.0909 |
| X_1462560 | 3 | 0.3333 | 1 | 0 | 3 | 0 | 3 | 0 | 10 | 0.1 | NA | NA | 1 | 0 | 2 | 0 | 4 | 0.5 | 7 | 0.2857 |
| X_1491921 | 2 | 0.5 | NA | NA | 1 | 0 | 1 | 0 | 4 | 0.25 | 1 | 0 | 2 | 1 | NA | NA | 1 | 1 | 4 | 0.75 |
| X_1491923 | 3 | 0.6667 | NA | NA | 1 | 1 | 1 | 1 | 5 | 0.8 | 2 | 0.5 | 4 | 0.75 | 2 | 1 | 1 | 1 | 9 | 0.7778 |
| X_1491924 | 2 | 0.5 | NA | NA | 1 | 0 | 2 | 0.5 | 5 | 0.4 | 2 | 0.5 | 4 | 0 | 1 | 1 | NA | NA | 7 | 0.2857 |
| X_1491925 | 2 | 0.5 | NA | NA | 1 | 0 | 2 | 1 | 5 | 0.6 | 2 | 0 | 4 | 0.25 | 2 | 0 | 1 | 0 | 9 | 0.1111 |
| X_1493549 | 1 | 1 | 1 | 1 | NA | NA | 1 | 1 | 3 | 1 | 1 | 1 | NA | NA | NA | NA | NA | NA | 1 | 1 |
| X_1493552 | 1 | 1 | 1 | 1 | 2 | 0 | 1 | 0 | 5 | 0.4 | 1 | 1 | NA | NA | NA | NA | NA | NA | 1 | 1 |
| X_1494279 | 2 | 0 | 3 | 0 | 2 | 0 | 2 | 0.5 | 9 | 0.1111 | 1 | 0 | 2 | 0 | 1 | 1 | 1 | 0 | 5 | 0.2 |
| X_1494287 | 12 | 0 | 14 | 0 | 13 | 0.0769 | 15 | 0.1333 | 54 | 0.0556 | 7 | 0 | 16 | 0.125 | 19 | 0.1053 | 17 | 0.1176 | 59 | 0.1017 |
| X_1494288 | 8 | 0.875 | 15 | 0.7333 | 12 | 0.75 | 13 | 0.7692 | 48 | 0.7708 | 8 | 0.875 | 16 | 0.6875 | 20 | 0.75 | 16 | 0.6875 | 60 | 0.7333 |
| X_1494288 | 13 | 0.9231 | 15 | 0.8667 | 12 | 0.9167 | 11 | 1 | 51 | 0.9216 | 12 | 1 | 16 | 0.9375 | 21 | 0.8095 | 16 | 0.875 | 65 | 0.8923 |
| X_1494289 | 12 | 0.25 | 15 | 0.0667 | 11 | 0 | 13 | 0.0769 | 51 | 0.098 | 11 | 0.0909 | 16 | 0.0625 | 23 | 0.087 | 14 | 0.0714 | 64 | 0.0781 |
| X_1494289 | 14 | 0.0714 | 16 | 0 | 13 | 0.1538 | 10 | 0.2 | 53 | 0.0943 | 10 | 0.1 | 16 | 0.125 | 26 | 0.2692 | 16 | 0.25 | 68 | 0.2059 |
| X_1494289 | 12 | 0.25 | 15 | 0.1333 | 13 | 0.3077 | 14 | 0.2857 | 54 | 0.2407 | 11 | 0.1818 | 16 | 0.1875 | 24 | 0.2083 | 14 | 0.2143 | 65 | 0.2 |
| X_1494290 | 12 | 0.1667 | 13 | 0.1538 | 15 | 0 | 13 | 0.0769 | 53 | 0.0943 | 9 | 0 | 14 | 0.0714 | 24 | 0.0833 | 17 | 0.1765 | 64 | 0.0938 |
| X_1494290 | 14 | 0.0714 | 15 | 0.0667 | 13 | 0.0769 | 14 | 0.1429 | 56 | 0.0893 | 9 | 0.2222 | 16 | 0.125 | 25 | 0.12 | 15 | 0.0667 | 65 | 0.1231 |
| X_1494290 | 12 | 0.0833 | 7 | 0.4286 | 12 | 0.25 | 13 | 0.1538 | 44 | 0.2045 | 11 | 0.1818 | 14 | 0.2143 | 20 | 0.2 | 13 | 0.1538 | 58 | 0.1897 |
| X_1494291 | 10 | 0.2 | 12 | 0.25 | 9 | 0.1111 | 12 | 0.4167 | 43 | 0.2558 | 10 | 0.2 | 15 | 0.4 | 20 | 0.3 | 14 | 0.4286 | 59 | 0.339 |
| X_1494291 | 12 | 0.9167 | 15 | 0.8667 | 13 | 0.8462 | 12 | 0.9167 | 52 | 0.8846 | 11 | 0.9091 | 15 | 0.8667 | 23 | 0.8696 | 14 | 0.9286 | 63 | 0.8889 |
| X_1494292 | 10 | 0.9 | 11 | 0.9091 | 11 | 0.9091 | 10 | 1 | 42 | 0.9286 | 10 | 1 | 14 | 0.8571 | 18 | 0.8889 | 13 | 0.8462 | 55 | 0.8909 |
| X_1494293 | 12 | 0.9167 | 13 | 0.7692 | 12 | 0.8333 | 10 | 1 | 47 | 0.8723 | 11 | 0.9091 | 15 | 1 | 23 | 0.8696 | 12 | 0.9167 | 61 | 0.918 |
| X_1494294 | 12 | 1 | 11 | 0.7273 | 11 | 1 | 11 | 0.9091 | 45 | 0.9111 | 9 | 0.8889 | 13 | 0.9231 | 19 | 0.9474 | 9 | 0.8889 | 50 | 0.92 |
| X_1494294 | 11 | 0.6364 | 7 | 0.5714 | 9 | 0.7778 | 8 | 0.625 | 35 | 0.6571 | 8 | 1 | 13 | 0.6154 | 16 | 0.5 | 10 | 0.6 | 47 | 0.6383 |
| X_1494294 | 12 | 0.9167 | 11 | 0.8182 | 10 | 0.9 | 10 | 1 | 43 | 0.907 | 10 | 0.9 | 14 | 0.9286 | 20 | 0.8 | 11 | 1 | 55 | 0.8909 |
| X_1494294 | 11 | 1 | 11 | 1 | 9 | 0.8889 | 12 | 1 | 43 | 0.9767 | 10 | 1 | 12 | 1 | 19 | 0.9474 | 9 | 0.7778 | 50 | 0.94 |
| X_1494295 | 10 | 0.9 | 13 | 0.9231 | 10 | 1 | 11 | 1 | 44 | 0.9545 | 9 | 0.8889 | 8 | 0.625 | 19 | 0.8421 | 10 | 0.8 | 46 | 0.8043 |
| X_1494295 | 6 | 0.5 | 10 | 0.4 | 7 | 0.7143 | 9 | 0.6667 | 32 | 0.5625 | 8 | 0.75 | 9 | 0.2222 | 15 | 0.4667 | 10 | 0.7 | 42 | 0.5238 |
| X_1494299 | 9 | 0.5556 | 11 | 0.2727 | 8 | 0.25 | 9 | 0.5556 | 37 | 0.4054 | 6 | 0.1667 | 9 | 0.6667 | 12 | 0.5 | 8 | 0.375 | 35 | 0.4571 |
| X_1494299 | 8 | 0.25 | 11 | 0.0909 | 9 | 0 | 10 | 0.2 | 38 | 0.1316 | 4 | 0 | 11 | 0.1818 | 14 | 0.0714 | 11 | 0.0909 | 40 | 0.1 |
| X_1494305 | 4 | 0.25 | 7 | 0.1429 | 6 | 0.1667 | 10 | 0.1 | 27 | 0.1481 | 4 | 0 | 6 | 0.1667 | 9 | 0 | 6 | 0.3333 | 25 | 0.12 |
| X_1494324 | 1 | 0 | 1 | 1 | 2 | 0 | NA | NA | 4 | 0.25 | 2 | 0 | 3 | 0 | 7 | 0.1429 | 3 | 0 | 15 | 0.0667 |
| X_1494325 | 2 | 1 | 3 | 0.6667 | 1 | 1 | 2 | 0.5 | 8 | 0.75 | 1 | 1 | 1 | 0 | 7 | 0.8571 | 1 | 1 | 10 | 0.8 |
| X_1494380 | NA | NA | 1 | 0 | 2 | 1 | 1 | 0 | 4 | 0.5 | NA | NA | 2 | 1 | 2 | 1 | NA | NA | 4 | 1 |
| X_1495940 | 3 | 0 | 2 | 0 | 6 | 0.1667 | 6 | 0.3333 | 17 | 0.1765 | 4 | 0.25 | 1 | 0 | 7 | 0.2857 | NA | NA | 12 | 0.25 |
| X_1495949 | 3 | 0 | 2 | 0.5 | 6 | 0 | 6 | 0 | 17 | 0.0588 | 3 | 0.3333 | 1 | 0 | 5 | 0 | NA | NA | 9 | 0.1111 |
| X_1495949 | 3 | 0.3333 | 1 | 0 | 5 | 0 | 5 | 0 | 14 | 0.0714 | 2 | 0 | 1 | 0 | 5 | 0.2 | NA | NA | 8 | 0.125 |
| X_1497663 | 24 | 0.125 | 23 | 0.3478 | 15 | 0.4 | 28 | 0.3571 | 90 | 0.3 | 27 | 0.2593 | 16 | 0.375 | 25 | 0.2 | 18 | 0.1667 | 86 | 0.2442 |
| X_1497671 | 18 | 0 | 18 | 0.1111 | 12 | 0 | 20 | 0.15 | 68 | 0.0735 | 24 | 0.0417 | 13 | 0.0769 | 19 | 0.1053 | 21 | 0.0476 | 77 | 0.0649 |
| X_1497686 | 11 | 0.0909 | 11 | 0.2727 | 7 | 0.1429 | 17 | 0.1176 | 46 | 0.1522 | 12 | 0.0833 | 7 | 0 | 12 | 0.1667 | 15 | 0.2 | 46 | 0.1304 |
| X_1497690 | 9 | 0.1111 | 6 | 0 | 3 | 0 | 14 | 0.0714 | 32 | 0.0625 | 8 | 0.125 | 6 | 0.1667 | 9 | 0 | 9 | 0.1111 | 32 | 0.0938 |
| X_1497691 | 6 | 0 | 7 | 0.1429 | 5 | 0.2 | 18 | 0.1111 | 36 | 0.1111 | 13 | 0.5385 | 7 | 0 | 12 | 0.0833 | 11 | 0.0909 | 43 | 0.2093 |
| X_1497691 | 6 | 0 | 8 | 0.125 | 4 | 0.25 | 17 | 0.2353 | 35 | 0.1714 | 11 | 0.0909 | 6 | 0 | 10 | 0 | 9 | 0.2222 | 36 | 0.0833 |
| X_1498306 | 1 | 1 | NA | NA | NA | NA | NA | NA | 1 | 1 | 1 | 1 | NA | NA | NA | NA | NA | NA | 1 | 1 |
| X_1498708 | 1 | 0 | 6 | 0.1667 | 4 | 0 | 4 | 0 | 15 | 0.0667 | 4 | 0.5 | 4 | 0.25 | 6 | 0.1667 | 2 | 0.5 | 16 | 0.3125 |
| X_1498714 | 1 | 0 | 4 | 0.25 | 3 | 0 | 2 | 0 | 10 | 0.1 | 2 | 0.5 | 4 | 0 | 2 | 0 | NA | NA | 8 | 0.125 |
| X_1498719 | 1 | 0 | 4 | 0.25 | 2 | 0 | 3 | 0 | 10 | 0.1 | 2 | 0.5 | 3 | 0 | 2 | 0 | NA | NA | 7 | 0.1429 |
| X_1498725 | 7 | 0 | 12 | 0.1667 | 9 | 0.1111 | 6 | 0.1667 | 34 | 0.1176 | 4 | 0 | 7 | 0.1429 | 6 | 0.1667 | 9 | 0.1111 | 26 | 0.1154 |
| X_1498727 | 7 | 0 | 11 | 0.0909 | 8 | 0.25 | 6 | 0.5 | 32 | 0.1875 | 4 | 0.5 | 8 | 0.375 | 6 | 0.1667 | 9 | 0.4444 | 27 | 0.3704 |
| X_1498732 | 5 | 0.2 | 10 | 0.3 | 7 | 0 | 4 | 0 | 26 | 0.1538 | 4 | 0.25 | 8 | 0.25 | 5 | 0.4 | 8 | 0.125 | 25 | 0.24 |
| X_1500785 | NA | NA | 1 | 1 | NA | NA | NA | NA | 1 | 1 | NA | NA | NA | NA | NA | NA | 1 | 1 | 1 | 1 |
| X_1500978 | NA | NA | 1 | 1 | NA | NA | NA | NA | 1 | 1 | NA | NA | NA | NA | NA | NA | 2 | 1 | 2 | 1 |
| X_1500979 | NA | NA | 1 | 1 | NA | NA | NA | NA | 1 | 1 | NA | NA | NA | NA | NA | NA | 1 | 1 | 1 | 1 |
| X_1500981 | NA | NA | 1 | 1 | NA | NA | 1 | 1 | 2 | 1 | NA | NA | NA | NA | 2 | 1 | 2 | 1 | 4 | 1 |
| X_1500982 | NA | NA | 1 | 1 | NA | NA | 1 | 1 | 2 | 1 | NA | NA | NA | NA | 2 | 1 | 2 | 0.5 | 4 | 0.75 |
| X_1500987 | NA | NA | 1 | 0 | 1 | 0 | 1 | 1 | 3 | 0.3333 | NA | NA | NA | NA | NA | NA | 1 | 1 | 1 | 1 |
| X_1501015 | NA | NA | 3 | 0.3333 | 1 | 0 | 1 | 1 | 5 | 0.4 | NA | NA | 1 | 0 | 3 | 0.3333 | 4 | 0.25 | 8 | 0.25 |
| X_1501018 | NA | NA | 3 | 0.3333 | 2 | 1 | 1 | 1 | 6 | 0.6667 | NA | NA | 2 | 0 | 3 | 0.3333 | 3 | 0.6667 | 8 | 0.375 |
| X_1501278 | NA | NA | 1 | 0 | 1 | 1 | 1 | 0 | 3 | 0.3333 | NA | NA | NA | NA | NA | NA | 1 | 1 | 1 | 1 |
| X_1502192 | NA | NA | 1 | 0 | 1 | 1 | 2 | 0 | 4 | 0.25 | 1 | 1 | NA | NA | 2 | 0 | 4 | 0.5 | 7 | 0.4286 |
| X_1502526 | NA | NA | 2 | 0.5 | 1 | 1 | 5 | 1 | 8 | 0.875 | 2 | 1 | NA | NA | 2 | 1 | 4 | 1 | 8 | 1 |
| X_1502535 | NA | NA | 1 | 0 | 1 | 0 | 5 | 0.4 | 7 | 0.2857 | 2 | 0 | 1 | 1 | 4 | 0 | 4 | 0 | 11 | 0.0909 |
| X_1502545 | NA | NA | 3 | 0 | 4 | 0.25 | 4 | 0 | 11 | 0.0909 | 2 | 0 | 1 | 0 | 3 | 0.3333 | 3 | 0.3333 | 9 | 0.2222 |

| | | | | | | | | | | | | | | | | | | | | |
|---|---|---|---|---|---|---|---|---|---|---|---|---|---|---|---|---|---|---|---|---|
| X_1502550 | 1 | 1 | 3 | 0 | 6 | 0.1667 | 8 | 0.125 | 18 | 0.1667 | 2 | 0 | NA | NA | 3 | 0.3333 | 4 | 0 | 9 | 0.1111 |
| X_1502554 | 1 | 1 | 4 | 0.5 | 5 | 0.2 | 9 | 0.3333 | 19 | 0.3684 | 2 | 0 | 2 | 0.5 | 4 | 0.25 | 5 | 0 | 13 | 0.1538 |
| X_1502556 | 1 | 1 | 5 | 0 | 6 | 0.3333 | 8 | 0.25 | 20 | 0.25 | 4 | 0 | 3 | 0.6667 | 5 | 0.6 | 5 | 0.4 | 17 | 0.4118 |
| X_1502569 | 4 | 0.25 | 5 | 0.2 | 8 | 0 | 11 | 0.0909 | 28 | 0.1071 | 7 | 0.1429 | 8 | 0.125 | 7 | 0 | 4 | 0 | 26 | 0.0769 |
| X_1504143 | NA | NA | 4 | 0.25 | NA | NA | 1 | 0 | 5 | 0.2 | NA | NA | 3 | 0 | 1 | 0 | 1 | 1 | 5 | 0.2 |
| X_1504145 | NA | NA | 3 | 1 | NA | NA | 1 | 1 | 4 | 1 | NA | NA | 2 | 0.5 | 1 | 1 | 1 | 1 | 4 | 0.75 |
| X_1504145 | NA | NA | 4 | 1 | NA | NA | 1 | 1 | 5 | 1 | NA | NA | 3 | 1 | 1 | 1 | 1 | 1 | 5 | 1 |
| X_1504146 | NA | NA | 3 | 0.3333 | 1 | 0 | 2 | 0.5 | 6 | 0.3333 | NA | NA | 3 | 0.3333 | 1 | 0 | 1 | 0 | 5 | 0.2 |
| X_1504150 | 1 | 0 | 4 | 0.25 | NA | NA | 1 | 0 | 6 | 0.1667 | NA | NA | 3 | 0.3333 | 2 | 0 | 1 | 0 | 6 | 0.1667 |
| X_1504150 | 1 | 0 | 4 | 0 | 1 | 1 | 2 | 0.5 | 8 | 0.25 | NA | NA | 3 | 0 | 2 | 1 | 1 | 1 | 6 | 0.5 |
| X_1504151 | 1 | 0 | 3 | 0.3333 | 1 | 0 | 2 | 0.5 | 7 | 0.2857 | NA | NA | 3 | 0.3333 | 2 | 0 | 1 | 0 | 6 | 0.1667 |
| X_1504154 | 1 | 0 | 4 | 0.25 | 1 | 0 | 1 | 0 | 7 | 0.1429 | NA | NA | 1 | 1 | 2 | 0 | 1 | 0 | 4 | 0.25 |
| X_1506545 | 1 | 0 | NA | NA | 1 | 1 | NA | NA | 2 | 0.5 | NA | NA | 1 | 1 | NA | NA | NA | NA | 1 | 1 |
| X_1507001 | 7 | 0 | 6 | 0.3333 | 5 | 0.2 | 9 | 0.1111 | 27 | 0.1481 | 4 | 0.5 | 9 | 0.2222 | 5 | 0 | 3 | 0 | 21 | 0.1905 |
| X_1507002 | 8 | 0 | 5 | 0.2 | 7 | 0.2857 | 9 | 0 | 29 | 0.1034 | 3 | 0 | 8 | 0.375 | 5 | 0 | 4 | 0 | 20 | 0.15 |
| X_1507003 | 8 | 0 | 5 | 0.2 | 6 | 0 | 8 | 0.125 | 27 | 0.0741 | 4 | 0 | 7 | 0.1429 | 4 | 0.25 | 4 | 0 | 19 | 0.1053 |
| X_1507017 | 8 | 0.625 | 7 | 0.5714 | 9 | 0.5556 | 11 | 0.8182 | 35 | 0.6571 | 4 | 0.5 | 9 | 0.7778 | 5 | 0.6 | 7 | 0.8571 | 25 | 0.72 |
| X_1507026 | 9 | 0.5556 | 8 | 0.375 | 10 | 0.5 | 8 | 0.25 | 35 | 0.4286 | 5 | 0.4 | 8 | 0.625 | 7 | 0.1429 | 8 | 0.375 | 28 | 0.3929 |
| X_1507027 | 9 | 0.7778 | 7 | 0.5714 | 8 | 0.875 | 9 | 1 | 33 | 0.8182 | 6 | 1 | 8 | 0.75 | 7 | 0.5714 | 8 | 1 | 29 | 0.8276 |
| X_1507027 | 9 | 0.2222 | 6 | 0.1667 | 10 | 0.1 | 10 | 0.1 | 35 | 0.1429 | 5 | 0.2 | 7 | 0.1429 | 6 | 0 | 6 | 0 | 24 | 0.0833 |
| X_1507027 | 10 | 0.4 | 6 | 0.1667 | 7 | 0.2857 | 11 | 0.5455 | 34 | 0.3824 | 4 | 0.25 | 7 | 0.5714 | 6 | 0.5 | 6 | 0.1667 | 23 | 0.3913 |
| X_1507027 | 9 | 0.2222 | 7 | 0.4286 | 9 | 0.5556 | 10 | 0.7 | 35 | 0.4857 | 6 | 0.3333 | 7 | 0.5714 | 7 | 0.2857 | 6 | 0.8333 | 26 | 0.5 |
| X_1507028 | 8 | 0.75 | 4 | 0.75 | 7 | 0.7143 | 8 | 0.75 | 27 | 0.7407 | 4 | 0.25 | 3 | 0.3333 | 6 | 0.5 | 5 | 0.6 | 18 | 0.4444 |
| X_1507029 | 11 | 0.4545 | 6 | 0.3333 | 7 | 0 | 8 | 0.125 | 32 | 0.25 | 6 | 0.1667 | 8 | 0.25 | 6 | 0.1667 | 9 | 0.3333 | 29 | 0.2414 |
| X_1507156 | NA | NA | NA | NA | NA | NA | 1 | 1 | 1 | 1 | 1 | 1 | NA | NA | NA | NA | NA | NA | 1 | 1 |
| X_1507174 | 19 | 0.3158 | 11 | 0.7273 | 14 | 0.1429 | 9 | 0.5556 | 53 | 0.3962 | 10 | 0.1 | 9 | 0.3333 | 12 | 0.4167 | 16 | 0.375 | 47 | 0.3191 |
| X_1507176 | 14 | 0 | 18 | 0.0556 | 16 | 0.0625 | 18 | 0.1111 | 66 | 0.0606 | 13 | 0.0769 | 14 | 0 | 15 | 0.0667 | 13 | 0.2308 | 55 | 0.0909 |
| X_1507853 | 12 | 0.1667 | 9 | 0 | 6 | 0 | 7 | 0 | 34 | 0.0588 | 9 | 0 | 11 | 0.0909 | 8 | 0.125 | 7 | 0.1429 | 35 | 0.0857 |
| X_1507971 | NA | NA | NA | NA | 1 | 1 | NA | NA | 1 | 1 | 1 | 1 | 1 | 1 | NA | NA | NA | NA | 2 | 1 |
| X_1507995 | 1 | 0 | 6 | 0 | 3 | 0.3333 | 3 | 0 | 13 | 0.0769 | NA | NA | 1 | 1 | 4 | 0 | 1 | 0 | 6 | 0.1667 |
| X_1508448 | 6 | 0 | 19 | 0.1053 | 8 | 0 | 11 | 0.0909 | 44 | 0.0682 | 5 | 0 | 7 | 0.1429 | 9 | 0 | 6 | 0.3333 | 27 | 0.1111 |
| X_1509276 | NA | NA | 2 | 0 | 3 | 0.3333 | 2 | 0 | 7 | 0.1429 | 3 | 0.3333 | 2 | 0 | 5 | 0 | 1 | 0 | 11 | 0.0909 |
| X_1509278 | NA | NA | 4 | 0.5 | 3 | 0.6667 | 2 | 0.5 | 9 | 0.5556 | 4 | 0.25 | 2 | 0.5 | 5 | 0.2 | NA | NA | 11 | 0.2727 |
| X_1509773 | 1 | 0 | 3 | 0.3333 | 1 | 0 | NA | NA | 5 | 0.2 | 3 | 0.3333 | 1 | 0 | 2 | 0 | NA | NA | 6 | 0.1667 |
| X_1509870 | 1 | 0 | 3 | 1 | 2 | 0.5 | 2 | 0.5 | 8 | 0.625 | 2 | 1 | 1 | 1 | 1 | 1 | NA | NA | 4 | 1 |
| X_1509871 | 1 | 0 | 5 | 0.2 | 3 | 0 | 3 | 0.3333 | 12 | 0.1667 | 2 | 0 | 2 | 0.5 | 4 | 0 | 3 | 0 | 11 | 0.0909 |
| X_1509873 | 1 | 0 | 5 | 0.2 | 4 | 0 | 3 | 0.6667 | 13 | 0.2308 | 2 | 0.5 | 2 | 0 | 2 | 0 | 3 | 0 | 9 | 0.1111 |
| X_1509920 | 1 | 1 | 6 | 0.5 | 4 | 0.5 | 3 | 0.3333 | 14 | 0.5 | 2 | 0.5 | 3 | 1 | 4 | 0.5 | 3 | 0.3333 | 12 | 0.5833 |
| X_1509921 | 1 | 1 | 4 | 0.75 | 4 | 0.25 | 4 | 0.75 | 13 | 0.6154 | 2 | 1 | 2 | 1 | 3 | 1 | 3 | 0.6667 | 10 | 0.9 |
| X_1509921 | 1 | 1 | 5 | 0 | 4 | 0 | 4 | 0 | 14 | 0.0714 | 2 | 0.5 | 3 | 0 | 1 | 0 | 2 | 0 | 8 | 0.125 |
| X_1509922 | 1 | 1 | 6 | 1 | 3 | 0.3333 | 4 | 0.5 | 14 | 0.7143 | 2 | 1 | 2 | 0.5 | 3 | 0.6667 | 3 | 0.6667 | 10 | 0.7 |
| X_1509924 | 1 | 0 | 6 | 0.6667 | 3 | 1 | 3 | 0.6667 | 13 | 0.6923 | 2 | 1 | 4 | 0.75 | 4 | 0.25 | 3 | 0.6667 | 13 | 0.6154 |
| X_1509924 | 1 | 0 | 5 | 0.2 | 4 | 0.25 | 3 | 0.3333 | 13 | 0.2308 | 1 | 0 | 4 | 0.25 | 4 | 0.75 | NA | NA | 9 | 0.4444 |
| X_1509925 | 1 | 0 | 6 | 0.1667 | 4 | 0 | 3 | 0 | 14 | 0.0714 | 2 | 0.5 | 4 | 0.25 | 5 | 0.4 | 4 | 0 | 15 | 0.2667 |
| X_1510062 | 2 | 0.5 | 8 | 0.25 | 5 | 0.2 | 6 | 0 | 21 | 0.1905 | 3 | 0 | 5 | 0.2 | 8 | 0.125 | 5 | 0.2 | 21 | 0.1429 |
| X_1510064 | 4 | 0 | 13 | 0.0769 | 6 | 0.3333 | 6 | 0.1667 | 29 | 0.1379 | 4 | 0.25 | 7 | 0.1429 | 11 | 0 | 7 | 0 | 29 | 0.069 |
| X_1510064 | 5 | 0.2 | 14 | 0.2143 | 7 | 0.1429 | 9 | 0 | 35 | 0.1429 | 4 | 0 | 10 | 0.2 | 11 | 0.1818 | 6 | 0.1667 | 31 | 0.1613 |
| X_1510541 | 2 | 1 | 2 | 1 | 5 | 1 | 1 | 1 | 10 | 1 | 3 | 0.6667 | NA | NA | 1 | 1 | NA | NA | 4 | 0.75 |
| X_1510542 | 2 | 0.5 | 2 | 0 | 6 | 0 | 2 | 0.5 | 12 | 0.1667 | 3 | 0.3333 | NA | NA | 1 | 0 | NA | NA | 4 | 0.25 |
| X_1510542 | 3 | 0.6667 | 2 | 0 | 5 | 0.2 | 2 | 0 | 12 | 0.25 | 3 | 0.3333 | NA | NA | 1 | 0 | NA | NA | 4 | 0.25 |
| X_1510543 | 2 | 0.5 | 2 | 0.5 | 5 | 0.6 | 2 | 1 | 11 | 0.6364 | 3 | 0.3333 | NA | NA | 1 | 1 | NA | NA | 4 | 0.5 |
| X_1510559 | 4 | 0.25 | 3 | 0 | 7 | 0.1429 | 3 | 0 | 17 | 0.1176 | 4 | 0.5 | NA | NA | 2 | 0.5 | 2 | 0 | 8 | 0.375 |
| X_1510559 | 4 | 0 | 4 | 0 | 7 | 0.1429 | 3 | 0 | 18 | 0.0556 | 3 | 0 | 1 | 0 | 2 | 0.5 | 2 | 0 | 8 | 0.125 |
| X_1510560 | 2 | 1 | 4 | 0 | 8 | 0.5 | 3 | 0.3333 | 17 | 0.4118 | 3 | 0.3333 | NA | NA | 2 | 0 | 1 | 0 | 6 | 0.1667 |
| X_1510560 | 3 | 0 | 3 | 0.6667 | 5 | 0.2 | 3 | 0.3333 | 14 | 0.2857 | 2 | 0.5 | NA | NA | 1 | 0 | 1 | 0 | 4 | 0.25 |
| X_1510560 | 4 | 0.25 | 4 | 0.25 | 8 | 0.125 | 3 | 0 | 19 | 0.1579 | 5 | 0.6 | 1 | 1 | 2 | 0.5 | 2 | 0.5 | 10 | 0.6 |
| X_1510561 | 4 | 0.5 | 4 | 0.25 | 8 | 0.25 | 2 | 0 | 18 | 0.2778 | 5 | 0 | 1 | 1 | 2 | 0.5 | 2 | 0 | 10 | 0.2 |
| X_1510562 | 3 | 1 | 4 | 0.75 | 7 | 0.8571 | 3 | 0 | 17 | 0.7059 | 4 | 0.75 | 1 | 1 | 1 | 0 | 1 | 0 | 7 | 0.5714 |
| X_1510613 | NA | NA | 3 | 0.6667 | 2 | 0 | 1 | 0 | 6 | 0.3333 | 3 | 0.3333 | 1 | 0 | 1 | 0 | 3 | 0 | 8 | 0.125 |
| X_1510615 | NA | NA | 4 | 0.25 | 2 | 0.5 | 1 | 0 | 7 | 0.2857 | 3 | 0 | 1 | 1 | 1 | 0 | 3 | 0.3333 | 8 | 0.25 |
| X_1510616 | NA | NA | 4 | 0.5 | 2 | 0.5 | 1 | 1 | 7 | 0.5714 | 3 | 0.3333 | 1 | 0 | NA | NA | 3 | 1 | 7 | 0.5714 |
| X_1510813 | 1 | 0 | 4 | 0.5 | 3 | 0 | 4 | 0.25 | 12 | 0.25 | NA | NA | NA | NA | 1 | 0 | 5 | 0.2 | 6 | 0.1667 |
| X_1510813 | 2 | 0 | 5 | 0.2 | 3 | 0 | 4 | 0 | 14 | 0.0714 | NA | NA | NA | NA | 2 | 0.5 | 6 | 0.1667 | 8 | 0.25 |
| X_1510814 | 2 | 1 | 5 | 1 | 3 | 1 | 4 | 1 | 14 | 1 | NA | NA | NA | NA | 1 | 1 | 6 | 0.8333 | 7 | 0.8571 |
| X_1510816 | 2 | 0 | 3 | 0 | 3 | 0 | 4 | 0.25 | 12 | 0.0833 | NA | NA | NA | NA | 1 | 0 | 6 | 0.1667 | 7 | 0.1429 |

| | | | | | | | | | | | | | | | | | | | | |
|---|---|---|---|---|---|---|---|---|---|---|---|---|---|---|---|---|---|---|---|---|
| X_1510816 | 2 | 1 | 4 | 0.75 | 3 | 1 | 3 | 1 | 12 | 0.9167 | NA | NA | NA | NA | 1 | 1 | 6 | 1 | 7 | 1 |
| X_1510816 | 2 | 1 | 3 | 0.3333 | 2 | 0.5 | 3 | 0.3333 | 10 | 0.5 | NA | NA | NA | NA | 1 | 0 | 6 | 0.3333 | 7 | 0.2857 |
| X_1510816 | 2 | 0.5 | 4 | 0.75 | 3 | 0.6667 | 4 | 0.5 | 13 | 0.6154 | NA | NA | NA | NA | NA | NA | 6 | 0.8333 | 6 | 0.8333 |
| X_1510817 | 2 | 0 | 1 | 1 | 2 | 0 | 3 | 0 | 8 | 0.125 | NA | NA | NA | NA | 1 | 0 | 3 | 0.3333 | 4 | 0.25 |
| X_1510818 | 2 | 0.5 | 4 | 0.75 | 2 | 0.5 | 3 | 0.3333 | 11 | 0.5455 | NA | NA | NA | NA | 1 | 0 | 3 | 0.3333 | 4 | 0.25 |
| X_1511200 | 9 | 0.3333 | 9 | 0 | 15 | 0.0667 | 10 | 0.2 | 43 | 0.1395 | 11 | 0.0909 | 14 | 0.5 | 16 | 0.25 | 7 | 0 | 48 | 0.25 |
| X_1511200 | 7 | 0.1429 | 7 | 0 | 15 | 0.0667 | 10 | 0.1 | 39 | 0.0769 | 10 | 0.2 | 12 | 0.1667 | 14 | 0.1429 | 6 | 0.1667 | 42 | 0.1667 |
| X_1511203 | 7 | 0.2857 | 6 | 0.1667 | 15 | 0.0667 | 12 | 0.0833 | 40 | 0.125 | 11 | 0 | 11 | 0.0909 | 17 | 0.1765 | 6 | 0 | 45 | 0.0889 |
| X_1511218 | 6 | 1 | 7 | 0.7143 | 14 | 0.8571 | 7 | 0.8571 | 34 | 0.8529 | 9 | 0.6667 | 10 | 0.7 | 13 | 0.6923 | 5 | 0.8 | 37 | 0.7027 |
| X_1511218 | 5 | 0.2 | 6 | 0.1667 | 14 | 0 | 5 | 0 | 30 | 0.0667 | 6 | 0.1667 | 6 | 0.1667 | 13 | 0.0769 | 4 | 0 | 29 | 0.1034 |
| X_1511219 | 4 | 0.75 | 7 | 0.2857 | 14 | 0.8571 | 6 | 1 | 31 | 0.7419 | 8 | 0.875 | 9 | 0.7778 | 15 | 0.8 | 3 | 1 | 35 | 0.8286 |
| X_1511219 | 5 | 0.4 | 8 | 0.375 | 12 | 0.0833 | 5 | 0.4 | 30 | 0.2667 | 9 | 0.2222 | 9 | 0.6667 | 12 | 0.3333 | 3 | 0 | 33 | 0.3636 |
| X_1511219 | 6 | 0.3333 | 6 | 0.5 | 9 | 0.4444 | 6 | 0.6667 | 27 | 0.4815 | 8 | 0.375 | 10 | 0.4 | 14 | 0.2857 | 3 | 0.3333 | 35 | 0.3429 |
| X_1511246 | 4 | 0.75 | 5 | 0.2 | 9 | 0.1111 | 6 | 0.1667 | 24 | 0.25 | 10 | 0.2 | 7 | 0.1429 | 10 | 0.2 | 3 | 0 | 30 | 0.1667 |
| X_1511249 | 4 | 0.25 | 3 | 0.3333 | 7 | 0.4286 | 3 | 0.6667 | 17 | 0.4118 | 7 | 0.4286 | 6 | 0.1667 | 6 | 0.5 | 2 | 1 | 21 | 0.4286 |
| X_1511253 | 3 | 0 | 2 | 0.5 | 3 | 0 | 3 | 0 | 11 | 0.0909 | 4 | 0.25 | 5 | 0.4 | 3 | 0 | 2 | 0 | 14 | 0.2143 |
| X_1511291 | 1 | 0 | NA | NA | NA | NA | 1 | 1 | 2 | 0.5 | NA | NA | 1 | 1 | NA | NA | NA | NA | 1 | 1 |
| X_1511294 | 1 | 1 | NA | NA | 1 | 1 | 1 | 1 | 3 | 1 | 2 | 1 | 1 | 1 | NA | NA | NA | NA | 3 | 1 |
| X_1511294 | 1 | 1 | NA | NA | 1 | 0 | 2 | 0 | 4 | 0.25 | 1 | 1 | 1 | 1 | NA | NA | NA | NA | 2 | 1 |
| X_1511297 | 1 | 1 | NA | NA | 1 | 1 | 1 | 1 | 3 | 1 | 1 | 1 | 1 | 1 | NA | NA | NA | NA | 2 | 1 |
| X_1511299 | 1 | 1 | NA | NA | 1 | 0 | 2 | 0 | 4 | 0.25 | 2 | 1 | 1 | 0 | NA | NA | NA | NA | 3 | 0.6667 |
| X_1511300 | 1 | 1 | NA | NA | 1 | 1 | 1 | 1 | 3 | 1 | 1 | 0 | 1 | 1 | NA | NA | NA | NA | 2 | 0.5 |
| X_1511300 | 1 | 1 | NA | NA | NA | NA | 1 | 0 | 2 | 0.5 | 2 | 1 | 1 | 1 | NA | NA | NA | NA | 3 | 1 |
| X_1511302 | 1 | 1 | NA | NA | NA | NA | 1 | 1 | 2 | 1 | 2 | 1 | 1 | 1 | NA | NA | NA | NA | 3 | 1 |
| X_1511302 | 1 | 1 | NA | NA | NA | NA | NA | NA | 1 | 1 | 2 | 0.5 | 1 | 1 | NA | NA | NA | NA | 3 | 0.6667 |
| X_1511303 | 1 | 1 | NA | NA | 1 | 1 | 1 | 1 | 3 | 1 | 2 | 1 | 1 | 1 | NA | NA | NA | NA | 3 | 1 |
| X_1511305 | NA | NA | NA | NA | 1 | 1 | NA | NA | 1 | 1 | NA | NA | 1 | 1 | NA | NA | NA | NA | 1 | 1 |
| X_1511514 | NA | NA | NA | NA | 2 | 0.5 | 1 | 0 | 3 | 0.3333 | 3 | 0.3333 | 1 | 0 | NA | NA | NA | NA | 4 | 0.25 |
| X_1511516 | NA | NA | NA | NA | 1 | 0 | 1 | 1 | 2 | 0.5 | 3 | 0.3333 | 1 | 0 | NA | NA | NA | NA | 4 | 0.25 |
| X_1513930 | 4 | 0 | 5 | 0.4 | 14 | 0.0714 | 14 | 0.0714 | 37 | 0.1081 | 12 | 0.1667 | 16 | 0.375 | 13 | 0 | 6 | 0.3333 | 47 | 0.2128 |
| X_1513931 | 3 | 0 | 8 | 0.5 | 15 | 0.2667 | 16 | 0.3125 | 42 | 0.3095 | 11 | 0.2727 | 12 | 0.4167 | 15 | 0.2667 | 4 | 0.25 | 42 | 0.3095 |
| X_1513934 | 4 | 0 | 6 | 0.1667 | 11 | 0 | 14 | 0.0714 | 35 | 0.0571 | 11 | 0.0909 | 15 | 0.1333 | 15 | 0.0667 | 7 | 0 | 48 | 0.0833 |
| X_1513935 | 5 | 0.4 | 5 | 0.4 | 12 | 0.4167 | 16 | 0.3125 | 38 | 0.3684 | 13 | 0.6154 | 17 | 0.5882 | 14 | 0.2857 | 8 | 0.375 | 52 | 0.4808 |
| X_1513935 | 5 | 0.4 | 5 | 0.6 | 11 | 0.0909 | 15 | 0.0667 | 36 | 0.1944 | 13 | 0.1538 | 12 | 0 | 13 | 0.0769 | 7 | 0.1429 | 45 | 0.0889 |
| X_1513936 | 4 | 0.5 | 4 | 0.75 | 10 | 0.4 | 13 | 0.3846 | 31 | 0.4516 | 12 | 0.4167 | 17 | 0.5882 | 13 | 0.5385 | 6 | 0.6667 | 48 | 0.5417 |
| X_1513938 | 4 | 0 | 6 | 0.3333 | 9 | 0.1111 | 13 | 0.2308 | 32 | 0.1875 | 12 | 0.3333 | 13 | 0.0769 | 14 | 0.2857 | 7 | 0.2857 | 46 | 0.2391 |
| X_1513939 | 3 | 0 | 5 | 0.2 | 9 | 0.2222 | 13 | 0.3846 | 30 | 0.2667 | 12 | 0.1667 | 16 | 0.0625 | 13 | 0.2308 | 6 | 0.1667 | 47 | 0.1489 |
| X_1513942 | 4 | 0.75 | 5 | 0.8 | 8 | 1 | 10 | 0.6 | 27 | 0.7778 | 9 | 0.5556 | 14 | 0.5714 | 10 | 0.6 | 5 | 0.6 | 38 | 0.5789 |
| X_1513942 | 5 | 0.8 | 5 | 0.4 | 9 | 0.1111 | 10 | 0.4 | 29 | 0.3793 | 10 | 0.6 | 14 | 0.5714 | 14 | 0.2143 | 6 | 0.1667 | 44 | 0.4091 |
| X_1513943 | 4 | 0.25 | 6 | 0.3333 | 7 | 0 | 9 | 0.1111 | 26 | 0.1538 | 8 | 0.125 | 16 | 0.0625 | 12 | 0.3333 | 6 | 0 | 42 | 0.1429 |
| X_1513945 | 4 | 0.25 | 5 | 0 | 9 | 0.1111 | 9 | 0.1111 | 27 | 0.1111 | 11 | 0.1818 | 15 | 0.1333 | 13 | 0 | 7 | 0 | 46 | 0.087 |
| X_1513946 | 4 | 0 | 5 | 0.2 | 8 | 0.25 | 10 | 0.1 | 27 | 0.1481 | 10 | 0.3 | 14 | 0.0714 | 13 | 0.0769 | 8 | 0.125 | 45 | 0.1333 |
| X_1513946 | 4 | 0 | 6 | 0.3333 | 7 | 0.1429 | 9 | 0.3333 | 26 | 0.2308 | 11 | 0.4545 | 13 | 0 | 13 | 0 | 6 | 0 | 43 | 0.1163 |
| X_1513950 | 2 | 0.5 | 2 | 0.5 | 3 | 0 | 5 | 0.4 | 12 | 0.3333 | 8 | 0.25 | 7 | 0 | 4 | 0 | 4 | 0.25 | 23 | 0.1304 |
| X_1513950 | 3 | 0 | 6 | 0.3333 | 7 | 0.2857 | 8 | 0.625 | 24 | 0.375 | 11 | 0.5455 | 10 | 0 | 11 | 0.2727 | 7 | 0.2857 | 39 | 0.2821 |
| X_1513950 | 2 | 0 | 5 | 0 | 4 | 0.25 | 7 | 0 | 18 | 0.0556 | 5 | 0.4 | 9 | 0 | 7 | 0.2857 | 5 | 0 | 26 | 0.1538 |
| X_1513950 | 3 | 0 | 6 | 0.1667 | 6 | 0 | 9 | 0.1111 | 24 | 0.0833 | 10 | 0.1 | 12 | 0.1667 | 10 | 0.1 | 8 | 0.125 | 40 | 0.125 |
| X_1513950 | 3 | 0 | 6 | 0.5 | 7 | 0.1429 | 9 | 0.1111 | 25 | 0.2 | 11 | 0.1818 | 12 | 0 | 11 | 0.1818 | 8 | 0 | 42 | 0.0952 |
| X_1513951 | 3 | 0 | 6 | 0.1667 | 7 | 0.1429 | 9 | 0 | 25 | 0.08 | 9 | 0.4444 | 13 | 0.2308 | 11 | 0.1818 | 8 | 0 | 41 | 0.2195 |
| X_1513953 | 2 | 0 | 5 | 0.2 | 7 | 0.1429 | 8 | 0 | 22 | 0.0909 | 9 | 0.1111 | 10 | 0.1 | 10 | 0 | 5 | 0 | 34 | 0.0588 |
| X_1513954 | 3 | 0.3333 | 6 | 0.3333 | 6 | 0.1667 | 9 | 0.5556 | 24 | 0.375 | 11 | 0.5455 | 10 | 0.5 | 10 | 0.4 | 7 | 0.1429 | 38 | 0.4211 |
| X_1513954 | 3 | 0.6667 | 5 | 0.2 | 5 | 0.4 | 7 | 0.1429 | 20 | 0.3 | 10 | 0.5 | 10 | 0.2 | 10 | 0 | 6 | 0.1667 | 36 | 0.2222 |
| X_1513954 | 2 | 0.5 | 5 | 1 | 6 | 1 | 9 | 1 | 22 | 0.9545 | 11 | 0.9091 | 8 | 1 | 10 | 1 | 7 | 1 | 36 | 0.9722 |
| X_1513954 | 3 | 0.3333 | 5 | 1 | 6 | 0.3333 | 9 | 0.5556 | 23 | 0.5652 | 9 | 0.5556 | 9 | 0.7778 | 10 | 0.3 | 7 | 0.4286 | 35 | 0.5143 |
| X_1513956 | 3 | 0 | 6 | 0.5 | 6 | 0 | 9 | 0 | 24 | 0.125 | 11 | 0.0909 | 9 | 0.1111 | 9 | 0 | 7 | 0.1429 | 36 | 0.0833 |
| X_1513957 | 2 | 0 | 5 | 0.2 | 5 | 0 | 8 | 0.375 | 20 | 0.2 | 9 | 0.2222 | 8 | 0.125 | 7 | 0.2857 | 6 | 0.1667 | 30 | 0.2 |
| X_1514020 | 3 | 0 | 3 | 0 | 4 | 0 | 6 | 0.1667 | 16 | 0.0625 | 7 | 0 | 8 | 0.125 | 8 | 0.125 | 6 | 0 | 29 | 0.069 |
| X_1514020 | 3 | 0.3333 | 4 | 0.75 | 4 | 0.25 | 6 | 0.1667 | 17 | 0.3529 | 6 | 0.6667 | 8 | 0.375 | 8 | 0.625 | 6 | 0.8333 | 28 | 0.6071 |
| X_1514022 | 3 | 0 | 4 | 0.25 | 3 | 0 | 5 | 0.2 | 15 | 0.1333 | 5 | 0 | 7 | 0.2857 | 8 | 0.25 | 5 | 0 | 25 | 0.16 |
| X_1518197 | 8 | 0.875 | 9 | 0.5556 | 2 | 1 | 5 | 0.8 | 24 | 0.75 | 10 | 0.6 | 5 | 0.2 | 8 | 0.375 | 5 | 0.8 | 28 | 0.5 |
| X_1518197 | 8 | 0.125 | 12 | 0.1667 | 2 | 0 | 3 | 0 | 25 | 0.12 | 9 | 0.2222 | 5 | 0.4 | 6 | 0 | 5 | 0.2 | 25 | 0.2 |
| X_1518197 | 9 | 1 | 13 | 1 | 2 | 1 | 4 | 1 | 28 | 1 | 10 | 0.9 | 5 | 1 | 7 | 1 | 4 | 1 | 26 | 0.9615 |
| X_1518202 | 8 | 0.5 | 11 | 0.0909 | 2 | 0.5 | 5 | 0.6 | 26 | 0.3462 | 12 | 0.4167 | 5 | 0.4 | 6 | 0.3333 | 4 | 0.25 | 27 | 0.3704 |
| X_1519231 | 3 | 0 | 4 | 0.25 | 6 | 0 | 3 | 0.3333 | 16 | 0.125 | 9 | 0.1111 | 7 | 0 | 1 | 0 | 2 | 0.5 | 19 | 0.1053 |
| X_1519232 | 4 | 0 | 6 | 0.1667 | 4 | 0 | 2 | 0.5 | 16 | 0.125 | 8 | 0.125 | 6 | 0.3333 | 1 | 0 | 2 | 0 | 17 | 0.1765 |

| | | | | | | | | | | | | | | | | | | | | |
|---|---|---|---|---|---|---|---|---|---|---|---|---|---|---|---|---|---|---|---|---|
| X_1519233 | 3 | 0 | 5 | 0.2 | 4 | 0 | 3 | 0.3333 | 15 | 0.1333 | 8 | 0.375 | 7 | 0.1429 | 1 | 0 | 2 | 1 | 18 | 0.3333 |
| X_1519233 | 3 | 0 | 4 | 0.25 | 2 | 0 | 5 | 0.4 | 14 | 0.2143 | 7 | 0.2857 | 5 | 0.2 | 1 | 0 | 2 | 0 | 15 | 0.2 |
| X_1519234 | 3 | 0 | 6 | 0.6667 | 5 | 0.4 | 7 | 0 | 21 | 0.2857 | 8 | 0.25 | 8 | 0.25 | 1 | 0 | 3 | 0 | 20 | 0.2 |
| X_1519362 | 6 | 0.1667 | 4 | 0 | 6 | 0 | 8 | 0.375 | 24 | 0.1667 | 8 | 0.375 | 9 | 0 | 3 | 0.3333 | 4 | 0.5 | 24 | 0.25 |
| X_1519542 | 7 | 0 | 6 | 0.1667 | 8 | 0.125 | 10 | 0.1 | 31 | 0.0968 | 10 | 0.1 | 12 | 0.0833 | 5 | 0.2 | 7 | 0.2857 | 34 | 0.1471 |
| X_1522817 | 17 | 0.8235 | 17 | 1 | 20 | 0.85 | 23 | 0.8261 | 77 | 0.8701 | 16 | 0.8125 | 23 | 0.913 | 24 | 0.9583 | 15 | 0.8 | 78 | 0.8846 |
| X_1522817 | 22 | 0.1818 | 17 | 0.1765 | 17 | 0.2941 | 20 | 0.35 | 76 | 0.25 | 14 | 0 | 19 | 0.2632 | 18 | 0.1111 | 15 | 0.3333 | 66 | 0.1818 |
| X_1522819 | 21 | 0.0952 | 20 | 0.05 | 21 | 0 | 28 | 0.0714 | 90 | 0.0556 | 21 | 0.0952 | 26 | 0.0769 | 24 | 0.1667 | 17 | 0.1176 | 88 | 0.1136 |
| X_1522823 | 25 | 0.64 | 21 | 0.6667 | 20 | 0.45 | 34 | 0.7353 | 100 | 0.64 | 20 | 0.7 | 29 | 0.6207 | 23 | 0.5652 | 18 | 0.8333 | 90 | 0.6667 |
| X_1522823 | 29 | 0.1724 | 24 | 0.3333 | 24 | 0.2917 | 37 | 0.4595 | 114 | 0.3246 | 20 | 0.4 | 30 | 0.3333 | 26 | 0.3462 | 23 | 0.3913 | 99 | 0.3636 |
| X_1522826 | 27 | 0.2222 | 14 | 0.2143 | 18 | 0.2222 | 30 | 0.2333 | 89 | 0.2247 | 19 | 0.2105 | 27 | 0.3333 | 29 | 0.2069 | 21 | 0.0952 | 96 | 0.2188 |
| X_1522828 | 30 | 1 | 26 | 1 | 32 | 0.9688 | 36 | 1 | 124 | 0.9919 | 24 | 0.9583 | 40 | 1 | 37 | 0.973 | 25 | 1 | 126 | 0.9841 |
| X_1522829 | 30 | 0.0333 | 23 | 0.087 | 30 | 0.0333 | 36 | 0.1389 | 119 | 0.0756 | 25 | 0.12 | 40 | 0.1 | 37 | 0.0811 | 22 | 0.1364 | 124 | 0.1048 |
| X_1522830 | 35 | 0.8571 | 29 | 0.8621 | 33 | 0.7576 | 41 | 0.9024 | 138 | 0.8478 | 27 | 0.7407 | 43 | 0.814 | 40 | 0.925 | 31 | 0.7419 | 141 | 0.8156 |
| X_1522831 | 36 | 0.1389 | 27 | 0 | 34 | 0 | 37 | 0.1622 | 134 | 0.0821 | 27 | 0.037 | 40 | 0.075 | 35 | 0.0857 | 30 | 0.1 | 132 | 0.0758 |
| X_1522831 | 37 | 0.9189 | 27 | 0.8519 | 33 | 0.9394 | 41 | 1 | 138 | 0.9348 | 29 | 0.8276 | 42 | 0.9048 | 41 | 0.878 | 30 | 0.9333 | 142 | 0.8873 |
| X_1522832 | 25 | 0.36 | 19 | 0.2105 | 26 | 0.1538 | 32 | 0.25 | 102 | 0.2451 | 25 | 0.2 | 32 | 0.3438 | 30 | 0.1667 | 21 | 0.2381 | 108 | 0.2407 |
| X_1522832 | 24 | 0.2917 | 26 | 0.3077 | 30 | 0.2 | 38 | 0.1579 | 118 | 0.2288 | 27 | 0.2593 | 37 | 0.1892 | 34 | 0.2647 | 29 | 0.1034 | 127 | 0.2047 |
| X_1522843 | 46 | 0.0652 | 44 | 0.1136 | 43 | 0.0233 | 56 | 0.1071 | 189 | 0.0794 | 36 | 0.0833 | 55 | 0.0909 | 54 | 0.0185 | 47 | 0.0851 | 192 | 0.0677 |
| X_1525068 | 8 | 0.125 | 7 | 0 | 5 | 0.2 | 9 | 0 | 29 | 0.069 | 5 | 0.2 | 4 | 0.25 | 8 | 0 | 6 | 0 | 23 | 0.087 |
| X_1525068 | 5 | 0.2 | 8 | 0 | 10 | 0.2 | 8 | 0 | 31 | 0.0968 | 8 | 0.125 | 11 | 0.1818 | 12 | 0.1667 | 8 | 0.375 | 39 | 0.2051 |
| X_1525241 | 9 | 0.2222 | 12 | 0.25 | 10 | 0 | 10 | 0.4 | 41 | 0.2195 | 7 | 0.2857 | 11 | 0.2727 | 12 | 0.4167 | 10 | 0.4 | 40 | 0.35 |
| X_1526163 | 1 | 1 | NA | NA | 1 | 0 | 1 | 1 | 3 | 0.6667 | 1 | 1 | 1 | 1 | NA | NA | 1 | 1 | 3 | 1 |
| X_1526164 | 1 | 1 | NA | NA | 1 | 1 | 1 | 1 | 3 | 1 | NA | NA | 1 | 1 | NA | NA | 1 | 0 | 2 | 0.5 |
| X_1526164 | 1 | 1 | NA | NA | 1 | 1 | NA | NA | 2 | 1 | NA | NA | 1 | 1 | NA | NA | 1 | 1 | 2 | 1 |
| X_1526169 | 1 | 1 | NA | NA | 1 | 1 | 1 | 1 | 3 | 1 | NA | NA | 1 | 1 | 1 | 1 | 2 | 1 | 4 | 1 |
| X_1541214 | 5 | 0.2 | 1 | 1 | 1 | 1 | 4 | 0.25 | 11 | 0.3636 | 2 | 1 | 3 | 0.6667 | 3 | 0.3333 | 3 | 0 | 11 | 0.4545 |
| X_1543528 | 1 | 0 | 3 | 0 | 3 | 0.3333 | 3 | 0 | 10 | 0.1 | 4 | 0.25 | 3 | 0 | 1 | 0 | 1 | 0 | 9 | 0.1111 |
| X_1543530 | 1 | 1 | 2 | 0.5 | 2 | 0.5 | 3 | 0.3333 | 8 | 0.5 | 4 | 0.75 | 3 | 0.6667 | 1 | 0 | 1 | 1 | 9 | 0.6667 |
| X_1543530 | 1 | 1 | 1 | 1 | 2 | 0.5 | 3 | 0.3333 | 7 | 0.5714 | 4 | 0.75 | 2 | 1 | 1 | 1 | 1 | 1 | 8 | 0.875 |
| X_1543534 | 1 | 0 | 1 | 1 | 2 | 0.5 | 2 | 0.5 | 6 | 0.5 | 2 | 1 | 3 | 0 | 1 | 0 | 1 | 1 | 7 | 0.4286 |
| X_1543539 | NA | NA | NA | NA | 2 | 0.5 | 2 | 0.5 | 4 | 0.5 | NA | NA | 2 | 0 | NA | NA | 1 | 1 | 3 | 0.3333 |
| X_1543541 | NA | NA | NA | NA | 2 | 0.5 | 2 | 0.5 | 4 | 0.5 | 2 | 0.5 | 3 | 0.3333 | NA | NA | 1 | 0 | 6 | 0.3333 |
| X_1543547 | NA | NA | NA | NA | 1 | 0 | 2 | 0.5 | 3 | 0.3333 | 1 | 0 | 2 | 0 | NA | NA | 1 | 1 | 4 | 0.25 |
| X_1543549 | NA | NA | NA | NA | 1 | 0 | 2 | 0.5 | 3 | 0.3333 | 1 | 1 | 2 | 0 | NA | NA | 1 | 0 | 4 | 0.25 |
| X_1543559 | NA | NA | NA | NA | NA | NA | 2 | 0.5 | 2 | 0.5 | NA | NA | 2 | 0.5 | NA | NA | NA | NA | 2 | 0.5 |
| X_1543568 | NA | NA | NA | NA | 1 | 1 | NA | NA | 1 | 1 | NA | NA | 1 | 1 | NA | NA | NA | NA | 1 | 1 |
| X_1543569 | NA | NA | NA | NA | 1 | 1 | NA | NA | 1 | 1 | NA | NA | 1 | 1 | NA | NA | NA | NA | 1 | 1 |
| X_1543572 | NA | NA | NA | NA | 1 | 1 | NA | NA | 1 | 1 | NA | NA | 1 | 1 | NA | NA | NA | NA | 1 | 1 |
| X_1562288 | 10 | 0 | 8 | 0 | 8 | 0 | 14 | 0.1429 | 40 | 0.05 | 4 | 0 | 7 | 0.2857 | 9 | 0 | 10 | 0 | 30 | 0.0667 |
| X_1562289 | 11 | 0.0909 | 10 | 0.2 | 11 | 0.1818 | 17 | 0.1765 | 49 | 0.1633 | 4 | 0.25 | 8 | 0 | 13 | 0 | 15 | 0.4 | 40 | 0.175 |
| X_1562289 | 10 | 0.3 | 8 | 0.125 | 10 | 0 | 17 | 0.0588 | 45 | 0.1111 | 4 | 0 | 9 | 0 | 11 | 0.0909 | 15 | 0.0667 | 39 | 0.0513 |
| X_1563310 | NA | NA | 4 | 0 | 1 | 0 | 6 | 0.1667 | 11 | 0.0909 | 3 | 0 | 2 | 0 | 1 | 1 | 2 | 0 | 8 | 0.125 |
| X_1563310 | NA | NA | 4 | 0.25 | 1 | 0 | 5 | 0 | 10 | 0.1 | 5 | 0 | 2 | 0 | 2 | 0 | 3 | 0.3333 | 12 | 0.0833 |
| X_1563337 | NA | NA | NA | NA | 1 | 1 | 1 | 1 | 2 | 1 | 1 | 0 | NA | NA | 1 | 0 | 1 | 1 | 3 | 0.3333 |
| X_1563339 | NA | NA | 2 | 1 | 1 | 1 | 2 | 1 | 5 | 1 | 2 | 0.5 | NA | NA | 2 | 1 | 3 | 0.6667 | 7 | 0.7143 |
| X_1563339 | 1 | 0 | 2 | 0 | 1 | 1 | 1 | 0 | 5 | 0.2 | 1 | 0 | NA | NA | 2 | 1 | 3 | 0.6667 | 6 | 0.6667 |
| X_1563340 | NA | NA | 2 | 0.5 | NA | NA | 1 | 0 | 3 | 0.3333 | 2 | 0.5 | NA | NA | 2 | 0 | 3 | 0.3333 | 7 | 0.2857 |
| X_1563499 | NA | NA | NA | NA | 1 | 1 | 2 | 0 | 3 | 0.3333 | NA | NA | NA | NA | 1 | 0 | 1 | 1 | 2 | 0.5 |
| X_1563499 | NA | NA | NA | NA | 1 | 1 | 2 | 1 | 3 | 1 | NA | NA | NA | NA | 2 | 1 | 1 | 1 | 3 | 1 |
| X_1563500 | NA | NA | NA | NA | 1 | 0 | 1 | 1 | 2 | 0.5 | NA | NA | NA | NA | 1 | 1 | 1 | 1 | 2 | 1 |
| X_1563501 | NA | NA | NA | NA | 1 | 1 | 2 | 0.5 | 3 | 0.6667 | NA | NA | NA | NA | 1 | 1 | 1 | 1 | 2 | 1 |
| X_1563502 | NA | NA | NA | NA | 1 | 1 | NA | NA | 1 | 1 | NA | NA | NA | NA | 1 | 1 | NA | NA | 1 | 1 |
| X_1563502 | NA | NA | NA | NA | 1 | 1 | 2 | 0.5 | 3 | 0.6667 | NA | NA | NA | NA | 2 | 1 | 1 | 1 | 3 | 1 |
| X_1563503 | NA | NA | NA | NA | 1 | 0 | 1 | 1 | 2 | 0.5 | NA | NA | NA | NA | 1 | 0 | 1 | 1 | 2 | 0.5 |
| X_1563503 | NA | NA | NA | NA | NA | NA | 2 | 1 | 2 | 1 | NA | NA | NA | NA | 2 | 0.5 | 1 | 1 | 3 | 0.6667 |
| X_1563506 | NA | NA | NA | NA | NA | NA | 1 | 1 | 1 | 1 | NA | NA | NA | NA | 1 | 1 | 1 | 0 | 2 | 0.5 |
| X_1563507 | NA | NA | NA | NA | 1 | 1 | 2 | 0.5 | 3 | 0.6667 | NA | NA | NA | NA | 1 | 0 | 1 | 1 | 2 | 0.5 |
| X_1563507 | NA | NA | NA | NA | 1 | 1 | 1 | 0 | 2 | 0.5 | NA | NA | NA | NA | 2 | 0.5 | 1 | 1 | 3 | 0.6667 |
| X_1563510 | NA | NA | NA | NA | 1 | 1 | 2 | 1 | 3 | 1 | NA | NA | NA | NA | 1 | 1 | 1 | 1 | 2 | 1 |
| X_1563514 | NA | NA | NA | NA | 1 | 1 | 2 | 0.5 | 3 | 0.6667 | NA | NA | NA | NA | 1 | 1 | 1 | 1 | 2 | 1 |
| X_1565985 | 16 | 0.0625 | 18 | 0.0556 | 12 | 0.0833 | 12 | 0.25 | 58 | 0.1034 | 11 | 0.0909 | 19 | 0.0526 | 13 | 0.1538 | 16 | 0.1875 | 59 | 0.1186 |
| X_1565985 | 22 | 0.4091 | 21 | 0.3333 | 13 | 0.3077 | 16 | 0.125 | 72 | 0.3056 | 12 | 0.3333 | 20 | 0.1 | 16 | 0.125 | 15 | 0.1333 | 63 | 0.1587 |
| X_1565986 | 16 | 0.25 | 21 | 0.2381 | 14 | 0.1429 | 16 | 0.1875 | 67 | 0.209 | 11 | 0.2727 | 14 | 0.0714 | 19 | 0.2632 | 13 | 0.1538 | 57 | 0.193 |
| X_1565986 | 19 | 0.1053 | 22 | 0.3636 | 13 | 0.2308 | 17 | 0.0588 | 71 | 0.1972 | 10 | 0.3 | 17 | 0.2353 | 16 | 0.375 | 16 | 0.3125 | 59 | 0.3051 |

| | | | | | | | | | | | | | | | | | | | | |
|---|---|---|---|---|---|---|---|---|---|---|---|---|---|---|---|---|---|---|---|---|
| X_1566033 | 4 | 0.25 | 1 | 0 | 4 | 0 | 5 | 0.6 | 14 | 0.2857 | 1 | 0 | 2 | 0 | 3 | 0.3333 | 4 | 0.25 | 10 | 0.2 |
| X_1566054 | 4 | 0 | 3 | 0.3333 | 2 | 0 | 8 | 0.125 | 17 | 0.1176 | 5 | 0 | 7 | 0 | 3 | 0.6667 | 8 | 0 | 23 | 0.087 |
| X_1566063 | 6 | 0.1667 | 3 | 0.3333 | 6 | 0 | 7 | 0.1429 | 22 | 0.1364 | 5 | 0.2 | 3 | 0 | 5 | 0 | 5 | 0.2 | 18 | 0.1111 |
| X_1566066 | 6 | 0.1667 | 2 | 0.5 | 5 | 0 | 7 | 0.1429 | 20 | 0.15 | 4 | 0 | 4 | 0 | 4 | 0 | 6 | 0.1667 | 18 | 0.0556 |
| X_1566067 | 6 | 0.1667 | 2 | 0 | 5 | 0 | 6 | 0.1667 | 19 | 0.1053 | 4 | 0.25 | 4 | 0.25 | 3 | 0 | 4 | 0.25 | 15 | 0.2 |
| X_1566165 | 14 | 0.9286 | 14 | 1 | 6 | 0.8333 | 11 | 1 | 45 | 0.9556 | 6 | 1 | 7 | 1 | 11 | 0.9091 | 11 | 1 | 35 | 0.9714 |
| X_1566166 | 7 | 0.7143 | 10 | 1 | 6 | 0.8333 | 5 | 1 | 28 | 0.8929 | 4 | 0.75 | 6 | 0.5 | 4 | 1 | 7 | 1 | 21 | 0.8095 |
| X_1566167 | 12 | 0.0833 | 14 | 0.2857 | 5 | 0.2 | 11 | 0.2727 | 42 | 0.2143 | 5 | 0.2 | 10 | 0.5 | 8 | 0.25 | 11 | 0.0909 | 34 | 0.2647 |
| X_1566167 | 12 | 0.8333 | 14 | 1 | 6 | 1 | 10 | 0.9 | 42 | 0.9286 | 5 | 1 | 9 | 1 | 7 | 1 | 10 | 1 | 31 | 1 |
| X_1566168 | 11 | 1 | 11 | 1 | 6 | 1 | 11 | 1 | 39 | 1 | 5 | 1 | 6 | 0.8333 | 7 | 1 | 8 | 1 | 26 | 0.9615 |
| X_1566169 | 12 | 0.1667 | 11 | 0.1818 | 6 | 0.1667 | 10 | 0.2 | 39 | 0.1795 | 6 | 0.3333 | 9 | 0.1111 | 8 | 0 | 10 | 0.4 | 33 | 0.2121 |
| X_1566362 | 13 | 0 | 10 | 0.2 | 8 | 0 | 11 | 0.0909 | 42 | 0.0714 | 9 | 0.1111 | 9 | 0 | 6 | 0 | 11 | 0.0909 | 35 | 0.0571 |
| X_1566364 | 15 | 0.4667 | 9 | 0.5556 | 8 | 0.75 | 14 | 0.7857 | 46 | 0.6304 | 6 | 0.8333 | 11 | 0.9091 | 8 | 0.5 | 9 | 0.4444 | 34 | 0.6765 |
| X_1568258 | 7 | 0.1429 | 6 | 0 | 4 | 0.25 | 6 | 0 | 23 | 0.087 | 2 | 0 | 4 | 0.25 | 1 | 0 | 7 | 0.1429 | 14 | 0.1429 |
| X_1581464 | 4 | 0.25 | 3 | 0.6667 | 4 | 0.25 | 1 | 1 | 12 | 0.4167 | 2 | 0 | 5 | 0.4 | 1 | 0 | 2 | 0.5 | 10 | 0.3 |
| X_1581465 | 4 | 0.75 | 4 | 0.75 | 3 | 0.6667 | 1 | 1 | 12 | 0.75 | 2 | 1 | 4 | 0.5 | 1 | 1 | 2 | 1 | 9 | 0.7778 |
| X_1581467 | 4 | 0.75 | 4 | 1 | 4 | 0.75 | NA | NA | 12 | 0.8333 | 1 | 1 | 5 | 0.8 | 1 | 1 | 1 | 0 | 8 | 0.75 |
| X_1581468 | 4 | 1 | 2 | 1 | 3 | 1 | NA | NA | 9 | 1 | 2 | 1 | 4 | 0.5 | 1 | 0 | 2 | 0.5 | 9 | 0.5556 |
| X_1581493 | 3 | 0.3333 | 3 | 0 | 3 | 0 | 4 | 0.25 | 13 | 0.1538 | 2 | 0 | NA | NA | 2 | 0.5 | 2 | 0.5 | 6 | 0.3333 |
| X_1581495 | 4 | 0.25 | 3 | 0 | 3 | 0 | 3 | 0 | 13 | 0.0769 | 2 | 0 | NA | NA | 2 | 0 | 1 | 1 | 5 | 0.2 |
| X_1581495 | 4 | 0.25 | 3 | 0 | 3 | 0.3333 | 3 | 0 | 13 | 0.1538 | 2 | 0 | NA | NA | 2 | 0 | 1 | 1 | 5 | 0.2 |
| X_1581497 | 4 | 0.5 | 3 | 0 | 3 | 0.3333 | 3 | 0 | 13 | 0.2308 | 2 | 0 | NA | NA | 2 | 0 | 2 | 1 | 6 | 0.3333 |
| X_1581498 | 4 | 0.75 | 3 | 1 | 3 | 1 | 5 | 0.6 | 15 | 0.8 | 2 | 1 | NA | NA | 2 | 0 | 1 | 1 | 5 | 0.6 |
| X_1581502 | 2 | 0.5 | 1 | 1 | 2 | 0.5 | 4 | 0.25 | 9 | 0.4444 | 1 | 1 | 1 | 0 | 2 | 0.5 | 1 | 1 | 5 | 0.6 |
| X_1581505 | 3 | 0 | 2 | 0.5 | 3 | 0 | 5 | 0.2 | 13 | 0.1538 | 3 | 0 | 1 | 0 | 3 | 0.6667 | 3 | 0.3333 | 10 | 0.3 |
| X_1581505 | 5 | 0.2 | 3 | 0.6667 | 3 | 1 | 6 | 0 | 17 | 0.3529 | 3 | 0.6667 | 1 | 0 | 3 | 0 | 3 | 0 | 10 | 0.2 |
| X_1581685 | 8 | 0 | 11 | 0.2727 | 9 | 0.2222 | 10 | 0.3 | 38 | 0.2105 | 9 | 0.2222 | 14 | 0.0714 | 13 | 0.3846 | 9 | 0.1111 | 45 | 0.2 |
| X_1581686 | 11 | 0.1818 | 12 | 0.0833 | 10 | 0.3 | 8 | 0.5 | 41 | 0.2439 | 8 | 0 | 15 | 0.1333 | 15 | 0.1333 | 10 | 0.2 | 48 | 0.125 |
| X_1581687 | 4 | 0 | 6 | 0.1667 | 2 | 0 | 8 | 0 | 20 | 0.05 | 3 | 0.3333 | 4 | 0 | 6 | 0.1667 | 2 | 0 | 15 | 0.1333 |
| X_1581700 | 4 | 0 | 9 | 0.1111 | 6 | 0.3333 | 8 | 0.125 | 27 | 0.1481 | 7 | 0.2857 | 10 | 0.1 | 15 | 0.1333 | 10 | 0.1 | 42 | 0.1429 |
| X_1581719 | 8 | 0 | 14 | 0.1429 | 5 | 0 | 8 | 0.25 | 35 | 0.1143 | 3 | 0.3333 | 4 | 0 | 11 | 0.2727 | 8 | 0.5 | 26 | 0.3077 |
| X_1581720 | 8 | 0 | 18 | 0 | 9 | 0.1111 | 9 | 0.2222 | 44 | 0.0682 | 8 | 0 | 6 | 0 | 19 | 0.2105 | 13 | 0.0769 | 46 | 0.1087 |
| X_1582791 | 12 | 0.25 | 12 | 0.25 | 10 | 0.3 | 7 | 0.2857 | 41 | 0.2683 | 8 | 0.375 | 13 | 0 | 15 | 0.1333 | 16 | 0.5 | 52 | 0.25 |
| X_1582792 | 14 | 0.1429 | 13 | 0.0769 | 11 | 0 | 10 | 0.2 | 48 | 0.1042 | 11 | 0.0909 | 12 | 0.3333 | 12 | 0 | 17 | 0.1176 | 52 | 0.1346 |
| X_1582814 | 16 | 0.125 | 17 | 0.0588 | 13 | 0.0769 | 11 | 0.0909 | 57 | 0.0877 | 17 | 0.1176 | 17 | 0.1176 | 19 | 0.1579 | 16 | 0.1875 | 69 | 0.1449 |
| X_1583348 | NA | NA | NA | NA | 2 | 0.5 | 1 | 1 | 3 | 0.6667 | NA | NA | 1 | 1 | NA | NA | 1 | 0 | 2 | 0.5 |
| X_1583365 | NA | NA | 1 | 1 | 2 | 1 | 2 | 0.5 | 5 | 0.8 | NA | NA | 1 | 1 | NA | NA | 1 | 1 | 2 | 1 |
| X_1583366 | NA | NA | 1 | 1 | 2 | 1 | 2 | 1 | 5 | 1 | NA | NA | 1 | 1 | NA | NA | 2 | 0.5 | 3 | 0.6667 |
| X_1583366 | NA | NA | 1 | 0 | 2 | 1 | 2 | 0.5 | 5 | 0.6 | NA | NA | 1 | 1 | NA | NA | 1 | 1 | 2 | 1 |
| X_1583368 | NA | NA | 1 | 0 | 2 | 0.5 | 2 | 0 | 5 | 0.2 | NA | NA | 2 | 0 | 1 | 0 | 2 | 0.5 | 5 | 0.2 |
| X_1583754 | 1 | 0 | NA | NA | 1 | 1 | 2 | 1 | 4 | 0.75 | NA | NA | 2 | 0.5 | 2 | 0 | 1 | 1 | 5 | 0.4 |
| X_1587580 | 1 | 0 | 3 | 0.3333 | 1 | 0 | 1 | 0 | 6 | 0.1667 | 2 | 0.5 | NA | NA | 2 | 0.5 | 1 | 0 | 5 | 0.4 |
| X_1587584 | 2 | 1 | 2 | 1 | 2 | 0 | 1 | 0 | 7 | 0.5714 | 2 | 0.5 | NA | NA | 2 | 1 | 3 | 0.3333 | 7 | 0.5714 |
| X_1587585 | 2 | 1 | 3 | 0.6667 | 2 | 0.5 | 1 | 1 | 8 | 0.75 | 2 | 0 | NA | NA | 2 | 0.5 | 3 | 0.6667 | 7 | 0.4286 |
| X_1589060 | 35 | 0.1143 | 28 | 0.1071 | 23 | 0.087 | 21 | 0 | 107 | 0.0841 | 23 | 0.087 | 22 | 0.0455 | 25 | 0.08 | 26 | 0.0385 | 96 | 0.0625 |
| X_1589067 | 29 | 0.1034 | 20 | 0.05 | 15 | 0.0667 | 18 | 0.1667 | 82 | 0.0976 | 16 | 0.1875 | 18 | 0.2222 | 21 | 0 | 23 | 0.1304 | 78 | 0.1282 |
| X_1589067 | 25 | 0.2 | 23 | 0.1739 | 20 | 0.45 | 17 | 0.3529 | 85 | 0.2824 | 18 | 0.5 | 20 | 0.15 | 23 | 0.5217 | 21 | 0.2857 | 82 | 0.3659 |
| X_1589067 | 28 | 0.1786 | 22 | 0.0909 | 16 | 0.1875 | 19 | 0.1579 | 85 | 0.1529 | 17 | 0.1765 | 17 | 0.0588 | 21 | 0.1429 | 20 | 0.2 | 75 | 0.1467 |
| X_1593740 | 1 | 1 | 1 | 0 | 1 | 0 | 1 | 0 | 4 | 0.25 | 1 | 1 | NA | NA | NA | NA | 1 | 0 | 2 | 0.5 |
| X_1593740 | 1 | 0 | 1 | 1 | 2 | 0 | 2 | 0.5 | 6 | 0.3333 | 1 | 1 | NA | NA | NA | NA | 1 | 0 | 2 | 0.5 |
| X_1593743 | NA | NA | 1 | 1 | NA | NA | NA | NA | 1 | 1 | 1 | 1 | NA | NA | NA | NA | NA | NA | 1 | 1 |
| X_1593744 | NA | NA | 1 | 1 | NA | NA | NA | NA | 1 | 1 | 1 | 1 | NA | NA | NA | NA | 1 | 0 | 2 | 0.5 |
| X_1593807 | NA | NA | 1 | 1 | 1 | 0 | NA | NA | 2 | 0.5 | NA | NA | NA | NA | 2 | 0.5 | 2 | 0 | 4 | 0.25 |
| X_1593813 | NA | NA | 1 | 1 | NA | NA | 1 | 0 | 2 | 0.5 | NA | NA | NA | NA | 1 | 0 | 2 | 0.5 | 3 | 0.3333 |
| X_1595850 | 13 | 0.0769 | 13 | 0.0769 | 16 | 0.0625 | 15 | 0.0667 | 57 | 0.0702 | 15 | 0.1333 | 15 | 0.0667 | 20 | 0.05 | 14 | 0.0714 | 64 | 0.0781 |
| X_1595850 | 10 | 0.1 | 17 | 0.1765 | 18 | 0.2222 | 14 | 0.0714 | 59 | 0.1525 | 12 | 0.25 | 15 | 0.2 | 18 | 0.1111 | 15 | 0.1333 | 60 | 0.1667 |
| X_1595898 | 1 | 0 | 2 | 0.5 | 5 | 0.2 | 4 | 0 | 12 | 0.1667 | 1 | 0 | 5 | 0.2 | 2 | 0 | 4 | 0.25 | 12 | 0.1667 |
| X_1595908 | 1 | 0 | 2 | 0 | 6 | 0.3333 | 3 | 0 | 12 | 0.1667 | NA | NA | 8 | 0.375 | 2 | 0 | 3 | 0.3333 | 13 | 0.3077 |
| X_1595909 | 1 | 0 | 2 | 0 | 2 | 0.5 | 1 | 1 | 6 | 0.3333 | NA | NA | 6 | 0.6667 | 2 | 0 | 1 | 0 | 9 | 0.4444 |
| X_1605605 | NA | NA | NA | NA | 1 | 1 | NA | NA | 1 | 1 | 1 | 1 | 1 | 1 | 1 | 1 | NA | NA | 3 | 1 |
| X_1605606 | NA | NA | 1 | 0 | 2 | 0.5 | NA | NA | 3 | 0.3333 | 1 | 1 | 1 | 0 | 1 | 1 | NA | NA | 3 | 0.6667 |
| X_1605610 | NA | NA | 1 | 1 | 2 | 1 | NA | NA | 3 | 1 | 2 | 0.5 | 1 | 1 | NA | NA | NA | NA | 3 | 0.6667 |
| X_1605610 | 1 | 0 | 1 | 0 | 2 | 1 | NA | NA | 4 | 0.5 | 2 | 0.5 | NA | NA | 1 | 1 | NA | NA | 3 | 0.6667 |
| X_1605614 | NA | NA | 1 | 1 | 1 | 0 | NA | NA | 2 | 0.5 | 2 | 1 | 2 | 1 | NA | NA | NA | NA | 4 | 1 |
| X_1605616 | 1 | 1 | 2 | 1 | 2 | 1 | NA | NA | 5 | 1 | 2 | 1 | 2 | 1 | 1 | 1 | NA | NA | 5 | 1 |

| | | | | | | | | | | | | | | | | | | | | |
|---|---|---|---|---|---|---|---|---|---|---|---|---|---|---|---|---|---|---|---|---|
| X_1605616 | 1 | 0 | 1 | 0 | 2 | 0.5 | NA | NA | 4 | 0.25 | 2 | 0 | 2 | 0 | 1 | 1 | NA | NA | 5 | 0.2 |
| X_1605618 | 1 | 1 | 2 | 0 | 2 | 0.5 | NA | NA | 5 | 0.4 | 2 | 0.5 | 2 | 0 | 1 | 1 | NA | NA | 5 | 0.4 |
| X_1605619 | 1 | 0 | 2 | 0 | 2 | 0.5 | NA | NA | 5 | 0.2 | 2 | 0.5 | 2 | 0.5 | NA | NA | NA | NA | 4 | 0.5 |
| X_1605619 | NA | NA | 2 | 0.5 | 2 | 0.5 | NA | NA | 4 | 0.5 | 2 | 1 | 2 | 0 | 1 | 0 | NA | NA | 5 | 0.4 |
| X_1605623 | 1 | 1 | 3 | 0 | 3 | 0 | NA | NA | 7 | 0.1429 | 2 | 0.5 | 2 | 0 | 1 | 0 | NA | NA | 5 | 0.2 |
| X_1606169 | NA | NA | 1 | 1 | NA | NA | NA | NA | 1 | 1 | 1 | 1 | NA | NA | 1 | 0 | NA | NA | 2 | 0.5 |
| X_1606170 | NA | NA | 1 | 1 | NA | NA | 1 | 0 | 2 | 0.5 | 1 | 1 | NA | NA | 1 | 0 | NA | NA | 2 | 0.5 |
| X_1612004 | 3 | 0 | 2 | 0 | 3 | 0.3333 | 13 | 0.1538 | 21 | 0.1429 | 2 | 0 | 6 | 0 | 7 | 0.2857 | 10 | 0 | 25 | 0.08 |
| X_1615616 | 26 | 0.0385 | 29 | 0.069 | 21 | 0.0476 | 23 | 0.0435 | 99 | 0.0505 | 13 | 0.2308 | 22 | 0.0455 | 24 | 0.0417 | 31 | 0.1935 | 90 | 0.1222 |
| X_1615628 | 4 | 0.25 | 4 | 0.5 | 2 | 1 | 3 | 0 | 13 | 0.3846 | 1 | 0 | 3 | 0.3333 | 2 | 0 | NA | NA | 6 | 0.1667 |
| X_1615642 | 3 | 0.6667 | 2 | 0.5 | 4 | 0.75 | 2 | 1 | 11 | 0.7273 | 1 | 0 | 2 | 0.5 | 1 | 1 | NA | NA | 4 | 0.5 |
| X_1615642 | 4 | 0.75 | 3 | 0.6667 | 4 | 1 | 2 | 1 | 13 | 0.8462 | 1 | 0 | 2 | 1 | 1 | 1 | NA | NA | 4 | 0.75 |
| X_1615643 | 2 | 0 | 3 | 0 | 4 | 0.25 | 2 | 0 | 11 | 0.0909 | 1 | 0 | 1 | 1 | 2 | 0 | NA | NA | 4 | 0.25 |
| X_1615644 | 4 | 0.25 | 3 | 0.6667 | 4 | 0.5 | 1 | 0 | 12 | 0.4167 | 1 | 0 | 1 | 1 | 1 | 1 | NA | NA | 3 | 0.6667 |
| X_1615721 | 3 | 0.3333 | 1 | 0 | NA | NA | 1 | 1 | 5 | 0.4 | 3 | 1 | 1 | 1 | 1 | 0 | 2 | 0 | 7 | 0.5714 |
| X_1615723 | 3 | 0 | 1 | 0 | NA | NA | 1 | 1 | 5 | 0.2 | 3 | 0 | 1 | 1 | 1 | 0 | 2 | 0 | 7 | 0.1429 |
| X_1615733 | 3 | 0.3333 | 1 | 1 | NA | NA | 1 | 1 | 5 | 0.6 | 2 | 0.5 | 1 | 1 | NA | NA | 1 | 0 | 4 | 0.5 |
| X_1615736 | 5 | 0.4 | 2 | 0 | 1 | 0 | 2 | 0 | 10 | 0.2 | 2 | 1 | 3 | 0.6667 | 1 | 0 | 1 | 0 | 7 | 0.5714 |
| X_1615944 | 4 | 0.75 | 3 | 0.6667 | NA | NA | 1 | 0 | 8 | 0.625 | 1 | 1 | 1 | 1 | 1 | 0 | 1 | 0 | 4 | 0.5 |
| X_1627151 | 14 | 0.0714 | 7 | 0.1429 | 9 | 0.1111 | 21 | 0.3333 | 51 | 0.1961 | 6 | 0.3333 | 17 | 0.3529 | 16 | 0.1875 | 11 | 0.3636 | 50 | 0.3 |
| X_1627359 | 3 | 0 | 7 | 0.1429 | 9 | 0 | 7 | 0.1429 | 26 | 0.0769 | 7 | 0.1429 | 2 | 0.5 | 5 | 0 | 3 | 0 | 17 | 0.1176 |
| X_1627362 | 1 | 0 | 7 | 0 | 9 | 0.1111 | 4 | 0.5 | 21 | 0.1429 | 7 | 0.1429 | 2 | 0 | 4 | 0 | 3 | 0 | 16 | 0.0625 |
| X_1627580 | NA | NA | 1 | 1 | 1 | 1 | NA | NA | 2 | 1 | NA | NA | NA | NA | NA | NA | 1 | 1 | 1 | 1 |
| X_1627580 | NA | NA | 1 | 1 | 1 | 1 | NA | NA | 2 | 1 | NA | NA | NA | NA | NA | NA | 1 | 1 | 1 | 1 |
| X_1627581 | NA | NA | 1 | 1 | 1 | 1 | NA | NA | 2 | 1 | NA | NA | NA | NA | NA | NA | 1 | 1 | 1 | 1 |
| X_1628549 | NA | NA | NA | NA | NA | NA | 1 | 1 | 1 | 1 | 1 | 0 | 1 | 1 | NA | NA | 1 | 1 | 3 | 0.6667 |
| X_1628549 | 2 | 1 | NA | NA | NA | NA | 1 | 0 | 3 | 0.6667 | NA | NA | 1 | 1 | NA | NA | 1 | 0 | 2 | 0.5 |
| X_1631555 | NA | NA | 1 | 1 | 1 | 1 | NA | NA | 2 | 1 | 1 | 1 | NA | NA | 1 | 1 | NA | NA | 2 | 1 |
| X_1631557 | NA | NA | 1 | 1 | 1 | 0 | 1 | 1 | 3 | 0.6667 | 1 | 1 | NA | NA | 1 | 0 | NA | NA | 2 | 0.5 |
| X_1632056 | NA | NA | 2 | 0.5 | 1 | 0 | 2 | 0 | 5 | 0.2 | NA | NA | 1 | 1 | NA | NA | 1 | 1 | 2 | 1 |
| X_1632057 | NA | NA | 2 | 1 | 1 | 1 | 1 | 1 | 4 | 1 | NA | NA | 1 | 1 | NA | NA | 1 | 1 | 2 | 1 |
| X_1632060 | NA | NA | 1 | 1 | NA | NA | 1 | 0 | 2 | 0.5 | NA | NA | 1 | 1 | NA | NA | 1 | 0 | 2 | 0.5 |
| X_1632060 | NA | NA | 2 | 0 | NA | NA | 1 | 1 | 3 | 0.3333 | NA | NA | 1 | 0 | NA | NA | 1 | 1 | 2 | 0.5 |
| X_1632146 | NA | NA | 2 | 1 | 1 | 0 | 2 | 1 | 5 | 0.8 | NA | NA | 1 | 1 | 1 | 1 | 1 | 1 | 3 | 1 |
| X_1632147 | NA | NA | 2 | 1 | 1 | 1 | 1 | 0 | 4 | 0.75 | NA | NA | 1 | 0 | 1 | 1 | 1 | 0 | 3 | 0.3333 |
| X_1632148 | NA | NA | 2 | 1 | 1 | 1 | 1 | 1 | 4 | 1 | NA | NA | 1 | 1 | 1 | 1 | 1 | 1 | 3 | 1 |
| X_1632152 | NA | NA | 1 | 0 | 1 | 1 | 2 | 0.5 | 4 | 0.5 | NA | NA | NA | NA | 1 | 0 | 2 | 0.5 | 3 | 0.3333 |
| X_1632154 | NA | NA | 1 | 1 | NA | NA | 2 | 0.5 | 3 | 0.6667 | NA | NA | 1 | 1 | 1 | 1 | 1 | 0 | 3 | 0.6667 |
| X_1632155 | NA | NA | 1 | 1 | 1 | 0 | 2 | 0 | 4 | 0.25 | NA | NA | 1 | 1 | 1 | 1 | 2 | 0.5 | 4 | 0.75 |
| X_1637654 | 1 | 1 | NA | NA | NA | NA | NA | NA | 1 | 1 | NA | NA | NA | NA | 1 | 0 | 1 | 1 | 2 | 0.5 |
| X_1637657 | 1 | 1 | NA | NA | NA | NA | NA | NA | 1 | 1 | NA | NA | NA | NA | 1 | 1 | NA | NA | 1 | 1 |
| X_1637657 | 1 | 1 | NA | NA | NA | NA | NA | NA | 1 | 1 | NA | NA | NA | NA | 1 | 1 | 1 | 0 | 2 | 0.5 |
| X_1637658 | 1 | 1 | NA | NA | NA | NA | NA | NA | 1 | 1 | NA | NA | NA | NA | 1 | 1 | 1 | 1 | 2 | 1 |
| X_1637786 | 1 | 0 | 1 | 0 | NA | NA | 1 | 1 | 3 | 0.3333 | NA | NA | NA | NA | 1 | 0 | 3 | 0.3333 | 4 | 0.25 |
| X_1637786 | 1 | 1 | 1 | 0 | NA | NA | 1 | 1 | 3 | 0.6667 | NA | NA | NA | NA | 1 | 1 | 3 | 0 | 4 | 0.25 |
| X_1637798 | NA | NA | 1 | 0 | 1 | 1 | 1 | 0 | 3 | 0.3333 | NA | NA | NA | NA | 1 | 0 | 2 | 0.5 | 3 | 0.3333 |
| X_1639132 | NA | NA | NA | NA | 2 | 0.5 | NA | NA | 2 | 0.5 | 1 | 1 | 1 | 1 | NA | NA | 1 | 1 | 3 | 1 |
| X_1649635 | 1 | 1 | NA | NA | 1 | 1 | 1 | 1 | 3 | 1 | NA | NA | 1 | 1 | 1 | 0 | NA | NA | 2 | 0.5 |
| X_1649724 | 1 | 0 | 1 | 1 | 2 | 0.5 | 1 | 0 | 5 | 0.4 | NA | NA | 1 | 1 | 1 | 0 | NA | NA | 2 | 0.5 |
| X_1649725 | 1 | 1 | 1 | 1 | 1 | 1 | 1 | 1 | 4 | 1 | NA | NA | 1 | 1 | 1 | 0 | NA | NA | 2 | 0.5 |
| X_1651278 | 5 | 0.4 | 8 | 0 | 7 | 0.2857 | 5 | 0 | 25 | 0.16 | 6 | 0.1667 | 8 | 0 | 6 | 0.1667 | 3 | 0 | 23 | 0.087 |
| X_1651686 | 21 | 0.1429 | 40 | 0.1 | 30 | 0.1333 | 29 | 0.1379 | 120 | 0.125 | 22 | 0 | 30 | 0.0333 | 23 | 0.087 | 23 | 0.1739 | 98 | 0.0714 |
| X_1651700 | 67 | 0.0597 | 93 | 0.0323 | 80 | 0.125 | 59 | 0.0169 | 299 | 0.0602 | 46 | 0.0652 | 75 | 0.0667 | 63 | 0.0635 | 55 | 0.0364 | 239 | 0.0586 |
| X_1651700 | 58 | 0.0172 | 73 | 0.0822 | 72 | 0.1528 | 51 | 0.0392 | 254 | 0.0787 | 39 | 0.0513 | 65 | 0.0615 | 41 | 0.122 | 49 | 0.102 | 194 | 0.0825 |
| X_1651707 | 70 | 0.2714 | 94 | 0.3511 | 78 | 0.3205 | 65 | 0.2923 | 307 | 0.3127 | 42 | 0.2857 | 78 | 0.3462 | 58 | 0.2931 | 54 | 0.3704 | 232 | 0.3276 |
| X_1652353 | 9 | 0 | 7 | 0 | 13 | 0.1538 | 6 | 0.1667 | 35 | 0.0857 | 11 | 0 | 8 | 0.25 | 6 | 0 | 7 | 0.1429 | 32 | 0.0938 |
| X_1652355 | 8 | 0 | 7 | 0.1429 | 16 | 0.0625 | 7 | 0 | 38 | 0.0526 | 11 | 0.0909 | 8 | 0 | 6 | 0.1667 | 9 | 0.1111 | 34 | 0.0882 |
| X_1652404 | 5 | 0.4 | 11 | 0.1818 | 10 | 0.1 | 8 | 0 | 34 | 0.1471 | 7 | 0.1429 | 11 | 0 | 8 | 0 | 8 | 0.125 | 34 | 0.0588 |
| X_1652404 | 5 | 0.6 | 12 | 0.25 | 11 | 0.0909 | 8 | 0.375 | 36 | 0.2778 | 7 | 0.4286 | 9 | 0 | 8 | 0.125 | 6 | 0.3333 | 30 | 0.2 |
| X_1652405 | 5 | 0 | 12 | 0.5 | 9 | 0.4444 | 7 | 0.1429 | 33 | 0.3333 | 5 | 0.2 | 10 | 0.2 | 8 | 0.25 | 8 | 0.5 | 31 | 0.2903 |
| X_1652407 | 5 | 0.4 | 11 | 0.1818 | 11 | 0.2727 | 8 | 0.125 | 35 | 0.2286 | 6 | 0 | 12 | 0 | 8 | 0.25 | 9 | 0.2222 | 35 | 0.1143 |
| X_1652407 | 5 | 0 | 13 | 0.0769 | 10 | 0.2 | 7 | 0.1429 | 35 | 0.1143 | 5 | 0 | 11 | 0.1818 | 9 | 0.3333 | 9 | 0 | 34 | 0.1471 |
| X_1652408 | 5 | 0.2 | 14 | 0.2143 | 10 | 0.2 | 8 | 0.25 | 37 | 0.2162 | 6 | 0.1667 | 12 | 0.1667 | 9 | 0 | 9 | 0.3333 | 36 | 0.1667 |
| X_1652409 | 4 | 0.75 | 11 | 0.7273 | 8 | 0.875 | 6 | 0.8333 | 29 | 0.7931 | 6 | 0.5 | 10 | 1 | 7 | 0.8571 | 5 | 1 | 28 | 0.8571 |
| X_1652410 | 5 | 0.6 | 9 | 0.5556 | 8 | 0.625 | 6 | 0.6667 | 28 | 0.6071 | 6 | 0.6667 | 10 | 0.3 | 7 | 0.5714 | 7 | 0.5714 | 30 | 0.5 |

| | | | | | | | | | | | | | | | | | | | | |
|---|---|---|---|---|---|---|---|---|---|---|---|---|---|---|---|---|---|---|---|---|
| X_1652411 | 5 | 0.2 | 12 | 0.25 | 10 | 0.1 | 6 | 0.3333 | 33 | 0.2121 | 5 | 0 | 8 | 0.25 | 6 | 0 | 8 | 0 | 27 | 0.0741 |
| X_1652411 | 3 | 1 | 9 | 1 | 11 | 0.9091 | 4 | 1 | 27 | 0.963 | 5 | 1 | 11 | 0.7273 | 7 | 1 | 8 | 0.75 | 31 | 0.8387 |
| X_1652412 | 4 | 0.5 | 9 | 0.7778 | 8 | 0.625 | 6 | 0.6667 | 27 | 0.6667 | 4 | 0.5 | 5 | 0.6 | 4 | 1 | 5 | 0.6 | 18 | 0.6667 |
| X_1652854 | NA | NA | 1 | 1 | NA | NA | 1 | 1 | 2 | 1 | NA | NA | NA | NA | 1 | 1 | NA | NA | 1 | 1 |
| X_1652936 | NA | NA | NA | NA | NA | NA | 2 | 1 | 2 | 1 | NA | NA | 2 | 1 | NA | NA | NA | NA | 2 | 1 |
| X_1652937 | NA | NA | NA | NA | NA | NA | 2 | 0.5 | 2 | 0.5 | NA | NA | 2 | 1 | NA | NA | NA | NA | 2 | 1 |
| X_1659902 | 299 | 0.1505 | 375 | 0.144 | 319 | 0.185 | 268 | 0.1493 | 1261 | 0.157 | 288 | 0.1562 | 275 | 0.1273 | 274 | 0.1715 | 287 | 0.1463 | 1124 | 0.1504 |
| X_1659903 | 301 | 0.289 | 378 | 0.3783 | 314 | 0.3599 | 272 | 0.2941 | 1265 | 0.3344 | 280 | 0.3786 | 265 | 0.3396 | 266 | 0.3308 | 288 | 0.3785 | 1099 | 0.3576 |
| X_1660879 | NA | NA | NA | NA | 1 | 0 | 1 | 1 | 2 | 0.5 | NA | NA | 1 | 0 | 1 | 0 | 1 | 1 | 3 | 0.3333 |
| X_1660881 | 2 | 1 | NA | NA | 1 | 1 | 1 | 1 | 4 | 1 | NA | NA | 1 | 0 | 1 | 1 | 1 | 1 | 3 | 0.6667 |
| X_1660884 | 1 | 1 | NA | NA | 1 | 1 | 1 | 1 | 3 | 1 | NA | NA | 2 | 1 | 1 | 1 | 1 | 1 | 4 | 1 |
| X_1660885 | 1 | 1 | NA | NA | 1 | 1 | 1 | 1 | 3 | 1 | NA | NA | 2 | 1 | 1 | 1 | NA | NA | 3 | 1 |
| X_1660886 | 1 | 1 | NA | NA | 1 | 1 | NA | NA | 2 | 1 | NA | NA | 1 | 0 | 1 | 1 | 1 | 0 | 3 | 0.3333 |
| X_1660886 | 1 | 1 | NA | NA | 1 | 1 | 1 | 0 | 3 | 0.6667 | NA | NA | 1 | 1 | 1 | 0 | 1 | 1 | 3 | 0.6667 |
| X_1660887 | 1 | 0 | NA | NA | 1 | 1 | 1 | 1 | 3 | 0.6667 | NA | NA | 1 | 0 | 1 | 0 | 1 | 1 | 3 | 0.3333 |
| X_1660890 | 1 | 1 | NA | NA | 1 | 1 | 1 | 1 | 3 | 1 | NA | NA | 1 | 1 | NA | NA | 1 | 1 | 2 | 1 |
| X_1660893 | 1 | 1 | NA | NA | 1 | 1 | 1 | 0 | 3 | 0.6667 | NA | NA | 2 | 0.5 | 1 | 0 | 1 | 1 | 4 | 0.5 |
| X_1660894 | NA | NA | NA | NA | 1 | 1 | 1 | 0 | 2 | 0.5 | NA | NA | 2 | 0.5 | 1 | 1 | NA | NA | 3 | 0.6667 |
| X_1660895 | NA | NA | NA | NA | 1 | 1 | 1 | 1 | 2 | 1 | NA | NA | 2 | 1 | 1 | 1 | 1 | 1 | 4 | 1 |
| X_1660896 | NA | NA | NA | NA | 1 | 0 | 1 | 1 | 2 | 0.5 | NA | NA | 2 | 0.5 | 1 | 1 | 1 | 0 | 4 | 0.5 |
| X_1660909 | NA | NA | NA | NA | NA | NA | 1 | 1 | 1 | 1 | NA | NA | 1 | 1 | 1 | 1 | 1 | 0 | 3 | 0.6667 |
| X_1660909 | NA | NA | NA | NA | 1 | 1 | NA | NA | 1 | 1 | NA | NA | NA | NA | 1 | 1 | NA | NA | 1 | 1 |
| X_1660911 | NA | NA | NA | NA | 1 | 1 | 1 | 1 | 2 | 1 | NA | NA | 1 | 1 | 1 | 1 | 1 | 1 | 3 | 1 |
| X_1660913 | NA | NA | NA | NA | NA | NA | 1 | 1 | 1 | 1 | NA | NA | NA | NA | 1 | 1 | 1 | 1 | 2 | 1 |
| X_1661302 | 4 | 0.25 | 2 | 0.5 | 2 | 0 | 1 | 0 | 9 | 0.2222 | 3 | 0.6667 | 6 | 0.1667 | 4 | 0.25 | 3 | 0 | 16 | 0.25 |
| X_1661313 | 1 | 1 | 3 | 0 | 2 | 1 | 1 | 0 | 7 | 0.4286 | 1 | 0 | 4 | 0.75 | 2 | 0.5 | 3 | 0.3333 | 10 | 0.5 |
| X_1661316 | NA | NA | 3 | 0.3333 | 1 | 1 | 1 | 1 | 5 | 0.6 | 1 | 0 | 3 | 0 | 2 | 0.5 | 4 | 0.5 | 10 | 0.3 |
| X_1661316 | 2 | 0.5 | 3 | 0.6667 | 2 | 0.5 | NA | NA | 7 | 0.5714 | 1 | 1 | 4 | 0.5 | 3 | 0.6667 | 4 | 0.5 | 12 | 0.5833 |
| X_1661318 | 2 | 1 | 3 | 1 | 2 | 1 | 1 | 1 | 8 | 1 | 1 | 1 | 3 | 1 | 2 | 0 | 4 | 0.75 | 10 | 0.7 |
| X_1662549 | 6 | 0.8333 | 4 | 1 | 10 | 0.9 | 6 | 1 | 26 | 0.9231 | 1 | 1 | 7 | 1 | 9 | 1 | 3 | 1 | 20 | 1 |
| X_1662551 | 5 | 0.4 | 4 | 0 | 10 | 0.3 | 5 | 0.8 | 24 | 0.375 | 1 | 0 | 5 | 0 | 8 | 0.25 | 3 | 0.6667 | 17 | 0.2353 |
| X_1662553 | 5 | 0.2 | 5 | 0.4 | 9 | 0.5556 | 8 | 0.25 | 27 | 0.3704 | 1 | 0 | 4 | 0 | 8 | 0.25 | 2 | 0 | 15 | 0.1333 |
| X_1662568 | 4 | 0.25 | 2 | 0 | 7 | 0.1429 | 5 | 0.2 | 18 | 0.1667 | 1 | 0 | 5 | 0.2 | 5 | 0.2 | 3 | 0.3333 | 14 | 0.2143 |
| X_1662569 | 1 | 1 | NA | NA | 5 | 1 | 5 | 1 | 11 | 1 | 1 | 1 | 4 | 1 | 4 | 1 | 3 | 1 | 12 | 1 |
| X_1662570 | 4 | 1 | 1 | 1 | 8 | 1 | 6 | 1 | 19 | 1 | 1 | 1 | 5 | 1 | 5 | 1 | 3 | 1 | 14 | 1 |
| X_1662570 | 3 | 0.3333 | 3 | 0 | 7 | 0.1429 | 6 | 0.1667 | 19 | 0.1579 | 1 | 0 | 5 | 0 | 4 | 0.25 | 3 | 0 | 13 | 0.0769 |
| X_1662571 | 3 | 0.6667 | 1 | 1 | 4 | 1 | 6 | 1 | 14 | 0.9286 | 1 | 1 | 4 | 1 | 4 | 1 | NA | NA | 9 | 1 |
| X_1662571 | 4 | 0.75 | 2 | 0.5 | 6 | 1 | 4 | 0.75 | 16 | 0.8125 | 1 | 1 | 4 | 1 | 4 | 1 | NA | NA | 9 | 1 |
| X_1662573 | 4 | 0.25 | 2 | 0.5 | 7 | 0.8571 | 5 | 0.6 | 18 | 0.6111 | 1 | 1 | 4 | 0.5 | 2 | 1 | 2 | 1 | 9 | 0.7778 |
| X_1662573 | 4 | 0 | 2 | 0.5 | 7 | 0.7143 | 6 | 0.3333 | 19 | 0.4211 | 1 | 1 | 3 | 0 | 5 | 0.2 | 3 | 0.6667 | 12 | 0.3333 |
| X_1662573 | 3 | 0 | 3 | 0 | 6 | 0 | 5 | 0.2 | 17 | 0.0588 | NA | NA | 3 | 0.3333 | 5 | 0 | 3 | 0 | 11 | 0.0909 |
| X_1662575 | 4 | 1 | 3 | 1 | 8 | 1 | 5 | 1 | 20 | 1 | 1 | 1 | 2 | 1 | 5 | 1 | 3 | 1 | 11 | 1 |
| X_1662575 | 2 | 0.5 | 1 | 1 | 4 | 0.5 | 4 | 0 | 11 | 0.3636 | 1 | 0 | 2 | 0.5 | 4 | 0.25 | 2 | 0.5 | 9 | 0.3333 |
| X_1662575 | 4 | 0 | 2 | 0 | 8 | 0.375 | 6 | 0 | 20 | 0.15 | 1 | 0 | 2 | 0.5 | 5 | 0.4 | 2 | 0.5 | 10 | 0.4 |
| X_1662575 | 4 | 1 | 3 | 0.6667 | 5 | 0.6 | 3 | 0.3333 | 15 | 0.6667 | NA | NA | 2 | 1 | 3 | 0.3333 | 2 | 0.5 | 7 | 0.5714 |
| X_1662575 | 2 | 0 | 1 | 0 | 6 | 0.5 | 3 | 0.3333 | 12 | 0.3333 | NA | NA | 2 | 0.5 | 4 | 0.25 | NA | NA | 6 | 0.3333 |
| X_1662576 | 3 | 1 | 3 | 1 | 7 | 1 | 5 | 1 | 18 | 1 | 1 | 1 | 1 | 1 | 4 | 1 | 3 | 1 | 9 | 1 |
| X_1662577 | 4 | 0.5 | 3 | 0.6667 | 6 | 0.5 | 4 | 0.5 | 17 | 0.5294 | 1 | 0 | 2 | 0.5 | 4 | 0.5 | 1 | 0 | 8 | 0.375 |
| X_1662577 | 4 | 0.5 | 3 | 0.3333 | 8 | 0.125 | 6 | 0 | 21 | 0.1905 | 1 | 1 | 2 | 1 | 5 | 0 | 3 | 0 | 11 | 0.2727 |
| X_1662578 | 4 | 0.5 | 3 | 0.6667 | 8 | 0.25 | 6 | 0.6667 | 21 | 0.4762 | NA | NA | 2 | 0.5 | 5 | 0.2 | 3 | 1 | 10 | 0.5 |
| X_1662578 | 4 | 1 | 3 | 1 | 8 | 1 | 6 | 1 | 21 | 1 | NA | NA | 2 | 1 | 5 | 1 | 3 | 1 | 10 | 1 |
| X_1662579 | 2 | 0 | 2 | 0 | 7 | 0.1429 | 5 | 0.2 | 16 | 0.125 | NA | NA | 2 | 0.5 | 5 | 0 | 2 | 0.5 | 9 | 0.2222 |
| X_1662580 | 2 | 0.5 | 3 | 0.6667 | 8 | 0.75 | 6 | 0.8333 | 19 | 0.7368 | NA | NA | 2 | 1 | 4 | 0.75 | 3 | 0.3333 | 9 | 0.6667 |
| X_1662580 | 3 | 0.3333 | 3 | 0.6667 | 7 | 0.1429 | 6 | 0 | 19 | 0.2105 | NA | NA | 2 | 0 | 5 | 0 | 3 | 0.6667 | 10 | 0.2 |
| X_1662580 | 2 | 0 | 2 | 0.5 | 7 | 0 | 6 | 0.3333 | 17 | 0.1765 | NA | NA | 2 | 0.5 | 5 | 0.2 | 3 | 0.3333 | 10 | 0.3 |
| X_1662600 | 2 | 0 | 1 | 0 | 6 | 0.6667 | 3 | 0.3333 | 12 | 0.4167 | NA | NA | NA | NA | 5 | 0.6 | 2 | 0 | 7 | 0.4286 |
| X_1662600 | 2 | 0.5 | 3 | 0.6667 | 7 | 0.4286 | 4 | 0 | 16 | 0.375 | NA | NA | NA | NA | 5 | 0.2 | 1 | 0 | 6 | 0.1667 |
| X_1662601 | 1 | 0 | 1 | 0 | 6 | 0.3333 | 3 | 0 | 11 | 0.1818 | NA | NA | NA | NA | 4 | 0.5 | 2 | 0 | 6 | 0.3333 |
| X_1662602 | 1 | 0 | 1 | 1 | 7 | 0.5714 | 2 | 1 | 11 | 0.6364 | NA | NA | NA | NA | 5 | 0.4 | 2 | 1 | 7 | 0.5714 |
| X_1662602 | 1 | 1 | 2 | 1 | 6 | 0.8333 | 3 | 0.6667 | 12 | 0.8333 | NA | NA | NA | NA | 5 | 1 | 2 | 1 | 7 | 1 |
| X_1662604 | 1 | 1 | NA | NA | 5 | 0.8 | 3 | 0.6667 | 9 | 0.7778 | NA | NA | NA | NA | 3 | 0.3333 | 1 | 0 | 4 | 0.25 |
| X_1662604 | 1 | 1 | NA | NA | 6 | 0.6667 | 3 | 0 | 10 | 0.5 | NA | NA | NA | NA | 4 | 0.75 | 1 | 1 | 5 | 0.8 |
| X_1662605 | 1 | 0 | NA | NA | 6 | 0.5 | 2 | 0 | 9 | 0.3333 | NA | NA | NA | NA | 4 | 0.25 | 2 | 0 | 6 | 0.1667 |
| X_1662605 | 1 | 0 | NA | NA | 5 | 0.2 | 3 | 0.3333 | 9 | 0.2222 | NA | NA | NA | NA | 5 | 0.4 | 2 | 0 | 7 | 0.2857 |
| X_1662606 | 1 | 1 | NA | NA | 6 | 0 | 3 | 0 | 10 | 0.1 | NA | NA | NA | NA | 4 | 0 | 1 | 1 | 5 | 0.2 |

| | | | | | | | | | | | | | | | | | | | | |
|---|---|---|---|---|---|---|---|---|---|---|---|---|---|---|---|---|---|---|---|---|
| X_1662607 | 1 | 1 | NA | NA | 4 | 0 | 2 | 0 | 7 | 0.1429 | NA | NA | NA | NA | 4 | 0.25 | 2 | 0 | 6 | 0.1667 |
| X_1662607 | 1 | 1 | NA | NA | 5 | 0.6 | 3 | 1 | 9 | 0.7778 | NA | NA | NA | NA | 3 | 0.3333 | 2 | 0.5 | 5 | 0.4 |
| X_1662615 | 1 | 0 | NA | NA | 4 | 0.75 | 2 | 0.5 | 7 | 0.5714 | NA | NA | NA | NA | 3 | 0.3333 | 1 | 0 | 4 | 0.25 |
| X_1662616 | 1 | 1 | NA | NA | 3 | 0.6667 | 1 | 1 | 5 | 0.8 | NA | NA | NA | NA | 2 | 1 | 1 | 0 | 3 | 0.6667 |
| X_1662619 | 1 | 1 | NA | NA | 3 | 1 | NA | NA | 4 | 1 | NA | NA | NA | NA | 1 | 1 | NA | NA | 1 | 1 |
| X_1662623 | 1 | 0 | NA | NA | 4 | 0 | 1 | 1 | 6 | 0.1667 | NA | NA | NA | NA | 1 | 1 | 1 | 1 | 2 | 1 |
| X_1674344 | 1 | 0 | NA | NA | 1 | 1 | NA | NA | 2 | 0.5 | NA | NA | 1 | 1 | 1 | 1 | 1 | 1 | 3 | 1 |
| X_1674346 | 1 | 0 | NA | NA | 1 | 1 | NA | NA | 2 | 0.5 | NA | NA | 1 | 0 | 1 | 0 | 1 | 1 | 3 | 0.3333 |
| X_1674346 | 1 | 1 | NA | NA | 1 | 1 | NA | NA | 2 | 1 | NA | NA | 1 | 0 | NA | NA | 1 | 1 | 2 | 0.5 |
| X_1674347 | 1 | 0 | NA | NA | 1 | 1 | NA | NA | 2 | 0.5 | NA | NA | 1 | 1 | 1 | 1 | 1 | 1 | 3 | 1 |
| X_1674350 | 1 | 1 | NA | NA | 2 | 0.5 | NA | NA | 3 | 0.6667 | NA | NA | 1 | 1 | 1 | 0 | NA | NA | 2 | 0.5 |
| X_1674360 | NA | NA | NA | NA | 1 | 1 | NA | NA | 1 | 1 | NA | NA | 1 | 0 | 1 | 1 | 1 | 0 | 3 | 0.3333 |
| X_1680854 | 3 | 0.6667 | 1 | 0 | 1 | 1 | 1 | 0 | 6 | 0.5 | NA | NA | 2 | 0.5 | 3 | 0 | 5 | 0.4 | 10 | 0.3 |
| X_1680854 | 5 | 1 | 1 | 1 | 1 | 1 | 2 | 0.5 | 9 | 0.8889 | NA | NA | 2 | 1 | 3 | 1 | 5 | 1 | 10 | 1 |
| X_1680856 | 6 | 0.5 | 1 | 0 | 1 | 1 | 2 | 1 | 10 | 0.6 | NA | NA | 2 | 1 | 3 | 0.6667 | 6 | 0.8333 | 11 | 0.8182 |
| X_1680858 | 6 | 0.1667 | 1 | 0 | NA | NA | 2 | 0 | 9 | 0.1111 | NA | NA | 2 | 1 | 2 | 0 | 4 | 0.25 | 8 | 0.375 |
| X_1680858 | 4 | 1 | NA | NA | 1 | 1 | 2 | 0.5 | 7 | 0.8571 | NA | NA | 2 | 1 | 3 | 1 | 4 | 0.75 | 9 | 0.8889 |
| X_1680858 | 4 | 0.25 | 1 | 1 | 1 | 1 | 2 | 0.5 | 8 | 0.5 | NA | NA | 2 | 0.5 | 2 | 1 | 5 | 0.6 | 9 | 0.6667 |
| X_1680859 | 6 | 0.6667 | NA | NA | 1 | 1 | 2 | 1 | 9 | 0.7778 | NA | NA | 2 | 1 | 3 | 0.6667 | 4 | 1 | 9 | 0.8889 |
| X_1680860 | 6 | 1 | NA | NA | 1 | 0 | 2 | 1 | 9 | 0.8889 | NA | NA | 2 | 1 | 2 | 1 | 4 | 0.75 | 8 | 0.875 |
| X_1680860 | 6 | 0.8333 | 1 | 0 | 1 | 1 | 2 | 0 | 10 | 0.6 | NA | NA | 2 | 1 | NA | NA | 6 | 1 | 8 | 1 |
| X_1680957 | 6 | 0 | 4 | 0.25 | 3 | 0 | 6 | 0 | 19 | 0.0526 | 1 | 0 | 3 | 0 | 4 | 0 | 6 | 0.1667 | 14 | 0.0714 |
| X_1680958 | 5 | 0.8 | 2 | 1 | 3 | 0.3333 | 5 | 0.2 | 15 | 0.5333 | NA | NA | 3 | 0.3333 | 3 | 0.6667 | 4 | 0.5 | 10 | 0.5 |
| X_1680958 | 5 | 0 | 5 | 0.2 | 2 | 0.5 | 4 | 0.5 | 16 | 0.25 | 1 | 0 | 3 | 0.3333 | 3 | 0.3333 | 5 | 0.6 | 12 | 0.4167 |
| X_1687594 | 2 | 0.5 | NA | NA | 1 | 1 | 1 | 1 | 4 | 0.75 | NA | NA | NA | NA | 2 | 0 | 1 | 1 | 3 | 0.3333 |
| X_1687594 | 2 | 0.5 | 1 | 1 | 1 | 1 | NA | NA | 4 | 0.75 | NA | NA | NA | NA | 1 | 1 | 1 | 1 | 2 | 1 |
| X_1687595 | 2 | 1 | 1 | 1 | 1 | 1 | 1 | 1 | 5 | 1 | NA | NA | NA | NA | 2 | 0.5 | 1 | 1 | 3 | 0.6667 |
| X_1687596 | 1 | 1 | 1 | 1 | 1 | 1 | 1 | 0 | 4 | 0.75 | NA | NA | NA | NA | 2 | 0 | 1 | 1 | 3 | 0.3333 |
| X_1687596 | 2 | 1 | NA | NA | 1 | 1 | 1 | 1 | 4 | 1 | NA | NA | NA | NA | 2 | 0.5 | 1 | 1 | 3 | 0.6667 |
| X_1687613 | 2 | 1 | NA | NA | NA | NA | NA | NA | 2 | 1 | NA | NA | NA | NA | 1 | 1 | 1 | 1 | 2 | 1 |
| X_1687615 | 2 | 0 | 1 | 1 | 1 | 1 | NA | NA | 4 | 0.5 | NA | NA | NA | NA | 2 | 0.5 | 1 | 1 | 3 | 0.6667 |
| X_1687720 | NA | NA | NA | NA | 1 | 1 | NA | NA | 1 | 1 | NA | NA | NA | NA | NA | NA | 2 | 0.5 | 2 | 0.5 |
| X_1687721 | NA | NA | 1 | 1 | 1 | 0 | NA | NA | 2 | 0.5 | NA | NA | NA | NA | NA | NA | 2 | 0.5 | 2 | 0.5 |
| X_1687721 | NA | NA | 1 | 1 | 1 | 1 | NA | NA | 2 | 1 | NA | NA | NA | NA | NA | NA | 2 | 1 | 2 | 1 |
| X_1687729 | 1 | 1 | NA | NA | 1 | 1 | NA | NA | 2 | 1 | NA | NA | NA | NA | NA | NA | 2 | 1 | 2 | 1 |
| X_1687729 | 1 | 1 | 1 | 1 | 1 | 1 | 1 | 1 | 4 | 1 | NA | NA | NA | NA | NA | NA | 2 | 1 | 2 | 1 |
| X_1687730 | 1 | 1 | 1 | 1 | 1 | 1 | NA | NA | 3 | 1 | NA | NA | NA | NA | NA | NA | 2 | 1 | 2 | 1 |
| X_1687732 | 2 | 1 | 1 | 1 | NA | NA | NA | NA | 3 | 1 | NA | NA | NA | NA | NA | NA | 1 | 1 | 1 | 1 |
| X_1687733 | 2 | 0.5 | 1 | 1 | 1 | 1 | 1 | 1 | 5 | 0.8 | NA | NA | NA | NA | NA | NA | 2 | 1 | 2 | 1 |
| X_1687733 | 2 | 1 | 1 | 1 | 1 | 1 | NA | NA | 4 | 1 | NA | NA | NA | NA | NA | NA | 2 | 0.5 | 2 | 0.5 |
| X_1688561 | NA | NA | 7 | 0.5714 | 2 | 1 | NA | NA | 9 | 0.6667 | 3 | 1 | 2 | 0.5 | 1 | 0 | 1 | 1 | 7 | 0.7143 |
| X_1688561 | 1 | 1 | 10 | 0.6 | 1 | 1 | 1 | 0 | 13 | 0.6154 | 1 | 1 | 4 | 0.5 | 3 | 0.3333 | 2 | 1 | 10 | 0.6 |
| X_1688562 | 1 | 1 | 8 | 0.625 | 1 | 1 | 1 | 1 | 11 | 0.7273 | 3 | 0.6667 | 4 | 0.5 | 3 | 0.3333 | 3 | 0.3333 | 13 | 0.4615 |
| X_1688563 | 1 | 0 | 9 | 0.2222 | NA | NA | 1 | 0 | 11 | 0.1818 | 3 | 0.3333 | 5 | 0.4 | 4 | 0 | 3 | 0 | 15 | 0.2 |
| X_1688564 | 1 | 0 | 6 | 0.1667 | 2 | 0 | 1 | 0 | 10 | 0.1 | 3 | 0 | 4 | 0.25 | 4 | 0 | 3 | 0 | 14 | 0.0714 |
| X_1688565 | 1 | 1 | 8 | 0.75 | 2 | 0.5 | 1 | 0 | 12 | 0.6667 | 2 | 1 | 4 | 0.75 | 2 | 0.5 | 3 | 0 | 11 | 0.5455 |
| X_1690669 | 5 | 1 | 2 | 1 | NA | NA | 2 | 1 | 9 | 1 | 2 | 1 | 7 | 0.8571 | 4 | 1 | 3 | 0.6667 | 16 | 0.875 |
| X_1690669 | 5 | 0.2 | 3 | 0.3333 | NA | NA | 2 | 0 | 10 | 0.2 | 1 | 1 | 9 | 0 | 2 | 0 | 3 | 0.3333 | 15 | 0.1333 |
| X_1690669 | 4 | 0 | 3 | 0 | 1 | 0 | 2 | 1 | 10 | 0.2 | 2 | 0.5 | 8 | 0.375 | 5 | 0.2 | 3 | 0.3333 | 18 | 0.3333 |
| X_1690669 | 2 | 0.5 | NA | NA | NA | NA | 1 | 1 | 3 | 0.6667 | NA | NA | 4 | 0.75 | 1 | 1 | NA | NA | 5 | 0.8 |
| X_1690670 | 4 | 1 | 3 | 1 | NA | NA | 2 | 0.5 | 9 | 0.8889 | 2 | 1 | 7 | 0.5714 | 3 | 0.6667 | 3 | 1 | 15 | 0.7333 |
| X_1690670 | 2 | 0 | 2 | 0.5 | 1 | 0 | 2 | 0 | 7 | 0.1429 | 1 | 0 | 6 | 0 | 5 | 0.2 | 2 | 0 | 14 | 0.0714 |
| X_1690670 | 5 | 0 | 3 | 0 | 1 | 1 | 3 | 0 | 12 | 0.0833 | 2 | 0 | 8 | 0.125 | 4 | 0 | 3 | 0.3333 | 17 | 0.1176 |
| X_1690670 | 2 | 0.5 | 3 | 0.3333 | 1 | 1 | 3 | 0.6667 | 9 | 0.5556 | 2 | 1 | 9 | 0.6667 | 4 | 0.75 | 3 | 0.6667 | 18 | 0.7222 |
| X_1690671 | 5 | 0.6 | 2 | 0 | 1 | 0 | 2 | 1 | 10 | 0.5 | 2 | 0.5 | 8 | 0.75 | 4 | 0.75 | 3 | 0.3333 | 17 | 0.6471 |
| X_1690672 | 4 | 0 | 2 | 0.5 | 1 | 0 | 3 | 0.3333 | 10 | 0.2 | 2 | 0 | 6 | 0.1667 | 4 | 0.25 | 1 | 0 | 13 | 0.1538 |
| X_1690672 | 5 | 0.2 | 2 | 0.5 | 1 | 1 | 3 | 0.3333 | 11 | 0.3636 | 1 | 0 | 6 | 0.1667 | 4 | 0.5 | 2 | 0.5 | 13 | 0.3077 |
| X_1690672 | 4 | 0 | 2 | 0 | 1 | 1 | 1 | 0 | 8 | 0.125 | 2 | 0.5 | 6 | 0.1667 | 3 | 0.6667 | 2 | 0.5 | 13 | 0.3846 |
| X_1690672 | 4 | 0.75 | 2 | 0.5 | 1 | 0 | 3 | 0.6667 | 10 | 0.6 | 2 | 1 | 4 | 0.75 | 3 | 1 | 2 | 0.5 | 11 | 0.8182 |
| X_1690673 | 3 | 0.3333 | 2 | 0.5 | 1 | 0 | 2 | 0.5 | 8 | 0.375 | 2 | 0 | 5 | 0.2 | 2 | 0.5 | 3 | 0.3333 | 12 | 0.25 |
| X_1690673 | 4 | 0.25 | 2 | 1 | 1 | 0 | 2 | 1 | 9 | 0.5556 | 2 | 1 | 6 | 0.5 | 3 | 0.6667 | 3 | 0.6667 | 14 | 0.6429 |
| X_1690673 | 4 | 0.75 | 1 | 1 | 1 | 1 | 3 | 0.3333 | 9 | 0.6667 | 2 | 1 | 6 | 0.8333 | 3 | 0.6667 | 3 | 1 | 14 | 0.8571 |
| X_1690675 | 4 | 1 | 2 | 1 | 1 | 0 | 3 | 1 | 10 | 0.9 | 2 | 1 | 6 | 1 | 2 | 1 | 3 | 1 | 13 | 1 |
| X_1690675 | 4 | 0.75 | 2 | 1 | 1 | 1 | 2 | 1 | 9 | 0.8889 | 1 | 1 | 4 | 1 | 1 | 1 | 1 | 1 | 7 | 1 |
| X_1690675 | 2 | 0 | 2 | 1 | 1 | 1 | 3 | 0.3333 | 8 | 0.5 | 2 | 0 | 4 | 0.75 | 2 | 0.5 | 3 | 1 | 11 | 0.6364 |

| | | | | | | | | | | | | | | | | | | | | |
|---|---|---|---|---|---|---|---|---|---|---|---|---|---|---|---|---|---|---|---|---|
| X_1690676 | 2 | 0 | 1 | 0 | 1 | 1 | 2 | 1 | 6 | 0.5 | 2 | 0.5 | 5 | 0.6 | 1 | 1 | 1 | 1 | 9 | 0.6667 |
| X_1690677 | 1 | 1 | 2 | 0.5 | 1 | 1 | 2 | 0.5 | 6 | 0.6667 | 1 | 1 | 3 | 0.6667 | 2 | 1 | 2 | 0.5 | 8 | 0.75 |
| X_1690678 | 1 | 1 | 2 | 1 | 1 | 0 | 2 | 1 | 6 | 0.8333 | 2 | 1 | 6 | 1 | 2 | 1 | 3 | 1 | 13 | 1 |
| X_1690678 | 1 | 1 | 2 | 1 | 1 | 1 | 2 | 1 | 6 | 1 | 2 | 1 | 4 | 1 | 2 | 1 | 3 | 0.6667 | 11 | 0.9091 |
| X_1690683 | 1 | 1 | 2 | 0 | NA | NA | 2 | 0 | 5 | 0.2 | 2 | 0.5 | 4 | 0.5 | 2 | 0.5 | 3 | 0.3333 | 11 | 0.4545 |
| X_1690873 | 1 | 1 | 1 | 1 | NA | NA | NA | NA | 2 | 1 | NA | NA | 1 | 1 | 1 | 1 | 1 | 1 | 3 | 1 |
| X_1690877 | 1 | 1 | 2 | 0 | NA | NA | NA | NA | 3 | 0.3333 | NA | NA | 1 | 1 | 2 | 0.5 | NA | NA | 3 | 0.6667 |
| X_1690878 | 1 | 0 | 2 | 0.5 | NA | NA | NA | NA | 3 | 0.3333 | NA | NA | 1 | 0 | 2 | 0 | 1 | 1 | 4 | 0.25 |
| X_1690946 | 1 | 1 | NA | NA | NA | NA | NA | NA | 1 | 1 | 1 | 1 | 3 | 1 | NA | NA | 2 | 0.5 | 6 | 0.8333 |
| X_1690946 | 1 | 1 | NA | NA | NA | NA | NA | NA | 1 | 1 | NA | NA | 2 | 1 | NA | NA | 2 | 0.5 | 4 | 0.75 |
| X_1690946 | NA | NA | NA | NA | NA | NA | 1 | 1 | 1 | 1 | NA | NA | NA | NA | 1 | 1 | NA | NA | 1 | 1 |
| X_1691000 | 1 | 0 | 3 | 0.3333 | 2 | 0 | NA | NA | 6 | 0.1667 | NA | NA | 1 | 1 | NA | NA | NA | NA | 1 | 1 |
| X_1691001 | 1 | 0 | 1 | 0 | NA | NA | 1 | 1 | 3 | 0.3333 | NA | NA | 1 | 1 | 1 | 0 | NA | NA | 2 | 0.5 |
| X_1691003 | 2 | 1 | 3 | 0.3333 | 3 | 0 | 3 | 0.6667 | 11 | 0.4545 | 1 | 1 | 1 | 0 | 2 | 1 | NA | NA | 4 | 0.75 |
| X_1691050 | 5 | 0 | 4 | 0 | 5 | 0 | 6 | 0.1667 | 20 | 0.05 | 3 | 0 | 4 | 0 | 6 | 0.1667 | NA | NA | 13 | 0.0769 |
| X_1691051 | 5 | 0 | 4 | 0.5 | 4 | 0.25 | 7 | 0.4286 | 20 | 0.3 | 3 | 0.6667 | 4 | 0.25 | 7 | 0.1429 | NA | NA | 14 | 0.2857 |
| X_1691052 | 5 | 0.2 | 3 | 1 | 2 | 0.5 | 6 | 0.3333 | 16 | 0.4375 | 2 | 1 | 3 | 0.3333 | 7 | 0.1429 | NA | NA | 12 | 0.3333 |
| X_1691052 | 6 | 1 | 4 | 1 | 5 | 1 | 9 | 1 | 24 | 1 | 5 | 1 | 3 | 1 | 8 | 1 | NA | NA | 16 | 1 |
| X_1691053 | 8 | 0.75 | 6 | 0.3333 | 3 | 0 | 8 | 0.125 | 25 | 0.36 | 3 | 0.3333 | 5 | 0.4 | 8 | 0.375 | 2 | 1 | 18 | 0.4444 |
| Y_941418 | NA | NA | 3 | 1 | NA | NA | NA | NA | 3 | 1 | NA | NA | 1 | 1 | NA | NA | 1 | 0 | 2 | 0.5 |
| Y_941428 | NA | NA | 4 | 0.75 | NA | NA | NA | NA | 4 | 0.75 | NA | NA | 1 | 1 | NA | NA | 1 | 0 | 2 | 0.5 |
| Y_941586 | NA | NA | 4 | 1 | NA | NA | NA | NA | 4 | 1 | NA | NA | 1 | 1 | NA | NA | 1 | 1 | 2 | 1 |
| Y_941689 | NA | NA | 3 | 0.6667 | NA | NA | NA | NA | 3 | 0.6667 | NA | NA | 1 | 1 | NA | NA | 1 | 0 | 2 | 0.5 |
| Y_944927 | NA | NA | 1 | 1 | NA | NA | NA | NA | 1 | 1 | NA | NA | NA | NA | NA | NA | 1 | 1 | 1 | 1 |
| Y_944938 | NA | NA | 1 | 1 | NA | NA | NA | NA | 1 | 1 | NA | NA | NA | NA | NA | NA | 1 | 1 | 1 | 1 |
| Y_945200 | NA | NA | 1 | 1 | NA | NA | NA | NA | 1 | 1 | NA | NA | NA | NA | NA | NA | 1 | 1 | 1 | 1 |
| Y_945352 | NA | NA | 1 | 1 | NA | NA | NA | NA | 1 | 1 | NA | NA | NA | NA | NA | NA | 1 | 1 | 1 | 1 |
| Y_945497 | NA | NA | 1 | 1 | NA | NA | NA | NA | 1 | 1 | NA | NA | NA | NA | NA | NA | 1 | 1 | 1 | 1 |
| Y_955969 | 1 | 0 | 1 | 1 | 1 | 0 | NA | NA | 3 | 0.3333 | NA | NA | 1 | 1 | 1 | 0 | 1 | 0 | 3 | 0.3333 |
| Y_1017109 | NA | NA | 1 | 1 | 1 | 0 | 2 | 0.5 | 4 | 0.5 | NA | NA | NA | NA | 1 | 0 | 2 | 0.5 | 3 | 0.3333 |
| Y_1028325 | 3 | 0 | 7 | 0.1429 | 6 | 0 | 6 | 0.1667 | 22 | 0.0909 | 2 | 0.5 | 2 | 0 | 5 | 0.2 | 3 | 0.3333 | 12 | 0.25 |
| Y_1049596 | NA | NA | NA | NA | 2 | 0.5 | NA | NA | 2 | 0.5 | NA | NA | 1 | 1 | NA | NA | NA | NA | 1 | 1 |
| Y_1098474 | NA | NA | 3 | 1 | NA | NA | 1 | 1 | 4 | 1 | 1 | 1 | 1 | 1 | 2 | 0.5 | 3 | 0.3333 | 7 | 0.5714 |
| Y_1099679 | NA | NA | 3 | 0.6667 | NA | NA | 1 | 0 | 4 | 0.5 | NA | NA | 1 | 0 | 2 | 0 | 1 | 1 | 4 | 0.25 |
| Y_1158208 | NA | NA | NA | NA | NA | NA | 1 | 1 | 1 | 1 | 1 | 1 | 1 | 1 | 1 | 0 | 2 | 0 | 5 | 0.4 |
| Y_1261762 | 3 | 0 | 4 | 0 | 7 | 0 | 6 | 0.1667 | 20 | 0.05 | 2 | 0.5 | 8 | 0.125 | 3 | 0 | 4 | 0 | 17 | 0.1176 |
| Y_1262747 | 2 | 0 | 3 | 0 | 5 | 0.2 | 7 | 0 | 17 | 0.0588 | NA | NA | 10 | 0 | 3 | 0.3333 | 3 | 0 | 16 | 0.0625 |
| Y_1262846 | 2 | 0 | 4 | 0 | 7 | 0.1429 | 8 | 0.125 | 21 | 0.0952 | 1 | 0 | 11 | 0 | 4 | 0.25 | 4 | 0.25 | 20 | 0.1 |
| Y_1263211 | 1 | 0 | 3 | 0 | 7 | 0.2857 | 7 | 0.1429 | 18 | 0.1667 | 1 | 0 | 11 | 0.2727 | 3 | 0.6667 | 4 | 0.25 | 19 | 0.3158 |
| Y_1263233 | 1 | 0 | 4 | 0.5 | 8 | 0.625 | 7 | 0.4286 | 20 | 0.5 | 1 | 1 | 8 | 0.625 | 3 | 0.6667 | 4 | 0 | 16 | 0.5 |
| Y_1263304 | 1 | 0 | 4 | 0.25 | 7 | 0.2857 | 7 | 0.2857 | 19 | 0.2632 | 1 | 0 | 10 | 0.3 | 3 | 0 | 3 | 0 | 17 | 0.1765 |
| Y_9076389 | 1 | 1 | 1 | 1 | NA | NA | 1 | 1 | 3 | 1 | NA | NA | 1 | 1 | NA | NA | NA | NA | 1 | 1 |
| Y_9076397 | 1 | 1 | 1 | 1 | NA | NA | 1 | 1 | 3 | 1 | NA | NA | 1 | 1 | 1 | 0 | NA | NA | 2 | 0.5 |
| Y_9076407 | NA | NA | 1 | 1 | NA | NA | NA | NA | 1 | 1 | NA | NA | 1 | 1 | 1 | 0 | NA | NA | 2 | 0.5 |
| Y_9076617 | 1 | 0 | 1 | 1 | NA | NA | 1 | 1 | 3 | 0.6667 | NA | NA | 1 | 1 | 1 | 0 | NA | NA | 2 | 0.5 |
| Y_9076625 | 1 | 1 | 1 | 1 | NA | NA | NA | NA | 2 | 1 | NA | NA | 1 | 1 | 1 | 0 | NA | NA | 2 | 0.5 |
| Y_9077473 | NA | NA | 1 | 1 | NA | NA | NA | NA | 1 | 1 | NA | NA | 1 | 0 | 1 | 1 | NA | NA | 2 | 0.5 |
| Y_9077475 | NA | NA | 1 | 1 | NA | NA | NA | NA | 1 | 1 | NA | NA | 1 | 0 | 1 | 1 | NA | NA | 2 | 0.5 |
| Y_9077552 | NA | NA | 1 | 1 | NA | NA | NA | NA | 1 | 1 | 2 | 0 | 2 | 0.5 | 1 | 1 | NA | NA | 5 | 0.4 |
| Y_9077553 | NA | NA | 1 | 1 | NA | NA | NA | NA | 1 | 1 | 1 | 1 | 2 | 1 | 1 | 1 | NA | NA | 4 | 1 |
| Y_9077560 | NA | NA | 1 | 1 | NA | NA | NA | NA | 1 | 1 | 1 | 1 | 2 | 0.5 | 1 | 1 | NA | NA | 4 | 0.75 |
| Y_9077582 | NA | NA | 1 | 1 | NA | NA | NA | NA | 1 | 1 | 2 | 0.5 | 1 | 1 | 1 | 1 | NA | NA | 4 | 0.75 |
| Y_9077594 | NA | NA | 1 | 1 | NA | NA | NA | NA | 1 | 1 | 1 | 1 | 2 | 1 | 1 | 1 | NA | NA | 4 | 1 |
| Y_9079703 | 11 | 0.0909 | 25 | 0.12 | 6 | 0.3333 | 17 | 0.0588 | 59 | 0.1186 | 5 | 0 | 12 | 0.25 | 20 | 0.15 | 9 | 0.1111 | 46 | 0.1522 |
| Y_9084834 | 1 | 0 | NA | NA | NA | NA | 1 | 1 | 2 | 0.5 | 1 | 1 | NA | NA | NA | NA | NA | NA | 1 | 1 |
| GL456233.2 | 2 | 0 | 7 | 0 | 6 | 0 | 4 | 0.25 | 19 | 0.0526 | 6 | 0.1667 | 3 | 0 | 4 | 0.25 | 8 | 0.125 | 21 | 0.1429 |
| GL456233.2 | 2 | 0 | 4 | 0.25 | 1 | 0 | NA | NA | 7 | 0.1429 | 1 | 0 | 5 | 0.4 | 7 | 0.1429 | 1 | 1 | 14 | 0.2857 |
| GL456233.2 | 2 | 0 | 4 | 0.5 | 2 | 0.5 | NA | NA | 8 | 0.375 | 1 | 1 | 4 | 0.25 | 4 | 0.25 | 1 | 0 | 10 | 0.3 |
| GL456233.2 | 2 | 0 | 3 | 0 | 2 | 0.5 | NA | NA | 7 | 0.1429 | 1 | 1 | 4 | 0.25 | 4 | 0 | 1 | 0 | 10 | 0.2 |
| GL456233.2 | 2 | 0.5 | 4 | 1 | 1 | 1 | NA | NA | 7 | 0.8571 | 1 | 1 | 4 | 0.75 | 5 | 0.8 | 1 | 1 | 11 | 0.8182 |
| GL456233.2 | 2 | 0 | 4 | 0.5 | 1 | 0 | NA | NA | 7 | 0.2857 | NA | NA | 4 | 0.25 | 2 | 0 | 1 | 1 | 7 | 0.2857 |
| GL456233.2 | 2 | 0 | 4 | 0.5 | 1 | 1 | NA | NA | 7 | 0.4286 | 1 | 1 | 3 | 1 | 3 | 0.3333 | 1 | 0 | 8 | 0.625 |
| GL456233.2 | 2 | 0.5 | 4 | 1 | 1 | 1 | NA | NA | 7 | 0.8571 | 1 | 1 | 4 | 1 | 2 | 1 | 1 | 1 | 8 | 1 |
| GL456233.2 | 2 | 0 | 2 | 0.5 | 1 | 1 | NA | NA | 5 | 0.4 | 1 | 1 | 3 | 0.3333 | 3 | 0.3333 | 1 | 1 | 8 | 0.5 |
| GL456233.2 | 2 | 0 | 4 | 0.5 | NA | NA | NA | NA | 6 | 0.3333 | 1 | 0 | 4 | 0.5 | 1 | 1 | NA | NA | 6 | 0.5 |

| | | | | | | | | | | | | | | | | | | | | |
|---|---|---|---|---|---|---|---|---|---|---|---|---|---|---|---|---|---|---|---|---|
| GL456233.2 | 1 | 0 | 3 | 1 | 1 | 0 | NA | NA | 5 | 0.6 | NA | NA | 4 | 0.5 | 3 | 0.3333 | 1 | 0 | 8 | 0.375 |
| GL456233.2 | 1 | 1 | 4 | 1 | 1 | 0 | NA | NA | 6 | 0.8333 | 1 | 1 | 2 | 1 | NA | NA | 1 | 1 | 4 | 1 |
| GL456233.2 | 1 | 0 | 3 | 0.3333 | 1 | 0 | NA | NA | 5 | 0.2 | 1 | 0 | 3 | 0.6667 | 1 | 0 | NA | NA | 5 | 0.4 |
| GL456233.2 | NA | NA | 3 | 0 | 1 | 1 | NA | NA | 4 | 0.25 | 1 | 1 | 3 | 0 | 1 | 0 | 1 | 0 | 6 | 0.1667 |
| GL456233.2 | 1 | 0 | 1 | 1 | 1 | 1 | NA | NA | 3 | 0.6667 | 1 | 1 | 4 | 0 | 1 | 1 | 1 | 0 | 7 | 0.2857 |
| GL456233.2 | 1 | 0 | 3 | 0.3333 | NA | NA | NA | NA | 4 | 0.25 | NA | NA | 4 | 0 | NA | NA | 1 | 1 | 5 | 0.2 |
| GL456233.2 | 1 | 0 | 3 | 0.3333 | 1 | 0 | NA | NA | 5 | 0.2 | NA | NA | 3 | 0.3333 | 1 | 0 | NA | NA | 4 | 0.25 |
| GL456233.2 | NA | NA | NA | NA | 1 | 1 | 1 | 0 | 2 | 0.5 | NA | NA | NA | NA | 1 | 1 | NA | NA | 1 | 1 |
| JH584304.1 | NA | NA | 1 | 0 | NA | NA | 1 | 1 | 2 | 0.5 | 1 | 1 | NA | NA | NA | NA | NA | NA | 1 | 1 |
| JH584304.1 | NA | NA | NA | NA | NA | NA | 1 | 1 | 1 | 1 | 1 | 1 | NA | NA | NA | NA | NA | NA | 1 | 1 |
| JH584304.1 | 4 | 0.25 | 3 | 0 | 3 | 0.3333 | 2 | 0 | 12 | 0.1667 | 3 | 0 | 3 | 0.3333 | 1 | 0 | 1 | 0 | 8 | 0.125 |
| JH584304.1 | 9 | 0.4444 | 5 | 0.2 | 5 | 0 | 4 | 0.25 | 23 | 0.2609 | 6 | 0.5 | 5 | 0 | 5 | 0.6 | 4 | 0.25 | 20 | 0.35 |
| JH584304.1 | 9 | 0.2222 | 3 | 0.3333 | 5 | 0.4 | 4 | 0 | 21 | 0.2381 | 6 | 0.6667 | 5 | 0.2 | 5 | 0.6 | 4 | 0.25 | 20 | 0.45 |
| JH584304.1 | 12 | 0.5 | 4 | 0.75 | 8 | 0.375 | 4 | 0.75 | 28 | 0.5357 | 8 | 0.5 | 5 | 0.4 | 6 | 0.6667 | 5 | 0.4 | 24 | 0.5 |
| JH584304.1 | 10 | 0.7 | 3 | 0.3333 | 8 | 0.25 | 2 | 0 | 23 | 0.4348 | 7 | 0.2857 | 6 | 0.5 | 6 | 0.5 | 5 | 0.6 | 24 | 0.4583 |
| JH584304.1 | 6 | 0 | 3 | 0 | 6 | 0.1667 | 2 | 0 | 17 | 0.0588 | 4 | 0 | 4 | 0 | 5 | 0.4 | 3 | 0 | 16 | 0.125 |
| JH584304.1 | 7 | 0.4286 | 3 | 0 | 7 | 0 | 2 | 0 | 19 | 0.1579 | 5 | 0.2 | 5 | 0 | 5 | 0.2 | 3 | 0 | 18 | 0.1111 |
| JH584304.1 | 3 | 0.6667 | 1 | 1 | 5 | 0.6 | NA | NA | 9 | 0.6667 | 1 | 0 | 1 | 0 | 2 | 0.5 | 1 | 1 | 5 | 0.4 |
| JH584304.1 | 5 | 0.2 | 2 | 0.5 | 5 | 0 | 2 | 0 | 14 | 0.1429 | 3 | 0 | 4 | 0.25 | 5 | 0.2 | 3 | 0.6667 | 15 | 0.2667 |
| JH584304.1 | 2 | 0.5 | 1 | 0 | 1 | 0 | NA | NA | 4 | 0.25 | 2 | 1 | 1 | 0 | 4 | 0.75 | 2 | 1 | 9 | 0.7778 |
| JH584304.1 | 3 | 0.3333 | 1 | 0 | 3 | 0 | 1 | 0 | 8 | 0.125 | 3 | 0 | 2 | 0 | 4 | 0.25 | 4 | 0 | 13 | 0.0769 |
| JH584304.1 | 4 | 0.5 | 3 | 0 | 6 | 0.1667 | 2 | 0.5 | 15 | 0.2667 | 2 | 0.5 | 4 | 0 | 3 | 0.6667 | 4 | 0.25 | 13 | 0.3077 |
| JH584304.1 | 1 | 1 | NA | NA | 1 | 0 | NA | NA | 2 | 0.5 | 1 | 0 | NA | NA | NA | NA | 1 | 1 | 2 | 0.5 |
| JH584304.1 | 1 | 1 | NA | NA | 1 | 0 | NA | NA | 2 | 0.5 | 2 | 0.5 | NA | NA | NA | NA | 1 | 1 | 3 | 0.6667 |
